# Loss of PRDM16 Drives Nasal Septal Deviation through Dysregulated TGFβ2 Signaling

**DOI:** 10.64898/2026.07.28.741259

**Authors:** Eliya Tazreena Tashbib, Victoria L. Hansen, Maeve O’Brien, Alexis Klee, Eloise Fadial, Karin Pryharski, Gulzada Kulzhanova, Kathryn Lambright, Lomeli Shull, Chia-Lung Wu

## Abstract

Nasal septal deviation affects approximiately 20% of newborns globally and 80% individuals in the United States. GWAS have linked PRDM16, a histone methyltransferase, to craniofacial abnormalities, yet its role in nasal septum development remains poorly understood. Global *Prdm16* knockout mice (*Prdm16*^-/-^) exhibit severe craniofacial defects resembling Pierre Robin Syndrome but are neonatally lethal, precluding their potential applications for postnatal study. To address this, we generated an osteochondral lineage-specific, Prdm16 conditional knockout (Col2a1-Cre; Prdm16^flox/flox^; cKO) mouse model. Both sexes of cKO mice display significantly shorter nasal bone length, with a sex-dependent increase in nasal bone volume fraction of 12 wk old males. Nasal septal deviation is detectable as early as postnatal day 15 and progresses with age. Single-cell RNA sequencing (scRNA-seq) of 4 wk old nasal septal cartilage revealed a marked shift in chondrocyte composition: *Mgp^+^* chondrocytes were substantially reduced, while *Col10a1^+^*/*Serpina3n^+^* hypertrophic chondrocytes were dramatically increased, indicating PRDM16 regulates chondrocyte phenotypes. Spatial transcriptomics localized *Mgp^+^* chondrocytes and *Col1a1^high^*/*Col3a1^+^* fibrotic cells to the septal cartilage-bone interface (the site of deviation in cKO mice). Intercellular communication analyses revealed a switch in dominant sender cells from the fibrotic population in WT to *Mgp^+^* chondrocytes in cKO. MultiNicheNet bioinformatic analyses identified elevated TGFβ2 signaling at the nasal septal deviation site. Specifically, TGFβ2 secreted by *Mgp^+^* chondrocytes was predicted to promote *Col1a1*/*Col1a2* expression, resulting in fibrotic extracellular matrix (ECM) deposition and osteogenesis; consistent with elevated RUNX2 in cKO mice. TGFβ2 immunohistochemical staining confirmed increased TGFβ2⁺ cells in the fibrous ECM and apical nasal cartilage of cKO, but not WT mice. Loss of PRDM16 also increased chondrocyte apoptosis at 4 and 12 wks of age. These findings demonstrate that loss of PRDM16 drives hypertrophic and fibrotic remodeling of nasal septal cartilage through dysregulation of TGFβ2 signaling, establishing a mechanistic basis for nasal septal deviation.

## Introduction

Nasal septal cartilage is critical for craniofacial structure and overall respiratory health. However, deviation of the nasal septum is among the most common congenital craniofacial disorders, affecting up to 80% of the general population in the United States and approximately 20% of newborns globally every year (128 out of 652 children).^1,2^ In congenital cases, nasal septum deviation occurs during rapid midfacial growth and can lead to disordered breathing and long-term complications, including chronic sinusitis, nosebleeds, and sleep apnea.^3–5^ Correction of craniofacial anomalies often requires extensive surgical intervention, increasing the risk of cerebrospinal fluid leak and pneumocephalon.^6^ Despite the prevalence of nasal septal deviation, the molecular mechanism underlying its pathogenesis remains largely unknown.

A recent genome-wide association study (GWAS) has linked PRDM16, a histone methyltransferase, to abnormalities in human craniofacial structure.^7–9^ PRDM16 possesses a PR domain with the capacity for histone methylation and a zinc finger domain for protein-DNA and protein-protein interactions.^10^ Our bulk RNA sequencing analysis across chondrogenesis in human induced pluripotent stem cells (hiPSCs) showed that *PRDM16* was significantly upregulated, along with key chondrogenic markers including *SOX6*, *ACAN*, *COL2A1*, and *RUNX1*, during chondrogenic differentiation. This result suggests that PRDM16 may play a crucial role in chondrocyte cell fate determination.^11^ PRDM16 also contributes to various developmental processes, including cardiac function, adipose metabolism, neurogenesis, hematopoiesis, and immune regulation.^12–15^ In our recent work, we have shown that loss of PRDM16 delays ossification in several knee joint compartments and increases susceptibility to OA. These findings suggest that PRDM16 is a positive regulator of knee articular cartilage development and homeostasis, as well as chondrocyte specification.^16^ Furthermore, deletion of PRDM16 in the neural crest cell lineage of *Wnt1-Cre* mice led to disrupted development of Meckel’s cartilage and mandibular morphogenesis, resulting in mandibular hypoplasia and cleft palate.^17^ These features are consistent with Pierre Robin Syndrome (PRS)-like phenotype observed in global *Prdm16* knockout mice (Prdm16^-/-^) and humans.^8,17^ However, the functional role of PRDM16 in normal and deviated nasal septal cartilage is unclear.

To address this gap, we examined the role of PRDM16 in the development of nasal septal cartilage using an osteochondral lineage-specific, conditional Prdm16 knockout (Col2a1-Cre; Prdm16^flox/flox^; *Prdm16* cKO) mouse model, as well as single-cell RNA sequencing (scRNA-seq) and spatial transcriptomics (spatial-seq) approaches. Here, we show that PRDM16 is a critical regulator of nasal septal cartilage development and homeostasis by regulating chondrocyte phenotypes and osteogenesis. These findings identify that TGFβ2 signaling is a potential mediator of deviated nasal septal cartilage, by promoting fibrosis and activating the osteogenic pathway.

## Methods

Note that the full methodology is available in the **Supplemental Methods and Materials.**

### Animals

Prdm16^flox/flox^ mice (B6.129-Prdm16^tm1.1Brsp^/J) were crossed with Col2a1-Cre mice (Col2a1-Cre mice, B6;SJL-Tg(Col2a1-cre)1Bhr/J; #003554, JAX) to generate Col2a1-Cre; Prdm16^flox/flox^ (heretofore referred to as *Prdm16* cKO) mice, while Cre-negative littermates served as wild type (WT) controls. Embryonic day 18.5 (E18.5), postnatal day 5 (P5), postnatal day 15 (P15), 4 wk, 12 wk, and 1 year old mice were collected. Wnt1-Cre; Prdm16^flox/+^ mice were generated by crossing Wnt1-Cre transgenic mice with Prdm16^flox/flox^ mice. Mice carrying the Wnt1-Cre transgene and a single floxed Prdm16 allele (Wnt1-Cre; Prdm16^flox/+^) and Cre-negative littermates (Prdm16^flox/+^) were used for phenotypic analyses. Wnt1-Cre; Prdm16^flox/+^ and their control counterparts Prdm16^flox/+^ were harvested at 4 wks of age.

### Micro-computed Tomography (*μ*CT)

Skulls from WT and *Prdm16* cKO mice were harvested at 4 wks, 12 wks, and 1 year of age and analyzed by micro-computed tomography (*μ*CT; VivaCT 40, Scanco Medical). Three-dimensional reconstructions were generated to assess craniofacial morphology. Nasal bone volume fraction (BV/TV) and bone mineral density (BMD) were quantified using Scanco evaluation software. Nasal and cranial lengths were measured from dorsal reconstructions using a landmark-based approach in Amira software.

### Alcian blue and alizarin red (whole-mount) staining

Whole-mount alcian blue and alizarin red staining was performed on E18.5 WT and *Prdm16* cKO embryos, as previously described.^18^ Embryos were fixed in 95% ethanol, stained with alcian blue and alizarin red, cleared in KOH, and stored in glycerol prior to imaging.

### Tissue Processing

*Prdm16* cKO and WT mice were fixed in 10% neutral buffered formalin at room temperature (RT) for 48 hours. Following fixation, mouse skulls were decalcified using Cal-Ex solution (Fisher CS510-1D) at 4°C for 48 hours for P5 and P15 samples, and for 72 hours for 4 wk and 12 wk old samples, as previously described.^19^ After decalcification, the skulls were processed for paraffin embedding following standard protocols. Skulls were embedded in a frontal orientation, and 10*µm* coronal sections were collected for subsequent histological or immunohistochemical analyses.^20^

### Histological and Immunohistochemistry Analyses

Histological evaluation of the nasal septum was performed on coronal sections from WT and *Prdm16* cKO mice. Safranin-O/Fast Green staining was conducted at P5, P15, 4, and 12 wks of age to assess cartilage morphology, following previously established protocols.^21^ Septal deviation severity and nasal septal cartilage ratio were quantified from histological sections. Immunohistochemistry was performed from 4 wk and 12 wk old WT and *Prdm16* cKO male mice. Sections were stained either with hypertrophic chondrocyte markers (anti-RUNX2, 1:100; anti-Collagen X, 1:100), proliferation marker (anti-MKi67, 1:150), apoptosis marker (anti-Cleaved Caspase 3, CASP3, 1:600), or mature chondrocyte marker (anti-Collagen 2A1, 1:200), chondrocyte degradation marker (MMP13, 1:200), and anti-TGF-β2 antibody (1:400). A complete list of primary antibodies, including catalog numbers and suppliers, is provided in the **Supplementary Table 1**. Images were acquired using an Axioscope 5 microscope (Zeiss) with an Axiocam 305 color camera (Zeiss).

### scRNA-seq, spatial-seq, and bioinformatic analyses

For scRNA-seq, nasal septal cartilage samples from four 4 wk old male mice per genotype were pooled, and cell suspensions with >75% viability were submitted to the University of Rochester Genomics Research Center. Spatial-seq profiling was performed using the Visium HD platform on 4 wk old male WT and *Prdm16* cKO tissues (n = 2/genotype). Final scRNA-seq and spatial-seq libraries were sequenced on the NovaSeq X Plus platform (Illumina).

### Statistics

Data were presented as the mean ± standard deviation or standard error of mean. Student’s t-test or one-way analysis of variance (ANOVA) followed by *post-hoc* tests was performed as appropriate, with significance reported at the 95% confidence level.

## Results

### Loss of *Prdm16* results in shorter nasal length, nasal septal cartilage, and nasal septum deviation

At E18.5, although no significant difference in nasal length between WT and *Prdm16* cKO mice was observed, there was a decreasing trend in cranial length of *Prdm16* cKO mice compared to WT mice (**Figure 1A**). Starting at 2 wks of age, *Prdm16* cKO mice displayed a noticeably rounder facial morphology compared to WT mice (**Figure S1A**).

**Figure 1.**
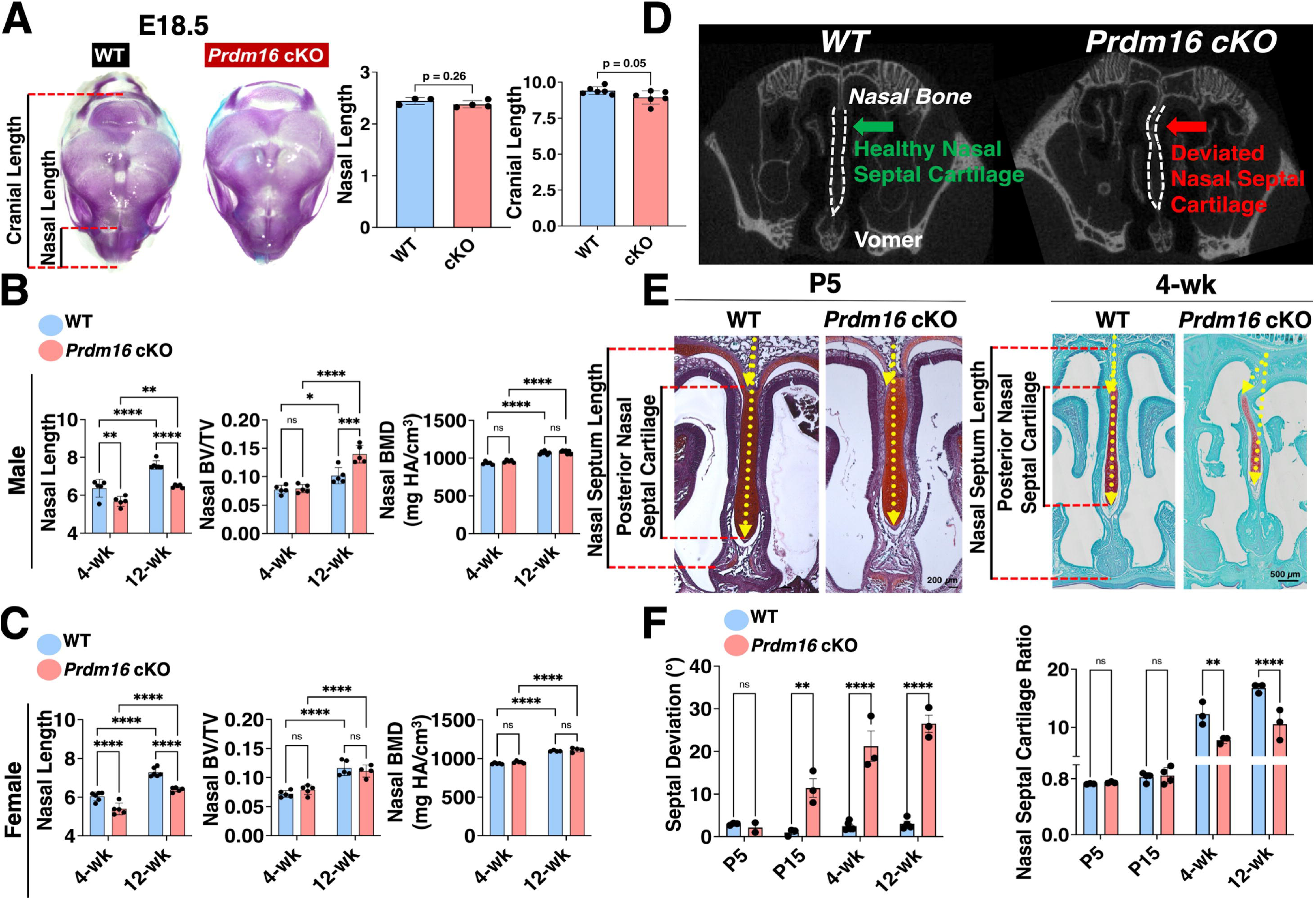
Loss of PRDM16 results in shorter nasal length, nasal septal cartilage, and deviated nasal septum. **(A)** Alizarin Red & Alcian Blue (whole-mount) staining of E18.5 embryos. *Prdm16* cKO shows a trend in decreased cranial length at E18.5 n ≥ 3/group. **(B, C)** Longitudinal morphometric measurement of nasal length in male and female WT and *Prdm16* cKO mice at 4 wks and 12 wks of age, indicating a shorter nasal length in the mutant mice. **(D)** *μ*CT of 4 wk old mice shows nasal septum deviation (red arrow). **(E)** Saf-O/Fast Green staining of nasal septal cartilage (coronal view) from P5 and 4 wk old male mice. **(F)** Measurements of septal deviation of P5, P15, 4 wk, and 12 wk males (left). The septal deviation angle is measured between the most deviated point of the septum and the midline. Quantification of nasal septal cartilage ratio (Posterior Nasal Septal Cartilage Length/Nasal Septum Length) in P5, P15, 4 wk, and 12 wk old male mice (right). Landmarks used for this measurement are depicted in red. All length measurements are reported in millimeters (mm). Data points represent individual mice and are presented as mean ± SEM. n ≥ 4/genotype/age. Student’s Unpaired t-test or two-way ANOVA with *post-hoc* Tukey’s test, accordingly. *p < 0.05, **p < 0.01, ***p < 0.001, ****p < 0.0001, and ns = not significant. Anatomical description of the landmarks is shown in **Figure S1B.**

*μ*CT analysis revealed that both female and male *Prdm16* cKO mice had significantly shorter nasal lengths compared to WT mice at 4 wks of age, with this reduced length maintained into adulthood at 12 wks of age (**Figure 1B, C**). Despite these differences in nasal length, no significant changes in nasal bone volume fraction (BV/TV) were detected between WT and *Prdm16* cKO mice of either sex at 4 wks of age (**Figure 1B, C**). However, at 12 wks, male *Prdm16* cKO mice showed a significantly higher nasal BV/TV compared with WT controls. Nasal bone mineral density (BMD) remained unchanged across genotypes, regardless of age. In contrast, female *Prdm16* cKO mice exhibited no significant differences in either nasal BV/TV or BMD relative to WT controls. A schematic illustrating the anatomical landmarks used for nasal length measurements is provided in **Figure S1B**.

Additionally, both male and female *Prdm16* cKO mice exhibited reduced cranial length compared with WT mice, while no significant differences in cranial BV/TV or BMD were observed (**Figure S1C**). No significant differences were detected between WT and cKO mice in other cranial bone measurements, including frontal and parietal bone length (**Figure S1D**). Interestingly, only 1 year old male cKO mice showed a significantly shorter nasal length than WT controls. Similarly, only aged male *Prdm16* cKO mice displayed significantly increased nasal BMD relative to WT controls. (**Figure S1E**)

The representative *μ*CT images of 4 wk male mice (**Figure 1D**) revealed apparent nasal septum deviation in *Prdm16* cKO mice, particularly at the junction of the nasal bone and the nasal septal cartilage. Histological analysis showed that at P5, there is no observable nasal septum deviation in *Prdm16* cKO mice. However, by P15, a marked deviation is evident, and this deviation persists postnatally (4 wk and 12 wk). (**Figure 1E, F**) Interestingly, *Prdm16* cKO mice have a comparable nasal septal cartilage ratio (NSC ratio; defined as Posterior Nasal Septal Cartilage Length/Nasal Septum Length) at P5 and P15, but significantly shorter septal cartilage (lower NSC ratio) at both juvenile (4 wk) and young adult (12 wk) stages compared to WT controls (**Figure 1E, F**). Additionally, nasal septal deviation was observed in 4 wk male and female neural crest cell-specific, Wnt1-Cre; Prdm16^flox/+^ mice (**Figure S1F**).

### *Prdm16* cKO mice show no changes in mandibular measurements

Anterior and posterior mandible length, effective mandible length, mandibular plane, and condylar axis did not differ significantly between cKO and WT mice, regardless of sex. **(Figure 2A, B, C**). Similarly, dorsal cranial measurements, including bitemporal distance, interzygomatic width, interorbital width, and the distance between the occipital point and the nasal tip, were comparable between cKO and WT mice in both sexes (**Figure S2A, B**).

**Figure 2.**
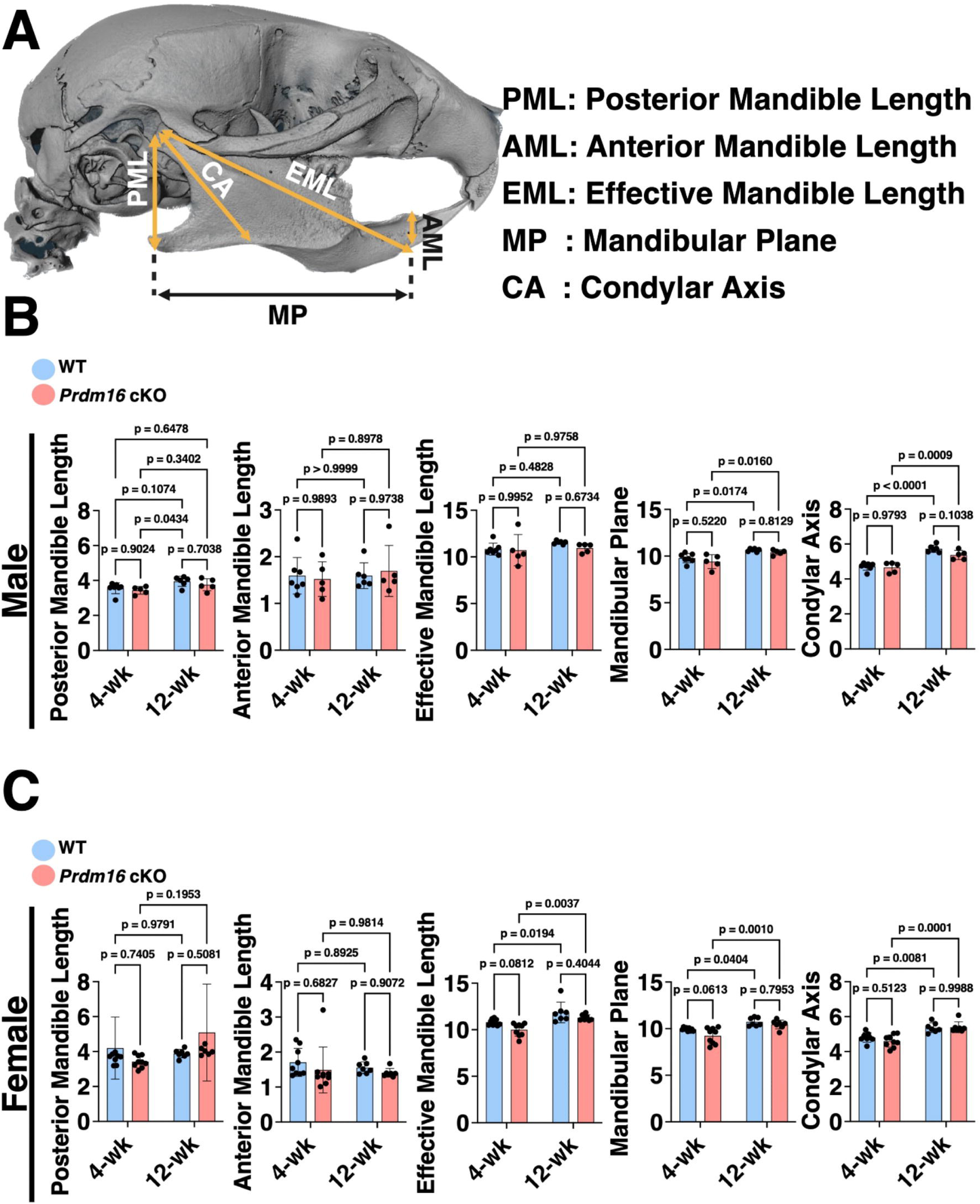
PRDM16 cKO does not significantly affect mandibular morphology in either male or female mice. PML: Posterior Mandible Length, AML: Anterior Mandible Length, EML: Effective Mandible Length, MP: Mandibular Plane, and CA: Condylar Axis. All length measurements are reported in millimeters (mm). Data points represent individual mice and are presented as mean ± SEM. n ≥ 4/genotype/age Two-way ANOVA with *post-hoc* Tukey’s test.

### *Prdm16* cKO mice show increased RUNX2 staining in the dorsal zone of nasal septal cartilage

To determine whether zonal differences in nasal septal chondrocytes contribute to the septal deviation observed in *Prdm16* cKO mice, the nasal septal cartilage was divided into three distinct regions: dorsal (D), middle (M), and ventral (V) zones (**Figure 3A**).^22^ Additionally, the nasal septal cartilage was organized into three distinct layers based on differences in chondrocyte morphology (**Figure 3A**).^23^ The perichondrium consists of flat and densely packed cells that surround the nasal septal cartilage matrix and are located adjacent to the epithelial layer (red dashed lines). The superficial zone is composed of chondrocytes with flattened morphology, and the central zone shows spheroidal-shaped chondrocytes.^24,25^ Immunohistochemistry (IHC) revealed that RUNX2 is significantly increased in *Prdm16* cKO mouse nasal cartilage, particularly in the dorsal zone (green arrow) near the nasal deviation (**Figure 3B, C**). Additionally, the percentage of RUNX2^+^ cells had an increasing trend in the perichondrium (yellow arrow) near the deviation, although no expression was observed at 12 wks of age (**Figure 3B, C; S2C)**. Male cKO mice had comparable COL10A1^+^ expression at 4 wks of age (**Figure 3D, E**).

**Figure 3.**
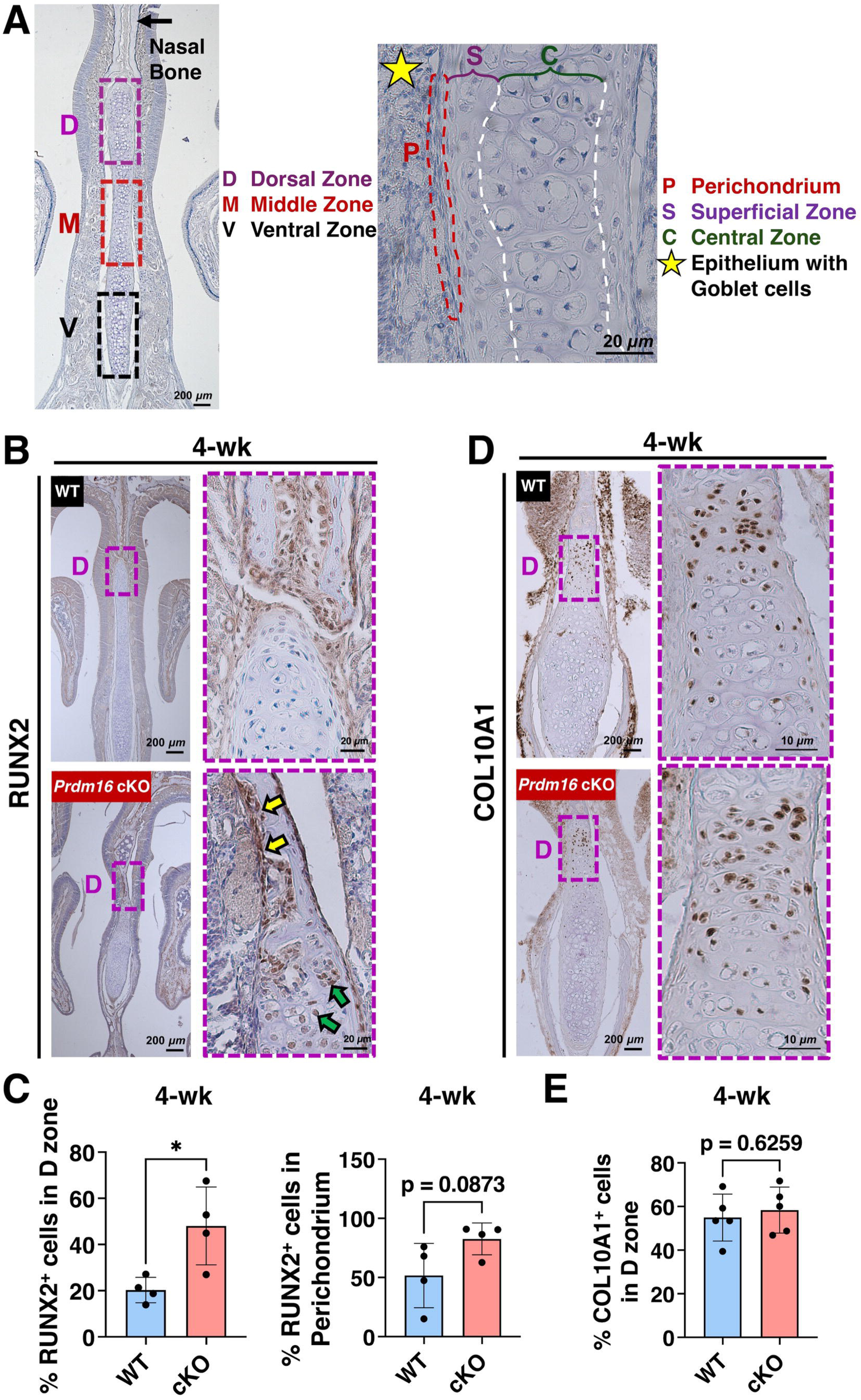
*Prdm16* cKO mice exhibited increased RUNX2 staining. **(A)** Anatomical and zonal organization of the nasal septal cartilage used as regions of interest for quantification. Zonal cartilage locations are defined as dorsal (D), middle (M), and ventral (V). Anatomical regions include the perichondrium (P), superficial (S), and central (C) cartilage. **(B, C)** Representative IHC images indicate significantly increased % RUNX2^+^ cells in the (D) dorsal zone (green arrows), but no significant difference in the perichondrium (yellow arrows) of the nasal cartilage in 4 wk old male mice. **(D, E)** Representative IHC images show no significant difference in %COL10A1^+^ cells in the (D) dorsal zone in 4 wk old male mice. Data are presented as mean ± SEM, with n ≥ 3 per group. Antibody staining appears in brown. Student’s unpaired t-test. *p < 0.05.

### Loss of PRDM16 reduces COL2A1 expression without altering MMP13 expression in the nasal septum

To determine whether loss of PRDM16 affects ECM composition in the nasal septum, we performed IHC for COL2A1 in 4 wk WT and *Prdm16* cKO mice (**Figure 4A, B**). In WT mice, COL2A1 was broadly distributed throughout both the pericellular and interterritorial matrix. In contrast, COL2A1 staining of cKO mice was concentrated in the pericellular matrix (yellow arrowheads). To assess whether PRDM16 deficiency promotes matrix degradation, we next examined MMP13 expression, a catabolic marker. However, MMP13 IHC staining revealed no significant differences between *Prdm16* WT and cKO mice (**Figure 4C, D**).

**Figure 4.**
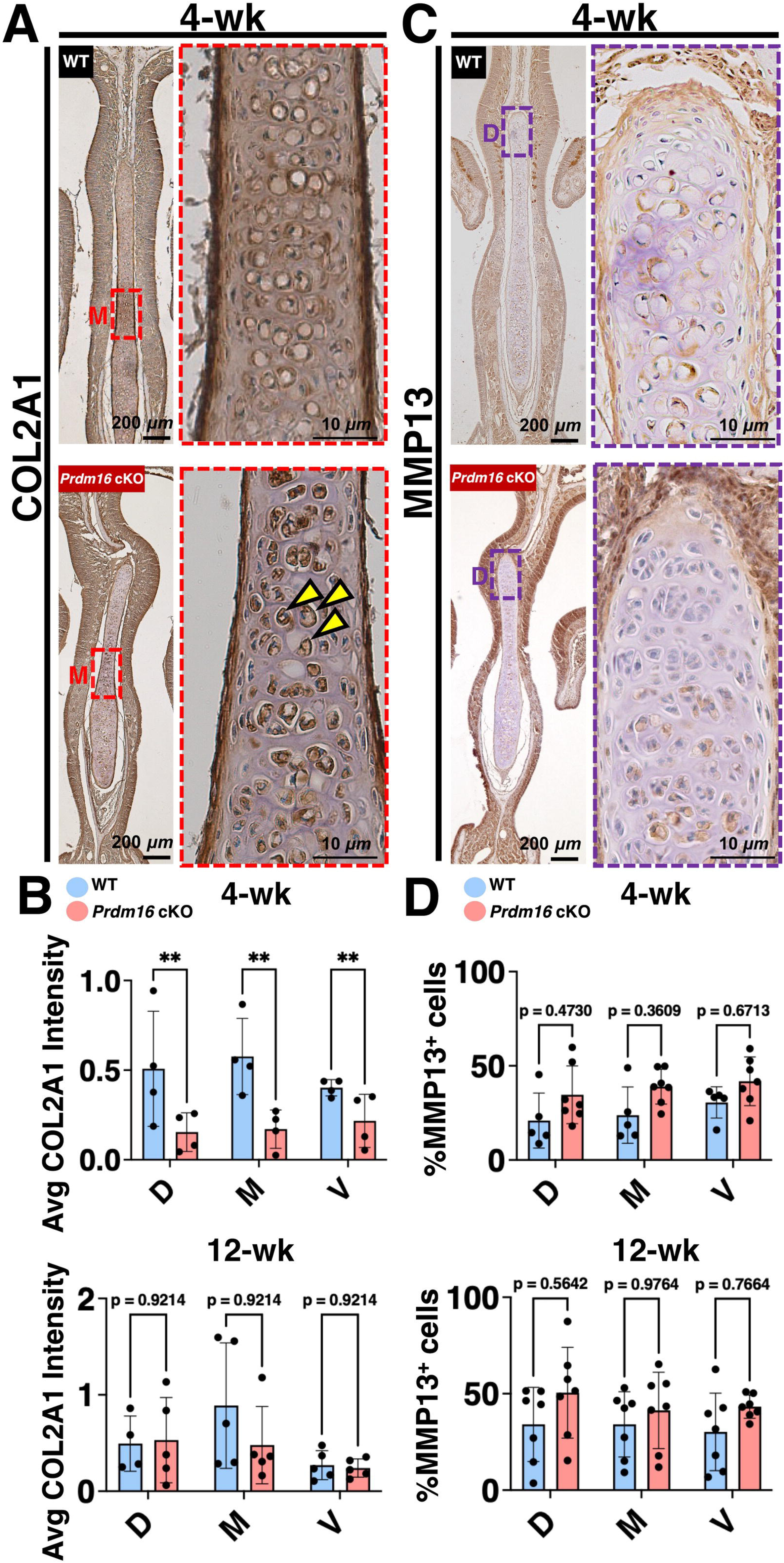
Loss of *Prdm16* leads to decreased COL2A1 expression across all nasal septal cartilage zones at 4 wks of age, while MMP13 expression remains unchanged. **(A, B)** Representative images and quantification of COL2A1 IHC staining in 4 wk old WT and *Prdm16* cKO mice. *Prdm16* cKO mice exhibited significantly reduced COL2A1 staining compared with WT controls. **(C, D)** Representative images and quantification of MMP13 IHC staining, showed no significant differences between WT and *Prdm16* cKO mice. D = dorsal zone, M = middle zone, V = ventral zone. Only cells in the central zone were counted, and not in the perichondrium. Data points represent individual mice and are presented as mean ± SEM. n ≥ 4/genotype/age. Antibody staining appears in brown, with arrowheads indicating positive signals. Two-way ANOVA with *post-hoc* Tukey’s test. **p < 0.01.

### Altered chondrocyte apoptosis may contribute to nasal septal deviation

We assessed the expression of chondrocyte proliferation and apoptosis markers of 4 wk and 12 wk old mice to determine if loss of PRDM16 results in abnormal cellular behavior. Expression of MKI67, a marker of chondrocyte proliferation, was comparable across all septal cartilage zones at both time points (**Figure 5A, B**). However, apoptosis was significantly increased in the middle zone of the septal cartilage at both 4 and 12 wks old. Notably, at 12 wks of age, CASP3 expression was also significantly elevated in the ventral zone. (**Figure 5C, D**).

**Figure 5.**
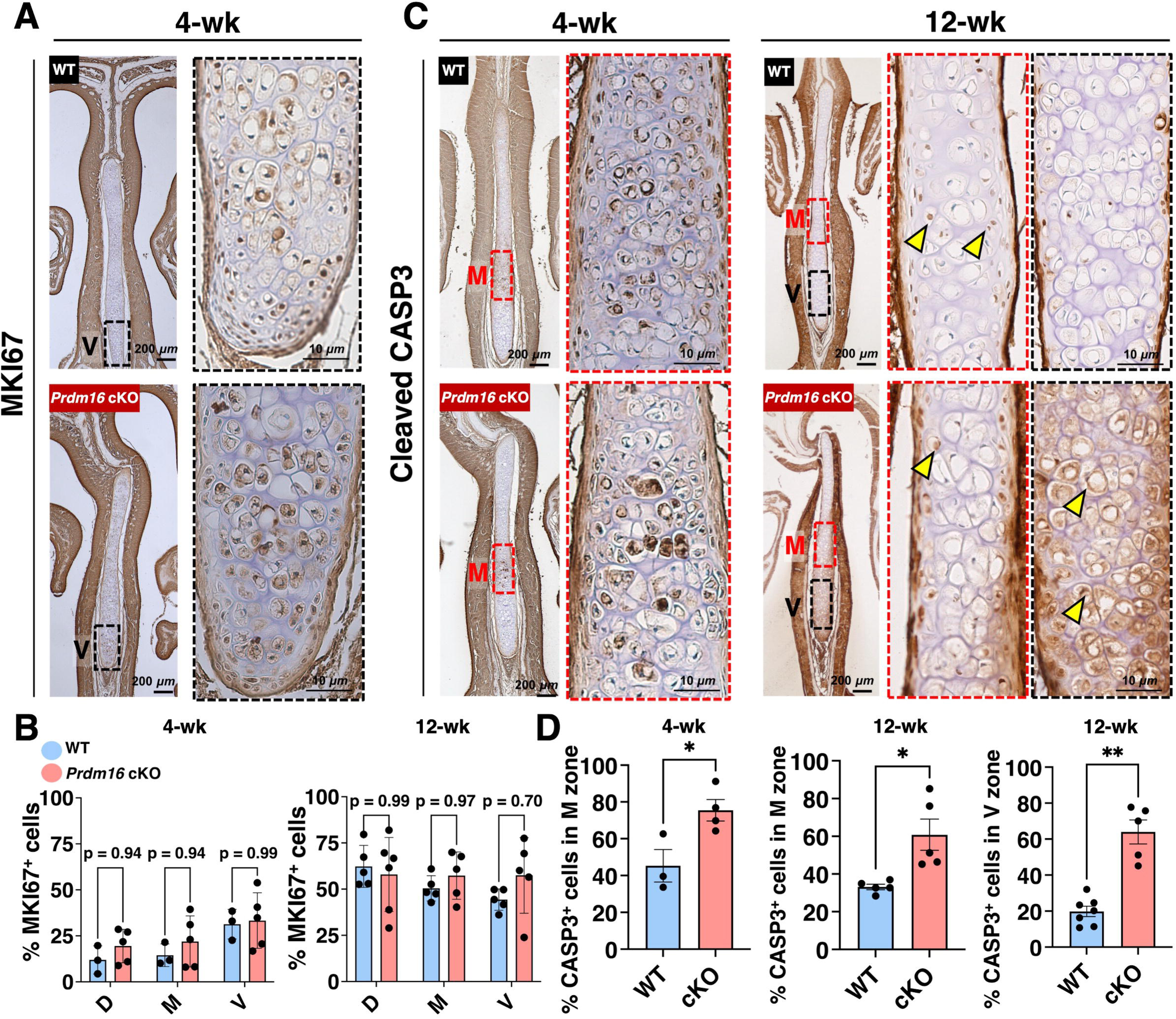
Altered chondrocyte apoptosis may lead to nasal septal deviation. **(A, B)** IHC staining of MKI67 revealed no difference in the nasal septal cartilage between WT and *Prdm16* cKO male mice at 4 or 12 wks of age. **(C, D)** Quantification of CASP3^+^ cells revealed significantly increased apoptosis in the middle zone (M) of the nasal septum in *Prdm16* cKO mice at 4 and 12 wks of age, and in the ventral zone (V) of 12 wk old male *Prdm16* cKO mice. Only cells in the central zone were counted (yellow arrowheads), and not in the perichondrium. Data are presented as mean ± SEM, with n ≥ 4 per group. Antibody staining appears in brown, with arrows indicating positive signals. Student’s Unpaired t-test or two-way ANOVA with *post-hoc* Tukey’s test, accordingly. *p < 0.05, and **p < 0.001.

### cKO of PRDM16 promotes chondrocyte hypertrophy while reducing *Mgp^+^* chondrocytes in the nasal septal cartilage

To elucidate how loss of PRDM16 may modulate chondrocyte phenotypes in nasal cartilage, we integrated our scRNA-seq datasets of 4 wk old male WT and cKO nasal septal cartilage samples. Unsupervised clustering identified 28 conserved cell populations across both genotypes (**Figure 6A**). To further focus on cartilage-specific populations, we next subset only the *Col2a1^+^* chondrocytes from the *Acan^+^*/*Runx2^high^* clusters to reveal four unique enriched chondrocyte populations shared between WT and cKO mice (n = 541 cells) (**Figure 6B, S3A–D**). Conserved marker genes and UMAP visualizations for each cluster across both genotypes are shown in **Figure S3A–F**. Cell cluster percentages of 4 wk old WT and *Prdm16* cKO cartilage datasets, further subsetted *Col2a1^+^* chondrocyte clusters, and the corresponding differentially expressed gene (DEG) lists are provided in **Supplementary File 1.** scRNA-seq analyses revealed that cKO mice had decreased septal *Mgp^+^* chondrocytes (WT: 71% vs. cKO: 33%) and increased *Col10a1^+^/Serpina3n^+^* hypertrophic chondrocytes (WT: 3% vs. cKO: 41%; **Figure 6C**). Gene Ontology (GO) analysis of genes upregulated in *Mgp^+^* chondrocytes (cluster 0) in cKO vs. WT mice revealed significant enrichment of biological processes related to connective tissue development, ECM organization, and osteoblast differentiation. Key DEGs associated with these pathways included *Mgp, Col9a1, Cnmd, Comp, Pkddc, Serpinh1, Snorc,* and *Pth1r*. Genes downregulated in cKO vs. WT mice were enriched for pathways associated with apoptosis, ECM organization, and cartilage development (**Figure 6D**). Furthermore, GO analysis of *Col10a1^+^/Serpina3n^+^* hypertrophic chondrocytes (cluster 1) from cKO vs. WT mice demonstrated enrichment of pathways related to ECM organization, connective tissue development, skeletal development, and bone mineralization. Representative DEGs contributing to these pathways included *Col10a1, Cytl1, Loxl1, Matn1, Bmpr2, Fgfr3, Slc26a2, Col6a1,* and *Slc39a14* (**Figure 6E**). DEGs and their associated GO terms for *Cdh13^+^*/*Dio2^+^* chondrocytes and *Foxp1^high^* chondrocytes are shown in **Figure S3G, H**.

**Figure 6.**
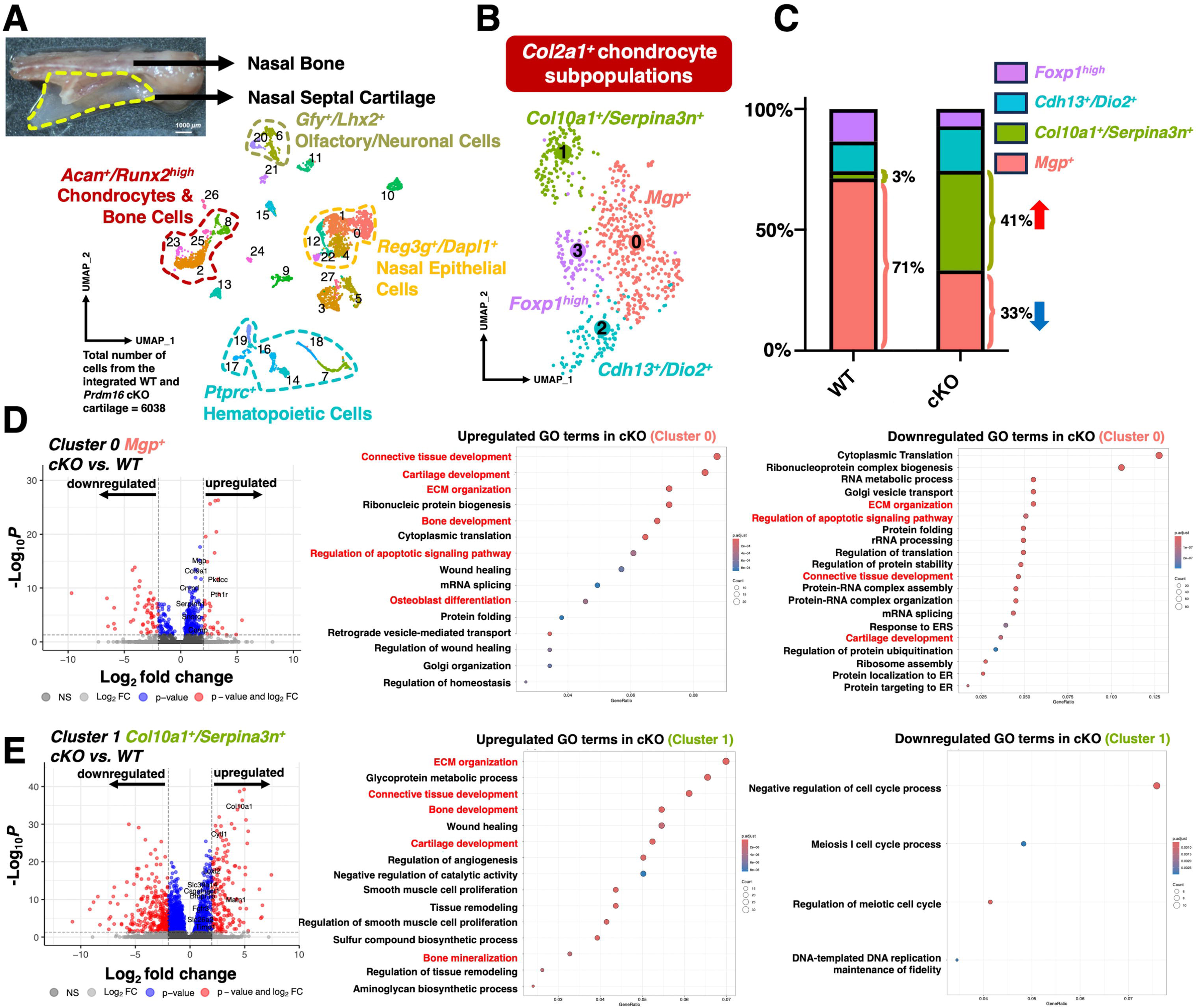
Loss of PRDM16 drives enrichment of hypertrophic chondrocytes and depletion of mature chondrocyte populations. **(A)** Integrated unsupervised clustering of WT and cKO nasal cartilage shows 28 conserved total cell populations (n = 6038 cells). **(B)** Reclustering of *Col2a1^+^* chondrocyte subpopulations (n = 541 cells) reveals 4 conserved chondrocyte populations between WT and cKO mice. **(C)** Percent cell profile showing significant reduction of *Mgp^+^* chondrocytes and enrichment of *Col10a1^+^/Serpina3n^+^* hypertrophic chondrocytes in cKO mice compared to 4 wk old WT mice. **(D)** Volcano plots showing differentially expressed genes (DEGs) in cluster 0 in cKO vs. WT mice. Gene Ontology (GO) biological process enrichment analysis of genes upregulated and downregulated in cKO cluster 0 compared to WT. **(E)** Volcano plots showing DEGs in cluster 1 in cKO vs. WT mice. GO biological process enrichment analysis of genes upregulated and downregulated in cKO cluster 1. Dot color reflects enrichment significance (red = higher enrichment; blue = lower enrichment), and dot size represents the proportion of genes associated with each GO term.

### Top ligand-receptor interactions between *Col1a1^high^*/*Col3a1^+^* fibrotic and *Mgp^+^* chondrocyte populations of 4 wk old male WT and cKO mice

To identify potential intercellular communication at the nasal sepal cartilage-nasal bone interface, we performed spatial-seq on 4 wk old male WT and cKO mice. From the spatial-seq dataset, we identified 14 distinct chondrocyte and bone cell populations within the selected region of interest from WT and cKO male mice, outlined by the white dashed line (**Figure 7A, B**). Conserved marker genes for each cluster, their relative proportions, and their spatial distribution within coronal sections are shown in **Figure S4A-C.** Cell cluster percentages and corresponding lists of DEGs for the complete 4 wk old WT and *Prdm16* cKO spatial transcriptomic datasets are provided in **Supplementary File 2**. Spatial-seq revealed that *Col1a1^high^/Col3a1^+^* fibrotic cells (cluster 1) and *Mgp^+^* chondrocytes (cluster 2) were located at the interface between nasal cartilage and bone (i.e., where deviation is present in the *Prdm16* cKO mice, **Figure 7C**).

**Figure 7.**
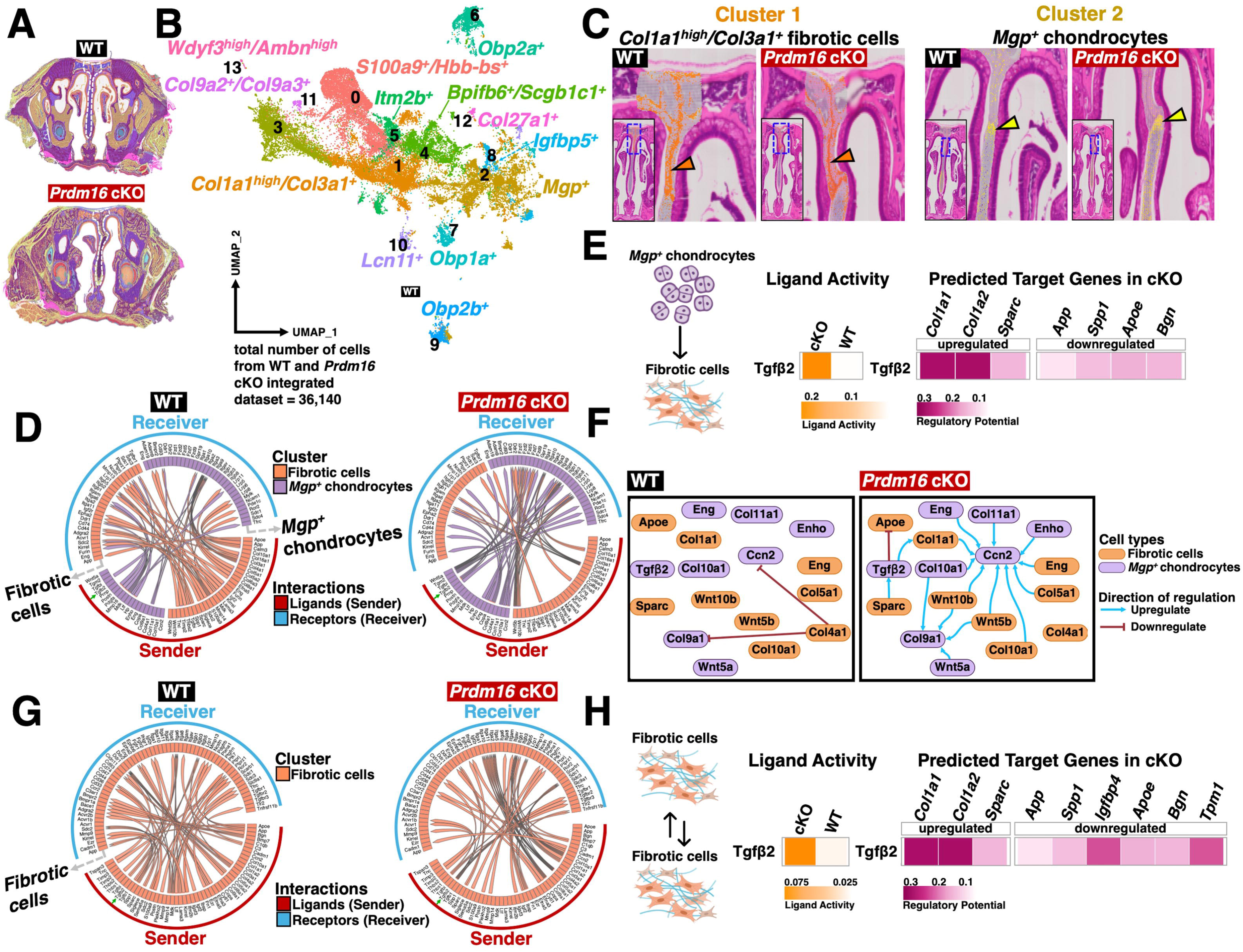
Top ligand-receptor interactions between fibrotic cells (cluster 1) and *Mgp^+^ chondrocyte* populations (cluster 2) of 4 wk old male WT and cKO mice. **(A)** Spatial capture spots (white dashed-lines) were selected for analysis. **(B)** Integrated unsupervised clustering identified 14 unique chondrocyte and bone populations across WT and cKO mice. **(C)** Fibrotic cells within the nasal bone (orange arrowhead) and *Mgp^+^* chondrocytes in nasal septal cartilage (yellow arrowhead) were both located near the deviation site. **(D)** Circos plot demonstrating top ligand-receptor interactions between *Mgp^+^* chondrocytes and fibrotic cells in WT and cKO mice. The green arrow indicates *Tgfβ2* as a ligand secreted by *Mgp^+^* chondrocytes. (E) *Tgfβ2,* along with its associated receptors, was the most prominent ligand-receptor pair along the *Mgp^+^* chondrocytes to fibrotic cells signaling axis. Increased orange indicates ligand upregulation in the respective genotype, while increased magenta represents higher predicted regulatory potential. **(F)** Visualization of inferred intercellular regulatory network. **(G)** Circos plot demonstrating top ligand-receptor interactions within fibrotic cells in WT and cKO mice. The green arrow indicates *Tgfβ2* as a ligand secreted by fibrotic cells. (H) *Tgfβ2,* along with its associated receptors, was the most prominent ligand-receptor pair within the fibrotic-to-fibrotic *cells* signaling axis. Increased orange indicates ligand upregulation in the respective genotype, while increased magenta represents higher predicted regulatory potential.

To identify potential intercellular communications at this interface, we compared predicted ligand-receptor interactions between two populations in WT and cKO mice^26^: fibrotic cells (cluster 1) and *Mgp^+^* chondrocytes (cluster 2) (**Figure 7D**). MultiNicheNet bioinformatic analysis revealed that in WT mice, signaling networks were predominantly driven by fibrotic cells functioning as senders; whereas in cKO, interaction activity shifted toward *Mgp*⁺ chondrocytes as sender cells (**Figure 7D**). We also observed that in cKO mice, *Tgfβ2* secreted by *Mgp*⁺ chondrocytes may signal through *Tgfβr1*, *Eng*, *Acvr1b*, and *Acvr1* receptors expressed in fibrotic cells (**Figure S4D**). This TGFβ2-mediated signaling pathway was predicted to promote upregulation of *Col1a1* and *Col1a2* expression, consistent with enhanced pro-fibrotic extracellular matrix deposition and accelerated osteogenesis (**Figure 7E**).^27^ Gene regulatory network analysis identified a marked shift in cell-cell communication following PRDM16 deletion. In WT mice, signaling was predominantly driven by *Col4a1* signaling from fibrotic cells and was predicted to repress *Ccn2* and *Col9a1* expression in *Mgp*⁺ chondrocytes (**Figure 7F**). In contrast, cKO mice displayed a coordinated pro-fibrotic signaling network in which multiple pathways converged to enhance *Ccn2* expression in *Mgp*⁺ chondrocytes. Furthermore, *Sparc*-derived signaling from fibrotic cells was predicted to induce TGFβ2 expression in *Mgp*⁺ chondrocytes, a regulatory interaction that was absent in the WT mice (**Figure 7F**).

Additionally, cell-cell interaction analysis among fibrotic cells revealed that the *Tgfβ2* signaling network, in which *Tgfβ2* engages multiple receptors, including *Tgfbr1*, *Tgfbr2*, *Tgfbr3*, *Eng*, *Acvr1b*, and *Acvr1* (**Figure S5A**). This signaling axis was uniquely enriched in cKO fibrotic-to-fibrotic cell interactions and was not detected in WT fibrotic cell communication networks, suggesting enhanced autocrine and paracrine *Tgfβ2* signaling in fibrotic cells following PRDM16 deletion **(Figure 7G)**. Notably, this autocrine-like signaling mirrors the response observed in *Mgp*⁺ chondrocytes to fibrotic cell interactions, resulting in upregulation of the extracellular matrix components *Col1a1*, *Col1a2*, and *Sparc* and downregulation of *App*, *Spp1*, *Igfbp4*, *Apoe*, *Bgn*, and *Tpm1* **(Figure 7H)**. Interestingly, gene regulatory network analysis revealed that *Tgfβ2* signaling was not significantly upregulated in WT samples. In *Prdm16* cKO samples, however, *Tgfβ2* was predicted to be a key upstream regulator, promoting the upregulation of *Col1a1*, *Col1a2*, and *Sparc*, while downregulating *Spp1* and *Apoe* (**Figure S5B**). The full repertoire of predicted cell-cell interactions between *Mgp*⁺ chondrocytes and fibrotic cells is presented in **Figure S4D and Figure S5**.

### Elevated TGFβ2 signaling is expressed across the nasal septal cartilage-bone interface in cKO mice

We next performed IHC to identify TGFβ2 protein expression levels on male 4 wk old WT and cKO mice. TGFβ2 expression was detected in osteoblasts (yellow arrowheads) adjacent to the nasal bone, and within the fibrous ECM (orange arrows) within the nasal bones in both WT and cKO samples (**Figure 8A**). Importantly, cKO mice exhibited a significantly increased percentage of TGFβ2^+^ cells within the fibrous ECM (orange arrows) and the apical region of the nasal septal cartilage matrix (dark green arrows) compared to WT mice (**Figure 8B, C**).

**Figure 8.**
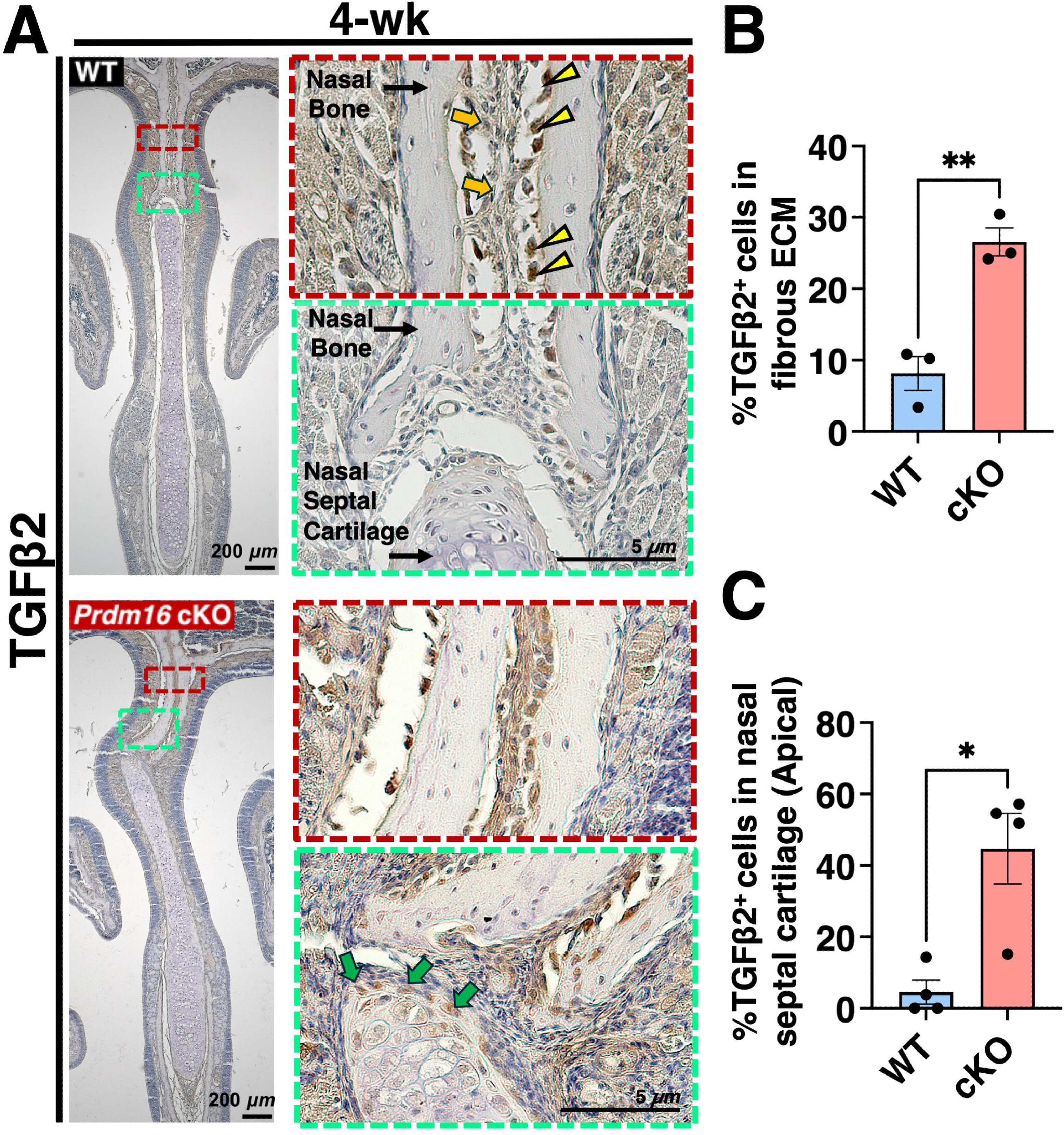
Elevated TGFβ2 signaling and matrix localization in 4 wk old male cKO mice. **(A)** IHC revealed increased TGFβ2 expression within the fibrous ECM and the apical region of the nasal septal cartilage in cKO vs. WT mice. **(B, C)** Quantification of %TGFβ2^+^ cells in the fibrous ECM and apical region of nasal septal cartilage. Orange arrows indicate TGFβ2^+^ cells within the fibrous ECM, yellow arrowheads indicate TGFβ2^+^ osteoblasts adjacent to the nasal bone, and green arrows indicate TGFβ2^+^ cells within the apical region of the nasal septal cartilage. Dark red box highlights fibrous ECM within the nasal bone, while the green box highlights the nasal bone-nasal septal cartilage interface at the site of septal deviation. Antibody staining is shown in brown. Data are presented as mean ± SEM, with n ≥ 3 per group. Student’s unpaired t-test. *p < 0.05 and **p < 0.01.

## Discussion

In this study, we demonstrate that loss of PRDM16 in chondrogenic cells results in shortened nasal septal cartilage and progressive nasal septal deviation. Integrative scRNA-seq analyses further revealed a marked shift in chondrocyte identity, characterized by depletion of *Mgp^+^* chondrocytes and expansion of *Col10a1^+^/Serpina3n^+^* hypertrophic chondrocytes. Importantly, spatial-seq identified a distinct nasal septal cartilage-bone interface niche composed of *Mgp^+^* chondrocytes and *Col1a1^high^/Col3a1^+^* fibrotic cells at the site of deviation in *Prdm16* cKO mice. Cell-cell communication analyses revealed extensive rewiring of signaling networks at this niche, with TGFβ2 emerging as a dominant altered signaling pathway. Together, these findings identify a previously unrecognized and novel role for PRDM16 in maintaining cartilage homeostasis and suppressing pathological cartilage-fibrotic cell communication during craniofacial development.

Our data also suggests that PRDM16 is required to maintain mature nasal septal chondrocyte identity. scRNA-seq analysis revealed a substantial reduction of *Mgp^+^* chondrocytes, accompanied by a significant expansion of *Col10a1^+^/Serpina3n^+^* hypertrophic chondrocytes following loss of PRDM16. These transcriptional changes were further supported by GO terms associated with upregulation of ECM organization, connective tissue development, osteoblast differentiation, and bone mineralization in cKO mice. Moreover, cKO mice exhibited higher nasal BV/TV and elevated RUNX2 expression within the dorsal septal cartilage adjacent to the deviation site, consistent with accelerated osteogenic differentiation. Interestingly, increased nasal BV/TV was primarily observed in male cKO mice, whereas both sexes exhibited shortened nasal lengths and septal deviation. These findings suggest that PRDM16-dependent regulation of osteogenic remodeling may contribute to sex-specific phenotypes, although the underlying mechanisms remain unclear. In addition, our data showed that cKO of PRDM16 increases chondrocyte apoptosis without significantly affecting proliferation, suggesting that the disruption of cartilage homeostasis arises from altered chondrocyte cell fate decisions and survival, rather than changes in cell proliferation. Indeed, these observations are consistent with our recent work demonstrating that PRDM16 regulates chondrocyte specification and endochondral ossification in the knee joint.^16^ Together, these findings support a model in which PRDM16 modulates chondrocyte phenotypes and osteogenic programs within the nasal septal cartilage. Disruptions of chondrocyte identity and survival due to loss of PRDM16 may represent one mechanism contributing to nasal septal deviation.

Most importantly, we identified that loss of PRDM16 leads to an altered TGFβ2-mediated communication axis between *Mgp*⁺ chondrocytes and adjacent fibrotic cells at the nasal cartilage-bone interface in cKO mice. In WT mice, signaling networks were largely dominated by fibrotic cell-derived signals that were predicted to suppress genes associated with matrix production and tissue remodeling. In contrast, loss of PRDM16 fundamentally altered these communication patterns, shifting signaling activity toward *Mgp*⁺ chondrocytes as dominant sender cells. MultiNicheNet and GRN analyses predicted that Tgfβ2, produced by *Mgp*⁺ chondrocytes, signals through *Tgfβr1*, *Eng, Acvr1b*, and *Acvr1* receptors expressed by fibrotic cells. This results in activation of profibrotic programs characterized by increased *Col1a1*, *Col1a2*, *Sparc,* and *Ccn2* expression, and decreased *Apoe* levels. A recent study reported that increased TGFβ signaling, with downregulation of *Apoe,* leads to vascular fibrosis by driving pericytes and vascular smooth muscle cells toward myofibroblast differentiation.^28^ Additionally, CCN2 (formerly CTGF) amplifies TGFβ signaling, promoting fibroblast differentiation into collagen-producing myofibroblasts.^29^ Our GRN analysis of cKO mice further indicated that SPARC-dependent signaling from fibrotic cells may reinforce TGFβ2 expression in *Mgp*⁺ chondrocytes, creating a reciprocal communication circuit that was absent in WT. Moreover, enhanced autocrine TGFβ2 signaling was predicted among fibrotic cells in cKO mice. Importantly, these computational predictions were supported by tissue-level validation. IHC demonstrated increased TGFβ2 localization within the fibrous ECM and in the apical region of nasal septal cartilage of cKO mice. Together, these findings lead us to propose that *Mgp*⁺ chondrocytes initiate TGFβ2 signaling in fibrotic cells, which subsequently upregulate autocrine TGFβ2 signaling, establishing a self-reinforcing feedback loop that drives progressive ECM remodeling and fibrosis within the fibrotic tissue between the nasal bones.

TGFβ2 has pleiotropic functions and regulates a wide range of biological processes, including ECM production, tissue homeostasis, and fibrosis.^30,31^ While TGFβ2 is essential for normal tissue function, excessive signaling is a major driver of fibrosis and pathological tissue remodeling.^32,33^ In knee cartilage, elevated TGFβ2 activity promotes synovial fibrosis and osteoarthritis-like progression.^34^ In neural crest-derived lineages, loss of TGFβ2 results in reduced proliferation in Meckel’s cartilage and the mandibular primordium, leading to a shortened mandible and cleft palate phenotype.^35–37^ Interestingly, PRDM16 suppresses profibrotic gene expression in adipocytes and inhibits myofibroblast activation in the kidney in both mice and humans.^38^ Furthermore, in mice with *Prdm16*-null cardiac cells, activation of the TGFβ/SMAD2 axis and upregulation of *Ccn2* result in severe cardiac fibrosis.^39^ While these prior studies imply a functional link between PRDM16 and TGFβ signaling, our findings extend these observations and suggest that PRDM16 may suppress fibrosis, at least in part, by inhibiting TGFβ2-mediated pathways.

In addition to fibrosis, TGFβ2 signaling has also been described for regulating chondrocyte survival and differentiation. Previous studies have demonstrated that TGFβ2 can modulate chondrocyte apoptosis by activating the TGFβ2/TAK1/p38 MAPK pathway and regulate hypertrophic differentiation through multiple downstream mechanisms.^40–43^ These observations may provide a potential explanation for the decreased COL2A1 IHC staining intensity and increased CASP3⁺ chondrocytes observed in *Prdm16* cKO mice. Despite the altered ECM organization in cKO mice, no significant differences were observed in the expression of the matrix degradation marker MMP13. Together, these findings suggest that the reduction in COL2A1 is likely due to impaired matrix synthesis rather than enhanced matrix degradation.

Nevertheless, the effects of TGFβ2 on cartilage are highly context-dependent and may vary according to developmental stage, tissue origin, and local signaling environment. Because the nasal septal cartilage is derived primarily from neural crest cells^44^, whereas many previous studies have focused on mesoderm-derived skeletal tissues, the mechanisms regulating PRDM16 may differ between these developmental lineages. Therefore, there is a need for further studies into how PRDM16 genetically and/or epigenetically modulates TGFβ2 expression in neural crest cell-derived lineages.

Several limitations should be considered when interpreting these findings. First, the ligand-receptor interactions and GRN identified in this study are computationally inferred and require direct experimental validation. Second, although elevated TGFβ2 signaling was strongly associated with septal deviation, causal relationships still need to be established through genetic and/or pharmacologic inhibition studies. Third, the relatively small spatial transcriptomic sample size may limit the detection of less abundant cell populations and signaling events. Future studies utilizing TGFβ2 pathway inhibition, cell-specific genetic models, and longitudinal analyses will be important for determining the precise contribution of these signaling pathways to disease progression.

Collectively, our findings establish PRDM16 as a critical regulator of nasal septal cartilage homeostasis and identify dysregulated TGFβ2 signaling as a central mechanism underlying nasal septal deviation following loss of PRDM16. We propose a model in which PRDM16 maintains chondrocyte identity while suppressing aberrant TGFβ2 signaling at the septal cartilage-bone interface. Loss of PRDM16 promotes chondrocyte hypertrophy, apoptosis, and activation of a TGFβ2-mediated signaling network between *Mgp*⁺ chondrocytes and fibrotic cells, resulting in excessive ECM deposition, fibrosis, and accelerated osteogenic remodeling. These molecular and cellular changes ultimately drive progressive nasal septal cartilage deviation and provide new mechanistic insights into previous human genetic studies linking PRDM16 to craniofacial morphology and developmental abnormalities.

## Acknowledgments

The authors thank UR Genomics Research Center for assisting with scRNA-seq and spatial-seq.

## Author contributions

Eliya Tazreena Tashbib (Conceptualization, Data curation, Formal analysis, Investigation, Methodology, Visualization, Writing—original draft, Writing—review & editing), Victoria L. Hansen (Methodology, Validation, Writing—review & editing), Maeve O’Brien (Methodology, Formal analysis, Validation), Alexis Klee (Methodology, Formal analysis, Validation), Eloise Fadial (Methodology, Validation), Karin Pryharski (Methodology, Validation), Gulzada Kulzhanova (Formal analysis, Writing—review & editing), Kathryn Lambright (Methodology), Lomeli Shull (Supervision, Writing—review & editing) and Chia-Lung Wu (Conceptualization, Funding acquisition, Methodology, Project administration, Resources, Supervision, Validation, Visualization, Writing— review & editing)

