## Supplementary figures and images for "Loss of PRDM16 Drives Nasal Septal Deviation through Dysregulated TGFβ2 Signaling"

### Supplemental Figure 1

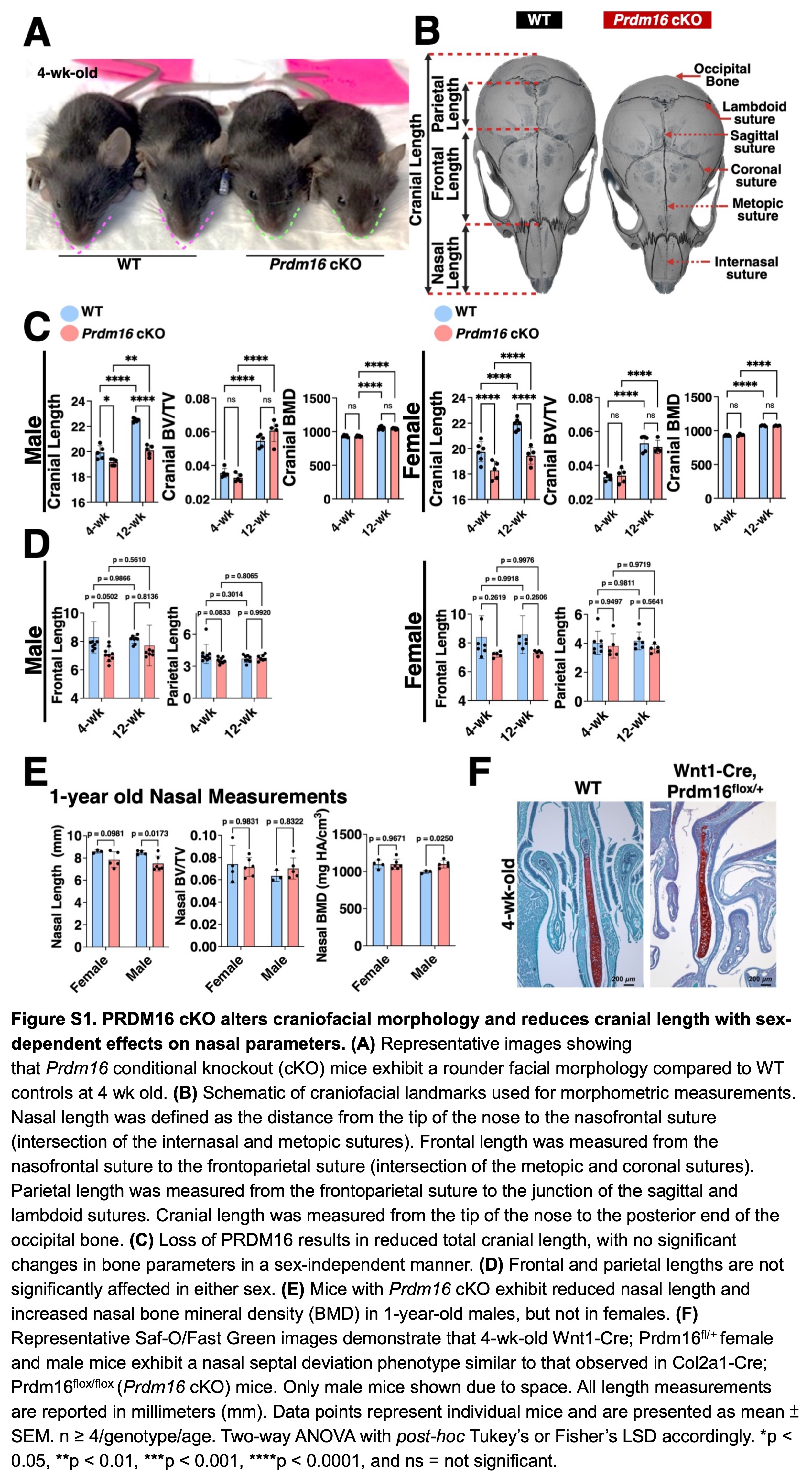

### Supplemental Figure 2

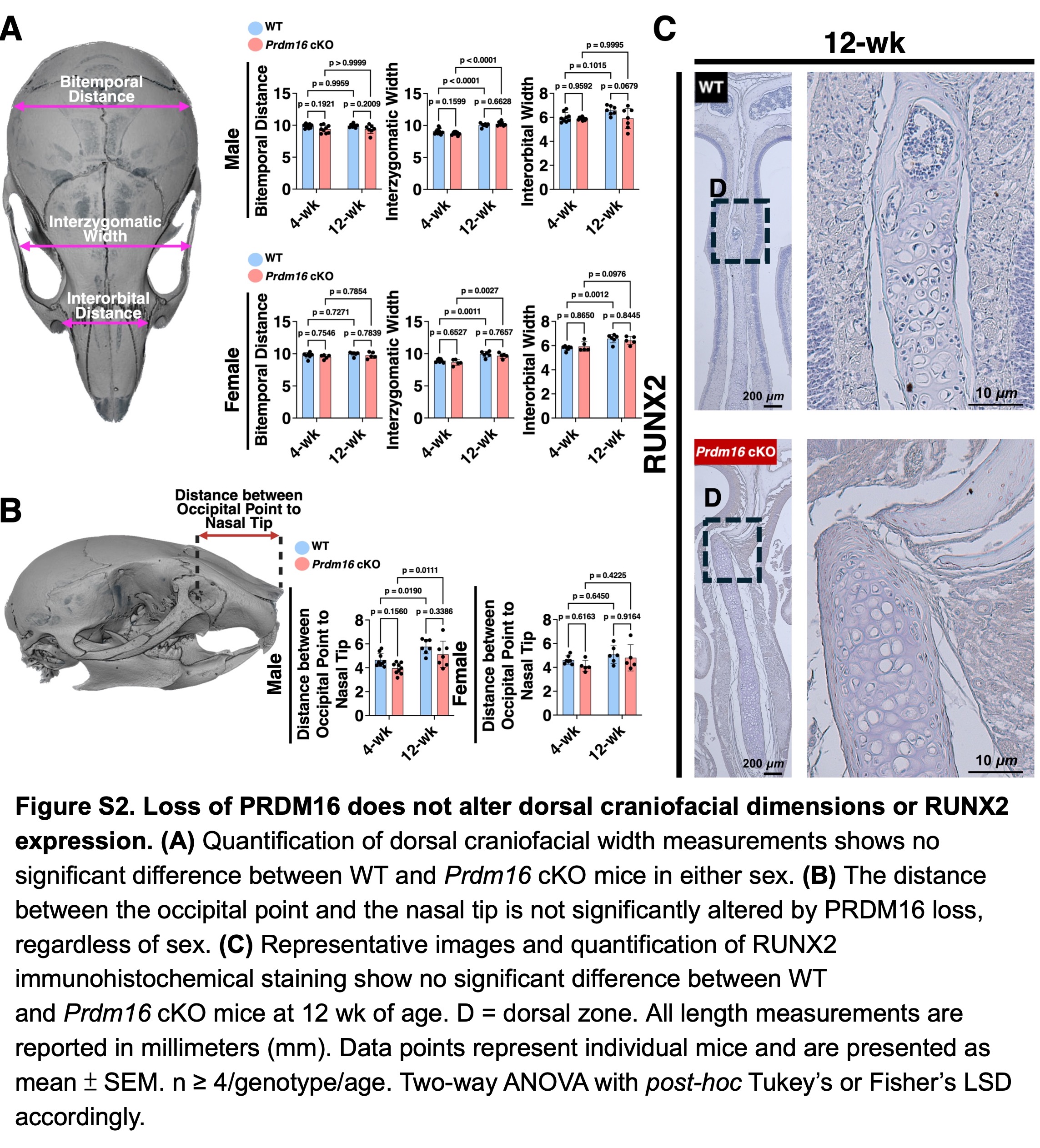

### Supplemental Figure 3

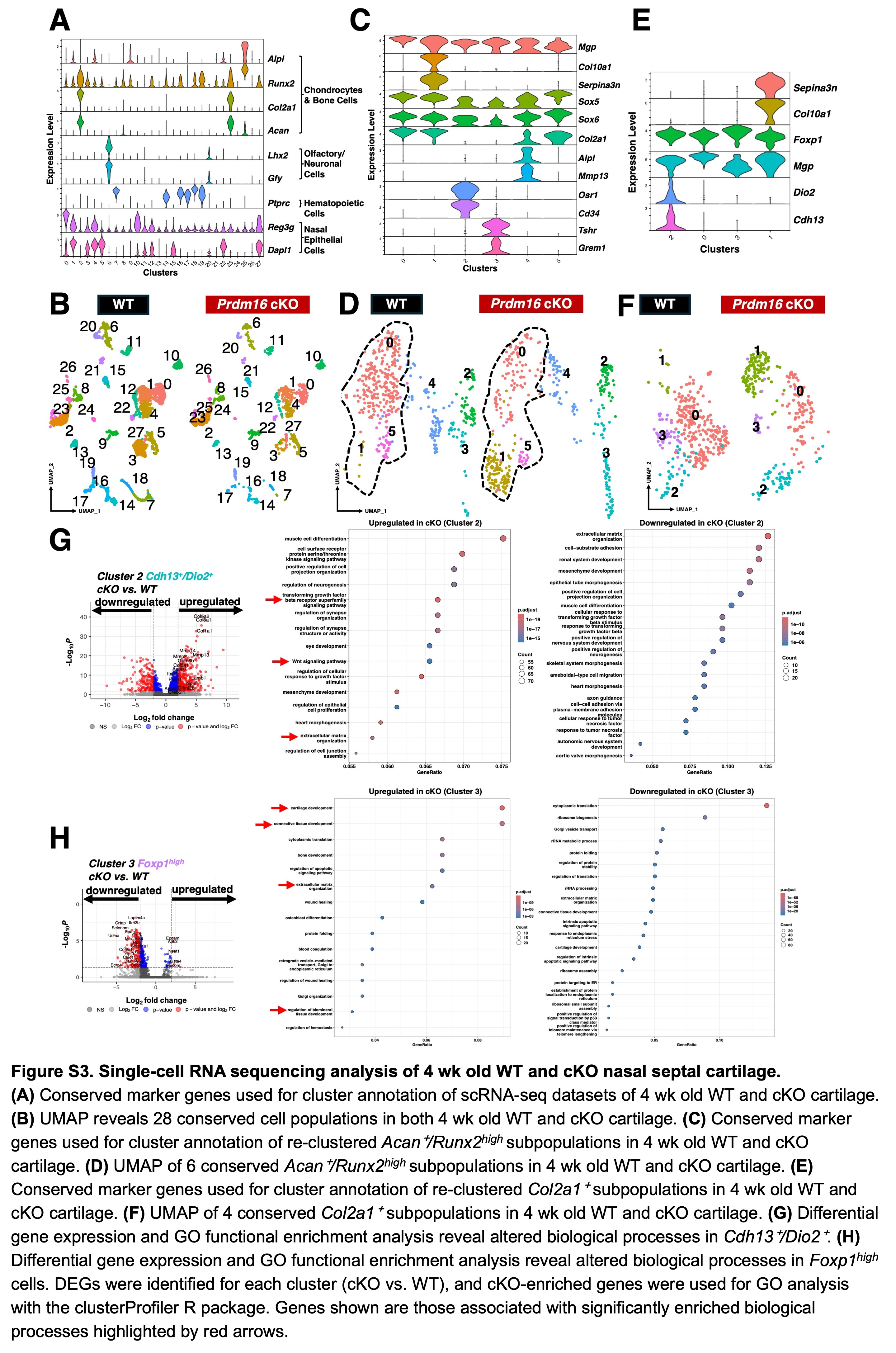

### Supplemental Figure 4

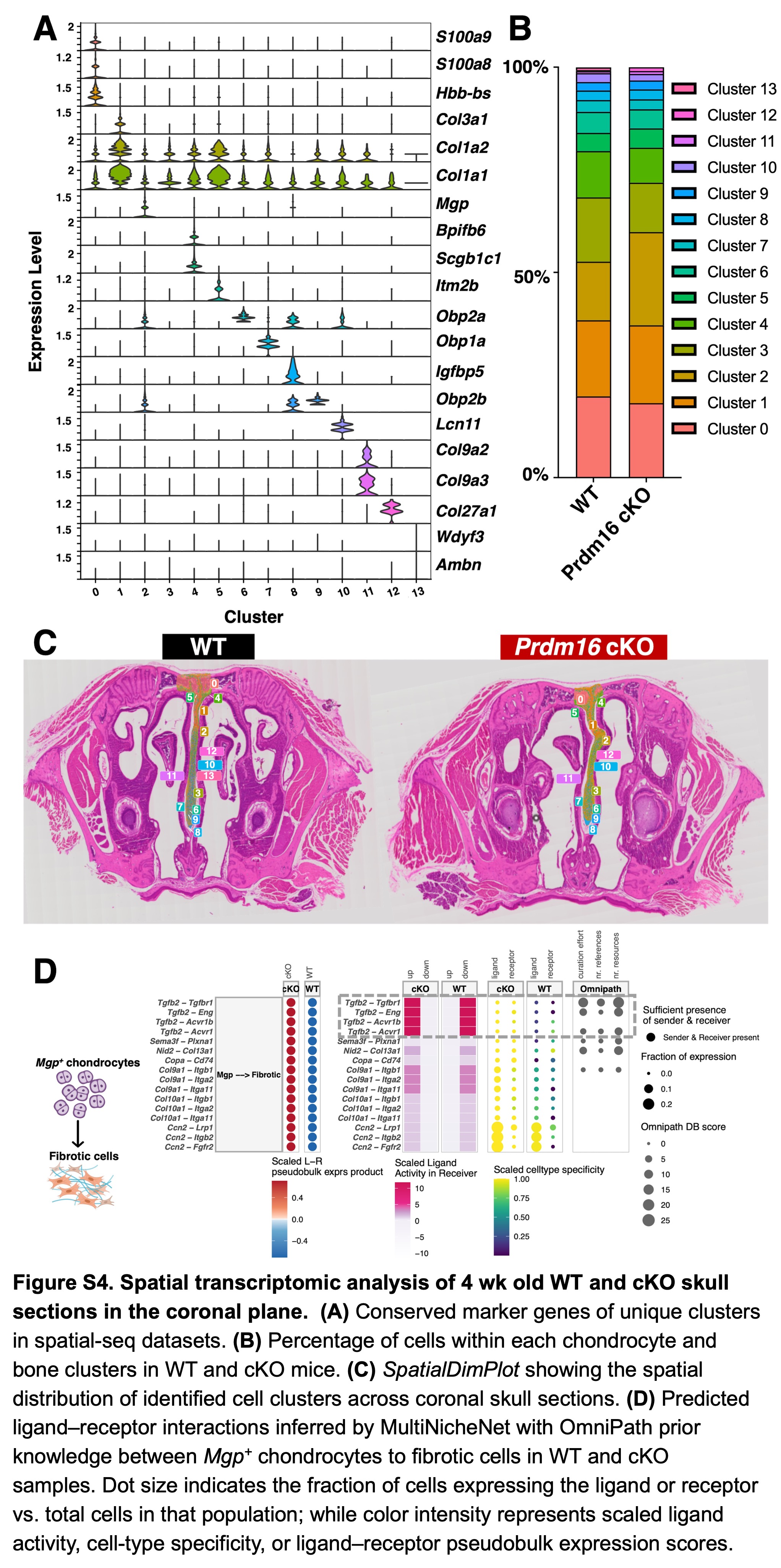

### Supplemental Figure 5

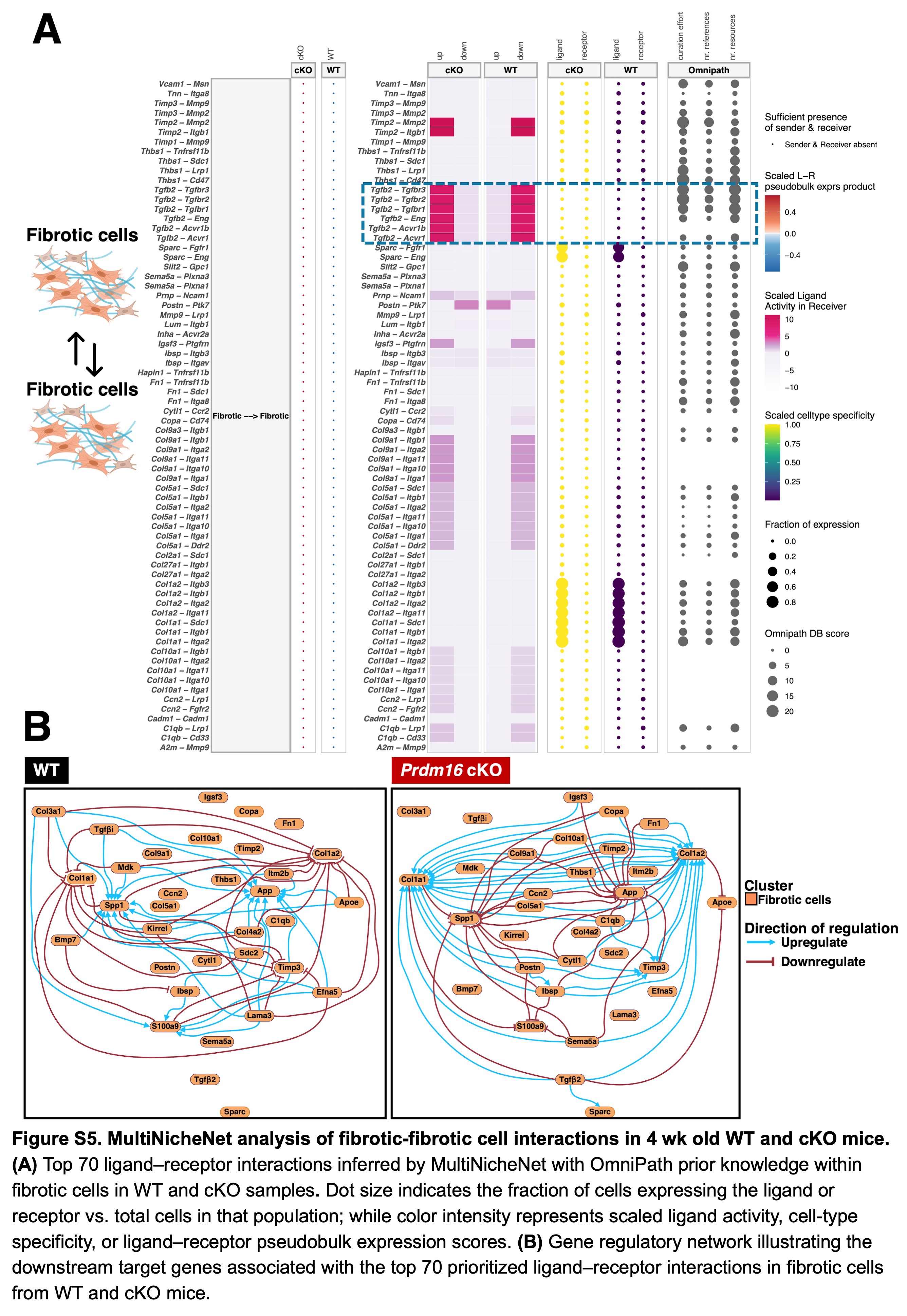
