## Supplemental Methods for "Loss of PRDM16 Drives Nasal Septal Deviation through Dysregulated TGFβ2 Signaling"

### Animal housing and developmental staging

All animal procedures were approved by the Health Sciences Animal Care and Use Committee at the University of Rochester and conducted in accordance with the Guide for the Care and Use of Laboratory Animals of the National Institutes of Health. For this study, *Prdm16$^{flox/flox}$* mice (B6.129-Prdm16$^{tm1.1Brsp}$/J) were crossed with Col2a1-Cre mice (Col2a1-Cre mice, B6;SJL-Tg(Col2a1-cre)1Bhr/J; #003554, the Jackson Laboratory) to generate Col2a1-Cre; *Prdm16$^{flox/flox}$* (referenced as *Prdm16* cKO) mice Cre-negative littermates were used as wild type (WT) controls. As previously described, nasal cartilage growth occurs in two phases: rapid growth from P0 to P4, and slower growth from P4 onward; with minimal growth between P4 and P14.[1,2] Key developmental stages (E18.5, P5, P15, 4 wk, 12 wk, and 1 year) were selected to capture craniofacial growth. The morning a vaginal plug was detected was considered E0.5. Wnt1-Cre; *Prdm16$^{flox/+}$* mice were generated by crossing Wnt1-Cre transgenic mice with *Prdm16$^{flox/flox}$* mice. Mice carrying the Wnt1-Cre transgene and one floxed Prdm16 allele (Wnt1-Cre; *Prdm16$^{flox/+}$*) were used for phenotypic analyses. Homozygous neural crest-specific deletion (Wnt1-Cre; *Prdm16$^{flox/flox}$*) resulted in embryonic lethality and was therefore not included in subsequent analyses. *Prdm16$^{flox/+}$* were used as WT controls. The number and sex of animals used in each experiment are detailed in the figure legends. Mice were housed under pathogen-free conditions (≤5 per cage) with standard light/dark cycles, controlled temperature and humidity, and *ad libitum* food and water.

### Alcian blue & alizarin red (whole-mount) skeletal staining for embryos

Alcian blue and alizarin red skeletal staining was performed on E18.5 WT and *Prdm16* cKO embryos, as previously described.[3] E18.5 embryos were collected in PBS, skinned, and eviscerated prior to fixation overnight in 95% EtOH at room temperature. Following fixation, embryos were stained with 0.01% Alcian blue 8GX (10 mg Alcian blue 8GX in 80 ml 95% EtOH and 20 mL glacial acetic acid) for 24 hours. Embryos were then rehydrated through a graded ethanol series (70%, 35%, and 17.5% EtOH) and cleared in 1% KOH until the tissues became transparent. After 24 hours, embryos were subsequently stained with 0.001% Alizarin Red S (1 mg Alizarin Red S in 100 mL 1% KOH) for 24 hours, with the staining solution refreshed twice during incubation. The next day, embryos were further cleared in 1% KOH and sequentially transferred through 20%, 50%, and 80% glycerol in 1% KOH (24 hours each), followed by incubation in 100% glycerol for 48 hours before imaging.

### Micro-computed tomography (*μ*CT) acquisition parameters

*Prdm16* cKO and WT skulls (n ≥ 4/genotype/sex) were assessed using *μ*CT (Scanco Medical VivaCT 40 cone-beam with 10.5 µm isotropic cubical voxels and 21.5mm OD).

1000 projections were generated over 180°, with a 300msec integration time, and reconstructed using proprietary Scanco algorithms. Scanco evaluation software V6.5 was used to analyze *μ*CT bone parameters. Skulls were scanned, and 3D reconstructions were generated for further analysis. The region used for nasal bone volume analysis begins at the first presence of bone frontally and proceeds in the posterior direction, ending just before the first presence of the maxilla. The first and last slices were for support only and were not analyzed. Contours are drawn close to the bone to minimize the size of the total contour and thus decrease the total volume measurement. Teeth embedded within or outside the skull bones were excluded from bone volume analyses. Bone volume fraction (BV/TV) and mean bone mineralization density (BMD) were quantified and plotted using GraphPad software. Statistical analyses were performed using Student's Unpaired t-test or Two-Way ANOVA, as appropriate. Nasal and cranial lengths of 4 wk, 12 wk, and 1 year old mice were obtained from dorsal views using Amira software with a landmark-based approach. The nasal length was defined as the distance from the tip of the nose to the nasofrontal suture (intersection of the internasal and metopic sutures) (**Figure S1B**).[4] All landmarks used for craniofacial measurements, including cranial length, frontal length, parietal length, anterior and posterior mandibular length, mandibular plane length, effective mandibular length, condylar axis length, bitemporal distance, interorbital width, interzygomatic width, and the distance from the occipital point to the nasal tip, are provided in the supplementary materials (**Figure S1B and S2A, B**). Frontal length was measured from the nasofrontal suture to the frontoparietal suture (intersection of the metopic and coronal sutures). Parietal length was measured from the frontoparietal suture to the junction of the sagittal and lambdoid sutures. Cranial length was measured from the tip of the nose to the posterior end of the occipital bone.

### Tissue processing

*Prdm16* cKO and WT, along with Wnt1-Cre; *Prdm16*$^{flox/+}$ mice and their littermate controls *Prdm16*$^{flox/+}$, were fixed in 10% neutral buffered formalin at room temperature (RT) for 48 hours. Following fixation, mouse skulls were decalcified using Cal-Ex solution (Fisher CS510-1D) at 4°C for 48 hours for P5 and P15 samples, and 72 hours for 4 wk and 12 wk old samples, as previously described.[5] After decalcification, the skulls were processed for paraffin embedding following standard protocol. Skulls were embedded in a frontal orientation, and 10*μm* thick coronal sections were collected for subsequent histological or immunohistochemical analyses.[6]

### Septal deviation and cartilage ratio quantification

Histological sections from the junction of the nasal septum with the vomer and from the mid-septal region were used for quantitative analysis. Septal deviation was assessed by drawing a reference straight line between anterior and posterior septal landmarks, followed by measurement of the angle between this line and the most deviated point of the septum. Greater deviation angles corresponded to increased severity of septal curvature. The nasal septal cartilage ratio was calculated as posterior septal length divided by total septal length.

### Immunohistochemistry methods

Immunohistochemistry was performed on coronal sections of paraffin-embedded nasal septa from 4 wk and 12 wk old male WT and *Prdm16* cKO (n ≥ 3/genotype/age). Antigen retrieval was conducted using a pepsin-based protocol (R2283-100ML, Sigma-Aldrich) for 10 min at RT. The sections were incubated either with hypertrophic chondrocyte markers (anti-RUNX2, 1:100; anti-Collagen X, 1:100), proliferation marker (anti-MKi67, 1:150), apoptosis marker (anti-Cleaved Caspase 3, CASP3, 1:600), or mature chondrocyte marker (anti-Collagen 2A1, 1:200), chondrocyte degradation marker (MMP13, 1:200), anti-TGF-β2 antibody (1:400), respectively overnight at 4°C. Details of the antibodies are provided in **Supplementary Table 1**. Target proteins were detected using the ImmPRESS polymer detection kit (MP-7401, Vector Laboratories) and visualized by colorimetric development with DAB (Vector Laboratories SK-4105) or SG (Vector Laboratories SK-4700) substrates. Hematoxylin was used to counterstain nuclei. Staining (percentage of number of stained cells over total cells) was quantified using ImageJ (https://imagej.net/).

### Nasal septal cartilage cell isolation for scRNA-seq analysis

Nasal septal cartilage was dissected from WT and *Prdm16* cKO mice and minced into small fragments using scalpels. Tissue fragments were digested in DMEM/F12 supplemented with 1% penicillin-streptomycin, 1.4 mg/mL pronase, and 0.1 μg/mL DNase I for 1 hour at 37°C with rotation. Digestion was terminated by the addition of an equal volume of prewarmed DMEM/F12 containing 20% FBS. Following centrifugation (140 ×g, 1 minute, 4°C), the supernatant was carefully removed to avoid disturbing the pellet. To the pellet, 1 mL of prewarmed neutralization media was added, and the tube was gently inverted 12 times to ensure thorough mixing and to facilitate the release of additional cartilage cells. The resulting suspension was filtered through a 40*μ*m nylon mesh to obtain single-cell preparations, which were then counted using a hemocytometer. The average yield from two nasal septa was approximately $3.5 \times 10^4$ cells. For scRNA-seq experiments, samples from four animals per genotype were pooled before library preparation.

Samples with greater than 75% viability were submitted to UR Genomics Research Center for scRNA-seq library preparation. Cell suspensions were processed to generate single-cell RNA-seq libraries using the GEM-X Universal 3' Gene Expression kit (10X Genomics) per the manufacturer's recommendations, as summarized below. Samples were loaded onto the Chromium X (10X Genomics, Pleasanton, CA, USA) to generate single-cell GEMs (Gel Bead-in-Emulsions). GEM reverse transcription (GEM-RT) was performed to produce a barcoded, full-length cDNA from polyadenylated mRNA. Subsequently, GEMs were broken to recover the pooled GEM-RT reaction mixtures, followed by cDNA purification and limited PCR amplification to generate sufficient material for library construction. Enzymatic fragmentation and size selection were used to optimize the cDNA fragment size with final library construction via end repair, A-tailing, adaptor ligation, and PCR with Illumina-compatible indexed primers. Libraries were sequenced using NovaSeq X Plus (Illumina, San Diego, CA), targeting 100,000 reads per cell.

### *Quality metrics analysis and initial subgrouping of major cell types*

Pre-processed aligned datasets of cells from Cell Ranger were loaded into RStudio to create Seurat objects (Seurat V5.0.1). A quality metrics analysis was performed to exclude low-quality cells or cells likely undergoing apoptosis. WT cartilage cells were retained if they contained >1,500 and <7,500 detected genes (nFeature_RNA) and <25% mitochondrial gene content, while cKO cells were retained if they contained >2,000 and <8,000 detected genes and <25% mitochondrial content for further downstream analyses. Individual datasets were later normalized using SCTransformation.[7]

### *Integration of WT and cKO scRNA-seq datasets*

To integrate scRNA-seq datasets from WT and *Prdm16* cKO cartilage samples, an anchor-based integration was performed following Seurat tutorials. Initially, the top variable features were selected using the *SelectIntegrationFeatures* function. Seurat objects were normalized with *SCTransform*, followed by anchor identification through *FindIntegrationAnchors*. These anchors were then used to integrate the datasets using the *IntegrateData* function. *RunPCA* was performed for linear dimensionality reduction, and an Elbow Plot was used to assess the variance explained by each principal component, helping to determine the optimal number of dimensions. Non-linear dimensionality reduction was then applied using UMAP to visualize the integrated dataset. To explore cell composition, *FindNeighbors* and *FindClusters* were run at a resolution of 0.7 to identify distinct clusters. Marker genes for these clusters were identified with *FindMarkers*, and the top 50 upregulated genes in both WT and *Prdm16* cKO groups were confirmed through VlnPlot. Marker genes of each cluster were selected based on unique (+) or increased (high) relative expression of the selected gene in its distinct cluster (**Figure S3A, B**). For further analysis of chondrocyte populations, $Acan^{+}$ and $Runx2^{high}$ clusters (2, 8, 23, 25) were selected, re-clustered at a resolution of 0.7, and annotated using *FindConservedMarkers*. Supervised clustering revealed six distinct clusters (**Figure S3C, D**). Subsequently, only the $Col2a1^{+}$ clusters (0, 1, 5; black dashed lines) were selected for reclustering and further downstream analyses. Only the top-upregulated genes were selected for cluster annotation (**Figure S3E, F**). To further characterize the biological processes associated with each conserved cell population, Gene Ontology (GO) enrichment analysis was performed on the top 70 upregulated genes from each cluster using the R packages *clusterProfiler* and *org.Mm.eg.db*. Biological Process (BP) terms were identified using the *enrichGO* function with Benjamini–Hochberg correction for multiple testing. Redundant GO terms were reduced using the *simplify* function, and enriched pathways were visualized using GO dot plots based on GeneRatio values. DEGs between WT and cKO mice were additionally visualized using volcano plots generated with the *EnhancedVolcano* R package.

**Spatial transcriptomics (Visium HD)**

Spatial Transcriptomics was conducted using the Visium HD Spatial Gene Expression System FFPE (formalin-fixed, paraffin-embedded) workflow (10x Genomics). 4-wk-old skulls *Prdm16* cKO and WT (n=2/genotype) were harvested and immediately fixed in 10% neutral-buffered formalin (Sigma HT501128) at 4°C for 16 hours with gentle rocking. Samples were thoroughly washed with Milli-Q water (10 min x 3 times, and 5 min x 3 times) at 4°C, and then decalcified with Formic Acid Bone Decalcifier (Immunocal, StatLab UN3412) for 16 hours at 4°C with gentle rocking. After decalcification, samples were thoroughly washed with Milli-Q water (10 min x 3 times, and 5 min x 3 times) at 4°C and stored in 70% EtOH at 4°C for less than 12 hours.[8] Samples were submitted to the URMC Histology, Biochemistry, and Molecular Imaging (HBMI) Core for paraffin processing. Samples were immediately embedded after processing. Tissue quality checks and gene expression assays were carried out according to the manufacturer's protocol.

This protocol generated an RNA integrity DV200 score > 75%. 5 $\mu$m FFPE sections were placed in the allowable area of Nexterion slide h 3-d slides according to manufacturer's instructions (10X Genomics, CG000548). Following tissue placement, slides were dried and placed in a desiccator at room temperature before deparaffinization and H&E staining according to the manufacturer's instructions (10X Genomics, CG000684). Slides were imaged using a VS120 Slide Scanner (Olympus) at 20X magnification. Sequence-ready libraries were then constructed according to the manufacturer's instructions (10X Genomics, CG000685). Final libraries were sequenced on the NovaSeq X Plus sequencer (Illumina, San Diego, CA) to obtain at least 275M reads per area of covered tissue. After sequence alignment and image mapping, this workflow detected 19,033 genes within the 250,617 binned squares (8 $\mu$m) under tissue.

***Selecting the region of interest and quality metric analysis***

To subset the nasal septal cartilage connected to the perpendicular plate of the ethmoid bone (PPE), regions of interest (ROI) in WT and *Prdm16* cKO mice were manually selected using the Loupe Browser version 9. Raw counts of selected spatial spots were loaded into Seurat using the *Load10X_Spatial* function, and quality control was performed based on mitochondrial gene percentage, using thresholds of <27% for both WT samples, <27% for cKO sample 1, and <32% for cKO sample 2, respectively. WT and cKO data were then normalized using the *SCTransform* function (v2, regularization). We integrated the WT and *Prdm16* cKO mice datasets using *PrepSCTIntegration* and *Seurat Wrappers* R packages. To explore cell composition, *FindNeighbors* and *FindClusters* were run at a resolution of 0.1 to identify distinct clusters and annotated using *FindConservedMarkers* function. Marker genes of each cluster were selected based on unique (+) or increased (high) relative expression of the selected gene in its distinct cluster. (**Figure S4A**) The top 50 upregulated genes in both WT and *Prdm16* cKO groups were confirmed through *VlnPlot*, and the locations of each cell population were visualized using *SpatialDimPlot*.

### *Intercellular communication analysis using MultiNicheNet*

We next used the MultiNicheNet R package to investigate how the loss of PRDM16 alters cell-cell interactions between nasal septal chondrocytes and fibrotic tissue located within the nasal bone.[9,10] The integrated Seurat object containing both WT and cKO datasets was loaded, and clusters corresponding to *Mgp*$^{+}$ chondrocytes (cluster 2) and *Col1a1*$^{high}$/ *Col3a1*$^{+}$ fibrotic cells (cluster 1) were selected for downstream MultiNicheNet analysis. Samples with fewer than 10 cells (*min_cells = 10*) were excluded from the analysis for that specific cell type.

### *Gene filtering and differential expression analysis*

Prior to performing differential expression (DE) analysis, genes with insufficient expression were removed using an adapted version of the *edgeR::filterByExpr* procedure tailored for single-cell datasets. Next, we performed gene filtering within each cell type to ensure that only biologically relevant and consistently detectable genes were retained. To determine whether a gene was considered “expressed” in a given cell type, we required that it be expressed in at least one condition, corresponding to a setting of *min_sample_prop* = 1 and *fraction_cutoff* = 0.0005. Only genes expressed in at least one cell type were retained for downstream analysis. We next calculated and normalized per-condition pseudobulk expression levels for each expressed gene in each cell type. The *process_abundance_expression_info* function then linked ligand expression in sender cell types to the corresponding receptor expression in receiver cell types, enabling downstream assessment of ligand-receptor specificity. Genome-wide differential expression (DE) analysis was performed to identify genes with altered expression between experimental conditions in both sender and receiver cell types. Differentially expressed (DE) genes were identified for each cell type across contrasts using *FindMarkers* (*logFC_threshold = 0.014, and p_val_threshold* = *1.5*). The resulting DE gene sets served as the "genesets of interest" for subsequent ligand activity prediction.

***Ligand activity analysis and ligand-target Inference (gene regulatory network)***

For ligand-target inference (i.e., predicted active genes downstream of a ligand-induced signaling pathway), the top 250 predicted target genes per ligand were considered (*top_n_target = 250*). Only ligands predicted to upregulate target genes were considered (*ligand_activity_down = FALSE)* to simplify interpretability. Next, differential expression and ligand activity were integrated to show ligand-receptor interactions which were ranked using a composite prioritization score. The score incorporated: (i) upregulation of ligands in sender cells and/or receptors in receiver cells, (ii) cell-type–specific expression of ligands and receptors, and (iii) NicheNet ligand activity scores. Prioritization was performed using *scenario = "no_frac_LR_expr"*. Top 50 prioritized interactions across all contrasts were visualized using ChordDiagram circos plots and bubble plots. Finally, the across-sample correlation between ligand-receptor pairs and inferred downstream target genes was calculated. The top 50 intercellular regulatory network (ligands of sender cell types, their ligand/receptor-annotated target genes in receiver cell types) was constructed.
