## Supplementary Table 1 for "Loss of PRDM16 Drives Nasal Septal Deviation through Dysregulated TGFβ2 Signaling"

**Supplementary Table 1: Immunohistochemistry antibody details**

| Antibody name | Dilution | Catalog number | Company |
|---|---|---|---|
| RUNX2 | 1:100 | MA5-41185 | Invitrogen |
| MKi67 | 1:150 | NB500-170 | R&D Systems |
| COL10A1 | 1:100 | PA5-115039 | Invitrogen |
| COL2A1 | 1:200 | PA126206 | Invitrogen |
| MMP13 | 1:200 | AB39012 | Abcam |
| Cleaved CASP3 | 1:600 | 9661S | Cell Signaling Technology |
| TGFβ2 | 1:400 | AB269279 | Abcam |
