## Supplementary File 1 for "Loss of PRDM16 Drives Nasal Septal Deviation through Dysregulated TGFβ2 Signaling"

**Cell cluster composition: scRNA seq 4-wk_WT & cKO cart_28clusters**

| Group | Cluster | Cell Count | Percentage |
|---|---|---|---|
| cKO cart | 0 | 139 | 8.162066941 |
| WT cart | 0 | 404 | 9.319492503 |
| cKO cart | 1 | 133 | 7.809747504 |
| WT cart | 1 | 388 | 8.950403691 |
| cKO cart | 2 | 244 | 14.32765708 |
| WT cart | 2 | 268 | 6.182237601 |
| cKO cart | 3 | 43 | 2.52495596 |
| WT cart | 3 | 415 | 9.573241061 |
| cKO cart | 4 | 115 | 6.752789196 |
| WT cart | 4 | 286 | 6.597462514 |
| cKO cart | 5 | 119 | 6.98766882 |
| WT cart | 5 | 203 | 4.682814302 |
| cKO cart | 6 | 75 | 4.403992954 |
| WT cart | 6 | 221 | 5.098039216 |
| cKO cart | 7 | 3 | 0.176159718 |
| WT cart | 7 | 262 | 6.043829296 |
| cKO cart | 8 | 128 | 7.516147974 |
| WT cart | 8 | 102 | 2.352941176 |
| cKO cart | 9 | 132 | 7.751027598 |
| WT cart | 9 | 74 | 1.707035755 |
| cKO cart | 10 | 61 | 3.581914269 |
| WT cart | 10 | 143 | 3.298731257 |
| cKO cart | 11 | 34 | 1.996476806 |
| WT cart | 11 | 152 | 3.506343714 |
| cKO cart | 12 | 38 | 2.23135643 |
| WT cart | 12 | 144 | 3.321799308 |
| cKO cart | 13 | 60 | 3.523194363 |
| WT cart | 13 | 114 | 2.629757785 |
| cKO cart | 14 | 34 | 1.996476806 |
| WT cart | 14 | 136 | 3.137254902 |
| cKO cart | 15 | 33 | 1.9377569 |
| WT cart | 15 | 137 | 3.160322953 |
| cKO cart | 16 | 50 | 2.935995302 |
| WT cart | 16 | 111 | 2.560553633 |
| cKO cart | 17 | 15 | 0.880798591 |
| WT cart | 17 | 119 | 2.745098039 |
| cKO cart | 18 | 28 | 1.644157369 |
| WT cart | 18 | 105 | 2.422145329 |
| cKO cart | 19 | 48 | 2.81855549 |
| WT cart | 19 | 81 | 1.868512111 |
| cKO cart | 20 | 20 | 1.174398121 |
| WT cart | 20 | 99 | 2.283737024 |
| cKO cart | 21 | 17 | 0.998238403 |

| | | | |
|---|---|---|---|
| WT cart | 21 | 76 | 1.753171857 |
| cKO cart | 22 | 15 | 0.880798591 |
| WT cart | 22 | 72 | 1.660899654 |
| cKO cart | 23 | 30 | 1.761597181 |
| WT cart | 23 | 51 | 1.176470588 |
| cKO cart | 24 | 10 | 0.58719906 |
| WT cart | 24 | 63 | 1.453287197 |
| cKO cart | 25 | 13 | 0.763358779 |
| WT cart | 25 | 54 | 1.24567474 |
| cKO cart | 26 | 28 | 1.644157369 |
| WT cart | 26 | 38 | 0.876585928 |
| cKO cart | 27 | 38 | 2.23135643 |
| WT cart | 27 | 17 | 0.392156863 |

**Cell cluster composition: scRNA seq 4-wk_WT & cKO cart_6clusters**

| Group | Cluster | Cell_Count | Percentage |
|---|---|---|---|
| cKO cart | 0 | 100 | 24.09638554 |
| WT cart | 0 | 252 | 53.05263158 |
| cKO cart | 1 | 117 | 28.19277108 |
| WT cart | 1 | 10 | 2.105263158 |
| cKO cart | 2 | 55 | 13.25301205 |
| WT cart | 2 | 54 | 11.36842105 |
| cKO cart | 3 | 67 | 16.14457831 |
| WT cart | 3 | 37 | 7.789473684 |
| cKO cart | 4 | 54 | 13.01204819 |
| WT cart | 4 | 82 | 17.26315789 |
| cKO cart | 5 | 22 | 5.301204819 |
| WT cart | 5 | 40 | 8.421052632 |

**Cell cluster composition: scRNA seq 4-wk_WT & cKO cart_4clusters**

| Group | Cluster | Cell Count | Percentage |
|---|---|---|---|
| WT cart | 0 | 238 | 71.257485 |
| cKO cart | 0 | 96 | 33.217993 |
| WT cart | 1 | 10 | 2.994012 |
| cKO cart | 1 | 119 | 41.176471 |
| WT cart | 2 | 41 | 12.275449 |
| cKO cart | 2 | 53 | 18.3391 |
| WT cart | 3 | 45 | 13.473054 |
| cKO cart | 3 | 21 | 7.266436 |

scRNA seq_Col2a1/Acan enriched cluster subset: cluster 0

| | cKO cart_p_val | cKO cart_avg_log2FC | cKO cart_pct.1 | cKO cart_pct.2 | cKO cart_p_val_adj | WT cart_p_val | WT cart_avg_log2FC | WT cart_pct.1 | WT cart_pct.2 | WT cart_p_val_adj | max_pval | minimump_p_val |
|---|---|---|---|---|---|---|---|---|---|---|---|---|
| Snorc | 6.6584E-10 | 0.95454321 | 0.99 | 0.736 | 0.000016175 | 1.2624E-32 | 2.66481037 | 0.979 | 0.427 | 3.0667E-28 | 6.6584E-10 | 2.5249E-32 |
| Plac9 | 2.09E-31 | 3.32440456 | 0.958 | 0.399 | 5.0771E-27 | 1.0673E-16 | 1.75623393 | 0.887 | 0.396 | 2.5927E-12 | 1.0673E-16 | 4.18E-31 |
| Fbln7 | 2.5718E-31 | 3.06813589 | 0.854 | 0.171 | 6.2474E-27 | 2.3972E-14 | 1.31008338 | 0.887 | 0.479 | 5.8232E-10 | 2.3972E-14 | 5.1436E-31 |
| Chrdl1 | 9.8831E-31 | 2.61438768 | 0.969 | 0.399 | 2.4008E-26 | 4.3971E-12 | 1.51230834 | 0.744 | 0.292 | 1.0682E-07 | 4.3971E-12 | 1.9766E-30 |
| Cnmd | 3.4301E-15 | 0.78450025 | 1 | 0.948 | 8.3324E-11 | 1.5202E-30 | 1.90801502 | 1 | 0.729 | 3.6928E-26 | 3.4301E-15 | 3.0403E-30 |
| Col9a2 | 3.1888E-07 | 0.61577256 | 1 | 0.922 | 0.00774613 | 4.6907E-28 | 1.86706228 | 1 | 0.729 | 1.1395E-23 | 3.1888E-07 | 9.3814E-28 |
| Mia | 0.000025555 | 0.45835473 | 1 | 0.891 | 0.62078073 | 9.2583E-28 | 1.94865852 | 0.987 | 0.51 | 2.249E-23 | 2.5555E-05 | 1.8517E-27 |
| Ucma | 0.10647911 | -0.529612 | 0.948 | 0.813 | 1 | 9.4502E-28 | 2.50647857 | 0.95 | 0.323 | 2.2956E-23 | 0.10647911 | 1.89E-27 |
| Col2a1 | 6.8163E-10 | 1.13600222 | 1 | 0.927 | 0.000016558 | 2.0798E-26 | 2.01959749 | 0.996 | 0.771 | 5.0523E-22 | 6.8163E-10 | 4.1596E-26 |
| Vit | 1.4847E-25 | 3.16503581 | 0.729 | 0.13 | 3.6067E-21 | 1.5248E-12 | 1.86232571 | 0.643 | 0.198 | 3.704E-08 | 1.5248E-12 | 2.9694E-25 |
| Rps3a1 | 0.00043159 | 0.38013184 | 1 | 0.99 | 1 | 1.813E-25 | 1.23187374 | 1 | 0.979 | 4.4043E-21 | 0.00043159 | 3.6261E-25 |
| Col9a1 | 2.5318E-18 | 1.38768877 | 1 | 0.798 | 6.1502E-14 | 4.1556E-25 | 1.6228757 | 1 | 0.76 | 1.0095E-20 | 2.5318E-18 | 8.3112E-25 |
| Susd5 | 0.00004742 | 0.47917562 | 0.99 | 0.896 | 1 | 5.9717E-25 | 1.64046804 | 0.979 | 0.615 | 1.4506E-20 | 0.00004742 | 1.1943E-24 |
| Ecrg4 | 1.1583E-24 | 2.22506672 | 0.969 | 0.611 | 2.8136E-20 | 5.0768E-20 | 2.05692707 | 0.912 | 0.385 | 1.2332E-15 | 5.0768E-20 | 2.3165E-24 |
| Maged2 | 2.9959E-06 | 0.64463976 | 0.969 | 0.772 | 0.07277534 | 4.4873E-24 | 1.91451752 | 0.954 | 0.417 | 1.0901E-19 | 2.9959E-06 | 8.9746E-24 |
| Col9a3 | 2.9009E-07 | 0.80164381 | 1 | 0.876 | 0.00704686 | 4.5738E-24 | 1.94214691 | 0.996 | 0.667 | 1.1111E-19 | 2.9009E-07 | 9.1477E-24 |
| Mgp | 2.7694E-20 | 1.66108062 | 1 | 0.969 | 6.7273E-16 | 5.4074E-24 | 1.83146656 | 1 | 0.969 | 1.3136E-19 | 2.7694E-20 | 1.0815E-23 |
| Eif1 | 0.0000279 | 0.37302854 | 1 | 0.995 | 0.67775208 | 1.3292E-23 | 1.08835272 | 1 | 0.99 | 3.2288E-19 | 0.0000279 | 2.6583E-23 |
| Fmod | 1.3665E-11 | 1.1072283 | 0.979 | 0.839 | 3.3195E-07 | 1.428E-23 | 1.8552275 | 0.954 | 0.604 | 3.4689E-19 | 1.3665E-11 | 2.856E-23 |
| Islr | 9.4913E-23 | 1.71907481 | 0.99 | 0.43 | 2.3056E-18 | 2.5568E-15 | 1.20998297 | 0.933 | 0.406 | 6.2109E-11 | 2.5568E-15 | 1.8983E-22 |
| Il17d | 4.6293E-10 | 1.16017799 | 0.948 | 0.694 | 0.000011245 | 1.1426E-22 | 1.98999033 | 0.954 | 0.417 | 2.7757E-18 | 4.6293E-10 | 2.2852E-22 |
| Omd | 2.8921E-06 | 1.10826561 | 0.99 | 0.948 | 0.07025575 | 1.4097E-22 | 1.93913058 | 0.979 | 0.75 | 3.4244E-18 | 2.8921E-06 | 2.8193E-22 |
| Rpl9 | 2.6985E-09 | 0.55734073 | 1 | 0.979 | 0.000065552 | 1.4272E-22 | 1.11263005 | 1 | 1 | 3.4669E-18 | 2.6985E-09 | 2.8544E-22 |
| Serpinh1 | 3.1818E-12 | 0.90177599 | 1 | 0.969 | 7.7291E-08 | 2.4099E-22 | 1.32799992 | 0.992 | 0.896 | 5.8541E-18 | 3.1818E-12 | 4.8198E-22 |
| Ostc | 8.0451E-13 | 0.9425052 | 0.979 | 0.891 | 1.9543E-08 | 5.7144E-22 | 1.38767202 | 0.983 | 0.771 | 1.3881E-17 | 8.0451E-13 | 1.1429E-21 |
| Tpt1 | 0.00028489 | 0.3269259 | 1 | 1 | 1 | 9.143E-22 | 1.13057539 | 1 | 1 | 2.221E-17 | 0.00028489 | 1.8286E-21 |
| Rpl19 | 0.01400763 | 0.25545321 | 1 | 0.984 | 1 | 1.0562E-21 | 0.98390293 | 1 | 0.99 | 2.5658E-17 | 0.01400763 | 2.1125E-21 |
| Gdf10 | 1.2007E-21 | 3.01102623 | 0.76 | 0.254 | 2.9167E-17 | 0.01846182 | -0.3430221 | 0.387 | 0.208 | 1 | 0.01846182 | 2.4014E-21 |
| Rpl34 | 7.1893E-06 | 0.44256247 | 1 | 0.969 | 0.1746417 | 2.6899E-21 | 1.1722663 | 1 | 0.948 | 6.5343E-17 | 7.1893E-06 | 5.3798E-21 |
| Tnfrsf12a | 0.00743701 | 0.47057147 | 0.854 | 0.819 | 1 | 4.6956E-21 | 1.64229527 | 0.962 | 0.854 | 1.1407E-16 | 0.00743701 | 9.3913E-21 |
| Rps7 | 0.000035319 | 0.40595837 | 1 | 0.979 | 0.85796178 | 5.0133E-21 | 1.06889474 | 1 | 0.969 | 1.2178E-16 | 3.5319E-05 | 1.0027E-20 |
| Rplp1 | 5.6645E-11 | 0.67717697 | 1 | 1 | 0.000001376 | 5.3597E-21 | 1.16244944 | 1 | 0.99 | 1.302E-16 | 5.6645E-11 | 1.0719E-20 |
| Smoc2 | 1.0598E-11 | 1.91096642 | 0.844 | 0.642 | 2.5745E-07 | 6.0233E-21 | 2.42812189 | 0.891 | 0.562 | 1.4632E-16 | 1.0598E-11 | 1.2047E-20 |
| Sparc | 0.35532404 | -0.616719 | 1 | 1 | 1 | 6.6972E-21 | 1.1993106 | 1 | 1 | 1.6269E-16 | 0.35532404 | 1.3394E-20 |
| Dcn | 4.6218E-10 | 0.88735451 | 0.99 | 0.746 | 0.000011227 | 8.0537E-21 | 2.21486782 | 0.954 | 0.76 | 1.9564E-16 | 4.6218E-10 | 1.6107E-20 |
| Chad | 0.000067036 | 0.13119409 | 0.99 | 0.87 | 1 | 9.1605E-21 | 1.34595319 | 0.996 | 0.625 | 2.2253E-16 | 6.7036E-05 | 1.8321E-20 |
| Rps3 | 0.00001114 | 0.49245074 | 1 | 0.964 | 0.2706168 | 3.7434E-20 | 1.0225333 | 1 | 0.969 | 9.0934E-16 | 0.00001114 | 7.4867E-20 |
| Srpx | 5.3171E-20 | 2.36868206 | 0.719 | 0.176 | 1.2916E-15 | 3.5608E-11 | 2.50435333 | 0.496 | 0.094 | 8.6498E-07 | 3.5608E-11 | 1.0634E-19 |
| Rps4x | 0.000022827 | 0.43597316 | 1 | 0.984 | 0.5545215 | 6.7714E-20 | 1.09494528 | 1 | 0.958 | 1.6449E-15 | 2.2827E-05 | 1.3543E-19 |
| Gcnt2 | 0.00122745 | -1.6537036 | 0.156 | 0.337 | 1 | 7.217E-20 | -2.6634922 | 0.088 | 0.542 | 1.7532E-15 | 0.00122745 | 1.4434E-19 |
| Eef1a1 | 0.00006211 | 0.39255891 | 1 | 1 | 1 | 1.1441E-19 | 0.99719744 | 1 | 0.99 | 2.7794E-15 | 0.00006211 | 2.2883E-19 |
| Col11a2 | 0.00011124 | 0.4285339 | 1 | 0.922 | 1 | 1.1622E-19 | 1.20734888 | 0.996 | 0.781 | 2.8231E-15 | 0.00011124 | 2.3243E-19 |
| Prnp | 1.1672E-09 | 0.77901101 | 0.99 | 0.922 | 0.000028353 | 1.2928E-19 | 1.60189699 | 0.966 | 0.781 | 3.1405E-15 | 1.1672E-09 | 2.5856E-19 |
| Rps21 | 0.00172825 | 0.30658266 | 1 | 0.99 | 1 | 1.4623E-19 | 1.01761102 | 0.996 | 0.979 | 3.5521E-15 | 0.00172825 | 2.9245E-19 |
| Rpl10 | 0.00060579 | 0.35660524 | 1 | 0.984 | 1 | 1.6265E-19 | 1.02610808 | 1 | 0.979 | 3.951E-15 | 0.00060579 | 3.2529E-19 |
| Rplp0 | 0.00223628 | 0.35860829 | 1 | 0.995 | 1 | 2.3618E-19 | 1.05222996 | 1 | 0.99 | 5.7374E-15 | 0.00223628 | 4.7237E-19 |
| Serf2 | 1.4958E-06 | 0.49466586 | 1 | 0.979 | 0.03633579 | 3.0098E-19 | 0.92469546 | 1 | 0.99 | 7.3115E-15 | 1.4958E-06 | 6.0197E-19 |
| Selenok | 0.00036182 | 0.32179045 | 0.99 | 0.979 | 1 | 3.1608E-19 | 1.06217144 | 0.996 | 0.938 | 7.6783E-15 | 0.00036182 | 6.3217E-19 |
| Tubb2a | 0.00152091 | 0.18788505 | 0.948 | 0.886 | 1 | 3.5276E-19 | 1.61094764 | 0.941 | 0.792 | 8.5693E-15 | 0.00152091 | 7.0553E-19 |
| Rpl5 | 0.00013721 | 0.38119764 | 1 | 0.979 | 1 | 3.9974E-19 | 0.97796891 | 1 | 0.979 | 9.7106E-15 | 0.00013721 | 7.9949E-19 |
| Rps6 | 3.5528E-06 | 0.44841067 | 1 | 0.979 | 0.08630491 | 4.1948E-19 | 0.97482579 | 1 | 0.99 | 1.019E-14 | 3.5528E-06 | 8.3895E-19 |
| mt-Cytb | 0.04790041 | -0.4711631 | 1 | 1 | 1 | 4.6004E-19 | -1.345906 | 1 | 1 | 1.1175E-14 | 0.04790041 | 9.2008E-19 |
| Rpl12 | 6.6326E-06 | 0.45655517 | 1 | 0.964 | 0.16111973 | 4.9741E-19 | 1.04533639 | 0.996 | 0.979 | 1.2083E-14 | 6.6326E-06 | 9.9482E-19 |
| Rps27a | 0.000043885 | 0.38209944 | 0.99 | 0.984 | 1 | 5.1155E-19 | 0.99400665 | 1 | 0.969 | 1.2426E-14 | 4.3885E-05 | 1.0231E-18 |
| Chadl | 0.00010345 | 0.45626767 | 0.958 | 0.762 | 1 | 5.7382E-19 | 1.80944367 | 0.861 | 0.302 | 1.3939E-14 | 0.00010345 | 1.1476E-18 |
| Gna14 | 6.0175E-19 | -4.0732019 | 0.208 | 0.705 | 1.4618E-14 | 0.00066794 | -1.2987147 | 0.37 | 0.562 | 1 | 0.00066794 | 1.2035E-18 |
| Col11a1 | 0.000057798 | 0.69794586 | 1 | 1 | 1 | 7.8511E-19 | 1.16751619 | 1 | 0.958 | 1.9072E-14 | 5.7798E-05 | 1.5702E-18 |
| Meis2 | 0.02599035 | -1.0320338 | 0.031 | 0.109 | 1 | 8.0565E-19 | -2.5706195 | 0.097 | 0.552 | 1.9571E-14 | 0.02599035 | 1.6113E-18 |
| Rpl30 | 0.00582072 | 0.28672529 | 1 | 0.974 | 1 | 8.3997E-19 | 0.99268248 | 0.996 | 0.99 | 2.0405E-14 | 0.00582072 | 1.6799E-18 |
| Cytl1 | 0.09039432 | -1.3416921 | 1 | 0.938 | 1 | 9.9531E-19 | 1.32832142 | 0.983 | 0.719 | 2.4178E-14 | 0.09039432 | 1.9906E-18 |
| Rpl28 | 0.000076454 | 0.40412679 | 1 | 0.959 | 1 | 9.9893E-19 | 0.94570115 | 1 | 0.969 | 2.4266E-14 | 7.6454E-05 | 1.9979E-18 |
| Cd63 | 1.6061E-09 | 0.59322138 | 1 | 0.99 | 0.000039016 | 1.0664E-18 | 1.02543705 | 1 | 0.979 | 2.5905E-14 | 1.6061E-09 | 2.1328E-18 |
| Rplp2 | 0.05141302 | 0.20395716 | 1 | 0.969 | 1 | 1.2079E-18 | 0.95317914 | 1 | 0.979 | 2.9342E-14 | 0.05141302 | 2.4158E-18 |
| Peg3 | 1.4267E-18 | 1.28483817 | 0.958 | 0.974 | 3.4658E-14 | 1.58E-13 | 0.83994519 | 1 | 0.979 | 3.838E-09 | 1.58E-13 | 2.8535E-18 |
| Rpl15 | 7.7761E-07 | 0.47519591 | 1 | 0.959 | 0.01888974 | 1.6443E-18 | 0.98341524 | 1 | 0.99 | 3.9944E-14 | 7.7761E-07 | 3.2886E-18 |
| Fndc1 | 1.945E-18 | -4.2805476 | 0.083 | 0.627 | 4.7248E-14 | 0.50473628 | -1.2045023 | 0.345 | 0.323 | 1 | 0.50473628 | 3.89E-18 |
| Pawr | 0.00397232 | -1.8150468 | 0.198 | 0.352 | 1 | 2.0011E-18 | -2.2587811 | 0.235 | 0.74 | 4.8612E-14 | 0.00397232 | 4.0023E-18 |
| Rps9 | 0.00059803 | 0.31103924 | 1 | 0.979 | 1 | 2.3125E-18 | 0.89383854 | 1 | 0.958 | 5.6176E-14 | 0.00059803 | 4.6251E-18 |
| Rack1 | 0.00136377 | 0.32631672 | 0.979 | 0.979 | 1 | 2.5164E-18 | 0.94268677 | 1 | 0.958 | 6.1128E-14 | 0.00136377 | 5.0328E-18 |
| Celf2 | 0.06157414 | -0.6929452 | 0.052 | 0.124 | 1 | 2.5696E-18 | -3.2139952 | 0.113 | 0.552 | 6.2421E-14 | 0.06157414 | 5.1392E-18 |
| Rps10 | 0.01487324 | 0.27221053 | 1 | 0.99 | 1 | 3.0018E-18 | 0.9618573 | 1 | 0.99 | 7.2919E-14 | 0.01487324 | 6.0035E-18 |
| Gng5 | 0.00068342 | 0.33260544 | 1 | 0.959 | 1 | 3.3336E-18 | 1.06646797 | 0.987 | 0.938 | 8.098E-14 | 0.00068342 | 6.6672E-18 |
| Comp | 2.0387E-07 | 1.54566783 | 0.979 | 0.855 | 0.00495242 | 3.6383E-18 | 1.22771518 | 0.971 | 0.635 | 8.8381E-14 | 2.0387E-07 | 7.2766E-18 |
| Bsg | 0.000016714 | 0.42918348 | 0.99 | 0.969 | 0.40602223 | 3.9476E-18 | 1.05835101 | 1 | 0.99 | 9.5894E-14 | 1.6714E-05 | 7.8951E-18 |
| Rpl6 | 0.000056707 | 0.3762556 | 1 | 0.974 | 1 | 4.5397E-18 | 0.92437449 | 1 | 0.969 | 1.1028E-13 | 5.6707E-05 | 9.0793E-18 |
| Mecom | 0.06470909 | -0.2809366 | 0.167 | 0.275 | 1 | 5.5803E-18 | -2.9797067 | 0.109 | 0.531 | 1.3556E-13 | 0.06470909 | 1.1161E-17 |
| Rpl21 | 0.00014026 | 0.36803239 | 1 | 0.979 | 1 | 6.4697E-18 | 0.88459677 | 1 | 0.969 | 1.5716E-13 | 0.00014026 | 1.2939E-17 |
| Malat1 | 1.2437E-11 | -1.0609544 | 1 | 1 | 3.0211E-07 | 9.5758E-18 | -1.1654628 | 1 | 1 | 2.3262E-13 | 1.2437E-11 | 1.9152E-17 |
| Vasn | 0.00515048 | 0.41686329 | 0.906 | 0.813 | 1 | 1.0458E-17 | 1.24285304 | 0.979 | 0.635 | 2.5406E-13 | 0.00515048 | 2.0917E-17 |
| Cst3 | 0.00145975 | -1.0618407 | 1 | 0.984 | 1 | 1.204E-17 | 0.99934617 | 1 | 0.979 | 2.9247E-13 | 0.00145975 | 2.4079E-17 |
| Pclo | 0.2050453 | -1.8993961 | 0.01 | 0.036 | 1 | 1.2087E-17 | -2.2818409 | 0.059 | 0.458 | 2.9361E-13 | 0.2050453 | 2.4173E-17 |
| Rps13 | 3.4865E-06 | 0.4462363 | 1 | 0.959 | 0.08469387 | 1.2812E-17 | 0.94191315 | 0.996 | 0.958 | 3.1122E-13 | 3.4865E-06 | 2.5623E-17 |
| Rpl35a | 0.00035076 | 0.35133088 | 1 | 0.953 | 1 | 1.4927E-17 | 0.92636501 | 1 | 0.969 | 3.6261E-13 | 0.00035076 | 2.9854E-17 |
| Thbs1 | 2.4655E-11 | 1.35921979 | 1 | 0.917 | 5.9893E-07 | 1.6009E-17 | 1.47708213 | 0.979 | 0.938 | 3.8889E-13 | 2.4655E-11 | 3.2018E-17 |
| Rpl11 | 0.000040954 | 0.39100795 | 1 | 0.974 | 0.99485379 | 1.7744E-17 | 0.9159635 | 1 | 0.969 | 4.3103E-13 | 4.0954E-05 | 3.5488E-17 |
| Rps18 | 4.2737E-07 | 0.49842965 | 1 | 0.974 | 0.01038162 | 2.0813E-17 | 0.88767181 | 1 | 0.969 | 5.0558E-13 | 4.2737E-07 | 4.1625E-17 |
| Rpl23 | 0.07902301 | 0.18391499 | 1 | 0.99 | 1 | 2.0916E-17 | 0.95338858 | 1 | 1 | 5.0809E-13 | 0.07902301 | 4.1832E-17 |
| Rps2 | 0.00002243 | 0.43483875 | 1 | 0.984 | 0.54487203 | 2.1165E-17 | 0.90202644 | 1 | 0.99 | 5.1413E-13 | 0.00002243 | 4.2329E-17 |
| Rpsa | 4.7142E-10 | 0.66917895 | 1 | 0.979 | 0.000011452 | 2.1574E-17 | 0.98009652 | 1 | 0.99 | 5.2408E-13 | 4.7142E-10 | 4.3148E-17 |
| Adgrl3 | 1.927E-11 | -5.903398 | 0.021 | 0.394 | 4.681E-07 | 2.4969E-17 | -2.8208172 | 0.059 | 0.448 | 6.0656E-13 | 1.927E-11 | 4.9939E-17 |

| | | | | | | | | | | | | |
|---|---|---|---|---|---|---|---|---|---|---|---|---|
| Rpl27a | 0.0199216 | 0.26739343 | 1 | 0.964 | 1 | 2.6885E-17 | 0.87959801 | 1 | 0.979 | 6.5309E-13 | 0.0199216 | 5.377E-17 |
| Eif1a | 0.00025124 | 0.44705097 | 0.917 | 0.969 | 1 | 3.1338E-17 | 1.02812077 | 0.996 | 0.958 | 7.6127E-13 | 0.00025124 | 6.2677E-17 |
| Rpl39 | 0.0457909 | 0.2078486 | 1 | 0.959 | 1 | 3.4096E-17 | 0.95735269 | 1 | 0.958 | 8.2826E-13 | 0.0457909 | 6.8192E-17 |
| Jund | 2.6674E-07 | 0.68780491 | 0.99 | 0.984 | 0.00647967 | 3.6452E-17 | 1.04513458 | 1 | 0.979 | 8.8549E-13 | 2.6674E-07 | 7.2904E-17 |
| Rps14 | 0.000014919 | 0.42354428 | 1 | 0.995 | 0.36242237 | 3.7058E-17 | 0.90985622 | 1 | 1 | 9.0022E-13 | 1.4919E-05 | 7.4116E-17 |
| Rps16 | 0.00191987 | 0.33624789 | 1 | 0.979 | 1 | 3.8395E-17 | 0.91177433 | 1 | 0.979 | 9.3268E-13 | 0.00191987 | 7.6789E-17 |
| Acan | 0.29721142 | -0.2217464 | 1 | 0.969 | 1 | 4.0665E-17 | 0.7706411 | 0.996 | 0.885 | 9.8783E-13 | 0.29721142 | 8.133E-17 |
| Prelp | 0.00724893 | -0.5164218 | 0.927 | 0.87 | 1 | 4.8939E-17 | 1.39171643 | 0.916 | 0.469 | 1.1888E-12 | 0.00724893 | 9.7878E-17 |
| Fxyd5 | 5.0262E-17 | 1.32382717 | 0.958 | 0.617 | 1.221E-12 | 2.5447E-06 | 0.83013245 | 0.84 | 0.656 | 0.06181615 | 2.5447E-06 | 1.0052E-16 |
| Eif4a1 | 0.000073202 | 0.41395475 | 0.979 | 0.984 | 1 | 5.7161E-17 | 0.92524821 | 1 | 0.979 | 1.3886E-12 | 7.3202E-05 | 1.1432E-16 |
| Rps23 | 0.00203545 | 0.32868449 | 1 | 0.99 | 1 | 6.795E-17 | 0.92198079 | 1 | 0.958 | 1.6507E-12 | 0.00203545 | 1.359E-16 |
| Rps11 | 0.01558282 | 0.23226961 | 1 | 0.969 | 1 | 7.0446E-17 | 0.8983889 | 1 | 0.969 | 1.7113E-12 | 0.01558282 | 1.4089E-16 |
| Maged1 | 1.5316E-06 | 0.53389314 | 1 | 0.917 | 0.03720623 | 7.676E-17 | 1.10719096 | 0.987 | 0.802 | 1.8647E-12 | 1.5316E-06 | 1.5352E-16 |
| Esrp1 | 0.20784723 | -1.1624305 | 0.01 | 0.036 | 1 | 7.8451E-17 | -2.9575535 | 0.055 | 0.427 | 1.9057E-12 | 0.20784723 | 1.569E-16 |
| Atf4 | 1.5478E-10 | 0.81816982 | 0.99 | 0.969 | 3.7599E-06 | 8.5749E-17 | 1.01106712 | 1 | 0.958 | 2.083E-12 | 1.5478E-10 | 1.715E-16 |
| Ncmap | 1.2862E-06 | 0.69852002 | 0.99 | 0.741 | 0.03124451 | 8.909E-17 | 1.55556872 | 0.878 | 0.375 | 2.1642E-12 | 1.2862E-06 | 1.7818E-16 |
| Rpl7 | 5.9058E-08 | 0.52076935 | 1 | 0.974 | 0.00143464 | 9.0025E-17 | 0.98069853 | 1 | 0.948 | 2.1869E-12 | 5.9058E-08 | 1.8005E-16 |
| Emilin1 | 9.1235E-17 | 1.80424136 | 0.844 | 0.394 | 2.2163E-12 | 1.1824E-10 | 1.04108392 | 0.882 | 0.448 | 2.8722E-06 | 1.1824E-10 | 1.8247E-16 |
| Pkdcc | 9.677E-17 | 3.30027629 | 0.583 | 0.14 | 2.3507E-12 | 0.0005371 | 1.25547843 | 0.412 | 0.208 | 1 | 0.0005371 | 1.9354E-16 |
| Cnbp | 9.1388E-08 | 0.50716598 | 0.99 | 0.974 | 0.00222001 | 1.0201E-16 | 0.77515749 | 1 | 0.979 | 2.478E-12 | 9.1388E-08 | 2.0402E-16 |
| 2900026A02Rik | 0.06422465 | -0.8993961 | 0.073 | 0.15 | 1 | 1.1591E-16 | -3.368749 | 0.038 | 0.385 | 2.8156E-12 | 0.06422465 | 2.3181E-16 |
| Sfr1 | 1.8396E-06 | 0.5006911 | 0.969 | 0.953 | 0.04468821 | 1.2804E-16 | 1.07976242 | 0.992 | 0.927 | 3.1103E-12 | 1.8396E-06 | 2.5607E-16 |
| Ddx5 | 7.0364E-06 | 0.43624934 | 1 | 0.995 | 0.17092869 | 1.3345E-16 | 0.71804081 | 1 | 1 | 3.2419E-12 | 7.0364E-06 | 2.6691E-16 |
| Rpl32 | 3.1576E-06 | 0.46421861 | 1 | 0.974 | 0.0767053 | 1.4571E-16 | 0.88184446 | 1 | 0.958 | 3.5395E-12 | 3.1576E-06 | 2.9141E-16 |
| Rpl26 | 0.00023183 | 0.36659676 | 1 | 0.964 | 1 | 1.6358E-16 | 0.91408015 | 1 | 0.958 | 3.9738E-12 | 0.00023183 | 3.2717E-16 |
| Morf4l2 | 7.3849E-11 | 0.5424415 | 0.99 | 0.99 | 1.7939E-06 | 2.0566E-16 | 0.79036529 | 0.996 | 0.99 | 4.9959E-12 | 7.3849E-11 | 4.1132E-16 |
| Col10a1 | 2.1672E-16 | -3.6324004 | 0.167 | 0.642 | 5.2647E-12 | 0.32096281 | -2.823801 | 0.084 | 0.115 | 1 | 0.32096281 | 4.3345E-16 |
| Hapln1 | 0.000074974 | 0.67493924 | 1 | 0.948 | 1 | 2.1836E-16 | 0.64117717 | 0.996 | 0.76 | 5.3044E-12 | 7.4974E-05 | 4.3672E-16 |
| Nfkbiz | 6.3351E-06 | -1.4282024 | 0.365 | 0.632 | 0.1538928 | 2.2871E-16 | -1.9298242 | 0.248 | 0.729 | 5.5558E-12 | 6.3351E-06 | 4.5742E-16 |
| Rps12 | 0.00082212 | 0.36056351 | 1 | 0.979 | 1 | 2.3855E-16 | 0.91384979 | 1 | 0.99 | 5.795E-12 | 0.00082212 | 4.7711E-16 |
| Fmo2 | 0.22187388 | -1.3144336 | 0 | 0.016 | 1 | 2.4407E-16 | -3.7692869 | 0.021 | 0.333 | 5.929E-12 | 0.22187388 | 4.8814E-16 |
| Wdr72 | 0.53395584 | -0.240433 | 0.01 | 0.021 | 1 | 2.6545E-16 | -4.3493836 | 0.034 | 0.365 | 6.4484E-12 | 0.53395584 | 5.3091E-16 |
| Rps19 | 0.000080803 | 0.42540924 | 1 | 0.979 | 1 | 2.9432E-16 | 0.88985629 | 0.996 | 0.969 | 7.1495E-12 | 8.0803E-05 | 5.8863E-16 |
| Meis1 | 0.70862459 | -1.1217885 | 0.062 | 0.073 | 1 | 3.4142E-16 | -2.7351611 | 0.088 | 0.479 | 8.2939E-12 | 0.70862459 | 6.8285E-16 |
| mt-Nd4 | 0.06891387 | -0.4950972 | 1 | 1 | 1 | 3.5696E-16 | -1.3371115 | 1 | 1 | 8.6712E-12 | 0.06891387 | 7.1392E-16 |
| Rpl22 | 0.01820753 | 0.2710614 | 1 | 0.959 | 1 | 3.7051E-16 | 0.86804524 | 0.992 | 0.948 | 9.0005E-12 | 0.01820753 | 7.4102E-16 |
| Epb41l5 | 0.08282186 | -1.6929452 | 0.073 | 0.14 | 1 | 4.4041E-16 | -2.6508922 | 0.084 | 0.479 | 1.0699E-11 | 0.08282186 | 8.8083E-16 |
| Rpl10a | 5.0049E-08 | 0.56311308 | 1 | 0.974 | 0.00121578 | 4.5449E-16 | 0.86328287 | 0.996 | 0.979 | 1.104E-11 | 5.0049E-08 | 9.0898E-16 |
| Kdelr2 | 5.3023E-11 | 0.74446013 | 1 | 0.969 | 0.000001288 | 5.9234E-16 | 0.97447209 | 0.996 | 0.917 | 1.4389E-11 | 5.3023E-11 | 1.1847E-15 |
| Rpl17 | 0.000019526 | 0.46453761 | 1 | 0.984 | 0.47432751 | 6.3122E-16 | 0.89274826 | 1 | 1 | 1.5334E-11 | 1.9526E-05 | 1.2624E-15 |
| Etl4 | 0.32750729 | 0.65957123 | 0.344 | 0.306 | 1 | 7.6363E-16 | -2.1649069 | 0.273 | 0.74 | 1.855E-11 | 0.32750729 | 1.5273E-15 |
| Rpl7a | 0.0008821 | 0.33077762 | 1 | 0.953 | 1 | 7.9217E-16 | 0.80020901 | 1 | 0.969 | 1.9243E-11 | 0.0008821 | 1.5843E-15 |
| Morf4l1 | 0.0001543 | 0.31243979 | 1 | 0.979 | 1 | 8.8835E-16 | 0.66403885 | 1 | 0.99 | 2.158E-11 | 0.0001543 | 1.7767E-15 |
| Fbxo32 | 0.07703983 | -1.10213 | 0.104 | 0.181 | 1 | 1.0651E-15 | -1.1432054 | 0.118 | 0.552 | 2.5873E-11 | 0.07703983 | 2.1302E-15 |
| mt-Nd1 | 0.11396548 | -0.4715987 | 1 | 1 | 1 | 1.1788E-15 | -1.2620113 | 1 | 1 | 2.8636E-11 | 0.11396548 | 2.3577E-15 |
| Itm2a | 1.2117E-07 | 0.31670271 | 1 | 0.902 | 0.00294337 | 1.347E-15 | 1.30723144 | 0.971 | 0.542 | 3.2721E-11 | 1.2117E-07 | 2.694E-15 |
| Ecm2 | 2.4381E-07 | 0.88664014 | 0.844 | 0.554 | 0.00592273 | 1.596E-15 | 1.49012013 | 0.924 | 0.531 | 3.877E-11 | 2.4381E-07 | 3.192E-15 |
| Tmed9 | 7.5249E-08 | 0.43232513 | 0.99 | 0.974 | 0.00182795 | 2.1703E-15 | 0.92587833 | 0.996 | 0.938 | 5.272E-11 | 7.5249E-08 | 4.3405E-15 |
| Vkorc1 | 4.7341E-07 | 0.6623201 | 0.969 | 0.86 | 0.01150014 | 2.3084E-15 | 1.19924278 | 0.95 | 0.667 | 5.6075E-11 | 4.7341E-07 | 4.6168E-15 |
| Rpl14 | 0.00030246 | 0.40743001 | 1 | 0.974 | 1 | 2.3508E-15 | 0.84294911 | 1 | 0.979 | 5.7105E-11 | 0.00030246 | 4.7015E-15 |
| Chst5 | 2.4089E-15 | -3.6949255 | 0.188 | 0.632 | 5.8517E-11 | 0.50917174 | -0.6538097 | 0.139 | 0.104 | 1 | 0.50917174 | 4.8178E-15 |
| Unc13b | 0.19908094 | -0.8225805 | 0.042 | 0.083 | 1 | 2.4621E-15 | -2.1678363 | 0.084 | 0.469 | 5.9809E-11 | 0.19908094 | 4.9242E-15 |
| Rps8 | 1.2019E-06 | 0.49337995 | 1 | 0.99 | 0.02919641 | 2.4727E-15 | 0.88644982 | 1 | 0.99 | 6.0068E-11 | 1.2019E-06 | 4.9455E-15 |
| C1d | 0.03750267 | 0.24775763 | 0.823 | 0.705 | 1 | 2.5237E-15 | 1.46817558 | 0.845 | 0.438 | 6.1306E-11 | 0.03750267 | 5.0474E-15 |
| Pebp1 | 0.03608257 | 0.22885091 | 0.969 | 0.907 | 1 | 2.8395E-15 | 0.95253951 | 0.962 | 0.781 | 6.8977E-11 | 0.03608257 | 5.679E-15 |
| Eef1b2 | 0.00061566 | 0.32669096 | 1 | 0.969 | 1 | 2.8733E-15 | 0.90429374 | 1 | 0.958 | 6.9797E-11 | 0.00061566 | 5.7465E-15 |
| Rpl13a | 0.00184167 | 0.32361526 | 1 | 0.99 | 1 | 3.1141E-15 | 0.83462304 | 1 | 1 | 7.5647E-11 | 0.00184167 | 6.2282E-15 |
| Rpl8 | 0.000024649 | 0.43479658 | 1 | 0.959 | 0.59878351 | 3.2749E-15 | 0.81248969 | 1 | 1 | 7.9555E-11 | 2.4649E-05 | 6.5499E-15 |
| Rps17 | 0.000092722 | 0.39834516 | 1 | 0.953 | 1 | 3.2771E-15 | 0.93913137 | 0.996 | 0.927 | 7.9608E-11 | 9.2722E-05 | 6.5542E-15 |
| Rps15 | 8.964E-08 | 0.51759423 | 1 | 0.969 | 0.00217753 | 3.2789E-15 | 0.86030511 | 1 | 0.979 | 7.965E-11 | 8.964E-08 | 6.5577E-15 |
| Bcl11a | 0.2723155 | -1.9925055 | 0.021 | 0.047 | 1 | 3.2975E-15 | -2.5322477 | 0.042 | 0.375 | 8.0103E-11 | 0.2723155 | 6.595E-15 |
| Rps24 | 0.00026581 | 0.42077586 | 1 | 0.99 | 1 | 3.7103E-15 | 0.85539103 | 1 | 0.99 | 9.0131E-11 | 0.00026581 | 7.4206E-15 |
| Matn4 | 4.2072E-15 | 1.39775976 | 0.927 | 0.523 | 1.022E-10 | 1.6888E-10 | 0.9476544 | 0.958 | 0.688 | 4.1023E-06 | 1.6888E-10 | 8.4144E-15 |
| Ifitm3 | 0.00523704 | 0.1921733 | 0.156 | 0.321 | 1 | 4.2845E-15 | -3.0728161 | 0.181 | 0.604 | 1.0408E-10 | 0.00523704 | 8.569E-15 |
| Spcs1 | 0.0000114 | 0.42462745 | 0.99 | 0.933 | 0.2769358 | 4.6445E-15 | 1.02434246 | 0.971 | 0.885 | 1.1282E-10 | 0.0000114 | 9.2889E-15 |
| Ston2 | 0.000036524 | -3.1354634 | 0.083 | 0.29 | 0.88723068 | 4.8732E-15 | -2.9084927 | 0.076 | 0.448 | 1.1838E-10 | 3.6524E-05 | 9.7464E-15 |
| Fkbp9 | 2.041E-10 | 0.70664824 | 0.99 | 0.907 | 0.000004958 | 5.4234E-15 | 0.95119059 | 0.983 | 0.75 | 1.3174E-10 | 2.041E-10 | 1.0847E-14 |
| Rps25 | 0.01040808 | 0.26170602 | 1 | 0.979 | 1 | 5.4856E-15 | 0.86754099 | 1 | 0.948 | 1.3326E-10 | 0.01040808 | 1.0971E-14 |
| Norad | 7.5295E-08 | 0.715447 | 0.927 | 0.824 | 0.00182906 | 5.9747E-15 | 1.06218977 | 0.971 | 0.823 | 1.4514E-10 | 7.5295E-08 | 1.1949E-14 |
| Serpina3n | 6.0767E-15 | -4.0799683 | 0.167 | 0.611 | 1.4761E-10 | 0.29686694 | -1.8102843 | 0.055 | 0.083 | 1 | 0.29686694 | 1.2153E-14 |
| Sar1a | 1.6454E-10 | 0.72811214 | 0.969 | 0.855 | 3.9969E-06 | 6.4025E-15 | 1.00048486 | 0.979 | 0.833 | 1.5553E-10 | 1.6454E-10 | 1.2805E-14 |
| Adgrg6 | 7.3689E-15 | 1.3571723 | 0.896 | 0.487 | 1.7901E-10 | 1.6231E-07 | 0.72404894 | 0.853 | 0.51 | 0.00394281 | 1.6231E-07 | 1.4738E-14 |
| Snhg18 | 0.00367614 | 0.43607857 | 0.875 | 0.777 | 1 | 8.0306E-15 | 1.52797201 | 0.84 | 0.406 | 1.9508E-10 | 0.00367614 | 1.6061E-14 |
| Pstpip2 | 0.15990294 | -1.3461424 | 0.052 | 0.098 | 1 | 8.2007E-15 | -2.7981417 | 0.084 | 0.458 | 1.9921E-10 | 0.15990294 | 1.6401E-14 |
| Aox3 | 0.08967201 | -2.1079827 | 0.021 | 0.067 | 1 | 8.9157E-15 | -3.3422767 | 0.029 | 0.333 | 2.1658E-10 | 0.08967201 | 1.7831E-14 |
| Tceal9 | 2.055E-13 | 1.00885621 | 0.99 | 0.896 | 4.9919E-09 | 9.2519E-15 | 1.02298861 | 0.987 | 0.885 | 2.2475E-10 | 2.055E-13 | 1.8504E-14 |
| mt-Co3 | 0.29304601 | -0.3556736 | 1 | 1 | 1 | 9.796E-15 | -1.1108562 | 1 | 1 | 2.3797E-10 | 0.29304601 | 1.9592E-14 |
| Oat | 0.00390897 | 0.35933821 | 0.979 | 0.907 | 1 | 9.8374E-15 | 1.0112821 | 0.987 | 0.854 | 2.3897E-10 | 0.00390897 | 1.9675E-14 |
| Rps15a | 0.0020129 | 0.35201249 | 1 | 0.959 | 1 | 1.0111E-14 | 0.82631305 | 1 | 0.948 | 2.4561E-10 | 0.0020129 | 2.0221E-14 |
| Tmem251 | 0.00096075 | 0.54652075 | 0.594 | 0.378 | 1 | 1.03E-14 | 1.66950497 | 0.697 | 0.208 | 2.502E-10 | 0.00096075 | 2.06E-14 |
| Rpl37a | 0.00417654 | 0.26925036 | 1 | 0.964 | 1 | 1.1357E-14 | 0.84719994 | 1 | 0.948 | 2.7588E-10 | 0.00417654 | 2.2714E-14 |
| Rpl24 | 0.00392421 | 0.2835356 | 1 | 0.974 | 1 | 1.1433E-14 | 0.77975198 | 1 | 1 | 2.7773E-10 | 0.00392421 | 2.2866E-14 |
| Htra1 | 1.2531E-14 | 1.16687838 | 0.99 | 0.503 | 3.044E-10 | 7.7679E-06 | 0.62741669 | 0.966 | 0.76 | 0.18869695 | 7.7679E-06 | 2.5062E-14 |
| Fap | 1.3348E-14 | 1.32062918 | 0.958 | 0.56 | 3.2424E-10 | 3.3036E-06 | 0.6549669 | 0.966 | 0.76 | 0.0802507 | 3.3036E-06 | 2.6695E-14 |
| Sec61g | 3.4087E-08 | 0.70041172 | 1 | 0.912 | 0.00082804 | 1.3926E-14 | 1.06014757 | 0.971 | 0.812 | 3.3829E-10 | 3.4087E-08 | 2.7852E-14 |
| mt-Atp6 | 0.02786025 | -0.571496 | 1 | 1 | 1 | 1.7322E-14 | -1.1167921 | 1 | 1 | 4.2078E-10 | 0.02786025 | 3.4643E-14 |
| Armcx3 | 0.00083367 | 0.70644052 | 0.75 | 0.627 | 1 | 1.769E-14 | 1.50212369 | 0.836 | 0.385 | 4.2974E-10 | 0.00083367 | 3.5381E-14 |
| Fam180a | 1.3265E-10 | 1.25460182 | 0.865 | 0.544 | 3.2223E-06 | 1.8291E-14 | 1.58607612 | 0.912 | 0.542 | 4.4432E-10 | 1.3265E-10 | 3.6581E-14 |
| Bgn | 1.9619E-14 | 0.79192306 | 1 | 0.912 | 4.7659E-10 | 1.0301E-10 | 0.74352862 | 1 | 0.802 | 2.5022E-06 | 1.0301E-10 | 3.9239E-14 |
| Cotl1 | 0.23212365 | 0.48812038 | 0.219 | 0.166 | 1 | 1.9921E-14 | -1.5873892 | 0.147 | 0.562 | 4.8393E-10 | 0.23212365 | 3.9843E-14 |
| Ppl | 0.78764899 | -0.4519371 | 0.021 | 0.026 | 1 | 2.0408E-14 | -3.0824448 | 0.046 | 0.365 | 4.9576E-10 | 0.78764899 | 4.0817E-14 |

| | | | | | | | | | | | | |
|---|---|---|---|---|---|---|---|---|---|---|---|---|
| Hnrnpa0 | 1.5621E-09 | 0.71455864 | 0.979 | 0.891 | 0.000037947 | 2.1594E-14 | 0.83828429 | 0.996 | 0.906 | 5.2456E-10 | 1.5621E-09 | 4.3188E-14 |
| Rpl13 | 0.00455555 | 0.28732201 | 1 | 0.99 | 1 | 2.182E-14 | 0.83116344 | 1 | 0.99 | 5.3005E-10 | 0.00455555 | 4.364E-14 |
| Car8 | 1.0571E-07 | 1.53017276 | 0.719 | 0.461 | 0.00256785 | 2.3366E-14 | 1.3213427 | 0.866 | 0.385 | 5.6761E-10 | 1.0571E-07 | 4.6732E-14 |
| Ift20 | 2.5113E-14 | 1.06989107 | 0.969 | 0.829 | 6.1006E-10 | 2.1767E-10 | 0.88258394 | 0.945 | 0.792 | 5.2875E-06 | 2.1767E-10 | 5.0227E-14 |
| Nme2 | 1.2843E-07 | 0.59611539 | 0.99 | 0.953 | 0.00311975 | 2.7372E-14 | 0.9385438 | 1 | 0.927 | 6.6493E-10 | 1.2843E-07 | 5.4745E-14 |
| Rpl23a | 3.1237E-06 | 0.4689847 | 1 | 0.974 | 0.07588058 | 2.8113E-14 | 0.79976923 | 1 | 0.99 | 6.8292E-10 | 3.1237E-06 | 5.6226E-14 |
| Tmed10 | 0.03322098 | 0.20039758 | 1 | 0.99 | 1 | 2.8582E-14 | 0.80680459 | 0.992 | 0.958 | 6.943E-10 | 0.03322098 | 5.7163E-14 |
| Rps26 | 0.00011735 | 0.38452319 | 1 | 0.953 | 1 | 2.8879E-14 | 0.86918708 | 1 | 0.948 | 7.0152E-10 | 0.00011735 | 5.7757E-14 |
| Rps5 | 0.00051925 | 0.36919019 | 1 | 0.964 | 1 | 2.9864E-14 | 0.82835607 | 1 | 0.958 | 7.2546E-10 | 0.00051925 | 5.9728E-14 |
| Krt8 | 0.21868423 | -0.8405024 | 0.115 | 0.171 | 1 | 3.0798E-14 | -2.4318458 | 0.122 | 0.5 | 7.4815E-10 | 0.21868423 | 6.1597E-14 |
| Sec22b | 4.1002E-07 | 0.60325135 | 0.948 | 0.896 | 0.00996023 | 3.2072E-14 | 1.05478974 | 0.971 | 0.688 | 7.791E-10 | 4.1002E-07 | 6.4144E-14 |
| Aqp4 | 0.47294388 | -0.9925055 | 0.021 | 0.036 | 1 | 3.2728E-14 | -2.4167705 | 0.076 | 0.427 | 7.9504E-10 | 0.47294388 | 6.5457E-14 |
| Timp3 | 3.3509E-14 | -2.691659 | 0.865 | 0.995 | 8.14E-10 | 0.58285075 | -0.5157312 | 0.924 | 0.948 | 1 | 0.58285075 | 6.7018E-14 |
| Ost4 | 0.000072782 | 0.37097685 | 0.99 | 0.948 | 1 | 3.6449E-14 | 0.99991522 | 0.966 | 0.854 | 8.8542E-10 | 7.2782E-05 | 7.2898E-14 |
| Hs6st3 | 3.6684E-14 | -9.6920779 | 0 | 0.44 | 8.9113E-10 | 0.00097129 | -6.1257722 | 0.013 | 0.083 | 1 | 0.00097129 | 7.3368E-14 |
| Rpl36a | 0.00731794 | 0.29581643 | 0.99 | 0.964 | 1 | 3.7914E-14 | 0.90666059 | 0.996 | 0.938 | 9.2101E-10 | 0.00731794 | 7.5829E-14 |
| Tuba1c | 0.15542704 | 0.14033022 | 0.958 | 0.891 | 1 | 4.5052E-14 | 0.87565363 | 0.983 | 0.865 | 1.0944E-09 | 0.15542704 | 9.0104E-14 |
| Myo5b | 0.0303018 | 2.00749454 | 0.052 | 0.01 | 1 | 4.5143E-14 | -2.7386986 | 0.042 | 0.354 | 1.0966E-09 | 0.0303018 | 9.0285E-14 |
| Rpl3 | 0.000032542 | 0.46672591 | 1 | 0.969 | 0.79051099 | 4.9782E-14 | 0.75328433 | 1 | 0.979 | 1.2093E-09 | 3.2542E-05 | 9.9565E-14 |
| Rpl41 | 1.9528E-06 | 0.47092352 | 1 | 0.979 | 0.04743805 | 5.7122E-14 | 0.78774085 | 0.996 | 0.979 | 1.3876E-09 | 1.9528E-06 | 1.1424E-13 |
| Adh7 | 0.31369561 | -1.3771693 | 0.062 | 0.098 | 1 | 5.7307E-14 | -3.2890967 | 0.122 | 0.5 | 1.3921E-09 | 0.31369561 | 1.1461E-13 |
| Kdelr1 | 1.3217E-06 | 0.55415899 | 0.99 | 0.907 | 0.03210688 | 5.9255E-14 | 0.89757489 | 0.975 | 0.885 | 1.4394E-09 | 1.3217E-06 | 1.1851E-13 |
| Pth1r | 6.055E-14 | 3.41611875 | 0.781 | 0.373 | 1.4709E-09 | 0.00014155 | 1.11189558 | 0.55 | 0.281 | 1 | 0.00014155 | 1.211E-13 |
| Mcu | 6.1537E-14 | 0.98012357 | 0.99 | 0.896 | 1.4948E-09 | 0.77760983 | -0.4217737 | 0.971 | 0.938 | 1 | 0.77760983 | 1.2307E-13 |
| Elf3 | 0.84326553 | -0.130009 | 0.042 | 0.047 | 1 | 6.1885E-14 | -2.3422767 | 0.076 | 0.427 | 1.5033E-09 | 0.84326553 | 1.2377E-13 |
| Rpl4 | 6.3305E-10 | 0.55611519 | 1 | 0.99 | 0.000015378 | 6.2597E-14 | 0.7745234 | 1 | 0.969 | 1.5206E-09 | 6.3305E-10 | 1.2519E-13 |
| Xkr4 | 0.61951934 | 0.24393473 | 0.094 | 0.078 | 1 | 6.2814E-14 | -1.6442743 | 0.088 | 0.458 | 1.5259E-09 | 0.61951934 | 1.2563E-13 |
| Atp1b1 | 0.00291413 | -1.7791018 | 0.083 | 0.223 | 1 | 6.8205E-14 | 0.11725507 | 0.197 | 0.604 | 1.6568E-09 | 0.00291413 | 1.3641E-13 |
| Xbp1 | 0.0207404 | 0.30197311 | 0.896 | 0.896 | 1 | 6.8363E-14 | 1.21110624 | 0.912 | 0.74 | 1.6607E-09 | 0.0207404 | 1.3673E-13 |
| Rps29 | 0.00481496 | 0.28447024 | 1 | 0.974 | 1 | 7.2744E-14 | 0.81341234 | 0.996 | 0.979 | 1.7671E-09 | 0.00481496 | 1.4549E-13 |
| Ssr3 | 0.00001472 | 0.53947839 | 0.99 | 0.959 | 0.35757249 | 7.3892E-14 | 0.81877921 | 0.996 | 0.938 | 1.795E-09 | 0.00001472 | 1.4778E-13 |
| mt-Nd2 | 0.07652858 | -0.4300045 | 1 | 1 | 1 | 7.8375E-14 | -1.1687937 | 1 | 1 | 1.9039E-09 | 0.07652858 | 1.5675E-13 |
| Ckap4 | 1.2853E-07 | 0.6345211 | 0.99 | 0.964 | 0.00312232 | 7.9851E-14 | 0.86006974 | 0.987 | 0.917 | 1.9397E-09 | 1.2853E-07 | 1.597E-13 |
| Rpl29 | 0.00838359 | 0.27819731 | 1 | 0.984 | 1 | 8.448E-14 | 0.72247814 | 1 | 0.969 | 2.0522E-09 | 0.00838359 | 1.6896E-13 |
| Atp8a1 | 0.14807125 | -2.0799683 | 0.062 | 0.114 | 1 | 8.5735E-14 | -2.1815312 | 0.122 | 0.521 | 2.0827E-09 | 0.14807125 | 1.7147E-13 |
| Mgat2 | 0.00646472 | 0.49033202 | 0.76 | 0.611 | 1 | 1.0047E-13 | 1.45011216 | 0.84 | 0.438 | 2.4407E-09 | 0.00646472 | 2.0095E-13 |
| mt-Co2 | 0.0102355 | -0.5695915 | 1 | 1 | 1 | 1.1068E-13 | -1.1270747 | 1 | 1 | 2.6887E-09 | 0.0102355 | 2.2137E-13 |
| Chchd10 | 0.70867244 | 0.37006462 | 0.042 | 0.052 | 1 | 1.1079E-13 | -0.9394868 | 0.055 | 0.385 | 2.6913E-09 | 0.70867244 | 2.2158E-13 |
| Sox2 | 0.00337433 | -1.6770036 | 0.052 | 0.181 | 1 | 1.124E-13 | -3.1723517 | 0.029 | 0.312 | 2.7305E-09 | 0.00337433 | 2.2481E-13 |
| Hsd11b1 | 1.1796E-13 | -3.8104815 | 0.188 | 0.596 | 2.8656E-09 | 0.09301855 | -2.2167459 | 0.025 | 0.062 | 1 | 0.09301855 | 2.3593E-13 |
| Ptpn13 | 0.01112006 | -0.9850497 | 0.385 | 0.539 | 1 | 1.2113E-13 | -2.0011697 | 0.475 | 0.844 | 2.9424E-09 | 0.01112006 | 2.4225E-13 |
| Dsp | 0.25775837 | 0.37522632 | 0.125 | 0.187 | 1 | 1.2163E-13 | -1.4618584 | 0.147 | 0.542 | 2.9546E-09 | 0.25775837 | 2.4325E-13 |
| Pmp22 | 0.000016035 | 0.61820712 | 0.948 | 0.834 | 0.38952345 | 1.2331E-13 | 1.32939739 | 0.908 | 0.469 | 2.9954E-09 | 1.6035E-05 | 2.4662E-13 |
| Paics | 0.00019549 | 0.43511833 | 0.948 | 0.922 | 1 | 1.7708E-13 | 0.97582507 | 0.975 | 0.885 | 4.3017E-09 | 0.00019549 | 3.5417E-13 |
| Fgfrl1 | 0.00037239 | 0.59330793 | 0.958 | 0.808 | 1 | 1.9243E-13 | 1.22930355 | 0.908 | 0.49 | 4.6745E-09 | 0.00037239 | 3.8486E-13 |
| Cyp2f2 | 0.43153541 | -1.5301622 | 0.188 | 0.223 | 1 | 2.0828E-13 | -0.4442643 | 0.218 | 0.625 | 5.0595E-09 | 0.43153541 | 4.1655E-13 |
| Npm1 | 6.0637E-11 | 0.65323842 | 1 | 0.974 | 0.000001473 | 2.379E-13 | 0.82618211 | 0.996 | 0.99 | 5.779E-09 | 6.0637E-11 | 4.758E-13 |
| Copb2 | 2.4534E-13 | 1.00284817 | 0.979 | 0.886 | 5.9599E-09 | 2.528E-12 | 0.91677033 | 0.983 | 0.875 | 6.141E-08 | 2.528E-12 | 4.9069E-13 |
| Igfbp7 | 2.7567E-13 | 1.19826963 | 0.948 | 0.653 | 6.6965E-09 | 4.5518E-13 | 0.98912166 | 0.983 | 0.875 | 1.1057E-08 | 4.5518E-13 | 5.5133E-13 |
| Tenm4 | 0.3702693 | -0.9185049 | 0.156 | 0.114 | 1 | 3.0918E-13 | -2.0134623 | 0.16 | 0.562 | 7.5105E-09 | 0.3702693 | 6.1836E-13 |
| Ece1 | 0.40729419 | -0.4379166 | 0.271 | 0.321 | 1 | 3.107E-13 | -1.8580592 | 0.294 | 0.698 | 7.5476E-09 | 0.40729419 | 6.2141E-13 |
| Ifitm1 | 0.22020496 | -0.8550019 | 0.073 | 0.119 | 1 | 3.1169E-13 | -2.3353904 | 0.101 | 0.458 | 7.5715E-09 | 0.22020496 | 6.2338E-13 |
| Rpl37 | 0.00053116 | 0.38111211 | 1 | 0.959 | 1 | 3.2909E-13 | 0.73032282 | 1 | 0.969 | 7.9943E-09 | 0.00053116 | 6.5818E-13 |
| Alox12 | 3.3971E-13 | -6.5413274 | 0.031 | 0.451 | 8.2522E-09 | 0.00251048 | -2.5952575 | 0.021 | 0.094 | 1 | 0.00251048 | 6.7942E-13 |
| Pcbp1 | 0.000011769 | 0.46323657 | 0.979 | 0.938 | 0.28589592 | 3.4763E-13 | 0.87218432 | 0.996 | 0.938 | 8.4446E-09 | 1.1769E-05 | 6.9526E-13 |
| Iqgap2 | 0.21012843 | -0.9665103 | 0.115 | 0.166 | 1 | 3.5357E-13 | -2.3483294 | 0.118 | 0.49 | 8.5888E-09 | 0.21012843 | 7.0713E-13 |
| Becn1 | 3.5488E-13 | -2.3887809 | 0.656 | 0.824 | 8.6207E-09 | 0.15615608 | -0.4126148 | 0.655 | 0.688 | 1 | 0.15615608 | 7.0976E-13 |
| Wwc1 | 0.15746505 | -0.4519371 | 0.042 | 0.088 | 1 | 3.6453E-13 | -2.0874628 | 0.097 | 0.448 | 8.8551E-09 | 0.15746505 | 7.2905E-13 |
| Ces1d | 0.11169194 | -3.1217885 | 0.01 | 0.047 | 1 | 3.752E-13 | -1.8143278 | 0.092 | 0.448 | 9.1145E-09 | 0.11169194 | 7.5041E-13 |
| Ccdc88c | 0.05377316 | -0.4009703 | 0.229 | 0.352 | 1 | 3.7928E-13 | -1.9077568 | 0.172 | 0.583 | 9.2134E-09 | 0.05377316 | 7.5855E-13 |
| Atp10d | 0.28924211 | 0.4088571 | 0.167 | 0.119 | 1 | 3.7931E-13 | -2.2766884 | 0.055 | 0.365 | 9.2142E-09 | 0.28924211 | 7.5862E-13 |
| U2af1 | 0.000013781 | 0.5853326 | 0.917 | 0.808 | 0.33476752 | 3.9045E-13 | 0.85459461 | 0.954 | 0.844 | 9.4848E-09 | 1.3781E-05 | 7.809E-13 |
| Tram1 | 4.5573E-09 | 0.70354249 | 0.979 | 0.927 | 0.00011071 | 4.0938E-13 | 0.89642568 | 0.962 | 0.875 | 9.9446E-09 | 4.5573E-09 | 8.1876E-13 |
| Ccdc3 | 0.01684266 | -3.1422526 | 0.031 | 0.114 | 1 | 4.2097E-13 | -1.8086611 | 0.101 | 0.458 | 1.0226E-08 | 0.01684266 | 8.4194E-13 |
| Clec11a | 0.00307442 | 0.27833731 | 0.865 | 0.736 | 1 | 4.3636E-13 | 1.02252719 | 0.941 | 0.49 | 1.06E-08 | 0.00307442 | 8.7271E-13 |
| 2610035D17Rik | 7.3141E-08 | -2.427275 | 0.354 | 0.637 | 0.00177674 | 5.0592E-13 | -1.9718588 | 0.311 | 0.719 | 1.229E-08 | 7.3141E-08 | 1.0118E-12 |
| Bag5 | 0.02982025 | 0.35629834 | 0.729 | 0.632 | 1 | 5.1261E-13 | 1.32170765 | 0.761 | 0.281 | 1.2452E-08 | 0.02982025 | 1.0252E-12 |
| Crtap | 0.08968687 | 0.16930525 | 0.969 | 0.834 | 1 | 5.1782E-13 | 1.07324363 | 0.92 | 0.51 | 1.2579E-08 | 0.08968687 | 1.0356E-12 |
| S100a4 | 5.7895E-13 | -2.8256511 | 0.24 | 0.668 | 1.4064E-08 | 0.69300345 | -0.8021183 | 0.408 | 0.385 | 1 | 0.69300345 | 1.1579E-12 |
| Shroom3 | 0.20778076 | -1.0968421 | 0.073 | 0.119 | 1 | 6.2833E-13 | -2.0665841 | 0.067 | 0.396 | 1.5263E-08 | 0.20778076 | 1.2567E-12 |
| Rabac1 | 0.00655177 | 0.31158789 | 0.979 | 0.953 | 1 | 6.4933E-13 | 0.80203762 | 0.979 | 0.896 | 1.5773E-08 | 0.00655177 | 1.2987E-12 |
| Selenos | 0.00065563 | 0.38522135 | 0.969 | 0.943 | 1 | 6.8568E-13 | 0.85787218 | 0.992 | 0.885 | 1.6656E-08 | 0.00065563 | 1.3714E-12 |
| Lmo7 | 0.000009802 | -0.8416593 | 0.625 | 0.85 | 0.23810924 | 6.9541E-13 | -1.3036233 | 0.563 | 0.896 | 1.6893E-08 | 9.802E-06 | 1.3908E-12 |
| Tmed2 | 1.0261E-09 | 0.55893608 | 0.979 | 0.964 | 0.000024925 | 7.0681E-13 | 0.71480679 | 0.996 | 0.99 | 1.717E-08 | 1.0261E-09 | 1.4136E-12 |
| Naca | 1.061E-08 | 0.49049763 | 1 | 0.969 | 0.00025774 | 7.1179E-13 | 0.71272802 | 1 | 0.948 | 1.7291E-08 | 1.061E-08 | 1.4236E-12 |
| Cp | 7.234E-13 | -2.4779323 | 0.344 | 0.751 | 1.7573E-08 | 0.38740193 | -1.2318528 | 0.176 | 0.208 | 1 | 0.38740193 | 1.4468E-12 |
| Arl1 | 0.000097038 | 0.52109587 | 0.969 | 0.865 | 1 | 7.4759E-13 | 1.01510689 | 0.941 | 0.75 | 1.816E-08 | 9.7038E-05 | 1.4952E-12 |
| Rps20 | 0.078487 | 0.22312546 | 1 | 0.979 | 1 | 7.7816E-13 | 0.73814145 | 1 | 0.969 | 1.8903E-08 | 0.078487 | 1.5563E-12 |
| Srpx2 | 7.8162E-13 | -2.868847 | 0.208 | 0.617 | 1.8987E-08 | 0.80971065 | -0.5502142 | 0.307 | 0.271 | 1 | 0.80971065 | 1.5632E-12 |
| Ephb2 | 8.2673E-13 | 1.60717854 | 0.677 | 0.212 | 2.0083E-08 | 0.64376421 | -0.4707465 | 0.672 | 0.667 | 1 | 0.64376421 | 1.6535E-12 |
| Herpud1 | 1.8789E-07 | 0.66503904 | 0.969 | 0.927 | 0.00456433 | 8.5476E-13 | 0.95212416 | 0.975 | 0.854 | 2.0764E-08 | 1.8789E-07 | 1.7095E-12 |
| Gsta4 | 0.0784391 | -1.1079827 | 0.073 | 0.145 | 1 | 9.0525E-13 | -1.4999581 | 0.168 | 0.552 | 2.199E-08 | 0.0784391 | 1.8105E-12 |
| Slc38a3 | 9.3756E-13 | 2.12533103 | 0.573 | 0.181 | 2.2775E-08 | 2.9208E-07 | 1.32676936 | 0.496 | 0.177 | 0.00709529 | 2.9208E-07 | 1.8751E-12 |
| Tubb4b | 0.000030867 | 0.80644702 | 0.885 | 0.772 | 0.74981175 | 1.1013E-12 | 1.25350498 | 0.958 | 0.885 | 2.6753E-08 | 3.0867E-05 | 2.2027E-12 |
| Sorbs2 | 1.1345E-12 | 1.40193513 | 1 | 0.87 | 2.7558E-08 | 3.7444E-09 | 0.91560278 | 0.992 | 0.865 | 0.00009096 | 3.7444E-09 | 2.2689E-12 |
| Casz1 | 0.86614761 | 0.22988696 | 0.042 | 0.047 | 1 | 1.1981E-12 | -2.1012686 | 0.067 | 0.385 | 2.9103E-08 | 0.86614761 | 2.3961E-12 |
| Eif3a | 1.5988E-06 | 0.49231595 | 1 | 0.99 | 0.0388373 | 1.2215E-12 | 0.70796824 | 1 | 0.969 | 2.9674E-08 | 1.5988E-06 | 2.4431E-12 |
| Six3 | 0.78979801 | -0.2148979 | 0.021 | 0.026 | 1 | 1.2684E-12 | -3.4797803 | 0.021 | 0.271 | 3.0812E-08 | 0.78979801 | 2.5368E-12 |
| Pik3r3 | 0.00342714 | -3.178372 | 0.042 | 0.155 | 1 | 1.3681E-12 | -2.4974823 | 0.101 | 0.448 | 3.3234E-08 | 0.00342714 | 2.7362E-12 |
| Gas5 | 1.2789E-08 | 0.58790697 | 1 | 0.969 | 0.00031067 | 1.3959E-12 | 0.82284194 | 0.992 | 0.948 | 3.391E-08 | 1.2789E-08 | 2.7919E-12 |
| Sdc4 | 0.01078166 | -0.4979268 | 1 | 1 | 1 | 1.3967E-12 | 0.90596574 | 1 | 1 | 3.3929E-08 | 0.01078166 | 2.7934E-12 |

| | | | | | | | | | | | | |
|---|---|---|---|---|---|---|---|---|---|---|---|---|
| Krt19 | 0.6135606 | -0.2148979 | 0.052 | 0.067 | 1 | 1.44E-12 | -2.7021727 | 0.092 | 0.427 | 3.4982E-08 | 0.6135606 | 2.8801E-12 |
| Cd9 | 0.44993952 | -0.1588019 | 1 | 1 | 1 | 1.492E-12 | 0.6483042 | 1 | 1 | 3.6244E-08 | 0.44993952 | 2.984E-12 |
| Efna1 | 0.0117801 | 0.38661246 | 0.854 | 0.782 | 1 | 1.5547E-12 | 1.08063712 | 0.941 | 0.688 | 3.7768E-08 | 0.0117801 | 3.1095E-12 |
| Sub1 | 0.00219632 | 0.32500697 | 0.979 | 0.948 | 1 | 1.5788E-12 | 0.74294326 | 0.992 | 0.948 | 3.8352E-08 | 0.00219632 | 3.1575E-12 |
| Chst12 | 0.00023277 | 0.47406234 | 0.885 | 0.679 | 1 | 1.6655E-12 | 1.32757466 | 0.773 | 0.292 | 4.0459E-08 | 0.00023277 | 3.3311E-12 |
| Cd36 | 0.36606794 | -1.9467018 | 0.021 | 0.041 | 1 | 1.7713E-12 | -3.1843244 | 0.029 | 0.292 | 4.3028E-08 | 0.36606794 | 3.5426E-12 |
| Epyc | 1.787E-12 | 1.54598547 | 0.885 | 0.627 | 4.341E-08 | 8.0506E-12 | 2.06938152 | 0.689 | 0.292 | 1.9557E-07 | 8.0506E-12 | 3.574E-12 |
| Cpxm2 | 0.00066798 | 0.89560166 | 0.469 | 0.259 | 1 | 1.857E-12 | 1.39780396 | 0.735 | 0.281 | 4.511E-08 | 0.00066798 | 3.714E-12 |
| Sec13 | 0.00154347 | 0.45495351 | 0.875 | 0.782 | 1 | 2.2041E-12 | 1.06611267 | 0.87 | 0.583 | 5.3541E-08 | 0.00154347 | 4.4081E-12 |
| Polr2d | 0.00010225 | 0.71798792 | 0.76 | 0.57 | 1 | 2.2557E-12 | 1.42286086 | 0.735 | 0.281 | 5.4796E-08 | 0.00010225 | 4.5115E-12 |
| Sik1 | 0.55686368 | -0.130009 | 0.208 | 0.244 | 1 | 2.598E-12 | -1.7793405 | 0.189 | 0.594 | 6.311E-08 | 0.55686368 | 5.196E-12 |
| BC006965 | 2.7265E-12 | -3.8216409 | 0.25 | 0.622 | 6.6231E-08 | 0.90929126 | -0.9496797 | 0.218 | 0.198 | 1 | 0.90929126 | 5.4529E-12 |
| Arpc1b | 0.01300749 | -0.4641879 | 0.938 | 0.964 | 1 | 2.9067E-12 | 0.78058967 | 0.992 | 0.958 | 7.061E-08 | 0.01300749 | 5.8135E-12 |
| Gsta3 | 0.99603711 | -0.1624305 | 0.031 | 0.031 | 1 | 2.9079E-12 | -2.1012686 | 0.092 | 0.427 | 7.064E-08 | 0.99603711 | 5.8159E-12 |
| Pcp4 | 2.961E-12 | 2.53800925 | 0.396 | 0.062 | 7.1929E-08 | 0.27154471 | 1.49749966 | 0.013 | 0 | 1 | 0.27154471 | 5.9221E-12 |
| Ssr4 | 0.0006252 | 0.37739712 | 0.979 | 0.964 | 1 | 3.2385E-12 | 0.86909491 | 0.962 | 0.865 | 7.8669E-08 | 0.0006252 | 6.477E-12 |
| Enpp2 | 3.2698E-12 | -1.9575371 | 0.917 | 0.896 | 7.943E-08 | 0.71491515 | -0.5581811 | 0.681 | 0.583 | 1 | 0.71491515 | 6.5396E-12 |
| Slc15a2 | 0.38484207 | -0.8225805 | 0.083 | 0.114 | 1 | 3.2759E-12 | -1.2598146 | 0.185 | 0.604 | 7.9579E-08 | 0.38484207 | 6.5518E-12 |
| Mcfd2 | 0.15597854 | 0.20566084 | 0.875 | 0.762 | 1 | 3.3036E-12 | 1.27206036 | 0.803 | 0.406 | 8.0251E-08 | 0.15597854 | 6.6072E-12 |
| Spon1 | 3.3247E-12 | -2.8253955 | 0.25 | 0.648 | 8.0764E-08 | 0.06340936 | -0.7581552 | 0.433 | 0.281 | 1 | 0.06340936 | 6.6495E-12 |
| Pglyrp1 | 0.60063579 | 0.27052894 | 0.146 | 0.171 | 1 | 3.3798E-12 | -1.6317834 | 0.105 | 0.448 | 8.2101E-08 | 0.60063579 | 6.7595E-12 |
| Sdc3 | 6.0197E-11 | 1.09966438 | 0.927 | 0.611 | 1.4623E-06 | 3.4439E-12 | 1.05937855 | 0.92 | 0.531 | 8.3658E-08 | 6.0197E-11 | 6.8877E-12 |
| Cdo1 | 3.4611E-12 | -2.1094816 | 0.292 | 0.689 | 8.4076E-08 | 0.34364284 | 0.10606166 | 0.639 | 0.75 | 1 | 0.34364284 | 6.9221E-12 |
| Rpl22l1 | 0.000035993 | 0.63260722 | 0.99 | 0.964 | 0.87434806 | 3.4845E-12 | 0.81942775 | 0.992 | 0.938 | 8.4645E-08 | 3.5993E-05 | 6.969E-12 |
| Purb | 3.5468E-12 | 0.81172336 | 0.99 | 0.86 | 8.616E-08 | 2.1344E-10 | 0.7014359 | 0.987 | 0.906 | 5.1848E-06 | 2.1344E-10 | 7.0937E-12 |
| Pgam1 | 0.03114213 | 0.21025093 | 0.969 | 0.876 | 1 | 3.6452E-12 | 0.91546174 | 0.929 | 0.771 | 8.855E-08 | 0.03114213 | 7.2904E-12 |
| Gpx8 | 3.2559E-08 | 0.68021321 | 0.969 | 0.834 | 0.00079093 | 3.7054E-12 | 1.0790421 | 0.92 | 0.594 | 9.0011E-08 | 3.2559E-08 | 7.4107E-12 |
| Igfbp3 | 0.31169919 | -1.7669458 | 0.062 | 0.098 | 1 | 3.7465E-12 | -1.0072925 | 0.126 | 0.5 | 9.1011E-08 | 0.31169919 | 7.4931E-12 |
| Rpl18a | 0.0014174 | 0.33242959 | 1 | 0.948 | 1 | 3.8351E-12 | 0.67902871 | 1 | 0.958 | 9.3161E-08 | 0.0014174 | 7.6701E-12 |
| Arsi | 3.8666E-12 | 2.19191911 | 0.573 | 0.187 | 9.3928E-08 | 1.6204E-07 | 1.32724004 | 0.643 | 0.312 | 0.0039364 | 1.6204E-07 | 7.7332E-12 |
| Rhoj | 4.1816E-12 | -1.7033119 | 0.719 | 0.891 | 1.0158E-07 | 0.09096336 | -0.4247501 | 0.824 | 0.562 | 1 | 0.09096336 | 8.3631E-12 |
| Pir | 4.3493E-12 | -1.4668488 | 0.188 | 0.617 | 1.0565E-07 | 0.03808581 | -0.2214173 | 0.399 | 0.573 | 1 | 0.03808581 | 8.6986E-12 |
| Kalrn | 0.21952024 | -1.2700394 | 0.208 | 0.264 | 1 | 4.4904E-12 | -2.225243 | 0.218 | 0.625 | 1.0908E-07 | 0.21952024 | 8.9809E-12 |
| Tspan13 | 0.13729503 | -0.3696391 | 0.958 | 0.922 | 1 | 4.5385E-12 | 0.92345845 | 0.966 | 0.906 | 1.1025E-07 | 0.13729503 | 9.0769E-12 |
| Pof1b | 0.00059979 | -3.0585947 | 0.031 | 0.171 | 1 | 4.7815E-12 | -2.9343461 | 0.046 | 0.323 | 1.1615E-07 | 0.00059979 | 9.563E-12 |
| B3gnt9 | 4.8206E-12 | 1.74446013 | 0.604 | 0.207 | 1.171E-07 | 5.0112E-07 | 1.10518224 | 0.542 | 0.219 | 0.01217323 | 5.0112E-07 | 9.6412E-12 |
| Ccdc80 | 0.000068444 | 0.33207369 | 1 | 0.969 | 1 | 5.0078E-12 | 0.71702515 | 0.996 | 0.865 | 1.2165E-07 | 6.8444E-05 | 1.0016E-11 |
| Grb10 | 5.2523E-12 | 0.80176881 | 1 | 0.855 | 1.2759E-07 | 0.0209754 | 0.18343368 | 0.979 | 0.906 | 1 | 0.0209754 | 1.0505E-11 |
| P4hb | 1.0318E-07 | 0.4954735 | 0.99 | 0.979 | 0.00250652 | 5.7623E-12 | 0.74593252 | 0.987 | 0.969 | 1.3998E-07 | 1.0318E-07 | 1.1525E-11 |
| Vamp3 | 0.000024662 | 0.7031011 | 0.76 | 0.56 | 0.59909738 | 5.9068E-12 | 1.27116005 | 0.798 | 0.354 | 1.4349E-07 | 2.4662E-05 | 1.1814E-11 |
| Rpl38 | 0.000040319 | 0.47234126 | 1 | 0.927 | 0.97944075 | 6.0797E-12 | 0.78351624 | 0.992 | 0.906 | 1.4769E-07 | 4.0319E-05 | 1.2159E-11 |
| Tspan7 | 6.1635E-12 | -3.0619906 | 0.156 | 0.56 | 1.4972E-07 | 0.01164092 | 0.33311284 | 0.445 | 0.26 | 1 | 0.01164092 | 1.2327E-11 |
| Pla2g5 | 5.3783E-08 | -1.6989136 | 0.385 | 0.663 | 0.0013065 | 6.2407E-12 | 1.28142101 | 0.693 | 0.229 | 1.516E-07 | 5.3783E-08 | 1.2481E-11 |
| Slc2a1 | 6.6887E-12 | -2.5311638 | 0.521 | 0.813 | 1.6248E-07 | 0.03198152 | -0.8181649 | 0.504 | 0.677 | 1 | 0.03198152 | 1.3377E-11 |
| Clic1 | 0.00240871 | 0.50305795 | 0.958 | 0.943 | 1 | 6.8412E-12 | 0.77010004 | 0.979 | 0.896 | 1.6619E-07 | 0.00240871 | 1.3682E-11 |
| Fxyd1 | 7.0032E-12 | 1.21360187 | 0.854 | 0.503 | 1.7012E-07 | 8.7407E-12 | 1.13308823 | 0.874 | 0.458 | 2.1233E-07 | 8.7407E-12 | 1.4006E-11 |
| Ywhah | 0.00012976 | 0.85173687 | 0.812 | 0.71 | 1 | 7.0807E-12 | 1.21852371 | 0.899 | 0.75 | 0.000000172 | 0.00012976 | 1.4161E-11 |
| Cct8 | 0.00160709 | 0.44067938 | 0.896 | 0.87 | 1 | 7.238E-12 | 0.85262931 | 0.966 | 0.865 | 1.7583E-07 | 0.00160709 | 1.4476E-11 |
| Nap1l1 | 7.2546E-12 | 0.7455868 | 1 | 0.938 | 1.7623E-07 | 1.3789E-10 | 0.78010811 | 0.992 | 0.948 | 3.3495E-06 | 1.3789E-10 | 1.4509E-11 |
| Chst9 | 0.62104899 | 0.42253204 | 0.01 | 0.005 | 1 | 7.4967E-12 | -2.9378865 | 0.038 | 0.302 | 1.8211E-07 | 0.62104899 | 1.4993E-11 |
| Krt5 | 0.11273737 | -2.240433 | 0.01 | 0.047 | 1 | 8.1867E-12 | -1.010295 | 0.126 | 0.479 | 1.9887E-07 | 0.11273737 | 1.6373E-11 |
| Blnk | 0.17179529 | 2.59245704 | 0.042 | 0.016 | 1 | 8.2134E-12 | -2.3923174 | 0.055 | 0.344 | 1.9952E-07 | 0.17179529 | 1.6427E-11 |
| Ptch1 | 8.304E-12 | 1.8955051 | 0.646 | 0.301 | 2.0172E-07 | 0.05850508 | 0.26588871 | 0.475 | 0.333 | 1 | 0.05850508 | 1.6608E-11 |
| Sec61b | 4.1825E-06 | 0.59808327 | 0.969 | 0.917 | 0.10160016 | 8.9725E-12 | 0.81383229 | 0.966 | 0.906 | 2.1796E-07 | 4.1825E-06 | 1.7945E-11 |
| Dph3 | 0.02603559 | 0.33430485 | 0.75 | 0.627 | 1 | 9.0444E-12 | 1.54812573 | 0.723 | 0.312 | 2.1971E-07 | 0.02603559 | 1.8089E-11 |
| Fau | 0.01475517 | 0.25667157 | 1 | 0.979 | 1 | 9.6044E-12 | 0.68035423 | 1 | 0.99 | 2.3331E-07 | 0.01475517 | 1.9209E-11 |
| Fcgrt | 0.03614855 | 0.29062036 | 0.948 | 0.829 | 1 | 1.0126E-11 | 0.94598364 | 0.954 | 0.531 | 2.4599E-07 | 0.03614855 | 2.0252E-11 |
| Runx3 | 1.0596E-11 | 2.65135073 | 0.417 | 0.083 | 2.5739E-07 | 0.000026612 | 0.67229674 | 0.424 | 0.167 | 0.64646367 | 2.6612E-05 | 2.1192E-11 |
| Il11ra1 | 7.9075E-06 | 0.77651265 | 0.823 | 0.663 | 0.1920887 | 1.1001E-11 | 0.81235067 | 0.908 | 0.448 | 2.6723E-07 | 7.9075E-06 | 2.2001E-11 |
| Slc12a2 | 1.1236E-11 | -1.6397522 | 0.406 | 0.767 | 2.7294E-07 | 5.5796E-07 | -1.6191833 | 0.576 | 0.812 | 0.01355395 | 5.5796E-07 | 2.2471E-11 |
| Loxl2 | 1.1367E-11 | -1.9588971 | 0.344 | 0.71 | 2.7612E-07 | 0.70079839 | -0.8608704 | 0.189 | 0.167 | 1 | 0.70079839 | 2.2733E-11 |
| Surf4 | 0.000019653 | 0.61644653 | 0.917 | 0.813 | 0.47741107 | 1.2861E-11 | 0.92997175 | 0.916 | 0.76 | 3.1241E-07 | 1.9653E-05 | 2.5721E-11 |
| Plod2 | 0.03746912 | 0.19520008 | 1 | 1 | 1 | 1.3648E-11 | 0.86281736 | 0.987 | 0.844 | 3.3153E-07 | 0.03746912 | 2.7295E-11 |
| Dpysl3 | 0.00078641 | -3.6267115 | 0.031 | 0.166 | 1 | 1.4428E-11 | -2.9957467 | 0.05 | 0.323 | 3.5048E-07 | 0.00078641 | 2.8855E-11 |
| Nbdy | 0.02982861 | 0.42855582 | 0.76 | 0.674 | 1 | 1.4479E-11 | 1.49269867 | 0.672 | 0.229 | 3.5173E-07 | 0.02982861 | 2.8959E-11 |
| Dlgap1 | 1.4704E-11 | -3.2166735 | 0.156 | 0.549 | 3.572E-07 | 0.97738083 | -0.1528116 | 0.311 | 0.302 | 1 | 0.97738083 | 2.9409E-11 |
| Rcn3 | 1.1815E-09 | 0.73391083 | 0.99 | 0.87 | 0.0000287 | 1.5343E-11 | 0.9868145 | 0.933 | 0.625 | 3.727E-07 | 1.1815E-09 | 3.0685E-11 |
| Rpl31 | 0.0003938 | 0.34655578 | 1 | 0.959 | 1 | 1.6264E-11 | 0.73241046 | 1 | 0.958 | 3.9508E-07 | 0.0003938 | 3.2527E-11 |
| Krt15 | 0.26929823 | -1.8449483 | 0.073 | 0.114 | 1 | 1.6279E-11 | -1.8634536 | 0.185 | 0.542 | 3.9546E-07 | 0.26929823 | 3.2559E-11 |
| Srp9 | 0.00075341 | 0.3816808 | 0.958 | 0.865 | 1 | 1.6579E-11 | 0.90798367 | 0.95 | 0.812 | 4.0273E-07 | 0.00075341 | 3.3157E-11 |
| Rpl18 | 0.00021302 | 0.38843933 | 1 | 0.969 | 1 | 1.6649E-11 | 0.6573548 | 1 | 0.969 | 4.0443E-07 | 0.00021302 | 3.3297E-11 |
| Calm1 | 0.13156021 | 0.19215085 | 1 | 0.979 | 1 | 1.7016E-11 | 0.80356312 | 1 | 1 | 4.1336E-07 | 0.13156021 | 3.4032E-11 |
| Filip1l | 0.00752891 | 0.12046471 | 0.979 | 0.902 | 1 | 1.707E-11 | 0.68756619 | 1 | 0.979 | 4.1467E-07 | 0.00752891 | 3.4141E-11 |
| Slc35e4 | 0.05903326 | 0.29967529 | 0.677 | 0.617 | 1 | 1.7524E-11 | 1.33046346 | 0.786 | 0.354 | 4.257E-07 | 0.05903326 | 3.5049E-11 |
| Gsto1 | 0.02733285 | -0.6247737 | 0.188 | 0.306 | 1 | 1.8099E-11 | -1.2651862 | 0.332 | 0.698 | 4.3966E-07 | 0.02733285 | 3.6198E-11 |
| Arcn1 | 1.826E-11 | 0.83700322 | 0.979 | 0.933 | 4.4356E-07 | 6.6045E-08 | 0.49143889 | 0.983 | 0.927 | 0.00160437 | 6.6045E-08 | 3.6519E-11 |
| Crim1 | 6.7631E-06 | 0.84995326 | 0.979 | 0.974 | 0.1642896 | 1.8394E-11 | 0.57503525 | 1 | 0.979 | 4.4682E-07 | 6.7631E-06 | 3.6787E-11 |
| Aldh3a1 | 0.90285512 | -0.3404288 | 0.146 | 0.135 | 1 | 1.8422E-11 | -1.3098553 | 0.143 | 0.5 | 4.4751E-07 | 0.90285512 | 3.6845E-11 |
| Gapdh | 0.80273492 | -0.2120641 | 1 | 0.969 | 1 | 1.8576E-11 | 0.69480396 | 1 | 0.969 | 4.5126E-07 | 0.80273492 | 3.7153E-11 |
| Col8a1 | 1.8748E-11 | -2.0701655 | 0.562 | 0.876 | 4.5543E-07 | 0.01490774 | 0.37478428 | 0.794 | 0.625 | 1 | 0.01490774 | 3.7497E-11 |
| Tceal8 | 0.000063496 | 0.59873548 | 0.958 | 0.793 | 1 | 2.0344E-11 | 1.01728583 | 0.941 | 0.76 | 4.9421E-07 | 6.3496E-05 | 4.0689E-11 |
| Cldn4 | 0.05519205 | -1.5160674 | 0.031 | 0.093 | 1 | 2.1658E-11 | -0.4276918 | 0.109 | 0.448 | 5.2611E-07 | 0.05519205 | 4.3316E-11 |
| Arhgef28 | 2.1703E-11 | -2.5353263 | 0.271 | 0.679 | 5.2722E-07 | 0.00055829 | -0.9640804 | 0.378 | 0.594 | 1 | 0.00055829 | 4.3407E-11 |
| Stox2 | 0.14420529 | -0.6929452 | 0.177 | 0.254 | 1 | 2.2432E-11 | -1.4557061 | 0.214 | 0.615 | 5.4493E-07 | 0.14420529 | 4.4865E-11 |
| Txnl4a | 0.65932026 | 0.28450188 | 0.479 | 0.508 | 1 | 2.3752E-11 | 1.01664556 | 0.706 | 0.25 | 5.7699E-07 | 0.65932026 | 4.7505E-11 |
| Atraid | 0.06560753 | 0.23040083 | 0.906 | 0.751 | 1 | 2.4255E-11 | 1.07185406 | 0.849 | 0.573 | 5.892E-07 | 0.06560753 | 4.851E-11 |
| Tagln2 | 6.0803E-08 | 0.87761425 | 0.896 | 0.658 | 0.00147702 | 2.637E-11 | 0.9599608 | 0.971 | 0.927 | 6.4058E-07 | 6.0803E-08 | 5.274E-11 |
| Rpl36al | 0.0689872 | 0.21614154 | 1 | 0.964 | 1 | 2.6441E-11 | 0.77127045 | 0.987 | 0.906 | 6.4231E-07 | 0.0689872 | 5.2882E-11 |
| Mrfap1 | 0.00011081 | 0.39338569 | 0.99 | 0.917 | 1 | 2.774E-11 | 0.77492516 | 0.992 | 0.833 | 6.7387E-07 | 0.00011081 | 5.5481E-11 |
| Polr1d | 0.1779756 | 0.11089603 | 0.979 | 0.917 | 1 | 2.8468E-11 | 0.71189012 | 0.954 | 0.885 | 6.9155E-07 | 0.1779756 | 5.6936E-11 |
| Cox4i2 | 2.859E-11 | 1.75503642 | 0.76 | 0.477 | 6.9452E-07 | 9.4368E-11 | 2.02956026 | 0.521 | 0.125 | 2.2924E-06 | 9.4368E-11 | 5.7181E-11 |

| | | | | | | | | | | | | |
|---|---|---|---|---|---|---|---|---|---|---|---|---|
| Krt18 | 0.247449 | -0.5564063 | 0.156 | 0.212 | 1 | 2.8909E-11 | -1.7297585 | 0.168 | 0.521 | 7.0226E-07 | 0.247449 | 5.7818E-11 |
| Slc3a2 | 3.0685E-11 | 1.02808381 | 0.927 | 0.824 | 7.454E-07 | 0.000046596 | 0.51101159 | 0.954 | 0.906 | 1 | 4.6596E-05 | 6.137E-11 |
| Fn1 | 0.32986575 | -0.181699 | 1 | 1 | 1 | 3.1219E-11 | 0.59723895 | 1 | 1 | 7.5838E-07 | 0.32986575 | 6.2439E-11 |
| Copb1 | 2.8153E-06 | 0.57394374 | 0.969 | 0.933 | 0.06838984 | 3.2159E-11 | 0.79690474 | 0.975 | 0.958 | 7.812E-07 | 2.8153E-06 | 6.4318E-11 |
| Aga | 0.00782912 | 0.3502072 | 0.844 | 0.668 | 1 | 3.2424E-11 | 1.27161825 | 0.777 | 0.333 | 7.8764E-07 | 0.00782912 | 6.4848E-11 |
| Grhl2 | 0.22187388 | -1.7998604 | 0 | 0.016 | 1 | 3.3868E-11 | -2.5508634 | 0.025 | 0.26 | 8.2271E-07 | 0.22187388 | 6.7735E-11 |
| Tsn | 5.7032E-06 | 0.62718164 | 0.969 | 0.845 | 0.1385414 | 3.5596E-11 | 0.76083403 | 0.95 | 0.833 | 8.647E-07 | 5.7032E-06 | 7.1192E-11 |
| Ptges3 | 3.593E-11 | 0.66780216 | 1 | 0.974 | 8.7281E-07 | 1.5339E-07 | 0.4372702 | 0.996 | 0.979 | 0.00372621 | 1.5339E-07 | 7.186E-11 |
| Nfib | 3.6595E-11 | -1.6266156 | 0.51 | 0.845 | 8.8896E-07 | 0.0049342 | -0.9730217 | 0.836 | 0.969 | 1 | 0.0049342 | 7.3189E-11 |
| A2m | 3.6758E-11 | -4.0237242 | 0.146 | 0.518 | 8.9291E-07 | 0.12465713 | -2.6424306 | 0.059 | 0.104 | 1 | 0.12465713 | 7.3515E-11 |
| Ppia | 0.00970724 | 0.20813541 | 1 | 0.979 | 1 | 3.7561E-11 | 0.72184828 | 1 | 0.99 | 9.1244E-07 | 0.00970724 | 7.5123E-11 |
| Rpl36 | 0.00542984 | 0.31208143 | 1 | 0.969 | 1 | 3.7609E-11 | 0.67598667 | 0.996 | 0.938 | 9.136E-07 | 0.00542984 | 7.5218E-11 |
| Aif1l | 0.0119804 | -0.8694946 | 0.344 | 0.466 | 1 | 3.8892E-11 | 3.04769674 | 0.441 | 0.062 | 9.4477E-07 | 0.0119804 | 7.7784E-11 |
| Nrk | 2.6981E-06 | 0.68872678 | 0.781 | 0.565 | 0.0655425 | 3.9881E-11 | 1.12410724 | 0.845 | 0.448 | 9.6879E-07 | 2.6981E-06 | 7.9762E-11 |
| Tmed3 | 0.000057759 | 0.51840294 | 0.906 | 0.767 | 1 | 4.046E-11 | 0.95753555 | 0.861 | 0.458 | 9.8285E-07 | 5.7759E-05 | 8.092E-11 |
| Cct6a | 0.000062206 | 0.5374678 | 0.979 | 0.896 | 1 | 4.0917E-11 | 0.84328978 | 0.958 | 0.844 | 9.9396E-07 | 6.2206E-05 | 8.1834E-11 |
| Ssr2 | 3.6317E-06 | 0.64156725 | 0.979 | 0.902 | 0.08822216 | 4.2629E-11 | 0.84780446 | 0.945 | 0.76 | 1.0356E-06 | 3.6317E-06 | 8.5259E-11 |
| Slc25a25 | 0.82564561 | -0.4157169 | 0.177 | 0.181 | 1 | 4.4318E-11 | -1.7137522 | 0.206 | 0.562 | 1.0766E-06 | 0.82564561 | 8.8636E-11 |
| Wif1 | 0.49363286 | -1.8014045 | 0.729 | 0.523 | 1 | 4.4654E-11 | 0.63275139 | 0.853 | 0.438 | 1.0847E-06 | 0.49363286 | 8.9309E-11 |
| Cldnd1 | 0.13735214 | 0.3803038 | 0.76 | 0.72 | 1 | 4.5014E-11 | 0.9186763 | 0.899 | 0.656 | 1.0935E-06 | 0.13735214 | 9.0027E-11 |
| Gng12 | 4.5168E-11 | -1.0784132 | 0.823 | 0.938 | 1.0972E-06 | 0.00222247 | -0.6569539 | 0.958 | 0.958 | 1 | 0.00222247 | 9.0335E-11 |
| Crispld1 | 4.6782E-11 | 1.24954569 | 0.99 | 0.803 | 1.1364E-06 | 7.9912E-07 | 0.80490115 | 0.899 | 0.604 | 0.01941221 | 7.9912E-07 | 9.3563E-11 |
| Ikbip | 6.0137E-06 | 0.54961894 | 0.969 | 0.865 | 0.14608441 | 4.7629E-11 | 0.89483516 | 0.945 | 0.583 | 0.000001157 | 6.0137E-06 | 9.5259E-11 |
| Rbm47 | 4.4141E-06 | -1.7379326 | 0.083 | 0.337 | 0.10722759 | 4.7834E-11 | -1.6898863 | 0.172 | 0.531 | 0.000001162 | 4.4141E-06 | 9.5668E-11 |
| AW112010 | 5.6955E-11 | -2.9387342 | 0.177 | 0.56 | 1.3836E-06 | 0.09493175 | -2.07539 | 0.025 | 0.062 | 1 | 0.09493175 | 1.1391E-10 |
| mt-Co1 | 0.17026874 | -0.3527271 | 1 | 1 | 1 | 5.8298E-11 | -1.0447154 | 1 | 1 | 1.4162E-06 | 0.17026874 | 1.166E-10 |
| Fam98a | 0.000023728 | 0.78207442 | 0.688 | 0.43 | 0.57639811 | 5.8553E-11 | 1.2422426 | 0.761 | 0.292 | 1.4224E-06 | 2.3728E-05 | 1.1711E-10 |
| Rps27 | 0.06943124 | 0.21847235 | 1 | 0.959 | 1 | 6.0997E-11 | 0.66276268 | 0.996 | 0.958 | 1.4817E-06 | 0.06943124 | 1.2199E-10 |
| Ran | 7.6562E-08 | 0.70915326 | 0.938 | 0.881 | 0.00185985 | 6.33E-11 | 0.81101338 | 0.966 | 0.792 | 1.5377E-06 | 7.6562E-08 | 1.266E-10 |
| Gda | 0.22187388 | -1.7998604 | 0 | 0.016 | 1 | 6.4595E-11 | -1.0128735 | 0.067 | 0.354 | 1.5691E-06 | 0.22187388 | 1.2919E-10 |
| Hnrnpf | 6.49E-11 | 0.7910385 | 0.979 | 0.943 | 1.5765E-06 | 5.6957E-09 | 0.53938857 | 0.992 | 0.979 | 0.00013836 | 5.6957E-09 | 1.298E-10 |
| Map9 | 6.4917E-11 | 2.55866872 | 0.562 | 0.212 | 0.000001577 | 1.4362E-07 | 1.94390133 | 0.483 | 0.177 | 0.00348881 | 1.4362E-07 | 1.2983E-10 |
| Ninj1 | 0.00016718 | -1.0480401 | 0.625 | 0.72 | 1 | 6.717E-11 | 1.11811447 | 0.79 | 0.354 | 1.6317E-06 | 0.00016718 | 1.3434E-10 |
| Map1lc3b | 0.36856368 | -0.135054 | 1 | 0.953 | 1 | 6.7463E-11 | 0.79051322 | 1 | 0.979 | 1.6388E-06 | 0.36856368 | 1.3493E-10 |
| Fibin | 0.00078685 | 0.92170983 | 0.802 | 0.663 | 1 | 6.7932E-11 | 1.61825182 | 0.66 | 0.219 | 1.6502E-06 | 0.00078685 | 1.3586E-10 |
| Mmp2 | 6.831E-11 | 0.71424552 | 0.896 | 0.342 | 1.6594E-06 | 0.00039736 | 0.1786733 | 0.933 | 0.573 | 1 | 0.00039736 | 1.3662E-10 |
| Pqbp1 | 0.1742454 | 0.24529165 | 0.635 | 0.518 | 1 | 6.9711E-11 | 1.48113621 | 0.622 | 0.188 | 1.6934E-06 | 0.1742454 | 1.3942E-10 |
| Cope | 0.00012181 | 0.40655518 | 0.969 | 0.902 | 1 | 7.047E-11 | 0.89285967 | 0.937 | 0.771 | 1.7119E-06 | 0.00012181 | 1.4094E-10 |
| Ppib | 1.7783E-08 | 0.49936067 | 0.99 | 0.974 | 0.00043198 | 7.5157E-11 | 0.75686308 | 1 | 0.958 | 1.8257E-06 | 1.7783E-08 | 1.5031E-10 |
| Ucp2 | 0.76791351 | 0.62898291 | 0.115 | 0.13 | 1 | 7.6059E-11 | 0.15869775 | 0.097 | 0.417 | 1.8476E-06 | 0.76791351 | 1.5212E-10 |
| Snx10 | 7.6289E-11 | -1.9502772 | 0.271 | 0.637 | 1.8532E-06 | 0.617791 | 1.17447488 | 0.282 | 0.25 | 1 | 0.617791 | 1.5258E-10 |
| Arf4 | 3.1242E-09 | 0.46265886 | 0.99 | 1 | 0.000075893 | 7.8611E-11 | 0.55860907 | 1 | 0.979 | 1.9096E-06 | 3.1242E-09 | 1.5722E-10 |
| Bod1 | 0.01463592 | 0.42879103 | 0.76 | 0.591 | 1 | 8.2727E-11 | 1.20422375 | 0.761 | 0.365 | 2.0096E-06 | 0.01463592 | 1.6545E-10 |
| Bcam | 0.52098413 | -1.6929452 | 0.01 | 0.021 | 1 | 8.7589E-11 | -3.1463565 | 0.021 | 0.24 | 2.1277E-06 | 0.52098413 | 1.7518E-10 |
| Aldh1a1 | 0.96743136 | 1.00749454 | 0.052 | 0.052 | 1 | 8.8263E-11 | -2.2273931 | 0.059 | 0.333 | 2.1441E-06 | 0.96743136 | 1.7653E-10 |
| Cep83 | 8.8421E-11 | 1.53667958 | 0.76 | 0.435 | 2.1479E-06 | 0.54650274 | -0.3958285 | 0.71 | 0.677 | 1 | 0.54650274 | 1.7684E-10 |
| Pdia6 | 0.27283054 | 0.12559914 | 0.99 | 0.969 | 1 | 8.8695E-11 | 0.79920026 | 0.992 | 0.875 | 2.1546E-06 | 0.27283054 | 1.7739E-10 |
| Rps27l | 0.0018321 | 0.346458 | 0.927 | 0.881 | 1 | 9.0459E-11 | 0.83248391 | 0.933 | 0.792 | 2.1974E-06 | 0.0018321 | 1.8092E-10 |
| Yif1b | 0.0004943 | 0.59517142 | 0.719 | 0.477 | 1 | 9.0716E-11 | 1.30987266 | 0.672 | 0.24 | 2.2037E-06 | 0.0004943 | 1.8143E-10 |
| Rarg | 9.0863E-11 | 0.96993112 | 0.938 | 0.777 | 2.2073E-06 | 0.00012642 | 0.53181163 | 0.954 | 0.896 | 1 | 0.00012642 | 1.8173E-10 |
| Gabarap | 0.03689771 | 0.22819744 | 0.99 | 0.964 | 1 | 9.2778E-11 | 0.68672738 | 0.996 | 0.927 | 2.2538E-06 | 0.03689771 | 1.8556E-10 |
| Tmem263 | 4.4386E-10 | 0.89598622 | 0.927 | 0.715 | 0.000010782 | 9.6339E-11 | 0.9514524 | 0.954 | 0.604 | 2.3403E-06 | 4.4386E-10 | 1.9268E-10 |
| Hspa9 | 2.4338E-06 | 0.61086907 | 0.938 | 0.902 | 0.05912278 | 9.7667E-11 | 0.80736856 | 0.962 | 0.906 | 2.3725E-06 | 2.4338E-06 | 1.9533E-10 |
| Copz2 | 4.6658E-09 | 0.81103561 | 0.938 | 0.793 | 0.00011334 | 1.0544E-10 | 1.13850231 | 0.824 | 0.406 | 2.5615E-06 | 4.6658E-09 | 2.1089E-10 |
| Rab25 | 0.80496186 | 0.42253204 | 0.021 | 0.026 | 1 | 1.0561E-10 | -2.5828738 | 0.042 | 0.292 | 2.5654E-06 | 0.80496186 | 2.1122E-10 |
| Hnrnpk | 7.6925E-07 | 0.50863277 | 1 | 0.969 | 0.01868658 | 1.0692E-10 | 0.62371738 | 0.996 | 0.979 | 2.5974E-06 | 7.6925E-07 | 2.1385E-10 |
| Slc25a3 | 0.00260139 | 0.27950071 | 0.99 | 0.974 | 1 | 1.1447E-10 | 0.68496184 | 1 | 0.969 | 2.7808E-06 | 0.00260139 | 2.2894E-10 |
| Cpe | 0.12042986 | -0.2220912 | 0.938 | 0.938 | 1 | 1.1947E-10 | 0.86961695 | 0.987 | 0.875 | 2.9022E-06 | 0.12042986 | 2.3894E-10 |
| S100a14 | 0.2825047 | -0.9925055 | 0.01 | 0.031 | 1 | 1.2024E-10 | -2.0874628 | 0.055 | 0.323 | 2.9209E-06 | 0.2825047 | 2.4048E-10 |
| Magoh | 5.7433E-08 | 0.70330281 | 0.958 | 0.762 | 0.00139516 | 1.253E-10 | 0.7553624 | 0.933 | 0.771 | 3.0439E-06 | 5.7433E-08 | 2.5061E-10 |
| Tubb6 | 0.00478081 | 0.59471301 | 0.688 | 0.539 | 1 | 1.2866E-10 | 1.20422375 | 0.714 | 0.292 | 3.1253E-06 | 0.00478081 | 2.5731E-10 |
| Zfp9 | 6.0736E-07 | 1.10650454 | 0.75 | 0.472 | 0.01475409 | 1.3305E-10 | 1.24311712 | 0.727 | 0.312 | 3.2322E-06 | 6.0736E-07 | 2.6611E-10 |
| S100b | 0.20407109 | -0.9710895 | 0.917 | 0.762 | 1 | 1.3314E-10 | 0.85600581 | 0.756 | 0.281 | 3.2343E-06 | 0.20407109 | 2.6628E-10 |
| Cystm1 | 1.336E-10 | -2.9874874 | 0.115 | 0.497 | 3.2455E-06 | 1.0415E-07 | -1.3284709 | 0.176 | 0.479 | 0.00253002 | 1.0415E-07 | 2.6721E-10 |
| Mbp | 0.19011488 | -0.536826 | 0.052 | 0.098 | 1 | 1.3702E-10 | -1.4117349 | 0.088 | 0.396 | 3.3285E-06 | 0.19011488 | 2.7404E-10 |
| Camk4 | 1.3756E-10 | 1.07964432 | 0.698 | 0.269 | 3.3417E-06 | 0.06340768 | -0.3243548 | 0.676 | 0.469 | 1 | 0.06340768 | 2.7513E-10 |
| Epcam | 0.00071358 | -2.1624305 | 0.062 | 0.218 | 1 | 1.4087E-10 | -0.90323 | 0.164 | 0.51 | 3.4221E-06 | 0.00071358 | 2.8174E-10 |
| Emp3 | 0.000029168 | 0.56033429 | 0.979 | 0.813 | 0.70855439 | 1.4302E-10 | 0.96124857 | 0.95 | 0.625 | 3.4743E-06 | 2.9168E-05 | 2.8605E-10 |
| Ckmt1 | 0.8061167 | -0.577468 | 0.031 | 0.036 | 1 | 1.4503E-10 | -1.8948178 | 0.05 | 0.312 | 0.000003523 | 0.8061167 | 2.9006E-10 |
| Slc6a8 | 0.02534505 | -0.7275715 | 0.479 | 0.565 | 1 | 1.4612E-10 | 1.55364274 | 0.592 | 0.167 | 3.5495E-06 | 0.02534505 | 2.9224E-10 |
| Cnpy2 | 2.1535E-09 | 0.69784023 | 0.958 | 0.891 | 0.000052313 | 1.4801E-10 | 0.79419461 | 0.945 | 0.771 | 3.5955E-06 | 2.1535E-09 | 2.9602E-10 |
| Slc7a1 | 1.4916E-10 | 1.21225211 | 0.833 | 0.513 | 3.6234E-06 | 5.7148E-07 | 0.62874419 | 0.853 | 0.625 | 0.01388233 | 5.7148E-07 | 2.9832E-10 |
| Aqp3 | 0.00230451 | -3.4187702 | 0.031 | 0.15 | 1 | 1.5239E-10 | -2.07539 | 0.193 | 0.531 | 3.7018E-06 | 0.00230451 | 3.0477E-10 |
| Strap | 6.1178E-08 | 0.70478006 | 0.927 | 0.731 | 0.00148615 | 1.6486E-10 | 0.79920521 | 0.899 | 0.74 | 4.0049E-06 | 6.1178E-08 | 3.2973E-10 |
| Lbp | 1.7359E-10 | -2.6728298 | 0.219 | 0.58 | 4.2168E-06 | 0.03652147 | -0.677587 | 0.206 | 0.312 | 1 | 0.03652147 | 3.4717E-10 |
| Eif2s2 | 9.912E-07 | 0.52048725 | 0.99 | 0.974 | 0.02407812 | 1.7842E-10 | 0.69779831 | 0.987 | 0.958 | 4.3343E-06 | 9.912E-07 | 3.5685E-10 |
| mt-Nd5 | 0.5323106 | -0.1667507 | 1 | 1 | 1 | 1.8202E-10 | -0.9603413 | 1 | 1 | 4.4215E-06 | 0.5323106 | 3.6403E-10 |
| Colgalt1 | 1.8259E-10 | 0.78149086 | 0.906 | 0.777 | 4.4355E-06 | 4.6426E-10 | 1.06996651 | 0.87 | 0.594 | 0.000011278 | 4.6426E-10 | 3.6518E-10 |
| Rimoc1 | 0.6079913 | 0.32028198 | 0.375 | 0.352 | 1 | 1.8285E-10 | 1.32676936 | 0.559 | 0.135 | 4.4417E-06 | 0.6079913 | 3.657E-10 |
| Slc16a4 | 9.5358E-06 | 0.80354943 | 0.854 | 0.653 | 0.23164416 | 1.898E-10 | 1.05859617 | 0.895 | 0.542 | 4.6107E-06 | 9.5358E-06 | 3.796E-10 |
| Gipc2 | 0.60556246 | -1.1338613 | 0.062 | 0.078 | 1 | 2.0161E-10 | -2.5974319 | 0.076 | 0.354 | 4.8976E-06 | 0.60556246 | 4.0323E-10 |
| Jkamp | 0.28277806 | 0.14043337 | 0.75 | 0.653 | 1 | 2.1382E-10 | 1.2538595 | 0.731 | 0.302 | 5.1941E-06 | 0.28277806 | 4.2764E-10 |
| Dsc2 | 0.03171128 | 2.32942263 | 0.062 | 0.016 | 1 | 2.1649E-10 | -2.0323213 | 0.063 | 0.333 | 5.2589E-06 | 0.03171128 | 4.3298E-10 |
| Snrpd1 | 2.1905E-10 | 0.88779947 | 0.917 | 0.777 | 5.3212E-06 | 0.000032768 | 0.40853823 | 0.924 | 0.802 | 0.79601212 | 3.2768E-05 | 4.381E-10 |
| F5 | 2.2385E-10 | -2.628589 | 0.188 | 0.565 | 5.4377E-06 | 0.04315156 | -2.9245651 | 0.071 | 0.135 | 1 | 0.04315156 | 4.477E-10 |
| Btf3 | 2.4139E-07 | 0.49204283 | 0.99 | 0.953 | 0.00586375 | 2.2469E-10 | 0.64620139 | 0.996 | 0.938 | 5.4583E-06 | 2.4139E-07 | 4.4939E-10 |
| Ctsh | 2.4217E-10 | -1.5338787 | 0.521 | 0.808 | 5.8829E-06 | 0.23225478 | -0.1200307 | 0.559 | 0.688 | 1 | 0.23225478 | 4.8435E-10 |
| Wnt4 | 0.05725783 | 2.04144187 | 0.104 | 0.047 | 1 | 2.4262E-10 | -1.4093909 | 0.118 | 0.438 | 5.8938E-06 | 0.05725783 | 4.8524E-10 |
| Zswim6 | 4.2446E-06 | -1.0219396 | 0.792 | 0.943 | 0.10311013 | 2.5109E-10 | -1.6016589 | 0.882 | 0.958 | 6.0994E-06 | 4.2446E-06 | 5.0218E-10 |
| F11r | 0.50502113 | -0.407543 | 0.052 | 0.073 | 1 | 2.5536E-10 | -1.8662486 | 0.101 | 0.406 | 6.2031E-06 | 0.50502113 | 5.1071E-10 |
| Col12a1 | 2.5803E-10 | 0.28211692 | 0.99 | 0.57 | 0.000006268 | 7.8956E-08 | 0.49185637 | 0.987 | 0.76 | 0.00191799 | 7.8956E-08 | 5.1605E-10 |

| | | | | | | | | | | | | |
|---|---|---|---|---|---|---|---|---|---|---|---|---|
| Spats2l | 2.6527E-10 | 0.77704743 | 0.969 | 0.86 | 0.000006444 | 0.02940012 | 0.21772771 | 0.933 | 0.906 | 1 | 0.02940012 | 5.3054E-10 |
| Pdia3 | 5.7121E-07 | 0.43161661 | 1 | 0.99 | 0.01387579 | 2.7071E-10 | 0.60401882 | 1 | 1 | 6.5761E-06 | 5.7121E-07 | 5.4142E-10 |
| Mcl1 | 0.000054769 | 0.63599234 | 0.938 | 0.948 | 1 | 2.7678E-10 | 0.74531333 | 0.987 | 0.969 | 6.7236E-06 | 5.4769E-05 | 5.5356E-10 |
| Stt3a | 2.9626E-10 | 0.72572357 | 0.958 | 0.829 | 7.1967E-06 | 1.0013E-06 | 0.41032517 | 0.958 | 0.823 | 0.02432359 | 1.0013E-06 | 5.9251E-10 |
| Rbp4 | 1.4597E-07 | 1.52999295 | 0.604 | 0.321 | 0.00354583 | 3.0195E-10 | 1.48694588 | 0.71 | 0.292 | 7.3349E-06 | 1.4597E-07 | 6.039E-10 |
| Il1r1 | 3.0609E-10 | -2.0080277 | 0.323 | 0.679 | 7.4355E-06 | 0.01064611 | 0.16762281 | 0.336 | 0.167 | 1 | 0.01064611 | 6.1218E-10 |
| Rraga | 0.00046121 | 0.57517905 | 0.771 | 0.611 | 1 | 3.0707E-10 | 1.12925637 | 0.786 | 0.406 | 7.4593E-06 | 0.00046121 | 6.1413E-10 |
| Hspa8 | 0.10435612 | 0.10339549 | 1 | 0.979 | 1 | 3.1352E-10 | 0.70342602 | 0.996 | 0.99 | 0.000007616 | 0.10435612 | 6.2704E-10 |
| Ldhb | 0.00483043 | 0.80104366 | 0.594 | 0.477 | 1 | 3.1491E-10 | 1.00134643 | 0.845 | 0.615 | 7.6497E-06 | 0.00483043 | 6.2982E-10 |
| Gstm2 | 0.11302678 | -2.4519371 | 0 | 0.026 | 1 | 3.1721E-10 | -2.3098553 | 0.05 | 0.302 | 7.7056E-06 | 0.11302678 | 6.3441E-10 |
| Dst | 3.2164E-10 | -1.4254063 | 0.969 | 0.959 | 7.8133E-06 | 0.89878152 | -0.4327523 | 0.983 | 0.948 | 1 | 0.89878152 | 6.4328E-10 |
| Irf6 | 0.32025586 | -1.3144336 | 0 | 0.01 | 1 | 3.2886E-10 | -2.8948178 | 0.034 | 0.26 | 7.9887E-06 | 0.32025586 | 6.5772E-10 |
| Nrip1 | 3.4539E-10 | -1.4819766 | 0.74 | 0.912 | 8.3902E-06 | 0.08831695 | -0.2696662 | 0.819 | 0.917 | 1 | 0.08831695 | 6.9078E-10 |
| Pdrg1 | 0.38653594 | -0.2884903 | 0.802 | 0.756 | 1 | 3.5506E-10 | 1.23138586 | 0.731 | 0.292 | 0.000008625 | 0.38653594 | 7.1011E-10 |
| Myl12a | 0.0049778 | 0.23645331 | 0.99 | 0.969 | 1 | 3.557E-10 | 0.57900141 | 0.996 | 0.969 | 8.6407E-06 | 0.0049778 | 7.114E-10 |
| Lrrc59 | 1.6143E-06 | 0.68191866 | 0.917 | 0.663 | 0.03921375 | 3.6936E-10 | 0.95190861 | 0.903 | 0.729 | 8.9724E-06 | 1.6143E-06 | 7.3871E-10 |
| Tspan3 | 0.0451845 | -0.2741573 | 0.99 | 0.979 | 1 | 3.8184E-10 | 0.69754699 | 0.992 | 0.938 | 9.2757E-06 | 0.0451845 | 7.6369E-10 |
| Camk1 | 0.000050313 | 0.80155264 | 0.802 | 0.596 | 1 | 3.8511E-10 | 1.12871775 | 0.765 | 0.323 | 9.3552E-06 | 5.0313E-05 | 7.7023E-10 |
| Pfn1 | 0.00052297 | 0.35347708 | 1 | 0.927 | 1 | 3.8752E-10 | 0.70320965 | 1 | 0.948 | 9.4137E-06 | 0.00052297 | 7.7504E-10 |
| Eif1ax | 0.01479914 | 0.21338504 | 0.917 | 0.845 | 1 | 3.9326E-10 | 0.81364969 | 0.962 | 0.792 | 9.5531E-06 | 0.01479914 | 7.8652E-10 |
| Syn3 | 4.6025E-10 | -2.577468 | 0.333 | 0.648 | 0.00001118 | 0.13962598 | -0.9082383 | 0.277 | 0.177 | 1 | 0.13962598 | 9.205E-10 |
| mt-Nd4l | 0.36356616 | -0.4339238 | 0.948 | 0.922 | 1 | 4.9003E-10 | -1.0773509 | 0.887 | 0.948 | 0.000011904 | 0.36356616 | 9.8006E-10 |
| Fscn1 | 5.3351E-10 | 1.28051303 | 0.719 | 0.321 | 0.00001296 | 0.000017358 | 0.55947579 | 0.739 | 0.375 | 0.42164908 | 1.7358E-05 | 1.067E-09 |
| Snrpf | 0.00203689 | 0.42426918 | 0.906 | 0.881 | 1 | 5.3548E-10 | 0.81388011 | 0.903 | 0.771 | 0.000013008 | 0.00203689 | 1.071E-09 |
| Rgs2 | 6.7379E-09 | -1.8426396 | 0.385 | 0.689 | 0.00016368 | 5.4623E-10 | -1.3932713 | 0.155 | 0.49 | 0.000013269 | 6.7379E-09 | 1.0925E-09 |
| Cd109 | 5.5792E-10 | 0.76089369 | 1 | 0.876 | 0.000013553 | 1.3013E-07 | 0.77772241 | 0.987 | 0.802 | 0.00316122 | 1.3013E-07 | 1.1158E-09 |
| Asap2 | 5.6636E-10 | -1.3655832 | 0.562 | 0.803 | 0.000013758 | 0.62983891 | -0.4526559 | 0.513 | 0.479 | 1 | 0.62983891 | 1.1327E-09 |
| Spint2 | 0.00425637 | -1.1010299 | 0.198 | 0.368 | 1 | 5.6831E-10 | -1.5006776 | 0.239 | 0.583 | 0.000013805 | 0.00425637 | 1.1366E-09 |
| Sorl1 | 0.00672156 | -1.5160674 | 0.083 | 0.207 | 1 | 5.6855E-10 | -1.5757493 | 0.206 | 0.562 | 0.000013811 | 0.00672156 | 1.1371E-09 |
| Lcp1 | 0.05939926 | 0.59245704 | 0.073 | 0.026 | 1 | 5.841E-10 | 0.54000439 | 0.101 | 0.406 | 0.000014189 | 0.05939926 | 1.1682E-09 |
| S100a10 | 0.00936576 | 0.38652897 | 0.979 | 0.969 | 1 | 5.9156E-10 | 0.91489066 | 0.987 | 0.948 | 0.00001437 | 0.00936576 | 1.1831E-09 |
| Tspan2 | 5.9176E-10 | 2.14951354 | 0.49 | 0.171 | 0.000014375 | 0.43508078 | -0.2377055 | 0.206 | 0.156 | 1 | 0.43508078 | 1.1835E-09 |
| Etv6 | 2.6612E-06 | -0.7454931 | 0.917 | 0.974 | 0.06464558 | 5.9599E-10 | -1.3542494 | 0.874 | 0.979 | 0.000014478 | 2.6612E-06 | 1.192E-09 |
| Ehf | 0.33643557 | -0.407543 | 0.156 | 0.207 | 1 | 6.0061E-10 | -1.7793405 | 0.16 | 0.49 | 0.00001459 | 0.33643557 | 1.2012E-09 |
| Slc16a11 | 0.28505098 | -1.2148979 | 0.01 | 0.031 | 1 | 6.7214E-10 | -2.1843244 | 0.059 | 0.312 | 0.000016328 | 0.28505098 | 1.3443E-09 |
| Tm2d3 | 0.68833432 | -0.130009 | 0.5 | 0.497 | 1 | 6.7499E-10 | 1.42923038 | 0.618 | 0.208 | 0.000016397 | 0.68833432 | 1.35E-09 |
| Tbrg1 | 6.7652E-10 | 0.75717118 | 1 | 0.891 | 0.000016434 | 8.7264E-10 | 0.83949953 | 0.975 | 0.823 | 0.000021198 | 8.7264E-10 | 1.353E-09 |
| Ube2q2 | 6.785E-10 | -1.7433746 | 0.76 | 0.876 | 0.000016482 | 0.37062157 | -0.2590594 | 0.794 | 0.802 | 1 | 0.37062157 | 1.357E-09 |
| Nrxn1 | 0.00199237 | -4.9232428 | 0.021 | 0.135 | 1 | 7.0144E-10 | -1.6577786 | 0.097 | 0.385 | 0.000017039 | 0.00199237 | 1.4029E-09 |
| Aqp1 | 7.1299E-10 | -2.4120444 | 0.031 | 0.378 | 0.00001732 | 0.00128948 | -3.4549056 | 0.059 | 0.167 | 1 | 0.00128948 | 1.426E-09 |
| Rnase4 | 0.00482413 | 0.40393482 | 0.823 | 0.689 | 1 | 7.5731E-10 | 0.65558752 | 0.95 | 0.75 | 0.000018397 | 0.00482413 | 1.5146E-09 |
| Ncbp2 | 3.1434E-06 | 0.79336973 | 0.792 | 0.534 | 0.07635854 | 7.7229E-10 | 1.09068267 | 0.769 | 0.417 | 0.00001876 | 3.1434E-06 | 1.5446E-09 |
| Gorasp2 | 7.8368E-10 | 0.74074936 | 0.958 | 0.896 | 0.000019037 | 1.0356E-06 | 0.40543443 | 0.958 | 0.865 | 0.02515692 | 1.0356E-06 | 1.5674E-09 |
| Six2 | 0.00662257 | 0.42495064 | 0.76 | 0.596 | 1 | 7.9391E-10 | 0.95220138 | 0.845 | 0.396 | 0.000019286 | 0.00662257 | 1.5878E-09 |
| Eif1b | 0.00038856 | 0.46534512 | 0.885 | 0.746 | 1 | 8.29E-10 | 0.96157803 | 0.908 | 0.677 | 0.000020138 | 0.00038856 | 1.658E-09 |
| Uqcrb | 0.00178298 | 0.38321911 | 0.958 | 0.886 | 1 | 8.5197E-10 | 0.71110436 | 0.954 | 0.771 | 0.000020696 | 0.00178298 | 1.7039E-09 |
| Sult1d1 | 0.57818601 | -0.577468 | 0.042 | 0.057 | 1 | 8.8537E-10 | -2.352924 | 0.122 | 0.427 | 0.000021507 | 0.57818601 | 1.7707E-09 |
| Mctp2 | 0.00010796 | -1.3348977 | 0.167 | 0.389 | 1 | 8.8737E-10 | -1.9669675 | 0.08 | 0.354 | 0.000021556 | 0.00010796 | 1.7747E-09 |
| Slc5a8 | 0.61737927 | 1.00749454 | 0.01 | 0.005 | 1 | 8.8837E-10 | -3.5728897 | 0.017 | 0.208 | 0.00002158 | 0.61737927 | 1.7767E-09 |
| Rgs3 | 9.0535E-10 | 1.36594851 | 0.781 | 0.461 | 0.000021993 | 0.000017925 | 0.18468016 | 0.79 | 0.469 | 0.43543357 | 1.7925E-05 | 1.8107E-09 |
| Eef1d | 0.01363263 | 0.2858362 | 0.979 | 0.891 | 1 | 9.345E-10 | 0.79019191 | 0.958 | 0.812 | 0.000022701 | 0.01363263 | 1.869E-09 |
| Prdx4 | 9.4921E-10 | 1.14971661 | 0.812 | 0.57 | 0.000023058 | 1.8595E-09 | 0.99680608 | 0.748 | 0.354 | 0.00004517 | 1.8595E-09 | 1.8984E-09 |
| Slco3a1 | 9.8016E-10 | -2.5647569 | 0.115 | 0.482 | 0.00002381 | 0.00131099 | -1.1843244 | 0.164 | 0.323 | 1 | 0.00131099 | 1.9603E-09 |
| Mfsd6 | 0.03579822 | -1.577468 | 0.01 | 0.067 | 1 | 9.9203E-10 | -1.9943534 | 0.092 | 0.375 | 0.000024098 | 0.03579822 | 1.9841E-09 |
| Ndn | 0.03549568 | 0.38061839 | 0.677 | 0.606 | 1 | 1.0398E-09 | 1.23203852 | 0.651 | 0.25 | 0.000025258 | 0.03549568 | 2.0795E-09 |
| Tsc22d3 | 0.01144338 | 0.40152154 | 0.948 | 0.891 | 1 | 1.0567E-09 | 1.01925043 | 0.945 | 0.812 | 0.00002567 | 0.01144338 | 2.1134E-09 |
| Nop10 | 0.01342992 | 0.28345144 | 0.979 | 0.845 | 1 | 1.0763E-09 | 0.83926842 | 0.899 | 0.698 | 0.000026146 | 0.01342992 | 2.1527E-09 |
| Plekha6 | 0.000025188 | -2.5208844 | 0.094 | 0.311 | 0.61186088 | 1.0833E-09 | -1.6454583 | 0.109 | 0.406 | 0.000026316 | 2.5188E-05 | 2.1667E-09 |
| Srp19 | 0.06374376 | 0.26591969 | 0.854 | 0.767 | 1 | 1.1562E-09 | 0.93231809 | 0.866 | 0.635 | 0.000028086 | 0.06374376 | 2.3124E-09 |
| Grcc10 | 0.08558926 | 0.19294012 | 0.885 | 0.824 | 1 | 1.1637E-09 | 0.92871709 | 0.866 | 0.75 | 0.000028269 | 0.08558926 | 2.3274E-09 |
| Lmnb1 | 0.84521296 | 0.53800925 | 0.135 | 0.15 | 1 | 1.1846E-09 | -1.5366261 | 0.109 | 0.417 | 0.000028777 | 0.84521296 | 2.3692E-09 |
| Cdc42 | 0.02673942 | 0.22286331 | 0.99 | 0.995 | 1 | 1.1971E-09 | 0.48693797 | 1 | 1 | 0.000029079 | 0.02673942 | 2.3941E-09 |
| Dapl1 | 0.06116328 | -2.5160674 | 0.01 | 0.057 | 1 | 1.2466E-09 | -1.484942 | 0.067 | 0.333 | 0.000030282 | 0.06116328 | 2.4931E-09 |
| Sult1e1 | 0.61553741 | -0.9467018 | 0.021 | 0.031 | 1 | 1.2564E-09 | -2.8123556 | 0.042 | 0.271 | 0.00003052 | 0.61553741 | 2.5127E-09 |
| Irf2bp1 | 0.33356682 | 0.47406234 | 0.271 | 0.223 | 1 | 1.2902E-09 | 1.74903843 | 0.492 | 0.125 | 0.000031341 | 0.33356682 | 2.5804E-09 |
| Plaur | 0.66997473 | 0.32497673 | 0.208 | 0.238 | 1 | 1.3005E-09 | -1.4353861 | 0.084 | 0.365 | 0.000031591 | 0.66997473 | 2.6009E-09 |
| Cadps2 | 0.43274423 | -0.3144336 | 0.031 | 0.052 | 1 | 1.3005E-09 | -1.525584 | 0.084 | 0.365 | 0.000031592 | 0.43274423 | 2.601E-09 |
| Penk | 1.3224E-09 | 1.82367331 | 0.604 | 0.223 | 0.000032124 | 9.9899E-09 | 1.54024931 | 0.592 | 0.24 | 0.00024268 | 9.9899E-09 | 2.6448E-09 |
| Bmp7 | 1.3252E-09 | 1.88791292 | 0.417 | 0.109 | 0.000032192 | 0.1281494 | 0.75725893 | 0.218 | 0.146 | 1 | 0.1281494 | 2.6504E-09 |
| Hspa5 | 0.02753099 | 0.21986504 | 1 | 0.984 | 1 | 1.339E-09 | 0.72275119 | 1 | 0.99 | 0.000032528 | 0.02753099 | 2.6781E-09 |
| Snu13 | 3.2526E-07 | 0.5950921 | 0.958 | 0.891 | 0.0079012 | 1.3615E-09 | 0.843391 | 0.929 | 0.812 | 0.000033074 | 3.2526E-07 | 2.7231E-09 |
| Scara5 | 1.3838E-09 | -2.6237616 | 0.167 | 0.523 | 0.000033616 | 0.0066844 | -3.4117349 | 0.084 | 0.177 | 1 | 0.0066844 | 2.7676E-09 |
| Ogn | 1.4011E-09 | 1.26773031 | 0.771 | 0.409 | 0.000034036 | 0.000091874 | 0.15875092 | 0.874 | 0.531 | 1 | 9.1874E-05 | 2.8023E-09 |
| Txnl1 | 1.3301E-06 | 0.69715442 | 0.917 | 0.87 | 0.03231162 | 1.4035E-09 | 0.71780877 | 0.975 | 0.875 | 0.000034094 | 1.3301E-06 | 2.807E-09 |
| Srbd1 | 1.4431E-09 | 0.92167517 | 0.906 | 0.611 | 0.000035056 | 0.000021686 | 0.50581493 | 0.773 | 0.417 | 0.52679329 | 2.1686E-05 | 2.8862E-09 |
| Trim24 | 0.07006691 | -0.561614 | 0.354 | 0.456 | 1 | 1.5009E-09 | -1.1636345 | 0.374 | 0.74 | 0.000036459 | 0.07006691 | 3.0017E-09 |
| Lncpint | 0.000020319 | -1.1782755 | 0.917 | 0.943 | 0.49359143 | 1.5265E-09 | -1.1039903 | 0.95 | 0.979 | 0.000037081 | 2.0319E-05 | 3.0529E-09 |
| Sdk1 | 0.68381741 | -0.590407 | 0.177 | 0.155 | 1 | 1.546E-09 | -1.6887214 | 0.239 | 0.594 | 0.000037555 | 0.68381741 | 3.092E-09 |
| Pmepa1 | 0.00038114 | 0.4014916 | 0.979 | 0.953 | 1 | 1.5595E-09 | 0.6145103 | 0.992 | 1 | 0.000037884 | 0.00038114 | 3.1191E-09 |
| Cox4i1 | 0.16741099 | 0.15284228 | 1 | 0.953 | 1 | 1.5781E-09 | 0.58462879 | 1 | 0.958 | 0.000038336 | 0.16741099 | 3.1563E-09 |
| Marchf3 | 0.37489417 | -0.5717992 | 0.292 | 0.358 | 1 | 1.5888E-09 | -1.7692869 | 0.244 | 0.615 | 0.000038595 | 0.37489417 | 3.1776E-09 |
| Mt2 | 0.00036528 | -1.1447718 | 0.281 | 0.518 | 1 | 1.6121E-09 | -0.4888254 | 0.206 | 0.583 | 0.00003916 | 0.00036528 | 3.2241E-09 |
| Cd55 | 0.04523461 | 1.69338595 | 0.229 | 0.145 | 1 | 1.6499E-09 | -1.1934458 | 0.193 | 0.542 | 0.00004008 | 0.04523461 | 3.2998E-09 |
| Aldoa | 0.2854541 | -0.272802 | 0.948 | 0.948 | 1 | 1.6734E-09 | 0.71934471 | 0.975 | 0.885 | 0.000040649 | 0.2854541 | 3.3467E-09 |
| Dad1 | 0.0008762 | 0.33181318 | 0.979 | 0.948 | 1 | 1.6848E-09 | 0.64578562 | 0.983 | 0.865 | 0.000040928 | 0.0008762 | 3.3697E-09 |
| Armh4 | 0.11606975 | 0.4544161 | 0.562 | 0.508 | 1 | 1.6867E-09 | 1.18121567 | 0.71 | 0.302 | 0.000040973 | 0.11606975 | 3.3733E-09 |
| Dap | 0.32412349 | -0.1631491 | 0.948 | 0.907 | 1 | 1.7729E-09 | 0.81250834 | 0.929 | 0.74 | 0.000043067 | 0.32412349 | 3.5458E-09 |
| Rpl27 | 0.03225249 | 0.2604122 | 0.99 | 0.959 | 1 | 1.8239E-09 | 0.62352117 | 1 | 0.958 | 0.000044306 | 0.03225249 | 3.6478E-09 |
| Tomm20 | 0.000515 | 0.41363564 | 0.958 | 0.938 | 1 | 1.8478E-09 | 0.61170338 | 0.975 | 0.896 | 0.000044886 | 0.000515 | 3.6956E-09 |
| Slc20a1 | 1.8651E-09 | -1.9245481 | 0.49 | 0.793 | 0.000045307 | 7.4539E-06 | -1.1786107 | 0.441 | 0.76 | 0.18107103 | 7.4539E-06 | 3.7302E-09 |
| Swi5 | 0.00099741 | 0.40652126 | 0.969 | 0.922 | 1 | 1.8678E-09 | 0.70977354 | 0.975 | 0.833 | 0.000045372 | 0.00099741 | 3.7355E-09 |
| Gar1 | 1.8692E-09 | 1.23356262 | 0.688 | 0.352 | 0.000045406 | 5.1414E-07 | 1.2488757 | 0.542 | 0.229 | 0.01248958 | 5.1414E-07 | 3.7384E-09 |

| | | | | | | | | | | | | |
|---|---|---|---|---|---|---|---|---|---|---|---|---|
| Gnai2 | 0.000090792 | 0.40742514 | 1 | 0.964 | 1 | 1.9177E-09 | 0.64151325 | 0.996 | 0.958 | 0.000046584 | 9.0792E-05 | 3.8353E-09 |
| Cd302 | 0.00117364 | -0.5455839 | 0.802 | 0.845 | 1 | 1.9443E-09 | 0.92209147 | 0.878 | 0.49 | 0.000047232 | 0.00117364 | 3.8887E-09 |
| Ndrg2 | 0.2713305 | -0.4399644 | 0.521 | 0.56 | 1 | 1.9918E-09 | 1.39796399 | 0.681 | 0.292 | 0.000048385 | 0.2713305 | 3.9836E-09 |
| Ssbp1 | 0.00142831 | 0.74446013 | 0.635 | 0.482 | 1 | 2.0453E-09 | 1.40286279 | 0.597 | 0.208 | 0.000049684 | 0.00142831 | 4.0905E-09 |
| Smim11 | 0.000076483 | 0.67207421 | 0.802 | 0.585 | 1 | 2.1176E-09 | 0.94456601 | 0.761 | 0.365 | 0.000051441 | 7.6483E-05 | 4.2352E-09 |
| Atxn7l3b | 0.00311373 | 0.39907306 | 0.948 | 0.896 | 1 | 2.2154E-09 | 0.65189727 | 0.958 | 0.844 | 0.000053817 | 0.00311373 | 4.4308E-09 |
| Tmem254 | 2.2441E-09 | 1.25111764 | 0.812 | 0.632 | 0.000054514 | 0.00086518 | 0.69330508 | 0.748 | 0.667 | 1 | 0.00086518 | 4.4882E-09 |
| Rpl35 | 0.000060777 | 0.44620208 | 1 | 0.922 | 1 | 2.246E-09 | 0.76670707 | 0.95 | 0.865 | 0.00005456 | 6.0777E-05 | 4.492E-09 |
| Mageh1 | 0.09059554 | 0.4134869 | 0.552 | 0.492 | 1 | 2.3483E-09 | 0.94887901 | 0.706 | 0.302 | 0.000057044 | 0.09059554 | 4.6965E-09 |
| Praf2 | 0.00219566 | 0.59838187 | 0.677 | 0.497 | 1 | 2.4221E-09 | 1.36601573 | 0.634 | 0.24 | 0.000058838 | 0.00219566 | 4.8443E-09 |
| Acbd3 | 1.7127E-06 | 0.59517142 | 0.938 | 0.907 | 0.04160492 | 2.4388E-09 | 0.68235806 | 0.987 | 0.917 | 0.000059243 | 1.7127E-06 | 4.8776E-09 |
| Cilp2 | 0.02757523 | 0.68901924 | 0.604 | 0.58 | 1 | 2.5007E-09 | 0.99826703 | 0.878 | 0.521 | 0.000060746 | 0.02757523 | 5.0013E-09 |
| Tacstd2 | 0.12782048 | -1.6363617 | 0.021 | 0.062 | 1 | 2.5186E-09 | -0.5799444 | 0.088 | 0.375 | 0.000061182 | 0.12782048 | 5.0372E-09 |
| Cdkn1c | 0.01326054 | 0.33825151 | 0.781 | 0.642 | 1 | 2.5313E-09 | 1.21162996 | 0.929 | 0.625 | 0.000061491 | 0.01326054 | 5.0627E-09 |
| F2r | 0.19775553 | -1.3144336 | 0.031 | 0.067 | 1 | 2.554E-09 | -2.2940879 | 0.118 | 0.406 | 0.000062041 | 0.19775553 | 5.108E-09 |
| Dnajc25 | 2.5745E-09 | 0.78510212 | 0.958 | 0.793 | 0.000062539 | 5.3547E-09 | 0.92201821 | 0.899 | 0.604 | 0.00013008 | 5.3547E-09 | 5.149E-09 |
| Itpr2 | 2.6272E-09 | -1.0521484 | 0.875 | 0.938 | 0.00006382 | 0.10719183 | -0.5479322 | 0.845 | 0.865 | 1 | 0.10719183 | 5.2544E-09 |
| Cibar1 | 0.27197183 | 0.15817345 | 0.698 | 0.617 | 1 | 2.6704E-09 | 1.02553509 | 0.697 | 0.281 | 0.000064869 | 0.27197183 | 5.3408E-09 |
| Gpsm2 | 1.7034E-08 | 0.94207146 | 0.75 | 0.383 | 0.00041379 | 2.798E-09 | 1.27510724 | 0.748 | 0.375 | 0.000067968 | 1.7034E-08 | 5.5959E-09 |
| Fkbp10 | 1.8168E-07 | 0.64985935 | 0.958 | 0.813 | 0.00441341 | 2.8186E-09 | 0.76297105 | 0.924 | 0.531 | 0.000068468 | 1.8168E-07 | 5.6371E-09 |
| Rnf11 | 0.00365563 | 0.30533082 | 0.969 | 0.891 | 1 | 2.9294E-09 | 0.67778224 | 0.992 | 0.885 | 0.00007116 | 0.00365563 | 5.8588E-09 |
| Srsf7 | 8.4848E-06 | 0.82391678 | 0.979 | 0.917 | 0.20611182 | 3.077E-09 | 0.733584 | 0.979 | 0.938 | 0.000074746 | 8.4848E-06 | 6.1539E-09 |
| Man2a1 | 3.1005E-09 | -1.6252437 | 0.427 | 0.725 | 0.000075317 | 1.5376E-06 | -1.2788284 | 0.441 | 0.698 | 0.03735074 | 1.5376E-06 | 6.201E-09 |
| Cox7c | 0.00434303 | 0.278043 | 0.99 | 0.969 | 1 | 3.1093E-09 | 0.59929661 | 0.996 | 0.917 | 0.000075531 | 0.00434303 | 6.2186E-09 |
| Zim1 | 3.1109E-09 | 2.04144187 | 0.5 | 0.181 | 0.000075569 | 0.000086754 | 1.09007534 | 0.403 | 0.167 | 1 | 8.6754E-05 | 6.2217E-09 |
| Btg1 | 0.05727029 | -0.3510698 | 0.99 | 1 | 1 | 3.3329E-09 | 0.54395996 | 1 | 1 | 0.000080963 | 0.05727029 | 6.6658E-09 |
| Trnt1 | 0.0002354 | 0.7716816 | 0.615 | 0.389 | 1 | 3.3501E-09 | 1.27510724 | 0.55 | 0.167 | 0.00008138 | 0.0002354 | 6.7002E-09 |
| Eif4ebp1 | 3.3877E-09 | 1.63910713 | 0.552 | 0.207 | 0.000082293 | 1.2763E-06 | 1.49176329 | 0.496 | 0.188 | 0.03100484 | 1.2763E-06 | 6.7753E-09 |
| Reck | 0.00010262 | 0.73619251 | 0.729 | 0.508 | 1 | 3.4594E-09 | 1.01544362 | 0.866 | 0.49 | 0.000084035 | 0.00010262 | 6.9188E-09 |
| Sbspon | 3.4683E-09 | -4.2019588 | 0.052 | 0.373 | 0.000084252 | 0.57448543 | -0.9754362 | 0.172 | 0.135 | 1 | 0.57448543 | 6.9366E-09 |
| Epas1 | 0.000079485 | -1.3078609 | 0.292 | 0.539 | 1 | 3.4738E-09 | -1.4782528 | 0.534 | 0.771 | 0.000084384 | 7.9485E-05 | 6.9475E-09 |
| Kif26b | 3.4982E-09 | 1.29048968 | 0.698 | 0.337 | 0.000084978 | 0.75869048 | -0.6396455 | 0.487 | 0.458 | 1 | 0.75869048 | 6.9964E-09 |
| AI506816 | 1.5199E-08 | 0.97547286 | 0.906 | 0.762 | 0.00036921 | 3.7186E-09 | 0.93807225 | 0.824 | 0.469 | 0.000090333 | 1.5199E-08 | 7.4372E-09 |
| Gm17225 | 3.8175E-09 | 5.43375929 | 0.188 | 0.005 | 0.000092735 | 0.08663267 | 2.01207283 | 0.05 | 0.01 | 1 | 0.08663267 | 7.635E-09 |
| Tbc1d8 | 0.29163323 | -1.130009 | 0.042 | 0.073 | 1 | 3.8981E-09 | -2.3345173 | 0.097 | 0.365 | 0.000094692 | 0.29163323 | 7.7961E-09 |
| Fkbp1a | 0.00030667 | 0.47726151 | 0.958 | 0.87 | 1 | 4.0637E-09 | 0.72797681 | 0.945 | 0.802 | 0.000098716 | 0.00030667 | 8.1275E-09 |
| Bcl7b | 0.19063089 | -0.2536303 | 0.438 | 0.523 | 1 | 4.0638E-09 | 1.39709776 | 0.571 | 0.208 | 0.000098718 | 0.19063089 | 8.1276E-09 |
| Lurap1l | 0.03210886 | -2.9925055 | 0 | 0.047 | 1 | 4.0878E-09 | -4.6317834 | 0.004 | 0.156 | 0.0000993 | 0.03210886 | 8.1755E-09 |
| Cpm | 4.1655E-09 | -1.5184236 | 0.594 | 0.808 | 0.00010119 | 0.75378158 | -0.5885713 | 0.303 | 0.333 | 1 | 0.75378158 | 8.3309E-09 |
| Irf1 | 0.06211919 | -1.1851505 | 0.26 | 0.342 | 1 | 4.1801E-09 | -1.5679234 | 0.193 | 0.521 | 0.00010154 | 0.06211919 | 8.3601E-09 |
| Cggbp1 | 0.00206571 | 0.32801444 | 0.896 | 0.808 | 1 | 4.2487E-09 | 0.73964992 | 0.958 | 0.875 | 0.00010321 | 0.00206571 | 8.4975E-09 |
| Slc39a8 | 4.2819E-09 | -1.8870123 | 0.25 | 0.606 | 0.00010402 | 0.79769791 | -0.1245002 | 0.248 | 0.219 | 1 | 0.79769791 | 8.5639E-09 |
| Cecr2 | 0.08034351 | 1.68556644 | 0.042 | 0.01 | 1 | 4.3581E-09 | -2.0468209 | 0.055 | 0.292 | 0.00010587 | 0.08034351 | 8.7163E-09 |
| Atp5g2 | 0.01806731 | 0.21135483 | 0.958 | 0.927 | 1 | 4.3945E-09 | 0.65914959 | 0.962 | 0.875 | 0.00010675 | 0.01806731 | 8.789E-09 |
| Cdk19 | 0.00077931 | -1.1335178 | 0.323 | 0.492 | 1 | 4.4061E-09 | -1.6714535 | 0.349 | 0.698 | 0.00010703 | 0.00077931 | 8.8122E-09 |
| Fkbp7 | 5.1984E-07 | 0.65708275 | 0.927 | 0.705 | 0.01262791 | 4.5737E-09 | 0.88041501 | 0.87 | 0.427 | 0.0001111 | 5.1984E-07 | 9.1474E-09 |
| Psmb6 | 0.000000918 | 0.65957123 | 0.927 | 0.829 | 0.02230013 | 4.6544E-09 | 0.81381911 | 0.903 | 0.708 | 0.00011306 | 9.18E-07 | 9.3087E-09 |
| Ncl | 4.6724E-09 | 0.62360574 | 1 | 0.984 | 0.0001135 | 2.291E-07 | 0.50183425 | 1 | 0.99 | 0.00556531 | 2.291E-07 | 9.3449E-09 |
| Olfml3 | 4.706E-09 | 1.02370476 | 0.896 | 0.596 | 0.00011432 | 0.06818565 | 0.18058964 | 0.643 | 0.469 | 1 | 0.06818565 | 9.412E-09 |
| Xpot | 3.0971E-06 | 0.80702995 | 0.781 | 0.528 | 0.07523527 | 4.7849E-09 | 1.04230503 | 0.782 | 0.448 | 0.00011623 | 3.0971E-06 | 9.5697E-09 |
| Abca13 | 0.32025586 | -1.3144336 | 0 | 0.01 | 1 | 4.9366E-09 | -2.8334172 | 0.029 | 0.229 | 0.00011992 | 0.32025586 | 9.8732E-09 |
| Gnl1 | 0.03017002 | 0.64071221 | 0.5 | 0.378 | 1 | 5.0649E-09 | 1.22619764 | 0.613 | 0.219 | 0.00012304 | 0.03017002 | 1.013E-08 |
| Tvp23b | 0.01169553 | 0.4810105 | 0.802 | 0.674 | 1 | 5.1254E-09 | 1.13349263 | 0.752 | 0.385 | 0.00012451 | 0.01169553 | 1.0251E-08 |
| Cox14 | 0.05334605 | 0.22018186 | 0.896 | 0.845 | 1 | 5.195E-09 | 0.82814497 | 0.882 | 0.698 | 0.0001262 | 0.05334605 | 1.039E-08 |
| Srp14 | 0.08011482 | 0.21212956 | 0.979 | 0.896 | 1 | 5.2626E-09 | 0.60768258 | 0.954 | 0.823 | 0.00012784 | 0.08011482 | 1.0525E-08 |
| Rsl1d1 | 0.00151072 | 0.42253204 | 0.927 | 0.824 | 1 | 5.4248E-09 | 0.8038309 | 0.941 | 0.812 | 0.00013178 | 0.00151072 | 1.085E-08 |
| Gk | 0.00029677 | -2.4843586 | 0.042 | 0.202 | 1 | 5.5867E-09 | -0.9337068 | 0.139 | 0.458 | 0.00013571 | 0.00029677 | 1.1173E-08 |
| Tspan5 | 5.6677E-09 | -1.7998604 | 0.448 | 0.736 | 0.00013768 | 0.00570577 | -0.7623675 | 0.601 | 0.75 | 1 | 0.00570577 | 1.1335E-08 |
| Csrp1 | 5.7107E-09 | -1.9298499 | 0.281 | 0.632 | 0.00013872 | 0.000014903 | -1.2692133 | 0.34 | 0.625 | 0.36202834 | 1.4903E-05 | 1.1421E-08 |
| 4930523C07Rik | 0.26085504 | 0.31943854 | 0.917 | 0.891 | 1 | 5.7384E-09 | 0.73351651 | 0.975 | 0.844 | 0.0001394 | 0.26085504 | 1.1477E-08 |
| Msmo1 | 0.59548508 | 0.17979435 | 0.667 | 0.637 | 1 | 5.8818E-09 | 1.56236632 | 0.651 | 0.292 | 0.00014288 | 0.59548508 | 1.1764E-08 |
| Ogfod3 | 0.54591122 | 0.33258991 | 0.385 | 0.368 | 1 | 5.9169E-09 | 1.9808219 | 0.441 | 0.094 | 0.00014373 | 0.54591122 | 1.1834E-08 |
| Cdkn1a | 0.41948193 | 0.14894633 | 0.969 | 0.964 | 1 | 5.92E-09 | 0.69820451 | 0.987 | 0.969 | 0.00014381 | 0.41948193 | 1.184E-08 |
| Scin | 0.07170011 | -0.4065757 | 0.677 | 0.731 | 1 | 5.9915E-09 | 0.55509478 | 0.782 | 0.312 | 0.00014555 | 0.07170011 | 1.1983E-08 |
| Dipk2a | 0.04457819 | 0.33313295 | 0.958 | 0.865 | 1 | 6.1555E-09 | 0.72338107 | 0.971 | 0.75 | 0.00014953 | 0.04457819 | 1.2311E-08 |
| Pef1 | 0.37721988 | 0.11099911 | 0.5 | 0.43 | 1 | 6.1625E-09 | 1.0671142 | 0.534 | 0.146 | 0.0001497 | 0.37721988 | 1.2325E-08 |
| Trip11 | 6.2129E-09 | 0.60894516 | 0.99 | 0.922 | 0.00015092 | 0.00033469 | 0.36584184 | 0.979 | 0.948 | 1 | 0.00033469 | 1.2426E-08 |
| Ddah1 | 0.000029085 | 0.65763551 | 0.74 | 0.503 | 0.70652787 | 6.2712E-09 | 0.78968041 | 0.693 | 0.26 | 0.00015234 | 2.9085E-05 | 1.2542E-08 |
| Rcn1 | 0.02221199 | 0.30250133 | 0.948 | 0.824 | 1 | 6.34E-09 | 0.9289316 | 0.857 | 0.458 | 0.00015401 | 0.02221199 | 1.268E-08 |
| Ykt6 | 0.00091763 | 0.65565086 | 0.667 | 0.446 | 1 | 6.371E-09 | 0.96698494 | 0.731 | 0.354 | 0.00015476 | 0.00091763 | 1.2742E-08 |
| Hsp90b1 | 0.000011412 | 0.42273101 | 1 | 0.984 | 0.27721577 | 6.4254E-09 | 0.62795468 | 1 | 0.979 | 0.00015608 | 1.1412E-05 | 1.2851E-08 |
| Sumo2 | 0.00204391 | 0.32136209 | 0.99 | 0.938 | 1 | 6.5325E-09 | 0.57623142 | 0.996 | 0.938 | 0.00015869 | 0.00204391 | 1.3065E-08 |
| Eef2 | 0.000011083 | 0.4276269 | 0.99 | 0.969 | 0.26922082 | 6.7394E-09 | 0.57143096 | 1 | 0.979 | 0.00016371 | 1.1083E-05 | 1.3479E-08 |
| Hs3st3b1 | 0.0345741 | -0.747149 | 0.719 | 0.725 | 1 | 6.9072E-09 | 0.78836827 | 0.727 | 0.323 | 0.00016779 | 0.0345741 | 1.3814E-08 |
| Glt8d1 | 0.00461634 | 0.43163424 | 0.781 | 0.679 | 1 | 7.2158E-09 | 1.04590643 | 0.689 | 0.323 | 0.00017529 | 0.00461634 | 1.4432E-08 |
| Ocln | 0.48064264 | 0.59245704 | 0.021 | 0.01 | 1 | 7.2939E-09 | -1.6842508 | 0.071 | 0.323 | 0.00017718 | 0.48064264 | 1.4588E-08 |
| Kdelr3 | 0.00098561 | 0.47286371 | 0.896 | 0.751 | 1 | 7.5975E-09 | 0.89811664 | 0.857 | 0.479 | 0.00018456 | 0.00098561 | 1.5195E-08 |
| Tesc | 7.774E-09 | -4.440966 | 0.042 | 0.347 | 0.00018885 | 0.55991918 | -0.3838558 | 0.076 | 0.094 | 1 | 0.55991918 | 1.5548E-08 |
| Ccdc90b | 0.34725877 | 0.31447229 | 0.604 | 0.56 | 1 | 7.7913E-09 | 1.34069558 | 0.609 | 0.24 | 0.00018927 | 0.34725877 | 1.5583E-08 |
| Tcim | 0.25620655 | -1.240433 | 0.031 | 0.062 | 1 | 7.8564E-09 | -2.34733 | 0.059 | 0.292 | 0.00019085 | 0.25620655 | 1.5713E-08 |
| Rgs10 | 8.0456E-09 | 1.13873907 | 0.625 | 0.29 | 0.00019544 | 0.00043452 | 1.80235424 | 0.315 | 0.135 | 1 | 0.00043452 | 1.6091E-08 |
| Spcs2 | 0.000043678 | 0.45855771 | 0.979 | 0.876 | 1 | 8.074E-09 | 0.75140803 | 0.971 | 0.823 | 0.00019613 | 4.3678E-05 | 1.6148E-08 |
| Selenof | 0.00067319 | 0.30619479 | 1 | 0.979 | 1 | 8.445E-09 | 0.5784005 | 0.996 | 0.958 | 0.00020515 | 0.00067319 | 1.689E-08 |
| Erp29 | 0.00453239 | 0.34064609 | 0.969 | 0.886 | 1 | 8.6059E-09 | 0.7054927 | 0.937 | 0.74 | 0.00020906 | 0.00453239 | 1.7212E-08 |
| Pantr1 | 8.9668E-09 | -2.9126016 | 0.208 | 0.528 | 0.00021782 | 0.0845515 | 0.24115991 | 0.319 | 0.219 | 1 | 0.0845515 | 1.7934E-08 |
| Tspan31 | 0.00060882 | 0.44637878 | 0.865 | 0.782 | 1 | 9.0127E-09 | 0.83439833 | 0.891 | 0.615 | 0.00021894 | 0.00060882 | 1.8025E-08 |
| Gng10 | 0.00945746 | 0.46932625 | 0.74 | 0.648 | 1 | 9.2025E-09 | 1.10787132 | 0.761 | 0.385 | 0.00022355 | 0.00945746 | 1.8405E-08 |
| Plk2 | 0.0087024 | -0.7797761 | 0.198 | 0.358 | 1 | 9.2544E-09 | -1.7167649 | 0.147 | 0.438 | 0.00022481 | 0.0087024 | 1.8509E-08 |
| Pcsk6 | 9.3978E-09 | -1.3448565 | 0.344 | 0.689 | 0.00022829 | 0.59227743 | -1.1519756 | 0.214 | 0.219 | 1 | 0.59227743 | 1.8796E-08 |
| Trmt10c | 0.04817678 | 0.61517711 | 0.427 | 0.311 | 1 | 9.634E-09 | 1.19476513 | 0.58 | 0.208 | 0.00023403 | 0.04817678 | 1.9268E-08 |
| Map1b | 0.14293034 | -2.0180406 | 0.146 | 0.218 | 1 | 9.7212E-09 | -2.0772958 | 0.269 | 0.635 | 0.00023615 | 0.14293034 | 1.9442E-08 |

| | | | | | | | | | | | | |
|---|---|---|---|---|---|---|---|---|---|---|---|---|
| Litaf | 9.7568E-09 | -1.1248128 | 0.708 | 0.896 | 0.00023701 | 0.16838473 | -0.2656703 | 0.681 | 0.812 | 1 | 0.16838473 | 1.9514E-08 |
| Trim27 | 0.00059406 | 0.83756954 | 0.49 | 0.29 | 1 | 1.0142E-08 | 1.32111964 | 0.58 | 0.219 | 0.00024636 | 0.00059406 | 2.0284E-08 |
| Gm16351 | 1.0492E-08 | 3.28433474 | 0.333 | 0.073 | 0.00025486 | 0.00124672 | 1.3950167 | 0.332 | 0.146 | 1 | 0.00124672 | 2.0983E-08 |
| Hspd1 | 0.00129629 | 0.31752642 | 0.948 | 0.891 | 1 | 1.0685E-08 | 0.7478805 | 0.983 | 0.906 | 0.00025955 | 0.00129629 | 2.1369E-08 |
| Ly6a | 1.0706E-08 | -2.5046174 | 0.219 | 0.565 | 0.00026006 | 0.93318327 | -0.4307592 | 0.622 | 0.698 | 1 | 0.93318327 | 2.1412E-08 |
| Macf1 | 0.000006369 | -0.8749282 | 0.979 | 0.979 | 0.15471549 | 1.0719E-08 | -0.7395631 | 0.983 | 1 | 0.00026037 | 6.369E-06 | 2.1437E-08 |
| Dleu2 | 0.84287363 | -0.2551073 | 0.552 | 0.549 | 1 | 1.0771E-08 | -1.2757877 | 0.735 | 0.906 | 0.00026165 | 0.84287363 | 2.1542E-08 |
| Osbpl3 | 0.44405127 | -0.6091768 | 0.115 | 0.083 | 1 | 1.1111E-08 | -1.9186645 | 0.13 | 0.417 | 0.0002699 | 0.44405127 | 2.2221E-08 |
| Gclc | 0.13805583 | -0.6579178 | 0.427 | 0.492 | 1 | 1.1118E-08 | -1.4154655 | 0.475 | 0.792 | 0.00027008 | 0.13805583 | 2.2236E-08 |
| Krtcap2 | 9.8192E-07 | 0.69387736 | 0.927 | 0.834 | 0.02385287 | 1.1454E-08 | 0.84371513 | 0.866 | 0.615 | 0.00027825 | 9.8192E-07 | 2.2909E-08 |
| Arhgdib | 0.01785159 | 0.32942263 | 0.865 | 0.741 | 1 | 1.1512E-08 | 0.8779599 | 0.882 | 0.771 | 0.00027964 | 0.01785159 | 2.3023E-08 |
| Eif5 | 0.00970686 | 0.31700337 | 0.99 | 0.974 | 1 | 1.1599E-08 | 0.56211135 | 1 | 1 | 0.00028175 | 0.00970686 | 2.3197E-08 |
| Ifi30 | 0.00073845 | 0.75115478 | 0.688 | 0.523 | 1 | 1.1772E-08 | 1.86407167 | 0.609 | 0.281 | 0.00028596 | 0.00073845 | 2.3544E-08 |
| Ranbp1 | 0.000002057 | 0.74386508 | 0.927 | 0.813 | 0.04996833 | 1.1944E-08 | 0.69014474 | 0.941 | 0.802 | 0.00029016 | 2.057E-06 | 2.3889E-08 |
| Itgb4 | 0.40224917 | -0.1851505 | 0.042 | 0.067 | 1 | 1.207E-08 | -1.5508634 | 0.084 | 0.344 | 0.0002932 | 0.40224917 | 2.414E-08 |
| Snx18 | 0.00145689 | 0.50443229 | 0.854 | 0.679 | 1 | 1.208E-08 | 0.8893984 | 0.899 | 0.729 | 0.00029345 | 0.00145689 | 2.416E-08 |
| Krt14 | 0.18018935 | -2.1624305 | 0.052 | 0.098 | 1 | 1.2169E-08 | -1.9669675 | 0.101 | 0.365 | 0.00029562 | 0.18018935 | 2.4339E-08 |
| Capns1 | 2.1357E-08 | -1.188977 | 0.948 | 0.969 | 0.0005188 | 1.2649E-08 | 0.57395172 | 0.954 | 0.906 | 0.00030728 | 2.1357E-08 | 2.5299E-08 |
| Tmx1 | 0.03315546 | 0.28718018 | 0.875 | 0.777 | 1 | 1.2667E-08 | 0.6947911 | 0.912 | 0.708 | 0.00030771 | 0.03315546 | 2.5334E-08 |
| Nenf | 0.00468881 | 0.37331906 | 0.969 | 0.834 | 1 | 1.2738E-08 | 0.62874419 | 0.908 | 0.625 | 0.00030943 | 0.00468881 | 2.5476E-08 |
| Fgf2 | 0.000087228 | -1.1349189 | 0.594 | 0.746 | 1 | 1.2801E-08 | 1.06269891 | 0.718 | 0.333 | 0.00031097 | 8.7228E-05 | 2.5602E-08 |
| Mtss1 | 0.01070098 | -1.6680705 | 0.073 | 0.181 | 1 | 1.2887E-08 | -1.384623 | 0.109 | 0.385 | 0.00031306 | 0.01070098 | 2.5775E-08 |
| Med31 | 0.00044931 | 0.6256239 | 0.646 | 0.446 | 1 | 1.2936E-08 | 1.1342447 | 0.634 | 0.25 | 0.00031424 | 0.00044931 | 2.5872E-08 |
| Akap13 | 0.04432445 | -0.6421054 | 0.823 | 0.85 | 1 | 1.319E-08 | -1.0391448 | 0.845 | 0.969 | 0.0003204 | 0.04432445 | 2.6379E-08 |
| Id3 | 0.40952712 | -0.4714956 | 0.583 | 0.611 | 1 | 1.3444E-08 | -1.7890903 | 0.374 | 0.698 | 0.00032657 | 0.40952712 | 2.6887E-08 |
| H19 | 1.3529E-08 | 1.20846288 | 0.719 | 0.383 | 0.00032866 | 0.000000179 | 1.25431877 | 0.681 | 0.375 | 0.00434823 | 1.79E-07 | 2.7059E-08 |
| Dennd2b | 0.02260316 | -0.6314068 | 0.125 | 0.244 | 1 | 1.3709E-08 | -1.4782528 | 0.273 | 0.604 | 0.00033303 | 0.02260316 | 2.7419E-08 |
| Ap1s3 | 0.03616125 | -1.3848229 | 0.01 | 0.067 | 1 | 1.443E-08 | -3.2358547 | 0.034 | 0.229 | 0.00035054 | 0.03616125 | 2.8861E-08 |
| Tmem39a | 0.00011624 | 0.54183096 | 0.833 | 0.622 | 1 | 1.4529E-08 | 0.79142807 | 0.811 | 0.417 | 0.00035293 | 0.00011624 | 2.9058E-08 |
| Bcl2l11 | 0.2211382 | -0.1702926 | 0.188 | 0.244 | 1 | 1.545E-08 | -2.5388808 | 0.189 | 0.49 | 0.00037532 | 0.2211382 | 3.0901E-08 |
| Hmgcs1 | 0.00072818 | 0.95155105 | 0.823 | 0.767 | 1 | 1.5552E-08 | 1.3536827 | 0.895 | 0.844 | 0.00037779 | 0.00072818 | 3.1104E-08 |
| Tmem167 | 0.00918267 | 0.32750543 | 0.927 | 0.907 | 1 | 1.5668E-08 | 0.71810975 | 0.962 | 0.833 | 0.00038061 | 0.00918267 | 3.1336E-08 |
| Ccdc127 | 0.17064937 | 0.27052894 | 0.75 | 0.71 | 1 | 1.5786E-08 | 0.86327217 | 0.811 | 0.385 | 0.00038347 | 0.17064937 | 3.1572E-08 |
| Cspg4 | 1.6187E-08 | 0.9574761 | 0.969 | 0.782 | 0.00039321 | 1.4085E-07 | 0.74786024 | 0.941 | 0.708 | 0.00342164 | 1.4085E-07 | 3.2373E-08 |
| Zeb2 | 4.0728E-06 | -1.8059094 | 0.042 | 0.269 | 0.09893551 | 1.6598E-08 | -2.2184322 | 0.084 | 0.333 | 0.00040319 | 4.0728E-06 | 3.3196E-08 |
| Dusp5 | 0.40146863 | -0.240433 | 0.24 | 0.285 | 1 | 1.6711E-08 | -1.5728897 | 0.202 | 0.531 | 0.00040596 | 0.40146863 | 3.3423E-08 |
| Arl14ep | 1.7002E-08 | 1.38936517 | 0.542 | 0.207 | 0.00041301 | 2.7127E-06 | 0.9994728 | 0.529 | 0.208 | 0.06589586 | 2.7127E-06 | 3.4004E-08 |
| Arl4a | 1.7024E-08 | 0.77735425 | 0.885 | 0.648 | 0.00041356 | 0.0030475 | 0.53032068 | 0.887 | 0.823 | 1 | 0.0030475 | 3.4049E-08 |
| Col4a4 | 1.7268E-08 | -5.9232428 | 0 | 0.275 | 0.00041947 | 0.64761161 | -0.9139266 | 0.08 | 0.062 | 1 | 0.64761161 | 3.4536E-08 |
| Muc16 | 0.15729403 | -1.577468 | 0 | 0.021 | 1 | 1.7302E-08 | -2.8948178 | 0.025 | 0.208 | 0.0004203 | 0.15729403 | 3.4604E-08 |
| Cited2 | 2.1985E-06 | -2.1976199 | 0.177 | 0.446 | 0.05340495 | 1.7639E-08 | -1.2554075 | 0.155 | 0.469 | 0.00042848 | 2.1985E-06 | 3.5277E-08 |
| Rps28 | 0.00036131 | 0.3705374 | 1 | 0.959 | 1 | 1.7661E-08 | 0.62312649 | 0.992 | 0.906 | 0.00042902 | 0.00036131 | 3.5322E-08 |
| Stk26 | 0.93528172 | -0.3959859 | 0.792 | 0.679 | 1 | 1.79E-08 | 1.40006551 | 0.651 | 0.292 | 0.00043482 | 0.93528172 | 3.5799E-08 |
| H3f3a | 0.00393902 | 0.23007974 | 1 | 0.969 | 1 | 1.884E-08 | 0.48860143 | 1 | 0.979 | 0.00045766 | 0.00393902 | 3.768E-08 |
| Psap | 0.0856752 | -0.1885706 | 1 | 0.979 | 1 | 1.9494E-08 | 0.62753276 | 1 | 0.969 | 0.00047355 | 0.0856752 | 3.8988E-08 |
| Stmn1 | 0.5859658 | 1.6765213 | 0.177 | 0.155 | 1 | 1.9804E-08 | -1.5993619 | 0.122 | 0.406 | 0.00048107 | 0.5859658 | 3.9608E-08 |
| Papss1 | 1.9989E-08 | 1.07358373 | 0.781 | 0.523 | 0.00048558 | 2.2255E-06 | 0.8548895 | 0.828 | 0.688 | 0.05406106 | 2.2255E-06 | 3.9978E-08 |
| Cthrc1 | 0.64973679 | -0.3961382 | 0.812 | 0.684 | 1 | 2.0005E-08 | 0.88335895 | 0.916 | 0.51 | 0.00048597 | 0.64973679 | 4.0011E-08 |
| Polr1h | 0.01368827 | 0.60150218 | 0.646 | 0.503 | 1 | 2.0087E-08 | 1.0167192 | 0.727 | 0.354 | 0.00048795 | 0.01368827 | 4.0174E-08 |
| Golt1b | 0.01922724 | 0.41491885 | 0.708 | 0.575 | 1 | 2.0351E-08 | 1.28432577 | 0.655 | 0.312 | 0.00049437 | 0.01922724 | 4.0703E-08 |
| Cdc42se1 | 0.00034735 | 0.52272632 | 0.896 | 0.777 | 1 | 2.0363E-08 | 0.8513314 | 0.878 | 0.802 | 0.00049466 | 0.00034735 | 4.0726E-08 |
| Ctxn3 | 2.0472E-08 | -3.3500575 | 0.052 | 0.358 | 0.0004973 | 0.51188611 | -0.7248928 | 0.004 | 0.01 | 1 | 0.51188611 | 4.0943E-08 |
| Plagl1 | 2.0593E-08 | 1.08578573 | 0.938 | 0.803 | 0.00050024 | 0.00413955 | 0.26654014 | 0.941 | 0.833 | 1 | 0.00413955 | 4.1186E-08 |
| Samd4b | 0.24329663 | 0.16562373 | 0.865 | 0.782 | 1 | 2.0699E-08 | 0.83350891 | 0.861 | 0.646 | 0.00050281 | 0.24329663 | 4.1397E-08 |
| Abhd14a | 0.03236355 | 0.45729745 | 0.552 | 0.461 | 1 | 2.2301E-08 | 1.11004799 | 0.58 | 0.208 | 0.00054174 | 0.03236355 | 4.4603E-08 |
| Rtl3 | 2.2686E-08 | 1.41795931 | 0.458 | 0.145 | 0.00055109 | 0.00598964 | 0.53814164 | 0.286 | 0.125 | 1 | 0.00598964 | 4.5372E-08 |
| P3h4 | 0.05228849 | 0.18784519 | 0.969 | 0.839 | 1 | 2.3021E-08 | 0.89625208 | 0.84 | 0.5 | 0.00055924 | 0.05228849 | 4.6043E-08 |
| Fxyd3 | 2.3042E-08 | -2.4954907 | 0.531 | 0.731 | 0.00055974 | 0.46237864 | 0.35046624 | 0.542 | 0.635 | 1 | 0.46237864 | 4.6084E-08 |
| Map2k3 | 0.50591315 | 0.14583319 | 0.896 | 0.917 | 1 | 2.3081E-08 | 0.77550197 | 0.92 | 0.823 | 0.00056069 | 0.50591315 | 4.6163E-08 |
| Snx17 | 0.0792628 | 0.36909278 | 0.635 | 0.523 | 1 | 2.3437E-08 | 1.03903988 | 0.681 | 0.333 | 0.00056934 | 0.0792628 | 4.6875E-08 |
| Ppp3ca | 2.4099E-08 | -1.2213945 | 1 | 1 | 0.00058541 | 0.00083789 | -0.7043502 | 1 | 0.979 | 1 | 0.00083789 | 4.8198E-08 |
| Pla2g7 | 0.00179542 | -1.0931995 | 0.354 | 0.508 | 1 | 2.41E-08 | 2.86006974 | 0.382 | 0.073 | 0.00058544 | 0.00179542 | 4.82E-08 |
| Eif3h | 0.00171051 | 0.36442908 | 0.969 | 0.917 | 1 | 2.4124E-08 | 0.50540486 | 0.983 | 0.948 | 0.00058602 | 0.00171051 | 4.8248E-08 |
| Abca1 | 0.00120446 | -1.9633591 | 0.115 | 0.275 | 1 | 2.5026E-08 | -1.6152683 | 0.239 | 0.562 | 0.00060792 | 0.00120446 | 5.0051E-08 |
| Tiprl | 0.02089752 | 0.28823094 | 0.802 | 0.674 | 1 | 2.512E-08 | 0.72604139 | 0.882 | 0.615 | 0.00061022 | 0.02089752 | 5.024E-08 |
| Gars | 2.5454E-08 | 1.01936864 | 0.76 | 0.513 | 0.00061834 | 0.000008256 | 0.62383539 | 0.739 | 0.448 | 0.20055541 | 8.256E-06 | 5.0909E-08 |
| Arih1 | 0.13089866 | -0.5137967 | 0.99 | 0.99 | 1 | 2.5633E-08 | -0.7054269 | 0.996 | 1 | 0.00062267 | 0.13089866 | 5.1266E-08 |
| Atrnl1 | 0.39300244 | -0.2467468 | 0.417 | 0.466 | 1 | 2.5837E-08 | -1.2295536 | 0.462 | 0.802 | 0.00062763 | 0.39300244 | 5.1674E-08 |
| Cyp2a5 | 0.12946822 | -0.1730777 | 0.083 | 0.15 | 1 | 2.5882E-08 | -1.5508634 | 0.235 | 0.562 | 0.00062873 | 0.12946822 | 5.1764E-08 |
| Pabpc1 | 2.6101E-08 | 0.53199802 | 1 | 0.99 | 0.00063404 | 0.00003061 | 0.38824128 | 0.996 | 0.99 | 0.74356651 | 0.00003061 | 5.2202E-08 |
| Mtch1 | 2.6619E-08 | 0.51241754 | 0.969 | 0.772 | 0.00064662 | 0.00879379 | 0.20355398 | 0.891 | 0.802 | 1 | 0.00879379 | 5.3237E-08 |
| Col16a1 | 2.5988E-06 | 0.73740537 | 0.896 | 0.689 | 0.06312934 | 2.6657E-08 | 0.79961927 | 0.941 | 0.667 | 0.00064755 | 2.5988E-06 | 5.3314E-08 |
| Dusp18 | 8.8408E-06 | 1.47856607 | 0.552 | 0.337 | 0.21476054 | 2.6716E-08 | 1.49345283 | 0.588 | 0.229 | 0.00064899 | 8.8408E-06 | 5.3432E-08 |
| Fundc2 | 0.00078613 | 0.46000674 | 0.802 | 0.674 | 1 | 2.6876E-08 | 0.9261301 | 0.777 | 0.438 | 0.00065288 | 0.00078613 | 5.3752E-08 |
| Calu | 0.000035885 | 0.34434831 | 1 | 0.99 | 0.87171613 | 2.7355E-08 | 0.52778223 | 1 | 0.958 | 0.0006645 | 3.5885E-05 | 5.4709E-08 |
| Car6 | 2.8878E-08 | -3.3600762 | 0.073 | 0.378 | 0.00070151 | 0.5818708 | -0.5728897 | 0.013 | 0.021 | 1 | 0.5818708 | 5.7757E-08 |
| Arf2 | 0.000013149 | 0.91332783 | 0.885 | 0.694 | 0.31940567 | 2.9333E-08 | 1.21807629 | 0.794 | 0.448 | 0.00071255 | 1.3149E-05 | 5.8665E-08 |
| Zfp260 | 0.00092676 | 0.48359356 | 0.833 | 0.596 | 1 | 3.006E-08 | 0.8799693 | 0.832 | 0.458 | 0.00073022 | 0.00092676 | 6.012E-08 |
| Ptprn2 | 3.0524E-08 | 1.18479907 | 0.292 | 0.057 | 0.00074149 | 0.23638791 | 0.31574922 | 0.105 | 0.062 | 1 | 0.23638791 | 6.1048E-08 |
| Rnf41 | 0.13200225 | 0.26688117 | 0.521 | 0.42 | 1 | 3.0699E-08 | 0.85477544 | 0.765 | 0.385 | 0.00074574 | 0.13200225 | 6.1398E-08 |
| Ptprj | 3.0754E-08 | -1.380695 | 0.51 | 0.829 | 0.00074709 | 0.00021521 | -0.9039612 | 0.534 | 0.74 | 1 | 0.00021521 | 6.1509E-08 |
| Polr1c | 0.03490638 | 0.42794553 | 0.583 | 0.451 | 1 | 3.0863E-08 | 1.21807629 | 0.605 | 0.25 | 0.00074973 | 0.03490638 | 6.1727E-08 |
| Ier3ip1 | 0.0006372 | 0.43450468 | 0.927 | 0.829 | 1 | 3.0911E-08 | 0.7306829 | 0.933 | 0.75 | 0.00075089 | 0.0006372 | 6.1822E-08 |
| Myo1b | 0.34603297 | -0.9925055 | 0.104 | 0.14 | 1 | 3.0969E-08 | -0.7383133 | 0.101 | 0.365 | 0.00075229 | 0.34603297 | 6.1937E-08 |
| Adgrg2 | 3.1318E-08 | 2.17741954 | 0.417 | 0.135 | 0.00076078 | 0.6139163 | 0.37151172 | 0.29 | 0.25 | 1 | 0.6139163 | 6.2636E-08 |
| Chil1 | 0.00013795 | -2.2675275 | 0.542 | 0.653 | 1 | 3.1497E-08 | 0.56224878 | 0.609 | 0.24 | 0.00076512 | 0.00013795 | 6.2993E-08 |
| Blzf1 | 0.0008221 | 0.80881054 | 0.667 | 0.508 | 1 | 3.1978E-08 | 0.92238494 | 0.697 | 0.281 | 0.0007768 | 0.0008221 | 6.3955E-08 |
| Jam2 | 3.3361E-08 | -1.7896212 | 0.562 | 0.736 | 0.0008104 | 0.00148299 | 0.20179056 | 0.441 | 0.24 | 1 | 0.00148299 | 6.6721E-08 |
| Pycr1 | 3.3772E-08 | 1.58335483 | 0.49 | 0.181 | 0.0008204 | 1.2861E-06 | 2.12310414 | 0.328 | 0.073 | 0.03124262 | 1.2861E-06 | 6.7545E-08 |
| Mphosph6 | 0.00801836 | 0.53059178 | 0.562 | 0.404 | 1 | 3.4688E-08 | 1.6260144 | 0.475 | 0.135 | 0.00084265 | 0.00801836 | 6.9377E-08 |
| Krt17 | 0.06120877 | -0.4872702 | 0.052 | 0.124 | 1 | 3.562E-08 | -1.8541758 | 0.088 | 0.333 | 0.00086529 | 0.06120877 | 7.124E-08 |
| Mtfr1l | 0.14780498 | 0.33465928 | 0.542 | 0.446 | 1 | 3.5883E-08 | 1.71834163 | 0.475 | 0.146 | 0.00087168 | 0.14780498 | 7.1767E-08 |

| | | | | | | | | | | | | |
|---|---|---|---|---|---|---|---|---|---|---|---|---|
| Tbc1d10a | 0.3246914 | 0.19191911 | 0.719 | 0.674 | 1 | 3.6876E-08 | 0.72707131 | 0.845 | 0.479 | 0.00089579 | 0.3246914 | 7.3752E-08 |
| Ubb | 0.01462715 | 0.21209242 | 1 | 0.99 | 1 | 3.8516E-08 | 0.60383885 | 1 | 0.99 | 0.00093563 | 0.01462715 | 7.7032E-08 |
| Srsf9 | 0.06295437 | 0.29477649 | 0.833 | 0.741 | 1 | 3.859E-08 | 0.77429738 | 0.903 | 0.729 | 0.00093742 | 0.06295437 | 7.718E-08 |
| Thra | 0.00041912 | 0.45697268 | 0.906 | 0.803 | 1 | 3.8952E-08 | 0.59479686 | 0.954 | 0.76 | 0.00094622 | 0.00041912 | 7.7904E-08 |
| Rab11fip4 | 0.12310078 | 0.14084492 | 0.938 | 0.839 | 1 | 4.0688E-08 | 0.63074774 | 0.975 | 0.729 | 0.00098838 | 0.12310078 | 8.1375E-08 |
| Tsr1 | 4.5656E-08 | 1.25721459 | 0.688 | 0.389 | 0.00110907 | 4.0706E-08 | 0.94110631 | 0.685 | 0.333 | 0.00098884 | 4.5656E-08 | 8.1413E-08 |
| Gpx7 | 0.000023332 | 0.71715278 | 0.802 | 0.591 | 0.5667814 | 4.1142E-08 | 1.05040095 | 0.66 | 0.281 | 0.00099941 | 2.3332E-05 | 8.2283E-08 |
| Tmem98 | 2.911E-07 | 0.93710521 | 0.688 | 0.363 | 0.0070713 | 4.251E-08 | 1.13308823 | 0.643 | 0.312 | 0.00103266 | 2.911E-07 | 8.5021E-08 |
| Celsr1 | 0.32025586 | -0.9925055 | 0 | 0.01 | 1 | 4.3037E-08 | -2.5508634 | 0.042 | 0.24 | 0.00104545 | 0.32025586 | 8.6073E-08 |
| Hdlbp | 2.9115E-06 | 0.40402359 | 1 | 1 | 0.07072518 | 4.3275E-08 | 0.47607558 | 1 | 1 | 0.00105123 | 2.9115E-06 | 8.6549E-08 |
| Matn1 | 0.01583891 | -2.9647673 | 0.406 | 0.492 | 1 | 4.3743E-08 | 0.32616335 | 0.496 | 0.156 | 0.0010626 | 0.01583891 | 8.7486E-08 |
| Pank1 | 4.38E-08 | -2.0232677 | 0.271 | 0.58 | 0.00106399 | 0.66812893 | -0.5175823 | 0.475 | 0.375 | 1 | 0.66812893 | 8.76E-08 |
| Irs1 | 4.4264E-08 | 1.8494927 | 0.719 | 0.461 | 0.00107526 | 0.59441163 | -0.4624435 | 0.584 | 0.542 | 1 | 0.59441163 | 8.8528E-08 |
| Sec31a | 4.4524E-08 | 0.53542609 | 1 | 0.959 | 0.00108157 | 0.00021955 | 0.30611787 | 0.992 | 0.979 | 1 | 0.00021955 | 8.9047E-08 |
| Mxi1 | 0.0013014 | -1.293675 | 0.146 | 0.321 | 1 | 4.4995E-08 | -0.9679357 | 0.34 | 0.729 | 0.00109301 | 0.0013014 | 8.999E-08 |
| Tspan18 | 4.5135E-08 | -1.3261629 | 0.562 | 0.803 | 0.00109643 | 0.23811934 | -0.2576591 | 0.63 | 0.5 | 1 | 0.23811934 | 9.0271E-08 |
| Pim3 | 0.03989192 | -0.6774094 | 0.5 | 0.632 | 1 | 4.5684E-08 | -1.1338249 | 0.361 | 0.698 | 0.00110977 | 0.03989192 | 9.1369E-08 |
| Siva1 | 4.6095E-08 | 1.32585602 | 0.583 | 0.264 | 0.00111974 | 0.00842061 | 0.61128932 | 0.504 | 0.354 | 1 | 0.00842061 | 9.219E-08 |
| Lypd2 | 0.375678 | -1.1354634 | 0.104 | 0.14 | 1 | 4.6323E-08 | -1.5142138 | 0.168 | 0.458 | 0.00112528 | 0.375678 | 9.2646E-08 |
| Armcx2 | 1.7567E-06 | 1.68556644 | 0.396 | 0.155 | 0.04267343 | 4.6731E-08 | 1.4821517 | 0.5 | 0.177 | 0.00113519 | 1.7567E-06 | 9.3462E-08 |
| Mrpl50 | 0.01151804 | 0.61845225 | 0.542 | 0.42 | 1 | 4.7685E-08 | 1.49749966 | 0.546 | 0.208 | 0.00115836 | 0.01151804 | 9.537E-08 |
| Sav1 | 4.7761E-08 | -1.1400147 | 0.781 | 0.855 | 0.00116022 | 0.75806248 | -0.2746658 | 0.912 | 0.781 | 1 | 0.75806248 | 9.5523E-08 |
| Slc44a1 | 4.7975E-08 | -1.0394358 | 0.781 | 0.933 | 0.0011654 | 0.84438217 | -0.2412204 | 0.903 | 0.927 | 1 | 0.84438217 | 9.5949E-08 |
| Ppp1r1b | 3.2434E-06 | -2.3408039 | 0.396 | 0.57 | 0.07878937 | 4.8261E-08 | 1.00242266 | 0.483 | 0.135 | 0.00117236 | 3.2434E-06 | 9.6522E-08 |
| Cadm1 | 4.8911E-08 | 0.83273286 | 0.854 | 0.549 | 0.00118814 | 0.54862992 | -0.3459527 | 0.962 | 0.938 | 1 | 0.54862992 | 9.7822E-08 |
| Ccn6 | 0.00712551 | 1.06889508 | 0.469 | 0.342 | 1 | 4.9848E-08 | 1.51531094 | 0.433 | 0.104 | 0.00121092 | 0.00712551 | 9.9697E-08 |
| Hlf | 0.90315176 | -0.7173982 | 0.188 | 0.176 | 1 | 5.0159E-08 | -1.0922638 | 0.223 | 0.562 | 0.00121845 | 0.90315176 | 1.0032E-07 |
| Tfrc | 3.4317E-07 | 1.24650167 | 0.76 | 0.451 | 0.00833622 | 5.0676E-08 | 0.83816012 | 0.857 | 0.656 | 0.00123103 | 3.4317E-07 | 1.0135E-07 |
| Tspan4 | 0.05512691 | 0.35367018 | 0.875 | 0.72 | 1 | 5.1249E-08 | 0.89022257 | 0.761 | 0.396 | 0.00124493 | 0.05512691 | 1.025E-07 |
| Agmo | 5.2244E-08 | -6.4437166 | 0 | 0.259 | 0.0012691 | 0.12731831 | -1.6724253 | 0.021 | 0.052 | 1 | 0.12731831 | 1.0449E-07 |
| Uqcrh | 0.33311425 | 0.11459228 | 0.99 | 0.953 | 1 | 5.2487E-08 | 0.54976485 | 0.987 | 0.979 | 0.00127502 | 0.33311425 | 1.0497E-07 |
| Prkd1 | 5.3203E-08 | -1.8146384 | 0.406 | 0.684 | 0.00129241 | 0.00162793 | -0.8647682 | 0.613 | 0.75 | 1 | 0.00162793 | 1.0641E-07 |
| Lpcat2 | 0.00093334 | 0.96567436 | 0.26 | 0.104 | 1 | 5.3722E-08 | -1.6030516 | 0.189 | 0.5 | 0.00130501 | 0.00093334 | 1.0744E-07 |
| Nectin1 | 0.77589184 | -1.0920411 | 0.125 | 0.13 | 1 | 5.413E-08 | -1.6645201 | 0.193 | 0.479 | 0.00131492 | 0.77589184 | 1.0826E-07 |
| Rab33b | 2.5787E-07 | 1.14499806 | 0.667 | 0.378 | 0.00626428 | 5.4818E-08 | 1.02041213 | 0.672 | 0.281 | 0.00133163 | 2.5787E-07 | 1.0964E-07 |
| Itm2c | 0.12598919 | -0.2057176 | 0.938 | 0.938 | 1 | 5.5484E-08 | 0.5710221 | 0.954 | 0.844 | 0.00134782 | 0.12598919 | 1.1097E-07 |
| Eya1 | 5.6052E-08 | -1.2093285 | 0.812 | 0.933 | 0.00136161 | 0.01211581 | -0.6135692 | 0.966 | 0.958 | 1 | 0.01211581 | 1.121E-07 |
| Nolc1 | 5.628E-08 | 0.93466166 | 0.906 | 0.679 | 0.00136715 | 0.0015139 | 0.44768669 | 0.828 | 0.781 | 1 | 0.0015139 | 1.1256E-07 |
| Myadm | 5.6888E-08 | 1.25916269 | 0.844 | 0.689 | 0.00138192 | 1.0702E-06 | 1.18295739 | 0.782 | 0.531 | 0.0259985 | 1.0702E-06 | 1.1378E-07 |
| P2ry10b | 5.8445E-08 | 1.58335483 | 0.406 | 0.124 | 0.00141975 | 0.000088428 | 1.18011072 | 0.399 | 0.167 | 1 | 8.8428E-05 | 1.1689E-07 |
| Cct2 | 0.00128299 | 0.37006462 | 0.958 | 0.917 | 1 | 5.8564E-08 | 0.54740457 | 0.983 | 0.885 | 0.00142264 | 0.00128299 | 1.1713E-07 |
| Esrrg | 5.9874E-08 | -2.2681399 | 0.125 | 0.456 | 0.00145446 | 1.8161E-07 | -2.4093909 | 0.118 | 0.375 | 0.0044117 | 1.8161E-07 | 1.1975E-07 |
| Shb | 0.480984 | 0.72276506 | 0.198 | 0.171 | 1 | 6.064E-08 | -1.2016024 | 0.218 | 0.531 | 0.00147308 | 0.480984 | 1.2128E-07 |
| Cldn7 | 0.08281325 | -0.4519371 | 0.042 | 0.104 | 1 | 6.6205E-08 | -0.870971 | 0.151 | 0.448 | 0.00160826 | 0.08281325 | 1.3241E-07 |
| Tango6 | 0.06224756 | -0.1368954 | 0.073 | 0.15 | 1 | 6.6727E-08 | -1.1672509 | 0.189 | 0.51 | 0.00162094 | 0.06224756 | 1.3345E-07 |
| Map3k1 | 0.04453747 | -0.3303751 | 0.385 | 0.492 | 1 | 6.6742E-08 | -1.2431291 | 0.441 | 0.75 | 0.00162129 | 0.04453747 | 1.3348E-07 |
| Kctd12 | 0.1030993 | -1.1675922 | 0.219 | 0.295 | 1 | 6.6784E-08 | -2.5232589 | 0.202 | 0.49 | 0.00162231 | 0.1030993 | 1.3357E-07 |
| Rab1a | 6.891E-08 | 0.57290647 | 0.99 | 0.964 | 0.00167396 | 0.00024229 | 0.29932348 | 0.992 | 0.99 | 1 | 0.00024229 | 1.3782E-07 |
| Atp5g1 | 0.000049511 | 0.57302366 | 0.927 | 0.876 | 1 | 6.9281E-08 | 0.80273656 | 0.882 | 0.792 | 0.00168297 | 4.9511E-05 | 1.3856E-07 |
| Mydgf | 0.0003407 | 0.55044951 | 0.896 | 0.741 | 1 | 6.9389E-08 | 0.95693128 | 0.752 | 0.427 | 0.0016856 | 0.0003407 | 1.3878E-07 |
| Efna5 | 7.0004E-08 | -1.2097062 | 0.948 | 0.969 | 0.00170054 | 0.57835556 | -0.2134693 | 0.975 | 0.979 | 1 | 0.57835556 | 1.4001E-07 |
| Col17a1 | 0.38357309 | -1.577468 | 0.01 | 0.026 | 1 | 7.0097E-08 | -0.8244284 | 0.059 | 0.281 | 0.00170279 | 0.38357309 | 1.4019E-07 |
| Rbbp7 | 7.1136E-08 | 0.6319192 | 0.958 | 0.865 | 0.00172804 | 6.5024E-06 | 0.46704908 | 0.983 | 0.906 | 0.15795701 | 6.5024E-06 | 1.4227E-07 |
| Rora | 7.2598E-08 | -0.936599 | 0.99 | 1 | 0.00176354 | 0.000035708 | -0.8087527 | 0.983 | 1 | 0.86741845 | 3.5708E-05 | 1.452E-07 |
| Tspyl1 | 0.28194657 | 0.20013961 | 0.615 | 0.57 | 1 | 7.2946E-08 | 0.96316323 | 0.744 | 0.365 | 0.00177201 | 0.28194657 | 1.4589E-07 |
| Slc41a1 | 0.13608873 | 0.37630902 | 0.792 | 0.762 | 1 | 7.5151E-08 | 0.80944367 | 0.903 | 0.542 | 0.00182558 | 0.13608873 | 1.503E-07 |
| Pdlim5 | 0.00893888 | -0.6615614 | 0.823 | 0.933 | 1 | 7.6699E-08 | -1.2369123 | 0.866 | 0.906 | 0.00186318 | 0.00893888 | 1.534E-07 |
| Gm29157 | 7.6789E-08 | 2.90057933 | 0.229 | 0.031 | 0.00186537 | 0.00570557 | 3.01207283 | 0.076 | 0 | 1 | 0.00570557 | 1.5358E-07 |
| Scrg1 | 0.07640555 | -0.2238797 | 0.854 | 0.601 | 1 | 7.6968E-08 | 1.63938162 | 0.45 | 0.125 | 0.0018697 | 0.07640555 | 1.5394E-07 |
| Lamb2 | 0.0002031 | 0.57486298 | 0.823 | 0.596 | 1 | 7.7161E-08 | 0.88325937 | 0.786 | 0.427 | 0.0018744 | 0.0002031 | 1.5432E-07 |
| Oaf | 7.7334E-08 | 0.95070851 | 0.948 | 0.767 | 0.0018786 | 0.000015181 | 0.83545318 | 0.849 | 0.635 | 0.36877521 | 1.5181E-05 | 1.5467E-07 |
| Dtymk | 0.00723169 | 0.6797686 | 0.656 | 0.523 | 1 | 7.911E-08 | 1.02484013 | 0.588 | 0.219 | 0.00192174 | 0.00723169 | 1.5822E-07 |
| C1qtnf3 | 7.9742E-08 | 2.63976275 | 0.208 | 0.021 | 0.00193709 | 0.40076941 | -1.3098553 | 0.017 | 0.031 | 1 | 0.40076941 | 1.5948E-07 |
| Serinc1 | 0.03641903 | 0.23650154 | 0.958 | 0.896 | 1 | 8.1059E-08 | 0.57051295 | 0.971 | 0.938 | 0.00196908 | 0.03641903 | 1.6212E-07 |
| Dnaja1 | 0.00362234 | 0.25271528 | 1 | 0.943 | 1 | 8.133E-08 | 0.53415564 | 1 | 0.958 | 0.00197566 | 0.00362234 | 1.6266E-07 |
| Nip7 | 0.00017893 | 0.81635305 | 0.646 | 0.43 | 1 | 8.2328E-08 | 1.01807157 | 0.597 | 0.26 | 0.00199991 | 0.00017893 | 1.6466E-07 |
| Hmgn1 | 0.00018352 | 0.32942263 | 0.99 | 0.943 | 1 | 8.3073E-08 | 0.53167215 | 0.983 | 0.844 | 0.00201801 | 0.00018352 | 1.6615E-07 |
| Slc26a2 | 0.00375226 | 0.23894405 | 0.927 | 0.767 | 1 | 8.5667E-08 | 1.0482325 | 0.857 | 0.656 | 0.00208102 | 0.00375226 | 1.7133E-07 |
| Snhg1 | 3.4469E-06 | 0.66141729 | 0.896 | 0.845 | 0.08373224 | 0.000000086 | 0.7583856 | 0.878 | 0.75 | 0.00208912 | 3.4469E-06 | 0.000000172 |
| Fam20c | 8.6553E-08 | 0.91958745 | 0.75 | 0.43 | 0.00210253 | 5.5434E-06 | 0.4822329 | 0.832 | 0.427 | 0.01346605 | 5.5434E-07 | 1.7311E-07 |
| Laptm4a | 0.24663799 | 0.13669874 | 1 | 0.995 | 1 | 8.7565E-08 | 0.48033911 | 1 | 1 | 0.00212714 | 0.24663799 | 1.7513E-07 |
| Tln1 | 0.01686519 | 0.28109363 | 0.979 | 0.938 | 1 | 8.7696E-08 | 0.53019052 | 0.996 | 0.896 | 0.00213031 | 0.01686519 | 1.7539E-07 |
| Gnas | 9.102E-08 | 0.4116605 | 1 | 0.99 | 0.00221106 | 5.5442E-06 | 0.36227703 | 1 | 1 | 0.13467888 | 5.5442E-06 | 1.8204E-07 |
| Matn3 | 9.1146E-08 | 1.58196466 | 0.812 | 0.591 | 0.00221411 | 0.000012096 | 0.64406235 | 0.563 | 0.271 | 0.29383961 | 1.2096E-05 | 1.8229E-07 |
| Snai2 | 9.1195E-08 | 1.28969635 | 0.635 | 0.352 | 0.00221531 | 1.1837E-06 | 1.40063812 | 0.42 | 0.135 | 0.02875416 | 1.1837E-06 | 1.8239E-07 |
| Ndufa13 | 0.2238994 | 0.1668503 | 0.969 | 0.876 | 1 | 9.2314E-08 | 0.55243272 | 0.912 | 0.823 | 0.0022425 | 0.2238994 | 1.8463E-07 |
| Runx1 | 9.2415E-08 | -0.851033 | 0.948 | 0.99 | 0.00224495 | 0.00942593 | -0.6552073 | 0.987 | 0.99 | 1 | 0.00942593 | 1.8483E-07 |
| Apoe | 0.00096175 | 1.55562027 | 0.417 | 0.244 | 1 | 9.2439E-08 | 1.52639064 | 0.807 | 0.781 | 0.00224552 | 0.00096175 | 1.8488E-07 |
| Rsl24d1 | 9.9937E-06 | 0.95913151 | 0.562 | 0.301 | 0.24276783 | 9.2492E-08 | 1.14957636 | 0.584 | 0.25 | 0.00224682 | 9.9937E-06 | 1.8498E-07 |
| Hnrnpa2b1 | 0.00488106 | 0.38666737 | 1 | 1 | 1 | 9.3889E-08 | 0.4279028 | 1 | 1 | 0.00228076 | 0.00488106 | 1.8778E-07 |
| Ubtf | 0.02818494 | 0.38477711 | 0.74 | 0.642 | 1 | 9.4853E-08 | 0.79824242 | 0.92 | 0.823 | 0.00230416 | 0.02818494 | 1.8971E-07 |
| Arf1 | 0.11568611 | 0.14217202 | 0.99 | 0.984 | 1 | 9.6594E-08 | 0.47206461 | 0.996 | 0.99 | 0.00234646 | 0.11568611 | 1.9319E-07 |
| Unc50 | 0.00578646 | 0.37522632 | 0.812 | 0.637 | 1 | 9.7442E-08 | 0.84579561 | 0.748 | 0.385 | 0.00236707 | 0.00578646 | 1.9488E-07 |
| Yipf3 | 0.00194024 | 0.53704392 | 0.781 | 0.674 | 1 | 9.7704E-08 | 0.85222899 | 0.765 | 0.406 | 0.00237343 | 0.00194024 | 1.9541E-07 |
| Yipf4 | 0.00029588 | 0.52129601 | 0.917 | 0.762 | 1 | 9.8696E-08 | 0.60575255 | 0.912 | 0.75 | 0.00239753 | 0.00029588 | 1.9739E-07 |
| Cox7b | 0.000016041 | 0.51781824 | 0.948 | 0.829 | 0.38967003 | 9.9833E-08 | 0.66367253 | 0.903 | 0.76 | 0.00242514 | 1.6041E-05 | 1.9967E-07 |
| Prkg1 | 1.0168E-07 | 0.73101787 | 0.906 | 0.653 | 0.00246996 | 0.00543299 | 0.32829295 | 0.966 | 0.885 | 1 | 0.00543299 | 2.0336E-07 |
| Ndufa4 | 0.3391897 | 0.13471509 | 0.979 | 0.948 | 1 | 1.0193E-07 | 0.61855614 | 0.971 | 0.885 | 0.00247601 | 0.3391897 | 2.0385E-07 |
| Sc5d | 0.06505427 | 0.35349039 | 0.76 | 0.668 | 1 | 1.029E-07 | 1.16569602 | 0.689 | 0.354 | 0.00249958 | 0.06505427 | 2.0579E-07 |
| Tmem117 | 1.039E-07 | -1.9925055 | 0.302 | 0.58 | 0.00252382 | 0.757075 | -0.8983237 | 0.277 | 0.229 | 1 | 0.757075 | 2.0779E-07 |
| Rab11fip1 | 5.2158E-06 | -1.889412 | 0.188 | 0.451 | 0.12670167 | 1.0402E-07 | -1.3287143 | 0.21 | 0.51 | 0.00252684 | 5.2158E-06 | 2.0804E-07 |

| | | | | | | | | | | | |
|---|---|---|---|---|---|---|---|---|---|---|---|
| Panx3 | 1.0499E-07 | 3.696408 | 0.281 | 0.062 | 0.00255041 | 0.33550266 | 4.28260177 | 0.042 | 0.021 | 1 | 0.33550266 | 2.0998E-07 |
| Psmd12 | 0.00016271 | 0.56487121 | 0.875 | 0.782 | 1 | 1.0878E-07 | 0.73295782 | 0.924 | 0.844 | 0.00264242 | 0.00016271 | 2.1755E-07 |
| Zfp330 | 0.01451719 | 0.61691008 | 0.635 | 0.518 | 1 | 1.0891E-07 | 0.94110631 | 0.618 | 0.25 | 0.00264575 | 0.01451719 | 2.1783E-07 |
| Efcab1 | 1.1097E-07 | 2.10703021 | 0.531 | 0.269 | 0.00269577 | 5.1647E-06 | 1.46152036 | 0.412 | 0.146 | 0.12546181 | 5.1647E-06 | 2.2195E-07 |
| Golim4 | 1.1197E-07 | 0.63841849 | 0.99 | 0.907 | 0.00271989 | 0.00163439 | 0.30273314 | 0.971 | 0.885 | 1 | 0.00163439 | 2.2393E-07 |
| Mogs | 1.128E-07 | 1.4612125 | 0.49 | 0.192 | 0.00274008 | 5.2179E-06 | 1.74459252 | 0.345 | 0.094 | 0.12675328 | 5.2179E-06 | 2.256E-07 |
| Calr | 0.000013057 | 0.44966357 | 0.99 | 0.953 | 0.31719159 | 1.1446E-07 | 0.55445399 | 0.996 | 0.99 | 0.00278044 | 1.3057E-05 | 2.2892E-07 |
| Timm23 | 0.73535512 | 0.10906552 | 0.844 | 0.845 | 1 | 1.1595E-07 | 0.67672422 | 0.891 | 0.688 | 0.00281659 | 0.73535512 | 2.3189E-07 |
| Emc7 | 0.2034854 | 0.18460992 | 0.917 | 0.834 | 1 | 1.1664E-07 | 0.64356948 | 0.916 | 0.812 | 0.0028334 | 0.2034854 | 2.3328E-07 |
| Cavin1 | 0.00020581 | 0.6320905 | 0.885 | 0.741 | 1 | 1.1732E-07 | 0.79618637 | 0.924 | 0.833 | 0.00284989 | 0.00020581 | 2.3464E-07 |
| Sf3b4 | 0.14681557 | 0.2741402 | 0.562 | 0.451 | 1 | 1.1841E-07 | 0.9766675 | 0.718 | 0.396 | 0.00287637 | 0.14681557 | 2.3682E-07 |
| Plrg1 | 0.00178375 | 0.81484946 | 0.354 | 0.181 | 1 | 1.1844E-07 | 2.16145046 | 0.387 | 0.094 | 0.0028772 | 0.00178375 | 2.3688E-07 |
| Ppp1cb | 1.2045E-07 | 0.47607491 | 0.99 | 0.943 | 0.00292591 | 0.04467668 | 0.13123946 | 0.996 | 0.99 | 1 | 0.04467668 | 2.4089E-07 |
| Lmna | 0.000010334 | 0.60585837 | 0.979 | 0.995 | 0.25102617 | 1.2061E-07 | 0.5179516 | 1 | 0.99 | 0.00292995 | 1.0334E-05 | 2.4123E-07 |
| Nsmce3 | 0.33543464 | 0.23377039 | 0.438 | 0.399 | 1 | 1.2148E-07 | 1.24473359 | 0.55 | 0.219 | 0.00295092 | 0.33543464 | 2.4295E-07 |
| Commd7 | 0.00458385 | 0.61703506 | 0.521 | 0.358 | 1 | 1.2168E-07 | 1.23129101 | 0.555 | 0.198 | 0.00295585 | 0.00458385 | 2.4336E-07 |
| Impdh2 | 2.9217E-06 | 0.60835068 | 0.906 | 0.751 | 0.07097506 | 1.2216E-07 | 0.90766812 | 0.824 | 0.677 | 0.00296742 | 2.9217E-06 | 2.4431E-07 |
| Nfic | 1.2256E-07 | 0.71333761 | 0.958 | 0.813 | 0.0029772 | 0.00018893 | 0.42179859 | 0.979 | 0.927 | 1 | 0.00018893 | 2.4512E-07 |
| Btg2 | 0.00546875 | -1.0314767 | 0.729 | 0.829 | 1 | 1.2386E-07 | -0.9613854 | 0.508 | 0.844 | 0.00300876 | 0.00546875 | 2.4772E-07 |
| Eef1g | 9.6075E-06 | 0.46000674 | 0.979 | 0.948 | 0.23338651 | 1.2427E-07 | 0.53611111 | 0.992 | 0.958 | 0.00301877 | 9.6075E-06 | 2.4854E-07 |
| Gosr2 | 0.02838414 | 0.50904103 | 0.469 | 0.337 | 1 | 1.2472E-07 | 1.44716798 | 0.475 | 0.146 | 0.00302971 | 0.02838414 | 2.4944E-07 |
| Pofut2 | 2.0496E-06 | 0.68687264 | 0.854 | 0.694 | 0.04978885 | 1.2583E-07 | 0.71661695 | 0.849 | 0.438 | 0.00305666 | 2.0496E-06 | 2.5166E-07 |
| Perp | 0.22569302 | -0.6649308 | 0.115 | 0.171 | 1 | 1.2774E-07 | -1.1578522 | 0.202 | 0.5 | 0.00310296 | 0.22569302 | 2.5547E-07 |
| Slc25a38 | 0.203782 | 0.32942263 | 0.302 | 0.228 | 1 | 1.3066E-07 | 1.82969609 | 0.424 | 0.115 | 0.00317408 | 0.203782 | 2.6133E-07 |
| Rlim | 1.3236E-07 | 0.68967474 | 0.958 | 0.902 | 0.00321522 | 9.7179E-07 | 0.61471078 | 0.958 | 0.844 | 0.02360678 | 9.7179E-07 | 2.6471E-07 |
| Pear1 | 0.00011775 | 1.52944724 | 0.344 | 0.15 | 1 | 1.324E-07 | 1.54258755 | 0.45 | 0.135 | 0.00321633 | 0.00011775 | 2.6481E-07 |
| Tmem106c | 0.6283664 | 0.24047171 | 0.406 | 0.383 | 1 | 1.3294E-07 | 1.98970502 | 0.391 | 0.094 | 0.00322933 | 0.6283664 | 2.6588E-07 |
| Rbpms | 0.00100893 | -1.6072153 | 0.156 | 0.332 | 1 | 1.3364E-07 | -1.2881602 | 0.286 | 0.604 | 0.00324629 | 0.00100893 | 2.6727E-07 |
| Selenom | 0.44396214 | -0.1482689 | 0.917 | 0.829 | 1 | 1.3488E-07 | 0.67917362 | 0.899 | 0.521 | 0.00327653 | 0.44396214 | 2.6976E-07 |
| Mdfi | 0.18382219 | -0.540315 | 0.865 | 0.793 | 1 | 1.3555E-07 | 0.8510104 | 0.849 | 0.49 | 0.00329284 | 0.18382219 | 2.711E-07 |
| Mmgt1 | 0.01707307 | 0.61892925 | 0.427 | 0.29 | 1 | 1.3565E-07 | 1.3635452 | 0.513 | 0.188 | 0.00329522 | 0.01707307 | 2.713E-07 |
| Ddx3y | 0.99061337 | 0.12931328 | 0.542 | 0.56 | 1 | 1.364E-07 | 0.58002473 | 0.966 | 0.896 | 0.00331339 | 0.99061337 | 2.728E-07 |
| Arf5 | 0.03984023 | 0.19658374 | 0.99 | 0.943 | 1 | 1.4193E-07 | 0.51669322 | 0.987 | 0.948 | 0.00344768 | 0.03984023 | 2.8385E-07 |
| Lysmd3 | 0.11029611 | 0.62524697 | 0.635 | 0.591 | 1 | 1.4381E-07 | 0.98340907 | 0.786 | 0.51 | 0.00349338 | 0.11029611 | 2.8762E-07 |
| Nav1 | 1.4435E-07 | -1.6746366 | 0.51 | 0.736 | 0.00350659 | 0.00377211 | -1.1243513 | 0.567 | 0.635 | 1 | 0.00377211 | 2.887E-07 |
| Zbtb38 | 1.4588E-07 | -1.0437437 | 0.812 | 0.912 | 0.00354384 | 0.34373947 | -0.2953158 | 0.861 | 0.781 | 1 | 0.34373947 | 2.9177E-07 |
| Adrb2 | 0.05017249 | -0.5118796 | 0.062 | 0.14 | 1 | 1.4715E-07 | -1.9537115 | 0.05 | 0.25 | 0.00357452 | 0.05017249 | 2.943E-07 |
| Prrc1 | 1.5062E-07 | 0.6942313 | 0.948 | 0.803 | 0.00365876 | 0.00232955 | 0.41469107 | 0.887 | 0.708 | 1 | 0.00232955 | 3.0123E-07 |
| Tpst1 | 0.5687415 | 0.23761432 | 0.458 | 0.43 | 1 | 1.5089E-07 | 1.122372 | 0.706 | 0.375 | 0.00366544 | 0.5687415 | 3.0178E-07 |
| Tmem30a | 0.00970556 | 0.23743454 | 0.979 | 0.948 | 1 | 1.5118E-07 | 0.54119589 | 0.983 | 0.906 | 0.00367237 | 0.00970556 | 3.0235E-07 |
| Col4a3 | 1.5228E-07 | -5.8379955 | 0 | 0.244 | 0.00369922 | 0.05139523 | -2.9728203 | 0.021 | 0.062 | 1 | 0.05139523 | 3.0456E-07 |
| B3gnt2 | 1.5565E-07 | -1.6562221 | 0.51 | 0.72 | 0.00378116 | 0.14329545 | -0.2886907 | 0.685 | 0.792 | 1 | 0.14329545 | 3.1131E-07 |
| Ik | 0.00155698 | 0.37240856 | 0.896 | 0.85 | 1 | 1.5574E-07 | 0.67402507 | 0.908 | 0.823 | 0.00378325 | 0.00155698 | 3.1148E-07 |
| Srgap1 | 1.5749E-07 | 0.79779719 | 0.906 | 0.637 | 0.00382579 | 0.07963748 | 0.18090165 | 0.962 | 0.865 | 1 | 0.07963748 | 3.1498E-07 |
| Sgta | 0.07988045 | 0.58778055 | 0.49 | 0.404 | 1 | 1.5777E-07 | 1.19448878 | 0.643 | 0.333 | 0.00383246 | 0.07988045 | 3.1553E-07 |
| Cers6 | 0.10643024 | -0.6299354 | 0.125 | 0.202 | 1 | 1.5856E-07 | -0.88497 | 0.168 | 0.479 | 0.00385178 | 0.10643024 | 3.1712E-07 |
| Ext1 | 0.01125787 | -0.8116568 | 0.969 | 0.964 | 1 | 1.5983E-07 | -1.0179903 | 0.941 | 0.99 | 0.00388256 | 0.01125787 | 3.1966E-07 |
| Rbmxl1 | 0.00043659 | 0.64622829 | 0.677 | 0.461 | 1 | 1.6015E-07 | 1.0952862 | 0.681 | 0.354 | 0.00389033 | 0.00043659 | 3.203E-07 |
| Tma7 | 0.03132311 | 0.2382992 | 0.938 | 0.891 | 1 | 1.6109E-07 | 0.70063711 | 0.891 | 0.74 | 0.00391319 | 0.03132311 | 3.2218E-07 |
| Ifi202b | 0.28001997 | -2.5160674 | 0.01 | 0.031 | 1 | 1.6491E-07 | 0.99757326 | 0.109 | 0.365 | 0.00400607 | 0.28001997 | 3.2983E-07 |
| Eif5a | 1.6778E-07 | 0.56599932 | 0.99 | 0.964 | 0.00407578 | 2.3402E-07 | 0.50209277 | 0.987 | 0.927 | 0.00568473 | 2.3402E-07 | 3.3557E-07 |
| 1110004F10Rik | 0.00143614 | 0.43149847 | 0.875 | 0.803 | 1 | 1.678E-07 | 0.67743369 | 0.933 | 0.802 | 0.00407613 | 0.00143614 | 3.3559E-07 |
| Slc38a2 | 1.6879E-07 | -0.8539446 | 1 | 0.995 | 0.00410034 | 0.54355206 | -0.5223999 | 0.992 | 1 | 1 | 0.54355206 | 3.3759E-07 |
| Canx | 0.000018138 | 0.38862986 | 1 | 0.99 | 0.44061396 | 1.723E-07 | 0.47978487 | 1 | 1 | 0.00418561 | 1.8138E-05 | 3.4461E-07 |
| Isg20l2 | 0.0021522 | 0.50225923 | 0.51 | 0.301 | 1 | 1.7785E-07 | 1.17292684 | 0.555 | 0.229 | 0.00432042 | 0.0021522 | 3.5571E-07 |
| Gdi2 | 1.7987E-07 | 0.59167488 | 0.979 | 0.912 | 0.00436947 | 0.01836734 | 0.13164115 | 0.979 | 0.958 | 1 | 0.01836734 | 3.5975E-07 |
| Myl6 | 0.09206547 | 0.13007119 | 1 | 0.969 | 1 | 1.799E-07 | 0.37317152 | 1 | 0.979 | 0.00437024 | 0.09206547 | 3.5981E-07 |
| Prkg2 | 1.8134E-07 | 2.90533999 | 0.365 | 0.114 | 0.00440506 | 0.13206532 | 3.15891422 | 0.076 | 0.031 | 1 | 0.13206532 | 3.6268E-07 |
| Meltf | 0.00264438 | -0.6924688 | 0.74 | 0.725 | 1 | 1.8607E-07 | 0.58431713 | 0.798 | 0.375 | 0.00451996 | 0.00264438 | 3.7214E-07 |
| Ppp2r2d | 0.01052863 | 0.35979628 | 0.802 | 0.663 | 1 | 1.891E-07 | 0.70380118 | 0.895 | 0.802 | 0.00459373 | 0.01052863 | 3.7821E-07 |
| Luzp2 | 1.8985E-07 | -2.9949404 | 0.156 | 0.461 | 0.00461176 | 0.57276199 | -0.8737561 | 0.239 | 0.24 | 1 | 0.57276199 | 3.7969E-07 |
| Gmppa | 0.02533123 | 0.58260925 | 0.479 | 0.352 | 1 | 1.9457E-07 | 1.55521516 | 0.462 | 0.156 | 0.0047266 | 0.02533123 | 3.8915E-07 |
| Sqle | 0.00889691 | 0.68623297 | 0.667 | 0.622 | 1 | 1.9777E-07 | 1.53697394 | 0.555 | 0.271 | 0.0048043 | 0.00889691 | 3.9555E-07 |
| Fkbp14 | 0.00140256 | 0.3706604 | 0.875 | 0.679 | 1 | 1.9901E-07 | 0.95416425 | 0.639 | 0.312 | 0.00483444 | 0.00140256 | 3.9803E-07 |
| Ddost | 0.000041021 | 0.48655496 | 0.938 | 0.927 | 0.99648121 | 1.9903E-07 | 0.60936208 | 0.937 | 0.823 | 0.00483489 | 4.1021E-05 | 3.9806E-07 |
| Pde4dip | 2.0481E-07 | 0.63841535 | 0.979 | 0.927 | 0.00497516 | 0.01547344 | -0.1148528 | 0.979 | 0.948 | 1 | 0.01547344 | 4.0961E-07 |
| Cmklr1 | 2.0636E-07 | 1.07234568 | 0.542 | 0.218 | 0.00501282 | 0.00092575 | 0.25691412 | 0.676 | 0.406 | 1 | 0.00092575 | 4.1271E-07 |
| Hsp90ab1 | 0.01551917 | 0.15866502 | 1 | 0.995 | 1 | 2.115E-07 | 0.42220425 | 1 | 0.99 | 0.00513781 | 0.01551917 | 0.000000423 |
| Ppid | 0.000015131 | 0.72535131 | 0.698 | 0.43 | 0.36757222 | 2.1223E-07 | 0.94146816 | 0.714 | 0.375 | 0.0051554 | 1.5131E-05 | 4.2445E-07 |
| Foxn3 | 0.73215229 | -0.3395047 | 0.979 | 0.959 | 1 | 2.1572E-07 | -0.7309658 | 0.987 | 1 | 0.00524026 | 0.73215229 | 4.3144E-07 |
| Hspa1a | 0.58618864 | -0.3017027 | 0.469 | 0.523 | 1 | 2.1993E-07 | -1.3996004 | 0.529 | 0.792 | 0.00534249 | 0.58618864 | 4.3986E-07 |
| Samd4 | 2.2042E-07 | -0.957587 | 0.917 | 0.984 | 0.00535456 | 0.00091336 | -0.8145194 | 0.979 | 1 | 1 | 0.00091336 | 4.4085E-07 |
| Slc29a1 | 0.27763922 | -0.3000481 | 0.896 | 0.876 | 1 | 2.2139E-07 | 0.66379341 | 0.899 | 0.542 | 0.00537791 | 0.27763922 | 4.4277E-07 |
| Snx15 | 0.16755635 | 0.30250133 | 0.521 | 0.425 | 1 | 2.2729E-07 | 1.33843317 | 0.462 | 0.146 | 0.00552133 | 0.16755635 | 4.5458E-07 |
| 2300009A05Rik | 0.01362899 | 0.64492446 | 0.396 | 0.254 | 1 | 2.3071E-07 | 1.87234807 | 0.366 | 0.083 | 0.00560442 | 0.01362899 | 4.6142E-07 |
| Zzz3 | 2.3121E-07 | 0.80246573 | 0.812 | 0.539 | 0.0056166 | 0.03520389 | -0.4567711 | 0.857 | 0.927 | 1 | 0.03520389 | 4.6242E-07 |
| Klf4 | 0.000017738 | 0.70194361 | 0.969 | 0.964 | 0.43088591 | 2.3148E-07 | 0.46976892 | 0.996 | 0.958 | 0.00562315 | 1.7738E-05 | 4.6296E-07 |
| Inppl1 | 0.99092706 | 0.13677755 | 0.365 | 0.358 | 1 | 2.3235E-07 | 0.93349081 | 0.697 | 0.312 | 0.00564433 | 0.99092706 | 4.6471E-07 |
| Glis3 | 2.3352E-07 | -1.7503351 | 0.698 | 0.793 | 0.00567271 | 0.05466175 | -0.9001917 | 0.861 | 0.844 | 1 | 0.05466175 | 4.6704E-07 |
| Adh1 | 0.00023538 | -4.4187702 | 0 | 0.13 | 1 | 2.3431E-07 | -1.3098553 | 0.122 | 0.375 | 0.00569174 | 0.00023538 | 4.6861E-07 |
| D8Ertd738e | 0.96612455 | 0.1051578 | 0.896 | 0.855 | 1 | 2.3445E-07 | 0.72600479 | 0.899 | 0.708 | 0.00569521 | 0.96612455 | 4.689E-07 |
| Rxylt1 | 0.32984345 | 0.10595718 | 0.74 | 0.663 | 1 | 2.3623E-07 | 0.79184013 | 0.739 | 0.375 | 0.00573855 | 0.32984345 | 4.7246E-07 |
| Kcnj16 | 0.38523254 | 0.74446013 | 0.031 | 0.016 | 1 | 2.383E-07 | -1.6634922 | 0.042 | 0.229 | 0.00578881 | 0.38523254 | 4.766E-07 |
| Armcx4 | 0.00736417 | 0.67944034 | 0.375 | 0.218 | 1 | 2.4028E-07 | 1.40006551 | 0.483 | 0.167 | 0.00583692 | 0.00736417 | 4.8056E-07 |
| Cdk14 | 0.04226422 | -1.0006794 | 0.469 | 0.565 | 1 | 2.4289E-07 | -1.2927817 | 0.706 | 0.885 | 0.00590029 | 0.04226422 | 4.8578E-07 |
| Slit2 | 0.00466131 | -2.2574777 | 0.094 | 0.233 | 1 | 2.485E-07 | -1.7322648 | 0.261 | 0.594 | 0.00603663 | 0.00466131 | 4.9701E-07 |
| Gmppb | 0.25501269 | 0.45647937 | 0.292 | 0.233 | 1 | 2.4932E-07 | 3.26000035 | 0.286 | 0.031 | 0.00605639 | 0.25501269 | 4.9863E-07 |
| Gsn | 0.00805016 | -0.5367216 | 0.969 | 0.99 | 1 | 2.5181E-07 | 0.46797089 | 1 | 0.99 | 0.00611686 | 0.00805016 | 5.0361E-07 |

| | | | | | | | | | | | |
|---|---|---|---|---|---|---|---|---|---|---|---|
| Zfp874b | 0.00505732 | 1.00749454 | 0.26 | 0.13 | 1 | 2.5428E-07 | 2.28707988 | 0.353 | 0.073 | 0.00617696 | 0.00505732 | 5.0856E-07 |
| Lrrc8a | 0.22144517 | 0.40676472 | 0.51 | 0.44 | 1 | 2.6159E-07 | 0.90392403 | 0.689 | 0.302 | 0.00635455 | 0.22144517 | 5.2318E-07 |
| Ssrp1 | 2.6163E-07 | 0.86034084 | 0.781 | 0.508 | 0.0063554 | 0.016206 | 0.36474445 | 0.71 | 0.615 | 1 | 0.016206 | 5.2325E-07 |
| Emc3 | 0.00042647 | 0.50108268 | 0.854 | 0.658 | 1 | 2.6437E-07 | 0.84361922 | 0.807 | 0.625 | 0.00642212 | 0.00042647 | 5.2874E-07 |
| Ctsc | 0.41200015 | -0.6843832 | 0.052 | 0.078 | 1 | 2.6795E-07 | -1.2713811 | 0.088 | 0.323 | 0.00650894 | 0.41200015 | 5.3589E-07 |
| Pabpc4 | 1.2374E-06 | 0.7008332 | 0.875 | 0.658 | 0.03005933 | 2.7062E-07 | 0.89572887 | 0.798 | 0.552 | 0.00657392 | 1.2374E-06 | 5.4124E-07 |
| Abcc4 | 0.0006262 | -1.3815478 | 0.104 | 0.285 | 1 | 2.712E-07 | -1.7047149 | 0.16 | 0.427 | 0.00658796 | 0.0006262 | 5.424E-07 |
| Nomo1 | 0.89566678 | 0.22988696 | 0.354 | 0.358 | 1 | 2.7441E-07 | 1.42711033 | 0.466 | 0.146 | 0.00666605 | 0.89566678 | 5.4883E-07 |
| Rbm3 | 0.00047359 | 0.4692803 | 0.99 | 0.938 | 1 | 2.7459E-07 | 0.46479294 | 1 | 0.969 | 0.00667026 | 0.00047359 | 5.4917E-07 |
| AI837181 | 0.00691337 | 0.69656544 | 0.521 | 0.373 | 1 | 2.8102E-07 | 1.13071733 | 0.555 | 0.229 | 0.00682646 | 0.00691337 | 5.6203E-07 |
| Dis3 | 2.8801E-07 | 1.43325844 | 0.552 | 0.259 | 0.00699638 | 0.00465339 | 0.49269867 | 0.408 | 0.219 | 1 | 0.00465339 | 5.7602E-07 |
| Taf11 | 0.00027353 | 0.70186825 | 0.625 | 0.399 | 1 | 2.9281E-07 | 1.02484013 | 0.576 | 0.229 | 0.00711301 | 0.00027353 | 5.8563E-07 |
| Rragc | 0.00129034 | 0.33372277 | 0.76 | 0.513 | 1 | 2.9351E-07 | 0.87054307 | 0.845 | 0.667 | 0.00712994 | 0.00129034 | 5.8702E-07 |
| Clstn2 | 2.9843E-07 | 1.79476521 | 0.448 | 0.171 | 0.00724951 | 0.48051784 | -0.7389679 | 0.525 | 0.5 | 1 | 0.48051784 | 5.9686E-07 |
| Magi3 | 0.1301905 | 0.5093753 | 0.417 | 0.352 | 1 | 2.9963E-07 | -1.3600896 | 0.454 | 0.74 | 0.00727849 | 0.1301905 | 5.9925E-07 |
| Castor2 | 0.17265276 | 0.46907462 | 0.26 | 0.192 | 1 | 3.023E-07 | 1.01207283 | 0.639 | 0.26 | 0.00734355 | 0.17265276 | 6.0461E-07 |
| Znrf1 | 2.7485E-06 | -1.1105027 | 0.49 | 0.731 | 0.06676595 | 3.0298E-07 | -0.9622873 | 0.639 | 0.875 | 0.00736009 | 2.7485E-06 | 6.0597E-07 |
| Anapc2 | 0.10785923 | 0.407102 | 0.448 | 0.352 | 1 | 3.0716E-07 | 1.2190732 | 0.555 | 0.24 | 0.00746142 | 0.10785923 | 6.1431E-07 |
| Ppargc1a | 0.15985611 | -0.4519371 | 0.01 | 0.041 | 1 | 3.0821E-07 | -0.6577786 | 0.134 | 0.406 | 0.00748701 | 0.15985611 | 6.1642E-07 |
| Brwd3 | 0.00314135 | 0.39451766 | 0.406 | 0.218 | 1 | 3.0918E-07 | -1.3451307 | 0.42 | 0.719 | 0.00751071 | 0.00314135 | 6.1837E-07 |
| Trir | 0.4876472 | 0.11596633 | 0.854 | 0.85 | 1 | 3.1584E-07 | 0.65452083 | 0.899 | 0.792 | 0.00767238 | 0.4876472 | 6.3168E-07 |
| G6pc3 | 0.021762 | 0.39981196 | 0.51 | 0.352 | 1 | 3.1707E-07 | 1.31073115 | 0.42 | 0.115 | 0.0077022 | 0.021762 | 6.3413E-07 |
| Lrrc58 | 3.1972E-07 | 0.60784007 | 0.99 | 0.86 | 0.00776656 | 0.00021566 | 0.38701549 | 0.992 | 0.938 | 1 | 0.00021566 | 6.3943E-07 |
| Ube2g2 | 0.43337939 | -0.2929004 | 0.417 | 0.44 | 1 | 3.222E-07 | 0.92345845 | 0.601 | 0.25 | 0.00782686 | 0.43337939 | 6.444E-07 |
| S100a1 | 0.72040793 | -0.3565835 | 0.896 | 0.87 | 1 | 3.2702E-07 | 0.62650224 | 0.853 | 0.698 | 0.00794392 | 0.72040793 | 6.5404E-07 |
| Hs2st1 | 0.00849251 | -0.9015803 | 0.656 | 0.725 | 1 | 3.324E-07 | -1.3341028 | 0.601 | 0.833 | 0.00807472 | 0.00849251 | 6.648E-07 |
| Prdx1 | 0.11302451 | 0.15580096 | 1 | 0.953 | 1 | 3.3564E-07 | 0.53732116 | 0.996 | 0.99 | 0.00815347 | 0.11302451 | 6.7129E-07 |
| Nars | 0.000094033 | 0.7620185 | 0.854 | 0.819 | 1 | 3.3878E-07 | 0.67558011 | 0.945 | 0.76 | 0.00822967 | 9.4033E-05 | 6.7756E-07 |
| Gm56743 | 0.00821396 | 1.0470229 | 0.292 | 0.161 | 1 | 3.4075E-07 | 1.65361886 | 0.408 | 0.115 | 0.0082775 | 0.00821396 | 6.815E-07 |
| Rab2a | 3.4403E-07 | 0.38356562 | 1 | 0.995 | 0.00835714 | 0.01033607 | 0.14023154 | 1 | 1 | 1 | 0.01033607 | 6.8806E-07 |
| Psmd7 | 0.00348661 | 0.4187504 | 0.917 | 0.855 | 1 | 3.5053E-07 | 0.61049759 | 0.945 | 0.844 | 0.00851499 | 0.00348661 | 7.0105E-07 |
| Snrpa1 | 0.05283068 | 0.43847358 | 0.573 | 0.456 | 1 | 3.616E-07 | 0.80988398 | 0.592 | 0.24 | 0.00878398 | 0.05283068 | 7.232E-07 |
| Gstm5 | 0.1275517 | -0.3354715 | 0.698 | 0.741 | 1 | 3.6176E-07 | 0.7218536 | 0.693 | 0.312 | 0.00878799 | 0.1275517 | 7.2353E-07 |
| Arf6 | 0.001922 | 0.49578102 | 0.812 | 0.684 | 1 | 3.6867E-07 | 0.64031358 | 0.899 | 0.729 | 0.00895565 | 0.001922 | 7.3733E-07 |
| Tcp1 | 7.5755E-07 | 0.57392109 | 0.948 | 0.902 | 0.01840249 | 0.000000373 | 0.56340456 | 0.954 | 0.854 | 0.009061 | 7.5755E-07 | 7.4601E-07 |
| S100a16 | 0.86597511 | 0.11440974 | 0.594 | 0.679 | 1 | 3.7449E-07 | 1.08315593 | 0.769 | 0.542 | 0.00909709 | 0.86597511 | 7.4898E-07 |
| Anp32b | 3.7533E-07 | 0.66881508 | 0.969 | 0.87 | 0.00911742 | 0.00013848 | 0.36349423 | 0.966 | 0.896 | 1 | 0.00013848 | 7.5065E-07 |
| Klhl9 | 3.7622E-07 | 1.01548733 | 0.781 | 0.518 | 0.00913909 | 4.4939E-07 | 0.83197743 | 0.828 | 0.646 | 0.0109167 | 4.4939E-07 | 7.5244E-07 |
| Tada2b | 0.11527734 | 0.47315811 | 0.271 | 0.187 | 1 | 3.7749E-07 | 1.74542717 | 0.395 | 0.104 | 0.00916988 | 0.11527734 | 7.5497E-07 |
| Trmt112 | 0.00041967 | 0.52206771 | 0.854 | 0.731 | 1 | 3.7815E-07 | 0.66005267 | 0.828 | 0.604 | 0.00918605 | 0.00041967 | 7.563E-07 |
| Rps19bp1 | 0.01813818 | 0.59902969 | 0.385 | 0.244 | 1 | 3.7898E-07 | 1.63916081 | 0.42 | 0.125 | 0.0092062 | 0.01813818 | 7.5796E-07 |
| Fam104a | 0.00382345 | 0.42002953 | 0.948 | 0.886 | 1 | 3.8133E-07 | 0.61105191 | 0.945 | 0.896 | 0.00926326 | 0.00382345 | 7.6266E-07 |
| Tes | 0.03099967 | -1.6929452 | 0.042 | 0.119 | 1 | 3.9021E-07 | -1.6659991 | 0.088 | 0.312 | 0.0094789 | 0.03099967 | 7.8041E-07 |
| Cd24a | 0.00139697 | -0.3232395 | 0.49 | 0.736 | 1 | 3.9502E-07 | -0.480474 | 0.307 | 0.667 | 0.00959586 | 0.00139697 | 7.9004E-07 |
| Hadh | 0.12271991 | 0.41426442 | 0.615 | 0.518 | 1 | 4.0752E-07 | 1.13841857 | 0.668 | 0.427 | 0.00989937 | 0.12271991 | 8.1503E-07 |
| Nkd1 | 0.00060021 | 0.50450934 | 0.698 | 0.487 | 1 | 4.1264E-07 | 1.06211351 | 0.639 | 0.323 | 0.01002389 | 0.00060021 | 8.2528E-07 |
| Slc8a1 | 0.79463142 | 0.85549144 | 0.031 | 0.026 | 1 | 4.1594E-07 | -1.5783441 | 0.08 | 0.302 | 0.01010407 | 0.79463142 | 8.3188E-07 |
| Washc3 | 0.18918227 | 0.21260897 | 0.594 | 0.487 | 1 | 4.1858E-07 | 0.88157081 | 0.513 | 0.177 | 0.01016812 | 0.18918227 | 8.3716E-07 |
| Tmbim6 | 4.2491E-07 | -0.5329737 | 0.99 | 0.979 | 0.01032197 | 0.00448394 | 0.17201375 | 1 | 0.979 | 1 | 0.00448394 | 8.4982E-07 |
| Gps1 | 0.06031986 | 0.54117653 | 0.5 | 0.394 | 1 | 4.311E-07 | 0.95984287 | 0.571 | 0.24 | 0.0104724 | 0.06031986 | 8.6221E-07 |
| Tgm2 | 4.3474E-07 | -1.5408376 | 0.677 | 0.803 | 0.01056071 | 0.25365532 | -0.1119159 | 0.517 | 0.625 | 1 | 0.25365532 | 8.6948E-07 |
| Adrm1 | 0.00779627 | 0.52660449 | 0.625 | 0.466 | 1 | 4.3478E-07 | 0.99564728 | 0.66 | 0.344 | 0.01056168 | 0.00779627 | 8.6956E-07 |
| Itprid2 | 0.06118515 | -0.5636622 | 0.219 | 0.316 | 1 | 4.3889E-07 | -1.1590753 | 0.374 | 0.656 | 0.01066152 | 0.06118515 | 8.7778E-07 |
| Ssr1 | 4.4651E-07 | 0.54680479 | 1 | 0.917 | 0.01084652 | 2.4305E-06 | 0.5056205 | 0.962 | 0.896 | 0.0590416 | 2.4305E-06 | 8.9301E-07 |
| Cdc16 | 4.4858E-07 | 0.82692229 | 0.844 | 0.596 | 0.01089679 | 8.7368E-07 | 0.90515763 | 0.744 | 0.406 | 0.02122346 | 8.7368E-07 | 8.9715E-07 |
| Lman1 | 0.000056191 | 0.40279582 | 0.979 | 0.912 | 1 | 4.5241E-07 | 0.60636113 | 0.975 | 0.812 | 0.01099 | 5.6191E-05 | 9.0482E-07 |
| Skap2 | 4.5518E-07 | -2.1245339 | 0.177 | 0.472 | 0.01105734 | 0.15973308 | -0.1609919 | 0.206 | 0.281 | 1 | 0.15973308 | 9.1037E-07 |
| Marveld1 | 0.69521214 | -0.455906 | 0.698 | 0.642 | 1 | 4.6289E-07 | 0.81567562 | 0.706 | 0.344 | 0.01124461 | 0.69521214 | 9.2579E-07 |
| Pfdn4 | 0.13711559 | 0.25747279 | 0.5 | 0.383 | 1 | 4.6501E-07 | 1.23820766 | 0.521 | 0.219 | 0.01129595 | 0.13711559 | 9.3001E-07 |
| Rassf1 | 4.7008E-07 | 1.00749454 | 0.74 | 0.472 | 0.01141921 | 0.04857287 | 0.10518224 | 0.744 | 0.635 | 1 | 0.04857287 | 9.4016E-07 |
| Ptov1 | 0.00014537 | 0.52206771 | 0.896 | 0.72 | 1 | 4.7498E-07 | 0.69485174 | 0.866 | 0.698 | 0.01153824 | 0.00014537 | 9.4996E-07 |
| Mthfd2 | 5.1254E-07 | 1.34493563 | 0.646 | 0.373 | 0.01245054 | 4.785E-07 | 1.56964945 | 0.504 | 0.198 | 0.01162381 | 5.1254E-07 | 9.5701E-07 |
| Rasa4 | 4.7923E-07 | 3.28051303 | 0.188 | 0.021 | 0.01164149 | 0.84728961 | -0.6931839 | 0.071 | 0.062 | 1 | 0.84728961 | 9.5846E-07 |
| Atp6v0e | 0.00014076 | -0.7800811 | 0.979 | 0.969 | 1 | 4.8009E-07 | 0.71961165 | 0.975 | 0.885 | 0.01166245 | 0.00014076 | 9.6019E-07 |
| Rab18 | 0.17350826 | 0.14013231 | 0.927 | 0.865 | 1 | 4.9956E-07 | 0.66426611 | 0.941 | 0.812 | 0.01213542 | 0.17350826 | 9.9913E-07 |
| Lifr | 5.0378E-07 | -1.8059094 | 0.24 | 0.534 | 0.01223775 | 0.09553874 | -1.067285 | 0.218 | 0.281 | 1 | 0.09553874 | 1.0076E-06 |
| Hipk2 | 5.0554E-07 | -1.1112294 | 0.667 | 0.886 | 0.01228066 | 0.00817024 | -0.6063029 | 0.731 | 0.885 | 1 | 0.00817024 | 1.0111E-06 |
| Dctn6 | 0.01919535 | 0.42720852 | 0.656 | 0.518 | 1 | 5.075E-07 | 0.89296162 | 0.597 | 0.281 | 0.01232811 | 0.01919535 | 0.000001015 |
| Cldn11 | 5.1031E-07 | -2.6657878 | 0.125 | 0.399 | 0.01239651 | 0.000029409 | -4.129283 | 0.042 | 0.177 | 0.71441142 | 2.9409E-05 | 1.0206E-06 |
| Vma21 | 0.01065203 | 0.29945055 | 0.771 | 0.591 | 1 | 5.1223E-07 | 0.87100759 | 0.727 | 0.438 | 0.01244297 | 0.01065203 | 1.0245E-06 |
| Rtraf | 0.00040095 | 0.42253204 | 0.969 | 0.902 | 1 | 5.169E-07 | 0.57196331 | 0.912 | 0.792 | 0.01255662 | 0.00040095 | 1.0338E-06 |
| H3f3b | 0.67392467 | 0.1732203 | 1 | 0.995 | 1 | 5.2483E-07 | 0.57525613 | 1 | 1 | 0.0127492 | 0.67392467 | 1.0497E-06 |
| Lgalsl | 0.35911727 | -0.7981838 | 0.583 | 0.575 | 1 | 5.2572E-07 | 1.27931949 | 0.605 | 0.292 | 0.0127708 | 0.35911727 | 1.0514E-06 |
| Tmem214 | 0.01139288 | 0.5239773 | 0.615 | 0.503 | 1 | 5.3493E-07 | 0.8548895 | 0.634 | 0.292 | 0.01299444 | 0.01139288 | 1.0699E-06 |
| Rnh1 | 0.51922151 | 0.10417056 | 0.781 | 0.751 | 1 | 5.3755E-07 | 0.72690329 | 0.861 | 0.719 | 0.01305811 | 0.51922151 | 1.0751E-06 |
| Ndufb11 | 0.01426831 | 0.32032042 | 0.948 | 0.896 | 1 | 5.3884E-07 | 0.52561972 | 0.966 | 0.844 | 0.01308939 | 0.01426831 | 1.0777E-06 |
| Stag1 | 5.5095E-07 | 0.56794289 | 1 | 0.891 | 0.01338364 | 0.4099449 | -0.300621 | 0.996 | 0.99 | 1 | 0.4099449 | 1.1019E-06 |
| Klf2 | 5.6519E-07 | 1.37621231 | 0.833 | 0.617 | 0.01372967 | 0.36820082 | 0.67365661 | 0.71 | 0.74 | 1 | 0.36820082 | 1.1304E-06 |
| Timm8a1 | 0.000016196 | 1.50792353 | 0.375 | 0.15 | 0.39342117 | 5.6745E-07 | 1.59060906 | 0.336 | 0.062 | 0.01378448 | 1.6196E-05 | 1.1349E-06 |
| Hspg2 | 5.7453E-07 | 0.69235088 | 1 | 0.912 | 0.01395647 | 0.01461496 | 0.11083886 | 0.941 | 0.812 | 1 | 0.01461496 | 1.1491E-06 |
| Ninj2 | 5.7548E-07 | 1.02229154 | 0.729 | 0.544 | 0.01397951 | 7.2574E-07 | 0.54049706 | 0.622 | 0.25 | 0.01762963 | 7.2574E-07 | 0.000001151 |
| Cct3 | 0.00034002 | 0.51610615 | 0.875 | 0.803 | 1 | 5.7732E-07 | 0.64719813 | 0.912 | 0.792 | 0.01402421 | 0.00034002 | 1.1546E-06 |
| Alyref | 2.3527E-06 | 0.77250855 | 0.885 | 0.736 | 0.05715215 | 5.7833E-07 | 0.67733696 | 0.933 | 0.792 | 0.01404879 | 2.3527E-06 | 1.1567E-06 |
| Tceal3 | 1.0288E-06 | 1.22988696 | 0.542 | 0.269 | 0.02499112 | 5.7912E-07 | 1.30846584 | 0.597 | 0.281 | 0.01406795 | 1.0288E-06 | 1.1582E-06 |
| Lsp1 | 5.8287E-07 | 1.00749454 | 0.573 | 0.254 | 0.0141592 | 0.58784802 | -0.6871378 | 0.538 | 0.479 | 1 | 0.58784802 | 1.1657E-06 |
| Atp5pb | 0.00421726 | 0.28941884 | 0.958 | 0.907 | 1 | 5.837E-07 | 0.58544736 | 0.971 | 0.854 | 0.01417923 | 0.00421726 | 1.1674E-06 |
| Slit3 | 0.20651797 | -3.1329867 | 0.062 | 0.104 | 1 | 5.9318E-07 | -3.3322231 | 0.13 | 0.344 | 0.01440959 | 0.20651797 | 1.1864E-06 |
| Ctnnb1 | 0.00414943 | 0.29827393 | 0.958 | 0.943 | 1 | 5.9665E-07 | 0.47458713 | 1 | 0.99 | 0.01449389 | 0.00414943 | 1.1933E-06 |
| Mrpl4 | 0.37928417 | -0.1260886 | 0.458 | 0.534 | 1 | 5.9818E-07 | 0.73040861 | 0.613 | 0.25 | 0.01453107 | 0.37928417 | 1.1964E-06 |
| Lyn | 0.00175223 | 2.00749454 | 0.188 | 0.067 | 1 | 6.0031E-07 | -0.718495 | 0.223 | 0.531 | 0.01458277 | 0.00175223 | 1.2006E-06 |
| Prdx5 | 6.0886E-07 | 0.69543669 | 0.948 | 0.845 | 0.01479032 | 0.000027082 | 0.52042189 | 0.92 | 0.833 | 0.6578775 | 2.7082E-05 | 1.2177E-06 |
| Tfcp2l1 | 0.8044339 | -0.9185049 | 0.042 | 0.047 | 1 | 6.1232E-07 | -1.612418 | 0.067 | 0.271 | 0.01487442 | 0.8044339 | 1.2246E-06 |

| | | | | | | | | | | | | |
|---|---|---|---|---|---|---|---|---|---|---|---|---|
| Cr1l | 0.00295369 | 0.4485699 | 0.833 | 0.736 | 1 | 6.1241E-07 | 0.73289438 | 0.836 | 0.688 | 0.01487656 | 0.00295369 | 1.2248E-06 |
| Hnrnpa3 | 6.1331E-07 | 0.5180322 | 1 | 0.979 | 0.01489861 | 0.00060735 | 0.28851217 | 0.996 | 0.99 | 1 | 0.00060735 | 1.2266E-06 |
| Pja1 | 0.0671396 | 0.94448474 | 0.458 | 0.368 | 1 | 6.1708E-07 | 1.23267197 | 0.542 | 0.219 | 0.01499011 | 0.0671396 | 1.2342E-06 |
| Tmx4 | 6.1727E-07 | 1.35199304 | 0.802 | 0.549 | 0.01499473 | 0.000004937 | 1.19448878 | 0.697 | 0.521 | 0.1199299 | 4.937E-06 | 1.2345E-06 |
| Septin9 | 6.2194E-07 | -1.2016469 | 0.781 | 0.886 | 0.01510823 | 0.0648193 | -0.7278281 | 0.853 | 0.885 | 1 | 0.0648193 | 1.2439E-06 |
| Chchd7 | 0.16271682 | 0.24643327 | 0.615 | 0.508 | 1 | 6.2353E-07 | 1.09326892 | 0.538 | 0.219 | 0.01514668 | 0.16271682 | 1.2471E-06 |
| Arhgap15 | 0.99034437 | 2.42253204 | 0.021 | 0.021 | 1 | 6.328E-07 | -0.6317834 | 0.013 | 0.146 | 0.01537188 | 0.99034437 | 1.2656E-06 |
| Ipo5 | 6.3791E-07 | 0.98658555 | 0.729 | 0.409 | 0.01549618 | 0.16078877 | 0.19427616 | 0.685 | 0.615 | 1 | 0.16078877 | 1.2758E-06 |
| Fam3d | 0.53178362 | -0.407543 | 0.01 | 0.021 | 1 | 0.000000649 | -3.07539 | 0.017 | 0.156 | 0.01576559 | 0.53178362 | 0.000001298 |
| Srsf6 | 6.4949E-07 | 0.69979108 | 0.906 | 0.72 | 0.01577729 | 4.4531E-06 | 0.64730854 | 0.899 | 0.76 | 0.10817426 | 4.4531E-06 | 0.000001299 |
| Gsto2 | 0.22591488 | 0.59245704 | 0.104 | 0.062 | 1 | 6.5187E-07 | -2.5577828 | 0.059 | 0.25 | 0.01583516 | 0.22591488 | 1.3037E-06 |
| Slirp | 0.11260856 | 0.26305382 | 0.552 | 0.44 | 1 | 6.5821E-07 | 1.42711033 | 0.45 | 0.146 | 0.01598928 | 0.11260856 | 1.3164E-06 |
| Rtl8a | 0.02080615 | 0.57496961 | 0.625 | 0.518 | 1 | 6.6059E-07 | 0.931623 | 0.647 | 0.333 | 0.01604708 | 0.02080615 | 1.3212E-06 |
| Pcdh17 | 0.00186363 | -1.6348011 | 0.167 | 0.326 | 1 | 6.6202E-07 | -0.9662683 | 0.164 | 0.448 | 0.01608168 | 0.00186363 | 0.000001324 |
| Fam107a | 0.41478371 | 1.39451766 | 0.062 | 0.041 | 1 | 6.7325E-07 | 2.42194863 | 0.328 | 0.062 | 0.01635464 | 0.41478371 | 1.3465E-06 |
| Cdhr4 | 6.7546E-07 | -3.5160674 | 0.031 | 0.275 | 0.01640833 | 0.08430124 | -1.3098553 | 0.038 | 0.083 | 1 | 0.08430124 | 1.3509E-06 |
| Gprc5c | 0.04008598 | 0.74097956 | 0.438 | 0.321 | 1 | 6.8087E-07 | 1.43207658 | 0.508 | 0.208 | 0.01653973 | 0.04008598 | 1.3617E-06 |
| Snrpb | 0.00062738 | 0.37801337 | 0.969 | 0.865 | 1 | 6.848E-07 | 0.62466532 | 0.979 | 0.854 | 0.01663514 | 0.00062738 | 1.3696E-06 |
| Set | 5.0653E-06 | 0.41021176 | 0.969 | 0.927 | 0.12304538 | 6.9863E-07 | 0.49749966 | 0.996 | 0.958 | 0.01697123 | 5.0653E-06 | 1.3973E-06 |
| Ctsl | 6.989E-07 | -0.7754924 | 0.833 | 0.927 | 0.01697773 | 0.01542928 | 0.21119547 | 0.966 | 0.948 | 1 | 0.01542928 | 1.3978E-06 |
| Rer1 | 0.00042051 | 0.42253204 | 0.958 | 0.855 | 1 | 6.9918E-07 | 0.5319969 | 0.933 | 0.677 | 0.01698445 | 0.00042051 | 1.3984E-06 |
| Pabpn1 | 0.00030248 | 0.36294324 | 0.948 | 0.896 | 1 | 7.0163E-07 | 0.53563479 | 0.929 | 0.833 | 0.01704396 | 0.00030248 | 1.4033E-06 |
| Eif3i | 0.00786168 | 0.29866038 | 0.906 | 0.845 | 1 | 7.021E-07 | 0.60423083 | 0.899 | 0.75 | 0.01705536 | 0.00786168 | 1.4042E-06 |
| Pdia4 | 0.07943656 | 0.2288075 | 0.938 | 0.855 | 1 | 7.1705E-07 | 0.59314841 | 0.933 | 0.76 | 0.01741862 | 0.07943656 | 1.4341E-06 |
| Sem1 | 0.0002397 | 0.35828745 | 1 | 0.943 | 1 | 7.1904E-07 | 0.48765187 | 0.996 | 0.979 | 0.01746703 | 0.0002397 | 1.4381E-06 |
| Mrps25 | 0.01651769 | 0.58122978 | 0.479 | 0.352 | 1 | 7.2015E-07 | 0.70682348 | 0.525 | 0.198 | 0.01749399 | 0.01651769 | 1.4403E-06 |
| Yipf5 | 0.000012939 | 0.55159461 | 0.948 | 0.839 | 0.31431038 | 7.314E-07 | 0.5929885 | 0.933 | 0.792 | 0.01776724 | 1.2939E-05 | 1.4628E-06 |
| Baalc | 7.3514E-07 | -2.8543082 | 0.083 | 0.347 | 0.01785813 | 0.65799427 | -1.368749 | 0.063 | 0.073 | 1 | 0.65799427 | 1.4703E-06 |
| Lum | 0.02651433 | -0.8572075 | 0.75 | 0.523 | 1 | 7.3617E-07 | 0.18266433 | 0.849 | 0.531 | 0.017883 | 0.02651433 | 1.4723E-06 |
| Ddx17 | 7.4473E-07 | 0.54457634 | 1 | 0.943 | 0.01809094 | 0.000019375 | 0.43169214 | 1 | 0.958 | 0.47066129 | 1.9375E-05 | 1.4895E-06 |
| Atox1 | 7.4888E-07 | -1.6402037 | 0.927 | 0.974 | 0.01819189 | 0.00155887 | 0.4651155 | 0.933 | 0.854 | 1 | 0.00155887 | 1.4978E-06 |
| Mal2 | 0.01388843 | 3.00749454 | 0.031 | 0 | 1 | 7.5152E-07 | -2.6724253 | 0.025 | 0.177 | 0.01825583 | 0.01388843 | 0.000001503 |
| Ptprd | 0.12858867 | -2.4591551 | 0.385 | 0.415 | 1 | 7.5219E-07 | -1.9195503 | 0.601 | 0.823 | 0.01827226 | 0.12858867 | 1.5044E-06 |
| Gtf2a2 | 0.14738068 | 0.30508616 | 0.76 | 0.679 | 1 | 7.5475E-07 | 0.64216471 | 0.811 | 0.573 | 0.01833451 | 0.14738068 | 1.5095E-06 |
| Itm2b | 0.00014621 | -0.5258762 | 1 | 0.995 | 1 | 7.6937E-07 | 0.32121357 | 1 | 0.99 | 0.01868949 | 0.00014621 | 1.5387E-06 |
| Vdr | 7.7163E-07 | 1.27428108 | 0.458 | 0.181 | 0.0187445 | 0.00039801 | 1.27510724 | 0.227 | 0.062 | 1 | 0.00039801 | 1.5433E-06 |
| Zfand3 | 0.10976692 | -0.3069866 | 0.969 | 0.99 | 1 | 7.7814E-07 | -0.7005126 | 1 | 1 | 0.0189026 | 0.10976692 | 1.5563E-06 |
| Dpm3 | 7.799E-07 | 0.90680054 | 0.792 | 0.585 | 0.01894535 | 0.00169505 | 0.30871563 | 0.706 | 0.531 | 1 | 0.00169505 | 1.5598E-06 |
| Capn6 | 2.6324E-06 | 0.80611465 | 0.719 | 0.378 | 0.06394667 | 7.8123E-07 | 0.61136762 | 0.832 | 0.427 | 0.01897753 | 2.6324E-06 | 1.5625E-06 |
| Usf2 | 0.29273443 | 0.24298262 | 0.562 | 0.477 | 1 | 7.8473E-07 | 0.99053967 | 0.66 | 0.365 | 0.01906262 | 0.29273443 | 1.5695E-06 |
| Nr3c2 | 0.02565122 | -1.1217885 | 0.188 | 0.306 | 1 | 7.9093E-07 | -1.1041802 | 0.239 | 0.521 | 0.01921328 | 0.02565122 | 1.5819E-06 |
| Nifk | 0.00130733 | 0.73773427 | 0.615 | 0.415 | 1 | 8.0277E-07 | 1.34786358 | 0.483 | 0.167 | 0.01950095 | 0.00130733 | 1.6055E-06 |
| Poglut3 | 8.0588E-07 | 1.03550891 | 0.646 | 0.368 | 0.01957641 | 0.000012037 | 0.62405555 | 0.597 | 0.292 | 0.29240501 | 1.2037E-05 | 1.6118E-06 |
| Arid1b | 0.00027213 | -0.8402279 | 0.896 | 0.927 | 1 | 8.3665E-07 | -0.787028 | 0.962 | 1 | 0.02032389 | 0.00027213 | 1.6733E-06 |
| Psmb1 | 0.01165828 | 0.31932204 | 0.938 | 0.938 | 1 | 8.379E-07 | 0.44442721 | 0.971 | 0.896 | 0.02035421 | 0.01165828 | 1.6758E-06 |
| Tmem167b | 0.00979742 | 0.45495351 | 0.625 | 0.482 | 1 | 8.4184E-07 | 0.95724881 | 0.66 | 0.344 | 0.02044986 | 0.00979742 | 1.6837E-06 |
| Naf1 | 0.01371212 | 0.64944702 | 0.49 | 0.342 | 1 | 8.5224E-07 | 1.40959257 | 0.487 | 0.188 | 0.0207026 | 0.01371212 | 1.7045E-06 |
| Cldn3 | 0.2801998 | -0.8669746 | 0.021 | 0.047 | 1 | 8.6006E-07 | -2.0161241 | 0.055 | 0.24 | 0.0208927 | 0.2801998 | 1.7201E-06 |
| Rictor | 0.00734609 | -0.4806064 | 0.656 | 0.788 | 1 | 8.7183E-07 | -0.9799659 | 0.815 | 0.906 | 0.02117856 | 0.00734609 | 1.7437E-06 |
| Trp53i13 | 0.00013273 | 0.68854414 | 0.677 | 0.466 | 1 | 8.7797E-07 | 1.02543122 | 0.58 | 0.26 | 0.02132757 | 0.00013273 | 1.7559E-06 |
| Slc43a3 | 0.0595329 | 0.34542311 | 0.781 | 0.642 | 1 | 8.8265E-07 | 1.0812564 | 0.605 | 0.271 | 0.02144125 | 0.0595329 | 1.7653E-06 |
| Scara3 | 0.06569776 | 0.14720153 | 0.844 | 0.813 | 1 | 8.8288E-07 | 0.66113365 | 0.979 | 0.875 | 0.0214469 | 0.06569776 | 1.7658E-06 |
| Rbis | 0.0229148 | 0.30705482 | 0.677 | 0.503 | 1 | 8.8506E-07 | 1.0671142 | 0.538 | 0.219 | 0.02149997 | 0.0229148 | 1.7701E-06 |
| Rbp1 | 0.13550178 | 0.34270301 | 0.625 | 0.575 | 1 | 8.8744E-07 | 0.68385349 | 0.878 | 0.573 | 0.02155769 | 0.13550178 | 1.7749E-06 |
| Serpine1 | 8.8993E-07 | 1.93116549 | 0.542 | 0.264 | 0.02161829 | 0.23392739 | 1.12310414 | 0.235 | 0.177 | 1 | 0.23392739 | 1.7799E-06 |
| Tubb2b | 0.3325346 | -0.7886766 | 0.521 | 0.508 | 1 | 8.9575E-07 | 1.28793128 | 0.492 | 0.188 | 0.0217595 | 0.3325346 | 1.7915E-06 |
| Srm | 0.000030662 | 0.866632 | 0.719 | 0.487 | 0.74483067 | 8.9598E-07 | 1.34327874 | 0.521 | 0.219 | 0.02176508 | 3.0662E-05 | 0.000001792 |
| Fnta | 0.16113221 | 0.15715939 | 0.76 | 0.648 | 1 | 8.9659E-07 | 0.85103593 | 0.769 | 0.479 | 0.02178001 | 0.16113221 | 1.7932E-06 |
| Gm20186 | 0.01977398 | -0.7123975 | 0.312 | 0.435 | 1 | 8.9705E-07 | -1.0050007 | 0.328 | 0.625 | 0.0217912 | 0.01977398 | 1.7941E-06 |
| Lgals3 | 0.6922421 | 0.11391851 | 0.906 | 0.87 | 1 | 8.9796E-07 | 1.08117284 | 0.929 | 0.896 | 0.02181329 | 0.6922421 | 1.7959E-06 |
| Dhx58os | 0.06251708 | -0.73478 | 0.74 | 0.731 | 1 | 9.0032E-07 | 0.5636964 | 0.714 | 0.323 | 0.02187065 | 0.06251708 | 1.8006E-06 |
| Parp8 | 0.83778906 | -0.3057687 | 0.51 | 0.503 | 1 | 9.0629E-07 | -1.2736351 | 0.534 | 0.771 | 0.02201562 | 0.83778906 | 1.8126E-06 |
| Sra1 | 0.0201576 | 0.29544266 | 0.875 | 0.782 | 1 | 9.0671E-07 | 0.7116779 | 0.84 | 0.646 | 0.0220258 | 0.0201576 | 1.8134E-06 |
| Slc5a3 | 9.099E-07 | -1.2105977 | 0.552 | 0.777 | 0.02210319 | 0.0349373 | -0.6070957 | 0.622 | 0.698 | 1 | 0.0349373 | 1.8198E-06 |
| Jup | 0.01483842 | -0.9027088 | 0.198 | 0.337 | 1 | 9.1664E-07 | -0.961932 | 0.286 | 0.594 | 0.0222669 | 0.01483842 | 1.8333E-06 |
| Pxdc1 | 9.2621E-07 | -1.3904694 | 0.552 | 0.751 | 0.02249957 | 0.00210608 | 0.39596163 | 0.601 | 0.354 | 1 | 0.00210608 | 1.8524E-06 |
| Hspe1 | 0.01379525 | 0.2317342 | 0.948 | 0.922 | 1 | 9.4024E-07 | 0.59666958 | 0.908 | 0.844 | 0.0228404 | 0.01379525 | 1.8805E-06 |
| Yrdc | 0.00267797 | 0.56961706 | 0.729 | 0.56 | 1 | 9.5054E-07 | 0.87071698 | 0.714 | 0.438 | 0.02309063 | 0.00267797 | 1.9011E-06 |
| Siah2 | 9.5406E-07 | 0.93087325 | 0.74 | 0.472 | 0.02317599 | 1.0589E-06 | 0.91921322 | 0.655 | 0.375 | 0.02572229 | 1.0589E-06 | 1.9081E-06 |
| Derl1 | 0.08291373 | 0.26354251 | 0.844 | 0.777 | 1 | 9.6302E-07 | 0.66908312 | 0.878 | 0.719 | 0.02339371 | 0.08291373 | 0.000001926 |
| Bdp1 | 9.8854E-07 | 0.6546478 | 0.979 | 0.845 | 0.02401359 | 0.05110197 | 0.14184071 | 0.937 | 0.906 | 1 | 0.05110197 | 1.9771E-06 |
| Tmem115 | 0.01344401 | 0.66354014 | 0.406 | 0.259 | 1 | 9.9515E-07 | 0.85160816 | 0.521 | 0.198 | 0.02417412 | 0.01344401 | 1.9903E-06 |
| Aldh18a1 | 1.0129E-06 | 0.9517851 | 0.708 | 0.435 | 0.02460544 | 0.00017826 | 0.38162756 | 0.655 | 0.344 | 1 | 0.00017826 | 2.0258E-06 |
| Slc6a12 | 1.0186E-06 | -2.9743581 | 0.146 | 0.399 | 0.02474266 | 0.26927256 | -2.1242996 | 0.071 | 0.104 | 1 | 0.26927256 | 2.0371E-06 |
| Calm2 | 0.34198066 | 0.12831013 | 1 | 0.995 | 1 | 1.0289E-06 | 0.47169842 | 1 | 0.99 | 0.02499315 | 0.34198066 | 2.0577E-06 |
| Hnrnpu | 1.0358E-06 | 0.51211948 | 0.99 | 0.984 | 0.02516246 | 8.8244E-06 | 0.29859399 | 1 | 0.99 | 0.21436235 | 8.8244E-06 | 2.0717E-06 |
| Cd164 | 1.0444E-06 | -0.7235095 | 0.938 | 0.959 | 0.02537097 | 0.000037183 | 0.50224069 | 0.929 | 0.854 | 0.90324663 | 3.7183E-05 | 2.0888E-06 |
| Ooep | 3.7828E-06 | 1.61670458 | 0.438 | 0.197 | 0.09189126 | 1.0452E-06 | 1.34725702 | 0.5 | 0.198 | 0.02539075 | 3.7828E-06 | 2.0905E-06 |
| Plce1 | 0.000001059 | -1.6485511 | 0.49 | 0.684 | 0.02572579 | 0.04537777 | -0.8644441 | 0.479 | 0.542 | 1 | 0.04537777 | 0.000002118 |
| Ankrd44 | 0.48627875 | -0.3375172 | 0.479 | 0.415 | 1 | 1.0673E-06 | -1.2928822 | 0.328 | 0.625 | 0.02592614 | 0.48627875 | 2.1345E-06 |
| Zfp503 | 1.0795E-06 | 1.78121868 | 0.438 | 0.176 | 0.02622225 | 0.99240137 | -0.1276519 | 0.357 | 0.375 | 1 | 0.99240137 | 2.1589E-06 |
| Chpf | 0.000035547 | 0.86026858 | 0.677 | 0.492 | 0.86350276 | 1.0826E-06 | 0.84017018 | 0.71 | 0.365 | 0.02629926 | 3.5547E-05 | 2.1653E-06 |
| Bmp2 | 0.90996074 | 0.22829335 | 0.479 | 0.482 | 1 | 1.0905E-06 | 1.2853965 | 0.714 | 0.385 | 0.02648978 | 0.90996074 | 2.1809E-06 |
| Hnrnpab | 0.000036347 | 0.46368936 | 0.99 | 0.917 | 0.8829295 | 1.1002E-06 | 0.47893605 | 0.996 | 1 | 0.02672691 | 3.6347E-05 | 2.2005E-06 |
| Bola3 | 0.00321296 | 0.53404035 | 0.698 | 0.513 | 1 | 1.1113E-06 | 1.11464257 | 0.563 | 0.26 | 0.0269946 | 0.00321296 | 2.2225E-06 |
| Guk1 | 0.34675307 | 0.15791138 | 0.625 | 0.528 | 1 | 1.1614E-06 | 0.87026551 | 0.555 | 0.25 | 0.02821381 | 0.34675307 | 2.3229E-06 |
| Aff3 | 1.1689E-06 | 0.86692162 | 0.938 | 0.777 | 0.02839529 | 0.47628016 | -0.2856271 | 0.828 | 0.844 | 1 | 0.47628016 | 2.3378E-06 |
| Pak3 | 1.1697E-06 | 1.88196365 | 0.406 | 0.155 | 0.02841421 | 0.05275817 | 1.73196491 | 0.189 | 0.104 | 1 | 0.05275817 | 2.3394E-06 |
| Snhg4 | 0.0008202 | 0.83756954 | 0.615 | 0.409 | 1 | 1.1813E-06 | 1.2488757 | 0.504 | 0.198 | 0.0286963 | 0.0008202 | 2.3626E-06 |
| P3h3 | 5.8437E-06 | 0.68556644 | 0.823 | 0.622 | 0.14195405 | 1.1813E-06 | 0.73872492 | 0.84 | 0.479 | 0.02869682 | 5.8437E-06 | 2.3627E-06 |
| Ralgapa2 | 0.00576765 | -1.0174518 | 0.24 | 0.399 | 1 | 1.1863E-06 | -1.2667865 | 0.416 | 0.708 | 0.02881766 | 0.00576765 | 2.3726E-06 |

| | | | | | | | | | | | | |
|---|---|---|---|---|---|---|---|---|---|---|---|---|
| Cfl1 | 0.46019677 | -0.1354329 | 0.99 | 0.953 | 1 | 1.2021E-06 | 0.44588014 | 1 | 0.958 | 0.02920256 | 0.46019677 | 2.4043E-06 |
| Ripor1 | 1.2068E-06 | 1.30915024 | 0.427 | 0.161 | 0.02931503 | 0.08146213 | 0.22317945 | 0.529 | 0.396 | 1 | 0.08146213 | 2.4136E-06 |
| Nxt1 | 0.0182785 | 0.95502712 | 0.198 | 0.098 | 1 | 1.2105E-06 | 2.11640949 | 0.29 | 0.042 | 0.02940616 | 0.0182785 | 2.4211E-06 |
| Slc39a7 | 0.00010338 | 0.54610672 | 0.906 | 0.741 | 1 | 1.2122E-06 | 0.61234259 | 0.924 | 0.75 | 0.02944647 | 0.00010338 | 2.4244E-06 |
| Srxn1 | 0.8584966 | -0.2115152 | 0.448 | 0.415 | 1 | 0.000001214 | 1.22619764 | 0.466 | 0.167 | 0.02949115 | 0.8584966 | 2.4281E-06 |
| Cmpk1 | 0.15624285 | 0.24767627 | 0.865 | 0.824 | 1 | 1.2154E-06 | 0.70951006 | 0.874 | 0.792 | 0.02952369 | 0.15624285 | 2.4307E-06 |
| Hmgn2 | 0.000001218 | 1.07558353 | 0.833 | 0.637 | 0.02958882 | 0.32827629 | 0.23116389 | 0.584 | 0.604 | 1 | 0.32827629 | 2.4361E-06 |
| Nsun2 | 1.2191E-06 | 0.99222778 | 0.583 | 0.269 | 0.02961458 | 0.000019434 | 0.84729113 | 0.613 | 0.323 | 0.47210039 | 1.9434E-05 | 2.4382E-06 |
| Nrep | 1.2192E-06 | -1.4598907 | 0.75 | 0.824 | 0.0296167 | 0.00091564 | 0.21172452 | 0.895 | 0.583 | 1 | 0.00091564 | 2.4384E-06 |
| Mrps31 | 0.19031783 | 0.37225883 | 0.344 | 0.264 | 1 | 1.2199E-06 | 1.64434105 | 0.378 | 0.104 | 0.02963437 | 0.19031783 | 2.4398E-06 |
| Mpzl1 | 1.2261E-06 | 0.87625 | 0.802 | 0.528 | 0.02978543 | 0.01010977 | 0.26530961 | 0.811 | 0.625 | 1 | 0.01010977 | 2.4523E-06 |
| Smim10l1 | 0.16044948 | 0.19120649 | 0.896 | 0.798 | 1 | 1.2336E-06 | 0.58189185 | 0.882 | 0.688 | 0.02996611 | 0.16044948 | 2.4672E-06 |
| Smn1 | 0.00026338 | 0.8612738 | 0.51 | 0.29 | 1 | 1.2369E-06 | 1.14559651 | 0.492 | 0.177 | 0.03004755 | 0.00026338 | 2.4739E-06 |
| Cdk4 | 0.000001248 | 0.6213119 | 0.844 | 0.528 | 0.03031623 | 0.00420609 | 0.34786358 | 0.739 | 0.604 | 1 | 0.00420609 | 0.000002496 |
| Sgpp1 | 0.21829189 | -1.009579 | 0.312 | 0.363 | 1 | 1.2488E-06 | 1.78259099 | 0.416 | 0.146 | 0.03033539 | 0.21829189 | 2.4976E-06 |
| Bnip3l | 1.2547E-06 | -0.7864537 | 0.677 | 0.845 | 0.03047939 | 0.06100624 | 0.16999053 | 0.819 | 0.792 | 1 | 0.06100624 | 2.5094E-06 |
| Snrpd3 | 0.000041304 | 0.62257703 | 0.875 | 0.725 | 1 | 1.2647E-06 | 0.71439228 | 0.874 | 0.74 | 0.03072212 | 4.1304E-05 | 2.5294E-06 |
| Larp7 | 7.8017E-06 | 0.92700462 | 0.646 | 0.378 | 0.18951805 | 1.2692E-06 | 0.84718845 | 0.538 | 0.219 | 0.03083181 | 7.8017E-06 | 2.5384E-06 |
| Vegfa | 0.000077903 | -1.084428 | 0.375 | 0.591 | 1 | 1.2731E-06 | -1.3807387 | 0.336 | 0.615 | 0.03092587 | 7.7903E-05 | 2.5462E-06 |
| Lingo1 | 1.2754E-06 | 3.38600616 | 0.177 | 0.021 | 0.03098154 | 0.35432319 | 0.10518224 | 0.042 | 0.021 | 1 | 0.35432319 | 2.5508E-06 |
| Kcna6 | 0.02944036 | 0.56003556 | 0.646 | 0.497 | 1 | 1.2834E-06 | 0.22483403 | 0.735 | 0.354 | 0.03117734 | 0.02944036 | 2.5669E-06 |
| Wdr75 | 0.01698898 | 0.47046651 | 0.562 | 0.404 | 1 | 1.3069E-06 | 0.62786046 | 0.613 | 0.26 | 0.03174636 | 0.01698898 | 2.6137E-06 |
| 1810055G02Rik | 0.38187642 | 0.1215663 | 0.75 | 0.658 | 1 | 1.3121E-06 | 0.76469169 | 0.714 | 0.365 | 0.03187432 | 0.38187642 | 2.6243E-06 |
| Slc22a23 | 1.3222E-06 | -1.4583993 | 0.292 | 0.591 | 0.03211916 | 0.13435981 | -0.5870939 | 0.458 | 0.594 | 1 | 0.13435981 | 2.6444E-06 |
| Pappa2 | 1.3241E-06 | 2.37427687 | 0.229 | 0.041 | 0.03216607 | 0.0763749 | 1.35310975 | 0.202 | 0.125 | 1 | 0.0763749 | 2.6483E-06 |
| Brwd1 | 0.14813802 | 0.1546691 | 0.802 | 0.741 | 1 | 1.3269E-06 | -0.9337068 | 0.891 | 0.948 | 0.03223189 | 0.14813802 | 2.6537E-06 |
| Thumpd1 | 0.0003919 | 0.88992494 | 0.583 | 0.415 | 1 | 1.3338E-06 | 1.05829585 | 0.55 | 0.229 | 0.03240122 | 0.0003919 | 2.6676E-06 |
| Pafah1b2 | 1.3412E-06 | 0.74322123 | 0.823 | 0.606 | 0.03258152 | 0.00359368 | 0.2676127 | 0.874 | 0.729 | 1 | 0.00359368 | 2.6825E-06 |
| Maf1 | 0.12399162 | 0.42253204 | 0.344 | 0.249 | 1 | 1.3524E-06 | 1.51457317 | 0.462 | 0.177 | 0.03285141 | 0.12399162 | 2.7047E-06 |
| Gna11 | 1.3541E-06 | 0.6485231 | 0.906 | 0.705 | 0.03289263 | 0.000040619 | 0.51348939 | 0.845 | 0.615 | 0.98670547 | 4.0619E-05 | 2.7081E-06 |
| Snrpg | 1.4829E-06 | 0.59541641 | 0.927 | 0.876 | 0.03602351 | 1.3544E-06 | 0.47417859 | 0.933 | 0.812 | 0.03290048 | 1.4829E-06 | 2.7087E-06 |
| Elof1 | 0.37085257 | 0.30004803 | 0.5 | 0.466 | 1 | 1.3564E-06 | 1.01207283 | 0.643 | 0.375 | 0.03295028 | 0.37085257 | 2.7128E-06 |
| Lekr1 | 1.3586E-06 | 1.64492446 | 0.25 | 0.052 | 0.03300265 | 0.09121009 | 0.92718393 | 0.084 | 0.031 | 1 | 0.09121009 | 2.7172E-06 |
| Amer2 | 1.3657E-06 | 2.74446013 | 0.167 | 0.016 | 0.03317577 | 0.8076363 | 0.27510724 | 0.025 | 0.021 | 1 | 0.8076363 | 2.7314E-06 |
| Fam114a1 | 1.3721E-06 | 0.56929105 | 0.948 | 0.865 | 0.03333139 | 0.00293552 | 0.36391651 | 0.903 | 0.719 | 1 | 0.00293552 | 2.7442E-06 |
| 2810001G20Rik | 0.00058321 | 0.52356196 | 0.875 | 0.663 | 1 | 1.3778E-06 | 0.85203242 | 0.634 | 0.302 | 0.03346949 | 0.00058321 | 2.7556E-06 |
| Nop58 | 0.000013948 | 0.65023337 | 0.938 | 0.943 | 0.33883401 | 1.3819E-06 | 0.69654819 | 0.933 | 0.865 | 0.03356871 | 1.3948E-05 | 2.7638E-06 |
| Hspb8 | 0.68260645 | 0.31088668 | 0.177 | 0.161 | 1 | 1.3899E-06 | -1.520073 | 0.155 | 0.406 | 0.03376277 | 0.68260645 | 2.7797E-06 |
| Ednra | 0.00390295 | -3.9925055 | 0.01 | 0.104 | 1 | 1.3986E-06 | -6.6317834 | 0.008 | 0.125 | 0.03397363 | 0.00390295 | 2.7971E-06 |
| Magi1 | 0.000063305 | -1.8701315 | 0.323 | 0.554 | 1 | 1.4017E-06 | -1.0992883 | 0.458 | 0.76 | 0.03405088 | 6.3305E-05 | 2.8035E-06 |
| Gfod1 | 1.4036E-06 | -1.8322391 | 0.385 | 0.622 | 0.03409631 | 0.02275082 | -0.7248928 | 0.218 | 0.344 | 1 | 0.02275082 | 2.8072E-06 |
| Loxl4 | 0.000020986 | -1.7095926 | 0.74 | 0.725 | 0.50978701 | 1.4056E-06 | 0.48913134 | 0.849 | 0.469 | 0.03414411 | 2.0986E-05 | 2.8111E-06 |
| Angptl2 | 1.4077E-06 | 1.15433592 | 0.552 | 0.264 | 0.03419613 | 0.0703808 | 0.30485458 | 0.416 | 0.302 | 1 | 0.0703808 | 2.8154E-06 |
| Psmb4 | 0.2675693 | 0.14024975 | 0.917 | 0.865 | 1 | 1.4099E-06 | 0.52625865 | 0.87 | 0.74 | 0.03424837 | 0.2675693 | 2.8197E-06 |
| Slc35a2 | 1.4169E-06 | 0.87354008 | 0.802 | 0.56 | 0.03441816 | 0.000026952 | 0.67340147 | 0.609 | 0.312 | 0.65470987 | 2.6952E-05 | 2.8337E-06 |
| Eml1 | 1.4212E-06 | 1.09603121 | 0.406 | 0.14 | 0.0345237 | 0.33731362 | -0.6477249 | 0.466 | 0.344 | 1 | 0.33731362 | 2.8424E-06 |
| Gm2a | 0.77445802 | -0.2604387 | 0.542 | 0.544 | 1 | 1.4374E-06 | 0.7685158 | 0.773 | 0.417 | 0.03491792 | 0.77445802 | 2.8748E-06 |
| Bex3 | 0.0005036 | 0.61305394 | 0.833 | 0.611 | 1 | 1.4395E-06 | 0.59382578 | 0.798 | 0.469 | 0.03496776 | 0.0005036 | 0.000002879 |
| Ring1 | 0.06795834 | 0.55818714 | 0.396 | 0.301 | 1 | 1.4452E-06 | 1.22810976 | 0.483 | 0.188 | 0.03510635 | 0.06795834 | 2.8904E-06 |
| Ahr | 0.62735349 | -0.3404288 | 0.115 | 0.135 | 1 | 1.4478E-06 | -1.2733294 | 0.101 | 0.323 | 0.03516961 | 0.62735349 | 2.8956E-06 |
| Sqstm1 | 0.00403256 | 0.23412146 | 0.958 | 0.964 | 1 | 1.4579E-06 | 0.59354071 | 0.996 | 0.948 | 0.03541499 | 0.00403256 | 2.9158E-06 |
| Eif4g2 | 0.0001416 | 0.33719998 | 1 | 0.969 | 1 | 1.4656E-06 | 0.39594275 | 1 | 0.99 | 0.03560146 | 0.0001416 | 2.9311E-06 |
| Uvrag | 0.20113301 | -0.2065465 | 0.792 | 0.829 | 1 | 1.4873E-06 | -1.1397234 | 0.857 | 0.948 | 0.03612869 | 0.20113301 | 2.9745E-06 |
| Ablim1 | 0.42141455 | -0.7810014 | 0.198 | 0.238 | 1 | 1.4877E-06 | -1.0785297 | 0.231 | 0.521 | 0.03613914 | 0.42141455 | 2.9754E-06 |
| Ctsb | 8.3724E-06 | -0.4766945 | 0.979 | 0.979 | 0.20338186 | 1.5026E-06 | 0.57848494 | 0.987 | 0.958 | 0.03650173 | 8.3724E-06 | 3.0052E-06 |
| Fgd6 | 0.99866777 | -0.3050957 | 0.427 | 0.42 | 1 | 1.5322E-06 | -0.9328858 | 0.546 | 0.812 | 0.03722033 | 0.99866777 | 3.0644E-06 |
| Txndc5 | 1.7246E-06 | 0.59978038 | 0.969 | 0.85 | 0.04189445 | 1.5478E-06 | 0.64986588 | 0.933 | 0.656 | 0.03760031 | 1.7246E-06 | 3.0957E-06 |
| Rel | 0.21113361 | 0.34006988 | 0.406 | 0.337 | 1 | 1.5574E-06 | -1.3438026 | 0.315 | 0.615 | 0.03783152 | 0.21113361 | 3.1147E-06 |
| Pecam1 | 0.05708131 | -1.8643491 | 0.062 | 0.135 | 1 | 0.000001604 | -2.5769593 | 0.059 | 0.24 | 0.03896356 | 0.05708131 | 3.2079E-06 |
| Copa | 1.6057E-06 | 0.44223745 | 0.99 | 0.979 | 0.03900464 | 0.00009343 | 0.28536514 | 0.992 | 0.99 | 1 | 0.00009343 | 3.2113E-06 |
| Slc50a1 | 0.04741782 | 0.41219009 | 0.698 | 0.56 | 1 | 1.6083E-06 | 0.83698613 | 0.609 | 0.292 | 0.03907003 | 0.04741782 | 3.2167E-06 |
| Lcn2 | 0.20411197 | -3.4843586 | 0.01 | 0.036 | 1 | 1.6097E-06 | -0.1678363 | 0.084 | 0.292 | 0.03910212 | 0.20411197 | 3.2193E-06 |
| Iars | 1.6108E-06 | 0.92916618 | 0.76 | 0.554 | 0.03912881 | 3.2991E-06 | 0.62254666 | 0.782 | 0.448 | 0.08014268 | 3.2991E-06 | 3.2215E-06 |
| Fam3a | 0.05267672 | 0.69338595 | 0.312 | 0.212 | 1 | 1.6275E-06 | 1.29680231 | 0.424 | 0.135 | 0.03953511 | 0.05267672 | 0.000003255 |
| Phpt1 | 0.000028306 | 0.7651337 | 0.635 | 0.368 | 0.68760454 | 1.6654E-06 | 1.14957636 | 0.525 | 0.24 | 0.04045705 | 2.8306E-05 | 3.3309E-06 |
| Gtf2i | 1.6766E-06 | 0.79429129 | 0.833 | 0.611 | 0.04072726 | 0.02530968 | 0.1629395 | 0.878 | 0.865 | 1 | 0.02530968 | 3.3531E-06 |
| Aldh6a1 | 0.59313034 | -0.2389434 | 0.354 | 0.378 | 1 | 1.6989E-06 | 1.16587817 | 0.479 | 0.188 | 0.0412685 | 0.59313034 | 3.3977E-06 |
| Nid2 | 1.7084E-06 | 0.90293861 | 0.823 | 0.611 | 0.04150116 | 0.00658841 | 0.48342447 | 0.571 | 0.375 | 1 | 0.00658841 | 3.4169E-06 |
| Cmip | 0.00032767 | -0.6686335 | 0.896 | 0.953 | 1 | 1.7115E-06 | -0.8370472 | 0.815 | 0.938 | 0.04157481 | 0.00032767 | 3.4229E-06 |
| Arsj | 1.7214E-06 | 4.91438513 | 0.115 | 0 | 0.04181615 | 0.83552748 | 0.83698613 | 0.046 | 0.042 | 1 | 0.83552748 | 3.4428E-06 |
| Tmem216 | 0.52849312 | 0.15433592 | 0.25 | 0.218 | 1 | 1.7237E-06 | 1.56929034 | 0.349 | 0.083 | 0.04187245 | 0.52849312 | 3.4474E-06 |
| Pam | 0.00014505 | 0.36998817 | 1 | 0.995 | 1 | 1.7238E-06 | 0.50432162 | 1 | 1 | 0.04187479 | 0.00014505 | 3.4476E-06 |
| Gnb2 | 0.02149208 | 0.19819659 | 0.958 | 0.927 | 1 | 1.7262E-06 | 0.50185168 | 0.971 | 0.938 | 0.04193332 | 0.02149208 | 3.4524E-06 |
| Osgin2 | 0.35800108 | -0.432205 | 0.396 | 0.477 | 1 | 0.000001727 | 0.81320313 | 0.525 | 0.208 | 0.04195324 | 0.35800108 | 3.4541E-06 |
| Fgfr1 | 1.7899E-06 | -0.7132608 | 0.958 | 1 | 0.04347956 | 0.00023144 | 0.39107698 | 0.979 | 0.99 | 1 | 0.00023144 | 3.5797E-06 |
| Bloc1s4 | 0.007005 | 0.60539609 | 0.469 | 0.301 | 1 | 1.7928E-06 | 1.00208874 | 0.466 | 0.167 | 0.04355135 | 0.007005 | 3.5857E-06 |
| Arid5b | 1.7929E-06 | -0.9748516 | 0.729 | 0.917 | 0.043554 | 0.0010786 | -0.8726613 | 0.836 | 0.927 | 1 | 0.0010786 | 3.5859E-06 |
| Rfxap | 0.0863415 | 0.58526154 | 0.333 | 0.238 | 1 | 1.8009E-06 | 0.90515763 | 0.496 | 0.188 | 0.04374651 | 0.0863415 | 3.6017E-06 |
| Sfpq | 1.8222E-06 | 0.49603642 | 1 | 0.959 | 0.04426406 | 0.00062196 | 0.17111651 | 0.996 | 1 | 1 | 0.00062196 | 3.6443E-06 |
| Ndufb2 | 0.10889647 | 0.15949763 | 0.844 | 0.777 | 1 | 0.000001828 | 0.87734773 | 0.777 | 0.604 | 0.0444065 | 0.10889647 | 3.6561E-06 |
| Clta | 0.04664513 | 0.23324477 | 0.938 | 0.948 | 1 | 1.8282E-06 | 0.36009892 | 0.987 | 0.927 | 0.04441149 | 0.04664513 | 3.6565E-06 |
| Col27a1 | 0.62023614 | -0.1674151 | 0.99 | 0.938 | 1 | 1.8305E-06 | 0.30307341 | 0.996 | 0.896 | 0.04446539 | 0.62023614 | 3.6609E-06 |
| Naa50 | 1.8476E-06 | 0.51121719 | 0.906 | 0.72 | 0.04488177 | 0.01649857 | 0.1658289 | 0.916 | 0.958 | 1 | 0.01649857 | 3.6952E-06 |
| Slc39a13 | 0.00376152 | 0.73155265 | 0.74 | 0.642 | 1 | 1.8683E-06 | 0.69977344 | 0.731 | 0.396 | 0.04538399 | 0.00376152 | 3.7365E-06 |
| Gid8 | 0.01059594 | 0.62898291 | 0.51 | 0.373 | 1 | 0.000001904 | 1.31597952 | 0.5 | 0.219 | 0.04625131 | 0.01059594 | 3.8079E-06 |
| Ybx1 | 0.000069538 | 0.44101867 | 1 | 0.959 | 1 | 1.9052E-06 | 0.47828503 | 0.996 | 0.99 | 0.04628001 | 6.9538E-05 | 3.8103E-06 |
| Uap1 | 1.9214E-06 | -1.3950347 | 0.896 | 0.927 | 0.04667544 | 0.01590854 | 0.31627298 | 0.916 | 0.865 | 1 | 0.01590854 | 3.8429E-06 |
| Diaph1 | 0.00190725 | -0.6083289 | 0.667 | 0.845 | 1 | 1.9669E-06 | -0.9678196 | 0.769 | 0.917 | 0.04777903 | 0.00190725 | 3.9337E-06 |
| Arhgap12 | 0.24841819 | -0.3203101 | 0.25 | 0.311 | 1 | 1.9697E-06 | -1.2314286 | 0.361 | 0.646 | 0.04784813 | 0.24841819 | 3.9394E-06 |

| | | | | | | | | | | | | |
|---|---|---|---|---|---|---|---|---|---|---|---|---|
| Dnajc10 | 0.13193378 | 0.22003917 | 0.833 | 0.756 | 1 | 1.9898E-06 | 0.65509779 | 0.815 | 0.542 | 0.04833606 | 0.13193378 | 3.9796E-06 |
| Foxa1 | 0.47606502 | 1.00749454 | 0.021 | 0.01 | 1 | 1.9938E-06 | -2.1172102 | 0.038 | 0.198 | 0.04843221 | 0.47606502 | 3.9875E-06 |
| Rhbdl2 | 1.9966E-06 | 4.06638823 | 0.146 | 0.01 | 0.04850109 | 0.08663289 | 1.77760758 | 0.05 | 0.01 | 1 | 0.08663289 | 3.9932E-06 |
| Tshz2 | 2.0007E-06 | -1.0882916 | 0.938 | 0.984 | 0.04860032 | 0.04947752 | -0.7130089 | 0.874 | 0.844 | 1 | 0.04947752 | 4.0013E-06 |
| Krr1 | 0.00217863 | 0.82588773 | 0.49 | 0.311 | 1 | 2.0149E-06 | 1.16492432 | 0.517 | 0.208 | 0.04894523 | 0.00217863 | 4.0297E-06 |
| Igfbp5 | 0.60885421 | 0.43271341 | 0.281 | 0.342 | 1 | 2.0159E-06 | -1.2941736 | 0.244 | 0.552 | 0.04897085 | 0.60885421 | 4.0318E-06 |
| Cox7a2l | 0.38576233 | 0.1036444 | 0.969 | 0.912 | 1 | 2.0273E-06 | 0.63043849 | 0.916 | 0.875 | 0.04924781 | 0.38576233 | 4.0546E-06 |
| Ntm | 2.0325E-06 | 1.74446013 | 0.188 | 0.026 | 0.04937244 | 0.97266904 | -0.6577786 | 0.042 | 0.042 | 1 | 0.97266904 | 4.0649E-06 |
| Adam9 | 0.10672434 | -0.3419143 | 0.5 | 0.601 | 1 | 2.0472E-06 | -1.0280843 | 0.513 | 0.771 | 0.04973171 | 0.10672434 | 4.0945E-06 |
| Mfap1a | 0.04472165 | 0.31312082 | 0.427 | 0.295 | 1 | 2.0499E-06 | 1.44221722 | 0.403 | 0.125 | 0.04979558 | 0.04472165 | 4.0997E-06 |
| Hdgf | 0.03134662 | 0.19541439 | 0.969 | 0.933 | 1 | 2.0548E-06 | 0.47710972 | 0.992 | 0.917 | 0.04991588 | 0.03134662 | 4.1097E-06 |
| Arpc2 | 0.0239214 | 0.23740216 | 0.99 | 0.964 | 1 | 2.0984E-06 | 0.40762826 | 1 | 1 | 0.05097415 | 0.0239214 | 4.1968E-06 |
| 2210408F21Rik | 2.1094E-06 | -1.523575 | 0.24 | 0.554 | 0.0512415 | 0.04941795 | -0.8279863 | 0.399 | 0.49 | 1 | 0.04941795 | 4.2188E-06 |
| Myo19 | 2.1267E-06 | 1.27052894 | 0.438 | 0.176 | 0.05166103 | 0.14438021 | 0.24814019 | 0.307 | 0.219 | 1 | 0.14438021 | 4.2533E-06 |
| Srrm2 | 0.01140456 | 0.16989923 | 1 | 1 | 1 | 2.1775E-06 | 0.24870627 | 1 | 1 | 0.05289682 | 0.01140456 | 4.3551E-06 |
| Rrs1 | 0.000035021 | 0.96169085 | 0.562 | 0.321 | 0.85073153 | 2.1904E-06 | 0.86673347 | 0.525 | 0.208 | 0.05320975 | 3.5021E-05 | 4.3808E-06 |
| Btbd3 | 2.1905E-06 | -1.4254649 | 0.542 | 0.741 | 0.0532108 | 0.00054157 | 0.51708003 | 0.597 | 0.344 | 1 | 0.00054157 | 4.3809E-06 |
| Edil3 | 2.1974E-06 | 0.73910166 | 0.865 | 0.658 | 0.05337994 | 0.08806998 | -0.2574559 | 0.878 | 0.688 | 1 | 0.08806998 | 4.3949E-06 |
| Acvr1 | 2.2065E-06 | -0.9454853 | 0.854 | 0.907 | 0.05359918 | 0.68461785 | -0.1661709 | 0.887 | 0.896 | 1 | 0.68461785 | 4.4129E-06 |
| Adgrl2 | 0.4223661 | -0.7580402 | 0.135 | 0.171 | 1 | 2.2157E-06 | -2.0377757 | 0.181 | 0.417 | 0.05382405 | 0.4223661 | 4.4314E-06 |
| Mcrip1 | 0.000018622 | 0.62209653 | 0.896 | 0.637 | 0.45235383 | 2.2259E-06 | 0.75280877 | 0.777 | 0.552 | 0.05407168 | 1.8622E-05 | 4.4518E-06 |
| Ifitm2 | 0.000025566 | 0.6423439 | 0.958 | 0.917 | 0.62103897 | 2.2529E-06 | 0.48776812 | 0.971 | 0.927 | 0.05472861 | 2.5566E-05 | 4.5059E-06 |
| Tcaf1 | 0.92430477 | -0.101319 | 0.792 | 0.777 | 1 | 2.2593E-06 | 0.69231584 | 0.866 | 0.531 | 0.05488283 | 0.92430477 | 4.5186E-06 |
| Wdr45b | 2.2655E-06 | 0.90733037 | 0.75 | 0.492 | 0.05503329 | 0.65118777 | -0.1303058 | 0.782 | 0.74 | 1 | 0.65118777 | 0.000004531 |
| Mgst3 | 0.32005309 | 0.59245704 | 0.125 | 0.088 | 1 | 2.3013E-06 | -0.8017084 | 0.151 | 0.406 | 0.05590345 | 0.32005309 | 4.6026E-06 |
| Pnn | 2.3408E-06 | 0.58452286 | 0.969 | 0.86 | 0.05686168 | 0.000054937 | 0.46120353 | 0.95 | 0.917 | 1 | 5.4937E-05 | 4.6815E-06 |
| Fbn2 | 0.02589639 | -2.7294711 | 0.01 | 0.073 | 1 | 2.3422E-06 | -3.5142138 | 0.017 | 0.146 | 0.05689757 | 0.02589639 | 4.6845E-06 |
| Aff1 | 2.3553E-06 | -0.8990303 | 0.885 | 0.953 | 0.05721433 | 0.11918041 | -0.3910991 | 0.903 | 0.958 | 1 | 0.11918041 | 4.7105E-06 |
| Cops9 | 0.36261827 | -0.2468703 | 0.792 | 0.731 | 1 | 2.3702E-06 | 0.81043897 | 0.748 | 0.51 | 0.05757643 | 0.36261827 | 4.7404E-06 |
| Mtmr3 | 0.49111511 | -0.1531689 | 0.729 | 0.777 | 1 | 2.3839E-06 | -0.8177113 | 0.882 | 0.979 | 0.05790952 | 0.49111511 | 4.7678E-06 |
| Gfra1 | 2.3876E-06 | -2.8152356 | 0.25 | 0.477 | 0.05800027 | 0.76889144 | -1.013965 | 0.429 | 0.344 | 1 | 0.76889144 | 4.7753E-06 |
| Pcolce | 2.4052E-06 | 0.73294614 | 0.688 | 0.373 | 0.05842624 | 0.000033317 | 0.59538181 | 0.828 | 0.604 | 0.80933471 | 3.3317E-05 | 4.8103E-06 |
| Dennd10 | 0.49376768 | 0.24850264 | 0.469 | 0.456 | 1 | 2.4094E-06 | 0.89561386 | 0.584 | 0.281 | 0.05852973 | 0.49376768 | 4.8188E-06 |
| Pdzd2 | 2.4104E-06 | -1.4324555 | 0.333 | 0.57 | 0.05855405 | 0.00622127 | -0.6566235 | 0.492 | 0.667 | 1 | 0.00622127 | 4.8208E-06 |
| Zfp830 | 0.74813156 | 0.16277276 | 0.344 | 0.326 | 1 | 2.4488E-06 | 1.27510724 | 0.479 | 0.198 | 0.05948677 | 0.74813156 | 4.8976E-06 |
| Ing3 | 0.15339982 | 0.38189005 | 0.427 | 0.347 | 1 | 2.4492E-06 | 0.88374874 | 0.517 | 0.198 | 0.05949689 | 0.15339982 | 4.8985E-06 |
| Csde1 | 0.000030035 | 0.37687332 | 1 | 0.964 | 0.72960224 | 2.4592E-06 | 0.35927701 | 0.996 | 1 | 0.0597385 | 3.0035E-05 | 4.9184E-06 |
| Ogfr | 0.0111588 | 0.58428311 | 0.573 | 0.415 | 1 | 0.000002468 | 1.10292626 | 0.563 | 0.25 | 0.05995266 | 0.0111588 | 0.000004936 |
| B2m | 0.00265868 | 0.25824688 | 0.958 | 0.912 | 1 | 2.4835E-06 | 0.21760742 | 1 | 0.938 | 0.06032984 | 0.00265868 | 0.000004967 |
| Foxa3 | 0.43594724 | 0.29222401 | 0.406 | 0.358 | 1 | 2.5094E-06 | 1.95001186 | 0.37 | 0.104 | 0.0609587 | 0.43594724 | 5.0188E-06 |
| Gaa | 0.00163879 | 0.4338919 | 0.771 | 0.637 | 1 | 2.5298E-06 | 0.87821709 | 0.718 | 0.417 | 0.06145414 | 0.00163879 | 5.0596E-06 |
| Hmgb1 | 0.0001802 | 0.3736013 | 1 | 0.953 | 1 | 2.5304E-06 | 0.42864757 | 1 | 0.979 | 0.06146907 | 0.0001802 | 5.0608E-06 |
| Sf1 | 0.00454929 | 0.45971696 | 0.854 | 0.834 | 1 | 2.5583E-06 | 0.51154743 | 0.937 | 0.885 | 0.062146 | 0.00454929 | 5.1166E-06 |
| Ncbp2as2 | 0.43537307 | 0.1835249 | 0.521 | 0.466 | 1 | 2.5904E-06 | 1.30064233 | 0.458 | 0.177 | 0.06292493 | 0.43537307 | 5.1807E-06 |
| Ctdsp1 | 0.84036472 | 0.13928441 | 0.427 | 0.446 | 1 | 2.6078E-06 | 0.94933708 | 0.63 | 0.302 | 0.06334952 | 0.84036472 | 5.2157E-06 |
| Hdhd2 | 0.20573063 | 0.22250743 | 0.479 | 0.404 | 1 | 2.6238E-06 | 1.3409093 | 0.45 | 0.167 | 0.06373714 | 0.20573063 | 5.2476E-06 |
| Mllt10 | 0.01790907 | 0.33622246 | 0.844 | 0.741 | 1 | 2.6645E-06 | -0.6922041 | 0.924 | 0.99 | 0.06472571 | 0.01790907 | 0.000005329 |
| Ak6 | 0.00057002 | 0.83756954 | 0.49 | 0.28 | 1 | 2.6866E-06 | 0.99131427 | 0.546 | 0.25 | 0.06526218 | 0.00057002 | 5.3731E-06 |
| Proser2 | 0.16757222 | -0.4761338 | 0.604 | 0.689 | 1 | 0.000002692 | 0.47558488 | 0.803 | 0.448 | 0.06539506 | 0.16757222 | 5.3841E-06 |
| Aldoc | 2.6944E-06 | -2.33565 | 0.188 | 0.44 | 0.0654524 | 0.16799254 | -0.3098553 | 0.189 | 0.115 | 1 | 0.16799254 | 5.3888E-06 |
| Gspt1 | 0.000002703 | 0.58928976 | 0.979 | 0.891 | 0.06566106 | 0.00403889 | 0.33749806 | 0.983 | 0.958 | 1 | 0.00403889 | 0.000005406 |
| Prkcg | 2.7061E-06 | 1.81348392 | 0.5 | 0.249 | 0.06573568 | 0.32007228 | 0.55687821 | 0.34 | 0.25 | 1 | 0.32007228 | 5.4121E-06 |
| Notum | 0.000032825 | 1.34724391 | 0.469 | 0.238 | 0.79738669 | 2.7109E-06 | 1.03266974 | 0.756 | 0.458 | 0.06585279 | 3.2825E-05 | 5.4218E-06 |
| Slc9a9 | 0.79376775 | -0.4872702 | 0.24 | 0.212 | 1 | 2.7539E-06 | -1.194378 | 0.319 | 0.615 | 0.0668988 | 0.79376775 | 5.5079E-06 |
| Gm4258 | 0.13790678 | -1.8778629 | 0.052 | 0.104 | 1 | 2.7816E-06 | -1.3229114 | 0.013 | 0.135 | 0.06757141 | 0.13790678 | 5.5633E-06 |
| Commd3 | 0.02512257 | 0.38996417 | 0.688 | 0.575 | 1 | 2.8049E-06 | 0.86232571 | 0.71 | 0.448 | 0.0681357 | 0.02512257 | 5.6097E-06 |
| Mxd1 | 0.83699238 | -0.7924809 | 0.333 | 0.321 | 1 | 2.8158E-06 | -1.1090363 | 0.374 | 0.646 | 0.06840121 | 0.83699238 | 5.6316E-06 |
| Eps8l2 | 0.52235405 | -0.2629573 | 0.854 | 0.731 | 1 | 2.8245E-06 | 0.58948852 | 0.929 | 0.76 | 0.06861317 | 0.52235405 | 0.000005649 |
| Mrps17 | 0.01037387 | 0.44561565 | 0.656 | 0.513 | 1 | 2.8403E-06 | 0.97459813 | 0.529 | 0.229 | 0.06899676 | 0.01037387 | 5.6806E-06 |
| Stc2 | 2.8652E-06 | -2.1094816 | 0.406 | 0.606 | 0.06960129 | 0.10700327 | -0.1696776 | 0.239 | 0.146 | 1 | 0.10700327 | 5.7304E-06 |
| Cnnm2 | 2.8913E-06 | -1.5130774 | 0.24 | 0.518 | 0.07023582 | 0.00708539 | -1.768146 | 0.29 | 0.396 | 1 | 0.00708539 | 5.7826E-06 |
| Ampd3 | 2.8951E-06 | -3.3588277 | 0.073 | 0.306 | 0.07032712 | 0.03292991 | 0.84542296 | 0.134 | 0.052 | 1 | 0.03292991 | 5.7901E-06 |
| Tgfb3 | 0.00156297 | 0.66780216 | 0.521 | 0.316 | 1 | 2.9137E-06 | 0.6193045 | 0.706 | 0.344 | 0.07077961 | 0.00156297 | 5.8274E-06 |
| Myo1d | 2.9173E-06 | -0.8649554 | 0.969 | 0.984 | 0.07086699 | 0.01876866 | -0.4748824 | 0.992 | 0.979 | 1 | 0.01876866 | 5.8346E-06 |
| Dusp10 | 0.05051311 | -0.6496177 | 0.146 | 0.249 | 1 | 2.9759E-06 | -1.4473588 | 0.113 | 0.333 | 0.07229024 | 0.05051311 | 5.9518E-06 |
| Sh3glb1 | 0.00639545 | 0.22478007 | 0.99 | 0.979 | 1 | 2.9847E-06 | 0.45030256 | 0.992 | 0.99 | 0.07250405 | 0.00639545 | 5.9694E-06 |
| Nop16 | 0.03422116 | 0.70915326 | 0.479 | 0.358 | 1 | 3.0164E-06 | 1.12755005 | 0.424 | 0.135 | 0.07327357 | 0.03422116 | 6.0327E-06 |
| Cd200 | 3.0438E-06 | -0.9229356 | 0.729 | 0.85 | 0.0739393 | 0.1142309 | -0.2159264 | 0.702 | 0.479 | 1 | 0.1142309 | 6.0875E-06 |
| Gpx1 | 0.00313668 | -0.6451179 | 0.917 | 0.959 | 1 | 0.000003068 | 0.34776178 | 0.975 | 0.927 | 0.07452785 | 0.00313668 | 0.000006136 |
| Nabp1 | 0.0453106 | -1.293675 | 0.062 | 0.14 | 1 | 3.0685E-06 | -1.5322477 | 0.101 | 0.312 | 0.0745394 | 0.0453106 | 6.1369E-06 |
| Dusp7 | 0.01611086 | 0.42253204 | 0.479 | 0.326 | 1 | 3.0866E-06 | 0.8206763 | 0.609 | 0.302 | 0.07497848 | 0.01611086 | 6.1731E-06 |
| Tmem208 | 0.00459209 | 0.48595742 | 0.823 | 0.648 | 1 | 0.000003105 | 0.80249246 | 0.718 | 0.438 | 0.07542602 | 0.00459209 | 6.2099E-06 |
| Vamp5 | 0.2575498 | -0.1815393 | 0.427 | 0.497 | 1 | 3.1159E-06 | 1.37762215 | 0.454 | 0.177 | 0.07569058 | 0.2575498 | 6.2317E-06 |
| Enah | 3.1576E-06 | -1.0870063 | 0.896 | 0.948 | 0.07670559 | 0.08207946 | -0.2953953 | 0.857 | 0.906 | 1 | 0.08207946 | 6.3153E-06 |
| Lsm5 | 0.000056125 | 0.76422339 | 0.635 | 0.394 | 1 | 3.1634E-06 | 1.14957636 | 0.5 | 0.208 | 0.07684457 | 5.6125E-05 | 6.3267E-06 |
| Tex261 | 0.00106998 | 0.59868899 | 0.771 | 0.632 | 1 | 3.2305E-06 | 0.78457954 | 0.752 | 0.51 | 0.07847606 | 0.00106998 | 6.4611E-06 |
| Tmed1 | 0.11843901 | 0.19825003 | 0.688 | 0.58 | 1 | 3.2408E-06 | 0.70296878 | 0.634 | 0.292 | 0.07872606 | 0.11843901 | 6.4816E-06 |
| Gnai3 | 0.11824084 | 0.21754266 | 0.958 | 0.912 | 1 | 3.2522E-06 | 0.43570339 | 0.983 | 0.958 | 0.07900291 | 0.11824084 | 6.5044E-06 |
| Ccnd2 | 0.04987876 | 0.55004147 | 0.75 | 0.803 | 1 | 3.3782E-06 | 0.46095599 | 0.92 | 0.854 | 0.08206421 | 0.04987876 | 6.7565E-06 |
| Tpbg | 3.4005E-06 | 1.79599043 | 0.49 | 0.233 | 0.08260508 | 0.00010587 | 1.38646835 | 0.387 | 0.167 | 1 | 0.00010587 | 0.000006801 |
| Ttll3 | 3.4196E-06 | 2.54938832 | 0.365 | 0.14 | 0.08306958 | 0.15781438 | -0.1208214 | 0.34 | 0.219 | 1 | 0.15781438 | 6.8392E-06 |
| Josd1 | 0.69698239 | -0.1851505 | 0.562 | 0.549 | 1 | 3.4516E-06 | 0.71145524 | 0.702 | 0.385 | 0.08384593 | 0.69698239 | 6.9032E-06 |
| Fuca1 | 0.05674609 | -0.4068798 | 0.896 | 0.87 | 1 | 3.4751E-06 | 0.59861119 | 0.87 | 0.74 | 0.0844175 | 0.05674609 | 6.9502E-06 |
| Cemip2 | 0.00114869 | -1.329843 | 0.635 | 0.736 | 1 | 3.5157E-06 | -1.7237288 | 0.555 | 0.802 | 0.08540397 | 0.00114869 | 7.0314E-06 |
| Rpl7l1 | 0.00388748 | 0.46999339 | 0.802 | 0.731 | 1 | 3.6179E-06 | 0.65422516 | 0.824 | 0.573 | 0.08788579 | 0.00388748 | 7.2358E-06 |
| Ldha | 0.30750456 | 0.12082648 | 0.969 | 0.969 | 1 | 3.6467E-06 | 0.50122515 | 0.983 | 0.938 | 0.08858575 | 0.30750456 | 7.2934E-06 |
| Slc2a10 | 0.000011816 | 1.37006462 | 0.438 | 0.207 | 0.28703698 | 3.6521E-06 | 1.64188957 | 0.399 | 0.135 | 0.08871771 | 1.1816E-05 | 7.3043E-06 |
| Acsl1 | 0.0718725 | -0.8270819 | 0.427 | 0.503 | 1 | 3.6552E-06 | -0.8980417 | 0.37 | 0.667 | 0.08879273 | 0.0718725 | 7.3104E-06 |
| Gm43154 | 3.6562E-06 | 4.07358373 | 0.188 | 0.031 | 0.08881528 | 0.39655804 | 0.95317914 | 0.025 | 0.01 | 1 | 0.39655804 | 7.3123E-06 |
| Ttll10 | 3.6596E-06 | 4.39981196 | 0.125 | 0.005 | 0.08889984 | 0.02194919 | 0.51611534 | 0.13 | 0.042 | 1 | 0.02194919 | 7.3193E-06 |
| Mtfr1 | 0.14092772 | 0.34123193 | 0.604 | 0.513 | 1 | 3.6834E-06 | 0.80741079 | 0.643 | 0.333 | 0.08947729 | 0.14092772 | 7.3668E-06 |

| | | | | | | | | | | | | |
|---|---|---|---|---|---|---|---|---|---|---|---|---|
| Mrto4 | 0.00257355 | 1.12115132 | 0.385 | 0.238 | 1 | 3.7247E-06 | 1.6026819 | 0.378 | 0.115 | 0.0904806 | 0.00257355 | 7.4494E-06 |
| Mrpl54 | 0.44620748 | 0.1624692 | 0.615 | 0.554 | 1 | 3.7453E-06 | 0.99826703 | 0.534 | 0.24 | 0.09098172 | 0.44620748 | 7.4907E-06 |
| Yif1a | 0.00812163 | 0.52919214 | 0.74 | 0.585 | 1 | 3.7609E-06 | 0.55264121 | 0.63 | 0.312 | 0.09135986 | 0.00812163 | 7.5218E-06 |
| Selenbp1 | 0.26280002 | 0.65957123 | 0.073 | 0.041 | 1 | 3.8175E-06 | -1.4962684 | 0.097 | 0.302 | 0.09273351 | 0.26280002 | 7.6349E-06 |
| Uhmk1 | 0.1473743 | 0.22988696 | 0.375 | 0.275 | 1 | 0.000003827 | 0.85520398 | 0.475 | 0.167 | 0.09296478 | 0.1473743 | 7.6539E-06 |
| Fam107b | 0.03675858 | -0.59517 | 0.146 | 0.254 | 1 | 3.9402E-06 | -0.9454438 | 0.294 | 0.573 | 0.0957158 | 0.03675858 | 7.8804E-06 |
| Mob3b | 0.65298542 | -0.2483444 | 0.208 | 0.181 | 1 | 3.9409E-06 | -2.2746658 | 0.185 | 0.417 | 0.0957321 | 0.65298542 | 7.8818E-06 |
| Lage3 | 0.00317594 | 0.61754802 | 0.594 | 0.409 | 1 | 3.9656E-06 | 1.09102417 | 0.487 | 0.208 | 0.09633245 | 0.00317594 | 7.9312E-06 |
| Abce1 | 0.00704759 | 0.38892364 | 0.802 | 0.71 | 1 | 3.9925E-06 | 0.62612824 | 0.845 | 0.677 | 0.09698492 | 0.00704759 | 7.9849E-06 |
| Hivep2 | 0.000035486 | -0.7329281 | 0.938 | 0.995 | 0.86203039 | 4.0333E-06 | -0.8677848 | 0.979 | 0.979 | 0.09797584 | 3.5486E-05 | 8.0665E-06 |
| Rpl9-ps6 | 0.00063102 | 0.68556644 | 0.594 | 0.394 | 1 | 4.0974E-06 | 1.24566346 | 0.546 | 0.281 | 0.09953289 | 0.00063102 | 8.1947E-06 |
| Tars | 0.000091372 | 0.76709633 | 0.708 | 0.523 | 1 | 4.1396E-06 | 0.78636005 | 0.718 | 0.406 | 0.10055866 | 9.1372E-05 | 8.2791E-06 |
| Cttnbp2 | 0.14415248 | -1.6247737 | 0.062 | 0.114 | 1 | 4.1451E-06 | -1.5670131 | 0.118 | 0.333 | 0.10069178 | 0.14415248 | 8.2901E-06 |
| Lmcd1 | 0.00011527 | -1.0188053 | 0.688 | 0.767 | 1 | 4.1847E-06 | 0.26720204 | 0.786 | 0.375 | 0.10165564 | 0.00011527 | 8.3695E-06 |
| Lsm6 | 4.3185E-06 | 0.88411212 | 0.802 | 0.565 | 0.10490389 | 0.00089778 | 0.43105752 | 0.643 | 0.427 | 1 | 0.00089778 | 8.6369E-06 |
| Ttc9c | 0.000004326 | 1.2049406 | 0.542 | 0.29 | 0.10508746 | 0.000024512 | 0.59872171 | 0.592 | 0.26 | 0.59543358 | 2.4512E-05 | 0.000008652 |
| Ergic3 | 0.01685113 | 0.29036985 | 0.896 | 0.782 | 1 | 4.3309E-06 | 0.58826512 | 0.882 | 0.719 | 0.10520702 | 0.01685113 | 8.6618E-06 |
| Mt1 | 0.000009045 | -1.5259621 | 0.49 | 0.72 | 0.21971993 | 4.3496E-06 | 0.3705268 | 0.357 | 0.688 | 0.10565965 | 9.045E-06 | 8.6991E-06 |
| Bet1l | 0.00060282 | 0.50773436 | 0.823 | 0.622 | 1 | 4.3606E-06 | 0.83216374 | 0.71 | 0.396 | 0.10592788 | 0.00060282 | 8.7212E-06 |
| Prps1 | 0.0234527 | 0.73541499 | 0.396 | 0.269 | 1 | 0.000004391 | 1.03409914 | 0.42 | 0.146 | 0.10666632 | 0.0234527 | 0.000008782 |
| Thrap3 | 4.4088E-06 | 0.43724777 | 1 | 0.907 | 0.10709907 | 0.95782218 | -0.161228 | 0.979 | 0.99 | 1 | 0.95782218 | 8.8176E-06 |
| Eif3c | 0.00070148 | 0.24218139 | 0.896 | 0.876 | 1 | 4.4119E-06 | 0.53893238 | 0.962 | 0.906 | 0.10717472 | 0.00070148 | 8.8239E-06 |
| Rab3ip | 0.03945782 | -1.6649308 | 0.073 | 0.155 | 1 | 4.4529E-06 | -1.5971372 | 0.164 | 0.406 | 0.10817103 | 0.03945782 | 8.9059E-06 |
| Ndufb4 | 0.5911614 | 0.1730806 | 0.552 | 0.528 | 1 | 4.4624E-06 | 0.89854389 | 0.517 | 0.229 | 0.10839954 | 0.5911614 | 8.9247E-06 |
| Tgfbr2 | 4.4815E-06 | 0.58676143 | 0.885 | 0.694 | 0.10886408 | 0.02708091 | 0.14635964 | 0.958 | 0.948 | 1 | 0.02708091 | 8.9629E-06 |
| Mall | 4.4881E-06 | 1.28760246 | 0.49 | 0.228 | 0.10902372 | 0.000085003 | 1.10968363 | 0.403 | 0.167 | 1 | 8.5003E-05 | 8.9761E-06 |
| Prpf38a | 0.0975651 | 0.39314823 | 0.531 | 0.43 | 1 | 4.4969E-06 | 0.79567915 | 0.609 | 0.292 | 0.1092398 | 0.0975651 | 8.9939E-06 |
| Pde1c | 0.000004519 | -1.3912674 | 0.26 | 0.565 | 0.10977589 | 0.24314823 | 1.1698352 | 0.273 | 0.24 | 1 | 0.24314823 | 0.000009038 |
| Kank1 | 4.5437E-06 | -0.8325972 | 0.781 | 0.855 | 0.11037548 | 0.00934816 | 0.10476376 | 0.853 | 0.667 | 1 | 0.00934816 | 9.0874E-06 |
| Atp5o | 0.18304134 | 0.14699395 | 0.875 | 0.808 | 1 | 4.5494E-06 | 0.61361089 | 0.895 | 0.771 | 0.11051406 | 0.18304134 | 9.0988E-06 |
| Naa60 | 0.90496611 | -0.1153622 | 0.344 | 0.347 | 1 | 4.7198E-06 | 1.38130664 | 0.429 | 0.156 | 0.11465417 | 0.90496611 | 9.4396E-06 |
| Itgb5 | 4.7388E-06 | -0.7734488 | 0.865 | 0.902 | 0.11511441 | 0.07538812 | -0.4637021 | 0.937 | 0.896 | 1 | 0.07538812 | 9.4775E-06 |
| Commd2 | 0.09354315 | 0.26465238 | 0.396 | 0.275 | 1 | 4.7674E-06 | 1.15354526 | 0.5 | 0.219 | 0.11580942 | 0.09354315 | 9.5348E-06 |
| Tm9sf3 | 0.00612148 | 0.19252643 | 0.979 | 0.969 | 1 | 0.000004771 | 0.386516 | 0.996 | 0.99 | 0.11589682 | 0.00612148 | 0.000009542 |
| Mypn | 4.7713E-06 | -2.1742559 | 0.125 | 0.383 | 0.11590446 | 0.87851834 | -1.4797803 | 0.013 | 0.01 | 1 | 0.87851834 | 9.5426E-06 |
| Tfdp1 | 4.8008E-06 | 0.82616477 | 0.677 | 0.389 | 0.11662198 | 0.00959818 | 0.39909596 | 0.597 | 0.438 | 1 | 0.00959818 | 9.6017E-06 |
| Rbbp9 | 0.0079054 | 0.57538352 | 0.448 | 0.285 | 1 | 4.8071E-06 | 1.06269891 | 0.454 | 0.167 | 0.11677436 | 0.0079054 | 9.6142E-06 |
| Maob | 4.8224E-06 | 1.75846699 | 0.479 | 0.238 | 0.11714468 | 0.00010629 | 1.28310003 | 0.395 | 0.167 | 1 | 0.00010629 | 9.6447E-06 |
| Rab35 | 0.00105649 | 0.83588316 | 0.552 | 0.368 | 1 | 4.8614E-06 | 0.93917229 | 0.681 | 0.385 | 0.11809328 | 0.00105649 | 9.7228E-06 |
| Cntnap2 | 4.9302E-06 | -1.780408 | 0.125 | 0.394 | 0.1197647 | 0.01906751 | 0.62303054 | 0.328 | 0.188 | 1 | 0.01906751 | 9.8604E-06 |
| Prkcz | 0.00377481 | 0.92107978 | 0.448 | 0.295 | 1 | 4.9437E-06 | 1.28432577 | 0.382 | 0.115 | 0.12009136 | 0.00377481 | 9.8873E-06 |
| Eloc | 0.00743643 | 0.35541784 | 0.906 | 0.834 | 1 | 4.9796E-06 | 0.52604787 | 0.933 | 0.875 | 0.12096477 | 0.00743643 | 9.9592E-06 |
| Eif4b | 5.0131E-06 | 0.74209312 | 0.823 | 0.642 | 0.1217791 | 0.000020861 | 0.50626395 | 0.945 | 0.812 | 0.50676413 | 2.0861E-05 | 0.000010026 |
| Wsb2 | 0.2083571 | 0.21017929 | 0.656 | 0.549 | 1 | 0.000005022 | 0.68276521 | 0.744 | 0.458 | 0.12199378 | 0.2083571 | 0.000010044 |
| Ckb | 5.0515E-06 | -1.7008504 | 0.656 | 0.746 | 0.12271211 | 0.00020792 | 2.230253 | 0.605 | 0.396 | 1 | 0.00020792 | 0.000010103 |
| Psme3 | 0.35295054 | -0.1624305 | 0.25 | 0.306 | 1 | 5.0583E-06 | 1.41451029 | 0.429 | 0.156 | 0.12287554 | 0.35295054 | 0.000010117 |
| Polr3f | 0.02438572 | 0.88463779 | 0.323 | 0.212 | 1 | 5.0957E-06 | 1.27510724 | 0.395 | 0.125 | 0.1237859 | 0.02438572 | 0.000010191 |
| Ruvbl2 | 5.1261E-06 | 1.64904057 | 0.333 | 0.114 | 0.12452248 | 0.05792092 | 0.49749966 | 0.218 | 0.125 | 1 | 0.05792092 | 0.000010252 |
| Sgk1 | 9.3522E-06 | -1.2729776 | 0.594 | 0.788 | 0.22718363 | 5.1467E-06 | -0.9726815 | 0.639 | 0.896 | 0.12502266 | 9.3522E-06 | 0.000010293 |
| Erg | 5.1562E-06 | 0.5808087 | 0.958 | 0.813 | 0.12525445 | 0.43673849 | -0.6701167 | 0.95 | 0.823 | 1 | 0.43673849 | 0.000010312 |
| Pcolce2 | 0.05012498 | 0.45716567 | 0.385 | 0.285 | 1 | 5.1988E-06 | 2.02474666 | 0.357 | 0.104 | 0.12628827 | 0.05012498 | 0.000010397 |
| Atp8b2 | 0.000033916 | 0.76855581 | 0.75 | 0.565 | 0.82389036 | 5.2174E-06 | 0.67528678 | 0.794 | 0.458 | 0.12674022 | 3.3916E-05 | 0.000010435 |
| Arl8a | 0.08203342 | 0.24474492 | 0.698 | 0.544 | 1 | 5.2681E-06 | 0.73312791 | 0.845 | 0.656 | 0.12797184 | 0.08203342 | 0.000010536 |
| Eif3m | 0.01548288 | 0.35698627 | 0.875 | 0.829 | 1 | 5.2849E-06 | 0.5196453 | 0.908 | 0.792 | 0.128381 | 0.01548288 | 0.00001057 |
| Bnip2 | 0.1170249 | 0.12248706 | 0.885 | 0.829 | 1 | 5.2904E-06 | 0.54113479 | 0.933 | 0.792 | 0.12851374 | 0.1170249 | 0.000010581 |
| Cdca4 | 0.00094243 | 0.80894886 | 0.469 | 0.264 | 1 | 5.3278E-06 | 0.82138927 | 0.508 | 0.208 | 0.1294223 | 0.00094243 | 0.000010656 |
| Sec62 | 0.00850872 | 0.29761512 | 1 | 0.984 | 1 | 5.3717E-06 | 0.42032837 | 1 | 0.99 | 0.13048875 | 0.00850872 | 0.000010743 |
| Cenpw | 5.5068E-06 | 2.69555053 | 0.188 | 0.031 | 0.13377167 | 0.09020646 | 1.19264508 | 0.067 | 0.021 | 1 | 0.09020646 | 0.000011014 |
| Zfp704 | 5.5081E-06 | 0.63608397 | 0.792 | 0.508 | 0.13380254 | 0.10813389 | -0.1576261 | 0.79 | 0.729 | 1 | 0.10813389 | 0.000011016 |
| Sec16b | 0.00406204 | 0.46317402 | 0.573 | 0.368 | 1 | 5.5526E-06 | 1.23151797 | 0.395 | 0.125 | 0.13488368 | 0.00406204 | 0.000011105 |
| Cisd2 | 0.00174831 | 0.38523349 | 0.917 | 0.798 | 1 | 5.5723E-06 | 0.47382683 | 0.937 | 0.792 | 0.13536143 | 0.00174831 | 0.000011144 |
| Psmd9 | 0.35623321 | 0.20819189 | 0.479 | 0.415 | 1 | 5.5838E-06 | 1.03303245 | 0.55 | 0.271 | 0.13564202 | 0.35623321 | 0.000011168 |
| Gnb4 | 5.6153E-06 | 0.76400032 | 0.896 | 0.731 | 0.13640734 | 0.000028432 | 0.58391441 | 0.929 | 0.656 | 0.69066038 | 2.8432E-05 | 0.000011231 |
| Spock2 | 0.12661802 | 0.44725912 | 0.604 | 0.554 | 1 | 0.000005865 | 0.9739377 | 0.462 | 0.167 | 0.14247157 | 0.12661802 | 0.00001173 |
| Gm10076 | 0.000051406 | 0.47635474 | 0.938 | 0.881 | 1 | 5.9117E-06 | 0.73683797 | 0.836 | 0.719 | 0.14360697 | 5.1406E-05 | 0.000011823 |
| Sri | 0.00115721 | 0.44637878 | 0.844 | 0.689 | 1 | 5.9301E-06 | 0.65151208 | 0.828 | 0.698 | 0.14405372 | 0.00115721 | 0.00001186 |
| Cav1 | 5.9339E-06 | -2.3461424 | 0.156 | 0.409 | 0.14414548 | 0.00184747 | 0.71509109 | 0.303 | 0.125 | 1 | 0.00184747 | 0.000011868 |
| Pnp | 0.7319865 | -0.2646565 | 0.906 | 0.912 | 1 | 6.0103E-06 | 0.51414879 | 0.945 | 0.854 | 0.14600183 | 0.7319865 | 0.000012021 |
| L3hypdh | 6.0261E-06 | 1.43375929 | 0.354 | 0.124 | 0.14638567 | 0.02234355 | 0.76543286 | 0.185 | 0.083 | 1 | 0.02234355 | 0.000012052 |
| Tmed4 | 0.34123812 | -0.2440442 | 0.479 | 0.508 | 1 | 6.1466E-06 | 0.82764826 | 0.626 | 0.323 | 0.14931224 | 0.34123812 | 0.000012293 |
| Stard4 | 0.07247787 | 0.44299614 | 0.594 | 0.487 | 1 | 6.1888E-06 | 1.18937736 | 0.504 | 0.229 | 0.1503394 | 0.07247787 | 0.000012378 |
| Surf1 | 0.33990482 | 0.39907306 | 0.406 | 0.373 | 1 | 6.2555E-06 | 1.02356847 | 0.433 | 0.146 | 0.15195923 | 0.33990482 | 0.000012511 |
| Zfp706 | 0.10647535 | 0.18299133 | 0.885 | 0.808 | 1 | 6.2572E-06 | 0.47164623 | 0.912 | 0.792 | 0.15199974 | 0.10647535 | 0.000012514 |
| Aldh2 | 0.06269661 | 0.18170891 | 0.885 | 0.886 | 1 | 6.2606E-06 | 0.5003357 | 0.958 | 0.917 | 0.15208237 | 0.06269661 | 0.000012521 |
| Dpf3 | 6.2781E-06 | -2.4864943 | 0.073 | 0.311 | 0.15250711 | 0.34703906 | -1.3098553 | 0.008 | 0.021 | 1 | 0.34703906 | 0.000012556 |
| Map7 | 0.8664813 | -0.264585 | 0.448 | 0.472 | 1 | 6.3281E-06 | -1.0060076 | 0.483 | 0.74 | 0.15372233 | 0.8664813 | 0.000012656 |
| Ttc4 | 0.30596857 | 0.27814213 | 0.292 | 0.233 | 1 | 0.000006366 | 1.31073115 | 0.382 | 0.125 | 0.15464366 | 0.30596857 | 0.000012732 |
| Cct4 | 0.00009239 | 0.55008209 | 0.865 | 0.772 | 1 | 0.000006374 | 0.67629476 | 0.874 | 0.812 | 0.15483681 | 0.00009239 | 0.000012748 |
| Mettl9 | 0.08940446 | 0.15366518 | 0.802 | 0.736 | 1 | 6.3911E-06 | 0.58801195 | 0.853 | 0.677 | 0.15525296 | 0.08940446 | 0.000012782 |
| Lsm2 | 0.00011476 | 1.16182268 | 0.458 | 0.254 | 1 | 6.4135E-06 | 1.13594449 | 0.458 | 0.177 | 0.15579583 | 0.00011476 | 0.000012827 |
| Nfyb | 0.15905113 | 0.33110901 | 0.562 | 0.508 | 1 | 6.4425E-06 | 1.03760792 | 0.58 | 0.302 | 0.15650092 | 0.15905113 | 0.000012885 |
| Erp44 | 0.01075471 | 0.33560709 | 0.948 | 0.824 | 1 | 6.4732E-06 | 0.49246407 | 0.933 | 0.833 | 0.15724783 | 0.01075471 | 0.000012946 |
| Asph | 6.4833E-06 | 0.68983688 | 0.823 | 0.57 | 0.15749143 | 0.81570915 | -0.1915353 | 0.803 | 0.802 | 1 | 0.81570915 | 0.000012966 |
| Cgref1 | 6.5609E-06 | 1.19697234 | 0.531 | 0.269 | 0.15937777 | 0.00244348 | 0.83698613 | 0.34 | 0.167 | 1 | 0.00244348 | 0.000013122 |
| Psmd2 | 0.00004178 | 0.55254832 | 0.833 | 0.679 | 1 | 6.5944E-06 | 0.5796682 | 0.887 | 0.75 | 0.16019024 | 0.00004178 | 0.000013189 |
| Gm38910 | 6.6062E-06 | 2.36113149 | 0.198 | 0.036 | 0.16047773 | 0.03525699 | 1.56461386 | 0.084 | 0.021 | 1 | 0.03525699 | 0.000013212 |
| Prkar1a | 0.00203831 | 0.31836744 | 0.958 | 0.865 | 1 | 0.000006702 | 0.49543029 | 0.966 | 0.927 | 0.16280522 | 0.00203831 | 0.000013404 |
| Foxp1 | 0.12031472 | -0.2894872 | 0.948 | 0.99 | 1 | 6.7354E-06 | -0.7460349 | 0.992 | 1 | 0.16361548 | 0.12031472 | 0.000013471 |
| Fbxo2 | 6.7613E-06 | -2.8375308 | 0.219 | 0.446 | 0.16424533 | 0.00037677 | 1.07620317 | 0.298 | 0.104 | 1 | 0.00037677 | 0.000013523 |
| Ctsa | 0.49352629 | 0.25471224 | 0.667 | 0.632 | 1 | 6.7682E-06 | 0.78184457 | 0.718 | 0.406 | 0.16441299 | 0.49352629 | 0.000013536 |
| Stk16 | 0.01296707 | 0.56937342 | 0.49 | 0.342 | 1 | 6.7989E-06 | 0.98191086 | 0.471 | 0.177 | 0.16515842 | 0.01296707 | 0.000013598 |
| Lats1 | 6.8432E-06 | 0.80396114 | 0.792 | 0.528 | 0.16623585 | 0.00317168 | 0.21010998 | 0.912 | 0.771 | 1 | 0.00317168 | 0.000013686 |

| | | | | | | | | | | | |
|---|---|---|---|---|---|---|---|---|---|---|---|
| M1ap | 6.8574E-06 | -2.747393 | 0.083 | 0.316 | 0.16658027 | 0.84153357 | -0.5025003 | 0.038 | 0.042 | 1 | 0.84153357 | 0.000013715 |
| Cdv3 | 0.00121487 | 0.49736727 | 0.948 | 0.876 | 1 | 6.8701E-06 | 0.50265726 | 0.975 | 0.938 | 0.16688936 | 0.00121487 | 0.00001374 |
| Ebp | 0.30356273 | 0.1671512 | 0.635 | 0.539 | 1 | 6.9249E-06 | 1.22448117 | 0.471 | 0.188 | 0.16821896 | 0.30356273 | 0.00001385 |
| Pdcd6 | 6.9458E-06 | 0.66856002 | 0.885 | 0.777 | 0.16872711 | 0.00103604 | 0.34705708 | 0.87 | 0.75 | 1 | 0.00103604 | 0.000013892 |
| Acot1 | 6.9917E-06 | 1.61892925 | 0.354 | 0.135 | 0.16984208 | 0.00033537 | 1.57767001 | 0.248 | 0.073 | 1 | 0.00033537 | 0.000013983 |
| Clcc1 | 0.10396506 | 0.30122574 | 0.531 | 0.44 | 1 | 0.000007004 | 0.73615714 | 0.555 | 0.24 | 0.17014091 | 0.10396506 | 0.000014008 |
| Mpp1 | 0.03404743 | 0.63754493 | 0.427 | 0.311 | 1 | 7.0773E-06 | 1.09124305 | 0.5 | 0.208 | 0.17192087 | 0.03404743 | 0.000014154 |
| Srsf1 | 0.000076663 | 0.71930665 | 0.812 | 0.653 | 1 | 7.0887E-06 | 0.58358068 | 0.84 | 0.74 | 0.1721981 | 7.6663E-05 | 0.000014177 |
| Nudt9 | 0.69004761 | -0.2121835 | 0.573 | 0.554 | 1 | 7.1072E-06 | 0.92830447 | 0.521 | 0.219 | 0.17264894 | 0.69004761 | 0.000014214 |
| Rpn1 | 7.1332E-06 | 0.65026149 | 0.885 | 0.813 | 0.17327943 | 0.00130837 | 0.32663754 | 0.912 | 0.823 | 1 | 0.00130837 | 0.000014266 |
| Steap3 | 0.03031231 | 0.31809463 | 0.573 | 0.435 | 1 | 7.1425E-06 | 1.01527526 | 0.731 | 0.438 | 0.17350549 | 0.03031231 | 0.000014285 |
| Ehbp1 | 0.83252705 | -0.1800072 | 0.75 | 0.694 | 1 | 7.1461E-06 | -1.0350879 | 0.857 | 0.906 | 0.17359408 | 0.83252705 | 0.000014292 |
| Fundc1 | 0.25458482 | 0.34911097 | 0.552 | 0.477 | 1 | 7.2123E-06 | 0.92209147 | 0.55 | 0.24 | 0.17520212 | 0.25458482 | 0.000014425 |
| Zfp275 | 7.2775E-06 | 1.34091827 | 0.458 | 0.212 | 0.17678548 | 0.00109812 | 0.50228472 | 0.424 | 0.208 | 1 | 0.00109812 | 0.000014555 |
| Wnk4 | 7.2794E-06 | -1.2388165 | 0.26 | 0.528 | 0.17683111 | 0.27518201 | -0.154577 | 0.282 | 0.208 | 1 | 0.27518201 | 0.000014559 |
| Prkch | 0.0018307 | -1.353693 | 0.125 | 0.285 | 1 | 0.000007302 | -0.8160406 | 0.101 | 0.312 | 0.1773809 | 0.0018307 | 0.000014604 |
| Slc35a1 | 0.07128391 | 0.40604391 | 0.438 | 0.321 | 1 | 7.3652E-06 | 1.0671142 | 0.496 | 0.229 | 0.17891552 | 0.07128391 | 0.00001473 |
| Timm9 | 0.0010346 | 0.80005246 | 0.521 | 0.332 | 1 | 7.3889E-06 | 1.3146356 | 0.437 | 0.167 | 0.17949124 | 0.0010346 | 0.000014778 |
| Cyp51 | 0.04333976 | 0.39282073 | 0.771 | 0.731 | 1 | 0.000007422 | 1.21758714 | 0.689 | 0.521 | 0.18029551 | 0.04333976 | 0.000014844 |
| Arpc4 | 7.4578E-06 | 0.75115478 | 0.885 | 0.756 | 0.18116375 | 0.00142967 | 0.40163964 | 0.849 | 0.781 | 1 | 0.00142967 | 0.000014915 |
| Ppp1ca | 0.00168365 | 0.44924937 | 0.833 | 0.746 | 1 | 7.4845E-06 | 0.56728799 | 0.878 | 0.75 | 0.18181443 | 0.00168365 | 0.000014969 |
| Hbegf | 9.8847E-06 | -2.3144336 | 0.281 | 0.534 | 0.24012002 | 7.5027E-06 | -1.0168785 | 0.286 | 0.583 | 0.18225534 | 9.8847E-06 | 0.000015005 |
| Crybg1 | 0.01307511 | -1.577468 | 0.115 | 0.228 | 1 | 7.5662E-06 | -1.2196575 | 0.151 | 0.385 | 0.18379816 | 0.01307511 | 0.000015132 |
| Majin | 7.5864E-06 | 1.83480406 | 0.625 | 0.394 | 0.18428961 | 0.00015591 | 1.11682364 | 0.613 | 0.354 | 1 | 0.00015591 | 0.000015173 |
| Bckdk | 0.12631503 | 0.35263002 | 0.396 | 0.295 | 1 | 0.000007588 | 1.59060906 | 0.361 | 0.115 | 0.18432691 | 0.12631503 | 0.000015176 |
| Pdia5 | 7.6101E-06 | 1.2169479 | 0.562 | 0.306 | 0.18486352 | 0.00048887 | 0.35614381 | 0.534 | 0.271 | 1 | 0.00048887 | 0.00001522 |
| Slc36a4 | 0.0875736 | 0.3897421 | 0.427 | 0.321 | 1 | 7.8559E-06 | 0.79866919 | 0.534 | 0.24 | 0.1908364 | 0.0875736 | 0.000015712 |
| Exosc3 | 7.8621E-06 | 0.92407853 | 0.615 | 0.337 | 0.19098727 | 0.00015953 | 0.67103591 | 0.454 | 0.208 | 1 | 0.00015953 | 0.000015724 |
| Gm525 | 0.84329792 | -0.6346705 | 0.427 | 0.415 | 1 | 7.9334E-06 | 0.81964583 | 0.563 | 0.25 | 0.19271732 | 0.84329792 | 0.000015867 |
| Dab2 | 0.00394986 | 0.60613197 | 0.792 | 0.622 | 1 | 7.9438E-06 | 0.47960391 | 0.874 | 0.604 | 0.19297193 | 0.00394986 | 0.000015888 |
| Fam98b | 0.00049406 | 0.7326347 | 0.531 | 0.326 | 1 | 7.9589E-06 | 1.04008721 | 0.588 | 0.302 | 0.19333689 | 0.00049406 | 0.000015918 |
| Med7 | 0.08247382 | 0.56388789 | 0.469 | 0.383 | 1 | 7.9729E-06 | 0.93045907 | 0.504 | 0.208 | 0.19367658 | 0.08247382 | 0.000015946 |
| Eif3d | 0.01310719 | 0.56350566 | 0.615 | 0.482 | 1 | 0.000007974 | 0.87234807 | 0.718 | 0.49 | 0.19370357 | 0.01310719 | 0.000015948 |
| Tm9sf2 | 0.00095255 | 0.3337805 | 0.979 | 0.938 | 1 | 7.9756E-06 | 0.45163265 | 0.983 | 0.917 | 0.19374333 | 0.00095255 | 0.000015951 |
| Vill | 0.00611863 | 0.64333821 | 0.635 | 0.477 | 1 | 8.0353E-06 | 0.74223325 | 0.639 | 0.312 | 0.19519382 | 0.00611863 | 0.000016071 |
| Dsg2 | 0.74827573 | 0.59245704 | 0.021 | 0.016 | 1 | 8.0492E-06 | -1.0930439 | 0.118 | 0.333 | 0.19553003 | 0.74827573 | 0.000016098 |
| Ptgds | 0.00653349 | -1.7040004 | 0.177 | 0.301 | 1 | 8.0551E-06 | 3.06518417 | 0.223 | 0.021 | 0.19567448 | 0.00653349 | 0.00001611 |
| Dohh | 0.07922149 | 0.27052894 | 0.5 | 0.383 | 1 | 8.1099E-06 | 1.50728068 | 0.437 | 0.188 | 0.19700469 | 0.07922149 | 0.00001622 |
| Cbx4 | 0.32768934 | 0.11440974 | 0.375 | 0.311 | 1 | 8.1772E-06 | -0.9059583 | 0.349 | 0.635 | 0.1986416 | 0.32768934 | 0.000016354 |
| Slc35b2 | 0.01217796 | 0.45110119 | 0.74 | 0.632 | 1 | 0.00000831 | 0.99465578 | 0.592 | 0.302 | 0.20186546 | 0.01217796 | 0.00001662 |
| Hint1 | 0.03073487 | 0.21608116 | 0.948 | 0.907 | 1 | 8.3118E-06 | 0.51982593 | 0.937 | 0.854 | 0.20190914 | 0.03073487 | 0.000016623 |
| Tmem238 | 0.10448656 | 0.60223806 | 0.281 | 0.197 | 1 | 0.000008329 | 0.81942775 | 0.336 | 0.083 | 0.20232841 | 0.10448656 | 0.000016658 |
| Septin8 | 0.00353194 | 0.84603111 | 0.479 | 0.316 | 1 | 8.3316E-06 | 0.81567562 | 0.601 | 0.281 | 0.20239222 | 0.00353194 | 0.000016663 |
| Lsmem1 | 0.65559529 | -0.4779323 | 0.177 | 0.15 | 1 | 8.3968E-06 | -2.0215098 | 0.185 | 0.417 | 0.20397399 | 0.65559529 | 0.000016793 |
| Cryzl2 | 0.66494458 | 0.10060394 | 0.365 | 0.332 | 1 | 8.4374E-06 | 1.67620555 | 0.324 | 0.083 | 0.20496222 | 0.66494458 | 0.000016875 |
| Ripk4 | 0.22187575 | -0.9925055 | 0 | 0.016 | 1 | 8.4413E-06 | -2.5025003 | 0.025 | 0.156 | 0.20505599 | 0.22187575 | 0.000016883 |
| Psmd6 | 0.000096795 | 0.6730755 | 0.729 | 0.513 | 1 | 8.4459E-06 | 0.81413345 | 0.731 | 0.5 | 0.20516683 | 9.6795E-05 | 0.000016892 |
| Rce1 | 0.07859324 | 0.32236787 | 0.438 | 0.326 | 1 | 8.4617E-06 | 1.18199783 | 0.445 | 0.177 | 0.20555177 | 0.07859324 | 0.000016923 |
| C1qtnf12 | 0.000008466 | 1.90057933 | 0.229 | 0.052 | 0.20565662 | 0.02732023 | 1.11640949 | 0.151 | 0.062 | 1 | 0.02732023 | 0.000016932 |
| Pdcl3 | 0.03845293 | 0.17642355 | 0.792 | 0.679 | 1 | 8.4689E-06 | 0.50534468 | 0.845 | 0.656 | 0.2057261 | 0.03845293 | 0.000016938 |
| Ets2 | 0.00312506 | 0.63416729 | 0.75 | 0.622 | 1 | 8.4746E-06 | 0.57849938 | 0.903 | 0.854 | 0.20586513 | 0.00312506 | 0.000016949 |
| Hars | 0.00195212 | 0.47604755 | 0.698 | 0.503 | 1 | 0.000008487 | 0.61614416 | 0.693 | 0.344 | 0.2061659 | 0.00195212 | 0.000016974 |
| Saal1 | 0.12257383 | 0.83756954 | 0.177 | 0.114 | 1 | 8.5696E-06 | 4.27510724 | 0.181 | 0 | 0.20817383 | 0.12257383 | 0.000017139 |
| Pigx | 0.30581995 | 0.22809368 | 0.458 | 0.404 | 1 | 0.000008577 | 1.09696173 | 0.508 | 0.24 | 0.20835331 | 0.30581995 | 0.000017154 |
| Vmp1 | 0.000557 | -0.4840234 | 0.906 | 0.953 | 1 | 8.5896E-06 | -0.9288696 | 0.803 | 0.917 | 0.20865926 | 0.000557 | 0.000017179 |
| Zhx1 | 0.72481059 | -0.1268066 | 0.312 | 0.285 | 1 | 8.6324E-06 | 1.08955158 | 0.445 | 0.177 | 0.20969901 | 0.72481059 | 0.000017265 |
| Poldip2 | 0.02507631 | 0.59902969 | 0.417 | 0.29 | 1 | 8.7329E-06 | 1.12826585 | 0.508 | 0.229 | 0.21214063 | 0.02507631 | 0.000017466 |
| Flywch2 | 0.07710807 | 0.80104366 | 0.188 | 0.114 | 1 | 8.8255E-06 | 2.49749966 | 0.239 | 0.031 | 0.21438784 | 0.07710807 | 0.000017651 |
| Ankrd17 | 0.19084463 | 0.13896924 | 0.958 | 0.953 | 1 | 0.00000884 | -0.6837793 | 0.996 | 0.99 | 0.21474218 | 0.19084463 | 0.00001768 |
| Cacna2d1 | 8.9285E-06 | 0.88196365 | 0.771 | 0.554 | 0.21689065 | 0.80877642 | -0.7051224 | 0.782 | 0.583 | 1 | 0.80877642 | 0.000017857 |
| Bcl11b | 0.38192865 | -1.3144336 | 0.01 | 0.026 | 1 | 0.000008976 | -3.2029401 | 0.017 | 0.135 | 0.21804474 | 0.38192865 | 0.000017952 |
| Timm17a | 0.08181604 | 0.21163525 | 0.792 | 0.658 | 1 | 9.1338E-06 | 0.66157559 | 0.815 | 0.646 | 0.2218782 | 0.08181604 | 0.000018268 |
| Smpd3 | 9.1382E-06 | 4.75792239 | 0.229 | 0.062 | 0.2219854 | 0.00604323 | 4.70509508 | 0.097 | 0.01 | 1 | 0.00604323 | 0.000018276 |
| Sqor | 0.01508095 | 0.93878179 | 0.302 | 0.181 | 1 | 9.1525E-06 | 1.47564023 | 0.374 | 0.125 | 0.22233358 | 0.01508095 | 0.000018305 |
| Zbtb21 | 0.42320383 | 0.14499806 | 0.417 | 0.368 | 1 | 9.1549E-06 | 0.75885749 | 0.487 | 0.198 | 0.22239033 | 0.42320383 | 0.00001831 |
| Prg4 | 0.69571592 | -1.410358 | 0.344 | 0.332 | 1 | 9.1762E-06 | 2.06679868 | 0.387 | 0.135 | 0.22290888 | 0.69571592 | 0.000018352 |
| Rcn2 | 0.51292744 | -0.2066303 | 0.99 | 0.948 | 1 | 0.000009276 | 0.5715786 | 0.933 | 0.729 | 0.22533374 | 0.51292744 | 0.000018552 |
| Taf7 | 0.00747476 | 0.54117653 | 0.646 | 0.487 | 1 | 9.3225E-06 | 0.95718107 | 0.643 | 0.365 | 0.22646262 | 0.00747476 | 0.000018645 |
| Hspa1b | 0.0321315 | -0.5482034 | 0.479 | 0.596 | 1 | 0.000009348 | -1.763688 | 0.542 | 0.802 | 0.22708111 | 0.0321315 | 0.000018696 |
| Itgav | 9.3553E-06 | -0.7765593 | 0.76 | 0.907 | 0.22725886 | 0.20798726 | -0.2899007 | 0.832 | 0.906 | 1 | 0.20798726 | 0.000018711 |
| Slc39a1 | 0.00031389 | 0.64028556 | 0.75 | 0.518 | 1 | 9.3636E-06 | 0.83913908 | 0.777 | 0.469 | 0.22746035 | 0.00031389 | 0.000018727 |
| Anxa3 | 9.3984E-06 | -1.3873651 | 0.417 | 0.648 | 0.22830486 | 0.00041544 | -0.6062483 | 0.336 | 0.573 | 1 | 0.00041544 | 0.000018797 |
| Sspn | 9.4335E-06 | 0.83176725 | 0.802 | 0.565 | 0.22915814 | 0.00042166 | 0.40268429 | 0.752 | 0.458 | 1 | 0.00042166 | 0.000018867 |
| Cpox | 0.000009553 | 1.12297175 | 0.479 | 0.218 | 0.23206104 | 0.00033561 | 0.82400048 | 0.42 | 0.188 | 1 | 0.00033561 | 0.000019106 |
| Tsnax | 0.0024815 | 0.57517905 | 0.604 | 0.404 | 1 | 9.6191E-06 | 1.06070761 | 0.517 | 0.229 | 0.23366772 | 0.0024815 | 0.000019238 |
| Dip2c | 0.32355077 | -0.2761897 | 0.74 | 0.767 | 1 | 9.7282E-06 | -0.7819603 | 0.87 | 0.958 | 0.23631628 | 0.32355077 | 0.000019456 |
| Cacybp | 0.00287598 | 0.42965647 | 0.781 | 0.658 | 1 | 9.7656E-06 | 0.64655547 | 0.866 | 0.729 | 0.23722601 | 0.00287598 | 0.000019531 |
| Kctd14 | 9.7666E-06 | 3.32942263 | 0.156 | 0.021 | 0.23725116 | 0.1430089 | -0.5936482 | 0.076 | 0.031 | 1 | 0.1430089 | 0.000019533 |
| Car5b | 9.7704E-06 | -1.6149427 | 0.146 | 0.399 | 0.23734234 | 0.6093411 | -0.2774338 | 0.139 | 0.115 | 1 | 0.6093411 | 0.000019541 |
| Ndufab1 | 0.35107001 | 0.15858713 | 0.792 | 0.746 | 1 | 9.7737E-06 | 0.55143847 | 0.807 | 0.656 | 0.23742243 | 0.35107001 | 0.000019547 |
| Aldh7a1 | 0.11148762 | 0.2590333 | 0.635 | 0.518 | 1 | 9.8419E-06 | 0.91047469 | 0.592 | 0.292 | 0.23907914 | 0.11148762 | 0.000019684 |
| Rcan1 | 0.00788506 | -0.8822179 | 0.812 | 0.855 | 1 | 9.9503E-06 | 1.17116322 | 0.878 | 0.812 | 0.2417138 | 0.00788506 | 0.000019901 |
| Rapgef2 | 9.9619E-06 | 0.63799174 | 0.938 | 0.845 | 0.24199347 | 0.01607853 | -0.5305979 | 0.979 | 0.99 | 1 | 0.01607853 | 0.000019924 |
| Tmem63a | 0.000010627 | 0.87504424 | 0.646 | 0.394 | 0.25815701 | 0.000010008 | 0.67019013 | 0.735 | 0.396 | 0.24312561 | 1.0627E-05 | 0.000020017 |
| Dgka | 0.000010012 | 0.93623585 | 0.5 | 0.233 | 0.24321396 | 0.06714493 | -0.4838847 | 0.437 | 0.562 | 1 | 0.06714493 | 0.000020024 |
| Eif3e | 0.01187985 | 0.34810719 | 0.958 | 0.891 | 1 | 0.000010038 | 0.45190279 | 0.958 | 0.854 | 0.24384352 | 0.01187985 | 0.000020076 |
| Lgi2 | 0.03702282 | 1.04311845 | 0.26 | 0.161 | 1 | 0.000010053 | 1.99270751 | 0.399 | 0.167 | 0.24421354 | 0.03702282 | 0.000020106 |
| Slc35d1 | 0.000010057 | 0.78121868 | 0.917 | 0.71 | 0.2443009 | 0.03646644 | 0.24434498 | 0.853 | 0.74 | 1 | 0.03646644 | 0.000020114 |
| Igsf8 | 0.09575595 | 0.16503581 | 0.583 | 0.466 | 1 | 0.00001017 | 0.80750169 | 0.609 | 0.323 | 0.24703815 | 0.09575595 | 0.000020339 |
| Ccnc | 0.02215004 | 0.42253204 | 0.573 | 0.44 | 1 | 0.000010194 | 0.74992597 | 0.613 | 0.292 | 0.24764384 | 0.02215004 | 0.000020389 |
| Sfn | 0.03510896 | -1.1654518 | 0.208 | 0.321 | 1 | 0.000010288 | -0.4996798 | 0.239 | 0.5 | 0.24992428 | 0.03510896 | 0.000020577 |
| Ppp1r15b | 0.000010291 | 0.97293731 | 0.76 | 0.523 | 0.24998458 | 0.00314421 | 0.3590085 | 0.832 | 0.688 | 1 | 0.00314421 | 0.000020582 |

| | | | | | | | | | | | | |
|---|---|---|---|---|---|---|---|---|---|---|---|---|
| Srprb | 0.00068144 | 0.6730755 | 0.688 | 0.492 | 1 | 0.000010306 | 0.77927063 | 0.58 | 0.271 | 0.25034219 | 0.00068144 | 0.000020611 |
| Pno1 | 0.05921037 | 0.56692195 | 0.469 | 0.363 | 1 | 0.000010528 | 1.10518224 | 0.424 | 0.146 | 0.25575017 | 0.05921037 | 0.000021056 |
| Tfe3 | 0.04433876 | 0.53800925 | 0.438 | 0.321 | 1 | 0.000010552 | 0.81567562 | 0.588 | 0.292 | 0.25633352 | 0.04433876 | 0.000021104 |
| mt-Nd3 | 0.52067404 | -0.2958557 | 0.875 | 0.902 | 1 | 0.000010593 | -0.9594805 | 0.832 | 0.927 | 0.25732097 | 0.52067404 | 0.000021186 |
| Ssb | 0.000010602 | 0.42684503 | 0.99 | 0.969 | 0.25754324 | 0.000024016 | 0.45729334 | 0.983 | 0.917 | 0.58338778 | 2.4016E-05 | 0.000021204 |
| Rbm17 | 0.00019915 | 0.61219326 | 0.792 | 0.611 | 1 | 0.000010618 | 0.65361886 | 0.815 | 0.615 | 0.25794056 | 0.00019915 | 0.000021237 |
| Pacsin3 | 0.4854884 | 0.37672835 | 0.229 | 0.197 | 1 | 0.000010744 | 0.77193883 | 0.403 | 0.125 | 0.26098707 | 0.4854884 | 0.000021487 |
| Mmadhc | 0.000020388 | 0.85277194 | 0.688 | 0.415 | 0.4952611 | 0.000010831 | 0.50239846 | 0.681 | 0.375 | 0.2631058 | 2.0388E-05 | 0.000021662 |
| Rhoc | 0.28398877 | 0.11540988 | 0.938 | 0.907 | 1 | 0.000010896 | 0.65747855 | 0.937 | 0.792 | 0.26468102 | 0.28398877 | 0.000021792 |
| H2-T23 | 0.000010921 | -2.706751 | 0.062 | 0.285 | 0.26528349 | 0.000060575 | -1.3098553 | 0.076 | 0.24 | 1 | 6.0575E-05 | 0.000021841 |
| Pfdn2 | 0.02375771 | 0.40999768 | 0.812 | 0.777 | 1 | 0.000010946 | 0.45459107 | 0.891 | 0.74 | 0.26588979 | 0.02375771 | 0.000021891 |
| Klf16 | 0.000010986 | 1.15949763 | 0.406 | 0.166 | 0.26688156 | 0.00274214 | 0.56929034 | 0.319 | 0.146 | 1 | 0.00274214 | 0.000021973 |
| Lepr | 0.01376964 | 0.39844642 | 0.427 | 0.28 | 1 | 0.000010995 | 1.0337679 | 0.546 | 0.25 | 0.26710245 | 0.01376964 | 0.00002199 |
| Golga7 | 0.01313341 | 0.53490601 | 0.729 | 0.606 | 1 | 0.000011115 | 0.69448367 | 0.718 | 0.417 | 0.26999953 | 0.01313341 | 0.000022229 |
| Golm2 | 0.00001113 | 0.50302195 | 0.948 | 0.793 | 0.27038157 | 0.000044626 | 0.45456729 | 0.908 | 0.729 | 1 | 4.4626E-05 | 0.000022261 |
| Polr3d | 0.02669181 | 0.28472282 | 0.729 | 0.56 | 1 | 0.000011132 | 0.88278982 | 0.58 | 0.271 | 0.27043031 | 0.02669181 | 0.000022265 |
| Celf5 | 0.00001119 | 1.84702986 | 0.25 | 0.067 | 0.27183677 | 0.03235477 | 1.16407593 | 0.147 | 0.062 | 1 | 0.03235477 | 0.000022381 |
| Mex3d | 0.00032873 | 1.25260703 | 0.292 | 0.119 | 1 | 0.000011249 | 1.27510724 | 0.391 | 0.135 | 0.27326663 | 0.00032873 | 0.000022498 |
| Usp16 | 0.000011332 | 0.67045955 | 0.917 | 0.819 | 0.27527939 | 0.03730159 | 0.17785974 | 0.87 | 0.865 | 1 | 0.03730159 | 0.000022664 |
| Srpk2 | 0.000011426 | -0.5922145 | 0.844 | 0.907 | 0.27756424 | 0.10476636 | -0.4266689 | 0.912 | 0.958 | 1 | 0.10476636 | 0.000022852 |
| Meaf6 | 0.21103335 | 0.28051303 | 0.406 | 0.337 | 1 | 0.000011557 | 0.8917786 | 0.45 | 0.156 | 0.28075133 | 0.21103335 | 0.000023115 |
| Bmp2k | 0.12377305 | -0.6686159 | 0.312 | 0.378 | 1 | 0.000011561 | -1.1686093 | 0.391 | 0.646 | 0.2808332 | 0.12377305 | 0.000023121 |
| Togaram1 | 0.000011652 | 0.73684695 | 0.802 | 0.503 | 0.28305963 | 0.74937976 | -0.2484547 | 0.769 | 0.781 | 1 | 0.74937976 | 0.000023305 |
| Tmsb4x | 0.000011677 | -1.563753 | 0.708 | 0.85 | 0.28365727 | 0.000014841 | -1.7530737 | 0.832 | 0.917 | 0.36051591 | 1.4841E-05 | 0.000023354 |
| C1qtnf4 | 0.28094469 | -0.3037072 | 0.188 | 0.249 | 1 | 0.000011706 | 1.77760758 | 0.315 | 0.083 | 0.28437296 | 0.28094469 | 0.000023413 |
| Plcl2 | 0.000011761 | 1.27052894 | 0.281 | 0.083 | 0.28570131 | 0.02184636 | 0.48984609 | 0.21 | 0.094 | 1 | 0.02184636 | 0.000023522 |
| Znrf3 | 0.96956057 | -0.3351425 | 0.615 | 0.585 | 1 | 0.000011792 | -1.1659067 | 0.651 | 0.833 | 0.28645788 | 0.96956057 | 0.000023584 |
| Septin7 | 0.00604421 | 0.22141544 | 0.99 | 0.979 | 1 | 0.000011813 | 0.34839031 | 1 | 1 | 0.2869672 | 0.00604421 | 0.000023626 |
| Cks2 | 0.15245818 | 1.57139542 | 0.188 | 0.13 | 1 | 0.000011818 | -1.8102843 | 0.126 | 0.323 | 0.28708508 | 0.15245818 | 0.000023636 |
| Golga4 | 0.000011833 | 0.55525842 | 0.938 | 0.855 | 0.28744137 | 0.01188763 | 0.21370669 | 0.958 | 0.938 | 1 | 0.01188763 | 0.000023665 |
| Cdc26 | 0.000011896 | 0.61479337 | 0.875 | 0.617 | 0.2889725 | 0.02795704 | 0.17781004 | 0.756 | 0.625 | 1 | 0.02795704 | 0.000023791 |
| Snrpb2 | 0.00154935 | 0.44677958 | 0.76 | 0.606 | 1 | 0.000011929 | 0.93433308 | 0.655 | 0.438 | 0.28978761 | 0.00154935 | 0.000023859 |
| Heatr1 | 0.000012025 | 1.27542774 | 0.448 | 0.197 | 0.29211546 | 0.04535053 | 0.46355233 | 0.336 | 0.219 | 1 | 0.04535053 | 0.00002405 |
| Orc6 | 0.00342526 | 0.9920645 | 0.49 | 0.326 | 1 | 0.000012049 | 0.98217135 | 0.471 | 0.188 | 0.29270625 | 0.00342526 | 0.000024099 |
| Smim19 | 0.5146508 | -1.3600146 | 0.521 | 0.565 | 1 | 0.000012081 | 0.93445194 | 0.559 | 0.281 | 0.29346519 | 0.5146508 | 0.000024161 |
| Tspan6 | 0.08631958 | -0.2930599 | 0.583 | 0.699 | 1 | 0.000012142 | 0.71120635 | 0.664 | 0.344 | 0.29494819 | 0.08631958 | 0.000024283 |
| Slain2 | 0.000012157 | 0.75325325 | 0.781 | 0.58 | 0.29530616 | 0.09324287 | 0.15084914 | 0.845 | 0.792 | 1 | 0.09324287 | 0.000024313 |
| Fancc | 0.000012181 | 1.27304016 | 0.677 | 0.456 | 0.29590666 | 0.78511277 | -0.4001504 | 0.676 | 0.562 | 1 | 0.78511277 | 0.000024362 |
| Arpc1a | 0.000012182 | -0.7672132 | 0.833 | 0.855 | 0.29592037 | 0.0016087 | 0.34704113 | 0.861 | 0.708 | 1 | 0.0016087 | 0.000024363 |
| Rtl8b | 0.00377375 | 0.69338595 | 0.521 | 0.352 | 1 | 0.000012251 | 0.99270751 | 0.475 | 0.208 | 0.29760229 | 0.00377375 | 0.000024502 |
| Agtpbp1 | 0.000012327 | 0.87679084 | 0.625 | 0.342 | 0.29944155 | 0.11171646 | -0.6326764 | 0.63 | 0.698 | 1 | 0.11171646 | 0.000024653 |
| Gnl3 | 0.000012344 | 0.68063655 | 0.917 | 0.829 | 0.29984973 | 0.02894909 | 0.25326285 | 0.849 | 0.771 | 1 | 0.02894909 | 0.000024687 |
| Dynlrb1 | 0.1734855 | 0.12702814 | 0.948 | 0.876 | 1 | 0.000012392 | 0.44947814 | 0.908 | 0.823 | 0.30102543 | 0.1734855 | 0.000024784 |
| Ptpn4 | 0.372806 | -0.4682389 | 0.323 | 0.352 | 1 | 0.000012466 | -1.2937356 | 0.353 | 0.625 | 0.30282701 | 0.372806 | 0.000024932 |
| Tm9sf1 | 0.000016791 | 0.75522083 | 0.781 | 0.549 | 0.40789205 | 0.000012475 | 0.54178022 | 0.815 | 0.51 | 0.30303771 | 1.6791E-05 | 0.000024949 |
| Gm32618 | 0.000012499 | 1.26658176 | 0.312 | 0.104 | 0.30362987 | 0.69608545 | -0.6291338 | 0.21 | 0.177 | 1 | 0.69608545 | 0.000024998 |
| Cars | 0.00002501 | 0.95626921 | 0.667 | 0.446 | 0.60753711 | 0.000012562 | 0.50794141 | 0.761 | 0.417 | 0.30515363 | 0.00002501 | 0.000025124 |
| Ppp1r12a | 0.28543519 | -0.2891808 | 0.917 | 0.938 | 1 | 0.00001259 | -0.5685637 | 0.992 | 0.99 | 0.30584073 | 0.28543519 | 0.00002518 |
| 4931406C07Rik | 0.00325508 | 0.81247855 | 0.469 | 0.306 | 1 | 0.000012717 | 1.30752872 | 0.399 | 0.146 | 0.30892077 | 0.00325508 | 0.000025434 |
| Arhgef12 | 0.000012746 | -0.6611139 | 0.969 | 0.99 | 0.30962807 | 0.00010644 | -0.5818835 | 0.996 | 0.99 | 1 | 0.00010644 | 0.000025492 |
| Crls1 | 0.00082357 | 0.90931414 | 0.469 | 0.285 | 1 | 0.000012977 | 0.94312548 | 0.445 | 0.167 | 0.31524923 | 0.00082357 | 0.000025955 |
| Dusp15 | 0.000013062 | -5.1010299 | 0 | 0.176 | 0.31729182 | 0.18292349 | 1.69014474 | 0.038 | 0.01 | 1 | 0.18292349 | 0.000026123 |
| Clptm1l | 0.00794944 | 0.38958161 | 0.875 | 0.731 | 1 | 0.000013174 | 0.5580412 | 0.811 | 0.646 | 0.32003341 | 0.00794944 | 0.000026349 |
| Srp68 | 0.000013201 | 0.75813507 | 0.698 | 0.435 | 0.32067481 | 0.03420816 | 0.12776552 | 0.672 | 0.427 | 1 | 0.03420816 | 0.000026402 |
| Arhgap42 | 0.000013282 | -1.3094502 | 0.562 | 0.751 | 0.32265523 | 0.20517531 | -0.319295 | 0.693 | 0.719 | 1 | 0.20517531 | 0.000026565 |
| Poglut2 | 0.000013382 | 0.88196365 | 0.677 | 0.435 | 0.32507503 | 0.00695044 | 0.2182449 | 0.55 | 0.323 | 1 | 0.00695044 | 0.000026764 |
| Snx16 | 0.01186747 | 0.79804117 | 0.229 | 0.114 | 1 | 0.000013457 | 2.26000035 | 0.265 | 0.052 | 0.32689276 | 0.01186747 | 0.000026913 |
| Fmc1 | 0.01592953 | 0.61892925 | 0.438 | 0.295 | 1 | 0.000013465 | 0.78530197 | 0.441 | 0.167 | 0.32709929 | 0.01592953 | 0.00002693 |
| Ruvbl1 | 0.000013685 | 0.92107978 | 0.573 | 0.301 | 0.33243553 | 0.00323089 | 0.3752714 | 0.462 | 0.25 | 1 | 0.00323089 | 0.00002737 |
| Hnrnph2 | 0.00739207 | 0.40604391 | 0.865 | 0.746 | 1 | 0.000013688 | 0.57648796 | 0.866 | 0.771 | 0.33250183 | 0.00739207 | 0.000027375 |
| Ap1s1 | 0.68149952 | 0.2091284 | 0.333 | 0.321 | 1 | 0.000013703 | 0.80261947 | 0.475 | 0.188 | 0.33287152 | 0.68149952 | 0.000027406 |
| Psmd4 | 0.02300011 | 0.29299922 | 0.875 | 0.808 | 1 | 0.000013791 | 0.64582241 | 0.878 | 0.74 | 0.33500217 | 0.02300011 | 0.000027581 |
| Plpp5 | 0.00511297 | 0.47986521 | 0.781 | 0.617 | 1 | 0.000014061 | 0.54429387 | 0.706 | 0.375 | 0.34156171 | 0.00511297 | 0.000028121 |
| Abraxas2 | 0.46398982 | -0.2749052 | 0.292 | 0.326 | 1 | 0.000014102 | 0.96812948 | 0.504 | 0.229 | 0.34255719 | 0.46398982 | 0.000028203 |
| Hsd17b10 | 0.06218639 | 0.35541784 | 0.688 | 0.575 | 1 | 0.000014201 | 0.9107895 | 0.647 | 0.365 | 0.34496541 | 0.06218639 | 0.000028401 |
| Wdr5 | 0.01147939 | 0.59245704 | 0.365 | 0.218 | 1 | 0.000014213 | 0.77760758 | 0.483 | 0.198 | 0.34526263 | 0.01147939 | 0.000028426 |
| Tcf20 | 0.63037073 | -0.1705583 | 0.573 | 0.539 | 1 | 0.000014241 | -0.8225036 | 0.66 | 0.927 | 0.34594906 | 0.63037073 | 0.000028482 |
| Hikeshi | 0.69024594 | -0.1534973 | 0.479 | 0.487 | 1 | 0.000014305 | 0.69014474 | 0.563 | 0.26 | 0.34750628 | 0.69024594 | 0.000028611 |
| Glul | 0.00001437 | -0.7565763 | 0.708 | 0.876 | 0.34908637 | 0.08526744 | 0.43027761 | 0.803 | 0.833 | 1 | 0.08526744 | 0.000028741 |
| Map2k7 | 0.00057484 | 1.14499806 | 0.333 | 0.155 | 1 | 0.00001441 | 1.25492936 | 0.416 | 0.167 | 0.35005061 | 0.00057484 | 0.00002882 |
| Eif3g | 0.000014445 | 0.75956702 | 0.76 | 0.585 | 0.3508951 | 0.00012827 | 0.51422675 | 0.79 | 0.562 | 1 | 0.00012827 | 0.00002889 |
| Ccpg1 | 0.03036239 | 0.33073956 | 0.865 | 0.834 | 1 | 0.000014506 | 0.51543329 | 0.908 | 0.812 | 0.35237546 | 0.03036239 | 0.000029011 |
| Hnrnpa1 | 0.000014587 | 0.35863842 | 0.979 | 0.948 | 0.3543362 | 0.01825693 | 0.11963028 | 0.958 | 0.958 | 1 | 0.01825693 | 0.000029173 |
| Ctdsp2 | 0.00047029 | 0.49499123 | 0.906 | 0.767 | 1 | 0.000014692 | 0.50516951 | 0.937 | 0.885 | 0.35688625 | 0.00047029 | 0.000029383 |
| Serf1 | 0.00857508 | 0.63274974 | 0.594 | 0.435 | 1 | 0.000014741 | 0.97359868 | 0.492 | 0.219 | 0.35809594 | 0.00857508 | 0.000029482 |
| Dbi | 0.79496545 | -0.1839622 | 0.969 | 0.927 | 1 | 0.00001482 | 0.3909606 | 0.899 | 0.854 | 0.35999946 | 0.79496545 | 0.000029639 |
| Pccb | 0.0177639 | 0.72021258 | 0.438 | 0.306 | 1 | 0.000014956 | 1.04769674 | 0.458 | 0.198 | 0.36329904 | 0.0177639 | 0.000029911 |
| Ltf | 0.62914985 | 0.65282966 | 0.323 | 0.306 | 1 | 0.00001499 | -1.3277772 | 0.244 | 0.469 | 0.36413562 | 0.62914985 | 0.00002998 |
| Rai14 | 0.44524455 | -0.8370509 | 0.469 | 0.368 | 1 | 0.000015092 | -1.2378003 | 0.609 | 0.771 | 0.36660451 | 0.44524455 | 0.000030183 |
| Uty | 0.49363396 | -0.4121596 | 0.479 | 0.472 | 1 | 0.000015128 | -0.8927929 | 0.832 | 0.927 | 0.36747941 | 0.49363396 | 0.000030255 |
| Isyna1 | 0.000015286 | -1.6649308 | 0.458 | 0.627 | 0.37132129 | 0.0009679 | 0.36601573 | 0.63 | 0.365 | 1 | 0.0009679 | 0.000030571 |
| Hif1a | 0.10914153 | -0.4377729 | 0.979 | 0.979 | 1 | 0.000015324 | 0.47416953 | 0.996 | 0.958 | 0.37224087 | 0.10914153 | 0.000030647 |
| Bcl7c | 0.19220594 | 0.17741954 | 0.719 | 0.591 | 1 | 0.00001546 | 0.63882596 | 0.777 | 0.448 | 0.37554482 | 0.19220594 | 0.000030919 |
| Usp34 | 0.00077773 | 0.31437634 | 0.979 | 0.953 | 1 | 0.000015537 | -0.5750271 | 0.996 | 1 | 0.37743693 | 0.00077773 | 0.000031075 |
| Trappc5 | 0.01391826 | 0.64136664 | 0.385 | 0.244 | 1 | 0.000015651 | 1.41806519 | 0.366 | 0.125 | 0.38019618 | 0.01391826 | 0.000031302 |
| Bhlhe40 | 0.29454042 | -0.1497285 | 0.896 | 0.777 | 1 | 0.000015653 | 0.68006553 | 0.895 | 0.875 | 0.38024762 | 0.29454042 | 0.000031306 |
| Cebpg | 0.000015805 | 0.88830518 | 0.792 | 0.591 | 0.38393058 | 0.00559018 | 0.34577244 | 0.803 | 0.729 | 1 | 0.00559018 | 0.000031609 |
| Smg6 | 0.87434429 | -0.1538685 | 0.906 | 0.845 | 1 | 0.000015898 | -0.8332206 | 0.912 | 0.979 | 0.38619165 | 0.87434429 | 0.000031796 |
| Mrpl9 | 0.91031556 | -0.1539689 | 0.312 | 0.29 | 1 | 0.000016006 | 1.10799725 | 0.458 | 0.198 | 0.38882902 | 0.91031556 | 0.000032013 |
| Srgn | 0.59113409 | 0.18176109 | 0.656 | 0.642 | 1 | 0.000016034 | 0.73949465 | 0.828 | 0.677 | 0.3895092 | 0.59113409 | 0.000032069 |
| Atp5g3 | 0.14981678 | 0.16152817 | 0.896 | 0.845 | 1 | 0.000016091 | 0.62242065 | 0.874 | 0.812 | 0.39088355 | 0.14981678 | 0.000032182 |

| | | | | | | | | | | | | |
|---|---|---|---|---|---|---|---|---|---|---|---|---|
| Atosa | 0.40125805 | -0.3524014 | 0.438 | 0.456 | 1 | 0.000016106 | -1.3897086 | 0.454 | 0.677 | 0.39125807 | 0.40125805 | 0.000032213 |
| Ddt | 0.66745976 | -0.2231184 | 0.542 | 0.518 | 1 | 0.000016185 | 0.82185042 | 0.492 | 0.219 | 0.39316417 | 0.66745976 | 0.00003237 |
| Dpf2 | 0.09935572 | 0.53105649 | 0.354 | 0.259 | 1 | 0.00001621 | 0.66099839 | 0.555 | 0.26 | 0.39376334 | 0.09935572 | 0.000032419 |
| Mas1 | 0.000016249 | 4.81484946 | 0.094 | 0 | 0.39472975 | 0.08763061 | 1.49749966 | 0.05 | 0.01 | 1 | 0.08763061 | 0.000032498 |
| Odc1 | 0.05258668 | -0.2286207 | 0.896 | 0.917 | 1 | 0.00001634 | 0.74558906 | 0.874 | 0.823 | 0.39692104 | 0.05258668 | 0.000032679 |
| Isca2 | 0.07861865 | 0.4134869 | 0.427 | 0.316 | 1 | 0.000016377 | 2.02404547 | 0.328 | 0.104 | 0.39783196 | 0.07861865 | 0.000032754 |
| Gnrh1 | 0.000016552 | 2.22988696 | 0.198 | 0.041 | 0.40207148 | 0.04119697 | 0.56461386 | 0.147 | 0.062 | 1 | 0.04119697 | 0.000033103 |
| Sparcl1 | 0.71493515 | -0.1633257 | 0.75 | 0.642 | 1 | 0.000016704 | 0.77630138 | 0.542 | 0.24 | 0.40576217 | 0.71493515 | 0.000033407 |
| Scd1 | 0.000016743 | -2.3044495 | 0.5 | 0.668 | 0.40671917 | 0.02675211 | -0.8597708 | 0.513 | 0.312 | 1 | 0.02675211 | 0.000033486 |
| Ap2m1 | 0.000016749 | 0.70263996 | 0.823 | 0.637 | 0.40687535 | 0.00135679 | 0.44341214 | 0.824 | 0.74 | 1 | 0.00135679 | 0.000033498 |
| Gprasp2 | 0.30265553 | 1.00749454 | 0.188 | 0.145 | 1 | 0.000016917 | 1.46458504 | 0.345 | 0.104 | 0.41095439 | 0.30265553 | 0.000033834 |
| Rpn2 | 0.17482697 | 0.1726313 | 0.927 | 0.85 | 1 | 0.000016974 | 0.52103922 | 0.878 | 0.792 | 0.41232068 | 0.17482697 | 0.000033947 |
| Rpp14 | 0.02023874 | 0.69338595 | 0.323 | 0.197 | 1 | 0.000016989 | 1.65697787 | 0.286 | 0.062 | 0.41268572 | 0.02023874 | 0.000033977 |
| Ndufa8 | 0.0642763 | -0.5543844 | 0.802 | 0.746 | 1 | 0.000017195 | 0.62504971 | 0.765 | 0.552 | 0.41771176 | 0.0642763 | 0.000034391 |
| Hexa | 0.0826724 | -0.3703319 | 0.802 | 0.777 | 1 | 0.000017417 | 0.8169595 | 0.756 | 0.427 | 0.42308465 | 0.0826724 | 0.000034833 |
| Tmem43 | 0.38114754 | 0.53356335 | 0.448 | 0.435 | 1 | 0.000017429 | 0.94770401 | 0.588 | 0.302 | 0.42339582 | 0.38114754 | 0.000034859 |
| St7l | 0.000017465 | 0.88943878 | 0.656 | 0.394 | 0.42425972 | 0.00898466 | 0.24006029 | 0.693 | 0.458 | 1 | 0.00898466 | 0.00003493 |
| Etfdh | 0.08035699 | 0.45815595 | 0.302 | 0.202 | 1 | 0.000017471 | 1.11516633 | 0.433 | 0.167 | 0.42439619 | 0.08035699 | 0.000034941 |
| Tmem165 | 0.000017568 | 0.710061 | 0.885 | 0.741 | 0.42675654 | 0.00019847 | 0.46942889 | 0.87 | 0.771 | 1 | 0.00019847 | 0.000035135 |
| Fbl | 0.00010079 | 0.70726151 | 0.708 | 0.466 | 1 | 0.000017664 | 0.61697832 | 0.752 | 0.531 | 0.4290962 | 0.00010079 | 0.000035328 |
| Col4a2 | 0.01769564 | -0.8696487 | 0.5 | 0.601 | 1 | 0.000017729 | 0.44243606 | 0.706 | 0.354 | 0.43066761 | 0.01769564 | 0.000035457 |
| Polr2g | 0.12788312 | 0.30800365 | 0.656 | 0.56 | 1 | 0.000017849 | 0.96698494 | 0.534 | 0.25 | 0.4335902 | 0.12788312 | 0.000035698 |
| Kif19b | 0.00001785 | 2.03606369 | 0.26 | 0.078 | 0.43360216 | 0.03640652 | 0.52303475 | 0.248 | 0.135 | 1 | 0.03640652 | 0.000035699 |
| Tfam | 0.00039396 | 1.04496924 | 0.469 | 0.28 | 1 | 0.000017859 | 1.03118166 | 0.408 | 0.146 | 0.43383789 | 0.00039396 | 0.000035718 |
| Egfr | 0.25729313 | -1.1458562 | 0.312 | 0.358 | 1 | 0.000017918 | -1.060077 | 0.769 | 0.885 | 0.43526076 | 0.25729313 | 0.000035835 |
| Gpr108 | 0.04589034 | 0.40391636 | 0.552 | 0.415 | 1 | 0.000018021 | 0.70450003 | 0.576 | 0.281 | 0.43777791 | 0.04589034 | 0.000036043 |
| Vbp1 | 0.13338266 | 0.22549519 | 0.604 | 0.497 | 1 | 0.000018122 | 0.71399148 | 0.639 | 0.354 | 0.44020972 | 0.13338266 | 0.000036243 |
| B3galnt2 | 0.000018183 | 1.23305424 | 0.469 | 0.228 | 0.4417041 | 0.000043599 | 1.07201537 | 0.429 | 0.177 | 1 | 4.3599E-05 | 0.000036366 |
| Ipo4 | 0.00005311 | 0.90964721 | 0.479 | 0.238 | 1 | 0.000018227 | 1.14265694 | 0.366 | 0.115 | 0.4427583 | 0.00005311 | 0.000036453 |
| Polr2h | 0.00001823 | 0.85863115 | 0.562 | 0.295 | 0.44283355 | 0.21037221 | 0.22518201 | 0.412 | 0.333 | 1 | 0.21037221 | 0.000036459 |
| B3gat3 | 0.00030796 | 0.75832278 | 0.698 | 0.508 | 1 | 0.000018351 | 0.68431566 | 0.613 | 0.302 | 0.44577622 | 0.00030796 | 0.000036701 |
| Pigp | 0.6798865 | -0.1239313 | 0.771 | 0.705 | 1 | 0.000018365 | 0.67820512 | 0.739 | 0.562 | 0.44611428 | 0.6798865 | 0.000036729 |
| Timm22 | 0.38694673 | 0.14499806 | 0.448 | 0.378 | 1 | 0.000018385 | 1.06269891 | 0.433 | 0.177 | 0.44659727 | 0.38694673 | 0.000036769 |
| Isy1 | 0.000018489 | 1.07691594 | 0.625 | 0.394 | 0.44913062 | 0.29889747 | 0.14957636 | 0.66 | 0.615 | 1 | 0.29889747 | 0.000036977 |
| Sfrp4 | 0.00280232 | -5.2213242 | 0 | 0.088 | 1 | 0.000018565 | -7.1172102 | 0.008 | 0.104 | 0.45099302 | 0.00280232 | 0.000037131 |
| Cdh13 | 0.05265221 | -1.5436797 | 0.062 | 0.14 | 1 | 0.00001859 | -4.2794816 | 0.025 | 0.146 | 0.45159618 | 0.05265221 | 0.00003718 |
| Ndufb3 | 0.01624908 | 0.33372277 | 0.917 | 0.772 | 1 | 0.000018629 | 0.51411437 | 0.803 | 0.615 | 0.45253602 | 0.01624908 | 0.000037258 |
| Prelid3b | 0.00281157 | 0.64071221 | 0.521 | 0.347 | 1 | 0.00001865 | 0.6302022 | 0.504 | 0.219 | 0.45304711 | 0.00281157 | 0.0000373 |
| Ang | 0.02746952 | 0.5698083 | 0.406 | 0.285 | 1 | 0.00001885 | 1.13945214 | 0.475 | 0.198 | 0.45791344 | 0.02746952 | 0.0000377 |
| Socs4 | 0.00290094 | 0.77227807 | 0.573 | 0.42 | 1 | 0.000018906 | 0.74299662 | 0.66 | 0.354 | 0.45926824 | 0.00290094 | 0.000037812 |
| Ece2 | 0.000018946 | 3.59245704 | 0.125 | 0.01 | 0.46022755 | 0.00502286 | 1.86006974 | 0.101 | 0.01 | 1 | 0.00502286 | 0.000037891 |
| Trpv4 | 0.00001897 | 0.78445211 | 0.833 | 0.674 | 0.46081869 | 0.00005221 | 0.54705778 | 0.798 | 0.5 | 1 | 0.00005221 | 0.00003794 |
| Afap1 | 0.7754689 | -0.2858644 | 0.344 | 0.301 | 1 | 0.000019035 | -0.8244284 | 0.424 | 0.708 | 0.46240561 | 0.7754689 | 0.00003807 |
| Tuba1b | 0.000019976 | 0.7076367 | 0.938 | 0.803 | 0.48525827 | 0.000019093 | 0.39291076 | 0.908 | 0.833 | 0.46380779 | 1.9976E-05 | 0.000038186 |
| Lrba | 0.000019181 | -1.1557358 | 0.708 | 0.855 | 0.46594248 | 0.12387499 | -0.5521223 | 0.815 | 0.833 | 1 | 0.12387499 | 0.000038361 |
| Baz2b | 0.000019299 | -0.5894605 | 0.969 | 0.984 | 0.46880801 | 0.000032277 | -0.6727649 | 0.966 | 0.969 | 0.78407449 | 3.2277E-05 | 0.000038597 |
| Tmem107 | 0.000019315 | 1.16422004 | 0.458 | 0.212 | 0.46920524 | 0.00020775 | 1.70509508 | 0.282 | 0.094 | 1 | 0.00020775 | 0.00003863 |
| Nob1 | 0.08039265 | 0.17860645 | 0.562 | 0.399 | 1 | 0.000019354 | 0.70868069 | 0.592 | 0.302 | 0.47015209 | 0.08039265 | 0.000038708 |
| Klhdc2 | 0.37864799 | 0.20935814 | 0.708 | 0.658 | 1 | 0.000019407 | 0.4930446 | 0.861 | 0.562 | 0.47142345 | 0.37864799 | 0.000038813 |
| Ltv1 | 0.00318234 | 0.9652663 | 0.552 | 0.394 | 1 | 0.000019422 | 0.78164309 | 0.487 | 0.198 | 0.47181101 | 0.00318234 | 0.000038845 |
| Ncam1 | 0.000019725 | 0.60347003 | 0.854 | 0.725 | 0.47915084 | 0.44252757 | -0.3060736 | 0.765 | 0.688 | 1 | 0.44252757 | 0.000039449 |
| Cops7a | 0.09065441 | 0.25631508 | 0.667 | 0.565 | 1 | 0.000019799 | 0.73162137 | 0.639 | 0.385 | 0.48095144 | 0.09065441 | 0.000039597 |
| Golph3 | 0.000061935 | 0.47395213 | 0.99 | 0.938 | 1 | 0.000019893 | 0.28712977 | 0.996 | 0.917 | 0.48324673 | 6.1935E-05 | 0.000039786 |
| Cuedc2 | 0.04493765 | 0.34291487 | 0.625 | 0.492 | 1 | 0.000020062 | 0.8670225 | 0.563 | 0.26 | 0.48735762 | 0.04493765 | 0.000040125 |
| Pfdn5 | 0.02732954 | 0.25655717 | 0.99 | 0.948 | 1 | 0.000020068 | 0.43142802 | 0.975 | 0.927 | 0.48748282 | 0.02732954 | 0.000040135 |
| Anxa11 | 0.96079927 | 0.24095583 | 0.781 | 0.793 | 1 | 0.000020144 | 0.49106702 | 0.895 | 0.76 | 0.48934425 | 0.96079927 | 0.000040288 |
| Ccndbp1 | 0.23841305 | 0.28867629 | 0.406 | 0.332 | 1 | 0.000020199 | 1.02312802 | 0.412 | 0.156 | 0.49067576 | 0.23841305 | 0.000040398 |
| Eif3b | 0.000020214 | 0.76070077 | 0.719 | 0.446 | 0.49104376 | 0.08745187 | 0.27510724 | 0.685 | 0.656 | 1 | 0.08745187 | 0.000040428 |
| Gal3st4 | 0.00002036 | 2.1132892 | 0.375 | 0.166 | 0.49458628 | 0.01468384 | 0.73321346 | 0.294 | 0.156 | 1 | 0.01468384 | 0.00004072 |
| Pgm3 | 0.0096141 | 0.45576837 | 0.75 | 0.611 | 1 | 0.000020442 | 0.68634317 | 0.689 | 0.385 | 0.49656556 | 0.0096141 | 0.000040883 |
| Ppan | 0.000020467 | 0.86851812 | 0.573 | 0.306 | 0.49719117 | 0.000096782 | 0.77106473 | 0.487 | 0.229 | 1 | 9.6782E-05 | 0.000040934 |
| U2af1l4 | 0.44046999 | 0.31561683 | 0.312 | 0.28 | 1 | 0.000020575 | 1.91253716 | 0.29 | 0.073 | 0.49981647 | 0.44046999 | 0.00004115 |
| Eif3l | 0.00002062 | 0.76422339 | 0.792 | 0.539 | 0.5009005 | 0.00429934 | 0.30838339 | 0.752 | 0.604 | 1 | 0.00429934 | 0.00004124 |
| Mbd3 | 0.11347187 | 0.20680334 | 0.635 | 0.492 | 1 | 0.000020674 | 0.43621254 | 0.672 | 0.354 | 0.50221125 | 0.11347187 | 0.000041347 |
| Elp3 | 0.00037852 | 0.88501053 | 0.521 | 0.321 | 1 | 0.000020692 | 0.89367813 | 0.441 | 0.177 | 0.50265832 | 0.00037852 | 0.000041384 |
| Dhx57 | 0.00002074 | 0.68556644 | 0.531 | 0.254 | 0.50381464 | 0.29063491 | 0.13157993 | 0.462 | 0.385 | 1 | 0.29063491 | 0.000041479 |
| Apof | 0.00002092 | -2.7825824 | 0.042 | 0.244 | 0.50818401 | 0.27154327 | 1.01207283 | 0.013 | 0 | 1 | 0.27154327 | 0.000041839 |
| Zfp800 | 0.0080793 | 0.44390569 | 0.677 | 0.523 | 1 | 0.000020929 | 0.87431166 | 0.756 | 0.552 | 0.50840358 | 0.0080793 | 0.000041857 |
| Nt5dc1 | 0.000020966 | -2.1954455 | 0.365 | 0.57 | 0.5093116 | 0.81360074 | -0.5936482 | 0.332 | 0.333 | 1 | 0.81360074 | 0.000041932 |
| Scn1b | 0.000051694 | 1.08278266 | 0.562 | 0.332 | 1 | 0.000021038 | 0.8995981 | 0.588 | 0.312 | 0.51104354 | 5.1694E-05 | 0.000042075 |
| Eif4g1 | 0.000055774 | 0.45530237 | 0.99 | 0.969 | 1 | 0.000021145 | 0.42529284 | 0.996 | 0.99 | 0.51365817 | 5.5774E-05 | 0.00004229 |
| Pgrmc2 | 0.000021177 | 0.55136202 | 0.823 | 0.601 | 0.5144276 | 0.000022525 | 0.49316941 | 0.861 | 0.615 | 0.5471685 | 2.2525E-05 | 0.000042353 |
| Itpr1 | 0.55433025 | -0.1474069 | 0.854 | 0.865 | 1 | 0.000021351 | -0.7844095 | 0.966 | 1 | 0.51866966 | 0.55433025 | 0.000042702 |
| 2610037D02Rik | 0.000021365 | 1.17741954 | 0.5 | 0.249 | 0.51899408 | 0.66882534 | -0.6263289 | 0.55 | 0.552 | 1 | 0.66882534 | 0.000042729 |
| Pygl | 0.000021378 | 1.15751998 | 0.49 | 0.254 | 0.51930592 | 0.1938164 | -0.1723517 | 0.311 | 0.219 | 1 | 0.1938164 | 0.000042755 |
| Ino80e | 0.0904873 | 0.37946331 | 0.354 | 0.254 | 1 | 0.000021484 | 1.28770727 | 0.395 | 0.146 | 0.52190008 | 0.0904873 | 0.000042968 |
| Borcs7 | 0.00656063 | 0.73447604 | 0.458 | 0.306 | 1 | 0.000021507 | 1.6361049 | 0.366 | 0.135 | 0.52244886 | 0.00656063 | 0.000043014 |
| Map7d1 | 0.27642083 | 0.18462222 | 0.844 | 0.767 | 1 | 0.000021687 | 0.50735866 | 0.924 | 0.823 | 0.5268235 | 0.27642083 | 0.000043374 |
| Arhgap18 | 0.26281046 | -0.9185049 | 0.031 | 0.062 | 1 | 0.000021835 | -1.3390016 | 0.105 | 0.302 | 0.53042753 | 0.26281046 | 0.00004367 |
| Ppp3r1 | 0.02049964 | 0.43740531 | 0.708 | 0.596 | 1 | 0.000021908 | 0.66438164 | 0.803 | 0.667 | 0.53218188 | 0.02049964 | 0.000043815 |
| Ptma | 0.01021182 | 0.18895909 | 1 | 0.984 | 1 | 0.000022027 | 0.46110407 | 1 | 0.99 | 0.53507801 | 0.01021182 | 0.000044053 |
| Serbp1 | 0.000022092 | 0.36067357 | 1 | 0.964 | 0.53665864 | 0.00059326 | 0.2307918 | 0.983 | 0.969 | 1 | 0.00059326 | 0.000044183 |
| Pold4 | 0.62194001 | -0.2104105 | 0.365 | 0.383 | 1 | 0.00002213 | 1.61071027 | 0.336 | 0.104 | 0.53757496 | 0.62194001 | 0.000044259 |
| Hp1bp3 | 0.03125115 | 0.24850264 | 0.958 | 0.87 | 1 | 0.000022227 | 0.40973838 | 0.983 | 0.865 | 0.53993675 | 0.03125115 | 0.000044453 |
| Ppp1r3c | 0.000022274 | -1.6107441 | 0.302 | 0.513 | 0.54108487 | 0.92539297 | -0.352924 | 0.231 | 0.208 | 1 | 0.92539297 | 0.000044548 |
| Rab11a | 0.05273991 | -0.2409714 | 0.688 | 0.788 | 1 | 0.000022297 | 0.51611534 | 0.861 | 0.771 | 0.54163296 | 0.05273991 | 0.000044593 |
| Bad | 0.77242576 | -0.2185735 | 0.438 | 0.43 | 1 | 0.000022368 | 1.07384903 | 0.475 | 0.198 | 0.5433591 | 0.77242576 | 0.000044735 |
| Nt5dc3 | 0.000022428 | 0.85076903 | 0.75 | 0.544 | 0.54481207 | 0.04237905 | -0.5777885 | 0.697 | 0.781 | 1 | 0.04237905 | 0.000044855 |
| Fem1b | 0.26089998 | 0.24565427 | 0.458 | 0.383 | 1 | 0.000022497 | 0.76814725 | 0.555 | 0.271 | 0.54650421 | 0.26089998 | 0.000044994 |
| Rbm14 | 0.32106722 | 0.6718915 | 0.281 | 0.238 | 1 | 0.0000225 | 0.84542296 | 0.454 | 0.177 | 0.54657171 | 0.32106722 | 0.000045 |
| Mak16 | 0.02101692 | 0.49292136 | 0.542 | 0.415 | 1 | 0.000022548 | 0.84593941 | 0.592 | 0.312 | 0.54774169 | 0.02101692 | 0.000045096 |

| | | | | | | | | | | | |
|---|---|---|---|---|---|---|---|---|---|---|---|
| Sash1 | 0.000022936 | -0.8512563 | 0.885 | 0.974 | 0.557169 | 0.56977427 | -0.2630395 | 0.992 | 0.979 | 1 | 0.56977427 | 0.000045872 |
| Zfp397 | 0.00060807 | 0.74446013 | 0.562 | 0.352 | 1 | 0.000023194 | 1.09822948 | 0.496 | 0.229 | 0.56342355 | 0.00060807 | 0.000046387 |
| Exosc4 | 0.13554605 | 0.60310428 | 0.281 | 0.207 | 1 | 0.000023347 | 2.22263982 | 0.269 | 0.062 | 0.56714665 | 0.13554605 | 0.000046694 |
| Reg3g | 0.4978519 | 0.58059688 | 0.302 | 0.358 | 1 | 0.000023657 | -0.7913882 | 0.387 | 0.594 | 0.57466448 | 0.4978519 | 0.000047313 |
| Armc1 | 0.05963075 | 0.28502851 | 0.583 | 0.446 | 1 | 0.000023721 | 0.82430998 | 0.643 | 0.333 | 0.57623753 | 0.05963075 | 0.000047442 |
| Pmpca | 0.000023912 | 1.0609338 | 0.469 | 0.223 | 0.5808691 | 0.00093351 | 0.44503224 | 0.366 | 0.156 | 1 | 0.00093351 | 0.000047823 |
| Fuz | 0.06423297 | 0.85549144 | 0.292 | 0.197 | 1 | 0.000023976 | 2.46273424 | 0.239 | 0.042 | 0.58243513 | 0.06423297 | 0.000047952 |
| Smim30 | 0.37119377 | 0.1243082 | 0.573 | 0.497 | 1 | 0.000024082 | 0.95863357 | 0.576 | 0.302 | 0.58500228 | 0.37119377 | 0.000048164 |
| Cfap36 | 0.07601594 | 0.2541345 | 0.76 | 0.699 | 1 | 0.000024087 | 0.68049457 | 0.643 | 0.365 | 0.58512737 | 0.07601594 | 0.000048174 |
| Kctd10 | 0.1197535 | 0.52206771 | 0.375 | 0.285 | 1 | 0.000024193 | 0.84814657 | 0.588 | 0.302 | 0.58770389 | 0.1197535 | 0.000048386 |
| Pes1 | 0.20866443 | 0.14314964 | 0.323 | 0.238 | 1 | 0.000024195 | 1.23267197 | 0.387 | 0.135 | 0.58774656 | 0.20866443 | 0.00004839 |
| Max | 0.29784261 | 0.19607224 | 0.844 | 0.86 | 1 | 0.00002433 | 0.41208814 | 0.954 | 0.812 | 0.59101923 | 0.29784261 | 0.000048659 |
| Nudt21 | 0.000024344 | 0.60764344 | 0.875 | 0.658 | 0.59137505 | 0.00035207 | 0.54429387 | 0.836 | 0.708 | 1 | 0.00035207 | 0.000048688 |
| Fkrp | 0.04310169 | 1.1118312 | 0.271 | 0.181 | 1 | 0.000024358 | 1.12077909 | 0.399 | 0.146 | 0.5916943 | 0.04310169 | 0.000048715 |
| Cdk5rap3 | 0.20539387 | 0.4904361 | 0.396 | 0.332 | 1 | 0.000024364 | 1.03591957 | 0.399 | 0.146 | 0.59185749 | 0.20539387 | 0.000048728 |
| Tsr2 | 0.15868472 | 0.56003556 | 0.396 | 0.337 | 1 | 0.000024407 | 0.9808219 | 0.433 | 0.177 | 0.59288967 | 0.15868472 | 0.000048813 |
| Sbds | 0.00002442 | -0.7824853 | 0.76 | 0.85 | 0.59320168 | 0.00329779 | 0.5038861 | 0.836 | 0.812 | 1 | 0.00329779 | 0.000048839 |
| Smoc1 | 0.000024514 | -3.6478573 | 0.031 | 0.223 | 0.59549611 | 0.00123071 | -3.0402482 | 0.046 | 0.146 | 1 | 0.00123071 | 0.000049028 |
| Nipal2 | 0.14200994 | 0.45495351 | 0.104 | 0.057 | 1 | 0.000024604 | -0.612418 | 0.13 | 0.354 | 0.59768697 | 0.14200994 | 0.000049208 |
| Bcap31 | 0.000071518 | 0.42066929 | 0.969 | 0.927 | 1 | 0.000024644 | 0.37578375 | 0.962 | 0.875 | 0.59864037 | 7.1518E-05 | 0.000049286 |
| Atp6v1c2 | 0.000024952 | -3.1010299 | 0.021 | 0.207 | 0.60614138 | 0.21001636 | -1.4618584 | 0.025 | 0.052 | 1 | 0.21001636 | 0.000049904 |
| Slc33a1 | 0.00726445 | 0.74762741 | 0.49 | 0.332 | 1 | 0.000025097 | 0.57676294 | 0.454 | 0.177 | 0.60965451 | 0.00726445 | 0.000050193 |
| Polr1g | 0.00035276 | 1.21031142 | 0.385 | 0.192 | 1 | 0.000025098 | 1.22619764 | 0.357 | 0.115 | 0.60967499 | 0.00035276 | 0.000050195 |
| Rfx2 | 0.000025099 | 1.65602217 | 0.312 | 0.114 | 0.60969279 | 0.60949697 | -0.7651566 | 0.349 | 0.271 | 1 | 0.60949697 | 0.000050196 |
| Nt5c3 | 0.00022895 | 0.79650688 | 0.646 | 0.399 | 1 | 0.000025342 | 0.70557478 | 0.702 | 0.458 | 0.61560175 | 0.00022895 | 0.000050683 |
| Zfpl1 | 0.17371484 | 0.28839536 | 0.469 | 0.373 | 1 | 0.000025698 | 1.31073115 | 0.387 | 0.146 | 0.62424681 | 0.17371484 | 0.000051395 |
| Ggcx | 0.25364491 | 0.33846777 | 0.594 | 0.544 | 1 | 0.000025743 | 0.85453156 | 0.622 | 0.354 | 0.62534094 | 0.25364491 | 0.000051485 |
| Mbnl2 | 0.60174319 | -0.2238463 | 1 | 1 | 1 | 0.000025761 | -0.6536677 | 1 | 1 | 0.62577675 | 0.60174319 | 0.000051521 |
| Epha3 | 0.0046872 | -3.7204259 | 0.021 | 0.119 | 1 | 0.00002584 | -2.6587504 | 0.046 | 0.188 | 0.62770654 | 0.0046872 | 0.000051679 |
| Sgce | 0.00017002 | 1.00749454 | 0.417 | 0.202 | 1 | 0.000026085 | 0.3466605 | 0.508 | 0.198 | 0.63366796 | 0.00017002 | 0.00005217 |
| Pole3 | 0.01836452 | 0.46405131 | 0.573 | 0.456 | 1 | 0.000026123 | 0.78700628 | 0.45 | 0.177 | 0.6345823 | 0.01836452 | 0.000052246 |
| Ihh | 0.000026155 | 4.90561492 | 0.135 | 0.016 | 0.63536934 | 0.20315061 | 3.59703533 | 0.017 | 0 | 1 | 0.20315061 | 0.00005231 |
| Rmdn2 | 0.000056472 | 1.53876917 | 0.417 | 0.228 | 1 | 0.000026172 | 1.77356075 | 0.357 | 0.125 | 0.63575991 | 5.6472E-05 | 0.000052342 |
| Zcrb1 | 0.41519698 | 0.14275437 | 0.875 | 0.876 | 1 | 0.000026371 | 0.47564023 | 0.895 | 0.812 | 0.64060787 | 0.41519698 | 0.000052742 |
| Casd1 | 0.01586231 | 0.49468182 | 0.5 | 0.342 | 1 | 0.000026493 | 0.66996686 | 0.508 | 0.229 | 0.64355945 | 0.01586231 | 0.000052985 |
| Arhgef3 | 0.06715492 | -0.7948248 | 0.229 | 0.326 | 1 | 0.00002654 | -1.0524674 | 0.282 | 0.552 | 0.64471623 | 0.06715492 | 0.00005308 |
| Igf2 | 0.000026545 | 0.90561492 | 0.271 | 0.078 | 0.64484097 | 0.27103085 | 0.11464257 | 0.143 | 0.094 | 1 | 0.27103085 | 0.00005309 |
| Txndc15 | 0.41096323 | -0.1873844 | 0.708 | 0.71 | 1 | 0.000026607 | 0.45762164 | 0.811 | 0.562 | 0.64634327 | 0.41096323 | 0.000053214 |
| Ppfibp2 | 0.000074197 | -3.747393 | 0.01 | 0.171 | 1 | 0.000026645 | -2.7248928 | 0.071 | 0.229 | 0.64725493 | 7.4197E-05 | 0.000053289 |
| Gga2 | 0.86210139 | -0.1981805 | 0.385 | 0.383 | 1 | 0.000026899 | 0.9781255 | 0.508 | 0.229 | 0.65341848 | 0.86210139 | 0.000053796 |
| Cdk6 | 0.000027039 | -1.265246 | 0.583 | 0.736 | 0.65682588 | 0.000068351 | -0.8662486 | 0.387 | 0.646 | 1 | 6.8351E-05 | 0.000054077 |
| Dars | 0.00082364 | 0.72620843 | 0.719 | 0.57 | 1 | 0.000027095 | 0.83824338 | 0.676 | 0.469 | 0.65819875 | 0.00082364 | 0.00005419 |
| Cachd1 | 0.60128082 | -0.3581549 | 0.177 | 0.145 | 1 | 0.000027279 | -1.4101602 | 0.282 | 0.51 | 0.66265672 | 0.60128082 | 0.000054557 |
| Smarca1 | 0.00280983 | 0.33764314 | 0.552 | 0.352 | 1 | 0.000027364 | 0.76372587 | 0.63 | 0.344 | 0.66471694 | 0.00280983 | 0.000054726 |
| Tst | 0.00361984 | 1.16503581 | 0.219 | 0.093 | 1 | 0.000027482 | -1.2345671 | 0.197 | 0.427 | 0.66759205 | 0.00361984 | 0.000054963 |
| Smim7 | 0.43708262 | 0.11835513 | 0.823 | 0.736 | 1 | 0.000027707 | 0.62240013 | 0.807 | 0.542 | 0.67306705 | 0.43708262 | 0.000055414 |
| Sdhaf4 | 0.30046602 | 0.28760246 | 0.49 | 0.435 | 1 | 0.000028046 | 0.51903282 | 0.42 | 0.156 | 0.6812966 | 0.30046602 | 0.000056091 |
| Lratd2 | 0.00742159 | 0.81045769 | 0.281 | 0.145 | 1 | 0.000028085 | 1.40384055 | 0.387 | 0.146 | 0.6822364 | 0.00742159 | 0.000056169 |
| Ppt2 | 0.4875096 | 0.30952307 | 0.292 | 0.254 | 1 | 0.000028106 | 0.89401707 | 0.42 | 0.167 | 0.68274882 | 0.4875096 | 0.000056211 |
| Stk11 | 0.000028117 | 0.66991318 | 0.719 | 0.425 | 0.68300643 | 0.01688324 | 0.34599072 | 0.794 | 0.708 | 1 | 0.01688324 | 0.000056232 |
| Rpa3 | 0.000028117 | 1.64273024 | 0.427 | 0.218 | 0.68302287 | 0.089922 | 0.35640734 | 0.252 | 0.156 | 1 | 0.089922 | 0.000056234 |
| Trak1 | 0.21721462 | -0.3509594 | 0.604 | 0.668 | 1 | 0.000028126 | -1.021179 | 0.735 | 0.865 | 0.68324101 | 0.21721462 | 0.000056252 |
| Polr1f | 0.00223357 | 0.70199199 | 0.562 | 0.404 | 1 | 0.000028167 | 1.07007844 | 0.504 | 0.26 | 0.68422844 | 0.00223357 | 0.000056333 |
| Hsbp1 | 0.07560919 | 0.24120227 | 0.875 | 0.839 | 1 | 0.000028212 | 0.52402054 | 0.866 | 0.729 | 0.68532111 | 0.07560919 | 0.000056423 |
| Isoc1 | 0.000028309 | 1.10877787 | 0.458 | 0.218 | 0.68768535 | 0.00553176 | 0.49749966 | 0.45 | 0.26 | 1 | 0.00553176 | 0.000056617 |
| Tmem9 | 0.00334416 | 0.74121447 | 0.5 | 0.337 | 1 | 0.000028717 | 0.76543286 | 0.546 | 0.26 | 0.69760036 | 0.00334416 | 0.000057434 |
| Zbtb33 | 0.000028804 | 1.57139542 | 0.417 | 0.197 | 0.69970739 | 0.00065958 | 1.02312802 | 0.357 | 0.167 | 1 | 0.00065958 | 0.000057607 |
| 2600014E21Rik | 0.000028841 | -1.5015191 | 0.031 | 0.228 | 0.70060538 | 0.82047853 | 0.74903843 | 0.025 | 0.021 | 1 | 0.82047853 | 0.000057681 |
| Tlr2 | 0.06306452 | 0.85549144 | 0.25 | 0.161 | 1 | 0.000029073 | 1.88278982 | 0.336 | 0.115 | 0.7062392 | 0.06306452 | 0.000058145 |
| Col8a2 | 0.000029081 | 0.59245704 | 0.719 | 0.425 | 0.70643701 | 0.00031082 | 0.32886514 | 0.929 | 0.625 | 1 | 0.00031082 | 0.000058161 |
| Slc22a17 | 0.76890688 | -0.2045559 | 0.427 | 0.43 | 1 | 0.000029101 | 0.74459252 | 0.479 | 0.208 | 0.70691584 | 0.76890688 | 0.000058201 |
| Nudcd2 | 0.000029193 | 1.00749454 | 0.646 | 0.425 | 0.70914774 | 0.00015824 | 0.73196491 | 0.517 | 0.25 | 1 | 0.00015824 | 0.000058384 |
| Pithd1 | 0.23296115 | 0.17967551 | 0.375 | 0.29 | 1 | 0.000029198 | 1.26239619 | 0.378 | 0.135 | 0.70927905 | 0.23296115 | 0.000058395 |
| Rnf187 | 0.000045398 | 0.55890448 | 0.833 | 0.617 | 1 | 0.000029232 | 0.60853097 | 0.777 | 0.552 | 0.71011052 | 4.5398E-05 | 0.000058464 |
| Vat1 | 0.16860995 | 0.26556269 | 0.385 | 0.301 | 1 | 0.000029374 | 0.95223958 | 0.592 | 0.323 | 0.71355532 | 0.16860995 | 0.000058747 |
| Bloc1s1 | 0.63915881 | -0.1329867 | 0.365 | 0.378 | 1 | 0.000029405 | 1 | 0.416 | 0.156 | 0.71431707 | 0.63915881 | 0.00005881 |
| Arl6ip5 | 0.12873975 | -0.206893 | 1 | 0.99 | 1 | 0.000029586 | 0.38590056 | 1 | 1 | 0.71871442 | 0.12873975 | 0.000059172 |
| Erp27 | 0.00002963 | 1.3885847 | 0.292 | 0.098 | 0.71976224 | 0.22117765 | -0.3368223 | 0.164 | 0.104 | 1 | 0.22117765 | 0.000059258 |
| Tomm7 | 0.00180525 | 0.42548535 | 0.938 | 0.87 | 1 | 0.000029813 | 0.58199593 | 0.866 | 0.76 | 0.72422307 | 0.00180525 | 0.000059626 |
| Npdc1 | 0.00188876 | 0.41105925 | 0.927 | 0.834 | 1 | 0.000029874 | 0.55554027 | 0.895 | 0.604 | 0.72569379 | 0.00188876 | 0.000059747 |
| Dbp | 0.03013568 | 0.6878766 | 0.531 | 0.383 | 1 | 0.00003002 | 1.21560623 | 0.412 | 0.156 | 0.72923517 | 0.03013568 | 0.000060038 |
| Hspa4l | 0.01005196 | -1.0033938 | 0.135 | 0.275 | 1 | 0.000030046 | -0.8314918 | 0.269 | 0.542 | 0.72988698 | 0.01005196 | 0.000060092 |
| Lemd2 | 0.02298443 | 0.6241659 | 0.479 | 0.342 | 1 | 0.000030189 | 0.65025201 | 0.576 | 0.281 | 0.73335504 | 0.02298443 | 0.000060377 |
| Tpm3 | 0.00191536 | 0.34294675 | 0.969 | 0.984 | 1 | 0.00003028 | 0.41087129 | 1 | 1 | 0.73555749 | 0.00191536 | 0.000060559 |
| Vdac3 | 0.00740711 | 0.44212276 | 0.802 | 0.663 | 1 | 0.00003029 | 0.59622687 | 0.845 | 0.708 | 0.73580908 | 0.00740711 | 0.000060579 |
| Vamp2 | 0.000030385 | 0.89878828 | 0.646 | 0.425 | 0.73810652 | 0.00016837 | 0.56864373 | 0.592 | 0.312 | 1 | 0.00016837 | 0.000060769 |
| Bex2 | 0.000030463 | 3.38600616 | 0.146 | 0.021 | 0.74000391 | 0.02895661 | 0.84542296 | 0.139 | 0.052 | 1 | 0.02895661 | 0.000060925 |
| Tpd52 | 0.01810279 | 1.13302542 | 0.281 | 0.171 | 1 | 0.000030591 | -1.171149 | 0.298 | 0.552 | 0.74312326 | 0.01810279 | 0.000061182 |
| Stk17b | 0.20329233 | -0.8639907 | 0.271 | 0.311 | 1 | 0.000030938 | -1.6079366 | 0.349 | 0.594 | 0.75155633 | 0.20329233 | 0.000061876 |
| Vps37b | 0.01475512 | 0.47884618 | 0.885 | 0.798 | 1 | 0.000031099 | 0.42435597 | 0.945 | 0.885 | 0.75546242 | 0.01475512 | 0.000062197 |
| Adam17 | 0.41602588 | 0.21281344 | 0.469 | 0.409 | 1 | 0.000031173 | -1.3098553 | 0.55 | 0.75 | 0.75724566 | 0.41602588 | 0.000062344 |
| Igsf10 | 0.000031613 | 0.81644624 | 0.469 | 0.228 | 0.76795165 | 0.000046238 | 1.20471791 | 0.454 | 0.198 | 1 | 4.6238E-05 | 0.000063226 |
| Galnt18 | 0.000031695 | -1.5003962 | 0.688 | 0.793 | 0.7699362 | 0.16880618 | -0.1278111 | 0.773 | 0.74 | 1 | 0.16880618 | 0.000063389 |
| Slc35f5 | 0.03990779 | 0.3450759 | 0.719 | 0.606 | 1 | 0.000031843 | 0.54862815 | 0.777 | 0.49 | 0.77354195 | 0.03990779 | 0.000063686 |
| Ywhaz | 0.00072619 | 0.35731438 | 1 | 1 | 1 | 0.00003203 | 0.37419445 | 1 | 1 | 0.77806569 | 0.00072619 | 0.000064058 |
| Dnttip2 | 0.000032153 | 0.64723832 | 0.833 | 0.606 | 0.78105609 | 0.00116737 | 0.34980461 | 0.773 | 0.615 | 1 | 0.00116737 | 0.000064305 |
| Sowahc | 0.00013507 | -1.3581549 | 0.385 | 0.565 | 1 | 0.000032634 | 1.28125947 | 0.399 | 0.156 | 0.7927332 | 0.00013507 | 0.000065266 |
| Itga8 | 0.0159605 | -2.6169963 | 0.021 | 0.098 | 1 | 0.000032677 | -3.2818409 | 0.038 | 0.167 | 0.79380173 | 0.0159605 | 0.000065354 |
| Mfsd5 | 0.03639427 | 0.67045955 | 0.323 | 0.207 | 1 | 0.000032938 | 1.53341923 | 0.353 | 0.125 | 0.80012411 | 0.03639427 | 0.000065874 |
| Ubl4a | 0.01935451 | 0.53800925 | 0.396 | 0.259 | 1 | 0.000033031 | 1.3146356 | 0.353 | 0.115 | 0.80238009 | 0.01935451 | 0.00006606 |
| Ppm1g | 0.00003306 | 1.04171025 | 0.667 | 0.425 | 0.80310149 | 0.1921702 | 0.1515927 | 0.71 | 0.667 | 1 | 0.1921702 | 0.00006612 |

| | | | | | | | | | | | | |
|---|---|---|---|---|---|---|---|---|---|---|---|---|
| Shfl | 0.0009865 | 1.40604391 | 0.229 | 0.088 | 1 | 0.000033256 | 2.66742466 | 0.202 | 0.021 | 0.80786278 | 0.0009865 | 0.000066512 |
| Hes1 | 0.000033305 | -1.702504 | 0.208 | 0.456 | 0.80905134 | 0.01714852 | -0.6168918 | 0.504 | 0.698 | 1 | 0.01714852 | 0.000066609 |
| Dyrk2 | 0.00018714 | 1.11440974 | 0.427 | 0.223 | 1 | 0.000033526 | 1.24957215 | 0.504 | 0.26 | 0.81440868 | 0.00018714 | 0.00006705 |
| Foxp2 | 0.00003421 | 1.1626136 | 0.49 | 0.269 | 0.8310386 | 0.38530005 | -0.6145323 | 0.412 | 0.438 | 1 | 0.38530005 | 0.00006842 |
| Rrp1 | 0.000074845 | 0.65352219 | 0.823 | 0.637 | 1 | 0.000034238 | 0.65757687 | 0.777 | 0.667 | 0.83171437 | 7.4845E-05 | 0.000068475 |
| Zfp395 | 0.000034344 | 1.28125535 | 0.417 | 0.192 | 0.83428618 | 0.0932535 | 0.29104878 | 0.378 | 0.26 | 1 | 0.0932535 | 0.000068687 |
| Srpr | 0.01343243 | 0.24128284 | 0.979 | 0.974 | 1 | 0.000034672 | 0.45150956 | 0.971 | 0.958 | 0.84225569 | 0.01343243 | 0.000069343 |
| Tada3 | 0.04639725 | 0.68556644 | 0.281 | 0.181 | 1 | 0.000034689 | 1.89525917 | 0.269 | 0.062 | 0.84267202 | 0.04639725 | 0.000069377 |
| Tmem97 | 0.00011413 | 0.9534547 | 0.615 | 0.409 | 1 | 0.000034703 | 1.33703499 | 0.336 | 0.104 | 0.84300365 | 0.00011413 | 0.000069405 |
| Scn2b | 0.00003492 | 3.91438513 | 0.104 | 0.005 | 0.84827091 | 0.36435013 | 0.59703533 | 0.055 | 0.031 | 1 | 0.36435013 | 0.000069838 |
| S100a6 | 0.000034928 | -0.8187898 | 0.875 | 0.969 | 0.84847941 | 0.04854688 | 0.23366768 | 0.933 | 0.938 | 1 | 0.04854688 | 0.000069855 |
| Mrpl12 | 0.0218964 | 0.50302195 | 0.562 | 0.43 | 1 | 0.00003517 | 0.93807225 | 0.542 | 0.281 | 0.85435541 | 0.0218964 | 0.000070339 |
| Dennd2c | 0.07989077 | 0.69715442 | 0.365 | 0.264 | 1 | 0.000035171 | -0.9857263 | 0.319 | 0.573 | 0.85437793 | 0.07989077 | 0.000070341 |
| Ccdc86 | 0.000035282 | 0.96018882 | 0.562 | 0.306 | 0.85706544 | 0.0000677 | 1.54258755 | 0.37 | 0.156 | 1 | 0.0000677 | 0.000070562 |
| Mob1a | 0.000036102 | 0.77045534 | 0.656 | 0.399 | 0.87699038 | 0.00071073 | 0.46656148 | 0.748 | 0.604 | 1 | 0.00071073 | 0.000072203 |
| Tmem213 | 0.32025586 | -0.9925055 | 0 | 0.01 | 1 | 0.000036229 | 1.30279897 | 0.408 | 0.156 | 0.8800628 | 0.32025586 | 0.000072456 |
| Nfkbia | 0.70443886 | -0.4262622 | 0.625 | 0.617 | 1 | 0.000036402 | -1.2472103 | 0.697 | 0.885 | 0.88426985 | 0.70443886 | 0.000072802 |
| Nbn | 0.00951193 | 0.66645762 | 0.354 | 0.202 | 1 | 0.000036459 | 1.00242266 | 0.399 | 0.146 | 0.88565629 | 0.00951193 | 0.000072916 |
| S1pr1 | 0.000036614 | -2.4616196 | 0.156 | 0.373 | 0.88943366 | 0.61913845 | -0.733663 | 0.193 | 0.198 | 1 | 0.61913845 | 0.000073227 |
| Ppp5c | 0.02084886 | 0.4850735 | 0.438 | 0.285 | 1 | 0.000036687 | 0.69014474 | 0.424 | 0.167 | 0.89119979 | 0.02084886 | 0.000073373 |
| Gm47237 | 0.000036691 | 2.81484946 | 0.156 | 0.026 | 0.89129763 | 0.01911531 | 0.95863357 | 0.16 | 0.062 | 1 | 0.01911531 | 0.000073381 |
| Arsb | 0.09692911 | 0.2466822 | 0.312 | 0.223 | 1 | 0.000036751 | 2.57911343 | 0.408 | 0.167 | 0.89276707 | 0.09692911 | 0.000073502 |
| Tbc1d10b | 0.13266828 | 0.36113149 | 0.188 | 0.119 | 1 | 0.000036794 | 1.20110666 | 0.378 | 0.135 | 0.89381081 | 0.13266828 | 0.000073588 |
| Cds1 | 0.01566408 | -0.2244522 | 0.323 | 0.477 | 1 | 0.000037115 | -0.8083088 | 0.256 | 0.51 | 0.90160172 | 0.01566408 | 0.000074229 |
| Mthfd2l | 0.02704174 | 0.39314823 | 0.49 | 0.352 | 1 | 0.000037158 | 0.83310269 | 0.504 | 0.219 | 0.90265244 | 0.02704174 | 0.000074315 |
| Eno1 | 0.000060259 | 0.5989586 | 0.906 | 0.725 | 1 | 0.000037245 | 0.60216557 | 0.824 | 0.729 | 0.90475318 | 6.0259E-05 | 0.000074488 |
| 1600012H06Rik | 0.00039663 | 1.05996196 | 0.438 | 0.244 | 1 | 0.000037335 | 0.95863357 | 0.374 | 0.125 | 0.90695291 | 0.00039663 | 0.00007467 |
| Sgcd | 0.000037414 | 0.24752021 | 0.896 | 0.705 | 0.90885938 | 0.35279923 | -0.5527587 | 0.924 | 0.875 | 1 | 0.35279923 | 0.000074826 |
| Mzt1 | 0.0000375 | 0.75551532 | 0.708 | 0.497 | 0.91095101 | 0.00015927 | 0.57310532 | 0.672 | 0.417 | 1 | 0.00015927 | 0.000074999 |
| Arsk | 0.0009464 | 1.27542774 | 0.375 | 0.207 | 1 | 0.000037751 | 0.69014474 | 0.399 | 0.146 | 0.91705164 | 0.0009464 | 0.000075501 |
| Pdha1 | 0.27027799 | 0.23837842 | 0.667 | 0.585 | 1 | 0.000037926 | 0.53753432 | 0.815 | 0.719 | 0.92130134 | 0.27027799 | 0.000075851 |
| Serinc5 | 0.10699825 | 0.34904097 | 0.875 | 0.751 | 1 | 0.000037987 | 0.32143629 | 0.945 | 0.74 | 0.92277386 | 0.10699825 | 0.000075972 |
| Serpinb6a | 0.03939168 | -0.4448218 | 0.896 | 0.964 | 1 | 0.000038103 | 0.45646023 | 0.937 | 0.792 | 0.92559286 | 0.03939168 | 0.000076204 |
| Etfa | 0.21648767 | 0.14520656 | 0.906 | 0.907 | 1 | 0.000038335 | 0.44367696 | 0.966 | 0.917 | 0.93123519 | 0.21648767 | 0.000076669 |
| Manf | 0.07824512 | 0.14816402 | 0.917 | 0.912 | 1 | 0.000038344 | 0.38404161 | 0.941 | 0.781 | 0.93145714 | 0.07824512 | 0.000076687 |
| Aars | 0.00087833 | 0.5683829 | 0.615 | 0.404 | 1 | 0.000038413 | 0.6944321 | 0.651 | 0.344 | 0.93313162 | 0.00087833 | 0.000076825 |
| Large1 | 0.57162122 | -0.2365405 | 0.771 | 0.736 | 1 | 0.000038712 | -0.7973694 | 0.836 | 0.948 | 0.94038779 | 0.57162122 | 0.000077422 |
| Snip1 | 0.000039042 | 1.21771224 | 0.438 | 0.218 | 0.94841783 | 0.001674 | 0.56461386 | 0.395 | 0.198 | 1 | 0.001674 | 0.000078083 |
| Rnf7 | 0.04552915 | 0.27879656 | 0.792 | 0.736 | 1 | 0.000039309 | 0.49213647 | 0.87 | 0.688 | 0.95488772 | 0.04552915 | 0.000078616 |
| Kctd4 | 0.44557618 | 0.97869727 | 0.542 | 0.472 | 1 | 0.000039661 | 0.41142871 | 0.492 | 0.208 | 0.96345584 | 0.44557618 | 0.000079321 |
| C1qtnf1 | 0.04489191 | 0.38234295 | 0.396 | 0.269 | 1 | 0.000039904 | 0.88190554 | 0.475 | 0.208 | 0.96934346 | 0.04489191 | 0.000079806 |
| Ghitm | 0.03806666 | 0.19576117 | 0.875 | 0.798 | 1 | 0.000039944 | 0.55956063 | 0.874 | 0.771 | 0.97031433 | 0.03806666 | 0.000079886 |
| Ppp1r11 | 0.000040033 | 0.73026625 | 0.719 | 0.425 | 0.97247672 | 0.00037151 | 0.72507853 | 0.559 | 0.333 | 1 | 0.00037151 | 0.000080064 |
| Abcc1 | 0.26577995 | -0.3144336 | 0.146 | 0.197 | 1 | 0.000040045 | -0.973988 | 0.361 | 0.615 | 0.97277562 | 0.26577995 | 0.000080089 |
| Dnm1 | 0.000040362 | 0.71697939 | 0.677 | 0.394 | 0.98047497 | 0.00014942 | 0.48689156 | 0.798 | 0.5 | 1 | 0.00014942 | 0.000080722 |
| Cgnl1 | 0.000040507 | -1.5852036 | 0.208 | 0.446 | 0.98400274 | 0.61880897 | -0.2010188 | 0.391 | 0.333 | 1 | 0.61880897 | 0.000081013 |
| Fbxo9 | 0.76404329 | -0.211674 | 0.5 | 0.503 | 1 | 0.00004086 | 0.70900376 | 0.576 | 0.271 | 0.99255964 | 0.76404329 | 0.000081717 |
| Dmac1 | 0.10595399 | 0.36837627 | 0.458 | 0.352 | 1 | 0.000041009 | 1.47304662 | 0.37 | 0.146 | 0.99619972 | 0.10595399 | 0.000082017 |
| Trmt61a | 0.000041399 | 2.00749454 | 0.208 | 0.052 | 1 | 0.44357873 | 0.27510724 | 0.088 | 0.062 | 1 | 0.44357873 | 0.000082796 |
| Nup35 | 0.00502072 | 0.89045512 | 0.469 | 0.311 | 1 | 0.000041616 | 0.87549978 | 0.403 | 0.156 | 1 | 0.00502072 | 0.000083231 |
| Utp20 | 0.000041659 | 1.29926066 | 0.438 | 0.212 | 1 | 0.00044517 | 0.75427507 | 0.387 | 0.177 | 1 | 0.00044517 | 0.000083317 |
| Luc7l2 | 0.000041691 | 0.34898606 | 1 | 0.99 | 1 | 0.278777 | -0.1891414 | 1 | 1 | 1 | 0.278777 | 0.000083379 |
| Skp1 | 0.55074851 | -0.1308424 | 0.979 | 0.933 | 1 | 0.000042019 | 0.40075327 | 0.983 | 0.927 | 1 | 0.55074851 | 0.000084036 |
| Tipin | 0.00639336 | 0.77801269 | 0.49 | 0.337 | 1 | 0.000042322 | 0.90587343 | 0.412 | 0.167 | 1 | 0.00639336 | 0.000084642 |
| Ube2d3 | 0.14625218 | 0.13212861 | 1 | 0.99 | 1 | 0.000042438 | 0.26087585 | 0.996 | 0.99 | 1 | 0.14625218 | 0.000084875 |
| Tmcc3 | 0.000042677 | 1.5138472 | 0.365 | 0.161 | 1 | 0.86352123 | -0.6965305 | 0.5 | 0.542 | 1 | 0.86352123 | 0.000085351 |
| Scnn1a | 0.78117751 | -0.577468 | 0.021 | 0.026 | 1 | 0.000042809 | -1.561394 | 0.076 | 0.24 | 1 | 0.78117751 | 0.000085615 |
| Anxa2 | 0.15504632 | -0.213658 | 0.99 | 1 | 1 | 0.000042811 | 0.37049039 | 1 | 0.99 | 1 | 0.15504632 | 0.000085619 |
| Mrpl3 | 0.03319565 | 0.51196712 | 0.49 | 0.363 | 1 | 0.000043142 | 0.92460999 | 0.5 | 0.24 | 1 | 0.03319565 | 0.000086282 |
| Alg6 | 0.000043197 | 1.39744105 | 0.333 | 0.13 | 1 | 0.00302991 | 1.8917786 | 0.16 | 0.042 | 1 | 0.00302991 | 0.000086393 |
| Fastk | 0.08232866 | 0.45647937 | 0.521 | 0.435 | 1 | 0.000043222 | 0.926033 | 0.508 | 0.24 | 1 | 0.08232866 | 0.000086442 |
| Nans | 0.00141162 | 0.75751628 | 0.51 | 0.332 | 1 | 0.000043271 | 0.96174116 | 0.471 | 0.198 | 1 | 0.00141162 | 0.000086541 |
| Rrbp1 | 0.00012751 | 0.29499395 | 1 | 1 | 1 | 0.000043287 | 0.29205364 | 1 | 1 | 1 | 0.00012751 | 0.000086572 |
| C1qbp | 0.00025939 | 0.61434964 | 0.781 | 0.601 | 1 | 0.000043379 | 0.64821406 | 0.777 | 0.635 | 1 | 0.00025939 | 0.000086755 |
| Malsu1 | 0.71663516 | 0.10370985 | 0.531 | 0.554 | 1 | 0.000043411 | 0.78466234 | 0.534 | 0.271 | 1 | 0.71663516 | 0.000086821 |
| Park7 | 0.00794485 | 0.27052894 | 0.938 | 0.845 | 1 | 0.000043416 | 0.4669506 | 0.874 | 0.781 | 1 | 0.00794485 | 0.000086831 |
| Gm16638 | 0.00737841 | 0.35232303 | 0.5 | 0.321 | 1 | 0.000043566 | 0.81872894 | 0.618 | 0.323 | 1 | 0.00737841 | 0.000087131 |
| Rmc1 | 0.21036528 | 0.30705482 | 0.271 | 0.202 | 1 | 0.000043851 | 1.02165888 | 0.416 | 0.167 | 1 | 0.21036528 | 0.000087699 |
| Chek1 | 0.000043991 | 1.78510212 | 0.26 | 0.083 | 1 | 0.49682635 | -0.1878647 | 0.223 | 0.177 | 1 | 0.49682635 | 0.00008798 |
| Nfil3 | 0.11601234 | -0.9429234 | 0.229 | 0.306 | 1 | 0.000044026 | -1.0996376 | 0.298 | 0.531 | 1 | 0.11601234 | 0.000088049 |
| Efnb2 | 0.06415719 | 0.40676472 | 0.375 | 0.269 | 1 | 0.00004403 | -1.2294366 | 0.324 | 0.583 | 1 | 0.06415719 | 0.000088057 |
| Ufc1 | 0.00613314 | 0.25284244 | 0.885 | 0.788 | 1 | 0.000044133 | 0.55978394 | 0.828 | 0.688 | 1 | 0.00613314 | 0.000088265 |
| Cxadr | 0.00019515 | -1.5351762 | 0.292 | 0.503 | 1 | 0.000044275 | -0.4954109 | 0.235 | 0.5 | 1 | 0.00019515 | 0.000088549 |
| Npas2 | 0.000044387 | -2.0264528 | 0.219 | 0.44 | 1 | 0.00122873 | -1.233234 | 0.282 | 0.458 | 1 | 0.00122873 | 0.000088771 |
| Lims2 | 0.42328333 | -0.2898914 | 0.427 | 0.446 | 1 | 0.000044861 | 0.48900487 | 0.55 | 0.25 | 1 | 0.42328333 | 0.00008972 |
| Nxn | 0.00072784 | -1.3144336 | 0.417 | 0.622 | 1 | 0.000044897 | -1.404665 | 0.542 | 0.76 | 1 | 0.00072784 | 0.000089791 |
| Mrpl41 | 0.49263375 | 0.18549284 | 0.417 | 0.378 | 1 | 0.000045249 | 1.18199783 | 0.399 | 0.156 | 1 | 0.49263375 | 0.000090496 |
| Eprs | 0.000045262 | 0.54092674 | 0.979 | 0.948 | 1 | 0.00047109 | 0.4317567 | 0.962 | 0.938 | 1 | 0.00047109 | 0.000090523 |
| Tor2a | 0.31598486 | 0.19120649 | 0.385 | 0.321 | 1 | 0.000045275 | 1.23446525 | 0.361 | 0.135 | 1 | 0.31598486 | 0.000090548 |
| Igdcc4 | 0.00761802 | 0.61606959 | 0.479 | 0.306 | 1 | 0.000045419 | 0.73347217 | 0.458 | 0.188 | 1 | 0.00761802 | 0.000090835 |
| Qdpr | 0.35443288 | -0.3450219 | 0.469 | 0.508 | 1 | 0.000045676 | 0.84934333 | 0.45 | 0.188 | 1 | 0.35443288 | 0.000091349 |
| Pde7b | 0.33139553 | -0.5290508 | 0.24 | 0.176 | 1 | 0.000045687 | -1.5032186 | 0.197 | 0.406 | 1 | 0.33139553 | 0.000091372 |
| Yars2 | 0.00700429 | 1.06638823 | 0.229 | 0.109 | 1 | 0.000046084 | 2.19264508 | 0.244 | 0.052 | 1 | 0.00700429 | 0.000092166 |
| Sulf2 | 0.000046239 | 1.13006118 | 0.573 | 0.321 | 1 | 0.00019692 | 0.53814164 | 0.437 | 0.198 | 1 | 0.00019692 | 0.000092476 |
| Tapbp | 0.000046371 | -0.6915032 | 0.708 | 0.86 | 1 | 0.2864092 | -0.320297 | 0.748 | 0.792 | 1 | 0.2864092 | 0.00009274 |
| Epb41l2 | 0.000046386 | 0.42379479 | 0.917 | 0.725 | 1 | 0.0616001 | -0.5330024 | 0.891 | 0.875 | 1 | 0.0616001 | 0.000092769 |
| Dnajb11 | 0.00064012 | 0.51564144 | 0.833 | 0.658 | 1 | 0.000046675 | 0.51505803 | 0.849 | 0.667 | 1 | 0.00064012 | 0.000093348 |
| Postn | 0.00004677 | -2.5971503 | 0.229 | 0.456 | 1 | 0.22657174 | -3.8017084 | 0.088 | 0.125 | 1 | 0.22657174 | 0.000093538 |
| Rufy3 | 0.02580141 | -0.8069498 | 0.354 | 0.461 | 1 | 0.000046827 | -0.7819237 | 0.487 | 0.75 | 1 | 0.02580141 | 0.000093652 |
| Gsdme | 0.000046956 | 1.08445652 | 0.5 | 0.264 | 1 | 0.18074732 | -0.5493212 | 0.403 | 0.281 | 1 | 0.18074732 | 0.00009391 |
| Tanc1 | 0.00352893 | -0.4830566 | 0.927 | 0.959 | 1 | 0.000047081 | -0.7468417 | 0.811 | 0.917 | 1 | 0.00352893 | 0.00009416 |

| | | | | | | | | | | | | |
|---|---|---|---|---|---|---|---|---|---|---|---|---|
| Cops8 | 0.00474339 | 0.54900678 | 0.719 | 0.57 | 1 | 0.000047283 | 0.48683668 | 0.752 | 0.427 | 1 | 0.00474339 | 0.000094563 |
| Zfp84 | 0.09222488 | 0.47315811 | 0.25 | 0.161 | 1 | 0.000047378 | 1.34910782 | 0.366 | 0.135 | 1 | 0.09222488 | 0.000094753 |
| Lypla2 | 0.00259161 | 0.57453513 | 0.521 | 0.311 | 1 | 0.000047438 | 0.77760758 | 0.483 | 0.198 | 1 | 0.00259161 | 0.000094874 |
| Irak2 | 0.0939938 | -0.9153376 | 0.177 | 0.254 | 1 | 0.000047647 | -1.1370187 | 0.244 | 0.479 | 1 | 0.0939938 | 0.000095291 |
| Lta4h | 0.000048069 | 0.84783787 | 0.573 | 0.316 | 1 | 0.00018544 | 0.54024931 | 0.655 | 0.427 | 1 | 0.00018544 | 0.000096135 |
| 2410004B18Rik | 0.02021009 | 0.56319244 | 0.458 | 0.311 | 1 | 0.000048445 | 0.64434105 | 0.492 | 0.219 | 1 | 0.02021009 | 0.000096887 |
| Emp1 | 0.14271712 | 0.28954132 | 0.917 | 0.979 | 1 | 0.000048586 | 0.47526195 | 0.992 | 0.979 | 1 | 0.14271712 | 0.00009717 |
| Mamld1 | 0.000048617 | -1.3644742 | 0.177 | 0.42 | 1 | 0.8514831 | -0.4272122 | 0.151 | 0.156 | 1 | 0.8514831 | 0.000097231 |
| Ddx39a | 0.52007246 | 0.49934763 | 0.26 | 0.238 | 1 | 0.000048904 | 1.18047036 | 0.387 | 0.146 | 1 | 0.52007246 | 0.000097805 |
| Eif2b5 | 0.23804479 | 0.30488045 | 0.396 | 0.321 | 1 | 0.000048982 | 0.84309666 | 0.496 | 0.229 | 1 | 0.23804479 | 0.000097961 |
| Ypel2 | 0.000049406 | -2.536826 | 0.292 | 0.487 | 1 | 0.03595609 | -0.4066403 | 0.584 | 0.688 | 1 | 0.03595609 | 0.00009881 |
| Rftn2 | 0.62857803 | -0.3915425 | 0.562 | 0.549 | 1 | 0.000049465 | 0.64606367 | 0.651 | 0.323 | 1 | 0.62857803 | 0.000098928 |
| Igf1r | 0.00076935 | -0.7660305 | 0.99 | 0.99 | 1 | 0.000049508 | -0.6559647 | 0.992 | 0.99 | 1 | 0.00076935 | 0.000099014 |
| Klf10 | 0.2091472 | 0.30311687 | 0.594 | 0.539 | 1 | 0.000049673 | 1.00926876 | 0.664 | 0.417 | 1 | 0.2091472 | 0.000099343 |
| Snx5 | 0.000050231 | 0.59983657 | 0.885 | 0.731 | 1 | 0.00135914 | 0.29782731 | 0.853 | 0.729 | 1 | 0.00135914 | 0.00010046 |
| Tmem51 | 0.000050395 | -2.2852872 | 0.115 | 0.321 | 1 | 0.05272453 | -1.352924 | 0.088 | 0.156 | 1 | 0.05272453 | 0.00010079 |
| Kazald1 | 0.04359813 | -0.4506632 | 0.385 | 0.513 | 1 | 0.000051035 | 0.63146883 | 0.529 | 0.24 | 1 | 0.04359813 | 0.00010207 |
| Prmt1 | 0.000051049 | 0.4450746 | 0.823 | 0.575 | 1 | 0.00082506 | 0.51095146 | 0.761 | 0.646 | 1 | 0.00082506 | 0.00010209 |
| Atp5b | 0.00041568 | 0.34625627 | 0.979 | 0.943 | 1 | 0.000051099 | 0.434463 | 0.987 | 0.979 | 1 | 0.00041568 | 0.00010219 |
| Cmas | 0.000051179 | 0.91689199 | 0.635 | 0.394 | 1 | 0.1045916 | 0.15293537 | 0.534 | 0.427 | 1 | 0.1045916 | 0.00010236 |
| Rnasek | 0.000051211 | -1.0841862 | 0.885 | 0.907 | 1 | 0.18517242 | 0.12022164 | 0.786 | 0.781 | 1 | 0.18517242 | 0.00010242 |
| Cpq | 0.1155003 | -0.183189 | 0.646 | 0.798 | 1 | 0.000051254 | 0.29190959 | 0.861 | 0.656 | 1 | 0.1155003 | 0.00010251 |
| Decr1 | 0.0842185 | 0.50345203 | 0.417 | 0.321 | 1 | 0.000051353 | 1.20110666 | 0.357 | 0.125 | 1 | 0.0842185 | 0.0001027 |
| Adprm | 0.04171538 | 0.96685255 | 0.24 | 0.15 | 1 | 0.00005145 | 1.39058446 | 0.265 | 0.062 | 1 | 0.04171538 | 0.0001029 |
| Nme5 | 0.000051563 | 2.00749454 | 0.177 | 0.036 | 1 | 0.80178076 | -0.8150906 | 0.109 | 0.094 | 1 | 0.80178076 | 0.00010312 |
| Snhg3 | 0.05478466 | 0.4635593 | 0.583 | 0.477 | 1 | 0.000051596 | 0.84472177 | 0.529 | 0.271 | 1 | 0.05478466 | 0.00010319 |
| P3h1 | 0.00693353 | 0.40110066 | 0.854 | 0.746 | 1 | 0.000051861 | 0.4496529 | 0.798 | 0.469 | 1 | 0.00693353 | 0.00010372 |
| Abt1 | 0.37149823 | 0.16314541 | 0.354 | 0.29 | 1 | 0.000051974 | 1.49749966 | 0.357 | 0.135 | 1 | 0.37149823 | 0.00010395 |
| Jpt1 | 0.85202959 | 0.23544648 | 0.365 | 0.332 | 1 | 0.000052042 | -1.0443941 | 0.357 | 0.604 | 1 | 0.85202959 | 0.00010408 |
| Cebpd | 0.000052373 | -0.9644911 | 0.688 | 0.876 | 1 | 0.01747452 | -0.4121281 | 0.601 | 0.823 | 1 | 0.01747452 | 0.00010474 |
| S100a11 | 0.01138931 | 0.24376655 | 0.979 | 0.938 | 1 | 0.000052548 | 0.32148188 | 0.983 | 0.927 | 1 | 0.01138931 | 0.00010509 |
| Or8b53 | 0.00005262 | 2.35799178 | 0.25 | 0.083 | 1 | 0.01175768 | 0.6026819 | 0.269 | 0.135 | 1 | 0.01175768 | 0.00010524 |
| Zbtb1 | 0.39869573 | 0.41294599 | 0.458 | 0.394 | 1 | 0.000052933 | 0.64707602 | 0.756 | 0.521 | 1 | 0.39869573 | 0.00010586 |
| Tssc4 | 0.11289857 | 0.27052894 | 0.406 | 0.301 | 1 | 0.000053026 | 0.82054137 | 0.466 | 0.198 | 1 | 0.11289857 | 0.00010605 |
| Tmem248 | 0.26561674 | 0.3213532 | 0.74 | 0.72 | 1 | 0.000053601 | 0.50739812 | 0.882 | 0.667 | 1 | 0.26561674 | 0.0001072 |
| Arhgap10 | 0.000054343 | 0.76912218 | 0.802 | 0.565 | 1 | 0.37686567 | -0.192974 | 0.643 | 0.708 | 1 | 0.37686567 | 0.00010868 |
| Caprin1 | 0.000054424 | 0.43748238 | 0.948 | 0.845 | 1 | 0.80946877 | -0.1487234 | 0.962 | 0.948 | 1 | 0.80946877 | 0.00010885 |
| Phax | 0.02299899 | 0.40781053 | 0.688 | 0.565 | 1 | 0.000054442 | 0.63708349 | 0.735 | 0.49 | 1 | 0.02299899 | 0.00010888 |
| Akr1a1 | 0.00504806 | 0.29595869 | 0.979 | 0.938 | 1 | 0.000054499 | 0.3983063 | 0.992 | 0.969 | 1 | 0.00504806 | 0.000109 |
| Adgra3 | 0.00694753 | 0.38095293 | 0.812 | 0.715 | 1 | 0.000054547 | 0.3430113 | 0.933 | 0.615 | 1 | 0.00694753 | 0.00010909 |
| Ctsd | 0.09008773 | -0.2283741 | 0.938 | 0.953 | 1 | 0.000054621 | 0.40209844 | 0.983 | 0.938 | 1 | 0.09008773 | 0.00010924 |
| Rab5if | 0.23641218 | -0.1977886 | 0.812 | 0.813 | 1 | 0.000054707 | 0.47117238 | 0.878 | 0.729 | 1 | 0.23641218 | 0.00010941 |
| Pepd | 0.0854627 | 0.24931246 | 0.5 | 0.363 | 1 | 0.000054718 | 0.71567983 | 0.588 | 0.323 | 1 | 0.0854627 | 0.00010943 |
| Mrpl35 | 0.11020323 | 0.62898291 | 0.312 | 0.233 | 1 | 0.000054987 | 1.24653809 | 0.328 | 0.104 | 1 | 0.11020323 | 0.00010997 |
| Serp1 | 0.00017133 | 0.56228056 | 0.885 | 0.788 | 1 | 0.00005507 | 0.6140905 | 0.882 | 0.771 | 1 | 0.00017133 | 0.00011014 |
| Ebpl | 0.01916298 | 0.36638896 | 0.594 | 0.446 | 1 | 0.000055088 | 0.54429387 | 0.534 | 0.25 | 1 | 0.01916298 | 0.00011017 |
| Cat | 0.31940466 | -0.1026884 | 0.375 | 0.44 | 1 | 0.000055249 | 0.60900797 | 0.58 | 0.292 | 1 | 0.31940466 | 0.0001105 |
| Rfx3 | 0.00033764 | 0.5049942 | 0.635 | 0.415 | 1 | 0.000055269 | -1.2092229 | 0.668 | 0.854 | 1 | 0.00033764 | 0.00011053 |
| Tuba1a | 0.000055526 | 0.82122695 | 0.927 | 0.808 | 1 | 0.00093538 | 0.36810477 | 0.895 | 0.896 | 1 | 0.00093538 | 0.00011105 |
| Tmf1 | 0.00822602 | 0.27845586 | 0.969 | 0.865 | 1 | 0.000055856 | 0.41825035 | 0.937 | 0.875 | 1 | 0.00822602 | 0.00011171 |
| Erf | 0.00481788 | 0.72930791 | 0.708 | 0.611 | 1 | 0.000055907 | 0.63767732 | 0.87 | 0.688 | 1 | 0.00481788 | 0.00011181 |
| Phospho2 | 0.47771024 | 0.3239682 | 0.25 | 0.218 | 1 | 0.000056234 | 1.16658278 | 0.391 | 0.156 | 1 | 0.47771024 | 0.00011247 |
| Rbx1 | 0.00043059 | 0.36974462 | 0.948 | 0.896 | 1 | 0.00005643 | 0.38273492 | 0.937 | 0.844 | 1 | 0.00043059 | 0.00011286 |
| Maip1 | 0.12498761 | 0.31717904 | 0.365 | 0.264 | 1 | 0.000056457 | 1.02642813 | 0.353 | 0.125 | 1 | 0.12498761 | 0.00011291 |
| Tbc1d20 | 0.14405185 | 0.34452952 | 0.573 | 0.472 | 1 | 0.000056461 | 0.55215373 | 0.664 | 0.375 | 1 | 0.14405185 | 0.00011292 |
| Dele1 | 0.000056615 | 1.02952084 | 0.51 | 0.275 | 1 | 0.15695406 | 0.11395245 | 0.34 | 0.24 | 1 | 0.15695406 | 0.00011323 |
| Sbno2 | 0.000056836 | -0.9516542 | 0.667 | 0.824 | 1 | 0.00033844 | -0.9427609 | 0.618 | 0.75 | 1 | 0.00033844 | 0.00011367 |
| Pkp4 | 0.0979838 | 0.34452952 | 0.562 | 0.492 | 1 | 0.000056865 | -1.1995155 | 0.71 | 0.823 | 1 | 0.0979838 | 0.00011373 |
| Ebna1bp2 | 0.00122206 | 0.57255748 | 0.76 | 0.627 | 1 | 0.00005688 | 0.68062196 | 0.676 | 0.438 | 1 | 0.00122206 | 0.00011376 |
| Scml2 | 0.000057007 | 3.50999488 | 0.115 | 0.01 | 1 | 0.0412923 | 2.08246216 | 0.063 | 0.01 | 1 | 0.0412923 | 0.00011401 |
| Cttnbp2nl | 0.04898257 | 0.32222713 | 0.646 | 0.528 | 1 | 0.00005742 | 0.48775534 | 0.866 | 0.594 | 1 | 0.04898257 | 0.00011484 |
| Arpc5 | 0.00336408 | 0.40147042 | 0.938 | 0.886 | 1 | 0.000057638 | 0.37159076 | 0.983 | 0.906 | 1 | 0.00336408 | 0.00011527 |
| Sf3b6 | 0.00691574 | 0.376951 | 0.885 | 0.834 | 1 | 0.000058063 | 0.4408235 | 0.908 | 0.833 | 1 | 0.00691574 | 0.00011612 |
| Ankrd54 | 0.16711153 | 0.43325844 | 0.354 | 0.28 | 1 | 0.000058254 | 1.14957636 | 0.408 | 0.177 | 1 | 0.16711153 | 0.0001165 |
| Fam162a | 0.000058302 | 0.56754447 | 0.885 | 0.699 | 1 | 0.04445052 | 0.21370669 | 0.765 | 0.625 | 1 | 0.04445052 | 0.0001166 |
| Trnau1ap | 0.04139428 | 0.36465112 | 0.479 | 0.332 | 1 | 0.000058418 | 0.72761944 | 0.441 | 0.167 | 1 | 0.04139428 | 0.00011683 |
| Cd14 | 0.00005843 | -2.4224933 | 0.198 | 0.415 | 1 | 0.04522626 | -0.5322477 | 0.21 | 0.323 | 1 | 0.04522626 | 0.00011686 |
| Smagp | 0.00005887 | -2.3848229 | 0.115 | 0.321 | 1 | 0.55701222 | -1.4618584 | 0.029 | 0.042 | 1 | 0.55701222 | 0.00011774 |
| Trf | 0.000058896 | -3.3112939 | 0.083 | 0.285 | 1 | 0.03043348 | 0.19264508 | 0.21 | 0.104 | 1 | 0.03043348 | 0.00011779 |
| Cnnm1 | 0.000059163 | -2.6013147 | 0.021 | 0.197 | 1 | 0.13293947 | -0.8948178 | 0.042 | 0.083 | 1 | 0.13293947 | 0.00011832 |
| Foxd2 | 0.000059431 | 2.10703021 | 0.219 | 0.062 | 1 | 0.00183263 | 1.19264508 | 0.172 | 0.042 | 1 | 0.00183263 | 0.00011886 |
| Sgms2 | 0.000059509 | 0.77606683 | 0.948 | 0.865 | 1 | 0.90772034 | -0.202227 | 0.975 | 0.927 | 1 | 0.90772034 | 0.00011902 |
| Gpr180 | 0.05074701 | 0.40342321 | 0.656 | 0.539 | 1 | 0.000059935 | 0.62534961 | 0.601 | 0.312 | 1 | 0.05074701 | 0.00011987 |
| Klf5 | 0.09287646 | -0.4242217 | 0.479 | 0.617 | 1 | 0.000060201 | -0.5745585 | 0.475 | 0.76 | 1 | 0.09287646 | 0.0001204 |
| Gm16759 | 0.000060345 | 2.28760246 | 0.188 | 0.041 | 1 | 0.0705706 | 1.24473359 | 0.088 | 0.031 | 1 | 0.0705706 | 0.00012069 |
| Spg20 | 0.000060381 | 0.91537633 | 0.573 | 0.337 | 1 | 0.0049481 | 0.34592858 | 0.437 | 0.24 | 1 | 0.0049481 | 0.00012076 |
| Mxra8 | 0.25709901 | 0.19268381 | 0.896 | 0.87 | 1 | 0.000060594 | 0.49050707 | 0.958 | 0.646 | 1 | 0.25709901 | 0.00012118 |
| Trip10 | 0.000061004 | 0.92383661 | 0.479 | 0.233 | 1 | 0.00053752 | 0.87071698 | 0.471 | 0.24 | 1 | 0.00053752 | 0.000122 |
| P4ha2 | 0.03659971 | -0.5498557 | 0.677 | 0.699 | 1 | 0.000061197 | 0.40333295 | 0.664 | 0.344 | 1 | 0.03659971 | 0.00012239 |
| Ctbp1 | 0.000061264 | 0.67757365 | 0.74 | 0.565 | 1 | 0.0152854 | 0.27187612 | 0.794 | 0.646 | 1 | 0.0152854 | 0.00012252 |
| Axin2 | 0.000061305 | 1.08445652 | 0.448 | 0.212 | 1 | 0.88632568 | -0.4500329 | 0.336 | 0.292 | 1 | 0.88632568 | 0.00012261 |
| Add3 | 0.02095016 | -0.9311049 | 0.26 | 0.383 | 1 | 0.000061315 | -1.2667865 | 0.269 | 0.51 | 1 | 0.02095016 | 0.00012263 |
| Prelid1 | 0.06364506 | 0.24282601 | 0.833 | 0.725 | 1 | 0.000061361 | 0.53066652 | 0.853 | 0.75 | 1 | 0.06364506 | 0.00012272 |
| Rint1 | 0.00522957 | 0.76870768 | 0.531 | 0.394 | 1 | 0.000061959 | 0.73030186 | 0.584 | 0.312 | 1 | 0.00522957 | 0.00012391 |
| Palmd | 0.000062292 | -3.8379955 | 0.021 | 0.192 | 1 | 0.08215562 | -1.443711 | 0.164 | 0.229 | 1 | 0.08215562 | 0.00012458 |
| Plcb1 | 0.000062329 | 0.57372047 | 0.948 | 0.777 | 1 | 0.0374764 | 0.15731108 | 0.971 | 0.844 | 1 | 0.0374764 | 0.00012465 |
| Gem | 0.00006242 | -1.2094975 | 0.521 | 0.725 | 1 | 0.79835835 | -0.4082142 | 0.773 | 0.76 | 1 | 0.79835835 | 0.00012484 |
| Cops4 | 0.02589506 | 0.30866407 | 0.74 | 0.596 | 1 | 0.000062426 | 0.5404518 | 0.773 | 0.573 | 1 | 0.02589506 | 0.00012485 |
| Pbrm1 | 0.000062491 | 0.47060678 | 0.917 | 0.902 | 1 | 0.00135763 | 0.34452055 | 0.954 | 0.938 | 1 | 0.00135763 | 0.00012498 |
| Mea1 | 0.2205382 | -0.3968957 | 0.427 | 0.482 | 1 | 0.000062672 | 0.69014474 | 0.45 | 0.198 | 1 | 0.2205382 | 0.00012534 |
| Eeig2 | 0.000062715 | -0.4127715 | 0.615 | 0.829 | 1 | 0.21121263 | 0.17820302 | 0.79 | 0.677 | 1 | 0.21121263 | 0.00012543 |

| | | | | | | | | | | | | |
|---|---|---|---|---|---|---|---|---|---|---|---|---|
| Arpc5l | 0.00471139 | 0.62707079 | 0.635 | 0.523 | 1 | 0.000062993 | 0.74077081 | 0.668 | 0.49 | 1 | 0.00471139 | 0.00012598 |
| Prorsd1 | 0.07478462 | 0.32711247 | 0.49 | 0.383 | 1 | 0.000063212 | 1.33923758 | 0.399 | 0.167 | 1 | 0.07478462 | 0.00012642 |
| 4921524J17Rik | 0.02253693 | 0.60788708 | 0.458 | 0.316 | 1 | 0.000063244 | 0.80039005 | 0.609 | 0.344 | 1 | 0.02253693 | 0.00012648 |
| Eif2s3x | 0.00949112 | 0.44258969 | 0.792 | 0.637 | 1 | 0.000063589 | 0.42448486 | 0.761 | 0.542 | 1 | 0.00949112 | 0.00012717 |
| Smyd2 | 0.000063641 | 1.05188866 | 0.49 | 0.264 | 1 | 0.00540441 | 0.67286675 | 0.286 | 0.135 | 1 | 0.00540441 | 0.00012728 |
| Mamstr | 0.00006393 | 2.70793425 | 0.115 | 0.01 | 1 | 0.14428792 | 1.14957636 | 0.042 | 0.01 | 1 | 0.14428792 | 0.00012786 |
| Mycbp | 0.000064103 | 1.23530045 | 0.51 | 0.301 | 1 | 0.15052306 | 0.26168672 | 0.294 | 0.208 | 1 | 0.15052306 | 0.0001282 |
| Gpn3 | 0.00084507 | 0.64071221 | 0.542 | 0.326 | 1 | 0.000064389 | 1.22412331 | 0.391 | 0.156 | 1 | 0.00084507 | 0.00012877 |
| Pdk4 | 0.000064585 | -2.6363617 | 0.104 | 0.306 | 1 | 0.44449937 | -1.8244284 | 0.265 | 0.198 | 1 | 0.44449937 | 0.00012917 |
| Rex1bd | 0.35474897 | 0.11952115 | 0.573 | 0.513 | 1 | 0.000065168 | 0.75903982 | 0.55 | 0.292 | 1 | 0.35474897 | 0.00013033 |
| Psmb5 | 0.45732244 | 0.13440665 | 0.823 | 0.777 | 1 | 0.000065176 | 0.41918342 | 0.853 | 0.719 | 1 | 0.45732244 | 0.00013035 |
| Rpl10-ps3 | 0.32477168 | 0.15472049 | 0.75 | 0.679 | 1 | 0.000065616 | 0.70252846 | 0.727 | 0.594 | 1 | 0.32477168 | 0.00013123 |
| Ssna1 | 0.0006102 | 0.79336973 | 0.521 | 0.316 | 1 | 0.000066061 | 0.98638619 | 0.458 | 0.219 | 1 | 0.0006102 | 0.00013212 |
| Tceal5 | 0.000066224 | 1.25542205 | 0.302 | 0.109 | 1 | 0.06031591 | 0.27510724 | 0.197 | 0.104 | 1 | 0.06031591 | 0.00013244 |
| 9530068E07Rik | 0.1313791 | 0.20438659 | 0.917 | 0.845 | 1 | 0.000066426 | 0.46714805 | 0.916 | 0.802 | 1 | 0.1313791 | 0.00013285 |
| Prpf3 | 0.01605848 | 0.79026382 | 0.458 | 0.347 | 1 | 0.000066646 | 0.55041997 | 0.58 | 0.281 | 1 | 0.01605848 | 0.00013329 |
| Sec61a1 | 0.00006686 | 0.56388789 | 0.865 | 0.72 | 1 | 0.02562922 | 0.12536012 | 0.857 | 0.781 | 1 | 0.02562922 | 0.00013371 |
| ENSMUSG00000121513 | 0.24031804 | -0.577468 | 0.469 | 0.528 | 1 | 0.000067331 | -0.9601665 | 0.782 | 0.854 | 1 | 0.24031804 | 0.00013466 |
| Enpp1 | 0.33979085 | -0.2486862 | 0.896 | 0.87 | 1 | 0.000067831 | 0.47291402 | 0.887 | 0.51 | 1 | 0.33979085 | 0.00013566 |
| Lgr5 | 0.74827573 | 0.59245704 | 0.021 | 0.016 | 1 | 0.000068013 | -6.0377757 | 0.008 | 0.094 | 1 | 0.74827573 | 0.00013602 |
| Plxna4 | 0.000068285 | 1.67818691 | 0.25 | 0.078 | 1 | 0.02523726 | -0.3984866 | 0.475 | 0.292 | 1 | 0.02523726 | 0.00013657 |
| Fam114a2 | 0.00831266 | 0.66999364 | 0.646 | 0.539 | 1 | 0.000069111 | 0.65782702 | 0.693 | 0.479 | 1 | 0.00831266 | 0.00013822 |
| Arntl | 0.000069628 | -1.6060371 | 0.146 | 0.368 | 1 | 0.00076142 | -1.294982 | 0.366 | 0.583 | 1 | 0.00076142 | 0.00013925 |
| Elmo1 | 0.000069945 | -1.4434897 | 0.458 | 0.632 | 1 | 0.00118158 | -0.5811573 | 0.282 | 0.51 | 1 | 0.00118158 | 0.00013988 |
| Pura | 0.00471199 | 0.30145004 | 0.938 | 0.922 | 1 | 0.000070306 | 0.36556219 | 0.983 | 0.885 | 1 | 0.00471199 | 0.00014061 |
| Gatad2a | 0.000070404 | 0.80104366 | 0.823 | 0.637 | 1 | 0.58268978 | -0.2725018 | 0.861 | 0.938 | 1 | 0.58268978 | 0.0001408 |
| Ddx47 | 0.01166716 | 0.53686471 | 0.51 | 0.347 | 1 | 0.000070533 | 0.65299174 | 0.58 | 0.312 | 1 | 0.01166716 | 0.00014106 |
| Disp1 | 0.000071227 | -1.2837368 | 0.229 | 0.461 | 1 | 0.90013279 | -0.8131793 | 0.256 | 0.219 | 1 | 0.90013279 | 0.00014245 |
| Fam89b | 0.28299376 | 0.33348085 | 0.51 | 0.435 | 1 | 0.000071326 | 0.82764826 | 0.559 | 0.292 | 1 | 0.28299376 | 0.00014265 |
| Nr2f2 | 0.000071876 | 0.44114771 | 0.698 | 0.389 | 1 | 0.42463813 | 0.17242157 | 0.559 | 0.562 | 1 | 0.42463813 | 0.00014375 |
| Ep400 | 0.53006302 | -0.5965768 | 0.76 | 0.767 | 1 | 0.0000719 | -1.226768 | 0.87 | 0.948 | 1 | 0.53006302 | 0.0001438 |
| Rb1 | 0.12921691 | -0.4689435 | 0.312 | 0.399 | 1 | 0.000071943 | -1.0243135 | 0.567 | 0.802 | 1 | 0.12921691 | 0.00014388 |
| Kctd3 | 0.000072146 | 1.086929 | 0.427 | 0.202 | 1 | 0.00130465 | 0.37005564 | 0.437 | 0.219 | 1 | 0.00130465 | 0.00014429 |
| Pole4 | 0.07949192 | 0.31842363 | 0.604 | 0.482 | 1 | 0.000072236 | 0.83698613 | 0.525 | 0.281 | 1 | 0.07949192 | 0.00014447 |
| Atp6v1f | 0.71191427 | -0.1502558 | 0.917 | 0.865 | 1 | 0.000072239 | 0.592336 | 0.891 | 0.792 | 1 | 0.71191427 | 0.00014447 |
| Psmb2 | 0.000072412 | 0.38478994 | 0.958 | 0.938 | 1 | 0.04616067 | 0.10968363 | 0.979 | 0.948 | 1 | 0.04616067 | 0.00014482 |
| Cep170b | 0.04013387 | 0.68556644 | 0.302 | 0.192 | 1 | 0.000072944 | 1.15963002 | 0.387 | 0.156 | 1 | 0.04013387 | 0.00014588 |
| Cox6a1 | 0.69618001 | -0.1210897 | 0.917 | 0.907 | 1 | 0.000073144 | 0.3216172 | 0.929 | 0.896 | 1 | 0.69618001 | 0.00014628 |
| Arl15 | 0.13290122 | -0.474474 | 0.531 | 0.642 | 1 | 0.000073298 | -1.1144958 | 0.622 | 0.833 | 1 | 0.13290122 | 0.00014659 |
| Srsf3 | 0.000073481 | 0.41916361 | 1 | 0.964 | 1 | 0.000098382 | 0.29044355 | 0.996 | 0.979 | 1 | 9.8382E-05 | 0.00014696 |
| Gas6 | 0.00400957 | 1.18479907 | 0.365 | 0.207 | 1 | 0.000073631 | -0.9562183 | 0.223 | 0.469 | 1 | 0.00400957 | 0.00014726 |
| Larp6 | 0.00409657 | 0.62790587 | 0.562 | 0.383 | 1 | 0.000073794 | 0.8485741 | 0.66 | 0.396 | 1 | 0.00409657 | 0.00014758 |
| Suclg1 | 0.1429344 | 0.25830264 | 0.583 | 0.508 | 1 | 0.000074191 | 0.6339619 | 0.685 | 0.479 | 1 | 0.1429344 | 0.00014838 |
| Sod3 | 0.000074192 | -1.2082342 | 0.26 | 0.508 | 1 | 0.10172183 | 0.1369448 | 0.387 | 0.271 | 1 | 0.10172183 | 0.00014838 |
| Triap1 | 0.28761313 | 0.47406234 | 0.271 | 0.223 | 1 | 0.00007438 | 1.02512899 | 0.366 | 0.135 | 1 | 0.28761313 | 0.00014875 |
| Shmt2 | 0.000074415 | 1.26465238 | 0.375 | 0.166 | 1 | 0.00148246 | 1.40063812 | 0.248 | 0.094 | 1 | 0.00148246 | 0.00014882 |
| Mapk1ip1l | 0.01854521 | 0.40438469 | 0.865 | 0.772 | 1 | 0.000074677 | 0.42125761 | 0.933 | 0.854 | 1 | 0.01854521 | 0.00014935 |
| Foxd1 | 0.000074771 | 1.77922555 | 0.302 | 0.119 | 1 | 0.09068647 | 0.31574922 | 0.269 | 0.167 | 1 | 0.09068647 | 0.00014954 |
| Rela | 0.1181929 | 0.40342321 | 0.667 | 0.591 | 1 | 0.000075088 | 0.72256622 | 0.723 | 0.458 | 1 | 0.1181929 | 0.00015017 |
| Itpripl2 | 0.57828726 | 0.15581728 | 0.646 | 0.617 | 1 | 0.000075495 | 0.84692436 | 0.697 | 0.469 | 1 | 0.57828726 | 0.00015098 |
| Mfap1b | 0.08121226 | 0.44189736 | 0.354 | 0.259 | 1 | 0.000075618 | 0.90313846 | 0.466 | 0.208 | 1 | 0.08121226 | 0.00015123 |
| Ciao2b | 0.02998353 | 0.33211674 | 0.542 | 0.389 | 1 | 0.000075752 | 0.78380384 | 0.508 | 0.26 | 1 | 0.02998353 | 0.0001515 |
| Ppp1r13b | 0.000076041 | 0.80396114 | 0.594 | 0.332 | 1 | 0.87107533 | -0.1874192 | 0.685 | 0.677 | 1 | 0.87107533 | 0.00015208 |
| Arl13b | 0.00007624 | 0.61156586 | 0.823 | 0.637 | 1 | 0.00018116 | 0.59108607 | 0.824 | 0.667 | 1 | 0.00018116 | 0.00015247 |
| Smg1 | 0.000076558 | -0.6716504 | 0.958 | 0.979 | 1 | 0.01360711 | -0.5025003 | 0.987 | 0.969 | 1 | 0.01360711 | 0.00015311 |
| Mafb | 0.70391298 | -1.2332968 | 0.344 | 0.326 | 1 | 0.00007669 | -1.7937878 | 0.34 | 0.521 | 1 | 0.70391298 | 0.00015337 |
| Arfrp1 | 0.18579405 | 0.32160313 | 0.375 | 0.295 | 1 | 0.000076767 | 0.80988398 | 0.382 | 0.146 | 1 | 0.18579405 | 0.00015353 |
| Bud31 | 0.23929425 | 0.17543917 | 0.823 | 0.725 | 1 | 0.00007713 | 0.49749966 | 0.811 | 0.615 | 1 | 0.23929425 | 0.00015425 |
| Fbxl7 | 0.09127678 | -2.0273642 | 0.229 | 0.295 | 1 | 0.000077731 | -1.8484921 | 0.256 | 0.438 | 1 | 0.09127678 | 0.00015546 |
| Timp1 | 0.67554536 | -0.6111739 | 0.812 | 0.793 | 1 | 0.000077759 | 0.85750267 | 0.651 | 0.385 | 1 | 0.67554536 | 0.00015551 |
| Six1 | 0.12025833 | -1.0513992 | 0.135 | 0.202 | 1 | 0.000077831 | -0.9248303 | 0.282 | 0.51 | 1 | 0.12025833 | 0.00015566 |
| Ginm1 | 0.65239749 | -0.1641909 | 0.833 | 0.803 | 1 | 0.000078139 | 0.38933699 | 0.878 | 0.698 | 1 | 0.65239749 | 0.00015627 |
| Brca1 | 0.97580506 | 0.65957123 | 0.062 | 0.062 | 1 | 0.000078552 | -3.3802446 | 0.008 | 0.094 | 1 | 0.97580506 | 0.0001571 |
| Med25 | 0.01139356 | 0.44270992 | 0.573 | 0.43 | 1 | 0.000080146 | 0.43973794 | 0.664 | 0.375 | 1 | 0.01139356 | 0.00016028 |
| Gnpnat1 | 0.00585271 | 0.75668643 | 0.5 | 0.347 | 1 | 0.000080211 | 0.83853658 | 0.412 | 0.167 | 1 | 0.00585271 | 0.00016041 |
| BC005537 | 0.00931049 | 0.24881346 | 0.917 | 0.943 | 1 | 0.000080388 | 0.35448966 | 1 | 0.969 | 1 | 0.00931049 | 0.00016077 |
| Taf12 | 0.54294212 | 0.1243082 | 0.333 | 0.295 | 1 | 0.000080728 | 0.6361049 | 0.429 | 0.177 | 1 | 0.54294212 | 0.00016145 |
| Wnt11 | 0.000080867 | 1.41275102 | 0.448 | 0.244 | 1 | 0.00063158 | 1.13890285 | 0.374 | 0.177 | 1 | 0.00063158 | 0.00016173 |
| Prrx1 | 0.000081218 | -1.6868792 | 0.542 | 0.715 | 1 | 0.00511608 | -0.7940463 | 0.853 | 0.833 | 1 | 0.00511608 | 0.00016243 |
| Grina | 0.000081386 | -1.5973675 | 0.281 | 0.477 | 1 | 0.62530792 | -0.1442692 | 0.529 | 0.51 | 1 | 0.62530792 | 0.00016277 |
| 1300002E11Rik | 0.0898392 | 0.47315811 | 0.25 | 0.166 | 1 | 0.000081506 | 2.36821664 | 0.218 | 0.042 | 1 | 0.0898392 | 0.00016301 |
| Zmynd19 | 0.1762959 | 0.346981 | 0.271 | 0.192 | 1 | 0.000081562 | 1.06865636 | 0.307 | 0.094 | 1 | 0.1762959 | 0.00016312 |
| Nfx1 | 0.000081766 | 0.62384746 | 0.74 | 0.487 | 1 | 0.55985555 | -0.2309998 | 0.744 | 0.771 | 1 | 0.55985555 | 0.00016352 |
| Ndufv2 | 0.27277406 | 0.13381099 | 0.792 | 0.741 | 1 | 0.000082094 | 0.63464962 | 0.803 | 0.688 | 1 | 0.27277406 | 0.00016418 |
| Mllt11 | 0.01503291 | 0.877434 | 0.365 | 0.238 | 1 | 0.000082578 | 0.89825293 | 0.382 | 0.146 | 1 | 0.01503291 | 0.00016515 |
| Vim | 0.51802844 | 0.1939654 | 0.99 | 0.995 | 1 | 0.00008288 | 0.35621368 | 0.975 | 0.969 | 1 | 0.51802844 | 0.00016575 |
| Pip4k2c | 0.11301869 | -0.4719748 | 0.438 | 0.497 | 1 | 0.000082886 | 0.94136984 | 0.55 | 0.312 | 1 | 0.11301869 | 0.00016577 |
| Blm | 0.000083345 | -1.8873232 | 0.198 | 0.409 | 1 | 0.17823831 | -1.3098553 | 0.046 | 0.083 | 1 | 0.17823831 | 0.00016668 |
| Myo5a | 0.000083788 | -0.7729547 | 0.719 | 0.87 | 1 | 0.3818839 | -0.5354812 | 0.693 | 0.729 | 1 | 0.3818839 | 0.00016757 |
| A430018G15Rik | 0.75238773 | 0.30122574 | 0.146 | 0.135 | 1 | 0.000084059 | 2.65977109 | 0.185 | 0.021 | 1 | 0.75238773 | 0.00016811 |
| Myzap | 0.44578943 | 1.29379872 | 0.146 | 0.124 | 1 | 0.000084599 | 0.86239105 | 0.37 | 0.135 | 1 | 0.44578943 | 0.00016919 |
| Ankrd40 | 0.00035716 | 1.00749454 | 0.479 | 0.285 | 1 | 0.000084855 | 0.67340147 | 0.613 | 0.365 | 1 | 0.00035716 | 0.0001697 |
| Loxl3 | 0.00621748 | -0.7136461 | 0.719 | 0.731 | 1 | 0.00008499 | 0.30566471 | 0.668 | 0.365 | 1 | 0.00621748 | 0.00016997 |
| Gpr155 | 0.11437939 | -1.2555399 | 0.042 | 0.093 | 1 | 0.000085319 | -1.4473588 | 0.092 | 0.26 | 1 | 0.11437939 | 0.00017063 |

| | | | | | | | | | | | | |
|---|---|---|---|---|---|---|---|---|---|---|---|---|
| Chpf2 | 0.18065628 | 0.31842363 | 0.375 | 0.301 | 1 | 0.000085401 | 0.7739131 | 0.496 | 0.229 | 1 | 0.18065628 | 0.0001708 |
| Dapk3 | 0.01111235 | 0.61263492 | 0.438 | 0.285 | 1 | 0.000085476 | 0.78074729 | 0.387 | 0.135 | 1 | 0.01111235 | 0.00017094 |
| Eno1b | 0.02967926 | 0.65385758 | 0.333 | 0.212 | 1 | 0.000085714 | 2.29895398 | 0.218 | 0.042 | 1 | 0.02967926 | 0.00017142 |
| Chpt1 | 0.09594489 | 0.23837842 | 0.76 | 0.611 | 1 | 0.000085963 | 0.56461386 | 0.798 | 0.656 | 1 | 0.09594489 | 0.00017192 |
| Tgfbr1 | 0.38607198 | -0.4978797 | 0.562 | 0.585 | 1 | 0.000086202 | -0.8454517 | 0.685 | 0.854 | 1 | 0.38607198 | 0.0001724 |
| Ubl5 | 0.13001739 | 0.14965944 | 0.948 | 0.896 | 1 | 0.000086213 | 0.3943204 | 0.891 | 0.771 | 1 | 0.13001739 | 0.00017242 |
| Rab13 | 0.0505797 | 0.16152817 | 0.552 | 0.394 | 1 | 0.000086367 | 0.94206311 | 0.462 | 0.229 | 1 | 0.0505797 | 0.00017273 |
| Ftl1-ps1 | 0.2584344 | 0.22727574 | 0.385 | 0.316 | 1 | 0.00008664 | 0.65977109 | 0.55 | 0.271 | 1 | 0.2584344 | 0.00017327 |
| Pld3 | 0.10580577 | 0.17350449 | 0.552 | 0.409 | 1 | 0.000086849 | 0.42462078 | 0.55 | 0.26 | 1 | 0.10580577 | 0.00017369 |
| 8030456M14Rik | 0.9927582 | 0.42253204 | 0.021 | 0.021 | 1 | 0.000087083 | -3.2167459 | 0.013 | 0.104 | 1 | 0.9927582 | 0.00017416 |
| Prkag2 | 0.000087109 | -1.2879613 | 0.312 | 0.549 | 1 | 0.01347485 | -1.0436291 | 0.508 | 0.625 | 1 | 0.01347485 | 0.00017421 |
| Tlnrd1 | 0.10272964 | 0.43325844 | 0.323 | 0.228 | 1 | 0.00008724 | 0.8714745 | 0.366 | 0.135 | 1 | 0.10272964 | 0.00017447 |
| Il17ra | 0.000087608 | -1.3081473 | 0.312 | 0.513 | 1 | 0.82823364 | -0.2680351 | 0.218 | 0.198 | 1 | 0.82823364 | 0.00017521 |
| Rbm8a | 0.06127848 | 0.25047129 | 0.812 | 0.731 | 1 | 0.000088249 | 0.56994167 | 0.874 | 0.792 | 1 | 0.06127848 | 0.00017649 |
| Smarca5 | 0.01545305 | 0.31871774 | 0.99 | 0.964 | 1 | 0.000088659 | 0.34211009 | 0.996 | 0.938 | 1 | 0.01545305 | 0.00017731 |
| Bvht | 0.1525942 | -1.4519371 | 0.01 | 0.041 | 1 | 0.000089077 | -3.2167459 | 0.013 | 0.104 | 1 | 0.1525942 | 0.00017815 |
| Smad3 | 0.000089424 | -0.9374641 | 0.646 | 0.813 | 1 | 0.05772521 | -0.4741012 | 0.819 | 0.885 | 1 | 0.05772521 | 0.00017884 |
| Adam28 | 0.22187388 | 0.27052894 | 0.021 | 0.005 | 1 | 0.000089643 | -1.7573142 | 0.029 | 0.146 | 1 | 0.22187388 | 0.00017928 |
| Alg5 | 0.00166148 | 0.64960094 | 0.635 | 0.42 | 1 | 0.000089987 | 0.81503599 | 0.546 | 0.281 | 1 | 0.00166148 | 0.00017997 |
| Hsf1 | 0.41984451 | 0.35097877 | 0.396 | 0.358 | 1 | 0.000090716 | 0.52126783 | 0.454 | 0.188 | 1 | 0.41984451 | 0.00018142 |
| Fhip2a | 0.04479955 | 0.30705482 | 0.5 | 0.363 | 1 | 0.000091068 | 0.62303054 | 0.739 | 0.438 | 1 | 0.04479955 | 0.00018213 |
| Rnf213 | 0.56740145 | -0.6507169 | 0.156 | 0.176 | 1 | 0.00009124 | -2.1197524 | 0.189 | 0.385 | 1 | 0.56740145 | 0.00018247 |
| Car2 | 0.000091397 | -5.9697854 | 0 | 0.145 | 1 | 0.78710954 | 0.76053407 | 0.025 | 0.031 | 1 | 0.78710954 | 0.00018279 |
| Hivep1 | 0.12235829 | -0.4779323 | 0.406 | 0.477 | 1 | 0.000091446 | -1.1103695 | 0.567 | 0.792 | 1 | 0.12235829 | 0.00018288 |
| Seh1l | 0.02622199 | 0.47499946 | 0.646 | 0.477 | 1 | 0.000091639 | 0.60489745 | 0.576 | 0.281 | 1 | 0.02622199 | 0.00018327 |
| Mrpl28 | 0.28673935 | 0.35541784 | 0.427 | 0.378 | 1 | 0.000091829 | 0.90856826 | 0.42 | 0.177 | 1 | 0.28673935 | 0.00018365 |
| B3galt5 | 0.000091842 | -3.577468 | 0.01 | 0.166 | 1 | 0.20315061 | 2.14957636 | 0.017 | 0 | 1 | 0.20315061 | 0.00018368 |
| Wdr77 | 0.00095711 | 1.0790478 | 0.427 | 0.238 | 1 | 0.00009223 | 0.7944814 | 0.391 | 0.146 | 1 | 0.00095711 | 0.00018445 |
| Fam25c | 0.00215168 | 1.91438513 | 0.115 | 0.026 | 1 | 0.000092318 | 3.71251255 | 0.164 | 0.01 | 1 | 0.00215168 | 0.00018463 |
| Has2os | 0.264047 | 0.64136664 | 0.531 | 0.482 | 1 | 0.000092515 | 0.69568292 | 0.391 | 0.156 | 1 | 0.264047 | 0.00018502 |
| Rbm7 | 0.00882195 | 0.42563795 | 0.792 | 0.653 | 1 | 0.000092643 | 0.61686031 | 0.769 | 0.521 | 1 | 0.00882195 | 0.00018528 |
| Carnmt1 | 0.000092648 | 0.58511506 | 0.75 | 0.513 | 1 | 0.00314499 | 0.46304019 | 0.71 | 0.542 | 1 | 0.00314499 | 0.00018529 |
| Osbpl8 | 0.38864211 | -0.1289018 | 0.823 | 0.756 | 1 | 0.000093142 | -0.81014 | 0.828 | 0.948 | 1 | 0.38864211 | 0.00018628 |
| Ptdss1 | 0.00053949 | 0.71429816 | 0.531 | 0.306 | 1 | 0.000093357 | 0.59014895 | 0.592 | 0.292 | 1 | 0.00053949 | 0.00018671 |
| Gprasp1 | 0.03712398 | 0.44525211 | 0.625 | 0.508 | 1 | 0.000093611 | 0.65324515 | 0.689 | 0.406 | 1 | 0.03712398 | 0.00018721 |
| Pbdc1 | 0.00013057 | 0.62100358 | 0.906 | 0.762 | 1 | 0.000094056 | 0.36781761 | 0.87 | 0.656 | 1 | 0.00013057 | 0.0001881 |
| Med21 | 0.9925502 | -0.34944 | 0.74 | 0.622 | 1 | 0.000095187 | 0.67136753 | 0.555 | 0.312 | 1 | 0.9925502 | 0.00019037 |
| Cdc34 | 0.42701415 | 0.15559318 | 0.333 | 0.285 | 1 | 0.000095417 | 1.21586803 | 0.462 | 0.25 | 1 | 0.42701415 | 0.00019082 |
| Tmcc1 | 0.000095545 | 0.63836674 | 0.875 | 0.731 | 1 | 0.46673551 | -0.3547095 | 0.937 | 0.938 | 1 | 0.46673551 | 0.00019108 |
| Eci2 | 0.00790782 | -0.5168416 | 0.521 | 0.674 | 1 | 0.000095618 | 0.79567915 | 0.576 | 0.323 | 1 | 0.00790782 | 0.00019123 |
| Kcnk1 | 0.08048319 | 0.88117809 | 0.469 | 0.42 | 1 | 0.000095961 | 1.49328741 | 0.525 | 0.344 | 1 | 0.08048319 | 0.00019191 |
| Abracl | 0.34925897 | -0.5044045 | 0.406 | 0.43 | 1 | 0.000096076 | -0.790827 | 0.361 | 0.635 | 1 | 0.34925897 | 0.00019214 |
| Arhgap26 | 0.00087746 | -1.4088124 | 0.354 | 0.539 | 1 | 0.00009652 | -1.1172102 | 0.37 | 0.604 | 1 | 0.00087746 | 0.00019303 |
| Soga1 | 0.000096803 | 0.60695661 | 0.594 | 0.321 | 1 | 0.01450772 | -0.8981768 | 0.55 | 0.562 | 1 | 0.01450772 | 0.0001936 |
| Cxcl16 | 0.05525492 | 0.74446013 | 0.146 | 0.073 | 1 | 0.000096945 | 4.01207283 | 0.143 | 0 | 1 | 0.05525492 | 0.00019388 |
| Chchd1 | 0.02098793 | 0.39744105 | 0.75 | 0.617 | 1 | 0.000097158 | 0.59093513 | 0.723 | 0.458 | 1 | 0.02098793 | 0.00019431 |
| Plp2 | 0.000097239 | 0.87240691 | 0.667 | 0.451 | 1 | 0.05958768 | 0.46878836 | 0.66 | 0.635 | 1 | 0.05958768 | 0.00019447 |
| Acap2 | 0.70845705 | -0.3166686 | 0.562 | 0.565 | 1 | 0.000097305 | -0.1935856 | 0.655 | 0.854 | 1 | 0.70845705 | 0.0001946 |
| Tmem147 | 0.09609781 | 0.33506919 | 0.688 | 0.601 | 1 | 0.000097484 | 0.66297454 | 0.639 | 0.385 | 1 | 0.09609781 | 0.00019496 |
| Smg8 | 0.06727773 | 0.60695661 | 0.219 | 0.135 | 1 | 0.000097557 | 2.19264508 | 0.218 | 0.042 | 1 | 0.06727773 | 0.0001951 |
| Ppa2 | 0.83784812 | -0.1445086 | 0.229 | 0.212 | 1 | 0.000097652 | 1.35852325 | 0.282 | 0.083 | 1 | 0.83784812 | 0.0001953 |
| Alg13 | 0.000098129 | 1.25142012 | 0.302 | 0.114 | 1 | 0.00506821 | 0.84542296 | 0.261 | 0.115 | 1 | 0.00506821 | 0.00019625 |
| Spin4 | 0.000098211 | 1.83061677 | 0.208 | 0.057 | 1 | 0.03943087 | 0.39796399 | 0.176 | 0.083 | 1 | 0.03943087 | 0.00019641 |
| Rps27rt | 0.40003446 | 0.11996927 | 0.562 | 0.482 | 1 | 0.000098669 | 1.01207283 | 0.454 | 0.219 | 1 | 0.40003446 | 0.00019733 |
| Nedd4 | 0.000099044 | 0.25887858 | 1 | 1 | 1 | 0.00039502 | 0.29134694 | 1 | 1 | 1 | 0.00039502 | 0.00019808 |
| Gm2296 | 0.39008096 | 0.45110119 | 0.365 | 0.326 | 1 | 0.000099102 | 1.32757466 | 0.34 | 0.125 | 1 | 0.39008096 | 0.00019819 |
| Atp6v1g1 | 0.00720392 | 0.19215085 | 0.969 | 0.876 | 1 | 0.000099104 | 0.72300937 | 0.908 | 0.865 | 1 | 0.00720392 | 0.0001982 |
| Mrpl17 | 0.17867693 | 0.22472525 | 0.75 | 0.627 | 1 | 0.000099987 | 0.76339372 | 0.647 | 0.406 | 1 | 0.17867693 | 0.00019996 |
| Pcdh7 | 0.31757286 | -0.1578432 | 0.312 | 0.259 | 1 | 0.00010081 | -0.9097681 | 0.374 | 0.625 | 1 | 0.31757286 | 0.00020161 |
| Stim2 | 0.00010136 | 0.52552603 | 0.76 | 0.528 | 1 | 0.27866804 | -0.4167705 | 0.761 | 0.812 | 1 | 0.27866804 | 0.00020272 |
| C1ra | 0.03728363 | 0.79336973 | 0.271 | 0.171 | 1 | 0.00010189 | 0.29029364 | 0.441 | 0.177 | 1 | 0.03728363 | 0.00020376 |
| Rnd3 | 0.00010192 | -1.4137057 | 0.458 | 0.674 | 1 | 0.00555465 | -0.797207 | 0.597 | 0.771 | 1 | 0.00555465 | 0.00020383 |
| Dcaf13 | 0.0115187 | 0.73788748 | 0.479 | 0.342 | 1 | 0.0001022 | 0.52021974 | 0.483 | 0.219 | 1 | 0.0115187 | 0.00020438 |
| Pam16 | 0.20464609 | 0.17395568 | 0.573 | 0.482 | 1 | 0.00010229 | 0.96144676 | 0.429 | 0.198 | 1 | 0.20464609 | 0.00020456 |
| Spsb2 | 0.46036597 | -0.4919499 | 0.573 | 0.565 | 1 | 0.00010231 | 0.86646751 | 0.55 | 0.312 | 1 | 0.46036597 | 0.00020462 |
| Rexo4 | 0.37997486 | 0.11672361 | 0.365 | 0.301 | 1 | 0.00010248 | 0.76255323 | 0.429 | 0.177 | 1 | 0.37997486 | 0.00020494 |
| Prkce | 0.0002399 | -0.9559796 | 0.573 | 0.731 | 1 | 0.00010253 | -1.057357 | 0.66 | 0.875 | 1 | 0.0002399 | 0.00020505 |
| Sclt1 | 0.00010254 | 1.01665454 | 0.594 | 0.404 | 1 | 0.04988941 | 0.18490943 | 0.5 | 0.354 | 1 | 0.04988941 | 0.00020506 |
| Mtss2 | 0.00401856 | 0.36060429 | 0.75 | 0.549 | 1 | 0.00010256 | 0.74459252 | 0.676 | 0.417 | 1 | 0.00401856 | 0.00020511 |
| Gpha2 | 0.00010279 | 1.01747534 | 0.625 | 0.409 | 1 | 0.00339028 | 0.46756025 | 0.58 | 0.365 | 1 | 0.00339028 | 0.00020556 |
| Uap1l1 | 0.82412005 | 0.13873907 | 0.302 | 0.29 | 1 | 0.00010291 | 1.14080615 | 0.429 | 0.198 | 1 | 0.82412005 | 0.00020581 |
| Mtnr1a | 0.15729403 | -1.7998604 | 0 | 0.021 | 1 | 0.00010308 | -4.4797803 | 0 | 0.062 | 1 | 0.15729403 | 0.00020615 |
| Gm48898 | 0.15729907 | -1.3144336 | 0 | 0.021 | 1 | 0.00010309 | -4.8948178 | 0 | 0.062 | 1 | 0.15729907 | 0.00020617 |
| Iigp1 | 0.4837638 | -0.4950058 | 0.021 | 0.036 | 1 | 0.0001031 | -7.1427453 | 0 | 0.062 | 1 | 0.4837638 | 0.0002062 |
| Crispld2 | 0.00010312 | -1.3093327 | 0.521 | 0.694 | 1 | 0.01167664 | -2.0563279 | 0.223 | 0.323 | 1 | 0.01167664 | 0.00020623 |
| Czib | 0.0291435 | 0.45729745 | 0.573 | 0.44 | 1 | 0.00010355 | 0.74401161 | 0.529 | 0.26 | 1 | 0.0291435 | 0.00020709 |
| Magohb | 0.00163411 | 0.81846071 | 0.417 | 0.233 | 1 | 0.0001043 | 1.41806519 | 0.357 | 0.146 | 1 | 0.00163411 | 0.00020859 |
| Pard3 | 0.88283155 | -0.2380056 | 0.906 | 0.927 | 1 | 0.00010446 | -0.6163386 | 0.996 | 1 | 1 | 0.88283155 | 0.00020891 |
| Ccng1 | 0.0650818 | 0.77922555 | 0.479 | 0.404 | 1 | 0.00010446 | 0.92044236 | 0.466 | 0.219 | 1 | 0.0650818 | 0.00020891 |
| Agpat4 | 0.0001053 | 0.71922284 | 0.885 | 0.798 | 1 | 0.15164438 | -0.2016736 | 0.849 | 0.646 | 1 | 0.15164438 | 0.00021059 |
| Xpo1 | 0.06404742 | 0.61956888 | 0.594 | 0.482 | 1 | 0.00010562 | -0.8881231 | 0.529 | 0.75 | 1 | 0.06404742 | 0.00021123 |
| Pdxk | 0.10417571 | 0.60695661 | 0.198 | 0.124 | 1 | 0.00010607 | 1.60768258 | 0.218 | 0.042 | 1 | 0.10417571 | 0.00021213 |
| Tmpo | 0.00010623 | 1.16065132 | 0.604 | 0.358 | 1 | 0.17014899 | 0.20762364 | 0.664 | 0.635 | 1 | 0.17014899 | 0.00021244 |
| Clba1 | 0.12366759 | 1.04931471 | 0.198 | 0.13 | 1 | 0.00010647 | 1.48270266 | 0.269 | 0.073 | 1 | 0.12366759 | 0.00021292 |
| Plin2 | 0.00010658 | -1.8520903 | 0.188 | 0.409 | 1 | 0.00023684 | 1.20355398 | 0.349 | 0.146 | 1 | 0.00023684 | 0.00021315 |
| C1s1 | 0.11317789 | 0.2541345 | 0.26 | 0.171 | 1 | 0.00010662 | 0.69719949 | 0.483 | 0.24 | 1 | 0.11317789 | 0.00021323 |
| Naa10 | 0.00010693 | 1.18969787 | 0.427 | 0.212 | 1 | 0.00964181 | 0.41712624 | 0.319 | 0.167 | 1 | 0.00964181 | 0.00021385 |
| Rrp15 | 0.00558089 | 0.89646322 | 0.448 | 0.311 | 1 | 0.000107 | 0.79123086 | 0.408 | 0.167 | 1 | 0.00558089 | 0.00021398 |
| Slc23a2 | 0.00010778 | -0.8225805 | 0.594 | 0.762 | 1 | 0.00052848 | -0.8022516 | 0.706 | 0.865 | 1 | 0.00052848 | 0.00021556 |
| Cep126 | 0.0001079 | 1.02417328 | 0.479 | 0.259 | 1 | 0.48189307 | -0.8138978 | 0.239 | 0.26 | 1 | 0.48189307 | 0.00021579 |
| Hspb1 | 0.000108 | -1.2352126 | 0.583 | 0.793 | 1 | 0.23491028 | -0.2282346 | 0.727 | 0.865 | 1 | 0.23491028 | 0.00021599 |

| | | | | | | | | | | | | |
|---|---|---|---|---|---|---|---|---|---|---|---|---|
| Paxx | 0.79533039 | -0.130009 | 0.271 | 0.285 | 1 | 0.00010913 | 1.13071733 | 0.265 | 0.062 | 1 | 0.79533039 | 0.00021825 |
| Etnk2 | 0.00010913 | -2.5070786 | 0.031 | 0.202 | 1 | 0.55415894 | -0.5322477 | 0.038 | 0.052 | 1 | 0.55415894 | 0.00021825 |
| Socs2 | 0.00010982 | -0.9512292 | 0.667 | 0.793 | 1 | 0.01549294 | -0.4598178 | 0.479 | 0.656 | 1 | 0.01549294 | 0.00021964 |
| Fgd4 | 0.22089278 | -0.3461424 | 0.104 | 0.161 | 1 | 0.00011134 | -0.8623963 | 0.235 | 0.469 | 1 | 0.22089278 | 0.00022266 |
| Mcm6 | 0.0029789 | 0.71241766 | 0.604 | 0.461 | 1 | 0.00011208 | 0.94816107 | 0.496 | 0.24 | 1 | 0.0029789 | 0.00022414 |
| 1810053B23Rik | 0.00011279 | -4.6535709 | 0.042 | 0.212 | 1 | 0.42143167 | 2.05937855 | 0.038 | 0.021 | 1 | 0.42143167 | 0.00022557 |
| Ifrd2 | 0.00011281 | 1.13035128 | 0.396 | 0.187 | 1 | 0.00016664 | 1.43737867 | 0.29 | 0.094 | 1 | 0.00016664 | 0.0002256 |
| Eif2b1 | 0.07099411 | 0.56692195 | 0.344 | 0.254 | 1 | 0.00011289 | 1.62303054 | 0.282 | 0.083 | 1 | 0.07099411 | 0.00022578 |
| Tns2 | 0.00704237 | 0.75636222 | 0.625 | 0.518 | 1 | 0.00011293 | 0.66938618 | 0.714 | 0.458 | 1 | 0.00704237 | 0.00022586 |
| Ccdc85b | 0.11776375 | -0.6878455 | 0.51 | 0.58 | 1 | 0.0001132 | 0.62245793 | 0.496 | 0.24 | 1 | 0.11776375 | 0.00022639 |
| Btg3 | 0.00011322 | 0.61956888 | 0.885 | 0.741 | 1 | 0.00218905 | 0.44535604 | 0.811 | 0.76 | 1 | 0.00218905 | 0.00022643 |
| Usp25 | 0.12433423 | -0.2282193 | 0.604 | 0.741 | 1 | 0.00011326 | -0.8125726 | 0.634 | 0.854 | 1 | 0.12433423 | 0.00022651 |
| Foxd2os | 0.00011373 | 1.31447229 | 0.479 | 0.269 | 1 | 0.05936111 | 0.84542296 | 0.227 | 0.135 | 1 | 0.05936111 | 0.00022746 |
| Alg14 | 0.58184134 | 0.24195979 | 0.667 | 0.663 | 1 | 0.00011421 | 0.30194885 | 0.563 | 0.281 | 1 | 0.58184134 | 0.00022841 |
| Sf3b1 | 0.23786314 | 0.1315358 | 0.99 | 0.995 | 1 | 0.0001154 | 0.22215836 | 1 | 1 | 1 | 0.23786314 | 0.00023078 |
| Pdcd7 | 0.26758759 | 0.27668117 | 0.333 | 0.269 | 1 | 0.0001165 | 0.91253716 | 0.42 | 0.177 | 1 | 0.26758759 | 0.00023299 |
| Nhsl1 | 0.00115383 | -3.3144336 | 0.01 | 0.124 | 1 | 0.00011788 | -1.8948178 | 0.059 | 0.198 | 1 | 0.00115383 | 0.00023574 |
| Dkk2 | 0.00011819 | 1.76356895 | 0.188 | 0.047 | 1 | 0.90187771 | -1.6659991 | 0.05 | 0.052 | 1 | 0.90187771 | 0.00023636 |
| Gm56517 | 0.31113195 | 0.4088571 | 0.271 | 0.223 | 1 | 0.00011874 | 1.26303441 | 0.361 | 0.146 | 1 | 0.31113195 | 0.00023748 |
| Phf10 | 0.00011969 | 0.80313604 | 0.688 | 0.451 | 1 | 0.00124381 | 0.34786358 | 0.731 | 0.51 | 1 | 0.00124381 | 0.00023937 |
| Fnbp1l | 0.7033885 | -0.2751444 | 0.323 | 0.337 | 1 | 0.0001199 | -0.7091254 | 0.5 | 0.75 | 1 | 0.7033885 | 0.00023979 |
| Tmem230 | 0.08288083 | 0.3540523 | 0.573 | 0.503 | 1 | 0.00011995 | 0.55264121 | 0.462 | 0.208 | 1 | 0.08288083 | 0.00023988 |
| Msl1 | 0.00012061 | 0.61922395 | 0.677 | 0.404 | 1 | 0.00142622 | 0.35706283 | 0.887 | 0.76 | 1 | 0.00142622 | 0.0002412 |
| 0610009B22Rik | 0.29419502 | 0.3584017 | 0.354 | 0.29 | 1 | 0.00012111 | 0.830075 | 0.399 | 0.167 | 1 | 0.29419502 | 0.0002422 |
| Atp5e | 0.01168909 | 0.32160313 | 0.969 | 0.886 | 1 | 0.00012159 | 0.43073065 | 0.916 | 0.906 | 1 | 0.01168909 | 0.00024316 |
| Fkbp11 | 0.00017891 | 0.81484946 | 0.531 | 0.301 | 1 | 0.00012166 | 0.90313846 | 0.424 | 0.188 | 1 | 0.00017891 | 0.0002433 |
| Babam1 | 0.10565561 | 0.24453373 | 0.521 | 0.399 | 1 | 0.00012184 | 0.58199593 | 0.466 | 0.208 | 1 | 0.10565561 | 0.00024366 |
| Sap30l | 0.15365174 | 0.24195979 | 0.427 | 0.326 | 1 | 0.00012212 | 0.57412094 | 0.517 | 0.26 | 1 | 0.15365174 | 0.00024422 |
| Pon1 | 0.15837881 | 2.00749454 | 0.01 | 0 | 1 | 0.00012261 | -1.7404896 | 0.071 | 0.219 | 1 | 0.15837881 | 0.00024521 |
| Zcchc24 | 0.00012343 | -1.240433 | 0.302 | 0.523 | 1 | 0.62805289 | -0.6491253 | 0.504 | 0.448 | 1 | 0.62805289 | 0.00024685 |
| Sike1 | 0.01846109 | 0.70263996 | 0.375 | 0.244 | 1 | 0.00012393 | 0.76053407 | 0.45 | 0.198 | 1 | 0.01846109 | 0.00024784 |
| Adamtsl1 | 0.03621343 | -4.0714568 | 0.042 | 0.114 | 1 | 0.00012394 | -3.3851434 | 0.05 | 0.177 | 1 | 0.03621343 | 0.00024787 |
| Asap1 | 0.00012428 | 0.28437615 | 0.885 | 0.591 | 1 | 0.06552129 | -0.3846553 | 0.857 | 0.948 | 1 | 0.06552129 | 0.00024855 |
| Mob3a | 0.00012476 | 1.15949763 | 0.406 | 0.197 | 1 | 0.2972129 | -0.1578522 | 0.324 | 0.24 | 1 | 0.2972129 | 0.0002495 |
| Rab9 | 0.69802655 | 0.10153803 | 0.625 | 0.565 | 1 | 0.00012495 | 0.39352773 | 0.752 | 0.458 | 1 | 0.69802655 | 0.00024988 |
| Tnc | 0.00012574 | -7.3008445 | 0 | 0.14 | 1 | 0.14549382 | -2.4014857 | 0.088 | 0.135 | 1 | 0.14549382 | 0.00025146 |
| Drg2 | 0.04568653 | 0.59245704 | 0.344 | 0.228 | 1 | 0.00012706 | 0.78074729 | 0.382 | 0.146 | 1 | 0.04568653 | 0.00025411 |
| Scfd1 | 0.00012758 | 0.58428311 | 0.865 | 0.767 | 1 | 0.00034587 | 0.44913664 | 0.891 | 0.771 | 1 | 0.00034587 | 0.00025515 |
| Rai1 | 0.00012784 | 0.49292136 | 0.562 | 0.285 | 1 | 0.08197909 | -0.8794554 | 0.815 | 0.792 | 1 | 0.08197909 | 0.00025566 |
| Csgalnact2 | 0.14383802 | 0.28695449 | 0.75 | 0.653 | 1 | 0.00012798 | 0.42504787 | 0.605 | 0.292 | 1 | 0.14383802 | 0.00025595 |
| Tmem202 | 0.00012826 | -3.3500575 | 0.01 | 0.161 | 1 | 0.2363731 | -1.4473588 | 0.059 | 0.094 | 1 | 0.2363731 | 0.0002565 |
| Prpf38b | 0.02512067 | 0.2303459 | 0.875 | 0.819 | 1 | 0.00012919 | 0.44859006 | 0.945 | 0.823 | 1 | 0.02512067 | 0.00025836 |
| Tob1 | 0.36038741 | -0.1064232 | 0.927 | 0.927 | 1 | 0.00012942 | 0.53917713 | 0.979 | 0.969 | 1 | 0.36038741 | 0.00025883 |
| Efemp1 | 0.52636912 | 0.66645762 | 0.292 | 0.332 | 1 | 0.00012952 | 1.67544823 | 0.416 | 0.208 | 1 | 0.52636912 | 0.00025902 |
| Tnfrsf1a | 0.07737213 | -0.2435672 | 0.833 | 0.845 | 1 | 0.00012967 | 0.5414189 | 0.857 | 0.74 | 1 | 0.07737213 | 0.00025932 |
| Appl1 | 0.00013029 | 0.68435967 | 0.833 | 0.699 | 1 | 0.13027487 | -0.5263045 | 0.849 | 0.844 | 1 | 0.13027487 | 0.00026056 |
| Exoc6b | 0.01810977 | -0.7149715 | 0.885 | 0.912 | 1 | 0.00013069 | -0.7519356 | 0.924 | 0.979 | 1 | 0.01810977 | 0.00026137 |
| Alg2 | 0.01980496 | 0.80894886 | 0.417 | 0.295 | 1 | 0.00013166 | 0.64547569 | 0.471 | 0.219 | 1 | 0.01980496 | 0.00026331 |
| Zw10 | 0.19562569 | 0.41188479 | 0.323 | 0.254 | 1 | 0.00013243 | 0.71060884 | 0.45 | 0.198 | 1 | 0.19562569 | 0.00026485 |
| H2ax | 0.7834883 | 0.60223806 | 0.354 | 0.337 | 1 | 0.00013298 | 1.34835622 | 0.294 | 0.094 | 1 | 0.7834883 | 0.00026593 |
| Cebpzos | 0.62926513 | 0.22322323 | 0.365 | 0.342 | 1 | 0.00013298 | 0.98137604 | 0.315 | 0.104 | 1 | 0.62926513 | 0.00026595 |
| Pdlim7 | 0.06990335 | -0.5970968 | 0.448 | 0.497 | 1 | 0.00013349 | -1.2009209 | 0.437 | 0.583 | 1 | 0.06990335 | 0.00026696 |
| Cyba | 0.00013417 | 0.56779498 | 0.51 | 0.254 | 1 | 0.18884152 | -0.3098553 | 0.462 | 0.573 | 1 | 0.18884152 | 0.00026833 |
| Rbm15 | 0.06784744 | 0.59740625 | 0.448 | 0.342 | 1 | 0.00013439 | 0.63159256 | 0.504 | 0.229 | 1 | 0.06784744 | 0.00026876 |
| Cyb5b | 0.00013457 | 0.64236194 | 0.719 | 0.466 | 1 | 0.43989699 | -0.3327557 | 0.571 | 0.594 | 1 | 0.43989699 | 0.00026913 |
| 3110082I17Rik | 0.0192218 | 1.0392034 | 0.323 | 0.202 | 1 | 0.0001355 | 1.34549657 | 0.218 | 0.042 | 1 | 0.0192218 | 0.00027098 |
| Appl2 | 0.04406388 | 0.50053455 | 0.594 | 0.565 | 1 | 0.00013557 | 0.53024824 | 0.735 | 0.458 | 1 | 0.04406388 | 0.00027113 |
| Cbx3 | 0.00013587 | 0.51679277 | 0.938 | 0.839 | 1 | 0.00093805 | 0.39668834 | 0.95 | 0.823 | 1 | 0.00093805 | 0.00027173 |
| Yae1d1 | 0.02151293 | 0.60223806 | 0.521 | 0.383 | 1 | 0.0001359 | 0.78206723 | 0.466 | 0.229 | 1 | 0.02151293 | 0.00027178 |
| Actr1b | 0.17192528 | 0.29379872 | 0.552 | 0.461 | 1 | 0.00013728 | 0.76372587 | 0.508 | 0.271 | 1 | 0.17192528 | 0.00027454 |
| Egln2 | 0.21493883 | 0.17522198 | 0.344 | 0.264 | 1 | 0.00013866 | 0.42711033 | 0.437 | 0.188 | 1 | 0.21493883 | 0.0002773 |
| Ormdl3 | 0.00013893 | 0.61123424 | 0.833 | 0.632 | 1 | 0.01170071 | 0.40397383 | 0.782 | 0.635 | 1 | 0.01170071 | 0.00027783 |
| Mgat1 | 0.95622908 | 0.16182268 | 0.406 | 0.425 | 1 | 0.00013938 | 0.52079031 | 0.643 | 0.354 | 1 | 0.95622908 | 0.00027875 |
| Lzts2 | 0.00412739 | 0.5371746 | 0.604 | 0.404 | 1 | 0.00013957 | 0.55156951 | 0.706 | 0.396 | 1 | 0.00412739 | 0.00027912 |
| Nol12 | 0.00013983 | 0.71798792 | 0.562 | 0.306 | 1 | 0.00030138 | 0.83824338 | 0.475 | 0.25 | 1 | 0.00030138 | 0.00027964 |
| Ift81 | 0.29281459 | 0.31943854 | 0.344 | 0.29 | 1 | 0.00013995 | 0.57099747 | 0.37 | 0.135 | 1 | 0.29281459 | 0.00027987 |
| Mtrr | 0.00014006 | 1.85549144 | 0.167 | 0.036 | 1 | 0.53857377 | -0.1723517 | 0.084 | 0.062 | 1 | 0.53857377 | 0.00028009 |
| Pdlim1 | 0.3839366 | -0.1889027 | 0.167 | 0.124 | 1 | 0.0001401 | -0.80462 | 0.239 | 0.469 | 1 | 0.3839366 | 0.00028018 |
| Etfrf1 | 0.39102239 | -0.3144336 | 0.188 | 0.233 | 1 | 0.00014022 | 1.44016648 | 0.252 | 0.062 | 1 | 0.39102239 | 0.00028043 |
| Cfdp1 | 0.08095024 | 0.16467787 | 0.906 | 0.902 | 1 | 0.00014053 | 0.40326359 | 0.958 | 0.865 | 1 | 0.08095024 | 0.00028104 |
| Abcg2 | 0.42484041 | -0.3577899 | 0.427 | 0.446 | 1 | 0.00014134 | 0.85308331 | 0.454 | 0.219 | 1 | 0.42484041 | 0.00028266 |
| Ppp1r8 | 0.97316638 | -0.1815393 | 0.219 | 0.207 | 1 | 0.00014148 | 0.97554696 | 0.382 | 0.156 | 1 | 0.97316638 | 0.00028294 |
| Atxn2l | 0.00014265 | 0.62830224 | 0.792 | 0.57 | 1 | 0.03870968 | 0.29460341 | 0.824 | 0.698 | 1 | 0.03870968 | 0.00028528 |
| Hdac3 | 0.10705319 | 0.39907306 | 0.438 | 0.332 | 1 | 0.00014306 | 1.09953567 | 0.462 | 0.24 | 1 | 0.10705319 | 0.0002861 |
| Ccdc50 | 0.00014474 | -0.8575759 | 0.708 | 0.813 | 1 | 0.88455427 | -0.1820997 | 0.84 | 0.885 | 1 | 0.88455427 | 0.00028945 |
| S100a13 | 0.06044755 | -0.5326981 | 0.896 | 0.808 | 1 | 0.00014477 | 0.4461223 | 0.84 | 0.729 | 1 | 0.06044755 | 0.00028952 |
| Tmem175 | 0.00014548 | 0.77211647 | 0.479 | 0.264 | 1 | 0.00060461 | 0.47972496 | 0.366 | 0.167 | 1 | 0.00060461 | 0.00029094 |
| Rap2c | 0.00014571 | 1.02326185 | 0.521 | 0.321 | 1 | 0.01374048 | 0.40635177 | 0.42 | 0.271 | 1 | 0.01374048 | 0.0002914 |
| Pcnx4 | 0.00014661 | 0.98911601 | 0.479 | 0.264 | 1 | 0.04206707 | -0.2115532 | 0.445 | 0.26 | 1 | 0.04206707 | 0.0002932 |
| Tmco3 | 0.00069172 | -0.8874594 | 0.573 | 0.694 | 1 | 0.00014673 | 0.70450003 | 0.525 | 0.26 | 1 | 0.00069172 | 0.00029344 |
| Atic | 0.00048724 | 0.77382219 | 0.562 | 0.352 | 1 | 0.00014705 | 0.78530197 | 0.479 | 0.25 | 1 | 0.00048724 | 0.00029408 |
| Ptges | 0.01544068 | -1.7533178 | 0.083 | 0.187 | 1 | 0.0001473 | -1.1399303 | 0.076 | 0.229 | 1 | 0.01544068 | 0.00029459 |
| Atxn10 | 0.00014731 | 0.4921375 | 0.99 | 0.845 | 1 | 0.00441032 | 0.19710473 | 0.958 | 0.854 | 1 | 0.00441032 | 0.00029459 |
| Mcee | 0.51032604 | 0.20363706 | 0.438 | 0.383 | 1 | 0.00014763 | 0.97838971 | 0.391 | 0.167 | 1 | 0.51032604 | 0.00029524 |
| Ddr1 | 0.19339552 | 0.74446013 | 0.229 | 0.171 | 1 | 0.00014832 | -0.8116044 | 0.324 | 0.562 | 1 | 0.19339552 | 0.00029661 |
| Mtif2 | 0.00111369 | 0.67251029 | 0.542 | 0.358 | 1 | 0.00014855 | 0.50830641 | 0.492 | 0.219 | 1 | 0.00111369 | 0.00029708 |
| Habp4 | 0.00014896 | 0.9182272 | 0.406 | 0.187 | 1 | 0.00028913 | 0.93525724 | 0.378 | 0.167 | 1 | 0.00028913 | 0.00029789 |
| Med4 | 0.14116241 | 0.38300367 | 0.354 | 0.259 | 1 | 0.00014903 | 0.77760758 | 0.387 | 0.156 | 1 | 0.14116241 | 0.00029804 |

| | | | | | | | | | | | | |
|---|---|---|---|---|---|---|---|---|---|---|---|---|
| Homer3 | 0.76623397 | 0.34853145 | 0.24 | 0.228 | 1 | 0.00014971 | 0.66099839 | 0.462 | 0.219 | 1 | 0.76623397 | 0.00029939 |
| Cd28 | 0.00015002 | -3.9621318 | 0.021 | 0.176 | 1 | 0.19870886 | 0.27510724 | 0.067 | 0.031 | 1 | 0.19870886 | 0.00030002 |
| Mars1 | 0.242073 | 0.48393258 | 0.26 | 0.202 | 1 | 0.0001501 | 1.31950136 | 0.307 | 0.104 | 1 | 0.242073 | 0.00030018 |
| Map3k4 | 0.77899535 | -0.4779323 | 0.427 | 0.42 | 1 | 0.00015103 | 0.87777174 | 0.424 | 0.188 | 1 | 0.77899535 | 0.00030204 |
| Pfdn6 | 0.06562886 | 0.39451766 | 0.49 | 0.368 | 1 | 0.00015154 | 0.75192093 | 0.462 | 0.229 | 1 | 0.06562886 | 0.00030306 |
| Col1a1 | 0.00169604 | -3.5092472 | 0.156 | 0.301 | 1 | 0.00015168 | -1.9658096 | 0.382 | 0.51 | 1 | 0.00169604 | 0.00030333 |
| Prrc2a | 0.0001519 | 0.50355653 | 0.854 | 0.684 | 1 | 0.03262364 | 0.14684139 | 0.891 | 0.833 | 1 | 0.03262364 | 0.00030377 |
| Hspbp1 | 0.7033658 | 0.25076569 | 0.24 | 0.223 | 1 | 0.00015266 | 1.61297688 | 0.298 | 0.104 | 1 | 0.7033658 | 0.0003053 |
| Cenpx | 0.00994626 | 0.81904945 | 0.5 | 0.358 | 1 | 0.00015272 | 0.40142369 | 0.496 | 0.24 | 1 | 0.00994626 | 0.00030541 |
| Atp5a1 | 0.00015301 | 0.40028963 | 0.979 | 0.938 | 1 | 0.00259619 | 0.22178283 | 0.987 | 0.969 | 1 | 0.00259619 | 0.000306 |
| Snrnp27 | 0.04921673 | 0.37108527 | 0.615 | 0.513 | 1 | 0.00015362 | 0.63569695 | 0.542 | 0.281 | 1 | 0.04921673 | 0.00030721 |
| Arxes2 | 0.00015376 | 1.48142572 | 0.24 | 0.078 | 1 | 0.25750267 | 0.34725702 | 0.139 | 0.094 | 1 | 0.25750267 | 0.0003075 |
| Mtmr6 | 0.02379827 | 0.43218221 | 0.635 | 0.482 | 1 | 0.00015417 | 0.56912934 | 0.693 | 0.438 | 1 | 0.02379827 | 0.00030831 |
| Cnksr3 | 0.00015547 | 0.76018448 | 0.562 | 0.337 | 1 | 0.08182525 | -0.7623675 | 0.597 | 0.667 | 1 | 0.08182525 | 0.00031092 |
| Utp18 | 0.0001562 | 0.78865993 | 0.635 | 0.404 | 1 | 0.49452495 | -0.1977968 | 0.613 | 0.552 | 1 | 0.49452495 | 0.00031239 |
| Nol10 | 0.28381417 | 0.11267677 | 0.458 | 0.368 | 1 | 0.00015632 | 0.76909608 | 0.454 | 0.208 | 1 | 0.28381417 | 0.00031262 |
| S1pr2 | 0.05786548 | 0.866632 | 0.271 | 0.181 | 1 | 0.00015635 | 1.02999474 | 0.391 | 0.167 | 1 | 0.05786548 | 0.00031268 |
| Snrpe | 0.00020472 | 0.43037279 | 0.906 | 0.777 | 1 | 0.00015701 | 0.40820637 | 0.887 | 0.812 | 1 | 0.00020472 | 0.000314 |
| Snrpd2 | 0.02920913 | 0.33482744 | 0.865 | 0.777 | 1 | 0.00015743 | 0.52264531 | 0.803 | 0.698 | 1 | 0.02920913 | 0.00031484 |
| Lsamp | 0.286318 | -0.9185049 | 0.01 | 0.031 | 1 | 0.00015767 | 0.71251255 | 0.433 | 0.188 | 1 | 0.286318 | 0.00031531 |
| Ywhae | 0.01666157 | 0.19054081 | 0.99 | 0.995 | 1 | 0.00015828 | 0.28091954 | 1 | 0.99 | 1 | 0.01666157 | 0.00031654 |
| Zbtb41 | 0.10135176 | 0.4850735 | 0.396 | 0.301 | 1 | 0.00015953 | 0.29131747 | 0.534 | 0.24 | 1 | 0.10135176 | 0.00031903 |
| Elp5 | 0.16771411 | 0.27929915 | 0.635 | 0.544 | 1 | 0.00016186 | 0.59540425 | 0.588 | 0.312 | 1 | 0.16771411 | 0.0003237 |
| Prdx3 | 0.02120241 | 0.45943162 | 0.667 | 0.585 | 1 | 0.00016235 | 0.57056312 | 0.584 | 0.333 | 1 | 0.02120241 | 0.00032468 |
| Taf13 | 0.32444129 | -0.1532119 | 0.677 | 0.715 | 1 | 0.00016252 | 0.58322953 | 0.735 | 0.573 | 1 | 0.32444129 | 0.00032502 |
| Cxcl17 | 0.3723613 | 0.10060394 | 0.021 | 0.041 | 1 | 0.00016285 | -1.4542452 | 0.059 | 0.198 | 1 | 0.3723613 | 0.00032568 |
| Lrrc41 | 0.00016351 | 0.64184506 | 0.677 | 0.409 | 1 | 0.00450174 | 0.23593664 | 0.702 | 0.479 | 1 | 0.00450174 | 0.00032698 |
| Alcam | 0.10597072 | -0.5238869 | 0.229 | 0.326 | 1 | 0.00016367 | -0.4612323 | 0.521 | 0.802 | 1 | 0.10597072 | 0.00032731 |
| Slc25a48 | 0.00016394 | -3.0585947 | 0.031 | 0.192 | 1 | 0.71071963 | -0.6577786 | 0.034 | 0.042 | 1 | 0.71071963 | 0.00032785 |
| Cetn2 | 0.74004951 | -0.1345245 | 0.521 | 0.534 | 1 | 0.00016423 | 0.56461386 | 0.487 | 0.26 | 1 | 0.74004951 | 0.00032843 |
| Snhg6 | 0.11677941 | 0.2816695 | 0.792 | 0.668 | 1 | 0.00016455 | 0.54812573 | 0.622 | 0.344 | 1 | 0.11677941 | 0.00032908 |
| Zeb1 | 0.77082116 | -0.1846833 | 0.875 | 0.772 | 1 | 0.00016457 | -0.9062705 | 0.895 | 0.885 | 1 | 0.77082116 | 0.00032912 |
| Tex264 | 0.28585595 | 0.31644478 | 0.531 | 0.461 | 1 | 0.00016537 | 0.67229674 | 0.609 | 0.365 | 1 | 0.28585595 | 0.00033072 |
| Tspan14 | 0.5049923 | 0.1124641 | 0.271 | 0.316 | 1 | 0.00016651 | -0.3986645 | 0.324 | 0.573 | 1 | 0.5049923 | 0.000333 |
| Phf13 | 0.00978134 | 0.68556644 | 0.427 | 0.28 | 1 | 0.000167 | 0.62768065 | 0.521 | 0.25 | 1 | 0.00978134 | 0.00033398 |
| Ldlrad4 | 0.09402227 | -0.133071 | 0.948 | 0.922 | 1 | 0.00016766 | -0.7815081 | 0.971 | 0.99 | 1 | 0.09402227 | 0.00033528 |
| Neo1 | 0.00016903 | 0.21860115 | 0.917 | 0.725 | 1 | 0.58836014 | -0.185992 | 0.924 | 0.948 | 1 | 0.58836014 | 0.00033803 |
| Tead1 | 0.00023965 | -0.6947567 | 0.938 | 0.979 | 1 | 0.00016913 | -0.6457947 | 0.962 | 1 | 1 | 0.00023965 | 0.00033824 |
| Dr1 | 0.00398333 | 0.64994253 | 0.448 | 0.269 | 1 | 0.00017025 | 0.61260814 | 0.479 | 0.229 | 1 | 0.00398333 | 0.00034047 |
| Ccdc34 | 0.00017088 | 1.0142837 | 0.615 | 0.409 | 1 | 0.0127005 | 0.63767732 | 0.378 | 0.229 | 1 | 0.0127005 | 0.00034173 |
| E330013P04Rik | 0.00017163 | 3.17741954 | 0.104 | 0.01 | 1 | 0.14520233 | -0.9343461 | 0.101 | 0.156 | 1 | 0.14520233 | 0.00034323 |
| Mafk | 0.00550221 | 0.63754493 | 0.594 | 0.461 | 1 | 0.00017191 | 0.80562195 | 0.748 | 0.625 | 1 | 0.00550221 | 0.00034378 |
| Minpp1 | 0.02800769 | 0.54806292 | 0.49 | 0.373 | 1 | 0.00017201 | 0.59136658 | 0.592 | 0.302 | 1 | 0.02800769 | 0.00034399 |
| Yeats4 | 0.00017208 | 0.79804117 | 0.625 | 0.394 | 1 | 0.00082997 | 0.59113473 | 0.483 | 0.281 | 1 | 0.00082997 | 0.00034413 |
| Vps13b | 0.13108534 | 0.18414988 | 0.865 | 0.834 | 1 | 0.00017241 | -0.7548393 | 0.853 | 0.906 | 1 | 0.13108534 | 0.00034479 |
| Fbh1 | 0.62372622 | 0.23882008 | 0.208 | 0.181 | 1 | 0.00017245 | 0.7554362 | 0.391 | 0.156 | 1 | 0.62372622 | 0.00034487 |
| Cdc37 | 0.00829664 | 0.35394495 | 0.917 | 0.782 | 1 | 0.00017246 | 0.3493254 | 0.929 | 0.802 | 1 | 0.00829664 | 0.00034489 |
| Nfix | 0.01100902 | 0.14499806 | 1 | 0.964 | 1 | 0.00017254 | 0.28722617 | 0.992 | 1 | 1 | 0.01100902 | 0.00034505 |
| Slc1a4 | 0.00215201 | 0.54452256 | 0.583 | 0.415 | 1 | 0.00017269 | 0.59602256 | 0.647 | 0.375 | 1 | 0.00215201 | 0.00034535 |
| Olfml2b | 0.8440978 | -0.6169963 | 0.104 | 0.093 | 1 | 0.00017463 | -3.2358547 | 0.029 | 0.135 | 1 | 0.8440978 | 0.00034923 |
| Dhrs7 | 0.00030038 | 0.52884094 | 0.74 | 0.513 | 1 | 0.00017477 | 0.5413873 | 0.634 | 0.448 | 1 | 0.00030038 | 0.00034951 |
| Usp11 | 0.00017618 | 1.0790478 | 0.438 | 0.223 | 1 | 0.0005919 | 0.78034255 | 0.378 | 0.177 | 1 | 0.0005919 | 0.00035233 |
| Tfpt | 0.25101258 | 0.4972998 | 0.25 | 0.192 | 1 | 0.00017624 | 1.56461386 | 0.298 | 0.104 | 1 | 0.25101258 | 0.00035246 |
| Setd1b | 0.00017671 | 0.79989912 | 0.542 | 0.311 | 1 | 0.05571776 | 0.23827047 | 0.672 | 0.583 | 1 | 0.05571776 | 0.00035339 |
| Tmem100 | 0.01198467 | 0.36290805 | 0.729 | 0.58 | 1 | 0.00017715 | 0.93590115 | 0.601 | 0.354 | 1 | 0.01198467 | 0.00035426 |
| Tfg | 0.75825188 | 0.13410085 | 0.875 | 0.829 | 1 | 0.00017724 | 0.53391706 | 0.887 | 0.802 | 1 | 0.75825188 | 0.00035444 |
| Zbtb46 | 0.00017741 | 1.01917634 | 0.49 | 0.28 | 1 | 0.65387812 | -0.2807089 | 0.412 | 0.406 | 1 | 0.65387812 | 0.00035479 |
| Dpm2 | 0.07639838 | 0.36442908 | 0.521 | 0.43 | 1 | 0.0001783 | 0.79357433 | 0.466 | 0.219 | 1 | 0.07639838 | 0.00035657 |
| Prkca | 0.00017884 | -1.0188128 | 0.948 | 0.964 | 1 | 0.16611708 | -0.5760798 | 0.971 | 0.917 | 1 | 0.16611708 | 0.00035765 |
| Mphosph10 | 0.1472161 | 0.36703692 | 0.49 | 0.404 | 1 | 0.00017898 | 0.48369386 | 0.508 | 0.24 | 1 | 0.1472161 | 0.00035792 |
| Lrig1 | 0.58402005 | 0.35541784 | 0.24 | 0.207 | 1 | 0.0001794 | -1.1994313 | 0.265 | 0.49 | 1 | 0.58402005 | 0.00035876 |
| Map3k8 | 0.00018008 | -1.298008 | 0.323 | 0.518 | 1 | 0.97162022 | -0.2182248 | 0.307 | 0.292 | 1 | 0.97162022 | 0.00036013 |
| Adcy3 | 0.0728049 | 0.52944724 | 0.24 | 0.15 | 1 | 0.00018034 | 1.21138164 | 0.332 | 0.125 | 1 | 0.0728049 | 0.00036065 |
| Mfsd10 | 0.00018206 | 0.80679719 | 0.521 | 0.311 | 1 | 0.4368069 | -0.1920188 | 0.336 | 0.26 | 1 | 0.4368069 | 0.0003641 |
| Amz2 | 0.31342389 | -0.2894872 | 0.281 | 0.347 | 1 | 0.00018299 | 0.82764826 | 0.311 | 0.104 | 1 | 0.31342389 | 0.00036595 |
| Aph1a | 0.00019215 | 0.70560419 | 0.698 | 0.472 | 1 | 0.000183 | 0.54347509 | 0.685 | 0.479 | 1 | 0.00019215 | 0.00036596 |
| Nhs | 0.56012528 | -0.3144336 | 0.302 | 0.269 | 1 | 0.00018425 | -0.9583829 | 0.429 | 0.708 | 1 | 0.56012528 | 0.00036847 |
| Rgs19 | 0.0038002 | 1.39082318 | 0.24 | 0.114 | 1 | 0.00018498 | 1.88654195 | 0.21 | 0.042 | 1 | 0.0038002 | 0.00036993 |
| Lipo3 | 0.02984871 | 0.78510212 | 0.229 | 0.13 | 1 | 0.00018523 | 1.14957636 | 0.248 | 0.062 | 1 | 0.02984871 | 0.00037042 |
| Kti12 | 0.00776417 | 0.87319344 | 0.312 | 0.176 | 1 | 0.0001863 | 0.93807225 | 0.319 | 0.115 | 1 | 0.00776417 | 0.00037257 |
| Tmem70 | 0.00945637 | 0.49848089 | 0.583 | 0.42 | 1 | 0.00018756 | 0.55687821 | 0.521 | 0.25 | 1 | 0.00945637 | 0.00037509 |
| Naglu | 0.51521916 | 0.32942263 | 0.292 | 0.259 | 1 | 0.00018759 | 1.70806665 | 0.269 | 0.083 | 1 | 0.51521916 | 0.00037513 |
| Slc20a2 | 0.00018764 | -1.2521493 | 0.417 | 0.596 | 1 | 0.77242543 | -0.9406215 | 0.353 | 0.292 | 1 | 0.77242543 | 0.00037525 |
| Ltbp4 | 0.06454335 | -1.7397394 | 0.052 | 0.119 | 1 | 0.0001884 | -1.0078267 | 0.206 | 0.417 | 1 | 0.06454335 | 0.00037677 |
| Nae1 | 0.16055432 | 0.28664861 | 0.385 | 0.301 | 1 | 0.00018928 | 0.70296878 | 0.555 | 0.323 | 1 | 0.16055432 | 0.00037853 |
| P2rx4 | 0.41065047 | -0.4266013 | 0.344 | 0.368 | 1 | 0.00018969 | 0.3287452 | 0.399 | 0.167 | 1 | 0.41065047 | 0.00037935 |
| Gusb | 0.72956264 | 0.19556688 | 0.333 | 0.316 | 1 | 0.00018994 | 1.04663005 | 0.403 | 0.177 | 1 | 0.72956264 | 0.00037985 |
| Ncoa5 | 0.00019287 | 0.73929843 | 0.583 | 0.347 | 1 | 0.00684244 | 0.38336013 | 0.597 | 0.385 | 1 | 0.00684244 | 0.00038571 |
| Gpr89 | 0.00951864 | 0.51768927 | 0.396 | 0.244 | 1 | 0.00019749 | 1.02717972 | 0.29 | 0.094 | 1 | 0.00951864 | 0.00039495 |
| Trappc13 | 0.20974776 | 0.50345203 | 0.375 | 0.321 | 1 | 0.00019789 | 0.56226991 | 0.513 | 0.26 | 1 | 0.20974776 | 0.00039573 |
| Zbtb10 | 0.00019798 | 0.74797461 | 0.646 | 0.399 | 1 | 0.49149046 | -0.477115 | 0.622 | 0.573 | 1 | 0.49149046 | 0.00039591 |
| Mical3 | 0.03135165 | 0.37792673 | 0.812 | 0.741 | 1 | 0.00019827 | -0.9511252 | 0.483 | 0.698 | 1 | 0.03135165 | 0.00039651 |
| Rae1 | 0.01898949 | 0.53686471 | 0.49 | 0.352 | 1 | 0.00019828 | 0.80284487 | 0.471 | 0.219 | 1 | 0.01898949 | 0.00039652 |
| Ndufaf4 | 0.22990649 | 0.33706269 | 0.427 | 0.373 | 1 | 0.00019911 | 0.8917786 | 0.513 | 0.271 | 1 | 0.22990649 | 0.00039818 |
| Upf3a | 0.02968528 | 0.34223041 | 0.594 | 0.461 | 1 | 0.00019914 | 0.59411785 | 0.613 | 0.333 | 1 | 0.02968528 | 0.00039824 |
| Ints12 | 0.05604253 | 0.66460682 | 0.302 | 0.207 | 1 | 0.00019956 | 1.38316698 | 0.315 | 0.115 | 1 | 0.05604253 | 0.00039908 |
| Resf1 | 0.00019995 | -1.326623 | 0.583 | 0.793 | 1 | 0.26243067 | 0.77724331 | 0.651 | 0.708 | 1 | 0.26243067 | 0.00039985 |
| Sdad1 | 0.0030017 | 0.59961683 | 0.562 | 0.399 | 1 | 0.00020017 | 0.52741524 | 0.613 | 0.323 | 1 | 0.0030017 | 0.00040031 |
| Bag6 | 0.0906559 | 0.28867629 | 0.417 | 0.301 | 1 | 0.00020178 | 0.70900376 | 0.462 | 0.219 | 1 | 0.0906559 | 0.00040352 |
| Nav2 | 0.00254086 | -1.1451525 | 0.615 | 0.699 | 1 | 0.00020183 | -1.2447602 | 0.462 | 0.667 | 1 | 0.00254086 | 0.00040362 |

| | | | | | | | | | | | | |
|---|---|---|---|---|---|---|---|---|---|---|---|---|
| Pdzrn3 | 0.83240075 | -0.2860905 | 0.99 | 0.979 | 1 | 0.00020258 | -0.7665333 | 0.992 | 0.917 | 1 | 0.83240075 | 0.00040513 |
| Akirin1 | 0.00020444 | 0.82543421 | 0.677 | 0.549 | 1 | 0.04197626 | 0.31488714 | 0.769 | 0.802 | 1 | 0.04197626 | 0.00040885 |
| Tceal1 | 0.02076057 | 0.75813507 | 0.375 | 0.249 | 1 | 0.0002045 | 0.67765379 | 0.357 | 0.135 | 1 | 0.02076057 | 0.00040895 |
| Rab14 | 0.03732154 | 0.25530821 | 0.927 | 0.974 | 1 | 0.00020489 | 0.29967385 | 0.987 | 0.938 | 1 | 0.03732154 | 0.00040974 |
| Araf | 0.00107829 | 0.51970069 | 0.781 | 0.658 | 1 | 0.0002054 | 0.53965311 | 0.773 | 0.531 | 1 | 0.00107829 | 0.00041077 |
| Agr2 | 0.06075142 | -1.9467018 | 0.052 | 0.119 | 1 | 0.00020553 | -0.9575535 | 0.105 | 0.271 | 1 | 0.06075142 | 0.00041101 |
| Spsb3 | 0.29602514 | 0.26026061 | 0.24 | 0.187 | 1 | 0.00020561 | 1.01207283 | 0.353 | 0.146 | 1 | 0.29602514 | 0.00041118 |
| Prkra | 0.03271161 | 0.60086928 | 0.354 | 0.228 | 1 | 0.00020584 | 1.12013258 | 0.307 | 0.104 | 1 | 0.03271161 | 0.00041164 |
| Ccdc58 | 0.24067171 | -0.6770036 | 0.271 | 0.311 | 1 | 0.00020614 | 0.59060906 | 0.315 | 0.104 | 1 | 0.24067171 | 0.00041223 |
| Cln8 | 0.03134936 | 0.80955516 | 0.26 | 0.161 | 1 | 0.00020637 | 0.81447287 | 0.328 | 0.115 | 1 | 0.03134936 | 0.00041269 |
| Dkc1 | 0.00020791 | 1.06638823 | 0.469 | 0.259 | 1 | 0.00059968 | 0.96698494 | 0.319 | 0.125 | 1 | 0.00059968 | 0.00041579 |
| Nes | 0.00020796 | -3.2295447 | 0.115 | 0.29 | 1 | 0.43136466 | -0.6094155 | 0.042 | 0.062 | 1 | 0.43136466 | 0.00041588 |
| Rab39b | 0.01198146 | 1.17350449 | 0.271 | 0.155 | 1 | 0.00020831 | 1.17013768 | 0.29 | 0.094 | 1 | 0.01198146 | 0.00041657 |
| Pdgfrl | 0.09816636 | 0.33764314 | 0.75 | 0.648 | 1 | 0.00020993 | 0.17270623 | 0.731 | 0.385 | 1 | 0.09816636 | 0.00041981 |
| Tmem185a | 0.82093134 | 0.14647095 | 0.385 | 0.363 | 1 | 0.00021053 | 0.82969609 | 0.454 | 0.219 | 1 | 0.82093134 | 0.00042101 |
| Cmtm7 | 0.20814921 | -0.1702926 | 0.375 | 0.456 | 1 | 0.00021063 | 0.77572047 | 0.529 | 0.292 | 1 | 0.20814921 | 0.00042121 |
| Tafazzin | 0.60283523 | 0.11133035 | 0.51 | 0.461 | 1 | 0.00021112 | 0.74599797 | 0.471 | 0.229 | 1 | 0.60283523 | 0.00042219 |
| Gm56524 | 0.58630804 | 0.42253204 | 0.073 | 0.057 | 1 | 0.00021118 | 3.73453886 | 0.13 | 0 | 1 | 0.58630804 | 0.00042231 |
| Mylip | 0.31870069 | -0.2148979 | 0.427 | 0.332 | 1 | 0.00021154 | -0.9721105 | 0.513 | 0.76 | 1 | 0.31870069 | 0.00042303 |
| Cxxc1 | 0.13770388 | 0.40604391 | 0.25 | 0.171 | 1 | 0.00021211 | 0.82259503 | 0.387 | 0.156 | 1 | 0.13770388 | 0.00042418 |
| Fbxl2 | 0.00021452 | 0.7633416 | 0.688 | 0.513 | 1 | 0.00180627 | 0.33613667 | 0.576 | 0.302 | 1 | 0.00180627 | 0.000429 |
| Gatd3a | 0.30505735 | 0.26383429 | 0.323 | 0.264 | 1 | 0.00021535 | 0.8889246 | 0.361 | 0.146 | 1 | 0.30505735 | 0.00043066 |
| Ppp1r21 | 0.00486925 | 0.866632 | 0.271 | 0.135 | 1 | 0.00021539 | 0.98292649 | 0.273 | 0.083 | 1 | 0.00486925 | 0.00043072 |
| Tenm2 | 0.00021582 | 1.24505533 | 0.417 | 0.233 | 1 | 0.20023121 | -0.679704 | 0.357 | 0.469 | 1 | 0.20023121 | 0.0004316 |
| Nr1d1 | 0.06718211 | 0.53483501 | 0.677 | 0.601 | 1 | 0.00021609 | 0.61170338 | 0.777 | 0.594 | 1 | 0.06718211 | 0.00043212 |
| Ttll1 | 0.023417 | 0.52206771 | 0.229 | 0.119 | 1 | 0.00021639 | 1.69014474 | 0.193 | 0.031 | 1 | 0.023417 | 0.00043274 |
| Csk | 0.05538159 | 0.80396114 | 0.25 | 0.161 | 1 | 0.0002164 | 0.8995981 | 0.349 | 0.135 | 1 | 0.05538159 | 0.00043275 |
| Zmynd8 | 0.13836687 | 0.1222062 | 0.708 | 0.565 | 1 | 0.00021642 | -0.8948178 | 0.588 | 0.781 | 1 | 0.13836687 | 0.0004328 |
| Map7d2 | 0.00021689 | -2.1718292 | 0.177 | 0.383 | 1 | 0.4884557 | -0.9728203 | 0.055 | 0.073 | 1 | 0.4884557 | 0.00043373 |
| Ercc1 | 0.0214748 | 0.6926212 | 0.344 | 0.218 | 1 | 0.00021694 | 1.0369475 | 0.374 | 0.156 | 1 | 0.0214748 | 0.00043383 |
| Slc25a46 | 0.44009817 | 0.31415587 | 0.312 | 0.285 | 1 | 0.00021698 | 0.55364274 | 0.403 | 0.167 | 1 | 0.44009817 | 0.00043391 |
| C430049B03Rik | 0.00021826 | 1.13460645 | 0.312 | 0.124 | 1 | 0.34596242 | -1.3098553 | 0.151 | 0.188 | 1 | 0.34596242 | 0.00043648 |
| Commd9 | 0.16742449 | 0.56692195 | 0.229 | 0.161 | 1 | 0.0002196 | 1.42711033 | 0.282 | 0.094 | 1 | 0.16742449 | 0.00043916 |
| Mink1 | 0.1344524 | 0.35541784 | 0.406 | 0.326 | 1 | 0.00021993 | 0.36415843 | 0.609 | 0.333 | 1 | 0.1344524 | 0.0004398 |
| Zfx | 0.00022009 | 0.70193551 | 0.74 | 0.523 | 1 | 0.87275086 | -0.1227697 | 0.845 | 0.823 | 1 | 0.87275086 | 0.00044014 |
| Med12 | 0.02066451 | 0.68556644 | 0.479 | 0.352 | 1 | 0.00022066 | 0.743584 | 0.55 | 0.312 | 1 | 0.02066451 | 0.00044127 |
| Nampt | 0.65524364 | -0.2820121 | 0.417 | 0.425 | 1 | 0.00022161 | -1.033843 | 0.538 | 0.729 | 1 | 0.65524364 | 0.00044317 |
| Atp2b4 | 0.00022314 | 0.8682776 | 0.656 | 0.497 | 1 | 0.77987927 | -0.5577828 | 0.525 | 0.49 | 1 | 0.77987927 | 0.00044623 |
| Mrps16 | 0.00031028 | 0.86267798 | 0.698 | 0.534 | 1 | 0.0002245 | 0.70344156 | 0.588 | 0.385 | 1 | 0.00031028 | 0.00044895 |
| Ppp4r2 | 0.00047261 | 0.47735606 | 0.906 | 0.845 | 1 | 0.00022511 | 0.38982932 | 0.966 | 0.938 | 1 | 0.00047261 | 0.00045017 |
| Twsg1 | 0.08279429 | 0.1939903 | 0.885 | 0.865 | 1 | 0.00022513 | 0.41322638 | 0.975 | 0.917 | 1 | 0.08279429 | 0.00045022 |
| Ubxn4 | 0.00075694 | 0.44096928 | 1 | 0.99 | 1 | 0.00022562 | 0.26962431 | 0.996 | 0.99 | 1 | 0.00075694 | 0.00045118 |
| Mmp13 | 0.04588873 | -2.7359196 | 0.094 | 0.176 | 1 | 0.00022586 | -6.8275247 | 0.021 | 0.115 | 1 | 0.04588873 | 0.00045167 |
| Bri3bp | 0.00080092 | 0.80104366 | 0.5 | 0.301 | 1 | 0.00022616 | 0.62874419 | 0.441 | 0.198 | 1 | 0.00080092 | 0.00045227 |
| Pdcd11 | 0.17631391 | 0.38600616 | 0.448 | 0.373 | 1 | 0.00022626 | 0.84395007 | 0.454 | 0.219 | 1 | 0.17631391 | 0.00045246 |
| Ipmk | 0.03480492 | 0.39367717 | 0.333 | 0.212 | 1 | 0.0002265 | -1.084563 | 0.235 | 0.438 | 1 | 0.03480492 | 0.00045296 |
| Hsd17b7 | 0.59991488 | -0.4157169 | 0.656 | 0.642 | 1 | 0.00022718 | 0.73047683 | 0.559 | 0.302 | 1 | 0.59991488 | 0.00045431 |
| Gm8797 | 0.63842645 | -0.1753695 | 0.312 | 0.337 | 1 | 0.00022741 | 1.12077909 | 0.378 | 0.167 | 1 | 0.63842645 | 0.00045476 |
| Spag7 | 0.08789729 | 0.28890747 | 0.552 | 0.43 | 1 | 0.00022764 | 0.66060046 | 0.433 | 0.198 | 1 | 0.08789729 | 0.00045523 |
| Igip | 0.0013641 | 1.04496924 | 0.312 | 0.15 | 1 | 0.00022841 | 1.21370669 | 0.366 | 0.156 | 1 | 0.0013641 | 0.00045676 |
| Hyi | 0.22109652 | 0.59868899 | 0.365 | 0.311 | 1 | 0.0002285 | 0.80165305 | 0.387 | 0.156 | 1 | 0.22109652 | 0.00045695 |
| Cd320 | 0.67918221 | -0.1753695 | 0.333 | 0.342 | 1 | 0.00022851 | 1.11464257 | 0.357 | 0.146 | 1 | 0.67918221 | 0.00045697 |
| Kirrel | 0.00022854 | 0.59596298 | 0.802 | 0.575 | 1 | 0.00488742 | 0.17874581 | 0.84 | 0.521 | 1 | 0.00488742 | 0.00045702 |
| Rbm15b | 0.00023104 | 1.08149512 | 0.344 | 0.15 | 1 | 0.0045298 | 0.57208898 | 0.361 | 0.188 | 1 | 0.0045298 | 0.00046202 |
| Tmem41b | 0.60625861 | 0.10235372 | 0.604 | 0.57 | 1 | 0.00023133 | 0.20070485 | 0.744 | 0.49 | 1 | 0.60625861 | 0.0004626 |
| Ascc3 | 0.00023212 | 0.38673131 | 0.906 | 0.798 | 1 | 0.0026001 | -0.7499713 | 0.924 | 0.948 | 1 | 0.0026001 | 0.00046419 |
| Snhg16 | 0.06239835 | 0.6241659 | 0.312 | 0.218 | 1 | 0.00023215 | 1.01207283 | 0.273 | 0.083 | 1 | 0.06239835 | 0.00046424 |
| Tmem140 | 0.03517329 | -0.4718278 | 0.469 | 0.565 | 1 | 0.00023216 | 0.62057489 | 0.584 | 0.312 | 1 | 0.03517329 | 0.00046427 |
| Dpyd | 0.15729907 | -2.577468 | 0 | 0.021 | 1 | 0.00023298 | -2.522849 | 0.05 | 0.177 | 1 | 0.15729907 | 0.00046591 |
| Pmpcb | 0.30665921 | 0.17022404 | 0.531 | 0.466 | 1 | 0.00023309 | 0.53542214 | 0.605 | 0.354 | 1 | 0.30665921 | 0.00046613 |
| Nsmce1 | 0.01175648 | 0.70139141 | 0.396 | 0.254 | 1 | 0.00023354 | 1.13071733 | 0.349 | 0.146 | 1 | 0.01175648 | 0.00046702 |
| Scube1 | 0.00023372 | -1.0180406 | 0.417 | 0.637 | 1 | 0.39629991 | -0.4563414 | 0.298 | 0.333 | 1 | 0.39629991 | 0.00046739 |
| Smtn | 0.0002357 | 0.89201732 | 0.604 | 0.415 | 1 | 0.00572646 | 0.31977629 | 0.786 | 0.552 | 1 | 0.00572646 | 0.00047134 |
| Tmem159 | 0.53758846 | 0.16877544 | 0.385 | 0.337 | 1 | 0.00023581 | 0.82969609 | 0.412 | 0.177 | 1 | 0.53758846 | 0.00047156 |
| Tbl1x | 0.00023623 | 0.44054987 | 0.948 | 0.824 | 1 | 0.33762403 | -0.347899 | 0.962 | 0.979 | 1 | 0.33762403 | 0.0004724 |
| Slc35a4 | 0.21092079 | 0.34756998 | 0.406 | 0.337 | 1 | 0.00023631 | 0.77760758 | 0.336 | 0.125 | 1 | 0.21092079 | 0.00047256 |
| Fnbp1 | 0.2992143 | -0.3010545 | 0.302 | 0.378 | 1 | 0.00023778 | -0.8123556 | 0.504 | 0.719 | 1 | 0.2992143 | 0.0004755 |
| Gm19325 | 0.76891231 | -0.2335136 | 0.083 | 0.093 | 1 | 0.00023871 | 2.62088207 | 0.185 | 0.031 | 1 | 0.76891231 | 0.00047736 |
| Ints6 | 0.08331909 | -0.1294864 | 0.75 | 0.627 | 1 | 0.00023887 | -1.4481233 | 0.798 | 0.885 | 1 | 0.08331909 | 0.00047769 |
| Gpat4 | 0.00210303 | 0.65730818 | 0.583 | 0.409 | 1 | 0.00023902 | 0.58784616 | 0.689 | 0.417 | 1 | 0.00210303 | 0.00047799 |
| Ppdpf | 0.60248922 | -0.2049551 | 0.615 | 0.611 | 1 | 0.00023911 | 0.72377552 | 0.571 | 0.344 | 1 | 0.60248922 | 0.00047816 |
| Aldh1l2 | 0.00023984 | 2.20543391 | 0.188 | 0.052 | 1 | 0.08620958 | 0.84768601 | 0.101 | 0.042 | 1 | 0.08620958 | 0.00047962 |
| Slc22a4 | 0.00024 | -1.3574634 | 0.469 | 0.642 | 1 | 0.32917377 | -0.3735139 | 0.332 | 0.25 | 1 | 0.32917377 | 0.00047994 |
| Bpifb5 | 0.55521849 | 0.346981 | 0.146 | 0.119 | 1 | 0.00024056 | 1.86006974 | 0.218 | 0.052 | 1 | 0.55521849 | 0.00048106 |
| Rpf1 | 0.04972466 | 0.47499946 | 0.417 | 0.301 | 1 | 0.00024242 | 0.70806665 | 0.475 | 0.229 | 1 | 0.04972466 | 0.00048478 |
| Hnrnpd | 0.00024254 | 0.48653304 | 0.906 | 0.824 | 1 | 0.60672909 | -0.1024754 | 0.95 | 0.927 | 1 | 0.60672909 | 0.00048503 |
| Osbpl5 | 0.00024313 | 0.92176466 | 0.406 | 0.197 | 1 | 0.00437415 | 0.68009107 | 0.399 | 0.229 | 1 | 0.00437415 | 0.0004862 |
| Dhx32 | 0.00282048 | -0.6247737 | 0.458 | 0.648 | 1 | 0.00024435 | 0.40317228 | 0.521 | 0.25 | 1 | 0.00282048 | 0.00048865 |
| Bpnt2 | 0.2548179 | 0.13793673 | 0.812 | 0.793 | 1 | 0.00024478 | 0.49191863 | 0.882 | 0.771 | 1 | 0.2548179 | 0.0004895 |
| Prc1 | 0.02458863 | 2.89332352 | 0.125 | 0.052 | 1 | 0.00024588 | -3.4797803 | 0.021 | 0.115 | 1 | 0.02458863 | 0.00049171 |
| Bptf | 0.0002464 | 0.46820961 | 0.927 | 0.85 | 1 | 0.01783485 | 0.17884175 | 1 | 0.979 | 1 | 0.01783485 | 0.00049274 |
| Ccdc157 | 0.05159036 | 0.41055939 | 0.542 | 0.409 | 1 | 0.00024704 | 0.57705096 | 0.542 | 0.281 | 1 | 0.05159036 | 0.00049402 |
| Bet1 | 0.1229138 | 0.3105636 | 0.604 | 0.528 | 1 | 0.00024783 | 0.63874558 | 0.538 | 0.281 | 1 | 0.1229138 | 0.0004956 |
| Rrm1 | 0.00024942 | 2.05188866 | 0.281 | 0.114 | 1 | 0.2047868 | -0.5025003 | 0.197 | 0.125 | 1 | 0.2047868 | 0.00049877 |
| Plod3 | 0.04862887 | 0.35202406 | 0.74 | 0.663 | 1 | 0.00024997 | 0.67827064 | 0.752 | 0.51 | 1 | 0.04862887 | 0.00049988 |
| Zfp560 | 0.06100096 | 0.44719409 | 0.458 | 0.337 | 1 | 0.00025029 | 0.46187575 | 0.429 | 0.177 | 1 | 0.06100096 | 0.00050052 |
| 4930404H11Rik | 0.00025034 | 2.44806713 | 0.115 | 0.016 | 1 | 0.54135468 | 0.49749966 | 0.034 | 0.021 | 1 | 0.54135468 | 0.00050061 |
| Hmgxb4 | 0.09297004 | 0.48666237 | 0.375 | 0.285 | 1 | 0.00025085 | 1.09224318 | 0.328 | 0.115 | 1 | 0.09297004 | 0.00050164 |
| Zbed3 | 0.0201976 | 0.86310463 | 0.292 | 0.176 | 1 | 0.0002511 | 1.24473359 | 0.294 | 0.104 | 1 | 0.0201976 | 0.00050214 |

| | | | | | | | | | | | | |
|---|---|---|---|---|---|---|---|---|---|---|---|---|
| Tnfrsf11b | 0.00025203 | -0.9567402 | 0.75 | 0.762 | 1 | 0.04502422 | 0.6240994 | 0.529 | 0.365 | 1 | 0.04502422 | 0.00050401 |
| Naa20 | 0.00025295 | 0.78193484 | 0.74 | 0.549 | 1 | 0.79720082 | -0.1408162 | 0.718 | 0.667 | 1 | 0.79720082 | 0.00050583 |
| Egr3 | 0.03825606 | -1.2335136 | 0.073 | 0.161 | 1 | 0.00025305 | -1.9313436 | 0.071 | 0.208 | 1 | 0.03825606 | 0.00050604 |
| Vps26a | 0.02291818 | 0.33372277 | 0.688 | 0.497 | 1 | 0.00025407 | 0.49415363 | 0.891 | 0.781 | 1 | 0.02291818 | 0.00050807 |
| Cidea | 0.00025433 | -2.8466546 | 0.104 | 0.285 | 1 | 0.29079175 | 0.2109769 | 0.151 | 0.104 | 1 | 0.29079175 | 0.0005086 |
| Mrpl27 | 0.1320586 | 0.30849679 | 0.51 | 0.409 | 1 | 0.00025512 | 0.63418833 | 0.508 | 0.26 | 1 | 0.1320586 | 0.00051018 |
| Arfgap3 | 0.00025523 | 0.57343221 | 0.938 | 0.803 | 1 | 0.00736093 | 0.26801781 | 0.908 | 0.896 | 1 | 0.00736093 | 0.00051039 |
| Nup62 | 0.00025552 | 1.03786819 | 0.396 | 0.197 | 1 | 0.00231041 | 0.63356121 | 0.336 | 0.156 | 1 | 0.00231041 | 0.00051098 |
| Baz1a | 0.14350159 | -0.6351835 | 0.396 | 0.477 | 1 | 0.00025596 | -0.6830726 | 0.517 | 0.74 | 1 | 0.14350159 | 0.00051185 |
| Ado | 0.51798634 | 0.39981196 | 0.177 | 0.15 | 1 | 0.00025601 | 1.12310414 | 0.261 | 0.073 | 1 | 0.51798634 | 0.00051195 |
| Rarb | 0.36357259 | -1.3144336 | 0.021 | 0.041 | 1 | 0.00025779 | -3.4353861 | 0.021 | 0.115 | 1 | 0.36357259 | 0.00051551 |
| Serpinf1 | 0.00025893 | 0.72484702 | 0.594 | 0.363 | 1 | 0.05262532 | -0.2949049 | 0.643 | 0.406 | 1 | 0.05262532 | 0.0005178 |
| Mrpl16 | 0.92800083 | -0.1249558 | 0.219 | 0.218 | 1 | 0.00025957 | 1.7296731 | 0.261 | 0.083 | 1 | 0.92800083 | 0.00051907 |
| Tor1aip1 | 0.2034497 | 0.12613903 | 0.958 | 0.917 | 1 | 0.00026121 | 0.30929592 | 0.983 | 0.958 | 1 | 0.2034497 | 0.00052235 |
| Nono | 0.01343832 | 0.37006462 | 0.792 | 0.627 | 1 | 0.00026141 | 0.5047897 | 0.857 | 0.75 | 1 | 0.01343832 | 0.00052275 |
| Ube2z | 0.0020165 | 0.63093119 | 0.635 | 0.466 | 1 | 0.00026181 | 0.61541386 | 0.735 | 0.562 | 1 | 0.0020165 | 0.00052356 |
| Cenpp | 0.00026214 | 1.61107272 | 0.354 | 0.166 | 1 | 0.00139745 | 0.95317914 | 0.277 | 0.104 | 1 | 0.00139745 | 0.00052421 |
| Tnfsf11 | 0.49312889 | -0.4950058 | 0.052 | 0.073 | 1 | 0.00026262 | -4.7692869 | 0.008 | 0.083 | 1 | 0.49312889 | 0.00052517 |
| Bglap | 0.00026553 | -3.9943828 | 0.146 | 0.337 | 1 | 0.72110706 | 3.38440588 | 0.05 | 0.042 | 1 | 0.72110706 | 0.000531 |
| Tmem126a | 0.08060092 | -0.3348977 | 0.51 | 0.601 | 1 | 0.00026724 | 0.37580124 | 0.63 | 0.344 | 1 | 0.08060092 | 0.00053442 |
| Tafa5 | 0.06536354 | 0.55254832 | 0.562 | 0.518 | 1 | 0.00026729 | 0.15018063 | 0.735 | 0.458 | 1 | 0.06536354 | 0.00053451 |
| Lmtk2 | 0.40053013 | 0.24565427 | 0.271 | 0.228 | 1 | 0.00026763 | -0.7803059 | 0.37 | 0.625 | 1 | 0.40053013 | 0.00053518 |
| Cdc23 | 0.00026811 | 0.91438513 | 0.469 | 0.249 | 1 | 0.0648304 | 0.28552385 | 0.391 | 0.271 | 1 | 0.0648304 | 0.00053615 |
| Polr2e | 0.01671528 | 0.48666237 | 0.583 | 0.446 | 1 | 0.00026823 | 0.51486082 | 0.571 | 0.333 | 1 | 0.01671528 | 0.0005364 |
| Radil | 0.0002692 | -1.5330738 | 0.156 | 0.358 | 1 | 0.00739894 | -1.355659 | 0.063 | 0.156 | 1 | 0.00739894 | 0.00053833 |
| Wrap73 | 0.1308751 | 0.68556644 | 0.115 | 0.062 | 1 | 0.00027187 | 3.08246216 | 0.147 | 0.01 | 1 | 0.1308751 | 0.00054366 |
| Nme1 | 0.00051184 | 0.56535181 | 0.865 | 0.751 | 1 | 0.00027352 | 0.42609399 | 0.828 | 0.646 | 1 | 0.00051184 | 0.00054697 |
| Clic5 | 0.00027357 | -2.9925055 | 0.01 | 0.15 | 1 | 0.43040711 | -0.7248928 | 0.034 | 0.052 | 1 | 0.43040711 | 0.00054707 |
| Jag1 | 0.60073082 | -0.1624305 | 0.188 | 0.212 | 1 | 0.00027421 | -1.3225663 | 0.261 | 0.458 | 1 | 0.60073082 | 0.00054834 |
| Hgh1 | 0.00027465 | 1.67045955 | 0.188 | 0.052 | 1 | 0.01543255 | 2.77760758 | 0.059 | 0 | 1 | 0.01543255 | 0.00054922 |
| Tnpo2 | 0.00027533 | 0.80104366 | 0.479 | 0.249 | 1 | 0.4060879 | -0.1705805 | 0.471 | 0.365 | 1 | 0.4060879 | 0.00055058 |
| Celf4 | 0.69367172 | -0.2019588 | 0.219 | 0.238 | 1 | 0.00027536 | 2.39058446 | 0.185 | 0.031 | 1 | 0.69367172 | 0.00055064 |
| Rdx | 0.00072204 | 0.33987699 | 0.979 | 0.964 | 1 | 0.00027604 | 0.35504775 | 0.987 | 0.979 | 1 | 0.00072204 | 0.000552 |
| Sprtn | 0.00133125 | 1.27052894 | 0.26 | 0.114 | 1 | 0.00027694 | 0.95693128 | 0.256 | 0.073 | 1 | 0.00133125 | 0.0005538 |
| Arhgdia | 0.01493679 | 0.22161934 | 0.885 | 0.756 | 1 | 0.00027716 | 0.44913664 | 0.861 | 0.76 | 1 | 0.01493679 | 0.00055425 |
| Dusp6 | 0.0002778 | -2.3404288 | 0.073 | 0.249 | 1 | 0.00252696 | -1.1172102 | 0.097 | 0.219 | 1 | 0.00252696 | 0.00055552 |
| Cftr | 0.72298091 | -0.9925055 | 0.01 | 0.016 | 1 | 0.00027813 | -1.129283 | 0.067 | 0.208 | 1 | 0.72298091 | 0.00055618 |
| Eny2 | 0.00027844 | 0.49912446 | 0.896 | 0.824 | 1 | 0.02028099 | 0.20745029 | 0.903 | 0.844 | 1 | 0.02028099 | 0.00055681 |
| Mir100hg | 0.0002793 | -1.0497408 | 0.729 | 0.803 | 1 | 0.05852448 | -0.4853905 | 0.798 | 0.75 | 1 | 0.05852448 | 0.00055852 |
| Tbc1d19 | 0.00028118 | 1.0899567 | 0.365 | 0.166 | 1 | 0.46148424 | -0.4084191 | 0.29 | 0.24 | 1 | 0.46148424 | 0.00056228 |
| Zxdc | 0.03186961 | 0.76648644 | 0.333 | 0.223 | 1 | 0.00028289 | 0.32320153 | 0.538 | 0.271 | 1 | 0.03186961 | 0.0005657 |
| Mrpl43 | 0.23145269 | 0.12917309 | 0.5 | 0.378 | 1 | 0.00028429 | 0.89342834 | 0.487 | 0.281 | 1 | 0.23145269 | 0.0005685 |
| Arxes1 | 0.0002848 | 1.72370157 | 0.167 | 0.041 | 1 | 0.11754502 | 1.10518224 | 0.105 | 0.052 | 1 | 0.11754502 | 0.00056951 |
| Slc7a6 | 0.38836218 | 0.40177348 | 0.375 | 0.332 | 1 | 0.00028491 | 0.90856826 | 0.328 | 0.125 | 1 | 0.38836218 | 0.00056975 |
| Cdr2 | 0.00563076 | 0.87948692 | 0.458 | 0.306 | 1 | 0.00028575 | 0.66770735 | 0.622 | 0.354 | 1 | 0.00563076 | 0.00057142 |
| Pmvk | 0.36742825 | -0.4062813 | 0.552 | 0.58 | 1 | 0.00028654 | 0.99313127 | 0.424 | 0.198 | 1 | 0.36742825 | 0.00057299 |
| Iftap | 0.57216595 | -0.1059635 | 0.385 | 0.425 | 1 | 0.00028744 | 0.74903843 | 0.466 | 0.24 | 1 | 0.57216595 | 0.0005748 |
| Zfyve21 | 0.01398537 | 0.49687938 | 0.5 | 0.347 | 1 | 0.00028826 | 0.90104152 | 0.37 | 0.146 | 1 | 0.01398537 | 0.00057643 |
| Slc19a1 | 0.00028845 | 1.25142012 | 0.365 | 0.176 | 1 | 0.00052595 | 1.08703489 | 0.269 | 0.094 | 1 | 0.00052595 | 0.00057683 |
| Pin1 | 0.00038554 | 0.92343027 | 0.615 | 0.415 | 1 | 0.00028861 | 0.55408319 | 0.559 | 0.302 | 1 | 0.00038554 | 0.00057714 |
| Mettl5 | 0.12552812 | 0.36363835 | 0.281 | 0.197 | 1 | 0.00028867 | 0.78206723 | 0.294 | 0.094 | 1 | 0.12552812 | 0.00057726 |
| Bpgm | 0.23864605 | -0.5595461 | 0.51 | 0.534 | 1 | 0.00028907 | 0.60401332 | 0.769 | 0.531 | 1 | 0.23864605 | 0.00057805 |
| Tet1 | 0.37874831 | 0.40604391 | 0.323 | 0.275 | 1 | 0.00028932 | 1.47150445 | 0.29 | 0.104 | 1 | 0.37874831 | 0.00057855 |
| Rere | 0.26652176 | -0.1712179 | 1 | 0.995 | 1 | 0.00028934 | -0.6175112 | 0.992 | 0.979 | 1 | 0.26652176 | 0.00057859 |
| Snapc4 | 0.00029008 | 1.26125113 | 0.281 | 0.109 | 1 | 0.61558255 | 0.20471791 | 0.134 | 0.115 | 1 | 0.61558255 | 0.00058008 |
| Blvra | 0.00960846 | -0.8307544 | 0.26 | 0.404 | 1 | 0.00029083 | 1.06652062 | 0.353 | 0.146 | 1 | 0.00960846 | 0.00058158 |
| Hpse2 | 0.49738504 | -0.5170321 | 0.219 | 0.176 | 1 | 0.00029285 | 2.06781538 | 0.277 | 0.094 | 1 | 0.49738504 | 0.00058562 |
| Vegfd | 0.00029435 | -2.2075184 | 0.031 | 0.187 | 1 | 0.01607009 | -1.9378865 | 0.025 | 0.083 | 1 | 0.01607009 | 0.00058861 |
| Cdh19 | 0.00029442 | -1.765095 | 0.052 | 0.223 | 1 | 0.00345564 | -3.010295 | 0.008 | 0.062 | 1 | 0.00345564 | 0.00058875 |
| Gstt3 | 0.68887885 | 0.33506919 | 0.24 | 0.223 | 1 | 0.00029449 | 2.10518224 | 0.214 | 0.052 | 1 | 0.68887885 | 0.00058889 |
| Idh2 | 0.0002946 | 0.47343293 | 0.688 | 0.42 | 1 | 0.40245296 | -0.3098553 | 0.592 | 0.656 | 1 | 0.40245296 | 0.00058911 |
| Manbal | 0.23227827 | 0.3393381 | 0.49 | 0.425 | 1 | 0.00029471 | 0.66877109 | 0.534 | 0.281 | 1 | 0.23227827 | 0.00058934 |
| Rbsn | 0.00142118 | 0.56388789 | 0.458 | 0.249 | 1 | 0.00029482 | 0.86006974 | 0.445 | 0.219 | 1 | 0.00142118 | 0.00058954 |
| Ssbp4 | 0.35915005 | -0.1596154 | 0.354 | 0.409 | 1 | 0.00029549 | 0.60227198 | 0.546 | 0.292 | 1 | 0.35915005 | 0.0005909 |
| Stard13 | 0.00029698 | -0.8977576 | 0.875 | 0.917 | 1 | 0.39551891 | -0.164835 | 0.937 | 0.802 | 1 | 0.39551891 | 0.00059387 |
| Hsp90aa1 | 0.05060335 | 0.21220827 | 1 | 0.974 | 1 | 0.00029704 | 0.23790744 | 1 | 0.99 | 1 | 0.05060335 | 0.000594 |
| Eva1b | 0.31869112 | 0.11656624 | 0.51 | 0.472 | 1 | 0.00029795 | 0.66213036 | 0.597 | 0.365 | 1 | 0.31869112 | 0.00059582 |
| Tmem203 | 0.29892422 | -0.3144336 | 0.208 | 0.264 | 1 | 0.00029817 | 1.49749966 | 0.248 | 0.073 | 1 | 0.29892422 | 0.00059625 |
| Tubgcp6 | 0.00029873 | 1.71798792 | 0.177 | 0.047 | 1 | 0.38550879 | 0.23071312 | 0.118 | 0.083 | 1 | 0.38550879 | 0.00059737 |
| 2210016F16Rik | 0.25689661 | 0.2590333 | 0.365 | 0.301 | 1 | 0.00030011 | 0.8548895 | 0.408 | 0.188 | 1 | 0.25689661 | 0.00060013 |
| Lpar4 | 0.01382988 | 0.66084401 | 0.406 | 0.275 | 1 | 0.00030097 | 1.17057665 | 0.37 | 0.167 | 1 | 0.01382988 | 0.00060186 |
| Dennd4a | 0.84587701 | -0.8975089 | 0.458 | 0.43 | 1 | 0.00030122 | -0.7419087 | 0.504 | 0.74 | 1 | 0.84587701 | 0.00060235 |
| Ccnk | 0.01601739 | 0.64106698 | 0.542 | 0.42 | 1 | 0.00030131 | 0.68319198 | 0.504 | 0.271 | 1 | 0.01601739 | 0.00060254 |
| Tbc1d8b | 0.16724019 | 0.35707897 | 0.635 | 0.57 | 1 | 0.00030209 | 0.4259464 | 0.731 | 0.417 | 1 | 0.16724019 | 0.00060409 |
| Cep20 | 0.00030268 | 1.05812061 | 0.427 | 0.228 | 1 | 0.34992942 | -0.1815312 | 0.303 | 0.24 | 1 | 0.34992942 | 0.00060527 |
| Tro | 0.09976439 | 0.48024753 | 0.24 | 0.155 | 1 | 0.00030288 | 1.28840406 | 0.294 | 0.104 | 1 | 0.09976439 | 0.00060567 |
| Upf1 | 0.00575848 | 0.74446013 | 0.542 | 0.394 | 1 | 0.0003034 | 0.46676653 | 0.777 | 0.573 | 1 | 0.00575848 | 0.00060671 |
| Gadd45b | 0.00731949 | 0.50680095 | 0.885 | 0.834 | 1 | 0.00030437 | 0.52503868 | 0.941 | 0.885 | 1 | 0.00731949 | 0.00060864 |
| Dusp16 | 0.05546347 | -0.5450465 | 0.385 | 0.513 | 1 | 0.00030468 | -0.7457263 | 0.538 | 0.781 | 1 | 0.05546347 | 0.00060926 |
| Map1a | 0.06449871 | 0.55683313 | 0.375 | 0.275 | 1 | 0.00030484 | 0.46845113 | 0.521 | 0.271 | 1 | 0.06449871 | 0.00060959 |
| Polr2c | 0.00835779 | 0.5809614 | 0.688 | 0.528 | 1 | 0.00030509 | 0.55628899 | 0.681 | 0.458 | 1 | 0.00835779 | 0.00061009 |
| Nsd1 | 0.00030532 | 0.42749829 | 0.896 | 0.715 | 1 | 0.4692249 | -0.3240412 | 0.954 | 0.948 | 1 | 0.4692249 | 0.00061054 |
| Nrg1 | 0.27427808 | -1.4950058 | 0.021 | 0.047 | 1 | 0.00030814 | -3.0647428 | 0.017 | 0.104 | 1 | 0.27427808 | 0.00061618 |
| Ccdc59 | 0.01227295 | 0.44806713 | 0.771 | 0.653 | 1 | 0.00030925 | 0.5174242 | 0.756 | 0.604 | 1 | 0.01227295 | 0.00061841 |
| Mrpl32 | 0.00796133 | 0.63134605 | 0.583 | 0.42 | 1 | 0.0003098 | 0.88921608 | 0.567 | 0.385 | 1 | 0.00796133 | 0.0006195 |
| Dctd | 0.00030996 | 1.70793425 | 0.219 | 0.073 | 1 | 0.31571378 | 0.49749966 | 0.071 | 0.042 | 1 | 0.31571378 | 0.00061981 |
| Eloa | 0.00031067 | 0.71203865 | 0.729 | 0.56 | 1 | 0.03504563 | 0.39294373 | 0.769 | 0.771 | 1 | 0.03504563 | 0.00062125 |
| Bub3 | 0.04619247 | 0.49782016 | 0.594 | 0.461 | 1 | 0.00031069 | 0.49749966 | 0.723 | 0.479 | 1 | 0.04619247 | 0.00062128 |
| Hs3st3a1 | 0.35309241 | -0.1099283 | 0.667 | 0.591 | 1 | 0.00031108 | 0.64434105 | 0.508 | 0.25 | 1 | 0.35309241 | 0.00062206 |
| Bhlhb9 | 0.09709714 | 0.67818691 | 0.292 | 0.212 | 1 | 0.00031132 | 1.03497323 | 0.382 | 0.167 | 1 | 0.09709714 | 0.00062254 |

| | | | | | | | | | | | | |
|---|---|---|---|---|---|---|---|---|---|---|---|---|
| 9330154J02Rik | 0.00031136 | 4.09495738 | 0.083 | 0.005 | 1 | 0.31168168 | 0.39058446 | 0.029 | 0.01 | 1 | 0.31168168 | 0.00062262 |
| Nrcam | 0.05973235 | -3.7998604 | 0 | 0.036 | 1 | 0.00031233 | -3.7248928 | 0.017 | 0.104 | 1 | 0.05973235 | 0.00062455 |
| Arl6ip6 | 0.01596063 | 0.68867235 | 0.521 | 0.383 | 1 | 0.00031289 | 0.39468885 | 0.479 | 0.24 | 1 | 0.01596063 | 0.00062568 |
| Pde4c | 0.4259442 | -1.577468 | 0.062 | 0.088 | 1 | 0.0003138 | -2.8948178 | 0.021 | 0.115 | 1 | 0.4259442 | 0.0006275 |
| Kat5 | 0.02735118 | 0.68556644 | 0.312 | 0.187 | 1 | 0.00031512 | 1.07918703 | 0.366 | 0.156 | 1 | 0.02735118 | 0.00063015 |
| Slc12a8 | 0.98145284 | -0.3550755 | 0.073 | 0.073 | 1 | 0.00031586 | -3.0260623 | 0.013 | 0.094 | 1 | 0.98145284 | 0.00063162 |
| Pomc | 0.00031724 | 3.59245704 | 0.083 | 0.005 | 1 | 0.75977578 | -0.5025003 | 0.025 | 0.031 | 1 | 0.75977578 | 0.00063438 |
| Cask | 0.38327619 | -0.2722641 | 0.771 | 0.762 | 1 | 0.00031835 | -0.7049932 | 0.849 | 0.906 | 1 | 0.38327619 | 0.0006366 |
| Myc | 0.00031922 | 0.68253238 | 0.677 | 0.435 | 1 | 0.00061408 | 1.21370669 | 0.5 | 0.333 | 1 | 0.00061408 | 0.00063833 |
| Crebl2 | 0.83112015 | 0.16361374 | 0.26 | 0.254 | 1 | 0.0003208 | 0.66589719 | 0.332 | 0.135 | 1 | 0.83112015 | 0.0006415 |
| Dtx3 | 0.02420228 | 0.35232303 | 0.688 | 0.617 | 1 | 0.0003214 | 0.44100477 | 0.702 | 0.406 | 1 | 0.02420228 | 0.0006427 |
| Oca2 | 0.00032233 | -4.4843586 | 0 | 0.124 | 1 | 0.37864212 | -0.6239639 | 0.155 | 0.188 | 1 | 0.37864212 | 0.00064456 |
| Yap1 | 0.00032328 | -0.7305237 | 0.583 | 0.808 | 1 | 0.14523829 | -0.4207159 | 0.824 | 0.896 | 1 | 0.14523829 | 0.00064645 |
| Tspan8 | 0.01978817 | -1.6445822 | 0.031 | 0.114 | 1 | 0.00032737 | -0.7662336 | 0.164 | 0.365 | 1 | 0.01978817 | 0.00065462 |
| Mpzl2 | 0.00032793 | 2.03991601 | 0.24 | 0.088 | 1 | 0.87291177 | 0.44016648 | 0.206 | 0.208 | 1 | 0.87291177 | 0.00065576 |
| Apobec3 | 0.00379504 | -1.0628948 | 0.135 | 0.29 | 1 | 0.00032978 | -1.1759995 | 0.231 | 0.427 | 1 | 0.00379504 | 0.00065945 |
| Rac1 | 0.03851873 | 0.1971772 | 0.99 | 0.974 | 1 | 0.00033024 | 0.30563211 | 0.996 | 0.99 | 1 | 0.03851873 | 0.00066037 |
| Emc6 | 0.23567403 | 0.24969544 | 0.635 | 0.56 | 1 | 0.00033083 | 0.79216567 | 0.639 | 0.51 | 1 | 0.23567403 | 0.00066154 |
| Gne | 0.00033094 | 0.91019734 | 0.333 | 0.14 | 1 | 0.00063819 | 0.60575255 | 0.277 | 0.094 | 1 | 0.00063819 | 0.00066177 |
| Blmh | 0.00048743 | 0.47278681 | 0.802 | 0.668 | 1 | 0.00033165 | 0.42711033 | 0.84 | 0.719 | 1 | 0.00048743 | 0.00066318 |
| Ppm1n | 0.08587075 | -2.3500575 | 0.01 | 0.052 | 1 | 0.00033277 | -2.7692869 | 0.013 | 0.094 | 1 | 0.08587075 | 0.00066543 |
| Fam193a | 0.00634745 | 0.21144971 | 0.896 | 0.767 | 1 | 0.00033281 | -0.6102502 | 0.987 | 0.979 | 1 | 0.00634745 | 0.00066551 |
| Extl2 | 0.25321298 | 0.28760246 | 0.375 | 0.311 | 1 | 0.00033355 | 0.98676174 | 0.349 | 0.146 | 1 | 0.25321298 | 0.00066699 |
| Mrpl20 | 0.25619633 | 0.18704396 | 0.823 | 0.767 | 1 | 0.00033553 | 0.45939139 | 0.84 | 0.677 | 1 | 0.25619633 | 0.00067094 |
| Trappc6b | 0.00033558 | 0.4959942 | 0.76 | 0.565 | 1 | 0.01070345 | 0.30137289 | 0.773 | 0.688 | 1 | 0.01070345 | 0.00067105 |
| Pxylp1 | 0.19536925 | -0.4716733 | 0.344 | 0.425 | 1 | 0.00033606 | 0.86417414 | 0.382 | 0.167 | 1 | 0.19536925 | 0.00067201 |
| Micos10 | 0.27067951 | 0.11921962 | 0.948 | 0.933 | 1 | 0.00033809 | 0.39943537 | 0.962 | 0.938 | 1 | 0.27067951 | 0.00067606 |
| Pttg1 | 0.70387154 | -0.2219873 | 0.375 | 0.389 | 1 | 0.00033814 | 0.97435317 | 0.412 | 0.198 | 1 | 0.70387154 | 0.00067617 |
| Thoc3 | 0.00299355 | 0.72851859 | 0.479 | 0.316 | 1 | 0.00033829 | 0.4152849 | 0.5 | 0.25 | 1 | 0.00299355 | 0.00067646 |
| Acsl6 | 0.38521705 | 0.68556644 | 0.031 | 0.016 | 1 | 0.00033921 | 2.59703533 | 0.164 | 0.021 | 1 | 0.38521705 | 0.0006783 |
| 2410002F23Rik | 0.00033965 | 0.81045769 | 0.594 | 0.358 | 1 | 0.00313135 | 0.52920852 | 0.454 | 0.271 | 1 | 0.00313135 | 0.00067918 |
| Ubc | 0.09057311 | -0.3857476 | 1 | 0.979 | 1 | 0.00034023 | 0.28794425 | 1 | 0.969 | 1 | 0.09057311 | 0.00068034 |
| Itch | 0.00034368 | 0.3597052 | 0.948 | 0.824 | 1 | 0.09165999 | 0.10484 | 0.966 | 0.927 | 1 | 0.09165999 | 0.00068725 |
| Myd88 | 0.85568777 | 0.15534 | 0.417 | 0.425 | 1 | 0.00034576 | 0.44716798 | 0.479 | 0.24 | 1 | 0.85568777 | 0.00069141 |
| Rnaset2a | 0.00416623 | 0.42035766 | 0.865 | 0.715 | 1 | 0.00034653 | 0.21947837 | 0.777 | 0.615 | 1 | 0.00416623 | 0.00069293 |
| Slc39a14 | 0.00145525 | -0.9990483 | 0.938 | 0.855 | 1 | 0.00034692 | 0.22357694 | 0.714 | 0.417 | 1 | 0.00145525 | 0.00069371 |
| Scnm1 | 0.00034863 | 1.35896691 | 0.281 | 0.114 | 1 | 0.56228852 | 0.10518224 | 0.151 | 0.125 | 1 | 0.56228852 | 0.00069714 |
| A430073D23Rik | 0.00034927 | 1.59245704 | 0.156 | 0.036 | 1 | 0.25857653 | 0.49749966 | 0.063 | 0.031 | 1 | 0.25857653 | 0.00069842 |
| Kbtbd2 | 0.01046439 | 0.73541499 | 0.396 | 0.249 | 1 | 0.00034962 | 0.43336932 | 0.588 | 0.312 | 1 | 0.01046439 | 0.00069911 |
| Uso1 | 0.02002031 | 0.2626607 | 0.896 | 0.834 | 1 | 0.00034974 | 0.42515281 | 0.945 | 0.885 | 1 | 0.02002031 | 0.00069936 |
| Mrps7 | 0.00113745 | 0.74446013 | 0.552 | 0.342 | 1 | 0.00035125 | 0.59479686 | 0.462 | 0.229 | 1 | 0.00113745 | 0.00070239 |
| Cdc14a | 0.01385484 | -0.8608615 | 0.26 | 0.404 | 1 | 0.0003517 | -0.7962315 | 0.429 | 0.656 | 1 | 0.01385484 | 0.00070329 |
| Trim41 | 0.00090526 | 0.71572841 | 0.458 | 0.244 | 1 | 0.00035616 | 0.69014474 | 0.475 | 0.24 | 1 | 0.00090526 | 0.00071219 |
| Phyhd1 | 0.00035653 | 1.47144164 | 0.281 | 0.114 | 1 | 0.01619917 | 0.58544736 | 0.265 | 0.135 | 1 | 0.01619917 | 0.00071293 |
| Slc24a3 | 0.63158798 | -1.3144336 | 0.073 | 0.088 | 1 | 0.00035695 | -2.2733294 | 0.055 | 0.177 | 1 | 0.63158798 | 0.00071376 |
| Pcm1 | 0.00035864 | 0.40258904 | 0.948 | 0.876 | 1 | 0.32502215 | -0.2556995 | 0.958 | 0.958 | 1 | 0.32502215 | 0.00071716 |
| Fam3c | 0.00244653 | -0.730782 | 0.677 | 0.741 | 1 | 0.00036082 | 0.59575237 | 0.651 | 0.458 | 1 | 0.00244653 | 0.00072152 |
| Mlph | 0.00036102 | -2.9925055 | 0.094 | 0.259 | 1 | 0.03696427 | -1.0583165 | 0.118 | 0.042 | 1 | 0.03696427 | 0.0007219 |
| Fads1 | 0.00036298 | 1.15188445 | 0.5 | 0.311 | 1 | 0.0755919 | 0.20799304 | 0.374 | 0.25 | 1 | 0.0755919 | 0.00072583 |
| Zmym1 | 0.00036327 | 1.13302542 | 0.365 | 0.176 | 1 | 0.0197965 | 0.52021974 | 0.273 | 0.146 | 1 | 0.0197965 | 0.00072641 |
| Gm26782 | 0.17894515 | 0.37314401 | 0.427 | 0.352 | 1 | 0.00036365 | 0.84309666 | 0.437 | 0.219 | 1 | 0.17894515 | 0.00072718 |
| Zfp410 | 0.3307514 | 0.20013961 | 0.333 | 0.28 | 1 | 0.00036783 | 0.57188186 | 0.521 | 0.26 | 1 | 0.3307514 | 0.00073552 |
| Kpna4 | 0.44846899 | 0.1073298 | 0.896 | 0.876 | 1 | 0.00036798 | -0.5241106 | 0.962 | 0.958 | 1 | 0.44846899 | 0.00073582 |
| Khk | 0.96340749 | 0.40342321 | 0.167 | 0.171 | 1 | 0.00036953 | 1.96316323 | 0.197 | 0.042 | 1 | 0.96340749 | 0.00073893 |
| Ncoa2 | 0.805579 | -0.2432196 | 0.75 | 0.71 | 1 | 0.00036992 | -0.7434224 | 0.84 | 0.948 | 1 | 0.805579 | 0.0007397 |
| Arnt | 0.00037006 | -0.8449483 | 0.667 | 0.788 | 1 | 0.03089815 | -0.5235236 | 0.819 | 0.812 | 1 | 0.03089815 | 0.00073998 |
| Rabggtb | 0.04199032 | 0.35059814 | 0.74 | 0.663 | 1 | 0.00037019 | 0.55858503 | 0.765 | 0.552 | 1 | 0.04199032 | 0.00074024 |
| Medag | 0.00037244 | -1.5511813 | 0.281 | 0.492 | 1 | 0.32637412 | 0.75623393 | 0.227 | 0.188 | 1 | 0.32637412 | 0.00074474 |
| Arl6ip4 | 0.27030256 | 0.24850264 | 0.438 | 0.363 | 1 | 0.00037365 | 0.70806665 | 0.462 | 0.24 | 1 | 0.27030256 | 0.00074716 |
| Gnptg | 0.15354233 | 0.30866407 | 0.51 | 0.409 | 1 | 0.0003741 | 0.70137199 | 0.424 | 0.198 | 1 | 0.15354233 | 0.00074807 |
| Ndufb6 | 0.08758176 | 0.19035859 | 0.823 | 0.694 | 1 | 0.00037521 | 0.64898741 | 0.718 | 0.604 | 1 | 0.08758176 | 0.00075027 |
| Ash2l | 0.17848771 | 0.20013961 | 0.365 | 0.28 | 1 | 0.00037976 | 0.62212631 | 0.429 | 0.198 | 1 | 0.17848771 | 0.00075938 |
| Tmem258 | 0.00037995 | 0.54806292 | 0.927 | 0.756 | 1 | 0.00415856 | 0.38599134 | 0.744 | 0.656 | 1 | 0.00415856 | 0.00075976 |
| Pdcl | 0.07723004 | 0.4134869 | 0.396 | 0.285 | 1 | 0.00038009 | 0.9406882 | 0.382 | 0.167 | 1 | 0.07723004 | 0.00076004 |
| Pag1 | 0.01311301 | 0.7008332 | 0.198 | 0.093 | 1 | 0.00038014 | -0.6195398 | 0.244 | 0.469 | 1 | 0.01311301 | 0.00076014 |
| Lsm1 | 0.3773315 | -0.3404288 | 0.406 | 0.466 | 1 | 0.00038053 | 0.46066289 | 0.563 | 0.323 | 1 | 0.3773315 | 0.00076092 |
| Htra2 | 0.28079692 | 0.23576352 | 0.365 | 0.295 | 1 | 0.00038233 | 1.02717972 | 0.319 | 0.125 | 1 | 0.28079692 | 0.00076451 |
| Slc37a3 | 0.02150633 | 0.48481631 | 0.49 | 0.342 | 1 | 0.00038299 | 0.59363676 | 0.634 | 0.427 | 1 | 0.02150633 | 0.00076584 |
| Vezf1 | 0.00075287 | 0.44693338 | 0.896 | 0.788 | 1 | 0.00038388 | 0.42064808 | 0.954 | 0.875 | 1 | 0.00075287 | 0.00076762 |
| Zbed5 | 0.00038445 | 1.53356335 | 0.25 | 0.093 | 1 | 0.05334374 | 0.71251255 | 0.181 | 0.094 | 1 | 0.05334374 | 0.00076876 |
| Banf1 | 0.00038526 | 0.56892152 | 0.823 | 0.575 | 1 | 0.00426297 | 0.5580412 | 0.651 | 0.562 | 1 | 0.00426297 | 0.00077038 |
| Vcl | 0.00038572 | -0.7593813 | 0.823 | 0.948 | 1 | 0.07075584 | -0.563649 | 0.908 | 0.969 | 1 | 0.07075584 | 0.00077128 |
| Cand1 | 0.00038703 | 0.57179757 | 0.781 | 0.554 | 1 | 0.0209723 | 0.27054174 | 0.866 | 0.833 | 1 | 0.0209723 | 0.00077392 |
| Nacc2 | 0.40204484 | 0.21102793 | 0.406 | 0.363 | 1 | 0.0003874 | 0.55687821 | 0.634 | 0.354 | 1 | 0.40204484 | 0.00077464 |
| Aida | 0.0246881 | 0.30295042 | 0.646 | 0.482 | 1 | 0.00039019 | 0.47150445 | 0.672 | 0.417 | 1 | 0.0246881 | 0.00078024 |
| Endou | 0.00039059 | -2.2779077 | 0.073 | 0.238 | 1 | 0.28456652 | -0.2893912 | 0.168 | 0.115 | 1 | 0.28456652 | 0.00078103 |
| Camta1 | 0.05649308 | -0.1332638 | 0.76 | 0.642 | 1 | 0.00039085 | -0.7330667 | 0.597 | 0.823 | 1 | 0.05649308 | 0.00078155 |
| Mynn | 0.00675225 | 0.64492446 | 0.417 | 0.249 | 1 | 0.00039186 | 0.59392942 | 0.374 | 0.156 | 1 | 0.00675225 | 0.00078357 |
| Yod1 | 0.00067802 | 1.14665728 | 0.417 | 0.223 | 1 | 0.00039249 | 1.28599555 | 0.399 | 0.177 | 1 | 0.00067802 | 0.00078482 |
| Reep3 | 0.24151737 | 0.1162008 | 0.958 | 0.953 | 1 | 0.00039261 | 0.31243649 | 0.996 | 0.979 | 1 | 0.24151737 | 0.00078507 |
| Gtpbp4 | 0.00073105 | 0.57896119 | 0.875 | 0.756 | 1 | 0.000395 | 0.39573694 | 0.845 | 0.75 | 1 | 0.00073105 | 0.00078985 |
| P3h2 | 0.54000084 | -0.7858785 | 0.854 | 0.731 | 1 | 0.00039758 | -0.3512608 | 0.718 | 0.396 | 1 | 0.54000084 | 0.00079499 |
| Fbxo17 | 0.04074882 | 0.55769162 | 0.302 | 0.192 | 1 | 0.00039765 | 1.48270266 | 0.261 | 0.083 | 1 | 0.04074882 | 0.00079515 |
| Cercam | 0.00039879 | 1.22430593 | 0.292 | 0.119 | 1 | 0.01432157 | 1.20799304 | 0.181 | 0.073 | 1 | 0.01432157 | 0.00079742 |
| Lamb3 | 0.0143715 | -1.7580402 | 0.031 | 0.119 | 1 | 0.00039991 | -2.1723517 | 0.038 | 0.146 | 1 | 0.0143715 | 0.00079966 |
| Lyar | 0.05950246 | 0.45167838 | 0.531 | 0.415 | 1 | 0.00040001 | 0.55194744 | 0.487 | 0.24 | 1 | 0.05950246 | 0.00079987 |
| Yipf2 | 0.17399122 | 0.43633784 | 0.271 | 0.197 | 1 | 0.00040032 | 1.03118166 | 0.269 | 0.083 | 1 | 0.17399122 | 0.00080047 |
| Ngrn | 0.25977167 | 0.48142572 | 0.219 | 0.166 | 1 | 0.00040106 | 1.36821664 | 0.248 | 0.073 | 1 | 0.25977167 | 0.00080196 |
| Igfbp6 | 0.11302853 | -2.3144336 | 0 | 0.026 | 1 | 0.0004016 | -6.7692869 | 0 | 0.052 | 1 | 0.11302853 | 0.00080304 |

| | | | | | | | | | | | | |
|---|---|---|---|---|---|---|---|---|---|---|---|---|
| Ifi47 | 0.62104899 | -0.3144336 | 0.01 | 0.005 | 1 | 0.00040163 | -5.1172102 | 0 | 0.052 | 1 | 0.62104899 | 0.0008031 |
| Palm3 | 1 | 0.42253204 | 0.01 | 0.01 | 1 | 0.00040163 | -4.4797803 | 0 | 0.052 | 1 | 1 | 0.0008031 |
| Mrps23 | 0.03295827 | 0.63134605 | 0.385 | 0.269 | 1 | 0.0004021 | 1.10518224 | 0.349 | 0.146 | 1 | 0.03295827 | 0.00080403 |
| Msrb2 | 0.00040301 | 1.56692195 | 0.24 | 0.088 | 1 | 0.01122226 | 0.80562195 | 0.244 | 0.115 | 1 | 0.01122226 | 0.00080587 |
| Mzt2 | 0.00485592 | 1.13959807 | 0.406 | 0.254 | 1 | 0.00040414 | 1.12871775 | 0.319 | 0.125 | 1 | 0.00485592 | 0.00080812 |
| Ncs1 | 0.00040421 | 0.82040898 | 0.385 | 0.181 | 1 | 0.85825309 | -0.3736704 | 0.433 | 0.375 | 1 | 0.85825309 | 0.00080826 |
| Ptpn11 | 0.10567435 | 0.19058531 | 0.833 | 0.725 | 1 | 0.00040662 | 0.35975506 | 0.891 | 0.792 | 1 | 0.10567435 | 0.00081308 |
| Hycc1 | 0.9564174 | -0.2858644 | 0.323 | 0.311 | 1 | 0.00040717 | -0.7826083 | 0.332 | 0.552 | 1 | 0.9564174 | 0.00081418 |
| Gm56698 | 0.00040815 | 1.59245704 | 0.135 | 0.026 | 1 | 0.34625029 | 0.80562195 | 0.042 | 0.021 | 1 | 0.34625029 | 0.00081614 |
| Sccpdh | 0.00040823 | 0.9800138 | 0.385 | 0.197 | 1 | 0.00086999 | 0.61071027 | 0.37 | 0.167 | 1 | 0.00086999 | 0.0008163 |
| Wnk3 | 0.00040847 | 1.22161934 | 0.229 | 0.078 | 1 | 0.00146101 | 0.86006974 | 0.277 | 0.104 | 1 | 0.00146101 | 0.00081678 |
| Tbc1d9 | 0.47002772 | -0.4779323 | 0.042 | 0.062 | 1 | 0.00041246 | -1.1399303 | 0.109 | 0.271 | 1 | 0.47002772 | 0.00082476 |
| Thoc6 | 0.29901331 | 0.23611891 | 0.385 | 0.316 | 1 | 0.00041283 | 0.39415991 | 0.454 | 0.208 | 1 | 0.29901331 | 0.00082548 |
| Bcr | 0.00041479 | -0.9467018 | 0.5 | 0.699 | 1 | 0.00795199 | -0.7631812 | 0.681 | 0.75 | 1 | 0.00795199 | 0.0008294 |
| Pim1 | 0.00041489 | 0.47604755 | 0.469 | 0.269 | 1 | 0.00387124 | -0.4555029 | 0.416 | 0.625 | 1 | 0.00387124 | 0.00082961 |
| 1110065P20Rik | 0.0775364 | 0.35707897 | 0.552 | 0.435 | 1 | 0.00041515 | 0.43017663 | 0.496 | 0.229 | 1 | 0.0775364 | 0.00083013 |
| Mfsd14b | 0.00041573 | 0.60850498 | 0.625 | 0.409 | 1 | 0.33425737 | 0.11898804 | 0.685 | 0.646 | 1 | 0.33425737 | 0.00083129 |
| Atp5h | 0.12005892 | 0.14427327 | 0.969 | 0.938 | 1 | 0.00041599 | 0.3656061 | 0.941 | 0.865 | 1 | 0.12005892 | 0.00083181 |
| Sim2 | 0.31219908 | 0.22613482 | 0.531 | 0.466 | 1 | 0.00041955 | 0.75956614 | 0.538 | 0.302 | 1 | 0.31219908 | 0.00083893 |
| Tug1 | 0.00042003 | 0.529793 | 0.76 | 0.497 | 1 | 0.025057 | 0.31029667 | 0.71 | 0.583 | 1 | 0.025057 | 0.00083989 |
| Ddx19a | 0.03696336 | 0.55111624 | 0.354 | 0.228 | 1 | 0.00042036 | 0.71183981 | 0.429 | 0.198 | 1 | 0.03696336 | 0.00084055 |
| Mthfs | 0.00046164 | 1.13087695 | 0.427 | 0.228 | 1 | 0.00042043 | 0.98432784 | 0.307 | 0.115 | 1 | 0.00046164 | 0.00084067 |
| B9d2 | 0.00042174 | 1.35541784 | 0.25 | 0.093 | 1 | 0.0080011 | 0.90515763 | 0.21 | 0.083 | 1 | 0.0080011 | 0.00084331 |
| Fasn | 0.00405065 | 0.47099352 | 0.615 | 0.435 | 1 | 0.00042203 | 0.81352715 | 0.534 | 0.312 | 1 | 0.00405065 | 0.00084388 |
| Mthfr | 0.00042489 | 1.03396675 | 0.375 | 0.187 | 1 | 0.53041223 | -0.2654611 | 0.311 | 0.25 | 1 | 0.53041223 | 0.0008496 |
| Basp1 | 0.35107844 | -1.1624305 | 0.062 | 0.093 | 1 | 0.00042563 | -1.63702 | 0.109 | 0.26 | 1 | 0.35107844 | 0.00085108 |
| Dnajc9 | 0.00042634 | 1.50148338 | 0.531 | 0.352 | 1 | 0.00138652 | 0.19567277 | 0.408 | 0.198 | 1 | 0.00138652 | 0.00085251 |
| Rcc1l | 0.1401927 | 0.42253204 | 0.271 | 0.192 | 1 | 0.00042675 | 0.70395054 | 0.319 | 0.115 | 1 | 0.1401927 | 0.00085333 |
| Rhob | 0.00042742 | 1.14122934 | 0.802 | 0.622 | 1 | 0.05982353 | 0.7590066 | 0.739 | 0.75 | 1 | 0.05982353 | 0.00085466 |
| Anp32e | 0.00080495 | 0.51716891 | 0.74 | 0.57 | 1 | 0.00042963 | 0.52343645 | 0.786 | 0.719 | 1 | 0.00080495 | 0.00085907 |
| Robo2 | 0.00043186 | -2.240433 | 0.115 | 0.295 | 1 | 0.06854142 | -1.0778127 | 0.311 | 0.375 | 1 | 0.06854142 | 0.00086352 |
| Aamdc | 0.41524567 | 0.31234912 | 0.427 | 0.394 | 1 | 0.000433 | 0.17889192 | 0.458 | 0.208 | 1 | 0.41524567 | 0.00086581 |
| Pde7a | 0.38322859 | 0.35541784 | 0.25 | 0.212 | 1 | 0.00043419 | -0.7942665 | 0.437 | 0.698 | 1 | 0.38322859 | 0.0008682 |
| Dhx30 | 0.000436 | 0.85897201 | 0.531 | 0.316 | 1 | 0.02339599 | 0.25951038 | 0.496 | 0.312 | 1 | 0.02339599 | 0.00087181 |
| Limch1 | 0.87739557 | 0.23490503 | 0.177 | 0.171 | 1 | 0.0004368 | 0.80771433 | 0.282 | 0.094 | 1 | 0.87739557 | 0.00087341 |
| 9130401M01Rik | 0.07043944 | 0.44677958 | 0.344 | 0.238 | 1 | 0.00043699 | 1.04378169 | 0.311 | 0.115 | 1 | 0.07043944 | 0.0008738 |
| Kat2b | 0.00420492 | -1.235362 | 0.188 | 0.347 | 1 | 0.00043785 | -0.7526378 | 0.37 | 0.646 | 1 | 0.00420492 | 0.0008755 |
| Eif3k | 0.23751439 | 0.17291815 | 0.917 | 0.907 | 1 | 0.00043827 | 0.39058446 | 0.933 | 0.833 | 1 | 0.23751439 | 0.00087636 |
| Adra2c | 0.00043931 | -4.0368996 | 0 | 0.119 | 1 | 0.00630389 | -4.3098553 | 0 | 0.031 | 1 | 0.00630389 | 0.00087843 |
| N6amt1 | 0.85686727 | -0.1588638 | 0.333 | 0.337 | 1 | 0.00044129 | 1.64434105 | 0.269 | 0.094 | 1 | 0.85686727 | 0.00088238 |
| Zfp36l2 | 0.17950572 | -0.1787243 | 0.667 | 0.534 | 1 | 0.00044247 | -0.7667129 | 0.647 | 0.833 | 1 | 0.17950572 | 0.00088475 |
| Ccn4 | 0.00044318 | 0.65538829 | 0.542 | 0.326 | 1 | 0.00908818 | 0.74077081 | 0.324 | 0.167 | 1 | 0.00908818 | 0.00088616 |
| Shc1 | 0.00044322 | 0.51963072 | 0.771 | 0.611 | 1 | 0.00562295 | 0.24109835 | 0.878 | 0.562 | 1 | 0.00562295 | 0.00088625 |
| Lman2 | 0.0004458 | 0.60747407 | 0.802 | 0.663 | 1 | 0.04041538 | 0.29091775 | 0.761 | 0.667 | 1 | 0.04041538 | 0.0008914 |
| Pogk | 0.00044754 | 0.97507306 | 0.365 | 0.171 | 1 | 0.02110208 | 0.11516633 | 0.361 | 0.208 | 1 | 0.02110208 | 0.00089488 |
| Pdpk1 | 0.0004478 | 0.55818714 | 0.771 | 0.679 | 1 | 0.93038503 | -0.2887376 | 0.903 | 0.896 | 1 | 0.93038503 | 0.00089541 |
| Nudc | 0.01095491 | 0.50454045 | 0.792 | 0.648 | 1 | 0.00044867 | 0.62439029 | 0.727 | 0.615 | 1 | 0.01095491 | 0.00089713 |
| Dda1 | 0.00835386 | 0.48734033 | 0.74 | 0.554 | 1 | 0.00044919 | 0.42304067 | 0.609 | 0.365 | 1 | 0.00835386 | 0.00089818 |
| Kpnb1 | 0.00386489 | 0.40103097 | 0.708 | 0.513 | 1 | 0.00045323 | 0.49246407 | 0.794 | 0.646 | 1 | 0.00386489 | 0.00090625 |
| Scoc | 0.1601303 | 0.2542883 | 0.75 | 0.642 | 1 | 0.00045408 | 0.391705 | 0.672 | 0.396 | 1 | 0.1601303 | 0.00090795 |
| Nfatc2 | 0.00045439 | 1.35855369 | 0.667 | 0.575 | 1 | 0.66152174 | -0.4840969 | 0.639 | 0.583 | 1 | 0.66152174 | 0.00090858 |
| Atp5c1 | 0.13623903 | 0.15949763 | 0.927 | 0.896 | 1 | 0.00045547 | 0.38232241 | 0.933 | 0.854 | 1 | 0.13623903 | 0.00091074 |
| Selenoh | 0.15456963 | 0.48393258 | 0.396 | 0.306 | 1 | 0.00045581 | 1.48270266 | 0.294 | 0.115 | 1 | 0.15456963 | 0.00091141 |
| Prpf4 | 0.00045583 | 1.1124641 | 0.396 | 0.207 | 1 | 0.28742209 | -0.2952085 | 0.319 | 0.229 | 1 | 0.28742209 | 0.00091146 |
| Hmgcr | 0.46340034 | 0.25233945 | 0.74 | 0.715 | 1 | 0.00045634 | 0.81472416 | 0.706 | 0.562 | 1 | 0.46340034 | 0.00091247 |
| Rbbp4 | 0.16320999 | 0.13622785 | 0.938 | 0.881 | 1 | 0.00045688 | 0.37464291 | 0.929 | 0.896 | 1 | 0.16320999 | 0.00091356 |
| 9930021J03Rik | 0.15359021 | -0.1439172 | 0.5 | 0.575 | 1 | 0.00045821 | -0.6297736 | 0.681 | 0.854 | 1 | 0.15359021 | 0.0009162 |
| Vamp4 | 0.41794294 | 0.16480654 | 0.635 | 0.606 | 1 | 0.00045987 | 0.42711033 | 0.613 | 0.344 | 1 | 0.41794294 | 0.00091952 |
| Denr | 0.00046029 | 0.61226552 | 0.76 | 0.617 | 1 | 0.01206303 | 0.35882906 | 0.681 | 0.583 | 1 | 0.01206303 | 0.00092036 |
| Eef1e1 | 0.00046081 | 1.04813652 | 0.458 | 0.249 | 1 | 0.01374324 | 0.81116014 | 0.282 | 0.156 | 1 | 0.01374324 | 0.00092142 |
| Ogfod1 | 0.00046103 | 0.74097956 | 0.458 | 0.259 | 1 | 0.21277786 | -0.1216515 | 0.403 | 0.302 | 1 | 0.21277786 | 0.00092184 |
| Htatsf1 | 0.00056691 | 0.53356335 | 0.771 | 0.596 | 1 | 0.00046398 | 0.52613971 | 0.765 | 0.604 | 1 | 0.00056691 | 0.00092775 |
| Fkbp1b | 0.00188415 | 1.15433592 | 0.271 | 0.124 | 1 | 0.00046428 | 2.24473359 | 0.16 | 0.021 | 1 | 0.00188415 | 0.00092834 |
| Nrn1l | 0.00046434 | 4.00749454 | 0.062 | 0 | 1 | 0.27154471 | 0.69014474 | 0.013 | 0 | 1 | 0.27154471 | 0.00092847 |
| Sh2b1 | 0.01592352 | 0.58122978 | 0.292 | 0.161 | 1 | 0.00046453 | 1.14265694 | 0.328 | 0.135 | 1 | 0.01592352 | 0.00092884 |
| Vwc2 | 0.00046454 | 4.91438513 | 0.062 | 0 | 1 | 0.86312513 | -0.7248928 | 0.008 | 0.01 | 1 | 0.86312513 | 0.00092887 |
| Gm45902 | 0.00046459 | 3.81484946 | 0.062 | 0 | 1 | 0.17039078 | 0.27510724 | 0.071 | 0.031 | 1 | 0.17039078 | 0.00092897 |
| 4933433G19Rik | 0.00046459 | 3.81484946 | 0.062 | 0 | 1 | 0.24392267 | 0.77760758 | 0.063 | 0.031 | 1 | 0.24392267 | 0.00092897 |
| Rtcb | 0.0181715 | 0.32099401 | 0.74 | 0.601 | 1 | 0.0004657 | 0.24527201 | 0.735 | 0.49 | 1 | 0.0181715 | 0.00093119 |
| Ankrd46 | 0.10653585 | 0.42253204 | 0.385 | 0.295 | 1 | 0.00046639 | 0.85453156 | 0.412 | 0.198 | 1 | 0.10653585 | 0.00093257 |
| Arl16 | 0.00046862 | 1.00749454 | 0.281 | 0.109 | 1 | 0.23876664 | 0.31163311 | 0.143 | 0.094 | 1 | 0.23876664 | 0.00093702 |
| Tpcn1 | 0.0004688 | 0.95966601 | 0.51 | 0.306 | 1 | 0.80292855 | -0.1507888 | 0.622 | 0.521 | 1 | 0.80292855 | 0.00093739 |
| Cdh11 | 0.94789658 | -2.1901755 | 0.385 | 0.301 | 1 | 0.00047018 | -3.098541 | 0.319 | 0.417 | 1 | 0.94789658 | 0.00094014 |
| Rrp36 | 0.11275205 | 0.32160313 | 0.354 | 0.254 | 1 | 0.00047059 | 0.80117605 | 0.332 | 0.125 | 1 | 0.11275205 | 0.00094095 |
| Mia3 | 0.00870165 | 0.25168288 | 0.927 | 0.86 | 1 | 0.00047198 | 0.36552253 | 0.933 | 0.917 | 1 | 0.00870165 | 0.00094373 |
| Ints6l | 0.00047333 | 0.70155847 | 0.458 | 0.259 | 1 | 0.44781661 | -0.8478543 | 0.559 | 0.583 | 1 | 0.44781661 | 0.00094644 |
| Adamtsl3 | 0.00047573 | -1.7313619 | 0.615 | 0.741 | 1 | 0.3003769 | -0.8008977 | 0.815 | 0.688 | 1 | 0.3003769 | 0.00095124 |
| Rnf25 | 0.01067936 | 0.71010863 | 0.531 | 0.373 | 1 | 0.0004761 | 0.67103591 | 0.445 | 0.219 | 1 | 0.01067936 | 0.00095198 |
| Atg16l2 | 0.00047656 | 0.90347147 | 0.562 | 0.373 | 1 | 0.25151443 | -0.1667015 | 0.618 | 0.479 | 1 | 0.25151443 | 0.00095289 |
| Cog6 | 0.00431781 | 0.95686846 | 0.375 | 0.223 | 1 | 0.00047898 | 0.90104152 | 0.374 | 0.167 | 1 | 0.00431781 | 0.00095774 |
| Timm10b | 0.01336487 | 0.37881066 | 0.708 | 0.528 | 1 | 0.00048122 | 0.68401863 | 0.609 | 0.396 | 1 | 0.01336487 | 0.00096221 |
| Pycr2 | 0.04988121 | 0.35541784 | 0.448 | 0.321 | 1 | 0.00048211 | 0.72356774 | 0.382 | 0.167 | 1 | 0.04988121 | 0.00096398 |
| Csnk2b | 0.00691126 | 0.41294599 | 0.833 | 0.746 | 1 | 0.00048342 | 0.33923758 | 0.853 | 0.729 | 1 | 0.00691126 | 0.0009666 |
| Stard6 | 0.00048361 | 3.12297175 | 0.094 | 0.01 | 1 | 0.14259715 | 1.59703533 | 0.042 | 0.01 | 1 | 0.14259715 | 0.00096698 |
| Nadk | 0.0004839 | 0.74446013 | 0.594 | 0.394 | 1 | 0.30127415 | -0.3252853 | 0.605 | 0.656 | 1 | 0.30127415 | 0.00096757 |
| Pla2g12a | 0.04111903 | -0.8225805 | 0.24 | 0.342 | 1 | 0.00048483 | 0.77193883 | 0.403 | 0.188 | 1 | 0.04111903 | 0.00096942 |
| Amy1 | 0.00048558 | 1.00749454 | 0.323 | 0.145 | 1 | 0.2853542 | -0.3396026 | 0.244 | 0.177 | 1 | 0.2853542 | 0.00097092 |
| Lpin2 | 0.00048694 | -1.3296015 | 0.688 | 0.772 | 1 | 0.88044466 | -0.3784014 | 0.87 | 0.865 | 1 | 0.88044466 | 0.00097364 |

| | | | | | | | | | | | | |
|---|---|---|---|---|---|---|---|---|---|---|---|---|
| Cryab | 0.00048881 | -1.1112051 | 0.312 | 0.497 | 1 | 0.30372477 | -0.8482752 | 0.248 | 0.177 | 1 | 0.30372477 | 0.00097737 |
| Sgip1 | 0.00049315 | -1.1529701 | 0.021 | 0.161 | 1 | 0.00794277 | -2.8948178 | 0.034 | 0.104 | 1 | 0.00794277 | 0.00098606 |
| Vamp8 | 0.32461752 | 0.20848159 | 0.802 | 0.803 | 1 | 0.00049683 | 0.60034807 | 0.803 | 0.719 | 1 | 0.32461752 | 0.00099341 |
| Usp36 | 0.0004979 | 0.55804701 | 0.729 | 0.503 | 1 | 0.73685578 | -0.3587649 | 0.79 | 0.698 | 1 | 0.73685578 | 0.00099556 |
| Ociad1 | 0.00049806 | 0.37640287 | 0.958 | 0.86 | 1 | 0.00197039 | 0.28332774 | 0.853 | 0.823 | 1 | 0.00197039 | 0.00099587 |
| Mrgprf | 0.31116261 | -0.6139938 | 0.104 | 0.145 | 1 | 0.0004992 | 1.78700628 | 0.223 | 0.062 | 1 | 0.31116261 | 0.00099815 |
| Pck2 | 0.00081389 | 0.75846699 | 0.417 | 0.223 | 1 | 0.00049923 | 0.75242902 | 0.378 | 0.167 | 1 | 0.00081389 | 0.00099821 |
| Plekhf1 | 0.18884242 | 0.22762035 | 0.354 | 0.264 | 1 | 0.00049968 | 0.34725702 | 0.5 | 0.26 | 1 | 0.18884242 | 0.00099911 |
| Npepl1 | 0.89508635 | -0.1529701 | 0.146 | 0.15 | 1 | 0.00050188 | 0.92718393 | 0.332 | 0.135 | 1 | 0.89508635 | 0.00100351 |
| Veph1 | 0.03588274 | -0.4008483 | 0.292 | 0.415 | 1 | 0.00050231 | 0.35658864 | 0.479 | 0.24 | 1 | 0.03588274 | 0.00100437 |
| Mn1 | 0.0005054 | 0.98731665 | 0.438 | 0.264 | 1 | 0.00171672 | 0.31977629 | 0.618 | 0.354 | 1 | 0.00171672 | 0.00101054 |
| Swap70 | 0.55420139 | -0.1551257 | 0.458 | 0.482 | 1 | 0.00050635 | -1.0368368 | 0.5 | 0.677 | 1 | 0.55420139 | 0.00101245 |
| Igf2r | 0.00050718 | 0.46823897 | 0.969 | 0.943 | 1 | 0.11693566 | 0.12987152 | 0.971 | 0.958 | 1 | 0.11693566 | 0.00101411 |
| Banp | 0.1247983 | 0.25542205 | 0.469 | 0.368 | 1 | 0.00050825 | -0.8048163 | 0.555 | 0.688 | 1 | 0.1247983 | 0.00101624 |
| Gstm1 | 0.00407376 | -1.0734255 | 0.219 | 0.378 | 1 | 0.00050851 | -0.8544857 | 0.382 | 0.615 | 1 | 0.00407376 | 0.00101676 |
| Gm29418 | 0.50498933 | 0.29967529 | 0.229 | 0.197 | 1 | 0.00050982 | 0.86646751 | 0.361 | 0.156 | 1 | 0.50498933 | 0.00101939 |
| Prune2 | 0.00051068 | -2.2216871 | 0.208 | 0.383 | 1 | 0.2229266 | -1.280311 | 0.181 | 0.219 | 1 | 0.2229266 | 0.00102111 |
| Nfu1 | 0.04519223 | 0.54806292 | 0.5 | 0.389 | 1 | 0.00051132 | 0.12049529 | 0.576 | 0.312 | 1 | 0.04519223 | 0.00102238 |
| Ctnnal1 | 0.13541937 | -0.3407677 | 0.552 | 0.648 | 1 | 0.00051196 | 0.66414953 | 0.676 | 0.469 | 1 | 0.13541937 | 0.00102366 |
| Ugp2 | 0.126781 | -0.744636 | 0.958 | 0.917 | 1 | 0.00051211 | 0.49090698 | 0.853 | 0.823 | 1 | 0.126781 | 0.00102397 |
| Rfwd3 | 0.5431503 | -0.4128373 | 0.24 | 0.264 | 1 | 0.00051261 | 0.30258797 | 0.445 | 0.208 | 1 | 0.5431503 | 0.00102497 |
| Rbmx | 0.07905186 | 0.34452952 | 0.552 | 0.44 | 1 | 0.00051291 | 0.5290823 | 0.571 | 0.323 | 1 | 0.07905186 | 0.00102556 |
| Tmem185b | 0.05844996 | 0.61156586 | 0.323 | 0.223 | 1 | 0.00051412 | 1.57767001 | 0.252 | 0.083 | 1 | 0.05844996 | 0.00102798 |
| Ppp2ca | 0.06026941 | 0.19748841 | 0.948 | 0.953 | 1 | 0.00051435 | 0.16716912 | 0.992 | 0.99 | 1 | 0.06026941 | 0.00102844 |
| Phb2 | 0.01296317 | 0.44299614 | 0.74 | 0.611 | 1 | 0.00051452 | 0.48069137 | 0.798 | 0.688 | 1 | 0.01296317 | 0.00102877 |
| Gm10516 | 0.00051563 | 1.21899864 | 0.208 | 0.067 | 1 | 0.81217226 | -0.9313436 | 0.181 | 0.156 | 1 | 0.81217226 | 0.001031 |
| Pkm | 0.00290993 | -0.4210664 | 0.979 | 0.979 | 1 | 0.00051581 | 0.31516964 | 0.987 | 0.969 | 1 | 0.00290993 | 0.00103136 |
| Immp1l | 0.23712196 | 0.21659014 | 0.688 | 0.632 | 1 | 0.00051585 | 0.4506788 | 0.676 | 0.448 | 1 | 0.23712196 | 0.00103144 |
| Chd6 | 0.89641735 | -0.1114008 | 0.781 | 0.751 | 1 | 0.0005162 | -0.6374299 | 0.853 | 0.938 | 1 | 0.89641735 | 0.00103213 |
| Tnrc6c | 0.01445283 | 0.26206736 | 0.833 | 0.674 | 1 | 0.00051675 | -0.6741157 | 0.916 | 0.948 | 1 | 0.01445283 | 0.00103323 |
| Mrps10 | 0.00051914 | 0.70263996 | 0.438 | 0.223 | 1 | 0.93417263 | -0.5133887 | 0.223 | 0.208 | 1 | 0.93417263 | 0.001038 |
| Tmem35b | 0.0005208 | 1.53800925 | 0.229 | 0.083 | 1 | 0.01129214 | 1.39058446 | 0.139 | 0.042 | 1 | 0.01129214 | 0.00104132 |
| Dimt1 | 0.00052246 | 1.00749454 | 0.427 | 0.233 | 1 | 0.00575426 | 0.43122644 | 0.345 | 0.177 | 1 | 0.00575426 | 0.00104465 |
| Arglu1 | 0.00438014 | 0.36605927 | 0.948 | 0.876 | 1 | 0.0005229 | 0.29308498 | 0.971 | 0.896 | 1 | 0.00438014 | 0.00104553 |
| Fermt2 | 0.01962988 | -0.2715631 | 0.958 | 0.974 | 1 | 0.00052399 | 0.36381658 | 0.95 | 0.906 | 1 | 0.01962988 | 0.00104771 |
| Odr4 | 0.00325149 | 0.46450431 | 0.812 | 0.648 | 1 | 0.00052531 | 0.4604304 | 0.689 | 0.479 | 1 | 0.00325149 | 0.00105034 |
| Ift22 | 0.00052643 | 1.31943854 | 0.292 | 0.124 | 1 | 0.06249208 | 0.48369386 | 0.181 | 0.094 | 1 | 0.06249208 | 0.00105257 |
| Nucks1 | 0.04474031 | 0.51820248 | 0.969 | 0.886 | 1 | 0.00052841 | 0.37315413 | 0.937 | 0.854 | 1 | 0.04474031 | 0.00105655 |
| Khsrp | 0.00052859 | 0.62047141 | 0.604 | 0.352 | 1 | 0.80298597 | -0.1713733 | 0.689 | 0.656 | 1 | 0.80298597 | 0.00105689 |
| Pprc1 | 0.00052938 | 0.53800925 | 0.594 | 0.363 | 1 | 0.0152754 | 0.12806613 | 0.592 | 0.375 | 1 | 0.0152754 | 0.00105848 |
| Eftud2 | 0.43377188 | 0.29700115 | 0.396 | 0.352 | 1 | 0.00053394 | 0.83150059 | 0.454 | 0.229 | 1 | 0.43377188 | 0.00106759 |
| Hspb11 | 0.00274137 | 0.6730755 | 0.479 | 0.295 | 1 | 0.00053424 | 0.74261216 | 0.311 | 0.115 | 1 | 0.00274137 | 0.0010682 |
| Pvr | 0.36434196 | -0.3066141 | 0.25 | 0.295 | 1 | 0.00053449 | 0.47703556 | 0.517 | 0.281 | 1 | 0.36434196 | 0.00106869 |
| Ctdnep1 | 0.00558342 | 0.38336144 | 0.698 | 0.513 | 1 | 0.00053482 | 0.58322953 | 0.714 | 0.542 | 1 | 0.00558342 | 0.00106936 |
| Obsl1 | 0.00103421 | 0.82970742 | 0.438 | 0.244 | 1 | 0.00053912 | 0.52968007 | 0.424 | 0.208 | 1 | 0.00103421 | 0.00107796 |
| Tesk1 | 0.08413776 | 0.59245704 | 0.24 | 0.155 | 1 | 0.00053976 | 0.50728068 | 0.269 | 0.083 | 1 | 0.08413776 | 0.00107923 |
| Cobll1 | 0.00054093 | -0.9471625 | 0.573 | 0.762 | 1 | 0.52559854 | -0.6328636 | 0.634 | 0.667 | 1 | 0.52559854 | 0.00108158 |
| Cnot1 | 0.00054107 | 0.54568027 | 0.865 | 0.72 | 1 | 0.18405224 | -0.465264 | 0.882 | 0.896 | 1 | 0.18405224 | 0.00108185 |
| Parva | 0.00054171 | -0.6989136 | 0.656 | 0.845 | 1 | 0.14376156 | -0.4994874 | 0.84 | 0.854 | 1 | 0.14376156 | 0.00108312 |
| Crtac1 | 0.00054212 | 1.28312898 | 0.26 | 0.104 | 1 | 0.30447714 | 0.7296731 | 0.071 | 0.042 | 1 | 0.30447714 | 0.00108394 |
| Wdr11 | 0.40985037 | -0.3144336 | 0.208 | 0.259 | 1 | 0.00054241 | 1.06140054 | 0.366 | 0.167 | 1 | 0.40985037 | 0.00108453 |
| Dek | 0.00054492 | 0.68837051 | 0.927 | 0.819 | 1 | 0.01653496 | 0.11371892 | 0.92 | 0.885 | 1 | 0.01653496 | 0.00108955 |
| Sult1c1 | 0.31899281 | 0.52206771 | 0.042 | 0.021 | 1 | 0.0005469 | -1.6317834 | 0.05 | 0.167 | 1 | 0.31899281 | 0.00109351 |
| Chac1 | 0.01698099 | 1.30815401 | 0.229 | 0.124 | 1 | 0.00054775 | 2.91896343 | 0.155 | 0.021 | 1 | 0.01698099 | 0.00109521 |
| Rab12 | 0.22990441 | 0.17796282 | 0.802 | 0.731 | 1 | 0.00055054 | 0.41757612 | 0.861 | 0.667 | 1 | 0.22990441 | 0.00110078 |
| Sec63 | 0.00055185 | 0.43336584 | 0.979 | 0.907 | 1 | 0.62798971 | -0.2380646 | 0.971 | 0.938 | 1 | 0.62798971 | 0.0011034 |
| Vps36 | 0.00055365 | 0.74446013 | 0.562 | 0.358 | 1 | 0.55065276 | -0.2730185 | 0.55 | 0.615 | 1 | 0.55065276 | 0.001107 |
| Dock9 | 0.06441649 | -0.7117691 | 0.219 | 0.321 | 1 | 0.00055555 | -1.0771945 | 0.345 | 0.562 | 1 | 0.06441649 | 0.0011108 |
| Eya4 | 0.04442774 | -0.6649308 | 0.625 | 0.746 | 1 | 0.00055574 | -0.7318388 | 0.735 | 0.875 | 1 | 0.04442774 | 0.00111116 |
| Mef2a | 0.00055648 | 0.33084331 | 0.979 | 0.902 | 1 | 0.36891135 | -0.3023103 | 0.996 | 0.969 | 1 | 0.36891135 | 0.00111264 |
| Crk | 0.00055803 | 0.50454045 | 0.823 | 0.648 | 1 | 0.0047814 | 0.29256821 | 0.895 | 0.875 | 1 | 0.0047814 | 0.00111576 |
| Ctnnbl1 | 0.02961824 | 0.42253204 | 0.542 | 0.399 | 1 | 0.00056126 | 0.56461386 | 0.517 | 0.25 | 1 | 0.02961824 | 0.00112221 |
| Tomm5 | 0.00094877 | 0.73879138 | 0.667 | 0.477 | 1 | 0.0005614 | 0.59528555 | 0.576 | 0.365 | 1 | 0.00094877 | 0.00112248 |
| Gid4 | 0.003468 | 0.54452256 | 0.552 | 0.352 | 1 | 0.00056154 | 0.57528078 | 0.479 | 0.25 | 1 | 0.003468 | 0.00112276 |
| Stk19 | 0.19759308 | 0.37328852 | 0.562 | 0.497 | 1 | 0.00056179 | 0.65097414 | 0.542 | 0.323 | 1 | 0.19759308 | 0.00112326 |
| Pi4k2a | 0.00056425 | -0.716532 | 0.75 | 0.808 | 1 | 0.10892738 | 0.1671919 | 0.735 | 0.677 | 1 | 0.10892738 | 0.00112819 |
| Nt5c3b | 0.00710865 | 0.5962991 | 0.583 | 0.42 | 1 | 0.00056473 | 0.62232935 | 0.487 | 0.25 | 1 | 0.00710865 | 0.00112914 |
| Selenot | 0.01136434 | 0.44534192 | 0.792 | 0.679 | 1 | 0.00056496 | 0.32108267 | 0.903 | 0.802 | 1 | 0.01136434 | 0.00112961 |
| Hebp1 | 0.00056658 | 1.13677755 | 0.271 | 0.109 | 1 | 0.04359433 | 1.73453886 | 0.097 | 0.031 | 1 | 0.04359433 | 0.00113284 |
| Pym1 | 0.00208126 | 0.87422801 | 0.292 | 0.135 | 1 | 0.00056671 | 0.74672827 | 0.366 | 0.156 | 1 | 0.00208126 | 0.00113309 |
| Srsf5 | 0.00056886 | 0.36246547 | 0.979 | 0.969 | 1 | 0.08055734 | 0.1447106 | 0.966 | 0.948 | 1 | 0.08055734 | 0.00113739 |
| Msl3 | 0.00056968 | 0.78278825 | 0.562 | 0.352 | 1 | 0.00109309 | 0.49749966 | 0.479 | 0.25 | 1 | 0.00109309 | 0.00113903 |
| Naa30 | 0.11809857 | 0.6730755 | 0.323 | 0.244 | 1 | 0.00057054 | 0.54494682 | 0.475 | 0.229 | 1 | 0.11809857 | 0.00114076 |
| Nr2f6 | 0.95217949 | 0.26125113 | 0.219 | 0.223 | 1 | 0.0005719 | 0.77760758 | 0.391 | 0.177 | 1 | 0.95217949 | 0.00114348 |
| Nucb2 | 0.00404775 | 0.45495351 | 0.76 | 0.585 | 1 | 0.000572 | 0.60316144 | 0.681 | 0.5 | 1 | 0.00404775 | 0.00114367 |
| Maml3 | 0.00057347 | -1.0104274 | 0.396 | 0.622 | 1 | 0.0251459 | -0.7469786 | 0.584 | 0.74 | 1 | 0.0251459 | 0.00114661 |
| Calml3 | 0.00057364 | 1.52206771 | 0.323 | 0.15 | 1 | 0.00380143 | 0.53965311 | 0.34 | 0.167 | 1 | 0.00380143 | 0.00114695 |
| Bcas2 | 0.00057444 | 0.45532197 | 0.854 | 0.663 | 1 | 0.00058607 | 0.52021974 | 0.819 | 0.677 | 1 | 0.00058607 | 0.00114855 |
| Hmgn5 | 0.07077363 | 0.50053455 | 0.458 | 0.352 | 1 | 0.00057643 | -1.0055982 | 0.34 | 0.542 | 1 | 0.07077363 | 0.00115253 |
| Hibch | 0.86878363 | 0.51564144 | 0.188 | 0.192 | 1 | 0.00057701 | 1.07480859 | 0.29 | 0.104 | 1 | 0.86878363 | 0.00115369 |
| Has2 | 0.79048865 | -0.7941767 | 0.594 | 0.554 | 1 | 0.00057878 | 1.05692707 | 0.458 | 0.24 | 1 | 0.79048865 | 0.00115722 |
| Mrps27 | 0.15980704 | 0.6749192 | 0.208 | 0.145 | 1 | 0.00057889 | 1.20799304 | 0.21 | 0.052 | 1 | 0.15980704 | 0.00115744 |
| Man2b2 | 0.84734902 | 0.18258124 | 0.281 | 0.275 | 1 | 0.00057954 | 0.76543286 | 0.403 | 0.188 | 1 | 0.84734902 | 0.00115875 |
| Gadd45gip1 | 0.7617917 | -0.2097362 | 0.292 | 0.301 | 1 | 0.00057984 | 0.76696033 | 0.403 | 0.198 | 1 | 0.7617917 | 0.00115935 |
| Pwp1 | 0.0005805 | 0.91438513 | 0.365 | 0.176 | 1 | 0.00094914 | 0.82138927 | 0.294 | 0.115 | 1 | 0.00094914 | 0.00116067 |
| Anapc4 | 0.00058554 | 0.68296933 | 0.646 | 0.42 | 1 | 0.02841076 | 0.18446187 | 0.66 | 0.458 | 1 | 0.02841076 | 0.00117074 |
| Gpr153 | 0.01099537 | 0.81846071 | 0.375 | 0.238 | 1 | 0.00058567 | 0.69014474 | 0.534 | 0.312 | 1 | 0.01099537 | 0.001171 |
| Dnajb2 | 0.42564806 | 0.19535455 | 0.542 | 0.487 | 1 | 0.00058621 | 0.89367813 | 0.408 | 0.198 | 1 | 0.42564806 | 0.00117208 |
| Cpeb3 | 0.90659186 | -0.7636868 | 0.396 | 0.378 | 1 | 0.00058781 | -0.7422958 | 0.353 | 0.583 | 1 | 0.90659186 | 0.00117527 |
| Psma4 | 0.0020948 | 0.40618404 | 0.875 | 0.731 | 1 | 0.00059065 | 0.32538044 | 0.891 | 0.802 | 1 | 0.0020948 | 0.00118094 |

| | | | | | | | | | | | | |
|---|---|---|---|---|---|---|---|---|---|---|---|---|
| Nubpl | 0.53454154 | -0.3984978 | 0.177 | 0.202 | 1 | 0.0005909 | 1.30157945 | 0.176 | 0.031 | 1 | 0.53454154 | 0.00118146 |
| Chkb | 0.69445291 | 0.12166556 | 0.333 | 0.316 | 1 | 0.00059313 | 0.74261216 | 0.366 | 0.167 | 1 | 0.69445291 | 0.00118591 |
| Dusp23 | 0.63828658 | 0.42253204 | 0.188 | 0.171 | 1 | 0.00059377 | 1.27510724 | 0.227 | 0.062 | 1 | 0.63828658 | 0.00118719 |
| Hmga1 | 0.00261324 | 1.52206771 | 0.167 | 0.057 | 1 | 0.00059444 | 3.69014474 | 0.113 | 0 | 1 | 0.00261324 | 0.00118852 |
| Phf14 | 0.00059497 | 0.48748748 | 0.719 | 0.487 | 1 | 0.21641231 | -0.3187884 | 0.727 | 0.781 | 1 | 0.21641231 | 0.00118958 |
| Spag9 | 0.10352998 | 0.1009391 | 1 | 0.984 | 1 | 0.00059506 | -0.5088326 | 0.996 | 0.979 | 1 | 0.10352998 | 0.00118977 |
| Alx1 | 0.23167256 | -0.2237038 | 0.844 | 0.798 | 1 | 0.00059594 | 0.31808431 | 0.912 | 0.656 | 1 | 0.23167256 | 0.00119152 |
| Smad2 | 0.05337817 | 0.23462799 | 0.615 | 0.508 | 1 | 0.00060064 | -0.7274128 | 0.622 | 0.802 | 1 | 0.05337817 | 0.00120092 |
| Smad4 | 0.00060121 | 0.37145785 | 0.948 | 0.865 | 1 | 0.82331065 | -0.1005783 | 0.971 | 0.979 | 1 | 0.82331065 | 0.00120206 |
| Airn | 0.0006034 | 0.32256319 | 0.99 | 0.979 | 1 | 0.99850092 | -0.3289182 | 0.992 | 0.99 | 1 | 0.99850092 | 0.00120644 |
| Rhoa | 0.0007493 | 0.31922953 | 0.99 | 0.979 | 1 | 0.00060645 | 0.29285006 | 1 | 0.979 | 1 | 0.0007493 | 0.00121253 |
| Plbd2 | 0.602989 | -0.1517041 | 0.448 | 0.477 | 1 | 0.00060663 | 0.50484163 | 0.546 | 0.292 | 1 | 0.602989 | 0.0012129 |
| Mettl1 | 0.00190301 | 0.48393258 | 0.646 | 0.461 | 1 | 0.00060696 | 0.60303407 | 0.534 | 0.281 | 1 | 0.00190301 | 0.00121355 |
| Anapc5 | 0.00060932 | 0.3602029 | 0.958 | 0.824 | 1 | 0.00289152 | 0.27102606 | 0.929 | 0.802 | 1 | 0.00289152 | 0.00121826 |
| Timm29 | 0.34060327 | 0.25747279 | 0.198 | 0.15 | 1 | 0.00061177 | 0.90737545 | 0.315 | 0.125 | 1 | 0.34060327 | 0.00122316 |
| Rnmt | 0.05062654 | 0.36916881 | 0.625 | 0.497 | 1 | 0.0006125 | 0.55992216 | 0.605 | 0.354 | 1 | 0.05062654 | 0.00122463 |
| C1qtnf6 | 0.03343366 | 0.22988696 | 0.542 | 0.368 | 1 | 0.00061271 | 0.28226703 | 0.525 | 0.26 | 1 | 0.03343366 | 0.00122505 |
| Capn2 | 0.00350661 | 0.4090865 | 0.875 | 0.788 | 1 | 0.00061391 | 0.3933007 | 0.966 | 0.938 | 1 | 0.00350661 | 0.00122745 |
| Ahnak | 0.00126237 | 0.44187376 | 0.979 | 0.974 | 1 | 0.00061693 | 0.44804865 | 1 | 1 | 1 | 0.00126237 | 0.00123347 |
| Itga5 | 0.67535753 | -0.1106869 | 0.406 | 0.451 | 1 | 0.00061854 | 0.49280796 | 0.525 | 0.281 | 1 | 0.67535753 | 0.00123669 |
| Eif2b2 | 0.22466846 | 0.35979628 | 0.26 | 0.197 | 1 | 0.0006205 | 0.8808283 | 0.345 | 0.146 | 1 | 0.22466846 | 0.00124062 |
| Supt7l | 0.27719351 | -0.6805615 | 0.083 | 0.124 | 1 | 0.00062073 | 1.59703533 | 0.206 | 0.052 | 1 | 0.27719351 | 0.00124107 |
| Nav3 | 0.17969755 | -0.6545327 | 0.51 | 0.591 | 1 | 0.00062154 | -1.7939771 | 0.223 | 0.365 | 1 | 0.17969755 | 0.0012427 |
| Sox4 | 0.36791102 | -0.6602084 | 0.594 | 0.44 | 1 | 0.00062175 | -1.2201879 | 0.559 | 0.75 | 1 | 0.36791102 | 0.00124311 |
| Arl4c | 0.04856989 | 0.56692195 | 0.094 | 0.036 | 1 | 0.00062363 | -1.3769695 | 0.05 | 0.167 | 1 | 0.04856989 | 0.00124688 |
| Hormad2 | 0.00062524 | -2.8169339 | 0.052 | 0.202 | 1 | 0.64472654 | -1.5193086 | 0.071 | 0.083 | 1 | 0.64472654 | 0.0012501 |
| Lgals3bp | 0.20267489 | -0.3867844 | 0.25 | 0.326 | 1 | 0.00062621 | 0.7554362 | 0.336 | 0.146 | 1 | 0.20267489 | 0.00125203 |
| Zfp354c | 0.00063038 | 0.98240356 | 0.531 | 0.306 | 1 | 0.06088658 | 0.2538595 | 0.605 | 0.427 | 1 | 0.06088658 | 0.00126037 |
| Snx29 | 0.71688722 | -0.1659883 | 0.469 | 0.44 | 1 | 0.00063071 | -0.9798599 | 0.55 | 0.677 | 1 | 0.71688722 | 0.00126103 |
| Pinx1 | 0.00063169 | 1.57453513 | 0.26 | 0.109 | 1 | 0.1524922 | -0.473354 | 0.239 | 0.156 | 1 | 0.1524922 | 0.00126299 |
| Ndufa12 | 0.00063329 | 0.52350968 | 0.76 | 0.611 | 1 | 0.02251941 | 0.30248368 | 0.655 | 0.583 | 1 | 0.02251941 | 0.00126619 |
| Car12 | 0.00063465 | -1.7133514 | 0.094 | 0.269 | 1 | 0.03470143 | 0.46066289 | 0.118 | 0.042 | 1 | 0.03470143 | 0.0012689 |
| Snhg8 | 0.00168782 | 0.40839775 | 0.875 | 0.788 | 1 | 0.00063568 | 0.44596936 | 0.744 | 0.562 | 1 | 0.00168782 | 0.00127095 |
| Sdccag8 | 0.00063593 | 0.41787066 | 0.604 | 0.352 | 1 | 0.26634277 | -0.5906644 | 0.5 | 0.5 | 1 | 0.26634277 | 0.00127146 |
| Gtf2b | 0.10298512 | 0.30807331 | 0.812 | 0.725 | 1 | 0.0006374 | 0.50076737 | 0.765 | 0.625 | 1 | 0.10298512 | 0.00127439 |
| Maz | 0.04795857 | 0.51564144 | 0.469 | 0.337 | 1 | 0.00064206 | 0.65911784 | 0.508 | 0.271 | 1 | 0.04795857 | 0.0012837 |
| Cyp2u1 | 0.00064318 | 2.34629645 | 0.188 | 0.062 | 1 | 0.08467132 | 0.6026819 | 0.143 | 0.073 | 1 | 0.08467132 | 0.00128594 |
| Hibadh | 0.00570824 | 0.45269401 | 0.76 | 0.632 | 1 | 0.00064479 | 0.50131127 | 0.744 | 0.583 | 1 | 0.00570824 | 0.00128916 |
| Acad9 | 0.00065095 | 1.59245704 | 0.24 | 0.093 | 1 | 0.07662591 | 0.83310269 | 0.181 | 0.104 | 1 | 0.07662591 | 0.00130148 |
| Rp1 | 0.00065221 | 1.54036853 | 0.281 | 0.124 | 1 | 0.00119703 | 0.60662138 | 0.307 | 0.125 | 1 | 0.00119703 | 0.001304 |
| Copz1 | 0.01070088 | 0.55014623 | 0.708 | 0.57 | 1 | 0.00065498 | 0.32486027 | 0.689 | 0.448 | 1 | 0.01070088 | 0.00130953 |
| Mef2c | 0.00065536 | 2.05024418 | 0.396 | 0.249 | 1 | 0.79195089 | 1.33148971 | 0.172 | 0.156 | 1 | 0.79195089 | 0.0013103 |
| Spata5 | 0.38876983 | -0.6498049 | 0.688 | 0.679 | 1 | 0.00065537 | -1.0587935 | 0.626 | 0.771 | 1 | 0.38876983 | 0.0013103 |
| Gm35162 | 0.00065628 | 1.37006462 | 0.229 | 0.083 | 1 | 0.00508637 | 2.10518224 | 0.118 | 0.021 | 1 | 0.00508637 | 0.00131213 |
| 2310033P09Rik | 0.70054024 | -0.127155 | 0.333 | 0.352 | 1 | 0.00065687 | 0.88278982 | 0.366 | 0.167 | 1 | 0.70054024 | 0.00131331 |
| Trp53inp2 | 0.08139649 | -0.5147322 | 0.344 | 0.43 | 1 | 0.00065957 | 0.32241295 | 0.492 | 0.26 | 1 | 0.08139649 | 0.0013187 |
| Enpp5 | 0.00788169 | -0.8962901 | 0.24 | 0.383 | 1 | 0.00065992 | 0.85364347 | 0.353 | 0.156 | 1 | 0.00788169 | 0.0013194 |
| Usp29 | 0.00066149 | 1.06889508 | 0.271 | 0.109 | 1 | 0.26595352 | -1.5840302 | 0.13 | 0.167 | 1 | 0.26595352 | 0.00132254 |
| Creld2 | 0.93640746 | 0.16294906 | 0.5 | 0.523 | 1 | 0.00066217 | 0.59872171 | 0.534 | 0.292 | 1 | 0.93640746 | 0.0013239 |
| Shld2 | 0.00066226 | 0.95304675 | 0.333 | 0.161 | 1 | 0.07653347 | 0.22492148 | 0.269 | 0.167 | 1 | 0.07653347 | 0.00132409 |
| Cry2 | 0.00066288 | 0.48434804 | 0.854 | 0.777 | 1 | 0.89254376 | -0.141601 | 0.861 | 0.76 | 1 | 0.89254376 | 0.00132531 |
| Patj | 0.51664277 | 0.17741954 | 0.333 | 0.383 | 1 | 0.00066307 | -0.7188182 | 0.42 | 0.656 | 1 | 0.51664277 | 0.0013257 |
| Mllt1 | 0.00066508 | 1.20680334 | 0.281 | 0.119 | 1 | 0.05899409 | 0.40286279 | 0.223 | 0.125 | 1 | 0.05899409 | 0.00132972 |
| Psmc2 | 0.14835012 | 0.16389609 | 0.771 | 0.684 | 1 | 0.00066617 | 0.62232935 | 0.748 | 0.698 | 1 | 0.14835012 | 0.00133191 |
| St6gal1 | 0.00066806 | -2.6929452 | 0.021 | 0.155 | 1 | 0.01170817 | -1.961932 | 0.08 | 0.167 | 1 | 0.01170817 | 0.00133568 |
| Inpp5b | 0.00066929 | 1.63552576 | 0.24 | 0.093 | 1 | 0.03102891 | 0.76414532 | 0.235 | 0.125 | 1 | 0.03102891 | 0.00133813 |
| Faap100 | 0.00066933 | 1.55498233 | 0.177 | 0.052 | 1 | 0.03016043 | 0.30157945 | 0.189 | 0.083 | 1 | 0.03016043 | 0.00133822 |
| Rest | 0.03889777 | 0.2892655 | 0.667 | 0.534 | 1 | 0.00067059 | 0.55605184 | 0.798 | 0.719 | 1 | 0.03889777 | 0.00134072 |
| Zfp825 | 0.62052403 | -0.2148979 | 0.104 | 0.124 | 1 | 0.00067068 | 2.53144699 | 0.134 | 0.01 | 1 | 0.62052403 | 0.0013409 |
| Psma5 | 0.00067078 | 0.44204019 | 0.906 | 0.813 | 1 | 0.00084309 | 0.33564878 | 0.908 | 0.833 | 1 | 0.00084309 | 0.0013411 |
| Mitf | 0.00174521 | -1.577468 | 0.219 | 0.389 | 1 | 0.00067134 | -1.8504236 | 0.176 | 0.323 | 1 | 0.00174521 | 0.00134223 |
| Cnot11 | 0.67295296 | -0.1815393 | 0.229 | 0.249 | 1 | 0.00067393 | 0.31574922 | 0.378 | 0.156 | 1 | 0.67295296 | 0.0013474 |
| Zfp146 | 0.29971011 | 0.31943854 | 0.396 | 0.332 | 1 | 0.00067734 | 0.42968428 | 0.542 | 0.281 | 1 | 0.29971011 | 0.00135422 |
| Kif4 | 0.00067934 | 2.15433592 | 0.104 | 0.016 | 1 | 0.27154327 | 1.27510724 | 0.013 | 0 | 1 | 0.27154327 | 0.00135822 |
| Asrgl1 | 0.28849243 | -0.2277219 | 0.302 | 0.373 | 1 | 0.00068322 | 1.09224318 | 0.324 | 0.135 | 1 | 0.28849243 | 0.00136597 |
| Gm49454 | 0.27927855 | -2.0281294 | 0.021 | 0.047 | 1 | 0.00068515 | -2.0161241 | 0.029 | 0.125 | 1 | 0.27927855 | 0.00136984 |
| Mrps6 | 0.00068643 | -0.7146131 | 0.76 | 0.839 | 1 | 0.18741648 | -0.4506529 | 0.782 | 0.844 | 1 | 0.18741648 | 0.00137239 |
| Ppm1m | 0.42510883 | -0.1998304 | 0.406 | 0.435 | 1 | 0.00068738 | 0.76053407 | 0.399 | 0.188 | 1 | 0.42510883 | 0.00137429 |
| Hcfc2 | 0.00280695 | 0.55181505 | 0.51 | 0.316 | 1 | 0.00068847 | 0.52886383 | 0.588 | 0.354 | 1 | 0.00280695 | 0.00137647 |
| Clcn4 | 0.15704761 | 0.32617697 | 0.656 | 0.549 | 1 | 0.00068851 | 0.48613085 | 0.71 | 0.458 | 1 | 0.15704761 | 0.00137654 |
| Timm21 | 0.00219031 | 0.82970742 | 0.448 | 0.264 | 1 | 0.00068936 | 1.10042871 | 0.332 | 0.146 | 1 | 0.00219031 | 0.00137825 |
| Pelo | 0.00518113 | 0.95686846 | 0.385 | 0.233 | 1 | 0.00069039 | 0.79168276 | 0.332 | 0.135 | 1 | 0.00518113 | 0.00138031 |
| Pigbos1 | 0.00069079 | 0.9062112 | 0.438 | 0.249 | 1 | 0.00205391 | 1.10518224 | 0.286 | 0.125 | 1 | 0.00205391 | 0.00138111 |
| Dynlt2b | 0.44391042 | 0.12997854 | 0.354 | 0.306 | 1 | 0.00069083 | 0.78775553 | 0.336 | 0.146 | 1 | 0.44391042 | 0.00138118 |
| Fryl | 0.54192995 | 0.26253615 | 0.698 | 0.699 | 1 | 0.00069098 | -0.7383133 | 0.702 | 0.854 | 1 | 0.54192995 | 0.00138148 |
| Mrpl13 | 0.01621172 | 0.49848089 | 0.562 | 0.409 | 1 | 0.00069263 | 0.80261947 | 0.45 | 0.25 | 1 | 0.01621172 | 0.00138478 |
| Ehd2 | 0.00540347 | 0.52884094 | 0.667 | 0.487 | 1 | 0.00069312 | 0.42312498 | 0.769 | 0.542 | 1 | 0.00540347 | 0.00138575 |
| Zfp787 | 0.3857306 | 0.28621056 | 0.292 | 0.244 | 1 | 0.00069581 | 0.64547569 | 0.458 | 0.229 | 1 | 0.3857306 | 0.00139113 |
| Uba52 | 0.43431755 | 0.10442841 | 0.917 | 0.86 | 1 | 0.00069851 | 0.34940102 | 0.87 | 0.802 | 1 | 0.43431755 | 0.00139653 |
| 4930579G24Rik | 0.00069853 | 2.14499806 | 0.146 | 0.036 | 1 | 0.30340561 | 0.59703533 | 0.109 | 0.073 | 1 | 0.30340561 | 0.00139658 |
| Thyn1 | 0.01597567 | 0.57139542 | 0.417 | 0.275 | 1 | 0.00070072 | 0.88516072 | 0.319 | 0.135 | 1 | 0.01597567 | 0.00140095 |
| Ube2i | 0.00070375 | 0.28713695 | 0.938 | 0.907 | 1 | 0.00878974 | 0.14133124 | 0.966 | 0.906 | 1 | 0.00878974 | 0.00140701 |
| Cyhr1 | 0.00570011 | 0.80818573 | 0.25 | 0.119 | 1 | 0.00070391 | 1.06865636 | 0.298 | 0.115 | 1 | 0.00570011 | 0.00140733 |
| Cdipt | 0.5232114 | 0.17624899 | 0.51 | 0.451 | 1 | 0.00070399 | 0.25624821 | 0.597 | 0.354 | 1 | 0.5232114 | 0.00140748 |
| Mrpl15 | 0.02075451 | 0.52788504 | 0.5 | 0.358 | 1 | 0.00070597 | 0.67917362 | 0.424 | 0.208 | 1 | 0.02075451 | 0.00141143 |
| Mybbp1a | 0.00070737 | 0.61922395 | 0.677 | 0.487 | 1 | 0.00404674 | 0.4370258 | 0.668 | 0.521 | 1 | 0.00404674 | 0.00141424 |
| Lrrc71 | 0.00070814 | 2.65957123 | 0.115 | 0.021 | 1 | 0.0888909 | -0.3098553 | 0.134 | 0.062 | 1 | 0.0888909 | 0.00141578 |
| Abcb1b | 0.00070835 | 1.59245704 | 0.24 | 0.093 | 1 | 0.35658506 | 0.92044236 | 0.118 | 0.083 | 1 | 0.35658506 | 0.00141619 |
| Ube2j2 | 0.1559373 | 0.28221075 | 0.719 | 0.653 | 1 | 0.00070966 | 0.30157945 | 0.828 | 0.729 | 1 | 0.1559373 | 0.00141882 |
| Neb | 0.00071007 | -3.1393469 | 0.062 | 0.212 | 1 | 0.58036856 | -0.194378 | 0.029 | 0.042 | 1 | 0.58036856 | 0.00141965 |

| | | | | | | | | | | | | |
|---|---|---|---|---|---|---|---|---|---|---|---|---|
| Mrpl18 | 0.18761344 | 0.35079628 | 0.5 | 0.425 | 1 | 0.00071041 | 0.57911343 | 0.496 | 0.271 | 1 | 0.18761344 | 0.00142032 |
| Mmp16 | 0.00071047 | 0.73503512 | 0.583 | 0.358 | 1 | 0.01175871 | 0.11615335 | 0.529 | 0.344 | 1 | 0.01175871 | 0.00142043 |
| Dbn1 | 0.3748988 | -0.9518635 | 0.146 | 0.176 | 1 | 0.00071622 | -1.6094155 | 0.126 | 0.271 | 1 | 0.3748988 | 0.00143192 |
| Myo18a | 0.00072041 | 0.43362974 | 0.708 | 0.503 | 1 | 0.11424913 | -0.1016993 | 0.908 | 0.823 | 1 | 0.11424913 | 0.00144031 |
| Sec23ip | 0.27874266 | 0.23850009 | 0.562 | 0.503 | 1 | 0.00072052 | 0.54056838 | 0.702 | 0.438 | 1 | 0.27874266 | 0.00144053 |
| Lamb1 | 0.02043045 | -3.3382803 | 0.021 | 0.093 | 1 | 0.00072084 | -2.5114891 | 0.025 | 0.115 | 1 | 0.02043045 | 0.00144116 |
| 0610040J01Rik | 0.04161436 | -1.577468 | 0.156 | 0.254 | 1 | 0.00072122 | 0.45748398 | 0.454 | 0.208 | 1 | 0.04161436 | 0.00144193 |
| Commd6 | 0.09812098 | 0.19350652 | 0.542 | 0.409 | 1 | 0.00072342 | 0.55687821 | 0.412 | 0.198 | 1 | 0.09812098 | 0.00144631 |
| Mlec | 0.01031767 | 0.33714054 | 0.885 | 0.819 | 1 | 0.00072408 | 0.30727056 | 0.87 | 0.708 | 1 | 0.01031767 | 0.00144764 |
| Lamtor5 | 0.00177331 | 0.39531057 | 0.792 | 0.622 | 1 | 0.00072664 | 0.53226508 | 0.693 | 0.5 | 1 | 0.00177331 | 0.00145274 |
| Asap3 | 0.00072919 | -0.9540313 | 0.24 | 0.451 | 1 | 0.68493945 | -1.4618584 | 0.223 | 0.219 | 1 | 0.68493945 | 0.00145786 |
| E2f6 | 0.04812883 | 0.60310428 | 0.26 | 0.161 | 1 | 0.00073213 | 0.62560449 | 0.454 | 0.229 | 1 | 0.04812883 | 0.00146371 |
| Gm15337 | 0.68932378 | -0.577468 | 0.042 | 0.052 | 1 | 0.00073419 | -2.5322477 | 0.025 | 0.115 | 1 | 0.68932378 | 0.00146783 |
| Mgat3 | 0.01937011 | -1.4705528 | 0.052 | 0.145 | 1 | 0.00073847 | -0.7041342 | 0.197 | 0.385 | 1 | 0.01937011 | 0.00147639 |
| Nop14 | 0.44116625 | 0.14129303 | 0.729 | 0.689 | 1 | 0.00074009 | 0.29764981 | 0.744 | 0.562 | 1 | 0.44116625 | 0.00147963 |
| Mid1 | 0.95429137 | -0.5646211 | 0.667 | 0.585 | 1 | 0.00074305 | -1.1153959 | 0.513 | 0.75 | 1 | 0.95429137 | 0.00148556 |
| Cybrd1 | 0.02717254 | 1.00749454 | 0.25 | 0.15 | 1 | 0.00074308 | 0.81942775 | 0.315 | 0.125 | 1 | 0.02717254 | 0.0014856 |
| Atf5 | 0.87513739 | -0.2686299 | 0.385 | 0.352 | 1 | 0.00074503 | 0.85060941 | 0.445 | 0.26 | 1 | 0.87513739 | 0.0014895 |
| Cwc15 | 0.00074503 | 0.44915456 | 0.875 | 0.767 | 1 | 0.00103776 | 0.38282106 | 0.878 | 0.802 | 1 | 0.00103776 | 0.0014895 |
| Hyal2 | 0.04451199 | 0.75373794 | 0.229 | 0.135 | 1 | 0.00074506 | 1.20471791 | 0.256 | 0.083 | 1 | 0.04451199 | 0.00148956 |
| Taf4b | 0.0764982 | -0.3617393 | 0.5 | 0.591 | 1 | 0.00074795 | -1.39342 | 0.462 | 0.635 | 1 | 0.0764982 | 0.00149534 |
| Sertad4 | 0.7285176 | 0.17004755 | 0.344 | 0.326 | 1 | 0.00074898 | 0.43336932 | 0.525 | 0.271 | 1 | 0.7285176 | 0.0014974 |
| Xpo5 | 0.00075401 | 0.98477446 | 0.396 | 0.202 | 1 | 0.00970339 | 0.41806519 | 0.433 | 0.26 | 1 | 0.00970339 | 0.00150746 |
| Cab39 | 0.00075414 | 0.62335105 | 0.875 | 0.834 | 1 | 0.32847087 | -0.2787066 | 0.945 | 0.979 | 1 | 0.32847087 | 0.0015077 |
| Slc48a1 | 0.61931349 | -0.2061009 | 0.375 | 0.389 | 1 | 0.00075431 | 0.61128932 | 0.492 | 0.25 | 1 | 0.61931349 | 0.00150805 |
| Scrn3 | 0.00183465 | 1.45110119 | 0.25 | 0.114 | 1 | 0.00075601 | 0.40286279 | 0.206 | 0.052 | 1 | 0.00183465 | 0.00151146 |
| Ccdc97 | 0.11663149 | 0.79804117 | 0.229 | 0.161 | 1 | 0.00075612 | 1.02894665 | 0.294 | 0.115 | 1 | 0.11663149 | 0.00151167 |
| Grik3 | 0.54758872 | -0.3946039 | 0.125 | 0.15 | 1 | 0.00075863 | 1.13945214 | 0.269 | 0.094 | 1 | 0.54758872 | 0.00151669 |
| Aasdhppt | 0.36946692 | 0.24047171 | 0.406 | 0.352 | 1 | 0.00076178 | 0.44221722 | 0.399 | 0.177 | 1 | 0.36946692 | 0.00152298 |
| Smu1 | 0.13034116 | 0.40263248 | 0.542 | 0.482 | 1 | 0.00076227 | 0.60236878 | 0.672 | 0.49 | 1 | 0.13034116 | 0.00152395 |
| Ska2 | 0.00076237 | 2.15188445 | 0.219 | 0.083 | 1 | 0.25601148 | 0.10518224 | 0.143 | 0.094 | 1 | 0.25601148 | 0.00152417 |
| B4galt7 | 0.36051968 | 0.26641283 | 0.51 | 0.435 | 1 | 0.00076245 | 0.86006974 | 0.366 | 0.167 | 1 | 0.36051968 | 0.00152432 |
| Gm36129 | 0.61737927 | 1.00749454 | 0.01 | 0.005 | 1 | 0.00076285 | -3.7692869 | 0.004 | 0.062 | 1 | 0.61737927 | 0.00152513 |
| Cnr1 | 0.66681594 | -0.5160674 | 0.031 | 0.041 | 1 | 0.00076285 | -5.6674073 | 0.004 | 0.062 | 1 | 0.66681594 | 0.00152513 |
| Zscan12 | 0.78274108 | 0.17099327 | 0.115 | 0.104 | 1 | 0.00076286 | 2.93807225 | 0.13 | 0.01 | 1 | 0.78274108 | 0.00152514 |
| Cd1d1 | 0.15329198 | -1.3144336 | 0.01 | 0.041 | 1 | 0.00076287 | -3.8948178 | 0.004 | 0.062 | 1 | 0.15329198 | 0.00152515 |
| Slc25a33 | 0.47363066 | -0.4519371 | 0.198 | 0.228 | 1 | 0.00076352 | 1.1883956 | 0.294 | 0.115 | 1 | 0.47363066 | 0.00152646 |
| Carmil1 | 0.00218878 | 0.28257985 | 0.865 | 0.777 | 1 | 0.00076363 | 0.20271457 | 0.958 | 0.875 | 1 | 0.00218878 | 0.00152667 |
| Rrp8 | 0.00085496 | 0.74001423 | 0.427 | 0.223 | 1 | 0.00076463 | 0.74542717 | 0.387 | 0.177 | 1 | 0.00085496 | 0.00152867 |
| Nol11 | 0.27032559 | 0.15763489 | 0.51 | 0.446 | 1 | 0.00076582 | 0.24353716 | 0.559 | 0.302 | 1 | 0.27032559 | 0.00153106 |
| Gm11973 | 0.00076628 | 1.24453373 | 0.271 | 0.114 | 1 | 0.09079792 | 0.42711033 | 0.282 | 0.188 | 1 | 0.09079792 | 0.00153197 |
| Pnrc1 | 0.0007663 | -0.6247737 | 0.906 | 0.933 | 1 | 0.04876459 | -0.4780473 | 0.866 | 0.906 | 1 | 0.04876459 | 0.00153201 |
| Palld | 0.00076721 | -1.2140733 | 0.5 | 0.648 | 1 | 0.06367375 | -0.9710533 | 0.567 | 0.74 | 1 | 0.06367375 | 0.00153383 |
| Cux1 | 0.00076751 | -0.6617615 | 0.844 | 0.938 | 1 | 0.30379211 | -0.1474639 | 0.95 | 0.958 | 1 | 0.30379211 | 0.00153444 |
| Tut7 | 0.01133301 | 0.28222746 | 0.917 | 0.917 | 1 | 0.00076798 | 0.23108261 | 0.983 | 0.927 | 1 | 0.01133301 | 0.00153537 |
| Srfbp1 | 0.00880029 | 0.72370157 | 0.448 | 0.301 | 1 | 0.00076852 | 0.72861889 | 0.429 | 0.208 | 1 | 0.00880029 | 0.00153645 |
| mt-Atp8 | 0.15549727 | -0.4014684 | 0.667 | 0.705 | 1 | 0.00076891 | -0.919455 | 0.676 | 0.823 | 1 | 0.15549727 | 0.00153722 |
| Mrps21 | 0.05768383 | 0.2590333 | 0.781 | 0.653 | 1 | 0.00076921 | 0.52021974 | 0.685 | 0.49 | 1 | 0.05768383 | 0.00153783 |
| Smndc1 | 0.01698273 | 0.40272355 | 0.802 | 0.71 | 1 | 0.00077071 | 0.49597219 | 0.819 | 0.688 | 1 | 0.01698273 | 0.00154083 |
| Cep19 | 0.19517989 | 0.75373794 | 0.167 | 0.114 | 1 | 0.00077124 | 1.78968041 | 0.202 | 0.052 | 1 | 0.19517989 | 0.00154188 |
| Ppp6r1 | 0.08206612 | 0.56003556 | 0.271 | 0.181 | 1 | 0.00077244 | 0.83519507 | 0.353 | 0.156 | 1 | 0.08206612 | 0.00154429 |
| Trip4 | 0.85027225 | 0.10213141 | 0.406 | 0.394 | 1 | 0.0007736 | 0.69668758 | 0.508 | 0.281 | 1 | 0.85027225 | 0.00154661 |
| Brpf1 | 0.0007747 | 0.74969678 | 0.385 | 0.197 | 1 | 0.06922951 | 0.34725702 | 0.445 | 0.323 | 1 | 0.06922951 | 0.0015488 |
| Dennd4c | 0.97587139 | -0.2030724 | 0.385 | 0.368 | 1 | 0.00077527 | -1.0056459 | 0.718 | 0.854 | 1 | 0.97587139 | 0.00154994 |
| Lrrc8d | 0.00077555 | -0.6233838 | 0.552 | 0.731 | 1 | 0.55525419 | -0.2857596 | 0.853 | 0.854 | 1 | 0.55525419 | 0.00155049 |
| Syt16 | 0.03210508 | -3.240433 | 0 | 0.047 | 1 | 0.00077704 | -3.5322477 | 0.004 | 0.062 | 1 | 0.03210508 | 0.00155348 |
| Atp6v0a4 | 0.17503003 | 0.53356335 | 0.312 | 0.254 | 1 | 0.0007772 | 0.3524742 | 0.882 | 0.667 | 1 | 0.17503003 | 0.0015538 |
| Pcdh9 | 0.0007784 | -1.1906671 | 0.646 | 0.715 | 1 | 0.05127593 | -0.2259375 | 0.723 | 0.604 | 1 | 0.05127593 | 0.00155619 |
| Rilpl2 | 0.00077894 | 0.52547834 | 0.656 | 0.44 | 1 | 0.00465492 | 0.59378967 | 0.626 | 0.458 | 1 | 0.00465492 | 0.00155727 |
| Ube2b | 0.29521171 | 0.11344784 | 0.979 | 0.99 | 1 | 0.00078043 | 0.23393443 | 1 | 0.99 | 1 | 0.29521171 | 0.00156026 |
| Grn | 0.0403509 | 0.42253204 | 0.729 | 0.653 | 1 | 0.00078107 | 0.57242278 | 0.815 | 0.656 | 1 | 0.0403509 | 0.00156152 |
| Cnpy4 | 0.00315182 | 0.4721628 | 0.531 | 0.337 | 1 | 0.00078112 | 0.7233116 | 0.508 | 0.281 | 1 | 0.00315182 | 0.00156163 |
| Rapgef6 | 0.59395111 | -0.1061363 | 0.854 | 0.917 | 1 | 0.00078576 | -0.811022 | 0.941 | 0.969 | 1 | 0.59395111 | 0.0015709 |
| Nanp | 0.04923328 | 0.82108141 | 0.219 | 0.13 | 1 | 0.00078669 | 1.91253716 | 0.185 | 0.042 | 1 | 0.04923328 | 0.00157276 |
| Nup50 | 0.00079244 | 0.6613189 | 0.688 | 0.492 | 1 | 0.60337893 | -0.1878647 | 0.727 | 0.635 | 1 | 0.60337893 | 0.00158426 |
| Tmem160 | 0.17100154 | -0.3980202 | 0.458 | 0.518 | 1 | 0.00079341 | 0.81276402 | 0.429 | 0.229 | 1 | 0.17100154 | 0.00158619 |
| Irx3 | 0.07746962 | 0.31140122 | 0.542 | 0.42 | 1 | 0.00079403 | 0.21237148 | 0.5 | 0.26 | 1 | 0.07746962 | 0.00158742 |
| Jade2 | 0.04553849 | 0.6241659 | 0.354 | 0.249 | 1 | 0.00079748 | 0.89401707 | 0.361 | 0.167 | 1 | 0.04553849 | 0.00159433 |
| Gm46620 | 0.00080539 | 2.00749454 | 0.125 | 0.026 | 1 | 0.22089679 | 0.39058446 | 0.13 | 0.083 | 1 | 0.22089679 | 0.00161013 |
| Sema3c | 0.00080591 | -0.9518635 | 0.344 | 0.539 | 1 | 0.13886788 | 0.2190732 | 0.534 | 0.49 | 1 | 0.13886788 | 0.00161118 |
| Dusp22 | 0.00080731 | -1.2609943 | 0.177 | 0.363 | 1 | 0.19151455 | -0.1172102 | 0.202 | 0.135 | 1 | 0.19151455 | 0.00161398 |
| Sfrp1 | 0.50458574 | 0.39278469 | 0.177 | 0.145 | 1 | 0.00081189 | -0.5213594 | 0.197 | 0.396 | 1 | 0.50458574 | 0.00162313 |
| Nmd3 | 0.0008137 | 0.60279898 | 0.635 | 0.43 | 1 | 0.00359648 | 0.23530623 | 0.643 | 0.448 | 1 | 0.00359648 | 0.00162674 |
| Mrpl2 | 0.16122929 | 0.27052894 | 0.448 | 0.358 | 1 | 0.00081584 | 0.83698613 | 0.37 | 0.177 | 1 | 0.16122929 | 0.00163101 |
| Glis1 | 0.00081773 | 0.68172438 | 0.302 | 0.135 | 1 | 0.03291856 | -0.4187896 | 0.332 | 0.188 | 1 | 0.03291856 | 0.00163479 |
| Fgf2os | 0.00081773 | -4.3848229 | 0 | 0.109 | 1 | 0.79600331 | -0.9879272 | 0.017 | 0.021 | 1 | 0.79600331 | 0.0016348 |
| Adam10 | 0.00081784 | -0.863519 | 0.396 | 0.575 | 1 | 0.00107174 | -0.7714353 | 0.555 | 0.792 | 1 | 0.00107174 | 0.001635 |
| Smim26 | 0.12432624 | 0.60788708 | 0.344 | 0.259 | 1 | 0.00081858 | 0.91253716 | 0.298 | 0.115 | 1 | 0.12432624 | 0.00163649 |
| Tmem276-zftraf1 | 0.13800907 | 0.39605982 | 0.323 | 0.238 | 1 | 0.00081911 | 0.76469169 | 0.424 | 0.219 | 1 | 0.13800907 | 0.00163756 |
| Sp1 | 0.00081931 | 0.70981399 | 0.688 | 0.539 | 1 | 0.80846476 | -0.1840113 | 0.773 | 0.802 | 1 | 0.80846476 | 0.00163795 |
| Clca2 | 0.00082013 | -2.4519371 | 0.021 | 0.15 | 1 | 0.00157775 | -4.3098553 | 0 | 0.042 | 1 | 0.00157775 | 0.00163958 |
| Skic8 | 0.06241569 | 0.367538 | 0.74 | 0.699 | 1 | 0.00082115 | 0.50849866 | 0.769 | 0.677 | 1 | 0.06241569 | 0.00164163 |
| Taok1 | 0.00082411 | 0.37427687 | 0.958 | 0.788 | 1 | 0.90380482 | -0.111847 | 0.975 | 0.99 | 1 | 0.90380482 | 0.00164754 |
| Sf3b5 | 0.38902291 | 0.13653648 | 0.854 | 0.71 | 1 | 0.00082518 | 0.27995664 | 0.723 | 0.542 | 1 | 0.38902291 | 0.00164967 |
| Srgap2 | 0.0008254 | 0.7144213 | 0.865 | 0.756 | 1 | 0.04763052 | -0.2634753 | 0.92 | 0.938 | 1 | 0.04763052 | 0.00165011 |
| Zfp804b | 0.00082558 | -5.0905375 | 0.01 | 0.13 | 1 | 0.11689951 | -2.8948178 | 0 | 0.01 | 1 | 0.11689951 | 0.00165048 |
| Vdac2 | 0.09311912 | 0.22223339 | 0.906 | 0.829 | 1 | 0.0008294 | 0.36474445 | 0.941 | 0.917 | 1 | 0.09311912 | 0.00165811 |
| Erh | 0.01028811 | 0.34336182 | 0.865 | 0.684 | 1 | 0.00082993 | 0.2966404 | 0.811 | 0.635 | 1 | 0.01028811 | 0.00165917 |
| Dusp3 | 0.53518773 | -0.3056633 | 0.219 | 0.244 | 1 | 0.00083062 | 0.49061327 | 0.462 | 0.25 | 1 | 0.53518773 | 0.00166055 |
| Med30 | 0.54054096 | -0.2061009 | 0.375 | 0.399 | 1 | 0.00083151 | 0.74261216 | 0.361 | 0.167 | 1 | 0.54054096 | 0.00166233 |

| | | | | | | | | | | | | |
|---|---|---|---|---|---|---|---|---|---|---|---|---|
| Prtn3 | 0.01095279 | -1.8993961 | 0.01 | 0.088 | 1 | 0.00083206 | -2.7793405 | 0.038 | 0.135 | 1 | 0.01095279 | 0.00166343 |
| Asl | 0.51326518 | -0.407543 | 0.104 | 0.13 | 1 | 0.0008322 | 1.74903843 | 0.168 | 0.031 | 1 | 0.51326518 | 0.00166371 |
| Slc35g2 | 0.00083263 | 1.38189005 | 0.26 | 0.109 | 1 | 0.94524707 | -0.4284998 | 0.109 | 0.104 | 1 | 0.94524707 | 0.00166457 |
| Eng | 0.02818108 | 0.74055568 | 0.479 | 0.383 | 1 | 0.00083739 | 0.58453455 | 0.58 | 0.333 | 1 | 0.02818108 | 0.00167408 |
| Cops5 | 0.25694632 | 0.15433592 | 0.688 | 0.606 | 1 | 0.00084064 | 0.43908297 | 0.756 | 0.583 | 1 | 0.25694632 | 0.00168058 |
| Il6ra | 0.00084165 | -1.334425 | 0.25 | 0.44 | 1 | 0.70401859 | -0.2723806 | 0.315 | 0.271 | 1 | 0.70401859 | 0.00168258 |
| Tpm1 | 0.00084181 | 0.41977521 | 0.99 | 0.964 | 1 | 0.91106245 | -0.2557387 | 0.954 | 0.938 | 1 | 0.91106245 | 0.00168292 |
| Fgfr2 | 0.60369825 | -0.161195 | 0.969 | 0.953 | 1 | 0.00084628 | -0.6626531 | 0.996 | 1 | 1 | 0.60369825 | 0.00169185 |
| Paip2b | 0.00483222 | 0.54591445 | 0.469 | 0.29 | 1 | 0.00085001 | 0.93221952 | 0.357 | 0.167 | 1 | 0.00483222 | 0.00169929 |
| Dock4 | 0.88256045 | -0.4626476 | 0.365 | 0.311 | 1 | 0.00085094 | -1.1370187 | 0.517 | 0.635 | 1 | 0.88256045 | 0.00170115 |
| Zdhhc8 | 0.00085482 | 0.68051321 | 0.656 | 0.446 | 1 | 0.01194301 | 0.32955502 | 0.584 | 0.385 | 1 | 0.01194301 | 0.00170892 |
| Cnpy3 | 0.05533113 | 0.35933821 | 0.531 | 0.404 | 1 | 0.00085501 | 0.79794803 | 0.399 | 0.198 | 1 | 0.05533113 | 0.00170928 |
| Slc39a3 | 0.02913129 | 0.93710521 | 0.177 | 0.088 | 1 | 0.00085585 | 1.32241295 | 0.218 | 0.062 | 1 | 0.02913129 | 0.00171097 |
| Josd2 | 0.24650814 | 0.20013961 | 0.521 | 0.44 | 1 | 0.00085602 | 0.8714745 | 0.391 | 0.198 | 1 | 0.24650814 | 0.00171131 |
| Thoc5 | 0.09868454 | 0.38300367 | 0.229 | 0.145 | 1 | 0.00085818 | 1.47441605 | 0.218 | 0.062 | 1 | 0.09868454 | 0.00171562 |
| Eif4h | 0.0008608 | 0.41547728 | 0.906 | 0.891 | 1 | 0.08176188 | 0.19083832 | 0.971 | 0.927 | 1 | 0.08176188 | 0.00172085 |
| Dctpp1 | 0.05305503 | 0.63403614 | 0.312 | 0.218 | 1 | 0.00086264 | 1.66742466 | 0.202 | 0.052 | 1 | 0.05305503 | 0.00172453 |
| Derl2 | 0.0240602 | 0.37362244 | 0.521 | 0.352 | 1 | 0.00086312 | 0.82764826 | 0.408 | 0.208 | 1 | 0.0240602 | 0.00172549 |
| Tmem60 | 0.71415081 | -0.2708066 | 0.396 | 0.389 | 1 | 0.00086459 | 0.97025266 | 0.378 | 0.177 | 1 | 0.71415081 | 0.00172844 |
| Mtmr11 | 0.24847004 | 0.27459861 | 0.438 | 0.352 | 1 | 0.00086516 | 0.49512876 | 0.605 | 0.365 | 1 | 0.24847004 | 0.00172957 |
| Etf1 | 0.00086639 | 0.37650777 | 0.948 | 0.943 | 1 | 0.00127524 | 0.29221272 | 0.987 | 0.99 | 1 | 0.00127524 | 0.00173203 |
| Cetn3 | 0.00086733 | 0.51955649 | 0.771 | 0.658 | 1 | 0.00258274 | 0.40201935 | 0.739 | 0.583 | 1 | 0.00258274 | 0.00173391 |
| Actl6a | 0.00138708 | 0.53424242 | 0.552 | 0.368 | 1 | 0.00086782 | 0.38773609 | 0.618 | 0.406 | 1 | 0.00138708 | 0.00173488 |
| Mrps22 | 0.06339443 | 0.42253204 | 0.302 | 0.202 | 1 | 0.00086899 | 1.19264508 | 0.265 | 0.094 | 1 | 0.06339443 | 0.00173723 |
| Cycs | 0.68146993 | -0.1618344 | 0.844 | 0.87 | 1 | 0.00087044 | 0.40989126 | 0.845 | 0.729 | 1 | 0.68146993 | 0.00174013 |
| Hmbox1 | 0.00087524 | 0.61347482 | 0.635 | 0.409 | 1 | 0.00915263 | -0.7356192 | 0.66 | 0.771 | 1 | 0.00915263 | 0.00174972 |
| Maoa | 0.00087527 | 1.0899567 | 0.344 | 0.166 | 1 | 0.16719638 | -0.4948872 | 0.239 | 0.156 | 1 | 0.16719638 | 0.00174978 |
| Imp4 | 0.24870444 | 0.44806713 | 0.271 | 0.218 | 1 | 0.00087754 | 1.27510724 | 0.332 | 0.156 | 1 | 0.24870444 | 0.00175431 |
| Sorbs1 | 0.00087999 | -0.6845101 | 0.844 | 0.907 | 1 | 0.29328195 | -0.4871662 | 0.903 | 0.896 | 1 | 0.29328195 | 0.0017592 |
| Rbpj | 0.69219581 | -0.1941393 | 0.698 | 0.642 | 1 | 0.00088309 | -0.8499362 | 0.689 | 0.823 | 1 | 0.69219581 | 0.00176541 |
| Lin7c | 0.01002006 | 0.57199293 | 0.615 | 0.472 | 1 | 0.00088514 | 0.43896005 | 0.714 | 0.615 | 1 | 0.01002006 | 0.0017695 |
| Auts2 | 0.18443719 | -0.3905456 | 1 | 0.995 | 1 | 0.00088707 | -0.6961781 | 1 | 1 | 1 | 0.18443719 | 0.00177335 |
| Slc25a36 | 0.05624854 | 0.15208016 | 0.906 | 0.756 | 1 | 0.00088787 | 0.35844714 | 0.882 | 0.771 | 1 | 0.05624854 | 0.00177495 |
| Anapc11 | 0.00480659 | 0.37946331 | 0.865 | 0.715 | 1 | 0.00088871 | 0.44146562 | 0.786 | 0.562 | 1 | 0.00480659 | 0.00177663 |
| Ywhaq | 0.00088965 | 0.54565106 | 0.99 | 0.969 | 1 | 0.00800063 | 0.20272536 | 1 | 0.979 | 1 | 0.00800063 | 0.00177851 |
| Cops6 | 0.05166682 | 0.25672314 | 0.677 | 0.539 | 1 | 0.00089164 | 0.48405413 | 0.664 | 0.5 | 1 | 0.05166682 | 0.00178248 |
| Utp3 | 0.08702621 | 0.36923381 | 0.625 | 0.518 | 1 | 0.00089857 | 0.62335765 | 0.71 | 0.594 | 1 | 0.08702621 | 0.00179633 |
| Arrdc3 | 0.85085716 | -0.1714756 | 0.323 | 0.332 | 1 | 0.00089922 | -1.0619277 | 0.37 | 0.562 | 1 | 0.85085716 | 0.00179762 |
| Rtl5 | 0.00090091 | 1.70793425 | 0.125 | 0.026 | 1 | 0.01913922 | 0.78968041 | 0.113 | 0.031 | 1 | 0.01913922 | 0.001801 |
| Smc4 | 0.00090138 | 1.1891685 | 0.656 | 0.534 | 1 | 0.47188249 | 0.26023396 | 0.576 | 0.646 | 1 | 0.47188249 | 0.00180194 |
| Ccdc68 | 0.03631244 | 1.17741954 | 0.135 | 0.062 | 1 | 0.0009056 | 0.96515178 | 0.286 | 0.104 | 1 | 0.03631244 | 0.00181037 |
| Plekhh2 | 0.0009062 | -0.9299015 | 0.635 | 0.782 | 1 | 0.30723072 | -0.4857571 | 0.739 | 0.552 | 1 | 0.30723072 | 0.00181157 |
| Tgs1 | 0.07562545 | 0.23400307 | 0.635 | 0.503 | 1 | 0.00090751 | 0.50803033 | 0.597 | 0.396 | 1 | 0.07562545 | 0.0018142 |
| Immt | 0.00603275 | 0.59245704 | 0.625 | 0.56 | 1 | 0.000908 | 0.56233111 | 0.739 | 0.635 | 1 | 0.00603275 | 0.00181517 |
| Polr2k | 0.26614004 | 0.14194474 | 0.74 | 0.627 | 1 | 0.0009088 | 0.55382327 | 0.609 | 0.417 | 1 | 0.26614004 | 0.00181677 |
| Pax9 | 0.03355985 | -2.577468 | 0.01 | 0.067 | 1 | 0.00091013 | -4.5114891 | 0.013 | 0.083 | 1 | 0.03355985 | 0.00181943 |
| G930009F23Rik | 0.0009118 | -1.6554705 | 0.021 | 0.15 | 1 | 0.02918112 | -2.4318458 | 0.05 | 0.115 | 1 | 0.02918112 | 0.00182277 |
| Ptrhd1 | 0.14147372 | 0.55498233 | 0.302 | 0.228 | 1 | 0.00091424 | 1.42711033 | 0.202 | 0.052 | 1 | 0.14147372 | 0.00182764 |
| Oasl2 | 0.15986872 | -0.3144336 | 0.01 | 0.041 | 1 | 0.00091826 | -5.0468209 | 0.008 | 0.073 | 1 | 0.15986872 | 0.00183568 |
| Gata3 | 0.35859791 | -1.407543 | 0.021 | 0.041 | 1 | 0.00091826 | -5.0468209 | 0.008 | 0.073 | 1 | 0.35859791 | 0.00183568 |
| Gm28153 | 0.00091961 | 1.15188445 | 0.292 | 0.13 | 1 | 0.00126782 | 1.24473359 | 0.239 | 0.083 | 1 | 0.00126782 | 0.00183837 |
| Aup1 | 0.00727889 | 0.7503367 | 0.521 | 0.394 | 1 | 0.00091962 | 0.586767 | 0.508 | 0.281 | 1 | 0.00727889 | 0.0018384 |
| Tspan1 | 0.39189053 | -0.407543 | 0.01 | 0.026 | 1 | 0.0009228 | -2.1843244 | 0.021 | 0.104 | 1 | 0.39189053 | 0.00184474 |
| Chsy3 | 0.00092345 | -1.029444 | 0.927 | 0.891 | 1 | 0.3522311 | -0.2343881 | 0.718 | 0.604 | 1 | 0.3522311 | 0.00184605 |
| Jagn1 | 0.04322448 | 0.46939864 | 0.51 | 0.394 | 1 | 0.00092866 | 0.52126783 | 0.462 | 0.24 | 1 | 0.04322448 | 0.00185647 |
| Trabd | 0.72102969 | -0.1880563 | 0.292 | 0.254 | 1 | 0.00092903 | 0.51699584 | 0.45 | 0.229 | 1 | 0.72102969 | 0.0018572 |
| Sf3a2 | 0.0032586 | 0.73447604 | 0.406 | 0.233 | 1 | 0.00093117 | 0.84214783 | 0.37 | 0.177 | 1 | 0.0032586 | 0.00186148 |
| Urod | 0.8277041 | 0.13755508 | 0.24 | 0.228 | 1 | 0.00093134 | 0.99680608 | 0.324 | 0.135 | 1 | 0.8277041 | 0.00186181 |
| Klhl26 | 0.19402153 | -0.4828311 | 0.167 | 0.233 | 1 | 0.00093153 | 0.80056873 | 0.315 | 0.125 | 1 | 0.19402153 | 0.0018622 |
| Per2 | 0.00093251 | 0.95199942 | 0.531 | 0.347 | 1 | 0.39074883 | 0.22263982 | 0.475 | 0.427 | 1 | 0.39074883 | 0.00186415 |
| Polr2b | 0.00093405 | 0.79659775 | 0.562 | 0.368 | 1 | 0.03146251 | 0.28159127 | 0.605 | 0.438 | 1 | 0.03146251 | 0.00186722 |
| Cnep1r1 | 0.00093461 | 0.60747407 | 0.771 | 0.658 | 1 | 0.01424876 | 0.23584781 | 0.811 | 0.552 | 1 | 0.01424876 | 0.00186834 |
| Arpc3 | 0.00093486 | 0.36518033 | 0.958 | 0.886 | 1 | 0.09478325 | 0.1383491 | 0.916 | 0.917 | 1 | 0.09478325 | 0.00186885 |
| Ss18l2 | 0.07609057 | -0.6492037 | 0.51 | 0.57 | 1 | 0.0009358 | 0.63767732 | 0.382 | 0.188 | 1 | 0.07609057 | 0.00187073 |
| Rida | 0.03252679 | 0.60150218 | 0.333 | 0.212 | 1 | 0.00093593 | 1.16587817 | 0.282 | 0.115 | 1 | 0.03252679 | 0.00187099 |
| Chd2 | 0.02967325 | 0.25402631 | 0.896 | 0.834 | 1 | 0.00093707 | -0.7029349 | 0.966 | 1 | 1 | 0.02967325 | 0.00187325 |
| Thoc7 | 0.00093757 | 0.36909278 | 0.875 | 0.699 | 1 | 0.00531182 | 0.36999519 | 0.819 | 0.719 | 1 | 0.00531182 | 0.00187426 |
| H1f10 | 0.00093907 | 2.37427687 | 0.26 | 0.114 | 1 | 0.04406416 | 0.25951038 | 0.248 | 0.135 | 1 | 0.04406416 | 0.00187725 |
| Slc16a1 | 0.00094256 | 0.8572519 | 0.458 | 0.28 | 1 | 0.09693138 | -0.2307837 | 0.399 | 0.26 | 1 | 0.09693138 | 0.00188424 |
| Ndst1 | 0.00094638 | -0.8730903 | 0.552 | 0.699 | 1 | 0.15941431 | -0.4813856 | 0.765 | 0.812 | 1 | 0.15941431 | 0.00189185 |
| Larp1 | 0.00095067 | 0.37875034 | 0.875 | 0.715 | 1 | 0.30504446 | -0.2928265 | 0.941 | 0.927 | 1 | 0.30504446 | 0.00190043 |
| Cct5 | 0.00095606 | 0.54452256 | 0.823 | 0.699 | 1 | 0.01364337 | 0.23229415 | 0.861 | 0.771 | 1 | 0.01364337 | 0.00191121 |
| Lsm12 | 0.02036632 | 0.29595154 | 0.854 | 0.782 | 1 | 0.00095677 | 0.38675259 | 0.891 | 0.833 | 1 | 0.02036632 | 0.00191262 |
| Gng2 | 0.00095679 | 1.23697638 | 0.188 | 0.062 | 1 | 0.33613293 | -0.473354 | 0.134 | 0.094 | 1 | 0.33613293 | 0.00191266 |
| Gm29055 | 0.00095833 | 3.46692616 | 0.073 | 0.005 | 1 | 0.3937346 | 0.69014474 | 0.025 | 0.01 | 1 | 0.3937346 | 0.00191573 |
| Cd200r1 | 0.00095833 | 4.17741954 | 0.073 | 0.005 | 1 | 0.14313773 | 1.49749966 | 0.042 | 0.01 | 1 | 0.14313773 | 0.00191573 |
| Kcnq4 | 0.32025586 | -1.3144336 | 0 | 0.01 | 1 | 0.00096403 | -3.4797803 | 0.013 | 0.083 | 1 | 0.32025586 | 0.00192713 |
| Pak1ip1 | 0.00096692 | 0.5331761 | 0.771 | 0.549 | 1 | 0.00199352 | 0.41062221 | 0.769 | 0.667 | 1 | 0.00199352 | 0.0019329 |
| Tmub2 | 0.00190287 | 0.92949202 | 0.406 | 0.238 | 1 | 0.00096711 | 0.61614416 | 0.349 | 0.156 | 1 | 0.00190287 | 0.00193328 |
| As3mt | 0.10640612 | 0.60894516 | 0.198 | 0.124 | 1 | 0.0009682 | 0.78968041 | 0.269 | 0.094 | 1 | 0.10640612 | 0.00193546 |
| Metap2 | 0.00408387 | 0.30000614 | 0.938 | 0.824 | 1 | 0.00096823 | 0.34433527 | 0.924 | 0.833 | 1 | 0.00408387 | 0.00193551 |
| Dnm3 | 0.00096857 | -1.8886697 | 0.125 | 0.29 | 1 | 0.00891424 | -1.2663536 | 0.176 | 0.292 | 1 | 0.00891424 | 0.0019362 |
| Ndufs8 | 0.00096906 | 0.75543125 | 0.625 | 0.446 | 1 | 0.36700287 | 0.22619764 | 0.492 | 0.469 | 1 | 0.36700287 | 0.00193719 |
| Scarf2 | 0.00096942 | 0.38655274 | 0.771 | 0.497 | 1 | 0.00857299 | 0.21406103 | 0.794 | 0.51 | 1 | 0.00857299 | 0.0019379 |
| Trappc6a | 0.11007578 | 0.54282627 | 0.375 | 0.29 | 1 | 0.00096998 | 0.40439026 | 0.353 | 0.146 | 1 | 0.11007578 | 0.00193901 |
| 2610307P16Rik | 0.01221036 | -2.2277219 | 0.062 | 0.166 | 1 | 0.00097185 | -3.4000531 | 0.059 | 0.167 | 1 | 0.01221036 | 0.00194275 |
| Rtkn | 0.08887635 | 0.44455834 | 0.344 | 0.249 | 1 | 0.00097274 | 0.53814164 | 0.454 | 0.229 | 1 | 0.08887635 | 0.00194453 |
| Apod | 0.60490035 | -1.0940435 | 0.052 | 0.067 | 1 | 0.00097472 | -5.2093284 | 0.046 | 0.146 | 1 | 0.60490035 | 0.00194849 |
| Gm9908 | 0.00097707 | 3.00749454 | 0.073 | 0.005 | 1 | 0.34704016 | -1.3098553 | 0.008 | 0.021 | 1 | 0.34704016 | 0.00195318 |
| Gm57411 | 0.00097707 | 3.00749454 | 0.073 | 0.005 | 1 | 0.80442048 | -0.5728897 | 0.017 | 0.021 | 1 | 0.80442048 | 0.00195318 |

| | | | | | | | | | | | | |
|---|---|---|---|---|---|---|---|---|---|---|---|---|
| Gm35612 | 0.00097707 | 3.00749454 | 0.073 | 0.005 | 1 | 0.12814986 | 1.21370669 | 0.076 | 0.031 | 1 | 0.12814986 | 0.00195318 |
| Dync1i1 | 0.00544843 | 0.58299671 | 0.5 | 0.326 | 1 | 0.00097917 | 0.6008774 | 0.496 | 0.271 | 1 | 0.00544843 | 0.00195738 |
| Emc4 | 0.01309601 | 0.49292136 | 0.667 | 0.539 | 1 | 0.0009799 | 0.62128288 | 0.584 | 0.396 | 1 | 0.01309601 | 0.00195885 |
| Kremen1 | 0.0025291 | 0.40575632 | 0.615 | 0.42 | 1 | 0.00098212 | -0.9987687 | 0.517 | 0.646 | 1 | 0.0025291 | 0.00196329 |
| Gfpt2 | 0.00098228 | -0.5397628 | 0.865 | 0.876 | 1 | 0.00624232 | -0.2883291 | 0.908 | 0.667 | 1 | 0.00624232 | 0.00196359 |
| Myh9 | 0.11544877 | -0.5103538 | 0.708 | 0.803 | 1 | 0.00098244 | -0.5230292 | 0.895 | 0.948 | 1 | 0.11544877 | 0.00196392 |
| Gm36198 | 0.54923784 | 0.13302542 | 0.052 | 0.036 | 1 | 0.00098495 | -3.6317834 | 0.013 | 0.083 | 1 | 0.54923784 | 0.00196892 |
| Ccn2 | 0.00523895 | -0.5278605 | 0.969 | 0.984 | 1 | 0.00098593 | 0.97227102 | 0.912 | 0.76 | 1 | 0.00523895 | 0.0019709 |
| Ctnnd1 | 0.95554173 | -0.1293722 | 0.792 | 0.824 | 1 | 0.00098695 | -0.5286274 | 0.899 | 0.938 | 1 | 0.95554173 | 0.00197292 |
| Cisd3 | 0.00098889 | -1.2512397 | 0.135 | 0.311 | 1 | 0.13161584 | 0.22357694 | 0.202 | 0.125 | 1 | 0.13161584 | 0.00197681 |
| Entpd5 | 0.04151357 | 0.59245704 | 0.208 | 0.114 | 1 | 0.0009892 | -0.9332919 | 0.235 | 0.417 | 1 | 0.04151357 | 0.00197741 |
| Fgfbp3 | 0.21149904 | 0.68556644 | 0.135 | 0.088 | 1 | 0.00099009 | 3.86006974 | 0.105 | 0 | 1 | 0.21149904 | 0.00197919 |
| D030045P18Rik | 0.70865945 | -0.577468 | 0.042 | 0.052 | 1 | 0.00099082 | 0.27510724 | 0.34 | 0.135 | 1 | 0.70865945 | 0.00198065 |
| Cks1b | 0.00796912 | 0.96070033 | 0.458 | 0.311 | 1 | 0.00099193 | 0.88808412 | 0.408 | 0.198 | 1 | 0.00796912 | 0.00198287 |
| Otos | 0.00099372 | -5.0477879 | 0.031 | 0.161 | 1 | 0.02605524 | -3.3098553 | 0 | 0.021 | 1 | 0.02605524 | 0.00198645 |
| Ppih | 0.00099504 | 1.28312898 | 0.198 | 0.067 | 1 | 0.27098567 | 0.34222143 | 0.151 | 0.104 | 1 | 0.27098567 | 0.00198908 |
| Znhit1 | 0.00099551 | 0.84603111 | 0.51 | 0.321 | 1 | 0.00829874 | 0.54812573 | 0.37 | 0.219 | 1 | 0.00829874 | 0.00199002 |
| Mmgt2 | 0.68046192 | 0.24565427 | 0.302 | 0.285 | 1 | 0.00099783 | 0.82259503 | 0.361 | 0.167 | 1 | 0.68046192 | 0.00199466 |
| Pigh | 0.00099856 | 1.65957123 | 0.188 | 0.062 | 1 | 0.19941631 | -0.5091641 | 0.181 | 0.115 | 1 | 0.19941631 | 0.00199612 |
| Gm57315 | 0.19777351 | 0.54806292 | 0.312 | 0.254 | 1 | 0.00099928 | 0.81725666 | 0.29 | 0.115 | 1 | 0.19777351 | 0.00199757 |
| Nudt5 | 0.00099969 | 0.75115478 | 0.51 | 0.306 | 1 | 0.01450355 | 0.55521516 | 0.336 | 0.198 | 1 | 0.01450355 | 0.00199838 |
| Aplp2 | 0.02847056 | 0.18813376 | 1 | 0.984 | 1 | 0.00100153 | 0.21632517 | 1 | 0.979 | 1 | 0.02847056 | 0.00200206 |
| Pink1 | 0.95986841 | -0.1229021 | 0.521 | 0.487 | 1 | 0.00100172 | 0.5600842 | 0.576 | 0.333 | 1 | 0.95986841 | 0.00200243 |
| Zc3h15 | 0.00100328 | 0.38564463 | 0.958 | 0.902 | 1 | 0.00223976 | 0.2827446 | 0.95 | 0.896 | 1 | 0.00223976 | 0.00200555 |
| Phyhipl | 0.00100467 | -2.6445822 | 0.042 | 0.181 | 1 | 0.01074636 | -3.3098553 | 0.004 | 0.042 | 1 | 0.01074636 | 0.00200834 |
| Hid1 | 0.28001913 | -1.4519371 | 0.01 | 0.031 | 1 | 0.00100618 | -3.3098553 | 0.013 | 0.083 | 1 | 0.28001913 | 0.00201134 |
| Nutf2-ps1 | 0.00148524 | 0.92503238 | 0.417 | 0.238 | 1 | 0.00100645 | 1.21370669 | 0.261 | 0.094 | 1 | 0.00148524 | 0.00201189 |
| Grip1 | 0.30554982 | 0.68556644 | 0.219 | 0.166 | 1 | 0.00100674 | -0.3358505 | 0.269 | 0.521 | 1 | 0.30554982 | 0.00201246 |
| Mettl26 | 0.9343451 | -0.2048091 | 0.333 | 0.326 | 1 | 0.00100914 | 0.69014474 | 0.366 | 0.167 | 1 | 0.9343451 | 0.00201727 |
| Zfp622 | 0.06752001 | 0.37714013 | 0.469 | 0.368 | 1 | 0.00101128 | 0.50413275 | 0.571 | 0.365 | 1 | 0.06752001 | 0.00202154 |
| Dapk2 | 0.01144841 | -2.6205367 | 0.062 | 0.166 | 1 | 0.00101336 | -1.4727938 | 0.176 | 0.333 | 1 | 0.01144841 | 0.0020257 |
| Cemip | 0.00102255 | -1.083786 | 0.708 | 0.803 | 1 | 0.58546363 | -0.4067018 | 0.84 | 0.646 | 1 | 0.58546363 | 0.00204406 |
| Stoml2 | 0.00527382 | 0.72509481 | 0.49 | 0.332 | 1 | 0.00102398 | 0.66213036 | 0.441 | 0.24 | 1 | 0.00527382 | 0.0020469 |
| Pdk1 | 0.01642472 | 0.78121868 | 0.365 | 0.233 | 1 | 0.00102861 | 1.24653809 | 0.286 | 0.115 | 1 | 0.01642472 | 0.00205617 |
| Anks1 | 0.90492803 | -0.1851505 | 0.542 | 0.508 | 1 | 0.00102944 | -0.6278253 | 0.58 | 0.802 | 1 | 0.90492803 | 0.00205782 |
| Spryd3 | 0.00103089 | 0.6579101 | 0.479 | 0.28 | 1 | 0.00118748 | 0.58176858 | 0.424 | 0.208 | 1 | 0.00118748 | 0.00206071 |
| Gucy2g | 0.00103519 | -3.1624305 | 0.021 | 0.145 | 1 | 0.23976577 | -1.9879272 | 0.013 | 0.031 | 1 | 0.23976577 | 0.00206931 |
| Ccdc137 | 0.00103916 | 1.10060394 | 0.354 | 0.181 | 1 | 0.12646634 | 0.27510724 | 0.248 | 0.167 | 1 | 0.12646634 | 0.00207725 |
| Ift122 | 0.00103947 | 1.47315811 | 0.229 | 0.088 | 1 | 0.15551535 | 0.47150445 | 0.181 | 0.115 | 1 | 0.15551535 | 0.00207786 |
| Eml3 | 0.08552744 | 0.32942263 | 0.302 | 0.207 | 1 | 0.00103967 | 0.65361886 | 0.357 | 0.167 | 1 | 0.08552744 | 0.00207825 |
| Rras | 0.56793982 | -0.2450475 | 0.667 | 0.648 | 1 | 0.00104852 | 0.36152199 | 0.609 | 0.344 | 1 | 0.56793982 | 0.00209593 |
| Tmem138 | 0.00134792 | 1.45815595 | 0.292 | 0.14 | 1 | 0.0010498 | 1.81942775 | 0.21 | 0.062 | 1 | 0.00134792 | 0.00209851 |
| Apmap | 0.00789036 | 0.70371379 | 0.469 | 0.321 | 1 | 0.00105049 | 0.75862448 | 0.454 | 0.24 | 1 | 0.00789036 | 0.00209988 |
| Traf7 | 0.10812772 | 0.48393258 | 0.365 | 0.28 | 1 | 0.00105197 | 0.50830641 | 0.458 | 0.25 | 1 | 0.10812772 | 0.00210282 |
| Uxs1 | 0.76035504 | -0.1319085 | 0.708 | 0.699 | 1 | 0.00105249 | 0.30133878 | 0.651 | 0.385 | 1 | 0.76035504 | 0.00210387 |
| Colgalt2 | 0.00105518 | -1.6910305 | 0.604 | 0.622 | 1 | 0.5252077 | -0.6924713 | 0.303 | 0.24 | 1 | 0.5252077 | 0.00210924 |
| Galnt7 | 0.00105596 | -1.2633594 | 0.312 | 0.487 | 1 | 0.01170761 | -0.9667814 | 0.395 | 0.573 | 1 | 0.01170761 | 0.00211081 |
| D16Ertd472e | 0.00105944 | 0.75956702 | 0.385 | 0.202 | 1 | 0.19912911 | 0.38282106 | 0.353 | 0.271 | 1 | 0.19912911 | 0.00211777 |
| Ralb | 0.00105991 | 0.52802516 | 0.677 | 0.446 | 1 | 0.00575668 | 0.47564023 | 0.597 | 0.438 | 1 | 0.00575668 | 0.0021187 |
| Slc27a1 | 0.0212172 | 0.66235905 | 0.292 | 0.171 | 1 | 0.00106141 | 0.86599457 | 0.353 | 0.167 | 1 | 0.0212172 | 0.00212169 |
| Pdzd11 | 0.02380513 | 0.46175335 | 0.583 | 0.42 | 1 | 0.00106299 | 0.47150445 | 0.441 | 0.219 | 1 | 0.02380513 | 0.00212485 |
| Bpnt1 | 0.87210841 | 0.13302542 | 0.198 | 0.192 | 1 | 0.00106482 | 0.91795065 | 0.298 | 0.115 | 1 | 0.87210841 | 0.0021285 |
| Abcc5 | 0.00106631 | 0.46692616 | 0.698 | 0.477 | 1 | 0.29263216 | 0.34592858 | 0.756 | 0.667 | 1 | 0.29263216 | 0.00213148 |
| Ndp | 0.00106842 | 1.83346514 | 0.208 | 0.078 | 1 | 0.17652578 | 0.80562195 | 0.139 | 0.083 | 1 | 0.17652578 | 0.00213571 |
| Sox6 | 0.00107242 | 0.2945023 | 1 | 0.99 | 1 | 0.3354143 | -0.4884901 | 0.996 | 0.906 | 1 | 0.3354143 | 0.00214369 |
| Mrps34 | 0.29286648 | 0.23697638 | 0.396 | 0.321 | 1 | 0.00107427 | 0.66505376 | 0.37 | 0.177 | 1 | 0.29286648 | 0.00214738 |
| Prkcsh | 0.20744366 | 0.24565427 | 0.719 | 0.653 | 1 | 0.00107678 | 0.36821664 | 0.672 | 0.417 | 1 | 0.20744366 | 0.0021524 |
| Ephb3 | 0.16634646 | 0.42253204 | 0.281 | 0.207 | 1 | 0.00107868 | 0.89525917 | 0.441 | 0.25 | 1 | 0.16634646 | 0.0021562 |
| Zc3h11a | 0.16009692 | 0.27598337 | 0.427 | 0.337 | 1 | 0.00108269 | 0.67051593 | 0.458 | 0.25 | 1 | 0.16009692 | 0.0021642 |
| Mterf3 | 0.02533333 | 0.42807022 | 0.594 | 0.456 | 1 | 0.00108411 | 0.46219279 | 0.571 | 0.312 | 1 | 0.02533333 | 0.00216704 |
| Tmsb15b1 | 0.62495317 | 0.19035859 | 0.177 | 0.155 | 1 | 0.0010878 | 1.7944814 | 0.164 | 0.031 | 1 | 0.62495317 | 0.00217442 |
| Tor1aip2 | 0.40684017 | -0.1003504 | 0.885 | 0.896 | 1 | 0.00108801 | 0.42073864 | 0.962 | 0.875 | 1 | 0.40684017 | 0.00217484 |
| Pmm2 | 0.16937536 | 0.51650818 | 0.292 | 0.223 | 1 | 0.0010881 | 0.85308331 | 0.332 | 0.146 | 1 | 0.16937536 | 0.00217502 |
| Ddx3x | 0.03386954 | 0.27137164 | 0.969 | 0.938 | 1 | 0.00109107 | 0.30451047 | 0.987 | 0.917 | 1 | 0.03386954 | 0.00218095 |
| Mest | 0.01712014 | 0.705466 | 0.531 | 0.389 | 1 | 0.00109213 | 0.75697625 | 0.492 | 0.271 | 1 | 0.01712014 | 0.00218307 |
| Chst2 | 0.00109227 | -1.6072153 | 0.167 | 0.347 | 1 | 0.1566913 | 0.36257008 | 0.143 | 0.083 | 1 | 0.1566913 | 0.00218335 |
| Tspan9 | 0.00109258 | 0.96377316 | 0.396 | 0.223 | 1 | 0.64965669 | -0.2567439 | 0.458 | 0.375 | 1 | 0.64965669 | 0.00218397 |
| Ptbp1 | 0.02772621 | 0.23719176 | 0.896 | 0.788 | 1 | 0.00109591 | 0.3390703 | 0.945 | 0.823 | 1 | 0.02772621 | 0.00219062 |
| Hnrnpdl | 0.00110256 | 0.31042167 | 0.979 | 0.938 | 1 | 0.00419561 | 0.19427616 | 0.983 | 0.958 | 1 | 0.00419561 | 0.00220391 |
| Ccdc107 | 0.00110883 | 0.89856016 | 0.406 | 0.223 | 1 | 0.05629113 | 0.42320588 | 0.382 | 0.271 | 1 | 0.05629113 | 0.00221644 |
| Map4 | 0.00111066 | 0.53356335 | 0.927 | 0.907 | 1 | 0.03033459 | -0.3928404 | 0.962 | 0.969 | 1 | 0.03033459 | 0.00222009 |
| Inhba | 0.00111382 | -1.0161874 | 0.094 | 0.264 | 1 | 0.07380725 | -0.681824 | 0.017 | 0.052 | 1 | 0.07380725 | 0.00222639 |
| Pla1a | 0.00111384 | -3.7998604 | 0 | 0.104 | 1 | 0.92095259 | -0.194378 | 0.034 | 0.031 | 1 | 0.92095259 | 0.00222644 |
| Snhg20 | 0.00561485 | 1.0915588 | 0.365 | 0.223 | 1 | 0.00111903 | 1.29680231 | 0.231 | 0.073 | 1 | 0.00561485 | 0.00223681 |
| Plekhb1 | 0.00396449 | -1.1079827 | 0.312 | 0.461 | 1 | 0.00112869 | 1.25782925 | 0.239 | 0.083 | 1 | 0.00396449 | 0.0022561 |
| Slc30a6 | 0.00113088 | 1.1843723 | 0.375 | 0.212 | 1 | 0.55320523 | -0.3098553 | 0.294 | 0.24 | 1 | 0.55320523 | 0.00226048 |
| Sdhd | 0.03384408 | 0.32631672 | 0.708 | 0.57 | 1 | 0.00113236 | 0.69014474 | 0.668 | 0.531 | 1 | 0.03384408 | 0.00226344 |
| Klhl28 | 0.06716522 | 0.3529911 | 0.417 | 0.301 | 1 | 0.0011351 | 0.44503224 | 0.433 | 0.208 | 1 | 0.06716522 | 0.00226891 |
| Dock2 | 0.21867159 | 2.59245704 | 0.021 | 0.005 | 1 | 0.00113557 | -1.4797803 | 0.013 | 0.083 | 1 | 0.21867159 | 0.00226984 |
| Ankrd50 | 0.01719113 | 0.92423903 | 0.448 | 0.326 | 1 | 0.00113963 | 0.31234014 | 0.63 | 0.354 | 1 | 0.01719113 | 0.00227797 |
| Psma1 | 0.25197023 | 0.18615539 | 0.812 | 0.788 | 1 | 0.00114508 | 0.39709776 | 0.878 | 0.865 | 1 | 0.25197023 | 0.00228884 |
| Ankfy1 | 0.00114736 | 0.6613189 | 0.667 | 0.456 | 1 | 0.00209941 | -0.9802871 | 0.647 | 0.75 | 1 | 0.00209941 | 0.0022934 |
| Oard1 | 0.93996589 | -0.1445086 | 0.271 | 0.264 | 1 | 0.00114975 | 0.89659561 | 0.214 | 0.062 | 1 | 0.93996589 | 0.00229818 |
| Slc10a7 | 0.00115606 | 0.83061677 | 0.521 | 0.358 | 1 | 0.00427204 | -0.738832 | 0.529 | 0.729 | 1 | 0.00427204 | 0.00231079 |
| Gnaq | 0.02306893 | -0.3802906 | 0.958 | 0.922 | 1 | 0.00115666 | -0.688965 | 0.992 | 0.979 | 1 | 0.02306893 | 0.00231199 |
| Zfp385b | 0.00115692 | -1.2402047 | 0.99 | 0.876 | 1 | 0.37845663 | -0.6394498 | 0.916 | 0.75 | 1 | 0.37845663 | 0.0023125 |
| Ccdc115 | 0.00115786 | 0.76463801 | 0.458 | 0.259 | 1 | 0.0023102 | 0.70344156 | 0.319 | 0.146 | 1 | 0.0023102 | 0.00231439 |
| Vsir | 0.11271713 | 0.58428311 | 0.281 | 0.197 | 1 | 0.00115907 | 0.9416835 | 0.282 | 0.104 | 1 | 0.11271713 | 0.00231679 |
| Gpn1 | 0.00154161 | 0.93154568 | 0.323 | 0.155 | 1 | 0.00116167 | 1.69014474 | 0.223 | 0.073 | 1 | 0.00154161 | 0.002322 |
| Unkl | 0.0393326 | 0.51845646 | 0.417 | 0.29 | 1 | 0.00116477 | 0.64434105 | 0.5 | 0.271 | 1 | 0.0393326 | 0.00232819 |

| | | | | | | | | | | | | |
|---|---|---|---|---|---|---|---|---|---|---|---|---|
| Lars2 | 0.00235915 | -1.1475738 | 1 | 1 | 1 | 0.00116484 | -1.2024249 | 1 | 1 | 1 | 0.00235915 | 0.00232832 |
| Blcap | 0.00116596 | 0.89201732 | 0.385 | 0.202 | 1 | 0.01664543 | 0.46592958 | 0.357 | 0.219 | 1 | 0.01664543 | 0.00233055 |
| Bmi1 | 0.33578007 | -0.3353932 | 0.312 | 0.358 | 1 | 0.00116638 | 0.60871714 | 0.429 | 0.219 | 1 | 0.33578007 | 0.0023314 |
| Ubr5 | 0.08226487 | -0.3021028 | 1 | 0.984 | 1 | 0.00116859 | -0.4549755 | 1 | 1 | 1 | 0.08226487 | 0.00233582 |
| Clcn3 | 0.00117302 | -0.7096626 | 0.344 | 0.565 | 1 | 0.02458682 | -0.4152083 | 0.571 | 0.708 | 1 | 0.02458682 | 0.00234467 |
| Tlcd4 | 0.36065274 | -0.4711591 | 0.375 | 0.409 | 1 | 0.00117399 | 0.41691456 | 0.513 | 0.292 | 1 | 0.36065274 | 0.0023466 |
| Zfp326 | 0.00117628 | -0.7024866 | 0.917 | 0.927 | 1 | 0.34597917 | -0.1320983 | 0.92 | 0.927 | 1 | 0.34597917 | 0.00235117 |
| Mtm1 | 0.001177 | -1.7076667 | 0.229 | 0.389 | 1 | 0.72341525 | -0.8312017 | 0.214 | 0.177 | 1 | 0.72341525 | 0.00235262 |
| Mpv17 | 0.44872381 | -0.4807919 | 0.323 | 0.347 | 1 | 0.00117766 | 0.70344156 | 0.324 | 0.135 | 1 | 0.44872381 | 0.00235393 |
| 2500004C02Rik | 0.00117934 | 1.94609399 | 0.167 | 0.052 | 1 | 0.07794965 | 0.63356121 | 0.168 | 0.094 | 1 | 0.07794965 | 0.00235729 |
| Golga2 | 0.00117977 | 0.67375714 | 0.729 | 0.596 | 1 | 0.00553665 | 0.48482583 | 0.706 | 0.542 | 1 | 0.00553665 | 0.00235815 |
| Etv5 | 0.00118138 | -1.407543 | 0.104 | 0.275 | 1 | 0.00226484 | 0.66668576 | 0.437 | 0.24 | 1 | 0.00226484 | 0.00236137 |
| Stxbp5 | 0.98633225 | -0.5498368 | 0.667 | 0.606 | 1 | 0.00119058 | -0.6727415 | 0.697 | 0.844 | 1 | 0.98633225 | 0.00237974 |
| Txn2 | 0.00119589 | 0.46092741 | 0.854 | 0.637 | 1 | 0.00681556 | 0.29104878 | 0.748 | 0.646 | 1 | 0.00681556 | 0.00239035 |
| Creb3l1 | 0.00119736 | 0.35811531 | 0.573 | 0.358 | 1 | 0.06162913 | -0.1839708 | 0.588 | 0.365 | 1 | 0.06162913 | 0.00239329 |
| Ppie | 0.01903023 | 0.65055999 | 0.406 | 0.28 | 1 | 0.00119992 | 0.70395054 | 0.366 | 0.167 | 1 | 0.01903023 | 0.00239841 |
| Galt | 0.04699592 | 0.61754802 | 0.271 | 0.171 | 1 | 0.0012001 | 1.53449387 | 0.261 | 0.104 | 1 | 0.04699592 | 0.00239876 |
| Nat9 | 0.63979594 | 0.22988696 | 0.188 | 0.166 | 1 | 0.00120104 | 2.14957636 | 0.16 | 0.031 | 1 | 0.63979594 | 0.00240063 |
| Dyrk1a | 0.16040926 | 0.16397995 | 0.948 | 0.855 | 1 | 0.00120155 | -0.598675 | 0.966 | 0.948 | 1 | 0.16040926 | 0.00240166 |
| Mkrn1 | 0.14792974 | -0.4330781 | 0.438 | 0.497 | 1 | 0.0012035 | 0.52021974 | 0.752 | 0.583 | 1 | 0.14792974 | 0.00240556 |
| Eif4a3 | 0.00120373 | 0.51306958 | 0.74 | 0.518 | 1 | 0.00246306 | 0.3439164 | 0.714 | 0.542 | 1 | 0.00246306 | 0.002406 |
| Anapc16 | 0.04762315 | 0.28701707 | 0.812 | 0.694 | 1 | 0.00120633 | 0.36952284 | 0.803 | 0.646 | 1 | 0.04762315 | 0.0024112 |
| Eif1ad | 0.12020294 | 0.35541784 | 0.354 | 0.259 | 1 | 0.00121242 | 0.55612739 | 0.37 | 0.167 | 1 | 0.12020294 | 0.00242336 |
| Hmgcll1 | 0.00122411 | -2.8604019 | 0.031 | 0.161 | 1 | 0.00875978 | 0.74077081 | 0.248 | 0.115 | 1 | 0.00875978 | 0.00244671 |
| Plscr1 | 0.16976953 | -0.614344 | 0.406 | 0.461 | 1 | 0.00122658 | -0.3184684 | 0.319 | 0.542 | 1 | 0.16976953 | 0.00245165 |
| Pa2g4 | 0.00122821 | 0.46807282 | 0.875 | 0.762 | 1 | 0.00132191 | 0.32267292 | 0.882 | 0.792 | 1 | 0.00132191 | 0.00245492 |
| Gemin2 | 0.0012311 | 0.90795886 | 0.24 | 0.093 | 1 | 0.30809332 | 0.27510724 | 0.147 | 0.104 | 1 | 0.30809332 | 0.00246069 |
| Ngef | 0.00137637 | 0.77156536 | 0.625 | 0.497 | 1 | 0.00123259 | 0.21939781 | 0.597 | 0.333 | 1 | 0.00137637 | 0.00246367 |
| Ddx50 | 0.00123501 | 0.42464278 | 0.969 | 0.927 | 1 | 0.73072731 | -0.1803224 | 0.966 | 0.948 | 1 | 0.73072731 | 0.0024685 |
| Dennd1b | 0.09953026 | 0.33192949 | 0.615 | 0.549 | 1 | 0.00123818 | -0.6640105 | 0.643 | 0.812 | 1 | 0.09953026 | 0.00247483 |
| Nsfl1c | 0.00217234 | 0.50958877 | 0.75 | 0.642 | 1 | 0.00124026 | 0.43579009 | 0.803 | 0.656 | 1 | 0.00217234 | 0.00247899 |
| Os9 | 0.1749097 | 0.11163799 | 0.917 | 0.876 | 1 | 0.0012416 | 0.23903798 | 0.929 | 0.875 | 1 | 0.1749097 | 0.00248167 |
| Ankrd28 | 0.00124242 | 0.42977495 | 1 | 0.969 | 1 | 0.00874521 | 0.14512278 | 0.996 | 1 | 1 | 0.00874521 | 0.00248329 |
| Itpa | 0.05347849 | 0.35768089 | 0.448 | 0.321 | 1 | 0.00124315 | 0.47150445 | 0.382 | 0.188 | 1 | 0.05347849 | 0.00248476 |
| Nrf1 | 0.00124322 | 0.56270969 | 0.688 | 0.446 | 1 | 0.19508177 | -0.4797803 | 0.761 | 0.771 | 1 | 0.19508177 | 0.00248489 |
| Pwwp2b | 0.52697614 | 0.46692616 | 0.094 | 0.073 | 1 | 0.0012505 | 1.45567948 | 0.21 | 0.062 | 1 | 0.52697614 | 0.00249943 |
| Fiz1 | 0.00196518 | 1.00749454 | 0.26 | 0.114 | 1 | 0.00126189 | 1.05004068 | 0.261 | 0.094 | 1 | 0.00196518 | 0.00252219 |
| Pycrl | 0.00514093 | 1.10937415 | 0.312 | 0.176 | 1 | 0.00126498 | 1.63767732 | 0.193 | 0.052 | 1 | 0.00514093 | 0.00252836 |
| Pvt1 | 0.00126687 | 1.08656611 | 0.521 | 0.342 | 1 | 0.76736642 | -0.4609968 | 0.387 | 0.354 | 1 | 0.76736642 | 0.00253214 |
| Odf2 | 0.00126767 | 0.7716816 | 0.573 | 0.373 | 1 | 0.00213359 | -0.9612451 | 0.454 | 0.594 | 1 | 0.00213359 | 0.00253373 |
| Chrna1os | 0.73831607 | 0.32942263 | 0.073 | 0.062 | 1 | 0.00127144 | 3.49749966 | 0.101 | 0 | 1 | 0.73831607 | 0.00254126 |
| 4930550C14Rik | 0.17712887 | 0.64492446 | 0.146 | 0.093 | 1 | 0.00127198 | 1.52303475 | 0.193 | 0.052 | 1 | 0.17712887 | 0.00254235 |
| Ei24 | 0.02901111 | 0.49504454 | 0.479 | 0.332 | 1 | 0.00127326 | 0.53981046 | 0.471 | 0.26 | 1 | 0.02901111 | 0.00254491 |
| Terf2ip | 0.28717104 | 0.25006484 | 0.312 | 0.249 | 1 | 0.00127349 | 0.17201375 | 0.445 | 0.229 | 1 | 0.28717104 | 0.00254536 |
| Osbpl9 | 0.30749704 | -0.1983662 | 0.906 | 0.907 | 1 | 0.00127453 | -0.48292 | 0.945 | 0.958 | 1 | 0.30749704 | 0.00254743 |
| Brix1 | 0.00163729 | 0.6241659 | 0.74 | 0.617 | 1 | 0.00127672 | 0.45116636 | 0.739 | 0.552 | 1 | 0.00163729 | 0.00255181 |
| Phf1 | 0.33905664 | 0.24744533 | 0.427 | 0.368 | 1 | 0.00127882 | 0.47946574 | 0.496 | 0.26 | 1 | 0.33905664 | 0.002556 |
| Alpl | 0.00131032 | 1.71603172 | 0.229 | 0.088 | 1 | 0.00128158 | -1.131857 | 0.034 | 0.125 | 1 | 0.00131032 | 0.00256153 |
| Nabp2 | 0.23776871 | 0.27542774 | 0.448 | 0.368 | 1 | 0.00128167 | 0.39796399 | 0.466 | 0.24 | 1 | 0.23776871 | 0.00256169 |
| 1700025G04Rik | 0.00128233 | -0.6209953 | 0.958 | 0.969 | 1 | 0.66032629 | -0.3676058 | 0.966 | 0.896 | 1 | 0.66032629 | 0.00256303 |
| Mtln | 0.24028426 | -0.4450177 | 0.312 | 0.368 | 1 | 0.00128295 | 1.01207283 | 0.286 | 0.115 | 1 | 0.24028426 | 0.00256426 |
| Lrrc47 | 0.00941256 | 0.80955516 | 0.312 | 0.176 | 1 | 0.00128592 | 0.53563479 | 0.366 | 0.167 | 1 | 0.00941256 | 0.00257018 |
| Cmbl | 0.77038317 | -0.1670032 | 0.271 | 0.285 | 1 | 0.00128638 | 0.60372999 | 0.399 | 0.198 | 1 | 0.77038317 | 0.00257111 |
| Dync2li1 | 0.67543659 | 0.18033113 | 0.26 | 0.238 | 1 | 0.00128904 | 0.96868024 | 0.349 | 0.167 | 1 | 0.67543659 | 0.00257643 |
| Psmc3 | 0.01196381 | 0.39538043 | 0.885 | 0.689 | 1 | 0.00129149 | 0.30413629 | 0.857 | 0.74 | 1 | 0.01196381 | 0.00258131 |
| Ambra1 | 0.00129541 | -0.5298226 | 0.771 | 0.907 | 1 | 0.06327091 | -0.4952869 | 0.895 | 0.885 | 1 | 0.06327091 | 0.00258915 |
| Farsa | 0.00129574 | 0.80755697 | 0.479 | 0.295 | 1 | 0.00359787 | 0.47540589 | 0.466 | 0.281 | 1 | 0.00359787 | 0.0025898 |
| Knl1 | 0.00129732 | 3.72996056 | 0.083 | 0.01 | 1 | 0.52073627 | -0.3098553 | 0.021 | 0.01 | 1 | 0.52073627 | 0.00259295 |
| Luc7l3 | 0.00480727 | 0.35768089 | 0.938 | 0.87 | 1 | 0.00129742 | 0.29430581 | 0.971 | 0.979 | 1 | 0.00480727 | 0.00259316 |
| Kdm3b | 0.14883735 | -0.4891814 | 0.427 | 0.487 | 1 | 0.00130266 | -0.7627361 | 0.609 | 0.75 | 1 | 0.14883735 | 0.00260363 |
| Naa15 | 0.00130526 | 0.37114976 | 0.917 | 0.829 | 1 | 0.63703598 | -0.1546539 | 0.929 | 0.938 | 1 | 0.63703598 | 0.00260881 |
| Sap30bp | 0.00130769 | 0.64994253 | 0.604 | 0.42 | 1 | 0.10240338 | 0.16587817 | 0.588 | 0.448 | 1 | 0.10240338 | 0.00261367 |
| Glb1l | 0.00131258 | 1.24453373 | 0.24 | 0.098 | 1 | 0.01949618 | 0.29680231 | 0.227 | 0.104 | 1 | 0.01949618 | 0.00262343 |
| Cby1 | 0.58583497 | 0.22988696 | 0.271 | 0.249 | 1 | 0.00131468 | 0.99499932 | 0.298 | 0.125 | 1 | 0.58583497 | 0.00262763 |
| Dvl1 | 0.05295889 | 0.43921078 | 0.458 | 0.347 | 1 | 0.00131823 | 0.51326698 | 0.571 | 0.344 | 1 | 0.05295889 | 0.00263473 |
| Tcf7l2 | 0.30426707 | 0.27052894 | 0.906 | 0.917 | 1 | 0.00132078 | -0.4743201 | 0.983 | 0.979 | 1 | 0.30426707 | 0.00263981 |
| Hmgcl | 0.04815774 | 0.55292867 | 0.406 | 0.285 | 1 | 0.00132677 | 0.5580412 | 0.458 | 0.24 | 1 | 0.04815774 | 0.00265179 |
| Gpc6 | 0.00133301 | -0.731676 | 1 | 0.995 | 1 | 0.22024512 | -0.1082692 | 1 | 0.969 | 1 | 0.22024512 | 0.00266425 |
| Mef2b | 0.43235571 | 0.17350449 | 0.219 | 0.176 | 1 | 0.00133345 | 1.23446525 | 0.239 | 0.083 | 1 | 0.43235571 | 0.00266512 |
| Coq10b | 0.41626347 | 0.22630855 | 0.854 | 0.86 | 1 | 0.00133445 | 0.34678934 | 0.912 | 0.875 | 1 | 0.41626347 | 0.00266712 |
| Mad1l1 | 0.00133872 | 0.79176585 | 0.385 | 0.197 | 1 | 0.07582058 | -0.8733806 | 0.475 | 0.51 | 1 | 0.07582058 | 0.00267566 |
| D130009I18Rik | 0.13470811 | 0.16774162 | 0.802 | 0.699 | 1 | 0.00133914 | 0.69994791 | 0.731 | 0.708 | 1 | 0.13470811 | 0.00267649 |
| Sec23a | 0.00165103 | 0.48543879 | 0.865 | 0.731 | 1 | 0.00133925 | 0.41705667 | 0.866 | 0.729 | 1 | 0.00165103 | 0.0026767 |
| Hectd2os | 0.0013399 | -2.8669746 | 0.104 | 0.254 | 1 | 0.69817746 | -1.3098553 | 0.105 | 0.115 | 1 | 0.69817746 | 0.00267801 |
| Mgat4a | 0.00134159 | -0.8008462 | 0.812 | 0.803 | 1 | 0.03510801 | 0.22991069 | 0.794 | 0.75 | 1 | 0.03510801 | 0.00268138 |
| Wdr74 | 0.02665028 | 0.59868899 | 0.417 | 0.28 | 1 | 0.00134919 | 0.70071398 | 0.441 | 0.24 | 1 | 0.02665028 | 0.00269655 |
| Timm50 | 0.15839852 | 0.36504654 | 0.354 | 0.264 | 1 | 0.00135143 | 1.02894665 | 0.282 | 0.115 | 1 | 0.15839852 | 0.00270103 |
| Gorasp1 | 0.00135714 | 1.43812889 | 0.25 | 0.109 | 1 | 0.03050441 | 0.87777174 | 0.134 | 0.052 | 1 | 0.03050441 | 0.00271243 |
| Nsf | 0.00135783 | 0.40803247 | 0.677 | 0.482 | 1 | 0.61123304 | -0.3618499 | 0.693 | 0.729 | 1 | 0.61123304 | 0.00271381 |
| Gm57390 | 0.16395262 | 0.55498233 | 0.167 | 0.109 | 1 | 0.00135811 | 1.55521516 | 0.176 | 0.042 | 1 | 0.16395262 | 0.00271438 |
| Agap1 | 0.06291209 | -0.3068867 | 0.479 | 0.606 | 1 | 0.00136013 | -0.5569481 | 0.723 | 0.885 | 1 | 0.06291209 | 0.00271842 |
| Sh3bgrl3 | 0.00136213 | 0.48094872 | 0.812 | 0.679 | 1 | 0.07607941 | 0.44687359 | 0.723 | 0.719 | 1 | 0.07607941 | 0.00272241 |
| Nedd9 | 0.00136343 | -0.9121117 | 0.792 | 0.927 | 1 | 0.45432169 | -0.345144 | 0.861 | 0.875 | 1 | 0.45432169 | 0.002725 |
| Dctn3 | 0.08158944 | 0.57908981 | 0.635 | 0.518 | 1 | 0.0013681 | 0.98292649 | 0.647 | 0.531 | 1 | 0.08158944 | 0.00273433 |
| Kcnj15 | 0.00581932 | -2.1158879 | 0.021 | 0.119 | 1 | 0.00137445 | -1.9375386 | 0.324 | 0.479 | 1 | 0.00581932 | 0.002747 |
| Nuak1 | 0.00137459 | 0.64492446 | 0.552 | 0.316 | 1 | 0.04162711 | -0.1046301 | 0.622 | 0.417 | 1 | 0.04162711 | 0.0027473 |
| Psmc1 | 0.29254624 | 0.20939758 | 0.719 | 0.637 | 1 | 0.00137595 | 0.47531942 | 0.714 | 0.562 | 1 | 0.29254624 | 0.00275 |
| Hint2 | 0.05580081 | 0.40438469 | 0.573 | 0.44 | 1 | 0.00138358 | 1.08869411 | 0.378 | 0.208 | 1 | 0.05580081 | 0.00276524 |

| | | | | | | | | | | | | |
|---|---|---|---|---|---|---|---|---|---|---|---|---|
| Sesn2 | 0.00854106 | 1.0470229 | 0.417 | 0.264 | 1 | 0.00138478 | 0.89401707 | 0.319 | 0.135 | 1 | 0.00854106 | 0.00276765 |
| Orai1 | 0.00347167 | -0.6179028 | 0.5 | 0.642 | 1 | 0.00138561 | 0.40707258 | 0.555 | 0.333 | 1 | 0.00347167 | 0.00276929 |
| Rps6ka2 | 0.00138902 | 0.77671433 | 0.438 | 0.249 | 1 | 0.84031625 | -0.4878536 | 0.189 | 0.177 | 1 | 0.84031625 | 0.00277611 |
| Anxa1 | 0.11077105 | -0.3665505 | 0.708 | 0.86 | 1 | 0.00139028 | -0.7012221 | 0.718 | 0.854 | 1 | 0.11077105 | 0.00277863 |
| Utp14a | 0.00139241 | 0.80033663 | 0.635 | 0.472 | 1 | 0.10777161 | 0.24922861 | 0.542 | 0.448 | 1 | 0.10777161 | 0.00278288 |
| Bbs7 | 0.1057117 | 0.88885004 | 0.24 | 0.166 | 1 | 0.00139364 | 1.06211351 | 0.235 | 0.083 | 1 | 0.1057117 | 0.00278535 |
| Idh3b | 0.02085481 | 0.47099352 | 0.667 | 0.528 | 1 | 0.00139467 | 0.63405018 | 0.605 | 0.438 | 1 | 0.02085481 | 0.0027874 |
| Trmt1 | 0.0021454 | 0.71110153 | 0.469 | 0.29 | 1 | 0.00139635 | 0.46775232 | 0.42 | 0.198 | 1 | 0.0021454 | 0.00279075 |
| Slco2a1 | 0.00139669 | 1.57139542 | 0.177 | 0.057 | 1 | 0.46446396 | 0.20471791 | 0.097 | 0.125 | 1 | 0.46446396 | 0.00279143 |
| Mapre3 | 0.00139757 | 1.37564565 | 0.677 | 0.523 | 1 | 0.01086817 | -0.1752855 | 0.697 | 0.479 | 1 | 0.01086817 | 0.00279318 |
| Tatdn2 | 0.12551988 | 0.45110119 | 0.271 | 0.192 | 1 | 0.00139802 | 0.80165305 | 0.395 | 0.198 | 1 | 0.12551988 | 0.00279409 |
| Notch2 | 0.0014032 | 0.36397985 | 0.948 | 0.891 | 1 | 0.00148341 | -0.5137675 | 0.971 | 0.969 | 1 | 0.00148341 | 0.00280443 |
| Setmar | 0.00140608 | 2.74446013 | 0.083 | 0.01 | 1 | 0.12815003 | 1.14957636 | 0.076 | 0.031 | 1 | 0.12815003 | 0.00281018 |
| Camkk1 | 0.00140608 | 2.74446013 | 0.083 | 0.01 | 1 | 0.50844268 | 0.69014474 | 0.021 | 0.01 | 1 | 0.50844268 | 0.00281018 |
| Ndufb8 | 0.22193708 | -0.1806351 | 0.729 | 0.772 | 1 | 0.00140735 | 0.4129976 | 0.807 | 0.74 | 1 | 0.22193708 | 0.00281272 |
| Ccna2 | 0.00140818 | 3.15433592 | 0.083 | 0.01 | 1 | 0.20315163 | 1.01207283 | 0.017 | 0 | 1 | 0.20315163 | 0.00281438 |
| Sf3a3 | 0.0014082 | 0.60058492 | 0.583 | 0.389 | 1 | 0.01162036 | 0.29616885 | 0.559 | 0.385 | 1 | 0.01162036 | 0.00281442 |
| Slc40a1 | 0.13820236 | -2.1696985 | 0.115 | 0.171 | 1 | 0.00141414 | 0.2785628 | 0.458 | 0.24 | 1 | 0.13820236 | 0.00282628 |
| Paqr6 | 0.29318604 | -0.3144336 | 0.073 | 0.114 | 1 | 0.00141628 | 1.94887901 | 0.189 | 0.052 | 1 | 0.29318604 | 0.00283055 |
| Fnbp4 | 0.94773972 | -0.2465109 | 0.771 | 0.736 | 1 | 0.00142276 | -0.5942852 | 0.912 | 0.938 | 1 | 0.94773972 | 0.00284349 |
| Med9 | 0.0568641 | 0.73331957 | 0.312 | 0.218 | 1 | 0.00142479 | 1.22619764 | 0.286 | 0.125 | 1 | 0.0568641 | 0.00284754 |
| Gm15893 | 0.00142586 | 3.81484946 | 0.052 | 0 | 1 | 0.57170879 | -1.3098553 | 0.013 | 0.021 | 1 | 0.57170879 | 0.00284968 |
| Rsph1 | 0.00142586 | 4.17741954 | 0.052 | 0 | 1 | 0.46472867 | -0.3098553 | 0.05 | 0.031 | 1 | 0.46472867 | 0.00284968 |
| N4bp3 | 0.00142624 | 3.59245704 | 0.052 | 0 | 1 | 0.22356137 | 0.80562195 | 0.05 | 0.021 | 1 | 0.22356137 | 0.00285044 |
| Jrk | 0.00142624 | 3.59245704 | 0.052 | 0 | 1 | 0.06885683 | 1.49749966 | 0.055 | 0.01 | 1 | 0.06885683 | 0.00285044 |
| Mff | 0.3704235 | 0.30952307 | 0.698 | 0.663 | 1 | 0.0014269 | 0.32996917 | 0.794 | 0.646 | 1 | 0.3704235 | 0.00285176 |
| Fam20b | 0.7585483 | 0.11569328 | 0.719 | 0.715 | 1 | 0.00142826 | 0.29325458 | 0.765 | 0.531 | 1 | 0.7585483 | 0.00285448 |
| Arhgap5 | 0.0044926 | 0.28208123 | 0.906 | 0.834 | 1 | 0.00142932 | 0.27137452 | 0.962 | 0.906 | 1 | 0.0044926 | 0.00285659 |
| Phlpp1 | 0.00473768 | 0.76859179 | 0.552 | 0.404 | 1 | 0.00142982 | -1.1961985 | 0.651 | 0.781 | 1 | 0.00473768 | 0.00285759 |
| Prmt6 | 0.00143672 | 1.77302928 | 0.156 | 0.047 | 1 | 0.04997656 | 1.22619764 | 0.109 | 0.042 | 1 | 0.04997656 | 0.00287137 |
| Cd52 | 0.61371908 | 1.22988696 | 0.01 | 0.005 | 1 | 0.00143777 | -0.6659991 | 0.038 | 0.135 | 1 | 0.61371908 | 0.00287347 |
| Klhl5 | 0.99929916 | -0.2027232 | 0.365 | 0.358 | 1 | 0.00144367 | 0.71399148 | 0.462 | 0.25 | 1 | 0.99929916 | 0.00288526 |
| Npnt | 0.16489838 | -2.4235553 | 0.115 | 0.166 | 1 | 0.00144681 | -1.9172352 | 0.277 | 0.448 | 1 | 0.16489838 | 0.00289154 |
| Mvb12a | 0.00144865 | 0.68239916 | 0.625 | 0.42 | 1 | 0.00239691 | 0.26649411 | 0.487 | 0.271 | 1 | 0.00239691 | 0.00289519 |
| Nfkb1 | 0.00145093 | -0.4532613 | 0.896 | 0.964 | 1 | 0.06903313 | -0.4389938 | 0.983 | 0.979 | 1 | 0.06903313 | 0.00289976 |
| Klc4 | 0.93245706 | 0.30122574 | 0.125 | 0.124 | 1 | 0.001454 | 0.96316323 | 0.298 | 0.125 | 1 | 0.93245706 | 0.00290589 |
| Bcl2l1 | 0.11935595 | -0.3453865 | 0.365 | 0.44 | 1 | 0.00145639 | -0.9565468 | 0.458 | 0.604 | 1 | 0.11935595 | 0.00291065 |
| Trp53 | 0.00145709 | 0.22322323 | 0.781 | 0.513 | 1 | 0.11390836 | 0.11356338 | 0.811 | 0.771 | 1 | 0.11390836 | 0.00291206 |
| Sirt5 | 0.00145903 | -2.0722327 | 0.104 | 0.254 | 1 | 0.06825606 | 1.93807225 | 0.055 | 0.01 | 1 | 0.06825606 | 0.00291593 |
| Fam118a | 0.00146132 | 0.98364779 | 0.354 | 0.192 | 1 | 0.28745151 | 0.154813 | 0.181 | 0.125 | 1 | 0.28745151 | 0.00292051 |
| Trp53rkb | 0.00146136 | 1.50999488 | 0.167 | 0.052 | 1 | 0.18218068 | 0.23071312 | 0.126 | 0.073 | 1 | 0.18218068 | 0.00292058 |
| Rnf111 | 0.25720427 | -0.2784767 | 0.667 | 0.679 | 1 | 0.0014623 | -0.6736346 | 0.819 | 0.896 | 1 | 0.25720427 | 0.00292247 |
| Adcy2 | 0.00146473 | 0.94939158 | 0.521 | 0.332 | 1 | 0.64821626 | -0.1493906 | 0.786 | 0.75 | 1 | 0.64821626 | 0.00292731 |
| Mrpl47 | 0.42385464 | -0.4372903 | 0.146 | 0.181 | 1 | 0.00146964 | 1.31817596 | 0.223 | 0.073 | 1 | 0.42385464 | 0.00293712 |
| Tob2 | 0.00147069 | 0.49021884 | 0.885 | 0.72 | 1 | 0.01881969 | 0.25694357 | 0.933 | 0.896 | 1 | 0.01881969 | 0.00293922 |
| Nsa2 | 0.18608054 | 0.20730214 | 0.917 | 0.876 | 1 | 0.00147103 | 0.32308727 | 0.874 | 0.844 | 1 | 0.18608054 | 0.00293989 |
| Vdac1 | 0.00147458 | 0.43355344 | 0.854 | 0.731 | 1 | 0.07584917 | 0.14888952 | 0.832 | 0.823 | 1 | 0.07584917 | 0.00294699 |
| Jpt2 | 0.00255044 | 0.93594128 | 0.396 | 0.233 | 1 | 0.00147692 | 0.77760758 | 0.349 | 0.156 | 1 | 0.00255044 | 0.00295166 |
| Lars | 0.0014784 | 0.71324805 | 0.708 | 0.627 | 1 | 0.07924728 | 0.1387348 | 0.756 | 0.667 | 1 | 0.07924728 | 0.00295461 |
| Uba1 | 0.00148208 | 0.51564144 | 0.729 | 0.523 | 1 | 0.12367276 | 0.12384608 | 0.777 | 0.719 | 1 | 0.12367276 | 0.00296196 |
| Ccdc12 | 0.00148208 | 0.56074818 | 0.698 | 0.513 | 1 | 0.00905344 | -0.514394 | 0.42 | 0.604 | 1 | 0.00905344 | 0.00296197 |
| Lpgat1 | 0.00148213 | 0.56123831 | 0.688 | 0.534 | 1 | 0.10168026 | 0.26000035 | 0.773 | 0.812 | 1 | 0.10168026 | 0.00296206 |
| Lcmt2 | 0.41514102 | 0.65957123 | 0.094 | 0.067 | 1 | 0.00148531 | 2.23446525 | 0.139 | 0.021 | 1 | 0.41514102 | 0.00296841 |
| Steap1 | 0.00148633 | -0.9974378 | 0.323 | 0.497 | 1 | 0.60813648 | 0.11756596 | 0.181 | 0.156 | 1 | 0.60813648 | 0.00297045 |
| Gramd2 | 0.08461511 | 0.38018575 | 0.688 | 0.611 | 1 | 0.00148648 | 1.04064198 | 0.378 | 0.198 | 1 | 0.08461511 | 0.00297075 |
| Atg16l1 | 0.03944462 | 0.30705482 | 0.531 | 0.368 | 1 | 0.0014878 | 0.66938618 | 0.559 | 0.354 | 1 | 0.03944462 | 0.00297338 |
| Apoa1 | 0.00148942 | 2.46692616 | 0.083 | 0.01 | 1 | 0.31492632 | -0.961932 | 0.046 | 0.073 | 1 | 0.31492632 | 0.00297662 |
| Atpsckmt | 0.03053272 | 0.66645762 | 0.271 | 0.161 | 1 | 0.00149096 | 0.69014474 | 0.261 | 0.094 | 1 | 0.03053272 | 0.00297969 |
| Qars | 0.14755761 | 0.34336182 | 0.448 | 0.363 | 1 | 0.00149723 | 0.63767732 | 0.454 | 0.26 | 1 | 0.14755761 | 0.00299223 |
| Aspscr1 | 0.00288974 | 0.92503238 | 0.448 | 0.295 | 1 | 0.00149786 | 0.32435076 | 0.613 | 0.354 | 1 | 0.00288974 | 0.00299348 |
| Zpr1 | 0.21265278 | 0.55377657 | 0.323 | 0.264 | 1 | 0.00149813 | 0.76696033 | 0.416 | 0.219 | 1 | 0.21265278 | 0.00299401 |
| Mpg | 0.46389299 | 0.25260703 | 0.271 | 0.233 | 1 | 0.00149982 | 1.18657056 | 0.239 | 0.083 | 1 | 0.46389299 | 0.00299739 |
| Copg1 | 0.00643263 | 0.35997372 | 0.792 | 0.72 | 1 | 0.00150059 | 0.45506439 | 0.815 | 0.604 | 1 | 0.00643263 | 0.00299893 |
| Slc26a7 | 0.00151367 | -3.577468 | 0 | 0.098 | 1 | 0.21295306 | -1.6317834 | 0.025 | 0.052 | 1 | 0.21295306 | 0.00302504 |
| Pclaf | 0.00151544 | 2.32942263 | 0.083 | 0.01 | 1 | 0.34026631 | -3.0468209 | 0.008 | 0.021 | 1 | 0.34026631 | 0.00302859 |
| Arfgap1 | 0.00752278 | 0.34756998 | 0.833 | 0.689 | 1 | 0.00151684 | 0.40179924 | 0.786 | 0.49 | 1 | 0.00752278 | 0.00303139 |
| Ankrd6 | 0.77687354 | -1.1079827 | 0.021 | 0.026 | 1 | 0.00151839 | -1.1172102 | 0.038 | 0.135 | 1 | 0.77687354 | 0.00303448 |
| Pde3b | 0.0015185 | 0.81340748 | 0.521 | 0.332 | 1 | 0.87087178 | -0.6931839 | 0.521 | 0.427 | 1 | 0.87087178 | 0.00303469 |
| Psma7 | 0.09909798 | 0.17116762 | 0.865 | 0.855 | 1 | 0.0015213 | 0.34450057 | 0.924 | 0.885 | 1 | 0.09909798 | 0.00304029 |
| Thop1 | 0.1793887 | 0.68556644 | 0.146 | 0.093 | 1 | 0.0015279 | 2.14957636 | 0.155 | 0.031 | 1 | 0.1793887 | 0.00305347 |
| Zcchc10 | 0.26519058 | 0.56003556 | 0.177 | 0.13 | 1 | 0.00152813 | 1.33923758 | 0.223 | 0.073 | 1 | 0.26519058 | 0.00305392 |
| BC029722 | 0.64309832 | -0.1263612 | 0.292 | 0.316 | 1 | 0.00152818 | 1.19264508 | 0.29 | 0.125 | 1 | 0.64309832 | 0.00305403 |
| Rwdd4a | 0.11199924 | 0.32677305 | 0.49 | 0.383 | 1 | 0.0015307 | 0.52413479 | 0.475 | 0.26 | 1 | 0.11199924 | 0.00305905 |
| Slc25a40 | 0.00153265 | 1.25260703 | 0.25 | 0.109 | 1 | 0.82511179 | -0.2742314 | 0.139 | 0.146 | 1 | 0.82511179 | 0.00306294 |
| Ehd3 | 0.03296488 | -0.9978389 | 0.615 | 0.642 | 1 | 0.001541 | 0.32686867 | 0.492 | 0.26 | 1 | 0.03296488 | 0.00307962 |
| 1110059G10Rik | 0.03915752 | 0.70139141 | 0.365 | 0.264 | 1 | 0.00154159 | 0.774209 | 0.328 | 0.146 | 1 | 0.03915752 | 0.00308079 |
| Dhx33 | 0.09088903 | 0.42253204 | 0.292 | 0.197 | 1 | 0.00154364 | 1.27510724 | 0.265 | 0.104 | 1 | 0.09088903 | 0.00308489 |
| Zfp53 | 0.15906818 | 0.68556644 | 0.229 | 0.161 | 1 | 0.00154893 | 1.55612739 | 0.303 | 0.135 | 1 | 0.15906818 | 0.00309545 |
| Col3a1 | 0.01743607 | -3.1501819 | 0.188 | 0.28 | 1 | 0.00155577 | -2.5094198 | 0.324 | 0.427 | 1 | 0.01743607 | 0.00310912 |
| Pcnx3 | 0.09462029 | 0.5268687 | 0.365 | 0.269 | 1 | 0.00155966 | 0.33547986 | 0.395 | 0.208 | 1 | 0.09462029 | 0.00311689 |
| Srp72 | 0.00604604 | 0.39996428 | 0.906 | 0.855 | 1 | 0.001563 | 0.41016759 | 0.878 | 0.802 | 1 | 0.00604604 | 0.00312356 |
| Zfp46 | 0.15917336 | 0.77045534 | 0.198 | 0.135 | 1 | 0.00156311 | 0.81213526 | 0.244 | 0.083 | 1 | 0.15917336 | 0.00312378 |
| Exosc2 | 0.00156438 | 1.2912875 | 0.229 | 0.093 | 1 | 0.00628321 | 0.86006974 | 0.21 | 0.083 | 1 | 0.00628321 | 0.00312631 |
| Ddx1 | 0.05236619 | 0.32099401 | 0.698 | 0.58 | 1 | 0.00156856 | 0.38371774 | 0.651 | 0.417 | 1 | 0.05236619 | 0.00313466 |
| Dpagt1 | 0.08891442 | 0.64492446 | 0.281 | 0.197 | 1 | 0.00156944 | 1.14957636 | 0.223 | 0.073 | 1 | 0.08891442 | 0.00313642 |
| Chmp2a | 0.4787692 | -0.1162408 | 0.812 | 0.824 | 1 | 0.00157238 | 0.3950167 | 0.84 | 0.781 | 1 | 0.4787692 | 0.00314228 |
| Slc25a1 | 0.97465166 | -0.2962862 | 0.281 | 0.264 | 1 | 0.00157349 | 1.69014474 | 0.189 | 0.052 | 1 | 0.97465166 | 0.0031445 |
| Ppt1 | 0.16651546 | 0.29945055 | 0.667 | 0.632 | 1 | 0.00157487 | 0.52992114 | 0.672 | 0.49 | 1 | 0.16651546 | 0.00314727 |
| Gm44448 | 0.11302151 | -1.7998604 | 0 | 0.026 | 1 | 0.00157775 | -3.8948178 | 0 | 0.042 | 1 | 0.11302151 | 0.00315301 |
| Caln1 | 0.15729781 | -1.7998604 | 0 | 0.021 | 1 | 0.00157775 | -5.6317834 | 0 | 0.042 | 1 | 0.15729781 | 0.00315301 |

| | | | | | | | | | | | | |
|---|---|---|---|---|---|---|---|---|---|---|---|---|
| Tonsl | 0.32498038 | 0.42253204 | 0.062 | 0.036 | 1 | 0.00157775 | -3.8948178 | 0 | 0.042 | 1 | 0.32498038 | 0.00315301 |
| Prima1 | 0.2825047 | -0.9925055 | 0.01 | 0.031 | 1 | 0.00157786 | -4.7692869 | 0 | 0.042 | 1 | 0.2825047 | 0.00315324 |
| Gm26583 | 0.38521705 | -0.7998604 | 0.01 | 0.026 | 1 | 0.00157786 | -4.3098553 | 0 | 0.042 | 1 | 0.38521705 | 0.00315324 |
| Gm47507 | 0.49468065 | -0.5230202 | 0.021 | 0.036 | 1 | 0.00157786 | -5.2167459 | 0 | 0.042 | 1 | 0.49468065 | 0.00315324 |
| Lrguk | 0.59199182 | -0.1624305 | 0.031 | 0.021 | 1 | 0.00157786 | -4.1172102 | 0 | 0.042 | 1 | 0.59199182 | 0.00315324 |
| 4933431K23Rik | 0.0819164 | -2.4519371 | 0 | 0.031 | 1 | 0.0015779 | -4.7692869 | 0 | 0.042 | 1 | 0.0819164 | 0.00315332 |
| Col18a1 | 0.52744059 | -0.577468 | 0.01 | 0.021 | 1 | 0.0015779 | -3.6317834 | 0 | 0.042 | 1 | 0.52744059 | 0.00315332 |
| Cntn1 | 0.61737927 | 1.00749454 | 0.01 | 0.005 | 1 | 0.0015779 | -4.6317834 | 0 | 0.042 | 1 | 0.61737927 | 0.00315332 |
| Tomm6 | 0.01882806 | 0.32099401 | 0.896 | 0.813 | 1 | 0.00158198 | 0.44503224 | 0.832 | 0.708 | 1 | 0.01882806 | 0.00316145 |
| Bscl2 | 0.02156421 | 0.42807022 | 0.583 | 0.415 | 1 | 0.00158454 | 0.26168672 | 0.592 | 0.344 | 1 | 0.02156421 | 0.00316657 |
| Trp53i11 | 0.00158551 | -2.3848229 | 0.021 | 0.14 | 1 | 0.79188071 | -0.7692869 | 0.067 | 0.073 | 1 | 0.79188071 | 0.0031685 |
| Itgb3bp | 0.16504834 | 0.77922555 | 0.24 | 0.176 | 1 | 0.00158558 | 1.19264508 | 0.193 | 0.052 | 1 | 0.16504834 | 0.00316865 |
| Adamts1 | 0.24220942 | 0.15672055 | 0.208 | 0.269 | 1 | 0.00158615 | -1.3098553 | 0.227 | 0.375 | 1 | 0.24220942 | 0.00316979 |
| Hnrnpc | 0.00159298 | 0.24460672 | 0.979 | 0.979 | 1 | 0.00512366 | 0.18286929 | 0.996 | 0.979 | 1 | 0.00512366 | 0.00318342 |
| Rac3 | 0.00159374 | 1.83756954 | 0.146 | 0.041 | 1 | 0.00747777 | 0.83698613 | 0.185 | 0.062 | 1 | 0.00747777 | 0.00318494 |
| Dcp2 | 0.21759893 | 0.38486162 | 0.385 | 0.316 | 1 | 0.00159487 | 0.42005557 | 0.395 | 0.198 | 1 | 0.21759893 | 0.0031872 |
| Cmya5 | 0.02654012 | 0.78510212 | 0.094 | 0.031 | 1 | 0.00159513 | -2.6790891 | 0.029 | 0.115 | 1 | 0.02654012 | 0.00318771 |
| Snx6 | 0.21917376 | 0.26746264 | 0.74 | 0.699 | 1 | 0.00160413 | 0.26377415 | 0.895 | 0.833 | 1 | 0.21917376 | 0.00320568 |
| Atp9a | 0.60950578 | -0.1747088 | 0.198 | 0.223 | 1 | 0.00160636 | 0.99595317 | 0.282 | 0.115 | 1 | 0.60950578 | 0.00321015 |
| Pias4 | 0.97205016 | -0.2148979 | 0.25 | 0.238 | 1 | 0.00160656 | 0.73905434 | 0.378 | 0.188 | 1 | 0.97205016 | 0.00321053 |
| Spata1 | 0.00160751 | 0.58173063 | 0.562 | 0.358 | 1 | 0.46318259 | -0.6027912 | 0.382 | 0.292 | 1 | 0.46318259 | 0.00321244 |
| Mknk2 | 0.00669015 | 0.51674738 | 0.646 | 0.487 | 1 | 0.00161165 | 0.56461386 | 0.752 | 0.688 | 1 | 0.00669015 | 0.00322071 |
| Aar2 | 0.14395349 | 0.79727683 | 0.271 | 0.202 | 1 | 0.00161171 | 1.13308823 | 0.282 | 0.115 | 1 | 0.14395349 | 0.00322083 |
| Ncln | 0.00161529 | 0.9735472 | 0.365 | 0.197 | 1 | 0.04922927 | 0.62303054 | 0.294 | 0.188 | 1 | 0.04922927 | 0.00322796 |
| Alkbh5 | 0.06573131 | 0.32003804 | 0.802 | 0.705 | 1 | 0.00161597 | 0.27675321 | 0.954 | 0.844 | 1 | 0.06573131 | 0.00322933 |
| Retreg2 | 0.00291972 | 0.74121447 | 0.531 | 0.368 | 1 | 0.00162263 | 0.65537932 | 0.487 | 0.292 | 1 | 0.00291972 | 0.00324263 |
| Ift27 | 0.54406255 | 0.13302542 | 0.396 | 0.363 | 1 | 0.00162625 | 0.75102698 | 0.412 | 0.219 | 1 | 0.54406255 | 0.00324986 |
| Slc30a5 | 0.00300268 | 0.64273024 | 0.646 | 0.513 | 1 | 0.00162866 | 0.47131014 | 0.786 | 0.604 | 1 | 0.00300268 | 0.00325466 |
| Setd5 | 0.00163042 | 0.39396288 | 0.958 | 0.927 | 1 | 0.37485963 | -0.2507486 | 0.992 | 0.99 | 1 | 0.37485963 | 0.00325818 |
| Pde12 | 0.80600118 | 0.2466822 | 0.26 | 0.249 | 1 | 0.00163364 | 0.78129263 | 0.315 | 0.135 | 1 | 0.80600118 | 0.00326462 |
| Hspbap1 | 0.57879146 | 0.14284639 | 0.344 | 0.316 | 1 | 0.00163617 | 0.27510724 | 0.466 | 0.229 | 1 | 0.57879146 | 0.00326967 |
| Ythdf1 | 0.00274417 | 0.56350566 | 0.635 | 0.43 | 1 | 0.00164086 | 0.50145767 | 0.731 | 0.583 | 1 | 0.00274417 | 0.00327902 |
| Ewsr1 | 0.00164343 | 0.38440849 | 0.948 | 0.87 | 1 | 0.95361194 | -0.1129228 | 0.992 | 0.979 | 1 | 0.95361194 | 0.00328416 |
| Dcaf15 | 0.00164456 | 1.62898291 | 0.156 | 0.047 | 1 | 0.54987021 | 0.39058446 | 0.092 | 0.073 | 1 | 0.54987021 | 0.00328642 |
| Ccm2 | 0.59705795 | 0.28260177 | 0.427 | 0.399 | 1 | 0.00165962 | 0.50550724 | 0.559 | 0.323 | 1 | 0.59705795 | 0.00331648 |
| Fbxo7 | 0.22709998 | -0.9352806 | 0.552 | 0.585 | 1 | 0.00166076 | 0.84542296 | 0.492 | 0.292 | 1 | 0.22709998 | 0.00331875 |
| ENSMUSG00000095041 | 0.50892195 | 0.20013961 | 0.281 | 0.249 | 1 | 0.00166327 | -0.5342868 | 0.252 | 0.448 | 1 | 0.50892195 | 0.00332376 |
| Cnn3 | 0.38276165 | -0.12375 | 0.969 | 0.974 | 1 | 0.00168985 | 0.3311733 | 0.992 | 0.948 | 1 | 0.38276165 | 0.00337684 |
| Fen1 | 0.00169159 | 1.93349396 | 0.135 | 0.036 | 1 | 0.16904025 | 1.01207283 | 0.084 | 0.042 | 1 | 0.16904025 | 0.00338033 |
| Yju2 | 0.15277221 | 0.42253204 | 0.24 | 0.166 | 1 | 0.00169207 | 1.13945214 | 0.231 | 0.083 | 1 | 0.15277221 | 0.00338127 |
| Wnt5b | 0.00169381 | 1.8316447 | 0.271 | 0.135 | 1 | 0.26789243 | -0.1222283 | 0.067 | 0.104 | 1 | 0.26789243 | 0.00338475 |
| 1700027I24Rik | 0.00169411 | 2.59245704 | 0.094 | 0.016 | 1 | 0.47306041 | 0.12310414 | 0.063 | 0.042 | 1 | 0.47306041 | 0.00338535 |
| Lamtor1 | 0.23691694 | 0.16304452 | 0.833 | 0.751 | 1 | 0.00169456 | 0.3794439 | 0.811 | 0.708 | 1 | 0.23691694 | 0.00338624 |
| Nkiras1 | 0.93330951 | -0.1127997 | 0.208 | 0.207 | 1 | 0.00169541 | 1.18657056 | 0.265 | 0.104 | 1 | 0.93330951 | 0.00338795 |
| Hagh | 0.75614032 | -0.2416772 | 0.406 | 0.42 | 1 | 0.0017013 | 0.89730265 | 0.42 | 0.24 | 1 | 0.75614032 | 0.00339972 |
| Mcm4 | 0.07321762 | 0.76648644 | 0.365 | 0.28 | 1 | 0.0017032 | 1.27510724 | 0.261 | 0.104 | 1 | 0.07321762 | 0.00340351 |
| Zkscan17 | 0.53184712 | -0.4664367 | 0.156 | 0.181 | 1 | 0.00170422 | 0.93115284 | 0.349 | 0.167 | 1 | 0.53184712 | 0.00340553 |
| Sult5a1 | 0.00170593 | -1.3710171 | 0.073 | 0.223 | 1 | 0.42767579 | 0.74077081 | 0.076 | 0.052 | 1 | 0.42767579 | 0.00340895 |
| Phtf2 | 0.00170908 | 0.57255748 | 0.74 | 0.549 | 1 | 0.9219451 | -0.2386765 | 0.845 | 0.896 | 1 | 0.9219451 | 0.00341524 |
| Nubp2 | 0.17272463 | 0.71203865 | 0.177 | 0.119 | 1 | 0.00171544 | 1.69014474 | 0.172 | 0.042 | 1 | 0.17272463 | 0.00342795 |
| Klf8 | 0.00172162 | 2.50999488 | 0.125 | 0.031 | 1 | 0.21919902 | -0.5133887 | 0.097 | 0.052 | 1 | 0.21919902 | 0.00344029 |
| Ror2 | 0.00172251 | 0.51014043 | 0.854 | 0.705 | 1 | 0.52288014 | -0.3195054 | 0.71 | 0.542 | 1 | 0.52288014 | 0.00344206 |
| Pgap4 | 0.14452356 | 0.47144164 | 0.177 | 0.114 | 1 | 0.00172499 | 1.83310269 | 0.185 | 0.052 | 1 | 0.14452356 | 0.003447 |
| Casp8 | 0.67424199 | -0.1683553 | 0.198 | 0.218 | 1 | 0.00172621 | 0.72256622 | 0.29 | 0.125 | 1 | 0.67424199 | 0.00344944 |
| Ube2c | 0.00172728 | 5.12297175 | 0.104 | 0.021 | 1 | 0.8259932 | -1.0874628 | 0.025 | 0.021 | 1 | 0.8259932 | 0.00345157 |
| Fam120b | 0.00338397 | 0.59702977 | 0.51 | 0.311 | 1 | 0.00172737 | 0.43430583 | 0.504 | 0.281 | 1 | 0.00338397 | 0.00345176 |
| Zbtb37 | 0.00172838 | 1.05480025 | 0.24 | 0.098 | 1 | 0.92803788 | -0.3791179 | 0.202 | 0.188 | 1 | 0.92803788 | 0.00345377 |
| Phlda1 | 0.00172908 | -0.821567 | 0.792 | 0.902 | 1 | 0.86773773 | -0.1282998 | 0.731 | 0.781 | 1 | 0.86773773 | 0.00345516 |
| Cog2 | 0.01001785 | 0.87214268 | 0.396 | 0.254 | 1 | 0.00173341 | 0.53449387 | 0.445 | 0.24 | 1 | 0.01001785 | 0.00346382 |
| Tollip | 0.37043442 | 0.13581863 | 0.375 | 0.311 | 1 | 0.00173464 | 0.41451029 | 0.605 | 0.375 | 1 | 0.37043442 | 0.00346627 |
| Ly6e | 0.57908299 | -0.114496 | 0.656 | 0.606 | 1 | 0.00173822 | 0.20680567 | 0.941 | 0.917 | 1 | 0.57908299 | 0.00347343 |
| Map10 | 0.00174607 | 2.32942263 | 0.094 | 0.016 | 1 | 0.43323624 | 0.42711033 | 0.038 | 0.021 | 1 | 0.43323624 | 0.0034891 |
| Mis18a | 0.00174658 | 1.6241659 | 0.188 | 0.067 | 1 | 0.02557009 | 2.01207283 | 0.071 | 0.01 | 1 | 0.02557009 | 0.00349011 |
| Art3 | 0.00174787 | -2.3144336 | 0.083 | 0.228 | 1 | 0.28801198 | -1.7248928 | 0.021 | 0.042 | 1 | 0.28801198 | 0.00349269 |
| Caml | 0.1726538 | 0.21979814 | 0.615 | 0.497 | 1 | 0.00174952 | 0.50066694 | 0.563 | 0.323 | 1 | 0.1726538 | 0.00349598 |
| Dsel | 0.00204905 | 0.65519279 | 0.5 | 0.306 | 1 | 0.00175075 | 0.31930704 | 0.542 | 0.302 | 1 | 0.00204905 | 0.00349844 |
| Nf1 | 0.62605164 | -0.1600502 | 0.844 | 0.912 | 1 | 0.00175159 | -0.4968345 | 0.966 | 0.958 | 1 | 0.62605164 | 0.00350011 |
| Tle3 | 0.45785952 | -0.3144336 | 0.156 | 0.192 | 1 | 0.00175341 | -1.0362902 | 0.311 | 0.51 | 1 | 0.45785952 | 0.00350375 |
| Hbs1l | 0.00175371 | 0.65816028 | 0.698 | 0.575 | 1 | 0.89154624 | -0.1147249 | 0.714 | 0.688 | 1 | 0.89154624 | 0.00350435 |
| Tmem223 | 0.0746945 | 0.37946331 | 0.604 | 0.513 | 1 | 0.00175568 | 0.39540147 | 0.563 | 0.344 | 1 | 0.0746945 | 0.00350828 |
| Rnps1 | 0.00175761 | 0.44307941 | 0.844 | 0.663 | 1 | 0.03537693 | 0.26239619 | 0.811 | 0.76 | 1 | 0.03537693 | 0.00351213 |
| Esyt3 | 0.0017582 | -2.0368996 | 0.062 | 0.202 | 1 | 0.34625029 | 0.69014474 | 0.042 | 0.021 | 1 | 0.34625029 | 0.0035133 |
| Avpi1 | 0.00176581 | -1.5719928 | 0.354 | 0.482 | 1 | 0.00491599 | -0.9378865 | 0.273 | 0.427 | 1 | 0.00491599 | 0.00352849 |
| Krit1 | 0.00176596 | 0.28355562 | 0.812 | 0.627 | 1 | 0.48155286 | -0.2378691 | 0.739 | 0.729 | 1 | 0.48155286 | 0.0035288 |
| Dgcr8 | 0.00193381 | 0.73541499 | 0.375 | 0.202 | 1 | 0.0017711 | 0.38130664 | 0.433 | 0.219 | 1 | 0.00193381 | 0.00353906 |
| Arhgef19 | 0.0017712 | 1.39744105 | 0.25 | 0.109 | 1 | 0.01125872 | 0.40940833 | 0.382 | 0.208 | 1 | 0.01125872 | 0.00353926 |
| Tent4b | 0.05127673 | 0.25947375 | 0.406 | 0.28 | 1 | 0.00177388 | -0.8948178 | 0.538 | 0.719 | 1 | 0.05127673 | 0.00354461 |
| Ssbp3 | 0.54059845 | -0.3016474 | 0.635 | 0.668 | 1 | 0.00177549 | -0.7109198 | 0.765 | 0.875 | 1 | 0.54059845 | 0.00354782 |
| Ubxn11 | 0.00177769 | 2.37006462 | 0.125 | 0.031 | 1 | 0.63349001 | 0.10518224 | 0.055 | 0.042 | 1 | 0.63349001 | 0.00355223 |
| Ephx3 | 0.00177769 | 2.20013961 | 0.125 | 0.031 | 1 | 0.32036883 | -0.9728203 | 0.071 | 0.104 | 1 | 0.32036883 | 0.00355223 |
| Gm41409 | 0.00177784 | 2.42253204 | 0.115 | 0.026 | 1 | 0.04657758 | 1.33400093 | 0.08 | 0.021 | 1 | 0.04657758 | 0.00355252 |
| Ltbr | 0.16881162 | 0.16140722 | 0.625 | 0.497 | 1 | 0.00178019 | 0.57188186 | 0.542 | 0.333 | 1 | 0.16881162 | 0.00355722 |
| Zfp770 | 0.00178525 | 1.2912875 | 0.229 | 0.093 | 1 | 0.01343994 | 0.72256622 | 0.155 | 0.052 | 1 | 0.01343994 | 0.00356732 |
| Fcf1 | 0.15528036 | 0.17395568 | 0.542 | 0.451 | 1 | 0.00178617 | 0.60681635 | 0.504 | 0.302 | 1 | 0.15528036 | 0.00356916 |
| Cops3 | 0.0017869 | 0.50015293 | 0.688 | 0.523 | 1 | 0.00739251 | 0.36900738 | 0.702 | 0.594 | 1 | 0.00739251 | 0.00357061 |
| Septin4 | 0.00178732 | 1.63552576 | 0.146 | 0.041 | 1 | 0.053372 | 1.01207283 | 0.109 | 0.042 | 1 | 0.053372 | 0.00357145 |
| Sdf2l1 | 0.2056664 | 0.39653683 | 0.354 | 0.29 | 1 | 0.00178828 | 0.91253716 | 0.336 | 0.156 | 1 | 0.2056664 | 0.00357336 |
| Ilk | 0.14487767 | 0.28973315 | 0.667 | 0.591 | 1 | 0.00178837 | 0.31677642 | 0.647 | 0.406 | 1 | 0.14487767 | 0.00357355 |

| | | | | | | | | | | | | |
|---|---|---|---|---|---|---|---|---|---|---|---|---|
| Fat1 | 0.0017891 | 0.57644472 | 0.708 | 0.513 | 1 | 0.00242598 | -0.8581593 | 0.634 | 0.823 | 1 | 0.00242598 | 0.003575 |
| Arl6ip1 | 0.00179091 | -0.2159128 | 0.625 | 0.824 | 1 | 0.08962783 | -0.2658481 | 0.748 | 0.917 | 1 | 0.08962783 | 0.00357861 |
| Fgfr1op2 | 0.23024542 | 0.23895886 | 0.865 | 0.819 | 1 | 0.00179189 | 0.23165698 | 0.861 | 0.812 | 1 | 0.23024542 | 0.00358057 |
| Clmp | 0.00179214 | 0.38202431 | 0.552 | 0.306 | 1 | 0.40104304 | -1.3937941 | 0.37 | 0.354 | 1 | 0.40104304 | 0.00358107 |
| Fndc4 | 0.34205573 | 0.43163424 | 0.354 | 0.301 | 1 | 0.00179675 | 0.89450324 | 0.319 | 0.146 | 1 | 0.34205573 | 0.00359028 |
| Gm41496 | 0.03087484 | -1.9067756 | 0.073 | 0.161 | 1 | 0.00179784 | -2.3524996 | 0.13 | 0.25 | 1 | 0.03087484 | 0.00359245 |
| Dchs2 | 0.10310885 | -0.6445822 | 0.031 | 0.083 | 1 | 0.00179864 | -3.9943534 | 0.025 | 0.104 | 1 | 0.10310885 | 0.00359405 |
| Fastkd3 | 0.00180075 | 1.59245704 | 0.146 | 0.041 | 1 | 0.99385845 | -0.9378865 | 0.088 | 0.083 | 1 | 0.99385845 | 0.00359825 |
| Abr | 0.001804 | 0.74244096 | 0.531 | 0.337 | 1 | 0.08673029 | -0.5125892 | 0.542 | 0.635 | 1 | 0.08673029 | 0.00360475 |
| 5930430L01Rik | 0.0018058 | -1.4950058 | 0.062 | 0.207 | 1 | 0.64528986 | -0.7913882 | 0.155 | 0.167 | 1 | 0.64528986 | 0.00360834 |
| Tmem234 | 0.11155866 | 0.16989658 | 0.885 | 0.891 | 1 | 0.00180618 | 0.42875818 | 0.861 | 0.844 | 1 | 0.11155866 | 0.0036091 |
| Gpc3 | 0.04812177 | -0.5135562 | 0.458 | 0.285 | 1 | 0.00180662 | 1.08785542 | 0.555 | 0.427 | 1 | 0.04812177 | 0.00360997 |
| Pfkfb1 | 0.00180814 | 2.27052894 | 0.094 | 0.016 | 1 | 0.74451943 | -0.8948178 | 0.025 | 0.031 | 1 | 0.74451943 | 0.003613 |
| Galnt5 | 0.00181338 | 2.25542205 | 0.125 | 0.031 | 1 | 0.00570893 | 2.73453886 | 0.097 | 0.01 | 1 | 0.00570893 | 0.00362348 |
| Stk10 | 0.00181798 | 0.24565427 | 0.583 | 0.332 | 1 | 0.23321918 | -0.1900608 | 0.618 | 0.49 | 1 | 0.23321918 | 0.00363265 |
| Gmfb | 0.09510795 | 0.41515251 | 0.667 | 0.57 | 1 | 0.00182047 | 0.41686023 | 0.773 | 0.573 | 1 | 0.09510795 | 0.00363763 |
| Urgcp | 0.04780977 | 0.24312279 | 0.458 | 0.321 | 1 | 0.00182163 | 0.14850967 | 0.639 | 0.375 | 1 | 0.04780977 | 0.00363995 |
| F2rl1 | 0.52528691 | -0.7998604 | 0.01 | 0.021 | 1 | 0.0018236 | -1.9472852 | 0.029 | 0.115 | 1 | 0.52528691 | 0.00364387 |
| Washc2 | 0.00182915 | 0.44189736 | 0.646 | 0.404 | 1 | 0.16576093 | -0.3475749 | 0.622 | 0.719 | 1 | 0.16576093 | 0.00365494 |
| Arhgef6 | 0.05258203 | 0.43375929 | 0.531 | 0.425 | 1 | 0.00183681 | 0.40474252 | 0.546 | 0.333 | 1 | 0.05258203 | 0.00367025 |
| Ccdc84 | 0.52279299 | 0.37522632 | 0.167 | 0.14 | 1 | 0.00184177 | 0.84214783 | 0.256 | 0.094 | 1 | 0.52279299 | 0.00368015 |
| Mthfsd | 0.0808441 | 0.72370157 | 0.208 | 0.13 | 1 | 0.00184237 | 0.64010405 | 0.29 | 0.115 | 1 | 0.0808441 | 0.00368134 |
| Nlrc3 | 0.00184551 | 1.63552576 | 0.146 | 0.041 | 1 | 0.36791201 | -0.8948178 | 0.109 | 0.073 | 1 | 0.36791201 | 0.00368762 |
| Mxra7 | 0.20963267 | 0.11440974 | 0.583 | 0.523 | 1 | 0.00184893 | 0.17733202 | 0.672 | 0.375 | 1 | 0.20963267 | 0.00369443 |
| Galnt1 | 0.0018498 | 0.52706262 | 0.927 | 0.845 | 1 | 0.14132643 | -0.4686316 | 0.887 | 0.927 | 1 | 0.14132643 | 0.00369618 |
| Ppa1 | 0.00415461 | 1.69284778 | 0.885 | 0.772 | 1 | 0.0018519 | 1.76404285 | 0.752 | 0.646 | 1 | 0.00415461 | 0.00370037 |
| Lamc1 | 0.89318055 | -0.1427481 | 0.385 | 0.404 | 1 | 0.00185369 | -0.7582615 | 0.609 | 0.76 | 1 | 0.89318055 | 0.00370394 |
| Scrib | 0.0018537 | 0.94860085 | 0.24 | 0.098 | 1 | 0.53807025 | -0.113458 | 0.181 | 0.146 | 1 | 0.53807025 | 0.00370397 |
| Nsmce4a | 0.35196327 | 0.17460452 | 0.542 | 0.472 | 1 | 0.0018556 | 0.44868741 | 0.634 | 0.448 | 1 | 0.35196327 | 0.00370776 |
| Il34 | 0.0018634 | -1.8800307 | 0.052 | 0.187 | 1 | 0.75613631 | -0.6613276 | 0.16 | 0.135 | 1 | 0.75613631 | 0.00372333 |
| Tmem242 | 0.00688518 | 0.78510212 | 0.49 | 0.358 | 1 | 0.00186792 | 0.40940833 | 0.424 | 0.219 | 1 | 0.00688518 | 0.00373235 |
| Isg20 | 0.00187162 | -1.4718062 | 0.354 | 0.477 | 1 | 0.00367108 | 1.27510724 | 0.235 | 0.094 | 1 | 0.00367108 | 0.00373973 |
| Mrpl10 | 0.00330407 | 0.56388789 | 0.448 | 0.259 | 1 | 0.00187171 | 0.51866427 | 0.349 | 0.156 | 1 | 0.00330407 | 0.00373992 |
| Tarbp1 | 0.07263029 | 0.54806292 | 0.125 | 0.062 | 1 | 0.00187311 | 2.53144699 | 0.134 | 0.021 | 1 | 0.07263029 | 0.00374272 |
| Leng8 | 0.05633969 | 0.30571837 | 0.573 | 0.44 | 1 | 0.00187468 | 0.35640734 | 0.563 | 0.323 | 1 | 0.05633969 | 0.00374584 |
| Dgki | 0.00187568 | -2.7533178 | 0.052 | 0.187 | 1 | 0.09225603 | -0.6988976 | 0.084 | 0.146 | 1 | 0.09225603 | 0.00374785 |
| Phactr2 | 0.00188236 | 0.48812038 | 0.688 | 0.513 | 1 | 0.15937053 | 0.11147398 | 0.71 | 0.719 | 1 | 0.15937053 | 0.00376118 |
| Mcm2 | 0.00188442 | 2.28760246 | 0.115 | 0.026 | 1 | 0.88210307 | -0.4618584 | 0.046 | 0.042 | 1 | 0.88210307 | 0.00376528 |
| Vcpip1 | 0.00189293 | 0.58368683 | 0.667 | 0.503 | 1 | 0.12053007 | -0.3417393 | 0.71 | 0.792 | 1 | 0.12053007 | 0.00378228 |
| Prpf19 | 0.00189492 | 0.71798792 | 0.583 | 0.389 | 1 | 0.02937384 | 0.28142101 | 0.576 | 0.438 | 1 | 0.02937384 | 0.00378625 |
| Dph6 | 0.02263725 | 0.69490431 | 0.427 | 0.29 | 1 | 0.00189576 | 0.53226508 | 0.357 | 0.167 | 1 | 0.02263725 | 0.00378793 |
| Sh3kbp1 | 0.00189871 | -0.5275427 | 0.479 | 0.71 | 1 | 0.0464627 | -1.3227943 | 0.248 | 0.333 | 1 | 0.0464627 | 0.00379382 |
| Lamtor4 | 0.05501705 | 0.29584003 | 0.781 | 0.637 | 1 | 0.00190818 | 0.52079031 | 0.668 | 0.542 | 1 | 0.05501705 | 0.00381273 |
| Rnf6 | 0.09250279 | 0.34377792 | 0.604 | 0.518 | 1 | 0.00190906 | 0.35869387 | 0.689 | 0.49 | 1 | 0.09250279 | 0.00381447 |
| Cped1 | 0.0019104 | -1.5499564 | 0.49 | 0.591 | 1 | 0.02016915 | -0.7743123 | 0.676 | 0.677 | 1 | 0.02016915 | 0.00381715 |
| Fam228b | 0.00191325 | 1.91438513 | 0.125 | 0.031 | 1 | 0.60448 | -0.84037 | 0.059 | 0.073 | 1 | 0.60448 | 0.00382284 |
| Lsr | 0.15046861 | 1.22988696 | 0.052 | 0.021 | 1 | 0.00191403 | -1.5728897 | 0.046 | 0.146 | 1 | 0.15046861 | 0.00382441 |
| Eif3f | 0.36030783 | -0.1869528 | 0.938 | 0.891 | 1 | 0.00191462 | 0.20592367 | 0.929 | 0.917 | 1 | 0.36030783 | 0.00382557 |
| Plxna3 | 0.00191474 | 2.14499806 | 0.104 | 0.021 | 1 | 0.21171395 | 0.76053407 | 0.08 | 0.042 | 1 | 0.21171395 | 0.0038258 |
| Sgpl1 | 0.10266641 | -0.4110068 | 0.323 | 0.409 | 1 | 0.00191502 | -0.7788316 | 0.605 | 0.792 | 1 | 0.10266641 | 0.00382637 |
| 3110040N11Rik | 0.00191544 | 1.00749454 | 0.312 | 0.155 | 1 | 0.03872928 | 0.41806519 | 0.176 | 0.083 | 1 | 0.03872928 | 0.00382721 |
| Dnah12 | 0.05620212 | -2.8933723 | 0.062 | 0.135 | 1 | 0.00191708 | -4.0934616 | 0.025 | 0.104 | 1 | 0.05620212 | 0.00383048 |
| Mgst2 | 0.00192051 | -1.2871262 | 0.177 | 0.342 | 1 | 0.00334047 | 0.98970502 | 0.181 | 0.052 | 1 | 0.00334047 | 0.00383733 |
| Mpi | 0.00192123 | 1.90057933 | 0.125 | 0.031 | 1 | 0.9655645 | 0.13071733 | 0.063 | 0.062 | 1 | 0.9655645 | 0.00383876 |
| Hebp2 | 0.02650115 | -1.1079827 | 0.188 | 0.295 | 1 | 0.00192172 | 1.83698613 | 0.197 | 0.062 | 1 | 0.02650115 | 0.00383975 |
| Mme | 0.00192342 | -4.18233 | 0.031 | 0.15 | 1 | 0.04571799 | -2.0128735 | 0.097 | 0.167 | 1 | 0.04571799 | 0.00384314 |
| Wdfy2 | 0.1473747 | 0.30488045 | 0.354 | 0.259 | 1 | 0.00192727 | -0.9802871 | 0.332 | 0.521 | 1 | 0.1473747 | 0.00385082 |
| Pcmtd2 | 0.87398333 | 0.17589207 | 0.354 | 0.352 | 1 | 0.00192748 | 0.61017643 | 0.487 | 0.281 | 1 | 0.87398333 | 0.00385125 |
| Agfg2 | 0.18836358 | -0.1753695 | 0.208 | 0.285 | 1 | 0.00192888 | 1.38316698 | 0.231 | 0.083 | 1 | 0.18836358 | 0.00385403 |
| BC024063 | 0.00193688 | 2.12297175 | 0.104 | 0.021 | 1 | 0.85030707 | -0.6577786 | 0.038 | 0.042 | 1 | 0.85030707 | 0.00387001 |
| Yars | 0.00494134 | 0.52066635 | 0.667 | 0.518 | 1 | 0.00194243 | 0.2190732 | 0.731 | 0.458 | 1 | 0.00494134 | 0.00388109 |
| Cfap300 | 0.00194351 | 1.42253204 | 0.219 | 0.088 | 1 | 0.15303378 | 0.69014474 | 0.101 | 0.052 | 1 | 0.15303378 | 0.00388324 |
| Pitpnm3 | 0.11885255 | 1.12297175 | 0.094 | 0.047 | 1 | 0.00194638 | 0.66996686 | 0.189 | 0.052 | 1 | 0.11885255 | 0.00388897 |
| Taok3 | 0.0019497 | 0.44455834 | 0.656 | 0.446 | 1 | 0.76429507 | -0.1091579 | 0.693 | 0.698 | 1 | 0.76429507 | 0.0038956 |
| Atosb | 0.3288206 | 0.51845646 | 0.333 | 0.29 | 1 | 0.0019514 | 0.36821664 | 0.534 | 0.312 | 1 | 0.3288206 | 0.00389899 |
| Rngtt | 0.24520242 | 0.20441005 | 0.635 | 0.56 | 1 | 0.00195323 | -0.8654911 | 0.622 | 0.771 | 1 | 0.24520242 | 0.00390264 |
| Cav2 | 0.00195728 | -1.2054992 | 0.312 | 0.472 | 1 | 0.01912808 | 0.60853097 | 0.37 | 0.229 | 1 | 0.01912808 | 0.00391072 |
| Pex13 | 0.24669827 | 0.36525723 | 0.594 | 0.518 | 1 | 0.00196014 | 0.53923917 | 0.622 | 0.427 | 1 | 0.24669827 | 0.00391644 |
| Mrpl22 | 0.01736591 | 0.57139542 | 0.302 | 0.176 | 1 | 0.00196343 | 1.07841533 | 0.261 | 0.104 | 1 | 0.01736591 | 0.003923 |
| Mical1 | 0.31712771 | -0.6783969 | 0.198 | 0.244 | 1 | 0.00197078 | 0.98676174 | 0.328 | 0.156 | 1 | 0.31712771 | 0.00393768 |
| Slc27a3 | 0.00197826 | 0.89201732 | 0.292 | 0.14 | 1 | 0.00997969 | 0.50643278 | 0.277 | 0.135 | 1 | 0.00997969 | 0.0039526 |
| Cept1 | 0.0019808 | 0.71924957 | 0.583 | 0.394 | 1 | 0.87925072 | -0.2060195 | 0.639 | 0.698 | 1 | 0.87925072 | 0.00395767 |
| Psmg3 | 0.02130478 | 0.70263996 | 0.302 | 0.181 | 1 | 0.00198184 | 1.36257008 | 0.172 | 0.042 | 1 | 0.02130478 | 0.00395975 |
| Potegl | 0.69494775 | -0.9467018 | 0.052 | 0.062 | 1 | 0.00198244 | -2.5062525 | 0.055 | 0.156 | 1 | 0.69494775 | 0.00396095 |
| 9130410C08Rik | 0.00198833 | 2.26125113 | 0.125 | 0.031 | 1 | 0.18670454 | 0.32241295 | 0.084 | 0.042 | 1 | 0.18670454 | 0.00397271 |
| Snph | 0.00198881 | 1.28760246 | 0.156 | 0.047 | 1 | 0.01346273 | 1.73453886 | 0.118 | 0.031 | 1 | 0.01346273 | 0.00397366 |
| Ahsa2 | 0.88528311 | 0.20389175 | 0.323 | 0.352 | 1 | 0.00198936 | 0.24520575 | 0.513 | 0.281 | 1 | 0.88528311 | 0.00397476 |
| Firre | 0.00199521 | 0.86453658 | 0.375 | 0.202 | 1 | 0.7561135 | -0.5486421 | 0.294 | 0.281 | 1 | 0.7561135 | 0.00398643 |
| Adamts17 | 0.00200389 | -2.1490099 | 0.094 | 0.238 | 1 | 0.02800986 | -1.1382439 | 0.164 | 0.25 | 1 | 0.02800986 | 0.00400377 |
| Med29 | 0.2485579 | 0.28002039 | 0.458 | 0.399 | 1 | 0.00200408 | 0.88452278 | 0.391 | 0.219 | 1 | 0.2485579 | 0.00400415 |
| Asb1 | 0.00201871 | 1.59245704 | 0.135 | 0.036 | 1 | 0.75164421 | -0.194378 | 0.05 | 0.042 | 1 | 0.75164421 | 0.00403334 |
| Hey1 | 0.00269949 | -3.2019588 | 0.031 | 0.145 | 1 | 0.00201982 | -1.9102478 | 0.084 | 0.198 | 1 | 0.00269949 | 0.00403555 |
| Rpia | 0.06143467 | 0.64071221 | 0.271 | 0.176 | 1 | 0.00202992 | 0.83698613 | 0.315 | 0.135 | 1 | 0.06143467 | 0.00405573 |
| Hook1 | 1 | 0.42253204 | 0.01 | 0.01 | 1 | 0.00203068 | -1.129283 | 0.046 | 0.146 | 1 | 1 | 0.00405723 |
| Aebp2 | 0.034716 | 0.33976458 | 0.719 | 0.642 | 1 | 0.00203396 | -0.5455877 | 0.84 | 0.906 | 1 | 0.034716 | 0.00406378 |
| Erg28 | 0.00203634 | 0.8235317 | 0.552 | 0.373 | 1 | 0.0096953 | 0.76053407 | 0.399 | 0.24 | 1 | 0.0096953 | 0.00406854 |
| Arl6 | 0.01210137 | 0.91438513 | 0.26 | 0.14 | 1 | 0.00203845 | 0.83989186 | 0.269 | 0.104 | 1 | 0.01210137 | 0.00407274 |
| Nudcd1 | 0.00203845 | 0.89013759 | 0.417 | 0.254 | 1 | 0.64876172 | -0.4381794 | 0.244 | 0.208 | 1 | 0.64876172 | 0.00407275 |

| | | | | | | | | | | | | |
|---|---|---|---|---|---|---|---|---|---|---|---|---|
| Gstcd | 0.04980527 | 0.60031916 | 0.323 | 0.218 | 1 | 0.00203896 | 0.92460999 | 0.248 | 0.094 | 1 | 0.04980527 | 0.00407376 |
| Flnc | 0.20607496 | -1.7397394 | 0.052 | 0.093 | 1 | 0.00204518 | -2.681824 | 0.063 | 0.167 | 1 | 0.20607496 | 0.00408618 |
| Ticam1 | 0.09983119 | 1.00749454 | 0.104 | 0.052 | 1 | 0.00204741 | 1.36257008 | 0.202 | 0.062 | 1 | 0.09983119 | 0.00409063 |
| Bloc1s6 | 0.05398351 | 0.72996056 | 0.271 | 0.176 | 1 | 0.00204786 | 0.71004429 | 0.265 | 0.104 | 1 | 0.05398351 | 0.00409153 |
| Hip1r | 0.02271722 | 0.42253204 | 0.729 | 0.575 | 1 | 0.0020485 | 0.163431 | 0.761 | 0.562 | 1 | 0.02271722 | 0.00409281 |
| Mbd2 | 0.02496016 | 0.12979068 | 0.979 | 0.964 | 1 | 0.0020529 | 0.17070253 | 0.992 | 0.948 | 1 | 0.02496016 | 0.00410158 |
| Plekhg3 | 0.00205556 | 0.93154568 | 0.365 | 0.192 | 1 | 0.15763037 | 0.16231547 | 0.609 | 0.573 | 1 | 0.15763037 | 0.0041069 |
| Eef1akmt1 | 0.00205825 | 1.02436836 | 0.49 | 0.342 | 1 | 0.00264524 | 0.72073306 | 0.387 | 0.208 | 1 | 0.00264524 | 0.00411226 |
| Sytl5 | 0.00206053 | -4.0799683 | 0 | 0.093 | 1 | 0.09092803 | -2.8948178 | 0.013 | 0.042 | 1 | 0.09092803 | 0.00411681 |
| Fhod3 | 0.00206431 | -6.4187702 | 0 | 0.093 | 1 | 0.05954021 | -2.129283 | 0.042 | 0.094 | 1 | 0.05954021 | 0.00412435 |
| Akt2 | 0.00700473 | 0.36016615 | 0.854 | 0.741 | 1 | 0.00206546 | 0.13105032 | 0.899 | 0.698 | 1 | 0.00700473 | 0.00412666 |
| Rangrf | 0.04209314 | 1.05188866 | 0.24 | 0.15 | 1 | 0.00206732 | 1.41806519 | 0.185 | 0.052 | 1 | 0.04209314 | 0.00413036 |
| Dusp12 | 0.00761776 | 1.00749454 | 0.365 | 0.228 | 1 | 0.00206998 | 0.43737867 | 0.294 | 0.125 | 1 | 0.00761776 | 0.00413568 |
| Smim8 | 0.06862933 | 0.45729745 | 0.417 | 0.316 | 1 | 0.00207548 | 1.22263982 | 0.244 | 0.094 | 1 | 0.06862933 | 0.00414665 |
| Gm56878 | 0.00207604 | -0.5957302 | 0.667 | 0.756 | 1 | 0.19575638 | -0.3229114 | 0.513 | 0.635 | 1 | 0.19575638 | 0.00414776 |
| Pigc | 0.72041759 | 0.15949763 | 0.156 | 0.14 | 1 | 0.00207829 | 1.35310975 | 0.256 | 0.104 | 1 | 0.72041759 | 0.00415226 |
| Dock8 | 0.00208224 | 0.45766026 | 0.844 | 0.71 | 1 | 0.77934848 | -0.2139308 | 0.735 | 0.75 | 1 | 0.77934848 | 0.00416014 |
| Tm2d1 | 0.51504392 | -0.1563915 | 0.74 | 0.741 | 1 | 0.00208781 | 0.26726648 | 0.782 | 0.51 | 1 | 0.51504392 | 0.00417127 |
| Tnfsf12 | 0.88769501 | 0.20013961 | 0.24 | 0.249 | 1 | 0.00209109 | 0.67620555 | 0.34 | 0.156 | 1 | 0.88769501 | 0.00417781 |
| 2310039H08Rik | 0.00362643 | 0.49903017 | 0.552 | 0.347 | 1 | 0.00210199 | 0.43488768 | 0.387 | 0.188 | 1 | 0.00362643 | 0.00419956 |
| Zfp383 | 0.22704217 | 0.49292136 | 0.167 | 0.114 | 1 | 0.00210579 | 1.52664601 | 0.185 | 0.052 | 1 | 0.22704217 | 0.00420715 |
| Rock2 | 0.00210895 | 0.29329447 | 1 | 0.99 | 1 | 0.05355732 | -0.2840153 | 1 | 1 | 1 | 0.05355732 | 0.00421345 |
| Ppfia2 | 0.11116816 | -3.3144336 | 0.01 | 0.047 | 1 | 0.00210939 | -4.0260623 | 0.021 | 0.094 | 1 | 0.11116816 | 0.00421434 |
| Pqlc3 | 0.34288551 | 0.22161934 | 0.229 | 0.181 | 1 | 0.00210962 | 1.27510724 | 0.227 | 0.083 | 1 | 0.34288551 | 0.00421478 |
| Ganab | 0.00211039 | 0.44359365 | 0.802 | 0.684 | 1 | 0.00627102 | 0.27166815 | 0.79 | 0.625 | 1 | 0.00627102 | 0.00421633 |
| Shisa4 | 0.00211121 | -0.5600334 | 0.438 | 0.611 | 1 | 0.95787371 | -0.2457249 | 0.332 | 0.312 | 1 | 0.95787371 | 0.00421797 |
| Ddx49 | 0.0367585 | 0.30705482 | 0.448 | 0.301 | 1 | 0.00211168 | 0.5849625 | 0.378 | 0.188 | 1 | 0.0367585 | 0.0042189 |
| Psmd11 | 0.00211221 | 0.30459078 | 0.969 | 0.912 | 1 | 0.00827344 | 0.17484738 | 0.983 | 0.938 | 1 | 0.00827344 | 0.00421996 |
| BC004004 | 0.02784035 | 0.41233627 | 0.604 | 0.477 | 1 | 0.00211457 | 0.36821664 | 0.454 | 0.25 | 1 | 0.02784035 | 0.00422467 |
| Ylpm1 | 0.00211981 | 0.58236116 | 0.76 | 0.57 | 1 | 0.37909391 | -0.2985842 | 0.765 | 0.833 | 1 | 0.37909391 | 0.00423514 |
| Prickle3 | 0.00212234 | 1.06638823 | 0.229 | 0.093 | 1 | 0.07648425 | 0.24268576 | 0.164 | 0.083 | 1 | 0.07648425 | 0.00424018 |
| Zfp91 | 0.03472388 | 0.33716205 | 0.833 | 0.777 | 1 | 0.00213603 | 0.30831428 | 0.92 | 0.896 | 1 | 0.03472388 | 0.00426749 |
| Stx2 | 0.00214349 | 1.24453373 | 0.24 | 0.104 | 1 | 0.20813986 | -0.4579539 | 0.244 | 0.156 | 1 | 0.20813986 | 0.00428239 |
| Rcbtb1 | 0.05879952 | 0.30331888 | 0.562 | 0.446 | 1 | 0.00214606 | 0.25599842 | 0.563 | 0.323 | 1 | 0.05879952 | 0.00428752 |
| Gm20257 | 0.0263166 | 0.95304675 | 0.219 | 0.119 | 1 | 0.0021469 | 1.74903843 | 0.168 | 0.042 | 1 | 0.0263166 | 0.00428918 |
| Wdr90 | 0.0021517 | 1.92503238 | 0.115 | 0.026 | 1 | 0.15810273 | 0.69014474 | 0.101 | 0.052 | 1 | 0.15810273 | 0.00429877 |
| Hps5 | 0.03082601 | 0.56548999 | 0.396 | 0.264 | 1 | 0.00215284 | 0.6145937 | 0.45 | 0.26 | 1 | 0.03082601 | 0.00430104 |
| Trip6 | 0.0607202 | 0.56820949 | 0.385 | 0.285 | 1 | 0.00215292 | 0.58176858 | 0.424 | 0.229 | 1 | 0.0607202 | 0.00430121 |
| Fam50a | 0.0463093 | 0.20875274 | 0.615 | 0.461 | 1 | 0.00216412 | 0.34852831 | 0.622 | 0.438 | 1 | 0.0463093 | 0.00432355 |
| Fam135a | 0.00216676 | -0.6779408 | 0.646 | 0.741 | 1 | 0.77848835 | -0.1104112 | 0.647 | 0.677 | 1 | 0.77848835 | 0.00432882 |
| Slc25a24 | 0.07150803 | 0.62315383 | 0.49 | 0.409 | 1 | 0.00217065 | 0.42943538 | 0.508 | 0.292 | 1 | 0.07150803 | 0.00433658 |
| Casp2 | 0.00217591 | 1.45495351 | 0.135 | 0.036 | 1 | 0.21538546 | 0.40063812 | 0.122 | 0.073 | 1 | 0.21538546 | 0.00434708 |
| Oit3 | 0.00217902 | 1.70793425 | 0.115 | 0.026 | 1 | 0.40992831 | -1.6883669 | 0.034 | 0.052 | 1 | 0.40992831 | 0.00435329 |
| Tmed5 | 0.88300092 | -0.1005652 | 0.688 | 0.715 | 1 | 0.00217905 | -0.6239639 | 0.634 | 0.823 | 1 | 0.88300092 | 0.00435336 |
| E2f4 | 0.00217937 | 0.9200317 | 0.375 | 0.202 | 1 | 0.00759689 | 0.49291239 | 0.441 | 0.271 | 1 | 0.00759689 | 0.00435399 |
| 1810037I17Rik | 0.20609101 | 0.14019707 | 0.938 | 0.881 | 1 | 0.00217992 | 0.24990668 | 0.887 | 0.844 | 1 | 0.20609101 | 0.00435508 |
| Junos | 0.00553191 | -2.1889027 | 0.146 | 0.275 | 1 | 0.00218097 | -2.2387722 | 0.118 | 0.25 | 1 | 0.00553191 | 0.00435719 |
| Cyb5a | 0.0021811 | -0.7600594 | 0.49 | 0.684 | 1 | 0.17835647 | -0.4537647 | 0.664 | 0.698 | 1 | 0.17835647 | 0.00435745 |
| Gm56906 | 0.00218141 | -1.6627532 | 0.24 | 0.409 | 1 | 0.04804356 | 0.42075484 | 0.483 | 0.354 | 1 | 0.04804356 | 0.00435806 |
| Cyth3 | 0.55266306 | -0.2127037 | 0.875 | 0.896 | 1 | 0.00218286 | -0.7758297 | 0.853 | 0.896 | 1 | 0.55266306 | 0.00436096 |
| Sec22a | 0.237903 | 0.14284639 | 0.458 | 0.363 | 1 | 0.00219012 | 0.55116832 | 0.445 | 0.24 | 1 | 0.237903 | 0.00437544 |
| Lrif1 | 0.00219336 | 0.84084467 | 0.354 | 0.187 | 1 | 0.36428129 | -0.2067618 | 0.319 | 0.24 | 1 | 0.36428129 | 0.00438191 |
| Cnot10 | 0.00219623 | 0.72418774 | 0.594 | 0.43 | 1 | 0.02226476 | 0.2190732 | 0.622 | 0.438 | 1 | 0.02226476 | 0.00438763 |
| Slc35e1 | 0.0199003 | 0.66869262 | 0.458 | 0.326 | 1 | 0.00219769 | 0.3557257 | 0.529 | 0.302 | 1 | 0.0199003 | 0.00439055 |
| Nkd2 | 0.02153171 | -2.1624305 | 0.021 | 0.093 | 1 | 0.00221351 | -3.3098553 | 0.021 | 0.094 | 1 | 0.02153171 | 0.00442212 |
| Thrb | 0.00221391 | -0.8641027 | 0.771 | 0.834 | 1 | 0.35418243 | -0.2938379 | 0.819 | 0.865 | 1 | 0.35418243 | 0.00442291 |
| Baiap2 | 0.00221422 | 0.53004209 | 0.729 | 0.627 | 1 | 0.17290108 | -0.5018816 | 0.731 | 0.823 | 1 | 0.17290108 | 0.00442355 |
| 2810403D21Rik | 0.70354789 | -0.3946039 | 0.292 | 0.311 | 1 | 0.00221829 | 0.51326698 | 0.261 | 0.104 | 1 | 0.70354789 | 0.00443167 |
| Twf1 | 0.00223636 | 0.63077397 | 0.667 | 0.513 | 1 | 0.03357366 | 0.35145812 | 0.664 | 0.583 | 1 | 0.03357366 | 0.00446772 |
| Dym | 0.00225518 | 0.52441165 | 0.51 | 0.321 | 1 | 0.67663521 | -0.3753083 | 0.546 | 0.521 | 1 | 0.67663521 | 0.00450527 |
| Dnaaf9 | 0.00226488 | 0.69816648 | 0.74 | 0.575 | 1 | 0.57974039 | -0.3203172 | 0.819 | 0.865 | 1 | 0.57974039 | 0.00452463 |
| Ryk | 0.00226523 | 0.35634176 | 0.979 | 0.756 | 1 | 0.05167597 | 0.14674291 | 0.933 | 0.865 | 1 | 0.05167597 | 0.00452533 |
| Fam149b | 0.15303594 | 0.61845225 | 0.281 | 0.212 | 1 | 0.00226566 | 0.81790028 | 0.345 | 0.167 | 1 | 0.15303594 | 0.00452618 |
| Appbp2 | 0.04111031 | 0.18152394 | 0.948 | 0.933 | 1 | 0.00226712 | 0.26727561 | 0.962 | 0.958 | 1 | 0.04111031 | 0.00452911 |
| Cul4b | 0.0022686 | 0.44876358 | 0.521 | 0.311 | 1 | 0.00267142 | 0.40837377 | 0.504 | 0.281 | 1 | 0.00267142 | 0.00453205 |
| Rab8b | 0.54847378 | -0.389841 | 0.448 | 0.461 | 1 | 0.00227831 | -0.8150906 | 0.525 | 0.677 | 1 | 0.54847378 | 0.00455143 |
| Champ1 | 0.04890865 | 0.44045394 | 0.26 | 0.155 | 1 | 0.00227868 | 0.85308331 | 0.29 | 0.125 | 1 | 0.04890865 | 0.00455217 |
| Tmem80 | 0.00284737 | 1.12297175 | 0.292 | 0.15 | 1 | 0.00228321 | 0.85160816 | 0.286 | 0.125 | 1 | 0.00284737 | 0.0045612 |
| Nek7 | 0.00228416 | -0.432615 | 0.667 | 0.834 | 1 | 0.05714548 | -0.3098553 | 0.84 | 0.896 | 1 | 0.05714548 | 0.00456311 |
| Plpp1 | 0.12399748 | 0.22132494 | 0.573 | 0.492 | 1 | 0.00228768 | 0.37100267 | 0.618 | 0.354 | 1 | 0.12399748 | 0.00457013 |
| Ccdc8 | 0.00229714 | 1.12778877 | 0.219 | 0.088 | 1 | 0.03370105 | 0.46775232 | 0.168 | 0.073 | 1 | 0.03370105 | 0.004589 |
| Fbxo4 | 0.64253291 | -0.1547769 | 0.333 | 0.358 | 1 | 0.00230091 | 0.59196434 | 0.395 | 0.208 | 1 | 0.64253291 | 0.00459653 |
| Sap18 | 0.00230722 | 0.49758274 | 0.812 | 0.756 | 1 | 0.01096284 | 0.21689262 | 0.849 | 0.771 | 1 | 0.01096284 | 0.00460911 |
| Abl1 | 0.00231735 | 0.43921078 | 0.51 | 0.301 | 1 | 0.01937299 | -0.4594316 | 0.504 | 0.656 | 1 | 0.01937299 | 0.00462933 |
| Lrrfip1 | 0.00231767 | 1.1764477 | 0.719 | 0.653 | 1 | 0.21217594 | -0.6677715 | 0.832 | 0.896 | 1 | 0.21217594 | 0.00462996 |
| Tmem250-ps | 0.14869927 | 0.53800925 | 0.219 | 0.15 | 1 | 0.00232857 | 1.36821664 | 0.227 | 0.083 | 1 | 0.14869927 | 0.00465172 |
| Macrod2 | 0.00232877 | -1.180397 | 0.333 | 0.513 | 1 | 0.07724766 | -1.6055644 | 0.277 | 0.354 | 1 | 0.07724766 | 0.00465212 |
| Erc2 | 0.00232996 | -3.6743295 | 0.021 | 0.13 | 1 | 0.99396047 | 0.17557156 | 0.021 | 0.021 | 1 | 0.99396047 | 0.00465449 |
| 2810001A02Rik | 0.00233047 | 0.76870768 | 0.396 | 0.223 | 1 | 0.00720806 | 1.08246216 | 0.282 | 0.146 | 1 | 0.00720806 | 0.00465552 |
| Eef2kmt | 0.01111094 | 0.7899031 | 0.302 | 0.166 | 1 | 0.00234732 | 1.60768258 | 0.21 | 0.073 | 1 | 0.01111094 | 0.00468913 |
| Il1rapl1 | 0.00236058 | -2.6929452 | 0.042 | 0.166 | 1 | 0.09458395 | -1.4904275 | 0.046 | 0.094 | 1 | 0.09458395 | 0.00471559 |
| Cdc42ep5 | 0.01828735 | 0.47144164 | 0.26 | 0.145 | 1 | 0.00236122 | 1.01207283 | 0.29 | 0.125 | 1 | 0.01828735 | 0.00471687 |
| Mapre2 | 0.00236311 | 0.39851515 | 0.854 | 0.705 | 1 | 0.00287811 | 0.16089167 | 0.811 | 0.688 | 1 | 0.00287811 | 0.00472064 |
| Med6 | 0.22262944 | 0.22322323 | 0.448 | 0.368 | 1 | 0.00236663 | 0.25404562 | 0.391 | 0.177 | 1 | 0.22262944 | 0.00472767 |
| Erbin | 0.00237427 | -0.5171675 | 0.833 | 0.922 | 1 | 0.10041546 | -0.3109553 | 0.966 | 0.969 | 1 | 0.10041546 | 0.00474289 |
| Trnp1 | 0.09530576 | -0.8993961 | 0.073 | 0.14 | 1 | 0.00237487 | 0.95863357 | 0.155 | 0.031 | 1 | 0.09530576 | 0.00474411 |

| | | | | | | | | | | | | |
|---|---|---|---|---|---|---|---|---|---|---|---|---|
| Creg1 | 0.0023789 | 0.5170189 | 0.812 | 0.632 | 1 | 0.00961666 | 0.40024899 | 0.714 | 0.635 | 1 | 0.00961666 | 0.00475213 |
| Ppig | 0.02587162 | 0.31985665 | 0.896 | 0.85 | 1 | 0.00238022 | 0.28459869 | 0.95 | 0.875 | 1 | 0.02587162 | 0.00475477 |
| Cd2bp2 | 0.09116553 | 0.43921078 | 0.26 | 0.171 | 1 | 0.00238412 | 0.54812573 | 0.294 | 0.125 | 1 | 0.09116553 | 0.00476255 |
| Cog3 | 0.00239697 | 0.61845225 | 0.552 | 0.383 | 1 | 0.68927006 | -0.1555271 | 0.597 | 0.51 | 1 | 0.68927006 | 0.0047882 |
| Gdap2 | 0.97232832 | 0.13302542 | 0.292 | 0.29 | 1 | 0.00240218 | 0.55521516 | 0.294 | 0.125 | 1 | 0.97232832 | 0.00479859 |
| Lcorl | 0.0174022 | 0.60447241 | 0.5 | 0.358 | 1 | 0.00240851 | -0.7349464 | 0.563 | 0.708 | 1 | 0.0174022 | 0.00481122 |
| Rnf26 | 0.21067239 | 0.59245704 | 0.167 | 0.114 | 1 | 0.00241833 | 0.54812573 | 0.223 | 0.073 | 1 | 0.21067239 | 0.00483081 |
| Sucla2 | 0.00242131 | 0.64661281 | 0.594 | 0.42 | 1 | 0.75878219 | -0.2756395 | 0.618 | 0.583 | 1 | 0.75878219 | 0.00483676 |
| Tnik | 0.00242434 | -3.1079827 | 0.01 | 0.114 | 1 | 0.11950464 | -2.1765887 | 0.042 | 0.083 | 1 | 0.11950464 | 0.0048428 |
| Raly | 0.94967062 | -0.1048084 | 0.833 | 0.839 | 1 | 0.00242488 | 0.32730345 | 0.887 | 0.781 | 1 | 0.94967062 | 0.00484387 |
| Nat14 | 0.72212321 | 0.13302542 | 0.042 | 0.052 | 1 | 0.00242983 | 1.14957636 | 0.113 | 0.01 | 1 | 0.72212321 | 0.00485376 |
| Rusc1 | 0.05511231 | 0.64904057 | 0.292 | 0.187 | 1 | 0.00244619 | 0.87601128 | 0.294 | 0.125 | 1 | 0.05511231 | 0.00488639 |
| Trim35 | 0.00244916 | 0.89501981 | 0.49 | 0.301 | 1 | 0.14739517 | -0.1193334 | 0.471 | 0.323 | 1 | 0.14739517 | 0.00489233 |
| Cracr2b | 0.44156269 | 0.12613903 | 0.125 | 0.093 | 1 | 0.00246468 | 0.91721365 | 0.282 | 0.125 | 1 | 0.44156269 | 0.00492329 |
| Stx12 | 0.33801534 | -0.1486247 | 0.708 | 0.762 | 1 | 0.002466 | 0.3146356 | 0.794 | 0.635 | 1 | 0.33801534 | 0.00492591 |
| Nbea | 0.28279447 | -0.7183305 | 0.521 | 0.565 | 1 | 0.00246905 | -0.6350961 | 0.773 | 0.885 | 1 | 0.28279447 | 0.004932 |
| Samhd1 | 0.00247783 | -1.8249514 | 0.573 | 0.658 | 1 | 0.00835932 | -0.748597 | 0.508 | 0.635 | 1 | 0.00835932 | 0.00494952 |
| Akip1 | 0.65084491 | 0.14076107 | 0.146 | 0.124 | 1 | 0.00248226 | 1.18490943 | 0.202 | 0.062 | 1 | 0.65084491 | 0.00495836 |
| Tsc22d2 | 0.06234184 | -0.1211399 | 0.823 | 0.907 | 1 | 0.00248386 | -0.663298 | 0.887 | 0.906 | 1 | 0.06234184 | 0.00496156 |
| Cgrrf1 | 0.17206184 | 0.35038225 | 0.583 | 0.503 | 1 | 0.00248602 | 0.19955621 | 0.643 | 0.365 | 1 | 0.17206184 | 0.00496585 |
| Prkdc | 0.00248797 | 1.11767745 | 0.333 | 0.181 | 1 | 0.10354732 | 0.23948333 | 0.256 | 0.156 | 1 | 0.10354732 | 0.00496976 |
| Ube2q1 | 0.07959686 | 0.30705482 | 0.625 | 0.497 | 1 | 0.00248928 | 0.44305188 | 0.752 | 0.656 | 1 | 0.07959686 | 0.00497237 |
| Upp2 | 0.33332563 | -0.4008976 | 0.51 | 0.554 | 1 | 0.0024929 | 0.18826398 | 0.521 | 0.271 | 1 | 0.33332563 | 0.00497959 |
| Irgm1 | 0.08412364 | -1.0628948 | 0.26 | 0.337 | 1 | 0.00249444 | 0.29730298 | 0.294 | 0.125 | 1 | 0.08412364 | 0.00498266 |
| Adamts2 | 0.37179645 | -0.5625947 | 0.24 | 0.275 | 1 | 0.00249608 | 0.37444126 | 0.664 | 0.438 | 1 | 0.37179645 | 0.00498592 |
| Ikzf5 | 0.17333086 | 0.27569065 | 0.375 | 0.29 | 1 | 0.00250513 | 0.45769865 | 0.412 | 0.219 | 1 | 0.17333086 | 0.00500399 |
| Pigyl | 0.00250563 | 0.73186009 | 0.344 | 0.176 | 1 | 0.14737861 | 0.40635177 | 0.218 | 0.146 | 1 | 0.14737861 | 0.00500499 |
| Nelfa | 0.00250929 | 0.78885425 | 0.406 | 0.228 | 1 | 0.00765595 | 0.33106364 | 0.542 | 0.323 | 1 | 0.00765595 | 0.00501228 |
| Klhl13 | 0.00649687 | 0.9920645 | 0.396 | 0.254 | 1 | 0.00251496 | 0.36601573 | 0.319 | 0.135 | 1 | 0.00649687 | 0.00502359 |
| Tsen15 | 0.20123954 | 0.36594851 | 0.49 | 0.43 | 1 | 0.00251704 | 0.5611583 | 0.429 | 0.229 | 1 | 0.20123954 | 0.00502774 |
| Gt(ROSA)26Sor | 0.05137575 | 0.38809854 | 0.469 | 0.337 | 1 | 0.0025183 | 0.17713208 | 0.538 | 0.312 | 1 | 0.05137575 | 0.00503027 |
| Psenen | 0.60446044 | -0.1285263 | 0.833 | 0.808 | 1 | 0.00251872 | 0.30291909 | 0.794 | 0.698 | 1 | 0.60446044 | 0.00503109 |
| Zfp667 | 0.19181974 | 0.39082318 | 0.135 | 0.083 | 1 | 0.00252126 | 1.82400048 | 0.206 | 0.073 | 1 | 0.19181974 | 0.00503617 |
| Gm49797 | 0.00252832 | 0.50053455 | 0.656 | 0.461 | 1 | 0.39046733 | -0.2397104 | 0.765 | 0.792 | 1 | 0.39046733 | 0.00505024 |
| Crlf3 | 0.00252973 | -0.6087189 | 0.583 | 0.725 | 1 | 0.00975559 | -0.7855887 | 0.492 | 0.583 | 1 | 0.00975559 | 0.00505307 |
| Cstf1 | 0.49890171 | 0.25542205 | 0.177 | 0.145 | 1 | 0.00253181 | 1.76053407 | 0.164 | 0.042 | 1 | 0.49890171 | 0.00505721 |
| Yipf6 | 0.2851984 | 0.30027629 | 0.365 | 0.306 | 1 | 0.0025329 | 0.60208856 | 0.403 | 0.208 | 1 | 0.2851984 | 0.00505939 |
| Rnf215 | 0.46623369 | 0.18334437 | 0.438 | 0.394 | 1 | 0.00253656 | 0.61799495 | 0.441 | 0.25 | 1 | 0.46623369 | 0.00506669 |
| Hdgfl2 | 0.00253718 | 0.77302928 | 0.469 | 0.285 | 1 | 0.37652208 | -0.3217294 | 0.378 | 0.271 | 1 | 0.37652208 | 0.00506791 |
| Nptn | 0.00701184 | -0.1233057 | 1 | 0.995 | 1 | 0.00253837 | -0.3540487 | 1 | 1 | 1 | 0.00701184 | 0.00507029 |
| Fdft1 | 0.53875726 | -0.1835102 | 0.635 | 0.658 | 1 | 0.00253999 | 0.94575494 | 0.605 | 0.479 | 1 | 0.53875726 | 0.00507352 |
| Ubqln4 | 0.00254039 | 1.13302542 | 0.292 | 0.145 | 1 | 0.00841455 | 0.34414888 | 0.328 | 0.167 | 1 | 0.00841455 | 0.00507433 |
| Pex12 | 0.00254367 | 1.55498233 | 0.167 | 0.057 | 1 | 0.00941042 | 1.49749966 | 0.126 | 0.031 | 1 | 0.00941042 | 0.00508087 |
| Gm14305 | 0.38158055 | 0.31561683 | 0.208 | 0.166 | 1 | 0.00254892 | 1.27510724 | 0.168 | 0.042 | 1 | 0.38158055 | 0.00509134 |
| Elk3 | 0.00255064 | 0.5803842 | 0.615 | 0.43 | 1 | 0.04709307 | -0.4115851 | 0.723 | 0.479 | 1 | 0.04709307 | 0.00509478 |
| Arid3b | 0.21065255 | 1.00749454 | 0.062 | 0.031 | 1 | 0.00255488 | -2.6790891 | 0.038 | 0.125 | 1 | 0.21065255 | 0.00510324 |
| Trerf1 | 0.77222448 | -0.1445086 | 0.146 | 0.155 | 1 | 0.00255561 | -1.3098553 | 0.202 | 0.354 | 1 | 0.77222448 | 0.00510469 |
| Fubp1 | 0.01510776 | 0.2935456 | 0.958 | 0.943 | 1 | 0.00256164 | 0.265807 | 0.971 | 0.948 | 1 | 0.01510776 | 0.00511671 |
| Dnpep | 0.52956543 | -0.1688282 | 0.375 | 0.394 | 1 | 0.00256234 | 0.56050946 | 0.391 | 0.198 | 1 | 0.52956543 | 0.00511811 |
| Lpp | 0.05780212 | -0.430416 | 0.979 | 0.974 | 1 | 0.00256385 | -0.5615113 | 0.992 | 0.99 | 1 | 0.05780212 | 0.00512112 |
| Syvn1 | 0.07020909 | 0.36363835 | 0.375 | 0.264 | 1 | 0.00256891 | 0.62695091 | 0.424 | 0.229 | 1 | 0.07020909 | 0.00513122 |
| Impa1 | 0.09811568 | 0.34073794 | 0.531 | 0.435 | 1 | 0.00257684 | 0.63592435 | 0.605 | 0.448 | 1 | 0.09811568 | 0.00514704 |
| Cacfd1 | 0.28520438 | 0.38776662 | 0.344 | 0.285 | 1 | 0.00257913 | 0.61859148 | 0.357 | 0.177 | 1 | 0.28520438 | 0.00515162 |
| Gm10561 | 0.21298864 | 0.26333344 | 0.323 | 0.249 | 1 | 0.00258013 | 1.2095189 | 0.252 | 0.104 | 1 | 0.21298864 | 0.00515361 |
| Ptp4a1 | 0.84736847 | 0.10060394 | 0.062 | 0.057 | 1 | 0.00258163 | -2.4253325 | 0.021 | 0.094 | 1 | 0.84736847 | 0.0051566 |
| Zfp637 | 0.60401858 | -0.1539689 | 0.271 | 0.301 | 1 | 0.00258274 | 0.69014474 | 0.42 | 0.229 | 1 | 0.60401858 | 0.00515881 |
| Zfhx3 | 0.03900125 | -0.5294818 | 0.885 | 0.922 | 1 | 0.0025893 | -0.6040787 | 0.945 | 0.958 | 1 | 0.03900125 | 0.0051719 |
| Sdk2 | 0.00258991 | 0.18075063 | 1 | 0.933 | 1 | 0.11872478 | -0.1377464 | 0.992 | 0.906 | 1 | 0.11872478 | 0.0051731 |
| Zfp189 | 0.00317183 | 1.29700115 | 0.198 | 0.078 | 1 | 0.0025902 | 1.31817596 | 0.214 | 0.073 | 1 | 0.00317183 | 0.0051737 |
| Commd4 | 0.77367949 | 0.19512154 | 0.333 | 0.321 | 1 | 0.00259281 | 0.56050946 | 0.378 | 0.188 | 1 | 0.77367949 | 0.0051789 |
| Ctf1 | 0.01333881 | 1.12948506 | 0.271 | 0.155 | 1 | 0.00259841 | 0.98137604 | 0.294 | 0.135 | 1 | 0.01333881 | 0.00519008 |
| Smpd2 | 0.10219366 | -1.0320338 | 0.167 | 0.244 | 1 | 0.00259984 | 1.14080615 | 0.256 | 0.104 | 1 | 0.10219366 | 0.00519291 |
| Fam83g | 0.00553111 | 1.53800925 | 0.104 | 0.026 | 1 | 0.00260872 | 2.18199783 | 0.13 | 0.021 | 1 | 0.00553111 | 0.00521063 |
| Blvrb | 0.02405165 | 0.40083696 | 0.354 | 0.212 | 1 | 0.00261155 | -0.8591939 | 0.282 | 0.458 | 1 | 0.02405165 | 0.00521629 |
| Mios | 0.02410289 | 0.60086928 | 0.323 | 0.197 | 1 | 0.00261575 | 0.82138927 | 0.269 | 0.115 | 1 | 0.02410289 | 0.00522466 |
| Zfp449 | 0.00261896 | 1.48142572 | 0.198 | 0.078 | 1 | 0.02164283 | 1.63767732 | 0.151 | 0.062 | 1 | 0.02164283 | 0.00523106 |
| Plekhm2 | 0.63803906 | 0.10577405 | 0.771 | 0.751 | 1 | 0.00262287 | 0.14188844 | 0.908 | 0.781 | 1 | 0.63803906 | 0.00523886 |
| Vangl2 | 0.00262634 | 1.51445453 | 0.198 | 0.078 | 1 | 0.01141742 | 1.01207283 | 0.235 | 0.115 | 1 | 0.01141742 | 0.00524578 |
| Gm26532 | 0.26778282 | 0.35038225 | 0.448 | 0.383 | 1 | 0.00262679 | 0.32935028 | 0.592 | 0.344 | 1 | 0.26778282 | 0.00524668 |
| Lrrfip2 | 0.0026274 | -0.4634435 | 0.729 | 0.886 | 1 | 0.85089286 | -0.1605111 | 0.861 | 0.865 | 1 | 0.85089286 | 0.00524789 |
| Polr2i | 0.1898682 | 0.37006462 | 0.354 | 0.28 | 1 | 0.00262744 | 0.34619034 | 0.353 | 0.167 | 1 | 0.1898682 | 0.00524798 |
| Mindy2 | 0.00262953 | -0.9446769 | 0.344 | 0.503 | 1 | 0.01068005 | -0.7552664 | 0.534 | 0.677 | 1 | 0.01068005 | 0.00525214 |
| Glrx | 0.27228325 | 0.37362244 | 0.208 | 0.155 | 1 | 0.00264075 | 1.02717972 | 0.269 | 0.115 | 1 | 0.27228325 | 0.00527452 |
| Zup1 | 0.36685002 | 0.12778877 | 0.438 | 0.368 | 1 | 0.0026413 | 0.14582422 | 0.471 | 0.24 | 1 | 0.36685002 | 0.00527562 |
| Vps13a | 0.4912353 | 0.25626237 | 0.51 | 0.596 | 1 | 0.00265632 | -0.7664937 | 0.605 | 0.781 | 1 | 0.4912353 | 0.00530559 |
| Dpp9 | 0.00847939 | 0.96567436 | 0.26 | 0.135 | 1 | 0.00265953 | 0.45567948 | 0.357 | 0.177 | 1 | 0.00847939 | 0.00531198 |
| Socs7 | 0.35535091 | 0.41595938 | 0.406 | 0.358 | 1 | 0.00268038 | 0.38906343 | 0.567 | 0.365 | 1 | 0.35535091 | 0.00535358 |
| Top3b | 0.04636234 | 0.38456419 | 0.365 | 0.249 | 1 | 0.00269067 | 0.42397062 | 0.441 | 0.24 | 1 | 0.04636234 | 0.0053741 |
| Gch1 | 0.07454547 | -0.4872702 | 0.125 | 0.218 | 1 | 0.00269111 | 0.96316323 | 0.248 | 0.094 | 1 | 0.07454547 | 0.00537499 |
| Ndufa5 | 0.05512288 | 0.25173135 | 0.719 | 0.596 | 1 | 0.00269501 | 0.64883776 | 0.609 | 0.49 | 1 | 0.05512288 | 0.00538275 |
| Rspo3 | 0.00270366 | -1.4098413 | 0.594 | 0.663 | 1 | 0.00535066 | 0.80032766 | 0.382 | 0.208 | 1 | 0.00535066 | 0.00540001 |
| Gm16364.1 | 0.15729907 | -1.3144336 | 0 | 0.021 | 1 | 0.00270825 | -2.4797803 | 0.021 | 0.094 | 1 | 0.15729907 | 0.00540917 |
| Bmyc | 0.32764241 | -0.4812526 | 0.271 | 0.316 | 1 | 0.00270918 | 0.61975541 | 0.294 | 0.125 | 1 | 0.32764241 | 0.00541103 |
| Gm57158 | 0.00271713 | 2.14499806 | 0.146 | 0.047 | 1 | 0.08011187 | 1.3146356 | 0.113 | 0.052 | 1 | 0.08011187 | 0.00542688 |
| Fdxacb1 | 0.28985223 | 1.00749454 | 0.083 | 0.052 | 1 | 0.00271921 | 2.77760758 | 0.109 | 0.01 | 1 | 0.28985223 | 0.00543102 |
| Traf6 | 0.18409278 | 0.13302542 | 0.281 | 0.202 | 1 | 0.00272022 | 0.34381999 | 0.378 | 0.188 | 1 | 0.18409278 | 0.00543304 |
| Cbr2 | 0.11487659 | -1.1212918 | 0.469 | 0.549 | 1 | 0.00272056 | -0.8364011 | 0.639 | 0.698 | 1 | 0.11487659 | 0.00543372 |
| Fbxw8 | 0.25688577 | -0.1434474 | 0.677 | 0.731 | 1 | 0.00272061 | 0.2124432 | 0.777 | 0.51 | 1 | 0.25688577 | 0.00543381 |
| Txndc17 | 0.16019388 | 0.18532662 | 0.875 | 0.803 | 1 | 0.00272178 | 0.35439615 | 0.874 | 0.792 | 1 | 0.16019388 | 0.00543616 |
| Epb41l1 | 0.00272256 | 0.51485422 | 0.792 | 0.684 | 1 | 0.00396859 | 0.29134788 | 0.811 | 0.625 | 1 | 0.00396859 | 0.0054377 |

| | | | | | | | | | | | | |
|---|---|---|---|---|---|---|---|---|---|---|---|---|
| Coa3 | 0.00300623 | 0.55292867 | 0.562 | 0.373 | 1 | 0.00273259 | 0.54304044 | 0.529 | 0.354 | 1 | 0.00300623 | 0.00545772 |
| Polr3gl | 0.69493681 | 0.16314541 | 0.344 | 0.326 | 1 | 0.0027337 | 0.4777364 | 0.433 | 0.219 | 1 | 0.69493681 | 0.00545994 |
| Ddit3 | 0.31596646 | -0.744578 | 0.396 | 0.415 | 1 | 0.00274098 | 0.63891941 | 0.429 | 0.229 | 1 | 0.31596646 | 0.00547445 |
| Cyp4a12b | 0.99320802 | 0.81484946 | 0.01 | 0.01 | 1 | 0.00274446 | -2.0468209 | 0.021 | 0.094 | 1 | 0.99320802 | 0.00548139 |
| Tmem161a | 0.07518582 | 0.56319244 | 0.438 | 0.352 | 1 | 0.00274767 | 0.68009107 | 0.42 | 0.24 | 1 | 0.07518582 | 0.00548779 |
| Ube2l3 | 0.00275055 | 0.36437523 | 0.906 | 0.834 | 1 | 0.00360369 | 0.25061268 | 0.92 | 0.854 | 1 | 0.00360369 | 0.00549353 |
| Ift74 | 0.00275109 | 0.63826073 | 0.615 | 0.477 | 1 | 0.0035432 | 0.1447106 | 0.651 | 0.406 | 1 | 0.0035432 | 0.00549461 |
| Ubr3 | 0.09782328 | 0.26556269 | 0.865 | 0.798 | 1 | 0.00276395 | -0.5820788 | 0.937 | 0.948 | 1 | 0.09782328 | 0.00552025 |
| Actr3 | 0.03536669 | 0.26103566 | 0.979 | 0.959 | 1 | 0.00277014 | 0.43597636 | 0.958 | 0.927 | 1 | 0.03536669 | 0.0055326 |
| Gm30025 | 0.47181153 | 0.29700115 | 0.104 | 0.078 | 1 | 0.00278038 | 1.73453886 | 0.13 | 0.021 | 1 | 0.47181153 | 0.00555304 |
| Vps9d1 | 0.61569272 | 0.13873907 | 0.177 | 0.15 | 1 | 0.00278229 | 1.10518224 | 0.214 | 0.073 | 1 | 0.61569272 | 0.00555685 |
| Gphn | 0.05873444 | -0.5416695 | 1 | 1 | 1 | 0.00278241 | -0.8095267 | 1 | 1 | 1 | 0.05873444 | 0.00555708 |
| Pkd2 | 0.00278869 | 0.5002496 | 0.865 | 0.824 | 1 | 0.08140128 | 0.18560909 | 0.937 | 0.771 | 1 | 0.08140128 | 0.0055696 |
| Fzd10 | 0.98102622 | -0.6705774 | 0.042 | 0.041 | 1 | 0.00279081 | 0.85595363 | 0.303 | 0.135 | 1 | 0.98102622 | 0.00557383 |
| Ctbs | 0.94999889 | 0.26275159 | 0.26 | 0.275 | 1 | 0.00279093 | 0.74903843 | 0.261 | 0.104 | 1 | 0.94999889 | 0.00557407 |
| Paqr7 | 0.00279362 | -3.4519371 | 0 | 0.088 | 1 | 0.63004663 | -0.4618584 | 0.088 | 0.104 | 1 | 0.63004663 | 0.00557944 |
| Prcc | 0.13058447 | 0.36465112 | 0.406 | 0.311 | 1 | 0.00279446 | 0.34129752 | 0.584 | 0.333 | 1 | 0.13058447 | 0.00558111 |
| Pced1a | 0.0683198 | 0.93710521 | 0.167 | 0.093 | 1 | 0.00279599 | 1.7944814 | 0.147 | 0.031 | 1 | 0.0683198 | 0.00558416 |
| St3gal2 | 0.38683362 | -0.4163132 | 0.302 | 0.352 | 1 | 0.00280165 | 0.19071505 | 0.58 | 0.312 | 1 | 0.38683362 | 0.00559544 |
| Brinp1 | 0.00280205 | -4.8253955 | 0 | 0.088 | 1 | 0.11689951 | -3.8948178 | 0 | 0.01 | 1 | 0.11689951 | 0.00559625 |
| Npr2 | 0.07938245 | 0.43163424 | 0.417 | 0.306 | 1 | 0.00280468 | 0.43685831 | 0.538 | 0.302 | 1 | 0.07938245 | 0.00560148 |
| Plet1 | 0.17261719 | 0.65385758 | 0.094 | 0.052 | 1 | 0.00280719 | -0.2394659 | 0.059 | 0.167 | 1 | 0.17261719 | 0.00560651 |
| Wipi1 | 0.04197761 | 0.32942263 | 0.729 | 0.617 | 1 | 0.00280956 | 0.28678904 | 0.588 | 0.333 | 1 | 0.04197761 | 0.00561123 |
| Dhrs3 | 0.01252016 | 0.5606483 | 0.625 | 0.503 | 1 | 0.00281767 | 0.37720543 | 0.744 | 0.635 | 1 | 0.01252016 | 0.0056274 |
| Alg3 | 0.00282422 | 1.05188866 | 0.292 | 0.145 | 1 | 0.00784641 | 0.93807225 | 0.21 | 0.083 | 1 | 0.00784641 | 0.00564047 |
| Bcas3 | 0.03801733 | -0.5217585 | 0.604 | 0.741 | 1 | 0.00282445 | -0.7730725 | 0.685 | 0.833 | 1 | 0.03801733 | 0.00564092 |
| Nr1h2 | 0.44899005 | -0.2421194 | 0.365 | 0.409 | 1 | 0.00282476 | 0.48851088 | 0.454 | 0.26 | 1 | 0.44899005 | 0.00564154 |
| Kctd11 | 0.08749415 | 1.00749454 | 0.167 | 0.098 | 1 | 0.00282557 | 0.93115284 | 0.185 | 0.052 | 1 | 0.08749415 | 0.00564315 |
| Prdm16os | 0.26266721 | -0.5427025 | 0.125 | 0.176 | 1 | 0.00282989 | 0.79866919 | 0.248 | 0.094 | 1 | 0.26266721 | 0.00565176 |
| Tk1 | 0.00283123 | 4.09495738 | 0.062 | 0.005 | 1 | 0.57370068 | -1.0468209 | 0.021 | 0.031 | 1 | 0.57370068 | 0.00565445 |
| Pgk1 | 0.00283162 | 0.46622463 | 0.885 | 0.772 | 1 | 0.0093188 | 0.40920013 | 0.803 | 0.76 | 1 | 0.0093188 | 0.00565523 |
| Otor | 0.0088726 | 1.21753857 | 0.5 | 0.378 | 1 | 0.00283201 | 1.67454788 | 0.353 | 0.177 | 1 | 0.0088726 | 0.005656 |
| Rpp21 | 0.00283293 | 0.81484946 | 0.354 | 0.187 | 1 | 0.51741176 | -0.2273931 | 0.235 | 0.188 | 1 | 0.51741176 | 0.00565783 |
| Ivd | 0.00283336 | 0.70941318 | 0.427 | 0.249 | 1 | 0.04831566 | 0.20110666 | 0.366 | 0.24 | 1 | 0.04831566 | 0.0056587 |
| Cfh | 0.18352047 | -1.536826 | 0.042 | 0.083 | 1 | 0.00283504 | -2.2933671 | 0.05 | 0.146 | 1 | 0.18352047 | 0.00566204 |
| Zbtb39 | 0.0110069 | 1.07460873 | 0.188 | 0.083 | 1 | 0.00284029 | 0.84214783 | 0.273 | 0.115 | 1 | 0.0110069 | 0.00567252 |
| Mindy1 | 0.2356034 | 0.28890747 | 0.5 | 0.415 | 1 | 0.00284592 | 0.60191755 | 0.538 | 0.354 | 1 | 0.2356034 | 0.00568375 |
| Krt10 | 0.00285015 | 1.13677755 | 0.292 | 0.15 | 1 | 0.09039905 | 0.13760371 | 0.181 | 0.104 | 1 | 0.09039905 | 0.00569218 |
| Mdh2 | 0.00956368 | 0.24798637 | 0.927 | 0.85 | 1 | 0.00285409 | 0.39289462 | 0.87 | 0.833 | 1 | 0.00956368 | 0.00570004 |
| Klhl40 | 0.00285688 | 3.81484946 | 0.062 | 0.005 | 1 | 0.07248287 | 0.86006974 | 0.055 | 0.01 | 1 | 0.07248287 | 0.0057056 |
| Lrrc3 | 0.00285688 | 3.46692616 | 0.062 | 0.005 | 1 | 0.86312513 | -0.7248928 | 0.008 | 0.01 | 1 | 0.86312513 | 0.0057056 |
| Gm2895 | 0.00285688 | 3.17741954 | 0.062 | 0.005 | 1 | 0.14439146 | -2.8948178 | 0.004 | 0.021 | 1 | 0.14439146 | 0.0057056 |
| Eif4e2 | 0.11978327 | 0.18960895 | 0.792 | 0.668 | 1 | 0.00286278 | 0.43497857 | 0.769 | 0.635 | 1 | 0.11978327 | 0.00571736 |
| Ahcyl2 | 0.03860639 | 0.51689572 | 0.677 | 0.632 | 1 | 0.00286326 | -0.7297367 | 0.849 | 0.906 | 1 | 0.03860639 | 0.00571832 |
| Gm57402 | 0.00715218 | 1.32942263 | 0.229 | 0.114 | 1 | 0.00286594 | 0.78530197 | 0.315 | 0.146 | 1 | 0.00715218 | 0.00572367 |
| Ints15 | 0.20321639 | 0.65385758 | 0.167 | 0.114 | 1 | 0.00287417 | 1.21370669 | 0.227 | 0.083 | 1 | 0.20321639 | 0.00574009 |
| Ndel1 | 0.00287628 | 0.20687595 | 0.958 | 0.886 | 1 | 0.00422921 | 0.22840697 | 0.954 | 0.99 | 1 | 0.00422921 | 0.00574429 |
| Bcl9l | 0.00287811 | 0.65699729 | 0.594 | 0.409 | 1 | 0.01443631 | 0.12915088 | 0.782 | 0.646 | 1 | 0.01443631 | 0.00574793 |
| Ywhab | 0.00287866 | 0.29471841 | 0.969 | 0.943 | 1 | 0.00311717 | 0.28726144 | 0.983 | 0.958 | 1 | 0.00311717 | 0.00574903 |
| 5033426O07Rik | 0.63709119 | -0.240433 | 0.062 | 0.078 | 1 | 0.0028788 | 1.09453499 | 0.244 | 0.094 | 1 | 0.63709119 | 0.00574932 |
| Gm11520 | 0.54639121 | -0.3550755 | 0.031 | 0.047 | 1 | 0.00288463 | -3.8948178 | 0.004 | 0.052 | 1 | 0.54639121 | 0.00576093 |
| Vcam1 | 0.75328966 | 0.27052894 | 0.021 | 0.016 | 1 | 0.00288463 | -4.1172102 | 0.004 | 0.052 | 1 | 0.75328966 | 0.00576093 |
| Wdr62 | 0.82258416 | -0.1624305 | 0.031 | 0.036 | 1 | 0.00288463 | -3.7692869 | 0.004 | 0.052 | 1 | 0.82258416 | 0.00576093 |
| Galc | 0.00288649 | 1.34853145 | 0.167 | 0.057 | 1 | 0.33791267 | 0.18199783 | 0.147 | 0.104 | 1 | 0.33791267 | 0.00576464 |
| St3gal6 | 0.00533685 | 0.49925783 | 0.823 | 0.663 | 1 | 0.00289507 | 0.36433701 | 0.87 | 0.708 | 1 | 0.00533685 | 0.00578175 |
| Rab26os | 0.00289592 | 1.10060394 | 0.271 | 0.13 | 1 | 0.04359433 | 1.33400093 | 0.097 | 0.031 | 1 | 0.04359433 | 0.00578345 |
| Wdr83 | 0.64521792 | 0.13302542 | 0.156 | 0.135 | 1 | 0.00289703 | 1.12310414 | 0.197 | 0.062 | 1 | 0.64521792 | 0.00578567 |
| Abcb9 | 0.52284897 | -0.1249558 | 0.177 | 0.212 | 1 | 0.00289795 | 0.69014474 | 0.311 | 0.146 | 1 | 0.52284897 | 0.0057875 |
| Hmgn2-ps | 0.00290048 | 1.37006462 | 0.167 | 0.057 | 1 | 0.00354159 | 2.44503224 | 0.105 | 0.01 | 1 | 0.00354159 | 0.00579254 |
| 4930481B07Rik | 0.37536054 | 1.32942263 | 0.031 | 0.016 | 1 | 0.00290993 | -3.3098553 | 0.004 | 0.052 | 1 | 0.37536054 | 0.00581138 |
| Frem1 | 0.52528691 | -1.577468 | 0.01 | 0.021 | 1 | 0.00290993 | -3.4797803 | 0.004 | 0.052 | 1 | 0.52528691 | 0.00581138 |
| Selenon | 0.05251853 | 0.32942263 | 0.333 | 0.212 | 1 | 0.00291065 | 0.80944367 | 0.378 | 0.198 | 1 | 0.05251853 | 0.00581284 |
| Ercc6l | 0.00291069 | 2.81484946 | 0.062 | 0.005 | 1 | 0.27154471 | 0.69014474 | 0.013 | 0 | 1 | 0.27154471 | 0.00581291 |
| Shisal1 | 0.15527232 | -2.1624305 | 0.021 | 0.057 | 1 | 0.00291366 | -2.4473588 | 0.017 | 0.083 | 1 | 0.15527232 | 0.00581884 |
| Aldh9a1 | 0.27695766 | 0.30122574 | 0.302 | 0.238 | 1 | 0.00291453 | 1.04064198 | 0.319 | 0.156 | 1 | 0.27695766 | 0.00582057 |
| Ensa | 0.14834841 | 0.18690379 | 0.698 | 0.617 | 1 | 0.00291748 | 0.26836566 | 0.832 | 0.646 | 1 | 0.14834841 | 0.00582644 |
| Cryz | 0.00552596 | 0.95091101 | 0.25 | 0.119 | 1 | 0.00292203 | 1.69014474 | 0.176 | 0.052 | 1 | 0.00552596 | 0.00583552 |
| Dstn | 0.89287222 | -0.1428845 | 1 | 0.984 | 1 | 0.00292714 | 0.25446293 | 0.992 | 0.99 | 1 | 0.89287222 | 0.0058457 |
| Pomgnt1 | 0.41032965 | 0.24850264 | 0.302 | 0.254 | 1 | 0.00293615 | 0.81942775 | 0.328 | 0.156 | 1 | 0.41032965 | 0.00586368 |
| Agbl2 | 1 | -0.1079827 | 0.042 | 0.041 | 1 | 0.00293664 | -3.4253325 | 0.004 | 0.052 | 1 | 1 | 0.00586466 |
| Chrm3 | 0.04512622 | 4.46692616 | 0.021 | 0 | 1 | 0.00293673 | -2.8948178 | 0.004 | 0.052 | 1 | 0.04512622 | 0.00586483 |
| Shcbp1 | 0.00293719 | 3.22988696 | 0.062 | 0.005 | 1 | 0.87066325 | -0.3098553 | 0.013 | 0.01 | 1 | 0.87066325 | 0.00586575 |
| Hexb | 0.08255474 | 0.49091901 | 0.656 | 0.611 | 1 | 0.00294031 | 0.90446386 | 0.769 | 0.615 | 1 | 0.08255474 | 0.00587197 |
| Hadha | 0.13765028 | 0.22717257 | 0.792 | 0.782 | 1 | 0.00294282 | 0.23171464 | 0.849 | 0.792 | 1 | 0.13765028 | 0.00587697 |
| Sidt2 | 0.00294332 | 0.61102914 | 0.771 | 0.715 | 1 | 0.00894505 | 0.43017663 | 0.786 | 0.698 | 1 | 0.00894505 | 0.00587798 |
| Gria1 | 0.00294469 | -2.5160674 | 0.021 | 0.13 | 1 | 0.52895176 | -1.3769695 | 0.046 | 0.062 | 1 | 0.52895176 | 0.00588071 |
| Mrpl57 | 0.85578641 | 0.21560273 | 0.427 | 0.415 | 1 | 0.00294485 | 0.41676839 | 0.412 | 0.219 | 1 | 0.85578641 | 0.00588102 |
| Cand2 | 0.15272256 | 0.64492446 | 0.125 | 0.073 | 1 | 0.00294598 | 1.57767001 | 0.147 | 0.031 | 1 | 0.15272256 | 0.00588327 |
| Dact1 | 0.01307418 | 0.7810666 | 0.531 | 0.389 | 1 | 0.00294815 | 0.42462078 | 0.559 | 0.323 | 1 | 0.01307418 | 0.00588762 |
| Spsb4 | 0.04816755 | 0.15433592 | 0.25 | 0.145 | 1 | 0.00296043 | 0.42005557 | 0.328 | 0.146 | 1 | 0.04816755 | 0.00591209 |
| Mta1 | 0.00297189 | 0.5371746 | 0.667 | 0.446 | 1 | 0.01570378 | 0.3834834 | 0.655 | 0.458 | 1 | 0.01570378 | 0.00593496 |
| Samd12 | 0.0029725 | -3.0657544 | 0.031 | 0.145 | 1 | 0.10346909 | -1.5927892 | 0.08 | 0.135 | 1 | 0.10346909 | 0.00593616 |
| Gm49969 | 0.00297418 | -2.3382803 | 0.052 | 0.176 | 1 | 0.71139303 | -0.198824 | 0.168 | 0.146 | 1 | 0.71139303 | 0.00593951 |
| Ccn5 | 0.00297941 | -1.4102321 | 0.24 | 0.394 | 1 | 0.08115468 | -0.6377408 | 0.223 | 0.125 | 1 | 0.08115468 | 0.00594994 |
| Gm15987 | 0.00298723 | -3.5160674 | 0.031 | 0.145 | 1 | 0.34363875 | -2.0468209 | 0.008 | 0.021 | 1 | 0.34363875 | 0.00596554 |
| Epc1 | 0.0837103 | 0.12469575 | 0.875 | 0.839 | 1 | 0.00299387 | -0.6193684 | 0.945 | 0.938 | 1 | 0.0837103 | 0.00597878 |
| Golga1 | 0.00299491 | 0.63552576 | 0.51 | 0.321 | 1 | 0.05749114 | 0.11210165 | 0.475 | 0.344 | 1 | 0.05749114 | 0.00598085 |
| Zcchc17 | 0.01577607 | 0.4940853 | 0.677 | 0.528 | 1 | 0.0029995 | 0.51952597 | 0.525 | 0.333 | 1 | 0.01577607 | 0.00599001 |
| Deptor | 0.00300455 | -1.2201062 | 0.333 | 0.497 | 1 | 0.04135146 | 0.15727075 | 0.416 | 0.26 | 1 | 0.04135146 | 0.00600007 |

| | | | | | | | | | | | | |
|---|---|---|---|---|---|---|---|---|---|---|---|---|
| Ankhd1 | 0.59105831 | -0.1970442 | 0.854 | 0.829 | 1 | 0.00300776 | -0.6343573 | 0.887 | 0.938 | 1 | 0.59105831 | 0.00600647 |
| Acat1 | 0.00301199 | 0.43965621 | 0.854 | 0.694 | 1 | 0.0054413 | 0.16308062 | 0.832 | 0.677 | 1 | 0.0054413 | 0.00601491 |
| Gale | 0.19632825 | 0.56692195 | 0.333 | 0.264 | 1 | 0.00302572 | 0.8799693 | 0.218 | 0.073 | 1 | 0.19632825 | 0.00604228 |
| Mcts1 | 0.00303814 | 0.5049942 | 0.646 | 0.446 | 1 | 0.03403954 | 0.23763253 | 0.542 | 0.375 | 1 | 0.03403954 | 0.00606705 |
| Acp2 | 0.04609647 | 0.51106871 | 0.417 | 0.295 | 1 | 0.0030392 | 0.56461386 | 0.412 | 0.219 | 1 | 0.04609647 | 0.00606916 |
| Prune1 | 0.24996886 | 0.33611728 | 0.219 | 0.161 | 1 | 0.00304181 | 0.90737545 | 0.265 | 0.115 | 1 | 0.24996886 | 0.00607437 |
| Rspo1 | 0.03210862 | -3.3848229 | 0 | 0.047 | 1 | 0.00304217 | -2.5322477 | 0.004 | 0.052 | 1 | 0.03210862 | 0.00607508 |
| Exo5 | 0.00304643 | 1.86999101 | 0.146 | 0.047 | 1 | 0.39033524 | 0.27510724 | 0.092 | 0.062 | 1 | 0.39033524 | 0.00608358 |
| Cep350 | 0.00304644 | 0.27052894 | 0.833 | 0.71 | 1 | 0.01436658 | -0.5950316 | 0.857 | 0.906 | 1 | 0.01436658 | 0.00608359 |
| Srpk1 | 0.00305449 | 0.61465278 | 0.656 | 0.523 | 1 | 0.42814417 | -0.2457249 | 0.655 | 0.615 | 1 | 0.42814417 | 0.00609965 |
| Klk10 | 0.00305496 | -3.0953163 | 0.031 | 0.145 | 1 | 0.82047853 | -0.1399303 | 0.025 | 0.021 | 1 | 0.82047853 | 0.00610058 |
| Olfml2a | 0.11302941 | -2.1624305 | 0 | 0.026 | 1 | 0.00306348 | -2.5322477 | 0.017 | 0.083 | 1 | 0.11302941 | 0.00611758 |
| Fzd7 | 0.00306771 | 1.11938742 | 0.375 | 0.218 | 1 | 0.20643459 | -0.1843244 | 0.265 | 0.177 | 1 | 0.20643459 | 0.00612601 |
| Gdpd4 | 0.47911845 | -1.0628948 | 0.021 | 0.036 | 1 | 0.0030694 | -2.3098553 | 0.004 | 0.052 | 1 | 0.47911845 | 0.00612937 |
| Rnf150 | 0.0030704 | -3.4018964 | 0.031 | 0.145 | 1 | 0.028553 | -2.4253325 | 0.029 | 0.083 | 1 | 0.028553 | 0.00613138 |
| Gm16152 | 0.00307277 | 2.22988696 | 0.062 | 0.005 | 1 | 0.18504625 | 1.01207283 | 0.038 | 0.01 | 1 | 0.18504625 | 0.00613611 |
| Irx5 | 0.98024889 | 0.19035859 | 0.125 | 0.124 | 1 | 0.00307618 | -1.0560987 | 0.038 | 0.125 | 1 | 0.98024889 | 0.0061429 |
| Orai3 | 0.18940153 | 0.49292136 | 0.24 | 0.176 | 1 | 0.00307842 | 0.95693128 | 0.269 | 0.115 | 1 | 0.18940153 | 0.00614736 |
| Acox1 | 0.03521291 | -0.7574667 | 0.688 | 0.767 | 1 | 0.00307902 | -0.7248928 | 0.71 | 0.823 | 1 | 0.03521291 | 0.00614857 |
| Tbp | 0.21431958 | 0.22988696 | 0.302 | 0.228 | 1 | 0.00308527 | 0.69014474 | 0.307 | 0.146 | 1 | 0.21431958 | 0.00616103 |
| Fam199x | 0.16154212 | 0.21608116 | 0.229 | 0.15 | 1 | 0.00308954 | 0.56461386 | 0.361 | 0.177 | 1 | 0.16154212 | 0.00616954 |
| Zer1 | 0.79989104 | 0.13755508 | 0.229 | 0.218 | 1 | 0.00309091 | 0.79976923 | 0.282 | 0.125 | 1 | 0.79989104 | 0.00617226 |
| Fam8a1 | 0.0032814 | 0.875391 | 0.458 | 0.306 | 1 | 0.00310045 | 0.31258194 | 0.58 | 0.354 | 1 | 0.0032814 | 0.00619129 |
| Anxa4 | 0.0031032 | 0.43582886 | 0.781 | 0.648 | 1 | 0.89878128 | -0.2397397 | 0.718 | 0.656 | 1 | 0.89878128 | 0.00619676 |
| Prr5 | 0.00484886 | 1.21394541 | 0.125 | 0.036 | 1 | 0.0031053 | 2.19264508 | 0.126 | 0.021 | 1 | 0.00484886 | 0.00620095 |
| Gm41724 | 0.00310678 | -2.0249269 | 0.052 | 0.181 | 1 | 0.91070565 | -0.3098553 | 0.067 | 0.062 | 1 | 0.91070565 | 0.00620391 |
| Otud5 | 0.02759472 | 0.31633263 | 0.625 | 0.472 | 1 | 0.00312504 | 0.53226508 | 0.676 | 0.521 | 1 | 0.02759472 | 0.00624032 |
| Elp2 | 0.05529291 | 0.64492446 | 0.354 | 0.254 | 1 | 0.00315753 | 0.69014474 | 0.357 | 0.177 | 1 | 0.05529291 | 0.0063051 |
| Rhno1 | 0.08659736 | -0.9925055 | 0.094 | 0.166 | 1 | 0.00315945 | 1.27510724 | 0.181 | 0.052 | 1 | 0.08659736 | 0.00630892 |
| Csnk2a1 | 0.00316727 | 0.33192949 | 0.885 | 0.762 | 1 | 0.08015052 | 0.12525903 | 0.903 | 0.833 | 1 | 0.08015052 | 0.00632451 |
| Abl2 | 0.00316762 | -0.6249769 | 0.802 | 0.85 | 1 | 0.00812393 | -0.5899632 | 0.916 | 0.917 | 1 | 0.00812393 | 0.0063252 |
| St8sia6 | 0.00317103 | -2.3984978 | 0.031 | 0.145 | 1 | 0.46048942 | -0.1578522 | 0.05 | 0.031 | 1 | 0.46048942 | 0.00633201 |
| Noc2l | 0.06478433 | 0.55377657 | 0.365 | 0.259 | 1 | 0.00317527 | 0.47083171 | 0.378 | 0.188 | 1 | 0.06478433 | 0.00634045 |
| Pdap1 | 0.00317712 | 0.38884891 | 0.917 | 0.741 | 1 | 0.09590444 | 0.14744796 | 0.824 | 0.76 | 1 | 0.09590444 | 0.00634415 |
| Hmbs | 0.98536976 | -0.1217885 | 0.146 | 0.145 | 1 | 0.00318561 | 1.76053407 | 0.16 | 0.042 | 1 | 0.98536976 | 0.00636108 |
| Rnf5 | 0.16707915 | 0.25652208 | 0.354 | 0.264 | 1 | 0.00318799 | 0.24374307 | 0.324 | 0.146 | 1 | 0.16707915 | 0.00636583 |
| Epb41 | 0.59908713 | -0.5230202 | 0.323 | 0.337 | 1 | 0.00319631 | -0.6171137 | 0.525 | 0.729 | 1 | 0.59908713 | 0.0063824 |
| Foxred2 | 0.00319868 | 1.37006462 | 0.156 | 0.052 | 1 | 0.45864643 | -0.8504236 | 0.126 | 0.094 | 1 | 0.45864643 | 0.00638713 |
| Mrpl58 | 0.04908821 | 0.4649673 | 0.375 | 0.259 | 1 | 0.00320597 | 0.76053407 | 0.273 | 0.115 | 1 | 0.04908821 | 0.00640165 |
| Emc8 | 0.02171533 | 0.55432191 | 0.531 | 0.383 | 1 | 0.0032066 | 0.22689815 | 0.601 | 0.365 | 1 | 0.02171533 | 0.00640291 |
| Extl1 | 0.13335131 | 0.21102793 | 0.333 | 0.244 | 1 | 0.00320883 | 0.60575255 | 0.277 | 0.115 | 1 | 0.13335131 | 0.00640737 |
| Cdkn2c | 0.49755523 | 0.57139542 | 0.167 | 0.207 | 1 | 0.00321306 | 1.26000035 | 0.261 | 0.115 | 1 | 0.49755523 | 0.00641579 |
| Elp4 | 0.00321428 | 0.6400381 | 0.562 | 0.399 | 1 | 0.39779633 | -0.5588828 | 0.395 | 0.417 | 1 | 0.39779633 | 0.00641822 |
| Upk1b | 0.00946917 | -4.4843586 | 0 | 0.067 | 1 | 0.00322313 | -3.3098553 | 0.013 | 0.073 | 1 | 0.00946917 | 0.00643588 |
| 1700016P03Rik | 0.94287593 | 1.00749454 | 0.052 | 0.057 | 1 | 0.00322316 | -3.6317834 | 0.013 | 0.073 | 1 | 0.94287593 | 0.00643593 |
| Spg21 | 0.03854343 | 0.43748238 | 0.594 | 0.487 | 1 | 0.00323194 | 0.3873485 | 0.689 | 0.5 | 1 | 0.03854343 | 0.00645344 |
| Ccl27a | 0.42175358 | 0.39451766 | 0.135 | 0.104 | 1 | 0.00323588 | 2.01207283 | 0.143 | 0.031 | 1 | 0.42175358 | 0.00646128 |
| Tle1 | 0.08642375 | -0.6440657 | 0.552 | 0.617 | 1 | 0.00323595 | -0.927102 | 0.697 | 0.812 | 1 | 0.08642375 | 0.00646142 |
| Zfp954 | 0.36631236 | 0.30705482 | 0.198 | 0.155 | 1 | 0.00324209 | 1.23071312 | 0.223 | 0.083 | 1 | 0.36631236 | 0.00647367 |
| Farsb | 0.00324278 | 0.55111624 | 0.531 | 0.347 | 1 | 0.00894718 | 0.63464962 | 0.508 | 0.333 | 1 | 0.00894718 | 0.00647505 |
| Gpalpp1 | 0.00325474 | 0.74446013 | 0.417 | 0.249 | 1 | 0.02102629 | 0.53144699 | 0.361 | 0.219 | 1 | 0.02102629 | 0.00649888 |
| Klc3 | 0.06986886 | -1.5160674 | 0.031 | 0.088 | 1 | 0.00325524 | 1.66414953 | 0.189 | 0.062 | 1 | 0.06986886 | 0.00649988 |
| D130020L05Rik | 0.0032607 | 1.68556644 | 0.146 | 0.047 | 1 | 0.54550196 | -0.1578522 | 0.071 | 0.052 | 1 | 0.54550196 | 0.00651078 |
| Cmc4 | 0.01131268 | 0.7503367 | 0.344 | 0.202 | 1 | 0.00326609 | 0.97554696 | 0.277 | 0.125 | 1 | 0.01131268 | 0.00652151 |
| Fra10ac1 | 0.04977307 | 0.66460682 | 0.281 | 0.181 | 1 | 0.00326886 | 0.97554696 | 0.227 | 0.083 | 1 | 0.04977307 | 0.00652703 |
| Kcnk6 | 0.07661478 | 1.30705482 | 0.115 | 0.057 | 1 | 0.00328832 | 0.95693128 | 0.227 | 0.083 | 1 | 0.07661478 | 0.00656584 |
| Dtd1 | 0.85998289 | 0.41233627 | 0.24 | 0.238 | 1 | 0.00328839 | 0.53814164 | 0.403 | 0.219 | 1 | 0.85998289 | 0.00656596 |
| Thbd | 0.18373245 | -0.9925055 | 0.062 | 0.109 | 1 | 0.00328947 | -3.2167459 | 0.013 | 0.073 | 1 | 0.18373245 | 0.00656811 |
| Ccser2 | 0.22784619 | 0.2219382 | 0.75 | 0.699 | 1 | 0.00329318 | 0.35979406 | 0.866 | 0.833 | 1 | 0.22784619 | 0.00657551 |
| Septin5 | 0.0887391 | 0.63309902 | 0.24 | 0.155 | 1 | 0.00329342 | 1.25308093 | 0.223 | 0.083 | 1 | 0.0887391 | 0.00657599 |
| Rasgef1c | 0.5321119 | -0.1989563 | 0.031 | 0.047 | 1 | 0.00330141 | -4.1172102 | 0.008 | 0.062 | 1 | 0.5321119 | 0.00659192 |
| D130043K22Rik | 0.01740615 | -3.8504865 | 0 | 0.057 | 1 | 0.00330507 | -2.4353861 | 0.017 | 0.083 | 1 | 0.01740615 | 0.00659922 |
| Dapp1 | 0.4350647 | 0.13873907 | 0.26 | 0.218 | 1 | 0.00332889 | -1.0439612 | 0.248 | 0.406 | 1 | 0.4350647 | 0.00664669 |
| Urm1 | 0.00787629 | 0.64492446 | 0.438 | 0.275 | 1 | 0.00333414 | 0.42711033 | 0.391 | 0.198 | 1 | 0.00787629 | 0.00665716 |
| Wdr3 | 0.14347727 | 0.34853145 | 0.281 | 0.202 | 1 | 0.00333732 | 0.9739377 | 0.328 | 0.167 | 1 | 0.14347727 | 0.00666351 |
| Cstf3 | 0.00334954 | 0.52502911 | 0.792 | 0.622 | 1 | 0.00582394 | -0.7209239 | 0.803 | 0.885 | 1 | 0.00582394 | 0.00668786 |
| Tpst2 | 0.00335186 | 0.48465375 | 0.635 | 0.425 | 1 | 0.00598483 | 0.25492936 | 0.437 | 0.229 | 1 | 0.00598483 | 0.00669248 |
| Inpp5k | 0.85210819 | -0.407543 | 0.312 | 0.311 | 1 | 0.00335187 | 0.1738239 | 0.471 | 0.25 | 1 | 0.85210819 | 0.0066925 |
| Socs6 | 0.08091947 | -0.6178257 | 0.354 | 0.451 | 1 | 0.00336777 | 0.27510724 | 0.521 | 0.302 | 1 | 0.08091947 | 0.00672419 |
| Idh3g | 0.00337301 | 0.59961683 | 0.646 | 0.492 | 1 | 0.1495642 | 0.23001935 | 0.525 | 0.438 | 1 | 0.1495642 | 0.00673464 |
| Birc6 | 0.00337431 | 0.23864488 | 1 | 0.933 | 1 | 0.0711646 | -0.2862941 | 0.996 | 0.99 | 1 | 0.0711646 | 0.00673723 |
| Psmg4 | 0.04941629 | 0.49528838 | 0.458 | 0.342 | 1 | 0.00338671 | 0.52508549 | 0.42 | 0.229 | 1 | 0.04941629 | 0.00676194 |
| Crip2 | 0.92963118 | 0.22205439 | 0.719 | 0.798 | 1 | 0.00339606 | 0.58491002 | 0.777 | 0.667 | 1 | 0.92963118 | 0.00678059 |
| Yy1 | 0.00339684 | 0.44139106 | 0.875 | 0.72 | 1 | 0.63528336 | -0.157225 | 0.92 | 0.938 | 1 | 0.63528336 | 0.00678213 |
| Ccdc47 | 0.00339698 | 0.35541784 | 0.917 | 0.746 | 1 | 0.02571214 | 0.18156647 | 0.92 | 0.896 | 1 | 0.02571214 | 0.00678243 |
| Ypel3 | 0.3925448 | -0.179504 | 0.812 | 0.829 | 1 | 0.00340139 | 0.35035313 | 0.857 | 0.792 | 1 | 0.3925448 | 0.00679121 |
| Gm43466 | 0.10192277 | 1.59245704 | 0.073 | 0.031 | 1 | 0.00340263 | -3.6317834 | 0.008 | 0.062 | 1 | 0.10192277 | 0.00679368 |
| Acss1 | 0.74827573 | 0.59245704 | 0.021 | 0.016 | 1 | 0.0034063 | -1.2029401 | 0.038 | 0.125 | 1 | 0.74827573 | 0.00680099 |
| Llgl1 | 0.1410058 | 0.20013961 | 0.208 | 0.135 | 1 | 0.00341316 | 0.61906164 | 0.332 | 0.156 | 1 | 0.1410058 | 0.00681467 |
| Chchd6 | 0.0034159 | 0.9200317 | 0.427 | 0.275 | 1 | 0.00784721 | 0.33995838 | 0.29 | 0.135 | 1 | 0.00784721 | 0.00682013 |
| Snx22 | 0.42362827 | 0.56003556 | 0.094 | 0.067 | 1 | 0.00341596 | 1.93807225 | 0.126 | 0.021 | 1 | 0.42362827 | 0.00682026 |
| Zfand6 | 0.00341724 | -0.4337976 | 0.823 | 0.907 | 1 | 0.14405821 | -0.3213509 | 0.924 | 0.958 | 1 | 0.14405821 | 0.0068228 |
| Rmnd5a | 0.93421146 | -0.1745401 | 0.688 | 0.627 | 1 | 0.0034174 | -0.6507563 | 0.777 | 0.875 | 1 | 0.93421146 | 0.00682313 |
| Ythdf2 | 0.00342039 | 0.42253204 | 0.802 | 0.648 | 1 | 0.78217508 | -0.1129001 | 0.769 | 0.792 | 1 | 0.78217508 | 0.00682907 |
| Sec11a | 0.00342414 | 0.39542019 | 0.865 | 0.808 | 1 | 0.01824702 | 0.21023257 | 0.92 | 0.792 | 1 | 0.01824702 | 0.00683655 |
| Slc31a2 | 0.17624154 | 0.80586068 | 0.156 | 0.104 | 1 | 0.00342494 | 1.43264852 | 0.256 | 0.115 | 1 | 0.17624154 | 0.00683815 |
| Acot13 | 0.29908864 | 0.34960051 | 0.333 | 0.28 | 1 | 0.00343317 | 0.45019394 | 0.328 | 0.167 | 1 | 0.29908864 | 0.00685455 |
| Pofut1 | 0.01677419 | 0.87504424 | 0.354 | 0.233 | 1 | 0.00343338 | 0.40940833 | 0.424 | 0.229 | 1 | 0.01677419 | 0.00685497 |
| Mthfsl | 0.00929872 | 0.71471279 | 0.354 | 0.207 | 1 | 0.0034421 | 0.69014474 | 0.311 | 0.146 | 1 | 0.00929872 | 0.00687235 |

| | | | | | | | | | | | | |
|---|---|---|---|---|---|---|---|---|---|---|---|---|
| Lcat | 0.28156155 | 0.39082318 | 0.125 | 0.083 | 1 | 0.00344353 | 1.74903843 | 0.16 | 0.042 | 1 | 0.28156155 | 0.00687521 |
| Elapor1 | 0.00392182 | -3.6929452 | 0.01 | 0.104 | 1 | 0.00344382 | -1.3098553 | 0.038 | 0.125 | 1 | 0.00392182 | 0.00687579 |
| Gm49267 | 0.15729907 | -1.9925055 | 0 | 0.021 | 1 | 0.00345396 | -2.8948178 | 0.008 | 0.062 | 1 | 0.15729907 | 0.00689599 |
| Pex2 | 0.15267098 | -0.577468 | 0.292 | 0.358 | 1 | 0.00345583 | 0.41746016 | 0.433 | 0.24 | 1 | 0.15267098 | 0.00689971 |
| Gm16193 | 0.81500687 | 0.17741954 | 0.219 | 0.207 | 1 | 0.00346721 | 0.33867237 | 0.315 | 0.135 | 1 | 0.81500687 | 0.0069224 |
| Zfp709 | 0.13975474 | 1.14499806 | 0.083 | 0.041 | 1 | 0.00347203 | 3.33400093 | 0.084 | 0 | 1 | 0.13975474 | 0.006932 |
| Milr1 | 0.72574709 | -0.3144336 | 0.01 | 0.016 | 1 | 0.0034741 | -2.8948178 | 0.013 | 0.073 | 1 | 0.72574709 | 0.00693613 |
| Ppp3cb | 0.00349161 | 0.52206771 | 0.708 | 0.575 | 1 | 0.54423208 | -0.2915544 | 0.803 | 0.823 | 1 | 0.54423208 | 0.00697103 |
| Txndc9 | 0.03091083 | 0.36363835 | 0.719 | 0.585 | 1 | 0.00349417 | 0.33726371 | 0.744 | 0.552 | 1 | 0.03091083 | 0.00697614 |
| Psmd3 | 0.00350382 | 0.53455865 | 0.698 | 0.513 | 1 | 0.23034636 | 0.13522106 | 0.697 | 0.635 | 1 | 0.23034636 | 0.00699536 |
| Map4k4 | 0.00351589 | -0.4180788 | 0.948 | 0.979 | 1 | 0.05364719 | -0.3721785 | 0.987 | 1 | 1 | 0.05364719 | 0.00701941 |
| Shisa5 | 0.10192315 | -0.2392011 | 0.823 | 0.839 | 1 | 0.00353156 | 0.44046871 | 0.761 | 0.646 | 1 | 0.10192315 | 0.00705064 |
| Cep63 | 0.48543428 | 0.19556688 | 0.312 | 0.275 | 1 | 0.00353228 | 0.99270751 | 0.357 | 0.198 | 1 | 0.48543428 | 0.00705208 |
| Glt28d2 | 0.01930839 | -1.0596197 | 0.156 | 0.275 | 1 | 0.00353298 | 0.52664601 | 0.408 | 0.229 | 1 | 0.01930839 | 0.00705347 |
| A730049H05Rik | 0.61182672 | -0.577468 | 0.021 | 0.031 | 1 | 0.0035348 | -2.8948178 | 0.008 | 0.062 | 1 | 0.61182672 | 0.00705711 |
| Pea15a | 0.99502694 | -0.1224641 | 0.677 | 0.596 | 1 | 0.00353651 | 0.27213567 | 0.752 | 0.51 | 1 | 0.99502694 | 0.00706051 |
| Gtf2h5 | 0.01984692 | 0.37466119 | 0.875 | 0.767 | 1 | 0.00354086 | 0.3575694 | 0.782 | 0.698 | 1 | 0.01984692 | 0.00706919 |
| Armcx5 | 0.59178248 | -0.17693 | 0.281 | 0.244 | 1 | 0.003542 | 0.87601128 | 0.303 | 0.146 | 1 | 0.59178248 | 0.00707146 |
| Gm13375 | 0.88244184 | 0.17350449 | 0.188 | 0.181 | 1 | 0.00354375 | 1.59703533 | 0.16 | 0.042 | 1 | 0.88244184 | 0.00707494 |
| Psen2 | 0.01809563 | -0.6039402 | 0.25 | 0.394 | 1 | 0.00354509 | 0.62024272 | 0.403 | 0.219 | 1 | 0.01809563 | 0.00707761 |
| Rdh14 | 0.37845485 | -0.4372903 | 0.167 | 0.207 | 1 | 0.00354605 | 0.72576865 | 0.29 | 0.125 | 1 | 0.37845485 | 0.00707952 |
| Dusp14 | 0.01348933 | -0.5909512 | 0.292 | 0.44 | 1 | 0.00354861 | 0.14957636 | 0.307 | 0.135 | 1 | 0.01348933 | 0.00708463 |
| Fmo3 | 0.15837881 | 2.00749454 | 0.01 | 0 | 1 | 0.00356905 | -2.3098553 | 0.013 | 0.073 | 1 | 0.15837881 | 0.00712536 |
| Gramd3 | 0.92434614 | 0.16389609 | 0.552 | 0.554 | 1 | 0.0035715 | -0.8658179 | 0.559 | 0.719 | 1 | 0.92434614 | 0.00713024 |
| Mrpl55 | 0.06807412 | 0.52636785 | 0.396 | 0.29 | 1 | 0.00357649 | 0.96316323 | 0.303 | 0.146 | 1 | 0.06807412 | 0.00714018 |
| Xxylt1 | 0.29779143 | 0.20013961 | 0.25 | 0.192 | 1 | 0.0035806 | 0.93115284 | 0.319 | 0.156 | 1 | 0.29779143 | 0.00714838 |
| Pigf | 0.1409881 | 0.56692195 | 0.333 | 0.264 | 1 | 0.00358783 | 0.53814164 | 0.37 | 0.188 | 1 | 0.1409881 | 0.00716278 |
| Commd5 | 0.24555928 | 0.45110119 | 0.146 | 0.098 | 1 | 0.0035893 | 2.39058446 | 0.105 | 0.01 | 1 | 0.24555928 | 0.00716572 |
| Tnfrsf21 | 0.00359528 | -0.9306662 | 0.458 | 0.585 | 1 | 0.09275069 | 0.11648489 | 0.811 | 0.688 | 1 | 0.09275069 | 0.00717763 |
| Ufm1 | 0.0083059 | 0.36562964 | 0.74 | 0.549 | 1 | 0.00359681 | 0.38238397 | 0.693 | 0.521 | 1 | 0.0083059 | 0.00718068 |
| Rtf1 | 0.00360402 | 0.48838458 | 0.76 | 0.731 | 1 | 0.0320691 | 0.18028369 | 0.891 | 0.896 | 1 | 0.0320691 | 0.00719506 |
| Thada | 0.26038122 | 0.19810047 | 0.5 | 0.42 | 1 | 0.00361315 | -0.8073549 | 0.496 | 0.604 | 1 | 0.26038122 | 0.00721324 |
| Mrpl44 | 0.20478419 | 0.60695661 | 0.208 | 0.15 | 1 | 0.00361661 | 1.27510724 | 0.193 | 0.062 | 1 | 0.20478419 | 0.00722014 |
| Coq10a | 0.12760127 | 0.56388789 | 0.229 | 0.155 | 1 | 0.00363807 | 0.7944814 | 0.273 | 0.115 | 1 | 0.12760127 | 0.0072629 |
| Cgn | 0.47150075 | 2.00749454 | 0.021 | 0.01 | 1 | 0.0036403 | -1.07539 | 0.063 | 0.167 | 1 | 0.47150075 | 0.00726736 |
| Itga9 | 0.00364837 | 0.84522311 | 0.51 | 0.332 | 1 | 0.02352036 | -0.9104994 | 0.487 | 0.51 | 1 | 0.02352036 | 0.00728342 |
| Cep170 | 0.00365361 | 0.59484758 | 0.646 | 0.466 | 1 | 0.87573545 | -0.2757474 | 0.676 | 0.635 | 1 | 0.87573545 | 0.00729388 |
| Bfar | 0.13637744 | 0.35896691 | 0.615 | 0.549 | 1 | 0.00367212 | 0.38046024 | 0.634 | 0.385 | 1 | 0.13637744 | 0.00733077 |
| Vps51 | 0.05802276 | 0.72509481 | 0.271 | 0.171 | 1 | 0.00367413 | 1.23763253 | 0.244 | 0.104 | 1 | 0.05802276 | 0.00733476 |
| Rreb1 | 0.30802928 | 0.15676007 | 0.49 | 0.44 | 1 | 0.00368162 | -0.7157328 | 0.441 | 0.635 | 1 | 0.30802928 | 0.00734969 |
| Nus1 | 0.00368171 | 0.68068422 | 0.615 | 0.477 | 1 | 0.01909431 | 0.28878218 | 0.714 | 0.573 | 1 | 0.01909431 | 0.00734987 |
| Gab2 | 0.00369764 | -1.4602844 | 0.167 | 0.337 | 1 | 0.00668191 | -1.0645229 | 0.483 | 0.677 | 1 | 0.00668191 | 0.00738161 |
| Col25a1 | 0.0594374 | -0.3430027 | 0.156 | 0.254 | 1 | 0.00369872 | -0.9162893 | 0.315 | 0.49 | 1 | 0.0594374 | 0.00738376 |
| Ube2v1 | 0.0192614 | 0.46086015 | 0.604 | 0.435 | 1 | 0.00371094 | 0.39198854 | 0.647 | 0.49 | 1 | 0.0192614 | 0.00740811 |
| Tgfa | 0.00671627 | -1.6705774 | 0.375 | 0.503 | 1 | 0.00371793 | 0.33995838 | 0.433 | 0.229 | 1 | 0.00671627 | 0.00742204 |
| Ubtd2 | 0.00372178 | 0.25024467 | 0.854 | 0.684 | 1 | 0.59193641 | -0.5835033 | 0.853 | 0.75 | 1 | 0.59193641 | 0.0074297 |
| Taf5l | 0.0353315 | 0.56937342 | 0.271 | 0.161 | 1 | 0.00372818 | 0.66317769 | 0.332 | 0.167 | 1 | 0.0353315 | 0.00744246 |
| Alg10b | 0.22555588 | -0.3659639 | 0.156 | 0.212 | 1 | 0.00373458 | 0.86006974 | 0.256 | 0.104 | 1 | 0.22555588 | 0.00745521 |
| Hapln3 | 0.45456728 | 0.40483003 | 0.219 | 0.187 | 1 | 0.00373698 | 1.23446525 | 0.218 | 0.083 | 1 | 0.45456728 | 0.00746 |
| Akt1s1 | 0.03472947 | 0.39828449 | 0.458 | 0.321 | 1 | 0.00375075 | 0.34965419 | 0.454 | 0.26 | 1 | 0.03472947 | 0.00748743 |
| Utp23 | 0.01134211 | 1.0899567 | 0.24 | 0.124 | 1 | 0.00375167 | 1.11047654 | 0.265 | 0.115 | 1 | 0.01134211 | 0.00748926 |
| Lig3 | 0.00376273 | 0.86999101 | 0.375 | 0.218 | 1 | 0.01689265 | 0.56883844 | 0.357 | 0.219 | 1 | 0.01689265 | 0.0075113 |
| Mul1 | 0.696452 | 0.16503581 | 0.25 | 0.233 | 1 | 0.00377474 | 0.99392549 | 0.277 | 0.125 | 1 | 0.696452 | 0.00753523 |
| Podxl2 | 0.00378649 | -2.6247737 | 0.042 | 0.155 | 1 | 0.67167764 | 0.18199783 | 0.164 | 0.146 | 1 | 0.67167764 | 0.00755864 |
| Fut2 | 0.00379222 | -3.3848229 | 0 | 0.083 | 1 | 0.34704016 | -1.3098553 | 0.008 | 0.021 | 1 | 0.34704016 | 0.00757006 |
| Gulo | 0.00379612 | -4.0368996 | 0 | 0.083 | 1 | 0.04098703 | -0.6317834 | 0.008 | 0.042 | 1 | 0.04098703 | 0.00757784 |
| 1700109H08Rik | 0.38362277 | 0.4343092 | 0.292 | 0.249 | 1 | 0.00379835 | 0.32904605 | 0.34 | 0.156 | 1 | 0.38362277 | 0.00758228 |
| Azgp1 | 0.0037987 | -3.7998604 | 0 | 0.083 | 1 | 0.51194506 | 0.27510724 | 0.021 | 0.01 | 1 | 0.51194506 | 0.00758297 |
| Cspp1 | 0.01168765 | 0.53849805 | 0.604 | 0.446 | 1 | 0.00379899 | -0.8420763 | 0.626 | 0.781 | 1 | 0.01168765 | 0.00758354 |
| Abca5 | 0.21064093 | 0.23611891 | 0.375 | 0.301 | 1 | 0.00380402 | 0.6588938 | 0.357 | 0.177 | 1 | 0.21064093 | 0.00759357 |
| Ints3 | 0.00380678 | 0.93154568 | 0.292 | 0.145 | 1 | 0.02080414 | 0.50511284 | 0.29 | 0.156 | 1 | 0.02080414 | 0.00759907 |
| Itsn1 | 0.17683472 | -0.7834969 | 0.417 | 0.477 | 1 | 0.00381149 | -0.8180022 | 0.71 | 0.74 | 1 | 0.17683472 | 0.00760846 |
| Rhog | 0.39666251 | -0.3348977 | 0.229 | 0.269 | 1 | 0.00381507 | 0.77356075 | 0.307 | 0.146 | 1 | 0.39666251 | 0.00761558 |
| Ttl | 0.00384399 | 0.78510212 | 0.365 | 0.207 | 1 | 0.00584867 | 0.3794439 | 0.374 | 0.198 | 1 | 0.00584867 | 0.0076732 |
| Ncapg | 0.00384535 | 3.32942263 | 0.073 | 0.01 | 1 | 0.50587544 | -1.8948178 | 0.004 | 0.01 | 1 | 0.50587544 | 0.00767591 |
| Ackr3 | 0.006324 | 0.29676818 | 0.688 | 0.477 | 1 | 0.00384733 | 0.4278315 | 0.559 | 0.375 | 1 | 0.006324 | 0.00767985 |
| Tsfm | 0.03368983 | 0.70793425 | 0.302 | 0.192 | 1 | 0.00385746 | 0.91253716 | 0.324 | 0.167 | 1 | 0.03368983 | 0.00770003 |
| Relch | 0.94829402 | 0.12297175 | 0.542 | 0.575 | 1 | 0.00386109 | -0.7601682 | 0.643 | 0.76 | 1 | 0.94829402 | 0.00770727 |
| Slc4a2 | 0.11880709 | 0.14499806 | 0.385 | 0.269 | 1 | 0.00386318 | 0.54812573 | 0.471 | 0.271 | 1 | 0.11880709 | 0.00771143 |
| Arl2 | 0.05908702 | 0.40449811 | 0.604 | 0.466 | 1 | 0.00386508 | 0.55750697 | 0.387 | 0.208 | 1 | 0.05908702 | 0.00771521 |
| Tent5a | 0.20512318 | -0.7220807 | 0.604 | 0.637 | 1 | 0.00386713 | 0.98560062 | 0.454 | 0.302 | 1 | 0.20512318 | 0.0077193 |
| Ndst2 | 0.13794802 | 0.78510212 | 0.188 | 0.124 | 1 | 0.00387134 | 0.99595317 | 0.29 | 0.135 | 1 | 0.13794802 | 0.00772769 |
| Obp1b | 0.0038726 | 1.05812061 | 0.26 | 0.124 | 1 | 0.47661479 | 0.61071027 | 0.214 | 0.177 | 1 | 0.47661479 | 0.00773021 |
| Ap1g1 | 0.00387567 | 0.27767101 | 0.927 | 0.824 | 1 | 0.70564113 | -0.1038964 | 0.941 | 0.917 | 1 | 0.70564113 | 0.00773631 |
| Zbed6 | 0.00387792 | 0.34354374 | 0.781 | 0.591 | 1 | 0.10838353 | -0.4455933 | 0.803 | 0.823 | 1 | 0.10838353 | 0.00774079 |
| H4c4 | 0.01444123 | 1.16503581 | 0.375 | 0.244 | 1 | 0.00388898 | 0.69014474 | 0.319 | 0.156 | 1 | 0.01444123 | 0.00776285 |
| Lag3 | 0.46544015 | -1.0628948 | 0.042 | 0.062 | 1 | 0.00390073 | -2.0356803 | 0.084 | 0.188 | 1 | 0.46544015 | 0.00778624 |
| Nrgn | 0.00390218 | 1.00749454 | 0.135 | 0.041 | 1 | 0.58036856 | 0.10518224 | 0.029 | 0.042 | 1 | 0.58036856 | 0.00778913 |
| Bub1b | 0.00390557 | 3.12297175 | 0.073 | 0.01 | 1 | 0.09634589 | -1.3098553 | 0.013 | 0.042 | 1 | 0.09634589 | 0.00779588 |
| Lpcat1 | 0.00392711 | 0.50792353 | 0.479 | 0.29 | 1 | 0.00652482 | 0.68036372 | 0.378 | 0.219 | 1 | 0.00652482 | 0.0078388 |
| Asb13 | 0.87550013 | 0.1843723 | 0.094 | 0.088 | 1 | 0.00393279 | 0.25404562 | 0.227 | 0.083 | 1 | 0.87550013 | 0.00785011 |
| Sntb1 | 0.00394251 | -4.6218621 | 0.01 | 0.104 | 1 | 0.19198508 | -1.3098553 | 0.055 | 0.094 | 1 | 0.19198508 | 0.00786948 |
| Ndufaf6 | 0.3418764 | -0.407543 | 0.104 | 0.145 | 1 | 0.00394902 | 1.52303475 | 0.16 | 0.042 | 1 | 0.3418764 | 0.00788245 |
| Ints7 | 0.00395147 | 1.04448874 | 0.406 | 0.264 | 1 | 0.28402187 | -0.2055186 | 0.345 | 0.25 | 1 | 0.28402187 | 0.00788733 |
| Steap2 | 0.00395402 | -1.1996634 | 0.458 | 0.575 | 1 | 0.10650726 | -0.2268659 | 0.391 | 0.26 | 1 | 0.10650726 | 0.0078924 |
| Tprkb | 0.47648466 | 0.22988696 | 0.333 | 0.295 | 1 | 0.00396197 | 0.94648449 | 0.349 | 0.188 | 1 | 0.47648466 | 0.00790824 |
| Terf2 | 0.02712356 | 0.60695661 | 0.469 | 0.342 | 1 | 0.00396306 | 0.42082407 | 0.555 | 0.344 | 1 | 0.02712356 | 0.00791042 |
| Adgrg1 | 0.00396625 | 3.12297175 | 0.073 | 0.01 | 1 | 0.14430862 | 1.01207283 | 0.101 | 0.052 | 1 | 0.14430862 | 0.00791676 |
| Hinfp | 0.17538442 | 0.34191358 | 0.229 | 0.161 | 1 | 0.0039739 | 0.74077081 | 0.265 | 0.115 | 1 | 0.17538442 | 0.007932 |
| Rasgef1b | 0.00397445 | -1.9665103 | 0.094 | 0.228 | 1 | 0.7238611 | -0.4473588 | 0.063 | 0.052 | 1 | 0.7238611 | 0.00793311 |

| | | | | | | | | | | | | |
|---|---|---|---|---|---|---|---|---|---|---|---|---|
| Rnf146 | 0.07626556 | 0.39451766 | 0.594 | 0.492 | 1 | 0.00397756 | 0.37217466 | 0.681 | 0.479 | 1 | 0.07626556 | 0.00793929 |
| Ier3 | 0.00397803 | -0.5635372 | 0.708 | 0.834 | 1 | 0.01310353 | -0.8471411 | 0.634 | 0.792 | 1 | 0.01310353 | 0.00794024 |
| Dixdc1 | 0.003992 | -0.7029988 | 0.625 | 0.731 | 1 | 0.35944022 | -0.1669564 | 0.777 | 0.594 | 1 | 0.35944022 | 0.00796806 |
| Gm19510 | 0.00399986 | -3.3848229 | 0.01 | 0.104 | 1 | 0.51188611 | -0.7248928 | 0.004 | 0.01 | 1 | 0.51188611 | 0.00798371 |
| Kbtbd7 | 0.11295954 | 0.77302928 | 0.156 | 0.093 | 1 | 0.00400771 | 1.10518224 | 0.16 | 0.042 | 1 | 0.11295954 | 0.00799936 |
| Fhl2 | 0.00401266 | -0.8682114 | 0.323 | 0.497 | 1 | 0.42922035 | -1.0229741 | 0.303 | 0.323 | 1 | 0.42922035 | 0.00800923 |
| Wdr89 | 0.31213369 | 0.27289596 | 0.667 | 0.632 | 1 | 0.00401349 | 0.62777886 | 0.664 | 0.583 | 1 | 0.31213369 | 0.00801088 |
| Ppp2r3d | 0.06229616 | 0.35263002 | 0.417 | 0.29 | 1 | 0.00401655 | 0.90737545 | 0.303 | 0.146 | 1 | 0.06229616 | 0.00801696 |
| Ddx56 | 0.00402281 | 0.78510212 | 0.385 | 0.223 | 1 | 0.03923418 | 0.40896298 | 0.256 | 0.146 | 1 | 0.03923418 | 0.00802943 |
| Tmem128 | 0.28250312 | 0.27052894 | 0.594 | 0.565 | 1 | 0.00402331 | 0.50066694 | 0.613 | 0.417 | 1 | 0.28250312 | 0.00803044 |
| Adipor2 | 0.00402914 | 0.41188479 | 0.698 | 0.528 | 1 | 0.00820476 | 0.36323815 | 0.731 | 0.677 | 1 | 0.00820476 | 0.00804204 |
| Wwp2 | 0.05064923 | 0.22599778 | 0.969 | 0.912 | 1 | 0.00403163 | 0.1370626 | 0.987 | 0.917 | 1 | 0.05064923 | 0.00804701 |
| 4921511C10Rik | 0.00403343 | 2.42253204 | 0.073 | 0.01 | 1 | 0.32304157 | -0.2103196 | 0.059 | 0.031 | 1 | 0.32304157 | 0.00805059 |
| Pank3 | 0.08662139 | 0.28851469 | 0.656 | 0.528 | 1 | 0.00403603 | 0.45407738 | 0.714 | 0.542 | 1 | 0.08662139 | 0.00805577 |
| Tmem237 | 0.58785675 | 0.19094068 | 0.438 | 0.42 | 1 | 0.00403677 | 0.27510724 | 0.429 | 0.229 | 1 | 0.58785675 | 0.00805725 |
| Adra2a | 0.00403734 | -2.6929452 | 0.01 | 0.104 | 1 | 0.44094848 | 0.74077081 | 0.063 | 0.042 | 1 | 0.44094848 | 0.00805838 |
| Pdgfra | 0.0040428 | -1.1517041 | 0.271 | 0.43 | 1 | 0.2997915 | -0.9826905 | 0.374 | 0.385 | 1 | 0.2997915 | 0.00806926 |
| Nsd3 | 0.87659025 | -0.2561714 | 0.979 | 0.927 | 1 | 0.00404311 | -0.4559475 | 0.996 | 0.979 | 1 | 0.87659025 | 0.00806988 |
| Mospd1 | 0.07736962 | 0.68556644 | 0.375 | 0.28 | 1 | 0.00404936 | 0.64231621 | 0.416 | 0.229 | 1 | 0.07736962 | 0.00808233 |
| 2310057M21Rik | 0.00405657 | 0.74446013 | 0.385 | 0.212 | 1 | 0.00696938 | 0.48022167 | 0.261 | 0.115 | 1 | 0.00696938 | 0.00809668 |
| Atp11a | 0.10931742 | 0.12778877 | 0.667 | 0.544 | 1 | 0.00406061 | -0.6740116 | 0.639 | 0.812 | 1 | 0.10931742 | 0.00810474 |
| Lct | 0.00406425 | 2.46692616 | 0.073 | 0.01 | 1 | 0.742831 | -1.4797803 | 0.025 | 0.031 | 1 | 0.742831 | 0.00811199 |
| Thap2 | 0.39243497 | 0.16090445 | 0.573 | 0.518 | 1 | 0.00407786 | 0.39484648 | 0.613 | 0.385 | 1 | 0.39243497 | 0.00813909 |
| Gtf3c4 | 0.01404869 | 0.59245704 | 0.333 | 0.197 | 1 | 0.00408129 | 0.52664601 | 0.441 | 0.26 | 1 | 0.01404869 | 0.00814592 |
| Lrp5 | 0.02690205 | 0.34853145 | 0.385 | 0.254 | 1 | 0.00408236 | 0.48888653 | 0.433 | 0.24 | 1 | 0.02690205 | 0.00814805 |
| Nup54 | 0.39227538 | 0.17637145 | 0.521 | 0.451 | 1 | 0.00410059 | 0.4290586 | 0.597 | 0.427 | 1 | 0.39227538 | 0.00818437 |
| Camk2n2 | 0.00410821 | 1.49292136 | 0.135 | 0.041 | 1 | 0.1457911 | 0.13071733 | 0.076 | 0.031 | 1 | 0.1457911 | 0.00819954 |
| Gm56915 | 0.00410821 | 1.49292136 | 0.135 | 0.041 | 1 | 0.00570903 | 3.08246216 | 0.076 | 0 | 1 | 0.00570903 | 0.00819954 |
| Stat6 | 0.52765898 | 0.13302542 | 0.375 | 0.347 | 1 | 0.00412453 | 0.26908345 | 0.58 | 0.365 | 1 | 0.52765898 | 0.00823205 |
| Mansc4 | 0.00412784 | 2.27052894 | 0.073 | 0.01 | 1 | 0.517206 | 0.27510724 | 0.021 | 0.01 | 1 | 0.517206 | 0.00823863 |
| Gm15675 | 0.00412856 | 2.27052894 | 0.073 | 0.01 | 1 | 0.14544666 | -3.3098553 | 0.004 | 0.021 | 1 | 0.14544666 | 0.00824007 |
| Ubxn8 | 0.41781617 | 0.22727574 | 0.375 | 0.332 | 1 | 0.0041366 | 0.43122644 | 0.399 | 0.208 | 1 | 0.41781617 | 0.00825608 |
| Zfyve16 | 0.58052139 | 0.19486467 | 0.469 | 0.42 | 1 | 0.00415663 | 0.55468132 | 0.462 | 0.25 | 1 | 0.58052139 | 0.00829599 |
| Fam120aos | 0.42704096 | 0.30952307 | 0.354 | 0.311 | 1 | 0.00415713 | 0.5451964 | 0.399 | 0.219 | 1 | 0.42704096 | 0.00829698 |
| Snx27 | 0.00416443 | 0.55579857 | 0.698 | 0.508 | 1 | 0.90929683 | -0.1320681 | 0.714 | 0.708 | 1 | 0.90929683 | 0.00831152 |
| Ap4e1 | 0.00416476 | 0.79599043 | 0.333 | 0.176 | 1 | 0.65180509 | -0.2647674 | 0.349 | 0.292 | 1 | 0.65180509 | 0.00831218 |
| Mien1 | 0.43671421 | -0.4339407 | 0.625 | 0.596 | 1 | 0.00416657 | 0.48578624 | 0.521 | 0.354 | 1 | 0.43671421 | 0.00831578 |
| Gm56829 | 0.09713663 | 0.93349396 | 0.146 | 0.083 | 1 | 0.0041762 | 1.35310975 | 0.143 | 0.031 | 1 | 0.09713663 | 0.00833496 |
| Syt2 | 0.06803991 | -2.4224933 | 0.052 | 0.119 | 1 | 0.00417967 | -2.150377 | 0.067 | 0.167 | 1 | 0.06803991 | 0.00834186 |
| Gm11508 | 0.00635907 | 0.83346514 | 0.26 | 0.13 | 1 | 0.00418078 | 0.12871775 | 0.282 | 0.125 | 1 | 0.00635907 | 0.00834408 |
| 1810009A15Rik | 0.10566405 | 0.46317402 | 0.49 | 0.409 | 1 | 0.00418217 | 0.67564517 | 0.332 | 0.167 | 1 | 0.10566405 | 0.00834685 |
| Slc25a44 | 0.23510797 | -0.577468 | 0.188 | 0.244 | 1 | 0.00418978 | 0.52303475 | 0.361 | 0.188 | 1 | 0.23510797 | 0.008362 |
| Rnaseh2c | 0.14443482 | 0.33289482 | 0.396 | 0.301 | 1 | 0.00419878 | 0.99680608 | 0.319 | 0.167 | 1 | 0.14443482 | 0.00837993 |
| Stip1 | 0.00420352 | 0.48838458 | 0.656 | 0.466 | 1 | 0.01658485 | 0.3889752 | 0.752 | 0.75 | 1 | 0.01658485 | 0.00838938 |
| Ercc6l2 | 0.00593724 | 0.73704266 | 0.521 | 0.347 | 1 | 0.00420693 | 0.61260814 | 0.361 | 0.188 | 1 | 0.00593724 | 0.00839617 |
| D430041D05Rik | 0.069103 | -3.1010299 | 0.031 | 0.088 | 1 | 0.00421098 | -3.5468945 | 0.029 | 0.104 | 1 | 0.069103 | 0.00840423 |
| Ube3a | 0.00421317 | 0.32213014 | 0.917 | 0.917 | 1 | 0.84774178 | -0.1006345 | 0.966 | 0.969 | 1 | 0.84774178 | 0.00840858 |
| Mrpl45 | 0.03353814 | 0.59245704 | 0.354 | 0.228 | 1 | 0.0042158 | 0.71399148 | 0.345 | 0.177 | 1 | 0.03353814 | 0.00841382 |
| Slc10a6 | 0.0042235 | -1.0148733 | 0.073 | 0.207 | 1 | 0.75297348 | -0.9472852 | 0.025 | 0.031 | 1 | 0.75297348 | 0.00842915 |
| Cdh20 | 0.00422355 | -3.4843586 | 0.01 | 0.104 | 1 | 0.01591403 | -0.2759079 | 0.025 | 0.083 | 1 | 0.01591403 | 0.00842925 |
| Aimp2 | 0.00422571 | 0.89201732 | 0.323 | 0.171 | 1 | 0.00547868 | 0.72117163 | 0.311 | 0.156 | 1 | 0.00547868 | 0.00843357 |
| Gm12743 | 0.26124804 | 0.78510212 | 0.125 | 0.083 | 1 | 0.00422887 | 1.98292649 | 0.139 | 0.031 | 1 | 0.26124804 | 0.00843986 |
| Pcbd2 | 0.00424035 | 0.5049942 | 0.542 | 0.352 | 1 | 0.00961717 | 0.52664601 | 0.374 | 0.208 | 1 | 0.00961717 | 0.00846272 |
| Tmem67 | 0.00952011 | 1.06194232 | 0.25 | 0.13 | 1 | 0.00424525 | 1.33400093 | 0.172 | 0.052 | 1 | 0.00952011 | 0.00847248 |
| Dnase1l1 | 0.00425193 | 1.42253204 | 0.271 | 0.14 | 1 | 0.01738805 | 0.42711033 | 0.239 | 0.115 | 1 | 0.01738805 | 0.00848577 |
| Gm31834 | 0.00425672 | 2.00749454 | 0.073 | 0.01 | 1 | 0.23592811 | 1.27510724 | 0.034 | 0.01 | 1 | 0.23592811 | 0.00849533 |
| Cpeb2 | 0.15576775 | 0.28105961 | 0.312 | 0.233 | 1 | 0.00427022 | -0.4154094 | 0.483 | 0.677 | 1 | 0.15576775 | 0.0085222 |
| Ssh3 | 0.06018511 | 0.78510212 | 0.177 | 0.098 | 1 | 0.00428526 | 0.88278982 | 0.218 | 0.083 | 1 | 0.06018511 | 0.00855216 |
| Chka | 0.00620359 | -0.5197525 | 0.771 | 0.865 | 1 | 0.00429484 | -0.4770848 | 0.912 | 0.927 | 1 | 0.00620359 | 0.00857123 |
| Mtbp | 0.00429715 | 1.15949763 | 0.177 | 0.067 | 1 | 0.08717907 | 1.61614416 | 0.097 | 0.042 | 1 | 0.08717907 | 0.00857584 |
| Gcnt1 | 0.00429761 | 0.91400791 | 0.542 | 0.389 | 1 | 0.0072031 | 0.15311671 | 0.366 | 0.188 | 1 | 0.0072031 | 0.00857675 |
| Usp48 | 0.57471076 | -0.2671278 | 0.552 | 0.492 | 1 | 0.00429768 | -0.4820362 | 0.626 | 0.792 | 1 | 0.57471076 | 0.00857688 |
| Naa12 | 0.00430026 | 0.43541333 | 0.771 | 0.632 | 1 | 0.03962835 | 0.16640098 | 0.773 | 0.656 | 1 | 0.03962835 | 0.00858203 |
| Cnot7 | 0.0043078 | 0.57762962 | 0.656 | 0.528 | 1 | 0.00983784 | 0.34012703 | 0.693 | 0.542 | 1 | 0.00983784 | 0.00859705 |
| Gstz1 | 0.82788177 | -0.1162408 | 0.479 | 0.451 | 1 | 0.00431135 | 0.60768258 | 0.412 | 0.229 | 1 | 0.82788177 | 0.00860411 |
| Zfp771 | 0.08149761 | 0.39981196 | 0.469 | 0.358 | 1 | 0.00432267 | 0.78259099 | 0.441 | 0.271 | 1 | 0.08149761 | 0.00862666 |
| Alg1 | 0.14194994 | 0.49934763 | 0.354 | 0.275 | 1 | 0.0043305 | 0.56461386 | 0.336 | 0.167 | 1 | 0.14194994 | 0.00864224 |
| Adamts6 | 0.00434446 | 0.43558819 | 0.271 | 0.13 | 1 | 0.40729644 | -0.7525239 | 0.231 | 0.26 | 1 | 0.40729644 | 0.00867004 |
| Pphln1 | 0.04717193 | 0.38823886 | 0.625 | 0.487 | 1 | 0.00434556 | 0.29895398 | 0.66 | 0.417 | 1 | 0.04717193 | 0.00867223 |
| Tcfl5 | 0.01911758 | 0.58603077 | 0.385 | 0.244 | 1 | 0.0043473 | 0.65361886 | 0.357 | 0.188 | 1 | 0.01911758 | 0.0086757 |
| Gss | 0.00434912 | 0.59245704 | 0.375 | 0.212 | 1 | 0.04549179 | 0.28552385 | 0.37 | 0.24 | 1 | 0.04549179 | 0.00867933 |
| Sart1 | 0.12509195 | 0.15663798 | 0.438 | 0.321 | 1 | 0.00435243 | 0.45421556 | 0.55 | 0.354 | 1 | 0.12509195 | 0.00868591 |
| Rad1 | 0.00436027 | 1.59245704 | 0.167 | 0.062 | 1 | 0.03073777 | 0.45567948 | 0.122 | 0.042 | 1 | 0.03073777 | 0.00870154 |
| Hsph1 | 0.00436373 | 0.57477476 | 0.812 | 0.699 | 1 | 0.39052425 | 0.21154211 | 0.79 | 0.885 | 1 | 0.39052425 | 0.00870842 |
| Noc4l | 0.10768637 | 0.50053455 | 0.188 | 0.114 | 1 | 0.00437158 | 1.30157945 | 0.202 | 0.073 | 1 | 0.10768637 | 0.00872405 |
| Pde5a | 0.5369835 | -1.0368996 | 0.083 | 0.104 | 1 | 0.00438729 | -3.0545982 | 0.08 | 0.177 | 1 | 0.5369835 | 0.00875533 |
| Rsbn1l | 0.09397806 | 0.41862758 | 0.531 | 0.456 | 1 | 0.0043986 | -0.6590048 | 0.664 | 0.802 | 1 | 0.09397806 | 0.00877786 |
| Phrf1 | 0.00441078 | 0.27052894 | 0.688 | 0.466 | 1 | 0.73912615 | -0.2063177 | 0.761 | 0.719 | 1 | 0.73912615 | 0.0088021 |
| Tfr2 | 0.00441438 | 3.59245704 | 0.042 | 0 | 1 | 0.50887612 | -1.3098553 | 0.004 | 0.01 | 1 | 0.50887612 | 0.00880927 |
| Gm10827 | 0.00441438 | 3.59245704 | 0.042 | 0 | 1 | 0.11689951 | -2.8948178 | 0 | 0.01 | 1 | 0.11689951 | 0.00880927 |
| Gm15764 | 0.00441438 | 3.59245704 | 0.042 | 0 | 1 | 0.75977578 | -0.5025003 | 0.025 | 0.031 | 1 | 0.75977578 | 0.00880927 |
| 8030442B05Rik | 0.00441486 | 4.00749454 | 0.042 | 0 | 1 | 0.06982867 | 2.39058446 | 0.034 | 0 | 1 | 0.06982867 | 0.00881022 |
| Ttpa | 0.00441486 | 3.32942263 | 0.042 | 0 | 1 | 0.11215403 | 1.49749966 | 0.046 | 0.01 | 1 | 0.11215403 | 0.00881022 |
| Ttc41 | 0.00441486 | 3.32942263 | 0.042 | 0 | 1 | 0.14876424 | 0.80562195 | 0.042 | 0.01 | 1 | 0.14876424 | 0.00881022 |
| Gm30655 | 0.00441486 | 3.32942263 | 0.042 | 0 | 1 | 0.85710544 | -1.7248928 | 0.008 | 0.01 | 1 | 0.85710544 | 0.00881022 |
| Map3k21 | 0.00441753 | 2.59245704 | 0.094 | 0.021 | 1 | 0.18744336 | -0.8073549 | 0.046 | 0.083 | 1 | 0.18744336 | 0.00881554 |

| | | | | | | | | | | | | |
|---|---|---|---|---|---|---|---|---|---|---|---|---|
| Rorc | 0.86173844 | 0.11440974 | 0.062 | 0.057 | 1 | 0.00442732 | 1.33400093 | 0.16 | 0.042 | 1 | 0.86173844 | 0.00883504 |
| C330024C12Rik | 0.25824304 | 0.78510212 | 0.052 | 0.026 | 1 | 0.00445029 | 3.14957636 | 0.08 | 0 | 1 | 0.25824304 | 0.00888077 |
| Hmcn1 | 0.00879301 | -2.4719748 | 0.062 | 0.171 | 1 | 0.00446971 | -3.0517871 | 0.063 | 0.156 | 1 | 0.00879301 | 0.00891944 |
| Naprt | 0.23923812 | 0.59245704 | 0.115 | 0.073 | 1 | 0.0044848 | 2.73453886 | 0.101 | 0.01 | 1 | 0.23923812 | 0.00894949 |
| Srsf10 | 0.00449088 | 0.33863806 | 0.927 | 0.902 | 1 | 0.03863434 | 0.13904569 | 0.979 | 0.938 | 1 | 0.03863434 | 0.00896159 |
| Pitpnc1 | 0.00449901 | -0.6862156 | 0.99 | 0.979 | 1 | 0.38532173 | -0.3547167 | 0.987 | 0.99 | 1 | 0.38532173 | 0.00897777 |
| Mapre1 | 0.08071234 | 0.14812804 | 0.896 | 0.891 | 1 | 0.00450847 | 0.21472132 | 0.962 | 0.885 | 1 | 0.08071234 | 0.00899662 |
| Dbnl | 0.00450895 | 0.72370157 | 0.375 | 0.212 | 1 | 0.4414109 | -0.1820997 | 0.37 | 0.312 | 1 | 0.4414109 | 0.00899756 |
| Dock3 | 0.04759503 | -1.7294711 | 0.01 | 0.062 | 1 | 0.00451685 | -2.3098553 | 0.042 | 0.125 | 1 | 0.04759503 | 0.00901331 |
| Fitm2 | 0.94872673 | 0.18806678 | 0.135 | 0.135 | 1 | 0.00452943 | 1.69014474 | 0.139 | 0.031 | 1 | 0.94872673 | 0.00903835 |
| Cdk1 | 0.00453701 | 3.27598337 | 0.094 | 0.021 | 1 | 0.20315061 | 1.49749966 | 0.017 | 0 | 1 | 0.20315061 | 0.00905343 |
| Mcrs1 | 0.46066518 | 0.20543391 | 0.333 | 0.29 | 1 | 0.00453768 | 0.69014474 | 0.294 | 0.135 | 1 | 0.46066518 | 0.00905477 |
| Ppp1r13l | 0.11375458 | 0.53800925 | 0.125 | 0.067 | 1 | 0.00454046 | 0.71399148 | 0.206 | 0.073 | 1 | 0.11375458 | 0.00906029 |
| Sppl2a | 0.02551137 | -0.3499505 | 0.906 | 0.902 | 1 | 0.00454675 | 0.24917877 | 0.87 | 0.854 | 1 | 0.02551137 | 0.00907283 |
| Pus3 | 0.00454934 | 1.00749454 | 0.219 | 0.093 | 1 | 0.19250854 | 0.56461386 | 0.097 | 0.052 | 1 | 0.19250854 | 0.00907798 |
| Sh3rf1 | 0.00455462 | 0.56852702 | 0.74 | 0.565 | 1 | 0.01730509 | -0.6357186 | 0.71 | 0.865 | 1 | 0.01730509 | 0.0090885 |
| Atp6v1g2 | 0.00455532 | 2.12297175 | 0.104 | 0.026 | 1 | 0.08157035 | 0.49749966 | 0.101 | 0.042 | 1 | 0.08157035 | 0.0090899 |
| Bax | 0.00533979 | 0.42800717 | 0.781 | 0.606 | 1 | 0.00456165 | 0.48337855 | 0.672 | 0.562 | 1 | 0.00533979 | 0.0091025 |
| Tmem222 | 0.00456841 | 0.59245704 | 0.51 | 0.347 | 1 | 0.02667931 | 0.27510724 | 0.353 | 0.208 | 1 | 0.02667931 | 0.00911594 |
| Ptpn12 | 0.34057793 | -0.2054992 | 0.656 | 0.684 | 1 | 0.00457909 | -0.8149716 | 0.773 | 0.823 | 1 | 0.34057793 | 0.0091372 |
| Usb1 | 0.11061452 | 0.56003556 | 0.188 | 0.119 | 1 | 0.00458594 | 1.34222143 | 0.155 | 0.042 | 1 | 0.11061452 | 0.00915085 |
| Zfp654 | 0.26227752 | -0.1718292 | 0.448 | 0.513 | 1 | 0.00460065 | -0.8995713 | 0.584 | 0.76 | 1 | 0.26227752 | 0.00918013 |
| Tbpl1 | 0.02814896 | 0.56820949 | 0.417 | 0.285 | 1 | 0.00460478 | 0.38404161 | 0.45 | 0.26 | 1 | 0.02814896 | 0.00918835 |
| Gsr | 0.2122603 | -1.5107419 | 0.438 | 0.451 | 1 | 0.00461706 | -0.885784 | 0.592 | 0.75 | 1 | 0.2122603 | 0.0092128 |
| Tmco1 | 0.00463639 | 0.39531057 | 0.865 | 0.715 | 1 | 0.0063103 | 0.35244933 | 0.739 | 0.625 | 1 | 0.0063103 | 0.00925129 |
| Phkb | 0.20102505 | 0.2904285 | 0.396 | 0.321 | 1 | 0.00464053 | -0.6955985 | 0.454 | 0.646 | 1 | 0.20102505 | 0.00925953 |
| Atp2c1 | 0.21250371 | 0.2635929 | 0.469 | 0.378 | 1 | 0.00464257 | -0.6317834 | 0.555 | 0.729 | 1 | 0.21250371 | 0.00926359 |
| Dnajc16 | 0.73053841 | 0.15949763 | 0.156 | 0.14 | 1 | 0.00464326 | 1.14957636 | 0.16 | 0.042 | 1 | 0.73053841 | 0.00926496 |
| Cnot8 | 0.17054371 | 0.4367459 | 0.458 | 0.389 | 1 | 0.00464531 | 0.50138309 | 0.5 | 0.292 | 1 | 0.17054371 | 0.00926904 |
| Rnf4 | 0.00464932 | 0.46189447 | 0.74 | 0.637 | 1 | 0.18165165 | 0.11768302 | 0.811 | 0.781 | 1 | 0.18165165 | 0.00927703 |
| Gm13293 | 0.00465026 | 2.27052894 | 0.094 | 0.021 | 1 | 0.16109008 | 0.93807225 | 0.071 | 0.031 | 1 | 0.16109008 | 0.00927889 |
| Arhgap28 | 0.00504453 | -3.3264062 | 0.042 | 0.15 | 1 | 0.00465396 | -1.8948178 | 0.084 | 0.188 | 1 | 0.00504453 | 0.00928626 |
| Fuom | 0.13967385 | 0.41085023 | 0.302 | 0.218 | 1 | 0.00465772 | 0.83150059 | 0.261 | 0.115 | 1 | 0.13967385 | 0.00929375 |
| Prmt3 | 0.51127561 | 0.15188445 | 0.323 | 0.28 | 1 | 0.00466004 | 0.36821664 | 0.303 | 0.135 | 1 | 0.51127561 | 0.00929837 |
| Tpm3-rs7 | 0.65371216 | -0.240433 | 0.073 | 0.088 | 1 | 0.00466434 | 1.57767001 | 0.139 | 0.031 | 1 | 0.65371216 | 0.00930693 |
| Cdk16 | 0.00466779 | 0.9800138 | 0.375 | 0.238 | 1 | 0.32730091 | -0.3300331 | 0.42 | 0.312 | 1 | 0.32730091 | 0.0093138 |
| Mrps18c | 0.25644481 | 0.39451766 | 0.385 | 0.332 | 1 | 0.00467884 | 0.40474252 | 0.399 | 0.219 | 1 | 0.25644481 | 0.0093358 |
| Zswim3 | 0.02257853 | -2.3500575 | 0.031 | 0.109 | 1 | 0.00468549 | -2.1399303 | 0.029 | 0.104 | 1 | 0.02257853 | 0.00934902 |
| Ccnl2 | 0.00468633 | 0.26820389 | 0.958 | 0.876 | 1 | 0.02523569 | -0.2790273 | 0.954 | 0.979 | 1 | 0.02523569 | 0.0093507 |
| Tspyl2 | 0.51258452 | 0.17967551 | 0.292 | 0.254 | 1 | 0.00472089 | 0.47150445 | 0.353 | 0.177 | 1 | 0.51258452 | 0.00941949 |
| Smarcal1 | 0.40849752 | 0.31561683 | 0.115 | 0.083 | 1 | 0.00472104 | 0.84768601 | 0.193 | 0.062 | 1 | 0.40849752 | 0.0094198 |
| Chmp1a | 0.09544733 | 0.58067987 | 0.417 | 0.326 | 1 | 0.00473072 | 0.80988398 | 0.366 | 0.198 | 1 | 0.09544733 | 0.00943907 |
| Incenp | 0.00473304 | 2.42253204 | 0.104 | 0.026 | 1 | 0.43835168 | -0.4029647 | 0.042 | 0.062 | 1 | 0.43835168 | 0.00944368 |
| Btf3l4 | 0.9104896 | -0.3208313 | 0.396 | 0.363 | 1 | 0.00473317 | 0.57550217 | 0.403 | 0.219 | 1 | 0.9104896 | 0.00944394 |
| Mns1 | 0.00474581 | 2.81484946 | 0.094 | 0.021 | 1 | 0.24217871 | 0.69014474 | 0.034 | 0.01 | 1 | 0.24217871 | 0.00946909 |
| Gcsh | 0.00621218 | 0.65658737 | 0.5 | 0.337 | 1 | 0.00475004 | 0.78968041 | 0.349 | 0.188 | 1 | 0.00621218 | 0.00947752 |
| Emd | 0.02142319 | 0.44841067 | 0.729 | 0.611 | 1 | 0.00475074 | 0.35244933 | 0.756 | 0.625 | 1 | 0.02142319 | 0.00947891 |
| Mgat5 | 0.83320521 | -0.148907 | 0.583 | 0.523 | 1 | 0.00475097 | -0.7050956 | 0.756 | 0.812 | 1 | 0.83320521 | 0.00947938 |
| E530011L22Rik | 0.00477003 | 2.14499806 | 0.094 | 0.021 | 1 | 0.56125608 | -0.5728897 | 0.059 | 0.042 | 1 | 0.56125608 | 0.0095173 |
| Gcc1 | 0.12875268 | 0.58122978 | 0.26 | 0.187 | 1 | 0.00477339 | 0.70450003 | 0.332 | 0.167 | 1 | 0.12875268 | 0.009524 |
| Abhd16a | 0.21783388 | 0.31234912 | 0.458 | 0.383 | 1 | 0.00477979 | 0.51796376 | 0.433 | 0.24 | 1 | 0.21783388 | 0.00953674 |
| Gm43445 | 0.63898271 | -0.4779323 | 0.094 | 0.109 | 1 | 0.00480625 | -2.2484547 | 0.042 | 0.125 | 1 | 0.63898271 | 0.00958941 |
| Alas1 | 0.30770132 | -0.7820857 | 0.25 | 0.295 | 1 | 0.00480702 | -0.6638633 | 0.416 | 0.615 | 1 | 0.30770132 | 0.00959093 |
| Kansl1l | 0.16536278 | -0.1294647 | 0.635 | 0.56 | 1 | 0.00481381 | -0.8656231 | 0.87 | 0.854 | 1 | 0.16536278 | 0.00960446 |
| Wnt9a | 0.01200338 | -2.577468 | 0.042 | 0.135 | 1 | 0.00481764 | -2.9287651 | 0.05 | 0.135 | 1 | 0.01200338 | 0.00961206 |
| Mlx | 0.45558099 | 0.34452952 | 0.344 | 0.306 | 1 | 0.00483363 | 0.77106473 | 0.349 | 0.188 | 1 | 0.45558099 | 0.00964389 |
| Tert | 0.00483403 | 2.00749454 | 0.083 | 0.016 | 1 | 0.92738405 | -0.7248928 | 0.034 | 0.031 | 1 | 0.92738405 | 0.00964469 |
| Ccdc88a | 0.0327481 | 0.30630007 | 1 | 0.943 | 1 | 0.00483561 | 0.23741495 | 0.983 | 0.948 | 1 | 0.0327481 | 0.00964783 |
| Ppp1r36 | 0.00485749 | 1.14499806 | 0.167 | 0.062 | 1 | 0.01159308 | 1.41261076 | 0.105 | 0.021 | 1 | 0.01159308 | 0.00969139 |
| Map3k20 | 0.00488003 | 0.97187063 | 0.469 | 0.321 | 1 | 0.13419804 | -0.1874586 | 0.441 | 0.583 | 1 | 0.13419804 | 0.00973625 |
| Crlf1 | 0.00489698 | -2.407543 | 0.021 | 0.119 | 1 | 0.13148267 | -1.0323213 | 0.126 | 0.188 | 1 | 0.13148267 | 0.00976998 |
| Kif18a | 0.004904 | 2.10703021 | 0.094 | 0.021 | 1 | 0.18361019 | 1.27510724 | 0.038 | 0.01 | 1 | 0.18361019 | 0.00978395 |
| Eps15l1 | 0.00491425 | 0.56003556 | 0.635 | 0.472 | 1 | 0.80283864 | -0.2528773 | 0.748 | 0.75 | 1 | 0.80283864 | 0.00980434 |
| Gtf2e2 | 0.00491455 | 0.66304901 | 0.562 | 0.394 | 1 | 0.13463957 | 0.10518224 | 0.555 | 0.417 | 1 | 0.13463957 | 0.00980496 |
| Septin10 | 0.64686864 | 0.20013961 | 0.323 | 0.368 | 1 | 0.00491537 | 0.9994728 | 0.29 | 0.146 | 1 | 0.64686864 | 0.00980658 |
| Asns | 0.04361469 | 0.71331143 | 0.448 | 0.347 | 1 | 0.00491916 | 0.54812573 | 0.244 | 0.104 | 1 | 0.04361469 | 0.00981412 |
| Abhd17a | 0.18326578 | 0.26383429 | 0.604 | 0.492 | 1 | 0.00492228 | 0.51681313 | 0.555 | 0.365 | 1 | 0.18326578 | 0.00982034 |
| Lmf2 | 0.0049238 | 0.82463048 | 0.333 | 0.181 | 1 | 0.0225528 | 1.01207283 | 0.244 | 0.135 | 1 | 0.0225528 | 0.00982336 |
| Zfp931 | 0.00492493 | 1.15949763 | 0.177 | 0.067 | 1 | 0.83443998 | -0.5728897 | 0.105 | 0.094 | 1 | 0.83443998 | 0.0098256 |
| Gm12829 | 0.00493463 | 1.85549144 | 0.083 | 0.016 | 1 | 0.15394118 | 1.49749966 | 0.021 | 0 | 1 | 0.15394118 | 0.0098449 |
| Pdgfa | 0.00493724 | 0.75875079 | 0.552 | 0.394 | 1 | 0.02010672 | 0.39590538 | 0.815 | 0.771 | 1 | 0.02010672 | 0.0098501 |
| Snhg15 | 0.004949 | 1.16503581 | 0.24 | 0.114 | 1 | 0.8705501 | -0.1399303 | 0.088 | 0.094 | 1 | 0.8705501 | 0.00987351 |
| Sema4c | 0.60767884 | 0.15949763 | 0.115 | 0.093 | 1 | 0.00495561 | -1.3454792 | 0.105 | 0.219 | 1 | 0.60767884 | 0.00988666 |
| Golga3 | 0.00589558 | 0.37381675 | 0.76 | 0.622 | 1 | 0.00498296 | 0.1730863 | 0.798 | 0.573 | 1 | 0.00589558 | 0.00994109 |
| Stx17 | 0.17633632 | 0.4088571 | 0.448 | 0.363 | 1 | 0.00500166 | 0.34414888 | 0.487 | 0.281 | 1 | 0.17633632 | 0.0099783 |
| Cdh2 | 0.00500328 | -3.706751 | 0.031 | 0.135 | 1 | 0.03854771 | -1.9537115 | 0.076 | 0.146 | 1 | 0.03854771 | 0.00998153 |
| Naalad2 | 0.00501097 | 0.85549144 | 0.385 | 0.228 | 1 | 0.01982146 | 1.27510724 | 0.172 | 0.073 | 1 | 0.01982146 | 0.00999683 |
| Pbx1 | 0.7437554 | -0.1083361 | 0.927 | 0.959 | 1 | 0.00501161 | -0.5777448 | 0.958 | 0.969 | 1 | 0.7437554 | 0.00999811 |
| Mtpap | 0.00502693 | 0.57453513 | 0.531 | 0.358 | 1 | 0.36520898 | -0.1143045 | 0.424 | 0.333 | 1 | 0.36520898 | 0.01002858 |
| Lzic | 0.09646626 | 0.36837627 | 0.406 | 0.311 | 1 | 0.00502948 | 0.44503224 | 0.324 | 0.167 | 1 | 0.09646626 | 0.01003367 |
| Med8 | 0.96770229 | 0.11440974 | 0.385 | 0.399 | 1 | 0.00503008 | 0.4150375 | 0.382 | 0.198 | 1 | 0.96770229 | 0.01003487 |
| Smim29 | 0.76940456 | -0.2462621 | 0.115 | 0.124 | 1 | 0.00503747 | 0.84768601 | 0.189 | 0.062 | 1 | 0.76940456 | 0.01004957 |
| Pdgfrb | 0.00504598 | -0.2555399 | 0.615 | 0.347 | 1 | 0.11438128 | -0.2544785 | 0.739 | 0.5 | 1 | 0.11438128 | 0.0100665 |
| Nod1 | 0.00504615 | -1.2625946 | 0.146 | 0.29 | 1 | 0.46123448 | -0.129283 | 0.21 | 0.167 | 1 | 0.46123448 | 0.01006685 |
| Fam131a | 0.00505278 | 0.89201732 | 0.344 | 0.192 | 1 | 0.01068258 | 0.78968041 | 0.273 | 0.135 | 1 | 0.01068258 | 0.01008003 |
| Rhod | 0.05206542 | 0.64994253 | 0.375 | 0.259 | 1 | 0.00505383 | 0.28575448 | 0.395 | 0.198 | 1 | 0.05206542 | 0.01008212 |
| Zfand2a | 0.81695381 | -0.178372 | 0.438 | 0.394 | 1 | 0.00505401 | 0.76180572 | 0.433 | 0.26 | 1 | 0.81695381 | 0.01008249 |
| Dna2 | 0.00505892 | 1.88196365 | 0.094 | 0.021 | 1 | 0.73377172 | -1.0874628 | 0.055 | 0.062 | 1 | 0.73377172 | 0.01009225 |
| Antxr1 | 0.01011548 | -0.2023076 | 0.875 | 0.611 | 1 | 0.00505967 | -1.3619096 | 0.92 | 0.865 | 1 | 0.01011548 | 0.01009374 |
| Sepsecs | 0.00506148 | 0.75751628 | 0.302 | 0.155 | 1 | 0.02001603 | 0.67286675 | 0.273 | 0.146 | 1 | 0.02001603 | 0.01009734 |

| | | | | | | | | | | | | |
|---|---|---|---|---|---|---|---|---|---|---|---|---|
| Prmt5 | 0.02518122 | 0.80313604 | 0.333 | 0.218 | 1 | 0.00506915 | 0.6008774 | 0.307 | 0.146 | 1 | 0.02518122 | 0.01011261 |
| Gtf3c3 | 0.05913889 | 0.56548999 | 0.396 | 0.285 | 1 | 0.00507259 | 0.70721825 | 0.261 | 0.115 | 1 | 0.05913889 | 0.01011944 |
| Med16 | 0.55006512 | 0.18152394 | 0.188 | 0.155 | 1 | 0.00507307 | 0.73453886 | 0.235 | 0.094 | 1 | 0.55006512 | 0.01012041 |
| Anxa6 | 0.00507585 | 0.29879667 | 0.729 | 0.513 | 1 | 0.42152947 | -0.194378 | 0.622 | 0.448 | 1 | 0.42152947 | 0.01012594 |
| Ints5 | 0.00508721 | 1.05996196 | 0.25 | 0.119 | 1 | 0.14173121 | 0.29782731 | 0.218 | 0.146 | 1 | 0.14173121 | 0.01014855 |
| Slc24a4 | 0.00509166 | 1.65957123 | 0.094 | 0.021 | 1 | 0.0928609 | -2.4797803 | 0.013 | 0.042 | 1 | 0.0928609 | 0.0101574 |
| Sars | 0.00554604 | 0.41897422 | 0.667 | 0.503 | 1 | 0.0050955 | 0.43409676 | 0.681 | 0.552 | 1 | 0.00554604 | 0.01016503 |
| Wdr6 | 0.00509558 | 1.05056326 | 0.281 | 0.145 | 1 | 0.07606247 | 0.27510724 | 0.332 | 0.219 | 1 | 0.07606247 | 0.01016519 |
| Gm3331 | 0.00511622 | 1.74446013 | 0.104 | 0.026 | 1 | 0.03964888 | -1.961932 | 0.025 | 0.073 | 1 | 0.03964888 | 0.01020627 |
| Dpy30 | 0.00511788 | 0.61208584 | 0.646 | 0.503 | 1 | 0.6312339 | -0.1214102 | 0.576 | 0.562 | 1 | 0.6312339 | 0.01020956 |
| Atxn7 | 0.11304466 | 0.30807331 | 0.771 | 0.694 | 1 | 0.00513034 | -0.6245518 | 0.845 | 0.896 | 1 | 0.11304466 | 0.01023436 |
| Otub2 | 0.00513834 | 1.65957123 | 0.104 | 0.026 | 1 | 0.40319345 | -0.8244284 | 0.055 | 0.031 | 1 | 0.40319345 | 0.01025029 |
| G6pdx | 0.01191237 | -1.2852872 | 0.083 | 0.197 | 1 | 0.00514013 | -1.2812861 | 0.168 | 0.302 | 1 | 0.01191237 | 0.01025384 |
| Clvs1 | 0.00514464 | -3.5160674 | 0 | 0.078 | 1 | 0.52072676 | -1.0468209 | 0.021 | 0.01 | 1 | 0.52072676 | 0.01026282 |
| Cmtm5 | 0.48825768 | 0.2662288 | 0.365 | 0.316 | 1 | 0.00514482 | 1.12755005 | 0.189 | 0.062 | 1 | 0.48825768 | 0.01026318 |
| Bcan | 0.00514701 | -3.3848229 | 0 | 0.078 | 1 | 0.5859632 | -0.3098553 | 0.013 | 0.021 | 1 | 0.5859632 | 0.01026752 |
| Fxyd2 | 0.00514793 | -1.4084819 | 0.188 | 0.347 | 1 | 0.68536235 | -0.6593044 | 0.084 | 0.094 | 1 | 0.68536235 | 0.01026936 |
| Gm49575 | 0.00514884 | -3.577468 | 0 | 0.078 | 1 | 0.0419804 | 3.01207283 | 0.042 | 0 | 1 | 0.0419804 | 0.01027118 |
| Stpg2 | 0.00515053 | -3.747393 | 0 | 0.078 | 1 | 0.8023126 | -0.5025003 | 0.017 | 0.021 | 1 | 0.8023126 | 0.01027454 |
| Fbxo45 | 0.00515344 | 0.59245704 | 0.333 | 0.181 | 1 | 0.00611063 | 0.4777364 | 0.429 | 0.25 | 1 | 0.00611063 | 0.01028033 |
| Akr1e1 | 0.45127053 | 0.21899864 | 0.271 | 0.228 | 1 | 0.00516156 | 0.92280549 | 0.256 | 0.115 | 1 | 0.45127053 | 0.01029648 |
| Nup88 | 0.00517187 | 0.59721057 | 0.521 | 0.342 | 1 | 0.2977057 | -0.1578522 | 0.462 | 0.344 | 1 | 0.2977057 | 0.01031699 |
| Atp23 | 0.00518819 | 1.74446013 | 0.094 | 0.021 | 1 | 0.58758726 | 0.27510724 | 0.08 | 0.062 | 1 | 0.58758726 | 0.01034945 |
| Cth | 0.005195 | -0.8119332 | 0.25 | 0.404 | 1 | 0.97086465 | -0.5274467 | 0.105 | 0.104 | 1 | 0.97086465 | 0.01036301 |
| Zfp868 | 0.9000458 | 0.10479174 | 0.188 | 0.181 | 1 | 0.00520121 | 0.92718393 | 0.214 | 0.083 | 1 | 0.9000458 | 0.01037536 |
| Ldb1 | 0.55396011 | 0.1481934 | 0.542 | 0.508 | 1 | 0.00520747 | 0.29206037 | 0.777 | 0.646 | 1 | 0.55396011 | 0.01038783 |
| Ppp1r18 | 0.61800007 | 0.25076569 | 0.385 | 0.363 | 1 | 0.00520843 | 0.35439615 | 0.496 | 0.292 | 1 | 0.61800007 | 0.01038973 |
| Smco4 | 0.03644992 | 0.86999101 | 0.323 | 0.212 | 1 | 0.00521564 | 0.43090191 | 0.387 | 0.198 | 1 | 0.03644992 | 0.01040408 |
| Dzip1l | 0.2341931 | -0.219916 | 0.25 | 0.326 | 1 | 0.00521994 | 0.72256622 | 0.357 | 0.198 | 1 | 0.2341931 | 0.01041262 |
| D830036C21Rik | 0.00522067 | 1.59245704 | 0.083 | 0.016 | 1 | 0.46087389 | -0.8948178 | 0.071 | 0.094 | 1 | 0.46087389 | 0.01041408 |
| Upk1a | 0.00522067 | 1.37006462 | 0.083 | 0.016 | 1 | 0.66139566 | -0.6883669 | 0.08 | 0.062 | 1 | 0.66139566 | 0.01041408 |
| Fyttd1 | 0.38223085 | -0.1126883 | 0.854 | 0.886 | 1 | 0.00522501 | 0.29176672 | 0.929 | 0.917 | 1 | 0.38223085 | 0.01042271 |
| Adsl | 0.20648395 | 0.18646468 | 0.26 | 0.187 | 1 | 0.00524442 | 0.27510724 | 0.311 | 0.146 | 1 | 0.20648395 | 0.01046134 |
| Mpped2 | 0.00526663 | 0.92503238 | 0.583 | 0.435 | 1 | 0.82819733 | -0.1787278 | 0.643 | 0.562 | 1 | 0.82819733 | 0.01050552 |
| Sipa1l3 | 0.4604701 | 0.17522198 | 0.229 | 0.187 | 1 | 0.00527469 | -0.3844941 | 0.487 | 0.708 | 1 | 0.4604701 | 0.01052156 |
| Ugdh | 0.02837243 | -0.4508207 | 0.958 | 0.969 | 1 | 0.00527796 | 0.57447533 | 0.954 | 0.885 | 1 | 0.02837243 | 0.01052805 |
| Pparg | 0.00528632 | -1.7091736 | 0.24 | 0.389 | 1 | 0.37924806 | 0.57597372 | 0.21 | 0.167 | 1 | 0.37924806 | 0.0105447 |
| Sfxn2 | 0.02682707 | 0.5757783 | 0.333 | 0.207 | 1 | 0.00529509 | 0.84664922 | 0.311 | 0.156 | 1 | 0.02682707 | 0.01056215 |
| Mlf1 | 0.02888669 | 1.73411819 | 0.312 | 0.218 | 1 | 0.0053194 | 0.73745045 | 0.239 | 0.094 | 1 | 0.02888669 | 0.0106105 |
| Gm15283 | 0.44531377 | -1.1696985 | 0.302 | 0.326 | 1 | 0.00535106 | 0.25684503 | 0.395 | 0.208 | 1 | 0.44531377 | 0.01067349 |
| Nup37 | 0.00535454 | 1.25542205 | 0.167 | 0.062 | 1 | 0.33719256 | 0.10518224 | 0.084 | 0.052 | 1 | 0.33719256 | 0.01068042 |
| Ntrk2 | 0.34543898 | -1.4828311 | 0.135 | 0.093 | 1 | 0.00535859 | 1.62874419 | 0.193 | 0.073 | 1 | 0.34543898 | 0.01068847 |
| Gramd1c | 0.45926943 | 0.38189005 | 0.208 | 0.176 | 1 | 0.00536056 | 0.46386888 | 0.286 | 0.125 | 1 | 0.45926943 | 0.01069239 |
| AW146154 | 0.00536301 | 1.65957123 | 0.094 | 0.021 | 1 | 0.00811651 | 2.23446525 | 0.109 | 0.021 | 1 | 0.00811651 | 0.01069726 |
| Zfp36l1 | 0.00536333 | -0.721943 | 0.823 | 0.881 | 1 | 0.01663832 | -0.4568246 | 0.807 | 0.917 | 1 | 0.01663832 | 0.0106979 |
| Rab27a | 0.40541969 | -0.2866886 | 0.302 | 0.342 | 1 | 0.00536711 | 1.13492958 | 0.265 | 0.125 | 1 | 0.40541969 | 0.01070541 |
| LTO1 | 0.00538827 | 0.96018882 | 0.26 | 0.13 | 1 | 0.24674662 | 0.11898804 | 0.13 | 0.083 | 1 | 0.24674662 | 0.01074751 |
| Plpp3 | 0.26112374 | -0.1955897 | 0.885 | 0.953 | 1 | 0.00538862 | 0.34213119 | 0.979 | 0.948 | 1 | 0.26112374 | 0.01074821 |
| Phf2 | 0.09394851 | 0.28867629 | 0.573 | 0.472 | 1 | 0.00539073 | 0.36884268 | 0.828 | 0.677 | 1 | 0.09394851 | 0.01075241 |
| Ptx3 | 0.31804443 | -1.282898 | 0.146 | 0.192 | 1 | 0.00539339 | 2.12013258 | 0.118 | 0.021 | 1 | 0.31804443 | 0.01075769 |
| Mapkapk3 | 0.84593295 | 0.30705482 | 0.062 | 0.057 | 1 | 0.00539819 | -0.9957467 | 0.155 | 0.292 | 1 | 0.84593295 | 0.01076723 |
| Rab4a | 0.18027504 | -0.3848229 | 0.125 | 0.187 | 1 | 0.00540261 | 0.34222143 | 0.235 | 0.094 | 1 | 0.18027504 | 0.01077603 |
| Lacc1 | 0.00541298 | 0.77922555 | 0.281 | 0.145 | 1 | 0.04531951 | 0.66213036 | 0.16 | 0.073 | 1 | 0.04531951 | 0.01079667 |
| Ap3d1 | 0.00541316 | 0.41962044 | 0.854 | 0.71 | 1 | 0.0152033 | 0.12077909 | 0.861 | 0.708 | 1 | 0.0152033 | 0.01079701 |
| Uqcc4 | 0.22046518 | 0.46692616 | 0.281 | 0.218 | 1 | 0.00542163 | 0.74325607 | 0.29 | 0.135 | 1 | 0.22046518 | 0.01081387 |
| Tomm22 | 0.0054222 | 0.43944542 | 0.781 | 0.642 | 1 | 0.03329843 | 0.35882906 | 0.727 | 0.656 | 1 | 0.03329843 | 0.01081499 |
| Rbms1 | 0.00635791 | -0.5214737 | 0.948 | 0.99 | 1 | 0.00542339 | -0.5942564 | 0.992 | 0.979 | 1 | 0.00635791 | 0.01081736 |
| Dnajc18 | 0.18098068 | 0.31415587 | 0.333 | 0.254 | 1 | 0.0054354 | 0.46647276 | 0.45 | 0.26 | 1 | 0.18098068 | 0.01084127 |
| Ebf1 | 0.00543881 | -0.3940045 | 0.99 | 0.959 | 1 | 0.74138633 | -0.2521624 | 0.958 | 0.99 | 1 | 0.74138633 | 0.01084804 |
| Zfp51 | 0.01427796 | 1.08149512 | 0.167 | 0.073 | 1 | 0.0054421 | 1.17127143 | 0.227 | 0.094 | 1 | 0.01427796 | 0.01085459 |
| Parm1 | 0.5253825 | -0.2625946 | 0.115 | 0.088 | 1 | 0.00545568 | -2.961932 | 0.042 | 0.125 | 1 | 0.5253825 | 0.0108816 |
| Sec24c | 0.53039704 | -0.3867844 | 0.302 | 0.326 | 1 | 0.00547196 | 0.33259273 | 0.391 | 0.208 | 1 | 0.53039704 | 0.01091397 |
| Tnr | 0.0054735 | -3.0368996 | 0.01 | 0.098 | 1 | 0.27154471 | 1.27510724 | 0.013 | 0 | 1 | 0.27154471 | 0.01091704 |
| Ccsap | 0.25156132 | -0.2879613 | 0.062 | 0.104 | 1 | 0.00547568 | 2.18199783 | 0.118 | 0.021 | 1 | 0.25156132 | 0.01092137 |
| Gm5617 | 0.59800285 | -0.2842716 | 0.25 | 0.269 | 1 | 0.00547683 | 1.10518224 | 0.202 | 0.073 | 1 | 0.59800285 | 0.01092367 |
| Ddx31 | 0.0710975 | 0.62047141 | 0.219 | 0.135 | 1 | 0.00549393 | 0.67103591 | 0.218 | 0.083 | 1 | 0.0710975 | 0.01095768 |
| H2-T22 | 0.93624292 | -0.1521621 | 0.312 | 0.311 | 1 | 0.00549496 | 0.19264508 | 0.416 | 0.229 | 1 | 0.93624292 | 0.01095972 |
| Dcaf5 | 0.0409935 | 0.20321901 | 0.406 | 0.269 | 1 | 0.00550025 | -0.5443205 | 0.58 | 0.781 | 1 | 0.0409935 | 0.01097025 |
| Phtf1 | 0.05165059 | 0.48105811 | 0.521 | 0.409 | 1 | 0.0055019 | -0.8510454 | 0.567 | 0.667 | 1 | 0.05165059 | 0.01097353 |
| Lmf1 | 0.00949396 | 0.80586068 | 0.333 | 0.197 | 1 | 0.00550514 | 1.10518224 | 0.227 | 0.094 | 1 | 0.00949396 | 0.01097997 |
| Sdr39u1 | 0.58035768 | 0.23882008 | 0.229 | 0.202 | 1 | 0.00551465 | 0.34222143 | 0.227 | 0.094 | 1 | 0.58035768 | 0.01099889 |
| Pigt | 0.70117083 | 0.22809368 | 0.458 | 0.456 | 1 | 0.00551607 | 0.4209581 | 0.487 | 0.292 | 1 | 0.70117083 | 0.01100171 |
| Pkp1 | 0.48064264 | 0.59245704 | 0.021 | 0.01 | 1 | 0.00553817 | -1.3098553 | 0.029 | 0.104 | 1 | 0.48064264 | 0.01104567 |
| Erbb3 | 0.45632632 | -0.8405024 | 0.042 | 0.062 | 1 | 0.00554254 | -1.8123556 | 0.042 | 0.125 | 1 | 0.45632632 | 0.01105437 |
| Asb4 | 0.59123637 | -0.774914 | 0.115 | 0.135 | 1 | 0.00554256 | 1.21586803 | 0.252 | 0.115 | 1 | 0.59123637 | 0.0110544 |
| Tma16 | 0.02626267 | 0.76422339 | 0.333 | 0.207 | 1 | 0.00554957 | 0.88278982 | 0.307 | 0.156 | 1 | 0.02626267 | 0.01106834 |
| Neto2 | 0.00555027 | -1.5015191 | 0.042 | 0.155 | 1 | 0.12199143 | -1.9472852 | 0.076 | 0.125 | 1 | 0.12199143 | 0.01106972 |
| Dnajc1 | 0.00555605 | 0.24494533 | 1 | 0.974 | 1 | 0.68363252 | -0.3742762 | 0.945 | 0.979 | 1 | 0.68363252 | 0.01108124 |
| Cd248 | 0.15729403 | -1.577468 | 0 | 0.021 | 1 | 0.00556087 | -2.5025003 | 0.025 | 0.094 | 1 | 0.15729403 | 0.01109081 |
| Acss3 | 0.2632671 | 0.42253204 | 0.052 | 0.026 | 1 | 0.00556123 | 0.6242568 | 0.231 | 0.094 | 1 | 0.2632671 | 0.01109153 |
| Amotl1 | 0.90500403 | 0.22089817 | 0.802 | 0.808 | 1 | 0.00556402 | 0.38051274 | 0.933 | 0.854 | 1 | 0.90500403 | 0.01109708 |
| Zdbf2 | 0.00556706 | 0.85671458 | 0.458 | 0.29 | 1 | 0.12201153 | -0.2120079 | 0.454 | 0.292 | 1 | 0.12201153 | 0.01110312 |
| Zfp866 | 0.43048793 | 0.11440974 | 0.177 | 0.14 | 1 | 0.00557721 | 0.13945214 | 0.252 | 0.104 | 1 | 0.43048793 | 0.01112332 |
| Stk3 | 0.25888003 | 0.13464551 | 0.802 | 0.741 | 1 | 0.00558317 | -0.7386986 | 0.773 | 0.833 | 1 | 0.25888003 | 0.01113518 |
| Atg4c | 0.4023371 | 0.31943854 | 0.156 | 0.119 | 1 | 0.00558914 | 1.29895398 | 0.21 | 0.083 | 1 | 0.4023371 | 0.01114704 |
| Gm49144 | 0.00560024 | -2.1624305 | 0.031 | 0.135 | 1 | 0.08779795 | -1.5163061 | 0.046 | 0.094 | 1 | 0.08779795 | 0.01116912 |
| Mib2 | 0.0056063 | 1.36113149 | 0.188 | 0.078 | 1 | 0.2041152 | -0.194378 | 0.181 | 0.115 | 1 | 0.2041152 | 0.01118118 |
| Hfe | 0.3278642 | -0.5160674 | 0.073 | 0.109 | 1 | 0.00560811 | 1.08869411 | 0.202 | 0.073 | 1 | 0.3278642 | 0.01118478 |
| Nudt13 | 0.00562885 | 1.086929 | 0.333 | 0.197 | 1 | 0.08699393 | 0.22215836 | 0.269 | 0.167 | 1 | 0.08699393 | 0.01122603 |
| Spout1 | 0.00563483 | 0.67628863 | 0.292 | 0.145 | 1 | 0.04325832 | 1.27510724 | 0.126 | 0.052 | 1 | 0.04325832 | 0.01123791 |

| | | | | | | | | | | | | |
|---|---|---|---|---|---|---|---|---|---|---|---|---|
| Cblb | 0.18971467 | -0.3439005 | 0.615 | 0.642 | 1 | 0.00564179 | -0.7733543 | 0.718 | 0.844 | 1 | 0.18971467 | 0.01125176 |
| Chchd4 | 0.00564366 | 1.33870045 | 0.271 | 0.145 | 1 | 0.01685979 | 0.2584285 | 0.282 | 0.146 | 1 | 0.01685979 | 0.01125547 |
| Eda | 0.4939942 | -0.2583995 | 0.198 | 0.161 | 1 | 0.00564704 | -0.4949667 | 0.29 | 0.5 | 1 | 0.4939942 | 0.01126218 |
| Polr2f | 0.26025548 | 0.18495321 | 0.677 | 0.606 | 1 | 0.00565167 | 0.25464314 | 0.588 | 0.427 | 1 | 0.26025548 | 0.0112714 |
| Lsm3 | 0.2063481 | 0.31234912 | 0.542 | 0.477 | 1 | 0.00565693 | 0.154813 | 0.454 | 0.25 | 1 | 0.2063481 | 0.01128187 |
| Fzr1 | 0.39565053 | 0.27052894 | 0.198 | 0.155 | 1 | 0.0056586 | 0.82764826 | 0.256 | 0.115 | 1 | 0.39565053 | 0.01128517 |
| Inpp4b | 0.0056607 | -1.7397394 | 0.094 | 0.223 | 1 | 0.055418 | 0.60768258 | 0.168 | 0.083 | 1 | 0.055418 | 0.01128936 |
| Grb14 | 0.00568282 | -1.9162699 | 0.115 | 0.249 | 1 | 0.56148913 | -0.5213594 | 0.038 | 0.052 | 1 | 0.56148913 | 0.01133334 |
| Rcc1 | 0.00568958 | 1.08344339 | 0.26 | 0.13 | 1 | 0.0393282 | 0.36821664 | 0.218 | 0.115 | 1 | 0.0393282 | 0.01134678 |
| Zdhhc9 | 0.59065482 | 0.19512154 | 0.271 | 0.244 | 1 | 0.00570017 | 0.46592958 | 0.34 | 0.167 | 1 | 0.59065482 | 0.01136784 |
| Ndufs3 | 0.0057091 | 0.40011457 | 0.781 | 0.663 | 1 | 0.03746966 | 0.3206219 | 0.706 | 0.604 | 1 | 0.03746966 | 0.0113856 |
| B230369F24Rik | 0.81499642 | -0.9384656 | 0.177 | 0.176 | 1 | 0.00571769 | -1.4714444 | 0.252 | 0.375 | 1 | 0.81499642 | 0.01140269 |
| Ntan1 | 0.15279708 | 0.15827449 | 0.719 | 0.58 | 1 | 0.00572204 | 0.32805612 | 0.702 | 0.51 | 1 | 0.15279708 | 0.01141135 |
| Wdr5b | 0.53207093 | 0.49292136 | 0.052 | 0.036 | 1 | 0.00572635 | 2.93807225 | 0.076 | 0 | 1 | 0.53207093 | 0.01141991 |
| Rbm22 | 0.00573325 | 0.48105811 | 0.646 | 0.461 | 1 | 0.04419723 | 0.18535259 | 0.681 | 0.542 | 1 | 0.04419723 | 0.01143363 |
| Vwa5a | 0.117494 | 0.13780256 | 0.615 | 0.456 | 1 | 0.00574402 | 0.39755214 | 0.676 | 0.448 | 1 | 0.117494 | 0.01145504 |
| Galnt10 | 0.00574566 | 0.6026528 | 0.448 | 0.285 | 1 | 0.35854041 | -0.1654654 | 0.454 | 0.344 | 1 | 0.35854041 | 0.01145831 |
| Fbxo22 | 0.00574577 | -1.070508 | 0.323 | 0.456 | 1 | 0.49072109 | -0.5340704 | 0.366 | 0.292 | 1 | 0.49072109 | 0.01145852 |
| 6030458C11Rik | 0.21269186 | 0.55769162 | 0.26 | 0.197 | 1 | 0.00575291 | 0.52021974 | 0.231 | 0.094 | 1 | 0.21269186 | 0.01147273 |
| Apbb2 | 0.25552021 | -0.1689423 | 0.979 | 0.984 | 1 | 0.00575337 | -0.5903296 | 0.996 | 0.99 | 1 | 0.25552021 | 0.01147363 |
| Zfp362 | 0.03881273 | 0.29477649 | 0.292 | 0.171 | 1 | 0.005754 | 0.5580412 | 0.387 | 0.208 | 1 | 0.03881273 | 0.0114749 |
| Hgs | 0.61643129 | 0.22613482 | 0.25 | 0.223 | 1 | 0.00576131 | 0.70296878 | 0.336 | 0.177 | 1 | 0.61643129 | 0.01148943 |
| 2810433D01Rik | 0.00576812 | 1.67045955 | 0.146 | 0.052 | 1 | 0.63276695 | -0.2318528 | 0.067 | 0.052 | 1 | 0.63276695 | 0.01150297 |
| Tubd1 | 0.00577101 | 1.44806713 | 0.156 | 0.057 | 1 | 0.88346689 | -0.6094155 | 0.046 | 0.042 | 1 | 0.88346689 | 0.01150871 |
| Myg1 | 0.04316734 | 0.87422801 | 0.25 | 0.155 | 1 | 0.00577208 | 0.50830641 | 0.252 | 0.104 | 1 | 0.04316734 | 0.01151085 |
| Has1 | 0.0484475 | -1.5664968 | 0.083 | 0.166 | 1 | 0.00578046 | 1.17781004 | 0.185 | 0.062 | 1 | 0.0484475 | 0.01152751 |
| Ubn2 | 0.00578391 | 0.25221706 | 0.958 | 0.938 | 1 | 0.18741003 | -0.2824491 | 0.954 | 0.948 | 1 | 0.18741003 | 0.01153436 |
| Tmem127 | 0.45748961 | 0.1221371 | 0.51 | 0.451 | 1 | 0.00579997 | 0.50859737 | 0.626 | 0.448 | 1 | 0.45748961 | 0.0115663 |
| Zfp523 | 0.00580439 | 1.0899567 | 0.292 | 0.155 | 1 | 0.15807244 | 0.10518224 | 0.21 | 0.135 | 1 | 0.15807244 | 0.01157508 |
| Osr2 | 0.30763643 | 1.22988696 | 0.042 | 0.021 | 1 | 0.0058171 | -1.6424306 | 0.042 | 0.125 | 1 | 0.30763643 | 0.01160036 |
| Gm2000 | 0.01077122 | 0.48481631 | 0.76 | 0.596 | 1 | 0.00581791 | 0.58066626 | 0.626 | 0.531 | 1 | 0.01077122 | 0.01160198 |
| Ahdc1 | 0.00581811 | 0.36618118 | 0.719 | 0.523 | 1 | 0.95117368 | -0.1716487 | 0.87 | 0.812 | 1 | 0.95117368 | 0.01160238 |
| Tor1b | 0.01952235 | 0.50114187 | 0.552 | 0.409 | 1 | 0.00583542 | 0.44621915 | 0.445 | 0.26 | 1 | 0.01952235 | 0.01163679 |
| Map3k7 | 0.00583603 | 0.48579693 | 0.646 | 0.44 | 1 | 0.25762255 | -0.456076 | 0.664 | 0.667 | 1 | 0.25762255 | 0.01163801 |
| Mrps15 | 0.19431301 | 0.26688117 | 0.51 | 0.44 | 1 | 0.00584317 | 0.49345283 | 0.513 | 0.323 | 1 | 0.19431301 | 0.0116522 |
| Uchl5 | 0.1256356 | 0.49009532 | 0.344 | 0.254 | 1 | 0.00584982 | 0.47333335 | 0.399 | 0.229 | 1 | 0.1256356 | 0.01166542 |
| Clmn | 0.02724126 | 1.07620729 | 0.375 | 0.275 | 1 | 0.00585897 | 0.50264296 | 0.71 | 0.615 | 1 | 0.02724126 | 0.01168361 |
| Bag1 | 0.03292944 | -0.5845227 | 0.885 | 0.886 | 1 | 0.00586162 | 0.35893883 | 0.87 | 0.802 | 1 | 0.03292944 | 0.01168889 |
| Pias2 | 0.66328888 | -0.1847425 | 0.885 | 0.819 | 1 | 0.00587809 | -0.7410541 | 0.811 | 0.885 | 1 | 0.66328888 | 0.01172164 |
| Corin | 0.00588215 | -1.0864816 | 0.083 | 0.207 | 1 | 0.15002914 | 0.25672871 | 0.265 | 0.177 | 1 | 0.15002914 | 0.01172969 |
| Smim13 | 0.01288809 | 0.64136664 | 0.375 | 0.228 | 1 | 0.00589578 | 0.31723271 | 0.387 | 0.208 | 1 | 0.01288809 | 0.01175679 |
| Ilf3 | 0.00589665 | 0.29316675 | 0.781 | 0.544 | 1 | 0.86997037 | -0.2552012 | 0.824 | 0.812 | 1 | 0.86997037 | 0.01175853 |
| Mbd6 | 0.27664161 | 0.11852585 | 0.25 | 0.187 | 1 | 0.00589873 | 0.43737867 | 0.332 | 0.167 | 1 | 0.27664161 | 0.01176266 |
| Zc3hav1l | 0.00591756 | 1.53105649 | 0.177 | 0.073 | 1 | 0.27817042 | -1.0297473 | 0.109 | 0.146 | 1 | 0.27817042 | 0.01180011 |
| Tmem198b | 0.00591874 | 1.63552576 | 0.146 | 0.052 | 1 | 0.26007486 | 0.69014474 | 0.088 | 0.052 | 1 | 0.26007486 | 0.01180245 |
| Mtcp1 | 0.00596079 | 1.16795921 | 0.167 | 0.062 | 1 | 0.2772181 | -0.1172102 | 0.168 | 0.115 | 1 | 0.2772181 | 0.01188605 |
| Zfp68 | 0.01455238 | 0.82108141 | 0.396 | 0.254 | 1 | 0.00597887 | 0.32757466 | 0.471 | 0.271 | 1 | 0.01455238 | 0.01192198 |
| Wdr1 | 0.97391432 | 0.15609103 | 0.802 | 0.819 | 1 | 0.0059806 | 0.29280924 | 0.878 | 0.833 | 1 | 0.97391432 | 0.01192544 |
| Hmgb2 | 0.00598799 | 1.25710843 | 0.729 | 0.611 | 1 | 0.01874375 | -0.2854711 | 0.597 | 0.771 | 1 | 0.01874375 | 0.01194011 |
| Tra2b | 0.00600211 | 0.21526234 | 1 | 0.984 | 1 | 0.41019254 | -0.1280951 | 0.996 | 1 | 1 | 0.41019254 | 0.0119682 |
| Zfp12 | 0.23518906 | 0.39082318 | 0.135 | 0.088 | 1 | 0.00600409 | 1.03409914 | 0.223 | 0.094 | 1 | 0.23518906 | 0.01197212 |
| Map3k6 | 0.00601523 | -1.0856149 | 0.083 | 0.218 | 1 | 0.31337347 | 0.49749966 | 0.122 | 0.083 | 1 | 0.31337347 | 0.01199427 |
| Gm14168 | 0.00601603 | 0.53800925 | 0.104 | 0.026 | 1 | 0.06983842 | 1.17557156 | 0.088 | 0.031 | 1 | 0.06983842 | 0.01199587 |
| Mrps33 | 0.00601626 | 0.37086992 | 0.729 | 0.549 | 1 | 0.27488119 | 0.15692581 | 0.584 | 0.542 | 1 | 0.27488119 | 0.01199632 |
| Smpdl3a | 0.00602945 | -0.5189419 | 0.51 | 0.679 | 1 | 0.00936579 | 0.3950167 | 0.786 | 0.667 | 1 | 0.00936579 | 0.01202255 |
| Kbtbd4 | 0.76423235 | -0.2555399 | 0.094 | 0.104 | 1 | 0.00603532 | 1.33400093 | 0.181 | 0.062 | 1 | 0.76423235 | 0.01203422 |
| Prorp | 0.00603737 | -1.9424648 | 0.083 | 0.207 | 1 | 0.18088738 | -0.8591939 | 0.097 | 0.146 | 1 | 0.18088738 | 0.01203829 |
| 2810004N23Rik | 0.00604456 | 0.6947071 | 0.5 | 0.332 | 1 | 0.03098603 | 0.40343134 | 0.399 | 0.25 | 1 | 0.03098603 | 0.01205258 |
| Ilkap | 0.18661973 | 0.2816695 | 0.552 | 0.446 | 1 | 0.00605318 | 0.23884369 | 0.66 | 0.448 | 1 | 0.18661973 | 0.01206973 |
| Nsmaf | 0.0060693 | -1.2555399 | 0.219 | 0.368 | 1 | 0.46760131 | -0.4845338 | 0.445 | 0.344 | 1 | 0.46760131 | 0.01210177 |
| B4galt6 | 0.00608072 | 0.83346514 | 0.312 | 0.166 | 1 | 0.01726006 | 0.51214644 | 0.315 | 0.177 | 1 | 0.01726006 | 0.01212446 |
| Clcn6 | 0.17976422 | 0.43325844 | 0.333 | 0.259 | 1 | 0.00608314 | 0.35105609 | 0.378 | 0.198 | 1 | 0.17976422 | 0.01212928 |
| Gm56582 | 0.5797009 | 0.35541784 | 0.125 | 0.104 | 1 | 0.00609206 | 1.53144699 | 0.164 | 0.052 | 1 | 0.5797009 | 0.01214701 |
| Asb8 | 0.3086933 | 0.42253204 | 0.219 | 0.171 | 1 | 0.00609368 | 0.80562195 | 0.244 | 0.104 | 1 | 0.3086933 | 0.01215023 |
| Dclre1a | 0.03812847 | 0.88196365 | 0.167 | 0.083 | 1 | 0.00610243 | 1.63767732 | 0.164 | 0.052 | 1 | 0.03812847 | 0.01216761 |
| Gm42984 | 0.58046959 | 0.37006462 | 0.188 | 0.161 | 1 | 0.00614004 | 1.0961371 | 0.185 | 0.062 | 1 | 0.58046959 | 0.01224238 |
| Zfp318 | 0.00614836 | 0.67045955 | 0.49 | 0.326 | 1 | 0.03999678 | 0.22448117 | 0.429 | 0.281 | 1 | 0.03999678 | 0.01225891 |
| Miga2 | 0.89747249 | 0.14499806 | 0.083 | 0.088 | 1 | 0.00615018 | 1.08869411 | 0.185 | 0.062 | 1 | 0.89747249 | 0.01226253 |
| Acaa1a | 0.38589465 | 0.10605837 | 0.604 | 0.565 | 1 | 0.00616225 | 0.28237524 | 0.546 | 0.354 | 1 | 0.38589465 | 0.01228653 |
| Epm2aip1 | 0.0061701 | 0.58691885 | 0.479 | 0.301 | 1 | 0.04558057 | 0.23763253 | 0.429 | 0.281 | 1 | 0.04558057 | 0.01230214 |
| Tinagl1 | 0.00617481 | -1.9621318 | 0.042 | 0.15 | 1 | 0.33103112 | -0.6577786 | 0.038 | 0.062 | 1 | 0.33103112 | 0.0123115 |
| Kif1c | 0.17023178 | 0.27052894 | 0.385 | 0.301 | 1 | 0.00621254 | 0.18011072 | 0.508 | 0.292 | 1 | 0.17023178 | 0.01238648 |
| Dld | 0.00623695 | 0.57277467 | 0.594 | 0.435 | 1 | 0.51771364 | -0.1093223 | 0.618 | 0.552 | 1 | 0.51771364 | 0.01243499 |
| Rcl1 | 0.25942396 | -0.1413688 | 0.354 | 0.275 | 1 | 0.00624905 | 0.17781004 | 0.378 | 0.198 | 1 | 0.25942396 | 0.01245906 |
| Smpd4 | 0.26992268 | 0.44189736 | 0.219 | 0.166 | 1 | 0.00625202 | 0.58322953 | 0.29 | 0.135 | 1 | 0.26992268 | 0.01246495 |
| Gtf3c6 | 0.01330501 | 0.65957123 | 0.365 | 0.223 | 1 | 0.0062577 | 0.58322953 | 0.349 | 0.188 | 1 | 0.01330501 | 0.01247625 |
| Inafm2 | 0.66163784 | 0.30705482 | 0.125 | 0.109 | 1 | 0.00626823 | 0.69014474 | 0.185 | 0.062 | 1 | 0.66163784 | 0.01249717 |
| Rpp30 | 0.89003205 | 0.10479174 | 0.188 | 0.197 | 1 | 0.00628341 | 1.13945214 | 0.223 | 0.094 | 1 | 0.89003205 | 0.01252735 |
| Pak4 | 0.96001588 | 0.14076107 | 0.146 | 0.145 | 1 | 0.00630071 | 0.8917786 | 0.168 | 0.052 | 1 | 0.96001588 | 0.01256171 |
| Necab1 | 0.03210036 | -2.9925055 | 0 | 0.047 | 1 | 0.00630374 | -5.1172102 | 0 | 0.031 | 1 | 0.03210036 | 0.01256774 |
| Thy1 | 0.0819164 | -2.1624305 | 0 | 0.031 | 1 | 0.00630374 | -6.2640516 | 0 | 0.031 | 1 | 0.0819164 | 0.01256774 |
| St8sia5 | 0.15729781 | -2.3144336 | 0 | 0.021 | 1 | 0.00630374 | -3.6317834 | 0 | 0.031 | 1 | 0.15729781 | 0.01256774 |
| 5730419F03Rik | 0.22187388 | -1.3144336 | 0 | 0.016 | 1 | 0.00630374 | -4.1172102 | 0 | 0.031 | 1 | 0.22187388 | 0.01256774 |
| 6430562O15Rik | 0.22187388 | -1.577468 | 0 | 0.016 | 1 | 0.00630374 | -3.6317834 | 0 | 0.031 | 1 | 0.22187388 | 0.01256774 |
| Gm48202 | 0.22187575 | -0.9925055 | 0 | 0.016 | 1 | 0.00630374 | -3.8948178 | 0 | 0.031 | 1 | 0.22187575 | 0.01256774 |

| | | | | | | | | | | | | |
|---|---|---|---|---|---|---|---|---|---|---|---|---|
| Gm37245 | 0.28735681 | -0.3144336 | 0.021 | 0.047 | 1 | 0.00630374 | -3.8948178 | 0 | 0.031 | 1 | 0.28735681 | 0.01256774 |
| Soat2 | 0.99482447 | -0.2555399 | 0.042 | 0.041 | 1 | 0.00630374 | -3.8948178 | 0 | 0.031 | 1 | 0.99482447 | 0.01256774 |
| Cenpu | 0.07100615 | 0.85549144 | 0.062 | 0.021 | 1 | 0.00630389 | -3.3098553 | 0 | 0.031 | 1 | 0.07100615 | 0.01256805 |
| Gm48089 | 0.11302151 | -1.7998604 | 0 | 0.026 | 1 | 0.00630389 | -5.010295 | 0 | 0.031 | 1 | 0.11302151 | 0.01256805 |
| Csnka2ip | 0.15729403 | -1.577468 | 0 | 0.021 | 1 | 0.00630389 | -6.010295 | 0 | 0.031 | 1 | 0.15729403 | 0.01256805 |
| F830016B08Rik | 0.15729781 | -2.577468 | 0 | 0.021 | 1 | 0.00630389 | -5.1172102 | 0 | 0.031 | 1 | 0.15729781 | 0.01256805 |
| Hhip | 0.20275693 | 0.78510212 | 0.031 | 0.01 | 1 | 0.00630389 | -6.010295 | 0 | 0.031 | 1 | 0.20275693 | 0.01256805 |
| Gm47200 | 0.20317827 | -3.2779077 | 0.01 | 0.036 | 1 | 0.00630389 | -3.3098553 | 0 | 0.031 | 1 | 0.20317827 | 0.01256805 |
| Reep6 | 0.22187575 | -0.9925055 | 0 | 0.016 | 1 | 0.00630389 | -3.3098553 | 0 | 0.031 | 1 | 0.22187575 | 0.01256805 |
| Plekhg4 | 0.28126256 | -1.1624305 | 0.01 | 0.031 | 1 | 0.00630389 | -3.3098553 | 0 | 0.031 | 1 | 0.28126256 | 0.01256805 |
| Selenov | 0.32025586 | -1.7998604 | 0 | 0.01 | 1 | 0.00630389 | -3.3098553 | 0 | 0.031 | 1 | 0.32025586 | 0.01256805 |
| Gm4951 | 0.32025586 | -1.3144336 | 0 | 0.01 | 1 | 0.00630389 | -4.6317834 | 0 | 0.031 | 1 | 0.32025586 | 0.01256805 |
| Prph | 0.38192865 | -1.3144336 | 0.01 | 0.026 | 1 | 0.00630389 | -4.1172102 | 0 | 0.031 | 1 | 0.38192865 | 0.01256805 |
| Gm43434 | 0.38357309 | -1.7998604 | 0.01 | 0.026 | 1 | 0.00630389 | -4.1172102 | 0 | 0.031 | 1 | 0.38357309 | 0.01256805 |
| Duoxa1 | 0.46686094 | -1.6554705 | 0.021 | 0.036 | 1 | 0.00630389 | -4.3098553 | 0 | 0.031 | 1 | 0.46686094 | 0.01256805 |
| 4930544I03Rik | 0.48064264 | 0.59245704 | 0.021 | 0.01 | 1 | 0.00630389 | -4.6317834 | 0 | 0.031 | 1 | 0.48064264 | 0.01256805 |
| 9230112J17Rik | 0.61737927 | 1.00749454 | 0.01 | 0.005 | 1 | 0.00630389 | -4.6317834 | 0 | 0.031 | 1 | 0.61737927 | 0.01256805 |
| Gm6980 | 0.73685701 | 0.59245704 | 0.01 | 0.016 | 1 | 0.00630389 | -5.1172102 | 0 | 0.031 | 1 | 0.73685701 | 0.01256805 |
| Gm48293 | 0.75328966 | 0.27052894 | 0.021 | 0.016 | 1 | 0.00630389 | -3.8948178 | 0 | 0.031 | 1 | 0.75328966 | 0.01256805 |
| Sec1 | 1 | 0.42253204 | 0.01 | 0.01 | 1 | 0.00630389 | -3.3098553 | 0 | 0.031 | 1 | 1 | 0.01256805 |
| Gm12185 | 1 | -0.7294711 | 0.01 | 0.01 | 1 | 0.00630389 | -4.3098553 | 0 | 0.031 | 1 | 1 | 0.01256805 |
| Gsc | 1 | 0.42253204 | 0.01 | 0.01 | 1 | 0.00630389 | -3.3098553 | 0 | 0.031 | 1 | 1 | 0.01256805 |
| 5530601H04Rik | 0.11997727 | 0.56455104 | 0.396 | 0.321 | 1 | 0.00631101 | 0.17396051 | 0.445 | 0.24 | 1 | 0.11997727 | 0.0125822 |
| Rps6ka1 | 0.03268802 | 0.60695661 | 0.219 | 0.119 | 1 | 0.00633467 | -1.0050007 | 0.143 | 0.271 | 1 | 0.03268802 | 0.01262922 |
| Cyp20a1 | 0.00636235 | 0.38905733 | 0.708 | 0.56 | 1 | 0.03732348 | -0.2588713 | 0.634 | 0.385 | 1 | 0.03732348 | 0.01268422 |
| 1600020E01Rik | 0.1972031 | 0.19521742 | 0.688 | 0.591 | 1 | 0.00638014 | 0.36875008 | 0.693 | 0.51 | 1 | 0.1972031 | 0.01271958 |
| Trim59 | 0.00638314 | 1.30122574 | 0.219 | 0.098 | 1 | 0.91414379 | -0.368749 | 0.143 | 0.135 | 1 | 0.91414379 | 0.01272553 |
| Eppk1 | 0.29910836 | -1.0431315 | 0.042 | 0.073 | 1 | 0.00642224 | -1.2273931 | 0.042 | 0.125 | 1 | 0.29910836 | 0.01280323 |
| Egr2 | 0.00642408 | -3.3382803 | 0.021 | 0.114 | 1 | 0.79918975 | -2.7952821 | 0.038 | 0.031 | 1 | 0.79918975 | 0.01280689 |
| Ovgp1 | 0.00878768 | 1.20013961 | 0.167 | 0.067 | 1 | 0.00645354 | 1.27510724 | 0.134 | 0.031 | 1 | 0.00878768 | 0.01286542 |
| Ube2j1 | 0.02320179 | 0.30849679 | 0.781 | 0.648 | 1 | 0.00645576 | 0.31051257 | 0.815 | 0.729 | 1 | 0.02320179 | 0.01286984 |
| Nsun5 | 0.09726931 | 1.00749454 | 0.125 | 0.067 | 1 | 0.00649683 | 2.05937855 | 0.113 | 0.021 | 1 | 0.09726931 | 0.01295146 |
| Csnk1g2 | 0.00650894 | 0.30620792 | 0.781 | 0.596 | 1 | 0.01437222 | 0.3705268 | 0.752 | 0.656 | 1 | 0.01437222 | 0.01297551 |
| Psmd5 | 0.87422001 | 0.24850264 | 0.25 | 0.249 | 1 | 0.00650931 | 0.45344448 | 0.399 | 0.219 | 1 | 0.87422001 | 0.01297625 |
| Pxmp4 | 0.07425183 | -0.2894872 | 0.25 | 0.363 | 1 | 0.00652934 | 1.06865636 | 0.197 | 0.073 | 1 | 0.07425183 | 0.01301605 |
| Nosip | 0.00654202 | 1.10703021 | 0.26 | 0.135 | 1 | 0.02638361 | 0.25308093 | 0.223 | 0.104 | 1 | 0.02638361 | 0.01304125 |
| Prkci | 0.10161148 | 0.15601706 | 0.375 | 0.259 | 1 | 0.00654334 | -0.670737 | 0.513 | 0.667 | 1 | 0.10161148 | 0.01304386 |
| Slc6a6 | 0.29169982 | -0.2595107 | 0.927 | 0.948 | 1 | 0.00656156 | -0.871548 | 0.912 | 0.948 | 1 | 0.29169982 | 0.01308007 |
| Wdr35 | 0.00657057 | 0.62237262 | 0.583 | 0.415 | 1 | 0.03895361 | 0.12755005 | 0.651 | 0.417 | 1 | 0.03895361 | 0.01309798 |
| Mkrn2 | 0.1351792 | 0.45647937 | 0.312 | 0.233 | 1 | 0.00657318 | 0.72256622 | 0.273 | 0.125 | 1 | 0.1351792 | 0.01310315 |
| Tor1a | 0.08078733 | 0.346981 | 0.469 | 0.358 | 1 | 0.00658374 | 0.30258797 | 0.454 | 0.26 | 1 | 0.08078733 | 0.01312413 |
| Nfatc2ip | 0.03617627 | 1.18806678 | 0.135 | 0.062 | 1 | 0.0065883 | 1.71989208 | 0.147 | 0.042 | 1 | 0.03617627 | 0.0131332 |
| Adamtsl4 | 0.00659422 | 0.80818573 | 0.229 | 0.104 | 1 | 0.50116032 | -0.247571 | 0.202 | 0.156 | 1 | 0.50116032 | 0.01314496 |
| Clip4 | 0.43155177 | -0.577468 | 0.031 | 0.052 | 1 | 0.00661604 | -2.0260623 | 0.05 | 0.135 | 1 | 0.43155177 | 0.01318832 |
| Ctla4 | 0.13010731 | 0.83756954 | 0.146 | 0.088 | 1 | 0.00661921 | 1.93807225 | 0.13 | 0.031 | 1 | 0.13010731 | 0.01319462 |
| Kcnma1 | 0.52577687 | -0.3591579 | 0.979 | 0.974 | 1 | 0.00662114 | -1.1557778 | 0.912 | 0.865 | 1 | 0.52577687 | 0.01319843 |
| Ddx39b | 0.00663681 | 0.27295568 | 0.875 | 0.813 | 1 | 0.0923658 | 0.15071638 | 0.891 | 0.854 | 1 | 0.0923658 | 0.01322958 |
| Abcf3 | 0.27772543 | 0.29529685 | 0.365 | 0.295 | 1 | 0.00664199 | 0.33259273 | 0.416 | 0.229 | 1 | 0.27772543 | 0.01323987 |
| Ust | 0.00665428 | 0.24214923 | 0.802 | 0.751 | 1 | 0.98600448 | -0.1765082 | 0.958 | 0.927 | 1 | 0.98600448 | 0.01326427 |
| Tmem164 | 0.01788681 | 0.14783761 | 0.625 | 0.43 | 1 | 0.00665449 | -0.6611169 | 0.777 | 0.865 | 1 | 0.01788681 | 0.0132647 |
| Nfkbib | 0.40991823 | -0.1263612 | 0.271 | 0.332 | 1 | 0.00665985 | 0.58322953 | 0.336 | 0.177 | 1 | 0.40991823 | 0.01327534 |
| Reps1 | 0.15514049 | 0.2662288 | 0.385 | 0.306 | 1 | 0.0066697 | -0.5752567 | 0.433 | 0.604 | 1 | 0.15514049 | 0.01329491 |
| Ngly1 | 0.6147671 | -0.1029295 | 0.365 | 0.316 | 1 | 0.00667568 | 0.19915838 | 0.513 | 0.323 | 1 | 0.6147671 | 0.01330679 |
| Tnfaip6 | 0.00668709 | -2.778001 | 0.115 | 0.244 | 1 | 0.21284229 | -2.0570892 | 0.088 | 0.125 | 1 | 0.21284229 | 0.01332947 |
| Cox6c | 0.58795726 | -0.2232857 | 0.979 | 0.948 | 1 | 0.00669184 | 0.16831575 | 0.962 | 0.917 | 1 | 0.58795726 | 0.0133389 |
| Rsf1 | 0.1724986 | 0.14591922 | 0.771 | 0.663 | 1 | 0.00670071 | -0.5337735 | 0.882 | 0.927 | 1 | 0.1724986 | 0.01335651 |
| Pald1 | 0.00671415 | -2.4950058 | 0.021 | 0.114 | 1 | 0.09009452 | -0.1276519 | 0.197 | 0.115 | 1 | 0.09009452 | 0.01338322 |
| Ccdc124 | 0.17738531 | 0.32711247 | 0.521 | 0.461 | 1 | 0.00671473 | 0.44621915 | 0.441 | 0.26 | 1 | 0.17738531 | 0.01338437 |
| Pold2 | 0.02469811 | 0.86999101 | 0.25 | 0.145 | 1 | 0.00672999 | 1.49749966 | 0.176 | 0.062 | 1 | 0.02469811 | 0.01341469 |
| Srrt | 0.01435139 | 0.33525171 | 0.75 | 0.611 | 1 | 0.00673035 | 0.29708561 | 0.815 | 0.76 | 1 | 0.01435139 | 0.0134154 |
| Utp11 | 0.00673325 | 0.646095 | 0.688 | 0.554 | 1 | 0.00956232 | 0.18917706 | 0.643 | 0.438 | 1 | 0.00956232 | 0.01342116 |
| Foxn2 | 0.00673731 | 0.36538813 | 0.448 | 0.264 | 1 | 0.0425372 | -0.6454583 | 0.521 | 0.615 | 1 | 0.0425372 | 0.01342924 |
| Mier1 | 0.18131763 | 0.27306778 | 0.938 | 0.933 | 1 | 0.00674066 | 0.30878288 | 0.975 | 0.979 | 1 | 0.18131763 | 0.01343589 |
| Gnb1 | 0.00674156 | 0.2853869 | 0.99 | 0.953 | 1 | 0.6110125 | -0.2039218 | 0.992 | 0.969 | 1 | 0.6110125 | 0.01343768 |
| Tmem69 | 0.14827403 | 0.70793425 | 0.125 | 0.073 | 1 | 0.00674842 | 1.77760758 | 0.13 | 0.031 | 1 | 0.14827403 | 0.0134513 |
| Ppil6 | 0.67856411 | -0.3946039 | 0.135 | 0.15 | 1 | 0.0067711 | 0.94390133 | 0.168 | 0.052 | 1 | 0.67856411 | 0.01349634 |
| 1700047M11Rik | 0.00677216 | -2.2148979 | 0.021 | 0.114 | 1 | 0.1465102 | -1.8948178 | 0.004 | 0.021 | 1 | 0.1465102 | 0.01349845 |
| Gm48727 | 0.61737927 | 1.00749454 | 0.01 | 0.005 | 1 | 0.00678267 | -3.0260623 | 0.021 | 0.083 | 1 | 0.61737927 | 0.01351933 |
| Opa3 | 0.74378836 | 0.18152394 | 0.292 | 0.28 | 1 | 0.00678746 | 0.24339838 | 0.374 | 0.198 | 1 | 0.74378836 | 0.01352885 |
| Cox6b1 | 0.25189681 | 0.15907696 | 0.938 | 0.902 | 1 | 0.00681092 | 0.33204813 | 0.929 | 0.875 | 1 | 0.25189681 | 0.01357545 |
| Gbf1 | 0.00681448 | 0.36048134 | 0.938 | 0.896 | 1 | 0.91647604 | -0.2266099 | 0.937 | 0.938 | 1 | 0.91647604 | 0.01358253 |
| Triobp | 0.93319163 | -0.18233 | 0.281 | 0.264 | 1 | 0.00681577 | 0.49191863 | 0.403 | 0.229 | 1 | 0.93319163 | 0.01358508 |
| Rnf34 | 0.46426589 | -0.2646805 | 0.594 | 0.622 | 1 | 0.00681589 | 0.15430909 | 0.576 | 0.365 | 1 | 0.46426589 | 0.01358532 |
| Pfkm | 0.56090507 | 0.32299636 | 0.229 | 0.202 | 1 | 0.00682191 | 0.69014474 | 0.298 | 0.146 | 1 | 0.56090507 | 0.01359729 |
| Etfbkmt | 0.16600729 | 1.25542205 | 0.125 | 0.078 | 1 | 0.00682278 | 1.69014474 | 0.147 | 0.042 | 1 | 0.16600729 | 0.01359901 |
| Micall1 | 0.00682534 | -0.5976458 | 0.385 | 0.549 | 1 | 0.1131343 | -0.273876 | 0.441 | 0.292 | 1 | 0.1131343 | 0.0136041 |
| Ranbp2 | 0.00682995 | 0.30732886 | 0.948 | 0.933 | 1 | 0.86208304 | -0.1602995 | 0.966 | 0.979 | 1 | 0.86208304 | 0.01361325 |
| Rnf168 | 0.36916111 | 0.22430593 | 0.479 | 0.43 | 1 | 0.00683785 | 0.22186522 | 0.58 | 0.354 | 1 | 0.36916111 | 0.01362894 |
| A430005L14Rik | 0.00684268 | 0.69030436 | 0.427 | 0.269 | 1 | 0.12026959 | 0.46273424 | 0.256 | 0.177 | 1 | 0.12026959 | 0.01363854 |
| Erbb4 | 0.12050853 | -2.5285584 | 0.021 | 0.062 | 1 | 0.00684712 | 0.84070441 | 0.193 | 0.073 | 1 | 0.12050853 | 0.01364735 |
| Sorbs3 | 0.16691127 | -0.4779323 | 0.385 | 0.487 | 1 | 0.00685243 | 0.34725702 | 0.521 | 0.333 | 1 | 0.16691127 | 0.0136579 |
| Spred2 | 0.00685679 | 0.51735558 | 0.604 | 0.425 | 1 | 0.92935369 | -0.3194201 | 0.756 | 0.792 | 1 | 0.92935369 | 0.01366656 |
| Hsd3b7 | 0.77183935 | 0.16976597 | 0.333 | 0.321 | 1 | 0.00685995 | 0.35454171 | 0.433 | 0.24 | 1 | 0.77183935 | 0.01367284 |
| Rhot2 | 0.00686002 | 1.22988696 | 0.177 | 0.073 | 1 | 0.37031807 | 0.18490943 | 0.118 | 0.083 | 1 | 0.37031807 | 0.01367299 |
| Dtd2 | 0.1137105 | 0.48240349 | 0.333 | 0.249 | 1 | 0.00686257 | 1.25085969 | 0.193 | 0.073 | 1 | 0.1137105 | 0.01367804 |

| | | | | | | | | | | | | |
|---|---|---|---|---|---|---|---|---|---|---|---|---|
| Caap1 | 0.16336666 | 0.16619228 | 0.354 | 0.259 | 1 | 0.00686368 | 0.58826512 | 0.324 | 0.167 | 1 | 0.16336666 | 0.01368025 |
| Uhrf1bp1 | 0.00686855 | 1.14499806 | 0.188 | 0.078 | 1 | 0.66978335 | -0.4208866 | 0.181 | 0.156 | 1 | 0.66978335 | 0.01368993 |
| Pcif1 | 0.04532898 | 0.45389621 | 0.438 | 0.311 | 1 | 0.00689817 | 0.25813416 | 0.471 | 0.292 | 1 | 0.04532898 | 0.01374876 |
| Rnf167 | 0.00690569 | 0.5367476 | 0.719 | 0.56 | 1 | 0.48469151 | -0.2587811 | 0.63 | 0.521 | 1 | 0.48469151 | 0.0137637 |
| Gm57383 | 0.60599256 | 0.37946331 | 0.177 | 0.155 | 1 | 0.00693232 | 1.69014474 | 0.147 | 0.042 | 1 | 0.60599256 | 0.01381659 |
| Ctsz | 0.47437361 | -0.1164942 | 0.75 | 0.725 | 1 | 0.00694067 | 0.67692104 | 0.786 | 0.625 | 1 | 0.47437361 | 0.01383316 |
| Prkar1b | 0.00694919 | 0.86310463 | 0.281 | 0.15 | 1 | 0.0138666 | 0.24473359 | 0.361 | 0.198 | 1 | 0.0138666 | 0.01385009 |
| Usf3 | 0.46868572 | -0.240433 | 0.354 | 0.29 | 1 | 0.00695149 | -0.7991173 | 0.408 | 0.583 | 1 | 0.46868572 | 0.01385466 |
| Tor3a | 0.00695983 | 0.55683313 | 0.448 | 0.269 | 1 | 0.37254855 | -0.3712558 | 0.345 | 0.271 | 1 | 0.37254855 | 0.01387122 |
| Tpp1 | 0.03702897 | -0.4431669 | 0.49 | 0.606 | 1 | 0.00696316 | 0.53188265 | 0.601 | 0.438 | 1 | 0.03702897 | 0.01387784 |
| Spp1 | 0.00696455 | -0.7188332 | 0.208 | 0.352 | 1 | 0.09940967 | 0.11187689 | 0.458 | 0.333 | 1 | 0.09940967 | 0.01388059 |
| Rsph9 | 0.74289161 | 0.30122574 | 0.146 | 0.135 | 1 | 0.00697308 | 0.78530197 | 0.168 | 0.052 | 1 | 0.74289161 | 0.01389754 |
| Zc3h7b | 0.00697694 | 0.78242798 | 0.469 | 0.332 | 1 | 0.13742702 | 0.11210165 | 0.538 | 0.385 | 1 | 0.13742702 | 0.01390521 |
| Rbm4b | 0.07998065 | 0.36037556 | 0.385 | 0.275 | 1 | 0.00698271 | 0.47864063 | 0.433 | 0.25 | 1 | 0.07998065 | 0.01391666 |
| Rtl4 | 0.00698618 | -5.6507169 | 0 | 0.073 | 1 | 0.34026741 | -3.2358547 | 0.008 | 0.021 | 1 | 0.34026741 | 0.01392356 |
| Irak1bp1 | 0.0342088 | 0.65755207 | 0.344 | 0.233 | 1 | 0.006994 | 0.82084843 | 0.336 | 0.188 | 1 | 0.0342088 | 0.01393909 |
| Hyou1 | 0.05710015 | 0.59245704 | 0.396 | 0.285 | 1 | 0.00701345 | 0.66996686 | 0.395 | 0.229 | 1 | 0.05710015 | 0.01397771 |
| Pkn2 | 0.00701386 | 0.20353819 | 0.917 | 0.907 | 1 | 0.01474292 | -0.4291367 | 0.983 | 0.99 | 1 | 0.01474292 | 0.01397852 |
| Lancl1 | 0.17026008 | 0.45001277 | 0.406 | 0.332 | 1 | 0.00702155 | 0.55143847 | 0.34 | 0.177 | 1 | 0.17026008 | 0.0139938 |
| Col1a2 | 0.06200346 | -0.3031154 | 0.844 | 0.741 | 1 | 0.00702433 | 0.23010691 | 0.983 | 0.823 | 1 | 0.06200346 | 0.01399933 |
| Zranb2 | 0.00703227 | 0.3776355 | 0.875 | 0.762 | 1 | 0.03898506 | 0.21820485 | 0.786 | 0.75 | 1 | 0.03898506 | 0.01401509 |
| Abcb8 | 0.71458965 | 0.18806678 | 0.156 | 0.14 | 1 | 0.00704179 | 1.04064198 | 0.181 | 0.062 | 1 | 0.71458965 | 0.01403399 |
| Hivep3 | 0.00704653 | 0.85173687 | 0.615 | 0.492 | 1 | 0.06867778 | -0.4655061 | 0.458 | 0.594 | 1 | 0.06867778 | 0.0140434 |
| Dguok | 0.10120309 | 0.46317402 | 0.365 | 0.269 | 1 | 0.00704717 | 0.60768258 | 0.412 | 0.24 | 1 | 0.10120309 | 0.01404467 |
| Birc2 | 0.0121793 | 0.345491 | 0.51 | 0.352 | 1 | 0.00707101 | 0.41596977 | 0.366 | 0.198 | 1 | 0.0121793 | 0.01409203 |
| Eeig1 | 0.09085941 | -0.7667383 | 0.5 | 0.554 | 1 | 0.00707843 | -0.3218279 | 0.395 | 0.604 | 1 | 0.09085941 | 0.01410676 |
| Magee1 | 0.05669636 | 0.86999101 | 0.177 | 0.098 | 1 | 0.00707955 | 1.27510724 | 0.206 | 0.083 | 1 | 0.05669636 | 0.01410898 |
| Nkap | 0.01033897 | 0.70793425 | 0.479 | 0.342 | 1 | 0.0070815 | 0.49749966 | 0.433 | 0.25 | 1 | 0.01033897 | 0.01411285 |
| Grk6 | 0.37059974 | -0.3910548 | 0.302 | 0.342 | 1 | 0.00708815 | 0.70721825 | 0.273 | 0.125 | 1 | 0.37059974 | 0.01412605 |
| Phykpl | 0.00709252 | 0.90057933 | 0.312 | 0.176 | 1 | 0.28922683 | -0.1063219 | 0.269 | 0.198 | 1 | 0.28922683 | 0.01413473 |
| Sprr1a | 0.00710522 | 1.00749454 | 0.156 | 0.057 | 1 | 0.50671238 | 0.98782529 | 0.055 | 0.073 | 1 | 0.50671238 | 0.01415995 |
| Slc36a1 | 0.00712467 | 1.20013961 | 0.146 | 0.052 | 1 | 0.03687858 | 1.34222143 | 0.143 | 0.062 | 1 | 0.03687858 | 0.01419857 |
| Zfp579 | 0.98129662 | 0.21899864 | 0.083 | 0.083 | 1 | 0.00714014 | 1.73453886 | 0.113 | 0.021 | 1 | 0.98129662 | 0.01422931 |
| Gpc1 | 0.03236158 | 0.3625895 | 0.844 | 0.71 | 1 | 0.00714491 | 0.27102606 | 0.782 | 0.542 | 1 | 0.03236158 | 0.01423876 |
| Ints14 | 0.36374726 | 0.22680756 | 0.292 | 0.238 | 1 | 0.00716754 | 0.62088207 | 0.332 | 0.177 | 1 | 0.36374726 | 0.01428371 |
| Ippk | 0.00924174 | 0.57139542 | 0.417 | 0.264 | 1 | 0.00717398 | 0.46414106 | 0.42 | 0.24 | 1 | 0.00924174 | 0.01429649 |
| Ssh1 | 0.75843288 | -0.6436582 | 0.385 | 0.368 | 1 | 0.00717673 | 0.24566346 | 0.634 | 0.385 | 1 | 0.75843288 | 0.01430195 |
| Parp1 | 0.00718031 | 0.5065963 | 0.406 | 0.254 | 1 | 0.03405153 | 0.41949715 | 0.319 | 0.198 | 1 | 0.03405153 | 0.01430907 |
| Aamp | 0.12739681 | 0.179407 | 0.885 | 0.798 | 1 | 0.00718667 | 0.2839446 | 0.878 | 0.792 | 1 | 0.12739681 | 0.01432169 |
| Tex30 | 0.10186806 | 0.72209232 | 0.208 | 0.135 | 1 | 0.00718964 | 0.67620555 | 0.282 | 0.135 | 1 | 0.10186806 | 0.01432759 |
| Fbxl17 | 0.79488905 | -0.1799641 | 0.917 | 0.886 | 1 | 0.00719477 | -0.6526687 | 0.954 | 0.969 | 1 | 0.79488905 | 0.01433777 |
| Gm15271 | 0.00722076 | -1.6805615 | 0.135 | 0.269 | 1 | 0.79600331 | -1.3098553 | 0.017 | 0.021 | 1 | 0.79600331 | 0.01438938 |
| Pip4p2 | 0.17446058 | 0.2466822 | 0.583 | 0.508 | 1 | 0.00722812 | 0.2107726 | 0.676 | 0.438 | 1 | 0.17446058 | 0.01440399 |
| Ttc19 | 0.00725794 | 0.45001277 | 0.708 | 0.617 | 1 | 0.98473399 | -0.2476605 | 0.752 | 0.729 | 1 | 0.98473399 | 0.01446321 |
| Gas8 | 0.02616869 | 0.89787005 | 0.26 | 0.155 | 1 | 0.00726255 | 0.76814725 | 0.252 | 0.115 | 1 | 0.02616869 | 0.01447235 |
| Dazap1 | 0.00726559 | 0.55259258 | 0.656 | 0.503 | 1 | 0.8023596 | -0.1715003 | 0.735 | 0.74 | 1 | 0.8023596 | 0.01447839 |
| Gm7694 | 0.63343064 | 0.27052894 | 0.083 | 0.067 | 1 | 0.00726668 | 0.69014474 | 0.202 | 0.073 | 1 | 0.63343064 | 0.01448055 |
| Cdca2 | 0.00728237 | 3.66460682 | 0.125 | 0.041 | 1 | 0.30382073 | 1.01207283 | 0.029 | 0.01 | 1 | 0.30382073 | 0.01451172 |
| Vps53 | 0.18205424 | 0.15123001 | 0.417 | 0.321 | 1 | 0.0072896 | 0.56163 | 0.445 | 0.281 | 1 | 0.18205424 | 0.01452605 |
| Bcl10 | 0.00729126 | 0.64385619 | 0.698 | 0.591 | 1 | 0.42069032 | -0.1109512 | 0.584 | 0.667 | 1 | 0.42069032 | 0.01452936 |
| Tyw1 | 0.87409525 | 0.20604022 | 0.198 | 0.207 | 1 | 0.00729395 | 0.89525917 | 0.235 | 0.104 | 1 | 0.87409525 | 0.01453471 |
| Traf5 | 0.00730406 | 0.60756393 | 0.5 | 0.332 | 1 | 0.32748368 | -0.3402289 | 0.315 | 0.24 | 1 | 0.32748368 | 0.01455477 |
| Gng3 | 0.00730574 | 1.09495738 | 0.146 | 0.052 | 1 | 0.09753843 | -1.3098553 | 0.134 | 0.198 | 1 | 0.09753843 | 0.0145581 |
| Snapc2 | 0.06993397 | 0.88196365 | 0.198 | 0.119 | 1 | 0.00732089 | 0.86006974 | 0.197 | 0.073 | 1 | 0.06993397 | 0.01458819 |
| Clip3 | 0.35901367 | 0.14499806 | 0.25 | 0.197 | 1 | 0.00732144 | 0.5492822 | 0.395 | 0.229 | 1 | 0.35901367 | 0.01458928 |
| Cfap70 | 0.25491248 | -0.7204259 | 0.083 | 0.13 | 1 | 0.00732164 | 1.32904605 | 0.282 | 0.135 | 1 | 0.25491248 | 0.01458968 |
| Gm14743 | 0.05999981 | 1.42253204 | 0.073 | 0.026 | 1 | 0.00732483 | 3.08246216 | 0.071 | 0 | 1 | 0.05999981 | 0.014596 |
| Mmp28 | 0.04092384 | -1.2609943 | 0.125 | 0.223 | 1 | 0.00733258 | -1.378568 | 0.08 | 0.177 | 1 | 0.04092384 | 0.01461139 |
| Sbf2 | 0.92889125 | -0.1875214 | 0.615 | 0.617 | 1 | 0.007347 | -0.5742048 | 0.828 | 0.885 | 1 | 0.92889125 | 0.01464002 |
| Mybl1 | 0.00736227 | 1.69102087 | 0.219 | 0.104 | 1 | 0.88415025 | -0.9228321 | 0.101 | 0.104 | 1 | 0.88415025 | 0.01467034 |
| Npas3 | 0.00737402 | 0.33295144 | 0.885 | 0.798 | 1 | 0.11544105 | -0.1822927 | 0.933 | 0.729 | 1 | 0.11544105 | 0.01469367 |
| Dusp4 | 0.02362383 | -3.3848229 | 0 | 0.052 | 1 | 0.00737411 | -0.3923174 | 0.042 | 0.125 | 1 | 0.02362383 | 0.01469384 |
| Tmub1 | 0.01360086 | 0.97507306 | 0.323 | 0.202 | 1 | 0.0073763 | 0.79976923 | 0.277 | 0.135 | 1 | 0.01360086 | 0.0146982 |
| Magi2 | 0.00738569 | -0.697619 | 0.677 | 0.767 | 1 | 0.11544442 | -0.525394 | 0.702 | 0.729 | 1 | 0.11544442 | 0.01471683 |
| Slc7a11 | 0.00738914 | 2.24850264 | 0.125 | 0.041 | 1 | 0.04191592 | 3.21370669 | 0.08 | 0.021 | 1 | 0.04191592 | 0.01472368 |
| Cmtm3 | 0.00739383 | 0.3225906 | 0.51 | 0.316 | 1 | 0.04367227 | 0.11283581 | 0.357 | 0.219 | 1 | 0.04367227 | 0.01473298 |
| Mon2 | 0.00741761 | 0.42790521 | 0.854 | 0.762 | 1 | 0.41409185 | -0.3400049 | 0.912 | 0.927 | 1 | 0.41409185 | 0.01478021 |
| Rfesd | 0.22795613 | 0.45495351 | 0.135 | 0.088 | 1 | 0.00742676 | 2.44503224 | 0.092 | 0.01 | 1 | 0.22795613 | 0.01479837 |
| Chac2 | 0.40005756 | 0.54806292 | 0.167 | 0.13 | 1 | 0.00743589 | 0.33923758 | 0.231 | 0.094 | 1 | 0.40005756 | 0.01481649 |
| Cwc22 | 0.00743704 | 0.62283069 | 0.469 | 0.306 | 1 | 0.01509884 | 0.31529633 | 0.483 | 0.302 | 1 | 0.01509884 | 0.01481877 |
| Rfc2 | 0.10686537 | 0.83756954 | 0.302 | 0.228 | 1 | 0.00745532 | 0.49749966 | 0.273 | 0.125 | 1 | 0.10686537 | 0.01485506 |
| Dcps | 0.95949697 | -0.1118044 | 0.25 | 0.244 | 1 | 0.00747089 | 0.78325414 | 0.21 | 0.083 | 1 | 0.95949697 | 0.01488596 |
| Mir99ahg | 0.08802295 | -0.6927313 | 0.927 | 0.933 | 1 | 0.00750613 | -0.5993619 | 0.807 | 0.896 | 1 | 0.08802295 | 0.01495593 |
| Gpx3 | 0.00750877 | -1.360316 | 0.521 | 0.663 | 1 | 0.7800597 | -1.2407309 | 0.223 | 0.219 | 1 | 0.7800597 | 0.01496115 |
| Col5a1 | 0.3683113 | -0.4218171 | 0.594 | 0.503 | 1 | 0.00751004 | -1.080259 | 0.613 | 0.583 | 1 | 0.3683113 | 0.01496367 |
| Chid1 | 0.0075133 | 1.03174208 | 0.385 | 0.249 | 1 | 0.03969259 | 0.14184071 | 0.336 | 0.208 | 1 | 0.03969259 | 0.01497015 |
| Rbm28 | 0.00751416 | 0.61983348 | 0.615 | 0.472 | 1 | 0.28286483 | -0.370462 | 0.588 | 0.646 | 1 | 0.28286483 | 0.01497186 |
| Slc39a10 | 0.15418504 | -0.5059147 | 0.302 | 0.373 | 1 | 0.00752312 | -0.8017084 | 0.429 | 0.573 | 1 | 0.15418504 | 0.01498963 |
| Cyp4f16 | 0.16397414 | 0.5049942 | 0.198 | 0.135 | 1 | 0.00752566 | 0.71762547 | 0.197 | 0.073 | 1 | 0.16397414 | 0.01499468 |
| Msx2 | 0.00752905 | 1.62898291 | 0.135 | 0.047 | 1 | 0.12914744 | 0.61975541 | 0.197 | 0.125 | 1 | 0.12914744 | 0.01500142 |
| Cfap43 | 0.04397655 | 0.54183096 | 0.292 | 0.187 | 1 | 0.00753119 | -1.8017084 | 0.097 | 0.198 | 1 | 0.04397655 | 0.01500566 |
| Baiap2l1 | 0.00753132 | -1.0543447 | 0.771 | 0.782 | 1 | 0.78644969 | -0.3888436 | 0.857 | 0.844 | 1 | 0.78644969 | 0.01500592 |
| Grk2 | 0.00753288 | 0.72738662 | 0.365 | 0.212 | 1 | 0.87812343 | -0.5163061 | 0.416 | 0.365 | 1 | 0.87812343 | 0.01500901 |
| Pou3f3 | 0.045145 | -1.0547897 | 0.146 | 0.244 | 1 | 0.00753671 | 0.45407738 | 0.399 | 0.219 | 1 | 0.045145 | 0.01501662 |
| Scap | 0.72786598 | 0.37744415 | 0.292 | 0.28 | 1 | 0.00754273 | 0.30312161 | 0.437 | 0.25 | 1 | 0.72786598 | 0.01502856 |
| Slc35b4 | 0.02179434 | 0.81986753 | 0.281 | 0.166 | 1 | 0.00754598 | 0.36530505 | 0.319 | 0.167 | 1 | 0.02179434 | 0.01503501 |
| Lrriq1 | 0.00756973 | -1.7910634 | 0.229 | 0.373 | 1 | 0.14168937 | 0.54480537 | 0.252 | 0.177 | 1 | 0.14168937 | 0.01508216 |
| Sobp | 0.30418105 | -0.2072143 | 0.875 | 0.907 | 1 | 0.00758017 | -0.4369672 | 0.958 | 0.938 | 1 | 0.30418105 | 0.01510288 |
| Taok2 | 0.01180167 | 0.38734261 | 0.438 | 0.269 | 1 | 0.00758298 | 0.44256398 | 0.529 | 0.333 | 1 | 0.01180167 | 0.01510847 |
| Ppp2r3a | 0.28013586 | -0.3293453 | 0.604 | 0.658 | 1 | 0.0075944 | -0.7687471 | 0.723 | 0.802 | 1 | 0.28013586 | 0.01513112 |
| St13 | 0.00759513 | 0.25608913 | 0.969 | 0.912 | 1 | 0.01069973 | 0.18382393 | 0.975 | 0.927 | 1 | 0.01069973 | 0.01513257 |
| Rassf2 | 0.00760043 | 1.12070515 | 0.323 | 0.192 | 1 | 0.01978253 | 0.62874419 | 0.286 | 0.156 | 1 | 0.01978253 | 0.0151431 |

| | | | | | | | | | | | | |
|---|---|---|---|---|---|---|---|---|---|---|---|---|
| Gtf2h3 | 0.01262504 | 0.91019734 | 0.323 | 0.197 | 1 | 0.00762144 | 0.97025266 | 0.248 | 0.115 | 1 | 0.01262504 | 0.0151848 |
| H13 | 0.00762194 | 0.3317291 | 0.917 | 0.881 | 1 | 0.25131579 | 0.14639121 | 0.945 | 0.927 | 1 | 0.25131579 | 0.01518579 |
| Mcm9 | 0.00762725 | 0.68556644 | 0.302 | 0.161 | 1 | 0.5348899 | -0.6759832 | 0.235 | 0.188 | 1 | 0.5348899 | 0.01519633 |
| Eif3j1 | 0.33631114 | 0.14984746 | 0.896 | 0.922 | 1 | 0.00764961 | 0.27723354 | 0.92 | 0.854 | 1 | 0.33631114 | 0.01524071 |
| Homer2 | 0.66589719 | 0.48393258 | 0.062 | 0.078 | 1 | 0.00766701 | 1.15829357 | 0.193 | 0.073 | 1 | 0.66589719 | 0.01527523 |
| Cdin1 | 0.61325945 | 0.25260703 | 0.156 | 0.135 | 1 | 0.00767906 | 0.55687821 | 0.147 | 0.042 | 1 | 0.61325945 | 0.01529915 |
| Skic2 | 0.08799462 | 0.407102 | 0.281 | 0.187 | 1 | 0.00768647 | 0.58544736 | 0.315 | 0.167 | 1 | 0.08799462 | 0.01531386 |
| Srf | 0.06102834 | 0.78121868 | 0.323 | 0.228 | 1 | 0.00769089 | 0.56461386 | 0.315 | 0.167 | 1 | 0.06102834 | 0.01532264 |
| Gatd1 | 0.34641534 | -0.407543 | 0.167 | 0.212 | 1 | 0.00769244 | 0.86006974 | 0.206 | 0.083 | 1 | 0.34641534 | 0.0153257 |
| Psma3 | 0.02368383 | 0.22272717 | 0.979 | 0.974 | 1 | 0.00769664 | 0.21597211 | 0.987 | 0.917 | 1 | 0.02368383 | 0.01533405 |
| Emc1 | 0.6748897 | -0.2820121 | 0.25 | 0.264 | 1 | 0.00769667 | 0.89525917 | 0.298 | 0.156 | 1 | 0.6748897 | 0.01533411 |
| Grk3 | 0.10349288 | 0.51564144 | 0.24 | 0.155 | 1 | 0.00769719 | 0.52446671 | 0.399 | 0.219 | 1 | 0.10349288 | 0.01533513 |
| Ramac | 0.01044284 | 0.35634176 | 0.802 | 0.622 | 1 | 0.00773665 | 0.45019394 | 0.794 | 0.719 | 1 | 0.01044284 | 0.01541345 |
| Gpcpd1 | 0.48479662 | 0.31601171 | 0.583 | 0.58 | 1 | 0.00773858 | -0.3314853 | 0.534 | 0.719 | 1 | 0.48479662 | 0.01541726 |
| Gskip | 0.02923658 | 0.56692195 | 0.333 | 0.207 | 1 | 0.0077438 | 0.31692741 | 0.319 | 0.156 | 1 | 0.02923658 | 0.01542763 |
| U2af2 | 0.04373909 | 0.29074216 | 0.708 | 0.554 | 1 | 0.00774825 | 0.45665461 | 0.693 | 0.615 | 1 | 0.04373909 | 0.01543646 |
| Chtop | 0.00776198 | 0.529793 | 0.646 | 0.492 | 1 | 0.14746708 | 0.18675136 | 0.689 | 0.615 | 1 | 0.14746708 | 0.01546371 |
| Hspa4 | 0.01151813 | 0.31594956 | 0.938 | 0.927 | 1 | 0.00777209 | 0.2437809 | 0.966 | 0.938 | 1 | 0.01151813 | 0.01548378 |
| Plagl2 | 0.08963087 | 0.21608116 | 0.25 | 0.155 | 1 | 0.00777726 | 0.60768258 | 0.315 | 0.167 | 1 | 0.08963087 | 0.01549403 |
| Mettl2 | 0.00778207 | 0.97001983 | 0.302 | 0.166 | 1 | 0.04460653 | 0.33650778 | 0.227 | 0.125 | 1 | 0.04460653 | 0.01550359 |
| Hacd2 | 0.00780172 | -0.6811466 | 0.521 | 0.637 | 1 | 0.06333489 | -0.4196434 | 0.55 | 0.688 | 1 | 0.06333489 | 0.01554257 |
| Smg9 | 0.10868585 | 0.75595577 | 0.177 | 0.109 | 1 | 0.0078073 | 0.73453886 | 0.214 | 0.083 | 1 | 0.10868585 | 0.01555365 |
| 1110019D14Rik | 0.90770067 | 0.15949763 | 0.135 | 0.145 | 1 | 0.00780827 | 1.23071312 | 0.164 | 0.052 | 1 | 0.90770067 | 0.01555557 |
| Crat | 0.05892356 | 1.07460873 | 0.167 | 0.093 | 1 | 0.00782396 | 1.7944814 | 0.143 | 0.042 | 1 | 0.05892356 | 0.0155867 |
| Tmem42 | 0.10862553 | 0.46387283 | 0.417 | 0.326 | 1 | 0.00782939 | 0.87549978 | 0.353 | 0.208 | 1 | 0.10862553 | 0.01559748 |
| Cdk5 | 0.03566166 | 0.46000674 | 0.427 | 0.29 | 1 | 0.00785992 | 0.67503785 | 0.34 | 0.188 | 1 | 0.03566166 | 0.01565805 |
| Dnm3os | 0.20787066 | -0.9697854 | 0.094 | 0.145 | 1 | 0.00786443 | -1.8017084 | 0.088 | 0.188 | 1 | 0.20787066 | 0.01566701 |
| Lamtor3 | 0.01704409 | 0.53800925 | 0.604 | 0.477 | 1 | 0.00786905 | 0.33752974 | 0.689 | 0.521 | 1 | 0.01704409 | 0.01567617 |
| Mcm10 | 0.00788627 | 1.17741954 | 0.146 | 0.052 | 1 | 0.22483655 | 0.36821664 | 0.118 | 0.073 | 1 | 0.22483655 | 0.01571035 |
| Acvr2a | 0.01032722 | 0.81484946 | 0.406 | 0.269 | 1 | 0.00789906 | -0.8048611 | 0.475 | 0.615 | 1 | 0.01032722 | 0.01573573 |
| Zbtb4 | 0.5508824 | 0.23882008 | 0.354 | 0.316 | 1 | 0.00790058 | 0.17292684 | 0.508 | 0.302 | 1 | 0.5508824 | 0.01573875 |
| Edc3 | 0.22481015 | -0.3144336 | 0.167 | 0.233 | 1 | 0.00792599 | 0.83698613 | 0.29 | 0.146 | 1 | 0.22481015 | 0.01578917 |
| Cacng4 | 0.00793222 | -1.2762984 | 0.135 | 0.269 | 1 | 0.07934909 | -1.1070384 | 0.109 | 0.177 | 1 | 0.07934909 | 0.01580152 |
| Pgm1 | 0.53492712 | -0.1547769 | 0.177 | 0.207 | 1 | 0.0079546 | 0.63767732 | 0.193 | 0.073 | 1 | 0.53492712 | 0.01584593 |
| Tra2a | 0.00795829 | -0.3509594 | 0.938 | 0.974 | 1 | 0.02383329 | -0.3954377 | 0.95 | 0.979 | 1 | 0.02383329 | 0.01585325 |
| Eva1a | 0.22228981 | 1.11852585 | 0.125 | 0.083 | 1 | 0.0079743 | 0.20861183 | 0.248 | 0.104 | 1 | 0.22228981 | 0.01588501 |
| Mboat7 | 0.11304618 | 0.38600616 | 0.323 | 0.228 | 1 | 0.00798908 | 0.44817195 | 0.357 | 0.188 | 1 | 0.11304618 | 0.01591433 |
| Aip | 0.01121494 | 0.53017276 | 0.427 | 0.269 | 1 | 0.00799482 | 0.55264121 | 0.378 | 0.208 | 1 | 0.01121494 | 0.01592573 |
| B930036N10Rik | 0.91142238 | -0.3346114 | 0.26 | 0.254 | 1 | 0.00799551 | 1.47864063 | 0.176 | 0.062 | 1 | 0.91142238 | 0.01592709 |
| Slc22a15 | 0.00800347 | -1.083108 | 0.229 | 0.373 | 1 | 0.77516051 | -0.2625495 | 0.256 | 0.229 | 1 | 0.77516051 | 0.01594289 |
| Afg3l2 | 0.00802307 | 0.67045955 | 0.51 | 0.368 | 1 | 0.31711987 | -0.1024132 | 0.555 | 0.448 | 1 | 0.31711987 | 0.01598177 |
| Rnaseh1 | 0.0474481 | 0.66354014 | 0.229 | 0.135 | 1 | 0.00803171 | 1.46775232 | 0.16 | 0.052 | 1 | 0.0474481 | 0.01599892 |
| Cluap1 | 0.00804766 | 0.50999488 | 0.385 | 0.228 | 1 | 0.03464636 | 0.21669055 | 0.353 | 0.219 | 1 | 0.03464636 | 0.01603055 |
| Lrrc23 | 0.0080547 | 1.54354744 | 0.188 | 0.083 | 1 | 0.75374902 | -0.4301495 | 0.16 | 0.135 | 1 | 0.75374902 | 0.01604452 |
| Tcof1 | 0.00805687 | 0.86310463 | 0.375 | 0.228 | 1 | 0.01436537 | 0.41884272 | 0.357 | 0.198 | 1 | 0.01436537 | 0.01604882 |
| Atp5j2 | 0.05408339 | 0.18024006 | 0.927 | 0.86 | 1 | 0.00806845 | 0.31048157 | 0.87 | 0.823 | 1 | 0.05408339 | 0.01607179 |
| Exph5 | 0.11302678 | -2.1624305 | 0 | 0.026 | 1 | 0.00807212 | -0.6931839 | 0.055 | 0.146 | 1 | 0.11302678 | 0.01607908 |
| Bcdin3d | 0.01130153 | 1.46692616 | 0.104 | 0.031 | 1 | 0.00808915 | 1.91253716 | 0.092 | 0.01 | 1 | 0.01130153 | 0.01611287 |
| Nlgn1 | 0.00810082 | -4.6267115 | 0.021 | 0.109 | 1 | 0.27709938 | -1.1566985 | 0.13 | 0.083 | 1 | 0.27709938 | 0.01613601 |
| H3c14 | 0.00811358 | -1.8405024 | 0.031 | 0.13 | 1 | 0.87673803 | -0.1723517 | 0.038 | 0.042 | 1 | 0.87673803 | 0.01616134 |
| Zfp874a | 0.07375904 | 1.07788386 | 0.156 | 0.088 | 1 | 0.00812081 | 0.66213036 | 0.181 | 0.062 | 1 | 0.07375904 | 0.01617568 |
| Zfp955a | 0.37661394 | 0.10060394 | 0.208 | 0.161 | 1 | 0.0081595 | 1.20110666 | 0.189 | 0.073 | 1 | 0.37661394 | 0.01625243 |
| Fkbp2 | 0.01553956 | 0.57838185 | 0.656 | 0.58 | 1 | 0.00816129 | 0.59822225 | 0.66 | 0.573 | 1 | 0.01553956 | 0.01625598 |
| Nek9 | 0.47091176 | 0.11564334 | 0.562 | 0.534 | 1 | 0.00818148 | 0.29782731 | 0.676 | 0.5 | 1 | 0.47091176 | 0.01629603 |
| Gpm6a | 0.61737927 | 1.00749454 | 0.01 | 0.005 | 1 | 0.00818671 | -1.6724253 | 0.021 | 0.083 | 1 | 0.61737927 | 0.01630639 |
| Rab8a | 0.30897572 | 0.20472688 | 0.562 | 0.523 | 1 | 0.00819229 | 0.52079031 | 0.643 | 0.552 | 1 | 0.30897572 | 0.01631746 |
| Fut4 | 0.00819309 | -1.6705774 | 0.073 | 0.192 | 1 | 0.58180179 | -0.7248928 | 0.021 | 0.031 | 1 | 0.58180179 | 0.01631905 |
| Mepce | 0.02556299 | 0.64904057 | 0.302 | 0.181 | 1 | 0.00819527 | 0.55687821 | 0.353 | 0.188 | 1 | 0.02556299 | 0.01632339 |
| Fbxl20 | 0.00819689 | -0.7363247 | 0.375 | 0.513 | 1 | 0.24867205 | -0.3599787 | 0.634 | 0.708 | 1 | 0.24867205 | 0.01632658 |
| Pex5 | 0.00820737 | 0.45167838 | 0.604 | 0.44 | 1 | 0.01768408 | 0.2357138 | 0.609 | 0.427 | 1 | 0.01768408 | 0.01634738 |
| Sox8 | 0.00824101 | 0.74446013 | 0.302 | 0.161 | 1 | 0.3194551 | 0.18047036 | 0.151 | 0.104 | 1 | 0.3194551 | 0.01641411 |
| Wdr36 | 0.13609339 | 0.39531057 | 0.448 | 0.373 | 1 | 0.00824914 | 0.28351948 | 0.429 | 0.24 | 1 | 0.13609339 | 0.01643024 |
| Megf9 | 0.61296218 | -0.4147385 | 0.312 | 0.332 | 1 | 0.00825273 | -0.7305615 | 0.319 | 0.479 | 1 | 0.61296218 | 0.01643736 |
| Mrpl39 | 0.00827185 | 0.97507306 | 0.333 | 0.197 | 1 | 0.02989699 | 0.50830641 | 0.235 | 0.125 | 1 | 0.02989699 | 0.01647528 |
| Mrps30 | 0.16341873 | 0.39605982 | 0.333 | 0.254 | 1 | 0.00828651 | 0.58544736 | 0.319 | 0.167 | 1 | 0.16341873 | 0.01650435 |
| Prkab1 | 0.0314318 | 0.91127922 | 0.229 | 0.13 | 1 | 0.0082942 | 0.76339372 | 0.324 | 0.177 | 1 | 0.0314318 | 0.0165196 |
| Arsa | 0.46993088 | 0.24195979 | 0.25 | 0.212 | 1 | 0.00829543 | 0.76053407 | 0.223 | 0.094 | 1 | 0.46993088 | 0.01652204 |
| Vopp1 | 0.91089542 | -0.17693 | 0.188 | 0.176 | 1 | 0.00829818 | 0.41806519 | 0.315 | 0.156 | 1 | 0.91089542 | 0.01652749 |
| Zfp511 | 0.19634073 | -0.5505009 | 0.167 | 0.228 | 1 | 0.00830547 | 0.69014474 | 0.265 | 0.125 | 1 | 0.19634073 | 0.01654196 |
| Rnf170 | 0.00830919 | 0.8439958 | 0.365 | 0.223 | 1 | 0.82085874 | -0.2244638 | 0.248 | 0.229 | 1 | 0.82085874 | 0.01654933 |
| Ptprs | 0.00834233 | 0.10370985 | 0.604 | 0.378 | 1 | 0.82994428 | -0.4177113 | 0.567 | 0.531 | 1 | 0.82994428 | 0.01661507 |
| 1700037H04Rik | 0.49931559 | -0.3404288 | 0.083 | 0.109 | 1 | 0.00834432 | 1.96316323 | 0.109 | 0.021 | 1 | 0.49931559 | 0.01661902 |
| N4bp2os | 0.00835424 | 2.00749454 | 0.115 | 0.036 | 1 | 0.05353127 | 1.39058446 | 0.092 | 0.031 | 1 | 0.05353127 | 0.01663869 |
| Gm16153 | 0.00835992 | 1.68556644 | 0.125 | 0.041 | 1 | 0.30443422 | 0.30485458 | 0.139 | 0.094 | 1 | 0.30443422 | 0.01664994 |
| Eif4ebp2 | 0.68960061 | -0.1290075 | 0.729 | 0.731 | 1 | 0.00836232 | 0.29314116 | 0.79 | 0.75 | 1 | 0.68960061 | 0.0166547 |
| Srd5a3 | 0.00836455 | 0.90440104 | 0.25 | 0.124 | 1 | 0.00945077 | 0.66996686 | 0.235 | 0.104 | 1 | 0.00945077 | 0.01665913 |
| Acat2 | 0.15345873 | 0.31561683 | 0.375 | 0.301 | 1 | 0.00836595 | 0.86006974 | 0.248 | 0.115 | 1 | 0.15345873 | 0.01666191 |
| Wtip | 0.029335 | 0.242711 | 0.667 | 0.477 | 1 | 0.008367 | 0.33547986 | 0.807 | 0.573 | 1 | 0.029335 | 0.016664 |
| Zfp445 | 0.08531378 | 0.37858869 | 0.583 | 0.497 | 1 | 0.00836805 | 0.37710695 | 0.647 | 0.438 | 1 | 0.08531378 | 0.01666607 |
| Med22 | 0.00837033 | 0.80955516 | 0.302 | 0.166 | 1 | 0.0186501 | 0.47150445 | 0.366 | 0.219 | 1 | 0.0186501 | 0.01667059 |
| Pla2g4a | 0.00838071 | -0.6517617 | 0.344 | 0.513 | 1 | 0.38514154 | -0.5301214 | 0.319 | 0.25 | 1 | 0.38514154 | 0.01669119 |
| Bpifa1 | 0.4321177 | 0.22161934 | 0.406 | 0.368 | 1 | 0.00838325 | -0.5686527 | 0.702 | 0.781 | 1 | 0.4321177 | 0.01669622 |
| Srp54a | 0.26474006 | 0.27052894 | 0.375 | 0.301 | 1 | 0.00838396 | 0.51866427 | 0.332 | 0.177 | 1 | 0.26474006 | 0.01669763 |
| Ubap2l | 0.10032041 | 0.14235939 | 0.969 | 0.938 | 1 | 0.00838921 | -0.4509332 | 0.975 | 0.979 | 1 | 0.10032041 | 0.01670803 |
| Tbc1d17 | 0.0394281 | -0.6186906 | 0.167 | 0.285 | 1 | 0.00839935 | 0.54812573 | 0.277 | 0.135 | 1 | 0.0394281 | 0.01672816 |
| Gm10131 | 0.00840024 | 1.28760246 | 0.167 | 0.067 | 1 | 0.43110109 | -0.7248928 | 0.118 | 0.083 | 1 | 0.43110109 | 0.01672992 |
| Gdpd5 | 0.70713506 | -0.1730777 | 0.115 | 0.098 | 1 | 0.00840105 | 0.29730298 | 0.34 | 0.177 | 1 | 0.70713506 | 0.01673152 |
| Fmnl2 | 0.02842862 | -0.6908353 | 0.708 | 0.829 | 1 | 0.00840436 | -1.2276144 | 0.794 | 0.917 | 1 | 0.02842862 | 0.01673808 |
| Etv4 | 0.00840976 | -0.9593386 | 0.052 | 0.166 | 1 | 0.67811349 | -0.4093909 | 0.101 | 0.115 | 1 | 0.67811349 | 0.01674879 |

| | | | | | | | | | | | | |
|---|---|---|---|---|---|---|---|---|---|---|---|---|
| Mdm2 | 0.00841015 | 0.49512004 | 0.781 | 0.642 | 1 | 0.67569378 | -0.2582635 | 0.723 | 0.792 | 1 | 0.67569378 | 0.01674958 |
| 4930435F18Rik | 0.61023023 | 1.68556644 | 0.042 | 0.031 | 1 | 0.00842413 | -2.8948178 | 0.017 | 0.073 | 1 | 0.61023023 | 0.0167773 |
| Fkbp4 | 0.00843341 | 0.39082318 | 0.531 | 0.352 | 1 | 0.33734431 | 0.13922381 | 0.584 | 0.583 | 1 | 0.33734431 | 0.0167957 |
| Orc5 | 0.47538207 | -0.2726134 | 0.219 | 0.254 | 1 | 0.00844133 | 0.64434105 | 0.223 | 0.094 | 1 | 0.47538207 | 0.0168114 |
| Ufl1 | 0.1036912 | 0.32028198 | 0.552 | 0.472 | 1 | 0.00844581 | 0.37464291 | 0.618 | 0.458 | 1 | 0.1036912 | 0.0168203 |
| Sae1 | 0.00847035 | 0.361778 | 0.719 | 0.523 | 1 | 0.35964184 | -0.1970354 | 0.769 | 0.698 | 1 | 0.35964184 | 0.01686896 |
| Melk | 0.0084847 | 3.17741954 | 0.052 | 0.005 | 1 | 0.02605524 | -2.8948178 | 0 | 0.021 | 1 | 0.02605524 | 0.0168974 |
| Clk1 | 0.42027301 | -0.117357 | 0.99 | 0.99 | 1 | 0.00848617 | -0.5498258 | 0.979 | 0.969 | 1 | 0.42027301 | 0.01690032 |
| Entrep3 | 0.00848746 | 0.87422801 | 0.26 | 0.13 | 1 | 0.18948783 | -0.2103196 | 0.265 | 0.177 | 1 | 0.18948783 | 0.01690289 |
| Dnaja2 | 0.36973761 | 0.11326216 | 0.875 | 0.907 | 1 | 0.00849458 | 0.15238121 | 0.958 | 0.896 | 1 | 0.36973761 | 0.01691699 |
| Gm37305 | 0.00849684 | 1.71798792 | 0.125 | 0.041 | 1 | 0.01843529 | 0.97025266 | 0.13 | 0.042 | 1 | 0.01843529 | 0.01692148 |
| 2900060B14Rik | 0.39689294 | -0.4274425 | 0.188 | 0.145 | 1 | 0.00850363 | -1.878139 | 0.084 | 0.177 | 1 | 0.39689294 | 0.01693494 |
| Ints13 | 0.32414573 | 0.20013961 | 0.385 | 0.326 | 1 | 0.00851004 | 0.32840546 | 0.496 | 0.292 | 1 | 0.32414573 | 0.01694765 |
| Ctxnd1 | 0.1992517 | 2.22988696 | 0.031 | 0.01 | 1 | 0.00852617 | -4.1843244 | 0.017 | 0.073 | 1 | 0.1992517 | 0.01697965 |
| Zdhhc24 | 0.00855788 | 1.52206771 | 0.156 | 0.062 | 1 | 0.63598532 | 0.27510724 | 0.088 | 0.073 | 1 | 0.63598532 | 0.01704253 |
| Hhipl2 | 0.00855921 | 2.81484946 | 0.052 | 0.005 | 1 | 0.4313809 | -0.6317834 | 0.025 | 0.042 | 1 | 0.4313809 | 0.01704516 |
| 1700019G24Rik | 0.00855921 | 4.76238204 | 0.052 | 0.005 | 1 | 0.0540843 | 2.01207283 | 0.038 | 0 | 1 | 0.0540843 | 0.01704516 |
| Rmi2 | 0.00855921 | 2.81484946 | 0.052 | 0.005 | 1 | 0.39796128 | -1.5322477 | 0.017 | 0.031 | 1 | 0.39796128 | 0.01704516 |
| Ffar4 | 0.00855921 | 3.00749454 | 0.052 | 0.005 | 1 | 0.11765402 | 1.49749966 | 0.025 | 0 | 1 | 0.11765402 | 0.01704516 |
| 2310002F09Rik | 0.00856171 | -2.9925055 | 0.021 | 0.109 | 1 | 0.91360311 | -1.1172102 | 0.05 | 0.052 | 1 | 0.91360311 | 0.01705012 |
| Carm1 | 0.00856203 | 0.46000674 | 0.521 | 0.342 | 1 | 0.01631555 | 0.17447488 | 0.626 | 0.438 | 1 | 0.01631555 | 0.01705076 |
| Gm17477 | 0.08192574 | -2.577468 | 0 | 0.031 | 1 | 0.00859577 | -2.1034044 | 0.046 | 0.125 | 1 | 0.08192574 | 0.01711765 |
| Ube2k | 0.00860324 | 0.25989953 | 0.958 | 0.979 | 1 | 0.6868352 | -0.1827433 | 0.996 | 0.979 | 1 | 0.6868352 | 0.01713246 |
| Pold3 | 0.010126 | 0.71798792 | 0.292 | 0.155 | 1 | 0.00861347 | 0.36570979 | 0.324 | 0.156 | 1 | 0.010126 | 0.01715275 |
| Psmg2 | 0.17883442 | 0.28760246 | 0.479 | 0.399 | 1 | 0.0086202 | 0.6523427 | 0.345 | 0.188 | 1 | 0.17883442 | 0.01716608 |
| Mgst1 | 0.21157963 | 0.85109298 | 0.385 | 0.378 | 1 | 0.00862165 | -0.5924425 | 0.437 | 0.625 | 1 | 0.21157963 | 0.01716897 |
| H2ac24 | 0.00863811 | 3.32942263 | 0.052 | 0.005 | 1 | 0.27154471 | 0.69014474 | 0.013 | 0 | 1 | 0.27154471 | 0.01720161 |
| Moap1 | 0.00863838 | 2.59245704 | 0.052 | 0.005 | 1 | 0.02542376 | 2.39058446 | 0.05 | 0 | 1 | 0.02542376 | 0.01720214 |
| Klc1 | 0.00863858 | 0.40629139 | 0.833 | 0.777 | 1 | 0.94233367 | -0.2448355 | 0.924 | 0.906 | 1 | 0.94233367 | 0.01720254 |
| Ddx23 | 0.59458442 | 0.16166547 | 0.479 | 0.451 | 1 | 0.0086466 | 0.24075573 | 0.597 | 0.385 | 1 | 0.59458442 | 0.01721843 |
| Sun2 | 0.0606555 | -0.4828311 | 0.417 | 0.534 | 1 | 0.00866172 | 0.39709776 | 0.517 | 0.344 | 1 | 0.0606555 | 0.01724842 |
| Dnajb14 | 0.00866226 | 0.19837916 | 0.646 | 0.461 | 1 | 0.50035774 | -0.468553 | 0.563 | 0.625 | 1 | 0.50035774 | 0.01724948 |
| Ube2d-ps | 0.02592328 | 0.89201732 | 0.344 | 0.228 | 1 | 0.00867944 | 0.63125105 | 0.298 | 0.156 | 1 | 0.02592328 | 0.01728355 |
| Alkbh6 | 0.20562925 | 0.40959298 | 0.323 | 0.249 | 1 | 0.00868909 | 0.17127143 | 0.227 | 0.094 | 1 | 0.20562925 | 0.01730268 |
| Gls | 0.00869833 | -0.4495346 | 0.958 | 0.99 | 1 | 0.01526356 | -0.4367601 | 0.992 | 1 | 1 | 0.01526356 | 0.017321 |
| Gm20732 | 0.43768031 | -0.3664803 | 0.292 | 0.332 | 1 | 0.00870053 | -1.3815567 | 0.357 | 0.448 | 1 | 0.43768031 | 0.01732536 |
| Polr2m | 0.02262914 | 0.34767804 | 0.792 | 0.668 | 1 | 0.00870249 | 0.40369144 | 0.752 | 0.656 | 1 | 0.02262914 | 0.01732925 |
| Zadh2 | 0.00871157 | 0.93878179 | 0.302 | 0.171 | 1 | 0.76732605 | -0.3246523 | 0.282 | 0.25 | 1 | 0.76732605 | 0.01734725 |
| Dus4l | 0.00873451 | 1.00749454 | 0.177 | 0.073 | 1 | 0.03741335 | 0.86006974 | 0.084 | 0.021 | 1 | 0.03741335 | 0.01739274 |
| Xpo7 | 0.00875677 | 0.407102 | 0.729 | 0.596 | 1 | 0.63185766 | -0.1882458 | 0.782 | 0.865 | 1 | 0.63185766 | 0.01743687 |
| Slc35e3 | 0.48030212 | 0.29074216 | 0.562 | 0.518 | 1 | 0.0087639 | 0.47792412 | 0.462 | 0.281 | 1 | 0.48030212 | 0.01745099 |
| Tent5c | 0.14068549 | -0.9925055 | 0.073 | 0.13 | 1 | 0.00876398 | -1.3483294 | 0.08 | 0.177 | 1 | 0.14068549 | 0.01745115 |
| Galntl6 | 0.00879248 | 2.59245704 | 0.052 | 0.005 | 1 | 0.15394868 | 1.27510724 | 0.021 | 0 | 1 | 0.15394868 | 0.01750766 |
| Hip1 | 0.00882651 | 0.35691924 | 0.833 | 0.762 | 1 | 0.01077781 | 0.3207859 | 0.895 | 0.729 | 1 | 0.01077781 | 0.01757512 |
| Col5a2 | 0.00883503 | -2.9467018 | 0.198 | 0.301 | 1 | 0.38056355 | -1.2767834 | 0.555 | 0.458 | 1 | 0.38056355 | 0.01759201 |
| Supt20 | 0.00886991 | 0.58007331 | 0.625 | 0.492 | 1 | 0.18242773 | -0.4178379 | 0.504 | 0.573 | 1 | 0.18242773 | 0.01766114 |
| Gm5444 | 0.0088701 | 3.12297175 | 0.052 | 0.005 | 1 | 0.56565874 | -2.3098553 | 0.013 | 0.021 | 1 | 0.56565874 | 0.01766153 |
| Cdkn3 | 0.00887093 | 2.59245704 | 0.052 | 0.005 | 1 | 0.67471359 | -2.4473588 | 0.017 | 0.01 | 1 | 0.67471359 | 0.01766318 |
| Dcun1d4 | 0.00887558 | -1.3550755 | 0.146 | 0.285 | 1 | 0.04255633 | -1.2592292 | 0.168 | 0.25 | 1 | 0.04255633 | 0.01767238 |
| Atp6v0a2 | 0.00887836 | 0.50609675 | 0.552 | 0.383 | 1 | 0.01317547 | 0.4370258 | 0.651 | 0.458 | 1 | 0.01317547 | 0.0176779 |
| Lnx1 | 0.57850636 | 1.00749454 | 0.031 | 0.021 | 1 | 0.00888019 | -2.0468209 | 0.034 | 0.104 | 1 | 0.57850636 | 0.01768152 |
| Rbck1 | 0.72165062 | 0.1118312 | 0.323 | 0.306 | 1 | 0.00890383 | 0.57278779 | 0.37 | 0.208 | 1 | 0.72165062 | 0.01772837 |
| Stimate | 0.00891536 | 0.80818573 | 0.354 | 0.212 | 1 | 0.01316079 | 0.78636005 | 0.311 | 0.177 | 1 | 0.01316079 | 0.01775123 |
| Fyco1 | 0.00895893 | 0.94140535 | 0.385 | 0.244 | 1 | 0.10749177 | 0.17808278 | 0.324 | 0.219 | 1 | 0.10749177 | 0.01783759 |
| Lztr1 | 0.21068683 | 0.33911603 | 0.438 | 0.352 | 1 | 0.00896818 | 0.38473173 | 0.399 | 0.229 | 1 | 0.21068683 | 0.01785593 |
| Gnpda1 | 0.00899118 | 1.10060394 | 0.229 | 0.114 | 1 | 0.24746714 | 0.13594449 | 0.235 | 0.167 | 1 | 0.24746714 | 0.01790152 |
| Mtap | 0.50594615 | 0.2912875 | 0.365 | 0.326 | 1 | 0.00900119 | 0.21138164 | 0.437 | 0.24 | 1 | 0.50594615 | 0.01792135 |
| Emb | 0.02163916 | -0.8575759 | 0.677 | 0.751 | 1 | 0.00901919 | -0.194726 | 0.626 | 0.833 | 1 | 0.02163916 | 0.01795704 |
| Gpr37l1 | 0.00902859 | 1.81484946 | 0.052 | 0.005 | 1 | 0.23679813 | 1.01207283 | 0.034 | 0.01 | 1 | 0.23679813 | 0.01797567 |
| Spag5 | 0.04511317 | 0.90795886 | 0.104 | 0.041 | 1 | 0.0090421 | -2.3098553 | 0.017 | 0.073 | 1 | 0.04511317 | 0.01800245 |
| B230307C23Rik | 0.00904393 | 0.77801269 | 0.24 | 0.114 | 1 | 0.46372607 | -0.2823745 | 0.164 | 0.125 | 1 | 0.46372607 | 0.01800607 |
| Trmt2a | 0.09128443 | 0.42253204 | 0.271 | 0.181 | 1 | 0.00904868 | 0.83989186 | 0.244 | 0.115 | 1 | 0.09128443 | 0.01801548 |
| Ncbp1 | 0.00905733 | 0.48094872 | 0.5 | 0.352 | 1 | 0.17327107 | 0.10518224 | 0.605 | 0.51 | 1 | 0.17327107 | 0.01803263 |
| Nup85 | 0.00905875 | 0.80955516 | 0.281 | 0.15 | 1 | 0.05052631 | 0.14957636 | 0.282 | 0.167 | 1 | 0.05052631 | 0.01803543 |
| Adam19 | 0.00905887 | -2.7738652 | 0.021 | 0.109 | 1 | 0.31246832 | -1.3947442 | 0.084 | 0.115 | 1 | 0.31246832 | 0.01803567 |
| Dcaf12l1 | 0.62638799 | -0.4330781 | 0.062 | 0.078 | 1 | 0.00906438 | 1.69014474 | 0.109 | 0.021 | 1 | 0.62638799 | 0.0180466 |
| Exosc7 | 0.01472625 | 0.7503367 | 0.323 | 0.192 | 1 | 0.00907698 | 0.73745045 | 0.193 | 0.073 | 1 | 0.01472625 | 0.01807158 |
| 2310010J17Rik | 0.89361339 | 0.22322323 | 0.344 | 0.342 | 1 | 0.00908031 | 0.86848198 | 0.273 | 0.135 | 1 | 0.89361339 | 0.01807816 |
| Gm20712 | 0.35950357 | 0.68556644 | 0.073 | 0.047 | 1 | 0.00908907 | 1.62874419 | 0.155 | 0.052 | 1 | 0.35950357 | 0.01809553 |
| Syce2 | 0.34858595 | 0.83756954 | 0.104 | 0.073 | 1 | 0.00909922 | 1.64434105 | 0.109 | 0.021 | 1 | 0.34858595 | 0.01811564 |
| Atg2a | 0.00910563 | -1.240433 | 0.229 | 0.368 | 1 | 0.31229154 | -0.8123556 | 0.269 | 0.302 | 1 | 0.31229154 | 0.01812836 |
| Hdac4 | 0.26942556 | -0.2538335 | 0.75 | 0.58 | 1 | 0.00913292 | -0.6945942 | 0.723 | 0.865 | 1 | 0.26942556 | 0.01818243 |
| Galk2 | 0.00913301 | 0.58480346 | 0.354 | 0.207 | 1 | 0.14879772 | 0.22619764 | 0.34 | 0.24 | 1 | 0.14879772 | 0.01818261 |
| Iqsec1 | 0.00913714 | 0.15655728 | 0.99 | 0.845 | 1 | 0.06888097 | -0.1781311 | 0.979 | 0.885 | 1 | 0.06888097 | 0.01819078 |
| Luzp1 | 0.00915764 | -0.4986741 | 0.969 | 0.953 | 1 | 0.20536633 | -0.4169148 | 0.992 | 0.969 | 1 | 0.20536633 | 0.01823142 |
| Aurkaip1 | 0.14227758 | 0.22250743 | 0.74 | 0.632 | 1 | 0.00916183 | 0.54039762 | 0.643 | 0.542 | 1 | 0.14227758 | 0.01823972 |
| Iars2 | 0.22383522 | 0.43280037 | 0.323 | 0.249 | 1 | 0.00916259 | 0.44175711 | 0.475 | 0.302 | 1 | 0.22383522 | 0.01824123 |
| Zfp992 | 0.00917336 | 0.94609399 | 0.219 | 0.104 | 1 | 0.32460617 | 0.33060235 | 0.181 | 0.135 | 1 | 0.32460617 | 0.01826257 |
| Ankrd9 | 0.00918471 | -2.0513992 | 0.021 | 0.109 | 1 | 0.62637284 | -0.3923174 | 0.059 | 0.073 | 1 | 0.62637284 | 0.01828507 |
| Epb41l4aos | 0.02268104 | 0.8291573 | 0.323 | 0.207 | 1 | 0.00918584 | 1.45095707 | 0.197 | 0.083 | 1 | 0.02268104 | 0.01828729 |
| Ttc5 | 0.00918941 | 0.56319244 | 0.438 | 0.285 | 1 | 0.05255738 | 0.34500925 | 0.403 | 0.281 | 1 | 0.05255738 | 0.01829438 |
| Fntb | 0.76129692 | 0.28502851 | 0.219 | 0.244 | 1 | 0.00919333 | 0.71567983 | 0.303 | 0.156 | 1 | 0.76129692 | 0.01830214 |
| Sncaip | 0.00919345 | -1.8810842 | 0.104 | 0.233 | 1 | 0.75465972 | -0.6655142 | 0.214 | 0.188 | 1 | 0.75465972 | 0.01830238 |
| Kdm7a | 0.34220839 | -0.2737916 | 0.562 | 0.632 | 1 | 0.00919467 | -0.4603469 | 0.731 | 0.906 | 1 | 0.34220839 | 0.0183048 |
| Smarcd3 | 0.03191237 | 1.05640414 | 0.219 | 0.119 | 1 | 0.00919672 | 1.01207283 | 0.202 | 0.083 | 1 | 0.03191237 | 0.01830887 |

| | | | | | | | | | | | | |
|---|---|---|---|---|---|---|---|---|---|---|---|---|
| Ttpal | 0.28985239 | 0.49292136 | 0.177 | 0.13 | 1 | 0.00920432 | 0.58091567 | 0.303 | 0.156 | 1 | 0.28985239 | 0.01832392 |
| Coq6 | 0.00920563 | 1.13302542 | 0.198 | 0.088 | 1 | 0.59139721 | 0.27510724 | 0.147 | 0.125 | 1 | 0.59139721 | 0.01832652 |
| Tbc1d25 | 0.03933033 | 0.90795886 | 0.135 | 0.062 | 1 | 0.00920977 | 0.81942775 | 0.109 | 0.021 | 1 | 0.03933033 | 0.01833471 |
| Ndor1 | 0.00921694 | 1.08549705 | 0.167 | 0.067 | 1 | 0.22813347 | 0.12624385 | 0.16 | 0.104 | 1 | 0.22813347 | 0.01834894 |
| Khnyn | 0.81190575 | -0.1208296 | 0.333 | 0.311 | 1 | 0.00922613 | 0.39058446 | 0.483 | 0.302 | 1 | 0.81190575 | 0.01836714 |
| Mtr | 0.00923318 | 0.64492446 | 0.375 | 0.233 | 1 | 0.17119937 | -0.1493906 | 0.391 | 0.281 | 1 | 0.17119937 | 0.01838112 |
| Col20a1 | 0.00925247 | 1.11555428 | 0.385 | 0.269 | 1 | 0.10106352 | 0.36364391 | 0.454 | 0.344 | 1 | 0.10106352 | 0.01841933 |
| Mpp4 | 0.0092613 | 1.08549705 | 0.125 | 0.041 | 1 | 0.6178988 | -0.2625495 | 0.092 | 0.073 | 1 | 0.6178988 | 0.01843683 |
| Cyb5r3 | 0.00926527 | -0.4874866 | 0.802 | 0.876 | 1 | 0.04015879 | 0.13114146 | 0.84 | 0.823 | 1 | 0.04015879 | 0.0184447 |
| Cherp | 0.00926981 | 0.41674968 | 0.531 | 0.363 | 1 | 0.61687628 | -0.2433599 | 0.513 | 0.406 | 1 | 0.61687628 | 0.01845369 |
| Sirt7 | 0.74279381 | -0.1079827 | 0.188 | 0.166 | 1 | 0.00927058 | -1.2812861 | 0.155 | 0.271 | 1 | 0.74279381 | 0.01845521 |
| Dhx38 | 0.16246708 | 0.28760246 | 0.271 | 0.192 | 1 | 0.00929163 | 0.49749966 | 0.395 | 0.229 | 1 | 0.16246708 | 0.01849693 |
| Tmem131l | 0.0093133 | 0.77302928 | 0.448 | 0.311 | 1 | 0.82983509 | -0.4287327 | 0.651 | 0.719 | 1 | 0.82983509 | 0.01853985 |
| Map3k19 | 0.32025586 | -0.577468 | 0 | 0.01 | 1 | 0.00931549 | -2.7248928 | 0.017 | 0.073 | 1 | 0.32025586 | 0.01854421 |
| Larp4 | 0.00932157 | 0.28673917 | 0.906 | 0.896 | 1 | 0.01574129 | 0.26559344 | 0.954 | 0.969 | 1 | 0.01574129 | 0.01855624 |
| Tjp2 | 0.00936345 | -0.5543844 | 0.219 | 0.378 | 1 | 0.0449733 | -0.4087703 | 0.496 | 0.677 | 1 | 0.0449733 | 0.01863922 |
| Zfp942 | 0.61678226 | -0.2066303 | 0.208 | 0.233 | 1 | 0.00937681 | 0.55521516 | 0.311 | 0.156 | 1 | 0.61678226 | 0.0186657 |
| Slc52a2 | 0.01105408 | 1.42253204 | 0.146 | 0.057 | 1 | 0.00938289 | 2.86006974 | 0.067 | 0 | 1 | 0.01105408 | 0.01867774 |
| 5730414N17Rik | 0.07296636 | -1.3144336 | 0.031 | 0.088 | 1 | 0.00938289 | 2.93807225 | 0.067 | 0 | 1 | 0.07296636 | 0.01867774 |
| Dnd1 | 0.38521705 | -0.7998604 | 0.01 | 0.026 | 1 | 0.00938289 | 2.86006974 | 0.067 | 0 | 1 | 0.38521705 | 0.01867774 |
| Tincr | 0.12701924 | 1.22988696 | 0.062 | 0.026 | 1 | 0.00938727 | 2.93807225 | 0.067 | 0 | 1 | 0.12701924 | 0.01868643 |
| Zfp354a | 0.20910685 | 1.00749454 | 0.062 | 0.031 | 1 | 0.00940649 | 2.54812573 | 0.088 | 0.01 | 1 | 0.20910685 | 0.01872449 |
| Macrod1 | 0.0094207 | 0.78072367 | 0.344 | 0.202 | 1 | 0.04109723 | 0.10909729 | 0.311 | 0.177 | 1 | 0.04109723 | 0.01875265 |
| Vapa | 0.00972459 | 0.30324691 | 0.969 | 0.964 | 1 | 0.00943937 | 0.15313593 | 0.996 | 0.969 | 1 | 0.00972459 | 0.01878963 |
| Usp3 | 0.01154005 | -0.7497553 | 0.552 | 0.653 | 1 | 0.00944584 | -0.6381811 | 0.794 | 0.875 | 1 | 0.01154005 | 0.01880246 |
| Gm17586 | 0.04841914 | 0.82692229 | 0.208 | 0.119 | 1 | 0.00945034 | 1.03806804 | 0.202 | 0.083 | 1 | 0.04841914 | 0.01881138 |
| Ranbp3l | 0.00945051 | -3.240433 | 0 | 0.067 | 1 | 0.85764357 | -0.6094155 | 0.038 | 0.042 | 1 | 0.85764357 | 0.01881172 |
| Il3ra | 0.00945884 | -3.0799683 | 0 | 0.067 | 1 | 0.14428792 | 1.14957636 | 0.042 | 0.01 | 1 | 0.14428792 | 0.01882822 |
| Slc16a6 | 0.00946194 | -3.1624305 | 0 | 0.067 | 1 | 0.07942247 | -1.9155763 | 0.038 | 0.083 | 1 | 0.07942247 | 0.01883435 |
| Gm12940 | 0.19579377 | -0.2027232 | 0.354 | 0.259 | 1 | 0.00946326 | -1.3997408 | 0.332 | 0.438 | 1 | 0.19579377 | 0.01883696 |
| Ung | 0.00946527 | 1.74446013 | 0.115 | 0.036 | 1 | 0.78322506 | -0.6883669 | 0.076 | 0.083 | 1 | 0.78322506 | 0.01884094 |
| Ccl19-ps2 | 0.00946798 | 2.25542205 | 0.104 | 0.031 | 1 | 0.1460986 | 0.76053407 | 0.059 | 0.021 | 1 | 0.1460986 | 0.01884631 |
| Zfp930 | 0.11885255 | 1.12297175 | 0.094 | 0.047 | 1 | 0.00947817 | 2.39058446 | 0.088 | 0.01 | 1 | 0.11885255 | 0.01886651 |
| Secisbp2 | 0.54100504 | -0.1158879 | 0.417 | 0.461 | 1 | 0.00947911 | 0.19264508 | 0.433 | 0.25 | 1 | 0.54100504 | 0.01886837 |
| Pex7 | 0.31642284 | 0.27052894 | 0.365 | 0.306 | 1 | 0.00947927 | 0.52886383 | 0.303 | 0.156 | 1 | 0.31642284 | 0.01886868 |
| Gm16159 | 0.20883718 | -0.9925055 | 0.01 | 0.036 | 1 | 0.00948558 | 1.23446525 | 0.126 | 0.031 | 1 | 0.20883718 | 0.01888117 |
| Atp5md | 0.02764423 | 0.29498199 | 0.854 | 0.808 | 1 | 0.0094934 | 0.32164982 | 0.769 | 0.677 | 1 | 0.02764423 | 0.01889668 |
| Atp8b1 | 0.12880204 | -1.770113 | 0.062 | 0.114 | 1 | 0.00951167 | -2.2125581 | 0.076 | 0.167 | 1 | 0.12880204 | 0.01893287 |
| Hook3 | 0.00952674 | 0.25024467 | 0.969 | 0.907 | 1 | 0.9623531 | -0.1298352 | 0.979 | 0.927 | 1 | 0.9623531 | 0.01896273 |
| R3hdml | 0.0095392 | 1.05403712 | 0.115 | 0.036 | 1 | 0.12945354 | -2.7084046 | 0.021 | 0.052 | 1 | 0.12945354 | 0.01898741 |
| Nudt11 | 0.20691434 | -1.4519371 | 0.01 | 0.036 | 1 | 0.00954747 | 2.33400093 | 0.088 | 0.01 | 1 | 0.20691434 | 0.01900379 |
| Tead3 | 0.95920329 | -0.1035368 | 0.208 | 0.202 | 1 | 0.00956031 | 0.44757443 | 0.239 | 0.104 | 1 | 0.95920329 | 0.01902922 |
| Tasp1 | 0.00956067 | 0.4141198 | 0.677 | 0.549 | 1 | 0.131825 | -0.2933671 | 0.689 | 0.469 | 1 | 0.131825 | 0.01902994 |
| Slc2a3 | 0.81729589 | 1.35541784 | 0.052 | 0.047 | 1 | 0.00959768 | -2.07539 | 0.017 | 0.073 | 1 | 0.81729589 | 0.01910324 |
| Coq5 | 0.46743318 | 0.23377039 | 0.229 | 0.197 | 1 | 0.00960005 | 0.57597372 | 0.235 | 0.104 | 1 | 0.46743318 | 0.01910793 |
| Pld2 | 0.06355752 | 0.64994253 | 0.208 | 0.124 | 1 | 0.00963066 | 0.59703533 | 0.256 | 0.125 | 1 | 0.06355752 | 0.01916858 |
| Pdik1l | 0.00963241 | 1.22988696 | 0.188 | 0.083 | 1 | 0.05392211 | 0.91253716 | 0.151 | 0.073 | 1 | 0.05392211 | 0.01917203 |
| Ddrgk1 | 0.00963678 | 0.4637547 | 0.698 | 0.585 | 1 | 0.03096177 | 0.27047579 | 0.697 | 0.646 | 1 | 0.03096177 | 0.01918069 |
| Slc13a5 | 0.56820519 | 0.11440974 | 0.302 | 0.254 | 1 | 0.00966122 | 1.71834163 | 0.189 | 0.073 | 1 | 0.56820519 | 0.0192291 |
| Naa35 | 0.00968159 | 0.50406192 | 0.51 | 0.352 | 1 | 0.37329377 | -0.3619438 | 0.618 | 0.677 | 1 | 0.37329377 | 0.01926944 |
| Ptgr2 | 0.18815209 | -0.6205367 | 0.104 | 0.161 | 1 | 0.00968392 | 0.57767001 | 0.227 | 0.094 | 1 | 0.18815209 | 0.01927407 |
| Cacna1a | 0.03383461 | -0.366901 | 0.104 | 0.207 | 1 | 0.00971224 | 0.29131747 | 0.336 | 0.177 | 1 | 0.03383461 | 0.01933016 |
| Tsacc | 0.2423773 | 0.61156586 | 0.156 | 0.109 | 1 | 0.00972646 | 0.54812573 | 0.16 | 0.052 | 1 | 0.2423773 | 0.01935831 |
| Crkl | 0.00974001 | 0.43460487 | 0.521 | 0.342 | 1 | 0.01898428 | 0.28816339 | 0.672 | 0.5 | 1 | 0.01898428 | 0.01938516 |
| Errfi1 | 0.00975013 | -0.3727197 | 0.708 | 0.829 | 1 | 0.66820956 | 0.26617411 | 0.773 | 0.896 | 1 | 0.66820956 | 0.0194052 |
| Slc30a7 | 0.00976338 | 0.42253204 | 0.531 | 0.347 | 1 | 0.03115505 | 0.17929145 | 0.55 | 0.396 | 1 | 0.03115505 | 0.01943144 |
| Nalcn | 0.00978874 | -3.8751485 | 0.01 | 0.088 | 1 | 0.25326197 | -0.677587 | 0.042 | 0.073 | 1 | 0.25326197 | 0.01948166 |
| Slc27a4 | 0.12941707 | 0.74446013 | 0.167 | 0.104 | 1 | 0.00978926 | 0.67103591 | 0.261 | 0.125 | 1 | 0.12941707 | 0.0194827 |
| N4bp2l1 | 0.70179876 | 0.22988696 | 0.146 | 0.13 | 1 | 0.00979923 | 0.18318475 | 0.223 | 0.094 | 1 | 0.70179876 | 0.01950244 |
| Gm45187 | 0.00980667 | -1.9185049 | 0.042 | 0.14 | 1 | 0.35045411 | -0.8948178 | 0.008 | 0.021 | 1 | 0.35045411 | 0.01951716 |
| Smad7 | 0.00982074 | 0.59427061 | 0.677 | 0.565 | 1 | 0.04484962 | -0.4740074 | 0.605 | 0.74 | 1 | 0.04484962 | 0.01954503 |
| Grpel2 | 0.03928962 | 1.22988696 | 0.188 | 0.104 | 1 | 0.00982345 | 0.8889246 | 0.303 | 0.167 | 1 | 0.03928962 | 0.01955039 |
| Spx | 0.00985954 | 1.53800925 | 0.115 | 0.036 | 1 | 0.06170946 | 0.63767732 | 0.092 | 0.031 | 1 | 0.06170946 | 0.01962187 |
| Tgfbr3 | 0.00986454 | -0.5677528 | 0.76 | 0.824 | 1 | 0.20359465 | 0.15037879 | 0.945 | 0.812 | 1 | 0.20359465 | 0.01963178 |
| Mkln1os | 0.00987825 | -2.240433 | 0.031 | 0.124 | 1 | 0.31795984 | -0.7623675 | 0.055 | 0.083 | 1 | 0.31795984 | 0.01965892 |
| Prex2 | 0.02831854 | -1.577468 | 0.156 | 0.254 | 1 | 0.00989463 | 0.23267197 | 0.357 | 0.188 | 1 | 0.02831854 | 0.01969135 |
| Eola1 | 0.08251339 | 0.50792353 | 0.271 | 0.176 | 1 | 0.00992215 | 0.66877109 | 0.244 | 0.115 | 1 | 0.08251339 | 0.01974585 |
| Gm11210 | 0.00992735 | 0.59245704 | 0.135 | 0.047 | 1 | 0.60747398 | -0.4272122 | 0.176 | 0.146 | 1 | 0.60747398 | 0.01975615 |
| Ranbp9 | 0.10182985 | -0.4657196 | 0.698 | 0.715 | 1 | 0.00993133 | -0.5385459 | 0.866 | 0.938 | 1 | 0.10182985 | 0.01976403 |
| Gdpd1 | 0.0099342 | -2.8504865 | 0.01 | 0.088 | 1 | 0.29802668 | -0.8359241 | 0.08 | 0.115 | 1 | 0.29802668 | 0.0197697 |
| Ifi35 | 0.68963106 | -0.3404288 | 0.365 | 0.363 | 1 | 0.00993993 | -0.1267892 | 0.454 | 0.271 | 1 | 0.68963106 | 0.01978106 |
| Dnttip1 | 0.20938628 | 0.2635929 | 0.562 | 0.477 | 1 | 0.00996044 | 0.31763185 | 0.613 | 0.385 | 1 | 0.20938628 | 0.01982167 |
| Cenpo | 0.00996269 | 1.20013961 | 0.125 | 0.041 | 1 | 0.44744586 | 0.27510724 | 0.063 | 0.042 | 1 | 0.44744586 | 0.01982612 |
| Atad2b | 0.28941283 | -0.4258283 | 0.615 | 0.658 | 1 | 0.00999239 | -0.4678763 | 0.773 | 0.906 | 1 | 0.28941283 | 0.01988493 |
| Slc25a15 | 0.01001054 | 1.00749454 | 0.156 | 0.062 | 1 | 0.17537296 | 0.16407593 | 0.168 | 0.104 | 1 | 0.17537296 | 0.01992087 |
| Gm53028 | 0.91035045 | -0.1624305 | 0.229 | 0.228 | 1 | 0.01001499 | 0.78206723 | 0.269 | 0.135 | 1 | 0.91035045 | 0.01992969 |
| Wdr66 | 0.10161121 | -0.9311049 | 0.031 | 0.083 | 1 | 0.0100282 | 2.04769674 | 0.122 | 0.031 | 1 | 0.10161121 | 0.01995583 |
| Fubp3 | 0.42743617 | 0.25968156 | 0.375 | 0.326 | 1 | 0.01004062 | 0.20259474 | 0.504 | 0.302 | 1 | 0.42743617 | 0.01998042 |
| Hus1 | 0.02830142 | 0.93349396 | 0.156 | 0.073 | 1 | 0.01004606 | 1.43737867 | 0.155 | 0.052 | 1 | 0.02830142 | 0.01999119 |
| Spry2 | 0.03235646 | -0.8024669 | 0.385 | 0.513 | 1 | 0.01005299 | 0.67544823 | 0.492 | 0.323 | 1 | 0.03235646 | 0.02000492 |
| 1810062O18Rik | 0.97633298 | 0.14951354 | 0.104 | 0.104 | 1 | 0.01006265 | 0.81942775 | 0.109 | 0.021 | 1 | 0.97633298 | 0.02002404 |
| Akt1 | 0.0100645 | 0.47697982 | 0.729 | 0.591 | 1 | 0.01226712 | 0.23338011 | 0.849 | 0.719 | 1 | 0.01226712 | 0.0200277 |
| Adnp | 0.01007059 | 0.41365388 | 0.854 | 0.731 | 1 | 0.60388082 | -0.2108773 | 0.929 | 0.927 | 1 | 0.60388082 | 0.02003976 |
| Psmg1 | 0.01007335 | 0.63403614 | 0.385 | 0.233 | 1 | 0.5987951 | -0.3864765 | 0.324 | 0.26 | 1 | 0.5987951 | 0.02004523 |
| Nova1 | 0.01007724 | -3.1624305 | 0.01 | 0.088 | 1 | 0.34184135 | -0.4618584 | 0.046 | 0.073 | 1 | 0.34184135 | 0.02005294 |
| Cfap74 | 0.0101099 | 0.90057933 | 0.26 | 0.13 | 1 | 0.14284295 | 0.3500693 | 0.197 | 0.125 | 1 | 0.14284295 | 0.0201176 |
| Gak | 0.0101217 | 0.43763893 | 0.656 | 0.508 | 1 | 0.11695534 | -0.3487909 | 0.693 | 0.792 | 1 | 0.11695534 | 0.02014096 |
| Cd38 | 0.01012367 | -2.577468 | 0.01 | 0.088 | 1 | 0.27154471 | 0.69014474 | 0.013 | 0 | 1 | 0.27154471 | 0.02014486 |
| Fam91a1 | 0.01013482 | 0.63342882 | 0.646 | 0.492 | 1 | 0.40460613 | -0.248016 | 0.643 | 0.688 | 1 | 0.40460613 | 0.02016692 |
| Ssh2 | 0.14509303 | -0.733473 | 0.406 | 0.472 | 1 | 0.01015838 | -0.6250225 | 0.714 | 0.854 | 1 | 0.14509303 | 0.02021357 |

| | | | | | | | | | | | | |
|---|---|---|---|---|---|---|---|---|---|---|---|---|
| Mlxip | 0.01018841 | 0.6132156 | 0.583 | 0.409 | 1 | 0.03451362 | -0.6577786 | 0.626 | 0.74 | 1 | 0.03451362 | 0.02027301 |
| Ubox5 | 0.01019298 | 1.36113149 | 0.156 | 0.062 | 1 | 0.82988265 | -0.6442743 | 0.147 | 0.135 | 1 | 0.82988265 | 0.02028207 |
| Sp2 | 0.27490791 | 0.32065242 | 0.333 | 0.275 | 1 | 0.01019325 | 0.2274105 | 0.361 | 0.188 | 1 | 0.27490791 | 0.0202826 |
| Wdtc1 | 0.18425036 | 0.43633784 | 0.281 | 0.212 | 1 | 0.01019847 | 0.71251255 | 0.361 | 0.208 | 1 | 0.18425036 | 0.02029293 |
| C130050O18Rik | 0.04336531 | 1.46692616 | 0.094 | 0.036 | 1 | 0.01019882 | 1.20471791 | 0.126 | 0.031 | 1 | 0.04336531 | 0.02029362 |
| Tmem132a | 0.12236045 | -1.6554705 | 0.021 | 0.062 | 1 | 0.01022507 | 0.87777174 | 0.294 | 0.156 | 1 | 0.12236045 | 0.02034559 |
| Dctn5 | 0.05099924 | 0.64492446 | 0.281 | 0.181 | 1 | 0.01022926 | 0.65843588 | 0.311 | 0.167 | 1 | 0.05099924 | 0.02035388 |
| Mycbp2 | 0.01025441 | 0.12234178 | 0.948 | 0.891 | 1 | 0.25143025 | -0.2531327 | 0.975 | 1 | 1 | 0.25143025 | 0.02040367 |
| Nyap1 | 0.01026921 | 1.81484946 | 0.104 | 0.031 | 1 | 0.19251009 | 0.49749966 | 0.097 | 0.052 | 1 | 0.19251009 | 0.02043296 |
| Rgl1 | 0.01028211 | -0.7569032 | 0.677 | 0.756 | 1 | 0.05328069 | -0.1549284 | 0.819 | 0.625 | 1 | 0.05328069 | 0.0204585 |
| Ndufa2 | 0.42148318 | -0.1413688 | 0.906 | 0.876 | 1 | 0.01028606 | 0.41847141 | 0.782 | 0.76 | 1 | 0.42148318 | 0.02046632 |
| Atp5k | 0.01401696 | 0.53542609 | 0.583 | 0.451 | 1 | 0.01030264 | 0.44221722 | 0.542 | 0.406 | 1 | 0.01401696 | 0.02049914 |
| Prdm10 | 0.01031513 | 1.10937415 | 0.26 | 0.14 | 1 | 0.99308242 | -0.8359241 | 0.223 | 0.208 | 1 | 0.99308242 | 0.02052386 |
| Chaserr | 0.68984478 | -0.1277163 | 0.802 | 0.834 | 1 | 0.01034387 | -0.3831919 | 0.891 | 0.917 | 1 | 0.68984478 | 0.02058074 |
| Hilpda | 0.83235021 | -0.9760173 | 0.417 | 0.378 | 1 | 0.01036115 | 0.83519507 | 0.261 | 0.125 | 1 | 0.83235021 | 0.02061494 |
| Zfp574 | 0.22232706 | 0.48142572 | 0.208 | 0.15 | 1 | 0.01037965 | 0.22263982 | 0.29 | 0.135 | 1 | 0.22232706 | 0.02065157 |
| Sh3glb2 | 0.58706432 | 0.11996927 | 0.188 | 0.161 | 1 | 0.01038271 | 0.95863357 | 0.185 | 0.073 | 1 | 0.58706432 | 0.02065763 |
| Otx1 | 0.72574709 | -0.3144336 | 0.01 | 0.016 | 1 | 0.01039027 | -1.4473588 | 0.034 | 0.104 | 1 | 0.72574709 | 0.02067258 |
| BC051537 | 0.35369407 | -2.1624305 | 0.021 | 0.041 | 1 | 0.01039832 | -2.8948178 | 0.013 | 0.062 | 1 | 0.35369407 | 0.02068852 |
| Lap3 | 0.01039852 | 0.80916582 | 0.521 | 0.373 | 1 | 0.01897398 | 0.37328763 | 0.517 | 0.354 | 1 | 0.01897398 | 0.02068892 |
| Gnptab | 0.61066772 | -0.3435799 | 0.333 | 0.342 | 1 | 0.01040732 | 0.59994693 | 0.437 | 0.271 | 1 | 0.61066772 | 0.02070632 |
| Ecsit | 0.01040905 | 1.00749454 | 0.208 | 0.098 | 1 | 0.07731 | 0.41806519 | 0.189 | 0.104 | 1 | 0.07731 | 0.02070974 |
| Pygm | 0.55708127 | -0.3144336 | 0.073 | 0.093 | 1 | 0.01042898 | -1.5381243 | 0.101 | 0.198 | 1 | 0.55708127 | 0.02074919 |
| Dap3 | 0.21467494 | 0.25076569 | 0.656 | 0.58 | 1 | 0.01042993 | 0.19264508 | 0.727 | 0.562 | 1 | 0.21467494 | 0.02075107 |
| Lsm10 | 0.04192575 | 1.00749454 | 0.188 | 0.104 | 1 | 0.01043089 | 0.40635177 | 0.176 | 0.062 | 1 | 0.04192575 | 0.02075298 |
| Stim1 | 0.01043614 | -0.743146 | 0.406 | 0.539 | 1 | 0.88087165 | -0.1343362 | 0.634 | 0.688 | 1 | 0.88087165 | 0.02076337 |
| Bag2 | 0.23440147 | 0.37006462 | 0.208 | 0.15 | 1 | 0.01048344 | 0.20720317 | 0.273 | 0.135 | 1 | 0.23440147 | 0.02085697 |
| Nit2 | 0.05905809 | 0.89201732 | 0.271 | 0.181 | 1 | 0.01051857 | 0.87234807 | 0.218 | 0.094 | 1 | 0.05905809 | 0.0209265 |
| Cyb5r1 | 0.65199919 | 0.10479174 | 0.375 | 0.342 | 1 | 0.01053688 | 0.70900376 | 0.235 | 0.104 | 1 | 0.65199919 | 0.02096273 |
| Rpf2 | 0.01054644 | 1.13035128 | 0.302 | 0.181 | 1 | 0.03171523 | 0.31307509 | 0.239 | 0.125 | 1 | 0.03171523 | 0.02098165 |
| Stk33 | 0.03131631 | 1.42253204 | 0.062 | 0.016 | 1 | 0.0105642 | -4.7021727 | 0.004 | 0.042 | 1 | 0.03131631 | 0.0210168 |
| Sh3bp4 | 0.97921362 | -0.3144336 | 0.198 | 0.192 | 1 | 0.01056632 | 0.13760371 | 0.353 | 0.188 | 1 | 0.97921362 | 0.02102099 |
| Snap47 | 0.01058574 | -0.6623569 | 0.417 | 0.539 | 1 | 0.3922437 | -0.1299462 | 0.471 | 0.365 | 1 | 0.3922437 | 0.02105942 |
| Man1b1 | 0.01059352 | 0.77302928 | 0.385 | 0.238 | 1 | 0.02096463 | 0.2095189 | 0.483 | 0.312 | 1 | 0.02096463 | 0.02107481 |
| Cdca8 | 0.0105997 | 3.67045955 | 0.062 | 0.01 | 1 | 0.39515719 | 0.49749966 | 0.025 | 0.01 | 1 | 0.39515719 | 0.02108705 |
| Ap1m1 | 0.01060653 | 0.73541499 | 0.406 | 0.259 | 1 | 0.01349055 | 0.51411437 | 0.382 | 0.229 | 1 | 0.01349055 | 0.02110057 |
| Ext2 | 0.02581822 | -0.4045216 | 0.875 | 0.855 | 1 | 0.01062667 | 0.2036721 | 0.861 | 0.625 | 1 | 0.02581822 | 0.0211404 |
| Uggt2 | 0.02406297 | 0.58759128 | 0.677 | 0.56 | 1 | 0.01062822 | 0.26205109 | 0.723 | 0.49 | 1 | 0.02406297 | 0.02114347 |
| Ctns | 0.14758859 | 0.39278469 | 0.25 | 0.176 | 1 | 0.01063115 | 0.82138927 | 0.29 | 0.156 | 1 | 0.14758859 | 0.02114928 |
| Nap1l4 | 0.01065303 | 0.35629834 | 0.76 | 0.611 | 1 | 0.67898045 | -0.245119 | 0.782 | 0.812 | 1 | 0.67898045 | 0.02119258 |
| Mab21l2 | 0.22187575 | -2.1624305 | 0 | 0.016 | 1 | 0.01065536 | -4.010295 | 0.004 | 0.042 | 1 | 0.22187575 | 0.02119719 |
| Ildr2 | 0.47606368 | 1.00749454 | 0.021 | 0.01 | 1 | 0.01065536 | -5.3098553 | 0.004 | 0.042 | 1 | 0.47606368 | 0.02119719 |
| Gins3 | 0.0106701 | 2.12297175 | 0.094 | 0.026 | 1 | 0.65691192 | 0.56461386 | 0.029 | 0.021 | 1 | 0.65691192 | 0.02122636 |
| Ptar1 | 0.01067472 | 0.6730755 | 0.531 | 0.399 | 1 | 0.91831576 | -0.3038565 | 0.542 | 0.469 | 1 | 0.91831576 | 0.02123549 |
| Fhip1b | 0.16472627 | -0.7721154 | 0.26 | 0.326 | 1 | 0.01068111 | 0.22901082 | 0.429 | 0.24 | 1 | 0.16472627 | 0.02124813 |
| Dock10 | 0.14767323 | -0.3771693 | 0.062 | 0.119 | 1 | 0.01068215 | 1.53449387 | 0.227 | 0.104 | 1 | 0.14767323 | 0.02125019 |
| Rfng | 0.0155297 | 0.9433642 | 0.208 | 0.104 | 1 | 0.01068637 | 1.04064198 | 0.185 | 0.073 | 1 | 0.0155297 | 0.02125855 |
| ENSMUSG00000121395 | 0.01068884 | 1.46692616 | 0.208 | 0.104 | 1 | 0.03496572 | 0.50728068 | 0.223 | 0.115 | 1 | 0.03496572 | 0.02126343 |
| Ciao2a | 0.11355286 | 0.26529229 | 0.458 | 0.337 | 1 | 0.01069247 | 0.66414953 | 0.462 | 0.323 | 1 | 0.11355286 | 0.02127061 |
| Ccdc6 | 0.01071277 | 0.23576352 | 0.562 | 0.383 | 1 | 0.61181537 | -0.3154148 | 0.626 | 0.615 | 1 | 0.61181537 | 0.02131077 |
| Tmsb10 | 0.01071727 | -0.7479116 | 0.823 | 0.658 | 1 | 0.01223059 | -0.881012 | 0.769 | 0.854 | 1 | 0.01223059 | 0.02131967 |
| Retreg1 | 0.01073133 | 0.52379704 | 0.938 | 0.865 | 1 | 0.0815648 | -0.3430221 | 0.882 | 0.854 | 1 | 0.0815648 | 0.02134751 |
| Stard3nl | 0.01338638 | 0.47697982 | 0.51 | 0.363 | 1 | 0.01074185 | 0.29159536 | 0.487 | 0.302 | 1 | 0.01338638 | 0.02136831 |
| 2700099C18Rik | 0.18228138 | 0.20013961 | 0.073 | 0.036 | 1 | 0.01074636 | -3.4797803 | 0.004 | 0.042 | 1 | 0.18228138 | 0.02137724 |
| Cabp1 | 0.28126256 | -2.240433 | 0.01 | 0.031 | 1 | 0.01074636 | -3.1172102 | 0.004 | 0.042 | 1 | 0.28126256 | 0.02137724 |
| Ofd1 | 0.42950248 | 0.21231433 | 0.281 | 0.233 | 1 | 0.01076477 | 0.43207658 | 0.256 | 0.115 | 1 | 0.42950248 | 0.02141367 |
| ENSMUSG00000121477 | 0.01077181 | 1.13302542 | 0.24 | 0.124 | 1 | 0.30440876 | -0.1353575 | 0.202 | 0.146 | 1 | 0.30440876 | 0.02142758 |
| Gpbp1l1 | 0.09277504 | -0.4308755 | 0.396 | 0.503 | 1 | 0.01077641 | -0.4272122 | 0.626 | 0.792 | 1 | 0.09277504 | 0.02143669 |
| Tmem50a | 0.08566764 | -0.338002 | 0.896 | 0.886 | 1 | 0.01079655 | 0.34526047 | 0.861 | 0.823 | 1 | 0.08566764 | 0.02147653 |
| Sema3f | 0.82258416 | -0.1624305 | 0.031 | 0.036 | 1 | 0.01080044 | -2.6317834 | 0.013 | 0.062 | 1 | 0.82258416 | 0.02148423 |
| Zfp280c | 0.0108159 | 0.39384928 | 0.51 | 0.337 | 1 | 0.26883213 | -0.1196714 | 0.521 | 0.396 | 1 | 0.26883213 | 0.02151482 |
| Azin2 | 0.57232326 | 0.27052894 | 0.208 | 0.181 | 1 | 0.01081874 | 0.94648449 | 0.155 | 0.052 | 1 | 0.57232326 | 0.02152043 |
| Ap1s2 | 0.14916931 | -0.5009698 | 0.344 | 0.425 | 1 | 0.0108256 | 0.75831624 | 0.303 | 0.167 | 1 | 0.14916931 | 0.02153402 |
| Ccdc83 | 0.08192823 | -2.4519371 | 0 | 0.031 | 1 | 0.01083762 | -3.4797803 | 0.004 | 0.042 | 1 | 0.08192823 | 0.02155778 |
| 2810408I11Rik | 0.36353452 | -0.8669746 | 0.021 | 0.041 | 1 | 0.01083762 | -3.1172102 | 0.004 | 0.042 | 1 | 0.36353452 | 0.02155778 |
| Tcam1 | 0.72574709 | -0.577468 | 0.01 | 0.016 | 1 | 0.01083762 | -2.8948178 | 0.004 | 0.042 | 1 | 0.72574709 | 0.02155778 |
| Cdca7l | 0.75328966 | 0.27052894 | 0.021 | 0.016 | 1 | 0.01083762 | -3.6317834 | 0.004 | 0.042 | 1 | 0.75328966 | 0.02155778 |
| C230035I16Rik | 0.01083873 | 2.59245704 | 0.062 | 0.01 | 1 | 0.43262492 | 0.69014474 | 0.063 | 0.042 | 1 | 0.43262492 | 0.02155998 |
| Serhl | 0.01084247 | 0.79068315 | 0.323 | 0.181 | 1 | 0.01330028 | 0.74077081 | 0.202 | 0.083 | 1 | 0.01330028 | 0.02156737 |
| Gsdmd | 0.15189495 | -1.577468 | 0.01 | 0.041 | 1 | 0.01085084 | 1.69014474 | 0.122 | 0.031 | 1 | 0.15189495 | 0.02158394 |
| Ss18 | 0.01085426 | 0.50287414 | 0.76 | 0.606 | 1 | 0.10355675 | 0.11411536 | 0.794 | 0.688 | 1 | 0.10355675 | 0.0215907 |
| Tmem179b | 0.3461803 | 0.2520326 | 0.552 | 0.549 | 1 | 0.01085669 | 0.24424627 | 0.525 | 0.344 | 1 | 0.3461803 | 0.02159551 |
| Dmwd | 0.21785516 | 0.33506919 | 0.281 | 0.212 | 1 | 0.01086295 | 0.32091093 | 0.307 | 0.156 | 1 | 0.21785516 | 0.02160789 |
| Unc119 | 0.1126008 | -0.4983964 | 0.208 | 0.295 | 1 | 0.01086411 | 0.49749966 | 0.235 | 0.104 | 1 | 0.1126008 | 0.0216102 |
| Bak1 | 0.53283035 | 0.22988696 | 0.135 | 0.109 | 1 | 0.01087913 | 0.82764826 | 0.16 | 0.052 | 1 | 0.53283035 | 0.02163991 |
| Ddx27 | 0.07795363 | 0.4141198 | 0.625 | 0.534 | 1 | 0.01088006 | 0.28125947 | 0.567 | 0.365 | 1 | 0.07795363 | 0.02164175 |
| Proser1 | 0.12436857 | 0.5649763 | 0.292 | 0.207 | 1 | 0.01088128 | 0.70296878 | 0.34 | 0.198 | 1 | 0.12436857 | 0.02164416 |
| Smad9 | 0.01088195 | 1.17099327 | 0.188 | 0.083 | 1 | 0.09309163 | 0.78968041 | 0.155 | 0.083 | 1 | 0.09309163 | 0.02164548 |
| Zdhhc5 | 0.11863937 | 0.23966798 | 0.667 | 0.56 | 1 | 0.01088305 | 0.29159536 | 0.769 | 0.656 | 1 | 0.11863937 | 0.02164767 |
| Ntn1 | 0.09021148 | -2.9925055 | 0.031 | 0.083 | 1 | 0.01088731 | -1.3098553 | 0.046 | 0.125 | 1 | 0.09021148 | 0.02165609 |
| Rnf181 | 0.34426711 | -0.337334 | 0.375 | 0.409 | 1 | 0.01090748 | 0.47864063 | 0.366 | 0.208 | 1 | 0.34426711 | 0.02169599 |

| | | | | | | | | | | | | |
|---|---|---|---|---|---|---|---|---|---|---|---|---|
| Elp1 | 0.01092032 | 0.46592463 | 0.562 | 0.378 | 1 | 0.01150717 | 0.36152199 | 0.588 | 0.385 | 1 | 0.01150717 | 0.02172139 |
| Tspyl5 | 0.01092599 | 2.00749454 | 0.094 | 0.026 | 1 | 0.45302102 | -0.5213594 | 0.063 | 0.042 | 1 | 0.45302102 | 0.02173261 |
| Cenph | 0.81731911 | 0.90057933 | 0.052 | 0.047 | 1 | 0.0109313 | -2.8948178 | 0.004 | 0.042 | 1 | 0.81731911 | 0.02174311 |
| Rasef | 0.04512622 | 3.00749454 | 0.021 | 0 | 1 | 0.01093188 | -2.6317834 | 0.004 | 0.042 | 1 | 0.04512622 | 0.02174426 |
| Egf | 0.1469607 | 1.49292136 | 0.052 | 0.021 | 1 | 0.01093188 | -2.6317834 | 0.004 | 0.042 | 1 | 0.1469607 | 0.02174426 |
| Gm48913 | 0.22187388 | 1.00749454 | 0.021 | 0.005 | 1 | 0.01093188 | -2.6317834 | 0.004 | 0.042 | 1 | 0.22187388 | 0.02174426 |
| Inhbb | 0.62104899 | 0.42253204 | 0.01 | 0.005 | 1 | 0.01093188 | -2.6317834 | 0.004 | 0.042 | 1 | 0.62104899 | 0.02174426 |
| Gm14286 | 0.01100062 | 2.59245704 | 0.062 | 0.01 | 1 | 0.26194382 | 0.27510724 | 0.063 | 0.031 | 1 | 0.26194382 | 0.02188022 |
| Gm49958 | 0.01100062 | 2.42253204 | 0.062 | 0.01 | 1 | 0.65785899 | -0.3098553 | 0.042 | 0.031 | 1 | 0.65785899 | 0.02188022 |
| Plekhj1 | 0.36680177 | 0.11767745 | 0.667 | 0.58 | 1 | 0.01101447 | 0.13185019 | 0.559 | 0.344 | 1 | 0.36680177 | 0.02190762 |
| Letm2 | 0.98614089 | -0.3095348 | 0.385 | 0.363 | 1 | 0.01102178 | 0.19808482 | 0.588 | 0.396 | 1 | 0.98614089 | 0.02192208 |
| Ddx10 | 0.01102325 | 0.5268687 | 0.635 | 0.508 | 1 | 0.20042591 | -0.535415 | 0.471 | 0.531 | 1 | 0.20042591 | 0.02192499 |
| Pde1a | 0.0110667 | -2.0940435 | 0.073 | 0.181 | 1 | 0.08837714 | 2.62088207 | 0.067 | 0.021 | 1 | 0.08837714 | 0.02201094 |
| Oip5os1 | 0.05667674 | 0.31487432 | 0.792 | 0.689 | 1 | 0.01108116 | 0.2365044 | 0.887 | 0.833 | 1 | 0.05667674 | 0.02203953 |
| Ntng1 | 0.52528691 | -1.1624305 | 0.01 | 0.021 | 1 | 0.01110369 | -1.8817616 | 0.029 | 0.094 | 1 | 0.52528691 | 0.0220841 |
| Gm4107 | 0.55648527 | 0.92949202 | 0.031 | 0.047 | 1 | 0.01111907 | -2.4797803 | 0.004 | 0.042 | 1 | 0.55648527 | 0.02211451 |
| Gm42209 | 0.14979364 | -3.747393 | 0.01 | 0.041 | 1 | 0.01111951 | -2.7952821 | 0.004 | 0.042 | 1 | 0.14979364 | 0.02211538 |
| Satb2 | 0.01113528 | -2.4682389 | 0.031 | 0.124 | 1 | 0.03126088 | -2.1172102 | 0.059 | 0.125 | 1 | 0.03126088 | 0.02214656 |
| Ttc33 | 0.4725334 | -0.235362 | 0.24 | 0.275 | 1 | 0.01114233 | 0.41099067 | 0.324 | 0.177 | 1 | 0.4725334 | 0.02216052 |
| Neat1 | 0.13084974 | -0.6450519 | 0.979 | 0.99 | 1 | 0.01115552 | -1.0056192 | 0.996 | 1 | 1 | 0.13084974 | 0.0221866 |
| Slmap | 0.15534152 | -0.2604387 | 0.708 | 0.808 | 1 | 0.01116305 | -0.6644206 | 0.882 | 0.927 | 1 | 0.15534152 | 0.02220148 |
| Esco2 | 0.01117376 | 2.72370157 | 0.062 | 0.01 | 1 | 0.27154471 | 0.69014474 | 0.013 | 0 | 1 | 0.27154471 | 0.02222266 |
| A430078I02Rik | 0.01117426 | 2.22988696 | 0.062 | 0.01 | 1 | 0.87066325 | -0.3098553 | 0.013 | 0.01 | 1 | 0.87066325 | 0.02222365 |
| Lrrc75b | 0.01117426 | 2.22988696 | 0.062 | 0.01 | 1 | 0.02542376 | 2.39058446 | 0.05 | 0 | 1 | 0.02542376 | 0.02222365 |
| Ccdc74a | 0.01117426 | 2.22988696 | 0.062 | 0.01 | 1 | 0.23772424 | 0.86006974 | 0.034 | 0.01 | 1 | 0.23772424 | 0.02222365 |
| Thsd1 | 0.01117643 | 2.00749454 | 0.094 | 0.026 | 1 | 0.09891444 | 0.91253716 | 0.097 | 0.042 | 1 | 0.09891444 | 0.02222794 |
| Adi1 | 0.19997636 | 0.94038034 | 0.146 | 0.098 | 1 | 0.01118207 | 0.87777174 | 0.16 | 0.052 | 1 | 0.19997636 | 0.02223911 |
| Sord | 0.01118406 | 0.78510212 | 0.24 | 0.119 | 1 | 0.07255477 | 0.25574191 | 0.214 | 0.125 | 1 | 0.07255477 | 0.02224304 |
| Arid2 | 0.03014171 | 0.26078097 | 0.667 | 0.523 | 1 | 0.01120091 | -0.6011574 | 0.845 | 0.885 | 1 | 0.03014171 | 0.02227636 |
| Gnl2 | 0.01121378 | 0.52722941 | 0.802 | 0.699 | 1 | 0.02987855 | 0.40881918 | 0.761 | 0.708 | 1 | 0.02987855 | 0.02230181 |
| Angptl1 | 0.0404543 | -1.5285584 | 0.125 | 0.218 | 1 | 0.01121439 | 0.94794249 | 0.244 | 0.115 | 1 | 0.0404543 | 0.02230302 |
| Pnkd | 0.19443421 | -0.4843586 | 0.521 | 0.575 | 1 | 0.01121574 | 0.41596977 | 0.492 | 0.312 | 1 | 0.19443421 | 0.02230568 |
| Spc25 | 0.011217 | 2.75751628 | 0.083 | 0.021 | 1 | 0.41074402 | -0.7248928 | 0.017 | 0.031 | 1 | 0.41074402 | 0.02230819 |
| Rbfox3 | 0.15837881 | 2.00749454 | 0.01 | 0 | 1 | 0.0112196 | -3.7248928 | 0.008 | 0.052 | 1 | 0.15837881 | 0.02231332 |
| Paxip1 | 0.57920877 | 0.2590333 | 0.208 | 0.181 | 1 | 0.01122016 | 0.45567948 | 0.315 | 0.167 | 1 | 0.57920877 | 0.02231443 |
| Med19 | 0.01122409 | 0.55732881 | 0.479 | 0.326 | 1 | 0.02180851 | 0.22966427 | 0.382 | 0.229 | 1 | 0.02180851 | 0.02232219 |
| Cds2 | 0.592588 | -0.1486247 | 0.312 | 0.264 | 1 | 0.01122641 | 0.21370669 | 0.403 | 0.229 | 1 | 0.592588 | 0.02232678 |
| Coil | 0.0575446 | 0.60894516 | 0.177 | 0.098 | 1 | 0.01123211 | 1.06360313 | 0.185 | 0.073 | 1 | 0.0575446 | 0.02233806 |
| Kdm2a | 0.01124867 | 0.18958821 | 0.885 | 0.725 | 1 | 0.10823849 | -0.3552472 | 0.962 | 0.969 | 1 | 0.10823849 | 0.02237082 |
| Senp1 | 0.0112587 | 0.61956888 | 0.531 | 0.394 | 1 | 0.09061024 | 0.14227156 | 0.504 | 0.375 | 1 | 0.09061024 | 0.02239064 |
| Smcr8 | 0.01129544 | 0.91438513 | 0.26 | 0.14 | 1 | 0.42810899 | 0.22619764 | 0.185 | 0.146 | 1 | 0.42810899 | 0.0224633 |
| Dhfr | 0.01129786 | 1.30705482 | 0.177 | 0.078 | 1 | 0.33587366 | -0.4353861 | 0.113 | 0.073 | 1 | 0.33587366 | 0.02246808 |
| Gsk3a | 0.18080193 | 0.29783629 | 0.542 | 0.456 | 1 | 0.01130308 | 0.26046046 | 0.681 | 0.5 | 1 | 0.18080193 | 0.0224784 |
| Aspa | 0.39189053 | -0.407543 | 0.01 | 0.026 | 1 | 0.0113097 | -1.8948178 | 0.004 | 0.042 | 1 | 0.39189053 | 0.02249149 |
| 4930447K03Rik | 0.08192263 | -2.577468 | 0 | 0.031 | 1 | 0.01131014 | -1.0203486 | 0.004 | 0.042 | 1 | 0.08192263 | 0.02249237 |
| Poglut1 | 0.30674207 | -0.3018335 | 0.365 | 0.42 | 1 | 0.0113156 | 0.44913664 | 0.403 | 0.24 | 1 | 0.30674207 | 0.02250315 |
| Azin1 | 0.01134331 | 0.24413744 | 0.896 | 0.902 | 1 | 0.89253658 | -0.3344892 | 0.929 | 0.906 | 1 | 0.89253658 | 0.02255795 |
| Nfyc | 0.01135378 | 0.43535608 | 0.656 | 0.466 | 1 | 0.61697877 | -0.3886094 | 0.618 | 0.573 | 1 | 0.61697877 | 0.02257865 |
| Sod2 | 0.01137845 | -0.6196645 | 0.771 | 0.824 | 1 | 0.9540518 | -0.2554075 | 0.723 | 0.719 | 1 | 0.9540518 | 0.02262743 |
| Rif1 | 0.0113812 | 0.59702977 | 0.656 | 0.528 | 1 | 0.38266542 | -0.2505893 | 0.584 | 0.688 | 1 | 0.38266542 | 0.02263286 |
| Tram2 | 0.01985507 | 0.39082318 | 0.708 | 0.627 | 1 | 0.01139211 | 0.27215392 | 0.773 | 0.542 | 1 | 0.01985507 | 0.02265444 |
| Cyp2r1 | 0.01142414 | 2.17741954 | 0.062 | 0.01 | 1 | 0.43079893 | 0.56461386 | 0.038 | 0.021 | 1 | 0.43079893 | 0.02271776 |
| Esr1 | 0.17322789 | 0.46560076 | 0.438 | 0.378 | 1 | 0.01142689 | 0.19686293 | 0.676 | 0.469 | 1 | 0.17322789 | 0.0227232 |
| Plcl1 | 0.34055665 | 0.19270941 | 0.26 | 0.207 | 1 | 0.01142746 | -1.6619615 | 0.181 | 0.292 | 1 | 0.34055665 | 0.02272433 |
| Fhl1 | 0.62355498 | -0.5259377 | 0.25 | 0.269 | 1 | 0.01143802 | 0.24851303 | 0.424 | 0.25 | 1 | 0.62355498 | 0.02274521 |
| Alkbh1 | 0.03752878 | 0.30469555 | 0.771 | 0.674 | 1 | 0.01147618 | 0.26323314 | 0.672 | 0.448 | 1 | 0.03752878 | 0.02282065 |
| Camkk2 | 0.02362909 | 0.71798792 | 0.531 | 0.415 | 1 | 0.01147861 | 0.39612264 | 0.588 | 0.396 | 1 | 0.02362909 | 0.02282547 |
| Rpp25l | 0.03195659 | 0.73619251 | 0.26 | 0.155 | 1 | 0.01148083 | 0.73850776 | 0.307 | 0.167 | 1 | 0.03195659 | 0.02282984 |
| Myl3 | 0.01148407 | 2.46692616 | 0.073 | 0.016 | 1 | 0.57576763 | -0.9879272 | 0.013 | 0.021 | 1 | 0.57576763 | 0.02283627 |
| Cox5b | 0.01148687 | 0.34605331 | 0.948 | 0.891 | 1 | 0.08485367 | 0.17835401 | 0.861 | 0.865 | 1 | 0.08485367 | 0.02284179 |
| Cul5 | 0.01149065 | 0.26547571 | 0.719 | 0.539 | 1 | 0.02179585 | 0.27748596 | 0.845 | 0.781 | 1 | 0.02179585 | 0.02284927 |
| Kif23 | 0.03482239 | 2.04144187 | 0.104 | 0.041 | 1 | 0.01151588 | -1.7692869 | 0.013 | 0.062 | 1 | 0.03482239 | 0.02289914 |
| Pakap.1 | 0.01152189 | -0.577468 | 0.573 | 0.674 | 1 | 0.20489152 | -1.0827864 | 0.475 | 0.552 | 1 | 0.20489152 | 0.02291102 |
| Mpv17l2 | 0.01152549 | -0.8993961 | 0.24 | 0.363 | 1 | 0.05552576 | 0.69014474 | 0.206 | 0.115 | 1 | 0.05552576 | 0.02291815 |
| 2700062C07Rik | 0.76917355 | 0.11852585 | 0.229 | 0.212 | 1 | 0.01153184 | 1.06865636 | 0.172 | 0.062 | 1 | 0.76917355 | 0.0229307 |
| Daxx | 0.38551178 | -0.366901 | 0.219 | 0.264 | 1 | 0.01153879 | 0.56929034 | 0.324 | 0.177 | 1 | 0.38551178 | 0.02294444 |
| Nrbp1 | 0.01154188 | 0.38189005 | 0.698 | 0.528 | 1 | 0.01265265 | 0.28316701 | 0.773 | 0.646 | 1 | 0.01265265 | 0.02295054 |
| 4930549C15Rik | 0.04512622 | 2.59245704 | 0.021 | 0 | 1 | 0.01154547 | -2.7248928 | 0.008 | 0.052 | 1 | 0.04512622 | 0.02295764 |
| A630014C17Rik | 0.28126256 | -1.1624305 | 0.01 | 0.031 | 1 | 0.01154547 | -3.0468209 | 0.008 | 0.052 | 1 | 0.28126256 | 0.02295764 |
| Fhl4 | 0.48064264 | 0.59245704 | 0.021 | 0.01 | 1 | 0.01154547 | -3.0468209 | 0.008 | 0.052 | 1 | 0.48064264 | 0.02295764 |
| Deaf1 | 0.37033685 | 0.20304535 | 0.417 | 0.358 | 1 | 0.01156103 | 0.19342344 | 0.567 | 0.354 | 1 | 0.37033685 | 0.0229884 |
| 2810013P06Rik | 0.22335794 | -0.5384738 | 0.208 | 0.264 | 1 | 0.01156735 | 0.69014474 | 0.231 | 0.104 | 1 | 0.22335794 | 0.02300089 |
| Eif5a2 | 0.25280242 | 0.65957123 | 0.094 | 0.057 | 1 | 0.01156902 | 1.3146356 | 0.139 | 0.042 | 1 | 0.25280242 | 0.02300419 |
| Crbn | 0.02943204 | 0.5049942 | 0.302 | 0.181 | 1 | 0.01159157 | 0.91896343 | 0.239 | 0.115 | 1 | 0.02943204 | 0.02304877 |
| Dcbld1 | 0.01159767 | -2.7998604 | 0.021 | 0.104 | 1 | 0.14168625 | 1.16819203 | 0.101 | 0.052 | 1 | 0.14168625 | 0.02306083 |
| Negr1 | 0.01160408 | -1.6623569 | 0.042 | 0.14 | 1 | 0.46231294 | 0.14533936 | 0.227 | 0.188 | 1 | 0.46231294 | 0.02307351 |
| Arhgef1 | 0.10263398 | 0.20962482 | 0.792 | 0.756 | 1 | 0.01160418 | 0.23763253 | 0.878 | 0.698 | 1 | 0.10263398 | 0.0230737 |
| Klf13 | 0.34817077 | 0.12007381 | 0.865 | 0.788 | 1 | 0.01161351 | 0.30320777 | 0.971 | 0.948 | 1 | 0.34817077 | 0.02309215 |
| Prkcd | 0.21162399 | 0.22508597 | 0.167 | 0.109 | 1 | 0.01161787 | -0.7395395 | 0.286 | 0.448 | 1 | 0.21162399 | 0.02310077 |
| Gdap1 | 0.01162616 | 2.38600616 | 0.083 | 0.021 | 1 | 0.20315163 | 1.01207283 | 0.017 | 0 | 1 | 0.20315163 | 0.02311715 |
| Zfp607a | 0.01162616 | 2.14499806 | 0.083 | 0.021 | 1 | 0.12128116 | 0.80562195 | 0.092 | 0.042 | 1 | 0.12128116 | 0.02311715 |
| Pds5a | 0.01163929 | 0.37048529 | 0.792 | 0.653 | 1 | 0.01680276 | -0.4587186 | 0.908 | 0.917 | 1 | 0.01680276 | 0.02314311 |
| Mrps35 | 0.09373804 | 0.60539609 | 0.24 | 0.155 | 1 | 0.01164782 | 0.7944814 | 0.239 | 0.115 | 1 | 0.09373804 | 0.02315997 |
| Cyp4f17 | 0.93661635 | 0.19390766 | 0.219 | 0.223 | 1 | 0.01165285 | 0.66877109 | 0.235 | 0.104 | 1 | 0.93661635 | 0.02316991 |
| Snx9 | 0.01166105 | -0.5120335 | 0.75 | 0.85 | 1 | 0.01618406 | -0.464607 | 0.891 | 0.938 | 1 | 0.01618406 | 0.02318612 |

| | | | | | | | | | | | | |
|---|---|---|---|---|---|---|---|---|---|---|---|---|
| 4930417H01Rik | 0.0116808 | 1.81484946 | 0.062 | 0.01 | 1 | 0.11764285 | 1.86006974 | 0.025 | 0 | 1 | 0.11764285 | 0.02322515 |
| Hsf5 | 0.0116808 | 1.81484946 | 0.062 | 0.01 | 1 | 1 | -0.8948178 | 0.021 | 0.021 | 1 | 1 | 0.02322515 |
| Vipr2 | 0.01168621 | 2.00749454 | 0.062 | 0.01 | 1 | 0.99643633 | -0.1630139 | 0.063 | 0.062 | 1 | 0.99643633 | 0.02323585 |
| Slc25a32 | 0.01169645 | 0.85549144 | 0.312 | 0.181 | 1 | 0.23099955 | -0.1469167 | 0.311 | 0.229 | 1 | 0.23099955 | 0.02325609 |
| Wwp1 | 0.01169852 | 0.53984241 | 0.708 | 0.544 | 1 | 0.44275618 | -0.40596 | 0.689 | 0.76 | 1 | 0.44275618 | 0.02326018 |
| H2bc6 | 0.01171005 | -1.6186906 | 0.146 | 0.275 | 1 | 0.70703889 | -2.2318528 | 0.097 | 0.083 | 1 | 0.70703889 | 0.02328297 |
| B430212C06Rik | 0.27624715 | -1.1079827 | 0.021 | 0.047 | 1 | 0.01171071 | -2.5322477 | 0.008 | 0.052 | 1 | 0.27624715 | 0.02328428 |
| Dpf1 | 0.47150075 | 1.74446013 | 0.021 | 0.01 | 1 | 0.01171071 | -2.5322477 | 0.008 | 0.052 | 1 | 0.47150075 | 0.02328428 |
| Lrrc43 | 0.8006267 | 0.32942263 | 0.021 | 0.026 | 1 | 0.01171071 | -3.0468209 | 0.008 | 0.052 | 1 | 0.8006267 | 0.02328428 |
| Tox2 | 0.0255382 | 2.32942263 | 0.042 | 0.005 | 1 | 0.01171368 | -3.1172102 | 0.008 | 0.052 | 1 | 0.0255382 | 0.02329014 |
| Lhfp | 0.01171802 | 0.25677251 | 0.771 | 0.539 | 1 | 0.27958957 | -0.2511359 | 0.857 | 0.615 | 1 | 0.27958957 | 0.02329873 |
| Dhx16 | 0.01175169 | 0.42253204 | 0.333 | 0.192 | 1 | 0.11670013 | -0.1110754 | 0.332 | 0.219 | 1 | 0.11670013 | 0.02336527 |
| Gm43111 | 0.608151 | -0.9925055 | 0.021 | 0.031 | 1 | 0.01175413 | -2.0468209 | 0.029 | 0.094 | 1 | 0.608151 | 0.02337011 |
| Pdcd2 | 0.52593547 | 0.31943854 | 0.146 | 0.119 | 1 | 0.01175528 | 1.14957636 | 0.155 | 0.052 | 1 | 0.52593547 | 0.02337237 |
| Nrbf2 | 0.7753219 | -0.1708427 | 0.531 | 0.456 | 1 | 0.01175584 | 0.50169965 | 0.416 | 0.25 | 1 | 0.7753219 | 0.02337348 |
| Ncapg2 | 0.01177033 | 1.16976597 | 0.188 | 0.083 | 1 | 0.03537908 | -1.8948178 | 0.092 | 0.167 | 1 | 0.03537908 | 0.02340213 |
| Myo1e | 0.01182012 | -0.4993317 | 0.99 | 0.995 | 1 | 0.05217657 | -0.3863986 | 1 | 1 | 1 | 0.05217657 | 0.02350053 |
| Fut8 | 0.01182256 | -0.6763214 | 0.635 | 0.715 | 1 | 0.75419532 | -0.1004865 | 0.903 | 0.917 | 1 | 0.75419532 | 0.02350534 |
| Lin37 | 0.15208105 | -0.7029988 | 0.104 | 0.166 | 1 | 0.01182891 | 1.11640949 | 0.155 | 0.052 | 1 | 0.15208105 | 0.02351789 |
| Actr1a | 0.01184121 | 0.4898873 | 0.573 | 0.42 | 1 | 0.09136822 | -0.4491301 | 0.668 | 0.698 | 1 | 0.09136822 | 0.02354221 |
| Pi4k2b | 0.02511659 | 0.59761874 | 0.417 | 0.301 | 1 | 0.01184276 | 0.3086598 | 0.58 | 0.427 | 1 | 0.02511659 | 0.02354527 |
| Cacnb2 | 0.11750611 | -0.3435799 | 0.583 | 0.466 | 1 | 0.01185257 | -1.2295811 | 0.744 | 0.76 | 1 | 0.11750611 | 0.02356466 |
| Ddc | 0.01186126 | 2.65957123 | 0.062 | 0.01 | 1 | 0.87066325 | -0.3098553 | 0.013 | 0.01 | 1 | 0.87066325 | 0.02358183 |
| Cpt1c | 0.01186863 | 0.67628863 | 0.26 | 0.135 | 1 | 0.17915591 | -1.2742314 | 0.147 | 0.198 | 1 | 0.17915591 | 0.02359639 |
| Fam117a | 0.01187003 | -1.8886697 | 0.073 | 0.181 | 1 | 0.0734973 | -0.8504236 | 0.122 | 0.198 | 1 | 0.0734973 | 0.02359916 |
| Kctd5 | 0.13580623 | 0.35214271 | 0.333 | 0.244 | 1 | 0.0118747 | 0.19403563 | 0.496 | 0.292 | 1 | 0.13580623 | 0.0236084 |
| Med24 | 0.09088477 | 0.60539609 | 0.25 | 0.166 | 1 | 0.01188331 | 0.97554696 | 0.252 | 0.125 | 1 | 0.09088477 | 0.02362541 |
| Zfp3 | 0.38689439 | -0.577468 | 0.01 | 0.026 | 1 | 0.0118836 | -2.3098553 | 0.008 | 0.052 | 1 | 0.38689439 | 0.02362599 |
| Gm2885 | 0.01189308 | 1.65957123 | 0.094 | 0.026 | 1 | 0.28991808 | -1.8948178 | 0.029 | 0.052 | 1 | 0.28991808 | 0.02364471 |
| Egfl7 | 0.0119104 | 2.00749454 | 0.083 | 0.021 | 1 | 0.15394868 | 1.27510724 | 0.021 | 0 | 1 | 0.15394868 | 0.02367895 |
| A630066F11Rik | 0.0119104 | 2.00749454 | 0.083 | 0.021 | 1 | 0.15394868 | 1.27510724 | 0.021 | 0 | 1 | 0.15394868 | 0.02367895 |
| Gm15581 | 0.0119114 | -2.7294711 | 0.021 | 0.104 | 1 | 0.40782973 | -1.4167705 | 0.034 | 0.052 | 1 | 0.40782973 | 0.02368093 |
| Flrt2 | 0.02937278 | 0.24437815 | 0.865 | 0.731 | 1 | 0.01194593 | 0.13633378 | 0.945 | 0.833 | 1 | 0.02937278 | 0.02374916 |
| Extl3 | 0.2186104 | -0.5924183 | 0.271 | 0.326 | 1 | 0.01194792 | 0.56461386 | 0.399 | 0.24 | 1 | 0.2186104 | 0.02375309 |
| E430024P14Rik | 0.01194881 | 2.17741954 | 0.073 | 0.016 | 1 | 0.04197017 | 2.39058446 | 0.042 | 0 | 1 | 0.04197017 | 0.02375485 |
| Wdr20 | 0.01195969 | 0.4681289 | 0.49 | 0.332 | 1 | 0.0627367 | -0.6468902 | 0.55 | 0.635 | 1 | 0.0627367 | 0.02377634 |
| Cenpm | 0.01196022 | 1.77302928 | 0.073 | 0.016 | 1 | 0.57170879 | -1.6317834 | 0.013 | 0.021 | 1 | 0.57170879 | 0.02377739 |
| Flrt1 | 0.01197527 | 0.84190847 | 0.396 | 0.264 | 1 | 0.02925794 | 0.44621915 | 0.345 | 0.198 | 1 | 0.02925794 | 0.02380713 |
| Bap1 | 0.01198485 | 0.92503238 | 0.281 | 0.155 | 1 | 0.04738078 | 0.16407593 | 0.366 | 0.229 | 1 | 0.04738078 | 0.02382606 |
| Nup153 | 0.01262374 | 0.25599285 | 0.854 | 0.746 | 1 | 0.01200596 | -0.3335242 | 0.874 | 0.927 | 1 | 0.01262374 | 0.02386777 |
| Osbpl7 | 0.01201133 | 1.00749454 | 0.104 | 0.031 | 1 | 0.23750863 | 0.52021974 | 0.105 | 0.062 | 1 | 0.23750863 | 0.02387838 |
| Ackr4 | 0.0120155 | -2.407543 | 0.021 | 0.104 | 1 | 0.04103067 | -1.5322477 | 0.038 | 0.094 | 1 | 0.04103067 | 0.02388663 |
| Polr3k | 0.21437159 | 0.14076107 | 0.292 | 0.212 | 1 | 0.01202822 | 0.77356075 | 0.282 | 0.146 | 1 | 0.21437159 | 0.02391177 |
| Arhgef26 | 0.62126082 | -0.3500575 | 0.094 | 0.114 | 1 | 0.012035 | -0.4711362 | 0.214 | 0.365 | 1 | 0.62126082 | 0.02392517 |
| 1700037F03Rik | 0.22347423 | 0.10703021 | 0.125 | 0.078 | 1 | 0.01203545 | 3.01207283 | 0.063 | 0 | 1 | 0.22347423 | 0.02392604 |
| Chmp7 | 0.44759919 | 0.62047141 | 0.104 | 0.078 | 1 | 0.01203553 | 0.67365661 | 0.286 | 0.146 | 1 | 0.44759919 | 0.02392621 |
| Zfp239 | 0.02667324 | 1.65957123 | 0.083 | 0.026 | 1 | 0.01204557 | 3.14957636 | 0.063 | 0 | 1 | 0.02667324 | 0.02394604 |
| Pcdhb4 | 0.38192865 | 1.22988696 | 0.031 | 0.016 | 1 | 0.01204589 | 3.27510724 | 0.063 | 0 | 1 | 0.38192865 | 0.02394668 |
| 3222401L13Rik | 0.01204606 | -1.4843586 | 0.031 | 0.124 | 1 | 0.38788407 | 0.34222143 | 0.067 | 0.042 | 1 | 0.38788407 | 0.02394702 |
| Kcnj6 | 0.17534439 | 1.32942263 | 0.042 | 0.016 | 1 | 0.01205257 | -2.8948178 | 0.008 | 0.052 | 1 | 0.17534439 | 0.02395988 |
| Pole | 0.37340263 | 0.17741954 | 0.073 | 0.047 | 1 | 0.01205257 | -2.3098553 | 0.008 | 0.052 | 1 | 0.37340263 | 0.02395988 |
| Gm29683 | 0.48218219 | 0.15949763 | 0.042 | 0.026 | 1 | 0.01205257 | -2.3098553 | 0.008 | 0.052 | 1 | 0.48218219 | 0.02395988 |
| Keap1 | 0.13626806 | 0.39183524 | 0.406 | 0.311 | 1 | 0.01205494 | 0.32091093 | 0.496 | 0.302 | 1 | 0.13626806 | 0.02396456 |
| Atp6v1c1 | 0.33708292 | -0.3264062 | 0.333 | 0.389 | 1 | 0.01205741 | 0.88549234 | 0.479 | 0.323 | 1 | 0.33708292 | 0.02396945 |
| 8430426J06Rik | 0.08261745 | -2.3144336 | 0.01 | 0.052 | 1 | 0.01208608 | -2.4797803 | 0.029 | 0.094 | 1 | 0.08261745 | 0.02402609 |
| Mpst | 0.28871744 | 0.57139542 | 0.146 | 0.104 | 1 | 0.01209278 | 0.61614416 | 0.189 | 0.073 | 1 | 0.28871744 | 0.02403933 |
| Orc4 | 0.01210502 | 0.30411154 | 0.656 | 0.466 | 1 | 0.08668385 | -0.5393371 | 0.668 | 0.719 | 1 | 0.08668385 | 0.0240635 |
| Pou2f2 | 0.32700752 | -1.6465096 | 0.031 | 0.057 | 1 | 0.01211077 | -2.8028953 | 0.042 | 0.115 | 1 | 0.32700752 | 0.02407487 |
| Fabp4 | 0.01211078 | 1.00749454 | 0.094 | 0.026 | 1 | 0.91453058 | 0.52664601 | 0.034 | 0.031 | 1 | 0.91453058 | 0.02407489 |
| Tmem50b | 0.49451216 | -0.3056633 | 0.281 | 0.316 | 1 | 0.01212523 | 0.63043849 | 0.429 | 0.281 | 1 | 0.49451216 | 0.02410345 |
| Patz1 | 0.01212596 | -1.3910548 | 0.094 | 0.207 | 1 | 0.01407617 | 0.38613855 | 0.273 | 0.135 | 1 | 0.01407617 | 0.02410488 |
| Il13ra1 | 0.07486254 | -0.601921 | 0.24 | 0.347 | 1 | 0.01213049 | -0.766412 | 0.357 | 0.51 | 1 | 0.07486254 | 0.02411382 |
| Ptp4a1.1 | 0.0122037 | 2.00749454 | 0.073 | 0.016 | 1 | 0.04198609 | 2.14957636 | 0.042 | 0 | 1 | 0.04198609 | 0.02425846 |
| Gm43062 | 0.0122037 | 2.00749454 | 0.073 | 0.016 | 1 | 0.69407793 | -0.8948178 | 0.063 | 0.073 | 1 | 0.69407793 | 0.02425846 |
| Nsun4 | 0.88193455 | -0.2148979 | 0.24 | 0.238 | 1 | 0.01220478 | 0.3068161 | 0.311 | 0.167 | 1 | 0.88193455 | 0.02426061 |
| Sema3b | 0.61550914 | -0.407543 | 0.021 | 0.031 | 1 | 0.01222249 | 2.27510724 | 0.084 | 0.01 | 1 | 0.61550914 | 0.0242956 |
| Gm11713 | 0.01222417 | -2.265524 | 0.021 | 0.104 | 1 | 0.53700593 | -0.2654611 | 0.109 | 0.083 | 1 | 0.53700593 | 0.02429891 |
| Ppat | 0.01599817 | 0.65080102 | 0.469 | 0.321 | 1 | 0.01222992 | 0.29837702 | 0.332 | 0.177 | 1 | 0.01599817 | 0.02431026 |
| Gga1 | 0.3299132 | 0.35541784 | 0.271 | 0.223 | 1 | 0.01223969 | 0.39058446 | 0.319 | 0.167 | 1 | 0.3299132 | 0.02432956 |
| Tm4sf1 | 0.01227572 | -1.5069181 | 0.396 | 0.534 | 1 | 0.88647014 | 0.66454491 | 0.366 | 0.333 | 1 | 0.88647014 | 0.02440074 |
| Rbm44 | 0.01227936 | 2.00749454 | 0.073 | 0.016 | 1 | 0.55024964 | -1.4353861 | 0.029 | 0.042 | 1 | 0.55024964 | 0.02440793 |
| Lypd3 | 0.01228151 | 1.70793425 | 0.073 | 0.016 | 1 | 0.67254361 | -0.3098553 | 0.017 | 0.01 | 1 | 0.67254361 | 0.02441219 |
| Dhcr7 | 0.78545318 | -0.1393469 | 0.198 | 0.181 | 1 | 0.0122923 | 1.61071027 | 0.164 | 0.062 | 1 | 0.78545318 | 0.02443351 |
| Nelfe | 0.01980488 | 0.70031303 | 0.406 | 0.285 | 1 | 0.01230223 | 0.10002053 | 0.315 | 0.156 | 1 | 0.01980488 | 0.02445312 |
| Cant1 | 0.08994954 | 0.62898291 | 0.271 | 0.187 | 1 | 0.0123163 | 0.92044236 | 0.197 | 0.083 | 1 | 0.08994954 | 0.02448091 |
| Gm43328 | 0.01232763 | 1.21394541 | 0.146 | 0.057 | 1 | 0.1073383 | 0.3163299 | 0.231 | 0.146 | 1 | 0.1073383 | 0.02450329 |
| Arhgef10 | 0.08812644 | 0.44455834 | 0.292 | 0.202 | 1 | 0.01233276 | 0.43933663 | 0.382 | 0.219 | 1 | 0.08812644 | 0.02451343 |
| Dnaaf2 | 0.03768094 | 0.87422801 | 0.208 | 0.114 | 1 | 0.01235979 | 1.27510724 | 0.193 | 0.083 | 1 | 0.03768094 | 0.02456682 |
| Slc22a5 | 0.07802063 | -1.1586977 | 0.25 | 0.342 | 1 | 0.01237279 | -1.4313188 | 0.345 | 0.438 | 1 | 0.07802063 | 0.0245925 |
| Mrpl36 | 0.206306 | 0.29854332 | 0.417 | 0.347 | 1 | 0.01239483 | 0.34910782 | 0.357 | 0.198 | 1 | 0.206306 | 0.02463603 |
| Arl10 | 0.48648442 | 0.27052894 | 0.188 | 0.155 | 1 | 0.01240276 | 0.87234807 | 0.214 | 0.094 | 1 | 0.48648442 | 0.0246517 |
| Rxrb | 0.09116553 | 0.37362244 | 0.26 | 0.171 | 1 | 0.01240693 | 0.57767001 | 0.235 | 0.104 | 1 | 0.09116553 | 0.02465993 |
| Zfp386 | 0.01597256 | 0.62283069 | 0.323 | 0.192 | 1 | 0.01241236 | 0.78968041 | 0.277 | 0.146 | 1 | 0.01597256 | 0.02467066 |
| Nr4a2 | 0.01245723 | 0.41787066 | 0.781 | 0.627 | 1 | 0.38956864 | -0.3626567 | 0.706 | 0.812 | 1 | 0.38956864 | 0.02475928 |
| Tmprss2 | 0.32025586 | -0.9925055 | 0 | 0.01 | 1 | 0.01249211 | -1.8948178 | 0.029 | 0.094 | 1 | 0.32025586 | 0.02482816 |

| | | | | | | | | | | | | |
|---|---|---|---|---|---|---|---|---|---|---|---|---|
| Fbln2 | 0.01251267 | 1.74446013 | 0.083 | 0.021 | 1 | 0.67144323 | -0.5322477 | 0.042 | 0.031 | 1 | 0.67144323 | 0.02486878 |
| Kntc1 | 0.01252267 | 1.78510212 | 0.083 | 0.021 | 1 | 0.25080911 | -0.9879272 | 0.013 | 0.031 | 1 | 0.25080911 | 0.02488853 |
| Dbndd2 | 0.97791667 | -0.1249558 | 0.208 | 0.202 | 1 | 0.01252335 | 0.86006974 | 0.307 | 0.167 | 1 | 0.97791667 | 0.02488987 |
| 4930428N03Rik | 0.01252421 | 1.85549144 | 0.073 | 0.016 | 1 | 0.50587544 | -1.8948178 | 0.004 | 0.01 | 1 | 0.50587544 | 0.02489157 |
| Ppil1 | 0.0125957 | 1.09495738 | 0.25 | 0.135 | 1 | 0.01552817 | 0.95863357 | 0.206 | 0.094 | 1 | 0.01552817 | 0.02503275 |
| Nudt18 | 0.01262323 | -1.599163 | 0.094 | 0.207 | 1 | 0.65386933 | -0.2625495 | 0.113 | 0.094 | 1 | 0.65386933 | 0.02508711 |
| Isg15 | 0.04003751 | -2.6445822 | 0.021 | 0.083 | 1 | 0.01265265 | -3.7992401 | 0.05 | 0.125 | 1 | 0.04003751 | 0.02514521 |
| Lrrc8c | 0.01267891 | 0.44114771 | 0.875 | 0.762 | 1 | 0.64280221 | -0.4375238 | 0.832 | 0.635 | 1 | 0.64280221 | 0.02519707 |
| B3galt6 | 0.04275994 | 0.62640437 | 0.344 | 0.233 | 1 | 0.01270937 | 0.45567948 | 0.248 | 0.115 | 1 | 0.04275994 | 0.02525721 |
| Rab43 | 0.01271817 | 0.73331957 | 0.521 | 0.383 | 1 | 0.18958022 | -0.6368963 | 0.643 | 0.688 | 1 | 0.18958022 | 0.02527458 |
| Gm13944 | 0.0127183 | -0.9603251 | 0.208 | 0.347 | 1 | 0.03082507 | 0.39641353 | 0.185 | 0.083 | 1 | 0.03082507 | 0.02527485 |
| Cep290 | 0.01273013 | 0.89969125 | 0.417 | 0.29 | 1 | 0.51888461 | 0.17557156 | 0.256 | 0.229 | 1 | 0.51888461 | 0.0252982 |
| Hccs | 0.21113314 | 0.27052894 | 0.271 | 0.202 | 1 | 0.01273051 | 0.17995001 | 0.349 | 0.198 | 1 | 0.21113314 | 0.02529896 |
| Fads3 | 0.01273085 | 0.54686017 | 0.615 | 0.466 | 1 | 0.17024846 | 0.16522962 | 0.521 | 0.406 | 1 | 0.17024846 | 0.02529962 |
| Pelp1 | 0.83611762 | -0.1477837 | 0.219 | 0.197 | 1 | 0.01273564 | 0.65537932 | 0.286 | 0.146 | 1 | 0.83611762 | 0.02530908 |
| Cdk2 | 0.07954675 | 0.866632 | 0.24 | 0.161 | 1 | 0.012764 | 0.70509508 | 0.307 | 0.167 | 1 | 0.07954675 | 0.02536509 |
| Scrn1 | 0.01277251 | 1.68556644 | 0.073 | 0.016 | 1 | 0.16736519 | 0.49749966 | 0.113 | 0.062 | 1 | 0.16736519 | 0.02538188 |
| Fbxo42 | 0.01279953 | 0.40342321 | 0.552 | 0.383 | 1 | 0.98403532 | -0.2414301 | 0.697 | 0.646 | 1 | 0.98403532 | 0.02543523 |
| Ap2b1 | 0.01280987 | 0.39140952 | 0.844 | 0.751 | 1 | 0.050459 | 0.10837051 | 0.849 | 0.802 | 1 | 0.050459 | 0.02545564 |
| 1700007L15Rik | 0.01281607 | 1.59245704 | 0.083 | 0.021 | 1 | 0.92228543 | -0.7248928 | 0.029 | 0.031 | 1 | 0.92228543 | 0.02546788 |
| Mocos | 0.01282541 | -2.9925055 | 0 | 0.062 | 1 | 0.98931086 | -0.9472852 | 0.063 | 0.062 | 1 | 0.98931086 | 0.02548632 |
| Gtf2ird2 | 0.01282666 | 1.00749454 | 0.177 | 0.078 | 1 | 0.18189466 | 0.75427507 | 0.134 | 0.083 | 1 | 0.18189466 | 0.02548879 |
| Sgcg | 0.0128284 | -3.1624305 | 0 | 0.062 | 1 | 0.11689951 | -2.3098553 | 0 | 0.01 | 1 | 0.11689951 | 0.02549223 |
| Slc18a2 | 0.0128284 | -3.8504865 | 0 | 0.062 | 1 | 0.77510918 | 0.14957636 | 0.025 | 0.031 | 1 | 0.77510918 | 0.02549223 |
| Cbr1 | 0.38487292 | 0.3897421 | 0.365 | 0.337 | 1 | 0.01283202 | 0.52787331 | 0.471 | 0.323 | 1 | 0.38487292 | 0.02549938 |
| Chd7 | 0.02698736 | 0.3205544 | 0.635 | 0.508 | 1 | 0.01283286 | -0.5964351 | 0.756 | 0.802 | 1 | 0.02698736 | 0.02550103 |
| Timp4 | 0.01283288 | -3.5160674 | 0 | 0.062 | 1 | 0.50887612 | -1.3098553 | 0.004 | 0.01 | 1 | 0.50887612 | 0.02550108 |
| Mmp9 | 0.04373406 | -3.6363617 | 0 | 0.041 | 1 | 0.01285022 | 0.98265634 | 0.034 | 0.104 | 1 | 0.04373406 | 0.02553531 |
| Qki | 0.01286247 | 0.23223809 | 0.99 | 0.995 | 1 | 0.01779059 | -0.3914258 | 1 | 1 | 1 | 0.01779059 | 0.02555949 |
| Ubap1l | 0.01287621 | 1.32942263 | 0.094 | 0.026 | 1 | 0.07666135 | 0.22619764 | 0.105 | 0.042 | 1 | 0.07666135 | 0.02558663 |
| Rexo1 | 0.8335606 | -0.114496 | 0.365 | 0.373 | 1 | 0.01288108 | 0.32091093 | 0.529 | 0.365 | 1 | 0.8335606 | 0.02559624 |
| Nol3 | 0.06357472 | 1.21394541 | 0.125 | 0.062 | 1 | 0.01288649 | 1.61614416 | 0.134 | 0.042 | 1 | 0.06357472 | 0.02560693 |
| Haus3 | 0.20771796 | 0.43199237 | 0.323 | 0.254 | 1 | 0.01292098 | 0.41393494 | 0.29 | 0.146 | 1 | 0.20771796 | 0.025675 |
| Synj1 | 0.10865982 | -0.7756941 | 0.302 | 0.383 | 1 | 0.01292622 | -0.713488 | 0.487 | 0.625 | 1 | 0.10865982 | 0.02568535 |
| Tnfaip8 | 0.01293114 | 0.33903772 | 0.708 | 0.57 | 1 | 0.51261421 | 0.11464257 | 0.769 | 0.792 | 1 | 0.51261421 | 0.02569506 |
| Mettl16 | 0.14857816 | 0.27052894 | 0.594 | 0.487 | 1 | 0.01293291 | 0.27510724 | 0.63 | 0.458 | 1 | 0.14857816 | 0.02569856 |
| Rras2 | 0.01294467 | -0.7663786 | 0.781 | 0.788 | 1 | 0.22735268 | -0.7823309 | 0.861 | 0.865 | 1 | 0.22735268 | 0.02572178 |
| Zfyve19 | 0.51508949 | 0.27052894 | 0.083 | 0.062 | 1 | 0.01298907 | 0.86006974 | 0.105 | 0.021 | 1 | 0.51508949 | 0.02580942 |
| Adssl1 | 1 | -0.3848229 | 0.062 | 0.062 | 1 | 0.01299004 | 2.21370669 | 0.084 | 0.01 | 1 | 1 | 0.02581133 |
| Bbip1 | 0.11667849 | 0.27052894 | 0.521 | 0.404 | 1 | 0.01300196 | 0.19603567 | 0.542 | 0.365 | 1 | 0.11667849 | 0.02583487 |
| Bach2 | 0.63265109 | 0.22848424 | 0.521 | 0.56 | 1 | 0.013056 | -0.5411808 | 0.672 | 0.865 | 1 | 0.63265109 | 0.02594153 |
| Sh2d4a | 0.51724187 | -0.4170033 | 0.094 | 0.119 | 1 | 0.0130635 | 1.86006974 | 0.147 | 0.052 | 1 | 0.51724187 | 0.02595635 |
| Oaz2 | 0.34854569 | -0.2666804 | 0.552 | 0.57 | 1 | 0.01307143 | 0.30339628 | 0.718 | 0.552 | 1 | 0.34854569 | 0.02597199 |
| Uqcrc2 | 0.08272526 | 0.21102793 | 0.833 | 0.782 | 1 | 0.01307651 | 0.22543985 | 0.908 | 0.833 | 1 | 0.08272526 | 0.02598202 |
| Uhrf2 | 0.1837223 | -0.4344497 | 0.729 | 0.741 | 1 | 0.01311466 | -0.7028665 | 0.824 | 0.844 | 1 | 0.1837223 | 0.02605732 |
| Rptor | 0.85134153 | -0.1497194 | 0.76 | 0.699 | 1 | 0.01312214 | -0.7758439 | 0.798 | 0.865 | 1 | 0.85134153 | 0.02607209 |
| Ldah | 0.01313758 | 0.46367073 | 0.531 | 0.358 | 1 | 0.02168177 | 0.10985872 | 0.508 | 0.312 | 1 | 0.02168177 | 0.02610256 |
| ENSMUSG00000121374 | 0.01314073 | 1.52206771 | 0.083 | 0.021 | 1 | 0.92416907 | -0.5728897 | 0.034 | 0.031 | 1 | 0.92416907 | 0.02610878 |
| Arhgef10l | 0.01314928 | -2.0799683 | 0.031 | 0.119 | 1 | 0.93693454 | -0.6270454 | 0.155 | 0.146 | 1 | 0.93693454 | 0.02612566 |
| Epop | 0.11788474 | -0.1989563 | 0.073 | 0.14 | 1 | 0.01319048 | 1.11640949 | 0.151 | 0.052 | 1 | 0.11788474 | 0.02620698 |
| Ano6 | 0.01323962 | -0.5399391 | 0.917 | 0.933 | 1 | 0.19297441 | -0.5106515 | 0.971 | 0.938 | 1 | 0.19297441 | 0.02630394 |
| Kmt5b | 0.346222 | 0.34573529 | 0.719 | 0.72 | 1 | 0.01325898 | 0.35105609 | 0.874 | 0.844 | 1 | 0.346222 | 0.02634215 |
| Gm10125 | 0.57275342 | 0.24565427 | 0.125 | 0.104 | 1 | 0.01327167 | -1.3838558 | 0.092 | 0.188 | 1 | 0.57275342 | 0.02636719 |
| Zfp30 | 0.44515784 | -0.7998604 | 0.073 | 0.098 | 1 | 0.01332243 | 0.87071698 | 0.214 | 0.094 | 1 | 0.44515784 | 0.02646736 |
| Ak7 | 0.01333505 | 1.24850264 | 0.094 | 0.026 | 1 | 0.2573472 | -0.1878647 | 0.092 | 0.052 | 1 | 0.2573472 | 0.02649227 |
| Pcgf6 | 0.05420597 | 0.78931437 | 0.292 | 0.197 | 1 | 0.01333714 | 0.56461386 | 0.244 | 0.115 | 1 | 0.05420597 | 0.0264964 |
| Slc25a5 | 0.01334594 | 0.30745551 | 0.844 | 0.782 | 1 | 0.17950727 | 0.23021578 | 0.912 | 0.875 | 1 | 0.17950727 | 0.02651376 |
| Prmt7 | 0.0133473 | 0.84148459 | 0.271 | 0.15 | 1 | 0.26803073 | -0.3754436 | 0.244 | 0.167 | 1 | 0.26803073 | 0.02651645 |
| Rad51b | 0.17786604 | 0.20543391 | 0.24 | 0.171 | 1 | 0.01336297 | -2.8523825 | 0.05 | 0.125 | 1 | 0.17786604 | 0.02654737 |
| Glrx3 | 0.05790663 | 0.28253896 | 0.781 | 0.725 | 1 | 0.01337948 | 0.24548207 | 0.819 | 0.771 | 1 | 0.05790663 | 0.02657994 |
| Pou5f1 | 0.02858131 | 1.32942263 | 0.083 | 0.026 | 1 | 0.01339411 | 1.49749966 | 0.084 | 0.01 | 1 | 0.02858131 | 0.02660881 |
| Fam168b | 0.01339744 | 0.50302195 | 0.688 | 0.554 | 1 | 0.03022214 | 0.18291354 | 0.798 | 0.635 | 1 | 0.03022214 | 0.02661539 |
| Pigo | 0.0134228 | 1.39451766 | 0.135 | 0.052 | 1 | 0.3464269 | 0.10518224 | 0.109 | 0.073 | 1 | 0.3464269 | 0.02666543 |
| Nrtn | 0.0134228 | 1.39451766 | 0.135 | 0.052 | 1 | 0.08910678 | 0.88278982 | 0.113 | 0.052 | 1 | 0.08910678 | 0.02666543 |
| Adra1b | 0.16021093 | 0.32160313 | 0.177 | 0.114 | 1 | 0.01343563 | 2.20110666 | 0.118 | 0.031 | 1 | 0.16021093 | 0.02669075 |
| Snap23 | 0.03410042 | 0.48008306 | 0.75 | 0.648 | 1 | 0.01343645 | 0.20301562 | 0.866 | 0.708 | 1 | 0.03410042 | 0.02669236 |
| Lmx1a | 0.48221225 | 0.27052894 | 0.042 | 0.026 | 1 | 0.01344434 | 0.71989208 | 0.084 | 0.01 | 1 | 0.48221225 | 0.02670793 |
| Nle1 | 0.88172641 | -0.2625946 | 0.146 | 0.15 | 1 | 0.01344922 | 0.56461386 | 0.122 | 0.031 | 1 | 0.88172641 | 0.02671757 |
| Fam216a | 0.08000533 | 0.47462055 | 0.344 | 0.244 | 1 | 0.0134694 | 0.37664526 | 0.353 | 0.198 | 1 | 0.08000533 | 0.02675737 |
| D830025C05Rik | 0.22798009 | 0.52562553 | 0.25 | 0.197 | 1 | 0.01347096 | 0.78896247 | 0.429 | 0.281 | 1 | 0.22798009 | 0.02676045 |
| Psrc1 | 0.01347867 | 1.27052894 | 0.073 | 0.016 | 1 | 0.5778028 | -0.8948178 | 0.013 | 0.021 | 1 | 0.5778028 | 0.02677567 |
| Agpat1 | 0.21963038 | 0.51153704 | 0.292 | 0.233 | 1 | 0.01348134 | 0.72117163 | 0.315 | 0.177 | 1 | 0.21963038 | 0.02678094 |
| Rsph3a | 0.01350616 | 1.33870045 | 0.24 | 0.13 | 1 | 0.57077059 | -0.2580163 | 0.227 | 0.188 | 1 | 0.57077059 | 0.02682991 |
| Tarbp2 | 0.39426955 | 0.56692195 | 0.208 | 0.171 | 1 | 0.01351798 | 1.20471791 | 0.151 | 0.052 | 1 | 0.39426955 | 0.02685322 |
| Pgp | 0.56357867 | 0.23697638 | 0.458 | 0.415 | 1 | 0.01353528 | 0.40755753 | 0.458 | 0.292 | 1 | 0.56357867 | 0.02688736 |
| Vcan | 0.30995495 | -1.8669746 | 0.115 | 0.155 | 1 | 0.01354659 | -1.9228321 | 0.042 | 0.115 | 1 | 0.30995495 | 0.02690967 |
| Megf6 | 0.01356017 | -2.577468 | 0.01 | 0.083 | 1 | 0.08751624 | -1.8573431 | 0.046 | 0.094 | 1 | 0.08751624 | 0.02693646 |
| Mtx1 | 0.04959444 | 0.64106698 | 0.469 | 0.373 | 1 | 0.01356187 | 0.12572961 | 0.471 | 0.292 | 1 | 0.04959444 | 0.02693982 |
| Nupr1 | 0.01356296 | -0.5368851 | 0.698 | 0.839 | 1 | 0.09212204 | 0.29963364 | 0.819 | 0.906 | 1 | 0.09212204 | 0.02694197 |
| Abtb3 | 0.14635658 | -0.7294711 | 0.229 | 0.311 | 1 | 0.01356871 | -1.229124 | 0.248 | 0.365 | 1 | 0.14635658 | 0.02695331 |
| Slc17a5 | 0.06242305 | 0.21394541 | 0.417 | 0.295 | 1 | 0.01358202 | 0.35999614 | 0.534 | 0.354 | 1 | 0.06242305 | 0.02697956 |
| Tfeb | 0.55675697 | -0.1529701 | 0.229 | 0.264 | 1 | 0.01358968 | 0.39685251 | 0.424 | 0.25 | 1 | 0.55675697 | 0.02699467 |
| Acvr2b | 0.80274368 | 0.20013961 | 0.031 | 0.026 | 1 | 0.01364078 | -1.5163061 | 0.042 | 0.115 | 1 | 0.80274368 | 0.02709549 |
| Pop5 | 0.97544178 | 0.23697638 | 0.323 | 0.326 | 1 | 0.01364455 | 0.37698685 | 0.408 | 0.25 | 1 | 0.97544178 | 0.02710293 |
| Ldb2 | 0.01366009 | 0.68253238 | 0.354 | 0.212 | 1 | 0.08244035 | -1.9357895 | 0.151 | 0.208 | 1 | 0.08244035 | 0.02713357 |
| Tmem212 | 0.01366787 | -0.8770282 | 0.042 | 0.14 | 1 | 0.09528326 | -0.5163061 | 0.168 | 0.094 | 1 | 0.09528326 | 0.02714893 |
| Ralgps2 | 0.01368279 | -0.2148979 | 0.208 | 0.368 | 1 | 0.64460934 | -0.7879026 | 0.16 | 0.177 | 1 | 0.64460934 | 0.02717835 |

| | | | | | | | | | | | | |
|---|---|---|---|---|---|---|---|---|---|---|---|---|
| Zfp597 | 0.0320148 | 0.71376333 | 0.25 | 0.145 | 1 | 0.01371374 | 0.13640097 | 0.328 | 0.177 | 1 | 0.0320148 | 0.02723942 |
| Lrch4 | 0.23312978 | 0.68556644 | 0.115 | 0.073 | 1 | 0.013723 | 1.64434105 | 0.118 | 0.031 | 1 | 0.23312978 | 0.02725768 |
| Abhd10 | 0.12184475 | 0.64994253 | 0.24 | 0.166 | 1 | 0.0137458 | 0.81942775 | 0.252 | 0.125 | 1 | 0.12184475 | 0.02730266 |
| Vti1a | 0.01377272 | 0.25593157 | 0.688 | 0.528 | 1 | 0.7391204 | -0.3159298 | 0.672 | 0.75 | 1 | 0.7391204 | 0.02735575 |
| Rnf126 | 0.03259188 | 0.63134605 | 0.354 | 0.244 | 1 | 0.01377436 | 0.49246407 | 0.408 | 0.26 | 1 | 0.03259188 | 0.02735899 |
| Nras | 0.50527743 | 0.10060394 | 0.771 | 0.767 | 1 | 0.01379063 | 0.21149114 | 0.819 | 0.802 | 1 | 0.50527743 | 0.02739107 |
| C1rl | 0.87043174 | 0.15949763 | 0.073 | 0.067 | 1 | 0.01380279 | 1.42711033 | 0.134 | 0.042 | 1 | 0.87043174 | 0.02741506 |
| Ipo7 | 0.01765423 | 0.32037749 | 0.896 | 0.845 | 1 | 0.01381315 | 0.21608672 | 0.933 | 0.875 | 1 | 0.01765423 | 0.02743551 |
| Lsm14a | 0.01381445 | 0.27428108 | 0.729 | 0.58 | 1 | 0.67352584 | -0.1586365 | 0.92 | 0.833 | 1 | 0.67352584 | 0.02743807 |
| Zbtb7a | 0.013842 | 0.45263407 | 0.76 | 0.642 | 1 | 0.26926417 | 0.20299939 | 0.916 | 0.927 | 1 | 0.26926417 | 0.02749239 |
| Ephb1 | 0.52528691 | -2.0799683 | 0.01 | 0.021 | 1 | 0.0138437 | -3.4167705 | 0.025 | 0.083 | 1 | 0.52528691 | 0.02749576 |
| Zfp934 | 0.01384674 | 0.87625 | 0.24 | 0.124 | 1 | 0.08192684 | 0.54812573 | 0.172 | 0.094 | 1 | 0.08192684 | 0.02750175 |
| Slc25a21 | 0.01385681 | -3.0799683 | 0.01 | 0.083 | 1 | 0.91751776 | -1.0323213 | 0.05 | 0.052 | 1 | 0.91751776 | 0.0275216 |
| Sfxn3 | 0.26224583 | -0.4157169 | 0.177 | 0.233 | 1 | 0.01387074 | 0.56883844 | 0.349 | 0.198 | 1 | 0.26224583 | 0.02754908 |
| 4833413G10Rik | 0.01388804 | 3.32942263 | 0.031 | 0 | 1 | 0.01981438 | 2.49749966 | 0.055 | 0 | 1 | 0.01981438 | 0.02758321 |
| Spink2 | 0.01388804 | 3.32942263 | 0.031 | 0 | 1 | 0.27154471 | 0.69014474 | 0.013 | 0 | 1 | 0.27154471 | 0.02758321 |
| Gm57357 | 0.01388804 | 3.32942263 | 0.031 | 0 | 1 | 0.02605524 | -3.6317834 | 0 | 0.021 | 1 | 0.02605524 | 0.02758321 |
| Klhl10 | 0.01388804 | 3.32942263 | 0.031 | 0 | 1 | 0.5818708 | -0.5728897 | 0.013 | 0.021 | 1 | 0.5818708 | 0.02758321 |
| B3gnt5 | 0.01388804 | 3.32942263 | 0.031 | 0 | 1 | 0.34533788 | -2.3098553 | 0.008 | 0.021 | 1 | 0.34533788 | 0.02758321 |
| Gm28151 | 0.01388843 | 3.00749454 | 0.031 | 0 | 1 | 0.80442048 | -0.5728897 | 0.017 | 0.021 | 1 | 0.80442048 | 0.02758397 |
| Gm15402 | 0.01388843 | 3.59245704 | 0.031 | 0 | 1 | 0.51194506 | 0.27510724 | 0.021 | 0.01 | 1 | 0.51194506 | 0.02758397 |
| Tpbgl | 0.01388843 | 3.00749454 | 0.031 | 0 | 1 | 0.86312513 | -0.7248928 | 0.008 | 0.01 | 1 | 0.86312513 | 0.02758397 |
| Snx20 | 0.01388843 | 3.00749454 | 0.031 | 0 | 1 | 0.27154327 | 1.01207283 | 0.013 | 0 | 1 | 0.27154327 | 0.02758397 |
| Adgrg3 | 0.01388843 | 3.00749454 | 0.031 | 0 | 1 | 0.91723954 | -1.0468209 | 0.029 | 0.031 | 1 | 0.91723954 | 0.02758397 |
| Or8b54 | 0.01388843 | 3.00749454 | 0.031 | 0 | 1 | 0.11689951 | -3.3098553 | 0 | 0.01 | 1 | 0.11689951 | 0.02758397 |
| Gm48968 | 0.01388843 | 3.00749454 | 0.031 | 0 | 1 | 0.51188611 | -0.7248928 | 0.004 | 0.01 | 1 | 0.51188611 | 0.02758397 |
| 4930592A05Rik | 0.01388843 | 5.91438513 | 0.031 | 0 | 1 | 0.15394868 | 1.27510724 | 0.021 | 0 | 1 | 0.15394868 | 0.02758397 |
| Gm49521 | 0.01388843 | 3.59245704 | 0.031 | 0 | 1 | 0.24396338 | 0.42711033 | 0.034 | 0.01 | 1 | 0.24396338 | 0.02758397 |
| Ly6g6c | 0.01388843 | 3.00749454 | 0.031 | 0 | 1 | 0.15394868 | 1.27510724 | 0.021 | 0 | 1 | 0.15394868 | 0.02758397 |
| Gm50203 | 0.01388843 | 3.00749454 | 0.031 | 0 | 1 | 0.86312513 | -0.7248928 | 0.008 | 0.01 | 1 | 0.86312513 | 0.02758397 |
| Dmrt3 | 0.01388843 | 3.00749454 | 0.031 | 0 | 1 | 0.50887612 | -1.3098553 | 0.004 | 0.01 | 1 | 0.50887612 | 0.02758397 |
| Zcchc18 | 0.01388843 | 3.00749454 | 0.031 | 0 | 1 | 0.80020906 | -0.7248928 | 0.017 | 0.021 | 1 | 0.80020906 | 0.02758397 |
| Dipk1a | 0.60972566 | -0.5133376 | 0.156 | 0.176 | 1 | 0.01390075 | 0.97218011 | 0.273 | 0.146 | 1 | 0.60972566 | 0.02760828 |
| Pid1 | 0.01391128 | -1.6499376 | 0.417 | 0.528 | 1 | 0.95607542 | -0.3959442 | 0.349 | 0.312 | 1 | 0.95607542 | 0.02762904 |
| Ufd1 | 0.05651697 | 0.27052894 | 0.688 | 0.57 | 1 | 0.01395481 | 0.27986077 | 0.71 | 0.542 | 1 | 0.05651697 | 0.02771488 |
| Grik4 | 0.01396238 | 0.74446013 | 0.25 | 0.135 | 1 | 0.30144066 | -0.8860476 | 0.101 | 0.135 | 1 | 0.30144066 | 0.02772982 |
| Gtf2h4 | 0.0139777 | 1.00749454 | 0.229 | 0.114 | 1 | 0.13540124 | 0.7296731 | 0.13 | 0.073 | 1 | 0.13540124 | 0.02776002 |
| Ppip5k2 | 0.12405583 | 0.63064023 | 0.333 | 0.269 | 1 | 0.01399385 | 0.60121101 | 0.382 | 0.229 | 1 | 0.12405583 | 0.02779187 |
| Srsf2 | 0.0593898 | 0.16820545 | 0.979 | 0.943 | 1 | 0.01400376 | 0.23289909 | 0.992 | 0.969 | 1 | 0.0593898 | 0.02781141 |
| Ift172 | 0.35878376 | 0.27052894 | 0.24 | 0.192 | 1 | 0.01400831 | 0.29280924 | 0.261 | 0.125 | 1 | 0.35878376 | 0.02782038 |
| Rundc1 | 0.01401183 | 1.27052894 | 0.188 | 0.088 | 1 | 0.02855932 | 0.54258755 | 0.214 | 0.104 | 1 | 0.02855932 | 0.02782734 |
| Bcar1 | 0.01403375 | 0.61703506 | 0.49 | 0.337 | 1 | 0.1107826 | -0.1074659 | 0.525 | 0.354 | 1 | 0.1107826 | 0.02787054 |
| Kars | 0.01405337 | 0.61156586 | 0.344 | 0.207 | 1 | 0.10975519 | -0.1254307 | 0.349 | 0.229 | 1 | 0.10975519 | 0.02790925 |
| Spire1 | 0.01406668 | -0.4116126 | 0.948 | 0.917 | 1 | 0.95843763 | -0.2817175 | 0.912 | 0.865 | 1 | 0.95843763 | 0.02793549 |
| Mrpl37 | 0.64325568 | -0.1624305 | 0.208 | 0.233 | 1 | 0.01411243 | 0.57597372 | 0.265 | 0.135 | 1 | 0.64325568 | 0.0280257 |
| Sdha | 0.3333118 | -0.2193714 | 0.708 | 0.725 | 1 | 0.01415206 | 0.27837495 | 0.782 | 0.729 | 1 | 0.3333118 | 0.02810385 |
| Ddit4 | 0.1352439 | 0.13994483 | 0.25 | 0.166 | 1 | 0.01415222 | -1.5101539 | 0.235 | 0.365 | 1 | 0.1352439 | 0.02810416 |
| Polr2j | 0.25199197 | 0.37522632 | 0.396 | 0.342 | 1 | 0.01415769 | 0.76543286 | 0.34 | 0.208 | 1 | 0.25199197 | 0.02811495 |
| Zfp729a | 0.01416647 | 0.75373794 | 0.229 | 0.114 | 1 | 0.01843178 | 0.64434105 | 0.206 | 0.094 | 1 | 0.01843178 | 0.02813225 |
| Fam185a | 0.02147828 | 0.60413884 | 0.198 | 0.098 | 1 | 0.01418962 | 0.77760758 | 0.185 | 0.073 | 1 | 0.02147828 | 0.0281779 |
| Ubn1 | 0.04706699 | 0.19603522 | 0.917 | 0.865 | 1 | 0.01419919 | 0.22172614 | 0.945 | 0.917 | 1 | 0.04706699 | 0.02819676 |
| E330011O21Rik | 0.01420637 | -1.8338077 | 0.042 | 0.135 | 1 | 0.50587544 | -2.8948178 | 0.004 | 0.01 | 1 | 0.50587544 | 0.02821091 |
| Fdps | 0.08936132 | 0.68556644 | 0.594 | 0.539 | 1 | 0.01420734 | 1.16751312 | 0.555 | 0.5 | 1 | 0.08936132 | 0.02821284 |
| Tomm70a | 0.0142087 | 0.39663017 | 0.802 | 0.637 | 1 | 0.1011638 | 0.15458448 | 0.786 | 0.688 | 1 | 0.1011638 | 0.02821551 |
| Abhd2 | 0.01423154 | -0.4309241 | 0.948 | 0.933 | 1 | 0.02018566 | 0.12345796 | 0.979 | 0.927 | 1 | 0.02018566 | 0.02826055 |
| Mdc1 | 0.01423769 | 0.82463048 | 0.271 | 0.15 | 1 | 0.04360486 | 0.37464291 | 0.244 | 0.135 | 1 | 0.04360486 | 0.02827267 |
| Dnajc12 | 0.57843831 | 0.28502851 | 0.146 | 0.124 | 1 | 0.01423871 | 1.30485458 | 0.164 | 0.062 | 1 | 0.57843831 | 0.02827468 |
| Fabp7 | 0.01426106 | -1.8993961 | 0.042 | 0.135 | 1 | 0.0713678 | 1.64434105 | 0.071 | 0.021 | 1 | 0.0713678 | 0.02831875 |
| Ak1 | 0.02651525 | 0.89897008 | 0.323 | 0.218 | 1 | 0.01428365 | 0.40286279 | 0.366 | 0.208 | 1 | 0.02651525 | 0.02836328 |
| Slc35g1 | 0.32181805 | -0.3504619 | 0.448 | 0.503 | 1 | 0.01429138 | 1.56929034 | 0.223 | 0.115 | 1 | 0.32181805 | 0.02837852 |
| Mthfd1 | 0.01429175 | 0.96567436 | 0.25 | 0.135 | 1 | 0.03235426 | 0.6026819 | 0.21 | 0.104 | 1 | 0.03235426 | 0.02837925 |
| Nedd4l | 0.01429522 | -0.3434821 | 0.802 | 0.922 | 1 | 0.45132569 | -0.6951454 | 0.933 | 0.896 | 1 | 0.45132569 | 0.0283861 |
| Susd6 | 0.01432165 | 0.21959198 | 0.854 | 0.767 | 1 | 0.49852032 | -0.4753885 | 0.857 | 0.865 | 1 | 0.49852032 | 0.02843819 |
| Zfp317 | 0.01433632 | 0.69629285 | 0.365 | 0.228 | 1 | 0.04518577 | 0.33784299 | 0.307 | 0.198 | 1 | 0.04518577 | 0.02846711 |
| Irf2bpl | 0.45291385 | 0.13378373 | 0.635 | 0.601 | 1 | 0.01435119 | 0.42495545 | 0.79 | 0.74 | 1 | 0.45291385 | 0.02849643 |
| Lrch1 | 0.01435587 | -0.2138618 | 0.51 | 0.694 | 1 | 0.04530145 | -0.5793737 | 0.71 | 0.833 | 1 | 0.04530145 | 0.02850564 |
| Dok4 | 0.01437049 | -1.6114153 | 0.052 | 0.15 | 1 | 0.01552236 | 2.27510724 | 0.08 | 0.01 | 1 | 0.01552236 | 0.02853446 |
| Csnk1d | 0.01437695 | 0.46630444 | 0.698 | 0.56 | 1 | 0.05886455 | -0.484353 | 0.756 | 0.823 | 1 | 0.05886455 | 0.02854721 |
| Spr | 0.43115684 | -0.341655 | 0.344 | 0.363 | 1 | 0.01439862 | 0.70211738 | 0.357 | 0.219 | 1 | 0.43115684 | 0.02858993 |
| Zc3h7a | 0.01440156 | 0.2362734 | 0.927 | 0.886 | 1 | 0.46191631 | -0.137724 | 0.908 | 0.938 | 1 | 0.46191631 | 0.02859572 |
| Elk4 | 0.01442899 | 0.31071803 | 0.823 | 0.668 | 1 | 0.0361484 | 0.22138938 | 0.853 | 0.854 | 1 | 0.0361484 | 0.02864979 |
| Cep44 | 0.45371877 | 0.13302542 | 0.177 | 0.14 | 1 | 0.01443635 | 0.54258755 | 0.21 | 0.094 | 1 | 0.45371877 | 0.02866429 |
| Ccny | 0.01445535 | 0.15219955 | 0.885 | 0.803 | 1 | 0.39698807 | -0.1305316 | 0.937 | 0.906 | 1 | 0.39698807 | 0.02870173 |
| Hbp1 | 0.01448134 | -0.5334128 | 0.667 | 0.762 | 1 | 0.49809408 | -0.1275521 | 0.832 | 0.875 | 1 | 0.49809408 | 0.02875297 |
| Krcc1 | 0.03395633 | 0.41055939 | 0.75 | 0.622 | 1 | 0.01450696 | 0.40216397 | 0.672 | 0.573 | 1 | 0.03395633 | 0.02880347 |
| Chd3 | 0.88449792 | -0.2634668 | 0.635 | 0.606 | 1 | 0.01451942 | -0.5719501 | 0.836 | 0.906 | 1 | 0.88449792 | 0.02882803 |
| Rnf166 | 0.02967122 | 0.43595255 | 0.51 | 0.373 | 1 | 0.01452007 | 0.15963002 | 0.504 | 0.302 | 1 | 0.02967122 | 0.02882931 |
| Plgrkt | 0.2231084 | 0.37613913 | 0.375 | 0.306 | 1 | 0.01453504 | 0.61071027 | 0.349 | 0.208 | 1 | 0.2231084 | 0.02885881 |
| Slfn9 | 0.54692828 | 0.12613903 | 0.104 | 0.083 | 1 | 0.014554 | -1.5163061 | 0.029 | 0.094 | 1 | 0.54692828 | 0.02889619 |
| Kmt2c | 0.18227461 | 0.11366009 | 0.781 | 0.699 | 1 | 0.01456993 | -0.3684924 | 0.983 | 0.99 | 1 | 0.18227461 | 0.02892757 |
| Vps39 | 0.26825867 | 0.35541784 | 0.229 | 0.171 | 1 | 0.01458167 | 0.70900376 | 0.265 | 0.135 | 1 | 0.26825867 | 0.02895072 |
| Faap20 | 0.82381811 | 0.16361374 | 0.271 | 0.29 | 1 | 0.01461602 | 0.83989186 | 0.227 | 0.104 | 1 | 0.82381811 | 0.02901841 |
| Dctn1 | 0.10522203 | 0.20304535 | 0.552 | 0.425 | 1 | 0.01464588 | 0.31307509 | 0.597 | 0.458 | 1 | 0.10522203 | 0.02907726 |
| 8030462N17Rik | 0.10524373 | 0.54237181 | 0.51 | 0.425 | 1 | 0.01466243 | -0.7643074 | 0.655 | 0.729 | 1 | 0.10524373 | 0.02910988 |
| Rp2 | 0.01466838 | 0.83756954 | 0.25 | 0.13 | 1 | 0.70358747 | -0.1759995 | 0.227 | 0.198 | 1 | 0.70358747 | 0.0291216 |
| Xdh | 0.66582201 | -0.6705774 | 0.042 | 0.031 | 1 | 0.01468289 | -0.8244284 | 0.029 | 0.094 | 1 | 0.66582201 | 0.02915018 |
| Klhl12 | 0.97825927 | 0.32352203 | 0.312 | 0.321 | 1 | 0.01471244 | 0.65619741 | 0.391 | 0.24 | 1 | 0.97825927 | 0.02920842 |
| Ptcd3 | 0.01471747 | 0.42951847 | 0.469 | 0.311 | 1 | 0.9735801 | -0.3830217 | 0.429 | 0.385 | 1 | 0.9735801 | 0.02921833 |

| | | | | | | | | | | | | |
|---|---|---|---|---|---|---|---|---|---|---|---|---|
| Sema4b | 0.34206645 | 0.2590333 | 0.094 | 0.135 | 1 | 0.01473338 | -1.07539 | 0.055 | 0.135 | 1 | 0.34206645 | 0.02924969 |
| Aff4 | 0.13837288 | 0.15902762 | 0.948 | 0.974 | 1 | 0.01473658 | 0.16875241 | 1 | 1 | 1 | 0.13837288 | 0.02925599 |
| L2hgdh | 0.75570141 | -0.1941393 | 0.177 | 0.155 | 1 | 0.01476959 | 1.08246216 | 0.181 | 0.073 | 1 | 0.75570141 | 0.02932105 |
| Tdg | 0.02374175 | 0.38189005 | 0.51 | 0.363 | 1 | 0.01478294 | 0.33650778 | 0.563 | 0.365 | 1 | 0.02374175 | 0.02934735 |
| Lss | 0.46107159 | 0.14647095 | 0.302 | 0.249 | 1 | 0.01479144 | 1.13760371 | 0.164 | 0.062 | 1 | 0.46107159 | 0.02936409 |
| Gm17066 | 0.12701924 | 1.22988696 | 0.062 | 0.026 | 1 | 0.01481143 | 1.44503224 | 0.101 | 0.021 | 1 | 0.12701924 | 0.02940349 |
| 9330179D12Rik | 0.01481259 | 1.63552576 | 0.125 | 0.047 | 1 | 0.55912552 | -1.2167459 | 0.059 | 0.042 | 1 | 0.55912552 | 0.02940576 |
| 3930402G23Rik | 0.01481259 | 1.79599043 | 0.125 | 0.047 | 1 | 0.78571043 | -0.3098553 | 0.038 | 0.031 | 1 | 0.78571043 | 0.02940576 |
| Fmo5 | 0.37417731 | -0.3144336 | 0.062 | 0.093 | 1 | 0.0148308 | -2.6053111 | 0.038 | 0.104 | 1 | 0.37417731 | 0.02944164 |
| R74862 | 0.09026672 | 1.06638823 | 0.167 | 0.098 | 1 | 0.01483896 | 1.34222143 | 0.134 | 0.042 | 1 | 0.09026672 | 0.02945773 |
| Cntrob | 0.01484501 | 0.67045955 | 0.146 | 0.057 | 1 | 0.32198397 | 0.10518224 | 0.122 | 0.083 | 1 | 0.32198397 | 0.02946964 |
| Bche | 0.50546978 | -0.4286046 | 0.135 | 0.161 | 1 | 0.0148516 | 2.01207283 | 0.13 | 0.042 | 1 | 0.50546978 | 0.02948264 |
| Vps29 | 0.0148538 | 0.32942263 | 0.708 | 0.534 | 1 | 0.06913764 | 0.22978325 | 0.693 | 0.583 | 1 | 0.06913764 | 0.02948696 |
| Trmt61b | 0.37456597 | 0.51153704 | 0.229 | 0.187 | 1 | 0.01487264 | 0.67286675 | 0.239 | 0.115 | 1 | 0.37456597 | 0.02952409 |
| Lrpap1 | 0.12997549 | 0.13767553 | 0.917 | 0.813 | 1 | 0.01488031 | 0.14449516 | 0.882 | 0.656 | 1 | 0.12997549 | 0.02953919 |
| Glt8d2 | 0.02389736 | 0.70941318 | 0.396 | 0.269 | 1 | 0.01490068 | 1.14957636 | 0.227 | 0.115 | 1 | 0.02389736 | 0.02957934 |
| Pnpla8 | 0.0149311 | -0.3946039 | 0.542 | 0.699 | 1 | 0.68404148 | -0.1513712 | 0.803 | 0.771 | 1 | 0.68404148 | 0.02963926 |
| AW209491 | 0.01493536 | 0.80955516 | 0.292 | 0.166 | 1 | 0.55403982 | 0.10518224 | 0.185 | 0.156 | 1 | 0.55403982 | 0.02964766 |
| Nde1 | 0.06096091 | 0.57043073 | 0.427 | 0.342 | 1 | 0.01497343 | 0.37698685 | 0.416 | 0.24 | 1 | 0.06096091 | 0.02972265 |
| Zfp740 | 0.10309481 | 0.35263002 | 0.354 | 0.244 | 1 | 0.01499128 | 0.28118175 | 0.576 | 0.396 | 1 | 0.10309481 | 0.02975782 |
| Gtpbp10 | 0.01500322 | 1.14499806 | 0.188 | 0.088 | 1 | 0.04646643 | 0.8917786 | 0.168 | 0.083 | 1 | 0.04646643 | 0.02978135 |
| Rbbp6 | 0.04762299 | 0.37765882 | 0.896 | 0.907 | 1 | 0.01501614 | 0.21219443 | 0.983 | 0.948 | 1 | 0.04762299 | 0.0298068 |
| Cyp7b1 | 0.08587166 | 0.95091101 | 0.135 | 0.073 | 1 | 0.01501974 | 1.13760371 | 0.164 | 0.062 | 1 | 0.08587166 | 0.02981388 |
| Actr10 | 0.08806437 | 0.27052894 | 0.76 | 0.668 | 1 | 0.01501977 | 0.11945637 | 0.819 | 0.75 | 1 | 0.08806437 | 0.02981394 |
| Stx3 | 0.4526755 | 0.15601706 | 0.333 | 0.275 | 1 | 0.01502829 | 0.17040986 | 0.597 | 0.438 | 1 | 0.4526755 | 0.02983073 |
| Utp15 | 0.01694012 | 0.65519279 | 0.385 | 0.249 | 1 | 0.01505554 | 0.36152199 | 0.273 | 0.135 | 1 | 0.01694012 | 0.02988441 |
| Ncapd2 | 0.03377865 | 1.75751628 | 0.156 | 0.078 | 1 | 0.01507813 | 1.45567948 | 0.118 | 0.031 | 1 | 0.03377865 | 0.02992891 |
| Ftsj3 | 0.04297251 | 0.40845685 | 0.5 | 0.352 | 1 | 0.01511412 | 0.20584458 | 0.487 | 0.312 | 1 | 0.04297251 | 0.02999981 |
| Fh1 | 0.15735692 | 0.50160361 | 0.375 | 0.301 | 1 | 0.0151227 | 0.28918242 | 0.34 | 0.188 | 1 | 0.15735692 | 0.0300167 |
| Sfxn1 | 0.01518246 | 0.92503238 | 0.448 | 0.332 | 1 | 0.01745303 | 0.12038464 | 0.429 | 0.24 | 1 | 0.01745303 | 0.03013442 |
| Pum3 | 0.01520749 | 0.61942408 | 0.5 | 0.347 | 1 | 0.46806121 | -0.3534445 | 0.445 | 0.365 | 1 | 0.46806121 | 0.03018371 |
| Acp5 | 0.48383385 | 0.52206771 | 0.073 | 0.052 | 1 | 0.01522604 | 7.43699492 | 0.13 | 0.042 | 1 | 0.48383385 | 0.03022025 |
| Bphl | 0.01523382 | 0.70793425 | 0.385 | 0.244 | 1 | 0.53396535 | -0.4473588 | 0.349 | 0.292 | 1 | 0.53396535 | 0.03023557 |
| Rnf128 | 0.01530199 | -1.7825824 | 0.094 | 0.202 | 1 | 0.63064977 | -0.1578522 | 0.067 | 0.052 | 1 | 0.63064977 | 0.03036982 |
| Nicn1 | 0.40819994 | 0.11852585 | 0.24 | 0.192 | 1 | 0.01531978 | 0.61214223 | 0.235 | 0.115 | 1 | 0.40819994 | 0.03040487 |
| Tusc1 | 0.3337434 | 0.47697982 | 0.135 | 0.098 | 1 | 0.01533612 | 1.0961371 | 0.176 | 0.073 | 1 | 0.3337434 | 0.03043704 |
| Gcfc2 | 0.01534443 | 0.87625 | 0.177 | 0.078 | 1 | 0.53988016 | 0.35999614 | 0.092 | 0.073 | 1 | 0.53988016 | 0.0304534 |
| Hmgn3 | 0.01537864 | 0.46822413 | 0.688 | 0.57 | 1 | 0.09792745 | 0.16548275 | 0.681 | 0.677 | 1 | 0.09792745 | 0.03052079 |
| Ufsp2 | 0.01540151 | 0.77227807 | 0.531 | 0.409 | 1 | 0.02450759 | 0.52303475 | 0.42 | 0.271 | 1 | 0.02450759 | 0.03056581 |
| Frmd8 | 0.29290346 | -0.2527412 | 0.448 | 0.508 | 1 | 0.01540604 | 0.24383578 | 0.664 | 0.469 | 1 | 0.29290346 | 0.03057473 |
| Cib1 | 0.16172164 | -0.5418441 | 0.375 | 0.425 | 1 | 0.01540722 | -0.6291338 | 0.328 | 0.469 | 1 | 0.16172164 | 0.03057706 |
| Sbsn | 0.11691811 | 1.28760246 | 0.115 | 0.062 | 1 | 0.01542718 | 2.69014474 | 0.059 | 0 | 1 | 0.11691811 | 0.03061637 |
| 2310034G01Rik | 0.25792709 | 0.83756954 | 0.073 | 0.041 | 1 | 0.01542718 | 2.69014474 | 0.059 | 0 | 1 | 0.25792709 | 0.03061637 |
| Gm32282 | 0.12591542 | -0.6623569 | 0.094 | 0.161 | 1 | 0.01543255 | 2.77760758 | 0.059 | 0 | 1 | 0.12591542 | 0.03062693 |
| Mrps18a | 0.03249235 | 0.54092674 | 0.458 | 0.337 | 1 | 0.01543823 | 0.48600402 | 0.399 | 0.24 | 1 | 0.03249235 | 0.03063811 |
| Gm34312 | 0.21708019 | 2.00749454 | 0.021 | 0.005 | 1 | 0.01544086 | 2.93807225 | 0.059 | 0 | 1 | 0.21708019 | 0.0306433 |
| Lfng | 0.46907898 | 0.37006462 | 0.115 | 0.088 | 1 | 0.01544086 | 2.93807225 | 0.059 | 0 | 1 | 0.46907898 | 0.0306433 |
| Gm13199 | 0.38521705 | 0.68556644 | 0.031 | 0.016 | 1 | 0.01545221 | 2.59703533 | 0.059 | 0 | 1 | 0.38521705 | 0.03066564 |
| Ogfrl1 | 0.27059095 | 1.05188866 | 0.156 | 0.114 | 1 | 0.01547395 | 0.38404161 | 0.185 | 0.073 | 1 | 0.27059095 | 0.03070846 |
| Slc7a6os | 0.01548553 | 0.42253204 | 0.469 | 0.316 | 1 | 0.73556381 | -0.5499893 | 0.37 | 0.302 | 1 | 0.73556381 | 0.03073125 |
| Cenpb | 0.01550366 | 0.2049406 | 0.594 | 0.399 | 1 | 0.14654844 | 0.16819203 | 0.718 | 0.677 | 1 | 0.14654844 | 0.03076696 |
| Med17 | 0.32005455 | 0.17967551 | 0.292 | 0.233 | 1 | 0.01550856 | 0.39641353 | 0.366 | 0.208 | 1 | 0.32005455 | 0.03077661 |
| Coq8a | 0.46336809 | 0.71798792 | 0.073 | 0.052 | 1 | 0.01552217 | 2.33400093 | 0.08 | 0.01 | 1 | 0.46336809 | 0.0308034 |
| Rxfp3 | 0.993208 | 1.00749454 | 0.01 | 0.01 | 1 | 0.01552236 | 2.27510724 | 0.08 | 0.01 | 1 | 0.993208 | 0.03080379 |
| Stk40 | 0.01554591 | 0.30069091 | 0.771 | 0.648 | 1 | 0.04973415 | 0.22512739 | 0.815 | 0.781 | 1 | 0.04973415 | 0.03085014 |
| Tfap2b | 0.1352222 | -1.1912853 | 0.094 | 0.155 | 1 | 0.01557053 | 0.68412095 | 0.311 | 0.177 | 1 | 0.1352222 | 0.03089862 |
| Zfp94 | 0.02267558 | 1.78510212 | 0.094 | 0.031 | 1 | 0.01558078 | 2.21370669 | 0.08 | 0.01 | 1 | 0.02267558 | 0.0309188 |
| Smpd1 | 0.09782887 | -0.4224933 | 0.281 | 0.378 | 1 | 0.01559972 | 0.48600402 | 0.424 | 0.26 | 1 | 0.09782887 | 0.03095609 |
| Zdhhc15 | 0.01560669 | 0.78510212 | 0.135 | 0.052 | 1 | 0.61774687 | -0.473354 | 0.092 | 0.073 | 1 | 0.61774687 | 0.03096982 |
| Sec24a | 0.01563351 | 0.33142036 | 0.844 | 0.731 | 1 | 0.48678998 | -0.2803781 | 0.887 | 0.927 | 1 | 0.48678998 | 0.03102262 |
| Shld3 | 0.64082014 | -0.1851505 | 0.062 | 0.078 | 1 | 0.01563855 | 2.14957636 | 0.08 | 0.01 | 1 | 0.64082014 | 0.03103254 |
| Hmgb3 | 0.0156389 | 0.63552576 | 0.26 | 0.135 | 1 | 0.02002791 | 0.93115284 | 0.261 | 0.146 | 1 | 0.02002791 | 0.03103323 |
| Rad17 | 0.39127239 | 0.30705482 | 0.448 | 0.389 | 1 | 0.01566019 | 0.21739174 | 0.483 | 0.302 | 1 | 0.39127239 | 0.03107513 |
| Trpm6 | 0.44738557 | -0.8770282 | 0.146 | 0.181 | 1 | 0.01567715 | 1.19915838 | 0.134 | 0.042 | 1 | 0.44738557 | 0.03110853 |
| Mtarc2 | 0.81734894 | -0.1639859 | 0.604 | 0.617 | 1 | 0.01571868 | 0.4252108 | 0.634 | 0.542 | 1 | 0.81734894 | 0.03119027 |
| Cib2 | 0.12305387 | 0.95091101 | 0.177 | 0.114 | 1 | 0.015741 | 1.22065945 | 0.164 | 0.062 | 1 | 0.12305387 | 0.03123423 |
| Apoo | 0.51602073 | 0.10060394 | 0.156 | 0.124 | 1 | 0.01584377 | 0.96316323 | 0.206 | 0.094 | 1 | 0.51602073 | 0.03143651 |
| Zfp827 | 0.86202465 | -0.1053995 | 0.396 | 0.394 | 1 | 0.01584847 | -1.1326623 | 0.429 | 0.51 | 1 | 0.86202465 | 0.03144576 |
| Pacs1 | 0.15851372 | 0.48094872 | 0.354 | 0.269 | 1 | 0.01586621 | -0.9897662 | 0.45 | 0.583 | 1 | 0.15851372 | 0.03148068 |
| 0610030E20Rik | 0.63419343 | 0.18583178 | 0.302 | 0.275 | 1 | 0.01587336 | 0.28770727 | 0.345 | 0.188 | 1 | 0.63419343 | 0.03149475 |
| Mturn | 0.59742017 | 0.35541784 | 0.156 | 0.135 | 1 | 0.01591098 | 0.9730787 | 0.218 | 0.104 | 1 | 0.59742017 | 0.03156881 |
| Nr2c2 | 0.01593186 | 0.39542019 | 0.594 | 0.435 | 1 | 0.03098493 | -0.536689 | 0.714 | 0.802 | 1 | 0.03098493 | 0.03160989 |
| Snx19 | 0.22108551 | 0.1030722 | 0.333 | 0.254 | 1 | 0.01593744 | 0.34910782 | 0.387 | 0.229 | 1 | 0.22108551 | 0.03162088 |
| Afdn | 0.70219481 | 0.1910086 | 0.594 | 0.668 | 1 | 0.01594503 | -0.5836468 | 0.739 | 0.885 | 1 | 0.70219481 | 0.03163582 |
| Gm3336 | 0.10030276 | -1.1941393 | 0.042 | 0.098 | 1 | 0.01596249 | -2.4797803 | 0.038 | 0.104 | 1 | 0.10030276 | 0.03167017 |
| Klhl22 | 0.0159699 | 0.84084467 | 0.333 | 0.202 | 1 | 0.19589842 | -0.2241254 | 0.328 | 0.229 | 1 | 0.19589842 | 0.03168477 |
| Prr14l | 0.01597597 | 0.39384928 | 0.729 | 0.56 | 1 | 0.76433659 | -0.307146 | 0.803 | 0.792 | 1 | 0.76433659 | 0.03169671 |
| Sec14l1 | 0.01598438 | -0.510699 | 0.719 | 0.834 | 1 | 0.10730973 | -0.4522271 | 0.815 | 0.906 | 1 | 0.10730973 | 0.03171326 |
| C2cd2l | 0.39014731 | 0.46000674 | 0.115 | 0.083 | 1 | 0.01598862 | 1.57767001 | 0.13 | 0.042 | 1 | 0.39014731 | 0.03172161 |
| Gosr1 | 0.18397754 | 0.26499076 | 0.396 | 0.316 | 1 | 0.0159943 | 0.20062902 | 0.42 | 0.25 | 1 | 0.18397754 | 0.03173278 |
| Ift88 | 0.04606621 | 0.41055939 | 0.49 | 0.363 | 1 | 0.01599707 | 0.41949715 | 0.42 | 0.25 | 1 | 0.04606621 | 0.03173823 |
| Bmp8a | 0.01600831 | 1.9800138 | 0.26 | 0.155 | 1 | 0.21533569 | 0.69014474 | 0.122 | 0.073 | 1 | 0.21533569 | 0.03176035 |
| Gm47283 | 0.01602421 | 0.48006954 | 0.948 | 0.907 | 1 | 0.26524456 | -0.2204909 | 0.912 | 0.948 | 1 | 0.26524456 | 0.03179165 |
| Gga3 | 0.0160244 | 1.24195979 | 0.156 | 0.067 | 1 | 0.32576312 | 0.36821664 | 0.109 | 0.073 | 1 | 0.32576312 | 0.03179202 |
| Tcp11 | 0.01603232 | 1.37006462 | 0.125 | 0.047 | 1 | 0.54837489 | -0.1399303 | 0.034 | 0.021 | 1 | 0.54837489 | 0.03180761 |
| Cpsf4 | 0.1509794 | 0.26026061 | 0.24 | 0.161 | 1 | 0.01604207 | 0.7726069 | 0.235 | 0.115 | 1 | 0.1509794 | 0.03182678 |
| Ago2 | 0.0160431 | 0.16891521 | 0.844 | 0.71 | 1 | 0.15583809 | -0.3301467 | 0.929 | 0.979 | 1 | 0.15583809 | 0.03182883 |

| | | | | | | | | | | | | |
|---|---|---|---|---|---|---|---|---|---|---|---|---|
| Cct7 | 0.07144376 | 0.31644478 | 0.771 | 0.725 | 1 | 0.01605283 | 0.27510724 | 0.832 | 0.76 | 1 | 0.07144376 | 0.03184796 |
| Hjurp | 0.44949595 | 0.28105961 | 0.281 | 0.358 | 1 | 0.01606547 | 0.42711033 | 0.345 | 0.208 | 1 | 0.44949595 | 0.03187283 |
| Nat10 | 0.01611719 | 0.75373794 | 0.333 | 0.202 | 1 | 0.73764671 | -0.4268947 | 0.37 | 0.312 | 1 | 0.73764671 | 0.03197462 |
| E130308A19Rik | 0.01612849 | -1.2962862 | 0.104 | 0.218 | 1 | 0.89469986 | -0.8487213 | 0.261 | 0.25 | 1 | 0.89469986 | 0.03199686 |
| Sap18b | 0.0280587 | 0.8593959 | 0.302 | 0.192 | 1 | 0.016131 | 0.81352715 | 0.223 | 0.104 | 1 | 0.0280587 | 0.0320018 |
| 2310014F06Rik | 0.4362101 | 0.36113149 | 0.156 | 0.124 | 1 | 0.01613326 | 0.90515763 | 0.193 | 0.083 | 1 | 0.4362101 | 0.03200624 |
| Per3 | 0.19961534 | 0.23544648 | 0.458 | 0.363 | 1 | 0.01614695 | 0.23688792 | 0.399 | 0.229 | 1 | 0.19961534 | 0.03203318 |
| Cdt1 | 0.01615088 | 0.96442581 | 0.219 | 0.114 | 1 | 0.0658382 | 0.24268576 | 0.139 | 0.062 | 1 | 0.0658382 | 0.0320409 |
| Scaf8 | 0.01619046 | 0.27124867 | 0.948 | 0.917 | 1 | 0.04188571 | -0.3224187 | 0.987 | 0.979 | 1 | 0.04188571 | 0.0321188 |
| Tcf7 | 0.35693072 | 0.15949763 | 0.156 | 0.114 | 1 | 0.01621163 | 0.62560449 | 0.315 | 0.177 | 1 | 0.35693072 | 0.03216045 |
| Cfap68 | 0.6472242 | 0.3529911 | 0.385 | 0.378 | 1 | 0.01624395 | 0.73850776 | 0.286 | 0.156 | 1 | 0.6472242 | 0.03222403 |
| Helb | 0.01626157 | 1.09769235 | 0.219 | 0.114 | 1 | 0.9594504 | -0.6545034 | 0.197 | 0.188 | 1 | 0.9594504 | 0.0322587 |
| Agpat2 | 0.0392924 | 0.47911556 | 0.385 | 0.269 | 1 | 0.01628734 | 0.4725533 | 0.265 | 0.135 | 1 | 0.0392924 | 0.03230941 |
| Ephx1 | 0.01628762 | -0.3183275 | 0.406 | 0.622 | 1 | 0.96229638 | 0.30129117 | 0.487 | 0.552 | 1 | 0.96229638 | 0.03230995 |
| E230029C05Rik | 0.99534689 | -0.2148979 | 0.052 | 0.052 | 1 | 0.0163002 | -1.9472852 | 0.025 | 0.083 | 1 | 0.99534689 | 0.03233471 |
| 1700029J07Rik | 0.01630432 | 0.84702986 | 0.271 | 0.15 | 1 | 0.14598024 | 0.56461386 | 0.118 | 0.062 | 1 | 0.14598024 | 0.03234281 |
| Mcoln2 | 0.46066594 | -0.4961679 | 0.646 | 0.534 | 1 | 0.01632179 | -0.622531 | 0.479 | 0.26 | 1 | 0.46066594 | 0.03237718 |
| ENSMUSG000001214 85 | 0.27530647 | -0.770113 | 0.052 | 0.088 | 1 | 0.01633897 | 2.16407593 | 0.147 | 0.052 | 1 | 0.27530647 | 0.03241098 |
| Uros | 0.39194494 | -0.4619907 | 0.125 | 0.161 | 1 | 0.01634237 | 1.37464291 | 0.16 | 0.062 | 1 | 0.39194494 | 0.03241768 |
| Enpp4 | 0.51265384 | -0.2820121 | 0.156 | 0.187 | 1 | 0.01636528 | 1.01207283 | 0.147 | 0.052 | 1 | 0.51265384 | 0.03246273 |
| Stard7 | 0.051298 | 0.56820949 | 0.365 | 0.259 | 1 | 0.01638246 | 0.25001626 | 0.441 | 0.271 | 1 | 0.051298 | 0.03249654 |
| C1qtnf2 | 0.1126127 | 1.2091284 | 0.104 | 0.052 | 1 | 0.01640404 | 1.01207283 | 0.176 | 0.073 | 1 | 0.1126127 | 0.03253898 |
| Ift46 | 0.01644191 | 0.54950489 | 0.562 | 0.415 | 1 | 0.0273374 | 0.25341217 | 0.542 | 0.385 | 1 | 0.0273374 | 0.03261348 |
| Tfb2m | 0.1687413 | 0.34309757 | 0.417 | 0.326 | 1 | 0.01647132 | 0.36257008 | 0.416 | 0.25 | 1 | 0.1687413 | 0.03267133 |
| Gm45025 | 0.11449695 | 0.81484946 | 0.156 | 0.093 | 1 | 0.01650708 | 0.87234807 | 0.193 | 0.083 | 1 | 0.11449695 | 0.03274168 |
| Midn | 0.41994903 | 0.1533454 | 0.719 | 0.699 | 1 | 0.01650761 | 0.47214409 | 0.807 | 0.792 | 1 | 0.41994903 | 0.03274271 |
| Zfp212 | 0.47455403 | -0.265524 | 0.188 | 0.223 | 1 | 0.01651148 | 0.2488757 | 0.324 | 0.167 | 1 | 0.47455403 | 0.03275033 |
| Snrnp25 | 0.75486869 | 0.1843723 | 0.208 | 0.192 | 1 | 0.01654906 | 1.01207283 | 0.164 | 0.062 | 1 | 0.75486869 | 0.03282425 |
| Fat4 | 0.01662371 | 0.29329792 | 0.635 | 0.461 | 1 | 0.24460338 | -0.5594113 | 0.643 | 0.458 | 1 | 0.24460338 | 0.03297108 |
| Nfrkb | 0.48235099 | 0.24015529 | 0.323 | 0.275 | 1 | 0.01662391 | 0.19710473 | 0.433 | 0.25 | 1 | 0.48235099 | 0.03297146 |
| Trappc9 | 0.0701198 | -0.3899846 | 0.458 | 0.549 | 1 | 0.01663134 | -0.9537115 | 0.567 | 0.667 | 1 | 0.0701198 | 0.03298608 |
| Klk8 | 0.01663988 | -3.2213242 | 0.031 | 0.114 | 1 | 0.16771559 | 1.23446525 | 0.071 | 0.031 | 1 | 0.16771559 | 0.03300287 |
| D630003M21Rik | 0.33833248 | 0.32942263 | 0.177 | 0.14 | 1 | 0.01664106 | 0.9602339 | 0.231 | 0.115 | 1 | 0.33833248 | 0.0330052 |
| Fyn | 0.01664758 | -0.9293116 | 0.49 | 0.591 | 1 | 0.12684508 | -0.774818 | 0.5 | 0.542 | 1 | 0.12684508 | 0.03301801 |
| Pip4p1 | 0.05330619 | -0.6445822 | 0.479 | 0.549 | 1 | 0.01665832 | 0.11756596 | 0.437 | 0.26 | 1 | 0.05330619 | 0.03303915 |
| Bms1 | 0.01668311 | 0.39384928 | 0.562 | 0.399 | 1 | 0.02889689 | 0.35409553 | 0.563 | 0.417 | 1 | 0.02889689 | 0.03308789 |
| Crlf2 | 0.51534464 | -0.3208313 | 0.365 | 0.389 | 1 | 0.01675818 | 0.58322953 | 0.336 | 0.198 | 1 | 0.51534464 | 0.03323552 |
| Gps2 | 0.01677787 | 0.35541784 | 0.542 | 0.368 | 1 | 0.20189237 | 0.2422426 | 0.555 | 0.51 | 1 | 0.20189237 | 0.03327424 |
| Rhbdf1 | 0.35687496 | 0.18033113 | 0.271 | 0.218 | 1 | 0.01678001 | 0.37698685 | 0.395 | 0.229 | 1 | 0.35687496 | 0.03327845 |
| Fam161a | 0.04549166 | 0.64492446 | 0.135 | 0.062 | 1 | 0.01679903 | -1.4029647 | 0.088 | 0.177 | 1 | 0.04549166 | 0.03331586 |
| Arc | 0.26367817 | -1.3848229 | 0.031 | 0.062 | 1 | 0.01680074 | -1.2223924 | 0.025 | 0.083 | 1 | 0.26367817 | 0.03331922 |
| Rinl | 0.11858078 | -1.7294711 | 0.021 | 0.062 | 1 | 0.01680787 | 1.69014474 | 0.08 | 0.01 | 1 | 0.11858078 | 0.03333323 |
| Plec | 0.01681384 | 0.43152082 | 0.938 | 0.943 | 1 | 0.06777361 | 0.32279333 | 0.975 | 0.969 | 1 | 0.06777361 | 0.03334497 |
| Pkd1 | 0.59157279 | -0.1790817 | 0.688 | 0.725 | 1 | 0.01682139 | 0.19829164 | 0.803 | 0.552 | 1 | 0.59157279 | 0.03335982 |
| Ism1 | 0.01683327 | -0.3566463 | 0.479 | 0.326 | 1 | 0.29823021 | 0.52711132 | 0.357 | 0.292 | 1 | 0.29823021 | 0.03338318 |
| Lsm11 | 0.25234841 | 1.20013961 | 0.052 | 0.026 | 1 | 0.01683887 | 2.01207283 | 0.097 | 0.021 | 1 | 0.25234841 | 0.03339418 |
| Net1 | 0.01684187 | 0.64994253 | 0.427 | 0.295 | 1 | 0.04254125 | -0.4052748 | 0.45 | 0.594 | 1 | 0.04254125 | 0.03340009 |
| Smc6 | 0.0376225 | 0.46194672 | 0.771 | 0.658 | 1 | 0.01685441 | 0.26232863 | 0.79 | 0.729 | 1 | 0.0376225 | 0.03342475 |
| L3mbtl1 | 0.78904319 | 0.22988696 | 0.073 | 0.083 | 1 | 0.01685446 | -1.5790419 | 0.097 | 0.188 | 1 | 0.78904319 | 0.03342484 |
| Ift43 | 0.01687847 | 0.47588146 | 0.625 | 0.466 | 1 | 0.21731059 | -0.2103196 | 0.592 | 0.438 | 1 | 0.21731059 | 0.03347205 |
| Scaf11 | 0.01689108 | 0.27945417 | 0.958 | 0.943 | 1 | 0.07411911 | 0.14491409 | 0.983 | 0.979 | 1 | 0.07411911 | 0.03349686 |
| Cse1l | 0.01691232 | 0.57453513 | 0.5 | 0.342 | 1 | 0.89752709 | -0.3533569 | 0.525 | 0.479 | 1 | 0.89752709 | 0.03353861 |
| Dennd3 | 0.31209779 | 0.31117093 | 0.438 | 0.404 | 1 | 0.01702344 | 0.31399625 | 0.416 | 0.26 | 1 | 0.31209779 | 0.03375708 |
| Cklf | 0.01703268 | 1.09238343 | 0.229 | 0.124 | 1 | 0.02721916 | 0.62303054 | 0.143 | 0.052 | 1 | 0.02721916 | 0.03377526 |
| Gm5544 | 0.0170373 | 1.16361374 | 0.208 | 0.109 | 1 | 0.59430737 | -0.8287286 | 0.164 | 0.177 | 1 | 0.59430737 | 0.03378432 |
| Ddx51 | 0.01704891 | 1.22988696 | 0.125 | 0.047 | 1 | 0.72941731 | 0.16407593 | 0.084 | 0.073 | 1 | 0.72941731 | 0.03380715 |
| Ilrun | 0.01708167 | 0.29229415 | 0.812 | 0.725 | 1 | 0.95247904 | -0.1237288 | 0.887 | 0.906 | 1 | 0.95247904 | 0.03387155 |
| Tmem14c | 0.32744094 | 0.15559318 | 0.802 | 0.736 | 1 | 0.01710926 | 0.28904643 | 0.714 | 0.604 | 1 | 0.32744094 | 0.0339258 |
| Cdc42se2 | 0.18858048 | -0.4994655 | 0.448 | 0.508 | 1 | 0.01713362 | -0.8935068 | 0.664 | 0.74 | 1 | 0.18858048 | 0.03397367 |
| Supt4a | 0.01718296 | -0.5623611 | 0.438 | 0.575 | 1 | 0.46075983 | -0.1864728 | 0.508 | 0.573 | 1 | 0.46075983 | 0.03407067 |
| Fpgt | 0.07008857 | 0.9200317 | 0.146 | 0.078 | 1 | 0.01719081 | 1.80562195 | 0.097 | 0.021 | 1 | 0.07008857 | 0.0340861 |
| Ahcy | 0.44824387 | -0.5160674 | 0.062 | 0.088 | 1 | 0.01719081 | 1.80562195 | 0.097 | 0.021 | 1 | 0.44824387 | 0.0340861 |
| Fbxw7 | 0.23616576 | -0.4427577 | 0.469 | 0.549 | 1 | 0.01719568 | -0.6308493 | 0.588 | 0.75 | 1 | 0.23616576 | 0.03409566 |
| Gm57198 | 0.01720049 | 0.63826073 | 0.219 | 0.109 | 1 | 0.33712554 | -0.364303 | 0.189 | 0.135 | 1 | 0.33712554 | 0.03410512 |
| Spast | 0.02703062 | 0.27052894 | 0.469 | 0.321 | 1 | 0.01722098 | -0.7278102 | 0.576 | 0.698 | 1 | 0.02703062 | 0.0341454 |
| Shoc2 | 0.01724197 | 0.24964158 | 0.917 | 0.808 | 1 | 0.12561027 | -0.5517781 | 0.941 | 0.927 | 1 | 0.12561027 | 0.03418665 |
| Fst | 0.01727558 | -2.2295447 | 0.042 | 0.13 | 1 | 0.03811958 | -5.2917079 | 0.008 | 0.042 | 1 | 0.03811958 | 0.03425271 |
| Nox4 | 0.01729033 | -3.6929452 | 0.031 | 0.114 | 1 | 0.02605524 | -6.5193086 | 0 | 0.021 | 1 | 0.02605524 | 0.03428171 |
| Efnb3 | 0.65896352 | 0.15949763 | 0.042 | 0.031 | 1 | 0.01734899 | -1.8504236 | 0.038 | 0.104 | 1 | 0.65896352 | 0.03439699 |
| Srek1ip1 | 0.23306152 | 0.31359766 | 0.26 | 0.197 | 1 | 0.01735927 | 0.87777174 | 0.218 | 0.104 | 1 | 0.23306152 | 0.0344172 |
| Zfp975 | 0.13663142 | 0.26125113 | 0.281 | 0.192 | 1 | 0.01736786 | 0.60575255 | 0.239 | 0.115 | 1 | 0.13663142 | 0.03443408 |
| Prps1l3 | 0.21807932 | 0.48666237 | 0.208 | 0.15 | 1 | 0.01737706 | 1.06360313 | 0.202 | 0.094 | 1 | 0.21807932 | 0.03445216 |
| Taf1a | 0.01738728 | 0.82193888 | 0.344 | 0.212 | 1 | 0.43771667 | -0.662157 | 0.311 | 0.229 | 1 | 0.43771667 | 0.03447225 |
| Gm39302 | 0.01739269 | -2.7998604 | 0 | 0.057 | 1 | 0.1465102 | -1.8948178 | 0.004 | 0.021 | 1 | 0.1465102 | 0.03448287 |
| Tmem44 | 0.01739269 | -2.7998604 | 0 | 0.057 | 1 | 0.93071546 | -0.3098553 | 0.029 | 0.031 | 1 | 0.93071546 | 0.03448287 |
| Gm15246 | 0.01739269 | -2.7998604 | 0 | 0.057 | 1 | 0.09942934 | -1.0468209 | 0.013 | 0.042 | 1 | 0.09942934 | 0.03448287 |
| Pax3 | 0.01740538 | -4.2779077 | 0 | 0.057 | 1 | 0.20778448 | -1.5298209 | 0.105 | 0.146 | 1 | 0.20778448 | 0.03450782 |
| Col6a3 | 0.01742737 | -0.3433927 | 0.958 | 0.948 | 1 | 0.21720258 | -0.3980629 | 0.895 | 0.875 | 1 | 0.21720258 | 0.03455103 |
| Tyw5 | 0.01743347 | 0.62047141 | 0.229 | 0.114 | 1 | 0.87761612 | -0.1399303 | 0.172 | 0.177 | 1 | 0.87761612 | 0.03456302 |
| Syne2 | 0.70584763 | -0.2382619 | 0.292 | 0.326 | 1 | 0.017442 | -0.4353861 | 0.538 | 0.719 | 1 | 0.70584763 | 0.03457978 |
| N4bp1 | 0.01744962 | 0.31150072 | 0.552 | 0.394 | 1 | 0.93096429 | -0.1857842 | 0.723 | 0.76 | 1 | 0.93096429 | 0.03459474 |
| Pycard | 0.5011795 | -0.3364599 | 0.094 | 0.119 | 1 | 0.01745599 | -0.9986536 | 0.231 | 0.354 | 1 | 0.5011795 | 0.03460727 |
| Hsdl1 | 0.74682952 | -0.240433 | 0.062 | 0.073 | 1 | 0.01746522 | 1.35310975 | 0.13 | 0.042 | 1 | 0.74682952 | 0.0346254 |
| Dvl3 | 0.14522853 | 0.50999488 | 0.333 | 0.259 | 1 | 0.01747582 | 0.38130664 | 0.42 | 0.271 | 1 | 0.14522853 | 0.03464624 |
| Zfp385c | 0.55609543 | 0.31717904 | 0.24 | 0.212 | 1 | 0.01749192 | 0.17952958 | 0.332 | 0.177 | 1 | 0.55609543 | 0.03467788 |

| | | | | | | | | | | | | |
|---|---|---|---|---|---|---|---|---|---|---|---|---|
| Elovl7 | 0.01749204 | -1.0299802 | 0.156 | 0.285 | 1 | 0.63676884 | -1.7773602 | 0.147 | 0.156 | 1 | 0.63676884 | 0.03467812 |
| Bclaf1 | 0.01750977 | 0.26638751 | 0.958 | 0.927 | 1 | 0.89794588 | -0.1041141 | 0.975 | 1 | 1 | 0.89794588 | 0.03471295 |
| Hspa14 | 0.31879956 | 0.40525404 | 0.49 | 0.44 | 1 | 0.01754277 | 0.26545707 | 0.433 | 0.26 | 1 | 0.31879956 | 0.03477779 |
| Gm48882 | 0.01754332 | -1.5160674 | 0.062 | 0.161 | 1 | 0.99798677 | -0.3098553 | 0.021 | 0.021 | 1 | 0.99798677 | 0.03477887 |
| Hacd4 | 0.34240257 | -0.3461424 | 0.188 | 0.244 | 1 | 0.01754404 | 0.19793938 | 0.261 | 0.125 | 1 | 0.34240257 | 0.03478029 |
| Wtap | 0.01754436 | 0.49888292 | 0.552 | 0.399 | 1 | 0.07683773 | 0.19185173 | 0.66 | 0.542 | 1 | 0.07683773 | 0.03478092 |
| Hspa13 | 0.1129186 | -0.3680146 | 0.448 | 0.549 | 1 | 0.01754913 | 0.19513893 | 0.525 | 0.323 | 1 | 0.1129186 | 0.03479029 |
| Trim56 | 0.41607091 | -0.2477625 | 0.208 | 0.254 | 1 | 0.01755095 | 0.27510724 | 0.387 | 0.219 | 1 | 0.41607091 | 0.03479386 |
| Toe1 | 0.05486331 | 0.77922555 | 0.271 | 0.176 | 1 | 0.01763532 | 0.76414532 | 0.269 | 0.146 | 1 | 0.05486331 | 0.03495964 |
| Krba1 | 0.01763828 | 1.00749454 | 0.156 | 0.067 | 1 | 0.10787394 | 0.15963002 | 0.126 | 0.062 | 1 | 0.10787394 | 0.03496544 |
| Bambi | 0.73172197 | 0.15715939 | 0.656 | 0.668 | 1 | 0.01765307 | -0.2076318 | 0.546 | 0.354 | 1 | 0.73172197 | 0.03499451 |
| Ppm1d | 0.01770952 | 0.74097956 | 0.396 | 0.259 | 1 | 0.84955902 | -0.1920188 | 0.496 | 0.49 | 1 | 0.84955902 | 0.03510541 |
| Kdm5a | 0.01772004 | 0.35656877 | 0.896 | 0.902 | 1 | 0.47487602 | -0.293099 | 0.954 | 0.938 | 1 | 0.47487602 | 0.03512609 |
| Gm43813 | 0.4888993 | 0.34006988 | 0.115 | 0.088 | 1 | 0.0177555 | -1.9343461 | 0.084 | 0.167 | 1 | 0.4888993 | 0.03519575 |
| Rnf144b | 0.02059887 | 0.75751628 | 0.25 | 0.135 | 1 | 0.01776526 | 0.13071733 | 0.244 | 0.115 | 1 | 0.02059887 | 0.03521492 |
| Gm16599 | 0.09302126 | -1.2820121 | 0.083 | 0.15 | 1 | 0.01779751 | 0.27510724 | 0.193 | 0.083 | 1 | 0.09302126 | 0.03527827 |
| Rab23 | 0.14457585 | 0.78008404 | 0.25 | 0.176 | 1 | 0.01780446 | 1.01207283 | 0.227 | 0.115 | 1 | 0.14457585 | 0.03529192 |
| Tmem205 | 0.01780722 | 0.4955955 | 0.427 | 0.28 | 1 | 0.06066284 | -0.3402289 | 0.277 | 0.406 | 1 | 0.06066284 | 0.03529734 |
| Zmat3 | 0.42734286 | 0.48902745 | 0.25 | 0.218 | 1 | 0.01781764 | 0.65977109 | 0.269 | 0.146 | 1 | 0.42734286 | 0.03531781 |
| Cnot6l | 0.76914015 | -0.2019588 | 0.656 | 0.596 | 1 | 0.01782962 | -0.458762 | 0.811 | 0.938 | 1 | 0.76914015 | 0.03534134 |
| Dnaja3 | 0.31857045 | -0.1941393 | 0.417 | 0.326 | 1 | 0.0178533 | -0.1979624 | 0.513 | 0.302 | 1 | 0.31857045 | 0.03538785 |
| Tafa2 | 0.01788279 | -1.6595037 | 0.156 | 0.269 | 1 | 0.76847012 | -1.1320681 | 0.05 | 0.042 | 1 | 0.76847012 | 0.03544579 |
| Phactr1 | 0.01788927 | -1.1329867 | 0.115 | 0.233 | 1 | 0.95094503 | -1.1481042 | 0.273 | 0.24 | 1 | 0.95094503 | 0.03545851 |
| Scyl1 | 0.41257117 | 0.33465928 | 0.458 | 0.43 | 1 | 0.01793445 | 0.26256198 | 0.592 | 0.406 | 1 | 0.41257117 | 0.03554725 |
| Tango2 | 0.26365599 | -0.8855903 | 0.125 | 0.171 | 1 | 0.01794788 | 1.3146356 | 0.172 | 0.073 | 1 | 0.26365599 | 0.03557364 |
| Preb | 0.27183369 | 0.27459861 | 0.458 | 0.389 | 1 | 0.01795083 | 0.38831785 | 0.45 | 0.281 | 1 | 0.27183369 | 0.03557942 |
| Mnd1 | 0.29904364 | 0.78510212 | 0.083 | 0.052 | 1 | 0.0179683 | 1.76053407 | 0.097 | 0.021 | 1 | 0.29904364 | 0.03561373 |
| Gm45051 | 0.63776216 | 0.10703021 | 0.146 | 0.124 | 1 | 0.01797137 | 1.18199783 | 0.16 | 0.062 | 1 | 0.63776216 | 0.03561977 |
| Rgs7bp | 0.8151386 | -2.1624305 | 0.052 | 0.057 | 1 | 0.01797992 | -1.3098553 | 0.038 | 0.104 | 1 | 0.8151386 | 0.03563657 |
| Ccnq | 0.51548688 | 0.16503581 | 0.24 | 0.202 | 1 | 0.01798001 | 1.05937855 | 0.214 | 0.104 | 1 | 0.51548688 | 0.03563674 |
| 3010001F23Rik | 0.4204199 | 0.27052894 | 0.083 | 0.057 | 1 | 0.01798372 | 0.69014474 | 0.147 | 0.052 | 1 | 0.4204199 | 0.03564403 |
| Fam76a | 0.02140079 | 0.22261896 | 0.823 | 0.705 | 1 | 0.01799059 | 0.29208032 | 0.824 | 0.688 | 1 | 0.02140079 | 0.03565752 |
| Lrrc42 | 0.06131604 | 0.63976275 | 0.26 | 0.166 | 1 | 0.01800198 | 0.38046024 | 0.382 | 0.219 | 1 | 0.06131604 | 0.0356799 |
| Ttll4 | 0.05227563 | 0.34629645 | 0.302 | 0.187 | 1 | 0.01802951 | -0.1417325 | 0.5 | 0.271 | 1 | 0.05227563 | 0.03573396 |
| Lpar6 | 0.01805285 | -2.6929452 | 0.01 | 0.078 | 1 | 0.30613085 | -1.3098553 | 0.029 | 0.052 | 1 | 0.30613085 | 0.03577979 |
| Tbl3 | 0.48743222 | 0.34452952 | 0.198 | 0.166 | 1 | 0.01812246 | 0.95317914 | 0.239 | 0.125 | 1 | 0.48743222 | 0.0359165 |
| Nmnat3 | 0.01814967 | -2.0431315 | 0.042 | 0.13 | 1 | 0.06548174 | -0.4200382 | 0.155 | 0.073 | 1 | 0.06548174 | 0.03596993 |
| Ppp4r1 | 0.14797755 | 0.40859284 | 0.531 | 0.43 | 1 | 0.01817963 | -0.6428137 | 0.756 | 0.792 | 1 | 0.14797755 | 0.03602876 |
| Ranbp3 | 0.20885299 | 0.42253204 | 0.406 | 0.332 | 1 | 0.01820283 | 0.32435076 | 0.454 | 0.281 | 1 | 0.20885299 | 0.03607431 |
| Camk1d | 0.0182171 | -0.1817517 | 1 | 1 | 1 | 0.90926268 | -0.6468902 | 1 | 1 | 1 | 0.90926268 | 0.03610234 |
| Traf4 | 0.33012191 | -0.577468 | 0.198 | 0.238 | 1 | 0.01829462 | 0.65697787 | 0.34 | 0.208 | 1 | 0.33012191 | 0.03625455 |
| St3gal1 | 0.01831558 | 0.4150375 | 0.458 | 0.316 | 1 | 0.49454822 | -0.7391066 | 0.244 | 0.26 | 1 | 0.49454822 | 0.03629569 |
| Fam53c | 0.01835939 | 0.79450081 | 0.344 | 0.218 | 1 | 0.03728398 | 0.27510724 | 0.378 | 0.229 | 1 | 0.03728398 | 0.03638172 |
| Eif3j2 | 0.62481887 | 0.17099327 | 0.135 | 0.114 | 1 | 0.01837685 | 1.01207283 | 0.13 | 0.042 | 1 | 0.62481887 | 0.03641599 |
| Cdc42ep1 | 0.01837892 | 0.80586068 | 0.188 | 0.088 | 1 | 0.0764287 | 0.18199783 | 0.168 | 0.083 | 1 | 0.0764287 | 0.03642005 |
| Akr7a5 | 0.25991665 | 0.32523483 | 0.365 | 0.29 | 1 | 0.01838882 | 0.39468885 | 0.315 | 0.177 | 1 | 0.25991665 | 0.03643948 |
| Zfp521 | 0.76549416 | -0.3751942 | 0.917 | 0.793 | 1 | 0.0183993 | -0.6888913 | 0.878 | 0.812 | 1 | 0.76549416 | 0.03646007 |
| Paf1 | 0.37655416 | -0.1365518 | 0.542 | 0.58 | 1 | 0.01840314 | 0.34099517 | 0.693 | 0.542 | 1 | 0.37655416 | 0.0364676 |
| Zfp292 | 0.01840718 | 0.31447229 | 0.542 | 0.378 | 1 | 0.04827489 | -0.2989669 | 0.63 | 0.792 | 1 | 0.04827489 | 0.03647554 |
| Tcap | 0.01840819 | -2.5160674 | 0.01 | 0.078 | 1 | 0.27154327 | 1.01207283 | 0.013 | 0 | 1 | 0.27154327 | 0.03647753 |
| Taldo1 | 0.22633666 | -0.2284346 | 0.802 | 0.839 | 1 | 0.01841198 | 0.47340398 | 0.798 | 0.75 | 1 | 0.22633666 | 0.03648495 |
| Mosmo | 0.39083811 | -0.2102062 | 0.729 | 0.715 | 1 | 0.01843254 | 0.18408306 | 0.836 | 0.625 | 1 | 0.39083811 | 0.03652532 |
| Hk1 | 0.98418558 | -0.1668036 | 0.823 | 0.756 | 1 | 0.01844399 | 0.13214928 | 0.828 | 0.75 | 1 | 0.98418558 | 0.03654779 |
| Whrn | 0.01845571 | 0.45887388 | 0.427 | 0.275 | 1 | 0.61091437 | -0.1585459 | 0.466 | 0.396 | 1 | 0.61091437 | 0.0365708 |
| Desi1 | 0.88545613 | 0.10417056 | 0.573 | 0.57 | 1 | 0.01847891 | 0.154813 | 0.655 | 0.438 | 1 | 0.88545613 | 0.03661635 |
| Slc6a9 | 0.01848175 | 1.28760246 | 0.115 | 0.041 | 1 | 0.03454856 | 1.74903843 | 0.084 | 0.021 | 1 | 0.03454856 | 0.03662192 |
| Vhl | 0.97395715 | 0.17099327 | 0.208 | 0.212 | 1 | 0.01848782 | 0.43638815 | 0.332 | 0.198 | 1 | 0.97395715 | 0.03663385 |
| Trim32 | 0.73779294 | -0.3550755 | 0.062 | 0.073 | 1 | 0.01851564 | 0.82764826 | 0.239 | 0.125 | 1 | 0.73779294 | 0.03668846 |
| Ank2 | 0.22221177 | -0.2127716 | 0.344 | 0.264 | 1 | 0.01851968 | -1.5246454 | 0.361 | 0.469 | 1 | 0.22221177 | 0.03669639 |
| Rnf130 | 0.55714786 | -0.1604822 | 0.917 | 0.845 | 1 | 0.01853483 | 0.12843759 | 0.937 | 0.885 | 1 | 0.55714786 | 0.03672613 |
| Nelfcd | 0.57475661 | -0.2148979 | 0.083 | 0.104 | 1 | 0.0185893 | 1.41261076 | 0.113 | 0.031 | 1 | 0.57475661 | 0.03683303 |
| 1700061N14Rik | 0.3058807 | 0.877434 | 0.208 | 0.171 | 1 | 0.01861267 | 0.88278982 | 0.256 | 0.135 | 1 | 0.3058807 | 0.03687891 |
| Fus | 0.01862539 | 0.15819143 | 0.99 | 0.969 | 1 | 0.07466873 | -0.3028441 | 1 | 1 | 1 | 0.07466873 | 0.03690387 |
| Usp42 | 0.26394012 | 0.16924561 | 0.281 | 0.212 | 1 | 0.01866274 | 0.45769865 | 0.403 | 0.25 | 1 | 0.26394012 | 0.03697718 |
| Znrd2 | 0.04604309 | 0.72209232 | 0.26 | 0.161 | 1 | 0.01871309 | 0.80562195 | 0.218 | 0.104 | 1 | 0.04604309 | 0.037076 |
| Mtmr1 | 0.43853716 | 0.71504364 | 0.365 | 0.347 | 1 | 0.01876318 | 0.16059536 | 0.521 | 0.344 | 1 | 0.43853716 | 0.03717431 |
| Gm31641 | 0.01877358 | -2.0180406 | 0.042 | 0.13 | 1 | 0.05412949 | -2.1113096 | 0.176 | 0.24 | 1 | 0.05412949 | 0.03719472 |
| Pgm2l1 | 0.83011444 | -0.7294711 | 0.073 | 0.078 | 1 | 0.01878295 | -0.9378865 | 0.164 | 0.281 | 1 | 0.83011444 | 0.0372131 |
| Usp40 | 0.01878638 | 0.52788504 | 0.417 | 0.28 | 1 | 0.05138378 | 0.10518224 | 0.399 | 0.26 | 1 | 0.05138378 | 0.03721983 |
| Avil | 0.0187882 | 0.8114294 | 0.469 | 0.332 | 1 | 0.35046394 | -0.189561 | 0.634 | 0.49 | 1 | 0.35046394 | 0.0372234 |
| Ash1l | 0.01879741 | 0.18999423 | 0.979 | 0.974 | 1 | 0.02116741 | -0.2521398 | 0.996 | 1 | 1 | 0.02116741 | 0.03724148 |
| Cyb5d1 | 0.01880133 | 1.28760246 | 0.146 | 0.062 | 1 | 0.76604614 | -0.6659991 | 0.097 | 0.083 | 1 | 0.76604614 | 0.03724917 |
| B430010I23Rik | 0.01881338 | 0.4681289 | 0.385 | 0.254 | 1 | 0.15051508 | 0.11590864 | 0.361 | 0.26 | 1 | 0.15051508 | 0.03727281 |
| Camta2 | 0.07619973 | 0.62898291 | 0.188 | 0.109 | 1 | 0.01883 | 0.39058446 | 0.239 | 0.115 | 1 | 0.07619973 | 0.03730543 |
| Bmp4 | 0.01883829 | 1.52206771 | 0.115 | 0.041 | 1 | 0.36247737 | -0.9406215 | 0.076 | 0.104 | 1 | 0.36247737 | 0.03732169 |
| Dnmt1 | 0.01883865 | 0.46560076 | 0.479 | 0.326 | 1 | 0.07977638 | 0.23011978 | 0.487 | 0.354 | 1 | 0.07977638 | 0.0373224 |
| Hpcal1 | 0.44736479 | -0.6514685 | 0.135 | 0.166 | 1 | 0.01884086 | -1.1358259 | 0.126 | 0.229 | 1 | 0.44736479 | 0.03732674 |
| Specc1l | 0.01885443 | 0.26509506 | 0.729 | 0.58 | 1 | 0.0560377 | -0.5379613 | 0.71 | 0.781 | 1 | 0.0560377 | 0.03735337 |
| Suv39h1 | 0.01885662 | 1.00749454 | 0.198 | 0.098 | 1 | 0.63709408 | -0.5025003 | 0.105 | 0.083 | 1 | 0.63709408 | 0.03735767 |
| 9330188P03Rik | 0.90818761 | -0.4231398 | 0.229 | 0.233 | 1 | 0.01887827 | 0.32293576 | 0.311 | 0.167 | 1 | 0.90818761 | 0.03740016 |
| Gapvd1 | 0.01889251 | 0.267955 | 0.833 | 0.741 | 1 | 0.85467359 | -0.1977264 | 0.891 | 0.906 | 1 | 0.85467359 | 0.03742809 |
| Repin1 | 0.46385443 | -0.4519371 | 0.073 | 0.098 | 1 | 0.01890326 | 1.17127143 | 0.21 | 0.104 | 1 | 0.46385443 | 0.03744919 |
| Mlycd | 0.19374243 | 0.64492446 | 0.125 | 0.078 | 1 | 0.01890752 | 1.10518224 | 0.113 | 0.031 | 1 | 0.19374243 | 0.03745754 |
| Rasal2 | 0.0814557 | -0.8473075 | 0.427 | 0.508 | 1 | 0.01891862 | -0.6538097 | 0.517 | 0.688 | 1 | 0.0814557 | 0.03747933 |
| Sdhaf2 | 0.18008271 | 0.36363835 | 0.458 | 0.389 | 1 | 0.01893807 | 0.54429387 | 0.387 | 0.24 | 1 | 0.18008271 | 0.0375175 |
| Fas | 0.01894049 | -0.4187702 | 0.323 | 0.461 | 1 | 0.23064379 | -0.2654611 | 0.29 | 0.198 | 1 | 0.23064379 | 0.03752224 |
| Gadd45a | 0.01894797 | -0.554009 | 0.365 | 0.508 | 1 | 0.02643357 | 0.24060904 | 0.706 | 0.583 | 1 | 0.02643357 | 0.03753692 |
| Usp22 | 0.01895781 | 0.26405945 | 0.583 | 0.42 | 1 | 0.5083195 | -0.127163 | 0.555 | 0.479 | 1 | 0.5083195 | 0.03755622 |

| | | | | | | | | | | | | |
|---|---|---|---|---|---|---|---|---|---|---|---|---|
| Smdt1 | 0.42186491 | -0.1851505 | 0.823 | 0.85 | 1 | 0.01896889 | 0.27510724 | 0.819 | 0.76 | 1 | 0.42186491 | 0.03757796 |
| Tpcn2 | 0.01899749 | 0.86065315 | 0.198 | 0.098 | 1 | 0.20414998 | -0.7175132 | 0.256 | 0.177 | 1 | 0.20414998 | 0.03763408 |
| Smim15 | 0.01901658 | 0.66907832 | 0.531 | 0.378 | 1 | 0.02524621 | 0.31383963 | 0.424 | 0.271 | 1 | 0.02524621 | 0.03767154 |
| Hmgxb3 | 0.88984957 | -0.5015191 | 0.333 | 0.337 | 1 | 0.01901667 | 0.23446525 | 0.42 | 0.24 | 1 | 0.88984957 | 0.03767171 |
| Galr2 | 0.74661777 | 0.42253204 | 0.115 | 0.104 | 1 | 0.01902171 | 0.95317914 | 0.214 | 0.104 | 1 | 0.74661777 | 0.03768159 |
| Hecw2 | 0.01902335 | -1.5894406 | 0.094 | 0.202 | 1 | 0.24638153 | -1.182476 | 0.202 | 0.24 | 1 | 0.24638153 | 0.03768482 |
| Elmod2 | 0.01905028 | 0.8872003 | 0.198 | 0.098 | 1 | 0.35523722 | -0.154577 | 0.172 | 0.125 | 1 | 0.35523722 | 0.03773765 |
| Dnal4 | 0.0632679 | 0.5049942 | 0.292 | 0.192 | 1 | 0.01906227 | 0.75623393 | 0.231 | 0.115 | 1 | 0.0632679 | 0.03776117 |
| Trappc14 | 0.74947105 | 0.46000674 | 0.104 | 0.093 | 1 | 0.01908302 | 0.76814725 | 0.147 | 0.052 | 1 | 0.74947105 | 0.03780189 |
| Supt16 | 0.0191356 | 0.23236795 | 0.792 | 0.689 | 1 | 0.68571737 | -0.206586 | 0.857 | 0.865 | 1 | 0.68571737 | 0.03790502 |
| Immp2l | 0.06393956 | -0.5554417 | 0.385 | 0.492 | 1 | 0.01913736 | -0.8301449 | 0.395 | 0.531 | 1 | 0.06393956 | 0.03790849 |
| a | 0.89411003 | -1.7119533 | 0.198 | 0.181 | 1 | 0.01917821 | -1.882434 | 0.193 | 0.281 | 1 | 0.89411003 | 0.03798861 |
| Kif11 | 0.01918547 | 2.71531379 | 0.104 | 0.036 | 1 | 0.66171832 | 1.14957636 | 0.017 | 0.01 | 1 | 0.66171832 | 0.03800286 |
| Slc25a13 | 0.30323466 | 0.11996927 | 0.26 | 0.212 | 1 | 0.0191918 | 0.71347724 | 0.374 | 0.229 | 1 | 0.30323466 | 0.03801527 |
| Smim27 | 0.08217882 | 0.65699729 | 0.323 | 0.238 | 1 | 0.01919259 | 0.66589719 | 0.21 | 0.094 | 1 | 0.08217882 | 0.03801683 |
| Mettl6 | 0.06045655 | 0.46387283 | 0.417 | 0.301 | 1 | 0.01920084 | 0.30846584 | 0.462 | 0.281 | 1 | 0.06045655 | 0.038033 |
| Jazf1 | 0.07736981 | 0.36363835 | 0.292 | 0.192 | 1 | 0.01921628 | -0.200444 | 0.429 | 0.281 | 1 | 0.07736981 | 0.0380633 |
| Zfp131 | 0.01926574 | 0.24158599 | 0.812 | 0.715 | 1 | 0.77467796 | -0.1809575 | 0.87 | 0.917 | 1 | 0.77467796 | 0.03816031 |
| Hpf1 | 0.01926889 | 0.56350566 | 0.615 | 0.503 | 1 | 0.16434557 | 0.17843122 | 0.492 | 0.417 | 1 | 0.16434557 | 0.03816648 |
| Pgs1 | 0.01927952 | -0.8637721 | 0.333 | 0.44 | 1 | 0.10509031 | 0.23446525 | 0.42 | 0.292 | 1 | 0.10509031 | 0.03818735 |
| Galk1 | 0.01928575 | 0.54559043 | 0.438 | 0.29 | 1 | 0.03932391 | -0.1830981 | 0.37 | 0.219 | 1 | 0.03932391 | 0.03819956 |
| Gm20559 | 0.1969519 | 2.00749454 | 0.031 | 0.01 | 1 | 0.01928846 | -2.5322477 | 0.021 | 0.073 | 1 | 0.1969519 | 0.03820488 |
| Uevld | 0.01929262 | 1.40342321 | 0.177 | 0.088 | 1 | 0.01998653 | 0.92718393 | 0.202 | 0.094 | 1 | 0.01998653 | 0.03821304 |
| Cfap97 | 0.53833134 | 0.3529911 | 0.333 | 0.306 | 1 | 0.01935787 | 0.70721825 | 0.256 | 0.135 | 1 | 0.53833134 | 0.03834101 |
| Rbm34 | 0.25329804 | 0.59868899 | 0.333 | 0.285 | 1 | 0.01936889 | 0.69014474 | 0.307 | 0.177 | 1 | 0.25329804 | 0.03836262 |
| Zfp758 | 0.69915495 | -0.1624305 | 0.167 | 0.187 | 1 | 0.01938112 | 0.46458504 | 0.21 | 0.094 | 1 | 0.69915495 | 0.03838661 |
| Reg4 | 0.09419385 | -1.7294711 | 0.021 | 0.067 | 1 | 0.01939461 | -3.1172102 | 0.021 | 0.073 | 1 | 0.09419385 | 0.03841308 |
| Crnkl1 | 0.01940992 | 0.55818714 | 0.375 | 0.233 | 1 | 0.11837219 | -0.1567759 | 0.416 | 0.292 | 1 | 0.11837219 | 0.0384431 |
| Wdr18 | 0.07838696 | 0.63712609 | 0.302 | 0.212 | 1 | 0.01942616 | 0.80056873 | 0.29 | 0.167 | 1 | 0.07838696 | 0.03847494 |
| Ndufb5 | 0.35387961 | 0.13171447 | 0.875 | 0.819 | 1 | 0.01942736 | 0.25770419 | 0.803 | 0.771 | 1 | 0.35387961 | 0.0384773 |
| Ybx3 | 0.0449398 | 0.16353401 | 0.958 | 0.948 | 1 | 0.01948583 | 0.2377054 | 0.975 | 0.969 | 1 | 0.0449398 | 0.03859197 |
| Pfkfb2 | 0.09409232 | 0.65135073 | 0.188 | 0.114 | 1 | 0.01951714 | -0.9957467 | 0.261 | 0.396 | 1 | 0.09409232 | 0.03865337 |
| Cfap410 | 0.26159873 | -0.5505009 | 0.167 | 0.218 | 1 | 0.01954462 | 0.52303475 | 0.206 | 0.094 | 1 | 0.26159873 | 0.03870725 |
| Smarcd2 | 0.01955849 | 0.80396114 | 0.26 | 0.145 | 1 | 0.24551694 | 0.32241295 | 0.315 | 0.25 | 1 | 0.24551694 | 0.03873445 |
| Ccdc149 | 0.36552793 | 1.00749454 | 0.083 | 0.057 | 1 | 0.01955917 | 1.11640949 | 0.113 | 0.031 | 1 | 0.36552793 | 0.03873578 |
| Cep104 | 0.06047914 | 0.33870045 | 0.271 | 0.176 | 1 | 0.01958187 | 0.1438627 | 0.378 | 0.219 | 1 | 0.06047914 | 0.03878029 |
| Lats2 | 0.01958375 | 0.60223806 | 0.542 | 0.415 | 1 | 0.34445307 | -0.1441548 | 0.706 | 0.552 | 1 | 0.34445307 | 0.03878398 |
| Dnajc7 | 0.01958788 | 0.53741567 | 0.823 | 0.834 | 1 | 0.03777612 | 0.16762281 | 0.899 | 0.896 | 1 | 0.03777612 | 0.03879207 |
| Adprs | 0.01960471 | 0.97096866 | 0.281 | 0.171 | 1 | 0.18817557 | 0.57278779 | 0.172 | 0.115 | 1 | 0.18817557 | 0.03882507 |
| Taf1b | 0.01961118 | 0.75695107 | 0.365 | 0.238 | 1 | 0.57343066 | -0.6239639 | 0.307 | 0.25 | 1 | 0.57343066 | 0.03883777 |
| Sun1 | 0.01963089 | 0.65957123 | 0.448 | 0.311 | 1 | 0.77602515 | -0.2573878 | 0.454 | 0.385 | 1 | 0.77602515 | 0.03887641 |
| Map4k3 | 0.34069391 | 0.21394541 | 0.573 | 0.513 | 1 | 0.01964322 | -0.5073013 | 0.727 | 0.833 | 1 | 0.34069391 | 0.03890057 |
| Ssx2ip | 0.14402663 | 0.54183096 | 0.292 | 0.212 | 1 | 0.0197938 | 0.66060046 | 0.361 | 0.24 | 1 | 0.14402663 | 0.0391958 |
| Cldn34c1 | 0.13676936 | 1.24850264 | 0.083 | 0.041 | 1 | 0.01979495 | 2.69014474 | 0.055 | 0 | 1 | 0.13676936 | 0.03919807 |
| Gm45507 | 0.15837881 | 2.00749454 | 0.01 | 0 | 1 | 0.01979495 | 2.69014474 | 0.055 | 0 | 1 | 0.15837881 | 0.03919807 |
| Phex | 0.03210532 | -5.5160674 | 0 | 0.047 | 1 | 0.01980688 | 3.14957636 | 0.055 | 0 | 1 | 0.03210532 | 0.03922144 |
| Alg11 | 0.1222681 | 0.48666237 | 0.208 | 0.135 | 1 | 0.01981188 | 0.73745045 | 0.206 | 0.094 | 1 | 0.1222681 | 0.03923125 |
| Slc18a1 | 0.28126256 | -1.4519371 | 0.01 | 0.031 | 1 | 0.01981218 | 4.04769674 | 0.055 | 0 | 1 | 0.28126256 | 0.03923183 |
| Amigo1 | 0.46992036 | 0.74446013 | 0.042 | 0.026 | 1 | 0.01981438 | 2.49749966 | 0.055 | 0 | 1 | 0.46992036 | 0.03923616 |
| Dut | 0.01981628 | 0.45495351 | 0.667 | 0.544 | 1 | 0.07822442 | 0.17808278 | 0.668 | 0.594 | 1 | 0.07822442 | 0.03923988 |
| Coprs | 0.23627371 | -0.5505009 | 0.167 | 0.223 | 1 | 0.01982686 | 1.11335617 | 0.185 | 0.083 | 1 | 0.23627371 | 0.03926061 |
| Fbxo31 | 0.34175529 | -0.236431 | 0.146 | 0.192 | 1 | 0.01982861 | 1.16407593 | 0.16 | 0.062 | 1 | 0.34175529 | 0.03926405 |
| Itfg2 | 0.01982932 | 0.89501981 | 0.292 | 0.176 | 1 | 0.06202037 | -0.4117349 | 0.286 | 0.167 | 1 | 0.06202037 | 0.03926543 |
| Gatc | 0.01983926 | 1.07162487 | 0.208 | 0.109 | 1 | 0.03745954 | 1.24034182 | 0.143 | 0.062 | 1 | 0.03745954 | 0.03928493 |
| Pafah1b3 | 0.97447202 | 0.10060394 | 0.188 | 0.187 | 1 | 0.01984546 | 0.95317914 | 0.172 | 0.073 | 1 | 0.97447202 | 0.03929708 |
| Plekhm3 | 0.01985444 | -1.3144336 | 0.125 | 0.233 | 1 | 0.19828117 | -0.7552664 | 0.118 | 0.167 | 1 | 0.19828117 | 0.03931468 |
| Mis12 | 0.01987223 | 0.22988696 | 0.302 | 0.176 | 1 | 0.04103304 | 0.3500693 | 0.256 | 0.146 | 1 | 0.04103304 | 0.03934955 |
| Mppe1 | 0.48607316 | 0.17460452 | 0.292 | 0.249 | 1 | 0.01989958 | 0.76814725 | 0.252 | 0.135 | 1 | 0.48607316 | 0.03940316 |
| Synj2 | 0.01993019 | -1.1539689 | 0.135 | 0.254 | 1 | 0.42167947 | -0.6179776 | 0.294 | 0.312 | 1 | 0.42167947 | 0.03946318 |
| Parp2 | 0.01993298 | 0.55498233 | 0.438 | 0.295 | 1 | 0.17705386 | 0.2584285 | 0.273 | 0.198 | 1 | 0.17705386 | 0.03946864 |
| Zfp598 | 0.53393636 | 0.11767745 | 0.302 | 0.259 | 1 | 0.01998294 | 0.43488768 | 0.395 | 0.24 | 1 | 0.53393636 | 0.03956655 |
| Taf1c | 0.50666259 | 0.47697982 | 0.083 | 0.062 | 1 | 0.02000041 | 2.08246216 | 0.076 | 0.01 | 1 | 0.50666259 | 0.03960081 |
| Rab5a | 0.17530487 | 0.16629365 | 0.948 | 0.912 | 1 | 0.02000333 | 0.19885213 | 0.979 | 0.948 | 1 | 0.17530487 | 0.03960653 |
| 1700036A12Rik | 0.02004241 | 1.00749454 | 0.198 | 0.098 | 1 | 0.0779552 | 0.3068161 | 0.252 | 0.156 | 1 | 0.0779552 | 0.03968313 |
| Gnai1 | 0.02004459 | 0.11440974 | 0.365 | 0.228 | 1 | 0.54976044 | -0.1341756 | 0.555 | 0.49 | 1 | 0.54976044 | 0.03968739 |
| Slc25a14 | 0.79514074 | -0.1079827 | 0.104 | 0.114 | 1 | 0.02005629 | 0.95317914 | 0.16 | 0.062 | 1 | 0.79514074 | 0.03971033 |
| Pak1 | 0.02006029 | 0.52009681 | 0.729 | 0.611 | 1 | 0.32890282 | -0.3098553 | 0.756 | 0.74 | 1 | 0.32890282 | 0.03971817 |
| Adamts9 | 0.02006076 | 0.34831387 | 0.448 | 0.285 | 1 | 0.10370979 | -1.4000531 | 0.303 | 0.344 | 1 | 0.10370979 | 0.03971908 |
| Txnrd2 | 0.02746621 | 1.00749454 | 0.177 | 0.088 | 1 | 0.02009682 | 0.95317914 | 0.172 | 0.073 | 1 | 0.02746621 | 0.03978975 |
| Anxa8 | 0.02010757 | -0.4857608 | 0.573 | 0.71 | 1 | 0.14957697 | 0.49442027 | 0.345 | 0.51 | 1 | 0.14957697 | 0.03981083 |
| Tceanc | 0.27720863 | 1.11440974 | 0.083 | 0.052 | 1 | 0.02010897 | 1.17557156 | 0.097 | 0.021 | 1 | 0.27720863 | 0.03981357 |
| Rbm45 | 0.02011047 | 0.83061677 | 0.219 | 0.114 | 1 | 0.03581686 | 0.71399148 | 0.176 | 0.083 | 1 | 0.03581686 | 0.03981651 |
| Folr2 | 0.02017634 | -0.6205367 | 0.021 | 0.098 | 1 | 0.31394819 | -0.7248928 | 0.029 | 0.01 | 1 | 0.31394819 | 0.03994559 |
| Sec16a | 0.02021791 | 0.3191543 | 0.792 | 0.694 | 1 | 0.74340188 | -0.2197428 | 0.84 | 0.792 | 1 | 0.74340188 | 0.04002705 |
| 2510046G10Rik | 0.38181636 | 0.78510212 | 0.052 | 0.031 | 1 | 0.02021829 | 1.93807225 | 0.076 | 0.01 | 1 | 0.38181636 | 0.0400278 |
| Foxo1 | 0.02021897 | -0.4155664 | 0.917 | 0.948 | 1 | 0.71277054 | -0.5682002 | 0.979 | 0.938 | 1 | 0.71277054 | 0.04002914 |
| Pou2af1 | 0.07531985 | -0.9990483 | 0.24 | 0.326 | 1 | 0.02026887 | 0.69014474 | 0.206 | 0.094 | 1 | 0.07531985 | 0.04012691 |
| Gm527 | 0.34934772 | -0.8290067 | 0.062 | 0.093 | 1 | 0.02027688 | 1.61614416 | 0.126 | 0.042 | 1 | 0.34934772 | 0.04014261 |
| Psmc5 | 0.03455636 | 0.43898256 | 0.729 | 0.617 | 1 | 0.02028257 | 0.35505539 | 0.727 | 0.646 | 1 | 0.03455636 | 0.04015375 |
| Farp2 | 0.02030459 | 0.43099361 | 0.562 | 0.389 | 1 | 0.39184775 | -0.5334385 | 0.651 | 0.552 | 1 | 0.39184775 | 0.04019691 |
| Mier3 | 0.83736445 | -0.1445086 | 0.469 | 0.44 | 1 | 0.02030472 | 0.17065607 | 0.634 | 0.417 | 1 | 0.83736445 | 0.04019717 |
| Cad | 0.02030835 | 1.59245704 | 0.135 | 0.057 | 1 | 0.10933142 | 0.93807225 | 0.122 | 0.062 | 1 | 0.10933142 | 0.04020427 |
| Psmd13 | 0.10198892 | 0.34670062 | 0.562 | 0.508 | 1 | 0.02031011 | 0.36917556 | 0.693 | 0.583 | 1 | 0.10198892 | 0.04020772 |
| Cryl1 | 0.02035603 | 0.53854014 | 0.635 | 0.518 | 1 | 0.87127347 | -0.1149763 | 0.58 | 0.594 | 1 | 0.87127347 | 0.0402977 |
| Bola1 | 0.74006342 | 0.14767219 | 0.24 | 0.223 | 1 | 0.02038169 | 0.97965135 | 0.16 | 0.062 | 1 | 0.74006342 | 0.04034797 |
| Tmem241 | 0.03903561 | 0.85221631 | 0.25 | 0.145 | 1 | 0.02040848 | 0.55521516 | 0.235 | 0.115 | 1 | 0.03903561 | 0.04040045 |
| Chmp6 | 0.03517281 | 0.86999101 | 0.24 | 0.145 | 1 | 0.02041011 | 0.62874419 | 0.261 | 0.135 | 1 | 0.03517281 | 0.04040366 |
| Kif26a | 0.37536054 | 1.32942263 | 0.031 | 0.016 | 1 | 0.02041447 | 0.78968041 | 0.097 | 0.021 | 1 | 0.37536054 | 0.04041218 |
| Sdhaf3 | 0.02042131 | 1.34452952 | 0.167 | 0.078 | 1 | 0.08184065 | 0.61614416 | 0.143 | 0.073 | 1 | 0.08184065 | 0.04042559 |
| Cxxc5 | 0.02360747 | 0.58148592 | 0.573 | 0.461 | 1 | 0.02042636 | 0.43850124 | 0.794 | 0.698 | 1 | 0.02360747 | 0.04043549 |

| | | | | | | | | | | | | |
|---|---|---|---|---|---|---|---|---|---|---|---|---|
| Igfals | 0.02044375 | 0.89201732 | 0.115 | 0.041 | 1 | 0.04158548 | 0.87777174 | 0.13 | 0.052 | 1 | 0.04158548 | 0.04046956 |
| Abcb10 | 0.02045309 | 0.89201732 | 0.198 | 0.098 | 1 | 0.07480265 | 0.3163299 | 0.231 | 0.135 | 1 | 0.07480265 | 0.04048784 |
| Rab28 | 0.02052447 | 0.49949402 | 0.719 | 0.642 | 1 | 0.31595709 | -0.247571 | 0.643 | 0.562 | 1 | 0.31595709 | 0.04062769 |
| Zfp948 | 0.02052663 | 0.40859284 | 0.562 | 0.42 | 1 | 0.10505898 | 0.32351187 | 0.592 | 0.469 | 1 | 0.10505898 | 0.04063192 |
| Msl2 | 0.02055549 | 0.42253204 | 0.562 | 0.389 | 1 | 0.3680902 | -0.2423484 | 0.735 | 0.75 | 1 | 0.3680902 | 0.04068844 |
| Cdk5r1 | 0.02059755 | 1.24195979 | 0.146 | 0.062 | 1 | 0.17657564 | 0.46775232 | 0.139 | 0.083 | 1 | 0.17657564 | 0.04077083 |
| Dnajb12 | 0.49143083 | -0.1708427 | 0.604 | 0.627 | 1 | 0.02059825 | 0.22669011 | 0.668 | 0.479 | 1 | 0.49143083 | 0.04077221 |
| Ctdspl2 | 0.02059913 | 0.30062537 | 0.719 | 0.554 | 1 | 0.12786672 | -0.5596841 | 0.798 | 0.812 | 1 | 0.12786672 | 0.04077395 |
| Mob3c | 0.02063164 | 0.78008404 | 0.302 | 0.187 | 1 | 0.03558117 | 0.79976923 | 0.231 | 0.125 | 1 | 0.03558117 | 0.04083762 |
| Nktr | 0.08041447 | 0.15033763 | 1 | 0.995 | 1 | 0.02067852 | -0.3133308 | 1 | 0.99 | 1 | 0.08041447 | 0.04092945 |
| Gm57354 | 0.05805424 | 0.31717904 | 0.365 | 0.249 | 1 | 0.02079972 | 0.25256467 | 0.382 | 0.219 | 1 | 0.05805424 | 0.0411668 |
| Gla | 0.52432969 | 0.35896691 | 0.25 | 0.223 | 1 | 0.02085707 | 0.37464291 | 0.345 | 0.198 | 1 | 0.52432969 | 0.04127912 |
| Stau2 | 0.02731996 | -0.7019153 | 0.208 | 0.347 | 1 | 0.02087015 | -0.810866 | 0.357 | 0.5 | 1 | 0.02731996 | 0.04130474 |
| Pcx | 0.07012737 | -1.1445086 | 0.073 | 0.145 | 1 | 0.02088838 | -1.9472852 | 0.034 | 0.094 | 1 | 0.07012737 | 0.04134043 |
| Trim8 | 0.02089307 | 0.36638896 | 0.542 | 0.383 | 1 | 0.02994406 | 0.17466734 | 0.651 | 0.542 | 1 | 0.02994406 | 0.04134962 |
| Tmem144 | 0.08353315 | 1.00749454 | 0.042 | 0.01 | 1 | 0.02092652 | 2.14957636 | 0.092 | 0.021 | 1 | 0.08353315 | 0.04141511 |
| Itgb1bp2 | 0.02092861 | 1.59245704 | 0.104 | 0.036 | 1 | 0.05670515 | 1.10518224 | 0.059 | 0.01 | 1 | 0.05670515 | 0.04141922 |
| Ccdc102a | 0.02093831 | 0.73186009 | 0.156 | 0.067 | 1 | 0.09851627 | 0.18971575 | 0.248 | 0.156 | 1 | 0.09851627 | 0.04143821 |
| Supv3l1 | 0.02098508 | 0.72738662 | 0.292 | 0.171 | 1 | 0.71858088 | -0.4423056 | 0.298 | 0.25 | 1 | 0.71858088 | 0.04152978 |
| Slc4a1ap | 0.47607507 | 0.18681824 | 0.458 | 0.404 | 1 | 0.0210008 | 0.18629797 | 0.471 | 0.281 | 1 | 0.47607507 | 0.04156057 |
| Gzmm | 0.82342138 | 0.32942263 | 0.042 | 0.036 | 1 | 0.02101442 | 1.77760758 | 0.109 | 0.031 | 1 | 0.82342138 | 0.04158724 |
| Tuba4a | 0.29814454 | 1.38600616 | 0.062 | 0.036 | 1 | 0.02102318 | -0.7623675 | 0.185 | 0.302 | 1 | 0.29814454 | 0.04160439 |
| Scarb2 | 0.02103146 | 0.38505733 | 0.708 | 0.606 | 1 | 0.03844779 | 0.22090496 | 0.849 | 0.75 | 1 | 0.03844779 | 0.0416206 |
| Adam15 | 0.02104794 | -0.6349535 | 0.323 | 0.461 | 1 | 0.02999452 | 0.49749966 | 0.303 | 0.177 | 1 | 0.02999452 | 0.04165287 |
| Mif4gd | 0.20310334 | -0.3144336 | 0.167 | 0.233 | 1 | 0.02112319 | 1.03118166 | 0.223 | 0.115 | 1 | 0.20310334 | 0.04180019 |
| Sh3bp2 | 0.16249064 | 0.38600616 | 0.104 | 0.057 | 1 | 0.02117512 | 0.48532495 | 0.193 | 0.083 | 1 | 0.16249064 | 0.04190185 |
| Gtf3c1 | 0.07624615 | -0.6912368 | 0.438 | 0.513 | 1 | 0.02118917 | -0.4433569 | 0.702 | 0.844 | 1 | 0.07624615 | 0.04192936 |
| Ocrl | 0.71212315 | -0.1679475 | 0.365 | 0.383 | 1 | 0.02119768 | 0.19603567 | 0.492 | 0.323 | 1 | 0.71212315 | 0.04194601 |
| E130307A14Rik | 0.21006167 | 0.64136664 | 0.24 | 0.181 | 1 | 0.0212032 | 0.64434105 | 0.231 | 0.115 | 1 | 0.21006167 | 0.04195681 |
| Ttc23 | 0.02122314 | -1.0799683 | 0.135 | 0.254 | 1 | 0.65765889 | -0.4742421 | 0.282 | 0.281 | 1 | 0.65765889 | 0.04199585 |
| Mkks | 0.02126412 | 1.46692616 | 0.104 | 0.036 | 1 | 0.05353127 | 1.39058446 | 0.092 | 0.031 | 1 | 0.05353127 | 0.04207607 |
| Hacd3 | 0.29609224 | -0.2440442 | 0.302 | 0.368 | 1 | 0.02127144 | 0.12755005 | 0.517 | 0.323 | 1 | 0.29609224 | 0.0420904 |
| Ppp2r1a | 0.16335601 | 0.13097759 | 0.781 | 0.668 | 1 | 0.02131517 | 0.25106176 | 0.748 | 0.667 | 1 | 0.16335601 | 0.04217601 |
| Zmym5 | 0.13669382 | 0.18737666 | 0.854 | 0.772 | 1 | 0.02131637 | 0.22399825 | 0.895 | 0.865 | 1 | 0.13669382 | 0.04217834 |
| Platr25 | 0.04767862 | 0.77922555 | 0.25 | 0.15 | 1 | 0.02135604 | 0.7296731 | 0.147 | 0.052 | 1 | 0.04767862 | 0.04225599 |
| Bnc2 | 0.64246413 | -0.6101721 | 0.552 | 0.466 | 1 | 0.0213637 | -0.8240374 | 0.702 | 0.656 | 1 | 0.64246413 | 0.042271 |
| Barx2 | 0.05278817 | 1.17741954 | 0.198 | 0.119 | 1 | 0.02137421 | 2.49749966 | 0.092 | 0.021 | 1 | 0.05278817 | 0.04229156 |
| Nap1l3 | 0.09968317 | 1.00749454 | 0.104 | 0.052 | 1 | 0.02137421 | 1.91253716 | 0.092 | 0.021 | 1 | 0.09968317 | 0.04229156 |
| Sh3bgrl2 | 0.83354272 | -0.770113 | 0.042 | 0.047 | 1 | 0.02137715 | -1.3098553 | 0.059 | 0.135 | 1 | 0.83354272 | 0.04229732 |
| Psmc4 | 0.32259798 | 0.16834855 | 0.625 | 0.565 | 1 | 0.02138845 | 0.33238204 | 0.718 | 0.656 | 1 | 0.32259798 | 0.04231943 |
| Usp28 | 0.06673699 | 0.73512227 | 0.354 | 0.259 | 1 | 0.0213994 | 0.23903798 | 0.374 | 0.219 | 1 | 0.06673699 | 0.04234086 |
| C2cd5 | 0.64397134 | 0.21307867 | 0.333 | 0.311 | 1 | 0.0214152 | 0.20910968 | 0.45 | 0.292 | 1 | 0.64397134 | 0.04237179 |
| Mdfic | 0.42052925 | -0.1885706 | 0.385 | 0.332 | 1 | 0.02143299 | 0.22263982 | 0.555 | 0.365 | 1 | 0.42052925 | 0.04240661 |
| Pcdh11x | 0.02146326 | 1.14499806 | 0.104 | 0.036 | 1 | 0.14198298 | 0.32241295 | 0.13 | 0.073 | 1 | 0.14198298 | 0.04246585 |
| Suox | 0.40691807 | -0.3018335 | 0.188 | 0.228 | 1 | 0.02147943 | 1.27510724 | 0.168 | 0.073 | 1 | 0.40691807 | 0.04249749 |
| Surf2 | 0.72289117 | -0.1393469 | 0.219 | 0.238 | 1 | 0.02149 | 0.79866919 | 0.227 | 0.115 | 1 | 0.72289117 | 0.04251818 |
| Agpat3 | 0.46297304 | -0.1529701 | 0.479 | 0.554 | 1 | 0.02152041 | 0.33400093 | 0.723 | 0.625 | 1 | 0.46297304 | 0.04257768 |
| Dlg3 | 0.90267644 | 0.10937415 | 0.156 | 0.161 | 1 | 0.02152324 | 0.24653809 | 0.307 | 0.167 | 1 | 0.90267644 | 0.04258322 |
| Hcfc1 | 0.02153633 | 0.5821887 | 0.469 | 0.342 | 1 | 0.05271743 | 0.32049914 | 0.534 | 0.406 | 1 | 0.05271743 | 0.04260885 |
| Mrpl51 | 0.11820147 | 0.28051303 | 0.573 | 0.456 | 1 | 0.02158867 | 0.61937166 | 0.483 | 0.354 | 1 | 0.11820147 | 0.04271127 |
| Ttc32 | 0.72273569 | 0.11440974 | 0.167 | 0.15 | 1 | 0.02160529 | 0.74459252 | 0.176 | 0.073 | 1 | 0.72273569 | 0.0427438 |
| Nectin3 | 0.02163218 | 0.11320398 | 0.531 | 0.358 | 1 | 0.502942 | -0.1705805 | 0.58 | 0.448 | 1 | 0.502942 | 0.0427964 |
| Ier5l | 0.80748142 | -0.4213488 | 0.188 | 0.192 | 1 | 0.02164524 | -1.3522905 | 0.151 | 0.25 | 1 | 0.80748142 | 0.04282197 |
| Atxn1 | 0.03280665 | -0.520224 | 0.938 | 0.938 | 1 | 0.02165061 | -0.7878694 | 0.95 | 0.938 | 1 | 0.03280665 | 0.04283248 |
| Ccdc192 | 0.02165939 | -2.8993961 | 0.021 | 0.093 | 1 | 0.11689951 | -2.3098553 | 0 | 0.01 | 1 | 0.11689951 | 0.04284965 |
| Vps11 | 0.53612519 | 0.34452952 | 0.208 | 0.181 | 1 | 0.02167792 | 0.27510724 | 0.315 | 0.167 | 1 | 0.53612519 | 0.04288592 |
| Xab2 | 0.54997501 | 0.61517711 | 0.125 | 0.104 | 1 | 0.02168684 | 0.19264508 | 0.193 | 0.083 | 1 | 0.54997501 | 0.04290337 |
| Tada2a | 0.02172976 | 0.52206771 | 0.417 | 0.275 | 1 | 0.18391367 | -0.1242996 | 0.328 | 0.219 | 1 | 0.18391367 | 0.04298734 |
| Gm15956 | 0.07511393 | 2.00749454 | 0.031 | 0.005 | 1 | 0.02173427 | 1.08246216 | 0.076 | 0.01 | 1 | 0.07511393 | 0.04299615 |
| Itprip | 0.02177868 | 0.80467765 | 0.344 | 0.218 | 1 | 0.15305412 | 0.37295456 | 0.193 | 0.125 | 1 | 0.15305412 | 0.04308306 |
| Foxc2 | 0.14753354 | 0.44561565 | 0.333 | 0.259 | 1 | 0.02178095 | 0.32150014 | 0.433 | 0.281 | 1 | 0.14753354 | 0.04308748 |
| Diaph3 | 0.02178229 | 0.99119272 | 0.104 | 0.036 | 1 | 0.40490417 | -1.4301495 | 0.05 | 0.073 | 1 | 0.40490417 | 0.04309011 |
| Fastkd5 | 0.42179602 | -0.6929452 | 0.031 | 0.052 | 1 | 0.02181944 | 1.80562195 | 0.092 | 0.021 | 1 | 0.42179602 | 0.04316279 |
| Fstl4 | 0.6727142 | -0.8290067 | 0.042 | 0.031 | 1 | 0.02184746 | -2.1073624 | 0.055 | 0.125 | 1 | 0.6727142 | 0.04321761 |
| Rrnad1 | 0.10898281 | 0.56270969 | 0.281 | 0.202 | 1 | 0.02190378 | 0.76814725 | 0.248 | 0.135 | 1 | 0.10898281 | 0.04332779 |
| Pstk | 0.09032037 | 0.46692616 | 0.219 | 0.135 | 1 | 0.02191697 | 0.90313846 | 0.172 | 0.073 | 1 | 0.09032037 | 0.0433536 |
| Shf | 0.0219631 | -0.5606433 | 0.583 | 0.658 | 1 | 0.18459239 | 0.30905457 | 0.277 | 0.198 | 1 | 0.18459239 | 0.04344381 |
| Ano10 | 0.42410838 | -0.2920657 | 0.271 | 0.316 | 1 | 0.02196691 | 0.44647566 | 0.382 | 0.25 | 1 | 0.42410838 | 0.04345128 |
| 1700008J07Rik | 0.58995453 | 0.52206771 | 0.073 | 0.057 | 1 | 0.02203697 | 1.49749966 | 0.109 | 0.031 | 1 | 0.58995453 | 0.04358832 |
| Wdr73 | 0.18199979 | 0.66645762 | 0.146 | 0.093 | 1 | 0.02205309 | 1.08246216 | 0.197 | 0.094 | 1 | 0.18199979 | 0.04361983 |
| 0610010K14Rik | 0.41147553 | -0.5427025 | 0.344 | 0.368 | 1 | 0.02205699 | 0.27510724 | 0.345 | 0.208 | 1 | 0.41147553 | 0.04362747 |
| Dip2a | 0.02205806 | 0.73331957 | 0.24 | 0.13 | 1 | 0.2909549 | -0.1578522 | 0.315 | 0.229 | 1 | 0.2909549 | 0.04362957 |
| Bin3 | 0.0220625 | -1.1941393 | 0.281 | 0.394 | 1 | 0.97322752 | -0.5823811 | 0.303 | 0.281 | 1 | 0.97322752 | 0.04363824 |
| Aldh1a3 | 0.02207031 | -0.5295335 | 0.188 | 0.321 | 1 | 0.17128345 | 0.59703533 | 0.168 | 0.104 | 1 | 0.17128345 | 0.04365351 |
| Prmt2 | 0.06559604 | 0.6718915 | 0.281 | 0.187 | 1 | 0.02212995 | 0.22263982 | 0.277 | 0.146 | 1 | 0.06559604 | 0.04377017 |
| Ino80b | 0.64009697 | -0.1693832 | 0.146 | 0.166 | 1 | 0.02214395 | 1.01207283 | 0.155 | 0.062 | 1 | 0.64009697 | 0.04379755 |
| Polr3e | 0.03826236 | 0.24850264 | 0.333 | 0.212 | 1 | 0.02217431 | 0.14265694 | 0.45 | 0.271 | 1 | 0.03826236 | 0.04385693 |
| Zfp111 | 0.12178827 | 0.90057933 | 0.115 | 0.062 | 1 | 0.0221993 | 1.09224318 | 0.126 | 0.042 | 1 | 0.12178827 | 0.0439058 |
| Lipa | 0.3960949 | -0.3050957 | 0.219 | 0.264 | 1 | 0.0222005 | 0.36108046 | 0.332 | 0.198 | 1 | 0.3960949 | 0.04390813 |
| Pms2 | 0.02221421 | 0.66354014 | 0.229 | 0.119 | 1 | 0.72984881 | -0.3098553 | 0.147 | 0.125 | 1 | 0.72984881 | 0.04393494 |
| Stambpl1 | 0.44295618 | 0.14951354 | 0.219 | 0.269 | 1 | 0.02222748 | 0.80562195 | 0.218 | 0.104 | 1 | 0.44295618 | 0.04396089 |
| Stk38l | 0.02228566 | -0.4370202 | 0.656 | 0.798 | 1 | 0.07349634 | -0.2644985 | 0.819 | 0.594 | 1 | 0.07349634 | 0.04407466 |
| Icmt | 0.99545999 | -0.2488452 | 0.177 | 0.171 | 1 | 0.0222913 | 0.75725893 | 0.265 | 0.146 | 1 | 0.99545999 | 0.04408569 |
| Brd9 | 0.2271037 | 0.13550215 | 0.396 | 0.311 | 1 | 0.02230808 | 0.3794439 | 0.378 | 0.229 | 1 | 0.2271037 | 0.04411851 |
| Inpp4a | 0.02230876 | 0.46249844 | 0.438 | 0.29 | 1 | 0.05038186 | -0.7608395 | 0.508 | 0.573 | 1 | 0.05038186 | 0.04411984 |
| Gfer | 0.10625621 | 0.83756954 | 0.198 | 0.13 | 1 | 0.02232868 | 0.48369386 | 0.176 | 0.073 | 1 | 0.10625621 | 0.04415879 |
| D17H6S56E-5 | 0.02234388 | 1.69629285 | 0.135 | 0.057 | 1 | 0.30657391 | -1.7837865 | 0.046 | 0.073 | 1 | 0.30657391 | 0.0441885 |
| Fbxo6 | 0.51265384 | -0.2820121 | 0.156 | 0.187 | 1 | 0.02235465 | 0.98782529 | 0.21 | 0.104 | 1 | 0.51265384 | 0.04420958 |

| | | | | | | | | | | | | |
|---|---|---|---|---|---|---|---|---|---|---|---|---|
| Ppfibp1 | 0.02235653 | -0.3589037 | 0.927 | 0.959 | 1 | 0.597337 | -0.3415641 | 0.916 | 0.969 | 1 | 0.597337 | 0.04421325 |
| Gm49735 | 0.02237832 | 1.38600616 | 0.104 | 0.036 | 1 | 0.62901183 | -0.6217993 | 0.08 | 0.062 | 1 | 0.62901183 | 0.04425585 |
| Slc44a3 | 0.10152598 | -1.0368996 | 0.062 | 0.124 | 1 | 0.0224002 | -1.8948178 | 0.034 | 0.094 | 1 | 0.10152598 | 0.04429863 |
| Ly6c1 | 0.02242126 | -1.053047 | 0.135 | 0.249 | 1 | 0.07274036 | 0.43949406 | 0.235 | 0.146 | 1 | 0.07274036 | 0.0443398 |
| Morn2 | 0.05837576 | 0.5821887 | 0.292 | 0.192 | 1 | 0.02242199 | 1.01207283 | 0.21 | 0.104 | 1 | 0.05837576 | 0.04434124 |
| Smpdl3b | 0.02246808 | 0.92503238 | 0.146 | 0.062 | 1 | 0.45513939 | 0.17557156 | 0.05 | 0.031 | 1 | 0.45513939 | 0.04443134 |
| Ddx18 | 0.29824246 | 0.30705482 | 0.448 | 0.389 | 1 | 0.02250345 | 0.41261076 | 0.433 | 0.281 | 1 | 0.29824246 | 0.0445005 |
| Ube3c | 0.02252077 | 0.47697982 | 0.688 | 0.58 | 1 | 0.44356996 | -0.1300993 | 0.748 | 0.75 | 1 | 0.44356996 | 0.04453436 |
| Sipa1l2 | 0.26976796 | -0.8669746 | 0.208 | 0.264 | 1 | 0.02252739 | -1.1323171 | 0.223 | 0.333 | 1 | 0.26976796 | 0.0445473 |
| Rell1 | 0.04451865 | 0.40449811 | 0.542 | 0.394 | 1 | 0.02255025 | -0.4641285 | 0.655 | 0.812 | 1 | 0.04451865 | 0.04459198 |
| Akap12 | 0.02256949 | 0.79968965 | 0.26 | 0.14 | 1 | 0.42694349 | -0.1918581 | 0.29 | 0.229 | 1 | 0.42694349 | 0.0446296 |
| Efnb1 | 0.33797899 | 0.11099911 | 0.396 | 0.321 | 1 | 0.02260417 | 0.30846584 | 0.416 | 0.26 | 1 | 0.33797899 | 0.0446974 |
| E130317F20Rik | 0.02264162 | 0.78510212 | 0.115 | 0.041 | 1 | 0.35731895 | -0.3098553 | 0.042 | 0.021 | 1 | 0.35731895 | 0.0447706 |
| Serpinb1a | 0.57848839 | 1.00749454 | 0.031 | 0.021 | 1 | 0.02266769 | -0.7952821 | 0.059 | 0.135 | 1 | 0.57848839 | 0.04482156 |
| Bcor | 0.02268416 | 0.64841344 | 0.448 | 0.321 | 1 | 0.18577996 | -0.4155647 | 0.563 | 0.635 | 1 | 0.18577996 | 0.04485376 |
| Cenpt | 0.02269217 | 1.00749454 | 0.188 | 0.093 | 1 | 0.02531638 | 1.03806804 | 0.193 | 0.094 | 1 | 0.02531638 | 0.04486941 |
| Map2k6 | 0.20733444 | 0.70793425 | 0.083 | 0.047 | 1 | 0.02269469 | 1.8995981 | 0.092 | 0.021 | 1 | 0.20733444 | 0.04487433 |
| Mcmdc2 | 0.88076569 | -0.1445086 | 0.094 | 0.098 | 1 | 0.02270482 | 0.48369386 | 0.218 | 0.104 | 1 | 0.88076569 | 0.04489412 |
| Map2k3os | 0.08034485 | 0.33506919 | 0.26 | 0.166 | 1 | 0.02274444 | 0.10518224 | 0.16 | 0.062 | 1 | 0.08034485 | 0.04497158 |
| Parp6 | 0.13305562 | 0.372779 | 0.292 | 0.207 | 1 | 0.02274633 | 0.23328706 | 0.298 | 0.156 | 1 | 0.13305562 | 0.04497526 |
| Ksr1 | 0.42682216 | -1.0968421 | 0.062 | 0.088 | 1 | 0.02276181 | -1.4253325 | 0.059 | 0.135 | 1 | 0.42682216 | 0.04500553 |
| Cx3cr1 | 0.02277603 | 0.25542205 | 0.198 | 0.098 | 1 | 0.33402815 | -0.9472852 | 0.155 | 0.188 | 1 | 0.33402815 | 0.04503332 |
| Zfp398 | 0.02287458 | 0.84190847 | 0.385 | 0.249 | 1 | 0.6055494 | -0.3098553 | 0.408 | 0.333 | 1 | 0.6055494 | 0.04522592 |
| Nup43 | 0.03041733 | 1.00749454 | 0.125 | 0.052 | 1 | 0.02287759 | 1.45567948 | 0.109 | 0.031 | 1 | 0.03041733 | 0.0452318 |
| Apex1 | 0.24529509 | -0.3269245 | 0.427 | 0.477 | 1 | 0.02292012 | 0.5103237 | 0.336 | 0.198 | 1 | 0.24529509 | 0.04531492 |
| Bsn | 0.02292394 | 2.09495738 | 0.094 | 0.031 | 1 | 0.12250846 | 0.16407593 | 0.147 | 0.083 | 1 | 0.12250846 | 0.04532237 |
| Nsdhl | 0.1753213 | -0.4414903 | 0.302 | 0.389 | 1 | 0.02297772 | 0.19538005 | 0.248 | 0.125 | 1 | 0.1753213 | 0.04542747 |
| Gm16310 | 0.17609384 | -0.940038 | 0.052 | 0.098 | 1 | 0.02299127 | -1.3712558 | 0.076 | 0.156 | 1 | 0.17609384 | 0.04545394 |
| C1galt1 | 0.18061631 | -0.3733272 | 0.281 | 0.363 | 1 | 0.02299452 | -0.5936482 | 0.382 | 0.531 | 1 | 0.18061631 | 0.04546029 |
| Med13 | 0.20197468 | -0.2277219 | 0.938 | 0.933 | 1 | 0.02301243 | -0.3699763 | 0.983 | 0.99 | 1 | 0.20197468 | 0.04549528 |
| Vnn1 | 0.82911542 | -0.5427025 | 0.115 | 0.104 | 1 | 0.02304311 | 1.49749966 | 0.109 | 0.031 | 1 | 0.82911542 | 0.04555524 |
| Limk2 | 0.15438279 | 0.37006462 | 0.292 | 0.212 | 1 | 0.02306083 | -0.43837 | 0.366 | 0.531 | 1 | 0.15438279 | 0.04558986 |
| Pibf1 | 0.8241703 | -0.1884257 | 0.573 | 0.523 | 1 | 0.02307226 | -0.6778592 | 0.513 | 0.646 | 1 | 0.8241703 | 0.04561219 |
| Dglucy | 0.02308172 | 0.78510212 | 0.177 | 0.083 | 1 | 0.35822308 | 0.69014474 | 0.092 | 0.062 | 1 | 0.35822308 | 0.04563067 |
| H2-K2 | 0.02311926 | -1.9925055 | 0.031 | 0.109 | 1 | 0.43079893 | 0.56461386 | 0.038 | 0.021 | 1 | 0.43079893 | 0.04570402 |
| Thap11 | 0.31008435 | 0.35979628 | 0.25 | 0.197 | 1 | 0.02314576 | 0.37820073 | 0.357 | 0.219 | 1 | 0.31008435 | 0.04575579 |
| Rmi1 | 0.04795028 | 1.00749454 | 0.146 | 0.073 | 1 | 0.02315459 | 0.59703533 | 0.16 | 0.062 | 1 | 0.04795028 | 0.04577304 |
| Mrps36 | 0.20526231 | 0.2816695 | 0.406 | 0.321 | 1 | 0.02316958 | 0.35310975 | 0.273 | 0.146 | 1 | 0.20526231 | 0.04580233 |
| Alg9 | 0.03957987 | 0.56937342 | 0.271 | 0.161 | 1 | 0.02317635 | 0.74077081 | 0.261 | 0.146 | 1 | 0.03957987 | 0.04581556 |
| Entpd3 | 0.52961488 | -0.3144336 | 0.01 | 0.021 | 1 | 0.02318022 | -1.6577786 | 0.034 | 0.094 | 1 | 0.52961488 | 0.04582312 |
| Cfap53 | 0.84451844 | -0.130009 | 0.042 | 0.036 | 1 | 0.02318756 | -1.7248928 | 0.034 | 0.094 | 1 | 0.84451844 | 0.04583745 |
| Tut1 | 0.37851643 | 0.43812889 | 0.229 | 0.187 | 1 | 0.02320595 | 0.15409184 | 0.269 | 0.135 | 1 | 0.37851643 | 0.04587338 |
| Mustn1 | 0.56264634 | -0.6779948 | 0.271 | 0.295 | 1 | 0.02320657 | 0.97025266 | 0.16 | 0.062 | 1 | 0.56264634 | 0.04587459 |
| Plekhg5 | 0.46422629 | -0.8823225 | 0.115 | 0.14 | 1 | 0.02323284 | 0.12186098 | 0.303 | 0.167 | 1 | 0.46422629 | 0.04592592 |
| Hdac1 | 0.02325593 | 0.32731497 | 0.823 | 0.705 | 1 | 0.03084374 | 0.16685435 | 0.832 | 0.74 | 1 | 0.03084374 | 0.04597102 |
| Frmd4a | 0.02326407 | -1.4983964 | 0.375 | 0.446 | 1 | 0.04247213 | -0.7142455 | 0.193 | 0.302 | 1 | 0.04247213 | 0.04598691 |
| Ctps2 | 0.11117407 | 0.14499806 | 0.396 | 0.275 | 1 | 0.02331315 | 0.18113109 | 0.487 | 0.302 | 1 | 0.11117407 | 0.04608279 |
| Fkbp15 | 0.74622619 | -0.1904448 | 0.375 | 0.378 | 1 | 0.0233291 | 0.23267197 | 0.5 | 0.333 | 1 | 0.74622619 | 0.04611395 |
| Gm47096 | 0.02336801 | 1.65957123 | 0.094 | 0.031 | 1 | 0.92663792 | -0.5025003 | 0.05 | 0.052 | 1 | 0.92663792 | 0.04618995 |
| Tspan11 | 0.27877443 | -1.6929452 | 0.01 | 0.031 | 1 | 0.02338666 | -2.5322477 | 0.042 | 0.104 | 1 | 0.27877443 | 0.04622638 |
| Ppp1cc | 0.18181256 | 0.20462701 | 0.812 | 0.777 | 1 | 0.02339391 | 0.22040135 | 0.866 | 0.812 | 1 | 0.18181256 | 0.04624054 |
| Pygb | 0.02532604 | 0.73740537 | 0.271 | 0.155 | 1 | 0.02342063 | -0.8890121 | 0.273 | 0.396 | 1 | 0.02532604 | 0.04629272 |
| Tmem87a | 0.97360133 | -0.1810832 | 0.604 | 0.565 | 1 | 0.02342466 | -0.5894726 | 0.718 | 0.823 | 1 | 0.97360133 | 0.04630061 |
| Stard5 | 0.4665903 | -0.366901 | 0.198 | 0.228 | 1 | 0.02345356 | 0.14957636 | 0.319 | 0.177 | 1 | 0.4665903 | 0.04635705 |
| Sass6 | 0.02346269 | 1.76648644 | 0.188 | 0.098 | 1 | 0.17239253 | 0.77106473 | 0.139 | 0.083 | 1 | 0.17239253 | 0.04637487 |
| Aasdh | 0.91250174 | -0.2920657 | 0.115 | 0.109 | 1 | 0.02348902 | 1.14957636 | 0.109 | 0.031 | 1 | 0.91250174 | 0.0464263 |
| Rffl | 0.41460036 | 0.21394541 | 0.115 | 0.083 | 1 | 0.02348906 | -1.4194798 | 0.105 | 0.198 | 1 | 0.41460036 | 0.04642638 |
| Creb5 | 0.02357134 | 0.60653222 | 0.312 | 0.197 | 1 | 0.12811477 | -0.920813 | 0.206 | 0.26 | 1 | 0.12811477 | 0.04658707 |
| Gm14066 | 0.16330661 | 1.85549144 | 0.073 | 0.036 | 1 | 0.02361196 | 0.83698613 | 0.092 | 0.021 | 1 | 0.16330661 | 0.0466664 |
| Trim38 | 0.02361796 | -3.1624305 | 0 | 0.052 | 1 | 0.14544666 | -2.3098553 | 0.004 | 0.021 | 1 | 0.14544666 | 0.0466781 |
| Gm34934 | 0.02361796 | -2.7998604 | 0 | 0.052 | 1 | 0.55295159 | -0.6577786 | 0.038 | 0.052 | 1 | 0.55295159 | 0.0466781 |
| Itga2 | 0.02362141 | -3.0799683 | 0 | 0.052 | 1 | 0.58072684 | 0.48155812 | 0.046 | 0.062 | 1 | 0.58072684 | 0.04668486 |
| Adcy5 | 0.02362141 | -3.1624305 | 0 | 0.052 | 1 | 0.16838072 | -1.6577786 | 0.038 | 0.073 | 1 | 0.16838072 | 0.04668486 |
| Dnah2 | 0.02362349 | -3.9925055 | 0 | 0.052 | 1 | 0.29725568 | -1.4353861 | 0.021 | 0.042 | 1 | 0.29725568 | 0.04668891 |
| Timm10 | 0.02362723 | 0.64841344 | 0.344 | 0.212 | 1 | 0.11773038 | 0.5111746 | 0.197 | 0.125 | 1 | 0.11773038 | 0.04669621 |
| Gtf3a | 0.11577389 | 0.53105649 | 0.198 | 0.124 | 1 | 0.02365356 | 0.59703533 | 0.214 | 0.104 | 1 | 0.11577389 | 0.04674762 |
| Ubiad1 | 0.50650971 | 0.46692616 | 0.083 | 0.062 | 1 | 0.02375307 | 0.86006974 | 0.172 | 0.073 | 1 | 0.50650971 | 0.04694192 |
| Ccp110 | 0.02378585 | 0.85549144 | 0.24 | 0.13 | 1 | 0.12792612 | 0.11160851 | 0.218 | 0.135 | 1 | 0.12792612 | 0.04700594 |
| Nhlrc2 | 0.02388843 | 0.47911556 | 0.594 | 0.472 | 1 | 0.53679394 | 0.17127143 | 0.613 | 0.625 | 1 | 0.53679394 | 0.04720621 |
| Med14 | 0.02389436 | 0.36241104 | 0.51 | 0.363 | 1 | 0.6177125 | -0.2388887 | 0.571 | 0.615 | 1 | 0.6177125 | 0.04721777 |
| Ddx20 | 0.02391208 | 0.70793425 | 0.229 | 0.124 | 1 | 0.03819125 | 0.45953181 | 0.261 | 0.146 | 1 | 0.03819125 | 0.04725238 |
| Abhd3 | 0.02393988 | -1.9467018 | 0.031 | 0.109 | 1 | 0.97919113 | -1.3769695 | 0.076 | 0.073 | 1 | 0.97919113 | 0.04730665 |
| Smg7 | 0.02395834 | 0.42951847 | 0.677 | 0.554 | 1 | 0.03499512 | -0.3672332 | 0.723 | 0.854 | 1 | 0.03499512 | 0.04734267 |
| Thtpa | 0.34404896 | -0.4779323 | 0.031 | 0.057 | 1 | 0.02398661 | 0.8917786 | 0.092 | 0.021 | 1 | 0.34404896 | 0.04739786 |
| Tspan12 | 0.15494106 | -1.5160674 | 0.031 | 0.073 | 1 | 0.02401544 | -1.4410998 | 0.076 | 0.156 | 1 | 0.15494106 | 0.04745414 |
| Mbtps1 | 0.07044503 | 0.3748656 | 0.677 | 0.585 | 1 | 0.02401926 | 0.16315231 | 0.761 | 0.552 | 1 | 0.07044503 | 0.04746159 |
| Sppl2b | 0.72673157 | -0.2213242 | 0.281 | 0.29 | 1 | 0.02403824 | 0.36391651 | 0.244 | 0.125 | 1 | 0.72673157 | 0.04749865 |
| Dancr | 0.02403874 | 0.56692195 | 0.24 | 0.13 | 1 | 0.05027557 | 0.36257008 | 0.172 | 0.083 | 1 | 0.05027557 | 0.04749962 |
| Slc25a22 | 0.02406522 | 1.00749454 | 0.208 | 0.109 | 1 | 0.62873253 | -0.1119159 | 0.126 | 0.104 | 1 | 0.62873253 | 0.04755131 |
| Nudt2 | 0.23652538 | 0.53618882 | 0.323 | 0.269 | 1 | 0.02406848 | 0.86006974 | 0.21 | 0.104 | 1 | 0.23652538 | 0.04755766 |
| Ptn | 0.02407519 | -2.7505327 | 0.292 | 0.368 | 1 | 0.06444899 | 0.35904 | 0.374 | 0.531 | 1 | 0.06444899 | 0.04757076 |
| A530072M11Rik | 0.02414381 | 1.52206771 | 0.094 | 0.031 | 1 | 0.53159922 | 0.27510724 | 0.059 | 0.042 | 1 | 0.53159922 | 0.0477047 |
| Pdhb | 0.14635631 | 0.31682259 | 0.656 | 0.591 | 1 | 0.02416893 | 0.42132798 | 0.702 | 0.646 | 1 | 0.14635631 | 0.04775373 |
| Ccdc174 | 0.43396755 | -0.2498933 | 0.479 | 0.503 | 1 | 0.02419956 | 0.142019 | 0.601 | 0.385 | 1 | 0.43396755 | 0.0478135 |
| Bmf | 0.6952916 | -0.4330781 | 0.042 | 0.052 | 1 | 0.02423175 | -0.9728203 | 0.067 | 0.146 | 1 | 0.6952916 | 0.04787631 |
| Tec | 0.0242615 | -0.4519371 | 0.021 | 0.093 | 1 | 0.03807653 | -0.1034044 | 0.055 | 0.125 | 1 | 0.03807653 | 0.04793439 |
| Endog | 0.92113269 | 0.15188445 | 0.167 | 0.176 | 1 | 0.02427625 | 1.10518224 | 0.168 | 0.073 | 1 | 0.92113269 | 0.04796316 |
| Col14a1 | 0.07765657 | -1.4889313 | 0.042 | 0.104 | 1 | 0.02431577 | -3.6790891 | 0.017 | 0.062 | 1 | 0.07765657 | 0.04804028 |
| Lman2l | 0.02432787 | 0.45290568 | 0.302 | 0.176 | 1 | 0.02804927 | 0.26046046 | 0.307 | 0.177 | 1 | 0.02804927 | 0.0480639 |
| Trim68 | 0.02434535 | 1.81484946 | 0.094 | 0.031 | 1 | 0.02939419 | 1.08246216 | 0.122 | 0.042 | 1 | 0.02939419 | 0.04809801 |

| | | | | | | | | | | | | |
|---|---|---|---|---|---|---|---|---|---|---|---|---|
| Cd99l2 | 0.52449917 | 0.10479174 | 0.333 | 0.301 | 1 | 0.02436551 | -0.1379026 | 0.466 | 0.281 | 1 | 0.52449917 | 0.04813735 |
| Stx4a | 0.20121619 | -0.4223183 | 0.562 | 0.575 | 1 | 0.02438548 | 0.29348577 | 0.647 | 0.479 | 1 | 0.20121619 | 0.04817631 |
| Cog8 | 0.04057131 | 0.50868868 | 0.396 | 0.275 | 1 | 0.02441366 | -0.1189125 | 0.403 | 0.229 | 1 | 0.04057131 | 0.04823129 |
| Jrkl | 0.07568321 | 0.78510212 | 0.333 | 0.244 | 1 | 0.02444399 | 0.58322953 | 0.223 | 0.115 | 1 | 0.07568321 | 0.04829047 |
| Ndnf | 0.0446515 | -3.1624305 | 0.01 | 0.062 | 1 | 0.02445007 | -3.3098553 | 0.017 | 0.062 | 1 | 0.0446515 | 0.04830233 |
| Smyd4 | 0.02447678 | 1.14499806 | 0.177 | 0.088 | 1 | 0.04709316 | 0.63767732 | 0.08 | 0.021 | 1 | 0.04709316 | 0.04835446 |
| Fgd1 | 0.25988469 | 0.13677755 | 0.271 | 0.202 | 1 | 0.02449142 | 0.6008774 | 0.273 | 0.156 | 1 | 0.25988469 | 0.04838301 |
| Insyn2a | 0.02449988 | -2.7998604 | 0.01 | 0.073 | 1 | 0.58187664 | -0.9313436 | 0.059 | 0.073 | 1 | 0.58187664 | 0.04839951 |
| Rdm1 | 0.53765399 | 0.14689759 | 0.365 | 0.326 | 1 | 0.02453462 | 0.59196434 | 0.378 | 0.24 | 1 | 0.53765399 | 0.0484673 |
| Smarcc2 | 0.02453858 | 0.20716688 | 0.75 | 0.596 | 1 | 0.06859999 | 0.1330049 | 0.87 | 0.802 | 1 | 0.06859999 | 0.04847502 |
| Arid5a | 0.08923094 | -0.4980335 | 0.365 | 0.451 | 1 | 0.02461408 | 0.21370669 | 0.357 | 0.198 | 1 | 0.08923094 | 0.04862231 |
| Gstt2 | 0.86528499 | -0.2054992 | 0.188 | 0.192 | 1 | 0.02463632 | 0.55074168 | 0.214 | 0.104 | 1 | 0.86528499 | 0.0486657 |
| Slc35c1 | 0.07537979 | 0.72738662 | 0.125 | 0.062 | 1 | 0.02465194 | 1.34222143 | 0.139 | 0.052 | 1 | 0.07537979 | 0.04869616 |
| Srebf1 | 0.31644696 | -0.4286046 | 0.344 | 0.389 | 1 | 0.02468187 | 0.30548089 | 0.378 | 0.24 | 1 | 0.31644696 | 0.04875455 |
| Serpine3 | 0.32025586 | -1.577468 | 0 | 0.01 | 1 | 0.02471727 | -3.4473588 | 0.017 | 0.062 | 1 | 0.32025586 | 0.0488236 |
| Synpo2 | 0.0247407 | -2.6929452 | 0.01 | 0.073 | 1 | 0.02684133 | -1.7119537 | 0.08 | 0.156 | 1 | 0.02684133 | 0.0488693 |
| Ptpn3 | 0.02474978 | -0.4060995 | 0.417 | 0.606 | 1 | 0.6839422 | -0.1631695 | 0.723 | 0.719 | 1 | 0.6839422 | 0.04888701 |
| Nup107 | 0.02481166 | 0.47499946 | 0.344 | 0.212 | 1 | 0.54586782 | -0.9228321 | 0.294 | 0.302 | 1 | 0.54586782 | 0.04900769 |
| Fam76b | 0.02485759 | 0.41674968 | 0.521 | 0.358 | 1 | 0.63545704 | -0.3560783 | 0.508 | 0.51 | 1 | 0.63545704 | 0.04909728 |
| Acot11 | 0.02485897 | 1.38600616 | 0.094 | 0.031 | 1 | 0.37365275 | -1.0560987 | 0.097 | 0.125 | 1 | 0.37365275 | 0.04909998 |
| Trpc4ap | 0.02487328 | 0.2042675 | 0.844 | 0.731 | 1 | 0.25019925 | -0.2516101 | 0.861 | 0.844 | 1 | 0.25019925 | 0.04912788 |
| Nek2 | 0.02489078 | 3.46692616 | 0.042 | 0.005 | 1 | 0.1465102 | -1.8948178 | 0.004 | 0.021 | 1 | 0.1465102 | 0.04916201 |
| Garnl3 | 0.02491117 | -1.577468 | 0.115 | 0.218 | 1 | 0.70797178 | -0.7826083 | 0.126 | 0.135 | 1 | 0.70797178 | 0.04920178 |
| Lix1l | 0.79225061 | -0.145357 | 0.604 | 0.528 | 1 | 0.0249317 | 0.11677021 | 0.605 | 0.385 | 1 | 0.79225061 | 0.04924181 |
| Ndrg4 | 0.02494369 | 1.07162487 | 0.177 | 0.088 | 1 | 0.39803439 | 0.42711033 | 0.113 | 0.083 | 1 | 0.39803439 | 0.04926519 |
| Cobl | 0.02495256 | -0.7904001 | 0.51 | 0.611 | 1 | 0.70441419 | -0.5787503 | 0.739 | 0.667 | 1 | 0.70441419 | 0.04928248 |
| Gtpbp1 | 0.20861458 | -0.5027002 | 0.208 | 0.269 | 1 | 0.02495368 | -0.152314 | 0.349 | 0.188 | 1 | 0.20861458 | 0.04928467 |
| Smad6 | 0.02495925 | 0.8620641 | 0.344 | 0.233 | 1 | 0.72459484 | -0.2602245 | 0.399 | 0.354 | 1 | 0.72459484 | 0.04929553 |
| Arrdc4 | 0.28594182 | 0.24195979 | 0.281 | 0.218 | 1 | 0.0249784 | 0.26720204 | 0.462 | 0.344 | 1 | 0.28594182 | 0.04933288 |
| 2510002D24Rik | 0.97291484 | 0.28401217 | 0.24 | 0.244 | 1 | 0.02499532 | 0.3068161 | 0.282 | 0.156 | 1 | 0.97291484 | 0.04936587 |
| Ddb1 | 0.02500308 | 0.23697638 | 0.854 | 0.808 | 1 | 0.02673806 | 0.25107997 | 0.929 | 0.896 | 1 | 0.02673806 | 0.04938101 |
| Abcc9 | 0.1281217 | 1.17741954 | 0.062 | 0.026 | 1 | 0.02501269 | 1.55521516 | 0.122 | 0.042 | 1 | 0.1281217 | 0.04939975 |
| Arhgap23 | 0.03322817 | 0.25542205 | 0.677 | 0.539 | 1 | 0.0250467 | -0.5825862 | 0.84 | 0.865 | 1 | 0.03322817 | 0.04946607 |
| Gm37459 | 0.02510381 | 3.46692616 | 0.042 | 0.005 | 1 | 0.39234595 | 0.86006974 | 0.025 | 0.01 | 1 | 0.39234595 | 0.04957742 |
| Muc5ac | 0.02510381 | 6.31127528 | 0.042 | 0.005 | 1 | 0.54349536 | -1.7692869 | 0.029 | 0.042 | 1 | 0.54349536 | 0.04957742 |
| Ino80d | 0.02511133 | 0.18837312 | 0.708 | 0.575 | 1 | 0.962984 | -0.1014245 | 0.849 | 0.865 | 1 | 0.962984 | 0.04959209 |
| Lrp2bp | 0.66582162 | -0.130009 | 0.042 | 0.031 | 1 | 0.02513171 | 1.13071733 | 0.109 | 0.031 | 1 | 0.66582162 | 0.04963182 |
| Lynx1 | 0.28896054 | 0.44045394 | 0.219 | 0.166 | 1 | 0.02513898 | 0.50830641 | 0.231 | 0.115 | 1 | 0.28896054 | 0.04964598 |
| Dcxr | 0.28997527 | 0.14076107 | 0.135 | 0.093 | 1 | 0.02517869 | 0.78530197 | 0.155 | 0.062 | 1 | 0.28997527 | 0.04972341 |
| Lrp10 | 0.31809992 | -0.3644742 | 0.385 | 0.435 | 1 | 0.02521042 | 0.46219279 | 0.584 | 0.458 | 1 | 0.31809992 | 0.04978528 |
| Slc25a47 | 0.02525548 | 0.59245704 | 0.167 | 0.078 | 1 | 0.28205851 | -0.8866438 | 0.16 | 0.198 | 1 | 0.28205851 | 0.04987311 |
| Mrpl42 | 0.02529969 | 0.34743058 | 0.771 | 0.585 | 1 | 0.09689961 | 0.22966427 | 0.622 | 0.542 | 1 | 0.09689961 | 0.04995931 |
| G730003C15Rik | 0.02531735 | 2.59245704 | 0.042 | 0.005 | 1 | 0.20315163 | 1.01207283 | 0.017 | 0 | 1 | 0.20315163 | 0.04999373 |
| Cd79a | 0.02531735 | 5.13677755 | 0.042 | 0.005 | 1 | 0.37446771 | -1.0050007 | 0.067 | 0.094 | 1 | 0.37446771 | 0.04999373 |
| Gm56533 | 0.02531735 | 2.59245704 | 0.042 | 0.005 | 1 | 0.27154471 | 0.69014474 | 0.013 | 0 | 1 | 0.27154471 | 0.04999373 |
| Rad51ap1 | 0.02532056 | 3.22988696 | 0.042 | 0.005 | 1 | 0.35045411 | -0.8948178 | 0.008 | 0.021 | 1 | 0.35045411 | 0.04999998 |
| Top1mt | 0.02534061 | 1.45495351 | 0.125 | 0.052 | 1 | 0.08743045 | 0.91253716 | 0.084 | 0.031 | 1 | 0.08743045 | 0.05003907 |
| Fam207a | 0.11421141 | 0.81484946 | 0.281 | 0.207 | 1 | 0.02536237 | 0.19264508 | 0.37 | 0.219 | 1 | 0.11421141 | 0.05008149 |
| Myo6 | 0.02537846 | -0.3406102 | 0.927 | 0.953 | 1 | 0.61132298 | -0.1583788 | 0.882 | 0.906 | 1 | 0.61132298 | 0.05011286 |
| Wdr83os | 0.63790149 | 0.13662316 | 0.75 | 0.689 | 1 | 0.02538829 | 0.3794439 | 0.626 | 0.531 | 1 | 0.63790149 | 0.05013201 |
| Fchsd1 | 0.07511393 | 2.00749454 | 0.031 | 0.005 | 1 | 0.02539875 | 2.49749966 | 0.05 | 0 | 1 | 0.07511393 | 0.05015241 |
| Gdpgp1 | 0.08339615 | -1.6929452 | 0.01 | 0.052 | 1 | 0.025405 | 2.59703533 | 0.05 | 0 | 1 | 0.08339615 | 0.05016459 |
| Zfp202 | 0.40680632 | -0.2148979 | 0.052 | 0.031 | 1 | 0.025405 | 2.59703533 | 0.05 | 0 | 1 | 0.40680632 | 0.05016459 |
| Mir155hg | 0.85517292 | -0.2820121 | 0.083 | 0.088 | 1 | 0.025405 | 2.77760758 | 0.05 | 0 | 1 | 0.85517292 | 0.05016459 |
| Nudt17 | 0.63977237 | 0.59245704 | 0.021 | 0.031 | 1 | 0.02541012 | 2.69014474 | 0.05 | 0 | 1 | 0.63977237 | 0.05017456 |
| Gm10734 | 0.37203266 | 1.17741954 | 0.052 | 0.031 | 1 | 0.02542376 | 2.39058446 | 0.05 | 0 | 1 | 0.37203266 | 0.05020114 |
| Gm10135 | 0.38028858 | 1.00749454 | 0.031 | 0.016 | 1 | 0.02542376 | 2.39058446 | 0.05 | 0 | 1 | 0.38028858 | 0.05020114 |
| Gm29491 | 0.38521705 | 0.68556644 | 0.031 | 0.016 | 1 | 0.02542376 | 2.39058446 | 0.05 | 0 | 1 | 0.38521705 | 0.05020114 |
| Atg12 | 0.2597438 | -0.2830206 | 0.417 | 0.482 | 1 | 0.02543879 | -0.6946781 | 0.462 | 0.573 | 1 | 0.2597438 | 0.05023046 |
| Morn4 | 0.49997471 | 0.46692616 | 0.083 | 0.062 | 1 | 0.02547394 | 2.08246216 | 0.071 | 0.01 | 1 | 0.49997471 | 0.05029895 |
| Sh3bgrl | 0.18955577 | -0.3223823 | 0.771 | 0.736 | 1 | 0.02547707 | 0.19754625 | 0.887 | 0.833 | 1 | 0.18955577 | 0.05030506 |
| Gm30489 | 0.02548561 | -1.407543 | 0.031 | 0.109 | 1 | 0.41179613 | -1.5025003 | 0.025 | 0.042 | 1 | 0.41179613 | 0.05032171 |
| Acsf2 | 0.50219858 | 0.178975 | 0.396 | 0.337 | 1 | 0.02550425 | 0.44146562 | 0.315 | 0.177 | 1 | 0.50219858 | 0.05035804 |
| Gm45669 | 0.02550584 | 0.74446013 | 0.219 | 0.119 | 1 | 0.08960365 | 0.79250646 | 0.202 | 0.125 | 1 | 0.08960365 | 0.05036114 |
| H2ac4 | 0.02553784 | 2.81484946 | 0.042 | 0.005 | 1 | 0.74957918 | -0.8244284 | 0.025 | 0.031 | 1 | 0.74957918 | 0.05042351 |
| Wdr38 | 0.0255382 | 2.32942263 | 0.042 | 0.005 | 1 | 0.85710544 | -1.3098553 | 0.008 | 0.01 | 1 | 0.85710544 | 0.0504242 |
| Cyb561d1 | 0.0255382 | 2.32942263 | 0.042 | 0.005 | 1 | 0.20315163 | 1.01207283 | 0.017 | 0 | 1 | 0.20315163 | 0.0504242 |
| Gm12758 | 0.0255382 | 2.32942263 | 0.042 | 0.005 | 1 | 0.52072676 | -0.7248928 | 0.021 | 0.01 | 1 | 0.52072676 | 0.0504242 |
| Neu3 | 0.0255382 | 2.32942263 | 0.042 | 0.005 | 1 | 0.04177162 | 1.86006974 | 0.063 | 0.01 | 1 | 0.04177162 | 0.0504242 |
| Pbx4 | 0.0255382 | 2.32942263 | 0.042 | 0.005 | 1 | 0.69586392 | -0.3098553 | 0.042 | 0.052 | 1 | 0.69586392 | 0.0504242 |
| Unc13a | 0.0255382 | 2.32942263 | 0.042 | 0.005 | 1 | 0.9959736 | -0.3098553 | 0.021 | 0.021 | 1 | 0.9959736 | 0.0504242 |
| Med9os | 0.0255382 | 2.32942263 | 0.042 | 0.005 | 1 | 0.40647274 | -0.9879272 | 0.017 | 0.031 | 1 | 0.40647274 | 0.0504242 |
| 2900041M22Rik | 0.0255382 | 2.32942263 | 0.042 | 0.005 | 1 | 0.11764843 | 1.86006974 | 0.025 | 0 | 1 | 0.11764843 | 0.0504242 |
| Gm32934 | 0.0255382 | 2.32942263 | 0.042 | 0.005 | 1 | 0.27154471 | 1.69014474 | 0.013 | 0 | 1 | 0.27154471 | 0.0504242 |
| Cyp46a1 | 0.0255382 | 2.32942263 | 0.042 | 0.005 | 1 | 0.43079893 | 0.56461386 | 0.038 | 0.021 | 1 | 0.43079893 | 0.0504242 |
| Nhlrc1 | 0.0255382 | 2.32942263 | 0.042 | 0.005 | 1 | 0.20315163 | 1.01207283 | 0.017 | 0 | 1 | 0.20315163 | 0.0504242 |
| Gm2237 | 0.0255382 | 2.32942263 | 0.042 | 0.005 | 1 | 0.15394568 | 1.69014474 | 0.021 | 0 | 1 | 0.15394568 | 0.0504242 |
| Gm15943 | 0.0255382 | 2.32942263 | 0.042 | 0.005 | 1 | 0.50887612 | -1.3098553 | 0.004 | 0.01 | 1 | 0.50887612 | 0.0504242 |
| Gm39556 | 0.0255382 | 2.32942263 | 0.042 | 0.005 | 1 | 0.34874556 | -0.8244284 | 0.008 | 0.021 | 1 | 0.34874556 | 0.0504242 |
| Fam186a | 0.0255382 | 2.32942263 | 0.042 | 0.005 | 1 | 0.86613737 | -0.3098553 | 0.008 | 0.01 | 1 | 0.86613737 | 0.0504242 |
| Spink5 | 0.0255382 | 2.32942263 | 0.042 | 0.005 | 1 | 0.83706547 | -0.9879272 | 0.025 | 0.021 | 1 | 0.83706547 | 0.0504242 |
| Nit1 | 0.05001992 | 0.48812038 | 0.292 | 0.181 | 1 | 0.0255427 | 0.37664526 | 0.307 | 0.177 | 1 | 0.05001992 | 0.05043298 |
| Ace | 0.61182672 | -0.7294711 | 0.021 | 0.031 | 1 | 0.02557009 | 2.01207283 | 0.071 | 0.01 | 1 | 0.61182672 | 0.05048635 |
| Pnrc2 | 0.41567182 | -0.2285728 | 0.323 | 0.373 | 1 | 0.02557584 | 0.62212631 | 0.403 | 0.281 | 1 | 0.41567182 | 0.05049755 |
| Chrac1 | 0.12196739 | 0.52534284 | 0.417 | 0.332 | 1 | 0.02560678 | 0.49749966 | 0.303 | 0.177 | 1 | 0.12196739 | 0.05055784 |
| Gse1 | 0.05204729 | -0.8380898 | 0.542 | 0.585 | 1 | 0.02563897 | -0.7293387 | 0.576 | 0.74 | 1 | 0.05204729 | 0.05062058 |
| Gm43330 | 0.81708725 | -0.3144336 | 0.031 | 0.036 | 1 | 0.02565719 | 0.74077081 | 0.092 | 0.021 | 1 | 0.81708725 | 0.05065608 |
| Poli | 0.24179987 | 0.90795886 | 0.094 | 0.057 | 1 | 0.02566532 | 1.93807225 | 0.071 | 0.01 | 1 | 0.24179987 | 0.05067194 |
| Acot8 | 0.44615298 | 0.15188445 | 0.188 | 0.15 | 1 | 0.02569811 | 0.71567983 | 0.185 | 0.083 | 1 | 0.44615298 | 0.05073584 |

| | | | | | | | | | | | | |
|---|---|---|---|---|---|---|---|---|---|---|---|---|
| Wbp11 | 0.02570467 | 0.40115839 | 0.802 | 0.736 | 1 | 0.02960177 | 0.27510724 | 0.798 | 0.76 | 1 | 0.02960177 | 0.05074861 |
| Pnpo | 0.02572105 | -1.0994207 | 0.115 | 0.218 | 1 | 0.11428227 | 1.06865636 | 0.092 | 0.042 | 1 | 0.11428227 | 0.05078052 |
| Abca8b | 0.29782773 | 0.60086928 | 0.24 | 0.192 | 1 | 0.02574186 | 0.88654195 | 0.155 | 0.062 | 1 | 0.29782773 | 0.05082108 |
| Rap1b | 0.11142978 | 0.18590301 | 0.99 | 0.979 | 1 | 0.02580106 | 0.20945206 | 0.992 | 0.958 | 1 | 0.11142978 | 0.05093643 |
| Hddc3 | 0.3390465 | -0.577468 | 0.031 | 0.057 | 1 | 0.02583513 | 1.86006974 | 0.071 | 0.01 | 1 | 0.3390465 | 0.0510028 |
| Wdr41 | 0.65079543 | -0.2800821 | 0.271 | 0.29 | 1 | 0.02583702 | -0.1229771 | 0.319 | 0.167 | 1 | 0.65079543 | 0.05100649 |
| Ulk1 | 0.02586453 | -0.8493805 | 0.396 | 0.466 | 1 | 0.71154433 | -0.2202181 | 0.597 | 0.604 | 1 | 0.71154433 | 0.05106009 |
| Pank4 | 0.02587795 | 0.74446013 | 0.188 | 0.093 | 1 | 0.19161468 | 0.30019822 | 0.189 | 0.125 | 1 | 0.19161468 | 0.05108623 |
| Cdk10 | 0.02593104 | 0.81484946 | 0.188 | 0.093 | 1 | 0.05794964 | 0.30019822 | 0.21 | 0.115 | 1 | 0.05794964 | 0.05118966 |
| Ak3 | 0.02598567 | 0.75292171 | 0.375 | 0.259 | 1 | 0.6205242 | -0.1712017 | 0.567 | 0.521 | 1 | 0.6205242 | 0.05129608 |
| Celf1 | 0.03823711 | 0.25199766 | 0.906 | 0.725 | 1 | 0.026055 | -0.4771801 | 0.954 | 0.969 | 1 | 0.03823711 | 0.05143114 |
| Tex29 | 0.02619831 | 2.00749454 | 0.042 | 0.005 | 1 | 0.02605524 | -2.8948178 | 0 | 0.021 | 1 | 0.02619831 | 0.0514316 |
| 5730420D15Rik | 0.03210862 | -2.9925055 | 0 | 0.047 | 1 | 0.02605524 | -3.6317834 | 0 | 0.021 | 1 | 0.03210862 | 0.0514316 |
| Tfap2c | 0.04371754 | -2.3144336 | 0 | 0.041 | 1 | 0.02605524 | -3.3098553 | 0 | 0.021 | 1 | 0.04371754 | 0.0514316 |
| Mybpc2 | 0.04512622 | 2.59245704 | 0.021 | 0 | 1 | 0.02605524 | -3.3098553 | 0 | 0.021 | 1 | 0.04512622 | 0.0514316 |
| Ybx2 | 0.04623052 | -1.6929452 | 0.01 | 0.062 | 1 | 0.02605524 | -2.8948178 | 0 | 0.021 | 1 | 0.04623052 | 0.0514316 |
| Trim5 | 0.05972787 | -2.4519371 | 0 | 0.036 | 1 | 0.02605524 | -4.4797803 | 0 | 0.021 | 1 | 0.05972787 | 0.0514316 |
| Dmrt1 | 0.05973548 | -3.0799683 | 0 | 0.036 | 1 | 0.02605524 | -2.8948178 | 0 | 0.021 | 1 | 0.05973548 | 0.0514316 |
| Cenpi | 0.06812307 | 1.49292136 | 0.062 | 0.021 | 1 | 0.02605524 | -3.3098553 | 0 | 0.021 | 1 | 0.06812307 | 0.0514316 |
| Tas1r1 | 0.07391189 | 3.32942263 | 0.031 | 0.005 | 1 | 0.02605524 | -2.8948178 | 0 | 0.021 | 1 | 0.07391189 | 0.0514316 |
| Armc3 | 0.07511393 | 2.00749454 | 0.031 | 0.005 | 1 | 0.02605524 | -3.8948178 | 0 | 0.021 | 1 | 0.07511393 | 0.0514316 |
| Aurkb | 0.0787868 | 2.59245704 | 0.042 | 0.01 | 1 | 0.02605524 | -3.8948178 | 0 | 0.021 | 1 | 0.0787868 | 0.0514316 |
| Gm38287 | 0.0819164 | -1.9925055 | 0 | 0.031 | 1 | 0.02605524 | -3.3098553 | 0 | 0.021 | 1 | 0.0819164 | 0.0514316 |
| Tgtp1 | 0.08192263 | -2.577468 | 0 | 0.031 | 1 | 0.02605524 | -3.8948178 | 0 | 0.021 | 1 | 0.08192263 | 0.0514316 |
| Gm12866 | 0.08192636 | -2.8993961 | 0 | 0.031 | 1 | 0.02605524 | -2.8948178 | 0 | 0.021 | 1 | 0.08192636 | 0.0514316 |
| Camk1g | 0.11302151 | -1.7998604 | 0 | 0.026 | 1 | 0.02605524 | -3.6317834 | 0 | 0.021 | 1 | 0.11302151 | 0.0514316 |
| Gm11659 | 0.11302151 | -2.1624305 | 0 | 0.026 | 1 | 0.02605524 | -3.6317834 | 0 | 0.021 | 1 | 0.11302151 | 0.0514316 |
| Gm14009 | 0.11302678 | -1.9925055 | 0 | 0.026 | 1 | 0.02605524 | -3.6317834 | 0 | 0.021 | 1 | 0.11302678 | 0.0514316 |
| Styk1 | 0.15729403 | -1.7998604 | 0 | 0.021 | 1 | 0.02605524 | -3.8948178 | 0 | 0.021 | 1 | 0.15729403 | 0.0514316 |
| Smco3 | 0.15729403 | -1.577468 | 0 | 0.021 | 1 | 0.02605524 | -3.3098553 | 0 | 0.021 | 1 | 0.15729403 | 0.0514316 |
| Gm44767 | 0.15729403 | -1.7998604 | 0 | 0.021 | 1 | 0.02605524 | -5.3098553 | 0 | 0.021 | 1 | 0.15729403 | 0.0514316 |
| Foxs1 | 0.15729781 | -1.9925055 | 0 | 0.021 | 1 | 0.02605524 | -2.8948178 | 0 | 0.021 | 1 | 0.15729781 | 0.0514316 |
| Gm41821 | 0.15729781 | -2.3144336 | 0 | 0.021 | 1 | 0.02605524 | -4.3098553 | 0 | 0.021 | 1 | 0.15729781 | 0.0514316 |
| Gabra4 | 0.15837881 | 2.00749454 | 0.01 | 0 | 1 | 0.02605524 | -3.3098553 | 0 | 0.021 | 1 | 0.15837881 | 0.0514316 |
| Tmem184a | 0.15837881 | 2.00749454 | 0.01 | 0 | 1 | 0.02605524 | -2.8948178 | 0 | 0.021 | 1 | 0.15837881 | 0.0514316 |
| Gm56577 | 0.15837881 | 2.00749454 | 0.01 | 0 | 1 | 0.02605524 | -2.8948178 | 0 | 0.021 | 1 | 0.15837881 | 0.0514316 |
| 4930572G02Rik | 0.15837881 | 2.00749454 | 0.01 | 0 | 1 | 0.02605524 | -2.8948178 | 0 | 0.021 | 1 | 0.15837881 | 0.0514316 |
| Gm56611 | 0.15837881 | 2.00749454 | 0.01 | 0 | 1 | 0.02605524 | -2.8948178 | 0 | 0.021 | 1 | 0.15837881 | 0.0514316 |
| Cd160 | 0.17265227 | 1.59245704 | 0.042 | 0.016 | 1 | 0.02605524 | -3.3098553 | 0 | 0.021 | 1 | 0.17265227 | 0.0514316 |
| Gm11592 | 0.20158871 | 1.42253204 | 0.031 | 0.01 | 1 | 0.02605524 | -3.3098553 | 0 | 0.021 | 1 | 0.20158871 | 0.0514316 |
| Gdnf | 0.21062121 | -2.265524 | 0.021 | 0.052 | 1 | 0.02605524 | -3.8948178 | 0 | 0.021 | 1 | 0.21062121 | 0.0514316 |
| Ctcflos | 0.21062169 | -1.2148979 | 0.021 | 0.052 | 1 | 0.02605524 | -4.3098553 | 0 | 0.021 | 1 | 0.21062169 | 0.0514316 |
| 1700057G04Rik | 0.21460233 | -0.4779323 | 0.01 | 0.036 | 1 | 0.02605524 | -2.8948178 | 0 | 0.021 | 1 | 0.21460233 | 0.0514316 |
| Gm40841 | 0.21621572 | -1.240433 | 0.021 | 0.052 | 1 | 0.02605524 | -3.3098553 | 0 | 0.021 | 1 | 0.21621572 | 0.0514316 |
| Bdkrb1 | 0.21708019 | 2.00749454 | 0.021 | 0.005 | 1 | 0.02605524 | -2.8948178 | 0 | 0.021 | 1 | 0.21708019 | 0.0514316 |
| Gm44036 | 0.21867159 | 1.59245704 | 0.021 | 0.005 | 1 | 0.02605524 | -3.6317834 | 0 | 0.021 | 1 | 0.21867159 | 0.0514316 |
| Adap1 | 0.22187388 | 1.00749454 | 0.021 | 0.005 | 1 | 0.02605524 | -3.3098553 | 0 | 0.021 | 1 | 0.22187388 | 0.0514316 |
| Cyp3a57 | 0.22187388 | -1.3144336 | 0 | 0.016 | 1 | 0.02605524 | -4.3098553 | 0 | 0.021 | 1 | 0.22187388 | 0.0514316 |
| Atp2c2 | 0.22187388 | -0.2148979 | 0.021 | 0.005 | 1 | 0.02605524 | -3.6317834 | 0 | 0.021 | 1 | 0.22187388 | 0.0514316 |
| Acer1 | 0.22187388 | -1.3144336 | 0 | 0.016 | 1 | 0.02605524 | -2.8948178 | 0 | 0.021 | 1 | 0.22187388 | 0.0514316 |
| Lad1 | 0.22187575 | -0.9925055 | 0 | 0.016 | 1 | 0.02605524 | -3.3098553 | 0 | 0.021 | 1 | 0.22187575 | 0.0514316 |
| Nnmt | 0.22187575 | -0.9925055 | 0 | 0.016 | 1 | 0.02605524 | -3.3098553 | 0 | 0.021 | 1 | 0.22187575 | 0.0514316 |
| Flt4 | 0.22187575 | -0.9925055 | 0 | 0.016 | 1 | 0.02605524 | -2.8948178 | 0 | 0.021 | 1 | 0.22187575 | 0.0514316 |
| Gm48860 | 0.22187575 | -0.9925055 | 0 | 0.016 | 1 | 0.02605524 | -2.8948178 | 0 | 0.021 | 1 | 0.22187575 | 0.0514316 |
| Cdh6 | 0.22187575 | -0.9925055 | 0 | 0.016 | 1 | 0.02605524 | -2.8948178 | 0 | 0.021 | 1 | 0.22187575 | 0.0514316 |
| Phtf1os | 0.2825047 | -1.3144336 | 0.01 | 0.031 | 1 | 0.02605524 | -3.3098553 | 0 | 0.021 | 1 | 0.2825047 | 0.0514316 |
| Gm5533 | 0.32025586 | -0.9925055 | 0 | 0.01 | 1 | 0.02605524 | -3.6317834 | 0 | 0.021 | 1 | 0.32025586 | 0.0514316 |
| Tm4sf4 | 0.32025586 | -1.577468 | 0 | 0.01 | 1 | 0.02605524 | -3.6317834 | 0 | 0.021 | 1 | 0.32025586 | 0.0514316 |
| Slc46a2 | 0.32025586 | -0.577468 | 0 | 0.01 | 1 | 0.02605524 | -2.8948178 | 0 | 0.021 | 1 | 0.32025586 | 0.0514316 |
| Slc5a9 | 0.32025586 | -0.577468 | 0 | 0.01 | 1 | 0.02605524 | -2.8948178 | 0 | 0.021 | 1 | 0.32025586 | 0.0514316 |
| Gm6559 | 0.32025586 | -0.9925055 | 0 | 0.01 | 1 | 0.02605524 | -2.8948178 | 0 | 0.021 | 1 | 0.32025586 | 0.0514316 |
| Slc16a13 | 0.32025586 | -0.9925055 | 0 | 0.01 | 1 | 0.02605524 | -2.8948178 | 0 | 0.021 | 1 | 0.32025586 | 0.0514316 |
| Osr1 | 0.32025586 | -0.577468 | 0 | 0.01 | 1 | 0.02605524 | -6.3542494 | 0 | 0.021 | 1 | 0.32025586 | 0.0514316 |
| Onecut2 | 0.32025586 | -1.3144336 | 0 | 0.01 | 1 | 0.02605524 | -3.6317834 | 0 | 0.021 | 1 | 0.32025586 | 0.0514316 |
| Pgm5 | 0.46593104 | -1.0281294 | 0.062 | 0.041 | 1 | 0.02605524 | -3.3098553 | 0 | 0.021 | 1 | 0.46593104 | 0.0514316 |
| E2f2 | 0.4692344 | 1.59245704 | 0.021 | 0.01 | 1 | 0.02605524 | -4.4797803 | 0 | 0.021 | 1 | 0.4692344 | 0.0514316 |
| Them7 | 0.47150075 | 1.74446013 | 0.021 | 0.01 | 1 | 0.02605524 | -2.8948178 | 0 | 0.021 | 1 | 0.47150075 | 0.0514316 |
| Gm56593 | 0.52961488 | -0.3144336 | 0.01 | 0.021 | 1 | 0.02605524 | -2.8948178 | 0 | 0.021 | 1 | 0.52961488 | 0.0514316 |
| Lzts1 | 0.61550914 | -0.407543 | 0.021 | 0.031 | 1 | 0.02605524 | -2.8948178 | 0 | 0.021 | 1 | 0.61550914 | 0.0514316 |
| Fgf22 | 0.61737927 | 1.00749454 | 0.01 | 0.005 | 1 | 0.02605524 | -3.6317834 | 0 | 0.021 | 1 | 0.61737927 | 0.0514316 |
| Dscc1 | 0.61737927 | 1.00749454 | 0.01 | 0.005 | 1 | 0.02605524 | -2.8948178 | 0 | 0.021 | 1 | 0.61737927 | 0.0514316 |
| En2 | 0.72574709 | -0.3144336 | 0.01 | 0.016 | 1 | 0.02605524 | -3.3098553 | 0 | 0.021 | 1 | 0.72574709 | 0.0514316 |
| Fbxo41 | 0.72574709 | -0.3144336 | 0.01 | 0.016 | 1 | 0.02605524 | -2.8948178 | 0 | 0.021 | 1 | 0.72574709 | 0.0514316 |
| Gm35549 | 0.72574709 | -0.3144336 | 0.01 | 0.016 | 1 | 0.02605524 | -4.1172102 | 0 | 0.021 | 1 | 0.72574709 | 0.0514316 |
| 4930556N13Rik | 0.72574709 | -0.3144336 | 0.01 | 0.016 | 1 | 0.02605524 | -2.8948178 | 0 | 0.021 | 1 | 0.72574709 | 0.0514316 |
| Tlr13 | 0.72574709 | -0.3144336 | 0.01 | 0.016 | 1 | 0.02605524 | -2.8948178 | 0 | 0.021 | 1 | 0.72574709 | 0.0514316 |
| Cdh15 | 0.73407659 | 0.27052894 | 0.01 | 0.016 | 1 | 0.02605524 | -3.3098553 | 0 | 0.021 | 1 | 0.73407659 | 0.0514316 |
| 4930565N06Rik | 0.99275828 | 0.52206771 | 0.021 | 0.021 | 1 | 0.02605524 | -3.3098553 | 0 | 0.021 | 1 | 0.99275828 | 0.0514316 |
| Ifi211 | 0.993208 | 2.00749454 | 0.01 | 0.01 | 1 | 0.02605524 | -3.3098553 | 0 | 0.021 | 1 | 0.993208 | 0.0514316 |
| Pcsk2 | 0.99320802 | 1.00749454 | 0.01 | 0.01 | 1 | 0.02605524 | -5.3973181 | 0 | 0.021 | 1 | 0.99320802 | 0.0514316 |
| 2310039L15Rik | 0.99320802 | 1.68556644 | 0.01 | 0.01 | 1 | 0.02605524 | -3.3098553 | 0 | 0.021 | 1 | 0.99320802 | 0.0514316 |
| 2810030D12Rik | 0.99758604 | -0.2148979 | 0.021 | 0.021 | 1 | 0.02605524 | -2.8948178 | 0 | 0.021 | 1 | 0.99758604 | 0.0514316 |
| Hgf | 1 | 0.42253204 | 0.01 | 0.01 | 1 | 0.02605524 | -2.8948178 | 0 | 0.021 | 1 | 1 | 0.0514316 |
| Npas1 | 1 | 0.42253204 | 0.01 | 0.01 | 1 | 0.02605524 | -3.3098553 | 0 | 0.021 | 1 | 1 | 0.0514316 |

| | | | | | | | | | | | | |
|---|---|---|---|---|---|---|---|---|---|---|---|---|
| Ntrk3 | 1 | -0.577468 | 0.01 | 0.01 | 1 | 0.02605524 | -3.8948178 | 0 | 0.021 | 1 | 1 | 0.0514316 |
| C130071C03Rik | 1 | -0.3144336 | 0.01 | 0.01 | 1 | 0.02605524 | -2.8948178 | 0 | 0.021 | 1 | 1 | 0.0514316 |
| Tymp | 1 | 0.42253204 | 0.01 | 0.01 | 1 | 0.02605524 | -3.3098553 | 0 | 0.021 | 1 | 1 | 0.0514316 |
| Zc2hc1a | 0.02609016 | 0.66354014 | 0.344 | 0.228 | 1 | 0.52449413 | -0.1457359 | 0.391 | 0.333 | 1 | 0.52449413 | 0.05149963 |
| Ngf | 0.02617208 | -1.684802 | 0.188 | 0.29 | 1 | 0.79209349 | -1.9350069 | 0.084 | 0.073 | 1 | 0.79209349 | 0.05165918 |
| Fam177a | 0.71031978 | 0.35541784 | 0.062 | 0.052 | 1 | 0.02618328 | 0.37820073 | 0.109 | 0.031 | 1 | 0.71031978 | 0.05168099 |
| Dusp28 | 0.02618575 | 1.15949763 | 0.156 | 0.073 | 1 | 0.04192925 | 1.77760758 | 0.063 | 0.01 | 1 | 0.04192925 | 0.0516858 |
| 2810407A14Rik | 0.02619831 | 2.00749454 | 0.042 | 0.005 | 1 | 0.11765123 | 2.59703533 | 0.025 | 0 | 1 | 0.11765123 | 0.05171027 |
| Kin | 0.43553299 | 0.25076569 | 0.469 | 0.425 | 1 | 0.02619917 | 0.14637392 | 0.504 | 0.344 | 1 | 0.43553299 | 0.05171194 |
| Tiam1 | 0.02620655 | -0.7055584 | 0.76 | 0.777 | 1 | 0.40491693 | -0.6676679 | 0.744 | 0.74 | 1 | 0.40491693 | 0.05172631 |
| Jmjd1c | 0.04140728 | -0.3043007 | 1 | 0.984 | 1 | 0.02621759 | -0.4819475 | 0.992 | 0.99 | 1 | 0.04140728 | 0.05174782 |
| Armcx1 | 0.09249769 | 0.16361374 | 0.302 | 0.207 | 1 | 0.02628421 | 0.64283902 | 0.332 | 0.208 | 1 | 0.09249769 | 0.05187756 |
| Mfn2 | 0.02629187 | 0.5275016 | 0.333 | 0.207 | 1 | 0.07393935 | -0.368749 | 0.328 | 0.198 | 1 | 0.07393935 | 0.05189249 |
| Gli2 | 0.02631955 | -1.190919 | 0.24 | 0.358 | 1 | 0.89404838 | -0.6094155 | 0.513 | 0.427 | 1 | 0.89404838 | 0.05194639 |
| Ezh2 | 0.12801241 | -0.5669756 | 0.458 | 0.518 | 1 | 0.02633701 | -0.8476358 | 0.504 | 0.656 | 1 | 0.12801241 | 0.05198038 |
| Borcs5 | 0.02637116 | 1.64492446 | 0.167 | 0.083 | 1 | 0.19554301 | 0.24034182 | 0.139 | 0.083 | 1 | 0.19554301 | 0.05204687 |
| Gm56507 | 0.0263832 | 1.15949763 | 0.094 | 0.031 | 1 | 0.63913175 | -1.2167459 | 0.05 | 0.062 | 1 | 0.63913175 | 0.05207033 |
| Arhgef9 | 0.02638448 | 1.68556644 | 0.177 | 0.093 | 1 | 0.10322222 | 0.17013768 | 0.168 | 0.094 | 1 | 0.10322222 | 0.05207283 |
| Pnisr | 0.02639798 | 0.25975749 | 0.938 | 0.953 | 1 | 0.04829964 | 0.15771129 | 0.966 | 0.969 | 1 | 0.04829964 | 0.05209912 |
| Cstdc2 | 0.42186046 | -1.2555399 | 0.031 | 0.052 | 1 | 0.02639898 | -1.961932 | 0.017 | 0.062 | 1 | 0.42186046 | 0.05210106 |
| Efhd2 | 0.78178628 | -0.4806416 | 0.688 | 0.611 | 1 | 0.02639921 | 0.54381039 | 0.676 | 0.646 | 1 | 0.78178628 | 0.0521015 |
| Reno1 | 0.02640642 | 1.35541784 | 0.125 | 0.052 | 1 | 0.07047002 | 1.12310414 | 0.088 | 0.031 | 1 | 0.07047002 | 0.05211554 |
| Gm12204 | 0.02641989 | 1.74446013 | 0.042 | 0.005 | 1 | 0.07095091 | -1.7952821 | 0.017 | 0.052 | 1 | 0.07095091 | 0.05214177 |
| Gm50462 | 0.02641989 | 1.74446013 | 0.042 | 0.005 | 1 | 0.24413474 | -1.6317834 | 0.013 | 0.031 | 1 | 0.24413474 | 0.05214177 |
| Awat2 | 0.02641989 | 0.52206771 | 0.042 | 0.005 | 1 | 0.27154471 | 0.69014474 | 0.013 | 0 | 1 | 0.27154471 | 0.05214177 |
| Lyrm2 | 0.10748801 | 0.54452256 | 0.292 | 0.207 | 1 | 0.02643285 | 0.53341923 | 0.214 | 0.104 | 1 | 0.10748801 | 0.05216701 |
| Tagln | 0.15299208 | -3.3456523 | 0.104 | 0.155 | 1 | 0.02645694 | -2.1172102 | 0.08 | 0.156 | 1 | 0.15299208 | 0.0522139 |
| Mtus2 | 0.03328727 | 2.27052894 | 0.104 | 0.041 | 1 | 0.02653152 | 2.10518224 | 0.088 | 0.021 | 1 | 0.03328727 | 0.05235912 |
| Zfp869 | 0.16301476 | -0.4291452 | 0.396 | 0.466 | 1 | 0.02655983 | 0.18629797 | 0.517 | 0.323 | 1 | 0.16301476 | 0.05241424 |
| Ube2h | 0.14489332 | -0.2992612 | 0.885 | 0.912 | 1 | 0.0266023 | -0.5596841 | 0.979 | 0.969 | 1 | 0.14489332 | 0.05249692 |
| Nsd2 | 0.02664105 | 0.5283267 | 0.583 | 0.472 | 1 | 0.25985536 | -0.4562447 | 0.63 | 0.677 | 1 | 0.25985536 | 0.05257236 |
| Epha5 | 0.4146983 | -1.7998604 | 0.031 | 0.052 | 1 | 0.02668736 | -1.8948178 | 0.017 | 0.062 | 1 | 0.4146983 | 0.0526625 |
| Faim | 0.21894804 | -0.7767768 | 0.073 | 0.119 | 1 | 0.02669492 | 1.64434105 | 0.105 | 0.031 | 1 | 0.21894804 | 0.05267721 |
| Lrrc40 | 0.26538179 | 0.2590333 | 0.219 | 0.161 | 1 | 0.02671082 | 0.55687821 | 0.197 | 0.094 | 1 | 0.26538179 | 0.05270817 |
| 2310009B15Rik | 0.85987135 | 0.10937415 | 0.323 | 0.321 | 1 | 0.02672663 | 0.34222143 | 0.286 | 0.156 | 1 | 0.85987135 | 0.05273895 |
| Morn1 | 0.0267786 | 1.08766489 | 0.25 | 0.145 | 1 | 0.60388654 | -0.5025003 | 0.218 | 0.177 | 1 | 0.60388654 | 0.0528401 |
| Atp11b | 0.13930211 | -0.3959859 | 0.667 | 0.694 | 1 | 0.02678543 | -0.5514764 | 0.807 | 0.865 | 1 | 0.13930211 | 0.05285339 |
| Sox12 | 0.02679972 | 1.14499806 | 0.094 | 0.031 | 1 | 0.07198422 | 0.52664601 | 0.088 | 0.031 | 1 | 0.07198422 | 0.05288122 |
| Gm11906 | 0.02680949 | -3.2148979 | 0.021 | 0.088 | 1 | 0.63288235 | -0.1399303 | 0.08 | 0.062 | 1 | 0.63288235 | 0.05290023 |
| Klc2 | 0.02681239 | 0.54354744 | 0.25 | 0.14 | 1 | 0.04299282 | 0.45953181 | 0.256 | 0.146 | 1 | 0.04299282 | 0.05290588 |
| Rcor2 | 0.02682642 | 1.70793425 | 0.083 | 0.026 | 1 | 0.79021828 | -0.6094155 | 0.038 | 0.031 | 1 | 0.79021828 | 0.05293319 |
| Il16 | 0.02683224 | 0.12773365 | 0.854 | 0.772 | 1 | 0.04769682 | -0.1033539 | 0.929 | 0.677 | 1 | 0.04769682 | 0.05294451 |
| Gm14636 | 0.52313476 | -0.9925055 | 0.01 | 0.021 | 1 | 0.02685374 | -0.9175378 | 0.046 | 0.115 | 1 | 0.52313476 | 0.05298635 |
| Ccdc191 | 0.02686565 | 1.12948506 | 0.24 | 0.14 | 1 | 0.17342964 | -0.131518 | 0.13 | 0.073 | 1 | 0.17342964 | 0.05300953 |
| Gnao1 | 0.11192229 | -1.3144336 | 0.073 | 0.135 | 1 | 0.02686635 | -1.2728611 | 0.172 | 0.26 | 1 | 0.11192229 | 0.05301091 |
| Trrap | 0.0268858 | 0.1604754 | 0.781 | 0.668 | 1 | 0.945083 | -0.166683 | 0.857 | 0.844 | 1 | 0.945083 | 0.05304875 |
| Tbcc | 0.62160822 | -0.2879613 | 0.177 | 0.202 | 1 | 0.02688912 | 1.14957636 | 0.151 | 0.062 | 1 | 0.62160822 | 0.05305522 |
| Prr13 | 0.02691007 | -0.7212034 | 0.417 | 0.539 | 1 | 0.3575182 | 0.13922381 | 0.563 | 0.552 | 1 | 0.3575182 | 0.05309598 |
| Brd8dc | 0.14594947 | 0.48554183 | 0.208 | 0.14 | 1 | 0.02692308 | 0.50957249 | 0.244 | 0.125 | 1 | 0.14594947 | 0.0531213 |
| Tspyl4 | 0.34483835 | 0.20543391 | 0.302 | 0.249 | 1 | 0.02692358 | 0.52021974 | 0.206 | 0.104 | 1 | 0.34483835 | 0.05312229 |
| Peak1 | 0.10441398 | 0.20576416 | 0.562 | 0.627 | 1 | 0.02693353 | -0.7573142 | 0.513 | 0.635 | 1 | 0.10441398 | 0.05314164 |
| Npm2 | 0.06202887 | 1.20013961 | 0.073 | 0.026 | 1 | 0.0269492 | 1.59703533 | 0.105 | 0.031 | 1 | 0.06202887 | 0.05317215 |
| Klhdc8b | 0.07187312 | 0.57686018 | 0.208 | 0.124 | 1 | 0.02697987 | 0.31163311 | 0.244 | 0.125 | 1 | 0.07187312 | 0.05323182 |
| Hcar2 | 0.22187388 | -1.3144336 | 0 | 0.016 | 1 | 0.02698127 | -1.7952821 | 0.017 | 0.062 | 1 | 0.22187388 | 0.05323456 |
| Bysl | 0.25353001 | 0.54806292 | 0.188 | 0.135 | 1 | 0.0269824 | 0.30019822 | 0.218 | 0.104 | 1 | 0.25353001 | 0.05323674 |
| H1f0 | 0.02701777 | -0.4564086 | 0.635 | 0.725 | 1 | 0.40361527 | 0.39246419 | 0.66 | 0.656 | 1 | 0.40361527 | 0.05330558 |
| Zkscan8 | 0.02706324 | 1.37006462 | 0.146 | 0.067 | 1 | 0.51392872 | 0.27510724 | 0.084 | 0.062 | 1 | 0.51392872 | 0.05339407 |
| Ccn3 | 0.11600757 | -1.577468 | 0.01 | 0.047 | 1 | 0.0270678 | -3.1578522 | 0.029 | 0.083 | 1 | 0.11600757 | 0.05340294 |
| Dlg1 | 0.54183289 | -0.1120871 | 0.875 | 0.927 | 1 | 0.02709887 | -0.5087976 | 0.941 | 0.958 | 1 | 0.54183289 | 0.05346339 |
| Ibsp | 0.02717959 | 1.78306297 | 0.073 | 0.021 | 1 | 0.79667672 | 3.48022167 | 0.025 | 0.021 | 1 | 0.79667672 | 0.05362046 |
| Prkacb | 0.02720725 | 0.25293488 | 0.812 | 0.642 | 1 | 0.98010352 | -0.2462634 | 0.811 | 0.771 | 1 | 0.98010352 | 0.05367427 |
| Pnpla7 | 0.02724521 | 0.77616899 | 0.188 | 0.093 | 1 | 0.33279859 | -0.1463565 | 0.164 | 0.115 | 1 | 0.33279859 | 0.05374812 |
| Itprid1 | 0.72852243 | -0.2148979 | 0.01 | 0.016 | 1 | 0.02726389 | -1.7248928 | 0.017 | 0.062 | 1 | 0.72852243 | 0.05378445 |
| Gm15417 | 0.02727221 | -1.1354634 | 0.115 | 0.218 | 1 | 0.03664029 | 0.9730787 | 0.223 | 0.125 | 1 | 0.03664029 | 0.05380064 |
| Mtrf1l | 0.46553195 | 0.30705482 | 0.219 | 0.181 | 1 | 0.0272745 | 0.83150059 | 0.223 | 0.115 | 1 | 0.46553195 | 0.0538051 |
| Syngap1 | 0.02728142 | 0.80818573 | 0.208 | 0.109 | 1 | 0.46312321 | -0.9966974 | 0.143 | 0.167 | 1 | 0.46312321 | 0.05381856 |
| Lrrc49 | 0.02732316 | 0.4850735 | 0.646 | 0.513 | 1 | 0.11421864 | -0.1740785 | 0.609 | 0.417 | 1 | 0.11421864 | 0.05389977 |
| C030013E06Rik | 0.02736929 | -1.8993961 | 0.052 | 0.135 | 1 | 0.06226322 | -1.8792209 | 0.092 | 0.156 | 1 | 0.06226322 | 0.05398951 |
| Gm54798 | 0.02738549 | 1.52206771 | 0.083 | 0.026 | 1 | 0.3036287 | 0.82764826 | 0.071 | 0.042 | 1 | 0.3036287 | 0.05402101 |
| Sik2 | 0.72173724 | -0.3144336 | 0.417 | 0.43 | 1 | 0.02743988 | -0.6873757 | 0.664 | 0.792 | 1 | 0.72173724 | 0.05412682 |
| Arhgef11 | 0.02744035 | 0.17673139 | 0.604 | 0.446 | 1 | 0.38267593 | -0.2080404 | 0.714 | 0.812 | 1 | 0.38267593 | 0.05412772 |
| Hdac8 | 0.02744728 | -0.496122 | 0.521 | 0.632 | 1 | 0.66670697 | -0.4402098 | 0.735 | 0.74 | 1 | 0.66670697 | 0.0541412 |
| Gm15270 | 0.02745727 | -1.3437279 | 0.229 | 0.342 | 1 | 0.03213537 | 0.44503224 | 0.168 | 0.073 | 1 | 0.03213537 | 0.05416065 |
| Alkbh7 | 0.02746621 | 1.00749454 | 0.177 | 0.088 | 1 | 0.65861142 | -0.2554075 | 0.088 | 0.104 | 1 | 0.65861142 | 0.05417802 |
| Pgd | 0.95373179 | -0.1583144 | 0.562 | 0.503 | 1 | 0.02746646 | -0.5819348 | 0.542 | 0.698 | 1 | 0.95373179 | 0.05417852 |
| Cep295 | 0.02749708 | 0.97948016 | 0.271 | 0.161 | 1 | 0.73945913 | -0.4071525 | 0.218 | 0.229 | 1 | 0.73945913 | 0.05423807 |
| Rrp12 | 0.0275648 | 0.9200317 | 0.25 | 0.145 | 1 | 0.38789294 | -0.3955851 | 0.185 | 0.135 | 1 | 0.38789294 | 0.05436979 |
| Cc2d1b | 0.02759584 | 0.76648644 | 0.208 | 0.109 | 1 | 0.07351348 | 0.29348577 | 0.256 | 0.156 | 1 | 0.07351348 | 0.05443016 |
| Dock11 | 0.23480161 | -0.4806064 | 0.26 | 0.347 | 1 | 0.02761578 | -0.3510126 | 0.412 | 0.573 | 1 | 0.23480161 | 0.05446893 |
| Gm15708 | 0.71418464 | -0.2148979 | 0.052 | 0.041 | 1 | 0.0276601 | 1.69014474 | 0.088 | 0.021 | 1 | 0.71418464 | 0.05455511 |
| Dcun1d2 | 0.9096107 | -0.4286046 | 0.146 | 0.145 | 1 | 0.02768321 | -0.2904899 | 0.231 | 0.115 | 1 | 0.9096107 | 0.05460006 |
| Lsm7 | 0.08983783 | 0.2167225 | 0.615 | 0.477 | 1 | 0.02775006 | 0.53240527 | 0.58 | 0.458 | 1 | 0.08983783 | 0.05473005 |
| Elac2 | 0.0389764 | 0.78510212 | 0.167 | 0.083 | 1 | 0.02775608 | 1.08246216 | 0.151 | 0.062 | 1 | 0.0389764 | 0.05474175 |
| Cep250 | 0.07046235 | 0.82692229 | 0.198 | 0.119 | 1 | 0.02775941 | 0.38046024 | 0.214 | 0.104 | 1 | 0.07046235 | 0.05474824 |
| Gm31323 | 0.35945034 | 0.68556644 | 0.073 | 0.047 | 1 | 0.02776265 | -1.198824 | 0.063 | 0.135 | 1 | 0.35945034 | 0.05475453 |
| C920006O11Rik | 0.02777911 | 1.22988696 | 0.125 | 0.052 | 1 | 0.87528207 | -0.5025003 | 0.038 | 0.042 | 1 | 0.87528207 | 0.05478655 |
| Amd1 | 0.0278652 | -0.5633928 | 0.625 | 0.684 | 1 | 0.74205886 | -0.2761346 | 0.689 | 0.698 | 1 | 0.74205886 | 0.05495393 |

| | | | | | | | | | | | | |
|---|---|---|---|---|---|---|---|---|---|---|---|---|
| Thegl | 0.83516282 | -1.0513992 | 0.042 | 0.047 | 1 | 0.02786521 | -1.7692869 | 0.017 | 0.062 | 1 | 0.83516282 | 0.05495396 |
| Endod1 | 0.02791948 | 0.83502734 | 0.375 | 0.259 | 1 | 0.9620396 | 0.23071312 | 0.412 | 0.438 | 1 | 0.9620396 | 0.05505947 |
| Dgke | 0.02792085 | -1.2726134 | 0.042 | 0.124 | 1 | 0.87248534 | -0.4473588 | 0.08 | 0.073 | 1 | 0.87248534 | 0.05506212 |
| B230216N24Rik | 0.02793234 | 1.12778877 | 0.188 | 0.098 | 1 | 0.7755884 | -0.2927817 | 0.252 | 0.219 | 1 | 0.7755884 | 0.05508446 |
| Wrnip1 | 0.02798146 | 0.59245704 | 0.292 | 0.171 | 1 | 0.40573724 | -0.2318528 | 0.298 | 0.229 | 1 | 0.40573724 | 0.05517996 |
| Pdpn | 0.26073383 | 0.75751628 | 0.177 | 0.13 | 1 | 0.02808185 | -0.6394234 | 0.29 | 0.417 | 1 | 0.26073383 | 0.05537511 |
| Mcts2 | 0.71030843 | -0.1079827 | 0.104 | 0.119 | 1 | 0.02808795 | 1.14957636 | 0.122 | 0.042 | 1 | 0.71030843 | 0.05538697 |
| Dph7 | 0.02810403 | 0.80818573 | 0.219 | 0.119 | 1 | 0.49072817 | -0.189561 | 0.16 | 0.125 | 1 | 0.49072817 | 0.05541822 |
| Tmem170 | 0.04196819 | -1.130009 | 0.083 | 0.171 | 1 | 0.02810669 | 0.77760758 | 0.122 | 0.042 | 1 | 0.04196819 | 0.05542339 |
| Zfp970 | 0.0281125 | 1.37006462 | 0.083 | 0.026 | 1 | 0.31828711 | 0.19264508 | 0.059 | 0.031 | 1 | 0.31828711 | 0.05543469 |
| Bfsp2 | 0.0281134 | 2.14499806 | 0.073 | 0.021 | 1 | 0.09044888 | 1.69014474 | 0.029 | 0 | 1 | 0.09044888 | 0.05543645 |
| Kif1b | 0.02813992 | 0.2570331 | 0.906 | 0.917 | 1 | 0.22906644 | -0.2892703 | 0.987 | 0.99 | 1 | 0.22906644 | 0.05548798 |
| Efs | 0.03805418 | 0.71376333 | 0.281 | 0.176 | 1 | 0.02814417 | 0.93807225 | 0.265 | 0.156 | 1 | 0.03805418 | 0.05549625 |
| Tesk2 | 0.86553108 | -0.8770282 | 0.125 | 0.135 | 1 | 0.02815888 | 0.54429387 | 0.126 | 0.042 | 1 | 0.86553108 | 0.05552485 |
| BC034090 | 0.73971793 | 0.45495351 | 0.104 | 0.093 | 1 | 0.02816306 | 0.82764826 | 0.168 | 0.073 | 1 | 0.73971793 | 0.05553296 |
| Rab39 | 0.56588891 | 0.31561683 | 0.115 | 0.093 | 1 | 0.02816708 | 1.56461386 | 0.088 | 0.021 | 1 | 0.56588891 | 0.05554078 |
| Tmem260 | 0.02821032 | 0.51716891 | 0.333 | 0.207 | 1 | 0.1220945 | -0.3579496 | 0.319 | 0.198 | 1 | 0.1220945 | 0.05562483 |
| Cenpf | 0.0282221 | 5.78008404 | 0.052 | 0.01 | 1 | 0.27154471 | 0.69014474 | 0.013 | 0 | 1 | 0.27154471 | 0.05564771 |
| Hmmr | 0.02822237 | 4.65135073 | 0.052 | 0.01 | 1 | 0.50887612 | -1.3098553 | 0.004 | 0.01 | 1 | 0.50887612 | 0.05564824 |
| Itpkc | 0.06144057 | 0.61754802 | 0.323 | 0.228 | 1 | 0.02824843 | 0.40837377 | 0.311 | 0.177 | 1 | 0.06144057 | 0.05569889 |
| Gm47761 | 0.02827673 | 1.10703021 | 0.125 | 0.052 | 1 | 0.79810697 | -1.3098553 | 0.017 | 0.021 | 1 | 0.79810697 | 0.0557539 |
| Usp47 | 0.02827749 | 0.22148452 | 0.948 | 0.886 | 1 | 0.25628765 | -0.2409152 | 0.992 | 0.979 | 1 | 0.25628765 | 0.05575536 |
| Hectd2 | 0.86152095 | -0.1289247 | 0.75 | 0.746 | 1 | 0.02828023 | -0.7122454 | 0.605 | 0.635 | 1 | 0.86152095 | 0.05576069 |
| Xaf1 | 0.57799365 | 0.58299671 | 0.146 | 0.124 | 1 | 0.0282864 | -1.681824 | 0.08 | 0.156 | 1 | 0.57799365 | 0.05577269 |
| Adamts5 | 0.12376107 | -1.6169963 | 0.052 | 0.104 | 1 | 0.02832002 | -1.6766376 | 0.088 | 0.167 | 1 | 0.12376107 | 0.05583802 |
| Ddx41 | 0.85030075 | -0.1127997 | 0.365 | 0.332 | 1 | 0.02836393 | 0.39058446 | 0.37 | 0.229 | 1 | 0.85030075 | 0.05592334 |
| Plin3 | 0.86639899 | 0.23611891 | 0.333 | 0.337 | 1 | 0.02838953 | 0.67503785 | 0.471 | 0.375 | 1 | 0.86639899 | 0.05597309 |
| Rnf139 | 0.24261771 | 0.37006462 | 0.312 | 0.249 | 1 | 0.02848208 | 0.12390143 | 0.496 | 0.323 | 1 | 0.24261771 | 0.05615293 |
| Lpcat4 | 0.95839398 | -0.2462621 | 0.115 | 0.109 | 1 | 0.02848457 | 0.86006974 | 0.218 | 0.115 | 1 | 0.95839398 | 0.05615777 |
| Flot1 | 0.11449173 | 0.25947375 | 0.521 | 0.404 | 1 | 0.02850829 | 0.13492958 | 0.42 | 0.26 | 1 | 0.11449173 | 0.05620386 |
| Plpp2 | 0.25211261 | -0.2555399 | 0.073 | 0.119 | 1 | 0.02851082 | 0.76053407 | 0.151 | 0.062 | 1 | 0.25211261 | 0.05620877 |
| Pde4d | 0.71608142 | -0.5876963 | 0.896 | 0.922 | 1 | 0.02853979 | -1.1150059 | 0.92 | 0.885 | 1 | 0.71608142 | 0.05626507 |
| H1f4 | 0.40284041 | 1.52534284 | 0.427 | 0.42 | 1 | 0.02855059 | 0.59024617 | 0.408 | 0.271 | 1 | 0.40284041 | 0.05628605 |
| Pmm1 | 0.86085424 | -0.1585154 | 0.333 | 0.337 | 1 | 0.02856777 | 0.49037523 | 0.311 | 0.188 | 1 | 0.86085424 | 0.05631943 |
| Rbm12b2 | 0.40543564 | -0.5330738 | 0.083 | 0.114 | 1 | 0.02865672 | 0.7944814 | 0.168 | 0.073 | 1 | 0.40543564 | 0.05649223 |
| Yipf1 | 0.20189698 | 0.26529229 | 0.438 | 0.363 | 1 | 0.02866503 | 0.35690133 | 0.395 | 0.24 | 1 | 0.20189698 | 0.05650837 |
| Adarb1 | 0.84094072 | -0.746487 | 0.292 | 0.29 | 1 | 0.02872964 | -1.7777446 | 0.357 | 0.406 | 1 | 0.84094072 | 0.05663389 |
| Zfp87 | 0.12905259 | 0.77616899 | 0.177 | 0.114 | 1 | 0.02874108 | 1.0961371 | 0.189 | 0.094 | 1 | 0.12905259 | 0.0566561 |
| Nol6 | 0.85931398 | -0.240433 | 0.073 | 0.078 | 1 | 0.02874519 | 1.05937855 | 0.122 | 0.042 | 1 | 0.85931398 | 0.05666408 |
| 2810032G03Rik | 0.02874785 | -2.1624305 | 0.021 | 0.088 | 1 | 0.0540843 | 2.01207283 | 0.038 | 0 | 1 | 0.0540843 | 0.05666926 |
| Ecd | 0.02875265 | 0.69715442 | 0.323 | 0.212 | 1 | 0.05334541 | 0.36257008 | 0.298 | 0.188 | 1 | 0.05334541 | 0.05667858 |
| Zfp82 | 0.02875754 | 1.85549144 | 0.073 | 0.021 | 1 | 0.37706954 | 0.56461386 | 0.067 | 0.042 | 1 | 0.37706954 | 0.05668809 |
| Phlda2 | 0.02875754 | 1.85549144 | 0.073 | 0.021 | 1 | 0.23679813 | 1.01207283 | 0.034 | 0.01 | 1 | 0.23679813 | 0.05668809 |
| Gm47469 | 0.02875754 | 2.00749454 | 0.073 | 0.021 | 1 | 0.11130697 | 1.59703533 | 0.046 | 0.01 | 1 | 0.11130697 | 0.05668809 |
| Gm56660 | 0.02875754 | 1.85549144 | 0.073 | 0.021 | 1 | 0.11764843 | 1.86006974 | 0.025 | 0 | 1 | 0.11764843 | 0.05668809 |
| Spin2c | 0.02875754 | 1.85549144 | 0.073 | 0.021 | 1 | 0.04177162 | 2.33400093 | 0.063 | 0.01 | 1 | 0.04177162 | 0.05668809 |
| Slco1c1 | 0.02876735 | 1.24453373 | 0.146 | 0.067 | 1 | 0.16972365 | 0.62695091 | 0.189 | 0.125 | 1 | 0.16972365 | 0.05670713 |
| Klhl23 | 0.02882995 | 1.32942263 | 0.167 | 0.083 | 1 | 0.05942961 | 1.10518224 | 0.122 | 0.052 | 1 | 0.05942961 | 0.05682874 |
| Unk | 0.02883672 | 0.65239958 | 0.354 | 0.233 | 1 | 0.98807171 | -0.5302453 | 0.294 | 0.271 | 1 | 0.98807171 | 0.05684189 |
| Qtrt2 | 0.02887322 | 0.72919337 | 0.333 | 0.218 | 1 | 0.1357867 | 0.15963002 | 0.332 | 0.229 | 1 | 0.1357867 | 0.05691277 |
| Tcta | 0.02887604 | 0.49292136 | 0.312 | 0.187 | 1 | 0.04940024 | 0.64434105 | 0.223 | 0.125 | 1 | 0.04940024 | 0.05691825 |
| Tyk2 | 0.35740444 | 0.24850264 | 0.104 | 0.073 | 1 | 0.0288805 | 0.46273424 | 0.155 | 0.062 | 1 | 0.35740444 | 0.05692691 |
| Dvl2 | 0.02891704 | 0.43491576 | 0.365 | 0.233 | 1 | 0.46132345 | -0.2818409 | 0.303 | 0.229 | 1 | 0.46132345 | 0.05699789 |
| Mnat1 | 0.02892855 | 0.16987678 | 0.771 | 0.58 | 1 | 0.03834879 | -0.4143494 | 0.685 | 0.771 | 1 | 0.03834879 | 0.05702023 |
| Abhd14b | 0.97873228 | -0.17693 | 0.208 | 0.202 | 1 | 0.02893956 | 0.40063812 | 0.248 | 0.135 | 1 | 0.97873228 | 0.05704162 |
| Kank2 | 0.71767665 | -0.3037072 | 0.385 | 0.399 | 1 | 0.0290114 | 0.32158167 | 0.63 | 0.427 | 1 | 0.71767665 | 0.05718114 |
| Psmc3ip | 0.02902605 | 1.71798792 | 0.115 | 0.047 | 1 | 0.77050453 | -0.3923174 | 0.055 | 0.062 | 1 | 0.77050453 | 0.05720959 |
| Rab2b | 0.98656049 | -0.1170506 | 0.427 | 0.42 | 1 | 0.02903156 | 0.39294373 | 0.42 | 0.271 | 1 | 0.98656049 | 0.05722029 |
| Cep55 | 0.02903834 | 3.00749454 | 0.052 | 0.01 | 1 | 0.86312513 | -0.7248928 | 0.008 | 0.01 | 1 | 0.86312513 | 0.05723346 |
| Cenpk | 0.02904464 | 2.06638823 | 0.115 | 0.047 | 1 | 0.05407544 | 2.49749966 | 0.038 | 0 | 1 | 0.05407544 | 0.05724569 |
| C030006K11Rik | 0.38401093 | -0.3144336 | 0.062 | 0.093 | 1 | 0.02904546 | 1.54812573 | 0.088 | 0.021 | 1 | 0.38401093 | 0.05724728 |
| Capn11 | 0.20902867 | 1.00749454 | 0.062 | 0.031 | 1 | 0.02905528 | 1.49749966 | 0.088 | 0.021 | 1 | 0.20902867 | 0.05726634 |
| Csgalnact1 | 0.02908596 | -1.2588493 | 0.865 | 0.793 | 1 | 0.25123035 | -0.1283886 | 0.66 | 0.469 | 1 | 0.25123035 | 0.05732592 |
| AV099323 | 0.02910304 | 1.88196365 | 0.073 | 0.021 | 1 | 0.81129592 | 0.10518224 | 0.025 | 0.021 | 1 | 0.81129592 | 0.05735909 |
| Hsd17b12 | 0.22211221 | -0.308874 | 0.792 | 0.85 | 1 | 0.0291428 | -0.7435084 | 0.714 | 0.812 | 1 | 0.22211221 | 0.05743629 |
| Sec14l2 | 0.36586035 | 0.35541784 | 0.135 | 0.098 | 1 | 0.02916008 | 0.65537932 | 0.261 | 0.146 | 1 | 0.36586035 | 0.05746985 |
| Ppard | 0.02926002 | -0.4615688 | 0.458 | 0.627 | 1 | 0.66971866 | -0.3190153 | 0.634 | 0.604 | 1 | 0.66971866 | 0.05766388 |
| Gm20033 | 0.63771059 | -0.6247737 | 0.083 | 0.098 | 1 | 0.0292981 | 1.27510724 | 0.105 | 0.031 | 1 | 0.63771059 | 0.05773781 |
| Steap4 | 0.02930585 | -2.265524 | 0.094 | 0.187 | 1 | 0.81171963 | -0.9472852 | 0.071 | 0.062 | 1 | 0.81171963 | 0.05775286 |
| Nt5c | 0.5877699 | 0.14835707 | 0.312 | 0.28 | 1 | 0.02940771 | 0.40063812 | 0.328 | 0.198 | 1 | 0.5877699 | 0.0579506 |
| Frem2 | 0.02945296 | 1.68556644 | 0.073 | 0.021 | 1 | 0.87066325 | -0.3098553 | 0.013 | 0.01 | 1 | 0.87066325 | 0.05803844 |
| Dnajc6 | 0.02947779 | -2.240433 | 0.031 | 0.104 | 1 | 0.27154327 | 1.01207283 | 0.013 | 0 | 1 | 0.27154327 | 0.05808664 |
| Akap8 | 0.02948275 | 0.21990287 | 0.531 | 0.368 | 1 | 0.64605506 | -0.1961985 | 0.618 | 0.562 | 1 | 0.64605506 | 0.05809627 |
| Gm56786 | 0.02948515 | -1.9311049 | 0.021 | 0.088 | 1 | 0.34704016 | -1.3098553 | 0.008 | 0.021 | 1 | 0.34704016 | 0.05810092 |
| Chaf1a | 0.02950778 | 1.14499806 | 0.083 | 0.026 | 1 | 0.30704993 | -1.1399303 | 0.029 | 0.052 | 1 | 0.30704993 | 0.05814485 |
| Dtx2 | 0.78863118 | -0.1851505 | 0.135 | 0.145 | 1 | 0.0295273 | 0.71004429 | 0.235 | 0.125 | 1 | 0.78863118 | 0.05818275 |
| Gm10549 | 0.06082017 | 0.86310463 | 0.156 | 0.083 | 1 | 0.0295534 | 1.39058446 | 0.088 | 0.021 | 1 | 0.06082017 | 0.05823341 |
| Usp38 | 0.02958156 | 0.36703692 | 0.427 | 0.275 | 1 | 0.85395784 | -0.3915391 | 0.559 | 0.51 | 1 | 0.85395784 | 0.05828805 |
| Gbe1 | 0.72885155 | -0.2309102 | 0.354 | 0.316 | 1 | 0.02962196 | -0.7598526 | 0.479 | 0.604 | 1 | 0.72885155 | 0.05836647 |
| Map1s | 0.02968787 | 0.50225923 | 0.281 | 0.166 | 1 | 0.15292257 | -0.1266334 | 0.336 | 0.219 | 1 | 0.15292257 | 0.05849436 |
| Esyt1 | 0.02972607 | -0.3650596 | 0.312 | 0.451 | 1 | 0.77447844 | -0.5322477 | 0.34 | 0.333 | 1 | 0.77447844 | 0.05856849 |
| Erbb2 | 0.09218657 | -0.6805615 | 0.062 | 0.13 | 1 | 0.0297352 | 0.66742466 | 0.218 | 0.115 | 1 | 0.09218657 | 0.05858621 |
| Ppp2r5b | 0.93473581 | 0.10060394 | 0.365 | 0.373 | 1 | 0.02974086 | 0.20471791 | 0.429 | 0.281 | 1 | 0.93473581 | 0.0585972 |
| 4833418N02Rik | 0.02974375 | -1.293675 | 0.125 | 0.223 | 1 | 0.70127829 | -0.3623227 | 0.193 | 0.167 | 1 | 0.70127829 | 0.05860281 |
| Bpifb1 | 0.02978338 | 2.59245704 | 0.062 | 0.016 | 1 | 0.18349795 | 0.71989208 | 0.084 | 0.042 | 1 | 0.18349795 | 0.05867971 |
| Pold1 | 0.02978338 | 2.32942263 | 0.062 | 0.016 | 1 | 0.30245777 | -1.6317834 | 0.029 | 0.052 | 1 | 0.30245777 | 0.05867971 |

| | | | | | | | | | | | | |
|---|---|---|---|---|---|---|---|---|---|---|---|---|
| Gm10644 | 0.02978338 | 2.17741954 | 0.062 | 0.016 | 1 | 0.39234595 | 1.01207283 | 0.025 | 0.01 | 1 | 0.39234595 | 0.05867971 |
| Gm26664 | 0.02978338 | 2.32942263 | 0.062 | 0.016 | 1 | 0.45367952 | -0.4353861 | 0.038 | 0.021 | 1 | 0.45367952 | 0.05867971 |
| 4930509G22Rik | 0.02978338 | 2.17741954 | 0.062 | 0.016 | 1 | 0.19069923 | 0.56461386 | 0.038 | 0.01 | 1 | 0.19069923 | 0.05867971 |
| Alkbh3 | 0.0297938 | 0.64492446 | 0.333 | 0.223 | 1 | 0.91676447 | -0.4693335 | 0.252 | 0.24 | 1 | 0.91676447 | 0.05869992 |
| Mcph1 | 0.02979881 | 0.91019734 | 0.24 | 0.135 | 1 | 0.98829351 | -0.4769652 | 0.164 | 0.156 | 1 | 0.98829351 | 0.05870966 |
| Nudt16l1 | 0.06935438 | 0.77302928 | 0.271 | 0.187 | 1 | 0.02981252 | 0.32955502 | 0.286 | 0.156 | 1 | 0.06935438 | 0.05873626 |
| Rpp40 | 0.02982012 | 1.15949763 | 0.083 | 0.026 | 1 | 0.56384362 | -0.4353861 | 0.038 | 0.052 | 1 | 0.56384362 | 0.058751 |
| 1700001K19Rik | 0.02982509 | -2.1624305 | 0.031 | 0.104 | 1 | 0.23067783 | -1.3802446 | 0.067 | 0.104 | 1 | 0.23067783 | 0.05876065 |
| Snrpc | 0.02984957 | 0.34614599 | 0.74 | 0.642 | 1 | 0.48724211 | -0.17125 | 0.66 | 0.594 | 1 | 0.48724211 | 0.05880815 |
| Rmdn3 | 0.27735312 | 0.39868529 | 0.312 | 0.254 | 1 | 0.02986368 | 0.21370669 | 0.294 | 0.167 | 1 | 0.27735312 | 0.05883552 |
| Gm16008 | 0.02986564 | 2.22988696 | 0.052 | 0.01 | 1 | 0.04198609 | 2.14957636 | 0.042 | 0 | 1 | 0.04198609 | 0.05883933 |
| Gls2 | 0.02986564 | 2.22988696 | 0.052 | 0.01 | 1 | 0.50887612 | -1.3098553 | 0.004 | 0.01 | 1 | 0.50887612 | 0.05883933 |
| Gm11963 | 0.02986564 | 2.22988696 | 0.052 | 0.01 | 1 | 0.66171832 | 0.49749966 | 0.017 | 0.01 | 1 | 0.66171832 | 0.05883933 |
| Rnf125 | 0.02986564 | 2.22988696 | 0.052 | 0.01 | 1 | 0.14795145 | 0.19264508 | 0.059 | 0.021 | 1 | 0.14795145 | 0.05883933 |
| Cdca3 | 0.02987569 | 2.75751628 | 0.052 | 0.01 | 1 | 0.87851725 | -0.8948178 | 0.013 | 0.01 | 1 | 0.87851725 | 0.05885883 |
| Osbpl11 | 0.10696262 | 0.50345203 | 0.521 | 0.435 | 1 | 0.0299479 | -0.9250756 | 0.676 | 0.698 | 1 | 0.10696262 | 0.05899893 |
| Syne3 | 0.02995574 | 0.23256109 | 0.333 | 0.202 | 1 | 0.81127668 | -0.5588828 | 0.349 | 0.302 | 1 | 0.81127668 | 0.05901414 |
| Mcm3ap | 0.98648912 | -0.1345245 | 0.25 | 0.244 | 1 | 0.02996208 | 0.15646274 | 0.294 | 0.167 | 1 | 0.98648912 | 0.05902644 |
| Rhobtb3 | 0.16948082 | 0.30932143 | 0.5 | 0.415 | 1 | 0.02996232 | 0.17167765 | 0.521 | 0.344 | 1 | 0.16948082 | 0.05902689 |
| Kat6a | 0.02997073 | 0.40911152 | 0.802 | 0.694 | 1 | 0.94058491 | -0.2068887 | 0.945 | 0.896 | 1 | 0.94058491 | 0.05904322 |
| Gm48027 | 0.63327961 | -0.3144336 | 0.115 | 0.135 | 1 | 0.02998805 | -1.3098553 | 0.151 | 0.24 | 1 | 0.63327961 | 0.05907681 |
| Tmod2 | 0.02999568 | 1.00749454 | 0.125 | 0.052 | 1 | 0.05598048 | 0.54812573 | 0.168 | 0.083 | 1 | 0.05598048 | 0.05909162 |
| Ids | 0.03890451 | 0.56937342 | 0.281 | 0.171 | 1 | 0.03002779 | 0.21739174 | 0.454 | 0.292 | 1 | 0.03890451 | 0.05915392 |
| Zfp689 | 0.14789493 | -1.0628948 | 0.042 | 0.088 | 1 | 0.03004769 | 0.91253716 | 0.105 | 0.031 | 1 | 0.14789493 | 0.05919252 |
| Peg13 | 0.73154007 | 0.37946331 | 0.083 | 0.073 | 1 | 0.03004769 | 0.49749966 | 0.105 | 0.031 | 1 | 0.73154007 | 0.05919252 |
| Gtf2ird1 | 0.64005438 | -0.1914865 | 0.427 | 0.352 | 1 | 0.03007858 | -0.5385892 | 0.63 | 0.812 | 1 | 0.64005438 | 0.05925243 |
| Abi1 | 0.03010184 | -0.3346924 | 0.938 | 0.927 | 1 | 0.03277785 | -0.6296123 | 0.966 | 0.969 | 1 | 0.03277785 | 0.05929757 |
| Cntln | 0.03011074 | 0.75058623 | 0.479 | 0.373 | 1 | 0.12063687 | 0.15764966 | 0.483 | 0.375 | 1 | 0.12063687 | 0.05931482 |
| Stpg4 | 0.14427852 | -1.6247737 | 0.042 | 0.088 | 1 | 0.03013091 | -2.4391383 | 0.05 | 0.115 | 1 | 0.14427852 | 0.05935394 |
| B4galt4 | 0.29451552 | -0.1889027 | 0.198 | 0.259 | 1 | 0.03016453 | 0.39468885 | 0.315 | 0.188 | 1 | 0.29451552 | 0.05941915 |
| Fxyd4 | 0.0301665 | 2.14499806 | 0.062 | 0.016 | 1 | 0.23005698 | -1.4253325 | 0.042 | 0.073 | 1 | 0.23005698 | 0.05942298 |
| Spdl1 | 0.0301665 | 2.38600616 | 0.062 | 0.016 | 1 | 0.50887612 | -1.3098553 | 0.004 | 0.01 | 1 | 0.50887612 | 0.05942298 |
| Utp6 | 0.25157462 | 0.41381482 | 0.333 | 0.285 | 1 | 0.03023837 | 0.31383963 | 0.42 | 0.271 | 1 | 0.25157462 | 0.05956239 |
| Sinhcaf | 0.03961015 | -1.6554705 | 0.021 | 0.083 | 1 | 0.03024089 | -1.6508922 | 0.042 | 0.104 | 1 | 0.03961015 | 0.05956727 |
| Fbxl14 | 0.31868838 | 0.50053455 | 0.167 | 0.124 | 1 | 0.03029146 | 0.48661134 | 0.252 | 0.135 | 1 | 0.31868838 | 0.05966535 |
| Creb3l4 | 0.0303018 | 2.00749454 | 0.052 | 0.01 | 1 | 0.14367653 | 1.27510724 | 0.042 | 0.01 | 1 | 0.14367653 | 0.05968541 |
| C230012O17Rik | 0.0303018 | 2.00749454 | 0.052 | 0.01 | 1 | 0.20315163 | 1.01207283 | 0.017 | 0 | 1 | 0.20315163 | 0.05968541 |
| Tmem151a | 0.0303018 | 2.00749454 | 0.052 | 0.01 | 1 | 0.62294577 | 0.27510724 | 0.055 | 0.042 | 1 | 0.62294577 | 0.05968541 |
| Slpi | 0.05925632 | -1.143749 | 0.406 | 0.259 | 1 | 0.03031617 | -0.5966671 | 0.458 | 0.271 | 1 | 0.05925632 | 0.05971327 |
| Ldlr | 0.18790515 | 0.15301819 | 0.75 | 0.679 | 1 | 0.03035533 | 0.32623573 | 0.63 | 0.531 | 1 | 0.18790515 | 0.05978922 |
| Vta1 | 0.03037721 | 0.47381254 | 0.604 | 0.492 | 1 | 0.07409601 | 0.26943849 | 0.618 | 0.531 | 1 | 0.07409601 | 0.05983164 |
| Gm1976 | 0.03339124 | 0.56937342 | 0.229 | 0.13 | 1 | 0.03042648 | 0.52021974 | 0.185 | 0.083 | 1 | 0.03339124 | 0.05992719 |
| Aspm | 0.03043201 | 1.85549144 | 0.135 | 0.062 | 1 | 0.1911727 | -0.6883669 | 0.046 | 0.083 | 1 | 0.1911727 | 0.0599379 |
| Epdr1 | 0.03047757 | -1.0884299 | 0.292 | 0.404 | 1 | 0.94373274 | -1.152314 | 0.239 | 0.208 | 1 | 0.94373274 | 0.06002626 |
| Tiam2 | 0.12104678 | 0.56020835 | 0.969 | 0.964 | 1 | 0.030492 | -0.5147825 | 0.945 | 0.948 | 1 | 0.12104678 | 0.06005424 |
| Zfp783 | 0.45033636 | 0.42253204 | 0.188 | 0.155 | 1 | 0.03049433 | 0.92280549 | 0.164 | 0.073 | 1 | 0.45033636 | 0.06005876 |
| Snx30 | 0.58040041 | -0.4157169 | 0.135 | 0.161 | 1 | 0.03050102 | -1.4311616 | 0.155 | 0.24 | 1 | 0.58040041 | 0.06007173 |
| Pde3a | 0.03053264 | 0.39682192 | 0.875 | 0.788 | 1 | 0.04026485 | 0.17714699 | 0.916 | 0.688 | 1 | 0.04026485 | 0.06013304 |
| Agps | 0.37458037 | 0.14369892 | 0.531 | 0.487 | 1 | 0.0305376 | -0.7588401 | 0.676 | 0.771 | 1 | 0.37458037 | 0.06014266 |
| Nup98 | 0.19899768 | -0.3358072 | 0.823 | 0.896 | 1 | 0.03056232 | -0.4709177 | 0.945 | 0.99 | 1 | 0.19899768 | 0.06019058 |
| Gm11808 | 0.15591166 | 0.26206736 | 0.656 | 0.57 | 1 | 0.03056846 | 0.43017663 | 0.483 | 0.365 | 1 | 0.15591166 | 0.0602025 |
| Gm26911 | 0.85039899 | -0.2148979 | 0.052 | 0.057 | 1 | 0.03057361 | -2.010295 | 0.05 | 0.115 | 1 | 0.85039899 | 0.06021247 |
| Impa2 | 0.1172464 | 0.54183096 | 0.448 | 0.358 | 1 | 0.03058938 | 0.47196457 | 0.458 | 0.302 | 1 | 0.1172464 | 0.06024305 |
| Slc35f2 | 0.03060579 | 1.00749454 | 0.083 | 0.026 | 1 | 0.11038855 | 1.56461386 | 0.063 | 0.021 | 1 | 0.11038855 | 0.06027487 |
| Kif16b | 0.03061028 | 0.59245704 | 0.562 | 0.42 | 1 | 0.60041098 | -0.202409 | 0.714 | 0.677 | 1 | 0.60041098 | 0.06028356 |
| Msh4 | 0.03061851 | 1.42253204 | 0.073 | 0.021 | 1 | 0.7555463 | -0.4797803 | 0.05 | 0.042 | 1 | 0.7555463 | 0.06029953 |
| Gpatch4 | 0.03063563 | 1.00749454 | 0.229 | 0.135 | 1 | 0.08589508 | 0.37603615 | 0.21 | 0.125 | 1 | 0.08589508 | 0.06033272 |
| Riok2 | 0.03067188 | 0.57554365 | 0.583 | 0.477 | 1 | 0.03310285 | -0.2072855 | 0.513 | 0.333 | 1 | 0.03310285 | 0.060403 |
| Cox10 | 0.68190252 | 0.78510212 | 0.135 | 0.124 | 1 | 0.03067437 | 0.45095707 | 0.185 | 0.083 | 1 | 0.68190252 | 0.06040781 |
| Gm33696 | 0.03072727 | 2.00749454 | 0.052 | 0.01 | 1 | 0.55833448 | -0.5728897 | 0.034 | 0.021 | 1 | 0.55833448 | 0.06051038 |
| Ccdc14 | 0.03072727 | 2.00749454 | 0.052 | 0.01 | 1 | 0.81074235 | -0.1399303 | 0.017 | 0.021 | 1 | 0.81074235 | 0.06051038 |
| A330033J07Rik | 0.03073529 | 2.14499806 | 0.062 | 0.016 | 1 | 0.8023126 | -0.7952821 | 0.017 | 0.021 | 1 | 0.8023126 | 0.06052593 |
| Lrig2 | 0.2042488 | 0.36113149 | 0.406 | 0.332 | 1 | 0.03073925 | 0.12806613 | 0.58 | 0.375 | 1 | 0.2042488 | 0.0605336 |
| Ric8b | 0.0307407 | 0.76870768 | 0.479 | 0.389 | 1 | 0.84431444 | -0.2282168 | 0.634 | 0.562 | 1 | 0.84431444 | 0.06053641 |
| Irgq | 0.09697537 | 0.68556644 | 0.198 | 0.124 | 1 | 0.03074567 | 0.66589719 | 0.206 | 0.104 | 1 | 0.09697537 | 0.06054604 |
| Ganc | 0.71984462 | 0.42253204 | 0.167 | 0.155 | 1 | 0.03078612 | 0.60768258 | 0.223 | 0.115 | 1 | 0.71984462 | 0.06062445 |
| Chn2 | 0.99482516 | -0.7087125 | 0.042 | 0.041 | 1 | 0.03080322 | -1.8459082 | 0.05 | 0.115 | 1 | 0.99482516 | 0.06065761 |
| Frk | 0.87074163 | -0.9153376 | 0.177 | 0.161 | 1 | 0.03081468 | -1.3649968 | 0.202 | 0.302 | 1 | 0.87074163 | 0.06067981 |
| Polr2a | 0.03083274 | -0.8583201 | 0.635 | 0.756 | 1 | 0.84571011 | -0.4520649 | 0.727 | 0.802 | 1 | 0.84571011 | 0.06071483 |
| Aprt | 0.03192821 | 0.38263931 | 0.583 | 0.44 | 1 | 0.03084801 | 0.39796399 | 0.592 | 0.479 | 1 | 0.03192821 | 0.06074442 |
| Itgbl1 | 0.03085679 | -1.8425047 | 0.135 | 0.233 | 1 | 0.9271614 | -0.9857429 | 0.374 | 0.312 | 1 | 0.9271614 | 0.06076145 |
| Nt5dc2 | 0.14397785 | 0.21185304 | 0.365 | 0.269 | 1 | 0.03086085 | 0.77760758 | 0.218 | 0.115 | 1 | 0.14397785 | 0.06076931 |
| Eef1akmt4 | 0.3472761 | 0.39744105 | 0.177 | 0.135 | 1 | 0.0309289 | 1.24034182 | 0.147 | 0.062 | 1 | 0.3472761 | 0.0609012 |
| Tax1bp1 | 0.03092921 | 0.288123 | 0.979 | 0.99 | 1 | 0.03332965 | 0.13577821 | 0.992 | 0.969 | 1 | 0.03332965 | 0.0609018 |
| Gm14443 | 0.03093614 | 1.81484946 | 0.062 | 0.016 | 1 | 0.14259748 | 1.77760758 | 0.042 | 0.01 | 1 | 0.14259748 | 0.06091524 |
| A530041M06Rik | 0.03093614 | 1.81484946 | 0.062 | 0.016 | 1 | 0.65471267 | -0.5728897 | 0.055 | 0.042 | 1 | 0.65471267 | 0.06091524 |
| Gm36640 | 0.03093614 | 1.81484946 | 0.062 | 0.016 | 1 | 0.32529413 | 0.40635177 | 0.071 | 0.042 | 1 | 0.32529413 | 0.06091524 |
| A930033M14Rik | 0.03093614 | 1.81484946 | 0.062 | 0.016 | 1 | 0.04197017 | 2.49749966 | 0.042 | 0 | 1 | 0.04197017 | 0.06091524 |
| A630052C17Rik | 0.03093614 | 1.81484946 | 0.062 | 0.016 | 1 | 0.6390466 | 0.39058446 | 0.042 | 0.031 | 1 | 0.6390466 | 0.06091524 |
| Gng13 | 0.03094487 | 1.81484946 | 0.052 | 0.01 | 1 | 0.57854956 | -0.8244284 | 0.021 | 0.031 | 1 | 0.57854956 | 0.06093216 |
| Col13a1 | 0.03094782 | 1.00749454 | 0.052 | 0.01 | 1 | 0.27154327 | 2.01207283 | 0.013 | 0 | 1 | 0.27154327 | 0.06093787 |
| Zdhhc4 | 0.03842104 | 0.70583884 | 0.344 | 0.233 | 1 | 0.03099229 | 0.3500693 | 0.277 | 0.156 | 1 | 0.03842104 | 0.06102405 |
| Senp5 | 0.03102375 | 0.32846371 | 0.781 | 0.642 | 1 | 0.3604142 | -0.2709539 | 0.874 | 0.833 | 1 | 0.3604142 | 0.06108502 |

| | | | | | | | | | | | | |
|---|---|---|---|---|---|---|---|---|---|---|---|---|
| AU015336 | 0.03109358 | -0.8290067 | 0.135 | 0.244 | 1 | 0.41163688 | 0.13760371 | 0.08 | 0.052 | 1 | 0.41163688 | 0.06122035 |
| Gm56933 | 0.03115911 | 1.37006462 | 0.073 | 0.021 | 1 | 0.03836766 | -2.5322477 | 0.008 | 0.042 | 1 | 0.03836766 | 0.06134732 |
| Cct6b | 0.0311638 | 1.49292136 | 0.052 | 0.01 | 1 | 0.03260032 | -2.3098553 | 0.013 | 0.052 | 1 | 0.03260032 | 0.06135642 |
| Mapk14 | 0.46642713 | -0.1973253 | 0.333 | 0.363 | 1 | 0.03117337 | -0.7442581 | 0.525 | 0.625 | 1 | 0.46642713 | 0.06137495 |
| A630089N07Rik | 0.03650999 | 0.70583884 | 0.26 | 0.155 | 1 | 0.03118939 | 0.47441605 | 0.197 | 0.094 | 1 | 0.03650999 | 0.061406 |
| Gzf1 | 0.12657012 | 0.6132156 | 0.271 | 0.197 | 1 | 0.03127811 | 0.74077081 | 0.269 | 0.156 | 1 | 0.12657012 | 0.0615779 |
| Psmb3 | 0.03604872 | 0.27052894 | 0.896 | 0.829 | 1 | 0.03129464 | 0.2621216 | 0.874 | 0.823 | 1 | 0.03604872 | 0.06160993 |
| Orm1 | 0.03131143 | 1.85549144 | 0.062 | 0.016 | 1 | 0.22356137 | 0.80562195 | 0.05 | 0.021 | 1 | 0.22356137 | 0.06164245 |
| Timeless | 0.03131143 | 1.85549144 | 0.062 | 0.016 | 1 | 0.51018266 | 0.49749966 | 0.021 | 0.01 | 1 | 0.51018266 | 0.06164245 |
| Tjp1 | 0.73569798 | -0.2206756 | 0.74 | 0.674 | 1 | 0.03134635 | -0.4369672 | 0.803 | 0.875 | 1 | 0.73569798 | 0.06171011 |
| Ankmy2 | 0.03134802 | 0.69338595 | 0.26 | 0.161 | 1 | 0.07268941 | 0.22856465 | 0.206 | 0.115 | 1 | 0.07268941 | 0.06171334 |
| Tsku | 0.0313796 | 1.59245704 | 0.052 | 0.01 | 1 | 0.46142849 | -0.4797803 | 0.063 | 0.042 | 1 | 0.46142849 | 0.06177451 |
| Pgbd1 | 0.0313796 | 1.59245704 | 0.052 | 0.01 | 1 | 0.30492205 | 0.86006974 | 0.029 | 0.01 | 1 | 0.30492205 | 0.06177451 |
| Camk2a | 0.0313796 | 1.59245704 | 0.052 | 0.01 | 1 | 0.3807744 | 0.20471791 | 0.055 | 0.031 | 1 | 0.3807744 | 0.06177451 |
| 4930517O19Rik | 0.26141035 | 0.13302542 | 0.188 | 0.135 | 1 | 0.03138686 | 0.11756596 | 0.248 | 0.125 | 1 | 0.26141035 | 0.06178859 |
| Coq7 | 0.60463657 | -0.3509594 | 0.344 | 0.347 | 1 | 0.03145176 | 0.61352346 | 0.332 | 0.208 | 1 | 0.60463657 | 0.06191431 |
| Ppp1r9a | 0.47911362 | -1.3144336 | 0.021 | 0.036 | 1 | 0.03149264 | 0.81725666 | 0.168 | 0.073 | 1 | 0.47911362 | 0.06199349 |
| Polrmt | 0.03150708 | 1.68556644 | 0.062 | 0.016 | 1 | 0.03268339 | 1.93807225 | 0.067 | 0.01 | 1 | 0.03268339 | 0.06202147 |
| Suco | 0.10738894 | 0.26692671 | 0.823 | 0.751 | 1 | 0.03151277 | -0.2972111 | 0.798 | 0.917 | 1 | 0.10738894 | 0.06203248 |
| Cyb561d2 | 0.91790533 | 0.12115132 | 0.24 | 0.244 | 1 | 0.0316041 | 0.90515763 | 0.193 | 0.094 | 1 | 0.91790533 | 0.06220939 |
| Nck2 | 0.03166067 | -0.4990969 | 0.604 | 0.689 | 1 | 0.41818628 | -0.4862176 | 0.794 | 0.792 | 1 | 0.41818628 | 0.06231894 |
| Gja1 | 0.03166435 | 0.11133035 | 0.667 | 0.487 | 1 | 0.21642949 | -0.6487898 | 0.475 | 0.5 | 1 | 0.21642949 | 0.06232607 |
| Syngr2 | 0.77207795 | -0.1851505 | 0.135 | 0.119 | 1 | 0.0316848 | -0.3838558 | 0.168 | 0.281 | 1 | 0.77207795 | 0.06236568 |
| Map3k10 | 0.99608337 | -0.130009 | 0.26 | 0.254 | 1 | 0.03170971 | 0.52126783 | 0.408 | 0.281 | 1 | 0.99608337 | 0.06241391 |
| Pip4k2b | 0.47380185 | 0.32942263 | 0.177 | 0.145 | 1 | 0.03176973 | 0.32757466 | 0.277 | 0.156 | 1 | 0.47380185 | 0.06253015 |
| Plekhf2 | 0.03181336 | 0.47315811 | 0.417 | 0.275 | 1 | 0.16083979 | 0.36694611 | 0.529 | 0.521 | 1 | 0.16083979 | 0.06261463 |
| Gm36220 | 0.07836231 | 0.42253204 | 0.385 | 0.28 | 1 | 0.03184525 | 0.24814019 | 0.21 | 0.104 | 1 | 0.07836231 | 0.06267638 |
| 6330403K07Rik | 0.04372434 | -2.577468 | 0 | 0.041 | 1 | 0.03184777 | -2.6883669 | 0.013 | 0.052 | 1 | 0.04372434 | 0.06268126 |
| Fgf7 | 0.38689439 | -0.577468 | 0.01 | 0.026 | 1 | 0.03184895 | -3.1172102 | 0.013 | 0.052 | 1 | 0.38689439 | 0.06268354 |
| Rnf216 | 0.03187601 | 0.1653742 | 0.906 | 0.746 | 1 | 0.79023114 | -0.1512388 | 0.912 | 0.927 | 1 | 0.79023114 | 0.06273593 |
| Ccdc117 | 0.5226876 | 0.13064277 | 0.354 | 0.301 | 1 | 0.03188395 | 0.15127865 | 0.534 | 0.365 | 1 | 0.5226876 | 0.06275131 |
| Ipp | 0.58068304 | -0.2555399 | 0.083 | 0.104 | 1 | 0.03193405 | 1.05271482 | 0.134 | 0.052 | 1 | 0.58068304 | 0.06284832 |
| C2 | 0.11302151 | -1.7998604 | 0 | 0.026 | 1 | 0.03203197 | -2.4797803 | 0.013 | 0.052 | 1 | 0.11302151 | 0.0630379 |
| Atp2a1 | 0.15729907 | -1.3144336 | 0 | 0.021 | 1 | 0.03203197 | -2.4797803 | 0.013 | 0.052 | 1 | 0.15729907 | 0.0630379 |
| Gm56772 | 0.32025586 | -1.9925055 | 0 | 0.01 | 1 | 0.03203197 | -2.6317834 | 0.013 | 0.052 | 1 | 0.32025586 | 0.0630379 |
| Trib3 | 0.17320258 | 0.77302928 | 0.115 | 0.067 | 1 | 0.03205483 | 2.39058446 | 0.067 | 0.01 | 1 | 0.17320258 | 0.06308214 |
| Chrna4 | 0.03205645 | 1.42253204 | 0.052 | 0.01 | 1 | 0.50887612 | -1.3098553 | 0.004 | 0.01 | 1 | 0.50887612 | 0.06308529 |
| Gm16268 | 0.06540799 | 0.2912875 | 0.104 | 0.047 | 1 | 0.03208355 | -1.4391383 | 0.08 | 0.156 | 1 | 0.06540799 | 0.06313775 |
| Fads6 | 0.03209374 | -2.577468 | 0 | 0.047 | 1 | 0.5778028 | -0.8948178 | 0.013 | 0.021 | 1 | 0.5778028 | 0.06315748 |
| Gm48321 | 0.03209374 | -2.4519371 | 0 | 0.047 | 1 | 0.77622115 | -0.9879272 | 0.046 | 0.052 | 1 | 0.77622115 | 0.06315748 |
| Gm29488 | 0.03209374 | 1.49292136 | 0.062 | 0.016 | 1 | 0.03836657 | 0.82764826 | 0.134 | 0.052 | 1 | 0.03836657 | 0.06315748 |
| Gm56783 | 0.03209374 | 1.49292136 | 0.062 | 0.016 | 1 | 0.14243143 | 1.01207283 | 0.059 | 0.021 | 1 | 0.14243143 | 0.06315748 |
| Mterf2 | 0.03209374 | 1.22988696 | 0.062 | 0.016 | 1 | 0.30324014 | -1.0983512 | 0.063 | 0.094 | 1 | 0.30324014 | 0.06315748 |
| Zc3h14 | 0.17193942 | 0.29515273 | 0.677 | 0.591 | 1 | 0.03209831 | 0.24166076 | 0.853 | 0.75 | 1 | 0.17193942 | 0.06316632 |
| Gm56999 | 0.03210036 | -2.7998604 | 0 | 0.047 | 1 | 0.45003757 | -0.194378 | 0.038 | 0.021 | 1 | 0.45003757 | 0.06317028 |
| Shank2 | 0.03210532 | -2.6929452 | 0 | 0.047 | 1 | 0.75977078 | -0.6577786 | 0.025 | 0.031 | 1 | 0.75977078 | 0.06317988 |
| Apoh | 0.03210532 | -2.6929452 | 0 | 0.047 | 1 | 0.81864396 | -0.1723517 | 0.025 | 0.021 | 1 | 0.81864396 | 0.06317988 |
| Dach1 | 0.03211098 | -3.577468 | 0 | 0.047 | 1 | 0.98557348 | 0.91253716 | 0.05 | 0.052 | 1 | 0.98557348 | 0.06319085 |
| Gm30371 | 0.03211169 | -4.6363617 | 0 | 0.047 | 1 | 0.11954511 | -2.2029401 | 0.008 | 0.031 | 1 | 0.11954511 | 0.06319222 |
| Ankrd37 | 0.57286036 | 0.42253204 | 0.062 | 0.047 | 1 | 0.03218256 | 2.33400093 | 0.067 | 0.01 | 1 | 0.57286036 | 0.0633294 |
| Tnfaip2 | 0.03219652 | -0.8669746 | 0.021 | 0.088 | 1 | 0.0533901 | 1.86006974 | 0.059 | 0.01 | 1 | 0.0533901 | 0.06335643 |
| Sephs1 | 0.03222148 | 0.77483378 | 0.292 | 0.181 | 1 | 0.13932388 | -0.1432054 | 0.345 | 0.229 | 1 | 0.13932388 | 0.06340473 |
| Bcl7a | 0.0322229 | 0.94609399 | 0.167 | 0.083 | 1 | 0.58337853 | -0.1654654 | 0.122 | 0.146 | 1 | 0.58337853 | 0.06340749 |
| Msra | 0.11607782 | -0.496978 | 0.229 | 0.316 | 1 | 0.03226348 | -0.3985595 | 0.361 | 0.5 | 1 | 0.11607782 | 0.06348602 |
| Pfas | 0.0639867 | 0.8872003 | 0.188 | 0.109 | 1 | 0.03227734 | 1.05271482 | 0.134 | 0.052 | 1 | 0.0639867 | 0.06351286 |
| Tmem88 | 0.15775364 | -1.4372903 | 0.073 | 0.124 | 1 | 0.03230965 | -1.9472852 | 0.059 | 0.125 | 1 | 0.15775364 | 0.06357538 |
| Scp2 | 0.16842054 | 0.142099 | 0.844 | 0.741 | 1 | 0.03234058 | 0.16675752 | 0.756 | 0.74 | 1 | 0.16842054 | 0.06363525 |
| Ddx21 | 0.03235626 | 0.31005731 | 0.771 | 0.762 | 1 | 0.05144924 | 0.1525406 | 0.87 | 0.771 | 1 | 0.05144924 | 0.06366558 |
| Colq | 0.746428 | -0.2820121 | 0.073 | 0.083 | 1 | 0.03243511 | 2.08246216 | 0.067 | 0.01 | 1 | 0.746428 | 0.06381818 |
| Podnl1 | 0.03244718 | -1.3430027 | 0.052 | 0.135 | 1 | 0.64563398 | -0.8244284 | 0.071 | 0.083 | 1 | 0.64563398 | 0.06384154 |
| Stab2 | 0.03250545 | 1.27052894 | 0.052 | 0.01 | 1 | 0.8259932 | -0.8244284 | 0.025 | 0.021 | 1 | 0.8259932 | 0.0639543 |
| Zfp354b | 0.07633009 | 1.74446013 | 0.031 | 0.005 | 1 | 0.03255962 | 2.14957636 | 0.067 | 0.01 | 1 | 0.07633009 | 0.06405911 |
| Dph1 | 0.20784723 | -1.1624305 | 0.01 | 0.036 | 1 | 0.03255972 | 2.01207283 | 0.067 | 0.01 | 1 | 0.20784723 | 0.0640593 |
| Tfap2a | 0.6598863 | -0.577468 | 0.198 | 0.212 | 1 | 0.03259195 | 0.57911343 | 0.269 | 0.156 | 1 | 0.6598863 | 0.06412166 |
| Zfp941 | 1 | 0.42253204 | 0.01 | 0.01 | 1 | 0.03260032 | -2.4797803 | 0.013 | 0.052 | 1 | 1 | 0.06413786 |
| 4930452G13Rik | 0.06812307 | 1.49292136 | 0.062 | 0.021 | 1 | 0.03262672 | 2.39058446 | 0.046 | 0 | 1 | 0.06812307 | 0.06418895 |
| Nagk | 0.04005093 | 0.51367992 | 0.375 | 0.249 | 1 | 0.03262937 | 0.23446525 | 0.34 | 0.198 | 1 | 0.04005093 | 0.06419406 |
| Rab6b | 0.38521705 | -1.4519371 | 0.01 | 0.026 | 1 | 0.03263332 | 2.49749966 | 0.046 | 0 | 1 | 0.38521705 | 0.0642017 |
| H2-Aa | 0.82244178 | -0.7528351 | 0.125 | 0.114 | 1 | 0.03263332 | 2.69014474 | 0.046 | 0 | 1 | 0.82244178 | 0.0642017 |
| Nudt16 | 0.85039899 | -0.2148979 | 0.052 | 0.057 | 1 | 0.03263332 | 2.49749966 | 0.046 | 0 | 1 | 0.85039899 | 0.0642017 |
| Akap6 | 0.80376323 | 0.18806678 | 0.083 | 0.093 | 1 | 0.03263441 | 0.93115284 | 0.105 | 0.031 | 1 | 0.80376323 | 0.06420381 |
| Gm34189 | 0.03444511 | -1.4705528 | 0.031 | 0.104 | 1 | 0.0326386 | 3.27510724 | 0.046 | 0 | 1 | 0.03444511 | 0.06421191 |
| Cited1 | 0.11096483 | -1.3144336 | 0.052 | 0.109 | 1 | 0.0326386 | 4.11640949 | 0.046 | 0 | 1 | 0.11096483 | 0.06421191 |
| Ccdc189 | 0.57678264 | 0.35541784 | 0.062 | 0.047 | 1 | 0.0326386 | 2.59703533 | 0.046 | 0 | 1 | 0.57678264 | 0.06421191 |
| Ccdc73 | 0.03263972 | 1.27052894 | 0.115 | 0.047 | 1 | 0.78571043 | -0.5728897 | 0.038 | 0.031 | 1 | 0.78571043 | 0.06421409 |
| Nmral1 | 0.03263972 | 1.27052894 | 0.115 | 0.047 | 1 | 0.31219265 | 0.52664601 | 0.071 | 0.042 | 1 | 0.31219265 | 0.06421409 |
| Gm12840 | 0.15405555 | -1.1624305 | 0.01 | 0.041 | 1 | 0.03265091 | 2.27510724 | 0.046 | 0 | 1 | 0.15405555 | 0.06423573 |
| 2210406O10Rik | 0.98291322 | 0.65957123 | 0.062 | 0.062 | 1 | 0.03265091 | 2.27510724 | 0.046 | 0 | 1 | 0.98291322 | 0.06423573 |
| Bicd1 | 0.03266201 | -1.8575759 | 0.042 | 0.119 | 1 | 0.2311252 | -0.1723517 | 0.092 | 0.052 | 1 | 0.2311252 | 0.06425722 |
| Fam221a | 0.03802463 | 1.08149512 | 0.156 | 0.078 | 1 | 0.03268339 | 1.93807225 | 0.067 | 0.01 | 1 | 0.03802463 | 0.06429858 |
| 1700066M21Rik | 0.11303029 | -1.577468 | 0 | 0.026 | 1 | 0.03268339 | 1.93807225 | 0.067 | 0.01 | 1 | 0.11303029 | 0.06429858 |
| Far1os | 0.0327082 | 1.42253204 | 0.062 | 0.016 | 1 | 0.32079052 | -0.8948178 | 0.038 | 0.062 | 1 | 0.32079052 | 0.06434657 |
| Selenok-ps1 | 0.0327082 | 1.42253204 | 0.062 | 0.016 | 1 | 0.23772424 | 0.86006974 | 0.034 | 0.01 | 1 | 0.23772424 | 0.06434657 |
| Zgrf1 | 0.03272719 | 1.56388789 | 0.115 | 0.047 | 1 | 0.29417559 | 0.93807225 | 0.059 | 0.031 | 1 | 0.29417559 | 0.06438332 |

| | | | | | | | | | | | | |
|---|---|---|---|---|---|---|---|---|---|---|---|---|
| Kcnb1 | 0.20795311 | -0.3237114 | 0.229 | 0.161 | 1 | 0.03278894 | 0.27510724 | 0.311 | 0.177 | 1 | 0.20795311 | 0.06450277 |
| Dusp29 | 0.7854853 | -0.407543 | 0.021 | 0.026 | 1 | 0.0328001 | -2.4473588 | 0.013 | 0.052 | 1 | 0.7854853 | 0.06452435 |
| Fbxo44 | 0.25881726 | -0.4330781 | 0.042 | 0.078 | 1 | 0.03280603 | 1.86006974 | 0.067 | 0.01 | 1 | 0.25881726 | 0.06453583 |
| Gm57269 | 0.84109777 | 1.00749454 | 0.073 | 0.067 | 1 | 0.03280603 | 1.86006974 | 0.067 | 0.01 | 1 | 0.84109777 | 0.06453583 |
| Atf6 | 0.03287398 | -0.5533555 | 0.74 | 0.845 | 1 | 0.32261486 | -0.4566967 | 0.853 | 0.906 | 1 | 0.32261486 | 0.06466727 |
| Smurf2 | 0.03292965 | 0.37740266 | 0.938 | 0.902 | 1 | 0.07621687 | -0.4622679 | 0.983 | 0.979 | 1 | 0.07621687 | 0.06477495 |
| Gm43259 | 0.61737927 | 1.00749454 | 0.01 | 0.005 | 1 | 0.03299407 | -1.9879272 | 0.013 | 0.052 | 1 | 0.61737927 | 0.06489953 |
| Mknk1 | 0.03299419 | 0.45110119 | 0.271 | 0.155 | 1 | 0.64079028 | -0.4093909 | 0.273 | 0.229 | 1 | 0.64079028 | 0.06489975 |
| Gm57100 | 0.34911874 | 0.68556644 | 0.104 | 0.073 | 1 | 0.03300688 | 1.77760758 | 0.067 | 0.01 | 1 | 0.34911874 | 0.06492431 |
| 2010204K13Rik | 0.59199182 | 0.42253204 | 0.031 | 0.021 | 1 | 0.03300688 | 1.77760758 | 0.067 | 0.01 | 1 | 0.59199182 | 0.06492431 |
| Rrp9 | 0.09569231 | 0.45355893 | 0.375 | 0.28 | 1 | 0.03301286 | 0.82400048 | 0.265 | 0.156 | 1 | 0.09569231 | 0.06493586 |
| H6pd | 0.9837892 | -0.130009 | 0.177 | 0.176 | 1 | 0.0330464 | 0.18490943 | 0.223 | 0.115 | 1 | 0.9837892 | 0.06500074 |
| Sgk3 | 0.03308004 | 0.15445198 | 0.823 | 0.741 | 1 | 0.10029212 | -0.1142488 | 0.866 | 0.76 | 1 | 0.10029212 | 0.06506579 |
| Sf3a1 | 0.13877798 | 0.178975 | 0.458 | 0.342 | 1 | 0.03310634 | 0.17557156 | 0.563 | 0.354 | 1 | 0.13877798 | 0.06511665 |
| Prpf8 | 0.03311408 | 0.4343092 | 0.677 | 0.596 | 1 | 0.03390767 | 0.15750327 | 0.744 | 0.646 | 1 | 0.03390767 | 0.06513161 |
| Il17rc | 0.0331201 | 1.22988696 | 0.115 | 0.047 | 1 | 0.28743694 | 0.17557156 | 0.101 | 0.062 | 1 | 0.28743694 | 0.06514326 |
| Cdan1 | 0.03312244 | 1.00749454 | 0.146 | 0.067 | 1 | 0.13554546 | 0.24034182 | 0.147 | 0.083 | 1 | 0.13554546 | 0.06514779 |
| Gas2l3 | 0.43819216 | 1.62898291 | 0.073 | 0.052 | 1 | 0.03312479 | 2.14957636 | 0.084 | 0.021 | 1 | 0.43819216 | 0.06515234 |
| Echdc2 | 0.0331854 | 1.0392034 | 0.292 | 0.192 | 1 | 0.08714784 | 0.36821664 | 0.244 | 0.156 | 1 | 0.08714784 | 0.06526954 |
| Gins1 | 0.13862725 | 0.47697982 | 0.062 | 0.026 | 1 | 0.03319286 | -1.8948178 | 0.013 | 0.052 | 1 | 0.13862725 | 0.06528395 |
| Gpr75 | 0.61737927 | 1.00749454 | 0.01 | 0.005 | 1 | 0.03319286 | -1.8948178 | 0.013 | 0.052 | 1 | 0.61737927 | 0.06528395 |
| Zfp300 | 0.8163872 | 0.64492446 | 0.042 | 0.036 | 1 | 0.03319286 | -1.8948178 | 0.013 | 0.052 | 1 | 0.8163872 | 0.06528395 |
| Tex49 | 1 | -2.5160674 | 0.01 | 0.01 | 1 | 0.03319286 | -1.8948178 | 0.013 | 0.052 | 1 | 1 | 0.06528395 |
| Selenoo | 0.60710823 | -0.3848229 | 0.146 | 0.166 | 1 | 0.03322075 | 0.87234807 | 0.202 | 0.104 | 1 | 0.60710823 | 0.06533787 |
| Mospd3 | 0.89366809 | 0.14179563 | 0.333 | 0.332 | 1 | 0.03323372 | 0.33400093 | 0.282 | 0.156 | 1 | 0.89366809 | 0.06536296 |
| Col4a6 | 0.03323765 | -2.6363617 | 0.01 | 0.067 | 1 | 0.34363875 | -2.7248928 | 0.008 | 0.021 | 1 | 0.34363875 | 0.06537057 |
| Tsen34 | 0.09176783 | 0.17134735 | 0.771 | 0.668 | 1 | 0.03329418 | 0.14875125 | 0.786 | 0.667 | 1 | 0.09176783 | 0.06547986 |
| Strip2 | 0.03331566 | 1.22988696 | 0.062 | 0.016 | 1 | 0.97985874 | 0.62874419 | 0.042 | 0.042 | 1 | 0.97985874 | 0.06552139 |
| Lck | 0.205046 | -1.577468 | 0.01 | 0.036 | 1 | 0.03338151 | -1.9879272 | 0.013 | 0.052 | 1 | 0.205046 | 0.0656487 |
| Ddx42 | 0.0334297 | 0.23043354 | 0.833 | 0.762 | 1 | 0.76153718 | -0.1528588 | 0.853 | 0.885 | 1 | 0.76153718 | 0.06574186 |
| Tubb3 | 0.03345127 | 1.00749454 | 0.146 | 0.067 | 1 | 0.77165756 | -0.4029647 | 0.055 | 0.062 | 1 | 0.77165756 | 0.06578355 |
| Exoc5 | 0.30893684 | 0.15464823 | 0.688 | 0.679 | 1 | 0.03347005 | -0.4230142 | 0.824 | 0.854 | 1 | 0.30893684 | 0.06581985 |
| Gm42743 | 0.03356001 | -3.0799683 | 0.01 | 0.067 | 1 | 0.30037034 | -0.7952821 | 0.021 | 0.042 | 1 | 0.30037034 | 0.06599374 |
| Fam186b | 0.62104899 | 0.27052894 | 0.01 | 0.005 | 1 | 0.03357822 | -1.7952821 | 0.013 | 0.052 | 1 | 0.62104899 | 0.06602894 |
| Dok6 | 0.32025586 | -2.3144336 | 0 | 0.01 | 1 | 0.03358331 | -1.9879272 | 0.013 | 0.052 | 1 | 0.32025586 | 0.06603878 |
| Sertad2 | 0.20954323 | -0.2457658 | 0.896 | 0.922 | 1 | 0.03361246 | -0.4503491 | 0.929 | 0.906 | 1 | 0.20954323 | 0.06609512 |
| Rnf169 | 0.03363598 | 0.53599009 | 0.51 | 0.389 | 1 | 0.23212028 | -0.5605764 | 0.597 | 0.688 | 1 | 0.23212028 | 0.06614059 |
| Zfp35 | 0.14405605 | 0.45110119 | 0.167 | 0.104 | 1 | 0.03364037 | 0.87777174 | 0.134 | 0.052 | 1 | 0.14405605 | 0.06614906 |
| Drp2 | 0.03369746 | -2.4736322 | 0.052 | 0.13 | 1 | 0.12829617 | 0.32757466 | 0.092 | 0.042 | 1 | 0.12829617 | 0.06625941 |
| Ing4 | 0.24680091 | 0.19556688 | 0.531 | 0.451 | 1 | 0.03376484 | 0.18999062 | 0.513 | 0.354 | 1 | 0.24680091 | 0.06638962 |
| Zfp938 | 0.25818065 | 0.69433665 | 0.219 | 0.166 | 1 | 0.03381031 | 0.87456931 | 0.176 | 0.083 | 1 | 0.25818065 | 0.06647747 |
| Vasp | 0.37593418 | -0.2585176 | 0.646 | 0.648 | 1 | 0.03381921 | 0.27777642 | 0.819 | 0.781 | 1 | 0.37593418 | 0.06649468 |
| Katnal1 | 0.15390071 | 0.39082318 | 0.26 | 0.187 | 1 | 0.03382218 | 0.32486027 | 0.172 | 0.073 | 1 | 0.15390071 | 0.06650042 |
| Zfp142 | 0.03385181 | 0.31783466 | 0.448 | 0.311 | 1 | 0.14332942 | 0.17702224 | 0.487 | 0.365 | 1 | 0.14332942 | 0.06655767 |
| Sin3a | 0.28482767 | 0.24330747 | 0.354 | 0.285 | 1 | 0.03393582 | -0.612216 | 0.445 | 0.552 | 1 | 0.28482767 | 0.06672001 |
| Pms1 | 0.14754637 | 1.00749454 | 0.083 | 0.041 | 1 | 0.03394783 | 1.86006974 | 0.067 | 0.01 | 1 | 0.14754637 | 0.06674321 |
| Plk4 | 0.03397927 | 1.34452952 | 0.135 | 0.062 | 1 | 0.70474274 | 0.53814164 | 0.084 | 0.073 | 1 | 0.70474274 | 0.06680395 |
| Tgds | 0.18453967 | 0.61517711 | 0.25 | 0.181 | 1 | 0.03398153 | 0.71120635 | 0.244 | 0.135 | 1 | 0.18453967 | 0.06680832 |
| Scmh1 | 0.4309446 | 0.24615201 | 0.688 | 0.637 | 1 | 0.03400962 | -0.5222405 | 0.849 | 0.854 | 1 | 0.4309446 | 0.06686259 |
| Fem1a | 0.03406026 | 1.13302542 | 0.115 | 0.047 | 1 | 0.18835053 | 0.15580831 | 0.202 | 0.135 | 1 | 0.18835053 | 0.06696043 |
| Pls3 | 0.03407155 | -0.821025 | 0.51 | 0.632 | 1 | 0.13659731 | -0.5258922 | 0.672 | 0.708 | 1 | 0.13659731 | 0.06698223 |
| Nxf1 | 0.03414632 | 0.34091827 | 0.688 | 0.534 | 1 | 0.67615268 | -0.1846547 | 0.761 | 0.708 | 1 | 0.67615268 | 0.06712667 |
| Dnajc13 | 0.03415451 | 0.1830661 | 0.719 | 0.591 | 1 | 0.46161011 | -0.2471195 | 0.761 | 0.792 | 1 | 0.46161011 | 0.06714249 |
| Card6 | 0.46101073 | 0.27052894 | 0.062 | 0.041 | 1 | 0.03416998 | -1.5728897 | 0.013 | 0.052 | 1 | 0.46101073 | 0.06717237 |
| Pgghg | 0.0546061 | 1.06889508 | 0.188 | 0.109 | 1 | 0.03417619 | 1.08246216 | 0.16 | 0.073 | 1 | 0.0546061 | 0.06718436 |
| Megf8 | 0.0342262 | 0.58299671 | 0.312 | 0.197 | 1 | 0.88052962 | -0.3516754 | 0.239 | 0.219 | 1 | 0.88052962 | 0.06728096 |
| Tnfrsf23 | 0.03423499 | -1.1529701 | 0.365 | 0.466 | 1 | 0.28962914 | -0.5728897 | 0.336 | 0.229 | 1 | 0.28962914 | 0.06729795 |
| Dhh | 0.0342624 | -1.7998604 | 0.01 | 0.067 | 1 | 0.18504625 | 1.01207283 | 0.038 | 0.01 | 1 | 0.18504625 | 0.06735088 |
| Echdc1 | 0.36628513 | -0.1683553 | 0.375 | 0.409 | 1 | 0.0342685 | 0.12902898 | 0.37 | 0.219 | 1 | 0.36628513 | 0.06736268 |
| Vps52 | 0.03439815 | 1.06889508 | 0.198 | 0.109 | 1 | 0.07385113 | 0.3068161 | 0.176 | 0.094 | 1 | 0.07385113 | 0.06761306 |
| Bcl6 | 0.05624218 | -0.2941612 | 0.646 | 0.746 | 1 | 0.03440608 | -0.4579539 | 0.836 | 0.927 | 1 | 0.05624218 | 0.06762838 |
| Aak1 | 0.03445955 | 0.11703738 | 0.969 | 0.912 | 1 | 0.94144927 | -0.2736203 | 0.983 | 0.969 | 1 | 0.94144927 | 0.06773163 |
| Rap1a | 0.62998672 | 0.15155649 | 0.938 | 0.959 | 1 | 0.03447467 | 0.12610808 | 0.975 | 0.99 | 1 | 0.62998672 | 0.06776084 |
| Hal | 0.03448769 | 0.4440652 | 0.073 | 0.161 | 1 | 0.04475792 | -1.9199087 | 0.055 | 0.115 | 1 | 0.04475792 | 0.06778598 |
| Armc9 | 0.03449385 | 0.50999488 | 0.271 | 0.161 | 1 | 0.69373361 | -0.3923174 | 0.294 | 0.25 | 1 | 0.69373361 | 0.06779788 |
| Rfc4 | 0.03451382 | 1.00749454 | 0.146 | 0.067 | 1 | 0.61832604 | 0.36821664 | 0.055 | 0.042 | 1 | 0.61832604 | 0.06783644 |
| Zfp414 | 0.20411833 | 0.53105649 | 0.198 | 0.14 | 1 | 0.03455015 | 1.42711033 | 0.13 | 0.052 | 1 | 0.20411833 | 0.06790659 |
| Pla2g15 | 0.03455511 | -1.0431315 | 0.042 | 0.119 | 1 | 0.10907373 | 0.31817596 | 0.126 | 0.062 | 1 | 0.10907373 | 0.06791616 |
| Ggct | 0.03456851 | -1.1079827 | 0.167 | 0.269 | 1 | 0.09134834 | 0.81942775 | 0.113 | 0.052 | 1 | 0.09134834 | 0.06794204 |
| Timm17b | 0.3026145 | 0.25260703 | 0.458 | 0.409 | 1 | 0.03462076 | 0.22930355 | 0.298 | 0.167 | 1 | 0.3026145 | 0.06804292 |
| Hao1 | 0.03464025 | -0.9925055 | 0.031 | 0.104 | 1 | 0.29614538 | -1.8948178 | 0.029 | 0.052 | 1 | 0.29614538 | 0.06808056 |
| Msantd2 | 0.0346427 | 0.38600616 | 0.615 | 0.456 | 1 | 0.45379935 | -0.384623 | 0.773 | 0.688 | 1 | 0.45379935 | 0.06808529 |
| Cacna1g | 0.05048655 | -1.940038 | 0.125 | 0.202 | 1 | 0.03465985 | -1.6075358 | 0.134 | 0.219 | 1 | 0.05048655 | 0.06811839 |
| Tstd2 | 0.07435966 | 0.61015904 | 0.312 | 0.218 | 1 | 0.03466593 | 0.11715488 | 0.378 | 0.219 | 1 | 0.07435966 | 0.06813012 |
| Gm42047 | 0.03468955 | 1.26875985 | 0.281 | 0.187 | 1 | 0.63462732 | 0.16221318 | 0.139 | 0.115 | 1 | 0.63462732 | 0.06817573 |
| Rusf1 | 0.03470164 | 0.67045955 | 0.177 | 0.088 | 1 | 0.11531866 | 0.42711033 | 0.109 | 0.052 | 1 | 0.11531866 | 0.06819908 |
| H2ac20 | 0.23288369 | 1.81484946 | 0.094 | 0.057 | 1 | 0.03478483 | 1.69014474 | 0.084 | 0.021 | 1 | 0.23288369 | 0.06835968 |
| Nqo1 | 0.07284125 | -0.8919786 | 0.344 | 0.409 | 1 | 0.03479544 | 0.61859148 | 0.311 | 0.198 | 1 | 0.07284125 | 0.06838016 |
| Dcaf1 | 0.0348144 | 0.19801644 | 0.521 | 0.368 | 1 | 0.11653672 | -0.6724253 | 0.559 | 0.604 | 1 | 0.11653672 | 0.06841675 |
| Prkx | 0.75147852 | 0.19556688 | 0.271 | 0.259 | 1 | 0.03481863 | -0.6070563 | 0.416 | 0.542 | 1 | 0.75147852 | 0.06842493 |
| Vps26b | 0.38004534 | -0.3910548 | 0.094 | 0.13 | 1 | 0.03486503 | 0.49749966 | 0.261 | 0.146 | 1 | 0.38004534 | 0.06851449 |
| Ilvbl | 0.34238963 | 0.46983775 | 0.25 | 0.207 | 1 | 0.0348727 | 0.76053407 | 0.265 | 0.156 | 1 | 0.34238963 | 0.0685293 |
| Mrpl19 | 0.03487918 | 1.00749454 | 0.24 | 0.145 | 1 | 0.15145304 | 0.30157945 | 0.181 | 0.115 | 1 | 0.15145304 | 0.06854181 |
| Aox1 | 0.51221958 | -0.577468 | 0.115 | 0.088 | 1 | 0.03497029 | -2.3493836 | 0.067 | 0.135 | 1 | 0.51221958 | 0.06871766 |
| 2610008E11Rik | 0.0851847 | 0.2816695 | 0.365 | 0.259 | 1 | 0.03501244 | 0.23304318 | 0.424 | 0.292 | 1 | 0.0851847 | 0.06879901 |
| Pigg | 0.26424825 | 0.37522632 | 0.188 | 0.135 | 1 | 0.03501796 | 0.74903843 | 0.164 | 0.073 | 1 | 0.26424825 | 0.06880965 |
| Calm3 | 0.28952192 | 0.31920279 | 0.781 | 0.731 | 1 | 0.03503157 | 0.15548732 | 0.811 | 0.74 | 1 | 0.28952192 | 0.06883593 |
| Vps13d | 0.45358556 | 0.14809287 | 0.656 | 0.56 | 1 | 0.03506586 | -0.6171541 | 0.832 | 0.885 | 1 | 0.45358556 | 0.06890211 |
| Gimap6 | 0.08831993 | 0.78510212 | 0.135 | 0.073 | 1 | 0.03508263 | 0.92718393 | 0.118 | 0.042 | 1 | 0.08831993 | 0.06893447 |
| Sema3e | 0.03512879 | -1.9626898 | 0.125 | 0.223 | 1 | 0.84548937 | -1.3098553 | 0.059 | 0.062 | 1 | 0.84548937 | 0.06902355 |

| | | | | | | | | | | | | |
|---|---|---|---|---|---|---|---|---|---|---|---|---|
| Klhdc10 | 0.03514714 | 0.58547061 | 0.531 | 0.404 | 1 | 0.11303799 | -0.4991977 | 0.626 | 0.719 | 1 | 0.11303799 | 0.06905895 |
| Hdac10 | 0.12701924 | 1.22988696 | 0.062 | 0.026 | 1 | 0.0351842 | 0.61614416 | 0.067 | 0.01 | 1 | 0.12701924 | 0.06913047 |
| Fam89a | 0.38253808 | -0.4320375 | 0.281 | 0.316 | 1 | 0.03520797 | 0.78968041 | 0.067 | 0.01 | 1 | 0.38253808 | 0.06917633 |
| Txlna | 0.03522448 | 0.29529685 | 0.729 | 0.653 | 1 | 0.21873123 | 0.24610737 | 0.727 | 0.667 | 1 | 0.21873123 | 0.0692082 |
| Gpat3 | 0.10315734 | -0.6363617 | 0.156 | 0.244 | 1 | 0.03523512 | 0.32019513 | 0.168 | 0.073 | 1 | 0.10315734 | 0.06922874 |
| Ica1 | 0.29307367 | -0.2159437 | 0.646 | 0.663 | 1 | 0.03525704 | -0.1922857 | 0.504 | 0.323 | 1 | 0.29307367 | 0.06927103 |
| Gdi1 | 0.38033042 | 0.10703021 | 0.562 | 0.503 | 1 | 0.03525868 | 0.47214409 | 0.622 | 0.542 | 1 | 0.38033042 | 0.06927419 |
| Wdr47 | 0.03528161 | -1.4606543 | 0.229 | 0.326 | 1 | 0.04228925 | -0.926151 | 0.361 | 0.427 | 1 | 0.04228925 | 0.06931843 |
| Gfod2 | 0.38714491 | 0.51716891 | 0.177 | 0.14 | 1 | 0.03529985 | 0.48270266 | 0.244 | 0.135 | 1 | 0.38714491 | 0.06935362 |
| Fmr1 | 0.15496057 | 0.15303538 | 0.75 | 0.705 | 1 | 0.03530906 | 0.13419332 | 0.861 | 0.833 | 1 | 0.15496057 | 0.06937139 |
| Garre1 | 0.03533517 | -0.3999298 | 0.833 | 0.829 | 1 | 0.22691378 | -0.304923 | 0.832 | 0.875 | 1 | 0.22691378 | 0.06942177 |
| Ercc8 | 0.03536133 | 0.84995326 | 0.188 | 0.098 | 1 | 0.06627814 | 0.57466752 | 0.164 | 0.083 | 1 | 0.06627814 | 0.06947224 |
| Cpne2 | 0.06844987 | -0.9241185 | 0.198 | 0.29 | 1 | 0.03539764 | 0.27510724 | 0.378 | 0.24 | 1 | 0.06844987 | 0.06954228 |
| Washc1 | 0.04537401 | 0.60695661 | 0.24 | 0.14 | 1 | 0.03539948 | 0.93807225 | 0.189 | 0.094 | 1 | 0.04537401 | 0.06954583 |
| Pop7 | 0.26012623 | 0.25260703 | 0.281 | 0.218 | 1 | 0.03545207 | 0.43557191 | 0.206 | 0.104 | 1 | 0.26012623 | 0.06964729 |
| Spata5l1 | 0.31785244 | 0.68556644 | 0.062 | 0.036 | 1 | 0.03548698 | 0.69014474 | 0.118 | 0.042 | 1 | 0.31785244 | 0.06971463 |
| Dusp19 | 0.10896908 | 0.76648644 | 0.188 | 0.119 | 1 | 0.03550948 | 1.3146356 | 0.13 | 0.052 | 1 | 0.10896908 | 0.06975804 |
| Tmem41a | 0.08160299 | 0.86999101 | 0.094 | 0.041 | 1 | 0.03550969 | 1.27510724 | 0.13 | 0.052 | 1 | 0.08160299 | 0.06975844 |
| Vldlr | 0.35947144 | 0.95858494 | 0.115 | 0.083 | 1 | 0.03552522 | -0.8948178 | 0.071 | 0.146 | 1 | 0.35947144 | 0.0697884 |
| Ten1 | 0.03554418 | 0.59245704 | 0.25 | 0.145 | 1 | 0.05688573 | 0.69014474 | 0.164 | 0.083 | 1 | 0.05688573 | 0.06982497 |
| Commd8 | 0.3292182 | 0.10060394 | 0.344 | 0.28 | 1 | 0.03561813 | 0.18368421 | 0.349 | 0.208 | 1 | 0.3292182 | 0.06996761 |
| 4930556M19Rik | 0.0356739 | 1.64492446 | 0.104 | 0.041 | 1 | 0.24435619 | -0.8948178 | 0.042 | 0.073 | 1 | 0.24435619 | 0.07007517 |
| Ldlrad2 | 0.03569091 | 0.89201732 | 0.115 | 0.047 | 1 | 0.08510873 | 0.78968041 | 0.101 | 0.042 | 1 | 0.08510873 | 0.07010798 |
| Gprc5a | 0.24614867 | 0.24195979 | 0.167 | 0.119 | 1 | 0.03572572 | 1.8995981 | 0.084 | 0.021 | 1 | 0.24614867 | 0.07017512 |
| Haus7 | 0.14653285 | 0.50225923 | 0.25 | 0.176 | 1 | 0.03576165 | 0.55687821 | 0.206 | 0.104 | 1 | 0.14653285 | 0.07024441 |
| Trim37 | 0.05253625 | 0.69298391 | 0.427 | 0.332 | 1 | 0.03576286 | 0.29218075 | 0.433 | 0.271 | 1 | 0.05253625 | 0.07024675 |
| Cmtm8 | 0.0357665 | 1.26465238 | 0.229 | 0.14 | 1 | 0.04814793 | 0.46893051 | 0.513 | 0.417 | 1 | 0.04814793 | 0.07025376 |
| Wdr46 | 0.0357667 | 0.74446013 | 0.167 | 0.083 | 1 | 0.24271058 | 0.3146356 | 0.13 | 0.083 | 1 | 0.24271058 | 0.07025415 |
| Lbh | 0.0357847 | -0.9644911 | 0.323 | 0.44 | 1 | 0.68564398 | -0.7248928 | 0.126 | 0.135 | 1 | 0.68564398 | 0.07028886 |
| Ppp1r37 | 0.37688977 | 0.14577391 | 0.583 | 0.528 | 1 | 0.03579278 | 0.11898804 | 0.739 | 0.604 | 1 | 0.37688977 | 0.07030444 |
| Sos1 | 0.03580169 | 0.17212524 | 0.573 | 0.409 | 1 | 0.50477081 | -0.4858329 | 0.664 | 0.698 | 1 | 0.50477081 | 0.07032161 |
| Zfp472 | 0.03585999 | 1.59245704 | 0.104 | 0.041 | 1 | 0.13750701 | 0.56461386 | 0.118 | 0.062 | 1 | 0.13750701 | 0.07043404 |
| Lipg | 0.03593697 | -2.7596713 | 0.094 | 0.181 | 1 | 0.79600331 | -0.9879272 | 0.017 | 0.021 | 1 | 0.79600331 | 0.07058247 |
| Wbp1 | 0.03598042 | -0.9311049 | 0.156 | 0.259 | 1 | 0.64594518 | -0.1668973 | 0.172 | 0.146 | 1 | 0.64594518 | 0.07066624 |
| Prim1 | 0.11158492 | 0.82108141 | 0.177 | 0.109 | 1 | 0.03606566 | 1.0961371 | 0.147 | 0.062 | 1 | 0.11158492 | 0.07083059 |
| Retsat | 0.44522649 | -0.5070786 | 0.062 | 0.088 | 1 | 0.0360843 | 0.96316323 | 0.118 | 0.042 | 1 | 0.44522649 | 0.07086652 |
| Traf3ip1 | 0.22459103 | 0.65957123 | 0.219 | 0.166 | 1 | 0.03611334 | 0.58630893 | 0.193 | 0.094 | 1 | 0.22459103 | 0.07092251 |
| Avpr1a | 0.63812871 | -0.8290067 | 0.177 | 0.192 | 1 | 0.03619723 | 1.01207283 | 0.101 | 0.031 | 1 | 0.63812871 | 0.07108421 |
| Gm12905 | 0.4567757 | -0.9185049 | 0.094 | 0.119 | 1 | 0.03623043 | 0.79705994 | 0.176 | 0.083 | 1 | 0.4567757 | 0.07114821 |
| Ints4 | 0.03623916 | 1.07964432 | 0.26 | 0.166 | 1 | 0.09033964 | -0.1004019 | 0.315 | 0.198 | 1 | 0.09033964 | 0.07116504 |
| Sesn1 | 0.11695759 | -0.2062122 | 0.562 | 0.425 | 1 | 0.0362533 | -0.8684836 | 0.534 | 0.646 | 1 | 0.11695759 | 0.07119229 |
| Wbp2 | 0.03626458 | -0.5391398 | 0.625 | 0.648 | 1 | 0.10762654 | 0.16635957 | 0.664 | 0.594 | 1 | 0.10762654 | 0.07121404 |
| Slc25a42 | 0.03632901 | 1.81484946 | 0.125 | 0.057 | 1 | 0.48195431 | -0.8623963 | 0.05 | 0.031 | 1 | 0.48195431 | 0.07133821 |
| Necap2 | 0.19896807 | 0.17848073 | 0.583 | 0.492 | 1 | 0.03639559 | 0.36935756 | 0.634 | 0.5 | 1 | 0.19896807 | 0.07146654 |
| Cfap298 | 0.25159546 | 0.14499806 | 0.271 | 0.202 | 1 | 0.03640994 | 0.59703533 | 0.256 | 0.146 | 1 | 0.25159546 | 0.07149419 |
| Mpp7 | 0.06231305 | 0.18646468 | 0.292 | 0.399 | 1 | 0.03644123 | -0.7589202 | 0.538 | 0.656 | 1 | 0.06231305 | 0.07155449 |
| Zfp62 | 0.03644347 | 0.65135073 | 0.354 | 0.238 | 1 | 0.70709708 | -0.4006207 | 0.445 | 0.365 | 1 | 0.70709708 | 0.07155881 |
| Tpmt | 0.67639218 | 0.36363835 | 0.125 | 0.109 | 1 | 0.03648465 | 1.19264508 | 0.13 | 0.052 | 1 | 0.67639218 | 0.07163816 |
| Hps4 | 0.03664802 | 0.67818691 | 0.333 | 0.218 | 1 | 0.19668661 | 0.20471791 | 0.21 | 0.146 | 1 | 0.19668661 | 0.07195296 |
| Ndc1 | 0.0367023 | 1.24850264 | 0.177 | 0.093 | 1 | 0.08938415 | 0.16407593 | 0.143 | 0.073 | 1 | 0.08938415 | 0.07205755 |
| Gpr173 | 0.03672822 | 0.68556644 | 0.188 | 0.098 | 1 | 0.67662179 | -0.6724253 | 0.139 | 0.115 | 1 | 0.67662179 | 0.07210748 |
| Gm56955 | 0.49654876 | -0.8504865 | 0.115 | 0.14 | 1 | 0.03674709 | 0.21370669 | 0.118 | 0.042 | 1 | 0.49654876 | 0.07214383 |
| Arhgap21 | 0.03674931 | 0.22345026 | 0.885 | 0.86 | 1 | 0.10376608 | -0.3444421 | 0.945 | 0.969 | 1 | 0.10376608 | 0.0721481 |
| Lgr6 | 0.15761659 | -0.582175 | 0.354 | 0.425 | 1 | 0.03678221 | -0.8055112 | 0.399 | 0.542 | 1 | 0.15761659 | 0.07221149 |
| Slc30a4 | 0.03680796 | 0.32942263 | 0.438 | 0.295 | 1 | 0.45182135 | -0.198824 | 0.424 | 0.344 | 1 | 0.45182135 | 0.07226108 |
| Herc1 | 0.036842 | 0.24479138 | 0.854 | 0.819 | 1 | 0.79834124 | -0.1396116 | 0.924 | 0.927 | 1 | 0.79834124 | 0.07232666 |
| Slc15a4 | 0.0432123 | 0.77922555 | 0.271 | 0.171 | 1 | 0.03686523 | 0.37464291 | 0.294 | 0.177 | 1 | 0.0432123 | 0.07237141 |
| Hexdc | 0.03692898 | 1.42253204 | 0.104 | 0.041 | 1 | 0.0785613 | 0.41596977 | 0.143 | 0.073 | 1 | 0.0785613 | 0.07249421 |
| Ggnbp1 | 0.03692898 | 1.42253204 | 0.104 | 0.041 | 1 | 0.60342377 | 0.27510724 | 0.067 | 0.052 | 1 | 0.60342377 | 0.07249421 |
| Hltf | 0.8968686 | 0.18528166 | 0.375 | 0.383 | 1 | 0.03694717 | 0.47564023 | 0.366 | 0.229 | 1 | 0.8968686 | 0.07252924 |
| Psma6 | 0.03700099 | 0.34036349 | 0.802 | 0.731 | 1 | 0.07231345 | 0.1519305 | 0.84 | 0.771 | 1 | 0.07231345 | 0.07263292 |
| Trpm7 | 0.03700257 | 0.11875129 | 0.948 | 0.876 | 1 | 0.82292007 | -0.134981 | 0.971 | 1 | 1 | 0.82292007 | 0.07263596 |
| Pet100 | 0.63676213 | -0.1789186 | 0.24 | 0.264 | 1 | 0.03701448 | 0.80562195 | 0.176 | 0.083 | 1 | 0.63676213 | 0.07265888 |
| A3galt2 | 0.03702426 | 0.27052894 | 0.208 | 0.114 | 1 | 0.49849487 | -0.4374053 | 0.328 | 0.26 | 1 | 0.49849487 | 0.07267773 |
| Vmac | 0.30232292 | 0.31561683 | 0.229 | 0.176 | 1 | 0.03709399 | 0.22856465 | 0.227 | 0.115 | 1 | 0.30232292 | 0.07281202 |
| Snx25 | 0.19668006 | -0.4632969 | 0.458 | 0.539 | 1 | 0.03712744 | 0.22518201 | 0.643 | 0.438 | 1 | 0.19668006 | 0.07287643 |
| Plekhm1 | 0.03714033 | -0.9593386 | 0.156 | 0.259 | 1 | 0.99506032 | -0.2370989 | 0.307 | 0.292 | 1 | 0.99506032 | 0.07290125 |
| Cep89 | 0.71117949 | 0.10703021 | 0.083 | 0.098 | 1 | 0.03715911 | 1.20471791 | 0.13 | 0.052 | 1 | 0.71117949 | 0.07293742 |
| Eef1akmt3 | 0.03716355 | 1.21899864 | 0.156 | 0.078 | 1 | 0.34913161 | -0.1630139 | 0.097 | 0.062 | 1 | 0.34913161 | 0.07294597 |
| Kctd17 | 0.49405817 | 0.30388754 | 0.271 | 0.233 | 1 | 0.03718654 | 0.54575483 | 0.252 | 0.146 | 1 | 0.49405817 | 0.07299024 |
| Klf3 | 0.03720588 | 0.80396114 | 0.802 | 0.834 | 1 | 0.04995663 | 0.64074793 | 0.937 | 0.927 | 1 | 0.04995663 | 0.07302748 |
| Camsap1 | 0.03727398 | 0.1473522 | 0.531 | 0.373 | 1 | 0.06681147 | 0.12013258 | 0.718 | 0.646 | 1 | 0.06681147 | 0.07315861 |
| Clstn1 | 0.41412459 | -0.1624305 | 0.604 | 0.679 | 1 | 0.03733784 | 0.14957636 | 0.819 | 0.812 | 1 | 0.41412459 | 0.07328156 |
| Rgs20 | 0.03734545 | -2.6554705 | 0.021 | 0.083 | 1 | 0.31054693 | 0.69014474 | 0.029 | 0.01 | 1 | 0.31054693 | 0.07329621 |
| Mettl13 | 0.48342698 | 0.1843723 | 0.104 | 0.078 | 1 | 0.03736583 | 0.69014474 | 0.147 | 0.062 | 1 | 0.48342698 | 0.07333546 |
| Kcnj2 | 0.05972787 | -4.6072153 | 0 | 0.036 | 1 | 0.03736625 | -3.3098553 | 0.008 | 0.042 | 1 | 0.05972787 | 0.07333626 |
| Gm49003 | 0.48525351 | -0.577468 | 0.021 | 0.01 | 1 | 0.03736625 | -3.3098553 | 0.008 | 0.042 | 1 | 0.48525351 | 0.07333626 |
| Ampd2 | 0.03737372 | 0.51650818 | 0.312 | 0.197 | 1 | 0.66478534 | -0.5361311 | 0.328 | 0.312 | 1 | 0.66478534 | 0.07335064 |
| Ctdp1 | 0.24207252 | 0.41055939 | 0.438 | 0.378 | 1 | 0.03737589 | 0.15248795 | 0.58 | 0.385 | 1 | 0.24207252 | 0.07335482 |
| 4930503L19Rik | 0.4006652 | 0.29700115 | 0.167 | 0.13 | 1 | 0.03739367 | 0.98292649 | 0.16 | 0.073 | 1 | 0.4006652 | 0.07338905 |
| Slc9a8 | 0.03744088 | 0.53971558 | 0.375 | 0.259 | 1 | 0.11972171 | -0.142958 | 0.349 | 0.24 | 1 | 0.11972171 | 0.07347993 |
| Mrpl49 | 0.22051937 | 0.48812038 | 0.25 | 0.192 | 1 | 0.03745224 | 0.59196434 | 0.244 | 0.135 | 1 | 0.22051937 | 0.07350181 |
| 1110038F14Rik | 0.13990857 | 0.47956298 | 0.312 | 0.233 | 1 | 0.03746216 | 0.32757466 | 0.244 | 0.135 | 1 | 0.13990857 | 0.0735209 |
| Rnpepl1 | 0.72559473 | 0.19191911 | 0.198 | 0.181 | 1 | 0.03748616 | 0.46458504 | 0.235 | 0.125 | 1 | 0.72559473 | 0.0735671 |
| Zbtb11 | 0.03751101 | 0.19246342 | 0.781 | 0.689 | 1 | 0.37536366 | -0.2006485 | 0.824 | 0.875 | 1 | 0.37536366 | 0.07361495 |
| Tmsb15b2 | 0.03751848 | 0.68556644 | 0.229 | 0.13 | 1 | 0.04930108 | 0.44913664 | 0.155 | 0.073 | 1 | 0.04930108 | 0.07362933 |
| Cebpa | 0.03754045 | 1.18806678 | 0.135 | 0.062 | 1 | 0.03932498 | 0.86006974 | 0.21 | 0.115 | 1 | 0.03932498 | 0.07367161 |
| Chmp1b2 | 0.07865608 | 0.60695661 | 0.219 | 0.135 | 1 | 0.03754811 | 0.36257008 | 0.168 | 0.073 | 1 | 0.07865608 | 0.07368636 |
| Atp1a2 | 0.39477962 | -1.9518635 | 0.052 | 0.078 | 1 | 0.03757149 | -1.0358616 | 0.197 | 0.094 | 1 | 0.39477962 | 0.07373136 |

| | | | | | | | | | | | |
|---|---|---|---|---|---|---|---|---|---|---|---|
| Ccdc130 | 0.037576 | 1.24195979 | 0.156 | 0.078 | 1 | 0.35749142 | 0.31817596 | 0.105 | 0.073 | 1 | 0.35749142 | 0.07374004 |
| Rgs12 | 0.56752121 | 0.72370157 | 0.135 | 0.114 | 1 | 0.0376383 | -0.9175378 | 0.139 | 0.229 | 1 | 0.56752121 | 0.07385996 |
| Aatf | 0.03765252 | 0.47315811 | 0.469 | 0.321 | 1 | 0.97514441 | -0.5251929 | 0.42 | 0.365 | 1 | 0.97514441 | 0.07388733 |
| Tnn | 0.03766528 | -2.765095 | 0.021 | 0.083 | 1 | 0.92223547 | 0.20471791 | 0.055 | 0.052 | 1 | 0.92223547 | 0.07391188 |
| Xylt2 | 0.37594369 | -0.8405024 | 0.042 | 0.067 | 1 | 0.03776915 | 0.80562195 | 0.101 | 0.031 | 1 | 0.37594369 | 0.0741118 |
| Aimp1 | 0.03779634 | 0.29839574 | 0.615 | 0.487 | 1 | 0.11966399 | -0.4711362 | 0.5 | 0.594 | 1 | 0.11966399 | 0.07416411 |
| Crtc2 | 0.82575396 | 0.10370985 | 0.25 | 0.244 | 1 | 0.03780991 | 0.20259474 | 0.328 | 0.198 | 1 | 0.82575396 | 0.07419023 |
| Dis3l | 0.20519383 | 0.78510212 | 0.104 | 0.062 | 1 | 0.03783356 | 0.22357694 | 0.164 | 0.073 | 1 | 0.20519383 | 0.07423574 |
| Gas1 | 0.04915512 | 0.64127206 | 0.49 | 0.389 | 1 | 0.03784243 | 0.48815423 | 0.529 | 0.385 | 1 | 0.04915512 | 0.0742528 |
| Gnb3 | 0.22187575 | -0.9925055 | 0 | 0.016 | 1 | 0.03786471 | -3.0468209 | 0.008 | 0.042 | 1 | 0.22187575 | 0.07429569 |
| Fsip1 | 0.21708019 | 2.00749454 | 0.021 | 0.005 | 1 | 0.03786705 | -3.4797803 | 0.008 | 0.042 | 1 | 0.21708019 | 0.07430019 |
| Fam169b | 0.39356588 | -0.1624305 | 0.01 | 0.026 | 1 | 0.03786734 | -3.3098553 | 0.008 | 0.042 | 1 | 0.39356588 | 0.07430075 |
| Spry1 | 0.03798868 | -2.265524 | 0.021 | 0.083 | 1 | 0.04830554 | -2.6634922 | 0.034 | 0.083 | 1 | 0.04830554 | 0.07453423 |
| Rcan2 | 0.03799571 | 1.88885004 | 0.125 | 0.057 | 1 | 0.3201338 | -0.7248928 | 0.029 | 0.052 | 1 | 0.3201338 | 0.07454775 |
| Cacng7 | 0.03806549 | 0.43921078 | 0.24 | 0.135 | 1 | 0.85495617 | -0.525584 | 0.269 | 0.26 | 1 | 0.85495617 | 0.074682 |
| Parp14 | 0.24505505 | -0.536826 | 0.052 | 0.093 | 1 | 0.03806599 | -1.9879272 | 0.076 | 0.146 | 1 | 0.24505505 | 0.07468296 |
| Atp13a3 | 0.03806793 | -0.2649245 | 0.635 | 0.798 | 1 | 0.17462409 | -0.2303478 | 0.87 | 0.938 | 1 | 0.17462409 | 0.0746867 |
| Slc39a11 | 0.0380971 | -0.9274104 | 0.385 | 0.497 | 1 | 0.61403106 | -1.4253325 | 0.298 | 0.292 | 1 | 0.61403106 | 0.0747428 |
| Ahrr | 0.99660398 | 0.59245704 | 0.01 | 0.01 | 1 | 0.03811841 | -3.010295 | 0.008 | 0.042 | 1 | 0.99660398 | 0.07478381 |
| Pip4k2a | 0.11529125 | 0.19625618 | 0.281 | 0.192 | 1 | 0.0381457 | -1.1193334 | 0.252 | 0.344 | 1 | 0.11529125 | 0.0748363 |
| Znfx1 | 0.7787952 | 0.2912875 | 0.188 | 0.176 | 1 | 0.03822046 | -0.2601022 | 0.366 | 0.219 | 1 | 0.7787952 | 0.07498011 |
| 9830144P21Rik | 0.03831159 | -2.0513992 | 0.021 | 0.083 | 1 | 0.17334368 | -2.0874628 | 0.017 | 0.042 | 1 | 0.17334368 | 0.07515541 |
| Eml2 | 0.32594768 | 0.3239682 | 0.26 | 0.207 | 1 | 0.03832947 | -0.2713811 | 0.269 | 0.146 | 1 | 0.32594768 | 0.0751898 |
| Mccc1os | 0.21867159 | 1.59245704 | 0.021 | 0.005 | 1 | 0.03836766 | -2.5322477 | 0.008 | 0.042 | 1 | 0.21867159 | 0.07526323 |
| Nuak2 | 0.32025586 | -0.9925055 | 0 | 0.01 | 1 | 0.03836766 | -2.3098553 | 0.008 | 0.042 | 1 | 0.32025586 | 0.07526323 |
| Slfn5 | 0.32025586 | -1.9925055 | 0 | 0.01 | 1 | 0.03836766 | -2.3098553 | 0.008 | 0.042 | 1 | 0.32025586 | 0.07526323 |
| Mfap5 | 1 | -1.1624305 | 0.01 | 0.01 | 1 | 0.03836766 | -3.1843244 | 0.008 | 0.042 | 1 | 1 | 0.07526323 |
| Mfsd13b | 0.61550914 | -0.8669746 | 0.021 | 0.031 | 1 | 0.03837237 | -2.8948178 | 0.008 | 0.042 | 1 | 0.61550914 | 0.07527229 |
| Ncaph | 0.47599549 | 0.52206771 | 0.042 | 0.026 | 1 | 0.03837325 | -2.3098553 | 0.008 | 0.042 | 1 | 0.47599549 | 0.07527399 |
| Nmrk1 | 0.08433383 | 0.407102 | 0.24 | 0.15 | 1 | 0.03841226 | 0.61071027 | 0.176 | 0.083 | 1 | 0.08433383 | 0.07534901 |
| Lrrc73 | 0.0384257 | 1.38600616 | 0.104 | 0.041 | 1 | 0.42714514 | 0.46775232 | 0.088 | 0.062 | 1 | 0.42714514 | 0.07537487 |
| 2810002D19Rik | 0.47586556 | 0.67045955 | 0.135 | 0.109 | 1 | 0.03849798 | 0.77760758 | 0.16 | 0.073 | 1 | 0.47586556 | 0.07551387 |
| Marcks | 0.03851048 | -0.8064231 | 0.604 | 0.389 | 1 | 0.07001541 | -0.7864851 | 0.634 | 0.74 | 1 | 0.07001541 | 0.07553791 |
| 9430091E24Rik | 0.03859304 | 1.27052894 | 0.104 | 0.041 | 1 | 0.34281303 | -0.2759079 | 0.139 | 0.094 | 1 | 0.34281303 | 0.07569666 |
| Calhm5 | 0.74160752 | -0.577468 | 0.073 | 0.083 | 1 | 0.03860619 | -2.5993619 | 0.025 | 0.073 | 1 | 0.74160752 | 0.07572194 |
| Kcnip3 | 0.52961488 | -0.3144336 | 0.01 | 0.021 | 1 | 0.03862717 | -2.4797803 | 0.008 | 0.042 | 1 | 0.52961488 | 0.07576228 |
| Plppr3 | 0.78979801 | -0.2148979 | 0.021 | 0.026 | 1 | 0.03862717 | -2.3098553 | 0.008 | 0.042 | 1 | 0.78979801 | 0.07576228 |
| Cldn12 | 0.38360516 | -0.2779077 | 0.135 | 0.176 | 1 | 0.03866177 | 0.59196434 | 0.244 | 0.135 | 1 | 0.38360516 | 0.0758288 |
| Nt5c2 | 0.14035216 | 0.31481822 | 0.5 | 0.389 | 1 | 0.03876345 | -0.5079824 | 0.433 | 0.562 | 1 | 0.14035216 | 0.0760243 |
| Impdh1 | 0.03879703 | 0.72370157 | 0.208 | 0.114 | 1 | 0.55813248 | -0.2625495 | 0.227 | 0.188 | 1 | 0.55813248 | 0.07608884 |
| Iqcg | 0.07877705 | 2.00749454 | 0.042 | 0.01 | 1 | 0.03888519 | -2.0468209 | 0.008 | 0.042 | 1 | 0.07877705 | 0.07625832 |
| Oas1b | 0.15729781 | -1.7998604 | 0 | 0.021 | 1 | 0.03888519 | -2.0468209 | 0.008 | 0.042 | 1 | 0.15729781 | 0.07625832 |
| Oscp1 | 0.03891202 | 0.77302928 | 0.167 | 0.083 | 1 | 0.93841483 | -0.6853644 | 0.126 | 0.125 | 1 | 0.93841483 | 0.07630989 |
| Megf11 | 0.2059713 | -2.407543 | 0.021 | 0.052 | 1 | 0.0389584 | -2.4618584 | 0.025 | 0.073 | 1 | 0.2059713 | 0.07639904 |
| D330050G23Rik | 0.03896076 | -1.7998604 | 0.021 | 0.083 | 1 | 0.30606883 | 0.69014474 | 0.029 | 0.01 | 1 | 0.30606883 | 0.07640359 |
| Arhgap31 | 0.03897855 | -0.5809112 | 0.792 | 0.798 | 1 | 0.86721356 | -0.1190839 | 0.861 | 0.833 | 1 | 0.86721356 | 0.07643776 |
| Igf2bp3 | 0.03905406 | -0.760332 | 0.156 | 0.264 | 1 | 0.40129965 | 0.35065827 | 0.181 | 0.135 | 1 | 0.40129965 | 0.0765829 |
| Stx18 | 0.03911766 | 0.15663798 | 0.656 | 0.508 | 1 | 0.90254755 | -0.4807881 | 0.668 | 0.5 | 1 | 0.90254755 | 0.07670514 |
| Sntb2 | 0.73998743 | -0.3215933 | 0.74 | 0.725 | 1 | 0.0391349 | -0.7581446 | 0.718 | 0.854 | 1 | 0.73998743 | 0.07673825 |
| Dscaml1 | 0.72574709 | -2.240433 | 0.01 | 0.016 | 1 | 0.03914019 | -2.4473588 | 0.008 | 0.042 | 1 | 0.72574709 | 0.07674843 |
| Tmem11 | 0.03914584 | 0.44159037 | 0.667 | 0.58 | 1 | 0.04069437 | 0.13577056 | 0.706 | 0.521 | 1 | 0.04069437 | 0.07675928 |
| Niban1 | 0.14353351 | -0.3637613 | 0.25 | 0.326 | 1 | 0.03915159 | -0.6833137 | 0.458 | 0.604 | 1 | 0.14353351 | 0.07677033 |
| Zfp101 | 0.03915688 | 0.58122978 | 0.271 | 0.161 | 1 | 0.33160976 | -0.1432054 | 0.307 | 0.229 | 1 | 0.33160976 | 0.07678049 |
| Tmem109 | 0.33283804 | 0.21163525 | 0.469 | 0.42 | 1 | 0.03926223 | 0.20014518 | 0.567 | 0.406 | 1 | 0.33283804 | 0.07698294 |
| Ctxn1 | 0.03928534 | -1.9311049 | 0.021 | 0.083 | 1 | 0.41007651 | -0.7248928 | 0.05 | 0.073 | 1 | 0.41007651 | 0.07702734 |
| Prss12 | 0.11302941 | -3.5160674 | 0 | 0.026 | 1 | 0.03931277 | -2.2273931 | 0.025 | 0.073 | 1 | 0.11302941 | 0.07708004 |
| Mmp14 | 0.03932788 | -0.9904191 | 0.812 | 0.549 | 1 | 0.63869221 | -0.8629689 | 0.92 | 0.729 | 1 | 0.63869221 | 0.07710908 |
| Sidt1 | 1 | 0.42253204 | 0.01 | 0.01 | 1 | 0.03938645 | -3.8948178 | 0.004 | 0.031 | 1 | 1 | 0.07722161 |
| Nrxn3 | 0.99758609 | 0.27052894 | 0.021 | 0.021 | 1 | 0.0393869 | -3.6317834 | 0.004 | 0.031 | 1 | 0.99758609 | 0.07722247 |
| Btbd9 | 0.93351427 | -0.2116302 | 0.896 | 0.886 | 1 | 0.0394095 | -0.5767317 | 0.962 | 0.938 | 1 | 0.93351427 | 0.07726589 |
| Dgat2 | 0.03941089 | -1.7366666 | 0.083 | 0.166 | 1 | 0.73298297 | -0.9754362 | 0.105 | 0.115 | 1 | 0.73298297 | 0.07726857 |
| Rnf149 | 0.03942304 | 0.34629645 | 0.365 | 0.233 | 1 | 0.57104019 | -0.6045985 | 0.244 | 0.26 | 1 | 0.57104019 | 0.0772919 |
| Hsf4 | 0.38181636 | 0.78510212 | 0.052 | 0.031 | 1 | 0.03946234 | 0.88278982 | 0.101 | 0.031 | 1 | 0.38181636 | 0.07736741 |
| Larp1b | 0.03947003 | -0.2607576 | 0.427 | 0.57 | 1 | 0.34363254 | -0.3444539 | 0.672 | 0.74 | 1 | 0.34363254 | 0.07738218 |
| Shank3 | 0.25620695 | -1.0799683 | 0.031 | 0.062 | 1 | 0.03948362 | -2.010295 | 0.025 | 0.073 | 1 | 0.25620695 | 0.07740829 |
| Slc37a2 | 0.48683788 | -0.1624305 | 0.021 | 0.036 | 1 | 0.03952932 | 2.56461386 | 0.113 | 0.042 | 1 | 0.48683788 | 0.07749607 |
| Fhdc1 | 0.43437044 | -0.3382803 | 0.188 | 0.223 | 1 | 0.03957981 | 0.59703533 | 0.202 | 0.104 | 1 | 0.43437044 | 0.07759307 |
| Nxt2 | 0.88135929 | 0.13302542 | 0.104 | 0.098 | 1 | 0.03958973 | 0.96316323 | 0.101 | 0.031 | 1 | 0.88135929 | 0.07761212 |
| Gm47802 | 0.72574709 | -0.3144336 | 0.01 | 0.016 | 1 | 0.03960125 | -1.4253325 | 0.038 | 0.094 | 1 | 0.72574709 | 0.07763424 |
| Clcn7 | 0.88653254 | -0.2852872 | 0.198 | 0.197 | 1 | 0.03964016 | 0.17557156 | 0.345 | 0.208 | 1 | 0.88653254 | 0.07770897 |
| Gm35330 | 0.26694365 | -0.9925055 | 0.031 | 0.062 | 1 | 0.03970511 | -3.6317834 | 0.004 | 0.031 | 1 | 0.26694365 | 0.07783373 |
| Gm56549 | 0.28379282 | -0.7998604 | 0.01 | 0.031 | 1 | 0.03970511 | -2.8948178 | 0.004 | 0.031 | 1 | 0.28379282 | 0.07783373 |
| Stox1 | 0.32025586 | -0.9925055 | 0 | 0.01 | 1 | 0.03970511 | -3.1172102 | 0.004 | 0.031 | 1 | 0.32025586 | 0.07783373 |
| Phf11d | 0.3819238 | 0.81484946 | 0.031 | 0.016 | 1 | 0.03970511 | -4.1172102 | 0.004 | 0.031 | 1 | 0.3819238 | 0.07783373 |
| Xrra1 | 0.47606502 | 1.00749454 | 0.021 | 0.01 | 1 | 0.03970511 | -3.3098553 | 0.004 | 0.031 | 1 | 0.47606502 | 0.07783373 |
| Tmem275 | 0.47835349 | 0.74446013 | 0.021 | 0.01 | 1 | 0.03970511 | -2.8948178 | 0.004 | 0.031 | 1 | 0.47835349 | 0.07783373 |
| Kcnj4 | 0.62104899 | 0.42253204 | 0.01 | 0.005 | 1 | 0.03970511 | -2.8948178 | 0.004 | 0.031 | 1 | 0.62104899 | 0.07783373 |
| D430001F17Rik | 0.75328966 | -0.2148979 | 0.021 | 0.016 | 1 | 0.03970511 | -2.8948178 | 0.004 | 0.031 | 1 | 0.75328966 | 0.07783373 |
| Gm50186 | 0.75328966 | 0.27052894 | 0.021 | 0.016 | 1 | 0.03970511 | -3.1172102 | 0.004 | 0.031 | 1 | 0.75328966 | 0.07783373 |
| Vamp9 | 0.79666623 | 0.42253204 | 0.031 | 0.026 | 1 | 0.03970511 | -2.8948178 | 0.004 | 0.031 | 1 | 0.79666623 | 0.07783373 |
| Slc9a7 | 0.9951721 | 0.32942263 | 0.021 | 0.021 | 1 | 0.03970511 | -4.3973181 | 0.004 | 0.031 | 1 | 0.9951721 | 0.07783373 |
| Relt | 0.03972473 | 1.00749454 | 0.135 | 0.062 | 1 | 0.08763061 | 1.49749966 | 0.05 | 0.01 | 1 | 0.08763061 | 0.0778714 |
| Atp6v1d | 0.09356755 | 0.37946331 | 0.76 | 0.674 | 1 | 0.03974449 | 0.43987763 | 0.727 | 0.708 | 1 | 0.09356755 | 0.07790935 |
| Cbx7 | 0.79103722 | 0.47697982 | 0.115 | 0.109 | 1 | 0.03979164 | 0.32757466 | 0.206 | 0.104 | 1 | 0.79103722 | 0.0779999 |
| Gm56768 | 0.16214989 | -0.9078932 | 0.552 | 0.394 | 1 | 0.03981337 | -0.6734936 | 0.748 | 0.448 | 1 | 0.16214989 | 0.07804164 |
| Mrps2 | 0.09362446 | 0.79599043 | 0.167 | 0.098 | 1 | 0.03984185 | 1.13071733 | 0.143 | 0.062 | 1 | 0.09362446 | 0.07809633 |
| Rbm43 | 0.03985806 | 0.79176585 | 0.25 | 0.15 | 1 | 0.33358856 | 0.17013768 | 0.181 | 0.135 | 1 | 0.33358856 | 0.07812746 |

| | | | | | | | | | | | | |
|---|---|---|---|---|---|---|---|---|---|---|---|---|
| Dcp1b | 0.03986629 | 0.39981196 | 0.167 | 0.083 | 1 | 0.11474556 | 0.47441605 | 0.109 | 0.052 | 1 | 0.11474556 | 0.07814326 |
| Zfp58 | 0.03989116 | 0.73447604 | 0.208 | 0.119 | 1 | 0.09409131 | 0.36821664 | 0.21 | 0.125 | 1 | 0.09409131 | 0.07819102 |
| Rgs16 | 0.41897458 | 0.42253204 | 0.042 | 0.067 | 1 | 0.03992437 | -1.7952821 | 0.008 | 0.042 | 1 | 0.41897458 | 0.07825478 |
| Dhrs1 | 0.53588554 | -0.2371278 | 0.375 | 0.409 | 1 | 0.03993643 | 0.39709776 | 0.349 | 0.219 | 1 | 0.53588554 | 0.07827794 |
| Nup210l | 0.04001981 | -0.1127997 | 0.542 | 0.409 | 1 | 0.28023077 | -0.2723806 | 0.5 | 0.604 | 1 | 0.28023077 | 0.07843803 |
| Cd4 | 0.04373115 | -3.5160674 | 0 | 0.041 | 1 | 0.04002461 | -2.8948178 | 0.004 | 0.031 | 1 | 0.04373115 | 0.07844725 |
| Gm42699 | 0.07511393 | 2.00749454 | 0.031 | 0.005 | 1 | 0.04002461 | -2.6317834 | 0.004 | 0.031 | 1 | 0.07511393 | 0.07844725 |
| Tnni1 | 0.11302678 | -2.3144336 | 0 | 0.026 | 1 | 0.04002461 | -3.6317834 | 0.004 | 0.031 | 1 | 0.11302678 | 0.07844725 |
| Tnfsf10 | 0.11302853 | -2.6929452 | 0 | 0.026 | 1 | 0.04002461 | -2.8948178 | 0.004 | 0.031 | 1 | 0.11302853 | 0.07844725 |
| Aknaos | 0.15729907 | -2.577468 | 0 | 0.021 | 1 | 0.04002461 | -2.6317834 | 0.004 | 0.031 | 1 | 0.15729907 | 0.07844725 |
| Gm6345 | 0.22187388 | -1.3144336 | 0 | 0.016 | 1 | 0.04002461 | -2.6317834 | 0.004 | 0.031 | 1 | 0.22187388 | 0.07844725 |
| Aldh3b3 | 0.22187388 | -1.3144336 | 0 | 0.016 | 1 | 0.04002461 | -2.8948178 | 0.004 | 0.031 | 1 | 0.22187388 | 0.07844725 |
| Pars2 | 0.32025586 | -0.577468 | 0 | 0.01 | 1 | 0.04002461 | -2.6317834 | 0.004 | 0.031 | 1 | 0.32025586 | 0.07844725 |
| 6030440G07Rik | 0.32025586 | -1.9925055 | 0 | 0.01 | 1 | 0.04002461 | -2.6317834 | 0.004 | 0.031 | 1 | 0.32025586 | 0.07844725 |
| 9930111J21Rik2 | 0.53395584 | -0.4519371 | 0.01 | 0.021 | 1 | 0.04002461 | -2.8948178 | 0.004 | 0.031 | 1 | 0.53395584 | 0.07844725 |
| Gm32816 | 0.60815045 | -1.8669746 | 0.021 | 0.031 | 1 | 0.04002461 | -3.7692869 | 0.004 | 0.031 | 1 | 0.60815045 | 0.07844725 |
| Tmeff2 | 0.73685932 | 0.42253204 | 0.01 | 0.016 | 1 | 0.04002461 | -4.3098553 | 0.004 | 0.031 | 1 | 0.73685932 | 0.07844725 |
| Sv2c | 0.84167733 | -0.4843586 | 0.042 | 0.047 | 1 | 0.04002461 | -3.7692869 | 0.004 | 0.031 | 1 | 0.84167733 | 0.07844725 |
| Gm49083 | 1 | -0.3144336 | 0.01 | 0.01 | 1 | 0.04002461 | -3.1172102 | 0.004 | 0.031 | 1 | 1 | 0.07844725 |
| Pkig | 0.04005492 | 0.1607408 | 0.656 | 0.497 | 1 | 0.95231398 | -0.4115507 | 0.681 | 0.646 | 1 | 0.95231398 | 0.07850545 |
| Csf2ra | 0.14277232 | -0.4664367 | 0.229 | 0.311 | 1 | 0.04006022 | 0.54429387 | 0.147 | 0.062 | 1 | 0.14277232 | 0.07851562 |
| Cald1 | 0.04013506 | -0.5437296 | 0.979 | 0.964 | 1 | 0.10418338 | -0.2972796 | 0.958 | 0.865 | 1 | 0.10418338 | 0.07865929 |
| Slc25a30 | 0.04014907 | -1.1363412 | 0.24 | 0.337 | 1 | 0.55866019 | -0.6201954 | 0.168 | 0.188 | 1 | 0.55866019 | 0.07868619 |
| Lnpep | 0.04025731 | 0.19755491 | 0.969 | 0.933 | 1 | 0.17087264 | -0.3017223 | 0.983 | 0.979 | 1 | 0.17087264 | 0.07889397 |
| Cnot9 | 0.09351697 | 0.44806713 | 0.312 | 0.223 | 1 | 0.04034292 | 0.33400093 | 0.311 | 0.188 | 1 | 0.09351697 | 0.0790583 |
| Gm56588 | 0.17891971 | 1.00749454 | 0.042 | 0.016 | 1 | 0.04034946 | -2.3098553 | 0.004 | 0.031 | 1 | 0.17891971 | 0.07907084 |
| Ccdc69 | 0.22026949 | 1.32942263 | 0.021 | 0.005 | 1 | 0.04034946 | -2.3098553 | 0.004 | 0.031 | 1 | 0.22026949 | 0.07907084 |
| Gm16364 | 0.32025586 | -0.577468 | 0 | 0.01 | 1 | 0.04034946 | -2.3098553 | 0.004 | 0.031 | 1 | 0.32025586 | 0.07907084 |
| Tbx21 | 0.52744059 | -0.577468 | 0.01 | 0.021 | 1 | 0.04034946 | -2.3098553 | 0.004 | 0.031 | 1 | 0.52744059 | 0.07907084 |
| Gm57220 | 0.62104899 | 0.42253204 | 0.01 | 0.005 | 1 | 0.04034946 | -2.3098553 | 0.004 | 0.031 | 1 | 0.62104899 | 0.07907084 |
| Etaa1 | 0.0404602 | 0.90057933 | 0.188 | 0.104 | 1 | 0.04854399 | 1.11640949 | 0.139 | 0.062 | 1 | 0.04854399 | 0.07928337 |
| Macroh2a2 | 0.04047372 | 0.29700115 | 0.365 | 0.233 | 1 | 0.2472276 | 0.3146356 | 0.303 | 0.24 | 1 | 0.2472276 | 0.07930932 |
| 4930558N01Rik | 0.04049195 | 0.78510212 | 0.104 | 0.041 | 1 | 0.96556448 | 0.20471791 | 0.063 | 0.062 | 1 | 0.96556448 | 0.0793443 |
| Dcaf11 | 0.86790625 | 0.28867629 | 0.323 | 0.326 | 1 | 0.04049352 | 0.18047036 | 0.387 | 0.25 | 1 | 0.86790625 | 0.07934731 |
| Zmym6 | 0.04049603 | -1.210929 | 0.125 | 0.223 | 1 | 0.77857182 | -0.4507178 | 0.176 | 0.156 | 1 | 0.77857182 | 0.07935214 |
| Brf1 | 0.40134307 | 0.17522198 | 0.312 | 0.259 | 1 | 0.04050646 | 0.15727075 | 0.508 | 0.333 | 1 | 0.40134307 | 0.07937215 |
| Ulk2 | 0.7308827 | -0.4199267 | 0.615 | 0.611 | 1 | 0.04052923 | -0.4402009 | 0.559 | 0.667 | 1 | 0.7308827 | 0.07941584 |
| 4933400C23Rik | 0.05972787 | -2.577468 | 0 | 0.036 | 1 | 0.04067426 | -1.9879272 | 0.004 | 0.031 | 1 | 0.05972787 | 0.07969412 |
| Gm47923 | 0.99654992 | -0.7998604 | 0.042 | 0.041 | 1 | 0.04067471 | -2.3098553 | 0.004 | 0.031 | 1 | 0.99654992 | 0.07969499 |
| Gm12089 | 0.11325766 | -1.4519371 | 0.01 | 0.047 | 1 | 0.04069356 | 0.80562195 | 0.101 | 0.031 | 1 | 0.11325766 | 0.07973116 |
| Rnf138 | 0.0605511 | 0.41515251 | 0.479 | 0.358 | 1 | 0.04072619 | 0.15646274 | 0.529 | 0.354 | 1 | 0.0605511 | 0.07979375 |
| Lpar1 | 0.04077128 | 0.40629139 | 0.917 | 0.813 | 1 | 0.68070447 | -0.3379736 | 0.903 | 0.792 | 1 | 0.68070447 | 0.07988026 |
| B4galnt3 | 0.04077174 | -0.5393328 | 0.344 | 0.461 | 1 | 0.27907509 | -0.2090546 | 0.303 | 0.219 | 1 | 0.27907509 | 0.07988114 |
| Mtf1 | 0.04086135 | 0.56388789 | 0.365 | 0.249 | 1 | 0.07087535 | -0.2769406 | 0.42 | 0.26 | 1 | 0.07087535 | 0.08005306 |
| Gm14326 | 0.04094277 | 0.72738662 | 0.25 | 0.15 | 1 | 0.99926157 | -0.5728897 | 0.189 | 0.177 | 1 | 0.99926157 | 0.08020922 |
| Porcn | 0.84105847 | -0.2820121 | 0.167 | 0.15 | 1 | 0.04095157 | 0.23071312 | 0.21 | 0.104 | 1 | 0.84105847 | 0.0802261 |
| Pyroxd2 | 0.26059398 | 0.52206771 | 0.094 | 0.057 | 1 | 0.04095476 | 0.22065945 | 0.101 | 0.031 | 1 | 0.26059398 | 0.08023223 |
| Lefty1 | 0.67531582 | -0.3144336 | 0.156 | 0.171 | 1 | 0.04097846 | 1.11640949 | 0.143 | 0.062 | 1 | 0.67531582 | 0.08027768 |
| Myh14 | 0.0884303 | 0.91438513 | 0.115 | 0.057 | 1 | 0.04098779 | -0.3802446 | 0.269 | 0.396 | 1 | 0.0884303 | 0.08029558 |
| Btla | 0.27728505 | -1.3550755 | 0.021 | 0.047 | 1 | 0.04100173 | -2.0468209 | 0.004 | 0.031 | 1 | 0.27728505 | 0.08032231 |
| Mir181a-1hg | 0.72574709 | -0.3144336 | 0.01 | 0.016 | 1 | 0.04100173 | -2.0468209 | 0.004 | 0.031 | 1 | 0.72574709 | 0.08032231 |
| H2az2 | 0.23981836 | 0.3245941 | 0.823 | 0.824 | 1 | 0.04112206 | 0.20602293 | 0.891 | 0.844 | 1 | 0.23981836 | 0.0805531 |
| Itgb8 | 0.0412019 | -2.0281294 | 0.031 | 0.098 | 1 | 0.04543441 | 1.04590643 | 0.349 | 0.24 | 1 | 0.04543441 | 0.0807062 |
| Lbhd2 | 0.048541 | -1.5160674 | 0.01 | 0.062 | 1 | 0.04129206 | 2.33400093 | 0.063 | 0.01 | 1 | 0.048541 | 0.08087909 |
| Flrt3 | 0.34814322 | -0.6843832 | 0.094 | 0.13 | 1 | 0.04130306 | 0.33400093 | 0.118 | 0.042 | 1 | 0.34814322 | 0.08090018 |
| 1300014J16Rik | 1 | 0.42253204 | 0.01 | 0.01 | 1 | 0.04133005 | -1.7248928 | 0.004 | 0.031 | 1 | 1 | 0.08095192 |
| Mapk11 | 1 | 0.42253204 | 0.01 | 0.01 | 1 | 0.04133005 | -1.7248928 | 0.004 | 0.031 | 1 | 1 | 0.08095192 |
| Fabp12 | 0.38521705 | -0.7998604 | 0.01 | 0.026 | 1 | 0.04133143 | -1.3098553 | 0.004 | 0.031 | 1 | 0.38521705 | 0.08095457 |
| Mettl21a | 0.10284817 | 0.54036853 | 0.281 | 0.197 | 1 | 0.04136038 | 0.48661134 | 0.244 | 0.135 | 1 | 0.10284817 | 0.08101007 |
| Mdm1 | 0.04139639 | 0.92503238 | 0.135 | 0.062 | 1 | 0.45619353 | -0.5025003 | 0.05 | 0.031 | 1 | 0.45619353 | 0.08107912 |
| Ifngr2 | 0.10468587 | 0.11564334 | 0.5 | 0.383 | 1 | 0.04148211 | 0.30752872 | 0.618 | 0.521 | 1 | 0.10468587 | 0.08124345 |
| St3gal4 | 0.04149117 | -0.5431536 | 0.656 | 0.72 | 1 | 0.45184674 | -0.3802446 | 0.63 | 0.604 | 1 | 0.45184674 | 0.08126082 |
| Rgs6 | 0.99275829 | 0.59245704 | 0.021 | 0.021 | 1 | 0.04150515 | 1.38700012 | 0.113 | 0.042 | 1 | 0.99275829 | 0.08128763 |
| Snhg9 | 0.0415079 | 0.91438513 | 0.135 | 0.062 | 1 | 0.30362892 | 0.76053407 | 0.071 | 0.042 | 1 | 0.30362892 | 0.0812929 |
| Rbm41 | 0.50433057 | -0.1422526 | 0.25 | 0.207 | 1 | 0.04153127 | 0.48923204 | 0.374 | 0.25 | 1 | 0.50433057 | 0.08133769 |
| Gm45397 | 0.04160335 | -1.4779323 | 0.021 | 0.083 | 1 | 0.40078481 | 0.27510724 | 0.025 | 0.01 | 1 | 0.40078481 | 0.08147586 |
| Gm32342 | 0.21458328 | -0.577468 | 0.01 | 0.036 | 1 | 0.0416255 | 1.77760758 | 0.097 | 0.031 | 1 | 0.21458328 | 0.08151832 |
| Mex3c | 0.04163389 | 0.48240349 | 0.615 | 0.497 | 1 | 0.06163401 | 0.23834869 | 0.782 | 0.729 | 1 | 0.06163401 | 0.0815344 |
| Anapc7 | 0.28266539 | 0.20680334 | 0.458 | 0.373 | 1 | 0.04165519 | 0.20584458 | 0.496 | 0.344 | 1 | 0.28266539 | 0.08157523 |
| A830082N09Rik | 0.04166097 | 0.82692229 | 0.135 | 0.062 | 1 | 0.50652381 | -1.3098553 | 0.046 | 0.062 | 1 | 0.50652381 | 0.08158631 |
| Gm39929 | 0.04171274 | 1.00749454 | 0.104 | 0.041 | 1 | 0.14990668 | 0.39058446 | 0.042 | 0.01 | 1 | 0.14990668 | 0.08168552 |
| Tubgcp5 | 0.21230386 | 0.54515132 | 0.281 | 0.223 | 1 | 0.04172194 | 0.27510724 | 0.349 | 0.219 | 1 | 0.21230386 | 0.08170316 |
| Atcayos | 0.83676741 | -0.2555399 | 0.042 | 0.047 | 1 | 0.041761 | -1.7692869 | 0.046 | 0.104 | 1 | 0.83676741 | 0.08177802 |
| Lrmda | 0.04180691 | -0.5551496 | 0.406 | 0.544 | 1 | 0.10655633 | -0.5095551 | 0.5 | 0.677 | 1 | 0.10655633 | 0.081866 |
| Luc7l | 0.04183435 | 0.3005998 | 0.896 | 0.782 | 1 | 0.40624645 | 0.12077909 | 0.857 | 0.865 | 1 | 0.40624645 | 0.08191859 |
| Rad51 | 0.04189481 | 1.37946331 | 0.094 | 0.036 | 1 | 0.12096572 | -1.7248928 | 0.008 | 0.031 | 1 | 0.12096572 | 0.08203445 |
| Acbd5 | 0.04192517 | 0.45343808 | 0.698 | 0.653 | 1 | 0.95345792 | -0.1229117 | 0.782 | 0.792 | 1 | 0.95345792 | 0.08209262 |
| Guca1b | 0.25332778 | 1.00749454 | 0.052 | 0.026 | 1 | 0.04197017 | 2.39058446 | 0.042 | 0 | 1 | 0.25332778 | 0.08217884 |
| Pcdhb3 | 0.61737927 | 1.00749454 | 0.01 | 0.005 | 1 | 0.04197017 | 2.39058446 | 0.042 | 0 | 1 | 0.61737927 | 0.08217884 |
| 4930434J08Rik | 0.6834121 | 0.42253204 | 0.052 | 0.041 | 1 | 0.04197017 | 2.39058446 | 0.042 | 0 | 1 | 0.6834121 | 0.08217884 |
| Gm17455 | 0.85158597 | -0.4779323 | 0.042 | 0.036 | 1 | 0.04197017 | 2.39058446 | 0.042 | 0 | 1 | 0.85158597 | 0.08217884 |
| Hopx | 0.61182672 | -0.577468 | 0.021 | 0.031 | 1 | 0.04197548 | 2.77760758 | 0.042 | 0 | 1 | 0.61182672 | 0.08218901 |
| Cd53 | 0.61371908 | 2.00749454 | 0.01 | 0.005 | 1 | 0.04198344 | 3.39058446 | 0.042 | 0 | 1 | 0.61371908 | 0.08220427 |

| | | | | | | | | | | | | |
|---|---|---|---|---|---|---|---|---|---|---|---|---|
| Hmga1b | 0.07024899 | 1.22988696 | 0.062 | 0.021 | 1 | 0.04198609 | 2.14957636 | 0.042 | 0 | 1 | 0.07024899 | 0.08220935 |
| Gm16091 | 0.14152876 | 2.59245704 | 0.052 | 0.021 | 1 | 0.04198609 | 2.14957636 | 0.042 | 0 | 1 | 0.14152876 | 0.08220935 |
| Plekha2 | 0.34980021 | -0.3144336 | 0.052 | 0.083 | 1 | 0.04198609 | 2.14957636 | 0.042 | 0 | 1 | 0.34980021 | 0.08220935 |
| Ovca2 | 0.38028858 | 1.00749454 | 0.031 | 0.016 | 1 | 0.04198609 | 2.14957636 | 0.042 | 0 | 1 | 0.38028858 | 0.08220935 |
| Ghrl | 0.48064264 | 0.59245704 | 0.021 | 0.01 | 1 | 0.04198609 | 2.14957636 | 0.042 | 0 | 1 | 0.48064264 | 0.08220935 |
| Ttc7 | 0.04202055 | -0.4838549 | 0.698 | 0.731 | 1 | 0.2515716 | -0.1828824 | 0.849 | 0.75 | 1 | 0.2515716 | 0.08227537 |
| Rpap1 | 0.04203141 | 1.59245704 | 0.094 | 0.036 | 1 | 0.41233494 | -0.3098553 | 0.105 | 0.073 | 1 | 0.41233494 | 0.08229618 |
| Gm57446 | 0.99758609 | 0.27052894 | 0.021 | 0.021 | 1 | 0.04204284 | -1.4797803 | 0.025 | 0.073 | 1 | 0.99758609 | 0.08231807 |
| Pex6 | 0.29377875 | 0.40342321 | 0.219 | 0.166 | 1 | 0.04208311 | 0.51326698 | 0.307 | 0.188 | 1 | 0.29377875 | 0.08239524 |
| Gm6710 | 0.3390465 | -0.577468 | 0.031 | 0.057 | 1 | 0.04216784 | 1.69014474 | 0.063 | 0.01 | 1 | 0.3390465 | 0.08255755 |
| Rsph3b | 0.04232794 | 0.57453513 | 0.302 | 0.192 | 1 | 0.09065363 | 0.33923758 | 0.21 | 0.125 | 1 | 0.09065363 | 0.08286423 |
| Nptx1 | 0.04236963 | -1.3144336 | 0.031 | 0.098 | 1 | 0.90489059 | 0.59703533 | 0.034 | 0.031 | 1 | 0.90489059 | 0.08294407 |
| Fbxl9 | 0.41813223 | 0.32942263 | 0.125 | 0.093 | 1 | 0.04238343 | 0.86006974 | 0.13 | 0.052 | 1 | 0.41813223 | 0.08297051 |
| Chmp3 | 0.32817995 | -0.2831826 | 0.667 | 0.699 | 1 | 0.04241584 | 0.20009388 | 0.727 | 0.656 | 1 | 0.32817995 | 0.08303257 |
| Mrps28 | 0.04245636 | 0.15165155 | 0.771 | 0.642 | 1 | 0.26078645 | -0.1210936 | 0.777 | 0.74 | 1 | 0.26078645 | 0.08311018 |
| Exd1 | 0.42175358 | 0.34006988 | 0.135 | 0.104 | 1 | 0.04250577 | 2.19264508 | 0.08 | 0.021 | 1 | 0.42175358 | 0.08320479 |
| Metrn | 0.0425376 | -1.2503032 | 0.219 | 0.306 | 1 | 0.38231883 | -0.1723517 | 0.172 | 0.125 | 1 | 0.38231883 | 0.08326574 |
| Peds1 | 0.04272695 | -0.4100112 | 0.479 | 0.58 | 1 | 0.19183101 | 0.21832121 | 0.58 | 0.51 | 1 | 0.19183101 | 0.08362831 |
| Hprt | 0.11968173 | 0.35014611 | 0.708 | 0.648 | 1 | 0.04274583 | 0.3723094 | 0.718 | 0.677 | 1 | 0.11968173 | 0.08366446 |
| Hmcn2 | 0.21622576 | 0.88196365 | 0.062 | 0.031 | 1 | 0.04276809 | 1.7944814 | 0.097 | 0.031 | 1 | 0.21622576 | 0.08370707 |
| Acot9 | 0.10827496 | 0.22322323 | 0.625 | 0.518 | 1 | 0.04279435 | 0.34160265 | 0.66 | 0.573 | 1 | 0.10827496 | 0.08375735 |
| Gfm2 | 0.715694 | 0.14499806 | 0.229 | 0.212 | 1 | 0.04289099 | 0.18490943 | 0.261 | 0.146 | 1 | 0.715694 | 0.08394233 |
| Ccnb2 | 0.04292511 | 1.84879679 | 0.167 | 0.088 | 1 | 0.19642305 | -0.7862933 | 0.155 | 0.208 | 1 | 0.19642305 | 0.08400766 |
| Zfp867 | 0.04293546 | -1.0628948 | 0.083 | 0.171 | 1 | 0.74043751 | 0.12310414 | 0.084 | 0.073 | 1 | 0.74043751 | 0.08402747 |
| Dnah17 | 0.25707611 | -1.6929452 | 0.031 | 0.062 | 1 | 0.04296346 | -1.8504236 | 0.063 | 0.125 | 1 | 0.25707611 | 0.08408105 |
| Thoc1 | 0.55851613 | 0.34591075 | 0.531 | 0.523 | 1 | 0.04298275 | 0.25612413 | 0.618 | 0.5 | 1 | 0.55851613 | 0.08411799 |
| Gm14296 | 0.24260071 | 0.56692195 | 0.115 | 0.073 | 1 | 0.04306311 | 1.14957636 | 0.113 | 0.042 | 1 | 0.24260071 | 0.08427179 |
| Magt1 | 0.04315804 | 0.28693055 | 0.781 | 0.741 | 1 | 0.0478713 | 0.16904218 | 0.786 | 0.688 | 1 | 0.0478713 | 0.08445346 |
| Nr2c2ap | 0.28628515 | 0.52206771 | 0.198 | 0.15 | 1 | 0.0431701 | 0.46458504 | 0.214 | 0.115 | 1 | 0.28628515 | 0.08447654 |
| Rragd | 0.52640963 | -0.8993961 | 0.031 | 0.047 | 1 | 0.04319799 | 1.44503224 | 0.097 | 0.031 | 1 | 0.52640963 | 0.08452992 |
| Snta1 | 0.04325596 | -0.628094 | 0.208 | 0.321 | 1 | 0.69793195 | -0.2884816 | 0.382 | 0.333 | 1 | 0.69793195 | 0.08464084 |
| Zfp105 | 0.41239321 | 0.37006462 | 0.083 | 0.057 | 1 | 0.04325832 | 1.27510724 | 0.126 | 0.052 | 1 | 0.41239321 | 0.08464537 |
| Usp1 | 0.04328451 | 0.79551873 | 0.531 | 0.451 | 1 | 0.2791729 | 0.10145915 | 0.546 | 0.51 | 1 | 0.2791729 | 0.08469547 |
| Tmem134 | 0.04336102 | 0.27273321 | 0.792 | 0.632 | 1 | 0.21995984 | 0.15293537 | 0.605 | 0.552 | 1 | 0.21995984 | 0.08484187 |
| Rchy1 | 0.32348456 | -0.1941393 | 0.344 | 0.394 | 1 | 0.04340036 | 0.2950068 | 0.433 | 0.292 | 1 | 0.32348456 | 0.08491713 |
| E2f5 | 0.13940625 | 0.69555053 | 0.365 | 0.29 | 1 | 0.04341662 | 0.24596089 | 0.399 | 0.25 | 1 | 0.13940625 | 0.08494824 |
| Rab31 | 0.04344855 | 0.32142984 | 0.781 | 0.668 | 1 | 0.35014363 | -0.3098553 | 0.84 | 0.802 | 1 | 0.35014363 | 0.08500933 |
| Ttll12 | 0.04355408 | 0.56003556 | 0.104 | 0.041 | 1 | 0.07260016 | 0.8917786 | 0.088 | 0.031 | 1 | 0.07260016 | 0.08521119 |
| BC024978 | 0.75630185 | -0.3144336 | 0.083 | 0.093 | 1 | 0.04357903 | 0.64434105 | 0.113 | 0.042 | 1 | 0.75630185 | 0.08525893 |
| Npr1 | 0.1525942 | -1.4519371 | 0.01 | 0.041 | 1 | 0.04359589 | -1.1843244 | 0.025 | 0.073 | 1 | 0.1525942 | 0.08529117 |
| Trabd2b | 0.04365545 | 0.28919815 | 0.792 | 0.751 | 1 | 0.09403157 | -0.1201599 | 0.933 | 0.802 | 1 | 0.09403157 | 0.0854051 |
| Nynrin | 0.04367619 | 0.50706555 | 0.271 | 0.161 | 1 | 0.06234325 | 0.18490943 | 0.34 | 0.219 | 1 | 0.06234325 | 0.08544477 |
| Rps6ka6 | 0.5978326 | 0.20013961 | 0.031 | 0.021 | 1 | 0.04369202 | 1.74903843 | 0.08 | 0.021 | 1 | 0.5978326 | 0.08547504 |
| 1700029H14Rik | 0.12302921 | 0.56692195 | 0.115 | 0.062 | 1 | 0.0436923 | 1.86006974 | 0.08 | 0.021 | 1 | 0.12302921 | 0.08547559 |
| Gm56730 | 0.04370111 | -0.5133376 | 0.208 | 0.316 | 1 | 0.50032294 | -0.2167459 | 0.185 | 0.146 | 1 | 0.50032294 | 0.08549244 |
| Nmnat2 | 0.04371754 | -2.4519371 | 0 | 0.041 | 1 | 0.11689951 | -2.3098553 | 0 | 0.01 | 1 | 0.11689951 | 0.08552385 |
| Crabp1 | 0.04371754 | -2.577468 | 0 | 0.041 | 1 | 0.11689951 | -3.3098553 | 0 | 0.01 | 1 | 0.11689951 | 0.08552385 |
| Best3 | 0.04371754 | -2.3144336 | 0 | 0.041 | 1 | 0.91219225 | -0.8504236 | 0.029 | 0.031 | 1 | 0.91219225 | 0.08552385 |
| F730043M19Rik | 0.04371754 | -2.3144336 | 0 | 0.041 | 1 | 0.86312513 | -0.7248928 | 0.008 | 0.01 | 1 | 0.86312513 | 0.08552385 |
| D930032P07Rik | 0.04371754 | -2.3144336 | 0 | 0.041 | 1 | 0.11764285 | 1.69014474 | 0.025 | 0 | 1 | 0.11764285 | 0.08552385 |
| Gdpd2 | 0.04372434 | -2.4519371 | 0 | 0.041 | 1 | 0.27154471 | 0.69014474 | 0.013 | 0 | 1 | 0.27154471 | 0.08553687 |
| Gm19466 | 0.04372888 | -2.6929452 | 0 | 0.041 | 1 | 0.14757845 | -1.7248928 | 0.004 | 0.021 | 1 | 0.14757845 | 0.08554554 |
| Gm15551 | 0.0437292 | -2.9925055 | 0 | 0.041 | 1 | 0.11689951 | -2.8948178 | 0 | 0.01 | 1 | 0.11689951 | 0.08554616 |
| Sh3gl3 | 0.0437292 | -2.577468 | 0 | 0.041 | 1 | 0.11689951 | -2.3098553 | 0 | 0.01 | 1 | 0.11689951 | 0.08554616 |
| Csf1r | 0.0437292 | -2.8993961 | 0 | 0.041 | 1 | 0.25822788 | 1.30157945 | 0.034 | 0.062 | 1 | 0.25822788 | 0.08554616 |
| Insc | 0.04373244 | -3.8504865 | 0 | 0.041 | 1 | 0.14544666 | -3.3098553 | 0.004 | 0.021 | 1 | 0.14544666 | 0.08555236 |
| Optc | 0.04373309 | -3.1624305 | 0 | 0.041 | 1 | 0.07096478 | -2.0468209 | 0.017 | 0.052 | 1 | 0.07096478 | 0.0855536 |
| Gm42397 | 0.04373374 | -3.5160674 | 0 | 0.041 | 1 | 0.11689951 | -2.3098553 | 0 | 0.01 | 1 | 0.11689951 | 0.08555484 |
| Zswim5 | 0.04373471 | -3.6363617 | 0 | 0.041 | 1 | 0.68770589 | -1.7248928 | 0.034 | 0.042 | 1 | 0.68770589 | 0.0855567 |
| Capn3 | 0.04373568 | -2.1624305 | 0 | 0.041 | 1 | 0.8086289 | -0.3098553 | 0.017 | 0.021 | 1 | 0.8086289 | 0.08555856 |
| Zfp647 | 0.04373568 | -2.1624305 | 0 | 0.041 | 1 | 0.93576656 | -0.1399303 | 0.029 | 0.031 | 1 | 0.93576656 | 0.08555856 |
| Cln3 | 0.45898697 | -0.4330781 | 0.125 | 0.155 | 1 | 0.04374766 | 0.96698494 | 0.206 | 0.115 | 1 | 0.45898697 | 0.08558146 |
| Pacc1 | 0.04379068 | 1.42253204 | 0.094 | 0.036 | 1 | 0.14465573 | 0.35310975 | 0.13 | 0.073 | 1 | 0.14465573 | 0.08566374 |
| Gm12992 | 0.04379068 | 1.42253204 | 0.094 | 0.036 | 1 | 0.31338711 | -1.3098553 | 0.046 | 0.073 | 1 | 0.31338711 | 0.08566374 |
| Paox | 0.04388706 | 1.00749454 | 0.125 | 0.057 | 1 | 0.0446763 | 0.91253716 | 0.113 | 0.042 | 1 | 0.0446763 | 0.08584805 |
| Mad2l1bp | 0.0439035 | 0.60894516 | 0.208 | 0.114 | 1 | 0.63786843 | -0.4542452 | 0.139 | 0.115 | 1 | 0.63786843 | 0.08587949 |
| Fam117b | 0.85484529 | -0.3768976 | 0.573 | 0.513 | 1 | 0.04393782 | -0.7520776 | 0.706 | 0.781 | 1 | 0.85484529 | 0.08594511 |
| Vps4a | 0.10058569 | 0.33870045 | 0.531 | 0.43 | 1 | 0.04395616 | 0.21168186 | 0.466 | 0.312 | 1 | 0.10058569 | 0.08598017 |
| Gm37083 | 0.15315468 | 0.56692195 | 0.125 | 0.073 | 1 | 0.04398918 | 1.69014474 | 0.08 | 0.021 | 1 | 0.15315468 | 0.08604331 |
| Mmab | 0.19507453 | 0.45110119 | 0.125 | 0.078 | 1 | 0.04398928 | 1.74903843 | 0.08 | 0.021 | 1 | 0.19507453 | 0.0860435 |
| Epc2 | 0.31010739 | 0.16602198 | 0.781 | 0.715 | 1 | 0.04403097 | -0.3832976 | 0.866 | 0.917 | 1 | 0.31010739 | 0.08612322 |
| Spop | 0.05362597 | -0.2872487 | 0.896 | 0.938 | 1 | 0.04405113 | -0.5595313 | 0.933 | 0.906 | 1 | 0.05362597 | 0.08616176 |
| Gm41764 | 0.04408154 | 0.68556644 | 0.135 | 0.062 | 1 | 0.74803035 | -0.9537115 | 0.113 | 0.094 | 1 | 0.74803035 | 0.0862199 |
| Xrn2 | 0.97644 | -0.1637769 | 0.854 | 0.824 | 1 | 0.04413727 | -0.4602721 | 0.895 | 0.906 | 1 | 0.97644 | 0.08632644 |
| Arntl2 | 0.04415463 | 0.78220222 | 0.302 | 0.197 | 1 | 0.15358693 | 0.1704737 | 0.361 | 0.26 | 1 | 0.15358693 | 0.08635963 |
| F3 | 0.04418332 | -0.2406204 | 0.677 | 0.617 | 1 | 0.79177813 | 0.18291354 | 0.672 | 0.74 | 1 | 0.79177813 | 0.08641447 |
| Gm29394 | 0.04421071 | 1.38600616 | 0.094 | 0.036 | 1 | 0.53156006 | 0.69014474 | 0.034 | 0.021 | 1 | 0.53156006 | 0.08646683 |
| Heatr5b | 0.27380499 | -0.8993961 | 0.156 | 0.202 | 1 | 0.04422551 | 0.66505376 | 0.311 | 0.198 | 1 | 0.27380499 | 0.08649513 |
| 3300002I08Rik | 0.04426843 | 0.74446013 | 0.219 | 0.124 | 1 | 0.99273933 | -0.7248928 | 0.151 | 0.146 | 1 | 0.99273933 | 0.08657716 |
| Siah1b | 0.3573566 | 0.71798792 | 0.073 | 0.047 | 1 | 0.04428616 | 1.56461386 | 0.08 | 0.021 | 1 | 0.3573566 | 0.08661105 |
| Tle5 | 0.04432162 | -0.7655403 | 0.49 | 0.549 | 1 | 0.36180953 | 0.2023509 | 0.651 | 0.646 | 1 | 0.36180953 | 0.08667884 |
| Man2a2 | 0.39091927 | -0.4682389 | 0.26 | 0.301 | 1 | 0.04440585 | 0.17808278 | 0.353 | 0.208 | 1 | 0.39091927 | 0.08683982 |
| Anapc1 | 0.04444913 | 0.49027664 | 0.604 | 0.472 | 1 | 0.76450128 | -0.175877 | 0.714 | 0.594 | 1 | 0.76450128 | 0.08692254 |
| Ints1 | 0.58366177 | 0.32942263 | 0.125 | 0.104 | 1 | 0.04445669 | 0.69014474 | 0.143 | 0.062 | 1 | 0.58366177 | 0.08693699 |
| Gstk1 | 0.70827863 | -0.3144336 | 0.146 | 0.161 | 1 | 0.04450644 | 0.94648449 | 0.168 | 0.083 | 1 | 0.70827863 | 0.08703205 |
| Bcl3 | 0.20076314 | -0.7580402 | 0.292 | 0.352 | 1 | 0.04456612 | -0.3967327 | 0.328 | 0.448 | 1 | 0.20076314 | 0.0871461 |
| Smim14 | 0.04457971 | -0.3134682 | 0.958 | 0.99 | 1 | 0.12198682 | 0.12630878 | 0.979 | 0.979 | 1 | 0.12198682 | 0.08717207 |

| | | | | | | | | | | | | |
|---|---|---|---|---|---|---|---|---|---|---|---|---|
| Gm56976 | 0.12330917 | -1.407543 | 0.021 | 0.062 | 1 | 0.04458293 | 1.49749966 | 0.08 | 0.021 | 1 | 0.12330917 | 0.08717822 |
| Myo3b | 0.04464821 | -2.6363617 | 0.01 | 0.062 | 1 | 0.14866047 | -1.9879272 | 0.029 | 0.062 | 1 | 0.14866047 | 0.08730297 |
| Gtf3c5 | 0.04467667 | 0.47697982 | 0.229 | 0.13 | 1 | 0.84775327 | -0.3446207 | 0.13 | 0.135 | 1 | 0.84775327 | 0.08735734 |
| Wdhd1 | 0.04469664 | 1.06407806 | 0.146 | 0.073 | 1 | 0.6750905 | -0.8504236 | 0.113 | 0.094 | 1 | 0.6750905 | 0.08739548 |
| Pkn3 | 0.0447066 | -1.3550755 | 0.125 | 0.218 | 1 | 0.19961617 | 0.17557156 | 0.097 | 0.052 | 1 | 0.19961617 | 0.08741452 |
| Rnf19b | 0.04471378 | -0.6505314 | 0.479 | 0.554 | 1 | 0.21940559 | -0.3560254 | 0.546 | 0.646 | 1 | 0.21940559 | 0.08742824 |
| Trp53inp1 | 0.44392797 | -0.3598765 | 0.438 | 0.352 | 1 | 0.04472286 | -0.5803551 | 0.416 | 0.552 | 1 | 0.44392797 | 0.08744559 |
| Glb1 | 0.83301767 | 0.14076107 | 0.229 | 0.249 | 1 | 0.04475513 | 1.08009126 | 0.29 | 0.177 | 1 | 0.83301767 | 0.08750725 |
| Chmp4c | 0.1827687 | -0.7149715 | 0.052 | 0.098 | 1 | 0.04476829 | 0.93807225 | 0.063 | 0.01 | 1 | 0.1827687 | 0.08753239 |
| Zfp956 | 0.04483004 | 1.21394541 | 0.125 | 0.057 | 1 | 0.96573487 | -0.5163061 | 0.097 | 0.094 | 1 | 0.96573487 | 0.08765035 |
| Zfp60 | 0.2767819 | 0.46317402 | 0.188 | 0.14 | 1 | 0.04483683 | 0.76543286 | 0.193 | 0.104 | 1 | 0.2767819 | 0.08766332 |
| Rdh11 | 0.38153751 | 0.18503272 | 0.479 | 0.425 | 1 | 0.04488075 | 0.3068161 | 0.366 | 0.24 | 1 | 0.38153751 | 0.08774722 |
| Relb | 0.04493446 | 0.64841344 | 0.427 | 0.326 | 1 | 0.47508681 | -0.6847037 | 0.433 | 0.427 | 1 | 0.47508681 | 0.08784982 |
| Apba1 | 0.04495177 | 0.15949763 | 0.26 | 0.155 | 1 | 0.79729559 | 0.12186098 | 0.307 | 0.281 | 1 | 0.79729559 | 0.08788287 |
| Stk32b | 0.0449894 | -0.5226195 | 0.562 | 0.71 | 1 | 0.28621867 | -0.3323393 | 0.609 | 0.469 | 1 | 0.28621867 | 0.08795476 |
| Gprc5b | 0.06087851 | -1.17693 | 0.094 | 0.176 | 1 | 0.04499082 | 1.27510724 | 0.097 | 0.031 | 1 | 0.06087851 | 0.08795746 |
| Llph | 0.04500479 | 0.55710176 | 0.688 | 0.57 | 1 | 0.72016118 | 0.16911654 | 0.525 | 0.562 | 1 | 0.72016118 | 0.08798414 |
| Gm10550 | 0.04512622 | 2.59245704 | 0.021 | 0 | 1 | 0.29209623 | -1.0874628 | 0.021 | 0.042 | 1 | 0.29209623 | 0.08821607 |
| Gm16701.1 | 0.04512622 | 2.59245704 | 0.021 | 0 | 1 | 0.27154471 | 0.69014474 | 0.013 | 0 | 1 | 0.27154471 | 0.08821607 |
| Clic3 | 0.04512622 | 2.59245704 | 0.021 | 0 | 1 | 0.27154471 | 0.69014474 | 0.013 | 0 | 1 | 0.27154471 | 0.08821607 |
| Erich2 | 0.04512622 | 3.00749454 | 0.021 | 0 | 1 | 0.51897182 | -0.1399303 | 0.021 | 0.01 | 1 | 0.51897182 | 0.08821607 |
| Lingo4 | 0.04512622 | 2.59245704 | 0.021 | 0 | 1 | 0.11765178 | 2.01207283 | 0.025 | 0 | 1 | 0.11765178 | 0.08821607 |
| Tnfaip8l2 | 0.04512622 | 2.59245704 | 0.021 | 0 | 1 | 0.27154471 | 0.69014474 | 0.013 | 0 | 1 | 0.27154471 | 0.08821607 |
| Gm6297 | 0.04512622 | 2.59245704 | 0.021 | 0 | 1 | 0.15394868 | 1.27510724 | 0.021 | 0 | 1 | 0.15394868 | 0.08821607 |
| Cyp2j5 | 0.04512622 | 3.00749454 | 0.021 | 0 | 1 | 0.11689951 | -2.3098553 | 0 | 0.01 | 1 | 0.11689951 | 0.08821607 |
| 1700121C08Rik | 0.04512622 | 3.00749454 | 0.021 | 0 | 1 | 0.27154471 | 0.69014474 | 0.013 | 0 | 1 | 0.27154471 | 0.08821607 |
| Kdf1 | 0.04512622 | 2.59245704 | 0.021 | 0 | 1 | 0.11689951 | -2.3098553 | 0 | 0.01 | 1 | 0.11689951 | 0.08821607 |
| Upk3bl | 0.04512622 | 2.59245704 | 0.021 | 0 | 1 | 0.86011448 | -0.8948178 | 0.008 | 0.01 | 1 | 0.86011448 | 0.08821607 |
| D5Ertd605e | 0.04512622 | 2.59245704 | 0.021 | 0 | 1 | 0.50887612 | -1.3098553 | 0.004 | 0.01 | 1 | 0.50887612 | 0.08821607 |
| Fscn3 | 0.04512622 | 2.59245704 | 0.021 | 0 | 1 | 0.51194506 | 0.27510724 | 0.021 | 0.01 | 1 | 0.51194506 | 0.08821607 |
| 4933427D06Rik | 0.04512622 | 3.00749454 | 0.021 | 0 | 1 | 0.34026631 | -3.6317834 | 0.008 | 0.021 | 1 | 0.34026631 | 0.08821607 |
| Pinlyp | 0.04512622 | 3.00749454 | 0.021 | 0 | 1 | 0.50887612 | -1.3098553 | 0.004 | 0.01 | 1 | 0.50887612 | 0.08821607 |
| Map4k1 | 0.04512622 | 3.00749454 | 0.021 | 0 | 1 | 0.11689951 | -2.3098553 | 0 | 0.01 | 1 | 0.11689951 | 0.08821607 |
| Gm21057 | 0.04512622 | 2.59245704 | 0.021 | 0 | 1 | 0.81129592 | 0.27510724 | 0.025 | 0.021 | 1 | 0.81129592 | 0.08821607 |
| 1700095J03Rik | 0.04512622 | 2.59245704 | 0.021 | 0 | 1 | 0.50587544 | -1.8948178 | 0.004 | 0.01 | 1 | 0.50587544 | 0.08821607 |
| 4933417D19Rik | 0.04512622 | 3.00749454 | 0.021 | 0 | 1 | 0.3937346 | 0.69014474 | 0.025 | 0.01 | 1 | 0.3937346 | 0.08821607 |
| Zfp558 | 0.04512622 | 2.59245704 | 0.021 | 0 | 1 | 0.34704016 | -1.3098553 | 0.008 | 0.021 | 1 | 0.34704016 | 0.08821607 |
| Scg3 | 0.04512622 | 2.59245704 | 0.021 | 0 | 1 | 0.11689951 | -3.6317834 | 0 | 0.01 | 1 | 0.11689951 | 0.08821607 |
| Col6a5 | 0.04512622 | 2.59245704 | 0.021 | 0 | 1 | 0.11765402 | 1.49749966 | 0.025 | 0 | 1 | 0.11765402 | 0.08821607 |
| Gm48249 | 0.04512622 | 2.59245704 | 0.021 | 0 | 1 | 0.67471359 | -0.5728897 | 0.017 | 0.01 | 1 | 0.67471359 | 0.08821607 |
| Pkib | 0.04512622 | 3.32942263 | 0.021 | 0 | 1 | 0.20315163 | 1.01207283 | 0.017 | 0 | 1 | 0.20315163 | 0.08821607 |
| Gm48086 | 0.04512622 | 2.59245704 | 0.021 | 0 | 1 | 0.87066325 | -0.3098553 | 0.013 | 0.01 | 1 | 0.87066325 | 0.08821607 |
| Gm12166 | 0.04512622 | 2.59245704 | 0.021 | 0 | 1 | 0.80442048 | -0.5728897 | 0.017 | 0.021 | 1 | 0.80442048 | 0.08821607 |
| H2bc27 | 0.04512622 | 2.59245704 | 0.021 | 0 | 1 | 0.11689951 | -2.3098553 | 0 | 0.01 | 1 | 0.11689951 | 0.08821607 |
| 2310002D06Rik | 0.04512622 | 2.59245704 | 0.021 | 0 | 1 | 0.87851725 | -0.8948178 | 0.013 | 0.01 | 1 | 0.87851725 | 0.08821607 |
| Pnma1 | 0.04512622 | 2.59245704 | 0.021 | 0 | 1 | 0.3937346 | 1.01207283 | 0.025 | 0.01 | 1 | 0.3937346 | 0.08821607 |
| Zfp759 | 0.04512622 | 3.32942263 | 0.021 | 0 | 1 | 0.1465102 | -1.8948178 | 0.004 | 0.021 | 1 | 0.1465102 | 0.08821607 |
| Gm48970 | 0.04512622 | 2.59245704 | 0.021 | 0 | 1 | 0.20314753 | 1.69014474 | 0.017 | 0 | 1 | 0.20314753 | 0.08821607 |
| Gm26766 | 0.04512622 | 2.59245704 | 0.021 | 0 | 1 | 0.86613737 | -0.3098553 | 0.008 | 0.01 | 1 | 0.86613737 | 0.08821607 |
| Phf21b | 0.04512622 | 2.59245704 | 0.021 | 0 | 1 | 0.5778028 | -0.8948178 | 0.013 | 0.021 | 1 | 0.5778028 | 0.08821607 |
| Cfap44 | 0.04512622 | 2.59245704 | 0.021 | 0 | 1 | 0.58179312 | -0.7248928 | 0.021 | 0.031 | 1 | 0.58179312 | 0.08821607 |
| 1700001C19Rik | 0.04512622 | 2.59245704 | 0.021 | 0 | 1 | 0.50587544 | -1.8948178 | 0.004 | 0.01 | 1 | 0.50587544 | 0.08821607 |
| Acsbg3 | 0.04512622 | 2.59245704 | 0.021 | 0 | 1 | 0.39515719 | 0.49749966 | 0.025 | 0.01 | 1 | 0.39515719 | 0.08821607 |
| Gfra3 | 0.04512622 | 2.59245704 | 0.021 | 0 | 1 | 0.11689951 | -2.8948178 | 0 | 0.01 | 1 | 0.11689951 | 0.08821607 |
| Gm56632 | 0.04512622 | 2.59245704 | 0.021 | 0 | 1 | 0.50587544 | -2.3098553 | 0.004 | 0.01 | 1 | 0.50587544 | 0.08821607 |
| Gm56636 | 0.04512622 | 2.59245704 | 0.021 | 0 | 1 | 0.11689951 | -2.3098553 | 0 | 0.01 | 1 | 0.11689951 | 0.08821607 |
| Gm14703 | 0.04512622 | 2.59245704 | 0.021 | 0 | 1 | 1 | -1.3098553 | 0.021 | 0.021 | 1 | 1 | 0.08821607 |
| 5330434G04Rik | 0.04512622 | 2.59245704 | 0.021 | 0 | 1 | 0.11689951 | -2.3098553 | 0 | 0.01 | 1 | 0.11689951 | 0.08821607 |
| Cldn2 | 0.04512622 | 2.59245704 | 0.021 | 0 | 1 | 0.27154327 | 1.01207283 | 0.013 | 0 | 1 | 0.27154327 | 0.08821607 |
| Rbak | 0.08433359 | 0.80818573 | 0.177 | 0.104 | 1 | 0.04516883 | 0.27510724 | 0.231 | 0.125 | 1 | 0.08433359 | 0.08829744 |
| Arpin | 0.0451715 | 0.60894516 | 0.219 | 0.124 | 1 | 0.04662789 | 0.66589719 | 0.197 | 0.104 | 1 | 0.04662789 | 0.08830254 |
| Otud1 | 0.04518114 | 1.20013961 | 0.146 | 0.073 | 1 | 0.10682111 | 0.59060906 | 0.176 | 0.104 | 1 | 0.10682111 | 0.08832095 |
| Nt5e | 0.04523214 | -0.7410823 | 0.604 | 0.622 | 1 | 0.06899579 | 0.19603567 | 0.517 | 0.354 | 1 | 0.06899579 | 0.08841833 |
| Cdk9 | 0.04524174 | 0.32065242 | 0.542 | 0.404 | 1 | 0.60513246 | -0.1353575 | 0.605 | 0.646 | 1 | 0.60513246 | 0.08843666 |
| Cfap69 | 0.04526562 | -1.0264528 | 0.198 | 0.311 | 1 | 0.07963146 | -1.4579539 | 0.155 | 0.219 | 1 | 0.07963146 | 0.08848227 |
| Gm5820 | 0.04530229 | -2.0799683 | 0.01 | 0.062 | 1 | 0.40078605 | -1.5025003 | 0.017 | 0.031 | 1 | 0.40078605 | 0.08855229 |
| Fbxo10 | 0.04531956 | -1.3144336 | 0.094 | 0.181 | 1 | 0.66880967 | -0.6424306 | 0.143 | 0.156 | 1 | 0.66880967 | 0.08858527 |
| Fam149a | 0.04835074 | -0.8855903 | 0.115 | 0.207 | 1 | 0.04532677 | 0.50830641 | 0.223 | 0.125 | 1 | 0.04835074 | 0.08859902 |
| Ctcf | 0.15602522 | 0.21899864 | 0.771 | 0.653 | 1 | 0.04538974 | 0.2990861 | 0.836 | 0.802 | 1 | 0.15602522 | 0.08871925 |
| Trio | 0.04539154 | -0.5282697 | 0.917 | 0.959 | 1 | 0.05931789 | -0.5439005 | 0.979 | 0.969 | 1 | 0.05931789 | 0.08872269 |
| Uri1 | 0.171563 | 0.27052894 | 0.438 | 0.358 | 1 | 0.04547769 | -0.5588828 | 0.483 | 0.615 | 1 | 0.171563 | 0.08888717 |
| Irf7 | 0.04549556 | 0.46000674 | 0.104 | 0.041 | 1 | 0.67878043 | -0.6577786 | 0.076 | 0.062 | 1 | 0.67878043 | 0.08892128 |
| Ppp1r16a | 0.04550504 | 1.17741954 | 0.146 | 0.073 | 1 | 0.09709059 | 0.96316323 | 0.097 | 0.042 | 1 | 0.09709059 | 0.08893937 |
| Gabbr2 | 0.04551726 | -2.240433 | 0.01 | 0.062 | 1 | 0.11883906 | -2.8948178 | 0.008 | 0.031 | 1 | 0.11883906 | 0.08896271 |
| Gm56904 | 0.04551726 | -1.9925055 | 0.01 | 0.062 | 1 | 0.05407052 | 2.27510724 | 0.038 | 0 | 1 | 0.05407052 | 0.08896271 |
| Rnf113a2 | 0.5839156 | 0.20680334 | 0.25 | 0.223 | 1 | 0.04553951 | 0.73196491 | 0.235 | 0.135 | 1 | 0.5839156 | 0.08900517 |
| A4galt | 0.12222256 | -0.4623088 | 0.562 | 0.627 | 1 | 0.04554385 | 0.24102624 | 0.752 | 0.594 | 1 | 0.12222256 | 0.08901346 |
| Ripk2 | 0.04570127 | -1.4231398 | 0.188 | 0.28 | 1 | 0.80365578 | -0.9562183 | 0.286 | 0.281 | 1 | 0.80365578 | 0.08931393 |
| Ntpcr | 0.0508866 | 1.04596868 | 0.25 | 0.161 | 1 | 0.0457017 | 1.16819203 | 0.126 | 0.052 | 1 | 0.0508866 | 0.08931476 |
| Noa1 | 0.28887705 | 0.47697982 | 0.146 | 0.104 | 1 | 0.0457017 | 1.16819203 | 0.126 | 0.052 | 1 | 0.28887705 | 0.08931476 |
| Mlh1 | 0.46591346 | -0.2920657 | 0.073 | 0.098 | 1 | 0.04577943 | 0.81942775 | 0.113 | 0.042 | 1 | 0.46591346 | 0.08946311 |
| Trim62 | 0.10575978 | 0.73186009 | 0.146 | 0.083 | 1 | 0.04578378 | 0.10518224 | 0.256 | 0.146 | 1 | 0.10575978 | 0.08947141 |
| Cinp | 0.0458739 | 0.84243529 | 0.26 | 0.166 | 1 | 0.05462487 | 0.32019513 | 0.223 | 0.125 | 1 | 0.05462487 | 0.08964338 |

| | | | | | | | | | | | | |
|---|---|---|---|---|---|---|---|---|---|---|---|---|
| Gm17024 | 0.04594335 | -1.7998604 | 0.01 | 0.062 | 1 | 0.50666075 | -0.7248928 | 0.055 | 0.073 | 1 | 0.50666075 | 0.08977591 |
| Gpm6b | 0.04598902 | -0.6957879 | 0.521 | 0.326 | 1 | 0.09722867 | 0.14080615 | 0.424 | 0.281 | 1 | 0.09722867 | 0.08986304 |
| Pdzd8 | 0.04599504 | 0.54131819 | 0.521 | 0.404 | 1 | 0.5824693 | -0.2929419 | 0.786 | 0.812 | 1 | 0.5824693 | 0.08987454 |
| Yme1l1 | 0.0460033 | 0.23030158 | 0.927 | 0.902 | 1 | 0.88212387 | -0.1933963 | 0.916 | 0.917 | 1 | 0.88212387 | 0.0898903 |
| Gm42917 | 0.04601819 | 1.13302542 | 0.094 | 0.036 | 1 | 0.98041816 | -0.9139266 | 0.097 | 0.094 | 1 | 0.98041816 | 0.0899187 |
| Tnfaip1 | 0.08537438 | 0.37006462 | 0.469 | 0.363 | 1 | 0.0460775 | 0.25782925 | 0.58 | 0.427 | 1 | 0.08537438 | 0.09003186 |
| Runx1t1 | 0.04611439 | -1.8225805 | 0.094 | 0.176 | 1 | 0.07441684 | -1.3563978 | 0.126 | 0.198 | 1 | 0.07441684 | 0.09010224 |
| Mfhas1 | 0.04620446 | 0.5686103 | 0.24 | 0.145 | 1 | 0.12950677 | -0.9424842 | 0.277 | 0.365 | 1 | 0.12950677 | 0.09027406 |
| Usp5 | 0.0462156 | 0.51367992 | 0.375 | 0.264 | 1 | 0.06382208 | 0.35134282 | 0.408 | 0.271 | 1 | 0.06382208 | 0.09029532 |
| Cdk15 | 0.04625369 | 1.28760246 | 0.094 | 0.036 | 1 | 0.65956279 | 0.86006974 | 0.017 | 0.01 | 1 | 0.65956279 | 0.09036799 |
| Katnip | 0.04626254 | -0.8216209 | 0.333 | 0.43 | 1 | 0.68913577 | -0.1045364 | 0.412 | 0.354 | 1 | 0.68913577 | 0.09038486 |
| Pmf1 | 0.32947103 | 0.29700115 | 0.198 | 0.15 | 1 | 0.04626976 | 0.39058446 | 0.227 | 0.125 | 1 | 0.32947103 | 0.09039863 |
| Cyb561 | 0.04627348 | -1.1624305 | 0.031 | 0.098 | 1 | 0.54691335 | -1.1578522 | 0.038 | 0.052 | 1 | 0.54691335 | 0.09040572 |
| Flii | 0.44428917 | 0.15045249 | 0.667 | 0.585 | 1 | 0.04631302 | 0.22731934 | 0.769 | 0.698 | 1 | 0.44428917 | 0.09048114 |
| Cep57l1 | 0.0463393 | 0.77801269 | 0.198 | 0.109 | 1 | 0.53215816 | 0.64434105 | 0.092 | 0.073 | 1 | 0.53215816 | 0.09053127 |
| Tshz1 | 0.04928159 | 0.32488347 | 0.594 | 0.456 | 1 | 0.04636089 | -0.6796635 | 0.525 | 0.667 | 1 | 0.04928159 | 0.09057244 |
| Mypop | 0.19475489 | 0.31561683 | 0.115 | 0.067 | 1 | 0.0463836 | 0.49749966 | 0.202 | 0.104 | 1 | 0.19475489 | 0.09061576 |
| Anks3 | 0.04638788 | 0.36113149 | 0.375 | 0.249 | 1 | 0.15346272 | -0.2812861 | 0.445 | 0.292 | 1 | 0.15346272 | 0.09062393 |
| Gm14418 | 0.3249356 | 0.35541784 | 0.062 | 0.036 | 1 | 0.04641449 | 1.44503224 | 0.08 | 0.021 | 1 | 0.3249356 | 0.09067468 |
| Rala | 0.04642805 | 0.46453956 | 0.698 | 0.596 | 1 | 0.9021015 | -0.2188816 | 0.693 | 0.729 | 1 | 0.9021015 | 0.09070054 |
| Ube2cbp | 0.04646984 | 0.61517711 | 0.125 | 0.057 | 1 | 0.09149841 | 0.49749966 | 0.084 | 0.031 | 1 | 0.09149841 | 0.09078023 |
| Leprotl1 | 0.10200694 | 0.41313334 | 0.604 | 0.523 | 1 | 0.04655078 | 0.23293778 | 0.605 | 0.5 | 1 | 0.10200694 | 0.09093459 |
| Polr3a | 0.77626261 | 0.38189005 | 0.208 | 0.202 | 1 | 0.04657695 | 0.23763253 | 0.261 | 0.146 | 1 | 0.77626261 | 0.09098449 |
| Tcn2 | 0.04658084 | -0.6783969 | 0.188 | 0.295 | 1 | 0.84082617 | -0.4353861 | 0.324 | 0.302 | 1 | 0.84082617 | 0.09099191 |
| Tns1 | 0.04658226 | -0.3214278 | 0.76 | 0.902 | 1 | 0.06191243 | 0.20207981 | 0.903 | 0.75 | 1 | 0.06191243 | 0.09099461 |
| Gabrr2 | 0.04659156 | 0.86999101 | 0.094 | 0.036 | 1 | 0.86012621 | -0.8573431 | 0.08 | 0.083 | 1 | 0.86012621 | 0.09101235 |
| Lrp12 | 0.4357062 | 0.45110119 | 0.385 | 0.326 | 1 | 0.04659558 | 0.18199783 | 0.475 | 0.312 | 1 | 0.4357062 | 0.09102 |
| Ints9 | 0.93855169 | -0.407543 | 0.188 | 0.187 | 1 | 0.0466832 | 0.46937606 | 0.303 | 0.188 | 1 | 0.93855169 | 0.09118707 |
| Ankrd27 | 0.04676801 | 0.71376333 | 0.26 | 0.161 | 1 | 0.30759154 | -0.2692133 | 0.244 | 0.177 | 1 | 0.30759154 | 0.09134877 |
| Gm31763 | 0.52106084 | 0.61517711 | 0.104 | 0.083 | 1 | 0.04683774 | 0.36391651 | 0.202 | 0.104 | 1 | 0.52106084 | 0.0914817 |
| Scrn2 | 0.04687998 | 1.00749454 | 0.125 | 0.057 | 1 | 0.25577784 | -0.1172102 | 0.063 | 0.031 | 1 | 0.25577784 | 0.09156222 |
| Smarca4 | 0.04691762 | 0.17327301 | 0.792 | 0.637 | 1 | 0.05503173 | -0.4637679 | 0.786 | 0.844 | 1 | 0.05503173 | 0.09163398 |
| B230354K17Rik | 0.06782517 | 0.73186009 | 0.271 | 0.181 | 1 | 0.04691947 | 0.49749966 | 0.239 | 0.135 | 1 | 0.06782517 | 0.09163749 |
| Cnst | 0.04696093 | 0.86065315 | 0.188 | 0.104 | 1 | 0.324635 | 0.13760371 | 0.256 | 0.198 | 1 | 0.324635 | 0.09171654 |
| 6330403L08Rik | 0.38245654 | 0.90795886 | 0.083 | 0.057 | 1 | 0.0470796 | 1.05271482 | 0.126 | 0.052 | 1 | 0.38245654 | 0.09194271 |
| Klhdc1 | 0.30023885 | -0.2779077 | 0.104 | 0.15 | 1 | 0.04710278 | 0.25308093 | 0.193 | 0.094 | 1 | 0.30023885 | 0.09198689 |
| Ankrd39 | 0.04712394 | 0.54354744 | 0.25 | 0.15 | 1 | 0.04865631 | 0.47150445 | 0.172 | 0.083 | 1 | 0.04865631 | 0.09202722 |
| Opa1 | 0.13597178 | 0.11440974 | 0.604 | 0.508 | 1 | 0.04712544 | 0.16468325 | 0.538 | 0.396 | 1 | 0.13597178 | 0.09203007 |
| Gm26533 | 0.04713723 | -1.9925055 | 0.01 | 0.062 | 1 | 0.29109279 | -1.1172102 | 0.021 | 0.042 | 1 | 0.29109279 | 0.09205254 |
| Lgmn | 0.0471951 | -0.577468 | 0.49 | 0.57 | 1 | 0.41800865 | -0.4232371 | 0.45 | 0.479 | 1 | 0.41800865 | 0.09216283 |
| Zfp617 | 0.08031873 | 0.60413884 | 0.271 | 0.181 | 1 | 0.04721051 | 0.36821664 | 0.218 | 0.115 | 1 | 0.08031873 | 0.09219219 |
| Ptges3l | 0.20297196 | 0.90795886 | 0.104 | 0.062 | 1 | 0.0472378 | 1.01207283 | 0.08 | 0.021 | 1 | 0.20297196 | 0.0922442 |
| Zfp385a | 0.04727085 | -0.6340515 | 0.167 | 0.28 | 1 | 0.88953326 | 0.16407593 | 0.126 | 0.135 | 1 | 0.88953326 | 0.09230717 |
| Haghl | 0.65309077 | 0.16795921 | 0.167 | 0.145 | 1 | 0.04733485 | 0.75427507 | 0.155 | 0.073 | 1 | 0.65309077 | 0.09242911 |
| Metap1d | 0.04734174 | -0.3550755 | 0.635 | 0.699 | 1 | 0.05464113 | -0.1007597 | 0.622 | 0.406 | 1 | 0.05464113 | 0.09244223 |
| Evpl | 0.84449794 | 0.62898291 | 0.062 | 0.057 | 1 | 0.04740333 | 0.95317914 | 0.08 | 0.021 | 1 | 0.84449794 | 0.09255958 |
| Crybg3 | 0.15214206 | -0.2421194 | 0.354 | 0.446 | 1 | 0.04750941 | -0.1351767 | 0.408 | 0.583 | 1 | 0.15214206 | 0.09276168 |
| Ptpn1 | 0.04751868 | -0.2464947 | 0.948 | 0.943 | 1 | 0.08231584 | 0.14229858 | 0.971 | 0.969 | 1 | 0.08231584 | 0.09277933 |
| Htt | 0.81600915 | -0.2305455 | 0.688 | 0.642 | 1 | 0.04756144 | -0.4718751 | 0.769 | 0.833 | 1 | 0.81600915 | 0.09286079 |
| 5031425E22Rik | 0.53274046 | -0.2196671 | 0.823 | 0.834 | 1 | 0.0475743 | -0.9459566 | 0.845 | 0.833 | 1 | 0.53274046 | 0.09288528 |
| Stxbp2 | 0.29961448 | 0.37946331 | 0.104 | 0.067 | 1 | 0.04759077 | 0.78968041 | 0.113 | 0.042 | 1 | 0.29961448 | 0.09291666 |
| Pecr | 0.05449592 | 0.5049942 | 0.219 | 0.13 | 1 | 0.0476173 | 0.59703533 | 0.113 | 0.042 | 1 | 0.05449592 | 0.0929672 |
| Spryd4 | 0.72900758 | 0.42253204 | 0.083 | 0.073 | 1 | 0.04769147 | 1.08246216 | 0.08 | 0.021 | 1 | 0.72900758 | 0.09310846 |
| Ank1 | 0.04775389 | -1.7738652 | 0.052 | 0.124 | 1 | 0.37667701 | -1.07539 | 0.059 | 0.083 | 1 | 0.37667701 | 0.09322735 |
| Atp6v1b2 | 0.33076338 | 0.23697638 | 0.5 | 0.435 | 1 | 0.04778515 | 0.4060013 | 0.689 | 0.667 | 1 | 0.33076338 | 0.09328687 |
| Rrp7a | 0.2026163 | 0.49934763 | 0.344 | 0.28 | 1 | 0.04782598 | 0.1883956 | 0.357 | 0.219 | 1 | 0.2026163 | 0.09336464 |
| Rusc2 | 0.0739068 | -0.5362453 | 0.375 | 0.472 | 1 | 0.04785566 | 0.26461486 | 0.546 | 0.375 | 1 | 0.0739068 | 0.09342116 |
| Lrrc57 | 0.68258451 | 0.18646468 | 0.25 | 0.233 | 1 | 0.04787682 | 0.25782925 | 0.256 | 0.156 | 1 | 0.68258451 | 0.09346146 |
| Nhej1 | 0.04791061 | 0.10060394 | 0.229 | 0.13 | 1 | 0.3486235 | -1.1172102 | 0.181 | 0.125 | 1 | 0.3486235 | 0.09352579 |
| Grwd1 | 0.39293764 | 0.12613903 | 0.177 | 0.135 | 1 | 0.04792619 | 0.84542296 | 0.181 | 0.094 | 1 | 0.39293764 | 0.09355546 |
| Ogfod2 | 0.04795028 | 1.00749454 | 0.146 | 0.073 | 1 | 0.61669206 | 0.44503224 | 0.088 | 0.073 | 1 | 0.61669206 | 0.09360133 |
| Raver1 | 0.24368051 | 0.21899864 | 0.344 | 0.269 | 1 | 0.04795958 | 0.2862478 | 0.391 | 0.24 | 1 | 0.24368051 | 0.09361903 |
| Snhg5 | 0.64650974 | -0.6542252 | 0.438 | 0.383 | 1 | 0.0480072 | 0.43488768 | 0.282 | 0.167 | 1 | 0.64650974 | 0.09370971 |
| 9630014M24Rik | 0.04805594 | -1.8669746 | 0.01 | 0.062 | 1 | 0.14151274 | 2.14957636 | 0.042 | 0.01 | 1 | 0.14151274 | 0.0938025 |
| Dhx8 | 0.04809309 | 0.35933821 | 0.458 | 0.337 | 1 | 0.19473837 | -0.1815312 | 0.487 | 0.365 | 1 | 0.19473837 | 0.09387323 |
| Stambp | 0.12330578 | 0.3584017 | 0.333 | 0.238 | 1 | 0.04809907 | 0.65977109 | 0.286 | 0.177 | 1 | 0.12330578 | 0.09388462 |
| Sstr2 | 0.04812464 | -1.8669746 | 0.052 | 0.124 | 1 | 0.91774154 | -0.3098553 | 0.034 | 0.031 | 1 | 0.91774154 | 0.09393329 |
| Mtmr12 | 0.04876079 | 0.22370309 | 0.615 | 0.461 | 1 | 0.04813603 | 0.20148437 | 0.693 | 0.562 | 1 | 0.04876079 | 0.09395499 |
| Tent2 | 0.04830316 | 0.3022378 | 0.74 | 0.611 | 1 | 0.72083788 | -0.2821103 | 0.748 | 0.74 | 1 | 0.72083788 | 0.09427312 |
| Gm56936 | 0.51658936 | -0.2681399 | 0.156 | 0.187 | 1 | 0.04834763 | 0.97025266 | 0.126 | 0.052 | 1 | 0.51658936 | 0.09435777 |
| Itfg1 | 0.30618753 | -0.2204574 | 0.635 | 0.668 | 1 | 0.04841421 | 0.17462585 | 0.634 | 0.542 | 1 | 0.30618753 | 0.09448449 |
| Gjc1 | 0.04845899 | 0.41331351 | 0.531 | 0.394 | 1 | 0.43734473 | -0.2890967 | 0.584 | 0.448 | 1 | 0.43734473 | 0.0945697 |
| Spg7 | 0.10676653 | -0.5330738 | 0.354 | 0.435 | 1 | 0.04846676 | 0.52126783 | 0.399 | 0.271 | 1 | 0.10676653 | 0.09458449 |
| Abhd12 | 0.85001501 | -0.1914865 | 0.521 | 0.477 | 1 | 0.04851555 | 0.29005758 | 0.517 | 0.385 | 1 | 0.85001501 | 0.09467733 |
| Mgmt | 0.04851714 | 0.89646322 | 0.167 | 0.088 | 1 | 0.04883824 | -0.5025003 | 0.147 | 0.062 | 1 | 0.04883824 | 0.09468037 |
| Fam171a2 | 0.53063347 | -0.577468 | 0.031 | 0.047 | 1 | 0.04856711 | -1.2029401 | 0.046 | 0.104 | 1 | 0.53063347 | 0.09477546 |
| Cyp26b1 | 0.31884964 | -0.367717 | 0.469 | 0.513 | 1 | 0.04857859 | 0.60681635 | 0.487 | 0.333 | 1 | 0.31884964 | 0.09479729 |
| Gpr137b | 0.58075814 | -0.7294711 | 0.104 | 0.124 | 1 | 0.04859332 | 0.97965135 | 0.181 | 0.094 | 1 | 0.58075814 | 0.09482532 |
| Tmie | 0.86749097 | 0.42253204 | 0.271 | 0.306 | 1 | 0.04860307 | 1.32241295 | 0.08 | 0.021 | 1 | 0.86749097 | 0.09484387 |
| Tufm | 0.75897543 | -0.1477837 | 0.188 | 0.202 | 1 | 0.04875302 | 0.16145046 | 0.214 | 0.115 | 1 | 0.75897543 | 0.09512918 |
| Pwwp2a | 0.10537777 | 0.27052894 | 0.458 | 0.352 | 1 | 0.04880835 | -0.7156149 | 0.525 | 0.615 | 1 | 0.10537777 | 0.09523445 |
| Ophn1 | 0.18866476 | -0.115487 | 0.458 | 0.534 | 1 | 0.04886938 | -0.6681557 | 0.492 | 0.594 | 1 | 0.18866476 | 0.09535054 |
| H2-Q10 | 0.04901267 | -0.4330781 | 0.01 | 0.062 | 1 | 0.20315061 | 1.69014474 | 0.017 | 0 | 1 | 0.20315061 | 0.09562309 |
| Syt11 | 0.04919691 | -1.9925055 | 0.042 | 0.109 | 1 | 0.41554104 | -0.5961594 | 0.097 | 0.125 | 1 | 0.41554104 | 0.09597349 |
| Runx2 | 0.82975693 | -0.473603 | 0.844 | 0.772 | 1 | 0.04928038 | -0.8676701 | 0.824 | 0.917 | 1 | 0.82975693 | 0.09613221 |
| Slc7a5 | 0.04933785 | 0.4339368 | 0.562 | 0.42 | 1 | 0.13069275 | 0.14957636 | 0.567 | 0.479 | 1 | 0.13069275 | 0.09624147 |
| Ddah2 | 0.21312172 | -0.6770036 | 0.208 | 0.264 | 1 | 0.0493415 | 0.53814164 | 0.29 | 0.177 | 1 | 0.21312172 | 0.09624841 |

| | | | | | | | | | | | | |
|---|---|---|---|---|---|---|---|---|---|---|---|---|
| Gm35546 | 0.85494807 | -0.1624305 | 0.156 | 0.166 | 1 | 0.04939046 | 0.54330335 | 0.227 | 0.125 | 1 | 0.85494807 | 0.09634151 |
| Zfp512b | 0.12781339 | 0.63826073 | 0.198 | 0.13 | 1 | 0.04942042 | 0.32401684 | 0.277 | 0.167 | 1 | 0.12781339 | 0.09639846 |
| Rftn1 | 0.22080787 | 0.42253204 | 0.135 | 0.088 | 1 | 0.04944996 | -1.6391629 | 0.097 | 0.167 | 1 | 0.22080787 | 0.09645461 |
| Cyth2 | 0.06619451 | 0.48142572 | 0.552 | 0.472 | 1 | 0.04946981 | 0.31117649 | 0.559 | 0.458 | 1 | 0.06619451 | 0.09649236 |
| D030056L22Rik | 0.51647536 | 0.22988696 | 0.219 | 0.187 | 1 | 0.04951416 | 0.5580412 | 0.21 | 0.115 | 1 | 0.51647536 | 0.09657667 |
| Abca4 | 0.27709143 | 0.92503238 | 0.198 | 0.15 | 1 | 0.04952941 | -2.5993619 | 0.08 | 0.146 | 1 | 0.27709143 | 0.09660566 |
| Slc18b1 | 0.04956707 | 0.95502712 | 0.208 | 0.124 | 1 | 0.13956516 | 0.13945214 | 0.214 | 0.135 | 1 | 0.13956516 | 0.09667725 |
| A630001G21Rik | 0.04964446 | 1.37006462 | 0.115 | 0.052 | 1 | 0.21548646 | -0.1723517 | 0.067 | 0.031 | 1 | 0.21548646 | 0.09682435 |
| Pcdh18 | 0.04976191 | -1.8669746 | 0.042 | 0.109 | 1 | 0.12180912 | -1.3699763 | 0.109 | 0.167 | 1 | 0.12180912 | 0.09704757 |
| Gm39090 | 0.04976714 | -1.0368996 | 0.115 | 0.207 | 1 | 0.14970708 | 0.15580831 | 0.185 | 0.115 | 1 | 0.14970708 | 0.09705751 |
| Trp53bp1 | 0.04984214 | 0.49292136 | 0.5 | 0.383 | 1 | 0.8130487 | -0.4588652 | 0.517 | 0.427 | 1 | 0.8130487 | 0.09720005 |
| 1700024B18Rik | 0.0498727 | -3.4018964 | 0.031 | 0.093 | 1 | 0.3137635 | -0.4043199 | 0.08 | 0.115 | 1 | 0.3137635 | 0.09725812 |
| Itga2b | 0.04992123 | 0.59245704 | 0.177 | 0.093 | 1 | 0.1800745 | -1.0535155 | 0.126 | 0.177 | 1 | 0.1800745 | 0.09735034 |
| Mboat1 | 0.09185381 | -1.6554705 | 0.021 | 0.067 | 1 | 0.04997164 | -2.3973181 | 0.021 | 0.062 | 1 | 0.09185381 | 0.09744611 |
| Tex10 | 0.43198901 | -0.5554417 | 0.344 | 0.358 | 1 | 0.04999593 | -0.6188055 | 0.378 | 0.479 | 1 | 0.43198901 | 0.09749227 |
| Arhgef18 | 0.2874494 | 0.27052894 | 0.25 | 0.192 | 1 | 0.05003958 | -0.5176601 | 0.345 | 0.448 | 1 | 0.2874494 | 0.0975752 |
| Ndufv3 | 0.46945849 | -0.1624305 | 0.656 | 0.663 | 1 | 0.05004075 | 0.35893883 | 0.609 | 0.521 | 1 | 0.46945849 | 0.09757742 |
| Ube2u | 0.31015453 | 0.83756954 | 0.042 | 0.021 | 1 | 0.05009411 | -1.8692827 | 0.034 | 0.083 | 1 | 0.31015453 | 0.09767881 |
| Tmem259 | 0.13378133 | 0.23305424 | 0.542 | 0.446 | 1 | 0.05017924 | 0.2007599 | 0.559 | 0.396 | 1 | 0.13378133 | 0.09784052 |
| Gm17057 | 0.31991037 | 0.15949763 | 0.062 | 0.036 | 1 | 0.05020135 | -2.5322477 | 0.021 | 0.062 | 1 | 0.31991037 | 0.09788252 |
| Foxk1 | 0.05029663 | 0.48554183 | 0.49 | 0.373 | 1 | 0.65328849 | -0.2628918 | 0.618 | 0.521 | 1 | 0.65328849 | 0.09806351 |
| Coa5 | 0.05032919 | 1.07162487 | 0.177 | 0.098 | 1 | 0.27490625 | 0.1876444 | 0.164 | 0.115 | 1 | 0.27490625 | 0.09812535 |
| Mprip | 0.19978404 | -0.4223021 | 0.958 | 0.953 | 1 | 0.0504234 | -0.3078389 | 0.979 | 0.99 | 1 | 0.19978404 | 0.09830428 |
| Egln1 | 0.41332098 | 0.10589824 | 0.583 | 0.523 | 1 | 0.05054396 | 0.19915838 | 0.689 | 0.552 | 1 | 0.41332098 | 0.09853323 |
| Fem1c | 0.07334245 | 0.38505733 | 0.49 | 0.368 | 1 | 0.05054982 | -0.406285 | 0.521 | 0.646 | 1 | 0.07334245 | 0.09854435 |
| Pus7l | 0.05059254 | 1.52206771 | 0.083 | 0.031 | 1 | 0.20030164 | 0.8917786 | 0.067 | 0.031 | 1 | 0.20030164 | 0.09862548 |
| Rassf3 | 0.42670753 | 0.31682259 | 0.302 | 0.264 | 1 | 0.05059978 | -0.3982111 | 0.454 | 0.604 | 1 | 0.42670753 | 0.09863922 |
| E4f1 | 0.07240653 | 0.37672835 | 0.271 | 0.171 | 1 | 0.05065312 | 0.16819203 | 0.248 | 0.146 | 1 | 0.07240653 | 0.09874049 |
| Smarcc1 | 0.05066731 | 0.13632452 | 0.76 | 0.642 | 1 | 0.2645014 | -0.3959442 | 0.874 | 0.885 | 1 | 0.2645014 | 0.09876745 |
| Cmc1 | 0.13883019 | -0.5309254 | 0.177 | 0.254 | 1 | 0.05067905 | 0.34222143 | 0.277 | 0.156 | 1 | 0.13883019 | 0.09878974 |
| Naa40 | 0.0553492 | 0.7105128 | 0.26 | 0.166 | 1 | 0.05075472 | 0.33650778 | 0.231 | 0.125 | 1 | 0.0553492 | 0.09893339 |
| Arhgap1 | 0.05078456 | 0.47499946 | 0.417 | 0.301 | 1 | 0.4660901 | -0.1308851 | 0.374 | 0.302 | 1 | 0.4660901 | 0.09899005 |
| Slc35c2 | 0.19216086 | 0.46692616 | 0.198 | 0.14 | 1 | 0.05082633 | 0.44503224 | 0.197 | 0.104 | 1 | 0.19216086 | 0.09906934 |
| Cox5a | 0.05084242 | 0.27562231 | 0.885 | 0.824 | 1 | 0.20277842 | 0.12067332 | 0.87 | 0.833 | 1 | 0.20277842 | 0.09909989 |
| Arid1a | 0.29184964 | -0.3358649 | 0.615 | 0.684 | 1 | 0.05091082 | -0.4283895 | 0.866 | 0.906 | 1 | 0.29184964 | 0.09922972 |
| Ppp2r5c | 0.20132133 | -0.2286642 | 0.781 | 0.813 | 1 | 0.050923 | -0.4432968 | 0.811 | 0.885 | 1 | 0.20132133 | 0.09925284 |
| Gpr176 | 0.15049214 | -2.3144336 | 0.01 | 0.041 | 1 | 0.05092672 | -2.8948178 | 0.021 | 0.062 | 1 | 0.15049214 | 0.0992599 |
| Csnk1e | 0.58465422 | 0.1213625 | 0.615 | 0.549 | 1 | 0.05096476 | 0.18522173 | 0.765 | 0.698 | 1 | 0.58465422 | 0.09933211 |
| Zfp446 | 0.38050235 | 0.20013961 | 0.073 | 0.047 | 1 | 0.05099335 | 0.56461386 | 0.08 | 0.021 | 1 | 0.38050235 | 0.09938637 |
| Pcyox1 | 0.56705728 | 0.10265177 | 0.365 | 0.321 | 1 | 0.05113112 | 0.17557156 | 0.37 | 0.229 | 1 | 0.56705728 | 0.09964786 |
| Invs | 0.13697298 | 0.17004755 | 0.375 | 0.28 | 1 | 0.05113235 | -0.1058414 | 0.378 | 0.229 | 1 | 0.13697298 | 0.09965019 |
| Tctn1 | 0.05117078 | 1.39451766 | 0.115 | 0.052 | 1 | 0.21177993 | 0.14957636 | 0.122 | 0.073 | 1 | 0.21177993 | 0.09972312 |
| Hcfc1r1 | 0.68044836 | -0.2015 | 0.76 | 0.756 | 1 | 0.05117124 | 0.33867237 | 0.693 | 0.635 | 1 | 0.68044836 | 0.09972398 |
| Trim28 | 0.15867644 | 0.32695438 | 0.448 | 0.363 | 1 | 0.05120432 | 0.17908035 | 0.634 | 0.49 | 1 | 0.15867644 | 0.09978675 |
| Uba2 | 0.06460211 | 0.34006988 | 0.552 | 0.435 | 1 | 0.05123532 | 0.27510724 | 0.676 | 0.594 | 1 | 0.06460211 | 0.09984559 |
| Herc2 | 0.13726449 | -0.3505028 | 0.667 | 0.762 | 1 | 0.05126071 | -0.3688219 | 0.887 | 0.927 | 1 | 0.13726449 | 0.09989376 |
| Cep152 | 0.14668388 | 0.86999101 | 0.104 | 0.057 | 1 | 0.05144001 | -1.7248928 | 0.034 | 0.083 | 1 | 0.14668388 | 0.10023394 |
| Gm38115 | 0.05147535 | 1.35541784 | 0.115 | 0.052 | 1 | 0.07406724 | -1.3098553 | 0.092 | 0.156 | 1 | 0.07406724 | 0.10030099 |
| Col23a1 | 0.0516136 | -3.1217885 | 0.031 | 0.093 | 1 | 0.45982689 | -1.73612 | 0.092 | 0.115 | 1 | 0.45982689 | 0.10056323 |
| Itih5 | 0.17037548 | -1.0051055 | 0.188 | 0.119 | 1 | 0.0516474 | -3.07539 | 0.021 | 0.062 | 1 | 0.17037548 | 0.10062734 |
| Nop56 | 0.05165784 | 0.27969726 | 0.844 | 0.658 | 1 | 0.92978421 | -0.1054968 | 0.681 | 0.719 | 1 | 0.92978421 | 0.10064715 |
| Gli1 | 0.05172658 | 1.22988696 | 0.115 | 0.052 | 1 | 0.45513939 | 0.17557156 | 0.05 | 0.031 | 1 | 0.45513939 | 0.10077753 |
| Msantd4 | 0.23983879 | -0.4779323 | 0.312 | 0.363 | 1 | 0.05173121 | -0.154577 | 0.433 | 0.271 | 1 | 0.23983879 | 0.10078631 |
| Snrnp200 | 0.27791892 | 0.14151188 | 0.458 | 0.373 | 1 | 0.05174311 | 0.29473604 | 0.601 | 0.448 | 1 | 0.27791892 | 0.10080887 |
| Scgb1c1 | 0.20910685 | 1.00749454 | 0.062 | 0.031 | 1 | 0.05176591 | -1.214698 | 0.168 | 0.26 | 1 | 0.20910685 | 0.1008521 |
| Tom1l2 | 0.05180392 | -0.4648504 | 0.75 | 0.782 | 1 | 0.54069848 | -0.3724072 | 0.882 | 0.917 | 1 | 0.54069848 | 0.10092419 |
| Ankrd24 | 0.05187422 | 1.21899864 | 0.156 | 0.083 | 1 | 0.20106769 | 0.41261076 | 0.109 | 0.062 | 1 | 0.20106769 | 0.10105751 |
| Marchf8 | 0.05198585 | 0.42879103 | 0.5 | 0.383 | 1 | 0.85271157 | -0.4120357 | 0.534 | 0.479 | 1 | 0.85271157 | 0.10126917 |
| Zfp74 | 0.96645835 | -0.240433 | 0.135 | 0.13 | 1 | 0.05205466 | 0.33650778 | 0.214 | 0.115 | 1 | 0.96645835 | 0.10139963 |
| Fam222a | 0.05210932 | 1.37006462 | 0.083 | 0.031 | 1 | 0.37505363 | -1.3098553 | 0.059 | 0.083 | 1 | 0.37505363 | 0.10150325 |
| Gm50069 | 0.05210932 | 1.37006462 | 0.083 | 0.031 | 1 | 0.14367653 | 1.27510724 | 0.042 | 0.01 | 1 | 0.14367653 | 0.10150325 |
| Vps4b | 0.44288893 | 0.11970404 | 0.698 | 0.694 | 1 | 0.05216258 | 0.10164188 | 0.782 | 0.698 | 1 | 0.44288893 | 0.10160422 |
| Furin | 0.11341817 | -0.379868 | 0.719 | 0.741 | 1 | 0.05218824 | 0.15566369 | 0.866 | 0.792 | 1 | 0.11341817 | 0.10165287 |
| Dclre1c | 0.62628741 | -0.4994655 | 0.115 | 0.135 | 1 | 0.05219124 | -1.5443205 | 0.063 | 0.125 | 1 | 0.62628741 | 0.10165856 |
| Atg101 | 0.08203292 | -0.3509594 | 0.281 | 0.383 | 1 | 0.0522016 | 0.30064233 | 0.475 | 0.354 | 1 | 0.08203292 | 0.1016782 |
| Amot | 0.05221902 | -1.4779323 | 0.115 | 0.197 | 1 | 0.87695866 | -0.9186645 | 0.235 | 0.208 | 1 | 0.87695866 | 0.10171121 |
| Ppp1r14c | 0.0522439 | -1.9311049 | 0.021 | 0.078 | 1 | 0.08772573 | 1.62874419 | 0.067 | 0.021 | 1 | 0.08772573 | 0.10175838 |
| Mmp3 | 0.05224904 | 0.23456345 | 0.052 | 0.13 | 1 | 0.81130676 | 1.69014474 | 0.025 | 0.021 | 1 | 0.81130676 | 0.10176812 |
| Slc49a4 | 0.0522579 | -0.6397522 | 0.479 | 0.554 | 1 | 0.41541241 | -0.384128 | 0.765 | 0.823 | 1 | 0.41541241 | 0.1017849 |
| Snrk | 0.05226188 | 0.64136664 | 0.323 | 0.223 | 1 | 0.08096967 | 0.20887354 | 0.412 | 0.281 | 1 | 0.08096967 | 0.10179245 |
| Pigs | 0.15377132 | -0.4163132 | 0.365 | 0.435 | 1 | 0.05229195 | 0.19915838 | 0.345 | 0.219 | 1 | 0.15377132 | 0.10184945 |
| Zc4h2 | 0.0522923 | 1.46692616 | 0.083 | 0.031 | 1 | 0.20577072 | 0.61614416 | 0.067 | 0.031 | 1 | 0.20577072 | 0.10185012 |
| Ncapd3 | 0.98091277 | 0.70793425 | 0.135 | 0.135 | 1 | 0.05239764 | 0.61214223 | 0.193 | 0.104 | 1 | 0.98091277 | 0.10204976 |
| Zranb3 | 0.05246569 | 1.27052894 | 0.135 | 0.067 | 1 | 0.55696387 | -0.6217993 | 0.097 | 0.073 | 1 | 0.55696387 | 0.10217874 |
| Gm42869 | 0.0525366 | 1.32942263 | 0.083 | 0.031 | 1 | 0.53717181 | 0.27510724 | 0.034 | 0.021 | 1 | 0.53717181 | 0.1023131 |
| A330041J22Rik | 0.0525366 | 1.32942263 | 0.083 | 0.031 | 1 | 0.18589831 | 0.49749966 | 0.055 | 0.021 | 1 | 0.18589831 | 0.1023131 |
| Tubg2 | 0.0525366 | 1.32942263 | 0.083 | 0.031 | 1 | 0.23679813 | 1.01207283 | 0.034 | 0.01 | 1 | 0.23679813 | 0.1023131 |
| 1600014C10Rik | 0.05255043 | 1.08149512 | 0.156 | 0.083 | 1 | 0.39726179 | 0.35310975 | 0.139 | 0.104 | 1 | 0.39726179 | 0.10233931 |
| Mfsd12 | 0.17265227 | 1.59245704 | 0.042 | 0.016 | 1 | 0.05256007 | 1.69014474 | 0.092 | 0.031 | 1 | 0.17265227 | 0.10235758 |
| Nhlrc3 | 0.50100142 | 0.48142572 | 0.177 | 0.15 | 1 | 0.05260378 | 0.22357694 | 0.185 | 0.094 | 1 | 0.50100142 | 0.10244041 |
| Gm56962 | 0.07421827 | 2.12297175 | 0.052 | 0.016 | 1 | 0.0526232 | -1.961932 | 0.021 | 0.062 | 1 | 0.07421827 | 0.1024772 |
| Tmem219 | 0.05268708 | 0.69338595 | 0.292 | 0.192 | 1 | 0.08945458 | 0.20259474 | 0.282 | 0.188 | 1 | 0.08945458 | 0.10259822 |
| B830012L14Rik | 0.2160902 | -0.4275529 | 0.406 | 0.311 | 1 | 0.05274612 | -0.5515022 | 0.517 | 0.333 | 1 | 0.2160902 | 0.10271009 |
| Cep43 | 0.0527562 | 0.44489985 | 0.427 | 0.311 | 1 | 0.79140242 | -0.1812711 | 0.357 | 0.333 | 1 | 0.79140242 | 0.10272919 |
| Pou2af3 | 0.37838273 | -0.8504865 | 0.229 | 0.264 | 1 | 0.05277246 | 0.45567948 | 0.197 | 0.104 | 1 | 0.37838273 | 0.10276 |
| Krt79 | 0.15837881 | 2.00749454 | 0.01 | 0 | 1 | 0.05277622 | 3.08246216 | 0.059 | 0.01 | 1 | 0.15837881 | 0.10276712 |

| | | | | | | | | | | | | |
|---|---|---|---|---|---|---|---|---|---|---|---|---|
| Gnasas1 | 0.05278184 | 1.00749454 | 0.083 | 0.031 | 1 | 0.39292627 | -0.1843244 | 0.067 | 0.042 | 1 | 0.39292627 | 0.10277776 |
| Thbs2 | 0.10651115 | -1.9559796 | 0.104 | 0.171 | 1 | 0.05285541 | -3.2213186 | 0.113 | 0.177 | 1 | 0.10651115 | 0.10291713 |
| Gm6712 | 0.05292341 | 0.54806292 | 0.146 | 0.073 | 1 | 0.56698272 | 0.44913664 | 0.126 | 0.104 | 1 | 0.56698272 | 0.10304593 |
| Gpr68 | 0.0530186 | 1.17741954 | 0.135 | 0.067 | 1 | 0.29652436 | 0.95317914 | 0.059 | 0.031 | 1 | 0.29652436 | 0.10322623 |
| Lsg1 | 0.1108341 | 0.28867629 | 0.573 | 0.477 | 1 | 0.05313759 | 0.41393494 | 0.534 | 0.438 | 1 | 0.1108341 | 0.10345157 |
| A430105J06Rik | 0.66676245 | -0.3144336 | 0.031 | 0.041 | 1 | 0.05318658 | 1.86006974 | 0.059 | 0.01 | 1 | 0.66676245 | 0.10354435 |
| Lsm14b | 0.5539099 | -0.2920657 | 0.115 | 0.14 | 1 | 0.05324912 | 0.41806519 | 0.172 | 0.083 | 1 | 0.5539099 | 0.10366277 |
| Ifih1 | 0.15746105 | 0.83240783 | 0.156 | 0.098 | 1 | 0.05328177 | -0.9045988 | 0.155 | 0.073 | 1 | 0.15746105 | 0.10372459 |
| Arl5c | 0.05338659 | -1.6363617 | 0.031 | 0.093 | 1 | 0.99287329 | -0.9472852 | 0.063 | 0.062 | 1 | 0.99287329 | 0.10392305 |
| Rph3al | 0.08192886 | -1.7998604 | 0 | 0.031 | 1 | 0.0533901 | 1.86006974 | 0.059 | 0.01 | 1 | 0.08192886 | 0.10392969 |
| Gp1bb | 0.69098575 | 0.27052894 | 0.052 | 0.041 | 1 | 0.0533901 | 1.77760758 | 0.059 | 0.01 | 1 | 0.69098575 | 0.10392969 |
| C4b | 0.88179872 | 0.44189736 | 0.083 | 0.088 | 1 | 0.05345922 | 0.79250646 | 0.244 | 0.146 | 1 | 0.88179872 | 0.10406056 |
| Gm16062 | 0.64722748 | -0.5285584 | 0.083 | 0.098 | 1 | 0.05347177 | 0.77760758 | 0.164 | 0.083 | 1 | 0.64722748 | 0.1040843 |
| Tle2 | 0.05350701 | 0.79599043 | 0.167 | 0.088 | 1 | 0.32415722 | -0.2654611 | 0.126 | 0.083 | 1 | 0.32415722 | 0.10415102 |
| Cpt2 | 0.09962026 | 1.00749454 | 0.104 | 0.052 | 1 | 0.05353118 | 1.44503224 | 0.092 | 0.031 | 1 | 0.09962026 | 0.10419677 |
| Rnf182 | 0.05355995 | -1.1624305 | 0.052 | 0.124 | 1 | 0.34923114 | -0.7504279 | 0.084 | 0.052 | 1 | 0.34923114 | 0.10425124 |
| D130062J10Rik | 0.47294388 | -0.7294711 | 0.021 | 0.036 | 1 | 0.05359234 | 1.69014474 | 0.059 | 0.01 | 1 | 0.47294388 | 0.10431253 |
| Gm3839 | 0.82342138 | 0.32942263 | 0.042 | 0.036 | 1 | 0.05359234 | 1.69014474 | 0.059 | 0.01 | 1 | 0.82342138 | 0.10431253 |
| Gm43138 | 0.98603793 | 0.22988696 | 0.052 | 0.052 | 1 | 0.05359234 | 1.69014474 | 0.059 | 0.01 | 1 | 0.98603793 | 0.10431253 |
| 6530402F18Rik | 0.9927582 | 0.42253204 | 0.021 | 0.021 | 1 | 0.05359234 | 1.69014474 | 0.059 | 0.01 | 1 | 0.9927582 | 0.10431253 |
| Prdm16 | 0.05361008 | -0.6698109 | 0.75 | 0.824 | 1 | 0.16441464 | -0.7332441 | 0.916 | 0.844 | 1 | 0.16441464 | 0.10434612 |
| Fam122a | 0.28316891 | -0.4950058 | 0.208 | 0.264 | 1 | 0.0536604 | 0.20471791 | 0.197 | 0.104 | 1 | 0.28316891 | 0.10444136 |
| Abraxas1 | 0.05369336 | 1.13302542 | 0.115 | 0.052 | 1 | 0.11130697 | 1.59703533 | 0.046 | 0.01 | 1 | 0.11130697 | 0.10450375 |
| Ddx28 | 0.05373189 | 1.12297175 | 0.115 | 0.052 | 1 | 0.5272316 | 0.36821664 | 0.059 | 0.042 | 1 | 0.5272316 | 0.10457666 |
| Mipep | 0.96467504 | -0.7998604 | 0.156 | 0.15 | 1 | 0.05378956 | 0.41003682 | 0.168 | 0.083 | 1 | 0.96467504 | 0.10468581 |
| Dio2 | 0.3153425 | -2.2962862 | 0.115 | 0.155 | 1 | 0.05383428 | -1.4156499 | 0.164 | 0.24 | 1 | 0.3153425 | 0.10477044 |
| Sema7a | 0.0538613 | -1.9925055 | 0.021 | 0.078 | 1 | 0.60764206 | -1.3098553 | 0.059 | 0.073 | 1 | 0.60764206 | 0.10482157 |
| Gm13832 | 0.05386794 | -1.7998604 | 0.021 | 0.078 | 1 | 0.77381753 | -0.5443205 | 0.038 | 0.031 | 1 | 0.77381753 | 0.10483412 |
| Adam30 | 0.05395408 | 0.75595577 | 0.167 | 0.088 | 1 | 0.73328508 | -0.2573878 | 0.151 | 0.167 | 1 | 0.73328508 | 0.10499712 |
| Zfp623 | 0.31880524 | 0.64492446 | 0.062 | 0.036 | 1 | 0.05397133 | 0.96316323 | 0.109 | 0.042 | 1 | 0.31880524 | 0.10502976 |
| Zfp748 | 0.05398613 | 1.10703021 | 0.115 | 0.052 | 1 | 0.32515764 | 0.24653809 | 0.181 | 0.135 | 1 | 0.32515764 | 0.10505777 |
| Srd5a1 | 0.05400563 | 1.17741954 | 0.083 | 0.031 | 1 | 0.28766211 | -0.194378 | 0.046 | 0.021 | 1 | 0.28766211 | 0.10509466 |
| Gm5817 | 0.05403672 | 1.38600616 | 0.083 | 0.031 | 1 | 0.23679813 | 1.01207283 | 0.034 | 0.01 | 1 | 0.23679813 | 0.10515347 |
| Tmem39b | 0.05405946 | 0.41025371 | 0.281 | 0.176 | 1 | 0.48957398 | -0.2499838 | 0.324 | 0.26 | 1 | 0.48957398 | 0.1051965 |
| 1110018N20Rik | 0.15837881 | 2.00749454 | 0.01 | 0 | 1 | 0.05406363 | 2.14957636 | 0.038 | 0 | 1 | 0.15837881 | 0.10520438 |
| K230015D01Rik | 0.30584838 | 1.00749454 | 0.062 | 0.036 | 1 | 0.05406363 | 2.14957636 | 0.038 | 0 | 1 | 0.30584838 | 0.10520438 |
| Gm56795 | 0.38689439 | -0.577468 | 0.01 | 0.026 | 1 | 0.05406363 | 2.39058446 | 0.038 | 0 | 1 | 0.38689439 | 0.10520438 |
| Adam11 | 0.48064264 | 0.59245704 | 0.021 | 0.01 | 1 | 0.05406363 | 2.14957636 | 0.038 | 0 | 1 | 0.48064264 | 0.10520438 |
| Gm29170 | 0.58426772 | 0.74446013 | 0.031 | 0.021 | 1 | 0.05406363 | 2.14957636 | 0.038 | 0 | 1 | 0.58426772 | 0.10520438 |
| Gm56791 | 0.5862099 | 0.68556644 | 0.031 | 0.021 | 1 | 0.05406363 | 2.14957636 | 0.038 | 0 | 1 | 0.5862099 | 0.10520438 |
| Mettl18 | 1 | -0.3144336 | 0.01 | 0.01 | 1 | 0.05406363 | 2.49749966 | 0.038 | 0 | 1 | 1 | 0.10520438 |
| Prss46 | 0.78979801 | -0.2148979 | 0.021 | 0.026 | 1 | 0.05407052 | 2.39058446 | 0.038 | 0 | 1 | 0.78979801 | 0.10521742 |
| Atp1a3 | 0.32025586 | -0.577468 | 0 | 0.01 | 1 | 0.05407913 | 2.86006974 | 0.038 | 0 | 1 | 0.32025586 | 0.10523371 |
| Mst1r | 0.21867159 | 1.59245704 | 0.021 | 0.005 | 1 | 0.05408282 | 3.64434105 | 0.038 | 0 | 1 | 0.21867159 | 0.10524069 |
| Pik3ap1 | 0.20393009 | 1.00749454 | 0.031 | 0.01 | 1 | 0.05408307 | 3.86006974 | 0.038 | 0 | 1 | 0.20393009 | 0.10524115 |
| Syk | 0.73685932 | 1.83756954 | 0.01 | 0.016 | 1 | 0.05408307 | 3.39058446 | 0.038 | 0 | 1 | 0.73685932 | 0.10524115 |
| Dnase1l2 | 0.07982472 | 1.74446013 | 0.042 | 0.01 | 1 | 0.0540843 | 2.01207283 | 0.038 | 0 | 1 | 0.07982472 | 0.10524348 |
| Pbld1 | 0.20750194 | 1.15949763 | 0.062 | 0.031 | 1 | 0.0540843 | 2.01207283 | 0.038 | 0 | 1 | 0.20750194 | 0.10524348 |
| Gm56779 | 0.32025586 | -0.577468 | 0 | 0.01 | 1 | 0.0540843 | 2.01207283 | 0.038 | 0 | 1 | 0.32025586 | 0.10524348 |
| B3galt4 | 0.47606502 | 1.00749454 | 0.021 | 0.01 | 1 | 0.0540843 | 2.01207283 | 0.038 | 0 | 1 | 0.47606502 | 0.10524348 |
| Thnsl1 | 0.57678264 | 0.35541784 | 0.062 | 0.047 | 1 | 0.0540843 | 2.01207283 | 0.038 | 0 | 1 | 0.57678264 | 0.10524348 |
| Samd15 | 0.59199182 | 0.42253204 | 0.031 | 0.021 | 1 | 0.0540843 | 2.01207283 | 0.038 | 0 | 1 | 0.59199182 | 0.10524348 |
| Gm32655 | 0.68489185 | 0.15949763 | 0.031 | 0.041 | 1 | 0.0540843 | 2.01207283 | 0.038 | 0 | 1 | 0.68489185 | 0.10524348 |
| A430046D13Rik | 0.69869193 | 0.13302542 | 0.052 | 0.041 | 1 | 0.0540843 | 2.01207283 | 0.038 | 0 | 1 | 0.69869193 | 0.10524348 |
| 4930458D05Rik | 0.69905423 | 0.47697982 | 0.062 | 0.052 | 1 | 0.0540843 | 2.01207283 | 0.038 | 0 | 1 | 0.69905423 | 0.10524348 |
| Efhd1 | 0.65211073 | -0.9925055 | 0.031 | 0.041 | 1 | 0.05411828 | -1.9728203 | 0.021 | 0.062 | 1 | 0.65211073 | 0.10530777 |
| Pemt | 0.09997022 | 0.89201732 | 0.104 | 0.052 | 1 | 0.0541542 | -1.5322477 | 0.034 | 0.083 | 1 | 0.09997022 | 0.10537573 |
| Rbm38 | 0.29324014 | 0.89201732 | 0.083 | 0.052 | 1 | 0.05424731 | -1.1493906 | 0.055 | 0.115 | 1 | 0.29324014 | 0.10555185 |
| A130014A01Rik | 0.05430727 | 1.27052894 | 0.083 | 0.031 | 1 | 0.89406385 | -1.3098553 | 0.05 | 0.052 | 1 | 0.89406385 | 0.10566525 |
| Zfp945 | 0.43721334 | 0.14499806 | 0.323 | 0.275 | 1 | 0.05442276 | 0.63951866 | 0.319 | 0.208 | 1 | 0.43721334 | 0.10588369 |
| Smarcb1 | 0.05442282 | 0.39197457 | 0.604 | 0.492 | 1 | 0.16859098 | 0.22563856 | 0.588 | 0.521 | 1 | 0.16859098 | 0.1058838 |
| 1700123M08Rik | 0.05443816 | 0.9200317 | 0.115 | 0.052 | 1 | 0.55294827 | -0.8504236 | 0.029 | 0.042 | 1 | 0.55294827 | 0.1059128 |
| Fpgs | 0.07925891 | 0.53356335 | 0.167 | 0.093 | 1 | 0.05448459 | 0.45567948 | 0.172 | 0.083 | 1 | 0.07925891 | 0.1060006 |
| Cep68 | 0.89248997 | -0.3592878 | 0.167 | 0.166 | 1 | 0.05449722 | -1.0957305 | 0.168 | 0.25 | 1 | 0.89248997 | 0.1060245 |
| Iqcn | 0.05450014 | 0.95091101 | 0.146 | 0.073 | 1 | 0.67194606 | -0.4318458 | 0.185 | 0.156 | 1 | 0.67194606 | 0.10603001 |
| Zmat2 | 0.2047498 | 0.23908589 | 0.667 | 0.585 | 1 | 0.0545096 | 0.28125947 | 0.597 | 0.521 | 1 | 0.2047498 | 0.1060479 |
| Stard8 | 0.0545672 | 0.31312082 | 0.406 | 0.269 | 1 | 0.14613725 | -0.6577786 | 0.336 | 0.219 | 1 | 0.14613725 | 0.10615681 |
| Jmjd4 | 0.05458752 | -1.5160674 | 0.031 | 0.093 | 1 | 0.43086562 | 0.62874419 | 0.076 | 0.052 | 1 | 0.43086562 | 0.10619524 |
| Sdhaf1 | 0.67557798 | -0.3710171 | 0.094 | 0.109 | 1 | 0.05459615 | 0.53144699 | 0.151 | 0.073 | 1 | 0.67557798 | 0.10621157 |
| Ska3 | 0.14387055 | 1.00749454 | 0.083 | 0.041 | 1 | 0.0546316 | -2.010295 | 0.021 | 0.062 | 1 | 0.14387055 | 0.10627859 |
| Gins2 | 0.05465486 | 0.82306997 | 0.177 | 0.098 | 1 | 0.87451868 | -0.6317834 | 0.092 | 0.083 | 1 | 0.87451868 | 0.10632256 |
| Gys1 | 0.05468339 | 0.75595577 | 0.146 | 0.073 | 1 | 0.71566797 | -0.7459544 | 0.092 | 0.104 | 1 | 0.71566797 | 0.10637651 |
| Tysnd1 | 0.36955012 | 0.23697638 | 0.167 | 0.124 | 1 | 0.05469128 | 0.97965135 | 0.151 | 0.073 | 1 | 0.36955012 | 0.10639142 |
| Inka1 | 0.48037805 | 0.17741954 | 0.073 | 0.052 | 1 | 0.05473341 | 0.49749966 | 0.109 | 0.042 | 1 | 0.48037805 | 0.10647107 |
| 1700001L05Rik | 0.05476704 | 0.77302928 | 0.146 | 0.073 | 1 | 0.22862884 | -0.4797803 | 0.147 | 0.094 | 1 | 0.22862884 | 0.10653465 |
| Nme7 | 0.05481208 | 0.33372277 | 0.323 | 0.202 | 1 | 0.19829938 | 0.21852371 | 0.315 | 0.229 | 1 | 0.19829938 | 0.10661979 |
| Gnb5 | 0.05482136 | -1.290186 | 0.104 | 0.187 | 1 | 0.60992751 | -0.1347686 | 0.223 | 0.188 | 1 | 0.60992751 | 0.10663733 |
| Pde6h | 0.10600283 | -1.4895203 | 0.219 | 0.285 | 1 | 0.05486983 | 1.90515763 | 0.092 | 0.031 | 1 | 0.10600283 | 0.10672896 |
| Tcf7l1 | 0.05493657 | 0.15464823 | 0.615 | 0.487 | 1 | 0.88209154 | -0.3200223 | 0.786 | 0.802 | 1 | 0.88209154 | 0.10685512 |
| 9930014A18Rik | 0.05499785 | -1.3848229 | 0.021 | 0.078 | 1 | 0.15976273 | 1.01207283 | 0.071 | 0.031 | 1 | 0.15976273 | 0.10697093 |
| Kmt2e | 0.10746575 | -0.2287257 | 1 | 1 | 1 | 0.05501632 | -0.3068355 | 0.996 | 1 | 1 | 0.10746575 | 0.10700584 |

| | | | | | | | | | | | | |
|---|---|---|---|---|---|---|---|---|---|---|---|---|
| Gab3 | 0.05506844 | 1.00749454 | 0.083 | 0.031 | 1 | 0.14151408 | 2.01207283 | 0.042 | 0.01 | 1 | 0.14151408 | 0.10710434 |
| Dnai3 | 0.05506865 | 1.14499806 | 0.083 | 0.031 | 1 | 0.85710544 | -2.0468209 | 0.008 | 0.01 | 1 | 0.85710544 | 0.10710475 |
| Epha4 | 0.48326414 | -0.3783272 | 0.333 | 0.363 | 1 | 0.0551081 | -0.8048611 | 0.319 | 0.438 | 1 | 0.48326414 | 0.10717929 |
| C78859 | 0.46989842 | -1.1079827 | 0.021 | 0.036 | 1 | 0.0551428 | -2.010295 | 0.021 | 0.062 | 1 | 0.46989842 | 0.10724487 |
| Cplx2 | 0.05523207 | -1.3144336 | 0.021 | 0.078 | 1 | 0.23893677 | 0.23763253 | 0.118 | 0.073 | 1 | 0.23893677 | 0.10741356 |
| Renbp | 0.49195304 | -0.1918143 | 0.26 | 0.207 | 1 | 0.05524954 | 0.61297688 | 0.256 | 0.156 | 1 | 0.49195304 | 0.10744657 |
| R3hcc1 | 0.25245918 | 0.34452952 | 0.208 | 0.15 | 1 | 0.05526418 | 0.29895398 | 0.21 | 0.115 | 1 | 0.25245918 | 0.10747424 |
| Meis3 | 0.76840302 | -0.4170033 | 0.167 | 0.176 | 1 | 0.05526737 | 0.61614416 | 0.193 | 0.104 | 1 | 0.76840302 | 0.10748026 |
| Zfp644 | 0.49917919 | 0.13581863 | 0.75 | 0.715 | 1 | 0.05527008 | -0.6511487 | 0.853 | 0.875 | 1 | 0.49917919 | 0.10748537 |
| Otoa | 0.05528655 | -1.7366666 | 0.083 | 0.161 | 1 | 0.52074229 | 0.27510724 | 0.071 | 0.052 | 1 | 0.52074229 | 0.10751649 |
| Mid1ip1 | 0.05532748 | -0.4736322 | 0.188 | 0.098 | 1 | 0.102303 | 0.32241295 | 0.248 | 0.167 | 1 | 0.102303 | 0.10759382 |
| Fut10 | 0.05535303 | 1.08549705 | 0.135 | 0.067 | 1 | 0.69283858 | -0.5819348 | 0.147 | 0.125 | 1 | 0.69283858 | 0.1076421 |
| Atxn1l | 0.5233479 | 0.21394541 | 0.229 | 0.192 | 1 | 0.05536488 | 0.10885789 | 0.387 | 0.24 | 1 | 0.5233479 | 0.10766449 |
| Fam229b | 0.36133361 | 0.64492446 | 0.156 | 0.119 | 1 | 0.05537379 | 1.27510724 | 0.134 | 0.062 | 1 | 0.36133361 | 0.10768133 |
| Bmp3 | 0.82511604 | 1.12297175 | 0.052 | 0.047 | 1 | 0.05545644 | 0.73196491 | 0.193 | 0.104 | 1 | 0.82511604 | 0.10783746 |
| Eid3 | 0.44158998 | 0.49292136 | 0.062 | 0.041 | 1 | 0.05556097 | 1.56461386 | 0.076 | 0.021 | 1 | 0.44158998 | 0.10803491 |
| Zfp651 | 0.18721174 | 0.49292136 | 0.24 | 0.171 | 1 | 0.05560761 | 0.26068017 | 0.483 | 0.333 | 1 | 0.18721174 | 0.10812302 |
| Ergic1 | 0.89814647 | 0.10552662 | 0.625 | 0.668 | 1 | 0.05562658 | 0.12384608 | 0.693 | 0.604 | 1 | 0.89814647 | 0.10815885 |
| Gm31718 | 0.47360738 | -0.4950058 | 0.042 | 0.062 | 1 | 0.05563675 | -1.961932 | 0.042 | 0.094 | 1 | 0.47360738 | 0.10817806 |
| Zpbp | 0.3270442 | 0.68556644 | 0.094 | 0.062 | 1 | 0.05566286 | 1.57767001 | 0.092 | 0.031 | 1 | 0.3270442 | 0.10822737 |
| Gm56587 | 0.97828543 | 0.30705482 | 0.052 | 0.052 | 1 | 0.05584596 | 0.86006974 | 0.109 | 0.042 | 1 | 0.97828543 | 0.10857315 |
| Fndc3a | 0.08350934 | -0.2344844 | 0.99 | 0.984 | 1 | 0.05589172 | -0.5322477 | 0.945 | 0.917 | 1 | 0.08350934 | 0.10865955 |
| Zfp746 | 0.79850413 | 0.18806678 | 0.26 | 0.249 | 1 | 0.05592333 | 0.14652303 | 0.332 | 0.208 | 1 | 0.79850413 | 0.10871925 |
| 9030025P20Rik | 0.17534439 | 1.32942263 | 0.042 | 0.016 | 1 | 0.05593307 | 1.49749966 | 0.076 | 0.021 | 1 | 0.17534439 | 0.10873764 |
| Slco2b1 | 0.31835113 | -0.6445822 | 0.125 | 0.166 | 1 | 0.0559866 | 0.97025266 | 0.109 | 0.042 | 1 | 0.31835113 | 0.10883869 |
| Gm31045 | 0.31389644 | 0.74446013 | 0.042 | 0.021 | 1 | 0.05605935 | 1.35310975 | 0.059 | 0.01 | 1 | 0.31389644 | 0.10897605 |
| Dzip1 | 0.05606259 | -1.4705528 | 0.042 | 0.109 | 1 | 0.30798538 | -0.9688183 | 0.055 | 0.083 | 1 | 0.30798538 | 0.10898216 |
| Mapk13 | 0.05606699 | 1.00749454 | 0.115 | 0.052 | 1 | 0.11360323 | 0.3146356 | 0.126 | 0.062 | 1 | 0.11360323 | 0.10899047 |
| Nexn | 0.29415385 | -1.1079827 | 0.042 | 0.073 | 1 | 0.05610871 | 0.90313846 | 0.151 | 0.073 | 1 | 0.29415385 | 0.10906923 |
| Gtdc1 | 0.05613625 | 0.34518995 | 0.469 | 0.347 | 1 | 0.35981914 | -0.543286 | 0.429 | 0.479 | 1 | 0.35981914 | 0.10912122 |
| Gm32592 | 0.38521705 | -0.577468 | 0.031 | 0.016 | 1 | 0.0562903 | 1.33400093 | 0.059 | 0.01 | 1 | 0.38521705 | 0.10941199 |
| Cradd | 0.85289896 | 0.52884094 | 0.229 | 0.223 | 1 | 0.05635657 | -0.5759301 | 0.265 | 0.375 | 1 | 0.85289896 | 0.10953707 |
| Bcl2l2 | 0.99659582 | -0.1180363 | 0.208 | 0.202 | 1 | 0.05636959 | 0.20033947 | 0.324 | 0.198 | 1 | 0.99659582 | 0.10956165 |
| Asnsd1 | 0.0564292 | 0.69338595 | 0.302 | 0.202 | 1 | 0.06758537 | 0.71567983 | 0.189 | 0.104 | 1 | 0.06758537 | 0.10967414 |
| Coro1a | 0.73685932 | 1.11440974 | 0.01 | 0.016 | 1 | 0.05648248 | -0.8948178 | 0.034 | 0.083 | 1 | 0.73685932 | 0.10977469 |
| Lima1 | 0.05653722 | -0.1840976 | 0.646 | 0.762 | 1 | 0.70140004 | -0.593075 | 0.845 | 0.865 | 1 | 0.70140004 | 0.10987798 |
| Sft2d1 | 0.09937074 | 0.31447229 | 0.469 | 0.342 | 1 | 0.0565678 | -0.5728897 | 0.437 | 0.542 | 1 | 0.09937074 | 0.10993568 |
| Svep1 | 0.05658294 | -0.9874166 | 0.26 | 0.352 | 1 | 0.29423416 | -0.2318528 | 0.634 | 0.5 | 1 | 0.29423416 | 0.10996426 |
| 1810030O07Rik | 0.05662041 | 0.55292867 | 0.385 | 0.28 | 1 | 0.38050635 | -0.2788284 | 0.399 | 0.323 | 1 | 0.38050635 | 0.11003494 |
| Tmem119 | 0.05662577 | -1.8504865 | 0.062 | 0.135 | 1 | 0.15357104 | -0.8737561 | 0.067 | 0.115 | 1 | 0.15357104 | 0.11004505 |
| Aifm3 | 0.38028858 | 1.00749454 | 0.031 | 0.016 | 1 | 0.05668038 | -1.3098553 | 0.021 | 0.062 | 1 | 0.38028858 | 0.11014809 |
| Sirt3 | 0.67248206 | 0.18806678 | 0.135 | 0.119 | 1 | 0.05668293 | 0.36821664 | 0.168 | 0.083 | 1 | 0.67248206 | 0.1101529 |
| Zfp213 | 0.05675355 | 0.90795886 | 0.115 | 0.052 | 1 | 0.14049288 | 1.27510724 | 0.059 | 0.021 | 1 | 0.14049288 | 0.11028613 |
| Vamp1 | 0.25784924 | -1.0513992 | 0.052 | 0.088 | 1 | 0.05676447 | -1.1172102 | 0.034 | 0.083 | 1 | 0.25784924 | 0.11030673 |
| Dnase2a | 0.12162479 | -1.2295447 | 0.062 | 0.119 | 1 | 0.05676607 | 0.59703533 | 0.155 | 0.073 | 1 | 0.12162479 | 0.11030976 |
| Gm12227 | 0.06165575 | 1.06889508 | 0.135 | 0.067 | 1 | 0.05680495 | 0.23446525 | 0.109 | 0.042 | 1 | 0.06165575 | 0.1103831 |
| Gmds | 0.05687649 | 0.19040597 | 0.938 | 0.938 | 1 | 0.0787734 | -0.40709 | 0.945 | 0.979 | 1 | 0.0787734 | 0.11051804 |
| Tmem268 | 0.08424511 | -1.4519371 | 0.01 | 0.052 | 1 | 0.05695383 | 0.53814164 | 0.059 | 0.01 | 1 | 0.08424511 | 0.11066393 |
| Ncaph2 | 0.06368665 | 0.39278469 | 0.448 | 0.332 | 1 | 0.05712779 | 0.19437597 | 0.45 | 0.323 | 1 | 0.06368665 | 0.110992 |
| Usp49 | 0.05713107 | 0.51106871 | 0.344 | 0.233 | 1 | 0.74968167 | -0.3902739 | 0.374 | 0.312 | 1 | 0.74968167 | 0.11099818 |
| Xpnpep1 | 0.0571587 | -0.6805615 | 0.354 | 0.435 | 1 | 0.34224727 | -1.0803734 | 0.336 | 0.344 | 1 | 0.34224727 | 0.11105028 |
| Fbxo36 | 0.37649317 | 0.61156586 | 0.24 | 0.197 | 1 | 0.0571772 | 0.29895398 | 0.181 | 0.094 | 1 | 0.37649317 | 0.11108517 |
| Urb2 | 0.19952883 | 0.42253204 | 0.146 | 0.093 | 1 | 0.05725859 | 0.62088207 | 0.202 | 0.115 | 1 | 0.19952883 | 0.11123863 |
| Fancl | 0.96374095 | -0.1345245 | 0.208 | 0.207 | 1 | 0.05732281 | 0.11898804 | 0.227 | 0.125 | 1 | 0.96374095 | 0.11135972 |
| 1500004A13Rik | 0.47477351 | -0.407543 | 0.042 | 0.062 | 1 | 0.057441 | 0.86006974 | 0.151 | 0.073 | 1 | 0.47477351 | 0.11158254 |
| 4933431E20Rik | 0.05762236 | -1.3144336 | 0.031 | 0.093 | 1 | 0.13074931 | 0.54812573 | 0.105 | 0.052 | 1 | 0.13074931 | 0.11192438 |
| Efr3b | 0.36377334 | -0.7860546 | 0.115 | 0.15 | 1 | 0.05762891 | -1.6094155 | 0.088 | 0.156 | 1 | 0.36377334 | 0.11193673 |
| Smap1 | 0.05770812 | 0.15433592 | 0.854 | 0.731 | 1 | 0.6879915 | -0.244152 | 0.836 | 0.865 | 1 | 0.6879915 | 0.11208602 |
| Cetn4 | 0.50844809 | 0.40083696 | 0.292 | 0.259 | 1 | 0.05773376 | 0.48155812 | 0.155 | 0.073 | 1 | 0.50844809 | 0.11213434 |
| AI987944 | 0.26808976 | 0.57139542 | 0.25 | 0.197 | 1 | 0.05773536 | 0.23267197 | 0.21 | 0.115 | 1 | 0.26808976 | 0.11213734 |
| Ttll5 | 0.24808335 | 0.31828207 | 0.521 | 0.487 | 1 | 0.05788265 | -0.7564037 | 0.605 | 0.604 | 1 | 0.24808335 | 0.1124149 |
| Endov | 0.23743184 | 0.42253204 | 0.167 | 0.114 | 1 | 0.05792274 | 1.01207283 | 0.092 | 0.031 | 1 | 0.23743184 | 0.11249044 |
| 1700102P08Rik | 0.05809483 | 0.81484946 | 0.156 | 0.083 | 1 | 0.82219909 | -0.2554075 | 0.172 | 0.156 | 1 | 0.82219909 | 0.11281465 |
| Gm10371 | 0.45760721 | -0.4147385 | 0.188 | 0.223 | 1 | 0.05810716 | 0.20204378 | 0.244 | 0.135 | 1 | 0.45760721 | 0.11283788 |
| Mrtfa | 0.65431877 | -0.1568923 | 0.583 | 0.523 | 1 | 0.05819411 | -0.6541898 | 0.668 | 0.76 | 1 | 0.65431877 | 0.11300166 |
| Gm10130 | 0.05821168 | 0.89201732 | 0.115 | 0.052 | 1 | 0.1007018 | 1.49749966 | 0.08 | 0.031 | 1 | 0.1007018 | 0.11303476 |
| Tmem168 | 0.4087659 | 0.23377039 | 0.25 | 0.202 | 1 | 0.05827042 | 0.10134017 | 0.349 | 0.229 | 1 | 0.4087659 | 0.1131454 |
| Cadm4 | 0.38357309 | -1.577468 | 0.01 | 0.026 | 1 | 0.05827302 | -1.0203486 | 0.034 | 0.083 | 1 | 0.38357309 | 0.11315029 |
| Rin3 | 0.86348914 | 0.1843723 | 0.146 | 0.161 | 1 | 0.05828477 | 0.48270266 | 0.231 | 0.135 | 1 | 0.86348914 | 0.11317243 |
| Gm6169 | 0.84081505 | 0.27052894 | 0.052 | 0.047 | 1 | 0.05828654 | 0.33400093 | 0.092 | 0.031 | 1 | 0.84081505 | 0.11317575 |
| Mbtps2 | 0.05832163 | 0.41440415 | 0.438 | 0.311 | 1 | 0.47877617 | -0.5432188 | 0.328 | 0.25 | 1 | 0.47877617 | 0.11324186 |
| Slc8b1 | 0.05833522 | -0.5982265 | 0.135 | 0.233 | 1 | 0.08742814 | 0.24596089 | 0.361 | 0.26 | 1 | 0.08742814 | 0.11326744 |
| Il33 | 0.05834761 | -0.9925055 | 0.062 | 0.14 | 1 | 0.24308145 | 0.77760758 | 0.034 | 0.01 | 1 | 0.24308145 | 0.11329077 |
| Coq4 | 0.08541042 | 1.00749454 | 0.115 | 0.057 | 1 | 0.05842045 | 0.32241295 | 0.109 | 0.042 | 1 | 0.08541042 | 0.11342795 |
| Hyal1 | 0.42126413 | -0.3710171 | 0.083 | 0.114 | 1 | 0.0584446 | 0.95317914 | 0.092 | 0.031 | 1 | 0.42126413 | 0.11347342 |
| Gm17354 | 0.42914542 | -0.4519371 | 0.031 | 0.052 | 1 | 0.0584446 | 0.95317914 | 0.092 | 0.031 | 1 | 0.42914542 | 0.11347342 |
| Yes1 | 0.81783901 | -0.3573421 | 0.562 | 0.518 | 1 | 0.05848044 | -0.5742323 | 0.643 | 0.729 | 1 | 0.81783901 | 0.11354091 |
| Dnajc22 | 0.24199603 | 0.93349396 | 0.135 | 0.093 | 1 | 0.05848203 | 0.97025266 | 0.122 | 0.052 | 1 | 0.24199603 | 0.11354391 |
| 9330160F10Rik | 0.05848602 | 1.59245704 | 0.073 | 0.026 | 1 | 0.11751579 | 0.86006974 | 0.063 | 0.021 | 1 | 0.11751579 | 0.11355142 |
| Cdc45 | 0.05848602 | 1.88196365 | 0.073 | 0.026 | 1 | 0.87066325 | -0.3098553 | 0.013 | 0.01 | 1 | 0.87066325 | 0.11355142 |
| Ephx2 | 0.05853384 | -1.2469267 | 0.26 | 0.352 | 1 | 0.79071329 | -0.2457249 | 0.071 | 0.062 | 1 | 0.79071329 | 0.11364147 |
| Myl12b | 0.05857784 | 0.31783466 | 0.823 | 0.819 | 1 | 0.75576606 | -0.1096906 | 0.903 | 0.906 | 1 | 0.75576606 | 0.11372432 |
| Hpn | 0.33113577 | 0.22988696 | 0.062 | 0.036 | 1 | 0.05862224 | 0.25085969 | 0.168 | 0.083 | 1 | 0.33113577 | 0.11380792 |
| Nprl2 | 0.05865428 | 1.21899864 | 0.167 | 0.093 | 1 | 0.29861824 | 0.60768258 | 0.122 | 0.083 | 1 | 0.29861824 | 0.11386823 |
| Cpne1 | 0.05873185 | 0.30269227 | 0.677 | 0.57 | 1 | 0.55283305 | -0.1432054 | 0.84 | 0.844 | 1 | 0.55283305 | 0.11401427 |
| Eps8l1 | 0.05882922 | -0.8669746 | 0.021 | 0.078 | 1 | 0.20577072 | 0.61614416 | 0.067 | 0.031 | 1 | 0.20577072 | 0.11419756 |

| | | | | | | | | | | | | |
|---|---|---|---|---|---|---|---|---|---|---|---|---|
| Ptgfr | 0.62414763 | -1.5907648 | 0.104 | 0.119 | 1 | 0.05893431 | -0.7983549 | 0.071 | 0.135 | 1 | 0.62414763 | 0.11439536 |
| Gm49207 | 0.10680756 | 0.76648644 | 0.104 | 0.052 | 1 | 0.05896819 | 0.62874419 | 0.092 | 0.031 | 1 | 0.10680756 | 0.11445914 |
| Dync2i1 | 0.3223662 | -0.5285584 | 0.333 | 0.373 | 1 | 0.05897381 | -0.1172102 | 0.391 | 0.25 | 1 | 0.3223662 | 0.11446971 |
| Zfp821 | 0.05898863 | 0.27052894 | 0.271 | 0.166 | 1 | 0.06905474 | 0.1883956 | 0.319 | 0.198 | 1 | 0.06905474 | 0.1144976 |
| AI467606 | 0.57286036 | 0.42253204 | 0.062 | 0.047 | 1 | 0.05900138 | 1.06865636 | 0.076 | 0.021 | 1 | 0.57286036 | 0.11452159 |
| Topors | 0.05905929 | 0.19894878 | 0.625 | 0.482 | 1 | 0.61186276 | -0.1591764 | 0.744 | 0.76 | 1 | 0.61186276 | 0.11463059 |
| Slc45a1 | 0.05910519 | 1.59245704 | 0.073 | 0.026 | 1 | 0.23185002 | -1.0203486 | 0.034 | 0.062 | 1 | 0.23185002 | 0.11471696 |
| Stoml1 | 0.05921631 | 1.00749454 | 0.188 | 0.109 | 1 | 0.94074032 | -0.1578522 | 0.13 | 0.125 | 1 | 0.94074032 | 0.11492604 |
| Nbr1 | 0.05927494 | 0.23792428 | 0.635 | 0.492 | 1 | 0.19534671 | 0.10640021 | 0.697 | 0.667 | 1 | 0.19534671 | 0.11503637 |
| Galnt11 | 0.1857433 | 0.3540523 | 0.594 | 0.518 | 1 | 0.05929536 | -0.1332107 | 0.626 | 0.427 | 1 | 0.1857433 | 0.11507477 |
| Foxm1 | 0.05938205 | 1.46692616 | 0.073 | 0.026 | 1 | 0.5207331 | -0.8244284 | 0.021 | 0.01 | 1 | 0.5207331 | 0.11523788 |
| Stn1 | 0.05938267 | 0.27598337 | 0.417 | 0.29 | 1 | 0.62772302 | -0.1973805 | 0.252 | 0.208 | 1 | 0.62772302 | 0.11523904 |
| Rab29 | 0.05946178 | 1.77302928 | 0.104 | 0.047 | 1 | 0.37063341 | 0.63767732 | 0.08 | 0.052 | 1 | 0.37063341 | 0.11538785 |
| Mecp2 | 0.05952189 | 0.35449125 | 0.76 | 0.71 | 1 | 0.15855424 | -0.5785362 | 0.832 | 0.865 | 1 | 0.15855424 | 0.11550092 |
| Gca | 0.05972116 | -2.1624305 | 0 | 0.036 | 1 | 0.86312513 | -0.7248928 | 0.008 | 0.01 | 1 | 0.86312513 | 0.11587571 |
| Bsx | 0.05972116 | -2.1624305 | 0 | 0.036 | 1 | 0.50587544 | -1.8948178 | 0.004 | 0.01 | 1 | 0.50587544 | 0.11587571 |
| 9430065F17Rik | 0.05972116 | -2.1624305 | 0 | 0.036 | 1 | 0.20315061 | 1.49749966 | 0.017 | 0 | 1 | 0.20315061 | 0.11587571 |
| Slc14a1 | 0.05972116 | -2.1624305 | 0 | 0.036 | 1 | 0.85710544 | -1.7248928 | 0.008 | 0.01 | 1 | 0.85710544 | 0.11587571 |
| Tmem131 | 0.30534177 | -0.2148979 | 0.729 | 0.746 | 1 | 0.05972289 | -0.410522 | 0.87 | 0.885 | 1 | 0.30534177 | 0.11587896 |
| 2200002D01Rik | 0.05972787 | -2.3144336 | 0 | 0.036 | 1 | 0.62504414 | -0.1843244 | 0.08 | 0.062 | 1 | 0.62504414 | 0.11588833 |
| Gm45894 | 0.05972787 | -2.3144336 | 0 | 0.036 | 1 | 0.3106474 | -1.3098553 | 0.038 | 0.062 | 1 | 0.3106474 | 0.11588833 |
| Gm47400 | 0.05972787 | -2.4519371 | 0 | 0.036 | 1 | 0.11689951 | -2.3098553 | 0 | 0.01 | 1 | 0.11689951 | 0.11588833 |
| Fgf14 | 0.0597319 | -2.6929452 | 0 | 0.036 | 1 | 0.54889202 | -1.1399303 | 0.029 | 0.042 | 1 | 0.54889202 | 0.1158959 |
| Ppp1r1c | 0.05973235 | -4.0368996 | 0 | 0.036 | 1 | 0.11954511 | -2.3098553 | 0.008 | 0.031 | 1 | 0.11954511 | 0.11589674 |
| Lrrc75a | 0.05973235 | -2.7998604 | 0 | 0.036 | 1 | 0.06851509 | 1.59703533 | 0.055 | 0.01 | 1 | 0.06851509 | 0.11589674 |
| 9130015A21Rik | 0.05973458 | -3.3144336 | 0 | 0.036 | 1 | 0.11689951 | -2.3098553 | 0 | 0.01 | 1 | 0.11689951 | 0.11590095 |
| Igsf11 | 0.05973458 | -2.9925055 | 0 | 0.036 | 1 | 0.57576763 | -0.9879272 | 0.013 | 0.021 | 1 | 0.57576763 | 0.11590095 |
| Cdr1os | 0.05973458 | -3.9925055 | 0 | 0.036 | 1 | 0.27154471 | 0.69014474 | 0.013 | 0 | 1 | 0.27154471 | 0.11590095 |
| Gm13710 | 0.05973503 | -3.6363617 | 0 | 0.036 | 1 | 0.11689951 | -3.3098553 | 0 | 0.01 | 1 | 0.11689951 | 0.11590179 |
| Ifit1 | 0.05973503 | -5.679006 | 0 | 0.036 | 1 | 0.27508759 | -3.1172102 | 0.021 | 0.042 | 1 | 0.27508759 | 0.11590179 |
| Gpnmb | 0.05973548 | -3.1624305 | 0 | 0.036 | 1 | 0.99396042 | -0.1399303 | 0.021 | 0.021 | 1 | 0.99396042 | 0.11590263 |
| Itgam | 0.05973593 | -4.4843586 | 0 | 0.036 | 1 | 0.07781087 | -1.7103932 | 0.029 | 0.073 | 1 | 0.07781087 | 0.11590347 |
| Gm28501 | 0.05973682 | -1.9925055 | 0 | 0.036 | 1 | 0.79170562 | -0.3098553 | 0.038 | 0.031 | 1 | 0.79170562 | 0.11590515 |
| 4933423P22Rik | 0.05973682 | -1.9925055 | 0 | 0.036 | 1 | 0.14865142 | -1.3098553 | 0.004 | 0.021 | 1 | 0.14865142 | 0.11590515 |
| Gm36608 | 0.05973682 | -1.9925055 | 0 | 0.036 | 1 | 0.27154471 | 1.49749966 | 0.013 | 0 | 1 | 0.27154471 | 0.11590515 |
| Twist2 | 0.78977807 | -0.130009 | 0.083 | 0.073 | 1 | 0.05977093 | 0.82764826 | 0.076 | 0.021 | 1 | 0.78977807 | 0.1159693 |
| Ptbp2 | 0.06617159 | 0.28890747 | 0.833 | 0.762 | 1 | 0.05987211 | -0.4407027 | 0.882 | 0.927 | 1 | 0.06617159 | 0.11615955 |
| Clpp | 0.24943025 | 0.14853842 | 0.635 | 0.513 | 1 | 0.0599556 | 0.15331876 | 0.525 | 0.344 | 1 | 0.24943025 | 0.11631653 |
| 4930545L23Rik | 0.05999981 | 1.42253204 | 0.073 | 0.026 | 1 | 0.37605042 | -0.5728897 | 0.097 | 0.062 | 1 | 0.37605042 | 0.11639965 |
| 6330408M09Rik | 0.05999981 | 1.42253204 | 0.073 | 0.026 | 1 | 0.34704016 | -1.3098553 | 0.008 | 0.021 | 1 | 0.34704016 | 0.11639965 |
| Ighmbp2 | 0.05999981 | 1.42253204 | 0.073 | 0.026 | 1 | 0.3011615 | 0.59703533 | 0.059 | 0.031 | 1 | 0.3011615 | 0.11639965 |
| Rmnd1 | 0.06004019 | 0.74446013 | 0.177 | 0.098 | 1 | 0.07988196 | 0.29895398 | 0.189 | 0.104 | 1 | 0.07988196 | 0.11647556 |
| Nek10 | 0.0600921 | 0.44561565 | 0.146 | 0.073 | 1 | 0.95652137 | -0.3275573 | 0.113 | 0.115 | 1 | 0.95652137 | 0.11657314 |
| Tap2 | 0.42122717 | 0.11852585 | 0.208 | 0.166 | 1 | 0.06015671 | -0.1399303 | 0.256 | 0.146 | 1 | 0.42122717 | 0.1166946 |
| Pcsk5 | 0.3694985 | -1.830075 | 0.271 | 0.301 | 1 | 0.06026296 | -1.6546838 | 0.277 | 0.333 | 1 | 0.3694985 | 0.1168943 |
| Tmem236 | 0.06027858 | 1.00749454 | 0.073 | 0.026 | 1 | 0.09535395 | -0.84037 | 0.013 | 0.042 | 1 | 0.09535395 | 0.11692365 |
| Polr1b | 0.06041537 | 0.74446013 | 0.135 | 0.067 | 1 | 0.61131311 | -0.5114891 | 0.143 | 0.115 | 1 | 0.61131311 | 0.11718073 |
| Tardbp | 0.16367825 | 0.32859564 | 0.844 | 0.777 | 1 | 0.06043125 | 0.12811905 | 0.895 | 0.844 | 1 | 0.16367825 | 0.11721057 |
| Ip6k3 | 0.99750658 | 0.27052894 | 0.083 | 0.083 | 1 | 0.06049912 | 0.65977109 | 0.139 | 0.062 | 1 | 0.99750658 | 0.11733809 |
| Dedd | 0.24752677 | 0.10060394 | 0.375 | 0.29 | 1 | 0.06050876 | -0.1082214 | 0.357 | 0.219 | 1 | 0.24752677 | 0.11735621 |
| Tpgs1 | 0.06051917 | 0.72738662 | 0.323 | 0.228 | 1 | 0.32480884 | 0.23267197 | 0.239 | 0.188 | 1 | 0.32480884 | 0.11737578 |
| Cpt1b | 0.0605464 | 1.37006462 | 0.073 | 0.026 | 1 | 0.65065939 | -0.8073549 | 0.05 | 0.062 | 1 | 0.65065939 | 0.11742693 |
| Adad1 | 0.06058014 | -2.9467018 | 0.01 | 0.057 | 1 | 0.12312067 | -1.3098553 | 0.008 | 0.031 | 1 | 0.12312067 | 0.11749033 |
| Fjx1 | 0.10941605 | 0.56388789 | 0.281 | 0.192 | 1 | 0.06069827 | 0.78775553 | 0.261 | 0.167 | 1 | 0.10941605 | 0.11771226 |
| Eif5b | 0.06074951 | 0.20766892 | 0.969 | 0.953 | 1 | 0.06685214 | 0.12793489 | 0.992 | 0.99 | 1 | 0.06685214 | 0.11780851 |
| Lgals1 | 0.06075292 | -2.9177746 | 0.219 | 0.285 | 1 | 0.67545944 | -1.5266667 | 0.441 | 0.354 | 1 | 0.67545944 | 0.11781493 |
| Ripk3 | 0.0608291 | 0.77616899 | 0.208 | 0.124 | 1 | 0.33914679 | 0.22065945 | 0.097 | 0.062 | 1 | 0.33914679 | 0.11795802 |
| Micu2 | 0.06101221 | 0.36657563 | 0.635 | 0.523 | 1 | 0.91240853 | -0.2694975 | 0.576 | 0.615 | 1 | 0.91240853 | 0.11830193 |
| Myl4 | 0.0610325 | 0.80104366 | 0.156 | 0.083 | 1 | 0.1332159 | 0.20471791 | 0.185 | 0.115 | 1 | 0.1332159 | 0.11834002 |
| Gm43150 | 0.06116328 | -2.240433 | 0.01 | 0.057 | 1 | 0.56568674 | -1.7248928 | 0.021 | 0.031 | 1 | 0.56568674 | 0.11858562 |
| Gtf2a1 | 0.06120652 | -0.8405024 | 0.312 | 0.409 | 1 | 0.25729355 | -0.4441564 | 0.382 | 0.448 | 1 | 0.25729355 | 0.11866681 |
| Il11 | 0.19125169 | 1.81484946 | 0.104 | 0.062 | 1 | 0.06124859 | 1.77760758 | 0.118 | 0.052 | 1 | 0.19125169 | 0.1187458 |
| Zfyve27 | 0.32777843 | -0.4828311 | 0.188 | 0.233 | 1 | 0.06127802 | 0.47036353 | 0.256 | 0.156 | 1 | 0.32777843 | 0.11880104 |
| Matk | 0.22026949 | 1.42253204 | 0.021 | 0.005 | 1 | 0.06128508 | 1.12310414 | 0.076 | 0.021 | 1 | 0.22026949 | 0.1188143 |
| Slc35e2 | 0.86117769 | -0.17693 | 0.281 | 0.259 | 1 | 0.06129211 | -0.2202181 | 0.282 | 0.167 | 1 | 0.86117769 | 0.11882749 |
| Dolk | 0.20245052 | 0.46983775 | 0.26 | 0.197 | 1 | 0.06131033 | 0.7726069 | 0.235 | 0.146 | 1 | 0.20245052 | 0.11886171 |
| Acads | 0.53283035 | 0.22988696 | 0.135 | 0.109 | 1 | 0.06133208 | 0.51326698 | 0.164 | 0.083 | 1 | 0.53283035 | 0.11890254 |
| Cdon | 0.06134681 | -0.8156277 | 0.365 | 0.456 | 1 | 0.17721223 | -0.603968 | 0.429 | 0.469 | 1 | 0.17721223 | 0.11893018 |
| Lat2 | 0.88652237 | 0.10060394 | 0.062 | 0.067 | 1 | 0.06139381 | 0.71762547 | 0.139 | 0.062 | 1 | 0.88652237 | 0.11901842 |
| Telo2 | 0.06139683 | 0.65385758 | 0.156 | 0.083 | 1 | 0.13022901 | -0.2167459 | 0.109 | 0.052 | 1 | 0.13022901 | 0.1190241 |
| Mtmr9 | 0.45017604 | 0.42253204 | 0.177 | 0.145 | 1 | 0.06142614 | 0.58826512 | 0.282 | 0.177 | 1 | 0.45017604 | 0.11907911 |
| Zfp341 | 0.0614531 | -2.240433 | 0.01 | 0.057 | 1 | 0.44369039 | 0.61614416 | 0.05 | 0.031 | 1 | 0.44369039 | 0.11912971 |
| 1700030N03Rik | 0.0614531 | -2.0799683 | 0.01 | 0.057 | 1 | 0.66387437 | 0.27510724 | 0.017 | 0.01 | 1 | 0.66387437 | 0.11912971 |
| Lix1 | 0.06169559 | 1.55498233 | 0.104 | 0.047 | 1 | 0.34459445 | -0.3878578 | 0.097 | 0.062 | 1 | 0.34459445 | 0.11958484 |
| Fhl3 | 0.72628313 | -0.3550755 | 0.052 | 0.062 | 1 | 0.06170946 | 0.63767732 | 0.092 | 0.031 | 1 | 0.72628313 | 0.11961085 |
| Podn | 0.0617419 | -1.8993961 | 0.01 | 0.057 | 1 | 0.37828151 | -1.6468902 | 0.059 | 0.083 | 1 | 0.37828151 | 0.11967175 |
| Pacsin1 | 0.0617419 | -1.7998604 | 0.01 | 0.057 | 1 | 0.97712406 | -0.4253325 | 0.084 | 0.083 | 1 | 0.97712406 | 0.11967175 |
| Tslp | 0.0617419 | -1.9925055 | 0.01 | 0.057 | 1 | 0.39515719 | 0.49749966 | 0.025 | 0.01 | 1 | 0.39515719 | 0.11967175 |
| Il20rb | 0.0617442 | 0.85137533 | 0.229 | 0.14 | 1 | 0.15892438 | 0.27510724 | 0.256 | 0.177 | 1 | 0.15892438 | 0.11967605 |
| Tmem17 | 0.34454333 | 0.52206771 | 0.104 | 0.073 | 1 | 0.06184896 | 0.97554696 | 0.147 | 0.073 | 1 | 0.34454333 | 0.11987263 |
| Fbxl15 | 0.67809385 | 0.25260703 | 0.135 | 0.119 | 1 | 0.06186187 | 0.97554696 | 0.134 | 0.062 | 1 | 0.67809385 | 0.11989685 |
| Myef2 | 0.06187591 | -0.4477001 | 0.396 | 0.497 | 1 | 0.26537166 | -0.4040706 | 0.403 | 0.281 | 1 | 0.26537166 | 0.11992319 |
| Ccdc60 | 0.25337721 | 1.00749454 | 0.052 | 0.026 | 1 | 0.06191918 | 0.54812573 | 0.092 | 0.031 | 1 | 0.25337721 | 0.12000438 |
| Lpl | 0.09647728 | -1.9925055 | 0.052 | 0.109 | 1 | 0.06193478 | -3.2598146 | 0.05 | 0.104 | 1 | 0.09647728 | 0.12003364 |
| Gm28375 | 0.06202887 | 1.20013961 | 0.073 | 0.026 | 1 | 0.66391569 | -1.3098553 | 0.042 | 0.052 | 1 | 0.66391569 | 0.12021016 |

| | | | | | | | | | | | | |
|---|---|---|---|---|---|---|---|---|---|---|---|---|
| Enkd1 | 0.06202887 | 1.20013961 | 0.073 | 0.026 | 1 | 0.43323624 | 0.42711033 | 0.038 | 0.021 | 1 | 0.43323624 | 0.12021016 |
| Lst1 | 0.06202887 | 1.20013961 | 0.073 | 0.026 | 1 | 0.8076363 | 0.27510724 | 0.025 | 0.021 | 1 | 0.8076363 | 0.12021016 |
| Pisd | 0.0621654 | 0.65957123 | 0.24 | 0.15 | 1 | 0.11500171 | -0.1438453 | 0.311 | 0.198 | 1 | 0.11500171 | 0.12046627 |
| Ltbp2 | 0.06222146 | -0.1999475 | 0.354 | 0.233 | 1 | 0.4035353 | -1.316169 | 0.357 | 0.344 | 1 | 0.4035353 | 0.1205714 |
| Iffo2 | 0.18699976 | -0.130009 | 0.615 | 0.482 | 1 | 0.06237081 | -0.297816 | 0.571 | 0.76 | 1 | 0.18699976 | 0.1208515 |
| Aff2 | 0.06238183 | 1.54354744 | 0.146 | 0.078 | 1 | 0.06848933 | -1.3552982 | 0.164 | 0.24 | 1 | 0.06848933 | 0.12087216 |
| Parp9 | 0.53455989 | -0.2983139 | 0.177 | 0.207 | 1 | 0.06239788 | 0.20720317 | 0.239 | 0.135 | 1 | 0.53455989 | 0.12090227 |
| Epn1 | 0.06241015 | 0.43717881 | 0.479 | 0.368 | 1 | 0.10801837 | 0.26679197 | 0.496 | 0.385 | 1 | 0.10801837 | 0.12092527 |
| AW554918 | 0.19891541 | 0.41164372 | 0.406 | 0.347 | 1 | 0.06242422 | -0.8527537 | 0.437 | 0.51 | 1 | 0.19891541 | 0.12095165 |
| Pot1a | 0.3285454 | 0.27052894 | 0.177 | 0.13 | 1 | 0.06245897 | 0.65977109 | 0.151 | 0.073 | 1 | 0.3285454 | 0.12101682 |
| Dchs1 | 0.53965482 | 0.15949763 | 0.156 | 0.13 | 1 | 0.06247426 | 0.22357694 | 0.345 | 0.219 | 1 | 0.53965482 | 0.12104549 |
| Slf1 | 0.06248984 | 0.67541849 | 0.396 | 0.301 | 1 | 0.71190495 | -0.8598223 | 0.349 | 0.323 | 1 | 0.71190495 | 0.1210747 |
| Ppp2r5d | 0.06252278 | 0.47697982 | 0.25 | 0.155 | 1 | 0.3747833 | -0.1276519 | 0.218 | 0.167 | 1 | 0.3747833 | 0.12113646 |
| Cox5b-ps | 0.84144398 | 0.27052894 | 0.146 | 0.14 | 1 | 0.06255647 | 1.12310414 | 0.105 | 0.042 | 1 | 0.84144398 | 0.12119963 |
| Ttc8 | 0.06267214 | 0.83061677 | 0.302 | 0.207 | 1 | 0.06439478 | 0.47864063 | 0.231 | 0.135 | 1 | 0.06439478 | 0.12141647 |
| Insig1 | 0.52659608 | -0.8496373 | 0.771 | 0.725 | 1 | 0.06267663 | 0.56363476 | 0.689 | 0.646 | 1 | 0.52659608 | 0.12142491 |
| Dbr1 | 0.75393206 | 0.11852585 | 0.219 | 0.202 | 1 | 0.06269067 | 0.50572017 | 0.277 | 0.177 | 1 | 0.75393206 | 0.12145122 |
| Gm28154 | 0.06275339 | 0.68556644 | 0.135 | 0.067 | 1 | 0.2329196 | 0.16819203 | 0.122 | 0.073 | 1 | 0.2329196 | 0.1215688 |
| Nr2f1 | 0.3884793 | 0.26125113 | 0.177 | 0.135 | 1 | 0.06286815 | 0.1876444 | 0.143 | 0.062 | 1 | 0.3884793 | 0.12178389 |
| Bcar3 | 0.06287125 | -2.0628948 | 0.042 | 0.104 | 1 | 0.80283227 | -0.510768 | 0.139 | 0.146 | 1 | 0.80283227 | 0.1217897 |
| Erlin1 | 0.06287641 | 0.8291573 | 0.26 | 0.171 | 1 | 0.66346151 | -0.2640516 | 0.328 | 0.271 | 1 | 0.66346151 | 0.12179939 |
| Smim20 | 0.4065306 | -0.130009 | 0.156 | 0.202 | 1 | 0.06296235 | 0.61614416 | 0.151 | 0.073 | 1 | 0.4065306 | 0.12196043 |
| Slc38a1 | 0.06296784 | -0.4810516 | 0.635 | 0.71 | 1 | 0.43189905 | -0.2205879 | 0.223 | 0.177 | 1 | 0.43189905 | 0.12197073 |
| Msh5 | 0.06296894 | 1.00749454 | 0.073 | 0.026 | 1 | 0.20314753 | 1.27510724 | 0.017 | 0 | 1 | 0.20314753 | 0.12197279 |
| Rdh12 | 0.25280242 | 0.65957123 | 0.094 | 0.057 | 1 | 0.06296895 | 0.80562195 | 0.076 | 0.021 | 1 | 0.25280242 | 0.12197281 |
| Grk4 | 0.18598943 | 0.39744105 | 0.26 | 0.192 | 1 | 0.06302416 | -0.3217294 | 0.277 | 0.156 | 1 | 0.18598943 | 0.12207627 |
| Lgals4 | 0.06310025 | -0.9185049 | 0.073 | 0.15 | 1 | 0.9090921 | -0.8504236 | 0.151 | 0.146 | 1 | 0.9090921 | 0.12221886 |
| Chchd3 | 0.06311458 | 0.21799328 | 0.688 | 0.534 | 1 | 0.73791022 | -0.2982206 | 0.727 | 0.76 | 1 | 0.73791022 | 0.12224571 |
| Ankrd12 | 0.44329995 | -0.2331601 | 0.948 | 0.948 | 1 | 0.06313194 | -0.3617979 | 0.971 | 0.99 | 1 | 0.44329995 | 0.12227824 |
| Rnf24 | 0.95659185 | -0.1026884 | 0.385 | 0.373 | 1 | 0.06327699 | -0.7893246 | 0.521 | 0.594 | 1 | 0.95659185 | 0.12255 |
| Cables2 | 0.06333784 | 0.71798792 | 0.177 | 0.098 | 1 | 0.0758289 | 0.76543286 | 0.193 | 0.115 | 1 | 0.0758289 | 0.122664 |
| Fmn1 | 0.06337761 | -0.186043 | 0.896 | 0.865 | 1 | 0.75101595 | -0.6355322 | 0.945 | 0.906 | 1 | 0.75101595 | 0.1227385 |
| Micu3 | 0.06339207 | 0.23093277 | 0.583 | 0.451 | 1 | 0.54362423 | -0.2318528 | 0.71 | 0.625 | 1 | 0.54362423 | 0.12276559 |
| Mrpl1 | 0.06353619 | 0.37006462 | 0.344 | 0.233 | 1 | 0.18892865 | -0.1830981 | 0.311 | 0.208 | 1 | 0.18892865 | 0.12303554 |
| Sfswap | 0.1910767 | 0.15949763 | 0.792 | 0.725 | 1 | 0.06361654 | -0.5297624 | 0.807 | 0.875 | 1 | 0.1910767 | 0.12318602 |
| Mocs3 | 0.14209227 | 0.91438513 | 0.125 | 0.073 | 1 | 0.06363451 | 1.10518224 | 0.105 | 0.042 | 1 | 0.14209227 | 0.12321967 |
| Tsc2 | 0.06373664 | 0.59245704 | 0.271 | 0.176 | 1 | 0.83637743 | -0.4954109 | 0.185 | 0.167 | 1 | 0.83637743 | 0.12341092 |
| 4921531C22Rik | 0.06381611 | -1.4519371 | 0.052 | 0.119 | 1 | 0.68967768 | -0.4353861 | 0.113 | 0.094 | 1 | 0.68967768 | 0.12355973 |
| Mdn1 | 0.06386156 | 0.4752735 | 0.521 | 0.42 | 1 | 0.41039887 | 0.18490943 | 0.655 | 0.635 | 1 | 0.41039887 | 0.12364482 |
| Hoga1 | 0.06386474 | 1.22988696 | 0.104 | 0.047 | 1 | 0.65746468 | 0.33400093 | 0.076 | 0.062 | 1 | 0.65746468 | 0.12365077 |
| Gigyf2 | 0.06389466 | 0.26595621 | 0.812 | 0.71 | 1 | 0.88937012 | -0.3138572 | 0.87 | 0.802 | 1 | 0.88937012 | 0.1237068 |
| Sgo1 | 0.0639341 | 2.00749454 | 0.062 | 0.021 | 1 | 0.24217871 | 0.69014474 | 0.034 | 0.01 | 1 | 0.24217871 | 0.12378064 |
| Ube2w | 0.09625402 | 0.12368755 | 0.781 | 0.674 | 1 | 0.06403677 | 0.1145201 | 0.887 | 0.875 | 1 | 0.09625402 | 0.12397283 |
| Lrrc1 | 0.06413434 | 0.30097006 | 0.729 | 0.565 | 1 | 0.30170028 | -0.2692949 | 0.845 | 0.771 | 1 | 0.30170028 | 0.12415546 |
| Neu2 | 0.13649883 | 0.35214271 | 0.333 | 0.244 | 1 | 0.06417146 | -0.1833229 | 0.273 | 0.156 | 1 | 0.13649883 | 0.12422495 |
| Camk2d | 0.07097211 | -0.5528841 | 1 | 0.964 | 1 | 0.06417867 | -0.2017396 | 0.996 | 0.969 | 1 | 0.07097211 | 0.12423844 |
| Inka2 | 0.06419156 | 1.00749454 | 0.073 | 0.026 | 1 | 0.49984105 | -0.6424306 | 0.063 | 0.083 | 1 | 0.49984105 | 0.12426256 |
| Gmip | 0.06425668 | 1.37006462 | 0.156 | 0.088 | 1 | 0.53742004 | 0.16407593 | 0.151 | 0.125 | 1 | 0.53742004 | 0.12438444 |
| Zfp420 | 0.06431973 | 1.00749454 | 0.104 | 0.047 | 1 | 0.10559793 | 0.88278982 | 0.109 | 0.052 | 1 | 0.10559793 | 0.12450243 |
| Stk11ip | 0.06436575 | 1.12297175 | 0.104 | 0.047 | 1 | 0.23628393 | -1.010295 | 0.059 | 0.094 | 1 | 0.23628393 | 0.12458856 |
| Med18 | 0.52497972 | 0.64492446 | 0.052 | 0.036 | 1 | 0.06442523 | 1.59703533 | 0.088 | 0.031 | 1 | 0.52497972 | 0.12469985 |
| Ntn4 | 0.0644354 | -1.4170033 | 0.094 | 0.171 | 1 | 0.61199164 | -0.2103196 | 0.126 | 0.146 | 1 | 0.61199164 | 0.12471889 |
| E430024I08Rik | 0.33016965 | 0.56270969 | 0.24 | 0.197 | 1 | 0.0644897 | 0.25404562 | 0.223 | 0.125 | 1 | 0.33016965 | 0.12482048 |
| Aggf1 | 0.06449838 | 0.30952307 | 0.542 | 0.409 | 1 | 0.29402021 | -0.3637941 | 0.5 | 0.583 | 1 | 0.29402021 | 0.12483671 |
| Med1 | 0.46246571 | 0.18064344 | 0.5 | 0.461 | 1 | 0.06451831 | 0.1725375 | 0.66 | 0.552 | 1 | 0.46246571 | 0.12487402 |
| Tmem95 | 0.06453363 | -1.407543 | 0.01 | 0.057 | 1 | 0.22356137 | 0.80562195 | 0.05 | 0.021 | 1 | 0.22356137 | 0.12490267 |
| Mief2 | 0.29477093 | -0.770113 | 0.062 | 0.098 | 1 | 0.06453523 | 0.63356121 | 0.164 | 0.083 | 1 | 0.29477093 | 0.12490566 |
| Rnf123 | 0.06460875 | 1.2091284 | 0.167 | 0.093 | 1 | 0.1136764 | 0.2095189 | 0.155 | 0.083 | 1 | 0.1136764 | 0.12504322 |
| Qpct | 0.06465158 | 0.44806713 | 0.167 | 0.088 | 1 | 0.7474356 | -0.1082214 | 0.151 | 0.135 | 1 | 0.7474356 | 0.12512334 |
| Pcdhga10 | 0.35136973 | -0.4330781 | 0.115 | 0.155 | 1 | 0.06475868 | -1.7504279 | 0.05 | 0.104 | 1 | 0.35136973 | 0.12532367 |
| Skic3 | 0.06490503 | 0.11580635 | 0.656 | 0.585 | 1 | 0.49672245 | -0.3858041 | 0.601 | 0.49 | 1 | 0.49672245 | 0.1255974 |
| Tmem123 | 0.83564238 | -0.2088234 | 0.24 | 0.249 | 1 | 0.06498067 | -0.2847643 | 0.328 | 0.458 | 1 | 0.83564238 | 0.12573886 |
| Ehmt1 | 0.06499096 | 0.29255525 | 0.76 | 0.653 | 1 | 0.73047313 | -0.1662096 | 0.866 | 0.896 | 1 | 0.73047313 | 0.12575809 |
| Hes6 | 0.48727529 | 0.37006462 | 0.073 | 0.052 | 1 | 0.06502055 | 1.49749966 | 0.088 | 0.031 | 1 | 0.48727529 | 0.12581344 |
| Rgmb | 0.13872649 | 0.2091284 | 0.323 | 0.228 | 1 | 0.0650355 | 0.44913664 | 0.462 | 0.333 | 1 | 0.13872649 | 0.12584138 |
| Add1 | 0.06506278 | 0.38146938 | 0.635 | 0.544 | 1 | 0.56813922 | -0.2551646 | 0.748 | 0.833 | 1 | 0.56813922 | 0.12589239 |
| Ormdl1 | 0.08298863 | 0.71798792 | 0.177 | 0.104 | 1 | 0.06516512 | 0.94648449 | 0.16 | 0.083 | 1 | 0.08298863 | 0.12608375 |
| Thap1 | 0.06519184 | 0.8872003 | 0.198 | 0.119 | 1 | 0.28237122 | 0.27510724 | 0.164 | 0.115 | 1 | 0.28237122 | 0.12613371 |
| Hars2 | 0.06523655 | 0.61517711 | 0.271 | 0.176 | 1 | 0.14421062 | 0.29680231 | 0.223 | 0.146 | 1 | 0.14421062 | 0.12621729 |
| Sacm1l | 0.06533481 | 0.1404684 | 0.729 | 0.637 | 1 | 0.65458273 | -0.3453695 | 0.811 | 0.823 | 1 | 0.65458273 | 0.12640098 |
| AW011738 | 0.06537588 | 0.90057933 | 0.104 | 0.047 | 1 | 0.13572174 | -1.3098553 | 0.084 | 0.135 | 1 | 0.13572174 | 0.12647775 |
| Utp4 | 0.10362072 | 0.25947375 | 0.479 | 0.383 | 1 | 0.06543185 | 0.14125149 | 0.563 | 0.417 | 1 | 0.10362072 | 0.12658238 |
| Cnksr2 | 0.6453829 | 0.42253204 | 0.042 | 0.031 | 1 | 0.06545347 | -2.0803734 | 0.067 | 0.125 | 1 | 0.6453829 | 0.12662278 |
| Lsm8 | 0.06548815 | 0.42863224 | 0.521 | 0.404 | 1 | 0.08987404 | 0.19668059 | 0.521 | 0.417 | 1 | 0.08987404 | 0.1266876 |
| Brpf3 | 0.06560729 | 0.47406234 | 0.281 | 0.181 | 1 | 0.78903585 | -0.3682719 | 0.319 | 0.281 | 1 | 0.78903585 | 0.12691025 |
| Gbp6 | 0.25384445 | -0.5427025 | 0.042 | 0.078 | 1 | 0.06561645 | -2.4029647 | 0.029 | 0.073 | 1 | 0.25384445 | 0.12692737 |
| Agfg1 | 0.06574867 | -0.2990185 | 0.833 | 0.891 | 1 | 0.07600175 | 0.10212244 | 0.941 | 0.885 | 1 | 0.07600175 | 0.12717445 |
| Spaar | 0.06576295 | -0.3710171 | 0.01 | 0.057 | 1 | 0.20315163 | 1.01207283 | 0.017 | 0 | 1 | 0.20315163 | 0.12720113 |
| Bcs1l | 0.51547857 | 0.27052894 | 0.135 | 0.109 | 1 | 0.06581594 | 0.59703533 | 0.105 | 0.042 | 1 | 0.51547857 | 0.12730015 |
| Hacd1 | 0.06583397 | 0.2462814 | 0.406 | 0.29 | 1 | 0.07297911 | 0.11128244 | 0.441 | 0.292 | 1 | 0.07297911 | 0.12733383 |
| Sgsm2 | 0.34555054 | 0.13302542 | 0.531 | 0.466 | 1 | 0.06584716 | -1.0840433 | 0.609 | 0.552 | 1 | 0.34555054 | 0.12735847 |
| Rc3h1 | 0.17313141 | 0.14499806 | 0.885 | 0.855 | 1 | 0.06598853 | -0.4927193 | 0.962 | 0.958 | 1 | 0.17313141 | 0.12762257 |
| Ncoa1 | 0.45647742 | -0.1535679 | 0.521 | 0.451 | 1 | 0.0659905 | -0.7176612 | 0.803 | 0.833 | 1 | 0.45647742 | 0.12762625 |
| Ubqln1 | 0.06599349 | 0.42253204 | 0.615 | 0.549 | 1 | 0.12036415 | 0.16448807 | 0.786 | 0.729 | 1 | 0.12036415 | 0.12763184 |
| Cables1 | 0.06615131 | -1.5118796 | 0.115 | 0.197 | 1 | 0.14789907 | 0.35640734 | 0.168 | 0.104 | 1 | 0.14789907 | 0.12792662 |
| Suz12 | 0.06616768 | 0.32942263 | 0.656 | 0.591 | 1 | 0.2682276 | -0.2874874 | 0.689 | 0.74 | 1 | 0.2682276 | 0.1279572 |
| Gcg | 0.06619166 | 1.0899567 | 0.104 | 0.047 | 1 | 0.49926198 | -1.3923174 | 0.046 | 0.062 | 1 | 0.49926198 | 0.12800199 |
| Cog1 | 0.06623706 | 0.80104366 | 0.219 | 0.135 | 1 | 0.17479065 | -0.5133887 | 0.252 | 0.167 | 1 | 0.17479065 | 0.12808677 |

| | | | | | | | | | | | | |
|---|---|---|---|---|---|---|---|---|---|---|---|---|
| Kcnq3 | 0.06639583 | -1.9384656 | 0.073 | 0.145 | 1 | 0.39144669 | -1.3098553 | 0.088 | 0.115 | 1 | 0.39144669 | 0.12838326 |
| Gm56813 | 0.06666821 | 1.68556644 | 0.062 | 0.021 | 1 | 1 | -0.5025003 | 0.021 | 0.021 | 1 | 1 | 0.12889177 |
| ENSMUSG00000121319 | 0.06666821 | 1.68556644 | 0.062 | 0.021 | 1 | 0.86203455 | -0.6577786 | 0.038 | 0.042 | 1 | 0.86203455 | 0.12889177 |
| Gm15867 | 0.85736048 | -0.2285728 | 0.292 | 0.29 | 1 | 0.06670995 | 0.52472113 | 0.286 | 0.188 | 1 | 0.85736048 | 0.12896968 |
| Rbm26 | 0.06671014 | 0.13766183 | 0.875 | 0.829 | 1 | 0.53351761 | -0.2279377 | 0.908 | 0.927 | 1 | 0.53351761 | 0.12897004 |
| Gm17529 | 0.06672266 | 0.31561683 | 0.115 | 0.052 | 1 | 0.78112888 | -0.5961594 | 0.134 | 0.115 | 1 | 0.78112888 | 0.12899341 |
| CJ186046Rik | 0.49423765 | -0.240433 | 0.146 | 0.114 | 1 | 0.06674413 | 0.14080615 | 0.151 | 0.073 | 1 | 0.49423765 | 0.12903348 |
| Smim12 | 0.87024845 | -0.1624305 | 0.188 | 0.192 | 1 | 0.06674701 | 0.62874419 | 0.239 | 0.146 | 1 | 0.87024845 | 0.12903886 |
| 9930104L06Rik | 0.06679648 | 1.00749454 | 0.125 | 0.062 | 1 | 0.10386849 | 1.01207283 | 0.122 | 0.062 | 1 | 0.10386849 | 0.1291312 |
| Sirt2 | 0.23297572 | 0.17439185 | 0.646 | 0.58 | 1 | 0.06680006 | 0.14642522 | 0.706 | 0.542 | 1 | 0.23297572 | 0.12913787 |
| Ankrd42 | 0.0668426 | -0.3637613 | 0.271 | 0.166 | 1 | 0.17444992 | -1.4942798 | 0.214 | 0.26 | 1 | 0.17444992 | 0.12921726 |
| Obp2b | 0.36140575 | -0.3680146 | 0.208 | 0.244 | 1 | 0.06688336 | -4.8208172 | 0.017 | 0.052 | 1 | 0.36140575 | 0.12929334 |
| Zbtb16 | 0.06692101 | -0.2862367 | 0.26 | 0.368 | 1 | 0.88741931 | -0.8848337 | 0.357 | 0.302 | 1 | 0.88741931 | 0.12936359 |
| Ppcs | 0.06693532 | 1.00749454 | 0.104 | 0.047 | 1 | 0.08804305 | 1.39058446 | 0.05 | 0.01 | 1 | 0.08804305 | 0.1293903 |
| Ptk2b | 0.81159776 | -0.8993961 | 0.031 | 0.036 | 1 | 0.06694238 | 2.59703533 | 0.055 | 0.01 | 1 | 0.81159776 | 0.12940349 |
| Frzb | 0.25383203 | -0.6993511 | 0.344 | 0.394 | 1 | 0.0670295 | -0.662157 | 0.139 | 0.062 | 1 | 0.25383203 | 0.12956605 |
| Rasl12 | 0.26118486 | -0.9925055 | 0.031 | 0.062 | 1 | 0.06710373 | -2.1105465 | 0.076 | 0.135 | 1 | 0.26118486 | 0.12970455 |
| Rcor1 | 0.97092257 | -0.5666612 | 0.667 | 0.617 | 1 | 0.06712404 | -0.5183573 | 0.84 | 0.865 | 1 | 0.97092257 | 0.12974245 |
| Tead2 | 0.06719817 | -1.765095 | 0.042 | 0.104 | 1 | 0.07109529 | -1.3098553 | 0.092 | 0.156 | 1 | 0.07109529 | 0.12988075 |
| Mfn1 | 0.06724045 | 0.28621056 | 0.49 | 0.352 | 1 | 0.94855254 | -0.2246879 | 0.475 | 0.469 | 1 | 0.94855254 | 0.12995962 |
| Zcchc3 | 0.06727047 | 0.68556644 | 0.208 | 0.124 | 1 | 0.09602387 | 0.34910782 | 0.197 | 0.115 | 1 | 0.09602387 | 0.13001562 |
| Cfap61 | 0.3380241 | -1.577468 | 0.031 | 0.057 | 1 | 0.0673288 | -1.0203486 | 0.042 | 0.094 | 1 | 0.3380241 | 0.13012443 |
| Rnf14 | 0.52984807 | 0.10703021 | 0.583 | 0.528 | 1 | 0.06737085 | 0.20083448 | 0.643 | 0.49 | 1 | 0.52984807 | 0.13020288 |
| Otulin | 0.06749877 | -0.6070122 | 0.552 | 0.679 | 1 | 0.81037231 | -0.3271333 | 0.676 | 0.646 | 1 | 0.81037231 | 0.13044146 |
| Ercc2 | 0.6297019 | 0.10703021 | 0.135 | 0.114 | 1 | 0.06750162 | 0.54812573 | 0.202 | 0.115 | 1 | 0.6297019 | 0.13044678 |
| Casp4 | 0.71039151 | 0.35811531 | 0.333 | 0.394 | 1 | 0.06753014 | 0.49749966 | 0.294 | 0.188 | 1 | 0.71039151 | 0.13049997 |
| Uaca | 0.06758409 | -0.6126574 | 0.781 | 0.798 | 1 | 0.45469352 | -0.1609212 | 0.773 | 0.812 | 1 | 0.45469352 | 0.13060058 |
| Ccdc43 | 0.06759033 | 0.68556644 | 0.26 | 0.171 | 1 | 0.31923792 | 0.11098786 | 0.252 | 0.198 | 1 | 0.31923792 | 0.1306122 |
| Eif4enif1 | 0.24809714 | 0.32200516 | 0.49 | 0.409 | 1 | 0.06765062 | 0.1507791 | 0.66 | 0.552 | 1 | 0.24809714 | 0.13072464 |
| Cdkl4 | 0.06766457 | -1.8669746 | 0.021 | 0.073 | 1 | 0.08257549 | -1.6317834 | 0.038 | 0.083 | 1 | 0.08257549 | 0.13075065 |
| Cyria | 0.06766674 | -2.2148979 | 0.021 | 0.073 | 1 | 0.18361019 | 1.27510724 | 0.038 | 0.01 | 1 | 0.18361019 | 0.1307547 |
| Trmt2b | 0.42948723 | -0.4596315 | 0.146 | 0.181 | 1 | 0.06768367 | 1.20471791 | 0.118 | 0.052 | 1 | 0.42948723 | 0.13078626 |
| Nostrin | 0.39200989 | 0.31561683 | 0.083 | 0.057 | 1 | 0.06775726 | -0.5728897 | 0.172 | 0.083 | 1 | 0.39200989 | 0.13092347 |
| Gin1 | 0.8619318 | -0.3968957 | 0.125 | 0.13 | 1 | 0.0677808 | 0.51326698 | 0.164 | 0.083 | 1 | 0.8619318 | 0.13096737 |
| Dusp8 | 0.83932545 | 0.20013961 | 0.167 | 0.161 | 1 | 0.06788089 | 0.95693128 | 0.185 | 0.104 | 1 | 0.83932545 | 0.13115397 |
| Rnasel | 0.06789881 | -0.8550019 | 0.062 | 0.135 | 1 | 0.12272662 | 1.09224318 | 0.105 | 0.052 | 1 | 0.12272662 | 0.13118738 |
| Ankrd35 | 0.06791505 | 0.59245704 | 0.156 | 0.083 | 1 | 0.38371714 | -0.5349218 | 0.214 | 0.25 | 1 | 0.38371714 | 0.13121765 |
| Trmt44 | 0.06795392 | -1.1079827 | 0.052 | 0.119 | 1 | 0.12829616 | 0.23446525 | 0.122 | 0.062 | 1 | 0.12829616 | 0.1312901 |
| Csnk1g3 | 0.06795574 | 0.24482323 | 0.719 | 0.606 | 1 | 0.38537746 | -0.2294818 | 0.845 | 0.906 | 1 | 0.38537746 | 0.13129349 |
| Gna13 | 0.06800057 | -0.3062991 | 0.823 | 0.834 | 1 | 0.23939765 | 0.13841857 | 0.832 | 0.865 | 1 | 0.23939765 | 0.13137705 |
| Gm46563 | 0.13729324 | 0.69937224 | 0.177 | 0.114 | 1 | 0.06801495 | 1.21370669 | 0.088 | 0.031 | 1 | 0.13729324 | 0.13140387 |
| Zfp329 | 0.49297381 | 0.4088571 | 0.208 | 0.181 | 1 | 0.06806411 | 0.35999614 | 0.29 | 0.177 | 1 | 0.49297381 | 0.13149549 |
| 4930405A21Rik | 0.06810359 | 1.17741954 | 0.062 | 0.021 | 1 | 0.99396051 | -0.3098553 | 0.021 | 0.021 | 1 | 0.99396051 | 0.13156909 |
| Adat2 | 0.06812307 | 1.49292136 | 0.062 | 0.021 | 1 | 0.6322619 | -0.7793405 | 0.05 | 0.062 | 1 | 0.6322619 | 0.13160539 |
| Gm10184 | 0.06812307 | 1.49292136 | 0.062 | 0.021 | 1 | 0.08946279 | 1.27510724 | 0.067 | 0.021 | 1 | 0.08946279 | 0.13160539 |
| Mvd | 0.06816671 | 0.80586068 | 0.271 | 0.181 | 1 | 0.50719708 | 0.22619764 | 0.097 | 0.073 | 1 | 0.50719708 | 0.13168672 |
| St6galnac2 | 0.06822569 | -1.7294711 | 0.021 | 0.073 | 1 | 0.19241213 | 0.27510724 | 0.055 | 0.021 | 1 | 0.19241213 | 0.13179663 |
| Lrrcc1 | 0.06824597 | 0.5686103 | 0.281 | 0.187 | 1 | 0.90047856 | -0.714697 | 0.193 | 0.188 | 1 | 0.90047856 | 0.13183443 |
| Gm45159 | 0.33184333 | -0.577468 | 0.115 | 0.155 | 1 | 0.06825606 | 1.69014474 | 0.055 | 0.01 | 1 | 0.33184333 | 0.13185324 |
| Uba7 | 0.06826276 | -1.5285584 | 0.042 | 0.104 | 1 | 0.06884019 | 0.72256622 | 0.134 | 0.062 | 1 | 0.06884019 | 0.13186571 |
| Gm19522 | 0.06828306 | 1.65957123 | 0.156 | 0.088 | 1 | 0.41777167 | 0.27510724 | 0.139 | 0.104 | 1 | 0.41777167 | 0.13190353 |
| Msl3l2 | 0.49855877 | 0.16795921 | 0.156 | 0.124 | 1 | 0.06831491 | 0.7296731 | 0.134 | 0.062 | 1 | 0.49855877 | 0.1319629 |
| Rian | 0.71866889 | -0.5607892 | 0.594 | 0.518 | 1 | 0.06832742 | -0.4055024 | 0.714 | 0.542 | 1 | 0.71866889 | 0.1319862 |
| Hdac7 | 0.06834432 | -1.9925055 | 0.031 | 0.088 | 1 | 0.83671988 | -0.8504236 | 0.143 | 0.125 | 1 | 0.83671988 | 0.1320177 |
| 1110017D15Rik | 0.06848495 | 1.39451766 | 0.104 | 0.047 | 1 | 0.8788747 | -0.7248928 | 0.08 | 0.083 | 1 | 0.8788747 | 0.13227971 |
| Gm45238 | 0.61924305 | -0.2148979 | 0.021 | 0.031 | 1 | 0.06851509 | 1.59703533 | 0.055 | 0.01 | 1 | 0.61924305 | 0.13233586 |
| Selenop | 0.20181144 | -0.4823107 | 0.323 | 0.383 | 1 | 0.06851786 | 0.17057665 | 0.416 | 0.281 | 1 | 0.20181144 | 0.13234102 |
| Fgf1 | 0.24369012 | 0.35541784 | 0.115 | 0.166 | 1 | 0.06855187 | -1.9472852 | 0.029 | 0.073 | 1 | 0.24369012 | 0.13240438 |
| Man1a | 0.46965371 | 0.1895185 | 0.833 | 0.876 | 1 | 0.06858057 | -0.4728295 | 0.924 | 0.948 | 1 | 0.46965371 | 0.13245785 |
| Herc6 | 0.398797 | -0.2295447 | 0.146 | 0.192 | 1 | 0.06864876 | -2.3952468 | 0.122 | 0.188 | 1 | 0.398797 | 0.13258486 |
| Tcf3 | 0.06885693 | 0.19035859 | 0.354 | 0.233 | 1 | 0.10215808 | 0.10212244 | 0.466 | 0.344 | 1 | 0.10215808 | 0.13297258 |
| Fah | 0.1104186 | 1.00749454 | 0.073 | 0.031 | 1 | 0.06887658 | 1.14957636 | 0.088 | 0.031 | 1 | 0.1104186 | 0.13300917 |
| Gm20274 | 0.99205163 | 0.10060394 | 0.073 | 0.073 | 1 | 0.06887658 | 1.14957636 | 0.088 | 0.031 | 1 | 0.99205163 | 0.13300917 |
| Bend5 | 0.38689439 | -0.577468 | 0.01 | 0.026 | 1 | 0.0689015 | -2.3098553 | 0.017 | 0.052 | 1 | 0.38689439 | 0.13305558 |
| Dnajc5 | 0.0910468 | 0.12202292 | 0.833 | 0.736 | 1 | 0.06891049 | 0.2176381 | 0.924 | 0.865 | 1 | 0.0910468 | 0.13307232 |
| Ube2n | 0.14187636 | 0.12423517 | 0.948 | 0.86 | 1 | 0.06893826 | 0.13700496 | 0.916 | 0.875 | 1 | 0.14187636 | 0.13312403 |
| Lmnb2 | 0.22926276 | 0.66645762 | 0.115 | 0.073 | 1 | 0.06904378 | 0.23763253 | 0.151 | 0.073 | 1 | 0.22926276 | 0.13332052 |
| Prkd2 | 0.82255289 | -0.3535031 | 0.271 | 0.29 | 1 | 0.0691046 | 0.13455684 | 0.454 | 0.312 | 1 | 0.82255289 | 0.13343375 |
| Btbd8 | 0.06917097 | 1.00749454 | 0.062 | 0.021 | 1 | 0.69978947 | -0.2223924 | 0.042 | 0.052 | 1 | 0.69978947 | 0.13355731 |
| Ube2g1 | 0.11656129 | 0.24580651 | 0.781 | 0.777 | 1 | 0.06918602 | -0.6278612 | 0.912 | 0.948 | 1 | 0.11656129 | 0.13358534 |
| Shox2 | 0.38774279 | 0.10060394 | 0.073 | 0.047 | 1 | 0.06924238 | -2.4253325 | 0.017 | 0.052 | 1 | 0.38774279 | 0.13369026 |
| Zfp512 | 0.11196318 | -0.8225805 | 0.146 | 0.223 | 1 | 0.06932921 | 0.15829357 | 0.273 | 0.167 | 1 | 0.11196318 | 0.13385189 |
| Ankrd13b | 0.06934317 | -1.577468 | 0.021 | 0.073 | 1 | 0.66159589 | -0.5133887 | 0.109 | 0.125 | 1 | 0.66159589 | 0.13387787 |
| Tfcp2 | 0.63482518 | 0.1243082 | 0.312 | 0.28 | 1 | 0.06934869 | -0.3864765 | 0.336 | 0.208 | 1 | 0.63482518 | 0.13388814 |
| Gm47071 | 0.74827573 | 0.59245704 | 0.021 | 0.016 | 1 | 0.06958408 | -2.0468209 | 0.017 | 0.052 | 1 | 0.74827573 | 0.13432622 |
| Zdhhc19 | 0.99758607 | 0.37946331 | 0.021 | 0.021 | 1 | 0.06958408 | -2.3098553 | 0.017 | 0.052 | 1 | 0.99758607 | 0.13432622 |
| Gm15728 | 0.53063347 | -0.6929452 | 0.031 | 0.047 | 1 | 0.06958481 | -1.5322477 | 0.059 | 0.115 | 1 | 0.53063347 | 0.13432757 |
| Kif17 | 0.06969347 | 0.8872003 | 0.146 | 0.078 | 1 | 0.79239775 | -0.7366699 | 0.13 | 0.135 | 1 | 0.79239775 | 0.13452977 |
| Gm56599 | 0.06969366 | -2.0799683 | 0.021 | 0.073 | 1 | 0.12096301 | -2.7692869 | 0.008 | 0.031 | 1 | 0.12096301 | 0.13453012 |
| A630072M18Rik | 0.14679137 | 0.12613903 | 0.188 | 0.119 | 1 | 0.06971952 | 0.5580412 | 0.214 | 0.125 | 1 | 0.14679137 | 0.13457823 |
| Sphk1 | 0.0697438 | -0.9266175 | 0.198 | 0.285 | 1 | 0.12218442 | 0.40635177 | 0.21 | 0.135 | 1 | 0.12218442 | 0.13462341 |
| Rmnd5b | 0.06977846 | 0.70793425 | 0.219 | 0.135 | 1 | 0.29824862 | -0.152314 | 0.256 | 0.188 | 1 | 0.29824862 | 0.13468788 |
| Tigd5 | 0.14979643 | 1.27052894 | 0.052 | 0.021 | 1 | 0.0698139 | 2.01207283 | 0.034 | 0 | 1 | 0.14979643 | 0.13475383 |
| Tmem59l | 0.70575267 | -0.4950058 | 0.052 | 0.062 | 1 | 0.0698139 | 2.01207283 | 0.034 | 0 | 1 | 0.70575267 | 0.13475383 |

| | | | | | | | | | | | | |
|---|---|---|---|---|---|---|---|---|---|---|---|---|
| Gm48529 | 0.74827573 | 0.59245704 | 0.021 | 0.016 | 1 | 0.0698139 | 2.01207283 | 0.034 | 0 | 1 | 0.74827573 | 0.13475383 |
| Cxcr4 | 0.75832339 | -0.407543 | 0.021 | 0.016 | 1 | 0.0698139 | 2.01207283 | 0.034 | 0 | 1 | 0.75832339 | 0.13475383 |
| Ppp1r3b | 0.10943834 | 0.79599043 | 0.115 | 0.062 | 1 | 0.06982064 | 2.27510724 | 0.034 | 0 | 1 | 0.10943834 | 0.13476637 |
| Gm48008 | 0.21867159 | 1.59245704 | 0.021 | 0.005 | 1 | 0.06982064 | 2.14957636 | 0.034 | 0 | 1 | 0.21867159 | 0.13476637 |
| Mdfic2 | 0.32025586 | -1.3144336 | 0 | 0.01 | 1 | 0.06982064 | 2.49749966 | 0.034 | 0 | 1 | 0.32025586 | 0.13476637 |
| Kbtbd8 | 0.72574709 | -0.3144336 | 0.01 | 0.016 | 1 | 0.06982064 | 2.27510724 | 0.034 | 0 | 1 | 0.72574709 | 0.13476637 |
| Fbp2 | 0.74576744 | 0.68556644 | 0.021 | 0.016 | 1 | 0.06982064 | 2.49749966 | 0.034 | 0 | 1 | 0.74576744 | 0.13476637 |
| Tbc1d2 | 0.99758604 | -0.2148979 | 0.021 | 0.021 | 1 | 0.06982064 | 2.27510724 | 0.034 | 0 | 1 | 0.99758604 | 0.13476637 |
| Gm30978 | 0.99758609 | 0.27052894 | 0.021 | 0.021 | 1 | 0.06982064 | 2.14957636 | 0.034 | 0 | 1 | 0.99758609 | 0.13476637 |
| Arhgef4 | 0.29656877 | 1.90057933 | 0.042 | 0.021 | 1 | 0.06982546 | 2.27510724 | 0.034 | 0 | 1 | 0.29656877 | 0.13477532 |
| Ablim3 | 0.52744059 | -0.577468 | 0.01 | 0.021 | 1 | 0.06982546 | 2.27510724 | 0.034 | 0 | 1 | 0.52744059 | 0.13477532 |
| Smco2 | 0.52744059 | -0.577468 | 0.01 | 0.021 | 1 | 0.06982835 | 2.59703533 | 0.034 | 0 | 1 | 0.52744059 | 0.1347807 |
| Inpp5d | 0.61371908 | 2.00749454 | 0.01 | 0.005 | 1 | 0.06982995 | 3.01207283 | 0.034 | 0 | 1 | 0.61371908 | 0.13478368 |
| Zfp953 | 0.14976512 | 1.59245704 | 0.052 | 0.021 | 1 | 0.06983188 | 1.86006974 | 0.034 | 0 | 1 | 0.14976512 | 0.13478727 |
| Arhgap9 | 0.32025586 | -0.9925055 | 0 | 0.01 | 1 | 0.06983188 | 1.86006974 | 0.034 | 0 | 1 | 0.32025586 | 0.13478727 |
| B230322F03Rik | 0.61737927 | 1.00749454 | 0.01 | 0.005 | 1 | 0.06983188 | 1.86006974 | 0.034 | 0 | 1 | 0.61737927 | 0.13478727 |
| Gm42595 | 0.74827573 | 0.59245704 | 0.021 | 0.016 | 1 | 0.06983188 | 1.86006974 | 0.034 | 0 | 1 | 0.74827573 | 0.13478727 |
| Mtrfr | 0.75417491 | -0.130009 | 0.198 | 0.181 | 1 | 0.06990552 | 0.42711033 | 0.176 | 0.094 | 1 | 0.75417491 | 0.13492427 |
| Mrps11 | 0.49785692 | -0.1547769 | 0.156 | 0.192 | 1 | 0.06990606 | 0.76053407 | 0.147 | 0.073 | 1 | 0.49785692 | 0.13492526 |
| Cpa6 | 0.06991025 | 1.13677755 | 0.104 | 0.047 | 1 | 0.214483 | -2.0297473 | 0.025 | 0.052 | 1 | 0.214483 | 0.13493305 |
| Gpsm3 | 0.91314039 | -0.1079827 | 0.167 | 0.171 | 1 | 0.0699219 | 0.36821664 | 0.105 | 0.042 | 1 | 0.91314039 | 0.13495472 |
| Taf1d | 0.59538987 | -0.3095348 | 0.833 | 0.829 | 1 | 0.06999231 | -0.4291083 | 0.765 | 0.865 | 1 | 0.59538987 | 0.1350857 |
| Dxo | 0.83177979 | -0.1624305 | 0.177 | 0.187 | 1 | 0.07002044 | 0.74459252 | 0.197 | 0.115 | 1 | 0.83177979 | 0.13513802 |
| Adgrv1 | 0.07002147 | -1.577468 | 0.042 | 0.104 | 1 | 0.34533788 | -2.1578522 | 0.008 | 0.021 | 1 | 0.34533788 | 0.13513993 |
| Nipal3 | 0.15156236 | -0.9185049 | 0.042 | 0.088 | 1 | 0.07008935 | 1.49749966 | 0.071 | 0.021 | 1 | 0.15156236 | 0.13526619 |
| Pafah2 | 0.24492634 | 1.15949763 | 0.073 | 0.041 | 1 | 0.07008935 | 1.49749966 | 0.071 | 0.021 | 1 | 0.24492634 | 0.13526619 |
| Mettl4 | 0.07022532 | 0.42253204 | 0.177 | 0.098 | 1 | 0.1119944 | 0.42711033 | 0.172 | 0.104 | 1 | 0.1119944 | 0.13551904 |
| Cdkl5 | 0.24985218 | -0.2013195 | 0.396 | 0.306 | 1 | 0.07023563 | 0.2950068 | 0.437 | 0.312 | 1 | 0.24985218 | 0.13553822 |
| Ryr3 | 0.07024899 | 1.22988696 | 0.062 | 0.021 | 1 | 0.41674525 | -0.3664388 | 0.092 | 0.062 | 1 | 0.41674525 | 0.13556306 |
| Spef1 | 0.07024899 | 1.22988696 | 0.062 | 0.021 | 1 | 0.94078718 | -1.3098553 | 0.067 | 0.062 | 1 | 0.94078718 | 0.13556306 |
| Sap25 | 0.07024899 | 1.22988696 | 0.062 | 0.021 | 1 | 0.77159127 | -0.3098553 | 0.084 | 0.073 | 1 | 0.77159127 | 0.13556306 |
| St8sia1 | 0.07024899 | 1.22988696 | 0.062 | 0.021 | 1 | 0.14367653 | 1.27510724 | 0.042 | 0.01 | 1 | 0.14367653 | 0.13556306 |
| 2310068J16Rik | 0.07024899 | 1.22988696 | 0.062 | 0.021 | 1 | 0.18504625 | 1.01207283 | 0.038 | 0.01 | 1 | 0.18504625 | 0.13556306 |
| Abhd15 | 0.11302151 | -2.577468 | 0 | 0.026 | 1 | 0.0702574 | -1.4703199 | 0.05 | 0.104 | 1 | 0.11302151 | 0.13557869 |
| Napb | 0.07030327 | -0.6599301 | 0.104 | 0.192 | 1 | 0.66085178 | -0.1493906 | 0.113 | 0.094 | 1 | 0.66085178 | 0.13566399 |
| Fchsd2 | 0.40770887 | -0.2363776 | 0.917 | 0.933 | 1 | 0.07031133 | -0.8188491 | 0.937 | 0.958 | 1 | 0.40770887 | 0.13567897 |
| Setd4 | 0.26940715 | 0.42253204 | 0.073 | 0.041 | 1 | 0.07040371 | -1.4093909 | 0.05 | 0.104 | 1 | 0.26940715 | 0.13585073 |
| 4933406I18Rik | 0.07045602 | 0.60788708 | 0.333 | 0.233 | 1 | 0.2603617 | -0.3361665 | 0.265 | 0.188 | 1 | 0.2603617 | 0.135948 |
| Kif14 | 0.07045948 | 2.59245704 | 0.052 | 0.016 | 1 | 0.87066325 | -0.3098553 | 0.013 | 0.01 | 1 | 0.87066325 | 0.13595442 |
| Bmx | 0.07046432 | -1.7294711 | 0.021 | 0.073 | 1 | 0.24304534 | -1.7952821 | 0.013 | 0.031 | 1 | 0.24304534 | 0.13596343 |
| Nudt3 | 0.07050594 | 0.43180985 | 0.594 | 0.497 | 1 | 0.1906538 | 0.16606448 | 0.626 | 0.542 | 1 | 0.1906538 | 0.1360408 |
| Lysmd2 | 0.09847834 | 0.84702986 | 0.146 | 0.083 | 1 | 0.07052511 | 0.46775232 | 0.176 | 0.094 | 1 | 0.09847834 | 0.13607643 |
| Kxd1 | 0.07055253 | 0.60894516 | 0.333 | 0.238 | 1 | 0.24713139 | -0.1128184 | 0.303 | 0.229 | 1 | 0.24713139 | 0.13612739 |
| Zfp940 | 0.08780579 | 1.42253204 | 0.083 | 0.036 | 1 | 0.0705543 | 1.56461386 | 0.071 | 0.021 | 1 | 0.08780579 | 0.1361307 |
| Ptpru | 0.22063631 | -0.6929452 | 0.052 | 0.093 | 1 | 0.0705543 | 1.56461386 | 0.071 | 0.021 | 1 | 0.22063631 | 0.1361307 |
| 2010004M13Rik | 0.28992179 | 0.47144164 | 0.188 | 0.14 | 1 | 0.07057037 | 0.97025266 | 0.118 | 0.052 | 1 | 0.28992179 | 0.13616056 |
| Dmac2 | 0.29621645 | 0.64492446 | 0.167 | 0.124 | 1 | 0.07064066 | 0.88278982 | 0.105 | 0.042 | 1 | 0.29621645 | 0.13629121 |
| Ubash3b | 0.66315589 | -0.4481849 | 0.167 | 0.192 | 1 | 0.0706541 | -1.2318528 | 0.088 | 0.156 | 1 | 0.66315589 | 0.13631619 |
| Uqcrfs1 | 0.32265037 | 0.12905652 | 0.615 | 0.596 | 1 | 0.07069023 | 0.42711033 | 0.697 | 0.677 | 1 | 0.32265037 | 0.13638335 |
| Dtl | 0.42013973 | 1.2590333 | 0.094 | 0.067 | 1 | 0.07078047 | -1.6577786 | 0.029 | 0.073 | 1 | 0.42013973 | 0.13655107 |
| Zdhhc13 | 0.93279157 | -0.2950682 | 0.146 | 0.145 | 1 | 0.07080029 | 1.08246216 | 0.13 | 0.062 | 1 | 0.93279157 | 0.13658789 |
| Lama5 | 1 | 0.42253204 | 0.01 | 0.01 | 1 | 0.07085741 | -1.2103196 | 0.05 | 0.104 | 1 | 1 | 0.13669405 |
| Abhd17b | 0.1270948 | 0.38379964 | 0.5 | 0.409 | 1 | 0.07086506 | -0.5145173 | 0.567 | 0.667 | 1 | 0.1270948 | 0.13670826 |
| Ttc6 | 0.32025586 | -0.577468 | 0 | 0.01 | 1 | 0.07095091 | -2.1578522 | 0.017 | 0.052 | 1 | 0.32025586 | 0.13686779 |
| Lonrf2 | 0.45476301 | 1.22988696 | 0.042 | 0.026 | 1 | 0.07095091 | -1.9879272 | 0.017 | 0.052 | 1 | 0.45476301 | 0.13686779 |
| Spata21 | 0.52528691 | -0.7998604 | 0.01 | 0.021 | 1 | 0.07095091 | -1.7952821 | 0.017 | 0.052 | 1 | 0.52528691 | 0.13686779 |
| Gm44759 | 0.74827573 | 0.59245704 | 0.021 | 0.016 | 1 | 0.07095091 | -1.7952821 | 0.017 | 0.052 | 1 | 0.74827573 | 0.13686779 |
| Btbd19 | 0.81708725 | -0.3144336 | 0.031 | 0.036 | 1 | 0.07095091 | -1.7952821 | 0.017 | 0.052 | 1 | 0.81708725 | 0.13686779 |
| Gm53065 | 0.07100615 | 1.20013961 | 0.062 | 0.021 | 1 | 0.5493805 | 0.13071733 | 0.046 | 0.031 | 1 | 0.5493805 | 0.13697043 |
| Pcyox1l | 0.17327047 | 1.00749454 | 0.073 | 0.036 | 1 | 0.07101945 | 1.35310975 | 0.071 | 0.021 | 1 | 0.17327047 | 0.13699513 |
| Tnfrsf18 | 0.13572533 | 1.29700115 | 0.083 | 0.041 | 1 | 0.07106948 | 0.27510724 | 0.105 | 0.042 | 1 | 0.13572533 | 0.13708808 |
| Fabp5 | 0.99793929 | 0.22288357 | 0.365 | 0.383 | 1 | 0.07112742 | 0.36821664 | 0.361 | 0.271 | 1 | 0.99793929 | 0.13719573 |
| Mansc1 | 0.07117776 | -1.3144336 | 0.021 | 0.073 | 1 | 0.37884342 | -0.3098553 | 0.055 | 0.031 | 1 | 0.37884342 | 0.13728925 |
| Tlcd1 | 0.78720646 | -0.4485702 | 0.354 | 0.347 | 1 | 0.07118449 | 0.54812573 | 0.269 | 0.177 | 1 | 0.78720646 | 0.13730174 |
| Atr | 0.57511673 | 0.12880083 | 0.198 | 0.171 | 1 | 0.0712223 | 0.57648796 | 0.265 | 0.167 | 1 | 0.57511673 | 0.13737198 |
| Taco1 | 0.07126488 | 1.42253204 | 0.094 | 0.041 | 1 | 0.207422 | 0.53814164 | 0.067 | 0.031 | 1 | 0.207422 | 0.13745108 |
| Cmc2 | 0.52350869 | -0.2488452 | 0.156 | 0.187 | 1 | 0.0712938 | 0.97025266 | 0.118 | 0.052 | 1 | 0.52350869 | 0.1375048 |
| Gipc1 | 0.07135874 | 0.24195979 | 0.323 | 0.207 | 1 | 0.70437546 | -0.1662644 | 0.307 | 0.333 | 1 | 0.70437546 | 0.13762542 |
| Ppox | 0.41827871 | 0.28760246 | 0.167 | 0.13 | 1 | 0.07146605 | 0.74903843 | 0.172 | 0.094 | 1 | 0.41827871 | 0.13782471 |
| Zfp1 | 0.53368243 | -0.2742764 | 0.406 | 0.415 | 1 | 0.07154075 | 0.10195834 | 0.395 | 0.26 | 1 | 0.53368243 | 0.13796342 |
| Gm49692 | 0.27195804 | -0.9185049 | 0.073 | 0.114 | 1 | 0.07156386 | 0.80562195 | 0.088 | 0.031 | 1 | 0.27195804 | 0.13800633 |
| Zfp217 | 0.95726896 | 0.15388402 | 0.333 | 0.342 | 1 | 0.0715952 | 0.40755753 | 0.454 | 0.354 | 1 | 0.95726896 | 0.13806452 |
| Gpatch3 | 0.30584838 | 1.00749454 | 0.062 | 0.036 | 1 | 0.07161994 | 1.01207283 | 0.055 | 0.01 | 1 | 0.30584838 | 0.13811047 |
| Pkia | 0.79846236 | 0.15949763 | 0.021 | 0.026 | 1 | 0.071637 | -1.4167705 | 0.029 | 0.073 | 1 | 0.79846236 | 0.13814214 |
| Ip6k2 | 0.24846762 | 0.42994951 | 0.365 | 0.306 | 1 | 0.0716818 | -0.9406215 | 0.395 | 0.458 | 1 | 0.24846762 | 0.13822533 |
| Kif22 | 0.07169074 | 3.00749454 | 0.052 | 0.016 | 1 | 0.87851725 | -0.8948178 | 0.013 | 0.01 | 1 | 0.87851725 | 0.13824193 |
| Gm56812 | 0.07169274 | 0.65957123 | 0.104 | 0.047 | 1 | 0.22699337 | 0.53814164 | 0.105 | 0.062 | 1 | 0.22699337 | 0.13824563 |
| Zfp119b | 0.07169293 | 0.46692616 | 0.104 | 0.047 | 1 | 0.20997488 | -1.3098553 | 0.025 | 0.052 | 1 | 0.20997488 | 0.13824599 |
| Trpm4 | 0.19503551 | -1.1624305 | 0.062 | 0.109 | 1 | 0.07183049 | -1.3664388 | 0.092 | 0.156 | 1 | 0.19503551 | 0.13850136 |
| Bpifc | 0.76363769 | -0.2148979 | 0.188 | 0.171 | 1 | 0.07191235 | 1.08246216 | 0.055 | 0.01 | 1 | 0.76363769 | 0.13865331 |
| Slc25a23 | 0.38521705 | -0.7998604 | 0.01 | 0.026 | 1 | 0.07196065 | 0.95317914 | 0.088 | 0.031 | 1 | 0.38521705 | 0.13874297 |
| A330048O09Rik | 0.07218619 | -1.3144336 | 0.031 | 0.088 | 1 | 0.54837489 | -0.4618584 | 0.034 | 0.021 | 1 | 0.54837489 | 0.13916154 |
| Pcmtd1 | 0.07220466 | 0.32670654 | 0.812 | 0.674 | 1 | 0.99543606 | -0.2035464 | 0.807 | 0.854 | 1 | 0.99543606 | 0.1391958 |
| Htatip2 | 0.66660752 | -0.1624305 | 0.25 | 0.269 | 1 | 0.07220638 | 0.69014474 | 0.147 | 0.073 | 1 | 0.66660752 | 0.13919899 |
| Cacnb4 | 0.07232615 | -1.6169963 | 0.031 | 0.088 | 1 | 0.60412275 | -1.8334172 | 0.08 | 0.094 | 1 | 0.60412275 | 0.13942123 |
| Rbm6 | 0.07234958 | 0.23437994 | 0.844 | 0.803 | 1 | 0.11697539 | -0.4750423 | 0.916 | 0.948 | 1 | 0.11697539 | 0.13946469 |
| Trim12c | 0.5955065 | 0.27052894 | 0.167 | 0.145 | 1 | 0.07235057 | 0.27510724 | 0.134 | 0.062 | 1 | 0.5955065 | 0.13946654 |

| | | | | | | | | | | | | |
|---|---|---|---|---|---|---|---|---|---|---|---|---|
| Spindoc | 0.07235349 | -0.4567865 | 0.333 | 0.43 | 1 | 0.66300544 | -0.4167705 | 0.542 | 0.542 | 1 | 0.66300544 | 0.13947195 |
| P2ry1 | 0.20691434 | -1.3144336 | 0.01 | 0.036 | 1 | 0.07236555 | -1.8948178 | 0.017 | 0.052 | 1 | 0.20691434 | 0.13949433 |
| Ints10 | 0.94980451 | -0.1217885 | 0.146 | 0.14 | 1 | 0.07237882 | 0.53144699 | 0.147 | 0.073 | 1 | 0.94980451 | 0.13951894 |
| Tcf19 | 0.0856767 | 1.71798792 | 0.083 | 0.036 | 1 | 0.0723819 | -1.5728897 | 0.017 | 0.052 | 1 | 0.0856767 | 0.13952467 |
| Gm50321 | 0.52528691 | -0.7998604 | 0.01 | 0.021 | 1 | 0.0723819 | -1.5728897 | 0.017 | 0.052 | 1 | 0.52528691 | 0.13952467 |
| Mbnl3 | 0.12804036 | 2.00749454 | 0.083 | 0.041 | 1 | 0.0724531 | 0.59703533 | 0.055 | 0.01 | 1 | 0.12804036 | 0.13965676 |
| Rasl11b | 0.11432369 | -1.4596315 | 0.125 | 0.192 | 1 | 0.07245561 | -1.0608277 | 0.16 | 0.229 | 1 | 0.11432369 | 0.1396614 |
| Gm17494 | 0.07248493 | 1.00749454 | 0.062 | 0.021 | 1 | 0.57370068 | -1.3098553 | 0.021 | 0.031 | 1 | 0.57370068 | 0.1397158 |
| Eldr | 0.07250193 | 1.00749454 | 0.062 | 0.021 | 1 | 0.34704016 | -1.3098553 | 0.008 | 0.021 | 1 | 0.34704016 | 0.13974734 |
| Ezh1 | 0.35644373 | 0.26465238 | 0.354 | 0.295 | 1 | 0.07258452 | 0.41062221 | 0.391 | 0.271 | 1 | 0.35644373 | 0.13990052 |
| H2-M3 | 0.42126413 | 0.35541784 | 0.125 | 0.093 | 1 | 0.07259139 | 1.01207283 | 0.13 | 0.062 | 1 | 0.42126413 | 0.13991327 |
| Zfp961 | 0.31164318 | 0.32942263 | 0.083 | 0.052 | 1 | 0.07261441 | 1.32241295 | 0.071 | 0.021 | 1 | 0.31164318 | 0.13995598 |
| Ddx60 | 0.07265011 | -0.7767768 | 0.021 | 0.073 | 1 | 0.2386824 | -2.6724253 | 0.013 | 0.031 | 1 | 0.2386824 | 0.14002218 |
| Usp14 | 0.07265788 | 0.18395969 | 0.729 | 0.642 | 1 | 0.77773976 | -0.1043064 | 0.723 | 0.688 | 1 | 0.77773976 | 0.14003659 |
| Bglap2 | 0.07270071 | -2.7998604 | 0.083 | 0.155 | 1 | 0.85368393 | 1.76543286 | 0.046 | 0.052 | 1 | 0.85368393 | 0.14011602 |
| Snx14 | 0.07270724 | 0.13076591 | 0.562 | 0.456 | 1 | 0.72068647 | -0.3696885 | 0.588 | 0.604 | 1 | 0.72068647 | 0.14012814 |
| Nup133 | 0.56685686 | 0.19191911 | 0.177 | 0.15 | 1 | 0.0727279 | 1.19915838 | 0.189 | 0.115 | 1 | 0.56685686 | 0.14016645 |
| Kptn | 0.13102699 | 0.15505172 | 0.281 | 0.192 | 1 | 0.07282322 | -0.44958 | 0.324 | 0.208 | 1 | 0.13102699 | 0.14034321 |
| Riok1 | 0.07288047 | 0.46413294 | 0.604 | 0.523 | 1 | 0.5116324 | -0.3042742 | 0.584 | 0.573 | 1 | 0.5116324 | 0.14044938 |
| Socs3 | 0.64932287 | 0.10153803 | 0.562 | 0.508 | 1 | 0.07289952 | -0.5675808 | 0.483 | 0.615 | 1 | 0.64932287 | 0.1404847 |
| Atm | 0.71186554 | -0.1118044 | 0.219 | 0.192 | 1 | 0.07292513 | 0.39796399 | 0.269 | 0.167 | 1 | 0.71186554 | 0.14053218 |
| B130024G19Rik | 0.07293652 | 2.46692616 | 0.052 | 0.016 | 1 | 0.11764285 | 1.69014474 | 0.025 | 0 | 1 | 0.11764285 | 0.14055331 |
| Tifa | 0.19165086 | -0.5450465 | 0.156 | 0.218 | 1 | 0.07322118 | 0.51796376 | 0.21 | 0.125 | 1 | 0.19165086 | 0.14108102 |
| Map3k11 | 0.07330044 | 0.83756954 | 0.146 | 0.078 | 1 | 0.56731623 | -0.3390016 | 0.181 | 0.146 | 1 | 0.56731623 | 0.14122792 |
| 5730471H19Rik | 0.07332067 | 0.87625 | 0.167 | 0.093 | 1 | 0.2694899 | -0.3098553 | 0.16 | 0.104 | 1 | 0.2694899 | 0.14126543 |
| Gask1b | 0.80496186 | 1.00749454 | 0.021 | 0.026 | 1 | 0.07332267 | 1.75315453 | 0.143 | 0.073 | 1 | 0.80496186 | 0.14126912 |
| Cenpe | 0.07337238 | 2.31234912 | 0.052 | 0.016 | 1 | 0.80020906 | -0.7248928 | 0.017 | 0.021 | 1 | 0.80020906 | 0.14136125 |
| H3c3 | 0.07337238 | 2.69199271 | 0.052 | 0.016 | 1 | 0.11689951 | -2.3098553 | 0 | 0.01 | 1 | 0.11689951 | 0.14136125 |
| Chrd | 0.07341132 | 0.80104366 | 0.104 | 0.047 | 1 | 0.2277352 | 0.45567948 | 0.118 | 0.073 | 1 | 0.2277352 | 0.14143342 |
| 6330418K02Rik | 0.07356948 | 1.42253204 | 0.094 | 0.041 | 1 | 0.11129992 | 0.76053407 | 0.08 | 0.031 | 1 | 0.11129992 | 0.1417265 |
| Dhx37 | 0.47608023 | 0.35541784 | 0.115 | 0.088 | 1 | 0.07359773 | 0.30905457 | 0.134 | 0.062 | 1 | 0.47608023 | 0.14177883 |
| Txndc2 | 0.07363413 | 0.83756954 | 0.062 | 0.021 | 1 | 0.2518463 | 0.53814164 | 0.063 | 0.031 | 1 | 0.2518463 | 0.14184628 |
| Zfand4 | 0.07365223 | 1.35541784 | 0.094 | 0.041 | 1 | 0.12042191 | -0.7793405 | 0.034 | 0.073 | 1 | 0.12042191 | 0.14187981 |
| Bbs5 | 0.07368807 | 0.44806713 | 0.177 | 0.098 | 1 | 0.50359742 | 0.51326698 | 0.139 | 0.115 | 1 | 0.50359742 | 0.14194621 |
| Gm48678 | 0.98603885 | -0.2148979 | 0.052 | 0.052 | 1 | 0.07372548 | 0.32241295 | 0.118 | 0.052 | 1 | 0.98603885 | 0.14201551 |
| Dffb | 0.28535634 | -0.3144336 | 0.052 | 0.088 | 1 | 0.0738297 | 1.01207283 | 0.088 | 0.031 | 1 | 0.28535634 | 0.14220858 |
| Umps | 0.0738348 | 0.76648644 | 0.167 | 0.093 | 1 | 0.08132272 | 0.52968007 | 0.21 | 0.125 | 1 | 0.08132272 | 0.14221803 |
| Gripap1 | 0.07384774 | 0.36113149 | 0.583 | 0.497 | 1 | 0.85094192 | -0.2452023 | 0.601 | 0.625 | 1 | 0.85094192 | 0.14224199 |
| Frmd4b | 0.8874389 | 0.10775866 | 0.677 | 0.653 | 1 | 0.07386607 | -0.764496 | 0.672 | 0.719 | 1 | 0.8874389 | 0.14227595 |
| Niban3 | 0.07391189 | 3.32942263 | 0.031 | 0.005 | 1 | 0.17708318 | -1.9879272 | 0.017 | 0.042 | 1 | 0.17708318 | 0.14236082 |
| Atp6v0e2 | 0.0739968 | -1.1624305 | 0.031 | 0.088 | 1 | 0.40076941 | -1.3098553 | 0.017 | 0.031 | 1 | 0.40076941 | 0.14251807 |
| Tbrg4 | 0.14957389 | 0.49292136 | 0.25 | 0.176 | 1 | 0.0740262 | 0.21739174 | 0.164 | 0.083 | 1 | 0.14957389 | 0.14257252 |
| Cdk7 | 0.0740556 | 0.26259112 | 0.667 | 0.57 | 1 | 0.09440011 | 0.10002053 | 0.735 | 0.562 | 1 | 0.09440011 | 0.14262696 |
| Hpse | 0.07419689 | 2.00749454 | 0.052 | 0.016 | 1 | 0.5859632 | -0.3098553 | 0.013 | 0.021 | 1 | 0.5859632 | 0.14288861 |
| Gm56848 | 0.07419689 | 1.81484946 | 0.052 | 0.016 | 1 | 0.31280025 | 0.27510724 | 0.029 | 0.01 | 1 | 0.31280025 | 0.14288861 |
| Cfap91 | 0.07419689 | 2.00749454 | 0.052 | 0.016 | 1 | 0.15394118 | 1.49749966 | 0.021 | 0 | 1 | 0.15394118 | 0.14288861 |
| Cyp4f13 | 0.07421271 | -0.6623569 | 0.083 | 0.161 | 1 | 0.49217868 | -1.1234421 | 0.113 | 0.135 | 1 | 0.49217868 | 0.1429179 |
| Birc3 | 0.07424876 | 0.1533454 | 0.323 | 0.212 | 1 | 0.92806011 | -0.7523735 | 0.252 | 0.219 | 1 | 0.92806011 | 0.14298464 |
| Vps26c | 0.2167444 | 0.66354014 | 0.198 | 0.145 | 1 | 0.07432434 | -0.2394659 | 0.294 | 0.177 | 1 | 0.2167444 | 0.14312458 |
| Moxd1 | 0.50525915 | 0.10749033 | 0.396 | 0.337 | 1 | 0.07444517 | 0.42132798 | 0.277 | 0.177 | 1 | 0.50525915 | 0.14334826 |
| Cilk1 | 0.07446115 | 0.53105649 | 0.521 | 0.42 | 1 | 0.42681373 | -0.1239887 | 0.613 | 0.5 | 1 | 0.42681373 | 0.14337784 |
| ENSMUSG000001213 46 | 0.07450785 | 2.32942263 | 0.031 | 0.005 | 1 | 0.51194506 | 0.27510724 | 0.021 | 0.01 | 1 | 0.51194506 | 0.14346428 |
| 4933427D14Rik | 0.07450785 | 2.32942263 | 0.031 | 0.005 | 1 | 0.30606883 | 0.69014474 | 0.029 | 0.01 | 1 | 0.30606883 | 0.14346428 |
| 9430024F10Rik | 0.07450785 | 2.32942263 | 0.031 | 0.005 | 1 | 0.57576763 | -1.0468209 | 0.013 | 0.021 | 1 | 0.57576763 | 0.14346428 |
| H3c1 | 0.07450785 | 2.32942263 | 0.031 | 0.005 | 1 | 0.27154471 | 0.69014474 | 0.013 | 0 | 1 | 0.27154471 | 0.14346428 |
| Taf9 | 0.07450785 | 2.32942263 | 0.031 | 0.005 | 1 | 0.53433287 | 0.42711033 | 0.034 | 0.021 | 1 | 0.53433287 | 0.14346428 |
| Depdc1b | 0.07450785 | 3.00749454 | 0.031 | 0.005 | 1 | 0.27154471 | 0.69014474 | 0.013 | 0 | 1 | 0.27154471 | 0.14346428 |
| Gm20939 | 0.07450785 | 2.32942263 | 0.031 | 0.005 | 1 | 0.30606883 | 0.69014474 | 0.029 | 0.01 | 1 | 0.30606883 | 0.14346428 |
| Ncor1 | 0.07451317 | 0.22109001 | 0.99 | 0.984 | 1 | 0.5388419 | -0.1372281 | 1 | 1 | 1 | 0.5388419 | 0.14347413 |
| Sema4d | 0.22331513 | -0.7294711 | 0.104 | 0.155 | 1 | 0.0745635 | -0.3749503 | 0.143 | 0.229 | 1 | 0.22331513 | 0.14356729 |
| Mitd1 | 0.4246061 | 0.24393473 | 0.292 | 0.254 | 1 | 0.07457195 | 0.4767411 | 0.214 | 0.125 | 1 | 0.4246061 | 0.14358292 |
| Necab3 | 0.07463974 | 2.38600616 | 0.052 | 0.016 | 1 | 0.79600331 | -0.9879272 | 0.017 | 0.021 | 1 | 0.79600331 | 0.14370839 |
| Tnrc6a | 0.13847519 | -0.2543431 | 0.667 | 0.731 | 1 | 0.07466051 | -0.4490679 | 0.958 | 0.979 | 1 | 0.13847519 | 0.14374683 |
| Gm5577 | 0.83566413 | -0.577468 | 0.052 | 0.057 | 1 | 0.07474274 | -2.0468209 | 0.038 | 0.083 | 1 | 0.83566413 | 0.14389901 |
| Ccdc136 | 0.07477803 | 0.9433642 | 0.156 | 0.088 | 1 | 0.11085303 | 0.42132798 | 0.256 | 0.167 | 1 | 0.11085303 | 0.1439643 |
| Otud4 | 0.07478299 | 0.36916881 | 0.604 | 0.503 | 1 | 0.34771603 | -0.2728083 | 0.739 | 0.771 | 1 | 0.34771603 | 0.14397348 |
| Bicral | 0.19537092 | 0.30323311 | 0.583 | 0.508 | 1 | 0.07479294 | 0.10679149 | 0.63 | 0.458 | 1 | 0.19537092 | 0.14399191 |
| Apeh | 0.07490845 | 1.00749454 | 0.156 | 0.088 | 1 | 0.76843821 | -0.2273931 | 0.134 | 0.146 | 1 | 0.76843821 | 0.14420562 |
| Cacna1d | 0.62894915 | -0.5564063 | 0.135 | 0.155 | 1 | 0.07502574 | -0.9403324 | 0.345 | 0.396 | 1 | 0.62894915 | 0.14442261 |
| Tamm41 | 0.07505438 | 0.80104366 | 0.125 | 0.062 | 1 | 0.33144774 | 0.19264508 | 0.122 | 0.083 | 1 | 0.33144774 | 0.1444756 |
| Tsen2 | 0.07505438 | 0.70793425 | 0.125 | 0.062 | 1 | 0.07663795 | 0.64950275 | 0.118 | 0.052 | 1 | 0.07663795 | 0.1444756 |
| Gm17655 | 0.07506945 | 1.68556644 | 0.052 | 0.016 | 1 | 0.27154327 | 1.01207283 | 0.013 | 0 | 1 | 0.27154327 | 0.14450348 |
| Gm15287 | 0.07506945 | 1.68556644 | 0.052 | 0.016 | 1 | 0.56565662 | -3.1172102 | 0.013 | 0.021 | 1 | 0.56565662 | 0.14450348 |
| Dlgap5 | 0.07511305 | 2.59245704 | 0.031 | 0.005 | 1 | 0.20315163 | 1.01207283 | 0.017 | 0 | 1 | 0.20315163 | 0.14458412 |
| Gm15852 | 0.07511393 | 2.00749454 | 0.031 | 0.005 | 1 | 0.67474842 | -0.3098553 | 0.029 | 0.021 | 1 | 0.67474842 | 0.14458576 |
| Fcgr3 | 0.07511393 | 2.00749454 | 0.031 | 0.005 | 1 | 0.15394118 | 2.14957636 | 0.021 | 0 | 1 | 0.15394118 | 0.14458576 |
| 1700016C15Rik | 0.07511393 | 2.00749454 | 0.031 | 0.005 | 1 | 0.27154327 | 1.01207283 | 0.013 | 0 | 1 | 0.27154327 | 0.14458576 |
| Nsl1 | 0.07511393 | 2.00749454 | 0.031 | 0.005 | 1 | 0.27154327 | 1.01207283 | 0.013 | 0 | 1 | 0.27154327 | 0.14458576 |
| Gm13421 | 0.07511393 | 2.00749454 | 0.031 | 0.005 | 1 | 0.24413474 | -2.3098553 | 0.013 | 0.031 | 1 | 0.24413474 | 0.14458576 |
| Frmd5 | 0.07511393 | 2.00749454 | 0.031 | 0.005 | 1 | 0.87851834 | -2.1172102 | 0.013 | 0.01 | 1 | 0.87851834 | 0.14458576 |
| Lor | 0.07511393 | 2.00749454 | 0.031 | 0.005 | 1 | 0.27759424 | 0.80562195 | 0.046 | 0.021 | 1 | 0.27759424 | 0.14458576 |
| Gm12473 | 0.07511393 | 2.00749454 | 0.031 | 0.005 | 1 | 0.26467561 | -0.4618584 | 0.063 | 0.031 | 1 | 0.26467561 | 0.14458576 |
| Gm12503 | 0.07511393 | 2.00749454 | 0.031 | 0.005 | 1 | 0.5818708 | -0.5728897 | 0.013 | 0.021 | 1 | 0.5818708 | 0.14458576 |
| Gm57291 | 0.07511393 | 2.00749454 | 0.031 | 0.005 | 1 | 0.52072676 | -0.7248928 | 0.021 | 0.01 | 1 | 0.52072676 | 0.14458576 |

| | | | | | | | | | | | | |
|---|---|---|---|---|---|---|---|---|---|---|---|---|
| C3ar1 | 0.07511393 | 2.00749454 | 0.031 | 0.005 | 1 | 0.20314753 | 1.27510724 | 0.017 | 0 | 1 | 0.20314753 | 0.14458576 |
| Clec4a1 | 0.07511393 | 2.00749454 | 0.031 | 0.005 | 1 | 0.34363875 | -1.7248928 | 0.008 | 0.021 | 1 | 0.34363875 | 0.14458576 |
| Ttll13 | 0.07511393 | 2.00749454 | 0.031 | 0.005 | 1 | 0.36858645 | -0.1723517 | 0.042 | 0.021 | 1 | 0.36858645 | 0.14458576 |
| Nmb | 0.07511393 | 2.00749454 | 0.031 | 0.005 | 1 | 0.77529594 | 0.45567948 | 0.063 | 0.073 | 1 | 0.77529594 | 0.14458576 |
| Gm31816 | 0.07511393 | 2.00749454 | 0.031 | 0.005 | 1 | 0.14544666 | -2.6317834 | 0.004 | 0.021 | 1 | 0.14544666 | 0.14458576 |
| Gm32742 | 0.07511393 | 2.00749454 | 0.031 | 0.005 | 1 | 0.1465102 | -1.8948178 | 0.004 | 0.021 | 1 | 0.1465102 | 0.14458576 |
| Gm56948 | 0.07511393 | 2.00749454 | 0.031 | 0.005 | 1 | 0.23178796 | 0.39058446 | 0.05 | 0.021 | 1 | 0.23178796 | 0.14458576 |
| 1700085D07Rik | 0.07511393 | 2.00749454 | 0.031 | 0.005 | 1 | 0.99280559 | -0.194378 | 0.042 | 0.042 | 1 | 0.99280559 | 0.14458576 |
| Panx2 | 0.07511393 | 2.00749454 | 0.031 | 0.005 | 1 | 0.27935024 | 0.69014474 | 0.046 | 0.021 | 1 | 0.27935024 | 0.14458576 |
| 4930483P17Rik | 0.07511393 | 2.00749454 | 0.031 | 0.005 | 1 | 0.86312513 | -0.7248928 | 0.008 | 0.01 | 1 | 0.86312513 | 0.14458576 |
| Prr18 | 0.07511393 | 2.00749454 | 0.031 | 0.005 | 1 | 0.94026906 | -0.9472852 | 0.034 | 0.031 | 1 | 0.94026906 | 0.14458576 |
| Pdxk-ps | 0.07511393 | 2.00749454 | 0.031 | 0.005 | 1 | 0.34363875 | -1.7248928 | 0.008 | 0.021 | 1 | 0.34363875 | 0.14458576 |
| Dnph1 | 0.07511393 | 2.00749454 | 0.031 | 0.005 | 1 | 0.11689951 | -2.3098553 | 0 | 0.01 | 1 | 0.11689951 | 0.14458576 |
| Gm41604 | 0.07511393 | 2.00749454 | 0.031 | 0.005 | 1 | 0.21450688 | -1.6577786 | 0.042 | 0.073 | 1 | 0.21450688 | 0.14458576 |
| Gm11110 | 0.07511393 | 2.00749454 | 0.031 | 0.005 | 1 | 0.57170879 | -1.6317834 | 0.013 | 0.021 | 1 | 0.57170879 | 0.14458576 |
| Abcg5 | 0.07511393 | 2.00749454 | 0.031 | 0.005 | 1 | 0.14544666 | -2.3098553 | 0.004 | 0.021 | 1 | 0.14544666 | 0.14458576 |
| Lrrtm2 | 0.07511393 | 2.00749454 | 0.031 | 0.005 | 1 | 0.15394868 | 1.27510724 | 0.021 | 0 | 1 | 0.15394868 | 0.14458576 |
| Chd4 | 0.07517919 | 0.14715484 | 0.958 | 0.902 | 1 | 0.82724954 | -0.2128139 | 0.992 | 0.99 | 1 | 0.82724954 | 0.14470647 |
| S1pr3 | 0.2151309 | -1.3910548 | 0.094 | 0.14 | 1 | 0.07525669 | -1.6317834 | 0.017 | 0.052 | 1 | 0.2151309 | 0.14484982 |
| Ghdc | 0.5055902 | -0.1801325 | 0.125 | 0.155 | 1 | 0.07533219 | 0.24814019 | 0.181 | 0.094 | 1 | 0.5055902 | 0.14498945 |
| Tgfbrap1 | 0.60537799 | 0.21394541 | 0.135 | 0.114 | 1 | 0.0753734 | 0.55074168 | 0.223 | 0.135 | 1 | 0.60537799 | 0.14506565 |
| Stam2 | 0.07541236 | 0.31819538 | 0.479 | 0.368 | 1 | 0.81387506 | -0.1172102 | 0.576 | 0.573 | 1 | 0.81387506 | 0.14513769 |
| Ctc1 | 0.15070685 | 0.80104366 | 0.104 | 0.057 | 1 | 0.07542533 | 0.54812573 | 0.088 | 0.031 | 1 | 0.15070685 | 0.14516168 |
| Acsl5 | 0.54490418 | 0.54354744 | 0.177 | 0.15 | 1 | 0.07545632 | 0.11237774 | 0.218 | 0.125 | 1 | 0.54490418 | 0.14521898 |
| Lrfn3 | 0.07550972 | 1.59245704 | 0.052 | 0.016 | 1 | 0.17858821 | 0.91253716 | 0.055 | 0.021 | 1 | 0.17858821 | 0.14531773 |
| Gm56727 | 0.07550972 | 1.59245704 | 0.052 | 0.016 | 1 | 0.27154471 | 0.69014474 | 0.013 | 0 | 1 | 0.27154471 | 0.14531773 |
| Gm26724 | 0.0776711 | 1.27052894 | 0.052 | 0.016 | 1 | 0.07554679 | 0.61614416 | 0.071 | 0.021 | 1 | 0.0776711 | 0.14538627 |
| Gm16845 | 0.29653056 | 0.65957123 | 0.083 | 0.052 | 1 | 0.07554679 | -0.2318528 | 0.071 | 0.021 | 1 | 0.29653056 | 0.14538627 |
| AU023762 | 0.07571957 | 2.00749454 | 0.031 | 0.005 | 1 | 0.50887612 | -1.3098553 | 0.004 | 0.01 | 1 | 0.50887612 | 0.14570569 |
| Ltb | 0.07571957 | 3.12297175 | 0.031 | 0.005 | 1 | 0.87066325 | -0.3098553 | 0.013 | 0.01 | 1 | 0.87066325 | 0.14570569 |
| Ttk | 0.07572046 | 2.77302928 | 0.031 | 0.005 | 1 | 0.50887612 | -1.3098553 | 0.004 | 0.01 | 1 | 0.50887612 | 0.14570733 |
| AI480526 | 0.07579117 | 0.70583884 | 0.271 | 0.181 | 1 | 0.77848185 | -0.5840302 | 0.261 | 0.25 | 1 | 0.77848185 | 0.14583804 |
| Gm57093 | 0.28929471 | 0.87625 | 0.125 | 0.088 | 1 | 0.07584609 | 1.30157945 | 0.13 | 0.062 | 1 | 0.28929471 | 0.14593955 |
| Nsmf | 0.07588669 | 0.48393258 | 0.385 | 0.285 | 1 | 0.27433651 | -0.2433599 | 0.357 | 0.271 | 1 | 0.27433651 | 0.1460146 |
| Mafg | 0.08571015 | 0.2614696 | 0.823 | 0.788 | 1 | 0.07593266 | 0.17306954 | 0.954 | 0.875 | 1 | 0.08571015 | 0.14609956 |
| Sfmbt2 | 0.0759666 | -0.2148979 | 0.104 | 0.047 | 1 | 0.23825987 | 0.10518224 | 0.16 | 0.104 | 1 | 0.23825987 | 0.14616228 |
| Heatr6 | 0.43094234 | 0.33764314 | 0.292 | 0.254 | 1 | 0.07599617 | 0.46775232 | 0.294 | 0.188 | 1 | 0.43094234 | 0.14621692 |
| Gm8883 | 0.07603075 | 1.22988696 | 0.094 | 0.041 | 1 | 0.14243143 | 1.01207283 | 0.059 | 0.021 | 1 | 0.14243143 | 0.14628083 |
| Alyref2 | 0.15954569 | 1.00749454 | 0.115 | 0.067 | 1 | 0.07608034 | 0.74903843 | 0.088 | 0.031 | 1 | 0.15954569 | 0.14637247 |
| Stat2 | 0.07629637 | -0.6914242 | 0.385 | 0.477 | 1 | 0.12363027 | -1.2995133 | 0.29 | 0.323 | 1 | 0.12363027 | 0.1467716 |
| Adamts10 | 0.07630584 | 0.37006462 | 0.469 | 0.368 | 1 | 0.51760592 | -0.129283 | 0.563 | 0.458 | 1 | 0.51760592 | 0.1467891 |
| 2900052L18Rik | 0.07634515 | 1.39451766 | 0.115 | 0.057 | 1 | 0.48091084 | -1.0619277 | 0.063 | 0.083 | 1 | 0.48091084 | 0.14686172 |
| BB218582 | 0.07636256 | 1.49292136 | 0.052 | 0.016 | 1 | 0.80442048 | -0.5728897 | 0.017 | 0.021 | 1 | 0.80442048 | 0.14689388 |
| Hdac11 | 0.07636256 | 1.49292136 | 0.052 | 0.016 | 1 | 0.14458909 | 0.42711033 | 0.076 | 0.031 | 1 | 0.14458909 | 0.14689388 |
| Gm15910 | 0.07636256 | 1.49292136 | 0.052 | 0.016 | 1 | 0.66339728 | 0.10518224 | 0.029 | 0.021 | 1 | 0.66339728 | 0.14689388 |
| Tedc1 | 0.07636256 | 1.49292136 | 0.052 | 0.016 | 1 | 0.56793117 | -0.7248928 | 0.029 | 0.042 | 1 | 0.56793117 | 0.14689388 |
| Krt76 | 0.07636256 | 1.49292136 | 0.052 | 0.016 | 1 | 0.20314753 | 1.49749966 | 0.017 | 0 | 1 | 0.20314753 | 0.14689388 |
| Rasgrp2 | 0.07636256 | 1.49292136 | 0.052 | 0.016 | 1 | 0.84175461 | -0.1082214 | 0.046 | 0.052 | 1 | 0.84175461 | 0.14689388 |
| Plp1 | 0.07636256 | 1.49292136 | 0.052 | 0.016 | 1 | 0.11765402 | 1.49749966 | 0.025 | 0 | 1 | 0.11765402 | 0.14689388 |
| Letmd1 | 0.76422355 | -0.2148979 | 0.083 | 0.093 | 1 | 0.07637856 | 0.86006974 | 0.13 | 0.062 | 1 | 0.76422355 | 0.14692344 |
| Pop4 | 0.58565039 | 0.18152394 | 0.354 | 0.326 | 1 | 0.07639842 | 0.29348577 | 0.269 | 0.167 | 1 | 0.58565039 | 0.14696013 |
| Trmt10a | 0.38317436 | 0.31943854 | 0.281 | 0.233 | 1 | 0.07644317 | 0.3575694 | 0.189 | 0.104 | 1 | 0.38317436 | 0.14704279 |
| B3galt1 | 0.07651883 | -0.6860405 | 0.406 | 0.492 | 1 | 0.20091264 | -0.5245941 | 0.336 | 0.385 | 1 | 0.20091264 | 0.14718253 |
| Cage1 | 0.07659567 | 1.00749454 | 0.094 | 0.041 | 1 | 0.37342246 | 0.36821664 | 0.055 | 0.031 | 1 | 0.37342246 | 0.14732444 |
| Sms | 0.14353086 | 0.18646468 | 0.427 | 0.326 | 1 | 0.07670157 | -0.3664388 | 0.349 | 0.479 | 1 | 0.14353086 | 0.14752 |
| Gm33302 | 0.69158293 | 0.38600616 | 0.031 | 0.041 | 1 | 0.07672624 | 2.5580412 | 0.084 | 0.031 | 1 | 0.69158293 | 0.14756557 |
| Cntrl | 0.43579768 | 0.3951556 | 0.438 | 0.399 | 1 | 0.07673424 | -0.7389679 | 0.391 | 0.5 | 1 | 0.43579768 | 0.14758033 |
| Zfp639 | 0.66969747 | -0.2019588 | 0.26 | 0.285 | 1 | 0.07673475 | 0.31574922 | 0.324 | 0.219 | 1 | 0.66969747 | 0.14758127 |
| Vps28 | 0.41652719 | 0.10534186 | 0.719 | 0.658 | 1 | 0.07673654 | 0.22681168 | 0.601 | 0.542 | 1 | 0.41652719 | 0.14758459 |
| Surf6 | 0.19885672 | 0.33506919 | 0.271 | 0.202 | 1 | 0.07680021 | -0.3446207 | 0.298 | 0.177 | 1 | 0.19885672 | 0.14770215 |
| Idua | 0.13520581 | 0.75956702 | 0.146 | 0.088 | 1 | 0.07681848 | 0.69014474 | 0.172 | 0.094 | 1 | 0.13520581 | 0.14773589 |
| Lrrc14 | 0.45747574 | 0.19035859 | 0.188 | 0.15 | 1 | 0.07687291 | 0.8917786 | 0.155 | 0.083 | 1 | 0.45747574 | 0.14783637 |
| A930005H10Rik | 0.07688324 | 1.07788386 | 0.167 | 0.098 | 1 | 0.65077572 | -0.1630139 | 0.101 | 0.083 | 1 | 0.65077572 | 0.14785545 |
| Gm20536 | 0.07691413 | -0.9925055 | 0.031 | 0.088 | 1 | 0.24392267 | 0.77760758 | 0.063 | 0.031 | 1 | 0.24392267 | 0.14791247 |
| Zfp33b | 0.0769413 | 0.39981196 | 0.188 | 0.109 | 1 | 0.13445261 | -0.2045023 | 0.231 | 0.146 | 1 | 0.13445261 | 0.14796263 |
| Oip5 | 0.07694193 | 1.42253204 | 0.031 | 0.005 | 1 | 0.27154327 | 1.01207283 | 0.013 | 0 | 1 | 0.27154327 | 0.1479638 |
| Tnfrsf14 | 0.07694193 | 1.42253204 | 0.031 | 0.005 | 1 | 0.08763061 | 1.69014474 | 0.05 | 0.01 | 1 | 0.08763061 | 0.1479638 |
| Syt3 | 0.07694193 | 1.42253204 | 0.031 | 0.005 | 1 | 0.15394568 | 1.69014474 | 0.021 | 0 | 1 | 0.15394568 | 0.1479638 |
| Kif7 | 0.12601863 | 0.77302928 | 0.135 | 0.078 | 1 | 0.07695295 | 0.7296731 | 0.13 | 0.062 | 1 | 0.12601863 | 0.14798414 |
| Cdkn1b | 0.07701988 | 0.18997259 | 0.573 | 0.446 | 1 | 0.24746416 | -0.1554266 | 0.773 | 0.708 | 1 | 0.24746416 | 0.14810769 |
| Ttyh2 | 0.07716092 | 0.52206771 | 0.188 | 0.109 | 1 | 0.29348794 | 0.11545057 | 0.185 | 0.135 | 1 | 0.29348794 | 0.14836803 |
| Kcnip2 | 0.41466006 | -0.9925055 | 0.031 | 0.052 | 1 | 0.0772259 | 1.13071733 | 0.101 | 0.042 | 1 | 0.41466006 | 0.14848795 |
| Jpx | 0.76079244 | 0.15949763 | 0.104 | 0.093 | 1 | 0.07726034 | 0.40635177 | 0.172 | 0.094 | 1 | 0.76079244 | 0.14855153 |
| 6720427I07Rik | 0.077288 | 0.4134869 | 0.354 | 0.249 | 1 | 0.55912681 | -0.4618584 | 0.357 | 0.281 | 1 | 0.55912681 | 0.14860256 |
| Cacna1c | 0.27717596 | -0.556769 | 0.406 | 0.311 | 1 | 0.07730775 | -0.7125242 | 0.433 | 0.469 | 1 | 0.27717596 | 0.14863901 |
| Ifit3 | 0.93861547 | -1.4491439 | 0.156 | 0.155 | 1 | 0.07735606 | -1.7638869 | 0.134 | 0.062 | 1 | 0.93861547 | 0.14872815 |
| Hook2 | 0.84342372 | 0.34452952 | 0.083 | 0.078 | 1 | 0.07737018 | -1.2103196 | 0.088 | 0.156 | 1 | 0.84342372 | 0.14875421 |
| Tlr3 | 0.07748436 | -0.6705774 | 0.094 | 0.171 | 1 | 0.27382746 | -0.9995151 | 0.113 | 0.073 | 1 | 0.27382746 | 0.1489649 |
| Unc5c | 0.85406907 | 0.11760806 | 0.958 | 0.907 | 1 | 0.0775626 | -0.164327 | 0.95 | 0.719 | 1 | 0.85406907 | 0.14910924 |
| Maco1 | 0.64056808 | 0.11627088 | 0.667 | 0.658 | 1 | 0.07766756 | -0.5299811 | 0.735 | 0.812 | 1 | 0.64056808 | 0.14930287 |
| Cutal | 0.0776711 | 1.27052894 | 0.052 | 0.016 | 1 | 0.86312513 | -0.7248928 | 0.008 | 0.01 | 1 | 0.86312513 | 0.1493094 |
| BB557941 | 0.0776711 | 1.27052894 | 0.052 | 0.016 | 1 | 0.85710544 | -1.3098553 | 0.008 | 0.01 | 1 | 0.85710544 | 0.1493094 |
| 4930534D22Rik | 0.0776711 | 1.27052894 | 0.052 | 0.016 | 1 | 0.75466895 | -0.6317834 | 0.025 | 0.031 | 1 | 0.75466895 | 0.1493094 |
| Rab3b | 0.0776711 | 1.00749454 | 0.052 | 0.016 | 1 | 0.86312513 | -0.7248928 | 0.008 | 0.01 | 1 | 0.86312513 | 0.1493094 |

| | | | | | | | | | | | | |
|---|---|---|---|---|---|---|---|---|---|---|---|---|
| 6530437J22Rik | 0.0776711 | 1.27052894 | 0.052 | 0.016 | 1 | 0.98590808 | 0.10518224 | 0.021 | 0.021 | 1 | 0.98590808 | 0.1493094 |
| Gm16794 | 0.0776711 | 1.27052894 | 0.052 | 0.016 | 1 | 0.77878694 | -1.4253325 | 0.046 | 0.052 | 1 | 0.77878694 | 0.1493094 |
| 1700001P01Rik | 0.0776711 | 1.27052894 | 0.052 | 0.016 | 1 | 0.09375669 | -1.961932 | 0.025 | 0.062 | 1 | 0.09375669 | 0.1493094 |
| Gm26660 | 0.0776711 | 1.00749454 | 0.052 | 0.016 | 1 | 0.52072676 | -0.3098553 | 0.021 | 0.01 | 1 | 0.52072676 | 0.1493094 |
| Batf2 | 0.0776711 | 1.00749454 | 0.052 | 0.016 | 1 | 0.54697586 | -0.3098553 | 0.034 | 0.021 | 1 | 0.54697586 | 0.1493094 |
| Notch1 | 0.33288321 | -0.2085424 | 0.427 | 0.332 | 1 | 0.07774856 | -0.5709414 | 0.542 | 0.667 | 1 | 0.33288321 | 0.14945228 |
| Tbx2 | 0.38943692 | -0.5470943 | 0.094 | 0.13 | 1 | 0.07775417 | -2.1578522 | 0.038 | 0.083 | 1 | 0.38943692 | 0.14946262 |
| Tgfb1 | 0.07775709 | 0.24876497 | 0.812 | 0.741 | 1 | 0.86243513 | -0.1804514 | 0.87 | 0.792 | 1 | 0.86243513 | 0.14946802 |
| Gpr161 | 0.07775867 | 2.22988696 | 0.042 | 0.01 | 1 | 0.81798678 | -0.5025003 | 0.046 | 0.052 | 1 | 0.81798678 | 0.14947092 |
| Bbox1 | 0.07775867 | 2.22988696 | 0.042 | 0.01 | 1 | 0.14544666 | -2.3098553 | 0.004 | 0.021 | 1 | 0.14544666 | 0.14947092 |
| Slc47a1 | 0.07775867 | 2.42253204 | 0.042 | 0.01 | 1 | 0.87851725 | -1.6317834 | 0.013 | 0.01 | 1 | 0.87851725 | 0.14947092 |
| Tmem132e | 0.07775867 | 2.22988696 | 0.042 | 0.01 | 1 | 0.10534401 | 1.22619764 | 0.08 | 0.031 | 1 | 0.10534401 | 0.14947092 |
| Cep72 | 0.07775867 | 2.22988696 | 0.042 | 0.01 | 1 | 0.86312513 | -0.7248928 | 0.008 | 0.01 | 1 | 0.86312513 | 0.14947092 |
| Ccnb1 | 0.07775867 | 3.67045955 | 0.042 | 0.01 | 1 | 0.11689951 | -2.3098553 | 0 | 0.01 | 1 | 0.11689951 | 0.14947092 |
| Gm41541 | 0.07775867 | 2.42253204 | 0.042 | 0.01 | 1 | 0.86312513 | -0.7248928 | 0.008 | 0.01 | 1 | 0.86312513 | 0.14947092 |
| Bub1 | 0.07776351 | 3.07788386 | 0.042 | 0.01 | 1 | 0.34704016 | -1.3098553 | 0.008 | 0.021 | 1 | 0.34704016 | 0.14947985 |
| Ndufa3 | 0.11175427 | -0.4254649 | 0.531 | 0.585 | 1 | 0.07777916 | 0.57944127 | 0.466 | 0.396 | 1 | 0.11175427 | 0.14950872 |
| Ppp1r7 | 0.07787656 | 0.64731897 | 0.406 | 0.316 | 1 | 0.14074083 | 0.16289773 | 0.357 | 0.25 | 1 | 0.14074083 | 0.14968836 |
| Nudt10 | 0.0781338 | 1.20013961 | 0.052 | 0.016 | 1 | 0.34363875 | -1.7248928 | 0.008 | 0.021 | 1 | 0.34363875 | 0.1501627 |
| Thap4 | 0.07827328 | 0.80104366 | 0.208 | 0.13 | 1 | 0.29378397 | -0.4455104 | 0.252 | 0.177 | 1 | 0.29378397 | 0.15041986 |
| Cdkn2aip | 0.07835816 | 0.18082614 | 0.312 | 0.207 | 1 | 0.73013346 | -0.4938181 | 0.366 | 0.354 | 1 | 0.73013346 | 0.15057632 |
| Pdhx | 0.49259029 | -0.3364599 | 0.323 | 0.352 | 1 | 0.07836867 | -0.194378 | 0.387 | 0.24 | 1 | 0.49259029 | 0.15059569 |
| Adamts4 | 0.07838142 | -1.6445822 | 0.052 | 0.114 | 1 | 0.32421961 | -0.961932 | 0.038 | 0.062 | 1 | 0.32421961 | 0.15061919 |
| 2010315B03Rik | 0.19842758 | 0.63826073 | 0.188 | 0.13 | 1 | 0.07840192 | 0.24268576 | 0.147 | 0.073 | 1 | 0.19842758 | 0.15065698 |
| Dph5 | 0.15432708 | 0.66354014 | 0.385 | 0.29 | 1 | 0.07842054 | 1.40635177 | 0.214 | 0.125 | 1 | 0.15432708 | 0.1506913 |
| Tmem231 | 0.30571589 | 0.42253204 | 0.156 | 0.114 | 1 | 0.07844413 | 1.01207283 | 0.101 | 0.042 | 1 | 0.30571589 | 0.15073477 |
| Mob2 | 0.36473703 | -0.1624305 | 0.365 | 0.42 | 1 | 0.07853309 | -0.561394 | 0.588 | 0.677 | 1 | 0.36473703 | 0.15089874 |
| Ubxn2a | 0.07855013 | 0.37701738 | 0.604 | 0.503 | 1 | 0.53258918 | 0.15632109 | 0.597 | 0.604 | 1 | 0.53258918 | 0.15093013 |
| Smug1 | 0.63033469 | -0.5160674 | 0.073 | 0.088 | 1 | 0.0785613 | 0.62303054 | 0.143 | 0.073 | 1 | 0.63033469 | 0.15095072 |
| Cbl | 0.83580874 | -0.1934182 | 0.479 | 0.456 | 1 | 0.07861913 | -0.4864256 | 0.676 | 0.781 | 1 | 0.83580874 | 0.15105729 |
| Trappc12 | 0.72041227 | -0.196864 | 0.177 | 0.197 | 1 | 0.07862492 | 0.46775232 | 0.235 | 0.146 | 1 | 0.72041227 | 0.15106797 |
| Ccnd3 | 0.07862733 | 0.41962044 | 0.615 | 0.528 | 1 | 0.26498741 | -0.5304063 | 0.731 | 0.729 | 1 | 0.26498741 | 0.15107241 |
| Mcc | 0.24657097 | 0.50341374 | 0.729 | 0.756 | 1 | 0.07864301 | -0.6916272 | 0.702 | 0.771 | 1 | 0.24657097 | 0.1511013 |
| Qrsl1 | 0.87043183 | 0.14499806 | 0.073 | 0.067 | 1 | 0.07870116 | 1.23446525 | 0.101 | 0.042 | 1 | 0.87043183 | 0.15120844 |
| Sco2 | 0.1206108 | 0.64492446 | 0.188 | 0.119 | 1 | 0.07870555 | 0.84214783 | 0.143 | 0.073 | 1 | 0.1206108 | 0.15121653 |
| Glmn | 0.30166872 | -0.4638112 | 0.156 | 0.202 | 1 | 0.07871183 | -0.3769695 | 0.21 | 0.115 | 1 | 0.30166872 | 0.15122811 |
| Zdhhc12 | 0.24537345 | 0.62898291 | 0.156 | 0.109 | 1 | 0.07871591 | 0.76053407 | 0.071 | 0.021 | 1 | 0.24537345 | 0.15123562 |
| Armcx6 | 0.07872315 | 0.86310463 | 0.156 | 0.088 | 1 | 0.13232644 | 0.27510724 | 0.134 | 0.073 | 1 | 0.13232644 | 0.15124897 |
| Gm56847 | 0.07877705 | 2.00749454 | 0.042 | 0.01 | 1 | 0.86312513 | -0.7248928 | 0.008 | 0.01 | 1 | 0.86312513 | 0.15134827 |
| Zfp660 | 0.07877705 | 2.22988696 | 0.042 | 0.01 | 1 | 0.40503552 | -1.0468209 | 0.017 | 0.031 | 1 | 0.40503552 | 0.15134827 |
| Ccdc40 | 0.07877705 | 2.00749454 | 0.042 | 0.01 | 1 | 0.57069113 | -1.7952821 | 0.029 | 0.042 | 1 | 0.57069113 | 0.15134827 |
| H2bc15 | 0.07877705 | 2.42253204 | 0.042 | 0.01 | 1 | 0.50887612 | -1.3098553 | 0.004 | 0.01 | 1 | 0.50887612 | 0.15134827 |
| Ap1g2 | 0.07877705 | 2.00749454 | 0.042 | 0.01 | 1 | 0.15394118 | 1.49749966 | 0.021 | 0 | 1 | 0.15394118 | 0.15134827 |
| Gm20522 | 0.07877705 | 2.00749454 | 0.042 | 0.01 | 1 | 0.27154471 | 0.69014474 | 0.013 | 0 | 1 | 0.27154471 | 0.15134827 |
| Bltp2 | 0.07881702 | 0.18334437 | 0.698 | 0.56 | 1 | 0.19129079 | -0.2519743 | 0.689 | 0.833 | 1 | 0.19129079 | 0.15142192 |
| Acbd4 | 0.07885959 | 0.71798792 | 0.156 | 0.088 | 1 | 0.20117399 | 0.27510724 | 0.109 | 0.062 | 1 | 0.20117399 | 0.15150034 |
| Nmnat1 | 0.76661095 | -0.3404288 | 0.083 | 0.093 | 1 | 0.07909042 | 1.23446525 | 0.113 | 0.052 | 1 | 0.76661095 | 0.15192554 |
| Serpinb9 | 0.2035607 | 0.59245704 | 0.156 | 0.104 | 1 | 0.07911698 | 0.69014474 | 0.168 | 0.094 | 1 | 0.2035607 | 0.15197446 |
| Enpep | 0.1275299 | 1.20013961 | 0.062 | 0.026 | 1 | 0.07913339 | -1.8244284 | 0.038 | 0.083 | 1 | 0.1275299 | 0.15200469 |
| Gm56635 | 0.07930255 | 2.00749454 | 0.042 | 0.01 | 1 | 0.91891525 | -0.6317834 | 0.029 | 0.031 | 1 | 0.91891525 | 0.1523162 |
| Depdc5 | 0.07944881 | 0.54183096 | 0.26 | 0.176 | 1 | 0.82546115 | -0.1883918 | 0.34 | 0.333 | 1 | 0.82546115 | 0.15258551 |
| Fastkd1 | 0.07951885 | 0.11440974 | 0.177 | 0.098 | 1 | 0.13471623 | 0.32757466 | 0.172 | 0.104 | 1 | 0.13471623 | 0.15271446 |
| Gbp7 | 0.31925004 | -1.0398112 | 0.073 | 0.109 | 1 | 0.07964904 | -2.9879272 | 0.038 | 0.083 | 1 | 0.31925004 | 0.15295411 |
| Olfml1 | 0.8006267 | 0.20013961 | 0.021 | 0.026 | 1 | 0.07964904 | -2.07539 | 0.038 | 0.083 | 1 | 0.8006267 | 0.15295411 |
| Zfp729b | 0.20036415 | 0.63826073 | 0.188 | 0.13 | 1 | 0.07965375 | 0.77760758 | 0.206 | 0.125 | 1 | 0.20036415 | 0.15296278 |
| Fbxo8 | 0.07967763 | 0.26426995 | 0.365 | 0.249 | 1 | 0.94862063 | -0.5976576 | 0.239 | 0.219 | 1 | 0.94862063 | 0.15300674 |
| Borcs8 | 0.34778548 | 0.27052894 | 0.365 | 0.311 | 1 | 0.07976488 | 0.22448117 | 0.294 | 0.188 | 1 | 0.34778548 | 0.15316732 |
| Zfp112 | 0.08338697 | 0.83756954 | 0.115 | 0.057 | 1 | 0.07976757 | 0.87456931 | 0.13 | 0.062 | 1 | 0.08338697 | 0.15317227 |
| Ostf1 | 0.7643043 | -0.1119407 | 0.781 | 0.772 | 1 | 0.07976763 | 0.22098202 | 0.824 | 0.781 | 1 | 0.7643043 | 0.15317239 |
| Nudt8 | 0.96926163 | 0.34452952 | 0.094 | 0.093 | 1 | 0.07981498 | 0.18490943 | 0.118 | 0.052 | 1 | 0.96926163 | 0.15325954 |
| Gm42067 | 0.07981716 | 1.12297175 | 0.115 | 0.057 | 1 | 0.18504625 | 1.01207283 | 0.038 | 0.01 | 1 | 0.18504625 | 0.15326354 |
| Rgs5 | 0.07982472 | 1.74446013 | 0.042 | 0.01 | 1 | 0.65410681 | -0.5993619 | 0.067 | 0.052 | 1 | 0.65410681 | 0.15327745 |
| 9130604C24Rik | 0.07982472 | 1.74446013 | 0.042 | 0.01 | 1 | 0.15394868 | 1.27510724 | 0.021 | 0 | 1 | 0.15394868 | 0.15327745 |
| Zfp59 | 0.07982472 | 1.74446013 | 0.042 | 0.01 | 1 | 0.24241449 | -0.9313436 | 0.042 | 0.073 | 1 | 0.24241449 | 0.15327745 |
| Myo7a | 0.07982472 | 1.74446013 | 0.042 | 0.01 | 1 | 0.82231716 | -0.3098553 | 0.025 | 0.021 | 1 | 0.82231716 | 0.15327745 |
| Stk32c | 0.07982472 | 1.74446013 | 0.042 | 0.01 | 1 | 0.92416907 | -0.9879272 | 0.034 | 0.031 | 1 | 0.92416907 | 0.15327745 |
| Gm2814 | 0.07982472 | 1.74446013 | 0.042 | 0.01 | 1 | 0.39515719 | 0.49749966 | 0.025 | 0.01 | 1 | 0.39515719 | 0.15327745 |
| Vstm5 | 0.07982472 | 1.74446013 | 0.042 | 0.01 | 1 | 0.27154471 | 0.69014474 | 0.013 | 0 | 1 | 0.27154471 | 0.15327745 |
| Smim6 | 0.07982472 | 1.74446013 | 0.042 | 0.01 | 1 | 0.50587544 | -2.6317834 | 0.004 | 0.01 | 1 | 0.50587544 | 0.15327745 |
| Mtif3 | 0.98148359 | 0.12613903 | 0.146 | 0.145 | 1 | 0.07986686 | 0.27510724 | 0.172 | 0.094 | 1 | 0.98148359 | 0.15335501 |
| Pard3b | 0.77533067 | -0.3754818 | 0.677 | 0.705 | 1 | 0.07990311 | -0.4936765 | 0.828 | 0.917 | 1 | 0.77533067 | 0.15342172 |
| Tmem82 | 0.07991872 | 1.00749454 | 0.052 | 0.016 | 1 | 0.66492598 | -0.2625495 | 0.101 | 0.083 | 1 | 0.66492598 | 0.15345044 |
| Ccdc120 | 0.73724235 | 0.35541784 | 0.042 | 0.052 | 1 | 0.08003417 | 0.35310975 | 0.071 | 0.021 | 1 | 0.73724235 | 0.15366287 |
| Nr4a3 | 0.08003926 | 1.10703021 | 0.354 | 0.259 | 1 | 0.62685124 | -0.3970398 | 0.479 | 0.448 | 1 | 0.62685124 | 0.15367224 |
| A230057D06Rik | 0.08004143 | -0.3281085 | 0.229 | 0.332 | 1 | 0.30672147 | -0.1604776 | 0.231 | 0.167 | 1 | 0.30672147 | 0.15367622 |
| Chmp2b | 0.08009079 | 0.18646468 | 0.729 | 0.591 | 1 | 0.09728986 | 0.19754625 | 0.748 | 0.667 | 1 | 0.09728986 | 0.15376704 |
| Braf | 0.08016366 | 0.21427476 | 0.875 | 0.87 | 1 | 0.17255738 | -0.3402289 | 0.971 | 0.969 | 1 | 0.17255738 | 0.15390111 |
| Isoc2b | 0.53937361 | 0.56003556 | 0.156 | 0.13 | 1 | 0.08019278 | 0.76814725 | 0.13 | 0.062 | 1 | 0.53937361 | 0.15395469 |
| Pcbp4 | 0.13780669 | 0.41188479 | 0.333 | 0.249 | 1 | 0.0802158 | 0.31383963 | 0.42 | 0.292 | 1 | 0.13780669 | 0.15399702 |
| Zfp335 | 0.36829333 | 0.42253204 | 0.135 | 0.098 | 1 | 0.08025862 | 0.48661134 | 0.227 | 0.135 | 1 | 0.36829333 | 0.15407579 |
| Hint3 | 0.32087229 | 0.22762035 | 0.375 | 0.316 | 1 | 0.08027369 | 0.54162221 | 0.252 | 0.167 | 1 | 0.32087229 | 0.15410351 |
| Sipa1 | 0.42847613 | -0.6445822 | 0.062 | 0.088 | 1 | 0.08029868 | 0.58322953 | 0.172 | 0.094 | 1 | 0.42847613 | 0.15414949 |
| Impg2 | 0.08034413 | 1.59245704 | 0.042 | 0.01 | 1 | 0.64256011 | -1.4253325 | 0.042 | 0.052 | 1 | 0.64256011 | 0.15423307 |
| Ndufv1 | 0.08036324 | 0.33764314 | 0.479 | 0.358 | 1 | 0.19029113 | -0.232779 | 0.492 | 0.573 | 1 | 0.19029113 | 0.15426824 |
| Prrg2 | 0.08042007 | 0.35541784 | 0.188 | 0.109 | 1 | 0.43880462 | -0.3402289 | 0.181 | 0.135 | 1 | 0.43880462 | 0.15437275 |
| Tomm40 | 0.08061691 | 0.32942263 | 0.5 | 0.399 | 1 | 0.6536008 | -0.1259911 | 0.588 | 0.542 | 1 | 0.6536008 | 0.15473474 |

| | | | | | | | | | | | | |
|---|---|---|---|---|---|---|---|---|---|---|---|---|
| Oxr1 | 0.08062005 | -0.5070786 | 0.667 | 0.746 | 1 | 0.19404393 | -0.2497343 | 0.714 | 0.792 | 1 | 0.19404393 | 0.15474051 |
| 1110046J04Rik | 0.08064901 | 0.78510212 | 0.167 | 0.093 | 1 | 0.99503555 | -0.4942798 | 0.143 | 0.135 | 1 | 0.99503555 | 0.15479377 |
| Zbtb34 | 0.85084749 | -0.3910548 | 0.104 | 0.109 | 1 | 0.08073986 | 0.41261076 | 0.202 | 0.115 | 1 | 0.85084749 | 0.1549608 |
| H2ac25 | 0.10604553 | 0.86999101 | 0.156 | 0.093 | 1 | 0.0809792 | 0.75427507 | 0.143 | 0.073 | 1 | 0.10604553 | 0.15540078 |
| Ndufs6 | 0.08101285 | 0.32488347 | 0.719 | 0.637 | 1 | 0.10997029 | 0.26950453 | 0.626 | 0.573 | 1 | 0.10997029 | 0.15546261 |
| Sdhb | 0.149854 | 0.22776066 | 0.708 | 0.606 | 1 | 0.08102706 | 0.19684018 | 0.71 | 0.635 | 1 | 0.149854 | 0.15548873 |
| Ap5m1 | 0.08110735 | -0.3144336 | 0.115 | 0.207 | 1 | 0.60398119 | -0.2609457 | 0.202 | 0.167 | 1 | 0.60398119 | 0.1556363 |
| Lyrm4 | 0.69377148 | 0.25260703 | 0.146 | 0.13 | 1 | 0.0812317 | 0.55264121 | 0.118 | 0.052 | 1 | 0.69377148 | 0.15586482 |
| Kif28 | 0.37204076 | -1.130009 | 0.042 | 0.067 | 1 | 0.08124359 | -2.001733 | 0.046 | 0.094 | 1 | 0.37204076 | 0.15588666 |
| Hnrnpul1 | 0.08125162 | 0.16765176 | 0.708 | 0.617 | 1 | 0.92549881 | -0.2606513 | 0.866 | 0.875 | 1 | 0.92549881 | 0.15590142 |
| Plekha7 | 0.08130647 | -1.1180363 | 0.042 | 0.104 | 1 | 0.55320093 | -0.3991226 | 0.21 | 0.24 | 1 | 0.55320093 | 0.15600219 |
| Zbtb25 | 0.08131753 | 0.65385758 | 0.135 | 0.073 | 1 | 0.94012481 | -0.7386986 | 0.176 | 0.167 | 1 | 0.94012481 | 0.15602252 |
| R3hdm2 | 0.08138016 | 0.31905592 | 0.854 | 0.808 | 1 | 0.8685845 | -0.126733 | 0.866 | 0.896 | 1 | 0.8685845 | 0.15613758 |
| Fbxl12os | 0.08139244 | 1.00749454 | 0.042 | 0.01 | 1 | 0.67474842 | -0.3098553 | 0.029 | 0.021 | 1 | 0.67474842 | 0.15616016 |
| Pcdh19 | 0.08144362 | -3.0799683 | 0.01 | 0.052 | 1 | 0.57872844 | -0.5537808 | 0.202 | 0.167 | 1 | 0.57872844 | 0.15625417 |
| Apc | 0.08144542 | 0.21193429 | 0.729 | 0.663 | 1 | 0.50046548 | -0.2500573 | 0.849 | 0.844 | 1 | 0.50046548 | 0.15625748 |
| Rpap3 | 0.08148327 | 0.44806713 | 0.312 | 0.218 | 1 | 0.23406495 | -0.2781464 | 0.269 | 0.188 | 1 | 0.23406495 | 0.15632701 |
| Snrpn | 0.08154734 | -0.9925055 | 0.104 | 0.181 | 1 | 0.21877221 | 0.31163311 | 0.134 | 0.083 | 1 | 0.21877221 | 0.15644472 |
| A630023A22Rik | 0.08154986 | -1.577468 | 0.052 | 0.114 | 1 | 0.92738405 | -0.9472852 | 0.034 | 0.031 | 1 | 0.92738405 | 0.15644935 |
| Btbd16 | 0.08168434 | 0.92503238 | 0.115 | 0.057 | 1 | 0.41094077 | -1.6931839 | 0.067 | 0.042 | 1 | 0.41094077 | 0.15669634 |
| Atg13 | 0.08171955 | -0.6427594 | 0.229 | 0.326 | 1 | 0.37286288 | 0.21370669 | 0.336 | 0.281 | 1 | 0.37286288 | 0.15676102 |
| Lasp1 | 0.0906876 | -0.517347 | 0.333 | 0.425 | 1 | 0.0817715 | -0.2015226 | 0.5 | 0.656 | 1 | 0.0906876 | 0.15685643 |
| Slc38a7 | 0.53548476 | -0.5160674 | 0.073 | 0.093 | 1 | 0.08177688 | 1.01207283 | 0.168 | 0.094 | 1 | 0.53548476 | 0.1568663 |
| Unc80 | 0.08183406 | -2.3848229 | 0.01 | 0.052 | 1 | 0.86312513 | -0.7248928 | 0.008 | 0.01 | 1 | 0.86312513 | 0.15697131 |
| Snx31 | 0.08183441 | -2.8993961 | 0.01 | 0.052 | 1 | 0.24857751 | -1.0468209 | 0.013 | 0.031 | 1 | 0.24857751 | 0.15697194 |
| Mast1 | 0.0818351 | -3.1217885 | 0.01 | 0.052 | 1 | 0.41219986 | -0.5728897 | 0.017 | 0.031 | 1 | 0.41219986 | 0.15697321 |
| A230072E10Rik | 0.5663165 | 0.46692616 | 0.062 | 0.047 | 1 | 0.08187221 | -0.3878578 | 0.088 | 0.031 | 1 | 0.5663165 | 0.15704137 |
| Armh3 | 0.0819132 | 0.33764314 | 0.385 | 0.28 | 1 | 0.71792245 | -0.3988603 | 0.471 | 0.417 | 1 | 0.71792245 | 0.15711662 |
| Gm28411 | 0.0819164 | 1.32942263 | 0.042 | 0.01 | 1 | 0.20315163 | 1.01207283 | 0.017 | 0 | 1 | 0.20315163 | 0.1571225 |
| Exo1 | 0.0819164 | 1.00749454 | 0.042 | 0.01 | 1 | 0.11689951 | -2.3098553 | 0 | 0.01 | 1 | 0.11689951 | 0.1571225 |
| 4933404O12Rik | 0.0819164 | 1.32942263 | 0.042 | 0.01 | 1 | 0.91387303 | -1.1578522 | 0.029 | 0.031 | 1 | 0.91387303 | 0.1571225 |
| Opn1sw | 0.0819164 | 1.32942263 | 0.042 | 0.01 | 1 | 0.11765402 | 1.49749966 | 0.025 | 0 | 1 | 0.11765402 | 0.1571225 |
| 2310040G24Rik | 0.0819164 | 1.32942263 | 0.042 | 0.01 | 1 | 0.8259932 | -1.0874628 | 0.025 | 0.021 | 1 | 0.8259932 | 0.1571225 |
| Fgd5 | 0.0819164 | 0.74446013 | 0.042 | 0.01 | 1 | 0.14259748 | 1.69014474 | 0.042 | 0.01 | 1 | 0.14259748 | 0.1571225 |
| Itgae | 0.0819164 | 0.15949763 | 0.042 | 0.01 | 1 | 0.20314753 | 1.49749966 | 0.017 | 0 | 1 | 0.20314753 | 0.1571225 |
| Slc26a8 | 0.0819164 | 1.32942263 | 0.042 | 0.01 | 1 | 1 | -0.5728897 | 0.021 | 0.021 | 1 | 1 | 0.1571225 |
| Vwa3b | 0.0819164 | -2.1624305 | 0 | 0.031 | 1 | 0.58664899 | -0.5025003 | 0.021 | 0.031 | 1 | 0.58664899 | 0.1571225 |
| Tgm5 | 0.0819164 | -1.9925055 | 0 | 0.031 | 1 | 0.5778028 | -0.8948178 | 0.013 | 0.021 | 1 | 0.5778028 | 0.1571225 |
| Gm56773 | 0.0819164 | -1.9925055 | 0 | 0.031 | 1 | 0.87066325 | -0.3098553 | 0.013 | 0.01 | 1 | 0.87066325 | 0.1571225 |
| Atp8b5 | 0.0819164 | -1.9925055 | 0 | 0.031 | 1 | 0.15394868 | 1.27510724 | 0.021 | 0 | 1 | 0.15394868 | 0.1571225 |
| Pianp | 0.0819164 | -1.9925055 | 0 | 0.031 | 1 | 0.24747056 | -1.3098553 | 0.013 | 0.031 | 1 | 0.24747056 | 0.1571225 |
| Gm30934 | 0.0819164 | -1.9925055 | 0 | 0.031 | 1 | 0.3937346 | 0.86006974 | 0.025 | 0.01 | 1 | 0.3937346 | 0.1571225 |
| Rwdd2a | 0.0819164 | -2.1624305 | 0 | 0.031 | 1 | 0.65691192 | 0.42711033 | 0.029 | 0.021 | 1 | 0.65691192 | 0.1571225 |
| Gm34388 | 0.0819164 | -1.9925055 | 0 | 0.031 | 1 | 0.50887612 | -1.3098553 | 0.004 | 0.01 | 1 | 0.50887612 | 0.1571225 |
| Adcy4 | 0.0819164 | -1.9925055 | 0 | 0.031 | 1 | 0.23890421 | -0.7952821 | 0.034 | 0.062 | 1 | 0.23890421 | 0.1571225 |
| Dner | 0.08192263 | -2.3144336 | 0 | 0.031 | 1 | 0.14865142 | -1.3098553 | 0.004 | 0.021 | 1 | 0.14865142 | 0.15713394 |
| Ccdc158 | 0.08192263 | -2.1624305 | 0 | 0.031 | 1 | 0.20314753 | 1.27510724 | 0.017 | 0 | 1 | 0.20314753 | 0.15713394 |
| Ckm | 0.08192263 | -2.1624305 | 0 | 0.031 | 1 | 0.86613737 | -0.3098553 | 0.008 | 0.01 | 1 | 0.86613737 | 0.15713394 |
| Aplp1 | 0.08192263 | -2.3144336 | 0 | 0.031 | 1 | 0.24413474 | -1.6317834 | 0.013 | 0.031 | 1 | 0.24413474 | 0.15713394 |
| Zscan2 | 0.08192263 | -2.3144336 | 0 | 0.031 | 1 | 0.99396051 | -0.3098553 | 0.021 | 0.021 | 1 | 0.99396051 | 0.15713394 |
| Gm45356 | 0.08192263 | -3.577468 | 0 | 0.031 | 1 | 0.50887612 | -1.3098553 | 0.004 | 0.01 | 1 | 0.50887612 | 0.15713394 |
| Brme1 | 0.08192263 | -2.1624305 | 0 | 0.031 | 1 | 0.15394793 | 2.14957636 | 0.021 | 0 | 1 | 0.15394793 | 0.15713394 |
| Gm56858 | 0.08192263 | -2.1624305 | 0 | 0.031 | 1 | 0.40076941 | -1.3098553 | 0.017 | 0.031 | 1 | 0.40076941 | 0.15713394 |
| Kcnh4 | 0.08192263 | -2.1624305 | 0 | 0.031 | 1 | 0.34703906 | -1.3098553 | 0.008 | 0.021 | 1 | 0.34703906 | 0.15713394 |
| Six3os1 | 0.08192263 | -2.3144336 | 0 | 0.031 | 1 | 0.56565662 | -2.1172102 | 0.013 | 0.021 | 1 | 0.56565662 | 0.15713394 |
| Prr16 | 0.08192263 | -3.8993961 | 0 | 0.031 | 1 | 0.14023007 | -1.7862933 | 0.088 | 0.135 | 1 | 0.14023007 | 0.15713394 |
| Fer1l5 | 0.08192574 | -3.8504865 | 0 | 0.031 | 1 | 0.11813493 | -2.5322477 | 0.008 | 0.031 | 1 | 0.11813493 | 0.15713966 |
| Tnfrsf26 | 0.08192574 | -2.4519371 | 0 | 0.031 | 1 | 0.50587544 | -2.3098553 | 0.004 | 0.01 | 1 | 0.50587544 | 0.15713966 |
| Gm41031 | 0.08192574 | -2.9925055 | 0 | 0.031 | 1 | 0.14757845 | -1.7248928 | 0.004 | 0.021 | 1 | 0.14757845 | 0.15713966 |
| 0610033M10Rik | 0.08192636 | -2.3144336 | 0 | 0.031 | 1 | 0.50887612 | -1.3098553 | 0.004 | 0.01 | 1 | 0.50887612 | 0.1571408 |
| Clic6 | 0.08192636 | -3.0799683 | 0 | 0.031 | 1 | 0.31544539 | -0.8948178 | 0.029 | 0.052 | 1 | 0.31544539 | 0.1571408 |
| Lrrn1 | 0.08192761 | -3.3144336 | 0 | 0.031 | 1 | 0.57576763 | -0.9879272 | 0.013 | 0.021 | 1 | 0.57576763 | 0.15714309 |
| Myrfl | 0.08192823 | -3.3848229 | 0 | 0.031 | 1 | 1 | -0.8948178 | 0.021 | 0.021 | 1 | 1 | 0.15714423 |
| Gm13008 | 0.08192886 | -1.7998604 | 0 | 0.031 | 1 | 0.6390466 | 0.39058446 | 0.042 | 0.031 | 1 | 0.6390466 | 0.15714537 |
| Gm42456 | 0.08192886 | -1.7998604 | 0 | 0.031 | 1 | 0.11689951 | -3.3098553 | 0 | 0.01 | 1 | 0.11689951 | 0.15714537 |
| ENSMUSG00000121482 | 0.08192886 | -1.7998604 | 0 | 0.031 | 1 | 0.27154327 | 1.01207283 | 0.013 | 0 | 1 | 0.27154327 | 0.15714537 |
| Angpt2 | 0.08192886 | -1.7998604 | 0 | 0.031 | 1 | 0.50887612 | -1.3098553 | 0.004 | 0.01 | 1 | 0.50887612 | 0.15714537 |
| Gm12714 | 0.08192886 | -1.7998604 | 0 | 0.031 | 1 | 0.5818708 | -0.3098553 | 0.013 | 0.021 | 1 | 0.5818708 | 0.15714537 |
| Gm48420 | 0.08192886 | -1.7998604 | 0 | 0.031 | 1 | 0.20315163 | 1.01207283 | 0.017 | 0 | 1 | 0.20315163 | 0.15714537 |
| Plaat1 | 0.08192886 | -1.7998604 | 0 | 0.031 | 1 | 0.11689951 | -2.3098553 | 0 | 0.01 | 1 | 0.11689951 | 0.15714537 |
| Reep2 | 0.08192886 | -1.7998604 | 0 | 0.031 | 1 | 0.57854956 | -0.8244284 | 0.021 | 0.031 | 1 | 0.57854956 | 0.15714537 |
| Gm9456 | 0.08196938 | 0.88196365 | 0.094 | 0.041 | 1 | 0.69916441 | -1.3098553 | 0.034 | 0.042 | 1 | 0.69916441 | 0.15721979 |
| Snap29 | 0.77610836 | -0.1580915 | 0.312 | 0.321 | 1 | 0.08204171 | 0.13236707 | 0.319 | 0.208 | 1 | 0.77610836 | 0.15735258 |
| Atoh8 | 0.19108902 | 0.47144164 | 0.177 | 0.119 | 1 | 0.08210094 | 1.21370669 | 0.139 | 0.073 | 1 | 0.19108902 | 0.15746132 |
| Fam20a | 0.0821301 | -1.7183305 | 0.083 | 0.155 | 1 | 0.64348905 | -0.7623675 | 0.067 | 0.052 | 1 | 0.64348905 | 0.15751484 |
| Rexo5 | 0.08216947 | 0.68556644 | 0.156 | 0.088 | 1 | 0.26970644 | -1.2609457 | 0.109 | 0.146 | 1 | 0.26970644 | 0.15758712 |
| Car11 | 0.08221594 | 0.78510212 | 0.052 | 0.016 | 1 | 0.97841817 | 0.17557156 | 0.042 | 0.042 | 1 | 0.97841817 | 0.15767242 |
| Mtarc1 | 0.08222361 | -2.240433 | 0.01 | 0.052 | 1 | 0.34363875 | -2.0468209 | 0.008 | 0.021 | 1 | 0.34363875 | 0.15768649 |
| Aipl1 | 0.08222707 | -3.0368996 | 0.01 | 0.052 | 1 | 0.86312513 | -0.7248928 | 0.008 | 0.01 | 1 | 0.86312513 | 0.15769285 |
| Rabgap1 | 0.82143813 | -0.2221367 | 0.688 | 0.674 | 1 | 0.08223663 | -0.5443205 | 0.84 | 0.833 | 1 | 0.82143813 | 0.1577104 |
| Usp46 | 0.08224673 | -0.7183305 | 0.073 | 0.145 | 1 | 0.18121981 | -0.7057839 | 0.105 | 0.156 | 1 | 0.18121981 | 0.15772893 |
| Prxl2b | 0.37664525 | 0.69555053 | 0.167 | 0.13 | 1 | 0.08246855 | 1.39058446 | 0.084 | 0.031 | 1 | 0.37664525 | 0.15813604 |
| Trappc2 | 0.92470872 | -0.1035368 | 0.25 | 0.249 | 1 | 0.08261071 | 0.39058446 | 0.269 | 0.177 | 1 | 0.92470872 | 0.15839689 |
| Gm56530 | 0.08261641 | -2.0799683 | 0.01 | 0.052 | 1 | 0.12096301 | -1.8948178 | 0.008 | 0.031 | 1 | 0.12096301 | 0.15840735 |

| | | | | | | | | | | | | |
|---|---|---|---|---|---|---|---|---|---|---|---|---|
| Fam53b | 0.64491077 | 0.52206771 | 0.115 | 0.098 | 1 | 0.08285679 | 0.69014474 | 0.155 | 0.083 | 1 | 0.64491077 | 0.15884833 |
| Zrsr2 | 0.16173542 | 0.24251457 | 0.688 | 0.658 | 1 | 0.08289844 | 0.28616243 | 0.714 | 0.656 | 1 | 0.16173542 | 0.15892473 |
| Dcun1d3 | 0.63153063 | -0.2671278 | 0.229 | 0.197 | 1 | 0.08292104 | -0.3098553 | 0.336 | 0.219 | 1 | 0.63153063 | 0.15896618 |
| Stau1 | 0.08299812 | 0.15445198 | 0.854 | 0.756 | 1 | 0.96206449 | -0.273667 | 0.962 | 0.958 | 1 | 0.96206449 | 0.15910754 |
| Acox2 | 0.08300727 | -1.8993961 | 0.01 | 0.052 | 1 | 1 | -0.7248928 | 0.021 | 0.021 | 1 | 1 | 0.15912433 |
| Pllp | 0.08300761 | -1.7998604 | 0.01 | 0.052 | 1 | 0.39515719 | 0.49749966 | 0.025 | 0.01 | 1 | 0.39515719 | 0.15912496 |
| Rita1 | 0.24416069 | 0.22988696 | 0.135 | 0.088 | 1 | 0.08311598 | 0.77760758 | 0.168 | 0.094 | 1 | 0.24416069 | 0.1593237 |
| C230013L11Rik | 0.2610669 | -0.8993961 | 0.031 | 0.062 | 1 | 0.08320344 | 1.33400093 | 0.084 | 0.031 | 1 | 0.2610669 | 0.15948408 |
| Mdp1 | 0.12413099 | -0.7294711 | 0.135 | 0.207 | 1 | 0.08326874 | 0.54812573 | 0.21 | 0.125 | 1 | 0.12413099 | 0.1596038 |
| Mtfr2 | 0.0833792 | 0.34006988 | 0.146 | 0.078 | 1 | 0.89443992 | -0.5840302 | 0.126 | 0.125 | 1 | 0.89443992 | 0.15980632 |
| Vps33b | 0.08338325 | 0.81484946 | 0.188 | 0.114 | 1 | 0.74721825 | -0.8073549 | 0.118 | 0.125 | 1 | 0.74721825 | 0.15981373 |
| Tbx3os1 | 0.08342447 | -1.6705774 | 0.042 | 0.098 | 1 | 0.45003757 | -0.3098553 | 0.038 | 0.021 | 1 | 0.45003757 | 0.1598893 |
| Rab15 | 0.48726009 | 0.13302542 | 0.104 | 0.078 | 1 | 0.08347627 | 0.93807225 | 0.113 | 0.052 | 1 | 0.48726009 | 0.15998425 |
| Mre11a | 0.08350075 | 0.69555053 | 0.208 | 0.13 | 1 | 0.11272911 | 0.39058446 | 0.214 | 0.135 | 1 | 0.11272911 | 0.16002912 |
| 4833420G17Rik | 0.08361294 | 0.23810746 | 0.552 | 0.425 | 1 | 0.79746722 | -0.2198739 | 0.546 | 0.479 | 1 | 0.79746722 | 0.16023475 |
| B230217O12Rik | 0.08376606 | 0.52206771 | 0.125 | 0.062 | 1 | 0.10822561 | 0.78968041 | 0.109 | 0.052 | 1 | 0.10822561 | 0.16051537 |
| Gmpr2 | 0.16389611 | 0.39278469 | 0.281 | 0.202 | 1 | 0.08379267 | 0.39058446 | 0.223 | 0.135 | 1 | 0.16389611 | 0.16056413 |
| Pigq | 0.0838117 | 0.54806292 | 0.229 | 0.145 | 1 | 0.11106878 | 0.14957636 | 0.218 | 0.135 | 1 | 0.11106878 | 0.16059899 |
| Gm14410 | 0.08387128 | 0.73447604 | 0.177 | 0.104 | 1 | 0.42056088 | -0.6263289 | 0.181 | 0.135 | 1 | 0.42056088 | 0.16070818 |
| Gstm4 | 0.08401626 | 1.00749454 | 0.135 | 0.073 | 1 | 0.15777023 | 0.17557156 | 0.101 | 0.052 | 1 | 0.15777023 | 0.16097379 |
| Gm45518 | 0.08407144 | 0.74446013 | 0.042 | 0.01 | 1 | 0.24304534 | -2.1578522 | 0.013 | 0.031 | 1 | 0.24304534 | 0.16107487 |
| Gm47438 | 0.08407144 | 0.74446013 | 0.042 | 0.01 | 1 | 0.34363875 | -2.3098553 | 0.008 | 0.021 | 1 | 0.34363875 | 0.16107487 |
| AW822252 | 0.08407144 | 1.00749454 | 0.042 | 0.01 | 1 | 0.66171832 | 1.14957636 | 0.017 | 0.01 | 1 | 0.66171832 | 0.16107487 |
| 1700028E10Rik | 0.08407649 | 0.49292136 | 0.042 | 0.01 | 1 | 0.93704875 | -0.9879272 | 0.034 | 0.031 | 1 | 0.93704875 | 0.16108413 |
| Ampd1 | 0.08420034 | 0.96902039 | 0.167 | 0.098 | 1 | 0.73893876 | -0.177405 | 0.155 | 0.135 | 1 | 0.73893876 | 0.16131098 |
| Gm36862 | 0.08423669 | -1.9925055 | 0.01 | 0.052 | 1 | 0.40503552 | -1.0468209 | 0.017 | 0.031 | 1 | 0.40503552 | 0.16137756 |
| Pfkfb3 | 0.08431622 | 0.49292136 | 0.375 | 0.285 | 1 | 0.54989061 | -0.1935682 | 0.445 | 0.344 | 1 | 0.54989061 | 0.16152321 |
| Col7a1 | 0.22350374 | 0.11440974 | 0.062 | 0.031 | 1 | 0.08433711 | 0.84214783 | 0.113 | 0.052 | 1 | 0.22350374 | 0.16156148 |
| Trp53rka | 0.93294253 | -0.130009 | 0.135 | 0.13 | 1 | 0.0845074 | 0.90515763 | 0.202 | 0.125 | 1 | 0.93294253 | 0.16187331 |
| Nemp1 | 0.16626335 | 0.62898291 | 0.229 | 0.161 | 1 | 0.08463864 | 0.39641353 | 0.185 | 0.104 | 1 | 0.16626335 | 0.16211358 |
| Efl1 | 0.84039049 | -0.5133376 | 0.323 | 0.285 | 1 | 0.08472051 | -1.2219825 | 0.445 | 0.5 | 1 | 0.84039049 | 0.16226346 |
| Carhsp1 | 0.48722959 | 0.2545874 | 0.344 | 0.306 | 1 | 0.08488123 | 0.22301873 | 0.374 | 0.26 | 1 | 0.48722959 | 0.16255764 |
| 4930581F22Rik | 0.37629772 | 0.14076107 | 0.26 | 0.207 | 1 | 0.08492509 | 0.72576865 | 0.143 | 0.073 | 1 | 0.37629772 | 0.16263792 |
| Eid2 | 0.20175821 | -0.1560042 | 0.156 | 0.228 | 1 | 0.0849668 | 0.18490943 | 0.319 | 0.208 | 1 | 0.20175821 | 0.16271424 |
| Tpm2 | 0.08500955 | -0.1173119 | 0.365 | 0.238 | 1 | 0.12798849 | -0.1811219 | 0.382 | 0.26 | 1 | 0.12798849 | 0.16279248 |
| Tgfb2 | 0.08514736 | 0.31608625 | 0.823 | 0.808 | 1 | 0.09266298 | -0.1897215 | 0.941 | 0.792 | 1 | 0.09266298 | 0.16304464 |
| Spns1 | 0.89900015 | -0.3144336 | 0.344 | 0.326 | 1 | 0.0851683 | 0.26089338 | 0.336 | 0.229 | 1 | 0.89900015 | 0.16308296 |
| Uck1 | 0.88436552 | -0.1249558 | 0.229 | 0.212 | 1 | 0.08526099 | 0.21852371 | 0.277 | 0.177 | 1 | 0.88436552 | 0.16325254 |
| Pspc1 | 0.21069994 | 0.23731716 | 0.625 | 0.565 | 1 | 0.08537539 | -0.5600084 | 0.672 | 0.74 | 1 | 0.21069994 | 0.16346183 |
| Astn2 | 0.08542904 | -1.9723276 | 0.083 | 0.15 | 1 | 0.90595176 | -0.3415641 | 0.113 | 0.115 | 1 | 0.90595176 | 0.16355995 |
| Ndufaf1 | 0.85277561 | -0.1217885 | 0.135 | 0.124 | 1 | 0.08558347 | 1.01207283 | 0.113 | 0.052 | 1 | 0.85277561 | 0.1638424 |
| Vps18 | 0.48626213 | 0.32942263 | 0.208 | 0.176 | 1 | 0.08563344 | 0.23763253 | 0.244 | 0.146 | 1 | 0.48626213 | 0.1639338 |
| Qsox1 | 0.08566854 | 0.29358418 | 0.667 | 0.575 | 1 | 0.91034569 | -0.3356439 | 0.756 | 0.76 | 1 | 0.91034569 | 0.16399798 |
| Tfpi | 0.08586437 | -1.577468 | 0.01 | 0.052 | 1 | 0.14544666 | -2.3098553 | 0.004 | 0.021 | 1 | 0.14544666 | 0.16435605 |
| Slurp1 | 0.86948427 | -0.9650993 | 0.135 | 0.14 | 1 | 0.08596226 | 2.86006974 | 0.05 | 0.01 | 1 | 0.86948427 | 0.16453502 |
| Plekhb2 | 0.4379042 | -0.577468 | 0.062 | 0.088 | 1 | 0.08602115 | 1.07480859 | 0.126 | 0.062 | 1 | 0.4379042 | 0.16464267 |
| Mill2 | 0.08620218 | 0.83061677 | 0.24 | 0.166 | 1 | 0.16271522 | 0.27510724 | 0.143 | 0.083 | 1 | 0.16271522 | 0.16497354 |
| Ndufaf7 | 0.08622304 | 0.33506919 | 0.271 | 0.176 | 1 | 0.10887842 | 0.36821664 | 0.218 | 0.135 | 1 | 0.10887842 | 0.16501167 |
| Gm17106 | 0.08624203 | 0.65385758 | 0.156 | 0.088 | 1 | 0.78495218 | -0.7094627 | 0.168 | 0.167 | 1 | 0.78495218 | 0.16504638 |
| Kdm1b | 0.17327047 | 1.00749454 | 0.073 | 0.036 | 1 | 0.0862562 | 0.8995981 | 0.113 | 0.052 | 1 | 0.17327047 | 0.16507226 |
| Tafa3 | 0.08627445 | -1.4950058 | 0.01 | 0.052 | 1 | 0.66387437 | 0.27510724 | 0.017 | 0.01 | 1 | 0.66387437 | 0.16510561 |
| Ndufc1 | 0.24855218 | 0.17317257 | 0.854 | 0.725 | 1 | 0.08628388 | 0.19584396 | 0.689 | 0.604 | 1 | 0.24855218 | 0.16512284 |
| Cenpa | 0.70704608 | 1.43199237 | 0.104 | 0.093 | 1 | 0.08630261 | -0.525584 | 0.118 | 0.052 | 1 | 0.70704608 | 0.16515709 |
| Slc23a3 | 0.34635599 | -0.6363617 | 0.052 | 0.083 | 1 | 0.08633246 | -1.1463565 | 0.092 | 0.156 | 1 | 0.34635599 | 0.16521162 |
| Cep128 | 0.08639118 | 0.35214271 | 0.167 | 0.098 | 1 | 0.92491322 | -0.7248928 | 0.105 | 0.104 | 1 | 0.92491322 | 0.16531892 |
| Zfp219 | 0.96756563 | 0.11440974 | 0.115 | 0.114 | 1 | 0.0864403 | 0.52413479 | 0.261 | 0.167 | 1 | 0.96756563 | 0.16540867 |
| Stat5b | 0.08659286 | -0.824778 | 0.354 | 0.446 | 1 | 0.53716079 | -0.6805839 | 0.744 | 0.719 | 1 | 0.53716079 | 0.1656874 |
| Ube2e2 | 0.43611128 | 0.11208344 | 0.833 | 0.824 | 1 | 0.08680588 | -0.7894523 | 0.832 | 0.906 | 1 | 0.43611128 | 0.1660765 |
| Gdf15 | 0.93637696 | -3.1149021 | 0.073 | 0.067 | 1 | 0.08696653 | 2.14957636 | 0.05 | 0.01 | 1 | 0.93637696 | 0.16636988 |
| Mfsd3 | 0.25411632 | 0.62898291 | 0.094 | 0.057 | 1 | 0.08696675 | 1.69014474 | 0.05 | 0.01 | 1 | 0.25411632 | 0.16637029 |
| Ano7 | 0.30888487 | 1.00749454 | 0.042 | 0.021 | 1 | 0.08696675 | 1.86006974 | 0.05 | 0.01 | 1 | 0.30888487 | 0.16637029 |
| 4933427G23Rik | 0.64538863 | 0.52206771 | 0.042 | 0.031 | 1 | 0.08696675 | 1.69014474 | 0.05 | 0.01 | 1 | 0.64538863 | 0.16637029 |
| Aoc2 | 0.78979801 | -0.2148979 | 0.021 | 0.026 | 1 | 0.08696675 | 1.93807225 | 0.05 | 0.01 | 1 | 0.78979801 | 0.16637029 |
| Polk | 0.08698195 | 0.99100641 | 0.344 | 0.269 | 1 | 0.36630157 | 0.1438627 | 0.298 | 0.24 | 1 | 0.36630157 | 0.16639804 |
| Ptger4 | 0.08708935 | -1.2148979 | 0.01 | 0.052 | 1 | 0.89589007 | -0.6468902 | 0.046 | 0.042 | 1 | 0.89589007 | 0.16659415 |
| Zmat5 | 0.08712323 | 0.39605982 | 0.312 | 0.212 | 1 | 0.36231358 | -0.1063219 | 0.269 | 0.208 | 1 | 0.36231358 | 0.166656 |
| Nos1ap | 0.08714023 | -1.6585664 | 0.604 | 0.57 | 1 | 0.61365442 | -0.6246751 | 0.756 | 0.74 | 1 | 0.61365442 | 0.16668704 |
| Rab40b | 0.31392325 | 0.78510212 | 0.042 | 0.021 | 1 | 0.08729941 | 1.69014474 | 0.05 | 0.01 | 1 | 0.31392325 | 0.16697763 |
| Gm49463 | 0.72622123 | -0.2148979 | 0.052 | 0.062 | 1 | 0.08729941 | 1.59703533 | 0.05 | 0.01 | 1 | 0.72622123 | 0.16697763 |
| Aard | 0.74827573 | 0.59245704 | 0.021 | 0.016 | 1 | 0.08729941 | 1.59703533 | 0.05 | 0.01 | 1 | 0.74827573 | 0.16697763 |
| Mob1b | 0.09441269 | 0.11087228 | 0.552 | 0.425 | 1 | 0.08731174 | -0.4970583 | 0.693 | 0.812 | 1 | 0.09441269 | 0.16700013 |
| Pts | 0.32748954 | 0.29529685 | 0.344 | 0.29 | 1 | 0.08732944 | 0.54162221 | 0.265 | 0.177 | 1 | 0.32748954 | 0.16703246 |
| Nmi | 0.8078038 | -0.240433 | 0.188 | 0.197 | 1 | 0.08736763 | 0.56461386 | 0.218 | 0.135 | 1 | 0.8078038 | 0.16710215 |
| Btbd2 | 0.57512257 | -0.1730777 | 0.229 | 0.192 | 1 | 0.08746338 | -0.6045985 | 0.252 | 0.146 | 1 | 0.57512257 | 0.16727691 |
| Zbtb45 | 0.08750668 | -1.1079827 | 0.01 | 0.052 | 1 | 0.86695837 | -0.194378 | 0.046 | 0.042 | 1 | 0.86695837 | 0.16735595 |
| H4c8 | 0.12585255 | -1.1624305 | 0.021 | 0.062 | 1 | 0.08763061 | 1.59703533 | 0.05 | 0.01 | 1 | 0.12585255 | 0.1675821 |
| Fam234b | 0.15227405 | 0.71798792 | 0.083 | 0.041 | 1 | 0.08763061 | 1.59703533 | 0.05 | 0.01 | 1 | 0.15227405 | 0.1675821 |
| Gm20635 | 0.16643489 | 0.37946331 | 0.104 | 0.057 | 1 | 0.08763061 | 1.49749966 | 0.05 | 0.01 | 1 | 0.16643489 | 0.1675821 |
| Inca1 | 0.56103105 | 0.62898291 | 0.062 | 0.047 | 1 | 0.08763061 | 1.69014474 | 0.05 | 0.01 | 1 | 0.56103105 | 0.1675821 |
| Pcdhb2 | 0.75328966 | 0.27052894 | 0.021 | 0.016 | 1 | 0.08763061 | 1.59703533 | 0.05 | 0.01 | 1 | 0.75328966 | 0.1675821 |
| ENSMUSG000001214 95 | 0.08776094 | 1.59245704 | 0.083 | 0.036 | 1 | 0.37896119 | 0.46775232 | 0.055 | 0.031 | 1 | 0.37896119 | 0.1678199 |
| Lpar2 | 0.32025586 | -1.3144336 | 0 | 0.01 | 1 | 0.08804305 | 1.39058446 | 0.05 | 0.01 | 1 | 0.32025586 | 0.16833452 |
| Pih1d1 | 0.20359936 | 0.4972998 | 0.25 | 0.192 | 1 | 0.08809978 | 0.38529016 | 0.223 | 0.135 | 1 | 0.20359936 | 0.16843799 |

| | | | | | | | | | | | | |
|---|---|---|---|---|---|---|---|---|---|---|---|---|
| Ptprm | 0.3560804 | -0.1941393 | 0.906 | 0.876 | 1 | 0.08810524 | -0.569475 | 0.966 | 0.958 | 1 | 0.3560804 | 0.16844794 |
| Nfe2l3 | 0.3387285 | -0.577468 | 0.052 | 0.083 | 1 | 0.08810815 | -1.1269912 | 0.13 | 0.198 | 1 | 0.3387285 | 0.16845326 |
| Kpna1 | 0.08811078 | 0.30062537 | 0.708 | 0.58 | 1 | 0.56454749 | -0.1615488 | 0.803 | 0.812 | 1 | 0.56454749 | 0.16845805 |
| Brf2 | 0.08812608 | 0.74446013 | 0.135 | 0.073 | 1 | 0.37288878 | 0.22619764 | 0.105 | 0.073 | 1 | 0.37288878 | 0.16848595 |
| Capn10 | 0.3360916 | 0.27052894 | 0.188 | 0.14 | 1 | 0.088176 | 0.10518224 | 0.176 | 0.094 | 1 | 0.3360916 | 0.16857698 |
| Cenpv | 0.85409803 | -0.3500575 | 0.146 | 0.15 | 1 | 0.08821909 | 0.73453886 | 0.168 | 0.094 | 1 | 0.85409803 | 0.16865557 |
| Msi1 | 0.0882324 | 0.72738662 | 0.219 | 0.14 | 1 | 0.12644506 | -1.2079756 | 0.139 | 0.198 | 1 | 0.12644506 | 0.16867984 |
| Tmem220 | 0.43214328 | 0.68556644 | 0.062 | 0.041 | 1 | 0.08830431 | 1.42711033 | 0.067 | 0.021 | 1 | 0.43214328 | 0.16881097 |
| Plekhg1 | 0.39533168 | 0.7383079 | 0.125 | 0.093 | 1 | 0.08835106 | -0.1335325 | 0.206 | 0.302 | 1 | 0.39533168 | 0.1688962 |
| Dqx1 | 0.08835518 | 2.17741954 | 0.01 | 0.052 | 1 | 0.99194729 | -1.0874628 | 0.021 | 0.021 | 1 | 0.99194729 | 0.16890372 |
| Tmem232 | 0.0883613 | -0.9925055 | 0.01 | 0.052 | 1 | 0.1711102 | -3.4473588 | 0.017 | 0.042 | 1 | 0.1711102 | 0.16891487 |
| 4930533K18Rik | 0.20883718 | -0.9925055 | 0.01 | 0.036 | 1 | 0.08858031 | -2.1172102 | 0.025 | 0.062 | 1 | 0.20883718 | 0.16931414 |
| Rasa3 | 0.20756506 | 0.50171587 | 0.594 | 0.534 | 1 | 0.08878512 | -0.1541195 | 0.613 | 0.438 | 1 | 0.20756506 | 0.16968744 |
| Klhl42 | 0.25522315 | 0.62898291 | 0.177 | 0.13 | 1 | 0.08880708 | 0.52021974 | 0.206 | 0.125 | 1 | 0.25522315 | 0.16972746 |
| Acsf3 | 0.41822206 | -0.7998604 | 0.031 | 0.052 | 1 | 0.08888333 | 1.35310975 | 0.067 | 0.021 | 1 | 0.41822206 | 0.16986642 |
| Rtel1 | 0.77671342 | -0.240433 | 0.073 | 0.062 | 1 | 0.08888333 | 1.42711033 | 0.067 | 0.021 | 1 | 0.77671342 | 0.16986642 |
| Cdkal1 | 0.18557759 | -0.1458951 | 0.698 | 0.617 | 1 | 0.08890674 | -0.6485894 | 0.769 | 0.823 | 1 | 0.18557759 | 0.16990908 |
| Cul2 | 0.0889951 | 0.30557589 | 0.458 | 0.352 | 1 | 0.90831924 | -0.3035415 | 0.534 | 0.458 | 1 | 0.90831924 | 0.17007008 |
| Apba3 | 0.08901602 | 0.58007331 | 0.25 | 0.166 | 1 | 0.10379456 | 0.16819203 | 0.265 | 0.177 | 1 | 0.10379456 | 0.17010818 |
| Scamp5 | 0.86085369 | 0.24195979 | 0.083 | 0.078 | 1 | 0.08921147 | 0.8917786 | 0.084 | 0.031 | 1 | 0.86085369 | 0.17046426 |
| Armc7 | 0.33637275 | 0.27052894 | 0.115 | 0.078 | 1 | 0.08923637 | 0.46775232 | 0.084 | 0.031 | 1 | 0.33637275 | 0.1705096 |
| Gadd45g | 0.34864413 | -0.2065182 | 0.552 | 0.482 | 1 | 0.08929797 | -1.3973181 | 0.282 | 0.354 | 1 | 0.34864413 | 0.17062182 |
| Plxnc1 | 0.48064264 | 0.27052894 | 0.021 | 0.01 | 1 | 0.08929965 | -2.6724253 | 0.025 | 0.062 | 1 | 0.48064264 | 0.17062486 |
| Xrcc6 | 0.58312823 | 0.14951354 | 0.26 | 0.228 | 1 | 0.08949727 | 0.27510724 | 0.21 | 0.125 | 1 | 0.58312823 | 0.17098478 |
| Cyp4v3 | 0.2521778 | -0.9067756 | 0.104 | 0.15 | 1 | 0.08951232 | 0.59060906 | 0.151 | 0.083 | 1 | 0.2521778 | 0.17101219 |
| Fam168a | 0.08962002 | -0.3167775 | 0.792 | 0.855 | 1 | 0.64263444 | -0.2813729 | 0.857 | 0.854 | 1 | 0.64263444 | 0.17120828 |
| Mettl7a1 | 0.08963126 | -0.6134473 | 0.427 | 0.497 | 1 | 0.40531234 | -0.2366063 | 0.508 | 0.615 | 1 | 0.40531234 | 0.17122876 |
| Slc16a7 | 0.08967201 | -2.265524 | 0.021 | 0.067 | 1 | 0.49356036 | -0.8441917 | 0.063 | 0.083 | 1 | 0.49356036 | 0.17130294 |
| Fbxo3 | 0.10611326 | 0.31415587 | 0.583 | 0.487 | 1 | 0.08974094 | 0.15753764 | 0.702 | 0.604 | 1 | 0.10611326 | 0.17142845 |
| D330023K18Rik | 0.09822859 | 0.57139542 | 0.156 | 0.088 | 1 | 0.08980323 | 0.44016648 | 0.113 | 0.052 | 1 | 0.09822859 | 0.17154184 |
| Arfgef1 | 0.08980747 | 0.26097464 | 0.875 | 0.834 | 1 | 0.3941839 | -0.282787 | 0.95 | 0.948 | 1 | 0.3941839 | 0.17154957 |
| Dtna | 0.29658808 | 0.62047141 | 0.083 | 0.052 | 1 | 0.08989191 | -0.600904 | 0.155 | 0.083 | 1 | 0.29658808 | 0.17170327 |
| Slc2a9 | 0.25139341 | -3.1010299 | 0.031 | 0.062 | 1 | 0.08996546 | -4.3973181 | 0.013 | 0.042 | 1 | 0.25139341 | 0.17183714 |
| Serpind1 | 0.61924305 | -0.2148979 | 0.021 | 0.031 | 1 | 0.08996546 | -2.7692869 | 0.013 | 0.042 | 1 | 0.61924305 | 0.17183714 |
| Mcm7 | 0.0899847 | 0.54452256 | 0.24 | 0.155 | 1 | 0.33970769 | 0.32486027 | 0.189 | 0.146 | 1 | 0.33970769 | 0.17187216 |
| Cfap46 | 0.74827573 | 0.59245704 | 0.021 | 0.016 | 1 | 0.08999201 | 0.82764826 | 0.084 | 0.031 | 1 | 0.74827573 | 0.17188546 |
| Auh | 0.09003748 | -0.4610585 | 0.302 | 0.404 | 1 | 0.20214134 | -0.9354597 | 0.315 | 0.344 | 1 | 0.20214134 | 0.17196821 |
| Snx33 | 0.09012908 | 0.47697982 | 0.156 | 0.088 | 1 | 0.7070656 | -0.6062483 | 0.21 | 0.177 | 1 | 0.7070656 | 0.17213491 |
| Gm43560 | 0.09015816 | -1.6363617 | 0.031 | 0.083 | 1 | 0.57370068 | -1.3098553 | 0.021 | 0.031 | 1 | 0.57370068 | 0.17218783 |
| Vegfc | 0.09015816 | -1.6929452 | 0.031 | 0.083 | 1 | 0.96415509 | -0.9064996 | 0.118 | 0.115 | 1 | 0.96415509 | 0.17218783 |
| Rwdd1 | 0.09016241 | 0.21619947 | 0.885 | 0.803 | 1 | 0.10462356 | 0.20471791 | 0.79 | 0.698 | 1 | 0.10462356 | 0.17219556 |
| 2900093K20Rik | 0.45429213 | 0.35541784 | 0.104 | 0.078 | 1 | 0.09035793 | -1.8948178 | 0.046 | 0.094 | 1 | 0.45429213 | 0.1725513 |
| Wipf2 | 0.17221289 | 0.48142572 | 0.333 | 0.264 | 1 | 0.09036598 | -0.5961594 | 0.408 | 0.49 | 1 | 0.17221289 | 0.17256594 |
| Macc1 | 0.1992517 | 2.59245704 | 0.031 | 0.01 | 1 | 0.09043413 | 1.86006974 | 0.029 | 0 | 1 | 0.1992517 | 0.17268993 |
| L1cam | 0.20598035 | -1.4519371 | 0.01 | 0.036 | 1 | 0.09043413 | 1.86006974 | 0.029 | 0 | 1 | 0.20598035 | 0.17268993 |
| 3830408C21Rik | 0.2060181 | 1.00749454 | 0.083 | 0.047 | 1 | 0.09043413 | 1.86006974 | 0.029 | 0 | 1 | 0.2060181 | 0.17268993 |
| Aif1 | 0.22187575 | -0.9925055 | 0 | 0.016 | 1 | 0.09043413 | 1.86006974 | 0.029 | 0 | 1 | 0.22187575 | 0.17268993 |
| 9130008F23Rik | 0.22187575 | -0.9925055 | 0 | 0.016 | 1 | 0.09043413 | 2.01207283 | 0.029 | 0 | 1 | 0.22187575 | 0.17268993 |
| Gm31105 | 0.24365282 | 1.42253204 | 0.052 | 0.026 | 1 | 0.09043413 | 1.86006974 | 0.029 | 0 | 1 | 0.24365282 | 0.17268993 |
| Pard6b | 0.32025586 | -0.577468 | 0 | 0.01 | 1 | 0.09043413 | 1.86006974 | 0.029 | 0 | 1 | 0.32025586 | 0.17268993 |
| Gm13134 | 0.38689439 | -0.577468 | 0.01 | 0.026 | 1 | 0.09043413 | 1.86006974 | 0.029 | 0 | 1 | 0.38689439 | 0.17268993 |
| Gm41396 | 0.72574709 | -0.3144336 | 0.01 | 0.016 | 1 | 0.09043413 | 1.86006974 | 0.029 | 0 | 1 | 0.72574709 | 0.17268993 |
| F630206G17Rik | 0.99603723 | 0.15949763 | 0.031 | 0.031 | 1 | 0.09043413 | 2.01207283 | 0.029 | 0 | 1 | 0.99603723 | 0.17268993 |
| Gm16573 | 0.99758609 | 0.27052894 | 0.021 | 0.021 | 1 | 0.09043413 | 1.86006974 | 0.029 | 0 | 1 | 0.99758609 | 0.17268993 |
| Agtr1a | 0.15119482 | -1.6929452 | 0.01 | 0.041 | 1 | 0.09044045 | 2.14957636 | 0.029 | 0 | 1 | 0.15119482 | 0.17270143 |
| Gm43661 | 0.22187388 | 1.00749454 | 0.021 | 0.005 | 1 | 0.09044045 | 2.01207283 | 0.029 | 0 | 1 | 0.22187388 | 0.17270143 |
| H4c14 | 0.3445565 | 0.46692616 | 0.094 | 0.062 | 1 | 0.09044045 | 2.27510724 | 0.029 | 0 | 1 | 0.3445565 | 0.17270143 |
| Ankrd61 | 0.38028858 | 1.00749454 | 0.031 | 0.016 | 1 | 0.09044045 | 2.01207283 | 0.029 | 0 | 1 | 0.38028858 | 0.17270143 |
| Prodh | 0.47606502 | 1.00749454 | 0.021 | 0.01 | 1 | 0.09044045 | 2.01207283 | 0.029 | 0 | 1 | 0.47606502 | 0.17270143 |
| Kif9 | 0.74420148 | 0.37006462 | 0.052 | 0.062 | 1 | 0.09044045 | 2.01207283 | 0.029 | 0 | 1 | 0.74420148 | 0.17270143 |
| Lrrc3b | 0.15837881 | 2.00749454 | 0.01 | 0 | 1 | 0.09044466 | 2.39058446 | 0.029 | 0 | 1 | 0.15837881 | 0.17270909 |
| Sptssb | 0.47606502 | 1.00749454 | 0.021 | 0.01 | 1 | 0.09044466 | 2.49749966 | 0.029 | 0 | 1 | 0.47606502 | 0.17270909 |
| Trim66 | 0.72574709 | -0.3144336 | 0.01 | 0.016 | 1 | 0.09044466 | 2.14957636 | 0.029 | 0 | 1 | 0.72574709 | 0.17270909 |
| Pcdhb10 | 1 | 0.42253204 | 0.01 | 0.01 | 1 | 0.09044466 | 2.59703533 | 0.029 | 0 | 1 | 1 | 0.17270909 |
| Cyth4 | 0.15837881 | 2.59245704 | 0.01 | 0 | 1 | 0.09044677 | 2.86006974 | 0.029 | 0 | 1 | 0.15837881 | 0.17271293 |
| Gm29865 | 0.5881353 | 0.68556644 | 0.031 | 0.021 | 1 | 0.09044677 | 2.69014474 | 0.029 | 0 | 1 | 0.5881353 | 0.17271293 |
| Gm5608 | 0.15117462 | 0.81484946 | 0.052 | 0.021 | 1 | 0.09044888 | 1.69014474 | 0.029 | 0 | 1 | 0.15117462 | 0.17271676 |
| BC026513 | 0.15837881 | 2.00749454 | 0.01 | 0 | 1 | 0.09044888 | 1.69014474 | 0.029 | 0 | 1 | 0.15837881 | 0.17271676 |
| 9130221H12Rik | 0.21867159 | 1.59245704 | 0.021 | 0.005 | 1 | 0.09044888 | 1.69014474 | 0.029 | 0 | 1 | 0.21867159 | 0.17271676 |
| P2ry2 | 0.25350784 | 0.56003556 | 0.094 | 0.057 | 1 | 0.09044888 | 1.69014474 | 0.029 | 0 | 1 | 0.25350784 | 0.17271676 |
| 9330136K24Rik | 0.2632671 | 0.13302542 | 0.052 | 0.026 | 1 | 0.09044888 | 1.69014474 | 0.029 | 0 | 1 | 0.2632671 | 0.17271676 |
| Gm56728 | 0.30891093 | 1.00749454 | 0.042 | 0.021 | 1 | 0.09044888 | 1.69014474 | 0.029 | 0 | 1 | 0.30891093 | 0.17271676 |
| Gm57453 | 0.36601088 | -0.7294711 | 0.021 | 0.041 | 1 | 0.09044888 | 1.69014474 | 0.029 | 0 | 1 | 0.36601088 | 0.17271676 |
| Gm43260 | 0.38521705 | 0.68556644 | 0.031 | 0.016 | 1 | 0.09044888 | 1.69014474 | 0.029 | 0 | 1 | 0.38521705 | 0.17271676 |
| Ctsw | 0.5948306 | 0.47697982 | 0.073 | 0.057 | 1 | 0.09044888 | 1.69014474 | 0.029 | 0 | 1 | 0.5948306 | 0.17271676 |
| 4930500A05Rik | 0.60815045 | -1.1079827 | 0.021 | 0.031 | 1 | 0.09044888 | 1.69014474 | 0.029 | 0 | 1 | 0.60815045 | 0.17271676 |
| Apoc1 | 0.62104899 | 0.42253204 | 0.01 | 0.005 | 1 | 0.09044888 | 1.69014474 | 0.029 | 0 | 1 | 0.62104899 | 0.17271676 |
| Gm30455 | 0.72298091 | -0.577468 | 0.01 | 0.016 | 1 | 0.09044888 | 1.69014474 | 0.029 | 0 | 1 | 0.72298091 | 0.17271676 |
| Spata2l | 0.72574709 | -0.3144336 | 0.01 | 0.016 | 1 | 0.09044888 | 1.69014474 | 0.029 | 0 | 1 | 0.72574709 | 0.17271676 |
| Arhgap27os2 | 0.75328966 | 0.27052894 | 0.021 | 0.016 | 1 | 0.09044888 | 1.69014474 | 0.029 | 0 | 1 | 0.75328966 | 0.17271676 |
| Cerox1 | 0.78979801 | -0.407543 | 0.021 | 0.026 | 1 | 0.09044888 | 1.69014474 | 0.029 | 0 | 1 | 0.78979801 | 0.17271676 |

| | | | | | | | | | | | | |
|---|---|---|---|---|---|---|---|---|---|---|---|---|
| Atp6v0d2 | 0.8007213 | -0.577468 | 0.031 | 0.026 | 1 | 0.09044888 | 6.36962484 | 0.029 | 0 | 1 | 0.8007213 | 0.17271676 |
| Sorbs2os | 0.0905394 | 0.96310042 | 0.198 | 0.124 | 1 | 0.095119 | 0.39641353 | 0.21 | 0.125 | 1 | 0.095119 | 0.17288141 |
| Upf3b | 0.09065809 | 0.22698123 | 0.625 | 0.518 | 1 | 0.69509633 | -0.1272638 | 0.613 | 0.583 | 1 | 0.69509633 | 0.17309728 |
| Qsox2 | 0.314976 | 0.32942263 | 0.219 | 0.166 | 1 | 0.09066374 | 0.20471791 | 0.21 | 0.125 | 1 | 0.314976 | 0.17310757 |
| Zfp316 | 0.0907141 | 1.15949763 | 0.083 | 0.036 | 1 | 0.34625029 | 0.69014474 | 0.042 | 0.021 | 1 | 0.34625029 | 0.17319916 |
| Slc66a2 | 0.98750192 | 0.16277276 | 0.562 | 0.596 | 1 | 0.09076131 | -0.7326284 | 0.55 | 0.646 | 1 | 0.98750192 | 0.17328501 |
| 2700069I18Rik | 0.79259943 | 0.52206771 | 0.031 | 0.026 | 1 | 0.09077507 | 1.39058446 | 0.067 | 0.021 | 1 | 0.79259943 | 0.17331003 |
| Akr1b10 | 0.28794709 | 0.42253204 | 0.219 | 0.166 | 1 | 0.09079084 | 0.2095189 | 0.16 | 0.083 | 1 | 0.28794709 | 0.17333871 |
| Trmt9b | 0.97438 | -0.3848229 | 0.062 | 0.062 | 1 | 0.09093827 | 0.93807225 | 0.084 | 0.031 | 1 | 0.97438 | 0.17360677 |
| Ttc13 | 0.25099532 | 0.29700115 | 0.198 | 0.14 | 1 | 0.09110366 | 0.39058446 | 0.185 | 0.104 | 1 | 0.25099532 | 0.17390744 |
| Gm9975 | 0.09112487 | -1.577468 | 0.021 | 0.067 | 1 | 0.23969244 | -1.0203486 | 0.042 | 0.073 | 1 | 0.23969244 | 0.173946 |
| 0610040F04Rik | 0.09113532 | -2.5470943 | 0.031 | 0.083 | 1 | 0.3731428 | -2.9688183 | 0.05 | 0.073 | 1 | 0.3731428 | 0.17396498 |
| Gm16066 | 0.15401115 | 0.53105649 | 0.188 | 0.124 | 1 | 0.0913918 | 0.15580831 | 0.214 | 0.125 | 1 | 0.15401115 | 0.17443114 |
| Icam2 | 0.22187575 | -1.577468 | 0 | 0.016 | 1 | 0.091405 | -2.4797803 | 0.013 | 0.042 | 1 | 0.22187575 | 0.17445512 |
| Mtor | 0.09142366 | 0.14076107 | 0.406 | 0.285 | 1 | 0.41129679 | -0.1630139 | 0.403 | 0.302 | 1 | 0.41129679 | 0.17448904 |
| A430090L17Rik | 0.22187575 | -0.9925055 | 0 | 0.016 | 1 | 0.09150755 | -2.0874628 | 0.025 | 0.062 | 1 | 0.22187575 | 0.17464146 |
| Acad10 | 0.24262368 | 1.38600616 | 0.073 | 0.041 | 1 | 0.09155048 | 1.01207283 | 0.05 | 0.01 | 1 | 0.24262368 | 0.17471948 |
| Fcho2 | 0.09155241 | -0.2105061 | 0.906 | 0.907 | 1 | 0.10635795 | -0.4384599 | 0.971 | 0.979 | 1 | 0.10635795 | 0.17472298 |
| AB124611 | 0.09161032 | 0.63552576 | 0.135 | 0.073 | 1 | 0.41417623 | -0.7395395 | 0.118 | 0.146 | 1 | 0.41417623 | 0.17482818 |
| Usp21 | 0.09161525 | 0.71203865 | 0.188 | 0.114 | 1 | 0.16411704 | 0.33060235 | 0.181 | 0.115 | 1 | 0.16411704 | 0.17483715 |
| Specc1 | 0.67520466 | -0.5889636 | 0.438 | 0.383 | 1 | 0.09161609 | 0.20805191 | 0.79 | 0.719 | 1 | 0.67520466 | 0.17483868 |
| Prickle2 | 0.09161658 | -1.0140708 | 0.26 | 0.337 | 1 | 0.67114966 | -0.2863963 | 0.42 | 0.354 | 1 | 0.67114966 | 0.17483957 |
| Arfgap2 | 0.50922191 | 0.1624692 | 0.396 | 0.352 | 1 | 0.09169041 | 0.26205109 | 0.353 | 0.24 | 1 | 0.50922191 | 0.17497368 |
| Abitram | 0.09176639 | 0.80104366 | 0.208 | 0.135 | 1 | 0.74876911 | -0.3463811 | 0.122 | 0.104 | 1 | 0.74876911 | 0.17511171 |
| Dus3l | 0.18375613 | 0.42253204 | 0.24 | 0.171 | 1 | 0.09184365 | 0.42711033 | 0.181 | 0.104 | 1 | 0.18375613 | 0.17525204 |
| Mterf4 | 0.80201301 | 0.14076107 | 0.146 | 0.135 | 1 | 0.09185329 | 0.46273424 | 0.155 | 0.083 | 1 | 0.80201301 | 0.17526954 |
| Stt3b | 0.12912601 | -0.2223281 | 0.844 | 0.948 | 1 | 0.09187167 | -0.1619218 | 0.975 | 0.938 | 1 | 0.12912601 | 0.17530293 |
| Catsperg1 | 1 | 0.42253204 | 0.01 | 0.01 | 1 | 0.0918871 | 0.91253716 | 0.05 | 0.01 | 1 | 1 | 0.17533096 |
| Zfp358 | 0.70779422 | -0.1180363 | 0.104 | 0.119 | 1 | 0.09194167 | 0.34910782 | 0.21 | 0.125 | 1 | 0.70779422 | 0.17543007 |
| Irf2 | 0.92446986 | -0.1611873 | 0.604 | 0.58 | 1 | 0.09202732 | -0.3944675 | 0.676 | 0.823 | 1 | 0.92446986 | 0.17558562 |
| Hdhd5 | 0.09211581 | -1.5160674 | 0.031 | 0.083 | 1 | 0.1164329 | 0.37820073 | 0.109 | 0.052 | 1 | 0.1164329 | 0.1757463 |
| Siah1a | 0.09217928 | 0.20013961 | 0.521 | 0.404 | 1 | 0.28039566 | -0.4353861 | 0.605 | 0.635 | 1 | 0.28039566 | 0.17586153 |
| Rarres1 | 0.73111155 | -0.3542738 | 0.25 | 0.218 | 1 | 0.09218513 | 0.18657056 | 0.265 | 0.167 | 1 | 0.73111155 | 0.17587217 |
| Itpkb | 0.42691842 | -0.3861693 | 0.49 | 0.534 | 1 | 0.09224003 | -1.2370989 | 0.387 | 0.438 | 1 | 0.42691842 | 0.17597184 |
| Nupr1l | 0.21218051 | -1.1079827 | 0.021 | 0.052 | 1 | 0.09226049 | 0.59703533 | 0.05 | 0.01 | 1 | 0.21218051 | 0.17600897 |
| AU040972 | 0.75328966 | 0.27052894 | 0.021 | 0.016 | 1 | 0.092271 | -0.8073549 | 0.05 | 0.01 | 1 | 0.75328966 | 0.17602805 |
| Gm56488 | 0.11302151 | -1.7998604 | 0 | 0.026 | 1 | 0.09238206 | -2.4473588 | 0.013 | 0.042 | 1 | 0.11302151 | 0.17622967 |
| Ptch2 | 0.52744059 | -0.577468 | 0.01 | 0.021 | 1 | 0.09238206 | -2.4473588 | 0.013 | 0.042 | 1 | 0.52744059 | 0.17622967 |
| Got2 | 0.91144322 | -0.113969 | 0.531 | 0.508 | 1 | 0.09239423 | 0.33481348 | 0.559 | 0.51 | 1 | 0.91144322 | 0.17625176 |
| Lactb2 | 0.38733284 | -0.5611662 | 0.458 | 0.477 | 1 | 0.0924039 | 0.21370669 | 0.29 | 0.188 | 1 | 0.38733284 | 0.17626932 |
| Npr3 | 0.09244562 | -1.2359595 | 0.177 | 0.254 | 1 | 0.46471582 | -0.1172102 | 0.05 | 0.031 | 1 | 0.46471582 | 0.17634505 |
| Klrg2 | 0.09247508 | 0.68556644 | 0.115 | 0.057 | 1 | 0.14552074 | -0.2788284 | 0.147 | 0.083 | 1 | 0.14552074 | 0.17639852 |
| Sned1 | 0.092477 | -0.4974997 | 0.552 | 0.601 | 1 | 0.10703279 | 0.21333104 | 0.756 | 0.615 | 1 | 0.10703279 | 0.176402 |
| Tfdp2 | 0.13321027 | -0.3597144 | 0.375 | 0.477 | 1 | 0.09254195 | -0.718495 | 0.601 | 0.656 | 1 | 0.13321027 | 0.17651989 |
| Ptger3 | 0.09315171 | 0.90964721 | 0.208 | 0.135 | 1 | 0.09256932 | 0.59703533 | 0.084 | 0.031 | 1 | 0.09315171 | 0.17656956 |
| Parg | 0.09262968 | 0.10937415 | 0.625 | 0.492 | 1 | 0.21092998 | -0.4311616 | 0.744 | 0.75 | 1 | 0.21092998 | 0.1766791 |
| Erc1 | 0.2490483 | 0.27798678 | 0.917 | 0.974 | 1 | 0.09267709 | -0.509803 | 0.983 | 0.979 | 1 | 0.2490483 | 0.17676514 |
| Gata2 | 0.99517218 | -0.9925055 | 0.021 | 0.021 | 1 | 0.09271918 | -1.9343461 | 0.071 | 0.125 | 1 | 0.99517218 | 0.17684152 |
| Gm17135 | 0.11303029 | -1.577468 | 0 | 0.026 | 1 | 0.0928609 | -2.1172102 | 0.013 | 0.042 | 1 | 0.11303029 | 0.17709864 |
| Jag2 | 0.15729781 | -1.7998604 | 0 | 0.021 | 1 | 0.0928609 | -2.1172102 | 0.013 | 0.042 | 1 | 0.15729781 | 0.17709864 |
| Gm57065 | 0.47911121 | -0.9925055 | 0.021 | 0.036 | 1 | 0.0928609 | -1.8948178 | 0.013 | 0.042 | 1 | 0.47911121 | 0.17709864 |
| Psmb9 | 0.7609195 | 0.65957123 | 0.052 | 0.062 | 1 | 0.0928609 | -1.8948178 | 0.013 | 0.042 | 1 | 0.7609195 | 0.17709864 |
| Htr6 | 0.78979801 | -0.577468 | 0.021 | 0.026 | 1 | 0.0928609 | -2.3098553 | 0.013 | 0.042 | 1 | 0.78979801 | 0.17709864 |
| Xrcc5 | 0.11326696 | 0.58738605 | 0.365 | 0.269 | 1 | 0.09287774 | -0.1154772 | 0.395 | 0.26 | 1 | 0.11326696 | 0.1771292 |
| Ndrg3 | 0.18745111 | 0.19390766 | 0.542 | 0.472 | 1 | 0.09290001 | -0.6787808 | 0.597 | 0.615 | 1 | 0.18745111 | 0.1771696 |
| Cpeb1 | 0.09292115 | 0.75373794 | 0.24 | 0.161 | 1 | 0.65844239 | -0.1058414 | 0.303 | 0.26 | 1 | 0.65844239 | 0.17720796 |
| Snx11 | 0.80685636 | 0.13302542 | 0.156 | 0.145 | 1 | 0.09292162 | 0.57466752 | 0.181 | 0.104 | 1 | 0.80685636 | 0.17720881 |
| E2f1 | 0.09299877 | 0.59245704 | 0.115 | 0.057 | 1 | 0.59693489 | -0.5728897 | 0.067 | 0.083 | 1 | 0.59693489 | 0.17734877 |
| Ipo13 | 0.09299896 | 0.5049942 | 0.115 | 0.057 | 1 | 0.60349739 | -0.2394659 | 0.151 | 0.125 | 1 | 0.60349739 | 0.17734912 |
| Paqr8 | 0.15315302 | -0.9925055 | 0.042 | 0.088 | 1 | 0.09301373 | -1.4029647 | 0.046 | 0.094 | 1 | 0.15315302 | 0.1773759 |
| Ncald | 0.09310293 | -1.3144336 | 0.031 | 0.083 | 1 | 0.38484315 | -1.010295 | 0.076 | 0.104 | 1 | 0.38484315 | 0.1775377 |
| Trim25 | 0.09325964 | 0.54183096 | 0.271 | 0.181 | 1 | 0.9131803 | -0.2437661 | 0.324 | 0.333 | 1 | 0.9131803 | 0.17782192 |
| Gm41077 | 0.09326355 | 0.63309902 | 0.208 | 0.135 | 1 | 0.92148942 | -0.6984206 | 0.16 | 0.156 | 1 | 0.92148942 | 0.177829 |
| Hemk1 | 0.09331579 | 1.00749454 | 0.083 | 0.036 | 1 | 0.43079893 | 0.69014474 | 0.038 | 0.021 | 1 | 0.43079893 | 0.17792374 |
| Gm15614 | 0.09333117 | 0.77045534 | 0.198 | 0.124 | 1 | 0.38526909 | -0.5025003 | 0.172 | 0.125 | 1 | 0.38526909 | 0.17795163 |
| Mbnl1 | 0.09355929 | -0.2984666 | 1 | 1 | 1 | 0.11351663 | -0.283967 | 1 | 0.99 | 1 | 0.11351663 | 0.17836524 |
| Fam210a | 0.15815587 | 0.25340476 | 0.479 | 0.383 | 1 | 0.09356431 | -0.1200307 | 0.55 | 0.375 | 1 | 0.15815587 | 0.17837434 |
| Tox3 | 0.09369842 | 1.7693348 | 0.135 | 0.078 | 1 | 0.66519574 | -0.7504279 | 0.16 | 0.177 | 1 | 0.66519574 | 0.17861745 |
| Elf4 | 0.14332737 | 0.33738392 | 0.625 | 0.544 | 1 | 0.09374021 | -0.2608479 | 0.697 | 0.469 | 1 | 0.14332737 | 0.17869319 |
| Cab39l | 0.09377054 | -0.5018037 | 0.656 | 0.653 | 1 | 0.84604079 | -0.3742819 | 0.718 | 0.729 | 1 | 0.84604079 | 0.17874816 |
| Gm13919 | 0.27538452 | 0.42253204 | 0.073 | 0.041 | 1 | 0.09393539 | 0.56461386 | 0.113 | 0.052 | 1 | 0.27538452 | 0.17904693 |
| Dlst | 0.12050737 | 0.2407816 | 0.719 | 0.627 | 1 | 0.09393706 | -0.4347647 | 0.744 | 0.802 | 1 | 0.12050737 | 0.17904995 |
| Cars2 | 0.70899546 | -0.4330781 | 0.167 | 0.181 | 1 | 0.09402477 | 0.22065945 | 0.189 | 0.104 | 1 | 0.70899546 | 0.17920888 |
| Efna4 | 0.56320982 | -0.3710171 | 0.073 | 0.093 | 1 | 0.09406332 | 1.06865636 | 0.097 | 0.042 | 1 | 0.56320982 | 0.17927873 |
| Coasy | 0.30295194 | 0.32942263 | 0.177 | 0.13 | 1 | 0.0941322 | 0.80562195 | 0.151 | 0.083 | 1 | 0.30295194 | 0.17940352 |
| Slc39a6 | 0.09414259 | 0.23126951 | 0.615 | 0.487 | 1 | 0.13346827 | 0.24268576 | 0.55 | 0.469 | 1 | 0.13346827 | 0.17942236 |
| 9530077C05Rik | 0.09415758 | 1.00749454 | 0.083 | 0.036 | 1 | 0.10045901 | 1.05271482 | 0.109 | 0.052 | 1 | 0.10045901 | 0.17944951 |
| Catspere2 | 0.09416912 | 1.06889508 | 0.156 | 0.093 | 1 | 0.96767817 | -1.2055186 | 0.134 | 0.125 | 1 | 0.96767817 | 0.17947041 |
| Ntmt1 | 0.52243883 | 0.13302542 | 0.385 | 0.358 | 1 | 0.09419448 | -0.188549 | 0.403 | 0.271 | 1 | 0.52243883 | 0.17951636 |
| Poc1b | 0.09421226 | 0.33506919 | 0.229 | 0.145 | 1 | 0.11156262 | 0.11004799 | 0.307 | 0.198 | 1 | 0.11156262 | 0.17954858 |
| Gm44686 | 0.15405555 | -1.1624305 | 0.01 | 0.041 | 1 | 0.09421884 | -1.3098553 | 0.071 | 0.125 | 1 | 0.15405555 | 0.1795605 |
| Plcxd3 | 0.2069367 | -2.1079827 | 0.01 | 0.036 | 1 | 0.09434833 | -2.3098553 | 0.013 | 0.042 | 1 | 0.2069367 | 0.17979505 |
| Cxcl1 | 0.26188554 | -0.9785663 | 0.115 | 0.161 | 1 | 0.09434833 | -2.1578522 | 0.013 | 0.042 | 1 | 0.26188554 | 0.17979505 |
| Gm26737 | 0.38523157 | 2.90057933 | 0.031 | 0.016 | 1 | 0.09434833 | -1.7952821 | 0.013 | 0.042 | 1 | 0.38523157 | 0.17979505 |
| Gm32191 | 0.48064264 | 0.59245704 | 0.021 | 0.01 | 1 | 0.09434833 | -1.7952821 | 0.013 | 0.042 | 1 | 0.48064264 | 0.17979505 |
| Gm35550 | 1 | 0.42253204 | 0.01 | 0.01 | 1 | 0.09434833 | -1.9879272 | 0.013 | 0.042 | 1 | 1 | 0.17979505 |
| Gm5165 | 0.5867163 | 0.25260703 | 0.115 | 0.093 | 1 | 0.0946026 | 0.44503224 | 0.155 | 0.083 | 1 | 0.5867163 | 0.18025555 |

| | | | | | | | | | | | | |
|---|---|---|---|---|---|---|---|---|---|---|---|---|
| Camk2n1 | 0.09460423 | 0.40935665 | 0.49 | 0.373 | 1 | 0.14339407 | 0.17702224 | 0.71 | 0.656 | 1 | 0.14339407 | 0.18025851 |
| Muc1 | 0.13153526 | 0.83756954 | 0.062 | 0.026 | 1 | 0.09476388 | 0.10518224 | 0.084 | 0.031 | 1 | 0.13153526 | 0.18054757 |
| Acot2 | 0.28687627 | 0.57139542 | 0.115 | 0.078 | 1 | 0.09476388 | 0.46775232 | 0.084 | 0.031 | 1 | 0.28687627 | 0.18054757 |
| Prag1 | 0.09479132 | -0.4664367 | 0.094 | 0.171 | 1 | 0.7589818 | 0.49749966 | 0.038 | 0.031 | 1 | 0.7589818 | 0.18059724 |
| Obp1a | 0.0948373 | 0.45290568 | 0.271 | 0.181 | 1 | 0.70814964 | -0.3349462 | 0.231 | 0.198 | 1 | 0.70814964 | 0.18068048 |
| Golph3l | 0.09485155 | 0.43558819 | 0.281 | 0.187 | 1 | 0.34469203 | -0.1054968 | 0.252 | 0.188 | 1 | 0.34469203 | 0.18070628 |
| Ndufs1 | 0.09502709 | 0.30557589 | 0.521 | 0.409 | 1 | 0.76317307 | -0.270015 | 0.613 | 0.552 | 1 | 0.76317307 | 0.18102403 |
| Plpp7 | 0.72298091 | -0.7998604 | 0.01 | 0.016 | 1 | 0.09503269 | 0.45567948 | 0.067 | 0.021 | 1 | 0.72298091 | 0.18103416 |
| Alms1 | 0.17545312 | 0.80396114 | 0.198 | 0.135 | 1 | 0.09504725 | -0.7379489 | 0.126 | 0.198 | 1 | 0.17545312 | 0.18106052 |
| Tirap | 0.09522876 | 0.50792353 | 0.25 | 0.166 | 1 | 0.13494873 | 0.64707602 | 0.206 | 0.135 | 1 | 0.13494873 | 0.181389 |
| Rad23a | 0.09529444 | 0.35282306 | 0.604 | 0.523 | 1 | 0.16570201 | 0.16407593 | 0.58 | 0.479 | 1 | 0.16570201 | 0.18150785 |
| Bag4 | 0.46386328 | 0.12778877 | 0.188 | 0.15 | 1 | 0.0953133 | 0.42711033 | 0.197 | 0.115 | 1 | 0.46386328 | 0.18154198 |
| Cep70 | 0.09540524 | 0.7693348 | 0.24 | 0.155 | 1 | 0.441568 | -0.6468902 | 0.109 | 0.135 | 1 | 0.441568 | 0.18170833 |
| Plxnb1 | 0.09545652 | 0.47697982 | 0.229 | 0.145 | 1 | 0.20123518 | 0.24155468 | 0.286 | 0.208 | 1 | 0.20123518 | 0.18180109 |
| Puf60 | 0.09545837 | 0.23153481 | 0.76 | 0.668 | 1 | 0.17654552 | 0.15561696 | 0.723 | 0.698 | 1 | 0.17654552 | 0.18180444 |
| Mtrf1 | 0.09552912 | 1.32942263 | 0.104 | 0.052 | 1 | 0.22902882 | 0.59703533 | 0.05 | 0.021 | 1 | 0.22902882 | 0.18193243 |
| Tapbpl | 0.71658253 | -0.4779323 | 0.146 | 0.155 | 1 | 0.09555903 | 0.52021974 | 0.113 | 0.052 | 1 | 0.71658253 | 0.18198653 |
| Mtrex | 0.09559174 | 0.14665728 | 0.594 | 0.446 | 1 | 0.97915116 | -0.101914 | 0.651 | 0.615 | 1 | 0.97915116 | 0.1820457 |
| Gm28055 | 0.11302941 | -2.1624305 | 0 | 0.026 | 1 | 0.09568573 | -1.6317834 | 0.025 | 0.062 | 1 | 0.11302941 | 0.1822157 |
| Bbs10 | 0.44158998 | 0.49292136 | 0.062 | 0.041 | 1 | 0.09568573 | -1.6317834 | 0.025 | 0.062 | 1 | 0.44158998 | 0.1822157 |
| Tmem141 | 0.32587032 | -0.577468 | 0.125 | 0.166 | 1 | 0.09584216 | 0.23071312 | 0.113 | 0.052 | 1 | 0.32587032 | 0.18249859 |
| Tnks | 0.09586621 | 0.2134692 | 0.885 | 0.746 | 1 | 0.56835875 | -0.2641147 | 0.924 | 0.917 | 1 | 0.56835875 | 0.18254209 |
| Hsd17b8 | 0.09588773 | 0.31359766 | 0.302 | 0.202 | 1 | 0.11168903 | 0.52303475 | 0.202 | 0.125 | 1 | 0.11168903 | 0.182581 |
| H4c16 | 0.09591527 | -1.5285584 | 0.031 | 0.083 | 1 | 0.18638441 | -0.7952821 | 0.017 | 0.042 | 1 | 0.18638441 | 0.18263079 |
| Pgap3 | 0.12532804 | 1.00749454 | 0.115 | 0.062 | 1 | 0.09601973 | 0.3146356 | 0.13 | 0.062 | 1 | 0.12532804 | 0.18281967 |
| Mthfd1l | 0.09603957 | 0.12015234 | 0.625 | 0.528 | 1 | 0.61997405 | -0.5034963 | 0.555 | 0.448 | 1 | 0.61997405 | 0.18285554 |
| Trim65 | 0.33113624 | 0.35541784 | 0.062 | 0.036 | 1 | 0.09604755 | -1.6724253 | 0.025 | 0.062 | 1 | 0.33113624 | 0.18286996 |
| Cdc6 | 0.09626076 | 1.20013961 | 0.104 | 0.052 | 1 | 0.47476851 | -0.8948178 | 0.063 | 0.083 | 1 | 0.47476851 | 0.18325539 |
| Trafd1 | 0.09642069 | 0.3951556 | 0.688 | 0.611 | 1 | 0.58466705 | -0.3157318 | 0.613 | 0.583 | 1 | 0.58466705 | 0.18354443 |
| Tlr4 | 0.09644365 | -0.6072153 | 0.156 | 0.238 | 1 | 0.55333614 | -0.5091641 | 0.134 | 0.156 | 1 | 0.55333614 | 0.18358592 |
| Rab32 | 0.30389555 | 1.27052894 | 0.042 | 0.021 | 1 | 0.09646187 | -1.4353861 | 0.025 | 0.062 | 1 | 0.30389555 | 0.18361884 |
| Pfdn1 | 0.26722428 | -0.1821079 | 0.5 | 0.57 | 1 | 0.09653259 | 0.17238094 | 0.622 | 0.552 | 1 | 0.26722428 | 0.18374663 |
| 4732440D04Rik | 0.09660556 | 0.73373372 | 0.312 | 0.233 | 1 | 0.37872054 | -0.1054968 | 0.378 | 0.302 | 1 | 0.37872054 | 0.18387848 |
| Rab11fip2 | 0.39783417 | 0.19746548 | 0.458 | 0.389 | 1 | 0.09664778 | -0.1115265 | 0.475 | 0.312 | 1 | 0.39783417 | 0.18395476 |
| Itpk1 | 0.09666673 | 0.92503238 | 0.219 | 0.15 | 1 | 0.4483727 | -1.2742314 | 0.151 | 0.177 | 1 | 0.4483727 | 0.18398901 |
| Desi2 | 0.13309281 | 0.42589889 | 0.938 | 0.87 | 1 | 0.09677303 | 0.14366547 | 0.975 | 0.958 | 1 | 0.13309281 | 0.18418104 |
| Rab20 | 0.0968048 | -0.9925055 | 0.094 | 0.166 | 1 | 0.54258249 | -0.9562183 | 0.155 | 0.177 | 1 | 0.54258249 | 0.18423843 |
| Trpm3 | 0.285137 | -1.7350092 | 0.094 | 0.135 | 1 | 0.09683786 | 0.77846454 | 0.265 | 0.177 | 1 | 0.285137 | 0.18429816 |
| Tmem145 | 0.99758604 | -0.2148979 | 0.021 | 0.021 | 1 | 0.09686309 | -1.3098553 | 0.013 | 0.042 | 1 | 0.99758604 | 0.18434373 |
| Nasp | 0.0969326 | 0.12948506 | 0.708 | 0.668 | 1 | 0.35202437 | -0.1788859 | 0.861 | 0.865 | 1 | 0.35202437 | 0.18446928 |
| Fuca2 | 0.45528036 | 0.12613903 | 0.167 | 0.13 | 1 | 0.0970424 | 0.20471791 | 0.273 | 0.177 | 1 | 0.45528036 | 0.18466757 |
| Pnpla2 | 0.0970844 | -0.8135353 | 0.135 | 0.212 | 1 | 0.20750389 | 0.70721825 | 0.244 | 0.188 | 1 | 0.20750389 | 0.18474341 |
| Osgepl1 | 0.3577459 | 0.38600616 | 0.177 | 0.135 | 1 | 0.09708715 | 0.41596977 | 0.143 | 0.073 | 1 | 0.3577459 | 0.18474839 |
| Gtf2h1 | 0.09714503 | 0.15663798 | 0.646 | 0.518 | 1 | 0.32593824 | -0.40108 | 0.731 | 0.719 | 1 | 0.32593824 | 0.18485291 |
| Hmces | 0.15824254 | 0.32299636 | 0.385 | 0.29 | 1 | 0.09716337 | -0.1130546 | 0.42 | 0.281 | 1 | 0.15824254 | 0.18488602 |
| Itga7 | 0.28631176 | -1.3144336 | 0.021 | 0.047 | 1 | 0.09725677 | -1.0468209 | 0.025 | 0.062 | 1 | 0.28631176 | 0.18505465 |
| Umad1 | 0.09734637 | 0.67734593 | 0.312 | 0.223 | 1 | 0.90256201 | -0.5120441 | 0.416 | 0.406 | 1 | 0.90256201 | 0.18521642 |
| Casp3 | 0.09736512 | 0.44708914 | 0.573 | 0.456 | 1 | 0.32922493 | 0.25157172 | 0.546 | 0.5 | 1 | 0.32922493 | 0.18525027 |
| 9330133O14Rik | 0.09750585 | -0.8669746 | 0.042 | 0.098 | 1 | 0.28848731 | 0.27510724 | 0.046 | 0.021 | 1 | 0.28848731 | 0.1855043 |
| Kat14 | 0.10121416 | 0.59245704 | 0.198 | 0.124 | 1 | 0.097525 | 0.33400093 | 0.214 | 0.135 | 1 | 0.10121416 | 0.18553887 |
| Gm56974 | 0.09761362 | -0.9925055 | 0.021 | 0.067 | 1 | 0.23772424 | 0.86006974 | 0.034 | 0.01 | 1 | 0.23772424 | 0.18569881 |
| Axdnd1 | 0.39417399 | 0.11440974 | 0.052 | 0.031 | 1 | 0.09765173 | -1.3098553 | 0.025 | 0.062 | 1 | 0.39417399 | 0.18576761 |
| Gm46412 | 0.28678131 | 0.83756954 | 0.115 | 0.078 | 1 | 0.09768856 | 0.62874419 | 0.067 | 0.021 | 1 | 0.28678131 | 0.18583406 |
| Pdgfc | 0.0977803 | -0.337334 | 0.177 | 0.275 | 1 | 0.47592879 | -0.4952103 | 0.462 | 0.375 | 1 | 0.47592879 | 0.18599961 |
| Dab2ip | 0.09781342 | 0.3358204 | 0.594 | 0.461 | 1 | 0.71220819 | -0.1446995 | 0.71 | 0.781 | 1 | 0.71220819 | 0.18605938 |
| 1110002L01Rik | 0.47969243 | 0.37672835 | 0.24 | 0.202 | 1 | 0.09782188 | 0.29421606 | 0.235 | 0.146 | 1 | 0.47969243 | 0.18607464 |
| Timmdc1 | 0.11404528 | -0.4519371 | 0.229 | 0.316 | 1 | 0.09782976 | 0.16407593 | 0.256 | 0.167 | 1 | 0.11404528 | 0.18608885 |
| Mri1 | 0.50575263 | -0.1118044 | 0.24 | 0.28 | 1 | 0.09802565 | 0.3146356 | 0.223 | 0.135 | 1 | 0.50575263 | 0.18644227 |
| Il18 | 0.58534847 | -0.183647 | 0.375 | 0.399 | 1 | 0.09803083 | -0.1276519 | 0.328 | 0.208 | 1 | 0.58534847 | 0.18645161 |
| Irf3 | 0.09807277 | 0.39744105 | 0.312 | 0.218 | 1 | 0.21487021 | -0.2935535 | 0.307 | 0.219 | 1 | 0.21487021 | 0.18652727 |
| Fcer1g | 0.75832339 | -1.6554705 | 0.021 | 0.016 | 1 | 0.0981272 | 0.86006974 | 0.059 | 0.115 | 1 | 0.75832339 | 0.18662545 |
| G2e3 | 0.14889926 | 0.43237982 | 0.344 | 0.264 | 1 | 0.09814764 | -0.3594373 | 0.399 | 0.26 | 1 | 0.14889926 | 0.18666233 |
| Dcaf7 | 0.09817416 | 0.18731557 | 0.448 | 0.332 | 1 | 0.26442325 | -0.1185566 | 0.416 | 0.333 | 1 | 0.26442325 | 0.18671016 |
| Gm38551 | 0.14082853 | 1.32942263 | 0.083 | 0.041 | 1 | 0.09828549 | 0.35999614 | 0.084 | 0.031 | 1 | 0.14082853 | 0.18691093 |
| Ldhc | 0.46686094 | -1.577468 | 0.021 | 0.036 | 1 | 0.09841186 | -1.3098553 | 0.025 | 0.062 | 1 | 0.46686094 | 0.18713883 |
| Tmem186 | 0.09847834 | 0.84702986 | 0.146 | 0.083 | 1 | 0.61290487 | -0.1805722 | 0.113 | 0.135 | 1 | 0.61290487 | 0.1872587 |
| Zfp606 | 0.37952001 | 0.46000674 | 0.188 | 0.15 | 1 | 0.09848758 | 0.84214783 | 0.21 | 0.135 | 1 | 0.37952001 | 0.18727535 |
| Tcerg1 | 0.09851039 | 0.19368459 | 0.865 | 0.772 | 1 | 0.71407323 | -0.1049965 | 0.882 | 0.885 | 1 | 0.71407323 | 0.18731648 |
| Peli2 | 0.09853573 | -1.0602501 | 0.146 | 0.228 | 1 | 0.22681213 | -0.7044866 | 0.303 | 0.344 | 1 | 0.22681213 | 0.18736216 |
| Efcab14 | 0.10083885 | 0.20726405 | 0.792 | 0.715 | 1 | 0.09855718 | 0.13583245 | 0.807 | 0.75 | 1 | 0.10083885 | 0.18740084 |
| Kansl1 | 0.45617932 | -0.3460339 | 0.958 | 0.974 | 1 | 0.09875331 | -0.3123513 | 0.996 | 0.99 | 1 | 0.45617932 | 0.1877544 |
| Ahctf1 | 0.09879199 | 0.1843723 | 0.792 | 0.725 | 1 | 0.3563933 | -0.2500573 | 0.899 | 0.865 | 1 | 0.3563933 | 0.18782412 |
| Gm16740 | 0.09880928 | 0.47144164 | 0.167 | 0.098 | 1 | 0.52179784 | 0.27510724 | 0.118 | 0.094 | 1 | 0.52179784 | 0.18785528 |
| Pik3c3 | 0.31228125 | -0.4950058 | 0.167 | 0.212 | 1 | 0.09881467 | -1.1966447 | 0.176 | 0.24 | 1 | 0.31228125 | 0.18786501 |
| Usp6nl | 0.22619664 | -0.1455171 | 0.635 | 0.679 | 1 | 0.09882345 | -0.3129814 | 0.718 | 0.812 | 1 | 0.22619664 | 0.18788082 |
| Armc5 | 0.09894621 | 0.83756954 | 0.146 | 0.083 | 1 | 0.14596345 | 0.31163311 | 0.16 | 0.094 | 1 | 0.14596345 | 0.18810207 |
| Clec14a | 0.15837881 | 2.00749454 | 0.01 | 0 | 1 | 0.09902427 | 1.10518224 | 0.109 | 0.052 | 1 | 0.15837881 | 0.18824273 |
| Eno3 | 0.90192914 | -0.4157169 | 0.156 | 0.145 | 1 | 0.09902459 | 1.14957636 | 0.109 | 0.052 | 1 | 0.90192914 | 0.18824331 |
| Trappc10 | 0.78430873 | -0.1063254 | 0.438 | 0.44 | 1 | 0.09907474 | -0.5880419 | 0.542 | 0.635 | 1 | 0.78430873 | 0.18833368 |
| Gabpa | 0.09918314 | 0.48393258 | 0.344 | 0.249 | 1 | 0.38758057 | -0.2374468 | 0.412 | 0.333 | 1 | 0.38758057 | 0.18852899 |
| Exog | 0.25696442 | 0.85549144 | 0.073 | 0.041 | 1 | 0.09918625 | 0.86006974 | 0.097 | 0.042 | 1 | 0.25696442 | 0.18853458 |
| Bbln | 0.36347296 | -0.3482672 | 0.458 | 0.472 | 1 | 0.09938934 | 0.43430583 | 0.382 | 0.271 | 1 | 0.36347296 | 0.18890043 |
| Gm16287 | 0.09959647 | 0.71798792 | 0.083 | 0.036 | 1 | 0.1382474 | -0.9472852 | 0.021 | 0.052 | 1 | 0.1382474 | 0.18927348 |
| Zfp39 | 0.09962026 | 1.00749454 | 0.104 | 0.052 | 1 | 0.33151861 | 0.32757466 | 0.097 | 0.062 | 1 | 0.33151861 | 0.18931632 |
| Spata13 | 0.35801112 | -0.3550755 | 0.562 | 0.627 | 1 | 0.09966727 | -0.169534 | 0.647 | 0.448 | 1 | 0.35801112 | 0.18940098 |
| D330041H03Rik | 0.09968317 | 1.00749454 | 0.104 | 0.052 | 1 | 0.26765236 | 0.64950275 | 0.113 | 0.073 | 1 | 0.26765236 | 0.18942961 |
| Nop53 | 0.09970137 | 0.33184529 | 0.76 | 0.674 | 1 | 0.11442473 | 0.26641628 | 0.702 | 0.719 | 1 | 0.11442473 | 0.18946237 |
| Slc25a27 | 0.09972729 | 0.79890791 | 0.24 | 0.161 | 1 | 0.11563485 | -0.2223924 | 0.315 | 0.198 | 1 | 0.11563485 | 0.18950904 |

| | | | | | | | | | | | | |
|---|---|---|---|---|---|---|---|---|---|---|---|---|
| Fam13a | 0.09972997 | 1.00749454 | 0.104 | 0.052 | 1 | 0.2910923 | -0.7793405 | 0.021 | 0.042 | 1 | 0.2910923 | 0.18951388 |
| Cyld | 0.94161874 | 0.23882008 | 0.292 | 0.321 | 1 | 0.09984185 | -0.3263434 | 0.445 | 0.583 | 1 | 0.94161874 | 0.18971531 |
| Sh3d21 | 0.94799745 | -0.7900126 | 0.156 | 0.155 | 1 | 0.09987715 | -0.8094263 | 0.176 | 0.25 | 1 | 0.94799745 | 0.18977886 |
| Ankrd26 | 0.13182091 | 0.93710521 | 0.219 | 0.155 | 1 | 0.09988312 | 0.34414888 | 0.248 | 0.156 | 1 | 0.13182091 | 0.18978961 |
| Dtx3l | 0.09989378 | 0.79804117 | 0.167 | 0.098 | 1 | 0.72717476 | -1.2692133 | 0.118 | 0.125 | 1 | 0.72717476 | 0.18980879 |
| Ctps | 0.09995164 | 0.38936517 | 0.302 | 0.207 | 1 | 0.21367273 | 0.10518224 | 0.269 | 0.198 | 1 | 0.21367273 | 0.18991295 |
| Igfbp4 | 0.10001088 | -1.6618602 | 0.302 | 0.352 | 1 | 0.18639234 | -1.9580116 | 0.424 | 0.406 | 1 | 0.18639234 | 0.19001958 |
| Kank3 | 0.10003141 | 0.74446013 | 0.083 | 0.036 | 1 | 0.34428571 | 0.91253716 | 0.042 | 0.021 | 1 | 0.34428571 | 0.19005655 |
| Mtfmt | 0.32090252 | 0.39744105 | 0.156 | 0.114 | 1 | 0.10018458 | 0.12624385 | 0.168 | 0.094 | 1 | 0.32090252 | 0.19033222 |
| Strn3 | 0.31583326 | 0.15035129 | 0.969 | 0.969 | 1 | 0.10019287 | -0.324401 | 0.992 | 0.99 | 1 | 0.31583326 | 0.19034714 |
| Gm57463 | 0.22944575 | 0.2912875 | 0.135 | 0.088 | 1 | 0.10026427 | 1.14957636 | 0.109 | 0.052 | 1 | 0.22944575 | 0.19047561 |
| Alkbh4 | 0.59996055 | 0.42253204 | 0.073 | 0.057 | 1 | 0.10026437 | 1.01207283 | 0.109 | 0.052 | 1 | 0.59996055 | 0.1904758 |
| Ccdc171 | 0.10036159 | -0.770113 | 0.083 | 0.15 | 1 | 0.47319043 | -1.2592292 | 0.084 | 0.104 | 1 | 0.47319043 | 0.19065073 |
| Gm10138 | 0.10042457 | -0.5330738 | 0.115 | 0.192 | 1 | 0.64589072 | -1.0128735 | 0.13 | 0.146 | 1 | 0.64589072 | 0.19076404 |
| Agpat5 | 0.10063648 | 0.32435164 | 0.479 | 0.373 | 1 | 0.24751602 | -0.1506567 | 0.374 | 0.292 | 1 | 0.24751602 | 0.19114525 |
| C87436 | 0.10075787 | 0.73447604 | 0.292 | 0.212 | 1 | 0.77418747 | -0.7927968 | 0.227 | 0.219 | 1 | 0.77418747 | 0.19136359 |
| Pierce2 | 0.73629989 | 0.42253204 | 0.146 | 0.135 | 1 | 0.10079261 | 0.58322953 | 0.097 | 0.042 | 1 | 0.73629989 | 0.19142608 |
| Isl1 | 1 | -0.3144336 | 0.01 | 0.01 | 1 | 0.10079319 | -1.1172102 | 0.025 | 0.062 | 1 | 1 | 0.19142712 |
| Ppwd1 | 0.10079474 | 0.40263248 | 0.365 | 0.264 | 1 | 0.34877994 | -0.2358547 | 0.336 | 0.26 | 1 | 0.34877994 | 0.19142989 |
| Ccdc162 | 0.12633776 | -1.3144336 | 0.021 | 0.062 | 1 | 0.10083542 | -1.0688472 | 0.025 | 0.062 | 1 | 0.12633776 | 0.19150306 |
| Fibp | 0.93211277 | -0.1564812 | 0.479 | 0.451 | 1 | 0.10085295 | -0.258325 | 0.391 | 0.25 | 1 | 0.93211277 | 0.19153458 |
| Ckap2 | 0.10107584 | 2.85549144 | 0.073 | 0.031 | 1 | 0.80021055 | -0.8244284 | 0.017 | 0.021 | 1 | 0.80021055 | 0.19193536 |
| Plpp6 | 0.10108162 | 0.91438513 | 0.125 | 0.067 | 1 | 0.27547969 | 0.44503224 | 0.101 | 0.062 | 1 | 0.27547969 | 0.19194575 |
| Gm15787 | 0.17795165 | 0.20013961 | 0.167 | 0.109 | 1 | 0.10130306 | 0.53144699 | 0.139 | 0.073 | 1 | 0.17795165 | 0.19234381 |
| Ak2 | 0.11062667 | 0.39304527 | 0.594 | 0.492 | 1 | 0.10132688 | 0.324225 | 0.576 | 0.5 | 1 | 0.11062667 | 0.19238662 |
| Nop9 | 0.10916306 | 0.46000674 | 0.25 | 0.166 | 1 | 0.10136877 | 0.38529016 | 0.244 | 0.156 | 1 | 0.10916306 | 0.19246191 |
| Sema4a | 0.3369865 | -0.747393 | 0.083 | 0.119 | 1 | 0.10162908 | -1.7992401 | 0.08 | 0.135 | 1 | 0.3369865 | 0.19292968 |
| Med11 | 0.3366862 | 0.44806713 | 0.177 | 0.135 | 1 | 0.10163425 | 1.10518224 | 0.147 | 0.083 | 1 | 0.3366862 | 0.19293898 |
| Mapk8ip3 | 0.10178911 | 0.25260703 | 0.615 | 0.539 | 1 | 0.59891827 | -0.3098553 | 0.672 | 0.573 | 1 | 0.59891827 | 0.1932172 |
| Klhl2 | 0.97040126 | -0.2488452 | 0.229 | 0.218 | 1 | 0.10181365 | -0.6044252 | 0.357 | 0.469 | 1 | 0.97040126 | 0.19326129 |
| Mcm5 | 0.10184562 | 0.56692195 | 0.104 | 0.052 | 1 | 0.74198247 | -0.3098553 | 0.063 | 0.073 | 1 | 0.74198247 | 0.19331872 |
| Itgb6 | 0.55846748 | -1.0431315 | 0.042 | 0.057 | 1 | 0.10197859 | -0.1578522 | 0.046 | 0.094 | 1 | 0.55846748 | 0.19355755 |
| Gm16124 | 0.10201089 | 0.75956702 | 0.208 | 0.135 | 1 | 0.1338136 | -0.1329775 | 0.168 | 0.094 | 1 | 0.1338136 | 0.19361557 |
| Pou2f1 | 0.50750294 | 0.14951354 | 0.448 | 0.399 | 1 | 0.10217272 | -0.3020358 | 0.58 | 0.708 | 1 | 0.50750294 | 0.19390618 |
| Atp13a2 | 0.64197016 | 0.59245704 | 0.042 | 0.031 | 1 | 0.10231956 | 0.78968041 | 0.097 | 0.042 | 1 | 0.64197016 | 0.19416983 |
| Zdhhc1 | 0.10243438 | 0.43633784 | 0.458 | 0.363 | 1 | 0.85639362 | -0.3739856 | 0.378 | 0.323 | 1 | 0.85639362 | 0.19437596 |
| Adgre5 | 0.10244509 | -0.6864023 | 0.094 | 0.166 | 1 | 0.79197991 | -0.5062525 | 0.202 | 0.208 | 1 | 0.79197991 | 0.19439519 |
| Rhpn2 | 0.83363279 | 0.49292136 | 0.031 | 0.036 | 1 | 0.10251031 | -1.3098553 | 0.055 | 0.104 | 1 | 0.83363279 | 0.19451226 |
| Amn1 | 0.16612495 | -0.4633962 | 0.354 | 0.43 | 1 | 0.10252791 | 0.34549657 | 0.319 | 0.219 | 1 | 0.16612495 | 0.19454385 |
| Ergic2 | 0.10256446 | 0.20462701 | 0.823 | 0.751 | 1 | 0.52228361 | -0.1040851 | 0.849 | 0.792 | 1 | 0.52228361 | 0.19460945 |
| Nmt2 | 0.66444996 | 0.19145737 | 0.344 | 0.321 | 1 | 0.10275472 | -0.5082688 | 0.555 | 0.635 | 1 | 0.66444996 | 0.1949509 |
| Gm48742 | 0.10279988 | -0.6623569 | 0.073 | 0.14 | 1 | 0.37884342 | 0.17557156 | 0.055 | 0.031 | 1 | 0.37884342 | 0.19503195 |
| Alpk1 | 0.10282111 | -0.3221075 | 0.438 | 0.539 | 1 | 0.49173494 | -0.8971617 | 0.399 | 0.292 | 1 | 0.49173494 | 0.19507004 |
| Dnajc24 | 0.36136886 | 0.53215653 | 0.25 | 0.207 | 1 | 0.10287172 | 0.72256622 | 0.223 | 0.146 | 1 | 0.36136886 | 0.19516084 |
| Tmem132b | 0.32896838 | -1.6705774 | 0.031 | 0.057 | 1 | 0.103203 | -2.8623963 | 0.034 | 0.073 | 1 | 0.32896838 | 0.19575513 |
| Rp9 | 0.23527372 | 0.41567836 | 0.604 | 0.627 | 1 | 0.10323351 | 0.20838112 | 0.693 | 0.656 | 1 | 0.23527372 | 0.19580987 |
| Kmt5c | 0.1033086 | 0.59245704 | 0.167 | 0.098 | 1 | 0.36528174 | -0.1723517 | 0.147 | 0.104 | 1 | 0.36528174 | 0.19594453 |
| Msh6 | 0.66854343 | -0.3910548 | 0.094 | 0.109 | 1 | 0.10336188 | 0.24034182 | 0.139 | 0.073 | 1 | 0.66854343 | 0.19604009 |
| Gm56496 | 0.71864235 | 0.22988696 | 0.062 | 0.052 | 1 | 0.10338402 | 1.21370669 | 0.08 | 0.031 | 1 | 0.71864235 | 0.19607978 |
| Gpn2 | 0.55824217 | -0.2295447 | 0.125 | 0.15 | 1 | 0.10339832 | 0.94648449 | 0.147 | 0.083 | 1 | 0.55824217 | 0.19610543 |
| Rab10os | 0.24597172 | 0.39451766 | 0.375 | 0.311 | 1 | 0.10339834 | 0.35852325 | 0.29 | 0.198 | 1 | 0.24597172 | 0.19610546 |
| Arl5a | 0.10355725 | 0.33176657 | 0.51 | 0.394 | 1 | 0.85638138 | -0.1860267 | 0.571 | 0.562 | 1 | 0.85638138 | 0.1963904 |
| Map2k2 | 0.44767173 | -0.1653421 | 0.802 | 0.798 | 1 | 0.10362022 | 0.1939915 | 0.861 | 0.812 | 1 | 0.44767173 | 0.19650328 |
| Slc12a7 | 0.29645488 | 0.74446013 | 0.083 | 0.052 | 1 | 0.10369391 | 0.64434105 | 0.097 | 0.042 | 1 | 0.29645488 | 0.1966354 |
| 2810029C07Rik | 0.10373091 | -1.265524 | 0.052 | 0.109 | 1 | 0.92228543 | -0.7248928 | 0.029 | 0.031 | 1 | 0.92228543 | 0.19670172 |
| Casc1 | 0.10385756 | 0.59245704 | 0.156 | 0.088 | 1 | 0.37006305 | -0.6486572 | 0.185 | 0.135 | 1 | 0.37006305 | 0.19692873 |
| Sh3bp5 | 0.86162339 | 0.22988696 | 0.125 | 0.119 | 1 | 0.10386426 | -1.1630139 | 0.097 | 0.156 | 1 | 0.86162339 | 0.19694074 |
| Tead4 | 0.79960536 | -0.2751444 | 0.302 | 0.321 | 1 | 0.10393176 | -0.3302231 | 0.307 | 0.198 | 1 | 0.79960536 | 0.19706172 |
| Bin1 | 0.18317657 | 0.2342654 | 0.406 | 0.311 | 1 | 0.10399994 | 0.30987266 | 0.324 | 0.219 | 1 | 0.18317657 | 0.19718388 |
| Ptbp3 | 0.10402086 | -0.2314238 | 0.896 | 0.938 | 1 | 0.11603229 | -0.4441325 | 0.954 | 0.948 | 1 | 0.11603229 | 0.19722137 |
| Dgcr2 | 0.10447719 | 0.37006462 | 0.448 | 0.352 | 1 | 0.27638133 | -0.1485744 | 0.576 | 0.448 | 1 | 0.27638133 | 0.1980389 |
| 4930412C18Rik | 0.10451019 | 1.52206771 | 0.073 | 0.031 | 1 | 0.20314753 | 1.27510724 | 0.017 | 0 | 1 | 0.20314753 | 0.19809801 |
| Zhx3 | 0.10451416 | -0.3441338 | 0.615 | 0.679 | 1 | 0.93498646 | -0.2662505 | 0.786 | 0.667 | 1 | 0.93498646 | 0.19810512 |
| Gjb6 | 0.10452455 | 0.94860085 | 0.125 | 0.067 | 1 | 0.27154471 | 1.86006974 | 0.013 | 0 | 1 | 0.27154471 | 0.19812371 |
| Rangap1 | 0.66602656 | 0.1312299 | 0.438 | 0.415 | 1 | 0.10460159 | -0.7066536 | 0.529 | 0.594 | 1 | 0.66602656 | 0.19826168 |
| Angptl7 | 0.10477021 | 0.17522198 | 0.052 | 0.114 | 1 | 0.30577763 | -0.5133887 | 0.038 | 0.062 | 1 | 0.30577763 | 0.19856363 |
| Zc3h12c | 0.85432986 | -0.1158879 | 0.188 | 0.197 | 1 | 0.10521514 | -0.247571 | 0.378 | 0.25 | 1 | 0.85432986 | 0.19936005 |
| Stard3 | 0.73773053 | -0.223831 | 0.208 | 0.218 | 1 | 0.10521913 | 0.96698494 | 0.193 | 0.125 | 1 | 0.73773053 | 0.1993672 |
| Noc3l | 0.11548731 | 0.30915024 | 0.385 | 0.285 | 1 | 0.10523354 | 0.22492148 | 0.332 | 0.229 | 1 | 0.11548731 | 0.19939299 |
| Zfp148 | 0.10539931 | 0.23526784 | 0.885 | 0.855 | 1 | 0.37156269 | -0.2110877 | 0.962 | 0.938 | 1 | 0.37156269 | 0.19968961 |
| Sfxn5 | 0.10563261 | 0.45227938 | 0.302 | 0.202 | 1 | 0.1269574 | 0.10125653 | 0.546 | 0.406 | 1 | 0.1269574 | 0.20010697 |
| Agl | 0.69673667 | 0.29700115 | 0.135 | 0.119 | 1 | 0.10568981 | 0.66414953 | 0.164 | 0.094 | 1 | 0.69673667 | 0.20020928 |
| Ttll11 | 0.10576788 | 0.90057933 | 0.104 | 0.052 | 1 | 0.80620952 | 0.35999614 | 0.08 | 0.073 | 1 | 0.80620952 | 0.20034892 |
| Bod1l | 0.2698587 | 0.17569486 | 0.781 | 0.746 | 1 | 0.10586944 | 0.2541603 | 0.761 | 0.802 | 1 | 0.2698587 | 0.20053054 |
| C630043F03Rik | 0.65196801 | -0.130009 | 0.094 | 0.114 | 1 | 0.10611175 | 0.47441605 | 0.202 | 0.125 | 1 | 0.65196801 | 0.2009638 |
| Rab7b | 0.10624662 | 1.56692195 | 0.146 | 0.088 | 1 | 0.56729373 | -1.5025003 | 0.021 | 0.031 | 1 | 0.56729373 | 0.20120489 |
| Insyn1 | 0.11384978 | -1.3144336 | 0.01 | 0.047 | 1 | 0.10636452 | 1.01207283 | 0.08 | 0.031 | 1 | 0.11384978 | 0.20141562 |
| Cyp26a1 | 0.5702909 | -1.5945415 | 0.042 | 0.057 | 1 | 0.10641533 | 0.7219936 | 0.239 | 0.156 | 1 | 0.5702909 | 0.20150643 |
| Mphosph9 | 0.10658026 | -0.5259377 | 0.115 | 0.192 | 1 | 0.42997865 | -0.423512 | 0.181 | 0.135 | 1 | 0.42997865 | 0.20180117 |
| Fblim1 | 0.10660245 | 0.48393258 | 0.104 | 0.052 | 1 | 0.86493058 | -0.484942 | 0.109 | 0.115 | 1 | 0.86493058 | 0.20184082 |
| Frg1 | 0.10662397 | 0.38537904 | 0.583 | 0.539 | 1 | 0.59863101 | -0.4671673 | 0.538 | 0.542 | 1 | 0.59863101 | 0.20187927 |
| Mettl8 | 0.75515781 | -0.4254649 | 0.083 | 0.093 | 1 | 0.10694542 | 0.47441605 | 0.109 | 0.052 | 1 | 0.75515781 | 0.20245352 |
| Samd8 | 0.42165157 | 0.17161392 | 0.531 | 0.487 | 1 | 0.1072264 | -0.2575608 | 0.622 | 0.802 | 1 | 0.42165157 | 0.20295529 |
| Atp8b3 | 0.52528691 | -0.9925055 | 0.01 | 0.021 | 1 | 0.10729406 | -1.9879272 | 0.034 | 0.073 | 1 | 0.52528691 | 0.20307611 |
| Dhtkd1 | 0.10730202 | 1.20013961 | 0.073 | 0.031 | 1 | 0.44394565 | -0.3098553 | 0.038 | 0.021 | 1 | 0.44394565 | 0.20309032 |
| Dram1 | 0.10742238 | -0.6537036 | 0.26 | 0.342 | 1 | 0.21925556 | -0.1843244 | 0.227 | 0.156 | 1 | 0.21925556 | 0.20330519 |
| Gsg1l | 0.10768205 | 1.17741954 | 0.073 | 0.031 | 1 | 0.42482884 | -1.5728897 | 0.071 | 0.094 | 1 | 0.42482884 | 0.20376867 |
| Pheta2 | 0.10768205 | 1.17741954 | 0.073 | 0.031 | 1 | 0.37126881 | -0.3098553 | 0.097 | 0.062 | 1 | 0.37126881 | 0.20376867 |

| | | | | | | | | | | | | |
|---|---|---|---|---|---|---|---|---|---|---|---|---|
| Dsn1 | 0.52071264 | 0.24195979 | 0.083 | 0.062 | 1 | 0.10785672 | -1.010295 | 0.063 | 0.115 | 1 | 0.52071264 | 0.20408037 |
| Gm43331 | 0.44880848 | 0.35541784 | 0.062 | 0.041 | 1 | 0.1078823 | -1.5468945 | 0.105 | 0.156 | 1 | 0.44880848 | 0.20412601 |
| Zfp382 | 0.10792984 | 0.52206771 | 0.146 | 0.083 | 1 | 0.67846585 | -0.3623227 | 0.185 | 0.156 | 1 | 0.67846585 | 0.20421083 |
| Bend6 | 0.10806037 | -0.2638075 | 0.135 | 0.218 | 1 | 0.75427934 | -0.1723517 | 0.05 | 0.042 | 1 | 0.75427934 | 0.20444369 |
| Bnip1 | 0.10816911 | 0.56003556 | 0.26 | 0.181 | 1 | 0.66969861 | -0.2416838 | 0.197 | 0.167 | 1 | 0.66969861 | 0.20463766 |
| 1700034P13Rik | 0.10817209 | 0.1843723 | 0.073 | 0.031 | 1 | 0.66015411 | 0.69014474 | 0.029 | 0.021 | 1 | 0.66015411 | 0.20464298 |
| B3gntl1 | 0.10832925 | 1.00749454 | 0.135 | 0.078 | 1 | 0.36639797 | -0.4253325 | 0.084 | 0.052 | 1 | 0.36639797 | 0.20492327 |
| D3Ertd751e | 0.11875313 | 1.09864242 | 0.25 | 0.181 | 1 | 0.10833073 | -0.2592292 | 0.185 | 0.104 | 1 | 0.11875313 | 0.20492592 |
| Plekha8 | 0.10842622 | -1.0566358 | 0.198 | 0.269 | 1 | 0.39546535 | 0.52968007 | 0.181 | 0.146 | 1 | 0.39546535 | 0.20509619 |
| Brdt | 0.10858195 | 0.42253204 | 0.208 | 0.135 | 1 | 0.5714434 | 0.10518224 | 0.172 | 0.146 | 1 | 0.5714434 | 0.20537385 |
| Cnppd1 | 0.28464307 | 0.30488045 | 0.354 | 0.29 | 1 | 0.10862996 | -0.4105493 | 0.433 | 0.531 | 1 | 0.28464307 | 0.20545944 |
| Tank | 0.10875678 | -0.6538189 | 0.448 | 0.539 | 1 | 0.81092488 | -0.1094282 | 0.5 | 0.521 | 1 | 0.81092488 | 0.20568553 |
| Rpe | 0.28144903 | 0.54452256 | 0.271 | 0.218 | 1 | 0.10880448 | -0.1843244 | 0.252 | 0.156 | 1 | 0.28144903 | 0.20577054 |
| Prpf31 | 0.4978322 | 0.27631129 | 0.354 | 0.321 | 1 | 0.10881136 | 0.19710473 | 0.34 | 0.229 | 1 | 0.4978322 | 0.2057828 |
| Ostm1 | 0.10890698 | -0.6169963 | 0.073 | 0.14 | 1 | 0.23533636 | 0.3575694 | 0.235 | 0.167 | 1 | 0.23533636 | 0.20595323 |
| BC049715 | 0.10902985 | 1.49292136 | 0.115 | 0.062 | 1 | 0.89588146 | -0.961932 | 0.046 | 0.042 | 1 | 0.89588146 | 0.2061722 |
| Kif20b | 0.10910243 | 2.17741954 | 0.094 | 0.047 | 1 | 0.58017355 | -0.9472852 | 0.021 | 0.031 | 1 | 0.58017355 | 0.20630151 |
| Arpp19 | 0.10921902 | 0.15581728 | 0.823 | 0.731 | 1 | 0.13447496 | 0.10614628 | 0.84 | 0.802 | 1 | 0.13447496 | 0.20650925 |
| Sp3os | 0.22803824 | -0.1989563 | 0.167 | 0.233 | 1 | 0.10930076 | 0.41261076 | 0.218 | 0.135 | 1 | 0.22803824 | 0.20665486 |
| Abhd5 | 0.10937197 | 0.43130225 | 0.375 | 0.28 | 1 | 0.58459451 | -0.4514094 | 0.5 | 0.5 | 1 | 0.58459451 | 0.20678171 |
| Aktip | 0.23312567 | -0.321454 | 0.292 | 0.368 | 1 | 0.10939737 | -0.1569033 | 0.466 | 0.333 | 1 | 0.23312567 | 0.20682695 |
| Kcns1 | 0.10950331 | 0.44359365 | 0.188 | 0.114 | 1 | 0.53642442 | -0.2823745 | 0.176 | 0.146 | 1 | 0.53642442 | 0.20701565 |
| Morc2a | 0.10955868 | 0.33273537 | 0.542 | 0.446 | 1 | 0.21766025 | -0.5025003 | 0.634 | 0.583 | 1 | 0.21766025 | 0.20711425 |
| Ric8a | 0.37601644 | 0.39082318 | 0.25 | 0.207 | 1 | 0.10963635 | 0.27510724 | 0.261 | 0.167 | 1 | 0.37601644 | 0.20725257 |
| Lbr | 0.29977084 | 0.38234295 | 0.396 | 0.332 | 1 | 0.10975864 | -0.5452333 | 0.437 | 0.531 | 1 | 0.29977084 | 0.20747032 |
| Elovl5 | 0.77285589 | -0.1079827 | 0.219 | 0.228 | 1 | 0.10980299 | -1.261761 | 0.193 | 0.26 | 1 | 0.77285589 | 0.20754928 |
| Zscan25 | 0.18188912 | 0.74446013 | 0.094 | 0.052 | 1 | 0.10983053 | 0.69014474 | 0.109 | 0.052 | 1 | 0.18188912 | 0.20759831 |
| Abhd4 | 0.10986335 | 0.47315811 | 0.229 | 0.15 | 1 | 0.11625621 | 0.42711033 | 0.235 | 0.156 | 1 | 0.11625621 | 0.20765674 |
| Mecr | 0.11519367 | 0.55181505 | 0.271 | 0.192 | 1 | 0.11002905 | 0.29348577 | 0.252 | 0.167 | 1 | 0.11519367 | 0.20795171 |
| Ppp1r12b | 0.44811535 | 0.17586239 | 0.812 | 0.756 | 1 | 0.11008536 | -0.6051154 | 0.798 | 0.875 | 1 | 0.44811535 | 0.20805193 |
| Ralgds | 0.28726897 | -0.8307544 | 0.208 | 0.259 | 1 | 0.11010094 | -1.2944252 | 0.265 | 0.323 | 1 | 0.28726897 | 0.20807966 |
| 9530052E02Rik | 0.11022867 | 0.59245704 | 0.177 | 0.109 | 1 | 0.37270021 | -0.7415715 | 0.109 | 0.073 | 1 | 0.37270021 | 0.20830698 |
| Ccdc112 | 0.24831038 | 0.22322323 | 0.281 | 0.207 | 1 | 0.11028808 | 0.33703499 | 0.286 | 0.188 | 1 | 0.24831038 | 0.20841269 |
| Eri2 | 0.11031971 | -1.2726134 | 0.052 | 0.109 | 1 | 0.92310537 | -0.8244284 | 0.071 | 0.073 | 1 | 0.92310537 | 0.20846898 |
| Ttc16 | 0.1104186 | 1.00749454 | 0.073 | 0.031 | 1 | 0.69413423 | -0.2318528 | 0.076 | 0.062 | 1 | 0.69413423 | 0.20864493 |
| Mylpf | 0.1104186 | 1.00749454 | 0.073 | 0.031 | 1 | 0.71069052 | -0.7248928 | 0.034 | 0.042 | 1 | 0.71069052 | 0.20864493 |
| Pabpc4l | 0.72080211 | -0.3550755 | 0.052 | 0.062 | 1 | 0.11045515 | 1.77760758 | 0.046 | 0.01 | 1 | 0.72080211 | 0.20870996 |
| Ctnna3 | 0.11064456 | -3.2019588 | 0.01 | 0.047 | 1 | 0.86804649 | 0.27510724 | 0.013 | 0.01 | 1 | 0.86804649 | 0.20904691 |
| R3hdm1 | 0.16726264 | 0.12613903 | 0.771 | 0.725 | 1 | 0.11078242 | -0.4854268 | 0.819 | 0.938 | 1 | 0.16726264 | 0.2092921 |
| Adck1 | 0.11079121 | 0.49292136 | 0.188 | 0.114 | 1 | 0.16170485 | 0.42711033 | 0.139 | 0.083 | 1 | 0.16170485 | 0.20930773 |
| Proca1 | 0.11085528 | 0.75595577 | 0.156 | 0.093 | 1 | 0.38132457 | -0.9228321 | 0.113 | 0.146 | 1 | 0.38132457 | 0.20942166 |
| Gm27253 | 0.99482517 | 0.11440974 | 0.042 | 0.041 | 1 | 0.11088151 | 1.69014474 | 0.046 | 0.01 | 1 | 0.99482517 | 0.20946831 |
| Zfp37 | 0.15163858 | 0.30705482 | 0.24 | 0.166 | 1 | 0.11092623 | 0.27510724 | 0.303 | 0.208 | 1 | 0.15163858 | 0.20954784 |
| Ammecr1 | 0.96154114 | -0.3037072 | 0.302 | 0.295 | 1 | 0.110968 | 0.3500693 | 0.345 | 0.25 | 1 | 0.96154114 | 0.20962211 |
| Ltbp1 | 0.11107878 | -0.7522424 | 0.135 | 0.212 | 1 | 0.34752452 | -0.9826905 | 0.21 | 0.25 | 1 | 0.34752452 | 0.20981907 |
| Rasl10a | 0.25696442 | 0.85549144 | 0.073 | 0.041 | 1 | 0.11110684 | 1.35310975 | 0.063 | 0.021 | 1 | 0.25696442 | 0.20986895 |
| Garin5b | 0.11116771 | -2.240433 | 0.01 | 0.047 | 1 | 0.17409084 | -2.0874628 | 0.017 | 0.042 | 1 | 0.17409084 | 0.20997717 |
| Cyp3a13 | 0.4755186 | -0.5103538 | 0.146 | 0.176 | 1 | 0.11117827 | 1.81942775 | 0.063 | 0.021 | 1 | 0.4755186 | 0.20999593 |
| Lysmd1 | 0.25792709 | 0.83756954 | 0.073 | 0.041 | 1 | 0.11129992 | 0.76053407 | 0.08 | 0.031 | 1 | 0.25792709 | 0.21021216 |
| Map7d3 | 0.47795675 | 0.31359766 | 0.333 | 0.295 | 1 | 0.11132949 | 0.58705124 | 0.227 | 0.146 | 1 | 0.47795675 | 0.21026473 |
| Rps6kb2 | 0.47318297 | -0.3946039 | 0.104 | 0.135 | 1 | 0.11139324 | 0.27510724 | 0.168 | 0.094 | 1 | 0.47318297 | 0.21037803 |
| Wfdc18 | 0.45584703 | 0.50225923 | 0.333 | 0.301 | 1 | 0.11160373 | -1.0795576 | 0.168 | 0.229 | 1 | 0.45584703 | 0.21075206 |
| Kdm4d | 0.11169238 | -1.7998604 | 0.01 | 0.047 | 1 | 0.98188267 | 0.27510724 | 0.021 | 0.021 | 1 | 0.98188267 | 0.21090958 |
| Panx1 | 0.11169238 | -1.7998604 | 0.01 | 0.047 | 1 | 0.30934712 | 0.69014474 | 0.071 | 0.042 | 1 | 0.30934712 | 0.21090958 |
| Aldh3b1 | 0.1525942 | -1.4519371 | 0.01 | 0.041 | 1 | 0.11173124 | 1.49749966 | 0.046 | 0.01 | 1 | 0.1525942 | 0.21097861 |
| Tyw3 | 0.36625316 | 0.54806292 | 0.073 | 0.047 | 1 | 0.11173124 | 1.49749966 | 0.046 | 0.01 | 1 | 0.36625316 | 0.21097861 |
| Garin5a | 0.56492112 | 0.54806292 | 0.062 | 0.047 | 1 | 0.11173124 | 1.49749966 | 0.046 | 0.01 | 1 | 0.56492112 | 0.21097861 |
| Sytl4 | 0.80274368 | 0.20013961 | 0.031 | 0.026 | 1 | 0.11173124 | 1.49749966 | 0.046 | 0.01 | 1 | 0.80274368 | 0.21097861 |
| Zbtb7c | 0.15513191 | -0.4384916 | 0.521 | 0.606 | 1 | 0.11178308 | -0.984455 | 0.378 | 0.427 | 1 | 0.15513191 | 0.21107071 |
| Carns1 | 0.38521705 | 0.68556644 | 0.031 | 0.016 | 1 | 0.1118253 | 1.35310975 | 0.063 | 0.021 | 1 | 0.38521705 | 0.2111457 |
| Gm43707 | 0.15768412 | -1.577468 | 0.021 | 0.057 | 1 | 0.11205581 | -1.9378865 | 0.042 | 0.083 | 1 | 0.15768412 | 0.21155512 |
| Pih1d2 | 0.1469607 | 1.49292136 | 0.052 | 0.021 | 1 | 0.11215403 | 1.39058446 | 0.046 | 0.01 | 1 | 0.1469607 | 0.21172954 |
| Faim2 | 0.22187575 | -1.7998604 | 0 | 0.016 | 1 | 0.11215403 | 1.59703533 | 0.046 | 0.01 | 1 | 0.22187575 | 0.21172954 |
| Wdr24 | 0.25332778 | 1.00749454 | 0.052 | 0.026 | 1 | 0.11215403 | 1.39058446 | 0.046 | 0.01 | 1 | 0.25332778 | 0.21172954 |
| A830008E24Rik | 0.45166454 | -1.0513992 | 0.094 | 0.067 | 1 | 0.11215403 | 1.39058446 | 0.046 | 0.01 | 1 | 0.45166454 | 0.21172954 |
| Pcdhgc5 | 0.61492853 | -0.2148979 | 0.052 | 0.067 | 1 | 0.11215403 | 1.39058446 | 0.046 | 0.01 | 1 | 0.61492853 | 0.21172954 |
| Gm15510 | 0.6229455 | -0.1624305 | 0.021 | 0.031 | 1 | 0.11215403 | 1.39058446 | 0.046 | 0.01 | 1 | 0.6229455 | 0.21172954 |
| Asgr1 | 0.63038178 | 0.52206771 | 0.021 | 0.031 | 1 | 0.11215403 | 1.39058446 | 0.046 | 0.01 | 1 | 0.63038178 | 0.21172954 |
| Stx1a | 0.11218465 | 0.54806292 | 0.146 | 0.083 | 1 | 0.34611611 | 0.37464291 | 0.143 | 0.104 | 1 | 0.34611611 | 0.2117839 |
| Stx11 | 0.11218465 | 0.48393258 | 0.146 | 0.083 | 1 | 0.12498936 | -1.1723517 | 0.101 | 0.156 | 1 | 0.12498936 | 0.2117839 |
| Obp2a | 0.49083063 | -0.4564526 | 0.333 | 0.358 | 1 | 0.11219779 | -1.4767526 | 0.193 | 0.115 | 1 | 0.49083063 | 0.21180724 |
| Iws1 | 0.11220261 | 0.32008477 | 0.698 | 0.622 | 1 | 0.16624352 | 0.11147398 | 0.735 | 0.708 | 1 | 0.16624352 | 0.21181579 |
| Il12rb1 | 0.11221502 | -2.3144336 | 0.01 | 0.047 | 1 | 0.39796128 | -1.8948178 | 0.017 | 0.031 | 1 | 0.39796128 | 0.21183782 |
| Pla2g12b | 0.11221502 | -1.9925055 | 0.01 | 0.047 | 1 | 0.57370068 | -1.3098553 | 0.021 | 0.031 | 1 | 0.57370068 | 0.21183782 |
| Dmp1 | 0.11221546 | -4.6649308 | 0.01 | 0.047 | 1 | 0.27154327 | 1.69014474 | 0.013 | 0 | 1 | 0.27154327 | 0.21183861 |
| Plekha4 | 0.19123352 | 0.6241659 | 0.167 | 0.109 | 1 | 0.1122158 | -1.4208866 | 0.084 | 0.135 | 1 | 0.19123352 | 0.21183922 |
| Tox | 0.12065921 | -0.6659059 | 0.531 | 0.368 | 1 | 0.1122343 | -0.2345671 | 0.601 | 0.427 | 1 | 0.12065921 | 0.21187205 |
| Pola1 | 0.11227017 | 0.866632 | 0.281 | 0.202 | 1 | 0.84245319 | -0.3098553 | 0.298 | 0.281 | 1 | 0.84245319 | 0.21193574 |
| Cstf2 | 0.71684228 | 0.23882008 | 0.198 | 0.181 | 1 | 0.11250984 | 0.12826585 | 0.282 | 0.188 | 1 | 0.71684228 | 0.21236122 |
| Chst11 | 0.76036489 | -0.2300181 | 1 | 0.964 | 1 | 0.11264178 | -0.3210782 | 0.987 | 0.854 | 1 | 0.76036489 | 0.21259539 |
| Arhgef33 | 0.22187388 | -1.3144336 | 0 | 0.016 | 1 | 0.11265473 | 1.27510724 | 0.046 | 0.01 | 1 | 0.22187388 | 0.21261837 |
| Lef1os1 | 0.4622729 | 1.00749454 | 0.042 | 0.026 | 1 | 0.11265473 | 1.27510724 | 0.046 | 0.01 | 1 | 0.4622729 | 0.21261837 |
| 1810021B22Rik | 0.53924132 | -0.3144336 | 0.031 | 0.047 | 1 | 0.11265473 | 1.27510724 | 0.046 | 0.01 | 1 | 0.53924132 | 0.21261837 |
| S100a7a | 0.69098575 | 0.27052894 | 0.052 | 0.041 | 1 | 0.11265473 | 1.27510724 | 0.046 | 0.01 | 1 | 0.69098575 | 0.21261837 |
| 2500002B13Rik | 0.11273737 | -1.577468 | 0.01 | 0.047 | 1 | 0.37612872 | 0.27510724 | 0.055 | 0.031 | 1 | 0.37612872 | 0.21276503 |

| | | | | | | | | | | | | |
|---|---|---|---|---|---|---|---|---|---|---|---|---|
| Gm10687 | 0.11273737 | -1.6929452 | 0.01 | 0.047 | 1 | 0.50887612 | -1.3098553 | 0.004 | 0.01 | 1 | 0.50887612 | 0.21276503 |
| Kcnj12 | 0.11273737 | -1.577468 | 0.01 | 0.047 | 1 | 0.11689951 | -2.3098553 | 0 | 0.01 | 1 | 0.11689951 | 0.21276503 |
| Bicc1 | 0.40191439 | -0.5162238 | 0.812 | 0.736 | 1 | 0.11275615 | -1.6583416 | 0.857 | 0.729 | 1 | 0.40191439 | 0.21279836 |
| Pde4b | 0.76125617 | -0.1804063 | 0.76 | 0.731 | 1 | 0.11283464 | 0.45668942 | 0.79 | 0.75 | 1 | 0.76125617 | 0.21293762 |
| Zbtb6 | 0.15292629 | 0.52206771 | 0.24 | 0.171 | 1 | 0.11284481 | 0.11098786 | 0.277 | 0.177 | 1 | 0.15292629 | 0.21295566 |
| Tdp2 | 0.11295598 | 0.16697275 | 0.354 | 0.249 | 1 | 0.21199703 | -0.9680667 | 0.345 | 0.375 | 1 | 0.21199703 | 0.21315292 |
| Gckr | 0.11302151 | -1.9925055 | 0 | 0.026 | 1 | 0.50587544 | -1.8948178 | 0.004 | 0.01 | 1 | 0.50587544 | 0.21326916 |
| Sgsm1 | 0.11302151 | -1.7998604 | 0 | 0.026 | 1 | 0.27154327 | 1.01207283 | 0.013 | 0 | 1 | 0.27154327 | 0.21326916 |
| Fbxo24 | 0.11302151 | -1.9925055 | 0 | 0.026 | 1 | 0.80020906 | -0.7248928 | 0.017 | 0.021 | 1 | 0.80020906 | 0.21326916 |
| Gm29443 | 0.11302151 | -2.1624305 | 0 | 0.026 | 1 | 0.24857751 | -0.6317834 | 0.013 | 0.031 | 1 | 0.24857751 | 0.21326916 |
| St8sia2 | 0.11302151 | -2.1624305 | 0 | 0.026 | 1 | 0.66339728 | 0.10518224 | 0.029 | 0.021 | 1 | 0.66339728 | 0.21326916 |
| Slc6a15 | 0.11302151 | -1.7998604 | 0 | 0.026 | 1 | 0.14049288 | 1.35310975 | 0.059 | 0.021 | 1 | 0.14049288 | 0.21326916 |
| D630024D03Rik | 0.11302151 | -1.7998604 | 0 | 0.026 | 1 | 0.24485439 | -0.1399303 | 0.034 | 0.01 | 1 | 0.24485439 | 0.21326916 |
| Kcnip1 | 0.11302151 | -1.7998604 | 0 | 0.026 | 1 | 0.53618537 | -1.1578522 | 0.038 | 0.052 | 1 | 0.53618537 | 0.21326916 |
| Sox30 | 0.11302151 | -1.7998604 | 0 | 0.026 | 1 | 0.20315163 | 1.01207283 | 0.017 | 0 | 1 | 0.20315163 | 0.21326916 |
| Tex56 | 0.11302151 | -1.7998604 | 0 | 0.026 | 1 | 0.11689951 | -2.8948178 | 0 | 0.01 | 1 | 0.11689951 | 0.21326916 |
| 4930425P05Rik | 0.11302151 | -1.7998604 | 0 | 0.026 | 1 | 0.11689951 | -2.3098553 | 0 | 0.01 | 1 | 0.11689951 | 0.21326916 |
| Pnp2 | 0.11302151 | -1.7998604 | 0 | 0.026 | 1 | 0.40076941 | -1.3098553 | 0.017 | 0.031 | 1 | 0.40076941 | 0.21326916 |
| Sox7 | 0.11302151 | -1.7998604 | 0 | 0.026 | 1 | 0.40220446 | -1.3098553 | 0.017 | 0.031 | 1 | 0.40220446 | 0.21326916 |
| Rgs22 | 0.11302151 | -1.9925055 | 0 | 0.026 | 1 | 0.86312513 | -0.5728897 | 0.008 | 0.01 | 1 | 0.86312513 | 0.21326916 |
| Zfp7 | 0.11302151 | -1.7998604 | 0 | 0.026 | 1 | 0.66013325 | 0.27510724 | 0.029 | 0.021 | 1 | 0.66013325 | 0.21326916 |
| Dscam | 0.11302151 | -1.7998604 | 0 | 0.026 | 1 | 0.14439146 | -2.8948178 | 0.004 | 0.021 | 1 | 0.14439146 | 0.21326916 |
| Trim36 | 0.11302151 | -1.7998604 | 0 | 0.026 | 1 | 0.57372311 | -0.9879272 | 0.021 | 0.031 | 1 | 0.57372311 | 0.21326916 |
| Gm39526 | 0.11302151 | -1.7998604 | 0 | 0.026 | 1 | 0.52072676 | -0.3098553 | 0.021 | 0.01 | 1 | 0.52072676 | 0.21326916 |
| S100a2 | 0.11302678 | -2.1624305 | 0 | 0.026 | 1 | 0.50887612 | -1.3098553 | 0.004 | 0.01 | 1 | 0.50887612 | 0.2132785 |
| Chd5 | 0.11302678 | -1.9925055 | 0 | 0.026 | 1 | 0.95151634 | -0.3769695 | 0.055 | 0.052 | 1 | 0.95151634 | 0.2132785 |
| Gm42531 | 0.11302678 | -2.4519371 | 0 | 0.026 | 1 | 0.80442048 | -0.5728897 | 0.017 | 0.021 | 1 | 0.80442048 | 0.2132785 |
| Actg2 | 0.11302678 | -1.9925055 | 0 | 0.026 | 1 | 0.11689951 | -3.3098553 | 0 | 0.01 | 1 | 0.11689951 | 0.2132785 |
| Lgals7 | 0.11302678 | -2.4519371 | 0 | 0.026 | 1 | 0.9206048 | -0.3098553 | 0.029 | 0.031 | 1 | 0.9206048 | 0.2132785 |
| Sult2b1 | 0.11302678 | -2.4519371 | 0 | 0.026 | 1 | 0.80442048 | -0.5728897 | 0.017 | 0.021 | 1 | 0.80442048 | 0.2132785 |
| Gm45847 | 0.11302678 | -2.1624305 | 0 | 0.026 | 1 | 0.72991918 | -0.2742314 | 0.109 | 0.094 | 1 | 0.72991918 | 0.2132785 |
| Mmp12 | 0.11302678 | -1.9925055 | 0 | 0.026 | 1 | 0.27154327 | 1.27510724 | 0.013 | 0 | 1 | 0.27154327 | 0.2132785 |
| Gpr15lg | 0.11302678 | -2.3144336 | 0 | 0.026 | 1 | 0.57170879 | -1.6317834 | 0.013 | 0.021 | 1 | 0.57170879 | 0.2132785 |
| Oacyl | 0.11302678 | -1.9925055 | 0 | 0.026 | 1 | 0.1465102 | -1.8948178 | 0.004 | 0.021 | 1 | 0.1465102 | 0.2132785 |
| Gm12524 | 0.11302853 | -2.6929452 | 0 | 0.026 | 1 | 0.11689951 | -2.3098553 | 0 | 0.01 | 1 | 0.11689951 | 0.21328162 |
| Ksr2 | 0.11302853 | -3.1624305 | 0 | 0.026 | 1 | 0.68307776 | -1.2394659 | 0.042 | 0.052 | 1 | 0.68307776 | 0.21328162 |
| Iqub | 0.11302853 | -2.3144336 | 0 | 0.026 | 1 | 0.27154327 | 1.01207283 | 0.013 | 0 | 1 | 0.27154327 | 0.21328162 |
| C330002G04Rik | 0.11302853 | -2.577468 | 0 | 0.026 | 1 | 0.1465102 | -2.3098553 | 0.004 | 0.021 | 1 | 0.1465102 | 0.21328162 |
| Gm39185 | 0.11302941 | -2.1624305 | 0 | 0.026 | 1 | 0.12311454 | -1.3098553 | 0.008 | 0.031 | 1 | 0.12311454 | 0.21328317 |
| 1010001B22Rik | 0.11303029 | -1.577468 | 0 | 0.026 | 1 | 0.51194506 | 0.27510724 | 0.021 | 0.01 | 1 | 0.51194506 | 0.21328473 |
| Mlkl | 0.11303029 | -1.577468 | 0 | 0.026 | 1 | 0.86312513 | -0.7248928 | 0.008 | 0.01 | 1 | 0.86312513 | 0.21328473 |
| Osgin1 | 0.11303029 | -2.8993961 | 0 | 0.026 | 1 | 0.37381387 | 0.62874419 | 0.067 | 0.042 | 1 | 0.37381387 | 0.21328473 |
| Fat2 | 0.11303029 | -3.7998604 | 0 | 0.026 | 1 | 0.53855674 | 0.45567948 | 0.034 | 0.021 | 1 | 0.53855674 | 0.21328473 |
| Pebp4 | 0.11303029 | -1.577468 | 0 | 0.026 | 1 | 0.86804649 | 0.27510724 | 0.013 | 0.01 | 1 | 0.86804649 | 0.21328473 |
| Gm49171 | 0.11303029 | -1.577468 | 0 | 0.026 | 1 | 0.50587544 | -1.8948178 | 0.004 | 0.01 | 1 | 0.50587544 | 0.21328473 |
| Dgkg | 0.11303029 | -1.577468 | 0 | 0.026 | 1 | 0.2408519 | -2.1172102 | 0.013 | 0.031 | 1 | 0.2408519 | 0.21328473 |
| Rps6ka4 | 0.11303029 | -1.577468 | 0 | 0.026 | 1 | 0.43323624 | 0.42711033 | 0.038 | 0.021 | 1 | 0.43323624 | 0.21328473 |
| Gm30541 | 0.11303029 | -1.577468 | 0 | 0.026 | 1 | 0.14544666 | -2.8948178 | 0.004 | 0.021 | 1 | 0.14544666 | 0.21328473 |
| Coq8b | 0.11306027 | 0.63884994 | 0.427 | 0.321 | 1 | 0.17255582 | 0.10518224 | 0.282 | 0.188 | 1 | 0.17255582 | 0.21333792 |
| Tspan32 | 0.66539403 | 0.25260703 | 0.104 | 0.088 | 1 | 0.1130847 | 1.12310414 | 0.092 | 0.042 | 1 | 0.66539403 | 0.21338125 |
| Hr | 0.11321325 | -0.6507169 | 0.208 | 0.295 | 1 | 0.33954225 | -0.1187138 | 0.34 | 0.26 | 1 | 0.33954225 | 0.21360925 |
| Stum | 0.11325766 | -1.4519371 | 0.01 | 0.047 | 1 | 0.86312513 | -0.7248928 | 0.008 | 0.01 | 1 | 0.86312513 | 0.21368802 |
| D230022J07Rik | 0.11325766 | -1.577468 | 0.01 | 0.047 | 1 | 0.75297348 | -1.1172102 | 0.025 | 0.031 | 1 | 0.75297348 | 0.21368802 |
| Tmem79 | 0.11325766 | -2.4519371 | 0.01 | 0.047 | 1 | 0.41906801 | -1.0874628 | 0.025 | 0.042 | 1 | 0.41906801 | 0.21368802 |
| Nadsyn1 | 0.11325766 | -1.4519371 | 0.01 | 0.047 | 1 | 0.65517034 | -0.2103196 | 0.042 | 0.031 | 1 | 0.65517034 | 0.21368802 |
| Gm9922 | 0.11325766 | -1.4519371 | 0.01 | 0.047 | 1 | 0.30606883 | 0.69014474 | 0.029 | 0.01 | 1 | 0.30606883 | 0.21368802 |
| Rragb | 0.11329045 | 0.86999101 | 0.073 | 0.031 | 1 | 0.19376259 | 1.14957636 | 0.067 | 0.031 | 1 | 0.19376259 | 0.21374617 |
| Gal3st1 | 0.11337051 | -0.913071 | 0.094 | 0.161 | 1 | 0.42979096 | -0.5025003 | 0.05 | 0.073 | 1 | 0.42979096 | 0.21388814 |
| Pml | 0.11350742 | 0.82692229 | 0.094 | 0.047 | 1 | 0.71784168 | -0.6468902 | 0.134 | 0.146 | 1 | 0.71784168 | 0.21413091 |
| Pik3r2 | 0.70025803 | -0.1989563 | 0.115 | 0.13 | 1 | 0.11351472 | 0.45567948 | 0.176 | 0.104 | 1 | 0.70025803 | 0.21414386 |
| Cd82 | 0.1137051 | -0.4317905 | 0.677 | 0.663 | 1 | 0.39411082 | -0.2350875 | 0.475 | 0.542 | 1 | 0.39411082 | 0.21448135 |
| Tmc4 | 0.11373061 | 0.83756954 | 0.073 | 0.031 | 1 | 0.80935439 | -0.9139266 | 0.088 | 0.094 | 1 | 0.80935439 | 0.21452656 |
| Piga | 0.42614575 | 0.66645762 | 0.104 | 0.078 | 1 | 0.11374896 | 0.41261076 | 0.122 | 0.062 | 1 | 0.42614575 | 0.2145591 |
| Agbl4 | 0.11380221 | -1.940038 | 0.042 | 0.093 | 1 | 0.9632026 | -0.5899632 | 0.063 | 0.062 | 1 | 0.9632026 | 0.21465348 |
| Bloc1s5 | 0.11383634 | 0.56692195 | 0.125 | 0.067 | 1 | 0.64139385 | -0.1172102 | 0.113 | 0.094 | 1 | 0.64139385 | 0.21471397 |
| Gm31728 | 0.11384978 | -1.3144336 | 0.01 | 0.047 | 1 | 0.40078481 | -1.3098553 | 0.017 | 0.031 | 1 | 0.40078481 | 0.21473779 |
| Anxa9 | 0.11384978 | -1.3144336 | 0.01 | 0.047 | 1 | 0.74957918 | -1.0874628 | 0.025 | 0.031 | 1 | 0.74957918 | 0.21473779 |
| Elovl4 | 0.11384978 | -1.3144336 | 0.01 | 0.047 | 1 | 0.65854006 | -0.5508634 | 0.08 | 0.094 | 1 | 0.65854006 | 0.21473779 |
| Gm31814 | 0.11384978 | -1.3144336 | 0.01 | 0.047 | 1 | 0.27154327 | 1.01207283 | 0.013 | 0 | 1 | 0.27154327 | 0.21473779 |
| Fhod1 | 0.11391609 | 0.73186009 | 0.146 | 0.083 | 1 | 0.4975016 | 0.46775232 | 0.084 | 0.062 | 1 | 0.4975016 | 0.2148553 |
| Elk1 | 0.81871306 | 0.11996927 | 0.146 | 0.161 | 1 | 0.11398313 | 0.48155812 | 0.176 | 0.104 | 1 | 0.81871306 | 0.21497411 |
| Pxmp2 | 0.11399719 | -0.8855903 | 0.052 | 0.109 | 1 | 0.57693422 | -1.3098553 | 0.021 | 0.031 | 1 | 0.57693422 | 0.21499903 |
| A530046M15Rik | 0.11403374 | -0.4779323 | 0.052 | 0.109 | 1 | 0.67254361 | -0.3098553 | 0.017 | 0.01 | 1 | 0.67254361 | 0.21506379 |
| 1700030J22Rik | 0.11424165 | 0.85549144 | 0.073 | 0.031 | 1 | 0.51194506 | 0.27510724 | 0.021 | 0.01 | 1 | 0.51194506 | 0.21543214 |
| 2410006H16Rik | 0.30730615 | 0.19447828 | 0.792 | 0.803 | 1 | 0.11429996 | 0.13728308 | 0.727 | 0.656 | 1 | 0.30730615 | 0.21553545 |
| Cdkn2d | 0.23085649 | -0.3144336 | 0.479 | 0.544 | 1 | 0.11443751 | 0.43234698 | 0.387 | 0.292 | 1 | 0.23085649 | 0.21577907 |
| Tmeff1 | 0.71305088 | 0.24195979 | 0.062 | 0.052 | 1 | 0.11465165 | -0.9313436 | 0.08 | 0.135 | 1 | 0.71305088 | 0.21615829 |
| Atf1 | 0.11469117 | 0.50406192 | 0.5 | 0.435 | 1 | 0.23939752 | -0.5889279 | 0.681 | 0.698 | 1 | 0.23939752 | 0.21622828 |
| Scd2 | 0.11497478 | 0.29815319 | 0.854 | 0.741 | 1 | 0.23434585 | 0.29976929 | 0.664 | 0.677 | 1 | 0.23434585 | 0.21673036 |
| 1190005I06Rik | 0.11512046 | -1.070508 | 0.073 | 0.135 | 1 | 0.14250995 | 0.45567948 | 0.118 | 0.062 | 1 | 0.14250995 | 0.2169882 |
| Gm17749 | 0.91389901 | -0.4330781 | 0.062 | 0.057 | 1 | 0.11526849 | 1.06865636 | 0.063 | 0.021 | 1 | 0.91389901 | 0.21725016 |
| Bora | 0.11533447 | 0.77302928 | 0.125 | 0.067 | 1 | 0.17903342 | -0.8504236 | 0.038 | 0.073 | 1 | 0.17903342 | 0.21736691 |
| Spring1 | 0.14562784 | 0.34452952 | 0.219 | 0.145 | 1 | 0.1154251 | 0.46273424 | 0.151 | 0.083 | 1 | 0.14562784 | 0.21752725 |

| | | | | | | | | | | | | |
|---|---|---|---|---|---|---|---|---|---|---|---|---|
| Rerg | 0.60385447 | -0.4282744 | 0.281 | 0.249 | 1 | 0.11546365 | -0.7196561 | 0.433 | 0.469 | 1 | 0.60385447 | 0.21759545 |
| Hmox1 | 0.18503688 | -0.2098126 | 0.552 | 0.596 | 1 | 0.11551413 | 0.19053947 | 0.714 | 0.698 | 1 | 0.18503688 | 0.21768474 |
| Hk2 | 0.11551633 | 0.63552576 | 0.698 | 0.694 | 1 | 0.46187679 | -0.3065041 | 0.651 | 0.708 | 1 | 0.46187679 | 0.21768864 |
| Cxcl12 | 0.17320371 | 0.21771224 | 0.208 | 0.135 | 1 | 0.11570487 | -1.9378865 | 0.092 | 0.146 | 1 | 0.17320371 | 0.21802213 |
| 1700085D22Rik | 0.25203593 | 0.35541784 | 0.115 | 0.073 | 1 | 0.11585357 | 0.51611534 | 0.122 | 0.062 | 1 | 0.25203593 | 0.21828509 |
| Zglp1 | 0.11590147 | 1.13302542 | 0.094 | 0.047 | 1 | 0.29743566 | -1.4029647 | 0.055 | 0.083 | 1 | 0.29743566 | 0.21836978 |
| Prox2 | 0.11590147 | 1.13302542 | 0.094 | 0.047 | 1 | 0.66054955 | -0.4353861 | 0.042 | 0.031 | 1 | 0.66054955 | 0.21836978 |
| Trmt11 | 0.11597205 | 0.80104366 | 0.24 | 0.166 | 1 | 0.87154963 | -0.4550532 | 0.286 | 0.25 | 1 | 0.87154963 | 0.21849459 |
| Entrep1 | 0.11598828 | -2.0368996 | 0.031 | 0.078 | 1 | 0.25298758 | 0.97025266 | 0.088 | 0.052 | 1 | 0.25298758 | 0.21852329 |
| Abhd11 | 0.48192761 | 0.13302542 | 0.219 | 0.176 | 1 | 0.1160156 | 0.75427507 | 0.16 | 0.094 | 1 | 0.48192761 | 0.21857158 |
| Ahsa1 | 0.11618161 | 0.22988696 | 0.594 | 0.461 | 1 | 0.14153843 | 0.37196878 | 0.571 | 0.5 | 1 | 0.14153843 | 0.21886506 |
| Gm2011 | 0.15329198 | 0.78510212 | 0.052 | 0.021 | 1 | 0.11619623 | 1.01207283 | 0.046 | 0.01 | 1 | 0.15329198 | 0.2188909 |
| Prdm5 | 0.11620628 | 0.39114162 | 0.677 | 0.596 | 1 | 0.11702208 | -0.1329775 | 0.672 | 0.51 | 1 | 0.11702208 | 0.21890865 |
| Psd3 | 0.1162903 | -0.265716 | 0.927 | 0.964 | 1 | 0.44803159 | -0.3436307 | 0.983 | 0.958 | 1 | 0.44803159 | 0.21905716 |
| Gm57271 | 0.48684984 | 0.27052894 | 0.021 | 0.036 | 1 | 0.1163744 | 0.10518224 | 0.08 | 0.031 | 1 | 0.48684984 | 0.21920579 |
| Pon2 | 0.6408988 | 0.22322323 | 0.542 | 0.528 | 1 | 0.11641897 | -0.3583991 | 0.559 | 0.677 | 1 | 0.6408988 | 0.21928456 |
| Actr2 | 0.11647098 | 0.20013961 | 0.844 | 0.829 | 1 | 0.33103585 | -0.2131595 | 0.929 | 0.927 | 1 | 0.33103585 | 0.21937648 |
| Adamts18 | 0.11656608 | -2.0368996 | 0.01 | 0.047 | 1 | 0.8131384 | 0.14957636 | 0.025 | 0.021 | 1 | 0.8131384 | 0.21954451 |
| Tacc2 | 0.11659344 | -0.6485095 | 0.927 | 0.912 | 1 | 0.90051705 | -0.1816353 | 0.962 | 0.938 | 1 | 0.90051705 | 0.21959284 |
| Prss53 | 0.97579945 | -0.2726134 | 0.062 | 0.062 | 1 | 0.11662507 | 0.91253716 | 0.046 | 0.01 | 1 | 0.97579945 | 0.21964874 |
| Gm29266 | 0.1166931 | 2.66645762 | 0.062 | 0.026 | 1 | 0.87851807 | -2.5728897 | 0.013 | 0.01 | 1 | 0.87851807 | 0.21976892 |
| Crocc | 0.99482447 | -0.4779323 | 0.042 | 0.041 | 1 | 0.11669799 | -1.3098553 | 0.034 | 0.073 | 1 | 0.99482447 | 0.21977757 |
| Gm48024 | 0.57025555 | -0.8669746 | 0.042 | 0.057 | 1 | 0.11674247 | -2.8948178 | 0.008 | 0.031 | 1 | 0.57025555 | 0.21985614 |
| Gdf6 | 0.61737927 | 1.00749454 | 0.01 | 0.005 | 1 | 0.11674247 | -3.5322477 | 0.008 | 0.031 | 1 | 0.61737927 | 0.21985614 |
| Acta2 | 0.11675581 | -0.8445043 | 0.271 | 0.342 | 1 | 0.6565613 | -3.1367064 | 0.08 | 0.062 | 1 | 0.6565613 | 0.2198797 |
| Atxn3 | 0.11688529 | 0.63712609 | 0.292 | 0.212 | 1 | 0.23571372 | 0.30232871 | 0.277 | 0.208 | 1 | 0.23571372 | 0.22010841 |
| Gm47865 | 0.12141323 | -1.4950058 | 0.021 | 0.062 | 1 | 0.11689951 | -2.3098553 | 0 | 0.01 | 1 | 0.12141323 | 0.22013352 |
| Gm266 | 0.15729403 | -1.577468 | 0 | 0.021 | 1 | 0.11689951 | -2.3098553 | 0 | 0.01 | 1 | 0.15729403 | 0.22013352 |
| Gm38563 | 0.15729403 | -1.577468 | 0 | 0.021 | 1 | 0.11689951 | -2.3098553 | 0 | 0.01 | 1 | 0.15729403 | 0.22013352 |
| Fam124b | 0.15729781 | -2.577468 | 0 | 0.021 | 1 | 0.11689951 | -2.3098553 | 0 | 0.01 | 1 | 0.15729781 | 0.22013352 |
| Wnt16 | 0.15729781 | -1.9925055 | 0 | 0.021 | 1 | 0.11689951 | -2.3098553 | 0 | 0.01 | 1 | 0.15729781 | 0.22013352 |
| Gm10649 | 0.15729781 | -2.1624305 | 0 | 0.021 | 1 | 0.11689951 | -2.3098553 | 0 | 0.01 | 1 | 0.15729781 | 0.22013352 |
| Gm39323 | 0.15729907 | -1.3144336 | 0 | 0.021 | 1 | 0.11689951 | -2.8948178 | 0 | 0.01 | 1 | 0.15729907 | 0.22013352 |
| Gm16998 | 0.15729907 | -1.3144336 | 0 | 0.021 | 1 | 0.11689951 | -2.8948178 | 0 | 0.01 | 1 | 0.15729907 | 0.22013352 |
| Gm8206 | 0.15729907 | -1.3144336 | 0 | 0.021 | 1 | 0.11689951 | -2.3098553 | 0 | 0.01 | 1 | 0.15729907 | 0.22013352 |
| Gm26834 | 0.15729907 | -1.3144336 | 0 | 0.021 | 1 | 0.11689951 | -2.3098553 | 0 | 0.01 | 1 | 0.15729907 | 0.22013352 |
| 4930445N18Rik | 0.15729907 | -1.3144336 | 0 | 0.021 | 1 | 0.11689951 | -2.8948178 | 0 | 0.01 | 1 | 0.15729907 | 0.22013352 |
| Rgs8 | 0.15837881 | 2.59245704 | 0.01 | 0 | 1 | 0.11689951 | -2.3098553 | 0 | 0.01 | 1 | 0.15837881 | 0.22013352 |
| Grin1os | 0.15837881 | 2.00749454 | 0.01 | 0 | 1 | 0.11689951 | -2.3098553 | 0 | 0.01 | 1 | 0.15837881 | 0.22013352 |
| Gatm | 0.15837881 | 2.00749454 | 0.01 | 0 | 1 | 0.11689951 | -2.3098553 | 0 | 0.01 | 1 | 0.15837881 | 0.22013352 |
| Gm15472 | 0.15837881 | 2.00749454 | 0.01 | 0 | 1 | 0.11689951 | -2.8948178 | 0 | 0.01 | 1 | 0.15837881 | 0.22013352 |
| Gm35986 | 0.15837881 | 2.59245704 | 0.01 | 0 | 1 | 0.11689951 | -2.3098553 | 0 | 0.01 | 1 | 0.15837881 | 0.22013352 |
| Gm57078 | 0.15837881 | 2.00749454 | 0.01 | 0 | 1 | 0.11689951 | -3.3098553 | 0 | 0.01 | 1 | 0.15837881 | 0.22013352 |
| Insl5 | 0.15837881 | 2.00749454 | 0.01 | 0 | 1 | 0.11689951 | -2.3098553 | 0 | 0.01 | 1 | 0.15837881 | 0.22013352 |
| Gm3510 | 0.15837881 | 3.59245704 | 0.01 | 0 | 1 | 0.11689951 | -2.8948178 | 0 | 0.01 | 1 | 0.15837881 | 0.22013352 |
| Gm43016 | 0.15837881 | 2.00749454 | 0.01 | 0 | 1 | 0.11689951 | -2.3098553 | 0 | 0.01 | 1 | 0.15837881 | 0.22013352 |
| Gm57304 | 0.15837881 | 2.00749454 | 0.01 | 0 | 1 | 0.11689951 | -2.3098553 | 0 | 0.01 | 1 | 0.15837881 | 0.22013352 |
| Gm38883 | 0.15837881 | 2.59245704 | 0.01 | 0 | 1 | 0.11689951 | -2.3098553 | 0 | 0.01 | 1 | 0.15837881 | 0.22013352 |
| Ceacam19 | 0.15837881 | 2.00749454 | 0.01 | 0 | 1 | 0.11689951 | -3.8948178 | 0 | 0.01 | 1 | 0.15837881 | 0.22013352 |
| Gm10605 | 0.15837881 | 2.00749454 | 0.01 | 0 | 1 | 0.11689951 | -3.3098553 | 0 | 0.01 | 1 | 0.15837881 | 0.22013352 |
| Cmtm1 | 0.15837881 | 2.00749454 | 0.01 | 0 | 1 | 0.11689951 | -2.3098553 | 0 | 0.01 | 1 | 0.15837881 | 0.22013352 |
| Tmem45b | 0.15837881 | 2.00749454 | 0.01 | 0 | 1 | 0.11689951 | -2.3098553 | 0 | 0.01 | 1 | 0.15837881 | 0.22013352 |
| Paqr5 | 0.15837881 | 2.00749454 | 0.01 | 0 | 1 | 0.11689951 | -3.3098553 | 0 | 0.01 | 1 | 0.15837881 | 0.22013352 |
| Gnat1 | 0.15837881 | 2.00749454 | 0.01 | 0 | 1 | 0.11689951 | -2.8948178 | 0 | 0.01 | 1 | 0.15837881 | 0.22013352 |
| Pirt | 0.15837881 | 2.59245704 | 0.01 | 0 | 1 | 0.11689951 | -2.8948178 | 0 | 0.01 | 1 | 0.15837881 | 0.22013352 |
| Gm11629 | 0.15837881 | 2.00749454 | 0.01 | 0 | 1 | 0.11689951 | -3.8948178 | 0 | 0.01 | 1 | 0.15837881 | 0.22013352 |
| 4921501I09Rik | 0.15837881 | 2.00749454 | 0.01 | 0 | 1 | 0.11689951 | -2.3098553 | 0 | 0.01 | 1 | 0.15837881 | 0.22013352 |
| Gm56532 | 0.15837881 | 2.00749454 | 0.01 | 0 | 1 | 0.11689951 | -3.3098553 | 0 | 0.01 | 1 | 0.15837881 | 0.22013352 |
| Gm56536 | 0.15837881 | 2.59245704 | 0.01 | 0 | 1 | 0.11689951 | -2.3098553 | 0 | 0.01 | 1 | 0.15837881 | 0.22013352 |
| A530001N23Rik | 0.15837881 | 2.00749454 | 0.01 | 0 | 1 | 0.11689951 | -2.8948178 | 0 | 0.01 | 1 | 0.15837881 | 0.22013352 |
| A230107N01Rik | 0.15837881 | 2.00749454 | 0.01 | 0 | 1 | 0.11689951 | -2.3098553 | 0 | 0.01 | 1 | 0.15837881 | 0.22013352 |
| Tnfrsf13c | 0.15837881 | 3.81484946 | 0.01 | 0 | 1 | 0.11689951 | -2.3098553 | 0 | 0.01 | 1 | 0.15837881 | 0.22013352 |
| Gm56736 | 0.15837881 | 2.00749454 | 0.01 | 0 | 1 | 0.11689951 | -2.8948178 | 0 | 0.01 | 1 | 0.15837881 | 0.22013352 |
| Ostn | 0.15837881 | 2.00749454 | 0.01 | 0 | 1 | 0.11689951 | -2.3098553 | 0 | 0.01 | 1 | 0.15837881 | 0.22013352 |
| Rnf39 | 0.15837881 | 2.00749454 | 0.01 | 0 | 1 | 0.11689951 | -2.3098553 | 0 | 0.01 | 1 | 0.15837881 | 0.22013352 |
| Ppp1r32 | 0.15837881 | 2.00749454 | 0.01 | 0 | 1 | 0.11689951 | -2.3098553 | 0 | 0.01 | 1 | 0.15837881 | 0.22013352 |
| Gm56664 | 0.15837881 | 2.00749454 | 0.01 | 0 | 1 | 0.11689951 | -2.3098553 | 0 | 0.01 | 1 | 0.15837881 | 0.22013352 |
| Gm56673 | 0.15837881 | 2.00749454 | 0.01 | 0 | 1 | 0.11689951 | -2.3098553 | 0 | 0.01 | 1 | 0.15837881 | 0.22013352 |
| Cnga2 | 0.15837881 | 2.00749454 | 0.01 | 0 | 1 | 0.11689951 | -2.8948178 | 0 | 0.01 | 1 | 0.15837881 | 0.22013352 |
| Gm57476 | 0.20158871 | 1.42253204 | 0.031 | 0.01 | 1 | 0.11689951 | -2.8948178 | 0 | 0.01 | 1 | 0.20158871 | 0.22013352 |
| Fmr1nb | 0.20275473 | 1.32942263 | 0.031 | 0.01 | 1 | 0.11689951 | -2.3098553 | 0 | 0.01 | 1 | 0.20275473 | 0.22013352 |
| Gm30439 | 0.20510378 | 1.00749454 | 0.031 | 0.01 | 1 | 0.11689951 | -3.3098553 | 0 | 0.01 | 1 | 0.20510378 | 0.22013352 |
| Tmem125 | 0.21708019 | 2.00749454 | 0.021 | 0.005 | 1 | 0.11689951 | -2.3098553 | 0 | 0.01 | 1 | 0.21708019 | 0.22013352 |
| H2bc8 | 0.21708019 | 2.59245704 | 0.021 | 0.005 | 1 | 0.11689951 | -2.3098553 | 0 | 0.01 | 1 | 0.21708019 | 0.22013352 |
| Rbp7 | 0.21867159 | 1.59245704 | 0.021 | 0.005 | 1 | 0.11689951 | -2.3098553 | 0 | 0.01 | 1 | 0.21867159 | 0.22013352 |
| Gm15737 | 0.21867159 | 1.59245704 | 0.021 | 0.005 | 1 | 0.11689951 | -2.3098553 | 0 | 0.01 | 1 | 0.21867159 | 0.22013352 |
| Fam187b | 0.21867159 | 1.59245704 | 0.021 | 0.005 | 1 | 0.11689951 | -2.8948178 | 0 | 0.01 | 1 | 0.21867159 | 0.22013352 |
| Il27ra | 0.21867159 | 1.59245704 | 0.021 | 0.005 | 1 | 0.11689951 | -3.3098553 | 0 | 0.01 | 1 | 0.21867159 | 0.22013352 |
| Gm56535 | 0.21867159 | 1.59245704 | 0.021 | 0.005 | 1 | 0.11689951 | -2.3098553 | 0 | 0.01 | 1 | 0.21867159 | 0.22013352 |
| 4930586N03Rik | 0.21867159 | 1.59245704 | 0.021 | 0.005 | 1 | 0.11689951 | -2.8948178 | 0 | 0.01 | 1 | 0.21867159 | 0.22013352 |
| Phf11b | 0.21867159 | 1.59245704 | 0.021 | 0.005 | 1 | 0.11689951 | -2.8948178 | 0 | 0.01 | 1 | 0.21867159 | 0.22013352 |
| Lrfn2 | 0.21867159 | 1.59245704 | 0.021 | 0.005 | 1 | 0.11689951 | -3.6317834 | 0 | 0.01 | 1 | 0.21867159 | 0.22013352 |
| Cyp2c65 | 0.21867159 | 1.59245704 | 0.021 | 0.005 | 1 | 0.11689951 | -3.3098553 | 0 | 0.01 | 1 | 0.21867159 | 0.22013352 |
| 5830432E09Rik | 0.22026949 | 1.42253204 | 0.021 | 0.005 | 1 | 0.11689951 | -2.3098553 | 0 | 0.01 | 1 | 0.22026949 | 0.22013352 |

| | | | | | | | | | | | | |
|---|---|---|---|---|---|---|---|---|---|---|---|---|
| Gm57162 | 0.22187388 | -1.3144336 | 0 | 0.016 | 1 | 0.11689951 | -2.3098553 | 0 | 0.01 | 1 | 0.22187388 | 0.22013352 |
| Kcne3 | 0.22187388 | -1.3144336 | 0 | 0.016 | 1 | 0.11689951 | -2.3098553 | 0 | 0.01 | 1 | 0.22187388 | 0.22013352 |
| 1700047A11Rik | 0.22187388 | 1.00749454 | 0.021 | 0.005 | 1 | 0.11689951 | -2.3098553 | 0 | 0.01 | 1 | 0.22187388 | 0.22013352 |
| 4930517E14Rik | 0.22187388 | -1.3144336 | 0 | 0.016 | 1 | 0.11689951 | -2.3098553 | 0 | 0.01 | 1 | 0.22187388 | 0.22013352 |
| Gm36278 | 0.22187388 | -1.3144336 | 0 | 0.016 | 1 | 0.11689951 | -2.8948178 | 0 | 0.01 | 1 | 0.22187388 | 0.22013352 |
| Gast | 0.22187388 | -1.3144336 | 0 | 0.016 | 1 | 0.11689951 | -2.3098553 | 0 | 0.01 | 1 | 0.22187388 | 0.22013352 |
| Gm2245 | 0.22187388 | -1.3144336 | 0 | 0.016 | 1 | 0.11689951 | -2.3098553 | 0 | 0.01 | 1 | 0.22187388 | 0.22013352 |
| Sec14l5 | 0.22187388 | -1.3144336 | 0 | 0.016 | 1 | 0.11689951 | -2.3098553 | 0 | 0.01 | 1 | 0.22187388 | 0.22013352 |
| Hormad1 | 0.22187575 | -2.6929452 | 0 | 0.016 | 1 | 0.11689951 | -3.8948178 | 0 | 0.01 | 1 | 0.22187575 | 0.22013352 |
| Gm16253 | 0.22187575 | -0.9925055 | 0 | 0.016 | 1 | 0.11689951 | -2.3098553 | 0 | 0.01 | 1 | 0.22187575 | 0.22013352 |
| Gm43251 | 0.22187575 | -2.4519371 | 0 | 0.016 | 1 | 0.11689951 | -2.3098553 | 0 | 0.01 | 1 | 0.22187575 | 0.22013352 |
| Adam1b | 0.22187575 | -0.9925055 | 0 | 0.016 | 1 | 0.11689951 | -2.3098553 | 0 | 0.01 | 1 | 0.22187575 | 0.22013352 |
| Enpp6 | 0.22187575 | -0.9925055 | 0 | 0.016 | 1 | 0.11689951 | -2.3098553 | 0 | 0.01 | 1 | 0.22187575 | 0.22013352 |
| Capn9 | 0.22187575 | -0.9925055 | 0 | 0.016 | 1 | 0.11689951 | -2.8948178 | 0 | 0.01 | 1 | 0.22187575 | 0.22013352 |
| Gna15 | 0.22187575 | -0.9925055 | 0 | 0.016 | 1 | 0.11689951 | -2.8948178 | 0 | 0.01 | 1 | 0.22187575 | 0.22013352 |
| Slc17a3 | 0.22187575 | -2.4519371 | 0 | 0.016 | 1 | 0.11689951 | -3.6317834 | 0 | 0.01 | 1 | 0.22187575 | 0.22013352 |
| Gm56612 | 0.22187575 | -0.9925055 | 0 | 0.016 | 1 | 0.11689951 | -2.3098553 | 0 | 0.01 | 1 | 0.22187575 | 0.22013352 |
| Kcnk4 | 0.22187575 | -0.9925055 | 0 | 0.016 | 1 | 0.11689951 | -2.3098553 | 0 | 0.01 | 1 | 0.22187575 | 0.22013352 |
| Cxcr3 | 0.22187575 | -0.9925055 | 0 | 0.016 | 1 | 0.11689951 | -2.3098553 | 0 | 0.01 | 1 | 0.22187575 | 0.22013352 |
| Mog | 0.23737755 | -0.9185049 | 0.042 | 0.078 | 1 | 0.11689951 | -3.3098553 | 0 | 0.01 | 1 | 0.23737755 | 0.22013352 |
| Gm41555 | 0.28505098 | -1.9311049 | 0.01 | 0.031 | 1 | 0.11689951 | -2.8948178 | 0 | 0.01 | 1 | 0.28505098 | 0.22013352 |
| Aox4 | 0.32025586 | -1.577468 | 0 | 0.01 | 1 | 0.11689951 | -2.3098553 | 0 | 0.01 | 1 | 0.32025586 | 0.22013352 |
| Adam23 | 0.32025586 | -1.3144336 | 0 | 0.01 | 1 | 0.11689951 | -3.3098553 | 0 | 0.01 | 1 | 0.32025586 | 0.22013352 |
| B3gnt7 | 0.32025586 | -1.577468 | 0 | 0.01 | 1 | 0.11689951 | -2.3098553 | 0 | 0.01 | 1 | 0.32025586 | 0.22013352 |
| Gm4793 | 0.32025586 | -0.577468 | 0 | 0.01 | 1 | 0.11689951 | -2.8948178 | 0 | 0.01 | 1 | 0.32025586 | 0.22013352 |
| Tex50 | 0.32025586 | -1.3144336 | 0 | 0.01 | 1 | 0.11689951 | -2.3098553 | 0 | 0.01 | 1 | 0.32025586 | 0.22013352 |
| Lmx1b | 0.32025586 | -1.9925055 | 0 | 0.01 | 1 | 0.11689951 | -3.3098553 | 0 | 0.01 | 1 | 0.32025586 | 0.22013352 |
| 4930402F06Rik | 0.32025586 | -0.9925055 | 0 | 0.01 | 1 | 0.11689951 | -2.8948178 | 0 | 0.01 | 1 | 0.32025586 | 0.22013352 |
| Gssos1 | 0.32025586 | -0.577468 | 0 | 0.01 | 1 | 0.11689951 | -2.3098553 | 0 | 0.01 | 1 | 0.32025586 | 0.22013352 |
| Ctcfl | 0.32025586 | -0.577468 | 0 | 0.01 | 1 | 0.11689951 | -2.8948178 | 0 | 0.01 | 1 | 0.32025586 | 0.22013352 |
| Gm32950 | 0.32025586 | -0.9925055 | 0 | 0.01 | 1 | 0.11689951 | -2.3098553 | 0 | 0.01 | 1 | 0.32025586 | 0.22013352 |
| Gm33206 | 0.32025586 | -0.577468 | 0 | 0.01 | 1 | 0.11689951 | -2.3098553 | 0 | 0.01 | 1 | 0.32025586 | 0.22013352 |
| C030032O16Rik | 0.32025586 | -0.577468 | 0 | 0.01 | 1 | 0.11689951 | -2.3098553 | 0 | 0.01 | 1 | 0.32025586 | 0.22013352 |
| 4930447N08Rik | 0.32025586 | -0.9925055 | 0 | 0.01 | 1 | 0.11689951 | -2.8948178 | 0 | 0.01 | 1 | 0.32025586 | 0.22013352 |
| Clcnkb | 0.32025586 | -0.577468 | 0 | 0.01 | 1 | 0.11689951 | -2.3098553 | 0 | 0.01 | 1 | 0.32025586 | 0.22013352 |
| Clnk | 0.32025586 | -0.577468 | 0 | 0.01 | 1 | 0.11689951 | -2.3098553 | 0 | 0.01 | 1 | 0.32025586 | 0.22013352 |
| Cxcl5 | 0.32025586 | -1.577468 | 0 | 0.01 | 1 | 0.11689951 | -2.3098553 | 0 | 0.01 | 1 | 0.32025586 | 0.22013352 |
| Tcaf3 | 0.32025586 | -0.9925055 | 0 | 0.01 | 1 | 0.11689951 | -3.8948178 | 0 | 0.01 | 1 | 0.32025586 | 0.22013352 |
| Gm38832 | 0.32025586 | -0.577468 | 0 | 0.01 | 1 | 0.11689951 | -3.3098553 | 0 | 0.01 | 1 | 0.32025586 | 0.22013352 |
| Slc6a1 | 0.32025586 | -1.577468 | 0 | 0.01 | 1 | 0.11689951 | -2.3098553 | 0 | 0.01 | 1 | 0.32025586 | 0.22013352 |
| Rimklb | 0.32025586 | -0.577468 | 0 | 0.01 | 1 | 0.11689951 | -2.3098553 | 0 | 0.01 | 1 | 0.32025586 | 0.22013352 |
| Mir9-3hg | 0.32025586 | -0.9925055 | 0 | 0.01 | 1 | 0.11689951 | -2.8948178 | 0 | 0.01 | 1 | 0.32025586 | 0.22013352 |
| Itgax | 0.32025586 | -0.577468 | 0 | 0.01 | 1 | 0.11689951 | -4.4797803 | 0 | 0.01 | 1 | 0.32025586 | 0.22013352 |
| Gm15347 | 0.32025586 | -0.577468 | 0 | 0.01 | 1 | 0.11689951 | -2.3098553 | 0 | 0.01 | 1 | 0.32025586 | 0.22013352 |
| Atp7b | 0.32025586 | -1.3144336 | 0 | 0.01 | 1 | 0.11689951 | -2.3098553 | 0 | 0.01 | 1 | 0.32025586 | 0.22013352 |
| Ces2g | 0.32025586 | -0.577468 | 0 | 0.01 | 1 | 0.11689951 | -2.3098553 | 0 | 0.01 | 1 | 0.32025586 | 0.22013352 |
| Gm10701 | 0.32025586 | -0.577468 | 0 | 0.01 | 1 | 0.11689951 | -2.3098553 | 0 | 0.01 | 1 | 0.32025586 | 0.22013352 |
| 4930434F21Rik | 0.32025586 | -0.577468 | 0 | 0.01 | 1 | 0.11689951 | -2.3098553 | 0 | 0.01 | 1 | 0.32025586 | 0.22013352 |
| Gm30373 | 0.32025586 | -1.3144336 | 0 | 0.01 | 1 | 0.11689951 | -2.3098553 | 0 | 0.01 | 1 | 0.32025586 | 0.22013352 |
| Gm57059 | 0.32025586 | -0.577468 | 0 | 0.01 | 1 | 0.11689951 | -2.3098553 | 0 | 0.01 | 1 | 0.32025586 | 0.22013352 |
| Gm47818 | 0.32025586 | -0.577468 | 0 | 0.01 | 1 | 0.11689951 | -2.3098553 | 0 | 0.01 | 1 | 0.32025586 | 0.22013352 |
| Raet1d | 0.32025586 | -0.577468 | 0 | 0.01 | 1 | 0.11689951 | -2.3098553 | 0 | 0.01 | 1 | 0.32025586 | 0.22013352 |
| Samd3 | 0.32025586 | -0.577468 | 0 | 0.01 | 1 | 0.11689951 | -2.3098553 | 0 | 0.01 | 1 | 0.32025586 | 0.22013352 |
| Esrrb | 0.32025586 | -1.577468 | 0 | 0.01 | 1 | 0.11689951 | -2.3098553 | 0 | 0.01 | 1 | 0.32025586 | 0.22013352 |
| Gm20658 | 0.32025586 | -0.577468 | 0 | 0.01 | 1 | 0.11689951 | -2.8948178 | 0 | 0.01 | 1 | 0.32025586 | 0.22013352 |
| Gm31683 | 0.32025586 | -0.577468 | 0 | 0.01 | 1 | 0.11689951 | -2.8948178 | 0 | 0.01 | 1 | 0.32025586 | 0.22013352 |
| Gm48975 | 0.32025586 | -0.577468 | 0 | 0.01 | 1 | 0.11689951 | -2.3098553 | 0 | 0.01 | 1 | 0.32025586 | 0.22013352 |
| Jph4 | 0.32025586 | -0.577468 | 0 | 0.01 | 1 | 0.11689951 | -2.3098553 | 0 | 0.01 | 1 | 0.32025586 | 0.22013352 |
| Defb30 | 0.32025586 | -0.577468 | 0 | 0.01 | 1 | 0.11689951 | -3.3098553 | 0 | 0.01 | 1 | 0.32025586 | 0.22013352 |
| Gm5464 | 0.32025586 | -0.577468 | 0 | 0.01 | 1 | 0.11689951 | -2.8948178 | 0 | 0.01 | 1 | 0.32025586 | 0.22013352 |
| Rspo2 | 0.32025586 | -1.7998604 | 0 | 0.01 | 1 | 0.11689951 | -2.3098553 | 0 | 0.01 | 1 | 0.32025586 | 0.22013352 |
| Nell2 | 0.32025586 | -0.9925055 | 0 | 0.01 | 1 | 0.11689951 | -2.8948178 | 0 | 0.01 | 1 | 0.32025586 | 0.22013352 |
| Gm10791 | 0.32025586 | -0.577468 | 0 | 0.01 | 1 | 0.11689951 | -3.3098553 | 0 | 0.01 | 1 | 0.32025586 | 0.22013352 |
| Bnip5 | 0.32025586 | -1.3144336 | 0 | 0.01 | 1 | 0.11689951 | -2.3098553 | 0 | 0.01 | 1 | 0.32025586 | 0.22013352 |
| Ly6g6f | 0.32025586 | -0.577468 | 0 | 0.01 | 1 | 0.11689951 | -2.3098553 | 0 | 0.01 | 1 | 0.32025586 | 0.22013352 |
| Crip3 | 0.32025586 | -0.577468 | 0 | 0.01 | 1 | 0.11689951 | -2.3098553 | 0 | 0.01 | 1 | 0.32025586 | 0.22013352 |
| Gm41804 | 0.32025586 | -0.9925055 | 0 | 0.01 | 1 | 0.11689951 | -2.3098553 | 0 | 0.01 | 1 | 0.32025586 | 0.22013352 |
| Vwa2 | 0.32025586 | -0.577468 | 0 | 0.01 | 1 | 0.11689951 | -2.3098553 | 0 | 0.01 | 1 | 0.32025586 | 0.22013352 |
| Tac1 | 0.39524459 | 0.27052894 | 0.01 | 0.026 | 1 | 0.11689951 | -5.1172102 | 0 | 0.01 | 1 | 0.39524459 | 0.22013352 |
| Kifc1 | 0.4638161 | 0.64492446 | 0.042 | 0.026 | 1 | 0.11689951 | -2.3098553 | 0 | 0.01 | 1 | 0.4638161 | 0.22013352 |
| 4933430I17Rik | 0.47150075 | 1.42253204 | 0.021 | 0.01 | 1 | 0.11689951 | -2.3098553 | 0 | 0.01 | 1 | 0.47150075 | 0.22013352 |
| Gm34907 | 0.47150075 | 2.00749454 | 0.021 | 0.01 | 1 | 0.11689951 | -3.3098553 | 0 | 0.01 | 1 | 0.47150075 | 0.22013352 |
| Npl | 0.47606502 | 1.00749454 | 0.021 | 0.01 | 1 | 0.11689951 | -2.3098553 | 0 | 0.01 | 1 | 0.47606502 | 0.22013352 |
| A930004D18Rik | 0.48064264 | 0.27052894 | 0.021 | 0.01 | 1 | 0.11689951 | -2.8948178 | 0 | 0.01 | 1 | 0.48064264 | 0.22013352 |
| Ttn | 0.48064264 | 0.59245704 | 0.021 | 0.01 | 1 | 0.11689951 | -2.8948178 | 0 | 0.01 | 1 | 0.48064264 | 0.22013352 |
| ENSMUSG00000121511 | 0.48064264 | 0.59245704 | 0.021 | 0.01 | 1 | 0.11689951 | -2.3098553 | 0 | 0.01 | 1 | 0.48064264 | 0.22013352 |
| Mx1 | 0.48525351 | -1.1079827 | 0.021 | 0.01 | 1 | 0.11689951 | -6.3542494 | 0 | 0.01 | 1 | 0.48525351 | 0.22013352 |
| Aldh3b2 | 0.52313476 | -0.9925055 | 0.01 | 0.021 | 1 | 0.11689951 | -3.3098553 | 0 | 0.01 | 1 | 0.52313476 | 0.22013352 |
| Fgf16 | 0.52528691 | -0.7998604 | 0.01 | 0.021 | 1 | 0.11689951 | -2.3098553 | 0 | 0.01 | 1 | 0.52528691 | 0.22013352 |
| Il1rapl2 | 0.52528691 | -1.577468 | 0.01 | 0.021 | 1 | 0.11689951 | -5.3098553 | 0 | 0.01 | 1 | 0.52528691 | 0.22013352 |
| Gm55759 | 0.52744059 | -0.9925055 | 0.01 | 0.021 | 1 | 0.11689951 | -2.8948178 | 0 | 0.01 | 1 | 0.52744059 | 0.22013352 |
| Gm39816 | 0.52961488 | -0.3144336 | 0.01 | 0.021 | 1 | 0.11689951 | -2.3098553 | 0 | 0.01 | 1 | 0.52961488 | 0.22013352 |

| | | | | | | | | | | | | |
|---|---|---|---|---|---|---|---|---|---|---|---|---|
| 4833417C18Rik | 0.52961488 | -0.3144336 | 0.01 | 0.021 | 1 | 0.11689951 | -2.3098553 | 0 | 0.01 | 1 | 0.52961488 | 0.22013352 |
| Dgat2l6 | 0.5383068 | 0.27052894 | 0.01 | 0.021 | 1 | 0.11689951 | -3.6317834 | 0 | 0.01 | 1 | 0.5383068 | 0.22013352 |
| 1200007C13Rik | 0.61371908 | 2.32942263 | 0.01 | 0.005 | 1 | 0.11689951 | -2.3098553 | 0 | 0.01 | 1 | 0.61371908 | 0.22013352 |
| Dnah3 | 0.61371908 | 1.42253204 | 0.01 | 0.005 | 1 | 0.11689951 | -2.3098553 | 0 | 0.01 | 1 | 0.61371908 | 0.22013352 |
| Tnfrsf13b | 0.61371908 | 1.59245704 | 0.01 | 0.005 | 1 | 0.11689951 | -3.6317834 | 0 | 0.01 | 1 | 0.61371908 | 0.22013352 |
| Scn4a | 0.61371908 | 1.59245704 | 0.01 | 0.005 | 1 | 0.11689951 | -2.8948178 | 0 | 0.01 | 1 | 0.61371908 | 0.22013352 |
| Tagap | 0.61371908 | 1.59245704 | 0.01 | 0.005 | 1 | 0.11689951 | -2.3098553 | 0 | 0.01 | 1 | 0.61371908 | 0.22013352 |
| Gm16283 | 0.61550914 | -0.407543 | 0.021 | 0.031 | 1 | 0.11689951 | -2.3098553 | 0 | 0.01 | 1 | 0.61550914 | 0.22013352 |
| Gm13184 | 0.61737927 | 1.00749454 | 0.01 | 0.005 | 1 | 0.11689951 | -3.3098553 | 0 | 0.01 | 1 | 0.61737927 | 0.22013352 |
| Gm57460 | 0.61737927 | 1.00749454 | 0.01 | 0.005 | 1 | 0.11689951 | -2.3098553 | 0 | 0.01 | 1 | 0.61737927 | 0.22013352 |
| 4731419I09Rik | 0.61737927 | 1.00749454 | 0.01 | 0.005 | 1 | 0.11689951 | -2.3098553 | 0 | 0.01 | 1 | 0.61737927 | 0.22013352 |
| Gm42463 | 0.61737927 | 1.00749454 | 0.01 | 0.005 | 1 | 0.11689951 | -2.3098553 | 0 | 0.01 | 1 | 0.61737927 | 0.22013352 |
| Gm15527 | 0.61737927 | 1.00749454 | 0.01 | 0.005 | 1 | 0.11689951 | -2.3098553 | 0 | 0.01 | 1 | 0.61737927 | 0.22013352 |
| Peak1os | 0.61737927 | 1.00749454 | 0.01 | 0.005 | 1 | 0.11689951 | -2.3098553 | 0 | 0.01 | 1 | 0.61737927 | 0.22013352 |
| Vstm2a | 0.61737927 | 1.00749454 | 0.01 | 0.005 | 1 | 0.11689951 | -2.3098553 | 0 | 0.01 | 1 | 0.61737927 | 0.22013352 |
| Cd300lg | 0.61737927 | 1.00749454 | 0.01 | 0.005 | 1 | 0.11689951 | -3.6317834 | 0 | 0.01 | 1 | 0.61737927 | 0.22013352 |
| Vsx2 | 0.61737927 | 1.00749454 | 0.01 | 0.005 | 1 | 0.11689951 | -2.3098553 | 0 | 0.01 | 1 | 0.61737927 | 0.22013352 |
| Rhpn1 | 0.61737927 | 1.00749454 | 0.01 | 0.005 | 1 | 0.11689951 | -2.8948178 | 0 | 0.01 | 1 | 0.61737927 | 0.22013352 |
| Tulp1 | 0.61737927 | 1.00749454 | 0.01 | 0.005 | 1 | 0.11689951 | -2.3098553 | 0 | 0.01 | 1 | 0.61737927 | 0.22013352 |
| Gm36914 | 0.61737927 | 1.00749454 | 0.01 | 0.005 | 1 | 0.11689951 | -2.3098553 | 0 | 0.01 | 1 | 0.61737927 | 0.22013352 |
| Gm44284 | 0.62104899 | 0.42253204 | 0.01 | 0.005 | 1 | 0.11689951 | -2.3098553 | 0 | 0.01 | 1 | 0.62104899 | 0.22013352 |
| Exoc3l | 0.62104899 | 0.42253204 | 0.01 | 0.005 | 1 | 0.11689951 | -2.3098553 | 0 | 0.01 | 1 | 0.62104899 | 0.22013352 |
| Gm46336 | 0.62104899 | 0.42253204 | 0.01 | 0.005 | 1 | 0.11689951 | -2.3098553 | 0 | 0.01 | 1 | 0.62104899 | 0.22013352 |
| Marchf11 | 0.62104899 | 0.42253204 | 0.01 | 0.005 | 1 | 0.11689951 | -2.3098553 | 0 | 0.01 | 1 | 0.62104899 | 0.22013352 |
| Gm4651 | 0.72574709 | -0.3144336 | 0.01 | 0.016 | 1 | 0.11689951 | -2.8948178 | 0 | 0.01 | 1 | 0.72574709 | 0.22013352 |
| 5730435O14Rik | 0.72574709 | -0.7998604 | 0.01 | 0.016 | 1 | 0.11689951 | -2.3098553 | 0 | 0.01 | 1 | 0.72574709 | 0.22013352 |
| Trappc3l | 0.74576744 | 0.68556644 | 0.021 | 0.016 | 1 | 0.11689951 | -3.3098553 | 0 | 0.01 | 1 | 0.74576744 | 0.22013352 |
| Cnr2 | 0.74827573 | 0.59245704 | 0.021 | 0.016 | 1 | 0.11689951 | -2.3098553 | 0 | 0.01 | 1 | 0.74827573 | 0.22013352 |
| Cxcl10 | 0.7558087 | -1.7123975 | 0.021 | 0.016 | 1 | 0.11689951 | -8.3322231 | 0 | 0.01 | 1 | 0.7558087 | 0.22013352 |
| Gm29282 | 0.8006267 | 0.20013961 | 0.021 | 0.026 | 1 | 0.11689951 | -3.3098553 | 0 | 0.01 | 1 | 0.8006267 | 0.22013352 |
| Tff1 | 0.993208 | 2.22988696 | 0.01 | 0.01 | 1 | 0.11689951 | -3.3098553 | 0 | 0.01 | 1 | 0.993208 | 0.22013352 |
| Gm14936 | 0.993208 | 1.00749454 | 0.01 | 0.01 | 1 | 0.11689951 | -3.3098553 | 0 | 0.01 | 1 | 0.993208 | 0.22013352 |
| Lhfpl3 | 0.9951721 | 0.32942263 | 0.021 | 0.021 | 1 | 0.11689951 | -2.3098553 | 0 | 0.01 | 1 | 0.9951721 | 0.22013352 |
| Slco5a1 | 0.99758609 | 0.27052894 | 0.021 | 0.021 | 1 | 0.11689951 | -2.8948178 | 0 | 0.01 | 1 | 0.99758609 | 0.22013352 |
| Wnt10a | 1 | 0.42253204 | 0.01 | 0.01 | 1 | 0.11689951 | -2.3098553 | 0 | 0.01 | 1 | 1 | 0.22013352 |
| Sp110 | 1 | 0.42253204 | 0.01 | 0.01 | 1 | 0.11689951 | -3.3098553 | 0 | 0.01 | 1 | 1 | 0.22013352 |
| Lexm | 1 | 0.42253204 | 0.01 | 0.01 | 1 | 0.11689951 | -2.3098553 | 0 | 0.01 | 1 | 1 | 0.22013352 |
| Gm43698 | 1 | -0.3144336 | 0.01 | 0.01 | 1 | 0.11689951 | -2.3098553 | 0 | 0.01 | 1 | 1 | 0.22013352 |
| Gm15541 | 1 | 0.42253204 | 0.01 | 0.01 | 1 | 0.11689951 | -2.3098553 | 0 | 0.01 | 1 | 1 | 0.22013352 |
| Dpep1 | 1 | -0.130009 | 0.031 | 0.031 | 1 | 0.11689951 | -2.3098553 | 0 | 0.01 | 1 | 1 | 0.22013352 |
| Ptprb | 1 | 0.42253204 | 0.01 | 0.01 | 1 | 0.11689951 | -4.6317834 | 0 | 0.01 | 1 | 1 | 0.22013352 |
| Kcnk13 | 1 | -1.577468 | 0.01 | 0.01 | 1 | 0.11689951 | -2.3098553 | 0 | 0.01 | 1 | 1 | 0.22013352 |
| Ly6k | 1 | -0.3144336 | 0.01 | 0.01 | 1 | 0.11689951 | -2.3098553 | 0 | 0.01 | 1 | 1 | 0.22013352 |
| Grep1 | 1 | 0.42253204 | 0.01 | 0.01 | 1 | 0.11689951 | -2.8948178 | 0 | 0.01 | 1 | 1 | 0.22013352 |
| Gldc | 1 | 0.42253204 | 0.01 | 0.01 | 1 | 0.11689951 | -3.3098553 | 0 | 0.01 | 1 | 1 | 0.22013352 |
| Mtmr14 | 0.11700459 | 0.52206771 | 0.146 | 0.083 | 1 | 0.17553612 | -0.4838847 | 0.147 | 0.083 | 1 | 0.17553612 | 0.2203191 |
| Rrp1b | 0.11704866 | -0.6225559 | 0.156 | 0.238 | 1 | 0.13167923 | 0.20720317 | 0.261 | 0.177 | 1 | 0.13167923 | 0.22039693 |
| Morrbid | 0.11710119 | -2.0222528 | 0.042 | 0.093 | 1 | 0.30329364 | -1.8573431 | 0.038 | 0.062 | 1 | 0.30329364 | 0.2204897 |
| Ebf2 | 0.61737927 | 1.00749454 | 0.01 | 0.005 | 1 | 0.11710883 | 0.61614416 | 0.046 | 0.01 | 1 | 0.61737927 | 0.22050318 |
| Usp18 | 0.11714866 | 0.10060394 | 0.125 | 0.067 | 1 | 0.67674034 | 0.15727075 | 0.088 | 0.073 | 1 | 0.67674034 | 0.22057351 |
| Nfasc | 0.44805679 | -0.2518921 | 0.229 | 0.181 | 1 | 0.11719523 | -0.3757432 | 0.29 | 0.177 | 1 | 0.44805679 | 0.22065574 |
| Gm15559 | 0.11721645 | -1.577468 | 0.031 | 0.078 | 1 | 0.1610839 | -0.9879272 | 0.029 | 0.062 | 1 | 0.1610839 | 0.22069321 |
| Xpo6 | 0.11731037 | 0.33149696 | 0.531 | 0.425 | 1 | 0.68667242 | -0.4272122 | 0.727 | 0.656 | 1 | 0.68667242 | 0.22085902 |
| Tmem8b | 0.77406769 | -0.1941393 | 0.094 | 0.104 | 1 | 0.11732485 | 1.19264508 | 0.143 | 0.083 | 1 | 0.77406769 | 0.22088459 |
| Zbtb8a | 0.99603742 | 0.20013961 | 0.031 | 0.031 | 1 | 0.11751579 | 0.86006974 | 0.063 | 0.021 | 1 | 0.99603742 | 0.22122161 |
| Gm27010 | 0.72298091 | -0.577468 | 0.01 | 0.016 | 1 | 0.11751705 | 0.27510724 | 0.046 | 0.01 | 1 | 0.72298091 | 0.22122384 |
| Inafm1 | 0.78673961 | 0.22430593 | 0.292 | 0.285 | 1 | 0.11751756 | 0.16819203 | 0.265 | 0.177 | 1 | 0.78673961 | 0.22122474 |
| Dand5 | 0.11759217 | -0.8770282 | 0.094 | 0.161 | 1 | 0.42908791 | -1.4881925 | 0.105 | 0.125 | 1 | 0.42908791 | 0.22135641 |
| Pdgfd | 0.59158055 | -0.2295447 | 0.156 | 0.181 | 1 | 0.11763615 | -1.5080814 | 0.181 | 0.24 | 1 | 0.59158055 | 0.22143404 |
| 1500009L16Rik | 0.16393187 | -0.9925055 | 0.021 | 0.057 | 1 | 0.11764285 | 1.69014474 | 0.025 | 0 | 1 | 0.16393187 | 0.22144586 |
| Myef2l | 0.17534439 | 1.32942263 | 0.042 | 0.016 | 1 | 0.11764285 | 1.86006974 | 0.025 | 0 | 1 | 0.17534439 | 0.22144586 |
| Spef2 | 0.22187575 | -1.577468 | 0 | 0.016 | 1 | 0.11764285 | 2.01207283 | 0.025 | 0 | 1 | 0.22187575 | 0.22144586 |
| Akr1b8 | 0.32025586 | -0.577468 | 0 | 0.01 | 1 | 0.11764285 | 1.69014474 | 0.025 | 0 | 1 | 0.32025586 | 0.22144586 |
| Wfikkn1 | 0.32025586 | -0.9925055 | 0 | 0.01 | 1 | 0.11764285 | 1.69014474 | 0.025 | 0 | 1 | 0.32025586 | 0.22144586 |
| 9230116N13Rik | 0.38181636 | 0.78510212 | 0.052 | 0.031 | 1 | 0.11764285 | 1.69014474 | 0.025 | 0 | 1 | 0.38181636 | 0.22144586 |
| Slc35f1 | 0.39021373 | -0.8993961 | 0.031 | 0.016 | 1 | 0.11764285 | 1.69014474 | 0.025 | 0 | 1 | 0.39021373 | 0.22144586 |
| Pla2g10os | 0.39524555 | 0.42253204 | 0.01 | 0.026 | 1 | 0.11764285 | 1.69014474 | 0.025 | 0 | 1 | 0.39524555 | 0.22144586 |
| Gm43714 | 0.47606502 | 1.00749454 | 0.021 | 0.01 | 1 | 0.11764285 | 1.69014474 | 0.025 | 0 | 1 | 0.47606502 | 0.22144586 |
| Mir142hg | 0.61371908 | 2.46692616 | 0.01 | 0.005 | 1 | 0.11764285 | 2.59703533 | 0.025 | 0 | 1 | 0.61371908 | 0.22144586 |
| 9630002D21Rik | 0.61371908 | 1.42253204 | 0.01 | 0.005 | 1 | 0.11764285 | 1.86006974 | 0.025 | 0 | 1 | 0.61371908 | 0.22144586 |
| Gm12915 | 0.61737927 | 1.00749454 | 0.01 | 0.005 | 1 | 0.11764285 | 1.69014474 | 0.025 | 0 | 1 | 0.61737927 | 0.22144586 |
| Gm6787 | 0.61737927 | 1.00749454 | 0.01 | 0.005 | 1 | 0.11764285 | 1.69014474 | 0.025 | 0 | 1 | 0.61737927 | 0.22144586 |
| Lcp2 | 0.72574709 | -1.4519371 | 0.01 | 0.016 | 1 | 0.11764285 | 2.01207283 | 0.025 | 0 | 1 | 0.72574709 | 0.22144586 |
| Gm41760 | 0.8006267 | 0.20013961 | 0.021 | 0.026 | 1 | 0.11764285 | 1.69014474 | 0.025 | 0 | 1 | 0.8006267 | 0.22144586 |
| Fcrla | 0.98414882 | 1.78510212 | 0.031 | 0.031 | 1 | 0.11764285 | 1.69014474 | 0.025 | 0 | 1 | 0.98414882 | 0.22144586 |
| Cd83 | 1 | 0.42253204 | 0.01 | 0.01 | 1 | 0.11764285 | 1.69014474 | 0.025 | 0 | 1 | 1 | 0.22144586 |
| Scn8a | 0.28379282 | -0.7998604 | 0.01 | 0.031 | 1 | 0.11764843 | 2.01207283 | 0.025 | 0 | 1 | 0.28379282 | 0.22145571 |
| Ccnf | 0.40677356 | 1.35541784 | 0.062 | 0.041 | 1 | 0.11764843 | 1.86006974 | 0.025 | 0 | 1 | 0.40677356 | 0.22145571 |
| Gm11767 | 0.5862099 | 0.68556644 | 0.031 | 0.021 | 1 | 0.11764843 | 1.86006974 | 0.025 | 0 | 1 | 0.5862099 | 0.22145571 |
| Gm3455 | 0.66189631 | -1.1941393 | 0.031 | 0.041 | 1 | 0.11764843 | 2.14957636 | 0.025 | 0 | 1 | 0.66189631 | 0.22145571 |
| Lipt2 | 0.84862675 | 0.13302542 | 0.052 | 0.047 | 1 | 0.11764843 | 2.01207283 | 0.025 | 0 | 1 | 0.84862675 | 0.22145571 |
| Kif6 | 0.98414882 | 0.78510212 | 0.031 | 0.031 | 1 | 0.11764843 | 1.86006974 | 0.025 | 0 | 1 | 0.98414882 | 0.22145571 |
| Pkp2 | 0.99034437 | 1.22988696 | 0.021 | 0.021 | 1 | 0.11764843 | 2.01207283 | 0.025 | 0 | 1 | 0.99034437 | 0.22145571 |
| Ppm1j | 1 | 0.42253204 | 0.01 | 0.01 | 1 | 0.11764843 | 1.86006974 | 0.025 | 0 | 1 | 1 | 0.22145571 |

| | | | | | | | | | | | | |
|---|---|---|---|---|---|---|---|---|---|---|---|---|
| Tmprss9 | 0.11765021 | -0.9925055 | 0.01 | 0.047 | 1 | 0.65527364 | -0.8073549 | 0.05 | 0.062 | 1 | 0.65527364 | 0.22145885 |
| Gpr183 | 0.22187388 | -1.3144336 | 0 | 0.016 | 1 | 0.11765178 | 3.33400093 | 0.025 | 0 | 1 | 0.22187388 | 0.22146163 |
| Oscar | 0.22187575 | -0.9925055 | 0 | 0.016 | 1 | 0.11765346 | 3.86006974 | 0.025 | 0 | 1 | 0.22187575 | 0.22146458 |
| Ccdc184 | 0.15837881 | 2.00749454 | 0.01 | 0 | 1 | 0.11765402 | 1.49749966 | 0.025 | 0 | 1 | 0.15837881 | 0.22146557 |
| ENSMUSG00000121487 | 0.16289169 | 0.50999488 | 0.156 | 0.098 | 1 | 0.11765402 | 1.49749966 | 0.025 | 0 | 1 | 0.16289169 | 0.22146557 |
| Gm10053 | 0.21867159 | 1.59245704 | 0.021 | 0.005 | 1 | 0.11765402 | 1.49749966 | 0.025 | 0 | 1 | 0.21867159 | 0.22146557 |
| Gm42982 | 0.22187388 | 1.00749454 | 0.021 | 0.005 | 1 | 0.11765402 | 1.49749966 | 0.025 | 0 | 1 | 0.22187388 | 0.22146557 |
| Syngr1 | 0.32025586 | -0.577468 | 0 | 0.01 | 1 | 0.11765402 | 1.49749966 | 0.025 | 0 | 1 | 0.32025586 | 0.22146557 |
| 4932443L11Rik | 0.38028858 | 1.00749454 | 0.031 | 0.016 | 1 | 0.11765402 | 1.49749966 | 0.025 | 0 | 1 | 0.38028858 | 0.22146557 |
| Fgl1 | 0.38028858 | 1.00749454 | 0.031 | 0.016 | 1 | 0.11765402 | 1.49749966 | 0.025 | 0 | 1 | 0.38028858 | 0.22146557 |
| Gm57397 | 0.61550914 | -0.407543 | 0.021 | 0.031 | 1 | 0.11765402 | 1.49749966 | 0.025 | 0 | 1 | 0.61550914 | 0.22146557 |
| Hhla1 | 0.61924305 | -0.2148979 | 0.021 | 0.031 | 1 | 0.11765402 | 1.49749966 | 0.025 | 0 | 1 | 0.61924305 | 0.22146557 |
| Mro | 0.87089632 | 0.15949763 | 0.083 | 0.078 | 1 | 0.11765402 | 1.49749966 | 0.025 | 0 | 1 | 0.87089632 | 0.22146557 |
| Gm43598 | 0.99034437 | 0.74446013 | 0.021 | 0.021 | 1 | 0.11765402 | 1.49749966 | 0.025 | 0 | 1 | 0.99034437 | 0.22146557 |
| Gm57475 | 0.99603742 | 0.20013961 | 0.031 | 0.031 | 1 | 0.11765402 | 1.49749966 | 0.025 | 0 | 1 | 0.99603742 | 0.22146557 |
| Ido2 | 1 | 0.42253204 | 0.01 | 0.01 | 1 | 0.11765402 | 1.49749966 | 0.025 | 0 | 1 | 1 | 0.22146557 |
| Nectin2 | 0.20622925 | -0.577468 | 0.115 | 0.171 | 1 | 0.11765701 | -0.6217993 | 0.197 | 0.271 | 1 | 0.20622925 | 0.22147085 |
| Gltp | 0.11781048 | -0.5965768 | 0.354 | 0.425 | 1 | 0.98935094 | 0.13760371 | 0.349 | 0.375 | 1 | 0.98935094 | 0.22174166 |
| 2310047K21Rik | 0.38578008 | 0.21899864 | 0.104 | 0.073 | 1 | 0.11790164 | 0.61614416 | 0.063 | 0.021 | 1 | 0.38578008 | 0.22190248 |
| Rpgr | 0.11793519 | 0.31943854 | 0.167 | 0.098 | 1 | 0.8684586 | -0.9139266 | 0.08 | 0.083 | 1 | 0.8684586 | 0.22196168 |
| Ldlrad1 | 0.15837881 | 2.00749454 | 0.01 | 0 | 1 | 0.11813493 | -2.5322477 | 0.008 | 0.031 | 1 | 0.15837881 | 0.222314 |
| Gm31812 | 0.20978548 | -1.3144336 | 0.01 | 0.036 | 1 | 0.11813493 | -3.5322477 | 0.008 | 0.031 | 1 | 0.20978548 | 0.222314 |
| Hap1 | 0.22187575 | -1.7998604 | 0 | 0.016 | 1 | 0.11813493 | -2.3098553 | 0.008 | 0.031 | 1 | 0.22187575 | 0.222314 |
| Unc5d | 0.32025586 | -0.9925055 | 0 | 0.01 | 1 | 0.11813493 | -2.7248928 | 0.008 | 0.031 | 1 | 0.32025586 | 0.222314 |
| Ccr9 | 0.32025586 | -0.577468 | 0 | 0.01 | 1 | 0.11813493 | -2.5322477 | 0.008 | 0.031 | 1 | 0.32025586 | 0.222314 |
| Gm5535 | 0.47602456 | 0.42253204 | 0.042 | 0.026 | 1 | 0.11813493 | -2.7248928 | 0.008 | 0.031 | 1 | 0.47602456 | 0.222314 |
| Tex12 | 0.52744059 | -0.577468 | 0.01 | 0.021 | 1 | 0.11813493 | -2.3098553 | 0.008 | 0.031 | 1 | 0.52744059 | 0.222314 |
| Gm57107 | 0.61737927 | 1.00749454 | 0.01 | 0.005 | 1 | 0.11813493 | -2.3098553 | 0.008 | 0.031 | 1 | 0.61737927 | 0.222314 |
| Tbc1d13 | 0.70125283 | -0.1801325 | 0.156 | 0.135 | 1 | 0.11815634 | 0.37820073 | 0.181 | 0.104 | 1 | 0.70125283 | 0.22235176 |
| Ndufa9 | 0.48904449 | 0.14620081 | 0.458 | 0.425 | 1 | 0.11816522 | 0.36625513 | 0.445 | 0.354 | 1 | 0.48904449 | 0.22236741 |
| Lca5 | 0.11821089 | 0.67045955 | 0.146 | 0.083 | 1 | 0.61818981 | -0.6909454 | 0.13 | 0.146 | 1 | 0.61818981 | 0.22244797 |
| Arrb2 | 0.27624715 | -1.1079827 | 0.021 | 0.047 | 1 | 0.11828828 | 0.53814164 | 0.063 | 0.021 | 1 | 0.27624715 | 0.22258444 |
| Ikzf4 | 0.1183916 | 0.54452256 | 0.385 | 0.306 | 1 | 0.42463443 | -0.2667865 | 0.332 | 0.26 | 1 | 0.42463443 | 0.22276663 |
| Gm17167 | 0.64725171 | -1.0799683 | 0.031 | 0.041 | 1 | 0.11839722 | -1.1172102 | 0.034 | 0.073 | 1 | 0.64725171 | 0.22277655 |
| Lrfn4 | 0.1184225 | 0.9200317 | 0.135 | 0.078 | 1 | 0.19242691 | 0.32486027 | 0.197 | 0.135 | 1 | 0.19242691 | 0.2228211 |
| 4930593A02Rik | 0.11845546 | -1.6929452 | 0.031 | 0.078 | 1 | 0.93071316 | -0.3098553 | 0.029 | 0.031 | 1 | 0.93071316 | 0.22287922 |
| Pi4kb | 0.1184832 | 0.40483003 | 0.375 | 0.28 | 1 | 0.13274973 | 0.10274319 | 0.471 | 0.333 | 1 | 0.13274973 | 0.22292814 |
| Cul7 | 0.1185575 | 0.1843723 | 0.323 | 0.223 | 1 | 0.14326949 | 0.19762849 | 0.403 | 0.312 | 1 | 0.14326949 | 0.22305911 |
| Mtx3 | 0.11856889 | 1.13873907 | 0.146 | 0.088 | 1 | 0.12133282 | 0.65361886 | 0.134 | 0.073 | 1 | 0.12133282 | 0.22307919 |
| Efhc1 | 0.21374303 | 0.81484946 | 0.062 | 0.031 | 1 | 0.11864487 | 0.45567948 | 0.109 | 0.052 | 1 | 0.21374303 | 0.22321313 |
| Brd3os | 0.41079295 | -0.6554705 | 0.052 | 0.078 | 1 | 0.11865483 | 0.19264508 | 0.063 | 0.021 | 1 | 0.41079295 | 0.22323069 |
| 0610009L18Rik | 0.98129896 | -0.2148979 | 0.083 | 0.083 | 1 | 0.11865483 | 0.45567948 | 0.063 | 0.021 | 1 | 0.98129896 | 0.22323069 |
| Gm35867 | 0.11874875 | -0.4519371 | 0.01 | 0.047 | 1 | 0.43587652 | -0.7248928 | 0.034 | 0.052 | 1 | 0.43587652 | 0.22339624 |
| Myo5c | 0.11876155 | 0.11440974 | 0.01 | 0.047 | 1 | 0.20472052 | -1.9537115 | 0.059 | 0.094 | 1 | 0.20472052 | 0.2234188 |
| Cntd1 | 0.11876932 | -0.6929452 | 0.01 | 0.047 | 1 | 0.24635272 | -1.5322477 | 0.013 | 0.031 | 1 | 0.24635272 | 0.22343249 |
| D530017H19Rik | 0.47942097 | -0.3144336 | 0.042 | 0.062 | 1 | 0.11879066 | -1.3098553 | 0.034 | 0.073 | 1 | 0.47942097 | 0.22347009 |
| Bcl9 | 0.11884338 | 0.48240349 | 0.25 | 0.166 | 1 | 0.16339765 | -0.2984505 | 0.336 | 0.24 | 1 | 0.16339765 | 0.22356302 |
| Rabl3 | 0.64911394 | 0.42253204 | 0.115 | 0.098 | 1 | 0.11886613 | 1.09224318 | 0.092 | 0.042 | 1 | 0.64911394 | 0.22360311 |
| Vrk3 | 0.26390049 | 0.14499806 | 0.385 | 0.301 | 1 | 0.11920614 | 0.29005758 | 0.303 | 0.208 | 1 | 0.26390049 | 0.22420218 |
| Pcdhgc3 | 0.50416281 | -0.4843586 | 0.146 | 0.171 | 1 | 0.11950886 | 0.64434105 | 0.218 | 0.146 | 1 | 0.50416281 | 0.22473534 |
| Cish | 0.12236045 | -1.3144336 | 0.021 | 0.062 | 1 | 0.11953838 | -2.0468209 | 0.008 | 0.031 | 1 | 0.12236045 | 0.22478734 |
| Aldh1a2 | 0.15729781 | -2.1624305 | 0 | 0.021 | 1 | 0.11953838 | -2.3098553 | 0.008 | 0.031 | 1 | 0.15729781 | 0.22478734 |
| Gm16174 | 0.18258775 | 0.74446013 | 0.042 | 0.016 | 1 | 0.11953838 | -2.3098553 | 0.008 | 0.031 | 1 | 0.18258775 | 0.22478734 |
| Zfp786 | 0.22187575 | 1.00749454 | 0.021 | 0.005 | 1 | 0.11953838 | -2.0468209 | 0.008 | 0.031 | 1 | 0.22187575 | 0.22478734 |
| G530011O06Rik | 0.24010913 | -0.8405024 | 0.042 | 0.078 | 1 | 0.11953838 | -2.3098553 | 0.008 | 0.031 | 1 | 0.24010913 | 0.22478734 |
| Oaz3 | 0.32025586 | -0.577468 | 0 | 0.01 | 1 | 0.11953838 | -2.5322477 | 0.008 | 0.031 | 1 | 0.32025586 | 0.22478734 |
| Plscr2 | 0.38722939 | -1.6554705 | 0.052 | 0.078 | 1 | 0.11953838 | -2.0468209 | 0.008 | 0.031 | 1 | 0.38722939 | 0.22478734 |
| Gm15545 | 0.46989842 | -0.9925055 | 0.021 | 0.036 | 1 | 0.11953838 | -2.3098553 | 0.008 | 0.031 | 1 | 0.46989842 | 0.22478734 |
| Gm6034 | 0.48064796 | 0.68556644 | 0.021 | 0.01 | 1 | 0.11953838 | -2.3098553 | 0.008 | 0.031 | 1 | 0.48064796 | 0.22478734 |
| Rpl3l | 0.52528691 | -0.7998604 | 0.01 | 0.021 | 1 | 0.11953838 | -2.3098553 | 0.008 | 0.031 | 1 | 0.52528691 | 0.22478734 |
| 1700041G16Rik | 0.61737927 | 1.00749454 | 0.01 | 0.005 | 1 | 0.11953838 | -2.0468209 | 0.008 | 0.031 | 1 | 0.61737927 | 0.22478734 |
| Asphd2 | 0.83676741 | -0.2555399 | 0.042 | 0.047 | 1 | 0.11953838 | -2.0468209 | 0.008 | 0.031 | 1 | 0.83676741 | 0.22478734 |
| Gm47350 | 0.32025586 | -1.3144336 | 0 | 0.01 | 1 | 0.11954377 | -2.1172102 | 0.008 | 0.031 | 1 | 0.32025586 | 0.22479682 |
| Blk | 0.73685932 | 2.27052894 | 0.01 | 0.016 | 1 | 0.11954377 | -3.4797803 | 0.008 | 0.031 | 1 | 0.73685932 | 0.22479682 |
| Crybb3 | 0.79666623 | 0.42253204 | 0.031 | 0.026 | 1 | 0.11954377 | -2.3098553 | 0.008 | 0.031 | 1 | 0.79666623 | 0.22479682 |
| Tom1l1 | 0.19310651 | 0.29700115 | 0.094 | 0.052 | 1 | 0.11961301 | 0.53814164 | 0.122 | 0.062 | 1 | 0.19310651 | 0.22491874 |
| Gstt1 | 0.52367115 | 0.27052894 | 0.135 | 0.109 | 1 | 0.1196488 | 0.47150445 | 0.189 | 0.115 | 1 | 0.52367115 | 0.22498177 |
| 0610043K17Rik | 0.17364247 | -0.223831 | 0.323 | 0.43 | 1 | 0.11969656 | -0.889952 | 0.269 | 0.323 | 1 | 0.17364247 | 0.22506585 |
| Zfp950 | 0.29269455 | 0.25830264 | 0.51 | 0.44 | 1 | 0.11973558 | -0.3234018 | 0.517 | 0.344 | 1 | 0.29269455 | 0.22513455 |
| B9d1 | 0.82175351 | -0.12375 | 0.188 | 0.197 | 1 | 0.11978207 | 0.51326698 | 0.172 | 0.104 | 1 | 0.82175351 | 0.2252164 |
| Zscan26 | 0.1198667 | 0.31392154 | 0.521 | 0.399 | 1 | 0.25911481 | -0.1254307 | 0.542 | 0.396 | 1 | 0.25911481 | 0.22536538 |
| Ypel1 | 0.3083669 | -0.7998604 | 0.073 | 0.109 | 1 | 0.1199364 | 0.40063812 | 0.122 | 0.062 | 1 | 0.3083669 | 0.22548806 |
| Gtf2h2 | 0.11994675 | 0.52206771 | 0.198 | 0.124 | 1 | 0.90254273 | -0.1723517 | 0.265 | 0.25 | 1 | 0.90254273 | 0.22550628 |
| Stx7 | 0.63677054 | -0.1575647 | 0.635 | 0.622 | 1 | 0.12006134 | 0.29804405 | 0.693 | 0.646 | 1 | 0.63677054 | 0.22570796 |
| H1f3 | 0.12013396 | 1.86310463 | 0.115 | 0.062 | 1 | 0.14167065 | -0.4029647 | 0.021 | 0.052 | 1 | 0.14167065 | 0.22583576 |
| Khdrbs3 | 0.94664576 | -0.2148979 | 0.167 | 0.166 | 1 | 0.12016765 | -0.4990114 | 0.462 | 0.573 | 1 | 0.94664576 | 0.22589504 |
| Csrnp2 | 0.83242622 | -0.6843832 | 0.104 | 0.109 | 1 | 0.12017444 | 0.35999614 | 0.122 | 0.062 | 1 | 0.83242622 | 0.22590699 |
| Plch2 | 0.12019803 | -1.6705774 | 0.031 | 0.078 | 1 | 0.32853124 | 0.27510724 | 0.122 | 0.083 | 1 | 0.32853124 | 0.22594849 |
| Hcn3 | 0.15837881 | 2.00749454 | 0.01 | 0 | 1 | 0.1202531 | -1.8948178 | 0.008 | 0.031 | 1 | 0.15837881 | 0.22604539 |
| Gm11630 | 0.29012577 | -0.407543 | 0.01 | 0.031 | 1 | 0.1202531 | -1.8948178 | 0.008 | 0.031 | 1 | 0.29012577 | 0.22604539 |
| Hdac9 | 0.43361119 | -0.382452 | 0.073 | 0.104 | 1 | 0.12034281 | -0.1422553 | 0.185 | 0.115 | 1 | 0.43361119 | 0.22620323 |
| Zfp280d | 0.12039058 | 0.14263061 | 0.844 | 0.756 | 1 | 0.92501611 | -0.2430682 | 0.929 | 0.906 | 1 | 0.92501611 | 0.22628727 |

| | | | | | | | | | | | | |
|---|---|---|---|---|---|---|---|---|---|---|---|---|
| Gm29994 | 0.12057613 | 0.54806292 | 0.073 | 0.031 | 1 | 0.23592811 | 1.39058446 | 0.034 | 0.01 | 1 | 0.23592811 | 0.22661366 |
| 1700093J21Rik | 0.70184804 | 0.37006462 | 0.062 | 0.052 | 1 | 0.12063447 | 0.91253716 | 0.092 | 0.042 | 1 | 0.70184804 | 0.22671626 |
| Aftph | 0.12068569 | 0.22567471 | 0.708 | 0.632 | 1 | 0.92271626 | -0.1853471 | 0.861 | 0.833 | 1 | 0.92271626 | 0.22680635 |
| Zfp64 | 0.1232727 | -0.2621125 | 0.365 | 0.44 | 1 | 0.12069559 | -0.134981 | 0.471 | 0.323 | 1 | 0.1232727 | 0.22682376 |
| Nudt1 | 0.12090375 | 0.67045955 | 0.156 | 0.093 | 1 | 0.23366437 | 0.52021974 | 0.105 | 0.062 | 1 | 0.23366437 | 0.22718978 |
| Kcne4 | 0.14873214 | -2.2779077 | 0.031 | 0.073 | 1 | 0.12096301 | -1.7952821 | 0.008 | 0.031 | 1 | 0.14873214 | 0.22729398 |
| 1110002J07Rik | 0.32025586 | -0.9925055 | 0 | 0.01 | 1 | 0.12096301 | -1.8948178 | 0.008 | 0.031 | 1 | 0.32025586 | 0.22729398 |
| Gper1 | 0.3793372 | 0.83756954 | 0.052 | 0.031 | 1 | 0.12096301 | -1.7952821 | 0.008 | 0.031 | 1 | 0.3793372 | 0.22729398 |
| Ccdc13 | 0.15837881 | 2.00749454 | 0.01 | 0 | 1 | 0.12096572 | -1.7248928 | 0.008 | 0.031 | 1 | 0.15837881 | 0.22729873 |
| Ppfia3 | 0.32025586 | -0.577468 | 0 | 0.01 | 1 | 0.12096572 | -1.7248928 | 0.008 | 0.031 | 1 | 0.32025586 | 0.22729873 |
| H1f1 | 0.39045207 | 0.42253204 | 0.052 | 0.031 | 1 | 0.12096572 | -1.7248928 | 0.008 | 0.031 | 1 | 0.39045207 | 0.22729873 |
| 2900092C05Rik | 0.52528691 | -0.9925055 | 0.01 | 0.021 | 1 | 0.12096572 | -1.7248928 | 0.008 | 0.031 | 1 | 0.52528691 | 0.22729873 |
| 1500002F19Rik | 1 | 0.42253204 | 0.01 | 0.01 | 1 | 0.12096572 | -1.7248928 | 0.008 | 0.031 | 1 | 1 | 0.22729873 |
| Myl6b | 1 | 0.42253204 | 0.01 | 0.01 | 1 | 0.12096572 | -1.7248928 | 0.008 | 0.031 | 1 | 1 | 0.22729873 |
| Arhgap32 | 0.24584495 | -0.2965671 | 0.552 | 0.622 | 1 | 0.12101049 | -0.4511493 | 0.811 | 0.875 | 1 | 0.24584495 | 0.22737744 |
| Nphp1 | 0.44436903 | 0.31561683 | 0.302 | 0.264 | 1 | 0.12103745 | 0.27510724 | 0.277 | 0.188 | 1 | 0.44436903 | 0.22742483 |
| Psat1 | 0.12120141 | 0.59245704 | 0.188 | 0.119 | 1 | 0.25159864 | 1.01207283 | 0.076 | 0.042 | 1 | 0.25159864 | 0.22771304 |
| Stac3 | 0.12145082 | 1.59245704 | 0.062 | 0.026 | 1 | 0.63798802 | -0.8244284 | 0.05 | 0.062 | 1 | 0.63798802 | 0.22815134 |
| Rslcan18 | 0.12145082 | 1.59245704 | 0.062 | 0.026 | 1 | 0.17515487 | 1.19264508 | 0.055 | 0.021 | 1 | 0.17515487 | 0.22815134 |
| Pdlim2 | 0.12145658 | -1.0469532 | 0.104 | 0.171 | 1 | 0.14760165 | -1.010295 | 0.193 | 0.25 | 1 | 0.14760165 | 0.22816146 |
| Ube2t | 0.12148866 | 0.91438513 | 0.115 | 0.062 | 1 | 0.27596631 | 0.91253716 | 0.046 | 0.021 | 1 | 0.27596631 | 0.22821782 |
| Gm42726 | 0.12159042 | 0.37006462 | 0.073 | 0.031 | 1 | 0.75684155 | -1.5025003 | 0.046 | 0.052 | 1 | 0.75684155 | 0.22839661 |
| Gm57047 | 0.38027888 | 1.00749454 | 0.031 | 0.016 | 1 | 0.12167825 | -1.6317834 | 0.008 | 0.031 | 1 | 0.38027888 | 0.2285509 |
| Sgo2a | 0.21541906 | 1.45495351 | 0.062 | 0.031 | 1 | 0.12168028 | -1.1172102 | 0.008 | 0.031 | 1 | 0.21541906 | 0.22855447 |
| Scml4 | 0.26542389 | 0.94860085 | 0.156 | 0.114 | 1 | 0.12172381 | 0.3068161 | 0.227 | 0.146 | 1 | 0.26542389 | 0.22863094 |
| Wdr48 | 0.12182286 | 0.37455201 | 0.469 | 0.373 | 1 | 0.84811973 | -0.1555271 | 0.5 | 0.469 | 1 | 0.84811973 | 0.22880491 |
| Usp13 | 0.95141749 | -0.407543 | 0.083 | 0.083 | 1 | 0.12183297 | 0.13071733 | 0.172 | 0.094 | 1 | 0.95141749 | 0.22882266 |
| 2310008N11Rik | 0.12185659 | 0.88196365 | 0.135 | 0.078 | 1 | 0.83427857 | -0.5114891 | 0.109 | 0.115 | 1 | 0.83427857 | 0.22886416 |
| Mapkap1 | 0.45667751 | 0.1540432 | 0.771 | 0.705 | 1 | 0.12190992 | -0.4552446 | 0.777 | 0.854 | 1 | 0.45667751 | 0.22895782 |
| A530017D24Rik | 0.12832438 | -0.8993961 | 0.021 | 0.062 | 1 | 0.12193029 | 0.63767732 | 0.092 | 0.042 | 1 | 0.12832438 | 0.22899358 |
| Gm1673 | 0.31433221 | 0.69715442 | 0.167 | 0.124 | 1 | 0.12193371 | 0.96316323 | 0.105 | 0.052 | 1 | 0.31433221 | 0.22899959 |
| Eid2b | 0.50183787 | -0.7087125 | 0.052 | 0.073 | 1 | 0.12225594 | 0.44503224 | 0.092 | 0.042 | 1 | 0.50183787 | 0.22956536 |
| Coa7 | 0.12236045 | -1.3144336 | 0.021 | 0.062 | 1 | 0.76711903 | -0.7793405 | 0.05 | 0.042 | 1 | 0.76711903 | 0.22974883 |
| Gm31645 | 0.12236045 | -1.4950058 | 0.021 | 0.062 | 1 | 0.13105689 | -0.8948178 | 0.021 | 0.052 | 1 | 0.13105689 | 0.22974883 |
| Gm57242 | 0.15729781 | -2.6929452 | 0 | 0.021 | 1 | 0.12239678 | -0.6317834 | 0.008 | 0.031 | 1 | 0.15729781 | 0.22981259 |
| Cit | 0.12256558 | 2.07162487 | 0.062 | 0.026 | 1 | 0.20314753 | 1.27510724 | 0.017 | 0 | 1 | 0.20314753 | 0.23010883 |
| Zbtb17 | 0.86951599 | 0.14076107 | 0.146 | 0.14 | 1 | 0.12266002 | 0.1876444 | 0.303 | 0.208 | 1 | 0.86951599 | 0.23027456 |
| Fdxr | 0.54775651 | -0.6247737 | 0.073 | 0.093 | 1 | 0.1226706 | -1.9313436 | 0.05 | 0.094 | 1 | 0.54775651 | 0.23029313 |
| Sergef | 0.12276101 | 0.63552576 | 0.115 | 0.062 | 1 | 0.1880543 | 0.13071733 | 0.113 | 0.062 | 1 | 0.1880543 | 0.23045176 |
| Swt1 | 0.12285021 | 0.142099 | 0.698 | 0.601 | 1 | 0.14323524 | -0.4481834 | 0.87 | 0.865 | 1 | 0.14323524 | 0.23060825 |
| Phkg2 | 0.17273179 | -0.6770036 | 0.24 | 0.301 | 1 | 0.1228636 | 0.2488757 | 0.34 | 0.24 | 1 | 0.17273179 | 0.23063174 |
| Nusap1 | 0.52704519 | 1.66460682 | 0.104 | 0.083 | 1 | 0.12288285 | -1.6634922 | 0.059 | 0.104 | 1 | 0.52704519 | 0.23066551 |
| Rc3h2 | 0.23897128 | 0.17244394 | 0.865 | 0.767 | 1 | 0.12289326 | -0.2576798 | 0.933 | 0.938 | 1 | 0.23897128 | 0.23068377 |
| Vstm2b | 0.15729403 | -1.577468 | 0 | 0.021 | 1 | 0.12311454 | -1.3098553 | 0.008 | 0.031 | 1 | 0.15729403 | 0.23107188 |
| Serpinb6c | 0.22187575 | -0.9925055 | 0 | 0.016 | 1 | 0.12311454 | -0.9879272 | 0.008 | 0.031 | 1 | 0.22187575 | 0.23107188 |
| Rims2 | 0.1231261 | 1.49292136 | 0.062 | 0.026 | 1 | 0.82724759 | -1.4703199 | 0.071 | 0.062 | 1 | 0.82724759 | 0.23109216 |
| Rnf141 | 0.75957905 | 0.35541784 | 0.104 | 0.093 | 1 | 0.12323558 | 0.47864063 | 0.134 | 0.073 | 1 | 0.75957905 | 0.23128415 |
| C430014B12Rik | 0.12330917 | -1.2148979 | 0.021 | 0.062 | 1 | 0.14700587 | 0.77760758 | 0.059 | 0.021 | 1 | 0.14700587 | 0.23141319 |
| Syde2 | 0.12330917 | -1.407543 | 0.021 | 0.062 | 1 | 0.24740671 | -1.3098553 | 0.076 | 0.115 | 1 | 0.24740671 | 0.23141319 |
| Bicdl1 | 0.12330917 | -1.2148979 | 0.021 | 0.062 | 1 | 0.84007282 | -1.4093909 | 0.038 | 0.042 | 1 | 0.84007282 | 0.23141319 |
| Eef2k | 0.12334914 | 0.6241659 | 0.188 | 0.119 | 1 | 0.57015249 | -0.7151117 | 0.214 | 0.229 | 1 | 0.57015249 | 0.23148328 |
| Rgcc | 0.89199861 | 0.17814941 | 0.708 | 0.699 | 1 | 0.12337078 | 0.37956679 | 0.723 | 0.677 | 1 | 0.89199861 | 0.23152121 |
| Zfp563 | 0.12353858 | 0.88196365 | 0.094 | 0.047 | 1 | 0.26964917 | -0.1034044 | 0.105 | 0.062 | 1 | 0.26964917 | 0.23181539 |
| Fbxw11 | 0.1235544 | -0.2735355 | 0.958 | 0.953 | 1 | 0.24488918 | -0.4253325 | 0.983 | 0.979 | 1 | 0.24488918 | 0.23184311 |
| 2810402E24Rik | 0.12359763 | 0.52944724 | 0.229 | 0.155 | 1 | 0.25826794 | -0.5322477 | 0.118 | 0.073 | 1 | 0.25826794 | 0.23191889 |
| Creld1 | 0.1236595 | -1.0920411 | 0.062 | 0.119 | 1 | 0.24923222 | 0.8917786 | 0.076 | 0.042 | 1 | 0.24923222 | 0.23202734 |
| Sharpin | 0.12368889 | 0.53542609 | 0.25 | 0.176 | 1 | 0.85597893 | -0.5322477 | 0.189 | 0.177 | 1 | 0.85597893 | 0.23207883 |
| Lppos | 0.12369747 | -0.2879613 | 0.135 | 0.212 | 1 | 0.6666593 | 0.13492958 | 0.155 | 0.135 | 1 | 0.6666593 | 0.23209388 |
| Uhrf1bp1l | 0.1238341 | -0.2892801 | 0.698 | 0.772 | 1 | 0.1583964 | -0.3515412 | 0.828 | 0.917 | 1 | 0.1583964 | 0.23233332 |
| Upf2 | 0.15140161 | 0.18226894 | 0.719 | 0.658 | 1 | 0.12384196 | -0.3279268 | 0.769 | 0.885 | 1 | 0.15140161 | 0.2323471 |
| Bfsp1 | 0.99527305 | -0.1079827 | 0.094 | 0.093 | 1 | 0.12393354 | -0.6508922 | 0.109 | 0.052 | 1 | 0.99527305 | 0.23250756 |
| Zfp276 | 0.95865296 | -0.4254649 | 0.094 | 0.093 | 1 | 0.12394481 | 0.91253716 | 0.168 | 0.104 | 1 | 0.95865296 | 0.23252731 |
| Wapl | 0.12403756 | 0.14499806 | 0.979 | 0.948 | 1 | 0.99749205 | -0.1389532 | 0.983 | 0.99 | 1 | 0.99749205 | 0.2326898 |
| mt-Nd6 | 0.12405834 | -0.5536212 | 0.385 | 0.472 | 1 | 0.46937753 | -0.4766744 | 0.412 | 0.302 | 1 | 0.46937753 | 0.2327262 |
| Gtf3c2 | 0.16617057 | 0.24015529 | 0.562 | 0.446 | 1 | 0.12407901 | -0.4299317 | 0.664 | 0.667 | 1 | 0.16617057 | 0.23276243 |
| Gm49539 | 0.529236 | 1.00749454 | 0.052 | 0.036 | 1 | 0.1241214 | -1.3098553 | 0.042 | 0.083 | 1 | 0.529236 | 0.23283668 |
| 4933433G15Rik | 0.12416619 | 1.42253204 | 0.062 | 0.026 | 1 | 0.40229719 | -1.0688472 | 0.042 | 0.062 | 1 | 0.40229719 | 0.23291513 |
| 1700110C19Rik | 0.12416619 | 1.59245704 | 0.062 | 0.026 | 1 | 0.31171996 | -0.9228321 | 0.063 | 0.094 | 1 | 0.31171996 | 0.23291513 |
| Polr1e | 0.12441324 | -1.1079827 | 0.021 | 0.062 | 1 | 0.52551869 | -0.5025003 | 0.055 | 0.073 | 1 | 0.52551869 | 0.23334782 |
| Tab3 | 0.94433625 | -0.577468 | 0.208 | 0.207 | 1 | 0.12444302 | 0.2584285 | 0.21 | 0.135 | 1 | 0.94433625 | 0.23339998 |
| A530013C23Rik | 0.66186725 | -0.577468 | 0.031 | 0.041 | 1 | 0.12454614 | -1.7692869 | 0.059 | 0.104 | 1 | 0.66186725 | 0.23358053 |
| Adam2 | 0.20443113 | -2.9621318 | 0.021 | 0.052 | 1 | 0.12457435 | -1.1172102 | 0.008 | 0.031 | 1 | 0.20443113 | 0.23362993 |
| Catsperd | 0.12461947 | 0.91438513 | 0.115 | 0.062 | 1 | 0.1608467 | -1.7692869 | 0.046 | 0.083 | 1 | 0.1608467 | 0.23370894 |
| Mat2b | 0.15901141 | 0.29872583 | 0.521 | 0.451 | 1 | 0.12467792 | -0.303429 | 0.496 | 0.635 | 1 | 0.15901141 | 0.23381125 |
| Zfp873 | 0.12475332 | 1.37006462 | 0.062 | 0.026 | 1 | 0.40076941 | -1.7952821 | 0.017 | 0.031 | 1 | 0.40076941 | 0.23394325 |
| Amdhd1 | 0.12475332 | 1.46692616 | 0.062 | 0.026 | 1 | 0.53433287 | 0.42711033 | 0.034 | 0.021 | 1 | 0.53433287 | 0.23394325 |
| Usp54 | 0.124813 | 0.10417056 | 0.406 | 0.285 | 1 | 0.55874014 | -0.1353575 | 0.496 | 0.458 | 1 | 0.55874014 | 0.23404772 |
| Tmtc4 | 0.75469571 | -0.130009 | 0.177 | 0.192 | 1 | 0.12485033 | 0.70806665 | 0.206 | 0.135 | 1 | 0.75469571 | 0.23411305 |
| Lypd6 | 0.12487099 | -1.240433 | 0.021 | 0.062 | 1 | 0.70927356 | -1.4253325 | 0.034 | 0.042 | 1 | 0.70927356 | 0.23414921 |
| Snapc3 | 0.12487905 | 0.39531057 | 0.406 | 0.301 | 1 | 0.93172104 | -0.6883669 | 0.517 | 0.417 | 1 | 0.93172104 | 0.23416332 |
| Gcat | 0.46547476 | 0.54806292 | 0.125 | 0.098 | 1 | 0.12491504 | 0.86006974 | 0.105 | 0.052 | 1 | 0.46547476 | 0.23422632 |

| | | | | | | | | | | | | |
|---|---|---|---|---|---|---|---|---|---|---|---|---|
| Spata6 | 0.12495498 | 0.1867191 | 0.562 | 0.466 | 1 | 0.95448694 | -0.2620267 | 0.651 | 0.583 | 1 | 0.95448694 | 0.23429621 |
| Gtpbp3 | 0.1249985 | -0.9925055 | 0.031 | 0.078 | 1 | 0.62773385 | 0.17557156 | 0.055 | 0.042 | 1 | 0.62773385 | 0.23437237 |
| Phf12 | 0.12500631 | 0.20214197 | 0.594 | 0.466 | 1 | 0.42504191 | -0.2460237 | 0.794 | 0.792 | 1 | 0.42504191 | 0.23438604 |
| H2-D1 | 0.12521521 | -0.2389996 | 0.646 | 0.798 | 1 | 0.3304817 | 0.11828023 | 0.798 | 0.875 | 1 | 0.3304817 | 0.23475158 |
| Nexmif | 0.21541668 | -1.1941393 | 0.021 | 0.052 | 1 | 0.12522065 | -2.3098553 | 0.021 | 0.052 | 1 | 0.21541668 | 0.2347611 |
| Psmd10 | 0.21652179 | -0.4900051 | 0.146 | 0.207 | 1 | 0.12552212 | 0.59703533 | 0.16 | 0.094 | 1 | 0.21652179 | 0.23528844 |
| Gm56531 | 0.20158871 | 1.42253204 | 0.031 | 0.01 | 1 | 0.12573859 | -2.0874628 | 0.021 | 0.052 | 1 | 0.20158871 | 0.23566698 |
| Rab3a | 0.12577815 | -1.130009 | 0.031 | 0.078 | 1 | 0.51654578 | -0.5728897 | 0.055 | 0.073 | 1 | 0.51654578 | 0.23573616 |
| Tdrd3 | 0.12580447 | -0.11115 | 0.458 | 0.337 | 1 | 0.52217363 | -0.5607734 | 0.58 | 0.552 | 1 | 0.52217363 | 0.23578217 |
| Kif15 | 0.12589692 | 1.92503238 | 0.062 | 0.026 | 1 | 0.66177135 | 0.59703533 | 0.029 | 0.021 | 1 | 0.66177135 | 0.23594381 |
| Uprt | 0.85437996 | 0.27052894 | 0.073 | 0.067 | 1 | 0.12602419 | -1.4253325 | 0.042 | 0.083 | 1 | 0.85437996 | 0.23616629 |
| Nelfb | 0.12602622 | 0.19625618 | 0.448 | 0.326 | 1 | 0.79876173 | -0.4749145 | 0.433 | 0.365 | 1 | 0.79876173 | 0.23616982 |
| Pitpnm1 | 0.62298353 | -0.3848229 | 0.062 | 0.078 | 1 | 0.1260673 | 0.92718393 | 0.105 | 0.052 | 1 | 0.62298353 | 0.23624164 |
| Mettl17 | 0.1261156 | 0.80104366 | 0.115 | 0.062 | 1 | 0.42490304 | 0.37820073 | 0.101 | 0.073 | 1 | 0.42490304 | 0.23632605 |
| Shroom2 | 0.12618841 | 0.69555053 | 0.271 | 0.197 | 1 | 0.17702612 | -0.6493417 | 0.294 | 0.385 | 1 | 0.17702612 | 0.23645331 |
| Top2a | 0.12623571 | 3.55181505 | 0.083 | 0.041 | 1 | 0.41181946 | -1.5508634 | 0.025 | 0.042 | 1 | 0.41181946 | 0.23653596 |
| Imp3 | 0.67300991 | -0.1891348 | 0.604 | 0.565 | 1 | 0.12626072 | 0.1835533 | 0.626 | 0.531 | 1 | 0.67300991 | 0.23657967 |
| Best2 | 0.62104899 | 0.42253204 | 0.01 | 0.005 | 1 | 0.12626311 | -1.961932 | 0.021 | 0.052 | 1 | 0.62104899 | 0.23658384 |
| Asxl3 | 0.12633776 | -1.5160674 | 0.021 | 0.062 | 1 | 0.98188264 | 0.56461386 | 0.021 | 0.021 | 1 | 0.98188264 | 0.2367143 |
| Sting1 | 0.12635343 | -1.0799683 | 0.021 | 0.062 | 1 | 0.53433287 | 0.42711033 | 0.034 | 0.021 | 1 | 0.53433287 | 0.23674167 |
| Exoc8 | 0.83292802 | 0.54806292 | 0.052 | 0.047 | 1 | 0.12644672 | 0.27510724 | 0.134 | 0.073 | 1 | 0.83292802 | 0.23690467 |
| Dennd4b | 0.17609384 | -1.3946039 | 0.052 | 0.098 | 1 | 0.12665736 | 0.54812573 | 0.092 | 0.042 | 1 | 0.17609384 | 0.23727263 |
| Stil | 0.12666526 | 2.40604391 | 0.083 | 0.041 | 1 | 0.40076941 | -1.7952821 | 0.017 | 0.031 | 1 | 0.40076941 | 0.23728643 |
| Gm56555 | 0.12672524 | -0.6876509 | 0.125 | 0.197 | 1 | 0.91565461 | -0.1004019 | 0.109 | 0.104 | 1 | 0.91565461 | 0.23739119 |
| Usp30 | 0.18299744 | 0.42253204 | 0.115 | 0.067 | 1 | 0.12679129 | 0.24653809 | 0.176 | 0.104 | 1 | 0.18299744 | 0.23750654 |
| Fam227a | 0.89789172 | -0.2269707 | 0.115 | 0.119 | 1 | 0.12685419 | -1.6577786 | 0.105 | 0.156 | 1 | 0.89789172 | 0.23761639 |
| Nudt6 | 0.12685728 | 1.27052894 | 0.125 | 0.073 | 1 | 0.18899612 | 0.33400093 | 0.084 | 0.042 | 1 | 0.18899612 | 0.23762178 |
| Gm43305 | 0.42851218 | 1.05188866 | 0.292 | 0.259 | 1 | 0.12689994 | 0.32757466 | 0.319 | 0.219 | 1 | 0.42851218 | 0.23769628 |
| Gm56867 | 0.12701924 | 1.22988696 | 0.062 | 0.026 | 1 | 0.17858821 | 0.91253716 | 0.055 | 0.021 | 1 | 0.17858821 | 0.2379046 |
| D030028A08Rik | 0.59740513 | -0.8290067 | 0.052 | 0.067 | 1 | 0.12705961 | 1.27510724 | 0.076 | 0.031 | 1 | 0.59740513 | 0.23797508 |
| Aifm2 | 0.66223816 | -0.3335424 | 0.146 | 0.119 | 1 | 0.12706752 | 0.18199783 | 0.16 | 0.094 | 1 | 0.66223816 | 0.23798888 |
| Rccd1 | 0.12710133 | 0.30705482 | 0.177 | 0.109 | 1 | 0.72002092 | -0.2457249 | 0.113 | 0.125 | 1 | 0.72002092 | 0.23804792 |
| Gm57292 | 0.22187388 | -1.9925055 | 0 | 0.016 | 1 | 0.12730109 | -1.8948178 | 0.021 | 0.052 | 1 | 0.22187388 | 0.23839661 |
| Gm43768 | 0.74827573 | 0.59245704 | 0.021 | 0.016 | 1 | 0.12730109 | -1.8948178 | 0.021 | 0.052 | 1 | 0.74827573 | 0.23839661 |
| Mapk12 | 0.74827573 | 0.59245704 | 0.021 | 0.016 | 1 | 0.12730109 | -1.8948178 | 0.021 | 0.052 | 1 | 0.74827573 | 0.23839661 |
| Trmt13 | 0.29917558 | -0.1208296 | 0.323 | 0.249 | 1 | 0.12735946 | 0.16931257 | 0.21 | 0.135 | 1 | 0.29917558 | 0.23849848 |
| Pkn1 | 0.12738084 | 0.71798792 | 0.24 | 0.166 | 1 | 0.9195709 | 0.12508179 | 0.387 | 0.396 | 1 | 0.9195709 | 0.23853579 |
| Far1 | 0.12739107 | 0.20013961 | 0.844 | 0.777 | 1 | 0.46392958 | -0.3291917 | 0.903 | 0.865 | 1 | 0.46392958 | 0.23855366 |
| Rev1 | 0.1275163 | 0.21022843 | 0.688 | 0.575 | 1 | 0.99746095 | -0.2301281 | 0.84 | 0.76 | 1 | 0.99746095 | 0.23877219 |
| Gm45494 | 0.1275299 | 1.20013961 | 0.062 | 0.026 | 1 | 1 | -0.8244284 | 0.021 | 0.021 | 1 | 1 | 0.23879593 |
| Zfhx2 | 0.30773829 | -0.577468 | 0.062 | 0.098 | 1 | 0.12758386 | 0.44913664 | 0.122 | 0.062 | 1 | 0.30773829 | 0.23889007 |
| Plekho2 | 0.80735328 | 0.1118312 | 0.302 | 0.295 | 1 | 0.12772658 | 0.28918242 | 0.324 | 0.229 | 1 | 0.80735328 | 0.23913908 |
| Scgn | 0.52744059 | -0.577468 | 0.01 | 0.021 | 1 | 0.1278557 | -1.6883669 | 0.021 | 0.052 | 1 | 0.52744059 | 0.23936432 |
| Hmga2 | 0.19846393 | -2.0628948 | 0.031 | 0.067 | 1 | 0.12788172 | 2.39058446 | 0.076 | 0.031 | 1 | 0.19846393 | 0.23940971 |
| Jak2 | 0.12794702 | -0.3740133 | 0.688 | 0.756 | 1 | 0.32508253 | -0.215453 | 0.714 | 0.854 | 1 | 0.32508253 | 0.23952359 |
| Rlbp1 | 0.49662775 | -0.6205367 | 0.094 | 0.119 | 1 | 0.12815003 | 1.14957636 | 0.076 | 0.031 | 1 | 0.49662775 | 0.23987764 |
| Inpp5e | 0.12815821 | 0.17741954 | 0.156 | 0.093 | 1 | 0.76859065 | -0.3802446 | 0.143 | 0.125 | 1 | 0.76859065 | 0.2398919 |
| Tbl2 | 0.34600385 | 0.1533454 | 0.25 | 0.197 | 1 | 0.12837138 | 0.39641353 | 0.223 | 0.146 | 1 | 0.34600385 | 0.24026356 |
| Ttc30b | 0.12842382 | 0.56003556 | 0.094 | 0.047 | 1 | 0.30087117 | 0.8917786 | 0.071 | 0.042 | 1 | 0.30087117 | 0.24035496 |
| Sf3b3 | 0.12843927 | -0.2779077 | 0.698 | 0.762 | 1 | 0.46250699 | -0.3098553 | 0.79 | 0.771 | 1 | 0.46250699 | 0.24038189 |
| Smc2 | 0.12851386 | 1.55249063 | 0.219 | 0.15 | 1 | 0.54413419 | -0.4797803 | 0.113 | 0.135 | 1 | 0.54413419 | 0.24051191 |
| Dcaf17 | 0.19297239 | 0.58368683 | 0.24 | 0.181 | 1 | 0.12852641 | 0.13760371 | 0.206 | 0.125 | 1 | 0.19297239 | 0.24053378 |
| Zmym2 | 0.74264638 | -0.2537032 | 0.688 | 0.632 | 1 | 0.12855899 | -0.3544454 | 0.845 | 0.927 | 1 | 0.74264638 | 0.24059056 |
| Phf6 | 0.19277272 | 0.39314823 | 0.375 | 0.301 | 1 | 0.12856038 | -0.4892645 | 0.462 | 0.521 | 1 | 0.19277272 | 0.240593 |
| Asf1b | 0.29234656 | 1.81484946 | 0.062 | 0.036 | 1 | 0.12890601 | -1.961932 | 0.021 | 0.052 | 1 | 0.29234656 | 0.24119527 |
| Ptcra | 0.81708725 | -0.577468 | 0.031 | 0.036 | 1 | 0.12891202 | -1.4797803 | 0.021 | 0.052 | 1 | 0.81708725 | 0.24120573 |
| Klhl24 | 0.29617625 | -0.1989563 | 0.594 | 0.658 | 1 | 0.12898651 | -0.1275055 | 0.622 | 0.812 | 1 | 0.29617625 | 0.24133549 |
| Cdyl2 | 0.12912831 | 0.73447604 | 0.167 | 0.104 | 1 | 0.18966091 | -0.3267291 | 0.202 | 0.125 | 1 | 0.18966091 | 0.2415825 |
| Mmd | 0.12931341 | -0.7998604 | 0.021 | 0.062 | 1 | 0.43040579 | -1.1034044 | 0.034 | 0.052 | 1 | 0.43040579 | 0.24190486 |
| Steep1 | 0.3611903 | 0.11267677 | 0.448 | 0.373 | 1 | 0.12948959 | -0.1442692 | 0.424 | 0.302 | 1 | 0.3611903 | 0.24221162 |
| Med20 | 0.12959096 | 0.53105649 | 0.344 | 0.254 | 1 | 0.4392822 | -0.3923174 | 0.408 | 0.302 | 1 | 0.4392822 | 0.24238811 |
| Vgll3 | 0.22187575 | -1.577468 | 0 | 0.016 | 1 | 0.12974993 | -2.4797803 | 0.059 | 0.104 | 1 | 0.22187575 | 0.24266482 |
| B4galt1 | 0.26665326 | -0.4435736 | 0.708 | 0.746 | 1 | 0.12977122 | -0.2973643 | 0.676 | 0.812 | 1 | 0.26665326 | 0.24270187 |
| Irak3 | 0.12981456 | -0.7171927 | 0.354 | 0.43 | 1 | 0.84026148 | -0.6787226 | 0.261 | 0.24 | 1 | 0.84026148 | 0.2427773 |
| Ppm1a | 0.1301328 | 0.24342371 | 0.792 | 0.865 | 1 | 0.60306113 | -0.2129937 | 0.912 | 0.896 | 1 | 0.60306113 | 0.24333105 |
| Slc25a28 | 0.32643252 | 0.22508597 | 0.365 | 0.311 | 1 | 0.1301629 | -0.1604776 | 0.361 | 0.24 | 1 | 0.32643252 | 0.24338343 |
| Tmem243 | 0.18434951 | 0.17004755 | 0.417 | 0.337 | 1 | 0.13026465 | -0.5420287 | 0.521 | 0.552 | 1 | 0.18434951 | 0.24356042 |
| 4930539J05Rik | 0.13035064 | 1.00749454 | 0.062 | 0.026 | 1 | 0.13847693 | 0.61614416 | 0.076 | 0.031 | 1 | 0.13847693 | 0.24370999 |
| Nuggc | 0.13035064 | 1.00749454 | 0.062 | 0.026 | 1 | 0.24304534 | -2.1578522 | 0.013 | 0.031 | 1 | 0.24304534 | 0.24370999 |
| 1700056N10Rik | 0.13035064 | 0.49292136 | 0.062 | 0.026 | 1 | 0.3103272 | 0.10518224 | 0.059 | 0.031 | 1 | 0.3103272 | 0.24370999 |
| Ankrd13d | 0.13035064 | 0.64492446 | 0.062 | 0.026 | 1 | 0.2967189 | -0.194378 | 0.088 | 0.052 | 1 | 0.2967189 | 0.24370999 |
| Socs5 | 0.13041387 | -0.5080031 | 0.458 | 0.528 | 1 | 0.47016151 | -0.1641136 | 0.588 | 0.479 | 1 | 0.47016151 | 0.24381996 |
| Dsc3 | 0.15837881 | 2.00749454 | 0.01 | 0 | 1 | 0.13041681 | -1.0688472 | 0.042 | 0.083 | 1 | 0.15837881 | 0.24382507 |
| Med23 | 0.33771648 | 0.28760246 | 0.26 | 0.207 | 1 | 0.13046004 | 0.20471791 | 0.332 | 0.229 | 1 | 0.33771648 | 0.24390025 |
| R3hcc1l | 0.80077743 | -0.1198848 | 0.406 | 0.425 | 1 | 0.13051632 | -0.6400039 | 0.521 | 0.583 | 1 | 0.80077743 | 0.24399812 |
| Gm15336 | 0.78117751 | -1.1079827 | 0.021 | 0.026 | 1 | 0.13052369 | -1.8948178 | 0.021 | 0.052 | 1 | 0.78117751 | 0.24401094 |
| Gm41505 | 0.51381802 | -1.3144336 | 0.031 | 0.047 | 1 | 0.13058883 | -1.3587649 | 0.059 | 0.104 | 1 | 0.51381802 | 0.24412422 |
| Rdh5 | 0.1308751 | 0.5049942 | 0.115 | 0.062 | 1 | 0.99561839 | -0.8948178 | 0.076 | 0.073 | 1 | 0.99561839 | 0.24462192 |
| Slc16a3 | 0.1308751 | 0.68556644 | 0.115 | 0.062 | 1 | 0.13871832 | 1.74903843 | 0.059 | 0.021 | 1 | 0.13871832 | 0.24462192 |
| Nphp4 | 0.13097941 | 1.00749454 | 0.062 | 0.026 | 1 | 0.3937346 | 0.69014474 | 0.025 | 0.01 | 1 | 0.3937346 | 0.24480322 |
| Emilin2 | 0.13131246 | -1.536826 | 0.052 | 0.104 | 1 | 0.97506785 | -0.6659991 | 0.063 | 0.062 | 1 | 0.97506785 | 0.24538195 |
| Ppp4r3b | 0.13131916 | 0.18091094 | 0.771 | 0.699 | 1 | 0.99540038 | -0.1493906 | 0.857 | 0.885 | 1 | 0.99540038 | 0.24539359 |
| Zfp119a | 0.13136876 | 0.38600616 | 0.115 | 0.062 | 1 | 0.98212878 | -0.355659 | 0.105 | 0.104 | 1 | 0.98212878 | 0.24547978 |
| Usf1 | 0.13139717 | 0.5049942 | 0.219 | 0.145 | 1 | 0.29060259 | 0.35852325 | 0.294 | 0.24 | 1 | 0.29060259 | 0.24552913 |
| Nebl | 0.22432715 | -0.3647461 | 0.844 | 0.798 | 1 | 0.13141902 | -0.2089807 | 0.794 | 0.615 | 1 | 0.22432715 | 0.24556709 |
| Ppp2r2b | 0.21783604 | -0.7810014 | 0.062 | 0.031 | 1 | 0.13149184 | -2.1227697 | 0.084 | 0.135 | 1 | 0.21783604 | 0.24569357 |
| Apol7d | 0.14697413 | 1.54806292 | 0.083 | 0.041 | 1 | 0.13160642 | -1.4618584 | 0.021 | 0.052 | 1 | 0.14697413 | 0.24589259 |
| Riiad1 | 0.5376551 | -0.9100433 | 0.083 | 0.104 | 1 | 0.13173855 | 0.2095189 | 0.147 | 0.083 | 1 | 0.5376551 | 0.24612206 |
| Ptp4a3 | 0.43687829 | 0.68556644 | 0.104 | 0.078 | 1 | 0.13199748 | 0.49749966 | 0.105 | 0.052 | 1 | 0.43687829 | 0.24657163 |

| | | | | | | | | | | | | |
|---|---|---|---|---|---|---|---|---|---|---|---|---|
| Gm56494 | 0.13211356 | 0.54806292 | 0.062 | 0.026 | 1 | 0.99168611 | -0.5993619 | 0.063 | 0.062 | 1 | 0.99168611 | 0.24677312 |
| Nr4a1 | 0.92092251 | -0.1568147 | 0.625 | 0.648 | 1 | 0.13213319 | -0.6267396 | 0.714 | 0.812 | 1 | 0.92092251 | 0.24680721 |
| Cep76 | 0.257851 | 0.46692616 | 0.188 | 0.135 | 1 | 0.13236836 | 0.24034182 | 0.265 | 0.177 | 1 | 0.257851 | 0.24721534 |
| Nubp1 | 0.13239687 | -0.6154358 | 0.219 | 0.29 | 1 | 0.57264212 | -0.1644248 | 0.256 | 0.219 | 1 | 0.57264212 | 0.2472648 |
| Zfp932 | 0.68770614 | -0.2950682 | 0.125 | 0.14 | 1 | 0.13246152 | 0.7944814 | 0.143 | 0.083 | 1 | 0.68770614 | 0.24737699 |
| Rnls | 0.13256645 | -0.8184761 | 0.062 | 0.119 | 1 | 0.46786812 | 0.55521516 | 0.088 | 0.062 | 1 | 0.46786812 | 0.24755904 |
| Qtrt1 | 0.13261813 | 0.15188445 | 0.344 | 0.238 | 1 | 0.29105607 | -0.1423985 | 0.395 | 0.292 | 1 | 0.29105607 | 0.2476487 |
| B130034C11Rik | 0.13266532 | 0.35541784 | 0.115 | 0.062 | 1 | 0.88072636 | -0.2318528 | 0.046 | 0.042 | 1 | 0.88072636 | 0.24773056 |
| Rad51c | 0.3583427 | 0.59245704 | 0.073 | 0.047 | 1 | 0.13267821 | -1.3098553 | 0.021 | 0.052 | 1 | 0.3583427 | 0.24775291 |
| Slc6a20b | 0.61737927 | 1.00749454 | 0.01 | 0.005 | 1 | 0.13267821 | -1.3098553 | 0.021 | 0.052 | 1 | 0.61737927 | 0.24775291 |
| Ptpn18 | 0.13270389 | 1.00749454 | 0.062 | 0.026 | 1 | 0.99424445 | -0.3098553 | 0.042 | 0.042 | 1 | 0.99424445 | 0.24779746 |
| Gm46440 | 0.13272539 | -0.9925055 | 0.042 | 0.093 | 1 | 0.90321158 | 0.33400093 | 0.055 | 0.052 | 1 | 0.90321158 | 0.24783476 |
| Psmb8 | 0.13272539 | -0.6554705 | 0.042 | 0.093 | 1 | 0.69916441 | -2.4618584 | 0.034 | 0.042 | 1 | 0.69916441 | 0.24783476 |
| Topbp1 | 0.16741633 | 0.76830687 | 0.302 | 0.228 | 1 | 0.13281217 | -0.1070384 | 0.332 | 0.24 | 1 | 0.16741633 | 0.24798527 |
| Gtpbp6 | 0.13283829 | 0.57865124 | 0.229 | 0.155 | 1 | 0.23750863 | 0.52021974 | 0.105 | 0.062 | 1 | 0.23750863 | 0.24803057 |
| Afap1l2 | 0.13283913 | -0.4564526 | 0.135 | 0.218 | 1 | 0.70818531 | -0.5979806 | 0.185 | 0.156 | 1 | 0.70818531 | 0.24803203 |
| Fmo1 | 0.73407659 | 0.27052894 | 0.01 | 0.016 | 1 | 0.13322545 | -1.1399303 | 0.021 | 0.052 | 1 | 0.73407659 | 0.24870188 |
| Fam110b | 0.13324196 | -0.7208321 | 0.427 | 0.528 | 1 | 0.93621433 | -0.6657493 | 0.424 | 0.385 | 1 | 0.93621433 | 0.2487305 |
| Cx3cl1 | 0.13332093 | -0.6753153 | 0.146 | 0.218 | 1 | 0.19943145 | -0.9228321 | 0.185 | 0.229 | 1 | 0.19943145 | 0.24886739 |
| Nrp1 | 0.13335382 | 0.16422004 | 0.271 | 0.187 | 1 | 0.93251752 | -0.6790891 | 0.319 | 0.271 | 1 | 0.93251752 | 0.2489244 |
| Slc4a4 | 0.87389428 | 0.10060394 | 0.583 | 0.575 | 1 | 0.13335857 | -0.2533883 | 0.639 | 0.74 | 1 | 0.87389428 | 0.24893263 |
| Map4k2 | 0.34310205 | 0.6730755 | 0.156 | 0.119 | 1 | 0.13352288 | 0.33400093 | 0.16 | 0.094 | 1 | 0.34310205 | 0.2492174 |
| Prrg1 | 0.16561317 | 0.81045769 | 0.229 | 0.166 | 1 | 0.13364295 | 0.46458504 | 0.172 | 0.104 | 1 | 0.16561317 | 0.24942546 |
| Pgpep1 | 0.13368369 | 0.19035859 | 0.188 | 0.119 | 1 | 0.15603734 | 0.30485458 | 0.155 | 0.094 | 1 | 0.15603734 | 0.24949605 |
| Dcdc2b | 0.13384309 | 0.35541784 | 0.062 | 0.026 | 1 | 0.90002388 | -1.194378 | 0.046 | 0.042 | 1 | 0.90002388 | 0.24977221 |
| B230303O12Rik | 0.13384309 | 0.81484946 | 0.062 | 0.026 | 1 | 0.27759424 | 0.91253716 | 0.046 | 0.021 | 1 | 0.27759424 | 0.24977221 |
| Ttc12 | 0.13384309 | 0.64492446 | 0.062 | 0.026 | 1 | 0.39386297 | -0.9175378 | 0.067 | 0.094 | 1 | 0.39386297 | 0.24977221 |
| 4833445I07Rik | 0.13384309 | 0.64492446 | 0.062 | 0.026 | 1 | 0.85710544 | -1.3098553 | 0.008 | 0.01 | 1 | 0.85710544 | 0.24977221 |
| Pp2d1 | 0.13384309 | 0.81484946 | 0.062 | 0.026 | 1 | 0.41807105 | -0.8244284 | 0.05 | 0.073 | 1 | 0.41807105 | 0.24977221 |
| Rpgrip1 | 0.57256131 | 0.53800925 | 0.188 | 0.166 | 1 | 0.13386201 | 0.42194863 | 0.202 | 0.125 | 1 | 0.57256131 | 0.24980497 |
| Etnk1 | 0.13391148 | -0.3130909 | 0.844 | 0.85 | 1 | 0.98025421 | -0.2590672 | 0.861 | 0.885 | 1 | 0.98025421 | 0.24989068 |
| Tet2 | 0.13392232 | -0.2037895 | 0.688 | 0.736 | 1 | 0.22378514 | -0.2203383 | 0.832 | 0.917 | 1 | 0.22378514 | 0.24990946 |
| Gm50012 | 0.57635618 | -0.3710171 | 0.094 | 0.114 | 1 | 0.13393825 | -1.2471195 | 0.176 | 0.229 | 1 | 0.57635618 | 0.24993705 |
| Srebf2 | 0.13408074 | 0.27702758 | 0.792 | 0.725 | 1 | 0.99898313 | -0.1077314 | 0.748 | 0.771 | 1 | 0.99898313 | 0.25018384 |
| Gm13274 | 0.29012577 | -0.407543 | 0.01 | 0.031 | 1 | 0.13433489 | -1.4797803 | 0.021 | 0.052 | 1 | 0.29012577 | 0.25062392 |
| Cdh4 | 0.74326422 | 2.49292136 | 0.021 | 0.016 | 1 | 0.13433489 | -2.1172102 | 0.021 | 0.052 | 1 | 0.74326422 | 0.25062392 |
| Fbln1 | 0.51778097 | -0.3910548 | 0.115 | 0.14 | 1 | 0.13437589 | -1.3098553 | 0.05 | 0.094 | 1 | 0.51778097 | 0.2506949 |
| Col24a1 | 0.1344162 | -2.67577 | 0.073 | 0.13 | 1 | 0.70524329 | -0.6239639 | 0.084 | 0.094 | 1 | 0.70524329 | 0.25076469 |
| Trub2 | 0.23595877 | 0.22508597 | 0.219 | 0.161 | 1 | 0.13446082 | 0.19603567 | 0.239 | 0.156 | 1 | 0.23595877 | 0.25084193 |
| Cd59b | 0.13462511 | 1.16795921 | 0.125 | 0.073 | 1 | 0.80856706 | -0.6998018 | 0.151 | 0.156 | 1 | 0.80856706 | 0.25112631 |
| Kansl2 | 0.98770313 | -0.1040138 | 0.406 | 0.389 | 1 | 0.13479844 | 0.16975424 | 0.525 | 0.396 | 1 | 0.98770313 | 0.25142626 |
| Heg1 | 0.37285973 | -0.1889027 | 0.208 | 0.254 | 1 | 0.13489851 | -1.5811573 | 0.118 | 0.177 | 1 | 0.37285973 | 0.25159942 |
| Eed | 0.1349037 | 0.40033629 | 0.479 | 0.373 | 1 | 0.13927516 | -0.3098553 | 0.521 | 0.635 | 1 | 0.13927516 | 0.25160839 |
| Sag | 0.66775813 | 0.20013961 | 0.125 | 0.109 | 1 | 0.13496504 | 0.76543286 | 0.155 | 0.094 | 1 | 0.66775813 | 0.25171452 |
| Ccrl2 | 0.13504298 | 0.42253204 | 0.062 | 0.026 | 1 | 0.3937346 | 0.69014474 | 0.025 | 0.01 | 1 | 0.3937346 | 0.25184935 |
| Smim31 | 0.30265323 | 0.6740708 | 0.26 | 0.207 | 1 | 0.13532315 | 0.24268576 | 0.147 | 0.083 | 1 | 0.30265323 | 0.25233395 |
| Pcid2 | 0.13535891 | 0.53542609 | 0.26 | 0.187 | 1 | 0.25118691 | 0.22065945 | 0.252 | 0.188 | 1 | 0.25118691 | 0.25239579 |
| Ficd | 0.13544923 | 0.42253204 | 0.115 | 0.062 | 1 | 0.18366232 | 0.16819203 | 0.151 | 0.094 | 1 | 0.18366232 | 0.25255196 |
| Lrrn4cl | 0.15060174 | 0.4088571 | 0.219 | 0.145 | 1 | 0.13556467 | -1.3838558 | 0.059 | 0.104 | 1 | 0.15060174 | 0.25275156 |
| Acap3 | 0.13562238 | 0.8439958 | 0.198 | 0.135 | 1 | 0.80647001 | -0.1578522 | 0.269 | 0.24 | 1 | 0.80647001 | 0.25285133 |
| Mrm2 | 0.13575269 | 0.32942263 | 0.094 | 0.047 | 1 | 0.41076656 | -0.1082214 | 0.08 | 0.052 | 1 | 0.41076656 | 0.25307659 |
| Mtcl1 | 0.13578653 | -1.1217885 | 0.052 | 0.104 | 1 | 0.94559773 | -0.2812861 | 0.147 | 0.146 | 1 | 0.94559773 | 0.25313508 |
| Mark1 | 0.50682337 | -0.6814189 | 0.344 | 0.368 | 1 | 0.13596535 | -0.7623675 | 0.349 | 0.229 | 1 | 0.50682337 | 0.25344412 |
| Cbr3 | 0.16525156 | -0.7580402 | 0.125 | 0.187 | 1 | 0.13605814 | 0.73745045 | 0.155 | 0.094 | 1 | 0.16525156 | 0.25360446 |
| Spats2 | 0.60509913 | -0.179591 | 0.458 | 0.404 | 1 | 0.13617366 | -0.2017955 | 0.479 | 0.333 | 1 | 0.60509913 | 0.25380406 |
| Crtc3 | 0.13639779 | -0.4420668 | 0.646 | 0.689 | 1 | 0.20395319 | -0.4949245 | 0.815 | 0.844 | 1 | 0.20395319 | 0.25419122 |
| Gpatch8 | 0.13649444 | 0.14457628 | 0.917 | 0.87 | 1 | 0.67040293 | -0.2897087 | 0.958 | 0.979 | 1 | 0.67040293 | 0.25435815 |
| Crebrf | 0.49075227 | -0.2121835 | 0.781 | 0.767 | 1 | 0.13676596 | -0.4535009 | 0.782 | 0.823 | 1 | 0.49075227 | 0.25482699 |
| Polr3h | 0.1951615 | -0.4128373 | 0.302 | 0.363 | 1 | 0.13693213 | 0.16407593 | 0.303 | 0.208 | 1 | 0.1951615 | 0.25511386 |
| Pla2g4b | 0.47912569 | -0.407543 | 0.021 | 0.036 | 1 | 0.13696237 | 0.49749966 | 0.076 | 0.031 | 1 | 0.47912569 | 0.25516605 |
| Epg5 | 0.13696849 | 0.28867629 | 0.354 | 0.264 | 1 | 0.61879139 | -0.6016214 | 0.336 | 0.281 | 1 | 0.61879139 | 0.25517661 |
| Ccdc138 | 0.13703514 | 0.50561378 | 0.406 | 0.332 | 1 | 0.6691686 | -1.0846423 | 0.45 | 0.406 | 1 | 0.6691686 | 0.25529164 |
| Lipe | 0.18120426 | -0.8993961 | 0.052 | 0.098 | 1 | 0.13705768 | 1.12310414 | 0.088 | 0.042 | 1 | 0.18120426 | 0.25533054 |
| Vwa8 | 0.70491183 | -0.130009 | 0.438 | 0.373 | 1 | 0.13717747 | -0.8084575 | 0.441 | 0.5 | 1 | 0.70491183 | 0.25553728 |
| Gramd4 | 0.25440662 | -0.1413688 | 0.135 | 0.197 | 1 | 0.1372415 | -0.1099177 | 0.231 | 0.146 | 1 | 0.25440662 | 0.25564777 |
| Dync2i2 | 0.13729338 | 0.68556644 | 0.177 | 0.114 | 1 | 0.21099146 | -0.4353861 | 0.122 | 0.073 | 1 | 0.21099146 | 0.25573729 |
| Zfp593 | 0.1373043 | 0.29289676 | 0.333 | 0.244 | 1 | 0.22370718 | 0.10518224 | 0.147 | 0.094 | 1 | 0.22370718 | 0.25575614 |
| 4930509H03Rik | 0.6834121 | 0.42253204 | 0.052 | 0.041 | 1 | 0.13732544 | 0.69014474 | 0.076 | 0.031 | 1 | 0.6834121 | 0.25579261 |
| Pard6a | 0.68484253 | 0.13302542 | 0.104 | 0.088 | 1 | 0.13732544 | 0.69014474 | 0.076 | 0.031 | 1 | 0.68484253 | 0.25579261 |
| Gm56844 | 0.13740312 | 0.64492446 | 0.062 | 0.026 | 1 | 0.54455416 | 0.17557156 | 0.046 | 0.031 | 1 | 0.54455416 | 0.25592663 |
| Aaas | 0.16010142 | 0.42253204 | 0.198 | 0.135 | 1 | 0.13768664 | -1.3878578 | 0.067 | 0.115 | 1 | 0.16010142 | 0.25641566 |
| Cacna1e | 0.85781033 | 0.40483003 | 0.062 | 0.067 | 1 | 0.13771449 | 0.48482583 | 0.185 | 0.115 | 1 | 0.85781033 | 0.2564637 |
| Smarca2 | 0.13785058 | 0.12087634 | 0.854 | 0.756 | 1 | 0.30667116 | -0.4542452 | 0.824 | 0.917 | 1 | 0.30667116 | 0.25669837 |
| Mr1 | 0.43642306 | -0.3144336 | 0.146 | 0.181 | 1 | 0.13788338 | 0.24957215 | 0.172 | 0.104 | 1 | 0.43642306 | 0.25675493 |
| Kdm4a | 0.23991022 | -0.5140426 | 0.229 | 0.29 | 1 | 0.13824986 | -0.2851932 | 0.391 | 0.271 | 1 | 0.23991022 | 0.25738669 |
| Gcn1 | 0.1383557 | 0.26078097 | 0.656 | 0.539 | 1 | 0.73690533 | -0.2995869 | 0.622 | 0.562 | 1 | 0.73690533 | 0.2575691 |
| Syde1 | 0.13837244 | 0.38300367 | 0.219 | 0.145 | 1 | 0.37584807 | 0.17127143 | 0.21 | 0.167 | 1 | 0.37584807 | 0.25759794 |
| Lpin3 | 0.84235519 | 0.20013961 | 0.052 | 0.047 | 1 | 0.13848166 | 1.17557156 | 0.088 | 0.042 | 1 | 0.84235519 | 0.25778615 |
| Lrrc45 | 0.13863934 | 0.60539609 | 0.208 | 0.14 | 1 | 0.43193612 | -0.4473588 | 0.261 | 0.198 | 1 | 0.43193612 | 0.25805781 |
| Nsun3 | 0.4895757 | 0.17741954 | 0.073 | 0.052 | 1 | 0.13864668 | -1.2273931 | 0.05 | 0.094 | 1 | 0.4895757 | 0.25807046 |
| Rbpjl | 0.13878792 | 1.24850264 | 0.083 | 0.041 | 1 | 0.95634731 | -0.6968784 | 0.084 | 0.083 | 1 | 0.95634731 | 0.25831375 |
| Cyp2j6 | 0.91395852 | -0.4807919 | 0.281 | 0.269 | 1 | 0.13900595 | 0.49749966 | 0.172 | 0.104 | 1 | 0.91395852 | 0.25868924 |
| G0s2 | 0.36686196 | -0.4779323 | 0.156 | 0.114 | 1 | 0.13924016 | 0.5111746 | 0.105 | 0.052 | 1 | 0.36686196 | 0.2590925 |
| Zcwpw1 | 0.13932393 | 1.20013961 | 0.104 | 0.057 | 1 | 0.75386896 | -0.7248928 | 0.055 | 0.062 | 1 | 0.75386896 | 0.25923671 |
| Rnf31 | 0.98438516 | 0.14951354 | 0.125 | 0.124 | 1 | 0.13942548 | -0.84037 | 0.118 | 0.177 | 1 | 0.98438516 | 0.2594115 |
| Mtch2 | 0.13956721 | 0.19109447 | 0.844 | 0.699 | 1 | 0.69753228 | -0.1369464 | 0.765 | 0.74 | 1 | 0.69753228 | 0.25965542 |
| Fos | 0.35473212 | 0.75589634 | 0.74 | 0.731 | 1 | 0.13959476 | -1.4100109 | 0.84 | 0.917 | 1 | 0.35473212 | 0.25970282 |
| Ric3 | 0.58224975 | -0.2555399 | 0.042 | 0.057 | 1 | 0.13960478 | 1.35310975 | 0.059 | 0.021 | 1 | 0.58224975 | 0.25972007 |

| | | | | | | | | | | | | |
|---|---|---|---|---|---|---|---|---|---|---|---|---|
| Dynlt1a | 0.64197016 | 0.59245704 | 0.042 | 0.031 | 1 | 0.13960478 | 1.35310975 | 0.059 | 0.021 | 1 | 0.64197016 | 0.25972007 |
| Fam83e | 0.13969328 | 1.15949763 | 0.083 | 0.041 | 1 | 0.66391569 | -1.1723517 | 0.042 | 0.052 | 1 | 0.66391569 | 0.25987235 |
| Malt1 | 0.15592285 | 1.00749454 | 0.167 | 0.109 | 1 | 0.1400134 | -0.9879272 | 0.126 | 0.188 | 1 | 0.15592285 | 0.26042305 |
| Parp12 | 0.14017979 | -0.265524 | 0.125 | 0.202 | 1 | 0.50358641 | -0.4353861 | 0.231 | 0.188 | 1 | 0.50358641 | 0.26070921 |
| Cd68 | 0.14019879 | 0.57139542 | 0.146 | 0.088 | 1 | 0.15201925 | 2.5103237 | 0.101 | 0.052 | 1 | 0.15201925 | 0.26074187 |
| Tmem184c | 0.31476701 | -0.5564063 | 0.188 | 0.233 | 1 | 0.14037697 | -0.1399303 | 0.319 | 0.208 | 1 | 0.31476701 | 0.26104824 |
| Gm17344 | 0.82258416 | -0.1624305 | 0.031 | 0.036 | 1 | 0.14049288 | 1.19264508 | 0.059 | 0.021 | 1 | 0.82258416 | 0.26124751 |
| Mlst8 | 0.82342138 | 0.32942263 | 0.042 | 0.036 | 1 | 0.14049288 | 1.19264508 | 0.059 | 0.021 | 1 | 0.82342138 | 0.26124751 |
| Scly | 0.14057131 | 0.66869262 | 0.198 | 0.135 | 1 | 0.38178643 | 0.21852371 | 0.189 | 0.146 | 1 | 0.38178643 | 0.26138233 |
| Cdr2l | 0.55747013 | -0.3550755 | 0.26 | 0.29 | 1 | 0.14057318 | 0.57767001 | 0.118 | 0.062 | 1 | 0.55747013 | 0.26138553 |
| Pheta1 | 0.50736458 | -0.5070786 | 0.052 | 0.073 | 1 | 0.14087265 | 1.27510724 | 0.113 | 0.062 | 1 | 0.50736458 | 0.26190019 |
| Zfp518a | 0.49558985 | 0.10479174 | 0.188 | 0.15 | 1 | 0.1409171 | 0.64434105 | 0.155 | 0.094 | 1 | 0.49558985 | 0.26197657 |
| 2900076A07Rik | 0.3761963 | 0.68556644 | 0.188 | 0.15 | 1 | 0.14093753 | -0.1399303 | 0.231 | 0.146 | 1 | 0.3761963 | 0.26201168 |
| Plaa | 0.14095421 | 0.30705482 | 0.583 | 0.472 | 1 | 0.18085083 | 0.21416447 | 0.664 | 0.635 | 1 | 0.18085083 | 0.26204032 |
| Hps3 | 0.14113062 | 1.00749454 | 0.135 | 0.083 | 1 | 0.71127334 | 0.60441486 | 0.134 | 0.125 | 1 | 0.71127334 | 0.26234338 |
| Zfp951 | 0.14114708 | 0.52206771 | 0.167 | 0.104 | 1 | 0.41084329 | -0.5549678 | 0.08 | 0.052 | 1 | 0.41084329 | 0.26237167 |
| Gm50071 | 1 | 0.42253204 | 0.01 | 0.01 | 1 | 0.1413819 | 1.10518224 | 0.059 | 0.021 | 1 | 1 | 0.26277495 |
| Serpinb8 | 0.14152931 | 1.85549144 | 0.052 | 0.021 | 1 | 0.37714057 | 0.21370669 | 0.055 | 0.031 | 1 | 0.37714057 | 0.26302808 |
| Kif20a | 0.14152931 | 2.59245704 | 0.052 | 0.021 | 1 | 0.57170879 | -1.8948178 | 0.013 | 0.021 | 1 | 0.57170879 | 0.26302808 |
| Ndufaf3 | 0.41497176 | -0.2054992 | 0.177 | 0.218 | 1 | 0.1417109 | 0.61614416 | 0.143 | 0.083 | 1 | 0.41497176 | 0.26333982 |
| Mmp23 | 0.14198699 | 0.35541784 | 0.115 | 0.062 | 1 | 0.54837489 | -0.1399303 | 0.034 | 0.021 | 1 | 0.54837489 | 0.26381367 |
| Pdlim3 | 0.22027047 | -1.4254649 | 0.021 | 0.052 | 1 | 0.14205582 | 2.27510724 | 0.042 | 0.01 | 1 | 0.22027047 | 0.26393178 |
| Parp3 | 0.14205626 | 0.21899864 | 0.323 | 0.238 | 1 | 0.81276202 | -0.4592329 | 0.303 | 0.271 | 1 | 0.81276202 | 0.26393253 |
| Bcorl1 | 0.14225967 | -0.8993961 | 0.125 | 0.187 | 1 | 0.20041988 | -0.7521657 | 0.303 | 0.354 | 1 | 0.20041988 | 0.26428152 |
| Gm47675 | 0.19012124 | -0.9185049 | 0.042 | 0.083 | 1 | 0.14243143 | 1.01207283 | 0.059 | 0.021 | 1 | 0.19012124 | 0.26457615 |
| Gm19605 | 0.20824604 | 1.17741954 | 0.062 | 0.031 | 1 | 0.14243143 | 1.01207283 | 0.059 | 0.021 | 1 | 0.20824604 | 0.26457615 |
| Shroom1 | 0.85959124 | 0.35541784 | 0.042 | 0.047 | 1 | 0.14243143 | 1.01207283 | 0.059 | 0.021 | 1 | 0.85959124 | 0.26457615 |
| Gm56489 | 0.86247167 | 0.20013961 | 0.073 | 0.067 | 1 | 0.14243143 | 1.01207283 | 0.059 | 0.021 | 1 | 0.86247167 | 0.26457615 |
| Rnf122 | 0.14248695 | -1.4779323 | 0.042 | 0.088 | 1 | 0.72611527 | 0.27510724 | 0.05 | 0.042 | 1 | 0.72611527 | 0.26467137 |
| Slc2a4 | 0.14248695 | -1.3710171 | 0.042 | 0.088 | 1 | 0.58664899 | -0.5025003 | 0.021 | 0.031 | 1 | 0.58664899 | 0.26467137 |
| Apbb1ip | 0.62104899 | 0.15949763 | 0.01 | 0.005 | 1 | 0.14259715 | 2.08246216 | 0.042 | 0.01 | 1 | 0.62104899 | 0.26486035 |
| Armc2 | 0.62852542 | 0.32942263 | 0.021 | 0.031 | 1 | 0.14259748 | 1.49749966 | 0.042 | 0.01 | 1 | 0.62852542 | 0.26486092 |
| Inpp5f | 0.44189842 | -0.4893371 | 0.312 | 0.342 | 1 | 0.14276719 | 0.26564691 | 0.378 | 0.281 | 1 | 0.44189842 | 0.2651519 |
| Lig1 | 0.29159647 | 0.82306997 | 0.177 | 0.135 | 1 | 0.14287924 | -0.1878647 | 0.118 | 0.062 | 1 | 0.29159647 | 0.265344 |
| Nipsnap1 | 0.14291286 | 0.72738662 | 0.125 | 0.073 | 1 | 0.2003234 | 0.49749966 | 0.134 | 0.083 | 1 | 0.2003234 | 0.26540163 |
| Gm47163 | 0.15729907 | -1.3144336 | 0 | 0.021 | 1 | 0.14313773 | 1.39058446 | 0.042 | 0.01 | 1 | 0.15729907 | 0.26578705 |
| Ankrd23 | 0.22104638 | -0.4779323 | 0.021 | 0.052 | 1 | 0.14313773 | 1.39058446 | 0.042 | 0.01 | 1 | 0.22104638 | 0.26578705 |
| Gm45237 | 0.33585318 | -0.6929452 | 0.031 | 0.057 | 1 | 0.14313773 | 1.39058446 | 0.042 | 0.01 | 1 | 0.33585318 | 0.26578705 |
| Gm47167 | 0.47294388 | -0.7294711 | 0.021 | 0.036 | 1 | 0.14313773 | 1.49749966 | 0.042 | 0.01 | 1 | 0.47294388 | 0.26578705 |
| Dph2 | 0.78651912 | 0.74446013 | 0.031 | 0.026 | 1 | 0.14313773 | 1.39058446 | 0.042 | 0.01 | 1 | 0.78651912 | 0.26578705 |
| Dhps | 0.19944874 | -0.6139938 | 0.312 | 0.363 | 1 | 0.14322984 | -0.3208264 | 0.416 | 0.281 | 1 | 0.19944874 | 0.26594489 |
| Cebpz | 0.30581452 | 0.29477649 | 0.542 | 0.492 | 1 | 0.14331746 | -0.2940303 | 0.576 | 0.656 | 1 | 0.30581452 | 0.26609503 |
| Def6 | 0.14337035 | 0.61156586 | 0.146 | 0.088 | 1 | 0.83493614 | -0.2103196 | 0.092 | 0.083 | 1 | 0.83493614 | 0.26618565 |
| Zfp280b | 0.83746085 | -0.130009 | 0.042 | 0.036 | 1 | 0.14338692 | 0.91253716 | 0.088 | 0.042 | 1 | 0.83746085 | 0.26621403 |
| Gpc4 | 0.1434552 | -1.2454862 | 0.26 | 0.316 | 1 | 0.99660209 | -0.8008416 | 0.433 | 0.365 | 1 | 0.99660209 | 0.26633101 |
| Spns2 | 0.14346805 | 0.63552576 | 0.104 | 0.057 | 1 | 0.71649639 | -0.5728897 | 0.034 | 0.042 | 1 | 0.71649639 | 0.26635302 |
| Kctd1 | 0.23068652 | 0.59686222 | 0.354 | 0.295 | 1 | 0.14355664 | -0.5941062 | 0.445 | 0.51 | 1 | 0.23068652 | 0.26650477 |
| Samm50 | 0.59922582 | -0.1208296 | 0.344 | 0.378 | 1 | 0.14359827 | 0.33650778 | 0.399 | 0.312 | 1 | 0.59922582 | 0.26657607 |
| Adam22 | 0.24655377 | 0.70793425 | 0.094 | 0.057 | 1 | 0.14367322 | 0.69014474 | 0.13 | 0.073 | 1 | 0.24655377 | 0.26670445 |
| 5830448L01Rik | 0.26351343 | 0.68556644 | 0.073 | 0.041 | 1 | 0.14367653 | 1.27510724 | 0.042 | 0.01 | 1 | 0.26351343 | 0.26671012 |
| Gm20404 | 0.38027888 | 1.00749454 | 0.031 | 0.016 | 1 | 0.14367653 | 1.27510724 | 0.042 | 0.01 | 1 | 0.38027888 | 0.26671012 |
| Catip | 0.72622123 | -0.2148979 | 0.052 | 0.062 | 1 | 0.14367653 | 1.77760758 | 0.042 | 0.01 | 1 | 0.72622123 | 0.26671012 |
| Mrm1 | 0.52066612 | -0.5070786 | 0.062 | 0.083 | 1 | 0.14372073 | -1.0468209 | 0.067 | 0.115 | 1 | 0.52066612 | 0.26678582 |
| Aig1 | 0.46087463 | -0.5783189 | 0.865 | 0.839 | 1 | 0.14391261 | -0.8997485 | 0.727 | 0.802 | 1 | 0.46087463 | 0.26711438 |
| Ccdc106 | 0.14422724 | 1.85549144 | 0.052 | 0.021 | 1 | 0.76784916 | 0.14957636 | 0.038 | 0.031 | 1 | 0.76784916 | 0.26765299 |
| Ttc21a | 0.14422724 | 1.68556644 | 0.052 | 0.021 | 1 | 0.39567521 | -1.7504279 | 0.042 | 0.062 | 1 | 0.39567521 | 0.26765299 |
| Adam4 | 0.14422724 | 1.68556644 | 0.052 | 0.021 | 1 | 0.87589833 | -0.5728897 | 0.013 | 0.01 | 1 | 0.87589833 | 0.26765299 |
| Gm48512 | 0.53920586 | 0.35541784 | 0.052 | 0.036 | 1 | 0.14422919 | 1.54812573 | 0.059 | 0.021 | 1 | 0.53920586 | 0.26765631 |
| Mtres1 | 0.14428635 | 0.55498233 | 0.281 | 0.207 | 1 | 0.3928803 | -0.5644281 | 0.202 | 0.146 | 1 | 0.3928803 | 0.26775414 |
| 9330111N05Rik | 0.2825047 | -1.3144336 | 0.01 | 0.031 | 1 | 0.14428792 | 1.14957636 | 0.042 | 0.01 | 1 | 0.2825047 | 0.26775684 |
| Slc19a3 | 0.58426772 | 0.78510212 | 0.031 | 0.021 | 1 | 0.14428792 | 1.14957636 | 0.042 | 0.01 | 1 | 0.58426772 | 0.26775684 |
| Rps6ka3 | 0.14436428 | -0.1575647 | 0.812 | 0.907 | 1 | 0.47290883 | -0.3780268 | 0.912 | 0.906 | 1 | 0.47290883 | 0.26788751 |
| Sephs2 | 0.21555806 | 0.46692616 | 0.198 | 0.14 | 1 | 0.14437057 | 1.08246216 | 0.126 | 0.073 | 1 | 0.21555806 | 0.26789828 |
| Il20ra | 0.15837881 | 2.00749454 | 0.01 | 0 | 1 | 0.14439146 | -3.1172102 | 0.004 | 0.021 | 1 | 0.15837881 | 0.26793403 |
| Msln | 0.15837881 | 2.59245704 | 0.01 | 0 | 1 | 0.14439146 | -2.8948178 | 0.004 | 0.021 | 1 | 0.15837881 | 0.26793403 |
| Ddx4 | 0.21867159 | 1.59245704 | 0.021 | 0.005 | 1 | 0.14439146 | -2.8948178 | 0.004 | 0.021 | 1 | 0.21867159 | 0.26793403 |
| Gm42912 | 0.22187575 | -1.577468 | 0 | 0.016 | 1 | 0.14439146 | -3.1172102 | 0.004 | 0.021 | 1 | 0.22187575 | 0.26793403 |
| 4930479D17Rik | 0.32025586 | -0.9925055 | 0 | 0.01 | 1 | 0.14439146 | -2.8948178 | 0.004 | 0.021 | 1 | 0.32025586 | 0.26793403 |
| Rasgrf2 | 0.32025586 | -0.9925055 | 0 | 0.01 | 1 | 0.14439146 | -3.3098553 | 0.004 | 0.021 | 1 | 0.32025586 | 0.26793403 |
| Stpg1 | 0.39524555 | -0.1624305 | 0.031 | 0.016 | 1 | 0.14439146 | -2.8948178 | 0.004 | 0.021 | 1 | 0.39524555 | 0.26793403 |
| Gm9195 | 0.5383068 | 0.27052894 | 0.01 | 0.021 | 1 | 0.14439146 | -3.3098553 | 0.004 | 0.021 | 1 | 0.5383068 | 0.26793403 |
| Srrm4 | 0.61737927 | 1.00749454 | 0.01 | 0.005 | 1 | 0.14439146 | -2.8948178 | 0.004 | 0.021 | 1 | 0.61737927 | 0.26793403 |
| 4930471E19Rik | 0.68932405 | -1.3144336 | 0.042 | 0.052 | 1 | 0.14439146 | -3.8123556 | 0.004 | 0.021 | 1 | 0.68932405 | 0.26793403 |
| Ucp3 | 0.8006267 | 0.20013961 | 0.021 | 0.026 | 1 | 0.14439146 | -3.3098553 | 0.004 | 0.021 | 1 | 0.8006267 | 0.26793403 |
| Zfp251 | 0.88847094 | 0.18806678 | 0.135 | 0.13 | 1 | 0.14445989 | -1.010295 | 0.139 | 0.198 | 1 | 0.88847094 | 0.26805112 |
| Hdx | 0.2084996 | -0.9467018 | 0.031 | 0.067 | 1 | 0.14450165 | -1.7573142 | 0.029 | 0.062 | 1 | 0.2084996 | 0.26812257 |
| Fbxl18 | 0.14476287 | -1.6770036 | 0.062 | 0.114 | 1 | 0.62618163 | -1.0161241 | 0.13 | 0.104 | 1 | 0.62618163 | 0.26856945 |
| Txndc16 | 0.14480776 | 0.60894516 | 0.271 | 0.192 | 1 | 0.79062194 | -0.7728272 | 0.244 | 0.208 | 1 | 0.79062194 | 0.26864623 |
| Nr6a1os | 0.14492772 | 1.12297175 | 0.083 | 0.041 | 1 | 0.5375503 | -0.5993619 | 0.046 | 0.062 | 1 | 0.5375503 | 0.26885139 |
| Krt1 | 0.24843624 | 1.22988696 | 0.052 | 0.026 | 1 | 0.14499914 | 0.20471791 | 0.076 | 0.031 | 1 | 0.24843624 | 0.26897354 |
| Plxna1 | 0.36986302 | 0.50225923 | 0.229 | 0.187 | 1 | 0.1451102 | 0.26000035 | 0.307 | 0.219 | 1 | 0.36986302 | 0.26916343 |
| Prkn | 0.14511716 | -0.5257919 | 0.531 | 0.627 | 1 | 0.98071977 | -0.7248928 | 0.588 | 0.583 | 1 | 0.98071977 | 0.26917534 |
| Map3k13 | 0.14516009 | 1.10703021 | 0.104 | 0.057 | 1 | 0.67037303 | 0.56461386 | 0.017 | 0.01 | 1 | 0.67037303 | 0.26924873 |
| Rhot1 | 0.14517333 | 0.33846777 | 0.552 | 0.451 | 1 | 0.79534202 | -0.2390049 | 0.672 | 0.656 | 1 | 0.79534202 | 0.26927136 |
| Mon1a | 0.51440407 | 0.28312898 | 0.188 | 0.155 | 1 | 0.14530932 | 0.2950068 | 0.231 | 0.156 | 1 | 0.51440407 | 0.26950384 |
| Marchf9 | 0.43759857 | -0.1079827 | 0.031 | 0.052 | 1 | 0.14541045 | -0.4473588 | 0.076 | 0.031 | 1 | 0.43759857 | 0.26967671 |

| | | | | | | | | | | | | |
|---|---|---|---|---|---|---|---|---|---|---|---|---|
| Hey2 | 0.15729907 | -1.3144336 | 0 | 0.021 | 1 | 0.14544666 | -2.8948178 | 0.004 | 0.021 | 1 | 0.15729907 | 0.2697386 |
| Atp2b2 | 0.15837881 | 2.00749454 | 0.01 | 0 | 1 | 0.14544666 | -2.3098553 | 0.004 | 0.021 | 1 | 0.15837881 | 0.2697386 |
| Gm45605 | 0.15837881 | 2.00749454 | 0.01 | 0 | 1 | 0.14544666 | -2.3098553 | 0.004 | 0.021 | 1 | 0.15837881 | 0.2697386 |
| Gm50368 | 0.15837881 | 2.59245704 | 0.01 | 0 | 1 | 0.14544666 | -2.6317834 | 0.004 | 0.021 | 1 | 0.15837881 | 0.2697386 |
| Xlr4b | 0.15837881 | 2.00749454 | 0.01 | 0 | 1 | 0.14544666 | -2.6317834 | 0.004 | 0.021 | 1 | 0.15837881 | 0.2697386 |
| Cdkn2a | 0.1969519 | 2.00749454 | 0.031 | 0.01 | 1 | 0.14544666 | -2.3098553 | 0.004 | 0.021 | 1 | 0.1969519 | 0.2697386 |
| Ankle1 | 0.20393009 | 1.00749454 | 0.031 | 0.01 | 1 | 0.14544666 | -2.6317834 | 0.004 | 0.021 | 1 | 0.20393009 | 0.2697386 |
| Slc39a12 | 0.21708019 | 2.32942263 | 0.021 | 0.005 | 1 | 0.14544666 | -3.1172102 | 0.004 | 0.021 | 1 | 0.21708019 | 0.2697386 |
| Dennd2d | 0.22187388 | -1.3144336 | 0 | 0.016 | 1 | 0.14544666 | -2.3098553 | 0.004 | 0.021 | 1 | 0.22187388 | 0.2697386 |
| Gm36329 | 0.22187575 | -0.9925055 | 0 | 0.016 | 1 | 0.14544666 | -2.3098553 | 0.004 | 0.021 | 1 | 0.22187575 | 0.2697386 |
| Trip13 | 0.24365282 | 2.12297175 | 0.052 | 0.026 | 1 | 0.14544666 | -2.3098553 | 0.004 | 0.021 | 1 | 0.24365282 | 0.2697386 |
| Gm57170 | 0.2825047 | -0.9925055 | 0.01 | 0.031 | 1 | 0.14544666 | -2.6317834 | 0.004 | 0.021 | 1 | 0.2825047 | 0.2697386 |
| Cd27 | 0.2825047 | -0.9925055 | 0.01 | 0.031 | 1 | 0.14544666 | -2.6317834 | 0.004 | 0.021 | 1 | 0.2825047 | 0.2697386 |
| Tcf15 | 0.32025586 | -0.577468 | 0 | 0.01 | 1 | 0.14544666 | -2.8948178 | 0.004 | 0.021 | 1 | 0.32025586 | 0.2697386 |
| Cyyr1 | 0.47150075 | 2.00749454 | 0.021 | 0.01 | 1 | 0.14544666 | -2.6317834 | 0.004 | 0.021 | 1 | 0.47150075 | 0.2697386 |
| Mfsd9 | 0.47606368 | 1.00749454 | 0.021 | 0.01 | 1 | 0.14544666 | -2.3098553 | 0.004 | 0.021 | 1 | 0.47606368 | 0.2697386 |
| Gm28731 | 0.47606502 | 1.00749454 | 0.021 | 0.01 | 1 | 0.14544666 | -2.6317834 | 0.004 | 0.021 | 1 | 0.47606502 | 0.2697386 |
| Gm17227 | 0.48064264 | 0.59245704 | 0.021 | 0.01 | 1 | 0.14544666 | -2.3098553 | 0.004 | 0.021 | 1 | 0.48064264 | 0.2697386 |
| 2610203C22Rik | 0.59199182 | 0.20013961 | 0.031 | 0.021 | 1 | 0.14544666 | -2.3098553 | 0.004 | 0.021 | 1 | 0.59199182 | 0.2697386 |
| Bco1 | 0.61737927 | 1.00749454 | 0.01 | 0.005 | 1 | 0.14544666 | -2.3098553 | 0.004 | 0.021 | 1 | 0.61737927 | 0.2697386 |
| Rps6kl1 | 0.61737927 | 1.00749454 | 0.01 | 0.005 | 1 | 0.14544666 | -2.6317834 | 0.004 | 0.021 | 1 | 0.61737927 | 0.2697386 |
| Tktl1 | 0.61737927 | 1.00749454 | 0.01 | 0.005 | 1 | 0.14544666 | -2.8948178 | 0.004 | 0.021 | 1 | 0.61737927 | 0.2697386 |
| Rsph6a | 0.62104899 | 0.42253204 | 0.01 | 0.005 | 1 | 0.14544666 | -2.3098553 | 0.004 | 0.021 | 1 | 0.62104899 | 0.2697386 |
| Tomt | 0.62104899 | 0.42253204 | 0.01 | 0.005 | 1 | 0.14544666 | -2.3098553 | 0.004 | 0.021 | 1 | 0.62104899 | 0.2697386 |
| Sntg1 | 0.72298091 | -0.577468 | 0.01 | 0.016 | 1 | 0.14544666 | -2.3098553 | 0.004 | 0.021 | 1 | 0.72298091 | 0.2697386 |
| Slc30a10 | 0.72298091 | -0.7998604 | 0.01 | 0.016 | 1 | 0.14544666 | -2.8948178 | 0.004 | 0.021 | 1 | 0.72298091 | 0.2697386 |
| Lypd6b | 0.73129736 | -0.577468 | 0.01 | 0.016 | 1 | 0.14544666 | -2.8948178 | 0.004 | 0.021 | 1 | 0.73129736 | 0.2697386 |
| Ptchd4 | 0.73129736 | -0.2148979 | 0.01 | 0.016 | 1 | 0.14544666 | -3.8948178 | 0.004 | 0.021 | 1 | 0.73129736 | 0.2697386 |
| Slc5a10 | 0.99275828 | -0.577468 | 0.021 | 0.021 | 1 | 0.14544666 | -2.6317834 | 0.004 | 0.021 | 1 | 0.99275828 | 0.2697386 |
| Plppr1 | 0.993208 | 1.42253204 | 0.01 | 0.01 | 1 | 0.14544666 | -2.3098553 | 0.004 | 0.021 | 1 | 0.993208 | 0.2697386 |
| Gm56633 | 0.99758609 | 0.27052894 | 0.021 | 0.021 | 1 | 0.14544666 | -2.3098553 | 0.004 | 0.021 | 1 | 0.99758609 | 0.2697386 |
| Pnma8b | 0.29908439 | -0.9925055 | 0.042 | 0.073 | 1 | 0.14564777 | 0.69014474 | 0.088 | 0.042 | 1 | 0.29908439 | 0.27008226 |
| Atad2 | 0.14585668 | 1.02199411 | 0.281 | 0.207 | 1 | 0.89035987 | -0.8073549 | 0.193 | 0.188 | 1 | 0.89035987 | 0.27043919 |
| 2210408I21Rik | 0.14603084 | 0.20893656 | 0.625 | 0.549 | 1 | 0.32946878 | -0.4468362 | 0.609 | 0.635 | 1 | 0.32946878 | 0.27073667 |
| Ssc4d | 0.1460806 | -1.130009 | 0.042 | 0.088 | 1 | 0.66013325 | 0.27510724 | 0.029 | 0.021 | 1 | 0.66013325 | 0.27082166 |
| Gm56874 | 0.14629689 | 1.88196365 | 0.052 | 0.021 | 1 | 0.79600331 | -0.9879272 | 0.017 | 0.021 | 1 | 0.79600331 | 0.27119101 |
| Ythdf3 | 0.14635007 | 0.2733886 | 0.833 | 0.762 | 1 | 0.57787352 | -0.2981418 | 0.937 | 0.938 | 1 | 0.57787352 | 0.27128181 |
| Gm4631 | 0.14646167 | 0.89201732 | 0.104 | 0.057 | 1 | 0.19098953 | 0.24268576 | 0.164 | 0.104 | 1 | 0.19098953 | 0.27147231 |
| Dclre1b | 0.14646167 | 0.89201732 | 0.104 | 0.057 | 1 | 0.26202048 | 0.86006974 | 0.076 | 0.042 | 1 | 0.26202048 | 0.27147231 |
| Csrp2 | 0.14646228 | -0.3890229 | 0.438 | 0.347 | 1 | 0.52955469 | -0.2584561 | 0.613 | 0.688 | 1 | 0.52955469 | 0.27147335 |
| Gm10484 | 0.15837881 | 2.00749454 | 0.01 | 0 | 1 | 0.1465102 | -1.8948178 | 0.004 | 0.021 | 1 | 0.15837881 | 0.27155516 |
| Adgrf3 | 0.15837881 | 2.00749454 | 0.01 | 0 | 1 | 0.1465102 | -1.8948178 | 0.004 | 0.021 | 1 | 0.15837881 | 0.27155516 |
| Myom2 | 0.15837881 | 2.00749454 | 0.01 | 0 | 1 | 0.1465102 | -1.8948178 | 0.004 | 0.021 | 1 | 0.15837881 | 0.27155516 |
| Ubxn10 | 0.20158871 | 1.42253204 | 0.031 | 0.01 | 1 | 0.1465102 | -1.8948178 | 0.004 | 0.021 | 1 | 0.20158871 | 0.27155516 |
| Spag6 | 0.20275473 | 1.32942263 | 0.031 | 0.01 | 1 | 0.1465102 | -1.8948178 | 0.004 | 0.021 | 1 | 0.20275473 | 0.27155516 |
| Rgsl1 | 0.22187388 | -1.577468 | 0 | 0.016 | 1 | 0.1465102 | -1.8948178 | 0.004 | 0.021 | 1 | 0.22187388 | 0.27155516 |
| Gm48796 | 0.22187388 | -1.3144336 | 0 | 0.016 | 1 | 0.1465102 | -1.8948178 | 0.004 | 0.021 | 1 | 0.22187388 | 0.27155516 |
| Sgk2 | 0.22187575 | -1.577468 | 0 | 0.016 | 1 | 0.1465102 | -1.8948178 | 0.004 | 0.021 | 1 | 0.22187575 | 0.27155516 |
| Gm15997 | 0.22187575 | -1.577468 | 0 | 0.016 | 1 | 0.1465102 | -1.8948178 | 0.004 | 0.021 | 1 | 0.22187575 | 0.27155516 |
| 8030423F21Rik | 0.32025586 | -0.577468 | 0 | 0.01 | 1 | 0.1465102 | -1.8948178 | 0.004 | 0.021 | 1 | 0.32025586 | 0.27155516 |
| Meiosin | 0.32025586 | -0.577468 | 0 | 0.01 | 1 | 0.1465102 | -2.0468209 | 0.004 | 0.021 | 1 | 0.32025586 | 0.27155516 |
| Dgkk | 0.32025586 | -1.3144336 | 0 | 0.01 | 1 | 0.1465102 | -1.8948178 | 0.004 | 0.021 | 1 | 0.32025586 | 0.27155516 |
| Gm16230 | 0.36601088 | -0.7294711 | 0.021 | 0.041 | 1 | 0.1465102 | -1.8948178 | 0.004 | 0.021 | 1 | 0.36601088 | 0.27155516 |
| Gm56868 | 0.38027888 | 1.00749454 | 0.031 | 0.016 | 1 | 0.1465102 | -1.8948178 | 0.004 | 0.021 | 1 | 0.38027888 | 0.27155516 |
| A730056A06Rik | 0.46989903 | -1.577468 | 0.021 | 0.036 | 1 | 0.1465102 | -1.8948178 | 0.004 | 0.021 | 1 | 0.46989903 | 0.27155516 |
| Gm36839 | 0.47150075 | 2.74446013 | 0.021 | 0.01 | 1 | 0.1465102 | -1.8948178 | 0.004 | 0.021 | 1 | 0.47150075 | 0.27155516 |
| Gm57180 | 0.52961488 | -0.3144336 | 0.01 | 0.021 | 1 | 0.1465102 | -2.0468209 | 0.004 | 0.021 | 1 | 0.52961488 | 0.27155516 |
| Gm40323 | 0.52961488 | -0.3144336 | 0.01 | 0.021 | 1 | 0.1465102 | -1.8948178 | 0.004 | 0.021 | 1 | 0.52961488 | 0.27155516 |
| Ddx43 | 0.608151 | -0.7294711 | 0.021 | 0.031 | 1 | 0.1465102 | -1.8948178 | 0.004 | 0.021 | 1 | 0.608151 | 0.27155516 |
| 9430024E24Rik | 0.61737927 | 1.00749454 | 0.01 | 0.005 | 1 | 0.1465102 | -1.8948178 | 0.004 | 0.021 | 1 | 0.61737927 | 0.27155516 |
| 1700063D05Rik | 0.61737927 | 1.00749454 | 0.01 | 0.005 | 1 | 0.1465102 | -1.8948178 | 0.004 | 0.021 | 1 | 0.61737927 | 0.27155516 |
| Gm16136 | 0.61737927 | 1.00749454 | 0.01 | 0.005 | 1 | 0.1465102 | -1.8948178 | 0.004 | 0.021 | 1 | 0.61737927 | 0.27155516 |
| Ptgir | 0.72298091 | -0.9925055 | 0.01 | 0.016 | 1 | 0.1465102 | -1.8948178 | 0.004 | 0.021 | 1 | 0.72298091 | 0.27155516 |
| Cartpt | 0.75328966 | 0.27052894 | 0.021 | 0.016 | 1 | 0.1465102 | -2.3098553 | 0.004 | 0.021 | 1 | 0.75328966 | 0.27155516 |
| E2f8 | 0.77216219 | 0.28760246 | 0.052 | 0.062 | 1 | 0.1465102 | -1.8948178 | 0.004 | 0.021 | 1 | 0.77216219 | 0.27155516 |
| Krt222 | 0.80274368 | 0.20013961 | 0.031 | 0.026 | 1 | 0.1465102 | -1.8948178 | 0.004 | 0.021 | 1 | 0.80274368 | 0.27155516 |
| Tmem108 | 0.83746085 | -0.2555399 | 0.042 | 0.036 | 1 | 0.1465102 | -1.8948178 | 0.004 | 0.021 | 1 | 0.83746085 | 0.27155516 |
| Trim71 | 0.99758607 | 0.20013961 | 0.021 | 0.021 | 1 | 0.1465102 | -2.0468209 | 0.004 | 0.021 | 1 | 0.99758607 | 0.27155516 |
| Gm57150 | 1 | 0.42253204 | 0.01 | 0.01 | 1 | 0.1465102 | -1.8948178 | 0.004 | 0.021 | 1 | 1 | 0.27155516 |
| Gm33318 | 1 | 0.42253204 | 0.01 | 0.01 | 1 | 0.1465102 | -1.8948178 | 0.004 | 0.021 | 1 | 1 | 0.27155516 |
| Zdhhc6 | 0.94423689 | 0.14835707 | 0.302 | 0.295 | 1 | 0.14651434 | 0.40774501 | 0.336 | 0.25 | 1 | 0.94423689 | 0.27156223 |
| 4930570G19Rik | 0.38689439 | -0.577468 | 0.01 | 0.026 | 1 | 0.14653446 | 0.93807225 | 0.059 | 0.021 | 1 | 0.38689439 | 0.27159658 |
| Sh3yl1 | 0.14654179 | 0.81484946 | 0.104 | 0.057 | 1 | 0.76776885 | -1.2394659 | 0.076 | 0.083 | 1 | 0.76776885 | 0.27160908 |
| Adhfe1 | 0.49274499 | -0.4689435 | 0.177 | 0.207 | 1 | 0.14669544 | 0.23071312 | 0.118 | 0.062 | 1 | 0.49274499 | 0.27187132 |
| Ptprk | 0.14671517 | -0.3424795 | 0.969 | 0.984 | 1 | 0.2246351 | -0.3918124 | 0.962 | 0.969 | 1 | 0.2246351 | 0.271905 |
| Dus1l | 0.14686774 | 0.52206771 | 0.146 | 0.088 | 1 | 0.28015598 | 0.27510724 | 0.088 | 0.052 | 1 | 0.28015598 | 0.27216534 |
| Gabpb2 | 0.14695835 | 0.34367662 | 0.49 | 0.389 | 1 | 0.16427089 | 0.28840406 | 0.597 | 0.552 | 1 | 0.16427089 | 0.27231994 |
| A730017L22Rik | 0.1469607 | 1.49292136 | 0.052 | 0.021 | 1 | 0.84173394 | 0.10518224 | 0.046 | 0.052 | 1 | 0.84173394 | 0.27232395 |
| Napepld | 0.1469607 | 1.49292136 | 0.052 | 0.021 | 1 | 1 | -0.5025003 | 0.021 | 0.021 | 1 | 1 | 0.27232395 |
| Il12rb2 | 0.1469607 | 1.68556644 | 0.052 | 0.021 | 1 | 0.31401394 | -1.4353861 | 0.038 | 0.062 | 1 | 0.31401394 | 0.27232395 |
| Gm43920 | 0.1469607 | 1.68556644 | 0.052 | 0.021 | 1 | 0.27935024 | 0.69014474 | 0.046 | 0.021 | 1 | 0.27935024 | 0.27232395 |
| Tha1 | 0.1469607 | 1.68556644 | 0.052 | 0.021 | 1 | 0.70493467 | -0.7952821 | 0.034 | 0.042 | 1 | 0.70493467 | 0.27232395 |
| Gm20508 | 0.1469607 | 1.49292136 | 0.052 | 0.021 | 1 | 0.93071182 | -0.1723517 | 0.029 | 0.031 | 1 | 0.93071182 | 0.27232395 |
| Cpne3 | 0.68300209 | 0.11913989 | 0.49 | 0.487 | 1 | 0.14696282 | 0.15311671 | 0.664 | 0.646 | 1 | 0.68300209 | 0.27232758 |

| | | | | | | | | | | | | |
|---|---|---|---|---|---|---|---|---|---|---|---|---|
| Pcgf5 | 0.14696752 | 0.25640783 | 0.625 | 0.58 | 1 | 0.56979205 | -0.433844 | 0.769 | 0.74 | 1 | 0.56979205 | 0.27233559 |
| Gm5914 | 0.14700359 | 0.80104366 | 0.104 | 0.057 | 1 | 0.58195575 | 0.33400093 | 0.08 | 0.062 | 1 | 0.58195575 | 0.27239712 |
| 2810457G06Rik | 0.6493646 | -0.2820121 | 0.073 | 0.088 | 1 | 0.14700587 | 0.77760758 | 0.059 | 0.021 | 1 | 0.6493646 | 0.27240102 |
| Uhrf1 | 0.14701926 | 1.21394541 | 0.083 | 0.041 | 1 | 0.71649639 | -0.5728897 | 0.034 | 0.042 | 1 | 0.71649639 | 0.27242385 |
| Gm39150 | 0.5158726 | -0.6987743 | 0.156 | 0.181 | 1 | 0.14702039 | -0.1463565 | 0.134 | 0.073 | 1 | 0.5158726 | 0.27242579 |
| Mmrn2 | 0.61182672 | -0.8669746 | 0.021 | 0.031 | 1 | 0.14705306 | -2.2167459 | 0.029 | 0.062 | 1 | 0.61182672 | 0.27248152 |
| Gm43727 | 0.52961488 | -0.3144336 | 0.01 | 0.021 | 1 | 0.14705335 | -1.6317834 | 0.029 | 0.062 | 1 | 0.52961488 | 0.27248202 |
| Gbp3 | 0.14713494 | -1.3848229 | 0.042 | 0.088 | 1 | 0.15287087 | -3.0468209 | 0.029 | 0.062 | 1 | 0.15287087 | 0.27262119 |
| Anapc15 | 0.30940974 | -0.1801325 | 0.135 | 0.187 | 1 | 0.14724291 | 0.11898804 | 0.134 | 0.073 | 1 | 0.30940974 | 0.27280534 |
| Ube2s | 0.14736222 | -0.1430651 | 0.344 | 0.244 | 1 | 0.89838221 | -0.6551838 | 0.416 | 0.365 | 1 | 0.89838221 | 0.27300881 |
| Zdhhc16 | 0.16733332 | 0.407102 | 0.271 | 0.197 | 1 | 0.14748892 | 0.46775232 | 0.239 | 0.167 | 1 | 0.16733332 | 0.27322487 |
| Xrcc4 | 0.14755853 | 0.15188445 | 0.427 | 0.332 | 1 | 0.2987998 | -0.1454684 | 0.328 | 0.25 | 1 | 0.2987998 | 0.27334353 |
| Gm15756 | 0.32025586 | -1.7998604 | 0 | 0.01 | 1 | 0.14757845 | -1.6317834 | 0.004 | 0.021 | 1 | 0.32025586 | 0.2733775 |
| Adam5 | 0.32025586 | -1.7998604 | 0 | 0.01 | 1 | 0.14757845 | -1.7248928 | 0.004 | 0.021 | 1 | 0.32025586 | 0.2733775 |
| Vpreb3 | 0.61371908 | 3.59245704 | 0.01 | 0.005 | 1 | 0.14757845 | -1.7248928 | 0.004 | 0.021 | 1 | 0.61371908 | 0.2733775 |
| Gm2093 | 0.61737927 | 1.00749454 | 0.01 | 0.005 | 1 | 0.14757845 | -1.7248928 | 0.004 | 0.021 | 1 | 0.61737927 | 0.2733775 |
| Pik3c2b | 0.73685932 | 1.42253204 | 0.01 | 0.016 | 1 | 0.14757845 | -1.7248928 | 0.004 | 0.021 | 1 | 0.73685932 | 0.2733775 |
| Phactr3 | 1 | 0.42253204 | 0.01 | 0.01 | 1 | 0.14757845 | -1.7248928 | 0.004 | 0.021 | 1 | 1 | 0.2733775 |
| Arhgap11a | 0.14768138 | 1.65957123 | 0.052 | 0.021 | 1 | 0.30492205 | 1.01207283 | 0.029 | 0.01 | 1 | 0.30492205 | 0.27355298 |
| Efna2 | 0.55414751 | -0.1127997 | 0.188 | 0.15 | 1 | 0.14779814 | -1.2828882 | 0.164 | 0.219 | 1 | 0.55414751 | 0.27375198 |
| Fam217b | 0.46544015 | -0.577468 | 0.042 | 0.062 | 1 | 0.14793 | 0.69014474 | 0.059 | 0.021 | 1 | 0.46544015 | 0.27397671 |
| Gm11464 | 0.65045504 | 0.27052894 | 0.042 | 0.031 | 1 | 0.14795145 | 0.19264508 | 0.059 | 0.021 | 1 | 0.65045504 | 0.27401328 |
| Npff | 0.14796579 | 0.85549144 | 0.083 | 0.041 | 1 | 0.56266331 | -0.4167705 | 0.046 | 0.031 | 1 | 0.56266331 | 0.2740377 |
| A330032P22Rik | 0.61737927 | 1.00749454 | 0.01 | 0.005 | 1 | 0.148195 | 0.91253716 | 0.042 | 0.01 | 1 | 0.61737927 | 0.27442825 |
| Zik1 | 0.69098575 | 0.27052894 | 0.052 | 0.041 | 1 | 0.14820491 | 0.91253716 | 0.042 | 0.01 | 1 | 0.69098575 | 0.27444513 |
| Gcdh | 0.97046361 | -0.1445086 | 0.167 | 0.166 | 1 | 0.14821165 | 0.46273424 | 0.143 | 0.083 | 1 | 0.97046361 | 0.27445661 |
| Nudt22 | 0.4096296 | 0.52206771 | 0.135 | 0.104 | 1 | 0.14826174 | 1.22065945 | 0.113 | 0.062 | 1 | 0.4096296 | 0.27454194 |
| Sptbn1 | 0.18172147 | -0.3910548 | 0.979 | 0.979 | 1 | 0.14847778 | -0.2734023 | 1 | 0.99 | 1 | 0.18172147 | 0.2749099 |
| Mpzl3 | 0.1484869 | 0.92503238 | 0.104 | 0.057 | 1 | 0.85170874 | -0.2318528 | 0.067 | 0.073 | 1 | 0.85170874 | 0.27492545 |
| Tmem201 | 0.58889921 | -0.2879613 | 0.094 | 0.114 | 1 | 0.14861401 | 0.53814164 | 0.13 | 0.073 | 1 | 0.58889921 | 0.2751419 |
| Gm42439 | 0.14864059 | -0.9467018 | 0.146 | 0.207 | 1 | 0.44113041 | -0.5644281 | 0.185 | 0.135 | 1 | 0.44113041 | 0.27518716 |
| Srrm3 | 0.15729403 | -1.577468 | 0 | 0.021 | 1 | 0.14865142 | -1.3098553 | 0.004 | 0.021 | 1 | 0.15729403 | 0.27520559 |
| Gm57409 | 0.1992517 | 1.74446013 | 0.031 | 0.01 | 1 | 0.14865142 | -1.3098553 | 0.004 | 0.021 | 1 | 0.1992517 | 0.27520559 |
| Wnt7b | 0.28126256 | -1.1624305 | 0.01 | 0.031 | 1 | 0.14865142 | -1.3098553 | 0.004 | 0.021 | 1 | 0.28126256 | 0.27520559 |
| Obscn | 0.32025586 | -0.577468 | 0 | 0.01 | 1 | 0.14865142 | -1.3098553 | 0.004 | 0.021 | 1 | 0.32025586 | 0.27520559 |
| Chit1 | 0.34200395 | -0.6554705 | 0.031 | 0.057 | 1 | 0.14865142 | 0.80562195 | 0.004 | 0.021 | 1 | 0.34200395 | 0.27520559 |
| Pcp4l1 | 0.38192865 | -1.3144336 | 0.01 | 0.026 | 1 | 0.14865142 | -0.8948178 | 0.004 | 0.021 | 1 | 0.38192865 | 0.27520559 |
| H2-DMb2 | 0.61371908 | 2.59245704 | 0.01 | 0.005 | 1 | 0.14865142 | -1.3098553 | 0.004 | 0.021 | 1 | 0.61371908 | 0.27520559 |
| Ch25h | 0.66353842 | -0.7998604 | 0.031 | 0.041 | 1 | 0.14865142 | 0.42711033 | 0.004 | 0.021 | 1 | 0.66353842 | 0.27520559 |
| Amph | 0.15729781 | -1.7998604 | 0 | 0.021 | 1 | 0.14865264 | -1.0874628 | 0.004 | 0.021 | 1 | 0.15729781 | 0.27520768 |
| 1700061H18Rik | 0.72574709 | -0.9925055 | 0.01 | 0.016 | 1 | 0.14865264 | -0.7248928 | 0.004 | 0.021 | 1 | 0.72574709 | 0.27520768 |
| Rasgef1a | 0.99758604 | -0.8669746 | 0.021 | 0.021 | 1 | 0.14865264 | -1.3098553 | 0.004 | 0.021 | 1 | 0.99758604 | 0.27520768 |
| Usp50 | 0.47421406 | -0.7252216 | 0.24 | 0.285 | 1 | 0.14885004 | -0.2366063 | 0.277 | 0.188 | 1 | 0.47421406 | 0.27554375 |
| Mvk | 0.14891131 | 1.14499806 | 0.146 | 0.093 | 1 | 0.2188555 | 0.11898804 | 0.122 | 0.073 | 1 | 0.2188555 | 0.27564805 |
| Plcg2 | 0.36434798 | -0.8405024 | 0.167 | 0.197 | 1 | 0.14896076 | -0.6391629 | 0.513 | 0.521 | 1 | 0.36434798 | 0.27573221 |
| Spaca1 | 0.43960598 | -0.5070786 | 0.062 | 0.088 | 1 | 0.14897757 | 0.74903843 | 0.088 | 0.042 | 1 | 0.43960598 | 0.27576083 |
| Rhou | 0.14901342 | -1.2295447 | 0.042 | 0.088 | 1 | 0.51302871 | -0.7826083 | 0.151 | 0.177 | 1 | 0.51302871 | 0.27582184 |
| Csf1 | 0.14903062 | 0.26333344 | 0.24 | 0.166 | 1 | 0.77749174 | -0.7415715 | 0.235 | 0.229 | 1 | 0.77749174 | 0.27585111 |
| Wars1 | 0.2402901 | 0.18152394 | 0.208 | 0.15 | 1 | 0.14917305 | 0.46066289 | 0.193 | 0.125 | 1 | 0.2402901 | 0.27609351 |
| Pmel | 0.17258946 | 0.88196365 | 0.073 | 0.036 | 1 | 0.14932246 | 0.69014474 | 0.042 | 0.01 | 1 | 0.17258946 | 0.27634772 |
| Serpina1d | 0.1493803 | 1.42253204 | 0.167 | 0.109 | 1 | 0.29520099 | 0.11098786 | 0.021 | 0.042 | 1 | 0.29520099 | 0.27644613 |
| Serinc4 | 0.14979643 | 1.27052894 | 0.052 | 0.021 | 1 | 0.57854956 | -0.8244284 | 0.021 | 0.031 | 1 | 0.57854956 | 0.2771539 |
| Gm43566 | 0.14979643 | 1.27052894 | 0.052 | 0.021 | 1 | 0.18429559 | 1.14957636 | 0.038 | 0.01 | 1 | 0.18429559 | 0.2771539 |
| Clstn3 | 0.14979643 | 1.27052894 | 0.052 | 0.021 | 1 | 0.2414379 | 0.42711033 | 0.105 | 0.062 | 1 | 0.2414379 | 0.2771539 |
| Tmem91 | 0.14979643 | 1.27052894 | 0.052 | 0.021 | 1 | 0.1685647 | 0.42711033 | 0.071 | 0.031 | 1 | 0.1685647 | 0.2771539 |
| Hsd17b14 | 0.14979643 | 1.27052894 | 0.052 | 0.021 | 1 | 0.50844268 | 2.93807225 | 0.021 | 0.01 | 1 | 0.50844268 | 0.2771539 |
| Gm34552 | 0.14979643 | 1.27052894 | 0.052 | 0.021 | 1 | 0.15394118 | 1.49749966 | 0.021 | 0 | 1 | 0.15394118 | 0.2771539 |
| Gm47920 | 0.14979643 | 1.27052894 | 0.052 | 0.021 | 1 | 0.39177935 | -1.1034044 | 0.05 | 0.073 | 1 | 0.39177935 | 0.2771539 |
| Pcdhga1 | 0.14979643 | 1.27052894 | 0.052 | 0.021 | 1 | 0.56267767 | -0.6724253 | 0.046 | 0.031 | 1 | 0.56267767 | 0.2771539 |
| Gm57182 | 0.3573566 | 0.71798792 | 0.073 | 0.047 | 1 | 0.14987922 | 0.14957636 | 0.042 | 0.01 | 1 | 0.3573566 | 0.27729466 |
| Lrrc19 | 0.47150075 | 2.00749454 | 0.021 | 0.01 | 1 | 0.14987922 | 0.56461386 | 0.042 | 0.01 | 1 | 0.47150075 | 0.27729466 |
| 9630013D21Rik | 0.82341254 | 0.22988696 | 0.042 | 0.036 | 1 | 0.14987922 | 0.56461386 | 0.042 | 0.01 | 1 | 0.82341254 | 0.27729466 |
| Ptgs2os | 0.52434216 | 0.14076107 | 0.052 | 0.073 | 1 | 0.14992316 | 0.27510724 | 0.042 | 0.01 | 1 | 0.52434216 | 0.27736936 |
| Stk4 | 0.53489954 | 0.37672835 | 0.344 | 0.301 | 1 | 0.14993332 | -0.2723806 | 0.378 | 0.479 | 1 | 0.53489954 | 0.27738664 |
| Tshz3 | 0.30541972 | -0.6340515 | 0.125 | 0.171 | 1 | 0.15032391 | -0.9762454 | 0.298 | 0.354 | 1 | 0.30541972 | 0.27805055 |
| 2010007H06Rik | 0.15038393 | -0.8225805 | 0.146 | 0.212 | 1 | 0.29392394 | -0.3838558 | 0.13 | 0.083 | 1 | 0.29392394 | 0.27815254 |
| Hspa12a | 0.15039809 | 0.37522632 | 0.146 | 0.088 | 1 | 0.65443361 | -0.4071525 | 0.101 | 0.083 | 1 | 0.65443361 | 0.27817659 |
| Acad12 | 0.15046861 | 1.22988696 | 0.052 | 0.021 | 1 | 0.67471359 | -0.5728897 | 0.017 | 0.01 | 1 | 0.67471359 | 0.27829642 |
| Rtn4ip1 | 0.15046861 | 1.22988696 | 0.052 | 0.021 | 1 | 0.2928226 | 0.27510724 | 0.046 | 0.021 | 1 | 0.2928226 | 0.27829642 |
| Paqr4 | 0.15046861 | 1.22988696 | 0.052 | 0.021 | 1 | 0.1822355 | 1.69014474 | 0.038 | 0.01 | 1 | 0.1822355 | 0.27829642 |
| Cadm2 | 0.15049382 | -2.6363617 | 0.01 | 0.041 | 1 | 0.41182766 | -0.1200307 | 0.092 | 0.062 | 1 | 0.41182766 | 0.27833925 |
| Naaa | 0.96846814 | -0.1708427 | 0.177 | 0.176 | 1 | 0.15052122 | 0.27510724 | 0.118 | 0.062 | 1 | 0.96846814 | 0.27838581 |
| Ethe1 | 0.86175967 | -0.5330738 | 0.104 | 0.109 | 1 | 0.15061899 | 0.39641353 | 0.218 | 0.146 | 1 | 0.86175967 | 0.27855191 |
| Bpifb3 | 0.15837881 | 2.00749454 | 0.01 | 0 | 1 | 0.15071937 | -1.4797803 | 0.029 | 0.062 | 1 | 0.15837881 | 0.27872241 |
| Sars2 | 0.41584221 | 1.00749454 | 0.062 | 0.041 | 1 | 0.15071937 | -1.6317834 | 0.029 | 0.062 | 1 | 0.41584221 | 0.27872241 |
| Egln3 | 0.47014725 | -0.4950058 | 0.042 | 0.062 | 1 | 0.15071937 | -1.6317834 | 0.029 | 0.062 | 1 | 0.47014725 | 0.27872241 |
| Rogdi | 0.87948399 | -0.130009 | 0.094 | 0.098 | 1 | 0.1508809 | 0.62874419 | 0.088 | 0.042 | 1 | 0.87948399 | 0.27899676 |
| Mxd4 | 0.15089464 | 0.29501861 | 0.729 | 0.658 | 1 | 0.81077372 | -0.1912108 | 0.853 | 0.854 | 1 | 0.81077372 | 0.27902009 |
| Zfp513 | 0.35913937 | -0.6205367 | 0.104 | 0.14 | 1 | 0.15098005 | -0.5025003 | 0.109 | 0.167 | 1 | 0.35913937 | 0.27916513 |
| Plekhg6 | 0.15117462 | 0.81484946 | 0.052 | 0.021 | 1 | 0.18429559 | 1.14957636 | 0.038 | 0.01 | 1 | 0.18429559 | 0.27949547 |
| Efcab11 | 0.15117462 | 0.35541784 | 0.052 | 0.021 | 1 | 0.18292349 | 1.69014474 | 0.038 | 0.01 | 1 | 0.18292349 | 0.27949547 |
| Ablim2 | 0.15119482 | -2.1624305 | 0.01 | 0.041 | 1 | 0.86312513 | -0.7248928 | 0.008 | 0.01 | 1 | 0.86312513 | 0.27952977 |
| Lrrc4 | 0.56809598 | -0.2228031 | 0.229 | 0.187 | 1 | 0.15125558 | -1.2223924 | 0.185 | 0.24 | 1 | 0.56809598 | 0.27963292 |
| Sbno1 | 0.15156671 | 0.11362289 | 0.948 | 0.839 | 1 | 0.18285554 | -0.3716143 | 0.958 | 0.958 | 1 | 0.18285554 | 0.28016094 |
| Gm36899 | 0.15172331 | 0.62047141 | 0.104 | 0.057 | 1 | 0.92228543 | -0.7248928 | 0.029 | 0.031 | 1 | 0.92228543 | 0.28042665 |
| AA467197 | 0.15189439 | -1.9925055 | 0.01 | 0.041 | 1 | 0.50587544 | -2.3098553 | 0.004 | 0.01 | 1 | 0.50587544 | 0.28071687 |

| | | | | | | | | | | | | |
|---|---|---|---|---|---|---|---|---|---|---|---|---|
| Otulinl | 0.15199371 | -1.2295447 | 0.062 | 0.114 | 1 | 0.45342825 | -1.0535155 | 0.113 | 0.135 | 1 | 0.45342825 | 0.28088533 |
| Ptpn22 | 0.15207986 | 2.65135073 | 0.073 | 0.036 | 1 | 0.8386124 | -1.0468209 | 0.038 | 0.042 | 1 | 0.8386124 | 0.28103143 |
| Nme6 | 0.152128 | 0.59245704 | 0.104 | 0.057 | 1 | 0.19035232 | -0.8073549 | 0.055 | 0.094 | 1 | 0.19035232 | 0.28111307 |
| Camk2g | 0.36886928 | -0.4537326 | 0.427 | 0.466 | 1 | 0.15224862 | -0.1130546 | 0.618 | 0.448 | 1 | 0.36886928 | 0.28131759 |
| Klhdc8a | 0.15227405 | 0.47697982 | 0.083 | 0.041 | 1 | 0.66339728 | 0.10518224 | 0.029 | 0.021 | 1 | 0.66339728 | 0.28136071 |
| Rufy4 | 0.57870246 | -0.9039688 | 0.156 | 0.171 | 1 | 0.15233386 | 0.14957636 | 0.13 | 0.073 | 1 | 0.57870246 | 0.28146211 |
| Gm10635 | 0.81741022 | 1.63976275 | 0.083 | 0.078 | 1 | 0.15247931 | 0.31950136 | 0.101 | 0.052 | 1 | 0.81741022 | 0.28170868 |
| Xpo4 | 0.30264686 | 0.14033022 | 0.552 | 0.446 | 1 | 0.1525648 | -0.131213 | 0.529 | 0.375 | 1 | 0.30264686 | 0.28185359 |
| Gm7160 | 0.1525942 | -1.4519371 | 0.01 | 0.041 | 1 | 0.88484452 | -0.9313436 | 0.046 | 0.042 | 1 | 0.88484452 | 0.2819034 |
| Gm13963 | 0.1525942 | -1.4519371 | 0.01 | 0.041 | 1 | 0.15394118 | 1.49749966 | 0.021 | 0 | 1 | 0.15394118 | 0.2819034 |
| Gm33694 | 0.1525942 | -1.4519371 | 0.01 | 0.041 | 1 | 0.87066325 | -0.3098553 | 0.013 | 0.01 | 1 | 0.87066325 | 0.2819034 |
| Gm57272 | 0.1525942 | -1.4519371 | 0.01 | 0.041 | 1 | 0.80442048 | -0.5728897 | 0.017 | 0.021 | 1 | 0.80442048 | 0.2819034 |
| Papolb | 0.1525942 | -1.8993961 | 0.01 | 0.041 | 1 | 0.87851725 | -1.3098553 | 0.013 | 0.01 | 1 | 0.87851725 | 0.2819034 |
| Cdip1 | 0.1526496 | 0.24195979 | 0.583 | 0.472 | 1 | 0.91907844 | -0.3038062 | 0.782 | 0.75 | 1 | 0.91907844 | 0.28199731 |
| Adss | 0.15265034 | 0.34314961 | 0.896 | 0.907 | 1 | 0.37677612 | -0.1226853 | 0.794 | 0.896 | 1 | 0.37677612 | 0.28199855 |
| Crym | 0.15273227 | -0.7998604 | 0.042 | 0.088 | 1 | 0.69003583 | -0.9472852 | 0.084 | 0.094 | 1 | 0.69003583 | 0.2821374 |
| Sirt4 | 0.99379628 | 0.13302542 | 0.052 | 0.052 | 1 | 0.15273891 | 0.83698613 | 0.113 | 0.062 | 1 | 0.99379628 | 0.28214865 |
| Muc2 | 0.61737927 | 1.00749454 | 0.01 | 0.005 | 1 | 0.15290592 | -1.010295 | 0.029 | 0.062 | 1 | 0.61737927 | 0.28243162 |
| Orc3 | 0.15297794 | 0.48024753 | 0.406 | 0.337 | 1 | 0.88844686 | -0.7738024 | 0.374 | 0.312 | 1 | 0.88844686 | 0.28255363 |
| Fer | 0.48121992 | -0.3359133 | 0.448 | 0.482 | 1 | 0.1530823 | -0.7060829 | 0.471 | 0.531 | 1 | 0.48121992 | 0.2827304 |
| Gm12002 | 0.15329198 | -1.3144336 | 0.01 | 0.041 | 1 | 0.53864238 | 0.27510724 | 0.046 | 0.031 | 1 | 0.53864238 | 0.28308553 |
| Gm49179 | 0.15329198 | -1.3144336 | 0.01 | 0.041 | 1 | 0.25080911 | -0.9879272 | 0.013 | 0.031 | 1 | 0.25080911 | 0.28308553 |
| Tspyl3 | 0.15329198 | 1.00749454 | 0.052 | 0.021 | 1 | 0.67471359 | -0.5728897 | 0.017 | 0.01 | 1 | 0.67471359 | 0.28308553 |
| Gm43378 | 0.15329198 | 1.00749454 | 0.052 | 0.021 | 1 | 0.2430612 | 0.56461386 | 0.034 | 0.01 | 1 | 0.2430612 | 0.28308553 |
| Gm57385 | 0.15329198 | 1.00749454 | 0.052 | 0.021 | 1 | 0.99396043 | -0.8244284 | 0.021 | 0.021 | 1 | 0.99396043 | 0.28308553 |
| Shpk | 0.15329198 | 1.00749454 | 0.052 | 0.021 | 1 | 0.67149861 | -0.1399303 | 0.029 | 0.021 | 1 | 0.67149861 | 0.28308553 |
| Gm36264 | 0.15329198 | 1.00749454 | 0.052 | 0.021 | 1 | 0.15394868 | 1.27510724 | 0.021 | 0 | 1 | 0.15394868 | 0.28308553 |
| Zfp120 | 0.66933774 | -0.1500467 | 0.125 | 0.104 | 1 | 0.1533168 | 0.8995981 | 0.113 | 0.062 | 1 | 0.66933774 | 0.28312756 |
| Mybph | 0.15337569 | -1.747393 | 0.031 | 0.073 | 1 | 0.54277206 | -0.7248928 | 0.034 | 0.021 | 1 | 0.54277206 | 0.28322727 |
| Slx1b | 0.15337569 | -1.577468 | 0.031 | 0.073 | 1 | 0.94385422 | -0.3098553 | 0.055 | 0.052 | 1 | 0.94385422 | 0.28322727 |
| Star | 0.15337661 | -1.8993961 | 0.031 | 0.073 | 1 | 0.39644749 | 0.56461386 | 0.113 | 0.083 | 1 | 0.39644749 | 0.28322884 |
| Zfp959 | 0.1534794 | 0.62047141 | 0.125 | 0.073 | 1 | 0.40341407 | 0.35999614 | 0.126 | 0.094 | 1 | 0.40341407 | 0.28340287 |
| Ess2 | 0.15362983 | -0.9311049 | 0.042 | 0.088 | 1 | 0.2088742 | 0.31817596 | 0.122 | 0.073 | 1 | 0.2088742 | 0.28365754 |
| Snx4 | 0.66314069 | -0.2422255 | 0.677 | 0.663 | 1 | 0.15363629 | -0.4434798 | 0.79 | 0.854 | 1 | 0.66314069 | 0.28366848 |
| 4930549G23Rik | 0.15365762 | 0.81484946 | 0.104 | 0.057 | 1 | 0.90722795 | 0.23446525 | 0.097 | 0.094 | 1 | 0.90722795 | 0.28370458 |
| Shld1 | 0.71031978 | 0.35541784 | 0.062 | 0.052 | 1 | 0.15377116 | 0.63767732 | 0.088 | 0.042 | 1 | 0.71031978 | 0.28389674 |
| Bcl2l12 | 0.59095266 | 0.28760246 | 0.135 | 0.114 | 1 | 0.15378623 | -0.7952821 | 0.067 | 0.115 | 1 | 0.59095266 | 0.28392225 |
| Nlrp10 | 0.15837881 | 2.00749454 | 0.01 | 0 | 1 | 0.15394118 | 1.49749966 | 0.021 | 0 | 1 | 0.15837881 | 0.28418448 |
| Gm57273 | 0.17265227 | 1.59245704 | 0.042 | 0.016 | 1 | 0.15394118 | 1.49749966 | 0.021 | 0 | 1 | 0.17265227 | 0.28418448 |
| Gm19391 | 0.1992517 | 1.74446013 | 0.031 | 0.01 | 1 | 0.15394118 | 1.49749966 | 0.021 | 0 | 1 | 0.1992517 | 0.28418448 |
| Gabrp | 0.20158431 | 1.49292136 | 0.031 | 0.01 | 1 | 0.15394118 | 1.49749966 | 0.021 | 0 | 1 | 0.20158431 | 0.28418448 |
| Il18r1 | 0.21860846 | -1.1624305 | 0.021 | 0.052 | 1 | 0.15394118 | 1.69014474 | 0.021 | 0 | 1 | 0.21860846 | 0.28418448 |
| Fam110a | 0.21867159 | 1.59245704 | 0.021 | 0.005 | 1 | 0.15394118 | 1.49749966 | 0.021 | 0 | 1 | 0.21867159 | 0.28418448 |
| Ppp1r18os | 0.25008035 | -0.4330781 | 0.042 | 0.078 | 1 | 0.15394118 | 1.49749966 | 0.021 | 0 | 1 | 0.25008035 | 0.28418448 |
| Scnn1b | 0.32025586 | -0.577468 | 0 | 0.01 | 1 | 0.15394118 | 1.49749966 | 0.021 | 0 | 1 | 0.32025586 | 0.28418448 |
| Lrrc25 | 0.32025586 | -0.577468 | 0 | 0.01 | 1 | 0.15394118 | 1.86006974 | 0.021 | 0 | 1 | 0.32025586 | 0.28418448 |
| Gm47922 | 0.32025586 | -0.9925055 | 0 | 0.01 | 1 | 0.15394118 | 1.49749966 | 0.021 | 0 | 1 | 0.32025586 | 0.28418448 |
| Lrrc56 | 0.37536054 | 1.32942263 | 0.031 | 0.016 | 1 | 0.15394118 | 1.49749966 | 0.021 | 0 | 1 | 0.37536054 | 0.28418448 |
| Cbln2 | 0.47150075 | 1.42253204 | 0.021 | 0.01 | 1 | 0.15394118 | 1.49749966 | 0.021 | 0 | 1 | 0.47150075 | 0.28418448 |
| Arhgap25 | 0.61371908 | 2.22988696 | 0.01 | 0.005 | 1 | 0.15394118 | 1.49749966 | 0.021 | 0 | 1 | 0.61371908 | 0.28418448 |
| Vwf | 0.61737927 | 1.00749454 | 0.01 | 0.005 | 1 | 0.15394118 | 1.49749966 | 0.021 | 0 | 1 | 0.61737927 | 0.28418448 |
| Jaml | 0.61737927 | 1.00749454 | 0.01 | 0.005 | 1 | 0.15394118 | 2.27510724 | 0.021 | 0 | 1 | 0.61737927 | 0.28418448 |
| Gm7854 | 0.62104899 | -0.9925055 | 0.01 | 0.005 | 1 | 0.15394118 | 1.69014474 | 0.021 | 0 | 1 | 0.62104899 | 0.28418448 |
| Lrrc46 | 0.62104899 | 0.42253204 | 0.01 | 0.005 | 1 | 0.15394118 | 1.49749966 | 0.021 | 0 | 1 | 0.62104899 | 0.28418448 |
| Gm56516 | 0.74827573 | 0.59245704 | 0.021 | 0.016 | 1 | 0.15394118 | 1.49749966 | 0.021 | 0 | 1 | 0.74827573 | 0.28418448 |
| A930001C03Rik | 0.7854853 | -0.577468 | 0.021 | 0.026 | 1 | 0.15394118 | 1.86006974 | 0.021 | 0 | 1 | 0.7854853 | 0.28418448 |
| Foxa2 | 0.97983704 | 0.42253204 | 0.052 | 0.052 | 1 | 0.15394118 | 1.69014474 | 0.021 | 0 | 1 | 0.97983704 | 0.28418448 |
| Raet1e | 1 | 0.42253204 | 0.01 | 0.01 | 1 | 0.15394118 | 1.49749966 | 0.021 | 0 | 1 | 1 | 0.28418448 |
| Ccdc38 | 1 | -0.3144336 | 0.01 | 0.01 | 1 | 0.15394118 | 1.69014474 | 0.021 | 0 | 1 | 1 | 0.28418448 |
| 5430400D12Rik | 0.15837881 | 2.00749454 | 0.01 | 0 | 1 | 0.15394568 | 1.69014474 | 0.021 | 0 | 1 | 0.15837881 | 0.28419208 |
| Gm45267 | 0.15837881 | 2.00749454 | 0.01 | 0 | 1 | 0.15394568 | 1.69014474 | 0.021 | 0 | 1 | 0.15837881 | 0.28419208 |
| Gm12472 | 0.21708019 | 2.00749454 | 0.021 | 0.005 | 1 | 0.15394568 | 2.01207283 | 0.021 | 0 | 1 | 0.21708019 | 0.28419208 |
| Tm6sf1 | 0.22187388 | -1.3144336 | 0 | 0.016 | 1 | 0.15394568 | 1.86006974 | 0.021 | 0 | 1 | 0.22187388 | 0.28419208 |
| Nfam1 | 0.32025586 | -0.577468 | 0 | 0.01 | 1 | 0.15394568 | 1.69014474 | 0.021 | 0 | 1 | 0.32025586 | 0.28419208 |
| Apobr | 1 | 0.42253204 | 0.01 | 0.01 | 1 | 0.15394568 | 1.69014474 | 0.021 | 0 | 1 | 1 | 0.28419208 |
| Nckap1l | 0.38689439 | -0.577468 | 0.01 | 0.026 | 1 | 0.15394718 | 2.01207283 | 0.021 | 0 | 1 | 0.38689439 | 0.28419462 |
| Gm48395 | 0.8006267 | 0.20013961 | 0.021 | 0.026 | 1 | 0.15394793 | 1.86006974 | 0.021 | 0 | 1 | 0.8006267 | 0.28419589 |
| Gm44145 | 0.15837881 | 2.00749454 | 0.01 | 0 | 1 | 0.15394868 | 1.27510724 | 0.021 | 0 | 1 | 0.15837881 | 0.28419716 |
| Pcsk4 | 0.15837881 | 2.00749454 | 0.01 | 0 | 1 | 0.15394868 | 1.27510724 | 0.021 | 0 | 1 | 0.15837881 | 0.28419716 |
| Gm54359 | 0.15837881 | 2.00749454 | 0.01 | 0 | 1 | 0.15394868 | 1.27510724 | 0.021 | 0 | 1 | 0.15837881 | 0.28419716 |
| Gm5148 | 0.21708019 | 2.32942263 | 0.021 | 0.005 | 1 | 0.15394868 | 1.27510724 | 0.021 | 0 | 1 | 0.21708019 | 0.28419716 |
| Gm15704 | 0.22187388 | -1.577468 | 0 | 0.016 | 1 | 0.15394868 | 1.27510724 | 0.021 | 0 | 1 | 0.22187388 | 0.28419716 |
| Gm19569 | 0.31389644 | 0.74446013 | 0.042 | 0.021 | 1 | 0.15394868 | 1.27510724 | 0.021 | 0 | 1 | 0.31389644 | 0.28419716 |
| 6720407P12Rik | 0.32025586 | -0.9925055 | 0 | 0.01 | 1 | 0.15394868 | 1.27510724 | 0.021 | 0 | 1 | 0.32025586 | 0.28419716 |
| Asb16 | 0.32025586 | -1.3144336 | 0 | 0.01 | 1 | 0.15394868 | 1.27510724 | 0.021 | 0 | 1 | 0.32025586 | 0.28419716 |
| 6330415G19Rik | 0.32025586 | -0.577468 | 0 | 0.01 | 1 | 0.15394868 | 1.27510724 | 0.021 | 0 | 1 | 0.32025586 | 0.28419716 |
| 4930513N10Rik | 0.38357309 | -1.1624305 | 0.01 | 0.026 | 1 | 0.15394868 | 1.27510724 | 0.021 | 0 | 1 | 0.38357309 | 0.28419716 |
| Fam131c | 0.38521705 | 0.68556644 | 0.031 | 0.016 | 1 | 0.15394868 | 1.27510724 | 0.021 | 0 | 1 | 0.38521705 | 0.28419716 |
| E230013L22Rik | 0.38689439 | -0.577468 | 0.01 | 0.026 | 1 | 0.15394868 | 1.27510724 | 0.021 | 0 | 1 | 0.38689439 | 0.28419716 |
| E130006D01Rik | 0.47150075 | 1.42253204 | 0.021 | 0.01 | 1 | 0.15394868 | 1.27510724 | 0.021 | 0 | 1 | 0.47150075 | 0.28419716 |
| Spata33 | 0.47150075 | 1.42253204 | 0.021 | 0.01 | 1 | 0.15394868 | 1.27510724 | 0.021 | 0 | 1 | 0.47150075 | 0.28419716 |
| Dyrk3 | 0.47294388 | -1.3144336 | 0.021 | 0.036 | 1 | 0.15394868 | 1.27510724 | 0.021 | 0 | 1 | 0.47294388 | 0.28419716 |
| Gm13091 | 0.47606502 | 1.00749454 | 0.021 | 0.01 | 1 | 0.15394868 | 1.27510724 | 0.021 | 0 | 1 | 0.47606502 | 0.28419716 |

| | | | | | | | | | | | | |
|---|---|---|---|---|---|---|---|---|---|---|---|---|
| Fbxo27 | 0.47606502 | 1.00749454 | 0.021 | 0.01 | 1 | 0.15394868 | 1.27510724 | 0.021 | 0 | 1 | 0.47606502 | 0.28419716 |
| Hoatz | 0.47606502 | 1.00749454 | 0.021 | 0.01 | 1 | 0.15394868 | 1.27510724 | 0.021 | 0 | 1 | 0.47606502 | 0.28419716 |
| Gm57266 | 0.51793219 | 0.81484946 | 0.052 | 0.036 | 1 | 0.15394868 | 1.27510724 | 0.021 | 0 | 1 | 0.51793219 | 0.28419716 |
| Gm20337 | 0.52528691 | -0.7998604 | 0.01 | 0.021 | 1 | 0.15394868 | 1.27510724 | 0.021 | 0 | 1 | 0.52528691 | 0.28419716 |
| Tmc7 | 0.52744059 | -0.577468 | 0.01 | 0.021 | 1 | 0.15394868 | 1.27510724 | 0.021 | 0 | 1 | 0.52744059 | 0.28419716 |
| Zfp605 | 0.52961488 | -0.3144336 | 0.01 | 0.021 | 1 | 0.15394868 | 1.27510724 | 0.021 | 0 | 1 | 0.52961488 | 0.28419716 |
| Gm57399 | 0.52961488 | -0.3144336 | 0.01 | 0.021 | 1 | 0.15394868 | 1.27510724 | 0.021 | 0 | 1 | 0.52961488 | 0.28419716 |
| Pnck | 0.52961488 | -0.3144336 | 0.01 | 0.021 | 1 | 0.15394868 | 1.27510724 | 0.021 | 0 | 1 | 0.52961488 | 0.28419716 |
| Gm56945 | 0.58426772 | 0.74446013 | 0.031 | 0.021 | 1 | 0.15394868 | 1.27510724 | 0.021 | 0 | 1 | 0.58426772 | 0.28419716 |
| Gm34006 | 0.61737927 | 1.00749454 | 0.01 | 0.005 | 1 | 0.15394868 | 1.27510724 | 0.021 | 0 | 1 | 0.61737927 | 0.28419716 |
| Gm38560 | 0.72298091 | -0.577468 | 0.01 | 0.016 | 1 | 0.15394868 | 1.27510724 | 0.021 | 0 | 1 | 0.72298091 | 0.28419716 |
| Gm56735 | 0.72574709 | -0.3144336 | 0.01 | 0.016 | 1 | 0.15394868 | 1.27510724 | 0.021 | 0 | 1 | 0.72574709 | 0.28419716 |
| Tmprss11b | 0.73686008 | 0.35541784 | 0.01 | 0.016 | 1 | 0.15394868 | 1.27510724 | 0.021 | 0 | 1 | 0.73686008 | 0.28419716 |
| 1810010H24Rik | 0.78117751 | -0.577468 | 0.021 | 0.026 | 1 | 0.15394868 | 1.27510724 | 0.021 | 0 | 1 | 0.78117751 | 0.28419716 |
| Gm826 | 0.78980953 | -0.3710171 | 0.021 | 0.026 | 1 | 0.15394868 | 1.27510724 | 0.021 | 0 | 1 | 0.78980953 | 0.28419716 |
| Gm56977 | 0.79666623 | 0.42253204 | 0.031 | 0.026 | 1 | 0.15394868 | 1.27510724 | 0.021 | 0 | 1 | 0.79666623 | 0.28419716 |
| Treh | 1 | 0.42253204 | 0.01 | 0.01 | 1 | 0.15394868 | 1.27510724 | 0.021 | 0 | 1 | 1 | 0.28419716 |
| Pde2a | 0.15404368 | -1.4950058 | 0.01 | 0.041 | 1 | 0.25659463 | 0.40635177 | 0.063 | 0.031 | 1 | 0.25659463 | 0.28435791 |
| Vps8 | 0.30617492 | -0.4431669 | 0.208 | 0.259 | 1 | 0.15405156 | 0.14184071 | 0.42 | 0.302 | 1 | 0.30617492 | 0.28437123 |
| Zc2hc1c | 0.15405555 | -1.1624305 | 0.01 | 0.041 | 1 | 0.33526548 | -0.6094155 | 0.046 | 0.073 | 1 | 0.33526548 | 0.28437799 |
| Dennd5b | 0.15412901 | -0.5175965 | 0.344 | 0.42 | 1 | 0.4850585 | -0.3511622 | 0.42 | 0.333 | 1 | 0.4850585 | 0.28450227 |
| Tbc1d12 | 0.15412903 | -0.8040604 | 0.312 | 0.373 | 1 | 0.2493044 | -0.4774552 | 0.492 | 0.542 | 1 | 0.2493044 | 0.2845023 |
| Mvb12b | 0.15420204 | -0.4872702 | 0.24 | 0.326 | 1 | 0.83014837 | -0.6189593 | 0.328 | 0.302 | 1 | 0.83014837 | 0.28462581 |
| Myadml2os | 0.81497862 | -0.1624305 | 0.031 | 0.026 | 1 | 0.15424568 | 0.27510724 | 0.059 | 0.021 | 1 | 0.81497862 | 0.28469963 |
| Dnah10 | 0.36239198 | 0.56003556 | 0.073 | 0.047 | 1 | 0.15426046 | -2.3739856 | 0.038 | 0.073 | 1 | 0.36239198 | 0.28472463 |
| Usp8 | 0.1798581 | 0.21172759 | 0.719 | 0.622 | 1 | 0.15431463 | 0.13860524 | 0.752 | 0.729 | 1 | 0.1798581 | 0.28481626 |
| Robo1 | 0.15465734 | -0.391495 | 0.281 | 0.368 | 1 | 0.21586469 | -0.2329938 | 0.773 | 0.844 | 1 | 0.21586469 | 0.28539579 |
| Rnpep | 0.37869062 | 0.62746349 | 0.365 | 0.332 | 1 | 0.15476561 | -0.4698957 | 0.437 | 0.531 | 1 | 0.37869062 | 0.28557882 |
| A730036I17Rik | 0.15476747 | -2.3144336 | 0.01 | 0.041 | 1 | 0.39937186 | -1.3098553 | 0.017 | 0.031 | 1 | 0.39937186 | 0.28558197 |
| Gm19951 | 0.50640077 | -0.4399644 | 0.979 | 0.964 | 1 | 0.15495782 | -0.7675495 | 0.987 | 0.99 | 1 | 0.50640077 | 0.28590372 |
| Il6st | 0.15502585 | -0.2373091 | 0.906 | 0.876 | 1 | 0.77610601 | -0.1159134 | 0.912 | 0.948 | 1 | 0.77610601 | 0.28601869 |
| Cep57 | 0.15568023 | -0.1534973 | 0.365 | 0.461 | 1 | 0.84057558 | -0.2708611 | 0.424 | 0.375 | 1 | 0.84057558 | 0.28712413 |
| Kifap3 | 0.15574272 | 0.16182268 | 0.646 | 0.534 | 1 | 0.78182948 | -0.5792452 | 0.639 | 0.531 | 1 | 0.78182948 | 0.28722965 |
| Dck | 0.15576919 | 0.81484946 | 0.135 | 0.083 | 1 | 0.78056544 | -0.3098553 | 0.13 | 0.115 | 1 | 0.78056544 | 0.28727434 |
| Rbm4 | 0.28517916 | 0.25542205 | 0.479 | 0.409 | 1 | 0.1558244 | -0.2355079 | 0.597 | 0.427 | 1 | 0.28517916 | 0.28736755 |
| Xpnpep3 | 0.15624338 | 0.40545852 | 0.375 | 0.301 | 1 | 0.8047277 | -0.5539599 | 0.395 | 0.323 | 1 | 0.8047277 | 0.28807476 |
| Ap5s1 | 0.88293873 | -0.1941393 | 0.083 | 0.088 | 1 | 0.15630704 | 0.54812573 | 0.101 | 0.052 | 1 | 0.88293873 | 0.28818219 |
| Apol8 | 0.15640917 | 0.21899864 | 0.104 | 0.057 | 1 | 0.58260294 | -0.3463811 | 0.105 | 0.125 | 1 | 0.58260294 | 0.2883545 |
| Pxn | 0.15648036 | 0.36160304 | 0.469 | 0.409 | 1 | 0.90482234 | -0.3727153 | 0.563 | 0.583 | 1 | 0.90482234 | 0.28847462 |
| Ccdc163 | 0.15649656 | 0.86999101 | 0.135 | 0.083 | 1 | 0.2890792 | -1.7605167 | 0.101 | 0.135 | 1 | 0.2890792 | 0.28850195 |
| Snhg7 | 0.15654887 | 0.30938488 | 0.594 | 0.549 | 1 | 0.16847167 | 0.13967061 | 0.538 | 0.417 | 1 | 0.16847167 | 0.2885902 |
| Tmem204 | 0.15691242 | -0.5350327 | 0.125 | 0.192 | 1 | 0.67203981 | -0.5025003 | 0.101 | 0.083 | 1 | 0.67203981 | 0.28920332 |
| Zfp263 | 0.39544485 | 0.10937415 | 0.208 | 0.161 | 1 | 0.15702789 | 0.22930355 | 0.256 | 0.177 | 1 | 0.39544485 | 0.28939802 |
| Per1 | 0.2359876 | 0.32942263 | 0.375 | 0.306 | 1 | 0.1571545 | -0.6508922 | 0.353 | 0.417 | 1 | 0.2359876 | 0.28961147 |
| Kcnmb4 | 0.68578606 | -0.2148979 | 0.094 | 0.078 | 1 | 0.15720663 | -0.7644211 | 0.139 | 0.198 | 1 | 0.68578606 | 0.28969934 |
| Gm16150 | 0.15729403 | -1.577468 | 0 | 0.021 | 1 | 0.80442048 | -0.5728897 | 0.017 | 0.021 | 1 | 0.80442048 | 0.28984665 |
| Creg2 | 0.15729403 | -1.7998604 | 0 | 0.021 | 1 | 0.86312513 | -0.7248928 | 0.008 | 0.01 | 1 | 0.86312513 | 0.28984665 |
| Mreg | 0.15729403 | -1.577468 | 0 | 0.021 | 1 | 0.86350341 | -0.5322477 | 0.038 | 0.042 | 1 | 0.86350341 | 0.28984665 |
| Acbd7 | 0.15729403 | -1.577468 | 0 | 0.021 | 1 | 0.86312513 | -0.7248928 | 0.008 | 0.01 | 1 | 0.86312513 | 0.28984665 |
| Large2 | 0.15729403 | -1.577468 | 0 | 0.021 | 1 | 0.57210759 | -1.1172102 | 0.021 | 0.031 | 1 | 0.57210759 | 0.28984665 |
| Pax6os1 | 0.15729403 | -1.577468 | 0 | 0.021 | 1 | 0.50587544 | -1.8948178 | 0.004 | 0.01 | 1 | 0.50587544 | 0.28984665 |
| Itpka | 0.15729403 | -1.577468 | 0 | 0.021 | 1 | 0.5207331 | -0.5025003 | 0.021 | 0.01 | 1 | 0.5207331 | 0.28984665 |
| Gm20632 | 0.15729403 | -1.577468 | 0 | 0.021 | 1 | 0.75977578 | -0.5025003 | 0.025 | 0.031 | 1 | 0.75977578 | 0.28984665 |
| Slc9b1 | 0.15729403 | -1.577468 | 0 | 0.021 | 1 | 0.92899987 | -0.4167705 | 0.034 | 0.031 | 1 | 0.92899987 | 0.28984665 |
| Gm42705 | 0.15729403 | -1.577468 | 0 | 0.021 | 1 | 0.86613737 | -0.3098553 | 0.008 | 0.01 | 1 | 0.86613737 | 0.28984665 |
| Gpr63 | 0.15729403 | -1.577468 | 0 | 0.021 | 1 | 0.24746336 | -1.3098553 | 0.013 | 0.031 | 1 | 0.24746336 | 0.28984665 |
| Pm20d2 | 0.15729403 | -1.577468 | 0 | 0.021 | 1 | 0.27154471 | 0.69014474 | 0.013 | 0 | 1 | 0.27154471 | 0.28984665 |
| Gm52698 | 0.15729403 | -1.577468 | 0 | 0.021 | 1 | 0.24746336 | -1.3098553 | 0.013 | 0.031 | 1 | 0.24746336 | 0.28984665 |
| Zmynd12 | 0.15729403 | -1.577468 | 0 | 0.021 | 1 | 0.80231436 | -0.6317834 | 0.017 | 0.021 | 1 | 0.80231436 | 0.28984665 |
| Gm12979 | 0.15729403 | -1.7998604 | 0 | 0.021 | 1 | 0.86543325 | 0.49749966 | 0.013 | 0.01 | 1 | 0.86543325 | 0.28984665 |
| Tnfrsf25 | 0.15729403 | -1.7998604 | 0 | 0.021 | 1 | 0.27759424 | 0.80562195 | 0.046 | 0.021 | 1 | 0.27759424 | 0.28984665 |
| Kcna1 | 0.15729403 | -1.577468 | 0 | 0.021 | 1 | 0.93576656 | -0.1399303 | 0.029 | 0.031 | 1 | 0.93576656 | 0.28984665 |
| Ptpro | 0.15729403 | -1.577468 | 0 | 0.021 | 1 | 0.27154471 | 0.69014474 | 0.013 | 0 | 1 | 0.27154471 | 0.28984665 |
| Evi2a | 0.15729403 | -1.577468 | 0 | 0.021 | 1 | 0.83706049 | -1.3098553 | 0.025 | 0.021 | 1 | 0.83706049 | 0.28984665 |
| Pctp | 0.15729403 | -1.577468 | 0 | 0.021 | 1 | 0.41074402 | -0.3098553 | 0.017 | 0.031 | 1 | 0.41074402 | 0.28984665 |
| Rundc3a | 0.15729403 | -1.577468 | 0 | 0.021 | 1 | 0.51194506 | 0.27510724 | 0.021 | 0.01 | 1 | 0.51194506 | 0.28984665 |
| 6030468B19Rik | 0.15729403 | -1.577468 | 0 | 0.021 | 1 | 0.57170879 | -1.3098553 | 0.013 | 0.021 | 1 | 0.57170879 | 0.28984665 |
| Alkal2 | 0.15729403 | -1.577468 | 0 | 0.021 | 1 | 0.50587544 | -2.6317834 | 0.004 | 0.01 | 1 | 0.50587544 | 0.28984665 |
| B930059L03Rik | 0.15729403 | -1.577468 | 0 | 0.021 | 1 | 0.20315163 | 1.01207283 | 0.017 | 0 | 1 | 0.20315163 | 0.28984665 |
| Nkapl | 0.15729403 | -1.577468 | 0 | 0.021 | 1 | 0.50887612 | -1.3098553 | 0.004 | 0.01 | 1 | 0.50887612 | 0.28984665 |
| Fam217a | 0.15729403 | -1.577468 | 0 | 0.021 | 1 | 0.87851725 | -0.8948178 | 0.013 | 0.01 | 1 | 0.87851725 | 0.28984665 |
| Gm49173 | 0.15729403 | -1.577468 | 0 | 0.021 | 1 | 0.58180179 | -0.7248928 | 0.021 | 0.031 | 1 | 0.58180179 | 0.28984665 |
| Gm49864 | 0.15729403 | -1.577468 | 0 | 0.021 | 1 | 0.34363875 | -2.0468209 | 0.008 | 0.021 | 1 | 0.34363875 | 0.28984665 |
| Gm17315 | 0.15729403 | -1.577468 | 0 | 0.021 | 1 | 1 | -0.7248928 | 0.021 | 0.021 | 1 | 1 | 0.28984665 |
| Acy3 | 0.15729403 | -1.577468 | 0 | 0.021 | 1 | 0.86312513 | -0.7248928 | 0.008 | 0.01 | 1 | 0.86312513 | 0.28984665 |
| Gm9895 | 0.15729403 | -1.577468 | 0 | 0.021 | 1 | 0.85710544 | -1.3098553 | 0.008 | 0.01 | 1 | 0.85710544 | 0.28984665 |
| Pi15 | 0.15729781 | -2.4519371 | 0 | 0.021 | 1 | 0.86804649 | 0.49749966 | 0.013 | 0.01 | 1 | 0.86804649 | 0.28985302 |
| Dnah7a | 0.15729781 | -2.4519371 | 0 | 0.021 | 1 | 0.78761875 | -1.3098553 | 0.017 | 0.021 | 1 | 0.78761875 | 0.28985302 |
| Sphkap | 0.15729781 | -1.7998604 | 0 | 0.021 | 1 | 0.27154471 | 1.86006974 | 0.013 | 0 | 1 | 0.27154471 | 0.28985302 |
| Rgs4 | 0.15729781 | -3.7998604 | 0 | 0.021 | 1 | 0.50887612 | -1.3098553 | 0.004 | 0.01 | 1 | 0.50887612 | 0.28985302 |
| Gm6403 | 0.15729781 | -2.1624305 | 0 | 0.021 | 1 | 0.80442048 | -0.5728897 | 0.017 | 0.021 | 1 | 0.80442048 | 0.28985302 |
| Gm13387 | 0.15729781 | -3.5160674 | 0 | 0.021 | 1 | 0.50887612 | -1.3098553 | 0.004 | 0.01 | 1 | 0.50887612 | 0.28985302 |
| Jph2 | 0.15729781 | -3.1624305 | 0 | 0.021 | 1 | 0.34704016 | -1.3098553 | 0.008 | 0.021 | 1 | 0.34704016 | 0.28985302 |
| Hmgcs2 | 0.15729781 | -2.3144336 | 0 | 0.021 | 1 | 0.85710586 | -2.1172102 | 0.008 | 0.01 | 1 | 0.85710586 | 0.28985302 |
| Lingo2 | 0.15729781 | -1.9925055 | 0 | 0.021 | 1 | 0.87851807 | -1.3098553 | 0.013 | 0.01 | 1 | 0.87851807 | 0.28985302 |

| | | | | | | | | | | | | |
|---|---|---|---|---|---|---|---|---|---|---|---|---|
| 9130230L23Rik | 0.15729781 | -1.7998604 | 0 | 0.021 | 1 | 0.56024564 | -0.8948178 | 0.046 | 0.031 | 1 | 0.56024564 | 0.28985302 |
| Kctd8 | 0.15729781 | -2.3144336 | 0 | 0.021 | 1 | 0.91613797 | 0.17557156 | 0.034 | 0.031 | 1 | 0.91613797 | 0.28985302 |
| Cpne9 | 0.15729781 | -2.4519371 | 0 | 0.021 | 1 | 0.55024997 | -1.3098553 | 0.029 | 0.042 | 1 | 0.55024997 | 0.28985302 |
| Lpar5 | 0.15729781 | -1.9925055 | 0 | 0.021 | 1 | 0.27154471 | 0.69014474 | 0.013 | 0 | 1 | 0.27154471 | 0.28985302 |
| Cblc | 0.15729781 | -1.7998604 | 0 | 0.021 | 1 | 0.20314753 | 1.49749966 | 0.017 | 0 | 1 | 0.20314753 | 0.28985302 |
| Gm16183 | 0.15729781 | -2.1624305 | 0 | 0.021 | 1 | 0.87851725 | -0.8948178 | 0.013 | 0.01 | 1 | 0.87851725 | 0.28985302 |
| Slc35f3 | 0.15729781 | -1.7998604 | 0 | 0.021 | 1 | 0.51188611 | -0.3098553 | 0.004 | 0.01 | 1 | 0.51188611 | 0.28985302 |
| Jhy | 0.15729781 | -2.1624305 | 0 | 0.021 | 1 | 0.50587544 | -1.7248928 | 0.004 | 0.01 | 1 | 0.50587544 | 0.28985302 |
| Unc13c | 0.15729781 | -3.577468 | 0 | 0.021 | 1 | 0.27154327 | 1.27510724 | 0.013 | 0 | 1 | 0.27154327 | 0.28985302 |
| Hs3st5 | 0.15729781 | -1.9925055 | 0 | 0.021 | 1 | 0.67966683 | -1.0560987 | 0.042 | 0.031 | 1 | 0.67966683 | 0.28985302 |
| Mmp11 | 0.15729781 | -1.7998604 | 0 | 0.021 | 1 | 0.24413474 | -1.6317834 | 0.013 | 0.031 | 1 | 0.24413474 | 0.28985302 |
| Asic2 | 0.15729781 | -1.7998604 | 0 | 0.021 | 1 | 0.66171832 | 2.73453886 | 0.017 | 0.01 | 1 | 0.66171832 | 0.28985302 |
| Gm36660 | 0.15729781 | -1.9925055 | 0 | 0.021 | 1 | 0.51188611 | -0.3098553 | 0.004 | 0.01 | 1 | 0.51188611 | 0.28985302 |
| Pde8b | 0.15729781 | -1.9925055 | 0 | 0.021 | 1 | 0.79600331 | -1.3098553 | 0.017 | 0.021 | 1 | 0.79600331 | 0.28985302 |
| Gpc5 | 0.15729781 | -3.5160674 | 0 | 0.021 | 1 | 0.51188611 | 1.69014474 | 0.004 | 0.01 | 1 | 0.51188611 | 0.28985302 |
| Krt75 | 0.15729781 | -1.9925055 | 0 | 0.021 | 1 | 0.50887612 | -1.3098553 | 0.004 | 0.01 | 1 | 0.50887612 | 0.28985302 |
| Tnfaip8l1 | 0.15729781 | -2.4519371 | 0 | 0.021 | 1 | 0.34026631 | -2.5322477 | 0.008 | 0.021 | 1 | 0.34026631 | 0.28985302 |
| Gm36201 | 0.15729781 | -1.9925055 | 0 | 0.021 | 1 | 0.85710544 | -1.3098553 | 0.008 | 0.01 | 1 | 0.85710544 | 0.28985302 |
| Gm36279 | 0.15729781 | -1.7998604 | 0 | 0.021 | 1 | 0.67474842 | -0.3098553 | 0.029 | 0.021 | 1 | 0.67474842 | 0.28985302 |
| Ren1 | 0.15729907 | -1.3144336 | 0 | 0.021 | 1 | 0.50887612 | -1.3098553 | 0.004 | 0.01 | 1 | 0.50887612 | 0.28985515 |
| Gm36988 | 0.15729907 | -1.3144336 | 0 | 0.021 | 1 | 0.50887612 | -1.3098553 | 0.004 | 0.01 | 1 | 0.50887612 | 0.28985515 |
| Gm14024 | 0.15729907 | -1.3144336 | 0 | 0.021 | 1 | 0.50887612 | -1.3098553 | 0.004 | 0.01 | 1 | 0.50887612 | 0.28985515 |
| Vtcn1 | 0.15729907 | -1.3144336 | 0 | 0.021 | 1 | 0.86312513 | -0.7248928 | 0.008 | 0.01 | 1 | 0.86312513 | 0.28985515 |
| Cfap276 | 0.15729907 | -1.3144336 | 0 | 0.021 | 1 | 0.87851807 | -1.3098553 | 0.013 | 0.01 | 1 | 0.87851807 | 0.28985515 |
| Oprd1 | 0.15729907 | -1.9925055 | 0 | 0.021 | 1 | 0.50887612 | -1.3098553 | 0.004 | 0.01 | 1 | 0.50887612 | 0.28985515 |
| Atad3aos | 0.15729907 | -1.3144336 | 0 | 0.021 | 1 | 0.27154471 | 0.69014474 | 0.013 | 0 | 1 | 0.27154471 | 0.28985515 |
| Rundc3b | 0.15729907 | -3.1624305 | 0 | 0.021 | 1 | 0.51188611 | 0.69014474 | 0.004 | 0.01 | 1 | 0.51188611 | 0.28985515 |
| Fkbp6 | 0.15729907 | -2.3144336 | 0 | 0.021 | 1 | 0.50887612 | -1.3098553 | 0.004 | 0.01 | 1 | 0.50887612 | 0.28985515 |
| Zfp108 | 0.15729907 | -1.3144336 | 0 | 0.021 | 1 | 0.5207331 | -0.5025003 | 0.021 | 0.01 | 1 | 0.5207331 | 0.28985515 |
| 1700028J19Rik | 0.15729907 | -1.3144336 | 0 | 0.021 | 1 | 0.24413474 | -1.8948178 | 0.013 | 0.031 | 1 | 0.24413474 | 0.28985515 |
| BC051019 | 0.15729907 | -1.3144336 | 0 | 0.021 | 1 | 0.57373861 | -1.7952821 | 0.013 | 0.021 | 1 | 0.57373861 | 0.28985515 |
| 5830408C22Rik | 0.15729907 | -1.3144336 | 0 | 0.021 | 1 | 0.85710544 | -1.7248928 | 0.008 | 0.01 | 1 | 0.85710544 | 0.28985515 |
| Fam241b | 0.15729907 | -1.3144336 | 0 | 0.021 | 1 | 0.75977578 | -0.5025003 | 0.025 | 0.031 | 1 | 0.75977578 | 0.28985515 |
| Hcn2 | 0.15729907 | -1.3144336 | 0 | 0.021 | 1 | 0.57170879 | -1.6317834 | 0.013 | 0.021 | 1 | 0.57170879 | 0.28985515 |
| Tjp3 | 0.15729907 | -1.3144336 | 0 | 0.021 | 1 | 1 | -1.3098553 | 0.021 | 0.021 | 1 | 1 | 0.28985515 |
| 2810408A11Rik | 0.15729907 | -1.3144336 | 0 | 0.021 | 1 | 0.66387437 | 0.49749966 | 0.017 | 0.01 | 1 | 0.66387437 | 0.28985515 |
| 5730507C01Rik | 0.15729907 | -1.3144336 | 0 | 0.021 | 1 | 0.3937346 | 0.69014474 | 0.025 | 0.01 | 1 | 0.3937346 | 0.28985515 |
| Pgf | 0.15729907 | -1.3144336 | 0 | 0.021 | 1 | 0.87066325 | -0.3098553 | 0.013 | 0.01 | 1 | 0.87066325 | 0.28985515 |
| Tshr | 0.15729907 | -1.3144336 | 0 | 0.021 | 1 | 0.86242829 | 0.34725702 | 0.067 | 0.062 | 1 | 0.86242829 | 0.28985515 |
| Nnt | 0.15729907 | -1.3144336 | 0 | 0.021 | 1 | 0.39515719 | 0.49749966 | 0.025 | 0.01 | 1 | 0.39515719 | 0.28985515 |
| Zfp957 | 0.15729907 | -2.6929452 | 0 | 0.021 | 1 | 0.66171832 | 0.49749966 | 0.017 | 0.01 | 1 | 0.66171832 | 0.28985515 |
| C9 | 0.15729907 | -2.6929452 | 0 | 0.021 | 1 | 0.40078851 | -1.84037 | 0.017 | 0.031 | 1 | 0.40078851 | 0.28985515 |
| Gm56725 | 0.15729907 | -1.3144336 | 0 | 0.021 | 1 | 0.50887612 | -1.3098553 | 0.004 | 0.01 | 1 | 0.50887612 | 0.28985515 |
| Ildr1 | 0.15729907 | -1.3144336 | 0 | 0.021 | 1 | 0.8076363 | 0.27510724 | 0.025 | 0.021 | 1 | 0.8076363 | 0.28985515 |
| Guca1a | 0.15729907 | -1.3144336 | 0 | 0.021 | 1 | 0.20315061 | 1.49749966 | 0.017 | 0 | 1 | 0.20315061 | 0.28985515 |
| Dsg3 | 0.15729907 | -1.3144336 | 0 | 0.021 | 1 | 0.27154471 | 0.69014474 | 0.013 | 0 | 1 | 0.27154471 | 0.28985515 |
| Kctd16 | 0.15729907 | -3.3144336 | 0 | 0.021 | 1 | 0.77340501 | 0.45567948 | 0.025 | 0.031 | 1 | 0.77340501 | 0.28985515 |
| Lrpprc | 0.15732138 | 0.40923521 | 0.417 | 0.321 | 1 | 0.46318592 | -0.1723517 | 0.538 | 0.479 | 1 | 0.46318592 | 0.28989275 |
| Itga3 | 0.15742602 | -0.9311049 | 0.042 | 0.088 | 1 | 0.56633327 | -0.3098553 | 0.038 | 0.052 | 1 | 0.56633327 | 0.29006909 |
| Rbfox1 | 0.15985156 | -1.7228984 | 0.104 | 0.161 | 1 | 0.15752121 | -1.2089264 | 0.097 | 0.146 | 1 | 0.15985156 | 0.29022949 |
| Aurka | 0.1576632 | 0.81484946 | 0.052 | 0.021 | 1 | 1 | -0.7248928 | 0.021 | 0.021 | 1 | 1 | 0.29046872 |
| Odad1 | 0.1578264 | 0.32299636 | 0.229 | 0.155 | 1 | 0.59771399 | -0.7347405 | 0.206 | 0.219 | 1 | 0.59771399 | 0.29074363 |
| Fign | 0.15804369 | -0.6974987 | 0.271 | 0.358 | 1 | 0.20971104 | -0.2318528 | 0.387 | 0.281 | 1 | 0.20971104 | 0.29110957 |
| Ptk2 | 0.15808544 | -0.1198098 | 0.865 | 0.777 | 1 | 0.19923233 | -0.3711316 | 0.937 | 0.969 | 1 | 0.19923233 | 0.29117988 |
| Aldh1l1 | 0.1580975 | -1.240433 | 0.031 | 0.073 | 1 | 0.40361357 | -1.3098553 | 0.017 | 0.031 | 1 | 0.40361357 | 0.29120018 |
| Bloc1s3 | 0.1580975 | -1.0799683 | 0.031 | 0.073 | 1 | 0.7311848 | 0.17557156 | 0.05 | 0.042 | 1 | 0.7311848 | 0.29120018 |
| Rnf220 | 0.15824944 | 0.34853145 | 0.292 | 0.212 | 1 | 0.4739604 | -0.3537986 | 0.29 | 0.24 | 1 | 0.4739604 | 0.291456 |
| Gm19938 | 0.15837881 | 2.00749454 | 0.01 | 0 | 1 | 0.50887612 | -1.3098553 | 0.004 | 0.01 | 1 | 0.50887612 | 0.29167377 |
| Gm30788 | 0.15837881 | 2.00749454 | 0.01 | 0 | 1 | 0.86312513 | -0.7248928 | 0.008 | 0.01 | 1 | 0.86312513 | 0.29167377 |
| Cfap65 | 0.15837881 | 2.00749454 | 0.01 | 0 | 1 | 0.86312513 | -0.7248928 | 0.008 | 0.01 | 1 | 0.86312513 | 0.29167377 |
| Nyap2 | 0.15837881 | 2.00749454 | 0.01 | 0 | 1 | 0.29941358 | 3.62088207 | 0.029 | 0.01 | 1 | 0.29941358 | 0.29167377 |
| Tmem177 | 0.15837881 | 2.00749454 | 0.01 | 0 | 1 | 0.51194506 | 0.27510724 | 0.021 | 0.01 | 1 | 0.51194506 | 0.29167377 |
| Il19 | 0.15837881 | 2.00749454 | 0.01 | 0 | 1 | 0.50887612 | -1.3098553 | 0.004 | 0.01 | 1 | 0.50887612 | 0.29167377 |
| Lrrn2 | 0.15837881 | 2.00749454 | 0.01 | 0 | 1 | 0.50887612 | -1.3098553 | 0.004 | 0.01 | 1 | 0.50887612 | 0.29167377 |
| Gm37571 | 0.15837881 | 2.00749454 | 0.01 | 0 | 1 | 0.67254361 | -0.3098553 | 0.017 | 0.01 | 1 | 0.67254361 | 0.29167377 |
| Gm38304 | 0.15837881 | 2.00749454 | 0.01 | 0 | 1 | 0.34026631 | -2.5322477 | 0.008 | 0.021 | 1 | 0.34026631 | 0.29167377 |
| Ifi209 | 0.15837881 | 4.00749454 | 0.01 | 0 | 1 | 0.27154471 | 0.69014474 | 0.013 | 0 | 1 | 0.27154471 | 0.29167377 |
| Tlr5 | 0.15837881 | 2.00749454 | 0.01 | 0 | 1 | 0.27154471 | 1.27510724 | 0.013 | 0 | 1 | 0.27154471 | 0.29167377 |
| Gm38155 | 0.15837881 | 2.00749454 | 0.01 | 0 | 1 | 0.86312513 | -0.7248928 | 0.008 | 0.01 | 1 | 0.86312513 | 0.29167377 |
| Ltk | 0.15837881 | 2.00749454 | 0.01 | 0 | 1 | 0.50887612 | -1.3098553 | 0.004 | 0.01 | 1 | 0.50887612 | 0.29167377 |
| Ralyl | 0.15837881 | 2.00749454 | 0.01 | 0 | 1 | 0.24194938 | -3.6317834 | 0.013 | 0.031 | 1 | 0.24194938 | 0.29167377 |
| Gm43024 | 0.15837881 | 2.00749454 | 0.01 | 0 | 1 | 0.87851725 | -0.8948178 | 0.013 | 0.01 | 1 | 0.87851725 | 0.29167377 |
| Gm42696 | 0.15837881 | 2.00749454 | 0.01 | 0 | 1 | 0.5818708 | -0.5728897 | 0.013 | 0.021 | 1 | 0.5818708 | 0.29167377 |
| Gm16958 | 0.15837881 | 2.00749454 | 0.01 | 0 | 1 | 0.86613737 | 0.27510724 | 0.008 | 0.01 | 1 | 0.86613737 | 0.29167377 |
| Gm19708 | 0.15837881 | 2.00749454 | 0.01 | 0 | 1 | 1 | -0.5025003 | 0.021 | 0.021 | 1 | 1 | 0.29167377 |
| Arhgef39 | 0.15837881 | 3.00749454 | 0.01 | 0 | 1 | 0.40076941 | -1.3098553 | 0.017 | 0.031 | 1 | 0.40076941 | 0.29167377 |
| Gm12454 | 0.15837881 | 2.59245704 | 0.01 | 0 | 1 | 0.86312513 | -0.7248928 | 0.008 | 0.01 | 1 | 0.86312513 | 0.29167377 |
| Gm44037 | 0.15837881 | 2.00749454 | 0.01 | 0 | 1 | 0.86312513 | -0.7248928 | 0.008 | 0.01 | 1 | 0.86312513 | 0.29167377 |
| Gm17114 | 0.15837881 | 2.00749454 | 0.01 | 0 | 1 | 0.50587544 | -2.3098553 | 0.004 | 0.01 | 1 | 0.50587544 | 0.29167377 |
| C1qa | 0.15837881 | 2.00749454 | 0.01 | 0 | 1 | 0.27154327 | 1.01207283 | 0.013 | 0 | 1 | 0.27154327 | 0.29167377 |
| Sh2d5 | 0.15837881 | 2.00749454 | 0.01 | 0 | 1 | 0.50887612 | -1.3098553 | 0.004 | 0.01 | 1 | 0.50887612 | 0.29167377 |
| Zfp268 | 0.15837881 | 2.59245704 | 0.01 | 0 | 1 | 0.20315163 | 1.01207283 | 0.017 | 0 | 1 | 0.20315163 | 0.29167377 |
| Gm13201 | 0.15837881 | 3.32942263 | 0.01 | 0 | 1 | 0.87851725 | -2.6317834 | 0.013 | 0.01 | 1 | 0.87851725 | 0.29167377 |
| 5830416I19Rik | 0.15837881 | 2.00749454 | 0.01 | 0 | 1 | 0.27154471 | 0.69014474 | 0.013 | 0 | 1 | 0.27154471 | 0.29167377 |

| | | | | | | | | | | | | |
|---|---|---|---|---|---|---|---|---|---|---|---|---|
| BC051076 | 0.15837881 | 2.00749454 | 0.01 | 0 | 1 | 1 | -0.3769695 | 0.063 | 0.062 | 1 | 1 | 0.29167377 |
| Epgn | 0.15837881 | 2.00749454 | 0.01 | 0 | 1 | 0.20315163 | 1.01207283 | 0.017 | 0 | 1 | 0.20315163 | 0.29167377 |
| Sdsl | 0.15837881 | 2.00749454 | 0.01 | 0 | 1 | 0.20315163 | 1.01207283 | 0.017 | 0 | 1 | 0.20315163 | 0.29167377 |
| Tspan33 | 0.15837881 | 2.00749454 | 0.01 | 0 | 1 | 0.34704016 | -1.3098553 | 0.008 | 0.021 | 1 | 0.34704016 | 0.29167377 |
| 1700111E14Rik | 0.15837881 | 2.00749454 | 0.01 | 0 | 1 | 0.50587544 | -1.8948178 | 0.004 | 0.01 | 1 | 0.50587544 | 0.29167377 |
| Snca | 0.15837881 | 2.00749454 | 0.01 | 0 | 1 | 0.86613737 | -0.3098553 | 0.008 | 0.01 | 1 | 0.86613737 | 0.29167377 |
| Igkc | 0.15837881 | 8.14704589 | 0.01 | 0 | 1 | 0.20315163 | 1.01207283 | 0.017 | 0 | 1 | 0.20315163 | 0.29167377 |
| Ctnna2 | 0.15837881 | 2.00749454 | 0.01 | 0 | 1 | 0.51391803 | -2.4618584 | 0.038 | 0.052 | 1 | 0.51391803 | 0.29167377 |
| Nrip2 | 0.15837881 | 2.00749454 | 0.01 | 0 | 1 | 0.20314753 | 1.27510724 | 0.017 | 0 | 1 | 0.20314753 | 0.29167377 |
| Clec12a | 0.15837881 | 2.00749454 | 0.01 | 0 | 1 | 0.30051742 | 3.16263251 | 0.029 | 0.01 | 1 | 0.30051742 | 0.29167377 |
| Sbk2 | 0.15837881 | 2.00749454 | 0.01 | 0 | 1 | 0.20315163 | 1.01207283 | 0.017 | 0 | 1 | 0.20315163 | 0.29167377 |
| Cyp2g1 | 0.15837881 | 2.00749454 | 0.01 | 0 | 1 | 0.45202946 | -0.5025003 | 0.076 | 0.052 | 1 | 0.45202946 | 0.29167377 |
| Tsks | 0.15837881 | 2.00749454 | 0.01 | 0 | 1 | 0.99194729 | -1.0874628 | 0.021 | 0.021 | 1 | 0.99194729 | 0.29167377 |
| Rasip1 | 0.15837881 | 2.00749454 | 0.01 | 0 | 1 | 0.50887612 | -1.3098553 | 0.004 | 0.01 | 1 | 0.50887612 | 0.29167377 |
| Acsm4 | 0.15837881 | 2.00749454 | 0.01 | 0 | 1 | 0.85710544 | -2.0468209 | 0.008 | 0.01 | 1 | 0.85710544 | 0.29167377 |
| Mrgprd | 0.15837881 | 2.00749454 | 0.01 | 0 | 1 | 0.86543325 | 0.49749966 | 0.013 | 0.01 | 1 | 0.86543325 | 0.29167377 |
| Tdrp | 0.15837881 | 2.59245704 | 0.01 | 0 | 1 | 0.27154471 | 0.69014474 | 0.013 | 0 | 1 | 0.27154471 | 0.29167377 |
| Gm16933 | 0.15837881 | 2.00749454 | 0.01 | 0 | 1 | 0.34704016 | -1.3098553 | 0.008 | 0.021 | 1 | 0.34704016 | 0.29167377 |
| Mmp8 | 0.15837881 | 2.00749454 | 0.01 | 0 | 1 | 0.79600816 | 0.46775232 | 0.017 | 0.021 | 1 | 0.79600816 | 0.29167377 |
| Cntn5 | 0.15837881 | 2.00749454 | 0.01 | 0 | 1 | 0.27154471 | 0.69014474 | 0.013 | 0 | 1 | 0.27154471 | 0.29167377 |
| Gpr83 | 0.15837881 | 2.00749454 | 0.01 | 0 | 1 | 0.87066325 | -0.3098553 | 0.013 | 0.01 | 1 | 0.87066325 | 0.29167377 |
| Scn3b | 0.15837881 | 2.00749454 | 0.01 | 0 | 1 | 0.27154327 | 1.49749966 | 0.013 | 0 | 1 | 0.27154327 | 0.29167377 |
| Gm10642 | 0.15837881 | 2.00749454 | 0.01 | 0 | 1 | 0.20314753 | 1.27510724 | 0.017 | 0 | 1 | 0.20314753 | 0.29167377 |
| Zmynd10 | 0.15837881 | 2.00749454 | 0.01 | 0 | 1 | 0.87066325 | -0.3098553 | 0.013 | 0.01 | 1 | 0.87066325 | 0.29167377 |
| 1700019L13Rik | 0.15837881 | 2.59245704 | 0.01 | 0 | 1 | 0.87851807 | -0.7248928 | 0.013 | 0.01 | 1 | 0.87851807 | 0.29167377 |
| A730085K08Rik | 0.15837881 | 2.00749454 | 0.01 | 0 | 1 | 0.20315163 | 1.01207283 | 0.017 | 0 | 1 | 0.20315163 | 0.29167377 |
| Gm34184 | 0.15837881 | 2.00749454 | 0.01 | 0 | 1 | 0.86312513 | -0.7248928 | 0.008 | 0.01 | 1 | 0.86312513 | 0.29167377 |
| Kcnab3 | 0.15837881 | 2.00749454 | 0.01 | 0 | 1 | 0.87066325 | -0.3098553 | 0.013 | 0.01 | 1 | 0.87066325 | 0.29167377 |
| Smtnl2 | 0.15837881 | 2.00749454 | 0.01 | 0 | 1 | 0.86613737 | -0.3098553 | 0.008 | 0.01 | 1 | 0.86613737 | 0.29167377 |
| 1700016K19Rik | 0.15837881 | 2.00749454 | 0.01 | 0 | 1 | 0.86312513 | -0.7248928 | 0.008 | 0.01 | 1 | 0.86312513 | 0.29167377 |
| AI662270 | 0.15837881 | 2.00749454 | 0.01 | 0 | 1 | 0.51018266 | 0.69014474 | 0.021 | 0.01 | 1 | 0.51018266 | 0.29167377 |
| Wfdc21 | 0.15837881 | 2.00749454 | 0.01 | 0 | 1 | 0.20314753 | 2.59703533 | 0.017 | 0 | 1 | 0.20314753 | 0.29167377 |
| Chct1 | 0.15837881 | 2.59245704 | 0.01 | 0 | 1 | 0.34704016 | -1.3098553 | 0.008 | 0.021 | 1 | 0.34704016 | 0.29167377 |
| Ccr7 | 0.15837881 | 2.00749454 | 0.01 | 0 | 1 | 0.50587544 | -1.8948178 | 0.004 | 0.01 | 1 | 0.50587544 | 0.29167377 |
| 4732490B19Rik | 0.15837881 | 2.00749454 | 0.01 | 0 | 1 | 0.34195151 | -1.8948178 | 0.008 | 0.021 | 1 | 0.34195151 | 0.29167377 |
| Gm47371 | 0.15837881 | 2.00749454 | 0.01 | 0 | 1 | 0.87851725 | -0.8948178 | 0.013 | 0.01 | 1 | 0.87851725 | 0.29167377 |
| Gm48383 | 0.15837881 | 2.00749454 | 0.01 | 0 | 1 | 0.51018266 | 0.49749966 | 0.021 | 0.01 | 1 | 0.51018266 | 0.29167377 |
| 4930408O17Rik | 0.15837881 | 2.59245704 | 0.01 | 0 | 1 | 0.50587544 | -2.3098553 | 0.004 | 0.01 | 1 | 0.50587544 | 0.29167377 |
| 1700014D04Rik | 0.15837881 | 2.00749454 | 0.01 | 0 | 1 | 0.50887612 | -1.3098553 | 0.004 | 0.01 | 1 | 0.50887612 | 0.29167377 |
| 4932411K12Rik | 0.15837881 | 2.00749454 | 0.01 | 0 | 1 | 0.27154327 | 1.01207283 | 0.013 | 0 | 1 | 0.27154327 | 0.29167377 |
| Cd180 | 0.15837881 | 3.81484946 | 0.01 | 0 | 1 | 0.66387437 | 0.27510724 | 0.017 | 0.01 | 1 | 0.66387437 | 0.29167377 |
| Asb14 | 0.15837881 | 2.00749454 | 0.01 | 0 | 1 | 0.40076941 | -3.1578522 | 0.017 | 0.031 | 1 | 0.40076941 | 0.29167377 |
| Wdfy4 | 0.15837881 | 4.46692616 | 0.01 | 0 | 1 | 0.50887612 | -1.3098553 | 0.004 | 0.01 | 1 | 0.50887612 | 0.29167377 |
| 2310043O21Rik | 0.15837881 | 2.00749454 | 0.01 | 0 | 1 | 0.50887612 | -1.3098553 | 0.004 | 0.01 | 1 | 0.50887612 | 0.29167377 |
| Slurp2 | 0.15837881 | 2.00749454 | 0.01 | 0 | 1 | 0.66387437 | 0.27510724 | 0.017 | 0.01 | 1 | 0.66387437 | 0.29167377 |
| Mapk15 | 0.15837881 | 2.00749454 | 0.01 | 0 | 1 | 0.24413474 | -2.1172102 | 0.013 | 0.031 | 1 | 0.24413474 | 0.29167377 |
| AU022754 | 0.15837881 | 2.00749454 | 0.01 | 0 | 1 | 0.27154471 | 0.69014474 | 0.013 | 0 | 1 | 0.27154471 | 0.29167377 |
| Muc19 | 0.15837881 | 2.00749454 | 0.01 | 0 | 1 | 0.27154471 | 0.69014474 | 0.013 | 0 | 1 | 0.27154471 | 0.29167377 |
| Krt87 | 0.15837881 | 2.00749454 | 0.01 | 0 | 1 | 0.20315163 | 1.01207283 | 0.017 | 0 | 1 | 0.20315163 | 0.29167377 |
| Klhl6 | 0.15837881 | 2.59245704 | 0.01 | 0 | 1 | 0.67471359 | -0.5728897 | 0.017 | 0.01 | 1 | 0.67471359 | 0.29167377 |
| Muc20 | 0.15837881 | 2.00749454 | 0.01 | 0 | 1 | 0.27154471 | 0.69014474 | 0.013 | 0 | 1 | 0.27154471 | 0.29167377 |
| 1700010I14Rik | 0.15837881 | 2.00749454 | 0.01 | 0 | 1 | 0.67471693 | -0.7248928 | 0.017 | 0.01 | 1 | 0.67471693 | 0.29167377 |
| Dcpp3 | 0.15837881 | 2.00749454 | 0.01 | 0 | 1 | 0.27154471 | 0.69014474 | 0.013 | 0 | 1 | 0.27154471 | 0.29167377 |
| Prss27 | 0.15837881 | 2.00749454 | 0.01 | 0 | 1 | 0.91934726 | -0.3098553 | 0.034 | 0.031 | 1 | 0.91934726 | 0.29167377 |
| Gm22146 | 0.15837881 | 2.00749454 | 0.01 | 0 | 1 | 0.86312513 | -0.7248928 | 0.008 | 0.01 | 1 | 0.86312513 | 0.29167377 |
| C4a | 0.15837881 | 2.00749454 | 0.01 | 0 | 1 | 0.51018266 | 0.49749966 | 0.021 | 0.01 | 1 | 0.51018266 | 0.29167377 |
| Gm50397 | 0.15837881 | 2.00749454 | 0.01 | 0 | 1 | 0.86312513 | -0.7248928 | 0.008 | 0.01 | 1 | 0.86312513 | 0.29167377 |
| Kcnn2 | 0.15837881 | 2.00749454 | 0.01 | 0 | 1 | 0.85710544 | -2.0468209 | 0.008 | 0.01 | 1 | 0.85710544 | 0.29167377 |
| Cd5 | 0.15837881 | 2.00749454 | 0.01 | 0 | 1 | 0.39515719 | 0.49749966 | 0.025 | 0.01 | 1 | 0.39515719 | 0.29167377 |
| Slit1 | 0.15837881 | 2.00749454 | 0.01 | 0 | 1 | 0.34363875 | -2.0468209 | 0.008 | 0.021 | 1 | 0.34363875 | 0.29167377 |
| Gm50100 | 0.15837881 | 2.00749454 | 0.01 | 0 | 1 | 0.82231716 | -0.4618584 | 0.025 | 0.021 | 1 | 0.82231716 | 0.29167377 |
| Pnliprp1 | 0.15837881 | 2.00749454 | 0.01 | 0 | 1 | 0.86312513 | -0.7248928 | 0.008 | 0.01 | 1 | 0.86312513 | 0.29167377 |
| Slc6a14 | 0.15837881 | 2.59245704 | 0.01 | 0 | 1 | 0.20315163 | 1.01207283 | 0.017 | 0 | 1 | 0.20315163 | 0.29167377 |
| Xlr4a | 0.15837881 | 3.00749454 | 0.01 | 0 | 1 | 0.50887612 | -1.3098553 | 0.004 | 0.01 | 1 | 0.50887612 | 0.29167377 |
| Magee2 | 0.15837881 | 2.00749454 | 0.01 | 0 | 1 | 0.86312513 | -0.7248928 | 0.008 | 0.01 | 1 | 0.86312513 | 0.29167377 |
| Numb | 0.43581633 | -0.407543 | 0.562 | 0.575 | 1 | 0.15858829 | -0.1696776 | 0.752 | 0.865 | 1 | 0.43581633 | 0.29202634 |
| Ccni | 0.15871007 | 0.15239074 | 0.896 | 0.845 | 1 | 0.73796945 | -0.2410164 | 0.891 | 0.885 | 1 | 0.73796945 | 0.29223125 |
| Tspo | 0.15871165 | -0.4247327 | 0.458 | 0.528 | 1 | 0.19740466 | 0.36285013 | 0.66 | 0.688 | 1 | 0.19740466 | 0.29223391 |
| Tbccd1 | 0.15881586 | 0.53800925 | 0.125 | 0.073 | 1 | 0.49807522 | -0.3838558 | 0.176 | 0.135 | 1 | 0.49807522 | 0.29240925 |
| 2610206C17Rik | 0.15890071 | -1.7294711 | 0.021 | 0.057 | 1 | 0.1885774 | 0.93807225 | 0.038 | 0.01 | 1 | 0.1885774 | 0.29255199 |
| Zfand2b | 0.18093892 | 0.35979628 | 0.281 | 0.207 | 1 | 0.15890897 | 0.77760758 | 0.172 | 0.115 | 1 | 0.18093892 | 0.29256589 |
| Arl4d | 0.15893404 | -1.4705528 | 0.052 | 0.098 | 1 | 0.35906112 | 0.77760758 | 0.055 | 0.031 | 1 | 0.35906112 | 0.29260805 |
| Kctd6 | 0.2878968 | -0.7029988 | 0.094 | 0.135 | 1 | 0.15893473 | 0.25649156 | 0.231 | 0.156 | 1 | 0.2878968 | 0.29260922 |
| Phldb2 | 0.15898206 | -0.7250252 | 0.271 | 0.347 | 1 | 0.2570995 | -0.2988563 | 0.55 | 0.448 | 1 | 0.2570995 | 0.29268882 |
| Gm42941 | 0.25012481 | -0.2726134 | 0.083 | 0.13 | 1 | 0.15898898 | -0.4029647 | 0.118 | 0.062 | 1 | 0.25012481 | 0.29270046 |
| 4930447C04Rik | 0.66829458 | -0.4779323 | 0.083 | 0.067 | 1 | 0.15901555 | 1.61614416 | 0.084 | 0.042 | 1 | 0.66829458 | 0.29274516 |
| Adpgk | 0.29985927 | 0.35541784 | 0.365 | 0.301 | 1 | 0.15902656 | 0.32955502 | 0.277 | 0.198 | 1 | 0.29985927 | 0.29276367 |
| Gm17092 | 0.15907563 | 0.53105649 | 0.188 | 0.124 | 1 | 0.73498356 | -0.1034044 | 0.097 | 0.083 | 1 | 0.73498356 | 0.29284621 |

| | | | | | | | | | | | | |
|---|---|---|---|---|---|---|---|---|---|---|---|---|
| Adora1 | 0.15909079 | 0.50999488 | 0.104 | 0.057 | 1 | 0.425508 | -0.681824 | 0.042 | 0.062 | 1 | 0.425508 | 0.29287171 |
| 4930599N23Rik | 0.56169566 | 0.44189736 | 0.167 | 0.14 | 1 | 0.15914668 | 0.13071733 | 0.181 | 0.115 | 1 | 0.56169566 | 0.2929657 |
| Mkx | 0.15922047 | -1.7258598 | 0.115 | 0.176 | 1 | 0.78273476 | -0.5936482 | 0.097 | 0.104 | 1 | 0.78273476 | 0.29308979 |
| Ciao3 | 0.41559247 | -0.4950058 | 0.052 | 0.078 | 1 | 0.15924025 | 0.32757466 | 0.101 | 0.052 | 1 | 0.41559247 | 0.29312304 |
| Chfr | 0.15933761 | -0.4675903 | 0.365 | 0.425 | 1 | 0.28361904 | -0.5314517 | 0.584 | 0.573 | 1 | 0.28361904 | 0.29328675 |
| Eif4e3 | 0.62373932 | -0.5070786 | 0.156 | 0.171 | 1 | 0.1594762 | 0.17889192 | 0.223 | 0.146 | 1 | 0.62373932 | 0.29351974 |
| Neil1 | 0.25337721 | 1.00749454 | 0.052 | 0.026 | 1 | 0.15976273 | 1.08246216 | 0.071 | 0.031 | 1 | 0.25337721 | 0.29400132 |
| Foxred1 | 0.1599705 | 0.52206771 | 0.26 | 0.192 | 1 | 0.24153971 | 0.16407593 | 0.168 | 0.115 | 1 | 0.24153971 | 0.29435043 |
| Gm13684 | 0.16011721 | -1.7294711 | 0.021 | 0.057 | 1 | 0.82054184 | -0.6883669 | 0.067 | 0.073 | 1 | 0.82054184 | 0.2945969 |
| Ybey | 0.16011721 | -1.407543 | 0.021 | 0.057 | 1 | 0.24607399 | 0.69014474 | 0.063 | 0.031 | 1 | 0.24607399 | 0.2945969 |
| Cyp2j9 | 0.16011761 | -1.3144336 | 0.021 | 0.057 | 1 | 0.87066325 | -0.3098553 | 0.013 | 0.01 | 1 | 0.87066325 | 0.29459757 |
| Prdm1 | 0.16011761 | -1.577468 | 0.021 | 0.057 | 1 | 0.64087546 | 2.86006974 | 0.029 | 0.021 | 1 | 0.64087546 | 0.29459757 |
| Trpv2 | 0.16016958 | -2.1624305 | 0.021 | 0.057 | 1 | 0.62710605 | 1.14957636 | 0.042 | 0.031 | 1 | 0.62710605 | 0.29468486 |
| Zfp612 | 0.16025112 | 1.59245704 | 0.073 | 0.036 | 1 | 0.64175181 | 0.45567948 | 0.042 | 0.031 | 1 | 0.64175181 | 0.29482182 |
| Anln | 0.16032059 | 1.90057933 | 0.073 | 0.036 | 1 | 0.50844268 | 0.86006974 | 0.021 | 0.01 | 1 | 0.50844268 | 0.29493849 |
| Nkiras2 | 0.42972185 | 0.45110119 | 0.146 | 0.114 | 1 | 0.16046501 | 0.61614416 | 0.126 | 0.073 | 1 | 0.42972185 | 0.295181 |
| Spata24 | 0.91299952 | 0.40177348 | 0.156 | 0.155 | 1 | 0.1605255 | 0.23446525 | 0.13 | 0.073 | 1 | 0.91299952 | 0.29528257 |
| Gde1 | 0.16072879 | 0.35541784 | 0.458 | 0.383 | 1 | 0.31157952 | 0.22619764 | 0.559 | 0.51 | 1 | 0.31157952 | 0.29562384 |
| Osbpl10 | 0.16074765 | 0.73619251 | 0.177 | 0.119 | 1 | 0.88533133 | -0.602637 | 0.345 | 0.312 | 1 | 0.88533133 | 0.2956555 |
| Sertad1 | 0.16084548 | 0.22596339 | 0.573 | 0.508 | 1 | 0.52926644 | -0.4906587 | 0.714 | 0.594 | 1 | 0.52926644 | 0.29581969 |
| Plekhh3 | 0.8423867 | 0.35541784 | 0.083 | 0.093 | 1 | 0.16097421 | 0.69014474 | 0.139 | 0.083 | 1 | 0.8423867 | 0.29603573 |
| Crnde | 0.16098562 | -0.9467018 | 0.062 | 0.114 | 1 | 0.5034108 | -0.194378 | 0.088 | 0.062 | 1 | 0.5034108 | 0.29605486 |
| Zbtb22 | 0.41614867 | -0.407543 | 0.083 | 0.114 | 1 | 0.16098729 | 0.59703533 | 0.164 | 0.104 | 1 | 0.41614867 | 0.29605768 |
| Exosc9 | 0.16122417 | 0.14499806 | 0.396 | 0.295 | 1 | 0.4592946 | -0.1143045 | 0.248 | 0.198 | 1 | 0.4592946 | 0.29645511 |
| Tmem161b | 0.16127483 | 0.31717904 | 0.333 | 0.249 | 1 | 0.30223659 | -0.4353861 | 0.328 | 0.24 | 1 | 0.30223659 | 0.29654009 |
| Dll1 | 0.1613372 | -1.4950058 | 0.021 | 0.057 | 1 | 0.5778028 | -0.8948178 | 0.013 | 0.021 | 1 | 0.5778028 | 0.29664471 |
| Tbc1d5 | 0.29111399 | 0.17882911 | 0.646 | 0.606 | 1 | 0.16167153 | -0.5008525 | 0.647 | 0.74 | 1 | 0.29111399 | 0.29720538 |
| Cnrip1 | 0.16178422 | 0.79336973 | 0.135 | 0.083 | 1 | 0.43250941 | 0.23763253 | 0.113 | 0.083 | 1 | 0.43250941 | 0.29739431 |
| Exoc7 | 0.16191739 | 0.64071221 | 0.292 | 0.223 | 1 | 0.84631921 | -0.1469167 | 0.294 | 0.271 | 1 | 0.84631921 | 0.29761754 |
| Zfp983 | 0.16212091 | 0.52206771 | 0.135 | 0.083 | 1 | 0.50193937 | 0.37820073 | 0.084 | 0.062 | 1 | 0.50193937 | 0.29795863 |
| Kank4 | 0.684893 | 0.42253204 | 0.052 | 0.041 | 1 | 0.16222678 | -0.9879272 | 0.038 | 0.073 | 1 | 0.684893 | 0.29813604 |
| 3110009E18Rik | 0.16249014 | 1.24195979 | 0.115 | 0.067 | 1 | 0.61679385 | -0.8123556 | 0.08 | 0.094 | 1 | 0.61679385 | 0.29857723 |
| AU020206 | 0.81336205 | -0.2148979 | 0.094 | 0.083 | 1 | 0.16252298 | -1.2067618 | 0.088 | 0.135 | 1 | 0.81336205 | 0.29863224 |
| Efcab2 | 0.16253486 | 0.37522632 | 0.177 | 0.114 | 1 | 0.78760813 | -0.7655347 | 0.118 | 0.104 | 1 | 0.78760813 | 0.29865214 |
| Rnft2 | 0.16255946 | -1.2148979 | 0.021 | 0.057 | 1 | 0.34232469 | 0.91253716 | 0.042 | 0.021 | 1 | 0.34232469 | 0.29869335 |
| Man2c1os | 0.86599256 | -0.4872702 | 0.062 | 0.067 | 1 | 0.16257073 | -0.7041342 | 0.13 | 0.073 | 1 | 0.86599256 | 0.29871221 |
| Tyro3 | 0.21867159 | 1.59245704 | 0.021 | 0.005 | 1 | 0.16268844 | 0.86006974 | 0.071 | 0.031 | 1 | 0.21867159 | 0.29890936 |
| Hexim2 | 0.31880524 | 0.49292136 | 0.062 | 0.036 | 1 | 0.16268844 | 0.86006974 | 0.071 | 0.031 | 1 | 0.31880524 | 0.29890936 |
| Pik3ip1 | 0.16292923 | -0.5399933 | 0.24 | 0.311 | 1 | 0.9087643 | -0.3098553 | 0.387 | 0.365 | 1 | 0.9087643 | 0.29931252 |
| Zfp369 | 0.25056486 | 0.67734593 | 0.208 | 0.155 | 1 | 0.16297022 | -0.1172102 | 0.29 | 0.188 | 1 | 0.25056486 | 0.29938114 |
| Mrrf | 0.17360469 | 0.4959942 | 0.365 | 0.29 | 1 | 0.16300045 | 0.16658278 | 0.21 | 0.135 | 1 | 0.17360469 | 0.29943176 |
| Runx2os1 | 0.24244498 | -0.6205367 | 0.083 | 0.13 | 1 | 0.16304221 | 0.35310975 | 0.143 | 0.083 | 1 | 0.24244498 | 0.29950166 |
| Rgp1 | 0.62973467 | 0.14951354 | 0.146 | 0.124 | 1 | 0.16311639 | 0.69014474 | 0.126 | 0.073 | 1 | 0.62973467 | 0.29962582 |
| Pcnt | 0.6657551 | 0.16221713 | 0.281 | 0.311 | 1 | 0.16322926 | -0.5383324 | 0.42 | 0.49 | 1 | 0.6657551 | 0.29981472 |
| Sulf1 | 0.16344257 | -0.2797174 | 0.177 | 0.249 | 1 | 0.39868276 | -1.361886 | 0.227 | 0.26 | 1 | 0.39868276 | 0.30017167 |
| Gm56815 | 0.33585318 | -0.6929452 | 0.031 | 0.057 | 1 | 0.16349292 | 0.32241295 | 0.113 | 0.062 | 1 | 0.33585318 | 0.30025591 |
| Mad2l2 | 0.86902654 | -0.1624305 | 0.156 | 0.161 | 1 | 0.16349292 | 0.47441605 | 0.113 | 0.062 | 1 | 0.86902654 | 0.30025591 |
| Plekhg2 | 0.16351036 | -0.4213488 | 0.25 | 0.332 | 1 | 0.22737548 | -1.1723517 | 0.319 | 0.333 | 1 | 0.22737548 | 0.30028509 |
| Mfsd11 | 0.20381599 | 0.12297175 | 0.375 | 0.275 | 1 | 0.16358168 | -0.1794586 | 0.445 | 0.302 | 1 | 0.20381599 | 0.30040439 |
| Abca2 | 0.66850407 | 0.21394541 | 0.115 | 0.098 | 1 | 0.16366162 | 0.59703533 | 0.164 | 0.104 | 1 | 0.66850407 | 0.30053812 |
| Gemin5 | 0.28408411 | 0.32523483 | 0.302 | 0.244 | 1 | 0.16388474 | -0.3349462 | 0.294 | 0.208 | 1 | 0.28408411 | 0.30091128 |
| Igf2bp1 | 0.163901 | -1.1624305 | 0.021 | 0.057 | 1 | 0.52037348 | -0.8948178 | 0.046 | 0.062 | 1 | 0.52037348 | 0.30093846 |
| Ppp6c | 0.16406125 | 0.23377039 | 0.594 | 0.503 | 1 | 0.36642581 | -0.152314 | 0.735 | 0.677 | 1 | 0.36642581 | 0.30120641 |
| Tamalin | 0.36754634 | -0.6576383 | 0.219 | 0.259 | 1 | 0.1641945 | -0.612418 | 0.273 | 0.344 | 1 | 0.36754634 | 0.30142917 |
| Erich6 | 0.16449998 | 1.29700115 | 0.073 | 0.036 | 1 | 0.27525593 | 0.34222143 | 0.076 | 0.042 | 1 | 0.27525593 | 0.30193971 |
| Pramel12 | 0.99127113 | -0.1445086 | 0.083 | 0.083 | 1 | 0.1645118 | 0.14957636 | 0.13 | 0.073 | 1 | 0.99127113 | 0.30195947 |
| Zbtb18 | 0.16476362 | -0.4330781 | 0.031 | 0.073 | 1 | 0.70249915 | -0.2394659 | 0.076 | 0.062 | 1 | 0.70249915 | 0.30238018 |
| Ireb2 | 0.16477544 | 0.1486706 | 0.646 | 0.544 | 1 | 0.54136225 | -0.2943423 | 0.723 | 0.708 | 1 | 0.54136225 | 0.30239993 |
| Gm38882 | 0.16487118 | 0.86310463 | 0.146 | 0.093 | 1 | 0.80526491 | 0.10518224 | 0.059 | 0.052 | 1 | 0.80526491 | 0.30255985 |
| Gm12703 | 0.16492424 | -1.7294711 | 0.083 | 0.135 | 1 | 0.2977194 | -0.9399057 | 0.231 | 0.167 | 1 | 0.2977194 | 0.30264847 |
| Bcl2l13 | 0.16493672 | 0.27814213 | 0.312 | 0.228 | 1 | 0.74380291 | -0.3778737 | 0.387 | 0.375 | 1 | 0.74380291 | 0.30266932 |
| Cenps | 0.16497353 | 1.22988696 | 0.094 | 0.052 | 1 | 0.27596631 | 0.91253716 | 0.046 | 0.021 | 1 | 0.27596631 | 0.30273079 |
| Arhgap27 | 0.72021678 | -0.7998604 | 0.01 | 0.016 | 1 | 0.16501219 | -0.7952821 | 0.029 | 0.062 | 1 | 0.72021678 | 0.30279536 |
| Larp4b | 0.16507593 | 0.22211181 | 0.875 | 0.839 | 1 | 0.19612429 | -0.393291 | 0.912 | 0.917 | 1 | 0.19612429 | 0.3029018 |
| Flt3l | 0.99482536 | 0.15949763 | 0.042 | 0.041 | 1 | 0.16518364 | 0.32241295 | 0.101 | 0.052 | 1 | 0.99482536 | 0.30308164 |
| Wac | 0.18803008 | 0.14890251 | 0.969 | 0.943 | 1 | 0.16523386 | -0.1866872 | 0.983 | 0.99 | 1 | 0.18803008 | 0.3031655 |
| Iqck | 0.16527087 | -0.8405024 | 0.031 | 0.073 | 1 | 0.33773619 | -0.4473588 | 0.055 | 0.083 | 1 | 0.33773619 | 0.30322728 |
| Zbtb49 | 0.18254469 | 0.29700115 | 0.094 | 0.052 | 1 | 0.16527501 | 0.42711033 | 0.113 | 0.062 | 1 | 0.18254469 | 0.30323419 |
| Mettl14 | 0.42972185 | 0.45110119 | 0.146 | 0.114 | 1 | 0.16534627 | 0.24268576 | 0.143 | 0.083 | 1 | 0.42972185 | 0.30335314 |
| Zfp141 | 0.1653635 | -1.1624305 | 0.052 | 0.098 | 1 | 0.680146 | -1.0785297 | 0.092 | 0.104 | 1 | 0.680146 | 0.30338191 |
| Knstrn | 0.16571594 | 2.88196365 | 0.042 | 0.016 | 1 | 0.86312513 | -0.7248928 | 0.008 | 0.01 | 1 | 0.86312513 | 0.30397011 |
| Fzd10os | 0.42281098 | 0.85549144 | 0.062 | 0.041 | 1 | 0.16573034 | -0.129283 | 0.202 | 0.125 | 1 | 0.42281098 | 0.30399413 |
| D730003I15Rik | 0.20433883 | 1.37006462 | 0.062 | 0.031 | 1 | 0.16589208 | 0.49749966 | 0.101 | 0.052 | 1 | 0.20433883 | 0.30426398 |
| Apool | 0.1659495 | -0.2609943 | 0.135 | 0.207 | 1 | 0.28156773 | 0.12454756 | 0.252 | 0.188 | 1 | 0.28156773 | 0.30435977 |
| Gm56830 | 0.16600671 | 0.47315811 | 0.302 | 0.228 | 1 | 0.90443355 | -0.2120079 | 0.286 | 0.26 | 1 | 0.90443355 | 0.3044552 |
| Vav2 | 0.87069473 | 0.10060394 | 0.146 | 0.14 | 1 | 0.16614939 | -1.188549 | 0.189 | 0.24 | 1 | 0.87069473 | 0.30469315 |
| H2ac8 | 0.16635301 | 3.56820949 | 0.073 | 0.036 | 1 | 0.87066325 | -0.3098553 | 0.013 | 0.01 | 1 | 0.87066325 | 0.30503269 |
| Vac14 | 0.16654257 | 0.75956702 | 0.135 | 0.083 | 1 | 0.43419136 | -1.0170735 | 0.168 | 0.188 | 1 | 0.43419136 | 0.30534872 |
| Fxn | 0.49177058 | 0.14951354 | 0.146 | 0.114 | 1 | 0.16659213 | 0.13492958 | 0.164 | 0.104 | 1 | 0.49177058 | 0.30543132 |
| Tet3 | 0.66590323 | 0.13796447 | 0.49 | 0.497 | 1 | 0.16688563 | -0.4418508 | 0.777 | 0.875 | 1 | 0.66590323 | 0.30592045 |
| F8a | 0.16695275 | -0.4950058 | 0.031 | 0.073 | 1 | 0.24012143 | 1.14957636 | 0.063 | 0.031 | 1 | 0.24012143 | 0.30603228 |
| Necap1 | 0.16695925 | 0.58368683 | 0.302 | 0.238 | 1 | 0.54969687 | -0.2062326 | 0.34 | 0.281 | 1 | 0.54969687 | 0.30604311 |
| P4htm | 0.61005857 | -0.6013147 | 0.177 | 0.192 | 1 | 0.16709856 | -1.1755542 | 0.155 | 0.208 | 1 | 0.61005857 | 0.3062752 |
| Zhx2 | 0.70818358 | -0.2769589 | 0.719 | 0.746 | 1 | 0.16717907 | -0.4017778 | 0.811 | 0.865 | 1 | 0.70818358 | 0.3064093 |
| Zgpat | 0.20471521 | 0.73619251 | 0.208 | 0.15 | 1 | 0.16742605 | -0.1560499 | 0.294 | 0.198 | 1 | 0.20471521 | 0.30682063 |
| Stk39 | 0.1674934 | -0.6053411 | 0.906 | 0.87 | 1 | 0.22169398 | -0.6102419 | 0.891 | 0.927 | 1 | 0.22169398 | 0.30693276 |
| Rars | 0.47985595 | 0.11996927 | 0.531 | 0.472 | 1 | 0.16757212 | 0.1462095 | 0.567 | 0.51 | 1 | 0.47985595 | 0.30706383 |

| | | | | | | | | | | | | |
|---|---|---|---|---|---|---|---|---|---|---|---|---|
| Cenpl | 0.16758426 | 1.27052894 | 0.073 | 0.036 | 1 | 0.18896559 | -0.8244284 | 0.046 | 0.083 | 1 | 0.18896559 | 0.30708403 |
| Gm11476 | 0.98007198 | 0.15949763 | 0.062 | 0.062 | 1 | 0.16765222 | 0.34222143 | 0.071 | 0.031 | 1 | 0.98007198 | 0.30719718 |
| Erich1 | 0.16772902 | 0.54183096 | 0.167 | 0.109 | 1 | 0.58882812 | -0.5025003 | 0.139 | 0.115 | 1 | 0.58882812 | 0.30732501 |
| Rnf103 | 0.1679089 | -0.393289 | 0.542 | 0.606 | 1 | 0.88136503 | -0.2248192 | 0.626 | 0.583 | 1 | 0.88136503 | 0.30762441 |
| Nudt14 | 0.16812445 | -0.577468 | 0.083 | 0.14 | 1 | 0.39018934 | 0.20471791 | 0.08 | 0.052 | 1 | 0.39018934 | 0.30798308 |
| Gm50333 | 0.16834881 | 0.59245704 | 0.021 | 0.057 | 1 | 0.90183801 | -0.1578522 | 0.059 | 0.062 | 1 | 0.90183801 | 0.3083563 |
| Tbk1 | 0.16844879 | 0.22322323 | 0.49 | 0.399 | 1 | 0.36895265 | -0.2457249 | 0.618 | 0.667 | 1 | 0.36895265 | 0.30852259 |
| Styx | 0.40290451 | 0.22120123 | 0.344 | 0.295 | 1 | 0.16847797 | -0.2587164 | 0.441 | 0.302 | 1 | 0.40290451 | 0.30857111 |
| Thg1l | 0.31806045 | 1.00749454 | 0.094 | 0.062 | 1 | 0.1685647 | 0.20471791 | 0.071 | 0.031 | 1 | 0.31806045 | 0.30871534 |
| Pcyt2 | 0.16861736 | -0.3241815 | 0.521 | 0.601 | 1 | 0.84611574 | -0.3597312 | 0.412 | 0.365 | 1 | 0.84611574 | 0.30880291 |
| Tmem40 | 0.1687351 | 1.17741954 | 0.073 | 0.036 | 1 | 0.30382073 | 1.14957636 | 0.029 | 0.01 | 1 | 0.30382073 | 0.30899866 |
| Cdc7 | 0.16879021 | 1.15949763 | 0.073 | 0.036 | 1 | 0.82344487 | -0.3098553 | 0.059 | 0.052 | 1 | 0.82344487 | 0.30909028 |
| Lrp11 | 0.16879021 | 1.15949763 | 0.073 | 0.036 | 1 | 0.2144911 | -1.0874628 | 0.025 | 0.052 | 1 | 0.2144911 | 0.30909028 |
| Grin2d | 1 | -0.265524 | 0.104 | 0.104 | 1 | 0.16887206 | 0.32757466 | 0.113 | 0.062 | 1 | 1 | 0.30922635 |
| Vwa1 | 0.95978223 | -0.2148979 | 0.135 | 0.135 | 1 | 0.16900531 | 0.61614416 | 0.071 | 0.031 | 1 | 0.95978223 | 0.30944783 |
| Slc2a13 | 0.16917176 | 0.8206164 | 0.344 | 0.28 | 1 | 0.95155817 | -0.5311451 | 0.311 | 0.292 | 1 | 0.95155817 | 0.30972444 |
| Pthlh | 0.25379393 | -2.0799683 | 0.031 | 0.062 | 1 | 0.16938307 | -2.4391383 | 0.046 | 0.083 | 1 | 0.25379393 | 0.31007552 |
| Wdr12 | 0.29186105 | 0.36282579 | 0.406 | 0.352 | 1 | 0.16940143 | -0.2299317 | 0.437 | 0.312 | 1 | 0.29186105 | 0.31010601 |
| Gan | 0.98518068 | 0.15663798 | 0.354 | 0.358 | 1 | 0.16956413 | -0.1644248 | 0.416 | 0.281 | 1 | 0.98518068 | 0.31037626 |
| Uchl1 | 0.7854853 | -0.8669746 | 0.021 | 0.026 | 1 | 0.16972763 | 2.46273424 | 0.055 | 0.021 | 1 | 0.7854853 | 0.3106478 |
| Gm15853 | 0.17002829 | 1.81484946 | 0.042 | 0.016 | 1 | 0.66387437 | 0.27510724 | 0.017 | 0.01 | 1 | 0.66387437 | 0.31114696 |
| Gatad2b | 0.17008694 | -0.2729888 | 0.875 | 0.839 | 1 | 0.28259179 | -0.3476345 | 0.962 | 0.969 | 1 | 0.28259179 | 0.31124432 |
| Zfp958 | 0.17019364 | 0.48393258 | 0.156 | 0.098 | 1 | 0.90024482 | -0.3838558 | 0.122 | 0.125 | 1 | 0.90024482 | 0.3114214 |
| Iah1 | 0.3684915 | -0.4779323 | 0.208 | 0.249 | 1 | 0.17029414 | -0.1082214 | 0.29 | 0.198 | 1 | 0.3684915 | 0.31158819 |
| Rnf114 | 0.17051032 | 0.4367459 | 0.417 | 0.352 | 1 | 0.18210909 | -0.266001 | 0.496 | 0.354 | 1 | 0.18210909 | 0.31194687 |
| Mettl25 | 0.17083556 | 0.94737354 | 0.25 | 0.192 | 1 | 0.41296957 | -0.6883669 | 0.235 | 0.177 | 1 | 0.41296957 | 0.31248634 |
| Zfp664 | 0.17123612 | 0.24610547 | 0.51 | 0.42 | 1 | 0.60261044 | -0.2416838 | 0.664 | 0.74 | 1 | 0.60261044 | 0.31315042 |
| Casc3 | 0.17136574 | 0.26242389 | 0.49 | 0.394 | 1 | 0.56182126 | -0.4403252 | 0.588 | 0.615 | 1 | 0.56182126 | 0.31336526 |
| Gm48877 | 0.17178928 | 1.88196365 | 0.042 | 0.016 | 1 | 0.86613737 | -0.3098553 | 0.008 | 0.01 | 1 | 0.86613737 | 0.314067 |
| Myo1f | 0.33113577 | -0.577468 | 0.062 | 0.036 | 1 | 0.17201499 | 0.27510724 | 0.071 | 0.031 | 1 | 0.33113577 | 0.31444082 |
| Gtpbp2 | 0.50932182 | -0.25785 | 0.375 | 0.409 | 1 | 0.17234004 | -0.2251929 | 0.5 | 0.365 | 1 | 0.50932182 | 0.31497898 |
| Akna | 0.17254227 | 0.91438513 | 0.094 | 0.052 | 1 | 0.34222591 | -0.152314 | 0.084 | 0.052 | 1 | 0.34222591 | 0.31531371 |
| Odf3l1 | 0.17257326 | 1.24850264 | 0.073 | 0.036 | 1 | 0.65163905 | -0.2223924 | 0.101 | 0.083 | 1 | 0.65163905 | 0.31536499 |
| Gm45250 | 0.17265227 | 1.59245704 | 0.042 | 0.016 | 1 | 0.24396338 | 0.42711033 | 0.034 | 0.01 | 1 | 0.24396338 | 0.31549574 |
| Gm16853 | 0.17265227 | 1.59245704 | 0.042 | 0.016 | 1 | 0.51532169 | -0.3098553 | 0.097 | 0.073 | 1 | 0.51532169 | 0.31549574 |
| BC037032 | 0.17265227 | 1.59245704 | 0.042 | 0.016 | 1 | 0.27154471 | 0.69014474 | 0.013 | 0 | 1 | 0.27154471 | 0.31549574 |
| Zfp677 | 0.17265227 | 1.59245704 | 0.042 | 0.016 | 1 | 0.93221765 | -0.5322477 | 0.034 | 0.031 | 1 | 0.93221765 | 0.31549574 |
| Slc7a8 | 0.47912569 | -0.407543 | 0.021 | 0.036 | 1 | 0.17272847 | -1.6094155 | 0.046 | 0.083 | 1 | 0.47912569 | 0.31562181 |
| 2610044O15Rik8 | 0.17302468 | 0.34452952 | 0.188 | 0.124 | 1 | 0.53176845 | -0.9728203 | 0.084 | 0.104 | 1 | 0.53176845 | 0.31611182 |
| Il31ra | 0.37343246 | -1.407543 | 0.052 | 0.078 | 1 | 0.17321968 | -1.3098553 | 0.046 | 0.083 | 1 | 0.37343246 | 0.31643431 |
| 6230426M11Rik | 0.17327047 | 1.00749454 | 0.073 | 0.036 | 1 | 0.909702 | 0.14957636 | 0.034 | 0.031 | 1 | 0.909702 | 0.31651828 |
| 2010009K17Rik | 0.20224701 | -3.0799683 | 0.01 | 0.036 | 1 | 0.17334171 | -2.4253325 | 0.017 | 0.042 | 1 | 0.20224701 | 0.31663607 |
| Tep1 | 0.36552019 | 0.21394541 | 0.125 | 0.088 | 1 | 0.17359081 | -0.1475838 | 0.155 | 0.094 | 1 | 0.36552019 | 0.31704784 |
| Upp1 | 0.17364852 | -0.1989563 | 0.083 | 0.145 | 1 | 0.17479874 | 0.62874419 | 0.071 | 0.031 | 1 | 0.17479874 | 0.31714322 |
| Tefm | 0.32272309 | 0.27052894 | 0.146 | 0.104 | 1 | 0.17364988 | 0.32241295 | 0.113 | 0.062 | 1 | 0.32272309 | 0.31714548 |
| 2810432F15Rik | 0.46385443 | -0.4519371 | 0.073 | 0.098 | 1 | 0.17378055 | -1.1654654 | 0.063 | 0.104 | 1 | 0.46385443 | 0.31736141 |
| Irf8 | 0.17390589 | 0.20013961 | 0.031 | 0.073 | 1 | 0.31444841 | -0.8504236 | 0.029 | 0.052 | 1 | 0.31444841 | 0.31756852 |
| Ap4m1 | 0.17400607 | 0.38600616 | 0.208 | 0.14 | 1 | 0.23106388 | 0.1438627 | 0.185 | 0.125 | 1 | 0.23106388 | 0.31773402 |
| Ppm1h | 0.17402871 | -1.1875214 | 0.135 | 0.192 | 1 | 0.21390547 | -0.3425215 | 0.239 | 0.323 | 1 | 0.21390547 | 0.31777143 |
| Slc30a1 | 0.43464748 | -0.577468 | 0.219 | 0.244 | 1 | 0.17403318 | 0.63076619 | 0.273 | 0.198 | 1 | 0.43464748 | 0.31777881 |
| Hvcn1 | 0.46836151 | 0.78510212 | 0.042 | 0.026 | 1 | 0.17407898 | -2.3098553 | 0.017 | 0.042 | 1 | 0.46836151 | 0.31785447 |
| Egr4 | 0.21708019 | 3.46692616 | 0.021 | 0.005 | 1 | 0.17409232 | -2.010295 | 0.017 | 0.042 | 1 | 0.21708019 | 0.31787651 |
| Sim1 | 0.27752846 | -1.6929452 | 0.01 | 0.031 | 1 | 0.17409232 | -2.0874628 | 0.017 | 0.042 | 1 | 0.27752846 | 0.31787651 |
| Ccne2 | 0.19174744 | 1.29700115 | 0.104 | 0.062 | 1 | 0.17418321 | -1.0203486 | 0.038 | 0.073 | 1 | 0.19174744 | 0.31802663 |
| Zfp747 | 0.58480395 | 0.22988696 | 0.062 | 0.047 | 1 | 0.17438514 | 1.10518224 | 0.097 | 0.052 | 1 | 0.58480395 | 0.31836011 |
| Gm49708 | 0.28021658 | -0.7998604 | 0.083 | 0.124 | 1 | 0.17445747 | -1.0983512 | 0.071 | 0.115 | 1 | 0.28021658 | 0.31847953 |
| Gm14744 | 0.17448902 | 0.44189736 | 0.229 | 0.161 | 1 | 0.74015141 | -0.5003772 | 0.349 | 0.333 | 1 | 0.74015141 | 0.31853162 |
| Itga11 | 0.17451139 | -0.9977998 | 0.417 | 0.295 | 1 | 0.53328932 | -0.5813019 | 0.546 | 0.5 | 1 | 0.53328932 | 0.31856855 |
| Gm15879 | 0.26717799 | 0.42253204 | 0.094 | 0.057 | 1 | 0.17465759 | 0.43737867 | 0.151 | 0.094 | 1 | 0.26717799 | 0.31880991 |
| Fosl2 | 0.2901552 | -0.2587002 | 0.698 | 0.772 | 1 | 0.17482515 | 0.28113103 | 0.811 | 0.833 | 1 | 0.2901552 | 0.31908647 |
| Pask | 0.79462678 | 0.32942263 | 0.031 | 0.026 | 1 | 0.17484387 | -2.07539 | 0.017 | 0.042 | 1 | 0.79462678 | 0.31911736 |
| Dido1 | 0.17487026 | 0.13230099 | 0.792 | 0.684 | 1 | 0.75944656 | -0.178948 | 0.874 | 0.823 | 1 | 0.75944656 | 0.3191609 |
| Il17re | 0.52528691 | -0.9925055 | 0.01 | 0.021 | 1 | 0.1751546 | 1.27510724 | 0.055 | 0.021 | 1 | 0.52528691 | 0.31963006 |
| Usp53 | 0.1752719 | 0.52128129 | 0.635 | 0.591 | 1 | 0.30087688 | -0.1182352 | 0.794 | 0.927 | 1 | 0.30087688 | 0.31982357 |
| Cc2d1a | 0.17709819 | 0.86999101 | 0.094 | 0.052 | 1 | 0.17530428 | 0.27510724 | 0.139 | 0.083 | 1 | 0.17709819 | 0.31987697 |
| Pcdhgb1 | 0.17532926 | 2.00749454 | 0.042 | 0.016 | 1 | 0.41906801 | -1.0874628 | 0.025 | 0.042 | 1 | 0.41906801 | 0.31991818 |
| Lrrc8dos | 0.17534439 | 1.32942263 | 0.042 | 0.016 | 1 | 0.22208005 | 0.91253716 | 0.05 | 0.021 | 1 | 0.22208005 | 0.31994313 |
| Gm54771 | 0.17534439 | 1.32942263 | 0.042 | 0.016 | 1 | 0.23976648 | -1.8948178 | 0.013 | 0.031 | 1 | 0.23976648 | 0.31994313 |
| Zfp712 | 0.17534439 | 1.32942263 | 0.042 | 0.016 | 1 | 0.55837322 | -0.9472852 | 0.029 | 0.042 | 1 | 0.55837322 | 0.31994313 |
| Gm31282 | 0.17534439 | 1.32942263 | 0.042 | 0.016 | 1 | 0.22695751 | 0.77760758 | 0.05 | 0.021 | 1 | 0.22695751 | 0.31994313 |
| Armc8 | 0.17541176 | 0.38189005 | 0.438 | 0.347 | 1 | 0.37414571 | -0.2592292 | 0.618 | 0.5 | 1 | 0.37414571 | 0.32005423 |
| 6230400D17Rik | 0.97172541 | -0.240433 | 0.146 | 0.145 | 1 | 0.17566314 | 0.38202244 | 0.151 | 0.094 | 1 | 0.97172541 | 0.32046874 |
| Atad3a | 0.24468438 | 0.44045394 | 0.354 | 0.301 | 1 | 0.17587101 | -0.1059829 | 0.395 | 0.292 | 1 | 0.24468438 | 0.3208114 |
| Il17rb | 0.66447229 | -0.3144336 | 0.094 | 0.109 | 1 | 0.17592206 | -1.3098553 | 0.109 | 0.156 | 1 | 0.66447229 | 0.32089554 |
| Gpi1 | 0.67241815 | -0.148907 | 0.677 | 0.705 | 1 | 0.1759757 | -0.2375044 | 0.748 | 0.865 | 1 | 0.67241815 | 0.32098395 |
| Smyd5 | 0.34290247 | -0.3461424 | 0.052 | 0.083 | 1 | 0.17597869 | 0.49749966 | 0.126 | 0.073 | 1 | 0.34290247 | 0.32098887 |
| Dnajc28 | 0.17622462 | 1.27052894 | 0.042 | 0.016 | 1 | 0.29798391 | -0.6724253 | 0.046 | 0.021 | 1 | 0.29798391 | 0.32139412 |
| Scgb1b27 | 0.37661787 | 0.20779319 | 0.229 | 0.181 | 1 | 0.17622787 | 0.27510724 | 0.071 | 0.031 | 1 | 0.37661787 | 0.32139948 |
| Gm45435 | 0.38028858 | 1.00749454 | 0.031 | 0.016 | 1 | 0.17624705 | 1.10518224 | 0.055 | 0.021 | 1 | 0.38028858 | 0.32143109 |
| Zfpm2 | 0.17625892 | -0.4006924 | 0.906 | 0.902 | 1 | 0.47259152 | -0.3505305 | 0.899 | 0.76 | 1 | 0.47259152 | 0.32145064 |
| Zkscan6 | 0.34330962 | -0.3461424 | 0.073 | 0.109 | 1 | 0.17631583 | 0.76053407 | 0.084 | 0.042 | 1 | 0.34330962 | 0.32154438 |
| Hs6st2 | 0.99207454 | 0.42253204 | 0.031 | 0.031 | 1 | 0.17643151 | -1.0050007 | 0.038 | 0.073 | 1 | 0.99207454 | 0.32173494 |
| Rabl2 | 0.17649161 | 0.65135073 | 0.188 | 0.13 | 1 | 0.44691177 | -0.3587649 | 0.202 | 0.156 | 1 | 0.44691177 | 0.32183393 |
| Gm19265 | 0.17693365 | 0.34452952 | 0.094 | 0.052 | 1 | 0.83550943 | 0.45567948 | 0.046 | 0.042 | 1 | 0.83550943 | 0.32256178 |
| Serac1 | 0.17702056 | 0.86999101 | 0.073 | 0.036 | 1 | 0.49515253 | 0.10518224 | 0.084 | 0.062 | 1 | 0.49515253 | 0.32270485 |

| | | | | | | | | | | | | |
|---|---|---|---|---|---|---|---|---|---|---|---|---|
| 0610040B10Rik | 0.35353601 | 0.78510212 | 0.073 | 0.047 | 1 | 0.17708318 | -1.5728897 | 0.017 | 0.042 | 1 | 0.35353601 | 0.32280791 |
| 1700003E16Rik | 0.17712338 | 0.59245704 | 0.042 | 0.016 | 1 | 0.31055303 | 0.56461386 | 0.029 | 0.01 | 1 | 0.31055303 | 0.32287407 |
| Gm19412 | 0.62760217 | 0.32942263 | 0.083 | 0.067 | 1 | 0.17734089 | 1.10518224 | 0.055 | 0.021 | 1 | 0.62760217 | 0.32323199 |
| Rabggta | 0.17734988 | -0.7998604 | 0.052 | 0.098 | 1 | 0.54069922 | -0.1082214 | 0.071 | 0.052 | 1 | 0.54069922 | 0.32324678 |
| Lrrc15 | 0.1774942 | -1.9559796 | 0.042 | 0.083 | 1 | 0.34704016 | -1.3098553 | 0.008 | 0.021 | 1 | 0.34704016 | 0.32348421 |
| Ahcyl | 0.17760975 | 0.83756954 | 0.073 | 0.036 | 1 | 0.43079893 | 0.56461386 | 0.038 | 0.021 | 1 | 0.43079893 | 0.32367427 |
| Ptges2 | 0.37721847 | 0.37946331 | 0.104 | 0.073 | 1 | 0.17763797 | -0.1082214 | 0.168 | 0.104 | 1 | 0.37721847 | 0.3237207 |
| Eme2 | 0.61924305 | -0.2148979 | 0.021 | 0.031 | 1 | 0.17769294 | 0.13071733 | 0.071 | 0.031 | 1 | 0.61924305 | 0.3238111 |
| Kazn | 0.27919911 | -0.6965206 | 0.177 | 0.223 | 1 | 0.17780203 | -0.5679234 | 0.185 | 0.25 | 1 | 0.27919911 | 0.3239905 |
| Camkmt | 0.21340457 | 0.4877162 | 0.323 | 0.269 | 1 | 0.17793864 | -0.326534 | 0.298 | 0.208 | 1 | 0.21340457 | 0.32421513 |
| Ttc28 | 0.60309996 | -0.3390292 | 0.917 | 0.891 | 1 | 0.17805796 | -0.3280637 | 0.912 | 0.938 | 1 | 0.60309996 | 0.32441129 |
| Crip1 | 0.92791738 | -0.2907635 | 0.656 | 0.653 | 1 | 0.178111 | 0.29998191 | 0.525 | 0.458 | 1 | 0.92791738 | 0.32449848 |
| Tns3 | 0.71116344 | -0.2978507 | 0.729 | 0.72 | 1 | 0.17820452 | -0.4960607 | 0.887 | 0.875 | 1 | 0.71116344 | 0.32465218 |
| Timm13 | 0.17823943 | 0.22840347 | 0.792 | 0.699 | 1 | 0.64416566 | -0.1640044 | 0.676 | 0.646 | 1 | 0.64416566 | 0.32470956 |
| Acox3 | 1 | -0.9925055 | 0.125 | 0.119 | 1 | 0.17831441 | -0.1034044 | 0.172 | 0.104 | 1 | 1 | 0.32483279 |
| Srr | 0.9023624 | -0.1809506 | 0.292 | 0.29 | 1 | 0.17852886 | -0.9568395 | 0.29 | 0.333 | 1 | 0.9023624 | 0.32518517 |
| Foxl1 | 0.99758609 | 0.27052894 | 0.021 | 0.021 | 1 | 0.1785364 | -0.4797803 | 0.038 | 0.073 | 1 | 0.99758609 | 0.32519756 |
| Actr8 | 0.17853759 | 0.81045769 | 0.26 | 0.192 | 1 | 0.26745618 | -0.3490259 | 0.345 | 0.25 | 1 | 0.26745618 | 0.32519951 |
| Tvp23a | 0.72212321 | 0.13302542 | 0.042 | 0.052 | 1 | 0.17858821 | 0.91253716 | 0.055 | 0.021 | 1 | 0.72212321 | 0.32528267 |
| Pstpip1 | 0.45455389 | 0.30705482 | 0.25 | 0.212 | 1 | 0.17863461 | 0.1579237 | 0.282 | 0.188 | 1 | 0.45455389 | 0.32535889 |
| 2810414N06Rik | 0.17891971 | 1.00749454 | 0.042 | 0.016 | 1 | 0.50887612 | -1.3098553 | 0.004 | 0.01 | 1 | 0.50887612 | 0.32582716 |
| Zfp973 | 0.17891971 | 1.00749454 | 0.042 | 0.016 | 1 | 0.39515719 | 0.49749966 | 0.025 | 0.01 | 1 | 0.39515719 | 0.32582716 |
| 4930522L14Rik | 0.17891971 | 0.74446013 | 0.042 | 0.016 | 1 | 0.18429559 | 1.14957636 | 0.038 | 0.01 | 1 | 0.18429559 | 0.32582716 |
| Kctd21 | 0.17891971 | 1.00749454 | 0.042 | 0.016 | 1 | 0.40319318 | -0.5025003 | 0.055 | 0.031 | 1 | 0.40319318 | 0.32582716 |
| Zfp964 | 0.17891971 | 1.00749454 | 0.042 | 0.016 | 1 | 0.41848998 | -1.0468209 | 0.034 | 0.052 | 1 | 0.41848998 | 0.32582716 |
| D430019H16Rik | 0.17891971 | 0.52206771 | 0.042 | 0.016 | 1 | 0.53715828 | 0.93807225 | 0.034 | 0.021 | 1 | 0.53715828 | 0.32582716 |
| Zfp174 | 0.17891971 | 1.00749454 | 0.042 | 0.016 | 1 | 0.51018266 | 0.49749966 | 0.021 | 0.01 | 1 | 0.51018266 | 0.32582716 |
| Mrpl21 | 0.34701896 | 0.10703021 | 0.552 | 0.466 | 1 | 0.17922488 | 0.22065945 | 0.412 | 0.323 | 1 | 0.34701896 | 0.3263282 |
| Ifi204 | 0.61550914 | -0.407543 | 0.021 | 0.031 | 1 | 0.17938 | -1.5322477 | 0.017 | 0.042 | 1 | 0.61550914 | 0.32658282 |
| Nqo2 | 0.17941911 | -0.8225805 | 0.073 | 0.124 | 1 | 0.69504727 | -0.5133887 | 0.122 | 0.135 | 1 | 0.69504727 | 0.32664701 |
| Zfp780b | 0.44876784 | 0.49292136 | 0.104 | 0.078 | 1 | 0.17947635 | -0.189561 | 0.185 | 0.115 | 1 | 0.44876784 | 0.32674093 |
| Slc46a1 | 0.26870509 | 0.78510212 | 0.104 | 0.067 | 1 | 0.17971949 | 0.52664601 | 0.084 | 0.042 | 1 | 0.26870509 | 0.32713989 |
| Cplane2 | 0.17984777 | 1.00749454 | 0.042 | 0.016 | 1 | 0.45828033 | -0.194378 | 0.05 | 0.031 | 1 | 0.45828033 | 0.32735032 |
| Gm9970 | 0.17984777 | 1.00749454 | 0.042 | 0.016 | 1 | 1 | -0.5025003 | 0.021 | 0.021 | 1 | 1 | 0.32735032 |
| Spaca6 | 0.83881317 | -0.1807092 | 0.375 | 0.363 | 1 | 0.17984948 | -0.2971814 | 0.622 | 0.469 | 1 | 0.83881317 | 0.32735313 |
| Soat1 | 0.33806058 | 0.16727498 | 0.615 | 0.534 | 1 | 0.17990177 | -0.187627 | 0.639 | 0.5 | 1 | 0.33806058 | 0.32743889 |
| B3gnt6 | 0.22187575 | -0.9925055 | 0 | 0.016 | 1 | 0.18016449 | -1.3098553 | 0.017 | 0.042 | 1 | 0.22187575 | 0.32786973 |
| Gm16861 | 0.37863831 | 1.27052894 | 0.031 | 0.016 | 1 | 0.18016449 | -1.3098553 | 0.017 | 0.042 | 1 | 0.37863831 | 0.32786973 |
| 2510017J16Rik | 0.18059951 | 0.49292136 | 0.135 | 0.083 | 1 | 0.44450654 | -0.9378865 | 0.08 | 0.104 | 1 | 0.44450654 | 0.32858283 |
| Impact | 0.18064484 | 0.32828171 | 0.688 | 0.606 | 1 | 0.74718771 | -0.2799316 | 0.693 | 0.625 | 1 | 0.74718771 | 0.32865712 |
| Phka1 | 0.18070076 | 0.27322809 | 0.448 | 0.389 | 1 | 0.48742106 | -0.2081599 | 0.609 | 0.469 | 1 | 0.48742106 | 0.32874875 |
| Rsad1 | 0.70233152 | -0.2555399 | 0.115 | 0.13 | 1 | 0.18073904 | 0.36821664 | 0.113 | 0.062 | 1 | 0.70233152 | 0.32881148 |
| Itpripl1 | 0.20158871 | 1.42253204 | 0.031 | 0.01 | 1 | 0.18091641 | -1.0874628 | 0.017 | 0.042 | 1 | 0.20158871 | 0.32910207 |
| Nsg1 | 0.18123919 | -1.1671375 | 0.302 | 0.368 | 1 | 0.67435706 | -0.8754524 | 0.185 | 0.146 | 1 | 0.67435706 | 0.32963073 |
| Pcgf2 | 0.18127083 | 0.33506919 | 0.271 | 0.197 | 1 | 0.54707512 | -0.2223924 | 0.353 | 0.292 | 1 | 0.54707512 | 0.32968255 |
| Xlr3b | 0.86971511 | 0.10703021 | 0.083 | 0.078 | 1 | 0.18138832 | 0.82764826 | 0.055 | 0.021 | 1 | 0.86971511 | 0.32987491 |
| D2hgdh | 0.18140516 | 0.76648644 | 0.094 | 0.052 | 1 | 0.48887385 | -0.681824 | 0.063 | 0.083 | 1 | 0.48887385 | 0.32990249 |
| Ccdc93 | 0.18154652 | 0.36113149 | 0.198 | 0.135 | 1 | 0.23157954 | 0.14957636 | 0.282 | 0.208 | 1 | 0.23157954 | 0.33013389 |
| Nfkbil1 | 0.19859658 | 0.5049942 | 0.188 | 0.13 | 1 | 0.18169279 | 0.19264508 | 0.181 | 0.115 | 1 | 0.19859658 | 0.33037332 |
| Gm15726 | 0.62768649 | 1.07162487 | 0.083 | 0.067 | 1 | 0.1817033 | -0.6883669 | 0.017 | 0.042 | 1 | 0.62768649 | 0.33039051 |
| Tspan17 | 0.18181441 | 0.44189736 | 0.208 | 0.145 | 1 | 0.78720275 | -0.2103196 | 0.055 | 0.062 | 1 | 0.78720275 | 0.33057234 |
| B3galnt1 | 0.46336809 | 0.71798792 | 0.073 | 0.052 | 1 | 0.18193074 | 1.08246216 | 0.055 | 0.021 | 1 | 0.46336809 | 0.33076269 |
| Cubn | 0.18199539 | 0.51153704 | 0.198 | 0.14 | 1 | 0.58186823 | -0.1475838 | 0.118 | 0.094 | 1 | 0.58186823 | 0.33086846 |
| Eme1 | 0.18220597 | 0.68556644 | 0.073 | 0.036 | 1 | 0.56388113 | -0.9313436 | 0.046 | 0.031 | 1 | 0.56388113 | 0.33121292 |
| Socs1 | 0.62291128 | -0.3144336 | 0.219 | 0.244 | 1 | 0.18240413 | 0.44687359 | 0.206 | 0.135 | 1 | 0.62291128 | 0.33153699 |
| Zfp143 | 0.85616519 | 0.10937415 | 0.146 | 0.14 | 1 | 0.18246883 | 0.22448117 | 0.239 | 0.167 | 1 | 0.85616519 | 0.33164278 |
| Ip6k1 | 0.18252877 | 0.22818265 | 0.667 | 0.601 | 1 | 0.73560769 | -0.1822074 | 0.815 | 0.792 | 1 | 0.73560769 | 0.33174079 |
| Sema3g | 0.18258775 | 0.52206771 | 0.042 | 0.016 | 1 | 0.27154327 | 1.01207283 | 0.013 | 0 | 1 | 0.27154327 | 0.33183722 |
| Gm5468.1 | 0.18258775 | 0.74446013 | 0.042 | 0.016 | 1 | 0.50887612 | -1.3098553 | 0.004 | 0.01 | 1 | 0.50887612 | 0.33183722 |
| Dnaaf5 | 0.18258804 | 0.53800925 | 0.115 | 0.067 | 1 | 0.48659739 | -0.3587649 | 0.101 | 0.073 | 1 | 0.48659739 | 0.33183769 |
| Fam13b | 0.18263085 | -0.4830524 | 0.771 | 0.782 | 1 | 0.68443259 | -0.2621414 | 0.853 | 0.823 | 1 | 0.68443259 | 0.33190768 |
| Gm13889 | 0.18272111 | -1.1427481 | 0.125 | 0.078 | 1 | 0.36182578 | 1.47441605 | 0.08 | 0.052 | 1 | 0.36182578 | 0.33205522 |
| Scube2 | 0.18274992 | -0.4804435 | 0.281 | 0.181 | 1 | 0.77027912 | -1.1560499 | 0.181 | 0.177 | 1 | 0.77027912 | 0.33210231 |
| Slc4a5 | 0.2050453 | -1.8993961 | 0.01 | 0.036 | 1 | 0.18292308 | 1.49749966 | 0.038 | 0.01 | 1 | 0.2050453 | 0.33238531 |
| Pfkl | 0.55089365 | -0.2025407 | 0.49 | 0.497 | 1 | 0.18298651 | -0.1432054 | 0.345 | 0.24 | 1 | 0.55089365 | 0.33248896 |
| Slitrk6 | 0.53755395 | 0.29700115 | 0.135 | 0.161 | 1 | 0.18307665 | 0.91253716 | 0.055 | 0.021 | 1 | 0.53755395 | 0.33263624 |
| Calhm6 | 0.205046 | -1.577468 | 0.01 | 0.036 | 1 | 0.18322874 | -0.8244284 | 0.017 | 0.042 | 1 | 0.205046 | 0.3328847 |
| 4931414P19Rik | 0.99827501 | -0.1851505 | 0.042 | 0.041 | 1 | 0.18327882 | -0.961932 | 0.046 | 0.083 | 1 | 0.99827501 | 0.33296652 |
| Chml | 0.73147402 | -0.4519371 | 0.062 | 0.073 | 1 | 0.18330291 | 0.81942775 | 0.097 | 0.052 | 1 | 0.73147402 | 0.33300587 |
| Sdhc | 0.18345273 | 0.21899864 | 0.656 | 0.58 | 1 | 0.23397892 | -0.2198739 | 0.555 | 0.656 | 1 | 0.23397892 | 0.33325056 |
| Rcor3 | 0.31165632 | 0.49137091 | 0.479 | 0.42 | 1 | 0.18351069 | -0.1924983 | 0.601 | 0.406 | 1 | 0.31165632 | 0.3333452 |
| Ajuba | 0.85795684 | 0.13302542 | 0.042 | 0.047 | 1 | 0.18353804 | -0.8244284 | 0.038 | 0.073 | 1 | 0.85795684 | 0.33338987 |
| Plau | 0.27231495 | -1.7294711 | 0.021 | 0.047 | 1 | 0.18361019 | 1.39058446 | 0.038 | 0.01 | 1 | 0.27231495 | 0.33350767 |
| Gm46212 | 0.30296503 | 1.29700115 | 0.062 | 0.036 | 1 | 0.18361019 | 1.27510724 | 0.038 | 0.01 | 1 | 0.30296503 | 0.33350767 |
| Gm13938 | 0.46992036 | 0.74446013 | 0.042 | 0.026 | 1 | 0.18361019 | 1.27510724 | 0.038 | 0.01 | 1 | 0.46992036 | 0.33350767 |
| Gm47093 | 0.75328966 | 0.27052894 | 0.021 | 0.016 | 1 | 0.18361019 | 1.49749966 | 0.038 | 0.01 | 1 | 0.75328966 | 0.33350767 |
| Bmp15 | 0.99288067 | 0.10060394 | 0.062 | 0.062 | 1 | 0.18361019 | 1.39058446 | 0.038 | 0.01 | 1 | 0.99288067 | 0.33350767 |
| Gm26549 | 0.4152556 | 0.40803247 | 0.24 | 0.202 | 1 | 0.18362709 | 0.34725702 | 0.214 | 0.146 | 1 | 0.4152556 | 0.33353528 |
| Gpr157 | 0.39356009 | -0.2148979 | 0.01 | 0.026 | 1 | 0.18363337 | 0.69014474 | 0.055 | 0.021 | 1 | 0.39356009 | 0.33354553 |
| Cers5 | 0.18384109 | 0.22885091 | 0.635 | 0.534 | 1 | 0.25732892 | -0.3415641 | 0.765 | 0.76 | 1 | 0.25732892 | 0.33388463 |
| Nrp2 | 0.18399191 | -0.5955584 | 0.656 | 0.487 | 1 | 0.53829924 | -0.7118729 | 0.853 | 0.688 | 1 | 0.53829924 | 0.3341308 |
| Strn4 | 0.18408161 | -0.5907648 | 0.26 | 0.326 | 1 | 0.2619618 | -0.1723517 | 0.466 | 0.333 | 1 | 0.2619618 | 0.33427718 |
| Gm46447 | 0.21867159 | 1.59245704 | 0.021 | 0.005 | 1 | 0.18422625 | 0.45567948 | 0.055 | 0.021 | 1 | 0.21867159 | 0.33451318 |
| Il10 | 0.47923898 | 0.47697982 | 0.073 | 0.052 | 1 | 0.18422707 | -0.5993619 | 0.055 | 0.021 | 1 | 0.47923898 | 0.33451453 |
| Upb1 | 0.21867159 | 1.59245704 | 0.021 | 0.005 | 1 | 0.18429559 | 1.14957636 | 0.038 | 0.01 | 1 | 0.21867159 | 0.33462631 |

| | | | | | | | | | | | | |
|---|---|---|---|---|---|---|---|---|---|---|---|---|
| Irx3os | 0.30891093 | 1.00749454 | 0.042 | 0.021 | 1 | 0.18429559 | 1.14957636 | 0.038 | 0.01 | 1 | 0.30891093 | 0.33462631 |
| Wdr54 | 0.38028858 | 1.00749454 | 0.031 | 0.016 | 1 | 0.18429559 | 1.14957636 | 0.038 | 0.01 | 1 | 0.38028858 | 0.33462631 |
| Nthl1 | 0.69098575 | 0.27052894 | 0.052 | 0.041 | 1 | 0.18429559 | 1.14957636 | 0.038 | 0.01 | 1 | 0.69098575 | 0.33462631 |
| Gm10638 | 0.72152305 | 0.20013961 | 0.062 | 0.052 | 1 | 0.18429559 | 1.14957636 | 0.038 | 0.01 | 1 | 0.72152305 | 0.33462631 |
| Faap24 | 0.74827573 | 0.59245704 | 0.021 | 0.016 | 1 | 0.18429559 | 1.14957636 | 0.038 | 0.01 | 1 | 0.74827573 | 0.33462631 |
| 1700022I11Rik | 0.78979801 | -0.2148979 | 0.021 | 0.026 | 1 | 0.18429559 | 1.14957636 | 0.038 | 0.01 | 1 | 0.78979801 | 0.33462631 |
| Hs6st1 | 0.8366448 | -0.1693832 | 0.198 | 0.181 | 1 | 0.18439089 | 0.18368421 | 0.328 | 0.25 | 1 | 0.8366448 | 0.33478179 |
| Man2b1 | 0.92132825 | -0.1217885 | 0.406 | 0.373 | 1 | 0.18466238 | 0.22930355 | 0.525 | 0.448 | 1 | 0.92132825 | 0.33522457 |
| Nif3l1 | 0.18472623 | 0.47144164 | 0.177 | 0.119 | 1 | 0.20197622 | 0.13492958 | 0.172 | 0.115 | 1 | 0.20197622 | 0.33532869 |
| Dnajc14 | 0.22066853 | 0.26125113 | 0.25 | 0.181 | 1 | 0.18484096 | -0.1853102 | 0.399 | 0.281 | 1 | 0.22066853 | 0.33551574 |
| Chst14 | 0.18488958 | 0.47697982 | 0.167 | 0.109 | 1 | 0.52030007 | 0.19264508 | 0.13 | 0.104 | 1 | 0.52030007 | 0.33559501 |
| Apex2 | 0.43965483 | 0.20819189 | 0.458 | 0.404 | 1 | 0.18491102 | -0.3205025 | 0.366 | 0.26 | 1 | 0.43965483 | 0.33562995 |
| Aldh5a1 | 0.20691434 | -1.3144336 | 0.01 | 0.036 | 1 | 0.18504625 | 1.01207283 | 0.038 | 0.01 | 1 | 0.20691434 | 0.33585038 |
| Exoc3l4 | 0.28677173 | -0.577468 | 0.052 | 0.088 | 1 | 0.18504625 | 1.01207283 | 0.038 | 0.01 | 1 | 0.28677173 | 0.33585038 |
| Gpr3 | 0.36859058 | -0.577468 | 0.021 | 0.041 | 1 | 0.18504625 | 1.01207283 | 0.038 | 0.01 | 1 | 0.36859058 | 0.33585038 |
| 9330151L19Rik | 0.61550914 | -0.407543 | 0.021 | 0.031 | 1 | 0.18504625 | 1.01207283 | 0.038 | 0.01 | 1 | 0.61550914 | 0.33585038 |
| A230028O05Rik | 0.74827573 | 0.59245704 | 0.021 | 0.016 | 1 | 0.18504625 | 1.01207283 | 0.038 | 0.01 | 1 | 0.74827573 | 0.33585038 |
| Galnt12 | 0.82258416 | -0.1624305 | 0.031 | 0.036 | 1 | 0.18504625 | 1.01207283 | 0.038 | 0.01 | 1 | 0.82258416 | 0.33585038 |
| Gm7628 | 0.98603793 | 0.22988696 | 0.052 | 0.052 | 1 | 0.18504625 | 1.01207283 | 0.038 | 0.01 | 1 | 0.98603793 | 0.33585038 |
| N4bp2l2 | 0.23837988 | 0.21926793 | 0.927 | 0.865 | 1 | 0.18531946 | -0.270666 | 0.895 | 0.896 | 1 | 0.23837988 | 0.33629562 |
| Tgfb1i1 | 0.18534955 | 0.18199227 | 0.469 | 0.389 | 1 | 0.6657236 | -0.265001 | 0.508 | 0.427 | 1 | 0.6657236 | 0.33634465 |
| H2ac6 | 0.75447492 | 0.11440974 | 0.156 | 0.171 | 1 | 0.18537071 | -1.2167459 | 0.063 | 0.104 | 1 | 0.75447492 | 0.33637912 |
| Sema5a | 0.18557432 | -0.9457113 | 0.292 | 0.347 | 1 | 0.80777772 | -0.4587937 | 0.45 | 0.396 | 1 | 0.80777772 | 0.33671081 |
| Tmem62 | 0.18564572 | -1.1941393 | 0.042 | 0.083 | 1 | 0.32043676 | 0.97554696 | 0.105 | 0.073 | 1 | 0.32043676 | 0.3368271 |
| Atg2b | 0.18591685 | 0.25076569 | 0.406 | 0.316 | 1 | 0.35328817 | -0.2854711 | 0.496 | 0.406 | 1 | 0.35328817 | 0.33726862 |
| Trp53bp2 | 0.35586548 | -0.2864967 | 0.417 | 0.456 | 1 | 0.1859967 | -0.3140069 | 0.651 | 0.729 | 1 | 0.35586548 | 0.33739862 |
| Zmat1 | 0.18610697 | -0.1918143 | 0.167 | 0.233 | 1 | 0.4167677 | -0.4797803 | 0.13 | 0.094 | 1 | 0.4167677 | 0.33757813 |
| Lama3 | 0.72574709 | -1.4519371 | 0.01 | 0.016 | 1 | 0.18634143 | -0.5728897 | 0.038 | 0.073 | 1 | 0.72574709 | 0.33795973 |
| Gm10785 | 0.1863585 | -0.7425272 | 0.052 | 0.098 | 1 | 0.66347477 | -0.9562183 | 0.071 | 0.083 | 1 | 0.66347477 | 0.33798751 |
| Nppc | 0.72574709 | -0.577468 | 0.01 | 0.016 | 1 | 0.18637076 | -0.4618584 | 0.017 | 0.042 | 1 | 0.72574709 | 0.33800746 |
| Atf6b | 0.55336775 | -0.2148979 | 0.25 | 0.28 | 1 | 0.18652646 | -0.2881602 | 0.227 | 0.156 | 1 | 0.55336775 | 0.33826079 |
| Dram2 | 0.56819429 | 0.15559318 | 0.271 | 0.238 | 1 | 0.18685404 | -0.4370905 | 0.395 | 0.479 | 1 | 0.56819429 | 0.33879365 |
| 2310058D17Rik | 0.1868844 | 0.20013961 | 0.073 | 0.036 | 1 | 0.86695837 | -0.194378 | 0.046 | 0.042 | 1 | 0.86695837 | 0.33884302 |
| Aph1b | 0.30116057 | -0.4779323 | 0.125 | 0.171 | 1 | 0.18700592 | 0.30312161 | 0.164 | 0.104 | 1 | 0.30116057 | 0.33904063 |
| Ccdc51 | 0.1871904 | 0.56003556 | 0.094 | 0.052 | 1 | 0.24139962 | -1.0342208 | 0.067 | 0.104 | 1 | 0.24139962 | 0.33934055 |
| Gm9929 | 0.21858689 | 0.49292136 | 0.062 | 0.031 | 1 | 0.1872521 | 0.81942775 | 0.097 | 0.052 | 1 | 0.21858689 | 0.33944085 |
| Rad51d | 0.53666344 | -0.3848229 | 0.062 | 0.083 | 1 | 0.18758869 | 0.44503224 | 0.084 | 0.042 | 1 | 0.53666344 | 0.33998786 |
| Cwc27 | 0.18759332 | 0.30915024 | 0.51 | 0.42 | 1 | 0.20027052 | -0.6568743 | 0.504 | 0.531 | 1 | 0.20027052 | 0.33999539 |
| Olfm1 | 0.35987448 | -0.2587855 | 0.26 | 0.311 | 1 | 0.18765783 | -0.2493137 | 0.218 | 0.146 | 1 | 0.35987448 | 0.3401002 |
| Fam81a | 0.30888255 | 1.00749454 | 0.062 | 0.036 | 1 | 0.18785938 | 1.01207283 | 0.038 | 0.01 | 1 | 0.30888255 | 0.34042762 |
| Lgals9 | 0.18814989 | 0.71798792 | 0.146 | 0.093 | 1 | 0.53745815 | -0.7248928 | 0.084 | 0.104 | 1 | 0.53745815 | 0.3408994 |
| Mgrn1 | 0.44145855 | 0.22988696 | 0.438 | 0.399 | 1 | 0.18816026 | 0.11160851 | 0.58 | 0.5 | 1 | 0.44145855 | 0.34091623 |
| 9230114K14Rik | 0.18849817 | 0.48142572 | 0.167 | 0.109 | 1 | 0.21298107 | -0.1399303 | 0.08 | 0.042 | 1 | 0.21298107 | 0.34146479 |
| Septin2 | 0.18856059 | 0.59245704 | 0.177 | 0.119 | 1 | 0.76830776 | -0.8359241 | 0.189 | 0.188 | 1 | 0.76830776 | 0.34156608 |
| Hadhb | 0.23708253 | 0.11535053 | 0.688 | 0.611 | 1 | 0.18861365 | 0.15380094 | 0.714 | 0.698 | 1 | 0.23708253 | 0.34165219 |
| Bbs9 | 0.18864333 | 0.4942335 | 0.469 | 0.409 | 1 | 0.5836074 | -0.7003147 | 0.643 | 0.625 | 1 | 0.5836074 | 0.34170035 |
| Col4a1 | 0.18880462 | -1.6013147 | 0.24 | 0.285 | 1 | 0.27109328 | -1.1493906 | 0.399 | 0.281 | 1 | 0.27109328 | 0.34196205 |
| Harbi1 | 0.18888658 | -0.9925055 | 0.062 | 0.109 | 1 | 0.29978152 | -0.473354 | 0.088 | 0.052 | 1 | 0.29978152 | 0.34209502 |
| Gm28376 | 0.59199182 | 0.42253204 | 0.031 | 0.021 | 1 | 0.18901513 | -0.6468902 | 0.046 | 0.083 | 1 | 0.59199182 | 0.34230353 |
| Gm29825 | 0.21781545 | -0.9925055 | 0.021 | 0.052 | 1 | 0.18927658 | 0.91253716 | 0.038 | 0.01 | 1 | 0.21781545 | 0.34272753 |
| Pfkp | 0.53832321 | -0.1717202 | 0.781 | 0.782 | 1 | 0.18931459 | -0.571053 | 0.651 | 0.74 | 1 | 0.53832321 | 0.34278916 |
| Rnf43 | 0.5920074 | 0.32942263 | 0.031 | 0.021 | 1 | 0.18954922 | -0.3098553 | 0.017 | 0.042 | 1 | 0.5920074 | 0.34316953 |
| Gm50295 | 0.63038178 | 0.20013961 | 0.021 | 0.031 | 1 | 0.18998139 | 0.80562195 | 0.038 | 0.01 | 1 | 0.63038178 | 0.34386985 |
| Zfp692 | 0.19017294 | 0.46317402 | 0.198 | 0.135 | 1 | 0.73916575 | -0.4117349 | 0.143 | 0.125 | 1 | 0.73916575 | 0.34418013 |
| Oplah | 0.7699026 | 0.27052894 | 0.115 | 0.104 | 1 | 0.19024823 | 0.39796399 | 0.164 | 0.104 | 1 | 0.7699026 | 0.34430206 |
| Fzd1 | 0.46234153 | 0.10802141 | 0.417 | 0.347 | 1 | 0.19047449 | -0.4063632 | 0.391 | 0.271 | 1 | 0.46234153 | 0.34466845 |
| Jph1 | 0.19068686 | 0.56270969 | 0.208 | 0.15 | 1 | 0.27663594 | 0.12976787 | 0.391 | 0.302 | 1 | 0.27663594 | 0.34501224 |
| D030016M11Rik | 0.42179602 | -0.6929452 | 0.031 | 0.052 | 1 | 0.1907107 | 0.39058446 | 0.038 | 0.01 | 1 | 0.42179602 | 0.34505084 |
| Vipas39 | 0.19073087 | 0.36363835 | 0.281 | 0.207 | 1 | 0.47580981 | -0.3838558 | 0.294 | 0.229 | 1 | 0.47580981 | 0.34508348 |
| Ccnyl1 | 0.97694169 | -0.6314068 | 0.25 | 0.238 | 1 | 0.19089199 | -1.0003109 | 0.361 | 0.396 | 1 | 0.97694169 | 0.34534422 |
| Rabgef1 | 0.1909816 | 0.37475631 | 0.76 | 0.684 | 1 | 0.39847825 | -0.3283912 | 0.731 | 0.771 | 1 | 0.39847825 | 0.34548922 |
| Nepro | 0.35795316 | 0.35541784 | 0.177 | 0.135 | 1 | 0.19114181 | 0.39641353 | 0.21 | 0.146 | 1 | 0.35795316 | 0.34574843 |
| Gm28379 | 0.22187388 | 1.00749454 | 0.021 | 0.005 | 1 | 0.1914043 | 0.42711033 | 0.038 | 0.01 | 1 | 0.22187388 | 0.34617299 |
| Gm34788 | 0.32025586 | -0.577468 | 0 | 0.01 | 1 | 0.1914043 | -0.7952821 | 0.038 | 0.01 | 1 | 0.32025586 | 0.34617299 |
| Zkscan7 | 0.89616978 | 0.24195979 | 0.062 | 0.067 | 1 | 0.1914043 | 0.42711033 | 0.038 | 0.01 | 1 | 0.89616978 | 0.34617299 |
| Gm39529 | 0.72021678 | -0.7998604 | 0.01 | 0.016 | 1 | 0.19141907 | 0.14957636 | 0.038 | 0.01 | 1 | 0.72021678 | 0.34619688 |
| Tmem135 | 0.19156814 | -0.5676202 | 0.333 | 0.383 | 1 | 0.55406087 | -0.3674428 | 0.492 | 0.51 | 1 | 0.55406087 | 0.34643793 |
| Casp9 | 0.71480946 | 0.56003556 | 0.073 | 0.062 | 1 | 0.19164077 | -0.7248928 | 0.088 | 0.135 | 1 | 0.71480946 | 0.34655536 |
| Zwilch | 0.19233242 | 1.37946331 | 0.104 | 0.062 | 1 | 0.29722453 | -0.6317834 | 0.021 | 0.042 | 1 | 0.29722453 | 0.34767307 |
| Btrc | 0.19240224 | 0.15650449 | 0.562 | 0.456 | 1 | 0.56049746 | -0.3098553 | 0.761 | 0.76 | 1 | 0.56049746 | 0.34778587 |
| Chic2 | 0.19253305 | -0.2191411 | 0.74 | 0.839 | 1 | 0.25514215 | 0.12921593 | 0.87 | 0.896 | 1 | 0.25514215 | 0.34799713 |
| Id4 | 0.36973472 | -0.6050802 | 0.396 | 0.42 | 1 | 0.19263035 | -0.3008665 | 0.37 | 0.281 | 1 | 0.36973472 | 0.34815424 |
| Gm26632 | 0.52528691 | -0.7998604 | 0.01 | 0.021 | 1 | 0.19288249 | -2.5025003 | 0.025 | 0.052 | 1 | 0.52528691 | 0.34856132 |
| Haus1 | 0.39128128 | 0.20013961 | 0.146 | 0.109 | 1 | 0.1929417 | 0.35999614 | 0.122 | 0.073 | 1 | 0.39128128 | 0.34865691 |
| Nol8 | 0.88162861 | 0.2541345 | 0.333 | 0.373 | 1 | 0.19297895 | -0.2838601 | 0.307 | 0.208 | 1 | 0.88162861 | 0.34871703 |
| Dpy19l3 | 0.6313066 | -0.1851505 | 0.156 | 0.181 | 1 | 0.19305134 | -0.2103196 | 0.244 | 0.167 | 1 | 0.6313066 | 0.34883387 |
| Mdga2 | 0.63862513 | 0.92732419 | 0.042 | 0.031 | 1 | 0.19308139 | -0.4029647 | 0.055 | 0.021 | 1 | 0.63862513 | 0.34888235 |
| Masp2 | 0.21257404 | 0.86999101 | 0.083 | 0.047 | 1 | 0.19315497 | 0.33400093 | 0.097 | 0.052 | 1 | 0.21257404 | 0.34900109 |
| Numbl | 0.30003631 | 0.12707615 | 0.323 | 0.254 | 1 | 0.19320163 | -0.4816216 | 0.487 | 0.333 | 1 | 0.30003631 | 0.34907638 |
| Dazap2 | 0.38057794 | -0.2312923 | 0.604 | 0.642 | 1 | 0.19338637 | -0.2159883 | 0.66 | 0.76 | 1 | 0.38057794 | 0.34937445 |
| Coq9 | 0.80723472 | -0.130009 | 0.229 | 0.244 | 1 | 0.19349019 | 0.42711033 | 0.172 | 0.115 | 1 | 0.80723472 | 0.34954192 |
| Ccno | 0.19355003 | -1.1079827 | 0.062 | 0.109 | 1 | 0.57756442 | -0.4618584 | 0.029 | 0.042 | 1 | 0.57756442 | 0.34963844 |
| Gm29677 | 0.2039323 | 1.27052894 | 0.031 | 0.01 | 1 | 0.19358645 | -2.9378865 | 0.025 | 0.052 | 1 | 0.2039323 | 0.34969718 |
| Dhdh | 0.84451844 | -0.3710171 | 0.042 | 0.036 | 1 | 0.19360556 | -0.5728897 | 0.055 | 0.021 | 1 | 0.84451844 | 0.349728 |
| Slbp | 0.19371954 | 0.12778877 | 0.729 | 0.617 | 1 | 0.42009647 | -0.1915712 | 0.664 | 0.688 | 1 | 0.42009647 | 0.34991182 |
| Vwce | 0.19387718 | -1.577468 | 0.031 | 0.067 | 1 | 0.52045354 | -0.1843244 | 0.055 | 0.073 | 1 | 0.52045354 | 0.350166 |

| | | | | | | | | | | | | |
|---|---|---|---|---|---|---|---|---|---|---|---|---|
| 2610021A01Rik | 0.48714006 | 0.53800925 | 0.083 | 0.062 | 1 | 0.19423885 | 0.44913664 | 0.134 | 0.083 | 1 | 0.48714006 | 0.35074897 |
| 2010109A12Rik | 0.19424251 | -1.4187702 | 0.073 | 0.119 | 1 | 0.90026869 | -0.5728897 | 0.113 | 0.104 | 1 | 0.90026869 | 0.35075487 |
| Tef | 0.19431217 | 0.37701738 | 0.531 | 0.497 | 1 | 0.90911101 | -0.2136399 | 0.651 | 0.594 | 1 | 0.90911101 | 0.35086712 |
| Ap4b1 | 0.52440614 | 0.15949763 | 0.188 | 0.155 | 1 | 0.19433625 | 0.61614416 | 0.122 | 0.073 | 1 | 0.52440614 | 0.35090593 |
| Zswim1 | 0.81112941 | 0.59245704 | 0.042 | 0.036 | 1 | 0.19457069 | 0.27510724 | 0.084 | 0.042 | 1 | 0.81112941 | 0.35128362 |
| Gm38825 | 0.19466731 | 2.42253204 | 0.031 | 0.01 | 1 | 0.99396051 | -0.3098553 | 0.021 | 0.021 | 1 | 0.99396051 | 0.35143926 |
| Zfp524 | 0.19487741 | -0.7998604 | 0.042 | 0.083 | 1 | 0.45862002 | -0.2223924 | 0.063 | 0.042 | 1 | 0.45862002 | 0.35177762 |
| ENSMUSG000001214 45 | 0.51189113 | 0.32942263 | 0.083 | 0.062 | 1 | 0.19489086 | 0.46775232 | 0.097 | 0.052 | 1 | 0.51189113 | 0.35179927 |
| Recql5 | 0.19494313 | -0.8405024 | 0.135 | 0.192 | 1 | 0.46417371 | -1.0468209 | 0.113 | 0.135 | 1 | 0.46417371 | 0.35188344 |
| Racgap1 | 0.19508428 | 2.15433592 | 0.083 | 0.047 | 1 | 0.4427584 | -0.1723517 | 0.038 | 0.021 | 1 | 0.4427584 | 0.35211068 |
| Rhbdd3 | 0.25942789 | 0.21899864 | 0.208 | 0.15 | 1 | 0.1950925 | -0.131518 | 0.164 | 0.104 | 1 | 0.25942789 | 0.35212391 |
| Prr12 | 0.19532051 | 0.35541784 | 0.167 | 0.109 | 1 | 0.67590386 | -0.5025003 | 0.197 | 0.208 | 1 | 0.67590386 | 0.35249092 |
| Chp1 | 0.1953291 | -0.1748355 | 0.76 | 0.803 | 1 | 0.98132288 | -0.1437981 | 0.882 | 0.906 | 1 | 0.98132288 | 0.35250475 |
| Lca5l | 0.24611112 | 0.34452952 | 0.115 | 0.073 | 1 | 0.19534661 | 1.01207283 | 0.067 | 0.031 | 1 | 0.24611112 | 0.35253292 |
| Zfp532 | 0.19554593 | 0.30250133 | 0.417 | 0.337 | 1 | 0.59691577 | -0.4113933 | 0.462 | 0.438 | 1 | 0.59691577 | 0.35285364 |
| Krt16 | 0.19580129 | -0.472115 | 0.062 | 0.109 | 1 | 0.78126103 | 2.65504094 | 0.038 | 0.031 | 1 | 0.78126103 | 0.35326443 |
| Zzef1 | 0.79313408 | -0.1309195 | 0.677 | 0.694 | 1 | 0.19641816 | -0.4224728 | 0.79 | 0.865 | 1 | 0.79313408 | 0.35425623 |
| Foxo3 | 0.19647166 | 0.28373982 | 0.833 | 0.803 | 1 | 0.26928431 | -0.168843 | 0.92 | 0.896 | 1 | 0.26928431 | 0.35434221 |
| D930015M05Rik | 0.19648774 | 0.20013961 | 0.073 | 0.036 | 1 | 0.97980798 | -0.4797803 | 0.063 | 0.062 | 1 | 0.97980798 | 0.35436805 |
| Casp12 | 0.19657158 | 0.30705482 | 0.25 | 0.176 | 1 | 0.75193807 | -0.4618584 | 0.294 | 0.25 | 1 | 0.75193807 | 0.35450277 |
| Slc25a11 | 0.19661576 | 0.25191326 | 0.51 | 0.435 | 1 | 0.44438317 | -0.202052 | 0.399 | 0.333 | 1 | 0.44438317 | 0.35457377 |
| Vegfb | 0.70639802 | 0.36215942 | 0.25 | 0.233 | 1 | 0.19677925 | 0.32955502 | 0.235 | 0.167 | 1 | 0.70639802 | 0.35483643 |
| Gm17251 | 0.48569952 | 0.48393258 | 0.135 | 0.109 | 1 | 0.19692799 | 0.32241295 | 0.097 | 0.052 | 1 | 0.48569952 | 0.35507534 |
| Mfsd4b4 | 0.37440936 | 1.20013961 | 0.052 | 0.031 | 1 | 0.19693582 | 0.93807225 | 0.067 | 0.031 | 1 | 0.37440936 | 0.35508792 |
| 9330198N18Rik | 0.1969519 | 2.22988696 | 0.031 | 0.01 | 1 | 0.5778028 | -0.8948178 | 0.013 | 0.021 | 1 | 0.5778028 | 0.35511375 |
| Galnt2 | 0.67127537 | -0.1624305 | 0.615 | 0.648 | 1 | 0.19711778 | -0.3795642 | 0.761 | 0.812 | 1 | 0.67127537 | 0.35538015 |
| Scaf1 | 0.1972911 | 0.1843723 | 0.49 | 0.389 | 1 | 0.47205847 | -0.2242795 | 0.508 | 0.396 | 1 | 0.47205847 | 0.35565842 |
| Wdfy1 | 0.19736744 | -0.6263776 | 0.417 | 0.461 | 1 | 0.57295456 | -0.4353861 | 0.546 | 0.531 | 1 | 0.57295456 | 0.35578098 |
| Ap2a1 | 0.27390821 | 0.42253204 | 0.479 | 0.43 | 1 | 0.19767904 | -0.2175968 | 0.576 | 0.406 | 1 | 0.27390821 | 0.35628107 |
| Pop1 | 0.21214348 | 0.25260703 | 0.156 | 0.104 | 1 | 0.1980821 | -0.1254307 | 0.164 | 0.104 | 1 | 0.21214348 | 0.35692768 |
| Acsl3 | 0.19823575 | -0.4005278 | 0.875 | 0.845 | 1 | 0.54752922 | -0.3209105 | 0.819 | 0.865 | 1 | 0.54752922 | 0.35717408 |
| Mkln1 | 0.38393369 | -0.1912018 | 0.917 | 0.933 | 1 | 0.19838625 | -0.3042395 | 0.971 | 0.958 | 1 | 0.38393369 | 0.35741539 |
| Atg10 | 0.21954927 | 0.48964623 | 0.25 | 0.187 | 1 | 0.19858941 | -0.9589481 | 0.235 | 0.281 | 1 | 0.21954927 | 0.35774107 |
| Btbd6 | 0.21257404 | 0.86999101 | 0.083 | 0.047 | 1 | 0.1986539 | 0.49749966 | 0.109 | 0.062 | 1 | 0.21257404 | 0.35784443 |
| Slc25a53 | 0.32853848 | 0.28502851 | 0.146 | 0.104 | 1 | 0.19869791 | 0.14957636 | 0.176 | 0.115 | 1 | 0.32853848 | 0.35791496 |
| Csnk1g1 | 0.19908978 | 0.20980271 | 0.688 | 0.622 | 1 | 0.93358875 | -0.3134665 | 0.832 | 0.833 | 1 | 0.93358875 | 0.35854282 |
| Slc9a1 | 0.46063807 | -0.1158879 | 0.521 | 0.461 | 1 | 0.19909188 | -0.6434652 | 0.584 | 0.625 | 1 | 0.46063807 | 0.35854618 |
| Ppcdc | 0.76209583 | -0.2726134 | 0.083 | 0.093 | 1 | 0.19912865 | 0.24473359 | 0.139 | 0.083 | 1 | 0.76209583 | 0.35860508 |
| Pus1 | 0.23987655 | 0.40483003 | 0.229 | 0.171 | 1 | 0.19918139 | -0.4301495 | 0.181 | 0.115 | 1 | 0.23987655 | 0.35868955 |
| B3galt2 | 0.1992517 | 2.00749454 | 0.031 | 0.01 | 1 | 0.69200558 | -1.1578522 | 0.034 | 0.042 | 1 | 0.69200558 | 0.35880216 |
| Cd72 | 0.1992517 | 3.00749454 | 0.031 | 0.01 | 1 | 0.75752191 | 0.69014474 | 0.038 | 0.031 | 1 | 0.75752191 | 0.35880216 |
| Rpusd3 | 0.1992517 | 1.74446013 | 0.031 | 0.01 | 1 | 0.58664899 | -0.5025003 | 0.021 | 0.031 | 1 | 0.58664899 | 0.35880216 |
| Gm47484 | 0.1992517 | 2.00749454 | 0.031 | 0.01 | 1 | 0.87066325 | -0.3098553 | 0.013 | 0.01 | 1 | 0.87066325 | 0.35880216 |
| Gpr18 | 0.1992517 | 2.22988696 | 0.031 | 0.01 | 1 | 0.57170879 | -1.6317834 | 0.013 | 0.021 | 1 | 0.57170879 | 0.35880216 |
| Gm53058 | 0.1992517 | 2.22988696 | 0.031 | 0.01 | 1 | 0.73189081 | -0.2509616 | 0.063 | 0.052 | 1 | 0.73189081 | 0.35880216 |
| A930007A09Rik | 0.1992517 | 1.74446013 | 0.031 | 0.01 | 1 | 0.34704016 | -1.3098553 | 0.008 | 0.021 | 1 | 0.34704016 | 0.35880216 |
| Tff2 | 0.1992517 | 2.00749454 | 0.031 | 0.01 | 1 | 0.28597111 | 0.49749966 | 0.046 | 0.021 | 1 | 0.28597111 | 0.35880216 |
| Zbp1 | 0.74287632 | 0.15949763 | 0.052 | 0.062 | 1 | 0.19944259 | -1.3098553 | 0.092 | 0.135 | 1 | 0.74287632 | 0.35910783 |
| Tbkbp1 | 0.26940715 | 0.54806292 | 0.073 | 0.041 | 1 | 0.19961325 | 0.41261076 | 0.122 | 0.073 | 1 | 0.26940715 | 0.35938104 |
| H3c6 | 0.19977696 | 1.78510212 | 0.062 | 0.031 | 1 | 0.66339728 | 0.10518224 | 0.029 | 0.021 | 1 | 0.66339728 | 0.35964308 |
| Tusc2 | 0.83753353 | -0.2440442 | 0.198 | 0.202 | 1 | 0.19988289 | 0.39796399 | 0.185 | 0.125 | 1 | 0.83753353 | 0.35981262 |
| Rfc5 | 0.20021956 | 0.70263996 | 0.125 | 0.078 | 1 | 0.77426989 | -0.1399303 | 0.105 | 0.094 | 1 | 0.77426989 | 0.36035124 |
| Utp14b | 0.84899953 | -0.4055153 | 0.51 | 0.497 | 1 | 0.20022827 | -0.5647846 | 0.475 | 0.562 | 1 | 0.84899953 | 0.36036517 |
| Dmpk | 0.20049747 | 0.42253204 | 0.125 | 0.078 | 1 | 0.39917122 | -0.5322477 | 0.197 | 0.146 | 1 | 0.39917122 | 0.36079571 |
| Tpd52l1 | 0.20066998 | 0.34336182 | 0.406 | 0.332 | 1 | 0.25771801 | -0.3280026 | 0.256 | 0.188 | 1 | 0.25771801 | 0.36107152 |
| Slc2a12 | 0.2007803 | -0.7860546 | 0.135 | 0.192 | 1 | 0.24392267 | 0.77760758 | 0.063 | 0.031 | 1 | 0.24392267 | 0.36124787 |
| Kctd9 | 0.20096383 | 0.15773717 | 0.521 | 0.425 | 1 | 0.96829042 | -0.4838847 | 0.479 | 0.417 | 1 | 0.96829042 | 0.36154119 |
| Nt5m | 0.20116671 | 0.54452256 | 0.271 | 0.202 | 1 | 0.61406271 | -0.1820997 | 0.197 | 0.167 | 1 | 0.61406271 | 0.36186538 |
| Mrpl34 | 0.78000106 | 0.14499806 | 0.344 | 0.337 | 1 | 0.20128788 | -0.1578522 | 0.261 | 0.177 | 1 | 0.78000106 | 0.36205895 |
| Zfp708 | 0.2013617 | 1.14499806 | 0.083 | 0.047 | 1 | 0.3116597 | 0.74077081 | 0.084 | 0.052 | 1 | 0.3116597 | 0.36217686 |
| Rassf5 | 0.20137256 | -0.10213 | 0.156 | 0.098 | 1 | 0.31794855 | -1.2442669 | 0.143 | 0.177 | 1 | 0.31794855 | 0.36219421 |
| Zfr2 | 0.47249996 | -0.7087125 | 0.042 | 0.062 | 1 | 0.20144982 | 0.12310414 | 0.097 | 0.052 | 1 | 0.47249996 | 0.36231761 |
| Spc24 | 0.20151944 | 1.53105649 | 0.083 | 0.047 | 1 | 1 | -1.0468209 | 0.021 | 0.021 | 1 | 1 | 0.36242879 |
| Serpinb6b | 0.20158431 | 1.59245704 | 0.031 | 0.01 | 1 | 0.40076941 | -1.7952821 | 0.017 | 0.031 | 1 | 0.40076941 | 0.36253239 |
| Emx2 | 0.20158431 | 2.00749454 | 0.031 | 0.01 | 1 | 0.5859632 | -0.3098553 | 0.013 | 0.021 | 1 | 0.5859632 | 0.36253239 |
| Gm9821 | 0.20158871 | 1.42253204 | 0.031 | 0.01 | 1 | 0.86312513 | -0.7248928 | 0.008 | 0.01 | 1 | 0.86312513 | 0.36253941 |
| Gm11457 | 0.20158871 | 1.42253204 | 0.031 | 0.01 | 1 | 0.28681577 | 0.39058446 | 0.046 | 0.021 | 1 | 0.28681577 | 0.36253941 |
| Gm42658 | 0.20158871 | 1.42253204 | 0.031 | 0.01 | 1 | 0.65825923 | -0.5025003 | 0.055 | 0.042 | 1 | 0.65825923 | 0.36253941 |
| Mterf1b | 0.20158871 | 1.42253204 | 0.031 | 0.01 | 1 | 0.42275543 | -1.4797803 | 0.025 | 0.042 | 1 | 0.42275543 | 0.36253941 |
| Gm15860 | 0.20158871 | 1.42253204 | 0.031 | 0.01 | 1 | 0.41074402 | -0.7248928 | 0.017 | 0.031 | 1 | 0.41074402 | 0.36253941 |
| Gm43915 | 0.20158871 | 1.42253204 | 0.031 | 0.01 | 1 | 0.80442048 | -0.5728897 | 0.017 | 0.021 | 1 | 0.80442048 | 0.36253941 |
| Gm57243 | 0.20158871 | 1.42253204 | 0.031 | 0.01 | 1 | 0.40076941 | -1.7952821 | 0.017 | 0.031 | 1 | 0.40076941 | 0.36253941 |
| Zbtb32 | 0.20158871 | 1.42253204 | 0.031 | 0.01 | 1 | 0.34026631 | -2.5322477 | 0.008 | 0.021 | 1 | 0.34026631 | 0.36253941 |
| Tmem147os | 0.20158871 | 1.42253204 | 0.031 | 0.01 | 1 | 0.40361357 | -1.3098553 | 0.017 | 0.031 | 1 | 0.40361357 | 0.36253941 |
| Fgf20 | 0.20158871 | 1.42253204 | 0.031 | 0.01 | 1 | 0.40647274 | -0.9879272 | 0.017 | 0.031 | 1 | 0.40647274 | 0.36253941 |
| Amt | 0.20158871 | 1.42253204 | 0.031 | 0.01 | 1 | 0.30606883 | 0.69014474 | 0.029 | 0.01 | 1 | 0.30606883 | 0.36253941 |
| Efcab9 | 0.20158871 | 1.42253204 | 0.031 | 0.01 | 1 | 0.8259932 | -0.8244284 | 0.025 | 0.021 | 1 | 0.8259932 | 0.36253941 |
| Gm12111 | 0.20158871 | 1.42253204 | 0.031 | 0.01 | 1 | 0.39515719 | 0.49749966 | 0.025 | 0.01 | 1 | 0.39515719 | 0.36253941 |
| Gm57367 | 0.20158871 | 1.42253204 | 0.031 | 0.01 | 1 | 0.34363875 | -1.7248928 | 0.008 | 0.021 | 1 | 0.34363875 | 0.36253941 |
| Gm57337 | 0.20158871 | 1.42253204 | 0.031 | 0.01 | 1 | 0.5778028 | -0.8948178 | 0.013 | 0.021 | 1 | 0.5778028 | 0.36253941 |
| C030010C08Rik | 0.20158871 | 1.42253204 | 0.031 | 0.01 | 1 | 0.39515719 | 0.49749966 | 0.025 | 0.01 | 1 | 0.39515719 | 0.36253941 |
| Gm41291 | 0.20158871 | 1.42253204 | 0.031 | 0.01 | 1 | 0.39515719 | 0.49749966 | 0.025 | 0.01 | 1 | 0.39515719 | 0.36253941 |
| Pdxp | 0.20158871 | 1.42253204 | 0.031 | 0.01 | 1 | 0.20315163 | 1.01207283 | 0.017 | 0 | 1 | 0.20315163 | 0.36253941 |

| | | | | | | | | | | | | |
|---|---|---|---|---|---|---|---|---|---|---|---|---|
| Gpd1 | 0.20158871 | 1.42253204 | 0.031 | 0.01 | 1 | 0.74957918 | -1.3098553 | 0.025 | 0.031 | 1 | 0.74957918 | 0.36253941 |
| Gp5 | 0.20158871 | 1.42253204 | 0.031 | 0.01 | 1 | 0.20315163 | 1.01207283 | 0.017 | 0 | 1 | 0.20315163 | 0.36253941 |
| Gm16278 | 0.20158871 | 1.42253204 | 0.031 | 0.01 | 1 | 0.31054693 | 1.01207283 | 0.029 | 0.01 | 1 | 0.31054693 | 0.36253941 |
| Gm48696 | 0.20158871 | 1.42253204 | 0.031 | 0.01 | 1 | 0.53717181 | 0.27510724 | 0.034 | 0.021 | 1 | 0.53717181 | 0.36253941 |
| Banf2 | 0.20167483 | -1.240433 | 0.031 | 0.067 | 1 | 0.40647274 | -0.9879272 | 0.017 | 0.031 | 1 | 0.40647274 | 0.36267692 |
| Exoc3l2 | 0.20167483 | -1.4519371 | 0.031 | 0.067 | 1 | 0.87589833 | -0.5728897 | 0.013 | 0.01 | 1 | 0.87589833 | 0.36267692 |
| Mok | 0.43930534 | 0.47697982 | 0.062 | 0.041 | 1 | 0.20206361 | 0.36821664 | 0.097 | 0.052 | 1 | 0.43930534 | 0.36329751 |
| Dnah7b | 0.20219675 | 0.57686018 | 0.135 | 0.088 | 1 | 0.57728121 | -1.5322477 | 0.113 | 0.125 | 1 | 0.57728121 | 0.36350998 |
| Gba | 0.20229135 | 0.2912875 | 0.354 | 0.275 | 1 | 0.86646083 | -0.4557061 | 0.185 | 0.167 | 1 | 0.86646083 | 0.36366091 |
| Ncoa3 | 0.20229259 | -0.577468 | 0.833 | 0.829 | 1 | 0.90318394 | -0.2795727 | 0.912 | 0.948 | 1 | 0.90318394 | 0.36366289 |
| Taf15 | 0.20245281 | 0.20013961 | 0.844 | 0.756 | 1 | 0.60027682 | -0.1893216 | 0.79 | 0.854 | 1 | 0.60027682 | 0.36391848 |
| Rhd | 0.20275473 | 1.32942263 | 0.031 | 0.01 | 1 | 0.79600331 | -1.7952821 | 0.017 | 0.021 | 1 | 0.79600331 | 0.36439998 |
| Gm44837 | 0.20275473 | 1.32942263 | 0.031 | 0.01 | 1 | 0.50844268 | 0.69014474 | 0.021 | 0.01 | 1 | 0.50844268 | 0.36439998 |
| Ano4 | 0.20275473 | 1.32942263 | 0.031 | 0.01 | 1 | 0.23505706 | 1.27510724 | 0.034 | 0.01 | 1 | 0.23505706 | 0.36439998 |
| Clcn2 | 0.23968724 | 0.77302928 | 0.125 | 0.083 | 1 | 0.20275849 | 0.40063812 | 0.109 | 0.062 | 1 | 0.23968724 | 0.36440598 |
| Gm26802 | 0.42203853 | 0.46692616 | 0.198 | 0.155 | 1 | 0.2030814 | -0.6683092 | 0.223 | 0.281 | 1 | 0.42203853 | 0.36492075 |
| Phgdh | 0.20310446 | 0.37522632 | 0.146 | 0.093 | 1 | 0.24475855 | 0.8995981 | 0.113 | 0.073 | 1 | 0.24475855 | 0.3649575 |
| Ankrd60 | 0.21867159 | 1.59245704 | 0.021 | 0.005 | 1 | 0.20314753 | 1.27510724 | 0.017 | 0 | 1 | 0.21867159 | 0.36502615 |
| Spag16 | 0.22187388 | 1.00749454 | 0.021 | 0.005 | 1 | 0.20314753 | 1.27510724 | 0.017 | 0 | 1 | 0.22187388 | 0.36502615 |
| Lat | 0.22187388 | -1.577468 | 0 | 0.016 | 1 | 0.20314753 | 1.27510724 | 0.017 | 0 | 1 | 0.22187388 | 0.36502615 |
| 1700084E18Rik | 0.22187575 | -0.9925055 | 0 | 0.016 | 1 | 0.20314753 | 1.27510724 | 0.017 | 0 | 1 | 0.22187575 | 0.36502615 |
| Cxcr2 | 0.32025586 | -2.1624305 | 0 | 0.01 | 1 | 0.20314753 | 2.01207283 | 0.017 | 0 | 1 | 0.32025586 | 0.36502615 |
| Rgs7 | 0.32025586 | -0.9925055 | 0 | 0.01 | 1 | 0.20314753 | 1.27510724 | 0.017 | 0 | 1 | 0.32025586 | 0.36502615 |
| Cnfn | 0.32025586 | -0.9925055 | 0 | 0.01 | 1 | 0.20314753 | 1.27510724 | 0.017 | 0 | 1 | 0.32025586 | 0.36502615 |
| Csf2rb2 | 0.32025586 | -0.9925055 | 0 | 0.01 | 1 | 0.20314753 | 1.27510724 | 0.017 | 0 | 1 | 0.32025586 | 0.36502615 |
| Gm16178 | 0.38357309 | -1.4519371 | 0.01 | 0.026 | 1 | 0.20314753 | 1.27510724 | 0.017 | 0 | 1 | 0.38357309 | 0.36502615 |
| Gm30835 | 0.44890627 | -1.8368546 | 0.062 | 0.041 | 1 | 0.20314753 | 1.49749966 | 0.017 | 0 | 1 | 0.44890627 | 0.36502615 |
| Serpinb11 | 0.47150075 | 2.74446013 | 0.021 | 0.01 | 1 | 0.20314753 | 1.27510724 | 0.017 | 0 | 1 | 0.47150075 | 0.36502615 |
| Wfdc12 | 0.48219422 | -0.577468 | 0.021 | 0.036 | 1 | 0.20314753 | 1.27510724 | 0.017 | 0 | 1 | 0.48219422 | 0.36502615 |
| Cd200r4 | 0.61371908 | 2.00749454 | 0.01 | 0.005 | 1 | 0.20314753 | 1.27510724 | 0.017 | 0 | 1 | 0.61371908 | 0.36502615 |
| Bloc1s6os | 0.61737927 | 1.00749454 | 0.01 | 0.005 | 1 | 0.20314753 | 1.49749966 | 0.017 | 0 | 1 | 0.61737927 | 0.36502615 |
| Gm15902 | 0.70642533 | -0.3144336 | 0.052 | 0.041 | 1 | 0.20314753 | 1.27510724 | 0.017 | 0 | 1 | 0.70642533 | 0.36502615 |
| Dnali1 | 0.74827573 | 0.59245704 | 0.021 | 0.016 | 1 | 0.20314753 | 1.27510724 | 0.017 | 0 | 1 | 0.74827573 | 0.36502615 |
| Gm20597 | 0.22187575 | -0.9925055 | 0 | 0.016 | 1 | 0.20315061 | 1.86006974 | 0.017 | 0 | 1 | 0.22187575 | 0.36503105 |
| C130060C02Rik | 0.47606502 | 1.00749454 | 0.021 | 0.01 | 1 | 0.20315061 | 1.49749966 | 0.017 | 0 | 1 | 0.47606502 | 0.36503105 |
| Rasgrp1 | 0.993208 | 2.22988696 | 0.01 | 0.01 | 1 | 0.20315061 | 1.69014474 | 0.017 | 0 | 1 | 0.993208 | 0.36503105 |
| Gm12724 | 0.20784723 | -1.1624305 | 0.01 | 0.036 | 1 | 0.20315163 | 1.01207283 | 0.017 | 0 | 1 | 0.20784723 | 0.36503268 |
| Gm26594 | 0.21708019 | 2.00749454 | 0.021 | 0.005 | 1 | 0.20315163 | 1.01207283 | 0.017 | 0 | 1 | 0.21708019 | 0.36503268 |
| Gm5083 | 0.21867159 | 1.59245704 | 0.021 | 0.005 | 1 | 0.20315163 | 1.01207283 | 0.017 | 0 | 1 | 0.21867159 | 0.36503268 |
| Mypopos | 0.22187388 | -1.3144336 | 0 | 0.016 | 1 | 0.20315163 | 1.01207283 | 0.017 | 0 | 1 | 0.22187388 | 0.36503268 |
| Gm94 | 0.22187388 | -1.3144336 | 0 | 0.016 | 1 | 0.20315163 | 1.01207283 | 0.017 | 0 | 1 | 0.22187388 | 0.36503268 |
| Otud7a | 0.22187575 | -0.9925055 | 0 | 0.016 | 1 | 0.20315163 | 1.01207283 | 0.017 | 0 | 1 | 0.22187575 | 0.36503268 |
| Gm15832 | 0.25824304 | 0.59245704 | 0.052 | 0.026 | 1 | 0.20315163 | 1.01207283 | 0.017 | 0 | 1 | 0.25824304 | 0.36503268 |
| Miip | 0.25863087 | -1.0799683 | 0.031 | 0.062 | 1 | 0.20315163 | 1.01207283 | 0.017 | 0 | 1 | 0.25863087 | 0.36503268 |
| Adgrg7 | 0.28001913 | -1.577468 | 0.01 | 0.031 | 1 | 0.20315163 | 1.01207283 | 0.017 | 0 | 1 | 0.28001913 | 0.36503268 |
| Gm31224 | 0.28126256 | -1.1624305 | 0.01 | 0.031 | 1 | 0.20315163 | 1.01207283 | 0.017 | 0 | 1 | 0.28126256 | 0.36503268 |
| Plppr4 | 0.32025586 | -1.3144336 | 0 | 0.01 | 1 | 0.20315163 | 2.14957636 | 0.017 | 0 | 1 | 0.32025586 | 0.36503268 |
| Rab42 | 0.47606502 | 1.00749454 | 0.021 | 0.01 | 1 | 0.20315163 | 1.01207283 | 0.017 | 0 | 1 | 0.47606502 | 0.36503268 |
| Abcc2 | 0.47606502 | 1.00749454 | 0.021 | 0.01 | 1 | 0.20315163 | 1.01207283 | 0.017 | 0 | 1 | 0.47606502 | 0.36503268 |
| Gm21060 | 0.47606502 | 1.00749454 | 0.021 | 0.01 | 1 | 0.20315163 | 1.01207283 | 0.017 | 0 | 1 | 0.47606502 | 0.36503268 |
| Gm57374 | 0.52744059 | -0.9925055 | 0.01 | 0.021 | 1 | 0.20315163 | 1.01207283 | 0.017 | 0 | 1 | 0.52744059 | 0.36503268 |
| Gm57175 | 0.61737927 | 1.00749454 | 0.01 | 0.005 | 1 | 0.20315163 | 1.01207283 | 0.017 | 0 | 1 | 0.61737927 | 0.36503268 |
| A930012O16Rik | 0.61737927 | 1.00749454 | 0.01 | 0.005 | 1 | 0.20315163 | 1.01207283 | 0.017 | 0 | 1 | 0.61737927 | 0.36503268 |
| Dcst2 | 0.61737927 | 1.00749454 | 0.01 | 0.005 | 1 | 0.20315163 | 1.01207283 | 0.017 | 0 | 1 | 0.61737927 | 0.36503268 |
| Prss22 | 0.61737927 | 1.00749454 | 0.01 | 0.005 | 1 | 0.20315163 | 1.01207283 | 0.017 | 0 | 1 | 0.61737927 | 0.36503268 |
| Calb2 | 0.62104899 | 0.42253204 | 0.01 | 0.005 | 1 | 0.20315163 | 1.01207283 | 0.017 | 0 | 1 | 0.62104899 | 0.36503268 |
| Ica1l | 0.67270388 | -0.3710171 | 0.042 | 0.031 | 1 | 0.20315163 | 1.01207283 | 0.017 | 0 | 1 | 0.67270388 | 0.36503268 |
| Gcnt7 | 0.72298091 | -0.577468 | 0.01 | 0.016 | 1 | 0.20315163 | 1.01207283 | 0.017 | 0 | 1 | 0.72298091 | 0.36503268 |
| Gm30881 | 0.73685701 | 0.59245704 | 0.01 | 0.016 | 1 | 0.20315163 | 1.01207283 | 0.017 | 0 | 1 | 0.73685701 | 0.36503268 |
| Gm36445 | 0.74827573 | 0.59245704 | 0.021 | 0.016 | 1 | 0.20315163 | 1.01207283 | 0.017 | 0 | 1 | 0.74827573 | 0.36503268 |
| AV039307 | 0.82258416 | -0.1624305 | 0.031 | 0.036 | 1 | 0.20315163 | 1.01207283 | 0.017 | 0 | 1 | 0.82258416 | 0.36503268 |
| Pcdh15 | 0.993208 | 1.00749454 | 0.01 | 0.01 | 1 | 0.20315163 | 1.01207283 | 0.017 | 0 | 1 | 0.993208 | 0.36503268 |
| Gm49291 | 0.993208 | 1.00749454 | 0.01 | 0.01 | 1 | 0.20315163 | 1.01207283 | 0.017 | 0 | 1 | 0.993208 | 0.36503268 |
| Gm46210 | 0.99603742 | 0.20013961 | 0.031 | 0.031 | 1 | 0.20315163 | 1.01207283 | 0.017 | 0 | 1 | 0.99603742 | 0.36503268 |
| Psma8 | 0.99758609 | 0.27052894 | 0.021 | 0.021 | 1 | 0.20315163 | 1.01207283 | 0.017 | 0 | 1 | 0.99758609 | 0.36503268 |
| Gm14302 | 1 | 0.42253204 | 0.01 | 0.01 | 1 | 0.20315163 | 1.01207283 | 0.017 | 0 | 1 | 1 | 0.36503268 |
| Zfp750 | 1 | 0.42253204 | 0.01 | 0.01 | 1 | 0.20315163 | 1.01207283 | 0.017 | 0 | 1 | 1 | 0.36503268 |
| Gm17949 | 1 | 0.42253204 | 0.01 | 0.01 | 1 | 0.20315163 | 1.01207283 | 0.017 | 0 | 1 | 1 | 0.36503268 |
| Prob1 | 1 | 0.42253204 | 0.01 | 0.01 | 1 | 0.20315163 | 1.01207283 | 0.017 | 0 | 1 | 1 | 0.36503268 |
| 1700084C06Rik | 0.20355024 | 0.91438513 | 0.104 | 0.062 | 1 | 0.81220604 | -0.5854897 | 0.105 | 0.094 | 1 | 0.81220604 | 0.36566778 |
| Kirrel3 | 0.52289048 | -0.4450177 | 0.125 | 0.098 | 1 | 0.2036299 | 0.58091567 | 0.126 | 0.073 | 1 | 0.52289048 | 0.36579467 |
| Bcl2 | 0.20370548 | -0.68255 | 0.333 | 0.425 | 1 | 0.44988321 | 0.49935522 | 0.458 | 0.406 | 1 | 0.44988321 | 0.36591504 |
| Papln | 0.80977653 | -0.6705774 | 0.031 | 0.036 | 1 | 0.20412561 | 0.69014474 | 0.067 | 0.031 | 1 | 0.80977653 | 0.36658396 |
| Gm10353 | 0.95503809 | 0.59245704 | 0.052 | 0.052 | 1 | 0.20413293 | -0.473354 | 0.097 | 0.052 | 1 | 0.95503809 | 0.36659561 |
| Cdk17 | 0.44474076 | -0.2335136 | 0.74 | 0.679 | 1 | 0.20416849 | -0.2744064 | 0.811 | 0.927 | 1 | 0.44474076 | 0.36665221 |
| Ccl19 | 0.20438593 | 1.17741954 | 0.062 | 0.031 | 1 | 0.81129592 | 0.10518224 | 0.025 | 0.021 | 1 | 0.81129592 | 0.36699824 |
| Adnp2 | 0.51630594 | 0.46000674 | 0.26 | 0.223 | 1 | 0.20441914 | -0.1200307 | 0.244 | 0.167 | 1 | 0.51630594 | 0.3670511 |
| Tmem37 | 0.20470878 | 0.74446013 | 0.104 | 0.062 | 1 | 0.98848918 | -0.7248928 | 0.042 | 0.042 | 1 | 0.98848918 | 0.36751188 |
| Dgcr6 | 0.80584331 | -0.2295447 | 0.406 | 0.347 | 1 | 0.20473044 | 0.27510724 | 0.324 | 0.24 | 1 | 0.80584331 | 0.36754632 |
| Nin | 0.35190374 | -0.6810906 | 0.396 | 0.456 | 1 | 0.20490515 | -0.5266667 | 0.496 | 0.573 | 1 | 0.35190374 | 0.36782418 |
| Gm6598 | 0.20510378 | 0.42253204 | 0.031 | 0.01 | 1 | 0.34704016 | -1.3098553 | 0.008 | 0.021 | 1 | 0.34704016 | 0.36814001 |
| BC050972 | 0.20510378 | 1.00749454 | 0.031 | 0.01 | 1 | 0.87066325 | -0.3098553 | 0.013 | 0.01 | 1 | 0.87066325 | 0.36814001 |
| Zfp85 | 0.20510378 | 1.00749454 | 0.031 | 0.01 | 1 | 0.81129592 | 0.10518224 | 0.025 | 0.021 | 1 | 0.81129592 | 0.36814001 |
| Gm10037 | 0.20510378 | 1.00749454 | 0.031 | 0.01 | 1 | 0.27154471 | 0.69014474 | 0.013 | 0 | 1 | 0.27154471 | 0.36814001 |
| Espl1 | 0.20510378 | 0.20013961 | 0.031 | 0.01 | 1 | 0.86312513 | -0.7248928 | 0.008 | 0.01 | 1 | 0.86312513 | 0.36814001 |
| Rnf138rt1 | 0.20510378 | 1.00749454 | 0.031 | 0.01 | 1 | 0.66387437 | 0.49749966 | 0.017 | 0.01 | 1 | 0.66387437 | 0.36814001 |
| Trib1 | 0.51644174 | -0.1065772 | 0.74 | 0.767 | 1 | 0.20548993 | 0.37026447 | 0.71 | 0.76 | 1 | 0.51644174 | 0.36875374 |

| | | | | | | | | | | | | |
|---|---|---|---|---|---|---|---|---|---|---|---|---|
| Hk1os | 0.20562978 | -0.8993961 | 0.031 | 0.067 | 1 | 0.51194506 | 0.27510724 | 0.021 | 0.01 | 1 | 0.51194506 | 0.36897595 |
| Rpa1 | 0.57201948 | -0.1079827 | 0.198 | 0.228 | 1 | 0.20564153 | 0.12077909 | 0.218 | 0.146 | 1 | 0.57201948 | 0.36899462 |
| 6330411D24Rik | 0.57291331 | 0.20013961 | 0.062 | 0.047 | 1 | 0.20564316 | -1.4029647 | 0.025 | 0.052 | 1 | 0.57291331 | 0.3689972 |
| Rnf180 | 0.20596752 | -2.1079827 | 0.021 | 0.052 | 1 | 0.23259197 | 1.21370669 | 0.05 | 0.021 | 1 | 0.23259197 | 0.36951242 |
| Gm57095 | 0.20597966 | -1.577468 | 0.01 | 0.036 | 1 | 0.40076941 | -1.3098553 | 0.017 | 0.031 | 1 | 0.40076941 | 0.36953169 |
| Rassf6 | 0.20598035 | -1.4519371 | 0.01 | 0.036 | 1 | 0.5778028 | -0.8948178 | 0.013 | 0.021 | 1 | 0.5778028 | 0.3695328 |
| Tafa4 | 0.20598035 | -1.4519371 | 0.01 | 0.036 | 1 | 0.57170879 | -1.6317834 | 0.013 | 0.021 | 1 | 0.57170879 | 0.3695328 |
| Ppp1r14a | 0.20598035 | -1.6929452 | 0.01 | 0.036 | 1 | 0.23592811 | 1.14957636 | 0.034 | 0.01 | 1 | 0.23592811 | 0.3695328 |
| Gm15563 | 0.20598035 | -1.4519371 | 0.01 | 0.036 | 1 | 0.64978199 | 0.36821664 | 0.042 | 0.031 | 1 | 0.64978199 | 0.3695328 |
| Gng4 | 0.20598035 | -1.4519371 | 0.01 | 0.036 | 1 | 0.35831374 | -0.1723517 | 0.042 | 0.021 | 1 | 0.35831374 | 0.3695328 |
| Map3k15 | 0.20598035 | -1.4519371 | 0.01 | 0.036 | 1 | 0.85710544 | -1.7248928 | 0.008 | 0.01 | 1 | 0.85710544 | 0.3695328 |
| Syf2 | 0.96561353 | -0.1696985 | 0.573 | 0.57 | 1 | 0.20604327 | -0.5482276 | 0.664 | 0.719 | 1 | 0.96561353 | 0.36963272 |
| Mccc1 | 0.38381121 | 0.21899864 | 0.177 | 0.135 | 1 | 0.20617702 | -0.357161 | 0.193 | 0.125 | 1 | 0.38381121 | 0.36984508 |
| Wasf3 | 0.20630717 | -0.6340515 | 0.208 | 0.269 | 1 | 0.54343435 | -0.1222283 | 0.239 | 0.208 | 1 | 0.54343435 | 0.37005169 |
| D130040H23Rik | 0.20653035 | 0.56692195 | 0.104 | 0.062 | 1 | 0.88318309 | -0.5163061 | 0.08 | 0.073 | 1 | 0.88318309 | 0.37040591 |
| Nsrp1 | 0.20657762 | 0.24792751 | 0.552 | 0.466 | 1 | 0.4644775 | -0.1543377 | 0.592 | 0.5 | 1 | 0.4644775 | 0.37048092 |
| Cplx1 | 0.20691434 | -1.4519371 | 0.01 | 0.036 | 1 | 0.57854956 | -0.8948178 | 0.021 | 0.031 | 1 | 0.57854956 | 0.37101513 |
| Gm15991 | 0.20691434 | -1.3144336 | 0.01 | 0.036 | 1 | 0.96042911 | -0.5025003 | 0.055 | 0.052 | 1 | 0.96042911 | 0.37101513 |
| Evi2b | 0.20691434 | -1.3144336 | 0.01 | 0.036 | 1 | 0.92228543 | -0.8504236 | 0.029 | 0.031 | 1 | 0.92228543 | 0.37101513 |
| Pmaip1 | 0.20691434 | -1.3144336 | 0.01 | 0.036 | 1 | 0.41665803 | -1.3098553 | 0.025 | 0.042 | 1 | 0.41665803 | 0.37101513 |
| Bckdhb | 0.33415687 | 0.10605837 | 0.396 | 0.332 | 1 | 0.20720018 | -0.3225663 | 0.345 | 0.25 | 1 | 0.33415687 | 0.37146844 |
| Mmachc | 0.20720192 | 1.00749454 | 0.083 | 0.047 | 1 | 0.59609426 | -0.1723517 | 0.08 | 0.062 | 1 | 0.59609426 | 0.37147121 |
| BC051226 | 0.20720192 | 1.00749454 | 0.083 | 0.047 | 1 | 0.66987161 | -0.1723517 | 0.029 | 0.021 | 1 | 0.66987161 | 0.37147121 |
| Eefsec | 0.50152342 | 0.346981 | 0.219 | 0.187 | 1 | 0.2072706 | 0.13308823 | 0.248 | 0.177 | 1 | 0.50152342 | 0.3715801 |
| Fer1l6 | 0.20751518 | -1.8669746 | 0.021 | 0.052 | 1 | 0.50887612 | -1.3098553 | 0.004 | 0.01 | 1 | 0.50887612 | 0.37196782 |
| Galnt17 | 0.2075166 | -2.7998604 | 0.021 | 0.052 | 1 | 0.77977619 | -1.5025003 | 0.038 | 0.031 | 1 | 0.77977619 | 0.37197007 |
| Dock7 | 0.20753592 | -0.3500575 | 0.823 | 0.85 | 1 | 0.97382094 | -0.234631 | 0.866 | 0.885 | 1 | 0.97382094 | 0.37200069 |
| Plcb3 | 0.20768699 | 0.39744105 | 0.177 | 0.119 | 1 | 0.38474453 | -0.3098553 | 0.202 | 0.146 | 1 | 0.38474453 | 0.3722401 |
| Rapgef5 | 0.2077826 | -1.0628948 | 0.031 | 0.067 | 1 | 0.64838591 | -1.357161 | 0.05 | 0.062 | 1 | 0.64838591 | 0.37239159 |
| Gm49085 | 0.44126404 | -0.130009 | 0.031 | 0.052 | 1 | 0.20780492 | -1.6317834 | 0.025 | 0.052 | 1 | 0.44126404 | 0.37242695 |
| Gm26810 | 0.20784723 | -1.1624305 | 0.01 | 0.036 | 1 | 0.56243824 | -0.8948178 | 0.029 | 0.042 | 1 | 0.56243824 | 0.37249399 |
| Abca7 | 0.20784723 | -1.1624305 | 0.01 | 0.036 | 1 | 0.36719871 | 0.49749966 | 0.055 | 0.031 | 1 | 0.36719871 | 0.37249399 |
| BC049762 | 0.20784723 | -1.4519371 | 0.01 | 0.036 | 1 | 0.82415676 | -0.8948178 | 0.025 | 0.021 | 1 | 0.82415676 | 0.37249399 |
| Gm32036 | 0.20784723 | -1.1624305 | 0.01 | 0.036 | 1 | 0.86312513 | -0.7248928 | 0.008 | 0.01 | 1 | 0.86312513 | 0.37249399 |
| Gm33732 | 0.20784723 | -1.1624305 | 0.01 | 0.036 | 1 | 0.34703906 | -1.3098553 | 0.008 | 0.021 | 1 | 0.34703906 | 0.37249399 |
| Nudt19 | 0.59045904 | -0.1989563 | 0.312 | 0.337 | 1 | 0.20795063 | -0.3098553 | 0.239 | 0.167 | 1 | 0.59045904 | 0.37265779 |
| Ino80 | 0.20805901 | 0.10060394 | 0.844 | 0.824 | 1 | 0.58760455 | -0.3328386 | 0.895 | 0.896 | 1 | 0.58760455 | 0.37282946 |
| Etv3 | 0.20821192 | 0.46692616 | 0.323 | 0.244 | 1 | 0.7581849 | -0.1926717 | 0.382 | 0.344 | 1 | 0.7581849 | 0.37307163 |
| Bzw2 | 0.94622952 | -0.182397 | 0.74 | 0.756 | 1 | 0.20822826 | 0.16471344 | 0.676 | 0.667 | 1 | 0.94622952 | 0.37309751 |
| D730045B01Rik | 0.48840769 | 0.15949763 | 0.042 | 0.026 | 1 | 0.20853636 | -1.6317834 | 0.025 | 0.052 | 1 | 0.48840769 | 0.37358531 |
| Gm15564 | 0.52744059 | -0.577468 | 0.01 | 0.021 | 1 | 0.20853636 | -1.4797803 | 0.025 | 0.052 | 1 | 0.52744059 | 0.37358531 |
| Gm10399 | 0.20868113 | 0.68556644 | 0.031 | 0.01 | 1 | 0.39234595 | 1.14957636 | 0.025 | 0.01 | 1 | 0.39234595 | 0.37381444 |
| Ankfn1 | 0.20868113 | 0.68556644 | 0.031 | 0.01 | 1 | 0.85710544 | -1.3098553 | 0.008 | 0.01 | 1 | 0.85710544 | 0.37381444 |
| 1810059H22Rik | 0.20883718 | -0.9925055 | 0.01 | 0.036 | 1 | 0.53433287 | 0.42711033 | 0.034 | 0.021 | 1 | 0.53433287 | 0.37406139 |
| Tril | 0.20883718 | -0.9925055 | 0.01 | 0.036 | 1 | 0.87066325 | -0.3098553 | 0.013 | 0.01 | 1 | 0.87066325 | 0.37406139 |
| Cracr2a | 0.20883718 | -0.9925055 | 0.01 | 0.036 | 1 | 0.45095839 | 0.45567948 | 0.05 | 0.031 | 1 | 0.45095839 | 0.37406139 |
| B930046K06Rik | 0.20902867 | 1.00749454 | 0.062 | 0.031 | 1 | 0.79619779 | -0.6577786 | 0.038 | 0.031 | 1 | 0.79619779 | 0.37436436 |
| Gm49077 | 0.20902867 | 1.00749454 | 0.062 | 0.031 | 1 | 0.67150494 | -0.5728897 | 0.029 | 0.021 | 1 | 0.67150494 | 0.37436436 |
| Gm48693 | 0.20906516 | -1.407543 | 0.021 | 0.052 | 1 | 0.31354397 | -0.6094155 | 0.029 | 0.052 | 1 | 0.31354397 | 0.37442207 |
| Gm46367 | 0.20906516 | -1.6554705 | 0.021 | 0.052 | 1 | 0.77674145 | -0.7793405 | 0.084 | 0.073 | 1 | 0.77674145 | 0.37442207 |
| Aknad1 | 0.20910685 | 1.00749454 | 0.062 | 0.031 | 1 | 0.52989515 | -0.7248928 | 0.046 | 0.062 | 1 | 0.52989515 | 0.37448802 |
| Lrfn1 | 0.20910685 | 1.00749454 | 0.062 | 0.031 | 1 | 0.93071546 | -0.3098553 | 0.029 | 0.031 | 1 | 0.93071546 | 0.37448802 |
| Meikin | 0.20910685 | 1.00749454 | 0.062 | 0.031 | 1 | 0.53497636 | 0.39058446 | 0.046 | 0.031 | 1 | 0.53497636 | 0.37448802 |
| Zfp97 | 0.20910685 | 1.00749454 | 0.062 | 0.031 | 1 | 0.43079893 | 0.56461386 | 0.038 | 0.021 | 1 | 0.43079893 | 0.37448802 |
| Gm8251 | 0.91699449 | -0.1345245 | 0.083 | 0.088 | 1 | 0.20924018 | -0.1269912 | 0.151 | 0.094 | 1 | 0.91699449 | 0.37469891 |
| Ptcd2 | 0.26059398 | 0.42253204 | 0.094 | 0.057 | 1 | 0.20925939 | -0.1034044 | 0.109 | 0.062 | 1 | 0.26059398 | 0.37472929 |
| Fam110c | 0.48064264 | 0.59245704 | 0.021 | 0.01 | 1 | 0.20927043 | -1.3098553 | 0.025 | 0.052 | 1 | 0.48064264 | 0.37474675 |
| Gm9968 | 0.69676098 | -0.407543 | 0.042 | 0.052 | 1 | 0.20947543 | -1.3098553 | 0.067 | 0.104 | 1 | 0.69676098 | 0.3750709 |
| Gm17435 | 0.25544528 | -0.3144336 | 0.115 | 0.166 | 1 | 0.20953355 | -0.2358547 | 0.151 | 0.094 | 1 | 0.25544528 | 0.37516279 |
| Pakap | 0.45720871 | -0.8756918 | 0.24 | 0.264 | 1 | 0.20991654 | -1.8608704 | 0.206 | 0.24 | 1 | 0.45720871 | 0.37576813 |
| Mzf1 | 0.38028858 | 1.00749454 | 0.031 | 0.016 | 1 | 0.20997488 | -1.5025003 | 0.025 | 0.052 | 1 | 0.38028858 | 0.37586031 |
| Mettl15 | 0.34954614 | 0.27052894 | 0.25 | 0.202 | 1 | 0.21052598 | -0.2609457 | 0.185 | 0.115 | 1 | 0.34954614 | 0.37673077 |
| Eri1 | 0.21053526 | 0.70493177 | 0.333 | 0.275 | 1 | 0.21103744 | -0.1195241 | 0.261 | 0.177 | 1 | 0.21103744 | 0.37674543 |
| Gm56935 | 0.21062121 | -1.4950058 | 0.021 | 0.052 | 1 | 0.30323558 | -1.6883669 | 0.038 | 0.062 | 1 | 0.30323558 | 0.37688113 |
| Slc4a11 | 0.21062169 | -1.2148979 | 0.021 | 0.052 | 1 | 0.642845 | -0.3098553 | 0.055 | 0.042 | 1 | 0.642845 | 0.37688188 |
| Noxred1 | 0.21062169 | -1.2148979 | 0.021 | 0.052 | 1 | 0.61588728 | -0.2318528 | 0.067 | 0.052 | 1 | 0.61588728 | 0.37688188 |
| Gm16001 | 0.21062881 | 1.00749454 | 0.083 | 0.047 | 1 | 0.74545301 | -0.5508634 | 0.084 | 0.094 | 1 | 0.74545301 | 0.37689312 |
| Gen1 | 0.21063688 | 0.47697982 | 0.062 | 0.031 | 1 | 0.68770566 | -1.3098553 | 0.034 | 0.042 | 1 | 0.68770566 | 0.37690587 |
| Slu7 | 0.21069165 | 0.21899864 | 0.469 | 0.394 | 1 | 0.37394197 | 0.11072042 | 0.441 | 0.396 | 1 | 0.37394197 | 0.37699234 |
| Tmem26 | 0.32933347 | 0.15949763 | 0.042 | 0.021 | 1 | 0.2107538 | -1.5728897 | 0.025 | 0.052 | 1 | 0.32933347 | 0.37709044 |
| 1810062G17Rik | 0.21091066 | -0.5070786 | 0.115 | 0.171 | 1 | 0.76553682 | -0.486733 | 0.084 | 0.073 | 1 | 0.76553682 | 0.37733802 |
| Zcchc14 | 0.21139954 | -0.3550755 | 0.76 | 0.782 | 1 | 0.99195704 | -0.1370959 | 0.945 | 0.917 | 1 | 0.99195704 | 0.37810932 |
| Rpgrip1l | 0.21155448 | 0.81109732 | 0.281 | 0.223 | 1 | 0.7157766 | -0.5783441 | 0.244 | 0.208 | 1 | 0.7157766 | 0.37835366 |
| Gm16341 | 0.21169852 | -1.3144336 | 0.01 | 0.036 | 1 | 0.50887612 | -1.3098553 | 0.004 | 0.01 | 1 | 0.50887612 | 0.37858078 |
| Usp27x | 0.76422355 | -0.2148979 | 0.083 | 0.093 | 1 | 0.21171395 | 0.76053407 | 0.08 | 0.042 | 1 | 0.76422355 | 0.3786051 |
| Antxr2 | 0.21191353 | 0.11440974 | 0.375 | 0.29 | 1 | 0.29159302 | -0.9511686 | 0.508 | 0.5 | 1 | 0.29159302 | 0.37891971 |
| Pcdhga8 | 0.21218051 | -1.3144336 | 0.021 | 0.052 | 1 | 0.54933729 | -0.194378 | 0.046 | 0.031 | 1 | 0.54933729 | 0.37934045 |
| C1qtnf9 | 0.21257404 | 0.86999101 | 0.083 | 0.047 | 1 | 0.99233097 | -0.8692827 | 0.076 | 0.073 | 1 | 0.99233097 | 0.37996035 |
| 4833438C02Rik | 0.21257404 | 0.86999101 | 0.083 | 0.047 | 1 | 0.81129592 | 0.10518224 | 0.025 | 0.021 | 1 | 0.81129592 | 0.37996035 |
| Pcdhga12 | 0.21258545 | -0.7810014 | 0.104 | 0.155 | 1 | 0.46452232 | -1.5025003 | 0.084 | 0.104 | 1 | 0.46452232 | 0.37997833 |
| Kpna2 | 0.30772405 | 0.49292136 | 0.115 | 0.078 | 1 | 0.2129155 | 0.32757466 | 0.172 | 0.115 | 1 | 0.30772405 | 0.38049799 |
| Trp53cor1 | 0.21296291 | -1.4705528 | 0.052 | 0.093 | 1 | 0.59419174 | 0.14957636 | 0.08 | 0.062 | 1 | 0.59419174 | 0.38057262 |
| Gucy1a2 | 0.2130338 | -1.4519371 | 0.021 | 0.052 | 1 | 0.29215666 | 0.3068161 | 0.126 | 0.083 | 1 | 0.29215666 | 0.3806842 |

| | | | | | | | | | | | | |
|---|---|---|---|---|---|---|---|---|---|---|---|---|
| Szt2 | 0.21318438 | -0.9100433 | 0.083 | 0.13 | 1 | 0.35816195 | 0.23446525 | 0.13 | 0.094 | 1 | 0.35816195 | 0.38092118 |
| Zc3h8 | 0.21331308 | 0.76648644 | 0.083 | 0.047 | 1 | 0.37360153 | 0.63767732 | 0.08 | 0.052 | 1 | 0.37360153 | 0.3811237 |
| Chmp1b | 0.74031406 | 0.11767745 | 0.208 | 0.233 | 1 | 0.21332438 | 0.49749966 | 0.277 | 0.219 | 1 | 0.74031406 | 0.38114146 |
| 4632427E13Rik | 0.21365408 | -0.4167146 | 0.479 | 0.539 | 1 | 0.3073766 | -0.3928758 | 0.563 | 0.594 | 1 | 0.3073766 | 0.38166009 |
| Gm29585 | 0.28328178 | -1.9925055 | 0.021 | 0.047 | 1 | 0.21369798 | -1.1172102 | 0.025 | 0.052 | 1 | 0.28328178 | 0.38172913 |
| Gm4681 | 0.52313476 | -0.9925055 | 0.01 | 0.021 | 1 | 0.21369798 | -1.1172102 | 0.025 | 0.052 | 1 | 0.52313476 | 0.38172913 |
| Ribc1 | 0.21374303 | 0.49292136 | 0.062 | 0.031 | 1 | 0.66387437 | 0.27510724 | 0.017 | 0.01 | 1 | 0.66387437 | 0.38179997 |
| Abca6 | 0.21374303 | -1.3144336 | 0.021 | 0.052 | 1 | 0.82415676 | -0.6317834 | 0.025 | 0.021 | 1 | 0.82415676 | 0.38179997 |
| C030034I22Rik | 0.21374303 | -0.9925055 | 0.021 | 0.052 | 1 | 0.46727722 | -0.2394659 | 0.063 | 0.042 | 1 | 0.46727722 | 0.38179997 |
| 4930481A15Rik | 0.21379069 | 0.83756954 | 0.062 | 0.031 | 1 | 0.57373861 | -1.3098553 | 0.013 | 0.021 | 1 | 0.57373861 | 0.38187492 |
| Fanci | 0.27403968 | -0.4330781 | 0.094 | 0.057 | 1 | 0.21410415 | -2.189561 | 0.059 | 0.094 | 1 | 0.27403968 | 0.38236771 |
| Ice2 | 0.21426025 | -0.3795286 | 0.156 | 0.223 | 1 | 0.25032545 | 0.16221318 | 0.239 | 0.177 | 1 | 0.25032545 | 0.38261305 |
| Prpf40b | 0.5781445 | 0.1843723 | 0.198 | 0.166 | 1 | 0.21437079 | 0.20204378 | 0.261 | 0.188 | 1 | 0.5781445 | 0.38278675 |
| Agk | 0.67685006 | 0.21741761 | 0.281 | 0.264 | 1 | 0.21459678 | 0.12013258 | 0.265 | 0.188 | 1 | 0.67685006 | 0.38314179 |
| Phkg1 | 0.21461929 | -1.240433 | 0.021 | 0.052 | 1 | 0.32744845 | -0.7248928 | 0.046 | 0.073 | 1 | 0.32744845 | 0.38317715 |
| Vars | 0.2147322 | 0.43717881 | 0.26 | 0.197 | 1 | 0.40039516 | -0.1119159 | 0.265 | 0.208 | 1 | 0.40039516 | 0.38335449 |
| Gas7 | 0.21498459 | -0.6834014 | 0.156 | 0.223 | 1 | 0.79712366 | -0.5881564 | 0.218 | 0.188 | 1 | 0.79712366 | 0.3837508 |
| Cmtm6 | 0.21507349 | 0.61156586 | 0.292 | 0.228 | 1 | 1 | -0.1242996 | 0.353 | 0.354 | 1 | 1 | 0.38389037 |
| Arf4os | 0.21535119 | 0.70793425 | 0.104 | 0.062 | 1 | 0.69948596 | -1.0297473 | 0.063 | 0.073 | 1 | 0.69948596 | 0.38432625 |
| Enkur | 0.21540263 | -0.7087125 | 0.031 | 0.067 | 1 | 0.93495836 | -0.1723517 | 0.055 | 0.052 | 1 | 0.93495836 | 0.38440697 |
| Trim7 | 0.21544532 | -0.8669746 | 0.021 | 0.052 | 1 | 0.28600429 | 0.27510724 | 0.046 | 0.021 | 1 | 0.28600429 | 0.38447396 |
| E2f7 | 0.21550082 | 3.00749454 | 0.021 | 0.005 | 1 | 0.34363875 | -2.5322477 | 0.008 | 0.021 | 1 | 0.34363875 | 0.38456103 |
| Rpl39l | 0.21550082 | 3.17741954 | 0.021 | 0.005 | 1 | 0.51188611 | -0.3098553 | 0.004 | 0.01 | 1 | 0.51188611 | 0.38456103 |
| Gm13166 | 0.21555612 | -0.1624305 | 0.01 | 0.036 | 1 | 0.23772424 | 0.86006974 | 0.034 | 0.01 | 1 | 0.23772424 | 0.38464781 |
| Dhx58 | 0.21570844 | -0.8669746 | 0.052 | 0.093 | 1 | 0.53977436 | -1.6577786 | 0.038 | 0.052 | 1 | 0.53977436 | 0.38488675 |
| Lamc2 | 0.2159763 | -1.5532204 | 0.052 | 0.093 | 1 | 0.3719752 | 1.62874419 | 0.046 | 0.073 | 1 | 0.3719752 | 0.38530684 |
| Bmpr2 | 0.51695841 | -0.2854499 | 0.875 | 0.881 | 1 | 0.21601492 | -0.1892763 | 0.987 | 0.99 | 1 | 0.51695841 | 0.38536739 |
| Zfp946 | 0.21655875 | 0.53800925 | 0.104 | 0.062 | 1 | 0.40930998 | 0.56461386 | 0.101 | 0.073 | 1 | 0.40930998 | 0.38621981 |
| Gng7 | 0.37571376 | 1.00749454 | 0.083 | 0.057 | 1 | 0.21660476 | 0.58322953 | 0.08 | 0.042 | 1 | 0.37571376 | 0.38629189 |
| Prr5l | 0.97404148 | -0.2335136 | 0.198 | 0.192 | 1 | 0.21663073 | -0.6336437 | 0.319 | 0.375 | 1 | 0.97404148 | 0.38633259 |
| Dpcd | 0.94555245 | 0.16361374 | 0.281 | 0.285 | 1 | 0.21683863 | -0.1560499 | 0.265 | 0.188 | 1 | 0.94555245 | 0.38665827 |
| Tcp11l2 | 0.44906585 | 0.12501784 | 0.625 | 0.606 | 1 | 0.21697706 | 0.29150168 | 0.756 | 0.75 | 1 | 0.44906585 | 0.38687508 |
| Gm44127 | 0.21701069 | -1.4519371 | 0.021 | 0.052 | 1 | 0.8065254 | -0.1399303 | 0.017 | 0.021 | 1 | 0.8065254 | 0.38692774 |
| Chm | 0.21702692 | 0.13873907 | 0.698 | 0.632 | 1 | 0.93158741 | -0.2017254 | 0.765 | 0.74 | 1 | 0.93158741 | 0.38695316 |
| Bank1 | 0.21708019 | 4.59245704 | 0.021 | 0.005 | 1 | 0.50587544 | -2.3098553 | 0.004 | 0.01 | 1 | 0.50587544 | 0.38703656 |
| Zfp981 | 0.21708019 | 2.32942263 | 0.021 | 0.005 | 1 | 0.50887612 | -1.3098553 | 0.004 | 0.01 | 1 | 0.50887612 | 0.38703656 |
| Rhoh | 0.21708019 | 2.59245704 | 0.021 | 0.005 | 1 | 0.51188611 | -0.7248928 | 0.004 | 0.01 | 1 | 0.51188611 | 0.38703656 |
| Syne4 | 0.21708019 | 2.00749454 | 0.021 | 0.005 | 1 | 0.86613737 | -0.3098553 | 0.008 | 0.01 | 1 | 0.86613737 | 0.38703656 |
| B4galnt4 | 0.21708019 | 2.00749454 | 0.021 | 0.005 | 1 | 0.27154471 | 0.69014474 | 0.013 | 0 | 1 | 0.27154471 | 0.38703656 |
| Gm16104 | 0.21708019 | 2.00749454 | 0.021 | 0.005 | 1 | 0.86613737 | -0.3098553 | 0.008 | 0.01 | 1 | 0.86613737 | 0.38703656 |
| H3c11 | 0.21708019 | 2.32942263 | 0.021 | 0.005 | 1 | 0.50587544 | -2.6317834 | 0.004 | 0.01 | 1 | 0.50587544 | 0.38703656 |
| Zfp85os | 0.21708019 | 2.00749454 | 0.021 | 0.005 | 1 | 0.27154471 | 0.69014474 | 0.013 | 0 | 1 | 0.27154471 | 0.38703656 |
| Emx2os | 0.21708019 | 2.32942263 | 0.021 | 0.005 | 1 | 0.51018266 | 0.49749966 | 0.021 | 0.01 | 1 | 0.51018266 | 0.38703656 |
| Gemin8 | 0.33777011 | 0.42253204 | 0.094 | 0.062 | 1 | 0.21712302 | 0.80562195 | 0.092 | 0.052 | 1 | 0.33777011 | 0.38710363 |
| Fktn | 0.40067837 | 0.2912875 | 0.219 | 0.176 | 1 | 0.21729524 | -0.1172102 | 0.231 | 0.156 | 1 | 0.40067837 | 0.38737325 |
| E130311K13Rik | 0.21750829 | 0.75373794 | 0.167 | 0.119 | 1 | 0.29563324 | 0.69014474 | 0.109 | 0.073 | 1 | 0.29563324 | 0.38770673 |
| Alg12 | 0.56110545 | 0.45495351 | 0.125 | 0.104 | 1 | 0.21763515 | 0.62874419 | 0.08 | 0.042 | 1 | 0.56110545 | 0.38790524 |
| Fat3 | 0.21776205 | -0.130009 | 0.177 | 0.119 | 1 | 0.35851409 | -0.9414679 | 0.155 | 0.188 | 1 | 0.35851409 | 0.38810379 |
| Gm46515 | 0.21781736 | -1.3144336 | 0.021 | 0.052 | 1 | 0.43798374 | 0.27510724 | 0.038 | 0.021 | 1 | 0.43798374 | 0.38819033 |
| Gm8098 | 0.21803987 | 0.74446013 | 0.083 | 0.047 | 1 | 0.77750728 | -1.1034044 | 0.05 | 0.042 | 1 | 0.77750728 | 0.38853835 |
| Lrrn3 | 0.47756351 | -0.1079827 | 0.042 | 0.026 | 1 | 0.2180708 | 1.42711033 | 0.05 | 0.021 | 1 | 0.47756351 | 0.38858673 |
| Gm29292 | 0.66663755 | 0.68556644 | 0.052 | 0.041 | 1 | 0.2180708 | 1.35310975 | 0.05 | 0.021 | 1 | 0.66663755 | 0.38858673 |
| Abhd6 | 0.98487152 | 0.13302542 | 0.115 | 0.114 | 1 | 0.21821934 | -0.2457249 | 0.16 | 0.104 | 1 | 0.98487152 | 0.388819 |
| Abhd8 | 0.28812295 | -0.407543 | 0.292 | 0.347 | 1 | 0.21823021 | 0.25782925 | 0.273 | 0.208 | 1 | 0.28812295 | 0.388836 |
| Nemp2 | 0.21858689 | 0.64492446 | 0.062 | 0.031 | 1 | 0.76325722 | -0.7248928 | 0.05 | 0.042 | 1 | 0.76325722 | 0.38939356 |
| Heph | 0.21860846 | -1.1624305 | 0.021 | 0.052 | 1 | 0.57170879 | -1.6317834 | 0.013 | 0.021 | 1 | 0.57170879 | 0.38942726 |
| Dolpp1 | 0.21864803 | 0.59245704 | 0.083 | 0.047 | 1 | 0.71285614 | -0.7623675 | 0.076 | 0.062 | 1 | 0.71285614 | 0.3894891 |
| Gm26901 | 0.21867159 | 1.59245704 | 0.021 | 0.005 | 1 | 0.87851725 | -0.8948178 | 0.013 | 0.01 | 1 | 0.87851725 | 0.38952592 |
| Brinp2 | 0.21867159 | 1.59245704 | 0.021 | 0.005 | 1 | 0.50587544 | -2.3098553 | 0.004 | 0.01 | 1 | 0.50587544 | 0.38952592 |
| Gm37748 | 0.21867159 | 1.59245704 | 0.021 | 0.005 | 1 | 0.27154327 | 1.01207283 | 0.013 | 0 | 1 | 0.27154327 | 0.38952592 |
| Gm32250 | 0.21867159 | 1.59245704 | 0.021 | 0.005 | 1 | 0.56243824 | -0.8948178 | 0.029 | 0.042 | 1 | 0.56243824 | 0.38952592 |
| Skida1 | 0.21867159 | 1.59245704 | 0.021 | 0.005 | 1 | 0.86312513 | -0.7248928 | 0.008 | 0.01 | 1 | 0.86312513 | 0.38952592 |
| 4930589L23Rik | 0.21867159 | 1.59245704 | 0.021 | 0.005 | 1 | 0.27154471 | 0.69014474 | 0.013 | 0 | 1 | 0.27154471 | 0.38952592 |
| I830077J02Rik | 0.21867159 | 1.59245704 | 0.021 | 0.005 | 1 | 0.41074402 | -0.7248928 | 0.017 | 0.031 | 1 | 0.41074402 | 0.38952592 |
| Gm43017 | 0.21867159 | 1.59245704 | 0.021 | 0.005 | 1 | 0.86312513 | -0.7248928 | 0.008 | 0.01 | 1 | 0.86312513 | 0.38952592 |
| Oas1a | 0.21867159 | 2.42253204 | 0.021 | 0.005 | 1 | 0.57170879 | -1.6317834 | 0.013 | 0.021 | 1 | 0.57170879 | 0.38952592 |
| Gm15747 | 0.21867159 | 1.59245704 | 0.021 | 0.005 | 1 | 0.51194506 | 0.27510724 | 0.021 | 0.01 | 1 | 0.51194506 | 0.38952592 |
| Gm57270 | 0.21867159 | 1.59245704 | 0.021 | 0.005 | 1 | 0.52072676 | -0.7248928 | 0.021 | 0.01 | 1 | 0.52072676 | 0.38952592 |
| Klk14 | 0.21867159 | 1.59245704 | 0.021 | 0.005 | 1 | 0.86312513 | -0.7248928 | 0.008 | 0.01 | 1 | 0.86312513 | 0.38952592 |
| 2310016G11Rik | 0.21867159 | 1.59245704 | 0.021 | 0.005 | 1 | 0.86312513 | -0.7248928 | 0.008 | 0.01 | 1 | 0.86312513 | 0.38952592 |
| Gm45442 | 0.21867159 | 1.59245704 | 0.021 | 0.005 | 1 | 0.5818708 | -0.5728897 | 0.013 | 0.021 | 1 | 0.5818708 | 0.38952592 |
| Gm10974 | 0.21867159 | 1.59245704 | 0.021 | 0.005 | 1 | 0.40220015 | -1.0874628 | 0.017 | 0.031 | 1 | 0.40220015 | 0.38952592 |
| Angptl6 | 0.21867159 | 1.59245704 | 0.021 | 0.005 | 1 | 0.27154471 | 0.69014474 | 0.013 | 0 | 1 | 0.27154471 | 0.38952592 |
| Gm47204 | 0.21867159 | 1.59245704 | 0.021 | 0.005 | 1 | 0.23679813 | 1.01207283 | 0.034 | 0.01 | 1 | 0.23679813 | 0.38952592 |
| Stra6 | 0.21867159 | 1.59245704 | 0.021 | 0.005 | 1 | 0.86312513 | -0.7248928 | 0.008 | 0.01 | 1 | 0.86312513 | 0.38952592 |
| Filip1 | 0.21867159 | 2.68556644 | 0.021 | 0.005 | 1 | 0.80397952 | 0.80562195 | 0.025 | 0.021 | 1 | 0.80397952 | 0.38952592 |
| Nbeal2 | 0.21867159 | 1.59245704 | 0.021 | 0.005 | 1 | 0.66387437 | 0.27510724 | 0.017 | 0.01 | 1 | 0.66387437 | 0.38952592 |
| Gm57003 | 0.21867159 | 1.59245704 | 0.021 | 0.005 | 1 | 0.87066325 | -0.3098553 | 0.013 | 0.01 | 1 | 0.87066325 | 0.38952592 |
| Mmp19 | 0.21867159 | 1.59245704 | 0.021 | 0.005 | 1 | 0.87066325 | -0.3098553 | 0.013 | 0.01 | 1 | 0.87066325 | 0.38952592 |
| Gm12059 | 0.21867159 | 1.74446013 | 0.021 | 0.005 | 1 | 0.87066325 | -0.3098553 | 0.013 | 0.01 | 1 | 0.87066325 | 0.38952592 |
| Vmo1 | 0.21867159 | 1.59245704 | 0.021 | 0.005 | 1 | 0.66054955 | -0.6577786 | 0.042 | 0.031 | 1 | 0.66054955 | 0.38952592 |
| Dtnbos | 0.21867159 | 1.59245704 | 0.021 | 0.005 | 1 | 0.67149861 | -0.1399303 | 0.029 | 0.021 | 1 | 0.67149861 | 0.38952592 |
| Myt1l | 0.21867159 | 1.59245704 | 0.021 | 0.005 | 1 | 0.42156153 | -0.7952821 | 0.025 | 0.042 | 1 | 0.42156153 | 0.38952592 |
| D830013O20Rik | 0.21867159 | 1.59245704 | 0.021 | 0.005 | 1 | 0.51188611 | -0.7248928 | 0.004 | 0.01 | 1 | 0.51188611 | 0.38952592 |

| | | | | | | | | | | | | |
|---|---|---|---|---|---|---|---|---|---|---|---|---|
| Sptb | 0.21867159 | 1.59245704 | 0.021 | 0.005 | 1 | 0.55975127 | -0.9472852 | 0.034 | 0.021 | 1 | 0.55975127 | 0.38952592 |
| Tmem63c | 0.21867159 | 1.59245704 | 0.021 | 0.005 | 1 | 0.27154471 | 0.69014474 | 0.013 | 0 | 1 | 0.27154471 | 0.38952592 |
| Gm5082 | 0.21867159 | 1.59245704 | 0.021 | 0.005 | 1 | 0.50887612 | -1.3098553 | 0.004 | 0.01 | 1 | 0.50887612 | 0.38952592 |
| Phf2os1 | 0.21867159 | 1.59245704 | 0.021 | 0.005 | 1 | 0.53433287 | 0.42711033 | 0.034 | 0.021 | 1 | 0.53433287 | 0.38952592 |
| Gm3604 | 0.21867159 | 1.59245704 | 0.021 | 0.005 | 1 | 0.51194506 | 0.27510724 | 0.021 | 0.01 | 1 | 0.51194506 | 0.38952592 |
| Dusp13 | 0.21867159 | 1.59245704 | 0.021 | 0.005 | 1 | 0.86613737 | -0.3098553 | 0.008 | 0.01 | 1 | 0.86613737 | 0.38952592 |
| Ltb4r1 | 0.21867159 | 1.59245704 | 0.021 | 0.005 | 1 | 0.67471359 | -0.9879272 | 0.017 | 0.01 | 1 | 0.67471359 | 0.38952592 |
| Nudt15 | 0.21867159 | 1.59245704 | 0.021 | 0.005 | 1 | 0.51194506 | 0.27510724 | 0.021 | 0.01 | 1 | 0.51194506 | 0.38952592 |
| Zfp41 | 0.21867159 | 1.59245704 | 0.021 | 0.005 | 1 | 1 | -1.0468209 | 0.021 | 0.021 | 1 | 1 | 0.38952592 |
| Sult4a1 | 0.21867159 | 1.59245704 | 0.021 | 0.005 | 1 | 0.34704016 | -1.3098553 | 0.008 | 0.021 | 1 | 0.34704016 | 0.38952592 |
| Ptx4 | 0.21867159 | 1.59245704 | 0.021 | 0.005 | 1 | 0.34703906 | -1.3098553 | 0.008 | 0.021 | 1 | 0.34703906 | 0.38952592 |
| Gm20468 | 0.21867159 | 1.59245704 | 0.021 | 0.005 | 1 | 0.8086289 | 0.69014474 | 0.017 | 0.021 | 1 | 0.8086289 | 0.38952592 |
| Gm34235 | 0.21867159 | 1.59245704 | 0.021 | 0.005 | 1 | 0.87851725 | -0.8948178 | 0.013 | 0.01 | 1 | 0.87851725 | 0.38952592 |
| Dkk1 | 0.21867159 | 1.59245704 | 0.021 | 0.005 | 1 | 0.50587544 | -2.3098553 | 0.004 | 0.01 | 1 | 0.50587544 | 0.38952592 |
| Gm1141 | 0.21867159 | 1.59245704 | 0.021 | 0.005 | 1 | 0.86312513 | -0.7248928 | 0.008 | 0.01 | 1 | 0.86312513 | 0.38952592 |
| Gabbr1 | 0.21879818 | 0.10335455 | 0.521 | 0.42 | 1 | 0.81160977 | -0.2662283 | 0.576 | 0.479 | 1 | 0.81160977 | 0.38972371 |
| Eif2s3y | 0.21897302 | -0.6218621 | 0.333 | 0.389 | 1 | 0.47901385 | -0.273836 | 0.66 | 0.719 | 1 | 0.47901385 | 0.38999685 |
| Mgat4b | 0.44816286 | 0.11440974 | 0.385 | 0.326 | 1 | 0.2192183 | 0.33225115 | 0.525 | 0.51 | 1 | 0.44816286 | 0.39037994 |
| Entpd6 | 0.51153206 | 0.70793425 | 0.094 | 0.073 | 1 | 0.21923599 | 0.21370669 | 0.08 | 0.042 | 1 | 0.51153206 | 0.39040756 |
| E130114P18Rik | 0.31891721 | 0.34452952 | 0.062 | 0.036 | 1 | 0.21940497 | 1.10518224 | 0.05 | 0.021 | 1 | 0.31891721 | 0.39067141 |
| Opn1mw | 0.44258367 | -0.2148979 | 0.052 | 0.078 | 1 | 0.21940497 | 1.42711033 | 0.05 | 0.021 | 1 | 0.44258367 | 0.39067141 |
| Unc45a | 0.21941056 | 0.41188479 | 0.323 | 0.254 | 1 | 0.66452186 | -0.2823745 | 0.479 | 0.427 | 1 | 0.66452186 | 0.39068013 |
| 9330159F19Rik | 0.21941847 | -0.7998604 | 0.021 | 0.052 | 1 | 0.56969078 | -1.3098553 | 0.013 | 0.021 | 1 | 0.56969078 | 0.39069247 |
| Scai | 0.21945609 | 0.1312299 | 0.438 | 0.352 | 1 | 0.70067372 | -0.2140963 | 0.592 | 0.583 | 1 | 0.70067372 | 0.3907512 |
| Gnal | 0.41466006 | -1.0799683 | 0.031 | 0.052 | 1 | 0.21946963 | -1.198824 | 0.067 | 0.104 | 1 | 0.41466006 | 0.39077234 |
| Ro60 | 0.29513396 | -0.577468 | 0.354 | 0.389 | 1 | 0.21955547 | -0.1004019 | 0.487 | 0.354 | 1 | 0.29513396 | 0.39090633 |
| Adal | 0.3888941 | 0.42253204 | 0.188 | 0.15 | 1 | 0.22003498 | 0.33400093 | 0.147 | 0.094 | 1 | 0.3888941 | 0.39165457 |
| Gm38537 | 0.22023159 | -0.6929452 | 0.021 | 0.052 | 1 | 0.67149861 | -0.1399303 | 0.029 | 0.021 | 1 | 0.67149861 | 0.39196122 |
| Megf10 | 0.22024839 | -0.5160674 | 0.062 | 0.031 | 1 | 0.24413474 | -1.6317834 | 0.013 | 0.031 | 1 | 0.24413474 | 0.39198743 |
| Cap2 | 0.22026135 | -0.4950058 | 0.021 | 0.052 | 1 | 0.57170879 | -1.6317834 | 0.013 | 0.021 | 1 | 0.57170879 | 0.39200764 |
| 3300002A11Rik | 0.22026949 | 1.42253204 | 0.021 | 0.005 | 1 | 0.8086289 | -0.3098553 | 0.017 | 0.021 | 1 | 0.8086289 | 0.39202033 |
| Stxbp5l | 0.22026949 | 1.42253204 | 0.021 | 0.005 | 1 | 0.39234595 | 1.01207283 | 0.025 | 0.01 | 1 | 0.39234595 | 0.39202033 |
| Nhp2 | 0.22035835 | 0.18806678 | 0.5 | 0.415 | 1 | 0.72963061 | -0.378568 | 0.286 | 0.25 | 1 | 0.72963061 | 0.3921589 |
| Flot2 | 0.33955465 | -0.6507169 | 0.25 | 0.295 | 1 | 0.22059219 | -0.513996 | 0.319 | 0.385 | 1 | 0.33955465 | 0.39252346 |
| Bcat2 | 0.42954374 | 0.34629645 | 0.292 | 0.249 | 1 | 0.22059756 | 0.22856465 | 0.21 | 0.146 | 1 | 0.42954374 | 0.39253184 |
| Ptpa | 0.22065304 | 0.2466822 | 0.646 | 0.549 | 1 | 0.44000746 | -0.2680351 | 0.634 | 0.646 | 1 | 0.44000746 | 0.39261832 |
| Accs | 0.34370807 | -0.577468 | 0.052 | 0.083 | 1 | 0.22074139 | 1.01207283 | 0.05 | 0.021 | 1 | 0.34370807 | 0.39275602 |
| Rbks | 0.43592269 | 0.42253204 | 0.094 | 0.067 | 1 | 0.22080994 | 0.14957636 | 0.08 | 0.042 | 1 | 0.43592269 | 0.39286285 |
| Kifc3 | 0.31393482 | 0.14151188 | 0.427 | 0.358 | 1 | 0.22085156 | -0.3098553 | 0.307 | 0.208 | 1 | 0.31393482 | 0.39292771 |
| Ranbp6 | 0.59327414 | 0.20013961 | 0.125 | 0.104 | 1 | 0.22113486 | 0.49749966 | 0.168 | 0.115 | 1 | 0.59327414 | 0.39336909 |
| Rhbdd2 | 0.88412974 | 0.13755508 | 0.24 | 0.233 | 1 | 0.22174964 | 0.19710473 | 0.185 | 0.125 | 1 | 0.88412974 | 0.39432637 |
| 4930444A19Rik | 0.22187388 | -1.3144336 | 0 | 0.016 | 1 | 0.66387437 | 0.27510724 | 0.017 | 0.01 | 1 | 0.66387437 | 0.39451975 |
| Inava | 0.22187388 | -1.577468 | 0 | 0.016 | 1 | 0.79600331 | -1.3098553 | 0.017 | 0.021 | 1 | 0.79600331 | 0.39451975 |
| Vash2 | 0.22187388 | -1.3144336 | 0 | 0.016 | 1 | 0.41906801 | -1.0874628 | 0.025 | 0.042 | 1 | 0.41906801 | 0.39451975 |
| Cacna1b | 0.22187388 | -1.9925055 | 0 | 0.016 | 1 | 0.57373861 | -1.3098553 | 0.013 | 0.021 | 1 | 0.57373861 | 0.39451975 |
| Klhl41 | 0.22187388 | -1.3144336 | 0 | 0.016 | 1 | 0.34704016 | -1.3098553 | 0.008 | 0.021 | 1 | 0.34704016 | 0.39451975 |
| Coro2a | 0.22187388 | -2.3144336 | 0 | 0.016 | 1 | 0.27154327 | 1.01207283 | 0.013 | 0 | 1 | 0.27154327 | 0.39451975 |
| 8030451A03Rik | 0.22187388 | -1.3144336 | 0 | 0.016 | 1 | 0.86312513 | -0.7248928 | 0.008 | 0.01 | 1 | 0.86312513 | 0.39451975 |
| Gm11266 | 0.22187388 | 1.00749454 | 0.021 | 0.005 | 1 | 0.57170879 | -1.8948178 | 0.013 | 0.021 | 1 | 0.57170879 | 0.39451975 |
| Gm12708 | 0.22187388 | -1.3144336 | 0 | 0.016 | 1 | 0.51188611 | -0.7248928 | 0.004 | 0.01 | 1 | 0.51188611 | 0.39451975 |
| Sema3a | 0.22187388 | -1.7998604 | 0 | 0.016 | 1 | 0.56565662 | -1.8948178 | 0.013 | 0.021 | 1 | 0.56565662 | 0.39451975 |
| Drc1 | 0.22187388 | -1.3144336 | 0 | 0.016 | 1 | 0.39514482 | -2.1578522 | 0.017 | 0.031 | 1 | 0.39514482 | 0.39451975 |
| Tfpi2 | 0.22187388 | -1.3144336 | 0 | 0.016 | 1 | 0.27154471 | 0.69014474 | 0.013 | 0 | 1 | 0.27154471 | 0.39451975 |
| Gm26826 | 0.22187388 | 1.00749454 | 0.021 | 0.005 | 1 | 0.51194506 | 0.27510724 | 0.021 | 0.01 | 1 | 0.51194506 | 0.39451975 |
| Lmo3 | 0.22187388 | 1.00749454 | 0.021 | 0.005 | 1 | 0.86312513 | -0.7248928 | 0.008 | 0.01 | 1 | 0.86312513 | 0.39451975 |
| Pla2g4c | 0.22187388 | 1.00749454 | 0.021 | 0.005 | 1 | 0.85710544 | -1.3098553 | 0.008 | 0.01 | 1 | 0.85710544 | 0.39451975 |
| Il4i1 | 0.22187388 | 1.00749454 | 0.021 | 0.005 | 1 | 0.36655012 | -0.194378 | 0.042 | 0.021 | 1 | 0.36655012 | 0.39451975 |
| Gm10610 | 0.22187388 | -1.9925055 | 0 | 0.016 | 1 | 0.34363875 | -1.7248928 | 0.008 | 0.021 | 1 | 0.34363875 | 0.39451975 |
| Hpx | 0.22187388 | -1.3144336 | 0 | 0.016 | 1 | 0.50887612 | -1.3098553 | 0.004 | 0.01 | 1 | 0.50887612 | 0.39451975 |
| Tmed6 | 0.22187388 | -1.3144336 | 0 | 0.016 | 1 | 0.50887612 | -1.3098553 | 0.004 | 0.01 | 1 | 0.50887612 | 0.39451975 |
| Mlip | 0.22187388 | -2.1624305 | 0 | 0.016 | 1 | 0.24413474 | -1.6317834 | 0.013 | 0.031 | 1 | 0.24413474 | 0.39451975 |
| Snap91 | 0.22187388 | -1.577468 | 0 | 0.016 | 1 | 0.99712231 | -0.5443205 | 0.042 | 0.042 | 1 | 0.99712231 | 0.39451975 |
| Cabcoco1 | 0.22187388 | 1.00749454 | 0.021 | 0.005 | 1 | 0.43079893 | 0.56461386 | 0.038 | 0.021 | 1 | 0.43079893 | 0.39451975 |
| Slc26a10 | 0.22187388 | -1.3144336 | 0 | 0.016 | 1 | 0.98590808 | 0.10518224 | 0.021 | 0.021 | 1 | 0.98590808 | 0.39451975 |
| 4930524B15Rik | 0.22187388 | -1.3144336 | 0 | 0.016 | 1 | 0.34874556 | -0.9879272 | 0.008 | 0.021 | 1 | 0.34874556 | 0.39451975 |
| Gm57328 | 0.22187388 | 1.00749454 | 0.021 | 0.005 | 1 | 0.50887612 | -1.3098553 | 0.004 | 0.01 | 1 | 0.50887612 | 0.39451975 |
| Nlrp3 | 0.22187388 | -1.3144336 | 0 | 0.016 | 1 | 0.50887612 | -1.3098553 | 0.004 | 0.01 | 1 | 0.50887612 | 0.39451975 |
| Zmynd15 | 0.22187388 | 1.00749454 | 0.021 | 0.005 | 1 | 0.30606883 | 0.69014474 | 0.029 | 0.01 | 1 | 0.30606883 | 0.39451975 |
| Gm40437 | 0.22187388 | -1.9925055 | 0 | 0.016 | 1 | 0.7411443 | -1.4797803 | 0.025 | 0.031 | 1 | 0.7411443 | 0.39451975 |
| Ryr2 | 0.22187388 | 1.00749454 | 0.021 | 0.005 | 1 | 0.5154503 | 0.42711033 | 0.021 | 0.01 | 1 | 0.5154503 | 0.39451975 |
| Gm15327 | 0.22187388 | -1.3144336 | 0 | 0.016 | 1 | 0.86312513 | -0.7248928 | 0.008 | 0.01 | 1 | 0.86312513 | 0.39451975 |
| Ogdhl | 0.22187388 | -1.3144336 | 0 | 0.016 | 1 | 0.87066325 | -0.3098553 | 0.013 | 0.01 | 1 | 0.87066325 | 0.39451975 |
| Rem2 | 0.22187388 | -1.3144336 | 0 | 0.016 | 1 | 0.8086289 | -0.3098553 | 0.017 | 0.021 | 1 | 0.8086289 | 0.39451975 |
| Pkhd1l1 | 0.22187388 | -1.7998604 | 0 | 0.016 | 1 | 0.51188611 | -0.7248928 | 0.004 | 0.01 | 1 | 0.51188611 | 0.39451975 |
| Csmd3 | 0.22187388 | -1.577468 | 0 | 0.016 | 1 | 0.55772463 | -2.0468209 | 0.021 | 0.031 | 1 | 0.55772463 | 0.39451975 |
| Syngr3 | 0.22187388 | -1.3144336 | 0 | 0.016 | 1 | 0.81129592 | 0.27510724 | 0.025 | 0.021 | 1 | 0.81129592 | 0.39451975 |
| Gm30984 | 0.22187388 | -1.3144336 | 0 | 0.016 | 1 | 0.27154471 | 0.69014474 | 0.013 | 0 | 1 | 0.27154471 | 0.39451975 |
| Pcdhb19 | 0.22187388 | 1.00749454 | 0.021 | 0.005 | 1 | 0.41910282 | -1.4618584 | 0.025 | 0.042 | 1 | 0.41910282 | 0.39451975 |
| 2010308F09Rik | 0.22187388 | -1.7998604 | 0 | 0.016 | 1 | 0.50887612 | -1.3098553 | 0.004 | 0.01 | 1 | 0.50887612 | 0.39451975 |
| Gm21860 | 0.22187388 | -1.3144336 | 0 | 0.016 | 1 | 0.34704016 | -1.3098553 | 0.008 | 0.021 | 1 | 0.34704016 | 0.39451975 |
| D430040D24Rik | 0.22187575 | -0.9925055 | 0 | 0.016 | 1 | 0.85710544 | -1.3098553 | 0.008 | 0.01 | 1 | 0.85710544 | 0.39452265 |
| Kcnj13 | 0.22187575 | -0.9925055 | 0 | 0.016 | 1 | 0.26649778 | 0.49749966 | 0.076 | 0.042 | 1 | 0.26649778 | 0.39452265 |
| Gm19461 | 0.22187575 | -0.9925055 | 0 | 0.016 | 1 | 0.50887612 | -1.3098553 | 0.004 | 0.01 | 1 | 0.50887612 | 0.39452265 |
| Fasl | 0.22187575 | -0.9925055 | 0 | 0.016 | 1 | 0.51188611 | -0.7248928 | 0.004 | 0.01 | 1 | 0.51188611 | 0.39452265 |

| | | | | | | | | | | | | |
|---|---|---|---|---|---|---|---|---|---|---|---|---|
| C8g | 0.22187575 | -0.9925055 | 0 | 0.016 | 1 | 0.51194506 | 0.27510724 | 0.021 | 0.01 | 1 | 0.51194506 | 0.39452265 |
| Cd59a | 0.22187575 | -0.9925055 | 0 | 0.016 | 1 | 0.81798678 | -0.6724253 | 0.046 | 0.052 | 1 | 0.81798678 | 0.39452265 |
| Zfp661 | 0.22187575 | -0.9925055 | 0 | 0.016 | 1 | 0.86312513 | -0.7248928 | 0.008 | 0.01 | 1 | 0.86312513 | 0.39452265 |
| AI847159 | 0.22187575 | -0.9925055 | 0 | 0.016 | 1 | 0.50887612 | -1.3098553 | 0.004 | 0.01 | 1 | 0.50887612 | 0.39452265 |
| 0610039K10Rik | 0.22187575 | -0.9925055 | 0 | 0.016 | 1 | 0.50887612 | -1.3098553 | 0.004 | 0.01 | 1 | 0.50887612 | 0.39452265 |
| Fam209 | 0.22187575 | -0.9925055 | 0 | 0.016 | 1 | 0.39515719 | 0.49749966 | 0.025 | 0.01 | 1 | 0.39515719 | 0.39452265 |
| Stmn3 | 0.22187575 | -0.9925055 | 0 | 0.016 | 1 | 0.78761875 | -1.5728897 | 0.017 | 0.021 | 1 | 0.78761875 | 0.39452265 |
| 1110032F04Rik | 0.22187575 | -0.9925055 | 0 | 0.016 | 1 | 0.50844268 | 0.69014474 | 0.021 | 0.01 | 1 | 0.50844268 | 0.39452265 |
| A730011C13Rik | 0.22187575 | -0.9925055 | 0 | 0.016 | 1 | 0.40647274 | -0.9879272 | 0.017 | 0.031 | 1 | 0.40647274 | 0.39452265 |
| Cdh17 | 0.22187575 | -3.3144336 | 0 | 0.016 | 1 | 0.50587544 | -1.8948178 | 0.004 | 0.01 | 1 | 0.50587544 | 0.39452265 |
| Ermap | 0.22187575 | -0.9925055 | 0 | 0.016 | 1 | 0.57170879 | -1.3098553 | 0.013 | 0.021 | 1 | 0.57170879 | 0.39452265 |
| Gm57092 | 0.22187575 | -0.9925055 | 0 | 0.016 | 1 | 0.98993409 | -1.3098553 | 0.021 | 0.021 | 1 | 0.98993409 | 0.39452265 |
| Abcb4 | 0.22187575 | -0.9925055 | 0 | 0.016 | 1 | 0.50844268 | 0.86006974 | 0.021 | 0.01 | 1 | 0.50844268 | 0.39452265 |
| Hs3st1 | 0.22187575 | -1.9925055 | 0 | 0.016 | 1 | 0.87851725 | -0.8948178 | 0.013 | 0.01 | 1 | 0.87851725 | 0.39452265 |
| Fam184b | 0.22187575 | -1.7998604 | 0 | 0.016 | 1 | 0.86312513 | -0.7248928 | 0.008 | 0.01 | 1 | 0.86312513 | 0.39452265 |
| D430040L24Rik | 0.22187575 | -0.9925055 | 0 | 0.016 | 1 | 0.34363875 | -2.0468209 | 0.008 | 0.021 | 1 | 0.34363875 | 0.39452265 |
| Gm29707 | 0.22187575 | 0.42253204 | 0.021 | 0.005 | 1 | 0.50587544 | -1.8948178 | 0.004 | 0.01 | 1 | 0.50587544 | 0.39452265 |
| Hoxa7 | 0.22187575 | -0.9925055 | 0 | 0.016 | 1 | 0.50887612 | -1.3098553 | 0.004 | 0.01 | 1 | 0.50887612 | 0.39452265 |
| Cfap100 | 0.22187575 | -0.9925055 | 0 | 0.016 | 1 | 0.50887612 | -1.3098553 | 0.004 | 0.01 | 1 | 0.50887612 | 0.39452265 |
| Gm17216 | 0.22187575 | -0.9925055 | 0 | 0.016 | 1 | 0.85710544 | -1.7248928 | 0.008 | 0.01 | 1 | 0.85710544 | 0.39452265 |
| Nfkbid | 0.22187575 | -0.9925055 | 0 | 0.016 | 1 | 0.27154471 | 0.69014474 | 0.013 | 0 | 1 | 0.27154471 | 0.39452265 |
| Gm15489 | 0.22187575 | -0.9925055 | 0 | 0.016 | 1 | 0.39234595 | 0.86006974 | 0.025 | 0.01 | 1 | 0.39234595 | 0.39452265 |
| Gm45502 | 0.22187575 | -0.9925055 | 0 | 0.016 | 1 | 0.27154471 | 0.69014474 | 0.013 | 0 | 1 | 0.27154471 | 0.39452265 |
| Rassf7 | 0.22187575 | -0.9925055 | 0 | 0.016 | 1 | 0.66387437 | 0.27510724 | 0.017 | 0.01 | 1 | 0.66387437 | 0.39452265 |
| Klf1 | 0.22187575 | -0.9925055 | 0 | 0.016 | 1 | 0.87851725 | -1.3098553 | 0.013 | 0.01 | 1 | 0.87851725 | 0.39452265 |
| Pkd1l3 | 0.22187575 | -0.9925055 | 0 | 0.016 | 1 | 0.94026906 | -0.9472852 | 0.034 | 0.031 | 1 | 0.94026906 | 0.39452265 |
| Trim67 | 0.22187575 | -0.9925055 | 0 | 0.016 | 1 | 0.50887612 | -1.3098553 | 0.004 | 0.01 | 1 | 0.50887612 | 0.39452265 |
| Odad3 | 0.22187575 | -0.9925055 | 0 | 0.016 | 1 | 0.87851725 | -1.3098553 | 0.013 | 0.01 | 1 | 0.87851725 | 0.39452265 |
| Tecta | 0.22187575 | -2.3144336 | 0 | 0.016 | 1 | 0.24746336 | -1.3098553 | 0.013 | 0.031 | 1 | 0.24746336 | 0.39452265 |
| Gm30015 | 0.22187575 | -0.9925055 | 0 | 0.016 | 1 | 0.50887612 | -1.3098553 | 0.004 | 0.01 | 1 | 0.50887612 | 0.39452265 |
| Tektip1 | 0.22187575 | -0.9925055 | 0 | 0.016 | 1 | 0.67144323 | -0.5322477 | 0.042 | 0.031 | 1 | 0.67144323 | 0.39452265 |
| 1500026H17Rik | 0.22187575 | -0.9925055 | 0 | 0.016 | 1 | 0.99396051 | -0.3098553 | 0.021 | 0.021 | 1 | 0.99396051 | 0.39452265 |
| Hic1 | 0.22187575 | -0.9925055 | 0 | 0.016 | 1 | 0.34704016 | -1.3098553 | 0.008 | 0.021 | 1 | 0.34704016 | 0.39452265 |
| Cntnap1 | 0.22187575 | -0.9925055 | 0 | 0.016 | 1 | 0.87851725 | -1.3098553 | 0.013 | 0.01 | 1 | 0.87851725 | 0.39452265 |
| Gm16045 | 0.22187575 | -0.9925055 | 0 | 0.016 | 1 | 1 | -0.7248928 | 0.021 | 0.021 | 1 | 1 | 0.39452265 |
| Kcns3 | 0.22187575 | -0.9925055 | 0 | 0.016 | 1 | 0.40076941 | -1.3098553 | 0.017 | 0.031 | 1 | 0.40076941 | 0.39452265 |
| Mycn | 0.22187575 | -0.9925055 | 0 | 0.016 | 1 | 0.25080911 | -0.9879272 | 0.013 | 0.031 | 1 | 0.25080911 | 0.39452265 |
| 6530401F13Rik | 0.22187575 | -1.577468 | 0 | 0.016 | 1 | 0.87066325 | -0.3098553 | 0.013 | 0.01 | 1 | 0.87066325 | 0.39452265 |
| Gm47882 | 0.22187575 | -0.9925055 | 0 | 0.016 | 1 | 0.87066325 | -0.3098553 | 0.013 | 0.01 | 1 | 0.87066325 | 0.39452265 |
| Cdc20b | 0.22187575 | -1.577468 | 0 | 0.016 | 1 | 0.66339728 | 0.10518224 | 0.029 | 0.021 | 1 | 0.66339728 | 0.39452265 |
| 4930562C15Rik | 0.22187575 | -0.9925055 | 0 | 0.016 | 1 | 0.50887612 | -1.3098553 | 0.004 | 0.01 | 1 | 0.50887612 | 0.39452265 |
| Gm42477 | 0.22187575 | -0.9925055 | 0 | 0.016 | 1 | 0.24413474 | -2.3098553 | 0.013 | 0.031 | 1 | 0.24413474 | 0.39452265 |
| Fgf12 | 0.22187575 | -1.7998604 | 0 | 0.016 | 1 | 0.79810697 | -1.0874628 | 0.017 | 0.021 | 1 | 0.79810697 | 0.39452265 |
| Egfl8 | 0.22187575 | -0.9925055 | 0 | 0.016 | 1 | 0.42648204 | -0.8244284 | 0.025 | 0.042 | 1 | 0.42648204 | 0.39452265 |
| Plin4 | 0.22187575 | -0.9925055 | 0 | 0.016 | 1 | 0.87066325 | -0.3098553 | 0.013 | 0.01 | 1 | 0.87066325 | 0.39452265 |
| Gm50020 | 0.22187575 | -0.9925055 | 0 | 0.016 | 1 | 0.85710544 | -1.7248928 | 0.008 | 0.01 | 1 | 0.85710544 | 0.39452265 |
| Gm19696 | 0.22187575 | -0.9925055 | 0 | 0.016 | 1 | 0.85710544 | -1.3098553 | 0.008 | 0.01 | 1 | 0.85710544 | 0.39452265 |
| Syt7 | 0.22187575 | -0.9925055 | 0 | 0.016 | 1 | 0.66171832 | 0.69014474 | 0.017 | 0.01 | 1 | 0.66171832 | 0.39452265 |
| Rorb | 0.22187575 | -0.9925055 | 0 | 0.016 | 1 | 0.57780211 | -0.8244284 | 0.013 | 0.021 | 1 | 0.57780211 | 0.39452265 |
| Sema4g | 0.22187575 | -0.9925055 | 0 | 0.016 | 1 | 0.79600331 | -0.9879272 | 0.017 | 0.021 | 1 | 0.79600331 | 0.39452265 |
| Bex1 | 0.22187575 | -0.9925055 | 0 | 0.016 | 1 | 0.80020906 | -0.7248928 | 0.017 | 0.021 | 1 | 0.80020906 | 0.39452265 |
| Gm29650 | 0.22187575 | -0.9925055 | 0 | 0.016 | 1 | 0.51194506 | 0.27510724 | 0.021 | 0.01 | 1 | 0.51194506 | 0.39452265 |
| Ada | 0.38027888 | 1.00749454 | 0.031 | 0.016 | 1 | 0.22208005 | 0.91253716 | 0.05 | 0.021 | 1 | 0.38027888 | 0.39484055 |
| Gm4013 | 0.43461987 | 0.64492446 | 0.062 | 0.041 | 1 | 0.22208005 | 1.01207283 | 0.05 | 0.021 | 1 | 0.43461987 | 0.39484055 |
| Nudt7 | 0.74382634 | 0.21899864 | 0.083 | 0.073 | 1 | 0.22208005 | 0.91253716 | 0.05 | 0.021 | 1 | 0.74382634 | 0.39484055 |
| Dmxl2 | 0.81043957 | 0.32942263 | 0.167 | 0.155 | 1 | 0.22209297 | -0.9949819 | 0.151 | 0.198 | 1 | 0.81043957 | 0.39486066 |
| Lmbr1l | 0.22247196 | -0.3461424 | 0.365 | 0.435 | 1 | 0.50501726 | -0.5070876 | 0.441 | 0.427 | 1 | 0.50501726 | 0.39545015 |
| Gm13269 | 0.7885569 | 0.59245704 | 0.031 | 0.026 | 1 | 0.22257857 | -1.5728897 | 0.05 | 0.083 | 1 | 0.7885569 | 0.39561592 |
| Fam43a | 0.22272435 | -0.2962862 | 0.115 | 0.176 | 1 | 0.59634196 | 0.15963002 | 0.147 | 0.125 | 1 | 0.59634196 | 0.39584256 |
| Dmrta1 | 0.52961488 | -0.3144336 | 0.01 | 0.021 | 1 | 0.22356137 | 0.80562195 | 0.05 | 0.021 | 1 | 0.52961488 | 0.39714305 |
| A230056P14Rik | 0.73724235 | 0.86999101 | 0.042 | 0.052 | 1 | 0.22356137 | 0.80562195 | 0.05 | 0.021 | 1 | 0.73724235 | 0.39714305 |
| Mrpl23 | 0.24116897 | -0.2073147 | 0.594 | 0.658 | 1 | 0.22357774 | 0.30157945 | 0.576 | 0.552 | 1 | 0.24116897 | 0.39716847 |
| Neurl1a | 0.22423045 | 0.59245704 | 0.083 | 0.047 | 1 | 0.66900598 | -1.0688472 | 0.042 | 0.052 | 1 | 0.66900598 | 0.39818161 |
| Gm15788 | 0.22429699 | 0.24195979 | 0.083 | 0.047 | 1 | 0.53974558 | -0.681824 | 0.046 | 0.062 | 1 | 0.53974558 | 0.39828484 |
| Sh2b2 | 0.22456227 | 0.27052894 | 0.104 | 0.062 | 1 | 0.79193958 | -0.7248928 | 0.097 | 0.083 | 1 | 0.79193958 | 0.39869632 |
| Ice1 | 0.22490796 | 0.22661183 | 0.573 | 0.523 | 1 | 0.65120771 | -0.183478 | 0.693 | 0.583 | 1 | 0.65120771 | 0.39923233 |
| Lonrf1 | 0.22544138 | 0.2541345 | 0.427 | 0.342 | 1 | 0.27182396 | -0.4788314 | 0.408 | 0.479 | 1 | 0.27182396 | 0.40005895 |
| Cracd | 0.22573176 | 0.51768927 | 0.24 | 0.181 | 1 | 0.77708751 | -0.4518743 | 0.256 | 0.26 | 1 | 0.77708751 | 0.40050869 |
| Ddb2 | 0.22591617 | -0.577468 | 0.146 | 0.202 | 1 | 0.34322603 | -0.2318528 | 0.185 | 0.135 | 1 | 0.34322603 | 0.40079422 |
| Dpysl2 | 0.22607917 | 0.13167395 | 0.396 | 0.487 | 1 | 0.27298081 | -0.3664388 | 0.412 | 0.292 | 1 | 0.27298081 | 0.40104656 |
| Gm27017 | 0.31889482 | 0.47697982 | 0.062 | 0.036 | 1 | 0.22626574 | 1.01207283 | 0.05 | 0.021 | 1 | 0.31889482 | 0.4013353 |
| Abtb2 | 0.65092495 | -0.2968002 | 0.844 | 0.922 | 1 | 0.22635032 | -0.2714985 | 0.912 | 0.875 | 1 | 0.65092495 | 0.40146617 |
| Zfp790 | 0.63343064 | 0.27052894 | 0.083 | 0.067 | 1 | 0.22658231 | 0.52303475 | 0.181 | 0.125 | 1 | 0.63343064 | 0.40182508 |
| Retreg3 | 0.22658322 | 0.39868529 | 0.302 | 0.233 | 1 | 0.84502217 | -0.3292206 | 0.429 | 0.417 | 1 | 0.84502217 | 0.40182649 |
| Mark3 | 0.22671064 | 0.1269097 | 0.927 | 0.896 | 1 | 0.86149079 | -0.1199195 | 0.987 | 0.948 | 1 | 0.86149079 | 0.40202357 |
| Slc41a3 | 0.47484645 | 0.54806292 | 0.073 | 0.052 | 1 | 0.22685764 | -1.3098553 | 0.042 | 0.073 | 1 | 0.47484645 | 0.40225089 |
| Pcdhga6 | 0.88154275 | 0.22988696 | 0.052 | 0.057 | 1 | 0.22700977 | -1.0468209 | 0.076 | 0.115 | 1 | 0.88154275 | 0.4024861 |
| Dhx36 | 0.28210301 | -0.1725405 | 0.698 | 0.798 | 1 | 0.22705899 | -0.3706805 | 0.744 | 0.823 | 1 | 0.28210301 | 0.40256219 |
| Cptp | 0.22745246 | 0.29700115 | 0.198 | 0.14 | 1 | 0.27171308 | 0.52664601 | 0.088 | 0.052 | 1 | 0.27171308 | 0.4031703 |
| Ccnj | 0.26351343 | -0.7998604 | 0.031 | 0.062 | 1 | 0.22765297 | 0.69014474 | 0.05 | 0.021 | 1 | 0.26351343 | 0.40348006 |
| Rad52 | 0.90397123 | -0.3430027 | 0.177 | 0.176 | 1 | 0.22793139 | -0.1340054 | 0.21 | 0.146 | 1 | 0.90397123 | 0.40391005 |
| Afg1l | 0.22832288 | -0.2462621 | 0.135 | 0.197 | 1 | 0.56400586 | -1.0260623 | 0.139 | 0.104 | 1 | 0.56400586 | 0.40451443 |
| Mrnip | 0.22842269 | -0.3461424 | 0.052 | 0.093 | 1 | 0.72873125 | -1.1347686 | 0.105 | 0.115 | 1 | 0.72873125 | 0.40466845 |
| C79798 | 0.22849436 | 0.35541784 | 0.062 | 0.031 | 1 | 0.49032564 | -1.3802446 | 0.055 | 0.073 | 1 | 0.49032564 | 0.40477904 |

| | | | | | | | | | | | | |
|---|---|---|---|---|---|---|---|---|---|---|---|---|
| Tnfsf13 | 0.5425293 | -0.5160674 | 0.073 | 0.093 | 1 | 0.22874493 | -0.8028953 | 0.092 | 0.135 | 1 | 0.5425293 | 0.40516562 |
| Ceacam1 | 0.22875926 | -0.3348977 | 0.125 | 0.181 | 1 | 0.52130139 | 0.82764826 | 0.155 | 0.135 | 1 | 0.52130139 | 0.40518773 |
| 6430548M08Rik | 0.33457423 | 0.53800925 | 0.094 | 0.062 | 1 | 0.22885291 | -0.177405 | 0.147 | 0.094 | 1 | 0.33457423 | 0.40533217 |
| 1700030K09Rik | 0.50788394 | 0.36363835 | 0.135 | 0.109 | 1 | 0.22902354 | 0.16407593 | 0.08 | 0.042 | 1 | 0.50788394 | 0.4055953 |
| Atp7a | 0.22903235 | -0.4035741 | 0.479 | 0.523 | 1 | 0.76986221 | -0.2495548 | 0.672 | 0.625 | 1 | 0.76986221 | 0.40560889 |
| Tmco4 | 0.22908171 | -0.479383 | 0.396 | 0.44 | 1 | 0.55056379 | -0.2680351 | 0.517 | 0.406 | 1 | 0.55056379 | 0.405685 |
| Psme3ip1 | 0.22913048 | 0.29967529 | 0.417 | 0.342 | 1 | 0.89558432 | -0.352924 | 0.433 | 0.396 | 1 | 0.89558432 | 0.40576019 |
| Prtg | 0.2294558 | -1.9743581 | 0.094 | 0.135 | 1 | 0.61599926 | 0.33060235 | 0.101 | 0.083 | 1 | 0.61599926 | 0.40626164 |
| Iqcd | 0.57848839 | 1.00749454 | 0.031 | 0.021 | 1 | 0.22956679 | -1.7692869 | 0.05 | 0.083 | 1 | 0.57848839 | 0.40643266 |
| Sfxn4 | 0.51508949 | 0.37006462 | 0.083 | 0.062 | 1 | 0.22960842 | -0.1082214 | 0.08 | 0.042 | 1 | 0.51508949 | 0.40649681 |
| Pbld2 | 0.46992036 | 0.74446013 | 0.042 | 0.026 | 1 | 0.22974343 | 0.36821664 | 0.05 | 0.021 | 1 | 0.46992036 | 0.40670482 |
| Zfp788 | 0.22987255 | -1.0920411 | 0.062 | 0.104 | 1 | 0.26700571 | 0.44016648 | 0.139 | 0.094 | 1 | 0.26700571 | 0.40690371 |
| Tnfsf13os | 0.22990776 | -0.4664367 | 0.083 | 0.047 | 1 | 0.64880499 | -0.3098553 | 0.055 | 0.042 | 1 | 0.64880499 | 0.40695794 |
| Zfp13 | 0.22990776 | 0.47697982 | 0.083 | 0.047 | 1 | 0.23754211 | -0.5728897 | 0.034 | 0.062 | 1 | 0.23754211 | 0.40695794 |
| Snx32 | 0.22990776 | 0.47697982 | 0.083 | 0.047 | 1 | 0.86929642 | -0.3098553 | 0.109 | 0.115 | 1 | 0.86929642 | 0.40695794 |
| Tax1bp3 | 0.23003081 | -0.1198848 | 0.49 | 0.389 | 1 | 0.28484489 | 0.21883902 | 0.471 | 0.438 | 1 | 0.28484489 | 0.40714745 |
| Gopc | 0.23032187 | 0.26529229 | 0.365 | 0.29 | 1 | 0.50690196 | -0.4604149 | 0.504 | 0.375 | 1 | 0.50690196 | 0.40759557 |
| Sgms1os1 | 0.23083851 | 0.70793425 | 0.115 | 0.073 | 1 | 0.23328779 | 0.40635177 | 0.092 | 0.052 | 1 | 0.23328779 | 0.4083906 |
| Sgsh | 0.82258416 | -0.1624305 | 0.031 | 0.036 | 1 | 0.23110901 | 0.17557156 | 0.05 | 0.021 | 1 | 0.82258416 | 0.40880665 |
| Gm56560 | 0.88651895 | -0.407543 | 0.073 | 0.067 | 1 | 0.23120366 | -1.2103196 | 0.059 | 0.094 | 1 | 0.88651895 | 0.40895219 |
| Pex16 | 0.87994131 | -0.1393469 | 0.135 | 0.14 | 1 | 0.23142179 | 0.49749966 | 0.13 | 0.083 | 1 | 0.87994131 | 0.40928754 |
| Rpusd1 | 0.24972389 | 1.14499806 | 0.073 | 0.041 | 1 | 0.23178796 | -0.4167705 | 0.05 | 0.021 | 1 | 0.24972389 | 0.40985026 |
| Tyrobp | 1 | -1.3144336 | 0.01 | 0.01 | 1 | 0.2318339 | 2.25085969 | 0.055 | 0.094 | 1 | 1 | 0.40992085 |
| Gm44577 | 0.28001997 | -2.3848229 | 0.01 | 0.031 | 1 | 0.23186374 | -1.6968784 | 0.034 | 0.062 | 1 | 0.28001997 | 0.40996669 |
| Gm56501 | 0.45180013 | -0.9925055 | 0.042 | 0.062 | 1 | 0.23191252 | -1.1082214 | 0.076 | 0.115 | 1 | 0.45180013 | 0.41004163 |
| Echdc3 | 0.99482536 | 0.15949763 | 0.042 | 0.041 | 1 | 0.23202053 | -1.2029401 | 0.042 | 0.073 | 1 | 0.99482536 | 0.41020754 |
| Usp24 | 0.23238924 | -0.2427016 | 0.844 | 0.891 | 1 | 0.56319648 | -0.1673587 | 0.95 | 0.917 | 1 | 0.56319648 | 0.41077372 |
| Klhl33 | 0.84651478 | 0.20013961 | 0.104 | 0.098 | 1 | 0.23257855 | 0.49749966 | 0.05 | 0.021 | 1 | 0.84651478 | 0.41106432 |
| Rbm20 | 0.56101748 | -0.3617393 | 0.083 | 0.104 | 1 | 0.23275704 | 0.39058446 | 0.105 | 0.062 | 1 | 0.56101748 | 0.41133824 |
| Ubr1 | 0.31726953 | 0.2358774 | 0.74 | 0.689 | 1 | 0.23289176 | -0.4176344 | 0.739 | 0.771 | 1 | 0.31726953 | 0.41154495 |
| Atxn7l1 | 0.41553702 | -0.1445086 | 0.406 | 0.446 | 1 | 0.23301923 | -0.4672116 | 0.475 | 0.562 | 1 | 0.41553702 | 0.41174049 |
| Zfp518b | 0.8397555 | 0.32942263 | 0.073 | 0.067 | 1 | 0.23311801 | 0.23071312 | 0.118 | 0.073 | 1 | 0.8397555 | 0.41189201 |
| Hyal3 | 0.30891093 | 1.00749454 | 0.042 | 0.021 | 1 | 0.23322339 | -0.9879272 | 0.034 | 0.062 | 1 | 0.30891093 | 0.41205363 |
| Rnf217 | 0.36770312 | -0.5977024 | 0.458 | 0.503 | 1 | 0.23324263 | -0.4643652 | 0.702 | 0.688 | 1 | 0.36770312 | 0.41208313 |
| Birc5 | 0.23335994 | 2.79890791 | 0.052 | 0.026 | 1 | 0.58180179 | -0.7248928 | 0.021 | 0.031 | 1 | 0.58180179 | 0.41226303 |
| Mrpl30 | 0.47943996 | -0.1537864 | 0.615 | 0.617 | 1 | 0.23349638 | 0.18895059 | 0.542 | 0.479 | 1 | 0.47943996 | 0.4124722 |
| Gm9967 | 0.85795684 | 0.13302542 | 0.042 | 0.047 | 1 | 0.23359962 | -1.2358547 | 0.059 | 0.094 | 1 | 0.85795684 | 0.41263045 |
| Lmln | 0.23365009 | 0.73447604 | 0.115 | 0.073 | 1 | 0.582846 | -1.1034044 | 0.097 | 0.115 | 1 | 0.582846 | 0.41270781 |
| Nlk | 0.234338 | -0.638397 | 0.469 | 0.513 | 1 | 0.23740985 | -0.368749 | 0.66 | 0.698 | 1 | 0.23740985 | 0.41376171 |
| Zmiz1os1 | 0.76463872 | -0.2040096 | 0.125 | 0.114 | 1 | 0.23438109 | -1.4673965 | 0.088 | 0.125 | 1 | 0.76463872 | 0.41382768 |
| Wdr76 | 0.23439611 | 1.09495738 | 0.094 | 0.057 | 1 | 0.63933649 | -0.2103196 | 0.055 | 0.042 | 1 | 0.63933649 | 0.41385068 |
| Zfp236 | 0.23455301 | 0.23544648 | 0.417 | 0.337 | 1 | 0.40035037 | -0.2607076 | 0.534 | 0.417 | 1 | 0.40035037 | 0.41409091 |
| Zfp626 | 0.46815208 | 0.27052894 | 0.104 | 0.078 | 1 | 0.23463395 | 0.2095189 | 0.143 | 0.094 | 1 | 0.46815208 | 0.41421481 |
| Fbxo25 | 0.23466331 | 0.27052894 | 0.167 | 0.114 | 1 | 0.79021106 | -0.42233 | 0.118 | 0.104 | 1 | 0.79021106 | 0.41425976 |
| Phc1 | 0.23467335 | 0.23882008 | 0.198 | 0.145 | 1 | 0.29963346 | 0.16221318 | 0.277 | 0.219 | 1 | 0.29963346 | 0.41427513 |
| 1500015L24Rik | 0.23470712 | -1.3144336 | 0.062 | 0.104 | 1 | 0.76149424 | -0.1254307 | 0.084 | 0.073 | 1 | 0.76149424 | 0.41432681 |
| Il27 | 0.28001913 | -1.4519371 | 0.01 | 0.031 | 1 | 0.23472999 | -0.9472852 | 0.034 | 0.062 | 1 | 0.28001913 | 0.41436181 |
| Lrch2 | 0.40127909 | -0.7998604 | 0.052 | 0.078 | 1 | 0.23482457 | -1.2592292 | 0.105 | 0.146 | 1 | 0.40127909 | 0.41450656 |
| Zfp65 | 0.82963384 | 0.11440974 | 0.094 | 0.104 | 1 | 0.23482457 | 0.38202244 | 0.13 | 0.083 | 1 | 0.82963384 | 0.41450656 |
| Gmnn | 0.23489651 | 2.27052894 | 0.073 | 0.041 | 1 | 0.36655012 | -0.194378 | 0.042 | 0.021 | 1 | 0.36655012 | 0.41461664 |
| Stbd1 | 0.53487059 | 0.42253204 | 0.052 | 0.036 | 1 | 0.23504554 | 0.49749966 | 0.092 | 0.052 | 1 | 0.53487059 | 0.41484467 |
| Insl6 | 0.59199182 | 0.42253204 | 0.031 | 0.021 | 1 | 0.23505706 | 1.27510724 | 0.034 | 0.01 | 1 | 0.59199182 | 0.4148623 |
| Gm16541 | 0.2354563 | -3.0037327 | 0.042 | 0.078 | 1 | 0.70056469 | -1.3923174 | 0.063 | 0.073 | 1 | 0.70056469 | 0.41547293 |
| Tmem25 | 0.52961488 | -0.3144336 | 0.01 | 0.021 | 1 | 0.23592811 | 1.14957636 | 0.034 | 0.01 | 1 | 0.52961488 | 0.41619415 |
| Hecw1 | 0.99275829 | -0.7294711 | 0.021 | 0.021 | 1 | 0.23592811 | 1.86006974 | 0.034 | 0.01 | 1 | 0.99275829 | 0.41619415 |
| Oxld1 | 0.38896453 | -0.407543 | 0.042 | 0.067 | 1 | 0.2360842 | -1.0203486 | 0.034 | 0.062 | 1 | 0.38896453 | 0.41643265 |
| Gm57189 | 0.98914258 | -0.3144336 | 0.052 | 0.052 | 1 | 0.23620694 | -1.484942 | 0.109 | 0.146 | 1 | 0.98914258 | 0.41662016 |
| Yjefn3 | 0.63792828 | 0.45495351 | 0.115 | 0.098 | 1 | 0.23654237 | 0.57767001 | 0.118 | 0.073 | 1 | 0.63792828 | 0.41713244 |
| Ghr | 0.23665455 | -0.4746858 | 0.771 | 0.777 | 1 | 0.29074803 | -0.5846806 | 0.735 | 0.708 | 1 | 0.29074803 | 0.41730372 |
| Cox18 | 0.236725 | 0.2775837 | 0.312 | 0.244 | 1 | 0.58061973 | -0.6804986 | 0.307 | 0.302 | 1 | 0.58061973 | 0.41741128 |
| Gm14455 | 0.47606502 | 1.00749454 | 0.021 | 0.01 | 1 | 0.23679813 | 1.01207283 | 0.034 | 0.01 | 1 | 0.47606502 | 0.4175229 |
| H2-Q7 | 0.5862099 | 0.68556644 | 0.031 | 0.021 | 1 | 0.23679813 | 1.49749966 | 0.034 | 0.01 | 1 | 0.5862099 | 0.4175229 |
| Zfp128 | 0.75328966 | 0.27052894 | 0.021 | 0.016 | 1 | 0.23679813 | 1.01207283 | 0.034 | 0.01 | 1 | 0.75328966 | 0.4175229 |
| Gm26672 | 0.86441475 | -0.1079827 | 0.052 | 0.047 | 1 | 0.23679813 | 1.01207283 | 0.034 | 0.01 | 1 | 0.86441475 | 0.4175229 |
| Bco2 | 0.99137461 | 0.35541784 | 0.042 | 0.041 | 1 | 0.23679813 | 1.01207283 | 0.034 | 0.01 | 1 | 0.99137461 | 0.4175229 |
| Il4ra | 0.99309965 | 0.22988696 | 0.042 | 0.041 | 1 | 0.23679813 | 1.01207283 | 0.034 | 0.01 | 1 | 0.99309965 | 0.4175229 |
| 2010320M18Rik | 0.23737723 | -1.130009 | 0.042 | 0.078 | 1 | 0.57497285 | 0.16407593 | 0.092 | 0.073 | 1 | 0.57497285 | 0.41840652 |
| Dstyk | 0.23740577 | 0.20013961 | 0.583 | 0.503 | 1 | 0.46408687 | -0.2934608 | 0.651 | 0.646 | 1 | 0.46408687 | 0.41845003 |
| Gfra4 | 0.38789252 | 0.59245704 | 0.052 | 0.031 | 1 | 0.23772424 | 0.86006974 | 0.034 | 0.01 | 1 | 0.38789252 | 0.41893566 |
| BC025920 | 0.39356009 | -0.2148979 | 0.01 | 0.026 | 1 | 0.23772424 | 0.86006974 | 0.034 | 0.01 | 1 | 0.39356009 | 0.41893566 |
| Etaa1os | 0.50217061 | 0.62047141 | 0.083 | 0.062 | 1 | 0.23772424 | 0.86006974 | 0.034 | 0.01 | 1 | 0.50217061 | 0.41893566 |
| 5033430I15Rik | 0.69098575 | 0.27052894 | 0.052 | 0.041 | 1 | 0.23772424 | 0.86006974 | 0.034 | 0.01 | 1 | 0.69098575 | 0.41893566 |
| Gm54310 | 0.80639868 | 0.71798792 | 0.052 | 0.047 | 1 | 0.23772424 | 0.86006974 | 0.034 | 0.01 | 1 | 0.80639868 | 0.41893566 |
| 1700031C06Rik | 0.88736279 | -0.2462621 | 0.094 | 0.088 | 1 | 0.23772424 | 0.86006974 | 0.034 | 0.01 | 1 | 0.88736279 | 0.41893566 |
| Ano8 | 0.80939866 | 0.22988696 | 0.042 | 0.036 | 1 | 0.23782391 | -1.0688472 | 0.05 | 0.083 | 1 | 0.80939866 | 0.4190876 |
| Synpo | 0.23785379 | -0.407543 | 0.76 | 0.782 | 1 | 0.31665413 | -0.1920775 | 0.773 | 0.583 | 1 | 0.31665413 | 0.41913315 |
| Twist1 | 0.48836827 | -1.104715 | 0.188 | 0.202 | 1 | 0.23811259 | -1.7306984 | 0.311 | 0.312 | 1 | 0.48836827 | 0.41952758 |
| Polg2 | 0.70496531 | -0.8375308 | 0.198 | 0.202 | 1 | 0.23814451 | -0.8729914 | 0.269 | 0.312 | 1 | 0.70496531 | 0.41957622 |
| Col26a1 | 0.81497835 | -1.0799683 | 0.031 | 0.026 | 1 | 0.23825911 | -1.010295 | 0.05 | 0.021 | 1 | 0.81497835 | 0.41975082 |
| Fzd8 | 0.23828158 | -0.9232428 | 0.125 | 0.171 | 1 | 0.64739851 | -0.6356254 | 0.193 | 0.208 | 1 | 0.64739851 | 0.41978505 |
| Rassf8 | 0.71894857 | -0.2462621 | 0.24 | 0.249 | 1 | 0.2386247 | -0.8244284 | 0.214 | 0.25 | 1 | 0.71894857 | 0.42030765 |
| Galnt14 | 0.32025586 | -0.9925055 | 0 | 0.01 | 1 | 0.23868169 | -2.3098553 | 0.013 | 0.031 | 1 | 0.32025586 | 0.42039443 |
| 2610507I01Rik | 0.23869864 | 0.52206771 | 0.146 | 0.098 | 1 | 0.93787312 | -0.6931839 | 0.16 | 0.156 | 1 | 0.93787312 | 0.42042023 |
| Hdac5 | 0.2387402 | -0.7580402 | 0.229 | 0.28 | 1 | 0.9953519 | -0.5885713 | 0.391 | 0.333 | 1 | 0.9953519 | 0.42048351 |

| | | | | | | | | | | | | |
|---|---|---|---|---|---|---|---|---|---|---|---|---|
| Nckap5l | 0.78800338 | -0.1880563 | 0.229 | 0.212 | 1 | 0.23887175 | -0.3361665 | 0.445 | 0.312 | 1 | 0.78800338 | 0.42068379 |
| Strn | 0.23903515 | 0.15949763 | 0.594 | 0.523 | 1 | 0.76182825 | -0.1012686 | 0.702 | 0.677 | 1 | 0.76182825 | 0.42093249 |
| E2f3 | 0.23918892 | 0.42253204 | 0.25 | 0.187 | 1 | 0.25247746 | 0.13214928 | 0.366 | 0.292 | 1 | 0.25247746 | 0.42116649 |
| Spty2d1 | 0.23920907 | 0.21732158 | 0.729 | 0.668 | 1 | 0.82270886 | -0.1596935 | 0.744 | 0.792 | 1 | 0.82270886 | 0.42119717 |
| Ccdc122 | 0.23924094 | -0.3281085 | 0.146 | 0.202 | 1 | 0.28899498 | 0.13375139 | 0.189 | 0.135 | 1 | 0.28899498 | 0.42124565 |
| Cdc14b | 0.23927887 | -0.6329631 | 0.094 | 0.14 | 1 | 0.40962831 | -0.4010032 | 0.214 | 0.167 | 1 | 0.40962831 | 0.42130336 |
| 2810036E18Rik | 0.37360617 | -0.4519371 | 0.021 | 0.041 | 1 | 0.23949613 | 1.01207283 | 0.034 | 0.01 | 1 | 0.37360617 | 0.42163386 |
| Fetub | 0.85444268 | 0.18806678 | 0.073 | 0.067 | 1 | 0.2395365 | -0.3098553 | 0.134 | 0.188 | 1 | 0.85444268 | 0.42169527 |
| Zbed5-ps | 0.2395632 | 0.45495351 | 0.115 | 0.073 | 1 | 0.3900186 | -1.144796 | 0.088 | 0.115 | 1 | 0.3900186 | 0.42173587 |
| Cmss1 | 0.73208244 | -0.2425807 | 1 | 1 | 1 | 0.23962003 | -0.3998059 | 1 | 1 | 1 | 0.73208244 | 0.42182231 |
| Mdm4 | 0.6444885 | 0.20819189 | 0.729 | 0.668 | 1 | 0.23981064 | -0.3689904 | 0.798 | 0.865 | 1 | 0.6444885 | 0.42211214 |
| Zfp36 | 0.24007413 | -0.3144336 | 0.74 | 0.808 | 1 | 0.56845302 | -0.1635658 | 0.803 | 0.906 | 1 | 0.56845302 | 0.42251268 |
| Lrrc75aos2 | 0.24010913 | -1.0628948 | 0.042 | 0.078 | 1 | 0.72001996 | -0.5322477 | 0.088 | 0.073 | 1 | 0.72001996 | 0.42256586 |
| Rfx5 | 0.73055047 | 0.30705482 | 0.073 | 0.062 | 1 | 0.24012143 | 0.93807225 | 0.063 | 0.031 | 1 | 0.73055047 | 0.42258455 |
| Morc3 | 0.24023072 | 0.2857957 | 0.385 | 0.311 | 1 | 0.51640935 | -0.2089264 | 0.366 | 0.406 | 1 | 0.51640935 | 0.42275064 |
| Dennd6b | 0.24023509 | 0.88196365 | 0.094 | 0.057 | 1 | 0.43079893 | 0.69014474 | 0.038 | 0.021 | 1 | 0.43079893 | 0.42275728 |
| Ass1 | 0.2402932 | 0.46152617 | 0.281 | 0.228 | 1 | 0.26874008 | 0.48155812 | 0.126 | 0.083 | 1 | 0.26874008 | 0.42284557 |
| Tepsin | 0.8006267 | 0.20013961 | 0.021 | 0.026 | 1 | 0.24037027 | -0.194378 | 0.05 | 0.021 | 1 | 0.8006267 | 0.42296268 |
| Mal | 0.74076005 | 1.00749454 | 0.021 | 0.016 | 1 | 0.24038606 | 1.01207283 | 0.034 | 0.01 | 1 | 0.74076005 | 0.42298666 |
| Tmem156 | 0.32025586 | -0.577468 | 0 | 0.01 | 1 | 0.2408519 | -2.1172102 | 0.013 | 0.031 | 1 | 0.32025586 | 0.42369416 |
| Gm42679 | 0.33961853 | -1.3144336 | 0.073 | 0.104 | 1 | 0.2408519 | -2.3098553 | 0.013 | 0.031 | 1 | 0.33961853 | 0.42369416 |
| Slc44a4 | 0.53178162 | -0.577468 | 0.01 | 0.021 | 1 | 0.2408519 | -2.1172102 | 0.013 | 0.031 | 1 | 0.53178162 | 0.42369416 |
| Frmpd4 | 0.62109304 | -0.4872702 | 0.021 | 0.031 | 1 | 0.2408519 | -3.6317834 | 0.013 | 0.031 | 1 | 0.62109304 | 0.42369416 |
| Opn3 | 0.79259943 | 0.52206771 | 0.031 | 0.026 | 1 | 0.2408519 | -2.6317834 | 0.013 | 0.031 | 1 | 0.79259943 | 0.42369416 |
| Coch | 0.24089524 | -2.6705774 | 0.073 | 0.114 | 1 | 0.31485306 | -1.8504236 | 0.038 | 0.062 | 1 | 0.31485306 | 0.42375997 |
| Tpx2 | 0.24115086 | 2.31005731 | 0.104 | 0.067 | 1 | 0.24304534 | -1.9879272 | 0.013 | 0.031 | 1 | 0.24304534 | 0.42414799 |
| Pappa | 0.61738685 | -0.7580402 | 0.021 | 0.031 | 1 | 0.24126679 | -1.1353575 | 0.126 | 0.167 | 1 | 0.61738685 | 0.42432392 |
| Stxbp1 | 0.24170132 | 0.37443775 | 0.562 | 0.523 | 1 | 0.68667205 | -0.1874586 | 0.66 | 0.542 | 1 | 0.68667205 | 0.42498312 |
| D430020J02Rik | 0.24179987 | 1.00749454 | 0.094 | 0.057 | 1 | 0.64304313 | 0.27510724 | 0.042 | 0.031 | 1 | 0.64304313 | 0.42513256 |
| Nagpa | 0.24185348 | 0.52206771 | 0.146 | 0.098 | 1 | 0.39364455 | -0.2029401 | 0.105 | 0.073 | 1 | 0.39364455 | 0.42521385 |
| Xrcc3 | 0.38521705 | -0.7998604 | 0.01 | 0.026 | 1 | 0.24201865 | 1.01207283 | 0.063 | 0.031 | 1 | 0.38521705 | 0.42546428 |
| 5730522E02Rik | 0.24206453 | 0.21231433 | 0.167 | 0.114 | 1 | 0.67716259 | -0.5873892 | 0.122 | 0.135 | 1 | 0.67716259 | 0.42553383 |
| Tnfsf9 | 0.87232592 | -0.577468 | 0.052 | 0.047 | 1 | 0.24224109 | -1.3098553 | 0.05 | 0.083 | 1 | 0.87232592 | 0.42580143 |
| Bbc3 | 0.78360215 | -0.4705528 | 0.115 | 0.124 | 1 | 0.2424192 | -0.3098553 | 0.134 | 0.083 | 1 | 0.78360215 | 0.42607134 |
| Fbxo33 | 0.77722731 | -0.5477206 | 0.375 | 0.363 | 1 | 0.24243621 | -0.480656 | 0.412 | 0.49 | 1 | 0.77722731 | 0.4260971 |
| Cdc27 | 0.24256727 | -0.3108669 | 0.469 | 0.549 | 1 | 0.38635207 | -0.1813405 | 0.681 | 0.708 | 1 | 0.38635207 | 0.42629566 |
| Ikbkb | 0.24260308 | 0.37613913 | 0.333 | 0.264 | 1 | 0.63395077 | -0.4182314 | 0.382 | 0.385 | 1 | 0.63395077 | 0.4263499 |
| Pik3c2g | 0.43140301 | -0.2019588 | 0.125 | 0.093 | 1 | 0.24277242 | 1.06865636 | 0.063 | 0.031 | 1 | 0.43140301 | 0.42660639 |
| Jcad | 0.24285005 | 0.39451766 | 0.26 | 0.197 | 1 | 0.57673784 | -0.1139351 | 0.29 | 0.24 | 1 | 0.57673784 | 0.42672396 |
| Fam222b | 0.24286512 | -0.3720941 | 0.573 | 0.601 | 1 | 0.56099625 | -0.25977 | 0.668 | 0.635 | 1 | 0.56099625 | 0.42674678 |
| Gpr135 | 0.67877064 | 0.49292136 | 0.052 | 0.041 | 1 | 0.24307133 | 0.56461386 | 0.034 | 0.01 | 1 | 0.67877064 | 0.42705898 |
| Fsd1 | 1 | -0.2148979 | 0.021 | 0.021 | 1 | 0.24307133 | 0.56461386 | 0.034 | 0.01 | 1 | 1 | 0.42705898 |
| Scn5a | 0.24312877 | -0.4872702 | 0.042 | 0.078 | 1 | 0.73379677 | 0.34222143 | 0.05 | 0.042 | 1 | 0.73379677 | 0.42714594 |
| Mamdc2 | 0.24324506 | 1.05640414 | 0.125 | 0.083 | 1 | 0.25742507 | 0.23763253 | 0.118 | 0.073 | 1 | 0.25742507 | 0.42732197 |
| Prim2 | 0.24360566 | 0.12533103 | 0.271 | 0.207 | 1 | 0.66729606 | -0.7573142 | 0.244 | 0.24 | 1 | 0.66729606 | 0.4278676 |
| Lrrc32 | 0.5425293 | -0.5160674 | 0.073 | 0.093 | 1 | 0.24366712 | -1.2223924 | 0.059 | 0.094 | 1 | 0.5425293 | 0.42796057 |
| Stard9 | 0.24837714 | -1.0320338 | 0.073 | 0.114 | 1 | 0.2436933 | 0.19264508 | 0.092 | 0.052 | 1 | 0.24837714 | 0.42800018 |
| Mis18bp1 | 0.24369666 | 2.13302542 | 0.052 | 0.026 | 1 | 0.57373861 | -1.3098553 | 0.013 | 0.021 | 1 | 0.57373861 | 0.42800526 |
| Gm57147 | 0.30891093 | 1.00749454 | 0.042 | 0.021 | 1 | 0.24392267 | 0.77760758 | 0.063 | 0.031 | 1 | 0.30891093 | 0.42834708 |
| Slc25a18 | 0.46697007 | 1.74446013 | 0.021 | 0.01 | 1 | 0.24396338 | 0.42711033 | 0.034 | 0.01 | 1 | 0.46697007 | 0.42840863 |
| Gm48855 | 0.52744059 | -0.577468 | 0.01 | 0.021 | 1 | 0.24396338 | 0.42711033 | 0.034 | 0.01 | 1 | 0.52744059 | 0.42840863 |
| B230208H11Rik | 0.55841286 | 0.74446013 | 0.062 | 0.047 | 1 | 0.24396338 | 0.42711033 | 0.034 | 0.01 | 1 | 0.55841286 | 0.42840863 |
| Gm715 | 0.78979801 | -0.407543 | 0.021 | 0.026 | 1 | 0.24396338 | 0.42711033 | 0.034 | 0.01 | 1 | 0.78979801 | 0.42840863 |
| Abcd3 | 0.98488366 | -0.159513 | 0.583 | 0.585 | 1 | 0.24398747 | -0.3585111 | 0.672 | 0.75 | 1 | 0.98488366 | 0.42844506 |
| Mgme1 | 0.4320165 | 0.22988696 | 0.125 | 0.093 | 1 | 0.24403618 | -0.2103196 | 0.164 | 0.104 | 1 | 0.4320165 | 0.4285187 |
| Ret | 0.31287735 | -0.2920657 | 0.094 | 0.135 | 1 | 0.24413474 | -1.6317834 | 0.013 | 0.031 | 1 | 0.31287735 | 0.42866771 |
| Zfp658 | 0.47599549 | 0.52206771 | 0.042 | 0.026 | 1 | 0.24413474 | -1.6317834 | 0.013 | 0.031 | 1 | 0.47599549 | 0.42866771 |
| Gm3055 | 0.57848839 | 1.00749454 | 0.031 | 0.021 | 1 | 0.24413474 | -1.6317834 | 0.013 | 0.031 | 1 | 0.57848839 | 0.42866771 |
| Hk3 | 0.59199182 | 0.42253204 | 0.031 | 0.021 | 1 | 0.24413474 | -1.8948178 | 0.013 | 0.031 | 1 | 0.59199182 | 0.42866771 |
| Dcst1 | 0.61737927 | 1.00749454 | 0.01 | 0.005 | 1 | 0.24413474 | -1.8948178 | 0.013 | 0.031 | 1 | 0.61737927 | 0.42866771 |
| Gm50455 | 0.61737927 | 1.00749454 | 0.01 | 0.005 | 1 | 0.24413474 | -1.8948178 | 0.013 | 0.031 | 1 | 0.61737927 | 0.42866771 |
| Myorg | 0.74827573 | 0.59245704 | 0.021 | 0.016 | 1 | 0.24413474 | -1.6317834 | 0.013 | 0.031 | 1 | 0.74827573 | 0.42866771 |
| 1700023H06Rik | 0.75328966 | 0.27052894 | 0.021 | 0.016 | 1 | 0.24413474 | -1.6317834 | 0.013 | 0.031 | 1 | 0.75328966 | 0.42866771 |
| Ifi44 | 0.8238181 | -0.7294711 | 0.042 | 0.047 | 1 | 0.24413474 | -4.3098553 | 0.013 | 0.031 | 1 | 0.8238181 | 0.42866771 |
| Gm3830 | 0.98414882 | 0.52206771 | 0.031 | 0.031 | 1 | 0.24413474 | -1.8948178 | 0.013 | 0.031 | 1 | 0.98414882 | 0.42866771 |
| Gm57069 | 0.993208 | 1.00749454 | 0.01 | 0.01 | 1 | 0.24413474 | -1.8948178 | 0.013 | 0.031 | 1 | 0.993208 | 0.42866771 |
| Tbc1d10c | 0.993208 | 1.00749454 | 0.01 | 0.01 | 1 | 0.24413474 | -1.6317834 | 0.013 | 0.031 | 1 | 0.993208 | 0.42866771 |
| Amigo2 | 1 | 0.42253204 | 0.01 | 0.01 | 1 | 0.24413474 | -1.6317834 | 0.013 | 0.031 | 1 | 1 | 0.42866771 |
| Mettl5os | 0.38521705 | -0.7998604 | 0.01 | 0.026 | 1 | 0.24414622 | -1.5728897 | 0.013 | 0.031 | 1 | 0.38521705 | 0.42868506 |
| Popdc3 | 0.49466999 | -0.2555399 | 0.042 | 0.026 | 1 | 0.24414622 | -2.1399303 | 0.013 | 0.031 | 1 | 0.49466999 | 0.42868506 |
| Arhgap45 | 0.52744059 | -0.577468 | 0.01 | 0.021 | 1 | 0.24485439 | 0.27510724 | 0.034 | 0.01 | 1 | 0.52744059 | 0.42975511 |
| Gm37982 | 0.57774092 | -1.265524 | 0.052 | 0.067 | 1 | 0.24485439 | 0.27510724 | 0.034 | 0.01 | 1 | 0.57774092 | 0.42975511 |
| Morn3 | 0.74827573 | 0.59245704 | 0.021 | 0.016 | 1 | 0.24485439 | -0.4618584 | 0.034 | 0.01 | 1 | 0.74827573 | 0.42975511 |
| Fank1 | 0.61182672 | -0.577468 | 0.021 | 0.031 | 1 | 0.24487757 | 0.14957636 | 0.034 | 0.01 | 1 | 0.61182672 | 0.42979012 |
| Nox1 | 0.70121365 | -0.7294711 | 0.042 | 0.052 | 1 | 0.24487806 | -1.4167705 | 0.034 | 0.01 | 1 | 0.70121365 | 0.42979085 |
| Gm14221 | 0.24492634 | 1.74446013 | 0.073 | 0.041 | 1 | 0.83565066 | -0.2509616 | 0.059 | 0.052 | 1 | 0.83565066 | 0.42986376 |
| Gm38190 | 0.2449825 | 0.46000674 | 0.115 | 0.073 | 1 | 0.44571648 | -0.7692869 | 0.118 | 0.083 | 1 | 0.44571648 | 0.42994858 |
| Acyp2 | 0.24503484 | -0.4179898 | 0.469 | 0.503 | 1 | 0.64613591 | -0.8355368 | 0.374 | 0.354 | 1 | 0.64613591 | 0.43002761 |
| Musk | 1 | 0.42253204 | 0.01 | 0.01 | 1 | 0.24524916 | -1.5322477 | 0.013 | 0.031 | 1 | 1 | 0.43035117 |
| Kifc2 | 0.61924305 | -0.2148979 | 0.021 | 0.031 | 1 | 0.24607399 | 0.69014474 | 0.063 | 0.031 | 1 | 0.61924305 | 0.43159557 |
| Rin1 | 0.99603711 | -0.3144336 | 0.031 | 0.031 | 1 | 0.24635272 | -1.7952821 | 0.013 | 0.031 | 1 | 0.99603711 | 0.43201577 |
| Hyls1 | 0.71693558 | 0.17099327 | 0.083 | 0.073 | 1 | 0.24635853 | -0.1723517 | 0.092 | 0.052 | 1 | 0.71693558 | 0.43202454 |
| Tradd | 0.72016246 | 0.14951354 | 0.146 | 0.13 | 1 | 0.24642391 | 0.17557156 | 0.105 | 0.062 | 1 | 0.72016246 | 0.43212308 |
| Gm11423 | 0.24646602 | 0.76648644 | 0.094 | 0.057 | 1 | 0.2986794 | 0.77760758 | 0.059 | 0.031 | 1 | 0.2986794 | 0.43218654 |
| 1700052K11Rik | 0.24655256 | 0.5049942 | 0.094 | 0.057 | 1 | 0.72614543 | -0.5025003 | 0.063 | 0.052 | 1 | 0.72614543 | 0.43231696 |

| | | | | | | | | | | | | |
|---|---|---|---|---|---|---|---|---|---|---|---|---|
| Daam2 | 0.54036789 | -0.2510385 | 0.51 | 0.456 | 1 | 0.24673784 | -0.2430238 | 0.437 | 0.323 | 1 | 0.54036789 | 0.43259611 |
| Limd2 | 0.48397605 | 0.27052894 | 0.167 | 0.135 | 1 | 0.24674662 | 0.23446525 | 0.13 | 0.083 | 1 | 0.48397605 | 0.43260934 |
| Met | 0.55702458 | -0.1329867 | 0.406 | 0.446 | 1 | 0.24701806 | -0.3646973 | 0.739 | 0.677 | 1 | 0.55702458 | 0.4330182 |
| Gtpbp8 | 0.24743492 | 0.67734593 | 0.24 | 0.192 | 1 | 0.36619065 | -0.2358547 | 0.248 | 0.188 | 1 | 0.36619065 | 0.4336458 |
| Pomt1 | 0.2474384 | 0.28502851 | 0.177 | 0.124 | 1 | 0.5870762 | -0.7395395 | 0.16 | 0.177 | 1 | 0.5870762 | 0.43365104 |
| Gm24474 | 0.32025586 | -0.577468 | 0 | 0.01 | 1 | 0.24746336 | -1.3098553 | 0.013 | 0.031 | 1 | 0.32025586 | 0.43368861 |
| Gm17058 | 0.52961488 | -0.3144336 | 0.01 | 0.021 | 1 | 0.24746336 | -1.3098553 | 0.013 | 0.031 | 1 | 0.52961488 | 0.43368861 |
| Gm26787 | 1 | 0.42253204 | 0.01 | 0.01 | 1 | 0.24746336 | -1.3098553 | 0.013 | 0.031 | 1 | 1 | 0.43368861 |
| Txndc8 | 0.52528691 | -1.1624305 | 0.01 | 0.021 | 1 | 0.24746984 | -1.0203486 | 0.013 | 0.031 | 1 | 0.52528691 | 0.43369835 |
| H3c7 | 0.370501 | 2.46692616 | 0.031 | 0.016 | 1 | 0.24747056 | -1.3098553 | 0.013 | 0.031 | 1 | 0.370501 | 0.43369944 |
| Gm57275 | 0.38521705 | -0.7998604 | 0.01 | 0.026 | 1 | 0.24747056 | -1.3098553 | 0.013 | 0.031 | 1 | 0.38521705 | 0.43369944 |
| Muc4 | 0.5383068 | 1.00749454 | 0.01 | 0.021 | 1 | 0.24747056 | -1.3098553 | 0.013 | 0.031 | 1 | 0.5383068 | 0.43369944 |
| Kif21b | 0.61737927 | 1.00749454 | 0.01 | 0.005 | 1 | 0.24747056 | -1.3098553 | 0.013 | 0.031 | 1 | 0.61737927 | 0.43369944 |
| C530020B09Rik | 0.75328966 | 0.27052894 | 0.021 | 0.016 | 1 | 0.24747056 | -1.3098553 | 0.013 | 0.031 | 1 | 0.75328966 | 0.43369944 |
| Elovl2 | 0.7854853 | -0.577468 | 0.021 | 0.026 | 1 | 0.24747056 | -1.3098553 | 0.013 | 0.031 | 1 | 0.7854853 | 0.43369944 |
| Znrf2 | 0.84678268 | -0.2257046 | 0.385 | 0.373 | 1 | 0.24752093 | -0.1399303 | 0.487 | 0.615 | 1 | 0.84678268 | 0.43377524 |
| Dpm1 | 0.24763847 | -0.1141349 | 0.885 | 0.881 | 1 | 0.90567518 | -0.1303058 | 0.857 | 0.885 | 1 | 0.90567518 | 0.43395213 |
| Adcy7 | 0.24782362 | -2.1624305 | 0.052 | 0.088 | 1 | 0.46251598 | -1.194378 | 0.071 | 0.094 | 1 | 0.46251598 | 0.4342307 |
| Rad18 | 0.7488862 | 0.56270969 | 0.177 | 0.171 | 1 | 0.24810444 | 1.01207283 | 0.126 | 0.083 | 1 | 0.7488862 | 0.43465307 |
| Gm14295 | 0.2481486 | 0.74446013 | 0.094 | 0.057 | 1 | 0.91035079 | 0.21370669 | 0.076 | 0.073 | 1 | 0.91035079 | 0.43471948 |
| Trim13 | 0.2481486 | 0.74446013 | 0.094 | 0.057 | 1 | 0.3190026 | 0.56461386 | 0.097 | 0.062 | 1 | 0.3190026 | 0.43471948 |
| Ccdc18 | 0.24819262 | 2.17741954 | 0.073 | 0.041 | 1 | 0.99568332 | -1.2103196 | 0.042 | 0.042 | 1 | 0.99568332 | 0.43478567 |
| Gm26698 | 0.24843624 | 1.22988696 | 0.052 | 0.026 | 1 | 0.50587544 | -2.3098553 | 0.004 | 0.01 | 1 | 0.50587544 | 0.43515191 |
| Ppp1r3e | 0.24843624 | 1.42253204 | 0.052 | 0.026 | 1 | 0.92738405 | -0.4618584 | 0.034 | 0.031 | 1 | 0.92738405 | 0.43515191 |
| Dbpht2 | 0.24847378 | 1.46692616 | 0.052 | 0.026 | 1 | 0.27154327 | 1.27510724 | 0.013 | 0 | 1 | 0.27154327 | 0.43520835 |
| Caskin1 | 0.24847378 | 1.20013961 | 0.052 | 0.026 | 1 | 0.65825923 | -0.6724253 | 0.055 | 0.042 | 1 | 0.65825923 | 0.43520835 |
| Nrg2 | 0.38689439 | -0.577468 | 0.01 | 0.026 | 1 | 0.24857751 | -1.0468209 | 0.013 | 0.031 | 1 | 0.38689439 | 0.43536424 |
| Abcc3 | 0.83916629 | 0.13302542 | 0.031 | 0.036 | 1 | 0.24857751 | -0.8948178 | 0.013 | 0.031 | 1 | 0.83916629 | 0.43536424 |
| 5430402E10Rik | 0.30389555 | 1.49292136 | 0.042 | 0.021 | 1 | 0.2490006 | 1.10518224 | 0.088 | 0.052 | 1 | 0.30389555 | 0.43599991 |
| Alkbh8 | 0.24941473 | 0.25542205 | 0.302 | 0.228 | 1 | 0.43569552 | -0.1095566 | 0.282 | 0.219 | 1 | 0.43569552 | 0.43662175 |
| Gm49519 | 0.24946138 | 1.15949763 | 0.052 | 0.026 | 1 | 0.99396043 | -0.8244284 | 0.021 | 0.021 | 1 | 0.99396043 | 0.43669179 |
| Dnhd1 | 0.2495285 | 0.68556644 | 0.177 | 0.13 | 1 | 0.30410997 | 0.81567562 | 0.143 | 0.104 | 1 | 0.30410997 | 0.43679253 |
| Pex11b | 0.2697982 | 0.33506919 | 0.281 | 0.218 | 1 | 0.24982408 | 0.25492936 | 0.218 | 0.156 | 1 | 0.2697982 | 0.43723609 |
| Cutc | 0.96905096 | 0.15949763 | 0.115 | 0.119 | 1 | 0.24990013 | 0.19264508 | 0.13 | 0.083 | 1 | 0.96905096 | 0.43735019 |
| Ttf2 | 0.250043 | -0.6554705 | 0.042 | 0.078 | 1 | 0.36895003 | 0.42711033 | 0.055 | 0.031 | 1 | 0.36895003 | 0.43756449 |
| Scube3 | 0.25008035 | -0.4330781 | 0.042 | 0.078 | 1 | 0.75977006 | -0.7952821 | 0.025 | 0.031 | 1 | 0.75977006 | 0.43762052 |
| Ttc39b | 0.25570358 | -0.3171789 | 0.792 | 0.772 | 1 | 0.25012795 | -0.3054094 | 0.811 | 0.833 | 1 | 0.25570358 | 0.43769191 |
| Pacs2 | 0.46773898 | -0.536826 | 0.188 | 0.218 | 1 | 0.25014976 | -0.3098553 | 0.286 | 0.198 | 1 | 0.46773898 | 0.43772461 |
| Serping1 | 0.60239618 | -0.9467018 | 0.052 | 0.067 | 1 | 0.25017935 | -3.6724253 | 0.071 | 0.104 | 1 | 0.60239618 | 0.437769 |
| Gm4924 | 0.25027908 | 0.65385758 | 0.125 | 0.083 | 1 | 0.80619008 | -0.7692869 | 0.046 | 0.052 | 1 | 0.80619008 | 0.43791854 |
| Gclm | 0.97899036 | -0.2032726 | 0.552 | 0.534 | 1 | 0.2503929 | -0.3889268 | 0.588 | 0.667 | 1 | 0.97899036 | 0.4380892 |
| Eral1 | 0.33727461 | 0.34006988 | 0.156 | 0.114 | 1 | 0.25067193 | 0.27510724 | 0.168 | 0.115 | 1 | 0.33727461 | 0.43850744 |
| Gm56596 | 0.38181636 | 0.78510212 | 0.052 | 0.031 | 1 | 0.25080911 | -0.9879272 | 0.013 | 0.031 | 1 | 0.38181636 | 0.43871302 |
| Dpysl5 | 0.47606368 | 1.00749454 | 0.021 | 0.01 | 1 | 0.25080911 | -0.7248928 | 0.013 | 0.031 | 1 | 0.47606368 | 0.43871302 |
| Kcnu1 | 0.99603711 | -0.1624305 | 0.031 | 0.031 | 1 | 0.25080911 | -0.9879272 | 0.013 | 0.031 | 1 | 0.99603711 | 0.43871302 |
| Itih2 | 0.27531804 | -0.536826 | 0.104 | 0.15 | 1 | 0.25108403 | 0.15727075 | 0.168 | 0.115 | 1 | 0.27531804 | 0.43912486 |
| Trmt5 | 0.86973886 | 0.17741954 | 0.083 | 0.078 | 1 | 0.25118438 | 0.82764826 | 0.063 | 0.031 | 1 | 0.86973886 | 0.43927516 |
| Ttc21b | 0.65869466 | 0.39451766 | 0.125 | 0.109 | 1 | 0.25142085 | 0.13492958 | 0.143 | 0.094 | 1 | 0.65869466 | 0.43962926 |
| Exoc6 | 0.25159393 | -0.6445822 | 0.042 | 0.078 | 1 | 0.52791748 | 0.59703533 | 0.046 | 0.031 | 1 | 0.52791748 | 0.43988836 |
| Tmem121 | 0.49880793 | 0.20013961 | 0.073 | 0.052 | 1 | 0.2518463 | 0.53814164 | 0.063 | 0.031 | 1 | 0.49880793 | 0.44026604 |
| Zfp719 | 0.5948299 | 0.15949763 | 0.177 | 0.15 | 1 | 0.2518843 | -0.3769695 | 0.147 | 0.094 | 1 | 0.5948299 | 0.44032291 |
| Rec114 | 0.25194879 | 0.31561683 | 0.115 | 0.073 | 1 | 0.77179334 | -0.5728897 | 0.143 | 0.125 | 1 | 0.77179334 | 0.44041939 |
| Gpd1l | 0.2523837 | 0.88196365 | 0.052 | 0.026 | 1 | 0.62366761 | 0.23763253 | 0.134 | 0.115 | 1 | 0.62366761 | 0.44106987 |
| Mrpl14 | 0.25238962 | 0.22250743 | 0.49 | 0.389 | 1 | 0.44617153 | -0.1454684 | 0.416 | 0.365 | 1 | 0.44617153 | 0.44107872 |
| Rin2 | 0.25246164 | -0.1579895 | 0.906 | 0.927 | 1 | 0.82389526 | -0.2866587 | 0.937 | 0.958 | 1 | 0.82389526 | 0.44118641 |
| Recql | 0.3450614 | -0.536826 | 0.104 | 0.145 | 1 | 0.25255109 | -0.131518 | 0.143 | 0.094 | 1 | 0.3450614 | 0.44132013 |
| 2900009J06Rik | 0.25280242 | 0.65957123 | 0.094 | 0.057 | 1 | 0.48467052 | -0.7573142 | 0.063 | 0.042 | 1 | 0.48467052 | 0.44169578 |
| Kcnrg | 0.5383068 | 0.27052894 | 0.01 | 0.021 | 1 | 0.25286107 | -0.8948178 | 0.067 | 0.104 | 1 | 0.5383068 | 0.44178343 |
| Fam111a | 0.25297529 | 0.75956702 | 0.094 | 0.057 | 1 | 0.58180179 | -0.7248928 | 0.021 | 0.031 | 1 | 0.58180179 | 0.44195408 |
| Zfp937 | 0.25332778 | 1.00749454 | 0.052 | 0.026 | 1 | 0.80020906 | -0.7248928 | 0.017 | 0.021 | 1 | 0.80020906 | 0.44248059 |
| Zfp764 | 0.25332778 | 1.00749454 | 0.052 | 0.026 | 1 | 0.97410634 | 0.19264508 | 0.042 | 0.042 | 1 | 0.97410634 | 0.44248059 |
| Abcb6 | 0.25337721 | 1.00749454 | 0.052 | 0.026 | 1 | 0.55484807 | -0.3923174 | 0.059 | 0.042 | 1 | 0.55484807 | 0.44255441 |
| Tedc2 | 0.25337721 | 1.00749454 | 0.052 | 0.026 | 1 | 1 | -0.7248928 | 0.021 | 0.021 | 1 | 1 | 0.44255441 |
| Pcdhga9 | 0.25337721 | 1.00749454 | 0.052 | 0.026 | 1 | 0.49922624 | -1.3098553 | 0.046 | 0.062 | 1 | 0.49922624 | 0.44255441 |
| H2aj | 0.64782404 | -0.1380713 | 0.708 | 0.715 | 1 | 0.25341992 | 0.30945874 | 0.634 | 0.646 | 1 | 0.64782404 | 0.44261818 |
| Trim46 | 0.25350784 | 0.56003556 | 0.094 | 0.057 | 1 | 0.73874009 | -0.8123556 | 0.084 | 0.094 | 1 | 0.73874009 | 0.44274946 |
| 1500002C15Rik | 0.25360034 | -0.4950058 | 0.042 | 0.078 | 1 | 0.7311848 | 0.17557156 | 0.05 | 0.042 | 1 | 0.7311848 | 0.44288755 |
| Mks1 | 0.25364147 | 0.64492446 | 0.125 | 0.083 | 1 | 0.91218966 | -0.7248928 | 0.029 | 0.031 | 1 | 0.91218966 | 0.44294895 |
| A330023F24Rik | 0.41810841 | -0.407543 | 0.125 | 0.161 | 1 | 0.25366878 | -1.0674944 | 0.239 | 0.271 | 1 | 0.41810841 | 0.44298971 |
| Gm33887 | 0.71031978 | 0.35541784 | 0.062 | 0.052 | 1 | 0.25380864 | 0.45567948 | 0.063 | 0.031 | 1 | 0.71031978 | 0.44319845 |
| Zfp2 | 0.82342138 | 0.32942263 | 0.042 | 0.036 | 1 | 0.25380864 | 0.45567948 | 0.063 | 0.031 | 1 | 0.82342138 | 0.44319845 |
| Pola2 | 0.51525002 | -0.3848229 | 0.052 | 0.073 | 1 | 0.2539771 | 0.8917786 | 0.076 | 0.042 | 1 | 0.51525002 | 0.44344983 |
| 9430037G07Rik | 0.28429278 | -0.7998604 | 0.021 | 0.047 | 1 | 0.25420086 | -0.5025003 | 0.013 | 0.031 | 1 | 0.28429278 | 0.44378365 |
| 5330439K02Rik | 0.25464564 | -1.7149715 | 0.031 | 0.062 | 1 | 0.30798358 | -1.4473588 | 0.029 | 0.052 | 1 | 0.30798358 | 0.44444688 |
| Yeats2 | 0.25524475 | 0.22199905 | 0.438 | 0.332 | 1 | 0.56608147 | -0.7430859 | 0.475 | 0.427 | 1 | 0.56608147 | 0.44533961 |
| Ikbke | 0.25563945 | 0.2912875 | 0.24 | 0.176 | 1 | 0.79413899 | -0.4381794 | 0.252 | 0.25 | 1 | 0.79413899 | 0.44592738 |
| Mocs1 | 0.96760572 | 0.11440974 | 0.083 | 0.083 | 1 | 0.25577784 | 0.10518224 | 0.063 | 0.031 | 1 | 0.96760572 | 0.44613337 |
| Ccnt2 | 0.25597251 | 0.18666338 | 0.688 | 0.622 | 1 | 0.2576406 | -0.3407871 | 0.693 | 0.75 | 1 | 0.2576406 | 0.44642309 |
| Gm1123 | 0.25620655 | 0.88196365 | 0.073 | 0.041 | 1 | 0.57731374 | -0.5531264 | 0.13 | 0.104 | 1 | 0.57731374 | 0.4467713 |
| Gm29966 | 0.25629766 | 1.17741954 | 0.052 | 0.026 | 1 | 0.86205365 | -0.7248928 | 0.038 | 0.042 | 1 | 0.86205365 | 0.44690683 |
| H1f2 | 0.88768312 | -0.4120444 | 0.615 | 0.585 | 1 | 0.25679658 | 0.15818949 | 0.55 | 0.51 | 1 | 0.88768312 | 0.44764868 |
| Gm13483 | 0.25682187 | 0.48240349 | 0.219 | 0.166 | 1 | 0.87537688 | -0.3256226 | 0.202 | 0.208 | 1 | 0.87537688 | 0.44768626 |
| Mybl2 | 0.25728821 | 0.83756954 | 0.052 | 0.026 | 1 | 0.57621614 | -0.5322477 | 0.029 | 0.042 | 1 | 0.57621614 | 0.44837919 |
| Mroh1 | 0.25772845 | 0.17350449 | 0.208 | 0.15 | 1 | 0.92638388 | -0.7031977 | 0.206 | 0.188 | 1 | 0.92638388 | 0.44903294 |
| Dnajb5 | 0.25792709 | 0.83756954 | 0.073 | 0.041 | 1 | 0.25913701 | 0.61614416 | 0.076 | 0.042 | 1 | 0.25913701 | 0.44932779 |

| | | | | | | | | | | | | |
|---|---|---|---|---|---|---|---|---|---|---|---|---|
| 6720489N17Rik | 0.25824304 | 0.78510212 | 0.052 | 0.026 | 1 | 0.39796314 | 0.56461386 | 0.025 | 0.01 | 1 | 0.39796314 | 0.44979661 |
| Cd200r3 | 0.25824304 | 0.78510212 | 0.052 | 0.026 | 1 | 0.34428571 | 0.80562195 | 0.042 | 0.021 | 1 | 0.34428571 | 0.44979661 |
| Gpr39 | 0.25830207 | 0.72738662 | 0.052 | 0.026 | 1 | 0.81707832 | 0.27510724 | 0.017 | 0.021 | 1 | 0.81707832 | 0.44988418 |
| Pign | 0.25847562 | 0.29529685 | 0.479 | 0.409 | 1 | 0.936417 | -0.4710101 | 0.45 | 0.417 | 1 | 0.936417 | 0.4501416 |
| Gm26771 | 0.25851703 | -0.9697854 | 0.115 | 0.073 | 1 | 0.79058742 | -0.3587649 | 0.076 | 0.083 | 1 | 0.79058742 | 0.45020301 |
| Tmod1 | 0.99758609 | 0.27052894 | 0.021 | 0.021 | 1 | 0.25859607 | 0.16407593 | 0.063 | 0.031 | 1 | 0.99758609 | 0.45032022 |
| Dnaaf4 | 0.25863087 | -1.3144336 | 0.031 | 0.062 | 1 | 0.37709681 | 0.34222143 | 0.067 | 0.042 | 1 | 0.37709681 | 0.45037181 |
| Dnajc21 | 0.57873894 | -0.1877618 | 0.385 | 0.399 | 1 | 0.25863269 | -0.2534728 | 0.584 | 0.688 | 1 | 0.57873894 | 0.45037451 |
| Primpol | 0.4320165 | 0.22988696 | 0.125 | 0.093 | 1 | 0.25913701 | 0.61614416 | 0.076 | 0.042 | 1 | 0.4320165 | 0.45112203 |
| Zcchc4 | 0.25925528 | 0.37006462 | 0.146 | 0.098 | 1 | 0.57084271 | -0.5819348 | 0.172 | 0.135 | 1 | 0.57084271 | 0.45129727 |
| B230217C12Rik | 0.25931175 | -0.6072153 | 0.146 | 0.197 | 1 | 0.6724304 | -0.9547603 | 0.181 | 0.188 | 1 | 0.6724304 | 0.45138092 |
| Mapt | 0.25952473 | -1.0628948 | 0.031 | 0.062 | 1 | 0.46606465 | -1.2457249 | 0.063 | 0.083 | 1 | 0.46606465 | 0.45169638 |
| Gm49774 | 0.2595505 | -0.3144336 | 0.042 | 0.078 | 1 | 0.54697586 | -0.4353861 | 0.034 | 0.021 | 1 | 0.54697586 | 0.45173454 |
| Lcn11 | 0.26059398 | 0.42253204 | 0.094 | 0.057 | 1 | 0.33170642 | -1.6683092 | 0.143 | 0.167 | 1 | 0.33170642 | 0.45327874 |
| Sirt6 | 0.26059398 | 0.52206771 | 0.094 | 0.057 | 1 | 0.82712037 | -0.1399303 | 0.059 | 0.052 | 1 | 0.82712037 | 0.45327874 |
| Fam220a.1 | 0.74423402 | -0.3550755 | 0.062 | 0.052 | 1 | 0.26084209 | 0.33400093 | 0.076 | 0.042 | 1 | 0.74423402 | 0.45364559 |
| Slc22a18 | 0.2610669 | -0.8993961 | 0.031 | 0.062 | 1 | 0.53618537 | -1.5728897 | 0.038 | 0.052 | 1 | 0.53618537 | 0.45397787 |
| Acd | 0.55511444 | 0.17460452 | 0.292 | 0.259 | 1 | 0.26117977 | 0.18657056 | 0.277 | 0.208 | 1 | 0.55511444 | 0.45414467 |
| Rnd1 | 0.26123564 | -0.6929452 | 0.115 | 0.161 | 1 | 0.77380679 | 0.17557156 | 0.038 | 0.031 | 1 | 0.77380679 | 0.45422722 |
| Spred3 | 0.26124898 | -0.9925055 | 0.052 | 0.088 | 1 | 0.62828011 | -0.5025003 | 0.088 | 0.104 | 1 | 0.62828011 | 0.45424693 |
| Rap1gap | 0.53661492 | -0.7294711 | 0.073 | 0.093 | 1 | 0.26129424 | -0.6263289 | 0.164 | 0.104 | 1 | 0.53661492 | 0.45431381 |
| Slc25a37 | 0.2615236 | 0.52206771 | 0.323 | 0.275 | 1 | 0.91204888 | -0.5854897 | 0.248 | 0.229 | 1 | 0.91204888 | 0.45465261 |
| Mapk7 | 0.26182248 | 0.47697982 | 0.156 | 0.109 | 1 | 0.76637108 | -0.5728897 | 0.147 | 0.156 | 1 | 0.76637108 | 0.45509395 |
| Gm44649 | 0.26186197 | -0.7294711 | 0.115 | 0.161 | 1 | 0.40648972 | -0.364303 | 0.197 | 0.146 | 1 | 0.40648972 | 0.45515225 |
| Dync1h1 | 0.26186556 | -0.2010257 | 0.812 | 0.824 | 1 | 0.40525822 | -0.1314405 | 0.958 | 0.948 | 1 | 0.40525822 | 0.45515755 |
| Gm11613 | 0.3941419 | -0.1079827 | 0.052 | 0.031 | 1 | 0.26200121 | 0.69014474 | 0.101 | 0.062 | 1 | 0.3941419 | 0.45535778 |
| C030037D09Rik | 0.81651401 | 0.47144164 | 0.135 | 0.13 | 1 | 0.26205229 | 0.46775232 | 0.076 | 0.042 | 1 | 0.81651401 | 0.45543318 |
| Tnrc6b | 0.74789087 | 0.10901732 | 0.979 | 0.969 | 1 | 0.26213012 | -0.3330195 | 0.975 | 0.99 | 1 | 0.74789087 | 0.45554804 |
| Six5 | 0.32199593 | 0.76648644 | 0.094 | 0.062 | 1 | 0.26254433 | 0.64434105 | 0.101 | 0.062 | 1 | 0.32199593 | 0.45615913 |
| Adcy9 | 0.2626097 | -0.1014398 | 0.292 | 0.218 | 1 | 0.91904911 | -0.5711525 | 0.479 | 0.406 | 1 | 0.91904911 | 0.45625554 |
| Gm56688 | 0.26275183 | 0.71798792 | 0.073 | 0.041 | 1 | 0.54631812 | 0.13071733 | 0.059 | 0.042 | 1 | 0.54631812 | 0.45646513 |
| Rpap2 | 0.29496067 | -0.7998604 | 0.104 | 0.145 | 1 | 0.26279151 | 0.21739174 | 0.143 | 0.094 | 1 | 0.29496067 | 0.45652365 |
| Dynll2 | 0.39466424 | 0.15635792 | 0.604 | 0.539 | 1 | 0.26322889 | 0.1414288 | 0.676 | 0.667 | 1 | 0.39466424 | 0.45716833 |
| Gm12811 | 0.2632671 | -0.1079827 | 0.052 | 0.026 | 1 | 0.70639085 | -0.4618584 | 0.034 | 0.042 | 1 | 0.70639085 | 0.45722463 |
| Arl9 | 0.2632671 | 0.59245704 | 0.052 | 0.026 | 1 | 0.56210775 | -0.3098553 | 0.071 | 0.052 | 1 | 0.56210775 | 0.45722463 |
| Gm30906 | 0.2632671 | 0.59245704 | 0.052 | 0.026 | 1 | 0.29444919 | 0.27510724 | 0.101 | 0.062 | 1 | 0.29444919 | 0.45722463 |
| Agap2 | 0.2632671 | 0.59245704 | 0.052 | 0.026 | 1 | 0.34703906 | -1.3098553 | 0.008 | 0.021 | 1 | 0.34703906 | 0.45722463 |
| 2810454H06Rik | 0.26331526 | -0.3461424 | 0.042 | 0.078 | 1 | 0.63004106 | -0.9879272 | 0.05 | 0.062 | 1 | 0.63004106 | 0.45729559 |
| Cyp2d22 | 0.26351343 | -0.8993961 | 0.031 | 0.062 | 1 | 0.72007043 | -0.3838558 | 0.063 | 0.073 | 1 | 0.72007043 | 0.45758754 |
| A930037H05Rik | 0.26358095 | 0.47697982 | 0.073 | 0.041 | 1 | 0.74620884 | -0.9472852 | 0.025 | 0.031 | 1 | 0.74620884 | 0.45768698 |
| Cd300a | 0.26362956 | -0.9925055 | 0.031 | 0.062 | 1 | 0.81167924 | -0.8123556 | 0.071 | 0.062 | 1 | 0.81167924 | 0.45775858 |
| Zfpm1 | 0.81529054 | -0.1035368 | 0.177 | 0.192 | 1 | 0.26364146 | -0.4167705 | 0.273 | 0.188 | 1 | 0.81529054 | 0.4577761 |
| B930095G15Rik | 0.28352896 | -0.4699835 | 0.469 | 0.503 | 1 | 0.26367342 | -0.453691 | 0.597 | 0.406 | 1 | 0.28352896 | 0.45782317 |
| Rfx1 | 0.26383465 | 0.40803247 | 0.26 | 0.202 | 1 | 0.86977287 | -0.3098553 | 0.349 | 0.312 | 1 | 0.86977287 | 0.45806058 |
| Dgkd | 0.65639365 | -0.2809366 | 0.375 | 0.404 | 1 | 0.26402045 | -0.364011 | 0.437 | 0.51 | 1 | 0.65639365 | 0.45833411 |
| Osbpl2 | 0.26408529 | 0.38936517 | 0.292 | 0.228 | 1 | 0.40637734 | -0.5681673 | 0.286 | 0.312 | 1 | 0.40637734 | 0.45842954 |
| Strip1 | 0.26424392 | 0.29700115 | 0.26 | 0.197 | 1 | 0.7583518 | -0.5340704 | 0.332 | 0.323 | 1 | 0.7583518 | 0.45866299 |
| Afg3l1 | 0.41453861 | -0.2991668 | 0.292 | 0.332 | 1 | 0.26430307 | -0.3608392 | 0.399 | 0.281 | 1 | 0.41453861 | 0.45875003 |
| Anks6 | 0.58084729 | -0.3144336 | 0.042 | 0.057 | 1 | 0.26447202 | 0.56461386 | 0.076 | 0.042 | 1 | 0.58084729 | 0.45899859 |
| Dcaf12 | 0.37911675 | -0.265524 | 0.417 | 0.461 | 1 | 0.26447716 | 0.10518224 | 0.571 | 0.51 | 1 | 0.37911675 | 0.45900615 |
| Kmt2d | 0.85161804 | -0.1500467 | 0.469 | 0.43 | 1 | 0.26491933 | -0.1151269 | 0.752 | 0.708 | 1 | 0.85161804 | 0.45965641 |
| Tmem38b | 0.26506937 | -0.5505009 | 0.156 | 0.207 | 1 | 0.45007658 | -0.5213594 | 0.252 | 0.292 | 1 | 0.45007658 | 0.45987697 |
| Zfp429 | 0.82258416 | -0.1624305 | 0.031 | 0.036 | 1 | 0.26509638 | 0.49749966 | 0.076 | 0.042 | 1 | 0.82258416 | 0.45991667 |
| Grid1 | 1 | 0.27052894 | 0.01 | 0.01 | 1 | 0.26545017 | -0.2592292 | 0.063 | 0.031 | 1 | 1 | 0.46043655 |
| Csnk2a2 | 0.2658512 | 0.17028969 | 0.75 | 0.684 | 1 | 0.766 | -0.1835195 | 0.819 | 0.823 | 1 | 0.766 | 0.46102553 |
| C130036L24Rik | 0.26619579 | -0.6929452 | 0.031 | 0.062 | 1 | 0.30492205 | 0.86006974 | 0.029 | 0.01 | 1 | 0.30492205 | 0.46153138 |
| Fbln5 | 0.26688454 | -0.3364599 | 0.25 | 0.181 | 1 | 0.32940398 | -1.9360404 | 0.126 | 0.156 | 1 | 0.32940398 | 0.46254173 |
| Mapkapk5 | 0.2670186 | 0.15292498 | 0.479 | 0.399 | 1 | 0.5353399 | -0.1818477 | 0.559 | 0.448 | 1 | 0.5353399 | 0.46273827 |
| 1700113A16Rik | 0.26717768 | 0.32942263 | 0.094 | 0.057 | 1 | 0.37605058 | -0.6659991 | 0.097 | 0.062 | 1 | 0.37605058 | 0.46297145 |
| Pomk | 0.26717768 | 0.32942263 | 0.094 | 0.057 | 1 | 0.53416796 | -0.1082214 | 0.084 | 0.062 | 1 | 0.53416796 | 0.46297145 |
| Rassf4 | 0.26781885 | -0.4330781 | 0.031 | 0.062 | 1 | 0.27154471 | 0.69014474 | 0.013 | 0 | 1 | 0.27154471 | 0.46391077 |
| Pus10 | 0.63339712 | -0.1268066 | 0.25 | 0.275 | 1 | 0.26810226 | -0.3353904 | 0.332 | 0.25 | 1 | 0.63339712 | 0.46432569 |
| Zfp990 | 0.26812519 | -0.9311049 | 0.052 | 0.088 | 1 | 0.76849153 | -0.3483294 | 0.118 | 0.104 | 1 | 0.76849153 | 0.46435926 |
| Slc16a10 | 0.26829236 | -0.8077656 | 0.208 | 0.264 | 1 | 0.35623851 | 0.1935197 | 0.252 | 0.198 | 1 | 0.35623851 | 0.46460393 |
| Il2 | 0.26870509 | 0.78510212 | 0.104 | 0.067 | 1 | 0.43323624 | 0.42711033 | 0.038 | 0.021 | 1 | 0.43323624 | 0.46520775 |
| Klhl20 | 0.60273397 | -0.1413688 | 0.188 | 0.161 | 1 | 0.26905112 | 0.23071312 | 0.202 | 0.146 | 1 | 0.60273397 | 0.46571374 |
| Gm34343 | 0.42215896 | 0.2912875 | 0.115 | 0.083 | 1 | 0.26911342 | -0.3098553 | 0.105 | 0.062 | 1 | 0.42215896 | 0.46580481 |
| Gm10658 | 0.26940715 | 0.54806292 | 0.073 | 0.041 | 1 | 0.38906484 | -0.2103196 | 0.055 | 0.031 | 1 | 0.38906484 | 0.46623409 |
| Gm30963 | 0.26946357 | 0.47697982 | 0.073 | 0.041 | 1 | 0.71649639 | -0.5728897 | 0.034 | 0.042 | 1 | 0.71649639 | 0.46631652 |
| Gm9887 | 0.26946357 | 0.47697982 | 0.073 | 0.041 | 1 | 0.34625029 | 0.69014474 | 0.042 | 0.021 | 1 | 0.34625029 | 0.46631652 |
| Vps45 | 0.26954959 | 0.14767219 | 0.219 | 0.161 | 1 | 0.81128588 | -0.5508634 | 0.214 | 0.188 | 1 | 0.81128588 | 0.4664422 |
| Atad5 | 0.26959969 | 0.37522632 | 0.094 | 0.057 | 1 | 0.32668435 | -1.0983512 | 0.046 | 0.073 | 1 | 0.32668435 | 0.46651539 |
| Gcnt4 | 0.27028216 | -0.6705774 | 0.031 | 0.062 | 1 | 0.27154471 | 1.27510724 | 0.013 | 0 | 1 | 0.27154471 | 0.46751188 |
| Thsd7a | 0.89344406 | 0.39082318 | 0.062 | 0.067 | 1 | 0.27038817 | -0.7161755 | 0.168 | 0.208 | 1 | 0.89344406 | 0.46766658 |
| Defb1 | 0.32025586 | -0.9925055 | 0 | 0.01 | 1 | 0.27067179 | 0.21370669 | 0.042 | 0.073 | 1 | 0.32025586 | 0.46808036 |
| Cdc37l1 | 0.27081198 | 0.22375217 | 0.771 | 0.679 | 1 | 0.60437537 | -0.1168466 | 0.853 | 0.833 | 1 | 0.60437537 | 0.46828483 |
| Gm12236 | 0.27096766 | 0.20013961 | 0.094 | 0.057 | 1 | 0.97098462 | -0.1723517 | 0.071 | 0.073 | 1 | 0.97098462 | 0.46851185 |
| Vmn2r29 | 0.27112836 | -0.577468 | 0.031 | 0.062 | 1 | 0.76973532 | -0.3923174 | 0.05 | 0.042 | 1 | 0.76973532 | 0.46874613 |
| Exd2 | 0.50114554 | -0.4664367 | 0.177 | 0.145 | 1 | 0.27115475 | 0.42711033 | 0.126 | 0.083 | 1 | 0.50114554 | 0.46878461 |
| Gm4285 | 0.67456678 | -0.5427025 | 0.115 | 0.13 | 1 | 0.27117011 | 0.16407593 | 0.088 | 0.052 | 1 | 0.67456678 | 0.46880699 |
| Ripk1 | 0.27117033 | 0.12334287 | 0.729 | 0.679 | 1 | 0.86116768 | -0.15383 | 0.761 | 0.75 | 1 | 0.86116768 | 0.46880731 |
| Mylk | 0.27151175 | -0.8290067 | 0.229 | 0.269 | 1 | 0.73649213 | -0.8273548 | 0.382 | 0.302 | 1 | 0.73649213 | 0.46930487 |
| Grid2 | 0.32025586 | -0.577468 | 0 | 0.01 | 1 | 0.27154327 | 2.14957636 | 0.013 | 0 | 1 | 0.32025586 | 0.46935079 |

| | | | | | | | | | | | | |
|---|---|---|---|---|---|---|---|---|---|---|---|---|
| 9330162012Rik | 0.32025586 | -1.3144336 | 0 | 0.01 | 1 | 0.27154327 | 1.27510724 | 0.013 | 0 | 1 | 0.32025586 | 0.46935079 |
| A630012P03Rik | 0.32025586 | -3.5160674 | 0 | 0.01 | 1 | 0.27154327 | 1.01207283 | 0.013 | 0 | 1 | 0.32025586 | 0.46935079 |
| Gm13936 | 0.36106354 | -1.2148979 | 0.021 | 0.041 | 1 | 0.27154327 | 1.49749966 | 0.013 | 0 | 1 | 0.36106354 | 0.46935079 |
| Togaram2 | 0.37863831 | 1.22988696 | 0.031 | 0.016 | 1 | 0.27154327 | 1.01207283 | 0.013 | 0 | 1 | 0.37863831 | 0.46935079 |
| Gm49785 | 0.38028858 | 1.00749454 | 0.031 | 0.016 | 1 | 0.27154327 | 1.27510724 | 0.013 | 0 | 1 | 0.38028858 | 0.46935079 |
| Plxna4os1 | 0.46992036 | 0.74446013 | 0.042 | 0.026 | 1 | 0.27154327 | 1.01207283 | 0.013 | 0 | 1 | 0.46992036 | 0.46935079 |
| Gm29571 | 0.48064264 | 0.59245704 | 0.021 | 0.01 | 1 | 0.27154327 | 1.01207283 | 0.013 | 0 | 1 | 0.48064264 | 0.46935079 |
| Sox5os4 | 0.52744059 | -0.577468 | 0.01 | 0.021 | 1 | 0.27154327 | 1.01207283 | 0.013 | 0 | 1 | 0.52744059 | 0.46935079 |
| 4930578G10Rik | 0.61371908 | 2.00749454 | 0.01 | 0.005 | 1 | 0.27154327 | 1.01207283 | 0.013 | 0 | 1 | 0.61371908 | 0.46935079 |
| Kcnmb2 | 0.61737927 | 1.00749454 | 0.01 | 0.005 | 1 | 0.27154327 | 1.27510724 | 0.013 | 0 | 1 | 0.61737927 | 0.46935079 |
| Mcoln3 | 0.61737927 | 1.00749454 | 0.01 | 0.005 | 1 | 0.27154327 | 1.01207283 | 0.013 | 0 | 1 | 0.61737927 | 0.46935079 |
| Gm38944 | 0.61737927 | 1.00749454 | 0.01 | 0.005 | 1 | 0.27154327 | 1.01207283 | 0.013 | 0 | 1 | 0.61737927 | 0.46935079 |
| Gm31698 | 0.61737927 | 1.00749454 | 0.01 | 0.005 | 1 | 0.27154327 | 1.49749966 | 0.013 | 0 | 1 | 0.61737927 | 0.46935079 |
| Krt13 | 0.61737927 | 1.00749454 | 0.01 | 0.005 | 1 | 0.27154327 | 1.01207283 | 0.013 | 0 | 1 | 0.61737927 | 0.46935079 |
| Stmn4 | 0.61737927 | 1.00749454 | 0.01 | 0.005 | 1 | 0.27154327 | 1.01207283 | 0.013 | 0 | 1 | 0.61737927 | 0.46935079 |
| Cytip | 0.62104899 | -0.1624305 | 0.01 | 0.005 | 1 | 0.27154327 | 1.27510724 | 0.013 | 0 | 1 | 0.62104899 | 0.46935079 |
| Samsn1 | 0.62104899 | 0.27052894 | 0.01 | 0.005 | 1 | 0.27154327 | 2.86006974 | 0.013 | 0 | 1 | 0.62104899 | 0.46935079 |
| F930017D23Rik | 0.62108713 | -0.3144336 | 0.021 | 0.031 | 1 | 0.27154327 | 1.86006974 | 0.013 | 0 | 1 | 0.62108713 | 0.46935079 |
| AA465934 | 0.63038178 | 0.20013961 | 0.021 | 0.031 | 1 | 0.27154327 | 1.01207283 | 0.013 | 0 | 1 | 0.63038178 | 0.46935079 |
| Pex5l | 0.72574709 | -0.3144336 | 0.01 | 0.016 | 1 | 0.27154327 | 1.27510724 | 0.013 | 0 | 1 | 0.72574709 | 0.46935079 |
| Gm14547 | 0.99482447 | -0.130009 | 0.042 | 0.041 | 1 | 0.27154327 | 1.01207283 | 0.013 | 0 | 1 | 0.99482447 | 0.46935079 |
| Gm20324 | 0.99758604 | -0.2148979 | 0.021 | 0.021 | 1 | 0.27154327 | 1.01207283 | 0.013 | 0 | 1 | 0.99758604 | 0.46935079 |
| Sh3tc2 | 0.27822105 | -1.1079827 | 0.021 | 0.047 | 1 | 0.27154471 | 1.27510724 | 0.013 | 0 | 1 | 0.27822105 | 0.46935289 |
| Ccl21a | 0.32025586 | -0.577468 | 0 | 0.01 | 1 | 0.27154471 | 0.69014474 | 0.013 | 0 | 1 | 0.32025586 | 0.46935289 |
| Bspry | 0.32025586 | -0.577468 | 0 | 0.01 | 1 | 0.27154471 | 0.69014474 | 0.013 | 0 | 1 | 0.32025586 | 0.46935289 |
| Gm830 | 0.32025586 | -0.9925055 | 0 | 0.01 | 1 | 0.27154471 | 0.69014474 | 0.013 | 0 | 1 | 0.32025586 | 0.46935289 |
| Sel1l3 | 0.32025586 | -0.9925055 | 0 | 0.01 | 1 | 0.27154471 | 1.49749966 | 0.013 | 0 | 1 | 0.32025586 | 0.46935289 |
| Tmprss11a | 0.32025586 | -1.3144336 | 0 | 0.01 | 1 | 0.27154471 | 1.49749966 | 0.013 | 0 | 1 | 0.32025586 | 0.46935289 |
| Tnni3 | 0.32025586 | -0.577468 | 0 | 0.01 | 1 | 0.27154471 | 0.69014474 | 0.013 | 0 | 1 | 0.32025586 | 0.46935289 |
| Klk13 | 0.32025586 | -0.577468 | 0 | 0.01 | 1 | 0.27154471 | 0.69014474 | 0.013 | 0 | 1 | 0.32025586 | 0.46935289 |
| Cdcp3 | 0.32025586 | -0.9925055 | 0 | 0.01 | 1 | 0.27154471 | 1.69014474 | 0.013 | 0 | 1 | 0.32025586 | 0.46935289 |
| G430095P16Rik | 0.32025586 | -0.577468 | 0 | 0.01 | 1 | 0.27154471 | 0.69014474 | 0.013 | 0 | 1 | 0.32025586 | 0.46935289 |
| Gm11205 | 0.32025586 | -0.577468 | 0 | 0.01 | 1 | 0.27154471 | 0.69014474 | 0.013 | 0 | 1 | 0.32025586 | 0.46935289 |
| Gm29968 | 0.32025586 | -0.577468 | 0 | 0.01 | 1 | 0.27154471 | 0.69014474 | 0.013 | 0 | 1 | 0.32025586 | 0.46935289 |
| Akap5 | 0.32025586 | -0.9925055 | 0 | 0.01 | 1 | 0.27154471 | 0.69014474 | 0.013 | 0 | 1 | 0.32025586 | 0.46935289 |
| Tc2n | 0.32025586 | -0.9925055 | 0 | 0.01 | 1 | 0.27154471 | 0.69014474 | 0.013 | 0 | 1 | 0.32025586 | 0.46935289 |
| Gm48807 | 0.32025586 | -0.577468 | 0 | 0.01 | 1 | 0.27154471 | 0.69014474 | 0.013 | 0 | 1 | 0.32025586 | 0.46935289 |
| Gm17546 | 0.32025586 | -0.577468 | 0 | 0.01 | 1 | 0.27154471 | 0.69014474 | 0.013 | 0 | 1 | 0.32025586 | 0.46935289 |
| Pfn4 | 0.38028858 | 1.00749454 | 0.031 | 0.016 | 1 | 0.27154471 | 1.49749966 | 0.013 | 0 | 1 | 0.38028858 | 0.46935289 |
| Cxcl2 | 0.38357309 | -1.9925055 | 0.01 | 0.026 | 1 | 0.27154471 | 0.69014474 | 0.013 | 0 | 1 | 0.38357309 | 0.46935289 |
| C530005A16Rik | 0.38521705 | -0.7998604 | 0.01 | 0.026 | 1 | 0.27154471 | 0.69014474 | 0.013 | 0 | 1 | 0.38521705 | 0.46935289 |
| Gm49719 | 0.38521705 | -0.7998604 | 0.01 | 0.026 | 1 | 0.27154471 | 0.69014474 | 0.013 | 0 | 1 | 0.38521705 | 0.46935289 |
| Gm15241 | 0.42644715 | -0.7900126 | 0.125 | 0.155 | 1 | 0.27154471 | 1.49749966 | 0.013 | 0 | 1 | 0.42644715 | 0.46935289 |
| Gm57313 | 0.46992036 | 0.74446013 | 0.042 | 0.026 | 1 | 0.27154471 | 0.69014474 | 0.013 | 0 | 1 | 0.46992036 | 0.46935289 |
| Ppef1 | 0.47150075 | 1.42253204 | 0.021 | 0.01 | 1 | 0.27154471 | 1.86006974 | 0.013 | 0 | 1 | 0.47150075 | 0.46935289 |
| Iqca | 0.47606368 | 1.00749454 | 0.021 | 0.01 | 1 | 0.27154471 | 0.69014474 | 0.013 | 0 | 1 | 0.47606368 | 0.46935289 |
| Gm57103 | 0.47606502 | 1.00749454 | 0.021 | 0.01 | 1 | 0.27154471 | 0.69014474 | 0.013 | 0 | 1 | 0.47606502 | 0.46935289 |
| Gm44957 | 0.47606502 | 1.00749454 | 0.021 | 0.01 | 1 | 0.27154471 | 0.69014474 | 0.013 | 0 | 1 | 0.47606502 | 0.46935289 |
| Gm26537 | 0.47606502 | 1.00749454 | 0.021 | 0.01 | 1 | 0.27154471 | 0.69014474 | 0.013 | 0 | 1 | 0.47606502 | 0.46935289 |
| Gm36738 | 0.47606502 | 1.00749454 | 0.021 | 0.01 | 1 | 0.27154471 | 0.69014474 | 0.013 | 0 | 1 | 0.47606502 | 0.46935289 |
| Gm36419 | 0.47606502 | 1.00749454 | 0.021 | 0.01 | 1 | 0.27154471 | 0.69014474 | 0.013 | 0 | 1 | 0.47606502 | 0.46935289 |
| Gm45820 | 0.48064264 | 0.27052894 | 0.021 | 0.01 | 1 | 0.27154471 | 0.69014474 | 0.013 | 0 | 1 | 0.48064264 | 0.46935289 |
| Itgal | 0.52528691 | -1.1624305 | 0.01 | 0.021 | 1 | 0.27154471 | 0.69014474 | 0.013 | 0 | 1 | 0.52528691 | 0.46935289 |
| Igsf6 | 0.52744059 | -0.577468 | 0.01 | 0.021 | 1 | 0.27154471 | 0.69014474 | 0.013 | 0 | 1 | 0.52744059 | 0.46935289 |
| Gm2447 | 0.52961388 | -1.577468 | 0.01 | 0.021 | 1 | 0.27154471 | 0.69014474 | 0.013 | 0 | 1 | 0.52961388 | 0.46935289 |
| Gm15915 | 0.52961488 | -0.3144336 | 0.01 | 0.021 | 1 | 0.27154471 | 0.69014474 | 0.013 | 0 | 1 | 0.52961488 | 0.46935289 |
| 1700093K21Rik | 0.52961488 | -0.3144336 | 0.01 | 0.021 | 1 | 0.27154471 | 0.69014474 | 0.013 | 0 | 1 | 0.52961488 | 0.46935289 |
| Irf5 | 0.5862099 | 0.68556644 | 0.031 | 0.021 | 1 | 0.27154471 | 0.69014474 | 0.013 | 0 | 1 | 0.5862099 | 0.46935289 |
| Gm13270 | 0.59199182 | 0.20013961 | 0.031 | 0.021 | 1 | 0.27154471 | 0.69014474 | 0.013 | 0 | 1 | 0.59199182 | 0.46935289 |
| A230083N12Rik | 0.59199182 | 0.42253204 | 0.031 | 0.021 | 1 | 0.27154471 | 0.69014474 | 0.013 | 0 | 1 | 0.59199182 | 0.46935289 |
| Fbxo16 | 0.61371908 | 1.59245704 | 0.01 | 0.005 | 1 | 0.27154471 | 0.69014474 | 0.013 | 0 | 1 | 0.61371908 | 0.46935289 |
| Cdh18 | 0.61371908 | 2.00749454 | 0.01 | 0.005 | 1 | 0.27154471 | 1.69014474 | 0.013 | 0 | 1 | 0.61371908 | 0.46935289 |
| 4930592C13Rik | 0.61737927 | 1.00749454 | 0.01 | 0.005 | 1 | 0.27154471 | 0.69014474 | 0.013 | 0 | 1 | 0.61737927 | 0.46935289 |
| 4930597O21Rik | 0.61737927 | 1.00749454 | 0.01 | 0.005 | 1 | 0.27154471 | 0.69014474 | 0.013 | 0 | 1 | 0.61737927 | 0.46935289 |
| Gm56852 | 0.61737927 | 1.00749454 | 0.01 | 0.005 | 1 | 0.27154471 | 0.69014474 | 0.013 | 0 | 1 | 0.61737927 | 0.46935289 |
| Marchf1 | 0.61737927 | 1.00749454 | 0.01 | 0.005 | 1 | 0.27154471 | 0.69014474 | 0.013 | 0 | 1 | 0.61737927 | 0.46935289 |
| Gm9917 | 0.61737927 | 1.00749454 | 0.01 | 0.005 | 1 | 0.27154471 | 0.69014474 | 0.013 | 0 | 1 | 0.61737927 | 0.46935289 |
| Gm26873 | 0.61737927 | 1.00749454 | 0.01 | 0.005 | 1 | 0.27154471 | 0.69014474 | 0.013 | 0 | 1 | 0.61737927 | 0.46935289 |
| Gm17080 | 0.61737927 | 1.00749454 | 0.01 | 0.005 | 1 | 0.27154471 | 0.69014474 | 0.013 | 0 | 1 | 0.61737927 | 0.46935289 |
| Gm4219 | 0.61737927 | 1.00749454 | 0.01 | 0.005 | 1 | 0.27154471 | 0.69014474 | 0.013 | 0 | 1 | 0.61737927 | 0.46935289 |
| Gm35113 | 0.62104899 | 0.42253204 | 0.01 | 0.005 | 1 | 0.27154471 | 0.69014474 | 0.013 | 0 | 1 | 0.62104899 | 0.46935289 |
| Gm12511 | 0.6229455 | -0.3710171 | 0.021 | 0.031 | 1 | 0.27154471 | 1.49749966 | 0.013 | 0 | 1 | 0.6229455 | 0.46935289 |
| Atp2a3 | 0.6402809 | 1.00749454 | 0.042 | 0.031 | 1 | 0.27154471 | 0.69014474 | 0.013 | 0 | 1 | 0.6402809 | 0.46935289 |
| Was | 0.72574709 | -0.3144336 | 0.01 | 0.016 | 1 | 0.27154471 | 0.69014474 | 0.013 | 0 | 1 | 0.72574709 | 0.46935289 |
| 1700105P06Rik | 0.74326422 | 1.00749454 | 0.021 | 0.016 | 1 | 0.27154471 | 0.69014474 | 0.013 | 0 | 1 | 0.74326422 | 0.46935289 |
| Gm26708 | 0.75328966 | 0.27052894 | 0.021 | 0.016 | 1 | 0.27154471 | 1.27510724 | 0.013 | 0 | 1 | 0.75328966 | 0.46935289 |
| Apobec2 | 0.75552329 | -1.2745408 | 0.073 | 0.062 | 1 | 0.27154471 | 0.69014474 | 0.013 | 0 | 1 | 0.75552329 | 0.46935289 |
| Cpn1 | 0.78248887 | 0.78510212 | 0.031 | 0.026 | 1 | 0.27154471 | 0.69014474 | 0.013 | 0 | 1 | 0.78248887 | 0.46935289 |
| Syt6 | 0.7854853 | -0.407543 | 0.021 | 0.026 | 1 | 0.27154471 | 2.14957636 | 0.013 | 0 | 1 | 0.7854853 | 0.46935289 |
| Gm26620 | 0.8006267 | 0.20013961 | 0.021 | 0.026 | 1 | 0.27154471 | 0.69014474 | 0.013 | 0 | 1 | 0.8006267 | 0.46935289 |

| | | | | | | | | | | | | |
|---|---|---|---|---|---|---|---|---|---|---|---|---|
| Gm32585 | 0.82111059 | -1.0799683 | 0.031 | 0.026 | 1 | 0.27154471 | 0.69014474 | 0.013 | 0 | 1 | 0.82111059 | 0.46935289 |
| Gm41611 | 0.83041404 | 0.15949763 | 0.042 | 0.036 | 1 | 0.27154471 | 0.69014474 | 0.013 | 0 | 1 | 0.83041404 | 0.46935289 |
| Polq | 0.99207455 | 1.81484946 | 0.031 | 0.031 | 1 | 0.27154471 | 0.69014474 | 0.013 | 0 | 1 | 0.99207455 | 0.46935289 |
| Gm57061 | 0.993208 | 0.68556644 | 0.01 | 0.01 | 1 | 0.27154471 | 0.69014474 | 0.013 | 0 | 1 | 0.993208 | 0.46935289 |
| Abi3 | 0.993208 | 1.00749454 | 0.01 | 0.01 | 1 | 0.27154471 | 0.69014474 | 0.013 | 0 | 1 | 0.993208 | 0.46935289 |
| Dio3os | 0.993208 | 1.00749454 | 0.01 | 0.01 | 1 | 0.27154471 | 0.69014474 | 0.013 | 0 | 1 | 0.993208 | 0.46935289 |
| Gm47467 | 0.993208 | 1.00749454 | 0.01 | 0.01 | 1 | 0.27154471 | 0.69014474 | 0.013 | 0 | 1 | 0.993208 | 0.46935289 |
| Tg | 0.99517215 | 1.81484946 | 0.021 | 0.021 | 1 | 0.27154471 | 1.27510724 | 0.013 | 0 | 1 | 0.99517215 | 0.46935289 |
| Gm15635 | 0.99758609 | 0.27052894 | 0.021 | 0.021 | 1 | 0.27154471 | 0.69014474 | 0.013 | 0 | 1 | 0.99758609 | 0.46935289 |
| Ttc30a1 | 1 | 0.42253204 | 0.01 | 0.01 | 1 | 0.27154471 | 0.69014474 | 0.013 | 0 | 1 | 1 | 0.46935289 |
| 4933429H19Rik | 1 | 0.42253204 | 0.01 | 0.01 | 1 | 0.27154471 | 0.69014474 | 0.013 | 0 | 1 | 1 | 0.46935289 |
| Gm42701 | 1 | 0.42253204 | 0.01 | 0.01 | 1 | 0.27154471 | 0.69014474 | 0.013 | 0 | 1 | 1 | 0.46935289 |
| Ccdc142os | 1 | 0.42253204 | 0.01 | 0.01 | 1 | 0.27154471 | 0.69014474 | 0.013 | 0 | 1 | 1 | 0.46935289 |
| Slco1a6 | 1 | 0.42253204 | 0.01 | 0.01 | 1 | 0.27154471 | 1.27510724 | 0.013 | 0 | 1 | 1 | 0.46935289 |
| Proscos | 1 | 0.42253204 | 0.01 | 0.01 | 1 | 0.27154471 | 0.69014474 | 0.013 | 0 | 1 | 1 | 0.46935289 |
| Gsta2 | 1 | -0.3144336 | 0.01 | 0.01 | 1 | 0.27154471 | 0.69014474 | 0.013 | 0 | 1 | 1 | 0.46935289 |
| 2210039B01Rik | 1 | 0.42253204 | 0.01 | 0.01 | 1 | 0.27154471 | 0.69014474 | 0.013 | 0 | 1 | 1 | 0.46935289 |
| D030051J21Rik | 1 | 0.42253204 | 0.01 | 0.01 | 1 | 0.27154471 | 0.69014474 | 0.013 | 0 | 1 | 1 | 0.46935289 |
| Scel | 1 | 0.27052894 | 0.01 | 0.01 | 1 | 0.27154471 | 0.69014474 | 0.013 | 0 | 1 | 1 | 0.46935289 |
| Muc13 | 1 | 0.42253204 | 0.01 | 0.01 | 1 | 0.27154471 | 0.69014474 | 0.013 | 0 | 1 | 1 | 0.46935289 |
| Trem3 | 1 | -0.9925055 | 0.01 | 0.01 | 1 | 0.27154471 | 0.69014474 | 0.013 | 0 | 1 | 1 | 0.46935289 |
| Prdm15 | 0.27186517 | 0.37946331 | 0.167 | 0.119 | 1 | 0.33902364 | -0.2828882 | 0.282 | 0.198 | 1 | 0.33902364 | 0.46981967 |
| Rpain | 0.32100648 | 0.10703021 | 0.281 | 0.218 | 1 | 0.27197532 | -0.3098553 | 0.315 | 0.229 | 1 | 0.32100648 | 0.46998007 |
| Ubl3 | 0.58368771 | 0.18522791 | 0.823 | 0.829 | 1 | 0.27199488 | -0.1615957 | 0.933 | 0.833 | 1 | 0.58368771 | 0.47000855 |
| Dzank1 | 0.27204938 | 0.27052894 | 0.073 | 0.041 | 1 | 0.89726528 | -0.961932 | 0.046 | 0.042 | 1 | 0.89726528 | 0.47008789 |
| Sacs | 0.27215053 | 0.74446013 | 0.177 | 0.13 | 1 | 0.93656048 | -0.5577828 | 0.113 | 0.115 | 1 | 0.93656048 | 0.47023515 |
| Smchd1 | 0.35730852 | 0.13422917 | 0.906 | 0.896 | 1 | 0.27250548 | -0.3549714 | 0.916 | 0.906 | 1 | 0.35730852 | 0.47075173 |
| Lcn3 | 0.27253308 | 0.50792353 | 0.188 | 0.14 | 1 | 0.40700073 | -3.84037 | 0.025 | 0.042 | 1 | 0.40700073 | 0.47079187 |
| Mss51 | 0.27268347 | 0.18806678 | 0.125 | 0.083 | 1 | 0.75138869 | -0.4838847 | 0.122 | 0.104 | 1 | 0.75138869 | 0.47101067 |
| Gm56816 | 0.70461002 | 0.42253204 | 0.062 | 0.052 | 1 | 0.27275949 | 0.27510724 | 0.076 | 0.042 | 1 | 0.70461002 | 0.47112124 |
| Dusp2 | 0.27292827 | -0.6072153 | 0.031 | 0.062 | 1 | 0.85710544 | -1.7248928 | 0.008 | 0.01 | 1 | 0.85710544 | 0.4713667 |
| Osmr | 0.45518269 | -0.2413938 | 0.771 | 0.762 | 1 | 0.27307309 | -0.1675614 | 0.643 | 0.469 | 1 | 0.45518269 | 0.47157726 |
| St6galnac4 | 0.33428704 | -0.3968957 | 0.25 | 0.295 | 1 | 0.27398559 | 0.43430583 | 0.218 | 0.167 | 1 | 0.33428704 | 0.47290307 |
| Trmt10b | 0.7519699 | 0.20013961 | 0.094 | 0.083 | 1 | 0.27400893 | 0.34222143 | 0.076 | 0.042 | 1 | 0.7519699 | 0.47293697 |
| Ccdc152 | 0.27427646 | -1.7998604 | 0.021 | 0.047 | 1 | 0.29023084 | -1.681824 | 0.046 | 0.073 | 1 | 0.29023084 | 0.47332534 |
| Rbpms2 | 0.27427863 | -1.2148979 | 0.021 | 0.047 | 1 | 0.34730609 | 1.01207283 | 0.042 | 0.021 | 1 | 0.34730609 | 0.47332849 |
| Pkd2l2 | 0.27449815 | -0.2148979 | 0.031 | 0.062 | 1 | 0.39798032 | -0.8692827 | 0.067 | 0.042 | 1 | 0.39798032 | 0.47364707 |
| Rufy2 | 0.83296408 | 0.20418645 | 0.281 | 0.301 | 1 | 0.27487418 | -0.1208214 | 0.513 | 0.375 | 1 | 0.83296408 | 0.47419254 |
| Ninl | 0.27495552 | 0.67045955 | 0.135 | 0.093 | 1 | 0.35729377 | -0.3098553 | 0.164 | 0.115 | 1 | 0.35729377 | 0.47431051 |
| Kri1 | 0.27498207 | -0.3710171 | 0.26 | 0.316 | 1 | 0.7055837 | -0.6724253 | 0.231 | 0.198 | 1 | 0.7055837 | 0.474349 |
| Prox1 | 0.27528135 | 0.45495351 | 0.104 | 0.067 | 1 | 0.33351829 | -0.2654611 | 0.113 | 0.073 | 1 | 0.33351829 | 0.47478288 |
| Gm43387 | 0.27538452 | 0.42253204 | 0.073 | 0.041 | 1 | 0.56110733 | -0.9879272 | 0.029 | 0.042 | 1 | 0.56110733 | 0.4749324 |
| Capn1 | 0.47777025 | 0.2912875 | 0.104 | 0.078 | 1 | 0.27548705 | -0.8948178 | 0.088 | 0.125 | 1 | 0.47777025 | 0.47508099 |
| Dipk1b | 0.71642465 | -0.1801325 | 0.146 | 0.161 | 1 | 0.27552902 | -0.7890231 | 0.118 | 0.073 | 1 | 0.71642465 | 0.4751418 |
| Gm3734 | 0.3941419 | -0.1079827 | 0.052 | 0.031 | 1 | 0.27580226 | 0.21370669 | 0.088 | 0.052 | 1 | 0.3941419 | 0.47553764 |
| Pdk2 | 0.74041673 | 0.46000674 | 0.094 | 0.083 | 1 | 0.27580226 | 0.21370669 | 0.088 | 0.052 | 1 | 0.74041673 | 0.47553764 |
| Acot7 | 0.34040364 | 0.14797576 | 0.562 | 0.482 | 1 | 0.27581773 | -0.1569033 | 0.487 | 0.365 | 1 | 0.34040364 | 0.47556004 |
| Ero1b | 0.27592245 | 0.42253204 | 0.125 | 0.083 | 1 | 0.75244233 | -1.152314 | 0.076 | 0.083 | 1 | 0.75244233 | 0.4757117 |
| Gm10432 | 0.2762466 | -1.3144336 | 0.021 | 0.047 | 1 | 0.27596631 | 0.91253716 | 0.046 | 0.021 | 1 | 0.2762466 | 0.47577522 |
| Chek2 | 0.51793219 | 0.81484946 | 0.052 | 0.036 | 1 | 0.27596631 | 0.91253716 | 0.046 | 0.021 | 1 | 0.51793219 | 0.47577522 |
| Bcat1 | 0.27599614 | 0.33145794 | 0.521 | 0.497 | 1 | 0.38310299 | -0.6540093 | 0.277 | 0.219 | 1 | 0.38310299 | 0.47581841 |
| Ccdc82 | 0.45224912 | 0.33149696 | 0.438 | 0.435 | 1 | 0.27610984 | -0.4518743 | 0.504 | 0.385 | 1 | 0.45224912 | 0.47598303 |
| Pdcd6ip | 0.27619371 | 0.18319372 | 0.906 | 0.927 | 1 | 0.83458021 | -0.1258793 | 0.962 | 0.948 | 1 | 0.83458021 | 0.47610445 |
| Dok3 | 0.2762466 | -1.2148979 | 0.021 | 0.047 | 1 | 0.51839323 | 0.31817596 | 0.084 | 0.062 | 1 | 0.51839323 | 0.47618102 |
| Mroh5 | 0.27624715 | -1.407543 | 0.021 | 0.047 | 1 | 0.54837489 | -0.1399303 | 0.034 | 0.021 | 1 | 0.54837489 | 0.47618181 |
| Sirt1 | 0.70620034 | -0.2643101 | 0.583 | 0.565 | 1 | 0.27631123 | -0.3826116 | 0.769 | 0.771 | 1 | 0.70620034 | 0.47627457 |
| Gm4876 | 0.53909249 | 0.17741954 | 0.177 | 0.15 | 1 | 0.27648746 | -0.34733 | 0.08 | 0.115 | 1 | 0.53909249 | 0.47652961 |
| AU040320 | 0.65517978 | -0.1468336 | 0.25 | 0.212 | 1 | 0.27657302 | -0.2264393 | 0.273 | 0.198 | 1 | 0.65517978 | 0.4766534 |
| Tmem53 | 0.85039706 | -0.5160674 | 0.073 | 0.078 | 1 | 0.27659908 | -0.1082214 | 0.076 | 0.042 | 1 | 0.85039706 | 0.47669111 |
| Wasf1 | 0.54923784 | -0.2148979 | 0.052 | 0.036 | 1 | 0.27759424 | 1.01207283 | 0.046 | 0.021 | 1 | 0.54923784 | 0.47812991 |
| Gm56573 | 0.65896352 | 0.15949763 | 0.042 | 0.031 | 1 | 0.27759424 | 0.80562195 | 0.046 | 0.021 | 1 | 0.65896352 | 0.47812991 |
| 4932422M17Rik | 0.40901384 | -0.7087125 | 0.052 | 0.078 | 1 | 0.27765434 | -1.3973181 | 0.063 | 0.094 | 1 | 0.40901384 | 0.47821675 |
| Arhgef17 | 0.27768419 | 0.5498127 | 0.375 | 0.326 | 1 | 0.75342741 | -0.3533569 | 0.437 | 0.365 | 1 | 0.75342741 | 0.47825987 |
| Nkx6-1 | 0.27822105 | -0.9925055 | 0.021 | 0.047 | 1 | 0.93865971 | -0.9879272 | 0.034 | 0.031 | 1 | 0.93865971 | 0.47903515 |
| 4930527F14Rik | 0.27822105 | -0.9925055 | 0.021 | 0.047 | 1 | 0.53856312 | 0.61614416 | 0.034 | 0.021 | 1 | 0.53856312 | 0.47903515 |
| Gm17501 | 0.27848454 | 0.38405789 | 0.208 | 0.166 | 1 | 0.71704849 | -0.3098553 | 0.063 | 0.052 | 1 | 0.71704849 | 0.47941544 |
| BC002059 | 0.32163326 | 0.75595577 | 0.125 | 0.088 | 1 | 0.27854126 | 0.23446525 | 0.101 | 0.062 | 1 | 0.32163326 | 0.47949729 |
| Abca8a | 0.27877696 | -1.9925055 | 0.01 | 0.031 | 1 | 0.8086289 | -0.3098553 | 0.017 | 0.021 | 1 | 0.8086289 | 0.47983733 |
| Pradc1 | 0.30941276 | 0.6241659 | 0.177 | 0.135 | 1 | 0.27895869 | 0.39058446 | 0.088 | 0.052 | 1 | 0.30941276 | 0.48009942 |
| Matcap1 | 0.67174581 | -0.1624305 | 0.031 | 0.041 | 1 | 0.27935024 | 0.69014474 | 0.046 | 0.021 | 1 | 0.67174581 | 0.48066392 |
| Fam220a | 0.85317634 | 0.22988696 | 0.062 | 0.057 | 1 | 0.27935024 | 0.69014474 | 0.046 | 0.021 | 1 | 0.85317634 | 0.48066392 |
| Nkrf | 0.78050508 | 0.1843723 | 0.115 | 0.104 | 1 | 0.27936907 | -0.1475838 | 0.155 | 0.104 | 1 | 0.78050508 | 0.48069106 |
| Gdf5 | 0.60784603 | -0.1547769 | 0.104 | 0.124 | 1 | 0.27938323 | 0.82054137 | 0.206 | 0.156 | 1 | 0.60784603 | 0.48071146 |
| Gpr27 | 0.27987287 | 0.69937224 | 0.156 | 0.114 | 1 | 0.78647652 | -0.2136399 | 0.181 | 0.167 | 1 | 0.78647652 | 0.48141692 |
| Pard3bos1 | 0.28001913 | -1.577468 | 0.01 | 0.031 | 1 | 0.29314751 | -1.0468209 | 0.021 | 0.042 | 1 | 0.29314751 | 0.48162755 |
| Rtn1 | 0.28001913 | -1.577468 | 0.01 | 0.031 | 1 | 0.99424465 | -0.6724253 | 0.042 | 0.042 | 1 | 0.99424465 | 0.48162755 |
| E130202H07Rik | 0.28001913 | -1.4519371 | 0.01 | 0.031 | 1 | 0.50887612 | -1.3098553 | 0.004 | 0.01 | 1 | 0.50887612 | 0.48162755 |
| Acer2 | 0.28001997 | -1.6929452 | 0.01 | 0.031 | 1 | 0.7078281 | -1.0203486 | 0.034 | 0.042 | 1 | 0.7078281 | 0.48162876 |
| Ednrb | 0.28001997 | -1.6929452 | 0.01 | 0.031 | 1 | 0.50887612 | -1.3098553 | 0.004 | 0.01 | 1 | 0.50887612 | 0.48162876 |
| Gm29358 | 0.2801998 | -0.8669746 | 0.021 | 0.047 | 1 | 0.54349536 | -1.6317834 | 0.029 | 0.042 | 1 | 0.54349536 | 0.48188768 |
| Gm13427 | 0.2801998 | -0.9925055 | 0.021 | 0.047 | 1 | 0.34232469 | 0.91253716 | 0.042 | 0.021 | 1 | 0.34232469 | 0.48188768 |

| | | | | | | | | | | | | |
|---|---|---|---|---|---|---|---|---|---|---|---|---|
| Cnksr1 | 0.2801998 | -0.8669746 | 0.021 | 0.047 | 1 | 0.72234088 | -0.4618584 | 0.034 | 0.042 | 1 | 0.72234088 | 0.48188768 |
| Ptprt | 0.34427845 | -0.8993961 | 0.135 | 0.176 | 1 | 0.28029575 | -0.2246879 | 0.172 | 0.115 | 1 | 0.34427845 | 0.4820258 |
| Polh | 0.28058729 | 0.71203865 | 0.146 | 0.104 | 1 | 0.64389882 | -0.5873892 | 0.113 | 0.094 | 1 | 0.64389882 | 0.48244536 |
| Myo15a | 0.64616609 | -0.4779323 | 0.083 | 0.098 | 1 | 0.28077598 | 0.33784299 | 0.126 | 0.083 | 1 | 0.64616609 | 0.48271681 |
| Prdm14 | 0.28126256 | -1.3144336 | 0.01 | 0.031 | 1 | 1 | -1.0468209 | 0.021 | 0.021 | 1 | 1 | 0.48341649 |
| Dlx6 | 0.28126256 | -1.1624305 | 0.01 | 0.031 | 1 | 0.35217453 | -0.5025003 | 0.008 | 0.021 | 1 | 0.35217453 | 0.48341649 |
| Apold1 | 0.28126256 | -2.4519371 | 0.01 | 0.031 | 1 | 0.50887612 | -1.3098553 | 0.004 | 0.01 | 1 | 0.50887612 | 0.48341649 |
| Gm4673 | 0.28126256 | -1.3144336 | 0.01 | 0.031 | 1 | 0.41738636 | -1.3098553 | 0.034 | 0.052 | 1 | 0.41738636 | 0.48341649 |
| Rsf1os2 | 0.28126256 | -1.1624305 | 0.01 | 0.031 | 1 | 0.67471359 | -0.5728897 | 0.017 | 0.01 | 1 | 0.67471359 | 0.48341649 |
| AV356131 | 0.28126256 | -1.4519371 | 0.01 | 0.031 | 1 | 0.39515719 | 0.49749966 | 0.025 | 0.01 | 1 | 0.39515719 | 0.48341649 |
| Gm17231 | 0.28126256 | -1.4519371 | 0.01 | 0.031 | 1 | 0.40700723 | -2.2273931 | 0.025 | 0.042 | 1 | 0.40700723 | 0.48341649 |
| Gm48768 | 0.28126256 | -1.3144336 | 0.01 | 0.031 | 1 | 0.4301671 | -0.5728897 | 0.025 | 0.042 | 1 | 0.4301671 | 0.48341649 |
| Gm26555 | 0.28126256 | -1.1624305 | 0.01 | 0.031 | 1 | 0.34232432 | 1.01207283 | 0.042 | 0.021 | 1 | 0.34232432 | 0.48341649 |
| Gm50334 | 0.28126256 | -1.3144336 | 0.01 | 0.031 | 1 | 0.55461992 | -1.3923174 | 0.059 | 0.073 | 1 | 0.55461992 | 0.48341649 |
| Calr3 | 0.64263218 | 0.27052894 | 0.094 | 0.078 | 1 | 0.28135676 | 0.22065945 | 0.088 | 0.052 | 1 | 0.64263218 | 0.4835519 |
| Tmem199 | 0.28136955 | 0.42253204 | 0.167 | 0.119 | 1 | 0.77356785 | -0.2223924 | 0.118 | 0.104 | 1 | 0.77356785 | 0.48357028 |
| Gm15411 | 0.28144609 | -0.3144336 | 0.031 | 0.062 | 1 | 0.67368778 | -0.2457249 | 0.076 | 0.062 | 1 | 0.67368778 | 0.48368028 |
| Zyg11b | 0.28183162 | 0.10974459 | 0.521 | 0.446 | 1 | 0.70473622 | -0.2322524 | 0.727 | 0.792 | 1 | 0.70473622 | 0.48423418 |
| Aldh16a1 | 0.28184182 | 1.00749454 | 0.083 | 0.052 | 1 | 0.55110577 | -0.9659009 | 0.118 | 0.135 | 1 | 0.55110577 | 0.48424883 |
| Pogz | 0.28193258 | -0.3644742 | 0.323 | 0.373 | 1 | 0.50363073 | -0.202052 | 0.563 | 0.469 | 1 | 0.50363073 | 0.48437918 |
| Abi2 | 0.28196653 | 0.13712982 | 0.604 | 0.492 | 1 | 0.45448156 | -0.2484547 | 0.761 | 0.625 | 1 | 0.45448156 | 0.48442794 |
| Mx2 | 0.52744059 | -0.9925055 | 0.01 | 0.021 | 1 | 0.28201681 | -2.1578522 | 0.021 | 0.042 | 1 | 0.52744059 | 0.48450014 |
| Ythdc2 | 0.35920159 | 0.3625895 | 0.344 | 0.295 | 1 | 0.28215606 | -0.3363275 | 0.307 | 0.208 | 1 | 0.35920159 | 0.48470008 |
| 9130019O22Rik | 0.28230263 | -0.7294711 | 0.021 | 0.047 | 1 | 0.84600255 | -0.3769695 | 0.08 | 0.073 | 1 | 0.84600255 | 0.48491048 |
| Gm49380 | 0.28230263 | -0.7294711 | 0.021 | 0.047 | 1 | 0.69328577 | 0.27510724 | 0.063 | 0.052 | 1 | 0.69328577 | 0.48491048 |
| Gm34655 | 0.28230263 | -0.7294711 | 0.021 | 0.047 | 1 | 0.52072676 | -1.0468209 | 0.021 | 0.01 | 1 | 0.52072676 | 0.48491048 |
| 4930556G22Rik | 0.2825047 | -0.9925055 | 0.01 | 0.031 | 1 | 0.87851725 | -0.8948178 | 0.013 | 0.01 | 1 | 0.87851725 | 0.48520049 |
| Cd247 | 0.2825047 | -1.4519371 | 0.01 | 0.031 | 1 | 1 | -1.0468209 | 0.021 | 0.021 | 1 | 1 | 0.48520049 |
| Ttc39aos1 | 0.2825047 | -0.9925055 | 0.01 | 0.031 | 1 | 0.87066325 | -0.3098553 | 0.013 | 0.01 | 1 | 0.87066325 | 0.48520049 |
| Ephx4 | 0.2825047 | -0.9925055 | 0.01 | 0.031 | 1 | 0.57373861 | -1.3098553 | 0.013 | 0.021 | 1 | 0.57373861 | 0.48520049 |
| Gm15929 | 0.2825047 | -0.9925055 | 0.01 | 0.031 | 1 | 0.57576763 | -0.9879272 | 0.013 | 0.021 | 1 | 0.57576763 | 0.48520049 |
| Carmil2 | 0.2825047 | -0.9925055 | 0.01 | 0.031 | 1 | 0.43916995 | 0.27510724 | 0.038 | 0.021 | 1 | 0.43916995 | 0.48520049 |
| Hspb2 | 0.2825047 | -0.9925055 | 0.01 | 0.031 | 1 | 0.5834148 | -0.6317834 | 0.021 | 0.031 | 1 | 0.5834148 | 0.48520049 |
| Igdcc3 | 0.2825047 | -0.9925055 | 0.01 | 0.031 | 1 | 0.85710586 | -1.5728897 | 0.008 | 0.01 | 1 | 0.85710586 | 0.48520049 |
| Gm57050 | 0.2825047 | -0.9925055 | 0.01 | 0.031 | 1 | 0.53433287 | 0.42711033 | 0.034 | 0.021 | 1 | 0.53433287 | 0.48520049 |
| Gm57330 | 0.2825047 | -0.9925055 | 0.01 | 0.031 | 1 | 0.34704016 | -1.3098553 | 0.008 | 0.021 | 1 | 0.34704016 | 0.48520049 |
| Gm28529 | 0.2825047 | -0.9925055 | 0.01 | 0.031 | 1 | 0.44394565 | -0.3098553 | 0.038 | 0.021 | 1 | 0.44394565 | 0.48520049 |
| Tesmin | 0.2825047 | -0.9925055 | 0.01 | 0.031 | 1 | 0.55901854 | -0.5322477 | 0.038 | 0.052 | 1 | 0.55901854 | 0.48520049 |
| Tsx | 0.2825047 | -0.9925055 | 0.01 | 0.031 | 1 | 0.66387437 | 0.27510724 | 0.017 | 0.01 | 1 | 0.66387437 | 0.48520049 |
| Susd4 | 0.99758604 | -0.2148979 | 0.021 | 0.021 | 1 | 0.28264244 | 0.77760758 | 0.046 | 0.021 | 1 | 0.99758604 | 0.48539813 |
| Cnot6 | 0.2827026 | 0.15798616 | 0.781 | 0.798 | 1 | 0.288917 | -0.1916235 | 0.908 | 0.906 | 1 | 0.288917 | 0.48548445 |
| 2610029K11Rik | 0.98414882 | 0.52206771 | 0.031 | 0.031 | 1 | 0.28299179 | -2.1843244 | 0.021 | 0.042 | 1 | 0.98414882 | 0.48589922 |
| Gm48302 | 0.28318646 | -0.1989563 | 0.115 | 0.073 | 1 | 0.76883646 | -0.9472852 | 0.063 | 0.052 | 1 | 0.76883646 | 0.48617835 |
| Pnldc1 | 0.67174581 | -0.1624305 | 0.031 | 0.041 | 1 | 0.28348176 | 0.59703533 | 0.046 | 0.021 | 1 | 0.67174581 | 0.48660161 |
| Rapgefl1 | 0.47599549 | -0.577468 | 0.021 | 0.036 | 1 | 0.28348512 | 0.23446525 | 0.113 | 0.073 | 1 | 0.47599549 | 0.48660642 |
| Acer3 | 0.28363492 | 0.35613081 | 0.667 | 0.627 | 1 | 0.84380032 | -0.3737489 | 0.761 | 0.792 | 1 | 0.84380032 | 0.48682107 |
| Gm26626 | 0.38357309 | -0.9925055 | 0.01 | 0.026 | 1 | 0.28377576 | -2.010295 | 0.029 | 0.052 | 1 | 0.38357309 | 0.48702284 |
| D630039A03Rik | 0.28379282 | -0.7998604 | 0.01 | 0.031 | 1 | 0.99396051 | -0.3098553 | 0.021 | 0.021 | 1 | 0.99396051 | 0.48704727 |
| 9130023H24Rik | 0.28379282 | -0.7998604 | 0.01 | 0.031 | 1 | 0.67471693 | -0.7248928 | 0.017 | 0.01 | 1 | 0.67471693 | 0.48704727 |
| Adap2 | 0.28379282 | -0.7998604 | 0.01 | 0.031 | 1 | 0.34704016 | -1.3098553 | 0.008 | 0.021 | 1 | 0.34704016 | 0.48704727 |
| Tomm20l | 0.28379282 | -0.7998604 | 0.01 | 0.031 | 1 | 0.87328024 | -0.3098553 | 0.013 | 0.01 | 1 | 0.87328024 | 0.48704727 |
| Mcam | 0.3749729 | -0.4619907 | 0.062 | 0.093 | 1 | 0.28386196 | -0.5728897 | 0.105 | 0.146 | 1 | 0.3749729 | 0.48714631 |
| Gulp1 | 0.28403019 | -0.4702787 | 0.438 | 0.316 | 1 | 0.9190919 | -0.5911491 | 0.748 | 0.625 | 1 | 0.9190919 | 0.48738723 |
| Usp20 | 0.28456394 | -0.4779323 | 0.073 | 0.114 | 1 | 0.93454797 | -0.352924 | 0.092 | 0.094 | 1 | 0.93454797 | 0.48815125 |
| Chst8 | 0.43152513 | -0.3461424 | 0.031 | 0.052 | 1 | 0.28484131 | -2.5828738 | 0.038 | 0.062 | 1 | 0.43152513 | 0.48854805 |
| Tomm40l | 0.2848923 | 0.56003556 | 0.104 | 0.067 | 1 | 0.53801774 | 0.18490943 | 0.118 | 0.094 | 1 | 0.53801774 | 0.48862098 |
| Gm11714 | 0.64538863 | 0.52206771 | 0.042 | 0.031 | 1 | 0.28490958 | 0.16407593 | 0.088 | 0.052 | 1 | 0.64538863 | 0.4886457 |
| Lrrc69 | 0.78117751 | -0.577468 | 0.021 | 0.026 | 1 | 0.28500854 | -1.6724253 | 0.021 | 0.042 | 1 | 0.78117751 | 0.48878722 |
| Sox11 | 0.28505098 | -1.407543 | 0.01 | 0.031 | 1 | 0.54938851 | -0.681824 | 0.038 | 0.052 | 1 | 0.54938851 | 0.4888479 |
| Gm29642 | 0.83546978 | 0.15949763 | 0.031 | 0.036 | 1 | 0.28514714 | 0.49749966 | 0.046 | 0.021 | 1 | 0.83546978 | 0.48898539 |
| Cacnb3 | 0.28531004 | -0.6929452 | 0.021 | 0.047 | 1 | 0.58180179 | -0.7248928 | 0.021 | 0.031 | 1 | 0.58180179 | 0.48921826 |
| Bbs4 | 0.28532464 | -0.1477837 | 0.198 | 0.145 | 1 | 0.76083918 | -0.1820997 | 0.176 | 0.156 | 1 | 0.76083918 | 0.48923913 |
| Cep83os | 0.42175358 | 0.39451766 | 0.135 | 0.104 | 1 | 0.28577117 | -0.1630139 | 0.101 | 0.062 | 1 | 0.42175358 | 0.48987718 |
| Gm57184 | 0.31073311 | -0.9153376 | 0.188 | 0.228 | 1 | 0.28582205 | 0.30905457 | 0.113 | 0.073 | 1 | 0.31073311 | 0.48994986 |
| Cep131 | 0.36601088 | -0.8669746 | 0.021 | 0.041 | 1 | 0.28597111 | 0.59703533 | 0.046 | 0.021 | 1 | 0.36601088 | 0.49016275 |
| Ttll7 | 0.7916499 | 0.17350449 | 0.25 | 0.238 | 1 | 0.28616375 | -0.4665808 | 0.504 | 0.354 | 1 | 0.7916499 | 0.4904378 |
| Wsb1 | 0.28630009 | -0.1388548 | 0.938 | 0.948 | 1 | 0.7672329 | -0.2456328 | 0.996 | 0.99 | 1 | 0.7672329 | 0.49063244 |
| Gm56614 | 0.73685932 | 1.27052894 | 0.01 | 0.016 | 1 | 0.2864176 | -2.2427411 | 0.029 | 0.052 | 1 | 0.73685932 | 0.49080016 |
| Glp2r | 0.51741252 | 0.24195979 | 0.052 | 0.073 | 1 | 0.28683444 | 0.17557156 | 0.046 | 0.021 | 1 | 0.51741252 | 0.49139488 |
| Gm11747 | 1 | 0.42253204 | 0.01 | 0.01 | 1 | 0.28683444 | 0.17557156 | 0.046 | 0.021 | 1 | 1 | 0.49139488 |
| Fam78b | 0.74076005 | 1.00749454 | 0.021 | 0.016 | 1 | 0.28702455 | -1.5993619 | 0.021 | 0.042 | 1 | 0.74076005 | 0.49166601 |
| Cfap54 | 0.52744059 | -0.577468 | 0.01 | 0.021 | 1 | 0.28703146 | -1.6317834 | 0.021 | 0.042 | 1 | 0.52744059 | 0.49167586 |
| Adm | 0.28733227 | -0.577468 | 0.021 | 0.047 | 1 | 0.42818028 | -0.7952821 | 0.034 | 0.052 | 1 | 0.42818028 | 0.49210471 |
| B530045E10Rik | 0.28733227 | -0.577468 | 0.021 | 0.047 | 1 | 0.86915235 | 0.27510724 | 0.008 | 0.01 | 1 | 0.86915235 | 0.49210471 |
| Angptl4 | 0.97731546 | -0.1588638 | 0.156 | 0.155 | 1 | 0.28757545 | -0.2545728 | 0.231 | 0.167 | 1 | 0.97731546 | 0.49245126 |
| Gm41556 | 0.8463648 | -0.4872702 | 0.094 | 0.098 | 1 | 0.28775544 | -1.8359241 | 0.076 | 0.104 | 1 | 0.8463648 | 0.49270768 |
| Dcaf8 | 0.28791985 | 0.12226472 | 0.76 | 0.762 | 1 | 0.65343624 | -0.1294379 | 0.849 | 0.854 | 1 | 0.65343624 | 0.49294186 |
| Gm26511 | 0.38521705 | -0.7998604 | 0.01 | 0.026 | 1 | 0.28801198 | -1.5322477 | 0.021 | 0.042 | 1 | 0.38521705 | 0.49307306 |
| Gm44956 | 0.82258416 | -0.1624305 | 0.031 | 0.036 | 1 | 0.28801198 | -1.7248928 | 0.021 | 0.042 | 1 | 0.82258416 | 0.49307306 |
| Igsf9b | 0.28835469 | -0.3710171 | 0.021 | 0.047 | 1 | 0.37620733 | 0.42711033 | 0.055 | 0.031 | 1 | 0.37620733 | 0.49356096 |
| Phf11c | 0.80274368 | 0.20013961 | 0.031 | 0.026 | 1 | 0.28848731 | 0.27510724 | 0.046 | 0.021 | 1 | 0.80274368 | 0.49374969 |
| Rars2 | 0.38375406 | 0.2091284 | 0.281 | 0.228 | 1 | 0.28851419 | -0.5549678 | 0.324 | 0.229 | 1 | 0.38375406 | 0.49378794 |
| Gm33906 | 0.28885667 | -0.407543 | 0.01 | 0.031 | 1 | 0.56565662 | -1.8948178 | 0.013 | 0.021 | 1 | 0.56565662 | 0.49427516 |
| R3hdm4 | 0.28895413 | 0.31127528 | 0.531 | 0.472 | 1 | 0.40041349 | -0.186707 | 0.559 | 0.646 | 1 | 0.40041349 | 0.49441377 |
| Kif1a | 0.28935992 | -0.4519371 | 0.021 | 0.047 | 1 | 0.40321914 | -1.0468209 | 0.059 | 0.083 | 1 | 0.40321914 | 0.49499068 |
| Olfm2 | 0.28937956 | 0.20013961 | 0.021 | 0.047 | 1 | 0.66976882 | -0.8073549 | 0.101 | 0.115 | 1 | 0.66976882 | 0.49501859 |

| | | | | | | | | | | | | |
|---|---|---|---|---|---|---|---|---|---|---|---|---|
| Lypd1 | 0.54395837 | 0.34006988 | 0.094 | 0.073 | 1 | 0.28942211 | 0.34222143 | 0.046 | 0.021 | 1 | 0.54395837 | 0.49507906 |
| Mfsd13a | 0.98414882 | 0.78510212 | 0.031 | 0.031 | 1 | 0.2899192 | -2.1172102 | 0.029 | 0.052 | 1 | 0.98414882 | 0.49578526 |
| Adat1 | 0.29011528 | 0.27052894 | 0.26 | 0.202 | 1 | 0.56630794 | -0.701046 | 0.172 | 0.188 | 1 | 0.56630794 | 0.49606368 |
| Zfp296 | 0.29012577 | -0.407543 | 0.01 | 0.031 | 1 | 0.50587544 | -2.3098553 | 0.004 | 0.01 | 1 | 0.50587544 | 0.49607858 |
| Slc26a6 | 0.29012577 | -0.407543 | 0.01 | 0.031 | 1 | 0.85030707 | -0.6577786 | 0.038 | 0.042 | 1 | 0.85030707 | 0.49607858 |
| Sema6d | 0.29017706 | -0.8531024 | 0.083 | 0.124 | 1 | 0.89652131 | -0.2860085 | 0.13 | 0.135 | 1 | 0.89652131 | 0.4961514 |
| Ppp1r9b | 0.29024673 | 0.32942263 | 0.208 | 0.155 | 1 | 0.53537983 | -0.3579496 | 0.34 | 0.281 | 1 | 0.53537983 | 0.4962503 |
| Gm16023 | 0.29028692 | -0.3930434 | 0.177 | 0.233 | 1 | 0.55441107 | -0.5834203 | 0.261 | 0.208 | 1 | 0.55441107 | 0.49630734 |
| Tcaim | 0.41591855 | 0.44359365 | 0.177 | 0.14 | 1 | 0.29077065 | 0.33784299 | 0.147 | 0.104 | 1 | 0.41591855 | 0.49699373 |
| Msx1 | 0.29079415 | -1.7052235 | 0.083 | 0.119 | 1 | 0.81717909 | -1.7366699 | 0.168 | 0.167 | 1 | 0.81717909 | 0.49702706 |
| 2310022A10Rik | 0.29098917 | 0.39082318 | 0.135 | 0.093 | 1 | 0.3070073 | 0.16407593 | 0.172 | 0.125 | 1 | 0.3070073 | 0.49730365 |
| Sanbr | 0.50646538 | 0.10534186 | 0.385 | 0.352 | 1 | 0.29102119 | -0.1680813 | 0.429 | 0.312 | 1 | 0.50646538 | 0.49734905 |
| Efhb | 0.29105112 | 0.53356335 | 0.104 | 0.067 | 1 | 0.93756367 | -0.4167705 | 0.071 | 0.073 | 1 | 0.93756367 | 0.49739149 |
| Slfn8 | 0.29141558 | 0.35541784 | 0.01 | 0.031 | 1 | 0.34703906 | -1.3098553 | 0.008 | 0.021 | 1 | 0.34703906 | 0.49790812 |
| Foxq1 | 0.29141981 | 1.24850264 | 0.01 | 0.031 | 1 | 0.66387437 | 0.49749966 | 0.017 | 0.01 | 1 | 0.66387437 | 0.49791412 |
| Tex15 | 0.29142235 | -0.2726134 | 0.01 | 0.031 | 1 | 0.50670329 | 1.27510724 | 0.021 | 0.01 | 1 | 0.50670329 | 0.49791771 |
| 1700096K18Rik | 0.29163528 | -1.130009 | 0.042 | 0.073 | 1 | 0.41231187 | -0.3098553 | 0.092 | 0.062 | 1 | 0.41231187 | 0.49821942 |
| C030034L19Rik | 0.41466006 | -0.9925055 | 0.031 | 0.052 | 1 | 0.29177769 | 0.30312161 | 0.139 | 0.094 | 1 | 0.41466006 | 0.49842115 |
| Iffo1 | 0.99473465 | -0.4705528 | 0.125 | 0.119 | 1 | 0.29186391 | -0.6094155 | 0.113 | 0.156 | 1 | 0.99473465 | 0.49854328 |
| Espn | 0.99758609 | 0.27052894 | 0.021 | 0.021 | 1 | 0.29195341 | -1.8504236 | 0.038 | 0.062 | 1 | 0.99758609 | 0.49867002 |
| Arhgap8 | 0.32025586 | -1.577468 | 0 | 0.01 | 1 | 0.29211352 | -0.9879272 | 0.021 | 0.042 | 1 | 0.32025586 | 0.49889673 |
| Map2k4 | 0.29241323 | -0.2207745 | 0.667 | 0.725 | 1 | 0.44073988 | -0.4322852 | 0.836 | 0.865 | 1 | 0.44073988 | 0.49932097 |
| Trap1 | 0.29243797 | 0.12070515 | 0.396 | 0.316 | 1 | 0.36751884 | -0.3788026 | 0.412 | 0.302 | 1 | 0.36751884 | 0.49935598 |
| Septin11 | 0.68264726 | -0.1970766 | 0.781 | 0.767 | 1 | 0.29247229 | -0.5489086 | 0.929 | 0.875 | 1 | 0.68264726 | 0.49940454 |
| Lonp2 | 0.67426291 | -0.1058469 | 0.552 | 0.58 | 1 | 0.29250959 | -0.2728611 | 0.601 | 0.688 | 1 | 0.67426291 | 0.49945733 |
| Dalrd3 | 0.29250967 | 0.32942263 | 0.219 | 0.166 | 1 | 0.3904524 | -0.3838558 | 0.21 | 0.156 | 1 | 0.3904524 | 0.49945744 |
| Fhip2b | 0.95141351 | 0.34452952 | 0.083 | 0.083 | 1 | 0.29262672 | 0.27510724 | 0.126 | 0.083 | 1 | 0.95141351 | 0.49962305 |
| Tsga10 | 0.52802234 | 0.65385758 | 0.104 | 0.083 | 1 | 0.29267339 | -0.6179776 | 0.088 | 0.125 | 1 | 0.52802234 | 0.49968907 |
| Mmaa | 0.3304875 | 0.30122574 | 0.156 | 0.114 | 1 | 0.29304605 | -0.2867716 | 0.193 | 0.135 | 1 | 0.3304875 | 0.50021611 |
| Acsm3 | 0.38521705 | 0.68556644 | 0.031 | 0.016 | 1 | 0.29313566 | -0.9472852 | 0.021 | 0.042 | 1 | 0.38521705 | 0.5003428 |
| BC065397 | 0.29321192 | 0.45495351 | 0.104 | 0.067 | 1 | 0.42460668 | -0.9093173 | 0.088 | 0.115 | 1 | 0.42460668 | 0.50045061 |
| Nfkb2 | 0.29363445 | -0.407543 | 0.26 | 0.311 | 1 | 0.31502649 | -0.2293653 | 0.37 | 0.271 | 1 | 0.31502649 | 0.50104771 |
| Fan1 | 0.29369234 | 0.74446013 | 0.146 | 0.104 | 1 | 0.44858855 | -0.8792209 | 0.101 | 0.125 | 1 | 0.44858855 | 0.5011295 |
| Mmp25 | 0.84470067 | 0.54806292 | 0.031 | 0.036 | 1 | 0.29386915 | -1.5899632 | 0.046 | 0.073 | 1 | 0.84470067 | 0.50137922 |
| Nedd1 | 0.57509684 | 0.13302542 | 0.229 | 0.197 | 1 | 0.29407188 | 0.17557156 | 0.252 | 0.188 | 1 | 0.57509684 | 0.50166548 |
| Cspg4b | 0.32466654 | 0.70793425 | 0.094 | 0.062 | 1 | 0.29410374 | 0.43737867 | 0.147 | 0.104 | 1 | 0.32466654 | 0.50171046 |
| Gm30938 | 0.63520923 | 0.78510212 | 0.042 | 0.031 | 1 | 0.29415625 | -1.1399303 | 0.021 | 0.042 | 1 | 0.63520923 | 0.5017846 |
| Tmc8 | 0.57678264 | 0.35541784 | 0.062 | 0.047 | 1 | 0.29435837 | -1.8123556 | 0.038 | 0.062 | 1 | 0.57678264 | 0.50206989 |
| Fes | 0.57177892 | -0.3710171 | 0.083 | 0.104 | 1 | 0.29471707 | -0.3415641 | 0.143 | 0.094 | 1 | 0.57177892 | 0.50257599 |
| Anpep | 0.29482302 | -0.577468 | 0.083 | 0.124 | 1 | 0.54624575 | -0.7248928 | 0.046 | 0.062 | 1 | 0.54624575 | 0.50272543 |
| Nxpe3 | 0.29488228 | -1.130009 | 0.042 | 0.073 | 1 | 0.62117966 | -0.6468902 | 0.059 | 0.073 | 1 | 0.62117966 | 0.502809 |
| Prkcb | 0.41785345 | -0.577468 | 0.052 | 0.078 | 1 | 0.29529199 | -1.5961594 | 0.08 | 0.115 | 1 | 0.41785345 | 0.50338662 |
| Pou6f1 | 0.29564443 | -0.9274104 | 0.146 | 0.187 | 1 | 0.58468359 | -0.6179776 | 0.181 | 0.198 | 1 | 0.58468359 | 0.50388323 |
| Sytl2 | 0.54089148 | 0.59245704 | 0.094 | 0.073 | 1 | 0.29577883 | 0.96316323 | 0.059 | 0.031 | 1 | 0.54089148 | 0.50407254 |
| Zfp566 | 0.29578659 | 0.68556644 | 0.115 | 0.078 | 1 | 0.991531 | -0.2029401 | 0.105 | 0.104 | 1 | 0.991531 | 0.50408347 |
| Rrm2 | 0.41707893 | 2.44359365 | 0.062 | 0.041 | 1 | 0.29593903 | -1.9879272 | 0.038 | 0.062 | 1 | 0.41707893 | 0.50429815 |
| Gm13562 | 0.75328966 | 0.27052894 | 0.021 | 0.016 | 1 | 0.29621551 | -0.8948178 | 0.021 | 0.042 | 1 | 0.75328966 | 0.5046874 |
| Gm49066 | 0.29645488 | 0.74446013 | 0.083 | 0.052 | 1 | 0.31321232 | -1.3098553 | 0.038 | 0.062 | 1 | 0.31321232 | 0.50502427 |
| Ppm1e | 0.29656877 | 2.42253204 | 0.042 | 0.021 | 1 | 0.75467989 | -0.6468902 | 0.025 | 0.031 | 1 | 0.75467989 | 0.5051845 |
| Gm26944 | 0.8692145 | -0.2335136 | 0.094 | 0.098 | 1 | 0.2965774 | -1.2167459 | 0.063 | 0.094 | 1 | 0.8692145 | 0.50519664 |
| Ctu2 | 0.55789144 | 0.27052894 | 0.177 | 0.15 | 1 | 0.29688589 | -0.194378 | 0.176 | 0.125 | 1 | 0.55789144 | 0.50563055 |
| Tcea2 | 0.57678264 | 0.35541784 | 0.062 | 0.047 | 1 | 0.29722453 | -0.8244284 | 0.021 | 0.042 | 1 | 0.57678264 | 0.50610664 |
| Gm43113 | 0.74827573 | 0.59245704 | 0.021 | 0.016 | 1 | 0.29722453 | -0.8244284 | 0.021 | 0.042 | 1 | 0.74827573 | 0.50610664 |
| Cbwd1 | 0.87925877 | -0.190919 | 0.458 | 0.466 | 1 | 0.29723724 | -0.129283 | 0.29 | 0.208 | 1 | 0.87925877 | 0.5061245 |
| Txlng | 0.29730994 | 0.16389609 | 0.49 | 0.399 | 1 | 0.3561832 | -0.2857428 | 0.45 | 0.375 | 1 | 0.3561832 | 0.50622667 |
| Abat | 0.4662281 | 0.12115132 | 0.281 | 0.238 | 1 | 0.29741934 | -0.3754436 | 0.277 | 0.198 | 1 | 0.4662281 | 0.50638041 |
| Chd9 | 0.29774936 | 0.14129303 | 0.969 | 0.943 | 1 | 0.98245703 | -0.2111799 | 0.962 | 0.969 | 1 | 0.98245703 | 0.50684404 |
| Ccdc39 | 0.29787678 | 0.59245704 | 0.115 | 0.078 | 1 | 0.64911825 | -1.129283 | 0.092 | 0.104 | 1 | 0.64911825 | 0.50702299 |
| Gm39326 | 0.61737927 | 1.00749454 | 0.01 | 0.005 | 1 | 0.29794961 | -0.7248928 | 0.046 | 0.021 | 1 | 0.61737927 | 0.50712524 |
| Tmem240 | 0.29834215 | 0.71798792 | 0.083 | 0.052 | 1 | 0.64256494 | -0.5728897 | 0.08 | 0.062 | 1 | 0.64256494 | 0.50767626 |
| Gm10643 | 0.29834215 | 0.71798792 | 0.083 | 0.052 | 1 | 0.94625326 | -0.7248928 | 0.05 | 0.052 | 1 | 0.94625326 | 0.50767626 |
| Ciao1 | 0.38827601 | 0.27814213 | 0.302 | 0.254 | 1 | 0.29853933 | 0.14265694 | 0.324 | 0.25 | 1 | 0.38827601 | 0.50795293 |
| B430105G09Rik | 0.57678264 | -0.3710171 | 0.042 | 0.057 | 1 | 0.2986794 | 0.69014474 | 0.059 | 0.031 | 1 | 0.57678264 | 0.50814942 |
| Comtd1 | 0.85778553 | -0.1079827 | 0.052 | 0.057 | 1 | 0.2986794 | 0.69014474 | 0.059 | 0.031 | 1 | 0.85778553 | 0.50814942 |
| Clk3 | 0.31903803 | 0.15433592 | 0.26 | 0.202 | 1 | 0.29875336 | -0.2244638 | 0.29 | 0.219 | 1 | 0.31903803 | 0.50825315 |
| Ppara | 0.47757197 | -1.1624305 | 0.021 | 0.036 | 1 | 0.29888275 | -1.6094155 | 0.029 | 0.052 | 1 | 0.47757197 | 0.5084346 |
| Gm12655 | 0.29899098 | 1.49292136 | 0.042 | 0.021 | 1 | 0.34026741 | -2.4797803 | 0.008 | 0.021 | 1 | 0.34026741 | 0.50858635 |
| Slc22a14 | 0.45632632 | -0.8405024 | 0.042 | 0.062 | 1 | 0.29901687 | -1.5163061 | 0.046 | 0.073 | 1 | 0.45632632 | 0.50862265 |
| Bend3 | 0.29918805 | -0.6929452 | 0.062 | 0.098 | 1 | 0.39479423 | 0.17013768 | 0.143 | 0.104 | 1 | 0.39479423 | 0.50886262 |
| Zfp52 | 0.45541133 | -0.407543 | 0.24 | 0.275 | 1 | 0.29968577 | -0.1399303 | 0.5 | 0.375 | 1 | 0.45541133 | 0.50955998 |
| Gm16201 | 0.29997632 | 1.10703021 | 0.094 | 0.062 | 1 | 0.75886834 | -0.1805722 | 0.118 | 0.104 | 1 | 0.75886834 | 0.50996685 |
| Pikfyve | 0.30005787 | -0.3213419 | 0.573 | 0.648 | 1 | 0.51614969 | -0.3173889 | 0.689 | 0.594 | 1 | 0.51614969 | 0.51008102 |
| Prkaca | 0.62925437 | -0.11115 | 0.312 | 0.264 | 1 | 0.30046558 | -0.355659 | 0.277 | 0.208 | 1 | 0.62925437 | 0.51065159 |
| Spi1 | 0.3819238 | 0.42253204 | 0.031 | 0.016 | 1 | 0.30051687 | 2.91896343 | 0.029 | 0.01 | 1 | 0.3819238 | 0.51072335 |
| Fkbpl | 0.30103475 | 1.00749454 | 0.062 | 0.036 | 1 | 0.43323624 | 0.42711033 | 0.038 | 0.021 | 1 | 0.43323624 | 0.51144758 |
| Pcdhb18 | 0.52744059 | -0.577468 | 0.01 | 0.021 | 1 | 0.30111454 | 0.62874419 | 0.059 | 0.031 | 1 | 0.52744059 | 0.51155911 |
| Foxo4 | 0.30116425 | 0.74446013 | 0.125 | 0.088 | 1 | 0.40567096 | 0.15727075 | 0.164 | 0.125 | 1 | 0.40567096 | 0.5116286 |
| Patl1 | 0.30117358 | 0.27052894 | 0.385 | 0.321 | 1 | 0.9716907 | -0.408259 | 0.462 | 0.396 | 1 | 0.9716907 | 0.51164163 |
| Nme4 | 0.66186725 | -0.4519371 | 0.031 | 0.041 | 1 | 0.30139541 | -0.6317834 | 0.021 | 0.042 | 1 | 0.66186725 | 0.51195162 |
| 4921536K21Rik | 0.30147738 | 0.89201732 | 0.083 | 0.052 | 1 | 0.45128837 | 0.33400093 | 0.076 | 0.052 | 1 | 0.45128837 | 0.51206615 |
| Phf20 | 0.72919907 | -0.2349991 | 0.625 | 0.539 | 1 | 0.30148401 | -0.2451772 | 0.693 | 0.729 | 1 | 0.72919907 | 0.51207541 |
| Ifi27l2a | 0.83451385 | 0.52206771 | 0.073 | 0.067 | 1 | 0.30153441 | -1.8244284 | 0.055 | 0.083 | 1 | 0.83451385 | 0.51214582 |
| 2410080I02Rik | 0.61182672 | -0.577468 | 0.021 | 0.031 | 1 | 0.30159499 | -1.5508634 | 0.029 | 0.052 | 1 | 0.61182672 | 0.51223044 |
| Galnt16 | 0.46838288 | 0.31561683 | 0.042 | 0.026 | 1 | 0.3016162 | 1.49749966 | 0.029 | 0.01 | 1 | 0.46838288 | 0.51226006 |
| Gm28198 | 0.3018119 | 0.37522632 | 0.177 | 0.13 | 1 | 0.49585846 | -0.3358505 | 0.197 | 0.156 | 1 | 0.49585846 | 0.51253338 |

| | | | | | | | | | | | | |
|---|---|---|---|---|---|---|---|---|---|---|---|---|
| C5ar2 | 0.74827573 | 0.59245704 | 0.021 | 0.016 | 1 | 0.30185121 | 0.69014474 | 0.059 | 0.031 | 1 | 0.74827573 | 0.51258827 |
| Gm35769 | 0.30185951 | 0.53800925 | 0.115 | 0.078 | 1 | 0.4384834 | 0.18490943 | 0.113 | 0.083 | 1 | 0.4384834 | 0.51259986 |
| Ankzf1 | 0.3019435 | -0.3946039 | 0.115 | 0.161 | 1 | 0.49374562 | 0.23071312 | 0.109 | 0.083 | 1 | 0.49374562 | 0.51271712 |
| Alkbh2 | 0.33568997 | 0.53356335 | 0.135 | 0.098 | 1 | 0.30221289 | -0.2654611 | 0.113 | 0.073 | 1 | 0.33568997 | 0.51309315 |
| Trim11 | 0.8847728 | -0.1941393 | 0.24 | 0.244 | 1 | 0.30225545 | -0.3275573 | 0.269 | 0.198 | 1 | 0.8847728 | 0.51315255 |
| 1700048O20Rik | 0.30241792 | 0.59245704 | 0.083 | 0.052 | 1 | 0.68204624 | 0.49749966 | 0.063 | 0.052 | 1 | 0.68204624 | 0.51337924 |
| Zfp994 | 0.30262202 | -0.5399933 | 0.094 | 0.135 | 1 | 0.48236411 | 0.69014474 | 0.084 | 0.062 | 1 | 0.48236411 | 0.51366395 |
| Gm10857 | 0.30268653 | 1.42253204 | 0.042 | 0.021 | 1 | 0.86312513 | -0.7248928 | 0.008 | 0.01 | 1 | 0.86312513 | 0.51375393 |
| Odf3l2 | 0.47150075 | 1.42253204 | 0.021 | 0.01 | 1 | 0.30271878 | 1.14957636 | 0.029 | 0.01 | 1 | 0.47150075 | 0.5137989 |
| Slc30a9 | 0.30274326 | 0.25365512 | 0.552 | 0.497 | 1 | 0.62466099 | -0.1023668 | 0.702 | 0.656 | 1 | 0.62466099 | 0.51383304 |
| Ccz1 | 0.30296268 | 0.1730806 | 0.552 | 0.472 | 1 | 0.51342319 | -0.1463565 | 0.521 | 0.573 | 1 | 0.51342319 | 0.51413897 |
| Efcab8 | 0.30298932 | -0.3144336 | 0.115 | 0.161 | 1 | 0.61024141 | -1.5592147 | 0.172 | 0.135 | 1 | 0.61024141 | 0.51417612 |
| Zfp846 | 0.30321437 | 0.56003556 | 0.083 | 0.052 | 1 | 0.6360176 | -0.1843244 | 0.067 | 0.052 | 1 | 0.6360176 | 0.51448978 |
| Gm20682 | 0.32693714 | -1.0799683 | 0.031 | 0.057 | 1 | 0.30323558 | -1.6883669 | 0.038 | 0.062 | 1 | 0.32693714 | 0.51451934 |
| Smyd3 | 0.97705745 | -0.1118044 | 0.615 | 0.591 | 1 | 0.30356719 | -0.5904302 | 0.676 | 0.719 | 1 | 0.97705745 | 0.51498134 |
| 2410018L13Rik | 0.30358994 | 0.5283267 | 0.229 | 0.181 | 1 | 0.90906703 | -0.4904275 | 0.16 | 0.146 | 1 | 0.90906703 | 0.51501304 |
| Nars2 | 0.84259086 | 0.13755508 | 0.188 | 0.202 | 1 | 0.30363049 | -1.0948424 | 0.176 | 0.219 | 1 | 0.84259086 | 0.5150695 |
| Ovol2 | 0.32025586 | -0.9925055 | 0 | 0.01 | 1 | 0.30382073 | 1.01207283 | 0.029 | 0.01 | 1 | 0.32025586 | 0.51533443 |
| Kcnj8 | 0.32025586 | -0.9925055 | 0 | 0.01 | 1 | 0.30382073 | 1.01207283 | 0.029 | 0.01 | 1 | 0.32025586 | 0.51533443 |
| Dmkn | 0.32025586 | -0.577468 | 0 | 0.01 | 1 | 0.30382073 | 1.01207283 | 0.029 | 0.01 | 1 | 0.32025586 | 0.51533443 |
| Myoc | 0.5862099 | 0.68556644 | 0.031 | 0.021 | 1 | 0.30382073 | 1.01207283 | 0.029 | 0.01 | 1 | 0.5862099 | 0.51533443 |
| Zfp641 | 0.61737927 | 1.00749454 | 0.01 | 0.005 | 1 | 0.30382073 | 1.01207283 | 0.029 | 0.01 | 1 | 0.61737927 | 0.51533443 |
| Cdc20 | 0.67121057 | 1.75956702 | 0.073 | 0.062 | 1 | 0.30382073 | 1.27510724 | 0.029 | 0.01 | 1 | 0.67121057 | 0.51533443 |
| C030005K06Rik | 0.72574709 | -0.3144336 | 0.01 | 0.016 | 1 | 0.30382073 | 1.01207283 | 0.029 | 0.01 | 1 | 0.72574709 | 0.51533443 |
| Hnmt | 0.30389555 | 1.27052894 | 0.042 | 0.021 | 1 | 0.30645059 | 0.53814164 | 0.059 | 0.031 | 1 | 0.30645059 | 0.5154386 |
| Fancf | 0.30389555 | 1.27052894 | 0.042 | 0.021 | 1 | 0.66339728 | 0.10518224 | 0.029 | 0.021 | 1 | 0.66339728 | 0.5154386 |
| Hunk | 0.30389555 | 2.00749454 | 0.042 | 0.021 | 1 | 0.31154055 | -1.6968784 | 0.038 | 0.062 | 1 | 0.31154055 | 0.5154386 |
| Kcnd1 | 0.30389555 | 1.27052894 | 0.042 | 0.021 | 1 | 0.46074961 | -0.2457249 | 0.076 | 0.052 | 1 | 0.46074961 | 0.5154386 |
| 4933417E11Rik | 0.30394239 | 1.42253204 | 0.062 | 0.036 | 1 | 0.86312513 | -0.7248928 | 0.008 | 0.01 | 1 | 0.86312513 | 0.51550381 |
| Gm41664 | 0.30398061 | 1.00749454 | 0.062 | 0.036 | 1 | 0.86645678 | -0.5728897 | 0.038 | 0.042 | 1 | 0.86645678 | 0.51555701 |
| Pcdhga5 | 0.30416949 | -0.577468 | 0.062 | 0.098 | 1 | 0.90786026 | -0.7248928 | 0.092 | 0.094 | 1 | 0.90786026 | 0.5158199 |
| Grin3a | 1 | -0.130009 | 0.042 | 0.041 | 1 | 0.30418557 | 0.34222143 | 0.059 | 0.031 | 1 | 1 | 0.51584228 |
| Arrdc2 | 0.79310976 | -0.3364599 | 0.104 | 0.114 | 1 | 0.30420964 | -1.8948178 | 0.046 | 0.073 | 1 | 0.79310976 | 0.51587578 |
| Lcmt1 | 0.3992432 | 0.11133035 | 0.323 | 0.269 | 1 | 0.30429797 | -0.4589987 | 0.328 | 0.24 | 1 | 0.3992432 | 0.51599869 |
| Cdc42ep3 | 0.80074924 | 0.14662302 | 0.792 | 0.798 | 1 | 0.30440711 | 0.20148437 | 0.903 | 0.896 | 1 | 0.80074924 | 0.51615053 |
| Prpf18 | 0.34650426 | 0.17099327 | 0.344 | 0.28 | 1 | 0.30443903 | -0.3960119 | 0.382 | 0.271 | 1 | 0.34650426 | 0.51619494 |
| Caskin2 | 0.33500979 | 0.81484946 | 0.104 | 0.073 | 1 | 0.3047235 | -0.4618584 | 0.097 | 0.135 | 1 | 0.33500979 | 0.51659059 |
| Gm34455 | 0.30477403 | -1.5304705 | 0.115 | 0.155 | 1 | 0.83387057 | -0.8035026 | 0.256 | 0.25 | 1 | 0.83387057 | 0.51666085 |
| Ubxn7 | 0.99752858 | -0.2030724 | 0.656 | 0.637 | 1 | 0.30487069 | -0.3926681 | 0.929 | 0.885 | 1 | 0.99752858 | 0.51679524 |
| Sugp1 | 0.30487425 | 0.39744105 | 0.25 | 0.202 | 1 | 0.48912076 | -0.2942584 | 0.277 | 0.219 | 1 | 0.48912076 | 0.51680019 |
| Sema6c | 0.30891093 | 1.00749454 | 0.042 | 0.021 | 1 | 0.30492205 | 0.86006974 | 0.029 | 0.01 | 1 | 0.30891093 | 0.51686665 |
| Gm15406 | 0.32025586 | -0.577468 | 0 | 0.01 | 1 | 0.30492205 | 0.86006974 | 0.029 | 0.01 | 1 | 0.32025586 | 0.51686665 |
| Gm42788 | 0.49952136 | -0.4950058 | 0.052 | 0.073 | 1 | 0.30492205 | 0.86006974 | 0.029 | 0.01 | 1 | 0.49952136 | 0.51686665 |
| Gm12963 | 0.52961488 | -0.3144336 | 0.01 | 0.021 | 1 | 0.30492205 | 0.86006974 | 0.029 | 0.01 | 1 | 0.52961488 | 0.51686665 |
| Gm20548 | 0.6229455 | -0.1624305 | 0.021 | 0.031 | 1 | 0.30492205 | 0.86006974 | 0.029 | 0.01 | 1 | 0.6229455 | 0.51686665 |
| Irag2 | 0.80279603 | 1.37006462 | 0.021 | 0.026 | 1 | 0.30492205 | 0.86006974 | 0.029 | 0.01 | 1 | 0.80279603 | 0.51686665 |
| Coa4 | 0.99758609 | 0.27052894 | 0.021 | 0.021 | 1 | 0.30492205 | 0.86006974 | 0.029 | 0.01 | 1 | 0.99758609 | 0.51686665 |
| Llgl2 | 0.99758609 | 0.27052894 | 0.021 | 0.021 | 1 | 0.30492205 | 0.86006974 | 0.029 | 0.01 | 1 | 0.99758609 | 0.51686665 |
| Rgs9 | 1 | -1.3144336 | 0.01 | 0.01 | 1 | 0.30492205 | 0.86006974 | 0.029 | 0.01 | 1 | 1 | 0.51686665 |
| Col19a1 | 0.99758607 | 0.20013961 | 0.021 | 0.021 | 1 | 0.30492236 | 0.56461386 | 0.059 | 0.031 | 1 | 0.99758607 | 0.51686707 |
| Gm10575 | 0.70270252 | -0.3144336 | 0.042 | 0.052 | 1 | 0.30502812 | -1.3098553 | 0.046 | 0.073 | 1 | 0.70270252 | 0.51701409 |
| Tmem9b | 0.63186731 | -0.1777203 | 0.51 | 0.497 | 1 | 0.30503066 | -0.1472351 | 0.555 | 0.479 | 1 | 0.63186731 | 0.51701762 |
| Tmed8 | 0.99062458 | 0.21608116 | 0.156 | 0.161 | 1 | 0.3051446 | -0.2345671 | 0.189 | 0.135 | 1 | 0.99062458 | 0.51717597 |
| Trim72 | 0.47294388 | -0.8669746 | 0.021 | 0.036 | 1 | 0.30521433 | -1.84037 | 0.029 | 0.052 | 1 | 0.47294388 | 0.51727288 |
| Atl1 | 0.30530459 | -0.6929452 | 0.25 | 0.301 | 1 | 0.89649502 | -0.3695615 | 0.202 | 0.198 | 1 | 0.89649502 | 0.51739829 |
| Inf2 | 0.72365031 | -0.3313074 | 0.344 | 0.358 | 1 | 0.30546165 | -0.2022145 | 0.429 | 0.323 | 1 | 0.72365031 | 0.51761648 |
| Akr1b3 | 0.78151122 | -0.2820121 | 0.406 | 0.378 | 1 | 0.3056021 | -0.4261794 | 0.366 | 0.427 | 1 | 0.78151122 | 0.51781156 |
| Il1rl2 | 0.30584838 | 1.00749454 | 0.062 | 0.036 | 1 | 0.57170879 | -1.3098553 | 0.013 | 0.021 | 1 | 0.57170879 | 0.51815352 |
| Ier5 | 0.42727086 | 0.27801052 | 0.854 | 0.876 | 1 | 0.30594226 | 0.23175087 | 0.786 | 0.833 | 1 | 0.42727086 | 0.51828385 |
| Gm50146 | 0.38521705 | 0.68556644 | 0.031 | 0.016 | 1 | 0.30606883 | 0.69014474 | 0.029 | 0.01 | 1 | 0.38521705 | 0.51845953 |
| Neil2 | 0.47599549 | -0.577468 | 0.021 | 0.036 | 1 | 0.30606883 | 0.69014474 | 0.029 | 0.01 | 1 | 0.47599549 | 0.51845953 |
| Aloxe3 | 0.52313375 | -1.7998604 | 0.01 | 0.021 | 1 | 0.30606883 | 0.69014474 | 0.029 | 0.01 | 1 | 0.52313375 | 0.51845953 |
| Glb1l2 | 0.61737927 | 1.00749454 | 0.01 | 0.005 | 1 | 0.30606883 | 0.69014474 | 0.029 | 0.01 | 1 | 0.61737927 | 0.51845953 |
| 1700064N11Rik | 0.99009299 | 0.32942263 | 0.031 | 0.031 | 1 | 0.30606883 | 0.69014474 | 0.029 | 0.01 | 1 | 0.99009299 | 0.51845953 |
| Riox2 | 0.76291062 | 0.17099327 | 0.104 | 0.093 | 1 | 0.30639896 | 0.69014474 | 0.071 | 0.042 | 1 | 0.76291062 | 0.51891759 |
| Enc1 | 0.30655151 | -0.3364599 | 0.062 | 0.098 | 1 | 0.40992831 | -1.5728897 | 0.034 | 0.052 | 1 | 0.40992831 | 0.51912919 |
| Gm26688 | 0.3066054 | -0.577468 | 0.042 | 0.073 | 1 | 0.66391569 | -0.8504236 | 0.042 | 0.052 | 1 | 0.66391569 | 0.51920393 |
| Zcchc2 | 0.57098251 | 0.31005731 | 0.25 | 0.223 | 1 | 0.30683682 | -0.6191833 | 0.248 | 0.292 | 1 | 0.57098251 | 0.5195248 |
| Mdga1 | 0.30686102 | -0.4828311 | 0.344 | 0.264 | 1 | 0.99738348 | -0.5907561 | 0.261 | 0.24 | 1 | 0.99738348 | 0.51955835 |
| Wnt2b | 0.53487059 | 0.27052894 | 0.052 | 0.036 | 1 | 0.30704993 | -1.1399303 | 0.029 | 0.052 | 1 | 0.53487059 | 0.5198202 |
| Pcbp3 | 0.30748011 | -0.8832764 | 0.177 | 0.223 | 1 | 0.40673802 | -0.1012686 | 0.197 | 0.156 | 1 | 0.40673802 | 0.5204162 |
| Clcf1 | 0.30758384 | 0.37946331 | 0.135 | 0.093 | 1 | 0.92181556 | -0.4916936 | 0.206 | 0.198 | 1 | 0.92181556 | 0.52055986 |
| Gm36287 | 0.30763643 | 1.52206771 | 0.042 | 0.021 | 1 | 0.50887612 | -1.3098553 | 0.004 | 0.01 | 1 | 0.50887612 | 0.52063269 |
| Ppargc1b | 0.81149164 | 0.52206771 | 0.021 | 0.026 | 1 | 0.30772698 | -0.2103196 | 0.021 | 0.042 | 1 | 0.81149164 | 0.52075806 |
| Rep15 | 0.53003414 | -0.7767768 | 0.083 | 0.104 | 1 | 0.30786051 | -0.8244284 | 0.063 | 0.094 | 1 | 0.53003414 | 0.52094292 |
| L3mbtl2 | 0.82342138 | 0.32942263 | 0.042 | 0.036 | 1 | 0.30786051 | -0.8244284 | 0.063 | 0.094 | 1 | 0.82342138 | 0.52094292 |
| Grem1 | 0.30791243 | 0.72738662 | 0.062 | 0.036 | 1 | 0.98417516 | -0.9728203 | 0.042 | 0.042 | 1 | 0.98417516 | 0.52101479 |
| 5430416N02Rik | 0.30794246 | 0.37522632 | 0.292 | 0.233 | 1 | 0.67713979 | -0.6013181 | 0.223 | 0.188 | 1 | 0.67713979 | 0.52105637 |
| Pdcd5 | 0.30802804 | 0.24453373 | 0.708 | 0.684 | 1 | 0.50361669 | -0.1516552 | 0.706 | 0.667 | 1 | 0.50361669 | 0.5211748 |
| Gm42997 | 0.53395584 | -0.2148979 | 0.01 | 0.021 | 1 | 0.30830219 | 0.80562195 | 0.029 | 0.01 | 1 | 0.53395584 | 0.52155414 |
| Naif1 | 0.30891093 | 1.00749454 | 0.042 | 0.021 | 1 | 0.53433287 | 0.42711033 | 0.034 | 0.021 | 1 | 0.53433287 | 0.5223959 |
| Gm14403 | 0.30891093 | 1.00749454 | 0.042 | 0.021 | 1 | 0.87673803 | -0.1723517 | 0.038 | 0.042 | 1 | 0.87673803 | 0.5223959 |
| Ccdc17 | 0.30891093 | 1.00749454 | 0.042 | 0.021 | 1 | 0.93071546 | -0.3098553 | 0.029 | 0.031 | 1 | 0.93071546 | 0.5223959 |
| 4930563E22Rik | 0.30891093 | 1.00749454 | 0.042 | 0.021 | 1 | 0.40647274 | -0.9879272 | 0.017 | 0.031 | 1 | 0.40647274 | 0.5223959 |
| Uimc1 | 0.30906879 | -0.2555399 | 0.698 | 0.762 | 1 | 0.43790727 | -0.1803445 | 0.731 | 0.646 | 1 | 0.43790727 | 0.52261406 |

| | | | | | | | | | | | | |
|---|---|---|---|---|---|---|---|---|---|---|---|---|
| Enoph1 | 0.30909701 | 0.19035859 | 0.198 | 0.145 | 1 | 0.65427074 | -0.4167705 | 0.193 | 0.167 | 1 | 0.65427074 | 0.52265305 |
| Gm4117 | 0.3094701 | -2.1217885 | 0.031 | 0.057 | 1 | 0.70493467 | -1.1843244 | 0.034 | 0.042 | 1 | 0.70493467 | 0.52316846 |
| Map3k3 | 0.3095114 | -0.1660202 | 0.833 | 0.674 | 1 | 0.39915258 | -0.3512392 | 0.866 | 0.896 | 1 | 0.39915258 | 0.5232255 |
| 9530026P05Rik | 0.30960395 | -0.1253173 | 0.365 | 0.399 | 1 | 0.94017412 | -0.3015559 | 0.437 | 0.406 | 1 | 0.94017412 | 0.5233533 |
| Usp32 | 0.74429017 | -0.1654903 | 0.646 | 0.606 | 1 | 0.30973249 | -0.3564305 | 0.815 | 0.938 | 1 | 0.74429017 | 0.52353077 |
| Kif13b | 0.30981507 | -0.3276259 | 0.958 | 0.99 | 1 | 0.52406538 | -0.2880387 | 0.971 | 0.969 | 1 | 0.52406538 | 0.52364476 |
| Tbc1d16 | 0.31004008 | 0.2099874 | 0.24 | 0.181 | 1 | 0.40944663 | -0.3623227 | 0.353 | 0.281 | 1 | 0.40944663 | 0.52395531 |
| Abcd2 | 0.69523544 | -0.7580402 | 0.042 | 0.052 | 1 | 0.31007669 | -0.7504279 | 0.088 | 0.052 | 1 | 0.69523544 | 0.52400583 |
| Gm30173 | 0.31013666 | 0.13302542 | 0.042 | 0.021 | 1 | 0.5207331 | -0.5025003 | 0.021 | 0.01 | 1 | 0.5207331 | 0.52408857 |
| Nek11 | 0.83878643 | 0.37006462 | 0.062 | 0.057 | 1 | 0.31037218 | -1.3098553 | 0.046 | 0.073 | 1 | 0.83878643 | 0.52441347 |
| 4933406C10Rik | 0.31052779 | 0.10060394 | 0.146 | 0.104 | 1 | 0.80542865 | -0.9995151 | 0.109 | 0.094 | 1 | 0.80542865 | 0.52462807 |
| Stap1 | 0.32025586 | -0.577468 | 0 | 0.01 | 1 | 0.31055303 | 0.56461386 | 0.029 | 0.01 | 1 | 0.32025586 | 0.52466288 |
| Rtn4rl1 | 0.97828543 | -0.577468 | 0.052 | 0.052 | 1 | 0.31072003 | -2.1329775 | 0.038 | 0.062 | 1 | 0.97828543 | 0.52489312 |
| Gm11638 | 0.31078953 | 0.32942263 | 0.083 | 0.052 | 1 | 0.66052743 | -0.8573431 | 0.071 | 0.083 | 1 | 0.66052743 | 0.52498893 |
| Ptprv | 0.85932205 | -0.1180363 | 0.052 | 0.057 | 1 | 0.31079262 | -0.9688183 | 0.055 | 0.083 | 1 | 0.85932205 | 0.52499319 |
| Celsr2 | 0.8181385 | 0.42253204 | 0.042 | 0.036 | 1 | 0.31102179 | 0.42711033 | 0.134 | 0.094 | 1 | 0.8181385 | 0.52530902 |
| Lama1 | 0.31156634 | 0.47697982 | 0.083 | 0.052 | 1 | 0.99136707 | -0.5728897 | 0.042 | 0.042 | 1 | 0.99136707 | 0.5260591 |
| Agrn | 0.31159675 | -0.4306266 | 0.25 | 0.306 | 1 | 0.9956342 | -0.2625495 | 0.303 | 0.281 | 1 | 0.9956342 | 0.52610096 |
| Gm16156 | 0.81159799 | -0.4519371 | 0.031 | 0.036 | 1 | 0.31167668 | 0.42711033 | 0.029 | 0.01 | 1 | 0.81159799 | 0.52621101 |
| Ptdss2 | 0.31169326 | 0.30388754 | 0.271 | 0.218 | 1 | 0.46590414 | -0.2457249 | 0.286 | 0.229 | 1 | 0.46590414 | 0.52623383 |
| Xrcc2 | 0.78979801 | -0.2148979 | 0.021 | 0.026 | 1 | 0.31280025 | 0.27510724 | 0.029 | 0.01 | 1 | 0.78979801 | 0.5277565 |
| Sycp3 | 1 | 0.42253204 | 0.01 | 0.01 | 1 | 0.31280025 | 0.27510724 | 0.029 | 0.01 | 1 | 1 | 0.5277565 |
| Trim69 | 0.46382346 | 1.00749454 | 0.042 | 0.026 | 1 | 0.31280858 | 0.14957636 | 0.029 | 0.01 | 1 | 0.46382346 | 0.52776795 |
| Cabyr | 0.31281193 | 0.68556644 | 0.062 | 0.036 | 1 | 0.42265322 | -0.5025003 | 0.05 | 0.073 | 1 | 0.42265322 | 0.52777256 |
| Dnah7c | 0.61737927 | 1.00749454 | 0.01 | 0.005 | 1 | 0.31281525 | 0.39058446 | 0.029 | 0.01 | 1 | 0.61737927 | 0.52777711 |
| Gm31305 | 0.46990573 | 0.64492446 | 0.042 | 0.026 | 1 | 0.31321262 | -1.0468209 | 0.038 | 0.062 | 1 | 0.46990573 | 0.5283231 |
| Actr5 | 0.31352693 | 0.39082318 | 0.115 | 0.078 | 1 | 0.77728561 | -0.2223924 | 0.105 | 0.115 | 1 | 0.77728561 | 0.52875472 |
| Msh3 | 0.31373206 | -0.913071 | 0.104 | 0.145 | 1 | 0.42125096 | -0.3483294 | 0.185 | 0.135 | 1 | 0.42125096 | 0.52903632 |
| Gm45464 | 0.31389644 | 0.52206771 | 0.042 | 0.021 | 1 | 0.57370068 | -1.0468209 | 0.021 | 0.031 | 1 | 0.57370068 | 0.5292619 |
| Gm26812 | 0.31389644 | 0.74446013 | 0.042 | 0.021 | 1 | 0.51018266 | 0.49749966 | 0.021 | 0.01 | 1 | 0.51018266 | 0.5292619 |
| Gm30786 | 0.31389644 | 0.52206771 | 0.042 | 0.021 | 1 | 0.79600331 | -0.9879272 | 0.017 | 0.021 | 1 | 0.79600331 | 0.5292619 |
| Rilp | 0.31389644 | 0.74446013 | 0.042 | 0.021 | 1 | 0.51255415 | -0.8073549 | 0.055 | 0.073 | 1 | 0.51255415 | 0.5292619 |
| Rtl6 | 0.31389644 | 0.74446013 | 0.042 | 0.021 | 1 | 0.85910786 | -0.4167705 | 0.038 | 0.042 | 1 | 0.85910786 | 0.5292619 |
| Krt6a | 0.31389644 | 0.74446013 | 0.042 | 0.021 | 1 | 0.47000784 | 1.89525917 | 0.084 | 0.062 | 1 | 0.47000784 | 0.5292619 |
| Zfp811 | 0.31389644 | 0.74446013 | 0.042 | 0.021 | 1 | 0.94994462 | -1.0203486 | 0.034 | 0.031 | 1 | 0.94994462 | 0.5292619 |
| Iqcb1 | 0.31389888 | 0.21635997 | 0.552 | 0.482 | 1 | 0.62387291 | -0.2097669 | 0.634 | 0.552 | 1 | 0.62387291 | 0.52926525 |
| Pgpep1l | 0.32025586 | -0.577468 | 0 | 0.01 | 1 | 0.31392319 | -0.6317834 | 0.029 | 0.01 | 1 | 0.32025586 | 0.52929861 |
| Stk35os1 | 0.46992036 | 0.74446013 | 0.042 | 0.026 | 1 | 0.31392319 | 0.10518224 | 0.029 | 0.01 | 1 | 0.46992036 | 0.52929861 |
| Sfrp5 | 0.47603364 | -2.1217885 | 0.021 | 0.036 | 1 | 0.31392319 | -1.8948178 | 0.029 | 0.01 | 1 | 0.47603364 | 0.52929861 |
| Gm47123 | 0.62666781 | 0.42253204 | 0.021 | 0.031 | 1 | 0.31392319 | 0.10518224 | 0.029 | 0.01 | 1 | 0.62666781 | 0.52929861 |
| 1700102H20Rik | 0.78979801 | -0.2148979 | 0.021 | 0.026 | 1 | 0.31392319 | 0.10518224 | 0.029 | 0.01 | 1 | 0.78979801 | 0.52929861 |
| Tmem253 | 0.32025586 | -0.577468 | 0 | 0.01 | 1 | 0.31394319 | -0.1723517 | 0.029 | 0.01 | 1 | 0.32025586 | 0.52932605 |
| Nipal1 | 0.78169785 | -0.1393469 | 0.083 | 0.093 | 1 | 0.31395152 | 0.10518224 | 0.029 | 0.01 | 1 | 0.78169785 | 0.52933748 |
| Cux2 | 0.81708725 | -0.3144336 | 0.031 | 0.036 | 1 | 0.31444841 | -0.7952821 | 0.029 | 0.052 | 1 | 0.81708725 | 0.53001902 |
| Bclaf3 | 0.31491493 | 0.18873485 | 0.406 | 0.352 | 1 | 0.74118139 | -0.1614634 | 0.508 | 0.448 | 1 | 0.74118139 | 0.53065844 |
| Adgb | 0.31518767 | 0.59245704 | 0.042 | 0.021 | 1 | 0.36245392 | 0.17557156 | 0.042 | 0.021 | 1 | 0.36245392 | 0.53103207 |
| Dnaaf3 | 0.39021373 | 0.20013961 | 0.031 | 0.016 | 1 | 0.31636136 | -0.8504236 | 0.029 | 0.052 | 1 | 0.39021373 | 0.5326382 |
| Setx | 0.34082159 | 0.20013961 | 0.448 | 0.378 | 1 | 0.31636515 | -0.3584354 | 0.576 | 0.615 | 1 | 0.34082159 | 0.53264339 |
| Aph1c | 0.33578218 | 0.48393258 | 0.125 | 0.088 | 1 | 0.31641463 | 0.37820073 | 0.097 | 0.062 | 1 | 0.33578218 | 0.53271104 |
| Afmid | 0.3167227 | -0.8225805 | 0.083 | 0.119 | 1 | 0.42502192 | -0.8073549 | 0.118 | 0.146 | 1 | 0.42502192 | 0.53313212 |
| Rapgef4 | 0.31674062 | -0.9612545 | 0.167 | 0.212 | 1 | 0.76658322 | -0.7837865 | 0.155 | 0.156 | 1 | 0.76658322 | 0.53315662 |
| Slc12a6 | 0.32848632 | 0.50764865 | 0.469 | 0.435 | 1 | 0.31685036 | 0.29747505 | 0.693 | 0.729 | 1 | 0.32848632 | 0.53330657 |
| Gm28229 | 0.61550914 | -0.407543 | 0.021 | 0.031 | 1 | 0.31719351 | -1.0688472 | 0.046 | 0.073 | 1 | 0.61550914 | 0.5337753 |
| Fam83d | 0.82650045 | -0.6929452 | 0.062 | 0.067 | 1 | 0.31730613 | -0.3098553 | 0.029 | 0.052 | 1 | 0.82650045 | 0.53392908 |
| Apobec1 | 1 | -0.4519371 | 0.031 | 0.031 | 1 | 0.31749023 | 0.58322953 | 0.059 | 0.031 | 1 | 1 | 0.53418041 |
| Pnpla3 | 0.31755188 | -1.0199862 | 0.094 | 0.13 | 1 | 0.57050612 | -0.8504236 | 0.021 | 0.031 | 1 | 0.57050612 | 0.53426457 |
| Gm35608 | 0.35590751 | -1.0368996 | 0.062 | 0.093 | 1 | 0.3178724 | -0.6577786 | 0.071 | 0.042 | 1 | 0.35590751 | 0.53470194 |
| Napa | 0.31814854 | 0.14582269 | 0.812 | 0.793 | 1 | 0.8663407 | -0.1567255 | 0.803 | 0.771 | 1 | 0.8663407 | 0.53507859 |
| Ccne1 | 0.85936059 | 0.12297175 | 0.073 | 0.078 | 1 | 0.31817461 | -0.9879272 | 0.038 | 0.062 | 1 | 0.85936059 | 0.53511413 |
| Rtn2 | 0.99482471 | 0.13302542 | 0.042 | 0.041 | 1 | 0.31817461 | -1.7952821 | 0.038 | 0.062 | 1 | 0.99482471 | 0.53511413 |
| Zfp69 | 0.71937765 | -0.3144336 | 0.052 | 0.062 | 1 | 0.31830141 | 0.42711033 | 0.134 | 0.094 | 1 | 0.71937765 | 0.53528704 |
| Gm45509 | 0.31848397 | 0.37006462 | 0.083 | 0.052 | 1 | 0.74536407 | -0.2457249 | 0.084 | 0.073 | 1 | 0.74536407 | 0.5355359 |
| Spidr | 0.31868674 | -0.4271337 | 0.365 | 0.415 | 1 | 0.51956809 | -0.8799903 | 0.361 | 0.344 | 1 | 0.51956809 | 0.53581224 |
| 9530056K15Rik | 0.31880524 | 0.64492446 | 0.062 | 0.036 | 1 | 0.53156006 | 0.56461386 | 0.034 | 0.021 | 1 | 0.53156006 | 0.5359737 |
| Gm15862 | 0.31880524 | 0.64492446 | 0.062 | 0.036 | 1 | 0.34428571 | 0.91253716 | 0.042 | 0.021 | 1 | 0.34428571 | 0.5359737 |
| Gm57395 | 0.31880524 | 0.64492446 | 0.062 | 0.036 | 1 | 0.66900598 | -1.0468209 | 0.042 | 0.052 | 1 | 0.66900598 | 0.5359737 |
| Gm57388 | 0.31880524 | 0.64492446 | 0.062 | 0.036 | 1 | 0.66339728 | 0.10518224 | 0.029 | 0.021 | 1 | 0.66339728 | 0.5359737 |
| Gm15927 | 0.31899281 | 0.52206771 | 0.042 | 0.021 | 1 | 0.66013325 | 0.27510724 | 0.029 | 0.021 | 1 | 0.66013325 | 0.53622921 |
| Ermard | 0.60899821 | 0.57865124 | 0.167 | 0.145 | 1 | 0.31927994 | -0.189561 | 0.176 | 0.125 | 1 | 0.60899821 | 0.5366202 |
| Nxpe4 | 0.31958317 | 0.46152617 | 0.281 | 0.228 | 1 | 0.40264357 | -0.3547095 | 0.227 | 0.177 | 1 | 0.40264357 | 0.53703294 |
| Gm17382 | 0.31987305 | 0.20013961 | 0.062 | 0.036 | 1 | 0.53386646 | 0.45567948 | 0.046 | 0.031 | 1 | 0.53386646 | 0.53742734 |
| Tmem86a | 0.89049734 | 0.30705482 | 0.052 | 0.057 | 1 | 0.31992331 | 0.44016648 | 0.109 | 0.073 | 1 | 0.89049734 | 0.5374957 |
| 4930558J18Rik | 0.32025586 | -0.577468 | 0 | 0.01 | 1 | 0.50587544 | -2.3098553 | 0.004 | 0.01 | 1 | 0.50587544 | 0.5379479 |
| Platr12 | 0.32025586 | -0.9925055 | 0 | 0.01 | 1 | 0.50887612 | -1.3098553 | 0.004 | 0.01 | 1 | 0.50887612 | 0.5379479 |
| Slc11a1 | 0.32025586 | -0.9925055 | 0 | 0.01 | 1 | 0.87066325 | -0.3098553 | 0.013 | 0.01 | 1 | 0.87066325 | 0.5379479 |
| Gm28294 | 0.32025586 | -0.577468 | 0 | 0.01 | 1 | 0.50887612 | -1.3098553 | 0.004 | 0.01 | 1 | 0.50887612 | 0.5379479 |
| Gm17634 | 0.32025586 | -0.577468 | 0 | 0.01 | 1 | 0.57170879 | -1.3098553 | 0.013 | 0.021 | 1 | 0.57170879 | 0.5379479 |
| 4933439K11Rik | 0.32025586 | -0.9925055 | 0 | 0.01 | 1 | 0.87851725 | -0.8948178 | 0.013 | 0.01 | 1 | 0.87851725 | 0.5379479 |
| Vsig8 | 0.32025586 | -0.9925055 | 0 | 0.01 | 1 | 0.98993389 | 0.27510724 | 0.021 | 0.021 | 1 | 0.98993389 | 0.5379479 |
| Dnah14 | 0.32025586 | -1.3144336 | 0 | 0.01 | 1 | 0.40820031 | -1.6883669 | 0.025 | 0.042 | 1 | 0.40820031 | 0.5379479 |
| Card9 | 0.32025586 | -0.577468 | 0 | 0.01 | 1 | 0.86312513 | -0.7248928 | 0.008 | 0.01 | 1 | 0.86312513 | 0.5379479 |
| Cel | 0.32025586 | -0.577468 | 0 | 0.01 | 1 | 0.86312513 | -0.7248928 | 0.008 | 0.01 | 1 | 0.86312513 | 0.5379479 |
| Cerkl | 0.32025586 | -0.577468 | 0 | 0.01 | 1 | 0.34704016 | -1.3098553 | 0.008 | 0.021 | 1 | 0.34704016 | 0.5379479 |
| 4930412B13Rik | 0.32025586 | -1.577468 | 0 | 0.01 | 1 | 0.50887612 | -1.3098553 | 0.004 | 0.01 | 1 | 0.50887612 | 0.5379479 |
| Ell3 | 0.32025586 | -0.577468 | 0 | 0.01 | 1 | 0.50887612 | -1.3098553 | 0.004 | 0.01 | 1 | 0.50887612 | 0.5379479 |

| | | | | | | | | | | | | |
|---|---|---|---|---|---|---|---|---|---|---|---|---|
| Atp8b4 | 0.32025586 | -1.577468 | 0 | 0.01 | 1 | 0.51188611 | 0.27510724 | 0.004 | 0.01 | 1 | 0.51188611 | 0.5379479 |
| Fermt1 | 0.32025586 | -0.9925055 | 0 | 0.01 | 1 | 0.51018266 | 0.49749966 | 0.021 | 0.01 | 1 | 0.51018266 | 0.5379479 |
| Gm57459 | 0.32025586 | -0.577468 | 0 | 0.01 | 1 | 0.5818708 | -0.3098553 | 0.013 | 0.021 | 1 | 0.5818708 | 0.5379479 |
| Slc2a2 | 0.32025586 | -0.9925055 | 0 | 0.01 | 1 | 0.50887612 | -1.3098553 | 0.004 | 0.01 | 1 | 0.50887612 | 0.5379479 |
| P2ry12 | 0.32025586 | -1.3144336 | 0 | 0.01 | 1 | 0.67471359 | -0.5728897 | 0.017 | 0.01 | 1 | 0.67471359 | 0.5379479 |
| Sit1 | 0.32025586 | -0.577468 | 0 | 0.01 | 1 | 0.50887612 | -1.3098553 | 0.004 | 0.01 | 1 | 0.50887612 | 0.5379479 |
| Car9 | 0.32025586 | -0.577468 | 0 | 0.01 | 1 | 0.50587544 | -1.8948178 | 0.004 | 0.01 | 1 | 0.50587544 | 0.5379479 |
| Gm12678 | 0.32025586 | -0.577468 | 0 | 0.01 | 1 | 0.51188611 | -0.7248928 | 0.004 | 0.01 | 1 | 0.51188611 | 0.5379479 |
| Wdr31 | 0.32025586 | -0.577468 | 0 | 0.01 | 1 | 0.86312513 | -0.7248928 | 0.008 | 0.01 | 1 | 0.86312513 | 0.5379479 |
| Gm42303 | 0.32025586 | -4.1217885 | 0 | 0.01 | 1 | 0.87851807 | -0.7248928 | 0.013 | 0.01 | 1 | 0.87851807 | 0.5379479 |
| Tek | 0.32025586 | -0.577468 | 0 | 0.01 | 1 | 0.34363875 | -2.0468209 | 0.008 | 0.021 | 1 | 0.34363875 | 0.5379479 |
| Tex46 | 0.32025586 | -0.577468 | 0 | 0.01 | 1 | 0.50887612 | -1.3098553 | 0.004 | 0.01 | 1 | 0.50887612 | 0.5379479 |
| Gm57064 | 0.32025586 | -1.7998604 | 0 | 0.01 | 1 | 1 | -0.8244284 | 0.021 | 0.021 | 1 | 1 | 0.5379479 |
| Trp73 | 0.32025586 | -1.577468 | 0 | 0.01 | 1 | 0.86312513 | -0.7248928 | 0.008 | 0.01 | 1 | 0.86312513 | 0.5379479 |
| 4932441J04Rik | 0.32025586 | -0.577468 | 0 | 0.01 | 1 | 0.66387437 | 0.27510724 | 0.017 | 0.01 | 1 | 0.66387437 | 0.5379479 |
| Chrna9 | 0.32025586 | -0.577468 | 0 | 0.01 | 1 | 0.40360435 | -0.8244284 | 0.025 | 0.01 | 1 | 0.40360435 | 0.5379479 |
| Tmprss11d | 0.32025586 | -1.577468 | 0 | 0.01 | 1 | 0.50887612 | -1.3098553 | 0.004 | 0.01 | 1 | 0.50887612 | 0.5379479 |
| Tmem150c | 0.32025586 | -0.9925055 | 0 | 0.01 | 1 | 0.41182291 | -1.4473588 | 0.025 | 0.042 | 1 | 0.41182291 | 0.5379479 |
| Gfi1 | 0.32025586 | -1.3144336 | 0 | 0.01 | 1 | 0.50887612 | -1.3098553 | 0.004 | 0.01 | 1 | 0.50887612 | 0.5379479 |
| Galnt9 | 0.32025586 | -0.9925055 | 0 | 0.01 | 1 | 0.50587544 | -3.3098553 | 0.004 | 0.01 | 1 | 0.50587544 | 0.5379479 |
| Dao | 0.32025586 | -0.577468 | 0 | 0.01 | 1 | 0.51194506 | 0.27510724 | 0.021 | 0.01 | 1 | 0.51194506 | 0.5379479 |
| Gm14508 | 0.32025586 | -0.9925055 | 0 | 0.01 | 1 | 0.87066325 | -0.3098553 | 0.013 | 0.01 | 1 | 0.87066325 | 0.5379479 |
| Upk3b | 0.32025586 | -0.577468 | 0 | 0.01 | 1 | 0.50887612 | -1.3098553 | 0.004 | 0.01 | 1 | 0.50887612 | 0.5379479 |
| Nptx2 | 0.32025586 | -0.9925055 | 0 | 0.01 | 1 | 0.5778028 | -0.8948178 | 0.013 | 0.021 | 1 | 0.5778028 | 0.5379479 |
| 4930540M05Rik | 0.32025586 | -0.577468 | 0 | 0.01 | 1 | 0.57170879 | -1.3098553 | 0.013 | 0.021 | 1 | 0.57170879 | 0.5379479 |
| Gm10010 | 0.32025586 | -1.3144336 | 0 | 0.01 | 1 | 0.86613737 | -0.3098553 | 0.008 | 0.01 | 1 | 0.86613737 | 0.5379479 |
| Tnnt1 | 0.32025586 | -0.577468 | 0 | 0.01 | 1 | 0.5778028 | -0.8948178 | 0.013 | 0.021 | 1 | 0.5778028 | 0.5379479 |
| Ptprh | 0.32025586 | -0.9925055 | 0 | 0.01 | 1 | 0.58664899 | -0.3098553 | 0.021 | 0.031 | 1 | 0.58664899 | 0.5379479 |
| Gm26852 | 0.32025586 | -0.577468 | 0 | 0.01 | 1 | 0.52072676 | -0.7248928 | 0.021 | 0.01 | 1 | 0.52072676 | 0.5379479 |
| Hipk4 | 0.32025586 | -0.9925055 | 0 | 0.01 | 1 | 0.56568674 | -1.8948178 | 0.021 | 0.031 | 1 | 0.56568674 | 0.5379479 |
| Gm44997 | 0.32025586 | -1.3144336 | 0 | 0.01 | 1 | 1 | -0.7248928 | 0.021 | 0.021 | 1 | 1 | 0.5379479 |
| Klk9 | 0.32025586 | -0.9925055 | 0 | 0.01 | 1 | 0.86543325 | 0.27510724 | 0.013 | 0.01 | 1 | 0.86543325 | 0.5379479 |
| 5430431A17Rik | 0.32025586 | -0.577468 | 0 | 0.01 | 1 | 0.87851725 | -1.3098553 | 0.013 | 0.01 | 1 | 0.87851725 | 0.5379479 |
| Gm57226 | 0.32025586 | -0.577468 | 0 | 0.01 | 1 | 0.86312513 | -0.7248928 | 0.008 | 0.01 | 1 | 0.86312513 | 0.5379479 |
| Tmc3 | 0.32025586 | -0.577468 | 0 | 0.01 | 1 | 0.50887612 | -1.3098553 | 0.004 | 0.01 | 1 | 0.50887612 | 0.5379479 |
| Gm44993 | 0.32025586 | -2.3144336 | 0 | 0.01 | 1 | 0.50887612 | -1.3098553 | 0.004 | 0.01 | 1 | 0.50887612 | 0.5379479 |
| Rab38 | 0.32025586 | -0.577468 | 0 | 0.01 | 1 | 0.38679371 | 2.21370669 | 0.025 | 0.01 | 1 | 0.38679371 | 0.5379479 |
| Ascl3 | 0.32025586 | -0.9925055 | 0 | 0.01 | 1 | 0.50587544 | -2.3098553 | 0.004 | 0.01 | 1 | 0.50587544 | 0.5379479 |
| Gm32884 | 0.32025586 | -2.4519371 | 0 | 0.01 | 1 | 0.86543325 | 1.39058446 | 0.013 | 0.01 | 1 | 0.86543325 | 0.5379479 |
| Brsk2 | 0.32025586 | -0.9925055 | 0 | 0.01 | 1 | 0.50887612 | -1.3098553 | 0.004 | 0.01 | 1 | 0.50887612 | 0.5379479 |
| Gm6410 | 0.32025586 | -0.9925055 | 0 | 0.01 | 1 | 0.86011448 | -0.9879272 | 0.008 | 0.01 | 1 | 0.86011448 | 0.5379479 |
| Dlgap2 | 0.32025586 | -1.3144336 | 0 | 0.01 | 1 | 0.50887612 | -1.3098553 | 0.004 | 0.01 | 1 | 0.50887612 | 0.5379479 |
| Gm45370 | 0.32025586 | -0.577468 | 0 | 0.01 | 1 | 0.42648204 | -0.8244284 | 0.025 | 0.042 | 1 | 0.42648204 | 0.5379479 |
| B230317F23Rik | 0.32025586 | -1.3144336 | 0 | 0.01 | 1 | 0.86613737 | -0.3098553 | 0.008 | 0.01 | 1 | 0.86613737 | 0.5379479 |
| Plvap | 0.32025586 | -0.577468 | 0 | 0.01 | 1 | 0.51188611 | -0.7248928 | 0.004 | 0.01 | 1 | 0.51188611 | 0.5379479 |
| Slc12a3 | 0.32025586 | -0.577468 | 0 | 0.01 | 1 | 0.34026631 | -2.8948178 | 0.008 | 0.021 | 1 | 0.34026631 | 0.5379479 |
| 1810019D21Rik | 0.32025586 | -0.577468 | 0 | 0.01 | 1 | 0.50887612 | -1.3098553 | 0.004 | 0.01 | 1 | 0.50887612 | 0.5379479 |
| Cdh3 | 0.32025586 | -0.577468 | 0 | 0.01 | 1 | 0.34363875 | -2.0468209 | 0.008 | 0.021 | 1 | 0.34363875 | 0.5379479 |
| Has3 | 0.32025586 | -0.577468 | 0 | 0.01 | 1 | 0.87851725 | -1.3098553 | 0.013 | 0.01 | 1 | 0.87851725 | 0.5379479 |
| Clec18a | 0.32025586 | -0.9925055 | 0 | 0.01 | 1 | 0.51188611 | -0.7248928 | 0.004 | 0.01 | 1 | 0.51188611 | 0.5379479 |
| Vat1l | 0.32025586 | -2.3144336 | 0 | 0.01 | 1 | 0.86312513 | -0.7248928 | 0.008 | 0.01 | 1 | 0.86312513 | 0.5379479 |
| Pgr | 0.32025586 | -0.577468 | 0 | 0.01 | 1 | 0.50587544 | -1.8948178 | 0.004 | 0.01 | 1 | 0.50587544 | 0.5379479 |
| Gm47557 | 0.32025586 | -0.577468 | 0 | 0.01 | 1 | 0.50887612 | -1.3098553 | 0.004 | 0.01 | 1 | 0.50887612 | 0.5379479 |
| Gm47326 | 0.32025586 | -0.577468 | 0 | 0.01 | 1 | 0.87066325 | -0.3098553 | 0.013 | 0.01 | 1 | 0.87066325 | 0.5379479 |
| Gm16754 | 0.32025586 | -0.577468 | 0 | 0.01 | 1 | 0.87851807 | -1.3098553 | 0.013 | 0.01 | 1 | 0.87851807 | 0.5379479 |
| Ap1m2 | 0.32025586 | -0.577468 | 0 | 0.01 | 1 | 0.50887612 | -1.3098553 | 0.004 | 0.01 | 1 | 0.50887612 | 0.5379479 |
| Zfp810 | 0.32025586 | -0.577468 | 0 | 0.01 | 1 | 0.35045411 | -0.8948178 | 0.008 | 0.021 | 1 | 0.35045411 | 0.5379479 |
| Crtam | 0.32025586 | -0.577468 | 0 | 0.01 | 1 | 0.86312513 | -0.7248928 | 0.008 | 0.01 | 1 | 0.86312513 | 0.5379479 |
| Tnfaip8l3 | 0.32025586 | -0.9925055 | 0 | 0.01 | 1 | 0.50587544 | -2.3098553 | 0.004 | 0.01 | 1 | 0.50587544 | 0.5379479 |
| Gldn | 0.32025586 | -2.3144336 | 0 | 0.01 | 1 | 0.85710544 | -2.3098553 | 0.008 | 0.01 | 1 | 0.85710544 | 0.5379479 |
| Nr2e3 | 0.32025586 | -0.9925055 | 0 | 0.01 | 1 | 0.42645769 | -1.1399303 | 0.025 | 0.042 | 1 | 0.42645769 | 0.5379479 |
| C2cd4a | 0.32025586 | -1.7998604 | 0 | 0.01 | 1 | 0.50587544 | -1.8948178 | 0.004 | 0.01 | 1 | 0.50587544 | 0.5379479 |
| 5730403I07Rik | 0.32025586 | -0.9925055 | 0 | 0.01 | 1 | 0.87851725 | -1.3098553 | 0.013 | 0.01 | 1 | 0.87851725 | 0.5379479 |
| Ngp | 0.32025586 | -0.9925055 | 0 | 0.01 | 1 | 0.95136123 | -0.5163061 | 0.092 | 0.094 | 1 | 0.95136123 | 0.5379479 |
| Clec3b | 0.32025586 | -0.577468 | 0 | 0.01 | 1 | 0.85710544 | -1.7248928 | 0.008 | 0.01 | 1 | 0.85710544 | 0.5379479 |
| Gm28905 | 0.32025586 | -0.9925055 | 0 | 0.01 | 1 | 0.87851807 | -0.7248928 | 0.013 | 0.01 | 1 | 0.87851807 | 0.5379479 |
| Gm48084 | 0.32025586 | -0.9925055 | 0 | 0.01 | 1 | 0.87851725 | -1.3098553 | 0.013 | 0.01 | 1 | 0.87851725 | 0.5379479 |
| Gm26535 | 0.32025586 | -0.577468 | 0 | 0.01 | 1 | 0.79600331 | -1.3098553 | 0.017 | 0.021 | 1 | 0.79600331 | 0.5379479 |
| 4930547M16Rik | 0.32025586 | -0.577468 | 0 | 0.01 | 1 | 0.51188611 | -0.7248928 | 0.004 | 0.01 | 1 | 0.51188611 | 0.5379479 |
| Mfsd4b1 | 0.32025586 | -1.7998604 | 0 | 0.01 | 1 | 0.81864396 | -0.1399303 | 0.025 | 0.021 | 1 | 0.81864396 | 0.5379479 |
| Lilrb4a | 0.32025586 | -1.577468 | 0 | 0.01 | 1 | 0.86613737 | 1.01207283 | 0.008 | 0.01 | 1 | 0.86613737 | 0.5379479 |
| B4galnt1 | 0.32025586 | -0.577468 | 0 | 0.01 | 1 | 0.67415878 | -0.6577786 | 0.042 | 0.031 | 1 | 0.67415878 | 0.5379479 |
| Havcr2 | 0.32025586 | -1.7998604 | 0 | 0.01 | 1 | 0.86312513 | -0.7248928 | 0.008 | 0.01 | 1 | 0.86312513 | 0.5379479 |
| Gm46280 | 0.32025586 | -0.577468 | 0 | 0.01 | 1 | 0.56565662 | -2.1172102 | 0.013 | 0.021 | 1 | 0.56565662 | 0.5379479 |
| Mpp3 | 0.32025586 | -0.9925055 | 0 | 0.01 | 1 | 0.51188611 | -0.7248928 | 0.004 | 0.01 | 1 | 0.51188611 | 0.5379479 |
| Prcd | 0.32025586 | -0.577468 | 0 | 0.01 | 1 | 0.50587544 | -2.8948178 | 0.004 | 0.01 | 1 | 0.50587544 | 0.5379479 |
| Dcdc2c | 0.32025586 | -1.3144336 | 0 | 0.01 | 1 | 0.87066325 | -0.3098553 | 0.013 | 0.01 | 1 | 0.87066325 | 0.5379479 |
| Gm48236 | 0.32025586 | -0.9925055 | 0 | 0.01 | 1 | 0.50887612 | -1.3098553 | 0.004 | 0.01 | 1 | 0.50887612 | 0.5379479 |
| Gm56554 | 0.32025586 | -0.577468 | 0 | 0.01 | 1 | 0.50587544 | -1.8948178 | 0.004 | 0.01 | 1 | 0.50587544 | 0.5379479 |
| Lrfn5 | 0.32025586 | -0.577468 | 0 | 0.01 | 1 | 0.86613737 | 0.49749966 | 0.008 | 0.01 | 1 | 0.86613737 | 0.5379479 |
| Tmem229b | 0.32025586 | -0.577468 | 0 | 0.01 | 1 | 0.82415676 | -1.1172102 | 0.025 | 0.021 | 1 | 0.82415676 | 0.5379479 |
| Ttc9 | 0.32025586 | -1.7998604 | 0 | 0.01 | 1 | 0.39515719 | 0.49749966 | 0.025 | 0.01 | 1 | 0.39515719 | 0.5379479 |

| | | | | | | | | | | | | |
|---|---|---|---|---|---|---|---|---|---|---|---|---|
| Akr1c18 | 0.32025586 | -0.9925055 | 0 | 0.01 | 1 | 0.86312513 | -0.7248928 | 0.008 | 0.01 | 1 | 0.86312513 | 0.5379479 |
| Rbm24 | 0.32025586 | -1.9925055 | 0 | 0.01 | 1 | 0.80442048 | -0.5728897 | 0.017 | 0.021 | 1 | 0.80442048 | 0.5379479 |
| Zfp459 | 0.32025586 | -0.577468 | 0 | 0.01 | 1 | 0.86613737 | -0.3098553 | 0.008 | 0.01 | 1 | 0.86613737 | 0.5379479 |
| Gm35161 | 0.32025586 | -0.577468 | 0 | 0.01 | 1 | 0.50844268 | 0.69014474 | 0.021 | 0.01 | 1 | 0.50844268 | 0.5379479 |
| Gm47849 | 0.32025586 | -0.577468 | 0 | 0.01 | 1 | 0.50887612 | -1.3098553 | 0.004 | 0.01 | 1 | 0.50887612 | 0.5379479 |
| 4921509O07Rik | 0.32025586 | -0.577468 | 0 | 0.01 | 1 | 0.50887612 | -1.3098553 | 0.004 | 0.01 | 1 | 0.50887612 | 0.5379479 |
| Gm17509 | 0.32025586 | -0.577468 | 0 | 0.01 | 1 | 0.87066325 | -0.3098553 | 0.013 | 0.01 | 1 | 0.87066325 | 0.5379479 |
| Trdv4 | 0.32025586 | -0.577468 | 0 | 0.01 | 1 | 0.86312513 | -0.7248928 | 0.008 | 0.01 | 1 | 0.86312513 | 0.5379479 |
| Gm16549 | 0.32025586 | -0.577468 | 0 | 0.01 | 1 | 0.52072676 | -0.7248928 | 0.021 | 0.01 | 1 | 0.52072676 | 0.5379479 |
| Fbxo43 | 0.32025586 | -0.9925055 | 0 | 0.01 | 1 | 0.34533788 | -2.1172102 | 0.008 | 0.021 | 1 | 0.34533788 | 0.5379479 |
| Gm10373 | 0.32025586 | -0.9925055 | 0 | 0.01 | 1 | 0.86312513 | -0.7248928 | 0.008 | 0.01 | 1 | 0.86312513 | 0.5379479 |
| Parp10 | 0.32025586 | -0.9925055 | 0 | 0.01 | 1 | 0.56110733 | -0.9879272 | 0.029 | 0.042 | 1 | 0.56110733 | 0.5379479 |
| Card10 | 0.32025586 | -0.577468 | 0 | 0.01 | 1 | 0.82231716 | -0.3098553 | 0.025 | 0.021 | 1 | 0.82231716 | 0.5379479 |
| Gm17753 | 0.32025586 | -0.577468 | 0 | 0.01 | 1 | 0.51720983 | 0.10518224 | 0.021 | 0.01 | 1 | 0.51720983 | 0.5379479 |
| Parvb | 0.32025586 | -0.9925055 | 0 | 0.01 | 1 | 0.75298375 | -0.6577786 | 0.025 | 0.031 | 1 | 0.75298375 | 0.5379479 |
| Gm15722 | 0.32025586 | -0.9925055 | 0 | 0.01 | 1 | 0.50887612 | -1.3098553 | 0.004 | 0.01 | 1 | 0.50887612 | 0.5379479 |
| Kcnh3 | 0.32025586 | -0.9925055 | 0 | 0.01 | 1 | 0.85710544 | -1.7248928 | 0.008 | 0.01 | 1 | 0.85710544 | 0.5379479 |
| Fignl2 | 0.32025586 | -0.577468 | 0 | 0.01 | 1 | 0.995619 | -0.9562183 | 0.076 | 0.073 | 1 | 0.995619 | 0.5379479 |
| Krt78 | 0.32025586 | -0.577468 | 0 | 0.01 | 1 | 0.50887612 | -1.3098553 | 0.004 | 0.01 | 1 | 0.50887612 | 0.5379479 |
| Septin12 | 0.32025586 | -0.577468 | 0 | 0.01 | 1 | 0.50887612 | -1.3098553 | 0.004 | 0.01 | 1 | 0.50887612 | 0.5379479 |
| Gm15558 | 0.32025586 | -0.9925055 | 0 | 0.01 | 1 | 0.50887612 | -1.3098553 | 0.004 | 0.01 | 1 | 0.50887612 | 0.5379479 |
| Tm4sf19 | 0.32025586 | -0.9925055 | 0 | 0.01 | 1 | 0.99194729 | -1.6724253 | 0.021 | 0.021 | 1 | 0.99194729 | 0.5379479 |
| Fbxo40 | 0.32025586 | -0.9925055 | 0 | 0.01 | 1 | 0.50587544 | -2.3098553 | 0.004 | 0.01 | 1 | 0.50587544 | 0.5379479 |
| Gm38477 | 0.32025586 | -0.9925055 | 0 | 0.01 | 1 | 0.51188611 | 0.69014474 | 0.004 | 0.01 | 1 | 0.51188611 | 0.5379479 |
| Gm15965 | 0.32025586 | -0.577468 | 0 | 0.01 | 1 | 0.34026631 | -2.3098553 | 0.008 | 0.021 | 1 | 0.34026631 | 0.5379479 |
| Dact2 | 0.32025586 | -1.3144336 | 0 | 0.01 | 1 | 0.50887612 | -1.3098553 | 0.004 | 0.01 | 1 | 0.50887612 | 0.5379479 |
| Fpr3 | 0.32025586 | -1.3144336 | 0 | 0.01 | 1 | 0.85710544 | -1.7248928 | 0.008 | 0.01 | 1 | 0.85710544 | 0.5379479 |
| Rab44 | 0.32025586 | -1.7998604 | 0 | 0.01 | 1 | 0.87066325 | -0.3098553 | 0.013 | 0.01 | 1 | 0.87066325 | 0.5379479 |
| Pi16 | 0.32025586 | -0.9925055 | 0 | 0.01 | 1 | 0.50887612 | -1.3098553 | 0.004 | 0.01 | 1 | 0.50887612 | 0.5379479 |
| Dnah8 | 0.32025586 | -1.577468 | 0 | 0.01 | 1 | 0.92060545 | -0.5322477 | 0.029 | 0.031 | 1 | 0.92060545 | 0.5379479 |
| Tnxb | 0.32025586 | -0.577468 | 0 | 0.01 | 1 | 0.86011448 | -0.8948178 | 0.008 | 0.01 | 1 | 0.86011448 | 0.5379479 |
| Adgre1 | 0.32025586 | -0.9925055 | 0 | 0.01 | 1 | 0.50887612 | -1.3098553 | 0.004 | 0.01 | 1 | 0.50887612 | 0.5379479 |
| Rasgrp3 | 0.32025586 | -0.577468 | 0 | 0.01 | 1 | 0.50887612 | -1.3098553 | 0.004 | 0.01 | 1 | 0.50887612 | 0.5379479 |
| Gm50034 | 0.32025586 | -0.577468 | 0 | 0.01 | 1 | 0.86312513 | -0.7248928 | 0.008 | 0.01 | 1 | 0.86312513 | 0.5379479 |
| Kcng3 | 0.32025586 | -0.9925055 | 0 | 0.01 | 1 | 0.86613737 | -0.3098553 | 0.008 | 0.01 | 1 | 0.86613737 | 0.5379479 |
| Nol4 | 0.32025586 | -1.3144336 | 0 | 0.01 | 1 | 0.78762065 | -2.6317834 | 0.017 | 0.021 | 1 | 0.78762065 | 0.5379479 |
| Sh3rf2 | 0.32025586 | -0.577468 | 0 | 0.01 | 1 | 0.99194729 | -2.0874628 | 0.021 | 0.021 | 1 | 0.99194729 | 0.5379479 |
| Sptbn2 | 0.32025586 | -0.577468 | 0 | 0.01 | 1 | 0.5207331 | -0.5025003 | 0.021 | 0.01 | 1 | 0.5207331 | 0.5379479 |
| Pdcd1lg2 | 0.32025586 | -0.9925055 | 0 | 0.01 | 1 | 0.50887612 | -1.3098553 | 0.004 | 0.01 | 1 | 0.50887612 | 0.5379479 |
| Zcchc12 | 0.32025586 | -0.577468 | 0 | 0.01 | 1 | 0.86312513 | -0.7248928 | 0.008 | 0.01 | 1 | 0.86312513 | 0.5379479 |
| Gm14827 | 0.32025586 | -0.577468 | 0 | 0.01 | 1 | 0.34363875 | -1.7248928 | 0.008 | 0.021 | 1 | 0.34363875 | 0.5379479 |
| Asb11 | 0.32025586 | -0.9925055 | 0 | 0.01 | 1 | 0.87066325 | -0.3098553 | 0.013 | 0.01 | 1 | 0.87066325 | 0.5379479 |
| Wdpcp | 0.42978675 | 0.19458009 | 0.271 | 0.228 | 1 | 0.32043531 | -1.1780654 | 0.239 | 0.281 | 1 | 0.42978675 | 0.53819184 |
| Lrsam1 | 0.95200833 | 0.25260703 | 0.115 | 0.119 | 1 | 0.32048005 | 0.20799304 | 0.206 | 0.156 | 1 | 0.95200833 | 0.53825263 |
| Shq1 | 0.65249135 | 0.42253204 | 0.104 | 0.088 | 1 | 0.32114612 | 0.23763253 | 0.109 | 0.073 | 1 | 0.65249135 | 0.53915741 |
| Dzip3 | 0.69304537 | 0.17431363 | 0.385 | 0.43 | 1 | 0.32118166 | -0.317804 | 0.408 | 0.312 | 1 | 0.69304537 | 0.53920566 |
| Garem1 | 0.3702202 | -0.6851818 | 0.521 | 0.544 | 1 | 0.32132618 | 0.30587959 | 0.588 | 0.552 | 1 | 0.3702202 | 0.53940185 |
| Ankmy1 | 0.41766676 | 0.6241659 | 0.24 | 0.202 | 1 | 0.32135362 | -0.4797803 | 0.218 | 0.156 | 1 | 0.41766676 | 0.5394391 |
| Nrxn2 | 0.7645474 | -0.130009 | 0.073 | 0.083 | 1 | 0.32135373 | -0.3721395 | 0.244 | 0.292 | 1 | 0.7645474 | 0.53943924 |
| Adam32 | 0.39385604 | -0.7998604 | 0.073 | 0.104 | 1 | 0.3218837 | -1.8244284 | 0.092 | 0.125 | 1 | 0.39385604 | 0.54015828 |
| Tbc1d7 | 0.97501874 | -0.2295447 | 0.125 | 0.124 | 1 | 0.32256427 | 0.37820073 | 0.109 | 0.073 | 1 | 0.97501874 | 0.54108083 |
| Osbp2 | 0.80611646 | -0.8993961 | 0.031 | 0.036 | 1 | 0.32304157 | -0.2103196 | 0.059 | 0.031 | 1 | 0.80611646 | 0.54172728 |
| Gart | 0.32331788 | 0.38936517 | 0.312 | 0.259 | 1 | 0.43749728 | -0.2083172 | 0.303 | 0.24 | 1 | 0.43749728 | 0.54210131 |
| Zfp438 | 0.69621191 | -0.3710171 | 0.156 | 0.171 | 1 | 0.32345926 | -0.6147098 | 0.08 | 0.115 | 1 | 0.69621191 | 0.54229262 |
| Dpysl4 | 0.47599549 | 0.32942263 | 0.042 | 0.026 | 1 | 0.3235078 | 0.40635177 | 0.084 | 0.052 | 1 | 0.47599549 | 0.54235831 |
| Adcy10 | 0.32399159 | -1.4519371 | 0.031 | 0.057 | 1 | 0.40705175 | -1.4093909 | 0.042 | 0.062 | 1 | 0.40705175 | 0.54301263 |
| 4833422C13Rik | 0.32399159 | -1.3144336 | 0.031 | 0.057 | 1 | 0.51194506 | 0.27510724 | 0.021 | 0.01 | 1 | 0.51194506 | 0.54301263 |
| Rims1 | 0.3239929 | -2.1217885 | 0.031 | 0.057 | 1 | 0.6934539 | -1.3973181 | 0.034 | 0.042 | 1 | 0.6934539 | 0.5430144 |
| Slc8a3 | 0.44753953 | 1.19715576 | 0.458 | 0.477 | 1 | 0.32457491 | -0.1795472 | 0.286 | 0.323 | 1 | 0.44753953 | 0.54380094 |
| Gm7072 | 0.32471207 | 0.40177348 | 0.323 | 0.275 | 1 | 0.81700513 | -0.5916262 | 0.151 | 0.156 | 1 | 0.81700513 | 0.5439862 |
| Irgm2 | 0.90945802 | -0.2148979 | 0.177 | 0.176 | 1 | 0.32479658 | -1.1214102 | 0.143 | 0.177 | 1 | 0.90945802 | 0.54410034 |
| Clec2l | 0.3249356 | 0.49292136 | 0.062 | 0.036 | 1 | 0.93576656 | -0.1399303 | 0.029 | 0.031 | 1 | 0.93576656 | 0.54428806 |
| ENSMUSG00000121476 | 0.32498038 | 0.20013961 | 0.062 | 0.036 | 1 | 0.79185213 | -0.7248928 | 0.088 | 0.094 | 1 | 0.79185213 | 0.54434851 |
| Syt17 | 0.55458034 | -0.3550755 | 0.094 | 0.073 | 1 | 0.32513368 | -1.7952821 | 0.067 | 0.094 | 1 | 0.55458034 | 0.54455546 |
| Cchcr1 | 0.37520507 | 0.51272984 | 0.156 | 0.119 | 1 | 0.32565455 | 0.22065945 | 0.097 | 0.062 | 1 | 0.37520507 | 0.54525821 |
| Agap3 | 0.32584712 | 0.11996927 | 0.302 | 0.238 | 1 | 0.56387966 | -0.7108176 | 0.542 | 0.49 | 1 | 0.56387966 | 0.54551789 |
| D6Wsu163e | 0.46738434 | -0.1624305 | 0.062 | 0.088 | 1 | 0.3262517 | -0.3623227 | 0.088 | 0.125 | 1 | 0.46738434 | 0.54606323 |
| Slc16a2 | 0.32633757 | 0.26198383 | 0.688 | 0.674 | 1 | 0.86807277 | -0.5490429 | 0.891 | 0.833 | 1 | 0.86807277 | 0.54617894 |
| Alg8 | 0.32680295 | -0.3144336 | 0.083 | 0.124 | 1 | 0.33336882 | -1.3098553 | 0.084 | 0.115 | 1 | 0.33336882 | 0.54680574 |
| Frrs1 | 0.32693714 | -1.0799683 | 0.031 | 0.057 | 1 | 0.66236078 | 1.06211351 | 0.113 | 0.135 | 1 | 0.66236078 | 0.54698638 |
| Gngt2 | 0.32693844 | -1.3848229 | 0.031 | 0.057 | 1 | 0.35231982 | 0.13071733 | 0.042 | 0.021 | 1 | 0.35231982 | 0.54698814 |
| Abcb1a | 0.32705737 | 0.37006462 | 0.062 | 0.036 | 1 | 0.87851807 | -1.5728897 | 0.013 | 0.01 | 1 | 0.87851807 | 0.54714821 |
| Cpsf4l | 0.32742803 | -0.7294711 | 0.073 | 0.109 | 1 | 0.42315514 | -0.8792209 | 0.088 | 0.115 | 1 | 0.42315514 | 0.54764695 |
| Slc3a1 | 0.79196846 | -0.3144336 | 0.021 | 0.026 | 1 | 0.32758926 | -0.5728897 | 0.071 | 0.104 | 1 | 0.79196846 | 0.54786379 |
| Them4 | 0.32799428 | 0.59245704 | 0.198 | 0.155 | 1 | 0.50104836 | 0.49749966 | 0.118 | 0.094 | 1 | 0.50104836 | 0.54840831 |
| Acadl | 0.52649921 | 0.18374518 | 0.562 | 0.518 | 1 | 0.32801107 | 0.1566484 | 0.765 | 0.74 | 1 | 0.52649921 | 0.54843088 |
| Bdkrb2 | 0.32807662 | 0.37006462 | 0.062 | 0.036 | 1 | 0.39796128 | -2.1843244 | 0.017 | 0.031 | 1 | 0.39796128 | 0.54851897 |
| Cys1 | 0.99145797 | -0.1624305 | 0.062 | 0.062 | 1 | 0.3280914 | -0.9472852 | 0.092 | 0.125 | 1 | 0.99145797 | 0.54853883 |
| Pnkp | 0.32877223 | 0.59245704 | 0.135 | 0.098 | 1 | 0.86537225 | -0.2887936 | 0.218 | 0.219 | 1 | 0.86537225 | 0.54945327 |
| Pxk | 0.32881923 | 0.22048826 | 0.312 | 0.254 | 1 | 0.54941071 | -0.1836782 | 0.471 | 0.521 | 1 | 0.54941071 | 0.54951637 |
| Akap1 | 0.3290243 | 0.44561565 | 0.167 | 0.124 | 1 | 0.65180719 | -0.4618584 | 0.143 | 0.156 | 1 | 0.65180719 | 0.5497916 |
| Brca2 | 0.97968795 | -1.0414151 | 0.135 | 0.13 | 1 | 0.32902926 | -0.4208866 | 0.063 | 0.094 | 1 | 0.97968795 | 0.54979827 |
| 4933424G05Rik | 0.3336854 | 0.66645762 | 0.094 | 0.062 | 1 | 0.32903516 | 0.39058446 | 0.084 | 0.052 | 1 | 0.3336854 | 0.54980618 |
| Glod4 | 0.32907457 | 0.22706091 | 0.49 | 0.451 | 1 | 0.48069675 | -0.183478 | 0.445 | 0.51 | 1 | 0.48069675 | 0.54985906 |
| Fam167a | 0.32924416 | 0.35394495 | 0.323 | 0.275 | 1 | 0.67293322 | -0.5794623 | 0.164 | 0.135 | 1 | 0.67293322 | 0.55008661 |
| Nek3 | 0.32941367 | 0.68556644 | 0.094 | 0.062 | 1 | 0.41318145 | -0.486733 | 0.067 | 0.094 | 1 | 0.41318145 | 0.55031397 |

| | | | | | | | | | | | | |
|---|---|---|---|---|---|---|---|---|---|---|---|---|
| Spry4 | 0.44494194 | -0.7998604 | 0.073 | 0.098 | 1 | 0.32942598 | -0.5936482 | 0.071 | 0.104 | 1 | 0.44494194 | 0.55033048 |
| Gm57457 | 0.61816868 | 0.42253204 | 0.083 | 0.067 | 1 | 0.32988376 | -0.7248928 | 0.046 | 0.073 | 1 | 0.61816868 | 0.55094422 |
| 4933412O06Rik | 0.32989765 | -1.0799683 | 0.031 | 0.057 | 1 | 0.85710544 | -1.3098553 | 0.008 | 0.01 | 1 | 0.85710544 | 0.55096284 |
| Zfp697 | 0.33019613 | 0.32942263 | 0.115 | 0.166 | 1 | 0.87549616 | -0.9313436 | 0.13 | 0.115 | 1 | 0.87549616 | 0.55136277 |
| A930015D03Rik | 0.33024094 | 0.22988696 | 0.542 | 0.477 | 1 | 0.84270124 | -0.2285552 | 0.592 | 0.552 | 1 | 0.84270124 | 0.5514228 |
| Clcn5 | 0.33040374 | -0.6599301 | 0.323 | 0.373 | 1 | 0.7828355 | -0.4484605 | 0.441 | 0.406 | 1 | 0.7828355 | 0.55164085 |
| Eif2ak3 | 0.3305196 | 0.41121672 | 0.5 | 0.461 | 1 | 0.64979115 | -0.1214877 | 0.672 | 0.635 | 1 | 0.64979115 | 0.55179599 |
| Hsf2 | 0.33074935 | 0.32468471 | 0.26 | 0.202 | 1 | 0.44713621 | -0.3001401 | 0.391 | 0.302 | 1 | 0.44713621 | 0.55210356 |
| Gm10602 | 0.99603723 | 0.15949763 | 0.031 | 0.031 | 1 | 0.33103112 | -0.6577786 | 0.038 | 0.062 | 1 | 0.99603723 | 0.55248064 |
| Slc25a16 | 0.33103733 | 0.10842345 | 0.312 | 0.254 | 1 | 0.88691081 | -0.3939195 | 0.294 | 0.281 | 1 | 0.88691081 | 0.55248895 |
| Gm17178 | 0.33118518 | 0.14499806 | 0.062 | 0.036 | 1 | 0.89995605 | -0.5728897 | 0.092 | 0.094 | 1 | 0.89995605 | 0.55268673 |
| Gpr137c | 0.33118518 | -0.2555399 | 0.062 | 0.036 | 1 | 0.75428046 | -0.1034044 | 0.05 | 0.042 | 1 | 0.75428046 | 0.55268673 |
| Exosc5 | 0.33123843 | 0.22508597 | 0.229 | 0.176 | 1 | 0.40008865 | -0.2187074 | 0.172 | 0.125 | 1 | 0.40008865 | 0.55275797 |
| Lmo1 | 0.75328966 | 0.27052894 | 0.021 | 0.016 | 1 | 0.33173572 | -0.6317834 | 0.055 | 0.083 | 1 | 0.75328966 | 0.55342285 |
| Stx1b | 0.54203294 | 0.27052894 | 0.052 | 0.036 | 1 | 0.33223756 | -0.7248928 | 0.092 | 0.125 | 1 | 0.54203294 | 0.55409333 |
| Zfp346 | 0.33224972 | 0.37946331 | 0.24 | 0.197 | 1 | 0.37299017 | -0.2273931 | 0.218 | 0.167 | 1 | 0.37299017 | 0.55410957 |
| Lrrc51 | 0.33271172 | 0.42253204 | 0.125 | 0.088 | 1 | 0.88407153 | -0.6179776 | 0.08 | 0.083 | 1 | 0.88407153 | 0.55472636 |
| Zfp949 | 0.83080497 | -0.293675 | 0.219 | 0.223 | 1 | 0.33314932 | -0.2249664 | 0.387 | 0.292 | 1 | 0.83080497 | 0.55531017 |
| Ankrd13c | 0.50125029 | -0.3522356 | 0.646 | 0.632 | 1 | 0.33324975 | -0.3175497 | 0.748 | 0.75 | 1 | 0.50125029 | 0.55544411 |
| 2900089D17Rik | 0.69363445 | -0.2991668 | 0.156 | 0.171 | 1 | 0.33327135 | -0.3878578 | 0.244 | 0.177 | 1 | 0.69363445 | 0.5554729 |
| 3110045C21Rik | 0.33357992 | 0.14076107 | 0.167 | 0.119 | 1 | 0.69882111 | 0.19264508 | 0.063 | 0.052 | 1 | 0.69882111 | 0.55588428 |
| Ppp2r2a | 0.3337405 | 0.15653217 | 0.875 | 0.87 | 1 | 0.69178638 | -0.1417325 | 0.929 | 0.938 | 1 | 0.69178638 | 0.55609827 |
| Nck1 | 0.33376556 | 0.15068917 | 0.625 | 0.596 | 1 | 0.45762488 | -0.1224125 | 0.727 | 0.729 | 1 | 0.45762488 | 0.55613167 |
| Hsd17b4 | 0.89942399 | -0.2259956 | 0.302 | 0.301 | 1 | 0.33418051 | 0.18629797 | 0.475 | 0.406 | 1 | 0.89942399 | 0.5566844 |
| Gm47990 | 0.33418721 | 0.39278469 | 0.167 | 0.124 | 1 | 0.76146733 | -0.3098553 | 0.025 | 0.031 | 1 | 0.76146733 | 0.55669332 |
| Kdm5c | 0.33495375 | 0.14676933 | 0.729 | 0.684 | 1 | 0.73727776 | -0.1204779 | 0.811 | 0.833 | 1 | 0.73727776 | 0.55771348 |
| Wdr27 | 0.33509934 | 0.66645762 | 0.104 | 0.073 | 1 | 0.45487568 | 0.13071733 | 0.063 | 0.042 | 1 | 0.45487568 | 0.55790711 |
| Zfp619 | 0.33523767 | 1.00749454 | 0.073 | 0.047 | 1 | 0.45899181 | -1.5443205 | 0.063 | 0.083 | 1 | 0.45899181 | 0.55809104 |
| Slc26a11 | 0.33578004 | 0.43921078 | 0.219 | 0.176 | 1 | 0.80927194 | -0.3854063 | 0.29 | 0.25 | 1 | 0.80927194 | 0.55881184 |
| Rybp | 0.39562632 | 0.267955 | 0.802 | 0.808 | 1 | 0.33583411 | -0.3721037 | 0.84 | 0.833 | 1 | 0.39562632 | 0.55888367 |
| Slc16a9 | 0.56492112 | 0.54806292 | 0.062 | 0.047 | 1 | 0.3358496 | 0.10518224 | 0.084 | 0.052 | 1 | 0.56492112 | 0.55890425 |
| Cpsf3 | 0.77801698 | -0.3710171 | 0.281 | 0.285 | 1 | 0.33648354 | -0.2766884 | 0.303 | 0.219 | 1 | 0.77801698 | 0.55974591 |
| 4930594M22Rik | 0.3369646 | -0.7580402 | 0.031 | 0.057 | 1 | 0.66172649 | -0.5936482 | 0.113 | 0.094 | 1 | 0.66172649 | 0.56038405 |
| Gm48226 | 0.33727461 | 0.34006988 | 0.156 | 0.114 | 1 | 0.64539908 | -0.2592292 | 0.101 | 0.083 | 1 | 0.64539908 | 0.56079506 |
| Smim5 | 0.33731339 | 1.13302542 | 0.115 | 0.083 | 1 | 0.55298341 | -0.3098553 | 0.038 | 0.052 | 1 | 0.55298341 | 0.56084646 |
| Dhx35 | 0.3380367 | 0.90795886 | 0.104 | 0.073 | 1 | 0.64544695 | -0.6931839 | 0.08 | 0.094 | 1 | 0.64544695 | 0.56180459 |
| Dok5 | 0.93635708 | -0.599163 | 0.115 | 0.114 | 1 | 0.33841444 | 1.74903843 | 0.042 | 0.021 | 1 | 0.93635708 | 0.56230454 |
| Map3k12 | 0.33859098 | -0.5160674 | 0.115 | 0.155 | 1 | 0.48147874 | -0.566195 | 0.248 | 0.188 | 1 | 0.48147874 | 0.56253811 |
| Dnai4 | 0.33863727 | 0.60894516 | 0.135 | 0.098 | 1 | 0.99163975 | -0.6508922 | 0.109 | 0.104 | 1 | 0.99163975 | 0.56259934 |
| Sppl3 | 0.33864763 | -0.1508711 | 0.708 | 0.813 | 1 | 0.74008398 | -0.1232515 | 0.933 | 0.917 | 1 | 0.74008398 | 0.56261305 |
| Man2c1 | 0.43314662 | -0.3592878 | 0.188 | 0.223 | 1 | 0.33877138 | 0.11640949 | 0.172 | 0.125 | 1 | 0.43314662 | 0.56277671 |
| Mms19 | 0.97294414 | 0.22161934 | 0.188 | 0.192 | 1 | 0.33882632 | -0.3587649 | 0.252 | 0.188 | 1 | 0.97294414 | 0.56284937 |
| Stxbp4 | 0.79722101 | -0.1268066 | 0.229 | 0.212 | 1 | 0.33904647 | -0.1375397 | 0.349 | 0.26 | 1 | 0.79722101 | 0.56314043 |
| Zmym3 | 0.33934046 | 0.52206771 | 0.094 | 0.062 | 1 | 0.53046209 | -0.4703199 | 0.122 | 0.146 | 1 | 0.53046209 | 0.56352897 |
| Gm50431 | 0.33934046 | -0.577468 | 0.052 | 0.083 | 1 | 0.56833085 | -0.1172102 | 0.046 | 0.062 | 1 | 0.56833085 | 0.56352897 |
| Knop1 | 0.68781715 | 0.10703021 | 0.385 | 0.352 | 1 | 0.33947095 | 0.1515927 | 0.433 | 0.406 | 1 | 0.68781715 | 0.56370137 |
| Tnfrsf1b | 0.64499356 | -0.5230202 | 0.083 | 0.098 | 1 | 0.33969074 | -0.4542452 | 0.134 | 0.094 | 1 | 0.64499356 | 0.56399168 |
| Lrriq3 | 0.38689439 | -0.577468 | 0.01 | 0.026 | 1 | 0.33988591 | -0.5443205 | 0.038 | 0.062 | 1 | 0.38689439 | 0.56424939 |
| Peg10 | 0.3400253 | 0.32942263 | 0.031 | 0.057 | 1 | 0.90063186 | -0.1843244 | 0.059 | 0.062 | 1 | 0.90063186 | 0.56443339 |
| Cspg5 | 0.72498737 | -0.2920657 | 0.115 | 0.098 | 1 | 0.34012534 | 0.31163311 | 0.13 | 0.094 | 1 | 0.72498737 | 0.56456544 |
| Bc1 | 0.34018073 | 1.15949763 | 0.146 | 0.109 | 1 | 0.54445141 | -0.4797803 | 0.084 | 0.104 | 1 | 0.54445141 | 0.56463854 |
| Ggt7 | 0.48064834 | -0.8993961 | 0.021 | 0.036 | 1 | 0.34026631 | -3.0468209 | 0.008 | 0.021 | 1 | 0.48064834 | 0.56475146 |
| Gm16120 | 0.61737927 | 1.00749454 | 0.01 | 0.005 | 1 | 0.34026631 | -2.3098553 | 0.008 | 0.021 | 1 | 0.61737927 | 0.56475146 |
| Gm50107 | 0.61737927 | 1.00749454 | 0.01 | 0.005 | 1 | 0.34026631 | -2.3098553 | 0.008 | 0.021 | 1 | 0.61737927 | 0.56475146 |
| Gm33424 | 0.66676245 | -0.3144336 | 0.031 | 0.041 | 1 | 0.34026631 | -2.3098553 | 0.008 | 0.021 | 1 | 0.66676245 | 0.56475146 |
| Zfp457 | 0.75328966 | 0.27052894 | 0.021 | 0.016 | 1 | 0.34026631 | -2.3098553 | 0.008 | 0.021 | 1 | 0.75328966 | 0.56475146 |
| Gm50373 | 0.79413116 | -0.7998604 | 0.021 | 0.026 | 1 | 0.34026631 | -2.5322477 | 0.008 | 0.021 | 1 | 0.79413116 | 0.56475146 |
| Nek5 | 0.8061167 | -0.577468 | 0.031 | 0.036 | 1 | 0.34026631 | -2.3098553 | 0.008 | 0.021 | 1 | 0.8061167 | 0.56475146 |
| Tsix | 0.84090392 | -0.407543 | 0.135 | 0.14 | 1 | 0.34026631 | -2.7248928 | 0.008 | 0.021 | 1 | 0.84090392 | 0.56475146 |
| Tprg | 0.99758609 | 0.27052894 | 0.021 | 0.021 | 1 | 0.34026631 | -2.7248928 | 0.008 | 0.021 | 1 | 0.99758609 | 0.56475146 |
| Gm57096 | 1 | 0.42253204 | 0.01 | 0.01 | 1 | 0.34026631 | -2.3098553 | 0.008 | 0.021 | 1 | 1 | 0.56475146 |
| Gm56752 | 1 | 0.42253204 | 0.01 | 0.01 | 1 | 0.34026631 | -2.3098553 | 0.008 | 0.021 | 1 | 1 | 0.56475146 |
| Zmiz2 | 0.34048945 | -0.8550019 | 0.094 | 0.13 | 1 | 0.41683946 | -0.1172102 | 0.261 | 0.208 | 1 | 0.41683946 | 0.56504584 |
| Ehmt2 | 0.93780305 | -0.407543 | 0.312 | 0.275 | 1 | 0.34052033 | -0.2447602 | 0.399 | 0.302 | 1 | 0.93780305 | 0.56508657 |
| Fancm | 0.90620255 | -0.2148979 | 0.167 | 0.171 | 1 | 0.34055181 | -0.7479764 | 0.218 | 0.25 | 1 | 0.90620255 | 0.56512808 |
| Ulk4 | 0.5356124 | 0.12297175 | 0.25 | 0.295 | 1 | 0.34067811 | -0.5690981 | 0.202 | 0.24 | 1 | 0.5356124 | 0.56529465 |
| 4930453N24Rik | 0.43534172 | 0.15292498 | 0.51 | 0.446 | 1 | 0.34087242 | -0.1440464 | 0.479 | 0.396 | 1 | 0.43534172 | 0.56555084 |
| Cbx8 | 0.34100823 | -0.577468 | 0.031 | 0.057 | 1 | 0.73632333 | 0.10518224 | 0.05 | 0.042 | 1 | 0.73632333 | 0.56572984 |
| Gm50232 | 0.34100823 | -0.577468 | 0.031 | 0.057 | 1 | 0.69948596 | -1.0297473 | 0.063 | 0.073 | 1 | 0.69948596 | 0.56572984 |
| Pdk3 | 0.72816182 | -0.1472281 | 0.385 | 0.404 | 1 | 0.34108853 | -0.2805113 | 0.437 | 0.323 | 1 | 0.72816182 | 0.56583567 |
| Gm56970 | 0.62104899 | 0.42253204 | 0.01 | 0.005 | 1 | 0.34195151 | -1.8948178 | 0.008 | 0.021 | 1 | 0.62104899 | 0.56697218 |
| Dclk1 | 0.5161904 | -0.5070786 | 0.115 | 0.14 | 1 | 0.34216425 | -1.154577 | 0.092 | 0.125 | 1 | 0.5161904 | 0.56725213 |
| Lpar3 | 0.34220578 | 0.15949763 | 0.094 | 0.062 | 1 | 0.86613737 | -0.3098553 | 0.008 | 0.01 | 1 | 0.86613737 | 0.56730677 |
| Mapk6 | 0.41443853 | -0.1159778 | 0.771 | 0.839 | 1 | 0.34225649 | -0.1663111 | 0.84 | 0.885 | 1 | 0.41443853 | 0.56737348 |
| Tfip11 | 0.41880012 | 0.25542205 | 0.167 | 0.13 | 1 | 0.34353601 | -0.3098553 | 0.134 | 0.094 | 1 | 0.41880012 | 0.56905503 |
| Fndc5 | 0.38689439 | -0.577468 | 0.01 | 0.026 | 1 | 0.34363875 | -1.7248928 | 0.008 | 0.021 | 1 | 0.38689439 | 0.56918991 |
| Gm16068 | 0.47606502 | 1.00749454 | 0.021 | 0.01 | 1 | 0.34363875 | -1.7248928 | 0.008 | 0.021 | 1 | 0.47606502 | 0.56918991 |
| Cdc25c | 0.47835216 | -0.6929452 | 0.021 | 0.01 | 1 | 0.34363875 | -1.7248928 | 0.008 | 0.021 | 1 | 0.47835216 | 0.56918991 |
| Stmn2 | 0.52961087 | -0.577468 | 0.01 | 0.021 | 1 | 0.34363875 | -1.7248928 | 0.008 | 0.021 | 1 | 0.52961087 | 0.56918991 |
| BC055324 | 0.57848839 | 1.00749454 | 0.031 | 0.021 | 1 | 0.34363875 | -1.7248928 | 0.008 | 0.021 | 1 | 0.57848839 | 0.56918991 |
| Slc5a11 | 0.61182672 | -0.577468 | 0.021 | 0.031 | 1 | 0.34363875 | -2.0468209 | 0.008 | 0.021 | 1 | 0.61182672 | 0.56918991 |
| Perm1 | 0.61371908 | 1.59245704 | 0.01 | 0.005 | 1 | 0.34363875 | -1.7248928 | 0.008 | 0.021 | 1 | 0.61371908 | 0.56918991 |
| Dnai2 | 0.61737927 | 1.00749454 | 0.01 | 0.005 | 1 | 0.34363875 | -2.0468209 | 0.008 | 0.021 | 1 | 0.61737927 | 0.56918991 |
| Uckl1os | 0.62104899 | 0.42253204 | 0.01 | 0.005 | 1 | 0.34363875 | -2.3098553 | 0.008 | 0.021 | 1 | 0.62104899 | 0.56918991 |

| | | | | | | | | | | | | |
|---|---|---|---|---|---|---|---|---|---|---|---|---|
| 1700019D03Rik | 0.62852542 | 0.32942263 | 0.021 | 0.031 | 1 | 0.34363875 | -2.0468209 | 0.008 | 0.021 | 1 | 0.62852542 | 0.56918991 |
| Gm48677 | 0.64527612 | 0.15949763 | 0.052 | 0.067 | 1 | 0.34363875 | -2.0468209 | 0.008 | 0.021 | 1 | 0.64527612 | 0.56918991 |
| Cntf | 0.75328966 | 0.27052894 | 0.021 | 0.016 | 1 | 0.34363875 | -1.7248928 | 0.008 | 0.021 | 1 | 0.75328966 | 0.56918991 |
| Hsd17b13 | 0.75832339 | -0.577468 | 0.021 | 0.016 | 1 | 0.34363875 | -1.7248928 | 0.008 | 0.021 | 1 | 0.75832339 | 0.56918991 |
| Flacc1 | 0.99320802 | 1.00749454 | 0.01 | 0.01 | 1 | 0.34363875 | -2.7248928 | 0.008 | 0.021 | 1 | 0.99320802 | 0.56918991 |
| Gm47270 | 0.99758609 | 0.27052894 | 0.021 | 0.021 | 1 | 0.34363875 | -2.3098553 | 0.008 | 0.021 | 1 | 0.99758609 | 0.56918991 |
| D630036H23Rik | 0.99758609 | 0.27052894 | 0.021 | 0.021 | 1 | 0.34363875 | -1.7248928 | 0.008 | 0.021 | 1 | 0.99758609 | 0.56918991 |
| Asic4 | 1 | 0.42253204 | 0.01 | 0.01 | 1 | 0.34363875 | -1.7248928 | 0.008 | 0.021 | 1 | 1 | 0.56918991 |
| Prnd | 1 | -0.577468 | 0.01 | 0.01 | 1 | 0.34363875 | -2.0468209 | 0.008 | 0.021 | 1 | 1 | 0.56918991 |
| Fam83c | 1 | 0.42253204 | 0.01 | 0.01 | 1 | 0.34363875 | -2.0468209 | 0.008 | 0.021 | 1 | 1 | 0.56918991 |
| Gm15873 | 1 | 0.42253204 | 0.01 | 0.01 | 1 | 0.34363875 | -1.7248928 | 0.008 | 0.021 | 1 | 1 | 0.56918991 |
| Gm6270 | 1 | 0.42253204 | 0.01 | 0.01 | 1 | 0.34363875 | -1.7248928 | 0.008 | 0.021 | 1 | 1 | 0.56918991 |
| Slc7a7 | 0.43293505 | 0.13302542 | 0.229 | 0.181 | 1 | 0.34366126 | -0.3329389 | 0.298 | 0.219 | 1 | 0.43293505 | 0.56921945 |
| Acaca | 0.34369607 | 0.1124641 | 0.667 | 0.648 | 1 | 0.60885538 | -0.6605837 | 0.647 | 0.542 | 1 | 0.60885538 | 0.56926515 |
| Gm9856 | 0.34422149 | 0.1843723 | 0.125 | 0.088 | 1 | 0.73324232 | -0.7573142 | 0.063 | 0.073 | 1 | 0.73324232 | 0.56995455 |
| Tle6 | 0.65212989 | 0.15949763 | 0.042 | 0.031 | 1 | 0.34428571 | 0.91253716 | 0.042 | 0.021 | 1 | 0.65212989 | 0.57003877 |
| Tchh | 0.79867625 | -0.7791018 | 0.146 | 0.155 | 1 | 0.34428571 | 1.01207283 | 0.042 | 0.021 | 1 | 0.79867625 | 0.57003877 |
| Tulp2 | 0.99009327 | -0.4519371 | 0.031 | 0.031 | 1 | 0.34428571 | 0.80562195 | 0.042 | 0.021 | 1 | 0.99009327 | 0.57003877 |
| Arhgap22 | 0.65211183 | -1.240433 | 0.031 | 0.041 | 1 | 0.34446225 | -0.8459082 | 0.084 | 0.115 | 1 | 0.65211183 | 0.57027026 |
| Lrrc28 | 0.34446833 | -0.5982265 | 0.135 | 0.176 | 1 | 0.76615469 | -1.0785297 | 0.151 | 0.156 | 1 | 0.76615469 | 0.57027823 |
| Rmdn1 | 0.43250906 | -0.1118044 | 0.438 | 0.352 | 1 | 0.3445564 | -0.2480791 | 0.399 | 0.302 | 1 | 0.43250906 | 0.57039369 |
| Mfsd2a | 0.34533981 | 0.56692195 | 0.104 | 0.073 | 1 | 0.39419235 | 0.49749966 | 0.08 | 0.052 | 1 | 0.39419235 | 0.57142003 |
| Stxbp6 | 0.92227068 | -0.1260291 | 0.427 | 0.42 | 1 | 0.34559847 | -0.4044921 | 0.382 | 0.438 | 1 | 0.92227068 | 0.57175864 |
| Gm16093 | 0.34566853 | -0.577468 | 0.125 | 0.166 | 1 | 0.53583247 | -0.1654654 | 0.13 | 0.104 | 1 | 0.53583247 | 0.57185032 |
| Flna | 0.39960241 | -0.3144336 | 0.635 | 0.534 | 1 | 0.34603286 | -0.3098553 | 0.832 | 0.823 | 1 | 0.39960241 | 0.57232698 |
| 2310015A10Rik | 0.34606218 | 0.73186009 | 0.135 | 0.098 | 1 | 0.78534892 | -0.8768959 | 0.097 | 0.104 | 1 | 0.78534892 | 0.57236533 |
| Poln | 0.92478986 | 0.44806713 | 0.135 | 0.135 | 1 | 0.3461902 | -1.4518743 | 0.134 | 0.167 | 1 | 0.92478986 | 0.57253275 |
| 4833411C07Rik | 0.50823881 | 0.31561683 | 0.073 | 0.098 | 1 | 0.34625029 | 0.69014474 | 0.042 | 0.021 | 1 | 0.50823881 | 0.57261131 |
| Zfp28 | 0.80274368 | 0.20013961 | 0.031 | 0.026 | 1 | 0.34625029 | 0.69014474 | 0.042 | 0.021 | 1 | 0.80274368 | 0.57261131 |
| Ppp1r26 | 0.83546978 | 0.15949763 | 0.031 | 0.036 | 1 | 0.34625029 | 0.69014474 | 0.042 | 0.021 | 1 | 0.83546978 | 0.57261131 |
| Sugp2 | 0.3466123 | -0.4170033 | 0.302 | 0.347 | 1 | 0.67256419 | -0.3098553 | 0.361 | 0.302 | 1 | 0.67256419 | 0.57308451 |
| Ncoa6 | 0.92904866 | -0.1641909 | 0.51 | 0.492 | 1 | 0.34697645 | -0.5227306 | 0.79 | 0.833 | 1 | 0.92904866 | 0.57356024 |
| Tent5d | 0.47150075 | 1.42253204 | 0.021 | 0.01 | 1 | 0.34703906 | -1.3098553 | 0.008 | 0.021 | 1 | 0.47150075 | 0.57364201 |
| Rnf165 | 0.72574709 | -0.3144336 | 0.01 | 0.016 | 1 | 0.34703906 | -1.3098553 | 0.008 | 0.021 | 1 | 0.72574709 | 0.57364201 |
| Grin1 | 0.36731724 | -0.7998604 | 0.021 | 0.041 | 1 | 0.34704016 | -1.3098553 | 0.008 | 0.021 | 1 | 0.36731724 | 0.57364344 |
| Gm15908 | 0.39523977 | 1.20013961 | 0.01 | 0.026 | 1 | 0.34704016 | -1.3098553 | 0.008 | 0.021 | 1 | 0.39523977 | 0.57364344 |
| Slc29a4 | 0.52313476 | -0.9925055 | 0.01 | 0.021 | 1 | 0.34704016 | -1.3098553 | 0.008 | 0.021 | 1 | 0.52313476 | 0.57364344 |
| Fam178b | 0.52744059 | -1.6929452 | 0.01 | 0.021 | 1 | 0.34704016 | -1.3098553 | 0.008 | 0.021 | 1 | 0.52744059 | 0.57364344 |
| 4931415C17Rik | 0.61737927 | 1.00749454 | 0.01 | 0.005 | 1 | 0.34704016 | -1.3098553 | 0.008 | 0.021 | 1 | 0.61737927 | 0.57364344 |
| 2010001K21Rik | 0.61737927 | 1.00749454 | 0.01 | 0.005 | 1 | 0.34704016 | -1.3098553 | 0.008 | 0.021 | 1 | 0.61737927 | 0.57364344 |
| 4930414N06Rik | 0.61737927 | 1.00749454 | 0.01 | 0.005 | 1 | 0.34704016 | -1.3098553 | 0.008 | 0.021 | 1 | 0.61737927 | 0.57364344 |
| Gm10863 | 0.62104899 | -0.3144336 | 0.01 | 0.005 | 1 | 0.34704016 | -1.3098553 | 0.008 | 0.021 | 1 | 0.62104899 | 0.57364344 |
| Ttc39d | 0.62104899 | 0.42253204 | 0.01 | 0.005 | 1 | 0.34704016 | -1.3098553 | 0.008 | 0.021 | 1 | 0.62104899 | 0.57364344 |
| Gm13594 | 0.70116305 | -0.2555399 | 0.042 | 0.052 | 1 | 0.34704016 | -1.3098553 | 0.008 | 0.021 | 1 | 0.70116305 | 0.57364344 |
| 1110006O24Rik | 0.73324151 | -0.1079827 | 0.052 | 0.062 | 1 | 0.34704016 | -1.3098553 | 0.008 | 0.021 | 1 | 0.73324151 | 0.57364344 |
| Chrnb2 | 0.73407659 | 0.27052894 | 0.01 | 0.016 | 1 | 0.34704016 | -1.3098553 | 0.008 | 0.021 | 1 | 0.73407659 | 0.57364344 |
| 1110035H17Rik | 0.74076005 | 1.00749454 | 0.021 | 0.016 | 1 | 0.34704016 | -1.3098553 | 0.008 | 0.021 | 1 | 0.74076005 | 0.57364344 |
| Gm32261 | 0.74076005 | 1.00749454 | 0.021 | 0.016 | 1 | 0.34704016 | -1.3098553 | 0.008 | 0.021 | 1 | 0.74076005 | 0.57364344 |
| Gm29481 | 0.74576744 | 0.68556644 | 0.021 | 0.016 | 1 | 0.34704016 | -1.3098553 | 0.008 | 0.021 | 1 | 0.74576744 | 0.57364344 |
| 5033406O09Rik | 0.78979801 | -0.2148979 | 0.021 | 0.026 | 1 | 0.34704016 | -1.3098553 | 0.008 | 0.021 | 1 | 0.78979801 | 0.57364344 |
| Gm38250 | 0.79666623 | 0.42253204 | 0.031 | 0.026 | 1 | 0.34704016 | -1.3098553 | 0.008 | 0.021 | 1 | 0.79666623 | 0.57364344 |
| 1810044D09Rik | 0.82342138 | 0.32942263 | 0.042 | 0.036 | 1 | 0.34704016 | -1.3098553 | 0.008 | 0.021 | 1 | 0.82342138 | 0.57364344 |
| Nos1 | 0.99603711 | -0.3144336 | 0.031 | 0.031 | 1 | 0.34704016 | -1.3098553 | 0.008 | 0.021 | 1 | 0.99603711 | 0.57364344 |
| Swsap1 | 0.99603742 | 0.20013961 | 0.031 | 0.031 | 1 | 0.34704016 | -1.3098553 | 0.008 | 0.021 | 1 | 0.99603742 | 0.57364344 |
| Gm19497 | 1 | 0.42253204 | 0.01 | 0.01 | 1 | 0.34704016 | -1.3098553 | 0.008 | 0.021 | 1 | 1 | 0.57364344 |
| Nacad | 1 | 0.42253204 | 0.01 | 0.01 | 1 | 0.34704016 | -1.3098553 | 0.008 | 0.021 | 1 | 1 | 0.57364344 |
| Ube2ql1 | 0.34710248 | -0.3710171 | 0.031 | 0.057 | 1 | 0.74567031 | -0.7623675 | 0.063 | 0.052 | 1 | 0.74567031 | 0.57372483 |
| Kdm4c | 0.34817517 | 0.19390766 | 0.583 | 0.528 | 1 | 0.78842459 | -0.2786794 | 0.739 | 0.781 | 1 | 0.78842459 | 0.57512439 |
| Slc25a17 | 0.34824118 | 0.11111717 | 0.646 | 0.575 | 1 | 0.43989222 | -0.3366222 | 0.655 | 0.667 | 1 | 0.43989222 | 0.57521044 |
| Eipr1 | 0.44091496 | -0.1445086 | 0.198 | 0.238 | 1 | 0.34830088 | -0.6179776 | 0.193 | 0.229 | 1 | 0.44091496 | 0.57528826 |
| Pcdhgb7 | 0.38052601 | 0.81484946 | 0.052 | 0.031 | 1 | 0.34832484 | 0.56461386 | 0.042 | 0.021 | 1 | 0.38052601 | 0.57531949 |
| 9530077C14Rik | 0.46336809 | 0.71798792 | 0.073 | 0.052 | 1 | 0.34832484 | 0.56461386 | 0.042 | 0.021 | 1 | 0.46336809 | 0.57531949 |
| Slc12a1 | 0.34844305 | -0.1010299 | 0.177 | 0.218 | 1 | 0.42845063 | 2.35310975 | 0.038 | 0.021 | 1 | 0.42845063 | 0.57547355 |
| Fam53a | 0.96721912 | -0.2054992 | 0.188 | 0.181 | 1 | 0.34852058 | -0.774818 | 0.252 | 0.177 | 1 | 0.96721912 | 0.57557456 |
| Kdm6a | 0.7681511 | -0.1177248 | 0.76 | 0.689 | 1 | 0.34864652 | -0.4892088 | 0.828 | 0.833 | 1 | 0.7681511 | 0.57573865 |
| Rfc3 | 0.34893863 | -0.3404288 | 0.073 | 0.109 | 1 | 0.64450353 | -0.3098553 | 0.109 | 0.125 | 1 | 0.64450353 | 0.57611909 |
| Gm57301 | 0.34925442 | -0.407543 | 0.031 | 0.057 | 1 | 0.57370068 | -1.0468209 | 0.021 | 0.031 | 1 | 0.57370068 | 0.57653019 |
| Traf3ip2 | 0.51107282 | 0.12297175 | 0.115 | 0.088 | 1 | 0.34998111 | -1.0161241 | 0.126 | 0.156 | 1 | 0.51107282 | 0.57747545 |
| Nfkbie | 0.35026965 | -0.407543 | 0.031 | 0.057 | 1 | 0.66391569 | -1.0203486 | 0.042 | 0.052 | 1 | 0.66391569 | 0.57785047 |
| Naaladl2 | 0.45113728 | -0.1104688 | 0.688 | 0.658 | 1 | 0.35040441 | -0.2972111 | 0.803 | 0.865 | 1 | 0.45113728 | 0.57802557 |
| Gm26671 | 0.52502715 | -0.8405024 | 0.031 | 0.047 | 1 | 0.35045411 | -0.3098553 | 0.008 | 0.021 | 1 | 0.52502715 | 0.57809014 |
| Tmem151b | 0.61737927 | 1.00749454 | 0.01 | 0.005 | 1 | 0.35045411 | -0.8948178 | 0.008 | 0.021 | 1 | 0.61737927 | 0.57809014 |
| Spag17 | 0.62104899 | 0.42253204 | 0.01 | 0.005 | 1 | 0.35045411 | -0.8948178 | 0.008 | 0.021 | 1 | 0.62104899 | 0.57809014 |
| Vmn1r4 | 0.7507837 | 0.42253204 | 0.021 | 0.016 | 1 | 0.35045411 | -0.8948178 | 0.008 | 0.021 | 1 | 0.7507837 | 0.57809014 |
| Gm47772 | 1 | -0.3144336 | 0.01 | 0.01 | 1 | 0.35045411 | -0.5728897 | 0.008 | 0.021 | 1 | 1 | 0.57809014 |
| Akap10 | 0.37837372 | 0.48056416 | 0.396 | 0.363 | 1 | 0.35091084 | -0.4505992 | 0.521 | 0.562 | 1 | 0.37837372 | 0.57868326 |
| Tmem87b | 0.35122152 | 0.13677755 | 0.656 | 0.575 | 1 | 0.38288663 | -0.2108888 | 0.752 | 0.635 | 1 | 0.38288663 | 0.57908648 |
| C730034F03Rik | 0.35132231 | -0.2726134 | 0.031 | 0.057 | 1 | 0.41845995 | -1.3098553 | 0.034 | 0.052 | 1 | 0.41845995 | 0.57921726 |

| | | | | | | | | | | | | |
|---|---|---|---|---|---|---|---|---|---|---|---|---|
| Fam171b | 0.35154197 | -1.0547897 | 0.115 | 0.15 | 1 | 0.54785445 | -0.4492583 | 0.185 | 0.146 | 1 | 0.54785445 | 0.57950218 |
| Xkr6 | 0.35162153 | -0.7839188 | 0.104 | 0.14 | 1 | 0.97201637 | -0.9228321 | 0.101 | 0.094 | 1 | 0.97201637 | 0.57960537 |
| Pnpla6 | 0.45193315 | 0.36113149 | 0.167 | 0.135 | 1 | 0.35187644 | -0.1864728 | 0.21 | 0.156 | 1 | 0.45193315 | 0.57993585 |
| Dexi | 0.35945036 | 0.11133035 | 0.312 | 0.254 | 1 | 0.35193935 | 0.13760371 | 0.408 | 0.344 | 1 | 0.35945036 | 0.5800174 |
| Cirbp | 0.35225652 | -0.3627291 | 0.552 | 0.575 | 1 | 0.37750923 | -0.3271333 | 0.605 | 0.688 | 1 | 0.37750923 | 0.58042838 |
| Dyrk1b | 0.69523544 | -0.6705774 | 0.042 | 0.052 | 1 | 0.35248867 | -0.3098553 | 0.046 | 0.073 | 1 | 0.69523544 | 0.58072908 |
| Cers4 | 0.35250927 | -1.1801325 | 0.073 | 0.104 | 1 | 0.62027061 | -0.7623675 | 0.13 | 0.146 | 1 | 0.62027061 | 0.58075575 |
| Rhbdd1 | 0.89565903 | 0.13755508 | 0.333 | 0.342 | 1 | 0.35257024 | -0.4200382 | 0.324 | 0.24 | 1 | 0.89565903 | 0.58083471 |
| Stat5a | 0.89440829 | -0.3376409 | 0.26 | 0.264 | 1 | 0.35286762 | -0.8043272 | 0.349 | 0.354 | 1 | 0.89440829 | 0.58121969 |
| Nog | 0.35289767 | 1.35960078 | 0.198 | 0.259 | 1 | 0.48179579 | 0.8917786 | 0.097 | 0.073 | 1 | 0.48179579 | 0.58125858 |
| Gpr146 | 0.35302608 | -0.577468 | 0.219 | 0.259 | 1 | 0.75588861 | -0.3098553 | 0.134 | 0.146 | 1 | 0.75588861 | 0.58142475 |
| Lancl2 | 0.37133953 | 0.32942263 | 0.188 | 0.145 | 1 | 0.35348431 | 0.11756596 | 0.231 | 0.188 | 1 | 0.37133953 | 0.58201746 |
| Gm26930 | 0.74576744 | 0.68556644 | 0.021 | 0.016 | 1 | 0.35389699 | 0.10518224 | 0.008 | 0.021 | 1 | 0.74576744 | 0.5825509 |
| Ect2 | 0.3542203 | 2.81484946 | 0.052 | 0.031 | 1 | 0.58180179 | -0.7248928 | 0.021 | 0.031 | 1 | 0.58180179 | 0.58296857 |
| Mllt3 | 0.35430562 | -0.1059877 | 0.938 | 0.896 | 1 | 0.67243983 | -0.2376166 | 0.866 | 0.875 | 1 | 0.67243983 | 0.58307877 |
| Nid1 | 0.35489564 | -0.5568577 | 0.302 | 0.223 | 1 | 0.41018344 | -1.5404682 | 0.21 | 0.229 | 1 | 0.41018344 | 0.58384036 |
| Muc5b | 0.993208 | 1.00749454 | 0.01 | 0.01 | 1 | 0.35532256 | 0.17557156 | 0.042 | 0.021 | 1 | 0.993208 | 0.584391 |
| Susd1 | 0.35614361 | -1.2148979 | 0.021 | 0.041 | 1 | 0.92228543 | -0.7248928 | 0.029 | 0.031 | 1 | 0.92228543 | 0.58544894 |
| Zfp54 | 0.47606502 | 1.00749454 | 0.021 | 0.01 | 1 | 0.3563055 | 0.27510724 | 0.042 | 0.021 | 1 | 0.47606502 | 0.58565739 |
| Kcnk2 | 0.51600699 | -0.4031378 | 0.229 | 0.192 | 1 | 0.3564758 | -0.4765051 | 0.378 | 0.281 | 1 | 0.51600699 | 0.5858766 |
| Rab10 | 0.35655913 | -0.2234822 | 0.917 | 0.964 | 1 | 0.6455043 | -0.1537361 | 0.987 | 0.979 | 1 | 0.6455043 | 0.58598384 |
| Gm21859 | 0.35663684 | -1.3144336 | 0.042 | 0.067 | 1 | 0.54576607 | -1.7248928 | 0.038 | 0.052 | 1 | 0.54576607 | 0.58608385 |
| Gm37768 | 0.67008258 | -0.407543 | 0.031 | 0.041 | 1 | 0.35698851 | -0.194378 | 0.097 | 0.062 | 1 | 0.67008258 | 0.58653622 |
| Fads2 | 0.45984478 | -0.2070995 | 0.354 | 0.399 | 1 | 0.3571419 | -0.3252853 | 0.273 | 0.198 | 1 | 0.45984478 | 0.58673346 |
| Bpifb4 | 0.61737927 | 1.00749454 | 0.01 | 0.005 | 1 | 0.35731895 | -0.5322477 | 0.042 | 0.021 | 1 | 0.61737927 | 0.58696107 |
| Gm36975 | 0.3573566 | 0.74446013 | 0.073 | 0.047 | 1 | 0.84663765 | -0.677587 | 0.105 | 0.094 | 1 | 0.84663765 | 0.58700946 |
| Ggta1 | 0.35739797 | -0.2920657 | 0.083 | 0.119 | 1 | 0.4452538 | -0.6317834 | 0.118 | 0.083 | 1 | 0.4452538 | 0.58706262 |
| Kdm3a | 0.35776975 | -0.1901512 | 0.531 | 0.627 | 1 | 0.72048226 | -0.2496683 | 0.727 | 0.75 | 1 | 0.72048226 | 0.5875403 |
| Zdhhc2 | 0.90493976 | -0.4372903 | 0.094 | 0.098 | 1 | 0.35800223 | -0.7386986 | 0.109 | 0.073 | 1 | 0.90493976 | 0.58783886 |
| Gm52988 | 0.35801946 | -0.1624305 | 0.125 | 0.088 | 1 | 0.46188023 | -1.154577 | 0.134 | 0.156 | 1 | 0.46188023 | 0.58786099 |
| Gm34728 | 0.47606502 | 1.00749454 | 0.021 | 0.01 | 1 | 0.35824447 | 0.8917786 | 0.055 | 0.031 | 1 | 0.47606502 | 0.58814984 |
| Nat8f1 | 0.86824888 | -0.240433 | 0.073 | 0.078 | 1 | 0.35831374 | 0.14957636 | 0.042 | 0.021 | 1 | 0.86824888 | 0.58823874 |
| Gm13708 | 0.99482471 | 0.13302542 | 0.042 | 0.041 | 1 | 0.35831374 | -0.1723517 | 0.042 | 0.021 | 1 | 0.99482471 | 0.58823874 |
| Cog7 | 0.35894861 | 0.45110119 | 0.25 | 0.207 | 1 | 0.60857693 | -0.7890231 | 0.206 | 0.219 | 1 | 0.60857693 | 0.58905311 |
| Evl | 0.35898047 | -0.7843973 | 0.344 | 0.363 | 1 | 0.71753311 | -0.7546401 | 0.471 | 0.469 | 1 | 0.71753311 | 0.58909396 |
| Nradd | 0.35898451 | -0.7087125 | 0.062 | 0.093 | 1 | 0.6634615 | -1.152314 | 0.113 | 0.125 | 1 | 0.6634615 | 0.58909914 |
| Ccdc181 | 0.35950357 | 0.68556644 | 0.073 | 0.047 | 1 | 0.91891525 | -0.6317834 | 0.029 | 0.031 | 1 | 0.91891525 | 0.58976432 |
| Rps6kc1 | 0.70354996 | -0.2001009 | 0.26 | 0.233 | 1 | 0.35983456 | -0.3838558 | 0.445 | 0.354 | 1 | 0.70354996 | 0.59018821 |
| Spata17 | 0.36106354 | -1.2148979 | 0.021 | 0.041 | 1 | 0.81129592 | 0.10518224 | 0.025 | 0.021 | 1 | 0.81129592 | 0.59176019 |
| Gm33508 | 0.36106354 | -1.1079827 | 0.021 | 0.041 | 1 | 0.86543325 | 0.27510724 | 0.013 | 0.01 | 1 | 0.86543325 | 0.59176019 |
| Gm20544 | 0.36106354 | -1.2148979 | 0.021 | 0.041 | 1 | 0.8076363 | 0.42711033 | 0.025 | 0.021 | 1 | 0.8076363 | 0.59176019 |
| Arhgap20 | 0.36146558 | 0.63552576 | 0.073 | 0.047 | 1 | 0.57373861 | -1.7952821 | 0.013 | 0.021 | 1 | 0.57373861 | 0.5922738 |
| Usp35 | 0.70184804 | 0.27052894 | 0.062 | 0.052 | 1 | 0.36164088 | 0.76053407 | 0.067 | 0.042 | 1 | 0.70184804 | 0.59249763 |
| Gm44777 | 0.38521705 | 0.68556644 | 0.031 | 0.016 | 1 | 0.36170139 | 0.69014474 | 0.055 | 0.031 | 1 | 0.38521705 | 0.59257488 |
| Gm9828 | 0.46992036 | 0.74446013 | 0.042 | 0.026 | 1 | 0.36170139 | 0.69014474 | 0.055 | 0.031 | 1 | 0.46992036 | 0.59257488 |
| A330069E16Rik | 0.61933998 | -0.1941393 | 0.083 | 0.067 | 1 | 0.36170139 | 1.08246216 | 0.055 | 0.031 | 1 | 0.61933998 | 0.59257488 |
| Cbfa2t2 | 0.36201358 | 0.32942263 | 0.479 | 0.446 | 1 | 0.47401007 | -0.5936482 | 0.655 | 0.646 | 1 | 0.47401007 | 0.59297332 |
| Iqgap3 | 0.36248497 | 1.62898291 | 0.052 | 0.031 | 1 | 0.87066325 | -0.3098553 | 0.013 | 0.01 | 1 | 0.87066325 | 0.59357459 |
| Batf | 0.36249246 | -0.3404288 | 0.219 | 0.259 | 1 | 0.64998943 | -1.194378 | 0.113 | 0.125 | 1 | 0.64998943 | 0.59358413 |
| Nupl2 | 0.3627271 | 0.39451766 | 0.135 | 0.098 | 1 | 0.52850666 | 0.16819203 | 0.118 | 0.094 | 1 | 0.52850666 | 0.59388325 |
| Rtp4 | 0.36280846 | 0.36113149 | 0.094 | 0.135 | 1 | 0.74083174 | -1.4071525 | 0.088 | 0.094 | 1 | 0.74083174 | 0.59398695 |
| Rpa2 | 0.36302 | 0.34853145 | 0.229 | 0.181 | 1 | 0.87827205 | -0.4208866 | 0.172 | 0.177 | 1 | 0.87827205 | 0.59425647 |
| Nploc4 | 0.50934872 | -0.1322302 | 0.385 | 0.425 | 1 | 0.36316276 | -0.2812861 | 0.634 | 0.646 | 1 | 0.50934872 | 0.59443833 |
| Piwil2 | 0.36336537 | 0.36113149 | 0.115 | 0.083 | 1 | 0.68726051 | -0.8662486 | 0.084 | 0.094 | 1 | 0.68726051 | 0.59469634 |
| Gm20275 | 0.36350693 | -1.2319714 | 0.146 | 0.104 | 1 | 0.69600426 | -1.7091254 | 0.197 | 0.198 | 1 | 0.69600426 | 0.59487658 |
| Cntn4 | 0.36353452 | -2.536826 | 0.021 | 0.041 | 1 | 0.85741658 | -1.4391383 | 0.059 | 0.062 | 1 | 0.85741658 | 0.5949117 |
| Gm35248 | 0.36353452 | -0.8669746 | 0.021 | 0.041 | 1 | 0.51194506 | 0.27510724 | 0.021 | 0.01 | 1 | 0.51194506 | 0.5949117 |
| Serp2 | 0.36401945 | 0.38600616 | 0.104 | 0.073 | 1 | 0.878874 | -0.486733 | 0.08 | 0.083 | 1 | 0.878874 | 0.59552874 |
| Ipo9 | 0.36411199 | 0.42253204 | 0.312 | 0.269 | 1 | 0.979044 | -0.379865 | 0.471 | 0.427 | 1 | 0.979044 | 0.59564644 |
| Tasor | 0.36427236 | 0.30469555 | 0.719 | 0.674 | 1 | 0.85789233 | -0.1822672 | 0.807 | 0.833 | 1 | 0.85789233 | 0.59585036 |
| Gm48956 | 0.36429625 | -1.4254649 | 0.042 | 0.067 | 1 | 0.39937124 | 0.69014474 | 0.025 | 0.01 | 1 | 0.39937124 | 0.59588074 |
| Gm47271 | 0.36432975 | 0.59245704 | 0.073 | 0.047 | 1 | 0.92533164 | -0.8623963 | 0.05 | 0.052 | 1 | 0.92533164 | 0.59592334 |
| Chaf1b | 0.36432975 | 0.52206771 | 0.073 | 0.047 | 1 | 0.79171114 | -0.6094155 | 0.038 | 0.031 | 1 | 0.79171114 | 0.59592334 |
| Gm13563 | 0.38789252 | 0.59245704 | 0.052 | 0.031 | 1 | 0.36435013 | 0.59703533 | 0.055 | 0.031 | 1 | 0.38789252 | 0.59594925 |
| Gm19531 | 0.78979801 | -0.2148979 | 0.021 | 0.026 | 1 | 0.36435013 | 0.59703533 | 0.055 | 0.031 | 1 | 0.78979801 | 0.59594925 |
| Ciart | 0.7920382 | -0.2335136 | 0.083 | 0.093 | 1 | 0.36435013 | 0.59703533 | 0.055 | 0.031 | 1 | 0.7920382 | 0.59594925 |
| Tkfc | 0.87443987 | -0.1445086 | 0.083 | 0.088 | 1 | 0.36435013 | 0.59703533 | 0.055 | 0.031 | 1 | 0.87443987 | 0.59594925 |
| Dpp3 | 0.36452197 | -0.2019588 | 0.125 | 0.166 | 1 | 0.54003776 | 0.12310414 | 0.176 | 0.146 | 1 | 0.54003776 | 0.59616768 |
| Mindy4 | 0.36494905 | 0.69629285 | 0.312 | 0.269 | 1 | 0.86631903 | -0.6043026 | 0.298 | 0.281 | 1 | 0.86631903 | 0.59671029 |
| 2900005J15Rik | 0.36557148 | 0.5049942 | 0.104 | 0.073 | 1 | 0.43147365 | -0.8948178 | 0.034 | 0.052 | 1 | 0.43147365 | 0.59750045 |
| Nlrp5-ps | 0.36567603 | 2.46692616 | 0.031 | 0.016 | 1 | 0.91387303 | -0.7952821 | 0.029 | 0.031 | 1 | 0.91387303 | 0.5976331 |
| Slx4ip | 0.55422934 | 0.21231433 | 0.24 | 0.207 | 1 | 0.3657095 | -0.194378 | 0.21 | 0.156 | 1 | 0.55422934 | 0.59767557 |
| Gm48045 | 0.36601088 | -0.7294711 | 0.021 | 0.041 | 1 | 0.39515719 | 0.49749966 | 0.025 | 0.01 | 1 | 0.39515719 | 0.59805779 |
| Gm47962 | 0.36601088 | -1.1079827 | 0.021 | 0.041 | 1 | 0.909702 | 0.27510724 | 0.034 | 0.031 | 1 | 0.909702 | 0.59805779 |
| Qrich2 | 0.36601088 | -0.7294711 | 0.021 | 0.041 | 1 | 0.99396051 | -0.3098553 | 0.021 | 0.021 | 1 | 0.99396051 | 0.59805779 |
| Kif27 | 0.36601088 | -0.7294711 | 0.021 | 0.041 | 1 | 0.69919544 | -1.2223924 | 0.034 | 0.042 | 1 | 0.69919544 | 0.59805779 |
| Spdya | 0.36601088 | -0.7294711 | 0.021 | 0.041 | 1 | 0.8076363 | 0.42711033 | 0.025 | 0.021 | 1 | 0.8076363 | 0.59805779 |
| Cnbd2 | 0.36623755 | 0.36113149 | 0.167 | 0.124 | 1 | 0.88955876 | -0.7248928 | 0.193 | 0.188 | 1 | 0.88955876 | 0.59834515 |
| Ppp1r3f | 0.36625316 | 0.54806292 | 0.073 | 0.047 | 1 | 0.53717181 | 0.27510724 | 0.034 | 0.021 | 1 | 0.53717181 | 0.59836494 |
| Vps37a | 0.36630476 | -0.2272419 | 0.698 | 0.679 | 1 | 0.91768849 | -0.1563247 | 0.819 | 0.833 | 1 | 0.91768849 | 0.59843035 |
| Gm30054 | 0.67877064 | 0.49292136 | 0.052 | 0.041 | 1 | 0.36655161 | -0.3098553 | 0.042 | 0.021 | 1 | 0.67877064 | 0.59874313 |
| Fry | 0.7068007 | -0.1344516 | 0.927 | 0.886 | 1 | 0.36717071 | -0.1471634 | 0.861 | 0.792 | 1 | 0.7068007 | 0.59952709 |
| Sv2a | 0.36720299 | 1.20013961 | 0.052 | 0.031 | 1 | 0.6519798 | -0.8692827 | 0.067 | 0.052 | 1 | 0.6519798 | 0.59956795 |
| Fbxo5 | 0.36724689 | 2.08549705 | 0.052 | 0.031 | 1 | 0.75466895 | -0.6317834 | 0.025 | 0.031 | 1 | 0.75466895 | 0.59962351 |
| Elp6 | 0.38774429 | 0.20013961 | 0.073 | 0.047 | 1 | 0.36802272 | 0.82764826 | 0.067 | 0.042 | 1 | 0.38774429 | 0.60060471 |
| Marf1 | 0.41733333 | 0.118198 | 0.531 | 0.466 | 1 | 0.36806955 | -0.2840922 | 0.567 | 0.594 | 1 | 0.41733333 | 0.60066391 |
| A330015K06Rik | 0.36858168 | -1.1941393 | 0.021 | 0.041 | 1 | 0.50844268 | 1.01207283 | 0.021 | 0.01 | 1 | 0.50844268 | 0.6013109 |
| Clhc1 | 0.36858346 | -0.7998604 | 0.021 | 0.041 | 1 | 0.67922439 | -1.0297473 | 0.042 | 0.052 | 1 | 0.67922439 | 0.60131315 |
| Slc37a4 | 0.36859058 | -0.577468 | 0.021 | 0.041 | 1 | 0.42275543 | -1.1172102 | 0.025 | 0.042 | 1 | 0.42275543 | 0.60132215 |

| | | | | | | | | | | | | |
|---|---|---|---|---|---|---|---|---|---|---|---|---|
| Eqtn | 0.39021373 | 0.42253204 | 0.031 | 0.016 | 1 | 0.36861613 | -0.6094155 | 0.042 | 0.021 | 1 | 0.39021373 | 0.60135441 |
| Actr6 | 0.36864403 | 0.35541784 | 0.208 | 0.166 | 1 | 0.56849288 | -0.468553 | 0.147 | 0.115 | 1 | 0.56849288 | 0.60138964 |
| Acly | 0.37236816 | -0.4808865 | 0.896 | 0.87 | 1 | 0.36880762 | -0.360098 | 0.786 | 0.802 | 1 | 0.37236816 | 0.60159618 |
| Dgkq | 0.40599271 | 0.48393258 | 0.125 | 0.093 | 1 | 0.36938296 | 0.19915838 | 0.118 | 0.083 | 1 | 0.40599271 | 0.60232215 |
| Gm35019 | 0.5128082 | 0.65957123 | 0.094 | 0.073 | 1 | 0.36970988 | -1.5936482 | 0.059 | 0.083 | 1 | 0.5128082 | 0.60273436 |
| Scarf1 | 0.74827573 | 0.59245704 | 0.021 | 0.016 | 1 | 0.36981206 | 0.78968041 | 0.067 | 0.042 | 1 | 0.74827573 | 0.60286316 |
| Sh3bgr | 0.36982563 | -0.6929452 | 0.021 | 0.041 | 1 | 0.87066325 | -0.3098553 | 0.013 | 0.01 | 1 | 0.87066325 | 0.60288027 |
| Gdpd3 | 0.74827573 | 0.59245704 | 0.021 | 0.016 | 1 | 0.36990606 | -1.0297473 | 0.067 | 0.094 | 1 | 0.74827573 | 0.60298163 |
| Hpgds | 0.370501 | 2.17741954 | 0.031 | 0.016 | 1 | 0.42645812 | -0.9879272 | 0.025 | 0.042 | 1 | 0.42645812 | 0.603731 |
| Ccdc113 | 0.370501 | 1.59245704 | 0.031 | 0.016 | 1 | 0.76485982 | -0.3098553 | 0.025 | 0.031 | 1 | 0.76485982 | 0.603731 |
| Gm10190 | 0.370501 | 1.59245704 | 0.031 | 0.016 | 1 | 0.40503552 | -1.0468209 | 0.017 | 0.031 | 1 | 0.40503552 | 0.603731 |
| Phf7 | 0.40418925 | -0.4664367 | 0.073 | 0.104 | 1 | 0.37051676 | 0.64434105 | 0.092 | 0.062 | 1 | 0.40418925 | 0.60375085 |
| Arhgap24 | 0.93008641 | 0.22089817 | 0.177 | 0.176 | 1 | 0.37059877 | -0.7049932 | 0.113 | 0.146 | 1 | 0.93008641 | 0.60385409 |
| Rbmx2 | 0.63709119 | -0.240433 | 0.062 | 0.078 | 1 | 0.37123439 | 0.53814164 | 0.067 | 0.042 | 1 | 0.63709119 | 0.60465381 |
| Gm26749 | 0.3713814 | 0.47697982 | 0.073 | 0.047 | 1 | 0.56568391 | -0.2654611 | 0.109 | 0.083 | 1 | 0.56568391 | 0.60483865 |
| Id2 | 0.37181146 | 0.23046094 | 0.677 | 0.637 | 1 | 0.61476572 | -0.8095764 | 0.664 | 0.74 | 1 | 0.61476572 | 0.60537915 |
| Cyb561a3 | 0.54764662 | 0.49292136 | 0.115 | 0.093 | 1 | 0.37193145 | 0.49749966 | 0.08 | 0.052 | 1 | 0.54764662 | 0.6055299 |
| Tnfrsf22 | 0.82543011 | -0.4187702 | 0.135 | 0.14 | 1 | 0.37197927 | -1.0297473 | 0.059 | 0.083 | 1 | 0.82543011 | 0.60558996 |
| Gm56952 | 0.3720404 | -0.8405024 | 0.042 | 0.067 | 1 | 0.67751452 | -0.1843244 | 0.088 | 0.073 | 1 | 0.67751452 | 0.60566674 |
| Gm26879 | 0.37204076 | -0.7580402 | 0.042 | 0.067 | 1 | 0.92363734 | -0.5163061 | 0.092 | 0.094 | 1 | 0.92363734 | 0.60566719 |
| Sco1 | 0.37227565 | 0.72738662 | 0.115 | 0.083 | 1 | 0.70507825 | 0.12310414 | 0.097 | 0.083 | 1 | 0.70507825 | 0.60596214 |
| Rskr | 0.37233937 | -0.6929452 | 0.021 | 0.041 | 1 | 0.82600342 | -0.3098553 | 0.025 | 0.021 | 1 | 0.82600342 | 0.60604213 |
| Avl9 | 0.7475367 | -0.2512397 | 0.448 | 0.492 | 1 | 0.37261944 | -0.4491543 | 0.693 | 0.771 | 1 | 0.7475367 | 0.60639364 |
| Snupn | 0.37262469 | -0.4872702 | 0.104 | 0.14 | 1 | 0.60762848 | -0.4904275 | 0.097 | 0.115 | 1 | 0.60762848 | 0.60640022 |
| Tube1 | 0.37362512 | -0.2148979 | 0.021 | 0.041 | 1 | 0.39796128 | -1.7248928 | 0.017 | 0.031 | 1 | 0.39796128 | 0.60765452 |
| Zbtb48 | 0.98433693 | 0.20013961 | 0.062 | 0.062 | 1 | 0.37398879 | 0.27510724 | 0.118 | 0.083 | 1 | 0.98433693 | 0.60810996 |
| Tmem158 | 0.53159592 | -0.577468 | 0.073 | 0.093 | 1 | 0.37405857 | -0.7692869 | 0.113 | 0.146 | 1 | 0.53159592 | 0.60819732 |
| Gm48604 | 0.65896352 | 0.15949763 | 0.042 | 0.031 | 1 | 0.37433169 | 0.19264508 | 0.055 | 0.031 | 1 | 0.65896352 | 0.60853917 |
| 4930430F08Rik | 0.40328624 | 0.27052894 | 0.115 | 0.083 | 1 | 0.3743522 | 0.52664601 | 0.08 | 0.052 | 1 | 0.40328624 | 0.60856483 |
| Slc25a29 | 0.37440936 | 1.00749454 | 0.052 | 0.031 | 1 | 0.82344487 | -0.5443205 | 0.059 | 0.052 | 1 | 0.82344487 | 0.60863635 |
| Il15ra | 0.88471632 | -0.3424479 | 0.302 | 0.285 | 1 | 0.3744765 | -0.3483294 | 0.29 | 0.219 | 1 | 0.88471632 | 0.60872034 |
| Fahd1 | 0.73877691 | 0.27052894 | 0.094 | 0.109 | 1 | 0.37476227 | 0.45567948 | 0.067 | 0.042 | 1 | 0.73877691 | 0.60907778 |
| Tctn3 | 0.84593295 | 0.30705482 | 0.062 | 0.057 | 1 | 0.37476227 | 0.45567948 | 0.067 | 0.042 | 1 | 0.84593295 | 0.60907778 |
| Gm10648 | 0.78793893 | -0.7183305 | 0.167 | 0.171 | 1 | 0.37476681 | -0.4012783 | 0.21 | 0.156 | 1 | 0.78793893 | 0.60908345 |
| Fbf1 | 0.90435931 | 0.35038225 | 0.167 | 0.166 | 1 | 0.37495422 | -1.001733 | 0.168 | 0.198 | 1 | 0.90435931 | 0.60931777 |
| Rnf115 | 0.37511046 | 0.12613903 | 0.562 | 0.513 | 1 | 0.47728581 | -0.2865586 | 0.761 | 0.781 | 1 | 0.47728581 | 0.60951306 |
| Trim17 | 0.83841427 | -0.2148979 | 0.042 | 0.047 | 1 | 0.37524202 | 0.10518224 | 0.055 | 0.031 | 1 | 0.83841427 | 0.60967746 |
| Pla2r1 | 0.37536054 | 2.81484946 | 0.031 | 0.016 | 1 | 0.86804649 | 0.27510724 | 0.013 | 0.01 | 1 | 0.86804649 | 0.60982554 |
| Dnah9 | 0.37536054 | 1.32942263 | 0.031 | 0.016 | 1 | 0.79600331 | -1.3098553 | 0.017 | 0.021 | 1 | 0.79600331 | 0.60982554 |
| Arhgap4 | 0.37536054 | 1.59245704 | 0.031 | 0.016 | 1 | 0.70782098 | -0.6577786 | 0.034 | 0.042 | 1 | 0.70782098 | 0.60982554 |
| Gm46404 | 0.37537611 | 1.27052894 | 0.031 | 0.016 | 1 | 0.5818708 | -0.5728897 | 0.013 | 0.021 | 1 | 0.5818708 | 0.60984499 |
| Gm2449 | 0.3756878 | 0.59245704 | 0.052 | 0.031 | 1 | 0.99396042 | -0.1399303 | 0.021 | 0.021 | 1 | 0.99396042 | 0.61023427 |
| Zfp607b | 0.37594369 | -0.6705774 | 0.042 | 0.067 | 1 | 0.42845063 | 0.69014474 | 0.038 | 0.021 | 1 | 0.42845063 | 0.61055372 |
| Mboat2 | 0.59571442 | -0.4470713 | 0.188 | 0.212 | 1 | 0.37598581 | -0.2825823 | 0.483 | 0.344 | 1 | 0.59571442 | 0.61060629 |
| Ctu1 | 0.71031978 | 0.35541784 | 0.062 | 0.052 | 1 | 0.37607926 | 0.42711033 | 0.08 | 0.052 | 1 | 0.71031978 | 0.61072291 |
| Slc45a3 | 0.53635189 | -0.9185049 | 0.031 | 0.047 | 1 | 0.37612872 | 0.27510724 | 0.055 | 0.031 | 1 | 0.53635189 | 0.61078463 |
| H4c1 | 0.37612998 | -0.3144336 | 0.021 | 0.041 | 1 | 0.86613737 | -0.3098553 | 0.008 | 0.01 | 1 | 0.86613737 | 0.6107862 |
| Ndufa4l2 | 0.70532717 | -0.1837033 | 0.5 | 0.435 | 1 | 0.37678014 | 0.69014474 | 0.08 | 0.052 | 1 | 0.70532717 | 0.611597 |
| Fgl2 | 0.37845308 | -0.8669746 | 0.073 | 0.104 | 1 | 0.90566757 | -0.2499838 | 0.101 | 0.094 | 1 | 0.90566757 | 0.61367943 |
| Arhgap40 | 0.37863831 | 1.27052894 | 0.031 | 0.016 | 1 | 0.8076363 | 0.27510724 | 0.025 | 0.021 | 1 | 0.8076363 | 0.61390965 |
| Samd13 | 0.37865975 | 0.15949763 | 0.021 | 0.041 | 1 | 0.75636561 | -0.4618584 | 0.025 | 0.031 | 1 | 0.75636561 | 0.61393629 |
| Zfp707 | 0.8409956 | 0.32942263 | 0.031 | 0.036 | 1 | 0.37884342 | -0.5025003 | 0.055 | 0.031 | 1 | 0.8409956 | 0.6141645 |
| Zbtb2 | 0.37887276 | 0.26602755 | 0.292 | 0.244 | 1 | 0.75203561 | -0.5222636 | 0.357 | 0.333 | 1 | 0.75203561 | 0.61420096 |
| Tgif2 | 0.37891908 | -0.2219873 | 0.24 | 0.181 | 1 | 0.74175373 | -0.2567439 | 0.231 | 0.208 | 1 | 0.74175373 | 0.6142585 |
| Gm51425 | 0.37901216 | -0.7294711 | 0.042 | 0.067 | 1 | 0.65350257 | -1.3739856 | 0.071 | 0.083 | 1 | 0.65350257 | 0.6143741 |
| Usp31 | 0.37921576 | -0.1539689 | 0.312 | 0.244 | 1 | 0.92002281 | -0.4426105 | 0.475 | 0.438 | 1 | 0.92002281 | 0.61462693 |
| Clspn | 0.37936415 | 1.06889508 | 0.052 | 0.031 | 1 | 0.6390466 | 0.59703533 | 0.042 | 0.031 | 1 | 0.6390466 | 0.61481115 |
| Sil1 | 0.37941049 | -0.3313074 | 0.365 | 0.404 | 1 | 0.47490835 | -0.2809039 | 0.391 | 0.417 | 1 | 0.47490835 | 0.61486866 |
| Vil1 | 0.37986707 | -0.577468 | 0.042 | 0.067 | 1 | 0.40077865 | 0.27510724 | 0.025 | 0.01 | 1 | 0.40077865 | 0.61543515 |
| Fahd2a | 0.38773728 | 0.59245704 | 0.125 | 0.093 | 1 | 0.38005143 | 0.27510724 | 0.105 | 0.073 | 1 | 0.38773728 | 0.61566377 |
| Gm56897 | 0.38027888 | 1.00749454 | 0.031 | 0.016 | 1 | 0.41507275 | -0.5025003 | 0.017 | 0.031 | 1 | 0.41507275 | 0.61594573 |
| Sptbn5 | 0.38028373 | -1.4519371 | 0.01 | 0.026 | 1 | 0.5778028 | -0.8948178 | 0.013 | 0.021 | 1 | 0.5778028 | 0.61595174 |
| Tmem169 | 0.38028858 | 1.00749454 | 0.031 | 0.016 | 1 | 0.93576656 | -0.1399303 | 0.029 | 0.031 | 1 | 0.93576656 | 0.61595776 |
| Pak6 | 0.38028858 | 1.00749454 | 0.031 | 0.016 | 1 | 0.43559383 | 0.69014474 | 0.038 | 0.021 | 1 | 0.43559383 | 0.61595776 |
| Gm16098 | 0.38028858 | 1.00749454 | 0.031 | 0.016 | 1 | 0.67471359 | -0.9879272 | 0.017 | 0.01 | 1 | 0.67471359 | 0.61595776 |
| 4930557J02Rik | 0.38028858 | 1.00749454 | 0.031 | 0.016 | 1 | 1 | -1.5322477 | 0.021 | 0.021 | 1 | 1 | 0.61595776 |
| Ssc5d | 0.38028858 | 1.00749454 | 0.031 | 0.016 | 1 | 0.98992944 | -0.1723517 | 0.042 | 0.042 | 1 | 0.98992944 | 0.61595776 |
| Gm57192 | 0.38028858 | 1.00749454 | 0.031 | 0.016 | 1 | 0.65509988 | -1.1843244 | 0.042 | 0.052 | 1 | 0.65509988 | 0.61595776 |
| Irag1 | 0.38028858 | 1.00749454 | 0.031 | 0.016 | 1 | 0.79181108 | -1.0874628 | 0.017 | 0.021 | 1 | 0.79181108 | 0.61595776 |
| D830035M03Rik | 0.38028858 | 1.00749454 | 0.031 | 0.016 | 1 | 0.57050081 | -1.0874628 | 0.021 | 0.031 | 1 | 0.57050081 | 0.61595776 |
| Tgm4 | 0.38028858 | 1.00749454 | 0.031 | 0.016 | 1 | 0.8086289 | -0.3098553 | 0.017 | 0.021 | 1 | 0.8086289 | 0.61595776 |
| Gm12043 | 0.38028858 | 1.00749454 | 0.031 | 0.016 | 1 | 0.50887612 | -1.3098553 | 0.004 | 0.01 | 1 | 0.50887612 | 0.61595776 |
| Tmem30b | 0.38028858 | 1.00749454 | 0.031 | 0.016 | 1 | 0.62796184 | -0.2457249 | 0.08 | 0.062 | 1 | 0.62796184 | 0.61595776 |
| Ager | 0.38028858 | 1.00749454 | 0.031 | 0.016 | 1 | 0.87851725 | -1.3098553 | 0.013 | 0.01 | 1 | 0.87851725 | 0.61595776 |
| Gm56689 | 0.38028858 | 1.00749454 | 0.031 | 0.016 | 1 | 0.5818708 | -0.5728897 | 0.013 | 0.021 | 1 | 0.5818708 | 0.61595776 |
| Ranbp17 | 0.38031799 | 1.04813652 | 0.146 | 0.114 | 1 | 0.38959509 | -1.2794816 | 0.197 | 0.219 | 1 | 0.38959509 | 0.61599421 |
| Tlcd3a | 0.81639944 | 0.22988696 | 0.042 | 0.036 | 1 | 0.38036886 | 0.20471791 | 0.067 | 0.042 | 1 | 0.81639944 | 0.61605725 |
| Arid3a | 0.38050273 | 0.30705482 | 0.073 | 0.047 | 1 | 0.72474647 | -0.2774338 | 0.134 | 0.115 | 1 | 0.72474647 | 0.61622313 |
| Gm49741 | 0.38052601 | 0.81484946 | 0.052 | 0.031 | 1 | 0.40361911 | -0.1399303 | 0.025 | 0.01 | 1 | 0.40361911 | 0.61625197 |
| 2010310C07Rik | 0.38055552 | 0.17741954 | 0.073 | 0.047 | 1 | 0.53252125 | -1.3769695 | 0.076 | 0.094 | 1 | 0.53252125 | 0.61628854 |
| Tnfrsf19 | 0.60027455 | -2.765095 | 0.083 | 0.098 | 1 | 0.38068664 | -2.7692869 | 0.05 | 0.073 | 1 | 0.60027455 | 0.61645097 |
| Dubr | 0.38084354 | 0.31561683 | 0.104 | 0.073 | 1 | 0.38381448 | 0.15963002 | 0.105 | 0.073 | 1 | 0.38381448 | 0.61664528 |
| Unc93b1 | 0.77633969 | -0.1624305 | 0.208 | 0.218 | 1 | 0.38103677 | -0.8948178 | 0.206 | 0.24 | 1 | 0.77633969 | 0.61688452 |
| Prpf39 | 0.38122805 | 0.15768747 | 0.74 | 0.772 | 1 | 0.38787384 | -0.307387 | 0.874 | 0.906 | 1 | 0.38787384 | 0.61712127 |
| 4930417O13Rik | 0.38127812 | -0.4950058 | 0.021 | 0.041 | 1 | 0.87851725 | -1.8948178 | 0.013 | 0.01 | 1 | 0.87851725 | 0.61718324 |
| Stk36 | 0.38175174 | 0.64492446 | 0.052 | 0.031 | 1 | 0.82077588 | -0.1082214 | 0.076 | 0.083 | 1 | 0.82077588 | 0.61776909 |
| Hells | 0.38176919 | -0.3771693 | 0.104 | 0.14 | 1 | 0.51669575 | -1.3422767 | 0.118 | 0.135 | 1 | 0.51669575 | 0.61779066 |

| | | | | | | | | | | | | |
|---|---|---|---|---|---|---|---|---|---|---|---|---|
| A730063M14Rik | 0.38181636 | 0.78510212 | 0.052 | 0.031 | 1 | 0.45202482 | 0.10518224 | 0.05 | 0.031 | 1 | 0.45202482 | 0.61784899 |
| Gm56548 | 0.38181636 | 0.78510212 | 0.052 | 0.031 | 1 | 0.54837489 | -0.1399303 | 0.034 | 0.021 | 1 | 0.54837489 | 0.61784899 |
| Gm33667 | 0.3819238 | 0.52206771 | 0.031 | 0.016 | 1 | 0.50887612 | -1.3098553 | 0.004 | 0.01 | 1 | 0.50887612 | 0.61798182 |
| Phf19 | 0.3819238 | 0.74446013 | 0.031 | 0.016 | 1 | 0.51194506 | 0.27510724 | 0.021 | 0.01 | 1 | 0.51194506 | 0.61798182 |
| Gm15492 | 0.38192865 | -1.3144336 | 0.01 | 0.026 | 1 | 0.57855872 | -0.5728897 | 0.021 | 0.031 | 1 | 0.57855872 | 0.61798781 |
| 9330104G04Rik | 0.38192962 | -1.6929452 | 0.01 | 0.026 | 1 | 0.57693422 | -1.5025003 | 0.021 | 0.031 | 1 | 0.57693422 | 0.61798901 |
| Gm16576 | 0.40263565 | 0.53356335 | 0.219 | 0.176 | 1 | 0.38197943 | -1.1034044 | 0.109 | 0.135 | 1 | 0.40263565 | 0.61805057 |
| Hic2 | 0.38207195 | -0.577468 | 0.042 | 0.067 | 1 | 0.48160788 | -1.0297473 | 0.122 | 0.146 | 1 | 0.48160788 | 0.61816493 |
| Flt1 | 0.38256999 | -0.240433 | 0.021 | 0.041 | 1 | 0.5540789 | -0.681824 | 0.034 | 0.021 | 1 | 0.5540789 | 0.61878018 |
| Amd2 | 0.3828348 | -0.577468 | 0.073 | 0.104 | 1 | 0.71085062 | -0.5322477 | 0.092 | 0.104 | 1 | 0.71085062 | 0.61910712 |
| Vps37d | 0.6556999 | 0.31561683 | 0.104 | 0.088 | 1 | 0.38313921 | -0.7248928 | 0.076 | 0.104 | 1 | 0.6556999 | 0.61948276 |
| Kif5a | 0.91003996 | -0.1180363 | 0.094 | 0.088 | 1 | 0.38326423 | 0.32241295 | 0.092 | 0.062 | 1 | 0.91003996 | 0.61963699 |
| Fech | 0.38338279 | -0.4187702 | 0.281 | 0.321 | 1 | 0.89105872 | -0.5179635 | 0.332 | 0.292 | 1 | 0.89105872 | 0.61978321 |
| Kcnh1 | 0.38357309 | -1.8993961 | 0.01 | 0.026 | 1 | 0.92738405 | -0.4618584 | 0.034 | 0.031 | 1 | 0.92738405 | 0.62001786 |
| Chn1os3 | 0.38357309 | -0.9925055 | 0.01 | 0.026 | 1 | 0.87851807 | -0.9879272 | 0.013 | 0.01 | 1 | 0.87851807 | 0.62001786 |
| Gm9889 | 0.38357309 | -0.9925055 | 0.01 | 0.026 | 1 | 0.50887612 | -1.3098553 | 0.004 | 0.01 | 1 | 0.50887612 | 0.62001786 |
| Gm15494 | 0.38357309 | -1.1624305 | 0.01 | 0.026 | 1 | 0.87673803 | -0.1723517 | 0.038 | 0.042 | 1 | 0.87673803 | 0.62001786 |
| Zfp850 | 0.38357309 | -0.9925055 | 0.01 | 0.026 | 1 | 0.8086289 | -0.3098553 | 0.017 | 0.021 | 1 | 0.8086289 | 0.62001786 |
| Zfp791 | 0.38357309 | -1.1624305 | 0.01 | 0.026 | 1 | 0.38724403 | -0.3098553 | 0.055 | 0.031 | 1 | 0.38724403 | 0.62001786 |
| Tbx4 | 0.38357309 | -1.1624305 | 0.01 | 0.026 | 1 | 1 | -0.5025003 | 0.021 | 0.021 | 1 | 1 | 0.62001786 |
| Cacng5 | 0.38357309 | -1.4519371 | 0.01 | 0.026 | 1 | 0.85710544 | -1.3098553 | 0.008 | 0.01 | 1 | 0.85710544 | 0.62001786 |
| Gm33373 | 0.38357309 | -1.1624305 | 0.01 | 0.026 | 1 | 0.50887612 | -1.3098553 | 0.004 | 0.01 | 1 | 0.50887612 | 0.62001786 |
| Gm20045 | 0.53988469 | -0.2148979 | 0.062 | 0.083 | 1 | 0.38384482 | -0.5443205 | 0.084 | 0.115 | 1 | 0.53988469 | 0.62035279 |
| Phlda3 | 0.38385088 | 0.4474067 | 0.302 | 0.259 | 1 | 0.44929229 | 0.82920878 | 0.261 | 0.229 | 1 | 0.44929229 | 0.62036026 |
| Ahnak2 | 0.50448989 | -1.259292 | 0.125 | 0.145 | 1 | 0.38403807 | -0.4518743 | 0.076 | 0.104 | 1 | 0.50448989 | 0.6205909 |
| Gm26728 | 0.38422929 | 1.00749454 | 0.052 | 0.031 | 1 | 0.69119208 | 0.22619764 | 0.076 | 0.062 | 1 | 0.69119208 | 0.62082644 |
| Sbk1 | 0.4320165 | 0.22988696 | 0.125 | 0.093 | 1 | 0.38445331 | 0.27510724 | 0.067 | 0.042 | 1 | 0.4320165 | 0.62110227 |
| Slc1a1 | 0.59588202 | 0.17985858 | 0.438 | 0.518 | 1 | 0.38457291 | -0.2998712 | 0.315 | 0.25 | 1 | 0.59588202 | 0.6212495 |
| Zfp160 | 0.39760912 | 0.11996927 | 0.312 | 0.264 | 1 | 0.38482564 | -0.1200307 | 0.382 | 0.312 | 1 | 0.39760912 | 0.6215605 |
| Adgrb2 | 0.38521705 | -0.7998604 | 0.01 | 0.026 | 1 | 0.52879169 | 1.01207283 | 0.034 | 0.021 | 1 | 0.52879169 | 0.62204192 |
| Gm21411 | 0.38521705 | -0.7998604 | 0.01 | 0.026 | 1 | 0.81129592 | 0.10518224 | 0.025 | 0.021 | 1 | 0.81129592 | 0.62204192 |
| Epo | 0.38521705 | -0.7998604 | 0.01 | 0.026 | 1 | 0.73412881 | -0.3923174 | 0.063 | 0.052 | 1 | 0.73412881 | 0.62204192 |
| C230038L03Rik | 0.38521705 | -0.7998604 | 0.01 | 0.026 | 1 | 0.40647274 | -0.9879272 | 0.017 | 0.031 | 1 | 0.40647274 | 0.62204192 |
| Xylb | 0.38521705 | -0.7998604 | 0.01 | 0.026 | 1 | 0.75297348 | -1.1172102 | 0.025 | 0.031 | 1 | 0.75297348 | 0.62204192 |
| Pkd1l1 | 0.38521705 | -0.7998604 | 0.01 | 0.026 | 1 | 0.87851725 | -0.8948178 | 0.013 | 0.01 | 1 | 0.87851725 | 0.62204192 |
| Engase | 0.38521705 | -0.7998604 | 0.01 | 0.026 | 1 | 0.47582317 | -0.6317834 | 0.063 | 0.042 | 1 | 0.47582317 | 0.62204192 |
| Etohd2 | 0.38521705 | -0.7998604 | 0.01 | 0.026 | 1 | 0.79600331 | -0.9879272 | 0.017 | 0.021 | 1 | 0.79600331 | 0.62204192 |
| Hcls1 | 0.38521705 | -0.7998604 | 0.01 | 0.026 | 1 | 0.63505799 | 0.69014474 | 0.042 | 0.031 | 1 | 0.63505799 | 0.62204192 |
| Gap43 | 0.38521705 | -1.4519371 | 0.01 | 0.026 | 1 | 0.55565349 | -1.7248928 | 0.029 | 0.042 | 1 | 0.55565349 | 0.62204192 |
| Cdc42ep2 | 0.38521705 | -0.7998604 | 0.01 | 0.026 | 1 | 0.3937346 | 0.69014474 | 0.025 | 0.01 | 1 | 0.3937346 | 0.62204192 |
| Rnf113a1 | 0.38521705 | -0.7998604 | 0.01 | 0.026 | 1 | 0.93576656 | -0.1399303 | 0.029 | 0.031 | 1 | 0.93576656 | 0.62204192 |
| Gm13524 | 0.38521705 | 0.68556644 | 0.031 | 0.016 | 1 | 0.54837489 | -0.7248928 | 0.034 | 0.021 | 1 | 0.54837489 | 0.62204192 |
| Gm6260 | 0.38521705 | 0.68556644 | 0.031 | 0.016 | 1 | 0.86613737 | 0.56461386 | 0.008 | 0.01 | 1 | 0.86613737 | 0.62204192 |
| Gm15672 | 0.38521705 | 0.68556644 | 0.031 | 0.016 | 1 | 0.87851725 | -1.3098553 | 0.013 | 0.01 | 1 | 0.87851725 | 0.62204192 |
| 1700126G02Rik | 0.38521705 | 0.68556644 | 0.031 | 0.016 | 1 | 1 | -0.7248928 | 0.021 | 0.021 | 1 | 1 | 0.62204192 |
| Zc3h12d | 0.38521705 | 0.42253204 | 0.031 | 0.016 | 1 | 0.79600331 | -0.9879272 | 0.017 | 0.021 | 1 | 0.79600331 | 0.62204192 |
| Taco1os | 0.38521705 | 0.68556644 | 0.031 | 0.016 | 1 | 0.66387437 | 0.49749966 | 0.017 | 0.01 | 1 | 0.66387437 | 0.62204192 |
| Plek2 | 0.38521705 | 0.42253204 | 0.031 | 0.016 | 1 | 0.66339728 | 0.10518224 | 0.029 | 0.021 | 1 | 0.66339728 | 0.62204192 |
| A930001M01Rik | 0.38521705 | 0.68556644 | 0.031 | 0.016 | 1 | 0.98848918 | -0.7248928 | 0.042 | 0.042 | 1 | 0.98848918 | 0.62204192 |
| Prrg3 | 0.38521705 | 0.68556644 | 0.031 | 0.016 | 1 | 0.39515719 | 0.49749966 | 0.025 | 0.01 | 1 | 0.39515719 | 0.62204192 |
| Rbm19 | 0.39720008 | 0.52206771 | 0.125 | 0.093 | 1 | 0.3852248 | 0.11640949 | 0.143 | 0.104 | 1 | 0.39720008 | 0.62205146 |
| Pcdhb11 | 0.38523254 | 0.59245704 | 0.031 | 0.016 | 1 | 0.66171832 | 0.49749966 | 0.017 | 0.01 | 1 | 0.66171832 | 0.62206097 |
| Gm34102 | 0.84328969 | -0.2148979 | 0.042 | 0.047 | 1 | 0.38537087 | 0.10518224 | 0.055 | 0.031 | 1 | 0.84328969 | 0.62223103 |
| Gm26590 | 0.3866982 | 0.64492446 | 0.052 | 0.031 | 1 | 0.57170879 | -1.3098553 | 0.013 | 0.021 | 1 | 0.57170879 | 0.6238609 |
| 4930403P22Rik | 0.38689439 | -0.577468 | 0.01 | 0.026 | 1 | 0.91218966 | -0.7248928 | 0.029 | 0.031 | 1 | 0.91218966 | 0.62410151 |
| Inha | 0.38689439 | -0.577468 | 0.01 | 0.026 | 1 | 0.92228543 | -0.7248928 | 0.029 | 0.031 | 1 | 0.92228543 | 0.62410151 |
| Gm13703 | 0.38689439 | -0.577468 | 0.01 | 0.026 | 1 | 0.42275543 | -1.1172102 | 0.025 | 0.042 | 1 | 0.42275543 | 0.62410151 |
| She | 0.38689439 | -0.577468 | 0.01 | 0.026 | 1 | 0.95096139 | 0.49749966 | 0.029 | 0.031 | 1 | 0.95096139 | 0.62410151 |
| Gm43063 | 0.38689439 | -0.577468 | 0.01 | 0.026 | 1 | 0.86312513 | -0.7248928 | 0.008 | 0.01 | 1 | 0.86312513 | 0.62410151 |
| Pex26 | 0.38689439 | -0.577468 | 0.01 | 0.026 | 1 | 0.63592616 | -0.1399303 | 0.067 | 0.052 | 1 | 0.63592616 | 0.62410151 |
| Gm56971 | 0.38689439 | -0.577468 | 0.01 | 0.026 | 1 | 0.5778028 | -0.8948178 | 0.013 | 0.021 | 1 | 0.5778028 | 0.62410151 |
| Nudt12 | 0.38689439 | -0.577468 | 0.01 | 0.026 | 1 | 0.51018266 | 0.49749966 | 0.021 | 0.01 | 1 | 0.51018266 | 0.62410151 |
| D330050I16Rik | 0.38689439 | -0.577468 | 0.01 | 0.026 | 1 | 0.79600331 | -0.9879272 | 0.017 | 0.021 | 1 | 0.79600331 | 0.62410151 |
| Ptpn6 | 0.38715463 | 0.90057933 | 0.083 | 0.057 | 1 | 0.75383252 | -1.1034044 | 0.055 | 0.062 | 1 | 0.75383252 | 0.62442055 |
| Dnaja4 | 0.38733316 | -0.4274425 | 0.198 | 0.238 | 1 | 0.55683869 | 0.14184071 | 0.282 | 0.25 | 1 | 0.55683869 | 0.62463934 |
| Mrm3 | 0.38773728 | 0.59245704 | 0.125 | 0.093 | 1 | 0.79094611 | -0.3098553 | 0.084 | 0.073 | 1 | 0.79094611 | 0.62513436 |
| Gm29681 | 0.38789252 | 0.27052894 | 0.052 | 0.031 | 1 | 0.94364294 | -0.3098553 | 0.05 | 0.052 | 1 | 0.94364294 | 0.62532443 |
| Zfp772 | 0.38789252 | 0.42253204 | 0.052 | 0.031 | 1 | 0.76485982 | -0.3098553 | 0.025 | 0.031 | 1 | 0.76485982 | 0.62532443 |
| Acad8 | 0.38812386 | -0.5488988 | 0.156 | 0.192 | 1 | 0.83635505 | -0.2249664 | 0.231 | 0.208 | 1 | 0.83635505 | 0.62560759 |
| Clock | 0.56828057 | -0.1365158 | 0.698 | 0.715 | 1 | 0.38823058 | -0.3878578 | 0.853 | 0.906 | 1 | 0.56828057 | 0.62573817 |
| Gm40438 | 0.38823428 | 1.07162487 | 0.083 | 0.057 | 1 | 0.54211203 | -0.2554075 | 0.059 | 0.042 | 1 | 0.54211203 | 0.62574271 |
| Gm47664 | 0.38896453 | -0.407543 | 0.042 | 0.067 | 1 | 0.8259932 | -0.5025003 | 0.025 | 0.021 | 1 | 0.8259932 | 0.62663565 |
| Tmem267 | 0.9906923 | 0.37006462 | 0.052 | 0.052 | 1 | 0.3890866 | -0.3098553 | 0.055 | 0.031 | 1 | 0.9906923 | 0.62678482 |
| Pex1 | 0.38924243 | -0.236431 | 0.125 | 0.166 | 1 | 0.6444234 | -0.2860085 | 0.155 | 0.135 | 1 | 0.6444234 | 0.62697519 |
| Edn2 | 0.67903939 | -1.8338077 | 0.042 | 0.052 | 1 | 0.38956906 | 4.27510724 | 0.025 | 0.01 | 1 | 0.67903939 | 0.62737407 |
| Arvcf | 0.80885634 | -0.4519371 | 0.031 | 0.026 | 1 | 0.38956906 | 1.14957636 | 0.025 | 0.01 | 1 | 0.80885634 | 0.62737407 |
| Acadm | 0.38978279 | 0.31234912 | 0.385 | 0.347 | 1 | 0.70825256 | 0.21001221 | 0.458 | 0.552 | 1 | 0.70825256 | 0.62763495 |
| Cfap157 | 0.39021373 | 0.42253204 | 0.031 | 0.016 | 1 | 0.72387422 | -0.3769695 | 0.063 | 0.052 | 1 | 0.72387422 | 0.6281607 |
| Spata32 | 0.39021373 | 0.20013961 | 0.031 | 0.016 | 1 | 0.70494229 | -1.4353861 | 0.034 | 0.042 | 1 | 0.70494229 | 0.6281607 |
| A330032B11Rik | 0.39021373 | -0.1624305 | 0.031 | 0.016 | 1 | 0.39515719 | 0.49749966 | 0.025 | 0.01 | 1 | 0.39515719 | 0.6281607 |
| Kcnb2 | 0.39022145 | -1.5285584 | 0.01 | 0.026 | 1 | 0.86915235 | 0.49749966 | 0.008 | 0.01 | 1 | 0.86915235 | 0.62817013 |
| Arhgap17 | 0.39033656 | 0.18106892 | 0.812 | 0.782 | 1 | 0.49445353 | -0.2297214 | 0.849 | 0.844 | 1 | 0.49445353 | 0.62831049 |
| Zfp820 | 0.84451844 | -0.130009 | 0.042 | 0.036 | 1 | 0.39095723 | 1.27510724 | 0.025 | 0.01 | 1 | 0.84451844 | 0.62906691 |
| Drg1 | 0.53690597 | 0.18845781 | 0.635 | 0.627 | 1 | 0.3912187 | -0.3459747 | 0.739 | 0.708 | 1 | 0.53690597 | 0.62938533 |

| | | | | | | | | | | | | |
|---|---|---|---|---|---|---|---|---|---|---|---|---|
| Fermt3 | 0.99275828 | 0.15949763 | 0.021 | 0.021 | 1 | 0.3913111 | -0.4431218 | 0.084 | 0.115 | 1 | 0.99275828 | 0.62949782 |
| Gm44700 | 0.66186725 | -0.577468 | 0.031 | 0.041 | 1 | 0.39170927 | -1.4167705 | 0.05 | 0.073 | 1 | 0.66186725 | 0.62998239 |
| Pcdhb21 | 0.39188474 | -0.407543 | 0.01 | 0.026 | 1 | 0.99194729 | -1.3098553 | 0.021 | 0.021 | 1 | 0.99194729 | 0.63019584 |
| Crabp2 | 0.39188511 | -2.1624305 | 0.052 | 0.078 | 1 | 0.65060587 | -1.84037 | 0.063 | 0.073 | 1 | 0.65060587 | 0.63019629 |
| Slc8a2 | 0.39194862 | 0.59245704 | 0.083 | 0.057 | 1 | 0.42509673 | -1.0728161 | 0.08 | 0.104 | 1 | 0.42509673 | 0.63027352 |
| Oma1 | 0.3923061 | 0.51445453 | 0.177 | 0.14 | 1 | 0.86473327 | -0.4447848 | 0.151 | 0.156 | 1 | 0.86473327 | 0.63070813 |
| Nat2 | 0.47599549 | 0.52206771 | 0.042 | 0.026 | 1 | 0.39234595 | 0.86006974 | 0.025 | 0.01 | 1 | 0.47599549 | 0.63075656 |
| A230108P19Rik | 0.48065738 | -0.4519371 | 0.021 | 0.036 | 1 | 0.39234595 | 0.86006974 | 0.025 | 0.01 | 1 | 0.48065738 | 0.63075656 |
| Tmem255a | 0.73685701 | 0.59245704 | 0.01 | 0.016 | 1 | 0.39234595 | 0.86006974 | 0.025 | 0.01 | 1 | 0.73685701 | 0.63075656 |
| Mex3b | 0.39285382 | -0.6623569 | 0.083 | 0.114 | 1 | 0.55504515 | -1.0203486 | 0.067 | 0.083 | 1 | 0.55504515 | 0.63137351 |
| Rttn | 0.86003944 | 0.13302542 | 0.104 | 0.098 | 1 | 0.39341357 | -0.5393371 | 0.076 | 0.104 | 1 | 0.86003944 | 0.6320529 |
| Plxnd1 | 0.39367055 | -0.765095 | 0.083 | 0.114 | 1 | 0.49162408 | -0.7504279 | 0.282 | 0.292 | 1 | 0.49162408 | 0.63236459 |
| Tex26 | 0.44164906 | -0.1624305 | 0.062 | 0.041 | 1 | 0.3937346 | 1.14957636 | 0.025 | 0.01 | 1 | 0.44164906 | 0.63244227 |
| Gm49801 | 0.48064264 | 0.59245704 | 0.021 | 0.01 | 1 | 0.3937346 | 1.01207283 | 0.025 | 0.01 | 1 | 0.48064264 | 0.63244227 |
| 1700034H15Rik | 0.61737927 | 1.00749454 | 0.01 | 0.005 | 1 | 0.3937346 | 0.69014474 | 0.025 | 0.01 | 1 | 0.61737927 | 0.63244227 |
| A330009N23Rik | 0.61737927 | 1.00749454 | 0.01 | 0.005 | 1 | 0.3937346 | 0.69014474 | 0.025 | 0.01 | 1 | 0.61737927 | 0.63244227 |
| A330102I10Rik | 0.72574709 | -0.3144336 | 0.01 | 0.016 | 1 | 0.3937346 | 0.69014474 | 0.025 | 0.01 | 1 | 0.72574709 | 0.63244227 |
| 6820408C15Rik | 0.81497862 | -0.1624305 | 0.031 | 0.026 | 1 | 0.3937346 | 0.69014474 | 0.025 | 0.01 | 1 | 0.81497862 | 0.63244227 |
| Ddo | 0.99034437 | 0.74446013 | 0.021 | 0.021 | 1 | 0.3937346 | 1.01207283 | 0.025 | 0.01 | 1 | 0.99034437 | 0.63244227 |
| Rnf135 | 0.39379496 | 0.62898291 | 0.083 | 0.057 | 1 | 0.51814236 | -0.8948178 | 0.046 | 0.062 | 1 | 0.51814236 | 0.63251545 |
| Gm57027 | 0.39417399 | 0.22988696 | 0.052 | 0.031 | 1 | 0.42648204 | -0.8244284 | 0.025 | 0.042 | 1 | 0.42648204 | 0.63297485 |
| Skap1 | 0.39417399 | 0.35541784 | 0.052 | 0.031 | 1 | 0.86312513 | -0.7248928 | 0.008 | 0.01 | 1 | 0.86312513 | 0.63297485 |
| Tmem192 | 0.49468518 | -0.4254649 | 0.094 | 0.119 | 1 | 0.39468526 | 0.14957636 | 0.118 | 0.083 | 1 | 0.49468518 | 0.63359407 |
| Gm42939 | 0.82258416 | -0.1624305 | 0.031 | 0.036 | 1 | 0.39475201 | -0.9113059 | 0.097 | 0.125 | 1 | 0.82258416 | 0.63367487 |
| Agbl3 | 0.99119559 | -1.1564812 | 0.323 | 0.295 | 1 | 0.39476467 | -0.6414768 | 0.227 | 0.26 | 1 | 0.99119559 | 0.6336902 |
| Garin2 | 0.39506346 | -0.3144336 | 0.073 | 0.047 | 1 | 0.76416309 | -0.7428147 | 0.118 | 0.125 | 1 | 0.76416309 | 0.63405178 |
| Rsbn1 | 0.395087 | 0.18609184 | 0.656 | 0.601 | 1 | 0.59049944 | -0.3692338 | 0.79 | 0.812 | 1 | 0.59049944 | 0.63408026 |
| Gm28874 | 0.99009299 | 0.32942263 | 0.031 | 0.031 | 1 | 0.39514482 | -1.7952821 | 0.017 | 0.031 | 1 | 0.99009299 | 0.63415022 |
| T2 | 0.99758607 | 0.20013961 | 0.021 | 0.021 | 1 | 0.39514482 | -1.9879272 | 0.017 | 0.031 | 1 | 0.99758607 | 0.63415022 |
| Gm57013 | 0.52961488 | -0.3144336 | 0.01 | 0.021 | 1 | 0.39515719 | 0.49749966 | 0.025 | 0.01 | 1 | 0.52961488 | 0.63416517 |
| Notch4 | 0.53487059 | -0.4519371 | 0.031 | 0.047 | 1 | 0.39515719 | 0.49749966 | 0.025 | 0.01 | 1 | 0.53487059 | 0.63416517 |
| Cmtr2 | 0.5862099 | 0.68556644 | 0.031 | 0.021 | 1 | 0.39515719 | 0.49749966 | 0.025 | 0.01 | 1 | 0.5862099 | 0.63416517 |
| Gm56859 | 0.5862099 | 0.68556644 | 0.031 | 0.021 | 1 | 0.39515719 | 0.49749966 | 0.025 | 0.01 | 1 | 0.5862099 | 0.63416517 |
| Cdk20 | 0.5862099 | 0.68556644 | 0.031 | 0.021 | 1 | 0.39515719 | 0.49749966 | 0.025 | 0.01 | 1 | 0.5862099 | 0.63416517 |
| Fadd | 0.61550914 | -0.407543 | 0.021 | 0.031 | 1 | 0.39515719 | 0.49749966 | 0.025 | 0.01 | 1 | 0.61550914 | 0.63416517 |
| Mrgpre | 0.61737927 | 1.00749454 | 0.01 | 0.005 | 1 | 0.39515719 | 0.49749966 | 0.025 | 0.01 | 1 | 0.61737927 | 0.63416517 |
| Cnih2 | 0.61737927 | 1.00749454 | 0.01 | 0.005 | 1 | 0.39515719 | 0.49749966 | 0.025 | 0.01 | 1 | 0.61737927 | 0.63416517 |
| Zfp984 | 0.62104899 | 0.42253204 | 0.01 | 0.005 | 1 | 0.39515719 | 0.49749966 | 0.025 | 0.01 | 1 | 0.62104899 | 0.63416517 |
| Gm50250 | 0.72298091 | -0.577468 | 0.01 | 0.016 | 1 | 0.39515719 | 0.49749966 | 0.025 | 0.01 | 1 | 0.72298091 | 0.63416517 |
| 9530062K07Rik | 0.74326422 | 1.00749454 | 0.021 | 0.016 | 1 | 0.39515719 | 0.49749966 | 0.025 | 0.01 | 1 | 0.74326422 | 0.63416517 |
| Slc26a1 | 0.75328966 | 0.27052894 | 0.021 | 0.016 | 1 | 0.39515719 | 0.49749966 | 0.025 | 0.01 | 1 | 0.75328966 | 0.63416517 |
| Odad4 | 0.86441475 | -0.1079827 | 0.052 | 0.047 | 1 | 0.39515719 | 0.49749966 | 0.025 | 0.01 | 1 | 0.86441475 | 0.63416517 |
| Pld4 | 0.9951721 | 0.42253204 | 0.021 | 0.021 | 1 | 0.39515719 | 0.49749966 | 0.025 | 0.01 | 1 | 0.9951721 | 0.63416517 |
| Mapk8ip2 | 1 | 0.42253204 | 0.01 | 0.01 | 1 | 0.39515719 | 0.49749966 | 0.025 | 0.01 | 1 | 1 | 0.63416517 |
| Tbc1d30 | 0.39523977 | 0.78510212 | 0.01 | 0.026 | 1 | 0.68622073 | -0.7952821 | 0.029 | 0.021 | 1 | 0.68622073 | 0.63426507 |
| Ncoa7 | 0.56828622 | -0.6247737 | 0.219 | 0.244 | 1 | 0.39539699 | 0.28804629 | 0.239 | 0.198 | 1 | 0.56828622 | 0.6344552 |
| Rabep1 | 0.45166747 | -0.1545016 | 0.812 | 0.803 | 1 | 0.3961782 | -0.2607076 | 0.878 | 0.854 | 1 | 0.45166747 | 0.63539923 |
| Trpc3 | 0.47163774 | 1.28312898 | 0.094 | 0.073 | 1 | 0.39627789 | -0.7573142 | 0.067 | 0.094 | 1 | 0.47163774 | 0.63551961 |
| Arhgap6 | 0.39630201 | -0.4347278 | 0.177 | 0.228 | 1 | 0.86470029 | -0.55369 | 0.441 | 0.375 | 1 | 0.86470029 | 0.63554873 |
| Rai2 | 0.83601253 | -1.0513992 | 0.062 | 0.067 | 1 | 0.39640349 | 0.19915838 | 0.067 | 0.042 | 1 | 0.83601253 | 0.63567126 |
| Doc2b | 0.52961488 | -0.3144336 | 0.01 | 0.021 | 1 | 0.39655495 | -1.8948178 | 0.017 | 0.031 | 1 | 0.52961488 | 0.63585407 |
| Amer1 | 0.55043116 | 0.47697982 | 0.125 | 0.104 | 1 | 0.39673919 | -0.5184419 | 0.122 | 0.156 | 1 | 0.55043116 | 0.6360764 |
| Def8 | 0.56684126 | 0.18806678 | 0.156 | 0.13 | 1 | 0.39685559 | -0.2532717 | 0.172 | 0.125 | 1 | 0.56684126 | 0.63621683 |
| Gpt2 | 0.92568168 | 0.35541784 | 0.094 | 0.098 | 1 | 0.39732751 | 0.57767001 | 0.113 | 0.083 | 1 | 0.92568168 | 0.63678587 |
| Tigit | 0.72298091 | -0.577468 | 0.01 | 0.016 | 1 | 0.39746984 | -2.2029401 | 0.025 | 0.042 | 1 | 0.72298091 | 0.63695741 |
| Zbtb9 | 0.39760751 | -0.6299354 | 0.073 | 0.104 | 1 | 0.87979871 | -0.2703269 | 0.101 | 0.094 | 1 | 0.87979871 | 0.63712329 |
| Pcbd1 | 0.61737927 | 1.00749454 | 0.01 | 0.005 | 1 | 0.39796314 | 0.56461386 | 0.025 | 0.01 | 1 | 0.61737927 | 0.63755161 |
| Sh3pxd2b | 0.39824117 | -0.6671052 | 0.323 | 0.244 | 1 | 0.4157956 | -0.9968524 | 0.475 | 0.458 | 1 | 0.4157956 | 0.63788631 |
| Sox2ot | 0.39833389 | -0.5095639 | 0.104 | 0.14 | 1 | 0.42236215 | 0.57278779 | 0.168 | 0.135 | 1 | 0.42236215 | 0.63799789 |
| Armc6 | 0.98792413 | 0.27052894 | 0.042 | 0.041 | 1 | 0.39880045 | -0.1399303 | 0.067 | 0.042 | 1 | 0.98792413 | 0.6385591 |
| Arfgef2 | 0.39906915 | -0.3828954 | 0.802 | 0.731 | 1 | 0.71770983 | -0.1413168 | 0.815 | 0.865 | 1 | 0.71770983 | 0.63888211 |
| Nlrc5 | 0.39932174 | -0.5683658 | 0.292 | 0.342 | 1 | 0.59091557 | -0.805624 | 0.231 | 0.24 | 1 | 0.59091557 | 0.63918563 |
| H1f5 | 0.91791135 | 2.99963242 | 0.083 | 0.083 | 1 | 0.39937124 | 0.80562195 | 0.025 | 0.01 | 1 | 0.91791135 | 0.63924509 |
| Gm57236 | 0.98793039 | 0.68556644 | 0.021 | 0.021 | 1 | 0.39937186 | -1.3098553 | 0.017 | 0.031 | 1 | 0.98793039 | 0.63924583 |
| Ccdc28a | 0.3994615 | -0.4399644 | 0.073 | 0.104 | 1 | 0.88738128 | -0.5508634 | 0.059 | 0.062 | 1 | 0.88738128 | 0.63935351 |
| Gm21781 | 0.43926834 | 0.54806292 | 0.062 | 0.041 | 1 | 0.39965247 | -0.5993619 | 0.067 | 0.042 | 1 | 0.43926834 | 0.63958284 |
| Mafa | 0.71255442 | -0.4950058 | 0.052 | 0.062 | 1 | 0.40013243 | 0.83698613 | 0.088 | 0.062 | 1 | 0.71255442 | 0.64015889 |
| Kcng2 | 0.417486 | 0.32942263 | 0.083 | 0.057 | 1 | 0.40038903 | 0.10518224 | 0.092 | 0.062 | 1 | 0.417486 | 0.64046669 |
| Gm50399 | 0.40046717 | 0.22988696 | 0.052 | 0.031 | 1 | 0.415442 | -1.3098553 | 0.025 | 0.042 | 1 | 0.415442 | 0.64056038 |
| Tcp11l1 | 0.40076296 | 0.24195979 | 0.24 | 0.197 | 1 | 0.67160077 | -0.1148393 | 0.265 | 0.229 | 1 | 0.67160077 | 0.64091497 |
| 4930506C21Rik | 0.45476301 | 1.22988696 | 0.042 | 0.026 | 1 | 0.40076941 | -1.5728897 | 0.017 | 0.031 | 1 | 0.45476301 | 0.6409227 |
| Kcnc4 | 0.52528691 | -0.7998604 | 0.01 | 0.021 | 1 | 0.40076941 | -1.3098553 | 0.017 | 0.031 | 1 | 0.52528691 | 0.6409227 |
| 4921515G04Rik | 0.99758609 | 0.27052894 | 0.021 | 0.021 | 1 | 0.40076941 | -1.3098553 | 0.017 | 0.031 | 1 | 0.99758609 | 0.6409227 |
| Gm26631 | 0.48064264 | 0.59245704 | 0.021 | 0.01 | 1 | 0.40078481 | 0.27510724 | 0.025 | 0.01 | 1 | 0.48064264 | 0.64094116 |
| Srl | 1 | 0.42253204 | 0.01 | 0.01 | 1 | 0.40078481 | -1.3098553 | 0.017 | 0.031 | 1 | 1 | 0.64094116 |
| St8sia4 | 0.74576864 | 1.00749454 | 0.021 | 0.016 | 1 | 0.40078851 | 0.27510724 | 0.025 | 0.01 | 1 | 0.74576864 | 0.64094559 |
| 1700112J16Rik | 0.4014134 | -0.747393 | 0.052 | 0.078 | 1 | 0.96201719 | -0.7862933 | 0.063 | 0.062 | 1 | 0.96201719 | 0.64169408 |
| Plekha5 | 0.92447469 | -0.250442 | 0.708 | 0.689 | 1 | 0.40169494 | -0.3979862 | 0.84 | 0.885 | 1 | 0.92447469 | 0.64203105 |
| Angel1 | 0.47150075 | 2.42253204 | 0.021 | 0.01 | 1 | 0.40219461 | -0.3098553 | 0.025 | 0.01 | 1 | 0.47150075 | 0.64262872 |
| Rnf225 | 0.64197016 | 0.59245704 | 0.042 | 0.031 | 1 | 0.40219461 | 0.10518224 | 0.025 | 0.01 | 1 | 0.64197016 | 0.64262872 |
| Slc24a5 | 0.40224917 | -0.1851505 | 0.042 | 0.067 | 1 | 0.44357873 | 0.27510724 | 0.088 | 0.062 | 1 | 0.44357873 | 0.64269394 |
| Rap1gap2 | 0.40225053 | -1.4187702 | 0.062 | 0.088 | 1 | 0.50531363 | -1.0260623 | 0.134 | 0.156 | 1 | 0.50531363 | 0.64269557 |

| | | | | | | | | | | | | |
|---|---|---|---|---|---|---|---|---|---|---|---|---|
| Tkt | 0.5148875 | -0.2195744 | 0.719 | 0.741 | 1 | 0.40321663 | -0.1449483 | 0.63 | 0.771 | 1 | 0.5148875 | 0.64384961 |
| Rasd1 | 0.40340164 | -1.3323555 | 0.115 | 0.15 | 1 | 0.72766729 | -1.0297473 | 0.122 | 0.104 | 1 | 0.72766729 | 0.6440704 |
| Gm9920 | 0.52744059 | -0.577468 | 0.01 | 0.021 | 1 | 0.40360435 | -0.5025003 | 0.025 | 0.01 | 1 | 0.52744059 | 0.64431222 |
| Reep1 | 0.51793219 | 1.00749454 | 0.052 | 0.036 | 1 | 0.40361357 | -0.7248928 | 0.025 | 0.01 | 1 | 0.51793219 | 0.64432323 |
| Cpne5 | 0.52313476 | -1.3144336 | 0.01 | 0.021 | 1 | 0.40361357 | -0.3098553 | 0.025 | 0.01 | 1 | 0.52313476 | 0.64432323 |
| Gm56649 | 0.75328966 | 0.27052894 | 0.021 | 0.016 | 1 | 0.40361357 | -0.6317834 | 0.025 | 0.01 | 1 | 0.75328966 | 0.64432323 |
| Fam161b | 0.81708725 | -0.3144336 | 0.031 | 0.036 | 1 | 0.40361357 | -0.3098553 | 0.025 | 0.01 | 1 | 0.81708725 | 0.64432323 |
| Tbxas1 | 0.82258416 | -0.3144336 | 0.031 | 0.036 | 1 | 0.40361357 | -0.3098553 | 0.025 | 0.01 | 1 | 0.82258416 | 0.64432323 |
| Ccdc187 | 1 | 0.42253204 | 0.01 | 0.01 | 1 | 0.40361357 | -0.6317834 | 0.025 | 0.01 | 1 | 1 | 0.64432323 |
| Ppp1r16b | 0.52744059 | -0.577468 | 0.01 | 0.021 | 1 | 0.40362095 | -0.3098553 | 0.025 | 0.01 | 1 | 0.52744059 | 0.64433203 |
| Mus81 | 0.68932285 | -0.265524 | 0.104 | 0.119 | 1 | 0.4037894 | -0.7248928 | 0.059 | 0.083 | 1 | 0.68932285 | 0.64453292 |
| Abhd18 | 0.40404234 | 0.59245704 | 0.135 | 0.104 | 1 | 0.81089703 | -0.5393371 | 0.143 | 0.146 | 1 | 0.81089703 | 0.64483447 |
| Ccdc30 | 0.84099824 | -0.2148979 | 0.042 | 0.036 | 1 | 0.40416743 | -1.4253325 | 0.042 | 0.062 | 1 | 0.84099824 | 0.64498355 |
| Adamts7 | 0.47150075 | 1.42253204 | 0.021 | 0.01 | 1 | 0.40503552 | -1.0468209 | 0.017 | 0.031 | 1 | 0.47150075 | 0.64601727 |
| 4931408D14Rik | 0.64538863 | 0.52206771 | 0.042 | 0.031 | 1 | 0.40568086 | -1.3098553 | 0.034 | 0.052 | 1 | 0.64538863 | 0.64678476 |
| A430033K04Rik | 0.40602211 | -0.7998604 | 0.052 | 0.078 | 1 | 0.79862134 | -0.7248928 | 0.088 | 0.094 | 1 | 0.79862134 | 0.64719027 |
| Ttbk1 | 0.61550914 | -0.577468 | 0.021 | 0.031 | 1 | 0.40646046 | -0.7248928 | 0.017 | 0.031 | 1 | 0.61550914 | 0.64771082 |
| Gja4 | 0.74576744 | 0.68556644 | 0.021 | 0.016 | 1 | 0.40646046 | -0.6317834 | 0.017 | 0.031 | 1 | 0.74576744 | 0.64771082 |
| Cxcr5 | 0.52961488 | -0.3144336 | 0.01 | 0.021 | 1 | 0.40647274 | -0.9879272 | 0.017 | 0.031 | 1 | 0.52961488 | 0.64772539 |
| Espnl | 0.58426772 | 0.74446013 | 0.031 | 0.021 | 1 | 0.40647274 | -0.9879272 | 0.017 | 0.031 | 1 | 0.58426772 | 0.64772539 |
| Gm37233 | 0.61550914 | -0.7294711 | 0.021 | 0.031 | 1 | 0.40647274 | -0.9879272 | 0.017 | 0.031 | 1 | 0.61550914 | 0.64772539 |
| Gm45767 | 0.40660753 | 0.46692616 | 0.083 | 0.057 | 1 | 0.4844697 | -0.6883669 | 0.063 | 0.083 | 1 | 0.4844697 | 0.64788537 |
| Grik5 | 0.69744678 | -0.1393469 | 0.104 | 0.119 | 1 | 0.40663 | -0.169374 | 0.273 | 0.208 | 1 | 0.69744678 | 0.64791204 |
| St6galnac3 | 0.40758417 | -1.9925055 | 0.031 | 0.052 | 1 | 0.7563772 | -1.4904275 | 0.025 | 0.031 | 1 | 0.7563772 | 0.64904349 |
| Klhl31 | 0.9927582 | 0.42253204 | 0.021 | 0.021 | 1 | 0.40789759 | -0.5322477 | 0.017 | 0.031 | 1 | 0.9927582 | 0.64941473 |
| Lockd | 0.67759869 | 1.71798792 | 0.073 | 0.062 | 1 | 0.40886845 | -1.4473588 | 0.034 | 0.052 | 1 | 0.67759869 | 0.65056349 |
| Prps2 | 0.4723989 | 0.47697982 | 0.073 | 0.052 | 1 | 0.4088951 | -0.2592292 | 0.105 | 0.073 | 1 | 0.4723989 | 0.650595 |
| Zfp26 | 0.41002197 | 0.43325844 | 0.292 | 0.244 | 1 | 0.50569114 | -0.4077026 | 0.345 | 0.271 | 1 | 0.50569114 | 0.65192593 |
| Tmem47 | 0.41005162 | 0.1843723 | 0.177 | 0.135 | 1 | 0.90511568 | -0.6670118 | 0.256 | 0.229 | 1 | 0.90511568 | 0.65196091 |
| Gm30085 | 0.97828541 | -0.577468 | 0.052 | 0.052 | 1 | 0.41075871 | -0.6317834 | 0.017 | 0.031 | 1 | 0.97828541 | 0.6527947 |
| Zfp839 | 0.50251363 | 0.42253204 | 0.146 | 0.119 | 1 | 0.41111052 | -0.9313436 | 0.088 | 0.115 | 1 | 0.50251363 | 0.65320919 |
| Gm16192 | 0.98792469 | -0.577468 | 0.042 | 0.041 | 1 | 0.41179613 | -1.3098553 | 0.025 | 0.042 | 1 | 0.98792469 | 0.65401621 |
| Mfsd2b | 0.9943045 | -0.1624305 | 0.062 | 0.062 | 1 | 0.41231187 | -0.3098553 | 0.092 | 0.062 | 1 | 0.9943045 | 0.65462267 |
| Zfp319 | 0.41239321 | 0.27052894 | 0.083 | 0.057 | 1 | 0.5671618 | -0.129283 | 0.13 | 0.104 | 1 | 0.5671618 | 0.65471826 |
| Wdr25 | 0.74547103 | -0.1851505 | 0.062 | 0.073 | 1 | 0.41345952 | -1.1399303 | 0.067 | 0.042 | 1 | 0.74547103 | 0.65597027 |
| Pik3r4 | 0.91671584 | -0.3848229 | 0.135 | 0.135 | 1 | 0.41400563 | -0.2532717 | 0.189 | 0.146 | 1 | 0.91671584 | 0.65661059 |
| Ttc38 | 0.66676245 | -0.4519371 | 0.031 | 0.041 | 1 | 0.41415595 | -1.1399303 | 0.034 | 0.052 | 1 | 0.66676245 | 0.65678675 |
| Rac2 | 0.61371908 | 2.00749454 | 0.01 | 0.005 | 1 | 0.41446037 | 2.96316323 | 0.038 | 0.021 | 1 | 0.61371908 | 0.65714334 |
| Pip5kl1 | 0.41466006 | -0.9925055 | 0.031 | 0.052 | 1 | 0.53156006 | 0.56461386 | 0.034 | 0.021 | 1 | 0.53156006 | 0.65737716 |
| Tmem94 | 0.41470975 | 0.31561683 | 0.062 | 0.093 | 1 | 0.43221293 | -0.9093173 | 0.109 | 0.135 | 1 | 0.43221293 | 0.65743533 |
| Ccdc91 | 0.41498717 | -0.2448926 | 0.198 | 0.244 | 1 | 0.94707913 | -0.5073013 | 0.202 | 0.198 | 1 | 0.94707913 | 0.65775999 |
| Gm43462 | 0.41537628 | 0.53800925 | 0.167 | 0.135 | 1 | 0.84824907 | -0.5531264 | 0.151 | 0.156 | 1 | 0.84824907 | 0.6582151 |
| F420014N23Rik | 0.41538946 | -0.3500575 | 0.115 | 0.15 | 1 | 0.72832583 | -0.4618584 | 0.155 | 0.167 | 1 | 0.72832583 | 0.65823052 |
| Gm36037 | 0.41588701 | -1.4779323 | 0.031 | 0.052 | 1 | 0.55025165 | -1.3973181 | 0.029 | 0.042 | 1 | 0.55025165 | 0.65881202 |
| Gm57071 | 0.41614867 | -0.407543 | 0.083 | 0.114 | 1 | 0.75164421 | -0.4167705 | 0.05 | 0.042 | 1 | 0.75164421 | 0.65911762 |
| Cmpk2 | 0.61737927 | 1.00749454 | 0.01 | 0.005 | 1 | 0.41666664 | -2.5322477 | 0.025 | 0.042 | 1 | 0.61737927 | 0.65972219 |
| Gm49602 | 0.84862675 | 0.13302542 | 0.052 | 0.047 | 1 | 0.41709762 | -0.2394659 | 0.08 | 0.052 | 1 | 0.84862675 | 0.66022481 |
| D630045J12Rik | 0.41715535 | -0.2481603 | 0.5 | 0.534 | 1 | 0.79282674 | -0.1204341 | 0.639 | 0.531 | 1 | 0.79282674 | 0.66029211 |
| Gm46430 | 0.417486 | 0.32942263 | 0.083 | 0.057 | 1 | 0.95595883 | -0.2029401 | 0.097 | 0.094 | 1 | 0.95595883 | 0.66067744 |
| Gm26881 | 0.41822161 | -0.8993961 | 0.031 | 0.052 | 1 | 0.7812432 | -0.1723517 | 0.038 | 0.031 | 1 | 0.7812432 | 0.6615339 |
| 4930533B01Rik | 0.41822206 | -0.7998604 | 0.031 | 0.052 | 1 | 0.57854956 | -0.8244284 | 0.021 | 0.031 | 1 | 0.57854956 | 0.66153443 |
| Kcnn3 | 0.98793039 | 0.68556644 | 0.021 | 0.021 | 1 | 0.41850233 | -1.4703199 | 0.034 | 0.052 | 1 | 0.98793039 | 0.66186046 |
| Vstm2l | 0.47606502 | 1.00749454 | 0.021 | 0.01 | 1 | 0.41906801 | -1.0874628 | 0.025 | 0.042 | 1 | 0.47606502 | 0.66251803 |
| Gm57038 | 0.48525351 | 0.27052894 | 0.021 | 0.01 | 1 | 0.41906801 | -1.0874628 | 0.025 | 0.042 | 1 | 0.48525351 | 0.66251803 |
| Aspg | 0.85781561 | -0.3550755 | 0.052 | 0.057 | 1 | 0.41906801 | -1.0874628 | 0.025 | 0.042 | 1 | 0.85781561 | 0.66251803 |
| Gm26759 | 0.84235519 | 0.22988696 | 0.052 | 0.047 | 1 | 0.41946234 | -0.9879272 | 0.088 | 0.115 | 1 | 0.84235519 | 0.66297602 |
| 2610027K06Rik | 0.85600312 | 0.10060394 | 0.062 | 0.057 | 1 | 0.41953582 | -0.9879272 | 0.034 | 0.052 | 1 | 0.85600312 | 0.66306134 |
| Alpk3 | 1 | -1.3144336 | 0.021 | 0.021 | 1 | 0.4195919 | -1.3098553 | 0.042 | 0.062 | 1 | 1 | 0.66312643 |
| B4galt5 | 0.41986572 | -0.3919123 | 0.375 | 0.399 | 1 | 0.87945161 | -1.0498872 | 0.437 | 0.385 | 1 | 0.87945161 | 0.66344422 |
| Lypla1 | 0.4198728 | 0.25968156 | 0.365 | 0.316 | 1 | 0.98537174 | -0.1442692 | 0.424 | 0.417 | 1 | 0.98537174 | 0.66345243 |
| Bok | 0.4204199 | -0.1445086 | 0.083 | 0.057 | 1 | 0.56384121 | -0.7952821 | 0.029 | 0.042 | 1 | 0.56384121 | 0.66408691 |
| Haus4 | 0.42059033 | 0.10060394 | 0.115 | 0.083 | 1 | 0.85661661 | -0.486733 | 0.08 | 0.073 | 1 | 0.85661661 | 0.66428443 |
| Wdyhv1 | 0.69377148 | 0.25260703 | 0.146 | 0.13 | 1 | 0.42060823 | -0.3446207 | 0.143 | 0.104 | 1 | 0.69377148 | 0.66430517 |
| Gm20470 | 0.608151 | -0.7294711 | 0.021 | 0.031 | 1 | 0.42063965 | -0.9313436 | 0.034 | 0.052 | 1 | 0.608151 | 0.66434158 |
| Igf2bp2 | 0.64919465 | -0.3998813 | 0.448 | 0.389 | 1 | 0.42066299 | -0.1540606 | 0.761 | 0.74 | 1 | 0.64919465 | 0.66436863 |
| Gm20646 | 0.52640963 | -0.9925055 | 0.031 | 0.047 | 1 | 0.42139335 | -0.3098553 | 0.067 | 0.042 | 1 | 0.52640963 | 0.66521434 |
| Fez1 | 0.42164902 | 1.00749454 | 0.062 | 0.041 | 1 | 0.56039585 | 0.36821664 | 0.092 | 0.073 | 1 | 0.56039585 | 0.66551015 |
| Gm10382 | 0.85781561 | -0.1851505 | 0.052 | 0.057 | 1 | 0.42169144 | -0.8244284 | 0.034 | 0.052 | 1 | 0.85781561 | 0.66555921 |
| Gm36447 | 0.42179602 | -0.6929452 | 0.031 | 0.052 | 1 | 0.79735475 | -0.3098553 | 0.071 | 0.062 | 1 | 0.79735475 | 0.66568016 |
| Gm12762 | 0.42179602 | -0.6929452 | 0.031 | 0.052 | 1 | 0.70349278 | -0.9879272 | 0.034 | 0.042 | 1 | 0.70349278 | 0.66568016 |
| Cdyl | 0.53747076 | -0.3404288 | 0.25 | 0.275 | 1 | 0.4219441 | -0.7941554 | 0.416 | 0.458 | 1 | 0.53747076 | 0.66585138 |
| Tfb1m | 0.42268673 | 0.68556644 | 0.094 | 0.067 | 1 | 0.79469975 | -0.4353861 | 0.038 | 0.031 | 1 | 0.79469975 | 0.66670939 |
| Eif4ebp3 | 0.42268673 | 0.68556644 | 0.094 | 0.067 | 1 | 0.9192876 | -0.2554075 | 0.088 | 0.083 | 1 | 0.9192876 | 0.66670939 |
| Aoc3 | 0.63520923 | 0.78510212 | 0.042 | 0.031 | 1 | 0.42273193 | -0.9472852 | 0.034 | 0.052 | 1 | 0.63520923 | 0.66676157 |
| Pcdhgc4 | 0.99288067 | 0.10060394 | 0.062 | 0.062 | 1 | 0.42273193 | -1.3098553 | 0.034 | 0.052 | 1 | 0.99288067 | 0.66676157 |
| Dnajc27 | 0.42286259 | 0.23882008 | 0.135 | 0.176 | 1 | 0.79241256 | -0.5025003 | 0.181 | 0.156 | 1 | 0.79241256 | 0.66691241 |
| Sdc1 | 0.71626837 | -0.5907648 | 0.271 | 0.269 | 1 | 0.42306815 | -0.3779443 | 0.576 | 0.615 | 1 | 0.71626837 | 0.66714964 |
| Pbx2 | 0.45801499 | 0.2816695 | 0.198 | 0.161 | 1 | 0.42311472 | -0.4000531 | 0.218 | 0.167 | 1 | 0.45801499 | 0.66720337 |
| Plxdc1 | 0.423668 | -0.4664367 | 0.052 | 0.078 | 1 | 0.43079893 | 0.56461386 | 0.038 | 0.021 | 1 | 0.43079893 | 0.66784142 |
| Gm56606 | 0.85039899 | -0.2148979 | 0.052 | 0.057 | 1 | 0.42376525 | 1.01207283 | 0.038 | 0.021 | 1 | 0.85039899 | 0.66795351 |
| Gm34250 | 0.61737927 | 1.00749454 | 0.01 | 0.005 | 1 | 0.42398359 | -0.8948178 | 0.025 | 0.042 | 1 | 0.61737927 | 0.66820509 |
| Ccdc173 | 0.93274922 | -0.6169963 | 0.094 | 0.093 | 1 | 0.42412725 | -1.1329775 | 0.071 | 0.094 | 1 | 0.93274922 | 0.66837058 |
| Rbl1 | 0.42424162 | 0.9200317 | 0.146 | 0.114 | 1 | 0.7631968 | -0.2509616 | 0.097 | 0.083 | 1 | 0.7631968 | 0.66850229 |
| Zfp711 | 0.42426413 | -1.4254649 | 0.031 | 0.052 | 1 | 0.91175866 | -0.2079756 | 0.134 | 0.125 | 1 | 0.91175866 | 0.66852821 |
| Zkscan14 | 0.42450517 | -0.4330781 | 0.052 | 0.078 | 1 | 0.86293192 | -0.2394659 | 0.067 | 0.073 | 1 | 0.86293192 | 0.6688057 |

| | | | | | | | | | | | | |
|---|---|---|---|---|---|---|---|---|---|---|---|---|
| Zdhhc23 | 0.76074084 | 0.11440974 | 0.104 | 0.093 | 1 | 0.42482896 | 0.15963002 | 0.113 | 0.083 | 1 | 0.76074084 | 0.66917827 |
| Pcdhb9 | 0.46080869 | -2.2148979 | 0.021 | 0.036 | 1 | 0.42497971 | -0.9472852 | 0.034 | 0.052 | 1 | 0.46080869 | 0.66935166 |
| Gm43149 | 0.42538331 | -0.577468 | 0.031 | 0.052 | 1 | 0.71794749 | -0.4353861 | 0.034 | 0.042 | 1 | 0.71794749 | 0.66981565 |
| Gm53610 | 0.94333953 | 0.82306997 | 0.094 | 0.093 | 1 | 0.42538708 | 0.82764826 | 0.063 | 0.042 | 1 | 0.94333953 | 0.66982 |
| Ccdc15 | 0.42545975 | 0.1843723 | 0.156 | 0.119 | 1 | 1 | -0.5936482 | 0.143 | 0.135 | 1 | 1 | 0.66990351 |
| Kctd13 | 0.42602756 | 0.31561683 | 0.125 | 0.093 | 1 | 0.85344216 | -0.2457249 | 0.08 | 0.073 | 1 | 0.85344216 | 0.67055564 |
| Abhd1 | 0.70116305 | -0.2555399 | 0.042 | 0.052 | 1 | 0.42610607 | 0.91253716 | 0.038 | 0.021 | 1 | 0.70116305 | 0.67064576 |
| Prrg4 | 0.66582162 | -0.130009 | 0.042 | 0.031 | 1 | 0.42610646 | 1.01207283 | 0.038 | 0.021 | 1 | 0.66582162 | 0.67064621 |
| 4933424G06Rik | 0.86441462 | -0.2148979 | 0.052 | 0.047 | 1 | 0.42610646 | 1.01207283 | 0.038 | 0.021 | 1 | 0.86441462 | 0.67064621 |
| A930012L18Rik | 0.99482536 | 0.15949763 | 0.042 | 0.041 | 1 | 0.42610646 | 0.80562195 | 0.038 | 0.021 | 1 | 0.99482536 | 0.67064621 |
| Tll2 | 0.79869472 | 0.30705482 | 0.031 | 0.026 | 1 | 0.42714283 | -1.5163061 | 0.034 | 0.052 | 1 | 0.79869472 | 0.67183466 |
| Wdr4 | 0.89142858 | 0.10703021 | 0.135 | 0.13 | 1 | 0.42721167 | -0.5025003 | 0.155 | 0.115 | 1 | 0.89142858 | 0.67191353 |
| Ptgs1 | 0.42730505 | 0.21644143 | 0.385 | 0.342 | 1 | 0.7395578 | -0.7952821 | 0.189 | 0.156 | 1 | 0.7395578 | 0.6720205 |
| Gm17036 | 0.85158597 | -0.3710171 | 0.042 | 0.036 | 1 | 0.4273327 | -0.2554075 | 0.092 | 0.062 | 1 | 0.85158597 | 0.67205216 |
| Idnk | 0.87464172 | -0.1693832 | 0.198 | 0.202 | 1 | 0.42759001 | -0.1555271 | 0.231 | 0.177 | 1 | 0.87464172 | 0.67234681 |
| Gria4 | 0.77277122 | -1.6139938 | 0.073 | 0.062 | 1 | 0.42809153 | -1.7655347 | 0.071 | 0.094 | 1 | 0.77277122 | 0.6729207 |
| Ctif | 0.95929417 | -0.366901 | 0.344 | 0.326 | 1 | 0.42828133 | -0.6833137 | 0.609 | 0.583 | 1 | 0.95929417 | 0.67313776 |
| Amhr2 | 0.82258416 | -0.1624305 | 0.031 | 0.036 | 1 | 0.42845063 | 0.80562195 | 0.038 | 0.021 | 1 | 0.82258416 | 0.67333132 |
| Gm43948 | 0.42847613 | -0.6445822 | 0.062 | 0.088 | 1 | 0.93268086 | -0.5728897 | 0.067 | 0.062 | 1 | 0.93268086 | 0.67336047 |
| Sgtb | 0.42855963 | -0.3848229 | 0.052 | 0.078 | 1 | 0.66252781 | 0.22065945 | 0.076 | 0.062 | 1 | 0.66252781 | 0.6734559 |
| Casp7 | 0.42890914 | 0.79599043 | 0.115 | 0.088 | 1 | 0.45889212 | 0.18490943 | 0.088 | 0.062 | 1 | 0.45889212 | 0.67385523 |
| Jade1 | 0.42896637 | 0.14284639 | 0.396 | 0.352 | 1 | 0.72070422 | -0.3293076 | 0.529 | 0.521 | 1 | 0.72070422 | 0.67392059 |
| 9330159M07Rik | 0.86924922 | -0.265524 | 0.094 | 0.098 | 1 | 0.42897434 | -0.6468902 | 0.05 | 0.073 | 1 | 0.86924922 | 0.67392969 |
| Stat3 | 0.42911115 | -0.1016508 | 0.99 | 0.984 | 1 | 0.81020429 | -0.2050679 | 0.992 | 0.99 | 1 | 0.81020429 | 0.67408593 |
| D130017N08Rik | 0.42914542 | -0.4519371 | 0.031 | 0.052 | 1 | 0.65354244 | -0.6317834 | 0.055 | 0.042 | 1 | 0.65354244 | 0.67412504 |
| Bach1 | 0.42930611 | 0.2585563 | 0.542 | 0.534 | 1 | 0.63064727 | -0.2348046 | 0.693 | 0.75 | 1 | 0.63064727 | 0.67430848 |
| Rabepk | 0.64463569 | 0.13302542 | 0.25 | 0.228 | 1 | 0.42952159 | -0.4721267 | 0.277 | 0.219 | 1 | 0.64463569 | 0.67455439 |
| Gm12592 | 0.51800362 | -0.8993961 | 0.031 | 0.047 | 1 | 0.42964041 | 0.44503224 | 0.088 | 0.062 | 1 | 0.51800362 | 0.67468993 |
| Rspry1 | 0.47433866 | -0.1463108 | 0.625 | 0.622 | 1 | 0.4297626 | -0.3915056 | 0.714 | 0.792 | 1 | 0.47433866 | 0.67482931 |
| Slc22a21 | 0.42999729 | -0.2228031 | 0.281 | 0.342 | 1 | 0.90853426 | -0.5349218 | 0.214 | 0.208 | 1 | 0.90853426 | 0.67509691 |
| Tinf2 | 0.43012656 | 0.39082318 | 0.135 | 0.104 | 1 | 0.81613978 | -0.1034044 | 0.113 | 0.104 | 1 | 0.81613978 | 0.67524426 |
| Rarres2 | 0.51800188 | -1.0799683 | 0.031 | 0.047 | 1 | 0.43040711 | -0.7248928 | 0.034 | 0.052 | 1 | 0.51800188 | 0.67556394 |
| BC028528 | 0.55793691 | 0.62898291 | 0.031 | 0.047 | 1 | 0.43079893 | 0.56461386 | 0.038 | 0.021 | 1 | 0.55793691 | 0.67601015 |
| Zfp960 | 0.64538863 | 0.52206771 | 0.042 | 0.031 | 1 | 0.43079893 | 0.56461386 | 0.038 | 0.021 | 1 | 0.64538863 | 0.67601015 |
| Cdnf | 0.82342138 | 0.32942263 | 0.042 | 0.036 | 1 | 0.43079893 | 0.56461386 | 0.038 | 0.021 | 1 | 0.82342138 | 0.67601015 |
| Batf3 | 0.43081766 | 0.55498233 | 0.115 | 0.083 | 1 | 0.88978657 | -0.3838558 | 0.059 | 0.062 | 1 | 0.88978657 | 0.67603146 |
| Cacna2d2 | 0.5383068 | 0.27052894 | 0.01 | 0.021 | 1 | 0.4313809 | -0.6317834 | 0.025 | 0.042 | 1 | 0.5383068 | 0.67667232 |
| Snn | 0.8006267 | 0.20013961 | 0.021 | 0.026 | 1 | 0.4313809 | -0.6317834 | 0.025 | 0.042 | 1 | 0.8006267 | 0.67667232 |
| Lyrm7 | 0.80274368 | 0.20013961 | 0.031 | 0.026 | 1 | 0.43147332 | -0.8504236 | 0.034 | 0.052 | 1 | 0.80274368 | 0.67677741 |
| H4c3 | 0.67877064 | 0.49292136 | 0.052 | 0.041 | 1 | 0.43152816 | -0.7248928 | 0.034 | 0.052 | 1 | 0.67877064 | 0.67683976 |
| Pcca | 0.81815532 | 0.1472193 | 0.333 | 0.316 | 1 | 0.43173856 | -0.7117174 | 0.403 | 0.417 | 1 | 0.81815532 | 0.67707894 |
| Pdp1 | 0.43179682 | 0.16795921 | 0.156 | 0.119 | 1 | 0.93514717 | -0.6968784 | 0.172 | 0.167 | 1 | 0.93514717 | 0.67714515 |
| Zfp688 | 0.43214328 | 0.68556644 | 0.062 | 0.041 | 1 | 0.64304313 | 0.27510724 | 0.042 | 0.031 | 1 | 0.64304313 | 0.67753874 |
| Col15a1 | 0.43219166 | 0.37006462 | 0.062 | 0.041 | 1 | 0.7394685 | -0.2249664 | 0.143 | 0.125 | 1 | 0.7394685 | 0.67759369 |
| Sh3d19 | 0.71300539 | -0.1324017 | 0.958 | 0.974 | 1 | 0.43314361 | -0.3117939 | 0.971 | 0.948 | 1 | 0.71300539 | 0.67867383 |
| Gm57238 | 0.61550914 | -0.407543 | 0.021 | 0.031 | 1 | 0.43323624 | 0.42711033 | 0.038 | 0.021 | 1 | 0.61550914 | 0.67877885 |
| Gm19426 | 0.83041404 | 0.15949763 | 0.042 | 0.036 | 1 | 0.43323624 | 0.42711033 | 0.038 | 0.021 | 1 | 0.83041404 | 0.67877885 |
| Gm6277 | 0.43350237 | -0.6363617 | 0.052 | 0.078 | 1 | 0.92738405 | -0.9472852 | 0.034 | 0.031 | 1 | 0.92738405 | 0.67908043 |
| Vcpkmt | 0.43370755 | 0.15949763 | 0.083 | 0.057 | 1 | 0.75960893 | -0.1630139 | 0.118 | 0.104 | 1 | 0.75960893 | 0.67931286 |
| Gm53 | 0.43396444 | -0.2148979 | 0.031 | 0.052 | 1 | 0.82231716 | -0.4618584 | 0.025 | 0.021 | 1 | 0.82231716 | 0.67960374 |
| Ell | 0.85387832 | -0.2028124 | 0.74 | 0.689 | 1 | 0.4340555 | -0.2798328 | 0.744 | 0.729 | 1 | 0.85387832 | 0.67970683 |
| Sfrp2 | 0.94629149 | -2.1863287 | 0.24 | 0.233 | 1 | 0.43440238 | -2.1172102 | 0.071 | 0.094 | 1 | 0.94629149 | 0.68009934 |
| Tsen54 | 0.43461987 | 0.64492446 | 0.062 | 0.041 | 1 | 0.55321261 | -0.1399303 | 0.071 | 0.052 | 1 | 0.55321261 | 0.68034531 |
| Ift140 | 0.92080419 | -0.3404288 | 0.094 | 0.088 | 1 | 0.4359825 | -0.4703199 | 0.105 | 0.073 | 1 | 0.92080419 | 0.68188426 |
| Slc35b1 | 0.43601461 | -0.1696332 | 0.615 | 0.679 | 1 | 0.68610544 | 0.15526747 | 0.58 | 0.594 | 1 | 0.68610544 | 0.68192048 |
| Rhof | 0.43638172 | 0.37006462 | 0.125 | 0.093 | 1 | 0.67037747 | -0.7248928 | 0.13 | 0.146 | 1 | 0.67037747 | 0.68233443 |
| Gm57286 | 0.43647072 | -0.5185743 | 0.385 | 0.404 | 1 | 0.78890793 | -0.7988933 | 0.345 | 0.281 | 1 | 0.78890793 | 0.68243475 |
| 6330562C20Rik | 0.76513302 | 0.1843723 | 0.104 | 0.093 | 1 | 0.43916995 | 0.27510724 | 0.038 | 0.021 | 1 | 0.76513302 | 0.68546965 |
| ENSMUSG00000121502 | 0.63353706 | 0.37946331 | 0.083 | 0.067 | 1 | 0.43929092 | -1.0468209 | 0.08 | 0.104 | 1 | 0.63353706 | 0.68560532 |
| Plxna2 | 0.53674886 | -0.8124238 | 0.594 | 0.56 | 1 | 0.43981506 | -0.6027382 | 0.79 | 0.75 | 1 | 0.53674886 | 0.68619283 |
| Tbc1d23 | 0.44015059 | -0.2130565 | 0.635 | 0.622 | 1 | 0.78196292 | -0.2195347 | 0.71 | 0.677 | 1 | 0.78196292 | 0.68656864 |
| Gm44148 | 0.44045737 | 0.42253204 | 0.062 | 0.041 | 1 | 0.5530041 | -0.3098553 | 0.046 | 0.031 | 1 | 0.5530041 | 0.68691205 |
| Sorcs2 | 0.60200118 | 0.22988696 | 0.125 | 0.104 | 1 | 0.44049477 | -1.567653 | 0.101 | 0.125 | 1 | 0.60200118 | 0.6869539 |
| Crtc1 | 0.91906971 | -0.1801325 | 0.146 | 0.15 | 1 | 0.44087619 | -0.3438026 | 0.265 | 0.208 | 1 | 0.91906971 | 0.68738057 |
| Trmo | 0.44124401 | 0.22988696 | 0.125 | 0.093 | 1 | 0.90637231 | -0.1172102 | 0.109 | 0.104 | 1 | 0.90637231 | 0.68779174 |
| 9130004C02Rik | 0.44126404 | 0.13302542 | 0.031 | 0.052 | 1 | 0.8259932 | -0.8244284 | 0.025 | 0.021 | 1 | 0.8259932 | 0.68781413 |
| D6Ertd527e | 0.44158998 | 0.49292136 | 0.062 | 0.041 | 1 | 0.61792593 | -0.5728897 | 0.059 | 0.073 | 1 | 0.61792593 | 0.68817825 |
| Gm56901 | 0.44158998 | 0.49292136 | 0.062 | 0.041 | 1 | 0.53497636 | 0.39058446 | 0.046 | 0.031 | 1 | 0.53497636 | 0.68817825 |
| Gm17690 | 0.44184522 | 0.29700115 | 0.094 | 0.067 | 1 | 0.87085967 | -0.194378 | 0.038 | 0.042 | 1 | 0.87085967 | 0.68846325 |
| Taf6l | 0.99758609 | 0.27052894 | 0.021 | 0.021 | 1 | 0.44247713 | -0.4353861 | 0.034 | 0.052 | 1 | 0.99758609 | 0.68916825 |
| Polr1has | 0.48364323 | -0.2148979 | 0.073 | 0.098 | 1 | 0.44257125 | 0.49749966 | 0.05 | 0.031 | 1 | 0.48364323 | 0.68927319 |
| Ephb6 | 0.61737927 | 1.00749454 | 0.01 | 0.005 | 1 | 0.44257125 | 0.49749966 | 0.05 | 0.031 | 1 | 0.61737927 | 0.68927319 |
| Fndc10 | 0.84862675 | 0.13302542 | 0.052 | 0.047 | 1 | 0.44257125 | 0.49749966 | 0.05 | 0.031 | 1 | 0.84862675 | 0.68927319 |
| Haus5 | 0.55702983 | 0.68556644 | 0.062 | 0.047 | 1 | 0.4427584 | -0.1723517 | 0.038 | 0.021 | 1 | 0.55702983 | 0.6894818 |
| Tbc1d22a | 0.44299004 | 0.2590333 | 0.323 | 0.275 | 1 | 0.92933356 | -0.5184419 | 0.37 | 0.344 | 1 | 0.92933356 | 0.6897399 |
| Ankrd29 | 0.44356932 | -0.3461424 | 0.062 | 0.088 | 1 | 0.62887086 | 0.54812573 | 0.055 | 0.042 | 1 | 0.62887086 | 0.6903849 |
| Rubcn | 0.67420956 | 0.30295042 | 0.323 | 0.301 | 1 | 0.44363694 | -0.4294369 | 0.408 | 0.302 | 1 | 0.67420956 | 0.69046015 |
| Snhg10 | 0.67877064 | 0.49292136 | 0.052 | 0.041 | 1 | 0.44394565 | -0.3098553 | 0.038 | 0.021 | 1 | 0.67877064 | 0.69080356 |
| Zfp11 | 1 | 0.42253204 | 0.01 | 0.01 | 1 | 0.44394565 | -0.3098553 | 0.038 | 0.021 | 1 | 1 | 0.69080356 |
| Trim26 | 0.44431676 | -0.2862367 | 0.25 | 0.29 | 1 | 0.51440793 | -0.3098553 | 0.395 | 0.312 | 1 | 0.51440793 | 0.69121614 |
| Wwox | 0.62853482 | 0.10265177 | 0.823 | 0.819 | 1 | 0.44459233 | -0.300965 | 0.765 | 0.823 | 1 | 0.62853482 | 0.69152232 |
| Ss18l1 | 0.44465052 | 0.32942263 | 0.094 | 0.067 | 1 | 0.95133938 | -0.9472852 | 0.063 | 0.062 | 1 | 0.95133938 | 0.69158696 |

| | | | | | | | | | | | | |
|---|---|---|---|---|---|---|---|---|---|---|---|---|
| Plag1 | 0.85324768 | -0.3364599 | 0.094 | 0.098 | 1 | 0.44466733 | -1.1172102 | 0.101 | 0.125 | 1 | 0.85324768 | 0.69160563 |
| Ralgps1 | 0.71621616 | -0.5160674 | 0.062 | 0.073 | 1 | 0.44491759 | -0.3368223 | 0.126 | 0.094 | 1 | 0.71621616 | 0.69188352 |
| Il1rap | 0.4468331 | -0.6649308 | 0.115 | 0.145 | 1 | 0.89868896 | -1.2723806 | 0.126 | 0.125 | 1 | 0.89868896 | 0.69400638 |
| Pabpc1l | 0.47294388 | -0.8669746 | 0.021 | 0.036 | 1 | 0.44699433 | -1.5443205 | 0.063 | 0.083 | 1 | 0.47294388 | 0.69418473 |
| Pigv | 0.44712589 | -0.4705528 | 0.135 | 0.166 | 1 | 0.80117516 | -0.7248928 | 0.143 | 0.146 | 1 | 0.80117516 | 0.69433022 |
| Xndc1 | 0.44726666 | -0.2066303 | 0.188 | 0.145 | 1 | 0.47337121 | 0.11237774 | 0.193 | 0.156 | 1 | 0.47337121 | 0.69448586 |
| Lmod1 | 0.44731854 | 1.42253204 | 0.042 | 0.026 | 1 | 1 | -0.5025003 | 0.021 | 0.021 | 1 | 1 | 0.69454321 |
| Cop1 | 0.47815992 | -0.1258558 | 0.917 | 0.819 | 1 | 0.44809772 | -0.3841804 | 0.945 | 0.927 | 1 | 0.47815992 | 0.69540387 |
| Pcdhgb5 | 0.72900758 | 0.42253204 | 0.083 | 0.073 | 1 | 0.44864033 | -0.1082214 | 0.076 | 0.052 | 1 | 0.72900758 | 0.69600251 |
| Drd3 | 0.67876537 | -1.6929452 | 0.052 | 0.062 | 1 | 0.44998568 | 0.77760758 | 0.05 | 0.031 | 1 | 0.67876537 | 0.69748425 |
| Mcf2l | 0.56103105 | -0.8405024 | 0.042 | 0.057 | 1 | 0.45005361 | -0.3098553 | 0.038 | 0.021 | 1 | 0.56103105 | 0.69755896 |
| Plcd4 | 0.99009356 | 0.11440974 | 0.031 | 0.031 | 1 | 0.45023182 | -0.1723517 | 0.063 | 0.042 | 1 | 0.99009356 | 0.69775495 |
| Pik3cb | 0.45174519 | -0.1871529 | 0.365 | 0.409 | 1 | 0.9251034 | -0.1161307 | 0.542 | 0.552 | 1 | 0.9251034 | 0.69941666 |
| Gjc3 | 0.49613627 | 0.15449695 | 0.583 | 0.513 | 1 | 0.452063 | 0.58322953 | 0.05 | 0.031 | 1 | 0.49613627 | 0.69976505 |
| Dlk1 | 0.58426772 | 0.74446013 | 0.031 | 0.021 | 1 | 0.45206414 | 0.87456931 | 0.05 | 0.031 | 1 | 0.58426772 | 0.69976629 |
| Thumpd3 | 0.45236422 | 0.37946331 | 0.281 | 0.249 | 1 | 0.47472344 | -0.2335044 | 0.277 | 0.219 | 1 | 0.47472344 | 0.70009505 |
| Nhsl2 | 0.51382153 | -2.3848229 | 0.031 | 0.047 | 1 | 0.45240969 | -0.484942 | 0.126 | 0.094 | 1 | 0.51382153 | 0.70014485 |
| Tmem191 | 0.64538863 | 0.52206771 | 0.042 | 0.031 | 1 | 0.4529977 | 0.14957636 | 0.088 | 0.062 | 1 | 0.64538863 | 0.70078849 |
| Rnaseh2a | 0.95529461 | -0.4187702 | 0.135 | 0.13 | 1 | 0.45356852 | -0.3098553 | 0.101 | 0.073 | 1 | 0.95529461 | 0.70141264 |
| Gm56764 | 0.45370345 | 0.30705482 | 0.062 | 0.041 | 1 | 0.52674111 | -0.961932 | 0.046 | 0.062 | 1 | 0.52674111 | 0.70156008 |
| Ptpdc1 | 0.4540137 | 0.19035859 | 0.167 | 0.13 | 1 | 0.48407181 | -0.4029647 | 0.164 | 0.125 | 1 | 0.48407181 | 0.70189897 |
| Amz1 | 0.45476301 | 1.59245704 | 0.042 | 0.026 | 1 | 0.92228543 | -0.9472852 | 0.029 | 0.031 | 1 | 0.92228543 | 0.70271662 |
| Camsap3 | 1 | 0.42253204 | 0.01 | 0.01 | 1 | 0.45488747 | -0.7952821 | 0.038 | 0.021 | 1 | 1 | 0.70285233 |
| Simc1 | 0.74366005 | -0.3410278 | 0.396 | 0.352 | 1 | 0.4558787 | -0.4554606 | 0.445 | 0.469 | 1 | 0.74366005 | 0.70393201 |
| A330076H08Rik | 0.56121485 | -0.2950682 | 0.219 | 0.238 | 1 | 0.45620312 | -0.6053111 | 0.286 | 0.219 | 1 | 0.56121485 | 0.70428496 |
| Ak8 | 0.46914383 | 0.52206771 | 0.073 | 0.052 | 1 | 0.45731547 | -1.1578522 | 0.063 | 0.083 | 1 | 0.46914383 | 0.7054935 |
| Ebag9 | 0.86319695 | -0.3231508 | 0.302 | 0.301 | 1 | 0.45742271 | -0.1621016 | 0.324 | 0.26 | 1 | 0.86319695 | 0.70560988 |
| Lnx2 | 0.71208157 | 0.62047141 | 0.083 | 0.073 | 1 | 0.45745397 | -0.9363969 | 0.16 | 0.188 | 1 | 0.71208157 | 0.7056438 |
| Cdh23 | 0.755406 | -0.2019588 | 0.125 | 0.109 | 1 | 0.45822138 | -1.561394 | 0.063 | 0.083 | 1 | 0.755406 | 0.70647593 |
| Pip5k1b | 0.97605526 | 0.14076107 | 0.125 | 0.124 | 1 | 0.45843018 | -0.3098553 | 0.176 | 0.208 | 1 | 0.97605526 | 0.70670213 |
| Jmjd7 | 0.45898384 | 0.88196365 | 0.073 | 0.052 | 1 | 0.95405618 | -0.4029647 | 0.055 | 0.052 | 1 | 0.95405618 | 0.70730151 |
| Zscan29 | 0.45898697 | -0.5427025 | 0.125 | 0.155 | 1 | 0.73817195 | 0.27510724 | 0.126 | 0.115 | 1 | 0.73817195 | 0.70730491 |
| Eya3 | 0.74476325 | -0.2769589 | 0.479 | 0.461 | 1 | 0.46011929 | -0.1754289 | 0.58 | 0.531 | 1 | 0.74476325 | 0.70852882 |
| Zfp808 | 0.91245523 | 0.27052894 | 0.104 | 0.109 | 1 | 0.46082759 | 0.17557156 | 0.076 | 0.052 | 1 | 0.91245523 | 0.70929311 |
| Zfp516 | 0.46114871 | -0.3754493 | 0.625 | 0.627 | 1 | 0.736411 | -0.1005387 | 0.832 | 0.844 | 1 | 0.736411 | 0.70963929 |
| Nipsnap3b | 0.46136252 | 0.31561683 | 0.219 | 0.181 | 1 | 0.906487 | -0.4431218 | 0.202 | 0.188 | 1 | 0.906487 | 0.70986967 |
| Entpd2 | 0.73436042 | 0.17741954 | 0.073 | 0.062 | 1 | 0.46142849 | -0.3098553 | 0.063 | 0.042 | 1 | 0.73436042 | 0.70994072 |
| Haus2 | 0.70374409 | 0.27879656 | 0.25 | 0.238 | 1 | 0.46150373 | -0.1200307 | 0.223 | 0.177 | 1 | 0.70374409 | 0.71002176 |
| C130013H08Rik | 0.4622729 | 1.00749454 | 0.042 | 0.026 | 1 | 0.99136676 | -0.6094155 | 0.042 | 0.042 | 1 | 0.99136676 | 0.71084957 |
| Zfp526 | 0.4622729 | 1.00749454 | 0.042 | 0.026 | 1 | 0.88759525 | -1.0468209 | 0.046 | 0.042 | 1 | 0.88759525 | 0.71084957 |
| Ifit2 | 0.4623376 | 0.59245704 | 0.125 | 0.098 | 1 | 0.55724522 | -0.7248928 | 0.076 | 0.094 | 1 | 0.55724522 | 0.71091914 |
| Senp8 | 0.46283638 | -0.1624305 | 0.135 | 0.104 | 1 | 0.54526995 | -0.8073549 | 0.105 | 0.125 | 1 | 0.54526995 | 0.71145524 |
| Pcsk7 | 0.79084172 | 0.20604022 | 0.229 | 0.218 | 1 | 0.46348687 | -0.2764323 | 0.374 | 0.312 | 1 | 0.79084172 | 0.71215366 |
| Rnf227 | 0.46355545 | 0.2912875 | 0.104 | 0.078 | 1 | 0.84171242 | -0.3623227 | 0.067 | 0.073 | 1 | 0.84171242 | 0.71222725 |
| Rabep2 | 0.75742279 | 0.11440974 | 0.229 | 0.212 | 1 | 0.4637051 | 0.154813 | 0.206 | 0.167 | 1 | 0.75742279 | 0.71238778 |
| Gm44710 | 0.5434853 | -0.2555399 | 0.031 | 0.047 | 1 | 0.46439596 | 0.13071733 | 0.063 | 0.042 | 1 | 0.5434853 | 0.71312832 |
| Gm41790 | 0.80606188 | -0.4638112 | 0.167 | 0.15 | 1 | 0.46500534 | -0.933001 | 0.21 | 0.229 | 1 | 0.80606188 | 0.71378072 |
| Brat1 | 0.46544015 | -0.577468 | 0.042 | 0.062 | 1 | 0.75602831 | -0.4797803 | 0.063 | 0.052 | 1 | 0.75602831 | 0.71424577 |
| Strada | 0.46592297 | 0.31150072 | 0.26 | 0.223 | 1 | 0.4954002 | -0.9448584 | 0.223 | 0.24 | 1 | 0.4954002 | 0.71476172 |
| Hspa2 | 0.46640287 | 0.12297175 | 0.219 | 0.259 | 1 | 0.75628834 | -0.169374 | 0.269 | 0.24 | 1 | 0.75628834 | 0.7152741 |
| Trim34a | 0.58616998 | 0.59245704 | 0.073 | 0.057 | 1 | 0.46651581 | -0.2592292 | 0.101 | 0.073 | 1 | 0.58616998 | 0.71539462 |
| Prx | 0.4666761 | -1.1079827 | 0.042 | 0.062 | 1 | 0.81538333 | -0.5213594 | 0.046 | 0.052 | 1 | 0.81538333 | 0.71556562 |
| Tprn | 0.53487059 | 0.42253204 | 0.052 | 0.036 | 1 | 0.46684912 | -0.2103196 | 0.05 | 0.031 | 1 | 0.53487059 | 0.71575013 |
| Gm53018 | 0.4668591 | -1.1079827 | 0.021 | 0.036 | 1 | 0.742831 | -1.9472852 | 0.025 | 0.031 | 1 | 0.742831 | 0.71576078 |
| Sdcbp2 | 0.46686094 | 0.83756954 | 0.042 | 0.026 | 1 | 0.66391569 | -1.1723517 | 0.042 | 0.052 | 1 | 0.66391569 | 0.71576274 |
| Nkain1 | 0.46686094 | -1.7294711 | 0.021 | 0.036 | 1 | 0.50844268 | 1.27510724 | 0.021 | 0.01 | 1 | 0.50844268 | 0.71576274 |
| A930038B10Rik | 0.46697007 | 1.74446013 | 0.021 | 0.01 | 1 | 0.50587544 | -2.6317834 | 0.004 | 0.01 | 1 | 0.50587544 | 0.71587909 |
| Tmem38a | 0.8053787 | -0.240433 | 0.115 | 0.104 | 1 | 0.46736951 | 0.17557156 | 0.147 | 0.115 | 1 | 0.8053787 | 0.71630476 |
| Gm46218 | 0.4679031 | -0.407543 | 0.073 | 0.098 | 1 | 0.7751094 | 0.39058446 | 0.025 | 0.031 | 1 | 0.7751094 | 0.71687289 |
| Gm27241 | 0.52219952 | -0.7998604 | 0.031 | 0.047 | 1 | 0.46793352 | -0.3098553 | 0.05 | 0.031 | 1 | 0.52219952 | 0.71690526 |
| Lrrc8b | 0.46846392 | -0.6445822 | 0.229 | 0.254 | 1 | 0.68107005 | -0.8429248 | 0.357 | 0.365 | 1 | 0.68107005 | 0.71746939 |
| Tlk2 | 0.46877726 | 0.10623774 | 0.865 | 0.896 | 1 | 0.60545655 | -0.2129937 | 0.975 | 0.969 | 1 | 0.60545655 | 0.7178024 |
| AI661453 | 0.4692344 | 1.81484946 | 0.021 | 0.01 | 1 | 0.67471359 | -1.5728897 | 0.017 | 0.01 | 1 | 0.67471359 | 0.71828788 |
| Tmem30c | 0.46989903 | -0.8669746 | 0.021 | 0.036 | 1 | 0.87066325 | -0.3098553 | 0.013 | 0.01 | 1 | 0.87066325 | 0.71899296 |
| Gm15345 | 0.46989903 | -0.8669746 | 0.021 | 0.036 | 1 | 0.82415676 | -0.7248928 | 0.025 | 0.021 | 1 | 0.82415676 | 0.71899296 |
| Mc1r | 0.46992036 | 0.74446013 | 0.042 | 0.026 | 1 | 0.57170879 | -1.6317834 | 0.013 | 0.021 | 1 | 0.57170879 | 0.71901558 |
| Dppa5a | 0.46992036 | 0.74446013 | 0.042 | 0.026 | 1 | 0.78571043 | -0.3098553 | 0.038 | 0.031 | 1 | 0.78571043 | 0.71901558 |
| Gm19967 | 0.46992036 | 0.74446013 | 0.042 | 0.026 | 1 | 0.67357745 | -0.9472852 | 0.067 | 0.052 | 1 | 0.67357745 | 0.71901558 |
| Pcdh1 | 0.46992036 | 0.74446013 | 0.042 | 0.026 | 1 | 0.66339728 | 0.10518224 | 0.029 | 0.021 | 1 | 0.66339728 | 0.71901558 |
| Spon2 | 0.47016587 | -0.3461424 | 0.042 | 0.062 | 1 | 0.53156006 | 0.56461386 | 0.034 | 0.021 | 1 | 0.53156006 | 0.7192758 |
| Il17rd | 0.97295748 | -0.6363617 | 0.062 | 0.062 | 1 | 0.47042786 | -1.1172102 | 0.063 | 0.083 | 1 | 0.97295748 | 0.71955335 |
| Sac3d1 | 0.4710237 | 0.33764314 | 0.167 | 0.135 | 1 | 0.63675357 | 0.15963002 | 0.122 | 0.104 | 1 | 0.63675357 | 0.72018408 |
| Gm57387 | 0.64875704 | 0.42253204 | 0.042 | 0.031 | 1 | 0.4710948 | -0.4167705 | 0.05 | 0.031 | 1 | 0.64875704 | 0.72025929 |
| Evi5 | 0.68129981 | -0.1696332 | 0.531 | 0.549 | 1 | 0.47125543 | -0.2753168 | 0.718 | 0.76 | 1 | 0.68129981 | 0.72042917 |
| Wdr81 | 0.47126984 | 0.59245704 | 0.073 | 0.052 | 1 | 0.59680643 | 0.37820073 | 0.101 | 0.083 | 1 | 0.59680643 | 0.72044442 |
| Pram1 | 0.47126984 | 0.59245704 | 0.073 | 0.052 | 1 | 0.52440062 | 0.77760758 | 0.046 | 0.031 | 1 | 0.52440062 | 0.72044442 |
| Aven | 0.47134883 | 0.15949763 | 0.458 | 0.404 | 1 | 0.8724761 | -0.6808376 | 0.588 | 0.448 | 1 | 0.8724761 | 0.72052794 |
| Foxo6 | 0.47150075 | 1.42253204 | 0.021 | 0.01 | 1 | 0.57492803 | -0.8623963 | 0.046 | 0.031 | 1 | 0.57492803 | 0.72068854 |
| Slc23a4 | 0.47150075 | 1.42253204 | 0.021 | 0.01 | 1 | 0.8086289 | -0.3098553 | 0.017 | 0.021 | 1 | 0.8086289 | 0.72068854 |
| Zfp78 | 0.47150075 | 1.42253204 | 0.021 | 0.01 | 1 | 0.85030707 | -0.6577786 | 0.038 | 0.042 | 1 | 0.85030707 | 0.72068854 |
| Folr1 | 0.47150075 | 1.42253204 | 0.021 | 0.01 | 1 | 0.50887612 | -1.3098553 | 0.004 | 0.01 | 1 | 0.50887612 | 0.72068854 |
| Pcdhb12 | 0.47150075 | 1.42253204 | 0.021 | 0.01 | 1 | 0.66825766 | -0.3098553 | 0.029 | 0.021 | 1 | 0.66825766 | 0.72068854 |
| Gm31356 | 0.47150075 | 1.42253204 | 0.021 | 0.01 | 1 | 0.56565662 | -2.6317834 | 0.013 | 0.021 | 1 | 0.56565662 | 0.72068854 |
| Ar | 0.47171229 | -0.747393 | 0.062 | 0.088 | 1 | 0.49732944 | -1.1614634 | 0.261 | 0.198 | 1 | 0.49732944 | 0.7209121 |
| Git1 | 0.54047546 | -0.6843832 | 0.125 | 0.145 | 1 | 0.4721566 | -0.4532194 | 0.202 | 0.156 | 1 | 0.54047546 | 0.72138134 |
| Il10rb | 0.47235004 | -0.3592878 | 0.146 | 0.181 | 1 | 0.79018868 | -0.2433599 | 0.29 | 0.302 | 1 | 0.79018868 | 0.72158552 |
| Gm35417 | 0.47244528 | -0.7294711 | 0.042 | 0.062 | 1 | 0.68525028 | -1.4566967 | 0.097 | 0.104 | 1 | 0.68525028 | 0.72168602 |
| Mcub | 0.5301942 | -0.8106671 | 0.146 | 0.171 | 1 | 0.47250717 | -0.7119537 | 0.151 | 0.177 | 1 | 0.5301942 | 0.72175132 |

| | | | | | | | | | | | | |
|---|---|---|---|---|---|---|---|---|---|---|---|---|
| Ypel4 | 0.47294388 | -1.1079827 | 0.021 | 0.036 | 1 | 0.98705046 | -0.8948178 | 0.042 | 0.042 | 1 | 0.98705046 | 0.72221184 |
| Slc7a14 | 0.47294388 | -1.2148979 | 0.021 | 0.036 | 1 | 0.80020906 | -0.7248928 | 0.017 | 0.021 | 1 | 0.80020906 | 0.72221184 |
| Mutyh | 0.47294388 | -0.9925055 | 0.021 | 0.036 | 1 | 0.51194506 | 0.27510724 | 0.021 | 0.01 | 1 | 0.51194506 | 0.72221184 |
| Serinc2 | 0.47294388 | -0.7294711 | 0.021 | 0.036 | 1 | 0.76485982 | -0.3098553 | 0.025 | 0.031 | 1 | 0.76485982 | 0.72221184 |
| 1700019J19Rik | 0.47294388 | -0.8669746 | 0.021 | 0.036 | 1 | 0.87066325 | -0.3098553 | 0.013 | 0.01 | 1 | 0.87066325 | 0.72221184 |
| Acot12 | 0.47294388 | -0.8669746 | 0.021 | 0.036 | 1 | 0.5778028 | -0.8948178 | 0.013 | 0.021 | 1 | 0.5778028 | 0.72221184 |
| Hsbp1l1 | 0.47294388 | -0.7294711 | 0.021 | 0.036 | 1 | 0.74957918 | -0.8244284 | 0.025 | 0.031 | 1 | 0.74957918 | 0.72221184 |
| Zfp367 | 0.47333238 | 0.78510212 | 0.24 | 0.202 | 1 | 0.9744834 | -1.3098553 | 0.227 | 0.208 | 1 | 0.9744834 | 0.72262122 |
| Pld1 | 0.47412045 | -0.1283889 | 0.375 | 0.404 | 1 | 0.55234078 | -0.56104 | 0.378 | 0.302 | 1 | 0.55234078 | 0.7234507 |
| Gm56721 | 0.5862099 | 0.68556644 | 0.031 | 0.021 | 1 | 0.47427047 | -0.4167705 | 0.088 | 0.062 | 1 | 0.5862099 | 0.72360846 |
| Zfp995 | 0.96422244 | 0.32942263 | 0.073 | 0.073 | 1 | 0.47427047 | -0.3098553 | 0.088 | 0.062 | 1 | 0.96422244 | 0.72360846 |
| Bbof1 | 0.47448877 | 0.42253204 | 0.115 | 0.088 | 1 | 0.97191498 | -0.7248928 | 0.055 | 0.052 | 1 | 0.97191498 | 0.72383794 |
| Gm43774 | 0.47448954 | 0.59245704 | 0.042 | 0.026 | 1 | 0.98992944 | -0.1723517 | 0.042 | 0.042 | 1 | 0.98992944 | 0.72383875 |
| Gm28791 | 0.47477351 | -0.407543 | 0.042 | 0.062 | 1 | 0.76964251 | -0.1254307 | 0.084 | 0.073 | 1 | 0.76964251 | 0.72413713 |
| Mcat | 0.47484645 | 0.54806292 | 0.073 | 0.052 | 1 | 0.67368778 | -0.1082214 | 0.076 | 0.062 | 1 | 0.67368778 | 0.72421375 |
| Pcdhgb6 | 0.4748735 | -0.3710171 | 0.042 | 0.062 | 1 | 0.93749089 | -0.2103196 | 0.055 | 0.052 | 1 | 0.93749089 | 0.72424215 |
| Gm26512 | 0.47553167 | 0.53800925 | 0.052 | 0.078 | 1 | 0.89915467 | -0.3098553 | 0.067 | 0.062 | 1 | 0.89915467 | 0.72493297 |
| Nlrc4 | 0.85317634 | 0.22988696 | 0.062 | 0.057 | 1 | 0.47582317 | -0.4797803 | 0.063 | 0.042 | 1 | 0.85317634 | 0.72523865 |
| Gm14023 | 0.47599549 | 0.52206771 | 0.042 | 0.026 | 1 | 0.86312513 | -0.7248928 | 0.008 | 0.01 | 1 | 0.86312513 | 0.72541927 |
| Gm26982 | 0.47599549 | 0.52206771 | 0.042 | 0.026 | 1 | 0.86695837 | -0.194378 | 0.046 | 0.042 | 1 | 0.86695837 | 0.72541927 |
| E130304I02Rik | 0.47599549 | 0.52206771 | 0.042 | 0.026 | 1 | 0.86312513 | -0.7248928 | 0.008 | 0.01 | 1 | 0.86312513 | 0.72541927 |
| Gm20457 | 0.47599549 | 0.52206771 | 0.042 | 0.026 | 1 | 0.67149861 | -0.1399303 | 0.029 | 0.021 | 1 | 0.67149861 | 0.72541927 |
| Kcnn1 | 0.47599549 | 0.52206771 | 0.042 | 0.026 | 1 | 0.79600331 | -0.9879272 | 0.017 | 0.021 | 1 | 0.79600331 | 0.72541927 |
| 1700022H01Rik | 0.47599549 | 0.52206771 | 0.042 | 0.026 | 1 | 0.98590808 | 0.10518224 | 0.021 | 0.021 | 1 | 0.98590808 | 0.72541927 |
| Spdef | 0.47599549 | 0.15949763 | 0.042 | 0.026 | 1 | 0.91228894 | -0.8623963 | 0.05 | 0.052 | 1 | 0.91228894 | 0.72541927 |
| Gm26885 | 0.47599549 | 0.52206771 | 0.042 | 0.026 | 1 | 0.63231936 | 0.42711033 | 0.055 | 0.042 | 1 | 0.63231936 | 0.72541927 |
| Prrt1 | 0.47599549 | 0.52206771 | 0.042 | 0.026 | 1 | 0.8076363 | 0.42711033 | 0.025 | 0.021 | 1 | 0.8076363 | 0.72541927 |
| Scd4 | 0.47599549 | 0.52206771 | 0.042 | 0.026 | 1 | 0.53532186 | -1.0688472 | 0.046 | 0.062 | 1 | 0.53532186 | 0.72541927 |
| Adam1a | 0.47599549 | -0.577468 | 0.021 | 0.036 | 1 | 0.52072676 | -0.3098553 | 0.021 | 0.01 | 1 | 0.52072676 | 0.72541927 |
| Gm3625 | 0.47599549 | -0.577468 | 0.021 | 0.036 | 1 | 0.51194506 | 0.27510724 | 0.021 | 0.01 | 1 | 0.51194506 | 0.72541927 |
| Esrra | 0.47599549 | -0.577468 | 0.021 | 0.036 | 1 | 0.56906762 | -0.9688183 | 0.067 | 0.083 | 1 | 0.56906762 | 0.72541927 |
| Nkain2 | 0.47603909 | -2.130009 | 0.021 | 0.036 | 1 | 0.50587544 | -2.7248928 | 0.004 | 0.01 | 1 | 0.50587544 | 0.72546496 |
| Gm16897 | 0.47606368 | 1.00749454 | 0.021 | 0.01 | 1 | 0.86543325 | 0.27510724 | 0.013 | 0.01 | 1 | 0.86543325 | 0.72549073 |
| Gm128 | 0.47606368 | 1.00749454 | 0.021 | 0.01 | 1 | 0.88208974 | -0.4167705 | 0.046 | 0.042 | 1 | 0.88208974 | 0.72549073 |
| Bcl2l14 | 0.47606368 | 1.00749454 | 0.021 | 0.01 | 1 | 0.85710544 | -1.7248928 | 0.008 | 0.01 | 1 | 0.85710544 | 0.72549073 |
| Gask1a | 0.47606368 | 1.00749454 | 0.021 | 0.01 | 1 | 0.98590808 | 0.42711033 | 0.021 | 0.021 | 1 | 0.98590808 | 0.72549073 |
| B3gat2 | 0.47606502 | 1.00749454 | 0.021 | 0.01 | 1 | 0.57170879 | -1.8948178 | 0.013 | 0.021 | 1 | 0.57170879 | 0.72549214 |
| Myocos | 0.47606502 | 1.00749454 | 0.021 | 0.01 | 1 | 0.70493467 | -0.7952821 | 0.034 | 0.042 | 1 | 0.70493467 | 0.72549214 |
| P2rx3 | 0.47606502 | 1.00749454 | 0.021 | 0.01 | 1 | 0.87066325 | -0.3098553 | 0.013 | 0.01 | 1 | 0.87066325 | 0.72549214 |
| Ptk6 | 0.47606502 | 1.00749454 | 0.021 | 0.01 | 1 | 0.66387437 | 0.27510724 | 0.017 | 0.01 | 1 | 0.66387437 | 0.72549214 |
| Gm42744 | 0.47606502 | 1.00749454 | 0.021 | 0.01 | 1 | 0.79600331 | -1.3098553 | 0.017 | 0.021 | 1 | 0.79600331 | 0.72549214 |
| Gm43112 | 0.47606502 | 1.00749454 | 0.021 | 0.01 | 1 | 0.78273445 | -0.6577786 | 0.038 | 0.031 | 1 | 0.78273445 | 0.72549214 |
| Ccdc146 | 0.47606502 | 1.00749454 | 0.021 | 0.01 | 1 | 0.50887612 | -1.3098553 | 0.004 | 0.01 | 1 | 0.50887612 | 0.72549214 |
| Gm43660 | 0.47606502 | 1.00749454 | 0.021 | 0.01 | 1 | 0.87851725 | -0.8948178 | 0.013 | 0.01 | 1 | 0.87851725 | 0.72549214 |
| Gm30003 | 0.47606502 | 1.00749454 | 0.021 | 0.01 | 1 | 0.50587544 | -1.8948178 | 0.004 | 0.01 | 1 | 0.50587544 | 0.72549214 |
| Gm56980 | 0.47606502 | 1.00749454 | 0.021 | 0.01 | 1 | 0.89707195 | -1.4618584 | 0.029 | 0.031 | 1 | 0.89707195 | 0.72549214 |
| Gm38973 | 0.47606502 | 1.00749454 | 0.021 | 0.01 | 1 | 0.66391569 | -1.1723517 | 0.042 | 0.052 | 1 | 0.66391569 | 0.72549214 |
| Icam5 | 0.47606502 | 1.00749454 | 0.021 | 0.01 | 1 | 0.87851725 | -1.3098553 | 0.013 | 0.01 | 1 | 0.87851725 | 0.72549214 |
| 5830454E08Rik | 0.47606502 | 1.00749454 | 0.021 | 0.01 | 1 | 0.87066325 | -0.3098553 | 0.013 | 0.01 | 1 | 0.87066325 | 0.72549214 |
| Mettl24 | 0.47606502 | 1.00749454 | 0.021 | 0.01 | 1 | 0.52072676 | -0.3098553 | 0.021 | 0.01 | 1 | 0.52072676 | 0.72549214 |
| Edar | 0.47606502 | 1.00749454 | 0.021 | 0.01 | 1 | 0.50764019 | -1.1034044 | 0.046 | 0.062 | 1 | 0.50764019 | 0.72549214 |
| Ramp3 | 0.47606502 | 1.00749454 | 0.021 | 0.01 | 1 | 0.50887612 | -1.3098553 | 0.004 | 0.01 | 1 | 0.50887612 | 0.72549214 |
| 2810021J22Rik | 0.47606502 | 1.00749454 | 0.021 | 0.01 | 1 | 0.51018266 | 0.49749966 | 0.021 | 0.01 | 1 | 0.51018266 | 0.72549214 |
| Lratd1 | 0.47606502 | 1.00749454 | 0.021 | 0.01 | 1 | 0.86312513 | -0.7248928 | 0.008 | 0.01 | 1 | 0.86312513 | 0.72549214 |
| 2410004P03Rik | 0.47606502 | 1.00749454 | 0.021 | 0.01 | 1 | 0.80442048 | -0.5728897 | 0.017 | 0.021 | 1 | 0.80442048 | 0.72549214 |
| Ighm | 0.47606502 | 4.15045249 | 0.021 | 0.01 | 1 | 0.69869669 | -0.4167705 | 0.071 | 0.083 | 1 | 0.69869669 | 0.72549214 |
| Ciita | 0.47606502 | 1.00749454 | 0.021 | 0.01 | 1 | 0.86312513 | -0.7248928 | 0.008 | 0.01 | 1 | 0.86312513 | 0.72549214 |
| Gm49584 | 0.47606502 | 1.00749454 | 0.021 | 0.01 | 1 | 0.51194506 | 0.27510724 | 0.021 | 0.01 | 1 | 0.51194506 | 0.72549214 |
| Gm15063 | 0.47606502 | 1.00749454 | 0.021 | 0.01 | 1 | 0.50887612 | -1.3098553 | 0.004 | 0.01 | 1 | 0.50887612 | 0.72549214 |
| Mbd4 | 0.47608023 | 0.22988696 | 0.115 | 0.088 | 1 | 0.68619675 | -0.194378 | 0.088 | 0.073 | 1 | 0.68619675 | 0.72550808 |
| Rab30 | 0.61289741 | -0.4872702 | 0.198 | 0.223 | 1 | 0.47771394 | -0.9636796 | 0.223 | 0.24 | 1 | 0.61289741 | 0.72721727 |
| Cpz | 0.47815602 | 0.14499806 | 0.073 | 0.052 | 1 | 0.58664899 | -0.3098553 | 0.021 | 0.031 | 1 | 0.58664899 | 0.72767886 |
| Car13 | 0.47912569 | -0.407543 | 0.021 | 0.036 | 1 | 0.89212057 | -0.9754362 | 0.092 | 0.094 | 1 | 0.89212057 | 0.72868995 |
| Gm39167 | 0.47912569 | -0.407543 | 0.021 | 0.036 | 1 | 0.5207331 | -0.5025003 | 0.021 | 0.01 | 1 | 0.5207331 | 0.72868995 |
| Ccdc159 | 0.47912569 | -0.407543 | 0.021 | 0.036 | 1 | 0.92396969 | -0.4618584 | 0.029 | 0.031 | 1 | 0.92396969 | 0.72868995 |
| Trim3 | 0.72689598 | -0.2962862 | 0.25 | 0.264 | 1 | 0.47916175 | -0.5583536 | 0.328 | 0.333 | 1 | 0.72689598 | 0.72872752 |
| Atg4a | 0.47931114 | -0.4350237 | 0.135 | 0.166 | 1 | 0.57606991 | -0.4987579 | 0.252 | 0.208 | 1 | 0.57606991 | 0.72888311 |
| Nfxl1 | 0.8619088 | 0.21659014 | 0.323 | 0.326 | 1 | 0.47932838 | -0.1931317 | 0.475 | 0.365 | 1 | 0.8619088 | 0.72890107 |
| Pde10a | 0.47977266 | -0.8906258 | 0.083 | 0.109 | 1 | 0.81741159 | -0.2275813 | 0.336 | 0.312 | 1 | 0.81741159 | 0.72936351 |
| Maml2 | 0.48050539 | -0.2134928 | 1 | 0.99 | 1 | 0.52143468 | -0.528918 | 1 | 1 | 1 | 0.52143468 | 0.73012535 |
| Peg12 | 0.48059356 | -0.1851505 | 0.042 | 0.062 | 1 | 0.66662869 | 0.14957636 | 0.029 | 0.021 | 1 | 0.66662869 | 0.73021695 |
| Tmem130 | 0.48064264 | -0.2148979 | 0.021 | 0.01 | 1 | 0.86613737 | -0.3098553 | 0.008 | 0.01 | 1 | 0.86613737 | 0.73026793 |
| Slc17a7 | 0.48064264 | 0.59245704 | 0.021 | 0.01 | 1 | 0.80020906 | -0.7248928 | 0.017 | 0.021 | 1 | 0.80020906 | 0.73026793 |
| Gm10457 | 0.48064264 | 0.59245704 | 0.021 | 0.01 | 1 | 0.5818708 | 0.42711033 | 0.013 | 0.021 | 1 | 0.5818708 | 0.73026793 |
| Gm48250 | 0.48064264 | 0.59245704 | 0.021 | 0.01 | 1 | 0.86011448 | -0.9879272 | 0.008 | 0.01 | 1 | 0.86011448 | 0.73026793 |
| Gm20621 | 0.48064264 | 0.59245704 | 0.021 | 0.01 | 1 | 0.50887612 | -1.3098553 | 0.004 | 0.01 | 1 | 0.50887612 | 0.73026793 |
| 4932412D23Rik | 0.48064264 | 0.59245704 | 0.021 | 0.01 | 1 | 0.50887612 | -1.3098553 | 0.004 | 0.01 | 1 | 0.50887612 | 0.73026793 |
| Gng5c | 0.48064264 | 0.59245704 | 0.021 | 0.01 | 1 | 0.66054955 | -0.1723517 | 0.042 | 0.031 | 1 | 0.66054955 | 0.73026793 |
| 9430015G10Rik | 0.87173216 | 0.10060394 | 0.073 | 0.067 | 1 | 0.48090937 | -0.9472852 | 0.063 | 0.083 | 1 | 0.87173216 | 0.73054492 |
| Lrrc61 | 0.48218219 | 0.32942263 | 0.042 | 0.026 | 1 | 0.74957918 | -0.8244284 | 0.025 | 0.031 | 1 | 0.74957918 | 0.73186471 |
| 4930579K19Rik | 0.48218219 | 0.32942263 | 0.042 | 0.026 | 1 | 0.98993409 | -1.0468209 | 0.021 | 0.021 | 1 | 0.98993409 | 0.73186471 |
| Pex11g | 0.4825879 | 0.42253204 | 0.073 | 0.052 | 1 | 0.96042911 | -0.961932 | 0.055 | 0.052 | 1 | 0.96042911 | 0.73228472 |
| Mau2 | 0.50971698 | -0.1422526 | 0.396 | 0.44 | 1 | 0.48385092 | -0.3000075 | 0.702 | 0.708 | 1 | 0.50971698 | 0.73359012 |

| | | | | | | | | | | | | |
|---|---|---|---|---|---|---|---|---|---|---|---|---|
| Pot1b | 0.48388092 | 0.27052894 | 0.073 | 0.052 | 1 | 0.95149247 | -1.0297473 | 0.05 | 0.052 | 1 | 0.95149247 | 0.7336211 |
| Zfp687 | 0.92336642 | 0.13302542 | 0.094 | 0.098 | 1 | 0.4841661 | -0.3991226 | 0.172 | 0.135 | 1 | 0.92336642 | 0.73391539 |
| 1700017B05Rik | 0.48437439 | 0.21394541 | 0.115 | 0.088 | 1 | 0.99134053 | -0.3259749 | 0.248 | 0.229 | 1 | 0.99134053 | 0.73413023 |
| Nlrx1 | 0.48501409 | 0.68556644 | 0.083 | 0.062 | 1 | 0.7185684 | -1.3098553 | 0.055 | 0.062 | 1 | 0.7185684 | 0.73478951 |
| Epm2a | 0.8032501 | -0.6770036 | 0.125 | 0.109 | 1 | 0.48524772 | -0.7793405 | 0.143 | 0.167 | 1 | 0.8032501 | 0.73503009 |
| Teddm2 | 0.48534895 | -0.3144336 | 0.042 | 0.062 | 1 | 0.54837489 | -0.4618584 | 0.034 | 0.021 | 1 | 0.54837489 | 0.73513429 |
| Coa8 | 0.48649088 | -0.1026884 | 0.51 | 0.404 | 1 | 0.62816702 | -0.2403143 | 0.479 | 0.396 | 1 | 0.62816702 | 0.73630839 |
| Rnf32 | 0.48652615 | -0.2726134 | 0.042 | 0.062 | 1 | 1 | -0.8244284 | 0.021 | 0.021 | 1 | 1 | 0.73634461 |
| Kbtbd11 | 0.48683788 | -0.1624305 | 0.021 | 0.036 | 1 | 0.92010413 | -0.7793405 | 0.05 | 0.052 | 1 | 0.92010413 | 0.73666464 |
| Elmo3 | 0.48683788 | -0.1624305 | 0.021 | 0.036 | 1 | 0.72150599 | -0.4797803 | 0.063 | 0.052 | 1 | 0.72150599 | 0.73666464 |
| Cenpc1 | 0.50106802 | -0.2955745 | 0.385 | 0.415 | 1 | 0.48762256 | -0.361902 | 0.546 | 0.594 | 1 | 0.50106802 | 0.73746935 |
| Ak5 | 0.48868542 | -0.9925055 | 0.052 | 0.073 | 1 | 0.4881277 | -0.7692869 | 0.13 | 0.156 | 1 | 0.48868542 | 0.73798675 |
| Hdc | 0.4884071 | -0.7580402 | 0.042 | 0.026 | 1 | 0.80021055 | 0.27510724 | 0.017 | 0.021 | 1 | 0.80021055 | 0.7382727 |
| Tab1 | 0.48860545 | -0.7294711 | 0.052 | 0.073 | 1 | 0.66532443 | -1.3923174 | 0.063 | 0.073 | 1 | 0.66532443 | 0.73847561 |
| AA986860 | 0.48943323 | 1.00749454 | 0.094 | 0.073 | 1 | 0.9617148 | -0.8244284 | 0.055 | 0.052 | 1 | 0.9617148 | 0.73932157 |
| Nup93 | 0.4898205 | -0.3033784 | 0.417 | 0.415 | 1 | 0.63681149 | -1.5381243 | 0.298 | 0.25 | 1 | 0.63681149 | 0.73971687 |
| Gm12536 | 0.49828179 | -0.1693832 | 0.125 | 0.155 | 1 | 0.4899184 | -0.3942474 | 0.197 | 0.156 | 1 | 0.49828179 | 0.73981676 |
| Mir22hg | 0.7516598 | -0.2356192 | 0.51 | 0.503 | 1 | 0.49015234 | -0.3788026 | 0.613 | 0.604 | 1 | 0.7516598 | 0.74005537 |
| Lpin1 | 0.93107913 | 0.15472049 | 0.375 | 0.358 | 1 | 0.49032264 | -0.5773356 | 0.265 | 0.281 | 1 | 0.93107913 | 0.74022899 |
| Gm26887 | 0.49035831 | 0.40342321 | 0.167 | 0.135 | 1 | 0.78801852 | -0.4410998 | 0.118 | 0.125 | 1 | 0.78801852 | 0.74026535 |
| Nrde2 | 0.49066501 | 0.30705482 | 0.073 | 0.052 | 1 | 0.65297498 | -0.8359241 | 0.08 | 0.094 | 1 | 0.65297498 | 0.74057787 |
| Pcdhgb4 | 0.49066501 | 0.30705482 | 0.073 | 0.052 | 1 | 0.56663686 | -1.4703199 | 0.059 | 0.073 | 1 | 0.56663686 | 0.74057787 |
| Fyb | 0.49086811 | -1.5070786 | 0.052 | 0.073 | 1 | 0.75390898 | -0.3463811 | 0.097 | 0.083 | 1 | 0.75390898 | 0.74078472 |
| Camk2b | 0.87558365 | 0.15949763 | 0.094 | 0.088 | 1 | 0.49119514 | -1.1082214 | 0.084 | 0.104 | 1 | 0.87558365 | 0.74111762 |
| Usp4 | 0.51999238 | -0.2230372 | 0.615 | 0.549 | 1 | 0.49213919 | -0.2356543 | 0.744 | 0.719 | 1 | 0.51999238 | 0.7420774 |
| 4921504A21Rik | 0.52395107 | 0.27052894 | 0.083 | 0.062 | 1 | 0.49254343 | -0.8623963 | 0.088 | 0.062 | 1 | 0.52395107 | 0.74248783 |
| Tubgcp4 | 0.8250914 | 0.37225883 | 0.271 | 0.264 | 1 | 0.49269007 | -0.2187074 | 0.273 | 0.219 | 1 | 0.8250914 | 0.74263664 |
| Synm | 0.62067229 | 0.54354744 | 0.146 | 0.13 | 1 | 0.49309434 | 0.20799304 | 0.235 | 0.198 | 1 | 0.62067229 | 0.74304665 |
| Pard6g | 0.6748651 | -0.3884341 | 0.125 | 0.14 | 1 | 0.49378306 | -0.9986536 | 0.202 | 0.219 | 1 | 0.6748651 | 0.7437444 |
| Gm16973 | 0.49404972 | -0.7294711 | 0.052 | 0.073 | 1 | 0.80634752 | -0.7248928 | 0.084 | 0.073 | 1 | 0.80634752 | 0.74401432 |
| Fbrsl1 | 0.59021504 | 0.14499806 | 0.469 | 0.461 | 1 | 0.49439914 | -0.2292898 | 0.634 | 0.688 | 1 | 0.59021504 | 0.74436777 |
| Fzd3 | 0.49525591 | -0.3550755 | 0.198 | 0.228 | 1 | 0.70049622 | -0.1696776 | 0.282 | 0.24 | 1 | 0.70049622 | 0.7452334 |
| Gpank1 | 0.90166162 | 0.34452952 | 0.073 | 0.078 | 1 | 0.49563246 | -0.2167459 | 0.122 | 0.094 | 1 | 0.90166162 | 0.74561338 |
| Rab3il1 | 0.69865596 | -0.3044495 | 0.167 | 0.181 | 1 | 0.49594179 | -0.1578522 | 0.21 | 0.167 | 1 | 0.69865596 | 0.74592532 |
| Ripor2 | 0.49624236 | 2.00749454 | 0.021 | 0.036 | 1 | 0.79600786 | 0.17557156 | 0.017 | 0.021 | 1 | 0.79600786 | 0.74622824 |
| Sptbn4 | 0.49780735 | 0.15949763 | 0.021 | 0.036 | 1 | 0.56767339 | -1.5728897 | 0.013 | 0.021 | 1 | 0.56767339 | 0.74780255 |
| Zfp81 | 1 | -0.1079827 | 0.104 | 0.104 | 1 | 0.4978811 | -0.3098553 | 0.113 | 0.083 | 1 | 1 | 0.74787661 |
| Nlgn3 | 0.75328966 | 0.27052894 | 0.021 | 0.016 | 1 | 0.49797056 | -0.4353861 | 0.076 | 0.052 | 1 | 0.75328966 | 0.74796644 |
| Tbcd | 0.49865266 | -0.1730777 | 0.24 | 0.197 | 1 | 0.63166911 | -0.5728897 | 0.227 | 0.25 | 1 | 0.63166911 | 0.74865085 |
| Plekho1 | 0.9393034 | 0.51445453 | 0.146 | 0.15 | 1 | 0.49936525 | -0.4208866 | 0.113 | 0.135 | 1 | 0.9393034 | 0.74936485 |
| Atg4d | 0.57652005 | -0.2820121 | 0.083 | 0.104 | 1 | 0.49940013 | -0.6317834 | 0.139 | 0.104 | 1 | 0.57652005 | 0.74939977 |
| Zfp442 | 0.49952136 | -0.4950058 | 0.052 | 0.073 | 1 | 0.83808753 | -0.3769695 | 0.059 | 0.052 | 1 | 0.83808753 | 0.74952113 |
| Spice1 | 0.50065366 | -0.2148979 | 0.167 | 0.202 | 1 | 0.75439663 | -0.6442743 | 0.151 | 0.135 | 1 | 0.75439663 | 0.75065323 |
| Pals1 | 0.80023751 | -0.132262 | 0.875 | 0.912 | 1 | 0.50176112 | -0.1525083 | 0.891 | 0.958 | 1 | 0.80023751 | 0.75175802 |
| Ttc39c | 0.84218034 | -0.3795286 | 0.115 | 0.104 | 1 | 0.50260896 | -0.9113059 | 0.092 | 0.115 | 1 | 0.84218034 | 0.75260215 |
| Zfp799 | 0.5027571 | -0.2681399 | 0.146 | 0.171 | 1 | 0.58012005 | -0.5549678 | 0.097 | 0.073 | 1 | 0.58012005 | 0.7527495 |
| Cpped1 | 0.67957772 | -0.3461424 | 0.167 | 0.181 | 1 | 0.50461434 | -0.1200307 | 0.218 | 0.177 | 1 | 0.67957772 | 0.75459305 |
| Gm12216 | 0.50474402 | -0.8290067 | 0.146 | 0.171 | 1 | 0.98233608 | -0.6634922 | 0.357 | 0.323 | 1 | 0.98233608 | 0.75472151 |
| 5330438D12Rik | 0.86718161 | -0.1379359 | 0.292 | 0.269 | 1 | 0.50490253 | -0.5154394 | 0.357 | 0.281 | 1 | 0.86718161 | 0.7548785 |
| Igtp | 0.67872781 | -0.240433 | 0.125 | 0.14 | 1 | 0.50516004 | -2.6883669 | 0.076 | 0.094 | 1 | 0.67872781 | 0.75513341 |
| Reps2 | 0.70581537 | -0.7998604 | 0.052 | 0.062 | 1 | 0.50533291 | -0.3587649 | 0.063 | 0.083 | 1 | 0.70581537 | 0.75530447 |
| Dmd | 0.89051405 | 0.34452952 | 0.052 | 0.057 | 1 | 0.50539642 | -1.3897086 | 0.189 | 0.219 | 1 | 0.89051405 | 0.7553673 |
| Gm14798 | 0.59199182 | 0.20013961 | 0.031 | 0.021 | 1 | 0.50552304 | -1.0874628 | 0.046 | 0.062 | 1 | 0.59199182 | 0.75549254 |
| Mycl | 0.52744059 | -0.577468 | 0.01 | 0.021 | 1 | 0.50587544 | -1.8948178 | 0.004 | 0.01 | 1 | 0.52744059 | 0.75584092 |
| Mrc1 | 0.52961488 | -0.3144336 | 0.01 | 0.021 | 1 | 0.50587544 | -1.7248928 | 0.004 | 0.01 | 1 | 0.52961488 | 0.75584092 |
| Slco1a5 | 0.61371908 | 1.74446013 | 0.01 | 0.005 | 1 | 0.50587544 | -2.6317834 | 0.004 | 0.01 | 1 | 0.61371908 | 0.75584092 |
| Napsa | 0.61371908 | 2.42253204 | 0.01 | 0.005 | 1 | 0.50587544 | -2.6317834 | 0.004 | 0.01 | 1 | 0.61371908 | 0.75584092 |
| Gm28068 | 0.61371908 | 1.59245704 | 0.01 | 0.005 | 1 | 0.50587544 | -2.3098553 | 0.004 | 0.01 | 1 | 0.61371908 | 0.75584092 |
| Gm57320 | 0.61737927 | 1.00749454 | 0.01 | 0.005 | 1 | 0.50587544 | -2.6317834 | 0.004 | 0.01 | 1 | 0.61737927 | 0.75584092 |
| Naip2 | 0.61737927 | 1.00749454 | 0.01 | 0.005 | 1 | 0.50587544 | -1.8948178 | 0.004 | 0.01 | 1 | 0.61737927 | 0.75584092 |
| H2-Ob | 0.61737927 | 1.00749454 | 0.01 | 0.005 | 1 | 0.50587544 | -1.8948178 | 0.004 | 0.01 | 1 | 0.61737927 | 0.75584092 |
| Arl13a | 0.61737927 | 1.00749454 | 0.01 | 0.005 | 1 | 0.50587544 | -1.8948178 | 0.004 | 0.01 | 1 | 0.61737927 | 0.75584092 |
| Rfx4 | 0.62104899 | 0.42253204 | 0.01 | 0.005 | 1 | 0.50587544 | -1.8948178 | 0.004 | 0.01 | 1 | 0.62104899 | 0.75584092 |
| Vwa5b2 | 0.62104899 | 0.42253204 | 0.01 | 0.005 | 1 | 0.50587544 | -2.3098553 | 0.004 | 0.01 | 1 | 0.62104899 | 0.75584092 |
| Lvrn | 0.72298091 | -0.577468 | 0.01 | 0.016 | 1 | 0.50587544 | -2.0468209 | 0.004 | 0.01 | 1 | 0.72298091 | 0.75584092 |
| Traf3ip3 | 0.72574709 | -0.3144336 | 0.01 | 0.016 | 1 | 0.50587544 | -2.3098553 | 0.004 | 0.01 | 1 | 0.72574709 | 0.75584092 |
| Hcn1 | 0.74827336 | 0.42253204 | 0.021 | 0.016 | 1 | 0.50587544 | -1.6317834 | 0.004 | 0.01 | 1 | 0.74827336 | 0.75584092 |
| 2010002M09Rik | 0.74827573 | 0.59245704 | 0.021 | 0.016 | 1 | 0.50587544 | -1.8948178 | 0.004 | 0.01 | 1 | 0.74827573 | 0.75584092 |
| 4931428L18Rik | 0.75328966 | 0.27052894 | 0.021 | 0.016 | 1 | 0.50587544 | -1.8948178 | 0.004 | 0.01 | 1 | 0.75328966 | 0.75584092 |
| Gm57468 | 0.993208 | 1.00749454 | 0.01 | 0.01 | 1 | 0.50587544 | -2.3098553 | 0.004 | 0.01 | 1 | 0.993208 | 0.75584092 |
| Gm49975 | 0.993208 | 1.00749454 | 0.01 | 0.01 | 1 | 0.50587544 | -1.8948178 | 0.004 | 0.01 | 1 | 0.993208 | 0.75584092 |
| Cdrt4 | 1 | 0.42253204 | 0.01 | 0.01 | 1 | 0.50587544 | -1.8948178 | 0.004 | 0.01 | 1 | 1 | 0.75584092 |
| Svip | 0.52744059 | -0.577468 | 0.01 | 0.021 | 1 | 0.50670329 | 0.86006974 | 0.021 | 0.01 | 1 | 0.52744059 | 0.75665835 |
| Gm11290 | 0.54205144 | -0.407543 | 0.031 | 0.047 | 1 | 0.50670329 | 1.93807225 | 0.021 | 0.01 | 1 | 0.54205144 | 0.75665835 |
| Tfap4 | 0.61550914 | -0.407543 | 0.021 | 0.031 | 1 | 0.50670329 | 0.86006974 | 0.021 | 0.01 | 1 | 0.61550914 | 0.75665835 |
| Gm38604 | 0.7854853 | -0.577468 | 0.021 | 0.026 | 1 | 0.50670329 | 0.86006974 | 0.021 | 0.01 | 1 | 0.7854853 | 0.75665835 |
| Raver2 | 0.67168595 | -0.8405024 | 0.042 | 0.052 | 1 | 0.50675003 | -1.3802446 | 0.055 | 0.073 | 1 | 0.67168595 | 0.75670447 |
| Orc1 | 0.50701639 | 0.10060394 | 0.073 | 0.052 | 1 | 0.9750676 | 0.27510724 | 0.063 | 0.062 | 1 | 0.9750676 | 0.75696716 |
| Ap3s1 | 0.50762851 | -0.2029792 | 0.812 | 0.86 | 1 | 0.64969812 | -0.1016767 | 0.895 | 0.865 | 1 | 0.64969812 | 0.75757031 |
| Them6 | 0.61550914 | -0.407543 | 0.021 | 0.031 | 1 | 0.50844268 | 0.69014474 | 0.021 | 0.01 | 1 | 0.61550914 | 0.7583714 |
| Fbxo47 | 1 | -0.3144336 | 0.031 | 0.031 | 1 | 0.50844268 | 1.27510724 | 0.021 | 0.01 | 1 | 1 | 0.7583714 |
| Tnni2 | 0.50874134 | 0.34006988 | 0.135 | 0.109 | 1 | 0.74891106 | -0.3838558 | 0.105 | 0.115 | 1 | 0.74891106 | 0.75866493 |
| Gm57030 | 0.52744059 | -0.577468 | 0.01 | 0.021 | 1 | 0.50887612 | -1.3098553 | 0.004 | 0.01 | 1 | 0.52744059 | 0.75879734 |
| Alk | 0.52744059 | -0.577468 | 0.01 | 0.021 | 1 | 0.50887612 | -1.3098553 | 0.004 | 0.01 | 1 | 0.52744059 | 0.75879734 |
| Lin7b | 0.52961488 | -0.3144336 | 0.01 | 0.021 | 1 | 0.50887612 | -1.3098553 | 0.004 | 0.01 | 1 | 0.52961488 | 0.75879734 |
| Slc39a4 | 0.52961488 | -0.3144336 | 0.01 | 0.021 | 1 | 0.50887612 | -1.3098553 | 0.004 | 0.01 | 1 | 0.52961488 | 0.75879734 |
| Ufsp1 | 0.5862099 | 0.68556644 | 0.031 | 0.021 | 1 | 0.50887612 | -1.3098553 | 0.004 | 0.01 | 1 | 0.5862099 | 0.75879734 |

| | | | | | | | | | | | | |
|---|---|---|---|---|---|---|---|---|---|---|---|---|
| Dmbt1 | 0.5978326 | -0.9925055 | 0.031 | 0.021 | 1 | 0.50887612 | -1.3098553 | 0.004 | 0.01 | 1 | 0.5978326 | 0.75879734 |
| Slc9a2 | 0.61371908 | 1.59245704 | 0.01 | 0.005 | 1 | 0.50887612 | -1.3098553 | 0.004 | 0.01 | 1 | 0.61371908 | 0.75879734 |
| Gm57144 | 0.61371908 | 1.59245704 | 0.01 | 0.005 | 1 | 0.50887612 | -1.3098553 | 0.004 | 0.01 | 1 | 0.61371908 | 0.75879734 |
| Cdcp2 | 0.61737927 | 1.00749454 | 0.01 | 0.005 | 1 | 0.50887612 | -1.3098553 | 0.004 | 0.01 | 1 | 0.61737927 | 0.75879734 |
| Gm15884 | 0.61737927 | 1.00749454 | 0.01 | 0.005 | 1 | 0.50887612 | -1.3098553 | 0.004 | 0.01 | 1 | 0.61737927 | 0.75879734 |
| Chrna10 | 0.61737927 | 1.00749454 | 0.01 | 0.005 | 1 | 0.50887612 | -1.3098553 | 0.004 | 0.01 | 1 | 0.61737927 | 0.75879734 |
| Jph3 | 0.61737927 | 1.00749454 | 0.01 | 0.005 | 1 | 0.50887612 | -1.3098553 | 0.004 | 0.01 | 1 | 0.61737927 | 0.75879734 |
| Gm30639 | 0.61737927 | 1.00749454 | 0.01 | 0.005 | 1 | 0.50887612 | -1.3098553 | 0.004 | 0.01 | 1 | 0.61737927 | 0.75879734 |
| Gm56958 | 0.61737927 | 1.00749454 | 0.01 | 0.005 | 1 | 0.50887612 | -1.3098553 | 0.004 | 0.01 | 1 | 0.61737927 | 0.75879734 |
| Gm56957 | 0.61737927 | 1.00749454 | 0.01 | 0.005 | 1 | 0.50887612 | -1.3098553 | 0.004 | 0.01 | 1 | 0.61737927 | 0.75879734 |
| 1700020G17Rik | 0.61737927 | 1.00749454 | 0.01 | 0.005 | 1 | 0.50887612 | -1.3098553 | 0.004 | 0.01 | 1 | 0.61737927 | 0.75879734 |
| 4930422I22Rik | 0.61737927 | 1.00749454 | 0.01 | 0.005 | 1 | 0.50887612 | -1.3098553 | 0.004 | 0.01 | 1 | 0.61737927 | 0.75879734 |
| Gm4129 | 0.61737927 | 1.00749454 | 0.01 | 0.005 | 1 | 0.50887612 | -1.3098553 | 0.004 | 0.01 | 1 | 0.61737927 | 0.75879734 |
| Ccdc103 | 0.61737927 | 1.00749454 | 0.01 | 0.005 | 1 | 0.50887612 | -1.3098553 | 0.004 | 0.01 | 1 | 0.61737927 | 0.75879734 |
| Gm11716 | 0.61737927 | 1.00749454 | 0.01 | 0.005 | 1 | 0.50887612 | -1.3098553 | 0.004 | 0.01 | 1 | 0.61737927 | 0.75879734 |
| Gm15829 | 0.61737927 | 1.00749454 | 0.01 | 0.005 | 1 | 0.50887612 | -1.3098553 | 0.004 | 0.01 | 1 | 0.61737927 | 0.75879734 |
| Gm7168 | 0.61737927 | 1.00749454 | 0.01 | 0.005 | 1 | 0.50887612 | -1.3098553 | 0.004 | 0.01 | 1 | 0.61737927 | 0.75879734 |
| Plac8l1 | 0.61737927 | 1.00749454 | 0.01 | 0.005 | 1 | 0.50887612 | -1.3098553 | 0.004 | 0.01 | 1 | 0.61737927 | 0.75879734 |
| Clec4n | 0.62104899 | 0.42253204 | 0.01 | 0.005 | 1 | 0.50887612 | -1.3098553 | 0.004 | 0.01 | 1 | 0.62104899 | 0.75879734 |
| Acta1 | 0.62104899 | -1.0408685 | 0.01 | 0.005 | 1 | 0.50887612 | -1.3098553 | 0.004 | 0.01 | 1 | 0.62104899 | 0.75879734 |
| Gpr132 | 0.62104899 | 0.59245704 | 0.01 | 0.005 | 1 | 0.50887612 | -1.3098553 | 0.004 | 0.01 | 1 | 0.62104899 | 0.75879734 |
| Zfp456 | 0.62104899 | 0.42253204 | 0.01 | 0.005 | 1 | 0.50887612 | -1.3098553 | 0.004 | 0.01 | 1 | 0.62104899 | 0.75879734 |
| Rnase1 | 0.62104899 | 0.42253204 | 0.01 | 0.005 | 1 | 0.50887612 | -1.3098553 | 0.004 | 0.01 | 1 | 0.62104899 | 0.75879734 |
| Gm21691 | 0.62104899 | 0.42253204 | 0.01 | 0.005 | 1 | 0.50887612 | -1.3098553 | 0.004 | 0.01 | 1 | 0.62104899 | 0.75879734 |
| Tbx18 | 0.63038178 | 0.52206771 | 0.021 | 0.031 | 1 | 0.50887612 | -1.3098553 | 0.004 | 0.01 | 1 | 0.63038178 | 0.75879734 |
| Fndc7 | 0.72574709 | -0.9925055 | 0.01 | 0.016 | 1 | 0.50887612 | -1.3098553 | 0.004 | 0.01 | 1 | 0.72574709 | 0.75879734 |
| Gm57347 | 0.72574709 | -0.7998604 | 0.01 | 0.016 | 1 | 0.50887612 | -1.3098553 | 0.004 | 0.01 | 1 | 0.72574709 | 0.75879734 |
| F12 | 0.74076005 | 2.17741954 | 0.021 | 0.016 | 1 | 0.50887612 | -1.3098553 | 0.004 | 0.01 | 1 | 0.74076005 | 0.75879734 |
| Gm45447 | 0.74827573 | 0.59245704 | 0.021 | 0.016 | 1 | 0.50887612 | -1.3098553 | 0.004 | 0.01 | 1 | 0.74827573 | 0.75879734 |
| Ddias | 0.74827573 | 0.59245704 | 0.021 | 0.016 | 1 | 0.50887612 | -1.3098553 | 0.004 | 0.01 | 1 | 0.74827573 | 0.75879734 |
| Fam166b | 0.75328966 | 0.27052894 | 0.021 | 0.016 | 1 | 0.50887612 | -1.3098553 | 0.004 | 0.01 | 1 | 0.75328966 | 0.75879734 |
| Gm43623 | 0.75328966 | 0.27052894 | 0.021 | 0.016 | 1 | 0.50887612 | -1.3098553 | 0.004 | 0.01 | 1 | 0.75328966 | 0.75879734 |
| Gm49518 | 0.78979801 | -0.2148979 | 0.021 | 0.026 | 1 | 0.50887612 | -1.3098553 | 0.004 | 0.01 | 1 | 0.78979801 | 0.75879734 |
| Adarb2 | 0.79196846 | -1.3144336 | 0.021 | 0.026 | 1 | 0.50887612 | -1.3098553 | 0.004 | 0.01 | 1 | 0.79196846 | 0.75879734 |
| A930029G22Rik | 0.79666623 | 0.42253204 | 0.031 | 0.026 | 1 | 0.50887612 | -1.3098553 | 0.004 | 0.01 | 1 | 0.79666623 | 0.75879734 |
| Tpte | 0.993208 | 1.74446013 | 0.01 | 0.01 | 1 | 0.50887612 | -1.3098553 | 0.004 | 0.01 | 1 | 0.993208 | 0.75879734 |
| Clcnka | 1 | -0.3144336 | 0.01 | 0.01 | 1 | 0.50887612 | -1.3098553 | 0.004 | 0.01 | 1 | 1 | 0.75879734 |
| Catsperb | 1 | 0.42253204 | 0.01 | 0.01 | 1 | 0.50887612 | -1.3098553 | 0.004 | 0.01 | 1 | 1 | 0.75879734 |
| Ms4a6c | 1 | -0.9925055 | 0.01 | 0.01 | 1 | 0.50887612 | -1.3098553 | 0.004 | 0.01 | 1 | 1 | 0.75879734 |
| Tln2 | 0.64396623 | 0.37672835 | 0.417 | 0.435 | 1 | 0.5098594 | -0.4542452 | 0.218 | 0.167 | 1 | 0.64396623 | 0.75976219 |
| Styxl1 | 0.61182672 | -0.577468 | 0.021 | 0.031 | 1 | 0.51018266 | 1.14957636 | 0.021 | 0.01 | 1 | 0.61182672 | 0.76007897 |
| Krtdap | 0.61737927 | 1.00749454 | 0.01 | 0.005 | 1 | 0.51018266 | 0.49749966 | 0.021 | 0.01 | 1 | 0.61737927 | 0.76007897 |
| Pde6a | 0.61737927 | 1.00749454 | 0.01 | 0.005 | 1 | 0.51018266 | 0.49749966 | 0.021 | 0.01 | 1 | 0.61737927 | 0.76007897 |
| H2bc9 | 0.70642533 | -0.407543 | 0.052 | 0.041 | 1 | 0.51018266 | 0.49749966 | 0.021 | 0.01 | 1 | 0.70642533 | 0.76007897 |
| Pgap6 | 0.72574709 | -0.3144336 | 0.01 | 0.016 | 1 | 0.51018266 | 0.49749966 | 0.021 | 0.01 | 1 | 0.72574709 | 0.76007897 |
| Gpc2 | 0.85778553 | -0.1079827 | 0.052 | 0.057 | 1 | 0.51018266 | 0.49749966 | 0.021 | 0.01 | 1 | 0.85778553 | 0.76007897 |
| Gm57349 | 0.99207432 | -0.130009 | 0.031 | 0.031 | 1 | 0.51018266 | 0.49749966 | 0.021 | 0.01 | 1 | 0.99207432 | 0.76007897 |
| Crb3 | 1 | 0.42253204 | 0.01 | 0.01 | 1 | 0.51018266 | 0.49749966 | 0.021 | 0.01 | 1 | 1 | 0.76007897 |
| Rad23b | 0.51034124 | -0.1448364 | 0.844 | 0.86 | 1 | 0.9210679 | -0.1576758 | 0.958 | 0.885 | 1 | 0.9210679 | 0.7602343 |
| Sec22c | 0.51079863 | 0.42253204 | 0.083 | 0.062 | 1 | 0.97088779 | -0.7248928 | 0.084 | 0.083 | 1 | 0.97088779 | 0.76068202 |
| Fbxo46 | 0.51094722 | -0.1941393 | 0.083 | 0.109 | 1 | 0.54645491 | -0.4029647 | 0.109 | 0.083 | 1 | 0.54645491 | 0.76082738 |
| Ccdc77 | 0.51107987 | 0.33430485 | 0.26 | 0.228 | 1 | 0.92623278 | -0.4093909 | 0.214 | 0.198 | 1 | 0.92623278 | 0.76095711 |
| Usp33 | 0.51112445 | 0.1867191 | 0.562 | 0.518 | 1 | 0.68207282 | -0.2853131 | 0.756 | 0.677 | 1 | 0.68207282 | 0.7610007 |
| Rwdd3 | 0.54510221 | 0.57686018 | 0.146 | 0.124 | 1 | 0.51128849 | -0.3587649 | 0.097 | 0.073 | 1 | 0.54510221 | 0.76116106 |
| Gm57314 | 0.53487059 | 0.27052894 | 0.052 | 0.036 | 1 | 0.51146864 | 0.14080615 | 0.097 | 0.073 | 1 | 0.53487059 | 0.76133711 |
| Calhm2 | 0.51150563 | 0.44045394 | 0.156 | 0.13 | 1 | 0.79369681 | -0.2370989 | 0.185 | 0.167 | 1 | 0.79369681 | 0.76137325 |
| Sox10 | 0.52528691 | -0.7998604 | 0.01 | 0.021 | 1 | 0.51188611 | -0.7248928 | 0.004 | 0.01 | 1 | 0.52528691 | 0.76174483 |
| Gm10603 | 0.61737927 | 1.00749454 | 0.01 | 0.005 | 1 | 0.51188611 | -0.7248928 | 0.004 | 0.01 | 1 | 0.61737927 | 0.76174483 |
| Tex22 | 0.61737927 | 1.00749454 | 0.01 | 0.005 | 1 | 0.51188611 | -0.7248928 | 0.004 | 0.01 | 1 | 0.61737927 | 0.76174483 |
| 1700029J03Rik | 0.85781561 | -0.1851505 | 0.052 | 0.057 | 1 | 0.51188611 | -0.7248928 | 0.004 | 0.01 | 1 | 0.85781561 | 0.76174483 |
| Gm26510 | 0.993208 | 1.00749454 | 0.01 | 0.01 | 1 | 0.51188611 | -0.7248928 | 0.004 | 0.01 | 1 | 0.993208 | 0.76174483 |
| Lancl3 | 0.99517219 | -2.1889027 | 0.021 | 0.021 | 1 | 0.51188611 | -0.3098553 | 0.004 | 0.01 | 1 | 0.99517219 | 0.76174483 |
| Plbd1 | 0.99660398 | 0.59245027 | 0.01 | 0.01 | 1 | 0.51188611 | -0.3098553 | 0.004 | 0.01 | 1 | 0.99660398 | 0.76174483 |
| 4930461C15Rik | 0.99758609 | 0.27052894 | 0.021 | 0.021 | 1 | 0.51188611 | -0.7248928 | 0.004 | 0.01 | 1 | 0.99758609 | 0.76174483 |
| Acsbg1 | 1 | 0.42253204 | 0.01 | 0.01 | 1 | 0.51188611 | -0.7248928 | 0.004 | 0.01 | 1 | 1 | 0.76174483 |
| Gm50215 | 1 | -0.3144336 | 0.01 | 0.01 | 1 | 0.51188611 | -0.7248928 | 0.004 | 0.01 | 1 | 1 | 0.76174483 |
| Man1c1 | 0.80624112 | -0.3586334 | 0.292 | 0.264 | 1 | 0.51189399 | -0.6317834 | 0.391 | 0.385 | 1 | 0.80624112 | 0.76175252 |
| Gm4756 | 0.61737927 | 1.00749454 | 0.01 | 0.005 | 1 | 0.51193864 | 0.56461386 | 0.021 | 0.01 | 1 | 0.61737927 | 0.76179611 |
| Gm43646 | 0.52744059 | -0.577468 | 0.01 | 0.021 | 1 | 0.51194506 | 0.27510724 | 0.021 | 0.01 | 1 | 0.52744059 | 0.76180238 |
| Cfap126 | 0.5862099 | 0.68556644 | 0.031 | 0.021 | 1 | 0.51194506 | 0.27510724 | 0.021 | 0.01 | 1 | 0.5862099 | 0.76180238 |
| A030001D20Rik | 0.59199182 | 0.42253204 | 0.031 | 0.021 | 1 | 0.51194506 | 0.27510724 | 0.021 | 0.01 | 1 | 0.59199182 | 0.76180238 |
| Il31 | 0.61737927 | 1.00749454 | 0.01 | 0.005 | 1 | 0.51194506 | 0.27510724 | 0.021 | 0.01 | 1 | 0.61737927 | 0.76180238 |
| Gm13056 | 0.64538863 | 0.52206771 | 0.042 | 0.031 | 1 | 0.51194506 | 0.27510724 | 0.021 | 0.01 | 1 | 0.64538863 | 0.76180238 |
| Cdca7 | 0.71305088 | 0.30705482 | 0.062 | 0.052 | 1 | 0.51194506 | 0.27510724 | 0.021 | 0.01 | 1 | 0.71305088 | 0.76180238 |
| 5930403N24Rik | 0.72574709 | -0.3144336 | 0.01 | 0.016 | 1 | 0.51194506 | 0.27510724 | 0.021 | 0.01 | 1 | 0.72574709 | 0.76180238 |
| Gpr87 | 0.75328966 | 0.27052894 | 0.021 | 0.016 | 1 | 0.51194506 | 0.27510724 | 0.021 | 0.01 | 1 | 0.75328966 | 0.76180238 |
| Stag3 | 0.80274368 | 0.20013961 | 0.031 | 0.026 | 1 | 0.51194506 | 0.27510724 | 0.021 | 0.01 | 1 | 0.80274368 | 0.76180238 |
| Agt | 0.89397428 | -0.4519371 | 0.188 | 0.187 | 1 | 0.51194506 | 0.27510724 | 0.021 | 0.01 | 1 | 0.89397428 | 0.76180238 |
| Prelid2 | 1 | -0.3710171 | 0.042 | 0.041 | 1 | 0.51194506 | 0.27510724 | 0.021 | 0.01 | 1 | 1 | 0.76180238 |
| Capg | 0.51314887 | 0.20543391 | 0.427 | 0.378 | 1 | 0.84003966 | 0.41320951 | 0.466 | 0.49 | 1 | 0.84003966 | 0.76297598 |
| Dync2h1 | 0.51348969 | -0.3566993 | 0.677 | 0.596 | 1 | 0.7730242 | -0.4912394 | 0.706 | 0.677 | 1 | 0.7730242 | 0.76330771 |
| Tnfsf13b | 0.51378473 | 1.00749454 | 0.052 | 0.036 | 1 | 0.91219225 | -1.0203486 | 0.029 | 0.031 | 1 | 0.91219225 | 0.76359471 |
| Zfp618 | 0.51398782 | -0.401311 | 0.177 | 0.207 | 1 | 0.93434992 | -0.4473588 | 0.361 | 0.333 | 1 | 0.93434992 | 0.76379216 |
| Unc13d | 0.5141755 | 0.34452952 | 0.083 | 0.062 | 1 | 0.89750221 | -0.5443205 | 0.113 | 0.104 | 1 | 0.89750221 | 0.76397455 |
| Cast | 0.51437697 | -0.1010651 | 0.906 | 0.974 | 1 | 0.77330611 | -0.2154102 | 0.958 | 0.917 | 1 | 0.77330611 | 0.76417027 |

| | | | | | | | | | | | | |
|---|---|---|---|---|---|---|---|---|---|---|---|---|
| Dcaf4 | 0.51472687 | 0.28760246 | 0.125 | 0.098 | 1 | 0.84562666 | -0.8028953 | 0.105 | 0.094 | 1 | 0.84562666 | 0.76450999 |
| C2cd3 | 0.7190031 | -0.2088234 | 0.25 | 0.223 | 1 | 0.51517926 | -0.427911 | 0.277 | 0.302 | 1 | 0.7190031 | 0.76494885 |
| Sntg2 | 0.51520848 | 0.86999101 | 0.052 | 0.036 | 1 | 0.97447092 | -1.2318528 | 0.055 | 0.052 | 1 | 0.97447092 | 0.76497719 |
| Cpt1a | 0.9489083 | -0.1368954 | 0.156 | 0.155 | 1 | 0.51538256 | -0.1329775 | 0.395 | 0.458 | 1 | 0.9489083 | 0.76514593 |
| Gm32389 | 0.78979801 | -0.577468 | 0.021 | 0.026 | 1 | 0.5154503 | 0.27510724 | 0.021 | 0.01 | 1 | 0.78979801 | 0.76521158 |
| Gm36371 | 0.61550914 | -0.407543 | 0.021 | 0.031 | 1 | 0.51720983 | 0.10518224 | 0.021 | 0.01 | 1 | 0.61550914 | 0.76691365 |
| Gm29340 | 0.79666623 | 0.42253204 | 0.031 | 0.026 | 1 | 0.51720983 | 0.10518224 | 0.021 | 0.01 | 1 | 0.79666623 | 0.76691365 |
| 4933412E12Rik | 0.80274368 | 0.20013961 | 0.031 | 0.026 | 1 | 0.51720983 | 0.10518224 | 0.021 | 0.01 | 1 | 0.80274368 | 0.76691365 |
| Zfp763 | 0.67008293 | -0.2555399 | 0.031 | 0.041 | 1 | 0.51788491 | 0.44503224 | 0.071 | 0.052 | 1 | 0.67008293 | 0.76756504 |
| Ago4 | 0.84981799 | 0.32942263 | 0.104 | 0.098 | 1 | 0.51867092 | 0.15727075 | 0.143 | 0.115 | 1 | 0.84981799 | 0.76832232 |
| Lzts3 | 0.53487059 | 0.13302542 | 0.052 | 0.036 | 1 | 0.51895598 | 0.53814164 | 0.059 | 0.042 | 1 | 0.53487059 | 0.76859665 |
| Pcdhb17 | 0.51940912 | 0.74446013 | 0.052 | 0.036 | 1 | 0.83987076 | -0.355659 | 0.092 | 0.083 | 1 | 0.83987076 | 0.76903241 |
| Zmat4 | 1 | -0.7998604 | 0.01 | 0.01 | 1 | 0.52010386 | 0.61614416 | 0.059 | 0.042 | 1 | 1 | 0.76969969 |
| Fli1 | 0.52071614 | 0.10395738 | 0.677 | 0.627 | 1 | 0.8567795 | -0.1976458 | 0.555 | 0.469 | 1 | 0.8567795 | 0.77028698 |
| Gm6566 | 0.5383068 | 0.27052894 | 0.01 | 0.021 | 1 | 0.52072676 | -0.7248928 | 0.021 | 0.01 | 1 | 0.5383068 | 0.77029716 |
| Begain | 0.59199182 | 0.42253204 | 0.031 | 0.021 | 1 | 0.52072676 | -0.3098553 | 0.021 | 0.01 | 1 | 0.59199182 | 0.77029716 |
| 6430573P05Rik | 0.61550914 | -0.407543 | 0.021 | 0.031 | 1 | 0.52072676 | -0.7248928 | 0.021 | 0.01 | 1 | 0.61550914 | 0.77029716 |
| Spats1 | 0.75329431 | -0.1624305 | 0.021 | 0.016 | 1 | 0.52072676 | -0.3098553 | 0.021 | 0.01 | 1 | 0.75329431 | 0.77029716 |
| Atf7ip2 | 0.7854853 | -0.9925055 | 0.021 | 0.026 | 1 | 0.52072676 | -0.3098553 | 0.021 | 0.01 | 1 | 0.7854853 | 0.77029716 |
| Car15 | 1 | 0.42253204 | 0.01 | 0.01 | 1 | 0.52072676 | -0.7248928 | 0.021 | 0.01 | 1 | 1 | 0.77029716 |
| Gm14320 | 0.993208 | 1.00749454 | 0.01 | 0.01 | 1 | 0.5207331 | -0.5025003 | 0.021 | 0.01 | 1 | 0.993208 | 0.77030324 |
| Gtf2a1l | 0.52528691 | -0.9925055 | 0.01 | 0.021 | 1 | 0.52073627 | -0.4618584 | 0.021 | 0.01 | 1 | 0.52528691 | 0.77030628 |
| Adgrf5 | 0.96775093 | -0.4431669 | 0.125 | 0.124 | 1 | 0.5216711 | -1.8188689 | 0.088 | 0.104 | 1 | 0.96775093 | 0.77120147 |
| A430010J10Rik | 0.99310033 | 0.20013961 | 0.042 | 0.041 | 1 | 0.52169993 | 0.53814164 | 0.071 | 0.052 | 1 | 0.99310033 | 0.77122904 |
| Gm10501 | 0.83444593 | 0.68556644 | 0.073 | 0.067 | 1 | 0.52346322 | 0.21370669 | 0.071 | 0.052 | 1 | 0.83444593 | 0.77291269 |
| Bmper | 0.53855341 | -0.4504288 | 0.385 | 0.326 | 1 | 0.52390775 | -0.5130182 | 0.273 | 0.219 | 1 | 0.53855341 | 0.77333617 |
| Coro2b | 0.96731785 | -0.1624305 | 0.104 | 0.104 | 1 | 0.52440062 | 0.77760758 | 0.046 | 0.031 | 1 | 0.96731785 | 0.77380523 |
| Smim24 | 0.52497972 | 0.64492446 | 0.052 | 0.036 | 1 | 0.67415878 | -0.8504236 | 0.042 | 0.031 | 1 | 0.67415878 | 0.77435573 |
| Gm56912 | 0.52497972 | 0.64492446 | 0.052 | 0.036 | 1 | 0.55786396 | -0.5728897 | 0.038 | 0.052 | 1 | 0.55786396 | 0.77435573 |
| Rem1 | 0.52528691 | -0.7998604 | 0.01 | 0.021 | 1 | 0.66387437 | 0.27510724 | 0.017 | 0.01 | 1 | 0.66387437 | 0.77464748 |
| Add2 | 0.52528691 | -1.577468 | 0.01 | 0.021 | 1 | 0.87851725 | -0.8948178 | 0.013 | 0.01 | 1 | 0.87851725 | 0.77464748 |
| A930016O22Rik | 0.52528691 | -0.7998604 | 0.01 | 0.021 | 1 | 0.92396969 | -0.4618584 | 0.029 | 0.031 | 1 | 0.92396969 | 0.77464748 |
| Ryr1 | 0.52528691 | -1.1624305 | 0.01 | 0.021 | 1 | 0.67471693 | -0.7248928 | 0.017 | 0.01 | 1 | 0.67471693 | 0.77464748 |
| Zfp473 | 0.52528691 | -1.3144336 | 0.01 | 0.021 | 1 | 0.67312714 | -0.4618584 | 0.029 | 0.021 | 1 | 0.67312714 | 0.77464748 |
| Apba2 | 0.52528691 | -1.8993961 | 0.01 | 0.021 | 1 | 0.57370068 | -1.5322477 | 0.021 | 0.031 | 1 | 0.57370068 | 0.77464748 |
| Gm40528 | 0.52528691 | -2.747393 | 0.01 | 0.021 | 1 | 0.98993389 | 0.27510724 | 0.021 | 0.021 | 1 | 0.98993389 | 0.77464748 |
| Gm26839 | 0.52528691 | -0.7998604 | 0.01 | 0.021 | 1 | 0.87851725 | -1.6317834 | 0.013 | 0.01 | 1 | 0.87851725 | 0.77464748 |
| Slc39a2 | 0.52528691 | -1.3144336 | 0.01 | 0.021 | 1 | 0.85710544 | -1.3098553 | 0.008 | 0.01 | 1 | 0.85710544 | 0.77464748 |
| Wnt10b | 0.52528691 | -0.9925055 | 0.01 | 0.021 | 1 | 0.87066325 | -0.3098553 | 0.013 | 0.01 | 1 | 0.87066325 | 0.77464748 |
| Mapk4 | 0.52528691 | -1.4519371 | 0.01 | 0.021 | 1 | 0.67555181 | -0.3098553 | 0.042 | 0.031 | 1 | 0.67555181 | 0.77464748 |
| Depdc7 | 0.52744059 | -0.9925055 | 0.01 | 0.021 | 1 | 0.52556214 | -1.7952821 | 0.038 | 0.052 | 1 | 0.52744059 | 0.77490872 |
| S100a5 | 0.68340344 | 0.53800925 | 0.052 | 0.041 | 1 | 0.52556214 | -1.8948178 | 0.038 | 0.052 | 1 | 0.68340344 | 0.77490872 |
| Vangl1 | 0.52620648 | 0.16619228 | 0.198 | 0.166 | 1 | 0.93131679 | -0.2909962 | 0.248 | 0.229 | 1 | 0.93131679 | 0.7755197 |
| Ifrd1 | 0.52677606 | -0.125824 | 0.969 | 0.979 | 1 | 0.69303836 | -0.112606 | 0.992 | 1 | 1 | 0.69303836 | 0.7760591 |
| Smurf1 | 0.94569563 | -0.3954701 | 0.333 | 0.311 | 1 | 0.52690157 | -0.8512285 | 0.483 | 0.448 | 1 | 0.94569563 | 0.77617788 |
| Slc19a2 | 0.52697336 | -0.4330781 | 0.062 | 0.083 | 1 | 0.84749752 | -0.5163061 | 0.101 | 0.104 | 1 | 0.84749752 | 0.7762458 |
| A830018L16Rik | 0.52744059 | -0.577468 | 0.01 | 0.021 | 1 | 0.55773096 | -2.1172102 | 0.021 | 0.031 | 1 | 0.55773096 | 0.77668761 |
| Gm28535 | 0.52744059 | -0.577468 | 0.01 | 0.021 | 1 | 0.86204866 | -0.5025003 | 0.038 | 0.042 | 1 | 0.86204866 | 0.77668761 |
| Dmrta2 | 0.52744059 | -0.7998604 | 0.01 | 0.021 | 1 | 0.87066325 | -0.3098553 | 0.013 | 0.01 | 1 | 0.87066325 | 0.77668761 |
| Krtcap3 | 0.52744059 | -0.577468 | 0.01 | 0.021 | 1 | 0.57854956 | -0.8244284 | 0.021 | 0.031 | 1 | 0.57854956 | 0.77668761 |
| Adam8 | 0.52744059 | -0.577468 | 0.01 | 0.021 | 1 | 0.72105597 | 0.45567948 | 0.05 | 0.042 | 1 | 0.72105597 | 0.77668761 |
| Gm37612 | 0.52744059 | -0.577468 | 0.01 | 0.021 | 1 | 0.87851725 | -0.8948178 | 0.013 | 0.01 | 1 | 0.87851725 | 0.77668761 |
| Gm47095 | 0.52744059 | -0.577468 | 0.01 | 0.021 | 1 | 0.9071436 | -1.1172102 | 0.029 | 0.031 | 1 | 0.9071436 | 0.77668761 |
| Cbarp | 0.52744059 | -0.577468 | 0.01 | 0.021 | 1 | 0.88484452 | -0.7793405 | 0.046 | 0.042 | 1 | 0.88484452 | 0.77668761 |
| Gm15608 | 0.52744059 | -0.577468 | 0.01 | 0.021 | 1 | 0.85710544 | -1.3098553 | 0.008 | 0.01 | 1 | 0.85710544 | 0.77668761 |
| 9130230N09Rik | 0.52744059 | -0.577468 | 0.01 | 0.021 | 1 | 0.66339728 | 0.10518224 | 0.029 | 0.021 | 1 | 0.66339728 | 0.77668761 |
| 9930004E17Rik | 0.52744059 | -0.577468 | 0.01 | 0.021 | 1 | 0.80442048 | -0.5728897 | 0.017 | 0.021 | 1 | 0.80442048 | 0.77668761 |
| 1700037C18Rik | 0.52744059 | -0.577468 | 0.01 | 0.021 | 1 | 0.71069052 | -0.9472852 | 0.034 | 0.042 | 1 | 0.71069052 | 0.77668761 |
| Pdzrn4 | 0.61561415 | -2.5022322 | 0.115 | 0.135 | 1 | 0.52753889 | -2.8394747 | 0.076 | 0.094 | 1 | 0.61561415 | 0.7767805 |
| Mkrn2os | 0.52785427 | -0.9925055 | 0.031 | 0.047 | 1 | 0.85710586 | -1.6317834 | 0.008 | 0.01 | 1 | 0.85710586 | 0.77707841 |
| Fancg | 0.52787359 | 0.59245704 | 0.052 | 0.036 | 1 | 0.78571043 | -0.3098553 | 0.038 | 0.031 | 1 | 0.78571043 | 0.77709665 |
| Dffa | 0.52787359 | 0.59245704 | 0.052 | 0.036 | 1 | 0.80279833 | -0.2318528 | 0.071 | 0.062 | 1 | 0.80279833 | 0.77709665 |
| Zfp784 | 0.52787359 | 0.59245704 | 0.052 | 0.036 | 1 | 0.73028844 | -0.8948178 | 0.055 | 0.062 | 1 | 0.73028844 | 0.77709665 |
| Gm44899 | 0.52819938 | -0.4330781 | 0.125 | 0.15 | 1 | 0.62237314 | -0.2055186 | 0.151 | 0.125 | 1 | 0.62237314 | 0.77740417 |
| Sh3rf3 | 0.78621232 | -0.2664282 | 0.229 | 0.244 | 1 | 0.52880157 | -1.1370187 | 0.387 | 0.344 | 1 | 0.78621232 | 0.77797204 |
| Synrg | 0.52916471 | 0.28312898 | 0.312 | 0.285 | 1 | 0.76384087 | -0.2860085 | 0.345 | 0.323 | 1 | 0.76384087 | 0.77831413 |
| Selplg | 0.52961488 | -0.3144336 | 0.01 | 0.021 | 1 | 0.54958085 | -0.1254307 | 0.055 | 0.073 | 1 | 0.54958085 | 0.77873784 |
| Limk1 | 0.52961488 | -0.3144336 | 0.01 | 0.021 | 1 | 0.86695837 | -0.194378 | 0.046 | 0.042 | 1 | 0.86695837 | 0.77873784 |
| Gm10244 | 0.52961488 | -0.3144336 | 0.01 | 0.021 | 1 | 0.86312513 | -0.7248928 | 0.008 | 0.01 | 1 | 0.86312513 | 0.77873784 |
| Proser3 | 0.52961488 | -0.3144336 | 0.01 | 0.021 | 1 | 0.65352625 | -0.4797803 | 0.055 | 0.042 | 1 | 0.65352625 | 0.77873784 |
| Abcg4 | 0.52961488 | -0.3144336 | 0.01 | 0.021 | 1 | 0.86312513 | -0.7248928 | 0.008 | 0.01 | 1 | 0.86312513 | 0.77873784 |
| Gm57017 | 0.52961488 | -0.3144336 | 0.01 | 0.021 | 1 | 0.54837489 | -0.4618584 | 0.034 | 0.021 | 1 | 0.54837489 | 0.77873784 |
| Gm5627 | 0.52961488 | -0.3144336 | 0.01 | 0.021 | 1 | 0.5818763 | -0.5025003 | 0.013 | 0.021 | 1 | 0.5818763 | 0.77873784 |
| Iho1 | 0.52961488 | -0.3144336 | 0.01 | 0.021 | 1 | 0.98993422 | -1.5025003 | 0.021 | 0.021 | 1 | 0.98993422 | 0.77873784 |
| Gm54621 | 0.52961488 | -0.3144336 | 0.01 | 0.021 | 1 | 0.71069052 | -0.9472852 | 0.034 | 0.042 | 1 | 0.71069052 | 0.77873784 |
| Gm48952 | 0.52961488 | -0.3144336 | 0.01 | 0.021 | 1 | 0.86312513 | -0.7248928 | 0.008 | 0.01 | 1 | 0.86312513 | 0.77873784 |
| Ly6g5b | 0.52961488 | -0.3144336 | 0.01 | 0.021 | 1 | 0.75297348 | -0.8948178 | 0.025 | 0.031 | 1 | 0.75297348 | 0.77873784 |
| Abcc10 | 0.52961488 | -0.3144336 | 0.01 | 0.021 | 1 | 0.87244608 | -0.3098553 | 0.046 | 0.042 | 1 | 0.87244608 | 0.77873784 |
| 1700018L02Rik | 0.52961488 | -0.3144336 | 0.01 | 0.021 | 1 | 0.58180179 | -0.7248928 | 0.021 | 0.031 | 1 | 0.58180179 | 0.77873784 |
| Eln | 0.52985774 | -1.1875214 | 0.25 | 0.275 | 1 | 0.91233732 | -2.7494349 | 0.252 | 0.219 | 1 | 0.91233732 | 0.77896626 |
| Mif | 0.61378233 | -0.2395983 | 0.729 | 0.762 | 1 | 0.53047343 | -0.2205879 | 0.643 | 0.646 | 1 | 0.61378233 | 0.7795448 |
| Gpr45 | 0.53063347 | -0.577468 | 0.031 | 0.047 | 1 | 0.99856096 | -0.6577786 | 0.042 | 0.042 | 1 | 0.99856096 | 0.77969506 |
| Cyb5rl | 0.53063347 | -0.577468 | 0.031 | 0.047 | 1 | 0.59306394 | -1.1172102 | 0.059 | 0.073 | 1 | 0.59306394 | 0.77969506 |

| | | | | | | | | | | | | |
|---|---|---|---|---|---|---|---|---|---|---|---|---|
| Pcdhga7 | 0.84644605 | 0.39082318 | 0.104 | 0.098 | 1 | 0.53117911 | 0.21739174 | 0.097 | 0.073 | 1 | 0.84644605 | 0.78020697 |
| Dkk3 | 0.59661048 | -0.4960039 | 0.323 | 0.332 | 1 | 0.53134852 | -0.115293 | 0.634 | 0.531 | 1 | 0.59661048 | 0.78036579 |
| Hrct1 | 0.6084646 | -0.3144336 | 0.052 | 0.067 | 1 | 0.53144246 | 0.69014474 | 0.046 | 0.031 | 1 | 0.6084646 | 0.78045383 |
| Cdkn2b | 0.61813213 | 0.82692229 | 0.094 | 0.078 | 1 | 0.53144246 | 0.69014474 | 0.046 | 0.031 | 1 | 0.61813213 | 0.78045383 |
| Rad54l | 0.79057349 | 0.78510212 | 0.031 | 0.026 | 1 | 0.53144246 | 0.49749966 | 0.046 | 0.031 | 1 | 0.79057349 | 0.78045383 |
| Stoml3 | 0.87907915 | -0.2726134 | 0.062 | 0.057 | 1 | 0.53144246 | 0.59703533 | 0.046 | 0.031 | 1 | 0.87907915 | 0.78045383 |
| Gm57455 | 0.58480395 | 0.22988696 | 0.062 | 0.047 | 1 | 0.53156006 | 0.56461386 | 0.034 | 0.021 | 1 | 0.58480395 | 0.78056402 |
| Trak2 | 0.94420901 | -0.1314819 | 0.531 | 0.513 | 1 | 0.53161864 | -0.2385379 | 0.676 | 0.708 | 1 | 0.94420901 | 0.7806189 |
| Gm15839 | 0.53207093 | 0.49292136 | 0.052 | 0.036 | 1 | 0.57576763 | -1.0468209 | 0.013 | 0.021 | 1 | 0.57576763 | 0.78104238 |
| Col5a3 | 0.53209653 | -1.2555399 | 0.031 | 0.047 | 1 | 0.92273508 | -0.9175378 | 0.05 | 0.052 | 1 | 0.92273508 | 0.78106635 |
| Lrp4 | 0.77084005 | -0.6025589 | 0.094 | 0.083 | 1 | 0.53232956 | -0.3923174 | 0.109 | 0.083 | 1 | 0.77084005 | 0.78128436 |
| Fbxw17 | 0.53288345 | 0.17741954 | 0.083 | 0.062 | 1 | 0.87839392 | -0.5577828 | 0.113 | 0.104 | 1 | 0.87839392 | 0.78180213 |
| Fgf13 | 0.9861312 | 0.47697982 | 0.031 | 0.031 | 1 | 0.53327159 | -0.681824 | 0.046 | 0.062 | 1 | 0.9861312 | 0.78216459 |
| Nckipsd | 0.53336991 | 0.34452952 | 0.094 | 0.073 | 1 | 0.72742262 | -0.152314 | 0.097 | 0.083 | 1 | 0.72742262 | 0.78225636 |
| Lrrc20 | 0.89148737 | -0.2693457 | 0.188 | 0.176 | 1 | 0.5334283 | -0.8573431 | 0.084 | 0.104 | 1 | 0.89148737 | 0.78231085 |
| Tnfaip3 | 0.53395584 | -0.2148979 | 0.01 | 0.021 | 1 | 0.66501153 | 0.14957636 | 0.029 | 0.021 | 1 | 0.66501153 | 0.78280285 |
| Vars2 | 0.53487059 | 0.27052894 | 0.052 | 0.036 | 1 | 0.53433287 | 0.42711033 | 0.034 | 0.021 | 1 | 0.53487059 | 0.78315412 |
| Ccdc116 | 0.75832339 | -0.2148979 | 0.021 | 0.016 | 1 | 0.53433287 | 0.42711033 | 0.034 | 0.021 | 1 | 0.75832339 | 0.78315412 |
| Kcnk3 | 0.58619677 | 0.49292136 | 0.031 | 0.021 | 1 | 0.53437063 | 1.17557156 | 0.034 | 0.021 | 1 | 0.58619677 | 0.78318929 |
| Pou2f3 | 0.53444729 | -0.3479861 | 0.104 | 0.13 | 1 | 0.92362645 | -0.2923678 | 0.151 | 0.146 | 1 | 0.92362645 | 0.78326067 |
| Mex3a | 0.93381354 | -0.7294711 | 0.177 | 0.161 | 1 | 0.53450033 | -0.3851434 | 0.282 | 0.219 | 1 | 0.93381354 | 0.78331006 |
| Zfp93 | 0.53487059 | -0.4519371 | 0.031 | 0.047 | 1 | 0.66335784 | -0.4797803 | 0.05 | 0.062 | 1 | 0.66335784 | 0.78365464 |
| Gm47863 | 0.53487059 | -0.4519371 | 0.031 | 0.047 | 1 | 0.84444682 | -0.9879272 | 0.038 | 0.042 | 1 | 0.84444682 | 0.78365464 |
| Zswim4 | 0.53489645 | 0.17460452 | 0.344 | 0.311 | 1 | 0.65370453 | -0.2929814 | 0.437 | 0.344 | 1 | 0.65370453 | 0.78367868 |
| Pth2r | 0.5349386 | -0.7294711 | 0.031 | 0.047 | 1 | 0.55058229 | -0.7793405 | 0.038 | 0.052 | 1 | 0.55058229 | 0.78371789 |
| Gm57404 | 0.85651485 | -0.2148979 | 0.052 | 0.047 | 1 | 0.53497636 | 0.39058446 | 0.046 | 0.031 | 1 | 0.85651485 | 0.78375302 |
| 6530409C15Rik | 0.80274368 | 0.20013961 | 0.031 | 0.026 | 1 | 0.53504184 | 0.53814164 | 0.046 | 0.031 | 1 | 0.80274368 | 0.78381391 |
| Arfgef3 | 0.96433507 | 0.52206771 | 0.052 | 0.052 | 1 | 0.53526374 | -0.3483294 | 0.084 | 0.062 | 1 | 0.96433507 | 0.78402021 |
| Tenm1 | 0.53613057 | 0.42253204 | 0.01 | 0.021 | 1 | 0.55431107 | -2.7879026 | 0.029 | 0.042 | 1 | 0.55431107 | 0.78482515 |
| Best1 | 0.74827573 | 0.59245704 | 0.021 | 0.016 | 1 | 0.53618537 | -1.3098553 | 0.038 | 0.052 | 1 | 0.74827573 | 0.78487599 |
| Cdk18 | 0.79666623 | 0.42253204 | 0.031 | 0.026 | 1 | 0.53716429 | 0.39058446 | 0.034 | 0.021 | 1 | 0.79666623 | 0.78578311 |
| Gm15441 | 0.99758609 | 0.27052894 | 0.021 | 0.021 | 1 | 0.53717181 | 0.27510724 | 0.034 | 0.021 | 1 | 0.99758609 | 0.78579007 |
| Prss36 | 0.53746156 | 0.21394541 | 0.135 | 0.109 | 1 | 0.55449877 | -0.7573142 | 0.076 | 0.094 | 1 | 0.55449877 | 0.78605819 |
| B630019K06Rik | 0.53765145 | -0.6299354 | 0.083 | 0.104 | 1 | 0.63698724 | 0.10518224 | 0.055 | 0.042 | 1 | 0.63698724 | 0.78623382 |
| Iqsec2 | 0.91779649 | 0.3256705 | 0.219 | 0.212 | 1 | 0.53766753 | -1.0859593 | 0.336 | 0.271 | 1 | 0.91779649 | 0.78624869 |
| Ccdc126 | 0.53783959 | 0.27052894 | 0.146 | 0.119 | 1 | 0.892195 | -0.42233 | 0.134 | 0.125 | 1 | 0.892195 | 0.78640776 |
| Greb1 | 0.5383068 | 0.27052894 | 0.01 | 0.021 | 1 | 0.86011448 | -0.8948178 | 0.008 | 0.01 | 1 | 0.86011448 | 0.78683939 |
| Sp100 | 0.53831472 | 1.90057933 | 0.01 | 0.021 | 1 | 0.87327995 | 0.27510724 | 0.013 | 0.01 | 1 | 0.87327995 | 0.78684671 |
| 1700109K24Rik | 0.65896352 | 0.15949763 | 0.042 | 0.031 | 1 | 0.53864238 | 0.27510724 | 0.046 | 0.031 | 1 | 0.65896352 | 0.78714915 |
| Srsf12 | 0.80274368 | 0.20013961 | 0.031 | 0.026 | 1 | 0.53864238 | 0.27510724 | 0.046 | 0.031 | 1 | 0.80274368 | 0.78714915 |
| Gm19710 | 0.53873156 | -0.6705774 | 0.083 | 0.104 | 1 | 0.56127529 | -0.2273931 | 0.13 | 0.104 | 1 | 0.56127529 | 0.78723143 |
| Rlf | 0.53924252 | 0.10789643 | 0.854 | 0.845 | 1 | 0.68750316 | -0.2254421 | 0.916 | 0.938 | 1 | 0.68750316 | 0.78770254 |
| Bid | 0.63038178 | 0.20013961 | 0.021 | 0.031 | 1 | 0.53932729 | -0.2223924 | 0.055 | 0.073 | 1 | 0.63038178 | 0.78778066 |
| Sh2d3c | 0.73305022 | 0.27052894 | 0.073 | 0.062 | 1 | 0.53980036 | 0.42711033 | 0.046 | 0.031 | 1 | 0.73305022 | 0.78821629 |
| Gjb2 | 0.69975621 | -1.8225805 | 0.042 | 0.052 | 1 | 0.54135468 | 0.76053407 | 0.034 | 0.021 | 1 | 0.69975621 | 0.78964447 |
| Cxcl14 | 0.54143944 | 0.9062112 | 0.083 | 0.109 | 1 | 0.54417085 | -0.5393371 | 0.122 | 0.094 | 1 | 0.54417085 | 0.78972222 |
| Cep120 | 0.94148043 | 0.1124641 | 0.323 | 0.342 | 1 | 0.5414395 | -0.1676332 | 0.542 | 0.438 | 1 | 0.94148043 | 0.78972226 |
| Gm4890 | 0.54168579 | 0.46000674 | 0.104 | 0.083 | 1 | 0.86203455 | -0.6577786 | 0.038 | 0.042 | 1 | 0.86203455 | 0.78994808 |
| Wbp1l | 0.67871133 | -0.4139692 | 0.333 | 0.347 | 1 | 0.54200332 | -0.1781926 | 0.676 | 0.688 | 1 | 0.67871133 | 0.79023905 |
| Ankrd2 | 0.54203294 | 0.27052894 | 0.052 | 0.036 | 1 | 0.73617146 | -0.9472852 | 0.055 | 0.062 | 1 | 0.73617146 | 0.79026618 |
| Gm26903 | 0.56626163 | -0.9925055 | 0.042 | 0.057 | 1 | 0.54216163 | -0.8504236 | 0.038 | 0.052 | 1 | 0.56626163 | 0.79038403 |
| Gm42937 | 0.5425293 | -0.6363617 | 0.073 | 0.093 | 1 | 0.91620427 | -0.7692869 | 0.05 | 0.052 | 1 | 0.91620427 | 0.79072056 |
| Tnnt3 | 0.64741551 | -1.4763212 | 0.135 | 0.15 | 1 | 0.54291115 | -1.0983512 | 0.067 | 0.083 | 1 | 0.64741551 | 0.79106978 |
| A330035P11Rik | 0.61182672 | -0.577468 | 0.021 | 0.031 | 1 | 0.54335697 | 0.27510724 | 0.046 | 0.031 | 1 | 0.61182672 | 0.79147714 |
| Lrrd1 | 0.62479773 | -0.577468 | 0.021 | 0.031 | 1 | 0.54337178 | -1.1843244 | 0.038 | 0.052 | 1 | 0.62479773 | 0.79149067 |
| Gm14963 | 0.81906688 | -0.265524 | 0.104 | 0.093 | 1 | 0.54460315 | 0.10518224 | 0.071 | 0.052 | 1 | 0.81906688 | 0.79261371 |
| Cops7b | 0.76323226 | -0.6205367 | 0.094 | 0.104 | 1 | 0.5446669 | -0.7071908 | 0.134 | 0.156 | 1 | 0.76323226 | 0.79267177 |
| Mtmr10 | 0.5450904 | -0.1679475 | 0.427 | 0.347 | 1 | 0.66122947 | -0.7996605 | 0.387 | 0.312 | 1 | 0.66122947 | 0.79305726 |
| Heyl | 0.54546054 | -1.3710171 | 0.042 | 0.057 | 1 | 0.81628353 | -1.352924 | 0.088 | 0.094 | 1 | 0.81628353 | 0.79339388 |
| Galns | 0.99054142 | -0.2555399 | 0.094 | 0.093 | 1 | 0.54695165 | -0.3098553 | 0.046 | 0.031 | 1 | 0.99054142 | 0.79474719 |
| 4933405D12Rik | 0.88604104 | 0.42253204 | 0.104 | 0.109 | 1 | 0.54719271 | -1.0229741 | 0.197 | 0.208 | 1 | 0.88604104 | 0.79496555 |
| Dhrs13 | 0.60329331 | -0.4619907 | 0.115 | 0.135 | 1 | 0.54738954 | -0.4167705 | 0.092 | 0.115 | 1 | 0.60329331 | 0.79514377 |
| Mast3 | 0.69636827 | 0.13302542 | 0.115 | 0.098 | 1 | 0.54756179 | 0.31817596 | 0.105 | 0.083 | 1 | 0.69636827 | 0.79529966 |
| Pdzd4 | 0.54837487 | 0.31561683 | 0.188 | 0.161 | 1 | 0.99668572 | -0.5925855 | 0.252 | 0.229 | 1 | 0.99668572 | 0.79603474 |
| Galnt13 | 0.61737927 | 1.00749454 | 0.01 | 0.005 | 1 | 0.54837489 | -0.9472852 | 0.034 | 0.021 | 1 | 0.61737927 | 0.79603476 |
| Zfp90 | 1 | -0.2148979 | 0.021 | 0.021 | 1 | 0.54953198 | -0.4029647 | 0.046 | 0.062 | 1 | 1 | 0.79707856 |
| Fmnl3 | 0.55051828 | -0.4889313 | 0.375 | 0.373 | 1 | 0.92476023 | -0.408259 | 0.613 | 0.49 | 1 | 0.92476023 | 0.79796618 |
| Gm43329 | 0.55119014 | 0.15949763 | 0.094 | 0.073 | 1 | 0.55851142 | -1.1347686 | 0.134 | 0.146 | 1 | 0.55851142 | 0.79856971 |
| Nptxr | 0.55119014 | 0.15949763 | 0.094 | 0.073 | 1 | 0.7793991 | -0.5322477 | 0.118 | 0.125 | 1 | 0.7793991 | 0.79856971 |
| Zbtb12 | 0.55119014 | 0.24195979 | 0.094 | 0.073 | 1 | 0.64519202 | -0.3664388 | 0.101 | 0.083 | 1 | 0.64519202 | 0.79856971 |
| Gm56765 | 0.5862099 | 0.68556644 | 0.031 | 0.021 | 1 | 0.55174553 | -0.8504236 | 0.038 | 0.052 | 1 | 0.5862099 | 0.79906793 |
| C130046K22Rik | 0.61182672 | -0.577468 | 0.021 | 0.031 | 1 | 0.55174553 | -1.0203486 | 0.038 | 0.052 | 1 | 0.61182672 | 0.79906793 |
| Tmem106a | 0.71255463 | -0.407543 | 0.052 | 0.062 | 1 | 0.55213501 | -1.1399303 | 0.067 | 0.083 | 1 | 0.71255463 | 0.79941695 |
| Efcab7 | 0.8313893 | 0.90057933 | 0.052 | 0.047 | 1 | 0.5529563 | -1.1723517 | 0.029 | 0.042 | 1 | 0.8313893 | 0.80015193 |
| Gm45871 | 0.55317563 | 0.71798792 | 0.062 | 0.047 | 1 | 0.81860312 | -0.3923174 | 0.059 | 0.052 | 1 | 0.81860312 | 0.80034798 |
| Prelid3a | 0.9243361 | 0.10060394 | 0.115 | 0.119 | 1 | 0.55457455 | -0.4542452 | 0.122 | 0.094 | 1 | 0.9243361 | 0.80159617 |
| Septin6 | 0.83286906 | -0.4689435 | 0.156 | 0.14 | 1 | 0.55463125 | -0.7769813 | 0.185 | 0.198 | 1 | 0.83286906 | 0.80164667 |
| H4c9 | 0.98855404 | 0.16795921 | 0.125 | 0.124 | 1 | 0.55484807 | -0.2223924 | 0.059 | 0.042 | 1 | 0.98855404 | 0.80183976 |
| Kdm8 | 0.59199182 | -0.1624305 | 0.031 | 0.021 | 1 | 0.55548698 | -0.4353861 | 0.034 | 0.021 | 1 | 0.59199182 | 0.80240818 |
| Adck5 | 0.8647959 | -0.2148979 | 0.083 | 0.088 | 1 | 0.55566579 | -0.1723517 | 0.084 | 0.062 | 1 | 0.8647959 | 0.80256711 |
| H3c15 | 0.55702983 | 1.14499806 | 0.062 | 0.047 | 1 | 0.66339728 | 0.10518224 | 0.029 | 0.021 | 1 | 0.66339728 | 0.80377743 |
| Orai2 | 0.55741182 | 0.32942263 | 0.167 | 0.145 | 1 | 0.6862207 | -0.6217993 | 0.088 | 0.073 | 1 | 0.6862207 | 0.80411571 |
| Eml6 | 0.66641437 | -0.536826 | 0.177 | 0.15 | 1 | 0.55768 | 0.13760371 | 0.252 | 0.219 | 1 | 0.66641437 | 0.80435302 |
| Slc4a8 | 0.99009327 | -0.6929452 | 0.031 | 0.031 | 1 | 0.55780609 | -0.6094155 | 0.038 | 0.052 | 1 | 0.99009327 | 0.80446454 |
| Micall2 | 0.79869018 | 0.32942263 | 0.031 | 0.026 | 1 | 0.55820636 | -0.2394659 | 0.071 | 0.052 | 1 | 0.79869018 | 0.80481838 |

| | | | | | | | | | | | | |
|---|---|---|---|---|---|---|---|---|---|---|---|---|
| Lysmd4 | 0.74304562 | -0.2148979 | 0.115 | 0.098 | 1 | 0.55822346 | -0.4272122 | 0.206 | 0.167 | 1 | 0.74304562 | 0.80483349 |
| Prkab2 | 0.75474405 | -0.3389559 | 0.229 | 0.238 | 1 | 0.55845905 | -0.3847093 | 0.395 | 0.312 | 1 | 0.75474405 | 0.80504159 |
| 5031415H12Rik | 0.82515969 | 0.27052894 | 0.042 | 0.036 | 1 | 0.55975127 | -0.4618584 | 0.034 | 0.021 | 1 | 0.82515969 | 0.80618105 |
| Kif24 | 0.59694884 | 0.29700115 | 0.042 | 0.057 | 1 | 0.56024564 | -0.6724253 | 0.046 | 0.031 | 1 | 0.59694884 | 0.80661611 |
| Kcnq1 | 1 | 0.56003556 | 0.042 | 0.041 | 1 | 0.56110733 | -1.3098553 | 0.029 | 0.042 | 1 | 1 | 0.80737322 |
| Dnmt3a | 0.7255155 | -0.1992857 | 0.677 | 0.632 | 1 | 0.56128711 | -0.3344925 | 0.95 | 0.938 | 1 | 0.7255155 | 0.807531 |
| Dnmt3b | 0.99224353 | 0.20013961 | 0.052 | 0.052 | 1 | 0.56144729 | -0.194378 | 0.046 | 0.031 | 1 | 0.99224353 | 0.80767152 |
| Hectd4 | 0.98512334 | -0.223831 | 0.354 | 0.342 | 1 | 0.56160036 | -0.3318816 | 0.571 | 0.583 | 1 | 0.98512334 | 0.80780576 |
| Reep4 | 0.6426548 | 0.27052894 | 0.052 | 0.067 | 1 | 0.56182467 | 0.27510724 | 0.092 | 0.073 | 1 | 0.6426548 | 0.80800238 |
| Katnbl1 | 0.91616056 | 0.20945337 | 0.5 | 0.518 | 1 | 0.56241852 | -0.3205552 | 0.71 | 0.719 | 1 | 0.91616056 | 0.80852245 |
| Epha1 | 1 | 0.42253204 | 0.01 | 0.01 | 1 | 0.56243824 | -1.3098553 | 0.029 | 0.042 | 1 | 1 | 0.80853971 |
| Fxyd7 | 0.67724727 | 0.68556644 | 0.052 | 0.041 | 1 | 0.56266331 | -0.4167705 | 0.046 | 0.031 | 1 | 0.67724727 | 0.80873662 |
| Hykk | 0.58217865 | 0.20013961 | 0.062 | 0.047 | 1 | 0.56309211 | -0.7248928 | 0.071 | 0.052 | 1 | 0.58217865 | 0.80911149 |
| Als2 | 0.77466398 | 0.29289676 | 0.219 | 0.218 | 1 | 0.56324431 | -0.1966447 | 0.256 | 0.219 | 1 | 0.77466398 | 0.80924447 |
| Kyat3 | 0.56492112 | 0.54806292 | 0.062 | 0.047 | 1 | 0.642845 | -0.3098553 | 0.055 | 0.042 | 1 | 0.642845 | 0.81070637 |
| ENSMUSG00000121596 | 0.56492112 | 0.54806292 | 0.062 | 0.047 | 1 | 0.92663792 | -0.5025003 | 0.05 | 0.052 | 1 | 0.92663792 | 0.81070637 |
| Adar | 0.56525651 | 0.18152394 | 0.302 | 0.269 | 1 | 0.70463419 | -0.3739856 | 0.361 | 0.354 | 1 | 0.70463419 | 0.8109981 |
| Zfp109 | 1 | 0.42253204 | 0.01 | 0.01 | 1 | 0.56565662 | -1.6317834 | 0.013 | 0.021 | 1 | 1 | 0.81134582 |
| Slc1a2 | 0.681597 | 0.27052894 | 0.031 | 0.041 | 1 | 0.56614149 | -0.4093909 | 0.071 | 0.052 | 1 | 0.681597 | 0.81176679 |
| Cul9 | 0.84597594 | -0.1079827 | 0.177 | 0.187 | 1 | 0.56714742 | -0.473354 | 0.345 | 0.271 | 1 | 0.84597594 | 0.81263864 |
| Fancd2 | 0.57467311 | 1.42253204 | 0.031 | 0.021 | 1 | 0.56729373 | -1.6724253 | 0.021 | 0.031 | 1 | 0.57467311 | 0.81276528 |
| Manea | 0.98497948 | 0.15949763 | 0.156 | 0.161 | 1 | 0.56732062 | -0.3098553 | 0.181 | 0.146 | 1 | 0.98497948 | 0.81278855 |
| Gprin3 | 0.56742138 | -1.1079827 | 0.073 | 0.093 | 1 | 0.66647237 | 0.36821664 | 0.21 | 0.188 | 1 | 0.66647237 | 0.81287574 |
| Wdr91 | 0.56789015 | -0.4254649 | 0.094 | 0.114 | 1 | 0.96963074 | 0.29680231 | 0.126 | 0.125 | 1 | 0.96963074 | 0.81328108 |
| Arnt2 | 0.56819899 | -0.6139938 | 0.104 | 0.124 | 1 | 0.90976726 | -0.9654097 | 0.286 | 0.25 | 1 | 0.90976726 | 0.81354789 |
| Miat | 0.72574709 | -0.577468 | 0.01 | 0.016 | 1 | 0.56889575 | -1.3098553 | 0.021 | 0.031 | 1 | 0.72574709 | 0.81414912 |
| Gm19744 | 0.56898992 | 0.49292136 | 0.062 | 0.047 | 1 | 0.97446729 | -1.1034044 | 0.055 | 0.052 | 1 | 0.97446729 | 0.81423031 |
| Trmt12 | 0.56898992 | 0.49292136 | 0.062 | 0.047 | 1 | 0.93576656 | -0.1399303 | 0.029 | 0.031 | 1 | 0.93576656 | 0.81423031 |
| Cdk5rap2 | 0.56914956 | 0.22259447 | 0.5 | 0.57 | 1 | 0.97107273 | -0.6984206 | 0.492 | 0.417 | 1 | 0.97107273 | 0.8143679 |
| Nub1 | 0.56958349 | -0.130009 | 0.615 | 0.622 | 1 | 0.66709685 | -0.2542629 | 0.634 | 0.635 | 1 | 0.66709685 | 0.81474163 |
| Cep192 | 0.81338784 | 0.46833573 | 0.26 | 0.29 | 1 | 0.57094773 | -0.5345615 | 0.252 | 0.271 | 1 | 0.81338784 | 0.81591415 |
| Abcc6 | 0.59199182 | 0.42253204 | 0.031 | 0.021 | 1 | 0.57170879 | -1.8948178 | 0.013 | 0.021 | 1 | 0.59199182 | 0.81656664 |
| Omp | 0.5978326 | -0.3144336 | 0.031 | 0.021 | 1 | 0.57170879 | -1.6317834 | 0.013 | 0.021 | 1 | 0.5978326 | 0.81656664 |
| Gm33037 | 0.64366127 | 0.49292136 | 0.042 | 0.031 | 1 | 0.57170879 | -1.3098553 | 0.013 | 0.021 | 1 | 0.64366127 | 0.81656664 |
| Marchf10 | 0.75328966 | 0.27052894 | 0.021 | 0.016 | 1 | 0.57170879 | -2.1172102 | 0.013 | 0.021 | 1 | 0.75328966 | 0.81656664 |
| Kcnd3 | 0.75832339 | -0.2148979 | 0.021 | 0.016 | 1 | 0.57170879 | -1.3098553 | 0.013 | 0.021 | 1 | 0.75832339 | 0.81656664 |
| Kiss1r | 0.7854853 | -0.407543 | 0.021 | 0.026 | 1 | 0.57170879 | -1.6317834 | 0.013 | 0.021 | 1 | 0.7854853 | 0.81656664 |
| S100a3 | 0.81159799 | -0.4519371 | 0.031 | 0.036 | 1 | 0.57170879 | -1.3098553 | 0.013 | 0.021 | 1 | 0.81159799 | 0.81656664 |
| 1500035N22Rik | 0.82260168 | -0.2555399 | 0.031 | 0.036 | 1 | 0.57170879 | -1.3098553 | 0.013 | 0.021 | 1 | 0.82260168 | 0.81656664 |
| Gm29724 | 0.85158597 | -0.3710171 | 0.042 | 0.036 | 1 | 0.57170879 | -1.3098553 | 0.013 | 0.021 | 1 | 0.85158597 | 0.81656664 |
| 4833419F23Rik | 0.87560284 | 0.30705482 | 0.052 | 0.057 | 1 | 0.57170879 | -2.1172102 | 0.013 | 0.021 | 1 | 0.87560284 | 0.81656664 |
| 4930505A04Rik | 0.993208 | 1.00749454 | 0.01 | 0.01 | 1 | 0.57170879 | -1.6317834 | 0.013 | 0.021 | 1 | 0.993208 | 0.81656664 |
| Hgd | 0.993208 | 1.00749454 | 0.01 | 0.01 | 1 | 0.57170879 | -1.3098553 | 0.013 | 0.021 | 1 | 0.993208 | 0.81656664 |
| Gm17090 | 1 | 0.42253204 | 0.01 | 0.01 | 1 | 0.57170879 | -1.6317834 | 0.013 | 0.021 | 1 | 1 | 0.81656664 |
| Cavin4 | 1 | 0.42253204 | 0.01 | 0.01 | 1 | 0.57170879 | -1.3098553 | 0.013 | 0.021 | 1 | 1 | 0.81656664 |
| Gm52829 | 1 | -0.577468 | 0.01 | 0.01 | 1 | 0.57170879 | -1.3098553 | 0.013 | 0.021 | 1 | 1 | 0.81656664 |
| 1600002D24Rik | 0.97580314 | 0.21899864 | 0.062 | 0.062 | 1 | 0.57171509 | -1.3098553 | 0.013 | 0.021 | 1 | 0.97580314 | 0.81657204 |
| Lin7a | 0.89648536 | -0.3910548 | 0.062 | 0.057 | 1 | 0.57192881 | 0.69014474 | 0.126 | 0.104 | 1 | 0.89648536 | 0.81675506 |
| Gm15261 | 0.9927582 | 0.42253204 | 0.021 | 0.021 | 1 | 0.57215593 | -0.8073549 | 0.059 | 0.042 | 1 | 0.9927582 | 0.81694945 |
| Rwdd2b | 0.57277177 | 0.74446013 | 0.073 | 0.057 | 1 | 0.62294577 | 0.27510724 | 0.055 | 0.042 | 1 | 0.62294577 | 0.81747604 |
| Cfap45 | 0.57291912 | 0.15949763 | 0.062 | 0.047 | 1 | 0.57395309 | -0.9472852 | 0.164 | 0.125 | 1 | 0.57395309 | 0.81760193 |
| Ifit3b | 0.81813838 | -1.3592878 | 0.094 | 0.083 | 1 | 0.57346666 | -2.0368368 | 0.029 | 0.042 | 1 | 0.81813838 | 0.81806931 |
| Tpi1 | 0.57356071 | -0.1058469 | 0.781 | 0.756 | 1 | 0.623254 | -0.2969162 | 0.807 | 0.812 | 1 | 0.623254 | 0.81814953 |
| Grhl1 | 0.57363044 | -0.5015191 | 0.104 | 0.124 | 1 | 0.78179247 | -0.1755542 | 0.231 | 0.24 | 1 | 0.78179247 | 0.818209 |
| Bcl2l15 | 0.57848839 | 1.27052894 | 0.031 | 0.021 | 1 | 0.57370068 | -1.3098553 | 0.021 | 0.031 | 1 | 0.57848839 | 0.81826889 |
| Gm7467 | 0.67174581 | -0.1624305 | 0.031 | 0.041 | 1 | 0.57370068 | -1.0468209 | 0.021 | 0.031 | 1 | 0.67174581 | 0.81826889 |
| Tigar | 1 | -0.3144336 | 0.01 | 0.01 | 1 | 0.57370068 | -1.3098553 | 0.021 | 0.031 | 1 | 1 | 0.81826889 |
| Zpld1 | 0.61737927 | 1.00749454 | 0.01 | 0.005 | 1 | 0.57373861 | -1.3098553 | 0.013 | 0.021 | 1 | 0.61737927 | 0.81830123 |
| Cxxc4 | 0.73407659 | 0.27052894 | 0.01 | 0.016 | 1 | 0.57373861 | -1.3098553 | 0.013 | 0.021 | 1 | 0.73407659 | 0.81830123 |
| Gm35188 | 0.83792894 | -0.6496177 | 0.115 | 0.119 | 1 | 0.57567959 | -1.2079756 | 0.16 | 0.167 | 1 | 0.83792894 | 0.81995219 |
| Gm35853 | 0.58888131 | -0.9518635 | 0.062 | 0.047 | 1 | 0.57576763 | -0.9879272 | 0.013 | 0.021 | 1 | 0.58888131 | 0.8200269 |
| Gm35082 | 0.61737927 | 1.00749454 | 0.01 | 0.005 | 1 | 0.57576763 | -0.9879272 | 0.013 | 0.021 | 1 | 0.61737927 | 0.8200269 |
| A030005L19Rik | 1 | 0.42253204 | 0.01 | 0.01 | 1 | 0.57576763 | -1.0468209 | 0.013 | 0.021 | 1 | 1 | 0.8200269 |
| Elovl6 | 0.57616219 | -0.3777273 | 0.812 | 0.736 | 1 | 0.58361198 | -0.2931765 | 0.55 | 0.583 | 1 | 0.58361198 | 0.82036151 |
| Igf1 | 0.59692703 | -0.6007377 | 0.271 | 0.223 | 1 | 0.57650566 | -1.681824 | 0.282 | 0.271 | 1 | 0.59692703 | 0.82065254 |
| Lclat1 | 0.57655915 | 0.2920621 | 0.458 | 0.435 | 1 | 0.79300803 | -0.6179776 | 0.265 | 0.25 | 1 | 0.79300803 | 0.82069785 |
| Gm11696 | 0.69067656 | 0.59245704 | 0.062 | 0.052 | 1 | 0.57693422 | -1.3098553 | 0.021 | 0.031 | 1 | 0.69067656 | 0.82101534 |
| Gm2350 | 0.80885634 | -0.577468 | 0.031 | 0.026 | 1 | 0.57693422 | -1.0874628 | 0.021 | 0.031 | 1 | 0.80885634 | 0.82101534 |
| Gm50163 | 1 | 0.42253204 | 0.01 | 0.01 | 1 | 0.57756442 | -0.4618584 | 0.029 | 0.042 | 1 | 1 | 0.82154818 |
| Gm56741 | 0.87953438 | -0.17693 | 0.094 | 0.098 | 1 | 0.57767012 | -0.6577786 | 0.067 | 0.083 | 1 | 0.87953438 | 0.82163747 |
| Ighd | 0.61371908 | 2.00749454 | 0.01 | 0.005 | 1 | 0.5778028 | -0.8948178 | 0.013 | 0.021 | 1 | 0.61371908 | 0.82174952 |
| Baiap3 | 0.61737927 | 1.00749454 | 0.01 | 0.005 | 1 | 0.5778028 | -0.8948178 | 0.013 | 0.021 | 1 | 0.61737927 | 0.82174952 |
| Gm9776 | 0.64538863 | 0.52206771 | 0.042 | 0.031 | 1 | 0.5778028 | -0.8948178 | 0.013 | 0.021 | 1 | 0.64538863 | 0.82174952 |
| Gm43672 | 0.78979801 | -0.407543 | 0.021 | 0.026 | 1 | 0.5778028 | -0.8948178 | 0.013 | 0.021 | 1 | 0.78979801 | 0.82174952 |
| Slc17a8 | 0.80611646 | -0.7998604 | 0.031 | 0.036 | 1 | 0.5778028 | -0.8948178 | 0.013 | 0.021 | 1 | 0.80611646 | 0.82174952 |
| Gm4419 | 0.86441462 | -0.2148979 | 0.052 | 0.047 | 1 | 0.5778028 | -0.8948178 | 0.013 | 0.021 | 1 | 0.86441462 | 0.82174952 |
| Gm57263 | 1 | -0.577468 | 0.021 | 0.021 | 1 | 0.5778028 | -0.8948178 | 0.013 | 0.021 | 1 | 1 | 0.82174952 |
| Tbxa2r | 1 | 0.42253204 | 0.01 | 0.01 | 1 | 0.5778028 | -0.8948178 | 0.013 | 0.021 | 1 | 1 | 0.82174952 |
| Hspa12b | 0.57848839 | 1.00749454 | 0.031 | 0.021 | 1 | 0.93705023 | -0.8504236 | 0.034 | 0.031 | 1 | 0.93705023 | 0.82232796 |
| Mfsd7a | 0.57848839 | 1.00749454 | 0.031 | 0.021 | 1 | 0.79600331 | -0.9879272 | 0.017 | 0.021 | 1 | 0.79600331 | 0.82232796 |
| Mdk | 0.61209726 | -1.4195156 | 0.333 | 0.249 | 1 | 0.57880327 | -1.0758981 | 0.508 | 0.385 | 1 | 0.61209726 | 0.82259331 |
| Laptm4b | 0.74349186 | 0.18806678 | 0.302 | 0.342 | 1 | 0.57999572 | -0.4645779 | 0.454 | 0.354 | 1 | 0.74349186 | 0.8235964 |
| Cfap299 | 0.76336555 | -1.3144336 | 0.021 | 0.016 | 1 | 0.58017355 | -1.5728897 | 0.021 | 0.031 | 1 | 0.76336555 | 0.82374575 |

| | | | | | | | | | | | | |
|---|---|---|---|---|---|---|---|---|---|---|---|---|
| 4930557K07Rik | 0.74827573 | 0.59245704 | 0.021 | 0.016 | 1 | 0.58180179 | -0.7248928 | 0.021 | 0.031 | 1 | 0.74827573 | 0.82511026 |
| Dynlt1b | 1 | 0.42253204 | 0.01 | 0.01 | 1 | 0.58180179 | -0.7248928 | 0.021 | 0.031 | 1 | 1 | 0.82511026 |
| Mslnl | 0.61737927 | 1.00749454 | 0.01 | 0.005 | 1 | 0.5818708 | -0.5728897 | 0.013 | 0.021 | 1 | 0.61737927 | 0.82516797 |
| Gm11542 | 0.75328966 | -0.2148979 | 0.021 | 0.016 | 1 | 0.5818708 | -0.5728897 | 0.013 | 0.021 | 1 | 0.75328966 | 0.82516797 |
| Syt9 | 0.5978326 | 0.20013961 | 0.031 | 0.021 | 1 | 0.5818763 | -0.5025003 | 0.013 | 0.021 | 1 | 0.5978326 | 0.82517257 |
| Cep112 | 0.58215805 | 0.12979863 | 0.542 | 0.534 | 1 | 0.93823274 | -0.7789326 | 0.521 | 0.417 | 1 | 0.93823274 | 0.8254081 |
| Sardh | 0.58224975 | -0.2555399 | 0.042 | 0.057 | 1 | 0.69464767 | -0.6317834 | 0.071 | 0.083 | 1 | 0.69464767 | 0.82548473 |
| Gm15890 | 0.58234163 | 1.00749454 | 0.031 | 0.021 | 1 | 0.73945321 | -1.3945321 | 0.025 | 0.031 | 1 | 0.73945321 | 0.82556149 |
| Isoc2a | 0.58268119 | -0.17693 | 0.083 | 0.104 | 1 | 0.66872948 | -0.4167705 | 0.042 | 0.031 | 1 | 0.66872948 | 0.82584501 |
| Htra4 | 0.97047514 | 1.35896691 | 0.094 | 0.093 | 1 | 0.58370606 | -0.3300331 | 0.143 | 0.115 | 1 | 0.97047514 | 0.82669935 |
| Dcbld2 | 0.5842315 | -0.2845321 | 0.5 | 0.503 | 1 | 0.6624212 | -0.2103196 | 0.45 | 0.385 | 1 | 0.6624212 | 0.82713655 |
| Abtb1 | 0.58480395 | 0.22988696 | 0.062 | 0.047 | 1 | 0.88737574 | -0.3098553 | 0.059 | 0.062 | 1 | 0.88737574 | 0.82761224 |
| Rigi | 0.58552271 | -0.7149715 | 0.208 | 0.228 | 1 | 0.79148053 | -0.9472852 | 0.298 | 0.24 | 1 | 0.79148053 | 0.82820858 |
| Ubr2 | 0.58582991 | -0.2580956 | 0.854 | 0.756 | 1 | 0.92116491 | -0.1198388 | 0.895 | 0.948 | 1 | 0.92116491 | 0.82846314 |
| Plekhn1 | 0.5862099 | 0.68556644 | 0.031 | 0.021 | 1 | 0.9338291 | -0.6577786 | 0.034 | 0.031 | 1 | 0.9338291 | 0.82877775 |
| Gm32856 | 0.5862099 | 0.68556644 | 0.031 | 0.021 | 1 | 0.86312513 | -0.7248928 | 0.008 | 0.01 | 1 | 0.86312513 | 0.82877775 |
| 4933408J17Rik | 0.5862099 | 0.68556644 | 0.031 | 0.021 | 1 | 0.87066325 | -0.3098553 | 0.013 | 0.01 | 1 | 0.87066325 | 0.82877775 |
| Hhipl1 | 0.5862099 | 0.68556644 | 0.031 | 0.021 | 1 | 0.99396051 | -0.3098553 | 0.021 | 0.021 | 1 | 0.99396051 | 0.82877775 |
| 5730424H11Rik | 0.5862099 | 0.68556644 | 0.031 | 0.021 | 1 | 0.87066325 | -0.3098553 | 0.013 | 0.01 | 1 | 0.87066325 | 0.82877775 |
| Amdhd2 | 0.58657435 | 0.24850264 | 0.115 | 0.093 | 1 | 0.72337008 | 0.21370669 | 0.084 | 0.073 | 1 | 0.72337008 | 0.82907923 |
| Gbx1 | 0.61371908 | 2.00749454 | 0.01 | 0.005 | 1 | 0.58664899 | -0.5025003 | 0.021 | 0.031 | 1 | 0.61371908 | 0.82914094 |
| Dnmbp | 0.84477495 | -0.3359133 | 0.448 | 0.409 | 1 | 0.58678406 | -0.2134869 | 0.584 | 0.531 | 1 | 0.84477495 | 0.82925259 |
| Slc12a9 | 0.58697932 | -0.3910548 | 0.094 | 0.114 | 1 | 0.61012529 | 0.14080615 | 0.134 | 0.115 | 1 | 0.61012529 | 0.82941392 |
| Gm56692 | 0.76170181 | 0.13302542 | 0.104 | 0.093 | 1 | 0.58758726 | 0.27510724 | 0.08 | 0.062 | 1 | 0.76170181 | 0.82991573 |
| Galm | 0.65020091 | -0.4872702 | 0.094 | 0.109 | 1 | 0.58801757 | -0.1723517 | 0.084 | 0.104 | 1 | 0.65020091 | 0.83027047 |
| Rab5b | 0.85069802 | 0.14399654 | 0.469 | 0.451 | 1 | 0.58907636 | -0.3781245 | 0.576 | 0.573 | 1 | 0.85069802 | 0.83114176 |
| Zfp933 | 0.97990512 | -0.1941393 | 0.094 | 0.093 | 1 | 0.58918013 | -0.1668973 | 0.176 | 0.146 | 1 | 0.97990512 | 0.83122704 |
| Zdhhc18 | 0.5894035 | -0.2216395 | 0.417 | 0.425 | 1 | 0.74375962 | -0.4211723 | 0.471 | 0.365 | 1 | 0.74375962 | 0.83141051 |
| Myo1g | 0.59007022 | 0.52206771 | 0.031 | 0.021 | 1 | 0.66387437 | 0.27510724 | 0.017 | 0.01 | 1 | 0.66387437 | 0.83195757 |
| Tmem104 | 0.86047304 | -0.1851505 | 0.062 | 0.067 | 1 | 0.59013107 | -0.7623675 | 0.067 | 0.083 | 1 | 0.86047304 | 0.83200746 |
| Dnajc17 | 0.9673047 | -0.2335136 | 0.104 | 0.104 | 1 | 0.59013107 | -0.6468902 | 0.067 | 0.083 | 1 | 0.9673047 | 0.83200746 |
| Zfp61 | 0.59021958 | 0.10060394 | 0.062 | 0.047 | 1 | 0.8034692 | -0.8948178 | 0.088 | 0.094 | 1 | 0.8034692 | 0.83208001 |
| Snx7 | 0.5907282 | -0.1971959 | 0.417 | 0.44 | 1 | 0.78833126 | -0.7479764 | 0.458 | 0.354 | 1 | 0.78833126 | 0.83249659 |
| Nr2c1 | 0.59082919 | -0.2779077 | 0.219 | 0.249 | 1 | 0.90631049 | -0.424314 | 0.294 | 0.26 | 1 | 0.90631049 | 0.83257925 |
| Mlh3 | 0.59116427 | 0.46983775 | 0.167 | 0.145 | 1 | 0.82868201 | -0.7826083 | 0.13 | 0.115 | 1 | 0.82868201 | 0.83285334 |
| Rdh13 | 0.65347736 | -0.5230202 | 0.083 | 0.098 | 1 | 0.59118313 | -0.5728897 | 0.105 | 0.125 | 1 | 0.65347736 | 0.83286876 |
| Ears2 | 0.59128081 | 0.36363835 | 0.125 | 0.104 | 1 | 0.93453836 | -0.4117349 | 0.134 | 0.125 | 1 | 0.93453836 | 0.83294862 |
| Adcy1 | 0.99654979 | -0.1079827 | 0.042 | 0.041 | 1 | 0.59154822 | -0.3098553 | 0.021 | 0.031 | 1 | 0.99654979 | 0.83316715 |
| Gm10848 | 0.59199182 | 0.42253204 | 0.031 | 0.021 | 1 | 0.65117141 | -0.3098553 | 0.055 | 0.042 | 1 | 0.65117141 | 0.83352933 |
| Gm10778 | 0.59199182 | 0.42253204 | 0.031 | 0.021 | 1 | 0.86312513 | -0.7248928 | 0.008 | 0.01 | 1 | 0.86312513 | 0.83352933 |
| 5830428M24Rik | 0.59199182 | 0.42253204 | 0.031 | 0.021 | 1 | 0.66387437 | 0.27510724 | 0.017 | 0.01 | 1 | 0.66387437 | 0.83352933 |
| Nme3 | 0.59199182 | 0.42253204 | 0.031 | 0.021 | 1 | 0.66339728 | 0.10518224 | 0.029 | 0.021 | 1 | 0.66339728 | 0.83352933 |
| Taf3 | 0.65659395 | -0.2528951 | 0.51 | 0.44 | 1 | 0.59267624 | -0.3189004 | 0.567 | 0.573 | 1 | 0.65659395 | 0.83408736 |
| Cipc | 0.77760837 | -0.3050957 | 0.24 | 0.249 | 1 | 0.59291307 | -0.4167705 | 0.176 | 0.146 | 1 | 0.77760837 | 0.83428023 |
| Mipol1 | 0.59417818 | 0.12166556 | 0.292 | 0.337 | 1 | 0.97775773 | -0.1340054 | 0.336 | 0.323 | 1 | 0.97775773 | 0.83530865 |
| Bdh2 | 0.89646712 | -0.3550755 | 0.062 | 0.057 | 1 | 0.59437106 | 0.23071312 | 0.08 | 0.062 | 1 | 0.89646712 | 0.83546516 |
| Cdkl1 | 0.9596895 | -0.6445822 | 0.052 | 0.052 | 1 | 0.59443273 | -0.6468902 | 0.088 | 0.104 | 1 | 0.9596895 | 0.83551519 |
| Clybl | 0.59459872 | -0.114496 | 0.24 | 0.207 | 1 | 0.81846691 | -0.7333543 | 0.202 | 0.198 | 1 | 0.81846691 | 0.8356498 |
| Lin9 | 0.59473928 | 0.17741954 | 0.115 | 0.093 | 1 | 0.69400069 | 0.44016648 | 0.084 | 0.073 | 1 | 0.69400069 | 0.83576375 |
| Arsg | 0.59478715 | -0.1180363 | 0.104 | 0.083 | 1 | 0.8187081 | 0.12310414 | 0.08 | 0.073 | 1 | 0.8187081 | 0.83580255 |
| Gm26797 | 0.59512028 | 0.75956702 | 0.083 | 0.067 | 1 | 0.85565192 | -0.4353861 | 0.143 | 0.146 | 1 | 0.85565192 | 0.83607241 |
| Arg2 | 0.59589633 | 0.32942263 | 0.031 | 0.021 | 1 | 0.91452613 | 0.17557156 | 0.034 | 0.031 | 1 | 0.91452613 | 0.83670023 |
| Zfp595 | 0.59614866 | 1.00749454 | 0.073 | 0.057 | 1 | 0.66277483 | -0.6179776 | 0.067 | 0.052 | 1 | 0.66277483 | 0.8369041 |
| Dars2 | 0.61608977 | 0.59245704 | 0.073 | 0.057 | 1 | 0.59713508 | 0.42711033 | 0.067 | 0.052 | 1 | 0.61608977 | 0.83769985 |
| Lhpp | 0.59735703 | 0.59245704 | 0.083 | 0.067 | 1 | 0.61573613 | 0.10518224 | 0.113 | 0.094 | 1 | 0.61573613 | 0.83787864 |
| Nxph4 | 0.72251146 | -0.577468 | 0.062 | 0.073 | 1 | 0.59767445 | -0.3483294 | 0.105 | 0.125 | 1 | 0.72251146 | 0.83813415 |
| Nceh1 | 0.70364706 | 0.46317402 | 0.146 | 0.13 | 1 | 0.59780724 | -1.0406686 | 0.134 | 0.104 | 1 | 0.70364706 | 0.83824098 |
| Gm1604b | 0.5978326 | 0.20013961 | 0.031 | 0.021 | 1 | 0.76936342 | 0.10518224 | 0.038 | 0.031 | 1 | 0.76936342 | 0.83826139 |
| 6430590A07Rik | 0.9106918 | 0.31943854 | 0.094 | 0.098 | 1 | 0.59827013 | -0.3923174 | 0.118 | 0.094 | 1 | 0.9106918 | 0.83861311 |
| Chn1 | 0.59853633 | -1.1393469 | 0.062 | 0.078 | 1 | 0.84083281 | 0.33400093 | 0.08 | 0.073 | 1 | 0.84083281 | 0.83882692 |
| Cyp2ab1 | 0.7507837 | 0.42253204 | 0.021 | 0.016 | 1 | 0.59929838 | 0.39058446 | 0.067 | 0.052 | 1 | 0.7507837 | 0.83943821 |
| Pcdhb22 | 0.72903586 | -0.3144336 | 0.052 | 0.062 | 1 | 0.59955518 | 0.82764826 | 0.088 | 0.073 | 1 | 0.72903586 | 0.83964395 |
| AU041133 | 0.59988664 | -0.4330781 | 0.052 | 0.067 | 1 | 1 | -0.5322477 | 0.042 | 0.042 | 1 | 1 | 0.8399093 |
| Tph2 | 0.60081969 | -2.3144336 | 0.021 | 0.031 | 1 | 0.66142291 | -1.0874628 | 0.042 | 0.052 | 1 | 0.66142291 | 0.84065508 |
| Tmem170b | 0.60287133 | -0.3430027 | 0.167 | 0.187 | 1 | 0.87704801 | -0.7655347 | 0.231 | 0.198 | 1 | 0.87704801 | 0.84228882 |
| Pptc7 | 0.63795134 | -0.2675125 | 0.354 | 0.295 | 1 | 0.60339881 | -0.3181229 | 0.655 | 0.646 | 1 | 0.63795134 | 0.8427075 |
| Dus2 | 0.60346498 | -0.577468 | 0.115 | 0.135 | 1 | 0.64046199 | -0.7248928 | 0.105 | 0.083 | 1 | 0.64046199 | 0.84275998 |
| Cd74 | 0.88496376 | -0.2465109 | 0.156 | 0.15 | 1 | 0.60454331 | 0.27510724 | 0.067 | 0.052 | 1 | 0.88496376 | 0.843614 |
| Tyms | 0.60480609 | 0.32160313 | 0.125 | 0.104 | 1 | 0.97650955 | -0.7248928 | 0.097 | 0.094 | 1 | 0.97650955 | 0.84382177 |
| Slc25a10 | 0.60565358 | 0.27052894 | 0.156 | 0.135 | 1 | 0.92129416 | -0.4253325 | 0.088 | 0.083 | 1 | 0.92129416 | 0.8444909 |
| Chst15 | 0.91243605 | -0.130009 | 0.104 | 0.109 | 1 | 0.60636983 | -0.1786107 | 0.164 | 0.135 | 1 | 0.91243605 | 0.84505529 |
| Cep295nl | 0.60660007 | 0.21394541 | 0.125 | 0.104 | 1 | 0.95120046 | -0.6909454 | 0.143 | 0.135 | 1 | 0.95120046 | 0.8452365 |
| Syn1 | 0.60790766 | -0.4157169 | 0.125 | 0.145 | 1 | 0.68423355 | -0.5961594 | 0.126 | 0.104 | 1 | 0.68423355 | 0.8462636 |
| Zfp169 | 0.60808531 | 0.25260703 | 0.083 | 0.104 | 1 | 0.6633864 | -0.5728897 | 0.143 | 0.115 | 1 | 0.6633864 | 0.84640287 |
| Zfp947 | 0.60815045 | -0.8669746 | 0.021 | 0.031 | 1 | 0.86312513 | -0.5728897 | 0.008 | 0.01 | 1 | 0.86312513 | 0.84645393 |
| A730090N16Rik | 0.608151 | -0.7294711 | 0.021 | 0.031 | 1 | 0.99396051 | -0.3098553 | 0.021 | 0.021 | 1 | 0.99396051 | 0.84645436 |
| Pigw | 0.6084646 | 0.30705482 | 0.073 | 0.057 | 1 | 0.64304313 | 0.27510724 | 0.042 | 0.031 | 1 | 0.64304313 | 0.84670003 |
| Gm14858 | 0.60855533 | 0.42253204 | 0.073 | 0.057 | 1 | 0.64711752 | -1.4129488 | 0.084 | 0.094 | 1 | 0.64711752 | 0.84677107 |
| Lins1 | 0.60857745 | 0.12297175 | 0.115 | 0.093 | 1 | 0.64569402 | -1.0468209 | 0.113 | 0.125 | 1 | 0.64569402 | 0.84678838 |
| 4930511M06Rik | 0.98102622 | -0.4779323 | 0.042 | 0.041 | 1 | 0.60961735 | -0.8948178 | 0.059 | 0.073 | 1 | 0.98102622 | 0.84760139 |
| Pknox1 | 0.60999564 | 0.20013961 | 0.281 | 0.259 | 1 | 0.80756496 | -0.5025003 | 0.311 | 0.302 | 1 | 0.80756496 | 0.8478966 |
| Klhl8 | 0.61015953 | 0.12297175 | 0.125 | 0.104 | 1 | 0.72092401 | -0.677587 | 0.109 | 0.094 | 1 | 0.72092401 | 0.84802441 |
| Trim2 | 0.79210777 | -0.2842716 | 0.427 | 0.409 | 1 | 0.61067419 | -0.2405072 | 0.538 | 0.562 | 1 | 0.79210777 | 0.84842541 |

| | | | | | | | | | | | | |
|---|---|---|---|---|---|---|---|---|---|---|---|---|
| Zfp335os | 0.81038593 | -0.2148979 | 0.156 | 0.166 | 1 | 0.61069846 | -0.4167705 | 0.126 | 0.104 | 1 | 0.81038593 | 0.84844431 |
| Mpc1 | 0.61091944 | -0.1158879 | 0.854 | 0.839 | 1 | 0.98513214 | -0.1602027 | 0.807 | 0.885 | 1 | 0.98513214 | 0.84861632 |
| Lmbr1 | 0.87527384 | -0.1079827 | 0.177 | 0.166 | 1 | 0.61132774 | -1.3353904 | 0.176 | 0.146 | 1 | 0.87527384 | 0.84893387 |
| 5031434O11Rik | 0.61182672 | -0.7294711 | 0.021 | 0.031 | 1 | 0.79918975 | -0.9879272 | 0.038 | 0.031 | 1 | 0.79918975 | 0.8493215 |
| Fbxl13 | 0.61182672 | -0.8669746 | 0.021 | 0.031 | 1 | 0.86613737 | -0.3098553 | 0.008 | 0.01 | 1 | 0.86613737 | 0.8493215 |
| Serpinf2 | 0.61182672 | -0.7294711 | 0.021 | 0.031 | 1 | 0.85710544 | -1.3098553 | 0.008 | 0.01 | 1 | 0.85710544 | 0.8493215 |
| Dab1 | 0.8006332 | 0.32942263 | 0.021 | 0.026 | 1 | 0.61184384 | -0.3814085 | 0.059 | 0.073 | 1 | 0.8006332 | 0.84933479 |
| Kif13a | 0.61184699 | -0.3187988 | 0.594 | 0.585 | 1 | 0.8787339 | -0.1459433 | 0.681 | 0.719 | 1 | 0.8787339 | 0.84933724 |
| Kcnc3 | 0.6127901 | -0.2148979 | 0.104 | 0.083 | 1 | 0.94831386 | -0.4797803 | 0.105 | 0.104 | 1 | 0.94831386 | 0.85006849 |
| Glipr2 | 0.61355959 | 0.17741954 | 0.073 | 0.057 | 1 | 0.67989501 | -0.152314 | 0.076 | 0.062 | 1 | 0.67989501 | 0.85066381 |
| Pate2 | 0.61368387 | -0.7998604 | 0.021 | 0.031 | 1 | 0.79810697 | -1.0874628 | 0.017 | 0.021 | 1 | 0.79810697 | 0.85075985 |
| Tmem163 | 0.61371908 | 1.74446013 | 0.01 | 0.005 | 1 | 0.87066325 | -0.3098553 | 0.013 | 0.01 | 1 | 0.87066325 | 0.85078705 |
| 6030442K20Rik | 0.61371908 | 1.59245704 | 0.01 | 0.005 | 1 | 0.66013325 | 0.27510724 | 0.029 | 0.021 | 1 | 0.66013325 | 0.85078705 |
| Mndal | 0.61371908 | 1.85549144 | 0.01 | 0.005 | 1 | 0.86312513 | -0.7248928 | 0.008 | 0.01 | 1 | 0.86312513 | 0.85078705 |
| Nup210 | 0.61371908 | 1.59245704 | 0.01 | 0.005 | 1 | 0.85710544 | -1.5728897 | 0.008 | 0.01 | 1 | 0.85710544 | 0.85078705 |
| Ak9 | 0.61371908 | 1.59245704 | 0.01 | 0.005 | 1 | 0.86312513 | -0.7248928 | 0.008 | 0.01 | 1 | 0.86312513 | 0.85078705 |
| Mettl7a3 | 0.61371908 | 1.59245704 | 0.01 | 0.005 | 1 | 0.85710544 | -1.3098553 | 0.008 | 0.01 | 1 | 0.85710544 | 0.85078705 |
| Atp13a5 | 0.61371908 | 1.59245704 | 0.01 | 0.005 | 1 | 0.80442048 | -0.5728897 | 0.017 | 0.021 | 1 | 0.80442048 | 0.85078705 |
| Cfb | 0.61371908 | 1.59245704 | 0.01 | 0.005 | 1 | 0.98993409 | -1.3098553 | 0.021 | 0.021 | 1 | 0.98993409 | 0.85078705 |
| Arhgef16 | 0.61550914 | -0.407543 | 0.021 | 0.031 | 1 | 0.93382932 | -0.9313436 | 0.034 | 0.031 | 1 | 0.93382932 | 0.85216678 |
| Mmp15 | 0.61550914 | -0.407543 | 0.021 | 0.031 | 1 | 0.742064 | -0.2554075 | 0.063 | 0.073 | 1 | 0.742064 | 0.85216678 |
| Gm57028 | 0.61550914 | -0.407543 | 0.021 | 0.031 | 1 | 0.66387437 | 0.27510724 | 0.017 | 0.01 | 1 | 0.66387437 | 0.85216678 |
| Nsg2 | 0.61550914 | -0.577468 | 0.021 | 0.031 | 1 | 0.87851725 | -1.3098553 | 0.013 | 0.01 | 1 | 0.87851725 | 0.85216678 |
| Pcdhb5 | 0.61550914 | -0.407543 | 0.021 | 0.031 | 1 | 0.78761875 | -1.7952821 | 0.017 | 0.021 | 1 | 0.78761875 | 0.85216678 |
| Tbc1d31 | 0.61597512 | 0.42253204 | 0.083 | 0.067 | 1 | 0.67342912 | -0.9175378 | 0.071 | 0.083 | 1 | 0.67342912 | 0.85252489 |
| Tlcd5 | 0.75443521 | -0.3144336 | 0.094 | 0.104 | 1 | 0.61626903 | -0.5508634 | 0.08 | 0.062 | 1 | 0.75443521 | 0.85275054 |
| Gm13391 | 0.83746085 | -0.2555399 | 0.042 | 0.036 | 1 | 0.61679385 | -0.8123556 | 0.08 | 0.094 | 1 | 0.83746085 | 0.85315305 |
| C2cd6 | 0.61737927 | 1.00749454 | 0.01 | 0.005 | 1 | 0.87851807 | -1.3098553 | 0.013 | 0.01 | 1 | 0.87851807 | 0.85360138 |
| ENSMUSG00000121413 | 0.61737927 | 1.00749454 | 0.01 | 0.005 | 1 | 0.67254361 | -0.3098553 | 0.017 | 0.01 | 1 | 0.67254361 | 0.85360138 |
| Acnat1 | 0.61737927 | 1.00749454 | 0.01 | 0.005 | 1 | 0.85710544 | -2.0468209 | 0.008 | 0.01 | 1 | 0.85710544 | 0.85360138 |
| Gm12802 | 0.61737927 | 1.00749454 | 0.01 | 0.005 | 1 | 0.66387437 | 0.27510724 | 0.017 | 0.01 | 1 | 0.66387437 | 0.85360138 |
| Htr1d | 0.61737927 | 1.00749454 | 0.01 | 0.005 | 1 | 0.87066325 | -0.3098553 | 0.013 | 0.01 | 1 | 0.87066325 | 0.85360138 |
| 6030445D17Rik | 0.61737927 | 1.00749454 | 0.01 | 0.005 | 1 | 0.87851725 | -0.8948178 | 0.013 | 0.01 | 1 | 0.87851725 | 0.85360138 |
| Zfp979 | 0.61737927 | 1.00749454 | 0.01 | 0.005 | 1 | 0.98188264 | 0.42711033 | 0.021 | 0.021 | 1 | 0.98188264 | 0.85360138 |
| Iqca1l | 0.61737927 | 1.00749454 | 0.01 | 0.005 | 1 | 0.79600331 | -0.9879272 | 0.017 | 0.021 | 1 | 0.79600331 | 0.85360138 |
| Gm43458 | 0.61737927 | 1.00749454 | 0.01 | 0.005 | 1 | 0.87851725 | -1.3098553 | 0.013 | 0.01 | 1 | 0.87851725 | 0.85360138 |
| Dnah6 | 0.61737927 | 1.00749454 | 0.01 | 0.005 | 1 | 0.87066325 | -0.3098553 | 0.013 | 0.01 | 1 | 0.87066325 | 0.85360138 |
| 1700101I11Rik | 0.61737927 | 1.00749454 | 0.01 | 0.005 | 1 | 0.66339728 | 0.10518224 | 0.029 | 0.021 | 1 | 0.66339728 | 0.85360138 |
| Cend1 | 0.61737927 | 1.00749454 | 0.01 | 0.005 | 1 | 0.86312513 | -0.7248928 | 0.008 | 0.01 | 1 | 0.86312513 | 0.85360138 |
| Gm16208 | 0.61737927 | 1.00749454 | 0.01 | 0.005 | 1 | 0.86312513 | -0.7248928 | 0.008 | 0.01 | 1 | 0.86312513 | 0.85360138 |
| Misp | 0.61737927 | 1.00749454 | 0.01 | 0.005 | 1 | 0.93576656 | -0.1399303 | 0.029 | 0.031 | 1 | 0.93576656 | 0.85360138 |
| Gm47101 | 0.61737927 | 1.00749454 | 0.01 | 0.005 | 1 | 0.87066325 | -0.3098553 | 0.013 | 0.01 | 1 | 0.87066325 | 0.85360138 |
| Glipr1 | 0.61737927 | 1.00749454 | 0.01 | 0.005 | 1 | 1 | -0.6317834 | 0.021 | 0.021 | 1 | 1 | 0.85360138 |
| Gm57355 | 0.61737927 | 1.00749454 | 0.01 | 0.005 | 1 | 0.85710544 | -1.3098553 | 0.008 | 0.01 | 1 | 0.85710544 | 0.85360138 |
| Gm26559 | 0.61737927 | 1.00749454 | 0.01 | 0.005 | 1 | 0.80442048 | -0.5728897 | 0.017 | 0.021 | 1 | 0.80442048 | 0.85360138 |
| Carmil3 | 0.61737927 | 1.00749454 | 0.01 | 0.005 | 1 | 0.86804649 | 0.27510724 | 0.013 | 0.01 | 1 | 0.86804649 | 0.85360138 |
| Sox21 | 0.61737927 | 1.00749454 | 0.01 | 0.005 | 1 | 0.85710544 | -1.3098553 | 0.008 | 0.01 | 1 | 0.85710544 | 0.85360138 |
| Baiap2l2 | 0.61737927 | 1.00749454 | 0.01 | 0.005 | 1 | 0.86312513 | -0.7248928 | 0.008 | 0.01 | 1 | 0.86312513 | 0.85360138 |
| Gm26760 | 0.61737927 | 1.00749454 | 0.01 | 0.005 | 1 | 0.99396051 | -0.3098553 | 0.021 | 0.021 | 1 | 0.99396051 | 0.85360138 |
| Ccdc65 | 0.61737927 | 1.00749454 | 0.01 | 0.005 | 1 | 0.86312513 | -0.7248928 | 0.008 | 0.01 | 1 | 0.86312513 | 0.85360138 |
| Sytl3 | 0.61737927 | 1.00749454 | 0.01 | 0.005 | 1 | 0.87851807 | -0.9879272 | 0.013 | 0.01 | 1 | 0.87851807 | 0.85360138 |
| Rhbdl1 | 0.61737927 | 1.00749454 | 0.01 | 0.005 | 1 | 0.99856096 | -0.8504236 | 0.042 | 0.042 | 1 | 0.99856096 | 0.85360138 |
| Vav1 | 0.61737927 | 1.00749454 | 0.01 | 0.005 | 1 | 0.67471693 | -0.7248928 | 0.017 | 0.01 | 1 | 0.67471693 | 0.85360138 |
| Gm19689 | 0.61737927 | 1.00749454 | 0.01 | 0.005 | 1 | 0.87851725 | -0.8948178 | 0.013 | 0.01 | 1 | 0.87851725 | 0.85360138 |
| Pcdhb15 | 0.61737927 | 1.00749454 | 0.01 | 0.005 | 1 | 0.67254361 | -0.3098553 | 0.017 | 0.01 | 1 | 0.67254361 | 0.85360138 |
| Plaat5 | 0.61737927 | 1.00749454 | 0.01 | 0.005 | 1 | 1 | -0.8948178 | 0.021 | 0.021 | 1 | 1 | 0.85360138 |
| Gm13830 | 0.99069229 | 0.30705482 | 0.052 | 0.052 | 1 | 0.61795633 | 0.13071733 | 0.067 | 0.052 | 1 | 0.99069229 | 0.85404264 |
| H2bc21 | 0.61924305 | -0.2148979 | 0.021 | 0.031 | 1 | 0.8076363 | 0.27510724 | 0.025 | 0.021 | 1 | 0.8076363 | 0.85502414 |
| Gm56817 | 0.61924305 | -0.2148979 | 0.021 | 0.031 | 1 | 0.81129592 | 0.10518224 | 0.025 | 0.021 | 1 | 0.81129592 | 0.85502414 |
| Trim21 | 0.61924305 | -0.2148979 | 0.021 | 0.031 | 1 | 0.64304313 | 0.27510724 | 0.042 | 0.031 | 1 | 0.64304313 | 0.85502414 |
| Egflam | 0.62065712 | -0.3710171 | 0.062 | 0.078 | 1 | 0.66338952 | 0.27510724 | 0.029 | 0.021 | 1 | 0.66338952 | 0.85609898 |
| Irs2 | 0.62092482 | 0.15570854 | 0.688 | 0.653 | 1 | 0.83657912 | -0.2161198 | 0.706 | 0.74 | 1 | 0.83657912 | 0.85630201 |
| Gm37168 | 0.62104899 | 0.42253204 | 0.01 | 0.005 | 1 | 0.87851725 | -0.8948178 | 0.013 | 0.01 | 1 | 0.87851725 | 0.85639613 |
| 9530097N15Rik | 0.62104899 | 0.42253204 | 0.01 | 0.005 | 1 | 0.86312513 | -0.7248928 | 0.008 | 0.01 | 1 | 0.86312513 | 0.85639613 |
| Gm43569 | 0.62104899 | 0.42253204 | 0.01 | 0.005 | 1 | 1 | -1.0468209 | 0.021 | 0.021 | 1 | 1 | 0.85639613 |
| Stk31 | 0.62104899 | -0.577468 | 0.01 | 0.005 | 1 | 0.87066325 | -0.3098553 | 0.013 | 0.01 | 1 | 0.87066325 | 0.85639613 |
| Ccdc81 | 0.62104899 | -0.577468 | 0.01 | 0.005 | 1 | 0.87851725 | -1.8948178 | 0.013 | 0.01 | 1 | 0.87851725 | 0.85639613 |
| Gm15133 | 0.62104899 | 0.42253204 | 0.01 | 0.005 | 1 | 0.86312513 | -0.7248928 | 0.008 | 0.01 | 1 | 0.86312513 | 0.85639613 |
| Tspoap1 | 0.62104899 | 0.42253204 | 0.01 | 0.005 | 1 | 0.85710544 | -1.3098553 | 0.008 | 0.01 | 1 | 0.85710544 | 0.85639613 |
| Ofcc1 | 0.62104899 | 0.42253204 | 0.01 | 0.005 | 1 | 0.86312513 | -0.7248928 | 0.008 | 0.01 | 1 | 0.86312513 | 0.85639613 |
| Mfap4 | 0.6556649 | -0.2871262 | 0.188 | 0.161 | 1 | 0.62257888 | -1.194378 | 0.193 | 0.198 | 1 | 0.6556649 | 0.8575533 |
| Il18bp | 0.62261737 | -0.577468 | 0.073 | 0.088 | 1 | 0.68274811 | -0.6968784 | 0.101 | 0.083 | 1 | 0.68274811 | 0.85758235 |
| 4732491K20Rik | 0.64419166 | -0.1624305 | 0.062 | 0.078 | 1 | 0.62294577 | 0.27510724 | 0.055 | 0.042 | 1 | 0.64419166 | 0.85783011 |
| Greb1l | 0.62304841 | 0.55498233 | 0.083 | 0.067 | 1 | 0.89534025 | -1.0280843 | 0.126 | 0.125 | 1 | 0.89534025 | 0.8579075 |
| Klf15 | 0.92459877 | 0.23697638 | 0.104 | 0.104 | 1 | 0.62314489 | -0.6317834 | 0.092 | 0.073 | 1 | 0.92459877 | 0.85798022 |
| Pcdhga3 | 0.63827665 | 0.15949763 | 0.083 | 0.067 | 1 | 0.6232833 | -0.5728897 | 0.059 | 0.073 | 1 | 0.63827665 | 0.85808453 |
| Garin4 | 0.62716478 | -1.1249558 | 0.094 | 0.109 | 1 | 0.67550311 | -1.3098553 | 0.071 | 0.083 | 1 | 0.67550311 | 0.8609939 |
| Klhdc4 | 0.6870345 | 0.42253204 | 0.188 | 0.171 | 1 | 0.6282713 | -0.6366656 | 0.189 | 0.208 | 1 | 0.6870345 | 0.86181777 |
| St3gal3 | 0.62841958 | -0.4299108 | 0.406 | 0.415 | 1 | 0.96602282 | -0.4716063 | 0.412 | 0.365 | 1 | 0.96602282 | 0.86192799 |
| Tagap1 | 0.78487956 | 0.11440974 | 0.115 | 0.104 | 1 | 0.62966226 | -1.1843244 | 0.092 | 0.104 | 1 | 0.78487956 | 0.86284996 |
| Mad2l1 | 0.67211985 | 1.59245704 | 0.094 | 0.083 | 1 | 0.62994205 | -0.5322477 | 0.088 | 0.104 | 1 | 0.67211985 | 0.86305711 |
| Poc1a | 0.99482536 | 0.15949763 | 0.042 | 0.041 | 1 | 0.63004106 | -0.8623963 | 0.05 | 0.062 | 1 | 0.99482536 | 0.86313038 |
| Chic1 | 0.63053352 | 0.36113149 | 0.271 | 0.311 | 1 | 0.76750815 | -0.1088682 | 0.399 | 0.344 | 1 | 0.76750815 | 0.86349452 |
| Snx8 | 0.71687935 | 0.14951354 | 0.229 | 0.212 | 1 | 0.63061982 | -0.1399303 | 0.374 | 0.323 | 1 | 0.71687935 | 0.86355828 |

| | | | | | | | | | | | | |
|---|---|---|---|---|---|---|---|---|---|---|---|---|
| Zc3h12a | 0.97728875 | -0.1079827 | 0.115 | 0.114 | 1 | 0.63311957 | -0.2223924 | 0.113 | 0.094 | 1 | 0.97728875 | 0.86539875 |
| Fam72a | 0.63709119 | 0.24195979 | 0.083 | 0.067 | 1 | 0.63349001 | 0.10518224 | 0.055 | 0.042 | 1 | 0.63709119 | 0.86567043 |
| Vps33a | 0.69378408 | -0.6013147 | 0.177 | 0.192 | 1 | 0.63464125 | -0.2048857 | 0.328 | 0.271 | 1 | 0.69378408 | 0.86651299 |
| Gm49700 | 0.83676741 | -0.3710171 | 0.042 | 0.047 | 1 | 0.63505823 | 0.49749966 | 0.042 | 0.031 | 1 | 0.83676741 | 0.8668175 |
| Palb2 | 0.63520923 | 0.78510212 | 0.042 | 0.031 | 1 | 0.98417243 | -0.8504236 | 0.042 | 0.042 | 1 | 0.98417243 | 0.8669277 |
| 1110020A21Rik | 0.63520923 | 1.00749454 | 0.042 | 0.031 | 1 | 0.80442048 | -0.5728897 | 0.017 | 0.021 | 1 | 0.80442048 | 0.8669277 |
| Wrap53 | 0.63590582 | 0.21899864 | 0.083 | 0.067 | 1 | 0.83851818 | 0.35999614 | 0.109 | 0.104 | 1 | 0.83851818 | 0.86743543 |
| Angpt1 | 0.63622814 | -0.7062013 | 0.115 | 0.093 | 1 | 0.96336256 | -0.7623675 | 0.164 | 0.156 | 1 | 0.96336256 | 0.86767003 |
| Gyg | 0.6380899 | -0.4519371 | 0.073 | 0.088 | 1 | 0.63700172 | -0.5728897 | 0.105 | 0.083 | 1 | 0.6380899 | 0.86823225 |
| 1810024B03Rik | 0.63824323 | 0.39082318 | 0.094 | 0.078 | 1 | 0.74258822 | -1.0468209 | 0.088 | 0.073 | 1 | 0.74258822 | 0.86913204 |
| Gamt | 0.64082014 | -0.1851505 | 0.062 | 0.078 | 1 | 0.87176622 | -0.4473588 | 0.059 | 0.062 | 1 | 0.87176622 | 0.87098983 |
| Zfp870 | 0.64197016 | 0.59245704 | 0.042 | 0.031 | 1 | 0.76784916 | 0.14957636 | 0.038 | 0.031 | 1 | 0.76784916 | 0.87181463 |
| Lpxn | 0.64198874 | 0.49292136 | 0.042 | 0.031 | 1 | 0.8315276 | -0.7248928 | 0.025 | 0.021 | 1 | 0.8315276 | 0.87182794 |
| Washc4 | 0.64358498 | 0.20331386 | 0.583 | 0.56 | 1 | 0.78620808 | -0.1231126 | 0.748 | 0.656 | 1 | 0.78620808 | 0.87296833 |
| Zfp72 | 0.64538863 | 0.52206771 | 0.042 | 0.031 | 1 | 0.97319112 | -0.8623963 | 0.055 | 0.052 | 1 | 0.97319112 | 0.87425078 |
| Xk | 0.64538863 | 0.52206771 | 0.042 | 0.031 | 1 | 0.86312513 | -0.7248928 | 0.008 | 0.01 | 1 | 0.86312513 | 0.87425078 |
| Aldh4a1 | 0.64541945 | -0.6554705 | 0.146 | 0.161 | 1 | 0.88326739 | -0.9064996 | 0.147 | 0.146 | 1 | 0.88326739 | 0.87427263 |
| Myo18b | 0.64553931 | 1.34629645 | 0.062 | 0.078 | 1 | 0.87066325 | -0.3098553 | 0.013 | 0.01 | 1 | 0.87066325 | 0.87435762 |
| Sh3bp5l | 0.64592553 | 0.42253204 | 0.115 | 0.098 | 1 | 0.95355557 | -0.468553 | 0.16 | 0.156 | 1 | 0.95355557 | 0.87463127 |
| Arhgef37 | 0.81989071 | 0.35541784 | 0.042 | 0.036 | 1 | 0.64710514 | 0.27510724 | 0.042 | 0.031 | 1 | 0.81989071 | 0.87546522 |
| Herc3 | 0.64725171 | -0.9925055 | 0.031 | 0.041 | 1 | 0.66852084 | -0.8073549 | 0.071 | 0.083 | 1 | 0.66852084 | 0.87556864 |
| A130010J15Rik | 0.70116305 | -0.2555399 | 0.042 | 0.052 | 1 | 0.64762622 | -0.2223924 | 0.059 | 0.073 | 1 | 0.70116305 | 0.87583272 |
| Ankdd1a | 0.64875704 | 0.42253204 | 0.042 | 0.031 | 1 | 0.8065254 | -0.3098553 | 0.017 | 0.021 | 1 | 0.8065254 | 0.87662838 |
| Ulk3 | 0.83449605 | 0.42253204 | 0.052 | 0.047 | 1 | 0.64880517 | -0.1172102 | 0.055 | 0.042 | 1 | 0.83449605 | 0.87666219 |
| Eepd1 | 0.6494173 | -0.9593386 | 0.083 | 0.098 | 1 | 0.89652272 | -0.4141919 | 0.155 | 0.156 | 1 | 0.89652272 | 0.87709177 |
| Eaf2 | 0.66270356 | -0.3617393 | 0.208 | 0.176 | 1 | 0.64978718 | -0.7421708 | 0.181 | 0.146 | 1 | 0.66270356 | 0.87735098 |
| Gm17103 | 0.8319199 | -0.3144336 | 0.042 | 0.047 | 1 | 0.65009158 | -1.0874628 | 0.042 | 0.052 | 1 | 0.8319199 | 0.8775641 |
| Mpp2 | 0.65161384 | -0.2555399 | 0.073 | 0.088 | 1 | 0.83587423 | -0.5728897 | 0.088 | 0.094 | 1 | 0.83587423 | 0.87862709 |
| Gmpr | 0.6834121 | 0.42253204 | 0.052 | 0.041 | 1 | 0.65200917 | -0.473354 | 0.08 | 0.094 | 1 | 0.6834121 | 0.87890238 |
| 4930556J24Rik | 0.65211073 | -0.8993961 | 0.031 | 0.041 | 1 | 0.75501778 | -0.7248928 | 0.055 | 0.062 | 1 | 0.75501778 | 0.87897305 |
| A2ml1 | 0.65211183 | -1.0799683 | 0.031 | 0.041 | 1 | 0.86613737 | 1.27510724 | 0.008 | 0.01 | 1 | 0.86613737 | 0.87897382 |
| E430018J23Rik | 0.65212989 | 0.32942263 | 0.042 | 0.031 | 1 | 0.98417243 | -1.1723517 | 0.042 | 0.042 | 1 | 0.98417243 | 0.87898639 |
| Gnmt | 0.69869193 | 0.13302542 | 0.052 | 0.041 | 1 | 0.65369483 | 0.56461386 | 0.029 | 0.021 | 1 | 0.69869193 | 0.88007273 |
| Ncdn | 0.65371216 | -0.3144336 | 0.073 | 0.088 | 1 | 0.93279087 | -0.7248928 | 0.147 | 0.135 | 1 | 0.93279087 | 0.88008473 |
| Aplf | 0.65371216 | -0.240433 | 0.073 | 0.088 | 1 | 0.77975275 | -0.194378 | 0.038 | 0.031 | 1 | 0.77975275 | 0.88008473 |
| A130051J06Rik | 0.65496193 | -0.407543 | 0.073 | 0.088 | 1 | 0.95405618 | -0.4029647 | 0.055 | 0.052 | 1 | 0.95405618 | 0.88094873 |
| Med15 | 0.65502431 | -0.1445086 | 0.562 | 0.585 | 1 | 0.66964426 | -0.1037187 | 0.685 | 0.625 | 1 | 0.66964426 | 0.88099177 |
| Aox2 | 0.70747628 | 1.92949202 | 0.062 | 0.052 | 1 | 0.65529568 | -0.6883669 | 0.05 | 0.062 | 1 | 0.70747628 | 0.88117893 |
| Fbxw10 | 0.78979801 | -0.2148979 | 0.021 | 0.026 | 1 | 0.65691192 | 0.56461386 | 0.029 | 0.021 | 1 | 0.78979801 | 0.88229057 |
| 6030407O03Rik | 0.6569134 | -0.6247737 | 0.135 | 0.15 | 1 | 0.92940164 | -0.9879272 | 0.084 | 0.083 | 1 | 0.92940164 | 0.88229158 |
| Actr3b | 0.65698333 | -0.8993961 | 0.031 | 0.041 | 1 | 0.66013325 | 0.27510724 | 0.029 | 0.021 | 1 | 0.66013325 | 0.88233957 |
| Gm30551 | 0.65698333 | -1.0799683 | 0.031 | 0.041 | 1 | 0.99194729 | -1.5025003 | 0.021 | 0.021 | 1 | 0.99194729 | 0.88233957 |
| Tekt2 | 0.65698369 | -0.577468 | 0.031 | 0.041 | 1 | 0.91453215 | -0.1399303 | 0.034 | 0.031 | 1 | 0.91453215 | 0.88233981 |
| Cd274 | 0.65698369 | -0.9925055 | 0.031 | 0.041 | 1 | 0.87675566 | -0.2223924 | 0.038 | 0.042 | 1 | 0.87675566 | 0.88233981 |
| Ccdc71l | 0.65717952 | 0.35541784 | 0.115 | 0.098 | 1 | 0.82983066 | -0.3446207 | 0.143 | 0.146 | 1 | 0.82983066 | 0.88247412 |
| Tubg1 | 0.86916492 | -0.1941393 | 0.094 | 0.098 | 1 | 0.65838396 | -0.7248928 | 0.071 | 0.083 | 1 | 0.86916492 | 0.88329848 |
| Gm14634 | 0.74076005 | 1.00749454 | 0.021 | 0.016 | 1 | 0.66013325 | 0.27510724 | 0.029 | 0.021 | 1 | 0.74076005 | 0.88449059 |
| Slc5a6 | 0.85180191 | -0.1851505 | 0.052 | 0.047 | 1 | 0.66013325 | 0.27510724 | 0.029 | 0.021 | 1 | 0.85180191 | 0.88449059 |
| Vrk1 | 0.85443189 | 0.5049942 | 0.135 | 0.13 | 1 | 0.66063382 | -0.3098553 | 0.185 | 0.156 | 1 | 0.85443189 | 0.8848306 |
| Eml5 | 0.67743079 | -0.770113 | 0.083 | 0.098 | 1 | 0.66145309 | -1.010295 | 0.168 | 0.177 | 1 | 0.67743079 | 0.88538599 |
| Cd164l2 | 0.73436042 | 0.17741954 | 0.073 | 0.062 | 1 | 0.66147833 | -0.5163061 | 0.08 | 0.094 | 1 | 0.73436042 | 0.88540308 |
| Zdhhc17 | 0.66154936 | -0.308533 | 0.354 | 0.352 | 1 | 0.78136765 | -0.3769695 | 0.471 | 0.385 | 1 | 0.78136765 | 0.88545117 |
| Gm34917 | 0.74576744 | 0.68556644 | 0.021 | 0.016 | 1 | 0.66171832 | 0.69014474 | 0.017 | 0.01 | 1 | 0.74576744 | 0.8855655 |
| Gas2 | 0.66186725 | -0.4519371 | 0.031 | 0.041 | 1 | 0.83550943 | 0.61614416 | 0.046 | 0.042 | 1 | 0.83550943 | 0.88566625 |
| Gm26827 | 0.66186725 | -0.4519371 | 0.031 | 0.041 | 1 | 0.74559899 | -0.7248928 | 0.055 | 0.062 | 1 | 0.74559899 | 0.88566625 |
| Hs1bp3 | 0.66186725 | -0.6929452 | 0.031 | 0.041 | 1 | 0.87934103 | -0.6724253 | 0.046 | 0.042 | 1 | 0.87934103 | 0.88566625 |
| 1810034E14Rik | 0.66186725 | -0.6929452 | 0.031 | 0.041 | 1 | 0.67471693 | -0.7248928 | 0.017 | 0.01 | 1 | 0.67471693 | 0.88566625 |
| B230206L02Rik | 0.66280209 | 0.13302542 | 0.125 | 0.109 | 1 | 0.68703298 | -0.1853102 | 0.185 | 0.198 | 1 | 0.68703298 | 0.88629757 |
| Mroh7 | 0.74576744 | 0.68556644 | 0.021 | 0.016 | 1 | 0.66339728 | 0.10518224 | 0.029 | 0.021 | 1 | 0.74576744 | 0.88669861 |
| Epn3 | 0.75328966 | 0.27052894 | 0.021 | 0.016 | 1 | 0.66339728 | 0.10518224 | 0.029 | 0.021 | 1 | 0.75328966 | 0.88669861 |
| 4930537H20Rik | 0.99758604 | -0.2148979 | 0.021 | 0.021 | 1 | 0.66339728 | 0.10518224 | 0.029 | 0.021 | 1 | 0.99758604 | 0.88669861 |
| Calml4 | 0.99758604 | -0.2148979 | 0.021 | 0.021 | 1 | 0.66339728 | 0.10518224 | 0.029 | 0.021 | 1 | 0.99758604 | 0.88669861 |
| Gm47814 | 1 | 0.42253204 | 0.01 | 0.01 | 1 | 0.66339728 | 0.10518224 | 0.029 | 0.021 | 1 | 1 | 0.88669861 |
| Wdr95 | 0.72298091 | -0.9925055 | 0.01 | 0.016 | 1 | 0.66387437 | 0.27510724 | 0.017 | 0.01 | 1 | 0.72298091 | 0.88701956 |
| Gm37634 | 0.75328966 | 0.27052894 | 0.021 | 0.016 | 1 | 0.66387437 | 0.27510724 | 0.017 | 0.01 | 1 | 0.75328966 | 0.88701956 |
| Gm960 | 0.78979801 | -0.407543 | 0.021 | 0.026 | 1 | 0.66387437 | 0.27510724 | 0.017 | 0.01 | 1 | 0.78979801 | 0.88701956 |
| Cfap20dc | 0.80274368 | 0.20013961 | 0.031 | 0.026 | 1 | 0.66387437 | 0.27510724 | 0.017 | 0.01 | 1 | 0.80274368 | 0.88701956 |
| Gm5432 | 0.82258416 | -0.1624305 | 0.031 | 0.036 | 1 | 0.66387437 | 0.69014474 | 0.017 | 0.01 | 1 | 0.82258416 | 0.88701956 |
| B930094E09Rik | 0.993208 | 1.00749454 | 0.01 | 0.01 | 1 | 0.66387437 | 0.27510724 | 0.017 | 0.01 | 1 | 0.993208 | 0.88701956 |
| Gm47819 | 1 | 0.42253204 | 0.01 | 0.01 | 1 | 0.66387437 | 0.27510724 | 0.017 | 0.01 | 1 | 1 | 0.88701956 |
| Nrros | 1 | -0.9925055 | 0.021 | 0.021 | 1 | 0.66387437 | 0.49749966 | 0.017 | 0.01 | 1 | 1 | 0.88701956 |
| Gm2629 | 1 | 0.42253204 | 0.01 | 0.01 | 1 | 0.66387437 | 0.49749966 | 0.017 | 0.01 | 1 | 1 | 0.88701956 |
| Ube3b | 0.97458756 | -0.1851505 | 0.25 | 0.244 | 1 | 0.66438336 | -0.1082214 | 0.345 | 0.312 | 1 | 0.97458756 | 0.88736147 |
| Rab26 | 0.8409956 | 0.32942263 | 0.031 | 0.036 | 1 | 0.66501153 | 0.14957636 | 0.029 | 0.021 | 1 | 0.8409956 | 0.88778273 |
| Gm28809 | 0.85341815 | 0.27052894 | 0.083 | 0.078 | 1 | 0.66552075 | -0.9537115 | 0.118 | 0.125 | 1 | 0.85341815 | 0.88812363 |
| Gm34894 | 1 | -0.577468 | 0.021 | 0.021 | 1 | 0.66601167 | -0.194378 | 0.042 | 0.031 | 1 | 1 | 0.8884518 |
| Ppp4r4 | 0.99009299 | 0.32942263 | 0.031 | 0.031 | 1 | 0.66649533 | -0.8244284 | 0.042 | 0.052 | 1 | 0.99009299 | 0.88877464 |
| Dmtn | 0.66676245 | -0.3144336 | 0.031 | 0.041 | 1 | 0.76485982 | -0.1399303 | 0.025 | 0.031 | 1 | 0.76485982 | 0.88895274 |
| Cflar | 0.66788115 | -0.1691501 | 0.667 | 0.58 | 1 | 0.76732481 | -0.2784228 | 0.773 | 0.76 | 1 | 0.76732481 | 0.88969707 |
| Wfs1 | 0.96926163 | 0.34452952 | 0.094 | 0.093 | 1 | 0.66803084 | -0.4253325 | 0.08 | 0.094 | 1 | 0.96926163 | 0.88979648 |
| Polr3g | 0.66816474 | 0.86999101 | 0.052 | 0.041 | 1 | 0.78571043 | -0.3098553 | 0.038 | 0.031 | 1 | 0.78571043 | 0.88988536 |

| | | | | | | | | | | | | |
|---|---|---|---|---|---|---|---|---|---|---|---|---|
| 2700054A10Rik | 0.80242802 | 0.71798792 | 0.042 | 0.036 | 1 | 0.66820632 | 0.10518224 | 0.017 | 0.01 | 1 | 0.80242802 | 0.88991296 |
| Zfp467 | 0.66859021 | -0.17693 | 0.083 | 0.098 | 1 | 0.91832538 | -0.4797803 | 0.139 | 0.135 | 1 | 0.91832538 | 0.89016755 |
| Fgd3 | 0.99405607 | -0.7294711 | 0.031 | 0.031 | 1 | 0.66988274 | -0.6094155 | 0.029 | 0.021 | 1 | 0.99405607 | 0.89102259 |
| Odf2l | 0.89612 | 0.38456419 | 0.271 | 0.269 | 1 | 0.67048016 | -0.4974823 | 0.218 | 0.188 | 1 | 0.89612 | 0.89141667 |
| Acacb | 0.67090536 | -0.2555399 | 0.083 | 0.098 | 1 | 0.71033773 | -0.8692827 | 0.063 | 0.073 | 1 | 0.71033773 | 0.89169672 |
| Gm17359 | 0.74827573 | 0.59245704 | 0.021 | 0.016 | 1 | 0.67149861 | -0.1399303 | 0.029 | 0.021 | 1 | 0.74827573 | 0.89208683 |
| Platr22 | 0.78979801 | -0.577468 | 0.021 | 0.026 | 1 | 0.67149861 | -0.1399303 | 0.029 | 0.021 | 1 | 0.78979801 | 0.89208683 |
| Esam | 0.67174581 | -0.1624305 | 0.031 | 0.041 | 1 | 0.85762418 | -0.7952821 | 0.038 | 0.042 | 1 | 0.85762418 | 0.89224919 |
| Ranbp10 | 0.67212505 | -0.1665349 | 0.438 | 0.482 | 1 | 0.90696759 | -0.1598298 | 0.626 | 0.604 | 1 | 0.90696759 | 0.89249802 |
| Nprl3 | 0.67228034 | -0.196864 | 0.198 | 0.171 | 1 | 0.7622978 | -0.4797803 | 0.202 | 0.177 | 1 | 0.7622978 | 0.89259982 |
| Cfap161 | 0.72298091 | -0.7998604 | 0.01 | 0.016 | 1 | 0.67254361 | -0.5025003 | 0.017 | 0.01 | 1 | 0.72298091 | 0.89277232 |
| Meig1 | 0.75328966 | 0.27052894 | 0.021 | 0.016 | 1 | 0.67254361 | -0.6317834 | 0.017 | 0.01 | 1 | 0.75328966 | 0.89277232 |
| 2600002D14Rik | 0.75328966 | 0.27052894 | 0.021 | 0.016 | 1 | 0.67254361 | -0.3098553 | 0.017 | 0.01 | 1 | 0.75328966 | 0.89277232 |
| Mastl | 0.67271453 | 0.89201732 | 0.052 | 0.041 | 1 | 0.80410674 | 0.13071733 | 0.059 | 0.052 | 1 | 0.80410674 | 0.89288422 |
| Cacna2d3 | 0.8219591 | 0.22613482 | 0.094 | 0.104 | 1 | 0.67300258 | -1.9742522 | 0.097 | 0.104 | 1 | 0.8219591 | 0.89307269 |
| Rhobtb1 | 0.67350048 | -0.407543 | 0.125 | 0.104 | 1 | 0.82217583 | -0.2723806 | 0.092 | 0.083 | 1 | 0.82217583 | 0.89339806 |
| Ifi203 | 0.72298091 | -0.577468 | 0.01 | 0.016 | 1 | 0.67471693 | -0.7248928 | 0.017 | 0.01 | 1 | 0.72298091 | 0.89419093 |
| Elapor2 | 0.72574709 | -0.577468 | 0.01 | 0.016 | 1 | 0.67471693 | -0.7248928 | 0.017 | 0.01 | 1 | 0.72574709 | 0.89419093 |
| Liph | 0.993208 | 1.42253204 | 0.01 | 0.01 | 1 | 0.67471693 | -0.7248928 | 0.017 | 0.01 | 1 | 0.993208 | 0.89419093 |
| D6Ertd474e | 1 | 0.42253204 | 0.01 | 0.01 | 1 | 0.67471693 | -0.8244284 | 0.017 | 0.01 | 1 | 1 | 0.89419093 |
| Dnajb3 | 1 | 0.42253204 | 0.01 | 0.01 | 1 | 0.67471805 | -0.5025003 | 0.017 | 0.01 | 1 | 1 | 0.89419165 |
| Gm43728 | 0.99758609 | 0.27052894 | 0.021 | 0.021 | 1 | 0.67474842 | -0.3098553 | 0.029 | 0.021 | 1 | 0.99758609 | 0.89421141 |
| A430102K17Rik | 0.99758609 | 0.27052894 | 0.021 | 0.021 | 1 | 0.67474842 | -0.6317834 | 0.029 | 0.021 | 1 | 0.99758609 | 0.89421141 |
| Gm3435 | 1 | 0.42253204 | 0.01 | 0.01 | 1 | 0.67474842 | -0.3098553 | 0.029 | 0.021 | 1 | 1 | 0.89421141 |
| Zdhhc3 | 0.67515119 | -0.1354634 | 0.542 | 0.534 | 1 | 0.82739389 | -0.1095566 | 0.571 | 0.542 | 1 | 0.82739389 | 0.89447325 |
| Fam118b | 0.6756012 | 0.21899864 | 0.292 | 0.275 | 1 | 0.84554524 | -0.6006347 | 0.319 | 0.271 | 1 | 0.84554524 | 0.89476542 |
| Cda | 0.67659216 | -0.2779077 | 0.104 | 0.119 | 1 | 0.86203455 | -1.1723517 | 0.038 | 0.042 | 1 | 0.86203455 | 0.89540737 |
| Il1r2 | 0.67755108 | -1.3710171 | 0.042 | 0.052 | 1 | 0.8023126 | -0.5025003 | 0.017 | 0.021 | 1 | 0.8023126 | 0.8960267 |
| Tulp4 | 0.97158645 | -0.1371637 | 0.896 | 0.881 | 1 | 0.67839868 | -0.1273302 | 0.95 | 0.99 | 1 | 0.97158645 | 0.89657259 |
| Mapk9 | 0.82641619 | 0.22988696 | 0.333 | 0.352 | 1 | 0.67858405 | -1.2442669 | 0.324 | 0.312 | 1 | 0.82641619 | 0.89669179 |
| Igflr1 | 0.67877064 | 0.49292136 | 0.052 | 0.041 | 1 | 0.88759525 | -0.8948178 | 0.046 | 0.042 | 1 | 0.88759525 | 0.8968117 |
| Slc35d2 | 1 | -0.1079827 | 0.052 | 0.052 | 1 | 0.68180609 | -0.3664388 | 0.042 | 0.052 | 1 | 1 | 0.89875263 |
| Ccng2 | 0.68306192 | 0.2091284 | 0.167 | 0.145 | 1 | 0.88239637 | -0.3368223 | 0.147 | 0.135 | 1 | 0.88239637 | 0.89955025 |
| Gm47889 | 0.72437359 | -0.407543 | 0.062 | 0.052 | 1 | 0.68307064 | -0.7623675 | 0.042 | 0.052 | 1 | 0.72437359 | 0.89955578 |
| E130102H24Rik | 0.68309634 | 0.17460452 | 0.229 | 0.212 | 1 | 0.88082153 | -0.5805029 | 0.202 | 0.198 | 1 | 0.88082153 | 0.89957207 |
| Foxj3 | 0.68480745 | -0.4790053 | 0.583 | 0.544 | 1 | 0.68723281 | -0.3203476 | 0.752 | 0.719 | 1 | 0.68723281 | 0.90065366 |
| Trhde | 1 | -0.7294711 | 0.021 | 0.021 | 1 | 0.6848734 | -1.3923174 | 0.034 | 0.042 | 1 | 1 | 0.90069522 |
| Gm42829 | 0.99758609 | 0.27052894 | 0.021 | 0.021 | 1 | 0.68562457 | -0.4029647 | 0.042 | 0.052 | 1 | 0.99758609 | 0.90116809 |
| Ap5z1 | 0.68571443 | -0.3144336 | 0.125 | 0.14 | 1 | 0.95207244 | -0.4253325 | 0.155 | 0.146 | 1 | 0.95207244 | 0.90122458 |
| Pcgf1 | 0.89735252 | -0.2335136 | 0.083 | 0.078 | 1 | 0.68770566 | -1.4618584 | 0.034 | 0.042 | 1 | 0.89735252 | 0.90247225 |
| Wdr7 | 0.83653961 | -0.293675 | 0.344 | 0.311 | 1 | 0.68883323 | -0.3610806 | 0.475 | 0.458 | 1 | 0.83653961 | 0.90317524 |
| Gm41231 | 0.68932405 | -0.4779323 | 0.042 | 0.052 | 1 | 0.85617402 | -0.6094155 | 0.038 | 0.042 | 1 | 0.85617402 | 0.90348045 |
| Mc5r | 0.68932405 | -0.4779323 | 0.042 | 0.052 | 1 | 0.90191534 | -1.3098553 | 0.08 | 0.083 | 1 | 0.90191534 | 0.90348045 |
| Sgsm3 | 0.69018889 | -0.3968957 | 0.323 | 0.269 | 1 | 0.76869069 | -0.2345671 | 0.307 | 0.312 | 1 | 0.76869069 | 0.90401708 |
| Gm13523 | 0.69098575 | 0.27052894 | 0.052 | 0.041 | 1 | 0.74908033 | -0.6724253 | 0.055 | 0.062 | 1 | 0.74908033 | 0.90451019 |
| Zfp628 | 0.69098575 | -0.1079827 | 0.052 | 0.041 | 1 | 0.76982931 | -0.7248928 | 0.076 | 0.083 | 1 | 0.76982931 | 0.90451019 |
| Npat | 0.69267607 | 0.37946331 | 0.49 | 0.482 | 1 | 0.9249783 | -0.2644123 | 0.592 | 0.521 | 1 | 0.9249783 | 0.905552 |
| Thap3 | 0.6930445 | -0.1730777 | 0.198 | 0.223 | 1 | 0.8971893 | -0.3695615 | 0.206 | 0.188 | 1 | 0.8971893 | 0.90577832 |
| Gm36033 | 0.69410364 | 0.22988696 | 0.052 | 0.041 | 1 | 0.85963481 | -0.5549678 | 0.092 | 0.083 | 1 | 0.85963481 | 0.90642742 |
| Myl9 | 0.86704544 | -0.9214224 | 0.167 | 0.145 | 1 | 0.69518226 | -1.4713187 | 0.214 | 0.219 | 1 | 0.86704544 | 0.90708615 |
| Enox1 | 0.69550509 | -0.1705978 | 0.344 | 0.311 | 1 | 0.98768416 | -0.7180391 | 0.429 | 0.375 | 1 | 0.98768416 | 0.90728285 |
| Tdp1 | 0.82735665 | 0.42253204 | 0.135 | 0.13 | 1 | 0.69576026 | -0.1843244 | 0.08 | 0.094 | 1 | 0.82735665 | 0.90743818 |
| Senp2 | 0.69594956 | -0.2168228 | 0.792 | 0.86 | 1 | 0.77421709 | -0.2279136 | 0.899 | 0.906 | 1 | 0.77421709 | 0.90755333 |
| P2ry14 | 0.69620097 | 0.54806292 | 0.062 | 0.052 | 1 | 0.74014199 | 0.10518224 | 0.05 | 0.042 | 1 | 0.74014199 | 0.90770615 |
| Cep97 | 0.69676216 | 0.23697638 | 0.146 | 0.13 | 1 | 0.92774543 | -0.5828738 | 0.147 | 0.146 | 1 | 0.92774543 | 0.90804681 |
| Gm26621 | 0.69869193 | 0.13302542 | 0.052 | 0.041 | 1 | 0.93579004 | -0.3098553 | 0.05 | 0.052 | 1 | 0.93579004 | 0.90921344 |
| Samd14 | 0.7586101 | 0.15949763 | 0.083 | 0.073 | 1 | 0.6991509 | -0.2002308 | 0.155 | 0.135 | 1 | 0.7586101 | 0.90948982 |
| Gm43597 | 0.99758609 | 0.27052894 | 0.021 | 0.021 | 1 | 0.69916441 | -1.1399303 | 0.034 | 0.042 | 1 | 0.99758609 | 0.90949795 |
| Saxo2 | 0.99758609 | 0.27052894 | 0.021 | 0.021 | 1 | 0.69916441 | -1.1399303 | 0.034 | 0.042 | 1 | 0.99758609 | 0.90949795 |
| Dlc1 | 0.85465356 | -0.2230231 | 0.938 | 0.87 | 1 | 0.69971419 | -0.2807742 | 0.895 | 0.802 | 1 | 0.85465356 | 0.90982843 |
| Sh3bp1 | 0.89052796 | -0.5160674 | 0.073 | 0.067 | 1 | 0.69974414 | -0.1723517 | 0.122 | 0.104 | 1 | 0.89052796 | 0.90984642 |
| Lax1 | 0.70045085 | 0.42253204 | 0.062 | 0.052 | 1 | 0.94026906 | -1.3098553 | 0.034 | 0.031 | 1 | 0.94026906 | 0.91027031 |
| Zfp40 | 0.72180054 | 0.31561683 | 0.083 | 0.073 | 1 | 0.70062719 | -1.0468209 | 0.034 | 0.042 | 1 | 0.72180054 | 0.91037592 |
| Madd | 0.70116305 | -0.2555399 | 0.042 | 0.052 | 1 | 0.70457616 | -0.3769695 | 0.076 | 0.062 | 1 | 0.70457616 | 0.91069648 |
| Pcdhga2 | 0.70116305 | -0.2555399 | 0.042 | 0.052 | 1 | 0.72812226 | -0.1723517 | 0.034 | 0.042 | 1 | 0.72812226 | 0.91069648 |
| Narf | 0.93041015 | -0.4418129 | 0.198 | 0.197 | 1 | 0.70120866 | -0.3955851 | 0.147 | 0.125 | 1 | 0.93041015 | 0.91072373 |
| Crot | 0.70308593 | -0.1079827 | 0.156 | 0.135 | 1 | 0.82005146 | 0.13945214 | 0.193 | 0.188 | 1 | 0.82005146 | 0.91184204 |
| Adamts12 | 0.70313488 | -0.9115855 | 0.208 | 0.218 | 1 | 0.94670562 | -0.5907561 | 0.378 | 0.344 | 1 | 0.94670562 | 0.9118711 |
| Supt5 | 0.76533637 | -0.1835538 | 0.604 | 0.606 | 1 | 0.70322934 | -0.2622896 | 0.731 | 0.74 | 1 | 0.76533637 | 0.91192718 |
| Capn5 | 0.83448168 | 0.5049942 | 0.073 | 0.067 | 1 | 0.7038912 | -1.0785297 | 0.063 | 0.073 | 1 | 0.83448168 | 0.91231958 |
| Wdr55 | 0.74547103 | -0.1851505 | 0.062 | 0.073 | 1 | 0.70415341 | 0.17557156 | 0.097 | 0.083 | 1 | 0.74547103 | 0.9124748 |
| 4732471J01Rik | 0.70431386 | -0.3364599 | 0.135 | 0.15 | 1 | 0.90296221 | -0.8948178 | 0.092 | 0.094 | 1 | 0.90296221 | 0.91256971 |
| Ddr2 | 0.70487491 | -0.2354004 | 0.938 | 0.891 | 1 | 0.78585612 | -0.1234138 | 0.912 | 0.875 | 1 | 0.78585612 | 0.91290118 |
| Gm47460 | 0.78979801 | -0.2148979 | 0.021 | 0.026 | 1 | 0.70494229 | -1.0203486 | 0.034 | 0.042 | 1 | 0.78979801 | 0.91294095 |
| Tnks2 | 0.91595507 | -0.1199952 | 0.958 | 0.933 | 1 | 0.70552995 | -0.1942049 | 0.983 | 0.979 | 1 | 0.91595507 | 0.91328739 |
| 6820445E23Rik | 0.70575267 | -0.4950058 | 0.052 | 0.062 | 1 | 0.91453215 | -0.1399303 | 0.034 | 0.031 | 1 | 0.91453215 | 0.91341851 |
| Gm16133 | 0.70587411 | 0.16795921 | 0.292 | 0.275 | 1 | 0.80418855 | -0.5366261 | 0.542 | 0.479 | 1 | 0.80418855 | 0.91348996 |
| Mettl22 | 0.71031978 | 0.35541784 | 0.062 | 0.052 | 1 | 0.92663792 | -0.5025003 | 0.05 | 0.052 | 1 | 0.92663792 | 0.91608537 |
| Cenpq | 0.75937862 | 0.32942263 | 0.219 | 0.249 | 1 | 0.71094938 | -0.3098553 | 0.071 | 0.083 | 1 | 0.75937862 | 0.91644974 |
| Frg2f1 | 0.71145337 | -0.2462621 | 0.104 | 0.119 | 1 | 1 | 0.33400093 | 0.071 | 0.073 | 1 | 1 | 0.91674085 |
| Adam12 | 0.71159075 | -0.1608425 | 0.292 | 0.316 | 1 | 0.85784879 | -0.4452564 | 0.391 | 0.344 | 1 | 0.85784879 | 0.9168201 |
| Slc9a5 | 0.71305088 | 0.17741954 | 0.062 | 0.052 | 1 | 0.9071436 | -0.8948178 | 0.029 | 0.031 | 1 | 0.9071436 | 0.9176602 |
| Ehhadh | 0.71314887 | 0.11440974 | 0.042 | 0.052 | 1 | 1 | -1.0468209 | 0.021 | 0.021 | 1 | 1 | 0.91771643 |
| Rad9a | 0.7141857 | -0.1079827 | 0.052 | 0.041 | 1 | 0.93979958 | -0.2394659 | 0.076 | 0.073 | 1 | 0.93979958 | 0.91831018 |
| Gm49890 | 0.74076005 | 1.00749454 | 0.021 | 0.016 | 1 | 0.71538478 | 0.78325414 | 0.042 | 0.052 | 1 | 0.74076005 | 0.91899418 |

| | | | | | | | | | | | | |
|---|---|---|---|---|---|---|---|---|---|---|---|---|
| Gm15446 | 0.98939785 | -0.2820121 | 0.073 | 0.073 | 1 | 0.71571496 | -1.0297473 | 0.063 | 0.073 | 1 | 0.98939785 | 0.91918202 |
| Stx16 | 0.71582125 | 0.2442974 | 0.562 | 0.585 | 1 | 0.95085028 | -0.1130546 | 0.605 | 0.615 | 1 | 0.95085028 | 0.91924244 |
| 9530046B11Rik | 0.81708725 | -0.3144336 | 0.031 | 0.036 | 1 | 0.71649639 | -0.5728897 | 0.034 | 0.042 | 1 | 0.81708725 | 0.9196257 |
| Gmeb2 | 0.8721136 | -0.2075184 | 0.427 | 0.394 | 1 | 0.71862867 | -0.225791 | 0.403 | 0.354 | 1 | 0.8721136 | 0.92083018 |
| Gm15738 | 0.71864235 | 0.22988696 | 0.062 | 0.052 | 1 | 0.95336432 | -0.4353861 | 0.092 | 0.094 | 1 | 0.95336432 | 0.92083787 |
| Dagla | 0.71864235 | 0.11440974 | 0.062 | 0.052 | 1 | 0.73761517 | -0.2394659 | 0.063 | 0.052 | 1 | 0.73761517 | 0.92083787 |
| A330040F15Rik | 0.71917366 | -0.747393 | 0.042 | 0.052 | 1 | 0.79181108 | -1.1399303 | 0.017 | 0.021 | 1 | 0.79181108 | 0.92113657 |
| Sphk2 | 0.71937765 | -0.3144336 | 0.052 | 0.062 | 1 | 0.74963743 | -0.2029401 | 0.092 | 0.104 | 1 | 0.74963743 | 0.92125109 |
| Nipa1 | 0.79574854 | -0.2726134 | 0.125 | 0.135 | 1 | 0.72029884 | -0.5728897 | 0.13 | 0.115 | 1 | 0.79574854 | 0.92176726 |
| Eda2r | 0.72042936 | -0.8504865 | 0.052 | 0.041 | 1 | 0.81707832 | 0.14957636 | 0.017 | 0.021 | 1 | 0.81707832 | 0.92184026 |
| Zfp433 | 0.72212321 | 0.13302542 | 0.042 | 0.052 | 1 | 0.92396969 | -0.4618584 | 0.029 | 0.031 | 1 | 0.92396969 | 0.92278449 |
| Acvrl1 | 0.78979801 | -0.2148979 | 0.021 | 0.026 | 1 | 0.72234088 | -0.4618584 | 0.034 | 0.042 | 1 | 0.78979801 | 0.92290541 |
| Gm42567 | 0.72298091 | -0.9925055 | 0.01 | 0.016 | 1 | 0.8679252 | -0.1172102 | 0.038 | 0.042 | 1 | 0.8679252 | 0.92326043 |
| Tent5b | 0.72298091 | -1.1624305 | 0.01 | 0.016 | 1 | 0.86938864 | -0.3098553 | 0.038 | 0.042 | 1 | 0.86938864 | 0.92326043 |
| Art4 | 0.72298091 | -0.577468 | 0.01 | 0.016 | 1 | 0.86312513 | -0.7248928 | 0.008 | 0.01 | 1 | 0.86312513 | 0.92326043 |
| Slc5a5 | 0.72298091 | -0.577468 | 0.01 | 0.016 | 1 | 0.87659345 | -0.5443205 | 0.046 | 0.042 | 1 | 0.87659345 | 0.92326043 |
| Dlec1 | 0.72298091 | -0.577468 | 0.01 | 0.016 | 1 | 0.98188267 | 0.27510724 | 0.021 | 0.021 | 1 | 0.98188267 | 0.92326043 |
| Gm47561 | 0.72298091 | -0.7998604 | 0.01 | 0.016 | 1 | 0.8065254 | -0.3098553 | 0.017 | 0.021 | 1 | 0.8065254 | 0.92326043 |
| Ppp1r42 | 0.72574709 | -0.577468 | 0.01 | 0.016 | 1 | 0.81285149 | 0.42711033 | 0.017 | 0.021 | 1 | 0.81285149 | 0.92478534 |
| Ankrd45 | 0.72574709 | -0.3144336 | 0.01 | 0.016 | 1 | 0.87851807 | -0.7248928 | 0.013 | 0.01 | 1 | 0.87851807 | 0.92478534 |
| Galnt3 | 0.72574709 | -1.7998604 | 0.01 | 0.016 | 1 | 0.88855723 | -0.7692869 | 0.059 | 0.062 | 1 | 0.88855723 | 0.92478534 |
| Gm2164 | 0.72574709 | -0.3144336 | 0.01 | 0.016 | 1 | 0.86312513 | -0.7248928 | 0.008 | 0.01 | 1 | 0.86312513 | 0.92478534 |
| Tgfbr3l | 0.72574709 | -0.577468 | 0.01 | 0.016 | 1 | 0.87066325 | -0.3098553 | 0.013 | 0.01 | 1 | 0.87066325 | 0.92478534 |
| Gm35721 | 0.72574709 | -0.3144336 | 0.01 | 0.016 | 1 | 0.86613737 | -0.3098553 | 0.008 | 0.01 | 1 | 0.86613737 | 0.92478534 |
| B4galnt2 | 0.72574709 | -0.3144336 | 0.01 | 0.016 | 1 | 0.86312513 | -0.7248928 | 0.008 | 0.01 | 1 | 0.86312513 | 0.92478534 |
| Gm11739 | 0.72574709 | -0.3144336 | 0.01 | 0.016 | 1 | 0.86312513 | -0.7248928 | 0.008 | 0.01 | 1 | 0.86312513 | 0.92478534 |
| Marveld2 | 0.72574709 | -0.9925055 | 0.01 | 0.016 | 1 | 0.80442048 | -0.5728897 | 0.017 | 0.021 | 1 | 0.80442048 | 0.92478534 |
| Gm49313 | 0.72574709 | -0.3144336 | 0.01 | 0.016 | 1 | 0.86312513 | -0.7248928 | 0.008 | 0.01 | 1 | 0.86312513 | 0.92478534 |
| Gm41335 | 0.72574709 | -0.3144336 | 0.01 | 0.016 | 1 | 0.85710586 | -3.010295 | 0.008 | 0.01 | 1 | 0.85710586 | 0.92478534 |
| Gm16189 | 0.72574709 | -0.577468 | 0.01 | 0.016 | 1 | 0.79600331 | -1.3098553 | 0.017 | 0.021 | 1 | 0.79600331 | 0.92478534 |
| 4631405J19Rik | 0.72575028 | -1.2148979 | 0.01 | 0.016 | 1 | 0.87066325 | -0.3098553 | 0.013 | 0.01 | 1 | 0.87066325 | 0.92478709 |
| Kcnab1 | 0.72765847 | -0.2726134 | 0.052 | 0.062 | 1 | 0.88484452 | -1.0688472 | 0.046 | 0.042 | 1 | 0.88484452 | 0.92583009 |
| Tcirg1 | 0.95905932 | -0.1801325 | 0.167 | 0.161 | 1 | 0.72892331 | 0.18657056 | 0.172 | 0.188 | 1 | 0.95905932 | 0.92651743 |
| Faxc | 0.72963938 | -0.6705774 | 0.073 | 0.083 | 1 | 0.99702154 | 0.23446525 | 0.092 | 0.094 | 1 | 0.99702154 | 0.92690513 |
| Zfp423 | 0.72963995 | 0.21899864 | 0.135 | 0.119 | 1 | 0.73928053 | -1.250149 | 0.231 | 0.219 | 1 | 0.73928053 | 0.92690544 |
| Ccdc62 | 0.72990919 | -0.1180363 | 0.198 | 0.218 | 1 | 0.81638217 | -0.3438026 | 0.202 | 0.177 | 1 | 0.81638217 | 0.92705095 |
| 4930512B01Rik | 0.7301396 | -0.7087125 | 0.062 | 0.073 | 1 | 0.79330564 | -1.0072925 | 0.134 | 0.135 | 1 | 0.79330564 | 0.92717537 |
| Slc22a1 | 0.7301396 | -0.3550755 | 0.062 | 0.073 | 1 | 0.79368278 | -0.3623227 | 0.076 | 0.083 | 1 | 0.79368278 | 0.92717537 |
| Sp4 | 0.73171553 | 0.19676489 | 0.302 | 0.28 | 1 | 0.87832306 | -0.4945951 | 0.399 | 0.344 | 1 | 0.87832306 | 0.92802344 |
| Vps37c | 0.94095222 | 0.23490503 | 0.156 | 0.161 | 1 | 0.73263889 | -0.6917259 | 0.223 | 0.219 | 1 | 0.94095222 | 0.92851803 |
| Gm11655 | 0.86291181 | -0.2148979 | 0.104 | 0.109 | 1 | 0.73412881 | -0.3923174 | 0.063 | 0.052 | 1 | 0.86291181 | 0.92931251 |
| Pdss1 | 0.7350927 | -0.2295447 | 0.188 | 0.202 | 1 | 0.83367486 | -0.8244284 | 0.155 | 0.135 | 1 | 0.83367486 | 0.92982412 |
| Trim16 | 0.73685701 | 1.00749454 | 0.01 | 0.016 | 1 | 0.79600331 | -1.5728897 | 0.017 | 0.021 | 1 | 0.79600331 | 0.93075577 |
| Leng9 | 0.73779294 | -0.2726134 | 0.062 | 0.073 | 1 | 0.76931152 | -0.4618584 | 0.055 | 0.062 | 1 | 0.76931152 | 0.93124746 |
| Nfia | 0.73937252 | -0.2416301 | 0.917 | 0.907 | 1 | 0.80957869 | -0.2867716 | 0.962 | 0.979 | 1 | 0.80957869 | 0.93207332 |
| Mm2pr | 0.74827573 | 0.59245704 | 0.021 | 0.016 | 1 | 0.73945321 | -1.5025003 | 0.025 | 0.031 | 1 | 0.74827573 | 0.93211537 |
| Gpx6 | 0.74827573 | 0.59245704 | 0.021 | 0.016 | 1 | 0.73945321 | -1.0874628 | 0.025 | 0.031 | 1 | 0.74827573 | 0.93211537 |
| Dock5 | 0.82870673 | 0.29967529 | 0.115 | 0.124 | 1 | 0.73974696 | -0.44958 | 0.239 | 0.208 | 1 | 0.82870673 | 0.93226836 |
| Armh1 | 0.74076005 | 1.00749454 | 0.021 | 0.016 | 1 | 0.87066325 | -0.3098553 | 0.013 | 0.01 | 1 | 0.87066325 | 0.93279465 |
| Pgbd5 | 0.74076005 | 1.32942263 | 0.021 | 0.016 | 1 | 0.8086289 | -0.3098553 | 0.017 | 0.021 | 1 | 0.8086289 | 0.93279465 |
| Slc6a20a | 0.74076005 | 1.32942263 | 0.021 | 0.016 | 1 | 0.79181108 | -1.0874628 | 0.017 | 0.021 | 1 | 0.79181108 | 0.93279465 |
| Gm56767 | 0.74076005 | 1.00749454 | 0.021 | 0.016 | 1 | 0.87066325 | -0.3098553 | 0.013 | 0.01 | 1 | 0.87066325 | 0.93279465 |
| Popdc2 | 0.74076005 | 1.00749454 | 0.021 | 0.016 | 1 | 0.86312513 | -0.7248928 | 0.008 | 0.01 | 1 | 0.86312513 | 0.93279465 |
| Gm17276 | 0.74076005 | 2.46692616 | 0.021 | 0.016 | 1 | 0.97842093 | -1.2029401 | 0.042 | 0.042 | 1 | 0.97842093 | 0.93279465 |
| Kdm2b | 0.84309839 | 0.11711903 | 0.271 | 0.264 | 1 | 0.74149257 | -0.6558511 | 0.315 | 0.302 | 1 | 0.84309839 | 0.93317391 |
| Elfn1 | 0.74326422 | 1.00749454 | 0.021 | 0.016 | 1 | 0.77529375 | 0.27510724 | 0.038 | 0.031 | 1 | 0.77529375 | 0.93408674 |
| Gm28050 | 0.84157317 | -0.4330781 | 0.052 | 0.057 | 1 | 0.74348215 | -0.9879272 | 0.076 | 0.062 | 1 | 0.84157317 | 0.93419859 |
| Rnf185 | 0.89411652 | -0.1014398 | 0.354 | 0.352 | 1 | 0.74424392 | -0.413233 | 0.429 | 0.375 | 1 | 0.89411652 | 0.93458883 |
| Gm20735 | 1 | 0.42253204 | 0.01 | 0.01 | 1 | 0.74572089 | 0.17557156 | 0.034 | 0.042 | 1 | 1 | 0.93534213 |
| Aatk | 0.74576744 | 0.68556644 | 0.021 | 0.016 | 1 | 0.82415676 | -0.8948178 | 0.025 | 0.021 | 1 | 0.82415676 | 0.93536581 |
| Ppfia4 | 0.74811732 | -0.1624305 | 0.062 | 0.073 | 1 | 0.85517682 | -0.681824 | 0.059 | 0.052 | 1 | 0.85517682 | 0.93655512 |
| Dnah5 | 0.74827336 | 0.42253204 | 0.021 | 0.016 | 1 | 0.85710544 | -2.0468209 | 0.008 | 0.01 | 1 | 0.85710544 | 0.9366337 |
| 2810459M11Rik | 0.74827573 | 0.59245704 | 0.021 | 0.016 | 1 | 0.87066325 | -0.3098553 | 0.013 | 0.01 | 1 | 0.87066325 | 0.93663489 |
| Gpr35 | 0.74827573 | 0.59245704 | 0.021 | 0.016 | 1 | 0.90489059 | 0.14957636 | 0.034 | 0.031 | 1 | 0.90489059 | 0.93663489 |
| Gstm6 | 0.74827573 | 0.59245704 | 0.021 | 0.016 | 1 | 0.8076363 | 0.56461386 | 0.025 | 0.021 | 1 | 0.8076363 | 0.93663489 |
| Gm34121 | 0.74827573 | 0.59245704 | 0.021 | 0.016 | 1 | 0.87851725 | -1.3098553 | 0.013 | 0.01 | 1 | 0.87851725 | 0.93663489 |
| Gm45570 | 0.74827573 | 0.59245704 | 0.021 | 0.016 | 1 | 0.93749089 | -0.2103196 | 0.055 | 0.052 | 1 | 0.93749089 | 0.93663489 |
| Gm56994 | 0.74827573 | 0.59245704 | 0.021 | 0.016 | 1 | 0.86312513 | -0.7248928 | 0.008 | 0.01 | 1 | 0.86312513 | 0.93663489 |
| Shmt1 | 0.74827573 | 0.59245704 | 0.021 | 0.016 | 1 | 0.84641179 | 0.36821664 | 0.046 | 0.042 | 1 | 0.84641179 | 0.93663489 |
| Gm51903 | 0.74827573 | 0.59245704 | 0.021 | 0.016 | 1 | 0.86312513 | -0.7248928 | 0.008 | 0.01 | 1 | 0.86312513 | 0.93663489 |
| Gm16618 | 0.74827573 | 0.59245704 | 0.021 | 0.016 | 1 | 0.86312513 | -0.7248928 | 0.008 | 0.01 | 1 | 0.86312513 | 0.93663489 |
| 4833408A19Rik | 0.74827573 | 0.59245704 | 0.021 | 0.016 | 1 | 0.87851725 | -1.3098553 | 0.013 | 0.01 | 1 | 0.87851725 | 0.93663489 |
| Mb | 0.97053455 | -0.577468 | 0.052 | 0.052 | 1 | 0.74914333 | -0.3098553 | 0.05 | 0.042 | 1 | 0.97053455 | 0.93707093 |
| Atg14 | 0.74947309 | -0.4286046 | 0.135 | 0.145 | 1 | 0.88895937 | -0.1399303 | 0.218 | 0.208 | 1 | 0.88895937 | 0.93723627 |
| Kcnk7 | 0.75328966 | 0.27052894 | 0.021 | 0.016 | 1 | 0.74957918 | -0.8244284 | 0.025 | 0.031 | 1 | 0.75328966 | 0.93728941 |
| Glra1 | 0.86441475 | -0.9311049 | 0.052 | 0.047 | 1 | 0.75164421 | -0.4167705 | 0.05 | 0.042 | 1 | 0.86441475 | 0.9383194 |
| 4930473A02Rik | 0.99603742 | 0.20013961 | 0.031 | 0.031 | 1 | 0.75164421 | -0.6094155 | 0.05 | 0.042 | 1 | 0.99603742 | 0.9383194 |
| Enpp3 | 0.75328966 | 0.27052894 | 0.021 | 0.016 | 1 | 0.85710544 | -1.3098553 | 0.008 | 0.01 | 1 | 0.85710544 | 0.93913401 |
| Cfap206 | 0.75832339 | -0.407543 | 0.021 | 0.016 | 1 | 0.87066325 | -0.3098553 | 0.013 | 0.01 | 1 | 0.87066325 | 0.94159241 |
| Gm36283 | 0.75832339 | -0.7294711 | 0.021 | 0.016 | 1 | 0.86312513 | -0.7248928 | 0.008 | 0.01 | 1 | 0.86312513 | 0.94159241 |
| Slc9b2 | 1 | 0.42253204 | 0.01 | 0.01 | 1 | 0.75900278 | 2.23858136 | 0.038 | 0.031 | 1 | 1 | 0.94192034 |
| Sec31b | 0.80274368 | 0.20013961 | 0.031 | 0.026 | 1 | 0.75977578 | -0.5025003 | 0.025 | 0.031 | 1 | 0.80274368 | 0.94229233 |
| Ube4bos1 | 0.81290066 | 0.49292136 | 0.042 | 0.036 | 1 | 0.76196741 | -0.3098553 | 0.05 | 0.042 | 1 | 0.81290066 | 0.94334048 |
| Gm9725 | 0.7620979 | -0.1445086 | 0.073 | 0.062 | 1 | 0.99280559 | -0.194378 | 0.042 | 0.042 | 1 | 0.99280559 | 0.94340259 |
| Gm56760 | 0.99758609 | 0.27052894 | 0.021 | 0.021 | 1 | 0.76485982 | -0.3098553 | 0.025 | 0.031 | 1 | 0.99758609 | 0.9447091 |

| | | | | | | | | | | | |
|---|---|---|---|---|---|---|---|---|---|---|---|
| Dennd2a | 0.89971162 | -0.4743745 | 0.198 | 0.197 | 1 | 0.76673737 | -0.1375397 | 0.332 | 0.292 | 1 | 0.89971162 | 0.94558854 |
| Gypc | 0.77556069 | -0.4330781 | 0.167 | 0.176 | 1 | 0.85691367 | -0.177405 | 0.155 | 0.146 | 1 | 0.85691367 | 0.949627 |
| Vps72 | 0.96689575 | -0.2555399 | 0.323 | 0.301 | 1 | 0.77563672 | -0.4518743 | 0.357 | 0.312 | 1 | 0.96689575 | 0.94966112 |
| Catsper2 | 0.77597231 | -0.3461424 | 0.083 | 0.073 | 1 | 0.85555831 | 0.27510724 | 0.076 | 0.083 | 1 | 0.85555831 | 0.94981159 |
| Ppif | 0.77627397 | -0.5197525 | 0.188 | 0.192 | 1 | 0.79179869 | -0.8359241 | 0.088 | 0.094 | 1 | 0.79179869 | 0.94994667 |
| Uck2 | 0.90819156 | -0.4869304 | 0.552 | 0.523 | 1 | 0.77704066 | -0.1230868 | 0.483 | 0.521 | 1 | 0.90819156 | 0.95028913 |
| Kyat1 | 0.84149913 | -0.4519371 | 0.073 | 0.078 | 1 | 0.77823334 | -0.7793405 | 0.088 | 0.094 | 1 | 0.84149913 | 0.95081955 |
| Tulp3 | 0.99438536 | -0.2718879 | 0.646 | 0.601 | 1 | 0.7785471 | -0.4804291 | 0.563 | 0.438 | 1 | 0.99438536 | 0.95095861 |
| Zc3hav1 | 0.97483749 | 0.15768747 | 0.656 | 0.684 | 1 | 0.77945542 | -0.2403143 | 0.618 | 0.698 | 1 | 0.97483749 | 0.95136009 |
| Slc13a4 | 0.99207488 | 0.24195979 | 0.031 | 0.031 | 1 | 0.78011554 | -0.7952821 | 0.05 | 0.042 | 1 | 0.99207488 | 0.95165082 |
| Plet1os | 0.78046984 | 1.00749454 | 0.031 | 0.026 | 1 | 0.86312513 | -0.7248928 | 0.008 | 0.01 | 1 | 0.86312513 | 0.95180651 |
| Nanos1 | 0.98603793 | 0.22988696 | 0.052 | 0.052 | 1 | 0.78177474 | -0.1723517 | 0.105 | 0.094 | 1 | 0.98603793 | 0.95237774 |
| Fbxw5 | 0.78289236 | -0.240433 | 0.156 | 0.166 | 1 | 0.85068738 | -0.355659 | 0.126 | 0.115 | 1 | 0.85068738 | 0.95286427 |
| Fam174b | 0.7854853 | -0.407543 | 0.021 | 0.026 | 1 | 0.82231716 | -0.3098553 | 0.025 | 0.021 | 1 | 0.82231716 | 0.95398344 |
| Grtp1 | 0.7854853 | -0.407543 | 0.021 | 0.026 | 1 | 0.86312513 | -0.7248928 | 0.008 | 0.01 | 1 | 0.86312513 | 0.95398344 |
| Gm39363 | 0.7854853 | -0.407543 | 0.021 | 0.026 | 1 | 0.84007904 | -0.9313436 | 0.038 | 0.042 | 1 | 0.84007904 | 0.95398344 |
| 4732465J04Rik | 0.7854853 | -0.407543 | 0.021 | 0.026 | 1 | 1 | -0.5025003 | 0.021 | 0.021 | 1 | 1 | 0.95398344 |
| Dlk2 | 0.7854853 | -0.407543 | 0.021 | 0.026 | 1 | 0.99396051 | -0.3098553 | 0.021 | 0.021 | 1 | 0.99396051 | 0.95398344 |
| Inhca | 0.88008001 | 0.10060394 | 0.052 | 0.057 | 1 | 0.78571043 | -0.3098553 | 0.038 | 0.031 | 1 | 0.88008001 | 0.95407998 |
| Trmu | 0.82527428 | -0.1708427 | 0.135 | 0.145 | 1 | 0.78689297 | -1.0368368 | 0.109 | 0.115 | 1 | 0.82527428 | 0.95458539 |
| Znhit2 | 0.78948058 | -0.2148979 | 0.115 | 0.124 | 1 | 0.9543359 | -0.194378 | 0.092 | 0.094 | 1 | 0.9543359 | 0.95568157 |
| Zfp853 | 0.78979801 | -0.407543 | 0.021 | 0.026 | 1 | 0.87066325 | -0.3098553 | 0.013 | 0.01 | 1 | 0.87066325 | 0.95581512 |
| Gm12781 | 0.78979801 | -0.2148979 | 0.021 | 0.026 | 1 | 0.99396051 | -0.3098553 | 0.021 | 0.021 | 1 | 0.99396051 | 0.95581512 |
| Pam16l | 0.78979801 | -0.2148979 | 0.021 | 0.026 | 1 | 0.91891525 | -0.6317834 | 0.029 | 0.031 | 1 | 0.91891525 | 0.95581512 |
| Ggt5 | 0.78979801 | -0.2148979 | 0.021 | 0.026 | 1 | 0.87589833 | -0.5728897 | 0.013 | 0.01 | 1 | 0.87589833 | 0.95581512 |
| Gm49492 | 0.78979801 | -0.2148979 | 0.021 | 0.026 | 1 | 0.87066325 | -0.3098553 | 0.013 | 0.01 | 1 | 0.87066325 | 0.95581512 |
| 1700109G14Rik | 1 | 0.42253204 | 0.01 | 0.01 | 1 | 0.79181139 | -1.3098553 | 0.017 | 0.021 | 1 | 1 | 0.9566575 |
| Sos2 | 0.93713392 | -0.1118044 | 0.583 | 0.591 | 1 | 0.79272376 | -0.1532975 | 0.803 | 0.812 | 1 | 0.93713392 | 0.95703656 |
| Disc1 | 0.79413116 | -0.4519371 | 0.021 | 0.026 | 1 | 0.85874064 | -0.1172102 | 0.046 | 0.042 | 1 | 0.85874064 | 0.95761802 |
| Ebf4 | 0.81733549 | -1.8405024 | 0.042 | 0.047 | 1 | 0.79448212 | -0.8948178 | 0.109 | 0.094 | 1 | 0.81733549 | 0.9577624 |
| Appbp2os | 0.79462678 | 0.32942263 | 0.031 | 0.026 | 1 | 0.91891525 | -0.8948178 | 0.029 | 0.031 | 1 | 0.91891525 | 0.95782184 |
| Cbs | 0.99660398 | 0.59245704 | 0.01 | 0.01 | 1 | 0.79600331 | -0.9879272 | 0.017 | 0.021 | 1 | 0.99660398 | 0.95838535 |
| 4930595D18Rik | 0.99758604 | -0.2148979 | 0.021 | 0.021 | 1 | 0.79600331 | -0.9879272 | 0.017 | 0.021 | 1 | 0.99758604 | 0.95838535 |
| Slc14a2 | 0.99009361 | -0.9925055 | 0.031 | 0.031 | 1 | 0.79600877 | -0.9472852 | 0.017 | 0.021 | 1 | 0.99009361 | 0.95838758 |
| Cldn15 | 0.79666623 | 0.42253204 | 0.031 | 0.026 | 1 | 0.90837471 | -0.961932 | 0.05 | 0.052 | 1 | 0.90837471 | 0.95865538 |
| Lmtk3 | 0.79666623 | 0.42253204 | 0.031 | 0.026 | 1 | 0.99396051 | -0.3098553 | 0.021 | 0.021 | 1 | 0.99396051 | 0.95865538 |
| Gm4566 | 0.79666623 | 0.42253204 | 0.031 | 0.026 | 1 | 0.79810697 | -1.5322477 | 0.017 | 0.021 | 1 | 0.79810697 | 0.95865538 |
| Btk | 0.79846236 | 0.13302542 | 0.021 | 0.026 | 1 | 0.99194729 | -2.0874628 | 0.021 | 0.021 | 1 | 0.99194729 | 0.95938258 |
| 6430550D23Rik | 0.79869018 | 0.32942263 | 0.031 | 0.026 | 1 | 0.82968402 | -0.4618584 | 0.025 | 0.021 | 1 | 0.82968402 | 0.95947435 |
| Nfatc4 | 0.97283264 | -0.3996808 | 0.177 | 0.171 | 1 | 0.79897118 | -0.4050125 | 0.277 | 0.24 | 1 | 0.97283264 | 0.95958741 |
| Gm52870 | 0.99758604 | -0.2148979 | 0.021 | 0.021 | 1 | 0.79918975 | -0.7952821 | 0.038 | 0.031 | 1 | 0.99758604 | 0.95967524 |
| Rrh | 0.80274368 | 0.20013961 | 0.031 | 0.026 | 1 | 0.80020906 | -0.7248928 | 0.017 | 0.021 | 1 | 0.80274368 | 0.96008358 |
| Gm19303 | 0.82167605 | 0.54806292 | 0.062 | 0.057 | 1 | 0.80020906 | -0.7248928 | 0.017 | 0.021 | 1 | 0.82167605 | 0.96008358 |
| Gm16833 | 0.99603742 | 0.20013961 | 0.031 | 0.031 | 1 | 0.80020906 | -0.7248928 | 0.017 | 0.021 | 1 | 0.99603742 | 0.96008358 |
| 4933428G20Rik | 1 | 0.42253204 | 0.01 | 0.01 | 1 | 0.80020906 | -0.7248928 | 0.017 | 0.021 | 1 | 1 | 0.96008358 |
| Ly75 | 0.8007213 | 0.15949763 | 0.031 | 0.026 | 1 | 0.87851725 | -0.8948178 | 0.013 | 0.01 | 1 | 0.87851725 | 0.960288 |
| Rfk | 0.80139438 | 0.1835249 | 0.469 | 0.472 | 1 | 0.87827876 | -0.2039218 | 0.563 | 0.604 | 1 | 0.87827876 | 0.96055581 |
| Dynlt2a2 | 0.84739935 | 0.27052894 | 0.062 | 0.057 | 1 | 0.80191518 | -0.5728897 | 0.076 | 0.083 | 1 | 0.84739935 | 0.9607624 |
| Gm35315 | 0.80274368 | 0.20013961 | 0.031 | 0.026 | 1 | 0.85710544 | -1.7248928 | 0.008 | 0.01 | 1 | 0.85710544 | 0.96108994 |
| 1700030C12Rik | 0.80274368 | 0.20013961 | 0.031 | 0.026 | 1 | 0.98992944 | -0.1723517 | 0.042 | 0.042 | 1 | 0.98992944 | 0.96108994 |
| Fancb | 0.80274368 | 0.20013961 | 0.031 | 0.026 | 1 | 0.86613737 | -0.3098553 | 0.008 | 0.01 | 1 | 0.86613737 | 0.96108994 |
| Gm47512 | 0.86441475 | -0.1079827 | 0.052 | 0.047 | 1 | 0.80442048 | -0.5728897 | 0.017 | 0.021 | 1 | 0.86441475 | 0.96174865 |
| Tbck | 0.98936713 | 0.20013961 | 0.271 | 0.28 | 1 | 0.80443272 | -0.4366853 | 0.437 | 0.375 | 1 | 0.98936713 | 0.96175344 |
| Gpr160 | 0.80479144 | 0.15949763 | 0.031 | 0.026 | 1 | 1 | -0.8244284 | 0.021 | 0.021 | 1 | 1 | 0.96189362 |
| Utrn | 0.81651622 | 0.16223001 | 0.979 | 0.984 | 1 | 0.80594156 | -0.28878 | 0.987 | 0.979 | 1 | 0.81651622 | 0.96234132 |
| Faah | 0.82705505 | -0.7087125 | 0.042 | 0.047 | 1 | 0.8086289 | -0.3098553 | 0.017 | 0.021 | 1 | 0.82705505 | 0.9633771 |
| D430018E03Rik | 0.9927582 | 0.42253204 | 0.021 | 0.021 | 1 | 0.8086289 | -0.3098553 | 0.017 | 0.021 | 1 | 0.9927582 | 0.9633771 |
| Gm16751 | 0.82691519 | 0.13302542 | 0.042 | 0.036 | 1 | 0.80972361 | -1.1843244 | 0.038 | 0.031 | 1 | 0.82691519 | 0.9637949 |
| Ercc3 | 0.92894729 | -0.4187702 | 0.146 | 0.145 | 1 | 0.81102138 | -0.6217993 | 0.214 | 0.188 | 1 | 0.92894729 | 0.96428708 |
| Arhgap35 | 0.81114094 | -0.243467 | 0.521 | 0.503 | 1 | 0.92897422 | -0.194378 | 0.765 | 0.792 | 1 | 0.92897422 | 0.96433226 |
| Serpinb5 | 0.99758609 | 0.27052894 | 0.021 | 0.021 | 1 | 0.81129592 | 0.10518224 | 0.025 | 0.021 | 1 | 0.99758609 | 0.96439077 |
| ENSMUSG00000121439 | 1 | 0.42253204 | 0.01 | 0.01 | 1 | 0.81129592 | 0.27510724 | 0.025 | 0.021 | 1 | 1 | 0.96439077 |
| Dhrs9 | 0.81149229 | 3.17741954 | 0.021 | 0.026 | 1 | 0.88263976 | 0.27510724 | 0.038 | 0.042 | 1 | 0.88263976 | 0.96446484 |
| AW046200 | 0.81210369 | -0.4254649 | 0.094 | 0.083 | 1 | 0.82828167 | -1.1399303 | 0.071 | 0.062 | 1 | 0.82828167 | 0.96469498 |
| Ptgfrn | 0.99110032 | -0.235362 | 0.24 | 0.228 | 1 | 0.81340372 | -0.6577786 | 0.168 | 0.167 | 1 | 0.99110032 | 0.96518183 |
| Zfp365 | 0.98414882 | 0.52206771 | 0.031 | 0.031 | 1 | 0.81403482 | -0.4167705 | 0.046 | 0.052 | 1 | 0.98414882 | 0.96541695 |
| Anks1b | 0.81533635 | -1.6096006 | 0.229 | 0.223 | 1 | 0.85255161 | -0.7248928 | 0.202 | 0.198 | 1 | 0.85255161 | 0.96589934 |
| Fras1 | 0.8163117 | 0.32942263 | 0.104 | 0.098 | 1 | 0.85591384 | -0.7504279 | 0.139 | 0.125 | 1 | 0.85591384 | 0.96625861 |
| Trac | 0.81708725 | -0.577468 | 0.031 | 0.036 | 1 | 0.91613473 | -0.194378 | 0.034 | 0.031 | 1 | 0.91613473 | 0.96654293 |
| Emc9 | 0.81708725 | -0.6929452 | 0.031 | 0.036 | 1 | 0.92416907 | -0.5728897 | 0.034 | 0.031 | 1 | 0.92416907 | 0.96654293 |
| Donson | 0.87554892 | -0.1624305 | 0.062 | 0.067 | 1 | 0.81759614 | 0.21370669 | 0.08 | 0.073 | 1 | 0.87554892 | 0.96672883 |
| Gpam | 0.88161007 | -0.7533178 | 0.146 | 0.145 | 1 | 0.81762383 | -0.1034044 | 0.105 | 0.115 | 1 | 0.88161007 | 0.96673893 |
| Zswim7 | 0.81777767 | -0.2762984 | 0.198 | 0.202 | 1 | 0.96409272 | -0.4954109 | 0.164 | 0.156 | 1 | 0.96409272 | 0.96679502 |
| B4galt3 | 0.84051124 | 0.2091284 | 0.188 | 0.181 | 1 | 0.81812021 | -0.1017471 | 0.231 | 0.208 | 1 | 0.84051124 | 0.96691974 |
| Gm36411 | 0.8181385 | 0.42253204 | 0.042 | 0.036 | 1 | 0.87066325 | -0.3098553 | 0.013 | 0.01 | 1 | 0.87066325 | 0.96692639 |
| Gdf9 | 0.81989071 | 0.13302542 | 0.042 | 0.036 | 1 | 0.85762418 | -0.5728897 | 0.038 | 0.042 | 1 | 0.85762418 | 0.96756064 |
| Ap3m2 | 0.83041404 | 0.15949763 | 0.042 | 0.036 | 1 | 0.82048279 | -0.5728897 | 0.025 | 0.021 | 1 | 0.83041404 | 0.96777357 |
| Pik3cd | 0.82165076 | 0.49292136 | 0.042 | 0.036 | 1 | 0.96979028 | -1.1723517 | 0.042 | 0.042 | 1 | 0.96979028 | 0.96819155 |
| Dhdds | 0.82189169 | 0.10060394 | 0.396 | 0.378 | 1 | 0.96107162 | -0.5936482 | 0.286 | 0.25 | 1 | 0.96107162 | 0.96827743 |
| Gm28729 | 0.99009359 | -0.577468 | 0.031 | 0.031 | 1 | 0.82231997 | -0.4618584 | 0.025 | 0.021 | 1 | 0.99009359 | 0.96842981 |
| Iqch | 0.8581186 | -1.0368996 | 0.052 | 0.047 | 1 | 0.82232217 | 0.42711033 | 0.025 | 0.021 | 1 | 0.8581186 | 0.96843059 |
| Ttc26 | 0.82258416 | -0.1624305 | 0.031 | 0.036 | 1 | 0.88545586 | -1.1082214 | 0.071 | 0.073 | 1 | 0.88545586 | 0.96852362 |

| | | | | | | | | | | | | |
|---|---|---|---|---|---|---|---|---|---|---|---|---|
| Gm56637 | 0.82258416 | -0.1624305 | 0.031 | 0.036 | 1 | 0.91994491 | -0.6468902 | 0.067 | 0.062 | 1 | 0.91994491 | 0.96852362 |
| Bmp1 | 0.83818225 | -0.2977548 | 0.51 | 0.492 | 1 | 0.82271991 | -0.5206224 | 0.504 | 0.375 | 1 | 0.83818225 | 0.96857177 |
| Pwwp3a | 0.82328074 | 0.11440974 | 0.198 | 0.187 | 1 | 0.95933035 | -0.2667865 | 0.214 | 0.208 | 1 | 0.95933035 | 0.9687703 |
| 9430038I01Rik | 0.8619039 | 0.36113149 | 0.146 | 0.14 | 1 | 0.82370294 | -1.2136399 | 0.113 | 0.115 | 1 | 0.8619039 | 0.96891935 |
| Cgas | 0.85044533 | -0.4872702 | 0.052 | 0.057 | 1 | 0.82415676 | -0.6317834 | 0.025 | 0.021 | 1 | 0.85044533 | 0.96907916 |
| Lurap1 | 0.82443929 | -0.130009 | 0.031 | 0.036 | 1 | 0.9959736 | -0.3098553 | 0.021 | 0.021 | 1 | 0.9959736 | 0.96917844 |
| Gm43625 | 0.99603742 | 0.20013961 | 0.031 | 0.031 | 1 | 0.8259932 | -0.8244284 | 0.025 | 0.021 | 1 | 0.99603742 | 0.96972163 |
| Ctss | 0.82627679 | -0.8855903 | 0.031 | 0.036 | 1 | 0.96398894 | -1.2318528 | 0.067 | 0.062 | 1 | 0.96398894 | 0.96982025 |
| Smarcad1 | 0.82809709 | 0.16946202 | 0.51 | 0.554 | 1 | 0.945381 | -0.1854449 | 0.702 | 0.635 | 1 | 0.945381 | 0.97044939 |
| Kics2 | 0.83041404 | 0.15949763 | 0.042 | 0.036 | 1 | 0.91453215 | -0.1399303 | 0.034 | 0.031 | 1 | 0.91453215 | 0.9712406 |
| Tsbp1 | 0.83041404 | 0.15949763 | 0.042 | 0.036 | 1 | 0.97554341 | -0.7248928 | 0.042 | 0.042 | 1 | 0.97554341 | 0.9712406 |
| Med26 | 0.87694944 | -0.2382619 | 0.479 | 0.446 | 1 | 0.83193037 | -0.315017 | 0.647 | 0.562 | 1 | 0.87694944 | 0.9717526 |
| Gm16675 | 0.83546978 | 0.15949763 | 0.031 | 0.036 | 1 | 0.87851725 | -1.6317834 | 0.013 | 0.01 | 1 | 0.87851725 | 0.97292981 |
| Gm16279 | 0.83566413 | -0.7998604 | 0.052 | 0.057 | 1 | 0.91460443 | 0.10518224 | 0.055 | 0.052 | 1 | 0.91460443 | 0.97299372 |
| Manba | 0.87717568 | 0.25652208 | 0.167 | 0.171 | 1 | 0.83645684 | -0.6006347 | 0.218 | 0.219 | 1 | 0.87717568 | 0.97325364 |
| Gm15458 | 0.83676741 | -0.2555399 | 0.042 | 0.047 | 1 | 0.96681282 | -0.6317834 | 0.055 | 0.052 | 1 | 0.96681282 | 0.97335512 |
| Noxo1 | 0.99379628 | 0.13302542 | 0.052 | 0.052 | 1 | 0.83706049 | -1.0874628 | 0.025 | 0.021 | 1 | 0.99379628 | 0.97345072 |
| Nnat | 0.99758604 | -0.577468 | 0.021 | 0.021 | 1 | 0.83706049 | -1.3098553 | 0.025 | 0.021 | 1 | 0.99758604 | 0.97345072 |
| Gm56492 | 0.83746085 | -0.130009 | 0.042 | 0.036 | 1 | 0.98848961 | -0.1172102 | 0.042 | 0.042 | 1 | 0.98848961 | 0.97358102 |
| Fam32a | 0.86176908 | -0.1624305 | 0.385 | 0.378 | 1 | 0.83809052 | -0.1998017 | 0.471 | 0.458 | 1 | 0.86176908 | 0.97378532 |
| Nlgn2 | 0.83934145 | -0.3807761 | 0.271 | 0.269 | 1 | 0.89406435 | -0.6430987 | 0.387 | 0.344 | 1 | 0.89406435 | 0.97418883 |
| Gm41349 | 0.84081505 | 0.27052894 | 0.052 | 0.047 | 1 | 0.85710544 | -1.3098553 | 0.008 | 0.01 | 1 | 0.85710544 | 0.97466015 |
| Pde11a | 0.98133981 | -0.1868835 | 0.208 | 0.202 | 1 | 0.84113126 | -0.4431218 | 0.101 | 0.104 | 1 | 0.98133981 | 0.97476072 |
| Klhl7 | 0.84264448 | -0.1904448 | 0.562 | 0.554 | 1 | 0.86297546 | -0.611236 | 0.576 | 0.573 | 1 | 0.86297546 | 0.97523924 |
| Gm33046 | 0.84285787 | 0.35541784 | 0.031 | 0.036 | 1 | 0.87109831 | -0.3838558 | 0.046 | 0.042 | 1 | 0.87109831 | 0.97530635 |
| Ak4 | 0.87822318 | 0.44900425 | 0.312 | 0.311 | 1 | 0.84325219 | -0.5443205 | 0.181 | 0.156 | 1 | 0.87822318 | 0.97543012 |
| Trpt1 | 0.98759716 | -0.2148979 | 0.104 | 0.104 | 1 | 0.84387055 | -0.3098553 | 0.067 | 0.073 | 1 | 0.98759716 | 0.9756236 |
| Efcab5 | 0.84451844 | -0.3710171 | 0.042 | 0.036 | 1 | 0.87086793 | -0.2103196 | 0.038 | 0.042 | 1 | 0.87086793 | 0.97582548 |
| 5730409E04Rik | 0.85795684 | 0.13302542 | 0.042 | 0.047 | 1 | 0.84641179 | 0.27510724 | 0.046 | 0.042 | 1 | 0.85795684 | 0.97641066 |
| Ttc23l | 0.97722938 | -0.2335136 | 0.062 | 0.062 | 1 | 0.8542058 | -0.8441917 | 0.101 | 0.104 | 1 | 0.97722938 | 0.97874405 |
| Rad51ap2 | 0.99758604 | -0.2148979 | 0.021 | 0.021 | 1 | 0.85710544 | -1.3098553 | 0.008 | 0.01 | 1 | 0.99758604 | 0.97958114 |
| Ccdc96 | 1 | 0.42253204 | 0.01 | 0.01 | 1 | 0.85710544 | -1.7248928 | 0.008 | 0.01 | 1 | 1 | 0.97958114 |
| Entrep2 | 1 | -0.3144336 | 0.01 | 0.01 | 1 | 0.85710544 | -1.3098553 | 0.008 | 0.01 | 1 | 1 | 0.97958114 |
| Gm40630 | 1 | -0.7998604 | 0.01 | 0.01 | 1 | 0.85710544 | -3.4253325 | 0.008 | 0.01 | 1 | 1 | 0.97958114 |
| Slc1a3 | 0.99801866 | -1.4519371 | 0.031 | 0.031 | 1 | 0.85911566 | -0.4618584 | 0.038 | 0.042 | 1 | 0.99801866 | 0.9801516 |
| Thumpd2 | 0.85960946 | 0.11440974 | 0.042 | 0.047 | 1 | 0.93762985 | 0.22619764 | 0.076 | 0.073 | 1 | 0.93762985 | 0.9802905 |
| Prcp | 0.85962858 | -0.4950058 | 0.104 | 0.109 | 1 | 0.88269663 | -1.0397661 | 0.139 | 0.135 | 1 | 0.88269663 | 0.98029587 |
| Kcnmb3 | 1 | 0.42253204 | 0.01 | 0.01 | 1 | 0.86011448 | -0.8948178 | 0.008 | 0.01 | 1 | 1 | 0.98043204 |
| Tmbim7 | 1 | 0.42253204 | 0.01 | 0.01 | 1 | 0.86011448 | -0.8244284 | 0.008 | 0.01 | 1 | 1 | 0.98043204 |
| A430035B10Rik | 0.99603736 | -0.130009 | 0.031 | 0.031 | 1 | 0.86203455 | -0.6577786 | 0.038 | 0.042 | 1 | 0.99603736 | 0.98096553 |
| Txlnb | 0.993208 | 1.00749454 | 0.01 | 0.01 | 1 | 0.86312513 | -0.3098553 | 0.008 | 0.01 | 1 | 0.993208 | 0.98126527 |
| Zfp474 | 0.99758609 | 0.27052894 | 0.021 | 0.021 | 1 | 0.86312513 | -0.7248928 | 0.008 | 0.01 | 1 | 0.99758609 | 0.98126527 |
| Gm29128 | 1 | 0.42253204 | 0.01 | 0.01 | 1 | 0.86312513 | -0.7248928 | 0.008 | 0.01 | 1 | 1 | 0.98126527 |
| Myh7b | 1 | 0.42253204 | 0.01 | 0.01 | 1 | 0.86312513 | -0.7248928 | 0.008 | 0.01 | 1 | 1 | 0.98126527 |
| Fmnl1 | 1 | 0.42253204 | 0.01 | 0.01 | 1 | 0.86312513 | -0.7248928 | 0.008 | 0.01 | 1 | 1 | 0.98126527 |
| Gm56602 | 1 | -0.3144336 | 0.01 | 0.01 | 1 | 0.86312513 | -0.7248928 | 0.008 | 0.01 | 1 | 1 | 0.98126527 |
| Gm50019 | 1 | 0.42253204 | 0.01 | 0.01 | 1 | 0.86312513 | -0.7248928 | 0.008 | 0.01 | 1 | 1 | 0.98126527 |
| Hspe1-rs1 | 1 | 0.42253204 | 0.01 | 0.01 | 1 | 0.86312513 | -0.7248928 | 0.008 | 0.01 | 1 | 1 | 0.98126527 |
| Cfap47 | 1 | 0.27052894 | 0.01 | 0.01 | 1 | 0.86543325 | 0.27510724 | 0.013 | 0.01 | 1 | 1 | 0.98189179 |
| Gm10101 | 1 | 0.42253204 | 0.01 | 0.01 | 1 | 0.86613737 | -0.3098553 | 0.008 | 0.01 | 1 | 1 | 0.98208079 |
| Cybb | 1 | -2.1079827 | 0.01 | 0.01 | 1 | 0.86613737 | 0.27510724 | 0.008 | 0.01 | 1 | 1 | 0.98208079 |
| Hps6 | 1 | 0.42253204 | 0.01 | 0.01 | 1 | 0.86695837 | -0.194378 | 0.046 | 0.042 | 1 | 1 | 0.98229993 |
| Serpini1 | 0.86729999 | -0.3364599 | 0.083 | 0.088 | 1 | 0.88346544 | -1.2103196 | 0.118 | 0.104 | 1 | 0.88346544 | 0.98239071 |
| Ugcg | 0.94714173 | -0.186678 | 0.406 | 0.404 | 1 | 0.86982185 | -0.2249664 | 0.525 | 0.469 | 1 | 0.94714173 | 0.98305365 |
| Ngb | 0.99275828 | -0.577468 | 0.021 | 0.021 | 1 | 0.87066325 | -0.3098553 | 0.013 | 0.01 | 1 | 0.99275828 | 0.98327201 |
| Jsrp1 | 0.99758609 | 0.27052894 | 0.021 | 0.021 | 1 | 0.87066325 | -0.3098553 | 0.013 | 0.01 | 1 | 0.99758609 | 0.98327201 |
| A230103J11Rik | 1 | 0.42253204 | 0.01 | 0.01 | 1 | 0.87066325 | -0.3098553 | 0.013 | 0.01 | 1 | 1 | 0.98327201 |
| Gm10629 | 1 | 0.42253204 | 0.01 | 0.01 | 1 | 0.87066325 | -0.3098553 | 0.013 | 0.01 | 1 | 1 | 0.98327201 |
| 1700003D09Rik | 1 | 0.42253204 | 0.01 | 0.01 | 1 | 0.87066325 | -0.3098553 | 0.013 | 0.01 | 1 | 1 | 0.98327201 |
| Gm26531 | 1 | 0.42253204 | 0.01 | 0.01 | 1 | 0.87066325 | -0.3098553 | 0.013 | 0.01 | 1 | 1 | 0.98327201 |
| Gm48342 | 1 | 0.42253204 | 0.01 | 0.01 | 1 | 0.87066325 | -0.3098553 | 0.013 | 0.01 | 1 | 1 | 0.98327201 |
| Myrip | 0.87089632 | 0.15949763 | 0.083 | 0.078 | 1 | 0.87328024 | 0.59703533 | 0.013 | 0.01 | 1 | 0.87328024 | 0.98333224 |
| Nrip3 | 0.88135348 | 0.17350449 | 0.146 | 0.14 | 1 | 0.87260874 | -0.5322477 | 0.092 | 0.083 | 1 | 0.88135348 | 0.98377147 |
| Gm57036 | 0.99758609 | 0.27052894 | 0.021 | 0.021 | 1 | 0.87589833 | -0.5728897 | 0.013 | 0.01 | 1 | 0.99758609 | 0.98459878 |
| Kcnab2 | 0.9927582 | 0.42253204 | 0.021 | 0.021 | 1 | 0.87851725 | -0.8948178 | 0.013 | 0.01 | 1 | 0.9927582 | 0.98524194 |
| Acot4 | 1 | 0.42253204 | 0.01 | 0.01 | 1 | 0.87851725 | -0.8948178 | 0.013 | 0.01 | 1 | 1 | 0.98524194 |
| Gm50209 | 1 | 0.42253204 | 0.01 | 0.01 | 1 | 0.87851725 | -1.6317834 | 0.013 | 0.01 | 1 | 1 | 0.98524194 |
| Fam124a | 0.99758609 | 0.27052894 | 0.021 | 0.021 | 1 | 0.87851807 | -0.9879272 | 0.013 | 0.01 | 1 | 0.99758609 | 0.98524214 |
| Zfp963 | 0.87907906 | -0.2726134 | 0.062 | 0.057 | 1 | 0.97220162 | -0.352924 | 0.118 | 0.115 | 1 | 0.97220162 | 0.98537813 |
| H2-Ab1 | 0.8831674 | -0.4269083 | 0.115 | 0.119 | 1 | 0.87934103 | -0.5025003 | 0.046 | 0.042 | 1 | 0.8831674 | 0.98544141 |
| Laptm5 | 1 | 0.59245704 | 0.01 | 0.01 | 1 | 0.88164483 | 2.83310269 | 0.046 | 0.052 | 1 | 1 | 0.98599205 |
| Foxj1 | 0.97622723 | 1.53800925 | 0.031 | 0.031 | 1 | 0.88485604 | -0.5443205 | 0.046 | 0.042 | 1 | 0.97622723 | 0.98674187 |
| Ankrd33b | 0.98792413 | 0.27052894 | 0.042 | 0.041 | 1 | 0.88485625 | -0.5213594 | 0.046 | 0.042 | 1 | 0.98792413 | 0.98674192 |
| Me3 | 0.89995565 | 0.30705482 | 0.083 | 0.088 | 1 | 0.88622972 | -0.5433454 | 0.223 | 0.219 | 1 | 0.89995565 | 0.98705632 |
| Mnt | 0.88667383 | -0.1217885 | 0.271 | 0.269 | 1 | 0.93034843 | -0.8174589 | 0.261 | 0.229 | 1 | 0.93034843 | 0.98715718 |
| 1600010M07Rik | 0.93530263 | -0.1670032 | 0.271 | 0.264 | 1 | 0.8877216 | -0.8114018 | 0.303 | 0.25 | 1 | 0.93530263 | 0.98739356 |
| Gm11944 | 0.88834602 | -0.1624305 | 0.156 | 0.145 | 1 | 0.98985757 | -0.6798049 | 0.197 | 0.188 | 1 | 0.98985757 | 0.98753339 |
| Wee1 | 0.95748366 | -0.4238448 | 0.271 | 0.259 | 1 | 0.88848876 | -0.9332919 | 0.214 | 0.198 | 1 | 0.95748366 | 0.98756524 |
| Dhrs11 | 0.98539043 | -0.2040096 | 0.146 | 0.145 | 1 | 0.88888504 | -0.9354597 | 0.126 | 0.115 | 1 | 0.98539043 | 0.98765346 |
| Cox11 | 0.89390184 | 0.11672361 | 0.323 | 0.321 | 1 | 0.88941041 | -0.5401529 | 0.345 | 0.292 | 1 | 0.89390184 | 0.98776994 |
| Nln | 0.890518 | -0.1624305 | 0.115 | 0.119 | 1 | 0.89052059 | -0.1617566 | 0.172 | 0.177 | 1 | 0.89052059 | 0.98801369 |
| Pamr1 | 0.94961253 | -0.1580915 | 0.219 | 0.218 | 1 | 0.89643922 | -0.2779124 | 0.185 | 0.177 | 1 | 0.94961253 | 0.98927517 |
| Camp | 0.99758609 | 0.27052894 | 0.021 | 0.021 | 1 | 0.90722351 | -0.7248928 | 0.067 | 0.062 | 1 | 0.99758609 | 0.99139252 |
| Lif | 0.9980186 | -0.3710171 | 0.031 | 0.031 | 1 | 0.90971018 | 1.49749966 | 0.034 | 0.031 | 1 | 0.9980186 | 0.99184775 |
| Gm32687 | 0.99009299 | 0.32942263 | 0.031 | 0.031 | 1 | 0.91453215 | -0.1399303 | 0.034 | 0.031 | 1 | 0.99009299 | 0.99269525 |
| Vsig10l | 1 | -0.3144336 | 0.01 | 0.01 | 1 | 0.91453215 | -0.1399303 | 0.034 | 0.031 | 1 | 1 | 0.99269525 |

| | | | | | | | | | | | | |
|---|---|---|---|---|---|---|---|---|---|---|---|---|
| Pcdhgb2 | 1 | -0.1624305 | 0.031 | 0.031 | 1 | 0.91891525 | -1.1172102 | 0.029 | 0.031 | 1 | 1 | 0.99342526 |
| Arid4b | 0.91934395 | -0.1278732 | 0.99 | 0.995 | 1 | 0.98898624 | -0.2026303 | 0.996 | 0.979 | 1 | 0.98898624 | 0.9934946 |
| Zfp286 | 0.97828113 | 0.35541784 | 0.052 | 0.052 | 1 | 0.92483664 | -0.2703269 | 0.101 | 0.104 | 1 | 0.97828113 | 0.99435047 |
| Gm57325 | 0.97518595 | 0.27052894 | 0.052 | 0.052 | 1 | 0.92738405 | -0.4618584 | 0.034 | 0.031 | 1 | 0.97518595 | 0.99472692 |
| Gm56546 | 0.99758609 | 0.27052894 | 0.021 | 0.021 | 1 | 0.92738405 | -0.4618584 | 0.034 | 0.031 | 1 | 0.99758609 | 0.99472692 |
| Syt1 | 1 | 0.42253204 | 0.01 | 0.01 | 1 | 0.92899987 | -0.2103196 | 0.034 | 0.031 | 1 | 1 | 0.99495898 |
| Mir670hg | 0.99758604 | -0.407543 | 0.021 | 0.021 | 1 | 0.92902634 | -0.3098553 | 0.029 | 0.031 | 1 | 0.99758604 | 0.99496274 |
| Gm52992 | 0.99758609 | 0.27052894 | 0.021 | 0.021 | 1 | 0.94385422 | -0.3098553 | 0.055 | 0.052 | 1 | 0.99758609 | 0.99684765 |
| Gm16835 | 0.98718788 | 0.24195979 | 0.062 | 0.062 | 1 | 0.94494948 | 0.21370669 | 0.05 | 0.052 | 1 | 0.98718788 | 0.99696944 |
| C330018D20Rik | 0.98792413 | 0.27052894 | 0.042 | 0.041 | 1 | 0.95673288 | -0.3098553 | 0.05 | 0.052 | 1 | 0.98792413 | 0.99812796 |
| Ccnl1 | 0.96824382 | 0.10310893 | 0.917 | 0.979 | 1 | 0.99949806 | -0.1603791 | 0.975 | 0.969 | 1 | 0.99949806 | 0.99899155 |
| Mtx2 | 0.99321121 | -0.1708427 | 0.292 | 0.28 | 1 | 0.97641972 | -0.6317834 | 0.403 | 0.354 | 1 | 0.99321121 | 0.99944397 |
| C330011M18Rik | 0.99603742 | 0.20013961 | 0.031 | 0.031 | 1 | 0.98992944 | -0.1723517 | 0.042 | 0.042 | 1 | 0.99603742 | 0.99989858 |
| Gm38505 | 1 | 0.42253204 | 0.01 | 0.01 | 1 | 0.99194729 | -1.0874628 | 0.021 | 0.021 | 1 | 1 | 0.99993515 |

**scRNA seq_Col2a1/Acan enriched cluster subset: cluster 1**

| | cKO cart_p_val | cKO cart_avg_log2FC | cKO cart_pct.1 | cKO cart_pct.2 | cKO cart_p_val_adj | WT cart_p_val | WT cart_avg_log2FC | WT cart_pct.1 | WT cart_pct.2 | WT cart_p_val_adj | max_pval | minimump_p_val |
|---|---|---|---|---|---|---|---|---|---|---|---|---|
| Serpina3n | 2.52852E-44 | 4.945245798 | 0.933 | 0.135 | 6.14228E-40 | 9.54688E-18 | 5.399012075 | 0.7 | 0.043 | 2.31913E-13 | 9.54688E-18 | 5.05704E-44 |
| Chst5 | 8.52783E-44 | 4.532899821 | 0.933 | 0.171 | 2.07158E-39 | 6.54057E-09 | 3.672147071 | 0.7 | 0.111 | 0.000158884 | 6.54057E-09 | 1.70557E-43 |
| Hsd11b1 | 1.79192E-41 | 4.775100723 | 0.899 | 0.153 | 4.35293E-37 | 6.4559E-16 | 5.655351829 | 0.5 | 0.022 | 1.56827E-11 | 6.4559E-16 | 3.58384E-41 |
| Col10a1 | 6.62453E-40 | 4.525934448 | 0.908 | 0.188 | 1.60923E-35 | 1.48822E-20 | 6.512572109 | 0.9 | 0.068 | 3.61519E-16 | 1.48822E-20 | 1.32491E-39 |
| AW112010 | 5.59846E-39 | 4.334608676 | 0.882 | 0.118 | 1.35998E-34 | 1.11789E-15 | 5.339850003 | 0.5 | 0.022 | 2.71558E-11 | 1.11789E-15 | 1.11969E-38 |
| Gna14 | 5.08248E-37 | 2.676460855 | 0.958 | 0.247 | 1.23463E-32 | 2.67911E-06 | 3.070723324 | 0.9 | 0.41 | 0.065080958 | 2.67911E-06 | 1.0165E-36 |
| A2m | 3.14333E-36 | 5.115972563 | 0.815 | 0.1 | 7.63578E-32 | 4.15709E-20 | 5.861196404 | 0.8 | 0.049 | 1.00984E-15 | 4.15709E-20 | 6.28666E-36 |
| Ppp1r1b | 3.39714E-35 | 3.802393413 | 0.899 | 0.241 | 8.25234E-31 | 9.41129E-05 | 2.361168724 | 0.9 | 0.367 | 1 | 9.41129E-05 | 6.79428E-35 |
| Mmp2 | 4.39569E-35 | -5.596040633 | 0.084 | 0.835 | 1.0678E-30 | 2.74833E-05 | -3.637429921 | 0.3 | 0.846 | 0.667623799 | 2.74833E-05 | 8.79138E-35 |
| F13a1 | 1.02614E-34 | 2.335801835 | 1 | 0.612 | 2.4927E-30 | 2.72178E-06 | 2.331489048 | 1 | 0.642 | 0.066117589 | 2.72178E-06 | 2.05228E-34 |
| Pla2g5 | 1.28169E-34 | 2.840637042 | 0.966 | 0.294 | 3.11349E-30 | 0.000515285 | 1.943497067 | 0.9 | 0.549 | 1 | 0.000515285 | 2.56339E-34 |
| Col12a1 | 3.34221E-34 | -4.701275601 | 0.387 | 0.935 | 8.11889E-30 | 5.94426E-07 | -6.095331899 | 0.2 | 0.944 | 0.014439795 | 5.94426E-07 | 6.68441E-34 |
| BC006965 | 5.623E-34 | 4.064488726 | 0.891 | 0.224 | 1.36594E-29 | 7.32524E-07 | 3.568118991 | 0.8 | 0.194 | 0.017794466 | 7.32524E-07 | 1.1246E-33 |
| Enpp2 | 2.37488E-33 | 2.461160246 | 0.992 | 0.841 | 5.76905E-29 | 9.65397E-08 | 3.261956906 | 1 | 0.642 | 0.002345143 | 9.65397E-08 | 4.74976E-33 |
| Alox12 | 3.27083E-33 | 5.319945009 | 0.706 | 0.035 | 7.94549E-29 | 1.0993E-19 | 5.713067326 | 0.6 | 0.025 | 2.67042E-15 | 1.0993E-19 | 6.54165E-33 |
| Cytl1 | 1.51595E-32 | 2.689832746 | 0.992 | 0.935 | 3.68255E-28 | 0.002565961 | 1.338346279 | 1 | 0.904 | 1 | 0.002565961 | 3.03191E-32 |
| Becn1 | 1.12988E-31 | 2.826417487 | 0.95 | 0.641 | 2.7447E-27 | 0.011664404 | 1.592517693 | 0.7 | 0.664 | 1 | 0.011664404 | 2.25976E-31 |
| Fstl1 | 5.92019E-31 | -4.092668 | 0.311 | 0.865 | 1.43813E-26 | 0.001112507 | -1.815784341 | 0.6 | 0.892 | 1 | 0.001112507 | 1.18404E-30 |
| F5 | 5.97598E-31 | 3.314614036 | 0.84 | 0.159 | 1.45168E-26 | 3.62358E-21 | 5.602884409 | 0.9 | 0.065 | 8.8024E-17 | 3.62358E-21 | 1.1952E-30 |
| Timp3 | 9.1579E-31 | 2.995465077 | 0.992 | 0.924 | 2.22464E-26 | 0.000170301 | 2.373977617 | 1 | 0.929 | 1 | 0.000170301 | 1.83158E-30 |
| Sdc4 | 1.55148E-30 | 1.449928533 | 1 | 1 | 3.76884E-26 | 0.001278233 | 0.659311131 | 1 | 1 | 1 | 0.001278233 | 3.10295E-30 |
| Syne1 | 1.84193E-30 | -2.635823287 | 0.647 | 0.971 | 4.47443E-26 | 0.000767486 | -1.91248691 | 0.8 | 0.988 | 1 | 0.000767486 | 3.68387E-30 |
| Hs6st3 | 5.51414E-30 | 5.230394545 | 0.664 | 0.035 | 1.3395E-25 | 8.52503E-25 | 8.142617655 | 0.6 | 0.015 | 2.0709E-20 | 8.52503E-25 | 1.10283E-29 |
| Fxyd3 | 8.78008E-30 | 3.527397213 | 0.908 | 0.494 | 2.13286E-25 | 4.3107E-06 | 2.348546202 | 1 | 0.556 | 0.104715615 | 4.3107E-06 | 1.75602E-29 |
| Slc2a1 | 2.3258E-29 | 3.165124015 | 0.941 | 0.559 | 5.64983E-25 | 0.000700161 | 3.096531743 | 0.8 | 0.546 | 1 | 0.000700161 | 4.6516E-29 |
| Loxl4 | 1.63202E-28 | 2.522102477 | 0.95 | 0.576 | 3.96449E-24 | 5.22831E-05 | 2.051356136 | 1 | 0.731 | 1 | 5.22831E-05 | 3.26403E-28 |
| Islr | 2.64854E-28 | -3.189294733 | 0.269 | 0.859 | 6.43384E-24 | 0.011081494 | -0.784271309 | 0.4 | 0.793 | 1 | 0.011081494 | 5.29709E-28 |
| Meltf | 5.16719E-28 | 1.807047125 | 1 | 0.541 | 1.25521E-23 | 3.4435E-06 | 2.068314403 | 1 | 0.667 | 0.08364959 | 3.4435E-06 | 1.03344E-27 |
| Scd1 | 5.84816E-28 | 3.445471391 | 0.924 | 0.394 | 1.42064E-23 | 1.96448E-08 | 3.952448071 | 1 | 0.438 | 0.000477211 | 1.96448E-08 | 1.16963E-27 |
| Syn3 | 1.35354E-27 | 3.202088325 | 0.866 | 0.318 | 3.28803E-23 | 0.001344555 | 3.391736745 | 0.6 | 0.238 | 1 | 0.001344555 | 2.70709E-27 |
| Atox1 | 2.34172E-27 | 2.432042128 | 0.992 | 0.935 | 5.6885E-23 | 0.01024047 | 1.124837112 | 1 | 0.907 | 1 | 0.01024047 | 4.68343E-27 |
| Aebp1 | 3.39742E-27 | -3.603424037 | 0.16 | 0.788 | 8.25302E-23 | 2.51884E-05 | -4.432258339 | 0.1 | 0.809 | 0.611876745 | 2.51884E-05 | 6.79484E-27 |
| Atp6v0e | 9.03541E-27 | 1.504377406 | 1 | 0.953 | 2.19488E-22 | 0.207174201 | 0.408362044 | 0.9 | 0.951 | 1 | 0.207174201 | 1.80708E-26 |
| Dlgap1 | 1.206E-26 | 3.307544177 | 0.782 | 0.165 | 2.92962E-22 | 0.000903618 | 1.171834591 | 0.8 | 0.293 | 1 | 0.000903618 | 2.412E-26 |
| Capns1 | 1.62549E-26 | 1.741499633 | 1 | 0.935 | 3.94865E-22 | 0.01917617 | 0.983851232 | 0.9 | 0.941 | 1 | 0.01917617 | 3.25098E-26 |
| Fap | 2.06868E-26 | -2.274895872 | 0.37 | 0.918 | 5.02525E-22 | 0.000800338 | -2.008569366 | 0.6 | 0.917 | 1 | 0.000800338 | 4.13737E-26 |
| Il1r1 | 2.95304E-26 | 2.243611851 | 0.916 | 0.312 | 7.17352E-22 | 0.004725457 | 2.276454922 | 0.6 | 0.278 | 1 | 0.004725457 | 5.90608E-26 |
| Rgs2 | 3.37605E-26 | 3.030472352 | 0.882 | 0.382 | 8.20109E-22 | 0.241383567 | -2.408342847 | 0.1 | 0.256 | 1 | 0.241383567 | 6.75209E-26 |
| Igfbp7 | 4.59905E-26 | -3.039339405 | 0.529 | 0.906 | 1.1172E-21 | 1.34868E-05 | -3.04943336 | 0.7 | 0.96 | 0.327620863 | 1.34868E-05 | 9.19809E-26 |
| Tgm2 | 5.55635E-26 | 1.899888515 | 0.992 | 0.6 | 1.34975E-21 | 7.01151E-06 | 2.367157349 | 1 | 0.534 | 0.170323702 | 7.01151E-06 | 1.11127E-25 |
| Tnfrsf11b | 5.59447E-26 | 1.973513494 | 0.983 | 0.6 | 1.35901E-21 | 0.000968186 | 1.569885044 | 1 | 0.466 | 1 | 0.000968186 | 1.11889E-25 |
| Lbp | 8.63406E-26 | 2.665439838 | 0.815 | 0.212 | 2.09739E-21 | 0.01018673 | 2.443930525 | 0.5 | 0.228 | 1 | 0.01018673 | 1.72681E-25 |
| Fndc1 | 9.97007E-26 | 2.533938498 | 0.815 | 0.188 | 2.42193E-21 | 7.92899E-05 | 2.592437621 | 0.8 | 0.324 | 1 | 7.92899E-05 | 1.99401E-25 |
| Stc2 | 2.13281E-25 | 3.055356225 | 0.857 | 0.318 | 5.18102E-21 | 0.000533648 | 2.936127817 | 0.6 | 0.201 | 1 | 0.000533648 | 4.26562E-25 |
| Jam2 | 3.06349E-25 | 2.204460828 | 0.924 | 0.506 | 7.44183E-21 | 0.002853865 | 1.438434537 | 0.9 | 0.367 | 1 | 0.002853865 | 6.12698E-25 |
| Chil1 | 5.3961E-25 | 3.375557261 | 0.899 | 0.418 | 1.31082E-20 | 0.000238696 | 2.853103797 | 0.9 | 0.491 | 1 | 0.000238696 | 1.07922E-24 |
| Rcan1 | 6.39584E-25 | 2.066899525 | 0.966 | 0.753 | 1.55368E-20 | 0.405090464 | 0.316910259 | 0.7 | 0.864 | 1 | 0.405090464 | 1.27917E-24 |
| Stag1 | 7.66896E-25 | -1.687261077 | 0.832 | 0.994 | 1.86294E-20 | 0.001145049 | -1.189040473 | 0.9 | 0.997 | 1 | 0.001145049 | 1.53379E-24 |
| S100b | 1.53364E-24 | 2.176280496 | 0.983 | 0.694 | 3.72553E-20 | 0.000155412 | 2.111031312 | 0.9 | 0.611 | 1 | 0.000155412 | 3.06729E-24 |
| Cst3 | 1.63811E-24 | 1.835062649 | 1 | 0.982 | 3.97929E-20 | 0.116471833 | 0.62235692 | 1 | 0.994 | 1 | 0.116471833 | 3.27622E-24 |
| Kif21a | 1.90362E-24 | 1.50697233 | 1 | 0.788 | 4.62428E-20 | 0.001996272 | 0.947041951 | 1 | 0.929 | 1 | 0.001996272 | 3.80725E-24 |
| Pir | 2.11808E-24 | 2.078656944 | 0.807 | 0.241 | 5.14524E-20 | 0.060349501 | 1.438695777 | 0.6 | 0.444 | 1 | 0.060349501 | 4.23616E-24 |
| Kcnq5 | 2.15421E-24 | -2.701511845 | 0.571 | 0.935 | 5.23301E-20 | 3.04042E-05 | -4.119737531 | 0.6 | 0.914 | 0.738578143 | 3.04042E-05 | 4.30843E-24 |
| Prkg1 | 2.27172E-24 | -3.076657696 | 0.496 | 0.906 | 5.51845E-20 | 8.80272E-06 | -3.32629612 | 0.3 | 0.963 | 0.213835607 | 8.80272E-06 | 4.54343E-24 |
| Uap1 | 5.16677E-24 | 1.791103421 | 0.992 | 0.865 | 1.25511E-19 | 0.229794581 | 0.230915631 | 0.9 | 0.901 | 1 | 0.229794581 | 1.03335E-23 |
| Fbxo2 | 5.2155E-24 | 4.145339363 | 0.689 | 0.147 | 1.26695E-19 | 0.000982467 | 2.293029146 | 0.6 | 0.231 | 1 | 0.000982467 | 1.0431E-23 |
| Luzp2 | 5.56008E-24 | 4.082257682 | 0.697 | 0.124 | 1.35065E-19 | 2.11212E-10 | 3.128104826 | 1 | 0.216 | 5.13076E-06 | 2.11212E-10 | 1.11202E-23 |
| Fscn1 | 6.01369E-24 | -3.980837743 | 0.101 | 0.7 | 1.46085E-19 | 0.001360613 | -3.094083118 | 0.1 | 0.651 | 1 | 0.001360613 | 1.20274E-23 |
| Grb10 | 7.98331E-24 | -1.597453445 | 0.807 | 0.971 | 1.93931E-19 | 4.2105E-05 | -2.156587712 | 0.7 | 0.966 | 1 | 4.2105E-05 | 1.59666E-23 |
| Syncrip | 1.15121E-23 | -1.490381709 | 0.857 | 0.971 | 2.79653E-19 | 0.00130169 | -1.107392207 | 0.9 | 0.988 | 1 | 0.00130169 | 2.30243E-23 |
| Sbspon | 1.33629E-23 | 5.049494991 | 0.571 | 0.053 | 3.24611E-19 | 3.65847E-07 | 3.715359138 | 0.7 | 0.145 | 0.008887156 | 3.65847E-07 | 2.67258E-23 |
| Car6 | 1.5073E-23 | 4.49946628 | 0.588 | 0.059 | 3.66152E-19 | 1.1674E-06 | 4.280956314 | 0.2 | 0.009 | 0.028358426 | 1.1674E-06 | 3.01459E-23 |
| Colgalt2 | 1.5324E-23 | 2.718229032 | 0.866 | 0.441 | 3.72251E-19 | 5.58714E-07 | 3.336451426 | 0.9 | 0.265 | 0.013572288 | 5.58714E-07 | 3.0648E-23 |
| Dst | 1.65259E-23 | 1.381733951 | 1 | 0.935 | 4.01447E-19 | 3.62809E-05 | 1.59025987 | 1 | 0.972 | 0.881336249 | 3.62809E-05 | 3.30518E-23 |
| Lmcd1 | 1.72717E-23 | 1.831274106 | 0.966 | 0.582 | 4.19564E-19 | 0.000442664 | 2.15317298 | 0.9 | 0.66 | 1 | 0.000442664 | 3.45434E-23 |
| Cdh19 | 1.61671E-11 | 2.659623506 | 0.345 | 0.041 | 3.92731E-07 | 2.38646E-23 | 5.532495081 | 0.5 | 0.009 | 5.79719E-19 | 1.61671E-11 | 4.77292E-23 |
| Marcks | 2.75774E-23 | -4.858727024 | 0.126 | 0.694 | 6.69911E-19 | 0.000145045 | -4.618702713 | 0 | 0.685 | 1 | 0.000145045 | 5.51549E-23 |
| Pcolce | 2.83619E-23 | -4.706530553 | 0.16 | 0.7 | 6.88967E-19 | 5.00547E-05 | -4.411484834 | 0.1 | 0.784 | 1 | 5.00547E-05 | 5.67238E-23 |
| Pla2g7 | 3.74077E-23 | 2.490050854 | 0.79 | 0.224 | 9.08708E-19 | 0.481805547 | 0.140177658 | 0.4 | 0.29 | 1 | 0.481805547 | 7.48154E-23 |
| Ucma | 3.82263E-23 | 1.534121866 | 1 | 0.759 | 9.28594E-19 | 0.102831798 | 0.428086733 | 1 | 0.762 | 1 | 0.102831798 | 7.64527E-23 |
| Lox | 4.74434E-23 | 2.663647299 | 0.924 | 0.559 | 1.15249E-18 | 0.01570019 | 0.340294982 | 1 | 0.719 | 1 | 0.01570019 | 9.48868E-23 |
| Pxdc1 | 5.05904E-23 | 2.206178234 | 0.924 | 0.518 | 1.22894E-18 | 0.003824252 | 1.39698011 | 0.8 | 0.522 | 1 | 0.003824252 | 1.01181E-22 |
| Loxl2 | 9.79064E-23 | 1.988721291 | 0.891 | 0.376 | 2.37834E-18 | 6.90669E-06 | 3.147557188 | 0.7 | 0.167 | 0.167777388 | 6.90669E-06 | 1.95813E-22 |
| Ccn2 | 1.01761E-22 | 1.462045972 | 1 | 0.965 | 2.47198E-18 | 0.000730314 | 1.166811894 | 1 | 0.864 | 1 | 0.000730314 | 2.03522E-22 |
| Aldoc | 1.09611E-22 | 3.487806263 | 0.672 | 0.135 | 2.66268E-18 | 0.157872685 | 1.744903414 | 0.3 | 0.164 | 1 | 0.157872685 | 2.19223E-22 |
| S100a13 | 1.74041E-22 | 1.551646606 | 0.941 | 0.765 | 4.2278E-18 | 0.053062494 | 1.20515349 | 0.9 | 0.806 | 1 | 0.053062494 | 3.48082E-22 |
| Usp34 | 2.21083E-22 | -1.299650648 | 0.924 | 0.988 | 5.37054E-18 | 0.000428733 | -1.07918614 | 1 | 0.997 | 1 | 0.000428733 | 4.42165E-22 |
| Pantr1 | 2.65148E-22 | 3.610669762 | 0.723 | 0.212 | 6.44098E-18 | 0.232056472 | 0.485426827 | 0.5 | 0.284 | 1 | 0.232056472 | 5.30296E-22 |
| Btg1 | 3.83476E-22 | 1.203917127 | 1 | 0.994 | 9.31541E-18 | 0.799132612 | 0.435613861 | 1 | 1 | 1 | 0.799132612 | 7.66953E-22 |
| Fgf2 | 4.42093E-22 | 1.865534085 | 0.958 | 0.512 | 1.07393E-17 | 0.024583096 | 0.698181372 | 1 | 0.596 | 1 | 0.024583096 | 8.84186E-22 |
| Peg3 | 4.6489E-22 | -1.608683301 | 0.958 | 0.976 | 1.12931E-17 | 0.000800525 | -1.352016859 | 1 | 0.994 | 1 | 0.000800525 | 9.2978E-22 |
| Ogn | 6.48209E-22 | -3.476602977 | 0.227 | 0.741 | 1.57463E-17 | 0.000415414 | -3.725229486 | 0.4 | 0.787 | 1 | 0.000415414 | 1.29642E-21 |
| Fbln7 | 9.23671E-22 | -4.866248611 | 0.076 | 0.624 | 2.24378E-17 | 0.004042892 | -1.833827133 | 0.4 | 0.781 | 1 | 0.004042892 | 1.84734E-21 |
| Tesc | 1.04765E-21 | 5.060766103 | 0.529 | 0.047 | 2.54496E-17 | 0.00029417 | 1.611929548 | 0.4 | 0.071 | 1 | 0.00029417 | 2.09531E-21 |
| Col4a4 | 1.24212E-21 | 7.44531051 | 0.445 | 0 | 3.01736E-17 | 0.005664693 | 3.058563892 | 0.3 | 0.068 | 1 | 0.005664693 | 2.48424E-21 |
| Prelp | 1.49953E-21 | 1.30790121 | 0.992 | 0.818 | 3.64266E-17 | 0.224562718 | 0.511662322 | 1 | 0.781 | 1 | 0.224562718 | 2.99906E-21 |
| Spon1 | 1.58968E-21 | 2.48681248 | 0.815 | 0.306 | 3.86166E-17 | 0.000705367 | 3.133887926 | 0.8 | 0.377 | 1 | 0.000705367 | 3.17937E-21 |
| Clmp | 3.16958E-21 | -4.848831558 | 0.067 | 0.612 | 7.69955E-17 | 0.018593753 | -3.782977992 | 0 | 0.377 | 1 | 0.018593753 | 6.33916E-21 |
| Sparc | 3.22974E-21 | 1.629443186 | 1 | 1 | 7.84569E-17 | 0.001942054 | 1.287618606 | 1 | 1 | 1 | 0.001942054 | 6.45948E-21 |
| Ninj1 | 4.42193E-21 | 1.928865286 | 0.908 | 0.535 | 1.07418E-16 | 0.007581347 | 1.265090992 | 1 | 0.654 | 1 | 0.007581347 | 8.84387E-21 |
| Snx10 | 4.55707E-21 | 1.983727221 | 0.815 | 0.306 | 1.107E-16 | 8.86631E-06 | 1.443013072 | 0.8 | 0.256 | 0.215380406 | 8.86631E-06 | 9.11414E-21 |
| Tmsb10 | 5.68389E-21 | -3.066925473 | 0.496 | 0.865 | 1.38073E-16 | 0.011238296 | -1.123852053 | 0.3 | 0.809 | 1 | 0.011238296 | 1.13678E-20 |
| Col8a1 | 6.4164E-21 | 1.933402251 | 0.95 | 0.647 | 1.55867E-16 | 0.028911948 | 0.85559145 | 0.8 | 0.744 | 1 | 0.028911948 | 1.28328E-20 |
| Rbfox2 | 6.72105E-21 | -1.701385438 | 0.815 | 0.976 | 1.63268E-16 | 0.002684216 | -1.265758718 | 0.9 | 0.975 | 1 | 0.002684216 | 1.34421E-20 |
| Ly6a | 7.08704E-21 | 3.532495081 | 0.731 | 0.253 | 1.72158E-16 | 0.241424157 | 1.759045984 | 0.6 | 0.645 | 1 | 0.241424157 | 1.41741E-20 |
| Neo1 | 7.91964E-21 | -2.068774672 | 0.613 | 0.912 | 1.92384E-16 | 0.020636023 | -1.125036046 | 0.7 | 0.938 | 1 | 0.020636023 | 1.58393E-20 |

| | | | | | | | | | | | | |
|---|---|---|---|---|---|---|---|---|---|---|---|---|
| Wnk4 | 8.25344E-21 | 2.142604395 | 0.765 | 0.212 | 2.00493E-16 | 8.06784E-06 | 2.720241359 | 0.8 | 0.244 | 0.195983986 | 8.06784E-06 | 1.65069E-20 |
| Sfpq | 1.2826E-20 | -1.112686349 | 0.933 | 1 | 3.11569E-16 | 0.000272221 | -1.022155531 | 1 | 0.997 | 1 | 0.000272221 | 2.5652E-20 |
| Camk4 | 1.51772E-20 | -3.797915226 | 0.092 | 0.635 | 3.68685E-16 | 0.003405859 | -3.196397213 | 0.2 | 0.63 | 1 | 0.003405859 | 3.03544E-20 |
| Sema3d | 1.60903E-20 | -4.452364806 | 0.193 | 0.7 | 3.90867E-16 | 0.008340389 | -3.08885911 | 0.1 | 0.556 | 1 | 0.008340389 | 3.21807E-20 |
| Ctxn3 | 1.73305E-20 | 3.210179737 | 0.546 | 0.053 | 4.20993E-16 | 0.000103743 | 4.432959407 | 0.1 | 0.003 | 1 | 0.000103743 | 3.4661E-20 |
| Steap1 | 1.81982E-20 | 2.284191984 | 0.756 | 0.218 | 4.4207E-16 | 5.9395E-06 | 2.510127268 | 0.7 | 0.157 | 0.144282412 | 5.9395E-06 | 3.63963E-20 |
| Thrap3 | 2.06823E-20 | -1.267554603 | 0.874 | 0.982 | 5.02414E-16 | 0.327208325 | -0.154313254 | 1 | 0.981 | 1 | 0.327208325 | 4.13645E-20 |
| Ppp1r3c | 2.29256E-20 | 2.52469739 | 0.739 | 0.235 | 5.56908E-16 | 4.65557E-05 | 2.399012075 | 0.7 | 0.21 | 1 | 4.65557E-05 | 4.58511E-20 |
| Slc6a12 | 2.48074E-20 | 3.529121481 | 0.605 | 0.112 | 6.02622E-16 | 5.71868E-10 | 3.70859385 | 0.6 | 0.065 | 1.38918E-05 | 5.71868E-10 | 4.96149E-20 |
| Cebpd | 2.78195E-20 | 1.987682246 | 0.958 | 0.712 | 6.75792E-16 | 0.002510087 | 1.886677375 | 0.8 | 0.66 | 1 | 0.002510087 | 5.5639E-20 |
| Galnt18 | 3.31557E-20 | 1.721744857 | 0.966 | 0.612 | 8.05419E-16 | 0.006156169 | 1.487060431 | 1 | 0.756 | 1 | 0.006156169 | 6.63115E-20 |
| Cdo1 | 4.06563E-20 | 2.004378441 | 0.824 | 0.371 | 9.87624E-16 | 0.127781763 | 0.938564116 | 0.8 | 0.667 | 1 | 0.127781763 | 8.13127E-20 |
| Matn4 | 4.87739E-20 | -3.202887833 | 0.454 | 0.8 | 1.18481E-15 | 0.044133985 | -1.232948214 | 0.8 | 0.883 | 1 | 0.044133985 | 9.75477E-20 |
| Gulo | 0.000116683 | 2.736965594 | 0.118 | 0.012 | 1 | 5.39474E-20 | 4.339850003 | 0.4 | 0.006 | 1.31049E-15 | 0.000116683 | 1.07895E-19 |
| Tenm3 | 5.52951E-20 | -1.428016039 | 0.992 | 1 | 1.34323E-15 | 0.007099068 | -0.788251753 | 1 | 1 | 1 | 0.007099068 | 1.1059E-19 |
| P4ha2 | 5.8578E-20 | 1.580101922 | 0.958 | 0.506 | 1.42298E-15 | 0.003942991 | 1.71726243 | 0.8 | 0.565 | 1 | 0.003942991 | 1.17156E-19 |
| Scara5 | 5.99969E-20 | 2.716207034 | 0.697 | 0.2 | 1.45744E-15 | 0.341014301 | 2.639410285 | 0.2 | 0.108 | 1 | 0.341014301 | 1.19994E-19 |
| Ctsh | 6.19065E-20 | 1.895177175 | 0.916 | 0.571 | 1.50383E-15 | 0.720693486 | 0.159940913 | 0.6 | 0.596 | 1 | 0.720693486 | 1.23813E-19 |
| Zfp385b | 6.45989E-20 | 1.312629332 | 0.992 | 0.859 | 1.56924E-15 | 0.000125682 | 1.57847167 | 1 | 0.864 | 1 | 0.000125682 | 1.29198E-19 |
| Btbd3 | 6.61341E-20 | 1.998252338 | 0.933 | 0.494 | 1.60653E-15 | 0.038871097 | 1.078342694 | 0.7 | 0.519 | 1 | 0.038871097 | 1.32268E-19 |
| Bptf | 8.26322E-20 | -1.634990875 | 0.773 | 0.947 | 2.0073E-15 | 0.001825908 | -1.171902651 | 0.8 | 1 | 1 | 0.001825908 | 1.65264E-19 |
| Htra1 | 8.98578E-20 | -1.281786427 | 0.345 | 0.888 | 2.18283E-15 | 0.126734451 | -0.322369624 | 0.5 | 0.92 | 1 | 0.126734451 | 1.79716E-19 |
| Col8a2 | 9.84579E-20 | -2.846883286 | 0.235 | 0.724 | 2.39174E-15 | 0.000419322 | -2.332414602 | 0.5 | 0.852 | 1 | 0.000419322 | 1.96916E-19 |
| Serpinf1 | 9.97853E-20 | -5.772741167 | 0.16 | 0.635 | 2.42399E-15 | 0.009876624 | -3.327483339 | 0.2 | 0.586 | 1 | 0.009876624 | 1.99571E-19 |
| Cpm | 1.24049E-19 | 1.625557531 | 0.958 | 0.582 | 3.01339E-15 | 0.003200997 | 2.799498389 | 0.7 | 0.299 | 1 | 0.003200997 | 2.48098E-19 |
| Col4a5 | 1.88101E-19 | 3.909635972 | 0.58 | 0.106 | 4.56936E-15 | 0.000911569 | 1.73588654 | 0.6 | 0.179 | 1 | 0.000911569 | 3.76203E-19 |
| Slc39a8 | 1.91807E-19 | 2.428416529 | 0.798 | 0.271 | 4.65939E-15 | 2.35038E-05 | 1.656465449 | 0.8 | 0.222 | 0.570954783 | 2.35038E-05 | 3.83615E-19 |
| Arhgef28 | 1.91984E-19 | 1.666576266 | 0.857 | 0.324 | 4.66367E-15 | 1.26954E-05 | 2.370223652 | 0.9 | 0.426 | 0.308397441 | 1.26954E-05 | 3.83968E-19 |
| Asap2 | 2.08058E-19 | 1.602358656 | 0.95 | 0.565 | 5.05414E-15 | 8.69295E-06 | 2.417226481 | 0.9 | 0.491 | 0.211169063 | 8.69295E-06 | 4.16115E-19 |
| Epb41l2 | 2.68652E-19 | -1.825617128 | 0.597 | 0.924 | 6.52609E-15 | 0.0123227 | -1.305132852 | 0.6 | 0.895 | 1 | 0.0123227 | 5.37304E-19 |
| Cav2 | 2.77543E-19 | 2.46419965 | 0.714 | 0.212 | 6.74208E-15 | 0.083314941 | 1.715802294 | 0.5 | 0.324 | 1 | 0.083314941 | 5.55087E-19 |
| Gulp1 | 3.37522E-19 | -5.132165525 | 0.059 | 0.565 | 8.19908E-15 | 0.000483387 | -3.731109474 | 0.2 | 0.728 | 1 | 0.000483387 | 6.75044E-19 |
| Cd164 | 3.5228E-19 | 1.089022447 | 0.992 | 0.924 | 8.55758E-15 | 0.532332513 | 0.176619654 | 0.9 | 0.907 | 1 | 0.532332513 | 7.0456E-19 |
| Esrrg | 3.5815E-19 | 2.272149047 | 0.664 | 0.124 | 8.70019E-15 | 0.000101599 | 2.847996907 | 0.6 | 0.179 | 1 | 0.000101599 | 7.16301E-19 |
| Cystm1 | 4.13413E-19 | 3.083047551 | 0.655 | 0.171 | 1.00426E-14 | 0.052400593 | 1.568614507 | 0.5 | 0.256 | 1 | 0.052400593 | 8.26825E-19 |
| Slc39a14 | 4.17056E-19 | 1.145735828 | 1 | 0.8 | 1.01311E-14 | 0.021322445 | 1.186521512 | 0.8 | 0.623 | 1 | 0.021322445 | 8.34111E-19 |
| Ckb | 5.42044E-19 | 2.273479906 | 0.882 | 0.6 | 1.31673E-14 | 0.015926163 | -3.667410918 | 0.2 | 0.556 | 1 | 0.015926163 | 1.08409E-18 |
| Kank1 | 6.20422E-19 | 1.30855658 | 0.966 | 0.735 | 1.50713E-14 | 0.001171336 | 1.426020977 | 1 | 0.793 | 1 | 0.001171336 | 1.24084E-18 |
| Ppp1cb | 6.94713E-19 | -1.378337121 | 0.916 | 0.988 | 1.6876E-14 | 0.027763735 | -0.624845044 | 1 | 0.994 | 1 | 0.027763735 | 1.38943E-18 |
| Rnasek | 8.35197E-19 | 1.683602369 | 0.966 | 0.853 | 2.02886E-14 | 0.002031458 | 1.242821185 | 1 | 0.778 | 1 | 0.002031458 | 1.67039E-18 |
| Nap1l1 | 8.53721E-19 | -1.211097952 | 0.916 | 0.988 | 2.07386E-14 | 0.025279247 | -0.89571552 | 0.7 | 0.988 | 1 | 0.025279247 | 1.70744E-18 |
| Setd2 | 1.05444E-18 | -1.509013647 | 0.765 | 0.953 | 2.56146E-14 | 0.033929407 | -0.634651499 | 0.9 | 0.978 | 1 | 0.033929407 | 2.10889E-18 |
| Tspan7 | 1.58864E-18 | 2.816109479 | 0.697 | 0.235 | 3.85913E-14 | 0.010682584 | 1.305847489 | 0.7 | 0.383 | 1 | 0.010682584 | 3.17728E-18 |
| Edil3 | 1.62258E-18 | -2.057867408 | 0.546 | 0.853 | 3.94158E-14 | 0.056696306 | -0.613966867 | 0.8 | 0.824 | 1 | 0.056696306 | 3.24517E-18 |
| Plac9 | 1.6264E-18 | -3.212222022 | 0.336 | 0.759 | 3.95086E-14 | 0.000692794 | -2.848584304 | 0.2 | 0.762 | 1 | 0.000692794 | 3.25281E-18 |
| Ephb2 | 1.8124E-18 | -3.86393845 | 0.067 | 0.576 | 4.40269E-14 | 0.0018602 | -3.093213762 | 0.2 | 0.685 | 1 | 0.0018602 | 3.62481E-18 |
| Prdx6 | 1.88098E-18 | 1.283858103 | 0.992 | 0.841 | 4.56927E-14 | 0.222342281 | 0.786003979 | 0.9 | 0.895 | 1 | 0.222342281 | 3.76196E-18 |
| S100a4 | 2.03507E-18 | 2.875839416 | 0.79 | 0.341 | 4.9436E-14 | 0.002074769 | 2.124485963 | 0.8 | 0.389 | 1 | 0.002074769 | 4.07015E-18 |
| Gm49144 | 0.00520595 | 1.569020957 | 0.16 | 0.059 | 1 | 2.11015E-18 | 4.602884409 | 0.7 | 0.04 | 5.12599E-14 | 0.00520595 | 4.22031E-18 |
| Glis3 | 2.28346E-18 | 1.435717752 | 0.95 | 0.629 | 5.54698E-14 | 0.014939145 | 1.049340894 | 0.9 | 0.855 | 1 | 0.014939145 | 4.56692E-18 |
| Cdh11 | 2.29497E-18 | -6.925603889 | 0.042 | 0.529 | 5.57495E-14 | 0.023217984 | -4.779739618 | 0 | 0.358 | 1 | 0.023217984 | 4.58995E-18 |
| Anxa8 | 2.77317E-18 | 1.633933825 | 0.933 | 0.476 | 6.73658E-14 | 0.001392648 | 1.388241487 | 0.8 | 0.38 | 1 | 0.001392648 | 5.54634E-18 |
| Sorbs2 | 2.89043E-18 | -1.87943682 | 0.84 | 0.965 | 7.02144E-14 | 0.00055603 | -2.358453971 | 0.9 | 0.957 | 1 | 0.00055603 | 5.78086E-18 |
| Antxr1 | 2.9423E-18 | -2.074027905 | 0.454 | 0.871 | 7.14744E-14 | 0.000800683 | -2.642192867 | 0.7 | 0.91 | 1 | 0.000800683 | 5.88461E-18 |
| B3gnt2 | 3.01245E-18 | 2.030273011 | 0.866 | 0.5 | 7.31784E-14 | 0.10244768 | 0.573137065 | 0.9 | 0.71 | 1 | 0.10244768 | 6.02489E-18 |
| Birc6 | 3.12883E-18 | -1.250961574 | 0.899 | 0.994 | 7.60056E-14 | 0.011592106 | -0.743895235 | 0.9 | 0.997 | 1 | 0.011592106 | 6.25766E-18 |
| Caprin1 | 3.69664E-18 | -1.428118144 | 0.782 | 0.947 | 8.97988E-14 | 0.00725676 | -1.010647244 | 0.8 | 0.963 | 1 | 0.00725676 | 7.39328E-18 |
| Mmp14 | 3.96453E-18 | -3.605030244 | 0.403 | 0.8 | 9.63064E-14 | 0.000370468 | -2.872038292 | 0.6 | 0.873 | 1 | 0.000370468 | 7.92906E-18 |
| Hs3st3b1 | 4.06999E-18 | 1.59208254 | 0.958 | 0.559 | 9.88683E-14 | 0.006661881 | 1.899672941 | 0.9 | 0.602 | 1 | 0.006661881 | 8.13999E-18 |
| Gpm6b | 4.5026E-18 | -3.99095486 | 0.101 | 0.594 | 1.09377E-13 | 0.014837606 | -3.786209113 | 0 | 0.395 | 1 | 0.014837606 | 9.00519E-18 |
| Rab2a | 4.85664E-18 | -0.834956147 | 1 | 0.994 | 1.17977E-13 | 0.016132506 | -0.670026866 | 1 | 1 | 1 | 0.016132506 | 9.71327E-18 |
| Srpx2 | 5.37007E-18 | 2.414815392 | 0.748 | 0.294 | 1.3045E-13 | 0.001893219 | 2.210566986 | 0.7 | 0.284 | 1 | 0.001893219 | 1.07401E-17 |
| Slc22a4 | 6.49066E-18 | 1.657400878 | 0.899 | 0.365 | 1.57671E-13 | 0.031246361 | 1.523157216 | 0.6 | 0.299 | 1 | 0.031246361 | 1.29813E-17 |
| Csrp1 | 7.58782E-18 | 2.656169451 | 0.765 | 0.341 | 1.84323E-13 | 1.18613E-05 | 3.432959407 | 0.9 | 0.407 | 0.288134799 | 1.18613E-05 | 1.51756E-17 |
| Col4a3 | 7.87297E-18 | 6.347463187 | 0.387 | 0.006 | 1.9125E-13 | 2.88544E-11 | 5.847996907 | 0.4 | 0.022 | 7.00932E-07 | 2.88544E-11 | 1.57459E-17 |
| Senp6 | 1.02906E-17 | -0.968709252 | 0.966 | 1 | 2.49978E-13 | 0.000255312 | -1.178475305 | 0.9 | 1 | 1 | 0.000255312 | 2.05811E-17 |
| Agmo | 1.08197E-17 | 4.906890596 | 0.403 | 0.012 | 2.62832E-13 | 3.07089E-07 | 4.280956314 | 0.3 | 0.022 | 0.007459807 | 3.07089E-07 | 2.16394E-17 |
| Ugdh | 1.27444E-17 | 1.288711562 | 1 | 0.941 | 3.09588E-13 | 0.006024268 | 0.715689614 | 1 | 0.932 | 1 | 0.006024268 | 2.54889E-17 |
| Khdrbs1 | 1.68189E-17 | -1.178269868 | 0.798 | 0.976 | 4.08564E-13 | 0.013471455 | -0.743186032 | 1 | 0.981 | 1 | 0.013471455 | 3.36377E-17 |
| Csgalnact1 | 1.70054E-17 | 1.092005189 | 0.992 | 0.694 | 4.13096E-13 | 0.00736214 | 1.159940913 | 0.9 | 0.596 | 1 | 0.00736214 | 3.40108E-17 |
| Cd24a | 1.75682E-17 | 1.506728156 | 0.899 | 0.482 | 4.26767E-13 | 1.90189E-05 | 2.510714565 | 0.9 | 0.395 | 0.46200658 | 1.90189E-05 | 3.51364E-17 |
| Maf | 1.79786E-17 | -2.586639079 | 0.361 | 0.771 | 4.36737E-13 | 0.000659956 | -2.205390306 | 0.5 | 0.864 | 1 | 0.000659956 | 3.59573E-17 |
| Adamts9 | 2.16746E-17 | -3.137503524 | 0.059 | 0.535 | 5.26519E-13 | 0.033920257 | -3.786209113 | 0 | 0.324 | 1 | 0.033920257 | 4.33492E-17 |
| Luc7l3 | 2.18454E-17 | -1.413873566 | 0.79 | 0.965 | 5.30669E-13 | 0.00128734 | -1.242890732 | 1 | 0.972 | 1 | 0.00128734 | 4.36909E-17 |
| Pole | 0.05857113 | -1.87774425 | 0.025 | 0.076 | 1 | 2.32841E-17 | 5.503348735 | 0.4 | 0.009 | 5.65616E-13 | 0.05857113 | 4.65681E-17 |
| Blm | 2.60909E-17 | 2.996224956 | 0.605 | 0.153 | 6.338E-13 | 4.58307E-14 | 4.280956314 | 0.6 | 0.04 | 1.11332E-09 | 4.58307E-14 | 5.21818E-17 |
| Ube2q2 | 2.61591E-17 | 1.968728931 | 0.933 | 0.771 | 6.35456E-13 | 0.006427718 | 1.184602967 | 0.9 | 0.793 | 1 | 0.006427718 | 5.23181E-17 |
| Lrguk | 0.935790323 | -0.655351829 | 0.025 | 0.024 | 1 | 2.62858E-17 | 5.754887502 | 0.3 | 0.003 | 6.38535E-13 | 0.935790323 | 5.25717E-17 |
| Clca2 | 4.87197E-08 | 2.836501268 | 0.227 | 0.024 | 0.0011835 | 2.83374E-17 | 4.695993813 | 0.3 | 0.003 | 6.88371E-13 | 4.87197E-08 | 5.66747E-17 |
| Hnrnpa0 | 2.89226E-17 | -1.223863809 | 0.874 | 0.953 | 7.02587E-13 | 0.003066264 | -1.020840343 | 0.9 | 0.972 | 1 | 0.003066264 | 5.78452E-17 |
| Tmem117 | 3.24009E-17 | 1.990442816 | 0.773 | 0.288 | 7.87083E-13 | 0.029226421 | 2.36882907 | 0.5 | 0.256 | 1 | 0.029226421 | 6.48019E-17 |
| Rapgef2 | 3.51371E-17 | -1.682193441 | 0.773 | 0.947 | 8.5355E-13 | 0.012519101 | -1.22202302 | 1 | 0.981 | 1 | 0.012519101 | 7.02742E-17 |
| Baalc | 3.74276E-17 | 2.317340825 | 0.521 | 0.076 | 9.09191E-13 | 5.75687E-09 | 4.358958826 | 0.5 | 0.052 | 0.000139846 | 5.75687E-09 | 7.48551E-17 |
| Hnrnpdl | 4.29252E-17 | -1.189617793 | 0.899 | 0.988 | 1.04274E-12 | 0.01434192 | -0.920922549 | 0.7 | 0.985 | 1 | 0.01434192 | 8.58504E-17 |
| Tmbim6 | 4.36371E-17 | 0.779412076 | 0.992 | 0.976 | 1.06003E-12 | 0.020603299 | 0.642309562 | 1 | 0.994 | 1 | 0.020603299 | 8.72742E-17 |
| Slc22a23 | 5.43589E-17 | 1.750640531 | 0.79 | 0.282 | 1.32049E-12 | 0.089989088 | 0.618847119 | 0.8 | 0.488 | 1 | 0.089989088 | 1.08718E-16 |
| Srpx | 5.46035E-17 | -3.50779464 | 0.076 | 0.553 | 1.32643E-12 | 0.014942083 | -2.795859283 | 0 | 0.392 | 1 | 0.014942083 | 1.09207E-16 |
| Crebbp | 5.52763E-17 | -1.120662534 | 0.95 | 0.994 | 1.34277E-12 | 0.00022926 | -1.10539481 | 1 | 1 | 1 | 0.00022926 | 1.10553E-16 |
| Rspo3 | 5.60304E-17 | 2.244005094 | 0.857 | 0.488 | 1.36109E-12 | 0.004196948 | 1.774747925 | 0.7 | 0.321 | 1 | 0.004196948 | 1.12061E-16 |
| Trp53 | 5.63889E-17 | -2.37721053 | 0.345 | 0.782 | 1.3698E-12 | 0.000748254 | -2.15033234 | 0.3 | 0.815 | 1 | 0.000748254 | 1.12778E-16 |
| Fn1 | 7.23587E-17 | 0.793761095 | 1 | 1 | 1.75774E-12 | 0.001866958 | 0.958914625 | 1 | 1 | 1 | 0.001866958 | 1.44717E-16 |
| Fbxo11 | 7.45183E-17 | -1.687060688 | 0.664 | 0.924 | 1.8102E-12 | 0.00259325 | -1.40974013 | 0.5 | 0.972 | 1 | 0.00259325 | 1.49037E-16 |
| Orai1 | 7.85319E-17 | 1.510468774 | 0.874 | 0.4 | 1.9077E-12 | 0.005026529 | 1.338181183 | 0.8 | 0.481 | 1 | 0.005026529 | 1.57064E-16 |
| Pdgfrb | 8.02513E-17 | -3.140778656 | 0.151 | 0.635 | 1.94946E-12 | 0.003929421 | -2.441509711 | 0.3 | 0.682 | 1 | 0.003929421 | 1.60503E-16 |
| Actn1 | 1.80189E-16 | -2.282774576 | 0.311 | 0.747 | 4.37715E-12 | 0.011733527 | -0.964256815 | 0.4 | 0.864 | 1 | 0.011733527 | 3.60378E-16 |
| Tgfbr2 | 2.02007E-16 | -1.754129983 | 0.613 | 0.859 | 4.90716E-12 | 0.005026392 | -1.153744427 | 0.8 | 0.96 | 1 | 0.005026392 | 4.04015E-16 |
| Cd14 | 2.31227E-16 | 3.115477217 | 0.605 | 0.159 | 5.61697E-12 | 0.825654724 | 0.236562194 | 0.2 | 0.244 | 1 | 0.825654724 | 4.62454E-16 |
| Mypn | 2.34059E-16 | 2.773765512 | 0.563 | 0.112 | 5.68575E-12 | 0.009005966 | 5.187846909 | 0.1 | 0.009 | 1 | 0.009005966 | 4.68117E-16 |

| | | | | | | | | | | | | |
|---|---|---|---|---|---|---|---|---|---|---|---|---|
| Cldn11 | 2.5336E-16 | 3.202221338 | 0.563 | 0.129 | 6.15462E-12 | 0.191871402 | -0.463204782 | 0.2 | 0.077 | 1 | 0.191871402 | 5.0672E-16 |
| Olfml3 | 2.68005E-16 | -2.144225209 | 0.521 | 0.818 | 6.51037E-12 | 0.019218575 | -2.260062839 | 0.3 | 0.602 | 1 | 0.019218575 | 5.3601E-16 |
| Pcsk6 | 2.82771E-16 | 1.4654677 | 0.84 | 0.388 | 6.86908E-12 | 2.83004E-11 | 3.260778432 | 1 | 0.191 | 6.87473E-07 | 2.83004E-11 | 5.65543E-16 |
| Emilin1 | 2.98839E-16 | -2.243225925 | 0.286 | 0.724 | 7.25939E-12 | 0.00136245 | -1.995168091 | 0.3 | 0.772 | 1 | 0.00136245 | 5.97677E-16 |
| Mtch1 | 3.29615E-16 | -1.680872349 | 0.739 | 0.906 | 8.00702E-12 | 0.000763118 | -1.699373706 | 0.3 | 0.883 | 1 | 0.000763118 | 6.59231E-16 |
| Mdk | 3.35022E-16 | -5.82527683 | 0.017 | 0.459 | 8.13835E-12 | 0.014706931 | -3.248864633 | 0.1 | 0.485 | 1 | 0.014706931 | 6.70043E-16 |
| Chd7 | 3.51786E-16 | -2.477079484 | 0.303 | 0.724 | 8.54559E-12 | 0.053895942 | -1.244534379 | 0.5 | 0.778 | 1 | 0.053895942 | 7.03572E-16 |
| Fus | 3.5352E-16 | -1.046235466 | 0.95 | 0.994 | 8.58771E-12 | 0.042350578 | -0.451529979 | 1 | 1 | 1 | 0.042350578 | 7.0704E-16 |
| Tnrc6c | 3.66222E-16 | -1.944858446 | 0.529 | 0.865 | 8.89626E-12 | 0.029154795 | -0.707903129 | 0.8 | 0.929 | 1 | 0.029154795 | 7.32444E-16 |
| Zfp354c | 3.8652E-16 | -3.226893814 | 0.109 | 0.571 | 9.38933E-12 | 0.060394841 | -1.461049897 | 0.3 | 0.562 | 1 | 0.060394841 | 7.73039E-16 |
| Bmpr1b | 4.19912E-16 | 1.156656952 | 0.975 | 0.553 | 1.02005E-11 | 0.017710106 | 0.901468115 | 0.9 | 0.802 | 1 | 0.017710106 | 8.39824E-16 |
| Cpxm1 | 4.86768E-16 | -2.7153766 | 0.202 | 0.665 | 1.18246E-11 | 0.074037705 | -1.672774536 | 0.5 | 0.654 | 1 | 0.074037705 | 9.73537E-16 |
| Sowahc | 4.93105E-16 | 2.067651613 | 0.756 | 0.329 | 1.19785E-11 | 0.784659837 | -0.235419102 | 0.4 | 0.327 | 1 | 0.784659837 | 9.86211E-16 |
| Pank1 | 5.10045E-16 | 2.028100536 | 0.739 | 0.294 | 1.239E-11 | 6.37828E-05 | 1.650905924 | 0.9 | 0.432 | 1 | 6.37828E-05 | 1.02009E-15 |
| Trpm7 | 5.35237E-16 | -1.26498005 | 0.824 | 0.953 | 1.3002E-11 | 0.079635792 | -0.571708434 | 1 | 0.978 | 1 | 0.079635792 | 1.07047E-15 |
| Slc38a2 | 5.55306E-16 | 1.083396915 | 0.992 | 1 | 1.34895E-11 | 0.00448793 | 1.541874189 | 1 | 0.994 | 1 | 0.00448793 | 1.11061E-15 |
| Ash1l | 5.67502E-16 | -0.968961595 | 0.966 | 0.982 | 1.37857E-11 | 0.01965728 | -0.606568957 | 1 | 0.997 | 1 | 0.01965728 | 1.135E-15 |
| Nsd1 | 5.88026E-16 | -1.750863723 | 0.63 | 0.876 | 1.42843E-11 | 0.128480191 | -0.66906153 | 0.8 | 0.957 | 1 | 0.128480191 | 1.17605E-15 |
| Aif1l | 5.98859E-16 | 1.967938791 | 0.697 | 0.235 | 1.45475E-11 | 0.726362806 | 0.227844977 | 0.4 | 0.33 | 1 | 0.726362806 | 1.19772E-15 |
| B3gnt9 | 6.43882E-16 | -3.109917692 | 0.067 | 0.529 | 1.56412E-11 | 0.305992915 | -0.579013234 | 0.3 | 0.454 | 1 | 0.305992915 | 1.28776E-15 |
| Cacna1c | 6.55244E-16 | -4.289557848 | 0.076 | 0.529 | 1.59172E-11 | 0.020050202 | -3.389345856 | 0.1 | 0.454 | 1 | 0.020050202 | 1.31049E-15 |
| Phlpp1 | 6.5731E-16 | -2.917701031 | 0.185 | 0.641 | 1.59674E-11 | 0.011025066 | -2.183385219 | 0.3 | 0.701 | 1 | 0.011025066 | 1.31462E-15 |
| Nid1 | 6.60305E-16 | -7.016808288 | 0 | 0.424 | 1.60401E-11 | 0.338657698 | -1.31299497 | 0.1 | 0.219 | 1 | 0.338657698 | 1.32061E-15 |
| H19 | 6.77743E-16 | -2.959749999 | 0.244 | 0.671 | 1.64637E-11 | 0.108103528 | -2.101921413 | 0.4 | 0.599 | 1 | 0.108103528 | 1.35549E-15 |
| Hnrnpd | 6.92514E-16 | -1.395094507 | 0.756 | 0.918 | 1.68226E-11 | 0.001824095 | -1.296561375 | 0.7 | 0.951 | 1 | 0.001824095 | 1.38503E-15 |
| Isg20 | 6.95383E-16 | 2.326713157 | 0.689 | 0.259 | 1.68922E-11 | 0.012334063 | 1.667424661 | 0.5 | 0.185 | 1 | 0.012334063 | 1.39077E-15 |
| Ptch1 | 6.99569E-16 | -3.888782521 | 0.16 | 0.594 | 1.69939E-11 | 0.036361027 | -2.677886361 | 0.1 | 0.444 | 1 | 0.036361027 | 1.39914E-15 |
| Mfap2 | 7.13922E-16 | -8.433794059 | 0 | 0.424 | 1.73426E-11 | 0.004451235 | -4.515407824 | 0 | 0.488 | 1 | 0.004451235 | 1.42784E-15 |
| Ybx2 | 0.00013176 | 3.215012891 | 0.101 | 0.006 | 1 | 8.31457E-16 | 6.602884409 | 0.2 | 0 | 2.01977E-11 | 0.00013176 | 1.66291E-15 |
| Gm12866 | 0.003173955 | 4.421463768 | 0.05 | 0 | 1 | 8.31457E-16 | 6.602884409 | 0.2 | 0 | 2.01977E-11 | 0.003173955 | 1.66291E-15 |
| Gm6559 | 0.091361251 | 2.514573173 | 0.017 | 0 | 1 | 8.31457E-16 | 6.602884409 | 0.2 | 0 | 2.01977E-11 | 0.091361251 | 1.66291E-15 |
| Gm13067 | 0.168156185 | 1.514573173 | 0.025 | 0.006 | 1 | 8.31457E-16 | 6.602884409 | 0.2 | 0 | 2.01977E-11 | 0.168156185 | 1.66291E-15 |
| Dscc1 | 0.237885589 | -1.070389328 | 0 | 0.012 | 1 | 8.31457E-16 | 6.602884409 | 0.2 | 0 | 2.01977E-11 | 0.237885589 | 1.66291E-15 |
| Cyp3a57 | 0.374548525 | 0.514573173 | 0.017 | 0.006 | 1 | 8.31457E-16 | 8.017921908 | 0.2 | 0 | 2.01977E-11 | 0.374548525 | 1.66291E-15 |
| Dmrt1 | 0.388707013 | 0.836501268 | 0.034 | 0.018 | 1 | 8.31457E-16 | 6.602884409 | 0.2 | 0 | 2.01977E-11 | 0.388707013 | 1.66291E-15 |
| 2310039L15Rik | 0.778665521 | -3.070389328 | 0.008 | 0.012 | 1 | 8.31457E-16 | 7.017921908 | 0.2 | 0 | 2.01977E-11 | 0.778665521 | 1.66291E-15 |
| Gtf2i | 9.22728E-16 | -1.939017567 | 0.504 | 0.812 | 2.24149E-11 | 0.00053556 | -1.875379623 | 0.6 | 0.883 | 1 | 0.00053556 | 1.84546E-15 |
| Mgat4a | 9.31074E-16 | 1.309895539 | 0.941 | 0.712 | 2.26177E-11 | 0.000829347 | 1.185031894 | 1 | 0.775 | 1 | 0.000829347 | 1.86215E-15 |
| Tbl1x | 9.496E-16 | -1.414944539 | 0.765 | 0.935 | 2.30677E-11 | 0.130513199 | -0.693572999 | 0.9 | 0.969 | 1 | 0.130513199 | 1.8992E-15 |
| Nrp2 | 9.63439E-16 | -3.245163729 | 0.303 | 0.712 | 2.34039E-11 | 0.008166707 | -1.953319099 | 0.5 | 0.815 | 1 | 0.008166707 | 1.92688E-15 |
| Mef2a | 1.00395E-15 | -1.211427054 | 0.857 | 0.976 | 2.43879E-11 | 0.001706994 | -1.18029574 | 0.8 | 0.994 | 1 | 0.001706994 | 2.0079E-15 |
| Cdc42bpa | 1.12077E-15 | -1.134815978 | 0.966 | 1 | 2.72257E-11 | 0.601016341 | -0.196298076 | 1 | 0.994 | 1 | 0.601016341 | 2.24154E-15 |
| Slc20a1 | 1.16053E-15 | 1.713118852 | 0.891 | 0.553 | 2.81915E-11 | 0.000561478 | 2.97412679 | 0.9 | 0.522 | 1 | 0.000561478 | 2.32105E-15 |
| Prkca | 1.20335E-15 | 0.968899201 | 1 | 0.929 | 2.92319E-11 | 0.033399976 | 0.873263665 | 1 | 0.954 | 1 | 0.033399976 | 2.40671E-15 |
| Gpc3 | 1.25652E-15 | -3.341416524 | 0.067 | 0.535 | 3.05235E-11 | 0.007696706 | -3.101019165 | 0.1 | 0.531 | 1 | 0.007696706 | 2.51305E-15 |
| Eps8l2 | 1.31202E-15 | 1.193707576 | 0.966 | 0.635 | 3.18715E-11 | 0.015242566 | 0.867474281 | 1 | 0.877 | 1 | 0.015242566 | 2.62403E-15 |
| Map7d2 | 1.38386E-15 | 3.274310075 | 0.571 | 0.135 | 3.36168E-11 | 1.57465E-09 | 3.967295835 | 0.5 | 0.046 | 3.82513E-05 | 1.57465E-09 | 2.76773E-15 |
| Scrg1 | 1.38829E-15 | 1.686482819 | 0.899 | 0.535 | 3.37243E-11 | 0.000188736 | 1.808468542 | 0.9 | 0.34 | 1 | 0.000188736 | 2.77658E-15 |
| Sat1 | 1.40753E-15 | 1.463233189 | 0.958 | 0.794 | 3.41918E-11 | 0.132211875 | 0.642478304 | 1 | 0.91 | 1 | 0.132211875 | 2.81507E-15 |
| Ccnl2 | 1.50603E-15 | -1.231852751 | 0.84 | 0.947 | 3.65844E-11 | 0.607119769 | -0.155163434 | 0.8 | 0.966 | 1 | 0.607119769 | 3.01206E-15 |
| Ehd3 | 1.52977E-15 | 1.914993576 | 0.84 | 0.488 | 3.71612E-11 | 0.003345354 | 2.001620096 | 0.7 | 0.417 | 1 | 0.003345354 | 3.05954E-15 |
| Chrdl1 | 2.00867E-15 | -2.537052516 | 0.37 | 0.741 | 4.87945E-11 | 0.013168696 | -2.672575007 | 0.3 | 0.623 | 1 | 0.013168696 | 4.01733E-15 |
| 2010009K17Rik | 0.20654211 | 2.099535674 | 0.042 | 0.018 | 1 | 2.11705E-15 | 6.133399125 | 0.4 | 0.012 | 5.14274E-11 | 0.20654211 | 4.2341E-15 |
| Ppp3ca | 2.11825E-15 | 0.857379997 | 1 | 1 | 5.14565E-11 | 0.046508662 | 0.795861943 | 1 | 0.994 | 1 | 0.046508662 | 4.2365E-15 |
| Vit | 2.1399E-15 | -4.452259963 | 0.076 | 0.506 | 5.19824E-11 | 0.007292555 | -3.451719909 | 0.1 | 0.528 | 1 | 0.007292555 | 4.27979E-15 |
| Dhx9 | 2.20024E-15 | -1.393112077 | 0.647 | 0.906 | 5.34481E-11 | 0.005683636 | -1.216803423 | 0.7 | 0.92 | 1 | 0.005683636 | 4.40047E-15 |
| Gm56768 | 2.48792E-15 | -3.889817082 | 0.202 | 0.618 | 6.04365E-11 | 0.002666554 | -2.921657306 | 0.2 | 0.676 | 1 | 0.002666554 | 4.97583E-15 |
| M1ap | 2.73043E-15 | 3.525800428 | 0.471 | 0.076 | 6.63277E-11 | 1.17928E-05 | 3.602884409 | 0.3 | 0.031 | 0.286469801 | 1.17928E-05 | 5.46087E-15 |
| Kmt2c | 3.13379E-15 | -1.907902658 | 0.571 | 0.835 | 7.61259E-11 | 0.145580642 | -0.598940677 | 0.9 | 0.988 | 1 | 0.145580642 | 6.26757E-15 |
| Cldn10 | 3.16806E-15 | 4.258734268 | 0.454 | 0.071 | 7.69586E-11 | 4.60862E-07 | 3.570462931 | 0.5 | 0.071 | 0.011195266 | 4.60862E-07 | 6.33612E-15 |
| Fxyd5 | 3.23786E-15 | -1.681683761 | 0.58 | 0.835 | 7.86542E-11 | 0.037686534 | -0.945295303 | 0.4 | 0.799 | 1 | 0.037686534 | 6.47573E-15 |
| Tgfa | 3.52737E-15 | 2.808607262 | 0.714 | 0.282 | 8.56868E-11 | 0.016108397 | 1.97689464 | 0.6 | 0.367 | 1 | 0.016108397 | 7.05473E-15 |
| Itga11 | 3.73963E-15 | -4.392317423 | 0.084 | 0.512 | 9.08432E-11 | 0.358044071 | -0.619508013 | 0.4 | 0.537 | 1 | 0.358044071 | 7.47927E-15 |
| Set | 3.79784E-15 | -1.009320628 | 0.908 | 0.965 | 9.2257E-11 | 0.000493698 | -1.434278141 | 0.7 | 0.994 | 1 | 0.000493698 | 7.59567E-15 |
| Dock4 | 3.83016E-15 | -4.478971805 | 0.076 | 0.506 | 9.30422E-11 | 0.018992082 | -2.652734341 | 0.2 | 0.562 | 1 | 0.018992082 | 7.66031E-15 |
| Matn1 | 3.91279E-15 | 4.211381641 | 0.697 | 0.3 | 9.50494E-11 | 0.017612825 | 3.261147833 | 0.7 | 0.389 | 1 | 0.017612825 | 7.82557E-15 |
| Dixdc1 | 3.9881E-15 | 1.228664301 | 0.933 | 0.529 | 9.6879E-11 | 0.008479727 | 1.136840499 | 1 | 0.716 | 1 | 0.008479727 | 7.97621E-15 |
| Capn6 | 4.00875E-15 | -3.064512762 | 0.244 | 0.665 | 9.73806E-11 | 0.013351112 | -1.923125698 | 0.4 | 0.725 | 1 | 0.013351112 | 8.0175E-15 |
| Asap1 | 4.31042E-15 | -1.795688861 | 0.471 | 0.841 | 1.04709E-10 | 0.080841518 | -0.739101339 | 0.6 | 0.892 | 1 | 0.080841518 | 8.62084E-15 |
| Mllt10 | 4.78098E-15 | -1.545041683 | 0.622 | 0.882 | 1.16139E-10 | 0.284843826 | -0.443330146 | 0.9 | 0.944 | 1 | 0.284843826 | 9.56195E-15 |
| P3h2 | 5.57008E-15 | 1.187074352 | 0.983 | 0.624 | 1.35308E-10 | 0.029976559 | 1.15508244 | 0.9 | 0.617 | 1 | 0.029976559 | 1.11402E-14 |
| Lhfp | 6.06283E-15 | -1.95535082 | 0.37 | 0.788 | 1.47278E-10 | 0.001459209 | -2.800837777 | 0.4 | 0.799 | 1 | 0.001459209 | 1.21257E-14 |
| Fam180a | 6.21892E-15 | -2.054658207 | 0.437 | 0.8 | 1.5107E-10 | 2.63422E-05 | -5.004908233 | 0.2 | 0.824 | 0.639903744 | 2.63422E-05 | 1.24378E-14 |
| Skap2 | 6.52267E-15 | 2.079906867 | 0.639 | 0.188 | 1.58449E-10 | 0.027922624 | 1.231325546 | 0.5 | 0.219 | 1 | 0.027922624 | 1.30453E-14 |
| Isyna1 | 6.676E-15 | 2.132325609 | 0.79 | 0.418 | 1.62173E-10 | 0.000514516 | 1.781545257 | 0.9 | 0.543 | 1 | 0.000514516 | 1.3352E-14 |
| Gpsm2 | 6.88469E-15 | -2.152604091 | 0.252 | 0.682 | 1.67243E-10 | 0.099444364 | -1.075891765 | 0.5 | 0.645 | 1 | 0.099444364 | 1.37694E-14 |
| Dhx58os | 6.97138E-15 | 1.60034898 | 0.908 | 0.612 | 1.69349E-10 | 0.029653589 | 1.122526951 | 0.8 | 0.596 | 1 | 0.029653589 | 1.39428E-14 |
| Acan | 7.09103E-15 | 1.342210966 | 1 | 0.965 | 1.72255E-10 | 4.89862E-05 | 1.637874933 | 1 | 0.963 | 1 | 4.89862E-05 | 1.41821E-14 |
| Fxyd1 | 7.95029E-15 | -1.747883114 | 0.387 | 0.782 | 1.93128E-10 | 0.042056367 | -1.062451508 | 0.7 | 0.756 | 1 | 0.042056367 | 1.59006E-14 |
| Itch | 8.67061E-15 | -1.218781168 | 0.748 | 0.947 | 2.10627E-10 | 0.016556145 | -0.873861795 | 0.7 | 0.963 | 1 | 0.016556145 | 1.73412E-14 |
| Tcf12 | 9.02155E-15 | -1.358219646 | 0.924 | 0.982 | 2.19151E-10 | 6.97618E-05 | -1.644239761 | 0.9 | 1 | 1 | 6.97618E-05 | 1.80431E-14 |
| N4bp2 | 1.28645E-14 | -2.38830436 | 0.277 | 0.665 | 3.12504E-10 | 0.000748397 | -2.321928095 | 0.2 | 0.769 | 1 | 0.000748397 | 2.5729E-14 |
| Srgap1 | 1.31315E-14 | -1.850999395 | 0.521 | 0.871 | 3.18991E-10 | 0.000150283 | -2.335855053 | 0.5 | 0.948 | 1 | 0.000150283 | 2.62631E-14 |
| Cobll1 | 1.41042E-14 | 1.370778893 | 0.908 | 0.553 | 3.42619E-10 | 0.006620445 | 1.178199391 | 1 | 0.633 | 1 | 0.006620445 | 2.82084E-14 |
| Arpc1a | 1.4257E-14 | 1.071743522 | 0.924 | 0.794 | 3.46331E-10 | 0.259298568 | 0.587190602 | 0.7 | 0.821 | 1 | 0.259298568 | 2.8514E-14 |
| Gfpt2 | 1.48851E-14 | 0.949815902 | 0.992 | 0.788 | 3.61588E-10 | 0.006437808 | 1.174843209 | 1 | 0.833 | 1 | 0.006437808 | 2.97701E-14 |
| Rhoj | 1.55031E-14 | 1.390837804 | 0.966 | 0.741 | 3.76601E-10 | 0.038098121 | 1.527137871 | 1 | 0.741 | 1 | 0.038098121 | 3.10062E-14 |
| Notch2 | 1.57961E-14 | -1.258734268 | 0.84 | 0.959 | 3.83719E-10 | 0.012696242 | -0.940257916 | 0.8 | 0.975 | 1 | 0.012696242 | 3.15922E-14 |
| Wls | 1.59376E-14 | -1.658949574 | 0.731 | 0.924 | 3.87155E-10 | 0.000173695 | -1.904910231 | 0.7 | 0.963 | 1 | 0.000173695 | 3.18751E-14 |
| Smo | 1.67845E-14 | -1.914270126 | 0.387 | 0.765 | 4.07728E-10 | 0.268416645 | -0.499022169 | 0.6 | 0.787 | 1 | 0.268416645 | 3.35689E-14 |
| Col1a2 | 1.80348E-14 | -3.943111577 | 0.697 | 0.829 | 4.38101E-10 | 0.0004312 | -2.729340514 | 0.8 | 0.941 | 1 | 0.0004312 | 3.60696E-14 |
| Atp6v1c2 | 1.81809E-14 | 4.62309763 | 0.336 | 0.012 | 4.41651E-10 | 0.00315387 | 2.602884409 | 0.2 | 0.028 | 1 | 0.00315387 | 3.63618E-14 |
| Loxl3 | 1.9395E-14 | 1.391139232 | 0.891 | 0.612 | 4.71142E-10 | 0.001100801 | 2.229087074 | 0.9 | 0.571 | 1 | 0.001100801 | 3.87899E-14 |
| Col5a1 | 1.9708E-14 | -2.809798098 | 0.328 | 0.676 | 4.78746E-10 | 0.004599841 | -3.046371769 | 0.2 | 0.617 | 1 | 0.004599841 | 3.94159E-14 |
| Rai14 | 2.00759E-14 | -3.432445535 | 0.151 | 0.576 | 4.87684E-10 | 0.000634415 | -3.550034167 | 0.1 | 0.673 | 1 | 0.000634415 | 4.01518E-14 |
| Lsp1 | 2.02526E-14 | -3.380957561 | 0.109 | 0.535 | 4.91975E-10 | 0.147105312 | -0.956082883 | 0.3 | 0.528 | 1 | 0.147105312 | 4.05051E-14 |
| Col3a1 | 2.15453E-14 | -8.828057126 | 0.017 | 0.412 | 5.23378E-10 | 0.021718601 | -5.795057563 | 0 | 0.364 | 1 | 0.021718601 | 4.30906E-14 |
| Il17ra | 2.16311E-14 | 1.851222773 | 0.689 | 0.276 | 5.25464E-10 | 0.008684508 | 1.938194716 | 0.5 | 0.204 | 1 | 0.008684508 | 4.32623E-14 |

| | | | | | | | | | | | | |
|---|---|---|---|---|---|---|---|---|---|---|---|---|
| Qki | 2.2804E-14 | -0.945066943 | 0.992 | 0.994 | 5.53955E-10 | 0.000195488 | -1.342082965 | 1 | 1 | 1 | 0.000195488 | 4.5608E-14 |
| Anxa3 | 2.38184E-14 | 1.711311005 | 0.815 | 0.4 | 5.78596E-10 | 0.025629475 | 1.48866884 | 0.7 | 0.395 | 1 | 0.025629475 | 4.76367E-14 |
| Vcp | 2.48618E-14 | -0.968208933 | 0.95 | 0.988 | 6.03943E-10 | 0.003753602 | -0.831743819 | 1 | 0.994 | 1 | 0.003753602 | 4.97236E-14 |
| Col5a2 | 2.91607E-14 | -7.288912203 | 0.034 | 0.429 | 7.08371E-10 | 0.001962159 | -5.428127499 | 0 | 0.543 | 1 | 0.001962159 | 5.83214E-14 |
| Serbp1 | 2.92204E-14 | -0.92003786 | 0.95 | 0.994 | 7.09821E-10 | 0.008543337 | -0.763437806 | 0.8 | 0.985 | 1 | 0.008543337 | 5.84407E-14 |
| Gm12349 | 2.9497E-14 | 2.96203215 | 0.412 | 0.047 | 7.16542E-10 | 0.288269199 | 2.795529487 | 0.1 | 0.037 | 1 | 0.288269199 | 5.89941E-14 |
| Lum | 3.51634E-14 | -3.106275808 | 0.387 | 0.747 | 8.5419E-10 | 0.02403527 | -2.092043803 | 0.6 | 0.762 | 1 | 0.02403527 | 7.03269E-14 |
| Zfp451 | 3.98093E-14 | -2.20109302 | 0.403 | 0.718 | 9.67048E-10 | 0.034756455 | -0.954442525 | 0.6 | 0.833 | 1 | 0.034756455 | 7.96186E-14 |
| Pcm1 | 4.19298E-14 | -1.164159156 | 0.832 | 0.947 | 1.01856E-09 | 0.584524133 | 0.229908708 | 0.9 | 0.96 | 1 | 0.584524133 | 8.38595E-14 |
| Svil | 4.34574E-14 | -2.361576429 | 0.487 | 0.782 | 1.05567E-09 | 0.002435017 | -2.225932111 | 0.2 | 0.756 | 1 | 0.002435017 | 8.69147E-14 |
| Tex12 | 0.396130756 | 0.514573173 | 0.025 | 0.012 | 1 | 4.40492E-14 | 6.017921908 | 0.3 | 0.006 | 1.07004E-09 | 0.396130756 | 8.80984E-14 |
| Zzz3 | 4.5692E-14 | -1.7797992 | 0.429 | 0.771 | 1.10995E-09 | 0.005960026 | -1.302502596 | 0.8 | 0.88 | 1 | 0.005960026 | 9.13839E-14 |
| Cp | 4.58893E-14 | 1.512039906 | 0.849 | 0.453 | 1.11474E-09 | 1.53736E-05 | 1.665711246 | 0.7 | 0.17 | 0.373456098 | 1.53736E-05 | 9.17785E-14 |
| Gabbr2 | 0.007133032 | 2.602036014 | 0.084 | 0.018 | 1 | 4.68181E-14 | 6.602884409 | 0.3 | 0.006 | 1.1373E-09 | 0.007133032 | 9.36362E-14 |
| Nt5dc1 | 4.95634E-14 | 2.857804781 | 0.731 | 0.341 | 1.20399E-09 | 0.019646367 | 2.039726278 | 0.6 | 0.324 | 1 | 0.019646367 | 9.91268E-14 |
| Plekhb1 | 5.13873E-14 | 2.156679581 | 0.647 | 0.247 | 1.2483E-09 | 0.097015139 | 1.728415291 | 0.4 | 0.188 | 1 | 0.097015139 | 1.02775E-13 |
| Ddx50 | 5.14488E-14 | -1.12322872 | 0.908 | 0.965 | 1.24979E-09 | 0.284050137 | -0.362275873 | 0.9 | 0.963 | 1 | 0.284050137 | 1.02898E-13 |
| Gm42209 | 0.366497963 | 1.142604395 | 0.042 | 0.024 | 1 | 5.28457E-14 | 6.133399125 | 0.3 | 0.006 | 1.28373E-09 | 0.366497963 | 1.05691E-13 |
| Zbtb38 | 5.33623E-14 | 0.98108689 | 1 | 0.794 | 1.29628E-09 | 0.00240923 | 1.191608828 | 1 | 0.833 | 1 | 0.00240923 | 1.06725E-13 |
| Sox4 | 5.43887E-14 | -1.661222417 | 0.227 | 0.676 | 1.32121E-09 | 0.008146053 | -0.782053484 | 0.1 | 0.63 | 1 | 0.008146053 | 1.08777E-13 |
| Gm4107 | 0.23091891 | -0.36994961 | 0.059 | 0.029 | 1 | 5.61218E-14 | 5.695993813 | 0.3 | 0.006 | 1.36331E-09 | 0.23091891 | 1.12244E-13 |
| Fgfr2 | 5.89141E-14 | -1.286629522 | 0.933 | 0.976 | 1.43114E-09 | 0.137603892 | -0.370267629 | 1 | 0.997 | 1 | 0.137603892 | 1.17828E-13 |
| Ubr3 | 5.9723E-14 | -1.394086516 | 0.689 | 0.912 | 1.45079E-09 | 0.063868293 | -0.704153724 | 0.9 | 0.941 | 1 | 0.063868293 | 1.19446E-13 |
| 2600014E21Rik | 5.99788E-14 | 2.258734268 | 0.361 | 0.024 | 1.457E-09 | 0.115118737 | 1.159940913 | 0.1 | 0.022 | 1 | 0.115118737 | 1.19958E-13 |
| Bnc2 | 6.08435E-14 | -2.612098631 | 0.269 | 0.653 | 1.47801E-09 | 0.132957095 | -1.362899876 | 0.6 | 0.691 | 1 | 0.132957095 | 1.21687E-13 |
| Ttc3 | 6.56362E-14 | -1.288142854 | 0.891 | 0.959 | 1.59443E-09 | 0.04508241 | -0.738706928 | 1 | 0.981 | 1 | 0.04508241 | 1.31272E-13 |
| Lgals1 | 6.60483E-14 | -6.607823459 | 0.034 | 0.424 | 1.60445E-09 | 0.009689611 | -4.108782565 | 0 | 0.429 | 1 | 0.009689611 | 1.32097E-13 |
| Fam193a | 6.72408E-14 | -1.363494588 | 0.681 | 0.9 | 1.63341E-09 | 0.01585323 | -0.819180357 | 0.8 | 0.991 | 1 | 0.01585323 | 1.34482E-13 |
| Foxn3 | 6.78396E-14 | -1.313193882 | 0.933 | 0.988 | 1.64796E-09 | 0.72525523 | -0.251727342 | 1 | 0.991 | 1 | 0.72525523 | 1.35679E-13 |
| Bgn | 6.88428E-14 | -1.221919639 | 0.941 | 0.941 | 1.67233E-09 | 0.59229649 | -0.119025771 | 1 | 0.941 | 1 | 0.59229649 | 1.37686E-13 |
| Igf2bp2 | 6.97995E-14 | -2.735205136 | 0.16 | 0.582 | 1.69557E-09 | 0.000539761 | -2.762125668 | 0.2 | 0.772 | 1 | 0.000539761 | 1.39599E-13 |
| Taok1 | 7.67768E-14 | -1.335531397 | 0.714 | 0.935 | 1.86506E-09 | 0.012107379 | -0.888968688 | 1 | 0.978 | 1 | 0.012107379 | 1.53554E-13 |
| Ephx2 | 8.11991E-14 | 2.624197664 | 0.563 | 0.153 | 1.97249E-09 | 1.79443E-05 | 3.54399072 | 0.4 | 0.059 | 0.435903141 | 1.79443E-05 | 1.62398E-13 |
| Axl | 8.43699E-14 | -3.521050737 | 0.042 | 0.447 | 2.04951E-09 | 0.000331218 | -4.239465935 | 0 | 0.645 | 1 | 0.000331218 | 1.6874E-13 |
| Stard13 | 9.72676E-14 | 1.255773057 | 0.983 | 0.847 | 2.36282E-09 | 0.003120217 | 1.126257787 | 1 | 0.895 | 1 | 0.003120217 | 1.94535E-13 |
| Lgals3 | 1.00211E-13 | 0.982393103 | 1 | 0.8 | 2.43431E-09 | 0.801506776 | -0.523110865 | 1 | 0.917 | 1 | 0.801506776 | 2.00421E-13 |
| Aqp1 | 1.00852E-13 | 1.893084796 | 0.504 | 0.094 | 2.44989E-09 | 4.29211E-06 | 2.658840815 | 0.5 | 0.077 | 0.104263947 | 4.29211E-06 | 2.01704E-13 |
| Arid2 | 1.10505E-13 | -2.11345805 | 0.361 | 0.718 | 2.68438E-09 | 0.040940016 | -1.008601535 | 0.6 | 0.864 | 1 | 0.040940016 | 2.21009E-13 |
| Zbtb7a | 1.16044E-13 | -1.680774425 | 0.504 | 0.806 | 2.81894E-09 | 0.048219471 | -0.855684096 | 0.9 | 0.92 | 1 | 0.048219471 | 2.32088E-13 |
| Tmem164 | 1.19178E-13 | -2.014065721 | 0.261 | 0.659 | 2.89508E-09 | 0.013771957 | -1.752666909 | 0.6 | 0.809 | 1 | 0.013771957 | 2.38356E-13 |
| Acin1 | 1.19752E-13 | -1.041592064 | 0.882 | 0.941 | 2.90901E-09 | 0.016029389 | -0.832385159 | 0.9 | 0.969 | 1 | 0.016029389 | 2.39503E-13 |
| Pds5a | 1.20351E-13 | -1.897729359 | 0.555 | 0.8 | 2.92357E-09 | 0.026434713 | -0.967350806 | 0.8 | 0.914 | 1 | 0.026434713 | 2.40702E-13 |
| Nqo1 | 1.34805E-13 | 2.178824583 | 0.622 | 0.224 | 3.27468E-09 | 0.060444996 | 1.748461233 | 0.5 | 0.272 | 1 | 0.060444996 | 2.6961E-13 |
| Mt1 | 1.39317E-13 | 2.475811138 | 0.79 | 0.541 | 3.38429E-09 | 0.861874361 | -0.106199404 | 0.4 | 0.454 | 1 | 0.861874361 | 2.78634E-13 |
| Etl4 | 1.41402E-13 | -4.831201664 | 0.084 | 0.482 | 3.43494E-09 | 0.129208793 | -1.963169629 | 0.2 | 0.414 | 1 | 0.129208793 | 2.82804E-13 |
| Snrnp70 | 1.429E-13 | -0.87244995 | 0.958 | 0.988 | 3.47134E-09 | 0.013013991 | -0.743819265 | 0.9 | 0.985 | 1 | 0.013013991 | 2.85801E-13 |
| Prkd1 | 1.43856E-13 | 1.196832875 | 0.866 | 0.4 | 3.49455E-09 | 0.024426957 | 0.558490289 | 1 | 0.642 | 1 | 0.024426957 | 2.87712E-13 |
| Dusp15 | 1.48692E-13 | 6.62309763 | 0.286 | 0 | 3.61202E-09 | 0.195226183 | 2.017921908 | 0.1 | 0.028 | 1 | 0.195226183 | 2.97384E-13 |
| Septin11 | 1.53924E-13 | -1.462795853 | 0.655 | 0.853 | 3.73913E-09 | 0.002281821 | -1.609131926 | 0.6 | 0.923 | 1 | 0.002281821 | 3.07849E-13 |
| Ago2 | 1.60245E-13 | -1.497108632 | 0.597 | 0.865 | 3.89268E-09 | 0.061577 | -0.593102889 | 0.9 | 0.944 | 1 | 0.061577 | 3.2049E-13 |
| Pafah1b1 | 1.62147E-13 | -0.736407923 | 1 | 0.988 | 3.93888E-09 | 0.008761303 | -0.697758413 | 1 | 1 | 1 | 0.008761303 | 3.24294E-13 |
| Atp1b3 | 1.62717E-13 | 0.850269423 | 1 | 0.906 | 3.95272E-09 | 0.035208812 | 0.609837169 | 1 | 0.96 | 1 | 0.035208812 | 3.25434E-13 |
| Optc | 0.007148548 | 2.929610672 | 0.059 | 0.006 | 1 | 1.6629E-13 | 5.017921908 | 0.4 | 0.015 | 4.03951E-09 | 0.007148548 | 3.32579E-13 |
| Plekhh2 | 1.70701E-13 | 1.179096052 | 0.941 | 0.588 | 4.14666E-09 | 0.000959556 | 1.291090691 | 1 | 0.676 | 1 | 0.000959556 | 3.41401E-13 |
| Shf | 1.7467E-13 | 1.278600663 | 0.84 | 0.488 | 4.24309E-09 | 0.039005349 | 1.473601392 | 0.5 | 0.247 | 1 | 0.039005349 | 3.4934E-13 |
| Ust | 1.80649E-13 | -1.769751201 | 0.639 | 0.859 | 4.38832E-09 | 2.99133E-05 | -2.575110023 | 0.6 | 0.96 | 0.726654882 | 2.99133E-05 | 3.61298E-13 |
| Rfx7 | 1.81252E-13 | -1.810706132 | 0.361 | 0.759 | 4.40298E-09 | 0.117470024 | -0.876895855 | 0.6 | 0.787 | 1 | 0.117470024 | 3.62504E-13 |
| Usp7 | 1.90127E-13 | -1.310440416 | 0.63 | 0.865 | 4.61857E-09 | 0.022499053 | -0.894302132 | 0.7 | 0.935 | 1 | 0.022499053 | 3.80254E-13 |
| Asxl1 | 1.95103E-13 | -1.683366205 | 0.555 | 0.847 | 4.73945E-09 | 0.295111005 | -0.442770257 | 0.7 | 0.873 | 1 | 0.295111005 | 3.90207E-13 |
| Nfic | 2.03783E-13 | -1.325696372 | 0.798 | 0.906 | 4.95029E-09 | 0.000337309 | -1.633782144 | 0.8 | 0.969 | 1 | 0.000337309 | 4.07566E-13 |
| Atp1a1 | 2.08838E-13 | 0.928510638 | 0.975 | 0.965 | 5.0731E-09 | 0.003280479 | 0.575899968 | 1 | 0.972 | 1 | 0.003280479 | 4.17677E-13 |
| Smim5 | 0.264155408 | 0.13606155 | 0.118 | 0.076 | 1 | 2.11638E-13 | 4.017921908 | 0.5 | 0.028 | 5.14111E-09 | 0.264155408 | 4.23276E-13 |
| Gsn | 2.17927E-13 | 1.056143078 | 1 | 0.971 | 5.29389E-09 | 0.089609293 | 0.601850737 | 1 | 0.997 | 1 | 0.089609293 | 4.35855E-13 |
| Gapvd1 | 2.32658E-13 | -1.390789953 | 0.639 | 0.865 | 5.65172E-09 | 0.09750768 | -0.667304716 | 0.7 | 0.901 | 1 | 0.09750768 | 4.65316E-13 |
| Pbrm1 | 2.50245E-13 | -0.992386816 | 0.866 | 0.935 | 6.07896E-09 | 0.035216067 | -0.842721706 | 0.8 | 0.954 | 1 | 0.035216067 | 5.0049E-13 |
| Setbp1 | 2.5151E-13 | -1.199943665 | 0.916 | 0.976 | 6.10967E-09 | 0.011544385 | -1.022155801 | 1 | 0.988 | 1 | 0.011544385 | 5.03019E-13 |
| Cav1 | 2.5886E-13 | 1.79180146 | 0.571 | 0.153 | 6.28824E-09 | 0.253611131 | 0.394578316 | 0.4 | 0.247 | 1 | 0.253611131 | 5.17721E-13 |
| Gm16364.1 | 0.093245792 | -1.807354922 | 0 | 0.024 | 1 | 2.61697E-13 | 3.847996907 | 0.5 | 0.028 | 6.35713E-09 | 0.093245792 | 5.23393E-13 |
| Ecrg4 | 2.66977E-13 | -2.490970018 | 0.639 | 0.794 | 6.4854E-09 | 0.934591749 | -0.539491407 | 0.9 | 0.756 | 1 | 0.934591749 | 5.33954E-13 |
| Grb14 | 2.79395E-09 | 2.78958022 | 0.37 | 0.088 | 6.78707E-05 | 2.72783E-13 | 4.229426013 | 0.5 | 0.028 | 6.62643E-09 | 2.79395E-09 | 5.45565E-13 |
| Tmco3 | 2.78104E-13 | 1.422780697 | 0.866 | 0.506 | 6.7557E-09 | 0.082683716 | 1.135278859 | 0.7 | 0.441 | 1 | 0.082683716 | 5.56208E-13 |
| Ptgds | 2.98394E-13 | 3.06636881 | 0.471 | 0.112 | 7.24858E-09 | 0.551327697 | -0.391469028 | 0.1 | 0.167 | 1 | 0.551327697 | 5.96787E-13 |
| Top2b | 3.01988E-13 | -1.617197488 | 0.731 | 0.882 | 7.33589E-09 | 0.001844925 | -1.313890776 | 0.8 | 0.972 | 1 | 0.001844925 | 6.03976E-13 |
| Cacnb2 | 3.20127E-13 | -2.713608426 | 0.277 | 0.665 | 7.77652E-09 | 0.008662616 | -2.762500686 | 0.6 | 0.753 | 1 | 0.008662616 | 6.40254E-13 |
| Ncl | 3.27416E-13 | -0.800272458 | 0.983 | 0.994 | 7.95358E-09 | 0.000537832 | -1.096145739 | 0.9 | 1 | 1 | 0.000537832 | 6.54832E-13 |
| Prrx2 | 3.29333E-13 | -3.105035471 | 0.118 | 0.518 | 8.00016E-09 | 0.277280626 | -0.950744885 | 0.3 | 0.472 | 1 | 0.277280626 | 6.58666E-13 |
| Copa | 3.52648E-13 | -0.804987908 | 0.966 | 0.994 | 8.56651E-09 | 0.003281223 | -0.927382571 | 1 | 0.991 | 1 | 0.003281223 | 7.05295E-13 |
| Srrm2 | 3.66378E-13 | -0.684308613 | 1 | 1 | 8.90006E-09 | 3.46937E-06 | -1.301343022 | 1 | 1 | 0.084278051 | 3.46937E-06 | 7.32756E-13 |
| Fdft1 | 3.75589E-13 | 1.140553899 | 0.899 | 0.476 | 9.1238E-09 | 0.196112818 | 0.341729736 | 0.8 | 0.562 | 1 | 0.196112818 | 7.51177E-13 |
| Mdfi | 3.86577E-13 | 1.273183761 | 0.958 | 0.718 | 9.39074E-09 | 0.21178078 | 0.379848071 | 1 | 0.738 | 1 | 0.21178078 | 7.73155E-13 |
| Itga6 | 4.09044E-13 | 1.080299455 | 0.882 | 0.571 | 9.93651E-09 | 0.08320663 | 0.294813115 | 0.9 | 0.577 | 1 | 0.08320663 | 8.18089E-13 |
| Ptprj | 4.10391E-13 | 0.903323981 | 0.924 | 0.582 | 9.96923E-09 | 0.303394425 | 1.248534836 | 0.6 | 0.593 | 1 | 0.303394425 | 8.20783E-13 |
| Tagln2 | 4.1218E-13 | -1.411678487 | 0.538 | 0.876 | 1.00127E-08 | 0.00178637 | -1.466727396 | 0.4 | 0.975 | 1 | 0.00178637 | 8.2436E-13 |
| Gfod1 | 4.18304E-13 | 1.514573173 | 0.773 | 0.382 | 1.01614E-08 | 0.065268743 | 2.176619654 | 0.5 | 0.247 | 1 | 0.065268743 | 8.36608E-13 |
| Zfp521 | 4.3064E-13 | -1.601267856 | 0.689 | 0.935 | 1.04611E-08 | 0.041508362 | -1.045772767 | 0.8 | 0.861 | 1 | 0.041508362 | 8.6128E-13 |
| Purb | 4.56562E-13 | -0.965315471 | 0.815 | 0.965 | 1.10908E-08 | 0.000128004 | -1.662799576 | 0.7 | 0.972 | 1 | 0.000128004 | 9.13123E-13 |
| Insig1 | 4.75443E-13 | 1.849574322 | 0.891 | 0.635 | 1.15495E-08 | 0.014610552 | 0.708066645 | 1 | 0.667 | 1 | 0.014610552 | 9.50885E-13 |
| Ints6l | 4.81952E-13 | -3.649998594 | 0.092 | 0.488 | 1.17076E-08 | 0.000920037 | -3.900941329 | 0 | 0.583 | 1 | 0.000920037 | 9.63905E-13 |
| Smad4 | 4.85253E-13 | -1.009953474 | 0.832 | 0.935 | 1.17878E-08 | 0.087543105 | -0.561016805 | 0.9 | 0.975 | 1 | 0.087543105 | 9.70507E-13 |
| Rsf1 | 5.04392E-13 | -1.504571876 | 0.504 | 0.835 | 1.22527E-08 | 0.201180268 | -0.635138109 | 0.8 | 0.898 | 1 | 0.201180268 | 1.00878E-12 |
| Slc5a3 | 5.30189E-13 | 1.329592757 | 0.908 | 0.559 | 1.28793E-08 | 0.036159401 | 1.287528968 | 0.8 | 0.639 | 1 | 0.036159401 | 1.06038E-12 |
| Smagp | 6.45398E-13 | 3.174082627 | 0.462 | 0.106 | 1.5678E-08 | 0.236765379 | 1.930459067 | 0.1 | 0.031 | 1 | 0.236765379 | 1.2908E-12 |
| Tnfrsf21 | 6.5825E-13 | 1.306366658 | 0.782 | 0.376 | 1.59902E-08 | 0.024845443 | 1.117582647 | 0.8 | 0.775 | 1 | 0.024845443 | 1.3165E-12 |
| Fat4 | 7.53093E-13 | -2.416714077 | 0.294 | 0.676 | 1.82941E-08 | 0.008553212 | -1.9226364 | 0.1 | 0.605 | 1 | 0.008553212 | 1.50619E-12 |
| Fzd9 | 7.5347E-13 | 1.240668032 | 0.882 | 0.506 | 1.83033E-08 | 0.014631212 | 1.291090691 | 0.9 | 0.691 | 1 | 0.014631212 | 1.50694E-12 |
| Nufip2 | 7.74326E-13 | -1.045282752 | 0.983 | 0.988 | 1.88099E-08 | 0.001104689 | -1.090950984 | 1 | 1 | 1 | 0.001104689 | 1.54865E-12 |
| Nt5e | 8.53946E-13 | 1.650456601 | 0.798 | 0.488 | 2.0744E-08 | 0.020510167 | 1.606988807 | 0.7 | 0.463 | 1 | 0.020510167 | 1.70789E-12 |
| Fgfr3 | 9.08168E-13 | 0.985123883 | 0.941 | 0.671 | 2.20612E-08 | 0.001046172 | 1.431545693 | 0.9 | 0.796 | 1 | 0.001046172 | 1.81634E-12 |

| | | | | | | | | | | | | |
|---|---|---|---|---|---|---|---|---|---|---|---|---|
| Smim3 | 9.08664E-13 | 1.360013242 | 0.84 | 0.594 | 2.20733E-08 | 0.006106612 | 1.293556351 | 1 | 0.614 | 1 | 0.006106612 | 1.81733E-12 |
| Sdc2 | 9.41267E-13 | -1.690602676 | 0.807 | 0.882 | 2.28653E-08 | 0.00139554 | -2.336427665 | 0.5 | 0.827 | 1 | 0.00139554 | 1.88253E-12 |
| Slc25a48 | 9.66679E-13 | 4.580662363 | 0.311 | 0.018 | 2.34826E-08 | 2.85385E-10 | 4.365845211 | 0.4 | 0.025 | 6.93258E-06 | 2.85385E-10 | 1.93336E-12 |
| Srsf6 | 9.94242E-13 | -1.17951105 | 0.63 | 0.888 | 2.41521E-08 | 0.020628773 | -0.994902132 | 0.5 | 0.87 | 1 | 0.020628773 | 1.98848E-12 |
| Smoc2 | 9.95622E-13 | -2.631145656 | 0.597 | 0.788 | 2.41857E-08 | 0.005238214 | -2.650861795 | 0.7 | 0.799 | 1 | 0.005238214 | 1.99124E-12 |
| Rcn2 | 1.01092E-12 | 0.82533196 | 0.992 | 0.941 | 2.45572E-08 | 0.01242642 | 0.874963954 | 1 | 0.87 | 1 | 0.01242642 | 2.02183E-12 |
| Ankrd11 | 1.02685E-12 | -0.973844349 | 0.983 | 0.994 | 2.49442E-08 | 0.013863163 | -0.71705158 | 1 | 1 | 1 | 0.013863163 | 2.0537E-12 |
| Degs1 | 1.0425E-12 | 1.192645078 | 0.95 | 0.753 | 2.53244E-08 | 0.615466325 | 0.784302231 | 0.7 | 0.765 | 1 | 0.615466325 | 2.085E-12 |
| Cyba | 1.07905E-12 | -2.694880193 | 0.101 | 0.506 | 2.62124E-08 | 0.043592118 | -1.5519337 | 0.2 | 0.503 | 1 | 0.043592118 | 2.15811E-12 |
| Hook3 | 1.0856E-12 | -1.089435084 | 0.874 | 0.965 | 2.63713E-08 | 0.077591163 | -0.645322237 | 0.9 | 0.966 | 1 | 0.077591163 | 2.1712E-12 |
| Il6ra | 1.09131E-12 | 2.106915204 | 0.613 | 0.212 | 2.65101E-08 | 0.131257954 | 1.3571919 | 0.5 | 0.296 | 1 | 0.131257954 | 2.18262E-12 |
| Safb | 1.13514E-12 | -1.31636857 | 0.555 | 0.835 | 2.75749E-08 | 0.098936811 | -0.644484678 | 0.5 | 0.864 | 1 | 0.098936811 | 2.27029E-12 |
| S100a1 | 1.16388E-12 | 1.418733976 | 0.958 | 0.824 | 2.82729E-08 | 0.729627143 | 1.090025454 | 0.6 | 0.815 | 1 | 0.729627143 | 2.32775E-12 |
| 1810053B23Rik | 1.17536E-12 | 5.146841388 | 0.336 | 0.029 | 2.85518E-08 | 0.003266043 | 1.648688098 | 0.2 | 0.028 | 1 | 0.003266043 | 2.35071E-12 |
| Cdk4 | 1.2043E-12 | -1.75595577 | 0.471 | 0.747 | 2.92547E-08 | 0.060881871 | -0.870821341 | 0.4 | 0.71 | 1 | 0.060881871 | 2.40859E-12 |
| Nasp | 1.22418E-12 | -1.857395605 | 0.546 | 0.776 | 2.97378E-08 | 0.040886515 | -1.052199036 | 0.7 | 0.867 | 1 | 0.040886515 | 2.44836E-12 |
| Slc3a2 | 1.31206E-12 | -1.16002654 | 0.756 | 0.929 | 3.18725E-08 | 0.005011448 | -1.280080348 | 0.6 | 0.951 | 1 | 0.005011448 | 2.62412E-12 |
| Sbds | 1.41929E-12 | 1.041009597 | 0.95 | 0.729 | 3.44774E-08 | 0.736628162 | 0.14550353 | 0.9 | 0.827 | 1 | 0.736628162 | 2.83858E-12 |
| Prune2 | 1.42822E-12 | 2.433954713 | 0.546 | 0.171 | 3.46944E-08 | 5.01658E-08 | 3.555131277 | 0.8 | 0.173 | 0.001218628 | 5.01658E-08 | 2.85644E-12 |
| Rtl3 | 1.50156E-12 | -4.145677455 | 0.034 | 0.4 | 3.64759E-08 | 0.075160725 | -2.080110175 | 0 | 0.247 | 1 | 0.075160725 | 3.00312E-12 |
| Tspan3 | 1.52654E-12 | 0.754296807 | 1 | 0.971 | 3.70827E-08 | 0.668276821 | 0.208115507 | 1 | 0.975 | 1 | 0.668276821 | 3.05308E-12 |
| Kdm2a | 1.55655E-12 | -1.406737939 | 0.613 | 0.894 | 3.78117E-08 | 0.006423873 | -1.135354742 | 0.8 | 0.969 | 1 | 0.006423873 | 3.1131E-12 |
| Naa50 | 1.68649E-12 | -1.275634443 | 0.655 | 0.871 | 4.09683E-08 | 0.005551709 | -1.244172937 | 0.8 | 0.932 | 1 | 0.005551709 | 3.37299E-12 |
| Itgbl1 | 1.72591E-12 | -9.164906927 | 0 | 0.341 | 4.19258E-08 | 0.020936039 | -5.196397213 | 0 | 0.367 | 1 | 0.020936039 | 3.45182E-12 |
| Tcf4 | 1.7437E-12 | -0.983436412 | 0.983 | 0.988 | 4.23579E-08 | 0.000116816 | -1.426642897 | 1 | 1 | 1 | 0.000116816 | 3.48739E-12 |
| Cadm1 | 1.80811E-12 | -1.519057605 | 0.403 | 0.824 | 4.39226E-08 | 0.012190555 | -0.976719176 | 0.7 | 0.963 | 1 | 0.012190555 | 3.61622E-12 |
| Prrc2c | 1.89046E-12 | -0.707819249 | 0.983 | 1 | 4.5923E-08 | 0.18710306 | -0.361644813 | 1 | 1 | 1 | 0.18710306 | 3.78092E-12 |
| Spag9 | 1.96255E-12 | -0.810620787 | 0.975 | 1 | 4.76743E-08 | 0.277603008 | -0.380698944 | 0.8 | 0.997 | 1 | 0.277603008 | 3.92511E-12 |
| Irs1 | 1.96639E-12 | -2.524420959 | 0.336 | 0.694 | 4.77676E-08 | 0.60503688 | -0.840059087 | 0.6 | 0.571 | 1 | 0.60503688 | 3.93278E-12 |
| Chsy3 | 1.9709E-12 | 0.973353063 | 0.975 | 0.853 | 4.78771E-08 | 7.06134E-05 | 2.409464392 | 1 | 0.676 | 1 | 7.06134E-05 | 3.9418E-12 |
| Rab1a | 1.97154E-12 | -0.839216021 | 0.958 | 0.982 | 4.78926E-08 | 0.008159696 | -0.846441135 | 0.9 | 0.994 | 1 | 0.008159696 | 3.94308E-12 |
| Cdhr4 | 1.98543E-12 | 3.189172886 | 0.387 | 0.059 | 4.823E-08 | 0.022929821 | 3.091922489 | 0.2 | 0.046 | 1 | 0.022929821 | 3.97085E-12 |
| Itpr2 | 2.10273E-12 | 1.046110352 | 0.983 | 0.871 | 5.10794E-08 | 0.003491265 | 1.745343729 | 1 | 0.846 | 1 | 0.003491265 | 4.20545E-12 |
| Tpm1 | 2.14716E-12 | -1.582473862 | 0.95 | 0.988 | 5.21587E-08 | 0.633653992 | 0.285974389 | 1 | 0.948 | 1 | 0.633653992 | 4.29431E-12 |
| Colec12 | 2.15762E-12 | -1.468839798 | 0.874 | 0.924 | 5.24129E-08 | 0.002858252 | -1.214531431 | 1 | 0.944 | 1 | 0.002858252 | 4.31524E-12 |
| Tubb2b | 2.15825E-12 | 1.96880856 | 0.723 | 0.365 | 5.24283E-08 | 0.265602785 | 0.45186787 | 0.6 | 0.398 | 1 | 0.265602785 | 4.31651E-12 |
| Baiap2l1 | 2.18521E-12 | 0.970051598 | 0.933 | 0.671 | 5.30831E-08 | 0.107582029 | 0.698138022 | 1 | 0.849 | 1 | 0.107582029 | 4.37042E-12 |
| Ptprd | 2.34113E-12 | -4.512907564 | 0.193 | 0.553 | 5.68707E-08 | 0.014567325 | -2.289175382 | 0.3 | 0.676 | 1 | 0.014567325 | 4.68226E-12 |
| Cidea | 2.4504E-12 | 3.141846478 | 0.429 | 0.082 | 5.95252E-08 | 0.000709804 | 2.191951308 | 0.5 | 0.127 | 1 | 0.000709804 | 4.90081E-12 |
| Flna | 2.48492E-12 | -2.141773961 | 0.37 | 0.706 | 6.03638E-08 | 0.051758248 | -0.85947179 | 0.5 | 0.84 | 1 | 0.051758248 | 4.96985E-12 |
| Phf21a | 2.50427E-12 | -1.555531066 | 0.731 | 0.9 | 6.08337E-08 | 0.162015981 | 0.304782328 | 1 | 0.932 | 1 | 0.162015981 | 5.00854E-12 |
| Ahdc1 | 2.55174E-12 | -1.777781807 | 0.387 | 0.729 | 6.19869E-08 | 0.405707936 | -0.589408406 | 0.8 | 0.855 | 1 | 0.405707936 | 5.10348E-12 |
| Sec31a | 2.56102E-12 | -0.808541837 | 0.95 | 0.988 | 6.22122E-08 | 0.115398623 | -0.463012512 | 1 | 0.988 | 1 | 0.115398623 | 5.12204E-12 |
| Rev3l | 2.66035E-12 | -1.666178859 | 0.504 | 0.806 | 6.46252E-08 | 0.424702899 | -0.453172628 | 0.6 | 0.867 | 1 | 0.424702899 | 5.3207E-12 |
| Eya2 | 2.66361E-12 | -2.449507253 | 0.202 | 0.588 | 6.47044E-08 | 0.031257131 | -1.58694015 | 0.6 | 0.843 | 1 | 0.031257131 | 5.32722E-12 |
| Col16a1 | 2.69927E-12 | -1.645298164 | 0.622 | 0.853 | 6.55708E-08 | 0.00029413 | -2.257426646 | 0.7 | 0.867 | 1 | 0.00029413 | 5.39855E-12 |
| Eif4g1 | 2.81101E-12 | -0.759056348 | 0.966 | 0.982 | 6.82849E-08 | 0.016575761 | -0.696018245 | 1 | 0.994 | 1 | 0.016575761 | 5.62201E-12 |
| Gfpt1 | 2.90573E-12 | -0.840269545 | 0.95 | 0.988 | 7.05859E-08 | 0.027973363 | -0.707576901 | 0.9 | 0.985 | 1 | 0.027973363 | 5.81146E-12 |
| Tspan18 | 3.08492E-12 | 1.335348611 | 0.899 | 0.6 | 7.49389E-08 | 0.018752582 | 1.11804321 | 0.9 | 0.583 | 1 | 0.018752582 | 6.16984E-12 |
| Sec24d | 3.09849E-12 | -1.046984724 | 0.874 | 0.929 | 7.52684E-08 | 0.139250944 | -0.588668372 | 1 | 0.948 | 1 | 0.139250944 | 6.19697E-12 |
| Ampd3 | 3.10245E-12 | 3.5334322 | 0.42 | 0.094 | 7.53648E-08 | 0.259865236 | -0.864721141 | 0 | 0.114 | 1 | 0.259865236 | 6.20491E-12 |
| Hsd17b7 | 3.17724E-12 | 1.536487949 | 0.857 | 0.5 | 7.71816E-08 | 0.25391278 | 0.899365298 | 0.6 | 0.481 | 1 | 0.25391278 | 6.35448E-12 |
| Gm56878 | 3.19389E-12 | 1.076815597 | 0.908 | 0.6 | 7.75859E-08 | 0.457297464 | -0.143069969 | 0.4 | 0.552 | 1 | 0.457297464 | 6.38777E-12 |
| Etnk2 | 3.26367E-12 | 3.459431619 | 0.319 | 0.024 | 7.9281E-08 | 4.22619E-05 | 3.111031312 | 0.3 | 0.034 | 1 | 4.22619E-05 | 6.52733E-12 |
| Ankrd17 | 3.29489E-12 | -1.115207832 | 0.933 | 0.971 | 8.00395E-08 | 0.054111874 | -0.648302095 | 1 | 0.994 | 1 | 0.054111874 | 6.58978E-12 |
| Sppl2a | 3.35986E-12 | 0.791154111 | 0.966 | 0.859 | 8.16176E-08 | 0.635054458 | 0.778782412 | 0.8 | 0.867 | 1 | 0.635054458 | 6.71971E-12 |
| Rab11fip1 | 3.48244E-12 | 2.139064038 | 0.58 | 0.212 | 8.45954E-08 | 0.91240568 | 1.017921908 | 0.3 | 0.296 | 1 | 0.91240568 | 6.96488E-12 |
| Sox9 | 3.49736E-12 | 1.075662178 | 0.992 | 0.929 | 8.49578E-08 | 0.170439084 | 0.638087569 | 1 | 0.948 | 1 | 0.170439084 | 6.99471E-12 |
| Rerg | 3.51204E-12 | -4.737566592 | 0.05 | 0.406 | 8.53144E-08 | 0.693962035 | -0.800127615 | 0.4 | 0.444 | 1 | 0.693962035 | 7.02407E-12 |
| Mon2 | 3.53499E-12 | -1.232660757 | 0.664 | 0.882 | 8.5872E-08 | 0.036469603 | -0.812555572 | 0.8 | 0.92 | 1 | 0.036469603 | 7.06998E-12 |
| Creb3l1 | 3.82448E-12 | -2.765534746 | 0.21 | 0.582 | 9.29043E-08 | 0.00200013 | -4.058893689 | 0 | 0.54 | 1 | 0.00200013 | 7.64897E-12 |
| Pum1 | 3.93114E-12 | -0.948398804 | 0.916 | 0.982 | 9.54953E-08 | 0.013894203 | -0.744115708 | 1 | 0.988 | 1 | 0.013894203 | 7.86228E-12 |
| Ep400 | 3.93124E-12 | -1.473552727 | 0.647 | 0.847 | 9.54977E-08 | 0.492367365 | -0.408342847 | 1 | 0.889 | 1 | 0.492367365 | 7.86248E-12 |
| Cnot1 | 4.32142E-12 | -1.361066489 | 0.655 | 0.847 | 1.04976E-07 | 0.008451359 | -1.197090983 | 0.6 | 0.895 | 1 | 0.008451359 | 8.64285E-12 |
| Mycbp2 | 4.32989E-12 | -1.01705406 | 0.84 | 0.959 | 1.05182E-07 | 0.369805161 | -0.254293974 | 1 | 0.981 | 1 | 0.369805161 | 8.65979E-12 |
| Tuba1c | 4.33036E-12 | 0.995784435 | 0.992 | 0.859 | 1.05193E-07 | 0.887404632 | 0.623555202 | 1 | 0.948 | 1 | 0.887404632 | 8.66072E-12 |
| Slc12a2 | 4.4953E-12 | 1.192645078 | 0.849 | 0.506 | 1.092E-07 | 0.034470487 | 0.823165054 | 0.9 | 0.636 | 1 | 0.034470487 | 8.99059E-12 |
| Fuca1 | 4.54114E-12 | 0.943125479 | 0.95 | 0.829 | 1.10313E-07 | 0.035080228 | 0.887762139 | 0.9 | 0.83 | 1 | 0.035080228 | 9.08228E-12 |
| Kirrel | 4.71627E-12 | -1.523295013 | 0.471 | 0.776 | 1.14568E-07 | 0.000399915 | -2.730270942 | 0.2 | 0.765 | 1 | 0.000399915 | 9.43254E-12 |
| Bicc1 | 4.74057E-12 | -2.025401873 | 0.655 | 0.835 | 1.15158E-07 | 0.205830342 | -1.473931188 | 0.8 | 0.821 | 1 | 0.205830342 | 9.48113E-12 |
| Ugp2 | 5.06053E-12 | 1.210374272 | 0.958 | 0.912 | 1.2293E-07 | 0.011534795 | 0.728015486 | 1 | 0.84 | 1 | 0.011534795 | 1.01211E-11 |
| Tmcc1 | 5.10246E-12 | -1.469347088 | 0.655 | 0.865 | 1.23949E-07 | 0.019133603 | -1.086795963 | 0.9 | 0.938 | 1 | 0.019133603 | 1.02049E-11 |
| Fnbp1l | 5.32103E-12 | -3.132673606 | 0.118 | 0.482 | 1.29259E-07 | 0.071018151 | -1.217294554 | 0.3 | 0.58 | 1 | 0.071018151 | 1.06421E-11 |
| Cnot6l | 5.4126E-12 | -1.93060949 | 0.429 | 0.747 | 1.31483E-07 | 0.151688314 | -0.694152511 | 0.7 | 0.852 | 1 | 0.151688314 | 1.08252E-11 |
| Zeb1 | 5.46701E-12 | -1.46136956 | 0.672 | 0.9 | 1.32805E-07 | 0.341137258 | -0.686055274 | 0.9 | 0.892 | 1 | 0.341137258 | 1.0934E-11 |
| Hnrnpa1 | 5.47044E-12 | -0.966253587 | 0.933 | 0.976 | 1.32888E-07 | 0.192177371 | -0.469788486 | 0.9 | 0.96 | 1 | 0.192177371 | 1.09409E-11 |
| Tceal9 | 5.56514E-12 | -1.106013238 | 0.899 | 0.947 | 1.35188E-07 | 0.012706531 | -0.98310237 | 0.7 | 0.966 | 1 | 0.012706531 | 1.11303E-11 |
| Gm13274 | 0.38869288 | 0.777607579 | 0.034 | 0.018 | 1 | 5.67901E-12 | 3.754887502 | 0.4 | 0.019 | 1.37955E-07 | 0.38869288 | 1.1358E-11 |
| Scin | 5.81095E-12 | 1.205005225 | 0.924 | 0.565 | 1.4116E-07 | 0.111190271 | 0.60749219 | 0.9 | 0.639 | 1 | 0.111190271 | 1.16219E-11 |
| Nebl | 6.00615E-12 | 0.999085745 | 0.975 | 0.7 | 1.45902E-07 | 0.081379212 | 0.376721393 | 1 | 0.735 | 1 | 0.081379212 | 1.20123E-11 |
| Tpm2 | 6.39433E-12 | -2.503962779 | 0.059 | 0.435 | 1.55331E-07 | 0.88438906 | 0.644463512 | 0.3 | 0.349 | 1 | 0.88438906 | 1.27887E-11 |
| Malat1 | 6.55304E-12 | 0.795380824 | 1 | 1 | 1.59186E-07 | 0.001113263 | 1.610722349 | 1 | 1 | 1 | 0.001113263 | 1.31061E-11 |
| Yy1 | 6.60444E-12 | -1.356693513 | 0.613 | 0.882 | 1.60435E-07 | 0.019328973 | -0.907550113 | 0.8 | 0.929 | 1 | 0.019328973 | 1.32089E-11 |
| Dhx32 | 6.67486E-12 | 1.220344145 | 0.84 | 0.406 | 1.62146E-07 | 0.009606149 | 1.432959407 | 0.8 | 0.432 | 1 | 0.009606149 | 1.33497E-11 |
| Ssbp2 | 6.72033E-12 | -2.035022285 | 0.42 | 0.741 | 1.6325E-07 | 0.000886176 | -2.754836758 | 0.6 | 0.873 | 1 | 0.000886176 | 1.34407E-11 |
| Cacna2d1 | 6.82965E-12 | -1.716625146 | 0.445 | 0.753 | 1.65906E-07 | 0.938153809 | 0.1378551 | 0.6 | 0.728 | 1 | 0.938153809 | 1.36593E-11 |
| Smc3 | 7.58651E-12 | -1.094501784 | 0.681 | 0.9 | 1.84292E-07 | 0.001336129 | -1.268712857 | 0.7 | 0.926 | 1 | 0.001336129 | 1.5173E-11 |
| Mamld1 | 7.96985E-12 | 1.576857451 | 0.571 | 0.176 | 1.93604E-07 | 0.019715327 | 2.299103661 | 0.4 | 0.145 | 1 | 0.019715327 | 1.59397E-11 |
| Psme4 | 8.13963E-12 | -1.279751801 | 0.84 | 0.941 | 1.97728E-07 | 0.052733795 | -0.717508087 | 0.9 | 0.991 | 1 | 0.052733795 | 1.62793E-11 |
| Plekha6 | 9.02666E-12 | 1.527869995 | 0.445 | 0.094 | 2.19276E-07 | 0.803619957 | 0.847996907 | 0.2 | 0.194 | 1 | 0.803619957 | 1.80533E-11 |
| ENSMUSG00000121322 | 9.17897E-12 | 1.440572591 | 0.639 | 0.224 | 2.22976E-07 | 0.005821461 | 1.787624289 | 0.6 | 0.253 | 1 | 0.005821461 | 1.83579E-11 |
| Stx5a | 9.87148E-12 | -1.346023771 | 0.588 | 0.818 | 2.39798E-07 | 0.212774322 | -0.671377253 | 0.8 | 0.836 | 1 | 0.212774322 | 1.9743E-11 |
| Hnrnpul2 | 9.95818E-12 | -1.161537216 | 0.639 | 0.853 | 2.41904E-07 | 0.003783006 | -1.185018151 | 0.8 | 0.91 | 1 | 0.003783006 | 1.99164E-11 |
| Sh3pxd2b | 1.02407E-11 | -3.719482166 | 0.059 | 0.418 | 2.48768E-07 | 0.004551037 | -3.970762779 | 0 | 0.485 | 1 | 0.004551037 | 2.04815E-11 |
| Mgst2 | 1.0332E-11 | 2.278132977 | 0.496 | 0.141 | 2.50985E-07 | 0.782867506 | 1.169925001 | 0.1 | 0.145 | 1 | 0.782867506 | 2.0664E-11 |
| Pabpc1 | 1.068E-11 | -0.7744403 | 0.992 | 0.994 | 2.59438E-07 | 0.02242363 | -0.681056853 | 0.9 | 0.997 | 1 | 0.02242363 | 2.136E-11 |
| Tubb5 | 1.07937E-11 | -1.451479495 | 0.681 | 0.871 | 2.622E-07 | 0.004107432 | -1.479530788 | 0.6 | 0.917 | 1 | 0.004107432 | 2.15874E-11 |
| Efnb2 | 1.13245E-11 | -3.426264755 | 0.092 | 0.453 | 2.75094E-07 | 0.011701768 | -3.31299497 | 0 | 0.41 | 1 | 0.011701768 | 2.26489E-11 |

| | | | | | | | | | | | | |
|---|---|---|---|---|---|---|---|---|---|---|---|---|
| Cep350 | 1.1342E-11 | -1.225743631 | 0.605 | 0.853 | 2.7552E-07 | 0.485219283 | -0.35767812 | 0.7 | 0.877 | 1 | 0.485219283 | 2.2684E-11 |
| Atl1 | 0.005992086 | 1.186518975 | 0.37 | 0.224 | 1 | 1.13958E-11 | 3.051088772 | 1 | 0.176 | 2.76826E-07 | 0.005992086 | 2.27915E-11 |
| Ltbp2 | 1.14972E-11 | -4.171927354 | 0.067 | 0.418 | 2.7929E-07 | 0.021553335 | -3.814968106 | 0 | 0.364 | 1 | 0.021553335 | 2.29944E-11 |
| Rbbp7 | 1.18267E-11 | -0.974234532 | 0.849 | 0.929 | 2.87293E-07 | 0.004479115 | -1.065678294 | 0.8 | 0.966 | 1 | 0.004479115 | 2.36533E-11 |
| Pltp | 1.27128E-11 | -4.353323291 | 0.025 | 0.365 | 3.08819E-07 | 0.070761269 | -2.391469028 | 0 | 0.253 | 1 | 0.070761269 | 2.54256E-11 |
| Slc35g1 | 1.29204E-11 | 1.479807755 | 0.723 | 0.318 | 3.13861E-07 | 0.023283965 | 1.043917117 | 0.5 | 0.182 | 1 | 0.023283965 | 2.58407E-11 |
| Rbbp6 | 1.42062E-11 | -1.322529093 | 0.866 | 0.929 | 3.45097E-07 | 0.000304305 | -1.652238606 | 0.8 | 0.978 | 1 | 0.000304305 | 2.84124E-11 |
| Tox | 1.42953E-11 | -2.485426827 | 0.193 | 0.582 | 3.47262E-07 | 0.013092334 | -1.99888638 | 0.1 | 0.565 | 1 | 0.013092334 | 2.85907E-11 |
| Sav1 | 1.5108E-11 | 1.052022212 | 0.95 | 0.747 | 3.67003E-07 | 0.327798954 | 1.198003524 | 0.8 | 0.877 | 1 | 0.327798954 | 3.0216E-11 |
| Phip | 1.52279E-11 | -1.881075341 | 0.412 | 0.759 | 3.69916E-07 | 0.047852625 | -1.352369208 | 0.6 | 0.756 | 1 | 0.047852625 | 3.04558E-11 |
| Dusp14 | 1.53811E-11 | 1.684498174 | 0.622 | 0.229 | 3.73637E-07 | 0.037624137 | 2.159940913 | 0.5 | 0.25 | 1 | 0.037624137 | 3.07622E-11 |
| Mlph | 1.59487E-11 | 3.592575685 | 0.387 | 0.076 | 3.87427E-07 | 0.025902593 | 2.414580878 | 0.3 | 0.09 | 1 | 0.025902593 | 3.18975E-11 |
| Brwd1 | 1.60585E-11 | -1.474371639 | 0.655 | 0.835 | 3.90092E-07 | 0.05666493 | -0.818856774 | 0.7 | 0.914 | 1 | 0.05666493 | 3.21169E-11 |
| Gdi2 | 1.70496E-11 | -0.89677928 | 0.908 | 0.953 | 4.14169E-07 | 0.000152073 | -1.483517237 | 0.8 | 0.978 | 1 | 0.000152073 | 3.40992E-11 |
| Tapbp | 1.70629E-11 | 0.977339403 | 0.941 | 0.718 | 4.14492E-07 | 0.016782827 | 0.767547074 | 1 | 0.753 | 1 | 0.016782827 | 3.41258E-11 |
| Srgap2 | 1.70701E-11 | -1.346993432 | 0.672 | 0.876 | 4.14667E-07 | 0.012935319 | -1.396890153 | 0.7 | 0.932 | 1 | 0.012935319 | 3.41402E-11 |
| Dnmt3a | 1.82569E-11 | -1.631745216 | 0.479 | 0.765 | 4.43495E-07 | 0.001170441 | -1.506619807 | 0.6 | 0.957 | 1 | 0.001170441 | 3.65137E-11 |
| Eif4g2 | 1.82885E-11 | -0.900464326 | 0.966 | 0.988 | 4.44263E-07 | 0.001972167 | -0.915844394 | 1 | 0.997 | 1 | 0.001972167 | 3.65769E-11 |
| Npm1 | 1.84066E-11 | -0.771905067 | 0.966 | 0.994 | 4.47132E-07 | 0.018892968 | -0.717016005 | 1 | 0.994 | 1 | 0.018892968 | 3.68131E-11 |
| Smarca4 | 1.88005E-11 | -1.372952098 | 0.513 | 0.812 | 4.56702E-07 | 0.060627734 | -0.961438326 | 0.7 | 0.806 | 1 | 0.060627734 | 3.7601E-11 |
| Eif3a | 1.91696E-11 | -0.787347374 | 0.983 | 1 | 4.65668E-07 | 7.45669E-05 | -1.40914146 | 0.8 | 0.997 | 1 | 7.45669E-05 | 3.83392E-11 |
| Cd9 | 2.03792E-11 | 0.67864856 | 1 | 1 | 4.95052E-07 | 0.196962161 | -0.259033304 | 1 | 1 | 1 | 0.196962161 | 4.07584E-11 |
| Atxn2 | 2.08022E-11 | -1.068456715 | 0.807 | 0.947 | 5.05327E-07 | 0.849176744 | -0.136263301 | 0.9 | 0.985 | 1 | 0.849176744 | 4.16044E-11 |
| Chd4 | 2.09131E-11 | -1.106639854 | 0.866 | 0.959 | 5.0802E-07 | 0.090660017 | -0.561647334 | 0.9 | 0.994 | 1 | 0.090660017 | 4.18261E-11 |
| Msl1 | 2.10215E-11 | -2.027954062 | 0.294 | 0.635 | 5.10655E-07 | 0.000887489 | -1.751915936 | 0.3 | 0.867 | 1 | 0.000887489 | 4.2043E-11 |
| Pth1r | 2.1335E-11 | -3.639855492 | 0.319 | 0.641 | 5.18269E-07 | 0.08867337 | -1.703177281 | 0.2 | 0.481 | 1 | 0.08867337 | 4.267E-11 |
| Ssh2 | 2.19558E-11 | -2.584962501 | 0.235 | 0.6 | 5.33349E-07 | 0.013033292 | -1.553101402 | 0.5 | 0.762 | 1 | 0.013033292 | 4.39115E-11 |
| Marcksl1 | 2.2235E-11 | -2.386648673 | 0.118 | 0.506 | 5.40133E-07 | 0.011940132 | -2.128434622 | 0.2 | 0.58 | 1 | 0.011940132 | 4.447E-11 |
| Naaladl2 | 2.27473E-11 | -2.164206688 | 0.529 | 0.765 | 5.52576E-07 | 0.002346263 | -2.130130913 | 0.4 | 0.833 | 1 | 0.002346263 | 4.54945E-11 |
| Hnrnpu | 2.30888E-11 | -0.796377383 | 0.975 | 0.994 | 5.60874E-07 | 0.000569816 | -1.08189118 | 0.9 | 1 | 1 | 0.000569816 | 4.61777E-11 |
| Itm2c | 2.36212E-11 | 0.70789267 | 0.992 | 0.9 | 5.73806E-07 | 0.05946404 | 0.704309081 | 0.9 | 0.923 | 1 | 0.05946404 | 4.72423E-11 |
| Gtf2ird1 | 2.4755E-11 | -2.364572432 | 0.151 | 0.535 | 6.0135E-07 | 0.009699464 | -2.141949429 | 0.4 | 0.691 | 1 | 0.009699464 | 4.95101E-11 |
| Pkm | 2.53041E-11 | 0.747946165 | 0.992 | 0.971 | 6.14688E-07 | 0.029389411 | 0.796095805 | 1 | 0.981 | 1 | 0.029389411 | 5.06082E-11 |
| Stim2 | 2.55429E-11 | -1.747648676 | 0.437 | 0.724 | 6.20488E-07 | 0.045113601 | -1.146527963 | 0.5 | 0.784 | 1 | 0.045113601 | 5.10858E-11 |
| Smarcc1 | 2.56362E-11 | -1.527434354 | 0.538 | 0.782 | 6.22754E-07 | 0.194419983 | -0.443070225 | 0.7 | 0.883 | 1 | 0.194419983 | 5.12724E-11 |
| Clstn2 | 2.62804E-11 | -3.816343705 | 0.059 | 0.406 | 6.38403E-07 | 0.028692105 | -2.976431529 | 0.2 | 0.528 | 1 | 0.028692105 | 5.25607E-11 |
| Acbd3 | 2.81741E-11 | -0.925089775 | 0.899 | 0.929 | 6.84406E-07 | 0.000583946 | -1.343846451 | 0.8 | 0.972 | 1 | 0.000583946 | 5.63482E-11 |
| Nrip1 | 2.91409E-11 | 1.099535674 | 0.916 | 0.812 | 7.07891E-07 | 0.838915523 | -0.164386818 | 0.8 | 0.849 | 1 | 0.838915523 | 5.82818E-11 |
| Eml1 | 2.95482E-11 | -4.46727948 | 0.034 | 0.365 | 7.17784E-07 | 0.208633159 | -1.846264237 | 0.3 | 0.435 | 1 | 0.208633159 | 5.90964E-11 |
| Lrrc58 | 2.98893E-11 | -0.863242505 | 0.849 | 0.941 | 7.2607E-07 | 0.029377389 | -0.73770753 | 0.9 | 0.978 | 1 | 0.029377389 | 5.97785E-11 |
| Luc7l2 | 3.17978E-11 | -0.708826717 | 0.983 | 1 | 7.72433E-07 | 0.00101088 | -0.942331913 | 1 | 1 | 1 | 0.00101088 | 6.35956E-11 |
| Mctp2 | 3.27279E-11 | 1.723520063 | 0.538 | 0.159 | 7.95026E-07 | 0.000936723 | 2.669998605 | 0.5 | 0.148 | 1 | 0.000936723 | 6.54558E-11 |
| Zfp36l2 | 3.32064E-11 | -1.965784285 | 0.387 | 0.712 | 8.06651E-07 | 0.014516582 | -2.292366271 | 0.4 | 0.71 | 1 | 0.014516582 | 6.64129E-11 |
| Zim1 | 3.33771E-11 | -3.392317423 | 0.084 | 0.429 | 8.10798E-07 | 0.976243004 | -0.448052557 | 0.4 | 0.333 | 1 | 0.976243004 | 6.67543E-11 |
| Gpx1 | 3.42745E-11 | 1.205255219 | 0.966 | 0.929 | 8.32597E-07 | 0.692969506 | 0.58329368 | 0.7 | 0.969 | 1 | 0.692969506 | 6.8549E-11 |
| Metrn | 3.53706E-11 | 2.555600441 | 0.471 | 0.141 | 8.59223E-07 | 0.001338401 | 2.36882907 | 0.5 | 0.148 | 1 | 0.001338401 | 7.07412E-11 |
| Slc10a6 | 3.74815E-11 | 2.536940986 | 0.336 | 0.041 | 9.105E-07 | 0.153152701 | 2.210566986 | 0.1 | 0.025 | 1 | 0.153152701 | 7.4963E-11 |
| Tlk1 | 3.82366E-11 | -1.602036014 | 0.487 | 0.788 | 9.28843E-07 | 0.454541892 | -0.28585884 | 0.7 | 0.855 | 1 | 0.454541892 | 7.64732E-11 |
| Prmt1 | 3.84376E-11 | -1.48966382 | 0.504 | 0.765 | 9.33727E-07 | 0.001726894 | -2.082740431 | 0.3 | 0.741 | 1 | 0.001726894 | 7.68753E-11 |
| Sema6a | 3.85766E-11 | -5.297925053 | 0.025 | 0.347 | 9.37103E-07 | 0.003520897 | -4.679045618 | 0 | 0.503 | 1 | 0.003520897 | 7.71532E-11 |
| Baz2a | 3.95416E-11 | -2.359895945 | 0.235 | 0.588 | 9.60546E-07 | 0.10962985 | -0.906350766 | 0.7 | 0.756 | 1 | 0.10962985 | 7.90833E-11 |
| Gm4681 | 0.389129094 | 1.099535674 | 0.025 | 0.012 | 1 | 3.99934E-11 | 4.432959407 | 0.4 | 0.022 | 9.71519E-07 | 0.389129094 | 7.99867E-11 |
| Gfra1 | 4.05818E-11 | 1.803334404 | 0.605 | 0.259 | 9.85812E-07 | 0.025614912 | 1.966874302 | 0.6 | 0.398 | 1 | 0.025614912 | 8.11635E-11 |
| Evl | 4.19484E-11 | -3.003752135 | 0.151 | 0.5 | 1.01901E-06 | 0.089940987 | -1.469918126 | 0.2 | 0.478 | 1 | 0.089940987 | 8.38968E-11 |
| Sos1 | 4.26371E-11 | -1.972051409 | 0.252 | 0.612 | 1.03574E-06 | 0.014031852 | -1.55950692 | 0.3 | 0.685 | 1 | 0.014031852 | 8.52743E-11 |
| Dnajc1 | 4.28183E-11 | -0.775201081 | 0.958 | 1 | 1.04014E-06 | 0.336835192 | -0.474572244 | 1 | 0.954 | 1 | 0.336835192 | 8.56365E-11 |
| Ptges3 | 4.43632E-11 | -0.71780068 | 0.983 | 0.982 | 1.07767E-06 | 0.037378701 | -0.699307468 | 0.8 | 0.997 | 1 | 0.037378701 | 8.87264E-11 |
| Ndrg2 | 4.44684E-11 | 1.410236513 | 0.773 | 0.388 | 1.08023E-06 | 0.053284777 | 0.796650027 | 0.8 | 0.562 | 1 | 0.053284777 | 8.89369E-11 |
| Cpa6 | 0.546771039 | 0.385290156 | 0.076 | 0.059 | 1 | 4.64032E-11 | 5.470434113 | 0.4 | 0.022 | 1.12723E-06 | 0.546771039 | 9.28065E-11 |
| Socs2 | 4.66929E-11 | 1.338606836 | 0.874 | 0.665 | 1.13426E-06 | 0.00330821 | 1.529635427 | 0.9 | 0.519 | 1 | 0.00330821 | 9.33859E-11 |
| Efhd1 | 0.028709407 | 2.514573173 | 0.067 | 0.018 | 1 | 4.97657E-11 | 3.494359952 | 0.4 | 0.022 | 1.20891E-06 | 0.028709407 | 9.95313E-11 |
| Pan3 | 5.03842E-11 | -1.202254776 | 0.798 | 0.924 | 1.22393E-06 | 0.479624374 | -0.410921391 | 0.9 | 0.941 | 1 | 0.479624374 | 1.00768E-10 |
| Ilf3 | 5.09845E-11 | -1.489519582 | 0.429 | 0.759 | 1.23852E-06 | 0.014452162 | -1.280369823 | 0.6 | 0.827 | 1 | 0.014452162 | 1.01969E-10 |
| Scube2 | 5.42703E-11 | -4.401903472 | 0.025 | 0.347 | 1.31834E-06 | 0.45439851 | -1.365782384 | 0.1 | 0.182 | 1 | 0.45439851 | 1.08541E-10 |
| Cyp26b1 | 5.44039E-11 | 1.46124367 | 0.731 | 0.335 | 1.32158E-06 | 0.049658927 | 0.870615209 | 0.7 | 0.435 | 1 | 0.049658927 | 1.08808E-10 |
| Cd28 | 5.60336E-11 | 3.80407979 | 0.277 | 0.018 | 1.36117E-06 | 0.044993069 | 3.252387162 | 0.2 | 0.052 | 1 | 0.044993069 | 1.12067E-10 |
| Rnf19a | 5.64609E-11 | -1.569027029 | 0.353 | 0.706 | 1.37155E-06 | 0.277059602 | -0.567040593 | 0.6 | 0.784 | 1 | 0.277059602 | 1.12922E-10 |
| Myo9a | 5.95144E-11 | -1.105395778 | 0.782 | 0.882 | 1.44572E-06 | 0.531019659 | -0.477215641 | 0.9 | 0.935 | 1 | 0.531019659 | 1.19029E-10 |
| Thbs3 | 6.04114E-11 | -2.710057721 | 0.126 | 0.471 | 1.46751E-06 | 0.703506566 | -0.304006187 | 0.3 | 0.34 | 1 | 0.703506566 | 1.20823E-10 |
| Rarg | 6.18077E-11 | -1.118598421 | 0.756 | 0.882 | 1.50143E-06 | 0.016154703 | -0.979474732 | 0.9 | 0.938 | 1 | 0.016154703 | 1.23615E-10 |
| Strn3 | 6.20899E-11 | -0.829195971 | 0.958 | 0.976 | 1.50829E-06 | 0.104124425 | -0.556638997 | 0.9 | 0.994 | 1 | 0.104124425 | 1.2418E-10 |
| Art3 | 6.349E-11 | 3.544320516 | 0.353 | 0.059 | 1.5423E-06 | 0.000653677 | 3.14345279 | 0.2 | 0.022 | 1 | 0.000653677 | 1.2698E-10 |
| Lcorl | 6.38549E-11 | -3.099340702 | 0.202 | 0.547 | 1.55116E-06 | 0.003652538 | -2.876895855 | 0.1 | 0.62 | 1 | 0.003652538 | 1.2771E-10 |
| Usp47 | 6.51254E-11 | -1.067091738 | 0.84 | 0.953 | 1.58203E-06 | 0.151615813 | -0.400469026 | 1 | 0.988 | 1 | 0.151615813 | 1.30251E-10 |
| Chd2 | 6.89442E-11 | -1.298134984 | 0.79 | 0.9 | 1.67479E-06 | 0.302924854 | -0.459184603 | 1 | 0.975 | 1 | 0.302924854 | 1.37888E-10 |
| Atrx | 6.89567E-11 | -1.029932174 | 0.882 | 0.994 | 1.6751E-06 | 0.01613447 | -0.846871151 | 0.8 | 0.994 | 1 | 0.01613447 | 1.37913E-10 |
| Mpzl1 | 6.99879E-11 | -1.583037624 | 0.454 | 0.735 | 1.70015E-06 | 0.638523568 | -0.165299916 | 0.8 | 0.756 | 1 | 0.638523568 | 1.39976E-10 |
| Gja1 | 7.13171E-11 | -1.943676393 | 0.353 | 0.682 | 1.73243E-06 | 0.136771524 | -0.840059087 | 0.2 | 0.491 | 1 | 0.136771524 | 1.42634E-10 |
| Lsm14a | 7.38171E-11 | -1.463534221 | 0.462 | 0.747 | 1.79316E-06 | 0.026833652 | -0.846676966 | 0.5 | 0.907 | 1 | 0.026833652 | 1.47634E-10 |
| Rbms3 | 7.92482E-11 | -2.022645228 | 0.412 | 0.729 | 1.9251E-06 | 0.000152235 | -5.455783842 | 0 | 0.688 | 1 | 0.000152235 | 1.58496E-10 |
| Smarce1 | 8.2406E-11 | -1.299013703 | 0.622 | 0.829 | 2.00181E-06 | 0.024229918 | -0.9715199 | 0.7 | 0.91 | 1 | 0.024229918 | 1.64812E-10 |
| Celf1 | 8.24493E-11 | -1.067669827 | 0.622 | 0.9 | 2.00286E-06 | 0.014657241 | -0.993752625 | 0.7 | 0.966 | 1 | 0.014657241 | 1.64899E-10 |
| Parp8 | 8.33185E-11 | -2.033137933 | 0.303 | 0.647 | 2.02397E-06 | 0.001856309 | -3.463877524 | 0.1 | 0.617 | 1 | 0.001856309 | 1.66637E-10 |
| Zfp292 | 8.44799E-11 | -2.027954062 | 0.218 | 0.582 | 2.05218E-06 | 0.058080145 | -1.379294314 | 0.5 | 0.682 | 1 | 0.058080145 | 1.6896E-10 |
| B3galt5 | 8.46983E-11 | 3.133483005 | 0.261 | 0.012 | 2.05749E-06 | 0.730750993 | 1.558490289 | 0 | 0.012 | 1 | 0.730750993 | 1.69397E-10 |
| Sez6l2 | 0.147029286 | -1.807354922 | 0 | 0.018 | 1 | 8.54368E-11 | 6.017921908 | 0.2 | 0.003 | 2.07543E-06 | 0.147029286 | 1.70874E-10 |
| Bbox1 | 0.217412808 | -1.485426827 | 0.008 | 0.029 | 1 | 8.54368E-11 | 6.017921908 | 0.2 | 0.003 | 2.07543E-06 | 0.217412808 | 1.70874E-10 |
| Cyyr1 | 0.508731273 | -1.292781749 | 0.008 | 0.018 | 1 | 8.54368E-11 | 6.339850003 | 0.2 | 0.003 | 2.07543E-06 | 0.508731273 | 1.70874E-10 |
| Trip13 | 0.926217033 | -0.970853654 | 0.034 | 0.035 | 1 | 8.54368E-11 | 6.017921908 | 0.2 | 0.003 | 2.07543E-06 | 0.926217033 | 1.70874E-10 |
| Cdkn2a | 0.949508638 | -0.485426827 | 0.017 | 0.018 | 1 | 8.54368E-11 | 6.017921908 | 0.2 | 0.003 | 2.07543E-06 | 0.949508638 | 1.70874E-10 |
| Adamts2 | 8.72481E-11 | -3.608809243 | 0.067 | 0.4 | 2.11943E-06 | 0.001791318 | -3.365782384 | 0.1 | 0.614 | 1 | 0.001791318 | 1.74496E-10 |
| Ints6 | 8.90472E-11 | -1.339399538 | 0.496 | 0.788 | 2.16314E-06 | 0.86402787 | -0.149946507 | 0.8 | 0.824 | 1 | 0.86402787 | 1.78094E-10 |
| Gnai1 | 8.91816E-11 | -3.515174171 | 0.076 | 0.412 | 2.1664E-06 | 0.001592721 | -3.802257054 | 0 | 0.552 | 1 | 0.001592721 | 1.78363E-10 |
| Rai1 | 9.03165E-11 | -2.137503524 | 0.16 | 0.529 | 2.19397E-06 | 0.019823934 | -1.369703819 | 0.5 | 0.818 | 1 | 0.019823934 | 1.80633E-10 |
| 1700047M11Rik | 1.42844E-06 | 3.269460675 | 0.176 | 0.018 | 0.03469971 | 9.14131E-11 | 5.602884409 | 0.2 | 0.003 | 2.22061E-06 | 1.42844E-06 | 1.82826E-10 |
| Gm15997 | 0.038142909 | 3.099535674 | 0.025 | 0 | 1 | 9.14131E-11 | 5.602884409 | 0.2 | 0.003 | 2.22061E-06 | 0.038142909 | 1.82826E-10 |
| Cenpu | 0.043234392 | -1.900464326 | 0.008 | 0.053 | 1 | 9.14131E-11 | 5.602884409 | 0.2 | 0.003 | 2.22061E-06 | 0.043234392 | 1.82826E-10 |
| E2f8 | 0.046722529 | 1.099535674 | 0.092 | 0.035 | 1 | 9.14131E-11 | 5.602884409 | 0.2 | 0.003 | 2.22061E-06 | 0.046722529 | 1.82826E-10 |

| | | | | | | | | | | | | |
|---|---|---|---|---|---|---|---|---|---|---|---|---|
| Spag6 | 0.060010312 | -2.485426827 | 0 | 0.029 | 1 | 9.14131E-11 | 5.602884409 | 0.2 | 0.003 | 2.22061E-06 | 0.060010312 | 1.82826E-10 |
| Gm32742 | 0.093245792 | -1.807354922 | 0 | 0.024 | 1 | 9.14131E-11 | 5.602884409 | 0.2 | 0.003 | 2.22061E-06 | 0.093245792 | 1.82826E-10 |
| Krt222 | 0.212902821 | 1.321928095 | 0.042 | 0.018 | 1 | 9.14131E-11 | 5.602884409 | 0.2 | 0.003 | 2.22061E-06 | 0.212902821 | 1.82826E-10 |
| Selenov | 0.237885589 | -2.292781749 | 0 | 0.012 | 1 | 9.14131E-11 | 5.602884409 | 0.2 | 0.003 | 2.22061E-06 | 0.237885589 | 1.82826E-10 |
| Rgsl1 | 0.368010882 | 1.836501268 | 0.017 | 0.006 | 1 | 9.14131E-11 | 5.602884409 | 0.2 | 0.003 | 2.22061E-06 | 0.368010882 | 1.82826E-10 |
| Sec1 | 0.370184089 | 1.099535674 | 0.017 | 0.006 | 1 | 9.14131E-11 | 5.602884409 | 0.2 | 0.003 | 2.22061E-06 | 0.370184089 | 1.82826E-10 |
| Gm47200 | 0.852288804 | 1.621488377 | 0.025 | 0.029 | 1 | 9.14131E-11 | 5.602884409 | 0.2 | 0.003 | 2.22061E-06 | 0.852288804 | 1.82826E-10 |
| Pdk4 | 9.2368E-11 | 3.393718778 | 0.42 | 0.112 | 2.2438E-06 | 3.57128E-07 | 4.921952849 | 0.8 | 0.228 | 0.008675352 | 3.57128E-07 | 1.84736E-10 |
| Smim14 | 9.32259E-11 | 0.763453484 | 0.992 | 0.971 | 2.26464E-06 | 0.03556503 | 0.655713996 | 1 | 0.978 | 1 | 0.03556503 | 1.86452E-10 |
| Phlda1 | 9.37541E-11 | 1.410179936 | 0.933 | 0.818 | 2.27748E-06 | 0.007577402 | 1.350218976 | 0.9 | 0.741 | 1 | 0.007577402 | 1.87508E-10 |
| Vezf1 | 9.44978E-11 | -1.197879063 | 0.731 | 0.888 | 2.29554E-06 | 0.002887792 | -1.310634925 | 0.6 | 0.941 | 1 | 0.002887792 | 1.88996E-10 |
| Gm41031 | 0.003173893 | 4.514573173 | 0.05 | 0 | 1 | 9.77849E-11 | 5.432959407 | 0.2 | 0.003 | 2.37539E-06 | 0.003173893 | 1.9557E-10 |
| Nrep | 9.80459E-11 | 1.535896942 | 0.899 | 0.729 | 2.38173E-06 | 0.413102634 | 0.673175731 | 0.8 | 0.806 | 1 | 0.413102634 | 1.96092E-10 |
| Smurf2 | 9.90613E-11 | -1.143289875 | 0.866 | 0.947 | 2.4064E-06 | 0.002405646 | -1.254207954 | 0.8 | 0.988 | 1 | 0.002405646 | 1.98123E-10 |
| Ctsk | 9.9601E-11 | -1.370752073 | 0.773 | 0.871 | 2.41951E-06 | 0.730655377 | -2.083708265 | 0.9 | 0.833 | 1 | 0.730655377 | 1.99202E-10 |
| Nfatc4 | 1.02686E-10 | -5.860466259 | 0 | 0.294 | 2.49444E-06 | 0.056301968 | -2.048167282 | 0 | 0.275 | 1 | 0.056301968 | 2.05371E-10 |
| Srrm3 | 0.167062889 | 1.836501268 | 0.025 | 0.006 | 1 | 1.04577E-10 | 5.017921908 | 0.2 | 0.003 | 2.54038E-06 | 0.167062889 | 2.09154E-10 |
| Rasgef1a | 0.661963295 | -0.333423734 | 0.025 | 0.018 | 1 | 1.0459E-10 | 5.017921908 | 0.2 | 0.003 | 2.54071E-06 | 0.661963295 | 2.09181E-10 |
| Pkp4 | 1.04794E-10 | -2.125430691 | 0.294 | 0.671 | 2.54566E-06 | 0.190260597 | -1.292366271 | 0.7 | 0.744 | 1 | 0.190260597 | 2.09588E-10 |
| Dbi | 1.04942E-10 | 1.127652423 | 0.966 | 0.924 | 2.54925E-06 | 0.002128556 | 1.489350589 | 1 | 0.883 | 1 | 0.002128556 | 2.09884E-10 |
| Pim1 | 1.06108E-10 | -3.169925001 | 0.134 | 0.476 | 2.57758E-06 | 0.015690478 | -2.703177281 | 0.1 | 0.488 | 1 | 0.015690478 | 2.12217E-10 |
| Kdm5b | 1.11052E-10 | -1.10134614 | 0.815 | 0.924 | 2.69767E-06 | 0.097135105 | -0.585100564 | 1 | 0.994 | 1 | 0.097135105 | 2.22104E-10 |
| Nbl1 | 1.14184E-10 | -1.782789776 | 0.345 | 0.694 | 2.77375E-06 | 0.001130675 | -2.455107314 | 0.4 | 0.802 | 1 | 0.001130675 | 2.28367E-10 |
| Hspb1 | 1.16028E-10 | 1.941790167 | 0.832 | 0.647 | 2.81855E-06 | 0.763073135 | -0.626835685 | 0.7 | 0.769 | 1 | 0.763073135 | 2.32056E-10 |
| Tgoln1 | 1.16303E-10 | 0.622587573 | 0.983 | 0.941 | 2.82523E-06 | 0.512950022 | 0.350218976 | 1 | 0.957 | 1 | 0.512950022 | 2.32606E-10 |
| Son | 1.22444E-10 | -0.520025489 | 1 | 1 | 2.9744E-06 | 0.235118003 | -0.235173474 | 1 | 1 | 1 | 0.235118003 | 2.44887E-10 |
| Mylip | 1.23267E-10 | -2.023595116 | 0.143 | 0.518 | 2.99441E-06 | 0.026756151 | -1.549400434 | 0.2 | 0.596 | 1 | 0.026756151 | 2.46535E-10 |
| Thoc2 | 1.2703E-10 | -0.913974613 | 0.916 | 0.941 | 3.0858E-06 | 0.29297674 | 0.238506528 | 0.9 | 0.944 | 1 | 0.29297674 | 2.54059E-10 |
| Cert1 | 1.29555E-10 | -1.494887156 | 0.529 | 0.806 | 3.14716E-06 | 0.063765001 | -0.957448425 | 0.6 | 0.793 | 1 | 0.063765001 | 2.59111E-10 |
| Usp9x | 1.30076E-10 | -1.047881928 | 0.899 | 0.976 | 3.15981E-06 | 0.008289359 | -1.02709662 | 0.7 | 0.991 | 1 | 0.008289359 | 2.60153E-10 |
| Tpm4 | 1.31342E-10 | -1.150048777 | 0.748 | 0.906 | 3.19057E-06 | 0.202590579 | -0.498303407 | 0.8 | 0.957 | 1 | 0.202590579 | 2.62685E-10 |
| Csnk1g1 | 1.317E-10 | -1.731864722 | 0.504 | 0.741 | 3.19926E-06 | 0.03968992 | -0.862273821 | 0.6 | 0.84 | 1 | 0.03968992 | 2.63401E-10 |
| Psmd1 | 1.33276E-10 | -1.039103639 | 0.798 | 0.918 | 3.23755E-06 | 0.112697969 | -0.574535129 | 0.9 | 0.917 | 1 | 0.112697969 | 2.66553E-10 |
| Ssb | 1.37198E-10 | -0.878787036 | 0.966 | 0.982 | 3.33281E-06 | 0.031223511 | -0.771215156 | 0.6 | 0.975 | 1 | 0.031223511 | 2.74396E-10 |
| Dnm1 | 1.38675E-10 | -1.777607579 | 0.277 | 0.635 | 3.3687E-06 | 0.001195244 | -2.420869945 | 0.2 | 0.728 | 1 | 0.001195244 | 2.77351E-10 |
| A130051J06Rik | 0.027938087 | 1.321928095 | 0.126 | 0.053 | 1 | 1.387E-10 | 4.111031312 | 0.5 | 0.04 | 3.36929E-06 | 0.027938087 | 2.77399E-10 |
| Apoe | 1.38952E-10 | -3.256255873 | 0.101 | 0.441 | 3.37543E-06 | 0.182588111 | -1.106597114 | 0.7 | 0.802 | 1 | 0.182588111 | 2.77905E-10 |
| Ppp2r5e | 1.39218E-10 | -1.521952703 | 0.471 | 0.729 | 3.38187E-06 | 0.099691836 | -0.938212207 | 0.8 | 0.809 | 1 | 0.099691836 | 2.78435E-10 |
| Cdk14 | 1.44568E-10 | -2.522516146 | 0.37 | 0.647 | 3.51184E-06 | 0.900987303 | -0.133471811 | 0.7 | 0.759 | 1 | 0.900987303 | 2.89135E-10 |
| Tspan5 | 1.45091E-10 | 1.324350757 | 0.824 | 0.512 | 3.52455E-06 | 0.186370681 | 0.983063151 | 0.7 | 0.642 | 1 | 0.186370681 | 2.90182E-10 |
| Stk26 | 1.5258E-10 | 0.711353666 | 0.891 | 0.594 | 3.70647E-06 | 0.040684927 | 0.494359952 | 1 | 0.534 | 1 | 0.040684927 | 3.0516E-10 |
| Sod3 | 1.52933E-10 | 1.380555825 | 0.647 | 0.271 | 3.71505E-06 | 0.822351923 | -0.720845929 | 0.4 | 0.352 | 1 | 0.822351923 | 3.05866E-10 |
| Tnpo1 | 1.55165E-10 | -0.757386463 | 0.849 | 0.965 | 3.76926E-06 | 0.006643734 | -1.066000576 | 1 | 0.96 | 1 | 0.006643734 | 3.1033E-10 |
| Ndrg1 | 1.61636E-10 | -2.733354341 | 0.109 | 0.453 | 3.92645E-06 | 0.000524774 | -4.045473173 | 0 | 0.614 | 1 | 0.000524774 | 3.23271E-10 |
| Ptk2 | 1.62462E-10 | -1.280987075 | 0.706 | 0.876 | 3.94652E-06 | 0.050588949 | -0.897957471 | 0.8 | 0.951 | 1 | 0.050588949 | 3.24924E-10 |
| Pi4k2a | 1.66449E-10 | 0.782730011 | 0.933 | 0.688 | 4.04337E-06 | 0.298989401 | 0.405889164 | 0.8 | 0.716 | 1 | 0.298989401 | 3.32898E-10 |
| Usp36 | 1.73006E-10 | -1.527862093 | 0.403 | 0.7 | 4.20267E-06 | 0.470480267 | 0.290001453 | 0.8 | 0.762 | 1 | 0.470480267 | 3.46013E-10 |
| Herc1 | 1.73485E-10 | -1.12076285 | 0.756 | 0.882 | 4.21429E-06 | 0.029821521 | -0.878242281 | 0.9 | 0.926 | 1 | 0.029821521 | 3.46969E-10 |
| Ubn2 | 1.73786E-10 | -0.909004998 | 0.908 | 0.971 | 4.22162E-06 | 0.338404119 | -0.362455901 | 0.9 | 0.954 | 1 | 0.338404119 | 3.47573E-10 |
| Serpinb6a | 1.77167E-10 | 0.954018747 | 0.975 | 0.918 | 4.30375E-06 | 0.920060315 | 0.165132336 | 0.8 | 0.898 | 1 | 0.920060315 | 3.54335E-10 |
| Mfap4 | 1.78055E-10 | -7.266786541 | 0 | 0.288 | 4.3253E-06 | 0.118240634 | -2.942080024 | 0 | 0.201 | 1 | 0.118240634 | 3.56109E-10 |
| Twist1 | 1.79558E-10 | -3.581351247 | 0.017 | 0.324 | 4.36181E-06 | 0.034783781 | -3.157003775 | 0 | 0.321 | 1 | 0.034783781 | 3.59115E-10 |
| Igsf10 | 1.8131E-10 | -2.957914599 | 0.109 | 0.447 | 4.40437E-06 | 0.015160567 | -3.116504412 | 0 | 0.392 | 1 | 0.015160567 | 3.62619E-10 |
| Cul1 | 1.85615E-10 | -1.096188797 | 0.924 | 0.976 | 4.50897E-06 | 0.000331393 | -1.294082694 | 0.9 | 0.997 | 1 | 0.000331393 | 3.71231E-10 |
| Ptprs | 1.92989E-10 | -1.948053785 | 0.244 | 0.6 | 4.68808E-06 | 0.227658212 | -0.767802998 | 0.4 | 0.562 | 1 | 0.227658212 | 3.85977E-10 |
| Otos | 1.93123E-10 | 6.569855608 | 0.261 | 0.018 | 4.69135E-06 | 0.000103743 | 4.432959407 | 0.1 | 0.003 | 1 | 0.000103743 | 3.86247E-10 |
| Chd1 | 1.97206E-10 | -1.138155884 | 0.807 | 0.941 | 4.79052E-06 | 0.154800149 | -0.564726727 | 0.7 | 0.904 | 1 | 0.154800149 | 3.94411E-10 |
| Nav1 | 1.97489E-10 | 1.377968245 | 0.857 | 0.524 | 4.79739E-06 | 0.016126581 | 0.728087443 | 0.9 | 0.577 | 1 | 0.016126581 | 3.94977E-10 |
| Cd302 | 2.01881E-10 | 0.871038213 | 0.95 | 0.747 | 4.90408E-06 | 0.923600226 | 0.175095412 | 0.8 | 0.765 | 1 | 0.923600226 | 4.03761E-10 |
| Hipk2 | 2.03402E-10 | 1.108013827 | 0.933 | 0.729 | 4.94104E-06 | 0.241523294 | 1.336523756 | 0.9 | 0.772 | 1 | 0.241523294 | 4.06804E-10 |
| Rfx3 | 2.06082E-10 | -1.917600239 | 0.294 | 0.624 | 5.00615E-06 | 0.037495763 | -1.736965594 | 0.5 | 0.728 | 1 | 0.037495763 | 4.12165E-10 |
| Scube1 | 2.20833E-10 | 1.133483005 | 0.765 | 0.424 | 5.36448E-06 | 0.025334013 | 2.009929117 | 0.6 | 0.299 | 1 | 0.025334013 | 4.41666E-10 |
| Cul3 | 2.20949E-10 | -0.919829651 | 0.908 | 0.947 | 5.36729E-06 | 0.00949248 | -0.787397704 | 1 | 0.981 | 1 | 0.00949248 | 4.41898E-10 |
| Frmd6 | 2.21166E-10 | -2.462219436 | 0.445 | 0.676 | 5.37257E-06 | 4.69817E-05 | -2.820581713 | 0.4 | 0.92 | 1 | 4.69817E-05 | 4.42332E-10 |
| Dsel | 2.35107E-10 | -2.470319935 | 0.176 | 0.506 | 5.71122E-06 | 0.29353776 | -0.898554736 | 0.4 | 0.475 | 1 | 0.29353776 | 4.70214E-10 |
| Egfr | 2.36504E-10 | -2.556370079 | 0.134 | 0.488 | 5.74514E-06 | 0.173397125 | -0.592692858 | 0.5 | 0.812 | 1 | 0.173397125 | 4.73007E-10 |
| Ube2i | 2.39908E-10 | -0.776475609 | 0.899 | 0.929 | 5.82784E-06 | 0.026666094 | -0.785915674 | 0.8 | 0.954 | 1 | 0.026666094 | 4.79816E-10 |
| Foxd1 | 2.41235E-10 | -5.253611152 | 0.008 | 0.3 | 5.86007E-06 | 0.075275291 | -2.131825212 | 0 | 0.247 | 1 | 0.075275291 | 4.82469E-10 |
| Hnrnpf | 2.41284E-10 | -0.828623539 | 0.924 | 0.976 | 5.86127E-06 | 0.000543987 | -1.039528364 | 1 | 0.988 | 1 | 0.000543987 | 4.82568E-10 |
| Spsb2 | 2.42645E-10 | 1.555445454 | 0.748 | 0.441 | 5.89434E-06 | 0.740380781 | 0.18142064 | 0.5 | 0.481 | 1 | 0.740380781 | 4.85291E-10 |
| Morf4l2 | 2.44989E-10 | -0.573182611 | 0.992 | 0.988 | 5.95126E-06 | 0.001599814 | -0.885416708 | 1 | 0.994 | 1 | 0.001599814 | 4.89977E-10 |
| Avil | 2.56266E-10 | -2.747521673 | 0.185 | 0.512 | 6.2252E-06 | 0.09272727 | -1.477933119 | 0.4 | 0.599 | 1 | 0.09272727 | 5.12531E-10 |
| Dhx15 | 2.58408E-10 | -1.235448574 | 0.681 | 0.824 | 6.27725E-06 | 0.002332111 | -1.389770741 | 0.4 | 0.895 | 1 | 0.002332111 | 5.16816E-10 |
| Mxra7 | 2.59947E-10 | -2.167855137 | 0.395 | 0.647 | 6.31464E-06 | 0.023607879 | -1.766712938 | 0.3 | 0.596 | 1 | 0.023607879 | 5.19895E-10 |
| Hnrnpa3 | 2.87874E-10 | -0.811888463 | 0.983 | 0.988 | 6.99304E-06 | 0.008837689 | -0.708296251 | 0.9 | 0.997 | 1 | 0.008837689 | 5.75748E-10 |
| Cspp1 | 2.88767E-10 | -1.785328616 | 0.294 | 0.641 | 7.01472E-06 | 0.013138001 | -1.5519337 | 0.2 | 0.685 | 1 | 0.013138001 | 5.77533E-10 |
| Man2a1 | 2.89214E-10 | 1.009023381 | 0.815 | 0.494 | 7.02558E-06 | 0.003828 | 0.95491211 | 0.9 | 0.503 | 1 | 0.003828 | 5.78427E-10 |
| Eed | 3.04411E-10 | -1.913663824 | 0.202 | 0.553 | 7.39476E-06 | 0.093774616 | -1.056754778 | 0.3 | 0.562 | 1 | 0.093774616 | 6.08822E-10 |
| Dnm3 | 7.76947E-05 | 1.374395515 | 0.353 | 0.153 | 1 | 3.07393E-10 | 2.839118755 | 1 | 0.185 | 7.4672E-06 | 7.76947E-05 | 6.14787E-10 |
| Pnn | 3.09017E-10 | -0.965527783 | 0.832 | 0.941 | 7.50663E-06 | 0.005211549 | -1.124635686 | 0.7 | 0.948 | 1 | 0.005211549 | 6.18033E-10 |
| Fat1 | 3.09821E-10 | -1.881011964 | 0.395 | 0.706 | 7.52616E-06 | 0.717599588 | -0.102171937 | 0.7 | 0.688 | 1 | 0.717599588 | 6.19641E-10 |
| Pkmyt1 | 0.725168649 | 0.12225575 | 0.101 | 0.088 | 1 | 3.13895E-10 | 5.503348735 | 0.3 | 0.012 | 7.62514E-06 | 0.725168649 | 6.2779E-10 |
| Gnb1 | 3.28204E-10 | -0.766209902 | 0.933 | 0.988 | 7.97274E-06 | 0.004040339 | -1.051330395 | 0.7 | 0.994 | 1 | 0.004040339 | 6.56409E-10 |
| Radil | 3.31449E-10 | 1.019808481 | 0.504 | 0.141 | 8.05155E-06 | 2.18471E-08 | 2.611929548 | 0.6 | 0.074 | 0.000530709 | 2.18471E-08 | 6.62897E-10 |
| Odc1 | 3.36851E-10 | 0.818460963 | 0.983 | 0.859 | 8.18278E-06 | 0.203630552 | 0.312052394 | 1 | 0.855 | 1 | 0.203630552 | 6.73702E-10 |
| Piezo2 | 3.4013E-10 | -1.488577964 | 0.529 | 0.824 | 8.26244E-06 | 0.001335633 | -1.733031327 | 0.6 | 0.904 | 1 | 0.001335633 | 6.8026E-10 |
| Dpf3 | 3.47969E-10 | 2.184424571 | 0.42 | 0.1 | 8.45286E-06 | 0.730750993 | 2.210566986 | 0 | 0.012 | 1 | 0.730750993 | 6.95938E-10 |
| Picalm | 3.53809E-10 | -0.978758852 | 0.899 | 0.935 | 8.59472E-06 | 0.002140389 | -1.269392432 | 0.9 | 0.994 | 1 | 0.002140389 | 7.07617E-10 |
| Sh3rf1 | 3.59937E-10 | -1.564911611 | 0.437 | 0.753 | 8.7436E-06 | 0.987900241 | -0.256477817 | 0.8 | 0.753 | 1 | 0.987900241 | 7.19875E-10 |
| Copb2 | 3.66522E-10 | -0.934184944 | 0.874 | 0.947 | 8.90355E-06 | 0.003253254 | -1.187464088 | 0.6 | 0.963 | 1 | 0.003253254 | 7.33043E-10 |
| Gjc3 | 3.69645E-10 | 0.843732837 | 0.765 | 0.376 | 8.97942E-06 | 0.0161889 | 2.532495081 | 0.2 | 0.04 | 1 | 0.0161889 | 7.3929E-10 |
| Col6a3 | 3.70482E-10 | 0.684148048 | 0.992 | 0.924 | 8.99975E-06 | 0.008343272 | 0.722609669 | 1 | 0.886 | 1 | 0.008343272 | 7.40964E-10 |
| Maoa | 3.71284E-10 | -3.733354341 | 0.042 | 0.353 | 9.01923E-06 | 0.095711339 | -2.191531458 | 0 | 0.222 | 1 | 0.095711339 | 7.42568E-10 |
| C1qtnf3 | 1.98037E-05 | -4.807354922 | 0 | 0.141 | 0.481070682 | 3.72642E-10 | 5.017921908 | 0.3 | 0.012 | 9.05221E-06 | 1.98037E-05 | 7.45284E-10 |
| Gm5820 | 0.037793735 | 1.292180751 | 0.076 | 0.024 | 1 | 3.73171E-10 | 5.210566986 | 0.3 | 0.012 | 9.06507E-06 | 0.037793735 | 7.46342E-10 |
| Fxyd2 | 3.83036E-10 | 2.770169515 | 0.487 | 0.159 | 9.30471E-06 | 7.14641E-07 | 3.847996907 | 0.5 | 0.074 | 0.017360061 | 7.14641E-07 | 7.66072E-10 |
| Gpc6 | 3.87597E-10 | 0.602760834 | 1 | 0.994 | 9.41551E-06 | 0.148049646 | 0.614993385 | 1 | 0.991 | 1 | 0.148049646 | 7.75194E-10 |
| Memo1 | 3.92713E-10 | -1.72080489 | 0.429 | 0.706 | 9.53978E-06 | 0.125703897 | -1.133855747 | 0.7 | 0.725 | 1 | 0.125703897 | 7.85426E-10 |

| | | | | | | | | | | | | |
|---|---|---|---|---|---|---|---|---|---|---|---|---|
| Car2 | 4.00704E-10 | 6.480357457 | 0.227 | 0.006 | 9.7339E-06 | 0.000847156 | 2.079322453 | 0.2 | 0.022 | 1 | 0.000847156 | 8.01408E-10 |
| Septin9 | 4.01673E-10 | 0.765572514 | 0.958 | 0.776 | 9.75745E-06 | 0.046500377 | 1.105384749 | 0.9 | 0.861 | 1 | 0.046500377 | 8.03347E-10 |
| Slc6a8 | 4.06238E-10 | 1.353029312 | 0.731 | 0.4 | 9.86833E-06 | 0.306011788 | -0.161987182 | 0.3 | 0.475 | 1 | 0.306011788 | 8.12476E-10 |
| Stk38l | 4.35599E-10 | 0.649813645 | 0.966 | 0.6 | 1.05816E-05 | 0.09818882 | 0.643969253 | 0.8 | 0.753 | 1 | 0.09818882 | 8.71197E-10 |
| Bmp7 | 4.42366E-10 | -4.070389328 | 0.034 | 0.335 | 1.0746E-05 | 0.4883384 | -0.125036046 | 0.1 | 0.201 | 1 | 0.4883384 | 8.84732E-10 |
| Banf2 | 0.004473483 | 2.362570079 | 0.101 | 0.024 | 1 | 4.43233E-10 | 4.695993813 | 0.3 | 0.012 | 1.0767E-05 | 0.004473483 | 8.86465E-10 |
| Ecpas | 4.43389E-10 | -1.144021358 | 0.849 | 0.947 | 1.07708E-05 | 0.034457071 | -0.841440318 | 0.9 | 0.972 | 1 | 0.034457071 | 8.86778E-10 |
| Abl1 | 4.44272E-10 | -2.292781749 | 0.168 | 0.512 | 1.07923E-05 | 0.538856969 | -0.308008117 | 0.4 | 0.552 | 1 | 0.538856969 | 8.88544E-10 |
| Pigs | 4.4927E-10 | 1.401362562 | 0.622 | 0.265 | 1.09137E-05 | 0.010924996 | 1.737813989 | 0.6 | 0.299 | 1 | 0.010924996 | 8.98539E-10 |
| Obsl1 | 4.61953E-10 | -2.318316841 | 0.109 | 0.447 | 1.12218E-05 | 0.236346631 | -0.814968106 | 0.2 | 0.367 | 1 | 0.236346631 | 9.23907E-10 |
| Zbtb11 | 4.91589E-10 | -1.366346844 | 0.588 | 0.812 | 1.19417E-05 | 0.131329692 | -0.842820227 | 0.7 | 0.843 | 1 | 0.131329692 | 9.83179E-10 |
| Srsf4 | 5.10828E-10 | -1.442358105 | 0.429 | 0.724 | 1.2409E-05 | 0.725334805 | -0.158001834 | 0.9 | 0.852 | 1 | 0.725334805 | 1.02166E-09 |
| Srek1 | 5.15743E-10 | -1.036347755 | 0.79 | 0.935 | 1.25284E-05 | 0.081394025 | -0.81691375 | 0.9 | 0.932 | 1 | 0.081394025 | 1.03149E-09 |
| Eif4ebp1 | 5.40502E-10 | -2.49441561 | 0.126 | 0.459 | 1.31299E-05 | 0.493874759 | -0.531541912 | 0.3 | 0.41 | 1 | 0.493874759 | 1.081E-09 |
| Supt16 | 5.4229E-10 | -0.976815213 | 0.588 | 0.818 | 1.31733E-05 | 0.065867356 | -0.755217299 | 0.6 | 0.867 | 1 | 0.065867356 | 1.08458E-09 |
| H2-T23 | 5.48864E-10 | 2.821234511 | 0.387 | 0.088 | 1.3333E-05 | 0.803962884 | 0.111031312 | 0.1 | 0.123 | 1 | 0.803962884 | 1.09773E-09 |
| Phf3 | 5.49377E-10 | -1.163026569 | 0.782 | 0.924 | 1.33455E-05 | 0.121920482 | -0.5519337 | 0.9 | 0.951 | 1 | 0.121920482 | 1.09875E-09 |
| Psmd11 | 5.54898E-10 | -0.8244791 | 0.882 | 0.965 | 1.34796E-05 | 0.012468497 | -0.92188331 | 0.7 | 0.978 | 1 | 0.012468497 | 1.1098E-09 |
| Sec61a1 | 5.66572E-10 | -1.252227651 | 0.689 | 0.824 | 1.37632E-05 | 0.012180304 | -1.126272818 | 0.5 | 0.846 | 1 | 0.012180304 | 1.13314E-09 |
| Eml4 | 5.78138E-10 | -1.565491216 | 0.563 | 0.812 | 1.40441E-05 | 0.003555973 | -2.152003093 | 0.6 | 0.873 | 1 | 0.003555973 | 1.15628E-09 |
| Dlg4 | 5.95946E-10 | -2.061776198 | 0.185 | 0.535 | 1.44767E-05 | 0.315640857 | 0.12217187 | 0.3 | 0.537 | 1 | 0.315640857 | 1.19189E-09 |
| Isoc1 | 5.98782E-10 | -2.321928095 | 0.101 | 0.435 | 1.45456E-05 | 0.461055475 | 0.558490289 | 0.5 | 0.392 | 1 | 0.461055475 | 1.19756E-09 |
| Col1a1 | 6.06695E-10 | -7.40449562 | 0.076 | 0.376 | 1.47378E-05 | 0.028899495 | -6.40177498 | 0.1 | 0.429 | 1 | 0.028899495 | 1.21339E-09 |
| Neb | 6.07795E-10 | 3.96203215 | 0.319 | 0.053 | 1.47645E-05 | 0.23891309 | 1.930459067 | 0.1 | 0.031 | 1 | 0.23891309 | 1.21559E-09 |
| Gng12 | 6.20615E-10 | 0.716395388 | 0.975 | 0.847 | 1.5076E-05 | 0.039172606 | 0.559964684 | 1 | 0.957 | 1 | 0.039172606 | 1.24123E-09 |
| Itm2a | 6.20995E-10 | -1.516925652 | 0.966 | 0.912 | 1.50852E-05 | 0.105728531 | -0.927862163 | 0.9 | 0.846 | 1 | 0.105728531 | 1.24199E-09 |
| Setd5 | 6.2525E-10 | -0.964359892 | 0.899 | 0.965 | 1.51886E-05 | 0.648663421 | -0.283500531 | 1 | 0.991 | 1 | 0.648663421 | 1.2505E-09 |
| Calu | 6.3696E-10 | -0.538735671 | 0.992 | 0.994 | 1.5473E-05 | 0.222569569 | -0.276821358 | 1 | 0.988 | 1 | 0.222569569 | 1.27392E-09 |
| Runx2 | 6.54584E-10 | -1.656925621 | 0.681 | 0.876 | 1.59012E-05 | 0.807583547 | -0.395101498 | 0.9 | 0.849 | 1 | 0.807583547 | 1.30917E-09 |
| Loxl1 | 6.76845E-10 | -2.488701959 | 0.311 | 0.582 | 1.64419E-05 | 0.910593777 | 0.191649773 | 0.8 | 0.784 | 1 | 0.910593777 | 1.35369E-09 |
| Abcc5 | 6.77885E-10 | -1.597774553 | 0.37 | 0.676 | 1.64672E-05 | 0.72113971 | -0.267480311 | 0.9 | 0.725 | 1 | 0.72113971 | 1.35577E-09 |
| Cpe | 6.8188E-10 | 0.80407979 | 0.983 | 0.906 | 1.65642E-05 | 0.203321152 | 0.317977706 | 1 | 0.954 | 1 | 0.203321152 | 1.36376E-09 |
| Slc26a2 | 6.84831E-10 | 0.965356333 | 0.95 | 0.729 | 1.66359E-05 | 0.091937051 | 0.485604949 | 1 | 0.793 | 1 | 0.091937051 | 1.36966E-09 |
| Gm13944 | 6.86718E-10 | 1.782506378 | 0.504 | 0.159 | 1.66818E-05 | 0.554456463 | 1.615823464 | 0.2 | 0.154 | 1 | 0.554456463 | 1.37344E-09 |
| Avpi1 | 7.16914E-10 | 2.221165118 | 0.605 | 0.324 | 1.74153E-05 | 0.878919908 | 0.808468542 | 0.3 | 0.318 | 1 | 0.878919908 | 1.43383E-09 |
| Serpine1 | 7.26617E-10 | -2.451406718 | 0.143 | 0.506 | 1.7651E-05 | 0.093529807 | -3.786209113 | 0 | 0.225 | 1 | 0.093529807 | 1.45323E-09 |
| Sec63 | 7.57467E-10 | -0.90908934 | 0.899 | 0.953 | 1.84004E-05 | 0.011736315 | -0.966386994 | 0.9 | 0.963 | 1 | 0.011736315 | 1.51493E-09 |
| Appl1 | 7.57756E-10 | -1.301415498 | 0.647 | 0.812 | 1.84074E-05 | 0.124065054 | -0.482343428 | 0.6 | 0.855 | 1 | 0.124065054 | 1.51551E-09 |
| Abca9 | 8.06725E-10 | -3.596935142 | 0.05 | 0.353 | 1.9597E-05 | 0.062304421 | -2.365782384 | 0 | 0.265 | 1 | 0.062304421 | 1.61345E-09 |
| Bdp1 | 8.16918E-10 | -0.921011704 | 0.824 | 0.935 | 1.98446E-05 | 0.576162751 | 0.205209776 | 0.9 | 0.929 | 1 | 0.576162751 | 1.63384E-09 |
| Wac | 8.33767E-10 | -0.927724099 | 0.924 | 0.971 | 2.02539E-05 | 0.090123117 | -0.313170662 | 1 | 0.985 | 1 | 0.090123117 | 1.66753E-09 |
| Itm2b | 8.4958E-10 | 0.717910095 | 1 | 0.994 | 2.0638E-05 | 0.853546205 | 0.398570682 | 1 | 0.997 | 1 | 0.853546205 | 1.69916E-09 |
| Tmem140 | 8.54987E-10 | 1.367825805 | 0.714 | 0.406 | 2.07693E-05 | 0.509638969 | 1.229236197 | 0.5 | 0.506 | 1 | 0.509638969 | 1.70997E-09 |
| Tmod3 | 8.57221E-10 | -1.416356434 | 0.647 | 0.806 | 2.08236E-05 | 0.01866354 | -0.939760578 | 0.7 | 0.91 | 1 | 0.01866354 | 1.71444E-09 |
| Ecm2 | 8.61773E-10 | -1.416517118 | 0.471 | 0.776 | 2.09342E-05 | 0.022418529 | -1.324737628 | 0.8 | 0.812 | 1 | 0.022418529 | 1.72355E-09 |
| 4931414P19Rik | 0.240743347 | -1.333423734 | 0.025 | 0.053 | 1 | 9.18156E-10 | 3.655351829 | 0.5 | 0.043 | 2.23038E-05 | 0.240743347 | 1.83631E-09 |
| Ly6c1 | 9.26317E-10 | 2.378918073 | 0.387 | 0.088 | 2.25021E-05 | 0.020368636 | 2.177702352 | 0.5 | 0.201 | 1 | 0.020368636 | 1.85263E-09 |
| Eif3b | 9.39999E-10 | -1.598884877 | 0.37 | 0.653 | 2.28345E-05 | 0.052676763 | -0.946212639 | 0.4 | 0.685 | 1 | 0.052676763 | 1.88E-09 |
| Gm15271 | 9.4172E-10 | 2.370837695 | 0.403 | 0.1 | 2.28763E-05 | 0.049344356 | 3.017921908 | 0.1 | 0.015 | 1 | 0.049344356 | 1.88344E-09 |
| Tanc2 | 9.56366E-10 | -1.481954635 | 0.429 | 0.735 | 2.3232E-05 | 0.843865901 | -0.398992879 | 0.8 | 0.713 | 1 | 0.843865901 | 1.91273E-09 |
| Upf2 | 9.80715E-10 | -1.268196111 | 0.529 | 0.782 | 2.38235E-05 | 0.666694831 | -0.105164843 | 0.8 | 0.802 | 1 | 0.666694831 | 1.96143E-09 |
| Tmem51 | 1.00691E-09 | 2.427110332 | 0.429 | 0.129 | 2.44599E-05 | 4.41759E-05 | 2.506959989 | 0.5 | 0.096 | 1 | 4.41759E-05 | 2.01382E-09 |
| Gal3st1 | 0.000248961 | 1.74723393 | 0.227 | 0.076 | 1 | 1.01076E-09 | 3.602884409 | 0.5 | 0.043 | 2.45533E-05 | 0.000248961 | 2.02151E-09 |
| Enpp5 | 1.01852E-09 | 1.558967292 | 0.538 | 0.194 | 2.4742E-05 | 0.000189836 | 1.994075166 | 0.8 | 0.281 | 1 | 0.000189836 | 2.03705E-09 |
| Tgfb3 | 1.02107E-09 | -2.192379852 | 0.185 | 0.524 | 2.48038E-05 | 0.013353153 | -1.820021334 | 0.2 | 0.614 | 1 | 0.013353153 | 2.04214E-09 |
| Papss2 | 1.06128E-09 | 0.78895946 | 1 | 0.912 | 2.57807E-05 | 0.954904014 | -0.297774233 | 1 | 0.96 | 1 | 0.954904014 | 2.12257E-09 |
| Kif1b | 1.10333E-09 | -1.058282725 | 0.891 | 0.929 | 2.68021E-05 | 0.011378196 | -0.9214774 | 1 | 0.988 | 1 | 0.011378196 | 2.20666E-09 |
| Wif1 | 1.11225E-09 | 3.099535674 | 0.706 | 0.512 | 2.70188E-05 | 0.06165365 | 1.765925774 | 0.8 | 0.731 | 1 | 0.06165365 | 2.22451E-09 |
| Nr2f2 | 1.11621E-09 | -2.063876021 | 0.311 | 0.618 | 2.71149E-05 | 0.007867864 | -2.345482823 | 0.1 | 0.574 | 1 | 0.007867864 | 2.23242E-09 |
| Tnrc6a | 1.13079E-09 | -1.402371087 | 0.597 | 0.788 | 2.74692E-05 | 0.009357477 | -1.031216362 | 0.9 | 0.966 | 1 | 0.009357477 | 2.26158E-09 |
| Cdk11b | 1.13467E-09 | -1.012037089 | 0.748 | 0.888 | 2.75633E-05 | 0.100310108 | -0.592364749 | 0.8 | 0.935 | 1 | 0.100310108 | 2.26933E-09 |
| Bag1 | 1.14047E-09 | 1.038135129 | 0.958 | 0.835 | 2.77043E-05 | 0.169157765 | 0.539874611 | 0.8 | 0.852 | 1 | 0.169157765 | 2.28094E-09 |
| Naca | 1.15428E-09 | -0.585072542 | 0.95 | 1 | 2.80398E-05 | 0.003949115 | -0.956082883 | 0.7 | 0.994 | 1 | 0.003949115 | 2.30856E-09 |
| Medag | 1.15787E-09 | 1.864706149 | 0.622 | 0.282 | 2.8127E-05 | 0.098704357 | -0.745843746 | 0.5 | 0.207 | 1 | 0.098704357 | 2.31575E-09 |
| Bglap | 1.21758E-09 | 4.486782107 | 0.454 | 0.147 | 2.95774E-05 | 0.463974022 | -1.888968688 | 0.1 | 0.046 | 1 | 0.463974022 | 2.43516E-09 |
| Frzb | 1.22518E-09 | 1.982595128 | 0.571 | 0.241 | 2.97622E-05 | 0.00201372 | 3.96133838 | 0.4 | 0.108 | 1 | 0.00201372 | 2.45037E-09 |
| Lpin2 | 1.23661E-09 | 1.232450444 | 0.857 | 0.665 | 3.00397E-05 | 0.582015552 | 0.267900161 | 0.7 | 0.873 | 1 | 0.582015552 | 2.47322E-09 |
| Glis2 | 1.25792E-09 | -2.737813989 | 0.126 | 0.447 | 3.05574E-05 | 0.019052082 | -2.020997081 | 0.2 | 0.571 | 1 | 0.019052082 | 2.51584E-09 |
| Camk1d | 1.26673E-09 | -1.173344625 | 1 | 1 | 3.07713E-05 | 0.013660207 | -0.139256577 | 1 | 1 | 1 | 0.013660207 | 2.53345E-09 |
| Foxn2 | 1.26705E-09 | -2.466318004 | 0.134 | 0.459 | 3.07791E-05 | 0.838916682 | -0.463204782 | 0.6 | 0.546 | 1 | 0.838916682 | 2.5341E-09 |
| Map4k3 | 1.28487E-09 | -1.644625422 | 0.345 | 0.665 | 3.12121E-05 | 0.08600571 | -0.840059087 | 0.5 | 0.765 | 1 | 0.08600571 | 2.56975E-09 |
| Mbtd1 | 1.32409E-09 | -1.867896464 | 0.504 | 0.729 | 3.21649E-05 | 0.142368258 | -0.816172862 | 0.8 | 0.864 | 1 | 0.142368258 | 2.64819E-09 |
| Ubtd2 | 1.40304E-09 | -1.239929867 | 0.63 | 0.818 | 3.40826E-05 | 0.019260589 | -1.248864633 | 0.5 | 0.833 | 1 | 0.019260589 | 2.80608E-09 |
| Rbm26 | 1.41514E-09 | -1.062396537 | 0.765 | 0.9 | 3.43766E-05 | 0.233327768 | -0.530257804 | 0.7 | 0.92 | 1 | 0.233327768 | 2.83028E-09 |
| Car12 | 1.42561E-09 | 1.621488377 | 0.387 | 0.088 | 3.46309E-05 | 0.23713486 | 2.894539492 | 0.2 | 0.093 | 1 | 0.23713486 | 2.85122E-09 |
| Syt8 | 1.45383E-09 | 1.446669108 | 0.521 | 0.188 | 3.53164E-05 | 9.99476E-06 | 1.855301707 | 0.9 | 0.265 | 0.242792647 | 9.99476E-06 | 2.90765E-09 |
| Arid4a | 1.46428E-09 | -1.692659441 | 0.471 | 0.724 | 3.55702E-05 | 0.02295175 | -1.472697588 | 0.6 | 0.818 | 1 | 0.02295175 | 2.92855E-09 |
| Mbd5 | 1.48324E-09 | -1.122383966 | 0.723 | 0.871 | 3.60309E-05 | 0.744293399 | -0.218674133 | 0.9 | 0.904 | 1 | 0.744293399 | 2.96649E-09 |
| Prrc2a | 1.49364E-09 | -1.05246742 | 0.622 | 0.824 | 3.62834E-05 | 0.060339944 | -0.534619115 | 0.6 | 0.883 | 1 | 0.060339944 | 2.98727E-09 |
| Tceal3 | 1.50784E-09 | -2.548929769 | 0.176 | 0.488 | 3.66284E-05 | 0.122503282 | -1.311201688 | 0.3 | 0.512 | 1 | 0.122503282 | 3.01568E-09 |
| Ube2k | 1.51738E-09 | -0.75205322 | 0.966 | 0.976 | 3.68601E-05 | 0.00100733 | -0.975773515 | 0.9 | 0.994 | 1 | 0.00100733 | 3.03475E-09 |
| Pprc1 | 1.57275E-09 | -1.86393845 | 0.252 | 0.571 | 3.82053E-05 | 0.279538484 | -0.687134438 | 0.4 | 0.534 | 1 | 0.279538484 | 3.1455E-09 |
| Mapk1 | 1.58122E-09 | -1.056681908 | 0.748 | 0.871 | 3.8411E-05 | 0.257816204 | -0.338276089 | 0.8 | 0.941 | 1 | 0.257816204 | 3.16244E-09 |
| Tiam1 | 1.5875E-09 | 0.914353413 | 0.933 | 0.659 | 3.85636E-05 | 0.019530374 | 1.406291457 | 0.9 | 0.738 | 1 | 0.019530374 | 3.175E-09 |
| Cep85l | 1.58912E-09 | 1.768505372 | 0.723 | 0.482 | 3.86028E-05 | 0.203398517 | 0.717367464 | 0.6 | 0.46 | 1 | 0.203398517 | 3.17823E-09 |
| Ppp1r12a | 1.63801E-09 | -0.968009886 | 0.908 | 0.947 | 3.97906E-05 | 0.185032221 | -0.283643569 | 1 | 0.991 | 1 | 0.185032221 | 3.27602E-09 |
| Naa15 | 1.64323E-09 | -0.915943059 | 0.782 | 0.912 | 3.99174E-05 | 0.236985574 | -0.413127909 | 0.9 | 0.932 | 1 | 0.236985574 | 3.28646E-09 |
| Mpped2 | 1.64706E-09 | -2.347923303 | 0.319 | 0.6 | 4.00104E-05 | 0.049948071 | -1.708296251 | 0.4 | 0.627 | 1 | 0.049948071 | 3.29412E-09 |
| Akap12 | 1.66495E-09 | -4.704595348 | 0.017 | 0.294 | 4.0445E-05 | 0.704015667 | 0.880418384 | 0.3 | 0.272 | 1 | 0.704015667 | 3.3299E-09 |
| Cnn2 | 1.73018E-09 | -2.451818435 | 0.185 | 0.506 | 4.20295E-05 | 0.041130155 | -0.587927959 | 0.2 | 0.583 | 1 | 0.041130155 | 3.46036E-09 |
| Spire1 | 1.76978E-09 | 0.649253418 | 0.992 | 0.882 | 4.29916E-05 | 0.054756592 | 0.558490289 | 1 | 0.895 | 1 | 0.054756592 | 3.53957E-09 |
| Mex3a | 1.78771E-09 | -5.330916878 | 0.008 | 0.276 | 4.34269E-05 | 0.058711159 | -2.391469028 | 0 | 0.272 | 1 | 0.058711159 | 3.57541E-09 |
| Ywhab | 1.81064E-09 | -0.772088292 | 0.933 | 0.965 | 4.3984E-05 | 0.041992799 | -0.630580647 | 0.8 | 0.981 | 1 | 0.041992799 | 3.62127E-09 |
| Norad | 1.86808E-09 | -0.891419187 | 0.807 | 0.894 | 4.53793E-05 | 0.002777484 | -1.251067221 | 0.6 | 0.938 | 1 | 0.002777484 | 3.73616E-09 |
| Tm4sf1 | 1.97306E-09 | 2.73812734 | 0.664 | 0.365 | 4.79296E-05 | 0.3526977 | 0.944413238 | 0.4 | 0.355 | 1 | 0.3526977 | 3.94612E-09 |
| Cds1 | 1.98214E-09 | 1 | 0.647 | 0.271 | 4.81502E-05 | 0.011031059 | 2.156119202 | 0.6 | 0.321 | 1 | 0.011031059 | 3.96428E-09 |
| Cab39l | 2.00436E-09 | 0.972822739 | 0.832 | 0.529 | 4.86899E-05 | 0.002913179 | 0.966391607 | 1 | 0.713 | 1 | 0.002913179 | 4.00872E-09 |
| Ehmt1 | 2.02988E-09 | -1.344727964 | 0.555 | 0.782 | 4.93097E-05 | 0.064079764 | -0.846676966 | 0.8 | 0.877 | 1 | 0.064079764 | 4.05975E-09 |

| | | | | | | | | | | | | |
|---|---|---|---|---|---|---|---|---|---|---|---|---|
| Pds5b | 2.11553E-09 | -1.242383897 | 0.63 | 0.8 | 5.13904E-05 | 0.073055929 | -0.803674937 | 0.6 | 0.87 | 1 | 0.073055929 | 4.23105E-09 |
| Pcf11 | 2.13266E-09 | -1.180572246 | 0.723 | 0.882 | 5.18065E-05 | 0.189817572 | -0.517908885 | 0.8 | 0.929 | 1 | 0.189817572 | 4.26531E-09 |
| Tmem263 | 2.23467E-09 | -1.056662345 | 0.739 | 0.818 | 5.42846E-05 | 0.06033196 | -0.738179473 | 0.6 | 0.861 | 1 | 0.06033196 | 4.46934E-09 |
| Arcn1 | 2.24021E-09 | -0.835172866 | 0.941 | 0.953 | 5.44191E-05 | 7.32845E-05 | -1.676665084 | 0.6 | 0.978 | 1 | 7.32845E-05 | 4.48042E-09 |
| Disp1 | 2.32904E-09 | 1.16966176 | 0.597 | 0.235 | 5.6577E-05 | 0.046515409 | 1.245332404 | 0.5 | 0.238 | 1 | 0.046515409 | 4.65808E-09 |
| Nfix | 2.37293E-09 | -0.707819249 | 0.958 | 0.988 | 5.76432E-05 | 0.006098546 | -0.790832466 | 1 | 0.994 | 1 | 0.006098546 | 4.74586E-09 |
| Hpgd | 2.38012E-09 | 1.968464427 | 0.748 | 0.465 | 5.78179E-05 | 0.725868115 | -0.43713127 | 0.4 | 0.454 | 1 | 0.725868115 | 4.76024E-09 |
| Chd6 | 2.39095E-09 | -1.351409479 | 0.655 | 0.835 | 5.80809E-05 | 0.56904371 | 0.131423101 | 0.9 | 0.877 | 1 | 0.56904371 | 4.7819E-09 |
| Pxylp1 | 2.4118E-09 | 1.523238034 | 0.605 | 0.253 | 5.85874E-05 | 1.65214E-05 | 2.156119202 | 0.9 | 0.302 | 0.40133703 | 1.65214E-05 | 4.8236E-09 |
| ENSMUSG00000121513 | 2.41438E-09 | -1.833350131 | 0.328 | 0.635 | 5.86501E-05 | 0.000625339 | -2.38782739 | 0.3 | 0.818 | 1 | 0.000625339 | 4.82876E-09 |
| Usp32 | 2.42222E-09 | -1.488958518 | 0.454 | 0.735 | 5.88405E-05 | 0.003719044 | -1.700439718 | 0.7 | 0.855 | 1 | 0.003719044 | 4.84444E-09 |
| Fkbp7 | 2.43219E-09 | -1.252449655 | 0.723 | 0.818 | 5.90829E-05 | 0.013622578 | -1.325880687 | 0.4 | 0.753 | 1 | 0.013622578 | 4.86439E-09 |
| Hnrnpm | 2.44443E-09 | -0.806297615 | 0.933 | 0.947 | 5.938E-05 | 0.024769369 | -0.733514169 | 0.9 | 0.988 | 1 | 0.024769369 | 4.88885E-09 |
| Golga4 | 2.4837E-09 | -0.874868011 | 0.815 | 0.929 | 6.0334E-05 | 0.188746663 | -0.512040973 | 0.9 | 0.954 | 1 | 0.188746663 | 4.9674E-09 |
| Trip11 | 2.48484E-09 | -0.736632946 | 0.933 | 0.953 | 6.03617E-05 | 0.329351022 | -0.225368202 | 0.9 | 0.972 | 1 | 0.329351022 | 4.96968E-09 |
| Dpy19l1 | 2.50122E-09 | -1.858885223 | 0.218 | 0.541 | 6.07595E-05 | 0.413379363 | -0.630735268 | 0.4 | 0.478 | 1 | 0.413379363 | 5.00243E-09 |
| Apaf1 | 2.54314E-09 | -2.233619677 | 0.16 | 0.488 | 6.1778E-05 | 0.396106122 | -0.625934282 | 0.4 | 0.5 | 1 | 0.396106122 | 5.08628E-09 |
| Cercam | 2.55469E-09 | -4.167250867 | 0.017 | 0.288 | 6.20586E-05 | 0.62153471 | 0.888638891 | 0.2 | 0.148 | 1 | 0.62153471 | 5.10939E-09 |
| Anp32b | 2.57211E-09 | -0.897268987 | 0.866 | 0.929 | 6.24816E-05 | 0.12296125 | -0.520662239 | 0.7 | 0.954 | 1 | 0.12296125 | 5.14421E-09 |
| Topors | 2.58517E-09 | -1.950238297 | 0.353 | 0.653 | 6.2799E-05 | 0.004767161 | -1.840059087 | 0.4 | 0.759 | 1 | 0.004767161 | 5.17034E-09 |
| Btg2 | 2.62872E-09 | 1.952080764 | 0.874 | 0.741 | 6.3857E-05 | 0.037356037 | 2.769484878 | 0.8 | 0.599 | 1 | 0.037356037 | 5.25745E-09 |
| Pgrmc1 | 2.64292E-09 | -0.911485733 | 0.647 | 0.865 | 6.42017E-05 | 0.018756336 | -0.92188331 | 0.6 | 0.895 | 1 | 0.018756336 | 5.28583E-09 |
| Micall1 | 2.66988E-09 | 1.116311386 | 0.706 | 0.347 | 6.48568E-05 | 0.906760948 | 0.759905577 | 0.4 | 0.398 | 1 | 0.906760948 | 5.33977E-09 |
| Polr2h | 2.68583E-09 | -1.82527683 | 0.193 | 0.518 | 6.52442E-05 | 0.172864348 | -0.951704443 | 0.2 | 0.395 | 1 | 0.172864348 | 5.37166E-09 |
| Tra2b | 2.73841E-09 | -0.801288921 | 0.975 | 1 | 6.65213E-05 | 0.162496437 | -0.378868991 | 1 | 0.997 | 1 | 0.162496437 | 5.47681E-09 |
| Pdxdc1 | 2.74387E-09 | -1.057542448 | 0.765 | 0.888 | 6.66542E-05 | 0.698710791 | -0.330520705 | 1 | 0.92 | 1 | 0.698710791 | 5.48775E-09 |
| Rexo2 | 2.9012E-09 | 0.726939167 | 0.992 | 0.953 | 7.04758E-05 | 0.494774116 | -0.25165245 | 0.8 | 0.951 | 1 | 0.494774116 | 5.80239E-09 |
| Apc | 2.90763E-09 | -1.250961574 | 0.563 | 0.771 | 7.06322E-05 | 0.261582425 | -0.501152657 | 0.6 | 0.855 | 1 | 0.261582425 | 5.81526E-09 |
| Ift20 | 3.06167E-09 | -0.911803043 | 0.84 | 0.9 | 7.43741E-05 | 0.059877197 | -0.749058666 | 0.7 | 0.907 | 1 | 0.059877197 | 6.12334E-09 |
| Dync1i2 | 3.0621E-09 | -1.130385293 | 0.689 | 0.871 | 7.43845E-05 | 0.012459578 | -1.201991229 | 0.6 | 0.883 | 1 | 0.012459578 | 6.12419E-09 |
| Hnrnpab | 3.15412E-09 | -0.845450224 | 0.908 | 0.965 | 7.66199E-05 | 0.023490731 | -0.724186989 | 1 | 0.997 | 1 | 0.023490731 | 6.30824E-09 |
| Cep83 | 3.17964E-09 | -1.589327693 | 0.37 | 0.665 | 7.72398E-05 | 0.000244287 | -3.425021588 | 0.1 | 0.719 | 1 | 0.000244287 | 6.35928E-09 |
| Rgs3 | 3.21109E-09 | -1.675251386 | 0.403 | 0.682 | 7.80038E-05 | 0.658908984 | -0.376302581 | 0.7 | 0.698 | 1 | 0.658908984 | 6.42218E-09 |
| Glul | 3.21516E-09 | 1.258583682 | 0.857 | 0.794 | 7.81026E-05 | 0.570115941 | -0.518130992 | 0.7 | 0.815 | 1 | 0.570115941 | 6.43031E-09 |
| Mtdh | 3.24174E-09 | -0.82303284 | 0.958 | 0.988 | 7.87483E-05 | 0.042705081 | -0.688055994 | 1 | 0.966 | 1 | 0.042705081 | 6.48348E-09 |
| Lifr | 3.25576E-09 | 1.060007309 | 0.647 | 0.288 | 7.90889E-05 | 0.000243896 | 2.089814826 | 0.7 | 0.222 | 1 | 0.000243896 | 6.51152E-09 |
| Slc20a2 | 3.41341E-09 | 1.359294947 | 0.723 | 0.406 | 8.29186E-05 | 0.034392009 | 1.421554644 | 0.6 | 0.327 | 1 | 0.034392009 | 6.82683E-09 |
| Mmp13 | 3.42821E-09 | -10.76677734 | 0 | 0.253 | 8.32781E-05 | 0.475017099 | -4.528972552 | 0 | 0.049 | 1 | 0.475017099 | 6.85642E-09 |
| Cnot4 | 3.46878E-09 | -0.805200037 | 0.849 | 0.918 | 8.42636E-05 | 0.500268968 | -0.313890776 | 0.9 | 0.969 | 1 | 0.500268968 | 6.93756E-09 |
| Zfp131 | 3.553E-09 | -1.023696092 | 0.63 | 0.829 | 8.63095E-05 | 0.869088042 | -0.133275885 | 1 | 0.88 | 1 | 0.869088042 | 7.106E-09 |
| Nktr | 3.64894E-09 | -0.606800084 | 0.992 | 1 | 8.864E-05 | 0.675817245 | -0.207806226 | 1 | 0.997 | 1 | 0.675817245 | 7.29788E-09 |
| Arpc1b | 3.66635E-09 | 0.827064891 | 0.975 | 0.941 | 8.90631E-05 | 0.325619487 | -0.276698841 | 1 | 0.981 | 1 | 0.325619487 | 7.33271E-09 |
| Zfp148 | 3.68335E-09 | -1.012383724 | 0.815 | 0.9 | 8.94759E-05 | 0.097827689 | -0.619404128 | 1 | 0.954 | 1 | 0.097827689 | 7.36669E-09 |
| Camk2d | 3.69746E-09 | 0.494764692 | 1 | 0.959 | 8.98187E-05 | 0.170642243 | 0.475233603 | 1 | 0.988 | 1 | 0.170642243 | 7.39492E-09 |
| Dek | 3.72913E-09 | -0.968317419 | 0.773 | 0.912 | 9.0588E-05 | 0.131478475 | -0.528378972 | 0.7 | 0.917 | 1 | 0.131478475 | 7.45826E-09 |
| Crkl | 3.73719E-09 | -1.979415668 | 0.218 | 0.529 | 9.07837E-05 | 0.004303391 | -2.178475305 | 0.2 | 0.636 | 1 | 0.004303391 | 7.47437E-09 |
| Steap2 | 3.84156E-09 | 1.571604118 | 0.706 | 0.418 | 9.33193E-05 | 0.267113452 | 0.789103218 | 0.5 | 0.349 | 1 | 0.267113452 | 7.68313E-09 |
| Eps15l1 | 3.85692E-09 | -1.497014801 | 0.345 | 0.653 | 9.36923E-05 | 0.1527141 | -0.714528205 | 0.6 | 0.753 | 1 | 0.1527141 | 7.71384E-09 |
| Klk10 | 3.87788E-09 | 4.617383978 | 0.235 | 0.018 | 9.42015E-05 | 0.000212566 | 3.847996907 | 0.2 | 0.019 | 1 | 0.000212566 | 7.75577E-09 |
| Sp7 | 3.90316E-09 | -4.500377169 | 0.017 | 0.282 | 9.48156E-05 | 0.25993628 | -1.004445905 | 0 | 0.114 | 1 | 0.25993628 | 7.80632E-09 |
| Eif4a3 | 3.94988E-09 | -1.352487995 | 0.437 | 0.7 | 9.59504E-05 | 0.064570473 | -1.063466419 | 0.4 | 0.673 | 1 | 0.064570473 | 7.89975E-09 |
| Lbh | 4.02422E-09 | 1.812914447 | 0.597 | 0.265 | 9.77564E-05 | 7.15646E-07 | 3.410239331 | 0.6 | 0.114 | 0.01738447 | 7.15646E-07 | 8.04844E-09 |
| Ndufa4l2 | 4.04468E-09 | 1.427321979 | 0.647 | 0.324 | 9.82534E-05 | 0.715898633 | 0.902444691 | 0.1 | 0.071 | 1 | 0.715898633 | 8.08936E-09 |
| St8sia6 | 4.11828E-09 | 3.920565533 | 0.235 | 0.018 | 0.000100041 | 0.016392512 | 2.374065718 | 0.2 | 0.04 | 1 | 0.016392512 | 8.23657E-09 |
| Arhgef11 | 4.11906E-09 | -1.85544921 | 0.328 | 0.618 | 0.00010006 | 0.272860125 | -0.684366235 | 0.7 | 0.744 | 1 | 0.272860125 | 8.23812E-09 |
| Setd1b | 4.13002E-09 | -1.997325866 | 0.202 | 0.518 | 0.000100326 | 0.04354304 | -1.245738532 | 0.4 | 0.654 | 1 | 0.04354304 | 8.26004E-09 |
| Gli3 | 4.15221E-09 | -1.310593025 | 0.546 | 0.753 | 0.000100866 | 0.288439292 | 0.44049308 | 0.8 | 0.586 | 1 | 0.288439292 | 8.30443E-09 |
| Hmgb1 | 4.18592E-09 | -0.753004825 | 0.95 | 0.982 | 0.000101684 | 0.035813225 | -0.677826115 | 0.9 | 0.997 | 1 | 0.035813225 | 8.37184E-09 |
| Zc3h18 | 4.1862E-09 | -1.327547167 | 0.479 | 0.724 | 0.000101691 | 0.176053278 | -0.75818208 | 0.8 | 0.75 | 1 | 0.176053278 | 8.3724E-09 |
| Gar1 | 4.25825E-09 | -1.744161096 | 0.294 | 0.582 | 0.000103441 | 0.366649541 | -0.12891948 | 0.3 | 0.457 | 1 | 0.366649541 | 8.51649E-09 |
| Trf | 4.47056E-09 | 3.941656014 | 0.378 | 0.106 | 0.000108599 | 4.77936E-06 | 3.215602576 | 0.7 | 0.164 | 0.116100279 | 4.77936E-06 | 8.94113E-09 |
| Cop1 | 4.5695E-09 | -0.976674893 | 0.765 | 0.912 | 0.000111002 | 0.022952214 | -0.777576656 | 0.6 | 0.951 | 1 | 0.022952214 | 9.139E-09 |
| Rasa1 | 4.59208E-09 | -1.517135687 | 0.462 | 0.706 | 0.000111551 | 0.012795186 | -1.252607034 | 0.4 | 0.836 | 1 | 0.012795186 | 9.18415E-09 |
| Sugt1 | 4.61216E-09 | -0.986779895 | 0.697 | 0.853 | 0.000112039 | 0.032356014 | -0.961118473 | 0.5 | 0.855 | 1 | 0.032356014 | 9.22431E-09 |
| Srsf1 | 4.73302E-09 | -1.185866545 | 0.588 | 0.788 | 0.000114975 | 0.120353996 | -0.688450766 | 0.6 | 0.818 | 1 | 0.120353996 | 9.46604E-09 |
| Rdx | 4.78774E-09 | -0.742480256 | 0.95 | 0.982 | 0.000116304 | 0.097542603 | -0.511204935 | 0.9 | 0.988 | 1 | 0.097542603 | 9.57549E-09 |
| Phtf2 | 4.84676E-09 | -1.631870395 | 0.462 | 0.718 | 0.000117738 | 0.006652721 | -1.843839312 | 0.5 | 0.87 | 1 | 0.006652721 | 9.69352E-09 |
| Pard3b | 4.88599E-09 | -1.588319709 | 0.605 | 0.759 | 0.00011869 | 0.023351212 | -1.253744102 | 0.7 | 0.858 | 1 | 0.023351212 | 9.77198E-09 |
| Ubr2 | 4.93343E-09 | -1.269698136 | 0.681 | 0.865 | 0.000119843 | 0.434373216 | -0.215834705 | 1 | 0.907 | 1 | 0.434373216 | 9.86685E-09 |
| Septin7 | 4.93611E-09 | -0.605008443 | 0.975 | 0.988 | 0.000119908 | 0.000911214 | -0.943869773 | 1 | 1 | 1 | 0.000911214 | 9.87223E-09 |
| Pon3 | 5.11468E-09 | -1.970853654 | 0.101 | 0.418 | 0.000124246 | 0.028644247 | -1.858595039 | 0.1 | 0.448 | 1 | 0.028644247 | 1.02294E-08 |
| Slc43a2 | 5.34842E-09 | -3.328701323 | 0.084 | 0.382 | 0.000129924 | 0.078579756 | -2.40274014 | 0.2 | 0.438 | 1 | 0.078579756 | 1.06968E-08 |
| Kdm5a | 5.36738E-09 | -1.033423002 | 0.874 | 0.918 | 0.000130384 | 0.147171856 | -0.534391399 | 0.8 | 0.954 | 1 | 0.147171856 | 1.07348E-08 |
| Ctdspl2 | 5.47222E-09 | -1.49872365 | 0.471 | 0.706 | 0.000132931 | 0.066681534 | -0.823380346 | 0.5 | 0.812 | 1 | 0.066681534 | 1.09444E-08 |
| Zc3h13 | 5.52473E-09 | -1.193246076 | 0.597 | 0.806 | 0.000134207 | 0.523539912 | -0.319947731 | 0.8 | 0.824 | 1 | 0.523539912 | 1.10495E-08 |
| Adh1 | 5.6831E-09 | 4.906890596 | 0.202 | 0.006 | 0.000138054 | 0.438613761 | -0.304006187 | 0.1 | 0.198 | 1 | 0.438613761 | 1.13662E-08 |
| Pnpla3 | 0.061541304 | 0.777607579 | 0.16 | 0.088 | 1 | 5.70669E-09 | 4.558490289 | 0.3 | 0.015 | 0.000138627 | 0.061541304 | 1.14134E-08 |
| Hnrnpk | 5.95852E-09 | -0.6856565 | 0.966 | 0.988 | 0.000144744 | 0.003244638 | -0.901702178 | 0.9 | 0.994 | 1 | 0.003244638 | 1.1917E-08 |
| Efhd2 | 5.98845E-09 | 1.514573173 | 0.773 | 0.541 | 0.000145471 | 0.936864864 | 0.118644496 | 0.7 | 0.667 | 1 | 0.936864864 | 1.19769E-08 |
| Magoh | 5.99282E-09 | -0.885277923 | 0.739 | 0.888 | 0.000145577 | 0.031835296 | -0.859822342 | 0.5 | 0.898 | 1 | 0.031835296 | 1.19856E-08 |
| Adcy2 | 6.02781E-09 | -2.259867127 | 0.21 | 0.524 | 0.000146428 | 0.283592154 | -0.679313007 | 0.6 | 0.781 | 1 | 0.283592154 | 1.20556E-08 |
| Grk5 | 6.09472E-09 | 0.675656298 | 0.891 | 0.594 | 0.000148053 | 0.695251473 | -0.493483986 | 0.9 | 0.728 | 1 | 0.695251473 | 1.21894E-08 |
| Arhgap10 | 6.1602E-09 | -1.63895739 | 0.504 | 0.741 | 0.000149643 | 0.596072886 | -0.598749452 | 0.6 | 0.664 | 1 | 0.596072886 | 1.23204E-08 |
| Diaph2 | 6.34829E-09 | -1.295355693 | 0.546 | 0.788 | 0.000154213 | 0.856585473 | -0.396506216 | 0.9 | 0.815 | 1 | 0.856585473 | 1.26966E-08 |
| Tfrc | 6.41149E-09 | -1.909924656 | 0.395 | 0.665 | 0.000155748 | 0.002945605 | -2.401616984 | 0.5 | 0.809 | 1 | 0.002945605 | 1.2823E-08 |
| Tmbim1 | 6.4221E-09 | 0.626238776 | 0.992 | 0.847 | 0.000156006 | 0.798572563 | 0.20469042 | 0.8 | 0.944 | 1 | 0.798572563 | 1.28442E-08 |
| Dleu2 | 6.4232E-09 | -2.394914534 | 0.387 | 0.665 | 0.000156032 | 0.156251338 | -1.102747979 | 0.6 | 0.79 | 1 | 0.156251338 | 1.28464E-08 |
| Raph1 | 6.6918E-09 | 0.869742089 | 0.975 | 0.906 | 0.000162557 | 0.558650269 | 0.554285054 | 1 | 0.929 | 1 | 0.558650269 | 1.33836E-08 |
| Nedd4 | 6.71004E-09 | -0.501216267 | 1 | 1 | 0.000163 | 0.043073634 | -0.446486506 | 1 | 1 | 1 | 0.043073634 | 1.34201E-08 |
| Gxylt2 | 6.72016E-09 | -1.624763509 | 0.571 | 0.759 | 0.000163246 | 0.257639733 | -0.665462646 | 0.7 | 0.694 | 1 | 0.257639733 | 1.34403E-08 |
| Fam89a | 6.74724E-09 | 1.694122593 | 0.487 | 0.176 | 0.000163904 | 0.434533558 | 2.918386234 | 0.1 | 0.049 | 1 | 0.434533558 | 1.34945E-08 |
| Stk10 | 6.75592E-09 | -1.89862298 | 0.21 | 0.559 | 0.000164115 | 0.073749064 | -1.678548908 | 0.4 | 0.586 | 1 | 0.073749064 | 1.35118E-08 |
| Igdcc4 | 6.81277E-09 | -2.032914622 | 0.176 | 0.494 | 0.000165496 | 0.01475524 | -2.689437224 | 0 | 0.392 | 1 | 0.01475524 | 1.36255E-08 |
| Fgf1 | 0.039482691 | 0.12755005 | 0.202 | 0.112 | 1 | 7.05308E-09 | 4.380491987 | 0.4 | 0.031 | 0.000171333 | 0.039482691 | 1.41062E-08 |
| Ube3a | 7.11273E-09 | -0.81259157 | 0.891 | 0.935 | 0.000172782 | 0.856727176 | -0.152893371 | 0.9 | 0.969 | 1 | 0.856727176 | 1.42255E-08 |
| Tent4b | 7.18988E-09 | -2.572889668 | 0.143 | 0.447 | 0.000174657 | 0.073973596 | -1.496200352 | 0.4 | 0.596 | 1 | 0.073973596 | 1.43798E-08 |
| Zfp609 | 7.2293E-09 | -1.374395515 | 0.487 | 0.741 | 0.000175614 | 0.134511654 | -0.871690696 | 0.6 | 0.769 | 1 | 0.134511654 | 1.44586E-08 |
| Ubap2l | 7.32827E-09 | -0.696137443 | 0.916 | 0.971 | 0.000178018 | 0.636604716 | -0.130674584 | 0.9 | 0.978 | 1 | 0.636604716 | 1.46565E-08 |

| | | | | | | | | | | | | |
|---|---|---|---|---|---|---|---|---|---|---|---|---|
| Alx1 | 7.36781E-09 | -1.361637805 | 0.773 | 0.841 | 0.000178979 | 0.002206527 | -2.051240117 | 0.6 | 0.846 | 1 | 0.002206527 | 1.47356E-08 |
| Cpsf6 | 7.40401E-09 | -1.167385654 | 0.639 | 0.824 | 0.000179858 | 0.004255563 | -1.128010238 | 0.8 | 0.935 | 1 | 0.004255563 | 1.4808E-08 |
| Rasgef1c | 0.968643239 | -0.5360529 | 0.042 | 0.041 | 1 | 7.41634E-09 | 4.532495081 | 0.3 | 0.015 | 0.000180158 | 0.968643239 | 1.48327E-08 |
| Fut4 | 1.83584E-08 | 2.883806982 | 0.294 | 0.053 | 0.000445963 | 7.41956E-09 | 4.432959407 | 0.3 | 0.015 | 0.000180236 | 1.83584E-08 | 1.48391E-08 |
| Ect2 | 0.110416608 | -2.944858446 | 0.017 | 0.053 | 1 | 7.41956E-09 | 4.432959407 | 0.3 | 0.015 | 0.000180236 | 0.110416608 | 1.48391E-08 |
| Smarca2 | 7.44795E-09 | -1.174340292 | 0.689 | 0.859 | 0.000180926 | 0.194419737 | -0.41670632 | 0.8 | 0.852 | 1 | 0.194419737 | 1.48959E-08 |
| Tmbim4 | 7.67791E-09 | 0.713506621 | 0.95 | 0.812 | 0.000186512 | 0.58236124 | -0.127302653 | 0.7 | 0.836 | 1 | 0.58236124 | 1.53558E-08 |
| Elk4 | 7.84533E-09 | -1.123300859 | 0.588 | 0.812 | 0.000190579 | 0.016460399 | -1.170254798 | 0.8 | 0.855 | 1 | 0.016460399 | 1.56907E-08 |
| Ncoa1 | 7.94426E-09 | -2.120142363 | 0.303 | 0.594 | 0.000192982 | 0.008271435 | -1.608517229 | 0.4 | 0.824 | 1 | 0.008271435 | 1.58885E-08 |
| Kalrn | 8.02461E-09 | -3.400310214 | 0.076 | 0.365 | 0.000194934 | 0.858719519 | -0.191531458 | 0.3 | 0.336 | 1 | 0.858719519 | 1.60492E-08 |
| Phka2 | 8.25693E-09 | 1.009131012 | 0.681 | 0.341 | 0.000200577 | 0.000306537 | 2.719263592 | 0.7 | 0.228 | 1 | 0.000306537 | 1.65139E-08 |
| Hpse2 | 8.49827E-09 | -5.111865964 | 0.034 | 0.3 | 0.00020644 | 0.362560714 | -1.195425374 | 0.1 | 0.228 | 1 | 0.362560714 | 1.69965E-08 |
| Cdc42ep5 | 8.52339E-09 | -1.448900951 | 0.025 | 0.294 | 0.00020705 | 0.263440752 | -0.936274402 | 0.1 | 0.247 | 1 | 0.263440752 | 1.70468E-08 |
| Ece1 | 8.6279E-09 | -2.522901533 | 0.126 | 0.429 | 0.000209589 | 0.396743628 | -0.660149997 | 0.3 | 0.414 | 1 | 0.396743628 | 1.72558E-08 |
| Lrrn4cl | 8.69695E-09 | -3.87774425 | 0.017 | 0.276 | 0.000211266 | 0.749384801 | 0.808468542 | 0.1 | 0.071 | 1 | 0.749384801 | 1.73939E-08 |
| Stk32b | 8.81685E-09 | 0.672798446 | 0.891 | 0.5 | 0.000214179 | 0.000242594 | 1.669623473 | 1 | 0.556 | 1 | 0.000242594 | 1.76337E-08 |
| Pmvk | 9.13205E-09 | 1.280982545 | 0.756 | 0.441 | 0.000221836 | 0.00406724 | 1.642882477 | 0.7 | 0.349 | 1 | 0.00406724 | 1.82641E-08 |
| Senp5 | 9.19507E-09 | -1.304186513 | 0.571 | 0.771 | 0.000223367 | 0.736230196 | -0.191531458 | 0.9 | 0.861 | 1 | 0.736230196 | 1.83901E-08 |
| Enho | 9.33766E-09 | 1.139970538 | 0.714 | 0.376 | 0.00022683 | 0.769101126 | 0.111031312 | 0.3 | 0.373 | 1 | 0.769101126 | 1.86753E-08 |
| Rbm3 | 9.49472E-09 | -0.818115617 | 0.924 | 0.976 | 0.000230646 | 0.184874263 | -0.405218438 | 1 | 0.991 | 1 | 0.184874263 | 1.89894E-08 |
| Kansl1l | 9.67146E-09 | -1.814101754 | 0.42 | 0.7 | 0.000234939 | 0.002060798 | -2.053127843 | 0.4 | 0.88 | 1 | 0.002060798 | 1.93429E-08 |
| Brd3 | 9.94097E-09 | -1.610957709 | 0.361 | 0.629 | 0.000241486 | 0.145810858 | -0.573466862 | 0.7 | 0.799 | 1 | 0.145810858 | 1.98819E-08 |
| Rasl12 | 0.008856755 | 2.230780207 | 0.092 | 0.024 | 1 | 1.01834E-08 | 4.573137065 | 0.6 | 0.077 | 0.000247375 | 0.008856755 | 2.03668E-08 |
| Zfp704 | 1.0276E-08 | -1.052600014 | 0.437 | 0.718 | 0.000249625 | 0.927470987 | 0.464795494 | 0.6 | 0.778 | 1 | 0.927470987 | 2.05521E-08 |
| Ewsr1 | 1.03863E-08 | -0.807354922 | 0.832 | 0.941 | 0.000252305 | 0.278801712 | -0.321419118 | 1 | 0.988 | 1 | 0.278801712 | 2.07727E-08 |
| Anp32e | 1.05522E-08 | -1.415037499 | 0.487 | 0.724 | 0.000256335 | 0.041563281 | -0.956811333 | 0.6 | 0.772 | 1 | 0.041563281 | 2.11045E-08 |
| Gucy2g | 1.05726E-08 | 3.602036014 | 0.227 | 0.018 | 0.00025683 | 0.047702188 | 3.280956314 | 0.1 | 0.015 | 1 | 0.047702188 | 2.11453E-08 |
| Psap | 1.07561E-08 | 0.542318163 | 0.992 | 0.982 | 0.000261286 | 0.610869963 | 0.152766285 | 1 | 0.991 | 1 | 0.610869963 | 2.15121E-08 |
| Nr6a1 | 1.07777E-08 | -2.382667253 | 0.202 | 0.506 | 0.000261811 | 0.003669166 | -2.756865152 | 0.2 | 0.651 | 1 | 0.003669166 | 2.15553E-08 |
| Vav3 | 1.0824E-08 | 0.876139464 | 0.933 | 0.594 | 0.000262937 | 0.162301754 | 1.08695497 | 1 | 0.753 | 1 | 0.162301754 | 2.1648E-08 |
| Arl4a | 1.13746E-08 | -1.737813989 | 0.622 | 0.8 | 0.000276313 | 0.049277216 | -1.207687226 | 0.8 | 0.87 | 1 | 0.049277216 | 2.27493E-08 |
| Csnk1g3 | 1.15886E-08 | -1.269698136 | 0.487 | 0.753 | 0.000281511 | 0.142661231 | -0.591872446 | 0.7 | 0.867 | 1 | 0.142661231 | 2.31772E-08 |
| Cryab | 1.17165E-08 | 2.182362105 | 0.597 | 0.324 | 0.000284618 | 0.86312411 | -0.546862711 | 0.2 | 0.228 | 1 | 0.86312411 | 2.34331E-08 |
| Runx1 | 1.22726E-08 | 0.664116674 | 1 | 0.959 | 0.000298125 | 0.682416516 | 0.267117406 | 1 | 0.988 | 1 | 0.682416516 | 2.45451E-08 |
| Lpgat1 | 1.23937E-08 | -1.382667253 | 0.437 | 0.688 | 0.000301068 | 0.192774265 | -0.830460282 | 0.7 | 0.787 | 1 | 0.192774265 | 2.47874E-08 |
| Cobl | 1.24186E-08 | 1.152387556 | 0.773 | 0.441 | 0.000301673 | 0.021228585 | 1.094004301 | 0.9 | 0.713 | 1 | 0.021228585 | 2.48372E-08 |
| Slc40a1 | 1.24501E-08 | -6.260213887 | 0.008 | 0.253 | 0.000302438 | 0.042521586 | -3.098422053 | 0.1 | 0.404 | 1 | 0.042521586 | 2.49002E-08 |
| Sox2 | 1.25705E-08 | 2.974004791 | 0.277 | 0.041 | 0.000305362 | 0.259359241 | -0.625934282 | 0 | 0.114 | 1 | 0.259359241 | 2.51409E-08 |
| Col27a1 | 1.26094E-08 | 0.794825658 | 1 | 0.924 | 0.000306308 | 0.029142192 | 0.756809468 | 1 | 0.966 | 1 | 0.029142192 | 2.52188E-08 |
| Myl9 | 1.27086E-08 | -5.427941333 | 0.008 | 0.253 | 0.000308718 | 0.975966169 | 0.760534065 | 0.2 | 0.216 | 1 | 0.975966169 | 2.54172E-08 |
| Zbed6 | 1.276E-08 | -1.178765489 | 0.521 | 0.747 | 0.000309966 | 0.040173537 | -0.93203741 | 0.6 | 0.815 | 1 | 0.040173537 | 2.552E-08 |
| Plxdc2 | 1.29315E-08 | -1.40019731 | 0.706 | 0.853 | 0.000314132 | 0.016721025 | -1.133428503 | 0.6 | 0.932 | 1 | 0.016721025 | 2.5863E-08 |
| Poglut3 | 1.30297E-08 | -1.494197037 | 0.277 | 0.588 | 0.000316517 | 0.009469108 | -2.308507579 | 0.1 | 0.522 | 1 | 0.009469108 | 2.60594E-08 |
| Tshz2 | 1.32739E-08 | 0.929174747 | 0.992 | 0.953 | 0.00032245 | 0.031757385 | 0.766240307 | 1 | 0.861 | 1 | 0.031757385 | 2.65478E-08 |
| Sec24a | 1.34625E-08 | -1.070389328 | 0.681 | 0.829 | 0.000327031 | 0.942607006 | -0.145308441 | 1 | 0.895 | 1 | 0.942607006 | 2.6925E-08 |
| Ccdc12 | 1.36687E-08 | -1.342686655 | 0.42 | 0.682 | 0.000332039 | 0.41181377 | -0.524336142 | 0.4 | 0.475 | 1 | 0.41181377 | 2.73373E-08 |
| Mtm1 | 1.37094E-08 | 2.057471614 | 0.504 | 0.218 | 0.000333029 | 0.006076725 | 2.444455046 | 0.5 | 0.194 | 1 | 0.006076725 | 2.74188E-08 |
| Kcns1 | 1.37238E-08 | -5.292781749 | 0 | 0.235 | 0.000333378 | 0.152227152 | -1.267480311 | 0 | 0.173 | 1 | 0.152227152 | 2.74476E-08 |
| Igfbp4 | 1.37812E-08 | -3.548079168 | 0.168 | 0.453 | 0.000334774 | 0.048137745 | -2.887765941 | 0.1 | 0.429 | 1 | 0.048137745 | 2.75625E-08 |
| Zmym4 | 1.38273E-08 | -1.324962155 | 0.496 | 0.747 | 0.000335893 | 0.033360872 | -1.07605424 | 0.7 | 0.833 | 1 | 0.033360872 | 2.76546E-08 |
| Endou | 1.3848E-08 | 2.432111013 | 0.336 | 0.076 | 0.000336395 | 0.528682188 | 2.526068812 | 0.2 | 0.151 | 1 | 0.528682188 | 2.7696E-08 |
| Sgcg | 2.45536E-05 | 4.684498174 | 0.101 | 0 | 0.596456832 | 1.39069E-08 | 6.017921908 | 0.1 | 0 | 0.000337826 | 2.45536E-05 | 2.78137E-08 |
| Kcnj12 | 0.011788405 | 1.836501268 | 0.067 | 0.012 | 1 | 1.39069E-08 | 6.017921908 | 0.1 | 0 | 0.000337826 | 0.011788405 | 2.78137E-08 |
| Gm47865 | 0.018806191 | 1.736965594 | 0.084 | 0.024 | 1 | 1.39069E-08 | 6.017921908 | 0.1 | 0 | 0.000337826 | 0.018806191 | 2.78137E-08 |
| Exo1 | 0.038974464 | -2.655351829 | 0 | 0.035 | 1 | 1.39069E-08 | 6.017921908 | 0.1 | 0 | 0.000337826 | 0.038974464 | 2.78137E-08 |
| Gm12524 | 0.074658411 | 3.099535674 | 0.034 | 0.006 | 1 | 1.39069E-08 | 6.017921908 | 0.1 | 0 | 0.000337826 | 0.074658411 | 2.78137E-08 |
| Gm55759 | 0.075690158 | 2.514573173 | 0.034 | 0.006 | 1 | 1.39069E-08 | 6.602884409 | 0.1 | 0 | 0.000337826 | 0.075690158 | 2.78137E-08 |
| 4833417C18Rik | 0.076220415 | 1.836501268 | 0.034 | 0.006 | 1 | 1.39069E-08 | 6.017921908 | 0.1 | 0 | 0.000337826 | 0.076220415 | 2.78137E-08 |
| B3gnt7 | 0.091361251 | 3.099535674 | 0.017 | 0 | 1 | 1.39069E-08 | 6.017921908 | 0.1 | 0 | 0.000337826 | 0.091361251 | 2.78137E-08 |
| Tex50 | 0.091361251 | 2.836501268 | 0.017 | 0 | 1 | 1.39069E-08 | 6.017921908 | 0.1 | 0 | 0.000337826 | 0.091361251 | 2.78137E-08 |
| Clnk | 0.091361251 | 2.099535674 | 0.017 | 0 | 1 | 1.39069E-08 | 6.017921908 | 0.1 | 0 | 0.000337826 | 0.091361251 | 2.78137E-08 |
| Raet1d | 0.091361251 | 2.099535674 | 0.017 | 0 | 1 | 1.39069E-08 | 6.017921908 | 0.1 | 0 | 0.000337826 | 0.091361251 | 2.78137E-08 |
| 1700047A11Rik | 0.147029286 | -1.807354922 | 0 | 0.018 | 1 | 1.39069E-08 | 6.017921908 | 0.1 | 0 | 0.000337826 | 0.147029286 | 2.78137E-08 |
| 5830432E09Rik | 0.147030916 | -2.070389328 | 0 | 0.018 | 1 | 1.39069E-08 | 6.017921908 | 0.1 | 0 | 0.000337826 | 0.147030916 | 2.78137E-08 |
| Gm16998 | 0.168156185 | 1.514573173 | 0.025 | 0.006 | 1 | 1.39069E-08 | 6.602884409 | 0.1 | 0 | 0.000337826 | 0.168156185 | 2.78137E-08 |
| Gm57460 | 0.237885589 | -1.070389328 | 0 | 0.012 | 1 | 1.39069E-08 | 6.017921908 | 0.1 | 0 | 0.000337826 | 0.237885589 | 2.78137E-08 |
| 4731419I09Rik | 0.237885589 | -1.070389328 | 0 | 0.012 | 1 | 1.39069E-08 | 6.017921908 | 0.1 | 0 | 0.000337826 | 0.237885589 | 2.78137E-08 |
| Gm42463 | 0.237885589 | -1.070389328 | 0 | 0.012 | 1 | 1.39069E-08 | 6.017921908 | 0.1 | 0 | 0.000337826 | 0.237885589 | 2.78137E-08 |
| Gm30373 | 0.237885589 | -1.807354922 | 0 | 0.012 | 1 | 1.39069E-08 | 6.017921908 | 0.1 | 0 | 0.000337826 | 0.237885589 | 2.78137E-08 |
| Vsx2 | 0.237885589 | -1.070389328 | 0 | 0.012 | 1 | 1.39069E-08 | 6.017921908 | 0.1 | 0 | 0.000337826 | 0.237885589 | 2.78137E-08 |
| Crip3 | 0.237885589 | -1.070389328 | 0 | 0.012 | 1 | 1.39069E-08 | 6.017921908 | 0.1 | 0 | 0.000337826 | 0.237885589 | 2.78137E-08 |
| Gfra3 | 0.237885589 | -1.070389328 | 0 | 0.012 | 1 | 1.39069E-08 | 6.602884409 | 0.1 | 0 | 0.000337826 | 0.237885589 | 2.78137E-08 |
| Gm56612 | 0.370184089 | 1.099535674 | 0.017 | 0.006 | 1 | 1.39069E-08 | 6.017921908 | 0.1 | 0 | 0.000337826 | 0.370184089 | 2.78137E-08 |
| Grep1 | 0.370184089 | 1.099535674 | 0.017 | 0.006 | 1 | 1.39069E-08 | 6.602884409 | 0.1 | 0 | 0.000337826 | 0.370184089 | 2.78137E-08 |
| Sec14l5 | 0.374548525 | 0.514573173 | 0.017 | 0.006 | 1 | 1.39069E-08 | 6.017921908 | 0.1 | 0 | 0.000337826 | 0.374548525 | 2.78137E-08 |
| 5730435O14Rik | 0.508731273 | -1.292781749 | 0.008 | 0.018 | 1 | 1.39069E-08 | 6.017921908 | 0.1 | 0 | 0.000337826 | 0.508731273 | 2.78137E-08 |
| Gm8206 | 0.511005543 | -0.485426827 | 0.008 | 0.018 | 1 | 1.39069E-08 | 6.017921908 | 0.1 | 0 | 0.000337826 | 0.511005543 | 2.78137E-08 |
| Dnph1 | 0.511005543 | -0.485426827 | 0.008 | 0.018 | 1 | 1.39069E-08 | 6.017921908 | 0.1 | 0 | 0.000337826 | 0.511005543 | 2.78137E-08 |
| Kifc1 | 0.614381021 | -1.070389328 | 0.025 | 0.035 | 1 | 1.39069E-08 | 6.017921908 | 0.1 | 0 | 0.000337826 | 0.614381021 | 2.78137E-08 |
| Sh3gl3 | 0.616555609 | 0.292180751 | 0.034 | 0.024 | 1 | 1.39069E-08 | 6.017921908 | 0.1 | 0 | 0.000337826 | 0.616555609 | 2.78137E-08 |
| Gm26834 | 0.721162042 | 0.514573173 | 0.017 | 0.012 | 1 | 1.39069E-08 | 6.017921908 | 0.1 | 0 | 0.000337826 | 0.721162042 | 2.78137E-08 |
| Fam124b | 0.729112716 | -0.655351829 | 0.017 | 0.012 | 1 | 1.39069E-08 | 6.017921908 | 0.1 | 0 | 0.000337826 | 0.729112716 | 2.78137E-08 |
| Kcne3 | 0.781790612 | -0.485426827 | 0.008 | 0.012 | 1 | 1.39069E-08 | 6.017921908 | 0.1 | 0 | 0.000337826 | 0.781790612 | 2.78137E-08 |
| Gm2245 | 0.781790612 | -0.485426827 | 0.008 | 0.012 | 1 | 1.39069E-08 | 6.017921908 | 0.1 | 0 | 0.000337826 | 0.781790612 | 2.78137E-08 |
| Kcnk13 | 0.784920947 | -1.359895945 | 0.008 | 0.012 | 1 | 1.39069E-08 | 6.017921908 | 0.1 | 0 | 0.000337826 | 0.784920947 | 2.78137E-08 |
| Fat3 | 1.42787E-08 | -6.936637939 | 0 | 0.235 | 0.000346857 | 0.850909231 | -0.936274402 | 0.2 | 0.164 | 1 | 0.850909231 | 2.85573E-08 |
| Stk17b | 1.43282E-08 | -2.61667136 | 0.126 | 0.418 | 0.000348061 | 0.107063075 | -2.058893689 | 0.2 | 0.426 | 1 | 0.107063075 | 2.86565E-08 |
| Prrc2b | 1.43306E-08 | -0.996083769 | 0.782 | 0.871 | 0.00034812 | 0.040104476 | -0.72647322 | 0.9 | 0.96 | 1 | 0.040104476 | 2.86613E-08 |
| Papola | 1.46453E-08 | -0.854860359 | 0.866 | 0.941 | 0.000355762 | 0.003813211 | -1.127541036 | 0.6 | 0.972 | 1 | 0.003813211 | 2.92905E-08 |
| Mob1b | 1.50717E-08 | -1.546538341 | 0.286 | 0.594 | 0.000366123 | 0.014893655 | -1.405544213 | 0.4 | 0.738 | 1 | 0.014893655 | 3.01435E-08 |
| Corin | 1.51124E-08 | 2.349149564 | 0.311 | 0.065 | 0.000367111 | 0.158187167 | 1.361574774 | 0.4 | 0.235 | 1 | 0.158187167 | 3.02249E-08 |
| Cacna1g | 1.55793E-08 | -4.033863452 | 0.025 | 0.282 | 0.000378453 | 0.166354453 | -1.709998547 | 0 | 0.164 | 1 | 0.166354453 | 3.11586E-08 |
| Spast | 1.57716E-08 | -2.062611977 | 0.193 | 0.494 | 0.000383123 | 0.03960987 | -1.329034981 | 0.3 | 0.62 | 1 | 0.03960987 | 3.15431E-08 |
| Rbm25 | 1.65202E-08 | -0.580681503 | 1 | 0.994 | 0.000401309 | 0.016771501 | -0.657586117 | 1 | 0.997 | 1 | 0.016771501 | 3.30404E-08 |
| Mt2 | 1.68326E-08 | 2.156834343 | 0.597 | 0.329 | 0.000408898 | 0.887019653 | -0.745843746 | 0.3 | 0.315 | 1 | 0.887019653 | 3.36652E-08 |
| Gna13 | 1.71357E-08 | 0.727707626 | 0.908 | 0.776 | 0.00041626 | 0.258174155 | -0.484020006 | 0.8 | 0.843 | 1 | 0.258174155 | 3.42713E-08 |
| Fndc3b | 1.72965E-08 | -0.51428758 | 1 | 1 | 0.000420167 | 0.225397264 | -0.381129226 | 1 | 1 | 1 | 0.225397264 | 3.45931E-08 |
| Mylk | 1.73432E-08 | -3.308961676 | 0.092 | 0.371 | 0.0004213 | 0.09751308 | -1.785132877 | 0.1 | 0.367 | 1 | 0.09751308 | 3.46864E-08 |

| | | | | | | | | | | | | |
|---|---|---|---|---|---|---|---|---|---|---|---|---|
| Naa16 | 1.74009E-08 | -1.660513534 | 0.185 | 0.5 | 0.000422702 | 0.207398382 | -1.314786026 | 0.4 | 0.494 | 1 | 0.207398382 | 3.48017E-08 |
| Resf1 | 1.76995E-08 | 1.919237084 | 0.832 | 0.647 | 0.000429956 | 0.447644507 | -0.345684218 | 0.7 | 0.667 | 1 | 0.447644507 | 3.5399E-08 |
| Map3k3 | 1.76996E-08 | -1.228427008 | 0.571 | 0.835 | 0.000429959 | 0.129048941 | -0.527686154 | 0.7 | 0.88 | 1 | 0.129048941 | 3.53993E-08 |
| Trip10 | 1.79994E-08 | -2.046141782 | 0.134 | 0.441 | 0.000437241 | 0.311498794 | -0.321928095 | 0.2 | 0.41 | 1 | 0.311498794 | 3.59988E-08 |
| Ccnd1 | 1.8006E-08 | -2.363052828 | 0.294 | 0.571 | 0.000437402 | 0.507471414 | -1.31836148 | 0.3 | 0.373 | 1 | 0.507471414 | 3.6012E-08 |
| Plcg1 | 1.81588E-08 | -2.628384781 | 0.084 | 0.371 | 0.000441113 | 0.279441406 | -0.974544419 | 0.2 | 0.346 | 1 | 0.279441406 | 3.63176E-08 |
| Atf7 | 1.82776E-08 | -1.13688474 | 0.739 | 0.9 | 0.000443999 | 0.017783867 | -1.020997081 | 0.7 | 0.957 | 1 | 0.017783867 | 3.65551E-08 |
| Ptk7 | 1.84987E-08 | -2.10433666 | 0.176 | 0.476 | 0.000449371 | 0.283103745 | -0.56034594 | 0.4 | 0.586 | 1 | 0.283103745 | 3.69974E-08 |
| Sbno1 | 1.86786E-08 | -0.88083552 | 0.79 | 0.935 | 0.000453741 | 0.05421381 | -0.781779442 | 0.8 | 0.963 | 1 | 0.05421381 | 3.73572E-08 |
| Brwd3 | 1.89322E-08 | -2.582288366 | 0.109 | 0.4 | 0.000459901 | 0.293735561 | -0.629536518 | 0.3 | 0.512 | 1 | 0.293735561 | 3.78644E-08 |
| Washc2 | 1.9337E-08 | -1.507341604 | 0.311 | 0.606 | 0.000469735 | 0.055953025 | -1.214099215 | 0.4 | 0.657 | 1 | 0.055953025 | 3.86741E-08 |
| 4833420G17Rik | 1.94804E-08 | -1.749926642 | 0.311 | 0.576 | 0.000473219 | 0.267156635 | -0.248864633 | 0.3 | 0.534 | 1 | 0.267156635 | 3.89609E-08 |
| Thbs2 | 1.97952E-08 | -6.253611152 | 0.008 | 0.247 | 0.000480864 | 0.214165394 | -2.618702713 | 0 | 0.136 | 1 | 0.214165394 | 3.95903E-08 |
| Trim41 | 1.98682E-08 | -1.931683057 | 0.134 | 0.441 | 0.000482638 | 0.326940453 | -0.668578619 | 0.3 | 0.41 | 1 | 0.326940453 | 3.97364E-08 |
| Ehmt2 | 1.99471E-08 | -2.14839184 | 0.109 | 0.412 | 0.000484555 | 0.199506212 | -1.026472211 | 0.2 | 0.377 | 1 | 0.199506212 | 3.98942E-08 |
| Ttc7 | 2.01749E-08 | 0.681223042 | 0.908 | 0.588 | 0.000490089 | 0.048919429 | 0.617991301 | 1 | 0.815 | 1 | 0.048919429 | 4.03498E-08 |
| 2610008E11Rik | 2.04044E-08 | -2.132317077 | 0.118 | 0.418 | 0.000495665 | 0.166704147 | -1.204470513 | 0.2 | 0.392 | 1 | 0.166704147 | 4.08089E-08 |
| Snx5 | 2.08017E-08 | -1.014679895 | 0.672 | 0.859 | 0.000505315 | 0.027908798 | -1.130261314 | 0.4 | 0.83 | 1 | 0.027908798 | 4.16034E-08 |
| Nr2f1 | 2.09765E-08 | -4.82527683 | 0.008 | 0.247 | 0.000509562 | 0.239050351 | -0.982078092 | 0 | 0.123 | 1 | 0.239050351 | 4.19531E-08 |
| 6820431F20Rik | 2.11054E-08 | -0.775396812 | 0.857 | 0.918 | 0.000512692 | 0.796426646 | -0.101272291 | 1 | 0.96 | 1 | 0.796426646 | 4.22108E-08 |
| Lars | 2.11077E-08 | -1.298443084 | 0.529 | 0.741 | 0.000512748 | 0.23678505 | -0.677886361 | 0.6 | 0.735 | 1 | 0.23678505 | 4.22154E-08 |
| Plcl2 | 2.11132E-08 | -5.240314329 | 0.008 | 0.247 | 0.000512881 | 0.480145801 | -0.876895855 | 0.1 | 0.179 | 1 | 0.480145801 | 4.22264E-08 |
| Camkk2 | 2.11186E-08 | -1.780882711 | 0.286 | 0.571 | 0.000513013 | 0.06779898 | -1.378682873 | 0.3 | 0.54 | 1 | 0.06779898 | 4.22372E-08 |
| Nfkbia | 2.11187E-08 | -1.223400946 | 0.454 | 0.735 | 0.000513015 | 0.153045803 | -1.309300487 | 0.6 | 0.756 | 1 | 0.153045803 | 4.22374E-08 |
| Scaf4 | 2.11192E-08 | -1.526903463 | 0.328 | 0.618 | 0.000513027 | 0.013766042 | -1.23237651 | 0.4 | 0.781 | 1 | 0.013766042 | 4.22384E-08 |
| Copb1 | 2.12731E-08 | -0.714296005 | 0.95 | 0.941 | 0.000516766 | 0.06278971 | -0.649502753 | 1 | 0.969 | 1 | 0.06278971 | 4.25462E-08 |
| Mafb | 2.1346E-08 | -1.104709401 | 0.143 | 0.465 | 0.000518536 | 0.30049608 | -2.344021866 | 0.3 | 0.395 | 1 | 0.30049608 | 4.26919E-08 |
| Eif3c | 2.17157E-08 | -0.783025593 | 0.832 | 0.918 | 0.000527517 | 0.017485768 | -0.949959318 | 0.5 | 0.96 | 1 | 0.017485768 | 4.34313E-08 |
| Ski | 2.21434E-08 | -0.941309379 | 0.866 | 0.894 | 0.000537908 | 0.000443353 | -1.362075243 | 0.9 | 0.981 | 1 | 0.000443353 | 4.42869E-08 |
| Cth | 2.2926E-08 | 1.486558797 | 0.538 | 0.224 | 0.000556918 | 0.002406492 | 2.111031312 | 0.4 | 0.096 | 1 | 0.002406492 | 4.58519E-08 |
| Alpl | 2.30316E-08 | -8.236970886 | 0 | 0.229 | 0.000559483 | 0.421072107 | -2.69632361 | 0 | 0.062 | 1 | 0.421072107 | 4.60632E-08 |
| Cul5 | 2.30467E-08 | -1.396319353 | 0.454 | 0.7 | 0.000559851 | 0.164723674 | -0.523971871 | 0.7 | 0.83 | 1 | 0.164723674 | 4.60934E-08 |
| Dennd2a | 2.32228E-08 | -3.364572432 | 0.042 | 0.306 | 0.000564128 | 0.310869919 | -1.145308441 | 0.2 | 0.324 | 1 | 0.310869919 | 4.64456E-08 |
| Pof1b | 2.38135E-08 | 3.707218251 | 0.252 | 0.035 | 0.000578478 | 0.00608363 | 2.432959407 | 0.4 | 0.117 | 1 | 0.00608363 | 4.76271E-08 |
| Sema5a | 2.42788E-08 | -2.648365398 | 0.16 | 0.447 | 0.00058978 | 0.620501305 | -0.984580599 | 0.4 | 0.435 | 1 | 0.620501305 | 4.85576E-08 |
| Cbfa2t2 | 2.45504E-08 | -1.714245518 | 0.294 | 0.571 | 0.000596379 | 0.028004268 | -1.49777793 | 0.3 | 0.664 | 1 | 0.028004268 | 4.91009E-08 |
| Tln1 | 2.5375E-08 | -0.758720147 | 0.924 | 0.971 | 0.000616409 | 0.002008639 | -1.181323667 | 0.9 | 0.969 | 1 | 0.002008639 | 5.075E-08 |
| Ino80d | 2.57123E-08 | -1.263034406 | 0.462 | 0.729 | 0.000624603 | 0.024637066 | -1.048167282 | 0.7 | 0.858 | 1 | 0.024637066 | 5.14246E-08 |
| Pclaf | 0.043247353 | -1.572889668 | 0.008 | 0.053 | 1 | 2.57985E-08 | 6.754887502 | 0.2 | 0.006 | 0.000626697 | 0.043247353 | 5.1597E-08 |
| Cgnl1 | 2.60442E-08 | 1.272755253 | 0.563 | 0.229 | 0.000632665 | 0.97552338 | -0.505640048 | 0.4 | 0.373 | 1 | 0.97552338 | 5.20883E-08 |
| Arhgap5 | 2.66514E-08 | -0.899699985 | 0.798 | 0.9 | 0.000647416 | 0.003819201 | -1.234743524 | 0.6 | 0.957 | 1 | 0.003819201 | 5.33028E-08 |
| 2610035D17Rik | 2.71738E-08 | 1.471319975 | 0.697 | 0.435 | 0.000660106 | 0.008441665 | 3.155425432 | 0.7 | 0.42 | 1 | 0.008441665 | 5.43476E-08 |
| Col4a6 | 0.003521814 | 2.652076697 | 0.092 | 0.018 | 1 | 2.85369E-08 | 6.432959407 | 0.2 | 0.006 | 0.000693219 | 0.003521814 | 5.70738E-08 |
| Tonsl | 0.051935865 | -1.987927168 | 0.017 | 0.065 | 1 | 2.85369E-08 | 5.432959407 | 0.2 | 0.006 | 0.000693219 | 0.051935865 | 5.70738E-08 |
| Fndc5 | 0.201658121 | 1.251538767 | 0.034 | 0.012 | 1 | 2.85369E-08 | 5.432959407 | 0.2 | 0.006 | 0.000693219 | 0.201658121 | 5.70738E-08 |
| Gm10610 | 0.374548525 | -0.485426827 | 0.017 | 0.006 | 1 | 2.85369E-08 | 5.432959407 | 0.2 | 0.006 | 0.000693219 | 0.374548525 | 5.70738E-08 |
| Cdc25c | 0.718517371 | 2.215012891 | 0.017 | 0.012 | 1 | 2.85369E-08 | 5.432959407 | 0.2 | 0.006 | 0.000693219 | 0.718517371 | 5.70738E-08 |
| Aknaos | 0.718518189 | 0.736965594 | 0.017 | 0.012 | 1 | 2.85369E-08 | 5.432959407 | 0.2 | 0.006 | 0.000693219 | 0.718518189 | 5.70738E-08 |
| Fancc | 2.91892E-08 | -1.763962327 | 0.37 | 0.641 | 0.000709065 | 0.096392275 | -1.391469028 | 0.5 | 0.648 | 1 | 0.096392275 | 5.83784E-08 |
| Rptor | 2.94923E-08 | -1.066474278 | 0.597 | 0.806 | 0.000716428 | 0.730578098 | -0.150093503 | 0.8 | 0.818 | 1 | 0.730578098 | 5.89847E-08 |
| Cmtm3 | 2.9907E-08 | -1.928370323 | 0.21 | 0.5 | 0.0007265 | 0.122488467 | -1.399930607 | 0.1 | 0.324 | 1 | 0.122488467 | 5.98139E-08 |
| Gm35330 | 0.001761967 | 2.514573173 | 0.101 | 0.018 | 1 | 3.00196E-08 | 6.017921908 | 0.2 | 0.006 | 0.000729236 | 0.001761967 | 6.00392E-08 |
| 1810055G02Rik | 3.06117E-08 | 0.817851211 | 0.857 | 0.571 | 0.00074362 | 0.026635225 | 0.948124941 | 0.9 | 0.605 | 1 | 0.026635225 | 6.12234E-08 |
| 1700025G04Rik | 3.10103E-08 | 0.734163814 | 1 | 0.941 | 0.000753303 | 0.107659402 | 0.913585248 | 1 | 0.944 | 1 | 0.107659402 | 6.20207E-08 |
| Pcdh9 | 3.12552E-08 | 1.363535416 | 0.824 | 0.6 | 0.00075925 | 0.386612029 | 0.576605084 | 0.7 | 0.688 | 1 | 0.386612029 | 6.25103E-08 |
| Rhoa | 3.13372E-08 | -0.574276721 | 0.983 | 0.982 | 0.000761242 | 0.005651618 | -0.814968106 | 0.9 | 0.997 | 1 | 0.005651618 | 6.26743E-08 |
| Grin1 | 0.058406048 | 1.893084796 | 0.059 | 0.018 | 1 | 3.15723E-08 | 5.017921908 | 0.2 | 0.006 | 0.000766955 | 0.058406048 | 6.31447E-08 |
| Gm16364 | 0.091361251 | 2.099535674 | 0.017 | 0 | 1 | 3.15723E-08 | 5.017921908 | 0.2 | 0.006 | 0.000766955 | 0.091361251 | 6.31447E-08 |
| Gm15908 | 0.209949382 | -0.485426827 | 0.034 | 0.012 | 1 | 3.15723E-08 | 5.017921908 | 0.2 | 0.006 | 0.000766955 | 0.209949382 | 6.31447E-08 |
| Bub1 | 0.21958736 | -1.944858446 | 0.008 | 0.029 | 1 | 3.15723E-08 | 5.017921908 | 0.2 | 0.006 | 0.000766955 | 0.21958736 | 6.31447E-08 |
| Prkd3 | 3.17272E-08 | -1.736965594 | 0.235 | 0.529 | 0.000770717 | 0.003963812 | -2.186649236 | 0.3 | 0.685 | 1 | 0.003963812 | 6.34544E-08 |
| Tsc22d1 | 3.18335E-08 | 0.779724759 | 1 | 0.988 | 0.000773298 | 0.581941558 | 0.322620497 | 1 | 0.997 | 1 | 0.581941558 | 6.36669E-08 |
| Dlx5 | 3.19916E-08 | -4.614709844 | 0.017 | 0.259 | 0.00077714 | 0.166405066 | -1.91281543 | 0 | 0.164 | 1 | 0.166405066 | 6.39832E-08 |
| Cyth1 | 3.21446E-08 | -1.342822432 | 0.345 | 0.635 | 0.000780857 | 0.638611006 | 0.556562841 | 0.7 | 0.735 | 1 | 0.638611006 | 6.42892E-08 |
| Klf13 | 3.22177E-08 | -1.011276234 | 0.714 | 0.882 | 0.000782633 | 0.010555653 | -1.065067457 | 0.8 | 0.969 | 1 | 0.010555653 | 6.44355E-08 |
| Adra2c | 3.22687E-08 | 4.514573173 | 0.185 | 0.006 | 0.000783872 | 0.001983182 | 4.695993813 | 0.1 | 0.006 | 1 | 0.001983182 | 6.45374E-08 |
| Cotl1 | 3.22958E-08 | -3.250961574 | 0.034 | 0.288 | 0.000784531 | 0.565804176 | -0.556986928 | 0.2 | 0.269 | 1 | 0.565804176 | 6.45917E-08 |
| Rev1 | 3.29977E-08 | -1.288653864 | 0.471 | 0.712 | 0.00080158 | 0.613319784 | 0.295455884 | 0.9 | 0.815 | 1 | 0.613319784 | 6.59954E-08 |
| Rcc2 | 3.37475E-08 | -1.935229745 | 0.294 | 0.565 | 0.000819793 | 0.04357772 | -1.233797185 | 0.3 | 0.623 | 1 | 0.04357772 | 6.74949E-08 |
| Fzd2 | 3.41566E-08 | -2.14363831 | 0.193 | 0.488 | 0.000829732 | 0.059623732 | -1.709998547 | 0.3 | 0.534 | 1 | 0.059623732 | 6.83132E-08 |
| Pura | 3.42632E-08 | -0.6927518 | 0.899 | 0.947 | 0.000832323 | 0.004612811 | -1.046115084 | 0.7 | 0.963 | 1 | 0.004612811 | 6.85265E-08 |
| Mxd4 | 3.45012E-08 | -1.428580967 | 0.571 | 0.759 | 0.000838104 | 0.329662256 | -0.152003093 | 0.7 | 0.858 | 1 | 0.329662256 | 6.90024E-08 |
| 2210408F21Rik | 3.45866E-08 | 0.574161969 | 0.681 | 0.288 | 0.000840177 | 0.002458601 | 0.851644684 | 0.9 | 0.41 | 1 | 0.002458601 | 6.91731E-08 |
| Hectd2os | 3.47619E-08 | 1.267933205 | 0.361 | 0.094 | 0.000844436 | 1.02627E-07 | 4.252387162 | 0.6 | 0.093 | 0.002493007 | 1.02627E-07 | 6.95238E-08 |
| Rad51ap1 | 0.332621252 | -1.707819249 | 0.008 | 0.024 | 1 | 3.49001E-08 | 4.602884409 | 0.2 | 0.006 | 0.000847793 | 0.332621252 | 6.98001E-08 |
| Gm47923 | 0.942269525 | 1.514573173 | 0.042 | 0.041 | 1 | 3.491E-08 | 4.695993813 | 0.2 | 0.006 | 0.000848034 | 0.942269525 | 6.982E-08 |
| Nr3c1 | 3.60553E-08 | -0.906072973 | 0.857 | 0.941 | 0.000875855 | 0.014520667 | -0.970131541 | 0.8 | 0.978 | 1 | 0.014520667 | 7.21106E-08 |
| Adnp | 3.61017E-08 | -1.077496214 | 0.681 | 0.835 | 0.000876983 | 0.050609478 | -0.715432433 | 0.8 | 0.932 | 1 | 0.050609478 | 7.22034E-08 |
| Pou2f1 | 3.64069E-08 | -2.146624914 | 0.252 | 0.529 | 0.000884397 | 0.198870197 | -0.607454861 | 0.4 | 0.623 | 1 | 0.198870197 | 7.28138E-08 |
| 4933400C23Rik | 0.01612589 | 2.251538767 | 0.05 | 0.006 | 1 | 3.85861E-08 | 3.280956314 | 0.2 | 0.006 | 0.000937333 | 0.01612589 | 7.71722E-08 |
| Fabp12 | 0.199602566 | 1.514573173 | 0.034 | 0.012 | 1 | 3.85861E-08 | 3.795529487 | 0.2 | 0.006 | 0.000937333 | 0.199602566 | 7.71722E-08 |
| Msmo1 | 3.91941E-08 | 1.049662885 | 0.807 | 0.535 | 0.000952104 | 0.868098256 | -0.677684656 | 0.6 | 0.546 | 1 | 0.868098256 | 7.83883E-08 |
| Rgs10 | 3.92203E-08 | -1.404808367 | 0.227 | 0.524 | 0.000952739 | 0.058231362 | -2.330806246 | 0 | 0.272 | 1 | 0.058231362 | 7.84406E-08 |
| Fbxl7 | 4.00662E-08 | -4.250961574 | 0.118 | 0.382 | 0.000973288 | 0.693327856 | -1.615073289 | 0.3 | 0.309 | 1 | 0.693327856 | 8.01324E-08 |
| Ggct | 4.02924E-08 | 2.099535674 | 0.395 | 0.124 | 0.000978783 | 0.01897765 | 2.393431043 | 0.3 | 0.09 | 1 | 0.01897765 | 8.05848E-08 |
| Gatad2a | 4.15712E-08 | -1.15758777 | 0.563 | 0.794 | 0.001009846 | 0.793738701 | -0.161589142 | 0.9 | 0.883 | 1 | 0.793738701 | 8.31423E-08 |
| Dhx57 | 4.17222E-08 | -1.987927168 | 0.168 | 0.471 | 0.001013516 | 0.659672212 | -0.125036046 | 0.4 | 0.441 | 1 | 0.659672212 | 8.34444E-08 |
| Mthfd2 | 4.18354E-08 | -1.539874611 | 0.286 | 0.588 | 0.001016267 | 0.193269763 | -0.986579484 | 0.2 | 0.423 | 1 | 0.193269763 | 8.36709E-08 |
| Myo5a | 4.19119E-08 | 0.898532361 | 0.941 | 0.735 | 0.001018123 | 0.02916162 | 1.09056721 | 1 | 0.694 | 1 | 0.02916162 | 8.38237E-08 |
| Srsf10 | 4.2754E-08 | -0.778208576 | 0.882 | 0.929 | 0.001038581 | 0.198931218 | -0.448448141 | 1 | 0.966 | 1 | 0.198931218 | 8.55081E-08 |
| Hdac2 | 4.37463E-08 | -1.572889668 | 0.361 | 0.612 | 0.001062686 | 0.183795926 | -0.668578619 | 0.5 | 0.688 | 1 | 0.183795926 | 8.74927E-08 |
| Magi3 | 4.37946E-08 | -2.250961574 | 0.202 | 0.494 | 0.001063858 | 0.521567363 | 0.139600465 | 0.4 | 0.54 | 1 | 0.521567363 | 8.75892E-08 |
| Fgd1 | 4.40915E-08 | -2.807354922 | 0.067 | 0.335 | 0.00107107 | 0.074726791 | -1.864721141 | 0 | 0.247 | 1 | 0.074726791 | 8.81829E-08 |
| Mrps6 | 4.45016E-08 | 0.826602807 | 0.916 | 0.741 | 0.001081033 | 0.037297522 | 0.766202815 | 1 | 0.793 | 1 | 0.037297522 | 8.90032E-08 |
| C1qtnf1 | 4.56153E-08 | -2.10433666 | 0.143 | 0.429 | 0.001108087 | 0.103175567 | 0.384596386 | 0.1 | 0.407 | 1 | 0.103175567 | 9.12306E-08 |
| Ttll4 | 4.59416E-08 | -2.821710215 | 0.067 | 0.335 | 0.001116014 | 0.028405264 | -2.146985019 | 0.1 | 0.444 | 1 | 0.028405264 | 9.18832E-08 |
| Tubb2a | 4.59575E-08 | 1.023018617 | 0.983 | 0.853 | 0.0011164 | 0.895229263 | 0.520181819 | 1 | 0.895 | 1 | 0.895229263 | 9.1915E-08 |
| Snrpb | 4.64839E-08 | -0.724765141 | 0.849 | 0.935 | 0.001129187 | 0.015950244 | -0.910185174 | 0.5 | 0.957 | 1 | 0.015950244 | 9.29678E-08 |

| | | | | | | | | | | | | |
|---|---|---|---|---|---|---|---|---|---|---|---|---|
| Nsd3 | 4.70234E-08 | -0.914113388 | 0.891 | 0.982 | 0.001142293 | 0.077514268 | -0.605470179 | 0.8 | 0.997 | 1 | 0.077514268 | 9.40468E-08 |
| Specc1 | 4.85947E-08 | -2.868345552 | 0.252 | 0.506 | 0.001180463 | 0.001768706 | -2.72161763 | 0.4 | 0.781 | 1 | 0.001768706 | 9.71895E-08 |
| Rapgef6 | 5.11079E-08 | -1.018859027 | 0.882 | 0.906 | 0.001241513 | 0.583975601 | -0.205985154 | 0.9 | 0.951 | 1 | 0.583975601 | 1.02216E-07 |
| Ctcf | 5.11401E-08 | -1.078951341 | 0.555 | 0.788 | 0.001242296 | 0.004107972 | -1.467907401 | 0.5 | 0.836 | 1 | 0.004107972 | 1.0228E-07 |
| Dis3 | 5.13412E-08 | -1.752907138 | 0.176 | 0.482 | 0.001247181 | 0.24141279 | -0.9669712 | 0.2 | 0.358 | 1 | 0.24141279 | 1.02682E-07 |
| Mapre1 | 5.19197E-08 | -0.690478049 | 0.874 | 0.906 | 0.001261233 | 0.100993073 | -0.594332284 | 0.7 | 0.948 | 1 | 0.100993073 | 1.03839E-07 |
| Pear1 | 5.34533E-08 | -2.74176658 | 0.059 | 0.324 | 0.001298487 | 0.214613514 | -1.171902651 | 0.2 | 0.364 | 1 | 0.214613514 | 1.06907E-07 |
| Chsy1 | 5.34949E-08 | -1.212050477 | 0.429 | 0.688 | 0.001299498 | 0.002290726 | -2.237578825 | 0.4 | 0.735 | 1 | 0.002290726 | 1.0699E-07 |
| Akt3 | 5.36357E-08 | -0.97174174 | 0.79 | 0.9 | 0.00130292 | 0.123093229 | -0.543733219 | 0.8 | 0.978 | 1 | 0.123093229 | 1.07271E-07 |
| Lpar4 | 5.38449E-08 | -2.365845211 | 0.151 | 0.435 | 0.001308 | 0.344252141 | -1.048167282 | 0.2 | 0.315 | 1 | 0.344252141 | 1.0769E-07 |
| Pnisr | 5.3966E-08 | -0.733130337 | 0.95 | 0.947 | 0.001310942 | 0.109909741 | -0.527335119 | 1 | 0.966 | 1 | 0.109909741 | 1.07932E-07 |
| Sirt5 | 5.40608E-08 | 2.497295181 | 0.353 | 0.1 | 0.001313246 | 0.360331175 | 1.769994395 | 0.1 | 0.04 | 1 | 0.360331175 | 1.08122E-07 |
| Msl2 | 5.53784E-08 | -1.524107295 | 0.269 | 0.571 | 0.001345253 | 0.003978249 | -1.660149997 | 0.3 | 0.753 | 1 | 0.003978249 | 1.10757E-07 |
| Ruvbl1 | 5.60649E-08 | -1.618693358 | 0.218 | 0.512 | 0.001361928 | 0.04513635 | -1.716787712 | 0.1 | 0.41 | 1 | 0.04513635 | 1.1213E-07 |
| Tmem39b | 5.6938E-08 | -2.977279923 | 0.059 | 0.318 | 0.001383138 | 0.13545437 | -1.473931188 | 0.1 | 0.312 | 1 | 0.13545437 | 1.13876E-07 |
| Pogk | 5.7384E-08 | -2.46727948 | 0.076 | 0.347 | 0.001393971 | 0.322826914 | -1.011825435 | 0.2 | 0.321 | 1 | 0.322826914 | 1.14768E-07 |
| Prkci | 5.74634E-08 | -2.018859027 | 0.126 | 0.418 | 0.0013959 | 0.864060984 | 0.101445264 | 0.6 | 0.556 | 1 | 0.864060984 | 1.14927E-07 |
| Ss18l2 | 5.75285E-08 | 1.381678902 | 0.697 | 0.447 | 0.001397482 | 0.0595773 | 1.017921908 | 0.6 | 0.318 | 1 | 0.0595773 | 1.15057E-07 |
| Naa20 | 5.78245E-08 | -1.31259023 | 0.479 | 0.706 | 0.001404673 | 0.043571891 | -1.161987182 | 0.5 | 0.71 | 1 | 0.043571891 | 1.15649E-07 |
| Ccdc85b | 5.79375E-08 | 1.542770065 | 0.723 | 0.441 | 0.001407419 | 0.672275429 | 1.34415014 | 0.5 | 0.42 | 1 | 0.672275429 | 1.15875E-07 |
| Zdhhc19 | 0.216340691 | -2.544320516 | 0.008 | 0.029 | 1 | 5.85027E-08 | 4.017921908 | 0.3 | 0.019 | 0.001421147 | 0.216340691 | 1.17005E-07 |
| Acsm3 | 0.653579641 | 0.836501268 | 0.025 | 0.018 | 1 | 5.85027E-08 | 4.280956314 | 0.3 | 0.019 | 0.001421147 | 0.653579641 | 1.17005E-07 |
| Wdr45b | 5.90793E-08 | -1.304511042 | 0.437 | 0.676 | 0.001435156 | 0.039385161 | -1.042774024 | 0.6 | 0.775 | 1 | 0.039385161 | 1.18159E-07 |
| Nono | 5.959E-08 | -1.128193779 | 0.546 | 0.776 | 0.00144756 | 0.071628347 | -0.567040593 | 0.5 | 0.836 | 1 | 0.071628347 | 1.1918E-07 |
| Zfx | 5.9662E-08 | -1.271596419 | 0.454 | 0.694 | 0.00144931 | 0.007458567 | -1.531541912 | 0.6 | 0.846 | 1 | 0.007458567 | 1.19324E-07 |
| Otud4 | 6.00199E-08 | -1.418875634 | 0.395 | 0.635 | 0.001458003 | 0.460346131 | -0.180347719 | 0.7 | 0.75 | 1 | 0.460346131 | 1.2004E-07 |
| Larp1 | 6.13857E-08 | -1.038819249 | 0.689 | 0.824 | 0.00149118 | 0.097364446 | -0.528972552 | 0.8 | 0.941 | 1 | 0.097364446 | 1.22771E-07 |
| Aebp2 | 6.21684E-08 | -1.12613188 | 0.538 | 0.759 | 0.001510196 | 0.163762997 | -0.776493958 | 0.7 | 0.864 | 1 | 0.163762997 | 1.24337E-07 |
| Gm30938 | 0.933394757 | -0.163498732 | 0.034 | 0.035 | 1 | 6.24903E-08 | 4.503348735 | 0.3 | 0.019 | 0.001518013 | 0.933394757 | 1.24981E-07 |
| P2ry10b | 6.28841E-08 | -2.243454037 | 0.059 | 0.329 | 0.001527581 | 0.100853492 | -1.782977992 | 0.1 | 0.34 | 1 | 0.100853492 | 1.25768E-07 |
| Mcoln2 | 6.30481E-08 | 1.003533854 | 0.756 | 0.441 | 0.001531563 | 0.000301882 | 2.833046565 | 0.8 | 0.404 | 1 | 0.000301882 | 1.26096E-07 |
| Fto | 6.44415E-08 | -0.870433605 | 0.899 | 0.971 | 0.001565413 | 0.2336037 | -0.481110414 | 0.9 | 0.985 | 1 | 0.2336037 | 1.28883E-07 |
| Adamts12 | 6.63148E-08 | -1.881011964 | 0.059 | 0.324 | 0.00161092 | 0.017727261 | -3.404142858 | 0 | 0.38 | 1 | 0.017727261 | 1.3263E-07 |
| Lactb2 | 6.63188E-08 | 1.275566038 | 0.647 | 0.347 | 0.001611016 | 0.012453639 | 1.685725478 | 0.6 | 0.25 | 1 | 0.012453639 | 1.32638E-07 |
| Snrpe | 6.64148E-08 | -0.774564987 | 0.739 | 0.876 | 0.001613349 | 0.008692319 | -0.874873858 | 0.3 | 0.883 | 1 | 0.008692319 | 1.3283E-07 |
| Srrt | 6.71078E-08 | -1.099244191 | 0.521 | 0.753 | 0.001630182 | 0.058810467 | -0.900941329 | 0.6 | 0.806 | 1 | 0.058810467 | 1.34216E-07 |
| Btbd7 | 6.71184E-08 | -1.085819368 | 0.706 | 0.859 | 0.001630439 | 0.230252202 | -0.324332988 | 0.9 | 0.957 | 1 | 0.230252202 | 1.34237E-07 |
| Gpha2 | 6.78146E-08 | -2.042032802 | 0.319 | 0.594 | 0.001647353 | 0.080048872 | -3.414619992 | 0.3 | 0.525 | 1 | 0.080048872 | 1.35629E-07 |
| Ptma | 6.80676E-08 | -0.764608473 | 0.992 | 0.988 | 0.001653498 | 0.01317141 | -0.821081867 | 1 | 0.997 | 1 | 0.01317141 | 1.36135E-07 |
| Fmnl3 | 6.8432E-08 | -1.696930932 | 0.202 | 0.494 | 0.001662351 | 0.002835263 | -3.004445905 | 0.1 | 0.593 | 1 | 0.002835263 | 1.36864E-07 |
| Bcorl1 | 6.87703E-08 | -3.655351829 | 0.025 | 0.265 | 0.001670569 | 0.684518297 | -0.736965594 | 0.3 | 0.318 | 1 | 0.684518297 | 1.37541E-07 |
| Col9a2 | 6.96326E-08 | 0.566489841 | 1 | 0.912 | 0.001691514 | 0.344005686 | 0.261859355 | 1 | 0.92 | 1 | 0.344005686 | 1.39265E-07 |
| Acap2 | 7.08897E-08 | -1.61667136 | 0.42 | 0.665 | 0.001722053 | 0.96046117 | -0.575365457 | 0.7 | 0.713 | 1 | 0.96046117 | 1.41779E-07 |
| Snrpd1 | 7.12402E-08 | -0.775682702 | 0.765 | 0.865 | 0.001730567 | 0.02838589 | -0.783475559 | 0.6 | 0.898 | 1 | 0.02838589 | 1.4248E-07 |
| Cacng4 | 3.47E-06 | 1.466744648 | 0.361 | 0.129 | 0.084293238 | 7.24125E-08 | 2.295455884 | 0.7 | 0.111 | 0.001759044 | 3.47E-06 | 1.44825E-07 |
| Anxa6 | 7.26457E-08 | -1.375549209 | 0.454 | 0.676 | 0.00176471 | 0.110483923 | -0.92291602 | 0.3 | 0.58 | 1 | 0.110483923 | 1.45291E-07 |
| Kif26b | 7.27154E-08 | -1.317510551 | 0.269 | 0.588 | 0.001766404 | 0.43548252 | -0.854906852 | 0.4 | 0.481 | 1 | 0.43548252 | 1.45431E-07 |
| Slc10a7 | 7.35668E-08 | -1.916061182 | 0.252 | 0.524 | 0.001787085 | 0.511276416 | -0.258918297 | 0.5 | 0.59 | 1 | 0.511276416 | 1.47134E-07 |
| Ahcyl2 | 7.49187E-08 | -1.672826795 | 0.521 | 0.735 | 0.001819924 | 0.015404308 | -1.506293295 | 0.5 | 0.877 | 1 | 0.015404308 | 1.49837E-07 |
| Enah | 7.55401E-08 | 0.998885519 | 0.983 | 0.894 | 0.001835019 | 0.297051607 | 1.211504105 | 0.9 | 0.87 | 1 | 0.297051607 | 1.5108E-07 |
| Trrap | 7.58393E-08 | -1.099876092 | 0.613 | 0.771 | 0.001842289 | 0.134773063 | -0.719043686 | 0.8 | 0.855 | 1 | 0.134773063 | 1.51679E-07 |
| Hmbox1 | 7.6017E-08 | -1.617877123 | 0.328 | 0.594 | 0.001846605 | 0.732004852 | -0.165299916 | 0.7 | 0.691 | 1 | 0.732004852 | 1.52034E-07 |
| Arglu1 | 7.60783E-08 | -0.801454321 | 0.849 | 0.935 | 0.001848094 | 0.014401837 | -0.896961478 | 0.7 | 0.957 | 1 | 0.014401837 | 1.52157E-07 |
| Ptbp2 | 7.68828E-08 | -1.091945471 | 0.714 | 0.835 | 0.001867637 | 0.054225017 | -0.803007872 | 0.7 | 0.901 | 1 | 0.054225017 | 1.53766E-07 |
| Rprd2 | 7.71029E-08 | -1.274344501 | 0.504 | 0.741 | 0.001872983 | 0.129726242 | -0.627993804 | 0.7 | 0.806 | 1 | 0.129726242 | 1.54206E-07 |
| Elf2 | 7.73868E-08 | -1.30016128 | 0.798 | 0.847 | 0.001879881 | 0.000838349 | -1.576195308 | 0.7 | 0.957 | 1 | 0.000838349 | 1.54774E-07 |
| Kdm6a | 7.78719E-08 | -1.064130337 | 0.571 | 0.812 | 0.001891665 | 0.011596254 | -1.445973959 | 0.7 | 0.833 | 1 | 0.011596254 | 1.55744E-07 |
| Atxn2l | 7.91346E-08 | -1.106523114 | 0.513 | 0.735 | 0.001922337 | 0.012458708 | -1.176834946 | 0.4 | 0.799 | 1 | 0.012458708 | 1.58269E-07 |
| Snrpg | 7.93693E-08 | -0.838279495 | 0.882 | 0.9 | 0.001928039 | 0.052427216 | -0.63270751 | 0.5 | 0.91 | 1 | 0.052427216 | 1.58739E-07 |
| St3gal1 | 7.9937E-08 | -2.562870913 | 0.202 | 0.476 | 0.00194183 | 0.068673277 | -2.374395515 | 0 | 0.256 | 1 | 0.068673277 | 1.59874E-07 |
| Eeig1 | 7.99799E-08 | 1.204551845 | 0.714 | 0.412 | 0.001942871 | 0.013486649 | 1.206279628 | 0.8 | 0.444 | 1 | 0.013486649 | 1.5996E-07 |
| Cmklr1 | 8.03008E-08 | -2.032914622 | 0.151 | 0.447 | 0.001950667 | 0.073120527 | -1.197090983 | 0.3 | 0.608 | 1 | 0.073120527 | 1.60602E-07 |
| Nadk2 | 8.04887E-08 | -4.070389328 | 0.017 | 0.247 | 0.001955232 | 0.079553364 | -1.91281543 | 0 | 0.241 | 1 | 0.079553364 | 1.60977E-07 |
| 4930523C07Rik | 8.05652E-08 | 0.515391723 | 0.983 | 0.841 | 0.001957089 | 0.112759584 | 0.38694701 | 1 | 0.935 | 1 | 0.112759584 | 1.6113E-07 |
| Hnrnpl | 8.1654E-08 | -0.754775007 | 0.941 | 0.947 | 0.001983538 | 0.357952813 | -0.31535116 | 0.9 | 0.981 | 1 | 0.357952813 | 1.63308E-07 |
| Timp1 | 8.23781E-08 | 1.435516444 | 0.891 | 0.735 | 0.002001128 | 0.39914398 | 1.99053911 | 0.7 | 0.571 | 1 | 0.39914398 | 1.64756E-07 |
| Mat2a | 8.39199E-08 | -0.865033198 | 0.824 | 0.935 | 0.002038582 | 0.024875153 | -1.030837404 | 0.8 | 0.926 | 1 | 0.024875153 | 1.6784E-07 |
| Cnot6 | 8.44814E-08 | -0.937415463 | 0.739 | 0.829 | 0.002052223 | 0.010708504 | -1.049602086 | 0.6 | 0.917 | 1 | 0.010708504 | 1.68963E-07 |
| Majin | 8.46834E-08 | -2.510032797 | 0.319 | 0.576 | 0.00205713 | 0.004684202 | -3.991906709 | 0.1 | 0.552 | 1 | 0.004684202 | 1.69367E-07 |
| Pparg | 8.58302E-08 | 1.85222562 | 0.513 | 0.218 | 0.002084986 | 0.016739477 | 1.558490289 | 0.5 | 0.188 | 1 | 0.016739477 | 1.7166E-07 |
| Ncmap | 8.6239E-08 | 0.581184775 | 0.983 | 0.712 | 0.002094919 | 0.082340162 | 0.716425713 | 0.8 | 0.731 | 1 | 0.082340162 | 1.72478E-07 |
| Gnas | 8.65115E-08 | -0.578666778 | 0.992 | 0.994 | 0.002101538 | 0.049706661 | -0.542989326 | 1 | 1 | 1 | 0.049706661 | 1.73023E-07 |
| Rnf128 | 8.80148E-08 | 2.929610672 | 0.303 | 0.071 | 0.002138055 | 0.638065809 | 1.263034406 | 0.1 | 0.062 | 1 | 0.638065809 | 1.7603E-07 |
| Lmo7 | 8.80841E-08 | 0.757290514 | 0.916 | 0.676 | 0.002139738 | 0.000377345 | 1.872071042 | 0.9 | 0.651 | 1 | 0.000377345 | 1.76168E-07 |
| Man1c1 | 8.86198E-08 | -2.387246433 | 0.109 | 0.388 | 0.002152751 | 0.178560417 | -1.574535129 | 0.2 | 0.395 | 1 | 0.178560417 | 1.7724E-07 |
| Chd3 | 8.87788E-08 | -1.498562229 | 0.479 | 0.712 | 0.002156615 | 0.284545649 | -0.542073677 | 0.8 | 0.858 | 1 | 0.284545649 | 1.77558E-07 |
| Dnm2 | 8.87821E-08 | -1.14839184 | 0.555 | 0.782 | 0.002156695 | 0.011989051 | -1.290417122 | 0.6 | 0.864 | 1 | 0.011989051 | 1.77564E-07 |
| Fabp7 | 8.98565E-08 | 3.421463768 | 0.218 | 0.024 | 0.002182794 | 0.048650655 | 2.063725598 | 0.2 | 0.052 | 1 | 0.048650655 | 1.79713E-07 |
| Eif4b | 8.99136E-08 | -1.135680788 | 0.613 | 0.765 | 0.002184181 | 0.025428844 | -0.855522205 | 0.6 | 0.917 | 1 | 0.025428844 | 1.79827E-07 |
| Tob2 | 9.01626E-08 | -0.977908412 | 0.655 | 0.859 | 0.002190231 | 0.184768056 | -0.479929929 | 0.8 | 0.926 | 1 | 0.184768056 | 1.80325E-07 |
| Itpr1 | 9.03405E-08 | -1.365797966 | 0.807 | 0.9 | 0.002194552 | 0.229008722 | -0.599291529 | 1 | 0.975 | 1 | 0.229008722 | 1.80681E-07 |
| Prkar1a | 9.05559E-08 | -0.803788307 | 0.857 | 0.924 | 0.002199783 | 0.006138847 | -1.030837404 | 0.7 | 0.963 | 1 | 0.006138847 | 1.81112E-07 |
| Atxn7l3 | 9.15017E-08 | -2.166148311 | 0.134 | 0.412 | 0.00222276 | 0.154928438 | -0.795859283 | 0.2 | 0.441 | 1 | 0.154928438 | 1.83003E-07 |
| Ctbp1 | 9.2959E-08 | -1.247517015 | 0.513 | 0.7 | 0.00225816 | 0.023783542 | -1.179294785 | 0.4 | 0.762 | 1 | 0.023783542 | 1.85918E-07 |
| Pmepa1 | 9.2994E-08 | -0.727953583 | 0.933 | 0.982 | 0.00225901 | 0.016741358 | -0.835404787 | 1 | 0.994 | 1 | 0.016741358 | 1.85988E-07 |
| Fnip1 | 9.38717E-08 | -0.918923151 | 0.899 | 0.947 | 0.002280332 | 0.003408655 | -1.146202858 | 0.9 | 0.985 | 1 | 0.003408655 | 1.87743E-07 |
| Ap1g1 | 1.00669E-07 | -0.835613182 | 0.807 | 0.894 | 0.002445449 | 0.027999597 | -0.834394788 | 0.7 | 0.941 | 1 | 0.027999597 | 2.01338E-07 |
| Ddx6 | 1.0153E-07 | -0.688668148 | 0.958 | 0.988 | 0.002466362 | 0.016184857 | -0.72691193 | 1 | 0.994 | 1 | 0.016184857 | 2.0306E-07 |
| Has2 | 1.02972E-07 | 1.817965316 | 0.731 | 0.453 | 0.002501393 | 0.028642617 | 0.60610831 | 0.8 | 0.383 | 1 | 0.028642617 | 2.05944E-07 |
| Bclaf1 | 1.03139E-07 | -0.75191905 | 0.899 | 0.965 | 0.002505447 | 0.221033224 | -0.403228197 | 1 | 0.981 | 1 | 0.221033224 | 2.06278E-07 |
| Cpxm2 | 1.05594E-07 | -2.205318908 | 0.168 | 0.441 | 0.00256509 | 0.006635327 | -2.410438265 | 0.2 | 0.617 | 1 | 0.006635327 | 2.11188E-07 |
| Zbtb20 | 1.07482E-07 | -0.531301645 | 1 | 1 | 0.002610947 | 0.236500881 | -0.191083207 | 1 | 1 | 1 | 0.236500881 | 2.14964E-07 |
| Grina | 1.07981E-07 | 1.591649396 | 0.58 | 0.294 | 0.002623078 | 0.027134806 | -1.876895855 | 0.2 | 0.534 | 1 | 0.027134806 | 2.15962E-07 |
| Gnb2 | 1.0961E-07 | -0.6732515 | 0.924 | 0.947 | 0.002662653 | 0.305031732 | -0.362570079 | 0.9 | 0.963 | 1 | 0.305031732 | 2.19221E-07 |
| Mmp16 | 1.10727E-07 | -1.742259894 | 0.261 | 0.553 | 0.002689777 | 0.526324079 | 0.326760003 | 0.5 | 0.475 | 1 | 0.526324079 | 2.21454E-07 |
| Nipbl | 1.11342E-07 | -0.789278632 | 0.966 | 0.988 | 0.00270471 | 0.199268239 | -0.363712433 | 0.9 | 1 | 1 | 0.199268239 | 2.22683E-07 |
| Rc3h1 | 1.11673E-07 | -1.008988783 | 0.815 | 0.9 | 0.002712761 | 0.002809568 | -1.287597027 | 0.7 | 0.969 | 1 | 0.002809568 | 2.23346E-07 |
| Csde1 | 1.11835E-07 | -0.548701002 | 0.966 | 0.982 | 0.002716694 | 0.025088134 | -0.567723525 | 1 | 0.997 | 1 | 0.025088134 | 2.2367E-07 |

| | | | | | | | | | | | | |
|---|---|---|---|---|---|---|---|---|---|---|---|---|
| Pdrg1 | 1.13439E-07 | 0.897108032 | 0.891 | 0.688 | 0.002755655 | 0.113986611 | 0.712481936 | 0.7 | 0.602 | 1 | 0.113986611 | 2.26878E-07 |
| Cd82 | 1.15574E-07 | 0.845397668 | 0.815 | 0.565 | 0.00280752 | 0.000618358 | 1.40053793 | 0.9 | 0.481 | 1 | 0.000618358 | 2.31148E-07 |
| Cbx3 | 1.15623E-07 | -0.84525153 | 0.832 | 0.9 | 0.002808718 | 0.051694216 | -0.789433014 | 0.7 | 0.92 | 1 | 0.051694216 | 2.31246E-07 |
| Pdss2 | 1.16111E-07 | -2.231670235 | 0.176 | 0.453 | 0.002820568 | 0.054119642 | -1.872849022 | 0.1 | 0.414 | 1 | 0.054119642 | 2.32222E-07 |
| Ehbp1 | 1.17052E-07 | -1.363220895 | 0.588 | 0.8 | 0.002843437 | 0.182726386 | -0.73613046 | 0.7 | 0.877 | 1 | 0.182726386 | 2.34105E-07 |
| Dok1 | 1.17145E-07 | 1.663095698 | 0.546 | 0.276 | 0.002845689 | 0.000518113 | 2.503348735 | 0.6 | 0.222 | 1 | 0.000518113 | 2.3429E-07 |
| Phtf1 | 1.20203E-07 | -1.561194335 | 0.286 | 0.559 | 0.002919964 | 0.276931414 | -0.754667596 | 0.4 | 0.602 | 1 | 0.276931414 | 2.40405E-07 |
| Idh2 | 1.20399E-07 | -1.37594835 | 0.336 | 0.629 | 0.002924742 | 0.469961298 | -0.198603506 | 0.4 | 0.617 | 1 | 0.469961298 | 2.40799E-07 |
| Macroh2a2 | 1.22699E-07 | -2.304186513 | 0.118 | 0.388 | 0.002980602 | 0.172484557 | -1.161987182 | 0.1 | 0.29 | 1 | 0.172484557 | 2.45398E-07 |
| Cnnm1 | 1.23122E-07 | 2.158429363 | 0.269 | 0.047 | 0.00299089 | 0.000614676 | 2.432959407 | 0.3 | 0.046 | 1 | 0.000614676 | 2.46245E-07 |
| Wdr82 | 1.25837E-07 | -1.517135687 | 0.294 | 0.565 | 0.003056833 | 0.023044106 | -1.81074352 | 0.2 | 0.543 | 1 | 0.023044106 | 2.51674E-07 |
| Tmem64 | 1.26016E-07 | -2.321928095 | 0.168 | 0.447 | 0.00306118 | 0.057001035 | -0.751732305 | 0.2 | 0.614 | 1 | 0.057001035 | 2.52032E-07 |
| Trib1 | 1.27249E-07 | 0.846892874 | 0.882 | 0.671 | 0.003091143 | 0.621890738 | 0.179978666 | 0.8 | 0.722 | 1 | 0.621890738 | 2.54499E-07 |
| Lrrc42 | 1.28263E-07 | -2.678071905 | 0.05 | 0.3 | 0.003115755 | 0.164463115 | -0.489872732 | 0.1 | 0.343 | 1 | 0.164463115 | 2.56525E-07 |
| Adgrl3 | 1.2836E-07 | 2.53447273 | 0.42 | 0.165 | 0.003118111 | 0.627299213 | 0.393431043 | 0.1 | 0.173 | 1 | 0.627299213 | 2.56719E-07 |
| Inka2 | 0.262060692 | -0.222392421 | 0.025 | 0.053 | 1 | 1.28707E-07 | 2.354956895 | 0.5 | 0.056 | 0.003126562 | 0.262060692 | 2.57415E-07 |
| Slc7a7 | 1.29862E-07 | -2.687060688 | 0.05 | 0.3 | 0.003154616 | 0.051545387 | -2.521236903 | 0 | 0.284 | 1 | 0.051545387 | 2.59725E-07 |
| Mxd1 | 1.30441E-07 | -2.865198997 | 0.168 | 0.435 | 0.003168663 | 0.072242977 | -1.754667596 | 0.2 | 0.46 | 1 | 0.072242977 | 2.60881E-07 |
| Scaf1 | 1.30828E-07 | -1.435844798 | 0.252 | 0.541 | 0.003178083 | 0.711712329 | 0.40053793 | 0.5 | 0.475 | 1 | 0.711712329 | 2.61657E-07 |
| Phf12 | 1.31207E-07 | -1.379436131 | 0.361 | 0.612 | 0.003187277 | 0.17418946 | -0.594521329 | 0.6 | 0.799 | 1 | 0.17418946 | 2.62414E-07 |
| Arfgef1 | 1.32583E-07 | -0.907856461 | 0.79 | 0.888 | 0.003220705 | 0.040994741 | -0.796774989 | 0.8 | 0.954 | 1 | 0.040994741 | 2.65166E-07 |
| Eif5b | 1.33527E-07 | -0.643442948 | 0.933 | 0.976 | 0.00324364 | 0.012153788 | -0.779548899 | 1 | 0.991 | 1 | 0.012153788 | 2.67054E-07 |
| Hmcn1 | 1.33734E-07 | -5.770829046 | 0.008 | 0.224 | 0.003248664 | 0.316168693 | -1.947862377 | 0 | 0.093 | 1 | 0.316168693 | 2.67468E-07 |
| Tut4 | 1.34564E-07 | -0.993803734 | 0.807 | 0.894 | 0.003268824 | 0.348928506 | -0.287625045 | 0.7 | 0.88 | 1 | 0.348928506 | 2.69128E-07 |
| Armcx4 | 1.3552E-07 | -2.086330872 | 0.109 | 0.382 | 0.003292046 | 0.139337567 | -1.425021588 | 0.2 | 0.398 | 1 | 0.139337567 | 2.7104E-07 |
| Gm32618 | 1.38197E-07 | -3.666756592 | 0.034 | 0.271 | 0.00335707 | 0.376147556 | -1.433289204 | 0.1 | 0.204 | 1 | 0.376147556 | 2.76393E-07 |
| Efr3b | 0.112807202 | 0.817135943 | 0.176 | 0.112 | 1 | 1.38517E-07 | 3.865918815 | 0.6 | 0.093 | 0.00336486 | 0.112807202 | 2.77034E-07 |
| Slc44a2 | 1.40527E-07 | 0.757902169 | 0.941 | 0.782 | 0.003413674 | 0.138544217 | 0.604071324 | 1 | 0.756 | 1 | 0.138544217 | 2.81053E-07 |
| Rbm6 | 1.4522E-07 | -1.002300338 | 0.765 | 0.853 | 0.003527681 | 0.06998253 | -0.734909008 | 1 | 0.923 | 1 | 0.06998253 | 2.9044E-07 |
| Tgfbr1 | 1.49437E-07 | -1.55818317 | 0.437 | 0.676 | 0.003630113 | 0.643417118 | -0.928497052 | 0.7 | 0.735 | 1 | 0.643417118 | 2.98873E-07 |
| Igf1 | 1.51041E-07 | -3.487736988 | 0.084 | 0.347 | 0.003669091 | 0.43198902 | -1.840059087 | 0.2 | 0.281 | 1 | 0.43198902 | 3.02082E-07 |
| Aftph | 1.51185E-07 | -1.263034406 | 0.529 | 0.747 | 0.003672574 | 0.248056516 | -0.573354367 | 0.8 | 0.855 | 1 | 0.248056516 | 3.02369E-07 |
| Smg7 | 1.52307E-07 | -1.207892852 | 0.462 | 0.688 | 0.003699845 | 0.956596623 | 0.14345279 | 0.7 | 0.762 | 1 | 0.956596623 | 3.04614E-07 |
| Srsf2 | 1.5251E-07 | -0.658604806 | 0.933 | 0.971 | 0.003704768 | 0.048698238 | -0.68251781 | 0.9 | 0.988 | 1 | 0.048698238 | 3.0502E-07 |
| Dync1li1 | 1.56213E-07 | -1.075096331 | 0.496 | 0.735 | 0.003794725 | 0.001747224 | -1.609351398 | 0.3 | 0.818 | 1 | 0.001747224 | 3.12426E-07 |
| Ube2o | 1.56752E-07 | -1.317029295 | 0.269 | 0.565 | 0.003807808 | 0.045374384 | -1.532824877 | 0.3 | 0.568 | 1 | 0.045374384 | 3.13503E-07 |
| Clip3 | 1.58919E-07 | -2.707819249 | 0.067 | 0.318 | 0.003860463 | 0.022450026 | -2.611434712 | 0 | 0.358 | 1 | 0.022450026 | 3.17838E-07 |
| Cdkn1a | 1.61044E-07 | 0.703599426 | 0.983 | 0.953 | 0.003912083 | 0.239598771 | 0.336376502 | 1 | 0.981 | 1 | 0.239598771 | 3.22088E-07 |
| Nuak1 | 1.6129E-07 | -2.096861539 | 0.218 | 0.518 | 0.003918065 | 0.031363457 | -1.990506714 | 0.3 | 0.571 | 1 | 0.031363457 | 3.22581E-07 |
| Nr2c2 | 1.62473E-07 | -1.550079786 | 0.336 | 0.594 | 0.003946788 | 0.02827953 | -1.160881245 | 0.4 | 0.75 | 1 | 0.02827953 | 3.24945E-07 |
| Eif4enif1 | 1.62848E-07 | -1.694013449 | 0.286 | 0.541 | 0.00395591 | 0.151805321 | -0.825774861 | 0.4 | 0.636 | 1 | 0.151805321 | 3.25696E-07 |
| St6gal1 | 0.000950879 | 1.514573173 | 0.185 | 0.059 | 1 | 1.63714E-07 | 3.116102302 | 0.6 | 0.09 | 0.003976942 | 0.000950879 | 3.27428E-07 |
| Mia | 1.64399E-07 | 0.70432201 | 1 | 0.876 | 0.003993573 | 0.300171403 | 0.595552384 | 1 | 0.846 | 1 | 0.300171403 | 3.28797E-07 |
| Ppig | 1.65406E-07 | -0.824737 | 0.815 | 0.9 | 0.004018051 | 0.157599169 | -0.55860023 | 0.9 | 0.929 | 1 | 0.157599169 | 3.30813E-07 |
| Slc22a15 | 1.68561E-07 | 1.133483005 | 0.504 | 0.2 | 0.004094673 | 0.007265104 | 1.48866884 | 0.6 | 0.238 | 1 | 0.007265104 | 3.37121E-07 |
| Phyhipl | 1.69983E-07 | 2.818353921 | 0.261 | 0.047 | 0.004129218 | 0.698532411 | 1.847996907 | 0 | 0.015 | 1 | 0.698532411 | 3.39965E-07 |
| Jmy | 1.70236E-07 | -1.583728901 | 0.445 | 0.641 | 0.004135362 | 0.078360556 | -0.739019253 | 0.7 | 0.799 | 1 | 0.078360556 | 3.40471E-07 |
| Dab2 | 1.70667E-07 | -0.873697417 | 0.521 | 0.788 | 0.004145845 | 0.010835739 | -1.496532617 | 0.6 | 0.802 | 1 | 0.010835739 | 3.41334E-07 |
| Dusp22 | 1.75933E-07 | 1.599964664 | 0.462 | 0.188 | 0.00427376 | 0.000225491 | 2.414580878 | 0.6 | 0.17 | 1 | 0.000225491 | 3.51866E-07 |
| Zfyve9 | 1.77854E-07 | -1.29061065 | 0.504 | 0.729 | 0.00432043 | 0.145717129 | -0.68251781 | 0.6 | 0.815 | 1 | 0.145717129 | 3.55708E-07 |
| Ncam1 | 1.78372E-07 | -1.242419476 | 0.731 | 0.794 | 0.004333018 | 0.619353771 | -0.149415299 | 0.9 | 0.738 | 1 | 0.619353771 | 3.56744E-07 |
| Igsf3 | 1.78972E-07 | -1.251686728 | 0.529 | 0.718 | 0.004347587 | 0.021869278 | -1.057705797 | 0.7 | 0.886 | 1 | 0.021869278 | 3.57944E-07 |
| Arsi | 1.80206E-07 | -2.292781749 | 0.151 | 0.429 | 0.004377565 | 0.569261877 | 0.346844249 | 0.5 | 0.549 | 1 | 0.569261877 | 3.60412E-07 |
| Zmynd11 | 1.80509E-07 | -0.763559804 | 0.874 | 0.953 | 0.004384914 | 0.009130381 | -0.960175638 | 0.8 | 0.975 | 1 | 0.009130381 | 3.61017E-07 |
| Ctsb | 1.82152E-07 | 0.446791835 | 1 | 0.965 | 0.004424845 | 0.065549291 | -0.623754984 | 0.8 | 0.985 | 1 | 0.065549291 | 3.64305E-07 |
| Prkn | 1.82615E-07 | 0.933212909 | 0.798 | 0.453 | 0.004436089 | 0.20983665 | 1.146246005 | 0.6 | 0.586 | 1 | 0.20983665 | 3.6523E-07 |
| Bcl9l | 1.82841E-07 | -1.572889668 | 0.319 | 0.576 | 0.004441572 | 0.004511469 | -1.876895855 | 0.4 | 0.753 | 1 | 0.004511469 | 3.65682E-07 |
| Hmgcr | 1.84673E-07 | 0.842147831 | 0.874 | 0.618 | 0.004486082 | 0.191449318 | 0.68342514 | 0.9 | 0.657 | 1 | 0.191449318 | 3.69346E-07 |
| Tmem202 | 1.85614E-07 | 2.64385619 | 0.227 | 0.029 | 0.004508939 | 0.10904137 | 1.31748219 | 0.2 | 0.065 | 1 | 0.10904137 | 3.71228E-07 |
| Ric1 | 1.85957E-07 | -1.340333679 | 0.462 | 0.688 | 0.004517273 | 0.099163375 | -0.664753531 | 0.4 | 0.707 | 1 | 0.099163375 | 3.71914E-07 |
| Txndc5 | 1.86859E-07 | -0.804059842 | 0.899 | 0.882 | 0.00453917 | 0.177890509 | -0.481923979 | 0.8 | 0.855 | 1 | 0.177890509 | 3.73717E-07 |
| Fbl | 1.88578E-07 | -1.333423734 | 0.403 | 0.647 | 0.004580949 | 0.003623182 | -1.740301307 | 0.2 | 0.704 | 1 | 0.003623182 | 3.77157E-07 |
| Elp1 | 1.92425E-07 | -1.576925182 | 0.277 | 0.553 | 0.004674392 | 0.368601369 | -0.549762601 | 0.4 | 0.534 | 1 | 0.368601369 | 3.8485E-07 |
| Dgka | 1.92926E-07 | -2.155278225 | 0.168 | 0.429 | 0.004686561 | 0.089478633 | -1.215697769 | 0.2 | 0.481 | 1 | 0.089478633 | 3.85852E-07 |
| Mia2 | 1.95643E-07 | -0.773816807 | 0.84 | 0.935 | 0.004752556 | 0.379318409 | -0.390162831 | 0.9 | 0.935 | 1 | 0.379318409 | 3.91286E-07 |
| Hmgn2 | 1.95946E-07 | -1.254352163 | 0.597 | 0.776 | 0.004759922 | 0.383974099 | -0.233797185 | 0.5 | 0.593 | 1 | 0.383974099 | 3.91892E-07 |
| Golim4 | 1.98901E-07 | -0.741514295 | 0.924 | 0.941 | 0.0048317 | 0.916301433 | -0.120898411 | 0.9 | 0.948 | 1 | 0.916301433 | 3.97802E-07 |
| Proser2 | 2.02554E-07 | 0.713728886 | 0.866 | 0.518 | 0.00492044 | 0.148582981 | 0.327606407 | 0.9 | 0.694 | 1 | 0.148582981 | 4.05108E-07 |
| Cdkn1c | 2.02826E-07 | -2.323370069 | 0.597 | 0.753 | 0.00492706 | 0.00428821 | -2.350469676 | 0.6 | 0.849 | 1 | 0.00428821 | 4.05653E-07 |
| Adgra2 | 2.02949E-07 | -2.778208576 | 0.067 | 0.318 | 0.004930036 | 0.272719195 | -0.647414009 | 0.2 | 0.38 | 1 | 0.272719195 | 4.05898E-07 |
| Tcf7l1 | 2.03668E-07 | -1.516790998 | 0.378 | 0.635 | 0.004947499 | 0.065809961 | -1.209522133 | 0.7 | 0.793 | 1 | 0.065809961 | 4.07336E-07 |
| Acsl3 | 2.03764E-07 | 0.913122549 | 0.941 | 0.794 | 0.004949826 | 0.155355848 | 0.498205342 | 0.9 | 0.83 | 1 | 0.155355848 | 4.07527E-07 |
| Lrrc59 | 2.04102E-07 | -0.841570637 | 0.622 | 0.835 | 0.004958054 | 0.147972494 | -0.309406175 | 0.7 | 0.858 | 1 | 0.147972494 | 4.08205E-07 |
| Zdbf2 | 2.06166E-07 | -2.097610797 | 0.185 | 0.459 | 0.005008193 | 0.010862157 | -3.461858356 | 0 | 0.42 | 1 | 0.010862157 | 4.12333E-07 |
| Peds1 | 2.07797E-07 | 0.817135943 | 0.739 | 0.412 | 0.00504781 | 0.39010386 | 0.242628195 | 0.8 | 0.552 | 1 | 0.39010386 | 4.15594E-07 |
| Nedd4l | 2.08468E-07 | 0.733058256 | 0.975 | 0.818 | 0.005064115 | 0.008393312 | 1.101215059 | 1 | 0.92 | 1 | 0.008393312 | 4.16937E-07 |
| Wipf1 | 2.09073E-07 | -1.649813645 | 0.294 | 0.559 | 0.005078807 | 0.284160162 | -0.507483748 | 0.4 | 0.608 | 1 | 0.284160162 | 4.18146E-07 |
| Csnk2a1 | 2.13129E-07 | -0.952758834 | 0.748 | 0.841 | 0.005177335 | 0.263865806 | -0.242605642 | 0.6 | 0.892 | 1 | 0.263865806 | 4.26258E-07 |
| Dnajb1 | 2.14457E-07 | 1.068729627 | 0.84 | 0.724 | 0.00520958 | 0.699244567 | 2.009021849 | 0.7 | 0.796 | 1 | 0.699244567 | 4.28913E-07 |
| Trim44 | 2.15026E-07 | -0.959358016 | 0.765 | 0.882 | 0.005223409 | 0.382791186 | -0.192923349 | 0.8 | 0.895 | 1 | 0.382791186 | 4.30052E-07 |
| Isg20l2 | 2.16914E-07 | -1.608809243 | 0.202 | 0.488 | 0.005269267 | 0.020095243 | -1.976431529 | 0.1 | 0.472 | 1 | 0.020095243 | 4.33827E-07 |
| Ndufa8 | 2.19633E-07 | 1.03155888 | 0.815 | 0.729 | 0.005335331 | 0.054246218 | -0.633129783 | 0.3 | 0.716 | 1 | 0.054246218 | 4.39266E-07 |
| Sod2 | 2.2006E-07 | 0.938072251 | 0.874 | 0.759 | 0.005345702 | 0.969829012 | 0.297182628 | 0.6 | 0.725 | 1 | 0.969829012 | 4.4012E-07 |
| Ralgapa1 | 2.23561E-07 | -0.916914847 | 0.782 | 0.888 | 0.005430745 | 0.257394205 | 0.413729936 | 1 | 0.935 | 1 | 0.257394205 | 4.47122E-07 |
| Dstn | 2.24085E-07 | 0.70046758 | 0.992 | 0.988 | 0.005443483 | 0.972105714 | 0.683393776 | 1 | 0.991 | 1 | 0.972105714 | 4.48171E-07 |
| Pkig | 2.24251E-07 | -1.566726929 | 0.412 | 0.647 | 0.0054475 | 0.046376048 | -1.476600375 | 0.5 | 0.676 | 1 | 0.046376048 | 4.48501E-07 |
| Sbno2 | 2.28981E-07 | 0.836501268 | 0.882 | 0.694 | 0.005562413 | 0.62279772 | 0.930459067 | 0.5 | 0.66 | 1 | 0.62279772 | 4.57963E-07 |
| Smap1 | 2.29304E-07 | -0.919522662 | 0.697 | 0.824 | 0.005570249 | 0.619834241 | 0.365348501 | 0.9 | 0.843 | 1 | 0.619834241 | 4.58608E-07 |
| Pcbp1 | 2.36806E-07 | -0.599168993 | 0.941 | 0.959 | 0.005752497 | 0.006009054 | -0.913406577 | 1 | 0.978 | 1 | 0.006009054 | 4.73612E-07 |
| Pcx | 0.005240582 | 1.173536255 | 0.185 | 0.076 | 1 | 2.40425E-07 | 3.82527683 | 0.4 | 0.04 | 0.005840406 | 0.005240582 | 4.8085E-07 |
| Alyref | 2.41367E-07 | -0.846098623 | 0.697 | 0.847 | 0.005863298 | 0.027401582 | -0.899450171 | 0.5 | 0.904 | 1 | 0.027401582 | 4.82735E-07 |
| Il11ra1 | 2.42099E-07 | -1.345450818 | 0.664 | 0.753 | 0.00588107 | 0.00038058 | -2.573039333 | 0.3 | 0.79 | 1 | 0.00038058 | 4.84198E-07 |
| Mir100hg | 2.42489E-07 | 0.882889941 | 0.857 | 0.724 | 0.005890541 | 0.725917188 | 0.120220484 | 0.8 | 0.784 | 1 | 0.725917188 | 4.84978E-07 |
| Ttc9c | 2.43726E-07 | -1.611959233 | 0.21 | 0.488 | 0.005920584 | 0.074786508 | -1.042774024 | 0.2 | 0.506 | 1 | 0.074786508 | 4.87451E-07 |
| Rbpj | 2.44643E-07 | -1.266566097 | 0.521 | 0.759 | 0.005942866 | 0.410894622 | -0.571657121 | 0.7 | 0.728 | 1 | 0.410894622 | 4.89286E-07 |
| Elovl6 | 2.4531E-07 | 0.747457968 | 0.882 | 0.676 | 0.00595906 | 0.178477413 | 0.597453445 | 0.7 | 0.556 | 1 | 0.178477413 | 4.90619E-07 |
| Tgif1 | 2.4886E-07 | 0.921482791 | 0.798 | 0.635 | 0.006045301 | 0.436962948 | -0.427388602 | 0.7 | 0.731 | 1 | 0.436962948 | 4.97719E-07 |
| Ncor1 | 2.49975E-07 | -0.614931043 | 0.983 | 0.988 | 0.006072383 | 0.2004538 | -0.358650403 | 1 | 1 | 1 | 0.2004538 | 4.99949E-07 |

| | | | | | | | | | | | | |
|---|---|---|---|---|---|---|---|---|---|---|---|---|
| Lrba | 2.51174E-07 | 0.784145863 | 0.924 | 0.724 | 0.00610151 | 0.073762121 | 0.815082284 | 1 | 0.815 | 1 | 0.073762121 | 5.02347E-07 |
| Mcu | 2.51561E-07 | -1.065591535 | 0.941 | 0.918 | 0.006110931 | 0.20826166 | -0.652805147 | 1 | 0.96 | 1 | 0.20826166 | 5.03123E-07 |
| Tgif2 | 2.53877E-07 | -2.961160258 | 0.059 | 0.3 | 0.00616718 | 0.252337557 | 0.457206954 | 0.4 | 0.219 | 1 | 0.252337557 | 5.07754E-07 |
| Xpo7 | 2.61508E-07 | -1.275007047 | 0.538 | 0.712 | 0.006352562 | 0.008236577 | -1.574535129 | 0.5 | 0.815 | 1 | 0.008236577 | 5.23017E-07 |
| Hnrnpll | 2.66491E-07 | -1.243805728 | 0.513 | 0.718 | 0.006473599 | 0.000326179 | -2.495805688 | 0.3 | 0.802 | 1 | 0.000326179 | 5.32982E-07 |
| Sec11a | 2.67257E-07 | -0.872943264 | 0.798 | 0.847 | 0.006492219 | 0.297564629 | -0.126124462 | 0.5 | 0.895 | 1 | 0.297564629 | 5.34515E-07 |
| Csnk1d | 2.68321E-07 | -1.203144537 | 0.454 | 0.712 | 0.006518043 | 0.084028087 | -0.858036828 | 0.6 | 0.781 | 1 | 0.084028087 | 5.36641E-07 |
| Vegfd | 2.73498E-07 | 1.893084796 | 0.261 | 0.047 | 0.006643803 | 0.010555306 | 3.432959407 | 0.2 | 0.037 | 1 | 0.010555306 | 5.46995E-07 |
| Me1 | 2.73872E-07 | 0.567040593 | 0.924 | 0.729 | 0.006652891 | 0.241354626 | 0.140656481 | 1 | 0.855 | 1 | 0.241354626 | 5.47743E-07 |
| Klf2 | 2.74222E-07 | -1.181761714 | 0.555 | 0.782 | 0.00666141 | 0.634152496 | -0.81082986 | 0.7 | 0.719 | 1 | 0.634152496 | 5.48445E-07 |
| Myh10 | 2.75777E-07 | -1.542760002 | 0.37 | 0.629 | 0.006699164 | 0.50181524 | -0.826173561 | 0.6 | 0.571 | 1 | 0.50181524 | 5.51553E-07 |
| Rhod | 2.80586E-07 | -2.226184 | 0.143 | 0.406 | 0.006815994 | 0.846927068 | 0.382333334 | 0.3 | 0.34 | 1 | 0.846927068 | 5.61172E-07 |
| Gm15987 | 2.81287E-07 | 4.215012891 | 0.218 | 0.029 | 0.006833012 | 0.009979004 | 3.432959407 | 0.1 | 0.009 | 1 | 0.009979004 | 5.62573E-07 |
| Chchd3 | 2.82977E-07 | -1.283792966 | 0.437 | 0.688 | 0.006874075 | 0.000949371 | -2.515407824 | 0.3 | 0.75 | 1 | 0.000949371 | 5.65954E-07 |
| Blmh | 2.86951E-07 | -1.040816213 | 0.63 | 0.771 | 0.006970616 | 0.110125295 | -0.684827971 | 0.7 | 0.809 | 1 | 0.110125295 | 5.73902E-07 |
| Gapdh | 2.88673E-07 | 0.75085254 | 0.975 | 0.982 | 0.007012453 | 0.129037614 | -0.241514345 | 0.9 | 0.994 | 1 | 0.129037614 | 5.77347E-07 |
| Col6a1 | 2.9067E-07 | 0.506448813 | 1 | 0.971 | 0.007060966 | 0.003407583 | 0.815707369 | 1 | 0.941 | 1 | 0.003407583 | 5.81341E-07 |
| Lipg | 2.93799E-07 | 4.058893689 | 0.277 | 0.065 | 0.007136957 | 0.044541052 | 4.017921908 | 0.1 | 0.015 | 1 | 0.044541052 | 5.87597E-07 |
| Ext2 | 2.94625E-07 | 0.656483798 | 0.941 | 0.806 | 0.007157037 | 0.001128822 | 1.290001453 | 1 | 0.787 | 1 | 0.001128822 | 5.8925E-07 |
| Chst2 | 2.96051E-07 | 1.956795501 | 0.445 | 0.176 | 0.007191679 | 0.226149833 | -1.026472211 | 0 | 0.13 | 1 | 0.226149833 | 5.92103E-07 |
| Hnrnpul1 | 2.96786E-07 | -1.177304532 | 0.538 | 0.724 | 0.007209532 | 0.059498523 | -0.811068928 | 0.8 | 0.87 | 1 | 0.059498523 | 5.93572E-07 |
| Tmpo | 2.99406E-07 | -1.547711105 | 0.269 | 0.559 | 0.00727316 | 0.106202248 | -1.057366219 | 0.5 | 0.66 | 1 | 0.106202248 | 5.98811E-07 |
| Selenoi | 3.03896E-07 | -1.563429339 | 0.311 | 0.571 | 0.007382251 | 0.94947232 | 0.137161044 | 0.5 | 0.562 | 1 | 0.94947232 | 6.07793E-07 |
| Tmem131l | 3.08046E-07 | -1.522263595 | 0.193 | 0.471 | 0.007483061 | 0.010248299 | -2.627376256 | 0.4 | 0.679 | 1 | 0.010248299 | 6.16093E-07 |
| Rbm27 | 3.09663E-07 | -1.167855137 | 0.555 | 0.776 | 0.007522326 | 0.755311609 | -0.139195317 | 0.8 | 0.827 | 1 | 0.755311609 | 6.19325E-07 |
| Cdv3 | 3.09665E-07 | -0.763688687 | 0.857 | 0.929 | 0.007522384 | 0.213551704 | -0.213642159 | 0.9 | 0.966 | 1 | 0.213551704 | 6.1933E-07 |
| Igf2 | 3.11575E-07 | -4.258016331 | 0.017 | 0.229 | 0.007568784 | 0.219844754 | -1.090602549 | 0 | 0.133 | 1 | 0.219844754 | 6.2315E-07 |
| Plce1 | 3.1334E-07 | 1.059734665 | 0.765 | 0.518 | 0.007611653 | 0.105036734 | 0.349543399 | 0.8 | 0.488 | 1 | 0.105036734 | 6.2668E-07 |
| Gm45187 | 3.13791E-07 | 2.877143252 | 0.218 | 0.029 | 0.007622604 | 0.009979004 | 3.695993813 | 0.1 | 0.009 | 1 | 0.009979004 | 6.27581E-07 |
| Kat6b | 3.16152E-07 | -1.211421304 | 0.664 | 0.788 | 0.007679961 | 0.199262964 | -0.78052196 | 0.9 | 0.907 | 1 | 0.199262964 | 6.32304E-07 |
| Brd4 | 3.20311E-07 | -0.813371532 | 0.874 | 0.929 | 0.007780998 | 0.024867279 | -0.728469517 | 1 | 0.997 | 1 | 0.024867279 | 6.40622E-07 |
| Chad | 3.23307E-07 | 1.138287434 | 0.95 | 0.882 | 0.007853764 | 0.066466669 | 1.055465727 | 1 | 0.886 | 1 | 0.066466669 | 6.46613E-07 |
| 4921524J17Rik | 3.2822E-07 | -1.921525942 | 0.202 | 0.476 | 0.007973123 | 0.01039962 | -2.207687226 | 0.1 | 0.546 | 1 | 0.01039962 | 6.5644E-07 |
| Plin2 | 3.32355E-07 | 1.952943176 | 0.496 | 0.224 | 0.008073563 | 0.708210725 | 0.988174565 | 0.2 | 0.293 | 1 | 0.708210725 | 6.64709E-07 |
| Rragc | 3.34775E-07 | -1.209042868 | 0.454 | 0.694 | 0.008132344 | 0.001850412 | -1.782977992 | 0.4 | 0.806 | 1 | 0.001850412 | 6.69549E-07 |
| Sec11c | 3.37166E-07 | 0.777607579 | 0.874 | 0.676 | 0.008190432 | 0.614264682 | 0.206279628 | 0.7 | 0.765 | 1 | 0.614264682 | 6.74331E-07 |
| Myo1b | 3.43461E-07 | -5.540709263 | 0.008 | 0.212 | 0.008343362 | 0.481675899 | -1.449683642 | 0.1 | 0.179 | 1 | 0.481675899 | 6.86922E-07 |
| Kremen1 | 3.4507E-07 | -1.661504056 | 0.345 | 0.582 | 0.008382448 | 0.700500105 | 0.699605067 | 0.4 | 0.559 | 1 | 0.700500105 | 6.9014E-07 |
| Arpc3 | 3.47426E-07 | -0.675326528 | 0.866 | 0.941 | 0.008439675 | 0.568968913 | -0.21012714 | 0.9 | 0.917 | 1 | 0.568968913 | 6.94852E-07 |
| Med14 | 3.48082E-07 | -1.729734025 | 0.261 | 0.518 | 0.00845561 | 0.009915789 | -1.91281543 | 0.2 | 0.596 | 1 | 0.009915789 | 6.96164E-07 |
| Rnf216 | 3.48157E-07 | -0.904316652 | 0.714 | 0.859 | 0.008457423 | 0.387520436 | -0.418249804 | 0.9 | 0.917 | 1 | 0.387520436 | 6.96313E-07 |
| Matr3 | 3.49255E-07 | -0.573076172 | 0.966 | 0.965 | 0.008484093 | 0.196868891 | -0.4090274 | 0.9 | 0.991 | 1 | 0.196868891 | 6.98509E-07 |
| Nolc1 | 3.50419E-07 | -0.957295549 | 0.664 | 0.818 | 0.008512384 | 0.229780532 | -0.58384888 | 0.8 | 0.815 | 1 | 0.229780532 | 7.00838E-07 |
| Pwwp2a | 3.50572E-07 | -1.773003417 | 0.235 | 0.494 | 0.008516099 | 0.045602409 | -1.570792728 | 0.3 | 0.559 | 1 | 0.045602409 | 7.01144E-07 |
| Zfp9 | 3.53692E-07 | -1.297680549 | 0.42 | 0.665 | 0.008591884 | 0.089337104 | -1.183151792 | 0.4 | 0.614 | 1 | 0.089337104 | 7.07384E-07 |
| Smarca1 | 3.53718E-07 | -1.718087584 | 0.269 | 0.524 | 0.008592528 | 0.212626517 | -1.185018151 | 0.5 | 0.549 | 1 | 0.212626517 | 7.07437E-07 |
| Sumo1 | 3.55234E-07 | -0.625950912 | 0.908 | 0.959 | 0.008629335 | 0.120579859 | -0.55024535 | 1 | 0.951 | 1 | 0.120579859 | 7.10467E-07 |
| Gm49797 | 3.56798E-07 | -1.475645806 | 0.387 | 0.624 | 0.008667329 | 0.10512861 | -0.917874564 | 0.7 | 0.775 | 1 | 0.10512861 | 7.13595E-07 |
| Mfge8 | 3.57796E-07 | 1.018449037 | 0.924 | 0.747 | 0.008691576 | 0.022263853 | 1.274132115 | 1 | 0.765 | 1 | 0.022263853 | 7.15591E-07 |
| D16Ertd472e | 3.58829E-07 | -1.987927168 | 0.109 | 0.371 | 0.008716674 | 0.398143289 | -0.457811523 | 0.2 | 0.333 | 1 | 0.398143289 | 7.17658E-07 |
| Ankrd50 | 3.58939E-07 | -2.111261609 | 0.218 | 0.471 | 0.008719356 | 0.008862331 | -2.087113563 | 0.1 | 0.565 | 1 | 0.008862331 | 7.17879E-07 |
| Pdgfa | 3.65275E-07 | -1.803105449 | 0.303 | 0.547 | 0.008873259 | 0.017169226 | -1.193030362 | 0.4 | 0.815 | 1 | 0.017169226 | 7.3055E-07 |
| Cdh13 | 3.69144E-07 | -7.427941333 | 0 | 0.194 | 0.008967255 | 0.421079405 | -1.69632361 | 0 | 0.062 | 1 | 0.421079405 | 7.38289E-07 |
| Nrbp1 | 3.70597E-07 | -1.342761598 | 0.462 | 0.671 | 0.009002536 | 0.011356863 | -1.279758641 | 0.3 | 0.75 | 1 | 0.011356863 | 7.41193E-07 |
| Eef2 | 3.71513E-07 | -0.586529022 | 0.95 | 0.994 | 0.009024794 | 0.051200232 | -0.488286481 | 1 | 0.994 | 1 | 0.051200232 | 7.43026E-07 |
| Mlxip | 3.78514E-07 | -1.652632327 | 0.319 | 0.571 | 0.009194856 | 0.635589991 | -0.46809031 | 0.7 | 0.657 | 1 | 0.635589991 | 7.57027E-07 |
| Jmjd6 | 3.7996E-07 | -1.546538341 | 0.303 | 0.553 | 0.009229996 | 0.667312808 | -0.270839323 | 0.6 | 0.642 | 1 | 0.667312808 | 7.5992E-07 |
| Rnf10 | 3.81571E-07 | -0.663167482 | 0.849 | 0.941 | 0.009269126 | 0.141213777 | -0.433573451 | 0.9 | 0.972 | 1 | 0.141213777 | 7.63142E-07 |
| Gorasp2 | 3.96783E-07 | -0.717469419 | 0.899 | 0.929 | 0.009638646 | 0.01751698 | -0.889970211 | 0.8 | 0.935 | 1 | 0.01751698 | 7.93565E-07 |
| Fbn1 | 3.98233E-07 | 0.822609703 | 0.924 | 0.782 | 0.009673867 | 0.267999376 | 0.844516338 | 1 | 0.895 | 1 | 0.267999376 | 7.96465E-07 |
| Dusp5 | 3.98536E-07 | -1.835011265 | 0.109 | 0.382 | 0.009681234 | 0.391834639 | 2.240314329 | 0.4 | 0.293 | 1 | 0.391834639 | 7.97072E-07 |
| Ncoa5 | 3.99946E-07 | -1.441837559 | 0.261 | 0.541 | 0.009715496 | 0.254779419 | -0.719043686 | 0.4 | 0.54 | 1 | 0.254779419 | 7.99893E-07 |
| Plp2 | 4.04047E-07 | -1.173482821 | 0.361 | 0.635 | 0.009815118 | 0.19889649 | -0.250305167 | 0.3 | 0.664 | 1 | 0.19889649 | 8.08094E-07 |
| Gpd2 | 4.04767E-07 | 0.523618312 | 0.824 | 0.565 | 0.009832605 | 0.488596272 | 0.146187758 | 0.9 | 0.809 | 1 | 0.488596272 | 8.09534E-07 |
| Sgce | 4.09201E-07 | -1.937122797 | 0.118 | 0.382 | 0.009940301 | 0.008760649 | -2.91281543 | 0 | 0.432 | 1 | 0.008760649 | 8.18401E-07 |
| Tinagl1 | 4.1401E-07 | 3.006426269 | 0.227 | 0.035 | 0.010057122 | 0.017003024 | 2.695993813 | 0.2 | 0.04 | 1 | 0.017003024 | 8.28019E-07 |
| Dcn | 4.15312E-07 | -0.566909269 | 0.697 | 0.918 | 0.01008877 | 0.014430845 | -2.013350625 | 0.7 | 0.904 | 1 | 0.014430845 | 8.30625E-07 |
| Zbtb2 | 4.18587E-07 | -2.485426827 | 0.109 | 0.365 | 0.010168318 | 0.297899174 | -0.795859283 | 0.2 | 0.355 | 1 | 0.297899174 | 8.37174E-07 |
| Pde1a | 4.19271E-07 | 2.762500686 | 0.269 | 0.059 | 0.01018492 | 0.040980916 | 1.08718457 | 0.2 | 0.049 | 1 | 0.040980916 | 8.38541E-07 |
| Kpnb1 | 4.21994E-07 | -1.12588444 | 0.445 | 0.671 | 0.010251082 | 0.040394194 | -0.896961478 | 0.5 | 0.759 | 1 | 0.040394194 | 8.43988E-07 |
| Idi1 | 4.27307E-07 | 1.291530742 | 0.756 | 0.518 | 0.01038013 | 0.261346118 | 0.169298968 | 0.7 | 0.448 | 1 | 0.261346118 | 8.54613E-07 |
| Hcfc2 | 4.29782E-07 | -1.337869639 | 0.21 | 0.5 | 0.010440253 | 0.685177564 | -0.255096586 | 0.5 | 0.522 | 1 | 0.685177564 | 8.59563E-07 |
| Ybx1 | 4.30284E-07 | -0.596117218 | 0.958 | 0.982 | 0.010452458 | 0.08864086 | -0.490561863 | 1 | 0.994 | 1 | 0.08864086 | 8.60568E-07 |
| Selenom | 4.32326E-07 | 0.654655241 | 0.933 | 0.806 | 0.010502057 | 0.472198617 | -0.304006187 | 0.9 | 0.787 | 1 | 0.472198617 | 8.64651E-07 |
| Phf13 | 4.36355E-07 | -2.031395196 | 0.176 | 0.435 | 0.010599938 | 0.355497268 | -0.293664243 | 0.3 | 0.448 | 1 | 0.355497268 | 8.7271E-07 |
| Lpp | 4.39683E-07 | -0.774613521 | 0.958 | 0.988 | 0.010680777 | 0.114485174 | -0.644793309 | 1 | 0.991 | 1 | 0.114485174 | 8.79366E-07 |
| Lpcat2 | 4.4078E-07 | -2.87774425 | 0.025 | 0.247 | 0.01070743 | 0.188430245 | -1.267480311 | 0.1 | 0.284 | 1 | 0.188430245 | 8.8156E-07 |
| Cbx6 | 4.40908E-07 | -1.777192952 | 0.185 | 0.453 | 0.010710528 | 0.0308641 | -1.083616118 | 0.3 | 0.667 | 1 | 0.0308641 | 8.81815E-07 |
| Furin | 4.42117E-07 | 0.88687479 | 0.866 | 0.641 | 0.010739914 | 0.058934049 | 0.604439875 | 0.9 | 0.843 | 1 | 0.058934049 | 8.84234E-07 |
| Ppp3cb | 4.42836E-07 | -1.285971217 | 0.496 | 0.706 | 0.010757384 | 0.161016443 | -0.619738016 | 0.7 | 0.812 | 1 | 0.161016443 | 8.85673E-07 |
| Ddx21 | 4.4317E-07 | -0.989364831 | 0.731 | 0.788 | 0.010765489 | 0.003135769 | -1.339630097 | 0.3 | 0.858 | 1 | 0.003135769 | 8.8634E-07 |
| Stag2 | 4.44876E-07 | -0.86393845 | 0.807 | 0.888 | 0.010806924 | 0.100401693 | -0.618952681 | 1 | 0.969 | 1 | 0.100401693 | 8.89751E-07 |
| Sspn | 4.48056E-07 | -1.259150971 | 0.538 | 0.718 | 0.010884178 | 0.929159725 | 0.337802174 | 0.6 | 0.67 | 1 | 0.929159725 | 8.96112E-07 |
| Aig1 | 4.507E-07 | 0.449478145 | 0.958 | 0.771 | 0.010948412 | 0.017234704 | 1.066995163 | 1 | 0.741 | 1 | 0.017234704 | 9.014E-07 |
| Nop56 | 4.63175E-07 | -1.002370904 | 0.588 | 0.812 | 0.011251457 | 0.241317742 | -0.191531458 | 0.5 | 0.698 | 1 | 0.241317742 | 9.26351E-07 |
| Irf2 | 4.6836E-07 | -1.502704819 | 0.471 | 0.671 | 0.011377409 | 0.042441769 | -1.210896782 | 0.5 | 0.725 | 1 | 0.042441769 | 9.3672E-07 |
| Usp53 | 4.69205E-07 | -1.692092375 | 0.496 | 0.682 | 0.011397933 | 0.134510995 | -0.613618959 | 0.7 | 0.836 | 1 | 0.134510995 | 9.3841E-07 |
| Tnks2 | 4.71874E-07 | -0.836781036 | 0.899 | 0.971 | 0.011462753 | 0.000378553 | -1.429341037 | 0.9 | 0.985 | 1 | 0.000378553 | 9.43747E-07 |
| Pdzd2 | 4.72845E-07 | 1.467897711 | 0.639 | 0.388 | 0.011486352 | 0.873656822 | -0.329541279 | 0.6 | 0.54 | 1 | 0.873656822 | 9.4569E-07 |
| 2510009E07Rik | 4.81668E-07 | -1.529820947 | 0.244 | 0.512 | 0.011700675 | 0.03604892 | -1.444784843 | 0.3 | 0.614 | 1 | 0.03604892 | 9.63335E-07 |
| C430049B03Rik | 4.83579E-07 | -3.544320516 | 0.05 | 0.282 | 0.011747095 | 0.161712708 | -2.141949429 | 0 | 0.167 | 1 | 0.161712708 | 9.67157E-07 |
| Trim33 | 4.8588E-07 | -1.386165196 | 0.504 | 0.706 | 0.011803001 | 0.932629194 | -0.161082387 | 0.9 | 0.833 | 1 | 0.932629194 | 9.7176E-07 |
| Pfn1 | 4.90812E-07 | -0.668764895 | 0.916 | 0.976 | 0.01192281 | 0.062041035 | -0.611434712 | 0.7 | 0.994 | 1 | 0.062041035 | 9.81624E-07 |
| Nes | 4.92105E-07 | 2.703606997 | 0.37 | 0.135 | 0.01195422 | 0.000184987 | 2.930459067 | 0.3 | 0.04 | 1 | 0.000184987 | 9.8421E-07 |
| Zfp91 | 4.92928E-07 | -0.996242436 | 0.714 | 0.853 | 0.011974208 | 0.028515348 | -0.900941329 | 0.8 | 0.917 | 1 | 0.028515348 | 9.85856E-07 |
| Scara3 | 4.96859E-07 | -0.871578877 | 0.79 | 0.847 | 0.012069689 | 0.036257182 | -0.825513834 | 0.9 | 0.951 | 1 | 0.036257182 | 9.93717E-07 |
| Axin1 | 4.98834E-07 | -1.766712938 | 0.218 | 0.482 | 0.012117667 | 0.000229307 | -3.808626579 | 0 | 0.657 | 1 | 0.000229307 | 9.97667E-07 |
| Slc39a7 | 4.99657E-07 | -0.943314631 | 0.756 | 0.824 | 0.012137661 | 0.120502773 | -0.609131926 | 0.9 | 0.873 | 1 | 0.120502773 | 9.99313E-07 |

| | | | | | | | | | | | | |
|---|---|---|---|---|---|---|---|---|---|---|---|---|
| Foxp4 | 5.03754E-07 | -1.262358829 | 0.437 | 0.653 | 0.012237185 | 0.317765635 | -0.528972552 | 0.7 | 0.806 | 1 | 0.317765635 | 1.00751E-06 |
| Wdr26 | 5.06761E-07 | -0.768579149 | 0.924 | 0.947 | 0.012310242 | 0.004005576 | -0.91281543 | 1 | 0.991 | 1 | 0.004005576 | 1.01352E-06 |
| Dcaf1 | 5.18515E-07 | -1.866516995 | 0.269 | 0.524 | 0.012595774 | 0.034761795 | -1.14865965 | 0.2 | 0.583 | 1 | 0.034761795 | 1.03703E-06 |
| Acly | 5.21927E-07 | 0.946221635 | 0.966 | 0.818 | 0.012678642 | 0.001519947 | 0.965742044 | 1 | 0.784 | 1 | 0.001519947 | 1.04385E-06 |
| Hip1 | 5.22049E-07 | -1.289480386 | 0.723 | 0.829 | 0.012681625 | 0.726035781 | -0.665296714 | 0.9 | 0.846 | 1 | 0.726035781 | 1.0441E-06 |
| Isy1 | 5.2764E-07 | -1.685853917 | 0.336 | 0.565 | 0.012817434 | 0.00997693 | -1.830701032 | 0.3 | 0.657 | 1 | 0.00997693 | 1.05528E-06 |
| Msantd2 | 5.3366E-07 | -1.437332539 | 0.37 | 0.606 | 0.012963662 | 0.032790711 | -0.954260148 | 0.4 | 0.759 | 1 | 0.032790711 | 1.06732E-06 |
| Rab3il1 | 5.37506E-07 | -2.839063782 | 0.042 | 0.271 | 0.013057104 | 0.114516103 | -1.827568143 | 0 | 0.204 | 1 | 0.114516103 | 1.07501E-06 |
| Kat6a | 5.42764E-07 | -1.006730691 | 0.63 | 0.8 | 0.013184825 | 0.000335516 | -1.652144161 | 0.7 | 0.938 | 1 | 0.000335516 | 1.08553E-06 |
| Zbtb10 | 5.45068E-07 | -1.181169759 | 0.319 | 0.594 | 0.0132408 | 0.083407829 | -1.029591553 | 0.3 | 0.617 | 1 | 0.083407829 | 1.09014E-06 |
| Chadl | 5.49208E-07 | 0.662579088 | 0.958 | 0.735 | 0.013341364 | 0.021698026 | 1.024454878 | 0.9 | 0.694 | 1 | 0.021698026 | 1.09842E-06 |
| Stxbp5 | 5.50747E-07 | -1.122420031 | 0.471 | 0.735 | 0.013378747 | 0.57009088 | -0.48280934 | 0.8 | 0.738 | 1 | 0.57009088 | 1.10149E-06 |
| Selenon | 5.51395E-07 | -1.944858446 | 0.101 | 0.359 | 0.013394483 | 0.884607971 | 0.111031312 | 0.3 | 0.327 | 1 | 0.884607971 | 1.10279E-06 |
| Ipo5 | 5.60868E-07 | -1.272697503 | 0.361 | 0.624 | 0.013624598 | 0.056675933 | -1.015501094 | 0.4 | 0.673 | 1 | 0.056675933 | 1.12174E-06 |
| Fbxo42 | 5.65051E-07 | -1.648365398 | 0.294 | 0.541 | 0.013726212 | 0.185168351 | -0.446964141 | 0.4 | 0.691 | 1 | 0.185168351 | 1.1301E-06 |
| Kansl1 | 5.69876E-07 | -0.832016976 | 0.958 | 0.976 | 0.013843428 | 0.760502484 | -0.147305843 | 1 | 0.994 | 1 | 0.760502484 | 1.13975E-06 |
| Tmem119 | 5.7355E-07 | -5.50779464 | 0 | 0.188 | 0.013932685 | 0.846549635 | 0.769994395 | 0.1 | 0.08 | 1 | 0.846549635 | 1.1471E-06 |
| Adcy6 | 5.74006E-07 | -2.222392421 | 0.092 | 0.341 | 0.013943763 | 0.398557787 | -0.633129783 | 0.3 | 0.395 | 1 | 0.398557787 | 1.14801E-06 |
| Enpp1 | 5.85061E-07 | 0.409603613 | 0.992 | 0.8 | 0.014212299 | 0.022430346 | 0.889254244 | 0.9 | 0.775 | 1 | 0.022430346 | 1.17012E-06 |
| Xylt1 | 5.86852E-07 | 0.422758307 | 0.992 | 0.906 | 0.014255801 | 0.024359908 | 0.507581106 | 1 | 0.963 | 1 | 0.024359908 | 1.1737E-06 |
| Hbegf | 5.93623E-07 | 2.521816845 | 0.571 | 0.365 | 0.014420289 | 0.016446089 | 1.381007328 | 0.7 | 0.361 | 1 | 0.016446089 | 1.18725E-06 |
| Alcam | 6.07612E-07 | -1.786596362 | 0.134 | 0.406 | 0.014760099 | 0.062057769 | -1.870821341 | 0.4 | 0.608 | 1 | 0.062057769 | 1.21522E-06 |
| Ylpm1 | 6.123E-07 | -1.246488097 | 0.513 | 0.718 | 0.014873982 | 0.051400037 | -0.843880798 | 0.4 | 0.796 | 1 | 0.051400037 | 1.2246E-06 |
| Acvr1 | 6.24074E-07 | 0.456885823 | 0.916 | 0.871 | 0.015160002 | 0.098158247 | -0.984893108 | 0.8 | 0.892 | 1 | 0.098158247 | 1.24815E-06 |
| Cavin1 | 6.34194E-07 | -0.86941589 | 0.739 | 0.824 | 0.015405831 | 0.006997936 | -1.347817281 | 0.8 | 0.901 | 1 | 0.006997936 | 1.26839E-06 |
| Gnb4 | 6.38011E-07 | -0.912707922 | 0.681 | 0.859 | 0.015498556 | 0.005477084 | -1.544320516 | 0.5 | 0.861 | 1 | 0.005477084 | 1.27602E-06 |
| Pabpn1 | 6.41185E-07 | -0.678071905 | 0.899 | 0.924 | 0.015575658 | 0.091500877 | -0.638860456 | 0.6 | 0.91 | 1 | 0.091500877 | 1.28237E-06 |
| Akap8 | 6.55023E-07 | -1.725741157 | 0.277 | 0.524 | 0.015911813 | 0.064131884 | -0.9669712 | 0.3 | 0.611 | 1 | 0.064131884 | 1.31005E-06 |
| Rcor1 | 6.57231E-07 | -1.478669461 | 0.513 | 0.718 | 0.015965464 | 0.142065028 | -0.441509711 | 0.6 | 0.855 | 1 | 0.142065028 | 1.31446E-06 |
| Smarcc2 | 6.59356E-07 | -1.065620084 | 0.521 | 0.735 | 0.016017079 | 0.05867892 | -0.787449928 | 0.6 | 0.858 | 1 | 0.05867892 | 1.31871E-06 |
| Khsrp | 6.62447E-07 | -1.442866365 | 0.277 | 0.547 | 0.016092159 | 0.096787157 | -0.895822359 | 0.5 | 0.685 | 1 | 0.096787157 | 1.32489E-06 |
| Trim47 | 6.65154E-07 | 0.938674755 | 0.824 | 0.594 | 0.016157919 | 0.881091355 | 0.11236776 | 0.8 | 0.759 | 1 | 0.881091355 | 1.33031E-06 |
| Sin3a | 6.7105E-07 | -2.021479727 | 0.16 | 0.412 | 0.016301142 | 0.290039837 | -0.230005605 | 0.3 | 0.481 | 1 | 0.290039837 | 1.3421E-06 |
| Siah1a | 6.73854E-07 | -1.564378169 | 0.303 | 0.541 | 0.016369271 | 0.074573951 | -0.939594761 | 0.3 | 0.623 | 1 | 0.074573951 | 1.34771E-06 |
| Rbm10 | 6.74415E-07 | -1.701155518 | 0.193 | 0.465 | 0.016382893 | 0.03695642 | -0.840059087 | 0.2 | 0.59 | 1 | 0.03695642 | 1.34883E-06 |
| Zfp275 | 6.8493E-07 | -1.885357434 | 0.143 | 0.4 | 0.016638321 | 0.073312902 | -1.567040593 | 0.1 | 0.37 | 1 | 0.073312902 | 1.36986E-06 |
| Insr | 6.90071E-07 | -1.49683159 | 0.395 | 0.612 | 0.0167632 | 0.005315349 | -2.056754778 | 0.3 | 0.71 | 1 | 0.005315349 | 1.38014E-06 |
| Lgr6 | 7.26469E-07 | 0.691161905 | 0.58 | 0.276 | 0.017647377 | 0.64378964 | -0.580337415 | 0.5 | 0.438 | 1 | 0.64378964 | 1.45294E-06 |
| S1pr1 | 7.29091E-07 | 1.568012432 | 0.454 | 0.194 | 0.017711089 | 0.930898309 | 0.569461407 | 0.2 | 0.194 | 1 | 0.930898309 | 1.45818E-06 |
| Tfap2b | 7.3296E-07 | -4.807354922 | 0.017 | 0.218 | 0.017805068 | 0.465558081 | -1.186649236 | 0.2 | 0.275 | 1 | 0.465558081 | 1.46592E-06 |
| B2m | 7.3768E-07 | -0.84669307 | 0.891 | 0.953 | 0.01791972 | 0.003957316 | -1.313684101 | 0.8 | 0.988 | 1 | 0.003957316 | 1.47536E-06 |
| Dnttip2 | 7.41872E-07 | -1.115285866 | 0.588 | 0.747 | 0.018021561 | 0.027158558 | -1.126736335 | 0.3 | 0.741 | 1 | 0.027158558 | 1.48374E-06 |
| Purg | 7.43729E-07 | -2.236747714 | 0.118 | 0.365 | 0.018066667 | 0.46970601 | -0.391469028 | 0.3 | 0.404 | 1 | 0.46970601 | 1.48746E-06 |
| Pafah1b2 | 7.52945E-07 | -0.994791709 | 0.605 | 0.729 | 0.018290534 | 0.009231718 | -1.088700512 | 0.4 | 0.846 | 1 | 0.009231718 | 1.50589E-06 |
| Kmt2a | 7.84662E-07 | -0.834283249 | 0.849 | 0.9 | 0.019061013 | 0.008364097 | -1.035350363 | 0.8 | 0.981 | 1 | 0.008364097 | 1.56932E-06 |
| Runx3 | 7.85854E-07 | -2.944858446 | 0.059 | 0.288 | 0.019089956 | 0.507712939 | -1.201246612 | 0.3 | 0.352 | 1 | 0.507712939 | 1.57171E-06 |
| Plagl1 | 7.91573E-07 | -0.924608701 | 0.79 | 0.888 | 0.019228887 | 0.00524532 | -1.512159241 | 0.6 | 0.92 | 1 | 0.00524532 | 1.58315E-06 |
| Erh | 7.9635E-07 | -0.779418046 | 0.63 | 0.824 | 0.019344945 | 0.032586685 | -0.859277003 | 0.4 | 0.772 | 1 | 0.032586685 | 1.5927E-06 |
| Eno3 | 8.03783E-07 | -3.411426246 | 0.025 | 0.235 | 0.019525502 | 0.248571837 | 1.432959407 | 0.2 | 0.09 | 1 | 0.248571837 | 1.60757E-06 |
| Gpbp1 | 8.05424E-07 | -0.713997823 | 0.849 | 0.918 | 0.019565356 | 0.002707115 | -1.21196505 | 0.8 | 0.957 | 1 | 0.002707115 | 1.61085E-06 |
| Tut7 | 8.05465E-07 | -0.764232557 | 0.908 | 0.924 | 0.019566355 | 0.085555814 | -0.577367731 | 0.9 | 0.969 | 1 | 0.085555814 | 1.61093E-06 |
| Crybg1 | 8.09804E-07 | 1.514573173 | 0.328 | 0.094 | 0.019671761 | 0.094507666 | 1.872871575 | 0.4 | 0.213 | 1 | 0.094507666 | 1.61961E-06 |
| Nsd2 | 8.11283E-07 | -1.508363638 | 0.395 | 0.588 | 0.019707692 | 0.029832618 | -1.28585884 | 0.3 | 0.654 | 1 | 0.029832618 | 1.62257E-06 |
| Elavl1 | 8.28071E-07 | -0.773003417 | 0.866 | 0.871 | 0.020115502 | 0.017680249 | -0.879779516 | 0.8 | 0.96 | 1 | 0.017680249 | 1.65614E-06 |
| Crk | 8.3191E-07 | -1.003079393 | 0.597 | 0.782 | 0.020208748 | 0.25539244 | -0.342977418 | 0.9 | 0.889 | 1 | 0.25539244 | 1.66382E-06 |
| Safb2 | 8.49653E-07 | -1.214530304 | 0.529 | 0.712 | 0.020639775 | 0.303147966 | -0.387219555 | 0.7 | 0.824 | 1 | 0.303147966 | 1.69931E-06 |
| Tcp1 | 8.49658E-07 | -0.652805147 | 0.891 | 0.935 | 0.020639887 | 0.021631785 | -0.853617966 | 0.8 | 0.929 | 1 | 0.021631785 | 1.69931E-06 |
| Faf1 | 8.50416E-07 | -0.984813596 | 0.731 | 0.835 | 0.020658308 | 0.118658801 | -0.56233666 | 0.8 | 0.923 | 1 | 0.118658801 | 1.70083E-06 |
| Brms1l | 8.5491E-07 | -1.650486073 | 0.227 | 0.476 | 0.02076748 | 0.734440421 | 0.199162223 | 0.3 | 0.438 | 1 | 0.734440421 | 1.70982E-06 |
| Bmpr2 | 8.60318E-07 | -0.915854635 | 0.824 | 0.918 | 0.020898848 | 0.329344875 | -0.437692045 | 0.9 | 0.991 | 1 | 0.329344875 | 1.72064E-06 |
| Rabgap1 | 8.72337E-07 | -0.986557737 | 0.555 | 0.765 | 0.021190816 | 0.377820734 | -0.207934369 | 0.7 | 0.843 | 1 | 0.377820734 | 1.74467E-06 |
| Rflnb | 8.92636E-07 | -1.231381205 | 0.538 | 0.735 | 0.021683923 | 0.034747523 | -1.654503434 | 0.4 | 0.657 | 1 | 0.034747523 | 1.78527E-06 |
| Dyrk2 | 8.93056E-07 | -1.694880193 | 0.134 | 0.4 | 0.021694107 | 0.237833447 | -0.993305347 | 0.3 | 0.438 | 1 | 0.237833447 | 1.78611E-06 |
| Podnl1 | 9.03763E-07 | -5.392317423 | 0 | 0.182 | 0.021954209 | 0.364073509 | -0.111361109 | 0 | 0.077 | 1 | 0.364073509 | 1.80753E-06 |
| Ppm1l | 9.06752E-07 | -2.752213368 | 0.025 | 0.235 | 0.022026809 | 0.05140472 | -0.979257573 | 0.1 | 0.463 | 1 | 0.05140472 | 1.8135E-06 |
| Tnn | 0.232015918 | -1.250961574 | 0.042 | 0.076 | 1 | 9.18509E-07 | 3.210566986 | 0.4 | 0.043 | 0.022312427 | 0.232015918 | 1.83702E-06 |
| Urb1 | 9.20693E-07 | -3.070389328 | 0.034 | 0.247 | 0.02236547 | 0.619856434 | -0.339630097 | 0.1 | 0.151 | 1 | 0.619856434 | 1.84138E-06 |
| Mtss1 | 9.23135E-07 | -4.496654083 | 0.025 | 0.229 | 0.022424802 | 0.428080086 | -1.55950692 | 0.1 | 0.191 | 1 | 0.428080086 | 1.84627E-06 |
| Map4 | 9.23209E-07 | -1.016734016 | 0.882 | 0.935 | 0.0224266 | 0.128882794 | -0.656746779 | 0.9 | 0.966 | 1 | 0.128882794 | 1.84642E-06 |
| Azgp1 | 9.26861E-07 | 5.321928095 | 0.134 | 0 | 0.02251531 | 0.048521704 | 3.432959407 | 0.1 | 0.015 | 1 | 0.048521704 | 1.85372E-06 |
| Epc2 | 9.30225E-07 | -0.967295835 | 0.63 | 0.812 | 0.022597025 | 0.013144821 | -1.126272818 | 0.7 | 0.886 | 1 | 0.013144821 | 1.86045E-06 |
| Tjp1 | 9.30808E-07 | -0.978101738 | 0.571 | 0.782 | 0.0226112 | 0.033789972 | -1.138063902 | 0.4 | 0.836 | 1 | 0.033789972 | 1.86162E-06 |
| Gem | 9.34043E-07 | 1.324089436 | 0.782 | 0.571 | 0.022689761 | 0.8403003 | -0.536666944 | 0.9 | 0.765 | 1 | 0.8403003 | 1.86808E-06 |
| Inhba | 9.44391E-07 | 1.402862785 | 0.345 | 0.112 | 0.022941146 | 0.160106345 | 1.31748219 | 0.1 | 0.025 | 1 | 0.160106345 | 1.88878E-06 |
| Tmed2 | 9.55209E-07 | -0.465703307 | 0.966 | 0.971 | 0.023203934 | 0.227071629 | -0.302823165 | 1 | 0.994 | 1 | 0.227071629 | 1.91042E-06 |
| Tcam1 | 0.167062889 | 2.099535674 | 0.025 | 0.006 | 1 | 9.55379E-07 | 5.017921908 | 0.2 | 0.009 | 0.023208062 | 0.167062889 | 1.91076E-06 |
| G530011O06Rik | 0.281966064 | 0.893084796 | 0.084 | 0.053 | 1 | 9.55379E-07 | 5.339850003 | 0.2 | 0.009 | 0.023208062 | 0.281966064 | 1.91076E-06 |
| Amot | 9.56911E-07 | -3.201633861 | 0.042 | 0.259 | 0.02324529 | 0.305620462 | -1.101019165 | 0.1 | 0.231 | 1 | 0.305620462 | 1.91382E-06 |
| Slc43a1 | 9.57298E-07 | -3.635173947 | 0.017 | 0.218 | 0.023254695 | 0.387373255 | -0.900941329 | 0.1 | 0.204 | 1 | 0.387373255 | 1.9146E-06 |
| Nynrin | 9.79017E-07 | -2.485426827 | 0.059 | 0.294 | 0.023782289 | 0.191347359 | -0.763437806 | 0.1 | 0.312 | 1 | 0.191347359 | 1.95803E-06 |
| Gpatch8 | 9.81651E-07 | -0.687919691 | 0.857 | 0.906 | 0.023846268 | 0.47801826 | 0.279926185 | 1 | 0.963 | 1 | 0.47801826 | 1.9633E-06 |
| Akirin1 | 9.89472E-07 | -1.205318908 | 0.471 | 0.676 | 0.02403626 | 0.061547763 | -0.94323835 | 0.7 | 0.781 | 1 | 0.061547763 | 1.97894E-06 |
| Fer1l5 | 0.203705868 | -0.707819249 | 0.034 | 0.012 | 1 | 9.94843E-07 | 5.017921908 | 0.2 | 0.009 | 0.024166729 | 0.203705868 | 1.98969E-06 |
| Luc7l | 1.01015E-06 | -0.803915838 | 0.748 | 0.871 | 0.024538579 | 0.375672281 | -0.432855245 | 0.8 | 0.861 | 1 | 0.375672281 | 2.0203E-06 |
| Atxn1 | 1.01142E-06 | 0.555382825 | 1 | 0.894 | 0.024569313 | 0.000192072 | 1.233696048 | 1 | 0.944 | 1 | 0.000192072 | 2.02283E-06 |
| Lncppara | 1.01657E-06 | -2.235448574 | 0.193 | 0.441 | 0.0246946 | 0.601342577 | -0.723545078 | 0.6 | 0.593 | 1 | 0.601342577 | 2.03315E-06 |
| P4ha1 | 1.01711E-06 | 0.479740601 | 1 | 0.953 | 0.024707689 | 0.405881468 | 0.226965151 | 1 | 0.991 | 1 | 0.405881468 | 2.03422E-06 |
| Arhgef2 | 1.02845E-06 | -2.017147307 | 0.176 | 0.424 | 0.024983182 | 0.812757738 | -0.304006187 | 0.4 | 0.423 | 1 | 0.812757738 | 2.05691E-06 |
| Wdr33 | 1.02906E-06 | -0.765534746 | 0.882 | 0.929 | 0.024997862 | 0.035555254 | -0.73050231 | 0.8 | 0.969 | 1 | 0.035555254 | 2.05811E-06 |
| Usp38 | 1.03273E-06 | -1.635173947 | 0.168 | 0.435 | 0.025087025 | 0.576584367 | -0.244172937 | 0.4 | 0.549 | 1 | 0.576584367 | 2.06545E-06 |
| Unc5d | 0.091361251 | 2.514573173 | 0.017 | 0 | 1 | 1.03559E-06 | 4.754887502 | 0.2 | 0.009 | 0.025156625 | 0.091361251 | 2.07118E-06 |
| Tcea1 | 1.05142E-06 | -0.689173496 | 0.941 | 0.929 | 0.025540996 | 0.137081765 | -0.510649411 | 0.8 | 0.935 | 1 | 0.137081765 | 2.10283E-06 |
| Simc1 | 1.05519E-06 | -1.906088876 | 0.218 | 0.471 | 0.025632704 | 0.169899557 | -1.420869945 | 0.3 | 0.457 | 1 | 0.169899557 | 2.11038E-06 |
| Sf3b2 | 1.05528E-06 | -0.677776241 | 0.866 | 0.953 | 0.025634848 | 0.220782863 | -0.374395515 | 0.8 | 0.957 | 1 | 0.220782863 | 2.11056E-06 |
| Gdf10 | 1.06494E-06 | -2.093578955 | 0.286 | 0.518 | 0.025869443 | 0.445143935 | -0.714528205 | 0.2 | 0.34 | 1 | 0.445143935 | 2.12987E-06 |
| Eea1 | 1.06925E-06 | -0.949312587 | 0.84 | 0.924 | 0.025974273 | 0.019638649 | -1.059289436 | 0.8 | 0.938 | 1 | 0.019638649 | 2.1385E-06 |
| Zfp277 | 1.07684E-06 | -2.040015679 | 0.176 | 0.424 | 0.026158541 | 0.177001904 | -1.111361109 | 0.3 | 0.463 | 1 | 0.177001904 | 2.15367E-06 |
| 2700099C18Rik | 0.037003642 | -1.970853654 | 0.017 | 0.071 | 1 | 1.07795E-06 | 4.210566986 | 0.2 | 0.009 | 0.026185487 | 0.037003642 | 2.15589E-06 |

| | | | | | | | | | | | | |
|---|---|---|---|---|---|---|---|---|---|---|---|---|
| Gm31812 | 0.595471914 | 1.166649869 | 0.034 | 0.024 | 1 | 1.07795E-06 | 4.503348735 | 0.2 | 0.009 | 0.026185487 | 0.595471914 | 2.15589E-06 |
| Cenph | 0.657006316 | -0.733354341 | 0.042 | 0.053 | 1 | 1.07795E-06 | 4.532495081 | 0.2 | 0.009 | 0.026185487 | 0.657006316 | 2.15589E-06 |
| Tgm5 | 0.034295975 | 2.321928095 | 0.042 | 0.006 | 1 | 1.07814E-06 | 4.602884409 | 0.2 | 0.009 | 0.026190198 | 0.034295975 | 2.15628E-06 |
| Rassf6 | 0.047669258 | 2.251538767 | 0.05 | 0.012 | 1 | 1.07814E-06 | 4.602884409 | 0.2 | 0.009 | 0.026190198 | 0.047669258 | 2.15628E-06 |
| Rad51 | 0.059329643 | -2.250961574 | 0.025 | 0.076 | 1 | 1.07814E-06 | 4.602884409 | 0.2 | 0.009 | 0.026190198 | 0.059329643 | 2.15628E-06 |
| Gm30371 | 0.390945734 | -0.887525271 | 0.042 | 0.024 | 1 | 1.07814E-06 | 4.432959407 | 0.2 | 0.009 | 0.026190198 | 0.390945734 | 2.15628E-06 |
| Gm43672 | 0.935790323 | -0.292781749 | 0.025 | 0.024 | 1 | 1.07814E-06 | 4.602884409 | 0.2 | 0.009 | 0.026190198 | 0.935790323 | 2.15628E-06 |
| Camsap1 | 1.0937E-06 | -1.4440069 | 0.269 | 0.535 | 0.026568226 | 0.014251744 | -1.436034581 | 0.3 | 0.71 | 1 | 0.014251744 | 2.1874E-06 |
| Shisa5 | 1.10129E-06 | 0.662671812 | 0.899 | 0.788 | 0.026752454 | 0.28425013 | 0.464668267 | 0.8 | 0.725 | 1 | 0.28425013 | 2.20257E-06 |
| Phyhd1 | 1.11465E-06 | -2.87774425 | 0.042 | 0.259 | 0.027077046 | 0.735121428 | -0.242605642 | 0.2 | 0.228 | 1 | 0.735121428 | 2.2293E-06 |
| Tacc2 | 1.12812E-06 | 0.603173836 | 0.966 | 0.882 | 0.027404257 | 0.036152706 | 0.512210582 | 1 | 0.954 | 1 | 0.036152706 | 2.25624E-06 |
| Lsm7 | 1.13353E-06 | -1.313245852 | 0.387 | 0.618 | 0.02753569 | 0.10267949 | -0.963930745 | 0.3 | 0.552 | 1 | 0.10267949 | 2.26706E-06 |
| Diaph3 | 0.040367597 | -2.918386234 | 0.025 | 0.082 | 1 | 1.13453E-06 | 4.532495081 | 0.4 | 0.046 | 0.027560109 | 0.040367597 | 2.26907E-06 |
| Foxf1 | 1.14052E-06 | 1.048686532 | 0.664 | 0.376 | 0.027705525 | 0.820199087 | -0.126007885 | 0.7 | 0.528 | 1 | 0.820199087 | 2.28104E-06 |
| Dapk2 | 1.14129E-06 | -5.070389328 | 0.017 | 0.212 | 0.027724213 | 0.31763018 | -1.574535129 | 0.1 | 0.225 | 1 | 0.31763018 | 2.28258E-06 |
| Xrn2 | 1.14561E-06 | -0.81780928 | 0.782 | 0.871 | 0.027829266 | 0.181466334 | -0.510857757 | 0.8 | 0.901 | 1 | 0.181466334 | 2.29123E-06 |
| Zfp503 | 1.16615E-06 | -2.292781749 | 0.118 | 0.365 | 0.028328208 | 0.018672997 | -2.970762779 | 0 | 0.373 | 1 | 0.018672997 | 2.33231E-06 |
| Serpinb6c | 0.038142909 | 2.514573173 | 0.025 | 0 | 1 | 1.1674E-06 | 4.017921908 | 0.2 | 0.009 | 0.028358426 | 0.038142909 | 2.33479E-06 |
| Mki67 | 0.063453471 | -2.924538461 | 0.008 | 0.047 | 1 | 1.1674E-06 | 3.14345279 | 0.2 | 0.009 | 0.028358426 | 0.063453471 | 2.33479E-06 |
| Gm31597 | 0.168156185 | 1.514573173 | 0.025 | 0.006 | 1 | 1.1674E-06 | 4.280956314 | 0.2 | 0.009 | 0.028358426 | 0.168156185 | 2.33479E-06 |
| Cdca7l | 0.33261065 | -1.070389328 | 0.008 | 0.024 | 1 | 1.1674E-06 | 3.432959407 | 0.2 | 0.009 | 0.028358426 | 0.33261065 | 2.33479E-06 |
| Gm56689 | 0.695893386 | -0.222392421 | 0.017 | 0.024 | 1 | 1.1674E-06 | 4.280956314 | 0.2 | 0.009 | 0.028358426 | 0.695893386 | 2.33479E-06 |
| Gm11542 | 0.954551343 | -0.707819249 | 0.017 | 0.018 | 1 | 1.1674E-06 | 4.280956314 | 0.2 | 0.009 | 0.028358426 | 0.954551343 | 2.33479E-06 |
| Zfp644 | 1.16949E-06 | -0.970853654 | 0.63 | 0.794 | 0.028409167 | 0.027051977 | -1.071055574 | 0.8 | 0.861 | 1 | 0.027051977 | 2.33897E-06 |
| Cul4b | 1.19947E-06 | -1.669851398 | 0.227 | 0.488 | 0.029137425 | 0.921912695 | 0.18222236 | 0.4 | 0.441 | 1 | 0.921912695 | 2.39893E-06 |
| Deptor | 1.19979E-06 | 1.325716355 | 0.613 | 0.324 | 0.029145257 | 0.013421592 | 1.635452271 | 0.7 | 0.361 | 1 | 0.013421592 | 2.39958E-06 |
| Lrrc8d | 1.20302E-06 | 0.425197365 | 0.849 | 0.547 | 0.029223757 | 0.088713957 | 0.389606569 | 1 | 0.849 | 1 | 0.088713957 | 2.40604E-06 |
| Kdelr2 | 1.20602E-06 | -0.605721061 | 0.983 | 0.976 | 0.029296643 | 0.080320788 | -0.535626038 | 0.8 | 0.978 | 1 | 0.080320788 | 2.41204E-06 |
| Arl14ep | 1.20957E-06 | -1.900464326 | 0.168 | 0.424 | 0.029382802 | 0.980826318 | 0.13799836 | 0.4 | 0.438 | 1 | 0.980826318 | 2.41913E-06 |
| Adam2 | 0.014382543 | 3.389042291 | 0.076 | 0.018 | 1 | 1.21541E-06 | 4.017921908 | 0.2 | 0.009 | 0.029524755 | 0.014382543 | 2.43082E-06 |
| Yeats2 | 1.2271E-06 | -1.733354341 | 0.21 | 0.476 | 0.02980862 | 0.529445243 | 0.290001453 | 0.6 | 0.457 | 1 | 0.529445243 | 2.45419E-06 |
| Pkn2 | 1.23913E-06 | -0.914661401 | 0.899 | 0.918 | 0.03010102 | 0.05695119 | -0.577267335 | 1 | 0.985 | 1 | 0.05695119 | 2.47826E-06 |
| Eif5 | 1.24781E-06 | -0.600322899 | 0.975 | 0.982 | 0.03031168 | 0.012967865 | -0.794903992 | 1 | 1 | 1 | 0.012967865 | 2.49561E-06 |
| Papss1 | 1.25209E-06 | -1.055742552 | 0.513 | 0.676 | 0.030415878 | 0.013374333 | -1.374395515 | 0.6 | 0.793 | 1 | 0.013374333 | 2.50419E-06 |
| Anapc5 | 1.25686E-06 | -0.721140627 | 0.84 | 0.888 | 0.030531758 | 0.232885185 | -0.436818015 | 0.7 | 0.898 | 1 | 0.232885185 | 2.51373E-06 |
| Hs3st3a1 | 1.25871E-06 | 0.976303908 | 0.79 | 0.494 | 0.030576473 | 0.026420914 | 1.582535763 | 0.7 | 0.426 | 1 | 0.026420914 | 2.51741E-06 |
| Mrps10 | 1.26198E-06 | -1.718087584 | 0.143 | 0.4 | 0.030656008 | 0.513502987 | 0.710493383 | 0.3 | 0.216 | 1 | 0.513502987 | 2.52396E-06 |
| Braf | 1.26225E-06 | -0.957701513 | 0.84 | 0.894 | 0.030662679 | 0.058922511 | -0.481367777 | 0.9 | 0.972 | 1 | 0.058922511 | 2.52451E-06 |
| Bcan | 2.10903E-06 | 4.906890596 | 0.126 | 0 | 0.051232463 | 1.26464E-06 | 4.017921908 | 0.2 | 0.009 | 0.030720743 | 2.10903E-06 | 2.52929E-06 |
| Hpse | 0.82670638 | -0.485426827 | 0.025 | 0.029 | 1 | 1.26464E-06 | 4.017921908 | 0.2 | 0.009 | 0.030720743 | 0.82670638 | 2.52929E-06 |
| Ttyh3 | 1.27443E-06 | -1.783107376 | 0.16 | 0.418 | 0.030958394 | 0.127407645 | -1.528972552 | 0.1 | 0.318 | 1 | 0.127407645 | 2.54885E-06 |
| Nup85 | 1.27685E-06 | -2.140778656 | 0.059 | 0.288 | 0.031017187 | 0.546066198 | 1.005097868 | 0.3 | 0.247 | 1 | 0.546066198 | 2.55369E-06 |
| Prdm10 | 1.2978E-06 | -2.752213368 | 0.05 | 0.271 | 0.03152604 | 0.093266706 | -2.399930607 | 0 | 0.225 | 1 | 0.093266706 | 2.59559E-06 |
| Ythdf2 | 1.30031E-06 | -1.033978688 | 0.605 | 0.765 | 0.031587046 | 0.03239738 | -1.064818523 | 0.5 | 0.784 | 1 | 0.03239738 | 2.60061E-06 |
| 4930447K03Rik | 0.204733108 | 0.292180751 | 0.034 | 0.012 | 1 | 1.31614E-06 | 3.017921908 | 0.2 | 0.009 | 0.031971769 | 0.204733108 | 2.63229E-06 |
| Thbs1 | 1.33051E-06 | -1.340330862 | 0.924 | 0.959 | 0.032320741 | 0.101400405 | -1.36710732 | 1 | 0.966 | 1 | 0.101400405 | 2.66102E-06 |
| N4bp1 | 1.33837E-06 | -1.641931313 | 0.311 | 0.541 | 0.032511652 | 0.004598164 | -1.739190259 | 0.3 | 0.747 | 1 | 0.004598164 | 2.67674E-06 |
| Frmd8 | 1.35208E-06 | 1.033947332 | 0.664 | 0.365 | 0.032844796 | 0.253192817 | 0.645688731 | 0.8 | 0.602 | 1 | 0.253192817 | 2.70416E-06 |
| Metap2 | 1.35653E-06 | -0.785875195 | 0.773 | 0.924 | 0.032952824 | 0.716879968 | -0.171902651 | 0.9 | 0.898 | 1 | 0.716879968 | 2.71306E-06 |
| Mib1 | 1.36492E-06 | -1.058084818 | 0.639 | 0.812 | 0.033156747 | 0.006365451 | -1.202322047 | 0.8 | 0.926 | 1 | 0.006365451 | 2.72985E-06 |
| Utp18 | 1.37029E-06 | -1.42161173 | 0.345 | 0.576 | 0.033287056 | 0.357041201 | -0.636714121 | 0.5 | 0.599 | 1 | 0.357041201 | 2.74058E-06 |
| Slc39a11 | 1.37863E-06 | 0.373710637 | 0.63 | 0.341 | 0.033489644 | 0.003760374 | 1.494359952 | 0.7 | 0.284 | 1 | 0.003760374 | 2.75726E-06 |
| Sema3c | 1.38259E-06 | 0.877143252 | 0.647 | 0.353 | 0.033585838 | 0.135215521 | -1.048167282 | 0.3 | 0.528 | 1 | 0.135215521 | 2.76517E-06 |
| Ubtf | 1.39087E-06 | -1.03191518 | 0.588 | 0.735 | 0.033787018 | 0.000467049 | -1.823380346 | 0.5 | 0.904 | 1 | 0.000467049 | 2.78174E-06 |
| Lars2 | 1.39402E-06 | -1.506737333 | 1 | 1 | 0.033863517 | 0.016356729 | -0.35226973 | 1 | 1 | 1 | 0.016356729 | 2.78804E-06 |
| Ahcyl1 | 1.40826E-06 | -0.732066795 | 0.756 | 0.871 | 0.034209433 | 0.016318689 | -0.918369604 | 0.7 | 0.914 | 1 | 0.016318689 | 2.81652E-06 |
| Pcbp2 | 1.40876E-06 | -0.514052668 | 0.966 | 0.982 | 0.034221556 | 0.016789619 | -0.620556102 | 0.9 | 0.997 | 1 | 0.016789619 | 2.81751E-06 |
| Rftn1 | 1.41071E-06 | -5.240314329 | 0 | 0.176 | 0.034268957 | 0.245888135 | -1.425021588 | 0 | 0.12 | 1 | 0.245888135 | 2.82142E-06 |
| Slc1a5 | 1.41264E-06 | 0.977307414 | 0.899 | 0.718 | 0.034315931 | 0.043908267 | 1.670622023 | 0.9 | 0.753 | 1 | 0.043908267 | 2.82528E-06 |
| Lamc3 | 1.41553E-06 | -5.292781749 | 0 | 0.176 | 0.034386175 | 0.282477202 | -0.596787936 | 0 | 0.105 | 1 | 0.282477202 | 2.83107E-06 |
| Pappa2 | 1.41573E-06 | -5.551516018 | 0 | 0.176 | 0.034390974 | 0.136671162 | -2.756865152 | 0 | 0.185 | 1 | 0.136671162 | 2.83146E-06 |
| Dennd1a | 1.43095E-06 | -0.706620499 | 0.866 | 0.935 | 0.034760713 | 0.19267868 | 0.188435816 | 1 | 0.938 | 1 | 0.19267868 | 2.8619E-06 |
| Six4 | 1.4342E-06 | -4.343407822 | 0.008 | 0.194 | 0.034839673 | 0.38607321 | -0.723545078 | 0.1 | 0.204 | 1 | 0.38607321 | 2.86841E-06 |
| Fgd6 | 1.43459E-06 | -1.905758626 | 0.294 | 0.512 | 0.034848956 | 0.605667259 | -0.623713042 | 0.6 | 0.623 | 1 | 0.605667259 | 2.86917E-06 |
| Psma5 | 1.45152E-06 | -0.780610067 | 0.782 | 0.888 | 0.035260209 | 0.002938777 | -1.26411346 | 0.5 | 0.898 | 1 | 0.002938777 | 2.90303E-06 |
| Gnl3 | 1.45324E-06 | -0.81832604 | 0.798 | 0.9 | 0.035301988 | 0.006901051 | -1.420491929 | 0.4 | 0.84 | 1 | 0.006901051 | 2.90647E-06 |
| Dmxl1 | 1.46879E-06 | -0.936533581 | 0.613 | 0.794 | 0.03567984 | 0.24500625 | -0.446964141 | 0.7 | 0.864 | 1 | 0.24500625 | 2.93758E-06 |
| Trip12 | 1.47917E-06 | -0.776077727 | 0.941 | 0.971 | 0.035932048 | 0.042037815 | -0.672331641 | 0.9 | 0.991 | 1 | 0.042037815 | 2.95834E-06 |
| Lgals3bp | 1.48818E-06 | 1.430180986 | 0.454 | 0.194 | 0.036150926 | 0.852688 | 0.536795218 | 0.3 | 0.281 | 1 | 0.852688 | 2.97636E-06 |
| Gtf2h1 | 1.48954E-06 | -1.082667658 | 0.429 | 0.653 | 0.036183929 | 0.469913885 | -0.235058833 | 0.6 | 0.731 | 1 | 0.469913885 | 2.97908E-06 |
| Map3k8 | 1.4905E-06 | 1.253981943 | 0.605 | 0.347 | 0.036207181 | 0.135411188 | 1.119068631 | 0.5 | 0.296 | 1 | 0.135411188 | 2.98099E-06 |
| Taf13 | 1.51585E-06 | 0.753674837 | 0.84 | 0.606 | 0.036823138 | 0.612623728 | 0.393431043 | 0.6 | 0.691 | 1 | 0.612623728 | 3.03171E-06 |
| Camk2n1 | 1.52407E-06 | -1.720643289 | 0.269 | 0.512 | 0.037022698 | 0.247562154 | -0.569770385 | 0.6 | 0.698 | 1 | 0.247562154 | 3.04814E-06 |
| Map1lc3b | 1.52461E-06 | 0.581372627 | 0.966 | 0.971 | 0.037035727 | 0.124353777 | -0.535789659 | 1 | 0.994 | 1 | 0.124353777 | 3.04921E-06 |
| Crtac1 | 1.52514E-06 | -3.021479727 | 0.034 | 0.241 | 0.037048625 | 0.652325951 | 0.558490289 | 0.1 | 0.062 | 1 | 0.652325951 | 3.05027E-06 |
| Setdb1 | 1.53873E-06 | -1.707819249 | 0.218 | 0.471 | 0.037378859 | 0.062998929 | -1.556986928 | 0.2 | 0.472 | 1 | 0.062998929 | 3.07746E-06 |
| Nudt18 | 1.55837E-06 | 2.357847669 | 0.294 | 0.082 | 0.037856044 | 0.040013389 | 2.280956314 | 0.3 | 0.102 | 1 | 0.040013389 | 3.11675E-06 |
| Uba5 | 1.55844E-06 | -1.008988783 | 0.513 | 0.706 | 0.037857576 | 0.489783144 | -0.31230551 | 0.7 | 0.698 | 1 | 0.489783144 | 3.11687E-06 |
| Itgb1 | 1.56197E-06 | -0.654804628 | 0.992 | 0.994 | 0.037943303 | 0.01274463 | -0.734588831 | 1 | 0.991 | 1 | 0.01274463 | 3.12393E-06 |
| H1f0 | 1.57505E-06 | 1.29514621 | 0.782 | 0.635 | 0.038261108 | 0.516742723 | 0.659364891 | 0.6 | 0.66 | 1 | 0.516742723 | 3.1501E-06 |
| Il1rapl1 | 1.58638E-06 | 2.755581272 | 0.235 | 0.047 | 0.03853625 | 0.000902344 | 3.091922489 | 0.3 | 0.052 | 1 | 0.000902344 | 3.17275E-06 |
| Cpne1 | 1.59398E-06 | -1.058893689 | 0.504 | 0.676 | 0.038720891 | 0.005271804 | -1.430538593 | 0.7 | 0.846 | 1 | 0.005271804 | 3.18795E-06 |
| Aggf1 | 1.59948E-06 | -1.417611443 | 0.311 | 0.553 | 0.038854672 | 0.313334798 | -0.687134438 | 0.4 | 0.528 | 1 | 0.313334798 | 3.19897E-06 |
| Usp48 | 1.59964E-06 | -0.848897184 | 0.361 | 0.618 | 0.03885837 | 0.643354144 | -0.152003093 | 0.7 | 0.673 | 1 | 0.643354144 | 3.19927E-06 |
| Dync1h1 | 1.6228E-06 | -0.908117899 | 0.748 | 0.871 | 0.039421089 | 0.017186263 | -0.899385304 | 0.8 | 0.96 | 1 | 0.017186263 | 3.2456E-06 |
| Pa2g4 | 1.62554E-06 | -0.861869081 | 0.765 | 0.824 | 0.039487736 | 0.003936364 | -1.313990275 | 0.5 | 0.867 | 1 | 0.003936364 | 3.25109E-06 |
| Ccdc115 | 1.63759E-06 | -1.641931313 | 0.176 | 0.429 | 0.039780283 | 0.05448564 | -2.058893689 | 0 | 0.278 | 1 | 0.05448564 | 3.27517E-06 |
| Gm30489 | 1.64197E-06 | 2.929610672 | 0.176 | 0.018 | 0.039886678 | 0.001520803 | 3.017921908 | 0.2 | 0.025 | 1 | 0.001520803 | 3.28393E-06 |
| Col6a2 | 1.64407E-06 | 0.529988876 | 0.992 | 0.947 | 0.03993783 | 0.102893813 | 0.449484381 | 1 | 0.935 | 1 | 0.102893813 | 3.28814E-06 |
| Nup88 | 1.64504E-06 | -1.50589093 | 0.252 | 0.506 | 0.039961201 | 0.510379754 | -0.217294554 | 0.3 | 0.432 | 1 | 0.510379754 | 3.29007E-06 |
| Klhl9 | 1.65278E-06 | -1.075722772 | 0.479 | 0.694 | 0.040149351 | 0.015953377 | -1.147543445 | 0.5 | 0.784 | 1 | 0.015953377 | 3.30556E-06 |
| Ostc | 1.66268E-06 | -0.674106969 | 0.924 | 0.918 | 0.040389814 | 0.223915931 | -0.38999237 | 0.8 | 0.926 | 1 | 0.223915931 | 3.32536E-06 |
| Krr1 | 1.67516E-06 | -1.645298164 | 0.218 | 0.476 | 0.040692962 | 0.106720561 | -1.178475305 | 0.2 | 0.435 | 1 | 0.106720561 | 3.35032E-06 |
| Aida | 1.69214E-06 | -1.180888869 | 0.403 | 0.629 | 0.041105581 | 0.233494632 | -0.885529604 | 0.6 | 0.599 | 1 | 0.233494632 | 3.38429E-06 |
| Kif5b | 1.69557E-06 | -0.667219627 | 0.966 | 0.971 | 0.041188684 | 0.014224673 | -0.707728373 | 1 | 0.997 | 1 | 0.014224673 | 3.39113E-06 |
| Lgalsl | 1.71277E-06 | 1.553886097 | 0.706 | 0.488 | 0.041606492 | 0.537851763 | -0.423177888 | 0.4 | 0.519 | 1 | 0.537851763 | 3.42553E-06 |
| Crlf3 | 2.20062E-06 | 0.800417462 | 0.832 | 0.571 | 0.053457533 | 1.71582E-06 | 2.140932798 | 1 | 0.503 | 0.041680777 | 2.20062E-06 | 3.43164E-06 |
| Snx4 | 1.72278E-06 | -1.074876719 | 0.597 | 0.718 | 0.041849892 | 0.102768352 | -0.892970618 | 0.7 | 0.812 | 1 | 0.102768352 | 3.44557E-06 |

| | | | | | | | | | | | | |
|---|---|---|---|---|---|---|---|---|---|---|---|---|
| Zfp618 | 1.72422E-06 | -2.748461233 | 0.067 | 0.288 | 0.041884807 | 0.21059782 | -1.505640048 | 0.2 | 0.358 | 1 | 0.21059782 | 3.44844E-06 |
| Spats2l | 1.73084E-06 | -0.658990406 | 0.874 | 0.912 | 0.042045619 | 0.007593075 | -1.286932674 | 0.9 | 0.926 | 1 | 0.007593075 | 3.46168E-06 |
| Glud1 | 1.73116E-06 | -0.755154251 | 0.975 | 0.982 | 0.042053421 | 0.535610086 | -0.455811637 | 1 | 0.988 | 1 | 0.535610086 | 3.46232E-06 |
| Peli2 | 1.77298E-06 | -2.715724447 | 0.067 | 0.294 | 0.043069321 | 0.900154465 | -0.344648171 | 0.3 | 0.315 | 1 | 0.900154465 | 3.54596E-06 |
| Nr4a2 | 1.78451E-06 | -1.115477217 | 0.529 | 0.782 | 0.043349248 | 0.805718516 | -0.231586641 | 0.7 | 0.738 | 1 | 0.805718516 | 3.56901E-06 |
| C1galt1c1 | 1.82954E-06 | 0.763252286 | 0.689 | 0.376 | 0.044443229 | 0.241638823 | -0.825056924 | 0.4 | 0.506 | 1 | 0.241638823 | 3.65908E-06 |
| Golm2 | 1.85356E-06 | -0.765534746 | 0.79 | 0.882 | 0.045026749 | 0.223331086 | -0.401030641 | 0.8 | 0.858 | 1 | 0.223331086 | 3.70712E-06 |
| Nectin3 | 1.85549E-06 | -1.840521786 | 0.277 | 0.512 | 0.045073471 | 0.046374752 | -1.858595039 | 0.3 | 0.549 | 1 | 0.046374752 | 3.71097E-06 |
| Tmie | 1.86151E-06 | 0.840559484 | 0.462 | 0.176 | 0.045219778 | 0.596379093 | 1.558490289 | 0.1 | 0.062 | 1 | 0.596379093 | 3.72301E-06 |
| Pms2 | 1.866E-06 | -2.628384781 | 0.034 | 0.241 | 0.045328856 | 0.555646585 | 0.995554095 | 0.2 | 0.139 | 1 | 0.555646585 | 3.732E-06 |
| Hmgn1 | 1.86845E-06 | -0.590105804 | 0.958 | 0.959 | 0.045388295 | 0.651003918 | -0.10919001 | 0.6 | 0.954 | 1 | 0.651003918 | 3.73689E-06 |
| Psmc3 | 1.87669E-06 | -0.872135603 | 0.63 | 0.841 | 0.04558861 | 0.19374837 | -0.42170123 | 0.6 | 0.83 | 1 | 0.19374837 | 3.75338E-06 |
| Bmp5 | 1.88237E-06 | 1.134618133 | 0.613 | 0.353 | 0.045726505 | 0.199848524 | 1.464437639 | 0.8 | 0.701 | 1 | 0.199848524 | 3.76473E-06 |
| Cacng7 | 1.89837E-06 | -2.117695043 | 0.042 | 0.259 | 0.046115077 | 0.551863198 | -0.567040593 | 0.2 | 0.269 | 1 | 0.551863198 | 3.79673E-06 |
| Togaram1 | 1.91088E-06 | -1.226893814 | 0.454 | 0.706 | 0.046419139 | 0.027838119 | -1.111361109 | 0.5 | 0.781 | 1 | 0.027838119 | 3.82176E-06 |
| Fam98b | 1.92524E-06 | -1.475373163 | 0.244 | 0.5 | 0.046767813 | 0.233418039 | -0.649781024 | 0.3 | 0.512 | 1 | 0.233418039 | 3.85047E-06 |
| Rgmb | 1.94233E-06 | -2.070389328 | 0.118 | 0.359 | 0.047183159 | 0.007988188 | -3.131825212 | 0 | 0.438 | 1 | 0.007988188 | 3.88466E-06 |
| Ncoa3 | 1.94506E-06 | 0.967938791 | 0.916 | 0.771 | 0.047249349 | 0.387798221 | 0.349970043 | 1 | 0.92 | 1 | 0.387798221 | 3.89011E-06 |
| Zcchc14 | 1.95045E-06 | -1.11345805 | 0.714 | 0.818 | 0.04738027 | 0.027563892 | -0.958362023 | 1 | 0.935 | 1 | 0.027563892 | 3.90089E-06 |
| Col9a3 | 1.98027E-06 | 0.383519019 | 1 | 0.859 | 0.048104726 | 0.290011417 | 0.153135343 | 1 | 0.898 | 1 | 0.290011417 | 3.96054E-06 |
| Ccdc34 | 1.98433E-06 | -1.731300682 | 0.345 | 0.571 | 0.048203406 | 0.912936874 | 0.580516596 | 0.3 | 0.336 | 1 | 0.912936874 | 3.96866E-06 |
| Cdc26 | 1.98527E-06 | -0.881355504 | 0.613 | 0.765 | 0.048226183 | 0.389242509 | -0.399930607 | 0.8 | 0.716 | 1 | 0.389242509 | 3.97054E-06 |
| Wbp1l | 2.00686E-06 | -1.676852898 | 0.193 | 0.447 | 0.048750611 | 0.276560259 | -0.437405312 | 0.6 | 0.682 | 1 | 0.276560259 | 4.01371E-06 |
| Nod1 | 2.03878E-06 | 1.075288127 | 0.387 | 0.141 | 0.049526068 | 0.000129739 | 1.712113478 | 0.7 | 0.182 | 1 | 0.000129739 | 4.07756E-06 |
| Ep300 | 2.03901E-06 | -1.154311811 | 0.639 | 0.735 | 0.049531724 | 0.439212849 | -0.272755253 | 0.9 | 0.914 | 1 | 0.439212849 | 4.07802E-06 |
| Nxf1 | 2.04268E-06 | -1.130013311 | 0.454 | 0.676 | 0.049620875 | 0.123148548 | -0.672393593 | 0.5 | 0.753 | 1 | 0.123148548 | 4.08536E-06 |
| Gphn | 2.10908E-06 | -1.250080359 | 1 | 1 | 0.051233667 | 0.270855538 | -0.274921694 | 1 | 1 | 1 | 0.270855538 | 4.21815E-06 |
| Stpg2 | 2.11294E-06 | 5.269460675 | 0.126 | 0 | 0.051327435 | 1.1147E-05 | 4.210566986 | 0.2 | 0.012 | 0.270783232 | 1.1147E-05 | 4.22587E-06 |
| Gsk3b | 2.11744E-06 | -0.738883784 | 0.882 | 0.988 | 0.051436803 | 0.511158307 | -0.306369324 | 1 | 0.981 | 1 | 0.511158307 | 4.23487E-06 |
| Crispld2 | 2.11774E-06 | 1.021914776 | 0.782 | 0.535 | 0.051444171 | 0.008403245 | 2.529343944 | 0.6 | 0.241 | 1 | 0.008403245 | 4.23548E-06 |
| Cpeb1 | 2.11904E-06 | -2.712495736 | 0.059 | 0.276 | 0.051475634 | 0.094821588 | 1.263034406 | 0.5 | 0.284 | 1 | 0.094821588 | 4.23807E-06 |
| Cdkal1 | 2.12933E-06 | -1.272578179 | 0.538 | 0.718 | 0.051725617 | 0.452848529 | 0.270423385 | 0.8 | 0.784 | 1 | 0.452848529 | 4.25865E-06 |
| Ccny | 2.17388E-06 | -0.822305264 | 0.765 | 0.876 | 0.052807898 | 0.025471647 | -0.890170433 | 0.8 | 0.932 | 1 | 0.025471647 | 4.34776E-06 |
| Cct6a | 2.18275E-06 | -0.686474346 | 0.891 | 0.947 | 0.053023455 | 0.120544269 | -0.541569905 | 0.8 | 0.929 | 1 | 0.120544269 | 4.3655E-06 |
| Cnrip1 | 2.21266E-06 | -5.213347282 | 0 | 0.171 | 0.053749939 | 0.987515054 | 1.017921908 | 0.1 | 0.105 | 1 | 0.987515054 | 4.42532E-06 |
| Osbp | 2.21348E-06 | -1.167530486 | 0.462 | 0.653 | 0.053769812 | 0.011940377 | -1.186649236 | 0.5 | 0.812 | 1 | 0.011940377 | 4.42695E-06 |
| Prpf38b | 2.2181E-06 | -0.828728573 | 0.79 | 0.871 | 0.053882006 | 0.000473162 | -1.685289559 | 0.4 | 0.926 | 1 | 0.000473162 | 4.43619E-06 |
| Sp3 | 2.21952E-06 | -1.030311889 | 0.689 | 0.788 | 0.053916632 | 0.243234224 | -0.495147674 | 0.7 | 0.914 | 1 | 0.243234224 | 4.43904E-06 |
| Ly6e | 2.23672E-06 | -0.766940654 | 0.429 | 0.759 | 0.054334309 | 0.02148137 | -0.372064828 | 0.6 | 0.944 | 1 | 0.02148137 | 4.47343E-06 |
| Sema3e | 2.24411E-06 | 1.819918384 | 0.319 | 0.1 | 0.054514029 | 0.565897789 | 1.432959407 | 0.1 | 0.059 | 1 | 0.565897789 | 4.48822E-06 |
| Trap1 | 2.25175E-06 | -1.684735635 | 0.193 | 0.447 | 0.054699608 | 0.180347672 | -1.145308441 | 0.2 | 0.386 | 1 | 0.180347672 | 4.5035E-06 |
| Lrrfip1 | 2.26066E-06 | -1.500912075 | 0.571 | 0.747 | 0.054915951 | 0.080335527 | -1.132748536 | 0.8 | 0.852 | 1 | 0.080335527 | 4.52131E-06 |
| Ccdc148 | 0.347964691 | -0.900464326 | 0.017 | 0.035 | 1 | 2.27915E-06 | 2.210566986 | 0.3 | 0.025 | 0.055365003 | 0.347964691 | 4.55829E-06 |
| Slain2 | 2.31428E-06 | -1.03858059 | 0.546 | 0.718 | 0.056218525 | 0.006308487 | -1.389581919 | 0.4 | 0.843 | 1 | 0.006308487 | 4.62856E-06 |
| Lzts2 | 2.34433E-06 | -1.288734916 | 0.336 | 0.565 | 0.056948546 | 0.279859833 | -0.654503434 | 0.5 | 0.62 | 1 | 0.279859833 | 4.68866E-06 |
| Slc30a7 | 2.35738E-06 | -1.350497247 | 0.261 | 0.512 | 0.05726545 | 0.010132828 | -2.276698841 | 0.1 | 0.519 | 1 | 0.010132828 | 4.71475E-06 |
| Arpc5 | 2.37877E-06 | -0.665999073 | 0.84 | 0.947 | 0.057785143 | 0.038992353 | -0.712013484 | 0.7 | 0.969 | 1 | 0.038992353 | 4.75754E-06 |
| Actr2 | 2.38953E-06 | -0.794264923 | 0.782 | 0.871 | 0.058046394 | 0.008621268 | -1.015501094 | 0.9 | 0.929 | 1 | 0.008621268 | 4.77905E-06 |
| Adgrg6 | 2.39627E-06 | -1.097897643 | 0.479 | 0.724 | 0.058210299 | 0.129498263 | -1.055573331 | 0.6 | 0.759 | 1 | 0.129498263 | 4.79254E-06 |
| Nlgn2 | 2.40226E-06 | -1.937939032 | 0.126 | 0.371 | 0.058355587 | 0.463949719 | -0.647414009 | 0.3 | 0.377 | 1 | 0.463949719 | 4.8045E-06 |
| Ank | 2.43298E-06 | 0.644109672 | 1 | 0.947 | 0.059101916 | 0.754544952 | -0.16403418 | 1 | 0.972 | 1 | 0.754544952 | 4.86595E-06 |
| Srsf5 | 2.43811E-06 | -0.557812753 | 0.975 | 0.971 | 0.059226651 | 0.011433145 | -0.916449424 | 0.9 | 0.963 | 1 | 0.011433145 | 4.87622E-06 |
| Frk | 2.44694E-06 | -1.121015401 | 0.042 | 0.253 | 0.059441061 | 0.294459208 | -1.258202497 | 0.1 | 0.235 | 1 | 0.294459208 | 4.89387E-06 |
| Ifitm2 | 2.45516E-06 | -0.928234615 | 0.933 | 0.929 | 0.059640801 | 0.010605482 | -1.212819095 | 0.7 | 0.966 | 1 | 0.010605482 | 4.91032E-06 |
| Ppm1g | 2.4901E-06 | -1.367163946 | 0.37 | 0.6 | 0.060489528 | 0.008662938 | -1.576029376 | 0.3 | 0.71 | 1 | 0.008662938 | 4.9802E-06 |
| Wapl | 2.50161E-06 | -0.642968104 | 0.95 | 0.965 | 0.060769228 | 0.051391829 | -0.518130992 | 0.9 | 0.988 | 1 | 0.051391829 | 5.00322E-06 |
| Ctsd | 2.53767E-06 | 0.562702344 | 0.992 | 0.918 | 0.061644962 | 0.053167462 | 0.858166539 | 0.9 | 0.972 | 1 | 0.053167462 | 5.07532E-06 |
| Qser1 | 2.55856E-06 | -1.169925001 | 0.353 | 0.594 | 0.06215244 | 0.075256449 | -1.104474723 | 0.7 | 0.753 | 1 | 0.075256449 | 5.11711E-06 |
| Tagln | 2.56072E-06 | -5.297069108 | 0.025 | 0.218 | 0.06220491 | 0.30757544 | 0.785261151 | 0.2 | 0.099 | 1 | 0.30757544 | 5.12143E-06 |
| Nrp1 | 2.57376E-06 | -1.30256277 | 0.076 | 0.312 | 0.062521866 | 0.037245303 | -2.864721141 | 0 | 0.315 | 1 | 0.037245303 | 5.14752E-06 |
| Fxr1 | 2.62096E-06 | -0.539703888 | 0.916 | 0.959 | 0.063668402 | 0.064288432 | -0.628916085 | 1 | 0.978 | 1 | 0.064288432 | 5.24192E-06 |
| Hif1a | 2.62274E-06 | 0.621907217 | 0.992 | 0.971 | 0.063711582 | 0.197090286 | -0.509862573 | 0.9 | 0.988 | 1 | 0.197090286 | 5.24547E-06 |
| Emc3 | 2.62352E-06 | -0.880911529 | 0.639 | 0.782 | 0.063730472 | 0.151747217 | -0.693572999 | 0.7 | 0.756 | 1 | 0.151747217 | 5.24703E-06 |
| Agtpbp1 | 2.63889E-06 | -1.362415937 | 0.277 | 0.547 | 0.064103965 | 0.89546202 | -0.281638374 | 0.7 | 0.648 | 1 | 0.89546202 | 5.27778E-06 |
| Osbpl5 | 2.70465E-06 | -1.595051318 | 0.118 | 0.371 | 0.065701347 | 0.556504512 | -0.433289204 | 0.3 | 0.352 | 1 | 0.556504512 | 5.40929E-06 |
| Gm49969 | 2.93503E-05 | 2.070966521 | 0.235 | 0.065 | 0.712976402 | 2.7106E-06 | 2.389890685 | 0.7 | 0.145 | 0.065845983 | 2.93503E-05 | 5.4212E-06 |
| Brd8dc | 2.71375E-06 | -3.137503524 | 0.042 | 0.247 | 0.065922488 | 0.930978251 | -0.210896782 | 0.2 | 0.21 | 1 | 0.930978251 | 5.4275E-06 |
| Rab27a | 2.71847E-06 | 1.182951682 | 0.479 | 0.224 | 0.066037162 | 0.45555799 | 1.057092505 | 0.3 | 0.222 | 1 | 0.45555799 | 5.43694E-06 |
| Dennd5a | 2.72599E-06 | -0.955829564 | 0.597 | 0.765 | 0.066219749 | 0.01936761 | -1.040547397 | 0.7 | 0.938 | 1 | 0.01936761 | 5.45197E-06 |
| Api5 | 2.7423E-06 | -0.940621453 | 0.521 | 0.735 | 0.066615921 | 0.000718479 | -1.991066875 | 0.4 | 0.806 | 1 | 0.000718479 | 5.48459E-06 |
| Git2 | 2.74723E-06 | -1.773003417 | 0.218 | 0.465 | 0.066735706 | 0.167277348 | -1.022093771 | 0.3 | 0.509 | 1 | 0.167277348 | 5.49445E-06 |
| Srpk1 | 2.75764E-06 | -1.193660703 | 0.462 | 0.641 | 0.066988686 | 0.241167501 | -0.743629324 | 0.5 | 0.648 | 1 | 0.241167501 | 5.51528E-06 |
| Runx1t1 | 2.76997E-06 | -4.029747343 | 0.034 | 0.229 | 0.067288039 | 0.186318034 | -1.924592597 | 0 | 0.151 | 1 | 0.186318034 | 5.53993E-06 |
| Cd63 | 2.78022E-06 | -0.485426827 | 1 | 0.988 | 0.067537139 | 0.196795169 | -0.385157127 | 1 | 0.994 | 1 | 0.196795169 | 5.56044E-06 |
| Nr3c2 | 2.79606E-06 | 1.327203037 | 0.42 | 0.159 | 0.067921837 | 0.590335783 | 0.176619654 | 0.2 | 0.324 | 1 | 0.590335783 | 5.59211E-06 |
| Clec3a | 2.85924E-06 | 0.998694984 | 0.765 | 0.524 | 0.069456668 | 0.05941628 | 1.29389537 | 0.5 | 0.265 | 1 | 0.05941628 | 5.71847E-06 |
| 2410002F23Rik | 2.86084E-06 | -1.298056691 | 0.294 | 0.535 | 0.069495542 | 0.14453085 | -1.062451508 | 0.2 | 0.407 | 1 | 0.14453085 | 5.72167E-06 |
| Prcc | 2.86317E-06 | -1.497810552 | 0.193 | 0.447 | 0.069552083 | 0.060285277 | -1.161987182 | 0.2 | 0.522 | 1 | 0.060285277 | 5.72633E-06 |
| Shroom4 | 2.87423E-06 | 0.577308928 | 0.639 | 0.371 | 0.069820882 | 0.041981611 | 0.73803971 | 0.8 | 0.509 | 1 | 0.041981611 | 5.74846E-06 |
| Lef1 | 2.87773E-06 | -1.726434927 | 0.109 | 0.353 | 0.069905915 | 0.691864117 | -0.533252279 | 0.4 | 0.426 | 1 | 0.691864117 | 5.75546E-06 |
| Stau1 | 2.87992E-06 | -0.818721177 | 0.697 | 0.853 | 0.069959044 | 0.007265885 | -1.029041571 | 0.8 | 0.966 | 1 | 0.007265885 | 5.75983E-06 |
| Kbtbd2 | 2.8856E-06 | -1.595850817 | 0.151 | 0.4 | 0.070097102 | 0.096678194 | -1.191531458 | 0.3 | 0.515 | 1 | 0.096678194 | 5.7712E-06 |
| Ldb1 | 2.90625E-06 | -1.390881191 | 0.412 | 0.594 | 0.070598691 | 0.10713845 | -0.78685447 | 0.6 | 0.744 | 1 | 0.10713845 | 5.8125E-06 |
| Cd200 | 2.91808E-06 | 0.522361019 | 0.916 | 0.735 | 0.070886105 | 0.032918695 | 0.869058522 | 0.9 | 0.63 | 1 | 0.032918695 | 5.83616E-06 |
| Fbln5 | 2.95192E-06 | -2.632268215 | 0.076 | 0.294 | 0.071707974 | 0.561764271 | 0.197742946 | 0.2 | 0.133 | 1 | 0.561764271 | 5.90383E-06 |
| Csrp2 | 2.95222E-06 | -2.387730153 | 0.244 | 0.471 | 0.071715367 | 0.155702479 | -1.174370906 | 0.5 | 0.639 | 1 | 0.155702479 | 5.90443E-06 |
| Prickle2 | 2.96611E-06 | -2.246239163 | 0.168 | 0.412 | 0.072052693 | 0.152717991 | -1.763437806 | 0.2 | 0.407 | 1 | 0.152717991 | 5.93221E-06 |
| Ipo4 | 2.97793E-06 | -1.531720479 | 0.168 | 0.424 | 0.072339885 | 0.463755808 | -0.304006187 | 0.2 | 0.296 | 1 | 0.463755808 | 5.95585E-06 |
| Wdr43 | 2.98779E-06 | -0.945815457 | 0.639 | 0.747 | 0.072579392 | 0.040019833 | -0.832385159 | 0.6 | 0.809 | 1 | 0.040019833 | 5.97557E-06 |
| Cacul1 | 2.995E-06 | -1.340691734 | 0.294 | 0.547 | 0.072754419 | 0.027334069 | -1.413923879 | 0.4 | 0.694 | 1 | 0.027334069 | 5.98998E-06 |
| Fkbp11 | 3.02003E-06 | -1.411426246 | 0.227 | 0.482 | 0.073362553 | 0.214083236 | -1.33375353 | 0.2 | 0.361 | 1 | 0.214083236 | 6.04005E-06 |
| Ror2 | 3.03281E-06 | -0.912436968 | 0.672 | 0.812 | 0.073672948 | 0.755242548 | -0.362899876 | 0.6 | 0.664 | 1 | 0.755242548 | 6.0656E-06 |
| Nsdhl | 3.06135E-06 | 1.228664301 | 0.521 | 0.247 | 0.074366391 | 0.099910323 | 1.861417422 | 0.4 | 0.207 | 1 | 0.099910323 | 6.1227E-06 |
| Trim27 | 3.06985E-06 | -1.64385619 | 0.218 | 0.453 | 0.074572875 | 0.016303292 | -1.987702641 | 0.1 | 0.488 | 1 | 0.016303292 | 6.1397E-06 |
| Trpc4ap | 3.09774E-06 | -0.896853073 | 0.714 | 0.806 | 0.075250362 | 0.243202315 | -0.38520237 | 0.7 | 0.861 | 1 | 0.243202315 | 6.19548E-06 |
| Pcyt2 | 3.14008E-06 | 0.870964677 | 0.723 | 0.471 | 0.076278816 | 0.320710985 | 0.543435944 | 0.5 | 0.395 | 1 | 0.320710985 | 6.28015E-06 |
| Myo10 | 3.15721E-06 | -0.903884387 | 0.807 | 0.888 | 0.076695047 | 0.012281854 | -1.143629912 | 1 | 0.972 | 1 | 0.012281854 | 6.31442E-06 |
| Iars | 3.16611E-06 | -1.283118652 | 0.546 | 0.676 | 0.076911236 | 0.497366629 | -0.562740455 | 0.8 | 0.682 | 1 | 0.497366629 | 6.33222E-06 |

| | | | | | | | | | | | | |
|---|---|---|---|---|---|---|---|---|---|---|---|---|
| Acta2 | 3.20076E-06 | 0.809770902 | 0.471 | 0.212 | 0.077752922 | 0.003194066 | 6.474779583 | 0.3 | 0.068 | 1 | 0.003194066 | 6.40151E-06 |
| Irf1 | 3.22298E-06 | -1.980686525 | 0.176 | 0.412 | 0.078292622 | 0.045004244 | -2.709998547 | 0 | 0.296 | 1 | 0.045004244 | 6.44595E-06 |
| Znrf3 | 3.22738E-06 | -1.133341234 | 0.462 | 0.688 | 0.078399424 | 0.508786608 | -0.365406732 | 0.7 | 0.704 | 1 | 0.508786608 | 6.45474E-06 |
| Cdkn1b | 3.2339E-06 | -1.322873814 | 0.345 | 0.588 | 0.07855788 | 0.000881473 | -2.363621043 | 0.3 | 0.769 | 1 | 0.000881473 | 6.46779E-06 |
| C1ra | 3.2505E-06 | -2.485426827 | 0.076 | 0.294 | 0.078961139 | 0.067491262 | -1.959358016 | 0.1 | 0.373 | 1 | 0.067491262 | 6.50099E-06 |
| Cyp51 | 3.25362E-06 | 0.698997744 | 0.882 | 0.647 | 0.07903698 | 0.775769784 | -0.315656547 | 0.6 | 0.642 | 1 | 0.775769784 | 6.50723E-06 |
| Ank2 | 3.2552E-06 | -1.802001688 | 0.143 | 0.394 | 0.079075375 | 0.954771029 | -1.127302653 | 0.5 | 0.389 | 1 | 0.954771029 | 6.51039E-06 |
| Nbea | 3.25774E-06 | -1.15758777 | 0.42 | 0.641 | 0.079137128 | 0.101645049 | -0.854388456 | 0.6 | 0.812 | 1 | 0.101645049 | 6.51548E-06 |
| Pgrmc2 | 3.26046E-06 | -0.942775131 | 0.588 | 0.735 | 0.079203101 | 0.046661541 | -0.824142668 | 0.5 | 0.799 | 1 | 0.046661541 | 6.52091E-06 |
| AU015336 | 3.26655E-06 | 1.251538767 | 0.345 | 0.112 | 0.079351151 | 0.110242015 | 1.808468542 | 0.2 | 0.068 | 1 | 0.110242015 | 6.5331E-06 |
| Ngf | 3.28037E-06 | 2.049675159 | 0.387 | 0.165 | 0.079686681 | 0.165780518 | 0.665405493 | 0.2 | 0.077 | 1 | 0.165780518 | 6.56072E-06 |
| Ddx42 | 3.29169E-06 | -0.708970182 | 0.697 | 0.847 | 0.079961694 | 0.18125182 | -0.492373793 | 0.8 | 0.864 | 1 | 0.18125182 | 6.58337E-06 |
| Mcm3 | 3.31104E-06 | -3.655351829 | 0.017 | 0.2 | 0.080431692 | 0.236586849 | 2.539874611 | 0.2 | 0.102 | 1 | 0.236586849 | 6.62206E-06 |
| Cfap74 | 3.31652E-06 | -2.675251386 | 0.05 | 0.259 | 0.080564896 | 0.858896277 | 0.374065718 | 0.2 | 0.176 | 1 | 0.858896277 | 6.63303E-06 |
| Cisd3 | 3.38914E-06 | 1.357031896 | 0.395 | 0.153 | 0.082329052 | 0.346970636 | 0.888638891 | 0.3 | 0.176 | 1 | 0.346970636 | 6.77827E-06 |
| ENSMUSG00000121477 | 3.39352E-06 | -2.632268215 | 0.042 | 0.247 | 0.082435383 | 0.443734742 | -0.789433014 | 0.1 | 0.188 | 1 | 0.443734742 | 6.78703E-06 |
| Cbl | 3.39922E-06 | -1.374868452 | 0.319 | 0.565 | 0.082573897 | 0.009102378 | -1.523820064 | 0.3 | 0.719 | 1 | 0.009102378 | 6.79843E-06 |
| Mme | 3.42193E-06 | -5.694880193 | 0.008 | 0.182 | 0.08312564 | 0.245983478 | -1.789433014 | 0 | 0.12 | 1 | 0.245983478 | 6.84386E-06 |
| Bnip3 | 3.44704E-06 | 1.136462243 | 0.739 | 0.465 | 0.083735449 | 0.507364436 | 0.18308625 | 0.7 | 0.54 | 1 | 0.507364436 | 6.89406E-06 |
| Ap2m1 | 3.45458E-06 | -0.814610738 | 0.605 | 0.765 | 0.08391858 | 0.538541079 | -0.113935053 | 0.7 | 0.802 | 1 | 0.538541079 | 6.90914E-06 |
| Phf20 | 3.47441E-06 | -0.955312803 | 0.429 | 0.665 | 0.08440044 | 0.344577798 | -0.41670632 | 0.5 | 0.71 | 1 | 0.344577798 | 6.94881E-06 |
| Abcc4 | 3.4843E-06 | 0.846416737 | 0.37 | 0.124 | 0.084640594 | 0.019698437 | 2.446379923 | 0.5 | 0.228 | 1 | 0.019698437 | 6.96859E-06 |
| Tlr4 | 3.4985E-06 | 1.432111013 | 0.345 | 0.118 | 0.084985632 | 0.017552122 | 1.539874611 | 0.4 | 0.133 | 1 | 0.017552122 | 6.99699E-06 |
| Amotl2 | 3.56964E-06 | -1.249996883 | 0.37 | 0.6 | 0.086713764 | 0.697292437 | 0.197742946 | 0.6 | 0.537 | 1 | 0.697292437 | 7.13927E-06 |
| Phgdh | 3.60369E-06 | -4.100136671 | 0.008 | 0.182 | 0.08754076 | 0.282342718 | -0.441509711 | 0 | 0.105 | 1 | 0.282342718 | 7.20736E-06 |
| Timm8a1 | 3.60406E-06 | -2.211251864 | 0.092 | 0.318 | 0.087549906 | 0.231668738 | -0.947862377 | 0.1 | 0.262 | 1 | 0.231668738 | 7.20811E-06 |
| Hacd1 | 3.60411E-06 | -1.840269545 | 0.185 | 0.429 | 0.087551047 | 0.315441224 | 0.646363045 | 0.5 | 0.395 | 1 | 0.315441224 | 7.20821E-06 |
| Rbm5 | 3.6199E-06 | -0.678770716 | 0.807 | 0.876 | 0.087934544 | 0.492012367 | 0.242275846 | 0.9 | 0.951 | 1 | 0.492012367 | 7.23978E-06 |
| Ap3d1 | 3.63134E-06 | -0.790281409 | 0.681 | 0.812 | 0.088212572 | 0.0589242 | -0.780819884 | 0.5 | 0.827 | 1 | 0.0589242 | 7.26267E-06 |
| C1qtnf12 | 3.63933E-06 | -3.733354341 | 0.008 | 0.182 | 0.088406554 | 0.225732244 | -0.625934282 | 0 | 0.13 | 1 | 0.225732244 | 7.27864E-06 |
| Snorc | 3.67382E-06 | 0.414761939 | 0.992 | 0.7 | 0.089244335 | 0.209192406 | 0.401901444 | 1 | 0.815 | 1 | 0.209192406 | 7.34762E-06 |
| Golga1 | 3.6776E-06 | -1.532120063 | 0.244 | 0.482 | 0.089336371 | 0.313038145 | -0.79970135 | 0.3 | 0.441 | 1 | 0.313038145 | 7.3552E-06 |
| Stmp1 | 3.75702E-06 | 0.86034801 | 0.899 | 0.753 | 0.091265496 | 0.382077234 | 0.359089511 | 0.7 | 0.722 | 1 | 0.382077234 | 7.51402E-06 |
| Shisa2 | 3.76421E-06 | -2.316827223 | 0.101 | 0.329 | 0.09144016 | 0.023241034 | -2.348400306 | 0.1 | 0.457 | 1 | 0.023241034 | 7.5284E-06 |
| Sik3 | 3.7801E-06 | -1.01678203 | 0.958 | 0.976 | 0.091826157 | 0.032151649 | -0.762688159 | 1 | 0.994 | 1 | 0.032151649 | 7.56018E-06 |
| Zdhhc8 | 3.78679E-06 | -1.108059746 | 0.37 | 0.618 | 0.091988701 | 0.611059538 | -0.272096939 | 0.5 | 0.528 | 1 | 0.611059538 | 7.57357E-06 |
| H1f2 | 3.79444E-06 | 1.514573173 | 0.714 | 0.512 | 0.092174646 | 0.155543505 | 0.796334787 | 0.7 | 0.534 | 1 | 0.155543505 | 7.58887E-06 |
| Sc5d | 3.80825E-06 | 0.729312022 | 0.84 | 0.6 | 0.092509914 | 0.892298975 | -0.131825212 | 0.6 | 0.593 | 1 | 0.892298975 | 7.61648E-06 |
| Specc1l | 3.8324E-06 | -1.093109404 | 0.538 | 0.694 | 0.093096771 | 0.35479126 | -0.564892126 | 0.6 | 0.735 | 1 | 0.35479126 | 7.66479E-06 |
| Uri1 | 3.83489E-06 | -1.713232745 | 0.252 | 0.476 | 0.093157108 | 0.635434279 | -0.31299497 | 0.5 | 0.522 | 1 | 0.635434279 | 7.66976E-06 |
| Tspan2 | 3.86709E-06 | -2.310593025 | 0.143 | 0.371 | 0.09393938 | 0.49420748 | -0.279758641 | 0.1 | 0.194 | 1 | 0.49420748 | 7.73417E-06 |
| Rtf1 | 3.8857E-06 | -1.014894215 | 0.706 | 0.765 | 0.094391482 | 0.005180223 | -1.342559428 | 0.7 | 0.898 | 1 | 0.005180223 | 7.77139E-06 |
| Kifap3 | 3.96496E-06 | -1.287980763 | 0.471 | 0.641 | 0.096316797 | 0.159017749 | -0.876895855 | 0.4 | 0.614 | 1 | 0.159017749 | 7.9299E-06 |
| St13 | 3.96975E-06 | -0.663401961 | 0.908 | 0.947 | 0.096433053 | 0.005484647 | -1.025478277 | 0.8 | 0.966 | 1 | 0.005484647 | 7.93947E-06 |
| Med13l | 4.01094E-06 | -0.480033568 | 0.941 | 0.982 | 0.097433844 | 0.000600946 | -1.288093268 | 1 | 0.997 | 1 | 0.000600946 | 8.02187E-06 |
| Arid4b | 4.02864E-06 | -0.614588246 | 1 | 0.988 | 0.097863769 | 0.009898831 | -0.995168091 | 0.9 | 0.994 | 1 | 0.009898831 | 8.05727E-06 |
| Ppid | 4.04187E-06 | -1.24961389 | 0.395 | 0.606 | 0.098185045 | 0.018675422 | -1.55950692 | 0.3 | 0.627 | 1 | 0.018675422 | 8.08372E-06 |
| Smim13 | 4.04474E-06 | -1.629816737 | 0.134 | 0.376 | 0.098254765 | 0.286587438 | -0.896961478 | 0.2 | 0.34 | 1 | 0.286587438 | 8.08946E-06 |
| Baiap2 | 4.06645E-06 | -0.988801793 | 0.529 | 0.753 | 0.098782181 | 0.108281982 | -0.890570792 | 0.5 | 0.765 | 1 | 0.108281982 | 8.13288E-06 |
| Ankrd42 | 4.10712E-06 | -3.032914622 | 0.076 | 0.288 | 0.099770244 | 0.646628622 | -0.175849835 | 0.3 | 0.225 | 1 | 0.646628622 | 8.21423E-06 |
| Rab3ip | 4.12029E-06 | 2.067114196 | 0.235 | 0.053 | 0.100090111 | 0.293859807 | -0.993305347 | 0.1 | 0.238 | 1 | 0.293859807 | 8.24057E-06 |
| Cdc73 | 4.12652E-06 | -0.721271048 | 0.739 | 0.859 | 0.100241514 | 0.30631886 | -0.447458977 | 0.8 | 0.886 | 1 | 0.30631886 | 8.25303E-06 |
| Palm | 4.13886E-06 | -2.783107376 | 0.034 | 0.229 | 0.100541067 | 0.504424364 | -0.28585884 | 0.1 | 0.179 | 1 | 0.504424364 | 8.27769E-06 |
| Rrbp1 | 4.14681E-06 | -0.311743107 | 1 | 1 | 0.100734385 | 0.267205612 | -0.12815936 | 1 | 1 | 1 | 0.267205612 | 8.29361E-06 |
| Creb1 | 4.16659E-06 | -1.458954616 | 0.286 | 0.529 | 0.101214913 | 0.150092596 | -1.164472445 | 0.4 | 0.611 | 1 | 0.150092596 | 8.33317E-06 |
| Nhs | 4.16964E-06 | -2.246239163 | 0.143 | 0.376 | 0.101288972 | 0.671876305 | -0.619260276 | 0.4 | 0.512 | 1 | 0.671876305 | 8.33927E-06 |
| Rab43 | 4.17877E-06 | -1.753360032 | 0.311 | 0.512 | 0.101510608 | 0.569074058 | -0.350416652 | 0.5 | 0.66 | 1 | 0.569074058 | 8.35752E-06 |
| Rassf1 | 4.20032E-06 | -1.161297824 | 0.437 | 0.647 | 0.102034119 | 0.128156413 | -0.596787936 | 0.4 | 0.722 | 1 | 0.128156413 | 8.40062E-06 |
| Pten | 4.22137E-06 | -0.835495402 | 0.874 | 0.929 | 0.102545533 | 0.098064959 | -0.544320516 | 1 | 0.966 | 1 | 0.098064959 | 8.44272E-06 |
| Ddx39b | 4.22846E-06 | -0.731737875 | 0.807 | 0.853 | 0.102717805 | 0.587959604 | 0.119644443 | 0.9 | 0.88 | 1 | 0.587959604 | 8.45691E-06 |
| Klhl22 | 4.25109E-06 | -1.944858446 | 0.109 | 0.341 | 0.10326743 | 0.148853806 | -1.220482831 | 0.1 | 0.306 | 1 | 0.148853806 | 8.50216E-06 |
| Snap47 | 4.27868E-06 | 1.064770255 | 0.647 | 0.394 | 0.103937646 | 0.000197666 | 1.425464871 | 1 | 0.423 | 1 | 0.000197666 | 8.55734E-06 |
| Epc1 | 4.37196E-06 | -0.824361347 | 0.798 | 0.888 | 0.106203753 | 0.049632574 | -0.855285561 | 0.7 | 0.951 | 1 | 0.049632574 | 8.74391E-06 |
| Ptprn2 | 4.38696E-06 | -4.754887502 | 0.025 | 0.212 | 0.10656812 | 0.994729443 | -1.121629444 | 0.1 | 0.093 | 1 | 0.994729443 | 8.77391E-06 |
| Pard6g | 4.39178E-06 | -3.50779464 | 0.025 | 0.212 | 0.106685122 | 0.104392046 | -1.888968688 | 0 | 0.213 | 1 | 0.104392046 | 8.78354E-06 |
| Agap3 | 4.46437E-06 | -2.100136671 | 0.126 | 0.353 | 0.108448408 | 0.007302431 | -2.391469028 | 0.1 | 0.54 | 1 | 0.007302431 | 8.92871E-06 |
| Cxcl12 | 4.4784E-06 | -3.74281467 | 0.042 | 0.241 | 0.108789349 | 0.267338584 | -1.357117523 | 0 | 0.111 | 1 | 0.267338584 | 8.95678E-06 |
| Slc25a39 | 4.49178E-06 | 0.734354381 | 0.756 | 0.535 | 0.109114259 | 0.452534995 | 0.601559011 | 0.7 | 0.633 | 1 | 0.452534995 | 8.98353E-06 |
| Prr7 | 4.49349E-06 | -2.313245852 | 0.067 | 0.282 | 0.10915586 | 0.021708428 | 1.744903414 | 0.4 | 0.139 | 1 | 0.021708428 | 8.98696E-06 |
| Pip5k1a | 4.54365E-06 | -1.237767647 | 0.454 | 0.659 | 0.11037433 | 0.493746706 | -0.220937725 | 0.8 | 0.858 | 1 | 0.493746706 | 9.08728E-06 |
| Tlk2 | 4.54835E-06 | -0.69751424 | 0.882 | 0.888 | 0.110488416 | 0.207783323 | -0.326607616 | 0.8 | 0.978 | 1 | 0.207783323 | 9.09667E-06 |
| Far1 | 4.54872E-06 | -0.778034348 | 0.706 | 0.865 | 0.110497559 | 0.663966873 | -0.216803423 | 0.8 | 0.895 | 1 | 0.663966873 | 9.09742E-06 |
| Cand1 | 4.55624E-06 | -1.023595116 | 0.529 | 0.7 | 0.110680248 | 0.037581105 | -0.868210127 | 0.7 | 0.861 | 1 | 0.037581105 | 9.11246E-06 |
| Lta4h | 4.60097E-06 | -1.431655571 | 0.261 | 0.5 | 0.111766653 | 0.083967027 | -1.090602549 | 0.4 | 0.596 | 1 | 0.083967027 | 9.20191E-06 |
| Afap1 | 4.60544E-06 | -1.887525271 | 0.176 | 0.412 | 0.111875239 | 0.484339716 | -0.709998547 | 0.5 | 0.506 | 1 | 0.484339716 | 9.21085E-06 |
| Srp68 | 4.64345E-06 | -1.070389328 | 0.387 | 0.618 | 0.112798664 | 0.337675209 | -0.528972552 | 0.5 | 0.605 | 1 | 0.337675209 | 9.28688E-06 |
| Sufu | 4.68176E-06 | -1.98550043 | 0.218 | 0.459 | 0.113729425 | 0.835172383 | -0.195857383 | 0.5 | 0.503 | 1 | 0.835172383 | 9.36351E-06 |
| H3f3a | 4.69608E-06 | -0.49908558 | 0.983 | 0.976 | 0.114077213 | 0.012177594 | -0.667518896 | 1 | 0.994 | 1 | 0.012177594 | 9.39214E-06 |
| Cped1 | 4.69932E-06 | -1.70104602 | 0.429 | 0.647 | 0.114155771 | 0.033889514 | -1.770415788 | 0.4 | 0.685 | 1 | 0.033889514 | 9.39861E-06 |
| Gm48882 | 4.74806E-06 | 2.321928095 | 0.235 | 0.053 | 0.115339907 | 0.643842184 | 1.558490289 | 0 | 0.022 | 1 | 0.643842184 | 9.4961E-06 |
| Uhrf1 | 0.174170701 | -1.359895945 | 0.034 | 0.071 | 1 | 4.78072E-06 | 3.880418384 | 0.3 | 0.028 | 0.116133171 | 0.174170701 | 9.56141E-06 |
| Krt16 | 0.000975356 | 1.939458458 | 0.16 | 0.047 | 1 | 4.78946E-06 | 1.053025709 | 0.3 | 0.028 | 0.116345588 | 0.000975356 | 9.5789E-06 |
| Zmym1 | 4.79503E-06 | -1.733354341 | 0.101 | 0.335 | 0.116480937 | 0.770943147 | 0.123104145 | 0.3 | 0.235 | 1 | 0.770943147 | 9.59004E-06 |
| Wnk1 | 4.80801E-06 | -0.725646245 | 0.992 | 0.982 | 0.116796233 | 0.026753343 | -0.776700249 | 0.9 | 0.997 | 1 | 0.026753343 | 9.616E-06 |
| Tial1 | 4.81853E-06 | -0.849117446 | 0.697 | 0.794 | 0.117051852 | 0.028668028 | -0.936274402 | 0.6 | 0.836 | 1 | 0.028668028 | 9.63705E-06 |
| Usp49 | 4.83109E-06 | -2.095480309 | 0.134 | 0.365 | 0.117356833 | 0.482176563 | 0.159940913 | 0.2 | 0.361 | 1 | 0.482176563 | 9.66216E-06 |
| Cdkn2d | 4.86437E-06 | 0.831046838 | 0.681 | 0.412 | 0.118165229 | 0.76578525 | 0.342356858 | 0.4 | 0.358 | 1 | 0.76578525 | 9.72871E-06 |
| Ptbp1 | 4.92124E-06 | -0.724345146 | 0.773 | 0.859 | 0.119546786 | 0.178869853 | -0.505640048 | 0.7 | 0.917 | 1 | 0.178869853 | 9.84246E-06 |
| Gltp | 4.93011E-06 | 1.274439169 | 0.546 | 0.3 | 0.119762156 | 0.778266859 | 0.299103661 | 0.4 | 0.355 | 1 | 0.778266859 | 9.86019E-06 |
| Sf3a1 | 4.9824E-06 | -1.441032708 | 0.235 | 0.482 | 0.121032575 | 0.010391418 | -2.220482831 | 0.1 | 0.515 | 1 | 0.010391418 | 9.96478E-06 |
| Setx | 5.0077E-06 | -1.6116906 | 0.269 | 0.494 | 0.121647043 | 0.127092297 | -1.168605061 | 0.4 | 0.593 | 1 | 0.127092297 | 1.00154E-05 |
| H13 | 5.04133E-06 | -0.681717838 | 0.891 | 0.894 | 0.122463879 | 0.062235419 | -0.694208221 | 0.9 | 0.941 | 1 | 0.062235419 | 1.00826E-05 |
| St3gal4 | 5.07929E-06 | 0.812164801 | 0.824 | 0.612 | 0.123386188 | 0.295292406 | 0.680396958 | 0.7 | 0.62 | 1 | 0.295292406 | 1.01586E-05 |
| Ano6 | 5.10913E-06 | 0.5901352 | 0.975 | 0.894 | 0.12411107 | 0.163277223 | 0.602189445 | 1 | 0.96 | 1 | 0.163277223 | 1.02182E-05 |
| Exoc5 | 5.12773E-06 | -1.016873819 | 0.605 | 0.735 | 0.124562937 | 0.049606406 | -0.923690067 | 0.6 | 0.84 | 1 | 0.049606406 | 1.02554E-05 |
| Bid | 0.345346582 | -0.900464326 | 0.017 | 0.035 | 1 | 5.1528E-06 | 3.017921908 | 0.4 | 0.049 | 0.125171702 | 0.345346582 | 1.03056E-05 |
| Rps6kb1 | 5.19956E-06 | -0.940926323 | 0.681 | 0.806 | 0.126307615 | 0.792536543 | -0.132577595 | 1 | 0.904 | 1 | 0.792536543 | 1.03991E-05 |
| Zfp398 | 5.27736E-06 | -1.885964757 | 0.151 | 0.394 | 0.128197645 | 0.603067295 | 0.210566986 | 0.5 | 0.383 | 1 | 0.603067295 | 1.05547E-05 |

| | | | | | | | | | | | | |
|---|---|---|---|---|---|---|---|---|---|---|---|---|
| Marveld1 | 5.28205E-06 | 1.019968416 | 0.765 | 0.588 | 0.128311583 | 0.006103251 | 1.375715455 | 0.8 | 0.596 | 1 | 0.006103251 | 1.05641E-05 |
| Acad9 | 0.000211513 | -1.87774425 | 0.05 | 0.206 | 1 | 5.30084E-06 | 2.185031894 | 0.7 | 0.142 | 0.128767931 | 0.000211513 | 1.06016E-05 |
| Sparcl1 | 1.32416E-05 | 0.789731138 | 0.807 | 0.588 | 0.321665221 | 5.30428E-06 | 2.244119546 | 1 | 0.438 | 0.128851565 | 1.32416E-05 | 1.06085E-05 |
| Kcnma1 | 5.32847E-06 | 0.258709882 | 1 | 0.959 | 0.129439295 | 0.000705678 | 1.096517316 | 1 | 0.895 | 1 | 0.000705678 | 1.06569E-05 |
| Cct5 | 5.33559E-06 | -0.886718319 | 0.664 | 0.794 | 0.129612256 | 0.0475157 | -0.756865152 | 0.5 | 0.846 | 1 | 0.0475157 | 1.06712E-05 |
| Foxd2os | 5.45619E-06 | -1.913663824 | 0.202 | 0.435 | 0.132541684 | 0.36640927 | 0.769994395 | 0.3 | 0.198 | 1 | 0.36640927 | 1.09123E-05 |
| Cdc42ep1 | 5.46948E-06 | -3.222392421 | 0.017 | 0.194 | 0.132864703 | 0.19086576 | -0.888968688 | 0 | 0.148 | 1 | 0.19086576 | 1.09389E-05 |
| Pxdn | 5.4928E-06 | -1.28838998 | 0.143 | 0.376 | 0.133431065 | 0.021983462 | -2.658035125 | 0.1 | 0.46 | 1 | 0.021983462 | 1.09856E-05 |
| Pdlim4 | 5.51096E-06 | -1.200024608 | 0.151 | 0.412 | 0.133872336 | 0.160429972 | -1.223664079 | 0.2 | 0.392 | 1 | 0.160429972 | 1.10219E-05 |
| Adamtsl4 | 5.51339E-06 | -2.891419187 | 0.034 | 0.224 | 0.133931172 | 0.124799693 | -1.68251781 | 0 | 0.194 | 1 | 0.124799693 | 1.10267E-05 |
| Zbed5 | 5.51556E-06 | -2.970853654 | 0.034 | 0.224 | 0.133983911 | 0.588658324 | -0.28585884 | 0.1 | 0.157 | 1 | 0.588658324 | 1.10311E-05 |
| Rcn3 | 5.53473E-06 | -0.631306355 | 0.941 | 0.888 | 0.134449748 | 0.168126093 | -0.577367731 | 1 | 0.84 | 1 | 0.168126093 | 1.10694E-05 |
| Cnksr1 | 0.031987759 | 1.362570079 | 0.067 | 0.018 | 1 | 5.55474E-06 | 3.695993813 | 0.3 | 0.028 | 0.134935761 | 0.031987759 | 1.11095E-05 |
| Tstd3 | 0.897527791 | 0.152003093 | 0.05 | 0.047 | 1 | 5.55474E-06 | 3.695993813 | 0.3 | 0.028 | 0.134935761 | 0.897527791 | 1.11095E-05 |
| Pdpk1 | 5.56859E-06 | -0.859069959 | 0.639 | 0.759 | 0.135272254 | 0.000878341 | -1.650640794 | 0.5 | 0.914 | 1 | 0.000878341 | 1.11372E-05 |
| Pum2 | 5.61399E-06 | -0.634701319 | 0.874 | 0.953 | 0.136375005 | 0.054344785 | -0.60498121 | 0.8 | 0.972 | 1 | 0.054344785 | 1.12279E-05 |
| Vezt | 5.65268E-06 | -1.351802263 | 0.361 | 0.582 | 0.137314874 | 0.121405284 | -1.05085637 | 0.6 | 0.698 | 1 | 0.121405284 | 1.13053E-05 |
| Gjc1 | 5.66833E-06 | -1.457946091 | 0.311 | 0.529 | 0.137695027 | 0.165160137 | -1.083616118 | 0.4 | 0.549 | 1 | 0.165160137 | 1.13366E-05 |
| Qtrt1 | 5.69294E-06 | -1.778208576 | 0.134 | 0.371 | 0.138292969 | 0.069054993 | -1.689437224 | 0.1 | 0.373 | 1 | 0.069054993 | 1.13859E-05 |
| Ampd1 | 5.69682E-06 | -3.635173947 | 0.017 | 0.194 | 0.138387058 | 0.642687779 | 0.71414116 | 0.2 | 0.148 | 1 | 0.642687779 | 1.13936E-05 |
| Angptl1 | 5.81129E-06 | 1.451379346 | 0.311 | 0.1 | 0.141167855 | 0.001981953 | 1.546616189 | 0.6 | 0.194 | 1 | 0.001981953 | 1.16225E-05 |
| Sptan1 | 5.85194E-06 | -0.866773132 | 0.857 | 0.935 | 0.142155213 | 0.328811154 | -0.299671597 | 0.9 | 0.972 | 1 | 0.328811154 | 1.17038E-05 |
| Kdm4c | 5.92239E-06 | -1.156225497 | 0.437 | 0.624 | 0.143866729 | 0.015701405 | -1.450847575 | 0.6 | 0.756 | 1 | 0.015701405 | 1.18447E-05 |
| Gramd2 | 5.96958E-06 | 0.514573173 | 0.84 | 0.494 | 0.145012985 | 0.043555137 | 1.260492211 | 0.6 | 0.318 | 1 | 0.043555137 | 1.19391E-05 |
| Etv4 | 5.98154E-06 | 1.893084796 | 0.235 | 0.053 | 0.14530347 | 0.339584249 | 1.280956314 | 0.2 | 0.102 | 1 | 0.339584249 | 1.1963E-05 |
| Ncoa4 | 5.98624E-06 | -2.152851488 | 0.109 | 0.335 | 0.145417751 | 0.045441276 | -1.419483404 | 0.2 | 0.506 | 1 | 0.045441276 | 1.19724E-05 |
| St7l | 6.0363E-06 | -1.285895364 | 0.345 | 0.576 | 0.146633739 | 0.01400435 | -1.909856054 | 0.3 | 0.636 | 1 | 0.01400435 | 1.20726E-05 |
| Lrpprc | 6.07409E-06 | -1.760049207 | 0.218 | 0.447 | 0.147551685 | 0.007994242 | -2.365782384 | 0.1 | 0.534 | 1 | 0.007994242 | 1.21481E-05 |
| Sptlc2 | 6.07438E-06 | 0.832347917 | 0.933 | 0.824 | 0.147558949 | 0.439645912 | 0.420490055 | 1 | 0.821 | 1 | 0.439645912 | 1.21487E-05 |
| Cggbp1 | 6.13845E-06 | -0.705756782 | 0.798 | 0.865 | 0.149115216 | 0.245731119 | -0.462343214 | 0.9 | 0.935 | 1 | 0.245731119 | 1.22769E-05 |
| Zbtb46 | 6.14094E-06 | -2.084064265 | 0.218 | 0.441 | 0.149175809 | 0.176391436 | 0.675529711 | 0.6 | 0.404 | 1 | 0.176391436 | 1.22819E-05 |
| Tcf3 | 6.1411E-06 | -1.748461233 | 0.134 | 0.371 | 0.149179524 | 0.291371317 | -0.709998547 | 0.3 | 0.435 | 1 | 0.291371317 | 1.22822E-05 |
| 0610040J01Rik | 6.20729E-06 | 1.707218251 | 0.353 | 0.129 | 0.150787535 | 0.841574516 | 0.381297287 | 0.3 | 0.386 | 1 | 0.841574516 | 1.24145E-05 |
| Ppp4r3a | 6.21043E-06 | -0.95491211 | 0.605 | 0.753 | 0.150863871 | 0.079330503 | -0.687405545 | 0.8 | 0.88 | 1 | 0.079330503 | 1.24208E-05 |
| Usp33 | 6.29149E-06 | -1.149460899 | 0.42 | 0.612 | 0.152832979 | 0.0033426 | -1.922244842 | 0.4 | 0.744 | 1 | 0.0033426 | 1.25829E-05 |
| Fbln1 | 6.35259E-06 | -3.733354341 | 0.025 | 0.206 | 0.154317077 | 0.408846153 | -0.473931188 | 0 | 0.065 | 1 | 0.408846153 | 1.27051E-05 |
| Usp11 | 6.36955E-06 | -1.821710215 | 0.16 | 0.388 | 0.154728992 | 0.448337064 | -0.294961047 | 0.2 | 0.324 | 1 | 0.448337064 | 1.2739E-05 |
| Actr1a | 6.37204E-06 | -1.341416524 | 0.353 | 0.553 | 0.154789569 | 0.124577535 | -0.625934282 | 0.4 | 0.685 | 1 | 0.124577535 | 1.2744E-05 |
| Ctnnal1 | 6.41658E-06 | 0.616326672 | 0.765 | 0.512 | 0.15587156 | 0.191215605 | -0.647414009 | 0.4 | 0.623 | 1 | 0.191215605 | 1.28331E-05 |
| Gspt1 | 6.44047E-06 | -0.681417933 | 0.899 | 0.935 | 0.156451829 | 0.109643432 | -0.548733044 | 0.9 | 0.978 | 1 | 0.109643432 | 1.28809E-05 |
| Rnmt | 6.46313E-06 | -1.163498732 | 0.429 | 0.618 | 0.157002251 | 0.336072479 | -0.563278674 | 0.4 | 0.537 | 1 | 0.336072479 | 1.29262E-05 |
| Uaca | 6.51831E-06 | 0.920786822 | 0.857 | 0.747 | 0.15834273 | 0.003389888 | 1.352132432 | 1 | 0.778 | 1 | 0.003389888 | 1.30366E-05 |
| Brd1 | 6.58369E-06 | -0.820510287 | 0.798 | 0.918 | 0.159931003 | 0.163110664 | -0.327157151 | 0.8 | 0.944 | 1 | 0.163110664 | 1.31673E-05 |
| Enox1 | 6.68562E-06 | -2.644625422 | 0.193 | 0.412 | 0.162407022 | 0.083459633 | -2.846264237 | 0.2 | 0.42 | 1 | 0.083459633 | 1.33712E-05 |
| Tnrc6b | 6.69644E-06 | -0.764710585 | 0.958 | 0.982 | 0.162670017 | 0.272790826 | -0.201938055 | 1 | 0.978 | 1 | 0.272790826 | 1.33928E-05 |
| Slmap | 6.75143E-06 | -1.002083315 | 0.739 | 0.8 | 0.164005838 | 0.295861981 | -0.55860023 | 0.8 | 0.898 | 1 | 0.295861981 | 1.35028E-05 |
| Krit1 | 6.76235E-06 | -1.005900601 | 0.597 | 0.753 | 0.164271027 | 0.457167522 | -0.328591825 | 0.6 | 0.741 | 1 | 0.457167522 | 1.35247E-05 |
| Rrp1 | 6.83441E-06 | -0.819229022 | 0.605 | 0.765 | 0.166021438 | 0.0533609 | -0.891637874 | 0.5 | 0.753 | 1 | 0.0533609 | 1.36688E-05 |
| Xpr1 | 6.92322E-06 | -1.447756409 | 0.487 | 0.671 | 0.168178821 | 0.620443795 | -0.342731556 | 0.9 | 0.852 | 1 | 0.620443795 | 1.38464E-05 |
| Nf1 | 6.94956E-06 | -0.922907376 | 0.891 | 0.888 | 0.168818751 | 0.897841448 | -0.110072413 | 1 | 0.963 | 1 | 0.897841448 | 1.38991E-05 |
| Hexa | 7.02481E-06 | 0.668143561 | 0.899 | 0.706 | 0.170646668 | 0.052853537 | 0.93531663 | 0.8 | 0.657 | 1 | 0.052853537 | 1.40496E-05 |
| Snrpn | 7.02978E-06 | 2.099535674 | 0.269 | 0.076 | 0.170767458 | 0.455031706 | 0.959028219 | 0.2 | 0.117 | 1 | 0.455031706 | 1.40595E-05 |
| Uqcrc1 | 7.05504E-06 | -1.123687554 | 0.42 | 0.618 | 0.171381085 | 0.668920223 | -0.187627003 | 0.6 | 0.636 | 1 | 0.668920223 | 1.411E-05 |
| Mta2 | 7.07593E-06 | -1.678071905 | 0.193 | 0.418 | 0.171888437 | 0.129331738 | -1.082214763 | 0.4 | 0.559 | 1 | 0.129331738 | 1.41518E-05 |
| Cachd1 | 7.1008E-06 | -2.538538164 | 0.042 | 0.235 | 0.172492533 | 0.085144393 | -1.987702641 | 0.1 | 0.355 | 1 | 0.085144393 | 1.42015E-05 |
| Zc3h14 | 7.11885E-06 | -0.933408425 | 0.504 | 0.7 | 0.172931059 | 0.055741053 | -0.548424915 | 0.4 | 0.836 | 1 | 0.055741053 | 1.42376E-05 |
| B4galnt3 | 7.15904E-06 | 0.418571353 | 0.588 | 0.306 | 0.173907344 | 0.01111567 | 1.920311111 | 0.6 | 0.269 | 1 | 0.01111567 | 1.4318E-05 |
| Pdap1 | 7.16696E-06 | -0.79639571 | 0.706 | 0.865 | 0.174099826 | 0.244960077 | -0.48032896 | 0.7 | 0.809 | 1 | 0.244960077 | 1.43339E-05 |
| Bace1 | 7.21046E-06 | -1.632268215 | 0.185 | 0.418 | 0.175156538 | 0.410129126 | -0.267480311 | 0.3 | 0.463 | 1 | 0.410129126 | 1.44209E-05 |
| Pdia3 | 7.23633E-06 | -0.459491918 | 1 | 0.988 | 0.175784952 | 0.100566514 | -0.386698975 | 1 | 1 | 1 | 0.100566514 | 1.44726E-05 |
| Btf3 | 7.27347E-06 | -0.446712562 | 0.95 | 0.976 | 0.176687044 | 0.005338797 | -0.960502557 | 0.6 | 0.991 | 1 | 0.005338797 | 1.45469E-05 |
| Canx | 7.29003E-06 | -0.454523015 | 1 | 0.988 | 0.177089518 | 0.035825822 | -0.602885792 | 1 | 1 | 1 | 0.035825822 | 1.458E-05 |
| Gm48521 | 7.37199E-06 | 1.836501268 | 0.235 | 0.053 | 0.179080301 | 0.338259829 | 1.111031312 | 0.1 | 0.037 | 1 | 0.338259829 | 1.47439E-05 |
| Ctdspl | 7.47173E-06 | -0.721867023 | 0.958 | 0.965 | 0.181503232 | 0.005156355 | -1.009842349 | 1 | 0.997 | 1 | 0.005156355 | 1.49434E-05 |
| Tlcd4 | 7.5465E-06 | 1.162271429 | 0.546 | 0.294 | 0.183319523 | 0.003094376 | 1.816288047 | 0.9 | 0.435 | 1 | 0.003094376 | 1.50929E-05 |
| Cdc5l | 7.58839E-06 | -0.879164925 | 0.613 | 0.782 | 0.184337252 | 0.002463059 | -1.659797734 | 0.5 | 0.861 | 1 | 0.002463059 | 1.51767E-05 |
| Zfp407 | 7.65827E-06 | -1.272211543 | 0.513 | 0.7 | 0.186034797 | 0.748860901 | 0.239278283 | 0.7 | 0.759 | 1 | 0.748860901 | 1.53165E-05 |
| Commd2 | 7.69101E-06 | -1.364572432 | 0.168 | 0.418 | 0.186830079 | 0.117086935 | -1.125036046 | 0.2 | 0.426 | 1 | 0.117086935 | 1.5382E-05 |
| Robo2 | 7.71144E-06 | -2.394914534 | 0.109 | 0.324 | 0.187326317 | 0.028703256 | -4.319699994 | 0 | 0.34 | 1 | 0.028703256 | 1.54228E-05 |
| Dock1 | 7.72813E-06 | -1.010579933 | 0.748 | 0.853 | 0.187731712 | 0.575168749 | -0.224934616 | 0.9 | 0.926 | 1 | 0.575168749 | 1.54562E-05 |
| Dgki | 7.76291E-06 | 2.745898719 | 0.252 | 0.065 | 0.188576705 | 0.001881053 | 1.614199722 | 0.4 | 0.093 | 1 | 0.001881053 | 1.55258E-05 |
| Podxl2 | 7.76291E-06 | -3.900464326 | 0.017 | 0.188 | 0.188576711 | 0.599773272 | 0.135278859 | 0.1 | 0.16 | 1 | 0.599773272 | 1.55258E-05 |
| Efr3a | 7.83015E-06 | -0.918885523 | 0.538 | 0.724 | 0.190210008 | 0.088607997 | -0.936274402 | 0.7 | 0.787 | 1 | 0.088607997 | 1.56602E-05 |
| Tbrg1 | 7.9301E-06 | -0.574802635 | 0.899 | 0.947 | 0.192637916 | 0.039713302 | -0.771807343 | 0.7 | 0.938 | 1 | 0.039713302 | 1.58601E-05 |
| Rlim | 7.93092E-06 | -0.717979115 | 0.916 | 0.924 | 0.192657889 | 0.360484308 | -0.402546555 | 0.9 | 0.926 | 1 | 0.360484308 | 1.58618E-05 |
| Cdc23 | 7.93492E-06 | -1.544320516 | 0.185 | 0.418 | 0.192755093 | 0.251233287 | 0.349543399 | 0.6 | 0.349 | 1 | 0.251233287 | 1.58698E-05 |
| Cep162 | 7.96129E-06 | -2.439623138 | 0.101 | 0.312 | 0.193395755 | 0.702070865 | -0.28585884 | 0.2 | 0.241 | 1 | 0.702070865 | 1.59225E-05 |
| Nav2 | 8.0428E-06 | 1.12322578 | 0.765 | 0.606 | 0.195375594 | 0.00349721 | 1.473601392 | 0.8 | 0.512 | 1 | 0.00349721 | 1.60855E-05 |
| Prkacb | 8.05414E-06 | -0.891419187 | 0.597 | 0.771 | 0.195651238 | 0.049054193 | -0.864721141 | 0.6 | 0.806 | 1 | 0.049054193 | 1.61082E-05 |
| Phactr1 | 8.05453E-06 | 1.283960245 | 0.319 | 0.106 | 0.195660604 | 0.22291428 | 0.708066645 | 0.4 | 0.259 | 1 | 0.22291428 | 1.6109E-05 |
| Irak3 | 8.10267E-06 | 1.161340958 | 0.555 | 0.3 | 0.196830097 | 0.203073999 | 0.980832589 | 0.4 | 0.25 | 1 | 0.203073999 | 1.62053E-05 |
| Atn1 | 8.1158E-06 | -1.09895848 | 0.303 | 0.553 | 0.197149039 | 0.027738705 | -1.350147969 | 0.4 | 0.66 | 1 | 0.027738705 | 1.62315E-05 |
| Dhx16 | 8.14103E-06 | -2.017921908 | 0.109 | 0.329 | 0.197762021 | 0.418873267 | -0.546862711 | 0.2 | 0.302 | 1 | 0.418873267 | 1.6282E-05 |
| Eps8 | 8.1697E-06 | -0.984997837 | 0.748 | 0.806 | 0.198458391 | 0.0733187 | -0.536666944 | 0.7 | 0.898 | 1 | 0.0733187 | 1.63393E-05 |
| Acvr2a | 8.29785E-06 | -1.987927168 | 0.185 | 0.406 | 0.201571366 | 0.556039959 | -0.134081185 | 0.4 | 0.519 | 1 | 0.556039959 | 1.65956E-05 |
| Itsn1 | 8.31073E-06 | -1.673630562 | 0.336 | 0.541 | 0.201884224 | 0.11881603 | -0.963169629 | 0.5 | 0.725 | 1 | 0.11881603 | 1.66214E-05 |
| Frs2 | 8.3573E-06 | -1.051178724 | 0.58 | 0.753 | 0.203015601 | 0.011751646 | -1.193030362 | 0.7 | 0.877 | 1 | 0.011751646 | 1.67145E-05 |
| Ythdf3 | 8.36962E-06 | -0.803602787 | 0.706 | 0.841 | 0.203314689 | 0.09403463 | -0.610109315 | 0.9 | 0.938 | 1 | 0.09403463 | 1.67392E-05 |
| Psmb6 | 8.39033E-06 | -0.693633342 | 0.807 | 0.9 | 0.203817974 | 0.000372565 | -1.924592597 | 0.4 | 0.861 | 1 | 0.000372565 | 1.67806E-05 |
| Zyg11b | 8.4903E-06 | -1.432959407 | 0.361 | 0.547 | 0.206246385 | 0.059077108 | -0.997028433 | 0.5 | 0.753 | 1 | 0.059077108 | 1.69805E-05 |
| Fam120a | 8.5364E-06 | -0.735564239 | 0.815 | 0.882 | 0.207366291 | 0.227839598 | -0.374395515 | 0.8 | 0.929 | 1 | 0.227839598 | 1.70727E-05 |
| Strap | 8.63001E-06 | -0.686339521 | 0.731 | 0.841 | 0.209640312 | 0.009187189 | -1.198603506 | 0.5 | 0.864 | 1 | 0.009187189 | 1.726E-05 |
| Scarf2 | 8.66214E-06 | -1.041820176 | 0.454 | 0.682 | 0.210420635 | 0.504886632 | -0.370095377 | 0.7 | 0.713 | 1 | 0.504886632 | 1.73242E-05 |
| Maz | 8.86387E-06 | -1.304854582 | 0.235 | 0.482 | 0.215321182 | 0.657701497 | -0.210896782 | 0.4 | 0.441 | 1 | 0.657701497 | 1.77277E-05 |
| Dennd4c | 8.88549E-06 | -1.705303651 | 0.244 | 0.465 | 0.215846403 | 0.1824008 | -1.156619722 | 0.8 | 0.756 | 1 | 0.1824008 | 1.77709E-05 |
| Lysmd2 | 8.90886E-06 | -3.572889668 | 0.008 | 0.171 | 0.216414073 | 0.175160723 | -0.91281543 | 0 | 0.157 | 1 | 0.175160723 | 1.78176E-05 |
| Ctsl | 8.96474E-06 | 0.763854815 | 0.941 | 0.865 | 0.217771392 | 0.179023518 | -0.401998531 | 0.8 | 0.966 | 1 | 0.179023518 | 1.79294E-05 |
| Adamts6 | 8.98557E-06 | -2.979415668 | 0.059 | 0.259 | 0.218277433 | 0.598336099 | -1.630735268 | 0.2 | 0.241 | 1 | 0.598336099 | 1.79711E-05 |

| | | | | | | | | | | | | |
|---|---|---|---|---|---|---|---|---|---|---|---|---|
| Angptl2 | 8.99191E-06 | -1.455053178 | 0.218 | 0.459 | 0.218431527 | 0.23735075 | -0.982078092 | 0.2 | 0.389 | 1 | 0.23735075 | 1.79837E-05 |
| Hsp90ab1 | 9.04331E-06 | -0.503883732 | 1 | 0.994 | 0.21968013 | 0.006996434 | -0.881938486 | 0.9 | 1 | 1 | 0.006996434 | 1.80865E-05 |
| Tubgcp3 | 9.06765E-06 | -2.027320606 | 0.118 | 0.335 | 0.220271441 | 0.403301257 | -0.673239997 | 0.2 | 0.302 | 1 | 0.403301257 | 1.81352E-05 |
| Taok2 | 9.10492E-06 | -1.511899039 | 0.185 | 0.424 | 0.221176624 | 0.051318828 | -1.531541912 | 0.2 | 0.481 | 1 | 0.051318828 | 1.82097E-05 |
| Tpx2 | 0.565575732 | -1.840907482 | 0.092 | 0.071 | 1 | 9.12378E-06 | 5.240314329 | 0.2 | 0.012 | 0.221634861 | 0.565575732 | 1.82475E-05 |
| Csf2ra | 9.26456E-06 | 1.087758506 | 0.429 | 0.182 | 0.225054802 | 0.803961834 | 0.159940913 | 0.1 | 0.123 | 1 | 0.803961834 | 1.8529E-05 |
| Lin54 | 9.30024E-06 | -2.555816155 | 0.076 | 0.276 | 0.225921397 | 0.144910611 | -1.661558192 | 0.1 | 0.302 | 1 | 0.144910611 | 1.86004E-05 |
| Cdon | 9.35977E-06 | 1.186687623 | 0.571 | 0.324 | 0.227367621 | 0.096239182 | 1.686078344 | 0.6 | 0.435 | 1 | 0.096239182 | 1.87195E-05 |
| Tmem170 | 9.36373E-06 | 1.636563697 | 0.252 | 0.065 | 0.227463845 | 1 | 0.695993813 | 0.1 | 0.099 | 1 | 1 | 1.87274E-05 |
| Sae1 | 9.44344E-06 | -0.986904342 | 0.471 | 0.671 | 0.229399995 | 0.008626253 | -1.527869995 | 0.4 | 0.759 | 1 | 0.008626253 | 1.88868E-05 |
| Cars | 9.47729E-06 | -1.246546283 | 0.403 | 0.6 | 0.230222332 | 0.26404831 | -0.848326703 | 0.7 | 0.66 | 1 | 0.26404831 | 1.89545E-05 |
| Mknk1 | 9.50363E-06 | -1.687060688 | 0.067 | 0.282 | 0.230862276 | 0.336157823 | 0.660369903 | 0.4 | 0.256 | 1 | 0.336157823 | 1.90072E-05 |
| Prep | 9.50565E-06 | -1.437171659 | 0.261 | 0.494 | 0.230911167 | 0.00663501 | -3.031926641 | 0 | 0.454 | 1 | 0.00663501 | 1.90112E-05 |
| Phrf1 | 9.53382E-06 | -0.900464326 | 0.395 | 0.641 | 0.231595625 | 0.235858191 | -0.411136062 | 0.6 | 0.753 | 1 | 0.235858191 | 1.90676E-05 |
| Ptov1 | 9.55746E-06 | -0.716471119 | 0.739 | 0.806 | 0.232169836 | 0.325541359 | -0.360974999 | 0.7 | 0.821 | 1 | 0.325541359 | 1.91148E-05 |
| Dnm3os | 9.56317E-06 | -3.714245518 | 0.025 | 0.2 | 0.23230851 | 0.245878587 | -1.230005605 | 0 | 0.12 | 1 | 0.245878587 | 1.91262E-05 |
| Pold3 | 9.68169E-06 | -1.987927168 | 0.076 | 0.288 | 0.235187725 | 0.617954506 | -0.181750437 | 0.2 | 0.278 | 1 | 0.617954506 | 1.93633E-05 |
| Ube2d2a | 9.71612E-06 | -0.624254532 | 0.933 | 0.929 | 0.236023944 | 0.001690931 | -1.109642592 | 0.8 | 0.972 | 1 | 0.001690931 | 1.94321E-05 |
| Hspa9 | 9.74155E-06 | -0.662965013 | 0.891 | 0.929 | 0.236641686 | 0.000364539 | -1.564892126 | 0.8 | 0.951 | 1 | 0.000364539 | 1.9483E-05 |
| Irf2bp2 | 9.75459E-06 | -1.177666064 | 0.42 | 0.629 | 0.236958382 | 0.005928394 | -1.530514717 | 0.4 | 0.821 | 1 | 0.005928394 | 1.95091E-05 |
| Pdia6 | 9.88082E-06 | 0.432554597 | 0.992 | 0.965 | 0.24002486 | 0.216662533 | 0.295846276 | 1 | 0.957 | 1 | 0.216662533 | 1.97615E-05 |
| Ier3 | 9.88396E-06 | 0.968635829 | 0.874 | 0.735 | 0.240101234 | 0.233213249 | -1.082214763 | 0.5 | 0.685 | 1 | 0.233213249 | 1.97678E-05 |
| Gcnt2 | 0.004104279 | 0.72562805 | 0.361 | 0.218 | 1 | 9.90977E-06 | 2.902444691 | 0.7 | 0.204 | 0.240728052 | 0.004104279 | 1.98194E-05 |
| Mex3d | 9.98596E-06 | -2.157852169 | 0.059 | 0.259 | 0.24257883 | 0.031813322 | -2.304006187 | 0 | 0.327 | 1 | 0.031813322 | 1.99718E-05 |
| Fgf2os | 2.10451E-05 | 1.565199246 | 0.151 | 0.018 | 0.511226362 | 1.00831E-05 | 4.695993813 | 0.2 | 0.012 | 0.2449396 | 2.10451E-05 | 2.01662E-05 |
| Necab3 | 0.349245302 | -1.292781749 | 0.017 | 0.035 | 1 | 1.00831E-05 | 4.695993813 | 0.2 | 0.012 | 0.2449396 | 0.349245302 | 2.01662E-05 |
| Ripor2 | 0.388317912 | -0.86393845 | 0.042 | 0.024 | 1 | 1.00878E-05 | 3.532495081 | 0.2 | 0.012 | 0.245053713 | 0.388317912 | 2.01756E-05 |
| Tbl1xr1 | 1.01173E-05 | -1.336283388 | 0.294 | 0.535 | 0.245769544 | 0.146231557 | -0.779739618 | 0.5 | 0.66 | 1 | 0.146231557 | 2.02345E-05 |
| Pygo1 | 1.02208E-05 | -2.273922722 | 0.076 | 0.282 | 0.248283925 | 0.15773774 | -1.374395515 | 0.1 | 0.299 | 1 | 0.15773774 | 2.04415E-05 |
| Kctd4 | 1.03644E-05 | 0.126377286 | 0.655 | 0.382 | 0.251771324 | 0.427773419 | 0.577349317 | 0.5 | 0.407 | 1 | 0.427773419 | 2.07286E-05 |
| Senp7 | 1.0395E-05 | -1.346023771 | 0.328 | 0.535 | 0.252515858 | 0.002524065 | -2.740301307 | 0.1 | 0.602 | 1 | 0.002524065 | 2.07899E-05 |
| Plscr3 | 1.0419E-05 | -1.852209158 | 0.151 | 0.371 | 0.253098904 | 0.166078878 | -0.970762779 | 0.1 | 0.299 | 1 | 0.166078878 | 2.08379E-05 |
| Cntd1 | 0.233061076 | 0.344648171 | 0.05 | 0.024 | 1 | 1.04287E-05 | 4.795529487 | 0.2 | 0.012 | 0.253332902 | 0.233061076 | 2.08572E-05 |
| Dscaml1 | 0.715874401 | 3.099535674 | 0.017 | 0.012 | 1 | 1.04287E-05 | 5.380491987 | 0.2 | 0.012 | 0.253332902 | 0.715874401 | 2.08572E-05 |
| Baz1b | 1.04353E-05 | -0.928916902 | 0.714 | 0.824 | 0.253494967 | 0.727159431 | -0.133179219 | 0.9 | 0.883 | 1 | 0.727159431 | 2.08705E-05 |
| Pou2af3 | 1.04921E-05 | 1.314274522 | 0.387 | 0.159 | 0.254873904 | 0.060052144 | 1.245332404 | 0.4 | 0.164 | 1 | 0.060052144 | 2.09841E-05 |
| Whrn | 1.05078E-05 | -1.378511623 | 0.185 | 0.424 | 0.255254821 | 0.02316385 | -2.361456459 | 0.1 | 0.457 | 1 | 0.02316385 | 2.10154E-05 |
| Ccar1 | 1.05356E-05 | -0.652455431 | 0.857 | 0.9 | 0.255930795 | 0.57889084 | -0.139195317 | 0.9 | 0.957 | 1 | 0.57889084 | 2.10711E-05 |
| Taf3 | 1.05666E-05 | -1.450661409 | 0.345 | 0.547 | 0.256684728 | 0.764812212 | -0.389925213 | 0.6 | 0.568 | 1 | 0.764812212 | 2.11332E-05 |
| 1810030O07Rik | 1.06238E-05 | -1.528071165 | 0.176 | 0.412 | 0.25807414 | 0.573804308 | -0.339630097 | 0.3 | 0.38 | 1 | 0.573804308 | 2.12476E-05 |
| Gas1 | 1.06249E-05 | -1.303588504 | 0.294 | 0.512 | 0.258099842 | 0.090295982 | -3.28585884 | 0.3 | 0.494 | 1 | 0.090295982 | 2.12497E-05 |
| Dr1 | 1.06392E-05 | -1.376720568 | 0.185 | 0.429 | 0.258446664 | 0.042356918 | -1.654503434 | 0.1 | 0.417 | 1 | 0.042356918 | 2.12782E-05 |
| Cpsf4 | 1.06472E-05 | -2.185866545 | 0.067 | 0.271 | 0.258642754 | 0.956187157 | 0.176619654 | 0.2 | 0.201 | 1 | 0.956187157 | 2.12944E-05 |
| Rhob | 1.06854E-05 | -1.074441845 | 0.538 | 0.782 | 0.259570795 | 0.169003424 | -1.247615005 | 0.5 | 0.75 | 1 | 0.169003424 | 2.13708E-05 |
| Rhot1 | 1.06927E-05 | -1.246639968 | 0.361 | 0.571 | 0.259747118 | 0.583031438 | -0.339630097 | 0.7 | 0.667 | 1 | 0.583031438 | 2.13853E-05 |
| Wbp2 | 1.073E-05 | 0.83836161 | 0.731 | 0.576 | 0.26065358 | 0.103826019 | -0.725469565 | 0.4 | 0.651 | 1 | 0.103826019 | 2.14599E-05 |
| H2-Q4 | 1.07873E-05 | 2.115655339 | 0.42 | 0.2 | 0.262044844 | 0.032509931 | 1.608530972 | 0.4 | 0.157 | 1 | 0.032509931 | 2.15745E-05 |
| Prr12 | 1.08292E-05 | -3.129283017 | 0.025 | 0.2 | 0.263062103 | 0.3012433 | 1.695993813 | 0.3 | 0.198 | 1 | 0.3012433 | 2.16582E-05 |
| Ranbp3l | 1.0834E-05 | 4.762500686 | 0.109 | 0 | 0.263178472 | 0.006949972 | 3.339850003 | 0.2 | 0.034 | 1 | 0.006949972 | 2.16678E-05 |
| Prr14l | 1.0867E-05 | -0.944858446 | 0.496 | 0.7 | 0.263980554 | 0.494848643 | 0.387155718 | 0.8 | 0.799 | 1 | 0.494848643 | 2.17338E-05 |
| Fkbp4 | 1.08795E-05 | -1.258834417 | 0.269 | 0.512 | 0.264285321 | 0.036184656 | -1.339630097 | 0.3 | 0.593 | 1 | 0.036184656 | 2.17589E-05 |
| Pde6h | 1.08981E-05 | 2.479807755 | 0.387 | 0.176 | 0.264736908 | 0.798645264 | -0.111361109 | 0.1 | 0.074 | 1 | 0.798645264 | 2.17961E-05 |
| Ldlr | 1.09452E-05 | 0.811384591 | 0.824 | 0.618 | 0.265880704 | 0.034537093 | 1.252387162 | 0.8 | 0.596 | 1 | 0.034537093 | 2.18903E-05 |
| Ebna1bp2 | 1.0979E-05 | -0.975996958 | 0.605 | 0.718 | 0.266700748 | 0.516429453 | -0.339630097 | 0.6 | 0.608 | 1 | 0.516429453 | 2.19578E-05 |
| Chd8 | 1.10082E-05 | -1.441987566 | 0.378 | 0.565 | 0.267411204 | 0.010622324 | -1.589408406 | 0.4 | 0.738 | 1 | 0.010622324 | 2.20163E-05 |
| Slc26a7 | 1.10182E-05 | 2.906890596 | 0.143 | 0.012 | 0.267653124 | 0.002939478 | 2.695993813 | 0.2 | 0.028 | 1 | 0.002939478 | 2.20362E-05 |
| Snx31 | 0.343641216 | -1.070389328 | 0.025 | 0.047 | 1 | 1.1147E-05 | 4.210566986 | 0.2 | 0.012 | 0.270783232 | 0.343641216 | 2.22939E-05 |
| Usf3 | 1.12484E-05 | -1.891419187 | 0.176 | 0.406 | 0.273245187 | 0.193814878 | -1.058893689 | 0.3 | 0.463 | 1 | 0.193814878 | 2.24966E-05 |
| Fut2 | 1.12687E-05 | 3.836501268 | 0.126 | 0.006 | 0.273739975 | 0.009727839 | 4.017921908 | 0.1 | 0.009 | 1 | 0.009727839 | 2.25373E-05 |
| Stxbp6 | 1.13134E-05 | 0.894537905 | 0.58 | 0.312 | 0.274825797 | 0.32527397 | -0.119581616 | 0.6 | 0.392 | 1 | 0.32527397 | 2.26267E-05 |
| Ppp2ca | 1.14634E-05 | -0.534624018 | 0.95 | 0.953 | 0.278470104 | 0.000233932 | -1.341513921 | 0.9 | 0.994 | 1 | 0.000233932 | 2.29268E-05 |
| Ddx5 | 1.15071E-05 | -0.423932378 | 1 | 0.994 | 0.279531089 | 0.001526681 | -0.770987111 | 1 | 1 | 1 | 0.001526681 | 2.30141E-05 |
| Gm57411 | 0.016715804 | -2.655351829 | 0 | 0.047 | 1 | 1.15272E-05 | 4.280956314 | 0.2 | 0.012 | 0.280018913 | 0.016715804 | 2.30543E-05 |
| Gm16150 | 0.171447336 | 0.929610672 | 0.025 | 0.006 | 1 | 1.15272E-05 | 4.280956314 | 0.2 | 0.012 | 0.280018913 | 0.171447336 | 2.30543E-05 |
| 1110020A21Rik | 0.225145359 | 0.514573173 | 0.05 | 0.024 | 1 | 1.15272E-05 | 4.280956314 | 0.2 | 0.012 | 0.280018913 | 0.225145359 | 2.30543E-05 |
| Calr4 | 0.368010882 | 1.514573173 | 0.017 | 0.006 | 1 | 1.15272E-05 | 4.280956314 | 0.2 | 0.012 | 0.280018913 | 0.368010882 | 2.30543E-05 |
| Iqcg | 0.691605844 | -0.485426827 | 0.017 | 0.024 | 1 | 1.15272E-05 | 4.280956314 | 0.2 | 0.012 | 0.280018913 | 0.691605844 | 2.30543E-05 |
| Hmgb3 | 1.16531E-05 | -2.250961574 | 0.059 | 0.259 | 0.28307652 | 0.752109176 | -0.339630097 | 0.2 | 0.228 | 1 | 0.752109176 | 2.3306E-05 |
| Stard4 | 1.17971E-05 | 0.667729961 | 0.689 | 0.406 | 0.286575191 | 0.256311661 | 0.554125097 | 0.6 | 0.42 | 1 | 0.256311661 | 2.35941E-05 |
| Marchf7 | 1.1804E-05 | -0.841991924 | 0.824 | 0.871 | 0.286742993 | 0.294911654 | -0.553101402 | 0.8 | 0.932 | 1 | 0.294911654 | 2.36079E-05 |
| Ankdd1a | 0.163453432 | -1.485426827 | 0.017 | 0.047 | 1 | 1.19143E-05 | 4.017921908 | 0.2 | 0.012 | 0.289421806 | 0.163453432 | 2.38284E-05 |
| Map2k5 | 1.19486E-05 | -1.336283388 | 0.286 | 0.524 | 0.290256227 | 0.878419294 | -0.114302384 | 0.8 | 0.66 | 1 | 0.878419294 | 2.38971E-05 |
| Pbx1 | 1.19661E-05 | -0.638364914 | 0.95 | 0.947 | 0.290679961 | 0.079238466 | -0.828658615 | 1 | 0.96 | 1 | 0.079238466 | 2.3932E-05 |
| Dip2c | 1.19874E-05 | -1.065666918 | 0.664 | 0.824 | 0.291198497 | 0.84756623 | -0.186768512 | 0.9 | 0.895 | 1 | 0.84756623 | 2.39747E-05 |
| Wdr20 | 1.2004E-05 | -1.379933698 | 0.252 | 0.476 | 0.291600757 | 0.406457603 | -0.489872732 | 0.5 | 0.577 | 1 | 0.406457603 | 2.40078E-05 |
| Rbbp4 | 1.22381E-05 | -0.612606908 | 0.874 | 0.918 | 0.297286712 | 0.109227462 | -0.479496075 | 0.6 | 0.929 | 1 | 0.109227462 | 2.4476E-05 |
| Ret | 0.000428393 | 1.514573173 | 0.202 | 0.065 | 1 | 1.231E-05 | 4.017921908 | 0.2 | 0.012 | 0.299034612 | 0.000428393 | 2.46199E-05 |
| Gm56933 | 0.343608466 | -0.622930351 | 0.025 | 0.047 | 1 | 1.231E-05 | 3.795529487 | 0.2 | 0.012 | 0.299034612 | 0.343608466 | 2.46199E-05 |
| Gm49179 | 0.381775243 | 0.514573173 | 0.042 | 0.024 | 1 | 1.231E-05 | 4.017921908 | 0.2 | 0.012 | 0.299034612 | 0.381775243 | 2.46199E-05 |
| Katnal2 | 0.602438591 | 0.777607579 | 0.034 | 0.024 | 1 | 1.231E-05 | 3.795529487 | 0.2 | 0.012 | 0.299034612 | 0.602438591 | 2.46199E-05 |
| Kcnu1 | 0.856663367 | -0.163498732 | 0.034 | 0.029 | 1 | 1.231E-05 | 4.017921908 | 0.2 | 0.012 | 0.299034612 | 0.856663367 | 2.46199E-05 |
| Rbak | 1.2351E-05 | -2.891419187 | 0.025 | 0.2 | 0.300031109 | 0.494726313 | 0.193493473 | 0.1 | 0.204 | 1 | 0.494726313 | 2.47019E-05 |
| Osmr | 1.25576E-05 | 0.522685819 | 0.899 | 0.671 | 0.305049125 | 0.007229528 | 1.132347544 | 0.9 | 0.583 | 1 | 0.007229528 | 2.5115E-05 |
| Izumo4 | 0.172045228 | 0.312939312 | 0.277 | 0.206 | 1 | 1.26325E-05 | 2.695993813 | 0.6 | 0.133 | 0.30686761 | 0.172045228 | 2.52648E-05 |
| Pbx3 | 1.26505E-05 | -1.799535418 | 0.303 | 0.524 | 0.307305934 | 0.004813646 | -2.965071667 | 0.2 | 0.648 | 1 | 0.004813646 | 2.53008E-05 |
| Oca2 | 1.26721E-05 | 2.762500686 | 0.168 | 0.024 | 0.307830872 | 0.004092787 | 2.017921908 | 0.5 | 0.154 | 1 | 0.004092787 | 2.53441E-05 |
| Ttc19 | 1.26863E-05 | -0.959912792 | 0.571 | 0.7 | 0.308175124 | 0.003538791 | -1.837984759 | 0.4 | 0.756 | 1 | 0.003538791 | 2.53724E-05 |
| Srxn1 | 1.26976E-05 | 1.209992322 | 0.571 | 0.324 | 0.30844954 | 0.60902453 | 0.272782382 | 0.3 | 0.383 | 1 | 0.60902453 | 2.5395E-05 |
| Hebp2 | 1.27528E-05 | 1.592575685 | 0.387 | 0.171 | 0.309791956 | 0.044960121 | 1.339850003 | 0.4 | 0.151 | 1 | 0.044960121 | 2.55055E-05 |
| Sdf4 | 1.27926E-05 | 0.45503985 | 0.966 | 0.888 | 0.310758749 | 0.007846955 | 0.730671265 | 1 | 0.883 | 1 | 0.007846955 | 2.55851E-05 |
| Pcdh18 | 1.28514E-05 | -5.100136671 | 0 | 0.147 | 0.312185793 | 0.226355142 | -1.5519337 | 0 | 0.13 | 1 | 0.226355142 | 2.57026E-05 |
| Gm4258 | 1.28747E-05 | -6.714245518 | 0 | 0.147 | 0.312752932 | 0.475021253 | -2.763437806 | 0 | 0.049 | 1 | 0.475021253 | 2.57493E-05 |
| Dock3 | 0.177603182 | -0.678071905 | 0.025 | 0.059 | 1 | 1.28883E-05 | 3.558490289 | 0.4 | 0.056 | 0.313082463 | 0.177603182 | 2.57764E-05 |
| Dop1b | 0.346260763 | 0.360767837 | 0.21 | 0.259 | 1 | 1.2942E-05 | 2.387871518 | 0.8 | 0.253 | 0.314387547 | 0.346260763 | 2.58839E-05 |
| Ythdf1 | 1.29941E-05 | -1.205741181 | 0.378 | 0.582 | 0.315651677 | 0.030770578 | -1.251983975 | 0.5 | 0.694 | 1 | 0.030770578 | 2.59879E-05 |
| Eif4h | 1.30736E-05 | -0.634023319 | 0.849 | 0.929 | 0.317582763 | 0.124212271 | -0.504275797 | 0.8 | 0.963 | 1 | 0.124212271 | 2.61469E-05 |
| Bglap3 | 0.093245792 | -1.807354922 | 0 | 0.024 | 1 | 1.31554E-05 | 3.280956314 | 0.2 | 0.012 | 0.319570254 | 0.093245792 | 2.63106E-05 |
| Frmd3 | 0.147029286 | -1.807354922 | 0 | 0.018 | 1 | 1.31554E-05 | 3.602884409 | 0.2 | 0.012 | 0.319570254 | 0.147029286 | 2.63106E-05 |

| | | | | | | | | | | | | |
|---|---|---|---|---|---|---|---|---|---|---|---|---|
| Lrp3 | 1.31906E-05 | -3.14839184 | 0.017 | 0.182 | 0.320426632 | 0.091789435 | -1.505640048 | 0 | 0.225 | 1 | 0.091789435 | 2.63811E-05 |
| Bvht | 0.619390946 | -0.655351829 | 0.025 | 0.035 | 1 | 1.31933E-05 | 3.902444691 | 0.3 | 0.031 | 0.320491019 | 0.619390946 | 2.63864E-05 |
| Dync2h1 | 1.32612E-05 | -1.254493265 | 0.504 | 0.706 | 0.322141024 | 0.809510769 | 0.237703116 | 0.7 | 0.698 | 1 | 0.809510769 | 2.65222E-05 |
| Anks3 | 1.32904E-05 | -1.693022247 | 0.16 | 0.382 | 0.32284957 | 0.347432904 | -0.789433014 | 0.3 | 0.404 | 1 | 0.347432904 | 2.65806E-05 |
| Kif19b | 1.33146E-05 | -3.029747343 | 0.034 | 0.212 | 0.323437203 | 0.782931319 | -0.556986928 | 0.2 | 0.216 | 1 | 0.782931319 | 2.66289E-05 |
| Ap3b1 | 1.34559E-05 | -0.779460916 | 0.723 | 0.841 | 0.326869713 | 0.547709608 | -0.149328959 | 0.8 | 0.904 | 1 | 0.547709608 | 2.69115E-05 |
| Sos2 | 1.35029E-05 | -1.26072054 | 0.513 | 0.641 | 0.328012307 | 0.098445451 | -0.932690065 | 0.7 | 0.809 | 1 | 0.098445451 | 2.70056E-05 |
| Cluap1 | 1.363E-05 | -1.765534746 | 0.151 | 0.371 | 0.331101053 | 0.184946888 | 0.921060369 | 0.5 | 0.309 | 1 | 0.184946888 | 2.72599E-05 |
| Prkcg | 1.37148E-05 | -2.353323291 | 0.202 | 0.424 | 0.333161069 | 0.154011806 | 0.354956895 | 0.5 | 0.309 | 1 | 0.154011806 | 2.74295E-05 |
| Ogt | 1.37599E-05 | -0.756196882 | 0.874 | 0.935 | 0.334255682 | 0.202767841 | -0.425021588 | 1 | 0.969 | 1 | 0.202767841 | 2.75196E-05 |
| Ubc | 1.37773E-05 | 0.713544584 | 0.992 | 0.982 | 0.334679008 | 0.939014713 | 0.658651659 | 0.9 | 0.994 | 1 | 0.939014713 | 2.75545E-05 |
| Npm3 | 1.38286E-05 | -1.359895945 | 0.244 | 0.476 | 0.33592394 | 0.013006544 | -2.716787712 | 0 | 0.401 | 1 | 0.013006544 | 2.7657E-05 |
| Plod2 | 1.38317E-05 | 0.375453787 | 1 | 1 | 0.336000436 | 0.15100644 | 0.19116574 | 1 | 0.944 | 1 | 0.15100644 | 2.76633E-05 |
| Srebf1 | 0.004446903 | 0.960829403 | 0.462 | 0.312 | 1 | 1.39132E-05 | 2.476548676 | 0.9 | 0.321 | 0.337980639 | 0.004446903 | 2.78263E-05 |
| Macrod2 | 1.39321E-05 | 0.999330691 | 0.622 | 0.335 | 0.338438917 | 0.053627769 | 1.742643906 | 0.5 | 0.293 | 1 | 0.053627769 | 2.7864E-05 |
| Mfsd13b | 0.602438591 | 1.362570079 | 0.034 | 0.024 | 1 | 1.40542E-05 | 2.902444691 | 0.2 | 0.012 | 0.341405324 | 0.602438591 | 2.81083E-05 |
| Mob3b | 1.40818E-05 | -3.412781525 | 0.076 | 0.271 | 0.342076223 | 0.066615787 | -2.91281543 | 0 | 0.259 | 1 | 0.066615787 | 2.81635E-05 |
| Abhd2 | 1.41008E-05 | 0.406749528 | 0.966 | 0.918 | 0.342537794 | 0.034808746 | 0.501885204 | 1 | 0.963 | 1 | 0.034808746 | 2.82015E-05 |
| Mtr | 1.41214E-05 | -1.918386234 | 0.151 | 0.371 | 0.343037239 | 0.626833288 | -0.138582578 | 0.5 | 0.355 | 1 | 0.626833288 | 2.82426E-05 |
| Hdac4 | 1.41234E-05 | -1.114907995 | 0.513 | 0.724 | 0.343084927 | 0.205564753 | -0.654503434 | 0.7 | 0.765 | 1 | 0.205564753 | 2.82465E-05 |
| Ccnyl1 | 1.42561E-05 | -1.736388401 | 0.118 | 0.329 | 0.346308604 | 0.748324617 | -0.169076607 | 0.3 | 0.373 | 1 | 0.748324617 | 2.85119E-05 |
| Atp2b4 | 1.43363E-05 | -1.46587403 | 0.454 | 0.618 | 0.348257429 | 0.678526144 | -0.852442812 | 0.5 | 0.515 | 1 | 0.678526144 | 2.86724E-05 |
| Cd55 | 1.44357E-05 | -2.794179533 | 0.059 | 0.253 | 0.350672886 | 0.541265388 | -0.315978829 | 0.4 | 0.29 | 1 | 0.541265388 | 2.88713E-05 |
| Stk38 | 1.44492E-05 | -0.9789663 | 0.403 | 0.659 | 0.350999885 | 0.662505208 | -0.31230551 | 0.8 | 0.704 | 1 | 0.662505208 | 2.88982E-05 |
| Zmym2 | 1.44514E-05 | -1.02322422 | 0.538 | 0.729 | 0.351053586 | 0.065341275 | -0.856064945 | 0.8 | 0.87 | 1 | 0.065341275 | 2.89026E-05 |
| Gm17690 | 0.670978617 | 0.215012891 | 0.084 | 0.071 | 1 | 1.44534E-05 | 3.532495081 | 0.3 | 0.031 | 0.351102424 | 0.670978617 | 2.89066E-05 |
| Boc | 1.45467E-05 | -1.58058406 | 0.319 | 0.529 | 0.353369184 | 0.006834782 | -2.53820091 | 0.3 | 0.67 | 1 | 0.006834782 | 2.90932E-05 |
| Vps13d | 1.47346E-05 | -0.927431374 | 0.462 | 0.682 | 0.357933329 | 0.087397152 | -0.963393529 | 0.7 | 0.852 | 1 | 0.087397152 | 2.9469E-05 |
| Cd109 | 1.47562E-05 | -0.801748133 | 0.874 | 0.947 | 0.358457024 | 0.438303056 | -0.428258252 | 1 | 0.932 | 1 | 0.438303056 | 2.95121E-05 |
| Rfxap | 1.48792E-05 | -1.852209158 | 0.143 | 0.359 | 0.361445408 | 0.615004899 | -0.518130992 | 0.4 | 0.407 | 1 | 0.615004899 | 2.97582E-05 |
| Angpt1 | 1.49831E-05 | -4.171927354 | 0.008 | 0.165 | 0.363968789 | 0.603715608 | -0.69173634 | 0.1 | 0.164 | 1 | 0.603715608 | 2.99659E-05 |
| Slc25a5 | 1.51846E-05 | -0.873294125 | 0.714 | 0.865 | 0.368864773 | 0.157563129 | -0.762056575 | 0.7 | 0.907 | 1 | 0.157563129 | 3.0369E-05 |
| Pik3ca | 1.5241E-05 | -0.874929234 | 0.697 | 0.782 | 0.370234491 | 0.014488215 | -1.156619722 | 0.6 | 0.87 | 1 | 0.014488215 | 3.04818E-05 |
| Slc8a3 | 0.008618117 | -0.638626681 | 0.613 | 0.371 | 1 | 1.52603E-05 | 1.966575028 | 0.9 | 0.278 | 0.370702826 | 0.008618117 | 3.05203E-05 |
| Foxo1 | 1.52942E-05 | 0.107972511 | 0.958 | 0.924 | 0.371526184 | 0.16893403 | -0.729312022 | 1 | 0.966 | 1 | 0.16893403 | 3.05881E-05 |
| Cct4 | 1.53123E-05 | -0.686725217 | 0.748 | 0.841 | 0.371966359 | 0.002983592 | -1.400867726 | 0.4 | 0.87 | 1 | 0.002983592 | 3.06244E-05 |
| Prkch | 1.53231E-05 | 1.10629304 | 0.361 | 0.141 | 0.372229293 | 0.706450078 | 0.795529487 | 0.2 | 0.16 | 1 | 0.706450078 | 3.0646E-05 |
| Sertad1 | 1.53592E-05 | -1.033630778 | 0.412 | 0.612 | 0.373105647 | 0.207410686 | -0.306580131 | 0.4 | 0.688 | 1 | 0.207410686 | 3.07182E-05 |
| Zbtb33 | 1.54161E-05 | -1.900464326 | 0.143 | 0.359 | 0.374486771 | 0.803446432 | -0.239465935 | 0.3 | 0.302 | 1 | 0.803446432 | 3.08319E-05 |
| Foxp2 | 1.54265E-05 | -2.096384536 | 0.218 | 0.429 | 0.374741666 | 0.14921654 | -1.448518769 | 0.2 | 0.426 | 1 | 0.14921654 | 3.08529E-05 |
| Slc7a1 | 1.54598E-05 | -1.52007297 | 0.529 | 0.682 | 0.375550588 | 0.155848266 | -1.082214763 | 0.8 | 0.787 | 1 | 0.155848266 | 3.09195E-05 |
| Arhgef37 | 0.767563352 | 0.514573173 | 0.042 | 0.035 | 1 | 1.54741E-05 | 3.432959407 | 0.3 | 0.031 | 0.375897429 | 0.767563352 | 3.0948E-05 |
| Ankhd1 | 1.55901E-05 | -0.879990856 | 0.773 | 0.882 | 0.378713828 | 0.013615298 | -1.223998539 | 0.7 | 0.907 | 1 | 0.013615298 | 3.11799E-05 |
| Cks2 | 1.56286E-05 | -3.029747343 | 0.042 | 0.224 | 0.379649341 | 0.324653928 | 0.785261151 | 0.3 | 0.179 | 1 | 0.324653928 | 3.12569E-05 |
| Gpn3 | 1.5726E-05 | -1.183157258 | 0.252 | 0.5 | 0.382015782 | 0.167307004 | -0.719043686 | 0.1 | 0.33 | 1 | 0.167307004 | 3.14517E-05 |
| Cenpb | 1.57629E-05 | -0.974174013 | 0.311 | 0.571 | 0.382912461 | 0.282392602 | -0.386832618 | 0.6 | 0.71 | 1 | 0.282392602 | 3.15256E-05 |
| Fnbp4 | 1.57974E-05 | -0.850711291 | 0.647 | 0.818 | 0.383750729 | 0.407617225 | -0.326906589 | 0.9 | 0.92 | 1 | 0.407617225 | 3.15946E-05 |
| Rsl24d1 | 1.58137E-05 | -1.282439805 | 0.252 | 0.482 | 0.384147497 | 0.002944427 | -2.924592597 | 0 | 0.503 | 1 | 0.002944427 | 3.16272E-05 |
| Sox6 | 1.58882E-05 | -0.632323486 | 0.992 | 0.994 | 0.385955853 | 0.171581343 | -0.560103116 | 0.9 | 0.972 | 1 | 0.171581343 | 3.17761E-05 |
| Akt1 | 1.6051E-05 | -1.005538184 | 0.555 | 0.694 | 0.389909918 | 0.004554106 | -1.374395515 | 0.4 | 0.824 | 1 | 0.004554106 | 3.21017E-05 |
| Fam3c | 1.60549E-05 | 0.714510743 | 0.824 | 0.647 | 0.390005674 | 0.864859848 | 0.29629686 | 0.6 | 0.596 | 1 | 0.864859848 | 3.21095E-05 |
| Ncs1 | 1.61273E-05 | -1.674460652 | 0.118 | 0.341 | 0.391764833 | 0.103875657 | -1.625934282 | 0.2 | 0.423 | 1 | 0.103875657 | 3.22544E-05 |
| Stx18 | 1.61653E-05 | -1.182864057 | 0.454 | 0.629 | 0.392686874 | 0.244431996 | -0.742890428 | 0.4 | 0.627 | 1 | 0.244431996 | 3.23303E-05 |
| Tfam | 1.62048E-05 | -1.416164165 | 0.21 | 0.435 | 0.393646665 | 0.382094567 | -0.457811523 | 0.2 | 0.336 | 1 | 0.382094567 | 3.24093E-05 |
| Hp1bp3 | 1.63169E-05 | -0.672304962 | 0.857 | 0.929 | 0.3963704 | 0.009933075 | -0.991330405 | 0.6 | 0.96 | 1 | 0.009933075 | 3.26336E-05 |
| E2f6 | 1.64258E-05 | -2.11345805 | 0.076 | 0.276 | 0.399015564 | 0.053741024 | -1.55950692 | 0.1 | 0.398 | 1 | 0.053741024 | 3.28513E-05 |
| Immp2l | 1.64333E-05 | 0.864515644 | 0.622 | 0.341 | 0.399197782 | 0.004628622 | 2.041229281 | 0.7 | 0.426 | 1 | 0.004628622 | 3.28663E-05 |
| Mnat1 | 1.66371E-05 | -1.017440448 | 0.538 | 0.718 | 0.404147842 | 0.316271393 | 0.421445419 | 0.8 | 0.707 | 1 | 0.316271393 | 3.32739E-05 |
| Lratd2 | 1.67605E-05 | -2.448900951 | 0.076 | 0.271 | 0.407145858 | 0.646190967 | 0.31748219 | 0.2 | 0.321 | 1 | 0.646190967 | 3.35207E-05 |
| Ssbp3 | 1.70722E-05 | -1.157100961 | 0.563 | 0.724 | 0.414718031 | 0.787295279 | -0.122167864 | 1 | 0.79 | 1 | 0.787295279 | 3.41441E-05 |
| Map1b | 1.71349E-05 | -3.138000234 | 0.076 | 0.276 | 0.416241872 | 0.402006175 | -1.879923548 | 0.3 | 0.377 | 1 | 0.402006175 | 3.42696E-05 |
| Cited1 | 1.71386E-05 | 2.836501268 | 0.176 | 0.029 | 0.416329807 | 0.557756607 | -0.408342847 | 0 | 0.034 | 1 | 0.557756607 | 3.42768E-05 |
| Kif1c | 1.72862E-05 | -1.49872365 | 0.202 | 0.418 | 0.419916913 | 0.910403513 | 0.222062625 | 0.5 | 0.444 | 1 | 0.910403513 | 3.45721E-05 |
| Polr2b | 1.72989E-05 | -1.357270476 | 0.303 | 0.524 | 0.420224133 | 0.516452181 | -0.35463226 | 0.5 | 0.559 | 1 | 0.516452181 | 3.45974E-05 |
| Sars | 1.73217E-05 | -1.104740833 | 0.445 | 0.635 | 0.42077842 | 0.132702429 | -0.763437806 | 0.4 | 0.651 | 1 | 0.132702429 | 3.46431E-05 |
| Bcl2l1 | 1.7429E-05 | 1.058893689 | 0.555 | 0.318 | 0.423385984 | 0.165414688 | 0.787355085 | 0.6 | 0.497 | 1 | 0.165414688 | 3.48578E-05 |
| Cdk13 | 1.74975E-05 | -0.888391494 | 0.782 | 0.865 | 0.425048502 | 0.042125223 | -0.603907975 | 0.9 | 0.969 | 1 | 0.042125223 | 3.49946E-05 |
| Tmem204 | 1.75472E-05 | -2.655351829 | 0.059 | 0.247 | 0.426257548 | 0.298574031 | -0.408342847 | 0 | 0.099 | 1 | 0.298574031 | 3.50942E-05 |
| Heg1 | 0.7489328 | -0.68182404 | 0.235 | 0.241 | 1 | 1.76577E-05 | 1.744903414 | 0.6 | 0.12 | 0.428940865 | 0.7489328 | 3.53151E-05 |
| Ctnnb1 | 1.80621E-05 | -0.594860068 | 0.95 | 0.947 | 0.438765552 | 0.07757074 | -0.518496741 | 0.9 | 1 | 1 | 0.07757074 | 3.6124E-05 |
| Samd12 | 1.82795E-05 | 1.434402824 | 0.202 | 0.041 | 0.444045038 | 0.001054794 | 2.147557188 | 0.4 | 0.086 | 1 | 0.001054794 | 3.65586E-05 |
| Atad2b | 1.83448E-05 | -1.171167098 | 0.546 | 0.712 | 0.445631088 | 0.540162408 | -0.28128611 | 0.8 | 0.812 | 1 | 0.540162408 | 3.66892E-05 |
| Dysf | 1.83453E-05 | 0.990672198 | 0.563 | 0.294 | 0.445643195 | 0.42303277 | 0.301714874 | 0.4 | 0.269 | 1 | 0.42303277 | 3.66902E-05 |
| Mbip | 1.86288E-05 | -1.622930351 | 0.151 | 0.371 | 0.452531012 | 0.517109925 | -0.339630097 | 0.2 | 0.299 | 1 | 0.517109925 | 3.72573E-05 |
| Efemp2 | 1.87204E-05 | 0.552666344 | 0.882 | 0.718 | 0.454756216 | 0.186869046 | 0.584962501 | 0.8 | 0.642 | 1 | 0.186869046 | 3.74405E-05 |
| Dcaf10 | 1.88075E-05 | -0.970853654 | 0.504 | 0.712 | 0.456871559 | 0.051806174 | -0.946812762 | 0.5 | 0.775 | 1 | 0.051806174 | 3.76146E-05 |
| Tmem45a | 1.88317E-05 | 0.465760923 | 0.95 | 0.824 | 0.457459906 | 0.046916499 | 0.791670291 | 1 | 0.806 | 1 | 0.046916499 | 3.76631E-05 |
| Smg5 | 1.88667E-05 | -1.519374159 | 0.134 | 0.359 | 0.458310278 | 0.151058611 | -1.125036046 | 0.2 | 0.404 | 1 | 0.151058611 | 3.77331E-05 |
| Rest | 1.88907E-05 | -1.195085075 | 0.479 | 0.647 | 0.458892668 | 0.006401804 | -1.778349515 | 0.6 | 0.781 | 1 | 0.006401804 | 3.7781E-05 |
| 2900005J15Rik | 0.370352969 | 0.415037499 | 0.101 | 0.071 | 1 | 1.89166E-05 | 2.930459067 | 0.3 | 0.031 | 0.459521148 | 0.370352969 | 3.78328E-05 |
| Ube2e3 | 1.90163E-05 | -0.818594673 | 0.773 | 0.841 | 0.461943536 | 0.013813747 | -1.202126573 | 0.6 | 0.858 | 1 | 0.013813747 | 3.80322E-05 |
| Zfp207 | 1.90638E-05 | -0.688394664 | 0.849 | 0.9 | 0.46309782 | 0.068212103 | -0.611434712 | 0.8 | 0.926 | 1 | 0.068212103 | 3.81272E-05 |
| Ppib | 1.91092E-05 | -0.408587846 | 0.983 | 0.976 | 0.464201146 | 0.230054267 | -0.389662271 | 0.9 | 0.991 | 1 | 0.230054267 | 3.82181E-05 |
| Dhfr | 1.91184E-05 | -3.68182404 | 0.017 | 0.176 | 0.464425256 | 0.024779198 | 2.591657153 | 0.3 | 0.096 | 1 | 0.024779198 | 3.82365E-05 |
| Zfp280d | 1.92441E-05 | -0.778734244 | 0.723 | 0.829 | 0.467478709 | 0.034989566 | -0.913997482 | 0.8 | 0.926 | 1 | 0.034989566 | 3.84879E-05 |
| A630089N07Rik | 1.92503E-05 | -2.292781749 | 0.076 | 0.271 | 0.467627733 | 0.152959385 | -1.876895855 | 0 | 0.173 | 1 | 0.152959385 | 3.85002E-05 |
| Tax1bp3 | 1.93681E-05 | -1.333423734 | 0.294 | 0.512 | 0.470489747 | 0.299216879 | -0.518130992 | 0.3 | 0.466 | 1 | 0.299216879 | 3.87358E-05 |
| Adamts10 | 1.9602E-05 | -1.547203025 | 0.277 | 0.488 | 0.476172712 | 0.895710491 | -0.237578825 | 0.7 | 0.528 | 1 | 0.895710491 | 3.92037E-05 |
| Aplp2 | 1.96828E-05 | -0.563548349 | 0.992 | 0.988 | 0.478135644 | 0.076279846 | -0.520884295 | 1 | 0.994 | 1 | 0.076279846 | 3.93653E-05 |
| Arxes1 | 1.97183E-05 | -4.655351829 | 0 | 0.141 | 0.478996697 | 0.927831145 | 0.847996907 | 0.1 | 0.09 | 1 | 0.927831145 | 3.94362E-05 |
| Txnl1 | 1.97242E-05 | -0.675758039 | 0.899 | 0.876 | 0.479141157 | 0.01870814 | -0.932390968 | 0.7 | 0.954 | 1 | 0.01870814 | 3.94481E-05 |
| Cygb | 1.97893E-05 | -4.770829046 | 0 | 0.141 | 0.480721195 | 0.489975126 | 0.159940913 | 0 | 0.046 | 1 | 0.489975126 | 3.95782E-05 |
| Tead2 | 1.98072E-05 | -5.008988783 | 0 | 0.141 | 0.481155587 | 0.259962028 | -0.864721141 | 0 | 0.114 | 1 | 0.259962028 | 3.96139E-05 |
| Mad1l1 | 1.98322E-05 | -1.996388746 | 0.134 | 0.347 | 0.481764938 | 0.637812205 | -0.767180207 | 0.5 | 0.485 | 1 | 0.637812205 | 3.96641E-05 |
| Tpr | 1.98659E-05 | -0.54946066 | 0.992 | 1 | 0.482581613 | 0.006395997 | -0.985921621 | 0.7 | 0.991 | 1 | 0.006395997 | 3.97313E-05 |
| Rock2 | 1.98713E-05 | -0.704951922 | 0.983 | 1 | 0.482713431 | 0.008931076 | -0.738910523 | 1 | 1 | 1 | 0.008931076 | 3.97422E-05 |
| Plscr1 | 1.99449E-05 | 1.468444002 | 0.563 | 0.359 | 0.484501394 | 0.990783811 | -0.33375353 | 0.4 | 0.383 | 1 | 0.990783811 | 3.98894E-05 |
| Tgfb1i1 | 1.99482E-05 | -1.396493099 | 0.286 | 0.506 | 0.484580784 | 0.329522482 | 0.34807051 | 0.7 | 0.478 | 1 | 0.329522482 | 3.98959E-05 |

| | | | | | | | | | | | | |
|---|---|---|---|---|---|---|---|---|---|---|---|---|
| Hdlbp | 1.99672E-05 | -0.355583649 | 1 | 1 | 0.485043552 | 0.198168592 | -0.350884879 | 1 | 1 | 1 | 0.198168592 | 3.9934E-05 |
| Nfrkb | 1.99699E-05 | -1.740240726 | 0.168 | 0.376 | 0.485109809 | 0.173633995 | -1.152003093 | 0.2 | 0.386 | 1 | 0.173633995 | 3.99395E-05 |
| Pdzd8 | 2.01543E-05 | -1.341416524 | 0.319 | 0.529 | 0.489589217 | 0.118620378 | -0.576259123 | 0.6 | 0.799 | 1 | 0.118620378 | 4.03083E-05 |
| Vcl | 2.02821E-05 | 0.923946373 | 0.958 | 0.871 | 0.492691832 | 0.000661738 | 2.088000344 | 1 | 0.923 | 1 | 0.000661738 | 4.05637E-05 |
| Apba1 | 2.03424E-05 | -2.655351829 | 0.076 | 0.271 | 0.494158299 | 0.442275763 | 0.465696194 | 0.4 | 0.296 | 1 | 0.442275763 | 4.06844E-05 |
| Sfxn5 | 2.04402E-05 | -1.736965594 | 0.109 | 0.324 | 0.496533404 | 0.061881043 | -1.360011597 | 0.2 | 0.515 | 1 | 0.061881043 | 4.088E-05 |
| Rnf2 | 2.04616E-05 | -0.971873587 | 0.538 | 0.706 | 0.497052146 | 0.607152239 | -0.117530876 | 0.7 | 0.781 | 1 | 0.607152239 | 4.09227E-05 |
| Fam20c | 2.0467E-05 | -1.22080617 | 0.412 | 0.624 | 0.497183622 | 0.046034407 | -1.370956431 | 0.5 | 0.722 | 1 | 0.046034407 | 4.09335E-05 |
| Hexim1 | 2.05471E-05 | -1.47720632 | 0.471 | 0.629 | 0.499130495 | 0.013741859 | -1.149997958 | 0.4 | 0.806 | 1 | 0.013741859 | 4.10938E-05 |
| Gm19510 | 2.07719E-05 | 1.929610672 | 0.151 | 0.018 | 0.504591683 | 0.000103743 | 4.432959407 | 0.1 | 0.003 | 1 | 0.000103743 | 4.15434E-05 |
| Plag1 | 2.08539E-05 | -3.944858446 | 0.008 | 0.159 | 0.506582866 | 0.267393657 | -1.191531458 | 0 | 0.111 | 1 | 0.267393657 | 4.17074E-05 |
| Tsr1 | 2.09267E-05 | -1.140436825 | 0.361 | 0.576 | 0.50835183 | 0.002556841 | -2.730270942 | 0.1 | 0.599 | 1 | 0.002556841 | 4.1853E-05 |
| Agps | 2.09823E-05 | -1.361852142 | 0.387 | 0.582 | 0.509701764 | 0.398496877 | -0.564726727 | 0.6 | 0.707 | 1 | 0.398496877 | 4.19641E-05 |
| Slc23a2 | 2.10123E-05 | 0.756424954 | 0.815 | 0.629 | 0.510430862 | 0.00186566 | 1.040641984 | 1 | 0.744 | 1 | 0.00186566 | 4.20242E-05 |
| Wnk2 | 2.11426E-05 | -3.14839184 | 0.042 | 0.218 | 0.513595832 | 0.510757291 | -0.827568143 | 0.3 | 0.355 | 1 | 0.510757291 | 4.22847E-05 |
| Psma3 | 2.1251E-05 | -0.468513444 | 0.966 | 0.982 | 0.516229765 | 0.047919509 | -0.652677237 | 0.8 | 0.972 | 1 | 0.047919509 | 4.25016E-05 |
| Gm41724 | 2.13071E-05 | 1.493199522 | 0.244 | 0.065 | 0.517592321 | 0.077136918 | 2.31748219 | 0.2 | 0.062 | 1 | 0.077136918 | 4.26138E-05 |
| Mapkapk5 | 2.14776E-05 | -1.315501826 | 0.303 | 0.512 | 0.521733156 | 0.026309818 | -1.654503434 | 0.2 | 0.537 | 1 | 0.026309818 | 4.29547E-05 |
| Sim2 | 2.15146E-05 | 0.591977677 | 0.664 | 0.365 | 0.522631522 | 0.010968744 | 0.967295835 | 0.9 | 0.457 | 1 | 0.010968744 | 4.30286E-05 |
| Dcaf5 | 2.16483E-05 | -1.626782676 | 0.185 | 0.406 | 0.525879576 | 0.176548594 | -0.470721632 | 0.4 | 0.645 | 1 | 0.176548594 | 4.32961E-05 |
| Shfl | 2.18161E-05 | -2.687060688 | 0.034 | 0.206 | 0.52995572 | 0.180369815 | -1.026472211 | 0 | 0.154 | 1 | 0.180369815 | 4.36316E-05 |
| Elovl7 | 6.82126E-05 | 1.470179053 | 0.361 | 0.159 | 1 | 2.19273E-05 | 2.792501794 | 0.6 | 0.136 | 0.53265789 | 6.82126E-05 | 4.38541E-05 |
| Zfp318 | 2.20104E-05 | -1.352789059 | 0.252 | 0.471 | 0.534675899 | 0.983246847 | 0.249737583 | 0.4 | 0.386 | 1 | 0.983246847 | 4.40203E-05 |
| Mid1 | 2.2268E-05 | -1.088471687 | 0.487 | 0.7 | 0.540933099 | 0.027369451 | -1.703177281 | 0.2 | 0.593 | 1 | 0.027369451 | 4.45354E-05 |
| Scnm1 | 2.23324E-05 | -2.343407822 | 0.059 | 0.247 | 0.542499015 | 0.681831366 | 0.290001453 | 0.1 | 0.145 | 1 | 0.681831366 | 4.46643E-05 |
| Chordc1 | 2.23656E-05 | -1.00270252 | 0.42 | 0.635 | 0.543304636 | 0.006634596 | -1.720845929 | 0.3 | 0.71 | 1 | 0.006634596 | 4.47307E-05 |
| Atp8b2 | 2.26933E-05 | -0.813114191 | 0.529 | 0.694 | 0.551265559 | 0.074463575 | -0.876895855 | 0.4 | 0.707 | 1 | 0.074463575 | 4.53861E-05 |
| Efna2 | 2.27627E-05 | -1.691877705 | 0.05 | 0.241 | 0.552951277 | 0.30625785 | 1.18142064 | 0.3 | 0.176 | 1 | 0.30625785 | 4.55249E-05 |
| Capn15 | 2.28423E-05 | -0.824737 | 0.319 | 0.559 | 0.554885253 | 0.494686289 | -0.38466385 | 0.6 | 0.617 | 1 | 0.494686289 | 4.56841E-05 |
| Tcof1 | 2.30008E-05 | -1.722466024 | 0.151 | 0.365 | 0.558735097 | 0.614164266 | 0.150025444 | 0.2 | 0.315 | 1 | 0.614164266 | 4.6001E-05 |
| Pwp1 | 2.30454E-05 | -1.86393845 | 0.118 | 0.324 | 0.559818282 | 0.271130687 | -0.776493958 | 0.1 | 0.247 | 1 | 0.271130687 | 4.60902E-05 |
| Smc5 | 2.31126E-05 | -0.921997488 | 0.63 | 0.741 | 0.561450695 | 0.67431411 | 0.189726643 | 0.7 | 0.781 | 1 | 0.67431411 | 4.62246E-05 |
| Zfp248 | 2.31277E-05 | -1.24726709 | 0.101 | 0.318 | 0.561819038 | 0.303001748 | -0.68251781 | 0.1 | 0.253 | 1 | 0.303001748 | 4.62549E-05 |
| Nos1ap | 2.31665E-05 | 0.514573173 | 0.681 | 0.512 | 0.562761462 | 0.058553402 | 0.546136862 | 0.9 | 0.747 | 1 | 0.058553402 | 4.63325E-05 |
| Usp15 | 2.32045E-05 | -0.984416105 | 0.513 | 0.706 | 0.563682716 | 0.099331174 | -0.897530701 | 0.7 | 0.83 | 1 | 0.099331174 | 4.64084E-05 |
| Zbtb1 | 2.3285E-05 | -1.680074258 | 0.303 | 0.494 | 0.565638762 | 0.009839014 | -1.759059365 | 0.3 | 0.701 | 1 | 0.009839014 | 4.65694E-05 |
| Prkag2 | 2.32868E-05 | 0.692360292 | 0.63 | 0.359 | 0.565684004 | 9.0057E-05 | 1.791509715 | 0.9 | 0.531 | 1 | 9.0057E-05 | 4.65731E-05 |
| Eif4e3 | 2.35223E-05 | -2.675251386 | 0.059 | 0.241 | 0.571404263 | 0.386101454 | -0.840059087 | 0.1 | 0.204 | 1 | 0.386101454 | 4.70441E-05 |
| Lmna | 2.3541E-05 | -0.770829046 | 0.992 | 0.988 | 0.57185749 | 0.07194947 | -0.636617364 | 0.9 | 1 | 1 | 0.07194947 | 4.70814E-05 |
| Rbm47 | 2.36921E-05 | 1.210566986 | 0.378 | 0.165 | 0.575528543 | 0.186524066 | -1.357117523 | 0.1 | 0.281 | 1 | 0.186524066 | 4.73836E-05 |
| Atpif1 | 2.38375E-05 | -0.700165676 | 0.706 | 0.853 | 0.579060938 | 0.121929963 | -0.394859617 | 0.4 | 0.719 | 1 | 0.121929963 | 4.76745E-05 |
| Ythdc1 | 2.40718E-05 | -0.499718915 | 0.975 | 0.982 | 0.584751692 | 0.246194345 | -0.322342246 | 1 | 0.981 | 1 | 0.246194345 | 4.8143E-05 |
| Cse1l | 2.43036E-05 | -1.327547167 | 0.261 | 0.488 | 0.590384249 | 0.992819241 | 0.174943076 | 0.5 | 0.512 | 1 | 0.992819241 | 4.86067E-05 |
| Zc3h15 | 2.43537E-05 | -0.560864962 | 0.908 | 0.929 | 0.591600518 | 0.242466906 | -0.442680254 | 1 | 0.932 | 1 | 0.242466906 | 4.87068E-05 |
| 9930021J03Rik | 2.43962E-05 | -1.299871174 | 0.429 | 0.635 | 0.592632461 | 0.598463564 | -0.495147674 | 0.8 | 0.728 | 1 | 0.598463564 | 4.87918E-05 |
| Gm9856 | 0.112938994 | -1.070389328 | 0.067 | 0.124 | 1 | 2.44643E-05 | 2.339850003 | 0.4 | 0.056 | 0.594287333 | 0.112938994 | 4.8928E-05 |
| Lats2 | 2.45267E-05 | -1.471487636 | 0.345 | 0.535 | 0.595802674 | 0.03645312 | -1.324893553 | 0.4 | 0.67 | 1 | 0.03645312 | 4.90528E-05 |
| Slc18a2 | 2.45536E-05 | 5.372554168 | 0.101 | 0 | 0.596456832 | 0.000713264 | 3.017921908 | 0.2 | 0.022 | 1 | 0.000713264 | 4.91067E-05 |
| Timp4 | 2.45766E-05 | 5.038135129 | 0.101 | 0 | 0.597014402 | 9.85609E-05 | 5.017921908 | 0.1 | 0.003 | 1 | 9.85609E-05 | 4.91526E-05 |
| Zfp592 | 2.46043E-05 | -1.242450074 | 0.311 | 0.529 | 0.59768715 | 0.164811911 | -0.586041555 | 0.4 | 0.685 | 1 | 0.164811911 | 4.9208E-05 |
| Bpgm | 2.46575E-05 | 0.894277599 | 0.664 | 0.429 | 0.598979586 | 0.408881437 | 0.469772073 | 0.8 | 0.698 | 1 | 0.408881437 | 4.93144E-05 |
| Mtrex | 2.48018E-05 | -0.985272714 | 0.353 | 0.594 | 0.602484973 | 0.607573462 | 0.135278859 | 0.7 | 0.639 | 1 | 0.607573462 | 4.9603E-05 |
| Cnksr3 | 2.48412E-05 | -1.380457268 | 0.294 | 0.494 | 0.603442392 | 0.024334245 | -1.896164189 | 0.3 | 0.627 | 1 | 0.024334245 | 4.96818E-05 |
| Tcf7l2 | 2.48661E-05 | -0.827613761 | 0.908 | 0.918 | 0.604046436 | 0.344769293 | -0.445394639 | 1 | 0.981 | 1 | 0.344769293 | 4.97315E-05 |
| Zbtb37 | 2.49631E-05 | -2.68182404 | 0.042 | 0.218 | 0.606404756 | 0.94455102 | 0.111031312 | 0.2 | 0.198 | 1 | 0.94455102 | 4.99257E-05 |
| Gm49575 | 2.50776E-05 | 4.038135129 | 0.118 | 0.006 | 0.609184466 | 0.577174234 | 0.695993813 | 0 | 0.031 | 1 | 0.577174234 | 5.01545E-05 |
| Marf1 | 2.51709E-05 | -1.318776958 | 0.378 | 0.565 | 0.611451752 | 0.039773397 | -1.361456459 | 0.3 | 0.583 | 1 | 0.039773397 | 5.03412E-05 |
| Dgcr8 | 2.5224E-05 | -1.763411574 | 0.134 | 0.347 | 0.612740221 | 0.064630341 | -1.633129783 | 0.1 | 0.38 | 1 | 0.064630341 | 5.04473E-05 |
| Fam20b | 2.52758E-05 | 0.479526226 | 0.849 | 0.624 | 0.614000602 | 0.011168321 | 0.86698001 | 0.9 | 0.691 | 1 | 0.011168321 | 5.0551E-05 |
| Ago1 | 2.53027E-05 | -1.911691582 | 0.109 | 0.312 | 0.614654188 | 0.066241272 | -2.101019165 | 0 | 0.259 | 1 | 0.066241272 | 5.06048E-05 |
| Phf6 | 2.53508E-05 | -1.529820947 | 0.193 | 0.418 | 0.61582091 | 0.660378291 | -0.567040593 | 0.5 | 0.478 | 1 | 0.660378291 | 5.07009E-05 |
| Axin2 | 2.53922E-05 | -1.485426827 | 0.16 | 0.382 | 0.616828392 | 0.109412612 | -1.802257054 | 0.1 | 0.33 | 1 | 0.109412612 | 5.07838E-05 |
| Clvs1 | 2.54495E-05 | 3.974004791 | 0.118 | 0.006 | 0.618218622 | 0.669843604 | 1.695993813 | 0 | 0.019 | 1 | 0.669843604 | 5.08983E-05 |
| Rsbn1 | 2.57534E-05 | -1.0138058 | 0.513 | 0.694 | 0.625602029 | 0.000658933 | -2.149997958 | 0.4 | 0.809 | 1 | 0.000658933 | 5.15062E-05 |
| Rnf144a | 2.57953E-05 | -1.525583954 | 0.353 | 0.535 | 0.626620445 | 0.446374373 | -1.004445905 | 0.4 | 0.451 | 1 | 0.446374373 | 5.159E-05 |
| Rad21 | 2.58775E-05 | -1.308901415 | 0.504 | 0.635 | 0.628616233 | 0.001463003 | -1.850386225 | 0.3 | 0.799 | 1 | 0.001463003 | 5.17543E-05 |
| Parg | 2.59715E-05 | -1.088767857 | 0.429 | 0.612 | 0.630899205 | 0.428432499 | -0.290066996 | 0.5 | 0.753 | 1 | 0.428432499 | 5.19423E-05 |
| Sh2b1 | 2.59822E-05 | -1.608809243 | 0.084 | 0.288 | 0.631158834 | 0.199769507 | -0.900941329 | 0.1 | 0.278 | 1 | 0.199769507 | 5.19637E-05 |
| Tamalin | 2.62752E-05 | -2.058893689 | 0.126 | 0.329 | 0.63827656 | 0.042062017 | -2.574535129 | 0 | 0.302 | 1 | 0.042062017 | 5.25497E-05 |
| Ccnd2 | 2.63236E-05 | -1.10831671 | 0.782 | 0.788 | 0.639452095 | 0.000348548 | -2.241128984 | 0.5 | 0.914 | 1 | 0.000348548 | 5.26464E-05 |
| Vgll4 | 2.6364E-05 | -1.094596188 | 0.647 | 0.765 | 0.640434669 | 0.936836759 | -0.31730301 | 0.9 | 0.809 | 1 | 0.936836759 | 5.27273E-05 |
| Sgcb | 2.6384E-05 | -1.845322772 | 0.101 | 0.306 | 0.640921222 | 0.750280499 | 0.210566986 | 0.3 | 0.235 | 1 | 0.750280499 | 5.27674E-05 |
| Fads3 | 2.65281E-05 | -1.122856748 | 0.403 | 0.594 | 0.644420168 | 0.305553452 | -0.936274402 | 0.4 | 0.491 | 1 | 0.305553452 | 5.30555E-05 |
| Tgfbi | 2.65526E-05 | -3.821710215 | 0.025 | 0.188 | 0.645015198 | 0.700618257 | -0.452398027 | 0.2 | 0.142 | 1 | 0.700618257 | 5.31044E-05 |
| Snai1 | 2.69257E-05 | -1.340228912 | 0.168 | 0.388 | 0.654080278 | 0.605674295 | 0.989725016 | 0.4 | 0.343 | 1 | 0.605674295 | 5.38508E-05 |
| Kdm1a | 2.69982E-05 | -0.61277204 | 0.891 | 0.924 | 0.655840623 | 0.000518957 | -1.451121305 | 0.6 | 0.972 | 1 | 0.000518957 | 5.39957E-05 |
| Abce1 | 2.70184E-05 | -0.757429697 | 0.689 | 0.776 | 0.656331804 | 0.003084394 | -1.600987925 | 0.6 | 0.802 | 1 | 0.003084394 | 5.40361E-05 |
| Dnm1l | 2.70322E-05 | -0.856070207 | 0.513 | 0.688 | 0.65666712 | 0.786956385 | 0.18733468 | 0.7 | 0.75 | 1 | 0.786956385 | 5.40637E-05 |
| Nacc1 | 2.71458E-05 | -1.414343729 | 0.168 | 0.394 | 0.65942692 | 0.680537216 | 0.451106754 | 0.5 | 0.435 | 1 | 0.680537216 | 5.4291E-05 |
| Tbc1d16 | 2.7393E-05 | -2.222392421 | 0.084 | 0.282 | 0.665429821 | 0.670284484 | -0.573039333 | 0.3 | 0.333 | 1 | 0.670284484 | 5.47852E-05 |
| Myo19 | 2.74226E-05 | -1.426533138 | 0.134 | 0.353 | 0.666148953 | 0.864362921 | 0.515421567 | 0.3 | 0.281 | 1 | 0.864362921 | 5.48444E-05 |
| Shc1 | 2.74708E-05 | -0.895582105 | 0.605 | 0.706 | 0.667320643 | 0.005368363 | -1.479929929 | 0.5 | 0.796 | 1 | 0.005368363 | 5.49408E-05 |
| Ube2z | 2.75628E-05 | -1.02666795 | 0.403 | 0.606 | 0.669556154 | 0.003535429 | -1.933362807 | 0.3 | 0.698 | 1 | 0.003535429 | 5.51249E-05 |
| Nudt21 | 2.77571E-05 | -0.706268797 | 0.647 | 0.788 | 0.674274926 | 0.208389879 | -0.478855067 | 0.6 | 0.806 | 1 | 0.208389879 | 5.55134E-05 |
| Acadm | 2.77806E-05 | -1.540709263 | 0.235 | 0.447 | 0.674847177 | 0.003213377 | -3.015501094 | 0 | 0.5 | 1 | 0.003213377 | 5.55605E-05 |
| Pde7b | 2.78101E-05 | -2.755753726 | 0.084 | 0.276 | 0.675562143 | 0.531741045 | -2.042774024 | 0.2 | 0.259 | 1 | 0.531741045 | 5.56194E-05 |
| Gstm5 | 2.79934E-05 | 0.606789544 | 0.857 | 0.635 | 0.680015891 | 0.521684727 | 0.638749861 | 0.6 | 0.583 | 1 | 0.521684727 | 5.5986E-05 |
| Sbf2 | 2.81746E-05 | -1.19626269 | 0.521 | 0.682 | 0.68441854 | 0.634768926 | -0.499353785 | 0.8 | 0.846 | 1 | 0.634768926 | 5.63485E-05 |
| Dbn1 | 2.83529E-05 | -2.448900951 | 0.059 | 0.241 | 0.688747719 | 0.839659257 | 0.210566986 | 0.2 | 0.167 | 1 | 0.839659257 | 5.67049E-05 |
| Clic5 | 2.84634E-05 | 2.377069649 | 0.193 | 0.041 | 0.691433963 | 0.522126681 | 0.769994395 | 0 | 0.04 | 1 | 0.522126681 | 5.69261E-05 |
| Shprh | 2.85412E-05 | -1.703850346 | 0.134 | 0.347 | 0.693323663 | 0.205963812 | -1.304006187 | 0.2 | 0.367 | 1 | 0.205963812 | 5.70817E-05 |
| Gm16351 | 2.9274E-05 | -2.533016286 | 0.05 | 0.235 | 0.711124144 | 0.204835486 | -1.940147412 | 0.1 | 0.284 | 1 | 0.204835486 | 5.85472E-05 |
| Ets2 | 2.92974E-05 | -0.737406044 | 0.529 | 0.759 | 0.711692893 | 0.067381208 | -0.475266399 | 0.6 | 0.898 | 1 | 0.067381208 | 5.8594E-05 |
| Sipa1l3 | 2.93201E-05 | -2.100136671 | 0.084 | 0.282 | 0.712242756 | 0.443162064 | -0.204470513 | 0.4 | 0.556 | 1 | 0.443162064 | 5.86392E-05 |
| Nup210l | 2.93894E-05 | -1.542570734 | 0.328 | 0.541 | 0.713927476 | 0.57890172 | 0.141112652 | 0.4 | 0.534 | 1 | 0.57890172 | 5.8778E-05 |
| Magohb | 2.94463E-05 | -1.519374159 | 0.168 | 0.382 | 0.715309158 | 0.593494668 | 0.135278859 | 0.2 | 0.299 | 1 | 0.593494668 | 5.88917E-05 |
| Myl4 | 2.95618E-05 | -3.628384781 | 0.017 | 0.171 | 0.718114192 | 0.156786818 | -1.304006187 | 0 | 0.17 | 1 | 0.156786818 | 5.91226E-05 |
| Btaf1 | 2.95816E-05 | -0.65972496 | 0.899 | 0.941 | 0.718596085 | 0.560547495 | -0.276450464 | 0.9 | 0.941 | 1 | 0.560547495 | 5.91623E-05 |

| | | | | | | | | | | | | |
|---|---|---|---|---|---|---|---|---|---|---|---|---|
| Frmd8os | 0.726024391 | -0.485426827 | 0.084 | 0.094 | 1 | 2.97789E-05 | 4.017921908 | 0.4 | 0.062 | 0.723388173 | 0.726024391 | 5.95568E-05 |
| Ppfia1 | 2.97816E-05 | -1.111031312 | 0.387 | 0.582 | 0.723453962 | 0.052834401 | -0.91281543 | 0.5 | 0.741 | 1 | 0.052834401 | 5.95623E-05 |
| Cct2 | 2.99732E-05 | -0.59812696 | 0.916 | 0.941 | 0.728109306 | 0.000366037 | -1.584277298 | 0.5 | 0.969 | 1 | 0.000366037 | 5.99455E-05 |
| Arl15 | 3.00783E-05 | -1.218712075 | 0.513 | 0.671 | 0.73066101 | 0.099773285 | -1.115220304 | 0.4 | 0.691 | 1 | 0.099773285 | 6.01556E-05 |
| Tnc | 3.02993E-05 | -6.789207575 | 0.008 | 0.153 | 0.736030383 | 0.282779044 | -2.5519337 | 0 | 0.105 | 1 | 0.282779044 | 6.05977E-05 |
| Bach1 | 3.03454E-05 | -1.466811149 | 0.462 | 0.588 | 0.737150529 | 0.169974475 | -0.464742017 | 0.5 | 0.716 | 1 | 0.169974475 | 6.06899E-05 |
| Cep135 | 0.049916561 | -1.93021167 | 0.067 | 0.135 | 1 | 3.04223E-05 | 2.233650599 | 0.6 | 0.127 | 0.739019679 | 0.049916561 | 6.08438E-05 |
| Mmp23 | 3.04353E-05 | -4.614709844 | 0 | 0.135 | 0.739334871 | 0.577142568 | 1.432959407 | 0 | 0.031 | 1 | 0.577142568 | 6.08697E-05 |
| Mia3 | 3.05106E-05 | -0.544320516 | 0.832 | 0.918 | 0.74116448 | 0.615677948 | -0.170584519 | 0.8 | 0.932 | 1 | 0.615677948 | 6.10204E-05 |
| Ntm | 3.05213E-05 | -5.635173947 | 0 | 0.135 | 0.74142273 | 0.345511791 | 1.558490289 | 0.1 | 0.04 | 1 | 0.345511791 | 6.10416E-05 |
| Adam19 | 3.05515E-05 | -5.343407822 | 0 | 0.135 | 0.742158023 | 0.30739492 | -1.048167282 | 0 | 0.096 | 1 | 0.30739492 | 6.11021E-05 |
| Ar | 3.05796E-05 | -5.593951284 | 0 | 0.135 | 0.742840213 | 0.073255924 | -2.959358016 | 0 | 0.25 | 1 | 0.073255924 | 6.11583E-05 |
| Haus3 | 3.0844E-05 | -1.586964854 | 0.151 | 0.365 | 0.749262802 | 0.244136011 | -1.111361109 | 0.1 | 0.253 | 1 | 0.244136011 | 6.16871E-05 |
| Sdc1 | 3.09222E-05 | -1.424026283 | 0.143 | 0.359 | 0.751163198 | 0.000566675 | -3.658035125 | 0 | 0.605 | 1 | 0.000566675 | 6.18435E-05 |
| Il33 | 3.11243E-05 | 0.845779081 | 0.21 | 0.047 | 0.756072426 | 0.597840427 | 0.695993813 | 0 | 0.028 | 1 | 0.597840427 | 6.22477E-05 |
| Fzd8 | 3.11319E-05 | -2.402964667 | 0.05 | 0.229 | 0.756255407 | 0.40042261 | -0.888968688 | 0.1 | 0.201 | 1 | 0.40042261 | 6.22628E-05 |
| Robo1 | 3.12267E-05 | -1.897687242 | 0.218 | 0.424 | 0.758558829 | 0.010040218 | -1.769478408 | 0.5 | 0.802 | 1 | 0.010040218 | 6.24524E-05 |
| Tram1 | 3.13725E-05 | -0.623987658 | 0.958 | 0.935 | 0.762101378 | 0.076002789 | -0.6333613 | 0.7 | 0.944 | 1 | 0.076002789 | 6.27441E-05 |
| Itih5 | 3.14762E-05 | -4.003752135 | 0.042 | 0.212 | 0.764618663 | 0.557772534 | -0.408342847 | 0 | 0.034 | 1 | 0.557772534 | 6.29513E-05 |
| Gm34934 | 0.218348362 | 1.099535674 | 0.05 | 0.024 | 1 | 3.15747E-05 | 3.639410285 | 0.3 | 0.034 | 0.767012892 | 0.218348362 | 6.31484E-05 |
| Tlcd1 | 3.18537E-05 | 1.283247627 | 0.479 | 0.259 | 0.773789705 | 0.221169562 | 1.515421567 | 0.4 | 0.238 | 1 | 0.221169562 | 6.37064E-05 |
| Nfia | 3.18577E-05 | -0.576359474 | 0.891 | 0.924 | 0.773886911 | 0.293026044 | -0.314894503 | 1 | 0.966 | 1 | 0.293026044 | 6.37144E-05 |
| BC005537 | 3.21607E-05 | -0.512179555 | 0.941 | 0.929 | 0.781246867 | 0.006077232 | -0.925658497 | 0.9 | 0.994 | 1 | 0.006077232 | 6.43203E-05 |
| Rrs1 | 3.2191E-05 | -1.382667253 | 0.286 | 0.482 | 0.78198369 | 0.084880258 | -1.430538593 | 0.2 | 0.441 | 1 | 0.084880258 | 6.4381E-05 |
| Htra4 | 3.22409E-05 | -4.485426827 | 0.008 | 0.153 | 0.783195421 | 0.208062724 | -1.709998547 | 0 | 0.139 | 1 | 0.208062724 | 6.44807E-05 |
| Polg2 | 3.28597E-05 | -1.951744831 | 0.084 | 0.282 | 0.79822828 | 0.048365077 | -2.567040593 | 0 | 0.29 | 1 | 0.048365077 | 6.57184E-05 |
| Bahcc1 | 3.2991E-05 | -2.08809133 | 0.193 | 0.4 | 0.801418415 | 0.676097019 | -0.368659145 | 0.5 | 0.352 | 1 | 0.676097019 | 6.5981E-05 |
| Stac2 | 3.30114E-05 | -3.614709844 | 0.008 | 0.153 | 0.801911761 | 0.209575853 | -1.210896782 | 0.1 | 0.269 | 1 | 0.209575853 | 6.60216E-05 |
| Mfsd10 | 3.30438E-05 | -1.235448574 | 0.252 | 0.471 | 0.802700314 | 0.366984998 | -0.58694015 | 0.2 | 0.318 | 1 | 0.366984998 | 6.60865E-05 |
| Nsun2 | 3.32544E-05 | -1.248986631 | 0.244 | 0.465 | 0.807816674 | 0.441614532 | -0.374395515 | 0.4 | 0.534 | 1 | 0.441614532 | 6.65078E-05 |
| Gm34189 | 3.32591E-05 | 2.184424571 | 0.16 | 0.024 | 0.807931005 | 0.557756607 | 0.432959407 | 0 | 0.034 | 1 | 0.557756607 | 6.65172E-05 |
| Galc | 3.34078E-05 | -3.485426827 | 0.008 | 0.153 | 0.811542707 | 0.207752534 | -0.888968688 | 0 | 0.139 | 1 | 0.207752534 | 6.68145E-05 |
| Tns1 | 3.34134E-05 | 0.176741736 | 0.966 | 0.776 | 0.811678193 | 0.376477763 | -0.269698136 | 0.7 | 0.864 | 1 | 0.376477763 | 6.68257E-05 |
| Nup107 | 3.34304E-05 | -2.033863452 | 0.134 | 0.341 | 0.812091965 | 0.834434802 | -0.441509711 | 0.3 | 0.296 | 1 | 0.834434802 | 6.68597E-05 |
| Gm12940 | 3.35099E-05 | -1.944858446 | 0.168 | 0.376 | 0.814021574 | 0.62326916 | -0.541773834 | 0.3 | 0.364 | 1 | 0.62326916 | 6.70186E-05 |
| Sap30l | 3.36174E-05 | -1.344828497 | 0.235 | 0.447 | 0.816633651 | 0.08040227 | -1.391469028 | 0.2 | 0.451 | 1 | 0.08040227 | 6.72337E-05 |
| Cdh20 | 0.0037881 | 2.140758336 | 0.126 | 0.035 | 1 | 3.37143E-05 | 2.210566986 | 0.3 | 0.034 | 0.818988264 | 0.0037881 | 6.74275E-05 |
| Pdzk1ip1 | 3.37288E-05 | 3.237039197 | 0.143 | 0.018 | 0.819338869 | 0.086046028 | 2.31748219 | 0.2 | 0.065 | 1 | 0.086046028 | 6.74564E-05 |
| Rock1 | 3.39105E-05 | -0.602692879 | 0.933 | 0.976 | 0.823752917 | 0.030346447 | -0.61165889 | 1 | 1 | 1 | 0.030346447 | 6.78198E-05 |
| Arfgef2 | 3.41268E-05 | 0.540635469 | 0.882 | 0.665 | 0.829009243 | 0.285012994 | 0.253713778 | 1 | 0.824 | 1 | 0.285012994 | 6.82525E-05 |
| Ctdsp2 | 3.43592E-05 | -0.618593279 | 0.773 | 0.841 | 0.834654686 | 0.007238552 | -0.953319099 | 0.8 | 0.926 | 1 | 0.007238552 | 6.87173E-05 |
| Lyn | 3.45373E-05 | -3.521050737 | 0.017 | 0.171 | 0.838979768 | 0.03376574 | -2.528972552 | 0 | 0.321 | 1 | 0.03376574 | 6.90734E-05 |
| Wdr75 | 3.46331E-05 | -1.311397427 | 0.336 | 0.541 | 0.841307184 | 0.00915031 | -2.395706021 | 0.1 | 0.525 | 1 | 0.00915031 | 6.9265E-05 |
| Spty2d1 | 3.47571E-05 | -0.955829564 | 0.605 | 0.747 | 0.844319472 | 0.120956007 | -0.798770879 | 0.6 | 0.762 | 1 | 0.120956007 | 6.9513E-05 |
| Serpinh1 | 3.47798E-05 | -0.59079773 | 0.992 | 0.971 | 0.844871129 | 0.064425488 | -0.677246372 | 1 | 0.963 | 1 | 0.064425488 | 6.95584E-05 |
| Mien1 | 3.4862E-05 | 1.04094102 | 0.689 | 0.547 | 0.846868366 | 0.819897548 | 0.789103218 | 0.4 | 0.475 | 1 | 0.819897548 | 6.97228E-05 |
| Hspa4 | 3.50568E-05 | -0.582724029 | 0.933 | 0.929 | 0.851600879 | 0.03134431 | -0.641182055 | 1 | 0.957 | 1 | 0.03134431 | 7.01125E-05 |
| Usp12 | 3.50572E-05 | -1.394453167 | 0.286 | 0.494 | 0.851610342 | 0.708137845 | -0.121629444 | 0.6 | 0.657 | 1 | 0.708137845 | 7.01132E-05 |
| Col9a1 | 3.52108E-05 | -0.836186105 | 0.832 | 0.888 | 0.85534194 | 0.006887016 | -1.300198884 | 0.8 | 0.935 | 1 | 0.006887016 | 7.04205E-05 |
| Fbxl19 | 3.55158E-05 | -2.521050737 | 0.034 | 0.2 | 0.862749213 | 0.408681027 | 0.574978412 | 0.3 | 0.182 | 1 | 0.408681027 | 7.10303E-05 |
| Npepps | 3.60104E-05 | -0.770703643 | 0.824 | 0.871 | 0.874764226 | 0.581866952 | -0.166572022 | 0.9 | 0.948 | 1 | 0.581866952 | 7.20195E-05 |
| Larp4 | 3.60113E-05 | -0.603499821 | 0.882 | 0.912 | 0.87478688 | 0.012894351 | -0.899385304 | 0.9 | 0.96 | 1 | 0.012894351 | 7.20213E-05 |
| Cstf3 | 3.60168E-05 | -0.809371268 | 0.538 | 0.776 | 0.874920924 | 0.886012819 | 0.538452536 | 0.8 | 0.827 | 1 | 0.886012819 | 7.20324E-05 |
| Rbm28 | 3.6026E-05 | -1.165966988 | 0.42 | 0.588 | 0.875144786 | 0.43170214 | -0.632332053 | 0.6 | 0.605 | 1 | 0.43170214 | 7.20508E-05 |
| Lamp1 | 3.62899E-05 | 0.362333978 | 1 | 0.982 | 0.881554331 | 0.206839673 | 0.275787594 | 1 | 0.991 | 1 | 0.206839673 | 7.25785E-05 |
| Kcnk2 | 3.65401E-05 | -2.883019192 | 0.092 | 0.282 | 0.887631149 | 0.215339933 | -1.640289575 | 0.2 | 0.355 | 1 | 0.215339933 | 7.30788E-05 |
| Gm5544 | 3.66054E-05 | -2.744161096 | 0.042 | 0.212 | 0.889219475 | 0.722211056 | 0.558490289 | 0.2 | 0.167 | 1 | 0.722211056 | 7.32096E-05 |
| A330048O09Rik | 3.66563E-05 | 2.836501268 | 0.143 | 0.018 | 0.890456001 | 0.19134412 | 2.432959407 | 0.1 | 0.028 | 1 | 0.19134412 | 7.33114E-05 |
| Ubn1 | 3.67509E-05 | -0.548554806 | 0.84 | 0.912 | 0.892752835 | 0.10211679 | -0.553754902 | 0.8 | 0.941 | 1 | 0.10211679 | 7.35004E-05 |
| Junos | 0.016123839 | 0.656592178 | 0.303 | 0.182 | 1 | 3.6987E-05 | 4.172896568 | 0.6 | 0.142 | 0.898488276 | 0.016123839 | 7.39726E-05 |
| Pfkp | 3.70474E-05 | 0.352533428 | 0.916 | 0.688 | 0.899954875 | 3.75986E-05 | 1.46985899 | 1 | 0.667 | 0.913345551 | 3.75986E-05 | 7.40934E-05 |
| Kctd5 | 3.71526E-05 | -1.365845211 | 0.143 | 0.365 | 0.902510395 | 0.028541575 | -1.840059087 | 0.1 | 0.448 | 1 | 0.028541575 | 7.43038E-05 |
| Arid1a | 3.71935E-05 | -1.145351386 | 0.597 | 0.706 | 0.903504574 | 0.000330711 | -2.025105376 | 0.6 | 0.886 | 1 | 0.000330711 | 7.43856E-05 |
| Dnajc14 | 3.71955E-05 | -2.050211446 | 0.092 | 0.282 | 0.903552615 | 0.2205646 | -0.974544419 | 0.2 | 0.37 | 1 | 0.2205646 | 7.43896E-05 |
| Ctnnd2 | 3.71978E-05 | 1.138197294 | 0.429 | 0.194 | 0.903608917 | 0.557787601 | 0.280956314 | 0.2 | 0.127 | 1 | 0.557787601 | 7.43942E-05 |
| Actl6a | 3.74157E-05 | -1.297354479 | 0.319 | 0.506 | 0.908901523 | 0.259219656 | -0.514573173 | 0.4 | 0.562 | 1 | 0.259219656 | 7.48299E-05 |
| Tspan8 | 3.742E-05 | 2.736965594 | 0.168 | 0.029 | 0.909005605 | 0.914143824 | 0.210566986 | 0.2 | 0.222 | 1 | 0.914143824 | 7.48385E-05 |
| Serp1 | 3.75041E-05 | -0.699347732 | 0.765 | 0.859 | 0.911049344 | 0.104502854 | -0.748607001 | 0.7 | 0.855 | 1 | 0.104502854 | 7.50068E-05 |
| Mmp28 | 0.001067183 | 0.37922132 | 0.286 | 0.124 | 1 | 3.77428E-05 | 2.568614507 | 0.5 | 0.096 | 0.916847476 | 0.001067183 | 7.54841E-05 |
| Kras | 3.78981E-05 | -0.841882605 | 0.824 | 0.888 | 0.920620497 | 0.001371086 | -1.664838643 | 0.7 | 0.932 | 1 | 0.001371086 | 7.57948E-05 |
| Ctdnep1 | 3.79859E-05 | -0.941309379 | 0.471 | 0.647 | 0.92275441 | 0.809755902 | -0.111361109 | 0.7 | 0.664 | 1 | 0.809755902 | 7.59704E-05 |
| Sucla2 | 3.82309E-05 | -1.079267479 | 0.361 | 0.559 | 0.928704424 | 0.563619964 | -0.195425374 | 0.6 | 0.608 | 1 | 0.563619964 | 7.64603E-05 |
| Mcrip1 | 3.83313E-05 | -0.759503404 | 0.639 | 0.782 | 0.931143465 | 0.099411199 | -0.90226551 | 0.6 | 0.716 | 1 | 0.099411199 | 7.66611E-05 |
| Obi1 | 3.83602E-05 | -2.070389328 | 0.059 | 0.241 | 0.931847154 | 0.465725671 | -0.640289575 | 0.1 | 0.185 | 1 | 0.465725671 | 7.6719E-05 |
| Tceal8 | 3.84252E-05 | -0.659797734 | 0.824 | 0.865 | 0.933423813 | 0.154956234 | -0.630366528 | 0.9 | 0.889 | 1 | 0.154956234 | 7.68488E-05 |
| Tmem87a | 3.85091E-05 | -0.987033581 | 0.445 | 0.671 | 0.935464192 | 0.278029333 | -0.484578433 | 0.6 | 0.753 | 1 | 0.278029333 | 7.70168E-05 |
| Abhd3 | 0.00040948 | 2.514573173 | 0.151 | 0.035 | 1 | 3.85914E-05 | 3.100384068 | 0.4 | 0.065 | 0.937461476 | 0.00040948 | 7.71812E-05 |
| Tnfrsf10b | 3.86214E-05 | 1.078925394 | 0.58 | 0.329 | 0.938190204 | 0.104155144 | 1.210566986 | 0.5 | 0.306 | 1 | 0.104155144 | 7.72412E-05 |
| Pde1b | 3.86705E-05 | 2.684498174 | 0.143 | 0.018 | 0.939383736 | 0.698534498 | 2.432959407 | 0 | 0.015 | 1 | 0.698534498 | 7.73395E-05 |
| Prr13 | 3.87202E-05 | 1.175589914 | 0.613 | 0.418 | 0.940590689 | 0.121836698 | 0.893226161 | 0.6 | 0.559 | 1 | 0.121836698 | 7.74389E-05 |
| Phf2 | 3.87281E-05 | -1.147146527 | 0.403 | 0.576 | 0.940781933 | 0.019657099 | -1.203665213 | 0.7 | 0.787 | 1 | 0.019657099 | 7.74546E-05 |
| Ccdc86 | 3.87735E-05 | -1.417408404 | 0.269 | 0.476 | 0.941885072 | 0.136580392 | -1.171902651 | 0.1 | 0.315 | 1 | 0.136580392 | 7.75454E-05 |
| Capza2 | 3.88208E-05 | -0.541352242 | 0.857 | 0.912 | 0.943033902 | 0.092342614 | -0.583134569 | 0.9 | 0.951 | 1 | 0.092342614 | 7.764E-05 |
| Dennd1b | 3.8944E-05 | -1.14332085 | 0.487 | 0.629 | 0.946028767 | 0.47095128 | -0.357117523 | 0.5 | 0.698 | 1 | 0.47095128 | 7.78866E-05 |
| Ica1 | 3.91672E-05 | 0.510181404 | 0.824 | 0.541 | 0.951449677 | 0.005095324 | 1.558490289 | 0.8 | 0.441 | 1 | 0.005095324 | 7.83329E-05 |
| Dync1i1 | 3.94836E-05 | -1.787989597 | 0.261 | 0.471 | 0.959136741 | 0.027709057 | -2.870821341 | 0.1 | 0.441 | 1 | 0.027709057 | 7.89657E-05 |
| Arhgap26 | 3.9538E-05 | 0.553047321 | 0.622 | 0.376 | 0.960457057 | 0.000356154 | 2.167065347 | 0.9 | 0.423 | 1 | 0.000356154 | 7.90744E-05 |
| Snx24 | 3.96639E-05 | -1.014355293 | 0.622 | 0.747 | 0.963516543 | 0.79039878 | -0.200170376 | 0.9 | 0.775 | 1 | 0.79039878 | 7.93263E-05 |
| Zfp101 | 3.99394E-05 | -1.87774425 | 0.084 | 0.276 | 0.970207998 | 0.484467241 | 0.345496566 | 0.4 | 0.281 | 1 | 0.484467241 | 7.98772E-05 |
| Syngap1 | 4.00746E-05 | -2.600904045 | 0.042 | 0.212 | 0.973493126 | 0.634770375 | -0.152003093 | 0.1 | 0.151 | 1 | 0.634770375 | 8.01477E-05 |
| Atg3 | 4.01729E-05 | -0.812788808 | 0.664 | 0.776 | 0.975880653 | 0.456985896 | -0.361456459 | 0.8 | 0.778 | 1 | 0.456985896 | 8.03442E-05 |
| Wdr70 | 4.02377E-05 | -0.861065509 | 0.613 | 0.729 | 0.977455357 | 0.412942768 | -0.443849849 | 0.6 | 0.738 | 1 | 0.412942768 | 8.04739E-05 |
| Etf1 | 4.02589E-05 | -0.518270084 | 0.933 | 0.953 | 0.977969986 | 0.282477057 | -0.31161572 | 1 | 0.988 | 1 | 0.282477057 | 8.05162E-05 |
| Ppp1ca | 4.03077E-05 | -0.703997538 | 0.731 | 0.806 | 0.979154096 | 0.237529146 | -0.372247048 | 0.6 | 0.849 | 1 | 0.237529146 | 8.06137E-05 |
| Stk11 | 4.03962E-05 | -0.868755467 | 0.387 | 0.618 | 0.981303391 | 0.041267259 | -0.96190021 | 0.6 | 0.775 | 1 | 0.041267259 | 8.07907E-05 |
| Pla1a | 4.04039E-05 | 2.453172628 | 0.143 | 0.018 | 0.981491379 | 0.214124936 | 3.111031312 | 0.1 | 0.031 | 1 | 0.214124936 | 8.08062E-05 |
| Trim28 | 4.05737E-05 | -1.438429348 | 0.277 | 0.471 | 0.985615131 | 0.020596526 | -1.352765499 | 0.2 | 0.605 | 1 | 0.020596526 | 8.11457E-05 |
| Comp | 4.05743E-05 | -0.271281766 | 0.992 | 0.829 | 0.985630766 | 0.007112086 | 1.372318056 | 1 | 0.87 | 1 | 0.007112086 | 8.11469E-05 |

| | | | | | | | | | | | | |
|---|---|---|---|---|---|---|---|---|---|---|---|---|
| Adipor1 | 4.08377E-05 | 0.459711238 | 0.958 | 0.935 | 0.99203011 | 0.749687537 | -0.164096165 | 0.9 | 0.966 | 1 | 0.749687537 | 8.16738E-05 |
| Cnpy4 | 4.10403E-05 | -1.222392421 | 0.277 | 0.488 | 0.996950422 | 0.247873259 | 0.632490871 | 0.6 | 0.438 | 1 | 0.247873259 | 8.20789E-05 |
| Cmip | 4.12863E-05 | 0.414525996 | 1 | 0.888 | 1 | 0.062474695 | 0.849650425 | 1 | 0.846 | 1 | 0.062474695 | 8.25709E-05 |
| Rmnd5a | 4.14815E-05 | -0.769219793 | 0.521 | 0.735 | 1 | 0.969041237 | 0.138542876 | 0.7 | 0.809 | 1 | 0.969041237 | 8.29614E-05 |
| Mcm9 | 4.14863E-05 | -2.266786541 | 0.092 | 0.288 | 1 | 0.54512878 | 1.068905837 | 0.3 | 0.219 | 1 | 0.54512878 | 8.29708E-05 |
| Slc13a5 | 4.4912E-05 | 0.857174836 | 0.395 | 0.182 | 1 | 4.17172E-05 | 1.902444691 | 0.6 | 0.142 | 1 | 4.4912E-05 | 8.34327E-05 |
| Arf4 | 4.20214E-05 | -0.317571347 | 1 | 0.994 | 1 | 0.014717055 | -0.647064449 | 0.9 | 0.997 | 1 | 0.014717055 | 8.40411E-05 |
| Sar1a | 4.20906E-05 | -0.506366165 | 0.899 | 0.888 | 1 | 0.104564148 | -0.431639467 | 0.9 | 0.938 | 1 | 0.104564148 | 8.41794E-05 |
| Higd1a | 4.22871E-05 | 0.753704392 | 0.874 | 0.794 | 1 | 0.967660217 | 0.194664491 | 0.6 | 0.849 | 1 | 0.967660217 | 8.45725E-05 |
| Gatad2b | 4.25043E-05 | -0.756910423 | 0.782 | 0.9 | 1 | 0.008266336 | -1.06777184 | 1 | 0.963 | 1 | 0.008266336 | 8.50069E-05 |
| Fhl2 | 4.26333E-05 | 0.765856573 | 0.58 | 0.341 | 1 | 0.186348001 | 1.049831156 | 0.5 | 0.302 | 1 | 0.186348001 | 8.52649E-05 |
| Ryk | 4.26911E-05 | -0.651785214 | 0.773 | 0.871 | 1 | 0.904877701 | 0.129536781 | 0.9 | 0.914 | 1 | 0.904877701 | 8.53803E-05 |
| Brca2 | 0.910758602 | 0.689070904 | 0.126 | 0.135 | 1 | 4.27259E-05 | 2.847996907 | 0.4 | 0.062 | 1 | 0.910758602 | 8.545E-05 |
| Ldlrad4 | 4.29108E-05 | -1.006894613 | 0.899 | 0.953 | 1 | 0.165936689 | -0.515857052 | 0.9 | 0.978 | 1 | 0.165936689 | 8.58197E-05 |
| 5730471H19Rik | 4.30089E-05 | -2.807354922 | 0.025 | 0.182 | 1 | 0.566399259 | 1.063725598 | 0.2 | 0.142 | 1 | 0.566399259 | 8.60159E-05 |
| Armcx2 | 4.31527E-05 | -1.714245518 | 0.118 | 0.318 | 1 | 0.32980634 | -0.69632361 | 0.3 | 0.41 | 1 | 0.32980634 | 8.63036E-05 |
| Fbxo32 | 4.32019E-05 | -1.790281409 | 0.05 | 0.229 | 1 | 0.072576549 | -2.67556505 | 0 | 0.25 | 1 | 0.072576549 | 8.64019E-05 |
| Cracr2b | 4.45805E-05 | -3.572889668 | 0.017 | 0.165 | 1 | 0.077391506 | -1.936274402 | 0 | 0.244 | 1 | 0.077391506 | 8.91591E-05 |
| Baz1a | 4.47432E-05 | -1.666289675 | 0.328 | 0.535 | 1 | 0.016479757 | -1.990506714 | 0.2 | 0.593 | 1 | 0.016479757 | 8.94844E-05 |
| Tenm4 | 4.49807E-05 | -4.008988783 | 0.034 | 0.194 | 1 | 0.190425001 | -1.489872732 | 0.1 | 0.281 | 1 | 0.190425001 | 8.99593E-05 |
| Lcor | 4.50699E-05 | -0.75461346 | 0.815 | 0.9 | 1 | 0.253825007 | -0.559904411 | 0.9 | 0.966 | 1 | 0.253825007 | 9.01378E-05 |
| Kdm4b | 4.51794E-05 | -2.185866545 | 0.076 | 0.259 | 1 | 0.228386291 | -1.161987182 | 0.1 | 0.259 | 1 | 0.228386291 | 9.03568E-05 |
| Sarnp | 4.5283E-05 | -0.563678273 | 0.882 | 0.953 | 1 | 0.0409151 | -0.706591945 | 0.9 | 0.944 | 1 | 0.0409151 | 9.05639E-05 |
| Ipo7 | 4.53131E-05 | -0.749446344 | 0.824 | 0.888 | 1 | 0.13588948 | -0.510457064 | 0.8 | 0.92 | 1 | 0.13588948 | 9.06242E-05 |
| Rabif | 4.55769E-05 | -1.690541257 | 0.134 | 0.335 | 1 | 0.203156011 | -1.181750437 | 0.1 | 0.275 | 1 | 0.203156011 | 9.11518E-05 |
| Tdp2 | 4.56057E-05 | -1.485426827 | 0.16 | 0.371 | 1 | 0.210680502 | 0.963474124 | 0.5 | 0.349 | 1 | 0.210680502 | 9.12093E-05 |
| Prex2 | 4.58356E-05 | 1.453172628 | 0.336 | 0.141 | 1 | 0.619403805 | 0.159940913 | 0.4 | 0.306 | 1 | 0.619403805 | 9.16691E-05 |
| Upf1 | 4.6018E-05 | -1.310593025 | 0.336 | 0.518 | 1 | 0.001917038 | -2.048167282 | 0.3 | 0.731 | 1 | 0.001917038 | 9.20339E-05 |
| Dnmt1 | 4.61153E-05 | -1.058893689 | 0.244 | 0.471 | 1 | 0.953093882 | 0.680618587 | 0.5 | 0.448 | 1 | 0.953093882 | 9.22285E-05 |
| Fiz1 | 4.61383E-05 | -2.439623138 | 0.059 | 0.235 | 1 | 0.446761382 | 1.176619654 | 0.3 | 0.21 | 1 | 0.446761382 | 9.22745E-05 |
| Ahctf1 | 4.62054E-05 | -0.805705185 | 0.697 | 0.782 | 1 | 0.035967387 | -0.728032469 | 0.6 | 0.898 | 1 | 0.035967387 | 9.24087E-05 |
| Foxo3 | 4.64175E-05 | -1.056806356 | 0.773 | 0.841 | 1 | 0.430313152 | -0.391983266 | 0.8 | 0.917 | 1 | 0.430313152 | 9.28328E-05 |
| Pgm2l1 | 4.68433E-05 | -4.733354341 | 0 | 0.129 | 1 | 0.114018944 | -1.840059087 | 0 | 0.204 | 1 | 0.114018944 | 9.36845E-05 |
| Ap2a2 | 4.68862E-05 | -1.173482821 | 0.37 | 0.571 | 1 | 0.173861431 | -0.785132877 | 0.6 | 0.756 | 1 | 0.173861431 | 9.37703E-05 |
| Ubp1 | 4.69109E-05 | -0.752365687 | 0.622 | 0.747 | 1 | 0.022754155 | -0.979822118 | 0.5 | 0.812 | 1 | 0.022754155 | 9.38196E-05 |
| Cxcl14 | 0.122764445 | -0.317699381 | 0.134 | 0.076 | 1 | 4.72566E-05 | 2.876566059 | 0.5 | 0.102 | 1 | 0.122764445 | 9.4511E-05 |
| Rbm15b | 4.72993E-05 | -1.807354922 | 0.101 | 0.294 | 1 | 0.358172971 | -0.728032469 | 0.2 | 0.315 | 1 | 0.358172971 | 9.45963E-05 |
| Kpna1 | 4.74881E-05 | -1.014355293 | 0.513 | 0.7 | 1 | 0.075301621 | -1.049960564 | 0.8 | 0.806 | 1 | 0.075301621 | 9.49739E-05 |
| Camta1 | 4.75788E-05 | -1.011344974 | 0.58 | 0.753 | 1 | 0.949358448 | -0.141949429 | 0.8 | 0.657 | 1 | 0.949358448 | 9.51554E-05 |
| Slc2a9 | 0.316865657 | 0.475044809 | 0.067 | 0.041 | 1 | 4.75934E-05 | 6.313377792 | 0.2 | 0.015 | 1 | 0.316865657 | 9.51844E-05 |
| Ube2w | 4.77728E-05 | -0.90600251 | 0.63 | 0.765 | 1 | 0.110758398 | -0.598518824 | 0.7 | 0.889 | 1 | 0.110758398 | 9.55433E-05 |
| R3hdm1 | 4.78654E-05 | -0.876067672 | 0.706 | 0.765 | 1 | 0.807339885 | -0.245390769 | 1 | 0.849 | 1 | 0.807339885 | 9.57285E-05 |
| Fam76b | 4.81437E-05 | -1.261754854 | 0.286 | 0.5 | 1 | 0.860497144 | -0.120492121 | 0.5 | 0.509 | 1 | 0.860497144 | 9.62851E-05 |
| Chek1 | 4.86234E-05 | -2.321928095 | 0.042 | 0.212 | 1 | 0.492325999 | 0.653349476 | 0.3 | 0.207 | 1 | 0.492325999 | 9.72445E-05 |
| Plcg2 | 4.8712E-05 | -0.635986504 | 0.067 | 0.271 | 1 | 0.040673723 | -1.959358016 | 0.2 | 0.525 | 1 | 0.040673723 | 9.74216E-05 |
| Pak1ip1 | 4.87143E-05 | -0.801686172 | 0.496 | 0.712 | 1 | 0.041796318 | -1.108782565 | 0.6 | 0.744 | 1 | 0.041796318 | 9.74263E-05 |
| Stxbp4 | 4.94255E-05 | -1.874469118 | 0.101 | 0.3 | 1 | 0.446510399 | -0.125036046 | 0.2 | 0.327 | 1 | 0.446510399 | 9.88486E-05 |
| Mcmbp | 4.96597E-05 | -1.057678276 | 0.403 | 0.588 | 1 | 0.022326828 | -1.272755253 | 0.5 | 0.747 | 1 | 0.022326828 | 9.93169E-05 |
| Ankrd40 | 5.00529E-05 | -1.351409479 | 0.227 | 0.435 | 1 | 0.324615045 | -0.700896339 | 0.5 | 0.543 | 1 | 0.324615045 | 0.000100103 |
| Gas6 | 5.00792E-05 | -1.776328026 | 0.134 | 0.347 | 1 | 0.419443896 | -1.055327074 | 0.2 | 0.296 | 1 | 0.419443896 | 0.000100156 |
| Setd3 | 5.01555E-05 | -0.950549558 | 0.521 | 0.665 | 1 | 0.109976689 | -0.780819884 | 0.5 | 0.719 | 1 | 0.109976689 | 0.000100308 |
| Smpdl3b | 5.09342E-05 | -3.529820947 | 0.008 | 0.147 | 1 | 0.489901827 | 0.847996907 | 0 | 0.046 | 1 | 0.489901827 | 0.000101866 |
| Ints7 | 5.13934E-05 | -1.616130519 | 0.193 | 0.394 | 1 | 0.838066928 | -0.183711953 | 0.3 | 0.318 | 1 | 0.838066928 | 0.000102784 |
| Wnt9a | 5.18134E-05 | -3.416164165 | 0.017 | 0.165 | 1 | 0.364146836 | -0.789433014 | 0 | 0.077 | 1 | 0.364146836 | 0.000103624 |
| Or8b53 | 5.2038E-05 | -2.922832139 | 0.042 | 0.206 | 1 | 0.802232813 | -0.28585884 | 0.3 | 0.228 | 1 | 0.802232813 | 0.000104073 |
| Pspc1 | 5.21071E-05 | -1.00775834 | 0.504 | 0.641 | 1 | 0.273291033 | -0.591872446 | 0.6 | 0.694 | 1 | 0.273291033 | 0.000104212 |
| Ralbp1 | 5.22933E-05 | -0.730792585 | 0.739 | 0.871 | 1 | 0.053883075 | -0.836501268 | 0.7 | 0.926 | 1 | 0.053883075 | 0.000104584 |
| Sart3 | 5.23459E-05 | -1.58058406 | 0.151 | 0.353 | 1 | 0.636084138 | 0.176619654 | 0.5 | 0.37 | 1 | 0.636084138 | 0.000104689 |
| Mcm7 | 5.24866E-05 | -2.14839184 | 0.076 | 0.259 | 1 | 0.794240883 | 0.808468542 | 0.2 | 0.176 | 1 | 0.794240883 | 0.00010497 |
| Jun | 5.27103E-05 | -0.944522974 | 0.958 | 0.941 | 1 | 0.033430529 | 0.385740521 | 1 | 0.985 | 1 | 0.033430529 | 0.000105418 |
| Taf5 | 5.29267E-05 | -1.847996907 | 0.101 | 0.294 | 1 | 0.368364572 | -0.864721141 | 0.1 | 0.21 | 1 | 0.368364572 | 0.000105851 |
| Rfc1 | 5.30695E-05 | -0.787989597 | 0.504 | 0.641 | 1 | 0.114979355 | 0.625604485 | 0.9 | 0.568 | 1 | 0.114979355 | 0.000106136 |
| Rnps1 | 5.34003E-05 | -0.702783659 | 0.647 | 0.776 | 1 | 0.145107248 | -0.606568957 | 0.6 | 0.802 | 1 | 0.145107248 | 0.000106798 |
| Nol12 | 5.34268E-05 | -1.070389328 | 0.261 | 0.482 | 1 | 0.342448346 | -0.604129911 | 0.3 | 0.414 | 1 | 0.342448346 | 0.000106851 |
| Arhgap28 | 5.35235E-05 | -4.427941333 | 0.025 | 0.176 | 1 | 0.252857058 | -1.090602549 | 0 | 0.117 | 1 | 0.252857058 | 0.000107044 |
| Nup62 | 5.36664E-05 | -1.582288366 | 0.143 | 0.347 | 1 | 0.179155586 | -0.936274402 | 0.1 | 0.29 | 1 | 0.179155586 | 0.00010733 |
| Snx2 | 5.37258E-05 | -0.787659121 | 0.63 | 0.753 | 1 | 0.151231994 | -0.596787936 | 0.7 | 0.849 | 1 | 0.151231994 | 0.000107449 |
| Ptgs2 | 5.37387E-05 | 0.602036014 | 0.445 | 0.224 | 1 | 0.967912423 | -1.41670632 | 0.1 | 0.09 | 1 | 0.967912423 | 0.000107475 |
| Zfp397 | 5.38962E-05 | -1.101416224 | 0.294 | 0.512 | 1 | 0.111933515 | -1.321928095 | 0.2 | 0.426 | 1 | 0.111933515 | 0.00010779 |
| Prpf4 | 5.38994E-05 | -1.655351829 | 0.151 | 0.353 | 1 | 0.430800066 | -0.577024681 | 0.2 | 0.296 | 1 | 0.430800066 | 0.000107796 |
| Btf3l4 | 5.4031E-05 | -1.194836699 | 0.244 | 0.465 | 1 | 0.092444355 | 1.026967048 | 0.6 | 0.343 | 1 | 0.092444355 | 0.000108059 |
| Itgb7 | 5.46189E-05 | -2.246239163 | 0.05 | 0.224 | 1 | 0.375492371 | -0.391469028 | 0.2 | 0.346 | 1 | 0.375492371 | 0.000109235 |
| ENSMUSG00000121307 | 5.46303E-05 | -1.510088881 | 0.193 | 0.4 | 1 | 0.988569236 | 0.119068631 | 0.3 | 0.306 | 1 | 0.988569236 | 0.000109258 |
| Taok3 | 5.48805E-05 | -1.111865964 | 0.403 | 0.594 | 1 | 0.276679252 | -0.743629324 | 0.6 | 0.698 | 1 | 0.276679252 | 0.000109758 |
| Tedc2 | 0.042093929 | -1.807354922 | 0.008 | 0.053 | 1 | 5.48887E-05 | 4.432959407 | 0.2 | 0.015 | 1 | 0.042093929 | 0.000109774 |
| Pnp2 | 0.33261065 | -1.070389328 | 0.008 | 0.024 | 1 | 5.48887E-05 | 4.432959407 | 0.2 | 0.015 | 1 | 0.33261065 | 0.000109774 |
| Zfp568 | 5.50631E-05 | -1.50779464 | 0.218 | 0.424 | 1 | 0.708894387 | -0.479264633 | 0.5 | 0.463 | 1 | 0.708894387 | 0.000110123 |
| Akap9 | 5.50804E-05 | -0.585493782 | 0.882 | 0.947 | 1 | 0.045726429 | -0.684827971 | 0.9 | 0.972 | 1 | 0.045726429 | 0.000110158 |
| Fzd1 | 5.51761E-05 | -1.565426149 | 0.244 | 0.459 | 1 | 0.076805376 | -1.900941329 | 0.1 | 0.364 | 1 | 0.076805376 | 0.000110349 |
| Aopep | 5.53309E-05 | -0.596087028 | 1 | 1 | 1 | 0.053719416 | -0.652616324 | 1 | 1 | 1 | 0.053719416 | 0.000110659 |
| Ccdc117 | 5.56301E-05 | -1.678071905 | 0.202 | 0.4 | 1 | 0.129959378 | -1.136896201 | 0.3 | 0.491 | 1 | 0.129959378 | 0.000111257 |
| Ranbp9 | 5.56799E-05 | -0.898289955 | 0.597 | 0.788 | 1 | 0.370556926 | -0.160742943 | 0.7 | 0.892 | 1 | 0.370556926 | 0.000111357 |
| Map3k7 | 5.58752E-05 | -0.962904901 | 0.378 | 0.6 | 1 | 0.503746542 | -0.493040011 | 0.7 | 0.664 | 1 | 0.503746542 | 0.000111747 |
| Ptgfr | 5.61258E-05 | -4.427941333 | 0.025 | 0.176 | 1 | 0.316140173 | -1.802257054 | 0 | 0.093 | 1 | 0.316140173 | 0.000112248 |
| Ptpn9 | 5.61615E-05 | -1.129283017 | 0.479 | 0.635 | 1 | 0.02343909 | -1.087382246 | 0.4 | 0.778 | 1 | 0.02343909 | 0.00011232 |
| Tbc1d10b | 5.6292E-05 | -2.157852169 | 0.042 | 0.212 | 1 | 0.524349353 | 0.602884409 | 0.4 | 0.306 | 1 | 0.524349353 | 0.000112581 |
| Atp5a1 | 5.64149E-05 | -0.490881257 | 0.924 | 0.971 | 1 | 0.026950717 | -0.73410026 | 0.7 | 0.991 | 1 | 0.026950717 | 0.000112827 |
| Smg6 | 5.64459E-05 | -0.646789815 | 0.807 | 0.906 | 1 | 0.434886293 | -0.559003274 | 1 | 0.929 | 1 | 0.434886293 | 0.000112889 |
| Gskip | 5.68426E-05 | -1.655351829 | 0.134 | 0.329 | 1 | 0.676042974 | 0.273760812 | 0.2 | 0.275 | 1 | 0.676042974 | 0.000113682 |
| Rilpl2 | 5.70107E-05 | -1.16480493 | 0.395 | 0.594 | 1 | 0.162393547 | -1.097555309 | 0.5 | 0.58 | 1 | 0.162393547 | 0.000114018 |
| Lrrc49 | 5.73932E-05 | -1.004155629 | 0.445 | 0.635 | 1 | 0.00429851 | -2.900941329 | 0.1 | 0.568 | 1 | 0.00429851 | 0.000114783 |
| Zfc3h1 | 5.74259E-05 | -0.594849656 | 0.832 | 0.924 | 1 | 0.163454251 | -0.641587546 | 0.9 | 0.966 | 1 | 0.163454251 | 0.000114848 |
| Tmem132b | 0.876265947 | -0.201633861 | 0.05 | 0.047 | 1 | 5.78377E-05 | 2.892391026 | 0.3 | 0.037 | 1 | 0.876265947 | 0.000115672 |
| Gm28153 | 5.80752E-05 | -2.134519665 | 0.076 | 0.259 | 1 | 0.948095714 | -0.138582578 | 0.2 | 0.194 | 1 | 0.948095714 | 0.000116147 |
| Cirbp | 5.80941E-05 | -1.076102979 | 0.471 | 0.635 | 1 | 0.417710441 | -0.501714345 | 0.6 | 0.63 | 1 | 0.417710441 | 0.000116185 |
| T2 | 0.666171662 | 0.251538767 | 0.025 | 0.018 | 1 | 5.81262E-05 | 4.339850003 | 0.2 | 0.015 | 1 | 0.666171662 | 0.000116249 |
| Nptx1 | 5.84528E-05 | 2.493199522 | 0.151 | 0.024 | 1 | 0.234627319 | 1.930459067 | 0.1 | 0.031 | 1 | 0.234627319 | 0.000116902 |
| Snx27 | 5.85616E-05 | -0.941805122 | 0.471 | 0.641 | 1 | 0.548547454 | -0.471462933 | 0.7 | 0.713 | 1 | 0.548547454 | 0.00011712 |

| | | | | | | | | | | | | |
|---|---|---|---|---|---|---|---|---|---|---|---|---|
| Stt3a | 5.85814E-05 | -0.50736938 | 0.832 | 0.9 | 1 | 0.963854554 | 0.184383054 | 0.9 | 0.92 | 1 | 0.963854554 | 0.000117159 |
| Tafa2 | 5.87198E-05 | 0.982122509 | 0.353 | 0.147 | 1 | 0.017685578 | 4.100384068 | 0.2 | 0.043 | 1 | 0.017685578 | 0.000117436 |
| Ppp4r4 | 0.062759149 | -1.944858446 | 0.008 | 0.047 | 1 | 5.89164E-05 | 3.737813989 | 0.3 | 0.037 | 1 | 0.062759149 | 0.000117829 |
| Esyt3 | 5.93936E-05 | 1.314274522 | 0.261 | 0.082 | 1 | 0.274256184 | 2.31748219 | 0.1 | 0.034 | 1 | 0.274256184 | 0.000118784 |
| Matn2 | 5.97964E-05 | -1.175471374 | 0.311 | 0.535 | 1 | 0.338137767 | -0.728032469 | 0.4 | 0.534 | 1 | 0.338137767 | 0.000119589 |
| Phf23 | 5.98643E-05 | -1.478047297 | 0.311 | 0.494 | 1 | 0.079467317 | -1.204470513 | 0.4 | 0.614 | 1 | 0.079467317 | 0.000119725 |
| Cul4a | 6.01308E-05 | -0.932226889 | 0.403 | 0.6 | 1 | 0.484392126 | 0.260224872 | 0.8 | 0.698 | 1 | 0.484392126 | 0.000120258 |
| Dock9 | 6.02393E-05 | 1.393266877 | 0.412 | 0.2 | 1 | 0.031215629 | 1.446379923 | 0.7 | 0.398 | 1 | 0.031215629 | 0.000120475 |
| Gpc1 | 6.04638E-05 | -1.011863251 | 0.697 | 0.794 | 1 | 0.190914567 | -1.14117316 | 0.7 | 0.713 | 1 | 0.190914567 | 0.000120924 |
| N4bp2l2 | 6.11135E-05 | -0.590863102 | 0.84 | 0.918 | 1 | 0.636598087 | 0.154266589 | 1 | 0.892 | 1 | 0.636598087 | 0.000122223 |
| Shisa4 | 6.11799E-05 | 0.76438829 | 0.664 | 0.476 | 1 | 0.002199318 | 1.362570079 | 0.8 | 0.312 | 1 | 0.002199318 | 0.000122356 |
| Dstyk | 6.13556E-05 | -1.15585867 | 0.437 | 0.594 | 1 | 0.266790861 | -0.539454761 | 0.5 | 0.654 | 1 | 0.266790861 | 0.000122707 |
| Tmem145 | 0.217412808 | -1.485426827 | 0.008 | 0.029 | 1 | 6.15371E-05 | 4.017921908 | 0.2 | 0.015 | 1 | 0.217412808 | 0.00012307 |
| Hlf | 6.16774E-05 | -2.817623258 | 0.076 | 0.253 | 1 | 0.462190159 | -0.159995884 | 0.2 | 0.324 | 1 | 0.462190159 | 0.000123351 |
| Usp37 | 6.17081E-05 | -0.963833227 | 0.471 | 0.659 | 1 | 0.574716651 | -0.26953397 | 0.8 | 0.821 | 1 | 0.574716651 | 0.000123412 |
| Cherp | 6.19131E-05 | -1.301714874 | 0.303 | 0.5 | 1 | 0.729918917 | -0.129565704 | 0.4 | 0.485 | 1 | 0.729918917 | 0.000123822 |
| Dhcr24 | 6.21006E-05 | 1.121230745 | 0.496 | 0.265 | 1 | 0.017175226 | 1.227844977 | 0.4 | 0.123 | 1 | 0.017175226 | 0.000124197 |
| Rsbn1l | 6.22227E-05 | -1.304854582 | 0.37 | 0.559 | 1 | 0.989006037 | -0.24081236 | 0.8 | 0.701 | 1 | 0.989006037 | 0.000124441 |
| Tceal5 | 6.22781E-05 | -2.070389328 | 0.067 | 0.247 | 1 | 0.880819654 | 0.263034406 | 0.2 | 0.17 | 1 | 0.880819654 | 0.000124552 |
| U2af2 | 6.23927E-05 | -0.898289955 | 0.496 | 0.682 | 1 | 0.138885615 | -0.532824877 | 0.4 | 0.679 | 1 | 0.138885615 | 0.000124782 |
| Rnf34 | 6.25121E-05 | 0.736965594 | 0.748 | 0.518 | 1 | 0.002201454 | 1.617991301 | 0.8 | 0.506 | 1 | 0.002201454 | 0.00012502 |
| Rmi2 | 0.218485013 | -1.292781749 | 0.008 | 0.029 | 1 | 6.33043E-05 | 3.847996907 | 0.2 | 0.015 | 1 | 0.218485013 | 0.000126605 |
| Gm2449 | 0.751159595 | 0.377069649 | 0.034 | 0.041 | 1 | 6.33043E-05 | 3.847996907 | 0.2 | 0.015 | 1 | 0.751159595 | 0.000126605 |
| A730011C13Rik | 0.038142909 | 2.514573173 | 0.025 | 0 | 1 | 6.33197E-05 | 4.017921908 | 0.2 | 0.015 | 1 | 0.038142909 | 0.000126635 |
| Zfp3 | 0.219589239 | -1.070389328 | 0.008 | 0.029 | 1 | 6.33197E-05 | 4.017921908 | 0.2 | 0.015 | 1 | 0.219589239 | 0.000126635 |
| 5033425G24Rik | 0.721162042 | 0.514573173 | 0.017 | 0.012 | 1 | 6.33197E-05 | 4.017921908 | 0.2 | 0.015 | 1 | 0.721162042 | 0.000126635 |
| Itga9 | 6.3392E-05 | -1.465248945 | 0.269 | 0.476 | 1 | 0.238864612 | -0.699754515 | 0.3 | 0.5 | 1 | 0.238864612 | 0.00012678 |
| Ypel5 | 6.39885E-05 | 0.590190219 | 0.782 | 0.612 | 1 | 0.041837507 | -1.203181817 | 0.2 | 0.59 | 1 | 0.041837507 | 0.000127973 |
| A230057D06Rik | 6.4909E-05 | 0.854059639 | 0.437 | 0.2 | 1 | 0.007688667 | 2.274697323 | 0.5 | 0.204 | 1 | 0.007688667 | 0.000129814 |
| Hps5 | 6.50257E-05 | -1.345249169 | 0.185 | 0.394 | 1 | 0.471628913 | -0.505640048 | 0.3 | 0.398 | 1 | 0.471628913 | 0.000130047 |
| Gm31728 | 0.220074704 | 1 | 0.05 | 0.024 | 1 | 6.51224E-05 | 3.432959407 | 0.2 | 0.015 | 1 | 0.220074704 | 0.00013024 |
| Panx3 | 6.51423E-05 | -5.409390936 | 0.042 | 0.2 | 1 | 0.539481436 | -2.596787936 | 0 | 0.037 | 1 | 0.539481436 | 0.00013028 |
| Sem1 | 6.58438E-05 | -0.536378015 | 0.95 | 0.971 | 1 | 0.014557705 | -0.800837777 | 0.8 | 0.997 | 1 | 0.014557705 | 0.000131683 |
| Slc35g2 | 0.465575803 | -0.555816155 | 0.143 | 0.171 | 1 | 6.59161E-05 | 2.270687978 | 0.5 | 0.096 | 1 | 0.465575803 | 0.000131828 |
| Tle2 | 6.60867E-05 | -2.694880193 | 0.025 | 0.176 | 1 | 0.252404124 | -0.567040593 | 0 | 0.117 | 1 | 0.252404124 | 0.000132169 |
| B430212C06Rik | 0.030981448 | 1.652076697 | 0.067 | 0.018 | 1 | 6.69515E-05 | 3.795529487 | 0.2 | 0.015 | 1 | 0.030981448 | 0.000133899 |
| Slc24a4 | 0.05288261 | -1.807354922 | 0.017 | 0.065 | 1 | 6.69515E-05 | 3.280956314 | 0.2 | 0.015 | 1 | 0.05288261 | 0.000133899 |
| Bub1b | 0.062759149 | -2.292781749 | 0.008 | 0.047 | 1 | 6.69515E-05 | 3.795529487 | 0.2 | 0.015 | 1 | 0.062759149 | 0.000133899 |
| Gm57065 | 0.113476214 | 2.099535674 | 0.05 | 0.018 | 1 | 6.69515E-05 | 3.795529487 | 0.2 | 0.015 | 1 | 0.113476214 | 0.000133899 |
| Gm56858 | 0.197589594 | 1.736965594 | 0.034 | 0.012 | 1 | 6.69515E-05 | 3.795529487 | 0.2 | 0.015 | 1 | 0.197589594 | 0.000133899 |
| Lrrc43 | 0.500300313 | -0.163498732 | 0.017 | 0.029 | 1 | 6.69515E-05 | 3.280956314 | 0.2 | 0.015 | 1 | 0.500300313 | 0.000133899 |
| 4921515G04Rik | 0.695893386 | -0.222392421 | 0.017 | 0.024 | 1 | 6.69515E-05 | 3.795529487 | 0.2 | 0.015 | 1 | 0.695893386 | 0.000133899 |
| Carnmt1 | 6.71033E-05 | -0.792708146 | 0.479 | 0.671 | 1 | 0.540888632 | -0.123090402 | 0.5 | 0.667 | 1 | 0.540888632 | 0.000134202 |
| Oxct1 | 6.71595E-05 | -0.713890567 | 0.908 | 0.865 | 1 | 0.72986296 | 0.280536987 | 0.7 | 0.889 | 1 | 0.72986296 | 0.000134315 |
| Zyx | 6.75367E-05 | 1.016322305 | 0.597 | 0.388 | 1 | 0.307495575 | 0.936533581 | 0.6 | 0.429 | 1 | 0.307495575 | 0.000135069 |
| Trappc6b | 6.78025E-05 | -0.970853654 | 0.571 | 0.671 | 1 | 0.020042965 | -1.163407857 | 0.5 | 0.756 | 1 | 0.020042965 | 0.0001356 |
| Dock10 | 6.86034E-05 | -3.854660637 | 0.017 | 0.159 | 1 | 0.121693352 | -2.465893869 | 0 | 0.198 | 1 | 0.121693352 | 0.000137202 |
| Fas | 6.89341E-05 | 0.883355081 | 0.546 | 0.324 | 1 | 0.021337576 | 2.549773072 | 0.5 | 0.256 | 1 | 0.021337576 | 0.000137863 |
| Tbc1d31 | 0.213899158 | -0.883976204 | 0.05 | 0.088 | 1 | 6.97237E-05 | 3.159940913 | 0.4 | 0.065 | 1 | 0.213899158 | 0.000139442 |
| Gm32282 | 7.01639E-05 | 1.707218251 | 0.235 | 0.071 | 1 | 0.505605027 | 0.930459067 | 0 | 0.043 | 1 | 0.505605027 | 0.000140323 |
| Aldh1l1 | 0.00037406 | 2.762500686 | 0.118 | 0.018 | 1 | 7.08575E-05 | 3.432959407 | 0.2 | 0.015 | 1 | 0.00037406 | 0.00014171 |
| Fhl4 | 0.508731273 | -0.807354922 | 0.008 | 0.018 | 1 | 7.08575E-05 | 3.280956314 | 0.2 | 0.015 | 1 | 0.508731273 | 0.00014171 |
| Kcnj6 | 0.929389703 | 0.192645078 | 0.025 | 0.024 | 1 | 7.08575E-05 | 2.902444691 | 0.2 | 0.015 | 1 | 0.929389703 | 0.00014171 |
| Klhl31 | 0.203701262 | 1.099535674 | 0.034 | 0.012 | 1 | 7.08807E-05 | 3.111031312 | 0.2 | 0.015 | 1 | 0.203701262 | 0.000141756 |
| Pole2 | 0.074132329 | -1.106915204 | 0.084 | 0.153 | 1 | 7.13495E-05 | 1.930459067 | 0.5 | 0.093 | 1 | 0.074132329 | 0.000142694 |
| Bicd2 | 7.13991E-05 | -1.82527683 | 0.126 | 0.318 | 1 | 0.009876428 | -2.802257054 | 0 | 0.423 | 1 | 0.009876428 | 0.000142793 |
| Gli1 | 7.14031E-05 | -4.129283017 | 0 | 0.124 | 1 | 0.489901827 | 0.847996907 | 0 | 0.046 | 1 | 0.489901827 | 0.000142801 |
| Gm48742 | 7.15537E-05 | 1.421463768 | 0.21 | 0.053 | 1 | 0.439528334 | 1.930459067 | 0.1 | 0.046 | 1 | 0.439528334 | 0.000143102 |
| Cdh24 | 7.18408E-05 | -4.733354341 | 0 | 0.124 | 1 | 0.822520652 | 0.135278859 | 0.1 | 0.12 | 1 | 0.822520652 | 0.000143676 |
| Rrad | 7.1896E-05 | -5.185866545 | 0 | 0.124 | 1 | 0.397023343 | -0.339630097 | 0 | 0.068 | 1 | 0.397023343 | 0.000143787 |
| 1700024B18Rik | 7.19887E-05 | -5.944858446 | 0 | 0.124 | 1 | 0.316192774 | -2.647414009 | 0 | 0.093 | 1 | 0.316192774 | 0.000143972 |
| Smn1 | 7.20698E-05 | -1.246239163 | 0.244 | 0.447 | 1 | 0.243919162 | -0.367509129 | 0.2 | 0.407 | 1 | 0.243919162 | 0.000144134 |
| Dchs1 | 7.21176E-05 | -2.292781749 | 0.042 | 0.206 | 1 | 0.135675165 | -1.171902651 | 0.1 | 0.315 | 1 | 0.135675165 | 0.00014423 |
| Srsf3 | 7.22546E-05 | -0.431206436 | 0.95 | 0.994 | 1 | 0.199211593 | -0.336834717 | 0.9 | 0.994 | 1 | 0.199211593 | 0.000144504 |
| Nr4a3 | 7.2445E-05 | -1.748461233 | 0.168 | 0.376 | 1 | 0.890393886 | -0.226838326 | 0.5 | 0.469 | 1 | 0.890393886 | 0.000144885 |
| Fbxl5 | 7.27264E-05 | 0.500633982 | 0.807 | 0.594 | 1 | 0.093300457 | 0.978057447 | 0.9 | 0.713 | 1 | 0.093300457 | 0.000145447 |
| Sema4c | 7.30176E-05 | -3.185866545 | 0.017 | 0.159 | 1 | 0.202435913 | -1.304006187 | 0 | 0.142 | 1 | 0.202435913 | 0.00014603 |
| Mrtfb | 7.32687E-05 | -1.187428749 | 0.353 | 0.547 | 1 | 0.109889331 | 0.965702225 | 0.8 | 0.691 | 1 | 0.109889331 | 0.000146532 |
| Mob1a | 7.33353E-05 | -1.039025157 | 0.37 | 0.565 | 1 | 0.012869967 | -1.181750437 | 0.3 | 0.719 | 1 | 0.012869967 | 0.000146665 |
| Hoga1 | 0.164390276 | -1.222392421 | 0.042 | 0.082 | 1 | 7.38885E-05 | 2.532495081 | 0.4 | 0.062 | 1 | 0.164390276 | 0.000147772 |
| Ccn4 | 7.3955E-05 | -1.652205273 | 0.277 | 0.482 | 1 | 0.174303109 | -1.457811523 | 0.1 | 0.284 | 1 | 0.174303109 | 0.000147904 |
| Sgk1 | 7.42325E-05 | 1.134854329 | 0.782 | 0.682 | 1 | 0.073668025 | 1.339532543 | 0.8 | 0.71 | 1 | 0.073668025 | 0.00014846 |
| Ttc13 | 7.42903E-05 | -2.129283017 | 0.059 | 0.229 | 1 | 0.780773173 | 0.580516596 | 0.2 | 0.16 | 1 | 0.780773173 | 0.000148575 |
| Gm30085 | 0.124328275 | 1.514573173 | 0.076 | 0.035 | 1 | 7.49685E-05 | 3.280956314 | 0.2 | 0.015 | 1 | 0.124328275 | 0.000149931 |
| Dlec1 | 0.721161231 | 0.514573173 | 0.017 | 0.012 | 1 | 7.49701E-05 | 3.432959407 | 0.2 | 0.015 | 1 | 0.721161231 | 0.000149935 |
| Drp2 | 0.000604395 | 2.234465254 | 0.176 | 0.053 | 1 | 7.49775E-05 | 3.380491987 | 0.4 | 0.068 | 1 | 0.000604395 | 0.000149949 |
| Snx13 | 7.51446E-05 | 0.956678915 | 0.697 | 0.471 | 1 | 0.267168013 | 0.237703116 | 0.9 | 0.679 | 1 | 0.267168013 | 0.000150284 |
| Spg20 | 7.58289E-05 | -1.19592021 | 0.294 | 0.5 | 1 | 0.585847612 | -0.138582578 | 0.3 | 0.383 | 1 | 0.585847612 | 0.000151652 |
| Bclaf3 | 7.59429E-05 | -1.521952703 | 0.252 | 0.453 | 1 | 0.659687793 | -0.214738849 | 0.4 | 0.494 | 1 | 0.659687793 | 0.00015188 |
| Zfp821 | 7.61E-05 | -2.03562391 | 0.092 | 0.276 | 1 | 0.682432631 | 0.130396637 | 0.2 | 0.287 | 1 | 0.682432631 | 0.000152194 |
| Zfp597 | 7.61224E-05 | -2.046141782 | 0.076 | 0.253 | 1 | 0.173621258 | -1.181750437 | 0.1 | 0.29 | 1 | 0.173621258 | 0.000152239 |
| Nog | 7.63171E-05 | -0.11345805 | 0.361 | 0.153 | 1 | 0.015042698 | 1.959028219 | 0.3 | 0.083 | 1 | 0.015042698 | 0.000152628 |
| Zfta | 7.64099E-05 | -2.070389328 | 0.092 | 0.271 | 1 | 0.058631931 | -1.441509711 | 0.1 | 0.392 | 1 | 0.058631931 | 0.000152814 |
| Snrpd3 | 7.6467E-05 | -0.72509724 | 0.731 | 0.806 | 1 | 0.05435859 | -0.591872446 | 0.6 | 0.843 | 1 | 0.05435859 | 0.000152928 |
| Tecr | 7.67007E-05 | 0.898532361 | 0.866 | 0.782 | 1 | 0.000672029 | 1.108205268 | 0.9 | 0.765 | 1 | 0.000672029 | 0.000153395 |
| Tex30 | 7.67487E-05 | -2.269698136 | 0.059 | 0.229 | 1 | 0.221305268 | 0.894539492 | 0.4 | 0.235 | 1 | 0.221305268 | 0.000153492 |
| Ifngr2 | 7.71989E-05 | -1.535057595 | 0.319 | 0.494 | 1 | 0.078787428 | -1.12367437 | 0.4 | 0.596 | 1 | 0.078787428 | 0.000154392 |
| Hsf2 | 7.74461E-05 | -1.655351829 | 0.109 | 0.3 | 1 | 0.018194133 | -2.776493958 | 0 | 0.377 | 1 | 0.018194133 | 0.000154886 |
| Gm8251 | 7.76761E-05 | -4.392317423 | 0.008 | 0.141 | 1 | 0.554364676 | 0.481869008 | 0.2 | 0.133 | 1 | 0.554364676 | 0.000155346 |
| Hmgn3 | 7.82657E-05 | -1.076035891 | 0.521 | 0.671 | 1 | 0.262010501 | -0.489872732 | 0.6 | 0.682 | 1 | 0.262010501 | 0.000156525 |
| Rnf227 | 7.84132E-05 | -3.614709844 | 0.008 | 0.141 | 1 | 0.385695028 | -0.304006187 | 0 | 0.071 | 1 | 0.385695028 | 0.00015682 |
| Grik5 | 7.94254E-05 | -2.185866545 | 0.025 | 0.176 | 1 | 0.063970357 | -2.111361109 | 0 | 0.262 | 1 | 0.063970357 | 0.000158845 |
| Cdk12 | 7.96123E-05 | -0.685683602 | 0.84 | 0.918 | 1 | 0.739109478 | -0.167944637 | 0.8 | 0.963 | 1 | 0.739109478 | 0.000159218 |
| Ccnl1 | 8.00081E-05 | -0.583202043 | 0.966 | 0.953 | 1 | 0.02912699 | -0.700135224 | 0.9 | 0.975 | 1 | 0.02912699 | 0.00016001 |
| Phlpp2 | 8.02912E-05 | -1.061619118 | 0.252 | 0.482 | 1 | 0.487789129 | 0.245332404 | 0.7 | 0.568 | 1 | 0.487789129 | 0.000160576 |
| Csf1 | 8.04263E-05 | -2.217230716 | 0.084 | 0.265 | 1 | 0.080076508 | -2.141949429 | 0 | 0.241 | 1 | 0.080076508 | 0.000160846 |
| Gm31305 | 0.62276256 | -0.622930351 | 0.025 | 0.035 | 1 | 8.04561E-05 | 3.252387162 | 0.3 | 0.037 | 1 | 0.62276256 | 0.000160906 |
| Ssr3 | 8.09348E-05 | -0.445406895 | 0.966 | 0.971 | 1 | 0.165357235 | -0.417632609 | 0.9 | 0.981 | 1 | 0.165357235 | 0.000161863 |

| | | | | | | | | | | | | |
|---|---|---|---|---|---|---|---|---|---|---|---|---|
| Cpsf2 | 8.10866E-05 | -1.819327564 | 0.168 | 0.359 | 1 | 0.011444697 | -2.723545078 | 0 | 0.41 | 1 | 0.011444697 | 0.000162167 |
| Tpp1 | 8.13302E-05 | 0.762500686 | 0.723 | 0.459 | 1 | 0.134524325 | 0.621317127 | 0.8 | 0.546 | 1 | 0.134524325 | 0.000162654 |
| Slc7a5 | 8.16516E-05 | -1.144389909 | 0.345 | 0.553 | 1 | 0.271247681 | -0.818128447 | 0.4 | 0.546 | 1 | 0.271247681 | 0.000163297 |
| Ndufv1 | 8.17143E-05 | -1.292781749 | 0.286 | 0.476 | 1 | 0.114961066 | -0.953621646 | 0.3 | 0.522 | 1 | 0.114961066 | 0.000163422 |
| Ptdss1 | 8.17257E-05 | -0.998239542 | 0.252 | 0.471 | 1 | 0.547160048 | 0.48866884 | 0.6 | 0.503 | 1 | 0.547160048 | 0.000163445 |
| Ccn5 | 8.18531E-05 | 1.213642159 | 0.462 | 0.259 | 1 | 0.085070965 | 2.292638119 | 0.4 | 0.188 | 1 | 0.085070965 | 0.0001637 |
| Cetn4 | 8.18807E-05 | 0.959984321 | 0.395 | 0.182 | 1 | 0.392281661 | 1.902444691 | 0.2 | 0.13 | 1 | 0.392281661 | 0.000163755 |
| Nfyc | 8.21055E-05 | -0.943108664 | 0.412 | 0.612 | 1 | 0.268140933 | -0.811800827 | 0.5 | 0.608 | 1 | 0.268140933 | 0.000164204 |
| Sec16a | 8.21499E-05 | -0.81620584 | 0.681 | 0.759 | 1 | 0.20829184 | -0.425021588 | 0.9 | 0.824 | 1 | 0.20829184 | 0.000164293 |
| Gadd45b | 8.2271E-05 | -0.73254805 | 0.782 | 0.9 | 1 | 0.02051705 | -1.163016086 | 0.7 | 0.932 | 1 | 0.02051705 | 0.000164535 |
| Ncor2 | 8.27729E-05 | -0.706335757 | 0.639 | 0.788 | 1 | 0.006770969 | -0.947862377 | 0.4 | 0.883 | 1 | 0.006770969 | 0.000165539 |
| Mdm2 | 8.29136E-05 | -0.517235565 | 0.588 | 0.759 | 1 | 0.046349859 | -1.097555309 | 0.6 | 0.747 | 1 | 0.046349859 | 0.00016582 |
| Bcat1 | 0.000125645 | 0.543803884 | 0.689 | 0.376 | 1 | 8.29518E-05 | 2.676563283 | 0.8 | 0.244 | 1 | 0.000125645 | 0.000165897 |
| Eln | 8.31751E-05 | -2.696137443 | 0.151 | 0.347 | 1 | 0.286737891 | 0.748461233 | 0.4 | 0.238 | 1 | 0.286737891 | 0.000166343 |
| Vti1a | 8.34567E-05 | -1.198805601 | 0.471 | 0.659 | 1 | 0.062521458 | -1.115477217 | 0.4 | 0.704 | 1 | 0.062521458 | 0.000166906 |
| Pcdh7 | 8.41223E-05 | -0.363171077 | 0.143 | 0.371 | 1 | 0.022650281 | -3.376540787 | 0.1 | 0.457 | 1 | 0.022650281 | 0.000168238 |
| Wdfy3 | 8.43153E-05 | -1.058893689 | 0.697 | 0.747 | 1 | 0.31810313 | -0.374395515 | 0.7 | 0.91 | 1 | 0.31810313 | 0.000168624 |
| Slit2 | 8.51903E-05 | -3.683366205 | 0.084 | 0.259 | 1 | 0.020574742 | -4.136896201 | 0 | 0.367 | 1 | 0.020574742 | 0.000170373 |
| Ilrun | 8.52907E-05 | -0.809399848 | 0.714 | 0.782 | 1 | 0.061659006 | -0.670788518 | 0.7 | 0.898 | 1 | 0.061659006 | 0.000170574 |
| Lsm2 | 8.57387E-05 | -1.299871174 | 0.202 | 0.406 | 1 | 0.278029605 | -0.49777793 | 0.2 | 0.383 | 1 | 0.278029605 | 0.00017147 |
| Tmem260 | 8.61384E-05 | -1.503348735 | 0.134 | 0.329 | 1 | 0.262192806 | -0.103093493 | 0.1 | 0.29 | 1 | 0.262192806 | 0.000172269 |
| Ankib1 | 8.73981E-05 | -0.932034405 | 0.563 | 0.706 | 1 | 0.612637063 | -0.359288622 | 0.8 | 0.778 | 1 | 0.612637063 | 0.000174788 |
| Ctr9 | 8.74488E-05 | -1.216368634 | 0.42 | 0.582 | 1 | 0.091274971 | -0.891971176 | 0.4 | 0.679 | 1 | 0.091274971 | 0.00017489 |
| Zc3h7b | 8.84877E-05 | -1.248726569 | 0.261 | 0.459 | 1 | 0.190584188 | -0.79970135 | 0.3 | 0.5 | 1 | 0.190584188 | 0.000176967 |
| Ywhaz | 0.00102457 | -0.320481663 | 1 | 1 | 1 | 8.9231E-05 | -1.291933355 | 1 | 1 | 1 | 0.00102457 | 0.000178454 |
| Ppp2r2a | 8.93353E-05 | -0.62792334 | 0.857 | 0.882 | 1 | 0.011181306 | -0.885167117 | 0.7 | 0.938 | 1 | 0.011181306 | 0.000178663 |
| Nelfa | 8.95098E-05 | -1.470319935 | 0.168 | 0.371 | 1 | 0.705124211 | 0.141404961 | 0.6 | 0.475 | 1 | 0.705124211 | 0.000179012 |
| Rdh10 | 8.96885E-05 | -2.014679895 | 0.118 | 0.306 | 1 | 0.118572968 | -0.702200176 | 0.2 | 0.478 | 1 | 0.118572968 | 0.000179369 |
| Tasor2 | 9.01949E-05 | -0.970853654 | 0.471 | 0.647 | 1 | 0.01134858 | -1.609003886 | 0.5 | 0.731 | 1 | 0.01134858 | 0.000180382 |
| Ttll3 | 9.11911E-05 | -2.160991877 | 0.101 | 0.294 | 1 | 0.707598386 | 0.530656081 | 0.2 | 0.309 | 1 | 0.707598386 | 0.000182374 |
| Tasor | 9.14253E-05 | -0.970853654 | 0.622 | 0.735 | 1 | 0.906043605 | -0.283474876 | 0.9 | 0.812 | 1 | 0.906043605 | 0.000182842 |
| Pygo2 | 9.16339E-05 | -1.561777713 | 0.168 | 0.365 | 1 | 0.542032144 | -0.230005605 | 0.3 | 0.395 | 1 | 0.542032144 | 0.000183259 |
| Anxa1 | 9.17217E-05 | 1.057236972 | 0.849 | 0.782 | 1 | 0.612121983 | 0.310102659 | 0.7 | 0.759 | 1 | 0.612121983 | 0.000183435 |
| Chst11 | 9.18475E-05 | 0.233166451 | 1 | 0.959 | 1 | 0.319941914 | 0.517831542 | 1 | 0.948 | 1 | 0.319941914 | 0.000183687 |
| P3h4 | 9.18809E-05 | 0.543428036 | 0.95 | 0.835 | 1 | 0.03160315 | 0.796818183 | 0.9 | 0.738 | 1 | 0.03160315 | 0.000183753 |
| Gsto1 | 9.21386E-05 | -1.117695043 | 0.143 | 0.353 | 1 | 0.027804761 | -2.308507579 | 0.1 | 0.448 | 1 | 0.027804761 | 0.000184269 |
| Cfap20 | 9.22098E-05 | -0.972778531 | 0.546 | 0.688 | 1 | 0.003076919 | -1.452398027 | 0.2 | 0.747 | 1 | 0.003076919 | 0.000184411 |
| Gramd1a | 9.2331E-05 | -1.191695624 | 0.261 | 0.453 | 1 | 0.116805611 | -1.080110175 | 0.3 | 0.506 | 1 | 0.116805611 | 0.000184654 |
| Tshz1 | 9.23634E-05 | -0.707819249 | 0.37 | 0.594 | 1 | 0.043753926 | -1.611434712 | 0.3 | 0.574 | 1 | 0.043753926 | 0.000184718 |
| Map3k6 | 0.001628711 | 0.918963428 | 0.261 | 0.112 | 1 | 9.27288E-05 | 2.31748219 | 0.5 | 0.099 | 1 | 0.001628711 | 0.000185449 |
| Smc1a | 9.27719E-05 | -0.424715413 | 0.874 | 0.929 | 1 | 0.050802083 | -0.826427222 | 0.9 | 0.948 | 1 | 0.050802083 | 0.000185535 |
| Notch1 | 9.33938E-05 | -1.5360529 | 0.244 | 0.447 | 1 | 0.024602349 | -1.593102889 | 0.2 | 0.59 | 1 | 0.024602349 | 0.000186779 |
| Trp53inp2 | 9.35102E-05 | 0.96203215 | 0.529 | 0.312 | 1 | 0.824167476 | 0.299103661 | 0.4 | 0.426 | 1 | 0.824167476 | 0.000187012 |
| Nkain2 | 0.023526018 | 3.652076697 | 0.059 | 0.012 | 1 | 9.36237E-05 | 6.432959407 | 0.1 | 0.003 | 1 | 0.023526018 | 0.000187239 |
| Traf3ip3 | 0.167062889 | 1.836501268 | 0.025 | 0.006 | 1 | 9.36237E-05 | 6.017921908 | 0.1 | 0.003 | 1 | 0.167062889 | 0.000187239 |
| Gm44767 | 0.167062889 | 2.099535674 | 0.025 | 0.006 | 1 | 9.36237E-05 | 7.240314329 | 0.1 | 0.003 | 1 | 0.167062889 | 0.000187239 |
| Gm41821 | 0.171451236 | 0.777607579 | 0.025 | 0.006 | 1 | 9.36237E-05 | 6.82527683 | 0.1 | 0.003 | 1 | 0.171451236 | 0.000187239 |
| Gckr | 0.396130756 | 0.192645078 | 0.025 | 0.012 | 1 | 9.36237E-05 | 5.602884409 | 0.1 | 0.003 | 1 | 0.396130756 | 0.000187239 |
| Smim6 | 0.695893386 | -0.222392421 | 0.017 | 0.024 | 1 | 9.36237E-05 | 6.339850003 | 0.1 | 0.003 | 1 | 0.695893386 | 0.000187239 |
| Bank1 | 0.781790612 | -3.070389328 | 0.008 | 0.012 | 1 | 9.36237E-05 | 6.017921908 | 0.1 | 0.003 | 1 | 0.781790612 | 0.000187239 |
| Wrn | 9.43049E-05 | -1.510961919 | 0.185 | 0.388 | 1 | 0.032236515 | -1.888968688 | 0.1 | 0.438 | 1 | 0.032236515 | 0.000188601 |
| Ppp4r2 | 9.47709E-05 | -0.597980649 | 0.84 | 0.882 | 1 | 0.138419713 | -0.48221916 | 1 | 0.957 | 1 | 0.138419713 | 0.000189533 |
| Lrmda | 9.50998E-05 | 0.750344624 | 0.655 | 0.388 | 1 | 0.006948674 | 1.424595934 | 0.9 | 0.54 | 1 | 0.006948674 | 0.000190191 |
| Tab2 | 9.51563E-05 | -0.665034382 | 0.824 | 0.888 | 1 | 0.419234329 | -0.282500697 | 0.9 | 0.975 | 1 | 0.419234329 | 0.000190303 |
| Ankrd52 | 9.51943E-05 | -1.351409479 | 0.244 | 0.441 | 1 | 0.249054354 | -0.773892163 | 0.3 | 0.472 | 1 | 0.249054354 | 0.00019038 |
| Xpot | 9.53897E-05 | -0.971985624 | 0.538 | 0.665 | 1 | 0.172200674 | -0.651671843 | 0.5 | 0.691 | 1 | 0.172200674 | 0.00019077 |
| Ccdc136 | 9.5488E-05 | -2.842978832 | 0.025 | 0.171 | 1 | 0.683450239 | 0.31748219 | 0.3 | 0.228 | 1 | 0.683450239 | 0.000190967 |
| Gmip | 9.55429E-05 | -2.944858446 | 0.025 | 0.171 | 1 | 0.660332477 | 0.580516596 | 0.2 | 0.142 | 1 | 0.660332477 | 0.000191077 |
| Amer1 | 9.55655E-05 | -2.807354922 | 0.025 | 0.171 | 1 | 0.213976815 | -1.111361109 | 0 | 0.136 | 1 | 0.213976815 | 0.000191122 |
| Itgb5 | 9.60655E-05 | 0.369924531 | 0.975 | 0.829 | 1 | 0.018396617 | 0.56096702 | 1 | 0.923 | 1 | 0.018396617 | 0.000192122 |
| Cops4 | 9.61501E-05 | -0.815734588 | 0.555 | 0.706 | 1 | 0.197124639 | -0.53359411 | 0.5 | 0.722 | 1 | 0.197124639 | 0.000192291 |
| 2810001G20Rik | 9.62658E-05 | 0.566882046 | 0.84 | 0.659 | 1 | 0.707600603 | 0.706335757 | 0.5 | 0.54 | 1 | 0.707600603 | 0.000192522 |
| Ppfibp1 | 9.63657E-05 | 0.419878762 | 0.983 | 0.924 | 1 | 0.078389582 | 0.488021908 | 1 | 0.929 | 1 | 0.078389582 | 0.000192722 |
| Ddah1 | 9.71367E-05 | -1.086330872 | 0.479 | 0.653 | 1 | 0.739177225 | 0.119068631 | 0.5 | 0.571 | 1 | 0.739177225 | 0.000194264 |
| Aatf | 9.73721E-05 | -1.228268988 | 0.235 | 0.465 | 1 | 0.553629672 | -0.28585884 | 0.3 | 0.407 | 1 | 0.553629672 | 0.000194735 |
| Cplane1 | 9.75229E-05 | -1.104822825 | 0.336 | 0.535 | 1 | 0.567731381 | -0.256052964 | 0.5 | 0.611 | 1 | 0.567731381 | 0.000195036 |
| mt-Atp6 | 0.000804569 | 0.336078869 | 1 | 1 | 1 | 9.78532E-05 | 1.173177012 | 1 | 1 | 1 | 0.000804569 | 0.000195697 |
| Zfp664 | 9.78806E-05 | -1.185866545 | 0.353 | 0.518 | 1 | 0.103823689 | -0.864721141 | 0.5 | 0.691 | 1 | 0.103823689 | 0.000195752 |
| Raver1 | 9.84974E-05 | -1.485426827 | 0.176 | 0.376 | 1 | 0.087781496 | -1.399930607 | 0.1 | 0.355 | 1 | 0.087781496 | 0.000196985 |
| Gm19461 | 0.038142909 | 2.514573173 | 0.025 | 0 | 1 | 9.85609E-05 | 5.017921908 | 0.1 | 0.003 | 1 | 0.038142909 | 0.000197112 |
| Hoxa7 | 0.038142909 | 2.514573173 | 0.025 | 0 | 1 | 9.85609E-05 | 5.017921908 | 0.1 | 0.003 | 1 | 0.038142909 | 0.000197112 |
| Cfap100 | 0.038142909 | 2.514573173 | 0.025 | 0 | 1 | 9.85609E-05 | 5.017921908 | 0.1 | 0.003 | 1 | 0.038142909 | 0.000197112 |
| Tex29 | 0.060010312 | -2.485426827 | 0 | 0.029 | 1 | 9.85609E-05 | 5.017921908 | 0.1 | 0.003 | 1 | 0.060010312 | 0.000197112 |
| Fer1l6 | 0.064654593 | 2.177538186 | 0.067 | 0.024 | 1 | 9.85609E-05 | 5.017921908 | 0.1 | 0.003 | 1 | 0.064654593 | 0.000197112 |
| Gm56725 | 0.093245792 | -1.807354922 | 0 | 0.024 | 1 | 9.85609E-05 | 5.017921908 | 0.1 | 0.003 | 1 | 0.093245792 | 0.000197112 |
| Bdkrb1 | 0.147029286 | -1.807354922 | 0 | 0.018 | 1 | 9.85609E-05 | 5.017921908 | 0.1 | 0.003 | 1 | 0.147029286 | 0.000197112 |
| Ttll9 | 0.168156185 | 1.514573173 | 0.025 | 0.006 | 1 | 9.85609E-05 | 5.017921908 | 0.1 | 0.003 | 1 | 0.168156185 | 0.000197112 |
| Gm15286 | 0.168156185 | 1.514573173 | 0.025 | 0.006 | 1 | 9.85609E-05 | 5.017921908 | 0.1 | 0.003 | 1 | 0.168156185 | 0.000197112 |
| Gm13387 | 0.169248346 | 1.794681092 | 0.025 | 0.006 | 1 | 9.85609E-05 | 5.017921908 | 0.1 | 0.003 | 1 | 0.169248346 | 0.000197112 |
| 0610033M10Rik | 0.201654447 | 1.321928095 | 0.034 | 0.012 | 1 | 9.85609E-05 | 5.017921908 | 0.1 | 0.003 | 1 | 0.201654447 | 0.000197112 |
| Gm45356 | 0.204733108 | -1.14839184 | 0.034 | 0.012 | 1 | 9.85609E-05 | 5.017921908 | 0.1 | 0.003 | 1 | 0.204733108 | 0.000197112 |
| Melk | 0.217412808 | -1.655351829 | 0.008 | 0.029 | 1 | 9.85609E-05 | 5.017921908 | 0.1 | 0.003 | 1 | 0.217412808 | 0.000197112 |
| Sit1 | 0.237885589 | -1.070389328 | 0 | 0.012 | 1 | 9.85609E-05 | 5.017921908 | 0.1 | 0.003 | 1 | 0.237885589 | 0.000197112 |
| Cdcp2 | 0.237885589 | -1.070389328 | 0 | 0.012 | 1 | 9.85609E-05 | 5.017921908 | 0.1 | 0.003 | 1 | 0.237885589 | 0.000197112 |
| D5Ertd605e | 0.237885589 | -1.070389328 | 0 | 0.012 | 1 | 9.85609E-05 | 5.017921908 | 0.1 | 0.003 | 1 | 0.237885589 | 0.000197112 |
| Clec4n | 0.237885589 | -1.485426827 | 0 | 0.012 | 1 | 9.85609E-05 | 5.017921908 | 0.1 | 0.003 | 1 | 0.237885589 | 0.000197112 |
| Pinlyp | 0.237885589 | -1.485426827 | 0 | 0.012 | 1 | 9.85609E-05 | 5.017921908 | 0.1 | 0.003 | 1 | 0.237885589 | 0.000197112 |
| Brsk2 | 0.237885589 | -1.485426827 | 0 | 0.012 | 1 | 9.85609E-05 | 5.017921908 | 0.1 | 0.003 | 1 | 0.237885589 | 0.000197112 |
| Gm56958 | 0.237885589 | -1.070389328 | 0 | 0.012 | 1 | 9.85609E-05 | 5.017921908 | 0.1 | 0.003 | 1 | 0.237885589 | 0.000197112 |
| Gpr132 | 0.237885589 | -2.070389328 | 0 | 0.012 | 1 | 9.85609E-05 | 5.017921908 | 0.1 | 0.003 | 1 | 0.237885589 | 0.000197112 |
| Gm47849 | 0.237885589 | -1.070389328 | 0 | 0.012 | 1 | 9.85609E-05 | 5.017921908 | 0.1 | 0.003 | 1 | 0.237885589 | 0.000197112 |
| A930029G22Rik | 0.347966916 | -0.707819249 | 0.017 | 0.035 | 1 | 9.85609E-05 | 5.017921908 | 0.1 | 0.003 | 1 | 0.347966916 | 0.000197112 |
| Acer1 | 0.368010882 | 1.514573173 | 0.017 | 0.006 | 1 | 9.85609E-05 | 5.017921908 | 0.1 | 0.003 | 1 | 0.368010882 | 0.000197112 |
| Catsperb | 0.370184089 | 1.099535674 | 0.017 | 0.006 | 1 | 9.85609E-05 | 5.017921908 | 0.1 | 0.003 | 1 | 0.370184089 | 0.000197112 |
| Hmmr | 0.47838547 | -3.129283017 | 0.017 | 0.029 | 1 | 9.85609E-05 | 5.017921908 | 0.1 | 0.003 | 1 | 0.47838547 | 0.000197112 |
| Oprd1 | 0.504214062 | -1.292781749 | 0.008 | 0.018 | 1 | 9.85609E-05 | 5.017921908 | 0.1 | 0.003 | 1 | 0.504214062 | 0.000197112 |
| Tas1r1 | 0.506471296 | -1.807354922 | 0.008 | 0.018 | 1 | 9.85609E-05 | 5.017921908 | 0.1 | 0.003 | 1 | 0.506471296 | 0.000197112 |
| Them7 | 0.508731273 | -1.070389328 | 0.008 | 0.018 | 1 | 9.85609E-05 | 5.017921908 | 0.1 | 0.003 | 1 | 0.508731273 | 0.000197112 |

| | | | | | | | | | | | | |
|---|---|---|---|---|---|---|---|---|---|---|---|---|
| Ramp3 | 0.511005543 | -0.485426827 | 0.008 | 0.018 | 1 | 9.85609E-05 | 5.017921908 | 0.1 | 0.003 | 1 | 0.511005543 | 0.000197112 |
| Gm15063 | 0.511005543 | -0.485426827 | 0.008 | 0.018 | 1 | 9.85609E-05 | 5.017921908 | 0.1 | 0.003 | 1 | 0.511005543 | 0.000197112 |
| Gm16341 | 0.618331095 | 0.836501268 | 0.034 | 0.024 | 1 | 9.85609E-05 | 5.017921908 | 0.1 | 0.003 | 1 | 0.618331095 | 0.000197112 |
| Gm9889 | 0.659865009 | 0.514573173 | 0.025 | 0.018 | 1 | 9.85609E-05 | 5.017921908 | 0.1 | 0.003 | 1 | 0.659865009 | 0.000197112 |
| Gm34388 | 0.666161437 | 0.192645078 | 0.025 | 0.018 | 1 | 9.85609E-05 | 5.017921908 | 0.1 | 0.003 | 1 | 0.666161437 | 0.000197112 |
| Gm33667 | 0.687345921 | -1.070389328 | 0.017 | 0.024 | 1 | 9.85609E-05 | 5.017921908 | 0.1 | 0.003 | 1 | 0.687345921 | 0.000197112 |
| Arhgap19 | 0.721162042 | 0.514573173 | 0.017 | 0.012 | 1 | 9.85609E-05 | 5.017921908 | 0.1 | 0.003 | 1 | 0.721162042 | 0.000197112 |
| Nlrp3 | 0.781790612 | -0.485426827 | 0.008 | 0.012 | 1 | 9.85609E-05 | 5.017921908 | 0.1 | 0.003 | 1 | 0.781790612 | 0.000197112 |
| Hpx | 0.791189474 | 0.514573173 | 0.008 | 0.012 | 1 | 9.85609E-05 | 5.017921908 | 0.1 | 0.003 | 1 | 0.791189474 | 0.000197112 |
| 1700057G04Rik | 0.814978101 | -0.970853654 | 0.025 | 0.029 | 1 | 9.85609E-05 | 5.017921908 | 0.1 | 0.003 | 1 | 0.814978101 | 0.000197112 |
| Gm36287 | 0.820836204 | -1.185866545 | 0.025 | 0.029 | 1 | 9.85609E-05 | 5.017921908 | 0.1 | 0.003 | 1 | 0.820836204 | 0.000197112 |
| Lzts1 | 0.842422009 | 0.251538767 | 0.025 | 0.029 | 1 | 9.85609E-05 | 5.017921908 | 0.1 | 0.003 | 1 | 0.842422009 | 0.000197112 |
| Cyp3a13 | 9.86801E-05 | 1.4639471 | 0.269 | 0.094 | 1 | 0.501413191 | 0.808468542 | 0.1 | 0.049 | 1 | 0.501413191 | 0.00019735 |
| Vps36 | 9.89225E-05 | -1.129283017 | 0.311 | 0.506 | 1 | 0.334569038 | -0.613515023 | 0.5 | 0.571 | 1 | 0.334569038 | 0.000197835 |
| Blvrb | 9.913E-05 | -1.501728639 | 0.143 | 0.341 | 1 | 0.025442562 | -2.121629444 | 0 | 0.343 | 1 | 0.025442562 | 0.00019825 |
| Rbm33 | 9.94723E-05 | -0.823595563 | 0.697 | 0.829 | 1 | 0.213222591 | -0.654503434 | 0.9 | 0.861 | 1 | 0.213222591 | 0.000198935 |
| Klhl20 | 9.96161E-05 | -2.003275132 | 0.067 | 0.241 | 1 | 0.41725734 | 0.660369903 | 0.3 | 0.182 | 1 | 0.41725734 | 0.000199222 |
| Rbm39 | 9.97739E-05 | -0.318428337 | 1 | 1 | 1 | 0.019742958 | -0.456910683 | 1 | 1 | 1 | 0.019742958 | 0.000199538 |
| Dclre1a | 9.97785E-05 | -2.87774425 | 0.025 | 0.171 | 1 | 0.5653083 | 0.744903414 | 0.2 | 0.13 | 1 | 0.5653083 | 0.000199547 |
| Fbh1 | 0.000100233 | -1.831201664 | 0.084 | 0.265 | 1 | 0.028958947 | -2.101019165 | 0 | 0.333 | 1 | 0.028958947 | 0.000200455 |
| Rcor3 | 0.000100672 | -1.264212597 | 0.336 | 0.512 | 1 | 0.017933571 | -1.932390968 | 0.2 | 0.556 | 1 | 0.017933571 | 0.000201334 |
| Sephs1 | 0.000100975 | -1.867896464 | 0.109 | 0.294 | 1 | 0.366301453 | -0.68251781 | 0.2 | 0.315 | 1 | 0.366301453 | 0.000201941 |
| Tnrc18 | 0.000101772 | -0.351802263 | 0.866 | 0.935 | 1 | 0.093191187 | -0.433723032 | 1 | 0.963 | 1 | 0.093191187 | 0.000203533 |
| Rab11fip4 | 0.000102054 | 0.387438048 | 0.941 | 0.824 | 1 | 0.648140161 | -0.257737037 | 1 | 0.901 | 1 | 0.648140161 | 0.000204098 |
| Reck | 0.000103319 | -0.895302621 | 0.487 | 0.647 | 1 | 0.254680865 | -0.368659145 | 0.6 | 0.762 | 1 | 0.254680865 | 0.000206627 |
| Slc22a17 | 0.000103339 | 1.027385888 | 0.555 | 0.341 | 1 | 0.100792186 | 1.27838237 | 0.6 | 0.395 | 1 | 0.100792186 | 0.000206668 |
| Mpv17l2 | 0.000103406 | 1.108580813 | 0.437 | 0.241 | 1 | 0.044808939 | 1.737813989 | 0.4 | 0.173 | 1 | 0.044808939 | 0.000206801 |
| Polr1g | 0.000103632 | -1.632268215 | 0.143 | 0.335 | 1 | 0.045110574 | -2.382957528 | 0 | 0.296 | 1 | 0.045110574 | 0.000207253 |
| Lancl3 | 0.038982035 | -5.392317423 | 0 | 0.035 | 1 | 0.000103743 | 4.017921908 | 0.1 | 0.003 | 1 | 0.038982035 | 0.000207475 |
| Adap1 | 0.147029286 | -1.807354922 | 0 | 0.018 | 1 | 0.000103743 | 4.432959407 | 0.1 | 0.003 | 1 | 0.147029286 | 0.000207475 |
| Atp2c2 | 0.147029286 | -2.655351829 | 0 | 0.018 | 1 | 0.000103743 | 4.017921908 | 0.1 | 0.003 | 1 | 0.147029286 | 0.000207475 |
| Cenpi | 0.167616515 | -1.070389328 | 0.017 | 0.047 | 1 | 0.000103743 | 4.432959407 | 0.1 | 0.003 | 1 | 0.167616515 | 0.000207475 |
| 4930461C15Rik | 0.201658121 | 1.251538767 | 0.034 | 0.012 | 1 | 0.000103743 | 4.432959407 | 0.1 | 0.003 | 1 | 0.201658121 | 0.000207475 |
| Tfap2c | 0.220389995 | 0.777607579 | 0.042 | 0.018 | 1 | 0.000103743 | 4.432959407 | 0.1 | 0.003 | 1 | 0.220389995 | 0.000207475 |
| Aurkb | 0.220674802 | -1.600904045 | 0.008 | 0.029 | 1 | 0.000103743 | 3.695993813 | 0.1 | 0.003 | 1 | 0.220674802 | 0.000207475 |
| Ctcflos | 0.235770388 | 0.344648171 | 0.059 | 0.029 | 1 | 0.000103743 | 4.017921908 | 0.1 | 0.003 | 1 | 0.235770388 | 0.000207475 |
| Mybpc2 | 0.237885589 | -1.070389328 | 0 | 0.012 | 1 | 0.000103743 | 4.432959407 | 0.1 | 0.003 | 1 | 0.237885589 | 0.000207475 |
| Gm10603 | 0.237885589 | -1.070389328 | 0 | 0.012 | 1 | 0.000103743 | 4.432959407 | 0.1 | 0.003 | 1 | 0.237885589 | 0.000207475 |
| Gm12708 | 0.374548525 | 0.514573173 | 0.017 | 0.006 | 1 | 0.000103743 | 4.432959407 | 0.1 | 0.003 | 1 | 0.374548525 | 0.000207475 |
| Pkhd1l1 | 0.374548525 | -0.222392421 | 0.017 | 0.006 | 1 | 0.000103743 | 4.432959407 | 0.1 | 0.003 | 1 | 0.374548525 | 0.000207475 |
| Gm38287 | 0.666161437 | 0.192645078 | 0.025 | 0.018 | 1 | 0.000103743 | 4.432959407 | 0.1 | 0.003 | 1 | 0.666161437 | 0.000207475 |
| Gm26510 | 0.781790612 | -0.485426827 | 0.008 | 0.012 | 1 | 0.000103743 | 4.432959407 | 0.1 | 0.003 | 1 | 0.781790612 | 0.000207475 |
| Sox10 | 0.962120409 | 0.192645078 | 0.017 | 0.018 | 1 | 0.000103743 | 4.432959407 | 0.1 | 0.003 | 1 | 0.962120409 | 0.000207475 |
| Eng | 0.000104026 | -1.362155452 | 0.303 | 0.494 | 1 | 0.777028701 | -0.271174794 | 0.5 | 0.509 | 1 | 0.777028701 | 0.000208041 |
| 2610307P16Rik | 0.000104201 | -3.79528209 | 0.042 | 0.194 | 1 | 0.316188655 | -2.321928095 | 0 | 0.093 | 1 | 0.316188655 | 0.00020839 |
| Ddit4l | 0.0001046 | 0.903068885 | 0.479 | 0.259 | 1 | 0.288702531 | -2.570727728 | 0.1 | 0.228 | 1 | 0.288702531 | 0.00020919 |
| Abi2 | 0.000104749 | -1.234508708 | 0.429 | 0.6 | 1 | 0.015718751 | -1.770912926 | 0.5 | 0.728 | 1 | 0.015718751 | 0.000209487 |
| Srsf7 | 0.000104803 | -0.595878278 | 0.891 | 0.971 | 1 | 0.012043128 | -0.945677184 | 0.9 | 0.969 | 1 | 0.012043128 | 0.000209595 |
| Man1a2 | 0.000105078 | -0.65358922 | 0.773 | 0.824 | 1 | 0.864784295 | -0.259416036 | 0.9 | 0.833 | 1 | 0.864784295 | 0.000210145 |
| Ppil2 | 0.000105124 | -1.374395515 | 0.202 | 0.4 | 1 | 0.085262631 | 0.888638891 | 0.7 | 0.414 | 1 | 0.085262631 | 0.000210238 |
| Mcl1 | 0.000105298 | -0.806907505 | 0.95 | 0.941 | 1 | 0.000298788 | -1.504792152 | 0.9 | 0.985 | 1 | 0.000298788 | 0.000210585 |
| Atm | 0.000105336 | -1.733354341 | 0.092 | 0.276 | 1 | 0.703289339 | 0.397335498 | 0.3 | 0.238 | 1 | 0.703289339 | 0.000210661 |
| Dnah10 | 0.066452457 | -0.655351829 | 0.025 | 0.076 | 1 | 0.000105518 | 4.67688499 | 0.3 | 0.04 | 1 | 0.066452457 | 0.000211025 |
| Taf1a | 0.000106293 | -1.639232163 | 0.143 | 0.335 | 1 | 0.913364669 | 0.393431043 | 0.3 | 0.287 | 1 | 0.913364669 | 0.000212574 |
| Ptpre | 0.000106724 | 0.787591667 | 0.429 | 0.194 | 1 | 0.007249241 | 1.930459067 | 0.5 | 0.194 | 1 | 0.007249241 | 0.000213436 |
| Klhl11 | 0.000106872 | -1.301714874 | 0.134 | 0.335 | 1 | 0.994021891 | -0.217294554 | 0.5 | 0.42 | 1 | 0.994021891 | 0.000213733 |
| Zfand5 | 0.000106896 | -0.585398294 | 0.992 | 0.971 | 1 | 0.003272273 | -1.179071337 | 0.9 | 0.994 | 1 | 0.003272273 | 0.000213781 |
| Car8 | 0.299221715 | -0.876664792 | 0.571 | 0.529 | 1 | 0.000107329 | -4.269790472 | 0.1 | 0.747 | 1 | 0.299221715 | 0.000214646 |
| Afg3l2 | 0.000107613 | -1.025995209 | 0.294 | 0.5 | 1 | 0.118843636 | 0.765256476 | 0.8 | 0.515 | 1 | 0.118843636 | 0.000215214 |
| Cited2 | 0.000108521 | 1.547631461 | 0.479 | 0.271 | 1 | 0.04683206 | 0.853733276 | 0.5 | 0.238 | 1 | 0.04683206 | 0.00021703 |
| Tmem184b | 0.000108545 | -1.392317423 | 0.328 | 0.506 | 1 | 0.30908088 | -1.011825435 | 0.5 | 0.537 | 1 | 0.30908088 | 0.000217078 |
| Srgn | 0.000108642 | 0.53792366 | 0.773 | 0.559 | 1 | 0.818072577 | 0.167369297 | 0.8 | 0.784 | 1 | 0.818072577 | 0.000217273 |
| Ints4 | 0.000108849 | -1.987927168 | 0.092 | 0.271 | 1 | 0.579745189 | -0.258202497 | 0.2 | 0.284 | 1 | 0.579745189 | 0.000217686 |
| Casp2 | 0.000109597 | -4.129283017 | 0 | 0.118 | 1 | 0.921182017 | 0.526068812 | 0.1 | 0.108 | 1 | 0.921182017 | 0.000219182 |
| Lztfl1 | 0.000109952 | -0.749926642 | 0.672 | 0.782 | 1 | 0.165252446 | -0.799307762 | 0.7 | 0.772 | 1 | 0.165252446 | 0.000219891 |
| C1qtnf2 | 0.000109998 | -4.87774425 | 0 | 0.118 | 1 | 0.185749549 | -1.248864633 | 0 | 0.151 | 1 | 0.185749549 | 0.000219985 |
| Mpp4 | 0.000110071 | -4.655351829 | 0 | 0.118 | 1 | 0.325051203 | -0.473931188 | 0 | 0.09 | 1 | 0.325051203 | 0.00022013 |
| Fnbp1 | 0.000110084 | -1.227930605 | 0.235 | 0.435 | 1 | 0.211936101 | -0.773240981 | 0.4 | 0.571 | 1 | 0.211936101 | 0.000220157 |
| Ddah2 | 0.000110216 | -1.693022247 | 0.134 | 0.324 | 1 | 0.062270312 | -2.304006187 | 0 | 0.265 | 1 | 0.062270312 | 0.000220419 |
| Prkcb | 0.000110246 | -5.157852169 | 0 | 0.118 | 1 | 0.316149521 | -1.473931188 | 0 | 0.093 | 1 | 0.316149521 | 0.00022048 |
| Mark1 | 0.000110285 | 1.558967292 | 0.479 | 0.276 | 1 | 0.632331397 | 2.089005006 | 0.3 | 0.315 | 1 | 0.632331397 | 0.000220558 |
| Mgp | 0.000110374 | -0.691155253 | 0.958 | 0.994 | 1 | 0.111976856 | -0.48703882 | 0.8 | 0.997 | 1 | 0.111976856 | 0.000220736 |
| Anxa7 | 0.00011047 | 0.408805546 | 0.983 | 0.876 | 1 | 0.03942369 | -0.736965594 | 0.8 | 0.963 | 1 | 0.03942369 | 0.000220928 |
| Ppp1r14b | 0.000110771 | -0.895212428 | 0.655 | 0.741 | 1 | 0.030691212 | -1.141031976 | 0.6 | 0.815 | 1 | 0.030691212 | 0.000221529 |
| Tstd2 | 0.000111064 | -1.485426827 | 0.134 | 0.329 | 1 | 0.403396316 | -0.321928095 | 0.2 | 0.336 | 1 | 0.403396316 | 0.000222117 |
| Xrn1 | 0.000111541 | -0.854660637 | 0.706 | 0.788 | 1 | 0.267609047 | -0.475213014 | 0.8 | 0.877 | 1 | 0.267609047 | 0.00022307 |
| Fars2 | 0.000111832 | -1.283792966 | 0.345 | 0.524 | 1 | 0.476610885 | -0.454565863 | 0.4 | 0.515 | 1 | 0.476610885 | 0.000223651 |
| Slc16a2 | 0.000111874 | -0.993146329 | 0.597 | 0.735 | 1 | 0.434241437 | -0.679993409 | 0.9 | 0.873 | 1 | 0.434241437 | 0.000223736 |
| Mansc1 | 0.000111941 | 2.836501268 | 0.118 | 0.012 | 1 | 0.474877085 | 0.695993813 | 0 | 0.049 | 1 | 0.474877085 | 0.00022387 |
| Iqcn | 0.000112014 | -3.485426827 | 0.017 | 0.153 | 1 | 0.688615584 | 1.922764675 | 0.2 | 0.176 | 1 | 0.688615584 | 0.000224015 |
| Rbbp9 | 0.000112663 | -1.401595236 | 0.227 | 0.418 | 1 | 0.305242651 | -0.382957528 | 0.2 | 0.377 | 1 | 0.305242651 | 0.000225313 |
| Dcp1b | 0.000112784 | -2.748461233 | 0.025 | 0.171 | 1 | 0.307032548 | -0.267480311 | 0 | 0.096 | 1 | 0.307032548 | 0.000225556 |
| Lims2 | 0.000112974 | 0.888638891 | 0.563 | 0.353 | 1 | 0.000766603 | 1.484741949 | 0.9 | 0.451 | 1 | 0.000766603 | 0.000225936 |
| Fam171b | 0.000113103 | -1.655351829 | 0.042 | 0.206 | 1 | 0.63551067 | 1.388565288 | 0.1 | 0.176 | 1 | 0.63551067 | 0.000226192 |
| Dnajc7 | 0.000113118 | -0.851554726 | 0.832 | 0.829 | 1 | 0.275508302 | -0.18552022 | 0.7 | 0.904 | 1 | 0.275508302 | 0.000226223 |
| Gipc1 | 0.000113516 | -1.622930351 | 0.134 | 0.324 | 1 | 0.978803798 | 0.284567567 | 0.3 | 0.315 | 1 | 0.978803798 | 0.000227019 |
| 2900060B14Rik | 0.527408075 | -0.315501826 | 0.143 | 0.171 | 1 | 0.000114004 | 2.017921908 | 0.5 | 0.099 | 1 | 0.527408075 | 0.000227994 |
| Psmc6 | 0.000114734 | -0.604512717 | 0.849 | 0.888 | 1 | 0.063625712 | -0.766712938 | 0.8 | 0.914 | 1 | 0.063625712 | 0.000229455 |
| Adss | 0.000115045 | -0.666974196 | 0.891 | 0.912 | 1 | 0.019890034 | -1.065678294 | 0.6 | 0.83 | 1 | 0.019890034 | 0.000230077 |
| Sumo2 | 0.000115046 | -0.445863493 | 0.941 | 0.965 | 1 | 0.371781237 | -0.270901453 | 0.8 | 0.985 | 1 | 0.371781237 | 0.000230078 |
| Tm9sf1 | 0.000115108 | -0.89662226 | 0.555 | 0.676 | 1 | 0.638507976 | 0.254156254 | 0.8 | 0.725 | 1 | 0.638507976 | 0.000230202 |
| Siah2 | 0.000115252 | -0.783606319 | 0.471 | 0.624 | 1 | 0.080392128 | -1.418373212 | 0.4 | 0.58 | 1 | 0.080392128 | 0.00023049 |
| Fmr1 | 0.000116393 | -0.805539069 | 0.655 | 0.765 | 1 | 0.016886263 | -1.1801138 | 0.7 | 0.858 | 1 | 0.016886263 | 0.000232772 |
| Nrf1 | 0.000117094 | -1.070389328 | 0.412 | 0.606 | 1 | 0.018656074 | -1.235173474 | 0.4 | 0.775 | 1 | 0.018656074 | 0.000234175 |
| Wdr46 | 0.00011729 | -2.448900951 | 0.025 | 0.171 | 1 | 0.428798435 | 1.079322453 | 0.2 | 0.114 | 1 | 0.428798435 | 0.000234566 |
| Zfp553 | 0.000117577 | -3.070389328 | 0.017 | 0.153 | 1 | 0.213668367 | -0.91281543 | 0 | 0.136 | 1 | 0.213668367 | 0.000235141 |
| Nf2 | 0.000117618 | -1.586512952 | 0.21 | 0.394 | 1 | 0.056389118 | -1.31299497 | 0.3 | 0.559 | 1 | 0.056389118 | 0.000235221 |
| Gm29157 | 0.00011866 | -2.807354922 | 0.017 | 0.153 | 1 | 0.446758355 | 0.695993813 | 0 | 0.056 | 1 | 0.446758355 | 0.000237306 |

| | | | | | | | | | | | | |
|---|---|---|---|---|---|---|---|---|---|---|---|---|
| Nr1d2 | 0.000119142 | -0.782762934 | 0.672 | 0.771 | 1 | 0.427925499 | -0.384812131 | 0.8 | 0.849 | 1 | 0.427925499 | 0.000238269 |
| Tle5 | 0.000119262 | -1.053901205 | 0.412 | 0.612 | 1 | 0.00441117 | -1.255096586 | 0.1 | 0.667 | 1 | 0.00441117 | 0.000238509 |
| Ksr2 | 0.07834103 | -0.263034406 | 0.034 | 0.006 | 1 | 0.000120284 | 1.808468542 | 0.3 | 0.037 | 1 | 0.07834103 | 0.000240555 |
| Gm15892 | 0.000120421 | -3.392317423 | 0.008 | 0.135 | 1 | 0.161506691 | -1.69632361 | 0 | 0.167 | 1 | 0.161506691 | 0.000240828 |
| Sppl3 | 0.000120653 | -0.737299127 | 0.731 | 0.812 | 1 | 0.115967334 | -0.632477128 | 0.6 | 0.938 | 1 | 0.115967334 | 0.000241292 |
| 2810433D01Rik | 0.000120982 | -3.343407822 | 0.008 | 0.135 | 1 | 0.408705208 | 0.263034406 | 0 | 0.065 | 1 | 0.408705208 | 0.000241948 |
| Dhx30 | 0.00012128 | -1.193964013 | 0.269 | 0.471 | 1 | 0.087236096 | -1.321928095 | 0.2 | 0.451 | 1 | 0.087236096 | 0.000242545 |
| Spock1 | 0.000121575 | -3.292781749 | 0.017 | 0.153 | 1 | 0.316175189 | -1.876895855 | 0 | 0.093 | 1 | 0.316175189 | 0.000243135 |
| Fam184a | 0.000121938 | -1.392317423 | 0.134 | 0.324 | 1 | 0.233477731 | -1.408342847 | 0.3 | 0.42 | 1 | 0.233477731 | 0.00024386 |
| Dab2ip | 0.000122125 | -0.824632913 | 0.378 | 0.594 | 1 | 0.554169389 | -0.498903717 | 0.6 | 0.735 | 1 | 0.554169389 | 0.000244235 |
| Gab1 | 0.000122151 | -0.905966303 | 0.664 | 0.788 | 1 | 0.049499128 | -1.187192522 | 0.5 | 0.824 | 1 | 0.049499128 | 0.000244287 |
| Adarb1 | 0.000122855 | -2.109917692 | 0.176 | 0.371 | 1 | 0.017215243 | -3.8829449 | 0 | 0.383 | 1 | 0.017215243 | 0.000245694 |
| Trpc3 | 0.864464472 | -0.485426827 | 0.084 | 0.076 | 1 | 0.000123909 | 2.339850003 | 0.4 | 0.065 | 1 | 0.864464472 | 0.000247802 |
| Tcf20 | 0.000123962 | -0.996388746 | 0.445 | 0.624 | 1 | 0.465267021 | 0.118746278 | 0.9 | 0.731 | 1 | 0.465267021 | 0.000247909 |
| Zfp458 | 0.203510547 | -1.201633861 | 0.05 | 0.088 | 1 | 0.00012407 | 2.695993813 | 0.4 | 0.068 | 1 | 0.203510547 | 0.000248124 |
| Birc3 | 0.000124828 | -1.560714954 | 0.134 | 0.329 | 1 | 0.336427815 | -0.397115591 | 0.1 | 0.247 | 1 | 0.336427815 | 0.00024964 |
| Ppm1a | 0.000126214 | -0.662015559 | 0.849 | 0.835 | 1 | 0.124030555 | -0.376030297 | 0.9 | 0.907 | 1 | 0.124030555 | 0.000252413 |
| Cntnap2 | 0.000126381 | 0.800624231 | 0.437 | 0.212 | 1 | 0.032718633 | 1.081737006 | 0.6 | 0.278 | 1 | 0.032718633 | 0.000252745 |
| Ero1b | 0.000126382 | -3.008988783 | 0.017 | 0.153 | 1 | 0.821582331 | 0.290001453 | 0.1 | 0.077 | 1 | 0.821582331 | 0.000252748 |
| Fign | 0.00012639 | 1.185111405 | 0.462 | 0.235 | 1 | 0.029414734 | 0.800691192 | 0.7 | 0.346 | 1 | 0.029414734 | 0.000252763 |
| Smad7 | 0.000126478 | -1.004800986 | 0.529 | 0.653 | 1 | 0.873310548 | -0.674569057 | 0.8 | 0.639 | 1 | 0.873310548 | 0.00025294 |
| Notum | 0.00012673 | -1.401448435 | 0.193 | 0.4 | 1 | 0.249626418 | -0.250060071 | 0.5 | 0.676 | 1 | 0.249626418 | 0.000253445 |
| Odf2 | 0.000127673 | -1.320796125 | 0.336 | 0.512 | 1 | 0.564921736 | 0.360809622 | 0.6 | 0.491 | 1 | 0.564921736 | 0.00025533 |
| Them4 | 0.947561834 | -0.333423734 | 0.176 | 0.165 | 1 | 0.000128111 | 2.017921908 | 0.5 | 0.099 | 1 | 0.947561834 | 0.000256206 |
| Atp5b | 0.000128126 | -0.449061082 | 0.933 | 0.971 | 1 | 0.123336415 | -0.507289785 | 0.9 | 0.988 | 1 | 0.123336415 | 0.000256235 |
| Olfml2b | 0.000128388 | -2.977279923 | 0.017 | 0.153 | 1 | 0.421047115 | -0.536666944 | 0 | 0.062 | 1 | 0.421047115 | 0.00025676 |
| Sdk1 | 0.000128942 | -3.134519665 | 0.067 | 0.229 | 1 | 0.884064648 | 0.494359952 | 0.3 | 0.343 | 1 | 0.884064648 | 0.000257867 |
| Map2k7 | 0.000129072 | -1.807354922 | 0.109 | 0.288 | 1 | 0.266960375 | -0.745843746 | 0.2 | 0.349 | 1 | 0.266960375 | 0.000258127 |
| Pde4c | 0.019094585 | -1.826463745 | 0.034 | 0.112 | 1 | 0.00012928 | 4.240314329 | 0.3 | 0.04 | 1 | 0.019094585 | 0.000258544 |
| Rnf187 | 0.000129553 | -0.718087584 | 0.63 | 0.729 | 1 | 0.505932432 | -0.14272528 | 0.6 | 0.716 | 1 | 0.505932432 | 0.000259089 |
| Ccdc47 | 0.000129828 | -0.596173893 | 0.697 | 0.876 | 1 | 0.124360873 | -0.432855245 | 0.8 | 0.917 | 1 | 0.124360873 | 0.000259639 |
| Eeig2 | 0.000129886 | 0.442912188 | 0.916 | 0.647 | 1 | 0.712763612 | -0.256339753 | 1 | 0.75 | 1 | 0.712763612 | 0.000259756 |
| Adcy3 | 0.000130754 | -1.68182404 | 0.076 | 0.253 | 1 | 0.24573788 | -0.880931369 | 0.1 | 0.278 | 1 | 0.24573788 | 0.000261491 |
| Ccdc102a | 0.000131131 | -2.733354341 | 0.017 | 0.153 | 1 | 0.901389987 | 0.276454922 | 0.2 | 0.222 | 1 | 0.901389987 | 0.000262246 |
| Atad3a | 0.000131819 | -1.472006311 | 0.21 | 0.394 | 1 | 0.882228175 | 0.119068631 | 0.4 | 0.364 | 1 | 0.882228175 | 0.00026362 |
| Rnf24 | 0.000132116 | -1.260360272 | 0.261 | 0.459 | 1 | 0.111438781 | -1.103093493 | 0.3 | 0.549 | 1 | 0.111438781 | 0.000264214 |
| Porcn | 0.000132121 | -2.04342228 | 0.059 | 0.224 | 1 | 0.481034913 | -0.374395515 | 0.1 | 0.182 | 1 | 0.481034913 | 0.000264224 |
| Tlcd2 | 0.000132492 | -1.350497247 | 0.168 | 0.365 | 1 | 0.054315846 | -2.080110175 | 0 | 0.278 | 1 | 0.054315846 | 0.000264966 |
| Rps6kc1 | 0.000132932 | -1.896359928 | 0.134 | 0.318 | 1 | 0.556030961 | -0.814968106 | 0.4 | 0.42 | 1 | 0.556030961 | 0.000265846 |
| Rp2 | 0.000133337 | -1.707819249 | 0.067 | 0.241 | 1 | 0.334922253 | -0.802257054 | 0.1 | 0.222 | 1 | 0.334922253 | 0.000266657 |
| Sdccag8 | 0.000133797 | -1.207892852 | 0.303 | 0.529 | 1 | 0.711156933 | 0.163772774 | 0.6 | 0.497 | 1 | 0.711156933 | 0.000267575 |
| Ruvbl2 | 0.000134267 | -1.753915663 | 0.084 | 0.259 | 1 | 0.994182647 | 0.31748219 | 0.2 | 0.191 | 1 | 0.994182647 | 0.000268516 |
| Pak3 | 0.00013512 | -2.451638514 | 0.134 | 0.312 | 1 | 0.780778291 | -0.239465935 | 0.2 | 0.164 | 1 | 0.780778291 | 0.000270221 |
| Tia1 | 0.000135454 | -1.598320884 | 0.269 | 0.459 | 1 | 0.352193125 | 0.272782382 | 0.7 | 0.481 | 1 | 0.352193125 | 0.00027089 |
| Add1 | 0.000135799 | -0.993976212 | 0.479 | 0.641 | 1 | 0.164567148 | -0.445874903 | 0.6 | 0.778 | 1 | 0.164567148 | 0.00027158 |
| Aldoa | 0.000136065 | 0.643688392 | 0.958 | 0.941 | 1 | 0.330431338 | -0.275076316 | 0.7 | 0.957 | 1 | 0.330431338 | 0.000272112 |
| Capn7 | 0.000136203 | -1.464053176 | 0.227 | 0.418 | 1 | 0.029777603 | -1.808626579 | 0.1 | 0.444 | 1 | 0.029777603 | 0.000272387 |
| Nat10 | 0.000136232 | -1.469659511 | 0.134 | 0.324 | 1 | 0.625643032 | -0.131825212 | 0.5 | 0.349 | 1 | 0.625643032 | 0.000272445 |
| Prc1 | 0.021508475 | -2.354842717 | 0.034 | 0.106 | 1 | 0.000136383 | 3.124837112 | 0.3 | 0.04 | 1 | 0.021508475 | 0.000272748 |
| Topbp1 | 0.000137206 | -1.743224585 | 0.143 | 0.329 | 1 | 0.389858196 | -0.823380346 | 0.2 | 0.309 | 1 | 0.389858196 | 0.000274393 |
| Kdm6b | 0.000138976 | -0.890683306 | 0.807 | 0.853 | 1 | 0.014429661 | -0.83507568 | 0.6 | 0.92 | 1 | 0.014429661 | 0.000277932 |
| Uggt2 | 0.000139536 | -0.950549558 | 0.513 | 0.659 | 1 | 0.545655332 | 0.524786986 | 0.8 | 0.651 | 1 | 0.545655332 | 0.000279052 |
| Stk39 | 0.000139886 | 0.291620808 | 0.95 | 0.835 | 1 | 0.025478529 | 1.025153477 | 1 | 0.898 | 1 | 0.025478529 | 0.000279752 |
| Aldh1a3 | 0.00014002 | 0.616111199 | 0.403 | 0.188 | 1 | 0.695461986 | -0.191531458 | 0.1 | 0.151 | 1 | 0.695461986 | 0.00028002 |
| Tpbg | 0.000140281 | -1.710601689 | 0.202 | 0.4 | 1 | 0.112427817 | -1.625934282 | 0.1 | 0.33 | 1 | 0.112427817 | 0.000280542 |
| Grip1 | 0.000140845 | -2.64689025 | 0.084 | 0.253 | 1 | 0.458453959 | -0.321928095 | 0.2 | 0.346 | 1 | 0.458453959 | 0.000281671 |
| H3c14 | 0.000140935 | 1.96203215 | 0.176 | 0.041 | 1 | 0.522066388 | 1.111031312 | 0 | 0.04 | 1 | 0.522066388 | 0.000281851 |
| 1700120C14Rik | 0.970184702 | 0.173536255 | 0.084 | 0.082 | 1 | 0.000141181 | 3.169925001 | 0.3 | 0.04 | 1 | 0.970184702 | 0.000282343 |
| Maf1 | 0.000141426 | -1.485426827 | 0.168 | 0.359 | 1 | 0.184758793 | -0.974544419 | 0.2 | 0.386 | 1 | 0.184758793 | 0.000282831 |
| Samd4 | 0.00014166 | 0.410348728 | 0.975 | 0.953 | 1 | 0.26590305 | 0.280483377 | 1 | 0.985 | 1 | 0.26590305 | 0.000283299 |
| Rnf38 | 0.00014219 | -1.17684027 | 0.378 | 0.571 | 1 | 0.260722543 | -0.971795119 | 0.7 | 0.704 | 1 | 0.260722543 | 0.00028436 |
| Mxra8 | 0.000142888 | -0.983196265 | 0.933 | 0.841 | 1 | 0.088244883 | -0.708990022 | 1 | 0.864 | 1 | 0.088244883 | 0.000285756 |
| Cfap69 | 0.000143485 | 0.676540652 | 0.403 | 0.182 | 1 | 0.002312566 | 2.411264336 | 0.5 | 0.164 | 1 | 0.002312566 | 0.000286949 |
| Ddx11 | 0.000143613 | -2.977279923 | 0.025 | 0.165 | 1 | 0.913227148 | 0.135278859 | 0.1 | 0.086 | 1 | 0.913227148 | 0.000287205 |
| Dnaaf2 | 0.000143844 | -2.122856748 | 0.05 | 0.212 | 1 | 0.686932588 | 0.973527789 | 0.2 | 0.16 | 1 | 0.686932588 | 0.000287666 |
| Dicer1 | 0.000144213 | -0.946560741 | 0.395 | 0.588 | 1 | 0.012256965 | -1.40354186 | 0.5 | 0.762 | 1 | 0.012256965 | 0.000288405 |
| Rell1 | 0.000144957 | -1.399171094 | 0.328 | 0.524 | 1 | 0.221002703 | -0.84750207 | 0.5 | 0.707 | 1 | 0.221002703 | 0.000289894 |
| Arfgap1 | 0.000145476 | -0.718486174 | 0.672 | 0.782 | 1 | 0.370961305 | 0.394278939 | 0.7 | 0.701 | 1 | 0.370961305 | 0.000290932 |
| Eif2s2 | 0.000147517 | -0.423636628 | 0.983 | 0.976 | 1 | 0.005466851 | -1.012745228 | 0.9 | 0.981 | 1 | 0.005466851 | 0.000295013 |
| 0610043K17Rik | 0.000149433 | 0.251538767 | 0.555 | 0.282 | 1 | 0.865156532 | 0.219180116 | 0.3 | 0.284 | 1 | 0.865156532 | 0.000298844 |
| Tuba1b | 0.000150097 | -0.440836676 | 0.79 | 0.888 | 1 | 0.224358916 | -0.239770076 | 0.6 | 0.895 | 1 | 0.224358916 | 0.000300171 |
| Rab5if | 0.000150159 | 0.578138302 | 0.849 | 0.788 | 1 | 0.160728882 | 0.767826558 | 0.7 | 0.84 | 1 | 0.160728882 | 0.000300296 |
| Ooep | 0.000150183 | -1.727501614 | 0.168 | 0.353 | 1 | 0.22839177 | -0.789433014 | 0.2 | 0.42 | 1 | 0.22839177 | 0.000300343 |
| Cttnbp2nl | 0.000151032 | -0.87621678 | 0.479 | 0.629 | 1 | 0.928823281 | -0.158955854 | 0.9 | 0.784 | 1 | 0.928823281 | 0.000302042 |
| Dpysl3 | 0.000152215 | -2.240314329 | 0.034 | 0.182 | 1 | 0.219939356 | -1.567040593 | 0 | 0.133 | 1 | 0.219939356 | 0.000304407 |
| Ckap5 | 0.00015267 | -1.103997721 | 0.513 | 0.641 | 1 | 0.132052557 | -0.849974556 | 0.5 | 0.71 | 1 | 0.132052557 | 0.000305316 |
| Nbr1 | 0.000152974 | -0.912127244 | 0.437 | 0.612 | 1 | 0.271223476 | -0.872849022 | 0.7 | 0.688 | 1 | 0.271223476 | 0.000305925 |
| Osbpl1a | 0.000153495 | 0.471799149 | 0.891 | 0.753 | 1 | 0.851478762 | 0.19994587 | 0.7 | 0.79 | 1 | 0.851478762 | 0.000306966 |
| Sirt1 | 0.000155378 | -1.182499694 | 0.471 | 0.641 | 1 | 0.175023609 | -0.613515023 | 0.5 | 0.778 | 1 | 0.175023609 | 0.000310732 |
| Ldb2 | 0.000155546 | -1.435386145 | 0.143 | 0.341 | 1 | 0.001346852 | 3.259627804 | 0.5 | 0.157 | 1 | 0.001346852 | 0.000311067 |
| Ing4 | 0.00015584 | -1.165026202 | 0.387 | 0.541 | 1 | 0.064344819 | -1.315978829 | 0.2 | 0.475 | 1 | 0.064344819 | 0.000311656 |
| Elp4 | 0.000156042 | -1.368069877 | 0.361 | 0.518 | 1 | 0.377816092 | 0.410022596 | 0.5 | 0.398 | 1 | 0.377816092 | 0.000312059 |
| Trim35 | 0.000156278 | -1.532732542 | 0.252 | 0.441 | 1 | 0.144686801 | -1.0853659 | 0.2 | 0.435 | 1 | 0.144686801 | 0.000312532 |
| Dclre1c | 0.567287312 | -0.992386816 | 0.118 | 0.135 | 1 | 0.000156962 | 2.918386234 | 0.4 | 0.071 | 1 | 0.567287312 | 0.000313899 |
| Rusc2 | 0.000158177 | 0.932638365 | 0.571 | 0.347 | 1 | 0.04346086 | 1.248150169 | 0.7 | 0.491 | 1 | 0.04346086 | 0.000316329 |
| Klk8 | 0.000158214 | 3.877143252 | 0.16 | 0.035 | 1 | 0.057899329 | 1.808468542 | 0.2 | 0.056 | 1 | 0.057899329 | 0.000316403 |
| Tpd52 | 0.00015986 | -1.626782676 | 0.101 | 0.282 | 1 | 0.627945749 | -0.198015491 | 0.3 | 0.373 | 1 | 0.627945749 | 0.000319695 |
| Adora1 | 0.497412033 | 0.868210127 | 0.084 | 0.065 | 1 | 0.000160578 | 3.14345279 | 0.3 | 0.04 | 1 | 0.497412033 | 0.000321131 |
| Sipa1l1 | 0.000161131 | -0.826785452 | 0.731 | 0.788 | 1 | 0.92804563 | -0.152003093 | 0.9 | 0.849 | 1 | 0.92804563 | 0.000322236 |
| Armc9 | 0.000161289 | -1.765534746 | 0.092 | 0.271 | 1 | 0.272978545 | 0.14345279 | 0.1 | 0.287 | 1 | 0.272978545 | 0.000322552 |
| Dcun1d1 | 0.000162312 | -1.051530301 | 0.487 | 0.624 | 1 | 0.520961964 | -0.256052964 | 0.7 | 0.741 | 1 | 0.520961964 | 0.000324597 |
| Morf4l1 | 0.000162536 | -0.27431697 | 0.983 | 0.988 | 1 | 0.050451412 | -0.49683159 | 0.9 | 1 | 1 | 0.050451412 | 0.000325046 |
| A830082K12Rik | 0.000162679 | -2.748461233 | 0.025 | 0.165 | 1 | 0.27486734 | -1.171902651 | 0 | 0.108 | 1 | 0.27486734 | 0.000325332 |
| Rfx2 | 0.000162713 | -2.418312631 | 0.084 | 0.247 | 1 | 0.279102332 | -1.446964141 | 0.2 | 0.33 | 1 | 0.279102332 | 0.0003254 |
| Lrp5 | 0.000162848 | -1.389211512 | 0.185 | 0.376 | 1 | 0.251115929 | -0.703177281 | 0.2 | 0.383 | 1 | 0.251115929 | 0.00032567 |
| Dok4 | 0.000162938 | 2.177538186 | 0.202 | 0.059 | 1 | 0.42093472 | 0.374065718 | 0 | 0.062 | 1 | 0.42093472 | 0.00032585 |
| Cdk6 | 0.000163145 | 0.398431578 | 0.79 | 0.612 | 1 | 0.026023628 | 1.492460419 | 0.7 | 0.454 | 1 | 0.026023628 | 0.000326264 |
| Atl2 | 0.000163731 | -0.967578522 | 0.37 | 0.559 | 1 | 0.02575604 | -1.099535674 | 0.3 | 0.716 | 1 | 0.02575604 | 0.000327435 |

| | | | | | | | | | | | | |
|---|---|---|---|---|---|---|---|---|---|---|---|---|
| Htatip2 | 0.000164896 | 1.126007885 | 0.378 | 0.182 | 1 | 0.08021098 | 1.847996907 | 0.3 | 0.12 | 1 | 0.08021098 | 0.000329765 |
| Qrich1 | 0.000165723 | -0.959358016 | 0.538 | 0.665 | 1 | 0.044573474 | -0.961265081 | 0.5 | 0.836 | 1 | 0.044573474 | 0.00033142 |
| Dcaf6 | 0.000166373 | -0.988925227 | 0.605 | 0.771 | 1 | 0.928034953 | -0.270999081 | 0.8 | 0.84 | 1 | 0.928034953 | 0.000332718 |
| Cdk17 | 0.000167145 | -0.9067781 | 0.622 | 0.753 | 1 | 0.595056461 | -0.454806533 | 0.9 | 0.843 | 1 | 0.595056461 | 0.000334262 |
| Dpp8 | 0.00016741 | -0.929838929 | 0.471 | 0.624 | 1 | 0.208946428 | -0.512859555 | 0.7 | 0.769 | 1 | 0.208946428 | 0.000334792 |
| Gdf11 | 0.000167426 | -4.070389328 | 0 | 0.112 | 1 | 0.364051785 | -0.111361109 | 0 | 0.077 | 1 | 0.364051785 | 0.000334823 |
| Prr5 | 0.000167793 | -4.240314329 | 0 | 0.112 | 1 | 0.298405825 | -0.152003093 | 0 | 0.099 | 1 | 0.298405825 | 0.000335558 |
| Tom1l1 | 0.000167803 | -4.292781749 | 0 | 0.112 | 1 | 0.274499721 | -0.473931188 | 0 | 0.108 | 1 | 0.274499721 | 0.000335578 |
| Gm46430 | 0.00016781 | -4.129283017 | 0 | 0.112 | 1 | 0.979200188 | 0.732519689 | 0.1 | 0.096 | 1 | 0.979200188 | 0.000335592 |
| Chrd | 0.000167931 | -4.240314329 | 0 | 0.112 | 1 | 0.947574013 | 0.625604485 | 0.1 | 0.105 | 1 | 0.947574013 | 0.000335834 |
| Crnkl1 | 0.000168966 | -1.384899951 | 0.168 | 0.359 | 1 | 0.181487482 | -1.131825212 | 0.2 | 0.386 | 1 | 0.181487482 | 0.000337904 |
| Tle4 | 0.000169332 | -0.917701031 | 0.521 | 0.682 | 1 | 0.241500576 | -1.068024701 | 0.7 | 0.747 | 1 | 0.241500576 | 0.000338635 |
| L3mbtl3 | 0.000169824 | -1.866248611 | 0.168 | 0.347 | 1 | 0.816726443 | -0.185018151 | 0.3 | 0.34 | 1 | 0.816726443 | 0.000339618 |
| Tpcn1 | 0.000169824 | -1.124027291 | 0.252 | 0.459 | 1 | 0.12300584 | -0.674170467 | 0.3 | 0.602 | 1 | 0.12300584 | 0.000339619 |
| Alg13 | 0.000170595 | -1.665999073 | 0.076 | 0.247 | 1 | 0.092902276 | -1.876895855 | 0 | 0.225 | 1 | 0.092902276 | 0.000341161 |
| Samhd1 | 0.0001706 | 1.949668324 | 0.655 | 0.612 | 1 | 0.02641873 | 1.325829533 | 0.7 | 0.54 | 1 | 0.02641873 | 0.000341171 |
| Poc1b | 0.00017092 | -2.050211446 | 0.076 | 0.241 | 1 | 0.051012227 | -2.058893689 | 0 | 0.284 | 1 | 0.051012227 | 0.000341811 |
| Nt5dc3 | 0.000171048 | -1.045642666 | 0.555 | 0.653 | 1 | 0.522997913 | -0.31535116 | 0.6 | 0.725 | 1 | 0.522997913 | 0.000342067 |
| Wdr74 | 0.000171127 | -1.292781749 | 0.21 | 0.406 | 1 | 0.19243242 | -0.789433014 | 0.2 | 0.389 | 1 | 0.19243242 | 0.000342225 |
| Xpo5 | 0.000171384 | -1.307428525 | 0.151 | 0.347 | 1 | 0.760284152 | 0.558490289 | 0.4 | 0.383 | 1 | 0.760284152 | 0.000342739 |
| Cfap43 | 0.451376069 | -0.442358105 | 0.202 | 0.235 | 1 | 0.000171469 | 2.718361626 | 0.5 | 0.114 | 1 | 0.451376069 | 0.000342908 |
| Rassf5 | 0.000172359 | -2.944858446 | 0.034 | 0.176 | 1 | 0.660732381 | 0.625604485 | 0.2 | 0.151 | 1 | 0.660732381 | 0.000344688 |
| Ccdc186 | 0.000172837 | -0.719892081 | 0.731 | 0.776 | 1 | 0.044773277 | -0.921237491 | 0.6 | 0.843 | 1 | 0.044773277 | 0.000345645 |
| Serpinc1 | 0.000173038 | -3.258016331 | 0.017 | 0.147 | 1 | 0.196968962 | -1.736965594 | 0 | 0.145 | 1 | 0.196968962 | 0.000346046 |
| Arhgdia | 0.000173569 | -0.59645814 | 0.748 | 0.835 | 1 | 0.027802727 | -0.906890596 | 0.6 | 0.84 | 1 | 0.027802727 | 0.000347107 |
| Lyst | 0.000174195 | -2.057678276 | 0.126 | 0.306 | 1 | 0.348254389 | -1.055327074 | 0.4 | 0.494 | 1 | 0.348254389 | 0.00034836 |
| 8430426J06Rik | 0.127754956 | 0.514573173 | 0.059 | 0.024 | 1 | 0.000175174 | 2.558490289 | 0.3 | 0.04 | 1 | 0.127754956 | 0.000350318 |
| Arpp19 | 0.000175631 | -0.657487573 | 0.706 | 0.8 | 1 | 0.020118599 | -1.061029433 | 0.6 | 0.836 | 1 | 0.020118599 | 0.000351231 |
| Mier3 | 0.000175906 | -1.137503524 | 0.345 | 0.524 | 1 | 0.18543749 | -0.785747967 | 0.4 | 0.577 | 1 | 0.18543749 | 0.000351781 |
| Cbfa2t3 | 0.000175974 | -3.050211446 | 0.034 | 0.176 | 1 | 0.186290958 | -1.802257054 | 0 | 0.151 | 1 | 0.186290958 | 0.000351916 |
| Cask | 0.000176541 | -0.95491211 | 0.697 | 0.812 | 1 | 0.658274947 | -0.231229785 | 0.8 | 0.867 | 1 | 0.658274947 | 0.00035305 |
| Gm10635 | 0.000176776 | -5.129283017 | 0.008 | 0.129 | 1 | 0.325140259 | -2.004445905 | 0 | 0.09 | 1 | 0.325140259 | 0.00035352 |
| Pla2g4a | 0.000176835 | 0.64157845 | 0.605 | 0.353 | 1 | 0.44163891 | -1.225252075 | 0.2 | 0.302 | 1 | 0.44163891 | 0.000353639 |
| Mrps22 | 0.000177692 | -1.587788545 | 0.126 | 0.312 | 1 | 0.095438066 | -1.611434712 | 0 | 0.222 | 1 | 0.095438066 | 0.000355353 |
| Vps37d | 0.0001782 | -2.854660637 | 0.017 | 0.147 | 1 | 0.189509455 | 1.602884409 | 0.2 | 0.08 | 1 | 0.189509455 | 0.000356369 |
| Psma4 | 0.000179513 | -0.612256849 | 0.714 | 0.824 | 1 | 0.008894363 | -0.9669712 | 0.4 | 0.88 | 1 | 0.008894363 | 0.000358994 |
| Ephx1 | 0.000180038 | 0.620251251 | 0.706 | 0.441 | 1 | 0.517380103 | 0.732519689 | 0.5 | 0.506 | 1 | 0.517380103 | 0.000360043 |
| Prdx4 | 0.000180385 | -0.89056829 | 0.613 | 0.676 | 1 | 0.491070693 | -0.307281319 | 0.6 | 0.636 | 1 | 0.491070693 | 0.000360738 |
| Ssna1 | 0.000180614 | -1.142539114 | 0.269 | 0.465 | 1 | 0.411595084 | -0.604129911 | 0.3 | 0.392 | 1 | 0.411595084 | 0.000361196 |
| Slc17a5 | 0.000180816 | -1.315501826 | 0.218 | 0.418 | 1 | 0.164907097 | -1.163975735 | 0.3 | 0.488 | 1 | 0.164907097 | 0.0003616 |
| Cmtm5 | 0.000181073 | 0.887382437 | 0.454 | 0.247 | 1 | 0.021391502 | 1.909397451 | 0.4 | 0.145 | 1 | 0.021391502 | 0.000362112 |
| Sh3kbp1 | 0.000181462 | 0.211326499 | 0.815 | 0.506 | 1 | 0.058401275 | 1.17199933 | 0.5 | 0.265 | 1 | 0.058401275 | 0.00036289 |
| Pfdn6 | 0.000181818 | -1.115477217 | 0.294 | 0.488 | 1 | 0.049163545 | -1.661558192 | 0.1 | 0.404 | 1 | 0.049163545 | 0.000363604 |
| Klhdc10 | 0.000181889 | -1.277433785 | 0.345 | 0.518 | 1 | 0.289433041 | -0.640289575 | 0.5 | 0.657 | 1 | 0.289433041 | 0.000363744 |
| Ptx3 | 0.000182185 | 2.457348304 | 0.277 | 0.106 | 1 | 0.316136211 | -1.69632361 | 0 | 0.093 | 1 | 0.316136211 | 0.000364338 |
| Rbfox1 | 0.000182645 | -2.835924074 | 0.05 | 0.206 | 1 | 0.857117169 | -1.121629444 | 0.1 | 0.111 | 1 | 0.857117169 | 0.000365257 |
| Usp10 | 0.000182773 | -1.316827223 | 0.235 | 0.424 | 1 | 0.25449933 | -0.68251781 | 0.5 | 0.614 | 1 | 0.25449933 | 0.000365513 |
| Wnt5b | 0.000182942 | -2.95419631 | 0.084 | 0.247 | 1 | 0.827168801 | 0.185031894 | 0.1 | 0.077 | 1 | 0.827168801 | 0.000365851 |
| Acox1 | 0.000183942 | 0.911528313 | 0.857 | 0.659 | 1 | 0.17207684 | 0.918386234 | 0.8 | 0.741 | 1 | 0.17207684 | 0.000367849 |
| Hdgfl2 | 0.000184701 | -1.286522759 | 0.235 | 0.424 | 1 | 0.086993662 | -1.473931188 | 0.1 | 0.355 | 1 | 0.086993662 | 0.000369367 |
| Rpl7 | 0.000185181 | -0.294071154 | 0.983 | 0.982 | 1 | 0.002046108 | -1.036831292 | 0.7 | 0.994 | 1 | 0.002046108 | 0.000370327 |
| Inppl1 | 0.000185663 | -1.559427409 | 0.261 | 0.429 | 1 | 0.027445113 | -1.273632538 | 0.2 | 0.599 | 1 | 0.027445113 | 0.000371292 |
| Sec14l1 | 0.000186864 | 0.555849448 | 0.882 | 0.735 | 1 | 0.209045414 | 0.298474075 | 1 | 0.836 | 1 | 0.209045414 | 0.000373694 |
| Stimate | 0.000187249 | -1.662965013 | 0.151 | 0.335 | 1 | 0.544061998 | 0.130396637 | 0.4 | 0.269 | 1 | 0.544061998 | 0.000374462 |
| Rbm22 | 0.000187653 | -0.860935962 | 0.42 | 0.594 | 1 | 0.118818158 | -0.796774989 | 0.4 | 0.648 | 1 | 0.118818158 | 0.000375271 |
| Zbtb12 | 0.000187889 | -3.129283017 | 0.008 | 0.129 | 1 | 0.29827118 | -0.191531458 | 0 | 0.099 | 1 | 0.29827118 | 0.000375743 |
| Stk4 | 0.000188836 | -1.638673087 | 0.21 | 0.388 | 1 | 0.4585584 | -0.486698484 | 0.3 | 0.41 | 1 | 0.4585584 | 0.000377637 |
| Gas5 | 0.000188901 | -0.436404543 | 0.966 | 0.988 | 1 | 0.024775562 | -0.817629906 | 0.9 | 0.981 | 1 | 0.024775562 | 0.000377766 |
| Brpf1 | 0.000189726 | -1.595051318 | 0.151 | 0.335 | 1 | 0.325562156 | -0.111361109 | 0.2 | 0.417 | 1 | 0.325562156 | 0.000379416 |
| Zfp24 | 0.000189818 | -1.09423607 | 0.403 | 0.547 | 1 | 0.420040352 | -0.242605642 | 0.5 | 0.583 | 1 | 0.420040352 | 0.0003796 |
| Bche | 0.000189974 | 1.726077278 | 0.244 | 0.088 | 1 | 0.053104141 | 1.345496566 | 0.3 | 0.099 | 1 | 0.053104141 | 0.000379912 |
| Nfat5 | 0.000190169 | -0.40344021 | 0.983 | 1 | 1 | 0.064040461 | -0.577860542 | 1 | 1 | 1 | 0.064040461 | 0.000380303 |
| Elk3 | 0.000263275 | -0.982102795 | 0.387 | 0.565 | 1 | 0.00019049 | -4.374395515 | 0 | 0.673 | 1 | 0.000263275 | 0.000380945 |
| Haus4 | 0.000191025 | -2.87774425 | 0.017 | 0.147 | 1 | 0.155164702 | 1.558490289 | 0.2 | 0.074 | 1 | 0.155164702 | 0.000382014 |
| Fibin | 0.000191096 | 0.211926911 | 0.874 | 0.594 | 1 | 0.002081019 | 0.998022351 | 1 | 0.519 | 1 | 0.002081019 | 0.000382155 |
| Ndel1 | 0.000191314 | -0.526372377 | 0.874 | 0.935 | 1 | 0.339913147 | -0.264955909 | 0.9 | 0.966 | 1 | 0.339913147 | 0.00038259 |
| Rsrc2 | 0.0001914 | -0.522292121 | 0.933 | 0.976 | 1 | 0.513221658 | -0.223509058 | 0.9 | 0.96 | 1 | 0.513221658 | 0.000382763 |
| Gypc | 0.000191706 | -1.807354922 | 0.076 | 0.241 | 1 | 0.609544212 | -0.321928095 | 0.1 | 0.154 | 1 | 0.609544212 | 0.000383376 |
| Gnb1l | 0.000192647 | -2.485426827 | 0.034 | 0.176 | 1 | 0.511794067 | 0.720241359 | 0.2 | 0.123 | 1 | 0.511794067 | 0.000385257 |
| Slc7a2 | 0.000192725 | -1.265036759 | 0.286 | 0.471 | 1 | 0.212276748 | -1.706591945 | 0.3 | 0.438 | 1 | 0.212276748 | 0.000385412 |
| Eif5a | 0.000193425 | -0.450077263 | 0.975 | 0.971 | 1 | 0.016494652 | -0.850991408 | 0.8 | 0.975 | 1 | 0.016494652 | 0.000386812 |
| Ikzf2 | 0.000193793 | -2.234365063 | 0.126 | 0.3 | 1 | 0.023926797 | -2.833827133 | 0 | 0.352 | 1 | 0.023926797 | 0.000387548 |
| Tet3 | 0.000194045 | -1.19592021 | 0.378 | 0.576 | 1 | 0.048242161 | -1.394295072 | 0.8 | 0.806 | 1 | 0.048242161 | 0.000388053 |
| Galnt10 | 0.000194447 | -1.378511623 | 0.227 | 0.418 | 1 | 0.704269995 | 0.135278859 | 0.5 | 0.42 | 1 | 0.704269995 | 0.000388856 |
| Rexo5 | 0.101308751 | -1.11345805 | 0.076 | 0.135 | 1 | 0.000195304 | 2.345496566 | 0.5 | 0.108 | 1 | 0.101308751 | 0.000390569 |
| Creb3l2 | 0.000195311 | 0.22659241 | 1 | 0.982 | 1 | 0.969472911 | -0.111062383 | 1 | 0.988 | 1 | 0.969472911 | 0.000390584 |
| Fpgs | 0.000196129 | -2.411426246 | 0.034 | 0.176 | 1 | 0.185734253 | -1.004445905 | 0 | 0.151 | 1 | 0.185734253 | 0.00039222 |
| Tgfb1 | 0.000196202 | -0.949978281 | 0.765 | 0.765 | 1 | 0.222132726 | -0.647752392 | 0.7 | 0.852 | 1 | 0.222132726 | 0.000392365 |
| Zfp941 | 0.147030916 | -1.485426827 | 0 | 0.018 | 1 | 0.000197248 | 4.795529487 | 0.2 | 0.019 | 1 | 0.147030916 | 0.000394456 |
| Atad2 | 0.000199075 | -1.816072139 | 0.126 | 0.306 | 1 | 0.121942114 | 0.821524695 | 0.4 | 0.185 | 1 | 0.121942114 | 0.00039811 |
| Nhej1 | 0.000199711 | -2.222392421 | 0.067 | 0.229 | 1 | 0.157210239 | -2.048167282 | 0 | 0.17 | 1 | 0.157210239 | 0.000399382 |
| Cyhr1 | 0.00020183 | -1.93021167 | 0.067 | 0.229 | 1 | 0.068849019 | -1.505640048 | 0 | 0.253 | 1 | 0.068849019 | 0.000403619 |
| Pxmp2 | 0.009021257 | 1.321928095 | 0.143 | 0.053 | 1 | 0.000202282 | 4.602884409 | 0.2 | 0.019 | 1 | 0.009021257 | 0.000404524 |
| Zfp606 | 0.000202919 | -1.918386234 | 0.067 | 0.229 | 1 | 0.139851256 | 0.800691192 | 0.4 | 0.182 | 1 | 0.139851256 | 0.000405796 |
| Phf10 | 0.000204432 | -1 | 0.437 | 0.594 | 1 | 0.079834662 | -1.066506873 | 0.5 | 0.673 | 1 | 0.079834662 | 0.000408822 |
| Cyp2f2 | 0.000204458 | 1.343461256 | 0.101 | 0.288 | 1 | 0.130429551 | -2.233165946 | 0.1 | 0.343 | 1 | 0.130429551 | 0.000408875 |
| Ncbp2 | 0.000205035 | -0.776328026 | 0.555 | 0.665 | 1 | 0.278805622 | -0.663902132 | 0.7 | 0.667 | 1 | 0.278805622 | 0.000410028 |
| Rpl4 | 0.000205305 | -0.322794318 | 0.992 | 0.994 | 1 | 0.170419913 | -0.453637189 | 0.9 | 0.994 | 1 | 0.170419913 | 0.000410568 |
| Zbtb44 | 0.00020599 | -1.119632847 | 0.294 | 0.476 | 1 | 0.152924955 | -0.94951023 | 0.5 | 0.623 | 1 | 0.152924955 | 0.000411938 |
| Reep2 | 0.219589239 | -1.070389328 | 0.008 | 0.029 | 1 | 0.000207364 | 4.017921908 | 0.2 | 0.019 | 1 | 0.219589239 | 0.000414686 |
| Kif20b | 0.821673064 | -1.021479727 | 0.059 | 0.065 | 1 | 0.000207417 | 4.280956314 | 0.2 | 0.019 | 1 | 0.821673064 | 0.000414791 |
| Scarb1 | 0.000207555 | 1.243271151 | 0.563 | 0.394 | 1 | 0.042913615 | 0.945972066 | 0.6 | 0.29 | 1 | 0.042913615 | 0.000415066 |
| Sec23a | 0.000208411 | -0.659631556 | 0.748 | 0.794 | 1 | 0.480185037 | -0.350782958 | 0.9 | 0.824 | 1 | 0.480185037 | 0.000416779 |
| Syf2 | 0.000208448 | -0.882085797 | 0.471 | 0.641 | 1 | 0.051851493 | -1.026472211 | 0.4 | 0.688 | 1 | 0.051851493 | 0.000416852 |
| Elavl3 | 0.00020881 | -3.185866545 | 0.025 | 0.159 | 1 | 0.719292164 | -0.763437806 | 0.1 | 0.133 | 1 | 0.719292164 | 0.000417577 |
| Rab33b | 0.000209395 | -1.029048533 | 0.37 | 0.547 | 1 | 0.083076533 | -1.076595691 | 0.3 | 0.568 | 1 | 0.083076533 | 0.000418746 |
| Acss2 | 0.119513809 | 0.790207615 | 0.118 | 0.065 | 1 | 0.000209456 | 2.18142064 | 0.5 | 0.108 | 1 | 0.119513809 | 0.000418869 |
| Ing1 | 0.000210685 | -1.218322757 | 0.412 | 0.582 | 1 | 0.115104771 | -1.214738849 | 0.4 | 0.565 | 1 | 0.115104771 | 0.000421326 |

| | | | | | | | | | | | | |
|---|---|---|---|---|---|---|---|---|---|---|---|---|
| Gpx4 | 0.000211226 | 0.506445292 | 0.933 | 0.9 | 1 | 0.383789801 | -0.171902651 | 0.5 | 0.935 | 1 | 0.383789801 | 0.000422408 |
| Oser1 | 0.000211634 | -0.8384426 | 0.58 | 0.688 | 1 | 0.00110161 | -2.33375353 | 0.2 | 0.707 | 1 | 0.00110161 | 0.000423223 |
| Gng2 | 0.000212054 | -3.368069877 | 0.025 | 0.159 | 1 | 0.91167625 | 1.150025444 | 0.1 | 0.123 | 1 | 0.91167625 | 0.000424063 |
| Ywhae | 0.000213669 | -0.298858957 | 0.992 | 0.994 | 1 | 0.001570917 | -0.851493982 | 0.9 | 1 | 1 | 0.001570917 | 0.000427293 |
| Cnpy2 | 0.000213706 | -0.460582756 | 0.941 | 0.894 | 1 | 0.407252399 | -0.220482831 | 0.9 | 0.895 | 1 | 0.407252399 | 0.000427366 |
| Qdpr | 0.000213872 | 0.878984599 | 0.613 | 0.412 | 1 | 0.643011478 | 0.732519689 | 0.4 | 0.373 | 1 | 0.643011478 | 0.000427698 |
| Tdrd3 | 0.000213946 | -1.584962501 | 0.269 | 0.453 | 1 | 0.453315801 | -0.580337415 | 0.5 | 0.574 | 1 | 0.453315801 | 0.000427846 |
| Gtf3c5 | 0.000214713 | -1.970853654 | 0.067 | 0.229 | 1 | 0.565312298 | 0.720241359 | 0.2 | 0.13 | 1 | 0.565312298 | 0.00042938 |
| Plxnd1 | 0.000215239 | -3.100136671 | 0.025 | 0.159 | 1 | 0.0468615 | -2.528972552 | 0 | 0.293 | 1 | 0.0468615 | 0.000430432 |
| Nop53 | 0.000215483 | -0.691877705 | 0.639 | 0.747 | 1 | 0.011033305 | -1.431639467 | 0.4 | 0.716 | 1 | 0.011033305 | 0.000430919 |
| Ccdc6 | 0.000216318 | -1.158385809 | 0.336 | 0.518 | 1 | 0.051003261 | -1.245112498 | 0.4 | 0.63 | 1 | 0.051003261 | 0.000432589 |
| Tbc1d1 | 0.000216514 | -0.604598528 | 0.647 | 0.788 | 1 | 0.905344297 | -0.191531458 | 0.7 | 0.883 | 1 | 0.905344297 | 0.000432982 |
| Cntfr | 0.000217218 | -1.139430972 | 0.151 | 0.353 | 1 | 0.623034875 | 0.123104145 | 0.3 | 0.213 | 1 | 0.623034875 | 0.000434389 |
| 2410006H16Rik | 0.014748643 | -0.565395132 | 0.815 | 0.788 | 1 | 0.000217545 | -3.103611609 | 0.1 | 0.725 | 1 | 0.014748643 | 0.000435042 |
| Dusp6 | 0.000217846 | 0.820676301 | 0.294 | 0.118 | 1 | 0.500466651 | 0.930459067 | 0.2 | 0.13 | 1 | 0.500466651 | 0.000435644 |
| Pask | 0.342795612 | -1.222392421 | 0.017 | 0.035 | 1 | 0.000217999 | 4.017921908 | 0.2 | 0.019 | 1 | 0.342795612 | 0.000435951 |
| Pfkl | 0.000218347 | 0.775224927 | 0.622 | 0.406 | 1 | 0.125223593 | 1.120681482 | 0.5 | 0.309 | 1 | 0.125223593 | 0.000436646 |
| Npr3 | 0.00022012 | 0.826517179 | 0.336 | 0.153 | 1 | 0.373655406 | 2.280956314 | 0.1 | 0.043 | 1 | 0.373655406 | 0.000440192 |
| Cct3 | 0.000220393 | -0.577685335 | 0.773 | 0.865 | 1 | 0.025987163 | -0.92291602 | 0.5 | 0.889 | 1 | 0.025987163 | 0.000440738 |
| Paxbp1 | 0.000222042 | -0.8372705 | 0.487 | 0.676 | 1 | 0.796734545 | -0.204470513 | 0.8 | 0.731 | 1 | 0.796734545 | 0.000444035 |
| Tasp1 | 0.000222265 | -0.925999419 | 0.513 | 0.647 | 1 | 0.587799786 | -0.17472317 | 0.5 | 0.63 | 1 | 0.587799786 | 0.000444481 |
| Cdc16 | 0.000222725 | -0.802099197 | 0.613 | 0.724 | 1 | 0.359208328 | -0.518130992 | 0.6 | 0.648 | 1 | 0.359208328 | 0.0004454 |
| Scand1 | 0.000223516 | -0.89862298 | 0.529 | 0.676 | 1 | 0.022384104 | -1.256339753 | 0.2 | 0.651 | 1 | 0.022384104 | 0.000446982 |
| Brpf3 | 0.000223553 | -1.645891499 | 0.109 | 0.288 | 1 | 0.131862943 | -1.489872732 | 0.1 | 0.315 | 1 | 0.131862943 | 0.000447056 |
| Ppp1r9b | 0.000223893 | -1.780882711 | 0.076 | 0.241 | 1 | 0.029722706 | -2.457811523 | 0 | 0.333 | 1 | 0.029722706 | 0.000447736 |
| Mzt1 | 0.000224897 | -0.758445322 | 0.487 | 0.624 | 1 | 0.318771632 | -0.445602465 | 0.5 | 0.602 | 1 | 0.318771632 | 0.000449743 |
| Rnf152 | 0.000225019 | 2.57870351 | 0.185 | 0.053 | 1 | 0.007718905 | 2.054447784 | 0.3 | 0.068 | 1 | 0.007718905 | 0.000449988 |
| Dnajc25 | 0.000225899 | -0.628031223 | 0.857 | 0.841 | 1 | 0.176486758 | -0.545978977 | 0.8 | 0.815 | 1 | 0.176486758 | 0.000451746 |
| Utp20 | 0.000226048 | -1.402964667 | 0.176 | 0.365 | 1 | 0.113089268 | -1.382957528 | 0.1 | 0.333 | 1 | 0.113089268 | 0.000452044 |
| Orc4 | 0.000227193 | -0.876067672 | 0.42 | 0.606 | 1 | 0.010246472 | -1.62881679 | 0.3 | 0.694 | 1 | 0.010246472 | 0.000454334 |
| Smim31 | 0.000520955 | 0.796344141 | 0.328 | 0.153 | 1 | 0.000228297 | 2.725140159 | 0.5 | 0.117 | 1 | 0.000520955 | 0.000456542 |
| Gpr75 | 0.237885589 | -1.070389328 | 0 | 0.012 | 1 | 0.000229049 | 3.795529487 | 0.2 | 0.019 | 1 | 0.237885589 | 0.000458046 |
| Slurp1 | 0.000230094 | 2.487167015 | 0.227 | 0.076 | 1 | 0.328344798 | 0.847996907 | 0.1 | 0.037 | 1 | 0.328344798 | 0.000460134 |
| Lrrc41 | 0.000231831 | -0.928370323 | 0.378 | 0.582 | 1 | 0.410422745 | -0.404983835 | 0.5 | 0.642 | 1 | 0.410422745 | 0.000463609 |
| Tars | 0.000232727 | -0.779488129 | 0.496 | 0.647 | 1 | 0.016440268 | -1.486698484 | 0.2 | 0.642 | 1 | 0.016440268 | 0.000465399 |
| Arhgap29 | 0.000233224 | -0.950043339 | 0.655 | 0.771 | 1 | 0.034762491 | -0.974941911 | 0.6 | 0.889 | 1 | 0.034762491 | 0.000466393 |
| Slco2b1 | 0.000233275 | 1.666576266 | 0.244 | 0.088 | 1 | 0.236382673 | 1.354956895 | 0.2 | 0.086 | 1 | 0.236382673 | 0.000466496 |
| Id3 | 0.000233638 | -1.141773961 | 0.496 | 0.676 | 1 | 0.264790874 | -0.68910848 | 0.3 | 0.472 | 1 | 0.264790874 | 0.000467222 |
| Fhip2a | 0.000234366 | -1.015941544 | 0.294 | 0.488 | 1 | 0.002318636 | -2.28585884 | 0.2 | 0.667 | 1 | 0.002318636 | 0.000468677 |
| Foxa3 | 0.000234405 | 0.628744193 | 0.496 | 0.288 | 1 | 0.319016434 | 0.762421175 | 0.4 | 0.29 | 1 | 0.319016434 | 0.000468755 |
| Lpl | 0.000234574 | -4.185866545 | 0.017 | 0.141 | 1 | 0.066822357 | 2.286118019 | 0.2 | 0.062 | 1 | 0.066822357 | 0.000469093 |
| Nup153 | 0.000234658 | -0.760049207 | 0.731 | 0.818 | 1 | 0.655476731 | -0.103981579 | 0.9 | 0.889 | 1 | 0.655476731 | 0.000469262 |
| Zfp931 | 0.000234907 | -2.572889668 | 0.025 | 0.159 | 1 | 0.317764748 | 1.111031312 | 0.2 | 0.099 | 1 | 0.317764748 | 0.000469759 |
| Ppp4r1 | 0.000235393 | -1.215160852 | 0.37 | 0.529 | 1 | 0.005189672 | -1.135883428 | 0.3 | 0.781 | 1 | 0.005189672 | 0.000470731 |
| Ube2s | 0.000235399 | -1.230169772 | 0.168 | 0.353 | 1 | 0.464596803 | -0.654503434 | 0.3 | 0.404 | 1 | 0.464596803 | 0.000470742 |
| Ctnnd1 | 0.000236736 | -0.810421225 | 0.798 | 0.824 | 1 | 0.538002305 | 0.351719559 | 0.9 | 0.91 | 1 | 0.538002305 | 0.000473416 |
| Arl5a | 0.000236963 | -1.070389328 | 0.319 | 0.512 | 1 | 0.044725052 | -0.541045384 | 0.2 | 0.58 | 1 | 0.044725052 | 0.00047387 |
| Stx2 | 0.000237105 | -2.213347282 | 0.059 | 0.212 | 1 | 0.805044113 | -0.204470513 | 0.2 | 0.219 | 1 | 0.805044113 | 0.000474153 |
| Ehd1 | 0.000238696 | 0.735988489 | 0.756 | 0.594 | 1 | 0.433978268 | -0.388070452 | 0.5 | 0.639 | 1 | 0.433978268 | 0.000477336 |
| Gpat4 | 0.000238871 | -0.987927168 | 0.361 | 0.541 | 1 | 0.217999378 | -0.412065933 | 0.4 | 0.617 | 1 | 0.217999378 | 0.000477685 |
| Tyw5 | 0.000239151 | -1.807354922 | 0.059 | 0.218 | 1 | 0.350315773 | 0.808468542 | 0.3 | 0.17 | 1 | 0.350315773 | 0.000478244 |
| Srbd1 | 0.000239326 | -0.802617003 | 0.639 | 0.759 | 1 | 0.026045899 | -1.113233523 | 0.2 | 0.685 | 1 | 0.026045899 | 0.000478595 |
| Mest | 0.000239673 | -0.797456461 | 0.319 | 0.518 | 1 | 0.43244844 | -1.616284112 | 0.4 | 0.429 | 1 | 0.43244844 | 0.000479288 |
| Slc2a4 | 0.000617478 | 2.573466862 | 0.134 | 0.029 | 1 | 0.000240381 | 3.602884409 | 0.2 | 0.019 | 1 | 0.000617478 | 0.000480704 |
| Cct6b | 0.025470389 | -2.944858446 | 0 | 0.041 | 1 | 0.000240381 | 3.432959407 | 0.2 | 0.019 | 1 | 0.025470389 | 0.000480704 |
| Gm56817 | 0.215728257 | 1.099535674 | 0.042 | 0.018 | 1 | 0.000240381 | 3.602884409 | 0.2 | 0.019 | 1 | 0.215728257 | 0.000480704 |
| Gbx1 | 0.237885589 | -1.807354922 | 0 | 0.012 | 1 | 0.000240381 | 3.602884409 | 0.2 | 0.019 | 1 | 0.237885589 | 0.000480704 |
| Bcl2l15 | 0.935790323 | -0.292781749 | 0.025 | 0.024 | 1 | 0.000240381 | 3.432959407 | 0.2 | 0.019 | 1 | 0.935790323 | 0.000480704 |
| Vamp2 | 0.0002411 | -0.90600251 | 0.403 | 0.565 | 1 | 0.030904427 | -1.630735268 | 0.2 | 0.522 | 1 | 0.030904427 | 0.000482141 |
| Pam | 0.00024217 | -0.412225301 | 1 | 0.994 | 1 | 0.021371134 | -0.663360071 | 1 | 1 | 1 | 0.021371134 | 0.000484281 |
| Itsn2 | 0.000242685 | -1.040015679 | 0.605 | 0.712 | 1 | 0.046318833 | -0.773069568 | 0.7 | 0.889 | 1 | 0.046318833 | 0.00048531 |
| Mdh2 | 0.000242699 | -0.533149868 | 0.84 | 0.9 | 1 | 0.014624059 | -0.722645923 | 0.6 | 0.867 | 1 | 0.014624059 | 0.000485338 |
| Slc2a10 | 0.000242972 | -1.359895945 | 0.176 | 0.359 | 1 | 0.114961157 | -1.339630097 | 0.1 | 0.33 | 1 | 0.114961157 | 0.000485885 |
| Glt8d1 | 0.000243489 | -0.635669463 | 0.664 | 0.747 | 1 | 0.658824141 | 0.27541813 | 0.7 | 0.58 | 1 | 0.658824141 | 0.000486919 |
| Kcnd2 | 0.128346244 | -0.371255807 | 0.101 | 0.053 | 1 | 0.000243763 | 1.609837169 | 0.7 | 0.213 | 1 | 0.128346244 | 0.000487467 |
| Sf3a2 | 0.00024452 | -1.18142064 | 0.176 | 0.371 | 1 | 0.373067504 | -0.567040593 | 0.2 | 0.318 | 1 | 0.373067504 | 0.000488981 |
| Gm15726 | 0.217405186 | -0.944858446 | 0.05 | 0.088 | 1 | 0.000246562 | 2.847996907 | 0.2 | 0.019 | 1 | 0.217405186 | 0.000493064 |
| Ddx10 | 0.000246598 | -0.978466838 | 0.462 | 0.612 | 1 | 0.717432225 | -0.215697769 | 0.4 | 0.491 | 1 | 0.717432225 | 0.000493135 |
| Ypel2 | 0.000248006 | 0.586052561 | 0.538 | 0.341 | 1 | 0.617459398 | -0.41621569 | 0.5 | 0.617 | 1 | 0.617459398 | 0.000495951 |
| Ufc1 | 0.000249186 | -0.510332169 | 0.79 | 0.841 | 1 | 0.052711379 | -0.67556505 | 0.5 | 0.796 | 1 | 0.052711379 | 0.00049831 |
| Mkln1os | 0.000249657 | 2.389042291 | 0.168 | 0.041 | 1 | 0.001868122 | 2.584962501 | 0.3 | 0.056 | 1 | 0.001868122 | 0.000499251 |
| Dera | 0.000251312 | 0.689070904 | 0.454 | 0.247 | 1 | 0.435582826 | -0.719043686 | 0.2 | 0.296 | 1 | 0.435582826 | 0.00050256 |
| Gsk3a | 0.000251912 | -1.13965199 | 0.387 | 0.553 | 1 | 0.230359093 | 0.148506018 | 0.4 | 0.636 | 1 | 0.230359093 | 0.000503761 |
| Anxa11 | 0.000251969 | 0.14919356 | 0.908 | 0.706 | 1 | 0.987950989 | -0.28585884 | 0.9 | 0.855 | 1 | 0.987950989 | 0.000503875 |
| Gars | 0.000252546 | -1.06248413 | 0.521 | 0.647 | 1 | 0.137124737 | -1.060080604 | 0.5 | 0.66 | 1 | 0.137124737 | 0.000505028 |
| Cplx1 | 0.611244073 | 0.292180751 | 0.034 | 0.024 | 1 | 0.000252557 | 3.14345279 | 0.2 | 0.019 | 1 | 0.611244073 | 0.00050505 |
| Snx3 | 0.000253673 | -0.464962725 | 0.882 | 0.924 | 1 | 0.075699085 | -0.592942456 | 0.8 | 0.935 | 1 | 0.075699085 | 0.000507281 |
| 2310002F09Rik | 0.000953373 | 1.777607579 | 0.143 | 0.035 | 1 | 0.000254046 | 3.718361626 | 0.3 | 0.043 | 1 | 0.000953373 | 0.000508028 |
| Tmem107 | 0.000254708 | -1.402964667 | 0.185 | 0.371 | 1 | 0.311485256 | -0.723545078 | 0.1 | 0.231 | 1 | 0.311485256 | 0.00050935 |
| Tlr3 | 0.000254946 | 1.514573173 | 0.235 | 0.082 | 1 | 0.311539133 | 0.874963954 | 0.2 | 0.099 | 1 | 0.311539133 | 0.000509828 |
| mt-Co2 | 0.00202361 | 0.272278074 | 1 | 1 | 1 | 0.000255417 | 0.972183804 | 1 | 1 | 1 | 0.00202361 | 0.000510769 |
| Tub | 0.000255659 | -4.070389328 | 0 | 0.106 | 1 | 0.364023002 | -0.111361109 | 0 | 0.077 | 1 | 0.364023002 | 0.000511252 |
| Fcho2 | 0.00025585 | -0.674283645 | 0.874 | 0.929 | 1 | 0.400034578 | -0.465240228 | 0.9 | 0.975 | 1 | 0.400034578 | 0.000511635 |
| Glipr2 | 0.000255922 | -4.070389328 | 0 | 0.106 | 1 | 0.374705201 | -0.339630097 | 0 | 0.074 | 1 | 0.374705201 | 0.000511779 |
| Rad23b | 0.000255964 | -0.571767676 | 0.849 | 0.859 | 1 | 0.005047928 | -1.077336294 | 0.4 | 0.954 | 1 | 0.005047928 | 0.000511863 |
| Cys1 | 0.000256142 | -4.129283017 | 0 | 0.106 | 1 | 0.28239418 | -0.536666944 | 0 | 0.105 | 1 | 0.28239418 | 0.000512219 |
| Eif1ad | 0.000256498 | -1.216610069 | 0.176 | 0.371 | 1 | 0.955998248 | 0.22090893 | 0.3 | 0.312 | 1 | 0.955998248 | 0.000512929 |
| Zfp462 | 0.00025658 | -1.053111336 | 0.319 | 0.5 | 1 | 0.386944779 | -0.441509711 | 0.2 | 0.367 | 1 | 0.386944779 | 0.000513094 |
| R3hdml | 0.000256914 | -6.427941333 | 0 | 0.106 | 1 | 0.577177401 | -0.267480311 | 0 | 0.031 | 1 | 0.577177401 | 0.000513762 |
| Cul2 | 0.000257044 | -1.241501244 | 0.277 | 0.465 | 1 | 0.948667157 | 0.102314095 | 0.5 | 0.512 | 1 | 0.948667157 | 0.000514022 |
| Smtn | 0.000258082 | -1.063402898 | 0.378 | 0.547 | 1 | 0.43884761 | -0.11821478 | 0.7 | 0.719 | 1 | 0.43884761 | 0.000516098 |
| Lrp6 | 0.000656303 | -0.625061363 | 0.941 | 0.959 | 1 | 0.000258198 | -1.349024714 | 0.9 | 0.988 | 1 | 0.000656303 | 0.00051633 |
| Smu1 | 0.000259829 | -1.118483616 | 0.42 | 0.559 | 1 | 0.060625184 | -1.061029433 | 0.4 | 0.627 | 1 | 0.060625184 | 0.000519591 |
| Tacc1 | 0.000260049 | -0.549769061 | 0.84 | 0.9 | 1 | 0.057929647 | -0.775311906 | 0.8 | 0.889 | 1 | 0.057929647 | 0.00052003 |
| Cdc37 | 0.000260738 | -0.590686615 | 0.782 | 0.859 | 1 | 0.051842968 | -0.711598874 | 0.7 | 0.898 | 1 | 0.051842968 | 0.000521408 |
| Gm56974 | 0.000261526 | 2.514573173 | 0.109 | 0.012 | 1 | 0.149795061 | 2.847996907 | 0.1 | 0.025 | 1 | 0.149795061 | 0.000522983 |
| Spry2 | 0.000262203 | 1.12321467 | 0.588 | 0.388 | 1 | 0.992673645 | 0.361752457 | 0.4 | 0.444 | 1 | 0.992673645 | 0.000524338 |
| mt-Co3 | 0.087001929 | 0.109124725 | 1 | 1 | 1 | 0.000262408 | 0.820078636 | 1 | 1 | 1 | 0.087001929 | 0.000524747 |

| | | | | | | | | | | | | |
|---|---|---|---|---|---|---|---|---|---|---|---|---|
| Vat1 | 0.000263271 | -1.485426827 | 0.227 | 0.4 | 1 | 0.259207096 | -0.908868245 | 0.4 | 0.519 | 1 | 0.259207096 | 0.000526473 |
| Med22 | 0.000264817 | -1.560714954 | 0.109 | 0.282 | 1 | 0.925761735 | 0.24181792 | 0.3 | 0.324 | 1 | 0.925761735 | 0.000529564 |
| Cfap299 | 0.97221701 | 1.514573173 | 0.017 | 0.018 | 1 | 0.000265283 | 2.354956895 | 0.2 | 0.019 | 1 | 0.97221701 | 0.000530495 |
| Cltc | 0.000265853 | -0.321646566 | 0.975 | 0.982 | 1 | 0.008103494 | -0.731502903 | 1 | 0.991 | 1 | 0.008103494 | 0.000531636 |
| Crispld1 | 0.000268538 | -1.255100974 | 0.891 | 0.847 | 1 | 0.110542372 | -1.158073098 | 0.8 | 0.815 | 1 | 0.110542372 | 0.000537003 |
| Zswim3 | 0.010583426 | -2.185866545 | 0.034 | 0.118 | 1 | 0.000271297 | 3.017921908 | 0.3 | 0.043 | 1 | 0.010583426 | 0.000542521 |
| Slit3 | 0.000271489 | -4.807354922 | 0.017 | 0.141 | 1 | 0.121847956 | -3.640289575 | 0 | 0.198 | 1 | 0.121847956 | 0.000542904 |
| Vegfa | 0.000272004 | 1.039664218 | 0.622 | 0.447 | 1 | 0.474263715 | 1.819652281 | 0.4 | 0.417 | 1 | 0.474263715 | 0.000543934 |
| Fhip1a | 0.000275019 | 0.630897291 | 0.571 | 0.341 | 1 | 0.539876972 | -0.294961047 | 0.5 | 0.593 | 1 | 0.539876972 | 0.000549962 |
| Susd6 | 0.000275709 | -0.71175835 | 0.756 | 0.824 | 1 | 0.517043727 | -0.367221482 | 0.8 | 0.861 | 1 | 0.517043727 | 0.000551341 |
| Sqle | 0.00027581 | 0.47902242 | 0.807 | 0.518 | 1 | 0.660366007 | -0.58635877 | 0.7 | 0.466 | 1 | 0.660366007 | 0.000551543 |
| Smad2 | 0.000276067 | -1.184495814 | 0.471 | 0.594 | 1 | 0.380603174 | -0.386832618 | 0.5 | 0.679 | 1 | 0.380603174 | 0.000552058 |
| Nars | 0.00027668 | -0.732408828 | 0.798 | 0.853 | 1 | 0.0151118 | -1.078793246 | 0.6 | 0.901 | 1 | 0.0151118 | 0.000553283 |
| Phactr4 | 0.00027687 | -0.892602209 | 0.504 | 0.641 | 1 | 0.466996426 | -0.493040011 | 0.5 | 0.627 | 1 | 0.466996426 | 0.000553663 |
| Rasl11b | 0.000277369 | -2.163498732 | 0.076 | 0.235 | 1 | 1 | -0.606568957 | 0.2 | 0.179 | 1 | 1 | 0.00055466 |
| Ttc28 | 0.000278877 | -0.910190872 | 0.857 | 0.929 | 1 | 0.109912096 | 0.590383625 | 1 | 0.917 | 1 | 0.109912096 | 0.000557677 |
| Igip | 0.000280271 | -1.532732542 | 0.101 | 0.276 | 1 | 0.144333227 | -1.037360528 | 0.1 | 0.312 | 1 | 0.144333227 | 0.000560464 |
| Twist2 | 0.00028058 | -3.392317423 | 0.008 | 0.124 | 1 | 0.420904619 | 0.31748219 | 0 | 0.062 | 1 | 0.420904619 | 0.000561081 |
| Rasa4 | 0.000280615 | -3.529820947 | 0.008 | 0.124 | 1 | 0.385678941 | -0.191531458 | 0 | 0.071 | 1 | 0.385678941 | 0.000561151 |
| Tmod2 | 0.000280638 | -3.485426827 | 0.008 | 0.124 | 1 | 0.191210788 | -1.171902651 | 0 | 0.148 | 1 | 0.191210788 | 0.000561197 |
| Cpd | 0.000280778 | -0.844113472 | 0.697 | 0.729 | 1 | 0.28557721 | 0.214388697 | 1 | 0.824 | 1 | 0.28557721 | 0.000561477 |
| Rab23 | 0.000281158 | -1.787989597 | 0.101 | 0.271 | 1 | 0.464551078 | 0.591657153 | 0.3 | 0.191 | 1 | 0.464551078 | 0.000562238 |
| Gm11210 | 0.000282055 | -3.529820947 | 0.008 | 0.124 | 1 | 0.325722325 | 0.574978412 | 0.3 | 0.164 | 1 | 0.325722325 | 0.00056403 |
| Id4 | 0.000282406 | 1.185950425 | 0.521 | 0.335 | 1 | 0.004483311 | 2.600513504 | 0.7 | 0.333 | 1 | 0.004483311 | 0.000564731 |
| Med21 | 0.000283292 | 0.94413109 | 0.748 | 0.6 | 1 | 0.50965859 | 1.497089745 | 0.5 | 0.485 | 1 | 0.50965859 | 0.000566504 |
| Net1 | 0.000283578 | -1.35710273 | 0.227 | 0.418 | 1 | 0.044192548 | -1.872849022 | 0.2 | 0.5 | 1 | 0.044192548 | 0.000567076 |
| Tor1b | 0.000283852 | -0.972692654 | 0.353 | 0.529 | 1 | 0.883077134 | 0.255421222 | 0.4 | 0.392 | 1 | 0.883077134 | 0.000567623 |
| Pabpc4 | 0.000284079 | -0.641077698 | 0.681 | 0.765 | 1 | 0.153118317 | -0.919893261 | 0.7 | 0.728 | 1 | 0.153118317 | 0.000568077 |
| Prss23 | 0.000284155 | -2.74669214 | 0.109 | 0.271 | 1 | 0.146276244 | -2.077475115 | 0.1 | 0.324 | 1 | 0.146276244 | 0.000568229 |
| Fam118a | 0.000284648 | -1.821710215 | 0.143 | 0.318 | 1 | 0.757107773 | 0.526068812 | 0.2 | 0.164 | 1 | 0.757107773 | 0.000569215 |
| Moxd1 | 0.001305557 | 0.739748035 | 0.462 | 0.282 | 1 | 0.000287427 | 2.385653692 | 0.7 | 0.235 | 1 | 0.001305557 | 0.000574771 |
| Arhgap23 | 0.000287713 | -0.887525271 | 0.504 | 0.641 | 1 | 0.094147464 | -0.750262417 | 0.8 | 0.849 | 1 | 0.094147464 | 0.000575344 |
| Cpox | 0.000288068 | -1.024585638 | 0.185 | 0.388 | 1 | 0.02087514 | -2.574535129 | 0 | 0.364 | 1 | 0.02087514 | 0.000576053 |
| Txlna | 0.000289405 | -0.79380839 | 0.639 | 0.706 | 1 | 0.651887159 | -0.290066996 | 0.6 | 0.713 | 1 | 0.651887159 | 0.000578727 |
| Wdr47 | 0.00029251 | 1.695145418 | 0.403 | 0.218 | 1 | 0.047077838 | 0.893800596 | 0.7 | 0.37 | 1 | 0.047077838 | 0.000584934 |
| Ran | 0.000292553 | -0.559136513 | 0.899 | 0.9 | 1 | 0.010370433 | -1.030110788 | 0.4 | 0.932 | 1 | 0.010370433 | 0.000585021 |
| Rpn2 | 0.000293059 | 0.385839859 | 0.958 | 0.818 | 1 | 0.32625551 | 0.526068812 | 1 | 0.849 | 1 | 0.32625551 | 0.000586032 |
| Dis3l2 | 0.000294453 | -1.21426454 | 0.437 | 0.588 | 1 | 0.855007992 | 0.144315904 | 0.5 | 0.586 | 1 | 0.855007992 | 0.00058882 |
| Ppp1r21 | 0.000295227 | -1.893511566 | 0.084 | 0.247 | 1 | 0.09273948 | -1.852442812 | 0 | 0.225 | 1 | 0.09273948 | 0.000590367 |
| Ppm1d | 0.000296805 | -1.569960337 | 0.202 | 0.376 | 1 | 0.427424789 | -0.58694015 | 0.4 | 0.497 | 1 | 0.427424789 | 0.000593522 |
| Plrg1 | 0.000297864 | -1.485426827 | 0.134 | 0.312 | 1 | 0.148876573 | -1.015501094 | 0.1 | 0.309 | 1 | 0.148876573 | 0.000595638 |
| Prpf40a | 0.000298522 | -0.578659072 | 0.933 | 0.924 | 1 | 0.55793132 | -0.231059822 | 1 | 0.951 | 1 | 0.55793132 | 0.000596955 |
| Myl12b | 0.000298691 | -0.669851398 | 0.765 | 0.859 | 1 | 0.373440248 | -0.369908124 | 0.7 | 0.91 | 1 | 0.373440248 | 0.000597293 |
| Tbc1d19 | 0.000299606 | -1.485426827 | 0.126 | 0.306 | 1 | 0.509034211 | -0.616284112 | 0.2 | 0.278 | 1 | 0.509034211 | 0.000599122 |
| Zfp68 | 0.000299671 | -1.470927257 | 0.193 | 0.376 | 1 | 0.127403097 | -1.171902651 | 0.2 | 0.42 | 1 | 0.127403097 | 0.000599252 |
| Pip4k2a | 0.000300139 | -1.691877705 | 0.118 | 0.294 | 1 | 0.476647874 | -0.880931369 | 0.2 | 0.281 | 1 | 0.476647874 | 0.000600187 |
| Appbp2 | 0.000300199 | -0.565480079 | 0.95 | 0.929 | 1 | 0.016426478 | -0.845196286 | 0.9 | 0.963 | 1 | 0.016426478 | 0.000600308 |
| Macir | 0.000301538 | -1.352160296 | 0.092 | 0.259 | 1 | 0.103614721 | -1.531541912 | 0.2 | 0.432 | 1 | 0.103614721 | 0.000602984 |
| Dctn6 | 0.000302325 | -0.874838519 | 0.471 | 0.629 | 1 | 0.101036672 | -1.176834946 | 0.3 | 0.512 | 1 | 0.101036672 | 0.000604558 |
| Bcl10 | 0.000304224 | -0.848831558 | 0.555 | 0.676 | 1 | 0.116017576 | -0.940147412 | 0.4 | 0.614 | 1 | 0.116017576 | 0.000608356 |
| Cdc25a | 0.000305224 | -2.510961919 | 0.05 | 0.194 | 1 | 0.084767767 | -1.864721141 | 0 | 0.235 | 1 | 0.084767767 | 0.000610354 |
| Ppp5c | 0.000305535 | -1.217731044 | 0.227 | 0.412 | 1 | 0.020665973 | -2.294961047 | 0 | 0.361 | 1 | 0.020665973 | 0.000610977 |
| Trim8 | 0.066783511 | -0.566726929 | 0.387 | 0.471 | 1 | 0.00030629 | -3.736965594 | 0 | 0.639 | 1 | 0.066783511 | 0.000612486 |
| Rbm45 | 0.000306774 | -1.842978832 | 0.059 | 0.212 | 1 | 0.169636372 | 1.473601392 | 0.3 | 0.145 | 1 | 0.169636372 | 0.000613453 |
| Trnt1 | 0.000306997 | -1.045727274 | 0.361 | 0.535 | 1 | 0.089184491 | -1.357117523 | 0.2 | 0.448 | 1 | 0.089184491 | 0.000613899 |
| Dido1 | 0.000309203 | -0.733354341 | 0.664 | 0.759 | 1 | 0.922053599 | 0.176619654 | 0.8 | 0.861 | 1 | 0.922053599 | 0.00061831 |
| Brsk1 | 0.00030947 | -2.485426827 | 0.017 | 0.141 | 1 | 0.48846329 | -0.267480311 | 0.1 | 0.182 | 1 | 0.48846329 | 0.000618844 |
| Ephb4 | 0.000309632 | -1.714245518 | 0.109 | 0.276 | 1 | 0.363069841 | -0.573039333 | 0.3 | 0.432 | 1 | 0.363069841 | 0.000619169 |
| Gmps | 0.000309801 | -0.848681784 | 0.571 | 0.682 | 1 | 0.240304108 | -0.642668298 | 0.6 | 0.756 | 1 | 0.240304108 | 0.000619506 |
| Casp12 | 0.000310684 | -1.663764068 | 0.101 | 0.271 | 1 | 0.558664206 | -0.567040593 | 0.2 | 0.284 | 1 | 0.558664206 | 0.000621271 |
| Pcna | 0.000310686 | -0.816485934 | 0.546 | 0.665 | 1 | 0.119736911 | 0.159940913 | 0.3 | 0.769 | 1 | 0.119736911 | 0.000621276 |
| Prkg2 | 0.000312005 | -2.020920652 | 0.101 | 0.265 | 1 | 0.618533069 | -0.709998547 | 0.1 | 0.062 | 1 | 0.618533069 | 0.000623912 |
| Zfp395 | 0.000312264 | -1.461179281 | 0.16 | 0.341 | 1 | 0.626767239 | -0.548893246 | 0.3 | 0.346 | 1 | 0.626767239 | 0.000624431 |
| Frmd4a | 0.000312674 | 1.250003168 | 0.513 | 0.359 | 1 | 0.171668569 | 1.653349476 | 0.4 | 0.219 | 1 | 0.171668569 | 0.00062525 |
| Arhgap35 | 0.000313774 | -1.151201373 | 0.42 | 0.571 | 1 | 0.105716706 | -0.748054773 | 0.5 | 0.781 | 1 | 0.105716706 | 0.000627449 |
| Nepro | 0.000313946 | -1.807354922 | 0.059 | 0.212 | 1 | 0.442388207 | -0.28585884 | 0.1 | 0.194 | 1 | 0.442388207 | 0.000627793 |
| Spaca6 | 0.00031456 | -1.353782821 | 0.269 | 0.435 | 1 | 0.022274578 | -1.648834684 | 0.2 | 0.59 | 1 | 0.022274578 | 0.000629021 |
| Pnkd | 0.000314639 | 0.789753009 | 0.681 | 0.471 | 1 | 0.703017795 | -0.165299916 | 0.4 | 0.441 | 1 | 0.703017795 | 0.00062918 |
| 4930556M19Rik | 0.089877159 | -1.333423734 | 0.034 | 0.082 | 1 | 0.000315335 | 2.930459067 | 0.3 | 0.043 | 1 | 0.089877159 | 0.000630571 |
| Wdr76 | 0.280380388 | -0.900464326 | 0.05 | 0.082 | 1 | 0.000315335 | 2.847996907 | 0.3 | 0.043 | 1 | 0.280380388 | 0.000630571 |
| Oxsm | 0.667442994 | -0.572889668 | 0.059 | 0.071 | 1 | 0.000315335 | 2.558490289 | 0.3 | 0.043 | 1 | 0.667442994 | 0.000630571 |
| Aff2 | 0.000316739 | -4.040015679 | 0.025 | 0.153 | 1 | 0.418424757 | -1.959358016 | 0.1 | 0.188 | 1 | 0.418424757 | 0.000633378 |
| Lrrc17 | 0.00031905 | -3.593951284 | 0.008 | 0.124 | 1 | 0.505605027 | 0.695993813 | 0 | 0.043 | 1 | 0.505605027 | 0.000637999 |
| Spaar | 0.000320148 | 1.893084796 | 0.092 | 0.006 | 1 | 0.73075158 | 2.695993813 | 0 | 0.012 | 1 | 0.73075158 | 0.000640193 |
| Mmadhc | 0.000320318 | -0.936088236 | 0.403 | 0.576 | 1 | 0.523403684 | -0.187627003 | 0.5 | 0.596 | 1 | 0.523403684 | 0.000640534 |
| Smc6 | 0.000320364 | -0.82136178 | 0.622 | 0.747 | 1 | 0.577062417 | 0.216524441 | 0.8 | 0.772 | 1 | 0.577062417 | 0.000640625 |
| Etv6 | 0.004201293 | 0.255838904 | 0.975 | 0.941 | 1 | 0.00032128 | 1.365326499 | 1 | 0.901 | 1 | 0.004201293 | 0.000642457 |
| Ubqln1 | 0.000321813 | -0.828314541 | 0.487 | 0.629 | 1 | 0.014147407 | -1.202629167 | 0.4 | 0.781 | 1 | 0.014147407 | 0.000643523 |
| Tmem47 | 0.000322783 | -1.807354922 | 0.059 | 0.212 | 1 | 0.617782368 | -0.556986928 | 0.2 | 0.25 | 1 | 0.617782368 | 0.000645461 |
| Nt5c3b | 0.000324877 | -0.891419187 | 0.37 | 0.547 | 1 | 0.273018381 | -0.840059087 | 0.3 | 0.423 | 1 | 0.273018381 | 0.000649648 |
| Gins2 | 0.152144555 | -0.87244995 | 0.092 | 0.147 | 1 | 0.000326737 | 2.82527683 | 0.4 | 0.08 | 1 | 0.152144555 | 0.000653367 |
| Gpx8 | 0.000328138 | -0.557396614 | 0.916 | 0.853 | 1 | 0.059415222 | -0.85049454 | 0.7 | 0.83 | 1 | 0.059415222 | 0.000656168 |
| Keap1 | 0.000330101 | -1.451211112 | 0.244 | 0.412 | 1 | 0.365960014 | -0.351311902 | 0.3 | 0.444 | 1 | 0.365960014 | 0.000660092 |
| E2f4 | 0.000330376 | -1.292781749 | 0.151 | 0.335 | 1 | 0.392498599 | -0.611434712 | 0.3 | 0.395 | 1 | 0.392498599 | 0.000660642 |
| Pycr1 | 0.000331061 | -1.250961574 | 0.176 | 0.359 | 1 | 0.239104235 | -0.840059087 | 0.1 | 0.259 | 1 | 0.239104235 | 0.000662011 |
| Marchf3 | 0.000331347 | -1.761872152 | 0.235 | 0.406 | 1 | 0.284446266 | 0.467724825 | 0.5 | 0.346 | 1 | 0.284446266 | 0.000662584 |
| Narf | 0.000332528 | -1.936088236 | 0.101 | 0.265 | 1 | 0.683596841 | -0.171902651 | 0.1 | 0.142 | 1 | 0.683596841 | 0.000664945 |
| Bcl2l11 | 0.000333085 | -2.219615337 | 0.126 | 0.294 | 1 | 0.180384651 | -2.25354112 | 0.1 | 0.281 | 1 | 0.180384651 | 0.000666059 |
| Uba2 | 0.000333999 | -0.900464326 | 0.378 | 0.541 | 1 | 0.076298709 | -0.864721141 | 0.4 | 0.66 | 1 | 0.076298709 | 0.000667887 |
| Pus1 | 0.000335721 | -1.566346823 | 0.092 | 0.259 | 1 | 0.160676288 | -1.111361109 | 0 | 0.167 | 1 | 0.160676288 | 0.000671328 |
| Zfp608 | 0.000336542 | 0.431075709 | 0.941 | 0.894 | 1 | 0.011692602 | 0.688915982 | 1 | 0.883 | 1 | 0.011692602 | 0.000672971 |
| Ccn6 | 0.000337239 | 0.360949984 | 0.529 | 0.282 | 1 | 0.244232021 | 1.026142415 | 0.5 | 0.333 | 1 | 0.244232021 | 0.000674364 |
| Flrt2 | 0.000339562 | -0.380743641 | 0.622 | 0.882 | 1 | 0.027020055 | -1.346411753 | 0.8 | 0.917 | 1 | 0.027020055 | 0.000679008 |
| Cilp2 | 0.000340265 | -1.077218666 | 0.504 | 0.647 | 1 | 0.025955604 | -1.570376305 | 0.7 | 0.778 | 1 | 0.025955604 | 0.000680414 |
| Stam2 | 0.000340297 | -1.079607858 | 0.294 | 0.482 | 1 | 0.012454397 | -1.997028433 | 0.2 | 0.586 | 1 | 0.012454397 | 0.000680479 |
| Dip2a | 0.000340487 | -2.275503758 | 0.076 | 0.229 | 1 | 0.960120217 | 0.210566986 | 0.3 | 0.29 | 1 | 0.960120217 | 0.000680857 |
| Ppp2r1b | 0.000341504 | -1.807354922 | 0.118 | 0.282 | 1 | 0.550929175 | 0.403212064 | 0.4 | 0.293 | 1 | 0.550929175 | 0.000682891 |
| Mkx | 0.000342096 | -3.175742328 | 0.067 | 0.218 | 1 | 0.29066254 | -1.171902651 | 0 | 0.102 | 1 | 0.29066254 | 0.000684074 |

| | | | | | | | | | | | | |
|---|---|---|---|---|---|---|---|---|---|---|---|---|
| Rassf3 | 0.000342202 | -1.866248611 | 0.176 | 0.347 | 1 | 0.042228982 | -1.567040593 | 0.2 | 0.506 | 1 | 0.042228982 | 0.000684286 |
| Gid4 | 0.000342227 | -0.900464326 | 0.294 | 0.506 | 1 | 0.308069177 | -0.802257054 | 0.3 | 0.417 | 1 | 0.308069177 | 0.000684336 |
| Rpl9 | 0.005610629 | -0.220395131 | 0.975 | 0.994 | 1 | 0.000342518 | -1.226419486 | 1 | 1 | 1 | 0.005610629 | 0.000684919 |
| Hmgxb4 | 0.000342626 | -1.359895945 | 0.202 | 0.394 | 1 | 0.595335224 | -0.408342847 | 0.2 | 0.269 | 1 | 0.595335224 | 0.000685134 |
| Rpl10a | 0.000342695 | -0.396330485 | 0.975 | 0.988 | 1 | 0.006944145 | -0.889413895 | 0.9 | 0.994 | 1 | 0.006944145 | 0.000685273 |
| Tm9sf4 | 0.000343703 | -0.537683064 | 0.84 | 0.876 | 1 | 0.301394913 | 0.33663329 | 1 | 0.886 | 1 | 0.301394913 | 0.000687288 |
| Acot1 | 0.000343935 | -1.607417352 | 0.109 | 0.276 | 1 | 0.383505032 | 1.036069255 | 0.3 | 0.194 | 1 | 0.383505032 | 0.000687751 |
| Septin8 | 0.00034483 | -1.28128611 | 0.269 | 0.441 | 1 | 0.575869793 | -0.146985019 | 0.4 | 0.512 | 1 | 0.575869793 | 0.000689542 |
| Arpc2 | 0.000346179 | -0.440673426 | 0.966 | 0.976 | 1 | 0.384599395 | -0.164058532 | 1 | 1 | 1 | 0.384599395 | 0.000692237 |
| Tmem254 | 0.000347429 | -0.857903864 | 0.655 | 0.718 | 1 | 0.479582483 | -0.296256687 | 0.5 | 0.731 | 1 | 0.479582483 | 0.000694737 |
| Nck1 | 0.00034818 | -0.928370323 | 0.546 | 0.647 | 1 | 0.316158306 | -0.541381638 | 0.7 | 0.728 | 1 | 0.316158306 | 0.000696239 |
| Gm50431 | 0.288105649 | 0.514573173 | 0.092 | 0.059 | 1 | 0.00034919 | 2.769994395 | 0.3 | 0.043 | 1 | 0.288105649 | 0.000698257 |
| Thra | 0.000350092 | -0.693268063 | 0.857 | 0.824 | 1 | 0.210874094 | -0.398724844 | 0.9 | 0.898 | 1 | 0.210874094 | 0.000700061 |
| Cep104 | 0.000350295 | -1.894817763 | 0.109 | 0.276 | 1 | 0.297823392 | -0.797995028 | 0.2 | 0.336 | 1 | 0.297823392 | 0.000700467 |
| Mapk8ip3 | 0.000350988 | -0.878571721 | 0.496 | 0.612 | 1 | 0.791486625 | 0.190102883 | 0.7 | 0.642 | 1 | 0.791486625 | 0.000701853 |
| Zfp667 | 0.00035109 | -2.572889668 | 0.025 | 0.153 | 1 | 0.614205493 | 0.193493473 | 0.1 | 0.17 | 1 | 0.614205493 | 0.000702057 |
| Septin6 | 0.000352571 | -2.430285273 | 0.059 | 0.206 | 1 | 0.436339157 | -0.802257054 | 0.1 | 0.191 | 1 | 0.436339157 | 0.000705017 |
| Dlg2 | 0.000352619 | 0.389269576 | 0.874 | 0.7 | 1 | 0.678674806 | 1.01523282 | 0.5 | 0.491 | 1 | 0.678674806 | 0.000705113 |
| Dnaja1 | 0.000352943 | -0.744252878 | 0.95 | 0.971 | 1 | 0.006054769 | -1.015501094 | 0.6 | 1 | 1 | 0.006054769 | 0.000705761 |
| Lats1 | 0.000353048 | -0.776206223 | 0.538 | 0.671 | 1 | 0.004796363 | -1.282534657 | 0.7 | 0.877 | 1 | 0.004796363 | 0.00070597 |
| Mybph | 0.000353175 | 3.269460675 | 0.118 | 0.018 | 1 | 0.199151937 | 1.494359952 | 0.1 | 0.028 | 1 | 0.199151937 | 0.000706226 |
| Pdlim1 | 0.000355145 | -1.829381228 | 0.05 | 0.2 | 1 | 0.145611571 | -1.191531458 | 0.1 | 0.312 | 1 | 0.145611571 | 0.000710164 |
| L3hypdh | 0.00035918 | -1.485426827 | 0.101 | 0.271 | 1 | 0.604704877 | -0.131825212 | 0.1 | 0.157 | 1 | 0.604704877 | 0.000718231 |
| Rbmx | 0.000359854 | -1.039528364 | 0.378 | 0.547 | 1 | 0.193282166 | -0.714528205 | 0.3 | 0.506 | 1 | 0.193282166 | 0.000719579 |
| Casp4 | 0.00036015 | 0.337380171 | 0.521 | 0.271 | 1 | 0.058541654 | 1.323041715 | 0.5 | 0.256 | 1 | 0.058541654 | 0.000720171 |
| Epha3 | 0.000360797 | -4.641931313 | 0.017 | 0.135 | 1 | 0.325201517 | -2.528972552 | 0 | 0.09 | 1 | 0.325201517 | 0.000721464 |
| Zmym3 | 0.220766719 | -0.765534746 | 0.05 | 0.088 | 1 | 0.000362005 | 2.301714874 | 0.5 | 0.117 | 1 | 0.220766719 | 0.000723878 |
| Srrm1 | 0.0003627 | -0.373467965 | 0.992 | 0.976 | 1 | 0.337800032 | -0.240390088 | 1 | 0.985 | 1 | 0.337800032 | 0.000725268 |
| Dym | 0.000364142 | -1.328187085 | 0.277 | 0.459 | 1 | 0.106492422 | -1.135883428 | 0.3 | 0.546 | 1 | 0.106492422 | 0.00072815 |
| Tbc1d15 | 0.000365838 | -0.840521786 | 0.597 | 0.724 | 1 | 0.121944282 | -0.678243155 | 0.8 | 0.796 | 1 | 0.121944282 | 0.000731542 |
| Nrk | 0.000367733 | -0.449751096 | 0.571 | 0.682 | 1 | 0.629015611 | -0.146577382 | 0.9 | 0.725 | 1 | 0.629015611 | 0.000735331 |
| Usp22 | 0.000367739 | -0.990662135 | 0.37 | 0.547 | 1 | 0.288500579 | -0.723545078 | 0.4 | 0.537 | 1 | 0.288500579 | 0.000735342 |
| Usf1 | 0.000370092 | -1.58058406 | 0.076 | 0.235 | 1 | 0.976290775 | 0.197742946 | 0.3 | 0.278 | 1 | 0.976290775 | 0.000740046 |
| Lsm12 | 0.000371447 | -0.609058176 | 0.773 | 0.829 | 1 | 0.041254846 | -0.79970135 | 0.7 | 0.88 | 1 | 0.041254846 | 0.000742755 |
| Usp14 | 0.000372489 | -0.712718048 | 0.613 | 0.712 | 1 | 0.958582109 | 0.190667429 | 0.6 | 0.716 | 1 | 0.958582109 | 0.000744839 |
| Rpsa | 0.000372827 | -0.393497485 | 0.983 | 0.988 | 1 | 0.018086667 | -0.783159368 | 1 | 0.997 | 1 | 0.018086667 | 0.000745515 |
| Pakap.1 | 0.000373326 | 0.76346641 | 0.731 | 0.576 | 1 | 0.223346665 | 0.563746015 | 0.7 | 0.491 | 1 | 0.223346665 | 0.000746513 |
| Lmf2 | 0.000373795 | -1.246239163 | 0.126 | 0.306 | 1 | 0.168370653 | 1.036069255 | 0.4 | 0.207 | 1 | 0.168370653 | 0.00074745 |
| Gria1 | 0.000375072 | 1.099535674 | 0.168 | 0.041 | 1 | 0.501412479 | 0.660369903 | 0.1 | 0.049 | 1 | 0.501412479 | 0.000750003 |
| Crtap | 0.000375713 | 0.416083309 | 0.95 | 0.829 | 1 | 0.288373437 | 0.578453392 | 0.9 | 0.799 | 1 | 0.288373437 | 0.000751285 |
| Kidins220 | 0.000377658 | -0.984659454 | 0.555 | 0.688 | 1 | 0.038345218 | -1.159271094 | 0.7 | 0.821 | 1 | 0.038345218 | 0.000755174 |
| Rasgef1b | 0.000378045 | 1.748063303 | 0.277 | 0.118 | 1 | 0.604714968 | 1.159940913 | 0.1 | 0.059 | 1 | 0.604714968 | 0.000755947 |
| Dnaja2 | 0.000379613 | -0.511610759 | 0.908 | 0.888 | 1 | 0.009951064 | -0.758660665 | 0.8 | 0.944 | 1 | 0.009951064 | 0.000759082 |
| Rps3 | 0.012273463 | -0.272740698 | 0.975 | 0.976 | 1 | 0.000380912 | -1.225465255 | 0.9 | 0.994 | 1 | 0.012273463 | 0.000761678 |
| Pck2 | 0.000380993 | -1.268328706 | 0.176 | 0.365 | 1 | 0.198789501 | 0.804142617 | 0.5 | 0.312 | 1 | 0.198789501 | 0.000761841 |
| Icam1 | 0.000381523 | -1.080373416 | 0.261 | 0.465 | 1 | 0.130514286 | -1.750262417 | 0.3 | 0.478 | 1 | 0.130514286 | 0.0007629 |
| Poln | 0.000383617 | -2.258016331 | 0.05 | 0.194 | 1 | 0.680398462 | -0.291933355 | 0.2 | 0.142 | 1 | 0.680398462 | 0.000767087 |
| Yipf5 | 0.00038652 | -0.568790712 | 0.899 | 0.859 | 1 | 0.432442844 | -0.278010978 | 0.8 | 0.895 | 1 | 0.432442844 | 0.000772891 |
| Mat2b | 0.000386865 | -0.985272714 | 0.37 | 0.547 | 1 | 0.249211582 | -0.767802998 | 0.4 | 0.54 | 1 | 0.249211582 | 0.00077358 |
| Heca | 0.000387106 | -1.314696526 | 0.336 | 0.512 | 1 | 0.415409812 | -0.685597203 | 0.6 | 0.617 | 1 | 0.415409812 | 0.000774062 |
| Zranb1 | 0.000387801 | -0.625908051 | 0.832 | 0.847 | 1 | 0.14337683 | -0.517353469 | 0.7 | 0.917 | 1 | 0.14337683 | 0.000775452 |
| Pid1 | 0.004288675 | 1 | 0.597 | 0.418 | 1 | 0.000388315 | 1.41635744 | 0.8 | 0.324 | 1 | 0.004288675 | 0.000776479 |
| 2310010J17Rik | 0.00038858 | 0.770412077 | 0.462 | 0.259 | 1 | 0.156287213 | 1.210566986 | 0.4 | 0.228 | 1 | 0.156287213 | 0.000777008 |
| Rpl10 | 0.021003609 | -0.235049705 | 0.983 | 0.994 | 1 | 0.000389719 | -1.213853126 | 0.9 | 0.997 | 1 | 0.021003609 | 0.000779287 |
| Itgb1bp2 | 0.000390365 | -4.070389328 | 0 | 0.1 | 1 | 0.363739469 | 2.602884409 | 0.1 | 0.043 | 1 | 0.363739469 | 0.000780577 |
| Cplx2 | 0.000390404 | -4.008988783 | 0 | 0.1 | 1 | 0.274607019 | -0.625934282 | 0 | 0.108 | 1 | 0.274607019 | 0.000780656 |
| Dclk2 | 0.00039084 | -4.292781749 | 0 | 0.1 | 1 | 0.260051931 | -1.374395515 | 0 | 0.114 | 1 | 0.260051931 | 0.000781528 |
| Kcne4 | 0.000391041 | -4.87774425 | 0 | 0.1 | 1 | 0.698532411 | 1.558490289 | 0 | 0.015 | 1 | 0.698532411 | 0.000781929 |
| Bcl7c | 0.000391156 | -0.68044281 | 0.529 | 0.706 | 1 | 0.001971573 | -1.789433014 | 0.1 | 0.701 | 1 | 0.001971573 | 0.000782158 |
| Sfrp4 | 0.000391407 | -5.714245518 | 0 | 0.1 | 1 | 0.539481121 | -2.391469028 | 0 | 0.037 | 1 | 0.539481121 | 0.000782661 |
| Taf11 | 0.000392472 | -0.910448415 | 0.37 | 0.547 | 1 | 0.053152637 | -1.468577954 | 0.2 | 0.485 | 1 | 0.053152637 | 0.000784789 |
| Klf11 | 0.000393017 | -1.600904045 | 0.143 | 0.312 | 1 | 0.53323585 | -0.304006187 | 0.2 | 0.296 | 1 | 0.53323585 | 0.000785879 |
| Lrp4 | 0.00039324 | -4.109917692 | 0.017 | 0.135 | 1 | 0.964594091 | 0.374065718 | 0.1 | 0.102 | 1 | 0.964594091 | 0.000786325 |
| Mob3c | 0.000395216 | -1.567888987 | 0.126 | 0.294 | 1 | 0.428647739 | 0.754887502 | 0.3 | 0.198 | 1 | 0.428647739 | 0.000790275 |
| Pepd | 0.000396136 | -1.123828587 | 0.294 | 0.488 | 1 | 0.127341399 | -1.111361109 | 0.3 | 0.519 | 1 | 0.127341399 | 0.000792116 |
| Vars | 0.000397016 | -1.529820947 | 0.118 | 0.288 | 1 | 0.675333027 | 0.598383016 | 0.3 | 0.247 | 1 | 0.675333027 | 0.000793874 |
| Evi5 | 0.000397314 | -1.167686529 | 0.471 | 0.594 | 1 | 0.060221991 | -1.073777926 | 0.5 | 0.738 | 1 | 0.060221991 | 0.000794469 |
| Pygm | 0.000397361 | -3.185866545 | 0.017 | 0.135 | 1 | 0.496225999 | 0.847996907 | 0.2 | 0.127 | 1 | 0.496225999 | 0.000794563 |
| Nup214 | 0.000397494 | -0.989193245 | 0.378 | 0.553 | 1 | 0.838591657 | -0.327852929 | 0.7 | 0.574 | 1 | 0.838591657 | 0.000794831 |
| U2af1 | 0.000398766 | -0.516058446 | 0.84 | 0.847 | 1 | 0.37702681 | -0.275166504 | 0.9 | 0.923 | 1 | 0.37702681 | 0.000797374 |
| Pdgfra | 0.00039962 | -1.796824255 | 0.286 | 0.441 | 1 | 0.015920965 | -3.517353469 | 0 | 0.389 | 1 | 0.015920965 | 0.000799081 |
| Prpf39 | 0.000403883 | -0.705050159 | 0.739 | 0.776 | 1 | 0.357622528 | -0.478984416 | 0.9 | 0.883 | 1 | 0.357622528 | 0.000807602 |
| Dcp2 | 0.000404333 | -1.375243909 | 0.244 | 0.406 | 1 | 0.099039944 | -1.433289204 | 0.1 | 0.346 | 1 | 0.099039944 | 0.000808503 |
| Cnbp | 0.000848726 | -0.311454601 | 0.975 | 0.982 | 1 | 0.000404508 | -1.195559459 | 0.9 | 0.997 | 1 | 0.000848726 | 0.000808852 |
| Utp25 | 0.00040462 | -1.568842835 | 0.134 | 0.306 | 1 | 0.762422017 | 0.469485283 | 0.2 | 0.259 | 1 | 0.762422017 | 0.000809077 |
| Gm15788 | 0.628300786 | 0.407657969 | 0.05 | 0.065 | 1 | 0.000406033 | 2.432959407 | 0.3 | 0.043 | 1 | 0.628300786 | 0.000811902 |
| Gm14296 | 0.000409521 | -2.807354922 | 0.017 | 0.135 | 1 | 0.306978493 | -0.230005605 | 0 | 0.096 | 1 | 0.306978493 | 0.000818875 |
| Nrgn | 0.000411465 | -4.529820947 | 0.008 | 0.118 | 1 | 0.557764571 | 0.625604485 | 0 | 0.034 | 1 | 0.557764571 | 0.00082276 |
| Pus7 | 0.000411763 | -1.179323699 | 0.21 | 0.394 | 1 | 0.223396143 | 0.769994395 | 0.5 | 0.321 | 1 | 0.223396143 | 0.000823357 |
| Ltf | 0.000414112 | -1.346720556 | 0.202 | 0.388 | 1 | 0.034857283 | -2.304006187 | 0 | 0.318 | 1 | 0.034857283 | 0.000828052 |
| Sppl2b | 0.00041441 | 1.088308418 | 0.395 | 0.212 | 1 | 0.319265358 | 1.884066161 | 0.3 | 0.207 | 1 | 0.319265358 | 0.000828648 |
| Large1 | 0.000414438 | -0.722466024 | 0.655 | 0.812 | 1 | 0.849159854 | -0.163828529 | 1 | 0.864 | 1 | 0.849159854 | 0.000828704 |
| Bcl7a | 0.000415176 | -2.555816155 | 0.034 | 0.165 | 1 | 0.219622256 | -0.888968688 | 0 | 0.133 | 1 | 0.219622256 | 0.00083018 |
| Hmces | 0.000416385 | -1.26121167 | 0.218 | 0.394 | 1 | 0.940252499 | 0.50836681 | 0.4 | 0.38 | 1 | 0.940252499 | 0.000832597 |
| Ednra | 0.000416558 | -4.485426827 | 0.008 | 0.118 | 1 | 0.505683207 | -2.654503434 | 0 | 0.043 | 1 | 0.505683207 | 0.000832943 |
| Chmp2b | 0.000417261 | -0.783107376 | 0.555 | 0.694 | 1 | 0.178283835 | -0.349194961 | 0.4 | 0.735 | 1 | 0.178283835 | 0.000834349 |
| Sfrp1 | 0.000417491 | -2.008988783 | 0.067 | 0.218 | 1 | 0.063792675 | -2.201246612 | 0 | 0.262 | 1 | 0.063792675 | 0.000834807 |
| Brox | 0.000418928 | -1.168236651 | 0.37 | 0.524 | 1 | 0.107912384 | -0.840059087 | 0.5 | 0.688 | 1 | 0.107912384 | 0.00083768 |
| Bcar1 | 0.000419611 | -1.10433666 | 0.277 | 0.465 | 1 | 0.74853994 | 0.13799836 | 0.4 | 0.478 | 1 | 0.74853994 | 0.000839045 |
| Prex1 | 0.000420329 | -1.819845866 | 0.151 | 0.324 | 1 | 0.911005402 | -0.294961047 | 0.5 | 0.42 | 1 | 0.911005402 | 0.000840481 |
| Gm15270 | 0.000420878 | 1.415658147 | 0.42 | 0.224 | 1 | 0.453779924 | 2.063725598 | 0.2 | 0.139 | 1 | 0.453779924 | 0.000841579 |
| Stk40 | 0.000423174 | -0.683898368 | 0.622 | 0.735 | 1 | 0.009979843 | -1.295968868 | 0.4 | 0.818 | 1 | 0.009979843 | 0.000846168 |
| Tent4a | 0.00042503 | -1.867896464 | 0.109 | 0.271 | 1 | 0.226264348 | -1.026472211 | 0.2 | 0.361 | 1 | 0.226264348 | 0.000849879 |
| Ppp2r1a | 0.000425344 | -0.668290884 | 0.647 | 0.747 | 1 | 0.021541428 | -1.05430642 | 0.3 | 0.738 | 1 | 0.021541428 | 0.000850507 |
| Dact1 | 0.000426129 | -0.9345059 | 0.328 | 0.512 | 1 | 0.011798181 | -2.947862377 | 0.1 | 0.503 | 1 | 0.011798181 | 0.000852077 |
| Usp16 | 0.00042618 | -0.603458844 | 0.832 | 0.865 | 1 | 0.125455992 | -0.443234391 | 0.7 | 0.873 | 1 | 0.125455992 | 0.000852179 |
| Rnf126 | 0.000428241 | -1.292781749 | 0.176 | 0.353 | 1 | 0.504445947 | -0.473931188 | 0.3 | 0.367 | 1 | 0.504445947 | 0.000856299 |
| Thap3 | 0.000428727 | -1.737813989 | 0.118 | 0.282 | 1 | 0.393628245 | -0.709998547 | 0.1 | 0.204 | 1 | 0.393628245 | 0.000857271 |

| | | | | | | | | | | | | |
|---|---|---|---|---|---|---|---|---|---|---|---|---|
| Cdk9 | 0.000431561 | -1.161729639 | 0.37 | 0.506 | 1 | 0.960690426 | 0.10666612 | 0.6 | 0.617 | 1 | 0.960690426 | 0.000862937 |
| Lingo1 | 0.0004325 | -3.343407822 | 0.008 | 0.118 | 1 | 0.539464158 | 0.625604485 | 0 | 0.037 | 1 | 0.539464158 | 0.000864813 |
| Sf1 | 0.000432681 | -0.57700343 | 0.84 | 0.841 | 1 | 0.42273564 | -0.25708514 | 1 | 0.92 | 1 | 0.42273564 | 0.000865175 |
| Ip6k2 | 0.000434462 | -1.142539114 | 0.21 | 0.406 | 1 | 0.009024362 | -2.756865152 | 0 | 0.426 | 1 | 0.009024362 | 0.000868735 |
| Alms1 | 0.000437486 | -2.09423607 | 0.067 | 0.218 | 1 | 0.64178071 | -0.473931188 | 0.1 | 0.148 | 1 | 0.64178071 | 0.00087478 |
| Maco1 | 0.000438968 | -0.97775835 | 0.597 | 0.706 | 1 | 0.610426658 | -0.441509711 | 0.8 | 0.756 | 1 | 0.610426658 | 0.000877743 |
| Gnai2 | 0.000439824 | -0.36201662 | 0.966 | 0.982 | 1 | 0.041805299 | -0.619669019 | 0.9 | 0.988 | 1 | 0.041805299 | 0.000879455 |
| Terf1 | 0.000440328 | -1.389211512 | 0.202 | 0.376 | 1 | 0.53188824 | 0.715802294 | 0.5 | 0.373 | 1 | 0.53188824 | 0.000880462 |
| Gtf3c4 | 0.000440356 | -1.505326385 | 0.143 | 0.312 | 1 | 0.170784855 | -0.959358016 | 0.2 | 0.395 | 1 | 0.170784855 | 0.000880519 |
| Tnfrsf23 | 0.000440792 | 0.607682577 | 0.563 | 0.341 | 1 | 0.066994201 | 1.703606997 | 0.5 | 0.299 | 1 | 0.066994201 | 0.00088139 |
| Osgin2 | 0.000441551 | 0.218748832 | 0.597 | 0.347 | 1 | 0.33663919 | 0.266856117 | 0.6 | 0.429 | 1 | 0.33663919 | 0.000882906 |
| Snrpf | 0.000441614 | -0.529434009 | 0.899 | 0.882 | 1 | 0.062703232 | -0.634154789 | 0.6 | 0.873 | 1 | 0.062703232 | 0.000883032 |
| Myh9 | 0.000445161 | -1.032914622 | 0.748 | 0.788 | 1 | 0.286790269 | -0.174893933 | 0.8 | 0.914 | 1 | 0.286790269 | 0.000890124 |
| Eps15 | 0.000445935 | -1.152003093 | 0.37 | 0.535 | 1 | 0.103049529 | -1.055327074 | 0.5 | 0.704 | 1 | 0.103049529 | 0.000891672 |
| Chmp4c | 0.000446228 | 2.055141554 | 0.151 | 0.035 | 1 | 0.474947236 | 0.558490289 | 0 | 0.049 | 1 | 0.474947236 | 0.000892256 |
| Hnrnpc | 0.000446587 | -0.280888071 | 0.975 | 0.982 | 1 | 0.000518166 | -1.014341371 | 0.9 | 0.994 | 1 | 0.000518166 | 0.000892975 |
| Atp5g1 | 0.00044705 | -0.533448342 | 0.882 | 0.9 | 1 | 0.031583525 | -0.633129783 | 0.5 | 0.867 | 1 | 0.031583525 | 0.0008939 |
| Ubfd1 | 0.000447211 | -0.875373345 | 0.353 | 0.559 | 1 | 0.084270566 | -0.389042291 | 0.2 | 0.525 | 1 | 0.084270566 | 0.000894221 |
| Drosha | 0.000448271 | -1.444784843 | 0.21 | 0.376 | 1 | 0.988946284 | -0.214738849 | 0.4 | 0.34 | 1 | 0.988946284 | 0.000896342 |
| Dhx29 | 0.00044935 | -1.402964667 | 0.193 | 0.365 | 1 | 0.127361767 | -1.465893869 | 0.1 | 0.318 | 1 | 0.127361767 | 0.000898498 |
| Slc27a6 | 0.000450682 | 0.96203215 | 0.403 | 0.212 | 1 | 0.294492914 | 0.255421222 | 0.5 | 0.287 | 1 | 0.294492914 | 0.00090116 |
| Casc1 | 0.000455298 | -2.326729081 | 0.034 | 0.165 | 1 | 0.515480678 | -0.750262417 | 0.1 | 0.173 | 1 | 0.515480678 | 0.000910388 |
| Scrib | 0.000455532 | -1.842978832 | 0.059 | 0.206 | 1 | 0.639857369 | 0.789103218 | 0.1 | 0.173 | 1 | 0.639857369 | 0.000910856 |
| Cacybp | 0.000457071 | -0.718697302 | 0.647 | 0.735 | 1 | 0.000622192 | -2.066000576 | 0.4 | 0.84 | 1 | 0.000622192 | 0.000913932 |
| Hace1 | 0.000458905 | -1.40103464 | 0.143 | 0.318 | 1 | 0.560676547 | -0.521236903 | 0.4 | 0.448 | 1 | 0.560676547 | 0.000917599 |
| Ncoa6 | 0.000459935 | -1.314808056 | 0.429 | 0.547 | 1 | 0.179624258 | -0.606568957 | 0.7 | 0.806 | 1 | 0.179624258 | 0.000919659 |
| Sap130 | 0.000460901 | -1.252592659 | 0.336 | 0.488 | 1 | 0.377258302 | -0.365782384 | 0.3 | 0.475 | 1 | 0.377258302 | 0.00092159 |
| Sqstm1 | 0.000461782 | -0.585878207 | 0.966 | 0.959 | 1 | 0.00258184 | -1.330961517 | 0.8 | 0.988 | 1 | 0.00258184 | 0.000923351 |
| Ash2l | 0.000462353 | -1.189442 | 0.202 | 0.382 | 1 | 0.228287492 | -0.888968688 | 0.2 | 0.367 | 1 | 0.228287492 | 0.000924492 |
| Snx14 | 0.000462394 | -1.029747343 | 0.395 | 0.559 | 1 | 0.610394544 | -0.355536488 | 0.6 | 0.593 | 1 | 0.610394544 | 0.000924573 |
| Psenen | 0.000463855 | 0.504377406 | 0.882 | 0.771 | 1 | 0.743721966 | 0.116384547 | 0.7 | 0.769 | 1 | 0.743721966 | 0.000927495 |
| Cnst | 0.000466966 | -2.378511623 | 0.05 | 0.188 | 1 | 0.273574766 | -0.900941329 | 0.1 | 0.244 | 1 | 0.273574766 | 0.000933714 |
| Lrrc20 | 0.07305434 | 1.072568626 | 0.227 | 0.147 | 1 | 0.000467281 | 2.808468542 | 0.4 | 0.08 | 1 | 0.07305434 | 0.000934344 |
| Postn | 0.000467536 | 0.326443576 | 0.504 | 0.294 | 1 | 1 | -1.69632361 | 0.1 | 0.099 | 1 | 1 | 0.000934854 |
| Glb1l | 0.000468447 | -1.515174171 | 0.059 | 0.206 | 1 | 0.526140694 | 0.176619654 | 0.1 | 0.194 | 1 | 0.526140694 | 0.000936674 |
| Ammecr1 | 0.000469834 | -1.466318004 | 0.193 | 0.371 | 1 | 0.328697944 | -1.076595691 | 0.2 | 0.321 | 1 | 0.328697944 | 0.000939447 |
| Poc1a | 0.079004402 | -1.359895945 | 0.017 | 0.059 | 1 | 0.000469982 | 2.94753258 | 0.3 | 0.046 | 1 | 0.079004402 | 0.000939743 |
| Lin7c | 0.000472005 | -0.881779763 | 0.429 | 0.582 | 1 | 0.022972788 | -0.875162888 | 0.3 | 0.698 | 1 | 0.022972788 | 0.000943788 |
| Iqsec2 | 0.000472366 | -1.634290213 | 0.118 | 0.282 | 1 | 0.113596799 | -2.186649236 | 0.1 | 0.324 | 1 | 0.113596799 | 0.000944509 |
| A3galt2 | 0.000472402 | -2.096861539 | 0.059 | 0.206 | 1 | 0.759316125 | -0.473931188 | 0.3 | 0.309 | 1 | 0.759316125 | 0.000944582 |
| Slf2 | 0.000472652 | -1.316240722 | 0.403 | 0.524 | 1 | 0.145172511 | -0.907550113 | 0.5 | 0.688 | 1 | 0.145172511 | 0.000945081 |
| Ccnb2 | 0.171562612 | -1.27897595 | 0.084 | 0.135 | 1 | 0.000475728 | 1.675529711 | 0.6 | 0.157 | 1 | 0.171562612 | 0.00095123 |
| Tent5a | 0.00047634 | 1.204020563 | 0.723 | 0.559 | 1 | 0.517996134 | -1.40274014 | 0.4 | 0.41 | 1 | 0.517996134 | 0.000952454 |
| Clasp1 | 0.000477532 | -0.827043257 | 0.588 | 0.718 | 1 | 0.702134968 | -0.178475305 | 0.8 | 0.83 | 1 | 0.702134968 | 0.000954836 |
| Psmd12 | 0.000477984 | -0.61127076 | 0.79 | 0.829 | 1 | 0.144457273 | -0.596787936 | 0.8 | 0.904 | 1 | 0.144457273 | 0.00095574 |
| Ptp4a3 | 0.000478009 | -2.117695043 | 0.017 | 0.135 | 1 | 0.919812465 | 0.930459067 | 0.1 | 0.09 | 1 | 0.919812465 | 0.000955789 |
| Glcci1 | 0.00047981 | -1.064440062 | 0.286 | 0.471 | 1 | 0.229581586 | -0.742299038 | 0.3 | 0.519 | 1 | 0.229581586 | 0.000959389 |
| Sfswap | 0.000481237 | -0.690541257 | 0.672 | 0.8 | 1 | 0.257471276 | -0.574535129 | 0.9 | 0.824 | 1 | 0.257471276 | 0.000962242 |
| Ccdc141 | 0.000483927 | -2.900464326 | 0.008 | 0.118 | 1 | 0.7558167 | 1.728415291 | 0.1 | 0.151 | 1 | 0.7558167 | 0.000967621 |
| Leo1 | 0.000484291 | -1.372952098 | 0.261 | 0.424 | 1 | 0.056771869 | -1.505640048 | 0.2 | 0.481 | 1 | 0.056771869 | 0.000968348 |
| B230307C23Rik | 0.000484767 | -1.726434927 | 0.067 | 0.218 | 1 | 0.723324402 | 0.354956895 | 0.2 | 0.151 | 1 | 0.723324402 | 0.000969299 |
| Mvd | 0.310738132 | 0.109316694 | 0.244 | 0.188 | 1 | 0.0004852 | 2.558490289 | 0.4 | 0.08 | 1 | 0.310738132 | 0.000970165 |
| Tifa | 0.007081544 | 1.160936218 | 0.269 | 0.147 | 1 | 0.000485463 | 2.299103661 | 0.6 | 0.173 | 1 | 0.007081544 | 0.000970691 |
| Krt18 | 0.00048632 | -0.368069877 | 0.092 | 0.265 | 1 | 0.198434965 | -1.399930607 | 0.1 | 0.275 | 1 | 0.198434965 | 0.000972403 |
| Shoc2 | 0.000488001 | -0.454818841 | 0.815 | 0.865 | 1 | 0.343508618 | 0.403767695 | 0.9 | 0.938 | 1 | 0.343508618 | 0.000975765 |
| mt-Co1 | 0.149954702 | 0.11390003 | 1 | 1 | 1 | 0.000489049 | 0.845355414 | 1 | 1 | 1 | 0.149954702 | 0.000977859 |
| Emp2 | 0.000489159 | 0.498690131 | 0.815 | 0.618 | 1 | 0.083326228 | 0.683680552 | 0.8 | 0.772 | 1 | 0.083326228 | 0.00097808 |
| Dpm3 | 0.000491179 | -0.711815636 | 0.58 | 0.706 | 1 | 0.031511071 | -1.030110788 | 0.3 | 0.667 | 1 | 0.031511071 | 0.000982117 |
| Tomm5 | 0.000491954 | -0.909924656 | 0.445 | 0.606 | 1 | 0.180692178 | -0.660149997 | 0.3 | 0.522 | 1 | 0.180692178 | 0.000983666 |
| Zmynd8 | 0.000493116 | -0.694013449 | 0.496 | 0.694 | 1 | 0.301693168 | 0.549962018 | 0.6 | 0.645 | 1 | 0.301693168 | 0.000985988 |
| Pim3 | 0.000493452 | 1.249228606 | 0.655 | 0.541 | 1 | 0.042949919 | 1.541483864 | 0.7 | 0.451 | 1 | 0.042949919 | 0.00098666 |
| Cox8a | 0.000496063 | 0.350377471 | 0.966 | 0.965 | 1 | 0.163004253 | -0.388283097 | 0.9 | 0.957 | 1 | 0.163004253 | 0.000991879 |
| Fig4 | 0.000500747 | -2.359895945 | 0.042 | 0.176 | 1 | 0.241824544 | 0.828097349 | 0.3 | 0.148 | 1 | 0.241824544 | 0.001001243 |
| Zfp740 | 0.000502382 | -1.354842717 | 0.176 | 0.353 | 1 | 0.195134702 | -0.945552216 | 0.4 | 0.528 | 1 | 0.195134702 | 0.001004512 |
| Cers5 | 0.000503365 | -0.898365861 | 0.487 | 0.624 | 1 | 0.467581265 | -0.201246612 | 0.6 | 0.769 | 1 | 0.467581265 | 0.001006476 |
| Grn | 0.000503404 | -0.861990179 | 0.655 | 0.694 | 1 | 0.164771123 | -0.791773153 | 0.6 | 0.775 | 1 | 0.164771123 | 0.001006555 |
| Ube2l3 | 0.000503929 | -0.449626106 | 0.832 | 0.876 | 1 | 0.281307634 | -0.309689825 | 0.8 | 0.904 | 1 | 0.281307634 | 0.001007603 |
| Cnot2 | 0.000505324 | -0.491853096 | 0.807 | 0.882 | 1 | 0.462691609 | 0.535794042 | 0.9 | 0.883 | 1 | 0.462691609 | 0.001010394 |
| Naxe | 0.000505859 | 0.472509113 | 0.773 | 0.571 | 1 | 0.95585596 | 0.231325546 | 0.5 | 0.543 | 1 | 0.95585596 | 0.001011463 |
| Fosb | 0.0005063 | -0.935679019 | 0.95 | 0.929 | 1 | 0.875752649 | 0.297957625 | 1 | 0.975 | 1 | 0.875752649 | 0.001012344 |
| Aff4 | 0.000506722 | -0.38681923 | 0.983 | 0.953 | 1 | 0.026392164 | -0.575482367 | 1 | 1 | 1 | 0.026392164 | 0.001013188 |
| Aldh18a1 | 0.000506958 | -0.918386234 | 0.437 | 0.588 | 1 | 0.563609161 | -0.299993124 | 0.5 | 0.568 | 1 | 0.563609161 | 0.001013659 |
| Map3k12 | 0.000507463 | -2.240314329 | 0.059 | 0.2 | 1 | 0.746226111 | -0.204470513 | 0.2 | 0.231 | 1 | 0.746226111 | 0.001014668 |
| Tst | 0.000510048 | -1.485426827 | 0.05 | 0.194 | 1 | 0.057011892 | -1.814968106 | 0 | 0.272 | 1 | 0.057011892 | 0.001019836 |
| Hcfc1 | 0.000510138 | -1.214779237 | 0.286 | 0.453 | 1 | 0.575617898 | -0.166953435 | 0.4 | 0.5 | 1 | 0.575617898 | 0.001020015 |
| Ubxn7 | 0.000511014 | -0.816072139 | 0.555 | 0.706 | 1 | 0.060404484 | -0.883732039 | 0.8 | 0.92 | 1 | 0.060404484 | 0.001021766 |
| Rbbp8 | 0.000511659 | -1.351409479 | 0.454 | 0.594 | 1 | 0.214084212 | 0.72683978 | 0.7 | 0.596 | 1 | 0.214084212 | 0.001023056 |
| Smarca5 | 0.000511954 | -0.464132049 | 0.966 | 0.976 | 1 | 0.003336542 | -1.043374271 | 0.7 | 0.988 | 1 | 0.003336542 | 0.001023645 |
| Ccdc88c | 0.000578146 | 0.817135943 | 0.429 | 0.229 | 1 | 0.000512633 | 2.396433531 | 0.7 | 0.278 | 1 | 0.000578146 | 0.001025003 |
| Spata13 | 0.000512816 | 0.583835835 | 0.723 | 0.524 | 1 | 0.034354577 | 1.296497165 | 0.8 | 0.583 | 1 | 0.034354577 | 0.001025369 |
| Mss51 | 0.000515307 | -2.911691582 | 0.025 | 0.147 | 1 | 0.836908848 | 0.135278859 | 0.1 | 0.117 | 1 | 0.836908848 | 0.001030349 |
| Rsrp1 | 0.000515397 | -0.628924937 | 0.958 | 0.959 | 1 | 0.011377585 | -0.911272165 | 0.9 | 0.994 | 1 | 0.011377585 | 0.001030529 |
| Spag1 | 0.000517064 | -2.266786541 | 0.05 | 0.188 | 1 | 0.502208324 | -0.68251781 | 0.1 | 0.176 | 1 | 0.502208324 | 0.001033861 |
| Cct8 | 0.000517166 | -0.573557737 | 0.891 | 0.871 | 1 | 0.089797017 | -0.575780442 | 0.8 | 0.941 | 1 | 0.089797017 | 0.001034064 |
| Reep5 | 0.000519136 | 0.42269488 | 0.992 | 0.947 | 1 | 0.304905306 | 0.277248265 | 1 | 0.985 | 1 | 0.304905306 | 0.001038003 |
| Nherf2 | 0.000520387 | -1.485426827 | 0.092 | 0.253 | 1 | 0.135445974 | -1.230005605 | 0 | 0.185 | 1 | 0.135445974 | 0.001040503 |
| Snip1 | 0.000520897 | -1.250961574 | 0.185 | 0.365 | 1 | 0.288755502 | -0.763437806 | 0.2 | 0.343 | 1 | 0.288755502 | 0.001041523 |
| Ipmk | 0.000520906 | -1.676568315 | 0.151 | 0.324 | 1 | 0.595226745 | 0.120681482 | 0.4 | 0.29 | 1 | 0.595226745 | 0.001041541 |
| Vps54 | 0.000521237 | -0.768959543 | 0.429 | 0.624 | 1 | 0.196287491 | -0.702200176 | 0.7 | 0.778 | 1 | 0.196287491 | 0.001042202 |
| Riok1 | 0.00052156 | -0.934016886 | 0.479 | 0.6 | 1 | 0.22008612 | 0.890603078 | 0.7 | 0.577 | 1 | 0.22008612 | 0.001042847 |
| 1700017B05Rik | 0.000522789 | -2.694880193 | 0.025 | 0.147 | 1 | 0.072883108 | -2.037360528 | 0 | 0.25 | 1 | 0.072883108 | 0.001045305 |
| Tnk2 | 0.000525307 | 1.230780207 | 0.471 | 0.271 | 1 | 0.026746562 | 1.584962501 | 0.6 | 0.302 | 1 | 0.026746562 | 0.001050337 |
| Fmc1 | 0.000527579 | -1.055282436 | 0.235 | 0.418 | 1 | 0.534422628 | -0.391469028 | 0.3 | 0.364 | 1 | 0.534422628 | 0.00105488 |
| Kti12 | 0.000529911 | -1.614709844 | 0.126 | 0.288 | 1 | 0.230031487 | -0.802257054 | 0.1 | 0.265 | 1 | 0.230031487 | 0.001059542 |
| Clcn2 | 0.000530422 | -2.529820947 | 0.025 | 0.147 | 1 | 0.981801623 | 0.526068812 | 0.1 | 0.096 | 1 | 0.981801623 | 0.001060562 |
| Gm16279 | 0.062022899 | -1.622930351 | 0.025 | 0.076 | 1 | 0.000532467 | 2.847996907 | 0.3 | 0.046 | 1 | 0.062022899 | 0.001064651 |
| Garin1a | 0.43719716 | 0.407657969 | 0.076 | 0.053 | 1 | 0.000532467 | 2.625604485 | 0.3 | 0.046 | 1 | 0.43719716 | 0.001064651 |
| Adcy9 | 0.000535506 | -1.929033479 | 0.143 | 0.312 | 1 | 0.089559002 | -1.760155222 | 0.2 | 0.466 | 1 | 0.089559002 | 0.001070725 |

| | | | | | | | | | | | | |
|---|---|---|---|---|---|---|---|---|---|---|---|---|
| Cdca4 | 0.00053578 | -0.997876828 | 0.218 | 0.412 | 1 | 0.114067947 | -1.198015491 | 0.2 | 0.429 | 1 | 0.114067947 | 0.001071274 |
| Dhh | 0.000536542 | 2.63005039 | 0.101 | 0.012 | 1 | 0.577184788 | 1.558490289 | 0 | 0.031 | 1 | 0.577184788 | 0.001072795 |
| Tmf1 | 0.000536763 | -0.41460549 | 0.874 | 0.918 | 1 | 0.172891769 | -0.450847575 | 0.9 | 0.92 | 1 | 0.172891769 | 0.001073237 |
| Gtf2ird2 | 0.000537484 | -2.10433666 | 0.034 | 0.165 | 1 | 0.238905555 | -0.789433014 | 0 | 0.123 | 1 | 0.238905555 | 0.00107468 |
| Zcchc8 | 0.000538646 | -1.329307625 | 0.277 | 0.435 | 1 | 0.00694772 | -2.258202497 | 0.1 | 0.543 | 1 | 0.00694772 | 0.001077001 |
| Gpr146 | 0.000542353 | 1.352822103 | 0.345 | 0.176 | 1 | 0.020893184 | 1.481869008 | 0.4 | 0.13 | 1 | 0.020893184 | 0.001084411 |
| Ak6 | 0.000543264 | -1.032914622 | 0.244 | 0.424 | 1 | 0.116012547 | -0.918716031 | 0.2 | 0.469 | 1 | 0.116012547 | 0.001086234 |
| Nup37 | 0.000545559 | -2.439623138 | 0.025 | 0.147 | 1 | 0.773317908 | 1.063725598 | 0.1 | 0.074 | 1 | 0.773317908 | 0.001090821 |
| Mybbp1a | 0.000545746 | -0.938599455 | 0.454 | 0.618 | 1 | 0.113619096 | -0.713881981 | 0.4 | 0.633 | 1 | 0.113619096 | 0.001091194 |
| Fam98a | 0.000545887 | -0.790281409 | 0.429 | 0.576 | 1 | 0.051949206 | -1.143389963 | 0.3 | 0.636 | 1 | 0.051949206 | 0.001091476 |
| Dio2 | 0.000546395 | -4.292781749 | 0.059 | 0.2 | 1 | 0.129027835 | -3.555725279 | 0 | 0.191 | 1 | 0.129027835 | 0.001092492 |
| Col14a1 | 0.000547468 | -4.392317423 | 0.017 | 0.129 | 1 | 0.577180918 | -0.152003093 | 0 | 0.031 | 1 | 0.577180918 | 0.001094636 |
| Xylb | 0.218485013 | -1.292781749 | 0.008 | 0.029 | 1 | 0.000549016 | 4.432959407 | 0.2 | 0.022 | 1 | 0.218485013 | 0.00109773 |
| Ddr2 | 0.000551215 | 0.5259139 | 0.941 | 0.882 | 1 | 0.996011535 | 0.136752651 | 0.9 | 0.901 | 1 | 0.996011535 | 0.001102126 |
| Ddhd1 | 0.000552527 | -1.431386987 | 0.21 | 0.388 | 1 | 0.85920879 | 0.170640802 | 0.5 | 0.537 | 1 | 0.85920879 | 0.001104748 |
| Sorbs1 | 0.000553111 | 0.372015578 | 0.958 | 0.835 | 1 | 0.015848598 | 1.033772376 | 1 | 0.898 | 1 | 0.015848598 | 0.001105915 |
| Steap4 | 0.00055507 | 0.755581272 | 0.244 | 0.094 | 1 | 0.38571429 | -0.959358016 | 0 | 0.071 | 1 | 0.38571429 | 0.001109832 |
| Bmp2k | 0.000555822 | -1.375802336 | 0.261 | 0.424 | 1 | 0.28446299 | -0.473931188 | 0.3 | 0.469 | 1 | 0.28446299 | 0.001111336 |
| Fam193b | 0.000556366 | -1.542010356 | 0.176 | 0.347 | 1 | 0.236863947 | -0.449683642 | 0.3 | 0.531 | 1 | 0.236863947 | 0.001112422 |
| 2810032G03Rik | 0.000557369 | 1.974004791 | 0.126 | 0.024 | 1 | 0.597845674 | 1.695993813 | 0 | 0.028 | 1 | 0.597845674 | 0.001114426 |
| Nop10 | 0.000558589 | 0.433696392 | 0.899 | 0.882 | 1 | 0.841195075 | 0.279829648 | 0.6 | 0.849 | 1 | 0.841195075 | 0.001116866 |
| Asxl2 | 0.000560228 | -1.167944637 | 0.429 | 0.576 | 1 | 0.054321142 | -1.105164843 | 0.4 | 0.698 | 1 | 0.054321142 | 0.001120143 |
| Ube2r2 | 0.00056342 | -0.448337508 | 0.899 | 0.953 | 1 | 0.112003952 | -0.506336834 | 0.8 | 0.991 | 1 | 0.112003952 | 0.001126523 |
| 1110038F14Rik | 0.000563423 | -1.40103464 | 0.16 | 0.329 | 1 | 0.354678169 | -0.750262417 | 0.1 | 0.216 | 1 | 0.354678169 | 0.001126529 |
| Man1b1 | 0.00056517 | -1.267835392 | 0.185 | 0.359 | 1 | 0.711024371 | 0.210566986 | 0.5 | 0.432 | 1 | 0.711024371 | 0.001130021 |
| Prkrip1 | 0.000566372 | -1.070389328 | 0.185 | 0.365 | 1 | 0.192821868 | -0.993305347 | 0.1 | 0.281 | 1 | 0.192821868 | 0.001132423 |
| Pnpo | 0.000567544 | 1.240398209 | 0.277 | 0.118 | 1 | 0.150972928 | 1.795529487 | 0.2 | 0.074 | 1 | 0.150972928 | 0.001134765 |
| Naa38 | 0.000570503 | 0.652076697 | 0.63 | 0.429 | 1 | 0.659683812 | 0.31748219 | 0.5 | 0.432 | 1 | 0.659683812 | 0.001140681 |
| Kptn | 0.000570943 | -1.690541257 | 0.126 | 0.288 | 1 | 0.163088983 | -1.481923979 | 0.1 | 0.296 | 1 | 0.163088983 | 0.001141561 |
| Npnt | 0.000573069 | -3.600904045 | 0.067 | 0.206 | 1 | 0.029401139 | -3.532824877 | 0 | 0.336 | 1 | 0.029401139 | 0.001145809 |
| Cep126 | 0.000574571 | -0.950549558 | 0.218 | 0.412 | 1 | 0.454789575 | -0.175849835 | 0.4 | 0.241 | 1 | 0.454789575 | 0.001148811 |
| Tor1aip1 | 0.000575183 | -0.478752821 | 0.924 | 0.935 | 1 | 0.186642476 | -0.392774044 | 1 | 0.975 | 1 | 0.186642476 | 0.001150035 |
| 2810029C07Rik | 0.000575549 | 1.836501268 | 0.16 | 0.041 | 1 | 0.193279696 | 2.31748219 | 0.1 | 0.028 | 1 | 0.193279696 | 0.001150767 |
| Cog2 | 0.000575624 | -1.388750808 | 0.202 | 0.371 | 1 | 0.309351125 | 0.418009066 | 0.6 | 0.38 | 1 | 0.309351125 | 0.001150916 |
| Hgh1 | 0.000576085 | -2.240314329 | 0.025 | 0.147 | 1 | 0.505605027 | 0.930459067 | 0 | 0.043 | 1 | 0.505605027 | 0.001151838 |
| Nup54 | 0.000576275 | -1.062352009 | 0.387 | 0.535 | 1 | 0.63181614 | -0.469343919 | 0.6 | 0.546 | 1 | 0.63181614 | 0.001152218 |
| Etv3 | 0.00057789 | -1.371255807 | 0.168 | 0.341 | 1 | 0.20446251 | -1.062451508 | 0.2 | 0.377 | 1 | 0.20446251 | 0.001155447 |
| Ube2g1 | 0.000582759 | -0.808897087 | 0.748 | 0.8 | 1 | 0.815961289 | -0.143759099 | 0.9 | 0.923 | 1 | 0.815961289 | 0.001165178 |
| Ppp6c | 0.000585686 | -0.921525942 | 0.462 | 0.582 | 1 | 0.005659064 | -1.623046002 | 0.3 | 0.731 | 1 | 0.005659064 | 0.001171029 |
| Dyrk1b | 0.332535457 | -0.807354922 | 0.034 | 0.059 | 1 | 0.000586113 | 2.695993813 | 0.3 | 0.046 | 1 | 0.332535457 | 0.001171883 |
| Ddx17 | 0.002309925 | -0.423587573 | 0.966 | 0.959 | 1 | 0.00058671 | -1.249533631 | 0.9 | 0.991 | 1 | 0.002309925 | 0.001173075 |
| Shmt2 | 0.000586736 | -1.292781749 | 0.134 | 0.306 | 1 | 0.397766863 | -0.321928095 | 0.1 | 0.207 | 1 | 0.397766863 | 0.001173128 |
| Tmem39a | 0.000587066 | -0.656654488 | 0.613 | 0.747 | 1 | 0.028414262 | -1.342141316 | 0.5 | 0.704 | 1 | 0.028414262 | 0.001173787 |
| Cept1 | 0.000587086 | -1.095480309 | 0.353 | 0.529 | 1 | 0.840823796 | -0.205641057 | 0.7 | 0.654 | 1 | 0.840823796 | 0.001173827 |
| Rtl4 | 0.000588366 | 0.514573173 | 0.101 | 0.012 | 1 | 0.73075158 | 0.494359952 | 0 | 0.012 | 1 | 0.73075158 | 0.001176386 |
| Msx1 | 0.000589308 | -3.140778656 | 0.034 | 0.159 | 1 | 0.152758215 | -2.131825212 | 0 | 0.173 | 1 | 0.152758215 | 0.001178268 |
| Ywhah | 0.000590821 | -0.911691582 | 0.689 | 0.782 | 1 | 0.101875681 | -0.787916782 | 0.7 | 0.861 | 1 | 0.101875681 | 0.001181293 |
| Meg3 | 0.006085275 | 0.828736954 | 0.798 | 0.753 | 1 | 0.000593823 | 1.509162577 | 1 | 0.852 | 1 | 0.006085275 | 0.001187294 |
| C87436 | 0.000594559 | -1.589763487 | 0.143 | 0.306 | 1 | 0.37612703 | -0.505640048 | 0.1 | 0.228 | 1 | 0.37612703 | 0.001188764 |
| Epb41l3 | 0.000594574 | -4.614709844 | 0 | 0.094 | 1 | 0.408781983 | -0.152003093 | 0 | 0.065 | 1 | 0.408781983 | 0.001188795 |
| Dram1 | 0.000596037 | 0.789433014 | 0.429 | 0.235 | 1 | 0.074396017 | 1.959028219 | 0.4 | 0.201 | 1 | 0.074396017 | 0.001191718 |
| Pcmt1 | 0.000596362 | -0.524955191 | 0.849 | 0.865 | 1 | 0.732345273 | -0.138582578 | 0.9 | 0.926 | 1 | 0.732345273 | 0.001192369 |
| Kmt5b | 0.000596625 | -0.756008042 | 0.689 | 0.741 | 1 | 0.175688902 | -0.635520331 | 0.9 | 0.864 | 1 | 0.175688902 | 0.001192895 |
| B4gat1 | 0.000600346 | -1.328701323 | 0.126 | 0.294 | 1 | 0.103950494 | -1.473931188 | 0 | 0.213 | 1 | 0.103950494 | 0.001200332 |
| Tenm2 | 0.000602387 | -2.398595869 | 0.202 | 0.359 | 1 | 0.013833697 | -5.111361109 | 0 | 0.401 | 1 | 0.013833697 | 0.001204411 |
| Gm3336 | 0.267893443 | -1.211251864 | 0.059 | 0.094 | 1 | 0.000604719 | 3.976101732 | 0.3 | 0.049 | 1 | 0.267893443 | 0.001209072 |
| Ptgfrn | 0.000606815 | -1.503348735 | 0.134 | 0.3 | 1 | 0.152677393 | -1.457811523 | 0 | 0.173 | 1 | 0.152677393 | 0.001213262 |
| Hhat | 0.000607392 | -1.450093754 | 0.21 | 0.376 | 1 | 0.281378076 | -0.288465336 | 0.2 | 0.389 | 1 | 0.281378076 | 0.001214415 |
| Basp1 | 0.000608862 | -3.258016331 | 0.017 | 0.129 | 1 | 0.176155845 | -1.924592597 | 0 | 0.157 | 1 | 0.176155845 | 0.001217353 |
| Scml4 | 0.00061134 | -2.977279923 | 0.05 | 0.182 | 1 | 0.996226686 | 0.135278859 | 0.2 | 0.204 | 1 | 0.996226686 | 0.001222305 |
| Nvl | 0.000613686 | -0.885357434 | 0.429 | 0.576 | 1 | 0.877399895 | -0.139930261 | 0.6 | 0.599 | 1 | 0.877399895 | 0.001226995 |
| Nova1 | 0.001317046 | 1.131244533 | 0.118 | 0.024 | 1 | 0.000614676 | 2.432959407 | 0.3 | 0.046 | 1 | 0.001317046 | 0.001228973 |
| Paip1 | 0.00061504 | -0.842643672 | 0.58 | 0.682 | 1 | 0.100360056 | -0.645773029 | 0.6 | 0.821 | 1 | 0.100360056 | 0.001229702 |
| Klf16 | 0.000615462 | -1.30256277 | 0.143 | 0.318 | 1 | 0.205097832 | -0.993305347 | 0.1 | 0.275 | 1 | 0.205097832 | 0.001230545 |
| Gmfb | 0.237966317 | -0.399957485 | 0.605 | 0.6 | 1 | 0.000615866 | -2.315978829 | 0.2 | 0.731 | 1 | 0.237966317 | 0.001231352 |
| Six2 | 0.000618401 | -0.586787542 | 0.555 | 0.718 | 1 | 0.015242281 | -1.443557539 | 0.4 | 0.725 | 1 | 0.015242281 | 0.00123642 |
| Rcc1 | 0.000621092 | -1.762267033 | 0.084 | 0.235 | 1 | 0.124041469 | -1.357117523 | 0 | 0.194 | 1 | 0.124041469 | 0.001241799 |
| Mypop | 0.000621198 | -2.854660637 | 0.017 | 0.129 | 1 | 0.865540859 | 0.494359952 | 0.2 | 0.173 | 1 | 0.865540859 | 0.001242009 |
| Numb | 0.000621684 | -0.807354922 | 0.462 | 0.647 | 1 | 0.253183388 | -0.810966176 | 0.7 | 0.787 | 1 | 0.253183388 | 0.001242982 |
| Kif3c | 0.000622799 | -2.485426827 | 0.034 | 0.159 | 1 | 0.701143615 | 0.558490289 | 0.2 | 0.148 | 1 | 0.701143615 | 0.001245211 |
| Ifitm3 | 0.000626296 | -0.497601541 | 0.151 | 0.347 | 1 | 0.141578123 | -2.152003093 | 0.1 | 0.309 | 1 | 0.141578123 | 0.001252199 |
| Tcf19 | 0.089019965 | -1.137503524 | 0.025 | 0.071 | 1 | 0.000626519 | 3.602884409 | 0.2 | 0.022 | 1 | 0.089019965 | 0.001252645 |
| Gm42670 | 0.147029286 | -1.807354922 | 0 | 0.018 | 1 | 0.000626519 | 3.602884409 | 0.2 | 0.022 | 1 | 0.147029286 | 0.001252645 |
| Gm50321 | 0.331040183 | -1.292781749 | 0.008 | 0.024 | 1 | 0.000626519 | 3.602884409 | 0.2 | 0.022 | 1 | 0.331040183 | 0.001252645 |
| Gng3 | 0.000627332 | -2.807354922 | 0.017 | 0.129 | 1 | 0.72332358 | 0.453137289 | 0.2 | 0.151 | 1 | 0.72332358 | 0.00125427 |
| Uba7 | 0.00062737 | -2.854660637 | 0.017 | 0.129 | 1 | 0.252713589 | -0.763437806 | 0 | 0.117 | 1 | 0.252713589 | 0.001254347 |
| Hal | 0.000627403 | 0.865647613 | 0.21 | 0.076 | 1 | 0.00318972 | 3.432959407 | 0.3 | 0.065 | 1 | 0.00318972 | 0.001254412 |
| Acat1 | 0.000631374 | -0.590893978 | 0.697 | 0.782 | 1 | 0.85676479 | 0.296822719 | 0.6 | 0.793 | 1 | 0.85676479 | 0.00126235 |
| 2700038G22Rik | 0.000631416 | -1.532732542 | 0.084 | 0.241 | 1 | 0.156529648 | -1.28585884 | 0 | 0.17 | 1 | 0.156529648 | 0.001262433 |
| Taldo1 | 0.000632455 | 0.580062254 | 0.866 | 0.8 | 1 | 0.037975254 | -1.05055783 | 0.5 | 0.793 | 1 | 0.037975254 | 0.00126451 |
| Zfp267 | 0.000633312 | -1.883976204 | 0.076 | 0.224 | 1 | 0.107270389 | -1.489872732 | 0 | 0.21 | 1 | 0.107270389 | 0.001266222 |
| Slc35c1 | 0.000633467 | -2.707819249 | 0.017 | 0.129 | 1 | 0.875234763 | 0.403212064 | 0.1 | 0.114 | 1 | 0.875234763 | 0.001266532 |
| Fbxl6 | 0.000633632 | -1.375243909 | 0.134 | 0.3 | 1 | 0.103870409 | -1.473931188 | 0 | 0.213 | 1 | 0.103870409 | 0.001266862 |
| Trmt11 | 0.000633966 | -1.807354922 | 0.101 | 0.253 | 1 | 0.678511704 | 0.859492545 | 0.3 | 0.275 | 1 | 0.678511704 | 0.00126753 |
| Crot | 0.000634681 | -1.900464326 | 0.059 | 0.2 | 1 | 0.427599374 | -0.5519337 | 0.1 | 0.194 | 1 | 0.427599374 | 0.00126896 |
| Dennd4a | 0.000636092 | -2.073869897 | 0.345 | 0.506 | 1 | 0.021526452 | -1.918716031 | 0.2 | 0.583 | 1 | 0.021526452 | 0.00127178 |
| Nrd1 | 0.000636814 | -0.59645814 | 0.84 | 0.847 | 1 | 0.191144919 | -0.531541912 | 0.7 | 0.892 | 1 | 0.191144919 | 0.001273223 |
| Sco2 | 0.000637083 | -1.807354922 | 0.059 | 0.2 | 1 | 0.81291432 | 0.463333056 | 0.1 | 0.123 | 1 | 0.81291432 | 0.00127376 |
| Pllp | 0.000637766 | 3.321928095 | 0.084 | 0.006 | 1 | 0.078038625 | 3.210566986 | 0.1 | 0.019 | 1 | 0.078038625 | 0.001275125 |
| Zfp236 | 0.000640959 | -1.070389328 | 0.261 | 0.435 | 1 | 0.528970175 | -0.645043105 | 0.5 | 0.5 | 1 | 0.528970175 | 0.001281507 |
| Ctsc | 0.000641285 | -4.213347282 | 0.008 | 0.112 | 1 | 0.170474125 | -1.191531458 | 0 | 0.16 | 1 | 0.170474125 | 0.001282159 |
| Gm5617 | 0.000644007 | 1.159908292 | 0.361 | 0.194 | 1 | 0.810319601 | 0.41305985 | 0.2 | 0.164 | 1 | 0.810319601 | 0.001287599 |
| Gxylt1 | 0.00064402 | -1.230169772 | 0.244 | 0.418 | 1 | 0.191541745 | -1.181750437 | 0.3 | 0.457 | 1 | 0.191541745 | 0.001287626 |
| Rab5a | 0.000644035 | -0.485426827 | 0.933 | 0.918 | 1 | 0.067379046 | -0.615950193 | 0.8 | 0.975 | 1 | 0.067379046 | 0.001287655 |
| Osbpl10 | 0.000645617 | -2.292781749 | 0.059 | 0.194 | 1 | 0.257204934 | -1.391469028 | 0.2 | 0.34 | 1 | 0.257204934 | 0.001290816 |
| Ninl | 0.00064594 | -2.411426246 | 0.034 | 0.159 | 1 | 0.181056121 | -1.723545078 | 0 | 0.154 | 1 | 0.181056121 | 0.001291462 |
| Prcp | 0.000645975 | -2.521050737 | 0.034 | 0.159 | 1 | 0.701355468 | -0.171902651 | 0.1 | 0.139 | 1 | 0.701355468 | 0.001291532 |
| Cxxc1 | 0.000646156 | -1.318316841 | 0.101 | 0.265 | 1 | 0.123330595 | -1.090602549 | 0.1 | 0.327 | 1 | 0.123330595 | 0.001291894 |

| | | | | | | | | | | | | |
|---|---|---|---|---|---|---|---|---|---|---|---|---|
| Meis1 | 0.000648275 | -3.842978832 | 0.008 | 0.112 | 1 | 0.110309335 | -1.982078092 | 0 | 0.207 | 1 | 0.110309335 | 0.00129613 |
| Slc38a6 | 0.000649065 | -2.056175469 | 0.118 | 0.271 | 1 | 0.406448672 | 0.140177658 | 0.4 | 0.247 | 1 | 0.406448672 | 0.001297709 |
| Emb | 0.000649216 | 1.236934354 | 0.748 | 0.712 | 1 | 0.028206226 | 1.2258015 | 0.9 | 0.679 | 1 | 0.028206226 | 0.001298011 |
| Mapk1ip1l | 0.000650789 | -0.618012467 | 0.756 | 0.835 | 1 | 0.318990751 | -0.423639034 | 1 | 0.907 | 1 | 0.318990751 | 0.001301155 |
| Vps13b | 0.000652609 | -0.632076686 | 0.824 | 0.859 | 1 | 0.11693254 | 0.442609577 | 1 | 0.864 | 1 | 0.11693254 | 0.001304792 |
| Ssc4d | 0.014871534 | 1.280107919 | 0.118 | 0.041 | 1 | 0.000653677 | 3.432959407 | 0.2 | 0.022 | 1 | 0.014871534 | 0.001306926 |
| Tcea2 | 0.086879909 | -1.292781749 | 0.025 | 0.071 | 1 | 0.000653677 | 3.432959407 | 0.2 | 0.022 | 1 | 0.086879909 | 0.001306926 |
| Ddx3x | 0.043717919 | -0.316736481 | 0.933 | 0.959 | 1 | 0.000654139 | -1.5146588 | 0.5 | 0.981 | 1 | 0.043717919 | 0.001307851 |
| Kcmf1 | 0.00065437 | -0.387542608 | 0.933 | 0.947 | 1 | 0.034197636 | -0.731454611 | 0.9 | 0.978 | 1 | 0.034197636 | 0.001308313 |
| Rmdn2 | 0.00065507 | -3.216610069 | 0.202 | 0.353 | 1 | 0.418288643 | -1.011825435 | 0.2 | 0.293 | 1 | 0.418288643 | 0.00130971 |
| H2-DMb1 | 0.000655917 | 1.010806755 | 0.546 | 0.329 | 1 | 0.167820526 | 0.836592143 | 0.5 | 0.284 | 1 | 0.167820526 | 0.001311404 |
| Lrch2 | 0.0006619 | -3.129283017 | 0.008 | 0.112 | 1 | 0.245621645 | -0.789433014 | 0 | 0.12 | 1 | 0.245621645 | 0.001323361 |
| Adam22 | 0.000661921 | -3.240314329 | 0.008 | 0.112 | 1 | 0.875234763 | 0.432959407 | 0.1 | 0.114 | 1 | 0.875234763 | 0.001323404 |
| Hars2 | 0.000666294 | -1.230854 | 0.109 | 0.276 | 1 | 0.941028406 | 0.176619654 | 0.2 | 0.201 | 1 | 0.941028406 | 0.001332144 |
| Telo2 | 0.000669449 | -2.333423734 | 0.034 | 0.159 | 1 | 0.307202183 | -0.505640048 | 0 | 0.096 | 1 | 0.307202183 | 0.001338449 |
| Ctif | 0.000670995 | -1.341036918 | 0.235 | 0.4 | 1 | 0.53571897 | 0.283534858 | 0.7 | 0.599 | 1 | 0.53571897 | 0.001341539 |
| Sprtn | 0.000672259 | -1.655351829 | 0.076 | 0.224 | 1 | 0.394164391 | -0.489872732 | 0.1 | 0.207 | 1 | 0.394164391 | 0.001344065 |
| Nme5 | 0.000676196 | -2.614709844 | 0.017 | 0.129 | 1 | 0.937632628 | 0.345496566 | 0.1 | 0.105 | 1 | 0.937632628 | 0.001351934 |
| Wipi2 | 0.000676841 | -1.023846742 | 0.462 | 0.606 | 1 | 0.14523057 | -0.709998547 | 0.5 | 0.698 | 1 | 0.14523057 | 0.001353223 |
| Zmat1 | 0.000680295 | 0.676844602 | 0.311 | 0.141 | 1 | 0.060633434 | 1.951832718 | 0.3 | 0.114 | 1 | 0.060633434 | 0.001360128 |
| S100a11 | 0.000681229 | -0.519696729 | 0.941 | 0.959 | 1 | 0.218722701 | -0.426462842 | 0.7 | 0.975 | 1 | 0.218722701 | 0.001361994 |
| Ccnd3 | 0.000682813 | -1.228454174 | 0.496 | 0.6 | 1 | 0.121938567 | -0.615592338 | 0.4 | 0.741 | 1 | 0.121938567 | 0.001365159 |
| 4833418N02Rik | 0.024544213 | 0.688602573 | 0.252 | 0.147 | 1 | 0.000685032 | 1.998022351 | 0.6 | 0.173 | 1 | 0.024544213 | 0.001369595 |
| Strip1 | 0.000686047 | -1.635173947 | 0.126 | 0.282 | 1 | 0.078734986 | 0.938970567 | 0.6 | 0.321 | 1 | 0.078734986 | 0.001371624 |
| Ubr7 | 0.000689124 | -1.028569152 | 0.21 | 0.4 | 1 | 0.092698446 | -1.399930607 | 0.1 | 0.352 | 1 | 0.092698446 | 0.001377774 |
| Gemin2 | 0.000689388 | -1.748461233 | 0.059 | 0.2 | 1 | 0.153399319 | 1.111031312 | 0.3 | 0.13 | 1 | 0.153399319 | 0.001378301 |
| Ankrd27 | 0.000689808 | -1.459431619 | 0.101 | 0.259 | 1 | 0.547620939 | 0.667424661 | 0.3 | 0.222 | 1 | 0.547620939 | 0.00137914 |
| Csrnp2 | 0.000691025 | -2.515174171 | 0.034 | 0.159 | 1 | 0.274499721 | -0.473931188 | 0 | 0.108 | 1 | 0.274499721 | 0.001381572 |
| Sass6 | 0.000692508 | -2.655351829 | 0.05 | 0.182 | 1 | 0.418619572 | 1.362570079 | 0.2 | 0.12 | 1 | 0.418619572 | 0.001384537 |
| Rcan3 | 0.000693377 | -2.117695043 | 0.034 | 0.159 | 1 | 0.196450798 | -0.936274402 | 0 | 0.145 | 1 | 0.196450798 | 0.001386274 |
| Zkscan17 | 0.000694139 | -1.432959407 | 0.084 | 0.235 | 1 | 0.160153678 | -0.936274402 | 0.1 | 0.302 | 1 | 0.160153678 | 0.001387795 |
| Mettl2 | 0.000695732 | -1.532732542 | 0.118 | 0.276 | 1 | 0.411933069 | -0.521236903 | 0.1 | 0.201 | 1 | 0.411933069 | 0.001390981 |
| Xxylt1 | 0.000695732 | -1.532732542 | 0.118 | 0.276 | 1 | 0.540031217 | -0.339630097 | 0.2 | 0.275 | 1 | 0.540031217 | 0.001390981 |
| Gm9801 | 0.952664701 | -0.177304532 | 0.176 | 0.176 | 1 | 0.00069576 | 2.210566986 | 0.5 | 0.123 | 1 | 0.952664701 | 0.001391035 |
| Ist1 | 0.000695814 | -0.631647568 | 0.739 | 0.8 | 1 | 0.002447881 | -1.334823779 | 0.6 | 0.914 | 1 | 0.002447881 | 0.001391143 |
| Pdcl | 0.000697039 | -1.14363831 | 0.218 | 0.394 | 1 | 0.030801099 | -2.357117523 | 0 | 0.33 | 1 | 0.030801099 | 0.001393591 |
| Gm39302 | 0.000700781 | 2.514573173 | 0.084 | 0.006 | 1 | 0.002052242 | 4.432959407 | 0.1 | 0.006 | 1 | 0.002052242 | 0.001401071 |
| Apobec3 | 0.000702353 | 0.96203215 | 0.345 | 0.165 | 1 | 0.946497847 | -0.181750437 | 0.3 | 0.287 | 1 | 0.946497847 | 0.001404213 |
| Hapstr1 | 0.000703133 | -0.497077187 | 0.714 | 0.847 | 1 | 0.094508123 | -0.702555563 | 0.8 | 0.926 | 1 | 0.094508123 | 0.001405771 |
| Ccser2 | 0.000703791 | -0.692523061 | 0.672 | 0.747 | 1 | 0.097843944 | -0.691161905 | 0.8 | 0.858 | 1 | 0.097843944 | 0.001407086 |
| Mthfd2l | 0.000704048 | -1.234618724 | 0.303 | 0.465 | 1 | 0.14969728 | -0.982078092 | 0.2 | 0.429 | 1 | 0.14969728 | 0.0014076 |
| Spin4 | 0.000705468 | -2.117695043 | 0.034 | 0.159 | 1 | 0.180293234 | -0.959358016 | 0 | 0.154 | 1 | 0.180293234 | 0.001410438 |
| S100a16 | 0.000707436 | 0.61497507 | 0.748 | 0.582 | 1 | 0.57827287 | -0.218092284 | 0.5 | 0.71 | 1 | 0.57827287 | 0.001414371 |
| Ncoa2 | 0.000709978 | -0.778208576 | 0.664 | 0.765 | 1 | 0.13422966 | -0.525792306 | 0.6 | 0.88 | 1 | 0.13422966 | 0.001419453 |
| Gm47990 | 0.056697405 | 0.908852112 | 0.185 | 0.106 | 1 | 0.000713264 | 3.017921908 | 0.2 | 0.022 | 1 | 0.056697405 | 0.001426019 |
| Cd80 | 0.213068927 | 0.755581272 | 0.076 | 0.041 | 1 | 0.000713264 | 2.795529487 | 0.2 | 0.022 | 1 | 0.213068927 | 0.001426019 |
| Gm46218 | 0.008787421 | 1.321928095 | 0.143 | 0.053 | 1 | 0.000713273 | 2.795529487 | 0.2 | 0.022 | 1 | 0.008787421 | 0.001426037 |
| Cenpq | 0.000714487 | 0.792874335 | 0.345 | 0.165 | 1 | 0.006693577 | 2.252387162 | 0.3 | 0.068 | 1 | 0.006693577 | 0.001428464 |
| Tmem80 | 0.007577123 | -1.368069877 | 0.126 | 0.247 | 1 | 0.000715671 | 2.079322453 | 0.7 | 0.225 | 1 | 0.007577123 | 0.001430831 |
| Dlst | 0.000720924 | -0.649998594 | 0.588 | 0.706 | 1 | 0.324890089 | -0.473931188 | 0.6 | 0.765 | 1 | 0.324890089 | 0.001441328 |
| Mtmr6 | 0.000723604 | -0.839959587 | 0.445 | 0.594 | 1 | 0.405858353 | -0.448777711 | 0.6 | 0.62 | 1 | 0.405858353 | 0.001446684 |
| Blzf1 | 0.000724957 | -0.839063782 | 0.479 | 0.618 | 1 | 0.002920434 | -2.67556505 | 0.1 | 0.593 | 1 | 0.002920434 | 0.001449389 |
| Hnrnph2 | 0.000731003 | -0.540839947 | 0.756 | 0.806 | 1 | 0.033031362 | -0.972654838 | 0.7 | 0.843 | 1 | 0.033031362 | 0.001461472 |
| Gm15738 | 0.752076132 | -0.263034406 | 0.05 | 0.059 | 1 | 0.000732824 | 2.591657153 | 0.4 | 0.083 | 1 | 0.752076132 | 0.001465111 |
| Rap1b | 0.000734661 | -0.422786108 | 0.992 | 0.976 | 1 | 0.117349838 | -0.314709856 | 0.9 | 0.985 | 1 | 0.117349838 | 0.001468783 |
| Psmc3ip | 0.524954938 | -0.87774425 | 0.059 | 0.076 | 1 | 0.000734982 | 2.94753258 | 0.3 | 0.049 | 1 | 0.524954938 | 0.001469424 |
| Rmnd5b | 0.000735696 | -1.58058406 | 0.076 | 0.224 | 1 | 0.077369843 | -1.947862377 | 0 | 0.244 | 1 | 0.077369843 | 0.00147085 |
| Cnnm2 | 0.00073727 | 0.647210941 | 0.563 | 0.329 | 1 | 0.00285247 | 1.945165566 | 0.7 | 0.309 | 1 | 0.00285247 | 0.001473996 |
| Relch | 0.000738758 | -1.053111336 | 0.513 | 0.6 | 1 | 0.400017423 | -0.357117523 | 0.5 | 0.682 | 1 | 0.400017423 | 0.00147697 |
| Cyp2j6 | 0.000738841 | 1.359294947 | 0.37 | 0.206 | 1 | 0.709403861 | 0.494359952 | 0.1 | 0.154 | 1 | 0.709403861 | 0.001477136 |
| Plxna2 | 0.000739088 | -0.911803043 | 0.471 | 0.641 | 1 | 0.027177656 | -1.593685365 | 0.4 | 0.79 | 1 | 0.027177656 | 0.00147763 |
| Cers4 | 0.000742323 | -3.008988783 | 0.025 | 0.141 | 1 | 0.20780632 | -0.959358016 | 0 | 0.139 | 1 | 0.20780632 | 0.001484095 |
| Ppp3cc | 0.017627459 | -0.130095574 | 0.538 | 0.371 | 1 | 0.000742965 | 1.597590109 | 0.7 | 0.238 | 1 | 0.017627459 | 0.001485378 |
| Hilpda | 0.007023579 | 1.635427568 | 0.454 | 0.347 | 1 | 0.000744013 | 2.151188439 | 0.6 | 0.21 | 1 | 0.007023579 | 0.001487473 |
| 2610037D02Rik | 0.000744052 | -1.317699381 | 0.235 | 0.4 | 1 | 0.039960298 | -1.391469028 | 0.2 | 0.562 | 1 | 0.039960298 | 0.001487551 |
| Gm47071 | 0.334204811 | -0.807354922 | 0.008 | 0.024 | 1 | 0.0007448 | 2.902444691 | 0.2 | 0.022 | 1 | 0.334204811 | 0.001489046 |
| Gm26533 | 0.335144931 | 1.280107919 | 0.059 | 0.035 | 1 | 0.000744838 | 3.017921908 | 0.2 | 0.022 | 1 | 0.335144931 | 0.00148912 |
| Zfp622 | 0.000746352 | -1.133125083 | 0.311 | 0.465 | 1 | 0.105644127 | -1.090602549 | 0.3 | 0.519 | 1 | 0.105644127 | 0.001492147 |
| Celf5 | 0.000747487 | -2.201633861 | 0.05 | 0.182 | 1 | 0.803962121 | 0.185031894 | 0.1 | 0.123 | 1 | 0.803962121 | 0.001494415 |
| Dimt1 | 0.000752449 | -1.269698136 | 0.202 | 0.365 | 1 | 0.422609337 | -0.616284112 | 0.2 | 0.299 | 1 | 0.422609337 | 0.001504331 |
| Rnf44 | 0.000752569 | -1.578536232 | 0.109 | 0.265 | 1 | 0.285576837 | -0.505640048 | 0.2 | 0.38 | 1 | 0.285576837 | 0.001504572 |
| Nol9 | 0.000756527 | -1.326729081 | 0.168 | 0.335 | 1 | 0.20359478 | 1.219180116 | 0.1 | 0.327 | 1 | 0.20359478 | 0.001512482 |
| Ppp1r15b | 0.00846654 | -0.753360032 | 0.563 | 0.629 | 1 | 0.000758859 | -1.831743819 | 0.2 | 0.809 | 1 | 0.00846654 | 0.001517143 |
| Scai | 0.000760978 | -0.916476644 | 0.277 | 0.453 | 1 | 0.262738666 | 0.494359952 | 0.7 | 0.586 | 1 | 0.262738666 | 0.001521376 |
| Met | 0.000763281 | 0.350687088 | 0.555 | 0.347 | 1 | 0.935488191 | 0.162657001 | 0.6 | 0.725 | 1 | 0.935488191 | 0.001525979 |
| Actg1 | 0.000765553 | 0.333654017 | 1 | 0.994 | 1 | 0.809475978 | 0.36564241 | 1 | 0.994 | 1 | 0.809475978 | 0.001530519 |
| Ston2 | 0.000765818 | -0.140515414 | 0.328 | 0.147 | 1 | 0.132048643 | -2.111361109 | 0 | 0.188 | 1 | 0.132048643 | 0.00153105 |
| Ahnak | 0.000767586 | -0.555192422 | 0.966 | 0.982 | 1 | 0.032580892 | -0.826951035 | 1 | 1 | 1 | 0.032580892 | 0.001534582 |
| Pdik1l | 0.000768709 | -2.169925001 | 0.042 | 0.171 | 1 | 0.772482885 | 0.403212064 | 0.1 | 0.13 | 1 | 0.772482885 | 0.001536827 |
| Utp6 | 0.000772026 | -1.155278225 | 0.202 | 0.371 | 1 | 0.429963432 | 0.432959407 | 0.5 | 0.373 | 1 | 0.429963432 | 0.001543456 |
| Alkbh5 | 0.000772757 | -0.707819249 | 0.689 | 0.771 | 1 | 0.00453943 | -1.348400306 | 0.8 | 0.926 | 1 | 0.00453943 | 0.001544918 |
| Myo6 | 0.000774821 | 0.422851643 | 0.975 | 0.924 | 1 | 0.011740298 | 0.788242164 | 1 | 0.886 | 1 | 0.011740298 | 0.001549042 |
| Usp28 | 0.000775115 | -1.432959407 | 0.193 | 0.359 | 1 | 0.101868442 | -1.69632361 | 0.1 | 0.336 | 1 | 0.101868442 | 0.00154963 |
| Ptpn21 | 0.000775678 | -1.683366205 | 0.118 | 0.271 | 1 | 0.355339587 | -0.965071667 | 0.3 | 0.389 | 1 | 0.355339587 | 0.001550755 |
| Slc39a3 | 0.000778367 | -2.029747343 | 0.042 | 0.171 | 1 | 0.143470666 | -1.131825212 | 0 | 0.179 | 1 | 0.143470666 | 0.001556128 |
| Pxmp4 | 0.000779228 | 0.729586064 | 0.437 | 0.247 | 1 | 0.760638394 | 0.720241359 | 0.2 | 0.16 | 1 | 0.760638394 | 0.001557849 |
| Orc6 | 0.000779554 | -0.97924144 | 0.269 | 0.459 | 1 | 0.276755719 | -0.204470513 | 0.2 | 0.395 | 1 | 0.276755719 | 0.001558501 |
| Ccdc137 | 0.00078483 | -1.446952679 | 0.143 | 0.306 | 1 | 0.19198446 | 0.988174565 | 0.4 | 0.219 | 1 | 0.19198446 | 0.001569043 |
| Pcsk7 | 0.000785437 | -1.504285854 | 0.126 | 0.288 | 1 | 0.325508409 | -0.55950692 | 0.2 | 0.361 | 1 | 0.325508409 | 0.001570257 |
| Dhx8 | 0.000789518 | -1.115477217 | 0.277 | 0.447 | 1 | 0.442570708 | -0.73161236 | 0.4 | 0.454 | 1 | 0.442570708 | 0.001578413 |
| Prkar1b | 0.000791941 | -1.485426827 | 0.101 | 0.259 | 1 | 0.525180425 | 0.518076021 | 0.4 | 0.312 | 1 | 0.525180425 | 0.001583255 |
| Rbm4b | 0.000792372 | -0.959358016 | 0.21 | 0.382 | 1 | 0.588215799 | -0.237578825 | 0.3 | 0.383 | 1 | 0.588215799 | 0.001584116 |
| Lrrc47 | 0.000792564 | -1.318316841 | 0.126 | 0.288 | 1 | 0.137853196 | -1.161987182 | 0.1 | 0.315 | 1 | 0.137853196 | 0.001584499 |
| Crym | 0.000793012 | 1.698997744 | 0.134 | 0.029 | 1 | 0.913227002 | 0.494359952 | 0.1 | 0.086 | 1 | 0.913227002 | 0.001585395 |
| Zmym5 | 0.000794302 | -0.72449799 | 0.748 | 0.835 | 1 | 0.183327933 | -0.480024946 | 0.7 | 0.892 | 1 | 0.183327933 | 0.001587973 |
| Scaf11 | 0.000794714 | -0.415993185 | 0.95 | 0.947 | 1 | 0.07387295 | -0.549880042 | 1 | 0.981 | 1 | 0.07387295 | 0.001588797 |
| Prkaca | 0.000797873 | -1.267835392 | 0.176 | 0.353 | 1 | 0.425033317 | 0.252387162 | 0.4 | 0.253 | 1 | 0.425033317 | 0.00159511 |
| Ssrp1 | 0.000798097 | -0.756280737 | 0.529 | 0.647 | 1 | 0.582582087 | -0.125036046 | 0.5 | 0.688 | 1 | 0.582582087 | 0.001595558 |

| | | | | | | | | | | | | |
|---|---|---|---|---|---|---|---|---|---|---|---|---|
| Ccdc134 | 0.000799493 | -1.359895945 | 0.076 | 0.229 | 1 | 0.617921771 | 0.847996907 | 0.2 | 0.139 | 1 | 0.617921771 | 0.001598346 |
| Rab11a | 0.000801709 | 0.44181683 | 0.849 | 0.688 | 1 | 0.026197703 | -0.842318594 | 0.5 | 0.846 | 1 | 0.026197703 | 0.001602776 |
| Yes1 | 0.000802422 | -0.684330896 | 0.42 | 0.612 | 1 | 0.099853627 | -1.124185149 | 0.5 | 0.673 | 1 | 0.099853627 | 0.001604201 |
| D830025C05Rik | 0.00081817 | -1.537894247 | 0.126 | 0.276 | 1 | 0.050102084 | -2.441509711 | 0.1 | 0.395 | 1 | 0.050102084 | 0.00163567 |
| Adamts4 | 0.000818964 | -2.687060688 | 0.025 | 0.141 | 1 | 0.489954768 | 0.432959407 | 0 | 0.046 | 1 | 0.489954768 | 0.001637258 |
| Rab14 | 0.000822015 | -0.4207574 | 0.983 | 0.941 | 1 | 0.381871662 | -0.276057596 | 0.9 | 0.975 | 1 | 0.381871662 | 0.001643355 |
| Prmt9 | 0.000823219 | -1.691877705 | 0.092 | 0.241 | 1 | 0.673082678 | 0.409112665 | 0.3 | 0.241 | 1 | 0.673082678 | 0.001645761 |
| Sdc3 | 0.0008267 | -0.644859455 | 0.639 | 0.771 | 1 | 0.146835175 | -0.412208937 | 0.7 | 0.812 | 1 | 0.146835175 | 0.001652716 |
| Entrep3 | 0.000826861 | -1.485426827 | 0.084 | 0.235 | 1 | 0.279683617 | -0.736965594 | 0.1 | 0.244 | 1 | 0.279683617 | 0.001653039 |
| Atp2c1 | 0.000827867 | -0.984997837 | 0.303 | 0.482 | 1 | 0.48805703 | -0.654503434 | 0.6 | 0.605 | 1 | 0.48805703 | 0.001655048 |
| Cast | 0.000830638 | 0.323779946 | 1 | 0.918 | 1 | 0.121173486 | 0.847370873 | 1 | 0.944 | 1 | 0.121173486 | 0.001660586 |
| Gigyf2 | 0.000830864 | -0.660513534 | 0.681 | 0.788 | 1 | 0.801152836 | -0.324665657 | 0.9 | 0.849 | 1 | 0.801152836 | 0.001661039 |
| Nudcd3 | 0.000832043 | -0.652721573 | 0.605 | 0.747 | 1 | 0.231817704 | -0.531936792 | 0.8 | 0.775 | 1 | 0.231817704 | 0.001663393 |
| Bnip3l | 0.00083282 | 0.463457857 | 0.866 | 0.735 | 1 | 0.146965007 | -0.707903129 | 0.7 | 0.815 | 1 | 0.146965007 | 0.001664947 |
| Polr3k | 0.00083547 | -1.411426246 | 0.143 | 0.306 | 1 | 0.680991518 | -0.138582578 | 0.2 | 0.244 | 1 | 0.680991518 | 0.001670243 |
| Fkbp1a | 0.000838956 | -0.49217366 | 0.882 | 0.912 | 1 | 0.029409802 | -0.833133029 | 0.7 | 0.91 | 1 | 0.029409802 | 0.001677209 |
| H2bc6 | 0.00084089 | 1.027023174 | 0.336 | 0.159 | 1 | 0.210383116 | 5.548436625 | 0.2 | 0.09 | 1 | 0.210383116 | 0.001681074 |
| Dmpk | 0.000844336 | -2.439623138 | 0.025 | 0.141 | 1 | 0.078167089 | 1.127150978 | 0.4 | 0.176 | 1 | 0.078167089 | 0.001687958 |
| Dhx38 | 0.000845485 | -1.587788545 | 0.126 | 0.282 | 1 | 0.259788158 | -0.789433014 | 0.2 | 0.352 | 1 | 0.259788158 | 0.001690254 |
| Cox6a1 | 0.000850249 | 0.473642909 | 0.95 | 0.882 | 1 | 0.722126811 | 0.495929852 | 0.9 | 0.92 | 1 | 0.722126811 | 0.001699775 |
| Mad2l1 | 0.530891315 | -1.292781749 | 0.076 | 0.094 | 1 | 0.000850398 | 2.295455884 | 0.4 | 0.083 | 1 | 0.530891315 | 0.001700073 |
| Alg8 | 0.38354194 | -0.642968104 | 0.092 | 0.124 | 1 | 0.000850411 | 2.130396637 | 0.4 | 0.083 | 1 | 0.38354194 | 0.001700099 |
| Rhobtb3 | 0.000850643 | -1.138560831 | 0.361 | 0.5 | 1 | 0.114655159 | -0.800432013 | 0.2 | 0.478 | 1 | 0.114655159 | 0.001700563 |
| Pdzd4 | 0.000850992 | -1.707819249 | 0.084 | 0.229 | 1 | 0.619625969 | -0.654503434 | 0.2 | 0.247 | 1 | 0.619625969 | 0.00170126 |
| Gm41077 | 0.000855722 | -1.86393845 | 0.076 | 0.218 | 1 | 0.573635741 | -0.457811523 | 0.1 | 0.16 | 1 | 0.573635741 | 0.001710712 |
| Tgds | 0.000856379 | -1.343407822 | 0.109 | 0.271 | 1 | 0.882935911 | 0.227844977 | 0.2 | 0.213 | 1 | 0.882935911 | 0.001712024 |
| Snx16 | 0.000861209 | -1.459431619 | 0.067 | 0.212 | 1 | 0.38933357 | -0.640289575 | 0.1 | 0.207 | 1 | 0.38933357 | 0.001721677 |
| Ptpn1 | 0.000863145 | 0.402283227 | 0.958 | 0.935 | 1 | 0.271497594 | 0.297377211 | 1 | 0.969 | 1 | 0.271497594 | 0.001725544 |
| Aco1 | 0.000863621 | -1.292781749 | 0.193 | 0.359 | 1 | 0.661252282 | -0.103093493 | 0.3 | 0.367 | 1 | 0.661252282 | 0.001726496 |
| P2rx4 | 0.000868482 | 1.017073513 | 0.462 | 0.288 | 1 | 0.692951695 | -0.449683642 | 0.3 | 0.333 | 1 | 0.692951695 | 0.00173621 |
| Cox7b | 0.000869963 | -0.474818728 | 0.824 | 0.9 | 1 | 0.27110793 | -0.291327053 | 0.6 | 0.87 | 1 | 0.27110793 | 0.00173917 |
| Bmp1 | 0.000872588 | -1.323675757 | 0.429 | 0.547 | 1 | 0.72740099 | 0.257700962 | 0.5 | 0.466 | 1 | 0.72740099 | 0.001744414 |
| Plcl1 | 0.00087667 | -2.0052943 | 0.134 | 0.288 | 1 | 0.848803355 | -0.763437806 | 0.2 | 0.213 | 1 | 0.848803355 | 0.001752571 |
| Tead1 | 0.000876938 | 0.457941353 | 0.983 | 0.953 | 1 | 0.019616084 | 0.894104802 | 1 | 0.972 | 1 | 0.019616084 | 0.001753107 |
| Mex3b | 0.000880083 | -2.521050737 | 0.034 | 0.153 | 1 | 0.374721324 | -0.267480311 | 0 | 0.074 | 1 | 0.374721324 | 0.001759391 |
| Mtbp | 0.000880922 | -2.86393845 | 0.034 | 0.153 | 1 | 0.343924324 | -0.374395515 | 0 | 0.083 | 1 | 0.343924324 | 0.001761069 |
| Mob3a | 0.000881622 | -1.215819767 | 0.168 | 0.335 | 1 | 0.998350482 | 0.210566986 | 0.3 | 0.299 | 1 | 0.998350482 | 0.001762468 |
| R3hdm2 | 0.00088281 | -0.644625422 | 0.79 | 0.847 | 1 | 0.605078385 | 0.159940913 | 1 | 0.87 | 1 | 0.605078385 | 0.00176484 |
| Plat | 0.000887781 | -3.309855263 | 0.042 | 0.165 | 1 | 0.252834158 | -1.321928095 | 0 | 0.117 | 1 | 0.252834158 | 0.001774773 |
| Hspd1 | 0.014036585 | -0.510270898 | 0.891 | 0.924 | 1 | 0.00089285 | -1.534241566 | 0.7 | 0.969 | 1 | 0.014036585 | 0.001784904 |
| Tspan13 | 0.00089313 | 0.343724793 | 0.958 | 0.918 | 1 | 0.109850926 | -0.683673353 | 0.8 | 0.954 | 1 | 0.109850926 | 0.001785462 |
| Egln2 | 0.000896881 | -1.244418728 | 0.193 | 0.359 | 1 | 0.071387359 | -1.473931188 | 0.1 | 0.373 | 1 | 0.071387359 | 0.001792958 |
| Dnah7b | 0.000897366 | -2.778208576 | 0.034 | 0.153 | 1 | 0.24572516 | -0.982078092 | 0 | 0.12 | 1 | 0.24572516 | 0.001793926 |
| Got1 | 0.000897682 | 1.251538767 | 0.277 | 0.124 | 1 | 0.540467099 | -0.391469028 | 0.1 | 0.17 | 1 | 0.540467099 | 0.001794558 |
| Pycrl | 0.000897723 | -1.381591016 | 0.126 | 0.288 | 1 | 0.625942698 | 0.135278859 | 0.1 | 0.154 | 1 | 0.625942698 | 0.001794639 |
| Rbl2 | 0.0008989 | -1.112700133 | 0.336 | 0.488 | 1 | 0.463968988 | -0.248864633 | 0.4 | 0.534 | 1 | 0.463968988 | 0.001796992 |
| Bmx | 0.000901177 | 1.152003093 | 0.109 | 0.018 | 1 | 0.051044361 | 2.695993813 | 0.1 | 0.015 | 1 | 0.051044361 | 0.001801542 |
| Ntn3 | 0.000901803 | -3.733354341 | 0 | 0.088 | 1 | 0.802052739 | 1.159940913 | 0.1 | 0.077 | 1 | 0.802052739 | 0.001802792 |
| Skil | 0.000901981 | -0.665872025 | 0.832 | 0.871 | 1 | 0.071197345 | -0.899787019 | 0.7 | 0.92 | 1 | 0.071197345 | 0.001803148 |
| Flnc | 0.000902367 | -3.572889668 | 0.017 | 0.124 | 1 | 0.963100513 | 0.135278859 | 0.1 | 0.093 | 1 | 0.963100513 | 0.00180392 |
| Chst1 | 0.000903121 | -4.008988783 | 0 | 0.088 | 1 | 0.132572648 | -2.131825212 | 0 | 0.188 | 1 | 0.132572648 | 0.001805425 |
| Swt1 | 0.000903425 | -0.8696392 | 0.571 | 0.676 | 1 | 0.159116039 | -0.731016328 | 0.7 | 0.873 | 1 | 0.159116039 | 0.001806034 |
| Trub2 | 0.000903476 | -1.707819249 | 0.092 | 0.241 | 1 | 0.204107633 | 0.832055363 | 0.4 | 0.21 | 1 | 0.204107633 | 0.001806135 |
| Ndufa7 | 0.000903505 | 0.394422142 | 0.891 | 0.912 | 1 | 0.602612028 | 0.178145508 | 0.6 | 0.836 | 1 | 0.602612028 | 0.001806194 |
| Lrrc71 | 0.000903527 | -4.292781749 | 0 | 0.088 | 1 | 0.863421541 | 0.159940913 | 0.1 | 0.114 | 1 | 0.863421541 | 0.001806238 |
| Cyp26a1 | 0.00090387 | -6.100136671 | 0 | 0.088 | 1 | 0.096372899 | -3.131825212 | 0 | 0.222 | 1 | 0.096372899 | 0.001806924 |
| Tspan9 | 0.00090675 | -1.512907564 | 0.185 | 0.347 | 1 | 0.150271033 | -0.900941329 | 0.2 | 0.441 | 1 | 0.150271033 | 0.001812678 |
| Thbd | 0.000906895 | -2.655351829 | 0.025 | 0.141 | 1 | 0.577165087 | 0.847996907 | 0 | 0.031 | 1 | 0.577165087 | 0.001812968 |
| Arhgap1 | 0.000909412 | -1.028569152 | 0.235 | 0.412 | 1 | 0.575079361 | -0.382957528 | 0.3 | 0.355 | 1 | 0.575079361 | 0.001817997 |
| Traf3 | 0.000909818 | -1.037222464 | 0.462 | 0.588 | 1 | 0.086790799 | -1.094083118 | 0.6 | 0.75 | 1 | 0.086790799 | 0.001818808 |
| Cep85 | 0.016693651 | 0.453172628 | 0.58 | 0.418 | 1 | 0.000910297 | 1.576486719 | 0.8 | 0.37 | 1 | 0.016693651 | 0.001819765 |
| Veph1 | 0.000919771 | 0.690252849 | 0.487 | 0.294 | 1 | 0.9151285 | -0.18827848 | 0.4 | 0.41 | 1 | 0.9151285 | 0.001838696 |
| Trmt13 | 0.000920601 | -1.44625623 | 0.176 | 0.341 | 1 | 0.986370577 | -0.327852929 | 0.2 | 0.188 | 1 | 0.986370577 | 0.001840355 |
| Fubp1 | 0.00122829 | -0.467772963 | 0.958 | 0.941 | 1 | 0.000924442 | -1.261301736 | 0.8 | 0.969 | 1 | 0.00122829 | 0.001848029 |
| Emp3 | 0.000929693 | -0.469814628 | 0.891 | 0.853 | 1 | 0.034800677 | -1.033740212 | 0.8 | 0.858 | 1 | 0.034800677 | 0.001858522 |
| Ranbp3 | 0.000933086 | -1.015941544 | 0.252 | 0.429 | 1 | 0.374700285 | -0.574535129 | 0.3 | 0.407 | 1 | 0.374700285 | 0.001865301 |
| Atxn7l3b | 0.000933369 | -0.45043655 | 0.899 | 0.924 | 1 | 0.168401399 | -0.445018496 | 0.7 | 0.932 | 1 | 0.168401399 | 0.001865867 |
| Gm15614 | 0.000936064 | -1.835924074 | 0.067 | 0.206 | 1 | 0.654137414 | 1.169925001 | 0.2 | 0.157 | 1 | 0.654137414 | 0.001871251 |
| Ier5l | 0.000937198 | -1.701744734 | 0.101 | 0.253 | 1 | 0.990081952 | -0.452398027 | 0.2 | 0.179 | 1 | 0.990081952 | 0.001873517 |
| Smcr8 | 0.000939311 | -1.544320516 | 0.092 | 0.241 | 1 | 0.267428625 | 1.150025444 | 0.3 | 0.17 | 1 | 0.267428625 | 0.001877739 |
| Gm50240 | 0.352905577 | -0.347923303 | 0.059 | 0.088 | 1 | 0.00093958 | 3.210566986 | 0.4 | 0.093 | 1 | 0.352905577 | 0.001878278 |
| Lig1 | 0.081058809 | -1.466318004 | 0.109 | 0.176 | 1 | 0.000941983 | 3.639410285 | 0.4 | 0.093 | 1 | 0.081058809 | 0.001883078 |
| Snx6 | 0.000942845 | -0.702349308 | 0.664 | 0.747 | 1 | 0.246171198 | -0.351893516 | 0.8 | 0.88 | 1 | 0.246171198 | 0.001884801 |
| Rrm1 | 0.000943239 | -1.953575663 | 0.084 | 0.229 | 1 | 0.671760586 | 1.245332404 | 0.2 | 0.176 | 1 | 0.671760586 | 0.001885588 |
| Zfp189 | 0.00094464 | -1.722466024 | 0.042 | 0.171 | 1 | 0.777306966 | 0.789103218 | 0.2 | 0.173 | 1 | 0.777306966 | 0.001888387 |
| Rpp14 | 0.000947533 | -1.237499314 | 0.143 | 0.306 | 1 | 0.335190361 | -0.5519337 | 0.1 | 0.225 | 1 | 0.335190361 | 0.001894168 |
| Jph1 | 0.000948109 | -1.707819249 | 0.084 | 0.229 | 1 | 0.120905568 | -0.849974556 | 0.1 | 0.373 | 1 | 0.120905568 | 0.001895318 |
| Exo5 | 0.000949123 | -2.707819249 | 0.017 | 0.124 | 1 | 0.334165224 | -0.111361109 | 0 | 0.086 | 1 | 0.334165224 | 0.001897346 |
| Fbxl8 | 0.000949161 | -2.854660637 | 0.017 | 0.124 | 1 | 0.32492745 | -0.339630097 | 0 | 0.09 | 1 | 0.32492745 | 0.001897422 |
| Cdkn2aip | 0.000951782 | -1.842978832 | 0.151 | 0.306 | 1 | 0.074870051 | -1.581990934 | 0.1 | 0.37 | 1 | 0.074870051 | 0.001902659 |
| Nsmf | 0.000953607 | -1.086877451 | 0.218 | 0.388 | 1 | 0.322322278 | -0.728032469 | 0.2 | 0.336 | 1 | 0.322322278 | 0.001906305 |
| Wtip | 0.000954941 | -0.833949132 | 0.454 | 0.6 | 1 | 0.057048307 | -0.968489027 | 0.5 | 0.747 | 1 | 0.057048307 | 0.00190897 |
| Urm1 | 0.00096016 | -1.070389328 | 0.227 | 0.4 | 1 | 0.378454304 | 0.504852326 | 0.5 | 0.33 | 1 | 0.378454304 | 0.001919399 |
| Dmac2 | 0.000962738 | -1.807354922 | 0.059 | 0.194 | 1 | 0.894290389 | 0.808468542 | 0.1 | 0.086 | 1 | 0.894290389 | 0.00192455 |
| Zswim8 | 0.000965391 | -0.810667656 | 0.479 | 0.635 | 1 | 0.400939337 | -0.417464237 | 0.6 | 0.738 | 1 | 0.400939337 | 0.001929851 |
| Rnf220 | 0.000965584 | -1.359895945 | 0.143 | 0.306 | 1 | 0.501113684 | -0.567040593 | 0.2 | 0.278 | 1 | 0.501113684 | 0.001930235 |
| Psma7 | 0.00096582 | -0.544513506 | 0.832 | 0.876 | 1 | 0.017321122 | -0.950168844 | 0.7 | 0.92 | 1 | 0.017321122 | 0.001930707 |
| Nckap5l | 0.000967656 | -1.707819249 | 0.126 | 0.282 | 1 | 0.969867592 | 0.123104145 | 0.4 | 0.407 | 1 | 0.969867592 | 0.001934376 |
| Sorbs2os | 0.00097169 | -1.780882711 | 0.067 | 0.206 | 1 | 0.404405797 | 0.751135367 | 0.3 | 0.182 | 1 | 0.404405797 | 0.001942435 |
| Gm17225 | 0.000973992 | -3.911691582 | 0.008 | 0.106 | 1 | 0.522115639 | 0.625604485 | 0 | 0.04 | 1 | 0.522115639 | 0.001947035 |
| Map3k21 | 0.168791026 | -1.485426827 | 0.025 | 0.059 | 1 | 0.000975552 | 2.374065718 | 0.3 | 0.049 | 1 | 0.168791026 | 0.001950153 |
| 2210408I21Rik | 0.000977252 | -0.892769935 | 0.504 | 0.624 | 1 | 0.147869567 | -0.302106654 | 0.4 | 0.623 | 1 | 0.147869567 | 0.001953548 |
| H1f10 | 0.000979705 | -1.858885223 | 0.076 | 0.224 | 1 | 0.968557568 | 0.457206954 | 0.2 | 0.216 | 1 | 0.968557568 | 0.00195845 |
| Mfn2 | 0.000981579 | -1.222392421 | 0.151 | 0.318 | 1 | 0.621915284 | 0.473601392 | 0.2 | 0.293 | 1 | 0.621915284 | 0.001962194 |
| Ptpn13 | 0.000984852 | -1.08236197 | 0.395 | 0.553 | 1 | 0.305636685 | -0.687134438 | 0.5 | 0.583 | 1 | 0.305636685 | 0.001968733 |
| Mynn | 0.000986403 | -1.070389328 | 0.202 | 0.376 | 1 | 0.134045622 | -1.161987182 | 0.1 | 0.318 | 1 | 0.134045622 | 0.001971833 |
| Apeh | 0.217832416 | -0.669851398 | 0.084 | 0.129 | 1 | 0.000986704 | 2.048295557 | 0.5 | 0.127 | 1 | 0.217832416 | 0.001972435 |
| Arl2bp | 0.000987778 | 0.514573173 | 0.849 | 0.712 | 1 | 0.475029303 | -0.210896782 | 0.6 | 0.744 | 1 | 0.475029303 | 0.001974581 |

| | | | | | | | | | | | | |
|---|---|---|---|---|---|---|---|---|---|---|---|---|
| Nfxl1 | 0.000987814 | -1.352789059 | 0.235 | 0.388 | 1 | 0.293738664 | -0.773892163 | 0.3 | 0.448 | 1 | 0.293738664 | 0.001974652 |
| Snapc3 | 0.000989416 | -1.308549065 | 0.235 | 0.406 | 1 | 0.933800047 | -0.138582578 | 0.5 | 0.488 | 1 | 0.933800047 | 0.001977852 |
| Vcpkmt | 0.000990599 | -3.185866545 | 0.008 | 0.106 | 1 | 0.882330095 | 0.591657153 | 0.1 | 0.114 | 1 | 0.882330095 | 0.001980217 |
| Rngtt | 0.000991792 | -1.060739158 | 0.538 | 0.618 | 1 | 0.238832399 | -0.739190259 | 0.5 | 0.67 | 1 | 0.238832399 | 0.0019826 |
| Nlgn1 | 0.000991816 | -3.655351829 | 0.017 | 0.124 | 1 | 0.24611316 | -3.020997081 | 0 | 0.12 | 1 | 0.24611316 | 0.001982648 |
| Antxr2 | 0.0009939 | -1.577349317 | 0.227 | 0.382 | 1 | 0.835965999 | -0.392774044 | 0.5 | 0.506 | 1 | 0.835965999 | 0.001986813 |
| Gale | 0.000994668 | 0.733582955 | 0.395 | 0.212 | 1 | 0.171869174 | 1.865918815 | 0.3 | 0.173 | 1 | 0.171869174 | 0.001988347 |
| Efhb | 0.000995151 | -2.770829046 | 0.017 | 0.124 | 1 | 0.749385093 | 0.769994395 | 0.1 | 0.071 | 1 | 0.749385093 | 0.001989312 |
| Syt12 | 0.00099583 | -3.392317423 | 0.008 | 0.106 | 1 | 0.259778249 | -0.840059087 | 0 | 0.114 | 1 | 0.259778249 | 0.001990667 |
| Bmp4 | 0.000995853 | -3.485426827 | 0.008 | 0.106 | 1 | 0.334427938 | -0.736965594 | 0 | 0.086 | 1 | 0.334427938 | 0.001990714 |
| Sfmbt2 | 0.000995861 | -3.733354341 | 0.008 | 0.106 | 1 | 0.191303296 | -1.230005605 | 0 | 0.148 | 1 | 0.191303296 | 0.00199073 |
| Ezh2 | 0.00099641 | -1.144936281 | 0.42 | 0.553 | 1 | 0.389478339 | -0.956082883 | 0.5 | 0.549 | 1 | 0.389478339 | 0.001991826 |
| Mapk8 | 0.000997047 | -0.992768431 | 0.361 | 0.524 | 1 | 0.945958611 | -0.123988717 | 0.6 | 0.651 | 1 | 0.945958611 | 0.0019931 |
| Stx1a | 0.000999994 | -2.207892852 | 0.034 | 0.153 | 1 | 0.553727093 | 0.795529487 | 0.2 | 0.13 | 1 | 0.553727093 | 0.001998988 |
| Washc1 | 0.001005959 | -1.276840205 | 0.084 | 0.235 | 1 | 0.772713406 | 0.558490289 | 0.2 | 0.16 | 1 | 0.772713406 | 0.002010907 |
| Gm19265 | 0.001006139 | -3.343407822 | 0.008 | 0.106 | 1 | 0.489939838 | 0.625604485 | 0 | 0.046 | 1 | 0.489939838 | 0.002011265 |
| Ccdc51 | 0.001006139 | -3.070389328 | 0.008 | 0.106 | 1 | 0.815998758 | 0.695993813 | 0.1 | 0.077 | 1 | 0.815998758 | 0.002011265 |
| Prxl2a | 0.001008936 | 1.251538767 | 0.37 | 0.182 | 1 | 0.003915824 | 2.678071905 | 0.5 | 0.194 | 1 | 0.003915824 | 0.002016855 |
| Ccdc61 | 0.001011184 | -2.944858446 | 0.008 | 0.106 | 1 | 0.374523929 | 0.210566986 | 0 | 0.074 | 1 | 0.374523929 | 0.002021346 |
| Aprt | 0.001012795 | -1.042238288 | 0.412 | 0.541 | 1 | 0.42602875 | -0.464742017 | 0.5 | 0.562 | 1 | 0.42602875 | 0.002024564 |
| Pbxip1 | 0.001012885 | 0.484702643 | 0.832 | 0.694 | 1 | 0.237093715 | -0.377035397 | 0.5 | 0.759 | 1 | 0.237093715 | 0.002024744 |
| Men1 | 0.00101771 | -2.163498732 | 0.034 | 0.153 | 1 | 0.081061831 | -1.567040593 | 0 | 0.238 | 1 | 0.081061831 | 0.002034384 |
| Wrnip1 | 0.001017842 | -1.304854582 | 0.118 | 0.276 | 1 | 0.187239963 | -1.090602549 | 0.1 | 0.284 | 1 | 0.187239963 | 0.002034649 |
| Slc4a3 | 0.001022653 | -1.642968104 | 0.101 | 0.247 | 1 | 0.857273827 | 1.036069255 | 0.2 | 0.191 | 1 | 0.857273827 | 0.00204426 |
| Cetn3 | 0.002288397 | -0.591104905 | 0.655 | 0.724 | 1 | 0.001024258 | -2.342559428 | 0.2 | 0.71 | 1 | 0.002288397 | 0.002047466 |
| Galnt1 | 0.001031236 | -0.55892464 | 0.849 | 0.888 | 1 | 0.481866055 | -0.468990447 | 0.8 | 0.901 | 1 | 0.481866055 | 0.002061408 |
| Ikzf5 | 0.00103393 | -1.243454037 | 0.218 | 0.388 | 1 | 0.019621823 | -2.505640048 | 0 | 0.367 | 1 | 0.019621823 | 0.00206679 |
| Wtap | 0.001036926 | -0.841912144 | 0.353 | 0.518 | 1 | 0.003114528 | -2.066000576 | 0.1 | 0.642 | 1 | 0.003114528 | 0.002072776 |
| Metap1d | 0.001038328 | 0.355716408 | 0.782 | 0.606 | 1 | 0.15620703 | 0.679119995 | 0.7 | 0.556 | 1 | 0.15620703 | 0.002075578 |
| Gopc | 0.00103939 | -1.273922722 | 0.218 | 0.382 | 1 | 0.940495647 | -0.225252075 | 0.5 | 0.466 | 1 | 0.940495647 | 0.002077699 |
| Slc12a1 | 0.001040724 | 1.004898799 | 0.294 | 0.141 | 1 | 0.557744661 | -0.304006187 | 0 | 0.034 | 1 | 0.557744661 | 0.002080366 |
| Adamtsl3 | 0.001041189 | 0.434437088 | 0.782 | 0.641 | 1 | 0.743820127 | 1.642412773 | 0.6 | 0.784 | 1 | 0.743820127 | 0.002081293 |
| Npat | 0.001045313 | -1.240314329 | 0.42 | 0.529 | 1 | 0.793025208 | 0.186980166 | 0.6 | 0.571 | 1 | 0.793025208 | 0.002089534 |
| Zdhhc20 | 0.001046891 | -0.64754924 | 0.706 | 0.841 | 1 | 0.319239369 | -0.177017301 | 1 | 0.901 | 1 | 0.319239369 | 0.002092686 |
| Rgcc | 0.001049993 | 0.801410603 | 0.782 | 0.647 | 1 | 0.014183244 | 1.043632027 | 0.9 | 0.704 | 1 | 0.014183244 | 0.002098883 |
| Cdc42 | 0.001061403 | -0.335974597 | 0.992 | 0.994 | 1 | 0.00815264 | -0.744345125 | 1 | 1 | 1 | 0.00815264 | 0.00212168 |
| Amy1 | 0.001066195 | -0.950095094 | 0.109 | 0.271 | 1 | 0.622285842 | 0.210566986 | 0.3 | 0.222 | 1 | 0.622285842 | 0.002131252 |
| Chtop | 0.001066554 | -0.72935241 | 0.462 | 0.6 | 1 | 0.158429066 | -0.740777208 | 0.6 | 0.67 | 1 | 0.158429066 | 0.002131971 |
| Optn | 0.001068835 | 0.477340267 | 0.714 | 0.465 | 1 | 0.91174475 | -0.115477217 | 0.6 | 0.503 | 1 | 0.91174475 | 0.002136528 |
| Kazald1 | 0.001069529 | 0.6492227 | 0.588 | 0.388 | 1 | 0.061316786 | -0.527869995 | 0.1 | 0.457 | 1 | 0.061316786 | 0.002137915 |
| Cycs | 0.001069771 | 0.646719954 | 0.899 | 0.835 | 1 | 0.544203282 | 0.140177658 | 0.6 | 0.818 | 1 | 0.544203282 | 0.002138398 |
| Scmh1 | 0.001072555 | -0.887288938 | 0.58 | 0.706 | 1 | 0.643375902 | -0.14068778 | 0.8 | 0.852 | 1 | 0.643375902 | 0.00214396 |
| Zfp800 | 0.001075409 | -0.89850281 | 0.513 | 0.618 | 1 | 0.034333472 | -0.775311906 | 0.3 | 0.71 | 1 | 0.034333472 | 0.002149662 |
| Mrpl1 | 0.001076304 | -1.333423734 | 0.176 | 0.335 | 1 | 0.470191848 | -0.654503434 | 0.2 | 0.284 | 1 | 0.470191848 | 0.002151449 |
| Nrtn | 0.132792179 | -0.707819249 | 0.05 | 0.1 | 1 | 0.001077288 | 2.252387162 | 0.4 | 0.086 | 1 | 0.132792179 | 0.002153415 |
| Bnip2 | 0.001077394 | -0.53475454 | 0.874 | 0.829 | 1 | 0.021273005 | -0.903919029 | 0.6 | 0.901 | 1 | 0.021273005 | 0.002153627 |
| Rpp21 | 0.001077443 | -1.11345805 | 0.143 | 0.312 | 1 | 0.910458605 | 0.345496566 | 0.2 | 0.222 | 1 | 0.910458605 | 0.002153725 |
| Vps45 | 0.001081593 | -1.509273569 | 0.092 | 0.241 | 1 | 0.375476121 | -0.668578619 | 0.1 | 0.21 | 1 | 0.375476121 | 0.002162016 |
| Zbed3 | 0.001082401 | -1.485426827 | 0.126 | 0.276 | 1 | 0.074469912 | -1.750262417 | 0 | 0.247 | 1 | 0.074469912 | 0.00216363 |
| Fam83d | 0.001082849 | -2.854660637 | 0.008 | 0.106 | 1 | 0.539470441 | 0.159940913 | 0 | 0.037 | 1 | 0.539470441 | 0.002164526 |
| Ing3 | 0.001085499 | -1.225906906 | 0.286 | 0.435 | 1 | 0.008042316 | -3.009984089 | 0 | 0.438 | 1 | 0.008042316 | 0.002169821 |
| Ireb2 | 0.001090864 | -0.638130088 | 0.487 | 0.641 | 1 | 0.012079799 | -1.508251262 | 0.4 | 0.728 | 1 | 0.012079799 | 0.002180537 |
| Zw10 | 0.001091432 | -1.301002256 | 0.185 | 0.341 | 1 | 0.933725845 | 0.185031894 | 0.4 | 0.377 | 1 | 0.933725845 | 0.002181673 |
| Lamc1 | 0.001098741 | -1.159198595 | 0.303 | 0.465 | 1 | 0.267586889 | -0.787284547 | 0.5 | 0.657 | 1 | 0.267586889 | 0.002196275 |
| Mocs2 | 0.001102416 | -1.061111514 | 0.294 | 0.459 | 1 | 0.133671256 | -0.869603363 | 0.3 | 0.531 | 1 | 0.133671256 | 0.002203617 |
| Wwox | 0.957571818 | -0.437598302 | 0.84 | 0.806 | 1 | 0.001103085 | 0.981879158 | 1 | 0.775 | 1 | 0.957571818 | 0.002204953 |
| Ak4 | 0.00110641 | 0.557008439 | 0.42 | 0.235 | 1 | 0.224642261 | 1.374065718 | 0.3 | 0.17 | 1 | 0.224642261 | 0.002211597 |
| Sstr2 | 0.00110996 | 2.340543773 | 0.168 | 0.053 | 1 | 0.23891309 | 2.017921908 | 0.1 | 0.031 | 1 | 0.23891309 | 0.002218687 |
| Rab13 | 0.001110509 | -1.010088818 | 0.353 | 0.512 | 1 | 0.705244629 | -0.401616984 | 0.4 | 0.395 | 1 | 0.705244629 | 0.002219786 |
| Bmpr1a | 0.00111173 | -0.597485334 | 0.739 | 0.853 | 1 | 0.021452312 | -1.024570174 | 0.8 | 0.941 | 1 | 0.021452312 | 0.002222225 |
| Spred2 | 0.001112472 | -1.040015679 | 0.387 | 0.553 | 1 | 0.229729528 | -0.549762601 | 0.6 | 0.772 | 1 | 0.229729528 | 0.002223706 |
| Rgma | 0.001113865 | -1.584962501 | 0.092 | 0.241 | 1 | 0.184128523 | 1.083809844 | 0.4 | 0.216 | 1 | 0.184128523 | 0.00222649 |
| Rnf214 | 0.001115422 | -1.027576244 | 0.361 | 0.506 | 1 | 0.239530695 | -0.526398608 | 0.3 | 0.494 | 1 | 0.239530695 | 0.002229601 |
| Arl13b | 0.001118681 | -0.812655706 | 0.664 | 0.724 | 1 | 0.204121743 | -0.731229225 | 0.7 | 0.781 | 1 | 0.204121743 | 0.002236111 |
| Amer2 | 0.001119669 | -2.424026283 | 0.008 | 0.106 | 1 | 0.619961444 | 1.558490289 | 0 | 0.025 | 1 | 0.619961444 | 0.002238084 |
| Nip7 | 0.00112011 | -0.96020641 | 0.429 | 0.553 | 1 | 0.038530318 | -1.556986928 | 0.2 | 0.509 | 1 | 0.038530318 | 0.002238965 |
| Dbndd2 | 0.00112051 | -1.665999073 | 0.118 | 0.265 | 1 | 0.818216401 | 0.351165316 | 0.3 | 0.265 | 1 | 0.818216401 | 0.002239764 |
| Eif2a | 0.001126872 | -0.735040717 | 0.487 | 0.647 | 1 | 0.094510383 | -0.864721141 | 0.5 | 0.676 | 1 | 0.094510383 | 0.002252474 |
| Brf2 | 0.003364551 | -1.970853654 | 0.034 | 0.135 | 1 | 0.001129626 | 2.252387162 | 0.4 | 0.086 | 1 | 0.003364551 | 0.002257975 |
| Dnah14 | 0.091361251 | 2.836501268 | 0.017 | 0 | 1 | 0.001133822 | 4.380491987 | 0.2 | 0.025 | 1 | 0.091361251 | 0.002266359 |
| Upk1b | 0.175720252 | -1.333423734 | 0.025 | 0.059 | 1 | 0.001133858 | 3.795529487 | 0.2 | 0.025 | 1 | 0.175720252 | 0.002266431 |
| Dcaf13 | 0.001134762 | -1.180013819 | 0.303 | 0.447 | 1 | 0.138253968 | -1.055327074 | 0.2 | 0.414 | 1 | 0.138253968 | 0.002268236 |
| Snhg4 | 0.001139566 | -0.986731399 | 0.378 | 0.547 | 1 | 0.108832023 | -1.374395515 | 0.2 | 0.423 | 1 | 0.108832023 | 0.002277834 |
| 3222401L13Rik | 0.022800608 | 0.897901812 | 0.143 | 0.059 | 1 | 0.001148476 | 2.754887502 | 0.3 | 0.052 | 1 | 0.022800608 | 0.002295632 |
| Bap1 | 0.001155939 | -1.485426827 | 0.109 | 0.259 | 1 | 0.943469072 | 0.306427001 | 0.3 | 0.327 | 1 | 0.943469072 | 0.002310541 |
| Cebpz | 0.001159224 | -1.131789873 | 0.437 | 0.559 | 1 | 0.108730738 | -0.832934653 | 0.3 | 0.608 | 1 | 0.108730738 | 0.002317105 |
| Rsl1d1 | 0.001161626 | -0.490644536 | 0.815 | 0.888 | 1 | 0.05874122 | -0.738449085 | 0.7 | 0.91 | 1 | 0.05874122 | 0.002321903 |
| Clock | 0.00116337 | -0.906159932 | 0.639 | 0.759 | 1 | 0.432707161 | -0.510113525 | 0.8 | 0.87 | 1 | 0.432707161 | 0.002325387 |
| Immt | 0.001168303 | -0.921525942 | 0.529 | 0.618 | 1 | 0.006893945 | -1.712718048 | 0.4 | 0.719 | 1 | 0.006893945 | 0.002335241 |
| Zfp532 | 0.001177074 | -1.411426246 | 0.277 | 0.424 | 1 | 0.189329085 | -1.166953435 | 0.3 | 0.46 | 1 | 0.189329085 | 0.002352763 |
| Cxxc5 | 0.001177736 | -1.014536093 | 0.437 | 0.541 | 1 | 0.888080849 | -0.257167936 | 0.9 | 0.762 | 1 | 0.888080849 | 0.002354085 |
| Sap30bp | 0.001182632 | -0.891419187 | 0.387 | 0.547 | 1 | 0.389377781 | -0.650456601 | 0.5 | 0.549 | 1 | 0.389377781 | 0.002363866 |
| Slc27a3 | 0.001185322 | -1.368069877 | 0.101 | 0.253 | 1 | 0.289357428 | -0.654503434 | 0.1 | 0.241 | 1 | 0.289357428 | 0.002369238 |
| Coil | 0.001187766 | -2.137503524 | 0.05 | 0.176 | 1 | 0.711343638 | 0.580516596 | 0.1 | 0.154 | 1 | 0.711343638 | 0.002374122 |
| Prkd2 | 0.001191253 | -0.473552727 | 0.185 | 0.353 | 1 | 0.007393611 | 1.079322453 | 0.8 | 0.401 | 1 | 0.007393611 | 0.002381087 |
| Dedd2 | 0.001194448 | -2.129283017 | 0.059 | 0.188 | 1 | 0.551691466 | 1.31748219 | 0.3 | 0.228 | 1 | 0.551691466 | 0.002387469 |
| Arhgef10 | 0.001195894 | -1.552541023 | 0.143 | 0.294 | 1 | 0.368482885 | -0.633129783 | 0.2 | 0.34 | 1 | 0.368482885 | 0.002390357 |
| Klf4 | 0.00279287 | -0.515295998 | 0.958 | 0.971 | 1 | 0.00119734 | -1.682684934 | 1 | 0.985 | 1 | 0.00279287 | 0.002393247 |
| Zfp638 | 0.001200236 | -0.588205424 | 0.857 | 0.865 | 1 | 0.379453759 | -0.345294355 | 0.9 | 0.917 | 1 | 0.379453759 | 0.002399032 |
| Rb1 | 0.001201308 | -1.077768858 | 0.277 | 0.435 | 1 | 0.357462797 | -0.237578825 | 0.5 | 0.639 | 1 | 0.357462797 | 0.002401174 |
| Aspg | 0.72759882 | -0.847996907 | 0.05 | 0.059 | 1 | 0.001202135 | 3.847996907 | 0.2 | 0.025 | 1 | 0.72759882 | 0.002402824 |
| Top2a | 0.016563331 | -2.776658125 | 0.017 | 0.082 | 1 | 0.00120327 | 3.737813989 | 0.2 | 0.025 | 1 | 0.016563331 | 0.002405092 |
| Malsu1 | 0.001206301 | 0.482151695 | 0.672 | 0.459 | 1 | 0.417849169 | -0.614346308 | 0.4 | 0.46 | 1 | 0.417849169 | 0.002411147 |
| Actr5 | 0.00121534 | -2.572889668 | 0.025 | 0.135 | 1 | 0.997533658 | 1.048295557 | 0.1 | 0.108 | 1 | 0.997533658 | 0.002429203 |
| Ncbp3 | 0.001215666 | -1.201633861 | 0.218 | 0.376 | 1 | 0.073694089 | -1.505640048 | 0.2 | 0.46 | 1 | 0.073694089 | 0.002429855 |
| Dvl2 | 0.001219996 | -1.090288885 | 0.176 | 0.347 | 1 | 0.843257554 | 0.383715888 | 0.3 | 0.281 | 1 | 0.843257554 | 0.002438504 |
| Fbxo22 | 0.001221067 | 1.074000581 | 0.513 | 0.341 | 1 | 0.897342165 | -0.103093493 | 0.4 | 0.343 | 1 | 0.897342165 | 0.002440643 |
| Kctd3 | 0.00122896 | -1.017921908 | 0.176 | 0.347 | 1 | 0.845394896 | -0.311201688 | 0.4 | 0.373 | 1 | 0.845394896 | 0.002456409 |

| | | | | | | | | | | | | |
|---|---|---|---|---|---|---|---|---|---|---|---|---|
| Fancm | 0.00122994 | -1.070389328 | 0.084 | 0.229 | 1 | 0.293710994 | -1.248864633 | 0.1 | 0.231 | 1 | 0.293710994 | 0.002458366 |
| Bivm | 0.001232444 | -2.439623138 | 0.025 | 0.135 | 1 | 0.972169102 | 0.625604485 | 0.1 | 0.102 | 1 | 0.972169102 | 0.002463369 |
| Pip4k2b | 0.001237551 | -1.780882711 | 0.076 | 0.212 | 1 | 0.680991518 | -0.152003093 | 0.2 | 0.244 | 1 | 0.680991518 | 0.00247357 |
| Tns2 | 0.001240328 | -0.94327741 | 0.479 | 0.606 | 1 | 0.921782585 | 0.376953998 | 0.7 | 0.639 | 1 | 0.921782585 | 0.002479118 |
| Taf4 | 0.001247401 | -2.070389328 | 0.05 | 0.176 | 1 | 0.282595287 | -0.936274402 | 0.1 | 0.238 | 1 | 0.282595287 | 0.002493246 |
| Hirip3 | 0.001250183 | -1.974811668 | 0.084 | 0.224 | 1 | 0.913033148 | 0.769994395 | 0.1 | 0.12 | 1 | 0.913033148 | 0.002498803 |
| Mast3 | 0.577205547 | -0.359895945 | 0.092 | 0.112 | 1 | 0.001254005 | 2.31748219 | 0.4 | 0.09 | 1 | 0.577205547 | 0.002506438 |
| Kctd1 | 0.001256488 | -1.494887156 | 0.227 | 0.376 | 1 | 0.28794742 | -0.646398384 | 0.3 | 0.469 | 1 | 0.28794742 | 0.002511396 |
| Fbxo30 | 0.001257593 | -0.669427014 | 0.588 | 0.688 | 1 | 0.083456627 | -0.905519864 | 0.7 | 0.812 | 1 | 0.083456627 | 0.002513604 |
| AI837181 | 0.001264967 | -0.870716983 | 0.328 | 0.488 | 1 | 0.201594309 | -0.441509711 | 0.2 | 0.469 | 1 | 0.201594309 | 0.002528334 |
| Naa12 | 0.001265056 | -0.615062108 | 0.63 | 0.712 | 1 | 0.369194505 | -0.361850261 | 0.7 | 0.741 | 1 | 0.369194505 | 0.002528511 |
| Rps18 | 0.029405882 | -0.215164336 | 0.992 | 0.976 | 1 | 0.001266091 | -1.195425374 | 0.8 | 0.997 | 1 | 0.029405882 | 0.002530579 |
| Pusl1 | 0.001266813 | -2.240314329 | 0.025 | 0.135 | 1 | 0.396884044 | 0.31748219 | 0 | 0.068 | 1 | 0.396884044 | 0.002532021 |
| Smoc1 | 0.001272174 | 1.859708659 | 0.244 | 0.1 | 1 | 0.146963474 | 0.510127268 | 0.2 | 0.071 | 1 | 0.146963474 | 0.002542729 |
| Med13 | 0.001275358 | -0.445287436 | 0.916 | 0.947 | 1 | 0.152379495 | -0.442116778 | 1 | 0.985 | 1 | 0.152379495 | 0.002549088 |
| Depdc5 | 0.001279275 | -1.773003417 | 0.118 | 0.265 | 1 | 0.454552796 | 0.191373421 | 0.5 | 0.333 | 1 | 0.454552796 | 0.002556914 |
| Igfbp5 | 0.00128654 | -1.813231488 | 0.218 | 0.394 | 1 | 0.650226797 | -1.489872732 | 0.3 | 0.333 | 1 | 0.650226797 | 0.002571426 |
| Stox2 | 0.001287731 | -1.432959407 | 0.134 | 0.294 | 1 | 0.027935816 | -3.191531458 | 0 | 0.34 | 1 | 0.027935816 | 0.002573803 |
| Atg16l1 | 0.00129003 | -0.883019192 | 0.319 | 0.494 | 1 | 0.532704182 | -0.208757872 | 0.4 | 0.503 | 1 | 0.532704182 | 0.002578396 |
| Penk | 0.001291091 | -0.994440475 | 0.235 | 0.429 | 1 | 0.998569768 | -0.922244842 | 0.6 | 0.488 | 1 | 0.998569768 | 0.002580515 |
| Sepsecs | 0.001291405 | -1.622930351 | 0.118 | 0.265 | 1 | 0.714373855 | 0.374065718 | 0.3 | 0.235 | 1 | 0.714373855 | 0.002581142 |
| Uhrf2 | 0.001291675 | -0.799180991 | 0.681 | 0.776 | 1 | 0.005759571 | -1.448942468 | 0.4 | 0.843 | 1 | 0.005759571 | 0.002581682 |
| Kansl2 | 0.001294891 | -1.125251263 | 0.311 | 0.453 | 1 | 0.272528679 | -0.208146171 | 0.3 | 0.494 | 1 | 0.272528679 | 0.002588106 |
| Amotl1 | 0.001296615 | -0.642230907 | 0.79 | 0.818 | 1 | 0.353078599 | 0.399536406 | 0.9 | 0.91 | 1 | 0.353078599 | 0.002591549 |
| Tubgcp6 | 0.001301372 | -2.129283017 | 0.025 | 0.135 | 1 | 0.355032511 | 1.280956314 | 0.2 | 0.105 | 1 | 0.355032511 | 0.002601051 |
| 4930403P22Rik | 0.659871503 | 0.514573173 | 0.025 | 0.018 | 1 | 0.001301471 | 3.558490289 | 0.2 | 0.025 | 1 | 0.659871503 | 0.002601248 |
| Tmem241 | 0.001302029 | -1.544320516 | 0.092 | 0.241 | 1 | 0.90330847 | -0.385800278 | 0.2 | 0.201 | 1 | 0.90330847 | 0.002602363 |
| Nab1 | 0.099718633 | -0.355791547 | 0.555 | 0.629 | 1 | 0.00130372 | -2.004445905 | 0.5 | 0.812 | 1 | 0.099718633 | 0.00260574 |
| Tm2d1 | 0.001308994 | 0.481098468 | 0.84 | 0.671 | 1 | 0.175815929 | -0.426528319 | 0.4 | 0.713 | 1 | 0.175815929 | 0.002616275 |
| Pank4 | 0.001309017 | -1.887525271 | 0.05 | 0.176 | 1 | 0.148024431 | -1.230005605 | 0 | 0.176 | 1 | 0.148024431 | 0.00261632 |
| Ift22 | 0.001309424 | -1.350497247 | 0.092 | 0.241 | 1 | 0.741300416 | 0.602884409 | 0.2 | 0.154 | 1 | 0.741300416 | 0.002617133 |
| Hbp1 | 0.001314922 | -0.681491966 | 0.689 | 0.759 | 1 | 0.093208833 | -0.693572999 | 0.6 | 0.852 | 1 | 0.093208833 | 0.002628116 |
| H2-Q10 | 0.001317623 | 1.432111013 | 0.092 | 0.012 | 1 | 0.730750993 | 2.017921908 | 0 | 0.012 | 1 | 0.730750993 | 0.00263351 |
| Rpl28 | 0.001317836 | -0.301124857 | 0.958 | 0.982 | 1 | 0.013769264 | -0.797890918 | 0.9 | 0.994 | 1 | 0.013769264 | 0.002633936 |
| Itpa | 0.001318252 | -1.083328384 | 0.269 | 0.429 | 1 | 0.116593896 | -1.111361109 | 0.1 | 0.333 | 1 | 0.116593896 | 0.002634766 |
| Clta | 0.001321578 | -0.398890077 | 0.958 | 0.935 | 1 | 0.11379531 | -0.412750718 | 0.7 | 0.978 | 1 | 0.11379531 | 0.00264141 |
| Apol7d | 0.179326482 | -1.292781749 | 0.034 | 0.071 | 1 | 0.001327528 | 3.532495081 | 0.2 | 0.025 | 1 | 0.179326482 | 0.002653295 |
| Gm45397 | 0.001327573 | 1.929610672 | 0.118 | 0.024 | 1 | 0.073613816 | 3.432959407 | 0.1 | 0.019 | 1 | 0.073613816 | 0.002653384 |
| Dmwd | 0.001329163 | -1.237499314 | 0.143 | 0.3 | 1 | 0.217310394 | -0.947862377 | 0.1 | 0.269 | 1 | 0.217310394 | 0.00265656 |
| Pde10a | 0.001333842 | -3.515174171 | 0.034 | 0.147 | 1 | 0.319596438 | -1.53447273 | 0.2 | 0.333 | 1 | 0.319596438 | 0.002665904 |
| Atp6ap2 | 0.002199517 | -0.479352313 | 0.824 | 0.853 | 1 | 0.001334247 | -1.600463594 | 0.7 | 0.938 | 1 | 0.002199517 | 0.002666714 |
| Clk2 | 0.001336351 | -0.998496409 | 0.294 | 0.459 | 1 | 0.317207154 | -0.345482823 | 0.3 | 0.472 | 1 | 0.317207154 | 0.002670917 |
| Dennd2b | 0.001338803 | -1.807354922 | 0.118 | 0.265 | 1 | 0.29077466 | -0.756865152 | 0.2 | 0.373 | 1 | 0.29077466 | 0.002675813 |
| Arf6 | 0.001578675 | -0.63837875 | 0.681 | 0.759 | 1 | 0.001339646 | -1.589408406 | 0.4 | 0.864 | 1 | 0.001578675 | 0.002677497 |
| Hpcal1 | 0.001341498 | -1.700439718 | 0.076 | 0.212 | 1 | 0.170730524 | -1.339630097 | 0 | 0.16 | 1 | 0.170730524 | 0.002681196 |
| Ttll7 | 0.001343977 | -1.457412451 | 0.151 | 0.306 | 1 | 0.328922345 | -0.71775499 | 0.3 | 0.466 | 1 | 0.328922345 | 0.002686149 |
| A430073D23Rik | 0.001346939 | -3.109917692 | 0.017 | 0.118 | 1 | 0.44686405 | 0.31748219 | 0 | 0.056 | 1 | 0.44686405 | 0.002692063 |
| 2610044O15Rik8 | 0.001350554 | -1.86393845 | 0.067 | 0.2 | 1 | 0.021952278 | 1.732519689 | 0.3 | 0.083 | 1 | 0.021952278 | 0.002699284 |
| Rps15a | 0.006454616 | -0.249154817 | 0.95 | 0.988 | 1 | 0.001354144 | -1.13207605 | 0.8 | 0.991 | 1 | 0.006454616 | 0.002706455 |
| Lbhd2 | 0.001354948 | 1.055141554 | 0.092 | 0.012 | 1 | 0.474964899 | 0.31748219 | 0 | 0.049 | 1 | 0.474964899 | 0.00270806 |
| Odf3l1 | 0.661155264 | -0.765534746 | 0.042 | 0.053 | 1 | 0.001361957 | 1.847996907 | 0.4 | 0.086 | 1 | 0.661155264 | 0.002722059 |
| H2az2 | 0.001362647 | -0.717936366 | 0.84 | 0.812 | 1 | 0.128061457 | -0.517785478 | 0.6 | 0.886 | 1 | 0.128061457 | 0.002723438 |
| Adam15 | 0.001364 | 0.599964664 | 0.538 | 0.329 | 1 | 0.333749526 | 0.808468542 | 0.4 | 0.262 | 1 | 0.333749526 | 0.00272614 |
| Klhl2 | 0.001365684 | -1.669851398 | 0.134 | 0.282 | 1 | 0.546671667 | -0.556986928 | 0.3 | 0.392 | 1 | 0.546671667 | 0.002729502 |
| Gm56901 | 0.00136738 | -3.485426827 | 0 | 0.082 | 1 | 0.505582845 | 1.017921908 | 0 | 0.043 | 1 | 0.505582845 | 0.002732891 |
| Utp23 | 0.001368972 | -1.908638258 | 0.084 | 0.218 | 1 | 0.55689788 | 0.653349476 | 0.3 | 0.219 | 1 | 0.55689788 | 0.00273607 |
| Bmp3 | 0.001370044 | -4.070389328 | 0 | 0.082 | 1 | 0.15308839 | -2.41670632 | 0 | 0.173 | 1 | 0.15308839 | 0.002738212 |
| Ttyh2 | 0.001370466 | -1.807354922 | 0.059 | 0.188 | 1 | 0.14864981 | -1.982078092 | 0 | 0.176 | 1 | 0.14864981 | 0.002739054 |
| Tspyl5 | 0.001370714 | -3.807354922 | 0 | 0.082 | 1 | 0.433635082 | 0.159940913 | 0 | 0.059 | 1 | 0.433635082 | 0.00273955 |
| Lypd6 | 0.001370933 | -3.944858446 | 0 | 0.082 | 1 | 0.00544582 | 2.14345279 | 0.2 | 0.031 | 1 | 0.00544582 | 0.002739987 |
| Rab30 | 0.001371158 | -1.891419187 | 0.126 | 0.276 | 1 | 0.305613919 | -1.026472211 | 0.1 | 0.231 | 1 | 0.305613919 | 0.002740437 |
| Slc19a2 | 0.077572724 | 0.96203215 | 0.109 | 0.053 | 1 | 0.001371672 | 2.473601392 | 0.4 | 0.093 | 1 | 0.077572724 | 0.002741463 |
| Supt5 | 0.001372068 | -0.807354922 | 0.546 | 0.647 | 1 | 0.930078236 | 1.567069274 | 0.6 | 0.738 | 1 | 0.930078236 | 0.002742253 |
| Styx | 0.001373338 | -1.070389328 | 0.21 | 0.382 | 1 | 0.993983905 | -0.161987182 | 0.4 | 0.401 | 1 | 0.993983905 | 0.00274479 |
| Ctnna3 | 0.00138393 | 4.724026538 | 0.076 | 0.006 | 1 | 0.009979004 | 3.432959407 | 0.1 | 0.009 | 1 | 0.009979004 | 0.002765945 |
| Larp7 | 0.001386688 | -0.972414168 | 0.387 | 0.524 | 1 | 0.283140553 | -0.726239188 | 0.3 | 0.451 | 1 | 0.283140553 | 0.002771453 |
| Saysd1 | 0.026501054 | 0.498453507 | 0.487 | 0.353 | 1 | 0.00138717 | 1.847996907 | 0.8 | 0.324 | 1 | 0.026501054 | 0.002772416 |
| B9d2 | 0.00139003 | -1.67305383 | 0.067 | 0.2 | 1 | 0.14387411 | -1.267480311 | 0 | 0.179 | 1 | 0.14387411 | 0.002778129 |
| Nob1 | 0.001390496 | -0.816940971 | 0.345 | 0.529 | 1 | 0.154134051 | -0.830074999 | 0.3 | 0.515 | 1 | 0.154134051 | 0.002779059 |
| Ube2v1 | 0.001391082 | -0.825851266 | 0.395 | 0.559 | 1 | 0.057270808 | -1.030837404 | 0.3 | 0.611 | 1 | 0.057270808 | 0.002780229 |
| Rnf111 | 0.001393136 | -0.663519149 | 0.605 | 0.724 | 1 | 0.410368745 | -0.123333751 | 0.7 | 0.846 | 1 | 0.410368745 | 0.002784332 |
| Mrpl9 | 0.001395241 | -1.117695043 | 0.202 | 0.365 | 1 | 0.057238519 | -1.633129783 | 0.1 | 0.392 | 1 | 0.057238519 | 0.002788536 |
| Prdx5 | 0.001398535 | -0.560137222 | 0.832 | 0.912 | 1 | 0.973169316 | 0.121268421 | 0.8 | 0.898 | 1 | 0.973169316 | 0.002795115 |
| Pcgf2 | 0.001400748 | -1.442358105 | 0.134 | 0.282 | 1 | 0.28312825 | -0.823380346 | 0.2 | 0.34 | 1 | 0.28312825 | 0.002799535 |
| Patl1 | 0.001401885 | -1.218487967 | 0.252 | 0.406 | 1 | 0.612642587 | -0.391469028 | 0.4 | 0.444 | 1 | 0.612642587 | 0.002801805 |
| Rnase4 | 0.001402249 | -0.266913663 | 0.613 | 0.818 | 1 | 0.032459769 | -0.649189634 | 0.6 | 0.901 | 1 | 0.032459769 | 0.002802531 |
| Gm10125 | 0.001403766 | -2.707819249 | 0.042 | 0.159 | 1 | 0.239132907 | -1.248864633 | 0 | 0.123 | 1 | 0.239132907 | 0.002805562 |
| Snrnp48 | 0.001405832 | -0.658832397 | 0.706 | 0.782 | 1 | 0.134700458 | -0.543229631 | 0.6 | 0.787 | 1 | 0.134700458 | 0.002809687 |
| Pcgf3 | 0.001407258 | -1.040641984 | 0.328 | 0.476 | 1 | 0.09598403 | -0.647414009 | 0.3 | 0.627 | 1 | 0.09598403 | 0.002812537 |
| Slc14a1 | 0.001408328 | 3.684498174 | 0.059 | 0 | 1 | 0.0019162 | 5.432959407 | 0.1 | 0.006 | 1 | 0.0019162 | 0.002814672 |
| Rgl1 | 0.001409102 | -0.972444277 | 0.714 | 0.741 | 1 | 0.009941764 | -1.668578619 | 0.5 | 0.772 | 1 | 0.009941764 | 0.002816219 |
| Ifit1 | 0.001409174 | 7.2010737 | 0.059 | 0 | 1 | 0.597844175 | -1.131825212 | 0 | 0.028 | 1 | 0.597844175 | 0.002816362 |
| Fastkd1 | 0.001409199 | -1.987927168 | 0.05 | 0.176 | 1 | 0.175683867 | -1.171902651 | 0 | 0.157 | 1 | 0.175683867 | 0.002816413 |
| Arpc4 | 0.001409904 | -0.608537369 | 0.773 | 0.818 | 1 | 0.27989834 | -0.461049897 | 0.7 | 0.833 | 1 | 0.27989834 | 0.00281782 |
| Pacsin2 | 0.001410337 | 0.22659241 | 0.95 | 0.753 | 1 | 0.47402214 | 0.148050502 | 0.9 | 0.861 | 1 | 0.47402214 | 0.002818684 |
| Rcsd1 | 0.001410758 | -2.589763487 | 0.034 | 0.147 | 1 | 0.275814486 | -1.048167282 | 0.1 | 0.241 | 1 | 0.275814486 | 0.002819526 |
| C1qbp | 0.001416459 | -0.693535023 | 0.605 | 0.7 | 1 | 0.183713741 | -0.657643142 | 0.6 | 0.741 | 1 | 0.183713741 | 0.002830912 |
| Cdh2 | 0.00141776 | -3.350497247 | 0.034 | 0.147 | 1 | 0.29582046 | 0.299103661 | 0.2 | 0.093 | 1 | 0.29582046 | 0.002833509 |
| Rpf1 | 0.001420354 | -1.084464513 | 0.244 | 0.406 | 1 | 0.502614503 | -0.282201817 | 0.3 | 0.407 | 1 | 0.502614503 | 0.00283869 |
| Atxn7 | 0.001421393 | -0.64353012 | 0.63 | 0.782 | 1 | 0.246217 | -0.322640361 | 0.7 | 0.864 | 1 | 0.246217 | 0.002840765 |
| Zscan18 | 0.00143212 | -2.758445322 | 0.017 | 0.118 | 1 | 0.721627425 | 0.135278859 | 0.1 | 0.136 | 1 | 0.721627425 | 0.002862189 |
| Vstm4 | 0.001432176 | -2.758445322 | 0.017 | 0.118 | 1 | 0.315694985 | -0.111361109 | 0 | 0.093 | 1 | 0.315694985 | 0.0028623 |
| Zfp334 | 0.001432213 | -2.600904045 | 0.017 | 0.118 | 1 | 0.139528548 | -1.28585884 | 0 | 0.182 | 1 | 0.139528548 | 0.002862374 |
| Mrto4 | 0.001435073 | -1.443198592 | 0.202 | 0.347 | 1 | 0.40631278 | -0.546862711 | 0.2 | 0.306 | 1 | 0.40631278 | 0.002868086 |
| Zfp516 | 0.001435161 | -0.852442812 | 0.571 | 0.665 | 1 | 0.026051886 | -1.035810076 | 0.5 | 0.846 | 1 | 0.026051886 | 0.002868262 |
| Wnt5a | 0.001435236 | -2.707819249 | 0.042 | 0.159 | 1 | 0.166140862 | -1.339630097 | 0 | 0.164 | 1 | 0.166140862 | 0.002868412 |
| Pip4p1 | 0.001445368 | 0.663950797 | 0.63 | 0.453 | 1 | 0.8151088 | 0.331421381 | 0.4 | 0.386 | 1 | 0.8151088 | 0.002888647 |
| Kcnk6 | 0.001446232 | -2.600904045 | 0.017 | 0.118 | 1 | 0.12829788 | -1.505640048 | 0 | 0.191 | 1 | 0.12829788 | 0.002890372 |

| | | | | | | | | | | | | |
|---|---|---|---|---|---|---|---|---|---|---|---|---|
| Septin4 | 0.001446269 | -2.544320516 | 0.017 | 0.118 | 1 | 0.19182373 | 2.017921908 | 0.2 | 0.086 | 1 | 0.19182373 | 0.002890447 |
| Wwp2 | 0.001448908 | 0.412510686 | 0.975 | 0.9 | 1 | 0.250260632 | 0.563037397 | 1 | 0.966 | 1 | 0.250260632 | 0.002895717 |
| Actb | 0.001453095 | 0.273440648 | 1 | 0.994 | 1 | 0.493332827 | 0.145432766 | 1 | 0.997 | 1 | 0.493332827 | 0.002904079 |
| Slc1a4 | 0.001454432 | -1.159198595 | 0.387 | 0.529 | 1 | 0.532031535 | -0.60005565 | 0.6 | 0.568 | 1 | 0.532031535 | 0.002906748 |
| Nisch | 0.001460738 | -0.501877348 | 0.79 | 0.871 | 1 | 0.568775893 | 0.178935377 | 0.8 | 0.904 | 1 | 0.568775893 | 0.002919342 |
| Pofut2 | 0.001460835 | -0.53359411 | 0.739 | 0.753 | 1 | 0.129207172 | -0.591424751 | 0.5 | 0.738 | 1 | 0.129207172 | 0.002919536 |
| Avpr1a | 0.00146107 | 1.868210127 | 0.269 | 0.129 | 1 | 0.849315046 | 0.695993813 | 0.1 | 0.08 | 1 | 0.849315046 | 0.002920005 |
| Mtfr2 | 0.001461824 | -2.448900951 | 0.034 | 0.147 | 1 | 0.083111983 | 1.625604485 | 0.3 | 0.12 | 1 | 0.083111983 | 0.00292151 |
| Zfp647 | 0.095769202 | -1.485426827 | 0.008 | 0.041 | 1 | 0.001462347 | 3.280956314 | 0.2 | 0.025 | 1 | 0.095769202 | 0.002922556 |
| Pcdhgb1 | 0.1433296 | -2.185866545 | 0.008 | 0.035 | 1 | 0.001462347 | 3.280956314 | 0.2 | 0.025 | 1 | 0.1433296 | 0.002922556 |
| Vstm2l | 0.511005543 | -0.485426827 | 0.008 | 0.018 | 1 | 0.001462347 | 3.280956314 | 0.2 | 0.025 | 1 | 0.511005543 | 0.002922556 |
| Hmox1 | 0.001465161 | 0.797604487 | 0.664 | 0.524 | 1 | 0.028542598 | -1.825475764 | 0.4 | 0.719 | 1 | 0.028542598 | 0.002928175 |
| Rgs19 | 0.001465273 | -1.589763487 | 0.076 | 0.212 | 1 | 0.584520763 | 0.135278859 | 0.1 | 0.164 | 1 | 0.584520763 | 0.0029284 |
| Acap3 | 0.001465273 | -1.589763487 | 0.076 | 0.212 | 1 | 0.882768943 | 0.210566986 | 0.3 | 0.259 | 1 | 0.882768943 | 0.0029284 |
| Pknox2 | 0.001466381 | -2.018461536 | 0.202 | 0.347 | 1 | 0.97409698 | -0.570376305 | 0.4 | 0.38 | 1 | 0.97409698 | 0.002930613 |
| Psmd4 | 0.001467451 | -0.448337508 | 0.798 | 0.853 | 1 | 0.031863649 | -0.955536305 | 0.5 | 0.849 | 1 | 0.031863649 | 0.002932749 |
| Zfp428 | 0.001474248 | -2.544320516 | 0.017 | 0.118 | 1 | 0.42093472 | 0.374065718 | 0 | 0.062 | 1 | 0.42093472 | 0.002946322 |
| Pde11a | 0.001475281 | 0.632754599 | 0.294 | 0.141 | 1 | 0.033767794 | 1.558490289 | 0.3 | 0.096 | 1 | 0.033767794 | 0.002948386 |
| Ndufs1 | 0.001475712 | -0.927431374 | 0.353 | 0.512 | 1 | 0.535379921 | 0.50222207 | 0.6 | 0.596 | 1 | 0.535379921 | 0.002949247 |
| Rab4a | 0.001476885 | 0.915111102 | 0.252 | 0.106 | 1 | 0.363687972 | 0.982297998 | 0.3 | 0.191 | 1 | 0.363687972 | 0.002951588 |
| Klf8 | 0.001477901 | -3.944858446 | 0.008 | 0.1 | 1 | 0.334379228 | -0.654503434 | 0 | 0.086 | 1 | 0.334379228 | 0.002953618 |
| Ptprt | 0.001480993 | -2.664750527 | 0.084 | 0.218 | 1 | 0.1714215 | -2.827568143 | 0 | 0.16 | 1 | 0.1714215 | 0.002959793 |
| Eif4ebp2 | 0.001481291 | 0.411233423 | 0.832 | 0.659 | 1 | 0.722343928 | -0.18232763 | 0.8 | 0.778 | 1 | 0.722343928 | 0.002960388 |
| Stip1 | 0.001482674 | -0.745816512 | 0.454 | 0.582 | 1 | 0.073354556 | -0.984580599 | 0.6 | 0.756 | 1 | 0.073354556 | 0.002963149 |
| Eno2 | 0.001483634 | -1.448900951 | 0.05 | 0.176 | 1 | 0.197803088 | -1.220482831 | 0.1 | 0.275 | 1 | 0.197803088 | 0.002965067 |
| Rab35 | 0.001485318 | -0.903739458 | 0.336 | 0.494 | 1 | 0.022760282 | -1.600463594 | 0.3 | 0.605 | 1 | 0.022760282 | 0.002968429 |
| Mrpl22 | 0.001493797 | -1.269698136 | 0.126 | 0.282 | 1 | 0.981493717 | 0.68163852 | 0.2 | 0.216 | 1 | 0.981493717 | 0.002985363 |
| Spag5 | 0.00149398 | -3.240314329 | 0.008 | 0.1 | 1 | 0.002939478 | 3.017921908 | 0.2 | 0.028 | 1 | 0.002939478 | 0.002985727 |
| Clptm1 | 0.001494141 | -0.691877705 | 0.496 | 0.624 | 1 | 0.289257704 | -0.237578825 | 0.5 | 0.685 | 1 | 0.289257704 | 0.002986049 |
| Zfp142 | 0.001494865 | -1.142539114 | 0.261 | 0.424 | 1 | 0.926761497 | -0.213642159 | 0.5 | 0.451 | 1 | 0.926761497 | 0.002987496 |
| Adcy7 | 0.001499385 | -3.292781749 | 0.017 | 0.118 | 1 | 0.353825447 | -0.596787936 | 0 | 0.08 | 1 | 0.353825447 | 0.002996521 |
| Ndrg3 | 0.001500338 | -0.99389119 | 0.429 | 0.541 | 1 | 0.606045682 | 0.585569823 | 0.6 | 0.602 | 1 | 0.606045682 | 0.002998424 |
| Gtpbp1 | 0.001507315 | -1.586964854 | 0.16 | 0.312 | 1 | 0.19977861 | -0.763437806 | 0.1 | 0.309 | 1 | 0.19977861 | 0.003012358 |
| Pml | 0.001510784 | -3.392317423 | 0.008 | 0.1 | 1 | 0.201730842 | -0.91281543 | 0 | 0.142 | 1 | 0.201730842 | 0.003019285 |
| Ddit4 | 0.001512176 | -0.472006311 | 0.101 | 0.259 | 1 | 0.828188484 | 0.595389872 | 0.2 | 0.275 | 1 | 0.828188484 | 0.003022066 |
| Mex3c | 0.002565065 | -0.753915663 | 0.462 | 0.588 | 1 | 0.001512828 | -1.933945596 | 0.3 | 0.781 | 1 | 0.002565065 | 0.003023367 |
| Uqcrc2 | 0.001518279 | -0.504384985 | 0.773 | 0.818 | 1 | 0.029657109 | -0.841971208 | 0.7 | 0.892 | 1 | 0.029657109 | 0.003034252 |
| Gm56862 | 0.001518776 | -3.129283017 | 0.008 | 0.1 | 1 | 0.266980603 | -0.473931188 | 0 | 0.111 | 1 | 0.266980603 | 0.003035244 |
| Dnajc8 | 0.00151981 | -0.432408445 | 0.723 | 0.829 | 1 | 0.10128275 | -0.680203944 | 0.5 | 0.821 | 1 | 0.10128275 | 0.00303731 |
| Gm57292 | 0.147029286 | -2.485426827 | 0 | 0.018 | 1 | 0.001520803 | 3.017921908 | 0.2 | 0.025 | 1 | 0.147029286 | 0.003039294 |
| Inha | 0.659871503 | 0.514573173 | 0.025 | 0.018 | 1 | 0.001520803 | 3.017921908 | 0.2 | 0.025 | 1 | 0.659871503 | 0.003039294 |
| Ube2h | 0.001521064 | -0.645673915 | 0.874 | 0.924 | 1 | 0.069316194 | -0.783152832 | 0.9 | 0.978 | 1 | 0.069316194 | 0.003039815 |
| Crybg3 | 0.001522515 | 0.628529362 | 0.529 | 0.335 | 1 | 0.176284284 | 0.989725016 | 0.6 | 0.454 | 1 | 0.176284284 | 0.003042711 |
| Copg1 | 0.001524582 | -0.695462042 | 0.748 | 0.741 | 1 | 0.691858895 | -0.189092412 | 0.8 | 0.753 | 1 | 0.691858895 | 0.00304684 |
| Speg | 0.001526725 | -3.008988783 | 0.008 | 0.1 | 1 | 0.659507303 | 1.393431043 | 0.1 | 0.17 | 1 | 0.659507303 | 0.003051118 |
| Zfp472 | 0.001526725 | -3.008988783 | 0.008 | 0.1 | 1 | 0.282110098 | -0.339630097 | 0 | 0.105 | 1 | 0.282110098 | 0.003051118 |
| Eif3e | 0.001528235 | -0.451016796 | 0.874 | 0.941 | 1 | 0.046456604 | -0.733994028 | 0.8 | 0.932 | 1 | 0.046456604 | 0.003054135 |
| Fos | 0.001530327 | -1.25640936 | 0.672 | 0.776 | 1 | 0.489675044 | 1.219681299 | 0.7 | 0.867 | 1 | 0.489675044 | 0.003058312 |
| Bri3bp | 0.001530558 | -1.070389328 | 0.277 | 0.429 | 1 | 0.506346092 | -0.391469028 | 0.3 | 0.373 | 1 | 0.506346092 | 0.003058774 |
| Gcfc2 | 0.001531968 | -2.14839184 | 0.042 | 0.159 | 1 | 0.325023254 | -0.473931188 | 0 | 0.09 | 1 | 0.325023254 | 0.00306159 |
| Ctxn1 | 0.001534523 | -3.070389328 | 0.008 | 0.1 | 1 | 0.433608183 | 0.432959407 | 0 | 0.059 | 1 | 0.433608183 | 0.003066691 |
| Rnf169 | 0.001539774 | -0.937939032 | 0.345 | 0.488 | 1 | 0.767583619 | 0.328090289 | 0.5 | 0.627 | 1 | 0.767583619 | 0.003077177 |
| Hspa1a | 0.001543098 | -1.802216199 | 0.403 | 0.576 | 1 | 0.067069994 | 0.728330787 | 0.3 | 0.614 | 1 | 0.067069994 | 0.003083814 |
| Limd1 | 0.041617116 | -0.45462078 | 0.563 | 0.618 | 1 | 0.001544467 | 1.350382094 | 1 | 0.707 | 1 | 0.041617116 | 0.003086548 |
| Dzip1 | 0.305710662 | -1.333423734 | 0.067 | 0.1 | 1 | 0.001547362 | 2.795529487 | 0.3 | 0.056 | 1 | 0.305710662 | 0.003092331 |
| Osbpl11 | 0.001548748 | -0.915414668 | 0.378 | 0.524 | 1 | 0.983584274 | 0.341811519 | 0.6 | 0.685 | 1 | 0.983584274 | 0.003095098 |
| Rora | 0.001549982 | 0.359556917 | 1 | 0.994 | 1 | 0.08443713 | 0.914256425 | 1 | 0.988 | 1 | 0.08443713 | 0.003097561 |
| Tox4 | 0.00155069 | -0.73405079 | 0.546 | 0.659 | 1 | 0.196889419 | -0.561565463 | 0.7 | 0.747 | 1 | 0.196889419 | 0.003098975 |
| Setd1a | 0.001550761 | -1.177304532 | 0.176 | 0.335 | 1 | 0.449829858 | -0.654503434 | 0.4 | 0.441 | 1 | 0.449829858 | 0.003099117 |
| Arpin | 0.001563295 | -1.521050737 | 0.076 | 0.212 | 1 | 0.25337182 | 1.231225546 | 0.3 | 0.167 | 1 | 0.25337182 | 0.003124146 |
| Gtf2f2 | 0.001564668 | 0.401191373 | 0.899 | 0.718 | 1 | 0.154074251 | 0.571238782 | 0.8 | 0.725 | 1 | 0.154074251 | 0.003126888 |
| Ddhd2 | 0.001566325 | -0.667367205 | 0.454 | 0.6 | 1 | 0.474181175 | -0.304006187 | 0.5 | 0.66 | 1 | 0.474181175 | 0.003130196 |
| Kpna4 | 0.001566362 | -0.589870675 | 0.857 | 0.9 | 1 | 0.037804806 | -0.869603363 | 0.8 | 0.966 | 1 | 0.037804806 | 0.00313027 |
| Fhod3 | 0.001568087 | -4.886306263 | 0.008 | 0.1 | 1 | 0.578877545 | 0.526068812 | 0.1 | 0.056 | 1 | 0.578877545 | 0.003133716 |
| Sap18 | 0.001571102 | -0.576925182 | 0.756 | 0.788 | 1 | 0.004021576 | -1.389345856 | 0.4 | 0.84 | 1 | 0.004021576 | 0.003139736 |
| H2az1 | 0.0015805 | -0.77347544 | 0.849 | 0.871 | 1 | 0.126784534 | -0.246989785 | 0.6 | 0.941 | 1 | 0.126784534 | 0.003158502 |
| Thegl | 0.426866601 | -1.544320516 | 0.034 | 0.053 | 1 | 0.001581274 | 2.602884409 | 0.2 | 0.025 | 1 | 0.426866601 | 0.003160048 |
| Kpna6 | 0.001582932 | -1.286522759 | 0.261 | 0.412 | 1 | 0.88552748 | 0.217022008 | 0.4 | 0.454 | 1 | 0.88552748 | 0.003163359 |
| Qsox1 | 0.005962799 | -0.802693545 | 0.555 | 0.641 | 1 | 0.001586405 | -2.207044457 | 0.3 | 0.772 | 1 | 0.005962799 | 0.003170294 |
| Ppp2r5d | 0.00158746 | -1.378511623 | 0.101 | 0.247 | 1 | 0.489318247 | 0.678071905 | 0.3 | 0.201 | 1 | 0.489318247 | 0.0031724 |
| Snx18 | 0.075110215 | -0.178289139 | 0.706 | 0.759 | 1 | 0.001588706 | -1.852442812 | 0.6 | 0.858 | 1 | 0.075110215 | 0.003174887 |
| Tmco4 | 0.001590041 | 0.664320292 | 0.538 | 0.347 | 1 | 0.013220354 | 1.256636635 | 0.9 | 0.472 | 1 | 0.013220354 | 0.003177553 |
| Cdc42ep4 | 0.001591254 | 0.632224761 | 0.706 | 0.482 | 1 | 0.778207666 | 0.173826439 | 0.8 | 0.657 | 1 | 0.778207666 | 0.003179977 |
| Nup188 | 0.001591937 | -1.307428525 | 0.193 | 0.353 | 1 | 0.403060762 | -0.536666944 | 0.3 | 0.423 | 1 | 0.403060762 | 0.003181341 |
| B3galnt2 | 0.001594611 | -1.084064265 | 0.21 | 0.376 | 1 | 0.622678338 | -0.304006187 | 0.3 | 0.358 | 1 | 0.622678338 | 0.003186679 |
| Mdm4 | 0.001596871 | -0.733961663 | 0.63 | 0.729 | 1 | 0.042974185 | -0.878915023 | 0.7 | 0.821 | 1 | 0.042974185 | 0.003191193 |
| Ube2e2 | 0.001598557 | -0.691618437 | 0.79 | 0.853 | 1 | 0.286295722 | -0.823873659 | 1 | 0.849 | 1 | 0.286295722 | 0.003194558 |
| Kat5 | 0.00160386 | -1.11345805 | 0.134 | 0.294 | 1 | 0.986857089 | 0.362570079 | 0.3 | 0.306 | 1 | 0.986857089 | 0.003205148 |
| Phkb | 0.001606112 | -1.001441974 | 0.252 | 0.412 | 1 | 0.471844899 | 0.630191755 | 0.6 | 0.506 | 1 | 0.471844899 | 0.003209644 |
| Bambi | 0.001608981 | 0.484515939 | 0.79 | 0.576 | 1 | 0.520316053 | 0.256370676 | 0.5 | 0.491 | 1 | 0.520316053 | 0.003215372 |
| Jazf1 | 0.001609151 | -1.96051171 | 0.143 | 0.282 | 1 | 0.553638954 | -0.657329478 | 0.6 | 0.38 | 1 | 0.553638954 | 0.003215713 |
| Kif23 | 0.001614293 | -3.392317423 | 0.008 | 0.1 | 1 | 0.154873683 | 1.930459067 | 0.1 | 0.025 | 1 | 0.154873683 | 0.00322598 |
| Slc25a32 | 0.001614379 | -1.469659511 | 0.134 | 0.288 | 1 | 0.051639626 | 0.878370556 | 0.6 | 0.278 | 1 | 0.051639626 | 0.003226152 |
| Usp1 | 0.001614388 | -0.896059147 | 0.395 | 0.535 | 1 | 0.338102787 | -0.844198817 | 0.5 | 0.537 | 1 | 0.338102787 | 0.003226169 |
| Nt5c3 | 0.001617209 | -0.788976393 | 0.37 | 0.559 | 1 | 0.05195154 | -1.251983975 | 0.4 | 0.639 | 1 | 0.05195154 | 0.003231803 |
| Iffo1 | 0.001617734 | -2.201633861 | 0.05 | 0.171 | 1 | 0.860133806 | 0.695993813 | 0.1 | 0.127 | 1 | 0.860133806 | 0.003232851 |
| Cald1 | 0.010739224 | -0.767967053 | 0.975 | 0.965 | 1 | 0.001618509 | -0.747423396 | 0.5 | 0.944 | 1 | 0.010739224 | 0.003234398 |
| Snupn | 0.086096876 | 0.790207615 | 0.168 | 0.1 | 1 | 0.001621915 | 2.245332404 | 0.4 | 0.093 | 1 | 0.086096876 | 0.003241199 |
| Arsb | 0.001628951 | -1.341036918 | 0.16 | 0.318 | 1 | 0.935064088 | -1.381249186 | 0.4 | 0.336 | 1 | 0.935064088 | 0.003255249 |
| Vps35 | 0.001631374 | -0.56250305 | 0.773 | 0.829 | 1 | 0.01335657 | -0.954063716 | 0.6 | 0.883 | 1 | 0.01335657 | 0.003260086 |
| Btg3 | 0.001633206 | -0.586355736 | 0.748 | 0.818 | 1 | 0.372811401 | -0.540753901 | 0.8 | 0.796 | 1 | 0.372811401 | 0.003263745 |
| Pigt | 0.174892633 | 0.228268988 | 0.504 | 0.424 | 1 | 0.001634898 | 1.566225938 | 0.8 | 0.42 | 1 | 0.174892633 | 0.003267124 |
| Tjap1 | 0.001635053 | -1.182034684 | 0.227 | 0.382 | 1 | 0.714346762 | -0.381249186 | 0.4 | 0.404 | 1 | 0.714346762 | 0.003267432 |
| Impact | 0.001635294 | -0.696137443 | 0.571 | 0.676 | 1 | 0.795208346 | 0.165805535 | 0.7 | 0.673 | 1 | 0.795208346 | 0.003267914 |
| Cda | 0.001643214 | 0.991011217 | 0.185 | 0.065 | 1 | 0.008171726 | 2.432959407 | 0.2 | 0.034 | 1 | 0.008171726 | 0.003283728 |
| Gm15336 | 0.929388469 | 0.514573173 | 0.025 | 0.024 | 1 | 0.001643818 | 2.515421567 | 0.2 | 0.025 | 1 | 0.929388469 | 0.003284934 |
| Gm16287 | 0.244853912 | -0.707819249 | 0.034 | 0.065 | 1 | 0.001643834 | 2.902444691 | 0.2 | 0.025 | 1 | 0.244853912 | 0.003284967 |
| H3f3b | 0.001645261 | 0.312446883 | 1 | 0.994 | 1 | 0.024230534 | -0.605868659 | 1 | 1 | 1 | 0.024230534 | 0.003287815 |

| | | | | | | | | | | | | |
|---|---|---|---|---|---|---|---|---|---|---|---|---|
| Trp53i11 | 0.001645973 | 1.421463768 | 0.168 | 0.053 | 1 | 0.385739706 | -0.709998547 | 0 | 0.071 | 1 | 0.385739706 | 0.003289237 |
| Evc2 | 0.001646287 | -1.070389328 | 0.176 | 0.341 | 1 | 0.150155848 | 0.732519689 | 0.6 | 0.349 | 1 | 0.150155848 | 0.003289864 |
| Il34 | 0.001647854 | -2.372952098 | 0.067 | 0.194 | 1 | 0.238589331 | 0.510127268 | 0.3 | 0.148 | 1 | 0.238589331 | 0.003292993 |
| Arf5 | 0.001652127 | -0.335613665 | 0.941 | 0.971 | 1 | 0.264126007 | -0.289676057 | 0.9 | 0.978 | 1 | 0.264126007 | 0.003301524 |
| Usp54 | 0.00165503 | -1.058005603 | 0.227 | 0.394 | 1 | 0.027255457 | 0.669193754 | 0.9 | 0.472 | 1 | 0.027255457 | 0.003307321 |
| Casd1 | 0.001655221 | -1.219252714 | 0.311 | 0.453 | 1 | 0.363104218 | -0.693572999 | 0.3 | 0.432 | 1 | 0.363104218 | 0.003307702 |
| Ppan | 0.001655985 | -1.060120992 | 0.303 | 0.459 | 1 | 0.199657318 | -0.625934282 | 0.2 | 0.42 | 1 | 0.199657318 | 0.003309228 |
| Sap30 | 0.001661302 | -1.780882711 | 0.059 | 0.188 | 1 | 0.021097199 | -2.441509711 | 0 | 0.361 | 1 | 0.021097199 | 0.003319844 |
| Htatsf1 | 0.001674251 | -0.723831566 | 0.605 | 0.688 | 1 | 0.148066102 | -0.51699991 | 0.5 | 0.725 | 1 | 0.148066102 | 0.0033457 |
| Eif2ak1 | 0.001676672 | -0.807354922 | 0.286 | 0.453 | 1 | 0.008383171 | -2.8829449 | 0 | 0.435 | 1 | 0.008383171 | 0.003350532 |
| Jpt2 | 0.001679083 | -1.217230716 | 0.193 | 0.353 | 1 | 0.708459393 | 0.695993813 | 0.3 | 0.293 | 1 | 0.708459393 | 0.003355347 |
| Gpx7 | 0.001683157 | -0.682318876 | 0.605 | 0.7 | 1 | 0.44646841 | -0.460125389 | 0.5 | 0.552 | 1 | 0.44646841 | 0.003363481 |
| Agr2 | 0.001685498 | 3.052007303 | 0.16 | 0.053 | 1 | 0.030329568 | 1.388565288 | 0.4 | 0.145 | 1 | 0.030329568 | 0.003368155 |
| Rbms2 | 0.001685631 | -1.109278341 | 0.412 | 0.529 | 1 | 0.715164952 | 0.5169378 | 0.5 | 0.481 | 1 | 0.715164952 | 0.00336842 |
| Gm11638 | 0.001686461 | -2.424026283 | 0.008 | 0.1 | 1 | 0.36414034 | -0.441509711 | 0 | 0.077 | 1 | 0.36414034 | 0.003370077 |
| Lman1 | 0.001688213 | -0.301150613 | 0.941 | 0.929 | 1 | 0.349544007 | 0.199567987 | 0.9 | 0.929 | 1 | 0.349544007 | 0.003373577 |
| Plxna4 | 0.001688956 | -1.513995979 | 0.059 | 0.188 | 1 | 0.652314887 | -0.907550113 | 0.4 | 0.423 | 1 | 0.652314887 | 0.003375059 |
| Sh3rf3 | 0.001690819 | -1.770829046 | 0.151 | 0.3 | 1 | 0.054896073 | -2.846264237 | 0.1 | 0.383 | 1 | 0.054896073 | 0.003378779 |
| Adgrl1 | 0.00169128 | -1.029747343 | 0.361 | 0.506 | 1 | 0.112074542 | -1.17978625 | 0.4 | 0.559 | 1 | 0.112074542 | 0.0033797 |
| Pfdn2 | 0.001691748 | -0.55359833 | 0.782 | 0.794 | 1 | 0.08639234 | -0.676101068 | 0.8 | 0.849 | 1 | 0.08639234 | 0.003380634 |
| Fam199x | 0.00169548 | -1.385891154 | 0.092 | 0.235 | 1 | 0.481528738 | -0.339630097 | 0.2 | 0.312 | 1 | 0.481528738 | 0.003388085 |
| Anapc13 | 0.001703242 | 0.4792401 | 0.689 | 0.465 | 1 | 0.788489254 | 0.558490289 | 0.4 | 0.358 | 1 | 0.788489254 | 0.003403583 |
| Fam167a | 0.001711867 | -1.907958863 | 0.21 | 0.347 | 1 | 0.171030379 | -1.840059087 | 0 | 0.16 | 1 | 0.171030379 | 0.003420804 |
| Zc3h11a | 0.001716028 | -0.923999841 | 0.277 | 0.429 | 1 | 0.421132486 | -0.465893869 | 0.3 | 0.401 | 1 | 0.421132486 | 0.003429111 |
| Adamts5 | 0.001716927 | -2.770829046 | 0.025 | 0.129 | 1 | 0.260094091 | -1.408342847 | 0 | 0.114 | 1 | 0.260094091 | 0.003430906 |
| Mafg | 0.001721264 | -0.617995947 | 0.798 | 0.8 | 1 | 0.040527279 | -0.831743819 | 0.6 | 0.941 | 1 | 0.040527279 | 0.003439565 |
| Rpl8 | 0.006585628 | -0.254052276 | 0.975 | 0.971 | 1 | 0.001723459 | -0.963462414 | 1 | 1 | 1 | 0.006585628 | 0.003443948 |
| Gm11508 | 0.001724005 | -1.786596362 | 0.092 | 0.229 | 1 | 0.277653948 | -1.101019165 | 0.1 | 0.241 | 1 | 0.277653948 | 0.003445037 |
| Esf1 | 0.001732584 | -0.722160272 | 0.647 | 0.747 | 1 | 0.8309914 | -0.161987182 | 0.6 | 0.704 | 1 | 0.8309914 | 0.003462166 |
| Nsun3 | 0.130089693 | -0.847996907 | 0.034 | 0.076 | 1 | 0.001747194 | 2.744903414 | 0.3 | 0.056 | 1 | 0.130089693 | 0.003491335 |
| Iffo2 | 0.001753214 | -0.589499281 | 0.412 | 0.606 | 1 | 0.522842715 | 1.167065347 | 0.6 | 0.627 | 1 | 0.522842715 | 0.003503354 |
| Mthfr | 0.001756351 | -1.309855263 | 0.16 | 0.312 | 1 | 0.973452266 | 0.190102883 | 0.3 | 0.293 | 1 | 0.973452266 | 0.003509617 |
| Cuedc1 | 0.001759264 | -0.898365861 | 0.496 | 0.606 | 1 | 0.07683701 | -0.940498988 | 0.6 | 0.793 | 1 | 0.07683701 | 0.003515433 |
| Mapre2 | 0.053631578 | -0.47618862 | 0.748 | 0.759 | 1 | 0.001765552 | -2.291933355 | 0.4 | 0.787 | 1 | 0.053631578 | 0.003527987 |
| Nme7 | 0.001766349 | -1.23388806 | 0.151 | 0.306 | 1 | 0.159354746 | -1.625934282 | 0.1 | 0.296 | 1 | 0.159354746 | 0.003529578 |
| Stard7 | 0.001770791 | -1.226893814 | 0.202 | 0.359 | 1 | 0.157824537 | -1.217294554 | 0.2 | 0.398 | 1 | 0.157824537 | 0.003538446 |
| Dgkh | 0.001776928 | -0.482098809 | 0.538 | 0.676 | 1 | 0.604011736 | 0.897292203 | 0.6 | 0.623 | 1 | 0.604011736 | 0.003550699 |
| Bbln | 0.001777523 | 0.817965316 | 0.546 | 0.412 | 1 | 0.024633142 | 0.982297998 | 0.7 | 0.34 | 1 | 0.024633142 | 0.003551886 |
| Xpo4 | 0.001787726 | -1.026573096 | 0.395 | 0.541 | 1 | 0.163613488 | -1.127755547 | 0.3 | 0.491 | 1 | 0.163613488 | 0.003572257 |
| Abcb10 | 0.001789229 | -1.87774425 | 0.059 | 0.182 | 1 | 0.10733274 | -1.521236903 | 0 | 0.21 | 1 | 0.10733274 | 0.003575258 |
| Efna4 | 0.001791801 | -2.485426827 | 0.025 | 0.129 | 1 | 0.829966406 | 1.159940913 | 0.1 | 0.08 | 1 | 0.829966406 | 0.003580392 |
| Arf1 | 0.030599718 | -0.224823248 | 0.992 | 0.982 | 1 | 0.001793531 | -1.031606363 | 0.9 | 0.997 | 1 | 0.030599718 | 0.003583844 |
| Pttg1 | 0.001793823 | 0.783062009 | 0.487 | 0.312 | 1 | 0.718553964 | -0.127755547 | 0.3 | 0.352 | 1 | 0.718553964 | 0.003584428 |
| Cep250 | 0.001793849 | -1.457412451 | 0.067 | 0.2 | 1 | 0.858673002 | 0.542188477 | 0.2 | 0.182 | 1 | 0.858673002 | 0.003584479 |
| Virma | 0.001795105 | -0.712062472 | 0.529 | 0.641 | 1 | 0.006753963 | -1.668578619 | 0.3 | 0.716 | 1 | 0.006753963 | 0.003586987 |
| Tom1l2 | 0.025802849 | 0.138662005 | 0.824 | 0.735 | 1 | 0.001808572 | 1.082945799 | 1 | 0.889 | 1 | 0.025802849 | 0.003613873 |
| Phc1 | 0.001809647 | -1.831201664 | 0.084 | 0.218 | 1 | 0.867268655 | 0.197742946 | 0.3 | 0.259 | 1 | 0.867268655 | 0.003616019 |
| Med25 | 0.001811383 | -1.02229504 | 0.412 | 0.524 | 1 | 0.018206477 | -1.600463594 | 0.2 | 0.593 | 1 | 0.018206477 | 0.003619486 |
| Mettl13 | 0.001817376 | -2.485426827 | 0.025 | 0.129 | 1 | 0.916271849 | 1.345496566 | 0.1 | 0.123 | 1 | 0.916271849 | 0.003631449 |
| Cisd1 | 0.001819096 | 0.651207865 | 0.672 | 0.5 | 1 | 0.071832606 | -0.80480824 | 0.2 | 0.531 | 1 | 0.071832606 | 0.003634882 |
| Dusp11 | 0.001819816 | -0.51249507 | 0.84 | 0.865 | 1 | 0.037212909 | -0.762056575 | 0.9 | 0.929 | 1 | 0.037212909 | 0.00363632 |
| Ormdl3 | 0.001827524 | -0.638583615 | 0.672 | 0.718 | 1 | 0.243311656 | -0.636388639 | 0.7 | 0.741 | 1 | 0.243311656 | 0.003651707 |
| Mtor | 0.001836067 | -1.235448574 | 0.235 | 0.388 | 1 | 0.676036613 | -0.171902651 | 0.3 | 0.377 | 1 | 0.676036613 | 0.003668764 |
| Bag4 | 0.001846535 | -1.642968104 | 0.084 | 0.218 | 1 | 0.881557161 | 0.374065718 | 0.2 | 0.173 | 1 | 0.881557161 | 0.003689661 |
| Gm16310 | 0.009995534 | -2.021479727 | 0.034 | 0.118 | 1 | 0.001848381 | 1.94753258 | 0.4 | 0.09 | 1 | 0.009995534 | 0.003693345 |
| Lncpint | 0.001852342 | 0.256564911 | 0.95 | 0.924 | 1 | 0.082729438 | 0.378303843 | 1 | 0.957 | 1 | 0.082729438 | 0.003701253 |
| Cpped1 | 0.001861385 | -1.193246076 | 0.092 | 0.235 | 1 | 0.37311131 | -0.668578619 | 0.1 | 0.21 | 1 | 0.37311131 | 0.003719305 |
| Rnf7 | 0.193568402 | -0.227358674 | 0.773 | 0.741 | 1 | 0.001861435 | -1.592575685 | 0.3 | 0.833 | 1 | 0.193568402 | 0.003719406 |
| Fam161a | 0.001868308 | -2.292781749 | 0.025 | 0.129 | 1 | 0.077488269 | 1.159940913 | 0.3 | 0.108 | 1 | 0.077488269 | 0.003733125 |
| Ap1ar | 0.001869882 | -1.216610069 | 0.176 | 0.335 | 1 | 0.364585578 | -0.220482831 | 0.2 | 0.361 | 1 | 0.364585578 | 0.003736269 |
| Srpk2 | 0.0018716 | 0.463086663 | 0.908 | 0.871 | 1 | 0.378536863 | 0.239479678 | 1 | 0.923 | 1 | 0.378536863 | 0.003739697 |
| Ap4e1 | 0.001871729 | -1.605721061 | 0.143 | 0.288 | 1 | 0.012193445 | 1.22650853 | 0.7 | 0.321 | 1 | 0.012193445 | 0.003739956 |
| Fanci | 0.342489739 | 0.292180751 | 0.05 | 0.082 | 1 | 0.001872539 | 3.956521363 | 0.3 | 0.062 | 1 | 0.342489739 | 0.003741571 |
| Farp2 | 0.001874518 | -0.990218979 | 0.353 | 0.512 | 1 | 0.670246826 | -0.345033455 | 0.6 | 0.623 | 1 | 0.670246826 | 0.003745523 |
| Foxp1 | 0.001880028 | -0.524955191 | 0.983 | 0.971 | 1 | 0.657038351 | -0.105205428 | 1 | 0.994 | 1 | 0.657038351 | 0.003756522 |
| Jcad | 0.001880804 | -1.025995209 | 0.126 | 0.282 | 1 | 0.051664508 | -2.473931188 | 0 | 0.284 | 1 | 0.051664508 | 0.00375807 |
| Zfp386 | 0.001882339 | -1.047305715 | 0.143 | 0.3 | 1 | 0.699336041 | -0.152003093 | 0.2 | 0.241 | 1 | 0.699336041 | 0.003761135 |
| Ndp | 0.001888667 | -2.015941544 | 0.05 | 0.171 | 1 | 0.232322773 | -0.959358016 | 0 | 0.127 | 1 | 0.232322773 | 0.003773766 |
| Trabd2b | 0.001890162 | -0.685011802 | 0.773 | 0.759 | 1 | 0.801370171 | -0.57832896 | 1 | 0.892 | 1 | 0.801370171 | 0.003776752 |
| Lmln | 0.001890192 | -2.455053178 | 0.025 | 0.129 | 1 | 0.964594122 | 0.263034406 | 0.1 | 0.102 | 1 | 0.964594122 | 0.00377681 |
| Ppdpf | 0.001891376 | 0.539664154 | 0.714 | 0.541 | 1 | 0.082827361 | 1.029237221 | 0.7 | 0.5 | 1 | 0.082827361 | 0.003779174 |
| Insig2 | 0.001894036 | 0.681223042 | 0.655 | 0.465 | 1 | 0.317253414 | 0.514457521 | 0.7 | 0.66 | 1 | 0.317253414 | 0.003784485 |
| Cfap298 | 0.001896776 | -1.263034406 | 0.134 | 0.288 | 1 | 0.314081684 | -0.789433014 | 0.1 | 0.228 | 1 | 0.314081684 | 0.003789953 |
| Rarres1 | 0.001897561 | 1.606495662 | 0.311 | 0.171 | 1 | 0.830124082 | 0.50836681 | 0.2 | 0.238 | 1 | 0.830124082 | 0.00379152 |
| Taf15 | 0.001898895 | -0.59893852 | 0.714 | 0.835 | 1 | 0.63792178 | -0.12891948 | 0.7 | 0.812 | 1 | 0.63792178 | 0.003794185 |
| Vps26c | 0.001899839 | -1.610957709 | 0.084 | 0.218 | 1 | 0.227225526 | -0.91281543 | 0.1 | 0.265 | 1 | 0.227225526 | 0.003796069 |
| Ifrd2 | 0.001901406 | -1.282933963 | 0.168 | 0.318 | 1 | 0.607593678 | 0.653349476 | 0.3 | 0.231 | 1 | 0.607593678 | 0.003799196 |
| mt-Nd1 | 0.817860835 | -0.16952252 | 1 | 1 | 1 | 0.001903529 | 0.576187837 | 1 | 1 | 1 | 0.817860835 | 0.003803435 |
| Dnaja3 | 0.001907307 | -1.01267383 | 0.252 | 0.429 | 1 | 0.664621816 | 0.332695284 | 0.4 | 0.454 | 1 | 0.664621816 | 0.003810976 |
| Mrps34 | 0.001907381 | -0.987927168 | 0.252 | 0.412 | 1 | 0.13104811 | -1.121629444 | 0.1 | 0.321 | 1 | 0.13104811 | 0.003811124 |
| Jund | 0.026658024 | -0.313670098 | 1 | 0.976 | 1 | 0.001907853 | -1.278025005 | 1 | 0.994 | 1 | 0.026658024 | 0.003812066 |
| Ints3 | 0.001914664 | -1.392317423 | 0.109 | 0.253 | 1 | 0.295524257 | 0.744903414 | 0.4 | 0.247 | 1 | 0.295524257 | 0.003825663 |
| Slc39a2 | 0.075174255 | 2.836501268 | 0.034 | 0.006 | 1 | 0.0019162 | 5.017921908 | 0.1 | 0.006 | 1 | 0.075174255 | 0.003828727 |
| Tcf15 | 0.091361251 | 2.099535674 | 0.017 | 0 | 1 | 0.0019162 | 5.754887502 | 0.1 | 0.006 | 1 | 0.091361251 | 0.003828727 |
| Dusp2 | 0.09262405 | 0.112474729 | 0.025 | 0.071 | 1 | 0.0019162 | 5.432959407 | 0.1 | 0.006 | 1 | 0.09262405 | 0.003828727 |
| Gm9895 | 0.171447336 | 0.929610672 | 0.025 | 0.006 | 1 | 0.0019162 | 5.017921908 | 0.1 | 0.006 | 1 | 0.171447336 | 0.003828727 |
| Gm36201 | 0.171451236 | 0.836501268 | 0.025 | 0.006 | 1 | 0.0019162 | 5.017921908 | 0.1 | 0.006 | 1 | 0.171451236 | 0.003828727 |
| Ankfn1 | 0.338976313 | -0.485426827 | 0.008 | 0.024 | 1 | 0.0019162 | 5.017921908 | 0.1 | 0.006 | 1 | 0.338976313 | 0.003828727 |
| St8sia5 | 0.513278666 | -0.900464326 | 0.008 | 0.018 | 1 | 0.0019162 | 5.017921908 | 0.1 | 0.006 | 1 | 0.513278666 | 0.003828727 |
| Slc30a10 | 0.51784324 | 0.192645078 | 0.008 | 0.018 | 1 | 0.0019162 | 5.754887502 | 0.1 | 0.006 | 1 | 0.51784324 | 0.003828727 |
| Trim38 | 0.561605963 | 0.362570079 | 0.042 | 0.029 | 1 | 0.0019162 | 5.017921908 | 0.1 | 0.006 | 1 | 0.561605963 | 0.003828727 |
| Gm35315 | 0.82670638 | -0.292781749 | 0.025 | 0.029 | 1 | 0.0019162 | 5.432959407 | 0.1 | 0.006 | 1 | 0.82670638 | 0.003828727 |
| BB557941 | 0.842422009 | 0.251538767 | 0.025 | 0.029 | 1 | 0.0019162 | 5.017921908 | 0.1 | 0.006 | 1 | 0.842422009 | 0.003828727 |
| C330002G04Rik | 0.00718634 | 4.099535674 | 0.042 | 0 | 1 | 0.00191626 | 5.339850003 | 0.1 | 0.006 | 1 | 0.00718634 | 0.003828848 |
| 4930471E19Rik | 0.202434389 | 2.230780207 | 0.067 | 0.035 | 1 | 0.00191626 | 5.602884409 | 0.1 | 0.006 | 1 | 0.202434389 | 0.003828848 |
| Gm41335 | 0.508731273 | -0.807354922 | 0.008 | 0.018 | 1 | 0.00191626 | 6.718361626 | 0.1 | 0.006 | 1 | 0.508731273 | 0.003828848 |
| Blvra | 0.001916275 | 0.763600721 | 0.462 | 0.282 | 1 | 0.513167293 | 0.558490289 | 0.4 | 0.29 | 1 | 0.513167293 | 0.003828879 |
| Phyh | 0.001920098 | 0.531316437 | 0.773 | 0.606 | 1 | 0.204750898 | 0.610074787 | 0.8 | 0.657 | 1 | 0.204750898 | 0.003836509 |

| | | | | | | | | | | | | |
|---|---|---|---|---|---|---|---|---|---|---|---|---|
| Pde1c | 0.001926251 | -0.124650922 | 0.613 | 0.359 | 1 | 0.059025796 | -3.186649236 | 0 | 0.272 | 1 | 0.059025796 | 0.003848791 |
| Ifi27 | 0.001927623 | 0.446468727 | 0.95 | 0.888 | 1 | 0.13572188 | -0.719390446 | 0.9 | 0.898 | 1 | 0.13572188 | 0.00385153 |
| Alg14 | 0.001930089 | 0.407657969 | 0.798 | 0.571 | 1 | 0.021836402 | 1.684938625 | 0.8 | 0.472 | 1 | 0.021836402 | 0.003856452 |
| Gm47283 | 0.001940606 | -0.727267011 | 0.882 | 0.947 | 1 | 0.090647037 | -0.880580902 | 0.9 | 0.923 | 1 | 0.090647037 | 0.003877445 |
| Zfp251 | 0.001941496 | -1.770829046 | 0.059 | 0.182 | 1 | 0.590451496 | -0.391469028 | 0.1 | 0.157 | 1 | 0.590451496 | 0.003879223 |
| Zfp280c | 0.001942644 | -1.015941544 | 0.303 | 0.459 | 1 | 0.692048224 | 0.490184276 | 0.5 | 0.485 | 1 | 0.692048224 | 0.003881514 |
| Golga2 | 0.001951513 | -0.625474787 | 0.571 | 0.688 | 1 | 0.418911972 | -0.401616984 | 0.6 | 0.66 | 1 | 0.418911972 | 0.003899218 |
| Inpp5b | 0.001952781 | -1.870090677 | 0.067 | 0.194 | 1 | 0.383212361 | -0.581990934 | 0.1 | 0.207 | 1 | 0.383212361 | 0.003901748 |
| Polk | 0.001967412 | -1.335531397 | 0.202 | 0.359 | 1 | 0.919539822 | 0.227844977 | 0.3 | 0.281 | 1 | 0.919539822 | 0.003930954 |
| Pptc7 | 0.001969812 | -1.10433666 | 0.218 | 0.382 | 1 | 0.147984474 | -0.394295072 | 0.3 | 0.664 | 1 | 0.147984474 | 0.003935745 |
| Gpalpp1 | 0.001974297 | -1.023083613 | 0.21 | 0.371 | 1 | 0.660772729 | 0.290001453 | 0.4 | 0.318 | 1 | 0.660772729 | 0.003944696 |
| Eif4a1 | 0.001974866 | -0.357644321 | 0.983 | 0.982 | 1 | 0.002006299 | -1.09818572 | 0.8 | 1 | 1 | 0.002006299 | 0.003945832 |
| Cox17 | 0.001978034 | 0.542770065 | 0.689 | 0.512 | 1 | 0.274401603 | -0.608517229 | 0.4 | 0.519 | 1 | 0.274401603 | 0.003952155 |
| Cisd2 | 0.00197888 | -0.440007702 | 0.824 | 0.847 | 1 | 0.502390458 | -0.176131364 | 0.6 | 0.904 | 1 | 0.502390458 | 0.003953844 |
| Tmbim7 | 0.370184089 | 1.099535674 | 0.017 | 0.006 | 1 | 0.001983182 | 4.532495081 | 0.1 | 0.006 | 1 | 0.370184089 | 0.00396243 |
| Bora | 0.001990118 | -2.163498732 | 0.025 | 0.129 | 1 | 0.474877085 | 0.847996907 | 0 | 0.049 | 1 | 0.474877085 | 0.003976275 |
| 4930558N01Rik | 0.028438641 | -1.944858446 | 0.025 | 0.088 | 1 | 0.001991835 | 2.494359952 | 0.3 | 0.056 | 1 | 0.028438641 | 0.003979702 |
| Rlbp1 | 0.00936653 | 1.099535674 | 0.168 | 0.071 | 1 | 0.001991844 | 2.558490289 | 0.3 | 0.056 | 1 | 0.00936653 | 0.003979721 |
| Smg9 | 0.001994316 | -1.733354341 | 0.059 | 0.182 | 1 | 0.495314046 | -0.304006187 | 0.1 | 0.179 | 1 | 0.495314046 | 0.003984654 |
| Rhobtb1 | 0.001997586 | -1.655351829 | 0.042 | 0.159 | 1 | 0.255685614 | 0.821524695 | 0.2 | 0.086 | 1 | 0.255685614 | 0.003991182 |
| Aimp2 | 0.001997727 | -1.255944981 | 0.134 | 0.282 | 1 | 0.902278685 | 0.197742946 | 0.3 | 0.265 | 1 | 0.902278685 | 0.003991463 |
| Cdk7 | 0.001999189 | -0.859822342 | 0.563 | 0.629 | 1 | 0.220127141 | 0.320405585 | 0.9 | 0.679 | 1 | 0.220127141 | 0.003994381 |
| Rapgef4 | 0.00200335 | 0.988504361 | 0.286 | 0.135 | 1 | 0.157621386 | 1.821524695 | 0.3 | 0.151 | 1 | 0.157621386 | 0.004002686 |
| Cenpp | 0.002011145 | -2.379717386 | 0.143 | 0.288 | 1 | 0.084664439 | -1.827568143 | 0 | 0.235 | 1 | 0.084664439 | 0.004018245 |
| Cdc42ep3 | 0.021289815 | -0.627686646 | 0.782 | 0.806 | 1 | 0.002017719 | -1.81074352 | 0.7 | 0.907 | 1 | 0.021289815 | 0.004031367 |
| Adgrl2 | 0.328602426 | -0.292781749 | 0.134 | 0.176 | 1 | 0.002018336 | 1.847996907 | 0.6 | 0.238 | 1 | 0.328602426 | 0.004032598 |
| Tomm70a | 0.002020815 | -0.555816155 | 0.613 | 0.747 | 1 | 0.372591793 | -0.374395515 | 0.6 | 0.762 | 1 | 0.372591793 | 0.004037546 |
| Gm26664 | 0.619390946 | -0.807354922 | 0.025 | 0.035 | 1 | 0.002022946 | 4.14345279 | 0.2 | 0.028 | 1 | 0.619390946 | 0.0040418 |
| mt-Nd5 | 0.105142787 | -0.20639424 | 1 | 1 | 1 | 0.002026937 | 0.787941783 | 1 | 1 | 1 | 0.105142787 | 0.004049766 |
| Heatr1 | 0.002030481 | -1.070389328 | 0.185 | 0.347 | 1 | 0.247131966 | 0.510127268 | 0.5 | 0.296 | 1 | 0.247131966 | 0.004056839 |
| Zxdc | 0.002033237 | -1.269698136 | 0.168 | 0.324 | 1 | 0.073301782 | -1.297982399 | 0.2 | 0.469 | 1 | 0.073301782 | 0.00406234 |
| Map9 | 0.002034208 | -1.67098248 | 0.244 | 0.388 | 1 | 0.881463246 | -0.463204782 | 0.4 | 0.395 | 1 | 0.881463246 | 0.004064279 |
| Sh3glb1 | 0.002035037 | -0.288983515 | 0.975 | 0.988 | 1 | 0.041637043 | -0.610109315 | 1 | 0.991 | 1 | 0.041637043 | 0.004065932 |
| Lrig2 | 0.002036146 | -1.125251263 | 0.269 | 0.418 | 1 | 0.073678736 | -0.687134438 | 0.2 | 0.531 | 1 | 0.073678736 | 0.004068146 |
| Phykpl | 0.033793469 | -0.648925559 | 0.16 | 0.265 | 1 | 0.002046009 | 2.002971567 | 0.6 | 0.238 | 1 | 0.033793469 | 0.004087832 |
| Csk | 0.002046264 | -1.510961919 | 0.109 | 0.247 | 1 | 0.474533034 | -0.441509711 | 0.2 | 0.29 | 1 | 0.474533034 | 0.00408834 |
| Adhfe1 | 0.002048615 | 0.948975997 | 0.286 | 0.135 | 1 | 0.040451025 | 1.732519689 | 0.3 | 0.096 | 1 | 0.040451025 | 0.004093033 |
| Ablim2 | 0.003112834 | 3.684498174 | 0.067 | 0.006 | 1 | 0.002052242 | 4.432959407 | 0.1 | 0.006 | 1 | 0.003112834 | 0.004100273 |
| Lrrc3 | 0.025470389 | -3.070389328 | 0 | 0.041 | 1 | 0.002052242 | 4.432959407 | 0.1 | 0.006 | 1 | 0.025470389 | 0.004100273 |
| Vat1l | 0.091361251 | 3.836501268 | 0.017 | 0 | 1 | 0.002052242 | 4.432959407 | 0.1 | 0.006 | 1 | 0.091361251 | 0.004100273 |
| Crtam | 0.091361251 | 2.099535674 | 0.017 | 0 | 1 | 0.002052242 | 4.432959407 | 0.1 | 0.006 | 1 | 0.091361251 | 0.004100273 |
| Gm10373 | 0.091361251 | 2.514573173 | 0.017 | 0 | 1 | 0.002052242 | 4.432959407 | 0.1 | 0.006 | 1 | 0.091361251 | 0.004100273 |
| Gm2164 | 0.093241817 | -2.070389328 | 0 | 0.024 | 1 | 0.002052242 | 4.432959407 | 0.1 | 0.006 | 1 | 0.093241817 | 0.004100273 |
| Rab3b | 0.095240327 | -1.807354922 | 0.008 | 0.041 | 1 | 0.002052242 | 4.432959407 | 0.1 | 0.006 | 1 | 0.095240327 | 0.004100273 |
| Fmnl1 | 0.147030916 | -1.485426827 | 0 | 0.018 | 1 | 0.002052242 | 4.432959407 | 0.1 | 0.006 | 1 | 0.147030916 | 0.004100273 |
| Hspe1-rs1 | 0.147030916 | -1.485426827 | 0 | 0.018 | 1 | 0.002052242 | 4.432959407 | 0.1 | 0.006 | 1 | 0.147030916 | 0.004100273 |
| Creg2 | 0.167062889 | 2.099535674 | 0.025 | 0.006 | 1 | 0.002052242 | 4.432959407 | 0.1 | 0.006 | 1 | 0.167062889 | 0.004100273 |
| Spaca7b | 0.168156185 | 1.514573173 | 0.025 | 0.006 | 1 | 0.002052242 | 4.432959407 | 0.1 | 0.006 | 1 | 0.168156185 | 0.004100273 |
| Zfp474 | 0.201658121 | 1.251538767 | 0.034 | 0.012 | 1 | 0.002052242 | 4.432959407 | 0.1 | 0.006 | 1 | 0.201658121 | 0.004100273 |
| 9530097N15Rik | 0.237885589 | -1.485426827 | 0 | 0.012 | 1 | 0.002052242 | 4.432959407 | 0.1 | 0.006 | 1 | 0.237885589 | 0.004100273 |
| 8030423F21Rik | 0.237885589 | -1.070389328 | 0 | 0.012 | 1 | 0.002052242 | 4.432959407 | 0.1 | 0.006 | 1 | 0.237885589 | 0.004100273 |
| Ofcc1 | 0.237885589 | -1.485426827 | 0 | 0.012 | 1 | 0.002052242 | 4.432959407 | 0.1 | 0.006 | 1 | 0.237885589 | 0.004100273 |
| Gm40323 | 0.334204811 | -0.807354922 | 0.008 | 0.024 | 1 | 0.002052242 | 4.432959407 | 0.1 | 0.006 | 1 | 0.334204811 | 0.004100273 |
| Gm51903 | 0.334204811 | -0.807354922 | 0.008 | 0.024 | 1 | 0.002052242 | 4.432959407 | 0.1 | 0.006 | 1 | 0.334204811 | 0.004100273 |
| Gm15327 | 0.374548525 | 0.514573173 | 0.017 | 0.006 | 1 | 0.002052242 | 4.432959407 | 0.1 | 0.006 | 1 | 0.374548525 | 0.004100273 |
| 4933417E11Rik | 0.423363634 | -1.163498732 | 0.034 | 0.053 | 1 | 0.002052242 | 4.432959407 | 0.1 | 0.006 | 1 | 0.423363634 | 0.004100273 |
| Xk | 0.467093859 | -0.485426827 | 0.025 | 0.041 | 1 | 0.002052242 | 4.432959407 | 0.1 | 0.006 | 1 | 0.467093859 | 0.004100273 |
| Gm10778 | 0.491791629 | -0.707819249 | 0.017 | 0.029 | 1 | 0.002052242 | 4.432959407 | 0.1 | 0.006 | 1 | 0.491791629 | 0.004100273 |
| Grtp1 | 0.500300313 | -0.292781749 | 0.017 | 0.029 | 1 | 0.002052242 | 4.432959407 | 0.1 | 0.006 | 1 | 0.500300313 | 0.004100273 |
| Gm49313 | 0.508731273 | -0.807354922 | 0.008 | 0.018 | 1 | 0.002052242 | 4.432959407 | 0.1 | 0.006 | 1 | 0.508731273 | 0.004100273 |
| Ptgir | 0.51784324 | 0.514573173 | 0.008 | 0.018 | 1 | 0.002052242 | 4.432959407 | 0.1 | 0.006 | 1 | 0.51784324 | 0.004100273 |
| Stpg1 | 0.706622275 | 0.292180751 | 0.017 | 0.024 | 1 | 0.002052242 | 4.280956314 | 0.1 | 0.006 | 1 | 0.706622275 | 0.004100273 |
| Gm11739 | 0.715874401 | 0.929610672 | 0.017 | 0.012 | 1 | 0.002052242 | 4.432959407 | 0.1 | 0.006 | 1 | 0.715874401 | 0.004100273 |
| Acaa1b | 0.721162042 | 0.514573173 | 0.017 | 0.012 | 1 | 0.002052242 | 4.432959407 | 0.1 | 0.006 | 1 | 0.721162042 | 0.004100273 |
| Unc80 | 0.772535179 | 0.13606155 | 0.042 | 0.035 | 1 | 0.002052242 | 4.432959407 | 0.1 | 0.006 | 1 | 0.772535179 | 0.004100273 |
| Gm48796 | 0.781790612 | -0.485426827 | 0.008 | 0.012 | 1 | 0.002052242 | 4.432959407 | 0.1 | 0.006 | 1 | 0.781790612 | 0.004100273 |
| Gm56602 | 0.781790612 | -0.807354922 | 0.008 | 0.012 | 1 | 0.002052242 | 4.432959407 | 0.1 | 0.006 | 1 | 0.781790612 | 0.004100273 |
| Fam184b | 0.784920947 | -0.222392421 | 0.008 | 0.012 | 1 | 0.002052242 | 4.432959407 | 0.1 | 0.006 | 1 | 0.784920947 | 0.004100273 |
| Zfp947 | 0.828676264 | -0.333423734 | 0.025 | 0.029 | 1 | 0.002052242 | 4.280956314 | 0.1 | 0.006 | 1 | 0.828676264 | 0.004100273 |
| Gm14023 | 0.832475776 | 0.514573173 | 0.034 | 0.029 | 1 | 0.002052242 | 4.432959407 | 0.1 | 0.006 | 1 | 0.832475776 | 0.004100273 |
| Cutal | 0.842422009 | 0.251538767 | 0.025 | 0.029 | 1 | 0.002052242 | 4.432959407 | 0.1 | 0.006 | 1 | 0.842422009 | 0.004100273 |
| Zfp266 | 0.002058027 | -0.501486678 | 0.521 | 0.641 | 1 | 0.009544979 | -1.283828305 | 0.6 | 0.827 | 1 | 0.009544979 | 0.004111818 |
| Cilk1 | 0.002059701 | -1.001264967 | 0.361 | 0.518 | 1 | 0.856856713 | -0.100075301 | 0.6 | 0.58 | 1 | 0.856856713 | 0.00411516 |
| Srp72 | 0.002069113 | -0.50284388 | 0.882 | 0.865 | 1 | 0.587020529 | -0.209147001 | 0.9 | 0.855 | 1 | 0.587020529 | 0.004133945 |
| Cdr2 | 0.002073385 | -1.137503524 | 0.269 | 0.418 | 1 | 0.722061525 | -0.126272818 | 0.5 | 0.546 | 1 | 0.722061525 | 0.004142471 |
| Taf2 | 0.002073947 | -1.122856748 | 0.244 | 0.406 | 1 | 0.298462689 | 0.32443495 | 0.6 | 0.377 | 1 | 0.298462689 | 0.004143592 |
| Iqck | 0.61095539 | -0.392317423 | 0.05 | 0.065 | 1 | 0.002074369 | 2.263034406 | 0.3 | 0.056 | 1 | 0.61095539 | 0.004144435 |
| Zfp970 | 0.002075029 | -3.392317423 | 0 | 0.076 | 1 | 0.46060033 | 0.558490289 | 0 | 0.052 | 1 | 0.46060033 | 0.004145752 |
| Fah | 0.002075029 | -3.392317423 | 0 | 0.076 | 1 | 0.729840207 | 1.432959407 | 0.1 | 0.071 | 1 | 0.729840207 | 0.004145752 |
| Mylpf | 0.002075029 | -3.392317423 | 0 | 0.076 | 1 | 0.539413885 | 1.210566986 | 0 | 0.037 | 1 | 0.539413885 | 0.004145752 |
| Asphd2 | 0.002076394 | -3.485426827 | 0 | 0.076 | 1 | 0.698529281 | 2.210566986 | 0 | 0.015 | 1 | 0.698529281 | 0.004148477 |
| Mamstr | 0.002076394 | -3.485426827 | 0 | 0.076 | 1 | 0.557784478 | 1.432959407 | 0 | 0.034 | 1 | 0.557784478 | 0.004148477 |
| Ajuba | 0.002076394 | -3.485426827 | 0 | 0.076 | 1 | 0.474962821 | 0.494359952 | 0 | 0.049 | 1 | 0.474962821 | 0.004148477 |
| Myo1f | 0.002077512 | -4.240314329 | 0 | 0.076 | 1 | 0.544188995 | 2.354956895 | 0.1 | 0.059 | 1 | 0.544188995 | 0.004150708 |
| Garem1 | 0.002078156 | -0.285313078 | 0.647 | 0.459 | 1 | 0.627905179 | 0.253851084 | 0.4 | 0.583 | 1 | 0.627905179 | 0.004151992 |
| Emc8 | 0.002078288 | -0.984659454 | 0.353 | 0.488 | 1 | 0.241385555 | 0.553035859 | 0.7 | 0.528 | 1 | 0.241385555 | 0.004152256 |
| Atrip | 0.158459275 | -0.409477974 | 0.244 | 0.318 | 1 | 0.002078395 | 1.342356858 | 0.7 | 0.241 | 1 | 0.158459275 | 0.00415247 |
| Rcor2 | 0.002079002 | -3.807354922 | 0 | 0.076 | 1 | 0.539462273 | 0.695993813 | 0 | 0.037 | 1 | 0.539462273 | 0.004153683 |
| Ebf4 | 0.002079189 | -4.807354922 | 0 | 0.076 | 1 | 0.274654967 | -0.763437806 | 0 | 0.108 | 1 | 0.274654967 | 0.004154055 |
| Ttll10 | 0.002079375 | -3.944858446 | 0 | 0.076 | 1 | 0.274689621 | -0.596787936 | 0 | 0.108 | 1 | 0.274689621 | 0.004154426 |
| Gm42726 | 0.002079375 | -3.944858446 | 0 | 0.076 | 1 | 0.474979184 | 0.159940913 | 0 | 0.049 | 1 | 0.474979184 | 0.004154426 |
| Pkd2 | 0.002079528 | -0.405125199 | 0.84 | 0.835 | 1 | 0.833162108 | -0.123090402 | 0.9 | 0.889 | 1 | 0.833162108 | 0.004154732 |
| Epha5 | 0.002080141 | -4.770829046 | 0 | 0.076 | 1 | 0.577179511 | 0.432959407 | 0 | 0.031 | 1 | 0.577179511 | 0.004155956 |
| Nfatc2ip | 0.002085576 | -1.918386234 | 0.025 | 0.129 | 1 | 0.245753697 | -0.736965594 | 0 | 0.12 | 1 | 0.245753697 | 0.004166803 |
| Mamdc2 | 0.002091076 | -2.918386234 | 0.034 | 0.141 | 1 | 0.274757409 | -0.625934282 | 0 | 0.108 | 1 | 0.274757409 | 0.004177778 |
| Trip6 | 0.002092051 | -1.226893814 | 0.235 | 0.376 | 1 | 0.614250457 | 0.168256181 | 0.5 | 0.364 | 1 | 0.614250457 | 0.004179726 |
| Kcnb1 | 0.00209887 | -1.823296466 | 0.101 | 0.241 | 1 | 0.461297092 | -1.310004928 | 0.2 | 0.275 | 1 | 0.461297092 | 0.004193335 |
| Ddx47 | 0.002105418 | -1.013358383 | 0.319 | 0.459 | 1 | 0.010723466 | -2.146985019 | 0.1 | 0.515 | 1 | 0.010723466 | 0.004206404 |

| | | | | | | | | | | | | |
|---|---|---|---|---|---|---|---|---|---|---|---|---|
| Zfp317 | 0.002107987 | -1.151007788 | 0.185 | 0.335 | 1 | 0.491961916 | 0.44503224 | 0.4 | 0.272 | 1 | 0.491961916 | 0.00421153 |
| Slc23a3 | 0.002112611 | -2.807354922 | 0.017 | 0.112 | 1 | 0.895499855 | 0.345496566 | 0.1 | 0.111 | 1 | 0.895499855 | 0.00422076 |
| Dcaf17 | 0.002121548 | -1.600904045 | 0.118 | 0.259 | 1 | 0.477284755 | -0.304006187 | 0.1 | 0.185 | 1 | 0.477284755 | 0.004238595 |
| Chit1 | 0.003718332 | 2.177538186 | 0.092 | 0.018 | 1 | 0.002123437 | 2.210566986 | 0.1 | 0.006 | 1 | 0.003718332 | 0.004242366 |
| Cd27 | 0.015539978 | 2.514573173 | 0.05 | 0.006 | 1 | 0.002123437 | 3.695993813 | 0.1 | 0.006 | 1 | 0.015539978 | 0.004242366 |
| Gm42809 | 0.023821715 | 1.929610672 | 0.059 | 0.012 | 1 | 0.002123437 | 3.695993813 | 0.1 | 0.006 | 1 | 0.023821715 | 0.004242366 |
| Ankle1 | 0.060010312 | -2.655351829 | 0 | 0.029 | 1 | 0.002123437 | 3.695993813 | 0.1 | 0.006 | 1 | 0.060010312 | 0.004242366 |
| Obscn | 0.091361251 | 2.099535674 | 0.017 | 0 | 1 | 0.002123437 | 4.017921908 | 0.1 | 0.006 | 1 | 0.091361251 | 0.004242366 |
| Tmem130 | 0.093241817 | -2.655351829 | 0 | 0.024 | 1 | 0.002123437 | 4.017921908 | 0.1 | 0.006 | 1 | 0.093241817 | 0.004242366 |
| Gm28731 | 0.093245792 | -1.807354922 | 0 | 0.024 | 1 | 0.002123437 | 3.695993813 | 0.1 | 0.006 | 1 | 0.093245792 | 0.004242366 |
| Gm37245 | 0.126151542 | 0.929610672 | 0.059 | 0.024 | 1 | 0.002123437 | 3.695993813 | 0.1 | 0.006 | 1 | 0.126151542 | 0.004242366 |
| Gm16104 | 0.147029286 | -1.807354922 | 0 | 0.018 | 1 | 0.002123437 | 4.017921908 | 0.1 | 0.006 | 1 | 0.147029286 | 0.004242366 |
| Trpm2 | 0.167062889 | 1.836501268 | 0.025 | 0.006 | 1 | 0.002123437 | 3.695993813 | 0.1 | 0.006 | 1 | 0.167062889 | 0.004242366 |
| Hey2 | 0.168156185 | 1.514573173 | 0.025 | 0.006 | 1 | 0.002123437 | 3.432959407 | 0.1 | 0.006 | 1 | 0.168156185 | 0.004242366 |
| Gm48877 | 0.485068998 | -1.707819249 | 0.017 | 0.029 | 1 | 0.002123437 | 4.017921908 | 0.1 | 0.006 | 1 | 0.485068998 | 0.004242366 |
| Gm57214 | 0.511005543 | -0.485426827 | 0.008 | 0.018 | 1 | 0.002123437 | 4.017921908 | 0.1 | 0.006 | 1 | 0.511005543 | 0.004242366 |
| Gm17163 | 0.511005543 | -0.485426827 | 0.008 | 0.018 | 1 | 0.002123437 | 4.017921908 | 0.1 | 0.006 | 1 | 0.511005543 | 0.004242366 |
| Soat2 | 0.563722678 | -0.748461233 | 0.034 | 0.047 | 1 | 0.002123437 | 3.695993813 | 0.1 | 0.006 | 1 | 0.563722678 | 0.004242366 |
| Ckm | 0.659865009 | 0.514573173 | 0.025 | 0.018 | 1 | 0.002123437 | 4.017921908 | 0.1 | 0.006 | 1 | 0.659865009 | 0.004242366 |
| Gm36246 | 0.721162042 | 0.514573173 | 0.017 | 0.012 | 1 | 0.002123437 | 4.017921908 | 0.1 | 0.006 | 1 | 0.721162042 | 0.004242366 |
| Cybb | 0.778665521 | -3.240314329 | 0.008 | 0.012 | 1 | 0.002123437 | 3.432959407 | 0.1 | 0.006 | 1 | 0.778665521 | 0.004242366 |
| Fancb | 0.842422009 | 0.251538767 | 0.025 | 0.029 | 1 | 0.002123437 | 4.017921908 | 0.1 | 0.006 | 1 | 0.842422009 | 0.004242366 |
| 4933423P22Rik | 0.929389703 | 0.192645078 | 0.025 | 0.024 | 1 | 0.002123437 | 4.017921908 | 0.1 | 0.006 | 1 | 0.929389703 | 0.004242366 |
| Gm57409 | 0.954551343 | -0.222392421 | 0.017 | 0.018 | 1 | 0.002123437 | 4.017921908 | 0.1 | 0.006 | 1 | 0.954551343 | 0.004242366 |
| Bet1l | 0.002124109 | -0.582943333 | 0.655 | 0.712 | 1 | 0.312211058 | -0.663902132 | 0.6 | 0.62 | 1 | 0.312211058 | 0.004243707 |
| Ddx46 | 0.002128598 | -0.616450478 | 0.689 | 0.741 | 1 | 0.022746052 | -0.921390165 | 0.6 | 0.818 | 1 | 0.022746052 | 0.004252665 |
| Gm16541 | 0.040097932 | 3.895394957 | 0.101 | 0.041 | 1 | 0.002130879 | 4.079322453 | 0.3 | 0.059 | 1 | 0.040097932 | 0.004257217 |
| Psma2 | 0.117399836 | -0.278357563 | 0.916 | 0.918 | 1 | 0.002133209 | -1.129283017 | 0.8 | 0.957 | 1 | 0.117399836 | 0.004261868 |
| Psmd7 | 0.003259221 | -0.512734173 | 0.849 | 0.894 | 1 | 0.002134168 | -1.41670632 | 0.7 | 0.923 | 1 | 0.003259221 | 0.004263781 |
| Rassf8 | 0.002139641 | -1.444784843 | 0.16 | 0.306 | 1 | 0.846384723 | -0.296774618 | 0.2 | 0.225 | 1 | 0.846384723 | 0.004274704 |
| Ccp110 | 0.002149485 | -1.280607035 | 0.084 | 0.224 | 1 | 0.117702899 | -1.654503434 | 0 | 0.201 | 1 | 0.117702899 | 0.00429435 |
| Rtf2 | 0.002151892 | -0.881355504 | 0.345 | 0.5 | 1 | 0.997062419 | 0.111031312 | 0.5 | 0.472 | 1 | 0.997062419 | 0.004299154 |
| Slco2a1 | 0.002156973 | -2.572889668 | 0.034 | 0.141 | 1 | 0.063154054 | -0.28585884 | 0.3 | 0.099 | 1 | 0.063154054 | 0.004309293 |
| Gas8 | 0.002161738 | -1.462706751 | 0.109 | 0.247 | 1 | 0.551288608 | 0.526068812 | 0.3 | 0.21 | 1 | 0.551288608 | 0.004318803 |
| Ptpn2 | 0.002162266 | 0.379523213 | 0.882 | 0.794 | 1 | 0.017605798 | 0.693419869 | 0.9 | 0.833 | 1 | 0.017605798 | 0.004319856 |
| Gm12743 | 0.002165988 | -2.250961574 | 0.034 | 0.141 | 1 | 0.267227073 | -0.709998547 | 0 | 0.111 | 1 | 0.267227073 | 0.004327284 |
| Gm20635 | 0.00217641 | -2.544320516 | 0.017 | 0.112 | 1 | 0.522066388 | 1.111031312 | 0 | 0.04 | 1 | 0.522066388 | 0.004348083 |
| Ino80dos | 0.002176564 | -1.485426827 | 0.118 | 0.259 | 1 | 0.018158181 | 0.937548492 | 0.7 | 0.312 | 1 | 0.018158181 | 0.004348392 |
| Mfsd11 | 0.002186813 | -0.869341809 | 0.21 | 0.376 | 1 | 0.200387749 | -0.840059087 | 0.2 | 0.41 | 1 | 0.200387749 | 0.004368844 |
| Tpst2 | 0.002190184 | -0.694880193 | 0.395 | 0.565 | 1 | 0.271557331 | 0.847996907 | 0.5 | 0.373 | 1 | 0.271557331 | 0.004375571 |
| Ktn1 | 0.002190424 | -0.487895083 | 0.899 | 0.882 | 1 | 0.605326929 | 0.125019651 | 0.8 | 0.904 | 1 | 0.605326929 | 0.00437605 |
| Vkorc1l1 | 0.002192294 | -0.745549638 | 0.588 | 0.688 | 1 | 0.043775599 | -1.075315571 | 0.6 | 0.778 | 1 | 0.043775599 | 0.004379782 |
| Dab1 | 0.491817436 | -0.925999419 | 0.017 | 0.029 | 1 | 0.002194432 | 1.847996907 | 0.3 | 0.056 | 1 | 0.491817436 | 0.004384048 |
| Cx3cl1 | 0.002195773 | 1.024767905 | 0.277 | 0.135 | 1 | 0.360330774 | 1.005097868 | 0.3 | 0.194 | 1 | 0.360330774 | 0.004386725 |
| 4930432B10Rik | 0.038142131 | 2.836501268 | 0.025 | 0 | 1 | 0.002197027 | 3.695993813 | 0.1 | 0.006 | 1 | 0.038142131 | 0.004389226 |
| Gm19466 | 0.213853217 | 1.362570079 | 0.042 | 0.018 | 1 | 0.002197027 | 3.695993813 | 0.1 | 0.006 | 1 | 0.213853217 | 0.004389226 |
| Vpreb3 | 0.237885589 | -3.185866545 | 0 | 0.012 | 1 | 0.002197027 | 3.695993813 | 0.1 | 0.006 | 1 | 0.237885589 | 0.004389226 |
| Trim71 | 0.666171662 | 0.251538767 | 0.025 | 0.018 | 1 | 0.002197027 | 3.432959407 | 0.1 | 0.006 | 1 | 0.666171662 | 0.004389226 |
| Mapk13 | 0.002197602 | -2.424026283 | 0.017 | 0.112 | 1 | 0.918736519 | 0.463333056 | 0.1 | 0.108 | 1 | 0.918736519 | 0.004390374 |
| Cbx4 | 0.002199886 | -1.269698136 | 0.244 | 0.394 | 1 | 0.138213322 | -1.201246612 | 0.2 | 0.438 | 1 | 0.138213322 | 0.004394932 |
| Btbd2 | 0.002200085 | -1.448900951 | 0.118 | 0.265 | 1 | 0.952384862 | 0.757394358 | 0.2 | 0.222 | 1 | 0.952384862 | 0.004395329 |
| Tatdn3 | 0.002203686 | -2.229587923 | 0.067 | 0.194 | 1 | 0.806285861 | 0.515421567 | 0.2 | 0.164 | 1 | 0.806285861 | 0.004402517 |
| Ppp1r14c | 0.002205756 | 2.584962501 | 0.109 | 0.024 | 1 | 0.526605007 | 1.432959407 | 0.1 | 0.052 | 1 | 0.526605007 | 0.004406646 |
| Pecam1 | 0.224107823 | -1.261754854 | 0.084 | 0.129 | 1 | 0.002207173 | 1.581822793 | 0.4 | 0.102 | 1 | 0.224107823 | 0.004409474 |
| Apol8 | 0.00221868 | -2.655351829 | 0.017 | 0.112 | 1 | 0.407995683 | 0.795529487 | 0.2 | 0.108 | 1 | 0.407995683 | 0.004432438 |
| Cdk2 | 0.002223233 | -1.765534746 | 0.109 | 0.241 | 1 | 0.055574572 | -1.888968688 | 0 | 0.275 | 1 | 0.055574572 | 0.004441523 |
| Syne3 | 0.002227696 | -1.591221491 | 0.16 | 0.306 | 1 | 0.595609862 | -0.736965594 | 0.3 | 0.336 | 1 | 0.595609862 | 0.004450429 |
| Jade3 | 0.002231094 | -2.572889668 | 0.017 | 0.112 | 1 | 0.186059268 | -1.267480311 | 0 | 0.151 | 1 | 0.186059268 | 0.00445721 |
| Pes1 | 0.002232136 | -1.177304532 | 0.176 | 0.329 | 1 | 0.183094121 | -0.663902132 | 0.1 | 0.321 | 1 | 0.183094121 | 0.00445929 |
| Ttc7b | 0.002234707 | -1.096861539 | 0.336 | 0.476 | 1 | 0.934604458 | -0.324470289 | 0.4 | 0.383 | 1 | 0.934604458 | 0.00446442 |
| Csnk1e | 0.172427279 | -0.391450679 | 0.513 | 0.612 | 1 | 0.002236664 | -1.534106703 | 0.2 | 0.762 | 1 | 0.172427279 | 0.004468326 |
| Pcdh11x | 0.002238149 | -3.807354922 | 0.008 | 0.094 | 1 | 0.830448747 | -0.900941329 | 0.1 | 0.114 | 1 | 0.830448747 | 0.004471289 |
| Napb | 0.005558232 | 1.133483005 | 0.235 | 0.112 | 1 | 0.002240066 | 2.240314329 | 0.4 | 0.099 | 1 | 0.005558232 | 0.004475115 |
| Mrps35 | 0.00224537 | -1.350497247 | 0.101 | 0.241 | 1 | 0.505647081 | 0.68163852 | 0.1 | 0.207 | 1 | 0.505647081 | 0.004485698 |
| Ltbp4 | 0.002246287 | -2.485426827 | 0.034 | 0.141 | 1 | 0.209164796 | -1.31299497 | 0.1 | 0.272 | 1 | 0.209164796 | 0.004487528 |
| Cep120 | 0.002249361 | -1.253253385 | 0.252 | 0.394 | 1 | 0.128567905 | -1.115477217 | 0.3 | 0.519 | 1 | 0.128567905 | 0.004493663 |
| Zfp362 | 0.002250772 | -1.30256277 | 0.126 | 0.271 | 1 | 0.419637406 | -0.159995884 | 0.2 | 0.34 | 1 | 0.419637406 | 0.004496477 |
| C630043F03Rik | 0.002252217 | -2.292781749 | 0.042 | 0.153 | 1 | 0.135522565 | -1.28585884 | 0 | 0.185 | 1 | 0.135522565 | 0.004499361 |
| Tmem38a | 0.002252573 | -2.485426827 | 0.042 | 0.153 | 1 | 0.11808318 | 1.639410285 | 0.3 | 0.133 | 1 | 0.11808318 | 0.004500071 |
| Rpp38 | 0.002253614 | 1.148445274 | 0.269 | 0.129 | 1 | 0.238854531 | -0.736965594 | 0 | 0.123 | 1 | 0.238854531 | 0.004502149 |
| Phldb2 | 0.002256144 | -1.336049203 | 0.227 | 0.388 | 1 | 0.580950575 | -0.456564056 | 0.4 | 0.525 | 1 | 0.580950575 | 0.004507198 |
| Tet1 | 0.00226219 | -1.26121167 | 0.202 | 0.353 | 1 | 0.685356495 | -0.339630097 | 0.2 | 0.238 | 1 | 0.685356495 | 0.004519262 |
| Zcchc3 | 0.002266878 | -1.452259963 | 0.076 | 0.206 | 1 | 0.51463874 | -0.248864633 | 0.1 | 0.176 | 1 | 0.51463874 | 0.004528617 |
| Zfp235 | 0.00226828 | -1.524955191 | 0.059 | 0.182 | 1 | 0.759498933 | 0.821524695 | 0.1 | 0.148 | 1 | 0.759498933 | 0.004531415 |
| Gpn1 | 0.002275085 | -1.280607035 | 0.126 | 0.271 | 1 | 0.92259318 | 0.31748219 | 0.2 | 0.179 | 1 | 0.92259318 | 0.004544994 |
| 2010007H06Rik | 0.00228085 | 0.868210127 | 0.277 | 0.129 | 1 | 0.372266122 | 1.611929548 | 0.2 | 0.114 | 1 | 0.372266122 | 0.004556498 |
| Aff3 | 0.002281393 | -0.779042745 | 0.807 | 0.847 | 1 | 0.014703254 | 0.703225382 | 1 | 0.827 | 1 | 0.014703254 | 0.004557581 |
| Dnaaf9 | 0.002285579 | -0.935323084 | 0.563 | 0.676 | 1 | 0.336395431 | 0.267373342 | 1 | 0.827 | 1 | 0.336395431 | 0.004565935 |
| Rab12 | 0.002286821 | -0.580375231 | 0.731 | 0.771 | 1 | 0.078579218 | -0.638247543 | 0.6 | 0.812 | 1 | 0.078579218 | 0.004568413 |
| Pot1b | 0.002288584 | -3.392317423 | 0.008 | 0.094 | 1 | 0.032761931 | 1.795529487 | 0.2 | 0.046 | 1 | 0.032761931 | 0.00457193 |
| Rcbtb2 | 0.002289646 | -1.265645619 | 0.176 | 0.324 | 1 | 0.52718393 | 0.103038522 | 0.5 | 0.336 | 1 | 0.52718393 | 0.00457405 |
| Sart1 | 0.002294198 | -0.892052087 | 0.269 | 0.424 | 1 | 0.012600094 | -1.965071667 | 0.1 | 0.506 | 1 | 0.012600094 | 0.004583132 |
| Hes6 | 0.002313001 | -2.87774425 | 0.008 | 0.094 | 1 | 0.746555869 | 1.111031312 | 0.1 | 0.071 | 1 | 0.746555869 | 0.004620652 |
| Eml3 | 0.002314896 | -1.222392421 | 0.151 | 0.3 | 1 | 0.851137552 | -0.141949429 | 0.3 | 0.302 | 1 | 0.851137552 | 0.004624433 |
| Mdn1 | 0.002320175 | -0.940222036 | 0.37 | 0.512 | 1 | 0.023701859 | -1.357117523 | 0.3 | 0.66 | 1 | 0.023701859 | 0.004634967 |
| Lrfn4 | 0.002320538 | -1.970853654 | 0.034 | 0.141 | 1 | 0.488460065 | -0.230005605 | 0.1 | 0.182 | 1 | 0.488460065 | 0.004635692 |
| Ice1 | 0.002321854 | -1.038399214 | 0.496 | 0.571 | 1 | 0.955918551 | 0.153323034 | 0.7 | 0.66 | 1 | 0.955918551 | 0.004638316 |
| Inka1 | 0.002324913 | -3.008988783 | 0.008 | 0.094 | 1 | 0.315635131 | -0.111361109 | 0 | 0.093 | 1 | 0.315635131 | 0.004644422 |
| Mafa | 0.002324913 | -2.87774425 | 0.008 | 0.094 | 1 | 0.343888998 | -0.191531458 | 0 | 0.083 | 1 | 0.343888998 | 0.004644422 |
| Gm49735 | 0.002324913 | -2.87774425 | 0.008 | 0.094 | 1 | 0.790190368 | 0.526068812 | 0.1 | 0.074 | 1 | 0.790190368 | 0.004644422 |
| Stoml3 | 0.002324931 | -2.944858446 | 0.008 | 0.094 | 1 | 0.505605027 | 0.847996907 | 0 | 0.043 | 1 | 0.505605027 | 0.004644457 |
| Snx32 | 0.002324931 | -2.807354922 | 0.008 | 0.094 | 1 | 0.044682096 | 2.31748219 | 0.3 | 0.105 | 1 | 0.044682096 | 0.004644457 |
| Npas2 | 0.002329333 | 0.244685096 | 0.479 | 0.288 | 1 | 0.275907553 | -1.198015491 | 0.2 | 0.336 | 1 | 0.275907553 | 0.00465324 |
| Noc3l | 0.002334026 | -1.168953162 | 0.227 | 0.382 | 1 | 0.397563396 | -0.616284112 | 0.2 | 0.306 | 1 | 0.397563396 | 0.004662604 |
| Hdgfl3 | 0.002334094 | -1.053710587 | 0.336 | 0.482 | 1 | 0.040596933 | -1.415037499 | 0.2 | 0.559 | 1 | 0.040596933 | 0.00466274 |
| Xylt2 | 0.002336671 | -2.87774425 | 0.008 | 0.094 | 1 | 0.829966406 | 1.111031312 | 0.1 | 0.08 | 1 | 0.829966406 | 0.004667883 |
| Taco1 | 0.002336671 | -2.733354341 | 0.008 | 0.094 | 1 | 0.433577142 | 0.558490289 | 0 | 0.059 | 1 | 0.433577142 | 0.004667883 |
| Gm46620 | 0.002336671 | -2.733354341 | 0.008 | 0.094 | 1 | 0.245560743 | -0.625934282 | 0 | 0.12 | 1 | 0.245560743 | 0.004667883 |

| | | | | | | | | | | | | |
|---|---|---|---|---|---|---|---|---|---|---|---|---|
| Cpeb2 | 0.002338928 | -1.096861539 | 0.168 | 0.324 | 1 | 0.173106734 | -1.017073513 | 0.3 | 0.546 | 1 | 0.173106734 | 0.004672386 |
| Sp4 | 0.002343565 | -1.517848305 | 0.202 | 0.347 | 1 | 0.26254438 | 0.36882907 | 0.6 | 0.377 | 1 | 0.26254438 | 0.004681638 |
| Gstt2 | 0.002351365 | -1.436517227 | 0.109 | 0.247 | 1 | 0.470055049 | -0.408342847 | 0.1 | 0.185 | 1 | 0.470055049 | 0.004697202 |
| Lamb3 | 0.002363077 | 1.684498174 | 0.151 | 0.047 | 1 | 0.087465478 | 2.263034406 | 0.2 | 0.065 | 1 | 0.087465478 | 0.00472057 |
| Atf4 | 0.002367375 | -0.420817746 | 0.966 | 0.982 | 1 | 0.003536974 | -1.125886441 | 0.9 | 0.991 | 1 | 0.003536974 | 0.004729145 |
| Proser1 | 0.002373813 | -1.427323873 | 0.151 | 0.294 | 1 | 0.151701884 | -1.101019165 | 0.1 | 0.306 | 1 | 0.151701884 | 0.004741991 |
| Rpl3 | 0.002375598 | -0.348981015 | 0.966 | 0.988 | 1 | 0.009355625 | -0.855801824 | 0.9 | 0.997 | 1 | 0.009355625 | 0.004745552 |
| Sort1 | 0.002385111 | 0.269460675 | 0.723 | 0.476 | 1 | 0.025097998 | 0.777607579 | 0.8 | 0.574 | 1 | 0.025097998 | 0.004764533 |
| Polr1has | 0.002385946 | 1.569020957 | 0.151 | 0.047 | 1 | 0.489877116 | 0.930459067 | 0 | 0.046 | 1 | 0.489877116 | 0.0047662 |
| Cd46 | 0.517848186 | -0.889817082 | 0.101 | 0.124 | 1 | 0.002386437 | 2.048295557 | 0.5 | 0.154 | 1 | 0.517848186 | 0.00476718 |
| Pitpnc1 | 0.002388477 | 0.263714231 | 0.992 | 0.976 | 1 | 0.126780175 | 0.399115207 | 1 | 0.988 | 1 | 0.126780175 | 0.00477125 |
| Ubxn1 | 0.005655075 | -0.642055006 | 0.571 | 0.676 | 1 | 0.002392041 | -1.973599938 | 0.3 | 0.731 | 1 | 0.005655075 | 0.00477836 |
| Bmper | 0.002393062 | -1.518849829 | 0.261 | 0.406 | 1 | 0.217558418 | -2.308507579 | 0.1 | 0.262 | 1 | 0.217558418 | 0.004780397 |
| Urah | 0.002395273 | 2.099535674 | 0.109 | 0.024 | 1 | 0.210167141 | 3.14345279 | 0.1 | 0.031 | 1 | 0.210167141 | 0.004784808 |
| Plaa | 0.002399011 | -0.807354922 | 0.437 | 0.559 | 1 | 0.254142522 | -0.685289559 | 0.5 | 0.66 | 1 | 0.254142522 | 0.004792267 |
| Ccdc3 | 0.002400204 | -2.025995209 | 0.025 | 0.129 | 1 | 0.106876031 | -1.763437806 | 0 | 0.21 | 1 | 0.106876031 | 0.004794646 |
| Slc25a30 | 0.005294274 | 0.824913293 | 0.387 | 0.247 | 1 | 0.002402353 | 2.34807051 | 0.5 | 0.164 | 1 | 0.005294274 | 0.004798934 |
| Cttnbp2 | 0.002404444 | -1.411426246 | 0.034 | 0.141 | 1 | 0.48402834 | -0.441509711 | 0.1 | 0.182 | 1 | 0.48402834 | 0.004803106 |
| Pou3f1 | 0.002427611 | -2.900464326 | 0.008 | 0.094 | 1 | 0.433680134 | -0.191531458 | 0 | 0.059 | 1 | 0.433680134 | 0.004849329 |
| Mrpl43 | 0.002429164 | -0.924538461 | 0.328 | 0.482 | 1 | 0.032659113 | -1.703177281 | 0.1 | 0.438 | 1 | 0.032659113 | 0.004852427 |
| Tent2 | 0.002430108 | -0.763130922 | 0.605 | 0.688 | 1 | 0.008909742 | -1.448518769 | 0.4 | 0.756 | 1 | 0.008909742 | 0.00485431 |
| Rae1 | 0.002433346 | -1.013358383 | 0.311 | 0.459 | 1 | 0.361960192 | -0.689437224 | 0.3 | 0.401 | 1 | 0.361960192 | 0.004860772 |
| Nupr1l | 0.002433805 | 2.63005039 | 0.084 | 0.012 | 1 | 0.522102209 | 0.847996907 | 0 | 0.04 | 1 | 0.522102209 | 0.004861686 |
| Rere | 0.002436969 | -0.377968406 | 1 | 0.994 | 1 | 0.475354312 | -0.379607272 | 1 | 0.988 | 1 | 0.475354312 | 0.004868 |
| Cklf | 0.002439966 | -1.858885223 | 0.084 | 0.212 | 1 | 0.245667588 | -0.68251781 | 0 | 0.12 | 1 | 0.245667588 | 0.004873979 |
| Pus10 | 0.002459962 | -1.430979043 | 0.185 | 0.324 | 1 | 0.137065135 | -1.365782384 | 0.1 | 0.315 | 1 | 0.137065135 | 0.004913872 |
| Rp9 | 0.002464204 | -0.894817763 | 0.571 | 0.653 | 1 | 0.04617206 | -0.789433014 | 0.3 | 0.694 | 1 | 0.04617206 | 0.004922335 |
| Tmem183a | 0.002465776 | -0.961864871 | 0.319 | 0.465 | 1 | 0.752375715 | 0.16492432 | 0.6 | 0.519 | 1 | 0.752375715 | 0.004925471 |
| Aldh1l2 | 0.002468535 | -2.070389328 | 0.034 | 0.141 | 1 | 0.334439903 | -1.111361109 | 0 | 0.086 | 1 | 0.334439903 | 0.004930976 |
| C030013E06Rik | 0.002473938 | -2.137503524 | 0.042 | 0.153 | 1 | 0.07666033 | 0.828097349 | 0.3 | 0.105 | 1 | 0.07666033 | 0.004941756 |
| Chst12 | 0.00247907 | 0.411109853 | 0.849 | 0.676 | 1 | 0.960192586 | -0.157716745 | 0.7 | 0.633 | 1 | 0.960192586 | 0.004951994 |
| Tnfrsf22 | 0.213995561 | 0.563482773 | 0.168 | 0.118 | 1 | 0.002491313 | 2.639410285 | 0.3 | 0.059 | 1 | 0.213995561 | 0.00497642 |
| Tgfb2 | 0.0024926 | -0.281502692 | 0.773 | 0.841 | 1 | 0.022453588 | -1.538087339 | 0.7 | 0.904 | 1 | 0.022453588 | 0.004978988 |
| Hsph1 | 0.002492684 | -0.997876828 | 0.689 | 0.771 | 1 | 0.036890014 | -0.766712938 | 0.5 | 0.827 | 1 | 0.036890014 | 0.004979154 |
| Ss18 | 0.002493852 | -0.709594873 | 0.622 | 0.682 | 1 | 0.628577117 | -0.154505601 | 0.6 | 0.769 | 1 | 0.628577117 | 0.004981485 |
| Zc3hav1l | 0.00250096 | -1.944858446 | 0.042 | 0.153 | 1 | 0.815687289 | 0.111031312 | 0.1 | 0.12 | 1 | 0.815687289 | 0.004995665 |
| Zbtb34 | 0.002501025 | -1.987927168 | 0.042 | 0.153 | 1 | 0.934105281 | 0.227844977 | 0.2 | 0.176 | 1 | 0.934105281 | 0.004995794 |
| Zfp422 | 0.002511517 | -1.304854582 | 0.092 | 0.229 | 1 | 0.061244687 | -1.840059087 | 0 | 0.265 | 1 | 0.061244687 | 0.005016727 |
| Aga | 0.002512335 | 0.391953886 | 0.849 | 0.641 | 1 | 0.693668078 | 0.153735763 | 0.7 | 0.648 | 1 | 0.693668078 | 0.005018358 |
| Megf8 | 0.586058604 | -0.179323699 | 0.218 | 0.247 | 1 | 0.002514486 | 1.800691192 | 0.6 | 0.222 | 1 | 0.586058604 | 0.005022649 |
| Rnh1 | 0.968593363 | 0.111744731 | 0.739 | 0.776 | 1 | 0.002516411 | -1.594946589 | 0.4 | 0.833 | 1 | 0.968593363 | 0.005026489 |
| Ints13 | 0.002523584 | -0.996388746 | 0.252 | 0.412 | 1 | 0.761475392 | 0.245332404 | 0.5 | 0.435 | 1 | 0.761475392 | 0.005040799 |
| Ska2 | 0.002524683 | -2.213347282 | 0.059 | 0.176 | 1 | 0.768173169 | 0.290001453 | 0.1 | 0.13 | 1 | 0.768173169 | 0.005042992 |
| Rheb | 0.00253038 | 0.328799343 | 0.966 | 0.965 | 1 | 0.731935335 | 0.173092281 | 1 | 0.951 | 1 | 0.731935335 | 0.005054356 |
| Gm43150 | 0.002537903 | 2.602036014 | 0.084 | 0.012 | 1 | 0.115118737 | 2.432959407 | 0.1 | 0.022 | 1 | 0.115118737 | 0.005069364 |
| Ccsap | 0.002545629 | 1.292180751 | 0.151 | 0.047 | 1 | 0.930503322 | 0.463333056 | 0.1 | 0.09 | 1 | 0.930503322 | 0.005084778 |
| Slc25a11 | 0.002546904 | -0.951994627 | 0.387 | 0.512 | 1 | 0.198341818 | -0.888968688 | 0.2 | 0.386 | 1 | 0.198341818 | 0.005087322 |
| Arl6 | 0.002546922 | -1.399696953 | 0.101 | 0.235 | 1 | 0.340265067 | -0.391469028 | 0.1 | 0.225 | 1 | 0.340265067 | 0.005087357 |
| Reps1 | 0.00254786 | -1.329307625 | 0.252 | 0.388 | 1 | 0.731620594 | -0.290417122 | 0.4 | 0.485 | 1 | 0.731620594 | 0.005089228 |
| Eva1a | 0.002548271 | -1.74176658 | 0.034 | 0.141 | 1 | 0.105054664 | -2.131825212 | 0 | 0.213 | 1 | 0.105054664 | 0.005090049 |
| Blcap | 0.002550426 | -0.920363884 | 0.168 | 0.329 | 1 | 0.779007085 | -0.131825212 | 0.3 | 0.318 | 1 | 0.779007085 | 0.005094348 |
| Mis18a | 0.002552815 | -1.900464326 | 0.042 | 0.153 | 1 | 0.446790883 | 0.625604485 | 0 | 0.056 | 1 | 0.446790883 | 0.005099113 |
| Prkab1 | 0.002565558 | -1.339575961 | 0.084 | 0.218 | 1 | 0.505585357 | -0.327852929 | 0.2 | 0.284 | 1 | 0.505585357 | 0.005124535 |
| Git1 | 0.002571499 | -1.839063782 | 0.067 | 0.188 | 1 | 0.124609705 | -1.668578619 | 0 | 0.194 | 1 | 0.124609705 | 0.005136385 |
| Il18r1 | 0.002571852 | 2.684498174 | 0.084 | 0.012 | 1 | 0.698529281 | 2.017921908 | 0 | 0.015 | 1 | 0.698529281 | 0.00513709 |
| Dynll1 | 0.01312259 | 0.288077582 | 0.966 | 0.982 | 1 | 0.002575121 | -1.233526688 | 0.9 | 0.994 | 1 | 0.01312259 | 0.005143611 |
| C9orf72 | 0.002584011 | 0.917928867 | 0.429 | 0.241 | 1 | 0.227234477 | -1.111361109 | 0.1 | 0.262 | 1 | 0.227234477 | 0.005161344 |
| Cep164 | 0.002591771 | -0.856264523 | 0.134 | 0.276 | 1 | 0.582679547 | -0.473931188 | 0.2 | 0.259 | 1 | 0.582679547 | 0.005176825 |
| Plekha4 | 0.002593886 | -1.977279923 | 0.059 | 0.176 | 1 | 0.020377921 | 2.333423734 | 0.3 | 0.093 | 1 | 0.020377921 | 0.005181043 |
| Psmd6 | 0.002597419 | -0.743224585 | 0.529 | 0.624 | 1 | 0.386795516 | -0.332575339 | 0.5 | 0.67 | 1 | 0.386795516 | 0.005188091 |
| Bmp6 | 0.002603303 | 0.218091302 | 0.84 | 0.682 | 1 | 0.479103792 | 2.135278859 | 0.5 | 0.364 | 1 | 0.479103792 | 0.005199829 |
| Akap6 | 0.002606798 | 1.334000927 | 0.151 | 0.047 | 1 | 0.334435522 | -0.936274402 | 0 | 0.086 | 1 | 0.334435522 | 0.005206801 |
| Sri | 0.023274218 | -0.372216217 | 0.697 | 0.771 | 1 | 0.002608351 | -1.616284112 | 0.3 | 0.806 | 1 | 0.023274218 | 0.005209898 |
| Egr3 | 0.063058752 | 1.02358682 | 0.176 | 0.1 | 1 | 0.00261212 | 2.847996907 | 0.4 | 0.102 | 1 | 0.063058752 | 0.005217416 |
| H2-Ab1 | 0.002615548 | 1.896002279 | 0.185 | 0.071 | 1 | 0.489939838 | 0.625604485 | 0 | 0.046 | 1 | 0.489939838 | 0.005224254 |
| Alpk1 | 0.002620447 | 0.629611681 | 0.622 | 0.424 | 1 | 0.063358887 | 2.251465017 | 0.6 | 0.361 | 1 | 0.063358887 | 0.005234027 |
| Otud5 | 0.002623114 | -0.743579012 | 0.454 | 0.571 | 1 | 0.237890083 | -0.702200176 | 0.6 | 0.633 | 1 | 0.237890083 | 0.005239347 |
| Tram2 | 0.002625874 | -0.762473323 | 0.63 | 0.671 | 1 | 0.361979037 | -0.599050988 | 0.7 | 0.707 | 1 | 0.361979037 | 0.005244852 |
| Zfr | 0.002627972 | -0.328737653 | 1 | 0.988 | 1 | 0.00802972 | -0.700724047 | 1 | 0.997 | 1 | 0.00802972 | 0.005249038 |
| Mocs1 | 0.002633284 | -2.655351829 | 0.025 | 0.124 | 1 | 0.446758355 | 0.625604485 | 0 | 0.056 | 1 | 0.446758355 | 0.005259634 |
| Fosl1 | 0.002633685 | 1.035405336 | 0.361 | 0.206 | 1 | 0.005028587 | 1.789103218 | 0.6 | 0.241 | 1 | 0.005028587 | 0.005260434 |
| Cflar | 0.00263916 | -0.768535673 | 0.513 | 0.676 | 1 | 0.120077977 | -0.804383356 | 0.6 | 0.775 | 1 | 0.120077977 | 0.005271354 |
| Gm56962 | 0.016714462 | -3.655351829 | 0 | 0.047 | 1 | 0.002646108 | 3.210566986 | 0.2 | 0.028 | 1 | 0.016714462 | 0.005285215 |
| Kcnq1 | 0.590614836 | -0.402964667 | 0.034 | 0.047 | 1 | 0.002646108 | 3.017921908 | 0.2 | 0.028 | 1 | 0.590614836 | 0.005285215 |
| Pak6 | 0.695893386 | -0.222392421 | 0.017 | 0.024 | 1 | 0.002646108 | 3.017921908 | 0.2 | 0.028 | 1 | 0.695893386 | 0.005285215 |
| Kitl | 0.002650853 | -2.359895945 | 0.042 | 0.153 | 1 | 0.0583578 | -2.640289575 | 0 | 0.272 | 1 | 0.0583578 | 0.00529468 |
| Mkln1 | 0.002651985 | -0.449094614 | 0.933 | 0.924 | 1 | 0.203735591 | -0.304773374 | 0.9 | 0.969 | 1 | 0.203735591 | 0.005296936 |
| Hells | 0.002662611 | -1.944858446 | 0.059 | 0.176 | 1 | 0.874768644 | 0.608530972 | 0.1 | 0.123 | 1 | 0.874768644 | 0.005318133 |
| Smyd4 | 0.002664468 | -1.807354922 | 0.05 | 0.165 | 1 | 0.646611563 | 1.017921908 | 0.1 | 0.062 | 1 | 0.646611563 | 0.005321837 |
| Zfp710 | 0.002670512 | -1.691877705 | 0.084 | 0.212 | 1 | 0.365528568 | -0.381249186 | 0.2 | 0.349 | 1 | 0.365528568 | 0.005333891 |
| Urgcp | 0.00267181 | -1.119632847 | 0.277 | 0.429 | 1 | 0.083912978 | -1.493040011 | 0.4 | 0.568 | 1 | 0.083912978 | 0.005336481 |
| Polr1a | 0.002681732 | -1.185866545 | 0.218 | 0.365 | 1 | 0.421171957 | -0.663902132 | 0.2 | 0.302 | 1 | 0.421171957 | 0.005356272 |
| Lekr1 | 0.002696698 | -1.592342031 | 0.05 | 0.165 | 1 | 0.718931935 | 0.769994395 | 0.1 | 0.068 | 1 | 0.718931935 | 0.005386124 |
| Usp13 | 0.002707773 | -2.343407822 | 0.025 | 0.124 | 1 | 0.180833345 | -1.668578619 | 0 | 0.154 | 1 | 0.180833345 | 0.005408214 |
| H2-K1 | 0.002710261 | 0.791090808 | 0.748 | 0.653 | 1 | 0.761781657 | 0.104177641 | 0.7 | 0.735 | 1 | 0.761781657 | 0.005413177 |
| Utp14b | 0.002710953 | 0.691813558 | 0.622 | 0.418 | 1 | 0.82498977 | 0.198725373 | 0.5 | 0.5 | 1 | 0.82498977 | 0.005414556 |
| Mis12 | 0.002711236 | -1.485426827 | 0.134 | 0.276 | 1 | 0.322155651 | -0.654503434 | 0.1 | 0.228 | 1 | 0.322155651 | 0.005415122 |
| Tmem212 | 0.002719566 | 0.467267458 | 0.176 | 0.059 | 1 | 0.649380916 | -0.304006187 | 0.1 | 0.148 | 1 | 0.649380916 | 0.005431736 |
| Il17rb | 0.002720758 | 1.514573173 | 0.168 | 0.059 | 1 | 0.085495512 | 1.409112665 | 0.3 | 0.117 | 1 | 0.085495512 | 0.005434114 |
| Rnf4 | 0.002722855 | -0.584962501 | 0.622 | 0.706 | 1 | 0.409533325 | -0.224934616 | 0.7 | 0.806 | 1 | 0.409533325 | 0.005438297 |
| Morc2a | 0.002723128 | -0.681213892 | 0.378 | 0.547 | 1 | 0.353451687 | 0.763406735 | 0.7 | 0.617 | 1 | 0.353451687 | 0.005438841 |
| Sdhaf3 | 0.002725591 | -1.847996907 | 0.042 | 0.153 | 1 | 0.466754113 | 1.111031312 | 0.2 | 0.12 | 1 | 0.466754113 | 0.005443753 |
| Ppic | 0.002725783 | -0.341192826 | 0.983 | 0.947 | 1 | 0.565913781 | -0.1682024 | 1 | 0.92 | 1 | 0.565913781 | 0.005444137 |
| Exosc2 | 0.002729393 | -1.707819249 | 0.067 | 0.188 | 1 | 0.514625925 | -0.210896782 | 0.1 | 0.176 | 1 | 0.514625925 | 0.005451336 |
| Phc2 | 0.002729528 | -0.610181858 | 0.723 | 0.747 | 1 | 0.215906542 | -0.479496075 | 0.8 | 0.907 | 1 | 0.215906542 | 0.005451606 |
| Gm33989 | 0.002732902 | -1.109917692 | 0.202 | 0.359 | 1 | 0.301632072 | 0.334225454 | 0.6 | 0.401 | 1 | 0.301632072 | 0.005458335 |
| Zzef1 | 0.002740939 | -0.551660526 | 0.63 | 0.729 | 1 | 0.340444492 | 0.274530045 | 0.9 | 0.809 | 1 | 0.340444492 | 0.005474366 |
| Enkd1 | 0.003147125 | -3.292781749 | 0 | 0.071 | 1 | 0.002742595 | 3.280956314 | 0.2 | 0.028 | 1 | 0.003147125 | 0.005477668 |

| | | | | | | | | | | | | |
|---|---|---|---|---|---|---|---|---|---|---|---|---|
| Rabl6 | 0.002744189 | -0.759865996 | 0.588 | 0.647 | 1 | 0.256887629 | 0.140177658 | 0.6 | 0.747 | 1 | 0.256887629 | 0.005480847 |
| Ipo11 | 0.002747368 | -1.388129626 | 0.134 | 0.276 | 1 | 0.179851926 | 0.669193754 | 0.5 | 0.293 | 1 | 0.179851926 | 0.005487189 |
| Pom121 | 0.00274767 | -0.750021747 | 0.429 | 0.571 | 1 | 0.045374793 | -1.326374 | 0.3 | 0.574 | 1 | 0.045374793 | 0.005487791 |
| Sesn2 | 0.002747881 | -1.318316841 | 0.218 | 0.382 | 1 | 0.202969163 | -1.258202497 | 0.1 | 0.272 | 1 | 0.202969163 | 0.00548821 |
| Llgl1 | 0.002761799 | -1.655351829 | 0.084 | 0.212 | 1 | 0.505600474 | -0.327852929 | 0.2 | 0.284 | 1 | 0.505600474 | 0.00551597 |
| Myof | 0.002763139 | -0.678646112 | 0.58 | 0.706 | 1 | 0.159811102 | -0.89068516 | 0.6 | 0.756 | 1 | 0.159811102 | 0.005518643 |
| Arhgap39 | 0.002765163 | -0.821029859 | 0.101 | 0.241 | 1 | 0.104450157 | -1.814968106 | 0 | 0.213 | 1 | 0.104450157 | 0.00552268 |
| Rps28 | 0.012779787 | -0.257032622 | 0.958 | 0.982 | 1 | 0.002771769 | -1.256509193 | 0.4 | 0.985 | 1 | 0.012779787 | 0.005535856 |
| Tnfrsf19 | 0.002772724 | -4.581351247 | 0.034 | 0.135 | 1 | 0.433698519 | -1.191531458 | 0 | 0.059 | 1 | 0.433698519 | 0.00553776 |
| P4hb | 0.002776547 | -0.291260721 | 0.983 | 0.982 | 1 | 0.13503355 | -0.463717375 | 1 | 0.981 | 1 | 0.13503355 | 0.005545384 |
| Sdad1 | 0.002788249 | -0.965858744 | 0.37 | 0.512 | 1 | 0.967379163 | -0.195857383 | 0.6 | 0.528 | 1 | 0.967379163 | 0.005568724 |
| Gm42951 | 0.947212699 | 0.514573173 | 0.042 | 0.041 | 1 | 0.002791087 | 2.902444691 | 0.2 | 0.028 | 1 | 0.947212699 | 0.005574383 |
| Tm9sf2 | 0.002791267 | -0.299226286 | 0.958 | 0.947 | 1 | 0.477386917 | -0.249399856 | 0.9 | 0.966 | 1 | 0.477386917 | 0.005574743 |
| Brcc3 | 0.002791492 | -1.15680408 | 0.202 | 0.347 | 1 | 0.749162638 | -0.394859617 | 0.4 | 0.383 | 1 | 0.749162638 | 0.005575192 |
| Xndc1 | 0.00279851 | -1.786596362 | 0.084 | 0.212 | 1 | 0.413080175 | 0.558490289 | 0.3 | 0.179 | 1 | 0.413080175 | 0.005589189 |
| Fzd7 | 0.002798829 | -1.336283388 | 0.185 | 0.329 | 1 | 0.698283415 | 0.263034406 | 0.3 | 0.238 | 1 | 0.698283415 | 0.005589825 |
| Nr4a1 | 0.002809567 | -0.649736037 | 0.571 | 0.688 | 1 | 0.78685208 | 0.357848483 | 0.7 | 0.744 | 1 | 0.78685208 | 0.00561124 |
| Wnk3 | 0.002819622 | -1.807354922 | 0.059 | 0.176 | 1 | 0.530996349 | 0.625604485 | 0.3 | 0.225 | 1 | 0.530996349 | 0.005631294 |
| Gpr173 | 0.002819622 | -1.807354922 | 0.059 | 0.176 | 1 | 0.213672862 | -1.069540933 | 0 | 0.136 | 1 | 0.213672862 | 0.005631294 |
| Mdfic | 0.00282185 | -1.45099333 | 0.261 | 0.412 | 1 | 0.330202826 | -0.919557007 | 0.4 | 0.503 | 1 | 0.330202826 | 0.005635737 |
| Mcm5 | 0.045705646 | -1.655351829 | 0.034 | 0.094 | 1 | 0.002822146 | 2.494359952 | 0.3 | 0.059 | 1 | 0.045705646 | 0.005636328 |
| Gstcd | 0.002825192 | -1.354182294 | 0.168 | 0.312 | 1 | 0.389328366 | -0.5519337 | 0.1 | 0.207 | 1 | 0.389328366 | 0.005642401 |
| Rpa3 | 0.002828446 | -1.269698136 | 0.202 | 0.347 | 1 | 0.92888978 | 0.494359952 | 0.2 | 0.225 | 1 | 0.92888978 | 0.005648892 |
| Vps4b | 0.002831498 | -0.67030217 | 0.672 | 0.712 | 1 | 0.090534166 | -0.623308812 | 0.4 | 0.769 | 1 | 0.090534166 | 0.005654978 |
| Ccno | 0.041422672 | 1.251538767 | 0.134 | 0.065 | 1 | 0.002837024 | 3.14345279 | 0.2 | 0.028 | 1 | 0.041422672 | 0.005666 |
| Gm26810 | 0.048798367 | 1.929610672 | 0.05 | 0.012 | 1 | 0.002837024 | 3.14345279 | 0.2 | 0.028 | 1 | 0.048798367 | 0.005666 |
| Zfp119b | 0.175759461 | -1 | 0.042 | 0.082 | 1 | 0.002837024 | 3.14345279 | 0.2 | 0.028 | 1 | 0.175759461 | 0.005666 |
| Gm47962 | 0.573820584 | -0.222392421 | 0.042 | 0.029 | 1 | 0.002837024 | 2.902444691 | 0.2 | 0.028 | 1 | 0.573820584 | 0.005666 |
| Gm56546 | 0.659871503 | 0.514573173 | 0.025 | 0.018 | 1 | 0.002837024 | 3.14345279 | 0.2 | 0.028 | 1 | 0.659871503 | 0.005666 |
| Cdnf | 0.770891247 | 0.292180751 | 0.042 | 0.035 | 1 | 0.002837024 | 3.14345279 | 0.2 | 0.028 | 1 | 0.770891247 | 0.005666 |
| Nrxn2 | 0.851241415 | 0.152003093 | 0.076 | 0.082 | 1 | 0.00284252 | 1.232819793 | 0.7 | 0.244 | 1 | 0.851241415 | 0.00567696 |
| Gm12703 | 0.002843478 | -1.333423734 | 0.05 | 0.165 | 1 | 0.828474811 | -0.58694015 | 0.2 | 0.213 | 1 | 0.828474811 | 0.005678871 |
| Dhx33 | 0.00284651 | -1.093109404 | 0.143 | 0.288 | 1 | 0.621259826 | 0.359710425 | 0.3 | 0.216 | 1 | 0.621259826 | 0.005684917 |
| Cd68 | 0.002852184 | -1.722466024 | 0.042 | 0.153 | 1 | 0.32503061 | -1.888968688 | 0 | 0.09 | 1 | 0.32503061 | 0.005696234 |
| Map3k2 | 0.002852877 | -0.511610759 | 0.899 | 0.906 | 1 | 0.126394093 | -0.527610448 | 0.9 | 0.938 | 1 | 0.126394093 | 0.005697614 |
| Pcdhgc3 | 0.002853319 | -1.232660757 | 0.084 | 0.218 | 1 | 0.450406084 | 0.808468542 | 0.3 | 0.194 | 1 | 0.450406084 | 0.005698497 |
| Leng1 | 0.002862474 | -1.025995209 | 0.126 | 0.276 | 1 | 0.768321712 | -0.217294554 | 0.2 | 0.225 | 1 | 0.768321712 | 0.005716754 |
| Aimp1 | 0.002874221 | -0.705552642 | 0.445 | 0.588 | 1 | 0.085472261 | -1.141949429 | 0.3 | 0.534 | 1 | 0.085472261 | 0.005740182 |
| Ypel3 | 0.002876094 | 0.451666421 | 0.874 | 0.788 | 1 | 0.614891357 | 0.116590838 | 0.8 | 0.84 | 1 | 0.614891357 | 0.005743916 |
| Mrps11 | 0.002889966 | -1.306456686 | 0.101 | 0.235 | 1 | 0.225799472 | -0.654503434 | 0 | 0.13 | 1 | 0.225799472 | 0.00577158 |
| Zadh2 | 0.002895114 | -1.464053176 | 0.134 | 0.271 | 1 | 0.984710668 | -0.131825212 | 0.3 | 0.272 | 1 | 0.984710668 | 0.005781847 |
| Cep20 | 0.002900783 | -1.036836768 | 0.202 | 0.359 | 1 | 0.045904074 | -2.069540933 | 0 | 0.293 | 1 | 0.045904074 | 0.005793151 |
| Snx29 | 0.002914685 | -0.999050546 | 0.378 | 0.5 | 1 | 0.425663679 | 0.628730586 | 0.6 | 0.586 | 1 | 0.425663679 | 0.005820875 |
| Tnni2 | 0.754770746 | -0.152851488 | 0.126 | 0.112 | 1 | 0.002916126 | 1.902444691 | 0.4 | 0.099 | 1 | 0.754770746 | 0.005823748 |
| Ccdc84 | 0.002923892 | -1.521050737 | 0.076 | 0.2 | 1 | 0.365500845 | -0.567040593 | 0.1 | 0.213 | 1 | 0.365500845 | 0.005839234 |
| Nek10 | 0.002924268 | -1.383547213 | 0.034 | 0.141 | 1 | 0.842208212 | -0.888968688 | 0.1 | 0.114 | 1 | 0.842208212 | 0.005839984 |
| Atp11a | 0.002927519 | -0.602884409 | 0.487 | 0.653 | 1 | 0.223871808 | 0.438983195 | 0.8 | 0.685 | 1 | 0.223871808 | 0.005846467 |
| Bag2 | 0.002936492 | -1.459431619 | 0.092 | 0.224 | 1 | 0.706568139 | 0.263034406 | 0.3 | 0.231 | 1 | 0.706568139 | 0.005864361 |
| Crip2 | 0.002941106 | 0.290532899 | 0.882 | 0.694 | 1 | 0.144678197 | -0.962049612 | 0.7 | 0.747 | 1 | 0.144678197 | 0.005873563 |
| Cinp | 0.002943202 | -1.359895945 | 0.118 | 0.253 | 1 | 0.986510726 | 0.227844977 | 0.2 | 0.194 | 1 | 0.986510726 | 0.005877741 |
| Golt1b | 0.002958848 | -0.554468472 | 0.546 | 0.671 | 1 | 0.215712747 | -0.662563204 | 0.4 | 0.562 | 1 | 0.215712747 | 0.005908942 |
| Xrcc1 | 0.002959511 | -1.411426246 | 0.101 | 0.235 | 1 | 0.733688892 | -0.204470513 | 0.2 | 0.235 | 1 | 0.733688892 | 0.005910264 |
| Usp19 | 0.002969933 | -0.970853654 | 0.345 | 0.476 | 1 | 0.713782293 | -0.384176536 | 0.6 | 0.534 | 1 | 0.713782293 | 0.005931046 |
| Kmt2d | 0.002970088 | -0.875373345 | 0.361 | 0.5 | 1 | 0.013589636 | -1.30914949 | 0.4 | 0.75 | 1 | 0.013589636 | 0.005931355 |
| Cmc1 | 0.002977855 | 0.868210127 | 0.319 | 0.165 | 1 | 0.272814359 | -0.814968106 | 0.1 | 0.247 | 1 | 0.272814359 | 0.005946842 |
| Umps | 0.002979149 | -1.600904045 | 0.05 | 0.165 | 1 | 0.457862439 | -0.391469028 | 0.1 | 0.188 | 1 | 0.457862439 | 0.005949423 |
| 4930594M22Rik | 0.67375286 | -0.292781749 | 0.042 | 0.053 | 1 | 0.002982027 | 1.973527789 | 0.4 | 0.099 | 1 | 0.67375286 | 0.005955162 |
| Mrps28 | 0.002985865 | -0.820611019 | 0.647 | 0.712 | 1 | 0.76236078 | -0.196397213 | 0.7 | 0.769 | 1 | 0.76236078 | 0.005962814 |
| Brip1 | 0.002997247 | -2.348924827 | 0.059 | 0.176 | 1 | 0.588685924 | -0.289506617 | 0.2 | 0.133 | 1 | 0.588685924 | 0.00598551 |
| Scube3 | 0.003003129 | 1.684498174 | 0.118 | 0.029 | 1 | 0.597836679 | 1.017921908 | 0 | 0.028 | 1 | 0.597836679 | 0.005997238 |
| Usp42 | 0.003003687 | -1.242856524 | 0.151 | 0.294 | 1 | 0.019148726 | -2.505640048 | 0 | 0.37 | 1 | 0.019148726 | 0.005998351 |
| AW046200 | 0.746775354 | 0.280107919 | 0.092 | 0.082 | 1 | 0.003005633 | 2.795529487 | 0.3 | 0.062 | 1 | 0.746775354 | 0.006002233 |
| Gm56786 | 0.00301714 | 1.955145764 | 0.118 | 0.029 | 1 | 0.73075158 | 2.695993813 | 0 | 0.012 | 1 | 0.73075158 | 0.006025178 |
| Pdha1 | 0.003021047 | -0.711399139 | 0.538 | 0.665 | 1 | 0.101644575 | -0.613861449 | 0.5 | 0.796 | 1 | 0.101644575 | 0.006032968 |
| Usp31 | 0.003024398 | -1.247927513 | 0.176 | 0.329 | 1 | 0.068973773 | -1.385800278 | 0.2 | 0.472 | 1 | 0.068973773 | 0.00603965 |
| 4632427E13Rik | 0.003031022 | 0.201633861 | 0.63 | 0.441 | 1 | 0.618526745 | -0.404983835 | 0.5 | 0.574 | 1 | 0.618526745 | 0.006052857 |
| Ptcd3 | 0.003034029 | -0.939144795 | 0.269 | 0.429 | 1 | 0.684702792 | -0.180347719 | 0.6 | 0.41 | 1 | 0.684702792 | 0.006058852 |
| Eva1b | 0.003045611 | -1.190683562 | 0.429 | 0.524 | 1 | 0.022045439 | -1.970762779 | 0.2 | 0.54 | 1 | 0.022045439 | 0.006081946 |
| Rtn3 | 0.003049441 | -0.388243161 | 0.983 | 0.971 | 1 | 0.94024901 | -0.166717334 | 1 | 0.985 | 1 | 0.94024901 | 0.006089583 |
| Ciz1 | 0.003051619 | -1.354182294 | 0.185 | 0.324 | 1 | 0.098317182 | 1.025455581 | 0.6 | 0.343 | 1 | 0.098317182 | 0.006093926 |
| Lgi2 | 0.003054884 | -0.849175098 | 0.109 | 0.253 | 1 | 0.027637584 | -3.91281543 | 0 | 0.343 | 1 | 0.027637584 | 0.006100436 |
| Rara | 0.003056318 | -1.436517227 | 0.092 | 0.224 | 1 | 0.30183514 | -0.806506527 | 0.2 | 0.34 | 1 | 0.30183514 | 0.006103294 |
| Fam13c | 0.003057671 | -1.762267033 | 0.067 | 0.188 | 1 | 0.144286345 | -1.68251781 | 0 | 0.179 | 1 | 0.144286345 | 0.006105992 |
| Zfp449 | 0.003058509 | -1.614709844 | 0.05 | 0.165 | 1 | 0.226093631 | -0.888968688 | 0 | 0.13 | 1 | 0.226093631 | 0.006107664 |
| Cpt1c | 0.003061152 | -1.617877123 | 0.101 | 0.229 | 1 | 0.0521042 | 1.091922489 | 0.4 | 0.154 | 1 | 0.0521042 | 0.006112932 |
| Vcpip1 | 0.003068748 | -0.86393845 | 0.504 | 0.594 | 1 | 0.245531964 | -0.444239503 | 0.6 | 0.738 | 1 | 0.245531964 | 0.006128078 |
| Trappc12 | 0.96657951 | -0.345249169 | 0.193 | 0.188 | 1 | 0.003073336 | 1.608530972 | 0.6 | 0.198 | 1 | 0.96657951 | 0.006137226 |
| Maff | 0.003077662 | 0.738436981 | 0.63 | 0.512 | 1 | 0.834483246 | -0.157716745 | 0.5 | 0.565 | 1 | 0.834483246 | 0.006145852 |
| Zfp84 | 0.003078354 | -1.152851488 | 0.109 | 0.247 | 1 | 0.152178561 | -1.080110175 | 0.1 | 0.306 | 1 | 0.152178561 | 0.006147231 |
| Foxj2 | 0.003086737 | -1.008988783 | 0.16 | 0.306 | 1 | 0.304462872 | 0.532495081 | 0.6 | 0.414 | 1 | 0.304462872 | 0.006163947 |
| Lurap1l | 0.003088931 | 3.421463768 | 0.067 | 0.006 | 1 | 0.474904369 | 0.625604485 | 0 | 0.049 | 1 | 0.474904369 | 0.00616832 |
| Bend6 | 0.003090356 | 0.543719518 | 0.277 | 0.129 | 1 | 0.023825722 | 2.210566986 | 0.2 | 0.043 | 1 | 0.023825722 | 0.006171162 |
| Maml3 | 0.003096326 | -1.301980686 | 0.513 | 0.571 | 1 | 0.246025945 | -0.719573991 | 0.4 | 0.636 | 1 | 0.246025945 | 0.006183064 |
| Txnl4a | 0.850309157 | -0.236399279 | 0.555 | 0.459 | 1 | 0.003098871 | -2.540498805 | 0.1 | 0.59 | 1 | 0.850309157 | 0.00618814 |
| Fblim1 | 0.00310754 | -3.070389328 | 0.017 | 0.106 | 1 | 0.367925426 | 0.816288047 | 0.2 | 0.108 | 1 | 0.367925426 | 0.006205423 |
| Nectin1 | 0.003127619 | -1.018859027 | 0.059 | 0.176 | 1 | 0.1970645 | -1.080110175 | 0.1 | 0.281 | 1 | 0.1970645 | 0.006245457 |
| Yipf4 | 0.003128435 | -0.4820521 | 0.79 | 0.829 | 1 | 0.008163654 | -1.262848862 | 0.7 | 0.87 | 1 | 0.008163654 | 0.006247082 |
| Exoc7 | 0.00313 | -1.137503524 | 0.16 | 0.306 | 1 | 0.496521601 | 0.443013072 | 0.4 | 0.284 | 1 | 0.496521601 | 0.006250204 |
| D17H6S56E-5 | 0.003133122 | -1.05941821 | 0.025 | 0.124 | 1 | 0.027737033 | 4.117457582 | 0.2 | 0.049 | 1 | 0.027737033 | 0.006256428 |
| Eif3d | 0.005997971 | -0.697345917 | 0.471 | 0.565 | 1 | 0.00313452 | -1.876895855 | 0.2 | 0.667 | 1 | 0.005997971 | 0.006259214 |
| Nrbp2 | 0.003135747 | 0.381306642 | 0.613 | 0.406 | 1 | 0.027114686 | 1.741797503 | 0.6 | 0.318 | 1 | 0.027114686 | 0.006261662 |
| Npm2 | 0.003147125 | -3.292781749 | 0 | 0.071 | 1 | 0.865301769 | 1.017921908 | 0.1 | 0.083 | 1 | 0.865301769 | 0.006284345 |
| Podn | 0.003148848 | -3.485426827 | 0 | 0.071 | 1 | 0.397032187 | -0.374395515 | 0 | 0.068 | 1 | 0.397032187 | 0.006287781 |
| Fam171a2 | 0.003148848 | -3.392317423 | 0 | 0.071 | 1 | 0.408706061 | 0.31748219 | 0 | 0.065 | 1 | 0.408706061 | 0.006287781 |
| Col7a1 | 0.003150259 | -3.733354341 | 0 | 0.071 | 1 | 0.989603182 | 0.463333056 | 0.1 | 0.096 | 1 | 0.989603182 | 0.006290594 |
| Slc45a1 | 0.003151952 | -3.733354341 | 0 | 0.071 | 1 | 0.365421586 | 1.847996907 | 0.1 | 0.04 | 1 | 0.365421586 | 0.00629397 |
| Rarres2 | 0.003152141 | -3.807354922 | 0 | 0.071 | 1 | 0.522126681 | 0.769994395 | 0 | 0.04 | 1 | 0.522126681 | 0.006294345 |
| E330013P04Rik | 0.003152297 | -3.87774425 | 0 | 0.071 | 1 | 0.47227707 | 0.245332404 | 0.2 | 0.114 | 1 | 0.47227707 | 0.006294658 |

| | | | | | | | | | | | | |
|---|---|---|---|---|---|---|---|---|---|---|---|---|
| Mms22l | 0.016396263 | -2.614709844 | 0.017 | 0.082 | 1 | 0.003153794 | 2.695993813 | 0.2 | 0.028 | 1 | 0.016396263 | 0.006297641 |
| Gm56606 | 0.199998077 | 1.099535674 | 0.076 | 0.041 | 1 | 0.003153794 | 2.695993813 | 0.2 | 0.028 | 1 | 0.199998077 | 0.006297641 |
| Mybl2 | 0.163453432 | -1.707819249 | 0.017 | 0.047 | 1 | 0.00315387 | 2.602884409 | 0.2 | 0.028 | 1 | 0.163453432 | 0.006297793 |
| Zfp354a | 0.003154023 | -3.185866545 | 0 | 0.071 | 1 | 0.396974588 | 0.111031312 | 0 | 0.068 | 1 | 0.396974588 | 0.006298098 |
| Egr1 | 0.003154261 | -0.868017704 | 0.496 | 0.671 | 1 | 0.918828891 | 1.774887153 | 0.7 | 0.682 | 1 | 0.918828891 | 0.006298573 |
| Gm13963 | 0.003159655 | 2.974004791 | 0.067 | 0.006 | 1 | 0.698529281 | 2.210566986 | 0 | 0.015 | 1 | 0.698529281 | 0.006309326 |
| Renbp | 0.003166179 | -1.272023189 | 0.143 | 0.282 | 1 | 0.304555047 | -0.802257054 | 0.1 | 0.231 | 1 | 0.304555047 | 0.006322334 |
| Osbpl3 | 0.003166267 | -2.68182404 | 0.034 | 0.135 | 1 | 0.097301008 | -1.625934282 | 0 | 0.219 | 1 | 0.097301008 | 0.006322508 |
| Atp8b5 | 0.003172967 | 3.514573173 | 0.05 | 0 | 1 | 0.698534498 | 2.432959407 | 0 | 0.015 | 1 | 0.698534498 | 0.006335866 |
| Snhg5 | 0.003176108 | -0.879705766 | 0.319 | 0.459 | 1 | 0.648447031 | -0.700896339 | 0.2 | 0.25 | 1 | 0.648447031 | 0.006342129 |
| Nifk | 0.003176449 | -0.880911529 | 0.403 | 0.535 | 1 | 0.948268035 | 0.522510992 | 0.3 | 0.395 | 1 | 0.948268035 | 0.006342808 |
| Gm12002 | 0.003182889 | 2.836501268 | 0.067 | 0.006 | 1 | 0.011622016 | 2.902444691 | 0.2 | 0.037 | 1 | 0.011622016 | 0.006355648 |
| Gm56686 | 0.003185767 | -1.424026283 | 0.084 | 0.212 | 1 | 0.098721018 | 1.299103661 | 0.4 | 0.188 | 1 | 0.098721018 | 0.006361385 |
| 5530601H04Rik | 0.003189693 | -0.909234536 | 0.261 | 0.406 | 1 | 0.151096295 | -1.304006187 | 0.2 | 0.392 | 1 | 0.151096295 | 0.006369212 |
| Akirin2 | 0.003190557 | -0.67030217 | 0.58 | 0.729 | 1 | 0.126949885 | -0.646398384 | 0.8 | 0.806 | 1 | 0.126949885 | 0.006370935 |
| Ube3c | 0.003194329 | -0.900464326 | 0.58 | 0.641 | 1 | 0.77925744 | -0.148514107 | 0.7 | 0.75 | 1 | 0.77925744 | 0.006378454 |
| Cib1 | 0.003199164 | 0.676036596 | 0.504 | 0.341 | 1 | 0.593446331 | 0.185031894 | 0.5 | 0.364 | 1 | 0.593446331 | 0.006388093 |
| Zgrf1 | 0.003203044 | -3.185866545 | 0.017 | 0.106 | 1 | 0.484801967 | 1.625604485 | 0.1 | 0.049 | 1 | 0.484801967 | 0.006395828 |
| Tipin | 0.003206745 | -0.844969214 | 0.303 | 0.447 | 1 | 0.932022243 | 1.378511623 | 0.3 | 0.343 | 1 | 0.932022243 | 0.006403207 |
| Gps2 | 0.003207409 | -0.974811668 | 0.345 | 0.482 | 1 | 0.07114248 | -1.146985019 | 0.3 | 0.549 | 1 | 0.07114248 | 0.00640453 |
| Timm8b | 0.003208496 | 0.456567569 | 0.756 | 0.641 | 1 | 0.877090052 | 0.253351852 | 0.6 | 0.559 | 1 | 0.877090052 | 0.006406698 |
| Mogs | 0.003209909 | -1.017921908 | 0.202 | 0.353 | 1 | 0.36277744 | 0.757394358 | 0.4 | 0.269 | 1 | 0.36277744 | 0.006409515 |
| Snhg1 | 0.003210418 | -0.478122027 | 0.866 | 0.859 | 1 | 0.006225968 | -1.341388409 | 0.5 | 0.852 | 1 | 0.006225968 | 0.006410528 |
| Smarcd1 | 0.003221366 | -1.006963948 | 0.202 | 0.347 | 1 | 0.514159881 | 0.151673297 | 0.5 | 0.336 | 1 | 0.514159881 | 0.006432356 |
| Gpr27 | 0.0032349 | -1.563429339 | 0.059 | 0.176 | 1 | 0.918149587 | 0.14345279 | 0.2 | 0.176 | 1 | 0.918149587 | 0.006459336 |
| Prkcd | 0.003251185 | -1.566346823 | 0.059 | 0.176 | 1 | 0.106924045 | -1.321928095 | 0.1 | 0.34 | 1 | 0.106924045 | 0.0064918 |
| Bach2 | 0.003267404 | -1.366920646 | 0.479 | 0.594 | 1 | 0.472538566 | -0.730270942 | 0.6 | 0.731 | 1 | 0.472538566 | 0.006524132 |
| Otor | 0.003281668 | 0.157391241 | 0.538 | 0.335 | 1 | 0.394595881 | -0.179071337 | 0.4 | 0.299 | 1 | 0.394595881 | 0.006552568 |
| Rhou | 0.003282494 | 2.099535674 | 0.126 | 0.035 | 1 | 0.165752854 | -1.339630097 | 0 | 0.164 | 1 | 0.165752854 | 0.006554213 |
| Mrpl17 | 0.00328452 | 0.525977936 | 0.756 | 0.606 | 1 | 0.757731918 | 0.432959407 | 0.5 | 0.58 | 1 | 0.757731918 | 0.006558252 |
| Zfp260 | 0.003288965 | -0.607142736 | 0.613 | 0.718 | 1 | 0.02163232 | -1.441509711 | 0.5 | 0.731 | 1 | 0.02163232 | 0.006567113 |
| Gm16153 | 0.003298307 | -2.424026283 | 0.017 | 0.106 | 1 | 0.22610108 | -0.982078092 | 0 | 0.13 | 1 | 0.22610108 | 0.006585736 |
| Asb1 | 0.003298327 | -2.359895945 | 0.017 | 0.106 | 1 | 0.439528334 | 1.930459067 | 0.1 | 0.046 | 1 | 0.439528334 | 0.006585776 |
| Slc22a1 | 0.003298327 | -2.359895945 | 0.017 | 0.106 | 1 | 0.353822752 | -0.304006187 | 0 | 0.08 | 1 | 0.353822752 | 0.006585776 |
| Rpl23 | 0.003300029 | 0.244126411 | 1 | 0.988 | 1 | 0.205019573 | -0.441332464 | 1 | 1 | 1 | 0.205019573 | 0.006589168 |
| Ift74 | 0.003314819 | -0.822076428 | 0.454 | 0.571 | 1 | 0.656740444 | 0.107336829 | 0.7 | 0.577 | 1 | 0.656740444 | 0.00661865 |
| Gm13483 | 0.00332193 | -1.67305383 | 0.109 | 0.235 | 1 | 0.371069464 | -1.058893689 | 0.1 | 0.207 | 1 | 0.371069464 | 0.006632824 |
| Ston1 | 0.003334165 | -1.017647866 | 0.387 | 0.506 | 1 | 0.410329527 | 0.133057619 | 0.7 | 0.472 | 1 | 0.410329527 | 0.006657213 |
| Sybu | 0.506298378 | 0.118644496 | 0.101 | 0.076 | 1 | 0.003338952 | 1.695993813 | 0.4 | 0.105 | 1 | 0.506298378 | 0.006666756 |
| Abca4 | 0.003347915 | -2.156362551 | 0.092 | 0.218 | 1 | 0.966698361 | -0.91281543 | 0.1 | 0.099 | 1 | 0.966698361 | 0.006684621 |
| Thap11 | 0.003354322 | -1.304854582 | 0.134 | 0.271 | 1 | 0.359618827 | -0.616284112 | 0.2 | 0.321 | 1 | 0.359618827 | 0.006697392 |
| Krt10 | 0.003359106 | -1.27897595 | 0.118 | 0.253 | 1 | 0.582344649 | -0.304006187 | 0.1 | 0.16 | 1 | 0.582344649 | 0.006706928 |
| Cdan1 | 0.003364566 | -1.918386234 | 0.034 | 0.135 | 1 | 0.76817204 | 0.290001453 | 0.1 | 0.13 | 1 | 0.76817204 | 0.006717812 |
| Pex12 | 0.003364566 | -1.918386234 | 0.034 | 0.135 | 1 | 0.290201571 | -0.267480311 | 0 | 0.102 | 1 | 0.290201571 | 0.006717812 |
| Faap100 | 0.003364566 | -1.918386234 | 0.034 | 0.135 | 1 | 0.165872588 | -1.152003093 | 0 | 0.164 | 1 | 0.165872588 | 0.006717812 |
| Rpl23a | 0.003365054 | -0.260561535 | 0.992 | 0.976 | 1 | 0.013801527 | -0.819637471 | 1 | 0.997 | 1 | 0.013801527 | 0.006718785 |
| Sult5a1 | 0.003382769 | 1.048005373 | 0.252 | 0.118 | 1 | 0.718929572 | 0.930459067 | 0.1 | 0.068 | 1 | 0.718929572 | 0.006754095 |
| Ihh | 0.003385677 | -5.920055055 | 0.008 | 0.088 | 1 | 0.730750993 | 0.111031312 | 0 | 0.012 | 1 | 0.730750993 | 0.006759891 |
| Prkce | 0.003387502 | 0.586559286 | 0.782 | 0.606 | 1 | 0.350263442 | 0.778616104 | 0.7 | 0.722 | 1 | 0.350263442 | 0.006763529 |
| Mprip | 0.003391824 | -0.400406243 | 0.924 | 0.976 | 1 | 0.230822873 | 0.307682186 | 1 | 0.981 | 1 | 0.230822873 | 0.006772144 |
| Mrfap1 | 0.003393477 | -0.347923303 | 0.933 | 0.947 | 1 | 0.515125493 | -0.203479311 | 0.7 | 0.954 | 1 | 0.515125493 | 0.006775439 |
| Socs3 | 0.003396851 | -0.837055156 | 0.429 | 0.594 | 1 | 0.165255907 | -1.337114499 | 0.3 | 0.528 | 1 | 0.165255907 | 0.006782164 |
| Gm26910 | 0.170716928 | -1.163498732 | 0.034 | 0.071 | 1 | 0.003421795 | 2.87108052 | 0.3 | 0.065 | 1 | 0.170716928 | 0.006831881 |
| Ctps | 0.003434839 | -1.08809133 | 0.151 | 0.3 | 1 | 0.387253618 | 0.231325546 | 0.4 | 0.244 | 1 | 0.387253618 | 0.006857881 |
| Gm20342 | 0.003435207 | -1.555816155 | 0.059 | 0.176 | 1 | 0.203125303 | -1.080110175 | 0.1 | 0.275 | 1 | 0.203125303 | 0.006858613 |
| Bcor | 0.003435898 | -1.485426827 | 0.294 | 0.412 | 1 | 0.085844524 | -1.419483404 | 0.4 | 0.59 | 1 | 0.085844524 | 0.00685999 |
| Pea15a | 0.00343591 | 0.40325615 | 0.706 | 0.565 | 1 | 0.112213035 | 0.604439875 | 1 | 0.673 | 1 | 0.112213035 | 0.006860014 |
| Mbd3 | 0.003436015 | -0.667630158 | 0.462 | 0.594 | 1 | 0.011257696 | -1.81074352 | 0.2 | 0.593 | 1 | 0.011257696 | 0.006860223 |
| Fhod1 | 0.003436832 | -2.070389328 | 0.042 | 0.147 | 1 | 0.799264655 | 1.159940913 | 0.1 | 0.077 | 1 | 0.799264655 | 0.006861851 |
| Sh3d21 | 0.003436884 | -2.258016331 | 0.084 | 0.206 | 1 | 0.150851686 | 0.785261151 | 0.4 | 0.191 | 1 | 0.150851686 | 0.006861955 |
| Trim62 | 0.003437133 | -2.070389328 | 0.042 | 0.147 | 1 | 0.086550376 | -1.596787936 | 0 | 0.231 | 1 | 0.086550376 | 0.006862452 |
| Gm31323 | 0.003442443 | -3.129283017 | 0.008 | 0.088 | 1 | 0.209131923 | 0.988174565 | 0.2 | 0.08 | 1 | 0.209131923 | 0.006873036 |
| Nmd3 | 0.003451375 | -0.713454783 | 0.42 | 0.553 | 1 | 0.858136103 | -0.207044457 | 0.6 | 0.586 | 1 | 0.858136103 | 0.006890838 |
| Slc30a6 | 0.003455508 | -1.190683562 | 0.185 | 0.324 | 1 | 0.976361955 | -0.111361109 | 0.3 | 0.278 | 1 | 0.976361955 | 0.006899075 |
| Egr2 | 0.329063979 | 1.868210127 | 0.101 | 0.071 | 1 | 0.003455944 | 6.313377792 | 0.2 | 0.031 | 1 | 0.329063979 | 0.006899945 |
| Cstpp1 | 0.003457291 | -1.183157258 | 0.227 | 0.365 | 1 | 0.72189321 | 0.494359952 | 0.3 | 0.383 | 1 | 0.72189321 | 0.006902629 |
| Mthfs | 0.00345866 | -1.222392421 | 0.21 | 0.353 | 1 | 0.24681559 | -0.91281543 | 0.1 | 0.256 | 1 | 0.24681559 | 0.006905357 |
| Mta1 | 0.003458687 | -0.728880864 | 0.437 | 0.576 | 1 | 0.265314971 | -0.589408406 | 0.5 | 0.602 | 1 | 0.265314971 | 0.00690541 |
| Rhbdl2 | 0.003461688 | -3.185866545 | 0.008 | 0.088 | 1 | 0.522115938 | 0.847996907 | 0 | 0.04 | 1 | 0.522115938 | 0.006911392 |
| Rbpjl | 0.003480255 | -2.944858446 | 0.008 | 0.088 | 1 | 0.873509911 | 0.660369903 | 0.1 | 0.083 | 1 | 0.873509911 | 0.006948398 |
| Cst6 | 0.00348041 | -3.008988783 | 0.008 | 0.088 | 1 | 0.730749233 | 2.432959407 | 0 | 0.012 | 1 | 0.730749233 | 0.006948707 |
| St6galnac2 | 0.003480514 | -2.87774425 | 0.008 | 0.088 | 1 | 0.489923006 | 0.769994395 | 0 | 0.046 | 1 | 0.489923006 | 0.006948914 |
| Tcfl5 | 0.003482347 | -0.941106311 | 0.202 | 0.353 | 1 | 0.745515086 | -0.567040593 | 0.3 | 0.309 | 1 | 0.745515086 | 0.006952567 |
| Ccdc126 | 0.003490759 | -1.411426246 | 0.059 | 0.176 | 1 | 0.748798999 | 0.236562194 | 0.1 | 0.133 | 1 | 0.748798999 | 0.006969332 |
| Dvl3 | 0.003491884 | -1.315501826 | 0.21 | 0.335 | 1 | 0.430045832 | -0.618702713 | 0.3 | 0.38 | 1 | 0.430045832 | 0.006971576 |
| Ska3 | 0.003498993 | -2.944858446 | 0.008 | 0.088 | 1 | 0.557765566 | 0.695993813 | 0 | 0.034 | 1 | 0.557765566 | 0.006985744 |
| Cldn34c1 | 0.003498993 | -2.944858446 | 0.008 | 0.088 | 1 | 0.52208609 | 1.017921908 | 0 | 0.04 | 1 | 0.52208609 | 0.006985744 |
| Mrps7 | 0.003509032 | -0.807354922 | 0.319 | 0.476 | 1 | 0.504108212 | -0.296774618 | 0.3 | 0.398 | 1 | 0.504108212 | 0.00700575 |
| Bri3 | 0.577726138 | -0.105531134 | 0.924 | 0.918 | 1 | 0.003513737 | -1.255984672 | 0.6 | 0.963 | 1 | 0.577726138 | 0.007015127 |
| Tmeff1 | 0.003517303 | -3.129283017 | 0.008 | 0.088 | 1 | 0.298704929 | -0.473931188 | 0 | 0.099 | 1 | 0.298704929 | 0.007022235 |
| Wdr90 | 0.003517303 | -3.070389328 | 0.008 | 0.088 | 1 | 0.206166426 | 1.602884409 | 0.2 | 0.083 | 1 | 0.206166426 | 0.007022235 |
| Cenpo | 0.00351944 | -2.185866545 | 0.017 | 0.106 | 1 | 0.433676456 | 0.263034406 | 0 | 0.059 | 1 | 0.433676456 | 0.007026494 |
| Klhl28 | 0.003528518 | -0.948398804 | 0.252 | 0.4 | 1 | 0.01743524 | -2.689437224 | 0 | 0.38 | 1 | 0.01743524 | 0.007044586 |
| Ttc5 | 0.003529248 | -0.86393845 | 0.252 | 0.394 | 1 | 0.893491983 | -0.230005605 | 0.4 | 0.367 | 1 | 0.893491983 | 0.007046041 |
| Heatr3 | 0.003529681 | -0.395229018 | 0.303 | 0.453 | 1 | 0.659979642 | 0.454985714 | 0.5 | 0.454 | 1 | 0.659979642 | 0.007046904 |
| Efna5 | 0.003537123 | 0.442448098 | 0.966 | 0.959 | 1 | 0.94958361 | 0.374065718 | 1 | 0.975 | 1 | 0.94958361 | 0.007061734 |
| Hormad2 | 0.003545165 | 1.126007885 | 0.227 | 0.1 | 1 | 0.364185996 | -1.069540933 | 0 | 0.077 | 1 | 0.364185996 | 0.007077762 |
| Nup205 | 0.003545483 | -1.124837112 | 0.143 | 0.288 | 1 | 0.677963529 | -0.121629444 | 0.2 | 0.262 | 1 | 0.677963529 | 0.007078396 |
| Rnaset2b | 0.003551361 | 0.529446449 | 0.765 | 0.612 | 1 | 0.807328589 | 0.107600407 | 0.7 | 0.667 | 1 | 0.807328589 | 0.00709011 |
| Fam83e | 0.003553087 | -2.572889668 | 0.008 | 0.088 | 1 | 0.401608067 | 1.769994395 | 0.1 | 0.043 | 1 | 0.401608067 | 0.00709355 |
| Tead4 | 0.018328593 | 0.1615574 | 0.412 | 0.247 | 1 | 0.003560564 | 1.120681482 | 0.7 | 0.262 | 1 | 0.018328593 | 0.00710845 |
| Rgs7bp | 0.003566299 | -4.167250867 | 0.008 | 0.088 | 1 | 0.433747232 | -0.763437806 | 0 | 0.059 | 1 | 0.433747232 | 0.00711988 |
| Cox14 | 0.003567046 | 0.366050648 | 0.924 | 0.818 | 1 | 0.83070559 | 0.294195337 | 0.7 | 0.833 | 1 | 0.83070559 | 0.007121368 |
| Bicdl1 | 0.003579395 | 2.215012891 | 0.092 | 0.018 | 1 | 0.522115639 | 0.210566986 | 0 | 0.04 | 1 | 0.522115639 | 0.007145977 |
| Snap23 | 0.003585976 | -0.709391668 | 0.655 | 0.7 | 1 | 0.048163991 | -0.932690065 | 0.6 | 0.827 | 1 | 0.048163991 | 0.007159093 |
| Kif2a | 0.003588587 | -0.723586564 | 0.706 | 0.759 | 1 | 0.519328184 | -0.383614045 | 0.9 | 0.87 | 1 | 0.519328184 | 0.007164295 |
| Eif3h | 0.034361417 | -0.265070673 | 0.924 | 0.941 | 1 | 0.003594698 | -0.961438326 | 1 | 0.972 | 1 | 0.034361417 | 0.007176474 |
| Gng11 | 0.003594834 | -1.196705778 | 0.286 | 0.418 | 1 | 0.021622408 | -2.152003093 | 0.1 | 0.466 | 1 | 0.021622408 | 0.007176745 |

| | | | | | | | | | | | | |
|---|---|---|---|---|---|---|---|---|---|---|---|---|
| Nsfl1c | 0.003597196 | -0.577349317 | 0.647 | 0.7 | 1 | 0.266036847 | -0.342559428 | 0.7 | 0.762 | 1 | 0.266036847 | 0.007181453 |
| Rreb1 | 0.003612579 | -0.779757363 | 0.37 | 0.518 | 1 | 0.901257984 | -0.245112498 | 0.5 | 0.497 | 1 | 0.901257984 | 0.007212108 |
| Fbrs | 0.003613779 | -0.948827348 | 0.235 | 0.388 | 1 | 0.492096892 | -0.357117523 | 0.4 | 0.485 | 1 | 0.492096892 | 0.007214498 |
| Herpud2 | 0.003617265 | -0.653880441 | 0.521 | 0.647 | 1 | 0.311978386 | -0.406104375 | 0.7 | 0.781 | 1 | 0.311978386 | 0.007221446 |
| 2310008N11Rik | 0.879222049 | -0.255944981 | 0.101 | 0.094 | 1 | 0.003623766 | 2.432959407 | 0.4 | 0.102 | 1 | 0.879222049 | 0.007234401 |
| Chaf1b | 0.059281554 | -1.807354922 | 0.025 | 0.076 | 1 | 0.003633737 | 4.017921908 | 0.2 | 0.031 | 1 | 0.059281554 | 0.007254269 |
| Tex261 | 0.006279886 | -0.533789849 | 0.647 | 0.7 | 1 | 0.003634608 | -1.776493958 | 0.2 | 0.698 | 1 | 0.006279886 | 0.007256005 |
| Cog3 | 0.003638274 | -0.897057725 | 0.361 | 0.494 | 1 | 0.090337404 | -1.28585884 | 0.4 | 0.577 | 1 | 0.090337404 | 0.007263311 |
| Krtcap2 | 0.003638426 | -0.442807193 | 0.899 | 0.841 | 1 | 0.844689047 | 0.155966544 | 0.7 | 0.796 | 1 | 0.844689047 | 0.007263614 |
| Prdm11 | 0.00363919 | -1.368069877 | 0.092 | 0.224 | 1 | 0.826476968 | 0.127150978 | 0.2 | 0.164 | 1 | 0.826476968 | 0.007265137 |
| B230354K17Rik | 0.003640573 | -1.464053176 | 0.134 | 0.265 | 1 | 0.979419435 | 0.785261151 | 0.2 | 0.21 | 1 | 0.979419435 | 0.007267891 |
| Folr2 | 0.003654891 | 1.372554168 | 0.126 | 0.035 | 1 | 0.619971082 | 0.769994395 | 0 | 0.025 | 1 | 0.619971082 | 0.007296423 |
| Tmem245 | 0.003657646 | -1.101416224 | 0.286 | 0.418 | 1 | 0.259047868 | -0.754667596 | 0.4 | 0.537 | 1 | 0.259047868 | 0.007301913 |
| Nudt14 | 0.047871697 | 0.634867407 | 0.168 | 0.088 | 1 | 0.003658775 | 2.880418384 | 0.3 | 0.065 | 1 | 0.047871697 | 0.007304164 |
| Cdip1 | 0.003672103 | -0.687060688 | 0.412 | 0.576 | 1 | 0.345471736 | -0.433289204 | 0.6 | 0.778 | 1 | 0.345471736 | 0.007330722 |
| Rap1gap2 | 0.003694514 | -3.040015679 | 0.025 | 0.118 | 1 | 0.196950202 | -1.640289575 | 0 | 0.145 | 1 | 0.196950202 | 0.007375378 |
| Cltb | 0.003702857 | 0.445088073 | 0.866 | 0.747 | 1 | 0.074069887 | -0.862681996 | 0.6 | 0.824 | 1 | 0.074069887 | 0.007392004 |
| Hes1 | 0.003707434 | -0.600328922 | 0.277 | 0.441 | 1 | 0.00401023 | -3.348400306 | 0.1 | 0.574 | 1 | 0.00401023 | 0.007401123 |
| Tcn2 | 0.003717169 | 0.783062009 | 0.353 | 0.194 | 1 | 0.427261706 | 0.838012818 | 0.4 | 0.315 | 1 | 0.427261706 | 0.00742052 |
| 4933431E20Rik | 0.003718981 | 1.736965594 | 0.126 | 0.035 | 1 | 0.228378513 | 1.515421567 | 0.2 | 0.086 | 1 | 0.228378513 | 0.007424131 |
| Abca1 | 0.003723114 | -1.296981738 | 0.143 | 0.276 | 1 | 0.357253242 | -1.064227133 | 0.2 | 0.336 | 1 | 0.357253242 | 0.007432367 |
| Rps13 | 0.083184046 | -0.144167254 | 0.958 | 0.982 | 1 | 0.003728671 | -1.062206423 | 0.7 | 0.994 | 1 | 0.083184046 | 0.007443438 |
| Zfp81 | 0.003730679 | -1.900464326 | 0.042 | 0.147 | 1 | 0.94260214 | 0.494359952 | 0.1 | 0.105 | 1 | 0.94260214 | 0.007447441 |
| Rbm19 | 0.003730741 | -1.807354922 | 0.042 | 0.147 | 1 | 0.213661944 | -0.840059087 | 0 | 0.136 | 1 | 0.213661944 | 0.007447563 |
| Timm29 | 0.003733738 | -1.455053178 | 0.092 | 0.218 | 1 | 0.606965189 | -0.178475305 | 0.2 | 0.262 | 1 | 0.606965189 | 0.007453535 |
| Ptges | 0.003734311 | 0.929610672 | 0.227 | 0.1 | 1 | 0.238666691 | -0.625934282 | 0 | 0.123 | 1 | 0.238666691 | 0.007454677 |
| Zfp423 | 0.003739497 | -2.185866545 | 0.059 | 0.171 | 1 | 0.306608161 | -1.141949429 | 0.1 | 0.231 | 1 | 0.306608161 | 0.00746501 |
| Tns3 | 0.003741565 | -0.648925559 | 0.681 | 0.753 | 1 | 0.126241756 | 0.192362391 | 1 | 0.88 | 1 | 0.126241756 | 0.00746913 |
| Sfi1 | 0.003742502 | -1.485426827 | 0.151 | 0.282 | 1 | 0.166247696 | -1.521236903 | 0.1 | 0.29 | 1 | 0.166247696 | 0.007470998 |
| Pqbp1 | 0.003757835 | -0.745489666 | 0.487 | 0.606 | 1 | 0.022258948 | -1.616284112 | 0.1 | 0.509 | 1 | 0.022258948 | 0.00750155 |
| Thoc3 | 0.003760868 | -1.093659107 | 0.294 | 0.424 | 1 | 0.18083845 | -0.67556505 | 0.2 | 0.435 | 1 | 0.18083845 | 0.007507592 |
| Erc2 | 0.003771023 | 2.004898799 | 0.151 | 0.053 | 1 | 0.643848687 | 0.847996907 | 0 | 0.022 | 1 | 0.643848687 | 0.007527825 |
| Cdkl4 | 0.003774448 | -2.359895945 | 0.008 | 0.088 | 1 | 0.460668048 | -0.111361109 | 0 | 0.052 | 1 | 0.460668048 | 0.00753465 |
| Eif2s1 | 0.003782664 | -0.577349317 | 0.622 | 0.688 | 1 | 0.740883489 | 0.196892049 | 0.7 | 0.676 | 1 | 0.740883489 | 0.007551019 |
| Ing2 | 0.003783364 | -0.857395605 | 0.328 | 0.482 | 1 | 0.098781212 | -1.58694015 | 0.2 | 0.432 | 1 | 0.098781212 | 0.007552413 |
| Leng8 | 0.003792466 | -0.720446857 | 0.412 | 0.535 | 1 | 0.551956919 | -0.536666944 | 0.5 | 0.494 | 1 | 0.551956919 | 0.00757055 |
| Gm9967 | 0.443210507 | -0.359895945 | 0.034 | 0.053 | 1 | 0.003801919 | 2.252387162 | 0.3 | 0.062 | 1 | 0.443210507 | 0.007589384 |
| Pigh | 0.003806539 | -1.807354922 | 0.042 | 0.147 | 1 | 0.796926992 | 0.245332404 | 0.2 | 0.16 | 1 | 0.796926992 | 0.007598588 |
| Mtss2 | 0.003809015 | -0.797012978 | 0.555 | 0.659 | 1 | 0.915918749 | -0.31230551 | 0.7 | 0.599 | 1 | 0.915918749 | 0.007603521 |
| Rabep1 | 0.003819055 | -0.582451282 | 0.756 | 0.841 | 1 | 0.460053527 | -0.52719108 | 0.9 | 0.87 | 1 | 0.460053527 | 0.007623525 |
| Dkk2 | 0.003821675 | -1.023083613 | 0.034 | 0.135 | 1 | 0.460672031 | -0.789433014 | 0 | 0.052 | 1 | 0.460672031 | 0.007628745 |
| Arhgap20 | 0.003821879 | -2.250961574 | 0.008 | 0.088 | 1 | 0.698532411 | 1.558490289 | 0 | 0.015 | 1 | 0.698532411 | 0.00762915 |
| Slc27a1 | 0.003822857 | -1.464053176 | 0.134 | 0.265 | 1 | 0.421157089 | -0.58694015 | 0.2 | 0.302 | 1 | 0.421157089 | 0.007631099 |
| Slc6a17 | 0.003823313 | -1.019763255 | 0.151 | 0.294 | 1 | 0.867237135 | -0.546862711 | 0.4 | 0.373 | 1 | 0.867237135 | 0.007632009 |
| Gm57286 | 0.038681536 | 0.514573173 | 0.471 | 0.347 | 1 | 0.003827336 | 1.896906507 | 0.7 | 0.315 | 1 | 0.038681536 | 0.007640024 |
| Ebf1 | 0.003844376 | 0.232359548 | 0.992 | 0.953 | 1 | 0.519860237 | -0.526602653 | 1 | 0.966 | 1 | 0.519860237 | 0.007673973 |
| Rps2 | 0.008048004 | -0.264690647 | 0.992 | 0.988 | 1 | 0.003844477 | -0.892147598 | 1 | 0.997 | 1 | 0.008048004 | 0.007674174 |
| Tmtc2 | 0.081152011 | -0.249251881 | 0.647 | 0.718 | 1 | 0.003855303 | -1.947862377 | 0.3 | 0.701 | 1 | 0.081152011 | 0.007695742 |
| Pdhb | 0.076496345 | -0.342468873 | 0.588 | 0.629 | 1 | 0.003861704 | -1.723545078 | 0.2 | 0.701 | 1 | 0.076496345 | 0.007708495 |
| Inip | 0.003863761 | -1.887525271 | 0.05 | 0.159 | 1 | 0.569198954 | -0.267480311 | 0.1 | 0.164 | 1 | 0.569198954 | 0.007712593 |
| Prr5l | 0.003866547 | -1.501728639 | 0.118 | 0.247 | 1 | 0.872193678 | -0.339630097 | 0.4 | 0.333 | 1 | 0.872193678 | 0.007718144 |
| Eif1 | 0.073056121 | 0.16529459 | 1 | 0.994 | 1 | 0.00386937 | -0.841316216 | 1 | 0.997 | 1 | 0.073056121 | 0.007723767 |
| Nemf | 0.003874805 | -0.488907396 | 0.748 | 0.812 | 1 | 0.054802928 | -0.895822359 | 0.6 | 0.827 | 1 | 0.054802928 | 0.007734597 |
| Sec22b | 0.003877158 | -0.332094673 | 0.95 | 0.888 | 1 | 0.297792416 | -0.40709968 | 0.8 | 0.892 | 1 | 0.297792416 | 0.007739283 |
| Jmjd4 | 0.013910304 | 1.514573173 | 0.118 | 0.041 | 1 | 0.003882137 | 2.374065718 | 0.3 | 0.062 | 1 | 0.013910304 | 0.007749203 |
| Gm56614 | 0.171451236 | 0.251538767 | 0.025 | 0.006 | 1 | 0.003882515 | 3.432959407 | 0.2 | 0.031 | 1 | 0.171451236 | 0.007749957 |
| Zfp382 | 0.003883361 | -1.807354922 | 0.042 | 0.147 | 1 | 0.906177881 | 0.299103661 | 0.2 | 0.176 | 1 | 0.906177881 | 0.007751642 |
| Fgfr1 | 0.916726167 | -0.155729594 | 1 | 0.976 | 1 | 0.003886824 | -1.165372638 | 0.9 | 0.985 | 1 | 0.916726167 | 0.007758541 |
| Smyd2 | 0.0038925 | -0.925999419 | 0.252 | 0.4 | 1 | 0.233001565 | 0.988174565 | 0.4 | 0.238 | 1 | 0.233001565 | 0.007769849 |
| Prrg2 | 0.003909205 | -1.807354922 | 0.067 | 0.182 | 1 | 0.657411233 | 0.951832718 | 0.1 | 0.17 | 1 | 0.657411233 | 0.007803128 |
| Prox1 | 0.003910203 | -2.529820947 | 0.025 | 0.118 | 1 | 0.962069915 | 0.463333056 | 0.1 | 0.102 | 1 | 0.962069915 | 0.007805117 |
| Dnajb2 | 0.003911381 | 0.502398459 | 0.613 | 0.429 | 1 | 0.553264439 | -0.481923979 | 0.3 | 0.349 | 1 | 0.553264439 | 0.007807463 |
| Sfr1 | 0.011379738 | -0.288896469 | 0.95 | 0.965 | 1 | 0.003929561 | -1.152544544 | 0.8 | 0.978 | 1 | 0.011379738 | 0.00784368 |
| Aldh6a1 | 0.003937427 | 0.85160816 | 0.462 | 0.306 | 1 | 0.28635054 | 0.695993813 | 0.6 | 0.389 | 1 | 0.28635054 | 0.007859351 |
| Gle1 | 0.00393974 | -0.900464326 | 0.244 | 0.394 | 1 | 0.324294697 | 0.966259788 | 0.5 | 0.358 | 1 | 0.324294697 | 0.007863957 |
| Ift80 | 0.00394562 | -0.894232373 | 0.328 | 0.482 | 1 | 0.170342063 | 0.775847121 | 0.6 | 0.423 | 1 | 0.170342063 | 0.007875672 |
| Mib2 | 0.003955 | -1.765534746 | 0.05 | 0.159 | 1 | 0.1610823 | -1.210896782 | 0 | 0.167 | 1 | 0.1610823 | 0.007894357 |
| Dpp9 | 0.003968935 | -1.333423734 | 0.101 | 0.229 | 1 | 0.616012769 | 0.351165316 | 0.4 | 0.302 | 1 | 0.616012769 | 0.007922117 |
| Psmd3 | 0.003973013 | -0.685566442 | 0.513 | 0.618 | 1 | 0.136110947 | -0.702200176 | 0.5 | 0.685 | 1 | 0.136110947 | 0.007930241 |
| Mfn1 | 0.003991992 | -0.807354922 | 0.311 | 0.459 | 1 | 0.415439146 | 0.217022008 | 0.7 | 0.466 | 1 | 0.415439146 | 0.007968047 |
| Hao1 | 0.003994598 | 1.514573173 | 0.134 | 0.041 | 1 | 0.004419961 | 3.017921908 | 0.2 | 0.031 | 1 | 0.004419961 | 0.007973239 |
| 1110059E24Rik | 0.004011017 | -1.057333175 | 0.202 | 0.353 | 1 | 0.254589345 | -0.663902132 | 0.1 | 0.278 | 1 | 0.254589345 | 0.008005946 |
| Ehd2 | 0.00401114 | -0.698420551 | 0.471 | 0.6 | 1 | 0.022548536 | -1.450847575 | 0.6 | 0.707 | 1 | 0.022548536 | 0.00800619 |
| Ppm1n | 0.342544753 | -1.292781749 | 0.025 | 0.047 | 1 | 0.00401142 | 4.017921908 | 0.2 | 0.031 | 1 | 0.342544753 | 0.008006748 |
| Rbm41 | 0.004015727 | -1.538538164 | 0.143 | 0.276 | 1 | 0.245376556 | -1.185018151 | 0.2 | 0.343 | 1 | 0.245376556 | 0.008015327 |
| Twf2 | 0.004016361 | -1.424026283 | 0.092 | 0.218 | 1 | 0.136093483 | -1.668578619 | 0 | 0.185 | 1 | 0.136093483 | 0.008016591 |
| Uqcc2 | 0.00402027 | 0.507078636 | 0.765 | 0.624 | 1 | 0.868815914 | 0.400949012 | 0.5 | 0.543 | 1 | 0.868815914 | 0.008024377 |
| Ube2cbp | 0.004020529 | -2.529820947 | 0.025 | 0.118 | 1 | 0.385680729 | -0.441509711 | 0 | 0.071 | 1 | 0.385680729 | 0.008024894 |
| Ap3s1 | 0.941187056 | -0.206611839 | 0.891 | 0.812 | 1 | 0.004021653 | -1.36434185 | 0.5 | 0.898 | 1 | 0.941187056 | 0.008027132 |
| H4c8 | 0.004028864 | 1.614108846 | 0.092 | 0.018 | 1 | 0.316137708 | 2.111031312 | 0.1 | 0.037 | 1 | 0.316137708 | 0.008041495 |
| Lix1l | 0.004029486 | -0.712837323 | 0.479 | 0.606 | 1 | 0.149134271 | -1.061029433 | 0.4 | 0.546 | 1 | 0.149134271 | 0.008042735 |
| Ndst2 | 0.004031911 | -1.555816155 | 0.076 | 0.194 | 1 | 0.805845331 | 0.331421381 | 0.2 | 0.247 | 1 | 0.805845331 | 0.008047566 |
| Asb4 | 0.033393866 | 1.633217669 | 0.176 | 0.094 | 1 | 0.004033658 | 1.31748219 | 0.6 | 0.201 | 1 | 0.033393866 | 0.008051046 |
| Cwc22 | 0.00403592 | -1.001676578 | 0.277 | 0.418 | 1 | 0.875903316 | -0.273632538 | 0.5 | 0.429 | 1 | 0.875903316 | 0.008055551 |
| Mapk7 | 0.004047516 | -1.655351829 | 0.059 | 0.171 | 1 | 0.701143376 | 0.494359952 | 0.2 | 0.148 | 1 | 0.701143376 | 0.008078649 |
| Rif1 | 0.004052911 | -0.6412215 | 0.487 | 0.629 | 1 | 0.152765774 | -0.858595039 | 0.4 | 0.62 | 1 | 0.152765774 | 0.008089396 |
| Vamp5 | 0.004055155 | 0.444183845 | 0.58 | 0.4 | 1 | 0.803004703 | -0.69173634 | 0.4 | 0.373 | 1 | 0.803004703 | 0.008093865 |
| Adgrg2 | 0.004055969 | -1.183599938 | 0.143 | 0.288 | 1 | 0.991596697 | -0.802257054 | 0.3 | 0.278 | 1 | 0.991596697 | 0.008095488 |
| Trim65 | 0.43851189 | -0.485426827 | 0.034 | 0.053 | 1 | 0.00407204 | 3.880418384 | 0.2 | 0.031 | 1 | 0.43851189 | 0.008127498 |
| Plekhm2 | 0.361315921 | -0.322748352 | 0.782 | 0.741 | 1 | 0.004074218 | -1.40354186 | 0.6 | 0.88 | 1 | 0.361315921 | 0.008131837 |
| Eny2 | 0.004074278 | -0.447265832 | 0.849 | 0.847 | 1 | 0.097544724 | -0.534860187 | 0.7 | 0.892 | 1 | 0.097544724 | 0.008131957 |
| Pigo | 0.00407618 | -2.185866545 | 0.025 | 0.118 | 1 | 0.255259517 | 1.695993813 | 0.2 | 0.096 | 1 | 0.255259517 | 0.008135746 |
| Tbc1d8b | 0.004080035 | -0.87774425 | 0.538 | 0.629 | 1 | 0.488214126 | -0.561565463 | 0.6 | 0.642 | 1 | 0.488214126 | 0.008143422 |
| Mitf | 0.004092238 | 1.113474865 | 0.429 | 0.265 | 1 | 0.253741353 | 0.117917698 | 0.4 | 0.213 | 1 | 0.253741353 | 0.00816773 |
| Katna1 | 0.004092689 | -0.980954244 | 0.345 | 0.494 | 1 | 0.517174468 | -0.489872732 | 0.5 | 0.515 | 1 | 0.517174468 | 0.008168628 |
| Gm43462 | 0.05772204 | -1.263034406 | 0.101 | 0.176 | 1 | 0.00409675 | 1.432959407 | 0.5 | 0.142 | 1 | 0.05772204 | 0.008176716 |
| Rnf115 | 0.004102224 | -0.953979836 | 0.471 | 0.571 | 1 | 0.09474047 | -0.719043686 | 0.6 | 0.772 | 1 | 0.09474047 | 0.00818762 |
| Gnl2 | 0.004105428 | -0.632580521 | 0.697 | 0.759 | 1 | 0.026802584 | -1.246185533 | 0.6 | 0.75 | 1 | 0.026802584 | 0.008194002 |

| | | | | | | | | | | | | |
|---|---|---|---|---|---|---|---|---|---|---|---|---|
| Zfas1 | 0.004111132 | -0.678071905 | 0.672 | 0.741 | 1 | 0.364756335 | -0.495147674 | 0.5 | 0.617 | 1 | 0.364756335 | 0.008205362 |
| Dlg5 | 0.004116623 | -0.98550043 | 0.185 | 0.329 | 1 | 0.623263829 | -0.239465935 | 0.4 | 0.478 | 1 | 0.623263829 | 0.008216299 |
| Ndufs8 | 0.004120565 | -0.803167125 | 0.445 | 0.547 | 1 | 0.546499391 | -0.230005605 | 0.4 | 0.488 | 1 | 0.546499391 | 0.008224152 |
| Pus3 | 0.004124533 | -1.622930351 | 0.067 | 0.182 | 1 | 0.199173461 | 1.31748219 | 0.2 | 0.08 | 1 | 0.199173461 | 0.008232053 |
| Sobp | 0.004129942 | -0.632091922 | 0.899 | 0.894 | 1 | 0.103737193 | 0.367667947 | 1 | 0.951 | 1 | 0.103737193 | 0.008242828 |
| Atf1 | 0.116475777 | -0.645891499 | 0.437 | 0.471 | 1 | 0.004130472 | -2.012745228 | 0.3 | 0.698 | 1 | 0.116475777 | 0.008243883 |
| Stx11 | 0.004141091 | -1.678071905 | 0.042 | 0.147 | 1 | 0.827648389 | -0.339630097 | 0.1 | 0.117 | 1 | 0.827648389 | 0.008265033 |
| Ftx | 0.1381491 | -0.789207575 | 0.227 | 0.282 | 1 | 0.004153641 | 1.258930008 | 0.7 | 0.272 | 1 | 0.1381491 | 0.008290029 |
| Slc35a3 | 0.004155732 | 0.382867494 | 0.672 | 0.476 | 1 | 0.40446759 | 0.31748219 | 0.8 | 0.627 | 1 | 0.40446759 | 0.008294194 |
| Hmgb2 | 0.004159031 | -0.964896199 | 0.571 | 0.706 | 1 | 0.111865937 | -1.020997081 | 0.4 | 0.654 | 1 | 0.111865937 | 0.008300764 |
| Ccn1 | 0.004159896 | 0.631008071 | 0.857 | 0.824 | 1 | 0.05885168 | 0.927295954 | 0.8 | 0.63 | 1 | 0.05885168 | 0.008302486 |
| Nt5c2 | 0.073496205 | -0.662304589 | 0.378 | 0.459 | 1 | 0.004163095 | -3.25354112 | 0 | 0.485 | 1 | 0.073496205 | 0.008308859 |
| 9430015G10Rik | 0.902908911 | -0.122856748 | 0.067 | 0.071 | 1 | 0.004177674 | 2.210566986 | 0.3 | 0.062 | 1 | 0.902908911 | 0.008337895 |
| Gm45767 | 0.064822935 | -1.292781749 | 0.034 | 0.088 | 1 | 0.004177769 | 2.159940913 | 0.3 | 0.062 | 1 | 0.064822935 | 0.008338084 |
| Klhl15 | 0.004179505 | -1.222392421 | 0.227 | 0.365 | 1 | 0.045706582 | -1.789433014 | 0.1 | 0.41 | 1 | 0.045706582 | 0.008341542 |
| Xpo6 | 0.00417959 | -0.856875058 | 0.378 | 0.518 | 1 | 0.021529743 | -1.473931188 | 0.5 | 0.713 | 1 | 0.021529743 | 0.00834171 |
| Tmem98 | 0.00418309 | -0.873697417 | 0.395 | 0.524 | 1 | 0.319417893 | -0.215697769 | 0.4 | 0.552 | 1 | 0.319417893 | 0.008348681 |
| Micos13 | 0.004201856 | 0.454108532 | 0.832 | 0.735 | 1 | 0.31674764 | -0.332575339 | 0.5 | 0.731 | 1 | 0.31674764 | 0.008386056 |
| Gen1 | 0.236604499 | -1.655351829 | 0.025 | 0.053 | 1 | 0.004209866 | 3.639410285 | 0.2 | 0.031 | 1 | 0.236604499 | 0.00840201 |
| Hivep3 | 0.004221449 | -1.014611871 | 0.471 | 0.576 | 1 | 0.279383637 | 0.530656081 | 0.6 | 0.494 | 1 | 0.279383637 | 0.008425077 |
| C030037D09Rik | 0.004221687 | 0.929610672 | 0.202 | 0.082 | 1 | 0.090971661 | 1.795529487 | 0.2 | 0.062 | 1 | 0.090971661 | 0.008425552 |
| Abhd4 | 0.004229376 | -1.266786541 | 0.101 | 0.229 | 1 | 0.363632391 | -0.457811523 | 0.1 | 0.216 | 1 | 0.363632391 | 0.008440864 |
| Wscd2 | 0.006950129 | -0.100603941 | 0.529 | 0.318 | 1 | 0.004232429 | 2.704505313 | 0.6 | 0.281 | 1 | 0.006950129 | 0.008446944 |
| Rad51b | 0.976000639 | 0.514573173 | 0.193 | 0.194 | 1 | 0.004241801 | 2.31748219 | 0.3 | 0.065 | 1 | 0.976000639 | 0.008465609 |
| Mtmr14 | 0.004243161 | -1.5360529 | 0.042 | 0.147 | 1 | 0.768174619 | 0.135278859 | 0.1 | 0.13 | 1 | 0.768174619 | 0.008468317 |
| Rdh5 | 0.004244338 | -2.129283017 | 0.025 | 0.118 | 1 | 0.776127749 | 0.973527789 | 0.1 | 0.074 | 1 | 0.776127749 | 0.008470661 |
| Caap1 | 0.004252794 | -0.93021167 | 0.202 | 0.353 | 1 | 0.605781322 | -0.230005605 | 0.2 | 0.281 | 1 | 0.605781322 | 0.008487502 |
| Cep97 | 0.004263022 | -1.524955191 | 0.067 | 0.182 | 1 | 0.185804252 | -1.230005605 | 0 | 0.151 | 1 | 0.185804252 | 0.00850787 |
| Pld2 | 0.004263173 | -1.980191519 | 0.084 | 0.2 | 1 | 0.025438645 | 1.473601392 | 0.5 | 0.21 | 1 | 0.025438645 | 0.008508172 |
| Vps26a | 0.056464669 | -0.505326385 | 0.513 | 0.594 | 1 | 0.004269459 | -1.495147674 | 0.6 | 0.867 | 1 | 0.056464669 | 0.008520689 |
| E2f1 | 0.006192956 | -2.070389328 | 0.025 | 0.112 | 1 | 0.004278578 | 2.584962501 | 0.3 | 0.065 | 1 | 0.006192956 | 0.008538849 |
| Pwwp3b | 0.004280135 | -1.272023189 | 0.109 | 0.241 | 1 | 0.262622204 | -1.080110175 | 0.1 | 0.247 | 1 | 0.262622204 | 0.00854195 |
| 1110038B12Rik | 0.004303713 | -0.617530363 | 0.605 | 0.7 | 1 | 0.238960295 | -0.600987925 | 0.4 | 0.611 | 1 | 0.238960295 | 0.008588904 |
| Zfp593 | 0.166240254 | 0.598637438 | 0.311 | 0.247 | 1 | 0.004315802 | 2.847996907 | 0.4 | 0.123 | 1 | 0.166240254 | 0.008612978 |
| Tulp4 | 0.004318462 | -0.438292852 | 0.882 | 0.888 | 1 | 0.354755414 | -0.27843628 | 1 | 0.96 | 1 | 0.354755414 | 0.008618274 |
| Ppp2r3d | 0.00432506 | -0.840907482 | 0.244 | 0.394 | 1 | 0.235274627 | -0.776493958 | 0.1 | 0.262 | 1 | 0.235274627 | 0.008631414 |
| Rnf181 | 0.004331739 | 0.694122593 | 0.487 | 0.335 | 1 | 0.737732946 | 0.190102883 | 0.4 | 0.318 | 1 | 0.737732946 | 0.008644714 |
| Pak1 | 0.004343434 | -0.716039755 | 0.58 | 0.7 | 1 | 0.255295495 | 0.604293979 | 0.8 | 0.75 | 1 | 0.255295495 | 0.008668003 |
| Rbx1 | 0.009259616 | -0.378896907 | 0.899 | 0.924 | 1 | 0.004349308 | -1.257486342 | 0.5 | 0.923 | 1 | 0.009259616 | 0.0086797 |
| Dctn1 | 0.004360035 | -0.76106127 | 0.378 | 0.529 | 1 | 0.769314079 | 0.130396637 | 0.5 | 0.559 | 1 | 0.769314079 | 0.00870106 |
| Prrg1 | 0.004374029 | -1.444784843 | 0.109 | 0.241 | 1 | 0.175648598 | -1.357117523 | 0 | 0.157 | 1 | 0.175648598 | 0.008728925 |
| Lrig1 | 0.004383441 | -1.807354922 | 0.143 | 0.271 | 1 | 0.412197541 | -0.465893869 | 0.2 | 0.333 | 1 | 0.412197541 | 0.008747668 |
| Tada1 | 0.004385065 | -1.432959407 | 0.109 | 0.235 | 1 | 0.911638078 | 0.245332404 | 0.2 | 0.207 | 1 | 0.911638078 | 0.008750902 |
| Inpp5e | 0.004393014 | -1.67305383 | 0.05 | 0.159 | 1 | 0.518884761 | 1.185031894 | 0.2 | 0.136 | 1 | 0.518884761 | 0.008766729 |
| Erlin1 | 0.004397289 | -1.50779464 | 0.126 | 0.253 | 1 | 0.721828686 | -0.521236903 | 0.3 | 0.312 | 1 | 0.721828686 | 0.008775242 |
| Gm49708 | 0.004397885 | -2.700439718 | 0.05 | 0.153 | 1 | 0.873509954 | 0.732519689 | 0.1 | 0.083 | 1 | 0.873509954 | 0.008776428 |
| Uhrf1bp1 | 0.00440529 | -1.572889668 | 0.05 | 0.159 | 1 | 0.319279272 | 0.828097349 | 0.3 | 0.17 | 1 | 0.319279272 | 0.008791174 |
| Zdhhc15 | 0.004405373 | -2.250961574 | 0.025 | 0.118 | 1 | 0.324842077 | -0.230005605 | 0 | 0.09 | 1 | 0.324842077 | 0.008791339 |
| Cnot3 | 0.004407682 | -0.929526792 | 0.294 | 0.435 | 1 | 0.693701657 | 0.57829877 | 0.4 | 0.401 | 1 | 0.693701657 | 0.008795936 |
| Ube2v2 | 0.004414578 | -0.566114673 | 0.555 | 0.682 | 1 | 0.074395273 | -1.042774024 | 0.6 | 0.728 | 1 | 0.074395273 | 0.008809668 |
| Mrpl40 | 0.004427352 | -0.889817082 | 0.218 | 0.365 | 1 | 0.468345891 | 0.695993813 | 0.3 | 0.194 | 1 | 0.468345891 | 0.008835102 |
| B230206L02Rik | 0.004427954 | -1.707819249 | 0.05 | 0.159 | 1 | 0.531100545 | -0.5519337 | 0.3 | 0.185 | 1 | 0.531100545 | 0.008836302 |
| Rhebl1 | 0.00443799 | -2.292781749 | 0.025 | 0.118 | 1 | 0.213439846 | 1.290001453 | 0.2 | 0.086 | 1 | 0.213439846 | 0.008856284 |
| Gigyf1 | 0.004439324 | -1.397963986 | 0.118 | 0.247 | 1 | 0.449659651 | 0.349543399 | 0.5 | 0.349 | 1 | 0.449659651 | 0.008858941 |
| Opa3 | 0.004441978 | -1.039025157 | 0.202 | 0.341 | 1 | 0.320838408 | -0.872849022 | 0.2 | 0.327 | 1 | 0.320838408 | 0.008864224 |
| Irak1bp1 | 0.004444792 | -1.349365278 | 0.193 | 0.324 | 1 | 0.568173374 | 0.374065718 | 0.4 | 0.29 | 1 | 0.568173374 | 0.008869827 |
| Nt5c | 0.004451722 | -1.086330872 | 0.21 | 0.347 | 1 | 0.043273705 | -2.111361109 | 0 | 0.299 | 1 | 0.043273705 | 0.008883626 |
| Zdhhc5 | 0.152527185 | -0.378511623 | 0.571 | 0.612 | 1 | 0.004462337 | -1.51501838 | 0.3 | 0.75 | 1 | 0.152527185 | 0.008904762 |
| Cables2 | 0.004487646 | -1.485426827 | 0.059 | 0.171 | 1 | 0.531373984 | -0.131825212 | 0.1 | 0.173 | 1 | 0.531373984 | 0.008955152 |
| Spata6l | 0.127331448 | 0.166649869 | 0.134 | 0.076 | 1 | 0.004492161 | 2.017921908 | 0.3 | 0.062 | 1 | 0.127331448 | 0.008964143 |
| Mcts1 | 0.00449581 | -0.755515991 | 0.429 | 0.571 | 1 | 0.465011894 | -0.357117523 | 0.4 | 0.497 | 1 | 0.465011894 | 0.008971408 |
| Plpp1 | 0.004497326 | -0.841570637 | 0.462 | 0.559 | 1 | 0.198583012 | -0.625934282 | 0.3 | 0.549 | 1 | 0.198583012 | 0.008974427 |
| Hmg20b | 0.004506594 | -0.67305383 | 0.605 | 0.688 | 1 | 0.234936221 | -0.916751844 | 0.5 | 0.571 | 1 | 0.234936221 | 0.008992878 |
| Zfp963 | 0.981806419 | 0.514573173 | 0.059 | 0.059 | 1 | 0.004528076 | 2.14345279 | 0.4 | 0.108 | 1 | 0.981806419 | 0.009035648 |
| Phka1 | 0.004528233 | -1.332130417 | 0.361 | 0.441 | 1 | 0.99583193 | -0.377035397 | 0.7 | 0.565 | 1 | 0.99583193 | 0.009035962 |
| Cdc40 | 0.004535779 | -0.733354341 | 0.437 | 0.571 | 1 | 0.365547377 | -0.795859283 | 0.6 | 0.62 | 1 | 0.365547377 | 0.009050986 |
| Eif3j1 | 0.004539425 | -0.40354982 | 0.924 | 0.906 | 1 | 0.197987316 | -0.449683642 | 0.8 | 0.904 | 1 | 0.197987316 | 0.009058243 |
| Micu3 | 0.004551258 | -0.856264523 | 0.429 | 0.541 | 1 | 0.145447791 | -1.057705797 | 0.7 | 0.685 | 1 | 0.145447791 | 0.009081802 |
| Rpl37 | 0.287589817 | -0.105218092 | 0.95 | 0.988 | 1 | 0.004560704 | -0.89749276 | 0.9 | 0.994 | 1 | 0.287589817 | 0.009100608 |
| Orai2 | 0.004568429 | -1.780882711 | 0.084 | 0.2 | 1 | 0.33428161 | -0.505640048 | 0 | 0.086 | 1 | 0.33428161 | 0.009115988 |
| Cnnm3 | 0.004568539 | -1.40599236 | 0.109 | 0.235 | 1 | 0.218067977 | 0.625604485 | 0.4 | 0.21 | 1 | 0.218067977 | 0.009116206 |
| Fbxo45 | 0.00457582 | -1.109917692 | 0.151 | 0.288 | 1 | 0.197333808 | -0.920677547 | 0.2 | 0.383 | 1 | 0.197333808 | 0.009130702 |
| Crbn | 0.004582056 | -1.229587923 | 0.143 | 0.276 | 1 | 0.163799944 | 0.913585248 | 0.4 | 0.198 | 1 | 0.163799944 | 0.009143116 |
| Nme2 | 0.004584798 | -0.352976531 | 0.958 | 0.971 | 1 | 0.045635008 | -0.342825436 | 0.8 | 0.985 | 1 | 0.045635008 | 0.009148575 |
| Gm10516 | 0.00459401 | -1.529820947 | 0.05 | 0.159 | 1 | 0.893579543 | 0.127150978 | 0.2 | 0.173 | 1 | 0.893579543 | 0.009166915 |
| Ccdc66 | 0.004597157 | -1.317304068 | 0.176 | 0.306 | 1 | 0.160807564 | 0.921060369 | 0.5 | 0.29 | 1 | 0.160807564 | 0.00917318 |
| Mrps16 | 0.004598245 | -0.804377222 | 0.538 | 0.624 | 1 | 0.217962142 | -0.720845929 | 0.4 | 0.534 | 1 | 0.217962142 | 0.009175346 |
| Slc25a1 | 0.004603482 | 1.067114196 | 0.353 | 0.212 | 1 | 0.022950418 | 1.769994395 | 0.4 | 0.142 | 1 | 0.022950418 | 0.009185771 |
| Ccdc93 | 0.004606757 | -1.455053178 | 0.084 | 0.206 | 1 | 0.596730924 | 1.005097868 | 0.3 | 0.259 | 1 | 0.596730924 | 0.009192291 |
| B230369F24Rik | 0.004619271 | -1.597319707 | 0.101 | 0.229 | 1 | 0.732414024 | 0.504852326 | 0.3 | 0.287 | 1 | 0.732414024 | 0.009217204 |
| Farsa | 0.004629232 | -1.058316496 | 0.277 | 0.412 | 1 | 0.663051988 | -0.332575339 | 0.4 | 0.414 | 1 | 0.663051988 | 0.009237034 |
| Gm20559 | 0.3342001 | -1.070389328 | 0.008 | 0.024 | 1 | 0.00464066 | 3.017921908 | 0.2 | 0.031 | 1 | 0.3342001 | 0.009259784 |
| Uchl5 | 0.004640669 | -1.19730144 | 0.202 | 0.341 | 1 | 0.652318859 | 0.50222207 | 0.4 | 0.349 | 1 | 0.652318859 | 0.009259803 |
| Tuba1a | 0.004643948 | -0.441837559 | 0.782 | 0.894 | 1 | 0.094622907 | -1.041692948 | 0.6 | 0.904 | 1 | 0.094622907 | 0.00926633 |
| Nusap1 | 0.112877814 | -1.996388746 | 0.059 | 0.112 | 1 | 0.004650049 | 2.473601392 | 0.3 | 0.065 | 1 | 0.112877814 | 0.009278476 |
| Sbf1 | 0.004655786 | -1.326729081 | 0.084 | 0.206 | 1 | 0.620866344 | -0.255096586 | 0.2 | 0.259 | 1 | 0.620866344 | 0.009289897 |
| Dus3l | 0.00467379 | -1.222392421 | 0.118 | 0.247 | 1 | 0.74702497 | 0.648688098 | 0.2 | 0.157 | 1 | 0.74702497 | 0.009325737 |
| Flrt1 | 0.004681146 | -1.082667658 | 0.227 | 0.365 | 1 | 0.87610696 | 0.455679484 | 0.3 | 0.302 | 1 | 0.87610696 | 0.00934038 |
| Med26 | 0.004694797 | -0.850423644 | 0.387 | 0.506 | 1 | 0.399371583 | -0.33610703 | 0.5 | 0.627 | 1 | 0.399371583 | 0.009367554 |
| Mettl8 | 0.004695053 | -2.163498732 | 0.034 | 0.129 | 1 | 0.952551372 | 0.769994395 | 0.1 | 0.093 | 1 | 0.952551372 | 0.009368062 |
| Kctd14 | 0.004698771 | -2.655351829 | 0.017 | 0.1 | 1 | 0.640912309 | 0.888638891 | 0.1 | 0.062 | 1 | 0.640912309 | 0.009375464 |
| Mad2l1bp | 0.004705903 | -1.347923303 | 0.076 | 0.194 | 1 | 0.750911865 | 0.185031894 | 0.1 | 0.133 | 1 | 0.750911865 | 0.009389661 |
| Casp3 | 0.004709513 | -0.609001512 | 0.395 | 0.565 | 1 | 0.108322683 | -1.311201688 | 0.3 | 0.54 | 1 | 0.108322683 | 0.009396846 |
| Zkscan6 | 0.520090423 | -0.418312631 | 0.084 | 0.106 | 1 | 0.004715812 | 2.558490289 | 0.3 | 0.065 | 1 | 0.520090423 | 0.009409385 |
| Zfp30 | 0.004731699 | -2.185866545 | 0.034 | 0.129 | 1 | 0.135595043 | -1.339630097 | 0 | 0.185 | 1 | 0.135595043 | 0.009441009 |
| Gm10371 | 0.004737878 | 0.957091409 | 0.294 | 0.153 | 1 | 0.006324009 | 2.333423734 | 0.5 | 0.204 | 1 | 0.006324009 | 0.009453308 |
| Lix1 | 0.004746593 | -2.758445322 | 0.017 | 0.1 | 1 | 0.910522355 | 0.290001453 | 0.1 | 0.086 | 1 | 0.910522355 | 0.009470655 |
| Srd5a3 | 0.004747086 | -1.211251864 | 0.092 | 0.218 | 1 | 0.413189184 | -0.425021588 | 0.1 | 0.201 | 1 | 0.413189184 | 0.009471638 |

| | | | | | | | | | | | | |
|---|---|---|---|---|---|---|---|---|---|---|---|---|
| Intu | 0.004747297 | -2.707819249 | 0.017 | 0.1 | 1 | 0.052020228 | 1.959028219 | 0.2 | 0.052 | 1 | 0.052020228 | 0.009472057 |
| Lemd2 | 0.004752275 | -0.871485259 | 0.303 | 0.447 | 1 | 0.275109066 | 0.165479096 | 0.3 | 0.497 | 1 | 0.275109066 | 0.009481965 |
| Tnfsf11 | 0.004753639 | -3.50779464 | 0.017 | 0.1 | 1 | 0.577177401 | -0.111361109 | 0 | 0.031 | 1 | 0.577177401 | 0.009484681 |
| Msh4 | 0.004772201 | -3.185866545 | 0 | 0.065 | 1 | 0.474962821 | 0.432959407 | 0 | 0.049 | 1 | 0.474962821 | 0.009521629 |
| Spin2c | 0.004772201 | -3.185866545 | 0 | 0.065 | 1 | 0.474904369 | 0.31748219 | 0 | 0.049 | 1 | 0.474904369 | 0.009521629 |
| Sv2a | 0.00477434 | -3.292781749 | 0 | 0.065 | 1 | 0.640912309 | 1.063725598 | 0.1 | 0.062 | 1 | 0.640912309 | 0.009525885 |
| Arsj | 0.00477434 | -3.392317423 | 0 | 0.065 | 1 | 0.393944866 | 1.473601392 | 0.1 | 0.043 | 1 | 0.393944866 | 0.009525885 |
| Upk1a | 0.00477434 | -3.392317423 | 0 | 0.065 | 1 | 0.773317908 | 1.063725598 | 0.1 | 0.074 | 1 | 0.773317908 | 0.009525885 |
| Gm36411 | 0.00477434 | -3.292781749 | 0 | 0.065 | 1 | 0.73075158 | 2.695993813 | 0 | 0.012 | 1 | 0.73075158 | 0.009525885 |
| Scn2b | 0.004776003 | -3.485426827 | 0 | 0.065 | 1 | 0.474877085 | 0.847996907 | 0 | 0.049 | 1 | 0.474877085 | 0.009529196 |
| Tekt2 | 0.004776051 | -3.392317423 | 0 | 0.065 | 1 | 0.557784478 | 1.432959407 | 0 | 0.034 | 1 | 0.557784478 | 0.009529291 |
| Arhgap22 | 0.004777334 | -3.944858446 | 0 | 0.065 | 1 | 0.307388615 | -1.069540933 | 0 | 0.096 | 1 | 0.307388615 | 0.009531846 |
| Gper1 | 0.004777382 | -3.485426827 | 0 | 0.065 | 1 | 0.698532411 | 1.558490289 | 0 | 0.015 | 1 | 0.698532411 | 0.00953194 |
| Cacnb3 | 0.004777382 | -3.485426827 | 0 | 0.065 | 1 | 0.6199755 | 1.847996907 | 0 | 0.025 | 1 | 0.6199755 | 0.00953194 |
| Zfp53 | 0.004778118 | -1.485426827 | 0.109 | 0.235 | 1 | 0.222195832 | -1.976431529 | 0.1 | 0.259 | 1 | 0.222195832 | 0.009533406 |
| Gm17167 | 0.00477819 | -3.807354922 | 0 | 0.065 | 1 | 0.489976483 | 0.159940913 | 0 | 0.046 | 1 | 0.489976483 | 0.009533549 |
| Nphp4 | 0.00477819 | -3.733354341 | 0 | 0.065 | 1 | 0.643835681 | 1.847996907 | 0 | 0.022 | 1 | 0.643835681 | 0.009533549 |
| Kctd12b | 0.00477838 | -3.572889668 | 0 | 0.065 | 1 | 0.374687157 | -0.191531458 | 0 | 0.074 | 1 | 0.374687157 | 0.009533927 |
| Pax3 | 0.004779093 | -4.770829046 | 0 | 0.065 | 1 | 0.923993469 | 0.558490289 | 0.1 | 0.117 | 1 | 0.923993469 | 0.009535347 |
| Ddt | 0.004784603 | 0.641105579 | 0.613 | 0.465 | 1 | 0.221291883 | -0.465893869 | 0.2 | 0.42 | 1 | 0.221291883 | 0.009546314 |
| Als2 | 0.004785613 | -1.450661409 | 0.143 | 0.271 | 1 | 0.786185503 | 0.111031312 | 0.3 | 0.244 | 1 | 0.786185503 | 0.009548324 |
| Cnpy3 | 0.00478661 | -0.772307975 | 0.37 | 0.5 | 1 | 0.113518292 | 0.828097349 | 0.6 | 0.333 | 1 | 0.113518292 | 0.009550308 |
| Utp3 | 0.782073684 | -0.123687554 | 0.563 | 0.547 | 1 | 0.004792949 | -1.953621646 | 0.3 | 0.688 | 1 | 0.782073684 | 0.009562926 |
| Gm36266 | 0.342795612 | -1.070389328 | 0.017 | 0.035 | 1 | 0.00479527 | 3.14345279 | 0.2 | 0.031 | 1 | 0.342795612 | 0.009567546 |
| 9530077C14Rik | 0.623249108 | 0.225066556 | 0.067 | 0.053 | 1 | 0.00479527 | 3.14345279 | 0.2 | 0.031 | 1 | 0.623249108 | 0.009567546 |
| Gm56743 | 0.004795762 | -1.205318908 | 0.126 | 0.259 | 1 | 0.716420584 | 0.351165316 | 0.4 | 0.321 | 1 | 0.716420584 | 0.009568524 |
| Kif17 | 0.004802237 | -2.292781749 | 0.042 | 0.141 | 1 | 0.812118963 | 0.403212064 | 0.1 | 0.133 | 1 | 0.812118963 | 0.009581413 |
| Pcolce2 | 0.00481814 | 0.241888597 | 0.42 | 0.247 | 1 | 0.422022867 | 0.522066881 | 0.4 | 0.281 | 1 | 0.422022867 | 0.009613065 |
| Ccdc43 | 0.004820874 | -1.292781749 | 0.126 | 0.253 | 1 | 0.708157708 | -0.191531458 | 0.2 | 0.238 | 1 | 0.708157708 | 0.009618508 |
| Ykt6 | 0.004830462 | -0.72799713 | 0.445 | 0.571 | 1 | 0.23476987 | -0.71775499 | 0.6 | 0.623 | 1 | 0.23476987 | 0.009637591 |
| Akna | 0.004842194 | -2.707819249 | 0.017 | 0.1 | 1 | 0.36412085 | -0.339630097 | 0 | 0.077 | 1 | 0.36412085 | 0.009660942 |
| Ppcs | 0.004842309 | -2.424026283 | 0.017 | 0.1 | 1 | 0.522151747 | 1.210566986 | 0 | 0.04 | 1 | 0.522151747 | 0.00966117 |
| Psme1 | 0.004844924 | -0.485426827 | 0.395 | 0.576 | 1 | 0.051151445 | -1.272096939 | 0.3 | 0.574 | 1 | 0.051151445 | 0.009666374 |
| Hmgcl | 0.004847261 | -0.839063782 | 0.235 | 0.388 | 1 | 0.394595079 | -0.489872732 | 0.3 | 0.398 | 1 | 0.394595079 | 0.009671026 |
| Cyb561a3 | 0.051318423 | -0.959358016 | 0.059 | 0.129 | 1 | 0.00484772 | 2.494359952 | 0.3 | 0.065 | 1 | 0.051318423 | 0.009671939 |
| Rnf20 | 0.004850608 | -0.621631211 | 0.546 | 0.647 | 1 | 0.152073531 | -0.775049096 | 0.5 | 0.71 | 1 | 0.152073531 | 0.009677687 |
| Bms1 | 0.004861794 | -0.77385801 | 0.37 | 0.512 | 1 | 0.083300727 | -1.290417122 | 0.3 | 0.528 | 1 | 0.083300727 | 0.009699952 |
| Adamts1 | 0.004863866 | -0.700743731 | 0.16 | 0.312 | 1 | 0.690347998 | -0.131825212 | 0.2 | 0.272 | 1 | 0.690347998 | 0.009704075 |
| Xpo1 | 0.004866813 | -1.06153841 | 0.445 | 0.571 | 1 | 0.407976831 | -0.760520322 | 0.6 | 0.593 | 1 | 0.407976831 | 0.009709941 |
| Chaserr | 0.004867459 | -0.47656685 | 0.807 | 0.835 | 1 | 0.150179753 | -0.589408406 | 0.7 | 0.904 | 1 | 0.150179753 | 0.009711226 |
| Sil1 | 0.004886532 | 0.192645078 | 0.504 | 0.312 | 1 | 0.639998706 | 0.256370676 | 0.5 | 0.395 | 1 | 0.639998706 | 0.009749185 |
| Rnf166 | 0.004897199 | -0.813001485 | 0.336 | 0.476 | 1 | 0.913173135 | 0.129178659 | 0.5 | 0.444 | 1 | 0.913173135 | 0.009770416 |
| Slc38a10 | 0.024481003 | 0.386416821 | 0.765 | 0.576 | 1 | 0.004921689 | 1.206279628 | 1 | 0.565 | 1 | 0.024481003 | 0.009819155 |
| Mirg | 0.186712935 | 0.397759508 | 0.403 | 0.335 | 1 | 0.00492778 | 2.031861099 | 0.7 | 0.377 | 1 | 0.186712935 | 0.009831277 |
| Ipo8 | 0.004936275 | -1.055742552 | 0.235 | 0.371 | 1 | 0.673066772 | 0.245332404 | 0.5 | 0.435 | 1 | 0.673066772 | 0.009848183 |
| Gm56812 | 0.00493711 | -2.359895945 | 0.017 | 0.1 | 1 | 0.271220423 | 1.176619654 | 0.2 | 0.09 | 1 | 0.271220423 | 0.009849844 |
| Zfp583 | 0.004937139 | -2.292781749 | 0.017 | 0.1 | 1 | 0.95518904 | 0.769994395 | 0.1 | 0.093 | 1 | 0.95518904 | 0.009849902 |
| Arhgap21 | 0.004939593 | -0.468009774 | 0.84 | 0.888 | 1 | 0.311165617 | -0.288828659 | 0.9 | 0.954 | 1 | 0.311165617 | 0.009854787 |
| Spg21 | 0.004943983 | -0.89224382 | 0.479 | 0.553 | 1 | 0.428013229 | 0.203713315 | 0.4 | 0.642 | 1 | 0.428013229 | 0.009863524 |
| 4833411C07Rik | 0.009793348 | 0.855610091 | 0.143 | 0.053 | 1 | 0.004944295 | 3.017921908 | 0.2 | 0.031 | 1 | 0.009793348 | 0.009864144 |
| Stoml1 | 0.004952132 | -1.452259963 | 0.067 | 0.182 | 1 | 0.763866289 | 0.210566986 | 0.1 | 0.13 | 1 | 0.763866289 | 0.00987974 |
| Arl16 | 0.004953694 | -1.133125083 | 0.092 | 0.218 | 1 | 0.44815723 | 1.403212064 | 0.2 | 0.127 | 1 | 0.44815723 | 0.00988285 |
| Lat2 | 0.004984503 | -2.222392421 | 0.017 | 0.1 | 1 | 0.245808312 | -1.004445905 | 0 | 0.12 | 1 | 0.245808312 | 0.00994416 |
| Rnf135 | 0.004984503 | -2.222392421 | 0.017 | 0.1 | 1 | 0.482051401 | 1.769994395 | 0.1 | 0.049 | 1 | 0.482051401 | 0.00994416 |
| Ovgp1 | 0.005004132 | -2.185866545 | 0.042 | 0.141 | 1 | 0.274689621 | -0.567040593 | 0 | 0.108 | 1 | 0.274689621 | 0.009983223 |
| Batf | 0.436866782 | 0.254929356 | 0.269 | 0.229 | 1 | 0.00500456 | 1.737813989 | 0.4 | 0.108 | 1 | 0.436866782 | 0.009984074 |
| Rps4x | 0.106190475 | -0.138581895 | 0.992 | 0.988 | 1 | 0.005014939 | -0.888776315 | 0.9 | 0.991 | 1 | 0.106190475 | 0.010004729 |
| Apcdd1 | 0.00502043 | -2.070389328 | 0.076 | 0.188 | 1 | 0.14815791 | -1.357117523 | 0 | 0.176 | 1 | 0.14815791 | 0.010015655 |
| F11r | 0.005031783 | -2.14839184 | 0.017 | 0.1 | 1 | 0.123676149 | -1.357117523 | 0 | 0.194 | 1 | 0.123676149 | 0.010038248 |
| Flcn | 0.005043206 | -1.359895945 | 0.134 | 0.265 | 1 | 0.207651257 | -0.577024681 | 0.1 | 0.309 | 1 | 0.207651257 | 0.010060978 |
| Sntb2 | 0.00505252 | -0.784545039 | 0.681 | 0.765 | 1 | 0.083108593 | -1.081613766 | 0.6 | 0.762 | 1 | 0.083108593 | 0.010079513 |
| Sec61g | 0.005056597 | -0.411180492 | 0.899 | 0.971 | 1 | 0.075699255 | -0.639796935 | 0.7 | 0.932 | 1 | 0.075699255 | 0.010087626 |
| Bin3 | 0.005070293 | 1.081157144 | 0.445 | 0.294 | 1 | 0.037444967 | 2.34807051 | 0.5 | 0.29 | 1 | 0.037444967 | 0.010114878 |
| Kif16b | 0.005076983 | -1.002485263 | 0.395 | 0.518 | 1 | 0.344072495 | -0.625934282 | 0.6 | 0.707 | 1 | 0.344072495 | 0.01012819 |
| Pkn3 | 0.005095194 | 0.177538186 | 0.269 | 0.129 | 1 | 0.868037657 | 0.847996907 | 0.1 | 0.083 | 1 | 0.868037657 | 0.010164427 |
| Gm9456 | 0.121845366 | -1.333423734 | 0.034 | 0.076 | 1 | 0.005106981 | 2.695993813 | 0.2 | 0.031 | 1 | 0.121845366 | 0.010187882 |
| Gm43112 | 0.511005543 | -0.485426827 | 0.008 | 0.018 | 1 | 0.005106981 | 2.695993813 | 0.2 | 0.031 | 1 | 0.511005543 | 0.010187882 |
| Gm30963 | 0.93771233 | 0.251538767 | 0.05 | 0.053 | 1 | 0.005106981 | 2.902444691 | 0.2 | 0.031 | 1 | 0.93771233 | 0.010187882 |
| Gm20705 | 0.962120409 | 0.192645078 | 0.017 | 0.018 | 1 | 0.005106981 | 2.902444691 | 0.2 | 0.031 | 1 | 0.962120409 | 0.010187882 |
| Lamtor5 | 0.005112169 | -0.540803584 | 0.63 | 0.712 | 1 | 0.668653066 | -0.331789343 | 0.7 | 0.636 | 1 | 0.668653066 | 0.010198203 |
| Cc2d1a | 0.005112926 | -2.070389328 | 0.017 | 0.1 | 1 | 0.810674074 | 0.374065718 | 0.1 | 0.123 | 1 | 0.810674074 | 0.01019971 |
| Suclg1 | 0.011942886 | -0.459740184 | 0.471 | 0.576 | 1 | 0.005125449 | -1.831743819 | 0.2 | 0.639 | 1 | 0.011942886 | 0.010224628 |
| Ddb2 | 0.977098774 | -0.137503524 | 0.185 | 0.182 | 1 | 0.005128359 | 1.819142044 | 0.5 | 0.16 | 1 | 0.977098774 | 0.010230418 |
| Gmds | 0.00513491 | -0.430550499 | 0.95 | 0.929 | 1 | 0.766897741 | -0.287779872 | 1 | 0.954 | 1 | 0.766897741 | 0.010243453 |
| Fut10 | 0.005139546 | -1.854660637 | 0.034 | 0.129 | 1 | 0.679626588 | -0.357117523 | 0.1 | 0.142 | 1 | 0.679626588 | 0.010252677 |
| Gm10432 | 0.005141447 | 2.836501268 | 0.076 | 0.012 | 1 | 0.318562315 | 2.111031312 | 0.1 | 0.037 | 1 | 0.318562315 | 0.010256461 |
| Psmc4 | 0.005149064 | -0.667367205 | 0.538 | 0.618 | 1 | 0.031096009 | -1.161987182 | 0.5 | 0.707 | 1 | 0.031096009 | 0.010271614 |
| Cdk8 | 0.005153603 | -0.469352064 | 0.958 | 0.941 | 1 | 0.490142459 | 0.21400607 | 1 | 0.948 | 1 | 0.490142459 | 0.010280647 |
| Fbn2 | 0.005171641 | -3.842978832 | 0.008 | 0.082 | 1 | 0.44694818 | -1.982078092 | 0 | 0.056 | 1 | 0.44694818 | 0.010316536 |
| Scd2 | 0.005181422 | 0.435889823 | 0.84 | 0.735 | 1 | 0.064718138 | 1.426295677 | 0.8 | 0.664 | 1 | 0.064718138 | 0.010335997 |
| Nbdy | 0.716310655 | -0.109669316 | 0.723 | 0.688 | 1 | 0.005196466 | -2.395706021 | 0.1 | 0.559 | 1 | 0.716310655 | 0.010365929 |
| Heyl | 0.005200809 | -3.343407822 | 0.008 | 0.082 | 1 | 0.316078209 | -1.026472211 | 0 | 0.093 | 1 | 0.316078209 | 0.010374569 |
| 1700110C19Rik | 0.736440063 | -0.485426827 | 0.034 | 0.041 | 1 | 0.005203772 | 2.31748219 | 0.3 | 0.065 | 1 | 0.736440063 | 0.010380466 |
| Szrd1 | 0.005220616 | -0.638278315 | 0.504 | 0.606 | 1 | 0.24824102 | -0.355536488 | 0.4 | 0.682 | 1 | 0.24824102 | 0.010413978 |
| Meis3 | 0.005221714 | -1.485426827 | 0.101 | 0.224 | 1 | 0.189663393 | 1.558490289 | 0.3 | 0.164 | 1 | 0.189663393 | 0.010416162 |
| Rbmxl1 | 0.005225631 | -0.666492386 | 0.462 | 0.582 | 1 | 0.062829953 | -1.030110788 | 0.3 | 0.596 | 1 | 0.062829953 | 0.010423955 |
| Gm57189 | 0.005229632 | -3.185866545 | 0.008 | 0.082 | 1 | 0.420494609 | 1.063725598 | 0.2 | 0.117 | 1 | 0.420494609 | 0.010431916 |
| Zfp512b | 0.005239178 | -1.485426827 | 0.084 | 0.2 | 1 | 0.66301577 | -0.255096586 | 0.2 | 0.247 | 1 | 0.66301577 | 0.010450906 |
| Pdp1 | 0.005253889 | -1.774933444 | 0.067 | 0.176 | 1 | 0.868755451 | 0.227844977 | 0.2 | 0.17 | 1 | 0.868755451 | 0.010480174 |
| Rpap2 | 0.005254331 | -1.807354922 | 0.067 | 0.176 | 1 | 0.557765089 | 0.625604485 | 0.2 | 0.127 | 1 | 0.557765089 | 0.010481053 |
| Cmya5 | 0.005258106 | -3.070389328 | 0.008 | 0.082 | 1 | 0.037348434 | 2.393431043 | 0.2 | 0.049 | 1 | 0.037348434 | 0.010488564 |
| Cd300a | 0.00525822 | -3.008988783 | 0.008 | 0.082 | 1 | 0.657567303 | 1.354956895 | 0.1 | 0.068 | 1 | 0.657567303 | 0.010488792 |
| Mrps5 | 0.005267519 | -1.163498732 | 0.218 | 0.347 | 1 | 0.041993032 | -2.339630097 | 0 | 0.302 | 1 | 0.041993032 | 0.010507291 |
| Iqank1 | 0.005286415 | -3.129283017 | 0.008 | 0.082 | 1 | 0.005274124 | 2.354956895 | 0.2 | 0.031 | 1 | 0.005286415 | 0.010520432 |
| Rnft2 | 0.037579775 | 1.029146346 | 0.076 | 0.024 | 1 | 0.005274124 | 2.795529487 | 0.2 | 0.031 | 1 | 0.037579775 | 0.010520432 |
| Ube2a | 0.005276312 | -0.664976248 | 0.479 | 0.606 | 1 | 0.021207 | -0.861661342 | 0.3 | 0.704 | 1 | 0.021207 | 0.010524784 |
| Morc3 | 0.005276366 | -1.108863476 | 0.261 | 0.388 | 1 | 0.184662273 | -1.138582578 | 0.2 | 0.383 | 1 | 0.184662273 | 0.010524893 |
| Cs | 0.00528049 | -0.953979836 | 0.252 | 0.394 | 1 | 0.31026868 | -0.505640048 | 0.4 | 0.583 | 1 | 0.31026868 | 0.010533097 |

| | | | | | | | | | | | | |
|---|---|---|---|---|---|---|---|---|---|---|---|---|
| Tubgcp5 | 0.005282416 | -1.354182294 | 0.168 | 0.294 | 1 | 0.726303262 | -0.408342847 | 0.3 | 0.312 | 1 | 0.726303262 | 0.010536928 |
| Znhit6 | 0.005285043 | -1.116782233 | 0.328 | 0.453 | 1 | 0.681175917 | 0.646363045 | 0.4 | 0.346 | 1 | 0.681175917 | 0.010542154 |
| Acot11 | 0.005286415 | -2.87774425 | 0.008 | 0.082 | 1 | 0.274802592 | -0.789433014 | 0 | 0.108 | 1 | 0.274802592 | 0.010544884 |
| Slc6a9 | 0.005286423 | -2.240314329 | 0.017 | 0.1 | 1 | 0.673292204 | 1.31748219 | 0.1 | 0.065 | 1 | 0.673292204 | 0.0105449 |
| Shank3 | 0.005286453 | -2.733354341 | 0.008 | 0.082 | 1 | 0.522115639 | 0.695993813 | 0 | 0.04 | 1 | 0.522115639 | 0.01054496 |
| Foxk1 | 0.005309805 | -0.970853654 | 0.336 | 0.465 | 1 | 0.870971411 | -0.187982391 | 0.6 | 0.59 | 1 | 0.870971411 | 0.010591416 |
| Zfp940 | 0.005314326 | -2.733354341 | 0.008 | 0.082 | 1 | 0.433577142 | 0.432959407 | 0 | 0.059 | 1 | 0.433577142 | 0.010600409 |
| Tmem87b | 0.005318148 | -0.641344971 | 0.538 | 0.647 | 1 | 0.611671143 | -0.18874364 | 0.6 | 0.722 | 1 | 0.611671143 | 0.010608013 |
| Rpf2 | 0.005318929 | -1.19592021 | 0.143 | 0.276 | 1 | 0.371884609 | -0.625934282 | 0.1 | 0.21 | 1 | 0.371884609 | 0.010609567 |
| Utp4 | 0.005324854 | -0.865850571 | 0.345 | 0.465 | 1 | 0.225059482 | -0.840059087 | 0.4 | 0.525 | 1 | 0.225059482 | 0.010621355 |
| Ube2q1 | 0.005325028 | -0.730539325 | 0.479 | 0.582 | 1 | 0.443319481 | -0.123988717 | 0.6 | 0.728 | 1 | 0.443319481 | 0.0106217 |
| Gm57352 | 0.276045493 | 0.277533976 | 0.185 | 0.135 | 1 | 0.005325693 | 1.785261151 | 0.4 | 0.105 | 1 | 0.276045493 | 0.010623024 |
| Kpna2 | 0.005340745 | -1.707819249 | 0.034 | 0.129 | 1 | 0.599357181 | -0.152003093 | 0.1 | 0.157 | 1 | 0.599357181 | 0.010652967 |
| Triap1 | 0.005355006 | -1.091147888 | 0.16 | 0.294 | 1 | 0.038627951 | -2.026472211 | 0 | 0.309 | 1 | 0.038627951 | 0.010681335 |
| Synpo | 0.005355972 | 0.640104055 | 0.857 | 0.718 | 1 | 0.398408498 | 0.50222207 | 0.8 | 0.716 | 1 | 0.398408498 | 0.010683258 |
| Arhgef9 | 0.00536034 | -1.944858446 | 0.059 | 0.165 | 1 | 0.682821373 | 0.494359952 | 0.2 | 0.145 | 1 | 0.682821373 | 0.010691948 |
| Pmf1 | 0.005364821 | -1.137503524 | 0.092 | 0.218 | 1 | 0.22436076 | 1.939919396 | 0.3 | 0.194 | 1 | 0.22436076 | 0.010700861 |
| Dcaf8 | 0.005365166 | -0.640312429 | 0.756 | 0.765 | 1 | 0.659819532 | 0.168256181 | 0.9 | 0.849 | 1 | 0.659819532 | 0.010701548 |
| Slc12a6 | 0.005366814 | -1.230497471 | 0.378 | 0.494 | 1 | 0.290463624 | -0.464470859 | 0.5 | 0.71 | 1 | 0.290463624 | 0.010704826 |
| Slc35d2 | 0.005369158 | -2.572889668 | 0.008 | 0.082 | 1 | 0.489989782 | -0.191531458 | 0 | 0.046 | 1 | 0.489989782 | 0.010709489 |
| Pogz | 0.005378159 | -0.977279923 | 0.277 | 0.412 | 1 | 0.704462983 | -0.135883428 | 0.5 | 0.537 | 1 | 0.704462983 | 0.010727394 |
| Sh3bp4 | 0.005380286 | -1.407424315 | 0.118 | 0.247 | 1 | 0.196709618 | -0.728032469 | 0.1 | 0.312 | 1 | 0.196709618 | 0.010731624 |
| Trim37 | 0.005385032 | -1.103556192 | 0.286 | 0.418 | 1 | 0.635825574 | 0.729160677 | 0.4 | 0.386 | 1 | 0.635825574 | 0.010741065 |
| Gm4956 | 0.005385504 | -3.14839184 | 0.008 | 0.082 | 1 | 0.408843595 | -1.026472211 | 0 | 0.065 | 1 | 0.408843595 | 0.010742004 |
| Adat1 | 0.263913569 | -0.661276663 | 0.193 | 0.241 | 1 | 0.005387972 | 1.664284953 | 0.5 | 0.167 | 1 | 0.263913569 | 0.010746915 |
| Fen1 | 0.014471136 | -1.424026283 | 0.025 | 0.1 | 1 | 0.005389893 | 2.31748219 | 0.3 | 0.065 | 1 | 0.014471136 | 0.010750734 |
| Lepr | 0.005398192 | -1.414343729 | 0.252 | 0.382 | 1 | 0.028978072 | -2.033740212 | 0.1 | 0.472 | 1 | 0.028978072 | 0.010767244 |
| Atp6v0a4 | 0.005419478 | -1.614709844 | 0.202 | 0.324 | 1 | 0.589456252 | 0.239563121 | 0.8 | 0.821 | 1 | 0.589456252 | 0.010809586 |
| Map2k3 | 0.005431645 | 0.245887099 | 0.933 | 0.894 | 1 | 0.247320746 | -0.505640048 | 0.9 | 0.892 | 1 | 0.247320746 | 0.010833787 |
| Atl3 | 0.005433245 | -0.432959407 | 0.655 | 0.718 | 1 | 0.291997375 | -0.443693958 | 0.7 | 0.824 | 1 | 0.291997375 | 0.010836971 |
| Nrip3 | 0.005435669 | 1.156119202 | 0.21 | 0.094 | 1 | 0.851068114 | 1.290001453 | 0.1 | 0.09 | 1 | 0.851068114 | 0.010841791 |
| Rttn | 0.005444769 | -2.198144875 | 0.042 | 0.141 | 1 | 0.206615906 | 1.176619654 | 0.2 | 0.08 | 1 | 0.206615906 | 0.010859893 |
| Gm20735 | 0.147030916 | -1.485426827 | 0 | 0.018 | 1 | 0.005445712 | 2.602884409 | 0.2 | 0.031 | 1 | 0.147030916 | 0.010861767 |
| Gm47460 | 0.099733028 | 1.736965594 | 0.042 | 0.012 | 1 | 0.00544582 | 2.515421567 | 0.2 | 0.031 | 1 | 0.099733028 | 0.010861983 |
| Klhl18 | 0.005451386 | -1.204245075 | 0.151 | 0.282 | 1 | 0.544571212 | -0.385800278 | 0.2 | 0.272 | 1 | 0.544571212 | 0.010873054 |
| Armc7 | 0.005455504 | -1.600904045 | 0.034 | 0.129 | 1 | 0.707564437 | 1.159940913 | 0.1 | 0.068 | 1 | 0.707564437 | 0.010881245 |
| Lyar | 0.00545927 | -0.911190733 | 0.37 | 0.512 | 1 | 0.395696183 | -0.542793046 | 0.3 | 0.42 | 1 | 0.395696183 | 0.010888737 |
| Trim2 | 0.024087495 | -0.617338503 | 0.336 | 0.471 | 1 | 0.005460401 | -2.650963076 | 0.1 | 0.559 | 1 | 0.024087495 | 0.010890986 |
| Ppme1 | 0.005472615 | 0.376599639 | 0.807 | 0.718 | 1 | 0.768289874 | 0.309935381 | 0.6 | 0.731 | 1 | 0.768289874 | 0.01091528 |
| Clk1 | 0.00548202 | -0.470078863 | 0.983 | 0.994 | 1 | 0.514612972 | -0.140167957 | 1 | 0.975 | 1 | 0.514612972 | 0.010933988 |
| Vangl1 | 0.00548618 | -0.93021167 | 0.101 | 0.229 | 1 | 0.681008559 | -0.204470513 | 0.2 | 0.244 | 1 | 0.681008559 | 0.010942262 |
| Banf1 | 0.005506551 | -0.491475872 | 0.58 | 0.712 | 1 | 0.016752538 | -1.505640048 | 0.3 | 0.636 | 1 | 0.016752538 | 0.01098278 |
| Slc38a3 | 0.005513689 | -1.135680788 | 0.235 | 0.365 | 1 | 0.675887665 | -0.48032896 | 0.4 | 0.404 | 1 | 0.675887665 | 0.010996977 |
| Arl8a | 0.005515884 | -0.639549972 | 0.529 | 0.641 | 1 | 0.005658481 | -1.469343919 | 0.4 | 0.802 | 1 | 0.005658481 | 0.011001343 |
| Dcp1a | 0.005517695 | -1.170791225 | 0.294 | 0.412 | 1 | 0.733820377 | 0.570462931 | 0.4 | 0.401 | 1 | 0.733820377 | 0.011004946 |
| Fgfr4 | 0.005519729 | -1.765534746 | 0.042 | 0.141 | 1 | 0.474962821 | 0.494359952 | 0 | 0.049 | 1 | 0.474962821 | 0.01100899 |
| Adprs | 0.005530405 | -1.352160296 | 0.134 | 0.259 | 1 | 0.170721584 | -1.191531458 | 0 | 0.16 | 1 | 0.170721584 | 0.011030224 |
| Lnx2 | 0.950373978 | -0.273922722 | 0.076 | 0.076 | 1 | 0.005533298 | 1.832055363 | 0.5 | 0.157 | 1 | 0.950373978 | 0.011035978 |
| Rps21 | 0.00553756 | 0.222704502 | 1 | 0.988 | 1 | 0.020239478 | -0.749424469 | 1 | 0.991 | 1 | 0.020239478 | 0.011044455 |
| Tm2d3 | 0.005542067 | 0.643470906 | 0.571 | 0.447 | 1 | 0.583521972 | -0.163407857 | 0.4 | 0.503 | 1 | 0.583521972 | 0.011053419 |
| Sf3a3 | 0.005543204 | -0.812237143 | 0.378 | 0.506 | 1 | 0.035652195 | -1.484578433 | 0.2 | 0.519 | 1 | 0.035652195 | 0.011055682 |
| Zfp329 | 0.005553434 | -1.547711105 | 0.118 | 0.241 | 1 | 0.293410341 | -0.473931188 | 0.1 | 0.262 | 1 | 0.293410341 | 0.011076027 |
| Dnajb14 | 0.005554489 | -1.034765418 | 0.462 | 0.565 | 1 | 0.770638261 | -0.133671271 | 0.5 | 0.583 | 1 | 0.770638261 | 0.011078125 |
| Lgals9 | 0.005563403 | -1.807354922 | 0.05 | 0.153 | 1 | 0.919812491 | 0.769994395 | 0.1 | 0.09 | 1 | 0.919812491 | 0.011095854 |
| Cdc42bpb | 0.005565676 | -0.693825976 | 0.672 | 0.759 | 1 | 0.457853494 | -0.199983122 | 0.7 | 0.861 | 1 | 0.457853494 | 0.011100374 |
| Ogg1 | 0.005573695 | -1.722466024 | 0.05 | 0.153 | 1 | 0.925158498 | 0.769994395 | 0.1 | 0.09 | 1 | 0.925158498 | 0.011116324 |
| Mcat | 0.613538034 | -0.263034406 | 0.05 | 0.065 | 1 | 0.005581377 | 2.159940913 | 0.3 | 0.065 | 1 | 0.613538034 | 0.011131602 |
| Zc3h8 | 0.628300786 | 0.273565073 | 0.067 | 0.053 | 1 | 0.005581492 | 2.111031312 | 0.3 | 0.065 | 1 | 0.628300786 | 0.01113183 |
| Adrb2 | 0.00558399 | 0.915111102 | 0.176 | 0.071 | 1 | 0.933420476 | 0.732519689 | 0.1 | 0.108 | 1 | 0.933420476 | 0.011136799 |
| Neurl4 | 0.005596002 | -1.207892852 | 0.059 | 0.171 | 1 | 0.175163934 | -0.959358016 | 0 | 0.157 | 1 | 0.175163934 | 0.011160688 |
| Sgpp1 | 0.00560583 | 1.504377406 | 0.429 | 0.288 | 1 | 0.288690389 | -0.709998547 | 0.2 | 0.343 | 1 | 0.288690389 | 0.011180234 |
| 9830144P21Rik | 0.005612081 | 1.836501268 | 0.109 | 0.029 | 1 | 0.110850287 | 2.602884409 | 0.1 | 0.022 | 1 | 0.110850287 | 0.011192666 |
| Sf3b1 | 0.005620501 | -0.246188081 | 0.992 | 0.994 | 1 | 0.155300156 | -0.276995234 | 1 | 1 | 1 | 0.155300156 | 0.011209411 |
| Ccdc162 | 0.217123465 | 1 | 0.067 | 0.035 | 1 | 0.005622165 | 2.210566986 | 0.2 | 0.031 | 1 | 0.217123465 | 0.011212722 |
| Dnaaf3 | 0.647331896 | 1.321928095 | 0.025 | 0.018 | 1 | 0.005622314 | 2.602884409 | 0.2 | 0.031 | 1 | 0.647331896 | 0.011213017 |
| Lrrk1 | 0.005626859 | -0.627337452 | 0.563 | 0.688 | 1 | 0.009739058 | -1.845576092 | 0.4 | 0.747 | 1 | 0.009739058 | 0.011222057 |
| Gm16740 | 0.005646728 | -1.842978832 | 0.059 | 0.165 | 1 | 0.890691321 | 0.374065718 | 0.1 | 0.111 | 1 | 0.890691321 | 0.01126157 |
| Rab22a | 0.005649913 | -0.726953415 | 0.58 | 0.659 | 1 | 0.01030334 | -1.302001051 | 0.3 | 0.802 | 1 | 0.01030334 | 0.011267905 |
| Banp | 0.005669895 | -0.959358016 | 0.328 | 0.453 | 1 | 0.007241508 | -1.625934282 | 0.1 | 0.608 | 1 | 0.007241508 | 0.011307642 |
| Rpl7a | 0.014731415 | -0.194745893 | 0.95 | 0.982 | 1 | 0.005679885 | -0.850628329 | 0.9 | 0.994 | 1 | 0.014731415 | 0.011327509 |
| Ppm1m | 0.005684364 | 0.482864313 | 0.521 | 0.359 | 1 | 0.310360276 | 0.60610831 | 0.5 | 0.333 | 1 | 0.310360276 | 0.011336416 |
| Atp1a2 | 0.005691401 | 3.301169535 | 0.118 | 0.035 | 1 | 0.589586793 | 3.662965013 | 0.2 | 0.167 | 1 | 0.589586793 | 0.011350409 |
| Arl6ip5 | 0.005692553 | 0.255838904 | 1 | 0.988 | 1 | 0.080207306 | -0.570257305 | 1 | 1 | 1 | 0.080207306 | 0.011352701 |
| Atxn10 | 0.005695458 | -0.507267876 | 0.857 | 0.918 | 1 | 0.055393069 | -0.708814554 | 0.7 | 0.935 | 1 | 0.055393069 | 0.011358477 |
| Abcd4 | 0.005702871 | -1.722466024 | 0.05 | 0.153 | 1 | 0.647478838 | -0.111361109 | 0.1 | 0.148 | 1 | 0.647478838 | 0.01137322 |
| Mical2 | 0.005707238 | -3.185866545 | 0.025 | 0.112 | 1 | 0.238734328 | -0.91281543 | 0 | 0.123 | 1 | 0.238734328 | 0.011381903 |
| Fst | 0.005709227 | -1.560714954 | 0.042 | 0.141 | 1 | 0.669852582 | -1.357117523 | 0 | 0.019 | 1 | 0.669852582 | 0.01138586 |
| Isoc2a | 0.031489901 | 0.836501268 | 0.143 | 0.065 | 1 | 0.00571334 | 3.795529487 | 0.2 | 0.034 | 1 | 0.031489901 | 0.011394038 |
| D130043K22Rik | 0.764297315 | 0.707218251 | 0.042 | 0.035 | 1 | 0.005713359 | 2.063725598 | 0.2 | 0.031 | 1 | 0.764297315 | 0.011394075 |
| Hlx | 0.005719854 | 1.062060968 | 0.202 | 0.088 | 1 | 0.323436159 | 1.063725598 | 0.1 | 0.037 | 1 | 0.323436159 | 0.011406991 |
| Golph3l | 0.005724107 | -1.242856524 | 0.143 | 0.271 | 1 | 0.340842072 | -0.616284112 | 0.1 | 0.238 | 1 | 0.340842072 | 0.011415448 |
| Grpel2 | 0.005727135 | -1.74176658 | 0.067 | 0.176 | 1 | 0.057665063 | -1.91281543 | 0 | 0.272 | 1 | 0.057665063 | 0.011421469 |
| Ube2n | 0.005736285 | -0.322165556 | 0.882 | 0.894 | 1 | 0.636665288 | -0.168605061 | 0.8 | 0.907 | 1 | 0.636665288 | 0.011439665 |
| Limd2 | 0.375566075 | -0.683366205 | 0.126 | 0.159 | 1 | 0.005739378 | 1.94753258 | 0.4 | 0.108 | 1 | 0.375566075 | 0.011445816 |
| Knop1 | 0.005771081 | -1.115477217 | 0.286 | 0.418 | 1 | 0.108339951 | -1.310004928 | 0.2 | 0.432 | 1 | 0.108339951 | 0.011508857 |
| Wdr1 | 0.005773261 | 0.237886666 | 0.891 | 0.759 | 1 | 0.462016032 | 0.438121112 | 0.8 | 0.867 | 1 | 0.462016032 | 0.011513191 |
| Nbas | 0.005779572 | -0.835233955 | 0.496 | 0.576 | 1 | 0.938440025 | -0.220482831 | 0.6 | 0.552 | 1 | 0.938440025 | 0.011525741 |
| D330050G23Rik | 0.005785604 | 1.794681092 | 0.109 | 0.029 | 1 | 0.112266525 | 3.017921908 | 0.1 | 0.022 | 1 | 0.112266525 | 0.011537735 |
| AI467606 | 0.005790342 | -1.987927168 | 0.008 | 0.082 | 1 | 0.421000218 | 0.111031312 | 0 | 0.062 | 1 | 0.421000218 | 0.011547155 |
| Ankrd37 | 0.005790342 | -1.987927168 | 0.008 | 0.082 | 1 | 0.460633694 | 0.31748219 | 0 | 0.052 | 1 | 0.460633694 | 0.011547155 |
| Btbd1 | 0.005798375 | -0.638673087 | 0.529 | 0.665 | 1 | 0.673327503 | 0.142141845 | 0.8 | 0.827 | 1 | 0.673327503 | 0.011563128 |
| 2410080I02Rik | 0.820836204 | -0.485426827 | 0.025 | 0.029 | 1 | 0.00580345 | 2.210566986 | 0.2 | 0.031 | 1 | 0.820836204 | 0.011573219 |
| Plxnb2 | 0.005815607 | -0.671839951 | 0.319 | 0.465 | 1 | 0.097594957 | -0.789433014 | 0.3 | 0.583 | 1 | 0.097594957 | 0.011597394 |
| Pcdhga7 | 0.005815972 | 1.131244533 | 0.16 | 0.059 | 1 | 0.316130348 | -1.004445905 | 0 | 0.093 | 1 | 0.316130348 | 0.011598119 |
| Ints5 | 0.005833693 | -1.326729081 | 0.092 | 0.212 | 1 | 0.967438895 | 0.245332404 | 0.2 | 0.198 | 1 | 0.967438895 | 0.011633355 |
| Fubp3 | 0.005846315 | -0.937939032 | 0.261 | 0.4 | 1 | 0.006059514 | -3.009984089 | 0 | 0.46 | 1 | 0.006059514 | 0.01165845 |
| Rspry1 | 0.005853829 | -0.691877705 | 0.571 | 0.659 | 1 | 0.774398996 | 0.222887258 | 0.9 | 0.731 | 1 | 0.774398996 | 0.011673391 |
| Atg5 | 0.005864734 | -0.606442228 | 0.412 | 0.559 | 1 | 0.095102821 | -1.167944637 | 0.4 | 0.586 | 1 | 0.095102821 | 0.011695074 |

| | | | | | | | | | | | | |
|---|---|---|---|---|---|---|---|---|---|---|---|---|
| Myadm | 0.005865313 | -0.638857625 | 0.714 | 0.759 | 1 | 0.942104051 | -0.147990031 | 0.8 | 0.707 | 1 | 0.942104051 | 0.011696224 |
| Gtf2e1 | 0.005903749 | -1.145351386 | 0.176 | 0.306 | 1 | 0.395755763 | 0.591657153 | 0.4 | 0.265 | 1 | 0.395755763 | 0.011772645 |
| Phf1 | 0.005904559 | 0.566412104 | 0.487 | 0.318 | 1 | 0.998502519 | 0.19592021 | 0.4 | 0.429 | 1 | 0.998502519 | 0.011774254 |
| Dpf2 | 0.005908988 | -1.03562391 | 0.21 | 0.347 | 1 | 0.790262737 | 0.299103661 | 0.5 | 0.469 | 1 | 0.790262737 | 0.011783061 |
| Acsl1 | 0.081627526 | 0.472447697 | 0.521 | 0.447 | 1 | 0.005915838 | 1.961090685 | 0.8 | 0.444 | 1 | 0.081627526 | 0.01179668 |
| Gm10131 | 0.005961065 | -1.544320516 | 0.042 | 0.141 | 1 | 0.926075469 | 0.432959407 | 0.1 | 0.108 | 1 | 0.926075469 | 0.011886595 |
| Rpia | 0.005964715 | -1.424026283 | 0.134 | 0.259 | 1 | 0.057694403 | -1.802257054 | 0 | 0.272 | 1 | 0.057694403 | 0.011893852 |
| Gm10648 | 0.204620047 | 0.772885168 | 0.202 | 0.147 | 1 | 0.005969929 | 2.169925001 | 0.5 | 0.185 | 1 | 0.204620047 | 0.011904218 |
| Fmod | 0.005973353 | -0.750595623 | 0.916 | 0.865 | 1 | 0.01120187 | -1.83268315 | 1 | 0.849 | 1 | 0.01120187 | 0.011911024 |
| Fam43a | 0.005980621 | 0.749038426 | 0.227 | 0.106 | 1 | 0.01407783 | 2.176619654 | 0.4 | 0.133 | 1 | 0.01407783 | 0.011925474 |
| Castor2 | 0.006014214 | -1.538538164 | 0.143 | 0.265 | 1 | 0.073954068 | -1.230005605 | 0.2 | 0.54 | 1 | 0.073954068 | 0.011992257 |
| Tfpt | 0.006016769 | -1.168236651 | 0.134 | 0.265 | 1 | 0.278615088 | -0.596787936 | 0.1 | 0.247 | 1 | 0.278615088 | 0.011997336 |
| Tmed4 | 0.380337021 | 0.146841388 | 0.538 | 0.471 | 1 | 0.006023138 | -2.186649236 | 0.1 | 0.552 | 1 | 0.380337021 | 0.012009998 |
| Cpeb3 | 0.057006524 | -0.150762744 | 0.328 | 0.424 | 1 | 0.006024026 | 1.276454922 | 0.8 | 0.407 | 1 | 0.057006524 | 0.012011763 |
| Tent5c | 0.0060284 | -1.694880193 | 0.05 | 0.153 | 1 | 0.894315581 | -0.171902651 | 0.1 | 0.108 | 1 | 0.894315581 | 0.012020459 |
| Psmd9 | 0.006040582 | -0.884833675 | 0.361 | 0.488 | 1 | 0.895169176 | 0.105032572 | 0.5 | 0.469 | 1 | 0.895169176 | 0.012044676 |
| Slc25a22 | 0.006068197 | -1.555816155 | 0.076 | 0.188 | 1 | 0.908353569 | 0.874963954 | 0.1 | 0.12 | 1 | 0.908353569 | 0.012099571 |
| Pou2f3 | 0.038442336 | -1.655351829 | 0.076 | 0.153 | 1 | 0.006068202 | 0.720241359 | 0.5 | 0.139 | 1 | 0.038442336 | 0.012099581 |
| Psmc5 | 0.065095054 | -0.409013712 | 0.605 | 0.688 | 1 | 0.006069628 | -1.576029376 | 0.3 | 0.716 | 1 | 0.065095054 | 0.012102416 |
| Ptpn22 | 0.637654067 | -1.655351829 | 0.042 | 0.053 | 1 | 0.006070672 | 3.602884409 | 0.2 | 0.034 | 1 | 0.637654067 | 0.012104491 |
| Prag1 | 0.006081639 | 0.59946207 | 0.218 | 0.094 | 1 | 0.539444993 | 0.930459067 | 0 | 0.037 | 1 | 0.539444993 | 0.012126292 |
| H2aj | 0.006090721 | 0.483810919 | 0.782 | 0.665 | 1 | 0.715638496 | 0.124206701 | 0.5 | 0.642 | 1 | 0.715638496 | 0.012144345 |
| Slc25a46 | 0.006097985 | -1.054792473 | 0.218 | 0.347 | 1 | 0.310457386 | 0.705038953 | 0.5 | 0.33 | 1 | 0.310457386 | 0.012158785 |
| Tmem11 | 0.006105666 | -0.675251386 | 0.58 | 0.629 | 1 | 0.244829094 | -0.563278674 | 0.5 | 0.657 | 1 | 0.244829094 | 0.012174053 |
| Arl5b | 0.006106328 | -0.81437535 | 0.412 | 0.535 | 1 | 0.474161185 | -0.539222649 | 0.5 | 0.664 | 1 | 0.474161185 | 0.01217537 |
| Dcps | 0.775691085 | 0.199699835 | 0.252 | 0.241 | 1 | 0.006125768 | 1.655351829 | 0.5 | 0.164 | 1 | 0.775691085 | 0.012214012 |
| Etv5 | 0.006153285 | 0.106915204 | 0.311 | 0.153 | 1 | 0.459292431 | -0.893769674 | 0.3 | 0.383 | 1 | 0.459292431 | 0.012268706 |
| Tusc3 | 0.006155374 | -0.623840856 | 0.672 | 0.706 | 1 | 0.041849854 | -1.283247627 | 0.6 | 0.71 | 1 | 0.041849854 | 0.012272859 |
| mt-Nd6 | 0.006171375 | 0.502600531 | 0.546 | 0.371 | 1 | 0.172363379 | 1.545434137 | 0.5 | 0.377 | 1 | 0.172363379 | 0.012304664 |
| Herpud1 | 0.125453485 | -0.201292301 | 0.924 | 0.953 | 1 | 0.00618059 | -1.193302722 | 0.8 | 0.944 | 1 | 0.125453485 | 0.01232298 |
| Exoc4 | 0.006190965 | -0.469414509 | 0.84 | 0.924 | 1 | 0.779044508 | -0.122038662 | 0.9 | 0.935 | 1 | 0.779044508 | 0.012343603 |
| Ipo13 | 0.006192987 | -2.129283017 | 0.025 | 0.112 | 1 | 0.674050765 | 0.111031312 | 0.1 | 0.145 | 1 | 0.674050765 | 0.01234762 |
| Aspm | 0.006193066 | -2.835924074 | 0.034 | 0.124 | 1 | 0.048178616 | 2.159940913 | 0.2 | 0.052 | 1 | 0.048178616 | 0.012347777 |
| 4933407K13Rik | 0.006205106 | -1.726434927 | 0.084 | 0.194 | 1 | 0.439805754 | 0.249737583 | 0.1 | 0.222 | 1 | 0.439805754 | 0.012371708 |
| Cep295 | 0.006242395 | -1.724892762 | 0.126 | 0.247 | 1 | 0.550899568 | 0.625604485 | 0.3 | 0.219 | 1 | 0.550899568 | 0.012445822 |
| Plagl2 | 0.006242956 | -0.736965594 | 0.109 | 0.241 | 1 | 0.199171249 | -0.970762779 | 0.1 | 0.278 | 1 | 0.199171249 | 0.012446938 |
| Nup155 | 0.006248812 | -1.142539114 | 0.168 | 0.306 | 1 | 0.848875292 | -0.201246612 | 0.3 | 0.296 | 1 | 0.848875292 | 0.012458577 |
| Phaf1 | 0.006266633 | -1.175742328 | 0.16 | 0.288 | 1 | 0.752932261 | -0.360589715 | 0.4 | 0.392 | 1 | 0.752932261 | 0.012493995 |
| Ercc4 | 0.202096105 | -0.385891154 | 0.084 | 0.135 | 1 | 0.006274905 | 1.558490289 | 0.5 | 0.154 | 1 | 0.202096105 | 0.012510435 |
| Gabpa | 0.006289118 | -1.021479727 | 0.202 | 0.335 | 1 | 0.165683015 | -1.097555309 | 0.2 | 0.395 | 1 | 0.165683015 | 0.012538683 |
| Slc35e3 | 0.006300317 | 0.457617288 | 0.63 | 0.465 | 1 | 0.053849934 | 0.917461726 | 0.7 | 0.401 | 1 | 0.053849934 | 0.01256094 |
| Ppih | 0.006311567 | -1.572889668 | 0.05 | 0.153 | 1 | 0.148092629 | 1.263034406 | 0.3 | 0.133 | 1 | 0.148092629 | 0.012583298 |
| Gal3st4 | 0.006313919 | -1.548436625 | 0.16 | 0.288 | 1 | 0.064522855 | -2.374395515 | 0 | 0.262 | 1 | 0.064522855 | 0.012587973 |
| Exosc8 | 0.006314669 | -1.318316841 | 0.109 | 0.229 | 1 | 0.974966647 | 0.227844977 | 0.2 | 0.194 | 1 | 0.974966647 | 0.012589463 |
| Mdc1 | 0.006333443 | -1.246239163 | 0.118 | 0.241 | 1 | 0.350177073 | -0.840059087 | 0.1 | 0.216 | 1 | 0.350177073 | 0.012626773 |
| Tubgcp2 | 0.0063531 | -1.139930261 | 0.193 | 0.329 | 1 | 0.593995307 | 0.695993813 | 0.4 | 0.34 | 1 | 0.593995307 | 0.012665837 |
| Llph | 0.006555969 | -0.620223605 | 0.529 | 0.665 | 1 | 0.006356407 | -2.215697769 | 0.1 | 0.549 | 1 | 0.006555969 | 0.012672411 |
| Sumf2 | 0.006357362 | -1.070389328 | 0.235 | 0.365 | 1 | 0.284391508 | 0.549152425 | 0.5 | 0.315 | 1 | 0.284391508 | 0.012674308 |
| Mbd6 | 0.00637171 | -1.304854582 | 0.134 | 0.259 | 1 | 0.988228485 | 0.135278859 | 0.3 | 0.284 | 1 | 0.988228485 | 0.012702822 |
| Hira | 0.006372384 | -1.080585095 | 0.269 | 0.388 | 1 | 0.084928709 | -1.017702002 | 0.2 | 0.5 | 1 | 0.084928709 | 0.01270416 |
| BC029722 | 0.006375047 | 0.640104055 | 0.403 | 0.241 | 1 | 0.839397754 | 0.432959407 | 0.2 | 0.244 | 1 | 0.839397754 | 0.012709452 |
| Mettl1 | 0.006385019 | -0.859551133 | 0.471 | 0.559 | 1 | 0.563614868 | 0.135278859 | 0.6 | 0.457 | 1 | 0.563614868 | 0.01272927 |
| Rab2b | 0.006395316 | -0.934734229 | 0.353 | 0.471 | 1 | 0.630393077 | 0.610418081 | 0.4 | 0.377 | 1 | 0.630393077 | 0.012749732 |
| Mllt1 | 0.0063965 | -1.137503524 | 0.101 | 0.224 | 1 | 0.433433693 | -0.210896782 | 0.1 | 0.198 | 1 | 0.433433693 | 0.012752085 |
| Mzt2 | 0.006405651 | -0.983677695 | 0.218 | 0.365 | 1 | 0.578997558 | -0.291933355 | 0.2 | 0.265 | 1 | 0.578997558 | 0.01277027 |
| Pgam1 | 0.006412478 | 0.406789346 | 0.924 | 0.894 | 1 | 0.556383116 | 0.434462998 | 0.8 | 0.886 | 1 | 0.556383116 | 0.012783837 |
| Psmb7 | 0.274390293 | -0.266007197 | 0.79 | 0.806 | 1 | 0.006414522 | -1.39389169 | 0.5 | 0.793 | 1 | 0.274390293 | 0.012787899 |
| Alg5 | 0.006415374 | -0.693231712 | 0.412 | 0.547 | 1 | 0.258271484 | -0.608517229 | 0.3 | 0.475 | 1 | 0.258271484 | 0.012789591 |
| Parn | 0.006424682 | -1.207892852 | 0.227 | 0.359 | 1 | 0.181713806 | 0.789103218 | 0.5 | 0.306 | 1 | 0.181713806 | 0.012808088 |
| Pcmtd1 | 0.006435314 | -0.793735313 | 0.672 | 0.753 | 1 | 0.64739521 | -0.133671271 | 0.7 | 0.824 | 1 | 0.64739521 | 0.012829215 |
| Aph1a | 0.006436661 | -0.595480373 | 0.479 | 0.594 | 1 | 0.191444133 | -0.579864633 | 0.4 | 0.633 | 1 | 0.191444133 | 0.012831892 |
| Mtln | 0.006440322 | 0.718106567 | 0.437 | 0.288 | 1 | 0.714732349 | -0.191531458 | 0.2 | 0.238 | 1 | 0.714732349 | 0.012839166 |
| G2e3 | 0.00644657 | -1.039025157 | 0.21 | 0.347 | 1 | 0.689301277 | -0.11821478 | 0.3 | 0.361 | 1 | 0.689301277 | 0.012851581 |
| Pag1 | 0.006447583 | -2.563429339 | 0.067 | 0.171 | 1 | 0.740945525 | -0.802257054 | 0.3 | 0.309 | 1 | 0.740945525 | 0.012853594 |
| B230219D22Rik | 0.006474957 | -0.4473222222 | 0.807 | 0.824 | 1 | 0.023787687 | -1.024649476 | 0.7 | 0.87 | 1 | 0.023787687 | 0.01290799 |
| Rufy4 | 0.006484303 | 0.80407979 | 0.235 | 0.118 | 1 | 0.063995139 | 1.737813989 | 0.3 | 0.108 | 1 | 0.063995139 | 0.012926561 |
| Igbp1 | 0.065858206 | -0.493000047 | 0.555 | 0.606 | 1 | 0.006485524 | -1.600463594 | 0.2 | 0.676 | 1 | 0.065858206 | 0.012928986 |
| Ptms | 0.068968627 | -0.211531429 | 0.933 | 0.976 | 1 | 0.006507947 | -0.936803153 | 1 | 0.988 | 1 | 0.068968627 | 0.01297354 |
| Slu7 | 0.006514264 | -0.944858446 | 0.353 | 0.465 | 1 | 0.104813505 | -1.279758641 | 0.2 | 0.435 | 1 | 0.104813505 | 0.012986093 |
| Sp1 | 0.006532144 | -0.714245518 | 0.563 | 0.606 | 1 | 0.505464131 | -0.396310291 | 0.8 | 0.781 | 1 | 0.505464131 | 0.013021619 |
| Snrpa | 0.00653894 | -0.739744016 | 0.496 | 0.6 | 1 | 0.218817592 | 0.591657153 | 0.7 | 0.54 | 1 | 0.218817592 | 0.013035123 |
| Prox2 | 0.224638472 | -0.987927168 | 0.042 | 0.076 | 1 | 0.006538984 | 3.754887502 | 0.2 | 0.034 | 1 | 0.224638472 | 0.01303521 |
| Rbbp5 | 0.006539824 | -1.214779237 | 0.134 | 0.259 | 1 | 0.063182632 | -1.888968688 | 0 | 0.262 | 1 | 0.063182632 | 0.013036878 |
| Ubl3 | 0.00654039 | -0.51752767 | 0.807 | 0.841 | 1 | 0.010730186 | -1.097555309 | 0.5 | 0.917 | 1 | 0.010730186 | 0.013038003 |
| Strn | 0.00654866 | -0.730539325 | 0.496 | 0.582 | 1 | 0.010663853 | -1.601686736 | 0.3 | 0.707 | 1 | 0.010663853 | 0.013054435 |
| Pde7a | 0.006569237 | -1.472935883 | 0.151 | 0.276 | 1 | 0.414744187 | -0.378682873 | 0.4 | 0.515 | 1 | 0.414744187 | 0.013095319 |
| Cfap410 | 0.006573005 | 0.882304957 | 0.277 | 0.147 | 1 | 0.52083926 | -0.131825212 | 0.1 | 0.176 | 1 | 0.52083926 | 0.013102807 |
| Airn | 0.006579891 | -0.403282306 | 0.983 | 0.982 | 1 | 0.334749891 | -0.469597584 | 1 | 0.991 | 1 | 0.334749891 | 0.013116487 |
| Gm17106 | 0.006586237 | -1.117695043 | 0.05 | 0.153 | 1 | 0.537183352 | -0.230005605 | 0.1 | 0.17 | 1 | 0.537183352 | 0.013129096 |
| Ggnbp2 | 0.006590263 | -0.429298815 | 0.958 | 0.935 | 1 | 0.257794038 | 0.181692873 | 1 | 0.978 | 1 | 0.257794038 | 0.013137094 |
| Szt2 | 0.587014071 | 0.312939312 | 0.126 | 0.106 | 1 | 0.006630985 | 1.94753258 | 0.4 | 0.111 | 1 | 0.587014071 | 0.013218001 |
| Ube2j2 | 0.062287029 | -0.459195286 | 0.672 | 0.676 | 1 | 0.006634271 | -1.347150679 | 0.6 | 0.806 | 1 | 0.062287029 | 0.013224528 |
| Gclm | 0.030319193 | -0.559427409 | 0.487 | 0.576 | 1 | 0.006638018 | -1.404983835 | 0.1 | 0.627 | 1 | 0.030319193 | 0.013231972 |
| Cbll1 | 0.006650856 | -0.868755467 | 0.311 | 0.447 | 1 | 0.305087715 | -0.457811523 | 0.3 | 0.469 | 1 | 0.305087715 | 0.013257478 |
| Eda | 0.006673691 | -1.398715194 | 0.101 | 0.224 | 1 | 0.07764858 | -3.018251705 | 0.1 | 0.358 | 1 | 0.07764858 | 0.013302844 |
| Phactr2 | 0.009580647 | -0.793549123 | 0.521 | 0.606 | 1 | 0.006677156 | -1.685289559 | 0.3 | 0.725 | 1 | 0.009580647 | 0.013309728 |
| Klhl23 | 0.00671385 | -1.485426827 | 0.05 | 0.153 | 1 | 0.282516096 | -0.789433014 | 0 | 0.105 | 1 | 0.282516096 | 0.013382625 |
| Surf4 | 0.006723065 | -0.404644172 | 0.849 | 0.847 | 1 | 0.889282581 | 0.206395335 | 0.9 | 0.87 | 1 | 0.889282581 | 0.013400931 |
| Rnf139 | 0.006737731 | -1.189970944 | 0.193 | 0.324 | 1 | 0.326929977 | -0.584962501 | 0.3 | 0.451 | 1 | 0.326929977 | 0.013430066 |
| Sirt2 | 0.00886519 | -0.714051202 | 0.555 | 0.635 | 1 | 0.006755852 | -1.840059087 | 0.3 | 0.67 | 1 | 0.00886519 | 0.013466062 |
| Zfp62 | 0.006756704 | -1.173482821 | 0.202 | 0.329 | 1 | 0.474517971 | 0.51967104 | 0.5 | 0.42 | 1 | 0.474517971 | 0.013467754 |
| Tmem106c | 0.006760246 | 0.514573173 | 0.487 | 0.324 | 1 | 0.190686181 | 0.962639472 | 0.5 | 0.299 | 1 | 0.190686181 | 0.013474791 |
| Acsl4 | 0.006763845 | 0.196211693 | 0.714 | 0.541 | 1 | 0.932160257 | 1.054447784 | 0.5 | 0.559 | 1 | 0.932160257 | 0.01348194 |
| Rbsn | 0.00676529 | -0.67305383 | 0.227 | 0.382 | 1 | 0.426172454 | -0.581990934 | 0.3 | 0.383 | 1 | 0.426172454 | 0.013484811 |
| Cd151 | 0.006776172 | 0.256775415 | 0.706 | 0.5 | 1 | 0.764985067 | 0.546616189 | 0.5 | 0.463 | 1 | 0.764985067 | 0.013506427 |
| Tcerg1 | 0.006812696 | -0.527862093 | 0.756 | 0.835 | 1 | 0.852594542 | -0.109711372 | 0.8 | 0.886 | 1 | 0.852594542 | 0.01357898 |
| Rps16 | 0.165164158 | -0.12202642 | 0.983 | 0.988 | 1 | 0.00683095 | -0.924113774 | 1 | 0.994 | 1 | 0.165164158 | 0.013615238 |
| Kansl3 | 0.006832567 | -0.734454375 | 0.437 | 0.565 | 1 | 0.251552144 | -0.401616984 | 0.5 | 0.691 | 1 | 0.251552144 | 0.01361845 |

| | | | | | | | | | | | | |
|---|---|---|---|---|---|---|---|---|---|---|---|---|
| Myoz3 | 0.006837602 | -0.826463745 | 0.092 | 0.218 | 1 | 0.052324913 | 0.732519689 | 0.5 | 0.21 | 1 | 0.052324913 | 0.013628452 |
| Tada2a | 0.006866438 | -0.900464326 | 0.235 | 0.382 | 1 | 0.412739333 | 0.884066161 | 0.4 | 0.293 | 1 | 0.412739333 | 0.013685727 |
| Csnk2b | 0.006870956 | -0.476813697 | 0.756 | 0.788 | 1 | 0.014326289 | -1.106773839 | 0.4 | 0.83 | 1 | 0.014326289 | 0.013694703 |
| Ppp1r13b | 0.006874102 | -0.835924074 | 0.336 | 0.476 | 1 | 0.272562776 | -0.256872211 | 0.4 | 0.691 | 1 | 0.272562776 | 0.013700952 |
| Patj | 0.573374621 | -0.253253385 | 0.403 | 0.341 | 1 | 0.006879664 | 1.35626665 | 0.8 | 0.478 | 1 | 0.573374621 | 0.013711999 |
| Vsir | 0.006887919 | -1.283792966 | 0.151 | 0.276 | 1 | 0.301137451 | -0.876895855 | 0.1 | 0.235 | 1 | 0.301137451 | 0.013728394 |
| Capn10 | 0.00690213 | -0.952552838 | 0.084 | 0.206 | 1 | 0.17569589 | -1.230005605 | 0 | 0.157 | 1 | 0.17569589 | 0.01375662 |
| Ptgs1 | 0.051590265 | 0.526447273 | 0.42 | 0.312 | 1 | 0.006920837 | 1.571665678 | 0.5 | 0.17 | 1 | 0.051590265 | 0.013793776 |
| Secisbp2l | 0.006928396 | 0.366960633 | 0.924 | 0.806 | 1 | 0.365096473 | 0.351509594 | 0.9 | 0.883 | 1 | 0.365096473 | 0.01380879 |
| Vdac1 | 0.006934672 | -0.469572876 | 0.739 | 0.794 | 1 | 0.119133285 | -0.739019253 | 0.8 | 0.83 | 1 | 0.119133285 | 0.013821255 |
| Dctd | 0.006945095 | -1.359895945 | 0.059 | 0.165 | 1 | 0.075916377 | 2.017921908 | 0.2 | 0.059 | 1 | 0.075916377 | 0.013841956 |
| Gpbp1l1 | 0.006956346 | -1.040487842 | 0.412 | 0.506 | 1 | 0.169432877 | -0.31706234 | 0.4 | 0.682 | 1 | 0.169432877 | 0.0138643 |
| Mif | 0.006972978 | 0.646245806 | 0.815 | 0.706 | 1 | 0.874274849 | 0.766547093 | 0.5 | 0.648 | 1 | 0.874274849 | 0.013897333 |
| Syde1 | 0.006974891 | -1.36994961 | 0.101 | 0.218 | 1 | 0.929250888 | 0.574978412 | 0.2 | 0.198 | 1 | 0.929250888 | 0.013901132 |
| Cyfip1 | 0.006979825 | -0.64948959 | 0.605 | 0.688 | 1 | 0.402672188 | -0.261032501 | 0.9 | 0.836 | 1 | 0.402672188 | 0.013910931 |
| Eef2k | 0.00698052 | -1.258016331 | 0.076 | 0.188 | 1 | 0.55657529 | 0.457206954 | 0.3 | 0.216 | 1 | 0.55657529 | 0.013912313 |
| Chac2 | 0.006995326 | -1.392317423 | 0.076 | 0.188 | 1 | 0.438381544 | -0.457811523 | 0.1 | 0.194 | 1 | 0.438381544 | 0.013941717 |
| Pfdn4 | 0.016723236 | -0.67305383 | 0.353 | 0.471 | 1 | 0.006996874 | -2.795859283 | 0 | 0.448 | 1 | 0.016723236 | 0.013944791 |
| Gm16133 | 0.00700477 | -1.310340121 | 0.21 | 0.329 | 1 | 0.472311316 | -0.290830798 | 0.4 | 0.528 | 1 | 0.472311316 | 0.013960474 |
| Lyrm1 | 0.925746628 | -0.10433666 | 0.185 | 0.176 | 1 | 0.007017053 | 2.87108052 | 0.3 | 0.071 | 1 | 0.925746628 | 0.013984866 |
| 1700001K19Rik | 0.826272069 | -0.539874611 | 0.076 | 0.082 | 1 | 0.00701718 | 2.973527789 | 0.3 | 0.071 | 1 | 0.826272069 | 0.013985119 |
| Hspb6 | 0.007026128 | 1.228268988 | 0.311 | 0.188 | 1 | 0.97216902 | 0.660369903 | 0.1 | 0.102 | 1 | 0.97216902 | 0.014002889 |
| Abhd5 | 0.0070305 | -0.763411574 | 0.227 | 0.371 | 1 | 0.215465678 | -0.601686736 | 0.3 | 0.506 | 1 | 0.215465678 | 0.014011572 |
| Gm12724 | 0.007037273 | 2.684498174 | 0.059 | 0.006 | 1 | 0.73075158 | 2.695993813 | 0 | 0.012 | 1 | 0.73075158 | 0.014025022 |
| D930032P07Rik | 0.007037273 | 2.684498174 | 0.059 | 0.006 | 1 | 0.669843604 | 2.017921908 | 0 | 0.019 | 1 | 0.669843604 | 0.014025022 |
| Erbin | 0.10878085 | -0.308706264 | 0.899 | 0.888 | 1 | 0.007050228 | -0.855056052 | 0.8 | 0.972 | 1 | 0.10878085 | 0.01405075 |
| Nek3 | 0.748064262 | -0.485426827 | 0.067 | 0.076 | 1 | 0.007050972 | 2.017921908 | 0.3 | 0.068 | 1 | 0.748064262 | 0.014052228 |
| Smad5 | 0.045673905 | -0.37506638 | 0.563 | 0.629 | 1 | 0.007073678 | -1.784271309 | 0.3 | 0.698 | 1 | 0.045673905 | 0.014097319 |
| Lin52 | 0.5706838 | -0.443362768 | 0.462 | 0.382 | 1 | 0.007075304 | 1.403212064 | 0.8 | 0.392 | 1 | 0.5706838 | 0.014100548 |
| Supt6 | 0.007078688 | -0.516790998 | 0.782 | 0.794 | 1 | 0.033249671 | -0.743629324 | 0.6 | 0.889 | 1 | 0.033249671 | 0.014107268 |
| Cdc25b | 0.007087433 | -2.544320516 | 0.017 | 0.094 | 1 | 0.902406098 | 0.732519689 | 0.1 | 0.086 | 1 | 0.902406098 | 0.014124633 |
| 9130410C08Rik | 0.007088041 | -3.258016331 | 0.017 | 0.094 | 1 | 0.374686965 | -0.304006187 | 0 | 0.074 | 1 | 0.374686965 | 0.014125842 |
| Il17rd | 0.007088041 | -2.807354922 | 0.017 | 0.094 | 1 | 0.385696815 | -0.441509711 | 0 | 0.071 | 1 | 0.385696815 | 0.014125842 |
| Prtn3 | 0.007088649 | -2.854660637 | 0.017 | 0.094 | 1 | 0.69612487 | 0.463333056 | 0.1 | 0.065 | 1 | 0.69612487 | 0.01412705 |
| Zfp445 | 0.007092627 | -0.738545764 | 0.471 | 0.565 | 1 | 0.028031314 | -1.5519337 | 0.3 | 0.596 | 1 | 0.028031314 | 0.014134949 |
| Dok7 | 0.007095537 | 2.514573173 | 0.059 | 0.006 | 1 | 0.557771538 | 0.769994395 | 0 | 0.034 | 1 | 0.557771538 | 0.014140727 |
| A830082N09Rik | 0.007101134 | -1.918386234 | 0.034 | 0.124 | 1 | 0.460577668 | 0.494359952 | 0 | 0.052 | 1 | 0.460577668 | 0.014151843 |
| ENSMUSG00000121500 | 0.007101225 | -2.021479727 | 0.034 | 0.124 | 1 | 0.135899886 | 1.744903414 | 0.2 | 0.071 | 1 | 0.135899886 | 0.014152022 |
| Helb | 0.007101789 | -1.753915663 | 0.084 | 0.194 | 1 | 0.105191083 | 1.240314329 | 0.4 | 0.188 | 1 | 0.105191083 | 0.014153142 |
| Fyco1 | 0.007102481 | -1.055742552 | 0.21 | 0.347 | 1 | 0.217873703 | -0.654503434 | 0.1 | 0.299 | 1 | 0.217873703 | 0.014154516 |
| Sec61b | 0.007110799 | -0.360278268 | 0.924 | 0.941 | 1 | 0.295692081 | -0.286304185 | 0.7 | 0.957 | 1 | 0.295692081 | 0.014171034 |
| Dbf4 | 0.007125569 | -2.070389328 | 0.05 | 0.147 | 1 | 0.143212177 | 2.339850003 | 0.2 | 0.077 | 1 | 0.143212177 | 0.014200365 |
| Klc3 | 0.00712693 | 1.514573173 | 0.118 | 0.035 | 1 | 0.627761381 | 0.111031312 | 0.1 | 0.154 | 1 | 0.627761381 | 0.014203067 |
| Gramd3 | 0.007129125 | -1.08931019 | 0.513 | 0.582 | 1 | 0.308504262 | 0.998022351 | 0.6 | 0.605 | 1 | 0.308504262 | 0.014207426 |
| Nbn | 0.007130571 | -0.887525271 | 0.168 | 0.312 | 1 | 0.108867111 | -1.449683642 | 0.1 | 0.333 | 1 | 0.108867111 | 0.014210296 |
| Cyp2j9 | 0.007133032 | 2.321928095 | 0.084 | 0.018 | 1 | 0.009727839 | 4.017921908 | 0.1 | 0.009 | 1 | 0.009727839 | 0.014215184 |
| Mtres1 | 0.007156302 | -1.292781749 | 0.16 | 0.282 | 1 | 0.340197816 | 0.913585248 | 0.3 | 0.182 | 1 | 0.340197816 | 0.014261392 |
| Cep43 | 0.007157346 | -0.851554726 | 0.261 | 0.412 | 1 | 0.992151339 | -0.175849835 | 0.4 | 0.349 | 1 | 0.992151339 | 0.014263465 |
| Myo3b | 0.417669916 | -1.292781749 | 0.034 | 0.053 | 1 | 0.007162426 | 3.280956314 | 0.2 | 0.034 | 1 | 0.417669916 | 0.014273551 |
| Stx6 | 0.007171994 | -0.967078611 | 0.42 | 0.518 | 1 | 0.136189143 | -1.175192721 | 0.3 | 0.528 | 1 | 0.136189143 | 0.01429255 |
| Jpt1 | 0.007172914 | -1.026996732 | 0.261 | 0.4 | 1 | 0.033124753 | -1.736965594 | 0.1 | 0.438 | 1 | 0.033124753 | 0.014294377 |
| Ids | 0.161083313 | -0.515174171 | 0.168 | 0.235 | 1 | 0.00717308 | 1.605538862 | 0.7 | 0.398 | 1 | 0.161083313 | 0.014294706 |
| Slc25a27 | 0.007183757 | -1.719892081 | 0.118 | 0.235 | 1 | 0.174800007 | -1.41670632 | 0.1 | 0.287 | 1 | 0.174800007 | 0.014315907 |
| Sox30 | 0.007185212 | 3.321928095 | 0.042 | 0 | 1 | 0.73075158 | 2.695993813 | 0 | 0.012 | 1 | 0.73075158 | 0.014318798 |
| Osgin1 | 0.007186622 | 4.421463768 | 0.042 | 0 | 1 | 0.054235621 | 2.374065718 | 0.2 | 0.056 | 1 | 0.054235621 | 0.014321596 |
| Pebp4 | 0.007186622 | 3.099535674 | 0.042 | 0 | 1 | 0.009979004 | 3.432959407 | 0.1 | 0.009 | 1 | 0.009979004 | 0.014321596 |
| Ankrd16 | 0.007190864 | -0.996388746 | 0.101 | 0.224 | 1 | 0.298611859 | 0.789103248 | 0.3 | 0.16 | 1 | 0.298611859 | 0.01433002 |
| Bex2 | 0.007229418 | -2.707819249 | 0.017 | 0.094 | 1 | 0.861061884 | 0.159940913 | 0.1 | 0.114 | 1 | 0.861061884 | 0.014406572 |
| Ryr3 | 0.007238297 | -3.070389328 | 0 | 0.059 | 1 | 0.868037793 | 0.847996907 | 0.1 | 0.083 | 1 | 0.868037793 | 0.014424202 |
| Wdr62 | 0.007238297 | -3.070389328 | 0 | 0.059 | 1 | 0.669848329 | 1.432959407 | 0 | 0.019 | 1 | 0.669848329 | 0.014424202 |
| Camkk1 | 0.007238297 | -3.070389328 | 0 | 0.059 | 1 | 0.669848329 | 1.847996907 | 0 | 0.019 | 1 | 0.669848329 | 0.014424202 |
| 6720489N17Rik | 0.007238297 | -3.070389328 | 0 | 0.059 | 1 | 0.643847387 | 1.31748219 | 0 | 0.022 | 1 | 0.643847387 | 0.014424202 |
| Ppp1r3e | 0.007238297 | -3.185866545 | 0 | 0.059 | 1 | 0.557729726 | 1.31748219 | 0 | 0.034 | 1 | 0.557729726 | 0.014424202 |
| Ccdc169 | 0.007240895 | -3.572889668 | 0 | 0.059 | 1 | 0.643835681 | 1.847996907 | 0 | 0.022 | 1 | 0.643835681 | 0.014429359 |
| Arl9 | 0.007240895 | -3.185866545 | 0 | 0.059 | 1 | 0.396948664 | 0.159940913 | 0 | 0.068 | 1 | 0.396948664 | 0.014429359 |
| Trac | 0.007240895 | -3.392317423 | 0 | 0.059 | 1 | 0.557765566 | 0.847996907 | 0 | 0.034 | 1 | 0.557765566 | 0.014429359 |
| Rbm44 | 0.007242915 | -3.572889668 | 0 | 0.059 | 1 | 0.557772534 | 0.558490289 | 0 | 0.034 | 1 | 0.557772534 | 0.01443337 |
| Gm16156 | 0.007242915 | -3.292781749 | 0 | 0.059 | 1 | 0.619967468 | 1.432959407 | 0 | 0.025 | 1 | 0.619967468 | 0.01443337 |
| Gm34045 | 0.007242915 | -3.292781749 | 0 | 0.059 | 1 | 0.362872338 | 1.847996907 | 0.1 | 0.04 | 1 | 0.362872338 | 0.01443337 |
| Gm57387 | 0.007242915 | -3.292781749 | 0 | 0.059 | 1 | 0.401608067 | 1.847996907 | 0.1 | 0.043 | 1 | 0.401608067 | 0.01443337 |
| Caskin1 | 0.007242915 | -3.292781749 | 0 | 0.059 | 1 | 0.460551517 | 0.558490289 | 0 | 0.052 | 1 | 0.460551517 | 0.01443337 |
| Ttc23l | 0.007242977 | -3.272023189 | 0.017 | 0.094 | 1 | 0.282750607 | -1.611434712 | 0 | 0.105 | 1 | 0.282750607 | 0.014433494 |
| Gm49519 | 0.007245225 | -3.655351829 | 0 | 0.059 | 1 | 0.64384652 | 1.558490289 | 0 | 0.022 | 1 | 0.64384652 | 0.014437956 |
| Adamts18 | 0.007245946 | -4.655351829 | 0 | 0.059 | 1 | 0.118051503 | 2.31748219 | 0.1 | 0.022 | 1 | 0.118051503 | 0.014439389 |
| Cnksr2 | 0.007245946 | -4.070389328 | 0 | 0.059 | 1 | 0.334429455 | -1.489872732 | 0 | 0.086 | 1 | 0.334429455 | 0.014439389 |
| Mdga2 | 0.00724638 | -5.635173947 | 0 | 0.059 | 1 | 0.489994396 | -1.069540933 | 0 | 0.046 | 1 | 0.489994396 | 0.014440249 |
| 4933427G23Rik | 0.007246957 | -2.944858446 | 0 | 0.059 | 1 | 0.320995017 | 2.017921908 | 0.1 | 0.037 | 1 | 0.320995017 | 0.014441396 |
| Rce1 | 0.007249729 | -0.807354922 | 0.277 | 0.424 | 1 | 0.232060474 | -0.68251781 | 0.2 | 0.373 | 1 | 0.232060474 | 0.014446899 |
| Plekhf2 | 0.032241693 | -0.640705053 | 0.252 | 0.371 | 1 | 0.007261898 | -2.191531458 | 0.1 | 0.54 | 1 | 0.032241693 | 0.014471061 |
| Prtg | 0.007275097 | -0.841570637 | 0.059 | 0.165 | 1 | 0.249021741 | 1.602884409 | 0.2 | 0.093 | 1 | 0.249021741 | 0.014497268 |
| Lipe | 0.381045639 | -0.970853654 | 0.067 | 0.094 | 1 | 0.007290591 | 2.210566986 | 0.3 | 0.068 | 1 | 0.381045639 | 0.014528029 |
| Rbm38 | 0.007299753 | -2.359895945 | 0.017 | 0.094 | 1 | 0.374632633 | -0.111361109 | 0 | 0.074 | 1 | 0.374632633 | 0.01454622 |
| Zfp961 | 0.007299878 | -2.424026283 | 0.017 | 0.094 | 1 | 0.433732528 | -0.111361109 | 0 | 0.059 | 1 | 0.433732528 | 0.014546468 |
| Gm26802 | 0.007302202 | -1.694880193 | 0.101 | 0.218 | 1 | 0.103823739 | 0.29389537 | 0.5 | 0.231 | 1 | 0.103823739 | 0.014551081 |
| Tpm3 | 0.349831366 | -0.116485742 | 0.992 | 0.971 | 1 | 0.007313058 | -0.853197557 | 1 | 1 | 1 | 0.349831366 | 0.014572634 |
| H2-K2 | 0.0073156 | 1.893084796 | 0.134 | 0.047 | 1 | 0.232498909 | 2.432959407 | 0.1 | 0.031 | 1 | 0.232498909 | 0.014577683 |
| Tmem175 | 0.007361474 | -1.207892852 | 0.261 | 0.388 | 1 | 0.331900255 | 0.374065718 | 0.5 | 0.302 | 1 | 0.331900255 | 0.014668756 |
| Fam228b | 0.007370418 | -2.222392421 | 0.017 | 0.094 | 1 | 0.643760887 | 1.159940913 | 0.1 | 0.062 | 1 | 0.643760887 | 0.014686514 |
| Asf1a | 0.007374452 | -0.869341809 | 0.244 | 0.371 | 1 | 0.357152029 | -0.528972552 | 0.2 | 0.346 | 1 | 0.357152029 | 0.014694521 |
| Cbr1 | 0.079109651 | -0.574694165 | 0.303 | 0.376 | 1 | 0.007388083 | -2.723545078 | 0 | 0.441 | 1 | 0.079109651 | 0.014721581 |
| Ilf2 | 0.00738917 | -0.437022195 | 0.739 | 0.812 | 1 | 0.03768433 | -1.08594869 | 0.4 | 0.775 | 1 | 0.03768433 | 0.014723741 |
| Tmem186 | 0.37271776 | -0.308549065 | 0.084 | 0.118 | 1 | 0.007402441 | 1.816288047 | 0.4 | 0.111 | 1 | 0.37271776 | 0.014750086 |
| Ddx55 | 0.007403084 | -1.715724447 | 0.101 | 0.212 | 1 | 0.459286191 | -0.441509711 | 0.1 | 0.188 | 1 | 0.459286191 | 0.014751361 |
| Sanbr | 0.00742033 | -0.930837975 | 0.286 | 0.418 | 1 | 0.77627051 | -0.572041274 | 0.4 | 0.395 | 1 | 0.77627051 | 0.014785599 |
| Coa3 | 0.007435894 | -0.693535023 | 0.353 | 0.494 | 1 | 0.175150551 | -0.776493958 | 0.3 | 0.485 | 1 | 0.175150551 | 0.014816496 |
| Gm9750 | 0.007443015 | 2.251538767 | 0.084 | 0.018 | 1 | 0.730749233 | 2.432959407 | 0 | 0.012 | 1 | 0.730749233 | 0.014830632 |
| Serpini1 | 0.00745388 | -1.963474124 | 0.034 | 0.124 | 1 | 0.918287749 | 0.210566986 | 0.1 | 0.114 | 1 | 0.918287749 | 0.014852199 |
| Pwwp3a | 0.007455752 | -1.168236651 | 0.118 | 0.241 | 1 | 0.362529271 | -0.611434712 | 0.1 | 0.216 | 1 | 0.362529271 | 0.014855917 |

| | | | | | | | | | | | | |
|---|---|---|---|---|---|---|---|---|---|---|---|---|
| Naprt | 0.007461588 | -1.687060688 | 0.034 | 0.124 | 1 | 0.776127749 | 0.973527789 | 0.1 | 0.074 | 1 | 0.776127749 | 0.0148675 |
| Slc52a2 | 0.007461588 | -1.687060688 | 0.034 | 0.124 | 1 | 0.474877085 | 0.847996907 | 0 | 0.049 | 1 | 0.474877085 | 0.0148675 |
| Il10rb | 0.0074646 | -1.306456686 | 0.101 | 0.218 | 1 | 0.430936117 | -0.728032469 | 0.2 | 0.296 | 1 | 0.430936117 | 0.014873479 |
| Daam2 | 0.007465557 | -1.044013912 | 0.403 | 0.524 | 1 | 0.9116727 | -0.263364202 | 0.4 | 0.404 | 1 | 0.9116727 | 0.01487538 |
| Dnah17 | 0.242675956 | -0.944858446 | 0.034 | 0.065 | 1 | 0.007508341 | 2.494359952 | 0.3 | 0.074 | 1 | 0.242675956 | 0.014960307 |
| Kat7 | 0.007510916 | -0.937939032 | 0.336 | 0.459 | 1 | 0.242188929 | -0.709998547 | 0.4 | 0.583 | 1 | 0.242188929 | 0.014965417 |
| Gabarapl2 | 0.007512452 | 0.335649722 | 0.924 | 0.912 | 1 | 0.283730817 | -0.281940246 | 0.8 | 0.951 | 1 | 0.283730817 | 0.014968466 |
| Prr14 | 0.007520755 | -0.796628515 | 0.328 | 0.459 | 1 | 0.465830059 | -0.573039333 | 0.4 | 0.451 | 1 | 0.465830059 | 0.014984948 |
| Usp4 | 0.007543893 | -0.695644535 | 0.496 | 0.624 | 1 | 0.662285849 | 0.475394674 | 0.7 | 0.738 | 1 | 0.662285849 | 0.015030875 |
| Riiad1 | 0.007544015 | -2.109917692 | 0.042 | 0.135 | 1 | 0.219672948 | -0.814968106 | 0 | 0.133 | 1 | 0.219672948 | 0.015031118 |
| Mfsd4a | 0.007544413 | -1.359895945 | 0.034 | 0.124 | 1 | 0.213410078 | -0.789433014 | 0 | 0.136 | 1 | 0.213410078 | 0.015031907 |
| Nol11 | 0.007549464 | -0.659677754 | 0.403 | 0.512 | 1 | 0.806969777 | -0.186649236 | 0.5 | 0.485 | 1 | 0.806969777 | 0.015041934 |
| Fxr2 | 0.194523442 | -0.415037499 | 0.403 | 0.447 | 1 | 0.007560921 | -1.860639965 | 0.2 | 0.614 | 1 | 0.194523442 | 0.015064675 |
| Rpl36 | 0.180447259 | -0.146580942 | 0.966 | 0.988 | 1 | 0.007572435 | -0.899787019 | 0.7 | 0.988 | 1 | 0.180447259 | 0.015087528 |
| Rbm17 | 0.007578117 | -0.546827372 | 0.639 | 0.694 | 1 | 0.180742718 | -0.339630097 | 0.4 | 0.769 | 1 | 0.180742718 | 0.015098806 |
| Cryz | 0.007587158 | -1.032914622 | 0.092 | 0.212 | 1 | 0.699481299 | 0.31748219 | 0.1 | 0.142 | 1 | 0.699481299 | 0.015116751 |
| Mdm1 | 0.007596537 | -1.758445322 | 0.034 | 0.124 | 1 | 0.401608067 | 1.695993813 | 0.1 | 0.043 | 1 | 0.401608067 | 0.015135367 |
| Gm42567 | 0.721161231 | 0.514573173 | 0.017 | 0.012 | 1 | 0.007597569 | 2.769994395 | 0.2 | 0.034 | 1 | 0.721161231 | 0.015137416 |
| Mpi | 0.007623139 | -1.987927168 | 0.017 | 0.094 | 1 | 0.074599625 | 2.079322453 | 0.2 | 0.059 | 1 | 0.074599625 | 0.015188166 |
| Lmnb2 | 0.007629482 | -1.707819249 | 0.034 | 0.124 | 1 | 0.432664857 | 1.816288047 | 0.2 | 0.127 | 1 | 0.432664857 | 0.015200756 |
| Ilkap | 0.232219866 | -0.361691459 | 0.454 | 0.5 | 1 | 0.007631496 | -1.91281543 | 0.2 | 0.611 | 1 | 0.232219866 | 0.015204751 |
| Pbx2 | 0.007632008 | -1.098403704 | 0.101 | 0.224 | 1 | 0.106964616 | -1.536666944 | 0 | 0.21 | 1 | 0.106964616 | 0.015205768 |
| Igsf8 | 0.007640699 | 0.669083122 | 0.597 | 0.441 | 1 | 0.383566376 | 0.479956887 | 0.4 | 0.531 | 1 | 0.383566376 | 0.015223017 |
| Rad54l2 | 0.007668905 | -0.733354341 | 0.328 | 0.465 | 1 | 0.156909934 | -1.17978625 | 0.4 | 0.534 | 1 | 0.156909934 | 0.015278998 |
| Lclat1 | 0.680467607 | -0.375549209 | 0.479 | 0.418 | 1 | 0.007673034 | 1.648688098 | 0.6 | 0.25 | 1 | 0.680467607 | 0.015287192 |
| Eloa | 0.007720722 | -0.685853917 | 0.58 | 0.641 | 1 | 0.093709137 | -0.724770297 | 0.5 | 0.778 | 1 | 0.093709137 | 0.015381834 |
| Vamp8 | 0.007722568 | 0.339630097 | 0.849 | 0.771 | 1 | 0.06720383 | -0.742299038 | 0.4 | 0.79 | 1 | 0.06720383 | 0.015385498 |
| Gm11476 | 0.00772909 | -2.070389328 | 0.017 | 0.094 | 1 | 0.601875604 | 1.263034406 | 0.1 | 0.059 | 1 | 0.601875604 | 0.015398441 |
| Morn1 | 0.007755854 | -1.292781749 | 0.109 | 0.229 | 1 | 0.863497756 | -0.279758641 | 0.2 | 0.207 | 1 | 0.863497756 | 0.015451555 |
| Gm47675 | 0.007761144 | 1.251538767 | 0.118 | 0.035 | 1 | 0.022740147 | 2.695993813 | 0.2 | 0.043 | 1 | 0.022740147 | 0.015462054 |
| Cyb5r3 | 0.007773361 | 0.313924871 | 0.924 | 0.8 | 1 | 0.697907131 | 0.339850003 | 0.8 | 0.836 | 1 | 0.697907131 | 0.015486297 |
| Sh3bgrl3 | 0.007803078 | -0.495996069 | 0.681 | 0.753 | 1 | 0.474967438 | 0.165805535 | 0.8 | 0.719 | 1 | 0.474967438 | 0.015545268 |
| Rab7b | 0.007815579 | -2 | 0.05 | 0.147 | 1 | 0.116579781 | 2.31748219 | 0.1 | 0.022 | 1 | 0.116579781 | 0.015570074 |
| Hebp1 | 0.007826605 | -1.27897595 | 0.092 | 0.212 | 1 | 0.796476766 | 0.888638891 | 0.1 | 0.077 | 1 | 0.796476766 | 0.015591954 |
| Card6 | 0.007847307 | -2.944858446 | 0.008 | 0.076 | 1 | 0.113669596 | 2.847996907 | 0.1 | 0.022 | 1 | 0.113669596 | 0.015633033 |
| Rad1 | 0.007847817 | -1.758445322 | 0.042 | 0.135 | 1 | 0.290313081 | -0.408342847 | 0 | 0.102 | 1 | 0.290313081 | 0.015634046 |
| Taf5l | 0.007863824 | -1.205318908 | 0.126 | 0.247 | 1 | 0.179177933 | -1.015501094 | 0.1 | 0.29 | 1 | 0.179177933 | 0.015665808 |
| AW554918 | 0.007865754 | -1.114243617 | 0.303 | 0.412 | 1 | 0.793061322 | 0.118448784 | 0.5 | 0.457 | 1 | 0.793061322 | 0.015669638 |
| Habp4 | 0.007874709 | -0.714908673 | 0.176 | 0.318 | 1 | 0.582244179 | 0.615823464 | 0.4 | 0.315 | 1 | 0.582244179 | 0.015687406 |
| Uvrag | 0.007880394 | -0.430896714 | 0.765 | 0.853 | 1 | 0.602685327 | -0.135883428 | 0.8 | 0.886 | 1 | 0.602685327 | 0.015698687 |
| Tfe3 | 0.007883479 | -0.774933444 | 0.277 | 0.418 | 1 | 0.733805563 | -0.146985019 | 0.5 | 0.503 | 1 | 0.733805563 | 0.015704809 |
| 2310057M21Rik | 0.007906129 | -0.827819025 | 0.185 | 0.329 | 1 | 0.334889884 | -0.69632361 | 0.1 | 0.222 | 1 | 0.334889884 | 0.015749751 |
| Dnajc5 | 0.007910463 | -0.429778798 | 0.714 | 0.806 | 1 | 0.622374443 | -0.217294554 | 0.9 | 0.907 | 1 | 0.622374443 | 0.01575835 |
| Gm56698 | 0.007922656 | -1.87774425 | 0.017 | 0.094 | 1 | 0.539413885 | 1.111031312 | 0 | 0.037 | 1 | 0.539413885 | 0.015782543 |
| Fam8a1 | 0.007924913 | -0.935229745 | 0.286 | 0.406 | 1 | 0.593346251 | -0.417897153 | 0.5 | 0.515 | 1 | 0.593346251 | 0.015787021 |
| Usp40 | 0.00792917 | -1.185866545 | 0.252 | 0.376 | 1 | 0.473643223 | -0.736965594 | 0.3 | 0.361 | 1 | 0.473643223 | 0.015795469 |
| D430020J02Rik | 0.900375516 | -0.273922722 | 0.067 | 0.071 | 1 | 0.007932225 | 2.902444691 | 0.2 | 0.034 | 1 | 0.900375516 | 0.015801529 |
| Syap1 | 0.007935186 | -0.517135687 | 0.655 | 0.741 | 1 | 0.226845408 | -0.567040593 | 0.7 | 0.815 | 1 | 0.226845408 | 0.015807405 |
| Cd2ap | 0.007962441 | -0.439927279 | 0.924 | 0.894 | 1 | 0.896013879 | -0.103093493 | 0.9 | 0.917 | 1 | 0.896013879 | 0.015861481 |
| Ino80e | 0.007988349 | -1.002485263 | 0.21 | 0.341 | 1 | 0.120407811 | -1.026472211 | 0.1 | 0.33 | 1 | 0.120407811 | 0.015912884 |
| Cep70 | 0.461196653 | -0.748461233 | 0.168 | 0.194 | 1 | 0.007998033 | 1.506959989 | 0.4 | 0.108 | 1 | 0.461196653 | 0.015932097 |
| Krba1 | 0.008002926 | -1.707819249 | 0.042 | 0.135 | 1 | 0.91873618 | 0.463333056 | 0.1 | 0.108 | 1 | 0.91873618 | 0.015941806 |
| Rab7 | 0.018475667 | -0.412576262 | 0.874 | 0.906 | 1 | 0.008003068 | -0.953319099 | 0.9 | 0.963 | 1 | 0.018475667 | 0.015942086 |
| Rps19bp1 | 0.008020009 | -1.131143359 | 0.218 | 0.341 | 1 | 0.305176835 | -0.663902132 | 0.2 | 0.34 | 1 | 0.305176835 | 0.015975698 |
| Ece2 | 0.008020866 | -2.733354341 | 0.008 | 0.076 | 1 | 0.363859515 | 0.159940913 | 0 | 0.077 | 1 | 0.363859515 | 0.015977397 |
| Cdc7 | 0.008020922 | -2.572889668 | 0.008 | 0.076 | 1 | 0.559081495 | 1.558490289 | 0.1 | 0.056 | 1 | 0.559081495 | 0.015977509 |
| Slc2a3 | 0.008020922 | -3.008988783 | 0.008 | 0.076 | 1 | 0.243243828 | 1.374065718 | 0.1 | 0.031 | 1 | 0.243243828 | 0.015977509 |
| BC024063 | 0.008020922 | -2.572889668 | 0.008 | 0.076 | 1 | 0.522102507 | 0.930459067 | 0 | 0.04 | 1 | 0.522102507 | 0.015977509 |
| Ufm1 | 0.029841346 | -0.529370175 | 0.571 | 0.641 | 1 | 0.008029148 | -1.618702713 | 0.2 | 0.657 | 1 | 0.029841346 | 0.015993829 |
| Kat8 | 0.008031298 | -1.193246076 | 0.109 | 0.229 | 1 | 0.267660189 | -0.611434712 | 0.1 | 0.25 | 1 | 0.267660189 | 0.015998094 |
| Fam131a | 0.008056892 | -1.070389328 | 0.168 | 0.294 | 1 | 0.711666697 | 0.276454922 | 0.3 | 0.231 | 1 | 0.711666697 | 0.01604887 |
| Ubap1l | 0.008062949 | -2.485426827 | 0.008 | 0.076 | 1 | 0.324842077 | -0.230005605 | 0 | 0.09 | 1 | 0.324842077 | 0.016060887 |
| Tmem132a | 0.008062949 | -2.807354922 | 0.008 | 0.076 | 1 | 0.228393429 | -1.191531458 | 0.1 | 0.259 | 1 | 0.228393429 | 0.016060887 |
| Zc4h2 | 0.008062949 | -2.572889668 | 0.008 | 0.076 | 1 | 0.433608413 | 0.494359952 | 0 | 0.059 | 1 | 0.433608413 | 0.016060887 |
| Exoc2 | 0.008067031 | -1.292781749 | 0.227 | 0.347 | 1 | 0.192494356 | -1.310004928 | 0.2 | 0.37 | 1 | 0.192494356 | 0.016068985 |
| Gm17586 | 0.008095636 | -1.600904045 | 0.084 | 0.194 | 1 | 0.151479157 | -1.026472211 | 0 | 0.173 | 1 | 0.151479157 | 0.016125733 |
| Syde2 | 0.008104493 | -2.572889668 | 0.008 | 0.076 | 1 | 0.226708798 | 0.874963954 | 0.2 | 0.083 | 1 | 0.226708798 | 0.016143304 |
| Gm49602 | 0.008104493 | -2.392317423 | 0.008 | 0.076 | 1 | 0.374563499 | 0.111031312 | 0 | 0.074 | 1 | 0.374563499 | 0.016143304 |
| Gm15867 | 0.008113367 | 0.957516669 | 0.37 | 0.235 | 1 | 0.445873364 | 0.295455884 | 0.4 | 0.253 | 1 | 0.445873364 | 0.016160908 |
| Vps9d1 | 0.008116667 | -1.350497247 | 0.092 | 0.206 | 1 | 0.513069853 | -0.191531458 | 0.1 | 0.176 | 1 | 0.513069853 | 0.016167453 |
| Abracl | 0.008118044 | -0.622930351 | 0.336 | 0.482 | 1 | 0.775829892 | 0.917461726 | 0.5 | 0.438 | 1 | 0.775829892 | 0.016170185 |
| AU020206 | 0.008127134 | -1.485426827 | 0.034 | 0.124 | 1 | 0.957023815 | 0.263034406 | 0.1 | 0.102 | 1 | 0.957023815 | 0.016188218 |
| Mospd1 | 0.008134147 | -1.292781749 | 0.244 | 0.359 | 1 | 0.213769059 | -1.185018151 | 0.2 | 0.367 | 1 | 0.213769059 | 0.016202129 |
| Gm39929 | 0.008152324 | -1.655351829 | 0.017 | 0.094 | 1 | 0.557756275 | 1.017921908 | 0 | 0.034 | 1 | 0.557756275 | 0.016238187 |
| Cbx5 | 0.008169654 | -0.643428661 | 0.79 | 0.8 | 1 | 0.428675494 | 0.228867803 | 0.9 | 0.836 | 1 | 0.428675494 | 0.016272566 |
| Fam222a | 0.008171567 | -2.292781749 | 0.008 | 0.076 | 1 | 0.676138216 | 1.111031312 | 0.1 | 0.065 | 1 | 0.676138216 | 0.016276359 |
| Gm50069 | 0.008171567 | -2.292781749 | 0.008 | 0.076 | 1 | 0.557729726 | 1.31748219 | 0 | 0.034 | 1 | 0.557729726 | 0.016276359 |
| Ube2t | 0.499402448 | 0.421463768 | 0.092 | 0.071 | 1 | 0.008171726 | 2.795529487 | 0.2 | 0.034 | 1 | 0.499402448 | 0.016276675 |
| Gm57446 | 0.695893386 | -0.222392421 | 0.017 | 0.024 | 1 | 0.008171726 | 2.795529487 | 0.2 | 0.034 | 1 | 0.695893386 | 0.016276675 |
| Psmb2 | 0.008180479 | -0.288481612 | 0.95 | 0.941 | 1 | 0.122013096 | -0.513459553 | 0.8 | 0.975 | 1 | 0.122013096 | 0.016294037 |
| C920021L13Rik | 0.008185767 | -1.722466024 | 0.034 | 0.124 | 1 | 0.324925979 | -0.304006187 | 0 | 0.09 | 1 | 0.324925979 | 0.016304528 |
| Gtf2e2 | 0.008193957 | -0.797370833 | 0.378 | 0.5 | 1 | 0.847672927 | 0.263034406 | 0.5 | 0.515 | 1 | 0.847672927 | 0.016320773 |
| Ramac | 0.008198923 | -0.515174171 | 0.639 | 0.712 | 1 | 0.52492021 | -0.224571719 | 0.6 | 0.778 | 1 | 0.52492021 | 0.016330625 |
| Calml3 | 0.908247369 | -0.203027096 | 0.21 | 0.206 | 1 | 0.008235689 | 2.070389328 | 0.6 | 0.281 | 1 | 0.908247369 | 0.016403551 |
| Fmn1 | 0.00823849 | -0.168051063 | 0.966 | 0.812 | 1 | 0.05218014 | 0.716059436 | 1 | 0.932 | 1 | 0.05218014 | 0.016409107 |
| Abhd12 | 0.008249363 | -0.957495271 | 0.437 | 0.529 | 1 | 0.893808407 | -0.146266724 | 0.6 | 0.475 | 1 | 0.893808407 | 0.016430674 |
| Afdn | 0.008251233 | -0.952785362 | 0.605 | 0.671 | 1 | 0.229954725 | -0.816549142 | 0.8 | 0.781 | 1 | 0.229954725 | 0.016434382 |
| Cntrob | 0.008266396 | -1.207892852 | 0.034 | 0.124 | 1 | 0.063718853 | 1.345496566 | 0.3 | 0.105 | 1 | 0.063718853 | 0.01646446 |
| Prickle3 | 0.008270864 | -1.448900951 | 0.076 | 0.182 | 1 | 0.693521278 | 0.185031894 | 0.1 | 0.142 | 1 | 0.693521278 | 0.01647332 |
| Fam227a | 0.008281326 | -1.900464326 | 0.059 | 0.159 | 1 | 0.398755273 | 1.494359952 | 0.2 | 0.117 | 1 | 0.398755273 | 0.016494072 |
| Selenow | 0.121622114 | -0.283792966 | 0.916 | 0.947 | 1 | 0.008286117 | -0.725469955 | 0.6 | 0.978 | 1 | 0.121622114 | 0.016503574 |
| Camsap2 | 0.00828881 | -0.603665939 | 0.866 | 0.871 | 1 | 0.08512691 | -0.669933475 | 0.8 | 0.948 | 1 | 0.08512691 | 0.016508916 |
| Eya1 | 0.224761818 | 0.145182676 | 0.924 | 0.871 | 1 | 0.008293579 | -1.262912407 | 0.7 | 0.972 | 1 | 0.224761818 | 0.016518374 |
| Rps10 | 0.149438749 | 0.117914203 | 0.983 | 1 | 1 | 0.008320618 | -0.849632088 | 1 | 0.997 | 1 | 0.149438749 | 0.016572002 |
| Gm41764 | 0.008327991 | -1.5360529 | 0.034 | 0.124 | 1 | 0.911403612 | 0.236562194 | 0.1 | 0.108 | 1 | 0.911403612 | 0.016586628 |
| Cog5 | 0.008339565 | -1.029250633 | 0.445 | 0.535 | 1 | 0.040903872 | 0.962639472 | 0.9 | 0.549 | 1 | 0.040903872 | 0.016609582 |
| H2-Eb1 | 0.008341855 | 1.828233652 | 0.134 | 0.047 | 1 | 0.446758355 | 0.695993813 | 0 | 0.056 | 1 | 0.446758355 | 0.016614123 |
| Pold4 | 0.008372414 | 0.684498174 | 0.462 | 0.318 | 1 | 0.791986753 | 0.494359952 | 0.3 | 0.269 | 1 | 0.791986753 | 0.01667473 |

| | | | | | | | | | | | | |
|---|---|---|---|---|---|---|---|---|---|---|---|---|
| Ugcg | 0.008382032 | -0.897057725 | 0.336 | 0.453 | 1 | 0.988521374 | -0.215697769 | 0.6 | 0.506 | 1 | 0.988521374 | 0.016693806 |
| Slc35e2 | 0.008409181 | -0.93122658 | 0.185 | 0.324 | 1 | 0.781261198 | 0.135278859 | 0.2 | 0.25 | 1 | 0.781261198 | 0.016747647 |
| C730034F03Rik | 0.663964809 | -0.485426827 | 0.042 | 0.053 | 1 | 0.008417093 | 2.515421567 | 0.2 | 0.034 | 1 | 0.663964809 | 0.016763338 |
| Stard8 | 0.008436385 | -0.965419768 | 0.235 | 0.371 | 1 | 0.038124936 | -2.604129911 | 0 | 0.312 | 1 | 0.038124936 | 0.016801598 |
| 4930404H11Rik | 0.008447175 | -2.424026283 | 0.008 | 0.076 | 1 | 0.577164735 | 0.930459067 | 0 | 0.031 | 1 | 0.577164735 | 0.016822995 |
| Kat14 | 0.008450544 | -1.359895945 | 0.084 | 0.194 | 1 | 0.426255271 | -0.581990934 | 0.1 | 0.194 | 1 | 0.426255271 | 0.016829676 |
| Crtc1 | 0.008450544 | -1.455053178 | 0.084 | 0.194 | 1 | 0.808719933 | 0.684021171 | 0.2 | 0.25 | 1 | 0.808719933 | 0.016829676 |
| Slc25a24 | 0.008463726 | -0.858470238 | 0.37 | 0.482 | 1 | 0.22110003 | 0.936127817 | 0.6 | 0.441 | 1 | 0.22110003 | 0.016855818 |
| Spata2 | 0.008467696 | -1.169925001 | 0.101 | 0.218 | 1 | 0.308680395 | -0.567040593 | 0.1 | 0.235 | 1 | 0.308680395 | 0.01686369 |
| Snta1 | 0.008922211 | 0.740849029 | 0.37 | 0.224 | 1 | 0.008468355 | 1.025455581 | 0.8 | 0.355 | 1 | 0.008922211 | 0.016864997 |
| Lactb | 0.008469866 | -1.119298928 | 0.16 | 0.288 | 1 | 0.238954587 | -1.141949429 | 0.1 | 0.256 | 1 | 0.238954587 | 0.016867994 |
| Srfbp1 | 0.008474014 | -0.983677695 | 0.277 | 0.4 | 1 | 0.486636843 | -0.513459553 | 0.3 | 0.367 | 1 | 0.486636843 | 0.016876218 |
| Uck2 | 0.008489641 | 1.19443836 | 0.605 | 0.482 | 1 | 0.991367087 | 0.421732152 | 0.5 | 0.494 | 1 | 0.991367087 | 0.016907208 |
| Tmem88 | 0.008489724 | -1.93021167 | 0.05 | 0.147 | 1 | 0.725741602 | 2.539874611 | 0.1 | 0.077 | 1 | 0.725741602 | 0.016907373 |
| Rrm2 | 0.008497363 | -2.485426827 | 0.008 | 0.076 | 1 | 0.015574879 | 2.558490289 | 0.2 | 0.04 | 1 | 0.015574879 | 0.016922521 |
| Mboat2 | 0.008502946 | -1.329775957 | 0.134 | 0.253 | 1 | 0.646364329 | 0.343611922 | 0.5 | 0.441 | 1 | 0.646364329 | 0.016933592 |
| Dock11 | 0.008533714 | -1.535179862 | 0.252 | 0.365 | 1 | 0.997084889 | 0.666781892 | 0.4 | 0.46 | 1 | 0.997084889 | 0.016994603 |
| Pdpn | 0.008536742 | -1.87244995 | 0.084 | 0.188 | 1 | 0.028634766 | -2.49777793 | 0 | 0.336 | 1 | 0.028634766 | 0.017000608 |
| Efna1 | 0.008555712 | 0.380717426 | 0.908 | 0.735 | 1 | 0.13584784 | -0.774933444 | 0.9 | 0.867 | 1 | 0.13584784 | 0.017038225 |
| 1700102P08Rik | 0.00856368 | -1.694880193 | 0.05 | 0.147 | 1 | 0.306831626 | 0.769994395 | 0.3 | 0.164 | 1 | 0.306831626 | 0.017054023 |
| Psma1 | 0.008568115 | -0.463265177 | 0.782 | 0.806 | 1 | 0.138967639 | -0.362220801 | 0.7 | 0.88 | 1 | 0.138967639 | 0.017062818 |
| Psmb1 | 0.126160263 | -0.201039181 | 0.95 | 0.929 | 1 | 0.008576316 | -0.968236167 | 0.7 | 0.957 | 1 | 0.126160263 | 0.017079078 |
| Sord | 0.00858659 | -1.100136671 | 0.092 | 0.206 | 1 | 0.441842537 | -0.581990934 | 0.1 | 0.191 | 1 | 0.441842537 | 0.01709945 |
| Smad9 | 0.008598866 | -1.589763487 | 0.059 | 0.159 | 1 | 0.717497446 | -0.152003093 | 0.1 | 0.136 | 1 | 0.717497446 | 0.017123791 |
| Id2 | 0.008611273 | -0.612976877 | 0.563 | 0.712 | 1 | 0.06086417 | -1.611434712 | 0.4 | 0.694 | 1 | 0.06086417 | 0.017148392 |
| Col25a1 | 0.008611826 | 1.027912746 | 0.294 | 0.171 | 1 | 0.451998482 | 1.386266172 | 0.4 | 0.364 | 1 | 0.451998482 | 0.017149489 |
| Ufd1 | 0.023689869 | -0.548314533 | 0.58 | 0.629 | 1 | 0.008612627 | -1.629536518 | 0.3 | 0.673 | 1 | 0.023689869 | 0.017151077 |
| Cmpk1 | 0.008628503 | -0.381975479 | 0.832 | 0.841 | 1 | 0.676888486 | -0.167944637 | 0.7 | 0.855 | 1 | 0.676888486 | 0.017182554 |
| Synj1 | 0.008636051 | -1.268615438 | 0.286 | 0.406 | 1 | 0.579620349 | -0.67556505 | 0.5 | 0.528 | 1 | 0.579620349 | 0.01719752 |
| Itga8 | 0.008647404 | -2.655351829 | 0.025 | 0.106 | 1 | 0.75674633 | 0.672147071 | 0.1 | 0.074 | 1 | 0.75674633 | 0.017220031 |
| Hdac1 | 0.008656261 | -0.490292582 | 0.697 | 0.776 | 1 | 0.056055708 | -0.791773153 | 0.5 | 0.815 | 1 | 0.056055708 | 0.017237591 |
| Wdr5 | 0.00866123 | -0.744813456 | 0.185 | 0.324 | 1 | 0.779179295 | -0.183711953 | 0.4 | 0.401 | 1 | 0.779179295 | 0.017247442 |
| B930095G15Rik | 0.008662993 | 0.514573173 | 0.588 | 0.424 | 1 | 0.416277388 | 0.490444902 | 0.6 | 0.54 | 1 | 0.416277388 | 0.017250938 |
| Sclt1 | 0.008667104 | -1.163498732 | 0.403 | 0.512 | 1 | 0.021998614 | 0.787972135 | 0.9 | 0.444 | 1 | 0.021998614 | 0.017259089 |
| Pdia5 | 0.008667633 | -0.916476644 | 0.311 | 0.447 | 1 | 0.650671757 | 0.238948787 | 0.3 | 0.463 | 1 | 0.650671757 | 0.017260137 |
| Oit3 | 0.189470318 | -0.485426827 | 0.034 | 0.071 | 1 | 0.008668433 | 2.280956314 | 0.2 | 0.034 | 1 | 0.189470318 | 0.017261724 |
| Gm35867 | 0.926217033 | -0.485426827 | 0.034 | 0.035 | 1 | 0.008668433 | 2.515421567 | 0.2 | 0.034 | 1 | 0.926217033 | 0.017261724 |
| Gm41231 | 0.12736459 | -0.970853654 | 0.025 | 0.065 | 1 | 0.008668577 | 2.432959407 | 0.2 | 0.034 | 1 | 0.12736459 | 0.017262009 |
| Krt19 | 0.008670485 | -1.201633861 | 0.017 | 0.094 | 1 | 0.12325073 | -1.408342847 | 0 | 0.194 | 1 | 0.12325073 | 0.017265793 |
| Alkbh7 | 0.008671316 | -1.485426827 | 0.059 | 0.159 | 1 | 0.256947236 | 1.393431043 | 0.2 | 0.09 | 1 | 0.256947236 | 0.017267441 |
| Rian | 0.325573495 | 0.328647693 | 0.571 | 0.524 | 1 | 0.008710761 | 1.930459067 | 1 | 0.654 | 1 | 0.325573495 | 0.017345645 |
| Slc35a1 | 0.15466335 | -0.470629825 | 0.319 | 0.388 | 1 | 0.008733973 | -2.776493958 | 0 | 0.432 | 1 | 0.15466335 | 0.017391664 |
| Dand5 | 0.008738041 | -1.36994961 | 0.076 | 0.182 | 1 | 0.260064347 | -1.304006187 | 0 | 0.114 | 1 | 0.260064347 | 0.017399728 |
| Zfp943 | 0.008739142 | -1.070389328 | 0.168 | 0.294 | 1 | 0.009763401 | 1.454985714 | 0.6 | 0.222 | 1 | 0.009763401 | 0.017401911 |
| Scly | 0.00873972 | -1.655351829 | 0.092 | 0.2 | 1 | 0.829401798 | 0.418009066 | 0.2 | 0.176 | 1 | 0.829401798 | 0.017403057 |
| H2-Aa | 0.00874564 | 2.162861609 | 0.176 | 0.076 | 1 | 0.557744661 | 1.017921908 | 0 | 0.034 | 1 | 0.557744661 | 0.017414793 |
| 2010310C07Rik | 0.834180595 | 0.166649869 | 0.059 | 0.053 | 1 | 0.00874905 | 3.477353527 | 0.3 | 0.074 | 1 | 0.834180595 | 0.017421554 |
| Paqr3 | 0.045016105 | 0.726077278 | 0.176 | 0.094 | 1 | 0.008753956 | 2.210566986 | 0.3 | 0.071 | 1 | 0.045016105 | 0.01743128 |
| Ung | 0.088617786 | -1.411426246 | 0.034 | 0.082 | 1 | 0.008753956 | 2.159940913 | 0.3 | 0.071 | 1 | 0.088617786 | 0.01743128 |
| Cspg4b | 0.00876515 | -2.129283017 | 0.025 | 0.106 | 1 | 0.584299096 | 0.744903414 | 0.2 | 0.133 | 1 | 0.584299096 | 0.017453472 |
| Pitpnm1 | 0.00876515 | -2.129283017 | 0.025 | 0.106 | 1 | 0.233699361 | 1.393431043 | 0.2 | 0.086 | 1 | 0.233699361 | 0.017453472 |
| Ddx20 | 0.008767491 | -1.276840205 | 0.092 | 0.206 | 1 | 0.319235496 | -0.5519337 | 0.1 | 0.231 | 1 | 0.319235496 | 0.017458114 |
| Il20rb | 0.00877008 | -1.122856748 | 0.101 | 0.218 | 1 | 0.717067935 | -0.165299916 | 0.2 | 0.235 | 1 | 0.717067935 | 0.017463246 |
| Hexdc | 0.231140974 | -0.807354922 | 0.042 | 0.076 | 1 | 0.008779968 | 1.667424661 | 0.4 | 0.114 | 1 | 0.231140974 | 0.017482848 |
| Afmid | 0.008793417 | -1.529820947 | 0.05 | 0.147 | 1 | 0.511794478 | 0.821524695 | 0.2 | 0.123 | 1 | 0.511794478 | 0.017509509 |
| Wdr3 | 0.008805404 | -1.359895945 | 0.16 | 0.276 | 1 | 0.047702132 | -2.048167282 | 0 | 0.29 | 1 | 0.047702132 | 0.017533273 |
| Ppp1r10 | 0.008813394 | -0.665634802 | 0.647 | 0.753 | 1 | 0.35175122 | 0.523018715 | 0.9 | 0.821 | 1 | 0.35175122 | 0.017549112 |
| Phf8 | 0.008833555 | -0.973220669 | 0.479 | 0.565 | 1 | 0.526495144 | -0.122311409 | 0.5 | 0.66 | 1 | 0.526495144 | 0.017589079 |
| Rack1 | 0.306558568 | -0.103451349 | 0.983 | 0.976 | 1 | 0.008847981 | -0.909974546 | 0.8 | 0.994 | 1 | 0.306558568 | 0.017617676 |
| Commd3 | 0.008852959 | -0.5088858 | 0.563 | 0.647 | 1 | 0.026615737 | -1.252607034 | 0.3 | 0.645 | 1 | 0.026615737 | 0.017627543 |
| Fbxw9 | 0.008864975 | -1.222392421 | 0.118 | 0.235 | 1 | 0.209039774 | 1.08718457 | 0.3 | 0.145 | 1 | 0.209039774 | 0.017651362 |
| Mtrf1l | 0.00887315 | -1.359895945 | 0.126 | 0.241 | 1 | 0.374302745 | 0.896906507 | 0.3 | 0.188 | 1 | 0.374302745 | 0.017667567 |
| Rin2 | 0.113752913 | -0.454771539 | 0.933 | 0.912 | 1 | 0.008874419 | 1.205890499 | 1 | 0.941 | 1 | 0.113752913 | 0.017670083 |
| 2700049A03Rik | 0.008879213 | -1.15168943 | 0.143 | 0.265 | 1 | 0.663282101 | -0.121629444 | 0.3 | 0.213 | 1 | 0.663282101 | 0.017679586 |
| Eef1e1 | 0.008881444 | -0.937939032 | 0.235 | 0.376 | 1 | 0.778480759 | 0.331421381 | 0.3 | 0.244 | 1 | 0.778480759 | 0.017684008 |
| Nup133 | 0.008882591 | -1.333423734 | 0.092 | 0.206 | 1 | 0.152528539 | -1.408342847 | 0 | 0.173 | 1 | 0.152528539 | 0.017686281 |
| Gm15446 | 0.008883337 | -2.240314329 | 0.025 | 0.106 | 1 | 0.670447518 | 1.159940913 | 0.1 | 0.065 | 1 | 0.670447518 | 0.017687761 |
| Rnf123 | 0.008889622 | -1.614709844 | 0.059 | 0.159 | 1 | 0.580398453 | 0.821524695 | 0.2 | 0.133 | 1 | 0.580398453 | 0.017700218 |
| Irx5 | 0.008892531 | -2.891419187 | 0.067 | 0.165 | 1 | 0.655185212 | 0.236562194 | 0.1 | 0.062 | 1 | 0.655185212 | 0.017705985 |
| Thrb | 0.008895648 | 0.283792966 | 0.882 | 0.765 | 1 | 0.715577675 | -0.217555086 | 0.9 | 0.83 | 1 | 0.715577675 | 0.017712164 |
| Cpne3 | 0.205840808 | -0.61667136 | 0.487 | 0.488 | 1 | 0.008902588 | -1.672076063 | 0.3 | 0.67 | 1 | 0.205840808 | 0.017725919 |
| Plekha1 | 0.008917381 | -1.070389328 | 0.294 | 0.418 | 1 | 0.013839976 | -2.034646143 | 0.3 | 0.633 | 1 | 0.013839976 | 0.017755243 |
| Borcs7 | 0.008923356 | -0.826463745 | 0.277 | 0.412 | 1 | 0.957153673 | 0.453137289 | 0.3 | 0.299 | 1 | 0.957153673 | 0.017767086 |
| Zhx1 | 0.008974951 | -1.055742552 | 0.218 | 0.347 | 1 | 0.016900353 | -2.481923979 | 0 | 0.38 | 1 | 0.016900353 | 0.017869353 |
| Metrnl | 0.008977078 | 0.77237093 | 0.672 | 0.535 | 1 | 0.995824241 | 0.393431043 | 0.5 | 0.568 | 1 | 0.995824241 | 0.017873569 |
| Mak16 | 0.008991316 | -0.780047576 | 0.395 | 0.5 | 1 | 0.522585949 | -0.225252075 | 0.4 | 0.515 | 1 | 0.522585949 | 0.017901789 |
| Fbxl17 | 0.008993341 | -0.536275969 | 0.866 | 0.918 | 1 | 0.589079427 | -0.47695254 | 0.9 | 0.96 | 1 | 0.589079427 | 0.017905801 |
| Fam72a | 0.009002626 | -2.008988783 | 0.025 | 0.106 | 1 | 0.460577668 | 0.625604485 | 0 | 0.052 | 1 | 0.460577668 | 0.017924205 |
| Gm15873 | 0.147030916 | -1.485426827 | 0 | 0.018 | 1 | 0.009005342 | 4.602884409 | 0.1 | 0.009 | 1 | 0.147030916 | 0.017929587 |
| Gm32816 | 0.34025168 | -2.359895945 | 0.017 | 0.035 | 1 | 0.009005342 | 6.187846909 | 0.1 | 0.009 | 1 | 0.34025168 | 0.017929587 |
| Tnni1 | 0.383916503 | 1.929610672 | 0.025 | 0.012 | 1 | 0.009005342 | 6.017921908 | 0.1 | 0.009 | 1 | 0.383916503 | 0.017929587 |
| Gm16675 | 0.458788392 | -0.807354922 | 0.025 | 0.041 | 1 | 0.009005342 | 5.339850003 | 0.1 | 0.009 | 1 | 0.458788392 | 0.017929587 |
| Pdxk-ps | 0.511005543 | -0.485426827 | 0.008 | 0.018 | 1 | 0.009005342 | 4.602884409 | 0.1 | 0.009 | 1 | 0.511005543 | 0.017929587 |
| Gm56646 | 0.511005543 | -0.485426827 | 0.008 | 0.018 | 1 | 0.00900581 | 5.017921908 | 0.1 | 0.009 | 1 | 0.511005543 | 0.017930515 |
| Nek5 | 0.566193127 | 0.514573173 | 0.042 | 0.029 | 1 | 0.00900581 | 4.695993813 | 0.1 | 0.009 | 1 | 0.566193127 | 0.017930515 |
| Ankrd45 | 0.715874401 | 0.929610672 | 0.017 | 0.012 | 1 | 0.00900581 | 4.432959407 | 0.1 | 0.009 | 1 | 0.715874401 | 0.017930515 |
| Mrpl15 | 0.009006475 | -0.873352481 | 0.336 | 0.453 | 1 | 0.541800416 | -0.304006187 | 0.3 | 0.364 | 1 | 0.541800416 | 0.017931834 |
| Ank3 | 0.009010537 | -1.698420551 | 0.067 | 0.171 | 1 | 0.101503696 | -2.004445905 | 0 | 0.216 | 1 | 0.101503696 | 0.017939883 |
| Tcf7 | 0.009012386 | -1.517848305 | 0.067 | 0.171 | 1 | 0.05165886 | -2.567040593 | 0 | 0.284 | 1 | 0.05165886 | 0.01794355 |
| Calr | 0.009053634 | -0.285513749 | 0.958 | 0.971 | 1 | 0.025189533 | -0.802442075 | 0.9 | 0.997 | 1 | 0.025189533 | 0.018025299 |
| Grem1 | 0.821958642 | -1.008988783 | 0.042 | 0.047 | 1 | 0.00906334 | 3.967295835 | 0.2 | 0.037 | 1 | 0.821958642 | 0.018044535 |
| Ern1 | 0.009066391 | -0.77535646 | 0.487 | 0.629 | 1 | 0.195761774 | 0.448556262 | 0.8 | 0.701 | 1 | 0.195761774 | 0.018050583 |
| Akr1a1 | 0.009070953 | -0.30157626 | 0.924 | 0.971 | 1 | 0.615108041 | -0.21595001 | 0.9 | 0.988 | 1 | 0.615108041 | 0.018059624 |
| Ankrd12 | 0.009078073 | -0.379112183 | 0.95 | 0.947 | 1 | 0.221390447 | -0.49284226 | 0.9 | 0.978 | 1 | 0.221390447 | 0.018073734 |
| C2cd2 | 0.009088426 | -1.424026283 | 0.109 | 0.224 | 1 | 0.482363774 | 0.782705446 | 0.3 | 0.207 | 1 | 0.482363774 | 0.018094252 |
| Bcap29 | 0.009098206 | -1.263034406 | 0.151 | 0.271 | 1 | 0.539650575 | -0.505640048 | 0.1 | 0.167 | 1 | 0.539650575 | 0.018113634 |
| Slc25a44 | 0.009099853 | -1.070389328 | 0.151 | 0.276 | 1 | 0.3028982 | 1.536795218 | 0.4 | 0.309 | 1 | 0.3028982 | 0.018116898 |
| Slc12a8 | 0.009112758 | -2.10433666 | 0.025 | 0.106 | 1 | 0.288140277 | 1.210566986 | 0.1 | 0.034 | 1 | 0.288140277 | 0.018142474 |

| | | | | | | | | | | | | |
|---|---|---|---|---|---|---|---|---|---|---|---|---|
| Amz2 | 0.00913098 | 0.347463187 | 0.429 | 0.253 | 1 | 0.79869498 | 0.31748219 | 0.3 | 0.25 | 1 | 0.79869498 | 0.018178586 |
| Rnf217 | 0.018723023 | -1.058316496 | 0.445 | 0.518 | 1 | 0.009131295 | -1.774868386 | 0.3 | 0.71 | 1 | 0.018723023 | 0.01817921 |
| Sf3b6 | 0.009145925 | -0.385891154 | 0.849 | 0.853 | 1 | 0.125515411 | -0.553830736 | 0.6 | 0.895 | 1 | 0.125515411 | 0.018208202 |
| Rps8 | 0.01833915 | -0.250185381 | 0.992 | 0.994 | 1 | 0.009163245 | -0.776032053 | 1 | 0.997 | 1 | 0.01833915 | 0.018242524 |
| Nipal1 | 0.009165426 | 1.372554168 | 0.143 | 0.053 | 1 | 0.102671865 | 3.602884409 | 0.1 | 0.022 | 1 | 0.102671865 | 0.018246848 |
| Map2k4 | 0.00918406 | -0.555361118 | 0.655 | 0.741 | 1 | 0.273752428 | -0.535131345 | 0.7 | 0.849 | 1 | 0.273752428 | 0.018283774 |
| Tll2 | 0.354463513 | -0.485426827 | 0.017 | 0.035 | 1 | 0.009189974 | 1.959028219 | 0.2 | 0.034 | 1 | 0.354463513 | 0.018295492 |
| Megf11 | 0.506600077 | 1.836501268 | 0.05 | 0.035 | 1 | 0.009190275 | 1.902444691 | 0.2 | 0.034 | 1 | 0.506600077 | 0.018296088 |
| Rhot2 | 0.00919639 | -1.439623138 | 0.05 | 0.147 | 1 | 0.266728707 | -0.339630097 | 0 | 0.111 | 1 | 0.266728707 | 0.018308206 |
| Lysmd3 | 0.009202119 | -0.809818962 | 0.58 | 0.624 | 1 | 0.014840608 | -1.652481436 | 0.4 | 0.716 | 1 | 0.014840608 | 0.018319559 |
| Tspan31 | 0.138224466 | -0.247927513 | 0.84 | 0.788 | 1 | 0.009211866 | -1.208757872 | 0.4 | 0.824 | 1 | 0.138224466 | 0.018338874 |
| Cnot7 | 0.009227544 | -0.688518693 | 0.538 | 0.594 | 1 | 0.048376504 | -0.978316173 | 0.3 | 0.66 | 1 | 0.048376504 | 0.01836994 |
| Thoc2l | 0.009229443 | -0.488286481 | 0.815 | 0.812 | 1 | 0.249842593 | -0.577522077 | 0.8 | 0.833 | 1 | 0.249842593 | 0.018373703 |
| Pum3 | 0.009230842 | -1.14332085 | 0.336 | 0.441 | 1 | 0.057578863 | -1.327852929 | 0.1 | 0.432 | 1 | 0.057578863 | 0.018376475 |
| Homer2 | 0.009232968 | -2.250961574 | 0.025 | 0.106 | 1 | 0.166449167 | -1.581990934 | 0 | 0.164 | 1 | 0.166449167 | 0.018380689 |
| Bdh1 | 0.009241906 | 0.893084796 | 0.244 | 0.124 | 1 | 0.872871184 | 0.374065718 | 0.1 | 0.114 | 1 | 0.872871184 | 0.0183984 |
| Pstpip1 | 0.00926123 | 0.790207615 | 0.303 | 0.171 | 1 | 0.756700708 | 0.185031894 | 0.2 | 0.256 | 1 | 0.756700708 | 0.01843669 |
| Aox3 | 0.009297822 | 1.906890596 | 0.092 | 0.024 | 1 | 0.245300203 | -0.763437806 | 0 | 0.12 | 1 | 0.245300203 | 0.018509195 |
| Arl8b | 0.213277242 | -0.361768359 | 0.714 | 0.7 | 1 | 0.00933014 | -1.380821784 | 0.5 | 0.802 | 1 | 0.213277242 | 0.018573228 |
| Rusf1 | 0.009334047 | -1.402964667 | 0.059 | 0.159 | 1 | 0.952551372 | 0.808468542 | 0.1 | 0.093 | 1 | 0.952551372 | 0.018580969 |
| Zbtb39 | 0.009334047 | -1.402964667 | 0.059 | 0.159 | 1 | 0.314392508 | -0.567040593 | 0.1 | 0.231 | 1 | 0.314392508 | 0.018580969 |
| Gfm1 | 0.009341099 | -1.027954062 | 0.168 | 0.294 | 1 | 0.664224071 | 0.403212064 | 0.4 | 0.327 | 1 | 0.664224071 | 0.018594941 |
| Ganab | 0.009345368 | -0.414807752 | 0.681 | 0.753 | 1 | 0.240774317 | 0.539489327 | 0.9 | 0.738 | 1 | 0.240774317 | 0.018603401 |
| Gm43154 | 0.009354696 | -3.008988783 | 0.034 | 0.118 | 1 | 0.643848687 | 0.210566986 | 0 | 0.022 | 1 | 0.643848687 | 0.018621882 |
| Armcx3 | 0.009362663 | -0.643763854 | 0.672 | 0.665 | 1 | 0.030721803 | -1.273632538 | 0.5 | 0.713 | 1 | 0.030721803 | 0.018637667 |
| Wdfy2 | 0.009366458 | -0.93021167 | 0.21 | 0.347 | 1 | 0.292399136 | 0.245332404 | 0.2 | 0.392 | 1 | 0.292399136 | 0.018645185 |
| Ddx18 | 0.009367441 | -0.844969214 | 0.336 | 0.459 | 1 | 0.872677603 | 0.32443495 | 0.4 | 0.389 | 1 | 0.872677603 | 0.018647133 |
| Suz12 | 0.009371763 | -0.815132273 | 0.58 | 0.635 | 1 | 0.061630868 | -0.820021334 | 0.3 | 0.716 | 1 | 0.061630868 | 0.018655696 |
| Tmem259 | 0.009371958 | -0.76836079 | 0.412 | 0.524 | 1 | 0.489399623 | -0.5004034 | 0.5 | 0.512 | 1 | 0.489399623 | 0.018656082 |
| Rps14 | 0.245669685 | -0.100442599 | 1 | 0.994 | 1 | 0.009375556 | -0.863302911 | 1 | 1 | 1 | 0.245669685 | 0.01866321 |
| Mbp | 0.00939064 | 1.202629167 | 0.134 | 0.047 | 1 | 0.508960956 | -0.357117523 | 0.1 | 0.179 | 1 | 0.508960956 | 0.018693095 |
| Hras | 0.009411417 | -0.880728108 | 0.294 | 0.424 | 1 | 0.164600857 | -0.763437806 | 0.3 | 0.522 | 1 | 0.164600857 | 0.01873426 |
| Slc25a38 | 0.009411476 | -0.966553517 | 0.176 | 0.306 | 1 | 0.411621608 | -0.239465935 | 0.2 | 0.34 | 1 | 0.411621608 | 0.018734376 |
| Brap | 0.009423408 | -0.907253492 | 0.319 | 0.441 | 1 | 0.494245985 | -0.391469028 | 0.5 | 0.614 | 1 | 0.494245985 | 0.018758016 |
| Cracd | 0.009424969 | -1.61667136 | 0.134 | 0.247 | 1 | 0.216030236 | -1.782977992 | 0.1 | 0.262 | 1 | 0.216030236 | 0.018761108 |
| Serinc3 | 0.009445499 | -0.6203309 | 0.992 | 0.941 | 1 | 0.07019213 | -0.56290933 | 1 | 0.994 | 1 | 0.07019213 | 0.01880178 |
| Qpctl | 0.617102321 | 0.204233052 | 0.168 | 0.147 | 1 | 0.009461673 | 2.290001453 | 0.4 | 0.139 | 1 | 0.617102321 | 0.018833822 |
| Slc2a12 | 0.00947294 | 0.730301864 | 0.244 | 0.124 | 1 | 0.03925842 | 2.432959407 | 0.2 | 0.049 | 1 | 0.03925842 | 0.018856144 |
| Gm33424 | 0.121879242 | 1.362570079 | 0.059 | 0.024 | 1 | 0.009481518 | 4.017921908 | 0.1 | 0.009 | 1 | 0.121879242 | 0.018873136 |
| Slc24a3 | 0.009504842 | -2.944858446 | 0.034 | 0.118 | 1 | 0.965239221 | -0.840059087 | 0.1 | 0.09 | 1 | 0.965239221 | 0.018919342 |
| Tmem150a | 0.009523175 | -1.051024003 | 0.185 | 0.312 | 1 | 0.293066927 | -0.339630097 | 0.1 | 0.269 | 1 | 0.293066927 | 0.018955658 |
| Ints8 | 0.009530197 | -0.932226889 | 0.437 | 0.512 | 1 | 0.080520712 | -0.86200164 | 0.5 | 0.701 | 1 | 0.080520712 | 0.018969569 |
| Cryzl1 | 0.009530346 | -0.744514048 | 0.37 | 0.506 | 1 | 0.336586056 | -0.575602606 | 0.4 | 0.568 | 1 | 0.336586056 | 0.018969864 |
| Nt5dc2 | 0.009539849 | -0.87774425 | 0.227 | 0.353 | 1 | 0.871869381 | 0.695993813 | 0.2 | 0.188 | 1 | 0.871869381 | 0.018988689 |
| Ubr1 | 0.009549685 | -0.570184445 | 0.672 | 0.729 | 1 | 0.451225988 | -0.21595001 | 0.7 | 0.75 | 1 | 0.451225988 | 0.019008173 |
| Pnpla6 | 0.009575563 | -1.610957709 | 0.084 | 0.188 | 1 | 0.978813346 | 0.176619654 | 0.2 | 0.194 | 1 | 0.978813346 | 0.019059434 |
| Uqcrb | 0.102025875 | -0.249218436 | 0.899 | 0.918 | 1 | 0.009581725 | -0.977845243 | 0.4 | 0.917 | 1 | 0.102025875 | 0.019071641 |
| Xkr4 | 0.009608411 | -2.600904045 | 0.034 | 0.118 | 1 | 0.434735961 | -0.654503434 | 0.1 | 0.198 | 1 | 0.434735961 | 0.0191245 |
| Rrp36 | 0.00961458 | -0.944858446 | 0.21 | 0.341 | 1 | 0.269951167 | -0.419483404 | 0.1 | 0.278 | 1 | 0.269951167 | 0.019136719 |
| Piwil2 | 0.009641426 | -2.628384781 | 0.042 | 0.129 | 1 | 0.17324519 | 1.874963954 | 0.2 | 0.083 | 1 | 0.17324519 | 0.019189894 |
| Vldlr | 0.009641461 | -2.628384781 | 0.042 | 0.129 | 1 | 0.247600602 | 0.695993813 | 0.2 | 0.09 | 1 | 0.247600602 | 0.019189964 |
| Sncaip | 0.009658302 | -2.418789634 | 0.126 | 0.235 | 1 | 0.105286248 | -3.230005605 | 0 | 0.213 | 1 | 0.105286248 | 0.019223321 |
| Stard10 | 0.009662354 | 0.763932642 | 0.37 | 0.224 | 1 | 0.01983669 | 1.236562194 | 0.7 | 0.373 | 1 | 0.01983669 | 0.019231346 |
| Ensa | 0.009678069 | -0.607056272 | 0.613 | 0.665 | 1 | 0.024928111 | -1.116504412 | 0.5 | 0.787 | 1 | 0.024928111 | 0.019262474 |
| Grk3 | 0.009701085 | -1.416164165 | 0.118 | 0.229 | 1 | 0.123212529 | -1.297982399 | 0.1 | 0.355 | 1 | 0.123212529 | 0.019308059 |
| Sh3bp2 | 0.009710346 | -1.655351829 | 0.025 | 0.106 | 1 | 0.208425713 | 1.930459067 | 0.3 | 0.157 | 1 | 0.208425713 | 0.019326401 |
| Fdx2 | 0.00971802 | -0.77915803 | 0.218 | 0.353 | 1 | 0.776005125 | 0.210566986 | 0.4 | 0.324 | 1 | 0.776005125 | 0.019341601 |
| Slc6a4 | 0.093245792 | -1.807354922 | 0 | 0.024 | 1 | 0.009727674 | 3.795529487 | 0.1 | 0.009 | 1 | 0.093245792 | 0.01936072 |
| Plin4 | 0.038142909 | 2.514573173 | 0.025 | 0 | 1 | 0.009727839 | 4.017921908 | 0.1 | 0.009 | 1 | 0.038142909 | 0.019361048 |
| Col18a1 | 0.06000657 | -2.292781749 | 0 | 0.029 | 1 | 0.009727839 | 4.017921908 | 0.1 | 0.009 | 1 | 0.06000657 | 0.019361048 |
| Cfap206 | 0.060010312 | -2.807354922 | 0 | 0.029 | 1 | 0.009727839 | 4.017921908 | 0.1 | 0.009 | 1 | 0.060010312 | 0.019361048 |
| Gm13594 | 0.075308142 | 1.029146346 | 0.076 | 0.029 | 1 | 0.009727839 | 4.017921908 | 0.1 | 0.009 | 1 | 0.075308142 | 0.019361048 |
| Cdc45 | 0.078232801 | -1.707819249 | 0.017 | 0.059 | 1 | 0.009727839 | 4.017921908 | 0.1 | 0.009 | 1 | 0.078232801 | 0.019361048 |
| Cerkl | 0.091361251 | 2.099535674 | 0.017 | 0 | 1 | 0.009727839 | 4.017921908 | 0.1 | 0.009 | 1 | 0.091361251 | 0.019361048 |
| Dcdc2c | 0.091361251 | 2.836501268 | 0.017 | 0 | 1 | 0.009727839 | 4.017921908 | 0.1 | 0.009 | 1 | 0.091361251 | 0.019361048 |
| Gm17509 | 0.091361251 | 2.099535674 | 0.017 | 0 | 1 | 0.009727839 | 4.017921908 | 0.1 | 0.009 | 1 | 0.091361251 | 0.019361048 |
| Jph2 | 0.093244798 | -3.655351829 | 0 | 0.024 | 1 | 0.009727839 | 4.017921908 | 0.1 | 0.009 | 1 | 0.093244798 | 0.019361048 |
| Shcbp1 | 0.145555939 | -1.707819249 | 0.008 | 0.035 | 1 | 0.009727839 | 4.017921908 | 0.1 | 0.009 | 1 | 0.145555939 | 0.019361048 |
| Mmp19 | 0.147030916 | -1.485426827 | 0 | 0.018 | 1 | 0.009727839 | 4.017921908 | 0.1 | 0.009 | 1 | 0.147030916 | 0.019361048 |
| Eldr | 0.16759755 | -1.292781749 | 0.017 | 0.047 | 1 | 0.009727839 | 4.017921908 | 0.1 | 0.009 | 1 | 0.16759755 | 0.019361048 |
| Tmem163 | 0.237885589 | -2.292781749 | 0 | 0.012 | 1 | 0.009727839 | 4.017921908 | 0.1 | 0.009 | 1 | 0.237885589 | 0.019361048 |
| Stk31 | 0.237885589 | -2.292781749 | 0 | 0.012 | 1 | 0.009727839 | 4.017921908 | 0.1 | 0.009 | 1 | 0.237885589 | 0.019361048 |
| Gm47326 | 0.237885589 | -1.070389328 | 0 | 0.012 | 1 | 0.009727839 | 4.017921908 | 0.1 | 0.009 | 1 | 0.237885589 | 0.019361048 |
| Gm47101 | 0.237885589 | -1.070389328 | 0 | 0.012 | 1 | 0.009727839 | 4.017921908 | 0.1 | 0.009 | 1 | 0.237885589 | 0.019361048 |
| Gm56767 | 0.33261065 | -1.070389328 | 0.008 | 0.024 | 1 | 0.009727839 | 4.017921908 | 0.1 | 0.009 | 1 | 0.33261065 | 0.019361048 |
| Dynlt2a3 | 0.368010882 | 1.514573173 | 0.017 | 0.006 | 1 | 0.009727839 | 4.017921908 | 0.1 | 0.009 | 1 | 0.368010882 | 0.019361048 |
| 1700003D09Rik | 0.370184089 | 1.099535674 | 0.017 | 0.006 | 1 | 0.009727839 | 4.017921908 | 0.1 | 0.009 | 1 | 0.370184089 | 0.019361048 |
| Gm48342 | 0.370184089 | 1.099535674 | 0.017 | 0.006 | 1 | 0.009727839 | 4.017921908 | 0.1 | 0.009 | 1 | 0.370184089 | 0.019361048 |
| Fam178b | 0.387377553 | 2.514573173 | 0.025 | 0.012 | 1 | 0.009727839 | 4.017921908 | 0.1 | 0.009 | 1 | 0.387377553 | 0.019361048 |
| Gm32261 | 0.396130756 | 0.514573173 | 0.025 | 0.012 | 1 | 0.009727839 | 4.017921908 | 0.1 | 0.009 | 1 | 0.396130756 | 0.019361048 |
| 5830454E08Rik | 0.511005543 | -0.485426827 | 0.008 | 0.018 | 1 | 0.009727839 | 4.017921908 | 0.1 | 0.009 | 1 | 0.511005543 | 0.019361048 |
| Ngb | 0.65150204 | 1 | 0.025 | 0.018 | 1 | 0.009727839 | 4.017921908 | 0.1 | 0.009 | 1 | 0.65150204 | 0.019361048 |
| Chrnb2 | 0.731769749 | -0.222392421 | 0.017 | 0.012 | 1 | 0.009727839 | 4.017921908 | 0.1 | 0.009 | 1 | 0.731769749 | 0.019361048 |
| Ogdhl | 0.781790612 | -0.485426827 | 0.008 | 0.012 | 1 | 0.009727839 | 4.017921908 | 0.1 | 0.009 | 1 | 0.781790612 | 0.019361048 |
| Iqgap3 | 0.810760968 | -1.378511623 | 0.042 | 0.035 | 1 | 0.009727839 | 4.017921908 | 0.1 | 0.009 | 1 | 0.810760968 | 0.019361048 |
| A930007A09Rik | 0.954551343 | -0.222392421 | 0.017 | 0.018 | 1 | 0.009727839 | 4.017921908 | 0.1 | 0.009 | 1 | 0.954551343 | 0.019361048 |
| Slc29a4 | 0.969692448 | 0.514573173 | 0.017 | 0.018 | 1 | 0.009727839 | 4.017921908 | 0.1 | 0.009 | 1 | 0.969692448 | 0.019361048 |
| Nxt1 | 0.009729606 | -0.645891499 | 0.067 | 0.176 | 1 | 0.092127669 | -1.521236903 | 0 | 0.225 | 1 | 0.092127669 | 0.019364547 |
| Rbm8a | 0.009730879 | -0.501546493 | 0.731 | 0.776 | 1 | 0.022267318 | -1.028292647 | 0.8 | 0.852 | 1 | 0.022267318 | 0.019367067 |
| Rpl13a | 0.198641176 | 0.104208374 | 0.992 | 0.994 | 1 | 0.009734331 | -0.804848066 | 1 | 1 | 1 | 0.198641176 | 0.019373906 |
| Gpr153 | 0.009739492 | -0.981852653 | 0.21 | 0.335 | 1 | 0.910721925 | 0.39994435 | 0.4 | 0.472 | 1 | 0.910721925 | 0.019384126 |
| Hey1 | 0.009744818 | -1.328701323 | 0.05 | 0.147 | 1 | 0.825331781 | -0.230005605 | 0.1 | 0.117 | 1 | 0.825331781 | 0.019394674 |
| Zfp213 | 0.009745566 | -1.748461233 | 0.025 | 0.106 | 1 | 0.474904369 | 0.695993813 | 0 | 0.049 | 1 | 0.474904369 | 0.019396156 |
| A230072C01Rik | 0.009747601 | 0.850176205 | 0.286 | 0.153 | 1 | 0.060595826 | 1.675529711 | 0.4 | 0.173 | 1 | 0.060595826 | 0.019400186 |
| Spcs2 | 0.07333957 | -0.234334094 | 0.899 | 0.918 | 1 | 0.00975047 | -1.018736563 | 0.6 | 0.938 | 1 | 0.07333957 | 0.019405868 |
| Fez2 | 0.009765807 | -0.97641318 | 0.244 | 0.376 | 1 | 0.483568617 | 0.64536774 | 0.4 | 0.315 | 1 | 0.483568617 | 0.019436244 |
| Abhd17b | 0.009774963 | -0.727184573 | 0.361 | 0.494 | 1 | 0.224391915 | -0.68251781 | 0.4 | 0.602 | 1 | 0.224391915 | 0.019454376 |
| Card19 | 0.009792588 | 0.505235308 | 0.664 | 0.506 | 1 | 0.624322711 | -0.232714893 | 0.6 | 0.667 | 1 | 0.624322711 | 0.01948928 |

| | | | | | | | | | | | | |
|---|---|---|---|---|---|---|---|---|---|---|---|---|
| Adar | 0.009802034 | -1.166604643 | 0.21 | 0.329 | 1 | 0.631292005 | 0.425464871 | 0.4 | 0.358 | 1 | 0.631292005 | 0.019507989 |
| Wwc1 | 0.009830054 | -1.807354922 | 0.025 | 0.106 | 1 | 0.425870342 | -0.191531458 | 0.1 | 0.201 | 1 | 0.425870342 | 0.019563478 |
| Trim59 | 0.009835735 | -1.333423734 | 0.076 | 0.182 | 1 | 0.649664235 | 0.473601392 | 0.2 | 0.139 | 1 | 0.649664235 | 0.019574728 |
| Otoa | 0.009839832 | 1.836501268 | 0.193 | 0.094 | 1 | 0.626078725 | 1.695993813 | 0.1 | 0.065 | 1 | 0.626078725 | 0.019582842 |
| Slc9a1 | 0.009876822 | -0.710393192 | 0.412 | 0.529 | 1 | 0.460068089 | -0.600463594 | 0.6 | 0.596 | 1 | 0.460068089 | 0.019656093 |
| Nif3l1 | 0.009901663 | -1.109917692 | 0.076 | 0.182 | 1 | 0.224071783 | 1.040641984 | 0.3 | 0.151 | 1 | 0.224071783 | 0.019705282 |
| Snrpd2 | 0.009920593 | -0.446384726 | 0.782 | 0.824 | 1 | 0.030449759 | -1.001977649 | 0.5 | 0.781 | 1 | 0.030449759 | 0.019742768 |
| Timm9 | 0.009926404 | -0.935459748 | 0.328 | 0.441 | 1 | 0.07443691 | -1.716787712 | 0.1 | 0.367 | 1 | 0.07443691 | 0.019754274 |
| A530046M15Rik | 0.009942469 | 0.819427754 | 0.143 | 0.053 | 1 | 0.698532411 | 2.017921908 | 0 | 0.015 | 1 | 0.698532411 | 0.019786086 |
| Mau2 | 0.321848766 | -0.474934453 | 0.412 | 0.435 | 1 | 0.009963655 | -1.425021588 | 0.3 | 0.716 | 1 | 0.321848766 | 0.019828035 |
| 4930500G05Rik | 0.009963853 | -1.608809243 | 0.092 | 0.2 | 1 | 0.445956184 | -0.374395515 | 0.1 | 0.191 | 1 | 0.445956184 | 0.019828427 |
| Ndufa4 | 0.009969562 | 0.246994 | 0.95 | 0.965 | 1 | 0.891017898 | 0.466044041 | 0.7 | 0.954 | 1 | 0.891017898 | 0.019839731 |
| Cyp2u1 | 0.009970184 | -2.185866545 | 0.05 | 0.141 | 1 | 0.460327488 | 0.751135367 | 0.2 | 0.12 | 1 | 0.460327488 | 0.019840963 |
| Nudt5 | 0.00997402 | -0.59645814 | 0.286 | 0.435 | 1 | 0.096952579 | 1.432959407 | 0.5 | 0.29 | 1 | 0.096952579 | 0.019848558 |
| Atpaf2 | 0.192018788 | -0.887525271 | 0.109 | 0.159 | 1 | 0.009976763 | 1.675529711 | 0.5 | 0.179 | 1 | 0.192018788 | 0.019853991 |
| Rbm46 | 0.034550051 | 2.099535674 | 0.042 | 0.006 | 1 | 0.009979004 | 3.695993813 | 0.1 | 0.009 | 1 | 0.034550051 | 0.019858427 |
| Tnfsf10 | 0.07674505 | 2.389042291 | 0.034 | 0.006 | 1 | 0.009979004 | 3.432959407 | 0.1 | 0.009 | 1 | 0.07674505 | 0.019858427 |
| Gm44079 | 0.091361251 | 2.099535674 | 0.017 | 0 | 1 | 0.009979004 | 3.695993813 | 0.1 | 0.009 | 1 | 0.091361251 | 0.019858427 |
| Gm14827 | 0.091361251 | 2.099535674 | 0.017 | 0 | 1 | 0.009979004 | 3.695993813 | 0.1 | 0.009 | 1 | 0.091361251 | 0.019858427 |
| Clec4a1 | 0.093245792 | -1.807354922 | 0 | 0.024 | 1 | 0.009979004 | 3.695993813 | 0.1 | 0.009 | 1 | 0.093245792 | 0.019858427 |
| Dnai2 | 0.237885589 | -1.070389328 | 0 | 0.012 | 1 | 0.009979004 | 3.432959407 | 0.1 | 0.009 | 1 | 0.237885589 | 0.019858427 |
| Carmil3 | 0.237885589 | -1.070389328 | 0 | 0.012 | 1 | 0.009979004 | 3.432959407 | 0.1 | 0.009 | 1 | 0.237885589 | 0.019858427 |
| Tmem151b | 0.237885589 | -1.070389328 | 0 | 0.012 | 1 | 0.009979004 | 3.695993813 | 0.1 | 0.009 | 1 | 0.237885589 | 0.019858427 |
| Mtarc1 | 0.335990222 | -1.163498732 | 0.025 | 0.047 | 1 | 0.009979004 | 3.432959407 | 0.1 | 0.009 | 1 | 0.335990222 | 0.019858427 |
| Sv2c | 0.345242044 | 0.943416472 | 0.059 | 0.035 | 1 | 0.009979004 | 2.558490289 | 0.1 | 0.009 | 1 | 0.345242044 | 0.019858427 |
| 1300014J16Rik | 0.370184089 | 1.099535674 | 0.017 | 0.006 | 1 | 0.009979004 | 3.695993813 | 0.1 | 0.009 | 1 | 0.370184089 | 0.019858427 |
| Aldh3b3 | 0.374548525 | 0.514573173 | 0.017 | 0.006 | 1 | 0.009979004 | 3.432959407 | 0.1 | 0.009 | 1 | 0.374548525 | 0.019858427 |
| P2ry6 | 0.396130756 | 0.514573173 | 0.025 | 0.012 | 1 | 0.009979004 | 3.695993813 | 0.1 | 0.009 | 1 | 0.396130756 | 0.019858427 |
| Cyp2c23 | 0.508731273 | -0.807354922 | 0.008 | 0.018 | 1 | 0.009979004 | 3.695993813 | 0.1 | 0.009 | 1 | 0.508731273 | 0.019858427 |
| Ydjc | 0.659865009 | 0.514573173 | 0.025 | 0.018 | 1 | 0.009979004 | 3.695993813 | 0.1 | 0.009 | 1 | 0.659865009 | 0.019858427 |
| Gm47270 | 0.659871503 | 0.514573173 | 0.025 | 0.018 | 1 | 0.009979004 | 3.210566986 | 0.1 | 0.009 | 1 | 0.659871503 | 0.019858427 |
| Gm57068 | 0.691605844 | -0.485426827 | 0.017 | 0.024 | 1 | 0.009979004 | 3.432959407 | 0.1 | 0.009 | 1 | 0.691605844 | 0.019858427 |
| Acbd5 | 0.009982352 | -0.645603852 | 0.622 | 0.7 | 1 | 0.117130473 | 0.432959407 | 0.9 | 0.781 | 1 | 0.117130473 | 0.019865057 |
| Tmsb15b2 | 0.166678069 | -0.485426827 | 0.126 | 0.188 | 1 | 0.009984992 | 1.930459067 | 0.4 | 0.123 | 1 | 0.166678069 | 0.019870283 |
| Atg4b | 0.010009612 | -0.944858446 | 0.345 | 0.453 | 1 | 0.348042253 | 0.489142243 | 0.7 | 0.525 | 1 | 0.348042253 | 0.019919033 |
| Myg1 | 0.010015898 | -1.100136671 | 0.118 | 0.235 | 1 | 0.834312687 | 1.280956314 | 0.2 | 0.21 | 1 | 0.834312687 | 0.019931477 |
| Slc35d1 | 0.023696644 | -0.532732542 | 0.765 | 0.788 | 1 | 0.010026296 | -1.510336835 | 0.6 | 0.827 | 1 | 0.023696644 | 0.019952066 |
| Sag | 0.010030092 | -2.040015679 | 0.059 | 0.153 | 1 | 0.630827111 | 0.473601392 | 0.2 | 0.136 | 1 | 0.630827111 | 0.01995958 |
| Hectd4 | 0.010032088 | -0.996388746 | 0.277 | 0.394 | 1 | 0.409217127 | -0.393353672 | 0.4 | 0.58 | 1 | 0.409217127 | 0.019963533 |
| Rffl | 0.010034441 | -2.029747343 | 0.042 | 0.129 | 1 | 0.21372809 | -1.267480311 | 0 | 0.136 | 1 | 0.21372809 | 0.019968191 |
| 1810062G17Rik | 0.010035197 | 0.945207527 | 0.218 | 0.106 | 1 | 0.170950984 | 1.558490289 | 0.2 | 0.077 | 1 | 0.170950984 | 0.019969689 |
| Ift172 | 0.010053227 | -1.070389328 | 0.134 | 0.259 | 1 | 0.51572366 | 0.916383882 | 0.3 | 0.219 | 1 | 0.51572366 | 0.020005387 |
| Atp5o | 0.010116376 | -0.450151352 | 0.773 | 0.871 | 1 | 0.734871969 | 0.281650084 | 0.7 | 0.864 | 1 | 0.734871969 | 0.020130411 |
| Hyal3 | 0.609500043 | 0.514573173 | 0.034 | 0.024 | 1 | 0.010117365 | 3.210566986 | 0.2 | 0.037 | 1 | 0.609500043 | 0.020132369 |
| E230029C05Rik | 0.22800107 | -1.485426827 | 0.034 | 0.065 | 1 | 0.010120629 | 3.515421567 | 0.2 | 0.037 | 1 | 0.22800107 | 0.020138831 |
| Cldn4 | 0.010136646 | -1.622930351 | 0.025 | 0.106 | 1 | 0.930290742 | -0.546862711 | 0.2 | 0.207 | 1 | 0.930290742 | 0.020170541 |
| Skic8 | 0.678988063 | -0.213173891 | 0.765 | 0.676 | 1 | 0.01015284 | -1.365782384 | 0.4 | 0.753 | 1 | 0.678988063 | 0.020202599 |
| Gm527 | 0.01016274 | -1.918386234 | 0.034 | 0.118 | 1 | 0.977222186 | 0.660369903 | 0.1 | 0.102 | 1 | 0.977222186 | 0.0202222 |
| Acvr1b | 0.010198058 | 0.546634382 | 0.471 | 0.318 | 1 | 0.118915645 | 1.054447784 | 0.5 | 0.278 | 1 | 0.118915645 | 0.020292116 |
| Nosip | 0.010202782 | -1.137503524 | 0.109 | 0.224 | 1 | 0.124271317 | -1.408342847 | 0 | 0.194 | 1 | 0.124271317 | 0.020301467 |
| Tmem59l | 0.010230705 | 2.017073513 | 0.101 | 0.029 | 1 | 0.619953009 | 1.695993813 | 0 | 0.025 | 1 | 0.619953009 | 0.020356742 |
| Zfp369 | 0.010232549 | -1.700439718 | 0.109 | 0.218 | 1 | 0.060030782 | -2.080110175 | 0 | 0.269 | 1 | 0.060030782 | 0.020360393 |
| Rab10 | 0.010235868 | -0.308278086 | 0.958 | 0.941 | 1 | 0.104662572 | -0.562973223 | 0.9 | 0.988 | 1 | 0.104662572 | 0.020366963 |
| Kcnh4 | 0.034054416 | 2.514573173 | 0.042 | 0.006 | 1 | 0.010237137 | 3.017921908 | 0.1 | 0.009 | 1 | 0.034054416 | 0.020369475 |
| Klk9 | 0.091361251 | 2.514573173 | 0.017 | 0 | 1 | 0.010237137 | 3.432959407 | 0.1 | 0.009 | 1 | 0.091361251 | 0.020369475 |
| Gm32884 | 0.091361251 | 3.974004791 | 0.017 | 0 | 1 | 0.010237137 | 2.31748219 | 0.1 | 0.009 | 1 | 0.091361251 | 0.020369475 |
| 4930524B15Rik | 0.147029286 | -1.807354922 | 0 | 0.018 | 1 | 0.010237137 | 3.210566986 | 0.1 | 0.009 | 1 | 0.147029286 | 0.020369475 |
| Gm26583 | 0.218485013 | -1.292781749 | 0.008 | 0.029 | 1 | 0.010237137 | 3.210566986 | 0.1 | 0.009 | 1 | 0.218485013 | 0.020369475 |
| Gm56549 | 0.495207728 | -0.485426827 | 0.017 | 0.029 | 1 | 0.010237137 | 3.432959407 | 0.1 | 0.009 | 1 | 0.495207728 | 0.020369475 |
| Gm16897 | 0.506471296 | -1.070389328 | 0.008 | 0.018 | 1 | 0.010237137 | 3.432959407 | 0.1 | 0.009 | 1 | 0.506471296 | 0.020369475 |
| Gm12979 | 0.508731273 | -1.070389328 | 0.008 | 0.018 | 1 | 0.010237137 | 3.210566986 | 0.1 | 0.009 | 1 | 0.508731273 | 0.020369475 |
| Vamp9 | 0.609500043 | 0.514573173 | 0.034 | 0.024 | 1 | 0.010237137 | 3.432959407 | 0.1 | 0.009 | 1 | 0.609500043 | 0.020369475 |
| Ip6k1 | 0.036719428 | -0.49674214 | 0.597 | 0.641 | 1 | 0.010276535 | -1.291933355 | 0.6 | 0.815 | 1 | 0.036719428 | 0.020447463 |
| Tomm22 | 0.01029464 | -0.491527027 | 0.647 | 0.718 | 1 | 0.036123896 | -0.942907495 | 0.4 | 0.716 | 1 | 0.036123896 | 0.0204833 |
| Orc3 | 0.010299394 | -1.001676578 | 0.294 | 0.406 | 1 | 0.074351264 | -1.906890596 | 0.1 | 0.364 | 1 | 0.074351264 | 0.02049271 |
| Prelid1 | 0.010324815 | -0.428524436 | 0.714 | 0.794 | 1 | 0.111710498 | -0.574535129 | 0.5 | 0.833 | 1 | 0.111710498 | 0.020543028 |
| Zfp513 | 0.895905609 | -0.205318908 | 0.126 | 0.129 | 1 | 0.010331431 | 1.695993813 | 0.4 | 0.117 | 1 | 0.895905609 | 0.020556124 |
| Tyk2 | 0.010332048 | -2.070389328 | 0.034 | 0.118 | 1 | 0.772481829 | 0.345496566 | 0.1 | 0.13 | 1 | 0.772481829 | 0.020557345 |
| Caskin2 | 0.01033209 | -1.807354922 | 0.034 | 0.118 | 1 | 0.351752874 | 1.31748219 | 0.2 | 0.105 | 1 | 0.351752874 | 0.020557428 |
| Rars2 | 0.010339151 | -1.040015679 | 0.168 | 0.3 | 1 | 0.770939211 | -0.374395515 | 0.4 | 0.293 | 1 | 0.770939211 | 0.020571405 |
| Snapc4 | 0.010362157 | -1.244418728 | 0.101 | 0.212 | 1 | 0.530895355 | 0.930459067 | 0.2 | 0.127 | 1 | 0.530895355 | 0.020616939 |
| Qars | 0.010385056 | -0.936088236 | 0.328 | 0.435 | 1 | 0.151976384 | -1.048167282 | 0.2 | 0.404 | 1 | 0.151976384 | 0.020662262 |
| Hacl1 | 0.202389891 | 0.514573173 | 0.218 | 0.159 | 1 | 0.010391806 | 2.804928185 | 0.4 | 0.133 | 1 | 0.202389891 | 0.020675623 |
| Rapgef3 | 0.507403698 | -0.584962501 | 0.143 | 0.165 | 1 | 0.01039646 | 0.888638891 | 0.7 | 0.259 | 1 | 0.507403698 | 0.020684834 |
| Ube4b | 0.072756769 | -0.464210341 | 0.824 | 0.776 | 1 | 0.010405626 | 0.655548287 | 1 | 0.861 | 1 | 0.072756769 | 0.020702974 |
| Gm50013 | 0.634825039 | -0.292781749 | 0.084 | 0.1 | 1 | 0.010435465 | 2.070389328 | 0.4 | 0.123 | 1 | 0.634825039 | 0.020762031 |
| Calm1 | 0.339846989 | 0.18219072 | 0.983 | 0.988 | 1 | 0.010450714 | -0.908918277 | 1 | 1 | 1 | 0.339846989 | 0.02079221 |
| Klhl13 | 0.010455704 | -1.060120992 | 0.227 | 0.353 | 1 | 0.613110917 | -0.374395515 | 0.2 | 0.269 | 1 | 0.613110917 | 0.020802086 |
| Agap2 | 0.472599012 | -0.333423734 | 0.025 | 0.041 | 1 | 0.010500667 | 3.017921908 | 0.1 | 0.009 | 1 | 0.472599012 | 0.020891071 |
| 2310047K21Rik | 0.010502677 | -1.807354922 | 0.034 | 0.118 | 1 | 0.460577668 | 0.494359952 | 0 | 0.052 | 1 | 0.460577668 | 0.020895047 |
| 9930104L06Rik | 0.010502677 | -1.807354922 | 0.034 | 0.118 | 1 | 0.333163571 | 1.31748219 | 0.2 | 0.102 | 1 | 0.333163571 | 0.020895047 |
| Cry1 | 0.010503527 | -0.907253492 | 0.303 | 0.424 | 1 | 0.015564414 | -2.080110175 | 0.2 | 0.568 | 1 | 0.015564414 | 0.020896729 |
| Epha4 | 0.010505406 | -0.706025967 | 0.277 | 0.406 | 1 | 0.220402603 | -1.745843746 | 0.2 | 0.358 | 1 | 0.220402603 | 0.020900448 |
| Myzap | 0.010523761 | -2.503348735 | 0.076 | 0.171 | 1 | 0.038860587 | -3.54813213 | 0 | 0.312 | 1 | 0.038860587 | 0.020936772 |
| Mfsd5 | 0.010533927 | -0.854660637 | 0.168 | 0.3 | 1 | 0.486864315 | -0.484578433 | 0.2 | 0.29 | 1 | 0.486864315 | 0.020956891 |
| Acbd6 | 0.010536806 | -0.427323873 | 0.605 | 0.735 | 1 | 0.032520715 | -1.349648852 | 0.3 | 0.667 | 1 | 0.032520715 | 0.020962588 |
| Nrcam | 0.387225194 | 0.415037499 | 0.034 | 0.018 | 1 | 0.010551263 | 2.602884409 | 0.2 | 0.037 | 1 | 0.387225194 | 0.020991197 |
| Pdcd7 | 0.01058755 | -1.024585638 | 0.218 | 0.341 | 1 | 0.084444397 | -1.513459553 | 0.1 | 0.358 | 1 | 0.084444397 | 0.021063004 |
| Gm17529 | 0.01059507 | -1.432959407 | 0.025 | 0.106 | 1 | 0.219873325 | -1.004445905 | 0 | 0.133 | 1 | 0.219873325 | 0.021077885 |
| Nphp3 | 0.317769646 | -0.715724447 | 0.185 | 0.224 | 1 | 0.010610038 | 1.92394576 | 0.5 | 0.194 | 1 | 0.317769646 | 0.021107503 |
| Retreg2 | 0.010615311 | -0.745954377 | 0.353 | 0.471 | 1 | 0.826897755 | 0.18865221 | 0.4 | 0.432 | 1 | 0.826897755 | 0.021117938 |
| Psmb3 | 0.010647086 | -0.338585439 | 0.824 | 0.871 | 1 | 0.267718867 | -0.30600854 | 0.5 | 0.87 | 1 | 0.267718867 | 0.021180813 |
| Tubb3 | 0.010651724 | -1.807354922 | 0.042 | 0.129 | 1 | 0.050020996 | 2.280956314 | 0.2 | 0.052 | 1 | 0.050020996 | 0.021189989 |
| Gabre | 0.010660677 | -2.854660637 | 0.017 | 0.088 | 1 | 0.964594132 | 0.31748219 | 0.1 | 0.102 | 1 | 0.964594132 | 0.021207704 |
| 2010109A12Rik | 0.010661882 | -2.137503524 | 0.05 | 0.141 | 1 | 0.25990199 | -0.959358016 | 0 | 0.114 | 1 | 0.25990199 | 0.021210088 |
| Zfp579 | 0.010674995 | -1.807354922 | 0.034 | 0.118 | 1 | 0.896995661 | 0.888638891 | 0.1 | 0.086 | 1 | 0.896995661 | 0.021236034 |
| Nol3 | 0.010674995 | -1.687060688 | 0.034 | 0.118 | 1 | 0.266798701 | -0.374395515 | 0 | 0.111 | 1 | 0.266798701 | 0.021236034 |
| Rab6a | 0.010679568 | -0.350497247 | 0.924 | 0.941 | 1 | 0.557224323 | -0.163407857 | 1 | 0.978 | 1 | 0.557224323 | 0.021245083 |

| | | | | | | | | | | | | |
|---|---|---|---|---|---|---|---|---|---|---|---|---|
| Plekhm3 | 0.188477577 | 0.192645078 | 0.235 | 0.171 | 1 | 0.010680368 | 1.868174788 | 0.4 | 0.123 | 1 | 0.188477577 | 0.021246665 |
| 3300002I08Rik | 0.01068755 | -1.378511623 | 0.092 | 0.2 | 1 | 0.689103442 | 0.432959407 | 0.2 | 0.148 | 1 | 0.689103442 | 0.021260877 |
| Tsbp1 | 0.354668871 | 0.929610672 | 0.05 | 0.029 | 1 | 0.010697473 | 3.017921908 | 0.2 | 0.037 | 1 | 0.354668871 | 0.02128051 |
| Rpl32 | 0.08283614 | -0.163662591 | 0.992 | 0.976 | 1 | 0.010721098 | -0.793666182 | 0.8 | 0.994 | 1 | 0.08283614 | 0.021327255 |
| Ncbp1 | 0.010728143 | -1.042374952 | 0.336 | 0.447 | 1 | 0.041372044 | -1.28128611 | 0.3 | 0.586 | 1 | 0.041372044 | 0.021341192 |
| Hyal2 | 0.01074482 | -1.244418728 | 0.101 | 0.212 | 1 | 0.383747025 | -0.505640048 | 0.1 | 0.21 | 1 | 0.383747025 | 0.021374189 |
| Cbr2 | 0.010761835 | 1.380717426 | 0.588 | 0.476 | 1 | 0.490502089 | -0.314274522 | 0.7 | 0.654 | 1 | 0.490502089 | 0.021407852 |
| Spa17 | 0.010763034 | 0.870716983 | 0.202 | 0.094 | 1 | 0.141230669 | 2.432959407 | 0.2 | 0.077 | 1 | 0.141230669 | 0.021410226 |
| Pabpc4l | 0.010764187 | -2.359895945 | 0.017 | 0.088 | 1 | 0.539465415 | 0.847996907 | 0 | 0.037 | 1 | 0.539465415 | 0.021412506 |
| Map2k6 | 0.01076454 | -2.600904045 | 0.017 | 0.088 | 1 | 0.374682358 | -0.304006187 | 0 | 0.074 | 1 | 0.374682358 | 0.021413205 |
| Cpz | 0.010764658 | -2.707819249 | 0.017 | 0.088 | 1 | 0.619953009 | 1.558490289 | 0 | 0.025 | 1 | 0.619953009 | 0.021413438 |
| Hspa1b | 0.010767623 | -1.166105751 | 0.462 | 0.624 | 1 | 0.086325244 | 0.295139121 | 0.3 | 0.627 | 1 | 0.086325244 | 0.021419305 |
| Ube2j1 | 0.010794001 | -0.593951284 | 0.655 | 0.718 | 1 | 0.341141043 | -0.304006187 | 0.7 | 0.793 | 1 | 0.341141043 | 0.021471492 |
| Nsmce4a | 0.010825626 | -0.583830531 | 0.42 | 0.547 | 1 | 0.07724194 | -0.959358016 | 0.3 | 0.59 | 1 | 0.07724194 | 0.021534058 |
| Itga3 | 0.044977942 | 1.440572591 | 0.109 | 0.047 | 1 | 0.010845217 | 2.930459067 | 0.2 | 0.037 | 1 | 0.044977942 | 0.021572814 |
| Cops9 | 0.010861497 | 0.492291725 | 0.765 | 0.741 | 1 | 0.043774046 | -0.900941329 | 0.3 | 0.691 | 1 | 0.043774046 | 0.021605022 |
| Acp5 | 0.010868719 | -2.292781749 | 0.017 | 0.088 | 1 | 0.349325625 | -4.467236536 | 0.2 | 0.102 | 1 | 0.349325625 | 0.021619309 |
| Dnhd1 | 0.010884508 | -1.572889668 | 0.084 | 0.188 | 1 | 0.128066841 | 1.236562194 | 0.3 | 0.127 | 1 | 0.128066841 | 0.021650544 |
| Spata17 | 0.010914179 | 2.736965594 | 0.067 | 0.012 | 1 | 0.619953009 | 1.695993813 | 0 | 0.025 | 1 | 0.619953009 | 0.021709238 |
| Ncaph2 | 0.010916485 | -1 | 0.303 | 0.418 | 1 | 0.504297652 | -0.175849835 | 0.3 | 0.417 | 1 | 0.504297652 | 0.021713801 |
| Nipa2 | 0.063795511 | 0.211181029 | 0.773 | 0.635 | 1 | 0.010942191 | 1.032539004 | 0.9 | 0.731 | 1 | 0.063795511 | 0.02176465 |
| Tsfm | 0.010943816 | -1.048363022 | 0.16 | 0.276 | 1 | 0.522990517 | -0.255096586 | 0.2 | 0.281 | 1 | 0.522990517 | 0.021767865 |
| Txn1 | 0.372289481 | 0.298928941 | 0.958 | 0.965 | 1 | 0.010951528 | -1.180347719 | 0.6 | 0.963 | 1 | 0.372289481 | 0.02178312 |
| Il10 | 0.010973268 | -2.14839184 | 0.017 | 0.088 | 1 | 0.406872117 | 1.263034406 | 0.1 | 0.043 | 1 | 0.406872117 | 0.021826123 |
| Kif28 | 0.010973268 | -2.485426827 | 0.017 | 0.088 | 1 | 0.057366432 | 2.111031312 | 0.2 | 0.056 | 1 | 0.057366432 | 0.021826123 |
| Gm29488 | 0.010987948 | -2.944858446 | 0 | 0.053 | 1 | 0.259848928 | -0.709998547 | 0 | 0.114 | 1 | 0.259848928 | 0.021855161 |
| Tspyl3 | 0.010987948 | -2.944858446 | 0 | 0.053 | 1 | 0.698529281 | 2.210566986 | 0 | 0.015 | 1 | 0.698529281 | 0.021855161 |
| Cd1d1 | 0.010987948 | -2.944858446 | 0 | 0.053 | 1 | 0.64384652 | 1.31748219 | 0 | 0.022 | 1 | 0.64384652 | 0.021855161 |
| Dpysl4 | 0.010987948 | -3.070389328 | 0 | 0.053 | 1 | 0.36390653 | 0.159940913 | 0 | 0.077 | 1 | 0.36390653 | 0.021855161 |
| Hsh2d | 0.010987948 | -3.292781749 | 0 | 0.053 | 1 | 0.557744661 | 1.017921908 | 0 | 0.034 | 1 | 0.557744661 | 0.021855161 |
| Gm57009 | 0.010987948 | -2.944858446 | 0 | 0.053 | 1 | 0.669843604 | 2.017921908 | 0 | 0.019 | 1 | 0.669843604 | 0.021855161 |
| Gm31834 | 0.010987948 | -2.944858446 | 0 | 0.053 | 1 | 0.597824685 | 1.31748219 | 0 | 0.028 | 1 | 0.597824685 | 0.021855161 |
| Prrt1 | 0.010987948 | -2.944858446 | 0 | 0.053 | 1 | 0.619961444 | 1.432959407 | 0 | 0.025 | 1 | 0.619961444 | 0.021855161 |
| Plekhg6 | 0.010991022 | -3.292781749 | 0 | 0.053 | 1 | 0.577142568 | 1.432959407 | 0 | 0.031 | 1 | 0.577142568 | 0.021861242 |
| Ccdc106 | 0.010991022 | -3.185866545 | 0 | 0.053 | 1 | 0.274256184 | 2.31748219 | 0.1 | 0.034 | 1 | 0.274256184 | 0.021861242 |
| Gm29683 | 0.010991022 | -3.185866545 | 0 | 0.053 | 1 | 0.080320981 | 2.695993813 | 0.1 | 0.019 | 1 | 0.080320981 | 0.021861242 |
| Polrmt | 0.010991022 | -3.070389328 | 0 | 0.053 | 1 | 0.460551517 | 0.695993813 | 0 | 0.052 | 1 | 0.460551517 | 0.021861242 |
| Ppargc1a | 0.010993219 | -3.292781749 | 0 | 0.053 | 1 | 0.098167853 | -2.31299497 | 0 | 0.219 | 1 | 0.098167853 | 0.021865587 |
| Mas1 | 0.010993219 | -3.292781749 | 0 | 0.053 | 1 | 0.316137708 | 2.210566986 | 0.1 | 0.037 | 1 | 0.316137708 | 0.021865587 |
| Arhgap11a | 0.010993329 | -3.572889668 | 0 | 0.053 | 1 | 0.619953009 | 1.558490289 | 0 | 0.025 | 1 | 0.619953009 | 0.021865804 |
| Selenok-ps1 | 0.010993329 | -3.185866545 | 0 | 0.053 | 1 | 0.597845674 | 1.695993813 | 0 | 0.028 | 1 | 0.597845674 | 0.021865804 |
| Fxyd4 | 0.010994537 | -3.392317423 | 0 | 0.053 | 1 | 0.487557552 | 1.494359952 | 0.1 | 0.049 | 1 | 0.487557552 | 0.021868194 |
| Popdc3 | 0.010994537 | -3.485426827 | 0 | 0.053 | 1 | 0.669851164 | 0.432959407 | 0 | 0.019 | 1 | 0.669851164 | 0.021868194 |
| Ldhc | 0.010994866 | -3.807354922 | 0 | 0.053 | 1 | 0.276546558 | 2.017921908 | 0.1 | 0.034 | 1 | 0.276546558 | 0.021868846 |
| Foxj1 | 0.010994866 | -3.87774425 | 0 | 0.053 | 1 | 0.489966712 | 0.31748219 | 0 | 0.046 | 1 | 0.489966712 | 0.021868846 |
| Aurka | 0.010994976 | -3.292781749 | 0 | 0.053 | 1 | 0.643835681 | 1.847996907 | 0 | 0.022 | 1 | 0.643835681 | 0.021869063 |
| 9330154J02Rik | 0.010995635 | -3.655351829 | 0 | 0.053 | 1 | 0.619971082 | 1.017921908 | 0 | 0.025 | 1 | 0.619971082 | 0.021870366 |
| Gm14443 | 0.010997173 | -2.807354922 | 0 | 0.053 | 1 | 0.557756607 | 0.847996907 | 0 | 0.034 | 1 | 0.557756607 | 0.021873408 |
| Zbtb8a | 0.010997173 | -2.807354922 | 0 | 0.053 | 1 | 0.460577668 | 0.625604485 | 0 | 0.052 | 1 | 0.460577668 | 0.021873408 |
| Sytl1 | 0.010997173 | -2.807354922 | 0 | 0.053 | 1 | 0.577142568 | 1.31748219 | 0 | 0.031 | 1 | 0.577142568 | 0.021873408 |
| Cog6 | 0.011003512 | -1.036836768 | 0.202 | 0.324 | 1 | 0.45701211 | 0.64536774 | 0.4 | 0.312 | 1 | 0.45701211 | 0.021885946 |
| 1700061N14Rik | 0.011056241 | -1.42129649 | 0.118 | 0.229 | 1 | 0.366950508 | -0.654503434 | 0.1 | 0.225 | 1 | 0.366950508 | 0.021990242 |
| Plekhg1 | 0.011074715 | -2.411426246 | 0.05 | 0.141 | 1 | 0.068748591 | 0.921060369 | 0.5 | 0.225 | 1 | 0.068748591 | 0.02202678 |
| Zfp276 | 0.011075715 | -1.854660637 | 0.042 | 0.129 | 1 | 0.69712267 | 0.580516596 | 0.2 | 0.148 | 1 | 0.69712267 | 0.022028759 |
| Vps4a | 0.011075958 | -0.797370833 | 0.412 | 0.5 | 1 | 0.741023396 | 0.159940913 | 0.5 | 0.42 | 1 | 0.741023396 | 0.022029238 |
| Zfp708 | 0.011077639 | -2.070389328 | 0.017 | 0.088 | 1 | 0.364023002 | -0.111361109 | 0 | 0.077 | 1 | 0.364023002 | 0.022032563 |
| C1qtnf9 | 0.011077639 | -2.070389328 | 0.017 | 0.088 | 1 | 0.133962235 | 1.695993813 | 0.2 | 0.071 | 1 | 0.133962235 | 0.022032563 |
| Neurl1a | 0.011077639 | -2.070389328 | 0.017 | 0.088 | 1 | 0.489922734 | 0.494359952 | 0 | 0.046 | 1 | 0.489922734 | 0.022032563 |
| Med27 | 0.131093921 | 0.163100802 | 0.555 | 0.424 | 1 | 0.011086317 | 1.178145508 | 1 | 0.485 | 1 | 0.131093921 | 0.022049727 |
| Fth1 | 0.011101898 | 0.327526453 | 1 | 0.988 | 1 | 0.363072768 | -0.288597856 | 1 | 1 | 1 | 0.363072768 | 0.022080544 |
| Ppp1r15a | 0.573387159 | -0.225202737 | 0.958 | 0.9 | 1 | 0.011104063 | -0.417401092 | 0.6 | 0.948 | 1 | 0.573387159 | 0.022084826 |
| Zc3h12b | 0.011127712 | -1.392317423 | 0.227 | 0.341 | 1 | 0.497748134 | -0.556986928 | 0.3 | 0.401 | 1 | 0.497748134 | 0.022131597 |
| Zfp944 | 0.011145839 | -1.333423734 | 0.118 | 0.229 | 1 | 0.388817677 | -0.339630097 | 0.1 | 0.225 | 1 | 0.388817677 | 0.022167448 |
| Azi2 | 0.01114814 | -0.608809243 | 0.479 | 0.594 | 1 | 0.060881228 | -0.67556505 | 0.4 | 0.698 | 1 | 0.060881228 | 0.022171999 |
| Emc1 | 0.011161113 | 0.794681092 | 0.336 | 0.206 | 1 | 0.259693173 | 1.0330288 | 0.4 | 0.253 | 1 | 0.259693173 | 0.022197655 |
| Dkc1 | 0.011267286 | -0.904965719 | 0.252 | 0.382 | 1 | 0.058678141 | -2.220482831 | 0 | 0.272 | 1 | 0.058678141 | 0.02240762 |
| Pde4d | 0.01128146 | 0.149669107 | 0.95 | 0.888 | 1 | 0.025762904 | 0.475541253 | 1 | 0.907 | 1 | 0.025762904 | 0.022435649 |
| Zbtb8os | 0.011290555 | 0.76611194 | 0.42 | 0.288 | 1 | 0.89720338 | 0.210566986 | 0.2 | 0.21 | 1 | 0.89720338 | 0.022453633 |
| Gm17382 | 0.423364081 | -1.250961574 | 0.034 | 0.053 | 1 | 0.011304686 | 2.847996907 | 0.2 | 0.037 | 1 | 0.423364081 | 0.022481577 |
| Gm12536 | 0.011316265 | 0.784662336 | 0.21 | 0.1 | 1 | 0.015000358 | 1.469485283 | 0.5 | 0.176 | 1 | 0.015000358 | 0.022504472 |
| Zmynd19 | 0.011324404 | -1.168236651 | 0.151 | 0.265 | 1 | 0.659886351 | -0.315978829 | 0.2 | 0.247 | 1 | 0.659886351 | 0.022520566 |
| Ddx56 | 0.011327772 | -0.655351829 | 0.193 | 0.335 | 1 | 0.657415668 | 0.388565288 | 0.3 | 0.222 | 1 | 0.657415668 | 0.022527226 |
| Gm42679 | 0.011344395 | 2.099535674 | 0.143 | 0.059 | 1 | 0.669848329 | 1.558490289 | 0 | 0.019 | 1 | 0.669848329 | 0.022560095 |
| Gpr3 | 0.011350971 | 2.099535674 | 0.067 | 0.012 | 1 | 0.577184788 | 1.558490289 | 0 | 0.031 | 1 | 0.577184788 | 0.022573097 |
| Kdm4a | 0.011351149 | -1.201633861 | 0.202 | 0.318 | 1 | 0.772442922 | 0.193493473 | 0.3 | 0.358 | 1 | 0.772442922 | 0.02257345 |
| 1500015L24Rik | 0.073658343 | 1.280107919 | 0.126 | 0.065 | 1 | 0.011385321 | 2.017921908 | 0.3 | 0.074 | 1 | 0.073658343 | 0.022641017 |
| Mthfd1 | 0.011390705 | -1.333423734 | 0.109 | 0.218 | 1 | 0.309609932 | 1.091922489 | 0.3 | 0.176 | 1 | 0.309609932 | 0.022651661 |
| Wdr19 | 0.011451058 | -1.485426827 | 0.034 | 0.118 | 1 | 0.225920219 | -0.840059087 | 0 | 0.13 | 1 | 0.225920219 | 0.02277099 |
| Ulk2 | 0.011452316 | -0.912848051 | 0.571 | 0.641 | 1 | 0.483715071 | 0.159940913 | 0.7 | 0.586 | 1 | 0.483715071 | 0.022773477 |
| Elmod3 | 0.106182313 | -0.900464326 | 0.101 | 0.165 | 1 | 0.011465985 | 2.978393544 | 0.3 | 0.086 | 1 | 0.106182313 | 0.0228005 |
| Tsc2 | 0.055570325 | -0.610957709 | 0.151 | 0.247 | 1 | 0.011476838 | 1.393431043 | 0.5 | 0.17 | 1 | 0.055570325 | 0.022821958 |
| Rpl15 | 0.011491163 | -0.21791312 | 0.975 | 0.971 | 1 | 0.02706798 | -0.702700173 | 1 | 0.997 | 1 | 0.02706798 | 0.022850279 |
| Tor2a | 0.011494472 | 0.628229955 | 0.429 | 0.282 | 1 | 0.922087723 | 0.123104145 | 0.3 | 0.296 | 1 | 0.922087723 | 0.022856821 |
| Tmem120b | 0.011498157 | -1.544320516 | 0.076 | 0.176 | 1 | 0.14411351 | -1.521236903 | 0 | 0.179 | 1 | 0.14411351 | 0.022864107 |
| Hmgn2-ps | 0.011511586 | -1.600904045 | 0.042 | 0.129 | 1 | 0.353533258 | 0.210566986 | 0 | 0.08 | 1 | 0.353533258 | 0.022890655 |
| Slc25a53 | 0.011526482 | -1.258016331 | 0.059 | 0.159 | 1 | 0.197143062 | 1.210566986 | 0.3 | 0.154 | 1 | 0.197143062 | 0.022920105 |
| Hnrnph1 | 0.011533454 | -0.269033146 | 0.95 | 0.982 | 1 | 0.041978236 | -0.686163743 | 0.7 | 0.997 | 1 | 0.041978236 | 0.022933888 |
| Zfp385c | 0.011547513 | 0.186998515 | 0.303 | 0.165 | 1 | 0.757453421 | 0.785261151 | 0.3 | 0.287 | 1 | 0.757453421 | 0.022961681 |
| Cib2 | 0.011549752 | -1.521050737 | 0.076 | 0.176 | 1 | 0.576513272 | 0.744903414 | 0.2 | 0.133 | 1 | 0.576513272 | 0.022966108 |
| Eif2s3x | 0.065801481 | -0.358122503 | 0.647 | 0.718 | 1 | 0.011560355 | -1.539733247 | 0.4 | 0.707 | 1 | 0.065801481 | 0.022987068 |
| Atp6v1d | 0.030120337 | -0.430564892 | 0.639 | 0.747 | 1 | 0.011570208 | -1.584467664 | 0.4 | 0.731 | 1 | 0.030120337 | 0.023006547 |
| Itgam | 0.381362131 | 0.263034406 | 0.034 | 0.018 | 1 | 0.011618673 | 1.611929548 | 0.2 | 0.037 | 1 | 0.381362131 | 0.023102353 |
| Mindy3 | 0.011638152 | -0.90945311 | 0.319 | 0.429 | 1 | 0.151536369 | 0.382333334 | 0.7 | 0.463 | 1 | 0.151536369 | 0.023140857 |
| Kcnmb4 | 0.011670652 | -1.722466024 | 0.034 | 0.118 | 1 | 0.036453611 | 1.432959407 | 0.4 | 0.148 | 1 | 0.036453611 | 0.0232051 |
| Qtrt2 | 0.011703169 | -0.997325866 | 0.185 | 0.306 | 1 | 0.399756096 | -0.567040593 | 0.2 | 0.306 | 1 | 0.399756096 | 0.023269374 |
| Cwc25 | 0.011710031 | -0.807354922 | 0.353 | 0.459 | 1 | 0.386360321 | -0.368659145 | 0.3 | 0.481 | 1 | 0.386360321 | 0.023282937 |
| Desi1 | 0.011729873 | 0.290740217 | 0.689 | 0.488 | 1 | 0.676194471 | 0.704031132 | 0.5 | 0.596 | 1 | 0.676194471 | 0.023322157 |
| Eif3l | 0.027484626 | -0.505699222 | 0.571 | 0.659 | 1 | 0.011739227 | -1.311201688 | 0.4 | 0.719 | 1 | 0.027484626 | 0.023340644 |

| | | | | | | | | | | | | |
|---|---|---|---|---|---|---|---|---|---|---|---|---|
| Hipk3 | 0.011747085 | -0.421650917 | 0.756 | 0.788 | 1 | 0.559097302 | 0.44378934 | 0.8 | 0.898 | 1 | 0.559097302 | 0.023356176 |
| Jak2 | 0.011783646 | 0.484290606 | 0.79 | 0.694 | 1 | 0.030629954 | 0.799498389 | 0.9 | 0.75 | 1 | 0.030629954 | 0.023428438 |
| Mrpl49 | 0.011790298 | -1.096861539 | 0.143 | 0.259 | 1 | 0.871979376 | 0.159940913 | 0.2 | 0.213 | 1 | 0.871979376 | 0.023441586 |
| Tsr3 | 0.011802944 | -1.003275132 | 0.151 | 0.276 | 1 | 0.356076235 | -0.391469028 | 0.1 | 0.219 | 1 | 0.356076235 | 0.023466578 |
| Snx15 | 0.011809456 | -0.622930351 | 0.387 | 0.506 | 1 | 0.206841491 | -0.936274402 | 0.2 | 0.377 | 1 | 0.206841491 | 0.023479448 |
| Epn1 | 0.011830371 | -0.758445322 | 0.336 | 0.453 | 1 | 0.446616612 | -0.48032896 | 0.4 | 0.466 | 1 | 0.446616612 | 0.023520784 |
| Orc1 | 0.011833697 | -1.733354341 | 0.017 | 0.088 | 1 | 0.070901818 | 1.847996907 | 0.2 | 0.059 | 1 | 0.070901818 | 0.023527357 |
| Mbd2 | 0.011988294 | -0.258776572 | 0.958 | 0.976 | 1 | 0.011858301 | -0.758328331 | 0.8 | 0.985 | 1 | 0.011988294 | 0.023575983 |
| Kif18a | 0.011899943 | -2.807354922 | 0.008 | 0.071 | 1 | 0.577155236 | 1.31748219 | 0 | 0.031 | 1 | 0.577155236 | 0.023658277 |
| Ckap2 | 0.011901416 | -3.944858446 | 0.008 | 0.071 | 1 | 0.051044361 | 2.695993813 | 0.1 | 0.015 | 1 | 0.051044361 | 0.023661187 |
| Bola3 | 0.011902541 | 0.364677742 | 0.681 | 0.5 | 1 | 0.299206485 | -0.279758641 | 0.3 | 0.481 | 1 | 0.299206485 | 0.023663412 |
| H2-M3 | 0.01191776 | -1.5360529 | 0.05 | 0.141 | 1 | 0.259731249 | -0.709998547 | 0 | 0.114 | 1 | 0.259731249 | 0.023693487 |
| Aspn | 0.01191843 | -1.660868834 | 0.244 | 0.371 | 1 | 0.03400731 | 1.102524375 | 0.9 | 0.574 | 1 | 0.03400731 | 0.02369481 |
| Cyb561 | 0.17792078 | 0.955145764 | 0.101 | 0.059 | 1 | 0.011921724 | 2.602884409 | 0.2 | 0.037 | 1 | 0.17792078 | 0.02370132 |
| Gm4013 | 0.327885716 | -0.622930351 | 0.034 | 0.059 | 1 | 0.011921724 | 2.695993813 | 0.2 | 0.037 | 1 | 0.327885716 | 0.02370132 |
| Eif2ak2 | 0.138334466 | -0.802909017 | 0.294 | 0.347 | 1 | 0.011942643 | -3.186649236 | 0 | 0.41 | 1 | 0.138334466 | 0.02374266 |
| Gnrh1 | 0.011958676 | -1.485426827 | 0.042 | 0.129 | 1 | 0.808436744 | 0.263034406 | 0.1 | 0.123 | 1 | 0.808436744 | 0.023774342 |
| Clec16a | 0.011985096 | -0.788818971 | 0.261 | 0.394 | 1 | 0.332529858 | -0.481923979 | 0.3 | 0.525 | 1 | 0.332529858 | 0.02382655 |
| E4f1 | 0.011997297 | -1.046141782 | 0.134 | 0.253 | 1 | 0.621260313 | 0.345496566 | 0.3 | 0.216 | 1 | 0.621260313 | 0.02385066 |
| Zfp869 | 0.012013835 | 0.447458977 | 0.529 | 0.382 | 1 | 0.142393439 | 1.111031312 | 0.6 | 0.457 | 1 | 0.142393439 | 0.023883338 |
| D630045J12Rik | 0.012027058 | 0.288145238 | 0.622 | 0.453 | 1 | 0.508484198 | 0.449234947 | 0.7 | 0.605 | 1 | 0.508484198 | 0.023909466 |
| Ddi2 | 0.44368515 | -0.143992418 | 0.773 | 0.747 | 1 | 0.012029286 | -1.10919001 | 0.7 | 0.944 | 1 | 0.44368515 | 0.023913868 |
| Gm43466 | 0.012031092 | -2.655351829 | 0.008 | 0.071 | 1 | 0.619961444 | 0.930459067 | 0 | 0.025 | 1 | 0.619961444 | 0.023917436 |
| Gm15708 | 0.012031092 | -2.655351829 | 0.008 | 0.071 | 1 | 0.385467319 | 0.31748219 | 0 | 0.071 | 1 | 0.385467319 | 0.023917436 |
| Polr3g | 0.012031339 | -2.733354341 | 0.008 | 0.071 | 1 | 0.539413885 | 1.210566986 | 0 | 0.037 | 1 | 0.539413885 | 0.023917925 |
| Rragb | 0.012031339 | -2.733354341 | 0.008 | 0.071 | 1 | 0.052020228 | 2.079322453 | 0.2 | 0.052 | 1 | 0.052020228 | 0.023917925 |
| Atg4a | 0.012049883 | -0.9789663 | 0.092 | 0.2 | 1 | 0.587576022 | 0.650351148 | 0.3 | 0.238 | 1 | 0.587576022 | 0.023954566 |
| Rtl5 | 0.012058872 | -1.572889668 | 0.017 | 0.088 | 1 | 0.315694985 | -0.152003093 | 0 | 0.093 | 1 | 0.315694985 | 0.023972327 |
| Selenot | 0.262495279 | -0.323837685 | 0.681 | 0.741 | 1 | 0.012064764 | -0.986170847 | 0.6 | 0.883 | 1 | 0.262495279 | 0.023983969 |
| Zeb2 | 0.012079529 | -1.163498732 | 0.277 | 0.135 | 1 | 0.700344404 | 0.530656081 | 0.1 | 0.157 | 1 | 0.700344404 | 0.024013143 |
| Slc35f2 | 0.012095741 | -2.485426827 | 0.008 | 0.071 | 1 | 0.460599334 | 0.432959407 | 0 | 0.052 | 1 | 0.460599334 | 0.024045176 |
| H2bc9 | 0.012095741 | -2.807354922 | 0.008 | 0.071 | 1 | 0.669843604 | 2.017921908 | 0 | 0.019 | 1 | 0.669843604 | 0.024045176 |
| Pou5f1 | 0.012095741 | -2.485426827 | 0.008 | 0.071 | 1 | 0.408596393 | 0.494359952 | 0 | 0.065 | 1 | 0.408596393 | 0.024045176 |
| Gm57198 | 0.012102529 | -1.326729081 | 0.084 | 0.188 | 1 | 0.873550019 | 0.354956895 | 0.2 | 0.173 | 1 | 0.873550019 | 0.024058586 |
| Bscl2 | 0.012116919 | -0.74281467 | 0.412 | 0.512 | 1 | 0.518950314 | 0.374065718 | 0.6 | 0.519 | 1 | 0.518950314 | 0.024087017 |
| Rars | 0.012137902 | -0.719892081 | 0.437 | 0.529 | 1 | 0.152536807 | -0.825056924 | 0.4 | 0.556 | 1 | 0.152536807 | 0.024128476 |
| Ipo9 | 0.012144644 | -1.04094555 | 0.21 | 0.335 | 1 | 0.614989714 | -0.246185533 | 0.4 | 0.46 | 1 | 0.614989714 | 0.024141795 |
| Gm13293 | 0.012159539 | -2.392317423 | 0.008 | 0.071 | 1 | 0.5933729 | 1.625604485 | 0.1 | 0.059 | 1 | 0.5933729 | 0.024171224 |
| 5730409E04Rik | 0.012159539 | -2.392317423 | 0.008 | 0.071 | 1 | 0.017442807 | 2.515421567 | 0.2 | 0.04 | 1 | 0.017442807 | 0.024171224 |
| Dna2 | 0.012159539 | -2.392317423 | 0.008 | 0.071 | 1 | 0.433608183 | 0.374065718 | 0 | 0.059 | 1 | 0.433608183 | 0.024171224 |
| Marchf9 | 0.012159539 | -2.572889668 | 0.008 | 0.071 | 1 | 0.408673423 | 0.111031312 | 0 | 0.065 | 1 | 0.408673423 | 0.024171224 |
| A430046D13Rik | 0.012159539 | -2.392317423 | 0.008 | 0.071 | 1 | 0.597845674 | 1.695993813 | 0 | 0.028 | 1 | 0.597845674 | 0.024171224 |
| Gm12248 | 0.012159539 | -2.392317423 | 0.008 | 0.071 | 1 | 0.232378687 | -0.91281543 | 0 | 0.127 | 1 | 0.232378687 | 0.024171224 |
| Pheta2 | 0.012159539 | -2.392317423 | 0.008 | 0.071 | 1 | 0.324884604 | -0.230005605 | 0 | 0.09 | 1 | 0.324884604 | 0.024171224 |
| Gak | 0.01216347 | -0.760049207 | 0.513 | 0.588 | 1 | 0.233785311 | 0.512393875 | 0.9 | 0.716 | 1 | 0.233785311 | 0.02417899 |
| 5430427G11Rik | 0.825035967 | -0.485426827 | 0.042 | 0.047 | 1 | 0.01218397 | 3.82527683 | 0.2 | 0.04 | 1 | 0.825035967 | 0.024219491 |
| Grsf1 | 0.012209578 | -0.604725756 | 0.496 | 0.588 | 1 | 0.177771181 | -0.621806396 | 0.4 | 0.602 | 1 | 0.177771181 | 0.024270083 |
| Elf3 | 0.012222723 | -2.292781749 | 0.008 | 0.071 | 1 | 0.138699708 | -1.026472211 | 0 | 0.182 | 1 | 0.138699708 | 0.024296052 |
| E530011L22Rik | 0.012222723 | -2.292781749 | 0.008 | 0.071 | 1 | 0.521021286 | 1.558490289 | 0.1 | 0.052 | 1 | 0.521021286 | 0.024296052 |
| Hrct1 | 0.022991853 | 1.514573173 | 0.101 | 0.035 | 1 | 0.012254024 | 2.515421567 | 0.2 | 0.037 | 1 | 0.022991853 | 0.024357888 |
| C130046K22Rik | 0.217575341 | 1 | 0.042 | 0.018 | 1 | 0.012254024 | 2.515421567 | 0.2 | 0.037 | 1 | 0.217575341 | 0.024357888 |
| Ptp4a1 | 0.303113451 | -0.987927168 | 0.042 | 0.071 | 1 | 0.012254024 | 2.602884409 | 0.2 | 0.037 | 1 | 0.303113451 | 0.024357888 |
| Rad54l | 0.351832008 | -0.655351829 | 0.017 | 0.035 | 1 | 0.012254024 | 2.695993813 | 0.2 | 0.037 | 1 | 0.351832008 | 0.024357888 |
| Agbl5 | 0.012254202 | -1.485426827 | 0.059 | 0.153 | 1 | 0.245823267 | 1.847996907 | 0.2 | 0.096 | 1 | 0.245823267 | 0.024358239 |
| Trim25 | 0.01226705 | -1.228651412 | 0.143 | 0.259 | 1 | 0.158113241 | -1.004445905 | 0.1 | 0.333 | 1 | 0.158113241 | 0.024383619 |
| Med31 | 0.01229055 | -0.630300457 | 0.454 | 0.553 | 1 | 0.090196433 | -1.074835233 | 0.3 | 0.531 | 1 | 0.090196433 | 0.024430043 |
| Naa10 | 0.012304999 | -0.984997837 | 0.21 | 0.335 | 1 | 0.532815472 | -0.327852929 | 0.2 | 0.278 | 1 | 0.532815472 | 0.024458585 |
| Dglucy | 0.101978448 | -0.402964667 | 0.076 | 0.141 | 1 | 0.012346625 | 2.210566986 | 0.3 | 0.077 | 1 | 0.101978448 | 0.024540811 |
| Magee1 | 0.012349598 | -1.359895945 | 0.067 | 0.165 | 1 | 0.147437462 | -1.090602549 | 0 | 0.176 | 1 | 0.147437462 | 0.024546683 |
| Smarcd2 | 0.012374827 | -1.266786541 | 0.118 | 0.229 | 1 | 0.039589461 | -1.91281543 | 0 | 0.306 | 1 | 0.039589461 | 0.024596518 |
| Zbtb16 | 0.426709875 | -0.465248945 | 0.378 | 0.3 | 1 | 0.012383517 | 1.552258336 | 0.7 | 0.33 | 1 | 0.426709875 | 0.024613683 |
| Ablim1 | 0.012416449 | 0.750640531 | 0.303 | 0.171 | 1 | 0.470847095 | -0.596787936 | 0.2 | 0.318 | 1 | 0.470847095 | 0.024678729 |
| Golga3 | 0.01242323 | -0.506037107 | 0.639 | 0.688 | 1 | 0.647341301 | -0.154358514 | 0.7 | 0.735 | 1 | 0.647341301 | 0.024692124 |
| Ncapd3 | 0.01244441 | -1.748461233 | 0.076 | 0.176 | 1 | 0.672849094 | 0.913585248 | 0.2 | 0.167 | 1 | 0.672849094 | 0.024733956 |
| Hccs | 0.012471134 | -1.070389328 | 0.151 | 0.276 | 1 | 0.929733977 | -0.103093493 | 0.3 | 0.306 | 1 | 0.929733977 | 0.024786739 |
| Zfp867 | 0.040179852 | 0.848992212 | 0.193 | 0.106 | 1 | 0.012476475 | 1.973527789 | 0.3 | 0.074 | 1 | 0.040179852 | 0.024797288 |
| Tacc3 | 0.150364336 | -1.485426827 | 0.084 | 0.135 | 1 | 0.012477236 | 1.808468542 | 0.3 | 0.074 | 1 | 0.150364336 | 0.024798791 |
| Foxd2 | 0.01248568 | -1.444784843 | 0.059 | 0.153 | 1 | 0.713367825 | -0.267480311 | 0.1 | 0.136 | 1 | 0.713367825 | 0.024815467 |
| Evc | 0.012556774 | -1.614709844 | 0.059 | 0.153 | 1 | 0.662822902 | 0.769994395 | 0.2 | 0.145 | 1 | 0.662822902 | 0.024955876 |
| Fam171a1 | 0.012580841 | -0.925999419 | 0.084 | 0.188 | 1 | 0.065975794 | 2.764165316 | 0.4 | 0.225 | 1 | 0.065975794 | 0.025003405 |
| Cfap418 | 0.012589763 | -1.204245075 | 0.168 | 0.282 | 1 | 0.526574705 | -0.441509711 | 0.2 | 0.275 | 1 | 0.526574705 | 0.025021025 |
| Oxld1 | 0.612185957 | -0.292781749 | 0.05 | 0.065 | 1 | 0.0125937 | 2.515421567 | 0.2 | 0.037 | 1 | 0.612185957 | 0.025028799 |
| Opn1mw | 0.088425833 | 0.707218251 | 0.101 | 0.047 | 1 | 0.012593759 | 2.280956314 | 0.2 | 0.037 | 1 | 0.088425833 | 0.025028915 |
| Polr2i | 0.01262564 | -0.737813989 | 0.227 | 0.359 | 1 | 0.154443936 | -1.058893689 | 0.1 | 0.306 | 1 | 0.154443936 | 0.025091872 |
| Tmem231 | 0.117781332 | -0.944858446 | 0.092 | 0.153 | 1 | 0.012632302 | 2.31748219 | 0.3 | 0.077 | 1 | 0.117781332 | 0.025105029 |
| Pikfyve | 0.012674122 | 0.336887555 | 0.731 | 0.547 | 1 | 0.455932966 | -0.201891969 | 0.6 | 0.664 | 1 | 0.455932966 | 0.025187611 |
| Dbt | 0.012682265 | -1.418312631 | 0.076 | 0.176 | 1 | 0.266865346 | -0.374395515 | 0 | 0.111 | 1 | 0.266865346 | 0.02520369 |
| Cdk1 | 0.012732835 | -2.292781749 | 0.008 | 0.071 | 1 | 0.730750993 | 2.210566986 | 0 | 0.012 | 1 | 0.730750993 | 0.025303544 |
| Pik3r4 | 0.012736738 | -1.424026283 | 0.076 | 0.176 | 1 | 0.139199287 | -1.230005605 | 0 | 0.182 | 1 | 0.139199287 | 0.025311252 |
| Slc25a10 | 0.017845629 | 0.877143252 | 0.202 | 0.1 | 1 | 0.012743029 | 2.648688098 | 0.3 | 0.08 | 1 | 0.017845629 | 0.025323673 |
| Ide | 0.012770291 | -0.669851398 | 0.639 | 0.718 | 1 | 0.584943893 | 0.180572246 | 0.9 | 0.713 | 1 | 0.584943893 | 0.025377501 |
| Mrtfa | 0.012775343 | -0.815422203 | 0.496 | 0.576 | 1 | 0.236938066 | -0.505640048 | 0.4 | 0.704 | 1 | 0.236938066 | 0.025387476 |
| CJ186046Rik | 0.012794314 | -1.622930351 | 0.067 | 0.165 | 1 | 0.770327644 | 0.263034406 | 0.1 | 0.13 | 1 | 0.770327644 | 0.025424933 |
| Chaf1a | 0.01279885 | -2.070389328 | 0.008 | 0.071 | 1 | 0.252293596 | 2.795529487 | 0.1 | 0.034 | 1 | 0.252293596 | 0.025433889 |
| Atp11c | 0.012807404 | -0.641931313 | 0.529 | 0.618 | 1 | 0.039236912 | -1.445148288 | 0.4 | 0.701 | 1 | 0.039236912 | 0.025450777 |
| Mrpl45 | 0.012809779 | -0.807354922 | 0.193 | 0.324 | 1 | 0.15395328 | -1.210896782 | 0.1 | 0.302 | 1 | 0.15395328 | 0.025455468 |
| Zbtb40 | 0.012828734 | -1.321928095 | 0.042 | 0.129 | 1 | 0.335993055 | 1.494359952 | 0.2 | 0.108 | 1 | 0.335993055 | 0.025492892 |
| Mical3 | 0.012837598 | -0.610413604 | 0.748 | 0.776 | 1 | 0.882163791 | -0.258202497 | 0.5 | 0.546 | 1 | 0.882163791 | 0.025510392 |
| Ptpn14 | 0.012839354 | -0.407774272 | 0.681 | 0.776 | 1 | 0.58434294 | 0.335662206 | 0.8 | 0.886 | 1 | 0.58434294 | 0.025513859 |
| Lacc1 | 0.012852193 | -1.392317423 | 0.126 | 0.235 | 1 | 0.207866038 | -0.959358016 | 0 | 0.139 | 1 | 0.207866038 | 0.025539207 |
| Chml | 0.012865105 | -2.185866545 | 0.025 | 0.1 | 1 | 0.3342467 | -0.374395515 | 0 | 0.086 | 1 | 0.3342467 | 0.025564698 |
| Fam13a | 0.012865683 | -2.185866545 | 0.025 | 0.1 | 1 | 0.141582823 | 2.847996907 | 0.1 | 0.025 | 1 | 0.141582823 | 0.025565839 |
| Socs6 | 0.01287059 | 0.424375364 | 0.513 | 0.353 | 1 | 0.114115241 | 0.896906507 | 0.7 | 0.451 | 1 | 0.114115241 | 0.025575528 |
| Tbk1 | 0.012906014 | -0.95491211 | 0.37 | 0.471 | 1 | 0.263021773 | -0.67943183 | 0.5 | 0.636 | 1 | 0.263021773 | 0.025645464 |
| Gm56936 | 0.012913371 | 0.916671616 | 0.244 | 0.129 | 1 | 0.274377708 | -0.304006187 | 0 | 0.108 | 1 | 0.274377708 | 0.025659988 |
| Nmt2 | 0.012932626 | -1.070389328 | 0.261 | 0.376 | 1 | 0.663918439 | 0.126138205 | 0.6 | 0.577 | 1 | 0.663918439 | 0.025697998 |
| Pram1 | 0.303113451 | -0.807354922 | 0.042 | 0.071 | 1 | 0.012940869 | 2.432959407 | 0.2 | 0.037 | 1 | 0.303113451 | 0.025714273 |
| Gm21859 | 0.603672523 | -0.485426827 | 0.05 | 0.065 | 1 | 0.012940869 | 1.959028219 | 0.2 | 0.037 | 1 | 0.603672523 | 0.025714273 |

| | | | | | | | | | | | | |
|---|---|---|---|---|---|---|---|---|---|---|---|---|
| Tob1 | 0.012966202 | 0.538906822 | 0.941 | 0.918 | 1 | 0.40767247 | -0.402653775 | 1 | 0.975 | 1 | 0.40767247 | 0.025764281 |
| Timm10 | 0.012973766 | -1.015941544 | 0.185 | 0.306 | 1 | 0.866283459 | 0.580516596 | 0.2 | 0.176 | 1 | 0.866283459 | 0.025779212 |
| Gba2 | 0.013006894 | -1.807354922 | 0.008 | 0.071 | 1 | 0.446819579 | 0.558490289 | 0 | 0.056 | 1 | 0.446819579 | 0.025844609 |
| Mcm2 | 0.062022899 | -1.411426246 | 0.025 | 0.076 | 1 | 0.013012886 | 3.432959407 | 0.2 | 0.04 | 1 | 0.062022899 | 0.025856437 |
| Rhof | 0.0130168 | -1.524955191 | 0.05 | 0.141 | 1 | 0.596057169 | 0.159940913 | 0.2 | 0.133 | 1 | 0.596057169 | 0.025864162 |
| Aamp | 0.013020529 | -0.383547213 | 0.798 | 0.847 | 1 | 0.266394001 | -0.394859617 | 0.8 | 0.855 | 1 | 0.266394001 | 0.025871524 |
| Gm56560 | 0.013037565 | -2.343407822 | 0.025 | 0.1 | 1 | 0.385508865 | 0.210566986 | 0 | 0.071 | 1 | 0.385508865 | 0.025905151 |
| Iqce | 0.013058554 | -1.517848305 | 0.076 | 0.176 | 1 | 0.66032688 | 0.515421567 | 0.2 | 0.142 | 1 | 0.66032688 | 0.025946581 |
| Map1s | 0.013071773 | -0.847996907 | 0.134 | 0.253 | 1 | 0.39325312 | -0.663902132 | 0.2 | 0.306 | 1 | 0.39325312 | 0.025972674 |
| Lpcat3 | 0.02496697 | 0.174233041 | 0.571 | 0.4 | 1 | 0.013075556 | 1.101445264 | 0.7 | 0.46 | 1 | 0.02496697 | 0.025980142 |
| Zc3h3 | 0.013111099 | -1.106915204 | 0.193 | 0.312 | 1 | 0.138235261 | -1.31299497 | 0.1 | 0.312 | 1 | 0.138235261 | 0.026050297 |
| Hat1 | 0.013115635 | -0.812077332 | 0.378 | 0.476 | 1 | 0.354791463 | 0.702017601 | 0.5 | 0.441 | 1 | 0.354791463 | 0.02605925 |
| Sra1 | 0.013123054 | 0.301399282 | 0.891 | 0.759 | 1 | 0.201920477 | -0.647960588 | 0.8 | 0.784 | 1 | 0.201920477 | 0.026073893 |
| Zfp384 | 0.013123759 | -1.089498151 | 0.185 | 0.306 | 1 | 0.135590653 | -1.040971781 | 0.2 | 0.414 | 1 | 0.135590653 | 0.026075284 |
| Tmem251 | 0.102101288 | -0.275973462 | 0.403 | 0.482 | 1 | 0.013143496 | -1.997028433 | 0.2 | 0.568 | 1 | 0.102101288 | 0.02611424 |
| Rnf8 | 0.013144074 | -0.714245518 | 0.387 | 0.494 | 1 | 0.363658807 | -0.241350579 | 0.4 | 0.586 | 1 | 0.363658807 | 0.026115381 |
| Pfas | 0.013146749 | -1.235448574 | 0.076 | 0.176 | 1 | 0.259359111 | -0.374395515 | 0 | 0.114 | 1 | 0.259359111 | 0.026120662 |
| Trp53i13 | 0.013173302 | -0.688960221 | 0.496 | 0.565 | 1 | 0.529417851 | -0.157716745 | 0.4 | 0.491 | 1 | 0.529417851 | 0.026173069 |
| Pdcd6 | 0.013186743 | -0.417963859 | 0.798 | 0.824 | 1 | 0.016166694 | -1.01449957 | 0.5 | 0.846 | 1 | 0.016166694 | 0.026199596 |
| Txnrd3 | 0.013208508 | -1.457412451 | 0.101 | 0.206 | 1 | 0.599366075 | -0.304006187 | 0.1 | 0.157 | 1 | 0.599366075 | 0.026242552 |
| Ttll11 | 0.013209904 | -2.008988783 | 0.025 | 0.1 | 1 | 0.824373597 | 0.558490289 | 0.1 | 0.077 | 1 | 0.824373597 | 0.026245306 |
| Dsn1 | 0.013209904 | -2.008988783 | 0.025 | 0.1 | 1 | 0.146852038 | 1.902444691 | 0.2 | 0.074 | 1 | 0.146852038 | 0.026245306 |
| D330041H03Rik | 0.01321014 | -2.292781749 | 0.025 | 0.1 | 1 | 0.040451025 | 1.695993813 | 0.3 | 0.096 | 1 | 0.040451025 | 0.026245772 |
| Psma6 | 0.016693641 | -0.51308317 | 0.739 | 0.765 | 1 | 0.013234349 | -1.083616118 | 0.5 | 0.83 | 1 | 0.016693641 | 0.026293551 |
| Rnf103 | 0.013234395 | 0.478949263 | 0.697 | 0.506 | 1 | 0.167546986 | 0.780386 | 0.7 | 0.611 | 1 | 0.167546986 | 0.026293641 |
| Castor1 | 0.013249238 | 0.472752997 | 0.454 | 0.294 | 1 | 0.300081682 | -0.719043686 | 0.2 | 0.34 | 1 | 0.300081682 | 0.026322933 |
| Fastkd3 | 0.022584338 | -1.485426827 | 0.034 | 0.106 | 1 | 0.013259545 | 2.558490289 | 0.3 | 0.08 | 1 | 0.022584338 | 0.026343275 |
| Zfp13 | 0.130956222 | -0.900464326 | 0.034 | 0.076 | 1 | 0.013295653 | 2.210566986 | 0.2 | 0.037 | 1 | 0.130956222 | 0.026414532 |
| Tm7sf3 | 0.013304484 | -0.750021747 | 0.479 | 0.553 | 1 | 0.224046624 | -0.651356879 | 0.5 | 0.623 | 1 | 0.224046624 | 0.026431958 |
| Ppip5k2 | 0.013306096 | -1.250961574 | 0.235 | 0.329 | 1 | 0.716617519 | -0.401616984 | 0.3 | 0.34 | 1 | 0.716617519 | 0.02643514 |
| mt-Nd4l | 0.82328187 | -0.127398807 | 0.924 | 0.935 | 1 | 0.01330846 | 0.744903414 | 1 | 0.901 | 1 | 0.82328187 | 0.026439805 |
| Tmem63a | 0.013321423 | -0.688960221 | 0.412 | 0.524 | 1 | 0.364026603 | 0.256370676 | 0.8 | 0.633 | 1 | 0.364026603 | 0.026465386 |
| Rexo1 | 0.01332401 | -1 | 0.311 | 0.412 | 1 | 0.356875953 | -0.620151929 | 0.4 | 0.485 | 1 | 0.356875953 | 0.026470491 |
| Rcan2 | 0.013327846 | -2.778208576 | 0.034 | 0.112 | 1 | 0.539470755 | 0.432959407 | 0 | 0.037 | 1 | 0.539470755 | 0.026478061 |
| Armh3 | 0.013336116 | -1.342062652 | 0.252 | 0.359 | 1 | 0.950369243 | -0.169705095 | 0.5 | 0.454 | 1 | 0.950369243 | 0.02649438 |
| Adam30 | 0.01335126 | -1.333423734 | 0.059 | 0.153 | 1 | 0.773682184 | 0.280956314 | 0.2 | 0.154 | 1 | 0.773682184 | 0.026524264 |
| Fuom | 0.013356576 | -1.126972856 | 0.176 | 0.294 | 1 | 0.345489484 | -0.473931188 | 0.1 | 0.222 | 1 | 0.345489484 | 0.026534753 |
| Gm47237 | 0.013383686 | -2.008988783 | 0.025 | 0.1 | 1 | 0.56531557 | 0.672147071 | 0.2 | 0.13 | 1 | 0.56531557 | 0.026588249 |
| Sh3bp1 | 0.013383686 | -2.240314329 | 0.025 | 0.1 | 1 | 0.245439637 | -0.536666944 | 0 | 0.12 | 1 | 0.245439637 | 0.026588249 |
| Eif5a2 | 0.013383746 | -1.87774425 | 0.025 | 0.1 | 1 | 0.259431972 | -0.374395515 | 0 | 0.114 | 1 | 0.259431972 | 0.026588367 |
| Asb6 | 0.013401976 | -1.137503524 | 0.109 | 0.218 | 1 | 0.744576209 | 0.135278859 | 0.1 | 0.133 | 1 | 0.744576209 | 0.026624339 |
| Ranbp17 | 0.013412479 | -1.783107376 | 0.067 | 0.165 | 1 | 0.363772218 | -1.536666944 | 0.1 | 0.207 | 1 | 0.363772218 | 0.026645064 |
| Ammecr1l | 0.013414594 | -0.862281133 | 0.429 | 0.506 | 1 | 0.260721915 | -0.709998547 | 0.6 | 0.71 | 1 | 0.260721915 | 0.026649237 |
| Srgap3 | 0.01342242 | -1 | 0.118 | 0.229 | 1 | 0.073287366 | -1.91281543 | 0.2 | 0.497 | 1 | 0.073287366 | 0.026664679 |
| Dhrs1 | 0.013439439 | 0.607682577 | 0.479 | 0.341 | 1 | 0.422318397 | 0.919889825 | 0.4 | 0.309 | 1 | 0.422318397 | 0.026698259 |
| Pigyl | 0.01347424 | -0.892602209 | 0.16 | 0.282 | 1 | 0.477424505 | 0.591657153 | 0.3 | 0.194 | 1 | 0.477424505 | 0.026766925 |
| Rps15 | 0.060962659 | -0.180798332 | 0.983 | 0.976 | 1 | 0.013474281 | -0.783174024 | 1 | 0.994 | 1 | 0.060962659 | 0.026767006 |
| Tsn | 0.013474488 | -0.373400209 | 0.891 | 0.882 | 1 | 0.297001496 | 0.380351981 | 0.8 | 0.92 | 1 | 0.297001496 | 0.026767414 |
| Samd1 | 0.013474751 | -1.359895945 | 0.092 | 0.194 | 1 | 0.101521568 | -1.736965594 | 0 | 0.216 | 1 | 0.101521568 | 0.026767934 |
| Relb | 0.013475579 | -0.841188517 | 0.294 | 0.406 | 1 | 0.721862992 | 0.432959407 | 0.5 | 0.429 | 1 | 0.721862992 | 0.026769566 |
| Mepce | 0.013484718 | -1.00625899 | 0.151 | 0.271 | 1 | 0.391958976 | -0.673239997 | 0.2 | 0.309 | 1 | 0.391958976 | 0.026787598 |
| Idh3b | 0.013498366 | -0.573864792 | 0.513 | 0.618 | 1 | 0.366666831 | -0.526398608 | 0.5 | 0.559 | 1 | 0.366666831 | 0.026814526 |
| 1110004F10Rik | 0.013532656 | -0.392730152 | 0.807 | 0.841 | 1 | 0.131210412 | -0.510857757 | 0.6 | 0.904 | 1 | 0.131210412 | 0.026882179 |
| Slc7a11 | 0.013536639 | -2.163498732 | 0.025 | 0.1 | 1 | 0.408830166 | -1.131825212 | 0 | 0.065 | 1 | 0.408830166 | 0.026890037 |
| 5033426O07Rik | 0.013553774 | 1.514573173 | 0.118 | 0.041 | 1 | 0.992396409 | 0.331421381 | 0.2 | 0.201 | 1 | 0.992396409 | 0.026923842 |
| Brd3os | 0.013558373 | -1.87774425 | 0.025 | 0.1 | 1 | 0.460551517 | 0.558490289 | 0 | 0.052 | 1 | 0.460551517 | 0.026932916 |
| 4921504A21Rik | 0.013558373 | -1.807354922 | 0.025 | 0.1 | 1 | 0.852081867 | 0.374065718 | 0.1 | 0.08 | 1 | 0.852081867 | 0.026932916 |
| Mrc2 | 0.013577397 | -0.594892807 | 0.773 | 0.729 | 1 | 0.810541508 | -0.21770634 | 0.8 | 0.799 | 1 | 0.810541508 | 0.026970449 |
| Ssbp1 | 0.013589607 | -0.687060688 | 0.496 | 0.559 | 1 | 0.034103521 | -1.048167282 | 0.1 | 0.497 | 1 | 0.034103521 | 0.026994537 |
| G0s2 | 0.037686869 | 0.929610672 | 0.076 | 0.165 | 1 | 0.013590086 | 2.644463512 | 0.3 | 0.083 | 1 | 0.037686869 | 0.026995481 |
| Dnaaf10 | 0.013599822 | -1.306456686 | 0.092 | 0.2 | 1 | 0.864524031 | 0.463333056 | 0.2 | 0.182 | 1 | 0.864524031 | 0.027014689 |
| Trmt61a | 0.013601698 | -1.292781749 | 0.05 | 0.141 | 1 | 0.832730137 | 1.111031312 | 0.1 | 0.08 | 1 | 0.832730137 | 0.02701839 |
| Acyp1 | 0.013620574 | 0.616452787 | 0.303 | 0.176 | 1 | 0.019174491 | 1.41305985 | 0.5 | 0.185 | 1 | 0.019174491 | 0.027055628 |
| Hnrnpr | 0.013646868 | -0.42246287 | 0.899 | 0.882 | 1 | 0.153192449 | -0.635520331 | 1 | 0.926 | 1 | 0.153192449 | 0.0271075 |
| Ppp2r2d | 0.039172994 | -0.429573592 | 0.681 | 0.729 | 1 | 0.013662646 | -1.234743524 | 0.9 | 0.867 | 1 | 0.039172994 | 0.027138624 |
| Emilin2 | 0.013682359 | 1.96203215 | 0.134 | 0.053 | 1 | 0.408816096 | -0.304006187 | 0 | 0.065 | 1 | 0.408816096 | 0.027177511 |
| 5930430L01Rik | 0.013696485 | 0.610788488 | 0.227 | 0.112 | 1 | 0.036963738 | 1.847996907 | 0.4 | 0.151 | 1 | 0.036963738 | 0.027205376 |
| Traf3ip2 | 0.834312469 | 0.280107919 | 0.101 | 0.094 | 1 | 0.013697398 | 1.87108052 | 0.4 | 0.127 | 1 | 0.834312469 | 0.027207178 |
| Gm15417 | 0.013701818 | 0.826517179 | 0.252 | 0.135 | 1 | 0.464540466 | 0.625604485 | 0.3 | 0.191 | 1 | 0.464540466 | 0.027215896 |
| Gtf2h3 | 0.013703554 | -1.00625899 | 0.168 | 0.288 | 1 | 0.378131303 | -0.304006187 | 0.1 | 0.213 | 1 | 0.378131303 | 0.027219321 |
| Polr2d | 0.013708103 | -0.491796352 | 0.588 | 0.665 | 1 | 0.1478469 | -0.755923752 | 0.4 | 0.611 | 1 | 0.1478469 | 0.027228293 |
| Sertad3 | 0.013710733 | -1.301714874 | 0.126 | 0.235 | 1 | 0.60805067 | -0.419483404 | 0.2 | 0.256 | 1 | 0.60805067 | 0.027233483 |
| Tnfaip2 | 0.013719811 | 1.142604395 | 0.109 | 0.035 | 1 | 0.489901827 | 0.769994395 | 0 | 0.046 | 1 | 0.489901827 | 0.027251389 |
| Gm56830 | 0.013735212 | -1.097610797 | 0.185 | 0.3 | 1 | 0.925979707 | -0.433289204 | 0.3 | 0.278 | 1 | 0.925979707 | 0.027281768 |
| Appl2 | 0.013735524 | -0.699295176 | 0.555 | 0.588 | 1 | 0.01621859 | -1.382957528 | 0.2 | 0.67 | 1 | 0.01621859 | 0.027282383 |
| Guk1 | 0.013757782 | 0.483984853 | 0.622 | 0.518 | 1 | 0.997079562 | 0.517351595 | 0.4 | 0.469 | 1 | 0.997079562 | 0.027326288 |
| Rhbdd1 | 0.01376955 | 0.196603092 | 0.437 | 0.271 | 1 | 0.323624312 | 1.095089769 | 0.4 | 0.296 | 1 | 0.323624312 | 0.027349499 |
| Ddb1 | 0.013789717 | -0.42433223 | 0.824 | 0.824 | 1 | 0.603673409 | 0.247604365 | 0.9 | 0.92 | 1 | 0.603673409 | 0.027389278 |
| Rps6kb2 | 0.013794766 | -1.347923303 | 0.067 | 0.165 | 1 | 0.590171457 | 1.017921908 | 0.2 | 0.145 | 1 | 0.590171457 | 0.027399236 |
| Crppa | 0.013795242 | -1.03562391 | 0.202 | 0.324 | 1 | 0.176489205 | 0.878370556 | 0.4 | 0.216 | 1 | 0.176489205 | 0.027400175 |
| Naa35 | 0.013816226 | -0.933700656 | 0.345 | 0.447 | 1 | 0.568007679 | 0.185031894 | 0.7 | 0.633 | 1 | 0.568007679 | 0.027441563 |
| Epha7 | 0.013820995 | 0.136550516 | 0.588 | 0.394 | 1 | 0.835741842 | -0.479748515 | 0.4 | 0.336 | 1 | 0.835741842 | 0.02745097 |
| Commd7 | 0.013829413 | -0.842978832 | 0.353 | 0.453 | 1 | 0.050896811 | -1.015501094 | 0.1 | 0.463 | 1 | 0.050896811 | 0.027467573 |
| Esd | 0.523971317 | -0.190298792 | 0.765 | 0.794 | 1 | 0.013836462 | -1.321928095 | 0.5 | 0.769 | 1 | 0.523971317 | 0.027481475 |
| Prim1 | 0.013837825 | -1.86393845 | 0.076 | 0.171 | 1 | 0.507581676 | 0.695993813 | 0.2 | 0.12 | 1 | 0.507581676 | 0.027484165 |
| Dnal4 | 0.01388055 | -1.137503524 | 0.16 | 0.271 | 1 | 0.463838723 | 0.71414116 | 0.3 | 0.194 | 1 | 0.463838723 | 0.02756843 |
| Gm46563 | 0.013884205 | -1.19592021 | 0.076 | 0.176 | 1 | 0.374430929 | 0.31748219 | 0 | 0.074 | 1 | 0.374430929 | 0.027575638 |
| Eaf1 | 0.013954983 | -0.614709844 | 0.252 | 0.394 | 1 | 0.27180039 | -0.831743819 | 0.2 | 0.349 | 1 | 0.27180039 | 0.027715224 |
| 4732491K20Rik | 0.013972097 | 1.432111013 | 0.118 | 0.041 | 1 | 0.460521625 | 0.769994395 | 0 | 0.052 | 1 | 0.460521625 | 0.027748975 |
| Nmral1 | 0.013992049 | -1.655351829 | 0.025 | 0.1 | 1 | 0.078598851 | 1.744903414 | 0.2 | 0.059 | 1 | 0.078598851 | 0.02778832 |
| Sergef | 0.014010424 | -2.250961574 | 0.034 | 0.112 | 1 | 0.290492439 | -0.654503434 | 0 | 0.102 | 1 | 0.290492439 | 0.027824556 |
| Ap4m1 | 0.014055806 | -1.378511623 | 0.101 | 0.206 | 1 | 0.152644918 | -1.391469028 | 0 | 0.173 | 1 | 0.152644918 | 0.027914046 |
| Brd7 | 0.014067829 | -0.807354922 | 0.513 | 0.6 | 1 | 0.026473881 | -1.357117523 | 0.4 | 0.694 | 1 | 0.026473881 | 0.027937754 |
| 2900076A07Rik | 0.014098893 | -1.292781749 | 0.101 | 0.206 | 1 | 0.374285054 | -0.654503434 | 0.1 | 0.21 | 1 | 0.374285054 | 0.027999008 |
| Mybl1 | 0.014132146 | -1.974811668 | 0.084 | 0.182 | 1 | 0.041414814 | 1.210566986 | 0.3 | 0.096 | 1 | 0.041414814 | 0.028064575 |
| Pgghg | 0.149561747 | -0.722466024 | 0.101 | 0.159 | 1 | 0.014183177 | 1.847996907 | 0.4 | 0.127 | 1 | 0.149561747 | 0.028165191 |
| 1300002E11Rik | 0.014199862 | -1.04342228 | 0.126 | 0.241 | 1 | 0.827576573 | 0.558490289 | 0.2 | 0.167 | 1 | 0.827576573 | 0.028198088 |
| Msh6 | 0.014218756 | -1.269698136 | 0.05 | 0.141 | 1 | 0.829358516 | 0.290001453 | 0.1 | 0.12 | 1 | 0.829358516 | 0.028235338 |

| | | | | | | | | | | | | |
|---|---|---|---|---|---|---|---|---|---|---|---|---|
| Itga10 | 0.014245739 | 0.205683078 | 0.891 | 0.706 | 1 | 0.217201665 | 0.489248538 | 0.9 | 0.846 | 1 | 0.217201665 | 0.028288536 |
| Pxk | 0.016218841 | -0.948398804 | 0.202 | 0.324 | 1 | 0.014250381 | -2.181750437 | 0.1 | 0.497 | 1 | 0.016218841 | 0.028297688 |
| Platr25 | 0.014276932 | -1.280607035 | 0.118 | 0.229 | 1 | 0.931729398 | 1.176619654 | 0.1 | 0.12 | 1 | 0.931729398 | 0.028350033 |
| Epm2aip1 | 0.014280655 | -0.780882711 | 0.286 | 0.412 | 1 | 0.560848551 | -0.296774618 | 0.3 | 0.389 | 1 | 0.560848551 | 0.028357372 |
| Tnnt3 | 0.014288898 | 2.397759508 | 0.202 | 0.106 | 1 | 0.374707504 | -0.304006187 | 0 | 0.074 | 1 | 0.374707504 | 0.028373624 |
| Washc4 | 0.014341011 | -0.807354922 | 0.538 | 0.588 | 1 | 0.021134105 | -1.50787852 | 0.5 | 0.728 | 1 | 0.021134105 | 0.028476357 |
| Samd9l | 0.014373632 | -2.14839184 | 0.059 | 0.147 | 1 | 0.059979965 | -2.171902651 | 0 | 0.269 | 1 | 0.059979965 | 0.028540663 |
| Impdh1 | 0.014380702 | -1.032914622 | 0.084 | 0.188 | 1 | 0.466867421 | 0.896906507 | 0.3 | 0.213 | 1 | 0.466867421 | 0.028554598 |
| Lap3 | 0.014393016 | -0.748461233 | 0.345 | 0.476 | 1 | 0.152319589 | -1.186649236 | 0.3 | 0.475 | 1 | 0.152319589 | 0.028578873 |
| Kcnq3 | 0.014397753 | 1.850176205 | 0.176 | 0.082 | 1 | 0.298995322 | -2.382957528 | 0 | 0.099 | 1 | 0.298995322 | 0.028588211 |
| Morn4 | 0.01439848 | -1.622930351 | 0.025 | 0.1 | 1 | 0.446819579 | 0.558490289 | 0 | 0.056 | 1 | 0.446819579 | 0.028589645 |
| Gprc5b | 0.014407935 | 1.340543773 | 0.21 | 0.106 | 1 | 0.159430078 | 1.31748219 | 0.2 | 0.074 | 1 | 0.159430078 | 0.028608282 |
| Csf1r | 0.830638286 | -0.222392421 | 0.025 | 0.029 | 1 | 0.014410499 | 0.672147071 | 0.2 | 0.037 | 1 | 0.830638286 | 0.028613335 |
| Rps6ka1 | 0.014417404 | -1.424026283 | 0.092 | 0.194 | 1 | 0.898760911 | 0.41305985 | 0.2 | 0.179 | 1 | 0.898760911 | 0.028626946 |
| Lnpep | 0.014441447 | -0.391056877 | 0.95 | 0.941 | 1 | 0.875399949 | 0.43483914 | 1 | 0.981 | 1 | 0.875399949 | 0.028674338 |
| Suox | 0.014441733 | 0.80407979 | 0.286 | 0.165 | 1 | 0.610048752 | 0.874963954 | 0.2 | 0.139 | 1 | 0.610048752 | 0.028674902 |
| Rpa2 | 0.014465045 | -0.735970289 | 0.126 | 0.247 | 1 | 0.266337253 | 1.169925001 | 0.3 | 0.17 | 1 | 0.266337253 | 0.028720853 |
| Stk35 | 0.014466421 | -1.215819767 | 0.151 | 0.265 | 1 | 0.196993785 | 0.695993813 | 0.5 | 0.287 | 1 | 0.196993785 | 0.028723565 |
| Agrp | 0.014479007 | -1.918386234 | 0.034 | 0.112 | 1 | 0.835493698 | 1.017921908 | 0.1 | 0.08 | 1 | 0.835493698 | 0.028748372 |
| Naa30 | 0.014494064 | -0.997325866 | 0.202 | 0.318 | 1 | 0.043793227 | -1.730270942 | 0.1 | 0.414 | 1 | 0.043793227 | 0.02877805 |
| Mustn1 | 0.014496717 | 1.409984272 | 0.353 | 0.241 | 1 | 0.21386954 | -1.321928095 | 0 | 0.136 | 1 | 0.21386954 | 0.02878328 |
| Timm13 | 0.014502668 | -0.46579802 | 0.689 | 0.759 | 1 | 0.32187941 | 0.504298189 | 0.7 | 0.667 | 1 | 0.32187941 | 0.028795008 |
| Cks1b | 0.014506095 | -0.918386234 | 0.294 | 0.406 | 1 | 0.12889006 | -0.944010051 | 0.1 | 0.355 | 1 | 0.12889006 | 0.028801764 |
| Glt28d2 | 0.014508688 | 0.541045384 | 0.311 | 0.182 | 1 | 0.539838162 | -0.536666944 | 0.3 | 0.358 | 1 | 0.539838162 | 0.028806874 |
| Fbxo44 | 0.014510952 | 1.192645078 | 0.109 | 0.035 | 1 | 0.479305119 | 1.847996907 | 0.1 | 0.049 | 1 | 0.479305119 | 0.028811337 |
| Wfs1 | 0.63771167 | -0.347923303 | 0.084 | 0.1 | 1 | 0.014532959 | 1.973527789 | 0.3 | 0.077 | 1 | 0.63771167 | 0.028854712 |
| Tdp1 | 0.341694523 | -0.485426827 | 0.109 | 0.147 | 1 | 0.014533097 | 2.063725598 | 0.3 | 0.077 | 1 | 0.341694523 | 0.028854982 |
| 1700030N03Rik | 0.014538593 | 2.421463768 | 0.076 | 0.018 | 1 | 0.698529281 | 2.210566986 | 0 | 0.015 | 1 | 0.698529281 | 0.028865815 |
| Tmem154 | 0.065494922 | 0.592575685 | 0.218 | 0.135 | 1 | 0.014555142 | 1.87108052 | 0.4 | 0.136 | 1 | 0.065494922 | 0.028898432 |
| Tmem138 | 0.014578778 | -1.266786541 | 0.126 | 0.235 | 1 | 0.542088078 | -0.230005605 | 0.1 | 0.17 | 1 | 0.542088078 | 0.028945014 |
| Phf20l1 | 0.014620209 | -0.532495081 | 0.773 | 0.788 | 1 | 0.340560807 | -0.386368156 | 0.9 | 0.793 | 1 | 0.340560807 | 0.029026667 |
| Crim1 | 0.014643018 | -0.55119107 | 0.975 | 0.976 | 1 | 0.144757517 | -0.509753885 | 1 | 0.994 | 1 | 0.144757517 | 0.029071618 |
| Spidr | 0.101659956 | 0.233672346 | 0.471 | 0.347 | 1 | 0.014661735 | 1.216000872 | 0.7 | 0.346 | 1 | 0.101659956 | 0.029108503 |
| Diaph1 | 0.01468389 | 0.159796198 | 0.891 | 0.712 | 1 | 0.902276446 | -0.14913776 | 0.7 | 0.815 | 1 | 0.902276446 | 0.029152164 |
| Slc25a42 | 0.014717621 | -1.970853654 | 0.034 | 0.112 | 1 | 0.489954768 | 0.374065718 | 0 | 0.046 | 1 | 0.489954768 | 0.029218633 |
| Tspyl2 | 0.014757591 | -1.122856748 | 0.202 | 0.312 | 1 | 0.521135257 | 0.515421567 | 0.4 | 0.299 | 1 | 0.521135257 | 0.029297396 |
| Bod1l | 0.014766643 | -0.617877123 | 0.748 | 0.765 | 1 | 0.030205327 | -1.156005024 | 0.5 | 0.781 | 1 | 0.030205327 | 0.029315232 |
| Bcl2 | 0.014780004 | 0.377938481 | 0.487 | 0.329 | 1 | 0.176056951 | 0.576486719 | 0.6 | 0.438 | 1 | 0.176056951 | 0.02934156 |
| Med12 | 0.01479199 | -0.776328026 | 0.328 | 0.441 | 1 | 0.61220064 | -0.457811523 | 0.5 | 0.481 | 1 | 0.61220064 | 0.029365177 |
| Ppa1 | 0.014810355 | -0.464865505 | 0.832 | 0.794 | 1 | 0.960601182 | -0.51724236 | 0.5 | 0.728 | 1 | 0.960601182 | 0.029401363 |
| Ltn1 | 0.014832566 | -0.885606368 | 0.378 | 0.476 | 1 | 0.432230233 | -0.505640048 | 0.5 | 0.546 | 1 | 0.432230233 | 0.029445127 |
| Ccdc59 | 0.042409068 | -0.494645357 | 0.655 | 0.718 | 1 | 0.014847421 | -1.385800278 | 0.4 | 0.722 | 1 | 0.042409068 | 0.029474397 |
| Terf2 | 0.014862896 | -0.715724447 | 0.311 | 0.435 | 1 | 0.042162748 | -1.601686736 | 0.2 | 0.503 | 1 | 0.042162748 | 0.029504886 |
| Sipa1 | 0.01495871 | -1.748461233 | 0.034 | 0.112 | 1 | 0.206064866 | 1.063725598 | 0.3 | 0.145 | 1 | 0.206064866 | 0.029693658 |
| Dhx35 | 0.205741212 | -0.87774425 | 0.059 | 0.1 | 1 | 0.014965258 | 1.930459067 | 0.3 | 0.077 | 1 | 0.205741212 | 0.029706558 |
| Gmpr | 0.711108838 | 0.321928095 | 0.05 | 0.041 | 1 | 0.014965258 | 1.888638891 | 0.3 | 0.077 | 1 | 0.711108838 | 0.029706558 |
| Med6 | 0.015023592 | -0.759601791 | 0.328 | 0.441 | 1 | 0.10615763 | -1.473931188 | 0.1 | 0.336 | 1 | 0.10615763 | 0.029821476 |
| L2hgdh | 0.015052748 | -1.129283017 | 0.101 | 0.206 | 1 | 0.631041736 | -0.171902651 | 0.1 | 0.151 | 1 | 0.631041736 | 0.02987891 |
| Tfip11 | 0.015053035 | -1.347923303 | 0.084 | 0.182 | 1 | 0.477766147 | 0.769994395 | 0.2 | 0.12 | 1 | 0.477766147 | 0.029879476 |
| Tll1 | 0.015069373 | -1.070389328 | 0.16 | 0.271 | 1 | 0.241027206 | 0.340789563 | 0.4 | 0.531 | 1 | 0.241027206 | 0.029911661 |
| Haus8 | 0.015115123 | -0.576192291 | 0.361 | 0.5 | 1 | 0.689224337 | 0.416657595 | 0.4 | 0.373 | 1 | 0.689224337 | 0.03000178 |
| Slc45a3 | 0.015136439 | 2.440572591 | 0.076 | 0.018 | 1 | 0.474904369 | 0.769994395 | 0 | 0.049 | 1 | 0.474904369 | 0.030043767 |
| Syngr2 | 0.015154352 | -0.987927168 | 0.067 | 0.165 | 1 | 0.952393575 | 0.210566986 | 0.2 | 0.201 | 1 | 0.952393575 | 0.03007905 |
| Katnbl1 | 0.015154668 | -0.827346397 | 0.471 | 0.541 | 1 | 0.289054915 | -0.611434712 | 0.6 | 0.716 | 1 | 0.289054915 | 0.030079672 |
| Pgap3 | 0.015201591 | -1.807354922 | 0.034 | 0.112 | 1 | 0.25967673 | -0.567040593 | 0 | 0.114 | 1 | 0.25967673 | 0.030172094 |
| Upp1 | 0.015221701 | 0.85160816 | 0.185 | 0.082 | 1 | 0.549475438 | 1.902444691 | 0.1 | 0.059 | 1 | 0.549475438 | 0.030211703 |
| Puf60 | 0.015235098 | -0.436517227 | 0.664 | 0.724 | 1 | 0.027785836 | -1.087612506 | 0.4 | 0.725 | 1 | 0.027785836 | 0.030238088 |
| Zfp945 | 0.015244609 | -0.889817082 | 0.218 | 0.341 | 1 | 0.462296303 | -0.505640048 | 0.2 | 0.29 | 1 | 0.462296303 | 0.030256821 |
| Ntan1 | 0.015245467 | 0.443800098 | 0.697 | 0.576 | 1 | 0.921325266 | 0.193493473 | 0.6 | 0.648 | 1 | 0.921325266 | 0.030258509 |
| Erf | 0.054682542 | -0.56584551 | 0.647 | 0.641 | 1 | 0.015250396 | -1.347605214 | 0.6 | 0.824 | 1 | 0.054682542 | 0.030268218 |
| Txnip | 0.064121816 | -0.271982111 | 0.697 | 0.788 | 1 | 0.01525562 | -0.577308928 | 0.6 | 0.849 | 1 | 0.064121816 | 0.030278506 |
| Ift140 | 0.015256209 | -1.944858446 | 0.042 | 0.124 | 1 | 0.298707021 | -0.68251781 | 0 | 0.099 | 1 | 0.298707021 | 0.030279667 |
| Mtf2 | 0.015305917 | -0.768598879 | 0.429 | 0.506 | 1 | 0.133811304 | -0.92291602 | 0.5 | 0.648 | 1 | 0.133811304 | 0.030377563 |
| Ncbp2as2 | 0.149104236 | -0.432959407 | 0.462 | 0.5 | 1 | 0.015323406 | -2.611434712 | 0 | 0.389 | 1 | 0.149104236 | 0.030412006 |
| Spryd3 | 0.015323876 | -0.673871917 | 0.269 | 0.4 | 1 | 0.967277866 | -0.103093493 | 0.4 | 0.361 | 1 | 0.967277866 | 0.03041293 |
| Tbc1d20 | 0.01533399 | -0.582525515 | 0.437 | 0.553 | 1 | 0.346088685 | -0.567040593 | 0.5 | 0.583 | 1 | 0.346088685 | 0.030432849 |
| Cr1l | 0.122682626 | -0.284962245 | 0.765 | 0.771 | 1 | 0.015343689 | -1.204470513 | 0.6 | 0.799 | 1 | 0.122682626 | 0.030451948 |
| Sorcs2 | 0.015436643 | -2.74176658 | 0.059 | 0.147 | 1 | 0.267367452 | -1.625934282 | 0 | 0.111 | 1 | 0.267367452 | 0.030634996 |
| Pkn1 | 0.015441594 | -1.168236651 | 0.126 | 0.235 | 1 | 0.247340083 | -0.596787936 | 0.2 | 0.395 | 1 | 0.247340083 | 0.030644745 |
| Wrap73 | 0.015446943 | -1.622930351 | 0.034 | 0.112 | 1 | 0.266864013 | -0.408342847 | 0 | 0.111 | 1 | 0.266864013 | 0.030655278 |
| Exoc1 | 0.015455181 | -1.424026283 | 0.084 | 0.182 | 1 | 0.778664744 | -0.111361109 | 0.2 | 0.225 | 1 | 0.778664744 | 0.0306715 |
| Cyren | 0.610898728 | -0.485426827 | 0.101 | 0.118 | 1 | 0.015458243 | 2.091922489 | 0.3 | 0.08 | 1 | 0.610898728 | 0.030677528 |
| Gstz1 | 0.032800269 | 0.528119705 | 0.529 | 0.412 | 1 | 0.015478088 | 1.280956314 | 0.7 | 0.349 | 1 | 0.032800269 | 0.030716604 |
| Bbc3 | 0.015478278 | -1.774933444 | 0.067 | 0.159 | 1 | 0.239052258 | -1.069540933 | 0 | 0.123 | 1 | 0.239052258 | 0.030716978 |
| Epha2 | 0.015479597 | -0.987927168 | 0.168 | 0.282 | 1 | 0.387778944 | -0.82128188 | 0.3 | 0.389 | 1 | 0.387778944 | 0.030719576 |
| Ubash3b | 0.015503344 | 0.316027493 | 0.252 | 0.135 | 1 | 0.042588371 | 3.272494735 | 0.3 | 0.102 | 1 | 0.042588371 | 0.030766335 |
| Skic2 | 0.015511113 | -1.021479727 | 0.151 | 0.265 | 1 | 0.279216991 | -0.242605642 | 0.1 | 0.278 | 1 | 0.279216991 | 0.030781631 |
| Atp6v0c | 0.015516516 | 0.1779859 | 0.992 | 0.982 | 1 | 0.410204347 | -0.500239448 | 0.8 | 0.997 | 1 | 0.410204347 | 0.03079227 |
| 4930556G22Rik | 0.015539978 | 2.514573173 | 0.05 | 0.006 | 1 | 0.730749233 | 2.432959407 | 0 | 0.012 | 1 | 0.730749233 | 0.030838466 |
| Gm57050 | 0.015539978 | 2.514573173 | 0.05 | 0.006 | 1 | 0.19134412 | 2.558490289 | 0.1 | 0.028 | 1 | 0.19134412 | 0.030838466 |
| 9330111N05Rik | 0.015539978 | 2.836501268 | 0.05 | 0.006 | 1 | 0.230466554 | 2.558490289 | 0.1 | 0.031 | 1 | 0.230466554 | 0.030838466 |
| Trappc5 | 0.015543277 | -0.836899198 | 0.218 | 0.341 | 1 | 0.160599909 | -0.888968688 | 0.1 | 0.302 | 1 | 0.160599909 | 0.030844961 |
| Gpnmb | 0.015545778 | 2.929610672 | 0.05 | 0.006 | 1 | 0.643846086 | 1.432959407 | 0 | 0.022 | 1 | 0.643846086 | 0.030849884 |
| Kif5a | 0.01555276 | -1.758445322 | 0.042 | 0.124 | 1 | 0.805895131 | 1.354956895 | 0.1 | 0.083 | 1 | 0.805895131 | 0.030863631 |
| Asb13 | 0.015553282 | -1.807354922 | 0.042 | 0.124 | 1 | 0.128105722 | -1.473931188 | 0 | 0.191 | 1 | 0.128105722 | 0.03086466 |
| Zfp37 | 0.015573612 | -1.259867127 | 0.126 | 0.235 | 1 | 0.503038407 | 0.351165316 | 0.4 | 0.272 | 1 | 0.503038407 | 0.030904687 |
| Arl6ip1 | 0.015612997 | 0.236685639 | 0.824 | 0.712 | 1 | 0.479933304 | -0.208757872 | 0.8 | 0.796 | 1 | 0.479933304 | 0.030982228 |
| Cdc6 | 0.015618502 | -1.201633861 | 0.025 | 0.1 | 1 | 0.716087628 | 0.973527789 | 0.1 | 0.068 | 1 | 0.716087628 | 0.030993066 |
| Nol4l | 0.015618932 | -1.807354922 | 0.059 | 0.147 | 1 | 0.117958402 | -2.101019165 | 0 | 0.201 | 1 | 0.117958402 | 0.030993912 |
| Clint1 | 0.015642263 | 0.158273447 | 0.992 | 1 | 1 | 0.936327987 | 0.116872191 | 1 | 1 | 1 | 0.936327987 | 0.031039846 |
| Aqr | 0.015649449 | -0.874993639 | 0.319 | 0.424 | 1 | 0.070724961 | -1.274859841 | 0.3 | 0.559 | 1 | 0.070724961 | 0.031053993 |
| Oaf | 0.015652811 | -0.454565863 | 0.773 | 0.865 | 1 | 0.110464599 | -0.930723433 | 0.7 | 0.79 | 1 | 0.110464599 | 0.031060612 |
| Svep1 | 0.083382591 | -0.114678604 | 0.252 | 0.371 | 1 | 0.015671358 | -2.782977992 | 0.3 | 0.605 | 1 | 0.083382591 | 0.031097124 |
| Ddx31 | 0.015685236 | -1.201633861 | 0.101 | 0.206 | 1 | 0.475116342 | -0.505640048 | 0.1 | 0.182 | 1 | 0.475116342 | 0.031124446 |
| Rel | 0.015693221 | -0.802767653 | 0.286 | 0.412 | 1 | 0.34749907 | -0.953621646 | 0.3 | 0.404 | 1 | 0.34749907 | 0.031140165 |
| Mios | 0.015693431 | -0.920363884 | 0.168 | 0.288 | 1 | 0.782135817 | -0.178475305 | 0.2 | 0.225 | 1 | 0.782135817 | 0.031140578 |
| Tomm40l | 0.015694536 | -1.555816155 | 0.034 | 0.112 | 1 | 0.907531274 | 0.695993813 | 0.1 | 0.111 | 1 | 0.907531274 | 0.031142754 |
| Scrn2 | 0.015694536 | -1.555816155 | 0.034 | 0.112 | 1 | 0.518235124 | 1.625604485 | 0.1 | 0.052 | 1 | 0.518235124 | 0.031142754 |

| | | | | | | | | | | | | |
|---|---|---|---|---|---|---|---|---|---|---|---|---|
| Nudt9 | 0.015711003 | 0.411887509 | 0.639 | 0.506 | 1 | 0.053000336 | -1.178475305 | 0.1 | 0.444 | 1 | 0.053000336 | 0.031175171 |
| Brms1 | 0.92377283 | -0.209792385 | 0.319 | 0.294 | 1 | 0.015714926 | 1.329865914 | 0.6 | 0.262 | 1 | 0.92377283 | 0.031182892 |
| Polr3b | 0.015734176 | -1.150559677 | 0.218 | 0.335 | 1 | 0.175882306 | -1.616284112 | 0.2 | 0.377 | 1 | 0.175882306 | 0.031220787 |
| Map2k2 | 0.015740348 | -0.377921202 | 0.79 | 0.806 | 1 | 0.63348327 | -0.121629444 | 0.7 | 0.852 | 1 | 0.63348327 | 0.031232938 |
| Lztr1 | 0.015776786 | -0.731587414 | 0.311 | 0.429 | 1 | 0.27065448 | -0.700896339 | 0.2 | 0.355 | 1 | 0.27065448 | 0.031304664 |
| Slc39a1 | 0.015782939 | -0.575408194 | 0.538 | 0.635 | 1 | 0.389326869 | -0.564533737 | 0.6 | 0.691 | 1 | 0.389326869 | 0.031316777 |
| Ubac2 | 0.015790599 | -1.046141782 | 0.395 | 0.488 | 1 | 0.058142449 | -1.107233223 | 0.3 | 0.605 | 1 | 0.058142449 | 0.031331854 |
| Hus1 | 0.015815691 | -1.678071905 | 0.05 | 0.135 | 1 | 0.508023035 | 0.930459067 | 0.2 | 0.123 | 1 | 0.508023035 | 0.031381247 |
| BC002059 | 0.01581689 | -1.963474124 | 0.05 | 0.135 | 1 | 0.241812663 | 1.14345279 | 0.2 | 0.086 | 1 | 0.241812663 | 0.031383606 |
| Spock2 | 0.395375356 | -0.267690768 | 0.63 | 0.529 | 1 | 0.015818945 | -3.053540455 | 0 | 0.389 | 1 | 0.395375356 | 0.031387652 |
| 1700016P03Rik | 0.015832096 | -2.424026283 | 0.017 | 0.082 | 1 | 0.577172475 | 0.494359952 | 0 | 0.031 | 1 | 0.577172475 | 0.031413536 |
| Iqgap2 | 0.015841309 | -0.301002256 | 0.084 | 0.194 | 1 | 0.422949675 | 0.249737583 | 0.1 | 0.228 | 1 | 0.422949675 | 0.031431671 |
| Gm5165 | 0.046253197 | -1.402964667 | 0.059 | 0.129 | 1 | 0.015883413 | 1.602884409 | 0.4 | 0.127 | 1 | 0.046253197 | 0.031514543 |
| Rbm34 | 0.01588443 | -1 | 0.235 | 0.347 | 1 | 0.054732499 | -2.152003093 | 0 | 0.278 | 1 | 0.054732499 | 0.031516545 |
| Tubgcp4 | 0.015888184 | -1.070389328 | 0.202 | 0.312 | 1 | 0.922160721 | -0.111361109 | 0.3 | 0.256 | 1 | 0.922160721 | 0.031523933 |
| Sash1 | 0.015922894 | 0.132986896 | 0.992 | 0.912 | 1 | 0.377784045 | 0.153436879 | 1 | 0.988 | 1 | 0.377784045 | 0.03159225 |
| Tsc1 | 0.015932347 | -0.903570237 | 0.353 | 0.453 | 1 | 0.616687907 | 0.304733697 | 0.6 | 0.519 | 1 | 0.616687907 | 0.031610854 |
| Drg2 | 0.015943735 | -0.807354922 | 0.193 | 0.318 | 1 | 0.131042127 | -1.111361109 | 0.1 | 0.321 | 1 | 0.131042127 | 0.031633267 |
| Gm56501 | 0.015984949 | -2.359895945 | 0.017 | 0.082 | 1 | 0.324927614 | -0.374395515 | 0 | 0.09 | 1 | 0.324927614 | 0.031714379 |
| Nr6a1os | 0.01598545 | -2.359895945 | 0.017 | 0.082 | 1 | 0.490317433 | 1.263034406 | 0.1 | 0.049 | 1 | 0.490317433 | 0.031715366 |
| 4930578M01Rik | 0.01598545 | -2.424026283 | 0.017 | 0.082 | 1 | 0.460521625 | 0.769994395 | 0 | 0.052 | 1 | 0.460521625 | 0.031715366 |
| Gm19409 | 0.015985617 | -2.655351829 | 0.017 | 0.082 | 1 | 0.629774271 | 0.602884409 | 0.2 | 0.139 | 1 | 0.629774271 | 0.031715695 |
| Gm44127 | 0.016022297 | 2.251538767 | 0.076 | 0.018 | 1 | 0.051044361 | 2.558490289 | 0.1 | 0.015 | 1 | 0.051044361 | 0.031787881 |
| Trim3 | 0.016087612 | -1.194378045 | 0.193 | 0.306 | 1 | 0.722238496 | -0.353636955 | 0.3 | 0.33 | 1 | 0.722238496 | 0.031916413 |
| Bpifc | 0.016093522 | 0.514573173 | 0.244 | 0.129 | 1 | 0.505651086 | 0.432959407 | 0 | 0.043 | 1 | 0.505651086 | 0.031928042 |
| Gm33906 | 0.016127751 | 1.929610672 | 0.05 | 0.006 | 1 | 0.698532411 | 1.847996907 | 0 | 0.015 | 1 | 0.698532411 | 0.031995398 |
| Shox2 | 0.016138691 | -2.222392421 | 0.017 | 0.082 | 1 | 0.597837803 | 0.847996907 | 0 | 0.028 | 1 | 0.597837803 | 0.032016924 |
| Gm10638 | 0.016139027 | -2.14839184 | 0.017 | 0.082 | 1 | 0.19134412 | 2.558490289 | 0.1 | 0.028 | 1 | 0.19134412 | 0.032017586 |
| Gm26749 | 0.016139027 | -2.14839184 | 0.017 | 0.082 | 1 | 0.282344833 | -0.567040593 | 0 | 0.105 | 1 | 0.282344833 | 0.032017586 |
| Vps16 | 0.016143693 | -0.807354922 | 0.143 | 0.259 | 1 | 0.347398522 | 0.625604485 | 0.4 | 0.256 | 1 | 0.347398522 | 0.032026766 |
| Bahd1 | 0.016157243 | -1.655351829 | 0.042 | 0.124 | 1 | 0.719291956 | 0.602884409 | 0.2 | 0.151 | 1 | 0.719291956 | 0.03205343 |
| Cacnb1 | 0.016157243 | -1.758445322 | 0.042 | 0.124 | 1 | 0.27473057 | -0.654503434 | 0 | 0.108 | 1 | 0.27473057 | 0.03205343 |
| Gm12227 | 0.01616264 | -1.770829046 | 0.042 | 0.124 | 1 | 0.315892224 | -0.505640048 | 0 | 0.093 | 1 | 0.315892224 | 0.032064049 |
| Ago4 | 0.016165547 | -1.678071905 | 0.05 | 0.135 | 1 | 0.207866038 | -0.959358016 | 0 | 0.139 | 1 | 0.207866038 | 0.032069769 |
| Auts2 | 0.016165896 | -0.503062247 | 1 | 0.994 | 1 | 0.750807888 | -0.106582733 | 1 | 1 | 1 | 0.750807888 | 0.032070455 |
| Zfp35 | 0.016170473 | -0.980191519 | 0.067 | 0.165 | 1 | 0.893094737 | 0.432959407 | 0.1 | 0.111 | 1 | 0.893094737 | 0.032079461 |
| Prdx2 | 0.016172972 | -0.490376043 | 0.824 | 0.853 | 1 | 0.087146439 | -0.160881245 | 0.7 | 0.914 | 1 | 0.087146439 | 0.032084378 |
| Zfp143 | 0.016176162 | -1.163498732 | 0.084 | 0.182 | 1 | 0.80150616 | -0.178475305 | 0.2 | 0.219 | 1 | 0.80150616 | 0.032090656 |
| Pcnx4 | 0.016176828 | -0.985855818 | 0.269 | 0.382 | 1 | 0.048353125 | -1.906890596 | 0.1 | 0.401 | 1 | 0.048353125 | 0.032091966 |
| Tph2 | 0.350551956 | -1.970853654 | 0.017 | 0.035 | 1 | 0.016183705 | 2.558490289 | 0.2 | 0.04 | 1 | 0.350551956 | 0.032105498 |
| Zfp385a | 0.208482772 | 0.261807102 | 0.286 | 0.212 | 1 | 0.016189907 | 1.362570079 | 0.4 | 0.12 | 1 | 0.208482772 | 0.032117701 |
| Elf1 | 0.016199458 | -0.565253458 | 0.958 | 0.9 | 1 | 0.066091566 | -0.57120903 | 0.8 | 0.975 | 1 | 0.066091566 | 0.032136494 |
| Gon7 | 0.016207468 | -0.892602209 | 0.176 | 0.294 | 1 | 0.224354449 | -0.864721141 | 0.1 | 0.265 | 1 | 0.224354449 | 0.032152255 |
| Rad9b | 0.016240802 | -0.765534746 | 0.092 | 0.2 | 1 | 0.39838031 | 0.880418384 | 0.3 | 0.194 | 1 | 0.39838031 | 0.032217841 |
| Ccm2 | 0.016248951 | -0.942284502 | 0.345 | 0.453 | 1 | 0.063446152 | -1.408342847 | 0.2 | 0.5 | 1 | 0.063446152 | 0.032233873 |
| Lman2l | 0.016257764 | -1.046141782 | 0.151 | 0.265 | 1 | 0.77751483 | 0.87108052 | 0.3 | 0.269 | 1 | 0.77751483 | 0.032251212 |
| Nabp2 | 0.016259002 | -0.583037624 | 0.319 | 0.447 | 1 | 0.393026101 | -0.536666944 | 0.3 | 0.404 | 1 | 0.393026101 | 0.032253649 |
| Tead3 | 0.016268322 | -0.769219793 | 0.134 | 0.253 | 1 | 0.404832074 | -0.473931188 | 0.1 | 0.204 | 1 | 0.404832074 | 0.032271985 |
| Eif1b | 0.116198306 | -0.253253385 | 0.807 | 0.782 | 1 | 0.016278696 | -1.098621997 | 0.4 | 0.855 | 1 | 0.116198306 | 0.032292397 |
| Gm41409 | 0.016292475 | -2.14839184 | 0.017 | 0.082 | 1 | 0.585423461 | 1.902444691 | 0.1 | 0.062 | 1 | 0.585423461 | 0.032319504 |
| Gm12992 | 0.016292559 | -2.070389328 | 0.017 | 0.082 | 1 | 0.52940302 | 1.210566986 | 0.1 | 0.052 | 1 | 0.52940302 | 0.032319671 |
| Tmem86a | 0.016292559 | -2.070389328 | 0.017 | 0.082 | 1 | 0.290449498 | -0.536666944 | 0 | 0.102 | 1 | 0.290449498 | 0.032319671 |
| Pglyrp1 | 0.01629426 | -1.269698136 | 0.101 | 0.206 | 1 | 0.40017468 | -0.581990934 | 0.1 | 0.207 | 1 | 0.40017468 | 0.032323018 |
| Sned1 | 0.016332785 | 0.503517984 | 0.664 | 0.529 | 1 | 0.323956808 | -0.283574287 | 0.9 | 0.71 | 1 | 0.323956808 | 0.032398809 |
| LTO1 | 0.016368075 | -0.857395605 | 0.109 | 0.218 | 1 | 0.857798779 | 0.494359952 | 0.1 | 0.117 | 1 | 0.857798779 | 0.032468236 |
| Dazap2 | 0.588949592 | -0.240741731 | 0.647 | 0.618 | 1 | 0.016384927 | -1.183711953 | 0.3 | 0.701 | 1 | 0.588949592 | 0.032501388 |
| Mrps9 | 0.016390534 | -0.631277694 | 0.261 | 0.394 | 1 | 0.308601221 | 0.762421175 | 0.5 | 0.383 | 1 | 0.308601221 | 0.032512419 |
| Snx33 | 0.016398057 | -1.655351829 | 0.059 | 0.147 | 1 | 0.406070607 | -0.473931188 | 0.1 | 0.204 | 1 | 0.406070607 | 0.032527218 |
| Borcs5 | 0.016398509 | -1.733354341 | 0.059 | 0.147 | 1 | 0.232319525 | -0.736965594 | 0 | 0.127 | 1 | 0.232319525 | 0.032528106 |
| Gm52698 | 0.016420704 | 3.099535674 | 0.034 | 0 | 1 | 0.050189208 | 3.017921908 | 0.1 | 0.015 | 1 | 0.050189208 | 0.032571768 |
| Kcna1 | 0.016420704 | 3.099535674 | 0.034 | 0 | 1 | 0.19134412 | 2.558490289 | 0.1 | 0.028 | 1 | 0.19134412 | 0.032571768 |
| Muc16 | 0.016420704 | 3.099535674 | 0.034 | 0 | 1 | 0.354000578 | 0.263034406 | 0 | 0.08 | 1 | 0.354000578 | 0.032571768 |
| Fam241b | 0.01642201 | 2.836501268 | 0.034 | 0 | 1 | 0.149795061 | 2.847996907 | 0.1 | 0.025 | 1 | 0.149795061 | 0.032574339 |
| Ildr1 | 0.01642201 | 2.836501268 | 0.034 | 0 | 1 | 0.619961444 | 1.558490289 | 0 | 0.025 | 1 | 0.619961444 | 0.032574339 |
| Med1 | 0.016424202 | -0.734454375 | 0.429 | 0.506 | 1 | 0.271325875 | -0.374395515 | 0.5 | 0.633 | 1 | 0.271325875 | 0.03257865 |
| Synj2bp | 0.056570884 | -0.508075566 | 0.739 | 0.718 | 1 | 0.016430899 | -1.214738849 | 0.5 | 0.812 | 1 | 0.056570884 | 0.032591823 |
| Lag3 | 0.016446283 | -2.424026283 | 0.017 | 0.082 | 1 | 0.386500816 | 0.995554095 | 0.2 | 0.111 | 1 | 0.386500816 | 0.032622085 |
| Fam234b | 0.016446283 | -1.987927168 | 0.017 | 0.082 | 1 | 0.316137708 | 2.111031312 | 0.1 | 0.037 | 1 | 0.316137708 | 0.032622085 |
| Sft2d1 | 0.016451898 | -0.668648651 | 0.303 | 0.441 | 1 | 0.073010053 | -1.131825212 | 0.2 | 0.475 | 1 | 0.073010053 | 0.03263313 |
| Tpt1 | 0.12788103 | 0.122228641 | 1 | 1 | 1 | 0.01650761 | -0.73760665 | 1 | 1 | 1 | 0.12788103 | 0.032742718 |
| A430005L14Rik | 0.016523836 | -0.56515402 | 0.244 | 0.376 | 1 | 0.862256628 | 0.345496566 | 0.2 | 0.235 | 1 | 0.862256628 | 0.032774634 |
| Heatr6 | 0.016524837 | -1.173482821 | 0.202 | 0.312 | 1 | 0.219378476 | -0.91281543 | 0.1 | 0.269 | 1 | 0.219378476 | 0.032776604 |
| Acaca | 0.016531282 | -0.406475486 | 0.63 | 0.671 | 1 | 0.023246418 | 1.002546682 | 0.9 | 0.608 | 1 | 0.023246418 | 0.032789281 |
| Abcb1b | 0.016533245 | -1.222392421 | 0.084 | 0.182 | 1 | 0.375007864 | 0.453137289 | 0.2 | 0.105 | 1 | 0.375007864 | 0.032793142 |
| Mysm1 | 0.188493222 | -0.198328716 | 0.555 | 0.6 | 1 | 0.016537297 | -1.602664502 | 0.4 | 0.682 | 1 | 0.188493222 | 0.032801112 |
| 1700036A12Rik | 0.016561266 | -1.383547213 | 0.076 | 0.171 | 1 | 0.929007499 | 0.469485283 | 0.2 | 0.225 | 1 | 0.929007499 | 0.032848256 |
| Srsf11 | 0.016567104 | -0.221560657 | 1 | 1 | 1 | 0.045571101 | -0.482422061 | 1 | 0.997 | 1 | 0.045571101 | 0.032859739 |
| Fbxw2 | 0.016585502 | -0.599498594 | 0.437 | 0.547 | 1 | 0.027461704 | -1.441509711 | 0.3 | 0.602 | 1 | 0.027461704 | 0.032895925 |
| Stx8 | 0.022888223 | -0.95836271 | 0.387 | 0.441 | 1 | 0.016592222 | -1.985830227 | 0.2 | 0.562 | 1 | 0.022888223 | 0.032909143 |
| C130050O18Rik | 0.016600098 | -1.900464326 | 0.017 | 0.082 | 1 | 0.290412643 | -0.505640048 | 0 | 0.102 | 1 | 0.290412643 | 0.032924633 |
| 1700087M22Rik | 0.511005543 | -0.485426827 | 0.008 | 0.018 | 1 | 0.016609983 | 2.795529487 | 0.2 | 0.04 | 1 | 0.511005543 | 0.032944074 |
| B830012L14Rik | 0.676154148 | 0.257122885 | 0.353 | 0.335 | 1 | 0.016642864 | 2.200596697 | 0.8 | 0.454 | 1 | 0.676154148 | 0.033008743 |
| Mapkap1 | 0.016652964 | -0.50738608 | 0.689 | 0.753 | 1 | 0.117952265 | -0.801274627 | 0.6 | 0.806 | 1 | 0.117952265 | 0.033028607 |
| Pdlim2 | 0.016673849 | -1.68182404 | 0.092 | 0.188 | 1 | 0.586685206 | 0.210566986 | 0.3 | 0.207 | 1 | 0.586685206 | 0.033069681 |
| Trmt10c | 0.036068499 | -0.821029859 | 0.294 | 0.388 | 1 | 0.016696233 | -2.031926641 | 0.1 | 0.485 | 1 | 0.036068499 | 0.033113702 |
| Gse1 | 0.016701424 | 0.642161228 | 0.647 | 0.518 | 1 | 0.11244261 | 1.472957475 | 0.8 | 0.617 | 1 | 0.11244261 | 0.03312391 |
| Gm37233 | 0.01670641 | -3.070389328 | 0 | 0.047 | 1 | 0.643850855 | 2.017921908 | 0 | 0.022 | 1 | 0.643850855 | 0.033133716 |
| Srsf12 | 0.01670641 | -2.807354922 | 0 | 0.047 | 1 | 0.355415506 | 2.210566986 | 0.1 | 0.04 | 1 | 0.355415506 | 0.033133716 |
| Dph2 | 0.01670641 | -2.807354922 | 0 | 0.047 | 1 | 0.557744661 | 1.210566986 | 0 | 0.034 | 1 | 0.557744661 | 0.033133716 |
| Zfp28 | 0.01670641 | -2.807354922 | 0 | 0.047 | 1 | 0.274256184 | 2.31748219 | 0.1 | 0.034 | 1 | 0.274256184 | 0.033133716 |
| 6530437J22Rik | 0.01670641 | -2.807354922 | 0 | 0.047 | 1 | 0.643842184 | 1.695993813 | 0 | 0.022 | 1 | 0.643842184 | 0.033133716 |
| Rab32 | 0.01670641 | -2.807354922 | 0 | 0.047 | 1 | 0.278846016 | 1.625604485 | 0.1 | 0.034 | 1 | 0.278846016 | 0.033133716 |
| Hsf5 | 0.01670641 | -2.807354922 | 0 | 0.047 | 1 | 0.643846086 | 1.31748219 | 0 | 0.022 | 1 | 0.643846086 | 0.033133716 |
| 1700001P01Rik | 0.01670641 | -2.807354922 | 0 | 0.047 | 1 | 0.276546558 | 2.017921908 | 0.1 | 0.034 | 1 | 0.276546558 | 0.033133716 |
| Tfap4 | 0.01670641 | -2.807354922 | 0 | 0.047 | 1 | 0.669851164 | 1.695993813 | 0 | 0.019 | 1 | 0.669851164 | 0.033133716 |
| Tcf24 | 0.016709933 | -2.944858446 | 0 | 0.047 | 1 | 0.573201119 | 1.159940913 | 0.1 | 0.056 | 1 | 0.573201119 | 0.033140643 |
| 1700019D03Rik | 0.016709933 | -3.070389328 | 0 | 0.047 | 1 | 0.730749233 | 2.210566986 | 0 | 0.012 | 1 | 0.730749233 | 0.033140643 |
| Gm29055 | 0.016709933 | -3.070389328 | 0 | 0.047 | 1 | 0.643835681 | 1.847996907 | 0 | 0.022 | 1 | 0.643835681 | 0.033140643 |

| | | | | | | | | | | | | |
|---|---|---|---|---|---|---|---|---|---|---|---|---|
| Opn3 | 0.016709933 | -2.944858446 | 0 | 0.047 | 1 | 0.669848329 | 1.31748219 | 0 | 0.019 | 1 | 0.669848329 | 0.033140643 |
| Pdzk1 | 0.016709933 | -2.944858446 | 0 | 0.047 | 1 | 0.619961444 | 1.432959407 | 0 | 0.025 | 1 | 0.619961444 | 0.033140643 |
| Zfp14 | 0.016709933 | -2.944858446 | 0 | 0.047 | 1 | 0.320995017 | 2.017921908 | 0.1 | 0.037 | 1 | 0.320995017 | 0.033140643 |
| 6330403K07Rik | 0.016709933 | -3.070389328 | 0 | 0.047 | 1 | 0.619971082 | 0.930459067 | 0 | 0.025 | 1 | 0.619971082 | 0.033140643 |
| Cd200r1 | 0.016709933 | -3.733354341 | 0 | 0.047 | 1 | 0.557744661 | 1.111031312 | 0 | 0.034 | 1 | 0.557744661 | 0.033140643 |
| Gm2350 | 0.016709933 | -3.392317423 | 0 | 0.047 | 1 | 0.619961444 | 1.432959407 | 0 | 0.025 | 1 | 0.619961444 | 0.033140643 |
| Gm32342 | 0.016709933 | -2.944858446 | 0 | 0.047 | 1 | 0.35375985 | -0.191531458 | 0 | 0.08 | 1 | 0.35375985 | 0.033140643 |
| Nudt11 | 0.016709933 | -3.070389328 | 0 | 0.047 | 1 | 0.396844932 | 0.31748219 | 0 | 0.068 | 1 | 0.396844932 | 0.033140643 |
| Col26a1 | 0.016712281 | -3.807354922 | 0 | 0.047 | 1 | 0.505649381 | 0.210566986 | 0 | 0.043 | 1 | 0.505649381 | 0.033145261 |
| Odad2 | 0.016712281 | -3.292781749 | 0 | 0.047 | 1 | 0.619961444 | 1.558490289 | 0 | 0.025 | 1 | 0.619961444 | 0.033145261 |
| Gm43111 | 0.016712449 | -3.292781749 | 0 | 0.047 | 1 | 0.025539196 | 2.210566986 | 0.2 | 0.043 | 1 | 0.025539196 | 0.033145591 |
| Car11 | 0.016712449 | -3.070389328 | 0 | 0.047 | 1 | 0.505605027 | 0.847996907 | 0 | 0.043 | 1 | 0.505605027 | 0.033145591 |
| Vipr2 | 0.016713455 | -3.292781749 | 0 | 0.047 | 1 | 0.073299054 | 1.210566986 | 0.2 | 0.059 | 1 | 0.073299054 | 0.033147571 |
| Nudt10 | 0.016713455 | -3.292781749 | 0 | 0.047 | 1 | 0.730749233 | 2.432959407 | 0 | 0.012 | 1 | 0.730749233 | 0.033147571 |
| Ppm1e | 0.016713958 | -3.87774425 | 0 | 0.047 | 1 | 0.597841551 | 0.111031312 | 0 | 0.028 | 1 | 0.597841551 | 0.03314856 |
| Tmem26 | 0.016714126 | -3.185866545 | 0 | 0.047 | 1 | 0.557765566 | 0.625604485 | 0 | 0.034 | 1 | 0.557765566 | 0.03314889 |
| Frmpd4 | 0.016714126 | -4.842978832 | 0 | 0.047 | 1 | 0.050189208 | 1.558490289 | 0.1 | 0.015 | 1 | 0.050189208 | 0.03314889 |
| Sema6c | 0.016715804 | -2.655351829 | 0 | 0.047 | 1 | 0.619953009 | 1.695993813 | 0 | 0.025 | 1 | 0.619953009 | 0.033152189 |
| Gm9908 | 0.016715804 | -2.655351829 | 0 | 0.047 | 1 | 0.73075158 | 2.695993813 | 0 | 0.012 | 1 | 0.73075158 | 0.033152189 |
| Gm35612 | 0.016715804 | -2.655351829 | 0 | 0.047 | 1 | 0.63522121 | 1.374065718 | 0.1 | 0.062 | 1 | 0.63522121 | 0.033152189 |
| Ppp1cc | 0.016735569 | -0.464053176 | 0.773 | 0.8 | 1 | 0.529432597 | -0.332575339 | 0.7 | 0.855 | 1 | 0.529432597 | 0.033191058 |
| Cchcr1 | 0.336029693 | -0.567888987 | 0.109 | 0.147 | 1 | 0.016754331 | 2.063725598 | 0.3 | 0.08 | 1 | 0.336029693 | 0.033227955 |
| Cdk5r1 | 0.016774092 | -1.584962501 | 0.042 | 0.124 | 1 | 0.88622897 | 0.720241359 | 0.1 | 0.123 | 1 | 0.88622897 | 0.033266815 |
| Nipsnap1 | 0.01677702 | -1.544320516 | 0.042 | 0.124 | 1 | 0.238442199 | -0.441509711 | 0 | 0.123 | 1 | 0.238442199 | 0.033272571 |
| Letmd1 | 0.01677702 | -1.485426827 | 0.042 | 0.124 | 1 | 0.259500407 | -0.408342847 | 0 | 0.114 | 1 | 0.259500407 | 0.033272571 |
| Pggt1b | 0.04914333 | -0.64689025 | 0.471 | 0.529 | 1 | 0.016791648 | -1.733622151 | 0.3 | 0.627 | 1 | 0.04914333 | 0.033301336 |
| Snw1 | 0.016807752 | -0.420789362 | 0.756 | 0.776 | 1 | 0.423402841 | -0.276698841 | 0.7 | 0.843 | 1 | 0.423402841 | 0.033333004 |
| Atp5h | 0.016821795 | -0.287228061 | 0.924 | 0.965 | 1 | 0.042543827 | -0.694152511 | 0.6 | 0.929 | 1 | 0.042543827 | 0.033360616 |
| Dact3 | 0.152006236 | -0.404808367 | 0.277 | 0.347 | 1 | 0.016824545 | -2.581990934 | 0 | 0.38 | 1 | 0.152006236 | 0.033366025 |
| Eif2b4 | 0.016844895 | -1.139930261 | 0.185 | 0.3 | 1 | 0.895916408 | 0.273760812 | 0.3 | 0.281 | 1 | 0.895916408 | 0.03340604 |
| Gm45669 | 0.016858079 | -1.01267383 | 0.092 | 0.194 | 1 | 0.946418123 | 0.159940913 | 0.2 | 0.179 | 1 | 0.946418123 | 0.033431963 |
| Mrpl3 | 0.016884308 | -0.665999073 | 0.336 | 0.453 | 1 | 0.950392401 | 0.27838237 | 0.4 | 0.426 | 1 | 0.950392401 | 0.033483537 |
| Dusp8 | 0.016908232 | 0.777607579 | 0.227 | 0.118 | 1 | 0.825295844 | 0.111031312 | 0.2 | 0.16 | 1 | 0.825295844 | 0.033530576 |
| C1qtnf6 | 0.01691354 | -0.600904045 | 0.345 | 0.482 | 1 | 0.564324445 | 0.466126271 | 0.5 | 0.448 | 1 | 0.564324445 | 0.033541012 |
| Nmt1 | 0.016934237 | -0.581004487 | 0.437 | 0.541 | 1 | 0.701174662 | 0.546616189 | 0.5 | 0.463 | 1 | 0.701174662 | 0.033581705 |
| Rxrb | 0.016939779 | -1.018859027 | 0.134 | 0.247 | 1 | 0.113927555 | -1.521236903 | 0 | 0.204 | 1 | 0.113927555 | 0.033592603 |
| Polr3a | 0.016955182 | -1.366782331 | 0.143 | 0.247 | 1 | 0.084215801 | -1.640289575 | 0 | 0.235 | 1 | 0.084215801 | 0.033622887 |
| Ephb6 | 0.237885589 | -1.070389328 | 0 | 0.012 | 1 | 0.017003024 | 2.695993813 | 0.2 | 0.04 | 1 | 0.237885589 | 0.033716946 |
| Tmem243 | 0.060276288 | -0.831201664 | 0.319 | 0.394 | 1 | 0.017004146 | -1.611434712 | 0.1 | 0.543 | 1 | 0.060276288 | 0.03371915 |
| Eif3i | 0.017021263 | -0.258764811 | 0.84 | 0.882 | 1 | 0.467761481 | -0.197368398 | 0.6 | 0.864 | 1 | 0.467761481 | 0.033752802 |
| Dennd11 | 0.017046044 | -1.87774425 | 0.017 | 0.082 | 1 | 0.539431168 | 0.695993813 | 0 | 0.037 | 1 | 0.539431168 | 0.033801521 |
| Zfp566 | 0.017096253 | -1.5360529 | 0.042 | 0.124 | 1 | 0.333164123 | 1.354956895 | 0.2 | 0.102 | 1 | 0.333164123 | 0.033900224 |
| Dhrs13 | 0.01711535 | 0.810029056 | 0.185 | 0.088 | 1 | 0.034126494 | 1.847996907 | 0.3 | 0.093 | 1 | 0.034126494 | 0.033937764 |
| Mtarc2 | 0.596623456 | -0.252329705 | 0.622 | 0.606 | 1 | 0.017144017 | -1.326374 | 0.2 | 0.62 | 1 | 0.596623456 | 0.033994117 |
| Arrdc4 | 0.017167368 | -1.216610069 | 0.176 | 0.282 | 1 | 0.031677821 | -1.894967428 | 0.1 | 0.438 | 1 | 0.031677821 | 0.034040018 |
| Thumpd1 | 0.017175436 | -0.844113472 | 0.437 | 0.494 | 1 | 0.441570377 | -0.185950425 | 0.3 | 0.463 | 1 | 0.441570377 | 0.034055876 |
| Xpnpep1 | 0.017184654 | 0.544946822 | 0.487 | 0.353 | 1 | 0.795651988 | -0.183711953 | 0.3 | 0.34 | 1 | 0.795651988 | 0.034073995 |
| Tnfrsf12a | 0.017186981 | 0.31301337 | 0.891 | 0.788 | 1 | 0.068178216 | -0.822801764 | 0.8 | 0.935 | 1 | 0.068178216 | 0.03407857 |
| Riok3 | 0.01719009 | -0.345344759 | 0.815 | 0.9 | 1 | 0.157792061 | -0.583429553 | 0.8 | 0.966 | 1 | 0.157792061 | 0.034084681 |
| F2r | 0.017210077 | -1.807354922 | 0.017 | 0.082 | 1 | 0.110543949 | -2.069540933 | 0 | 0.207 | 1 | 0.110543949 | 0.034123967 |
| Itga2b | 0.017214013 | -1.359895945 | 0.067 | 0.159 | 1 | 0.140029084 | 1.41305985 | 0.3 | 0.136 | 1 | 0.140029084 | 0.034131704 |
| Fam20a | 0.017236663 | -1.655351829 | 0.076 | 0.171 | 1 | 0.408830805 | -0.959358016 | 0 | 0.065 | 1 | 0.408830805 | 0.034176224 |
| Atp1b2 | 0.857206924 | -0.359895945 | 0.076 | 0.082 | 1 | 0.017239016 | 2.063725598 | 0.3 | 0.08 | 1 | 0.857206924 | 0.034180848 |
| Ppp2cb | 0.063594682 | -0.557576613 | 0.487 | 0.541 | 1 | 0.017250509 | -1.596787936 | 0.3 | 0.63 | 1 | 0.063594682 | 0.034203437 |
| A630023A22Rik | 0.017253371 | 1.070966521 | 0.143 | 0.059 | 1 | 0.232498909 | 2.210566986 | 0.1 | 0.031 | 1 | 0.232498909 | 0.034209063 |
| Dck | 0.017345623 | -1.485426827 | 0.05 | 0.135 | 1 | 0.515562449 | 0.847996907 | 0.2 | 0.123 | 1 | 0.515562449 | 0.034390375 |
| Tmtc4 | 0.716250379 | -0.126972856 | 0.176 | 0.194 | 1 | 0.017358482 | 1.270687978 | 0.5 | 0.176 | 1 | 0.716250379 | 0.034415647 |
| Slc26a11 | 0.017365816 | -1.070389328 | 0.126 | 0.235 | 1 | 0.494442096 | -0.516056664 | 0.2 | 0.281 | 1 | 0.494442096 | 0.03443006 |
| Dcun1d3 | 0.017374329 | -1.258016331 | 0.143 | 0.253 | 1 | 0.073507075 | 0.856034226 | 0.6 | 0.293 | 1 | 0.073507075 | 0.03444679 |
| Rabep2 | 0.017376531 | -0.975752454 | 0.151 | 0.265 | 1 | 0.308296208 | 1.14345279 | 0.3 | 0.191 | 1 | 0.308296208 | 0.034451117 |
| Gm11613 | 0.334866848 | -1.392317423 | 0.025 | 0.047 | 1 | 0.017396607 | 2.210566986 | 0.3 | 0.083 | 1 | 0.334866848 | 0.034490572 |
| Anapc2 | 0.01739885 | -0.787989597 | 0.319 | 0.429 | 1 | 0.421924268 | -0.518130992 | 0.4 | 0.466 | 1 | 0.421924268 | 0.034494981 |
| Ephb3 | 0.017399778 | -1.050211446 | 0.168 | 0.276 | 1 | 0.531511429 | -0.282201817 | 0.3 | 0.389 | 1 | 0.531511429 | 0.034496803 |
| Spopl | 0.017418727 | -0.642968104 | 0.303 | 0.424 | 1 | 0.929024096 | 0.190102883 | 0.4 | 0.472 | 1 | 0.929024096 | 0.034534042 |
| Myd88 | 0.017439202 | 0.407657969 | 0.521 | 0.353 | 1 | 0.677934436 | 0.296822719 | 0.3 | 0.414 | 1 | 0.677934436 | 0.034574278 |
| Gm26688 | 0.482805277 | -0.563429339 | 0.05 | 0.071 | 1 | 0.017442807 | 2.602884409 | 0.2 | 0.04 | 1 | 0.482805277 | 0.034581362 |
| Rnft1 | 0.017500685 | 0.253132025 | 0.714 | 0.535 | 1 | 0.847070014 | 0.222506599 | 0.6 | 0.58 | 1 | 0.847070014 | 0.034695097 |
| Slc9a9 | 0.017513352 | -1.572889668 | 0.16 | 0.265 | 1 | 0.403420904 | -0.924592597 | 0.3 | 0.407 | 1 | 0.403420904 | 0.034719986 |
| Zfp827 | 0.017547282 | -1.042238288 | 0.336 | 0.435 | 1 | 0.035301353 | 0.708066645 | 0.8 | 0.441 | 1 | 0.035301353 | 0.034786658 |
| Inpp5a | 0.140244224 | -0.634940369 | 0.697 | 0.682 | 1 | 0.017576119 | 1.154212136 | 0.9 | 0.765 | 1 | 0.140244224 | 0.034843318 |
| Pomt2 | 0.01758399 | -0.881355504 | 0.118 | 0.229 | 1 | 0.928011417 | 0.159940913 | 0.3 | 0.284 | 1 | 0.928011417 | 0.034858783 |
| Atr | 0.017605965 | -1.159198595 | 0.118 | 0.224 | 1 | 0.812597508 | -0.220482831 | 0.3 | 0.235 | 1 | 0.812597508 | 0.03490196 |
| Pacc1 | 0.017624353 | -1.572889668 | 0.017 | 0.082 | 1 | 0.25251941 | -0.596787936 | 0 | 0.117 | 1 | 0.25251941 | 0.034938088 |
| Tgfbrap1 | 0.017670847 | -1.315501826 | 0.067 | 0.159 | 1 | 0.415752776 | -0.408342847 | 0.1 | 0.201 | 1 | 0.415752776 | 0.035029434 |
| 4732471J01Rik | 0.372282342 | 0.389042291 | 0.168 | 0.129 | 1 | 0.017689225 | 2.930459067 | 0.3 | 0.086 | 1 | 0.372282342 | 0.035065542 |
| Jarid2 | 0.01769224 | -0.247383903 | 0.966 | 0.988 | 1 | 0.02802492 | -0.813478488 | 0.8 | 0.991 | 1 | 0.02802492 | 0.035071464 |
| Ppp2r2b | 0.017736905 | -4.040015679 | 0.008 | 0.065 | 1 | 0.290647173 | -1.640289575 | 0 | 0.102 | 1 | 0.290647173 | 0.035159211 |
| Col20a1 | 0.017765881 | -1.317699381 | 0.252 | 0.347 | 1 | 0.295421429 | -1.008878151 | 0.3 | 0.426 | 1 | 0.295421429 | 0.035216136 |
| Apbb1 | 0.017766057 | -1.240314329 | 0.042 | 0.124 | 1 | 0.057169256 | 1.130396637 | 0.4 | 0.157 | 1 | 0.057169256 | 0.035216481 |
| Krt17 | 0.017814231 | -1.572889668 | 0.05 | 0.135 | 1 | 0.601513634 | 0.185031894 | 0.1 | 0.16 | 1 | 0.601513634 | 0.035311116 |
| Nhsl1 | 0.017815567 | 1.070966521 | 0.134 | 0.053 | 1 | 1 | 0.591657153 | 0.1 | 0.099 | 1 | 1 | 0.035313739 |
| Dgat1 | 0.375511003 | -0.246239163 | 0.218 | 0.265 | 1 | 0.017822784 | 1.494359952 | 0.5 | 0.198 | 1 | 0.375511003 | 0.035327916 |
| Faxc | 0.042490194 | -2.070389328 | 0.042 | 0.106 | 1 | 0.017826654 | 2.245332404 | 0.3 | 0.086 | 1 | 0.042490194 | 0.035335518 |
| Arid3b | 0.01783525 | -2.807354922 | 0.008 | 0.065 | 1 | 0.649466592 | 0.660369903 | 0.1 | 0.062 | 1 | 0.649466592 | 0.035352405 |
| Dusp18 | 0.017845461 | -1.015046294 | 0.353 | 0.447 | 1 | 0.946950815 | -0.152003093 | 0.5 | 0.485 | 1 | 0.946950815 | 0.035372462 |
| Fryl | 0.017859177 | -0.748461233 | 0.655 | 0.729 | 1 | 0.7581188 | 0.168256181 | 0.7 | 0.747 | 1 | 0.7581188 | 0.035399403 |
| Hivep1 | 0.017868176 | -0.780882711 | 0.395 | 0.494 | 1 | 0.700780393 | 0.249737583 | 0.6 | 0.633 | 1 | 0.700780393 | 0.035417079 |
| Tubd1 | 0.121785453 | -0.900464326 | 0.059 | 0.112 | 1 | 0.017891477 | 2.432959407 | 0.2 | 0.04 | 1 | 0.121785453 | 0.035462849 |
| Coch | 0.610731388 | 2.554101537 | 0.109 | 0.094 | 1 | 0.017891477 | 2.017921908 | 0.2 | 0.04 | 1 | 0.610731388 | 0.035462849 |
| Gm20682 | 0.035283565 | -2.070389328 | 0.017 | 0.071 | 1 | 0.017891548 | 2.280956314 | 0.2 | 0.04 | 1 | 0.035283565 | 0.035462989 |
| Prmt5 | 0.017909173 | -1.051024003 | 0.193 | 0.3 | 1 | 0.321816622 | 0.782705446 | 0.4 | 0.256 | 1 | 0.321816622 | 0.035497607 |
| Ndfip2 | 0.92009743 | -0.194717463 | 0.849 | 0.806 | 1 | 0.017935979 | -1.051476217 | 0.5 | 0.886 | 1 | 0.92009743 | 0.035550259 |
| Gucy1a2 | 0.017937179 | -3.070389328 | 0.008 | 0.065 | 1 | 0.861061884 | 0.135278859 | 0.1 | 0.114 | 1 | 0.861061884 | 0.035552615 |
| Chn2 | 0.017937417 | -3.292781749 | 0.008 | 0.065 | 1 | 0.385705156 | -0.567040593 | 0 | 0.071 | 1 | 0.385705156 | 0.035553084 |
| Rnf170 | 0.018008586 | -0.894232373 | 0.202 | 0.318 | 1 | 0.839979749 | 0.906413593 | 0.2 | 0.244 | 1 | 0.839979749 | 0.035692863 |
| Faf2 | 0.018012221 | -0.417255325 | 0.664 | 0.724 | 1 | 0.519926834 | 0.152107255 | 0.9 | 0.769 | 1 | 0.519926834 | 0.035700002 |
| 1700027I24Rik | 0.018034461 | -2.655351829 | 0.008 | 0.065 | 1 | 0.433710928 | -0.152003093 | 0 | 0.059 | 1 | 0.433710928 | 0.03574368 |
| Klhdc8b | 0.308171942 | -0.366782331 | 0.126 | 0.171 | 1 | 0.018035862 | 1.754887502 | 0.5 | 0.201 | 1 | 0.308171942 | 0.035746431 |

| | | | | | | | | | | | | |
|---|---|---|---|---|---|---|---|---|---|---|---|---|
| Zfp729a | 0.018041713 | -1.10433666 | 0.092 | 0.194 | 1 | 0.517738808 | -0.230005605 | 0.1 | 0.176 | 1 | 0.517738808 | 0.035757923 |
| Med4 | 0.018045136 | -0.857395605 | 0.218 | 0.341 | 1 | 0.123002435 | -1.161987182 | 0.1 | 0.327 | 1 | 0.123002435 | 0.035764645 |
| Thop1 | 0.018046045 | -1.292781749 | 0.059 | 0.147 | 1 | 0.238521584 | -0.536666944 | 0 | 0.123 | 1 | 0.238521584 | 0.035766431 |
| Zswim7 | 0.018080992 | 0.770912926 | 0.269 | 0.153 | 1 | 0.279641193 | 0.769994395 | 0.3 | 0.157 | 1 | 0.279641193 | 0.035835061 |
| Naa25 | 0.018081889 | -0.756728849 | 0.361 | 0.459 | 1 | 0.715850584 | -0.441509711 | 0.5 | 0.472 | 1 | 0.715850584 | 0.035836824 |
| Dctpp1 | 0.0180848 | -0.728283351 | 0.185 | 0.294 | 1 | 0.576796795 | 1.295455884 | 0.2 | 0.157 | 1 | 0.576796795 | 0.035842539 |
| Sulf2 | 0.018098374 | -0.490533718 | 0.319 | 0.465 | 1 | 0.323407149 | 0.446898598 | 0.2 | 0.373 | 1 | 0.323407149 | 0.035869197 |
| BC028528 | 0.018133172 | -2.87774425 | 0.008 | 0.065 | 1 | 0.557729726 | 1.31748219 | 0 | 0.034 | 1 | 0.557729726 | 0.035937533 |
| C79798 | 0.018133293 | -2.485426827 | 0.008 | 0.065 | 1 | 0.61040405 | 0.732519689 | 0.1 | 0.059 | 1 | 0.61040405 | 0.035937769 |
| Tmem161b | 0.018154794 | -0.987927168 | 0.21 | 0.324 | 1 | 0.789333988 | -0.49777793 | 0.3 | 0.302 | 1 | 0.789333988 | 0.035979991 |
| Spata1 | 0.01819132 | -0.892052087 | 0.353 | 0.476 | 1 | 0.101682586 | 0.828097349 | 0.6 | 0.349 | 1 | 0.101682586 | 0.036051716 |
| Celf2 | 0.018200362 | -1.707819249 | 0.05 | 0.135 | 1 | 0.525082983 | 0.512393875 | 0.3 | 0.238 | 1 | 0.525082983 | 0.036069471 |
| ENSMUSG00000121374 | 0.018230429 | -2.392317423 | 0.008 | 0.065 | 1 | 0.234627319 | 2.210566986 | 0.1 | 0.031 | 1 | 0.234627319 | 0.036128509 |
| Heph | 0.018230429 | -2.807354922 | 0.008 | 0.065 | 1 | 0.698529281 | 2.017921908 | 0 | 0.015 | 1 | 0.698529281 | 0.036128509 |
| Cpsf7 | 0.032304751 | -0.61233894 | 0.538 | 0.594 | 1 | 0.018234082 | -1.090602549 | 0.4 | 0.75 | 1 | 0.032304751 | 0.036135681 |
| Pgap2 | 0.018263013 | -0.629816737 | 0.319 | 0.441 | 1 | 0.388448578 | -0.12891948 | 0.3 | 0.451 | 1 | 0.388448578 | 0.036192488 |
| Yeats4 | 0.018269142 | -0.645298164 | 0.412 | 0.512 | 1 | 0.283662902 | 0.689450967 | 0.6 | 0.42 | 1 | 0.283662902 | 0.036204523 |
| Unc45a | 0.018288067 | -0.953575663 | 0.21 | 0.324 | 1 | 0.628709312 | -0.248864633 | 0.4 | 0.466 | 1 | 0.628709312 | 0.036241681 |
| Csgalnact2 | 0.018289416 | 0.38426515 | 0.748 | 0.641 | 1 | 0.649409958 | 0.650351148 | 0.3 | 0.522 | 1 | 0.649409958 | 0.03624433 |
| Natd1 | 0.656008248 | -0.185866545 | 0.101 | 0.118 | 1 | 0.018294981 | 1.481869008 | 0.4 | 0.127 | 1 | 0.656008248 | 0.036255255 |
| Bltp2 | 0.018299752 | -0.421861698 | 0.529 | 0.659 | 1 | 0.570585236 | -0.287262923 | 0.7 | 0.731 | 1 | 0.570585236 | 0.036264622 |
| Ttc27 | 0.018300709 | -0.721055076 | 0.143 | 0.253 | 1 | 0.128743172 | -2.080110175 | 0 | 0.191 | 1 | 0.128743172 | 0.036266502 |
| Cpt1b | 0.018326936 | -2.292781749 | 0.008 | 0.065 | 1 | 0.043548461 | 1.902444691 | 0.2 | 0.049 | 1 | 0.043548461 | 0.036317995 |
| Ankrd2 | 0.018326936 | -2.292781749 | 0.008 | 0.065 | 1 | 0.573200928 | 1.111031312 | 0.1 | 0.056 | 1 | 0.573200928 | 0.036317995 |
| Nox1 | 0.018331825 | 2.251538767 | 0.084 | 0.024 | 1 | 0.597832556 | 0.31748219 | 0 | 0.028 | 1 | 0.597832556 | 0.036327593 |
| 4930532G15Rik | 0.038974464 | -2.485426827 | 0 | 0.035 | 1 | 0.018349159 | 2.280956314 | 0.2 | 0.04 | 1 | 0.038974464 | 0.036361627 |
| Sema4a | 0.132718412 | -1.359895945 | 0.076 | 0.129 | 1 | 0.018396401 | 4.035199899 | 0.3 | 0.09 | 1 | 0.132718412 | 0.036454374 |
| Pecr | 0.044750927 | -1.137503524 | 0.109 | 0.194 | 1 | 0.018411144 | 2.602884409 | 0.3 | 0.086 | 1 | 0.044750927 | 0.036483319 |
| Wnt2b | 0.018422681 | -2.292781749 | 0.008 | 0.065 | 1 | 0.539445308 | 1.017921908 | 0 | 0.037 | 1 | 0.539445308 | 0.036505967 |
| Sync | 0.018422681 | -2.185866545 | 0.008 | 0.065 | 1 | 0.318562315 | 2.017921908 | 0.1 | 0.037 | 1 | 0.318562315 | 0.036505967 |
| A630066F11Rik | 0.018422681 | -2.185866545 | 0.008 | 0.065 | 1 | 0.698534498 | 2.432959407 | 0 | 0.015 | 1 | 0.698534498 | 0.036505967 |
| Mir17hg | 0.018422681 | -2.485426827 | 0.008 | 0.065 | 1 | 0.489940652 | 0.494359952 | 0 | 0.046 | 1 | 0.489940652 | 0.036505967 |
| Lst1 | 0.018422681 | -2.185866545 | 0.008 | 0.065 | 1 | 0.619961444 | 1.558490289 | 0 | 0.025 | 1 | 0.619961444 | 0.036505967 |
| Lmbr1 | 0.018458146 | -1.280607035 | 0.109 | 0.212 | 1 | 0.843691781 | -0.178475305 | 0.2 | 0.167 | 1 | 0.843691781 | 0.036575589 |
| Slc4a4 | 0.018483081 | -0.155450204 | 0.513 | 0.624 | 1 | 0.111005567 | -1.902927145 | 0.6 | 0.67 | 1 | 0.111005567 | 0.036624539 |
| Cops5 | 0.018529268 | -0.61810931 | 0.597 | 0.659 | 1 | 0.122999139 | -0.663902132 | 0.4 | 0.716 | 1 | 0.122999139 | 0.036715202 |
| Ighmbp2 | 0.018552486 | -2.070389328 | 0.008 | 0.065 | 1 | 0.47676286 | 1.930459067 | 0.1 | 0.049 | 1 | 0.47676286 | 0.036760778 |
| Ak5 | 0.018563401 | -2.439623138 | 0.025 | 0.094 | 1 | 0.646822603 | -0.111361109 | 0.2 | 0.136 | 1 | 0.646822603 | 0.036782203 |
| Ivns1abp | 0.018599986 | -0.550610996 | 0.933 | 0.935 | 1 | 0.231552825 | -0.379992787 | 1 | 0.963 | 1 | 0.231552825 | 0.036854012 |
| Jtb | 0.018614596 | -0.602071746 | 0.529 | 0.618 | 1 | 0.569196104 | -0.203181817 | 0.5 | 0.605 | 1 | 0.569196104 | 0.036882688 |
| Med28 | 0.018661682 | 0.288184364 | 0.782 | 0.671 | 1 | 0.896277144 | 0.274957573 | 0.6 | 0.731 | 1 | 0.896277144 | 0.036975105 |
| Stxbp3 | 0.01866765 | -0.774933444 | 0.37 | 0.471 | 1 | 0.25185924 | -0.606568957 | 0.3 | 0.481 | 1 | 0.25185924 | 0.036986818 |
| Mycbp | 0.018701791 | -0.944858446 | 0.311 | 0.412 | 1 | 0.662602694 | 0.737813989 | 0.3 | 0.269 | 1 | 0.662602694 | 0.037053825 |
| Ftsj1 | 0.018777826 | -0.851076299 | 0.168 | 0.282 | 1 | 0.759743961 | -0.152003093 | 0.2 | 0.231 | 1 | 0.759743961 | 0.037203044 |
| Gnpat | 0.018786061 | -1.179323699 | 0.109 | 0.212 | 1 | 0.379605454 | 0.964810572 | 0.3 | 0.191 | 1 | 0.379605454 | 0.037219205 |
| Mpdz | 0.018788924 | -0.840907482 | 0.378 | 0.471 | 1 | 0.569666793 | 0.19088662 | 0.6 | 0.494 | 1 | 0.569666793 | 0.037224824 |
| Nup98 | 0.018810108 | -0.458119481 | 0.866 | 0.876 | 1 | 0.570756022 | 0.278073805 | 1 | 0.957 | 1 | 0.570756022 | 0.037266396 |
| Gngt2 | 0.018812055 | 1.344648171 | 0.084 | 0.024 | 1 | 0.248078127 | 3.280956314 | 0.1 | 0.034 | 1 | 0.248078127 | 0.037270217 |
| Btbd8 | 0.164808002 | -1.600904045 | 0.017 | 0.047 | 1 | 0.01881665 | 2.14345279 | 0.2 | 0.04 | 1 | 0.164808002 | 0.037279233 |
| Clhc1 | 0.933403655 | -0.250961574 | 0.034 | 0.035 | 1 | 0.01881665 | 1.795529487 | 0.2 | 0.04 | 1 | 0.933403655 | 0.037279233 |
| Potegl | 0.145997965 | -0.163498732 | 0.034 | 0.076 | 1 | 0.018819383 | 0.789103218 | 0.3 | 0.077 | 1 | 0.145997965 | 0.037284596 |
| Top3b | 0.018833386 | -0.767826558 | 0.218 | 0.335 | 1 | 0.182627069 | -1.011825435 | 0.2 | 0.389 | 1 | 0.182627069 | 0.037312075 |
| Pex14 | 0.140205127 | -0.485426827 | 0.437 | 0.471 | 1 | 0.018834723 | -1.792649727 | 0.3 | 0.623 | 1 | 0.140205127 | 0.037314699 |
| Dcakd | 0.018848684 | -1.131143359 | 0.227 | 0.335 | 1 | 0.494997863 | 0.383715888 | 0.2 | 0.327 | 1 | 0.494997863 | 0.037342095 |
| Eri3 | 0.018863583 | -0.375802336 | 0.84 | 0.824 | 1 | 0.434007178 | 0.452539776 | 1 | 0.806 | 1 | 0.434007178 | 0.037371331 |
| Plekhg5 | 0.018905476 | -0.707819249 | 0.076 | 0.171 | 1 | 0.759702778 | 1.858050571 | 0.3 | 0.262 | 1 | 0.759702778 | 0.037453535 |
| Ankrd46 | 0.01890808 | -0.964594664 | 0.261 | 0.371 | 1 | 0.086369767 | -1.276698841 | 0.1 | 0.358 | 1 | 0.086369767 | 0.037458645 |
| Fer | 0.018921082 | -0.905140814 | 0.42 | 0.506 | 1 | 0.024658807 | -1.705454621 | 0.1 | 0.5 | 1 | 0.024658807 | 0.037484157 |
| Ftl1 | 0.064861609 | -0.303892464 | 1 | 0.976 | 1 | 0.018954458 | -0.851434287 | 1 | 1 | 1 | 0.064861609 | 0.037549645 |
| Cap2 | 0.018962418 | -2.222392421 | 0.008 | 0.065 | 1 | 0.698529281 | 2.017921908 | 0 | 0.015 | 1 | 0.698529281 | 0.037565262 |
| Scyl3 | 0.598178241 | -0.5360529 | 0.21 | 0.224 | 1 | 0.019050237 | 1.135278859 | 0.6 | 0.244 | 1 | 0.598178241 | 0.037737562 |
| Ccni | 0.019095161 | -0.310748296 | 0.857 | 0.865 | 1 | 0.199608562 | -0.239016847 | 0.8 | 0.892 | 1 | 0.199608562 | 0.037825698 |
| Utp15 | 0.01909643 | -0.77915803 | 0.227 | 0.341 | 1 | 0.848214617 | 0.249737583 | 0.2 | 0.235 | 1 | 0.848214617 | 0.037828186 |
| Fbxo33 | 0.019154981 | -0.924538461 | 0.303 | 0.412 | 1 | 0.606206378 | -0.101019165 | 0.6 | 0.429 | 1 | 0.606206378 | 0.037943049 |
| Wdr73 | 0.019189021 | -1.185866545 | 0.059 | 0.147 | 1 | 0.299444619 | 0.868174788 | 0.3 | 0.164 | 1 | 0.299444619 | 0.038009824 |
| Map3k11 | 0.019201777 | -1.185866545 | 0.05 | 0.135 | 1 | 0.531371197 | -0.152003093 | 0.1 | 0.173 | 1 | 0.531371197 | 0.038034846 |
| Ier2 | 0.019245391 | -0.137197771 | 0.723 | 0.765 | 1 | 0.645153982 | 1.936204574 | 0.6 | 0.855 | 1 | 0.645153982 | 0.038120397 |
| Polr2c | 0.091917486 | -0.478047297 | 0.555 | 0.6 | 1 | 0.019245739 | -1.48590383 | 0.3 | 0.627 | 1 | 0.091917486 | 0.038121079 |
| Lrriq3 | 0.695893386 | -0.222392421 | 0.017 | 0.024 | 1 | 0.019293204 | 2.017921908 | 0.2 | 0.04 | 1 | 0.695893386 | 0.038214181 |
| Mark4 | 0.019299637 | -1.023083613 | 0.227 | 0.329 | 1 | 0.046068581 | -1.743629324 | 0.1 | 0.41 | 1 | 0.046068581 | 0.038226797 |
| 1110017D15Rik | 0.019302825 | -2.185866545 | 0.025 | 0.094 | 1 | 0.841021853 | 0.591657153 | 0.1 | 0.08 | 1 | 0.841021853 | 0.038233051 |
| AW011738 | 0.019303069 | -2.008988783 | 0.025 | 0.094 | 1 | 0.989745362 | 0.432959407 | 0.1 | 0.099 | 1 | 0.989745362 | 0.03823353 |
| Rad23a | 0.026382297 | -0.629392602 | 0.513 | 0.576 | 1 | 0.019325922 | -1.640289575 | 0.2 | 0.562 | 1 | 0.026382297 | 0.038278352 |
| Gtpbp10 | 0.019329466 | -1.163498732 | 0.067 | 0.159 | 1 | 0.679886964 | 0.263034406 | 0.1 | 0.145 | 1 | 0.679886964 | 0.038285303 |
| Atp6v1h | 0.139096162 | -0.304578177 | 0.706 | 0.771 | 1 | 0.019346722 | -1.396551928 | 0.5 | 0.818 | 1 | 0.139096162 | 0.038319149 |
| Zfp992 | 0.019354662 | -1.029747343 | 0.084 | 0.182 | 1 | 0.299452751 | 0.973527789 | 0.3 | 0.164 | 1 | 0.299452751 | 0.03833472 |
| Gm27241 | 0.019384336 | -1.807354922 | 0.008 | 0.065 | 1 | 0.489954768 | 0.494359952 | 0 | 0.046 | 1 | 0.489954768 | 0.038392919 |
| Hacd3 | 0.787286123 | -0.276840205 | 0.353 | 0.341 | 1 | 0.019417863 | -2.031926641 | 0.1 | 0.472 | 1 | 0.787286123 | 0.038458673 |
| Atg7 | 0.019429347 | -1.070389328 | 0.176 | 0.282 | 1 | 0.728653451 | 0.504852326 | 0.3 | 0.253 | 1 | 0.728653451 | 0.038481195 |
| Nfs1 | 0.019435117 | -1.269698136 | 0.252 | 0.359 | 1 | 0.82908151 | 0.217946516 | 0.3 | 0.367 | 1 | 0.82908151 | 0.038492509 |
| Gm4876 | 0.067470085 | 0.399095955 | 0.21 | 0.124 | 1 | 0.019454684 | 1.276454922 | 0.3 | 0.083 | 1 | 0.067470085 | 0.038530883 |
| Gm10138 | 0.054972787 | 0.380272081 | 0.218 | 0.129 | 1 | 0.019463358 | 1.150025444 | 0.4 | 0.127 | 1 | 0.054972787 | 0.038547894 |
| Arnt | 0.061409121 | 0.190410074 | 0.824 | 0.694 | 1 | 0.019498123 | 0.803161258 | 1 | 0.812 | 1 | 0.061409121 | 0.03861607 |
| A330040F15Rik | 0.019507445 | 2.269460675 | 0.084 | 0.024 | 1 | 0.669850692 | 1.017921908 | 0 | 0.019 | 1 | 0.669850692 | 0.038634349 |
| Ttf2 | 0.019552878 | -1.87774425 | 0.025 | 0.094 | 1 | 0.453044326 | 1.432959407 | 0.1 | 0.046 | 1 | 0.453044326 | 0.038723441 |
| Zfp420 | 0.169429949 | -1.070389328 | 0.042 | 0.082 | 1 | 0.01960384 | 2.252387162 | 0.3 | 0.086 | 1 | 0.169429949 | 0.038823369 |
| Ctbp2 | 0.019652912 | -0.900464326 | 0.244 | 0.353 | 1 | 0.182588179 | -0.936274402 | 0.2 | 0.389 | 1 | 0.182588179 | 0.038919587 |
| Ccdc28b | 0.436562618 | 0.192645078 | 0.361 | 0.312 | 1 | 0.019676072 | 1.432959407 | 0.6 | 0.269 | 1 | 0.436562618 | 0.038964996 |
| Grwd1 | 0.01968544 | -1.292781749 | 0.092 | 0.188 | 1 | 0.721183814 | 0.769994395 | 0.2 | 0.154 | 1 | 0.721183814 | 0.038983363 |
| Mrpl50 | 0.019688807 | -0.679599375 | 0.412 | 0.494 | 1 | 0.023298886 | -1.970762779 | 0.1 | 0.46 | 1 | 0.023298886 | 0.038989965 |
| Arhgap6 | 0.019701746 | -1.660513534 | 0.151 | 0.253 | 1 | 0.748346392 | -0.628554982 | 0.4 | 0.423 | 1 | 0.748346392 | 0.039015332 |
| Aip | 0.019708291 | -0.799087306 | 0.252 | 0.371 | 1 | 0.429818951 | -0.201246612 | 0.2 | 0.333 | 1 | 0.429818951 | 0.039028165 |
| a | 0.019764125 | -0.867572445 | 0.126 | 0.229 | 1 | 0.952260597 | 1.637779199 | 0.2 | 0.219 | 1 | 0.952260597 | 0.039137629 |
| Myl6 | 0.019773795 | -0.306651213 | 0.966 | 0.988 | 1 | 0.023772295 | -0.572397803 | 0.9 | 0.997 | 1 | 0.023772295 | 0.039156587 |
| Arxes2 | 0.019776414 | -1.025995209 | 0.076 | 0.171 | 1 | 0.500527417 | 0.930459067 | 0.2 | 0.123 | 1 | 0.500527417 | 0.039161721 |
| Cand2 | 0.019787519 | -1.070389328 | 0.042 | 0.124 | 1 | 0.887063664 | 0.695993813 | 0.1 | 0.114 | 1 | 0.887063664 | 0.039183493 |
| Pnpla8 | 0.019798413 | 0.302028537 | 0.731 | 0.588 | 1 | 0.063001583 | 0.417458314 | 1 | 0.787 | 1 | 0.063001583 | 0.039204849 |
| Cfap141 | 0.019804301 | -1.807354922 | 0.025 | 0.094 | 1 | 0.952551372 | 0.808468542 | 0.1 | 0.093 | 1 | 0.952551372 | 0.039216392 |

| | | | | | | | | | | | | |
|---|---|---|---|---|---|---|---|---|---|---|---|---|
| Gng7 | 0.019804301 | -1.807354922 | 0.025 | 0.094 | 1 | 0.105692031 | 1.695993813 | 0.2 | 0.065 | 1 | 0.105692031 | 0.039216392 |
| Pnrc1 | 0.019818162 | 0.288913038 | 0.933 | 0.918 | 1 | 0.731047879 | 0.543346803 | 0.7 | 0.883 | 1 | 0.731047879 | 0.039243565 |
| Pcdhga6 | 0.175202541 | -0.970853654 | 0.034 | 0.071 | 1 | 0.019833713 | 1.695993813 | 0.3 | 0.08 | 1 | 0.175202541 | 0.03927405 |
| Slc38a4 | 0.376913723 | 0.183927861 | 0.311 | 0.259 | 1 | 0.019834129 | 1.159940913 | 0.3 | 0.08 | 1 | 0.376913723 | 0.039274866 |
| Rgs12 | 0.019871289 | -1.185866545 | 0.067 | 0.159 | 1 | 0.156530452 | -1.171902651 | 0 | 0.17 | 1 | 0.156530452 | 0.03934771 |
| Ccdc125 | 0.019879144 | -2.333423734 | 0.034 | 0.106 | 1 | 1 | -0.171902651 | 0.1 | 0.096 | 1 | 1 | 0.039363108 |
| Nek1 | 0.019883643 | -0.929033479 | 0.471 | 0.535 | 1 | 0.285419835 | -0.370095377 | 0.6 | 0.673 | 1 | 0.285419835 | 0.039371927 |
| Slc30a5 | 0.028135416 | -0.902444691 | 0.521 | 0.582 | 1 | 0.019891396 | -1.384176536 | 0.5 | 0.741 | 1 | 0.028135416 | 0.039387124 |
| Ccdc107 | 0.01989166 | -0.630857267 | 0.21 | 0.335 | 1 | 0.484034477 | 0.336097868 | 0.5 | 0.346 | 1 | 0.484034477 | 0.039387642 |
| Pfdn5 | 0.115184625 | -0.1711274 | 0.958 | 0.965 | 1 | 0.019916036 | -0.787591667 | 0.8 | 0.966 | 1 | 0.115184625 | 0.039435423 |
| Tmem141 | 0.019931159 | 1.035405336 | 0.21 | 0.112 | 1 | 0.298522759 | -0.374395515 | 0 | 0.099 | 1 | 0.298522759 | 0.039465066 |
| Zfp507 | 0.019937791 | -0.829381228 | 0.269 | 0.382 | 1 | 0.295947974 | 1.494359952 | 0.4 | 0.302 | 1 | 0.295947974 | 0.039478066 |
| Tmx3 | 0.020002773 | -0.568991544 | 0.521 | 0.624 | 1 | 0.143436031 | -0.933362807 | 0.6 | 0.701 | 1 | 0.143436031 | 0.039605435 |
| Zfhx4 | 0.289084068 | -1.771210358 | 0.462 | 0.424 | 1 | 0.020006346 | -2.284553824 | 0.3 | 0.639 | 1 | 0.289084068 | 0.039612438 |
| Cblb | 0.020009273 | -0.665844202 | 0.571 | 0.676 | 1 | 0.783718009 | -0.822181596 | 0.8 | 0.753 | 1 | 0.783718009 | 0.039618176 |
| Gm973 | 0.350631965 | -0.204245075 | 0.345 | 0.271 | 1 | 0.020034031 | 2.466126271 | 0.5 | 0.204 | 1 | 0.350631965 | 0.039666701 |
| Bloc1s2 | 0.020037611 | -1 | 0.143 | 0.253 | 1 | 0.66270713 | 0.111031312 | 0.1 | 0.148 | 1 | 0.66270713 | 0.039673716 |
| Cmtm6 | 0.285699056 | -0.510088881 | 0.218 | 0.271 | 1 | 0.02004611 | -2.357117523 | 0 | 0.364 | 1 | 0.285699056 | 0.039690374 |
| Timp2 | 0.088461309 | -0.397215062 | 0.992 | 0.959 | 1 | 0.020067694 | -0.787688639 | 1 | 0.985 | 1 | 0.088461309 | 0.039732676 |
| Camkmt | 0.020071859 | -1.051406225 | 0.218 | 0.335 | 1 | 0.035014383 | 1.093109404 | 0.6 | 0.262 | 1 | 0.035014383 | 0.039740839 |
| Nfatc2 | 0.020095682 | -1.59059167 | 0.588 | 0.618 | 1 | 0.054927727 | 1.100384068 | 1 | 0.611 | 1 | 0.054927727 | 0.039787527 |
| Cd74 | 0.020121937 | 1.69273746 | 0.21 | 0.112 | 1 | 0.085604792 | 1.393431043 | 0.2 | 0.059 | 1 | 0.085604792 | 0.039838981 |
| Cacna1e | 0.020132716 | -2.198144875 | 0.025 | 0.094 | 1 | 0.157517356 | -2.750262417 | 0 | 0.17 | 1 | 0.157517356 | 0.039860105 |
| Spsb3 | 0.020145147 | -1.14839184 | 0.143 | 0.247 | 1 | 0.478119538 | -0.267480311 | 0.2 | 0.296 | 1 | 0.478119538 | 0.039884468 |
| Ltv1 | 0.020150299 | -0.861990179 | 0.387 | 0.488 | 1 | 0.043794151 | -1.756865152 | 0.1 | 0.414 | 1 | 0.043794151 | 0.039894564 |
| Zfand2a | 0.020153853 | 0.757429697 | 0.471 | 0.365 | 1 | 0.937599174 | 0.100384068 | 0.4 | 0.383 | 1 | 0.937599174 | 0.039901529 |
| Rcc1l | 0.020167785 | -0.847996907 | 0.151 | 0.265 | 1 | 0.223688594 | -0.993305347 | 0.1 | 0.265 | 1 | 0.223688594 | 0.039928831 |
| Mtmr3 | 0.020195578 | -0.412495305 | 0.723 | 0.788 | 1 | 0.136720729 | 0.643737576 | 1 | 0.907 | 1 | 0.136720729 | 0.039983294 |
| Stard9 | 0.020239461 | -1.155278225 | 0.05 | 0.135 | 1 | 0.85484857 | 0.558490289 | 0.1 | 0.08 | 1 | 0.85484857 | 0.040069286 |
| Zfat | 0.020271349 | -1.485426827 | 0.134 | 0.241 | 1 | 0.091999737 | 0.791853829 | 0.5 | 0.238 | 1 | 0.091999737 | 0.04013177 |
| Tap2 | 0.020290757 | -0.900464326 | 0.118 | 0.224 | 1 | 0.087075739 | -1.840059087 | 0 | 0.231 | 1 | 0.087075739 | 0.040169798 |
| Mtap | 0.020319711 | -0.968208933 | 0.277 | 0.382 | 1 | 0.862746838 | 0.167065347 | 0.3 | 0.383 | 1 | 0.862746838 | 0.040226532 |
| Osbpl7 | 0.184695148 | -0.563429339 | 0.034 | 0.071 | 1 | 0.02032022 | 2.169925001 | 0.3 | 0.086 | 1 | 0.184695148 | 0.040227529 |
| Irf2bp1 | 0.02033031 | -1.091147888 | 0.176 | 0.282 | 1 | 0.177173579 | -0.989572629 | 0.2 | 0.392 | 1 | 0.177173579 | 0.040247298 |
| Gm27003 | 0.020375995 | -1.637429921 | 0.067 | 0.153 | 1 | 0.245753697 | -0.763437806 | 0 | 0.12 | 1 | 0.245753697 | 0.040336809 |
| Clip2 | 0.020389907 | -1.512907564 | 0.084 | 0.176 | 1 | 0.335191553 | -0.625934282 | 0.1 | 0.225 | 1 | 0.335191553 | 0.040364066 |
| Coprs | 0.020397766 | 0.882304957 | 0.269 | 0.159 | 1 | 0.233532583 | 0.973527789 | 0.3 | 0.151 | 1 | 0.233532583 | 0.040379463 |
| Herc2 | 0.020399708 | -0.67018687 | 0.739 | 0.724 | 1 | 0.335267243 | 0.4906013 | 0.8 | 0.901 | 1 | 0.335267243 | 0.040383268 |
| Adrm1b | 0.020407691 | -1.572889668 | 0.025 | 0.094 | 1 | 0.482051401 | 1.769994395 | 0.1 | 0.049 | 1 | 0.482051401 | 0.040398909 |
| Ikbke | 0.020486524 | -0.918386234 | 0.134 | 0.241 | 1 | 0.28556221 | 0.847996907 | 0.4 | 0.247 | 1 | 0.28556221 | 0.04055335 |
| Mtmr12 | 0.020505518 | -0.534895503 | 0.445 | 0.559 | 1 | 0.752863181 | 0.30874738 | 0.7 | 0.654 | 1 | 0.752863181 | 0.040590559 |
| Tpcn2 | 0.02052106 | -1.169925001 | 0.076 | 0.171 | 1 | 0.730367103 | -0.719043686 | 0.2 | 0.235 | 1 | 0.730367103 | 0.040621007 |
| Ccne1 | 0.020534782 | -2.163498732 | 0.034 | 0.106 | 1 | 0.489877116 | 0.930459067 | 0 | 0.046 | 1 | 0.489877116 | 0.040647888 |
| Rhbdl3 | 0.020534857 | -2.292781749 | 0.034 | 0.106 | 1 | 0.156972122 | -2.048167282 | 0 | 0.17 | 1 | 0.156972122 | 0.040648033 |
| Mdga1 | 0.637635167 | 0.478503918 | 0.269 | 0.306 | 1 | 0.020534953 | 1 | 0.6 | 0.244 | 1 | 0.637635167 | 0.040648222 |
| Efemp1 | 0.124705155 | -0.549557165 | 0.378 | 0.276 | 1 | 0.020536435 | -3.344021866 | 0 | 0.367 | 1 | 0.124705155 | 0.040651124 |
| Yme1l1 | 0.020554074 | -0.349990202 | 0.941 | 0.888 | 1 | 0.83997471 | -0.103093493 | 1 | 0.914 | 1 | 0.83997471 | 0.040685678 |
| Vsig10 | 0.020608072 | -1.268328706 | 0.21 | 0.312 | 1 | 0.205952144 | 0.393431043 | 0.6 | 0.355 | 1 | 0.205952144 | 0.040791451 |
| Mfsd14b | 0.020609866 | -0.803502597 | 0.429 | 0.518 | 1 | 0.589957006 | -0.381249186 | 0.7 | 0.673 | 1 | 0.589957006 | 0.040794965 |
| Acp2 | 0.02061826 | -0.784987109 | 0.269 | 0.382 | 1 | 0.973429527 | -0.127755547 | 0.4 | 0.355 | 1 | 0.973429527 | 0.040811407 |
| Brix1 | 0.020641904 | -0.567888987 | 0.639 | 0.671 | 1 | 0.038046715 | -1.108413825 | 0.4 | 0.694 | 1 | 0.038046715 | 0.04085772 |
| Akt1s1 | 0.020696595 | -0.834576391 | 0.303 | 0.412 | 1 | 0.411898533 | -0.465893869 | 0.3 | 0.401 | 1 | 0.411898533 | 0.040964841 |
| Abcf2 | 0.020722261 | -0.649251629 | 0.479 | 0.547 | 1 | 0.459291562 | -0.469918126 | 0.7 | 0.623 | 1 | 0.459291562 | 0.04101511 |
| Kcng2 | 0.020732695 | -1.555816155 | 0.025 | 0.094 | 1 | 0.862565643 | 1.063725598 | 0.1 | 0.083 | 1 | 0.862565643 | 0.041035544 |
| Gna11 | 0.020738949 | -0.367933264 | 0.765 | 0.776 | 1 | 0.267503152 | -0.362899876 | 0.6 | 0.784 | 1 | 0.267503152 | 0.041047795 |
| Cep68 | 0.020799347 | -1.306456686 | 0.109 | 0.206 | 1 | 0.474238801 | 0.31748219 | 0.3 | 0.188 | 1 | 0.474238801 | 0.041166082 |
| Gm14410 | 0.020852521 | -1.589763487 | 0.076 | 0.165 | 1 | 0.835630645 | 0.227844977 | 0.2 | 0.167 | 1 | 0.835630645 | 0.041270214 |
| Zfp629 | 0.020868468 | -1.807354922 | 0.034 | 0.106 | 1 | 0.165669061 | -1.048167282 | 0 | 0.164 | 1 | 0.165669061 | 0.041301443 |
| Eif4e2 | 0.020873497 | -0.361620626 | 0.664 | 0.741 | 1 | 0.275900025 | -0.487228011 | 0.6 | 0.735 | 1 | 0.275900025 | 0.041311292 |
| Ccdc8 | 0.020884505 | -1.070389328 | 0.076 | 0.171 | 1 | 0.689549261 | 0.111031312 | 0.1 | 0.142 | 1 | 0.689549261 | 0.041332848 |
| AW209491 | 0.020885975 | -0.891419187 | 0.143 | 0.253 | 1 | 0.338895732 | 0.828097349 | 0.3 | 0.173 | 1 | 0.338895732 | 0.041335725 |
| A930007I19Rik | 0.020898126 | -2.070389328 | 0.05 | 0.129 | 1 | 0.089652851 | 2.339850003 | 0.2 | 0.065 | 1 | 0.089652851 | 0.04135952 |
| Zdhhc21 | 0.020899711 | -0.860935962 | 0.21 | 0.329 | 1 | 0.623275304 | -0.121629444 | 0.2 | 0.272 | 1 | 0.623275304 | 0.041362624 |
| Birc2 | 0.02090451 | -0.614709844 | 0.328 | 0.459 | 1 | 0.601237473 | 0.874963954 | 0.4 | 0.315 | 1 | 0.601237473 | 0.041372021 |
| Rpl6 | 0.044369173 | -0.156151188 | 0.983 | 0.982 | 1 | 0.0209062 | -0.681396399 | 0.9 | 0.994 | 1 | 0.044369173 | 0.041375331 |
| Gm26771 | 0.020922683 | -3.090288885 | 0.042 | 0.118 | 1 | 0.35387306 | -0.408342847 | 0 | 0.08 | 1 | 0.35387306 | 0.041407607 |
| Hk1os | 0.0209547 | 1.63005039 | 0.092 | 0.029 | 1 | 0.669853054 | 2.210566986 | 0 | 0.019 | 1 | 0.669853054 | 0.041470301 |
| Pik3r1 | 0.020974547 | -0.404722444 | 0.882 | 0.906 | 1 | 0.247044374 | -0.364851742 | 0.8 | 0.966 | 1 | 0.247044374 | 0.041509163 |
| Zfp948 | 0.020994256 | -0.753199152 | 0.412 | 0.506 | 1 | 0.887255205 | -0.270323061 | 0.7 | 0.552 | 1 | 0.887255205 | 0.041547754 |
| Nek6 | 0.021042245 | -0.667033634 | 0.218 | 0.329 | 1 | 0.159098932 | 0.552541023 | 0.6 | 0.352 | 1 | 0.159098932 | 0.041641713 |
| Gm5113 | 0.021083044 | 1.614108846 | 0.092 | 0.029 | 1 | 0.561898516 | 1.494359952 | 0.1 | 0.056 | 1 | 0.561898516 | 0.041721593 |
| Lpar6 | 0.021100066 | 1.592575685 | 0.092 | 0.029 | 1 | 0.539444993 | 0.930459067 | 0 | 0.037 | 1 | 0.539444993 | 0.041754919 |
| Dancr | 0.021136723 | -1.378511623 | 0.109 | 0.206 | 1 | 0.185734253 | -1.026472211 | 0 | 0.151 | 1 | 0.185734253 | 0.041826684 |
| Dtl | 0.021153696 | -2.222392421 | 0.034 | 0.106 | 1 | 0.367977634 | 1.494359952 | 0.1 | 0.04 | 1 | 0.367977634 | 0.041859914 |
| Nmnat3 | 0.021172942 | 0.112474729 | 0.151 | 0.065 | 1 | 0.475644714 | 2.678071905 | 0.2 | 0.13 | 1 | 0.475644714 | 0.041897591 |
| Arl6ip6 | 0.021188375 | -0.797704752 | 0.37 | 0.471 | 1 | 0.444962761 | -0.473931188 | 0.3 | 0.414 | 1 | 0.444962761 | 0.041927803 |
| Coq4 | 0.021204099 | -1.807354922 | 0.034 | 0.106 | 1 | 0.925158547 | 0.732519689 | 0.1 | 0.09 | 1 | 0.925158547 | 0.041958585 |
| Matn3 | 0.276766482 | -0.776624763 | 0.672 | 0.659 | 1 | 0.021207517 | 1.859774074 | 0.7 | 0.472 | 1 | 0.276766482 | 0.041965276 |
| Kctd20 | 0.021315444 | -0.807354922 | 0.319 | 0.424 | 1 | 0.384065877 | -0.200170376 | 0.4 | 0.562 | 1 | 0.384065877 | 0.04217654 |
| Mlec | 0.021325868 | 0.269576577 | 0.916 | 0.788 | 1 | 0.172115815 | 0.61479773 | 0.8 | 0.824 | 1 | 0.172115815 | 0.042196944 |
| 6430590A07Rik | 0.021356538 | -2 | 0.05 | 0.129 | 1 | 0.06238814 | 1.432959407 | 0.3 | 0.105 | 1 | 0.06238814 | 0.042256974 |
| Uba6 | 0.021429108 | -1.010088818 | 0.37 | 0.453 | 1 | 0.19205316 | -0.694359423 | 0.4 | 0.583 | 1 | 0.19205316 | 0.042399009 |
| Fads1 | 0.021437733 | -0.751016938 | 0.303 | 0.424 | 1 | 0.55832397 | 0.678071905 | 0.4 | 0.336 | 1 | 0.55832397 | 0.042415889 |
| Gabarapl1 | 0.692963797 | 0.123658468 | 0.664 | 0.647 | 1 | 0.021444858 | -1.196397213 | 0.4 | 0.75 | 1 | 0.692963797 | 0.042429834 |
| Polh | 0.021482782 | -1.637429921 | 0.067 | 0.153 | 1 | 0.266983933 | -0.68251781 | 0 | 0.111 | 1 | 0.266983933 | 0.042504054 |
| Tmem115 | 0.021487983 | -0.731183242 | 0.235 | 0.359 | 1 | 0.437563803 | -0.325485914 | 0.3 | 0.432 | 1 | 0.437563803 | 0.042514232 |
| Usp5 | 0.021502182 | -0.850423644 | 0.235 | 0.347 | 1 | 0.216712143 | -0.85654721 | 0.2 | 0.373 | 1 | 0.216712143 | 0.04254202 |
| Rnf13 | 0.021502324 | -0.560714954 | 0.723 | 0.735 | 1 | 0.027519138 | -1.073777926 | 0.6 | 0.815 | 1 | 0.027519138 | 0.042542297 |
| 2900052L18Rik | 0.021544634 | -1.807354922 | 0.034 | 0.106 | 1 | 0.71040436 | 0.973527789 | 0.1 | 0.068 | 1 | 0.71040436 | 0.042625098 |
| Samd14 | 0.021544711 | -1.687060688 | 0.034 | 0.106 | 1 | 0.180048469 | -0.864721141 | 0 | 0.154 | 1 | 0.180048469 | 0.042625248 |
| Rnf11 | 0.027802598 | -0.303412476 | 0.933 | 0.906 | 1 | 0.021570438 | -0.820021334 | 0.8 | 0.966 | 1 | 0.027802598 | 0.042675592 |
| Ube2d1 | 0.021577216 | 0.501862121 | 0.555 | 0.429 | 1 | 0.706337045 | -0.380109166 | 0.4 | 0.414 | 1 | 0.706337045 | 0.042688857 |
| Ifngr1 | 0.021578455 | -0.719892081 | 0.395 | 0.494 | 1 | 0.426361336 | 0.374065718 | 0.3 | 0.574 | 1 | 0.426361336 | 0.04269128 |
| Cdyl | 0.021629786 | -0.957495271 | 0.202 | 0.312 | 1 | 0.067367442 | -0.91753784 | 0.1 | 0.438 | 1 | 0.067367442 | 0.042791724 |
| Prkaa1 | 0.021651458 | -0.739183419 | 0.252 | 0.371 | 1 | 0.242822251 | -0.152003093 | 0.2 | 0.414 | 1 | 0.242822251 | 0.04283413 |
| 4933433G15Rik | 0.338096132 | -0.807354922 | 0.025 | 0.047 | 1 | 0.021677845 | 2.695993813 | 0.2 | 0.043 | 1 | 0.338096132 | 0.042885761 |
| Dchs2 | 0.063185079 | -2.350497247 | 0.034 | 0.088 | 1 | 0.021681011 | 2.558490289 | 0.2 | 0.043 | 1 | 0.063185079 | 0.042891955 |
| Yars2 | 0.021696523 | -1.109917692 | 0.092 | 0.188 | 1 | 0.123774183 | -1.171902651 | 0 | 0.194 | 1 | 0.123774183 | 0.042922307 |

| | | | | | | | | | | | | |
|---|---|---|---|---|---|---|---|---|---|---|---|---|
| B630019K06Rik | 0.021713923 | 1.573466862 | 0.143 | 0.065 | 1 | 0.482051401 | 1.695993813 | 0.1 | 0.049 | 1 | 0.482051401 | 0.042956352 |
| A4galt | 0.021734298 | 0.447741664 | 0.672 | 0.559 | 1 | 0.665275551 | -0.611434712 | 0.9 | 0.701 | 1 | 0.665275551 | 0.042996216 |
| Sec16b | 0.021767132 | -0.587525015 | 0.361 | 0.488 | 1 | 0.811447445 | -0.199308808 | 0.3 | 0.318 | 1 | 0.811447445 | 0.043060456 |
| Zbtb41 | 0.021809348 | -0.822461814 | 0.269 | 0.376 | 1 | 0.492269068 | -0.524336142 | 0.4 | 0.451 | 1 | 0.492269068 | 0.043143048 |
| Zfp438 | 0.745171527 | -0.137503524 | 0.176 | 0.159 | 1 | 0.021824112 | 1.695993813 | 0.3 | 0.083 | 1 | 0.745171527 | 0.043171933 |
| Ttc39c | 0.021835839 | -1.770829046 | 0.059 | 0.141 | 1 | 0.290411047 | -0.625934282 | 0 | 0.102 | 1 | 0.290411047 | 0.043194875 |
| Dctn5 | 0.021850677 | -0.975752454 | 0.151 | 0.259 | 1 | 0.70692541 | 0.249737583 | 0.2 | 0.272 | 1 | 0.70692541 | 0.043223903 |
| Gsta4 | 0.021886639 | -0.622930351 | 0.067 | 0.159 | 1 | 0.186011351 | -1.210896782 | 0.1 | 0.284 | 1 | 0.186011351 | 0.043294254 |
| Eloc | 0.197983961 | -0.256692879 | 0.857 | 0.859 | 1 | 0.021930073 | -0.740301307 | 0.7 | 0.923 | 1 | 0.197983961 | 0.043379218 |
| Rad50 | 0.021953055 | -0.678071905 | 0.294 | 0.429 | 1 | 0.610590737 | 0.808468542 | 0.4 | 0.312 | 1 | 0.610590737 | 0.043424173 |
| P3h1 | 0.888794161 | -0.111361109 | 0.849 | 0.735 | 1 | 0.021970361 | 0.933199229 | 1 | 0.694 | 1 | 0.888794161 | 0.043458025 |
| Prim2 | 0.022003117 | -1.548436625 | 0.168 | 0.271 | 1 | 0.036402467 | 1.515421567 | 0.5 | 0.235 | 1 | 0.036402467 | 0.043522096 |
| Negr1 | 0.022004619 | -1.744161096 | 0.059 | 0.141 | 1 | 0.987083603 | 0.210566986 | 0.2 | 0.216 | 1 | 0.987083603 | 0.043525034 |
| Slc4a2 | 0.022004765 | -0.774933444 | 0.235 | 0.359 | 1 | 0.718793276 | -0.380109166 | 0.4 | 0.414 | 1 | 0.718793276 | 0.04352532 |
| Phc3 | 0.022008095 | -0.550277972 | 0.454 | 0.535 | 1 | 0.725765357 | -0.114528388 | 0.6 | 0.648 | 1 | 0.725765357 | 0.043531833 |
| Pank3 | 0.996370618 | -0.163498732 | 0.597 | 0.553 | 1 | 0.02205314 | -1.242605642 | 0.3 | 0.676 | 1 | 0.996370618 | 0.043619939 |
| Armc8 | 0.022054187 | -0.762267033 | 0.311 | 0.424 | 1 | 0.12324338 | -1.125036046 | 0.4 | 0.59 | 1 | 0.12324338 | 0.043621988 |
| Timm44 | 0.022080069 | -0.888391494 | 0.311 | 0.412 | 1 | 0.074957987 | 0.826301835 | 0.7 | 0.386 | 1 | 0.074957987 | 0.043672609 |
| Zfp367 | 0.372921629 | -0.603608254 | 0.193 | 0.229 | 1 | 0.022092829 | 2.257109572 | 0.5 | 0.213 | 1 | 0.372921629 | 0.043697566 |
| Slc25a26 | 0.170852045 | -1.008988783 | 0.05 | 0.094 | 1 | 0.022150916 | 2.467724825 | 0.3 | 0.093 | 1 | 0.170852045 | 0.043811169 |
| 4833419F23Rik | 0.022198884 | 1.215012891 | 0.092 | 0.029 | 1 | 0.698529281 | 1.695993813 | 0 | 0.015 | 1 | 0.698529281 | 0.043904977 |
| Itgb3 | 0.022203365 | -2.292781749 | 0.034 | 0.106 | 1 | 0.433698059 | -1.171902651 | 0 | 0.059 | 1 | 0.433698059 | 0.043913741 |
| Pola1 | 0.022223894 | -1.474455709 | 0.168 | 0.271 | 1 | 0.138893994 | 0.930459067 | 0.5 | 0.287 | 1 | 0.138893994 | 0.043953886 |
| Top1mt | 0.022234523 | -1.555816155 | 0.034 | 0.106 | 1 | 0.716086493 | 1.017921908 | 0.1 | 0.068 | 1 | 0.716086493 | 0.043974672 |
| Eif4ebp3 | 0.022234523 | -1.622930351 | 0.034 | 0.106 | 1 | 0.902406098 | 0.769994395 | 0.1 | 0.086 | 1 | 0.902406098 | 0.043974672 |
| Dnal1 | 0.022241264 | -1.191695624 | 0.126 | 0.229 | 1 | 0.652407973 | 0.223506042 | 0.3 | 0.222 | 1 | 0.652407973 | 0.043987854 |
| Snx9 | 0.022243027 | 0.408419621 | 0.891 | 0.765 | 1 | 0.321462036 | 0.242275846 | 0.9 | 0.904 | 1 | 0.321462036 | 0.043991302 |
| Mxi1 | 0.159205165 | -1.047305715 | 0.235 | 0.282 | 1 | 0.02226412 | -2.581990934 | 0.1 | 0.463 | 1 | 0.159205165 | 0.044032549 |
| Rab40c | 0.022328892 | -1.010888316 | 0.227 | 0.329 | 1 | 0.685814564 | -0.428334322 | 0.3 | 0.346 | 1 | 0.685814564 | 0.044159204 |
| Fra10ac1 | 0.02234381 | -0.929033479 | 0.151 | 0.259 | 1 | 0.453637123 | -0.505640048 | 0.1 | 0.188 | 1 | 0.453637123 | 0.044188374 |
| Smad6 | 0.022394972 | -1.408805546 | 0.21 | 0.312 | 1 | 0.786711337 | -0.40354186 | 0.5 | 0.383 | 1 | 0.786711337 | 0.04428841 |
| Tiprl | 0.022402228 | -0.391680331 | 0.697 | 0.729 | 1 | 0.022897239 | -1 | 0.4 | 0.818 | 1 | 0.022897239 | 0.044302596 |
| Gm14167 | 0.908310465 | -0.301002256 | 0.193 | 0.182 | 1 | 0.022411796 | 1.868174788 | 0.4 | 0.139 | 1 | 0.908310465 | 0.044321304 |
| Klc2 | 0.022422677 | -0.987927168 | 0.118 | 0.218 | 1 | 0.521602513 | 0.816288047 | 0.3 | 0.222 | 1 | 0.521602513 | 0.044342577 |
| Lsm5 | 0.022437657 | -0.587788545 | 0.412 | 0.518 | 1 | 0.233403111 | 0.471027448 | 0.7 | 0.407 | 1 | 0.233403111 | 0.044371865 |
| Abcg2 | 0.022461628 | 0.637721404 | 0.504 | 0.394 | 1 | 0.921057697 | 0.263034406 | 0.4 | 0.386 | 1 | 0.921057697 | 0.044418731 |
| Snx25 | 0.022476126 | 0.506257905 | 0.605 | 0.447 | 1 | 0.969521097 | -0.103611609 | 0.6 | 0.583 | 1 | 0.969521097 | 0.044447076 |
| Rpl14 | 0.283403029 | -0.132166952 | 0.983 | 0.982 | 1 | 0.022492298 | -0.651838561 | 1 | 0.994 | 1 | 0.283403029 | 0.044478693 |
| Stard5 | 0.022502801 | 0.723159795 | 0.286 | 0.171 | 1 | 0.352554654 | 0.723301159 | 0.4 | 0.275 | 1 | 0.352554654 | 0.044499227 |
| Naaa | 0.022506868 | 1.080170349 | 0.235 | 0.135 | 1 | 0.967118498 | 0.494359952 | 0.1 | 0.102 | 1 | 0.967118498 | 0.044507177 |
| 4933405D12Rik | 0.02254713 | -2.185866545 | 0.059 | 0.141 | 1 | 0.111515095 | -2.769980651 | 0 | 0.207 | 1 | 0.111515095 | 0.044585887 |
| Zranb3 | 0.241040933 | -0.900464326 | 0.067 | 0.106 | 1 | 0.02255326 | 1.591657153 | 0.3 | 0.083 | 1 | 0.241040933 | 0.04459787 |
| Myo9b | 0.022553263 | -0.49966875 | 0.79 | 0.824 | 1 | 0.681900142 | -0.324470289 | 1 | 0.914 | 1 | 0.681900142 | 0.044597876 |
| Six5 | 0.092475087 | -1.070389328 | 0.042 | 0.094 | 1 | 0.022553481 | 1.888638891 | 0.3 | 0.083 | 1 | 0.092475087 | 0.044598303 |
| Rac3 | 0.022584338 | -1.485426827 | 0.034 | 0.106 | 1 | 0.715256804 | 0.41305985 | 0.1 | 0.151 | 1 | 0.715256804 | 0.044658624 |
| Traf3ip1 | 0.022620341 | -1.255944981 | 0.126 | 0.224 | 1 | 0.542977682 | -0.357117523 | 0.1 | 0.167 | 1 | 0.542977682 | 0.044729002 |
| Noa1 | 0.022632906 | -1.444784843 | 0.067 | 0.153 | 1 | 0.977523303 | 1.176619654 | 0.1 | 0.105 | 1 | 0.977523303 | 0.044753563 |
| Zfp770 | 0.022641391 | -1.185866545 | 0.084 | 0.176 | 1 | 0.785851498 | 0.263034406 | 0.1 | 0.127 | 1 | 0.785851498 | 0.044770149 |
| Impa2 | 0.250803171 | 0.225066556 | 0.429 | 0.359 | 1 | 0.022652883 | 0.811471031 | 0.8 | 0.401 | 1 | 0.250803171 | 0.044792612 |
| Ppm1k | 0.022696278 | -1.333423734 | 0.076 | 0.165 | 1 | 0.298744962 | -0.709998547 | 0 | 0.099 | 1 | 0.298744962 | 0.044877435 |
| Trdmt1 | 0.022745329 | 0.880701072 | 0.294 | 0.182 | 1 | 0.14140196 | 1.263034406 | 0.3 | 0.13 | 1 | 0.14140196 | 0.044973307 |
| Entr1 | 0.022789216 | -0.715724447 | 0.353 | 0.459 | 1 | 0.184348249 | -0.874469118 | 0.4 | 0.531 | 1 | 0.184348249 | 0.045059084 |
| Socs7 | 0.022815376 | -0.936806174 | 0.311 | 0.418 | 1 | 0.742238982 | 0.111031312 | 0.6 | 0.506 | 1 | 0.742238982 | 0.045110211 |
| Elmo1 | 0.977935311 | -0.2871572 | 0.588 | 0.565 | 1 | 0.022852335 | -3.517353469 | 0 | 0.358 | 1 | 0.977935311 | 0.04518244 |
| Sgk3 | 0.022908319 | -0.751086702 | 0.756 | 0.776 | 1 | 0.121454494 | -0.865943247 | 0.6 | 0.843 | 1 | 0.121454494 | 0.045291848 |
| Arap1 | 0.022926093 | -1.11345805 | 0.151 | 0.259 | 1 | 0.96035435 | -0.131825212 | 0.3 | 0.259 | 1 | 0.96035435 | 0.04532658 |
| Cpeb4 | 0.214331285 | -0.423516226 | 0.84 | 0.794 | 1 | 0.022951276 | -0.965784285 | 0.9 | 0.954 | 1 | 0.214331285 | 0.045375791 |
| Gm38190 | 0.022982681 | -1.600904045 | 0.042 | 0.118 | 1 | 0.918736721 | 0.31748219 | 0.1 | 0.108 | 1 | 0.918736721 | 0.045437158 |
| Plpp6 | 0.022982967 | -1.544320516 | 0.042 | 0.118 | 1 | 0.225744174 | 1.602884409 | 0.2 | 0.086 | 1 | 0.225744174 | 0.045437718 |
| E130317F20Rik | 0.023024864 | -0.925999419 | 0.025 | 0.094 | 1 | 0.539431168 | 0.930459067 | 0 | 0.037 | 1 | 0.539431168 | 0.045519584 |
| Tsnax | 0.023079802 | -0.452259963 | 0.387 | 0.529 | 1 | 0.871217262 | 0.117917698 | 0.5 | 0.432 | 1 | 0.871217262 | 0.045626927 |
| Eif6 | 0.087558755 | -0.310523332 | 0.605 | 0.688 | 1 | 0.023080567 | -1.030110788 | 0.2 | 0.673 | 1 | 0.087558755 | 0.045628422 |
| Stat5b | 0.023151371 | -0.999256151 | 0.37 | 0.447 | 1 | 0.59073481 | 0.853233555 | 0.6 | 0.741 | 1 | 0.59073481 | 0.045766756 |
| Psmd13 | 0.023161322 | -0.493419618 | 0.479 | 0.559 | 1 | 0.431489472 | -0.404610128 | 0.6 | 0.664 | 1 | 0.431489472 | 0.045786197 |
| Spr | 0.02319322 | 0.650228272 | 0.429 | 0.306 | 1 | 0.771647109 | 0.150025444 | 0.4 | 0.315 | 1 | 0.771647109 | 0.045848515 |
| Plk4 | 0.023220423 | -1.614709844 | 0.042 | 0.118 | 1 | 0.849315905 | 0.558490289 | 0.1 | 0.08 | 1 | 0.849315905 | 0.045901658 |
| 1700019J19Rik | 0.023227446 | 2.389042291 | 0.059 | 0.012 | 1 | 0.73075158 | 2.695993813 | 0 | 0.012 | 1 | 0.73075158 | 0.045915379 |
| Gm57071 | 0.026180687 | 1.13606155 | 0.151 | 0.071 | 1 | 0.023235847 | 2.515421567 | 0.2 | 0.043 | 1 | 0.026180687 | 0.045931789 |
| 1500002C15Rik | 0.045471556 | 1.152003093 | 0.101 | 0.041 | 1 | 0.023235847 | 2.602884409 | 0.2 | 0.043 | 1 | 0.045471556 | 0.045931789 |
| Ciart | 0.189968263 | 0.280107919 | 0.118 | 0.071 | 1 | 0.023235847 | 2.602884409 | 0.2 | 0.043 | 1 | 0.189968263 | 0.045931789 |
| 1700001L05Rik | 0.02326261 | -1.584962501 | 0.05 | 0.129 | 1 | 0.141655717 | 0.868174788 | 0.3 | 0.127 | 1 | 0.141655717 | 0.045984071 |
| Ubac1 | 0.023285552 | -0.984997837 | 0.193 | 0.306 | 1 | 0.512561792 | -0.495147674 | 0.2 | 0.281 | 1 | 0.512561792 | 0.046028888 |
| S100a3 | 0.057502101 | 1.974004791 | 0.059 | 0.018 | 1 | 0.023298155 | 4.280956314 | 0.1 | 0.012 | 1 | 0.057502101 | 0.046053505 |
| Il1rl2 | 0.432595587 | -0.622930351 | 0.034 | 0.053 | 1 | 0.023298155 | 4.280956314 | 0.1 | 0.012 | 1 | 0.432595587 | 0.046053505 |
| Capn1 | 0.023299011 | -1.584962501 | 0.042 | 0.118 | 1 | 0.296217728 | 1.31748219 | 0.2 | 0.096 | 1 | 0.296217728 | 0.046055178 |
| Gm5535 | 0.062431398 | -2.185866545 | 0.008 | 0.047 | 1 | 0.023299663 | 4.754887502 | 0.1 | 0.012 | 1 | 0.062431398 | 0.046056451 |
| Cacna1b | 0.368010882 | 1.514573173 | 0.017 | 0.006 | 1 | 0.023299663 | 4.432959407 | 0.1 | 0.012 | 1 | 0.368010882 | 0.046056451 |
| Gm57184 | 0.02335521 | 0.791413378 | 0.277 | 0.171 | 1 | 0.04544416 | 1.290001453 | 0.3 | 0.096 | 1 | 0.04544416 | 0.046164954 |
| Bsg | 0.154408826 | -0.204815648 | 0.992 | 0.965 | 1 | 0.023372439 | -0.736965594 | 1 | 0.997 | 1 | 0.154408826 | 0.046198608 |
| Tmem158 | 0.023386971 | -1.5360529 | 0.042 | 0.118 | 1 | 0.810675183 | 0.31748219 | 0.1 | 0.123 | 1 | 0.810675183 | 0.046226992 |
| Snx12 | 0.023388789 | -0.847996907 | 0.37 | 0.465 | 1 | 0.489709234 | -0.741411499 | 0.5 | 0.512 | 1 | 0.489709234 | 0.046230543 |
| Akap11 | 0.023395719 | 0.335603032 | 0.874 | 0.694 | 1 | 0.205653592 | 0.83432197 | 0.9 | 0.722 | 1 | 0.205653592 | 0.046244079 |
| Pink1 | 0.023425987 | 0.490120129 | 0.563 | 0.453 | 1 | 0.330171405 | 0.460545239 | 0.7 | 0.5 | 1 | 0.330171405 | 0.046303197 |
| Zfp263 | 0.023459119 | -1.040015679 | 0.118 | 0.218 | 1 | 0.310167581 | 0.432959407 | 0.4 | 0.228 | 1 | 0.310167581 | 0.046367907 |
| Zfp46 | 0.023497578 | -1.36994961 | 0.101 | 0.194 | 1 | 0.113798453 | -1.473931188 | 0 | 0.204 | 1 | 0.113798453 | 0.046443021 |
| Dock6 | 0.02349774 | -0.87244995 | 0.168 | 0.282 | 1 | 0.228956251 | -0.789433014 | 0.2 | 0.389 | 1 | 0.228956251 | 0.046443336 |
| Sfrp5 | 0.0235243 | 3.64385619 | 0.059 | 0.012 | 1 | 0.10265174 | 5.602884409 | 0.1 | 0.022 | 1 | 0.10265174 | 0.046495207 |
| Enkur | 0.023530699 | 0.901596296 | 0.092 | 0.029 | 1 | 0.037329331 | 2.210566986 | 0.2 | 0.049 | 1 | 0.037329331 | 0.046507703 |
| Impdh2 | 0.023535045 | -0.278194213 | 0.765 | 0.829 | 1 | 0.065893984 | -0.862273821 | 0.6 | 0.787 | 1 | 0.065893984 | 0.046516192 |
| Kcna6 | 0.023546964 | 0.180154134 | 0.63 | 0.488 | 1 | 0.817212068 | -0.188929662 | 0.7 | 0.623 | 1 | 0.817212068 | 0.046539468 |
| Ncln | 0.05817403 | -0.510961919 | 0.193 | 0.294 | 1 | 0.023602845 | 1.19592021 | 0.6 | 0.253 | 1 | 0.05817403 | 0.046648596 |
| Cfap68 | 0.023617764 | 0.285754482 | 0.471 | 0.318 | 1 | 0.282498369 | 0.861417422 | 0.4 | 0.244 | 1 | 0.282498369 | 0.046677729 |
| Lrrcc1 | 0.023627737 | -1.336904302 | 0.16 | 0.259 | 1 | 0.501929454 | 0.185031894 | 0.3 | 0.188 | 1 | 0.501929454 | 0.046697204 |
| Mid2 | 0.023629407 | -0.59946207 | 0.613 | 0.676 | 1 | 0.871920937 | -0.114411205 | 0.7 | 0.679 | 1 | 0.871920937 | 0.046700466 |
| Zfp61 | 0.023672427 | -2.222392421 | 0.017 | 0.076 | 1 | 0.927831027 | 0.695993813 | 0.1 | 0.09 | 1 | 0.927831027 | 0.04678447 |
| Gm48024 | 0.023673485 | -2.544320516 | 0.017 | 0.076 | 1 | 0.698533977 | 1.558490289 | 0 | 0.015 | 1 | 0.698533977 | 0.046786535 |
| Endod1 | 0.023728447 | -1.041820176 | 0.235 | 0.341 | 1 | 0.126050826 | -1.125036046 | 0.2 | 0.426 | 1 | 0.126050826 | 0.046893856 |
| 0610009E02Rik | 0.023757948 | -1.485426827 | 0.05 | 0.129 | 1 | 0.219557476 | -0.864721141 | 0 | 0.133 | 1 | 0.219557476 | 0.046951455 |

| | | | | | | | | | | | | |
|---|---|---|---|---|---|---|---|---|---|---|---|---|
| Tro | 0.023762322 | -1.237499314 | 0.126 | 0.224 | 1 | 0.271795412 | -0.947862377 | 0.1 | 0.244 | 1 | 0.271795412 | 0.046959996 |
| Lypd3 | 0.171101855 | -0.485426827 | 0.017 | 0.047 | 1 | 0.023785224 | 4.017921908 | 0.1 | 0.012 | 1 | 0.171101855 | 0.047004711 |
| Slc66a1 | 0.281631261 | 0.407657969 | 0.101 | 0.065 | 1 | 0.023802244 | 2.111031312 | 0.4 | 0.154 | 1 | 0.281631261 | 0.047037942 |
| Jmjd1c | 0.400824845 | -0.104518005 | 0.975 | 1 | 1 | 0.023816084 | -0.759974643 | 0.9 | 0.994 | 1 | 0.400824845 | 0.047064962 |
| Gm12840 | 0.023821715 | 1.929610672 | 0.059 | 0.012 | 1 | 0.557784478 | 1.432959407 | 0 | 0.034 | 1 | 0.557784478 | 0.047075956 |
| Gm39167 | 0.023821715 | 1.929610672 | 0.059 | 0.012 | 1 | 0.669848329 | 1.695993813 | 0 | 0.019 | 1 | 0.669848329 | 0.047075956 |
| Ccdc159 | 0.023821715 | 1.929610672 | 0.059 | 0.012 | 1 | 0.577155236 | 1.31748219 | 0 | 0.031 | 1 | 0.577155236 | 0.047075956 |
| Tmem256 | 0.023827125 | -0.367590337 | 0.613 | 0.712 | 1 | 0.071157635 | -0.880931369 | 0.3 | 0.605 | 1 | 0.071157635 | 0.047086518 |
| Trmt10b | 0.023845351 | -1.655351829 | 0.042 | 0.118 | 1 | 0.396950721 | 0.210566986 | 0 | 0.068 | 1 | 0.396950721 | 0.047122101 |
| Lca5l | 0.023845351 | -1.544320516 | 0.042 | 0.118 | 1 | 0.433608413 | 0.494359952 | 0 | 0.059 | 1 | 0.433608413 | 0.047122101 |
| AI506816 | 0.02385945 | -0.544539921 | 0.832 | 0.794 | 1 | 0.376050022 | 0.558490289 | 0.9 | 0.716 | 1 | 0.376050022 | 0.047149626 |
| Zfp469 | 0.135128647 | 0.414046297 | 0.471 | 0.394 | 1 | 0.023862171 | -1.640289575 | 0.4 | 0.67 | 1 | 0.135128647 | 0.047154939 |
| Bsn | 0.023897291 | -2.359895945 | 0.017 | 0.076 | 1 | 0.220191704 | -1.640289575 | 0 | 0.133 | 1 | 0.220191704 | 0.047223502 |
| Gm9887 | 0.023897291 | -2.14839184 | 0.017 | 0.076 | 1 | 0.274256184 | 2.31748219 | 0.1 | 0.034 | 1 | 0.274256184 | 0.047223502 |
| Gm15983 | 0.023897291 | -2.544320516 | 0.017 | 0.076 | 1 | 0.46060033 | 0.558490289 | 0 | 0.052 | 1 | 0.46060033 | 0.047223502 |
| Rtn4rl1 | 0.023897764 | -2.292781749 | 0.017 | 0.076 | 1 | 0.489966983 | -0.111361109 | 0 | 0.046 | 1 | 0.489966983 | 0.047224425 |
| Gm41556 | 0.023907736 | -1.572889668 | 0.05 | 0.129 | 1 | 0.334438555 | -0.888968688 | 0 | 0.086 | 1 | 0.334438555 | 0.047243893 |
| Otud1 | 0.023920427 | -1.485426827 | 0.05 | 0.129 | 1 | 0.170601261 | -1.111361109 | 0 | 0.16 | 1 | 0.170601261 | 0.047268667 |
| Cep57l1 | 0.023920917 | -1.163498732 | 0.084 | 0.176 | 1 | 0.213734676 | 1.432959407 | 0.2 | 0.083 | 1 | 0.213734676 | 0.047269624 |
| Zfp780b | 0.023922985 | -1.432959407 | 0.042 | 0.118 | 1 | 0.156452065 | -1.152003093 | 0 | 0.17 | 1 | 0.156452065 | 0.047273662 |
| Plekhf1 | 0.023933971 | -0.914270126 | 0.227 | 0.341 | 1 | 0.662246568 | -0.441509711 | 0.4 | 0.432 | 1 | 0.662246568 | 0.047295106 |
| Pam16 | 0.023935898 | 0.474680446 | 0.597 | 0.453 | 1 | 0.432870556 | 0.485426827 | 0.5 | 0.358 | 1 | 0.432870556 | 0.047298869 |
| Mrm1 | 0.023941304 | -1.292781749 | 0.034 | 0.106 | 1 | 0.832730078 | 1.063725998 | 0.1 | 0.08 | 1 | 0.832730078 | 0.047309422 |
| Tsga10 | 0.41463222 | -0.932885804 | 0.076 | 0.1 | 1 | 0.023947934 | 2.30520386 | 0.3 | 0.093 | 1 | 0.41463222 | 0.047322365 |
| Ldah | 0.023953043 | -0.582000087 | 0.353 | 0.459 | 1 | 0.819922767 | 0.247092862 | 0.4 | 0.454 | 1 | 0.819922767 | 0.047332337 |
| A230108P19Rik | 0.023965738 | 1.974004791 | 0.059 | 0.012 | 1 | 0.080320981 | 2.847996907 | 0.1 | 0.019 | 1 | 0.080320981 | 0.047357119 |
| Dmtf1 | 0.023988612 | -0.807354922 | 0.412 | 0.488 | 1 | 0.262165437 | -0.222392421 | 0.3 | 0.599 | 1 | 0.262165437 | 0.047401771 |
| Eppk1 | 0.023996415 | 1.698997744 | 0.101 | 0.035 | 1 | 0.073296931 | 3.017921908 | 0.2 | 0.062 | 1 | 0.073296931 | 0.047417003 |
| Ppp1r2 | 0.304695047 | -0.349584438 | 0.866 | 0.818 | 1 | 0.024001463 | -0.620514006 | 0.4 | 0.886 | 1 | 0.304695047 | 0.047426856 |
| Nfkb2 | 0.188220886 | -0.557576613 | 0.261 | 0.318 | 1 | 0.024064666 | -2.544320516 | 0 | 0.352 | 1 | 0.188220886 | 0.047550224 |
| Nckap1 | 0.097089314 | -0.272925155 | 0.966 | 0.965 | 1 | 0.024102354 | -0.790032016 | 1 | 0.978 | 1 | 0.097089314 | 0.047623785 |
| B4galt2 | 0.024122228 | -1.070389328 | 0.084 | 0.176 | 1 | 0.207416166 | -0.736965594 | 0 | 0.139 | 1 | 0.207416166 | 0.047662574 |
| Dnmt3b | 0.024122476 | -2.14839184 | 0.017 | 0.076 | 1 | 0.505623793 | 0.847996907 | 0 | 0.043 | 1 | 0.505623793 | 0.047663058 |
| Gm48678 | 0.024122476 | -1.987927168 | 0.017 | 0.076 | 1 | 0.290138826 | -0.304006187 | 0 | 0.102 | 1 | 0.290138826 | 0.047663058 |
| Csad | 0.024122975 | -0.822784963 | 0.479 | 0.559 | 1 | 0.453499088 | -0.674170467 | 0.7 | 0.667 | 1 | 0.453499088 | 0.047664031 |
| Capza1 | 0.063932098 | -0.426231035 | 0.849 | 0.806 | 1 | 0.024151239 | -0.85398333 | 0.8 | 0.898 | 1 | 0.063932098 | 0.047719195 |
| Hk1 | 0.02416708 | 0.291431494 | 0.849 | 0.729 | 1 | 0.093654976 | 0.455679484 | 0.9 | 0.802 | 1 | 0.093654976 | 0.047750112 |
| Macroh2a1 | 0.024181599 | -0.340478491 | 0.765 | 0.835 | 1 | 0.401001341 | 0.297799824 | 0.9 | 0.883 | 1 | 0.401001341 | 0.047778448 |
| Slc35b2 | 0.02418266 | 0.308122295 | 0.765 | 0.6 | 1 | 0.74338542 | 0.568614507 | 0.5 | 0.509 | 1 | 0.74338542 | 0.047780519 |
| Itih2 | 0.176836096 | 0.575973717 | 0.168 | 0.112 | 1 | 0.024203319 | 1.744903414 | 0.4 | 0.145 | 1 | 0.176836096 | 0.047820837 |
| Mrpl35 | 0.024250582 | -0.829381228 | 0.193 | 0.306 | 1 | 0.759025716 | 0.532495081 | 0.3 | 0.262 | 1 | 0.759025716 | 0.047913073 |
| Ppp1r16a | 0.024261129 | -1.432959407 | 0.05 | 0.129 | 1 | 0.84102208 | 0.973527789 | 0.1 | 0.08 | 1 | 0.84102208 | 0.047933656 |
| Cdkl1 | 0.02426277 | -2.129283017 | 0.017 | 0.076 | 1 | 0.280043603 | 0.720241359 | 0.2 | 0.09 | 1 | 0.280043603 | 0.047936859 |
| Cep112it | 0.741348152 | -0.222392421 | 0.034 | 0.041 | 1 | 0.024279539 | 3.795529487 | 0.1 | 0.012 | 1 | 0.741348152 | 0.047969582 |
| Lrp8os3 | 0.781790612 | -0.485426827 | 0.008 | 0.012 | 1 | 0.024279539 | 4.017921908 | 0.1 | 0.012 | 1 | 0.781790612 | 0.047969582 |
| Palm3 | 0.147030916 | -1.485426827 | 0 | 0.018 | 1 | 0.024280315 | 3.795529487 | 0.1 | 0.012 | 1 | 0.147030916 | 0.047971096 |
| Dnajb11 | 0.024312118 | -0.457412451 | 0.697 | 0.729 | 1 | 0.180571802 | -0.564726727 | 0.7 | 0.799 | 1 | 0.180571802 | 0.048033157 |
| Zfp316 | 0.024347604 | -1.900464326 | 0.017 | 0.076 | 1 | 0.539413885 | 1.210566986 | 0 | 0.037 | 1 | 0.539413885 | 0.048102402 |
| Zfp94 | 0.024347604 | -1.900464326 | 0.017 | 0.076 | 1 | 0.420904619 | 0.432959407 | 0 | 0.062 | 1 | 0.420904619 | 0.048102402 |
| Atp6v1g2 | 0.024347604 | -1.900464326 | 0.017 | 0.076 | 1 | 0.334119666 | -0.111361109 | 0 | 0.086 | 1 | 0.334119666 | 0.048102402 |
| Tmem168 | 0.024364017 | -0.807354922 | 0.151 | 0.265 | 1 | 0.351031852 | -0.797995028 | 0.2 | 0.318 | 1 | 0.351031852 | 0.048134428 |
| A730017L22Rik | 0.063085444 | -1.807354922 | 0.008 | 0.047 | 1 | 0.024366566 | 2.354956895 | 0.2 | 0.043 | 1 | 0.063085444 | 0.048139403 |
| Commd8 | 0.02437702 | -0.798584712 | 0.235 | 0.347 | 1 | 0.386220269 | -0.546862711 | 0.2 | 0.312 | 1 | 0.386220269 | 0.0481598 |
| Klf6 | 0.15401963 | -0.480735132 | 0.891 | 0.9 | 1 | 0.0243987 | -1.374395515 | 0.6 | 0.901 | 1 | 0.15401963 | 0.048202103 |
| 4930593A02Rik | 0.024407988 | 1.099535674 | 0.101 | 0.035 | 1 | 0.197183593 | 2.017921908 | 0.1 | 0.028 | 1 | 0.197183593 | 0.048220225 |
| H4c9 | 0.024427991 | 0.897901812 | 0.176 | 0.088 | 1 | 0.526604355 | 1.494359952 | 0.1 | 0.052 | 1 | 0.526604355 | 0.048259255 |
| Gin1 | 0.02444848 | -1.399696953 | 0.076 | 0.165 | 1 | 0.787106817 | 0.795529487 | 0.1 | 0.142 | 1 | 0.787106817 | 0.048299231 |
| Ppp1r36 | 0.024507956 | -1.572889668 | 0.05 | 0.129 | 1 | 0.343862196 | -0.191531458 | 0 | 0.083 | 1 | 0.343862196 | 0.048415272 |
| Tomm40 | 0.024556139 | -0.586512952 | 0.37 | 0.476 | 1 | 0.405569689 | -0.541045384 | 0.5 | 0.577 | 1 | 0.405569689 | 0.048509274 |
| Sgta | 0.024563787 | -0.784987109 | 0.387 | 0.465 | 1 | 0.290533934 | -0.40354186 | 0.4 | 0.559 | 1 | 0.290533934 | 0.048524195 |
| Abhd6 | 0.314489957 | -0.614709844 | 0.092 | 0.129 | 1 | 0.024570684 | 1.362570079 | 0.4 | 0.136 | 1 | 0.314489957 | 0.04853765 |
| Gm12258 | 0.024573011 | -1.807354922 | 0.017 | 0.076 | 1 | 0.153152701 | 2.31748219 | 0.1 | 0.025 | 1 | 0.153152701 | 0.04854219 |
| Tmc8 | 0.024573011 | -1.807354922 | 0.017 | 0.076 | 1 | 0.489966712 | 0.210566986 | 0 | 0.046 | 1 | 0.489966712 | 0.04854219 |
| Dgcr2 | 0.024619965 | -0.678071905 | 0.311 | 0.435 | 1 | 0.773227595 | 0.15497466 | 0.6 | 0.537 | 1 | 0.773227595 | 0.048633787 |
| Comt | 0.024702241 | -0.692584735 | 0.387 | 0.482 | 1 | 0.12261201 | -0.45600928 | 0.3 | 0.611 | 1 | 0.12261201 | 0.048794282 |
| Kif13b | 0.024708387 | 0.397033489 | 0.992 | 0.971 | 1 | 0.827488899 | 0.272478342 | 1 | 0.969 | 1 | 0.827488899 | 0.04880627 |
| Nat8f1 | 0.024712251 | 1.514573173 | 0.118 | 0.047 | 1 | 0.248008881 | 3.017921908 | 0.1 | 0.034 | 1 | 0.248008881 | 0.048813807 |
| Mkrn1 | 0.024716271 | 0.588573754 | 0.538 | 0.435 | 1 | 0.133380762 | -0.304006187 | 0.4 | 0.713 | 1 | 0.133380762 | 0.048821648 |
| 6330411D24Rik | 0.024732235 | -2.343407822 | 0.017 | 0.076 | 1 | 0.557776184 | 0.111031312 | 0 | 0.034 | 1 | 0.557776184 | 0.048852786 |
| Tmem100 | 0.024732899 | -0.509539283 | 0.563 | 0.676 | 1 | 0.44816163 | -1.153823529 | 0.5 | 0.531 | 1 | 0.44816163 | 0.048854082 |
| Azin1 | 0.024741014 | -0.325749806 | 0.908 | 0.894 | 1 | 0.79498556 | -0.173595624 | 0.9 | 0.923 | 1 | 0.79498556 | 0.048869911 |
| Fbxo31 | 0.024742194 | 0.877143252 | 0.235 | 0.135 | 1 | 0.213479811 | -0.840059087 | 0 | 0.136 | 1 | 0.213479811 | 0.048872212 |
| Pik3c2g | 0.024743268 | 1.724026538 | 0.151 | 0.071 | 1 | 0.037724104 | 2.263034406 | 0.2 | 0.049 | 1 | 0.037724104 | 0.048874307 |
| Ndufab1 | 0.141629585 | -0.245851394 | 0.756 | 0.765 | 1 | 0.024755074 | -1.080110175 | 0.4 | 0.775 | 1 | 0.141629585 | 0.048897333 |
| Wdr77 | 0.024781625 | -0.900464326 | 0.235 | 0.347 | 1 | 0.45277798 | -0.258202497 | 0.2 | 0.324 | 1 | 0.45277798 | 0.04894912 |
| Syt9 | 0.025470389 | -2.807354922 | 0 | 0.041 | 1 | 0.024783259 | 3.602884409 | 0.1 | 0.012 | 1 | 0.025470389 | 0.048952308 |
| Elf4 | 0.02478695 | -0.684549469 | 0.513 | 0.612 | 1 | 0.051940667 | -1.355184911 | 0.4 | 0.639 | 1 | 0.051940667 | 0.048959506 |
| Commd10 | 0.024792168 | -1.349771728 | 0.286 | 0.382 | 1 | 0.125010731 | 0.847996907 | 0.6 | 0.386 | 1 | 0.125010731 | 0.048969685 |
| Med20 | 0.024817034 | -0.835011265 | 0.218 | 0.329 | 1 | 0.433386442 | -0.654503434 | 0.3 | 0.38 | 1 | 0.433386442 | 0.049018184 |
| Clec11a | 0.522159071 | 0.169027973 | 0.765 | 0.788 | 1 | 0.024845793 | -1.200366115 | 0.7 | 0.815 | 1 | 0.522159071 | 0.049074272 |
| Def8 | 0.02485165 | -1.029747343 | 0.084 | 0.176 | 1 | 0.156485219 | 1.909397451 | 0.3 | 0.154 | 1 | 0.156485219 | 0.049085695 |
| Gm48321 | 0.024857109 | 1.514573173 | 0.059 | 0.012 | 1 | 0.455763535 | 1.374065718 | 0.1 | 0.046 | 1 | 0.455763535 | 0.049096341 |
| Sirt4 | 0.024866597 | -1.707819249 | 0.017 | 0.076 | 1 | 1 | 0.847996907 | 0.1 | 0.099 | 1 | 1 | 0.049114847 |
| Gm19744 | 0.024866597 | -1.707819249 | 0.017 | 0.076 | 1 | 0.526605007 | 1.31748219 | 0.1 | 0.052 | 1 | 0.526605007 | 0.049114847 |
| Ppm1h | 0.024933921 | -1.258016331 | 0.118 | 0.212 | 1 | 0.258959232 | -1.004445905 | 0.1 | 0.269 | 1 | 0.258959232 | 0.049246142 |
| Incenp | 0.085604716 | -1.970853654 | 0.025 | 0.071 | 1 | 0.024947578 | 2.280956314 | 0.2 | 0.043 | 1 | 0.085604716 | 0.049272774 |
| Pemt | 0.045284567 | -1.485426827 | 0.034 | 0.094 | 1 | 0.024947833 | 2.079322453 | 0.2 | 0.043 | 1 | 0.045284567 | 0.049273272 |
| Cbx8 | 0.663944693 | -0.378511623 | 0.042 | 0.053 | 1 | 0.024947919 | 2.354956895 | 0.2 | 0.043 | 1 | 0.663944693 | 0.049273438 |
| Zbtb49 | 0.063094072 | -1.748461233 | 0.034 | 0.088 | 1 | 0.025024198 | 2.558490289 | 0.3 | 0.093 | 1 | 0.063094072 | 0.049422186 |
| Ercc3 | 0.025029566 | -1.455053178 | 0.092 | 0.182 | 1 | 0.43462114 | 0.864116572 | 0.3 | 0.204 | 1 | 0.43462114 | 0.049432653 |
| Tmem101 | 0.025050916 | -0.93021167 | 0.143 | 0.247 | 1 | 0.673112449 | 0.795529487 | 0.2 | 0.148 | 1 | 0.673112449 | 0.049474283 |
| Ass1 | 0.025094018 | 0.584962501 | 0.319 | 0.194 | 1 | 0.868146068 | 0.210566986 | 0.1 | 0.114 | 1 | 0.868146068 | 0.049558326 |
| S1pr2 | 0.025104599 | -1.023846742 | 0.151 | 0.253 | 1 | 0.105832925 | -1.567040593 | 0.1 | 0.333 | 1 | 0.105832925 | 0.049578957 |
| Eef2kmt | 0.025150431 | -1.091147888 | 0.151 | 0.253 | 1 | 0.860713198 | 0.393431043 | 0.2 | 0.17 | 1 | 0.860713198 | 0.049668317 |
| Cebpa | 0.025188879 | -1.321928095 | 0.042 | 0.118 | 1 | 0.46569953 | -0.408342847 | 0.1 | 0.185 | 1 | 0.46569953 | 0.049743279 |
| Gm43813 | 0.025227181 | -0.900464326 | 0.05 | 0.129 | 1 | 0.267214296 | -0.888968688 | 0 | 0.111 | 1 | 0.267214296 | 0.049817952 |
| Meak7 | 0.060012807 | -2.070389328 | 0 | 0.029 | 1 | 0.025296002 | 3.695993813 | 0.1 | 0.012 | 1 | 0.060012807 | 0.049952116 |
| Gm8066 | 0.144054748 | -1.485426827 | 0.008 | 0.035 | 1 | 0.025296002 | 3.695993813 | 0.1 | 0.012 | 1 | 0.144054748 | 0.049952116 |

| | | | | | | | | | | | | |
|---|---|---|---|---|---|---|---|---|---|---|---|---|
| B430219N15Rik | 0.144054748 | -1.485426827 | 0.008 | 0.035 | 1 | 0.025296002 | 3.695993813 | 0.1 | 0.012 | 1 | 0.144054748 | 0.049952116 |
| Gm48913 | 0.147029286 | -2.655351829 | 0 | 0.018 | 1 | 0.025296002 | 3.695993813 | 0.1 | 0.012 | 1 | 0.147029286 | 0.049952116 |
| Psrc1 | 0.17252682 | -0.900464326 | 0.017 | 0.047 | 1 | 0.025296002 | 3.695993813 | 0.1 | 0.012 | 1 | 0.17252682 | 0.049952116 |
| Wdr86 | 0.218485013 | -1.292781749 | 0.008 | 0.029 | 1 | 0.025296002 | 3.695993813 | 0.1 | 0.012 | 1 | 0.218485013 | 0.049952116 |
| Gpld1 | 0.218515614 | 0.736965594 | 0.042 | 0.018 | 1 | 0.025296002 | 3.695993813 | 0.1 | 0.012 | 1 | 0.218515614 | 0.049952116 |
| Inhbb | 0.237885589 | -1.485426827 | 0 | 0.012 | 1 | 0.025296002 | 3.695993813 | 0.1 | 0.012 | 1 | 0.237885589 | 0.049952116 |
| Ppfia3 | 0.237885589 | -1.070389328 | 0 | 0.012 | 1 | 0.025296002 | 3.695993813 | 0.1 | 0.012 | 1 | 0.237885589 | 0.049952116 |
| Tmc5 | 0.237885589 | -1.070389328 | 0 | 0.012 | 1 | 0.025296002 | 3.695993813 | 0.1 | 0.012 | 1 | 0.237885589 | 0.049952116 |
| Egf | 0.240356677 | -1.070389328 | 0.017 | 0.041 | 1 | 0.025296002 | 3.695993813 | 0.1 | 0.012 | 1 | 0.240356677 | 0.049952116 |
| Gm4419 | 0.316514503 | -1.070389328 | 0.034 | 0.059 | 1 | 0.025296002 | 3.695993813 | 0.1 | 0.012 | 1 | 0.316514503 | 0.049952116 |
| 2900092C05Rik | 0.331040183 | -1.485426827 | 0.008 | 0.024 | 1 | 0.025296002 | 3.695993813 | 0.1 | 0.012 | 1 | 0.331040183 | 0.049952116 |
| Tdrkh | 0.33261065 | -1.292781749 | 0.008 | 0.024 | 1 | 0.025296002 | 3.695993813 | 0.1 | 0.012 | 1 | 0.33261065 | 0.049952116 |
| Gm57337 | 0.334204811 | -0.807354922 | 0.008 | 0.024 | 1 | 0.025296002 | 3.695993813 | 0.1 | 0.012 | 1 | 0.334204811 | 0.049952116 |
| H1f1 | 0.336002793 | -0.944858446 | 0.025 | 0.047 | 1 | 0.025296002 | 3.695993813 | 0.1 | 0.012 | 1 | 0.336002793 | 0.049952116 |
| 1500002F19Rik | 0.370184089 | 1.099535674 | 0.017 | 0.006 | 1 | 0.025296002 | 3.695993813 | 0.1 | 0.012 | 1 | 0.370184089 | 0.049952116 |
| 9930012K11Rik | 0.508731273 | -0.807354922 | 0.008 | 0.018 | 1 | 0.025296002 | 3.695993813 | 0.1 | 0.012 | 1 | 0.508731273 | 0.049952116 |
| Ttc36 | 0.635175693 | 1 | 0.059 | 0.047 | 1 | 0.025296002 | 3.695993813 | 0.1 | 0.012 | 1 | 0.635175693 | 0.049952116 |
| Sptbn5 | 0.655679748 | 0.736965594 | 0.025 | 0.018 | 1 | 0.025296002 | 3.695993813 | 0.1 | 0.012 | 1 | 0.655679748 | 0.049952116 |
| Dnai1 | 0.731769749 | -0.485426827 | 0.017 | 0.012 | 1 | 0.025296002 | 3.695993813 | 0.1 | 0.012 | 1 | 0.731769749 | 0.049952116 |
| Tardbp | 0.025308566 | -0.43986425 | 0.765 | 0.824 | 1 | 0.296067517 | -0.463771322 | 0.7 | 0.886 | 1 | 0.296067517 | 0.049976608 |
| Irgm1 | 0.025389607 | 0.859221344 | 0.37 | 0.271 | 1 | 0.24948225 | -1.408342847 | 0.1 | 0.25 | 1 | 0.24948225 | 0.050134582 |
| Casc3 | 0.025427689 | -0.780882711 | 0.37 | 0.465 | 1 | 0.681375824 | -0.175849835 | 0.5 | 0.599 | 1 | 0.681375824 | 0.05020881 |
| Bst2 | 0.025439338 | 0.597714408 | 0.21 | 0.353 | 1 | 0.144589478 | -2.453753306 | 0.1 | 0.299 | 1 | 0.144589478 | 0.050231515 |
| Mob2 | 0.025446259 | -0.655351829 | 0.345 | 0.441 | 1 | 0.059974443 | -1.293145194 | 0.4 | 0.62 | 1 | 0.059974443 | 0.050245006 |
| Kank3 | 0.025451659 | -1.655351829 | 0.017 | 0.076 | 1 | 0.539431168 | 1.017921908 | 0 | 0.037 | 1 | 0.539431168 | 0.050255531 |
| Hspa12b | 0.025466526 | -2.655351829 | 0 | 0.041 | 1 | 0.557765566 | 0.847996907 | 0 | 0.034 | 1 | 0.557765566 | 0.050284508 |
| Gask1b | 0.025466526 | -2.944858446 | 0 | 0.041 | 1 | 0.232763883 | -2.004445905 | 0 | 0.127 | 1 | 0.232763883 | 0.050284508 |
| Htr6 | 0.025466526 | -2.944858446 | 0 | 0.041 | 1 | 0.643835681 | 1.558490289 | 0 | 0.022 | 1 | 0.643835681 | 0.050284508 |
| 4930522L14Rik | 0.025466526 | -2.807354922 | 0 | 0.041 | 1 | 0.577142568 | 1.432959407 | 0 | 0.031 | 1 | 0.577142568 | 0.050284508 |
| Zfp853 | 0.025466526 | -2.807354922 | 0 | 0.041 | 1 | 0.73075158 | 2.695993813 | 0 | 0.012 | 1 | 0.73075158 | 0.050284508 |
| Abcc6 | 0.025466526 | -2.655351829 | 0 | 0.041 | 1 | 0.698529281 | 1.847996907 | 0 | 0.015 | 1 | 0.698529281 | 0.050284508 |
| Tsku | 0.025466526 | -2.655351829 | 0 | 0.041 | 1 | 0.556267414 | 1.494359952 | 0.1 | 0.056 | 1 | 0.556267414 | 0.050284508 |
| Tex45 | 0.025466526 | -2.655351829 | 0 | 0.041 | 1 | 0.619967066 | 1.31748219 | 0 | 0.025 | 1 | 0.619967066 | 0.050284508 |
| Zfp964 | 0.025466526 | -2.655351829 | 0 | 0.041 | 1 | 0.522129069 | 0.625604485 | 0 | 0.04 | 1 | 0.522129069 | 0.050284508 |
| Carmil2 | 0.025466526 | -2.655351829 | 0 | 0.041 | 1 | 0.557756607 | 1.111031312 | 0 | 0.034 | 1 | 0.557756607 | 0.050284508 |
| Ggt5 | 0.025466526 | -2.655351829 | 0 | 0.041 | 1 | 0.730750993 | 2.210566986 | 0 | 0.012 | 1 | 0.730750993 | 0.050284508 |
| Plppr3 | 0.025466526 | -2.655351829 | 0 | 0.041 | 1 | 0.669850692 | 1.558490289 | 0 | 0.019 | 1 | 0.669850692 | 0.050284508 |
| Samd15 | 0.025466526 | -2.655351829 | 0 | 0.041 | 1 | 0.597845674 | 1.695993813 | 0 | 0.028 | 1 | 0.597845674 | 0.050284508 |
| D430019H16Rik | 0.025466526 | -2.944858446 | 0 | 0.041 | 1 | 0.577172123 | 0.558490289 | 0 | 0.031 | 1 | 0.577172123 | 0.050284508 |
| Begain | 0.025466526 | -2.655351829 | 0 | 0.041 | 1 | 0.669843604 | 2.017921908 | 0 | 0.019 | 1 | 0.669843604 | 0.050284508 |
| Camk2a | 0.025466526 | -2.655351829 | 0 | 0.041 | 1 | 0.474964899 | 0.31748219 | 0 | 0.049 | 1 | 0.474964899 | 0.050284508 |
| Gm15853 | 0.025470389 | -2.807354922 | 0 | 0.041 | 1 | 0.698529281 | 2.210566986 | 0 | 0.015 | 1 | 0.698529281 | 0.050292037 |
| Uchl1 | 0.025470389 | -3.185866545 | 0 | 0.041 | 1 | 0.489984625 | -0.408342847 | 0 | 0.046 | 1 | 0.489984625 | 0.050292037 |
| Dlx6 | 0.025470389 | -2.807354922 | 0 | 0.041 | 1 | 0.730750993 | 1.695993813 | 0 | 0.012 | 1 | 0.730750993 | 0.050292037 |
| Gm16174 | 0.025470389 | -2.807354922 | 0 | 0.041 | 1 | 0.025814657 | 3.210566986 | 0.1 | 0.012 | 1 | 0.025814657 | 0.050292037 |
| Olfml1 | 0.025470389 | -2.807354922 | 0 | 0.041 | 1 | 0.460577668 | 0.31748219 | 0 | 0.052 | 1 | 0.460577668 | 0.050292037 |
| Klhl40 | 0.025470389 | -3.392317423 | 0 | 0.041 | 1 | 0.505649381 | 0.625604485 | 0 | 0.043 | 1 | 0.505649381 | 0.050292037 |
| Kcnj2 | 0.025470389 | -5.100136671 | 0 | 0.041 | 1 | 0.669851164 | 1.210566986 | 0 | 0.019 | 1 | 0.669851164 | 0.050292037 |
| Ccdc14 | 0.025470389 | -2.944858446 | 0 | 0.041 | 1 | 0.051909884 | 2.558490289 | 0.1 | 0.015 | 1 | 0.051909884 | 0.050292037 |
| Gng13 | 0.025470389 | -2.807354922 | 0 | 0.041 | 1 | 0.619961444 | 1.558490289 | 0 | 0.025 | 1 | 0.619961444 | 0.050292037 |
| Fgf14 | 0.025472707 | -3.185866545 | 0 | 0.041 | 1 | 0.557765898 | 1.017921908 | 0 | 0.034 | 1 | 0.557765898 | 0.050296555 |
| Pkia | 0.025472965 | -3.185866545 | 0 | 0.041 | 1 | 0.505664731 | 0.31748219 | 0 | 0.043 | 1 | 0.505664731 | 0.050297057 |
| Cplane2 | 0.025472965 | -2.944858446 | 0 | 0.041 | 1 | 0.489877116 | 0.847996907 | 0 | 0.046 | 1 | 0.489877116 | 0.050297057 |
| Gm9970 | 0.025472965 | -2.944858446 | 0 | 0.041 | 1 | 0.072537454 | 3.602884409 | 0.1 | 0.019 | 1 | 0.072537454 | 0.050297057 |
| Myo1g | 0.025472965 | -2.944858446 | 0 | 0.041 | 1 | 0.698529281 | 2.210566986 | 0 | 0.015 | 1 | 0.698529281 | 0.050297057 |
| Tspan11 | 0.02547348 | -3.292781749 | 0 | 0.041 | 1 | 0.421058395 | -0.596787936 | 0 | 0.062 | 1 | 0.421058395 | 0.050298061 |
| Dlx6os1 | 0.025474252 | -3.572889668 | 0 | 0.041 | 1 | 0.698534498 | 2.432959407 | 0 | 0.015 | 1 | 0.698534498 | 0.050299567 |
| Sox11 | 0.025474252 | -3.655351829 | 0 | 0.041 | 1 | 0.505659614 | 0.263034406 | 0 | 0.043 | 1 | 0.505659614 | 0.050299567 |
| Gm826 | 0.025474767 | -3.572889668 | 0 | 0.041 | 1 | 0.698534498 | 2.432959407 | 0 | 0.015 | 1 | 0.698534498 | 0.050300571 |
| Tex15 | 0.025475025 | -4.008988783 | 0 | 0.041 | 1 | 0.669851164 | 1.31748219 | 0 | 0.019 | 1 | 0.669851164 | 0.050301073 |
| C230012O17Rik | 0.02547554 | -2.485426827 | 0 | 0.041 | 1 | 0.73075158 | 2.695993813 | 0 | 0.012 | 1 | 0.73075158 | 0.050302077 |
| Txnl4b | 0.02547554 | -2.485426827 | 0 | 0.041 | 1 | 0.577155236 | 1.31748219 | 0 | 0.031 | 1 | 0.577155236 | 0.050302077 |
| Tomm20l | 0.02547554 | -2.485426827 | 0 | 0.041 | 1 | 0.73075158 | 2.017921908 | 0 | 0.012 | 1 | 0.73075158 | 0.050302077 |
| Zfp712 | 0.02547554 | -2.485426827 | 0 | 0.041 | 1 | 0.557756607 | 1.111031312 | 0 | 0.034 | 1 | 0.557756607 | 0.050302077 |
| Fbxl16 | 0.02547554 | -2.485426827 | 0 | 0.041 | 1 | 0.577142568 | 1.432959407 | 0 | 0.031 | 1 | 0.577142568 | 0.050302077 |
| H2-Q7 | 0.02547554 | -2.485426827 | 0 | 0.041 | 1 | 0.59781419 | 1.111031312 | 0 | 0.028 | 1 | 0.59781419 | 0.050302077 |
| Gm50214 | 0.02547554 | -2.485426827 | 0 | 0.041 | 1 | 0.597832556 | 1.31748219 | 0 | 0.028 | 1 | 0.597832556 | 0.050302077 |
| Khdrbs3 | 0.025498668 | -1.201633861 | 0.109 | 0.206 | 1 | 0.774081327 | 0.74645888 | 0.4 | 0.497 | 1 | 0.774081327 | 0.050347153 |
| Bpifa1 | 0.106481932 | -0.430979043 | 0.336 | 0.412 | 1 | 0.02553274 | -1.05085637 | 0.4 | 0.735 | 1 | 0.106481932 | 0.05041356 |
| Gm4924 | 0.78675278 | -0.592342031 | 0.092 | 0.1 | 1 | 0.025539888 | 2.017921908 | 0.2 | 0.043 | 1 | 0.78675278 | 0.050427491 |
| Adcy10 | 0.687848155 | -0.402964667 | 0.042 | 0.053 | 1 | 0.025539975 | 1.902444691 | 0.2 | 0.043 | 1 | 0.687848155 | 0.05042766 |
| Eral1 | 0.025560048 | -1.070389328 | 0.076 | 0.165 | 1 | 0.21051379 | 1.210566986 | 0.3 | 0.148 | 1 | 0.21051379 | 0.05046678 |
| Gm6277 | 0.025590742 | 1.716207034 | 0.109 | 0.041 | 1 | 0.557729726 | 1.111031312 | 0 | 0.034 | 1 | 0.557729726 | 0.050526597 |
| D130009I18Rik | 0.277298048 | -0.121404614 | 0.689 | 0.765 | 1 | 0.025593327 | -1.267480311 | 0.4 | 0.735 | 1 | 0.277298048 | 0.050531635 |
| Cdt1 | 0.025647412 | -0.718087584 | 0.092 | 0.188 | 1 | 0.843868 | 0.185031894 | 0.1 | 0.117 | 1 | 0.843868 | 0.050637035 |
| Ypel1 | 0.02565168 | -1.292781749 | 0.05 | 0.129 | 1 | 0.945087907 | 0.526068812 | 0.1 | 0.105 | 1 | 0.945087907 | 0.050645351 |
| Scyl2 | 0.025674194 | -0.674460652 | 0.277 | 0.394 | 1 | 0.588317889 | 0.141963172 | 0.4 | 0.599 | 1 | 0.588317889 | 0.050689224 |
| Crls1 | 0.025683107 | -0.807354922 | 0.286 | 0.388 | 1 | 0.22700909 | -0.806506527 | 0.2 | 0.37 | 1 | 0.22700909 | 0.050706592 |
| Efnb1 | 0.025685338 | -0.450661409 | 0.269 | 0.4 | 1 | 0.292416168 | -0.486698484 | 0.2 | 0.377 | 1 | 0.292416168 | 0.05071094 |
| Hnrnph3 | 0.025706609 | -0.37754207 | 0.874 | 0.824 | 1 | 0.150800201 | -0.536666944 | 0.7 | 0.877 | 1 | 0.150800201 | 0.050752388 |
| Cryzl2 | 0.025715063 | 0.514573173 | 0.42 | 0.288 | 1 | 0.305663684 | 0.821524695 | 0.4 | 0.25 | 1 | 0.305663684 | 0.050768861 |
| Ube2d3 | 0.065974849 | -0.20755738 | 0.983 | 1 | 1 | 0.025753972 | -0.590432441 | 1 | 0.994 | 1 | 0.065974849 | 0.050844677 |
| Tcp11l1 | 0.873656501 | -0.175086707 | 0.21 | 0.212 | 1 | 0.025759483 | 1.782705446 | 0.5 | 0.247 | 1 | 0.873656501 | 0.050855415 |
| Gm3331 | 0.025813827 | -1.411426246 | 0.017 | 0.076 | 1 | 0.318562315 | 2.017921908 | 0.1 | 0.037 | 1 | 0.318562315 | 0.050961301 |
| Gm29724 | 0.027323202 | -2.485426827 | 0.008 | 0.059 | 1 | 0.025814657 | 3.432959407 | 0.1 | 0.012 | 1 | 0.027323202 | 0.050962918 |
| Rnf138rt1 | 0.06000657 | -2.292781749 | 0 | 0.029 | 1 | 0.025814657 | 3.210566986 | 0.1 | 0.012 | 1 | 0.06000657 | 0.050962918 |
| Kif20a | 0.062431757 | -2.655351829 | 0.008 | 0.047 | 1 | 0.025814657 | 3.017921908 | 0.1 | 0.012 | 1 | 0.062431757 | 0.050962918 |
| Gm57459 | 0.091361251 | 2.099535674 | 0.017 | 0 | 1 | 0.025814657 | 3.210566986 | 0.1 | 0.012 | 1 | 0.091361251 | 0.050962918 |
| Gm11110 | 0.093245792 | -1.807354922 | 0 | 0.024 | 1 | 0.025814657 | 3.210566986 | 0.1 | 0.012 | 1 | 0.093245792 | 0.050962918 |
| Gm26590 | 0.112335908 | -1.600904045 | 0.017 | 0.053 | 1 | 0.025814657 | 3.432959407 | 0.1 | 0.012 | 1 | 0.112335908 | 0.050962918 |
| Tafa3 | 0.132134807 | 0.514573173 | 0.059 | 0.024 | 1 | 0.025814657 | 3.432959407 | 0.1 | 0.012 | 1 | 0.132134807 | 0.050962918 |
| Klhl10 | 0.147029286 | -1.807354922 | 0 | 0.018 | 1 | 0.025814657 | 3.432959407 | 0.1 | 0.012 | 1 | 0.147029286 | 0.050962918 |
| Zfp786 | 0.147030916 | -2.292781749 | 0 | 0.018 | 1 | 0.025814657 | 3.432959407 | 0.1 | 0.012 | 1 | 0.147030916 | 0.050962918 |
| Cenpm | 0.169706304 | -0.485426827 | 0.017 | 0.047 | 1 | 0.025814657 | 3.210566986 | 0.1 | 0.012 | 1 | 0.169706304 | 0.050962918 |
| Ccdc83 | 0.215269607 | -1.807354922 | 0.008 | 0.029 | 1 | 0.025814657 | 2.847996907 | 0.1 | 0.012 | 1 | 0.215269607 | 0.050962918 |
| Gm46404 | 0.216340691 | -1.655351829 | 0.008 | 0.029 | 1 | 0.025814657 | 3.432959407 | 0.1 | 0.012 | 1 | 0.216340691 | 0.050962918 |
| 1700041G16Rik | 0.237885589 | -1.070389328 | 0 | 0.012 | 1 | 0.025814657 | 3.432959407 | 0.1 | 0.012 | 1 | 0.237885589 | 0.050962918 |
| Mc1r | 0.242320046 | -0.900464326 | 0.017 | 0.041 | 1 | 0.025814657 | 3.210566986 | 0.1 | 0.012 | 1 | 0.242320046 | 0.050962918 |

| | | | | | | | | | | | | |
|---|---|---|---|---|---|---|---|---|---|---|---|---|
| Kcnd3 | 0.337379399 | -0.707819249 | 0.008 | 0.024 | 1 | 0.025814657 | 3.432959407 | 0.1 | 0.012 | 1 | 0.337379399 | 0.050962918 |
| Ermap | 0.370184089 | 1.099535674 | 0.017 | 0.006 | 1 | 0.025814657 | 3.432959407 | 0.1 | 0.012 | 1 | 0.370184089 | 0.050962918 |
| B930094E09Rik | 0.374548525 | 0.514573173 | 0.017 | 0.006 | 1 | 0.025814657 | 3.432959407 | 0.1 | 0.012 | 1 | 0.374548525 | 0.050962918 |
| Gm57465 | 0.511005543 | -0.485426827 | 0.008 | 0.018 | 1 | 0.025814657 | 3.432959407 | 0.1 | 0.012 | 1 | 0.511005543 | 0.050962918 |
| Ptk6 | 0.511005543 | -0.485426827 | 0.008 | 0.018 | 1 | 0.025814657 | 3.432959407 | 0.1 | 0.012 | 1 | 0.511005543 | 0.050962918 |
| Hcn2 | 0.511005543 | -0.485426827 | 0.008 | 0.018 | 1 | 0.025814657 | 3.210566986 | 0.1 | 0.012 | 1 | 0.511005543 | 0.050962918 |
| Wdr95 | 0.51784324 | 0.514573173 | 0.008 | 0.018 | 1 | 0.025814657 | 3.432959407 | 0.1 | 0.012 | 1 | 0.51784324 | 0.050962918 |
| Ribc1 | 0.519851852 | 1.029146346 | 0.05 | 0.035 | 1 | 0.025814657 | 3.432959407 | 0.1 | 0.012 | 1 | 0.519851852 | 0.050962918 |
| Gm6034 | 0.715877701 | 0.836501268 | 0.017 | 0.012 | 1 | 0.025814657 | 3.210566986 | 0.1 | 0.012 | 1 | 0.715877701 | 0.050962918 |
| Gm52829 | 0.781790612 | -1.070389328 | 0.008 | 0.012 | 1 | 0.025814657 | 3.432959407 | 0.1 | 0.012 | 1 | 0.781790612 | 0.050962918 |
| 4930505A04Rik | 0.781790612 | -0.485426827 | 0.008 | 0.012 | 1 | 0.025814657 | 3.210566986 | 0.1 | 0.012 | 1 | 0.781790612 | 0.050962918 |
| Cfap20dc | 0.842422009 | 0.251538767 | 0.025 | 0.029 | 1 | 0.025814657 | 3.432959407 | 0.1 | 0.012 | 1 | 0.842422009 | 0.050962918 |
| Gm33037 | 0.926217033 | -0.748461233 | 0.034 | 0.035 | 1 | 0.025814657 | 3.432959407 | 0.1 | 0.012 | 1 | 0.926217033 | 0.050962918 |
| Gm960 | 0.935790323 | -0.292781749 | 0.025 | 0.024 | 1 | 0.025814657 | 3.432959407 | 0.1 | 0.012 | 1 | 0.935790323 | 0.050962918 |
| 1500035N22Rik | 0.951374613 | 0.344648171 | 0.034 | 0.035 | 1 | 0.025814657 | 3.432959407 | 0.1 | 0.012 | 1 | 0.951374613 | 0.050962918 |
| Adad1 | 0.976891938 | -0.106915204 | 0.042 | 0.041 | 1 | 0.025817087 | 2.695993813 | 0.1 | 0.012 | 1 | 0.976891938 | 0.050967652 |
| Sgms1 | 0.025843903 | -0.516453723 | 0.773 | 0.818 | 1 | 0.044607562 | -0.818128447 | 0.6 | 0.886 | 1 | 0.044607562 | 0.051019898 |
| Arhgef40 | 0.025857812 | -0.4665678 | 0.353 | 0.488 | 1 | 0.615332002 | 0.751885014 | 0.6 | 0.583 | 1 | 0.615332002 | 0.051046998 |
| Ankrd54 | 0.02591197 | 0.566103473 | 0.378 | 0.253 | 1 | 0.385454775 | 0.504852326 | 0.5 | 0.336 | 1 | 0.385454775 | 0.051152509 |
| Zfp36l1 | 0.700639734 | 0.116917752 | 0.857 | 0.865 | 1 | 0.025954708 | -1.695464607 | 0.7 | 0.843 | 1 | 0.700639734 | 0.05123577 |
| Agap1 | 0.025966875 | -0.712962552 | 0.504 | 0.606 | 1 | 0.749402051 | -0.197674952 | 0.7 | 0.772 | 1 | 0.749402051 | 0.051259472 |
| Ak3 | 0.026015307 | -0.791235257 | 0.235 | 0.341 | 1 | 0.046804428 | -0.884955625 | 0.2 | 0.565 | 1 | 0.046804428 | 0.051353819 |
| Msantd3 | 0.026027277 | -1.211251864 | 0.101 | 0.194 | 1 | 0.229634278 | 0.754887502 | 0.4 | 0.219 | 1 | 0.229634278 | 0.051377135 |
| Acp6 | 0.026028926 | -1.03796785 | 0.109 | 0.206 | 1 | 0.124197723 | -1.357117523 | 0 | 0.194 | 1 | 0.124197723 | 0.051380347 |
| Sytl5 | 0.026087496 | 0.762500686 | 0.101 | 0.035 | 1 | 0.071483247 | 4.124837112 | 0.1 | 0.019 | 1 | 0.071483247 | 0.051494435 |
| Ier3ip1 | 0.098036207 | -0.267734331 | 0.857 | 0.865 | 1 | 0.026155767 | -0.816456188 | 0.5 | 0.892 | 1 | 0.098036207 | 0.05162741 |
| Zbtb26 | 0.026169515 | -1.392317423 | 0.017 | 0.076 | 1 | 0.913228316 | 0.695993813 | 0.1 | 0.086 | 1 | 0.913228316 | 0.051654186 |
| Gm10658 | 0.026169515 | -1.392317423 | 0.017 | 0.076 | 1 | 0.474927752 | 0.625604485 | 0 | 0.049 | 1 | 0.474927752 | 0.051654186 |
| Gm56507 | 0.026169515 | -1.392317423 | 0.017 | 0.076 | 1 | 0.446819579 | 0.111031312 | 0 | 0.056 | 1 | 0.446819579 | 0.051654186 |
| Rpl7l1 | 0.026224724 | -0.387938982 | 0.773 | 0.741 | 1 | 0.264443271 | -0.364547729 | 0.6 | 0.756 | 1 | 0.264443271 | 0.051761711 |
| Cic | 0.026266126 | -0.728697978 | 0.311 | 0.429 | 1 | 0.252602317 | -0.794576317 | 0.4 | 0.506 | 1 | 0.252602317 | 0.051842342 |
| Emx2 | 0.060010312 | -2.944858446 | 0 | 0.029 | 1 | 0.026346883 | 3.210566986 | 0.1 | 0.012 | 1 | 0.060010312 | 0.051999608 |
| Mis18bp1 | 0.164847278 | -2.070389328 | 0.017 | 0.047 | 1 | 0.026346883 | 2.695993813 | 0.1 | 0.012 | 1 | 0.164847278 | 0.051999608 |
| BC051019 | 0.168156185 | 1.514573173 | 0.025 | 0.006 | 1 | 0.026346883 | 2.695993813 | 0.1 | 0.012 | 1 | 0.168156185 | 0.051999608 |
| Gm34917 | 0.331040183 | -1.292781749 | 0.008 | 0.024 | 1 | 0.026346883 | 3.017921908 | 0.1 | 0.012 | 1 | 0.331040183 | 0.051999608 |
| Igfbp6 | 0.335789682 | -0.900464326 | 0.008 | 0.024 | 1 | 0.026346883 | 0.591657153 | 0.1 | 0.012 | 1 | 0.335789682 | 0.051999608 |
| Cxxc4 | 0.506471296 | -1.070389328 | 0.008 | 0.018 | 1 | 0.026346883 | 3.017921908 | 0.1 | 0.012 | 1 | 0.506471296 | 0.051999608 |
| Gm26608 | 0.721162042 | 0.514573173 | 0.017 | 0.012 | 1 | 0.026346883 | 3.210566986 | 0.1 | 0.012 | 1 | 0.721162042 | 0.051999608 |
| Gm35853 | 0.924405131 | -1.070389328 | 0.05 | 0.053 | 1 | 0.026346883 | 3.210566986 | 0.1 | 0.012 | 1 | 0.924405131 | 0.051999608 |
| Vps33b | 0.026363374 | -0.944858446 | 0.084 | 0.176 | 1 | 0.822519456 | 0.210566986 | 0.1 | 0.12 | 1 | 0.822519456 | 0.052031721 |
| Cnnm4 | 0.044105819 | 0.114332675 | 0.681 | 0.506 | 1 | 0.026446089 | 1.119068631 | 0.8 | 0.481 | 1 | 0.044105819 | 0.052192782 |
| Fhad1 | 0.026462866 | -0.567888987 | 0.109 | 0.041 | 1 | 0.139970154 | 3.14345279 | 0.1 | 0.025 | 1 | 0.139970154 | 0.052225448 |
| Hsd17b12 | 0.151620068 | 0.243521749 | 0.882 | 0.794 | 1 | 0.026489898 | 0.850356179 | 0.9 | 0.738 | 1 | 0.151620068 | 0.052278081 |
| Agfg2 | 0.026510149 | 0.538419915 | 0.328 | 0.212 | 1 | 0.434935356 | -0.763437806 | 0.1 | 0.191 | 1 | 0.434935356 | 0.05231751 |
| Gm47096 | 0.239542116 | -0.86393845 | 0.034 | 0.065 | 1 | 0.02655114 | 2.847996907 | 0.2 | 0.046 | 1 | 0.239542116 | 0.052397317 |
| AI597479 | 0.026569742 | -1.015941544 | 0.05 | 0.129 | 1 | 0.307031624 | -0.267480311 | 0 | 0.096 | 1 | 0.307031624 | 0.052433533 |
| Ptn | 0.026603005 | -0.213054557 | 0.244 | 0.412 | 1 | 0.959963519 | -0.588642847 | 0.4 | 0.42 | 1 | 0.959963519 | 0.052498291 |
| Ints12 | 0.02662553 | -0.851076299 | 0.176 | 0.282 | 1 | 0.236729579 | -0.763437806 | 0.1 | 0.262 | 1 | 0.236729579 | 0.052542141 |
| Snrnp200 | 0.026647438 | -0.847996907 | 0.345 | 0.441 | 1 | 0.109802443 | -0.840059087 | 0.3 | 0.565 | 1 | 0.109802443 | 0.05258479 |
| Ppp3r1 | 0.039343667 | -0.438776734 | 0.613 | 0.647 | 1 | 0.026669661 | -0.910185174 | 0.4 | 0.775 | 1 | 0.039343667 | 0.052628051 |
| Msi1 | 0.026723656 | -1.008988783 | 0.109 | 0.206 | 1 | 0.59401657 | -0.357117523 | 0.1 | 0.157 | 1 | 0.59401657 | 0.052733158 |
| Tmem68 | 0.060779905 | -0.752213368 | 0.235 | 0.324 | 1 | 0.026740553 | -2.473931188 | 0 | 0.343 | 1 | 0.060779905 | 0.052766049 |
| Gm41496 | 0.399956006 | 0.384512632 | 0.109 | 0.147 | 1 | 0.026776745 | 2.982297998 | 0.4 | 0.157 | 1 | 0.399956006 | 0.052836496 |
| 1600012H06Rik | 0.026809864 | -0.770829046 | 0.244 | 0.353 | 1 | 0.435183735 | -0.362899876 | 0.2 | 0.306 | 1 | 0.435183735 | 0.05290096 |
| Pcdh19 | 0.026882431 | -3.572889668 | 0.008 | 0.059 | 1 | 0.12122695 | -1.888968688 | 0 | 0.198 | 1 | 0.12122695 | 0.053042196 |
| Ibsp | 0.026884328 | -8.920749827 | 0.008 | 0.059 | 1 | 0.619973894 | -1.391469028 | 0 | 0.025 | 1 | 0.619973894 | 0.053045889 |
| Exoc3l4 | 0.026884747 | 1.362570079 | 0.118 | 0.047 | 1 | 0.189486514 | 2.695993813 | 0.1 | 0.028 | 1 | 0.189486514 | 0.053046705 |
| Sult6b1 | 0.093245792 | -1.807354922 | 0 | 0.024 | 1 | 0.026887441 | 2.558490289 | 0.1 | 0.012 | 1 | 0.093245792 | 0.053051948 |
| Sgo2a | 0.249990822 | -1.222392421 | 0.025 | 0.053 | 1 | 0.026887719 | 2.31748219 | 0.1 | 0.012 | 1 | 0.249990822 | 0.053052489 |
| Fastkd2 | 0.687545249 | -0.263034406 | 0.109 | 0.124 | 1 | 0.026899421 | 1.611929548 | 0.4 | 0.139 | 1 | 0.687545249 | 0.053075262 |
| Pex1 | 0.032018815 | -1.321928095 | 0.101 | 0.188 | 1 | 0.026932335 | 1.830294905 | 0.4 | 0.142 | 1 | 0.032018815 | 0.053139319 |
| Cct7 | 0.027000993 | -0.421076416 | 0.723 | 0.753 | 1 | 0.330367982 | -0.217294554 | 0.6 | 0.818 | 1 | 0.330367982 | 0.053272932 |
| Rims2 | 0.027029182 | -2.944858446 | 0.008 | 0.059 | 1 | 0.385726601 | -1.131825212 | 0 | 0.071 | 1 | 0.385726601 | 0.053327787 |
| Gins1 | 0.027029355 | -2.807354922 | 0.008 | 0.059 | 1 | 0.112266525 | 3.017921908 | 0.1 | 0.022 | 1 | 0.112266525 | 0.053328123 |
| Nus1 | 0.027043222 | -0.651436779 | 0.487 | 0.547 | 1 | 0.071384569 | -0.679045618 | 0.4 | 0.682 | 1 | 0.071384569 | 0.053355109 |
| Grk2 | 0.027063422 | -0.929033479 | 0.202 | 0.306 | 1 | 0.381390584 | -0.633129783 | 0.3 | 0.404 | 1 | 0.381390584 | 0.053394415 |
| Prmt7 | 0.02710612 | -1.324962155 | 0.134 | 0.229 | 1 | 0.572483592 | 0.362570079 | 0.3 | 0.219 | 1 | 0.572483592 | 0.053477498 |
| Atp8b1 | 0.895774964 | -0.292781749 | 0.101 | 0.094 | 1 | 0.027122622 | 2.159940913 | 0.3 | 0.096 | 1 | 0.895774964 | 0.053509607 |
| Abhd17c | 0.027165436 | -1.031395196 | 0.168 | 0.265 | 1 | 0.44122153 | 0.665405493 | 0.4 | 0.296 | 1 | 0.44122153 | 0.053592911 |
| Gm42937 | 0.260793067 | 0.251538767 | 0.109 | 0.071 | 1 | 0.027171466 | 2.880418384 | 0.2 | 0.046 | 1 | 0.260793067 | 0.053604644 |
| Gm50399 | 0.027175411 | -2.572889668 | 0.008 | 0.059 | 1 | 0.577164735 | 1.111031312 | 0 | 0.031 | 1 | 0.577164735 | 0.05361232 |
| Snu13 | 0.0271871 | -0.223998986 | 0.916 | 0.912 | 1 | 0.067081699 | -0.820286804 | 0.8 | 0.898 | 1 | 0.067081699 | 0.053635062 |
| Agbl4 | 0.027203272 | 1.23893873 | 0.118 | 0.047 | 1 | 0.061031695 | 2.31748219 | 0.2 | 0.059 | 1 | 0.061031695 | 0.053666526 |
| Ubxn6 | 0.290795234 | -0.384497918 | 0.58 | 0.594 | 1 | 0.027224716 | -1.290417122 | 0.3 | 0.614 | 1 | 0.290795234 | 0.053708248 |
| Rbl1 | 0.027236277 | -1.807354922 | 0.076 | 0.159 | 1 | 0.94991328 | 0.888638891 | 0.1 | 0.093 | 1 | 0.94991328 | 0.053730739 |
| Ccdc91 | 0.345206776 | -0.678071905 | 0.21 | 0.241 | 1 | 0.027242066 | 1.030860964 | 0.5 | 0.191 | 1 | 0.345206776 | 0.053742003 |
| Kctd18 | 0.027249921 | -1.485426827 | 0.084 | 0.171 | 1 | 0.796917492 | 0.515421567 | 0.2 | 0.16 | 1 | 0.796917492 | 0.053757284 |
| Bcs1l | 0.027261087 | -1.070389328 | 0.067 | 0.153 | 1 | 0.896995661 | 0.847996907 | 0.1 | 0.086 | 1 | 0.896995661 | 0.053779007 |
| Naga | 0.027289748 | 0.330148602 | 0.571 | 0.453 | 1 | 0.812112452 | 0.131789873 | 0.5 | 0.426 | 1 | 0.812112452 | 0.053834765 |
| Bloc1s4 | 0.02730295 | -0.603608254 | 0.286 | 0.406 | 1 | 0.061077784 | -1.399930607 | 0.1 | 0.389 | 1 | 0.061077784 | 0.053860449 |
| Gm17036 | 0.027323202 | -2.485426827 | 0.008 | 0.059 | 1 | 0.334207905 | -0.267480311 | 0 | 0.086 | 1 | 0.334207905 | 0.053899847 |
| Zfp202 | 0.0273239 | -2.655351829 | 0.008 | 0.059 | 1 | 0.539431168 | 1.111031312 | 0 | 0.037 | 1 | 0.539431168 | 0.053901205 |
| Rbpms2 | 0.0273239 | -2.392317423 | 0.008 | 0.059 | 1 | 0.539464786 | 0.494359952 | 0 | 0.037 | 1 | 0.539464786 | 0.053901205 |
| Thoc6 | 0.027342966 | -0.846174171 | 0.277 | 0.382 | 1 | 0.110433779 | -0.408342847 | 0.1 | 0.392 | 1 | 0.110433779 | 0.053938294 |
| Gm15559 | 0.027374334 | -2.070389328 | 0.025 | 0.088 | 1 | 0.320995017 | 2.017921908 | 0.1 | 0.037 | 1 | 0.320995017 | 0.053999314 |
| Arrdc2 | 0.072846557 | 0.893084796 | 0.151 | 0.082 | 1 | 0.027405989 | 4.432959407 | 0.2 | 0.049 | 1 | 0.072846557 | 0.054060889 |
| Mrgbp | 0.02741395 | -0.982926487 | 0.109 | 0.206 | 1 | 0.889909624 | 0.695993813 | 0.2 | 0.191 | 1 | 0.889909624 | 0.054076376 |
| Gm17092 | 0.027414662 | -1.177304532 | 0.092 | 0.182 | 1 | 0.221413195 | 1.732519689 | 0.2 | 0.09 | 1 | 0.221413195 | 0.054077761 |
| Steap3 | 0.027455333 | -0.111968432 | 0.395 | 0.541 | 1 | 0.391293201 | -0.50700301 | 0.5 | 0.651 | 1 | 0.391293201 | 0.05415687 |
| Kcnk5 | 0.084361132 | 0.437880416 | 0.739 | 0.618 | 1 | 0.027466774 | 1.14410871 | 0.9 | 0.627 | 1 | 0.084361132 | 0.054179124 |
| Actr3b | 0.02746994 | -2.572889668 | 0.008 | 0.059 | 1 | 0.59781419 | 1.558490289 | 0 | 0.028 | 1 | 0.59781419 | 0.054185283 |
| Itga7 | 0.02746994 | -2.944858446 | 0.008 | 0.059 | 1 | 0.258780271 | 2.354956895 | 0.1 | 0.034 | 1 | 0.258780271 | 0.054185283 |
| Dok3 | 0.02746994 | -2.392317423 | 0.008 | 0.059 | 1 | 0.810418519 | 0.591657153 | 0.1 | 0.077 | 1 | 0.810418519 | 0.054185283 |
| Cfap53 | 0.027470115 | -2.292781749 | 0.008 | 0.059 | 1 | 0.434539169 | 2.816288047 | 0.1 | 0.049 | 1 | 0.434539169 | 0.054185624 |
| Kifc3 | 0.027503422 | 0.49808505 | 0.462 | 0.324 | 1 | 0.06573387 | 1.306427001 | 0.5 | 0.272 | 1 | 0.06573387 | 0.054250407 |
| St7 | 0.027572422 | -0.755878705 | 0.504 | 0.571 | 1 | 0.463885711 | 0.808468542 | 0.6 | 0.568 | 1 | 0.463885711 | 0.054384605 |
| Twf1 | 0.16882684 | -0.485426827 | 0.571 | 0.559 | 1 | 0.027601398 | -1.222392421 | 0.3 | 0.651 | 1 | 0.16882684 | 0.054440959 |
| Nkx6-1 | 0.027615608 | -2.185866545 | 0.008 | 0.059 | 1 | 0.557756607 | 0.930459067 | 0 | 0.034 | 1 | 0.557756607 | 0.054468595 |

| | | | | | | | | | | | | |
|---|---|---|---|---|---|---|---|---|---|---|---|---|
| Ttc12 | 0.027615608 | -2.292781749 | 0.008 | 0.059 | 1 | 0.773317994 | 0.888638891 | 0.1 | 0.074 | 1 | 0.773317994 | 0.054468595 |
| Smim36 | 0.027615608 | -2.292781749 | 0.008 | 0.059 | 1 | 0.505605027 | 0.930459067 | 0 | 0.043 | 1 | 0.505605027 | 0.054468595 |
| Nme4 | 0.027615608 | -2.185866545 | 0.008 | 0.059 | 1 | 0.597824685 | 1.432959407 | 0 | 0.028 | 1 | 0.597824685 | 0.054468595 |
| Zfp300 | 0.027615608 | -2.292781749 | 0.008 | 0.059 | 1 | 0.112266525 | 3.017921908 | 0.1 | 0.022 | 1 | 0.112266525 | 0.054468595 |
| Mtpn | 0.027637551 | -0.475711672 | 0.697 | 0.753 | 1 | 0.206489137 | -0.47819952 | 0.6 | 0.861 | 1 | 0.206489137 | 0.054511268 |
| Pi4k2b | 0.027649763 | -0.920812972 | 0.286 | 0.376 | 1 | 0.332789729 | -0.382957528 | 0.4 | 0.54 | 1 | 0.332789729 | 0.054535016 |
| Sde2 | 0.027685359 | -0.570818318 | 0.714 | 0.706 | 1 | 0.360123789 | -0.431385493 | 0.6 | 0.738 | 1 | 0.360123789 | 0.054604239 |
| Wwp1 | 0.027698567 | -0.611875162 | 0.538 | 0.641 | 1 | 0.207499069 | -0.705454621 | 0.5 | 0.716 | 1 | 0.207499069 | 0.054629924 |
| Carmil1 | 0.02771771 | -0.712974247 | 0.782 | 0.824 | 1 | 0.275645138 | -0.514915652 | 1 | 0.932 | 1 | 0.275645138 | 0.054667148 |
| Snhg15 | 0.027741167 | -1.03562391 | 0.101 | 0.194 | 1 | 0.233697828 | 1.432959407 | 0.2 | 0.086 | 1 | 0.233697828 | 0.054712761 |
| Gm13563 | 0.027760189 | -2.070389328 | 0.008 | 0.059 | 1 | 0.474877085 | 0.847996907 | 0 | 0.049 | 1 | 0.474877085 | 0.054749751 |
| Gfra4 | 0.027760189 | -2.070389328 | 0.008 | 0.059 | 1 | 0.597845674 | 1.695993813 | 0 | 0.028 | 1 | 0.597845674 | 0.054749751 |
| Zfp82 | 0.027760189 | -2.070389328 | 0.008 | 0.059 | 1 | 0.601875084 | 1.31748219 | 0.1 | 0.059 | 1 | 0.601875084 | 0.054749751 |
| Polm | 0.027760189 | -2.185866545 | 0.008 | 0.059 | 1 | 0.355415506 | 2.210566986 | 0.1 | 0.04 | 1 | 0.355415506 | 0.054749751 |
| Gm47469 | 0.027760189 | -2.185866545 | 0.008 | 0.059 | 1 | 0.539445308 | 1.017921908 | 0 | 0.037 | 1 | 0.539445308 | 0.054749751 |
| 1700056N10Rik | 0.027760189 | -2.392317423 | 0.008 | 0.059 | 1 | 0.460551517 | 0.625604485 | 0 | 0.052 | 1 | 0.460551517 | 0.054749751 |
| Ankrd13d | 0.027760189 | -2.292781749 | 0.008 | 0.059 | 1 | 0.813207742 | 0.847996907 | 0.1 | 0.077 | 1 | 0.813207742 | 0.054749751 |
| Gm56660 | 0.027760189 | -2.070389328 | 0.008 | 0.059 | 1 | 0.669848329 | 1.847996907 | 0 | 0.019 | 1 | 0.669848329 | 0.054749751 |
| Nucb1 | 0.027787087 | 0.241700575 | 0.933 | 0.847 | 1 | 0.233580935 | 0.256572155 | 1 | 0.889 | 1 | 0.233580935 | 0.054802053 |
| Zfp783 | 0.027857234 | -0.944858446 | 0.109 | 0.206 | 1 | 0.717572959 | 0.236562194 | 0.1 | 0.139 | 1 | 0.717572959 | 0.054938443 |
| Gtpbp4 | 0.027884061 | -0.45432561 | 0.79 | 0.8 | 1 | 0.222780031 | -0.467907401 | 0.8 | 0.818 | 1 | 0.222780031 | 0.054990602 |
| Tbc1d4 | 0.988854633 | -1.007653573 | 0.269 | 0.253 | 1 | 0.027886633 | 1.248534836 | 0.5 | 0.213 | 1 | 0.988854633 | 0.054995602 |
| Frem2 | 0.027942624 | -1.944858446 | 0.008 | 0.059 | 1 | 0.73075158 | 2.695993813 | 0 | 0.012 | 1 | 0.73075158 | 0.055104458 |
| Pnldc1 | 0.027942624 | -1.944858446 | 0.008 | 0.059 | 1 | 0.522115938 | 0.847996907 | 0 | 0.04 | 1 | 0.522115938 | 0.055104458 |
| Tincr | 0.027942624 | -1.944858446 | 0.008 | 0.059 | 1 | 0.474904369 | 0.769994395 | 0 | 0.049 | 1 | 0.474904369 | 0.055104458 |
| Col4a1 | 0.027958407 | -0.736965594 | 0.202 | 0.318 | 1 | 0.161761506 | 0.363611361 | 0.6 | 0.358 | 1 | 0.161761506 | 0.055135142 |
| Nalcn | 0.028000593 | -2.432959407 | 0.025 | 0.088 | 1 | 0.460680742 | -0.625934282 | 0 | 0.052 | 1 | 0.460680742 | 0.055217152 |
| Sfmbt1 | 0.028007717 | -0.790772038 | 0.538 | 0.612 | 1 | 0.286166888 | 0.630798712 | 0.8 | 0.781 | 1 | 0.286166888 | 0.055231001 |
| Bak1 | 0.028044244 | -0.970853654 | 0.067 | 0.153 | 1 | 0.219552857 | -0.709998547 | 0 | 0.133 | 1 | 0.219552857 | 0.055302009 |
| Pcbp4 | 0.028060026 | -0.888782521 | 0.218 | 0.318 | 1 | 0.419170449 | -0.596787936 | 0.3 | 0.386 | 1 | 0.419170449 | 0.055332687 |
| Eef1a1 | 0.042325151 | -0.173643809 | 1 | 1 | 1 | 0.028064182 | -0.610383415 | 1 | 0.997 | 1 | 0.042325151 | 0.055340765 |
| Nudcd1 | 0.028083792 | -0.758445322 | 0.244 | 0.353 | 1 | 0.283536899 | -1.015501094 | 0.1 | 0.238 | 1 | 0.283536899 | 0.055378884 |
| Lig3 | 0.028094143 | -0.944858446 | 0.21 | 0.312 | 1 | 0.594326943 | 0.483943336 | 0.4 | 0.315 | 1 | 0.594326943 | 0.055399005 |
| Scyl1 | 0.028095759 | -0.62058641 | 0.378 | 0.482 | 1 | 0.125399859 | -0.893769674 | 0.3 | 0.546 | 1 | 0.125399859 | 0.055402146 |
| Dnajc4 | 0.028105444 | 0.437951891 | 0.37 | 0.241 | 1 | 0.568112634 | 0.832055363 | 0.3 | 0.231 | 1 | 0.568112634 | 0.055420972 |
| Rap1gds1 | 0.028119373 | -0.58185659 | 0.655 | 0.741 | 1 | 0.537205165 | -0.552541023 | 0.8 | 0.821 | 1 | 0.537205165 | 0.055448047 |
| Ubl4a | 0.028119556 | -0.450661409 | 0.235 | 0.353 | 1 | 0.647746441 | 0.385653692 | 0.2 | 0.287 | 1 | 0.647746441 | 0.055448402 |
| Zcchc4 | 0.028120754 | -1.524955191 | 0.067 | 0.147 | 1 | 0.570905211 | -0.357117523 | 0.1 | 0.164 | 1 | 0.570905211 | 0.05545073 |
| Rflna | 0.028181496 | 0.175367087 | 0.756 | 0.606 | 1 | 0.576278321 | -0.11037935 | 0.6 | 0.821 | 1 | 0.576278321 | 0.055568795 |
| Adam10 | 0.028187945 | -0.790281409 | 0.471 | 0.547 | 1 | 0.089781576 | -1.062451508 | 0.4 | 0.63 | 1 | 0.089781576 | 0.05558133 |
| Tapt1 | 0.028192433 | -0.599498594 | 0.42 | 0.512 | 1 | 0.28536439 | -0.471013705 | 0.6 | 0.716 | 1 | 0.28536439 | 0.055590054 |
| Zdhhc14 | 0.028196376 | -1.753915663 | 0.067 | 0.147 | 1 | 0.335500697 | -1.330806246 | 0.1 | 0.216 | 1 | 0.335500697 | 0.055597717 |
| Rps9 | 0.061390655 | -0.154612769 | 0.983 | 0.988 | 1 | 0.028206515 | -0.691161905 | 0.9 | 0.991 | 1 | 0.061390655 | 0.055617423 |
| Gm16062 | 0.629891029 | -0.485426827 | 0.084 | 0.1 | 1 | 0.028248765 | 1.036069255 | 0.4 | 0.133 | 1 | 0.629891029 | 0.055699538 |
| Irf2bpl | 0.028320381 | 0.408531535 | 0.689 | 0.559 | 1 | 0.561919143 | -0.184734295 | 0.8 | 0.775 | 1 | 0.561919143 | 0.055838717 |
| Map7 | 0.028365986 | -0.935362534 | 0.403 | 0.506 | 1 | 0.117729046 | 0.69350855 | 0.8 | 0.549 | 1 | 0.117729046 | 0.055927344 |
| Gm16835 | 0.454516499 | 0.192645078 | 0.076 | 0.053 | 1 | 0.028404366 | 2.263034406 | 0.2 | 0.046 | 1 | 0.454516499 | 0.056001924 |
| Reep3 | 0.028429164 | -0.257276056 | 0.958 | 0.953 | 1 | 0.557614544 | -0.226204035 | 0.9 | 0.994 | 1 | 0.557614544 | 0.056050111 |
| Kat2a | 0.028431481 | -1.109917692 | 0.092 | 0.182 | 1 | 0.156206274 | -1.026472211 | 0 | 0.17 | 1 | 0.156206274 | 0.056054614 |
| Tmcc2 | 0.028438641 | -1.87774425 | 0.025 | 0.088 | 1 | 0.90031049 | 0.403212064 | 0.1 | 0.111 | 1 | 0.90031049 | 0.056068526 |
| Zglp1 | 0.028438754 | -1.733354341 | 0.025 | 0.088 | 1 | 0.640911398 | 1.159940913 | 0.1 | 0.062 | 1 | 0.640911398 | 0.056068745 |
| Nme1 | 0.028463131 | -0.400814599 | 0.773 | 0.8 | 1 | 0.184290712 | -0.405944351 | 0.4 | 0.787 | 1 | 0.184290712 | 0.056116113 |
| Sacs | 0.028475505 | -1.378511623 | 0.092 | 0.182 | 1 | 0.875234763 | 0.403212064 | 0.1 | 0.114 | 1 | 0.875234763 | 0.056140156 |
| Fry | 0.053698331 | 0.227268808 | 0.95 | 0.865 | 1 | 0.028504426 | 1.453609917 | 0.9 | 0.84 | 1 | 0.053698331 | 0.056196349 |
| Sidt2 | 0.028515572 | -0.577685335 | 0.748 | 0.724 | 1 | 0.727175946 | -0.111361109 | 0.8 | 0.759 | 1 | 0.727175946 | 0.056218006 |
| Ctbs | 0.028525086 | 0.368722306 | 0.345 | 0.218 | 1 | 0.847158589 | 0.111031312 | 0.2 | 0.216 | 1 | 0.847158589 | 0.056236491 |
| Irf3 | 0.028560102 | -0.839063782 | 0.185 | 0.294 | 1 | 0.400139087 | 0.614199722 | 0.4 | 0.278 | 1 | 0.400139087 | 0.056304524 |
| Gm49454 | 0.028560541 | -2.87774425 | 0.008 | 0.059 | 1 | 0.584571279 | 0.432959407 | 0.1 | 0.056 | 1 | 0.584571279 | 0.056305378 |
| Rbm20 | 0.028589255 | 0.988504361 | 0.143 | 0.065 | 1 | 0.965736816 | 0.403212064 | 0.1 | 0.093 | 1 | 0.965736816 | 0.056361164 |
| Ptp4a2 | 0.028620625 | -0.315237765 | 0.992 | 0.988 | 1 | 0.358495713 | -0.244534379 | 1 | 0.994 | 1 | 0.358495713 | 0.05642211 |
| Hspa12a | 0.291329888 | -0.485426827 | 0.084 | 0.124 | 1 | 0.028639556 | 1.695993813 | 0.3 | 0.09 | 1 | 0.291329888 | 0.056458888 |
| Serinc5 | 0.028689022 | 0.33176938 | 0.866 | 0.741 | 1 | 0.109020903 | 0.664667479 | 1 | 0.883 | 1 | 0.109020903 | 0.056554985 |
| 1700031C06Rik | 0.02869942 | 1.181997834 | 0.134 | 0.059 | 1 | 0.149795061 | 2.847996907 | 0.1 | 0.025 | 1 | 0.149795061 | 0.056575184 |
| Prdm15 | 0.028706489 | -1.359895945 | 0.084 | 0.171 | 1 | 0.307152584 | -0.276698841 | 0.1 | 0.262 | 1 | 0.307152584 | 0.056588915 |
| Clcn3 | 0.207485501 | -0.225111923 | 0.571 | 0.435 | 1 | 0.028707535 | 0.669659665 | 0.9 | 0.602 | 1 | 0.207485501 | 0.056590948 |
| Npr2 | 0.028769864 | -0.687060688 | 0.277 | 0.388 | 1 | 0.747815698 | 0.233650599 | 0.5 | 0.469 | 1 | 0.747815698 | 0.056712023 |
| Msl3 | 0.028795092 | -0.668872968 | 0.37 | 0.459 | 1 | 0.800873176 | -0.171902651 | 0.4 | 0.414 | 1 | 0.800873176 | 0.056761027 |
| Ephx3 | 0.028797858 | -1.733354341 | 0.025 | 0.088 | 1 | 0.182375924 | 1.280956314 | 0.2 | 0.077 | 1 | 0.182375924 | 0.056766399 |
| Ndufa12 | 0.028851968 | -0.342592049 | 0.613 | 0.694 | 1 | 0.095727039 | -0.835907444 | 0.4 | 0.642 | 1 | 0.095727039 | 0.0568715 |
| Fnip2 | 0.18257309 | 0.112759369 | 0.782 | 0.588 | 1 | 0.028861244 | -1.601381047 | 0.6 | 0.83 | 1 | 0.18257309 | 0.056889517 |
| Plxdc1 | 0.028898683 | 0.684498174 | 0.109 | 0.041 | 1 | 0.232498909 | 2.432959407 | 0.1 | 0.031 | 1 | 0.232498909 | 0.056962231 |
| Ier5 | 0.1123391 | 0.106356939 | 0.891 | 0.853 | 1 | 0.028935102 | -1.481310719 | 0.5 | 0.809 | 1 | 0.1123391 | 0.057032964 |
| Rgl3 | 0.028978323 | -1.970853654 | 0.034 | 0.1 | 1 | 0.650890617 | 0.821524695 | 0.2 | 0.145 | 1 | 0.650890617 | 0.057116902 |
| Nck2 | 0.068690638 | 0.384980265 | 0.731 | 0.612 | 1 | 0.028996003 | -1.361456459 | 0.6 | 0.799 | 1 | 0.068690638 | 0.057151239 |
| Dedd | 0.028997986 | -0.737813989 | 0.252 | 0.365 | 1 | 0.462338094 | -0.31299497 | 0.2 | 0.321 | 1 | 0.462338094 | 0.057155089 |
| ENSMUSG00000121479 | 0.330649165 | -0.485426827 | 0.034 | 0.059 | 1 | 0.029046753 | 2.432959407 | 0.2 | 0.046 | 1 | 0.330649165 | 0.057249792 |
| Hbs1l | 0.029047661 | -0.577685335 | 0.597 | 0.629 | 1 | 0.454284628 | -0.258202497 | 0.6 | 0.71 | 1 | 0.454284628 | 0.057251555 |
| Fbxo8 | 0.029049451 | -0.702238216 | 0.218 | 0.335 | 1 | 0.590489099 | 0.720241359 | 0.3 | 0.231 | 1 | 0.590489099 | 0.057255032 |
| Wdtc1 | 0.029103788 | -1.048363022 | 0.176 | 0.276 | 1 | 0.338014134 | -0.709998547 | 0.2 | 0.321 | 1 | 0.338014134 | 0.057360545 |
| Mtx2 | 0.029156297 | -0.831877241 | 0.218 | 0.329 | 1 | 0.77985579 | 0.618750814 | 0.3 | 0.392 | 1 | 0.77985579 | 0.057462505 |
| Dcdc2b | 0.029173791 | -1.292781749 | 0.008 | 0.059 | 1 | 0.373655406 | 2.14345279 | 0.1 | 0.043 | 1 | 0.373655406 | 0.057496472 |
| Cars2 | 0.083107429 | -1.222392421 | 0.134 | 0.206 | 1 | 0.029197329 | 1.847996907 | 0.4 | 0.157 | 1 | 0.083107429 | 0.057542173 |
| Lonp1 | 0.02923108 | -0.920363884 | 0.193 | 0.294 | 1 | 0.179713238 | -1.004445905 | 0.1 | 0.29 | 1 | 0.179713238 | 0.057607705 |
| Slc39a13 | 0.029276933 | -0.577596671 | 0.647 | 0.694 | 1 | 0.323761117 | 0.528742946 | 0.8 | 0.63 | 1 | 0.323761117 | 0.057696727 |
| Tdrd7 | 0.029290906 | -1.137503524 | 0.151 | 0.253 | 1 | 0.05272486 | -2.161987182 | 0 | 0.281 | 1 | 0.05272486 | 0.057723856 |
| Elovl5 | 0.162742186 | 0.240398209 | 0.269 | 0.194 | 1 | 0.029316819 | 1.005097868 | 0.5 | 0.204 | 1 | 0.162742186 | 0.057774162 |
| Efcab1 | 0.496313191 | -0.967295835 | 0.361 | 0.353 | 1 | 0.029330584 | 0.854691559 | 0.7 | 0.324 | 1 | 0.496313191 | 0.057800885 |
| Gm49417 | 0.122416428 | 0.369142733 | 0.252 | 0.171 | 1 | 0.029332806 | 1.358958826 | 0.6 | 0.265 | 1 | 0.122416428 | 0.057805198 |
| D830036C21Rik | 0.029333692 | -1.485426827 | 0.008 | 0.059 | 1 | 0.815998758 | 0.695993813 | 0.1 | 0.077 | 1 | 0.815998758 | 0.057806919 |
| Ccdc88a | 0.029337462 | -0.292194452 | 0.95 | 0.971 | 1 | 0.558667476 | -0.234338026 | 1 | 0.972 | 1 | 0.558667476 | 0.057814237 |
| Pex5 | 0.029362054 | -0.600904045 | 0.445 | 0.529 | 1 | 0.586741214 | 0.167614841 | 0.7 | 0.552 | 1 | 0.586741214 | 0.057861978 |
| Vill | 0.029369843 | 0.175771259 | 0.63 | 0.459 | 1 | 0.267967076 | 0.726006382 | 0.7 | 0.54 | 1 | 0.267967076 | 0.057877098 |
| Echdc3 | 0.21910035 | 0.929610672 | 0.059 | 0.029 | 1 | 0.029372592 | 2.494359952 | 0.2 | 0.046 | 1 | 0.21910035 | 0.057882436 |
| Mrpl10 | 0.029413359 | -0.448900951 | 0.244 | 0.376 | 1 | 0.353639108 | 0.859492545 | 0.4 | 0.29 | 1 | 0.353639108 | 0.057961572 |
| Trp63 | 0.029513097 | -1.485426827 | 0.008 | 0.059 | 1 | 0.175080359 | -1.171902651 | 0 | 0.157 | 1 | 0.175080359 | 0.058155172 |
| Cdkn2b | 0.029557378 | 0.695145418 | 0.126 | 0.053 | 1 | 0.360331175 | 1.847996907 | 0.1 | 0.04 | 1 | 0.360331175 | 0.058241118 |
| Sac3d1 | 0.029564528 | -1.10433666 | 0.092 | 0.182 | 1 | 0.245440863 | -0.567040593 | 0 | 0.12 | 1 | 0.245440863 | 0.058254995 |
| Cep63 | 0.029571758 | -0.948827348 | 0.227 | 0.329 | 1 | 0.346126121 | -0.848326703 | 0.2 | 0.315 | 1 | 0.346126121 | 0.058269027 |

| | | | | | | | | | | | | |
|---|---|---|---|---|---|---|---|---|---|---|---|---|
| Nit2 | 0.029586066 | -0.944858446 | 0.151 | 0.253 | 1 | 0.131190612 | -1.131825212 | 0 | 0.188 | 1 | 0.131190612 | 0.058296797 |
| Tle1 | 0.029619031 | -0.906756072 | 0.58 | 0.606 | 1 | 0.204953021 | -0.963930745 | 0.6 | 0.735 | 1 | 0.204953021 | 0.058360776 |
| Stat6 | 0.066594722 | -0.887525271 | 0.319 | 0.382 | 1 | 0.029622449 | -1.700896339 | 0.2 | 0.528 | 1 | 0.066594722 | 0.058367408 |
| Limch1 | 0.029673238 | 0.470179053 | 0.235 | 0.129 | 1 | 0.085426346 | -3.121629444 | 0 | 0.235 | 1 | 0.085426346 | 0.058465976 |
| Gm20536 | 0.029691788 | 0.862496476 | 0.109 | 0.041 | 1 | 0.446758355 | 0.695993813 | 0 | 0.056 | 1 | 0.446758355 | 0.058501974 |
| Mcph1 | 0.035618338 | -1.465248945 | 0.118 | 0.206 | 1 | 0.029694412 | 2.036069255 | 0.4 | 0.154 | 1 | 0.035618338 | 0.058507065 |
| St3gal5 | 0.029708714 | -1.428843299 | 0.151 | 0.247 | 1 | 0.194791686 | 1.973527789 | 0.3 | 0.154 | 1 | 0.194791686 | 0.058534821 |
| Capn2 | 0.029757808 | -0.387376682 | 0.798 | 0.829 | 1 | 0.697308384 | 0.119588709 | 1 | 0.957 | 1 | 0.697308384 | 0.058630089 |
| Zfyve1 | 0.029776732 | -0.604071324 | 0.319 | 0.453 | 1 | 0.74399105 | 0.114040062 | 0.7 | 0.627 | 1 | 0.74399105 | 0.058666809 |
| Atp6v1a | 0.029790162 | -0.381997238 | 0.882 | 0.894 | 1 | 0.864679973 | -0.125591463 | 0.9 | 0.96 | 1 | 0.864679973 | 0.058692871 |
| Smim20 | 0.029795866 | 0.514573173 | 0.252 | 0.141 | 1 | 0.77894834 | 0.494359952 | 0.1 | 0.13 | 1 | 0.77894834 | 0.058703938 |
| Fam53a | 0.029803488 | -0.992386816 | 0.126 | 0.224 | 1 | 0.868778859 | 0.159940913 | 0.2 | 0.231 | 1 | 0.868778859 | 0.058718727 |
| Chrac1 | 0.029816003 | -0.676110389 | 0.294 | 0.406 | 1 | 0.843483622 | 0.22090893 | 0.3 | 0.265 | 1 | 0.843483622 | 0.058743012 |
| Mnd1 | 0.029847564 | -1.555816155 | 0.025 | 0.088 | 1 | 0.36410767 | -0.505640048 | 0 | 0.077 | 1 | 0.36410767 | 0.05880425 |
| 1700123M08Rik | 0.029911042 | -1.970853654 | 0.034 | 0.1 | 1 | 0.557764571 | 0.847996907 | 0 | 0.034 | 1 | 0.557764571 | 0.058927413 |
| Rpl39 | 0.029914208 | 0.216425913 | 0.966 | 0.976 | 1 | 0.761717745 | -0.148876992 | 0.9 | 0.991 | 1 | 0.761717745 | 0.058933557 |
| Erg | 0.029921245 | -0.696494333 | 0.857 | 0.865 | 1 | 0.48250349 | -0.650497535 | 1 | 0.91 | 1 | 0.48250349 | 0.058947209 |
| Cdc37l1 | 0.029938008 | -0.387124754 | 0.664 | 0.741 | 1 | 0.713776427 | 0.257517401 | 1 | 0.843 | 1 | 0.713776427 | 0.058979732 |
| Eepd1 | 0.029938948 | -2.232660757 | 0.05 | 0.124 | 1 | 0.671232313 | 0.123104145 | 0.1 | 0.157 | 1 | 0.671232313 | 0.058981555 |
| Gm11713 | 0.029953514 | 0.877143252 | 0.118 | 0.047 | 1 | 0.324050895 | 1.079322453 | 0.2 | 0.099 | 1 | 0.324050895 | 0.059009814 |
| Ccnk | 0.029954013 | -0.707819249 | 0.42 | 0.488 | 1 | 0.776976956 | -0.152003093 | 0.4 | 0.438 | 1 | 0.776976956 | 0.059010784 |
| Garnl3 | 0.030029487 | 0.824913293 | 0.244 | 0.141 | 1 | 0.816904597 | 0.280956314 | 0.1 | 0.13 | 1 | 0.816904597 | 0.059157204 |
| 4933406C10Rik | 0.694265848 | -0.532732542 | 0.109 | 0.124 | 1 | 0.0300857 | 2.432959407 | 0.3 | 0.099 | 1 | 0.694265848 | 0.05926625 |
| Ttc30b | 0.030096491 | -1.555816155 | 0.025 | 0.088 | 1 | 0.408705208 | 0.263034406 | 0 | 0.065 | 1 | 0.408705208 | 0.059287182 |
| Adam17 | 0.030104285 | -0.769880217 | 0.37 | 0.471 | 1 | 0.126904542 | 1.702566467 | 0.7 | 0.605 | 1 | 0.126904542 | 0.059302302 |
| Hadha | 0.030158067 | -0.396343284 | 0.79 | 0.782 | 1 | 0.264022613 | -0.31878154 | 0.7 | 0.836 | 1 | 0.264022613 | 0.059406625 |
| Slx4 | 0.030181785 | -1.372952098 | 0.076 | 0.159 | 1 | 0.492901201 | -0.111361109 | 0.1 | 0.182 | 1 | 0.492901201 | 0.059452629 |
| Spg11 | 0.030192099 | -1.004800986 | 0.101 | 0.194 | 1 | 0.936013976 | -0.201246612 | 0.3 | 0.278 | 1 | 0.936013976 | 0.059472634 |
| Nop9 | 0.030252889 | -0.839063782 | 0.134 | 0.235 | 1 | 0.9741693 | 0.608530972 | 0.2 | 0.219 | 1 | 0.9741693 | 0.059590541 |
| Ankrd28 | 0.274284585 | -0.316964551 | 0.975 | 0.982 | 1 | 0.03025648 | -0.865150068 | 1 | 0.997 | 1 | 0.274284585 | 0.059597505 |
| Idh3g | 0.0303039 | -0.550382269 | 0.504 | 0.571 | 1 | 0.078894524 | -0.947862377 | 0.2 | 0.509 | 1 | 0.078894524 | 0.059689473 |
| Actr3 | 0.03035121 | -0.335702616 | 0.95 | 0.976 | 1 | 0.928144993 | -0.138841845 | 1 | 0.948 | 1 | 0.928144993 | 0.059781225 |
| H2ac25 | 0.030373769 | -1.359895945 | 0.067 | 0.147 | 1 | 0.232373468 | -0.789433014 | 0 | 0.127 | 1 | 0.232373468 | 0.059824971 |
| 4930581F22Rik | 0.030396219 | 0.877143252 | 0.286 | 0.182 | 1 | 0.074738338 | 1.816288047 | 0.3 | 0.117 | 1 | 0.074738338 | 0.059868508 |
| Cdin1 | 0.030399722 | -1.847996907 | 0.092 | 0.176 | 1 | 0.245849206 | -1.267480311 | 0 | 0.12 | 1 | 0.245849206 | 0.059875301 |
| Thoc7 | 0.030419517 | -0.493419618 | 0.756 | 0.759 | 1 | 0.467769723 | -0.265166445 | 0.6 | 0.796 | 1 | 0.467769723 | 0.059913686 |
| Btd | 0.030464569 | 0.353292266 | 0.571 | 0.4 | 1 | 0.212038831 | -0.814968106 | 0.3 | 0.46 | 1 | 0.212038831 | 0.060001048 |
| Gm15581 | 0.030467361 | 0.581687369 | 0.118 | 0.047 | 1 | 0.328343757 | 1.374065718 | 0.1 | 0.037 | 1 | 0.328343757 | 0.060006461 |
| Wbp1 | 0.174155383 | 0.281912416 | 0.269 | 0.194 | 1 | 0.030468757 | 1.909397451 | 0.4 | 0.157 | 1 | 0.174155383 | 0.060009169 |
| Rbm4 | 0.0305205 | -0.714560826 | 0.378 | 0.471 | 1 | 0.059746376 | -1.299490706 | 0.3 | 0.556 | 1 | 0.059746376 | 0.0601095 |
| Rab11fip2 | 0.03053229 | -0.793094229 | 0.345 | 0.459 | 1 | 0.687006258 | 0.238948787 | 0.5 | 0.426 | 1 | 0.687006258 | 0.060132359 |
| Polr3e | 0.030537111 | -1.088767857 | 0.193 | 0.294 | 1 | 0.476715137 | -0.360589715 | 0.3 | 0.401 | 1 | 0.476715137 | 0.060141706 |
| Ubox5 | 0.030573768 | -1.765534746 | 0.05 | 0.124 | 1 | 0.668219851 | -0.171902651 | 0.1 | 0.145 | 1 | 0.668219851 | 0.060212781 |
| Dexi | 0.030635475 | -0.701155518 | 0.21 | 0.318 | 1 | 0.607856214 | 0.19592021 | 0.5 | 0.386 | 1 | 0.607856214 | 0.060332417 |
| Gm57084 | 0.123520686 | -1.622930351 | 0.034 | 0.076 | 1 | 0.030651873 | 2.515421567 | 0.2 | 0.046 | 1 | 0.123520686 | 0.060364208 |
| D830050J10Rik | 0.337704969 | 0.273565073 | 0.151 | 0.112 | 1 | 0.030651873 | 2.515421567 | 0.2 | 0.046 | 1 | 0.337704969 | 0.060364208 |
| Comtd1 | 0.403273565 | -0.600904045 | 0.042 | 0.065 | 1 | 0.030651873 | 2.515421567 | 0.2 | 0.046 | 1 | 0.403273565 | 0.060364208 |
| Tmem268 | 0.030662898 | 1.684498174 | 0.067 | 0.018 | 1 | 0.489939838 | 0.558490289 | 0 | 0.046 | 1 | 0.489939838 | 0.060385582 |
| Rtca | 0.0307003 | -0.769219793 | 0.176 | 0.282 | 1 | 0.188401719 | -1.026472211 | 0.1 | 0.284 | 1 | 0.188401719 | 0.060458092 |
| Tradd | 0.030705965 | -1.11345805 | 0.084 | 0.171 | 1 | 0.949913217 | 0.847996907 | 0.1 | 0.093 | 1 | 0.949913217 | 0.060469075 |
| Rnf167 | 0.181639099 | -0.379558192 | 0.597 | 0.624 | 1 | 0.03073674 | -1.378682873 | 0.3 | 0.608 | 1 | 0.181639099 | 0.060528732 |
| Adgrf5 | 0.030736742 | -1.517848305 | 0.076 | 0.159 | 1 | 0.905318082 | 0.695993813 | 0.1 | 0.093 | 1 | 0.905318082 | 0.060528737 |
| Arhgap42 | 0.030804178 | 0.389766717 | 0.756 | 0.641 | 1 | 0.33267123 | 0.100142996 | 0.9 | 0.694 | 1 | 0.33267123 | 0.060659459 |
| Tbc1d22a | 0.030816054 | -1.157852169 | 0.235 | 0.329 | 1 | 0.072426305 | -1.703177281 | 0.1 | 0.37 | 1 | 0.072426305 | 0.060682479 |
| Trir | 0.84514616 | -0.103918951 | 0.899 | 0.818 | 1 | 0.03084828 | -0.699068986 | 0.7 | 0.873 | 1 | 0.84514616 | 0.060744943 |
| Sh3yl1 | 0.030859647 | -1.807354922 | 0.034 | 0.1 | 1 | 0.155165164 | 1.354956895 | 0.2 | 0.074 | 1 | 0.155165164 | 0.060766977 |
| Uckl1 | 0.268033382 | 0.465663572 | 0.202 | 0.153 | 1 | 0.030886528 | 1.821524695 | 0.4 | 0.154 | 1 | 0.268033382 | 0.060819078 |
| Pnpla7 | 0.030928086 | -1.311397427 | 0.076 | 0.159 | 1 | 0.725349778 | 0.31748219 | 0.2 | 0.148 | 1 | 0.725349778 | 0.060899626 |
| Dusp3 | 0.030955736 | -1.189970944 | 0.176 | 0.276 | 1 | 0.312542418 | -1.031926641 | 0.3 | 0.404 | 1 | 0.312542418 | 0.060953214 |
| Kdm3b | 0.030966384 | -0.800123353 | 0.412 | 0.506 | 1 | 0.46246157 | 0.55551259 | 0.4 | 0.657 | 1 | 0.46246157 | 0.060973852 |
| Mrpl12 | 0.030976503 | -0.688055994 | 0.429 | 0.506 | 1 | 0.652145584 | 0.223506042 | 0.3 | 0.472 | 1 | 0.652145584 | 0.060993462 |
| Dnajc22 | 0.119486073 | 0.440572591 | 0.143 | 0.082 | 1 | 0.031008861 | 2.169925001 | 0.3 | 0.096 | 1 | 0.119486073 | 0.061056173 |
| Clk3 | 0.03106634 | -0.715724447 | 0.16 | 0.265 | 1 | 0.558328902 | -0.362899876 | 0.2 | 0.272 | 1 | 0.558328902 | 0.061167563 |
| B130055M24Rik | 0.031110297 | -1.411426246 | 0.118 | 0.206 | 1 | 0.384942007 | 0.769994395 | 0.3 | 0.185 | 1 | 0.384942007 | 0.061252743 |
| Gm8797 | 0.524548365 | 0.106488434 | 0.353 | 0.312 | 1 | 0.031172382 | -2.171902651 | 0 | 0.327 | 1 | 0.524548365 | 0.061373046 |
| Nt5m | 0.031200047 | -0.739183419 | 0.16 | 0.271 | 1 | 0.448725107 | -0.374395515 | 0.1 | 0.191 | 1 | 0.448725107 | 0.061426652 |
| 1700037F03Rik | 0.031218822 | -1.847996907 | 0.05 | 0.124 | 1 | 0.489901827 | 0.695993813 | 0 | 0.046 | 1 | 0.489901827 | 0.061463029 |
| Eif4e | 0.12457213 | 0.179224562 | 0.899 | 0.935 | 1 | 0.031258315 | -0.891453951 | 0.9 | 0.96 | 1 | 0.12457213 | 0.061539548 |
| Acox2 | 0.031318779 | 1.389042291 | 0.067 | 0.018 | 1 | 0.643835681 | 1.847996907 | 0 | 0.022 | 1 | 0.643835681 | 0.061656692 |
| Gemin8 | 0.0313406 | -1.807354922 | 0.034 | 0.1 | 1 | 0.829966406 | 1.111031312 | 0.1 | 0.08 | 1 | 0.829966406 | 0.061698967 |
| Rfc4 | 0.036384465 | -1.185866545 | 0.05 | 0.124 | 1 | 0.031343367 | 2.432959407 | 0.2 | 0.046 | 1 | 0.036384465 | 0.061704328 |
| Myo1h | 0.495207728 | -0.485426827 | 0.017 | 0.029 | 1 | 0.031343367 | 2.432959407 | 0.2 | 0.046 | 1 | 0.495207728 | 0.061704328 |
| P4ha3 | 0.031425878 | -1.111968432 | 0.277 | 0.371 | 1 | 0.143940154 | 1.058563892 | 0.4 | 0.219 | 1 | 0.143940154 | 0.061864171 |
| 4930527F14Rik | 0.031481996 | 1.514573173 | 0.067 | 0.018 | 1 | 0.577177049 | 0.494359952 | 0 | 0.031 | 1 | 0.577177049 | 0.061972876 |
| Sdhaf4 | 0.031488168 | 0.354108501 | 0.538 | 0.394 | 1 | 0.421792673 | 0.219180116 | 0.2 | 0.349 | 1 | 0.421792673 | 0.061984832 |
| Gm26759 | 0.316514938 | -0.970853654 | 0.034 | 0.059 | 1 | 0.03151788 | 1.660369903 | 0.3 | 0.09 | 1 | 0.316514938 | 0.062042383 |
| Grb2 | 0.031553849 | -0.430564892 | 0.773 | 0.788 | 1 | 0.638557375 | -0.119266307 | 0.7 | 0.901 | 1 | 0.638557375 | 0.062112053 |
| Slc30a9 | 0.03158617 | -0.814734452 | 0.487 | 0.535 | 1 | 0.108016226 | -0.473931188 | 0.4 | 0.698 | 1 | 0.108016226 | 0.062174653 |
| Siae | 0.840101566 | -0.275973462 | 0.303 | 0.294 | 1 | 0.031618263 | 1.135278859 | 0.6 | 0.272 | 1 | 0.840101566 | 0.062236812 |
| Ndufa2 | 0.045860885 | 0.217187262 | 0.891 | 0.882 | 1 | 0.031634859 | -0.964437933 | 0.4 | 0.787 | 1 | 0.045860885 | 0.062268954 |
| E2f3 | 0.031652018 | -1.105578756 | 0.151 | 0.247 | 1 | 0.700206744 | -0.245112498 | 0.3 | 0.346 | 1 | 0.700206744 | 0.062302186 |
| 2610507I01Rik | 0.163924323 | -0.914270126 | 0.084 | 0.135 | 1 | 0.031681111 | 2.121757719 | 0.4 | 0.151 | 1 | 0.163924323 | 0.06235853 |
| Gm56829 | 0.031759447 | -1.439623138 | 0.059 | 0.135 | 1 | 0.259431453 | -0.408342847 | 0 | 0.114 | 1 | 0.259431453 | 0.062510231 |
| Ddx28 | 0.031826273 | -1.485426827 | 0.034 | 0.1 | 1 | 0.518235124 | 1.769994395 | 0.1 | 0.052 | 1 | 0.518235124 | 0.062639635 |
| Ccdc22 | 0.031826273 | -1.485426827 | 0.034 | 0.1 | 1 | 0.750911357 | 0.210566986 | 0.1 | 0.133 | 1 | 0.750911357 | 0.062639635 |
| Nras | 0.031850241 | -0.481152172 | 0.739 | 0.788 | 1 | 0.148208747 | -0.614793724 | 0.8 | 0.815 | 1 | 0.148208747 | 0.062686044 |
| Ilk | 0.031860302 | -0.560477532 | 0.605 | 0.624 | 1 | 0.616807634 | -0.315978829 | 0.6 | 0.577 | 1 | 0.616807634 | 0.062705525 |
| Mettl14 | 0.031873461 | -1.163498732 | 0.076 | 0.159 | 1 | 0.226017437 | -0.840059087 | 0 | 0.13 | 1 | 0.226017437 | 0.062731004 |
| Sulf1 | 0.031887468 | -1.511222828 | 0.168 | 0.265 | 1 | 0.25214373 | -3.471926052 | 0.1 | 0.241 | 1 | 0.25214373 | 0.062758125 |
| Zcchc7 | 0.03190021 | -0.3707027 | 0.874 | 0.9 | 1 | 0.120616385 | 0.433870487 | 1 | 0.87 | 1 | 0.120616385 | 0.062782796 |
| Tarbp1 | 0.031905125 | -1.900464326 | 0.042 | 0.112 | 1 | 0.282175575 | -0.473931188 | 0 | 0.105 | 1 | 0.282175575 | 0.062792313 |
| Zfp938 | 0.344673806 | -0.544320516 | 0.16 | 0.2 | 1 | 0.031924861 | 1.532495081 | 0.4 | 0.142 | 1 | 0.344673806 | 0.062830526 |
| Rsph10b | 0.126323047 | -0.970853654 | 0.05 | 0.1 | 1 | 0.032046676 | 2.210566986 | 0.2 | 0.046 | 1 | 0.126323047 | 0.063066363 |
| Slc41a3 | 0.128864218 | -0.970853654 | 0.034 | 0.076 | 1 | 0.032046676 | 2.354956895 | 0.2 | 0.046 | 1 | 0.128864218 | 0.063066363 |
| Proser3 | 0.334204811 | -0.807354922 | 0.008 | 0.024 | 1 | 0.032046676 | 2.14345279 | 0.2 | 0.046 | 1 | 0.334204811 | 0.063066363 |
| Plekha7 | 0.032067463 | 0.759685671 | 0.126 | 0.053 | 1 | 0.897272471 | -0.111361109 | 0.2 | 0.219 | 1 | 0.897272471 | 0.063106605 |
| Srsf9 | 0.342121958 | -0.24132215 | 0.782 | 0.765 | 1 | 0.032122621 | -0.914633571 | 0.6 | 0.861 | 1 | 0.342121958 | 0.06321338 |
| Zfp33b | 0.032137858 | -0.963474124 | 0.084 | 0.171 | 1 | 0.376664725 | -0.654503434 | 0.1 | 0.21 | 1 | 0.376664725 | 0.063242973 |

| | | | | | | | | | | | | |
|---|---|---|---|---|---|---|---|---|---|---|---|---|
| Ssh1 | 0.589864721 | 0.494944366 | 0.387 | 0.365 | 1 | 0.032149773 | -1.993305347 | 0.3 | 0.571 | 1 | 0.589864721 | 0.063265939 |
| Jag1 | 0.780299203 | -0.464668267 | 0.202 | 0.206 | 1 | 0.032158202 | -2.802257054 | 0 | 0.327 | 1 | 0.780299203 | 0.063282255 |
| Shank1 | 0.032254834 | -1.163498732 | 0.109 | 0.2 | 1 | 0.170604907 | -1.131825212 | 0 | 0.16 | 1 | 0.170604907 | 0.063469293 |
| Camk2n2 | 0.032316362 | -1.411426246 | 0.034 | 0.1 | 1 | 0.408636939 | 0.374065718 | 0 | 0.065 | 1 | 0.408636939 | 0.063588377 |
| Gm56915 | 0.032316362 | -1.411426246 | 0.034 | 0.1 | 1 | 0.446790883 | 0.625604485 | 0 | 0.056 | 1 | 0.446790883 | 0.063588377 |
| Nrxn1 | 0.032317173 | -1.86393845 | 0.143 | 0.065 | 1 | 0.135193631 | -1.936274402 | 0 | 0.185 | 1 | 0.135193631 | 0.063589945 |
| Pmm2 | 0.032319941 | -0.762267033 | 0.185 | 0.288 | 1 | 0.515160503 | -0.339630097 | 0.2 | 0.281 | 1 | 0.515160503 | 0.063595303 |
| Vasn | 0.506404153 | -0.133001305 | 0.874 | 0.824 | 1 | 0.032331874 | -0.828113482 | 1 | 0.877 | 1 | 0.506404153 | 0.063618398 |
| Wsb1 | 0.032351794 | -0.391044162 | 0.933 | 0.953 | 1 | 0.564782268 | -0.285156286 | 1 | 0.994 | 1 | 0.564782268 | 0.06365695 |
| Niban2 | 0.032361907 | 0.215811063 | 0.924 | 0.841 | 1 | 0.562096365 | -0.268058198 | 0.8 | 0.941 | 1 | 0.562096365 | 0.063676521 |
| Denr | 0.03236991 | -0.440338938 | 0.639 | 0.682 | 1 | 0.505238575 | -0.277533976 | 0.6 | 0.654 | 1 | 0.505238575 | 0.06369201 |
| Htra3 | 0.032380215 | 0.387567896 | 0.58 | 0.471 | 1 | 0.198629124 | -0.165299916 | 0.2 | 0.429 | 1 | 0.198629124 | 0.063711951 |
| Rnf144b | 0.032470538 | -1.09423607 | 0.118 | 0.212 | 1 | 0.374275238 | -0.654503434 | 0.1 | 0.21 | 1 | 0.374275238 | 0.063886741 |
| Plbd2 | 0.032483341 | 0.395520501 | 0.555 | 0.406 | 1 | 0.307076169 | 0.689995072 | 0.6 | 0.469 | 1 | 0.307076169 | 0.063911514 |
| Dop1a | 0.643092063 | -0.357779004 | 0.588 | 0.553 | 1 | 0.032484543 | 0.717798183 | 0.9 | 0.54 | 1 | 0.643092063 | 0.06391384 |
| Tmem38b | 0.0324849 | 0.636563697 | 0.252 | 0.147 | 1 | 0.294011006 | 1.005097868 | 0.4 | 0.259 | 1 | 0.294011006 | 0.063914531 |
| Dnaaf5 | 0.032488801 | -1.544320516 | 0.042 | 0.112 | 1 | 0.957826589 | 0.625604485 | 0.1 | 0.093 | 1 | 0.957826589 | 0.06392208 |
| Ift122 | 0.032494601 | -1.14839184 | 0.084 | 0.171 | 1 | 0.219857668 | 1.295455884 | 0.3 | 0.157 | 1 | 0.219857668 | 0.063933302 |
| Pfdn1 | 0.032512363 | 0.256972653 | 0.639 | 0.482 | 1 | 0.898031257 | 0.127150978 | 0.6 | 0.602 | 1 | 0.898031257 | 0.063967673 |
| Wdyhv1 | 0.032541077 | -1.025995209 | 0.084 | 0.171 | 1 | 0.485902703 | 1.159940913 | 0.2 | 0.13 | 1 | 0.485902703 | 0.064023232 |
| Pelo | 0.032639341 | -0.780882711 | 0.218 | 0.329 | 1 | 0.843244971 | 0.409112665 | 0.3 | 0.275 | 1 | 0.843244971 | 0.064213355 |
| Srcin1 | 0.076383745 | 1.321928095 | 0.084 | 0.035 | 1 | 0.032649349 | 1.769994395 | 0.3 | 0.096 | 1 | 0.076383745 | 0.064232718 |
| Tkt | 0.032652826 | 0.44258706 | 0.773 | 0.706 | 1 | 0.841438065 | 0.280956314 | 0.7 | 0.67 | 1 | 0.841438065 | 0.064239444 |
| C1galt1 | 0.610203055 | -0.586710163 | 0.345 | 0.329 | 1 | 0.032660503 | -1.987702641 | 0.1 | 0.435 | 1 | 0.610203055 | 0.064254297 |
| Bbs5 | 0.032668357 | -1.19592021 | 0.076 | 0.159 | 1 | 0.134735583 | 1.236562194 | 0.3 | 0.127 | 1 | 0.134735583 | 0.064269493 |
| Tmem214 | 0.03277437 | -0.64731451 | 0.513 | 0.559 | 1 | 0.479371734 | 0.365214801 | 0.7 | 0.531 | 1 | 0.479371734 | 0.06447458 |
| Mfsd1 | 0.032799565 | -0.895302621 | 0.286 | 0.376 | 1 | 0.056289315 | -1.481923979 | 0.1 | 0.395 | 1 | 0.056289315 | 0.064523319 |
| Alkbh3 | 0.085194457 | -0.622930351 | 0.21 | 0.294 | 1 | 0.032814996 | 1.558490289 | 0.5 | 0.241 | 1 | 0.085194457 | 0.064553168 |
| Satb2 | 0.032904341 | -2.309855263 | 0.05 | 0.124 | 1 | 0.157443202 | 1.063725598 | 0.2 | 0.074 | 1 | 0.157443202 | 0.064725987 |
| Agbl3 | 0.03290634 | -1.230854 | 0.235 | 0.353 | 1 | 0.618499836 | -1.562022518 | 0.2 | 0.238 | 1 | 0.618499836 | 0.064729853 |
| Rab34 | 0.032935407 | -0.526454095 | 0.529 | 0.594 | 1 | 0.231587408 | -0.71775499 | 0.4 | 0.571 | 1 | 0.231587408 | 0.064786073 |
| Nudt1 | 0.032966937 | -1.207892852 | 0.067 | 0.147 | 1 | 0.307171866 | -0.304006187 | 0 | 0.096 | 1 | 0.307171866 | 0.064847056 |
| Tcta | 0.032982564 | -0.831201664 | 0.168 | 0.271 | 1 | 0.437821204 | 0.808468542 | 0.3 | 0.191 | 1 | 0.437821204 | 0.064877278 |
| Tpst1 | 0.032991672 | -0.750399037 | 0.387 | 0.476 | 1 | 0.884359201 | -0.226838326 | 0.7 | 0.608 | 1 | 0.884359201 | 0.064894894 |
| Cep44 | 0.033040696 | -1.070389328 | 0.101 | 0.188 | 1 | 0.499877607 | -0.321928095 | 0.1 | 0.179 | 1 | 0.499877607 | 0.064989704 |
| Klf9 | 0.0330478 | 0.388390562 | 0.95 | 0.924 | 1 | 0.069396753 | -0.744875355 | 0.9 | 0.969 | 1 | 0.069396753 | 0.065003442 |
| Flywch1 | 0.033050156 | -0.891419187 | 0.16 | 0.259 | 1 | 0.589533855 | -0.279758641 | 0.2 | 0.262 | 1 | 0.589533855 | 0.065007998 |
| Smim19 | 0.033056395 | 1.87313019 | 0.639 | 0.488 | 1 | 0.863224698 | 0.153735763 | 0.5 | 0.478 | 1 | 0.863224698 | 0.065020064 |
| Nradd | 0.033080292 | -1.485426827 | 0.042 | 0.112 | 1 | 0.245439637 | -0.736965594 | 0 | 0.12 | 1 | 0.245439637 | 0.065066278 |
| Slc36a1 | 0.033080292 | -1.485426827 | 0.042 | 0.112 | 1 | 0.238732419 | -0.625934282 | 0 | 0.123 | 1 | 0.238732419 | 0.065066278 |
| Cldn7 | 0.033080292 | -1.485426827 | 0.042 | 0.112 | 1 | 0.294660752 | -0.68251781 | 0.1 | 0.241 | 1 | 0.294660752 | 0.065066278 |
| Zfp839 | 0.372270612 | -0.632268215 | 0.109 | 0.141 | 1 | 0.033124855 | 1.625604485 | 0.3 | 0.09 | 1 | 0.372270612 | 0.065152454 |
| Gm11973 | 0.03325478 | -0.74176658 | 0.109 | 0.206 | 1 | 0.452185639 | 0.231325546 | 0.4 | 0.25 | 1 | 0.452185639 | 0.065403679 |
| 2610206C17Rik | 0.033265465 | 2.362570079 | 0.076 | 0.024 | 1 | 0.577177401 | 0.558490289 | 0 | 0.031 | 1 | 0.577177401 | 0.065424338 |
| Pygb | 0.03326997 | -0.847996907 | 0.134 | 0.235 | 1 | 0.034289637 | -2.152003093 | 0 | 0.318 | 1 | 0.034289637 | 0.065433049 |
| Myl12a | 0.033302883 | -0.254281 | 0.975 | 0.976 | 1 | 0.320035724 | -0.26211622 | 0.9 | 0.991 | 1 | 0.320035724 | 0.065496684 |
| Acot9 | 0.387947579 | -0.290854301 | 0.555 | 0.553 | 1 | 0.033353199 | -1.42170123 | 0.4 | 0.642 | 1 | 0.387947579 | 0.065593963 |
| Gm5914 | 0.033412874 | -1.359895945 | 0.034 | 0.1 | 1 | 0.776127243 | 1.063725598 | 0.1 | 0.074 | 1 | 0.776127243 | 0.065709328 |
| Ddx27 | 0.085615372 | -0.515800476 | 0.538 | 0.582 | 1 | 0.033444537 | -1.673239997 | 0.2 | 0.519 | 1 | 0.085615372 | 0.065770536 |
| Itfg1 | 0.033475051 | 0.115402079 | 0.748 | 0.594 | 1 | 0.895784876 | -0.177282502 | 0.7 | 0.605 | 1 | 0.895784876 | 0.065829523 |
| Bmp8a | 0.033549781 | -1.50851044 | 0.134 | 0.229 | 1 | 0.043011709 | 1.574978412 | 0.3 | 0.102 | 1 | 0.043011709 | 0.065973974 |
| Lmtk2 | 0.033554746 | -1.070389328 | 0.185 | 0.282 | 1 | 0.133539332 | -1.095820258 | 0.2 | 0.451 | 1 | 0.133539332 | 0.06598357 |
| Stpg4 | 0.033559964 | -0.669851398 | 0.034 | 0.1 | 1 | 0.718932948 | 0.403212064 | 0.1 | 0.068 | 1 | 0.718932948 | 0.065993657 |
| Pin1 | 0.033568695 | -0.635173947 | 0.412 | 0.529 | 1 | 0.854641495 | 0.210566986 | 0.5 | 0.485 | 1 | 0.854641495 | 0.066010532 |
| Twnk | 0.033581801 | -1.315501826 | 0.067 | 0.147 | 1 | 0.408732936 | 0.210566986 | 0 | 0.065 | 1 | 0.408732936 | 0.066035865 |
| Wdpcp | 0.033677514 | -1.659677754 | 0.185 | 0.282 | 1 | 0.580232512 | -1.291933355 | 0.2 | 0.253 | 1 | 0.580232512 | 0.066220853 |
| Tsen2 | 0.033678888 | -1.424026283 | 0.042 | 0.112 | 1 | 0.994871634 | 0.625604485 | 0.1 | 0.099 | 1 | 0.994871634 | 0.066223508 |
| Tmem198b | 0.033678888 | -1.359895945 | 0.042 | 0.112 | 1 | 0.148903646 | 1.847996907 | 0.2 | 0.074 | 1 | 0.148903646 | 0.066223508 |
| Tnks1bp1 | 0.638701638 | -0.222392421 | 0.37 | 0.382 | 1 | 0.033711626 | -1.317468447 | 0.2 | 0.565 | 1 | 0.638701638 | 0.066286778 |
| Sdhd | 0.033718831 | -0.521406124 | 0.597 | 0.629 | 1 | 0.034859887 | -1.152003093 | 0.3 | 0.639 | 1 | 0.034859887 | 0.066300702 |
| Fbxw7 | 0.033785801 | -0.714245518 | 0.462 | 0.565 | 1 | 0.161689442 | -0.602664502 | 0.4 | 0.642 | 1 | 0.161689442 | 0.066430121 |
| Kdm7a | 0.033926094 | -0.752600963 | 0.571 | 0.635 | 1 | 0.045744799 | 0.489283014 | 1 | 0.775 | 1 | 0.045744799 | 0.066701208 |
| Uxs1 | 0.03394608 | 0.318430651 | 0.79 | 0.641 | 1 | 0.777346536 | -0.145308441 | 0.8 | 0.568 | 1 | 0.777346536 | 0.066739824 |
| 4930549G23Rik | 0.757803678 | -0.273922722 | 0.067 | 0.076 | 1 | 0.033951608 | 1.591657153 | 0.3 | 0.09 | 1 | 0.757803678 | 0.066750504 |
| Crebl2 | 0.033959919 | 0.488100961 | 0.328 | 0.206 | 1 | 0.143277139 | 0.888638891 | 0.5 | 0.269 | 1 | 0.143277139 | 0.066766563 |
| Fam53b | 0.034012019 | -1.292781749 | 0.059 | 0.135 | 1 | 0.736119987 | 0.290001453 | 0.1 | 0.136 | 1 | 0.736119987 | 0.06686722 |
| Tmem60 | 0.034028136 | 0.532495081 | 0.454 | 0.347 | 1 | 0.975803952 | 0.299103661 | 0.3 | 0.321 | 1 | 0.975803952 | 0.066898358 |
| Srp9 | 0.156365597 | -0.192450106 | 0.908 | 0.888 | 1 | 0.034048202 | -0.881355504 | 0.8 | 0.914 | 1 | 0.156365597 | 0.066937123 |
| Atp6v1g1 | 0.034067527 | -0.260945662 | 0.874 | 0.929 | 1 | 0.289898595 | -0.475266399 | 0.8 | 0.898 | 1 | 0.289898595 | 0.066974458 |
| Foxk2 | 0.034071961 | -0.78958022 | 0.319 | 0.418 | 1 | 0.384932049 | -0.364547729 | 0.3 | 0.472 | 1 | 0.384932049 | 0.066983024 |
| Pdp2 | 0.197527188 | 0.316633795 | 0.235 | 0.171 | 1 | 0.034151353 | 2.097356375 | 0.3 | 0.099 | 1 | 0.197527188 | 0.067136391 |
| Ybx3 | 0.034166507 | -0.260325119 | 0.941 | 0.959 | 1 | 0.17898379 | -0.455287752 | 0.8 | 0.978 | 1 | 0.17898379 | 0.067165664 |
| Tmub2 | 0.034194796 | -0.789207575 | 0.235 | 0.335 | 1 | 0.968287663 | 0.159940913 | 0.3 | 0.293 | 1 | 0.968287663 | 0.067220307 |
| Stom | 0.034199374 | 0.769830228 | 0.218 | 0.124 | 1 | 0.339583559 | 1.079322453 | 0.2 | 0.102 | 1 | 0.339583559 | 0.06722915 |
| Vrk2 | 0.034218993 | -1.12588444 | 0.092 | 0.176 | 1 | 0.316595237 | 0.906413593 | 0.3 | 0.188 | 1 | 0.316595237 | 0.067267047 |
| Zbtb25 | 0.034222655 | -1.529820947 | 0.05 | 0.124 | 1 | 0.885565397 | 0.210566986 | 0.2 | 0.173 | 1 | 0.885565397 | 0.06727412 |
| Cenpw | 0.03425975 | -1.392317423 | 0.042 | 0.112 | 1 | 0.038538538 | 2.515421567 | 0.2 | 0.049 | 1 | 0.038538538 | 0.067345769 |
| Zfp282 | 0.034270799 | -1.325646383 | 0.21 | 0.306 | 1 | 0.909662447 | 0.176619654 | 0.3 | 0.318 | 1 | 0.909662447 | 0.06736711 |
| Nphp1 | 0.996349912 | -0.177304532 | 0.286 | 0.271 | 1 | 0.034281786 | 1.558490289 | 0.5 | 0.244 | 1 | 0.996349912 | 0.067388332 |
| 2210016F16Rik | 0.034330148 | -0.822461814 | 0.269 | 0.359 | 1 | 0.091255611 | -1.374395515 | 0.1 | 0.352 | 1 | 0.091255611 | 0.067481737 |
| Psen2 | 0.034350022 | 0.356311089 | 0.429 | 0.288 | 1 | 0.181152893 | 1.093649234 | 0.5 | 0.346 | 1 | 0.181152893 | 0.067520121 |
| Smim10l1 | 0.383902614 | 0.107484427 | 0.857 | 0.812 | 1 | 0.034374671 | -0.868478162 | 0.5 | 0.836 | 1 | 0.383902614 | 0.067567724 |
| Grik4 | 0.034447188 | -1.378511623 | 0.118 | 0.212 | 1 | 0.963481034 | 0.494359952 | 0.1 | 0.111 | 1 | 0.963481034 | 0.067707768 |
| Manbal | 0.034477442 | -0.770156304 | 0.412 | 0.471 | 1 | 0.181358519 | -0.936274402 | 0.3 | 0.466 | 1 | 0.181358519 | 0.06776619 |
| Arhgef17 | 0.034487655 | -1.048856166 | 0.294 | 0.376 | 1 | 0.888443267 | 0.174943076 | 0.4 | 0.417 | 1 | 0.888443267 | 0.067785912 |
| Tmem160 | 0.034533591 | 0.347923303 | 0.588 | 0.435 | 1 | 0.527879613 | -0.321928095 | 0.3 | 0.373 | 1 | 0.527879613 | 0.067874613 |
| Ano2 | 0.034550051 | 2.099535674 | 0.042 | 0.006 | 1 | 0.698533977 | 1.847996907 | 0 | 0.015 | 1 | 0.698533977 | 0.067906395 |
| C1qtnf4 | 0.034559595 | 0.514573173 | 0.294 | 0.182 | 1 | 0.818669855 | 0.571665678 | 0.2 | 0.25 | 1 | 0.818669855 | 0.067924824 |
| Smpd1 | 0.034578835 | 0.449478145 | 0.42 | 0.294 | 1 | 0.316070564 | 0.760534065 | 0.5 | 0.373 | 1 | 0.316070564 | 0.067961974 |
| Slc25a15 | 0.034597025 | -1.432959407 | 0.05 | 0.124 | 1 | 0.180218559 | -1.048167282 | 0 | 0.154 | 1 | 0.180218559 | 0.067997095 |
| Dcaf7 | 0.034623989 | -0.750771394 | 0.311 | 0.412 | 1 | 0.896885668 | -0.138582578 | 0.4 | 0.392 | 1 | 0.896885668 | 0.068049157 |
| Rinl | 0.034630271 | -2.222392421 | 0.017 | 0.071 | 1 | 0.420961276 | 0.31748219 | 0 | 0.062 | 1 | 0.420961276 | 0.068061286 |
| Efcab7 | 0.034631896 | -2.485426827 | 0.017 | 0.071 | 1 | 0.214157362 | 2.930459067 | 0.1 | 0.031 | 1 | 0.214157362 | 0.068064424 |
| Slc22a14 | 0.067243814 | -0.678071905 | 0.025 | 0.076 | 1 | 0.034688843 | 2.816288047 | 0.2 | 0.049 | 1 | 0.067243814 | 0.06817437 |
| Plgrkt | 0.037099222 | -0.707819249 | 0.269 | 0.371 | 1 | 0.034698973 | -2.037360528 | 0 | 0.318 | 1 | 0.037099222 | 0.068193928 |
| Pcnp | 0.034699213 | -0.287980763 | 0.95 | 0.894 | 1 | 0.441848096 | -0.31836148 | 0.9 | 0.957 | 1 | 0.441848096 | 0.068194391 |
| Rpl37a | 0.431084553 | 0.112711062 | 0.966 | 0.982 | 1 | 0.034714836 | -0.68251781 | 0.7 | 0.994 | 1 | 0.431084553 | 0.068224553 |
| Prkx | 0.034743499 | -1.070389328 | 0.21 | 0.3 | 1 | 0.082117481 | -1.327852929 | 0.2 | 0.46 | 1 | 0.082117481 | 0.068279888 |

| | | | | | | | | | | | | |
|---|---|---|---|---|---|---|---|---|---|---|---|---|
| Acat2 | 0.034743661 | 0.412693559 | 0.403 | 0.271 | 1 | 0.842210678 | -0.279758641 | 0.2 | 0.21 | 1 | 0.842210678 | 0.068280201 |
| Tmem62 | 0.532370314 | -0.807354922 | 0.059 | 0.076 | 1 | 0.034794136 | 1.591657153 | 0.3 | 0.09 | 1 | 0.532370314 | 0.06837764 |
| Cpsf4l | 0.183353927 | 0.142604395 | 0.126 | 0.076 | 1 | 0.034799186 | 1.374065718 | 0.3 | 0.09 | 1 | 0.183353927 | 0.068387388 |
| Ralgds | 0.034841836 | 1.195750989 | 0.303 | 0.2 | 1 | 0.750642137 | 1.017921908 | 0.3 | 0.281 | 1 | 0.750642137 | 0.068469718 |
| Wdr27 | 0.034898228 | -0.959358016 | 0.042 | 0.112 | 1 | 0.433608413 | 0.374065718 | 0 | 0.059 | 1 | 0.433608413 | 0.068578569 |
| Rilpl1 | 0.034935192 | -0.745954377 | 0.286 | 0.388 | 1 | 0.101794523 | -1.131825212 | 0.1 | 0.377 | 1 | 0.101794523 | 0.068649916 |
| Ercc1 | 0.03495315 | -0.907659828 | 0.202 | 0.3 | 1 | 0.13601231 | -1.090602549 | 0.1 | 0.318 | 1 | 0.13601231 | 0.068684578 |
| Mok | 0.034956669 | -2.14839184 | 0.017 | 0.071 | 1 | 0.334330677 | -0.339630097 | 0 | 0.086 | 1 | 0.334330677 | 0.068691369 |
| Gm37422 | 0.03495765 | -2.359895945 | 0.017 | 0.071 | 1 | 0.487556826 | 1.558490289 | 0.1 | 0.049 | 1 | 0.487556826 | 0.068693263 |
| H2ac8 | 0.03495765 | -3.944858446 | 0.017 | 0.071 | 1 | 0.73075158 | 2.695993813 | 0 | 0.012 | 1 | 0.73075158 | 0.068693263 |
| Gm56764 | 0.034958304 | -2.070389328 | 0.017 | 0.071 | 1 | 0.487557552 | 1.494359952 | 0.1 | 0.049 | 1 | 0.487557552 | 0.068694526 |
| Iqcc | 0.045625071 | -1.424026283 | 0.042 | 0.106 | 1 | 0.034972604 | 1.808468542 | 0.3 | 0.093 | 1 | 0.045625071 | 0.068722125 |
| 2310015A10Rik | 0.414325996 | -0.485426827 | 0.092 | 0.124 | 1 | 0.034973097 | 1.591657153 | 0.3 | 0.093 | 1 | 0.414325996 | 0.068723077 |
| Egln1 | 0.035059103 | -0.586120827 | 0.513 | 0.565 | 1 | 0.154067906 | -0.596787936 | 0.4 | 0.657 | 1 | 0.154067906 | 0.068889066 |
| Cyp7b1 | 0.035107509 | -1.521050737 | 0.05 | 0.124 | 1 | 0.502732924 | 1.295455884 | 0.2 | 0.133 | 1 | 0.502732924 | 0.068982481 |
| Pef1 | 0.081498757 | 0.328160049 | 0.521 | 0.406 | 1 | 0.035158691 | -1.769980651 | 0.1 | 0.432 | 1 | 0.081498757 | 0.069081249 |
| Col24a1 | 0.035190119 | -3.572889668 | 0.067 | 0.141 | 1 | 0.918638761 | 0.159940913 | 0.1 | 0.086 | 1 | 0.918638761 | 0.069141894 |
| Myh14 | 0.071114821 | -1.008988783 | 0.042 | 0.1 | 1 | 0.035212798 | -1.900941329 | 0 | 0.315 | 1 | 0.071114821 | 0.069185654 |
| Vmp1 | 0.035232763 | 0.191458163 | 0.966 | 0.918 | 1 | 0.049380902 | 0.516748184 | 0.9 | 0.833 | 1 | 0.049380902 | 0.069224178 |
| Lypla1 | 0.035233021 | -0.786086305 | 0.269 | 0.376 | 1 | 0.780226392 | 0.159940913 | 0.5 | 0.42 | 1 | 0.780226392 | 0.069224677 |
| 8030462N17Rik | 0.035236865 | -0.66726515 | 0.412 | 0.482 | 1 | 0.313040827 | -0.423745431 | 0.4 | 0.685 | 1 | 0.313040827 | 0.069232093 |
| Chchd10 | 0.035283565 | -2.222392421 | 0.017 | 0.071 | 1 | 0.18013682 | -1.521236903 | 0 | 0.154 | 1 | 0.18013682 | 0.069322201 |
| Asxl3 | 0.035284059 | -2.485426827 | 0.017 | 0.071 | 1 | 0.643846086 | 1.31748219 | 0 | 0.022 | 1 | 0.643846086 | 0.069323153 |
| D030056L22Rik | 0.03534042 | -1.070389328 | 0.143 | 0.235 | 1 | 0.13184901 | -1.505640048 | 0 | 0.188 | 1 | 0.13184901 | 0.069431895 |
| Plekha2 | 0.035356873 | -0.807354922 | 0.034 | 0.1 | 1 | 0.577184788 | 1.558490289 | 0 | 0.031 | 1 | 0.577184788 | 0.069463638 |
| Tpmt | 0.035414731 | -1.117695043 | 0.067 | 0.147 | 1 | 0.049438867 | 1.847996907 | 0.3 | 0.102 | 1 | 0.049438867 | 0.06957526 |
| Surf6 | 0.035545249 | -0.783107376 | 0.168 | 0.265 | 1 | 0.590427542 | -0.315978829 | 0.2 | 0.265 | 1 | 0.590427542 | 0.069827033 |
| Ccdc158 | 0.035548509 | 1.736965594 | 0.042 | 0.006 | 1 | 0.730749233 | 2.432959407 | 0 | 0.012 | 1 | 0.730749233 | 0.069833321 |
| Tln2 | 0.035581069 | -0.9510904 | 0.403 | 0.447 | 1 | 0.073388559 | 1.584962501 | 0.4 | 0.198 | 1 | 0.073388559 | 0.069896126 |
| Fez1 | 0.035610938 | -1.987927168 | 0.017 | 0.071 | 1 | 0.221461658 | 1.31748219 | 0.2 | 0.083 | 1 | 0.221461658 | 0.069953736 |
| Gins3 | 0.035611103 | -1.900464326 | 0.017 | 0.071 | 1 | 0.153152701 | 2.432959407 | 0.1 | 0.025 | 1 | 0.153152701 | 0.069954056 |
| Gjb2 | 0.035640086 | -3.593951284 | 0.017 | 0.071 | 1 | 0.577164735 | 0.374065718 | 0 | 0.031 | 1 | 0.577164735 | 0.070009956 |
| Grik3 | 0.035727768 | 1.029146346 | 0.193 | 0.106 | 1 | 0.349677934 | -0.339630097 | 0.1 | 0.222 | 1 | 0.349677934 | 0.070179062 |
| Sytl2 | 0.810909709 | -0.424026283 | 0.084 | 0.076 | 1 | 0.035747524 | 1.602884409 | 0.2 | 0.046 | 1 | 0.810909709 | 0.070217163 |
| Yju2 | 0.035758958 | -0.909924656 | 0.134 | 0.229 | 1 | 0.982449681 | 0.263034406 | 0.2 | 0.188 | 1 | 0.982449681 | 0.070239213 |
| Tarsl2 | 0.234168602 | -1.572889668 | 0.025 | 0.053 | 1 | 0.035788423 | 2.263034406 | 0.2 | 0.049 | 1 | 0.234168602 | 0.070296035 |
| Gnao1 | 0.035799034 | -1.339575961 | 0.067 | 0.147 | 1 | 0.386288477 | -1.248864633 | 0.1 | 0.201 | 1 | 0.386288477 | 0.070316498 |
| Vwa3b | 0.035799115 | 1.099535674 | 0.042 | 0.006 | 1 | 0.113669596 | 2.847996907 | 0.1 | 0.022 | 1 | 0.113669596 | 0.070316653 |
| Dolk | 0.303263572 | -0.600904045 | 0.193 | 0.235 | 1 | 0.035864725 | 1.227844977 | 0.5 | 0.201 | 1 | 0.303263572 | 0.070443171 |
| Pomt1 | 0.035907264 | -1.258016331 | 0.092 | 0.176 | 1 | 0.237367829 | 1.073063462 | 0.3 | 0.16 | 1 | 0.237367829 | 0.070525196 |
| Pcdhga10 | 0.035908207 | 0.96203215 | 0.193 | 0.106 | 1 | 0.612364586 | 2.159940913 | 0.1 | 0.065 | 1 | 0.612364586 | 0.070527015 |
| Wdfy1 | 0.035909239 | -0.807354922 | 0.387 | 0.488 | 1 | 0.125421119 | -0.852442812 | 0.3 | 0.549 | 1 | 0.125421119 | 0.070529005 |
| Gm42869 | 0.035938762 | -1.807354922 | 0.017 | 0.071 | 1 | 0.577184788 | 1.558490289 | 0 | 0.031 | 1 | 0.577184788 | 0.070585929 |
| Zswim9 | 0.035938762 | -1.987927168 | 0.017 | 0.071 | 1 | 0.967118498 | 0.494359952 | 0.1 | 0.102 | 1 | 0.967118498 | 0.070585929 |
| Tubg2 | 0.035938762 | -1.807354922 | 0.017 | 0.071 | 1 | 0.59781419 | 1.558490289 | 0 | 0.028 | 1 | 0.59781419 | 0.070585929 |
| Mrps27 | 0.035941568 | -0.74176658 | 0.109 | 0.206 | 1 | 0.15652788 | -1.171902651 | 0 | 0.17 | 1 | 0.15652788 | 0.07059134 |
| Mb21d2 | 0.508165242 | -0.49978212 | 0.319 | 0.335 | 1 | 0.035962274 | 1.432959407 | 0.6 | 0.278 | 1 | 0.508165242 | 0.070631262 |
| Tnip2 | 0.910164518 | -0.178765489 | 0.513 | 0.465 | 1 | 0.036000787 | 0.970958429 | 0.7 | 0.404 | 1 | 0.910164518 | 0.070705518 |
| Derl3 | 0.036015111 | 1.514573173 | 0.076 | 0.024 | 1 | 0.698534498 | 2.432959407 | 0 | 0.015 | 1 | 0.698534498 | 0.070733133 |
| Mall | 0.541050481 | 0.108580813 | 0.336 | 0.3 | 1 | 0.036091403 | 1.389332475 | 0.6 | 0.327 | 1 | 0.541050481 | 0.070880217 |
| C1s1 | 0.036106506 | -0.87244995 | 0.143 | 0.241 | 1 | 0.554907294 | -0.631693551 | 0.4 | 0.414 | 1 | 0.554907294 | 0.070909333 |
| Ube2c | 0.036142849 | -4.213347282 | 0.017 | 0.071 | 1 | 0.619953009 | 1.432959407 | 0 | 0.025 | 1 | 0.619953009 | 0.070979393 |
| Adgra3 | 0.036221482 | -0.591221491 | 0.723 | 0.765 | 1 | 0.163936386 | -0.797995028 | 0.8 | 0.843 | 1 | 0.163936386 | 0.071130968 |
| Eid3 | 0.036267015 | -1.707819249 | 0.017 | 0.071 | 1 | 0.599038816 | 1.494359952 | 0.1 | 0.059 | 1 | 0.599038816 | 0.071218733 |
| Srd5a1 | 0.036267015 | -1.707819249 | 0.017 | 0.071 | 1 | 0.52208609 | 0.847996907 | 0 | 0.04 | 1 | 0.52208609 | 0.071218733 |
| Smdt1 | 0.03627447 | 0.36389426 | 0.857 | 0.829 | 1 | 0.667005401 | -0.159057851 | 0.6 | 0.809 | 1 | 0.667005401 | 0.071233103 |
| Srp14 | 0.049736565 | 0.18905777 | 0.933 | 0.918 | 1 | 0.036348946 | -0.374395515 | 0.5 | 0.929 | 1 | 0.049736565 | 0.071376647 |
| Iftap | 0.036392174 | 0.381817964 | 0.496 | 0.353 | 1 | 0.733508943 | -0.159995884 | 0.4 | 0.401 | 1 | 0.733508943 | 0.071459958 |
| Nudc | 0.036400051 | -0.523900975 | 0.681 | 0.706 | 1 | 0.931571923 | 0.138338658 | 0.6 | 0.698 | 1 | 0.931571923 | 0.071475139 |
| Tmsb4x | 0.434035333 | -0.463027442 | 0.798 | 0.806 | 1 | 0.03644838 | -0.903252914 | 0.5 | 0.867 | 1 | 0.434035333 | 0.071568276 |
| Dld | 0.036471956 | -0.587788545 | 0.437 | 0.524 | 1 | 0.067482094 | -0.928497052 | 0.3 | 0.608 | 1 | 0.067482094 | 0.071613708 |
| Nfu1 | 0.03655362 | -0.707819249 | 0.37 | 0.465 | 1 | 0.041646296 | -1.625934282 | 0.2 | 0.509 | 1 | 0.041646296 | 0.071771073 |
| Foxred1 | 0.224979012 | 0.354108501 | 0.252 | 0.188 | 1 | 0.036598994 | 1.362570079 | 0.4 | 0.145 | 1 | 0.224979012 | 0.071858501 |
| Itpr3 | 0.036624987 | 0.130926093 | 0.664 | 0.512 | 1 | 0.588526551 | 0.133399125 | 0.9 | 0.725 | 1 | 0.588526551 | 0.071908585 |
| Plxna3 | 0.036672438 | -1.600904045 | 0.017 | 0.071 | 1 | 0.699053785 | 1.432959407 | 0.1 | 0.068 | 1 | 0.699053785 | 0.072000009 |
| Fam185a | 0.036751614 | -1.411426246 | 0.084 | 0.165 | 1 | 0.699916189 | 0.359710425 | 0.2 | 0.151 | 1 | 0.699916189 | 0.072152546 |
| 6030443J06Rik | 0.059091902 | -1.637429921 | 0.067 | 0.135 | 1 | 0.036797791 | 2.14345279 | 0.3 | 0.099 | 1 | 0.059091902 | 0.072241505 |
| Gm14066 | 0.036828068 | -2.070389328 | 0.017 | 0.071 | 1 | 0.746555377 | 0.847996907 | 0.1 | 0.071 | 1 | 0.746555377 | 0.072299829 |
| Ppie | 0.036853859 | -0.870090677 | 0.269 | 0.359 | 1 | 0.523111838 | 0.637100124 | 0.4 | 0.306 | 1 | 0.523111838 | 0.072349512 |
| Gng5 | 0.036867375 | 0.229573435 | 0.966 | 0.976 | 1 | 0.461354922 | -0.195071815 | 0.9 | 0.975 | 1 | 0.461354922 | 0.072375548 |
| Arhgef25 | 0.036872915 | -0.751501687 | 0.513 | 0.559 | 1 | 0.710031817 | -0.146985019 | 0.5 | 0.519 | 1 | 0.710031817 | 0.072386219 |
| Brd9 | 0.036888503 | -0.896059147 | 0.286 | 0.376 | 1 | 0.455491474 | 0.68700503 | 0.4 | 0.333 | 1 | 0.455491474 | 0.072416245 |
| Ldha | 0.036926832 | 0.221412382 | 0.975 | 0.965 | 1 | 0.134440262 | 0.40019545 | 1 | 0.969 | 1 | 0.134440262 | 0.072490073 |
| H1f4 | 0.037034574 | -0.405256478 | 0.504 | 0.365 | 1 | 0.932262805 | 0.957510391 | 0.4 | 0.367 | 1 | 0.932262805 | 0.072697587 |
| Fbxw4 | 0.037085222 | -1.04342228 | 0.126 | 0.218 | 1 | 0.718785945 | 0.549152425 | 0.3 | 0.265 | 1 | 0.718785945 | 0.072795131 |
| Tmem129 | 0.037095204 | -0.826463745 | 0.092 | 0.182 | 1 | 0.648370942 | 0.602884409 | 0.2 | 0.142 | 1 | 0.648370942 | 0.072814355 |
| Ankrd44 | 0.037106308 | -1.440656957 | 0.387 | 0.471 | 1 | 0.476041445 | 0.365214801 | 0.5 | 0.41 | 1 | 0.476041445 | 0.072835737 |
| Cox4i2 | 0.974988157 | -0.526145288 | 0.63 | 0.529 | 1 | 0.037116949 | -2.808626579 | 0.1 | 0.417 | 1 | 0.974988157 | 0.072856231 |
| Syvn1 | 0.037170145 | -0.538538164 | 0.235 | 0.347 | 1 | 0.070163237 | -1.365782384 | 0.1 | 0.377 | 1 | 0.070163237 | 0.07295867 |
| Tfdp2 | 0.037179963 | 0.126757142 | 0.538 | 0.376 | 1 | 0.079810211 | 1.100081013 | 0.7 | 0.614 | 1 | 0.079810211 | 0.072977576 |
| Cc2d2a | 0.355632483 | -0.724892762 | 0.252 | 0.276 | 1 | 0.037190732 | 1.22650853 | 0.6 | 0.306 | 1 | 0.355632483 | 0.072998314 |
| Pstpip2 | 0.037230819 | -1.177304532 | 0.042 | 0.112 | 1 | 0.121131306 | -2.069540933 | 0 | 0.198 | 1 | 0.121131306 | 0.073075504 |
| Snx30 | 0.037257443 | -1 | 0.101 | 0.188 | 1 | 0.136276961 | -1.864721141 | 0 | 0.185 | 1 | 0.136276961 | 0.073126769 |
| Pcmtd2 | 0.037269966 | -0.932885804 | 0.311 | 0.382 | 1 | 0.769483132 | 0.469485283 | 0.5 | 0.426 | 1 | 0.769483132 | 0.073150882 |
| Gucd1 | 0.037320161 | 1.131244533 | 0.16 | 0.082 | 1 | 0.204453065 | 0.828097349 | 0.3 | 0.142 | 1 | 0.204453065 | 0.073247528 |
| AA986860 | 0.109511739 | -1.807354922 | 0.05 | 0.1 | 1 | 0.037341032 | 2.385653692 | 0.2 | 0.049 | 1 | 0.109511739 | 0.073287712 |
| Pdpr | 0.037346415 | -1.168236651 | 0.134 | 0.224 | 1 | 0.668300563 | -0.279758641 | 0.2 | 0.244 | 1 | 0.668300563 | 0.073298074 |
| Serf1 | 0.037384983 | -0.74092756 | 0.445 | 0.518 | 1 | 0.432479965 | -0.428334322 | 0.3 | 0.417 | 1 | 0.432479965 | 0.073372329 |
| Coq6 | 0.037498274 | -0.987927168 | 0.076 | 0.159 | 1 | 0.195951444 | -0.763437806 | 0 | 0.145 | 1 | 0.195951444 | 0.073590427 |
| Synpo2 | 0.934755897 | 0.514573173 | 0.05 | 0.053 | 1 | 0.037617847 | 2.185031894 | 0.3 | 0.096 | 1 | 0.934755897 | 0.073820591 |
| Rit1 | 0.037627167 | -0.84450792 | 0.303 | 0.4 | 1 | 0.497634004 | -0.596787936 | 0.4 | 0.463 | 1 | 0.497634004 | 0.07383853 |
| 3110009E18Rik | 0.037680578 | -1.122856748 | 0.042 | 0.112 | 1 | 0.334306566 | -0.304006187 | 0 | 0.086 | 1 | 0.334306566 | 0.07394133 |
| Ak7 | 0.037706174 | -1.411426246 | 0.017 | 0.071 | 1 | 0.343976171 | -0.709998547 | 0 | 0.083 | 1 | 0.343976171 | 0.073990593 |
| Slc11a2 | 0.037733844 | 0.633217669 | 0.471 | 0.335 | 1 | 0.728128672 | -0.493040011 | 0.4 | 0.401 | 1 | 0.728128672 | 0.074043846 |
| Zfp787 | 0.037748418 | -0.807354922 | 0.202 | 0.3 | 1 | 0.162657858 | -1.019167411 | 0.2 | 0.398 | 1 | 0.162657858 | 0.074071893 |
| Scx | 0.037749822 | -1.235448574 | 0.042 | 0.112 | 1 | 0.082439996 | -2.090602549 | 0 | 0.238 | 1 | 0.082439996 | 0.074074596 |
| Dpysl2 | 0.037751194 | 0.230364744 | 0.546 | 0.394 | 1 | 0.413521143 | 1.190102883 | 0.4 | 0.377 | 1 | 0.413521143 | 0.074077236 |
| Tnfaip1 | 0.037779423 | -0.635986504 | 0.345 | 0.435 | 1 | 0.042312242 | -1.344021866 | 0.2 | 0.546 | 1 | 0.042312242 | 0.074131562 |

| | | | | | | | | | | | | |
|---|---|---|---|---|---|---|---|---|---|---|---|---|
| Bcl7b | 0.037897438 | 0.298439742 | 0.58 | 0.435 | 1 | 0.470301534 | 0.672147071 | 0.6 | 0.463 | 1 | 0.470301534 | 0.07435866 |
| Nkapd1 | 0.0379006 | -0.691101854 | 0.336 | 0.429 | 1 | 0.194251202 | -0.637429921 | 0.2 | 0.426 | 1 | 0.194251202 | 0.074364744 |
| Klhl8 | 0.415593984 | -0.36994961 | 0.092 | 0.124 | 1 | 0.037922276 | 1.816288047 | 0.3 | 0.099 | 1 | 0.415593984 | 0.074406452 |
| C2cd3 | 0.037927013 | -0.953575663 | 0.176 | 0.271 | 1 | 0.866910914 | 0.240314329 | 0.3 | 0.284 | 1 | 0.866910914 | 0.074415567 |
| Mindy2 | 0.194193975 | 0.334000927 | 0.496 | 0.418 | 1 | 0.03794493 | -1.593102889 | 0.3 | 0.583 | 1 | 0.194193975 | 0.074450041 |
| Fkbp9 | 0.448338981 | -0.14379527 | 0.975 | 0.906 | 1 | 0.038025664 | 0.738638151 | 1 | 0.914 | 1 | 0.448338981 | 0.074605378 |
| Slc43a3 | 0.08559849 | 0.217761755 | 0.748 | 0.647 | 1 | 0.038026049 | 1.093649234 | 0.8 | 0.5 | 1 | 0.08559849 | 0.074606117 |
| Lrp11 | 0.038039885 | -1.392317423 | 0.017 | 0.071 | 1 | 0.230466554 | 2.558490289 | 0.1 | 0.031 | 1 | 0.230466554 | 0.074632737 |
| Pisd | 0.038076198 | -0.847996907 | 0.126 | 0.218 | 1 | 0.690478535 | 0.847996907 | 0.3 | 0.278 | 1 | 0.690478535 | 0.074702599 |
| Mbnl1 | 1 | -0.277590242 | 1 | 1 | 1 | 0.038092423 | -0.660793914 | 0.9 | 1 | 1 | 1 | 0.074733813 |
| Polr1h | 0.038116039 | -0.65161911 | 0.513 | 0.576 | 1 | 0.142561859 | -0.493830746 | 0.4 | 0.627 | 1 | 0.142561859 | 0.074779245 |
| Mmp25 | 0.041399178 | -2.572889668 | 0.008 | 0.053 | 1 | 0.038118244 | 2.263034406 | 0.2 | 0.049 | 1 | 0.041399178 | 0.074783488 |
| Suclg2 | 0.960465998 | -0.121270497 | 0.395 | 0.376 | 1 | 0.038128074 | 1.484489708 | 0.6 | 0.46 | 1 | 0.960465998 | 0.074802398 |
| Fmo2 | 0.038142131 | 2.836501268 | 0.025 | 0 | 1 | 0.260206574 | -0.230005605 | 0 | 0.114 | 1 | 0.260206574 | 0.074829441 |
| Gm12500 | 0.038142131 | 2.836501268 | 0.025 | 0 | 1 | 0.730749233 | 2.432959407 | 0 | 0.012 | 1 | 0.730749233 | 0.074829441 |
| C8g | 0.038142909 | 2.514573173 | 0.025 | 0 | 1 | 0.669853054 | 2.210566986 | 0 | 0.019 | 1 | 0.669853054 | 0.074830937 |
| 1700084E18Rik | 0.038142909 | 2.514573173 | 0.025 | 0 | 1 | 0.730749233 | 2.432959407 | 0 | 0.012 | 1 | 0.730749233 | 0.074830937 |
| Rassf7 | 0.038142909 | 2.514573173 | 0.025 | 0 | 1 | 0.698529281 | 2.210566986 | 0 | 0.015 | 1 | 0.698529281 | 0.074830937 |
| Gm20597 | 0.038142909 | 2.514573173 | 0.025 | 0 | 1 | 0.730750993 | 1.847996907 | 0 | 0.012 | 1 | 0.730750993 | 0.074830937 |
| Syt7 | 0.038142909 | 2.514573173 | 0.025 | 0 | 1 | 0.698532411 | 1.847996907 | 0 | 0.015 | 1 | 0.698532411 | 0.074830937 |
| Rorb | 0.038142909 | 2.514573173 | 0.025 | 0 | 1 | 0.698533977 | 1.558490289 | 0 | 0.015 | 1 | 0.698533977 | 0.074830937 |
| Asb7 | 0.152677854 | -0.831201664 | 0.168 | 0.229 | 1 | 0.038152258 | -2.408342847 | 0 | 0.312 | 1 | 0.152677854 | 0.074848921 |
| Prdm1 | 0.038154432 | 1.215012891 | 0.076 | 0.024 | 1 | 0.597845299 | -0.789433014 | 0 | 0.028 | 1 | 0.597845299 | 0.074853104 |
| Sec22a | 0.038228449 | -0.807354922 | 0.345 | 0.429 | 1 | 0.734003254 | 0.419662585 | 0.4 | 0.386 | 1 | 0.734003254 | 0.074995483 |
| Tlcd5 | 0.038255952 | 1.273565073 | 0.143 | 0.071 | 1 | 0.135900341 | 1.602884409 | 0.2 | 0.071 | 1 | 0.135900341 | 0.075048386 |
| Bltp1 | 0.038294294 | -0.412383046 | 0.908 | 0.929 | 1 | 0.944015166 | 0.199760231 | 0.9 | 0.963 | 1 | 0.944015166 | 0.075122136 |
| Nsa2 | 0.038308625 | -0.322084476 | 0.866 | 0.906 | 1 | 0.038454775 | -0.880355557 | 0.5 | 0.877 | 1 | 0.038454775 | 0.075149699 |
| Galk2 | 0.038313979 | -0.835011265 | 0.193 | 0.3 | 1 | 0.168287028 | 0.808468542 | 0.5 | 0.306 | 1 | 0.168287028 | 0.075159996 |
| Dipk2a | 0.03831652 | 0.176533073 | 0.941 | 0.865 | 1 | 0.232181051 | 0.4304569 | 1 | 0.904 | 1 | 0.232181051 | 0.075164884 |
| Nars2 | 0.038372381 | -1.117695043 | 0.143 | 0.235 | 1 | 0.963038318 | -0.279758641 | 0.2 | 0.188 | 1 | 0.963038318 | 0.075272322 |
| Mmd | 0.038386616 | -1.292781749 | 0.017 | 0.071 | 1 | 0.522125488 | 0.263034406 | 0 | 0.04 | 1 | 0.522125488 | 0.0752997 |
| Seh1l | 0.623897227 | -0.127664133 | 0.513 | 0.547 | 1 | 0.038422749 | -0.5519337 | 0.1 | 0.503 | 1 | 0.623897227 | 0.075369191 |
| Edrf1 | 0.038486177 | -0.703850346 | 0.277 | 0.388 | 1 | 0.105242057 | -1.572041274 | 0.2 | 0.423 | 1 | 0.105242057 | 0.075491168 |
| Pex19 | 0.564736867 | -0.173482821 | 0.361 | 0.388 | 1 | 0.038544138 | -1.596787936 | 0.1 | 0.426 | 1 | 0.564736867 | 0.075602626 |
| Nup35 | 0.038593959 | -0.634290213 | 0.294 | 0.412 | 1 | 0.171733413 | -0.374395515 | 0.1 | 0.34 | 1 | 0.171733413 | 0.075698425 |
| Mcm10 | 0.039412462 | -0.959358016 | 0.042 | 0.112 | 1 | 0.038594921 | 2.611929548 | 0.3 | 0.099 | 1 | 0.039412462 | 0.075700273 |
| Gm20732 | 0.038601578 | -1.026996732 | 0.269 | 0.353 | 1 | 0.113760294 | 0.662717658 | 0.6 | 0.377 | 1 | 0.113760294 | 0.075713074 |
| Msh3 | 0.038603071 | -1.539874611 | 0.084 | 0.165 | 1 | 0.787785791 | 0.598383016 | 0.2 | 0.17 | 1 | 0.787785791 | 0.075715945 |
| Cables1 | 0.038700203 | 0.657531127 | 0.227 | 0.129 | 1 | 0.180880305 | 1.111031312 | 0.3 | 0.145 | 1 | 0.180880305 | 0.0759027 |
| Ccdc25 | 0.03871016 | -0.535550307 | 0.479 | 0.571 | 1 | 0.58063333 | -0.108186861 | 0.6 | 0.633 | 1 | 0.58063333 | 0.075921844 |
| Ccdc82 | 0.038785439 | -0.762960803 | 0.387 | 0.471 | 1 | 0.316867348 | -0.638247543 | 0.3 | 0.475 | 1 | 0.316867348 | 0.076066569 |
| Fli1 | 0.038854677 | -0.685267407 | 0.639 | 0.647 | 1 | 0.24671973 | -0.374395515 | 0.3 | 0.537 | 1 | 0.24671973 | 0.076199667 |
| Cstf2 | 0.038860349 | -1.096861539 | 0.134 | 0.224 | 1 | 0.757924023 | 0.223506042 | 0.2 | 0.256 | 1 | 0.757924023 | 0.076210571 |
| Gria3 | 0.810736537 | -0.194836699 | 0.454 | 0.412 | 1 | 0.038907025 | 0.984498906 | 0.7 | 0.407 | 1 | 0.810736537 | 0.076300293 |
| Gm13199 | 0.038974464 | -2.485426827 | 0 | 0.035 | 1 | 0.505686333 | 1.111031312 | 0 | 0.043 | 1 | 0.505686333 | 0.076429919 |
| Adgrb2 | 0.038974464 | -2.485426827 | 0 | 0.035 | 1 | 0.577165087 | 0.930459707 | 0 | 0.031 | 1 | 0.577165087 | 0.076429919 |
| Gm57263 | 0.038974464 | -2.944858446 | 0 | 0.035 | 1 | 0.698534498 | 2.432959407 | 0 | 0.015 | 1 | 0.698534498 | 0.076429919 |
| 1700019G24Rik | 0.038974464 | -4.292781749 | 0 | 0.035 | 1 | 0.597845674 | 1.695993813 | 0 | 0.028 | 1 | 0.597845674 | 0.076429919 |
| Opn1sw | 0.038974464 | -2.485426827 | 0 | 0.035 | 1 | 0.669853054 | 2.210566986 | 0 | 0.019 | 1 | 0.669853054 | 0.076429919 |
| Alpk3 | 0.038974464 | -3.572889668 | 0 | 0.035 | 1 | 0.474947236 | 0.374065718 | 0 | 0.049 | 1 | 0.474947236 | 0.076429919 |
| Zfp90 | 0.038974464 | -2.655351829 | 0 | 0.035 | 1 | 0.487557793 | 1.625604485 | 0.1 | 0.049 | 1 | 0.487557793 | 0.076429919 |
| Gm26511 | 0.038974464 | -2.485426827 | 0 | 0.035 | 1 | 0.59781419 | 1.432959407 | 0 | 0.028 | 1 | 0.59781419 | 0.076429919 |
| Itgae | 0.038974464 | -3.185866545 | 0 | 0.035 | 1 | 0.730749233 | 2.210566986 | 0 | 0.012 | 1 | 0.730749233 | 0.076429919 |
| Engase | 0.038974464 | -2.485426827 | 0 | 0.035 | 1 | 0.559081495 | 1.432959407 | 0.1 | 0.056 | 1 | 0.559081495 | 0.076429919 |
| Ccdc40 | 0.038974464 | -2.485426827 | 0 | 0.035 | 1 | 0.236765379 | 1.558490289 | 0.1 | 0.031 | 1 | 0.236765379 | 0.076429919 |
| Kif26a | 0.038974464 | -2.485426827 | 0 | 0.035 | 1 | 0.364022816 | -0.152003093 | 0 | 0.077 | 1 | 0.364022816 | 0.076429919 |
| Tes3-ps | 0.038974464 | -2.485426827 | 0 | 0.035 | 1 | 0.643835681 | 1.847996907 | 0 | 0.022 | 1 | 0.643835681 | 0.076429919 |
| Gap43 | 0.038974464 | -3.070389328 | 0 | 0.035 | 1 | 0.557756607 | 0.695993813 | 0 | 0.034 | 1 | 0.557756607 | 0.076429919 |
| Fsd1 | 0.038974464 | -2.655351829 | 0 | 0.035 | 1 | 0.597832556 | 1.31748219 | 0 | 0.028 | 1 | 0.597832556 | 0.076429919 |
| Cdc42ep2 | 0.038974464 | -2.485426827 | 0 | 0.035 | 1 | 0.643835681 | 1.847996907 | 0 | 0.022 | 1 | 0.643835681 | 0.076429919 |
| Ffar4 | 0.038974464 | -2.655351829 | 0 | 0.035 | 1 | 0.669853054 | 2.210566986 | 0 | 0.019 | 1 | 0.669853054 | 0.076429919 |
| Prrg3 | 0.038974464 | -2.485426827 | 0 | 0.035 | 1 | 0.643850855 | 2.017921908 | 0 | 0.022 | 1 | 0.643850855 | 0.076429919 |
| Arhgap4 | 0.038974464 | -2.655351829 | 0 | 0.035 | 1 | 0.254436177 | 2.695993813 | 0.1 | 0.034 | 1 | 0.254436177 | 0.076429919 |
| Gpr161 | 0.038978449 | -2.655351829 | 0 | 0.035 | 1 | 0.474927752 | 0.625604485 | 0 | 0.049 | 1 | 0.474927752 | 0.076437578 |
| Cfap157 | 0.038978449 | -2.655351829 | 0 | 0.035 | 1 | 0.607557782 | 1.111031312 | 0.1 | 0.059 | 1 | 0.607557782 | 0.076437578 |
| Phf19 | 0.038978449 | -2.807354922 | 0 | 0.035 | 1 | 0.669853054 | 2.210566986 | 0 | 0.019 | 1 | 0.669853054 | 0.076437578 |
| Arhgap15 | 0.038978449 | -4.911691582 | 0 | 0.035 | 1 | 0.460521625 | -0.230005605 | 0 | 0.052 | 1 | 0.460521625 | 0.076437578 |
| Arhgap40 | 0.038978449 | -2.807354922 | 0 | 0.035 | 1 | 0.619961444 | 1.558490289 | 0 | 0.025 | 1 | 0.619961444 | 0.076437578 |
| D430018E03Rik | 0.038978449 | -2.655351829 | 0 | 0.035 | 1 | 0.049344356 | 3.210566986 | 0.1 | 0.015 | 1 | 0.049344356 | 0.076437578 |
| Fbxl12os | 0.038978449 | -2.944858446 | 0 | 0.035 | 1 | 0.59781419 | 1.558490289 | 0 | 0.028 | 1 | 0.59781419 | 0.076437578 |
| Calml4 | 0.038978449 | -2.655351829 | 0 | 0.035 | 1 | 0.597845674 | 1.695993813 | 0 | 0.028 | 1 | 0.597845674 | 0.076437578 |
| AW822252 | 0.038978449 | -2.655351829 | 0 | 0.035 | 1 | 0.698532411 | 1.432959407 | 0 | 0.015 | 1 | 0.698532411 | 0.076437578 |
| Gm15261 | 0.038978449 | -2.655351829 | 0 | 0.035 | 1 | 0.446865484 | 0.210566986 | 0 | 0.056 | 1 | 0.446865484 | 0.076437578 |
| Tmem144 | 0.038980441 | -2.944858446 | 0 | 0.035 | 1 | 0.374682358 | -0.111361109 | 0 | 0.074 | 1 | 0.374682358 | 0.076441408 |
| Cecr2 | 0.038980441 | -3.070389328 | 0 | 0.035 | 1 | 0.474085213 | 0.874963954 | 0.2 | 0.12 | 1 | 0.474085213 | 0.076441408 |
| Nlrp5-ps | 0.038980441 | -3.292781749 | 0 | 0.035 | 1 | 0.197182351 | 2.111031312 | 0.1 | 0.028 | 1 | 0.197182351 | 0.076441408 |
| Gm56519 | 0.038980441 | -3.185866545 | 0 | 0.035 | 1 | 0.698533977 | 1.847996907 | 0 | 0.015 | 1 | 0.698533977 | 0.076441408 |
| Pcdhb4 | 0.038980441 | -3.070389328 | 0 | 0.035 | 1 | 0.489954768 | 0.432959407 | 0 | 0.046 | 1 | 0.489954768 | 0.076441408 |
| Gm56635 | 0.038980441 | -2.944858446 | 0 | 0.035 | 1 | 0.577142568 | 1.432959407 | 0 | 0.031 | 1 | 0.577142568 | 0.076441408 |
| Impg2 | 0.03898084 | -3.292781749 | 0 | 0.035 | 1 | 0.489954768 | 0.432959407 | 0 | 0.046 | 1 | 0.489954768 | 0.076442174 |
| Gata2 | 0.038981637 | -3.733354341 | 0 | 0.035 | 1 | 0.325029139 | -0.888968688 | 0 | 0.09 | 1 | 0.325029139 | 0.076443706 |
| Lrrn1 | 0.038981637 | -3.807354922 | 0 | 0.035 | 1 | 0.698532411 | 2.017921908 | 0 | 0.015 | 1 | 0.698532411 | 0.076443706 |
| Rgs5 | 0.038982434 | -2.292781749 | 0 | 0.035 | 1 | 0.63806535 | 1.263034406 | 0.1 | 0.062 | 1 | 0.63806535 | 0.076445237 |
| Gm45902 | 0.038982434 | -2.292781749 | 0 | 0.035 | 1 | 0.538649237 | 2.695993813 | 0.1 | 0.059 | 1 | 0.538649237 | 0.076445237 |
| Tmod1 | 0.038982434 | -2.292781749 | 0 | 0.035 | 1 | 0.532203993 | 1.017921908 | 0.1 | 0.052 | 1 | 0.532203993 | 0.076445237 |
| Ankrd61 | 0.038982434 | -2.292781749 | 0 | 0.035 | 1 | 0.643842184 | 1.695993813 | 0 | 0.022 | 1 | 0.643842184 | 0.076445237 |
| Ssc5d | 0.038982434 | -2.292781749 | 0 | 0.035 | 1 | 0.505686333 | 1.111031312 | 0 | 0.043 | 1 | 0.505686333 | 0.076445237 |
| Zfp59 | 0.038982434 | -2.292781749 | 0 | 0.035 | 1 | 0.442237182 | 2.280956314 | 0.1 | 0.049 | 1 | 0.442237182 | 0.076445237 |
| Moap1 | 0.038982434 | -2.292781749 | 0 | 0.035 | 1 | 0.539483006 | 1.31748219 | 0 | 0.037 | 1 | 0.539483006 | 0.076445237 |
| 4933433G19Rik | 0.038982434 | -2.292781749 | 0 | 0.035 | 1 | 0.518235124 | 1.769994395 | 0.1 | 0.052 | 1 | 0.518235124 | 0.076445237 |
| Gm41293 | 0.038982434 | -2.292781749 | 0 | 0.035 | 1 | 0.577142568 | 1.31748219 | 0 | 0.031 | 1 | 0.577142568 | 0.076445237 |
| Dnase1l2 | 0.038982434 | -2.292781749 | 0 | 0.035 | 1 | 0.597845674 | 1.695993813 | 0 | 0.028 | 1 | 0.597845674 | 0.076445237 |
| Ager | 0.038982434 | -2.292781749 | 0 | 0.035 | 1 | 0.730749233 | 2.210566986 | 0 | 0.012 | 1 | 0.730749233 | 0.076445237 |
| Mef2c | 0.039050322 | -2.103747925 | 0.252 | 0.329 | 1 | 0.235543461 | -0.152003093 | 0.3 | 0.164 | 1 | 0.235543461 | 0.076575716 |
| Elmod2 | 0.03909563 | -1.235448574 | 0.084 | 0.165 | 1 | 0.767180334 | 0.515421567 | 0.2 | 0.157 | 1 | 0.767180334 | 0.076662792 |
| Suv39h1 | 0.039097048 | -1.155278225 | 0.084 | 0.165 | 1 | 0.979498033 | 0.290001453 | 0.1 | 0.099 | 1 | 0.979498033 | 0.076665517 |
| 8430429K09Rik | 0.039156789 | -1.008988783 | 0.101 | 0.188 | 1 | 0.259767244 | -0.596787936 | 0 | 0.114 | 1 | 0.259767244 | 0.076780325 |
| Enoph1 | 0.039230485 | -0.87774425 | 0.109 | 0.2 | 1 | 0.094556159 | 1.111031312 | 0.4 | 0.179 | 1 | 0.094556159 | 0.07692194 |

| | | | | | | | | | | | | |
|---|---|---|---|---|---|---|---|---|---|---|---|---|
| Rbm7 | 0.606067569 | -0.17684027 | 0.723 | 0.682 | 1 | 0.039231653 | -0.777937375 | 0.4 | 0.707 | 1 | 0.606067569 | 0.076924183 |
| Svbp | 0.039253468 | -0.548162583 | 0.513 | 0.559 | 1 | 0.61558389 | -0.136896201 | 0.4 | 0.494 | 1 | 0.61558389 | 0.076966102 |
| Zcchc9 | 0.039343887 | -1.085339669 | 0.235 | 0.324 | 1 | 0.278836782 | -0.848326703 | 0.2 | 0.343 | 1 | 0.278836782 | 0.077139832 |
| Pla2g12a | 0.039363346 | 0.4911142 | 0.378 | 0.259 | 1 | 0.398817735 | -0.239465935 | 0.2 | 0.346 | 1 | 0.398817735 | 0.077177219 |
| Tnip1 | 0.039371699 | -1.255944981 | 0.126 | 0.212 | 1 | 0.268179984 | 0.457206954 | 0.4 | 0.219 | 1 | 0.268179984 | 0.077193266 |
| Asnsd1 | 0.707791333 | -0.231670235 | 0.227 | 0.241 | 1 | 0.039376364 | 1.580516596 | 0.4 | 0.157 | 1 | 0.707791333 | 0.077202231 |
| Hmgcll1 | 0.039415668 | 0.142604395 | 0.168 | 0.082 | 1 | 0.62636956 | -0.111361109 | 0.3 | 0.207 | 1 | 0.62636956 | 0.077277741 |
| Soat1 | 0.114651973 | 0.196397213 | 0.639 | 0.506 | 1 | 0.039448201 | 1.084231253 | 0.8 | 0.593 | 1 | 0.114651973 | 0.077340241 |
| Mto1 | 0.039461039 | -1.01267383 | 0.109 | 0.2 | 1 | 0.135746401 | -1.457811523 | 0 | 0.185 | 1 | 0.135746401 | 0.077364904 |
| Casp9 | 0.37567374 | -0.655351829 | 0.05 | 0.076 | 1 | 0.039504821 | 1.847996907 | 0.3 | 0.096 | 1 | 0.37567374 | 0.077449011 |
| Trappc3 | 0.039555342 | -0.491527027 | 0.655 | 0.635 | 1 | 0.822925659 | -0.105005614 | 0.7 | 0.673 | 1 | 0.822925659 | 0.077546058 |
| Zfp955a | 0.066383266 | -0.58058406 | 0.126 | 0.212 | 1 | 0.03958274 | 1.385653692 | 0.4 | 0.148 | 1 | 0.066383266 | 0.077598686 |
| Nid2 | 0.168471846 | -0.371027454 | 0.681 | 0.682 | 1 | 0.03962737 | 1.014586195 | 0.8 | 0.506 | 1 | 0.168471846 | 0.077684412 |
| Ino80b | 0.039645056 | -1.255944981 | 0.109 | 0.194 | 1 | 0.219557806 | -0.864721141 | 0 | 0.133 | 1 | 0.219557806 | 0.077718382 |
| Zfp74 | 0.039660965 | -1.292781749 | 0.084 | 0.165 | 1 | 0.452220282 | -0.521236903 | 0.1 | 0.188 | 1 | 0.452220282 | 0.077748938 |
| Arhgef18 | 0.03967159 | -0.622930351 | 0.151 | 0.253 | 1 | 0.976893376 | 0.598383016 | 0.3 | 0.377 | 1 | 0.976893376 | 0.077769346 |
| Nxpe4 | 0.774772407 | -0.538538164 | 0.244 | 0.247 | 1 | 0.039682578 | 0.727244747 | 0.5 | 0.204 | 1 | 0.774772407 | 0.07779045 |
| Ywhag | 0.952290449 | -0.131789873 | 0.807 | 0.759 | 1 | 0.039709986 | -0.930445324 | 0.6 | 0.91 | 1 | 0.952290449 | 0.07784309 |
| Ube4a | 0.03978416 | -0.680243004 | 0.504 | 0.576 | 1 | 0.078981938 | -0.9669712 | 0.4 | 0.642 | 1 | 0.078981938 | 0.077985541 |
| Bckdhb | 0.216875053 | -0.263034406 | 0.319 | 0.376 | 1 | 0.039824827 | 1.539874611 | 0.6 | 0.309 | 1 | 0.216875053 | 0.078063637 |
| Gtdc1 | 0.039936337 | -0.949830404 | 0.336 | 0.424 | 1 | 0.158839235 | 0.689140142 | 0.6 | 0.438 | 1 | 0.158839235 | 0.078277763 |
| Cep19 | 0.039975209 | -1.244418728 | 0.084 | 0.165 | 1 | 0.165439681 | -1.026472211 | 0 | 0.164 | 1 | 0.165439681 | 0.078352402 |
| Homer1 | 0.040008899 | -0.384228104 | 0.42 | 0.547 | 1 | 0.857099301 | 0.225363988 | 0.6 | 0.636 | 1 | 0.857099301 | 0.078417086 |
| Ifitm1 | 0.040049666 | -0.614709844 | 0.059 | 0.135 | 1 | 0.402563129 | -0.357117523 | 0.1 | 0.207 | 1 | 0.402563129 | 0.078495356 |
| Zfp64 | 0.188306859 | 0.27064759 | 0.454 | 0.388 | 1 | 0.040063913 | 0.968678389 | 0.7 | 0.42 | 1 | 0.188306859 | 0.078522709 |
| Hectd3 | 0.040081938 | -0.900464326 | 0.05 | 0.124 | 1 | 0.324887875 | -0.152003093 | 0 | 0.09 | 1 | 0.324887875 | 0.078557315 |
| Rtel1 | 0.061264795 | -1.622930351 | 0.034 | 0.088 | 1 | 0.040090028 | 2.280956314 | 0.2 | 0.049 | 1 | 0.061264795 | 0.078572845 |
| Adat2 | 0.167616515 | -1.070389328 | 0.017 | 0.047 | 1 | 0.040090028 | 2.354956895 | 0.2 | 0.049 | 1 | 0.167616515 | 0.078572845 |
| Fbxo3 | 0.204845151 | -0.432959407 | 0.504 | 0.529 | 1 | 0.040106562 | -1.135207851 | 0.5 | 0.679 | 1 | 0.204845151 | 0.078604588 |
| Bdh2 | 0.040156363 | -1.87774425 | 0.025 | 0.082 | 1 | 0.364139783 | -0.374395515 | 0 | 0.077 | 1 | 0.364139783 | 0.078700192 |
| 9930014A18Rik | 0.040156363 | -1.87774425 | 0.025 | 0.082 | 1 | 0.596204587 | 1.558490289 | 0.1 | 0.059 | 1 | 0.596204587 | 0.078700192 |
| Kif11 | 0.040186331 | -3.046141782 | 0.025 | 0.082 | 1 | 0.698532411 | 1.432959407 | 0 | 0.015 | 1 | 0.698532411 | 0.078757721 |
| Brk1 | 0.32613631 | 0.150209812 | 0.849 | 0.835 | 1 | 0.040209124 | -0.864721141 | 0.5 | 0.836 | 1 | 0.32613631 | 0.078801475 |
| Tank | 0.040283669 | 0.204505233 | 0.58 | 0.459 | 1 | 0.190745908 | -0.793720372 | 0.3 | 0.512 | 1 | 0.190745908 | 0.078944565 |
| Fam120b | 0.040317102 | -0.497203995 | 0.303 | 0.429 | 1 | 0.564160132 | 0.642882477 | 0.5 | 0.438 | 1 | 0.564160132 | 0.079008736 |
| Kin | 0.040387 | -0.569491092 | 0.387 | 0.476 | 1 | 0.771543969 | -0.40354186 | 0.5 | 0.457 | 1 | 0.771543969 | 0.07914289 |
| Zfyve27 | 0.043657197 | 0.658963082 | 0.277 | 0.176 | 1 | 0.040430238 | 1.463333056 | 0.5 | 0.219 | 1 | 0.043657197 | 0.079225872 |
| Gm49336 | 0.040435826 | -0.610122575 | 0.437 | 0.518 | 1 | 0.080892128 | -1.11821478 | 0.4 | 0.62 | 1 | 0.080892128 | 0.079236596 |
| Zc3h4 | 0.040457435 | -0.591221491 | 0.261 | 0.376 | 1 | 0.721680421 | -0.473931188 | 0.5 | 0.451 | 1 | 0.721680421 | 0.079278065 |
| Tusc1 | 0.040510487 | -1.444784843 | 0.067 | 0.141 | 1 | 0.185548958 | -0.936274402 | 0 | 0.151 | 1 | 0.185548958 | 0.079379874 |
| Fzd5 | 0.04054687 | -1.004800986 | 0.092 | 0.176 | 1 | 0.54208468 | -0.152003093 | 0.1 | 0.17 | 1 | 0.54208468 | 0.079449692 |
| Strada | 0.040554957 | -0.941106311 | 0.176 | 0.276 | 1 | 0.714920829 | -0.606568957 | 0.2 | 0.228 | 1 | 0.714920829 | 0.07946521 |
| Elob | 0.417524421 | 0.102544423 | 0.966 | 0.959 | 1 | 0.040555681 | -0.66757779 | 0.6 | 0.978 | 1 | 0.417524421 | 0.079466599 |
| Gpkow | 0.050974938 | -0.821029859 | 0.286 | 0.371 | 1 | 0.04055732 | -1.924592597 | 0.1 | 0.417 | 1 | 0.050974938 | 0.079469744 |
| 1700112J16Rik | 0.040699376 | -2.070389328 | 0.034 | 0.094 | 1 | 0.084166339 | 1.432959407 | 0.2 | 0.059 | 1 | 0.084166339 | 0.079742313 |
| Echdc1 | 0.041941861 | 0.448984831 | 0.462 | 0.353 | 1 | 0.040737626 | 1.191951308 | 0.6 | 0.318 | 1 | 0.041941861 | 0.079815697 |
| Knl1 | 0.040739512 | -3.029747343 | 0.008 | 0.053 | 1 | 0.044545914 | 4.017921908 | 0.1 | 0.015 | 1 | 0.044545914 | 0.079819317 |
| BC051537 | 0.040739762 | -3.292781749 | 0.008 | 0.053 | 1 | 0.158351333 | 1.847996907 | 0.1 | 0.025 | 1 | 0.158351333 | 0.079819795 |
| Mn1 | 0.040760861 | -0.655351829 | 0.269 | 0.359 | 1 | 0.043582502 | -1.730270942 | 0.3 | 0.549 | 1 | 0.043582502 | 0.079860275 |
| 4930447C04Rik | 0.040798104 | -0.428843299 | 0.034 | 0.1 | 1 | 0.757881238 | 0.660369903 | 0.1 | 0.071 | 1 | 0.757881238 | 0.079931722 |
| Cramp1 | 0.040837009 | -1.020464103 | 0.336 | 0.4 | 1 | 0.407727774 | -0.31706234 | 0.5 | 0.636 | 1 | 0.407727774 | 0.080006357 |
| Gm15706 | 0.04085784 | 0.679632419 | 0.235 | 0.141 | 1 | 0.282175716 | -0.267480311 | 0 | 0.105 | 1 | 0.282175716 | 0.080046317 |
| Poglut2 | 0.040869237 | -0.619282574 | 0.471 | 0.547 | 1 | 0.40201083 | 0.836592143 | 0.5 | 0.485 | 1 | 0.40201083 | 0.080068179 |
| Stac3 | 0.334866848 | -0.944858446 | 0.025 | 0.047 | 1 | 0.040934602 | 2.210566986 | 0.2 | 0.049 | 1 | 0.334866848 | 0.080193561 |
| Jmjd7 | 0.04116568 | -1.733354341 | 0.025 | 0.082 | 1 | 0.040934708 | 2.280956314 | 0.2 | 0.049 | 1 | 0.04116568 | 0.080193765 |
| Lzts3 | 0.981839274 | -0.359895945 | 0.042 | 0.041 | 1 | 0.040934708 | 2.280956314 | 0.2 | 0.049 | 1 | 0.981839274 | 0.080193765 |
| Gpd1l | 0.040959147 | -2.87774425 | 0.008 | 0.053 | 1 | 0.21925705 | -0.625934282 | 0 | 0.133 | 1 | 0.21925705 | 0.080240642 |
| Tube1 | 0.040959147 | -2.655351829 | 0.008 | 0.053 | 1 | 0.08148995 | 2.432959407 | 0.1 | 0.019 | 1 | 0.08148995 | 0.080240642 |
| Prr3 | 0.040995297 | -0.77915803 | 0.235 | 0.329 | 1 | 0.554624316 | -0.419483404 | 0.2 | 0.269 | 1 | 0.554624316 | 0.080309979 |
| Cops8 | 0.041067263 | -0.524955191 | 0.588 | 0.641 | 1 | 0.148266494 | -0.716787712 | 0.5 | 0.664 | 1 | 0.148266494 | 0.080448006 |
| Sema4b | 0.041150164 | 0.557641895 | 0.168 | 0.088 | 1 | 0.353792202 | -0.152003093 | 0 | 0.08 | 1 | 0.353792202 | 0.080606992 |
| Fbxl22 | 0.041165831 | -1.655351829 | 0.025 | 0.082 | 1 | 0.334280429 | -0.339630097 | 0 | 0.086 | 1 | 0.334280429 | 0.080637037 |
| Tmem121 | 0.041165831 | -1.655351829 | 0.025 | 0.082 | 1 | 0.523811244 | 1.625604485 | 0.1 | 0.052 | 1 | 0.523811244 | 0.080637037 |
| Gata3 | 0.041179037 | -2.572889668 | 0.008 | 0.053 | 1 | 0.5978438 | -0.374395515 | 0 | 0.028 | 1 | 0.5978438 | 0.080662361 |
| Lpxn | 0.041179037 | -2.392317423 | 0.008 | 0.053 | 1 | 0.619967468 | 1.210566986 | 0 | 0.025 | 1 | 0.619967468 | 0.080662361 |
| Slc17a8 | 0.041179791 | -2.485426827 | 0.008 | 0.053 | 1 | 0.698534498 | 2.432959407 | 0 | 0.015 | 1 | 0.698534498 | 0.080663806 |
| Plek | 0.041179791 | -2.807354922 | 0.008 | 0.053 | 1 | 0.23891309 | 1.847996907 | 0.1 | 0.031 | 1 | 0.23891309 | 0.080663806 |
| Adcy5 | 0.041179791 | -2.572889668 | 0.008 | 0.053 | 1 | 0.444918734 | 1.494359952 | 0.1 | 0.046 | 1 | 0.444918734 | 0.080663806 |
| Neurl1b | 0.041179791 | -2.485426827 | 0.008 | 0.053 | 1 | 0.698534498 | 2.432959407 | 0 | 0.015 | 1 | 0.698534498 | 0.080663806 |
| M6pr | 0.202714815 | 0.119344155 | 0.756 | 0.682 | 1 | 0.041277325 | 0.618390995 | 1 | 0.664 | 1 | 0.202714815 | 0.080850833 |
| Elovl1 | 0.041282578 | 0.797645332 | 0.597 | 0.488 | 1 | 0.545348099 | 0.667424661 | 0.5 | 0.386 | 1 | 0.545348099 | 0.080860905 |
| Chp1 | 0.041302959 | 0.197788381 | 0.857 | 0.741 | 1 | 0.463824127 | 0.244869483 | 0.9 | 0.889 | 1 | 0.463824127 | 0.080899983 |
| Pcyt1a | 0.041315066 | -0.394086516 | 0.672 | 0.718 | 1 | 0.7230261 | -0.22135114 | 0.8 | 0.802 | 1 | 0.7230261 | 0.080923197 |
| Lrp8 | 0.127977244 | 0.247092862 | 0.328 | 0.229 | 1 | 0.041392205 | 0.689450967 | 0.7 | 0.33 | 1 | 0.127977244 | 0.081071096 |
| Lsm10 | 0.041393562 | -1.070389328 | 0.084 | 0.165 | 1 | 0.190567652 | -0.864721141 | 0 | 0.148 | 1 | 0.190567652 | 0.081073698 |
| Rhpn2 | 0.041399178 | -2.392317423 | 0.008 | 0.053 | 1 | 0.38542139 | 0.263034406 | 0 | 0.071 | 1 | 0.38542139 | 0.081084464 |
| Myl3 | 0.04139943 | -2.185866545 | 0.008 | 0.053 | 1 | 0.698532411 | 2.017921908 | 0 | 0.015 | 1 | 0.698532411 | 0.081084948 |
| Spink10 | 0.04139943 | -2.572889668 | 0.008 | 0.053 | 1 | 0.328341676 | 1.374065718 | 0.1 | 0.037 | 1 | 0.328341676 | 0.081084948 |
| Pdzrn3 | 0.043431896 | -0.557111795 | 0.966 | 0.994 | 1 | 0.041415934 | -1.097712938 | 0.9 | 0.972 | 1 | 0.043431896 | 0.081116588 |
| Cep295nl | 0.041529697 | -1.315501826 | 0.067 | 0.141 | 1 | 0.629776542 | 0.515421567 | 0.2 | 0.139 | 1 | 0.629776542 | 0.081334677 |
| Ppp1r26 | 0.041618048 | -2.070389328 | 0.008 | 0.053 | 1 | 0.539413885 | 1.210566986 | 0 | 0.037 | 1 | 0.539413885 | 0.081504034 |
| Gm12811 | 0.041618048 | -2.572889668 | 0.008 | 0.053 | 1 | 0.539464786 | 0.263034406 | 0 | 0.037 | 1 | 0.539464786 | 0.081504034 |
| Btbd19 | 0.041618048 | -2.070389328 | 0.008 | 0.053 | 1 | 0.151443299 | 2.695993813 | 0.1 | 0.025 | 1 | 0.151443299 | 0.081504034 |
| Cux2 | 0.041618048 | -2.070389328 | 0.008 | 0.053 | 1 | 0.276546558 | 2.111031312 | 0.1 | 0.034 | 1 | 0.276546558 | 0.081504034 |
| Rnf225 | 0.041618048 | -2.070389328 | 0.008 | 0.053 | 1 | 0.643842184 | 1.695993813 | 0 | 0.022 | 1 | 0.643842184 | 0.081504034 |
| Apoa1 | 0.041618048 | -2.185866545 | 0.008 | 0.053 | 1 | 0.529403453 | 1.494359952 | 0.1 | 0.052 | 1 | 0.529403453 | 0.081504034 |
| Gm30906 | 0.041618048 | -2.070389328 | 0.008 | 0.053 | 1 | 0.315888259 | -0.267480311 | 0 | 0.093 | 1 | 0.315888259 | 0.081504034 |
| Ptcra | 0.041618048 | -2.292781749 | 0.008 | 0.053 | 1 | 0.577171068 | 1.017921908 | 0 | 0.031 | 1 | 0.577171068 | 0.081504034 |
| Foxa2 | 0.041623505 | 1.099535674 | 0.084 | 0.029 | 1 | 0.698529281 | 2.017921908 | 0 | 0.015 | 1 | 0.698529281 | 0.081514494 |
| Prpf8 | 0.041666674 | -0.549557165 | 0.63 | 0.618 | 1 | 0.043017071 | -1.111361109 | 0.5 | 0.722 | 1 | 0.043017071 | 0.081597236 |
| AB124611 | 0.041666916 | -0.833350131 | 0.05 | 0.124 | 1 | 0.828556619 | 0.299103661 | 0.1 | 0.127 | 1 | 0.828556619 | 0.081597701 |
| Btbd6 | 0.04167477 | -1.572889668 | 0.025 | 0.082 | 1 | 0.971392392 | 0.930459067 | 0.1 | 0.096 | 1 | 0.971392392 | 0.081612754 |
| Gm13832 | 0.041732062 | -1.918386234 | 0.025 | 0.082 | 1 | 0.283473298 | 1.374065718 | 0.1 | 0.034 | 1 | 0.283473298 | 0.081722559 |
| Zfp93 | 0.569304225 | -0.485426827 | 0.034 | 0.047 | 1 | 0.041792265 | 2.14345279 | 0.2 | 0.049 | 1 | 0.569304225 | 0.081837936 |
| Zfp580 | 0.041820386 | -1.10433666 | 0.109 | 0.194 | 1 | 0.742467401 | 0.111031312 | 0.1 | 0.133 | 1 | 0.742467401 | 0.081891828 |
| Gm15832 | 0.041835626 | -2.070389328 | 0.008 | 0.053 | 1 | 0.73075158 | 2.695993813 | 0 | 0.012 | 1 | 0.73075158 | 0.081921033 |
| Spef1 | 0.041835626 | -1.944858446 | 0.008 | 0.053 | 1 | 0.396976646 | -0.191531458 | 0 | 0.068 | 1 | 0.396976646 | 0.081921033 |
| Gm56813 | 0.041835626 | -1.944858446 | 0.008 | 0.053 | 1 | 0.080320981 | 2.847996907 | 0.1 | 0.019 | 1 | 0.080320981 | 0.081921033 |

| | | | | | | | | | | | | |
|---|---|---|---|---|---|---|---|---|---|---|---|---|
| Ttc26 | 0.041835626 | -1.944858446 | 0.008 | 0.053 | 1 | 0.757880107 | 0.660369903 | 0.1 | 0.071 | 1 | 0.757880107 | 0.081921033 |
| Scrn1 | 0.041835626 | -1.944858446 | 0.008 | 0.053 | 1 | 0.290409015 | -0.441509711 | 0 | 0.102 | 1 | 0.290409015 | 0.081921033 |
| Setmar | 0.041835626 | -1.944858446 | 0.008 | 0.053 | 1 | 0.408673423 | 0.374065718 | 0 | 0.065 | 1 | 0.408673423 | 0.081921033 |
| E430018J23Rik | 0.041835626 | -1.944858446 | 0.008 | 0.053 | 1 | 0.505605027 | 0.695993813 | 0 | 0.043 | 1 | 0.505605027 | 0.081921033 |
| Sema3f | 0.041835626 | -1.944858446 | 0.008 | 0.053 | 1 | 0.59781419 | 1.31748219 | 0 | 0.028 | 1 | 0.59781419 | 0.081921033 |
| Aoc3 | 0.041835626 | -1.944858446 | 0.008 | 0.053 | 1 | 0.316137708 | 2.210566986 | 0.1 | 0.037 | 1 | 0.316137708 | 0.081921033 |
| 2310068J16Rik | 0.041835626 | -1.944858446 | 0.008 | 0.053 | 1 | 0.577184788 | 1.558490289 | 0 | 0.031 | 1 | 0.577184788 | 0.081921033 |
| E430024P14Rik | 0.041835626 | -1.944858446 | 0.008 | 0.053 | 1 | 0.577155236 | 1.31748219 | 0 | 0.031 | 1 | 0.577155236 | 0.081921033 |
| C330024C12Rik | 0.041835626 | -1.944858446 | 0.008 | 0.053 | 1 | 0.433577142 | 0.558490289 | 0 | 0.059 | 1 | 0.433577142 | 0.081921033 |
| Gm57200 | 0.041882119 | -1.517848305 | 0.067 | 0.141 | 1 | 0.148279524 | -1.736965594 | 0 | 0.176 | 1 | 0.148279524 | 0.082010127 |
| Cdc27 | 0.041958389 | -0.657193175 | 0.487 | 0.547 | 1 | 0.184903913 | -0.789433014 | 0.5 | 0.694 | 1 | 0.184903913 | 0.082156271 |
| Vps33a | 0.041964879 | -1.292781749 | 0.134 | 0.224 | 1 | 0.583420503 | 0.503348735 | 0.4 | 0.309 | 1 | 0.583420503 | 0.082168706 |
| Anapc1 | 0.041998468 | -0.54297785 | 0.454 | 0.559 | 1 | 0.368614727 | -0.383733379 | 0.6 | 0.682 | 1 | 0.368614727 | 0.082233064 |
| Ripk3 | 0.042062657 | -0.847996907 | 0.101 | 0.188 | 1 | 0.894290425 | 0.973527789 | 0.1 | 0.086 | 1 | 0.894290425 | 0.082356047 |
| Peak1 | 0.042159533 | -0.841912144 | 0.546 | 0.647 | 1 | 0.042064401 | -1.397115591 | 0.2 | 0.559 | 1 | 0.042159533 | 0.082359389 |
| Abcb6 | 0.042093929 | -1.807354922 | 0.008 | 0.053 | 1 | 0.526604355 | 1.432959407 | 0.1 | 0.052 | 1 | 0.526604355 | 0.082415959 |
| Gm43062 | 0.042093929 | -1.807354922 | 0.008 | 0.053 | 1 | 0.396883838 | 0.263034406 | 0 | 0.068 | 1 | 0.396883838 | 0.082415959 |
| Neil1 | 0.042093929 | -1.807354922 | 0.008 | 0.053 | 1 | 0.42087075 | 0.432959407 | 0 | 0.062 | 1 | 0.42087075 | 0.082415959 |
| Insyn1 | 0.042093929 | -1.807354922 | 0.008 | 0.053 | 1 | 0.397118555 | 0.494359952 | 0 | 0.068 | 1 | 0.397118555 | 0.082415959 |
| Rab36 | 0.042093929 | -1.807354922 | 0.008 | 0.053 | 1 | 0.078596675 | 1.795529487 | 0.2 | 0.059 | 1 | 0.078596675 | 0.082415959 |
| Zxdb | 0.042093929 | -1.807354922 | 0.008 | 0.053 | 1 | 0.231844057 | -0.473931188 | 0 | 0.127 | 1 | 0.231844057 | 0.082415959 |
| Hgs | 0.042127324 | -0.87774425 | 0.176 | 0.271 | 1 | 0.451176512 | -0.505640048 | 0.2 | 0.293 | 1 | 0.451176512 | 0.082479937 |
| Eid2 | 0.042182429 | -1 | 0.151 | 0.241 | 1 | 0.889803747 | 0.420986766 | 0.3 | 0.287 | 1 | 0.889803747 | 0.082585501 |
| B3galnt1 | 0.042186797 | -1.485426827 | 0.025 | 0.082 | 1 | 0.371170836 | 2.14345279 | 0.1 | 0.043 | 1 | 0.371170836 | 0.082593867 |
| BC051226 | 0.042186797 | -1.485426827 | 0.025 | 0.082 | 1 | 0.153152701 | 2.210566986 | 0.1 | 0.025 | 1 | 0.153152701 | 0.082593867 |
| Ywhaq | 0.04220909 | -0.231334775 | 0.966 | 0.982 | 1 | 0.322926221 | -0.342038201 | 1 | 0.994 | 1 | 0.322926221 | 0.082636573 |
| Cfap300 | 0.04224566 | -1.109917692 | 0.084 | 0.165 | 1 | 0.902406098 | 0.769994395 | 0.1 | 0.086 | 1 | 0.902406098 | 0.082706624 |
| Lamc2 | 0.046897301 | 1.514573173 | 0.118 | 0.053 | 1 | 0.042264656 | 0.710493383 | 0.2 | 0.049 | 1 | 0.046897301 | 0.082743011 |
| Atp6v1f | 0.042326739 | 0.277657828 | 0.891 | 0.876 | 1 | 0.680042467 | -0.152003093 | 0.6 | 0.87 | 1 | 0.680042467 | 0.082861925 |
| Cdca2 | 0.042332223 | -3.376197757 | 0.034 | 0.094 | 1 | 0.619961444 | 1.558490289 | 0 | 0.025 | 1 | 0.619961444 | 0.082872429 |
| Gemin5 | 0.042369965 | -0.936088236 | 0.21 | 0.3 | 1 | 0.498650876 | -0.797995028 | 0.2 | 0.272 | 1 | 0.498650876 | 0.082944717 |
| Fbxo46 | 0.993424287 | -0.106915204 | 0.101 | 0.1 | 1 | 0.042394713 | 1.625604485 | 0.3 | 0.096 | 1 | 0.993424287 | 0.082992115 |
| Itpkc | 0.042395433 | -0.517848305 | 0.202 | 0.3 | 1 | 0.518506703 | -0.419483404 | 0.2 | 0.275 | 1 | 0.518506703 | 0.082993493 |
| Lsm1 | 0.419806842 | -0.655351829 | 0.454 | 0.441 | 1 | 0.042441363 | -1.611434712 | 0.2 | 0.503 | 1 | 0.419806842 | 0.083081457 |
| ENSMUSG00000121395 | 0.042453463 | -1.392317423 | 0.092 | 0.171 | 1 | 0.428961999 | -0.521236903 | 0.1 | 0.194 | 1 | 0.428961999 | 0.083104629 |
| Utp11 | 0.042496413 | -0.555135799 | 0.58 | 0.612 | 1 | 0.696611105 | -0.171902651 | 0.5 | 0.586 | 1 | 0.696611105 | 0.083186881 |
| Ctps2 | 0.042505904 | -0.866516995 | 0.252 | 0.359 | 1 | 0.185869305 | -0.278010978 | 0.2 | 0.441 | 1 | 0.185869305 | 0.083205057 |
| Alg9 | 0.042506078 | -0.891419187 | 0.143 | 0.235 | 1 | 0.065770133 | 0.930459067 | 0.5 | 0.219 | 1 | 0.065770133 | 0.08320539 |
| Gm39150 | 0.042517012 | 0.764551426 | 0.227 | 0.135 | 1 | 0.954804466 | 2.190102883 | 0.1 | 0.117 | 1 | 0.954804466 | 0.083226328 |
| Stoml2 | 0.042551884 | -0.888391494 | 0.345 | 0.412 | 1 | 0.435181301 | -0.536666944 | 0.3 | 0.386 | 1 | 0.435181301 | 0.083293105 |
| Sugp1 | 0.042621874 | -1.15168943 | 0.168 | 0.253 | 1 | 0.888161724 | 0.780882711 | 0.2 | 0.262 | 1 | 0.888161724 | 0.083427123 |
| Gm47802 | 0.167062889 | 1.836501268 | 0.025 | 0.006 | 1 | 0.042664232 | 2.14345279 | 0.2 | 0.049 | 1 | 0.167062889 | 0.083508227 |
| Rps5 | 0.042685378 | -0.198332341 | 0.975 | 0.976 | 1 | 0.120788251 | -0.5436766 | 0.9 | 0.991 | 1 | 0.120788251 | 0.083548714 |
| Ahsa1 | 0.04272265 | -0.513995979 | 0.445 | 0.547 | 1 | 0.694933564 | -0.186649236 | 0.5 | 0.552 | 1 | 0.694933564 | 0.083620076 |
| Crtc3 | 0.042797933 | 0.277214901 | 0.748 | 0.624 | 1 | 0.104953959 | 0.792820199 | 0.9 | 0.821 | 1 | 0.104953959 | 0.083764203 |
| Yaf2 | 0.042861054 | -0.48033795 | 0.622 | 0.694 | 1 | 0.451204817 | -0.467044803 | 0.7 | 0.769 | 1 | 0.451204817 | 0.083885038 |
| Spsb4 | 0.042897032 | -1.048363022 | 0.126 | 0.218 | 1 | 0.475366939 | -0.840059087 | 0.2 | 0.278 | 1 | 0.475366939 | 0.083953908 |
| Zfp180 | 0.042944674 | -1.311397427 | 0.076 | 0.153 | 1 | 0.114192598 | -1.709998547 | 0 | 0.204 | 1 | 0.114192598 | 0.084045104 |
| Nsmce3 | 0.306063767 | -0.259150971 | 0.387 | 0.429 | 1 | 0.042949281 | -1.158666824 | 0.1 | 0.466 | 1 | 0.306063767 | 0.084053922 |
| Lsmem1 | 0.042963197 | -1.347043343 | 0.109 | 0.194 | 1 | 0.655836326 | 0.753218682 | 0.3 | 0.25 | 1 | 0.655836326 | 0.084080557 |
| Star | 0.042963995 | -1.707819249 | 0.025 | 0.082 | 1 | 0.333163571 | 1.31748219 | 0.2 | 0.102 | 1 | 0.333163571 | 0.084082086 |
| Il31ra | 0.042987287 | -1.900464326 | 0.034 | 0.094 | 1 | 0.052020228 | 2.079322453 | 0.2 | 0.052 | 1 | 0.052020228 | 0.084126666 |
| Ccna2 | 0.043012953 | -1.900464326 | 0.008 | 0.053 | 1 | 0.73075158 | 2.695993813 | 0 | 0.012 | 1 | 0.73075158 | 0.084175791 |
| 2010001A14Rik | 0.043078443 | 0.647839704 | 0.252 | 0.153 | 1 | 0.766018887 | 0.263034406 | 0.1 | 0.13 | 1 | 0.766018887 | 0.084301133 |
| Tnfaip8 | 0.043109402 | -0.245406988 | 0.529 | 0.676 | 1 | 0.342060726 | -0.647104982 | 0.6 | 0.781 | 1 | 0.342060726 | 0.084360384 |
| S100a10 | 0.173492898 | 0.125011949 | 0.983 | 0.965 | 1 | 0.04312567 | -0.976677492 | 0.9 | 0.978 | 1 | 0.173492898 | 0.084391516 |
| St6galnac4 | 0.043146772 | 0.654297936 | 0.345 | 0.235 | 1 | 0.937522418 | 0.210566986 | 0.2 | 0.204 | 1 | 0.937522418 | 0.084431899 |
| Gm53028 | 0.043187347 | 0.736965594 | 0.286 | 0.188 | 1 | 0.893644023 | 0.432959407 | 0.2 | 0.231 | 1 | 0.893644023 | 0.084509546 |
| Mrps2 | 0.043202069 | -1.117695043 | 0.076 | 0.153 | 1 | 0.843049539 | 0.591657153 | 0.1 | 0.12 | 1 | 0.843049539 | 0.084537718 |
| Gm57383 | 0.043236653 | 0.514573173 | 0.218 | 0.124 | 1 | 0.850829949 | 0.31748219 | 0.1 | 0.117 | 1 | 0.850829949 | 0.084603899 |
| Ubiad1 | 0.043281206 | -1.622930351 | 0.034 | 0.094 | 1 | 0.533015027 | 1.345496566 | 0.2 | 0.142 | 1 | 0.533015027 | 0.084689149 |
| Taf4b | 0.148371843 | 0.152777695 | 0.605 | 0.529 | 1 | 0.043337222 | 0.739261267 | 0.8 | 0.503 | 1 | 0.148371843 | 0.084796329 |
| Gnpnat1 | 0.043347714 | -0.743738823 | 0.345 | 0.435 | 1 | 0.655665477 | -0.294961047 | 0.3 | 0.343 | 1 | 0.655665477 | 0.084816404 |
| Lin9 | 0.043385877 | -1.67305383 | 0.059 | 0.129 | 1 | 0.196699274 | 1.111031312 | 0.2 | 0.077 | 1 | 0.196699274 | 0.084889421 |
| Acot2 | 0.043415729 | -1.722466024 | 0.05 | 0.118 | 1 | 0.64934624 | 1.795529487 | 0.1 | 0.068 | 1 | 0.64934624 | 0.084946532 |
| Aqp3 | 0.255522311 | 1.736965594 | 0.084 | 0.129 | 1 | 0.043421235 | -2.536666944 | 0 | 0.299 | 1 | 0.255522311 | 0.084957067 |
| Epb41 | 0.043453876 | -0.847996907 | 0.277 | 0.371 | 1 | 0.501130389 | -0.325485914 | 0.5 | 0.586 | 1 | 0.501130389 | 0.085019513 |
| Armcx1 | 0.043458495 | -1.017921908 | 0.185 | 0.276 | 1 | 0.427156679 | -0.606568957 | 0.2 | 0.299 | 1 | 0.427156679 | 0.08502835 |
| Panx1 | 0.043473985 | -1.600904045 | 0.008 | 0.053 | 1 | 0.643759378 | 1.210566986 | 0.1 | 0.062 | 1 | 0.643759378 | 0.085057983 |
| Tmprss9 | 0.060533348 | 1.514573173 | 0.059 | 0.018 | 1 | 0.043547356 | 1.902444691 | 0.2 | 0.049 | 1 | 0.060533348 | 0.08519834 |
| Cdkn2aipnl | 0.043587568 | -1.029747343 | 0.084 | 0.165 | 1 | 0.259564031 | -0.567040593 | 0 | 0.114 | 1 | 0.259564031 | 0.08527526 |
| Rrnad1 | 0.884112809 | -0.213347282 | 0.227 | 0.229 | 1 | 0.043589373 | 1.14345279 | 0.5 | 0.207 | 1 | 0.884112809 | 0.085278713 |
| Sf3b4 | 0.043629069 | -0.609162196 | 0.445 | 0.518 | 1 | 0.103636604 | -0.974544419 | 0.5 | 0.63 | 1 | 0.103636604 | 0.085354643 |
| Usp46 | 0.043702164 | -1.517848305 | 0.076 | 0.153 | 1 | 0.238961217 | -0.936274402 | 0 | 0.123 | 1 | 0.238961217 | 0.085494448 |
| Lins1 | 0.043732783 | -1.343407822 | 0.059 | 0.129 | 1 | 0.850830868 | 0.290001453 | 0.1 | 0.117 | 1 | 0.850830868 | 0.085553009 |
| Tyms | 0.043749279 | -1.070389328 | 0.067 | 0.141 | 1 | 0.191393409 | 2.808468542 | 0.2 | 0.093 | 1 | 0.191393409 | 0.085584558 |
| Btbd10 | 0.366496021 | -0.580063701 | 0.336 | 0.353 | 1 | 0.043784922 | -1.04385429 | 0.2 | 0.565 | 1 | 0.366496021 | 0.085652725 |
| Rabggtb | 0.043826133 | -0.563429339 | 0.672 | 0.7 | 1 | 0.053889319 | -0.944974097 | 0.4 | 0.713 | 1 | 0.053889319 | 0.085731537 |
| Coro7 | 0.043937428 | -0.944858446 | 0.118 | 0.206 | 1 | 0.889361412 | -0.125036046 | 0.2 | 0.207 | 1 | 0.889361412 | 0.085944358 |
| Pgap1 | 0.043942609 | -1.555816155 | 0.034 | 0.094 | 1 | 0.238791096 | -0.654503434 | 0 | 0.123 | 1 | 0.238791096 | 0.085954265 |
| Il17rc | 0.043942609 | -1.555816155 | 0.034 | 0.094 | 1 | 0.925158351 | 0.847996907 | 0.1 | 0.09 | 1 | 0.925158351 | 0.085954265 |
| Smc4 | 0.043970354 | -1.089964866 | 0.571 | 0.576 | 1 | 0.368309539 | 0.385653692 | 0.7 | 0.593 | 1 | 0.368309539 | 0.086007317 |
| Kctd15 | 0.043985251 | -0.891419187 | 0.151 | 0.241 | 1 | 0.541332786 | 0.639410285 | 0.3 | 0.219 | 1 | 0.541332786 | 0.086035801 |
| Abcb9 | 0.044007316 | 0.479807755 | 0.261 | 0.159 | 1 | 0.962205691 | -0.171902651 | 0.3 | 0.262 | 1 | 0.962205691 | 0.086077988 |
| Eif2b2 | 0.332708598 | -0.542010356 | 0.193 | 0.235 | 1 | 0.04402622 | -1.970762779 | 0 | 0.296 | 1 | 0.332708598 | 0.086114132 |
| Arrdc3 | 0.044030924 | -0.950549558 | 0.277 | 0.365 | 1 | 0.179232481 | -0.936274402 | 0.2 | 0.432 | 1 | 0.179232481 | 0.086123126 |
| Sh3gl1 | 0.044052346 | -0.364572432 | 0.756 | 0.735 | 1 | 0.296753428 | -0.528972552 | 0.7 | 0.728 | 1 | 0.296753428 | 0.086164082 |
| Sirt6 | 0.045284567 | -1.411426246 | 0.034 | 0.094 | 1 | 0.044084765 | 2.558490289 | 0.2 | 0.052 | 1 | 0.045284567 | 0.086226063 |
| Sfn | 0.044124556 | 1.798669194 | 0.328 | 0.253 | 1 | 0.371073395 | -1.026472211 | 0.2 | 0.318 | 1 | 0.371073395 | 0.086302136 |
| Psmc2 | 0.044227608 | -0.37540404 | 0.697 | 0.724 | 1 | 0.592333836 | -0.242605642 | 0.8 | 0.731 | 1 | 0.592333836 | 0.086499135 |
| Nipal3 | 0.044255234 | 1.251538767 | 0.109 | 0.047 | 1 | 0.561898712 | 1.558490289 | 0.1 | 0.056 | 1 | 0.561898712 | 0.086551942 |
| Prkdc | 0.044324756 | -0.997876828 | 0.176 | 0.271 | 1 | 0.440863066 | 0.939919396 | 0.3 | 0.225 | 1 | 0.440863066 | 0.086684829 |
| Zfp383 | 0.044331192 | -1.032914622 | 0.084 | 0.165 | 1 | 0.65486424 | 0.847996907 | 0.2 | 0.145 | 1 | 0.65486424 | 0.08669713 |
| Plec | 0.044367676 | -0.51657021 | 0.933 | 0.947 | 1 | 0.742897195 | -0.27240419 | 1 | 0.972 | 1 | 0.742897195 | 0.086766861 |
| Ifitm10 | 0.780855378 | -0.350884704 | 0.681 | 0.641 | 1 | 0.044385495 | 0.53467383 | 1 | 0.753 | 1 | 0.780855378 | 0.086800917 |
| Perp | 0.365545818 | 0.471504451 | 0.126 | 0.171 | 1 | 0.044388659 | -2.391469028 | 0 | 0.296 | 1 | 0.365545818 | 0.086806966 |
| Acp1 | 0.044408369 | -0.521496082 | 0.445 | 0.535 | 1 | 0.856756616 | 0.456266781 | 0.6 | 0.571 | 1 | 0.856756616 | 0.086844634 |
| Nup93 | 0.108002058 | 0.375257597 | 0.454 | 0.388 | 1 | 0.044484823 | 2.028569152 | 0.6 | 0.275 | 1 | 0.108002058 | 0.086990747 |

| | | | | | | | | | | | | |
|---|---|---|---|---|---|---|---|---|---|---|---|---|
| Foxo4 | 0.044518735 | -1.444784843 | 0.059 | 0.129 | 1 | 0.721144327 | 0.672147071 | 0.1 | 0.154 | 1 | 0.721144327 | 0.087055553 |
| Tchp | 0.044540983 | -0.787989597 | 0.101 | 0.188 | 1 | 0.447801907 | 1.14345279 | 0.2 | 0.117 | 1 | 0.447801907 | 0.087098068 |
| Tnnc1 | 0.87097915 | -0.485426827 | 0.059 | 0.053 | 1 | 0.044541052 | 4.017921908 | 0.1 | 0.015 | 1 | 0.87097915 | 0.087098199 |
| Pate2 | 0.344092959 | -1.292781749 | 0.017 | 0.035 | 1 | 0.044544293 | 4.339850003 | 0.1 | 0.015 | 1 | 0.344092959 | 0.087104393 |
| Zdhhc16 | 0.048632639 | -0.833350131 | 0.168 | 0.259 | 1 | 0.044564563 | 1.159940913 | 0.5 | 0.21 | 1 | 0.048632639 | 0.087143125 |
| Ptgis | 0.044567287 | 0.432340358 | 0.891 | 0.812 | 1 | 0.473530101 | -0.398848559 | 0.8 | 0.836 | 1 | 0.473530101 | 0.08714833 |
| Glp2r | 0.044599673 | 0.836501268 | 0.101 | 0.041 | 1 | 0.522102209 | 0.847996907 | 0 | 0.04 | 1 | 0.522102209 | 0.087210216 |
| Trim46 | 0.044610298 | -1.555816155 | 0.034 | 0.094 | 1 | 0.213736224 | 1.354956895 | 0.2 | 0.083 | 1 | 0.213736224 | 0.087230517 |
| Gm15879 | 0.044610435 | -1.485426827 | 0.034 | 0.094 | 1 | 0.719552815 | 0.135278859 | 0.1 | 0.136 | 1 | 0.719552815 | 0.087230778 |
| Terf2ip | 0.044668266 | -0.641546029 | 0.21 | 0.312 | 1 | 0.248940572 | -0.736965594 | 0.2 | 0.389 | 1 | 0.248940572 | 0.087341278 |
| Zc3hc1 | 0.04469058 | -1.315501826 | 0.067 | 0.141 | 1 | 0.143394791 | -1.152003093 | 0 | 0.179 | 1 | 0.143394791 | 0.087383913 |
| 1700109H08Rik | 0.35406236 | -0.549557165 | 0.244 | 0.276 | 1 | 0.044707986 | -2.141949429 | 0 | 0.296 | 1 | 0.35406236 | 0.087417168 |
| Tafa5 | 0.518426865 | -0.392886421 | 0.563 | 0.512 | 1 | 0.044788232 | -1.831148033 | 0.5 | 0.66 | 1 | 0.518426865 | 0.087570479 |
| Ankrd10 | 0.044793834 | -0.597901556 | 0.521 | 0.576 | 1 | 0.146127704 | -0.293952522 | 0.7 | 0.784 | 1 | 0.146127704 | 0.08758118 |
| Phkg2 | 0.044817678 | -0.731183242 | 0.218 | 0.324 | 1 | 0.952744867 | 0.20029865 | 0.3 | 0.312 | 1 | 0.952744867 | 0.087626731 |
| Krt15 | 0.477811255 | 1.573466862 | 0.084 | 0.112 | 1 | 0.044843842 | -2.230005605 | 0 | 0.296 | 1 | 0.477811255 | 0.087676713 |
| L3mbtl1 | 0.044884686 | -1.632268215 | 0.042 | 0.106 | 1 | 0.121288073 | 0.660369903 | 0.3 | 0.117 | 1 | 0.121288073 | 0.087754737 |
| Zbp1 | 0.044997168 | -1.359895945 | 0.025 | 0.082 | 1 | 0.274832025 | -0.91281543 | 0 | 0.108 | 1 | 0.274832025 | 0.087969591 |
| Esyt1 | 0.656935071 | -0.23388806 | 0.445 | 0.376 | 1 | 0.04501865 | 1.473601392 | 0.6 | 0.33 | 1 | 0.656935071 | 0.08801062 |
| Ints15 | 0.045035579 | -0.932885804 | 0.084 | 0.165 | 1 | 0.127963676 | -1.304006187 | 0 | 0.191 | 1 | 0.127963676 | 0.088042955 |
| Tbpl1 | 0.045065465 | -0.843880798 | 0.277 | 0.365 | 1 | 0.150998416 | -1.131825212 | 0.2 | 0.401 | 1 | 0.150998416 | 0.088100034 |
| Arhgef10l | 0.951715279 | -0.428843299 | 0.092 | 0.088 | 1 | 0.04520551 | 0.785261151 | 0.4 | 0.145 | 1 | 0.951715279 | 0.088367482 |
| Trak1 | 0.045262937 | -0.595850817 | 0.622 | 0.665 | 1 | 0.40365765 | 0.374065718 | 0.9 | 0.769 | 1 | 0.40365765 | 0.088477141 |
| Maml1 | 0.148792681 | -0.622930351 | 0.37 | 0.435 | 1 | 0.045278694 | -1.101019165 | 0.3 | 0.623 | 1 | 0.148792681 | 0.088507228 |
| Ptgr1 | 0.045284567 | -1.411426246 | 0.034 | 0.094 | 1 | 0.979200149 | 0.732519689 | 0.1 | 0.096 | 1 | 0.979200149 | 0.088518441 |
| Ptcd2 | 0.045284567 | -1.485426827 | 0.034 | 0.094 | 1 | 0.298466542 | -0.374395515 | 0 | 0.099 | 1 | 0.298466542 | 0.088518441 |
| Gm49207 | 0.045284567 | -1.411426246 | 0.034 | 0.094 | 1 | 0.764891428 | 1.263034406 | 0.1 | 0.074 | 1 | 0.764891428 | 0.088518441 |
| Pnpla1 | 0.511005543 | -0.485426827 | 0.008 | 0.018 | 1 | 0.045316348 | 3.795529487 | 0.1 | 0.015 | 1 | 0.511005543 | 0.088579125 |
| Prickle1 | 0.045328396 | -0.911562035 | 0.63 | 0.665 | 1 | 0.907521466 | -0.242605642 | 0.6 | 0.528 | 1 | 0.907521466 | 0.088602129 |
| Gm46440 | 0.606651028 | -1.14839184 | 0.067 | 0.082 | 1 | 0.045360377 | 1.695993813 | 0.2 | 0.049 | 1 | 0.606651028 | 0.088663191 |
| Sgip1 | 0.226772745 | -0.214124805 | 0.143 | 0.094 | 1 | 0.045362305 | 1.245332404 | 0.2 | 0.049 | 1 | 0.226772745 | 0.08866687 |
| 0610010K14Rik | 0.04542403 | 0.74237909 | 0.42 | 0.318 | 1 | 0.993433891 | 0.179978666 | 0.3 | 0.306 | 1 | 0.993433891 | 0.088784717 |
| Cox15 | 0.045426378 | -1.392317423 | 0.05 | 0.118 | 1 | 0.046411016 | 1.732519689 | 0.3 | 0.099 | 1 | 0.046411016 | 0.088789201 |
| Ercc8 | 0.045481745 | -1.485426827 | 0.084 | 0.159 | 1 | 0.691535808 | 0.135278859 | 0.1 | 0.142 | 1 | 0.691535808 | 0.0888949 |
| Arl6ip4 | 0.04552559 | -0.610957709 | 0.336 | 0.424 | 1 | 0.15094648 | -1.033740212 | 0.2 | 0.404 | 1 | 0.15094648 | 0.0889786 |
| Rnf182 | 0.045528463 | 0.777607579 | 0.143 | 0.071 | 1 | 0.364105256 | -0.505640048 | 0 | 0.077 | 1 | 0.364105256 | 0.088984085 |
| Nfatc3 | 0.045529228 | -0.519696729 | 0.513 | 0.606 | 1 | 0.15407853 | -0.811800827 | 0.4 | 0.62 | 1 | 0.15407853 | 0.088985546 |
| Alg6 | 0.045569397 | -0.822461814 | 0.143 | 0.235 | 1 | 0.500529503 | 1.017921908 | 0.2 | 0.123 | 1 | 0.500529503 | 0.089062223 |
| Ahcy | 0.045624948 | -1.544320516 | 0.042 | 0.106 | 1 | 0.363859515 | 0.210566986 | 0 | 0.077 | 1 | 0.363859515 | 0.089168261 |
| 1700029H14Rik | 0.045624948 | -1.655351829 | 0.042 | 0.106 | 1 | 0.638065809 | 1.210566986 | 0.1 | 0.062 | 1 | 0.638065809 | 0.089168261 |
| Mboat1 | 0.045673764 | 0.514573173 | 0.084 | 0.029 | 1 | 0.557778506 | 0.432959407 | 0 | 0.034 | 1 | 0.557778506 | 0.089261436 |
| Mgst3 | 0.045721972 | -1.240314329 | 0.059 | 0.129 | 1 | 0.086640029 | -1.736965594 | 0 | 0.231 | 1 | 0.086640029 | 0.089353445 |
| Rab3a | 0.868440058 | 0.407657969 | 0.059 | 0.065 | 1 | 0.045764855 | 2.902444691 | 0.2 | 0.056 | 1 | 0.868440058 | 0.089435288 |
| Ndufaf5 | 0.045765307 | -0.847996907 | 0.084 | 0.165 | 1 | 0.361992708 | 1.345496566 | 0.2 | 0.111 | 1 | 0.361992708 | 0.089436151 |
| Acbd4 | 0.045784381 | -1.269698136 | 0.067 | 0.141 | 1 | 0.968788479 | 0.930459067 | 0.1 | 0.096 | 1 | 0.968788479 | 0.089472553 |
| Rpap3 | 0.045794327 | -0.87244995 | 0.193 | 0.288 | 1 | 0.659900789 | -0.463204782 | 0.2 | 0.247 | 1 | 0.659900789 | 0.089491533 |
| 2300009A05Rik | 0.636297996 | -0.333423734 | 0.294 | 0.306 | 1 | 0.045835891 | -2.004445905 | 0 | 0.293 | 1 | 0.636297996 | 0.089570853 |
| Gtf2a1 | 0.243873542 | -0.58058406 | 0.353 | 0.394 | 1 | 0.045866388 | -1.959358016 | 0.1 | 0.41 | 1 | 0.243873542 | 0.089629051 |
| Sp2 | 0.045887639 | -0.714245518 | 0.235 | 0.335 | 1 | 0.131619453 | -1.28585884 | 0.1 | 0.318 | 1 | 0.131619453 | 0.089669604 |
| Arhgap31 | 0.926357436 | -0.168953162 | 0.807 | 0.788 | 1 | 0.045896931 | -1.325485914 | 0.9 | 0.852 | 1 | 0.926357436 | 0.089687334 |
| Bhlhb9 | 0.045931391 | -0.900464326 | 0.185 | 0.276 | 1 | 0.951584775 | 0.41305985 | 0.3 | 0.321 | 1 | 0.951584775 | 0.089753088 |
| 1110019D14Rik | 0.045932872 | 0.617666666 | 0.193 | 0.106 | 1 | 0.734029927 | -0.171902651 | 0.1 | 0.133 | 1 | 0.734029927 | 0.089755915 |
| Fem1a | 0.045964883 | -1.333423734 | 0.034 | 0.094 | 1 | 0.938823795 | 0.354956895 | 0.2 | 0.182 | 1 | 0.938823795 | 0.089816995 |
| Slc25a45 | 0.045975884 | 1.177538186 | 0.118 | 0.053 | 1 | 0.342811714 | 1.210566986 | 0.2 | 0.102 | 1 | 0.342811714 | 0.089837985 |
| Dlc1 | 0.046019194 | -0.337480195 | 0.874 | 0.906 | 1 | 0.129849116 | 0.647937672 | 0.9 | 0.867 | 1 | 0.129849116 | 0.089920623 |
| Vps50 | 0.046059637 | -0.854660637 | 0.168 | 0.265 | 1 | 0.066112702 | -2.248864633 | 0 | 0.259 | 1 | 0.066112702 | 0.089997785 |
| Tmem219 | 0.046059637 | -0.783107376 | 0.168 | 0.265 | 1 | 0.39557319 | 0.362570079 | 0.4 | 0.25 | 1 | 0.39557319 | 0.089997785 |
| Ctnna1 | 0.046091895 | -0.238102504 | 0.975 | 0.994 | 1 | 0.115872172 | -0.202929547 | 1 | 0.988 | 1 | 0.115872172 | 0.090059327 |
| Gnb3 | 0.370184089 | 1.099535674 | 0.017 | 0.006 | 1 | 0.046099652 | 3.280956314 | 0.1 | 0.015 | 1 | 0.370184089 | 0.090074126 |
| Tef | 0.046231435 | -0.72080489 | 0.487 | 0.524 | 1 | 0.552388664 | -0.18827848 | 0.6 | 0.636 | 1 | 0.552388664 | 0.090325524 |
| Vps28 | 0.046252559 | 0.316406867 | 0.739 | 0.635 | 1 | 0.185790686 | -0.673239997 | 0.4 | 0.59 | 1 | 0.185790686 | 0.090365819 |
| Rsph3a | 0.046323276 | -1.240314329 | 0.118 | 0.2 | 1 | 0.793537679 | -0.339630097 | 0.2 | 0.216 | 1 | 0.793537679 | 0.090500706 |
| Rpl12 | 0.390970681 | -0.11106552 | 0.966 | 0.982 | 1 | 0.04634968 | -0.617252039 | 1 | 0.991 | 1 | 0.390970681 | 0.090551067 |
| Rufy1 | 0.052114883 | -0.581642142 | 0.151 | 0.247 | 1 | 0.046353666 | -2.201246612 | 0 | 0.293 | 1 | 0.052114883 | 0.09055867 |
| Cdc34 | 0.492479781 | -0.175742328 | 0.277 | 0.318 | 1 | 0.046363895 | -1.581990934 | 0.1 | 0.41 | 1 | 0.492479781 | 0.09057818 |
| Atp6v1b2 | 0.14221439 | -0.337328188 | 0.412 | 0.488 | 1 | 0.046415525 | -1.224299487 | 0.3 | 0.694 | 1 | 0.14221439 | 0.090676648 |
| Mettl17 | 0.046432979 | -1.359895945 | 0.042 | 0.106 | 1 | 0.952551523 | 0.847996907 | 0.1 | 0.093 | 1 | 0.952551523 | 0.090709937 |
| Prpf18 | 0.046450655 | -0.713695815 | 0.244 | 0.341 | 1 | 0.587624441 | 1.055989949 | 0.4 | 0.349 | 1 | 0.587624441 | 0.090743648 |
| Jak1 | 0.520193857 | 0.149731132 | 0.975 | 0.982 | 1 | 0.046458307 | -0.531936792 | 1 | 0.981 | 1 | 0.520193857 | 0.090758239 |
| Cyth2 | 0.053378612 | -0.629816737 | 0.479 | 0.512 | 1 | 0.046468981 | -1.294961047 | 0.2 | 0.54 | 1 | 0.053378612 | 0.090778596 |
| Morn2 | 0.732033407 | 0.160936218 | 0.235 | 0.218 | 1 | 0.046471835 | 1.675529711 | 0.4 | 0.173 | 1 | 0.732033407 | 0.090784039 |
| Rundc1 | 0.04647547 | -1.070389328 | 0.076 | 0.153 | 1 | 0.131703199 | -1.339630097 | 0 | 0.188 | 1 | 0.131703199 | 0.090790971 |
| Fam104a | 0.046539415 | -0.302226595 | 0.916 | 0.9 | 1 | 0.465249264 | -0.246925092 | 0.9 | 0.932 | 1 | 0.465249264 | 0.090912914 |
| Smim1 | 0.046552793 | 1.251538767 | 0.176 | 0.1 | 1 | 0.610048752 | 0.821524695 | 0.2 | 0.139 | 1 | 0.610048752 | 0.090938423 |
| Lcn2 | 0.046555568 | 4.389042291 | 0.05 | 0.012 | 1 | 0.685720281 | -0.489872732 | 0.1 | 0.145 | 1 | 0.685720281 | 0.090943716 |
| Ripk1 | 0.046560895 | -0.494825525 | 0.681 | 0.706 | 1 | 0.575693385 | -0.16841085 | 0.7 | 0.759 | 1 | 0.575693385 | 0.090953873 |
| Wfdc18 | 0.046616501 | -0.707819249 | 0.252 | 0.353 | 1 | 0.994123681 | -0.178475305 | 0.2 | 0.185 | 1 | 0.994123681 | 0.091059904 |
| Mob4 | 0.222318405 | -0.279842694 | 0.647 | 0.671 | 1 | 0.046688994 | -0.884590247 | 0.8 | 0.867 | 1 | 0.222318405 | 0.091198126 |
| Gm20300 | 0.046721892 | 1.04508789 | 0.101 | 0.041 | 1 | 0.812913407 | 0.403212064 | 0.1 | 0.123 | 1 | 0.812913407 | 0.09126085 |
| Pwp2 | 0.046741394 | -1.240314329 | 0.059 | 0.129 | 1 | 0.455028957 | 1.017921908 | 0.2 | 0.117 | 1 | 0.455028957 | 0.09129803 |
| Bax | 0.046749532 | -0.40240624 | 0.655 | 0.671 | 1 | 0.370992519 | -0.241350579 | 0.4 | 0.648 | 1 | 0.370992519 | 0.091313546 |
| Grk4 | 0.046819687 | -0.914270126 | 0.16 | 0.253 | 1 | 0.984096979 | 0.913585248 | 0.2 | 0.244 | 1 | 0.984096979 | 0.091447291 |
| Taf1c | 0.046839002 | -1.250961574 | 0.034 | 0.094 | 1 | 0.433577142 | 0.558490289 | 0 | 0.059 | 1 | 0.433577142 | 0.091484112 |
| Ticam1 | 0.046839002 | -1.250961574 | 0.034 | 0.094 | 1 | 0.752600125 | 0.795529487 | 0.2 | 0.16 | 1 | 0.752600125 | 0.091484112 |
| 6030458C11Rik | 0.046844295 | -1.254813899 | 0.168 | 0.253 | 1 | 0.443724895 | -0.304006187 | 0.1 | 0.194 | 1 | 0.443724895 | 0.091494202 |
| Gm14164 | 0.046853575 | 0.407657969 | 0.437 | 0.318 | 1 | 0.57313124 | -0.276698841 | 0.3 | 0.361 | 1 | 0.57313124 | 0.091511893 |
| Rnf138 | 0.046855561 | -0.6617496 | 0.345 | 0.435 | 1 | 0.583104539 | 0.231325546 | 0.6 | 0.475 | 1 | 0.583104539 | 0.091515678 |
| Pgp | 0.046884983 | 0.389042291 | 0.496 | 0.382 | 1 | 0.086663526 | 1.117917698 | 0.6 | 0.404 | 1 | 0.086663526 | 0.091571765 |
| Ggcx | 0.09773236 | 0.12225575 | 0.639 | 0.506 | 1 | 0.046889844 | 0.871765072 | 0.9 | 0.534 | 1 | 0.09773236 | 0.091581031 |
| Ociad2 | 0.046897263 | 0.836501268 | 0.092 | 0.035 | 1 | 0.632378953 | 1.494359952 | 0.1 | 0.062 | 1 | 0.632378953 | 0.091595172 |
| Vipas39 | 0.04695138 | -0.758445322 | 0.176 | 0.271 | 1 | 0.75338717 | 0.483943336 | 0.3 | 0.275 | 1 | 0.75338717 | 0.091698328 |
| Gm28809 | 0.047018669 | -1.321928095 | 0.042 | 0.106 | 1 | 0.238992918 | -0.789433014 | 0 | 0.123 | 1 | 0.238992918 | 0.091826582 |
| Ddost | 0.047019499 | -0.282497147 | 0.958 | 0.912 | 1 | 0.565880471 | -0.15815532 | 0.8 | 0.907 | 1 | 0.565880471 | 0.091828164 |
| Cyb5d1 | 0.047038131 | -1.263034406 | 0.05 | 0.118 | 1 | 0.248571837 | 1.354956895 | 0.2 | 0.09 | 1 | 0.248571837 | 0.091863677 |
| Ecd | 0.04704598 | -0.936088236 | 0.193 | 0.288 | 1 | 0.062186853 | 1.918386234 | 0.5 | 0.259 | 1 | 0.062186853 | 0.091878635 |
| Nxph4 | 0.047103235 | -1.359895945 | 0.034 | 0.094 | 1 | 0.407989827 | 0.874963954 | 0.2 | 0.108 | 1 | 0.407989827 | 0.091987755 |
| Hsf1 | 0.047165282 | -0.735040717 | 0.319 | 0.406 | 1 | 0.099000053 | -0.989572629 | 0.1 | 0.386 | 1 | 0.099000053 | 0.092105999 |
| Snrpa1 | 0.047165721 | -0.552990111 | 0.437 | 0.535 | 1 | 0.075850412 | -1.080110175 | 0.2 | 0.5 | 1 | 0.075850412 | 0.092106836 |

| | | | | | | | | | | | | |
|---|---|---|---|---|---|---|---|---|---|---|---|---|
| Rnpepl1 | 0.047168848 | -0.987927168 | 0.134 | 0.224 | 1 | 0.395355217 | -0.339630097 | 0.1 | 0.207 | 1 | 0.395355217 | 0.092112796 |
| Zfp111 | 0.047250572 | -1.292781749 | 0.042 | 0.106 | 1 | 0.282342718 | -0.408342847 | 0 | 0.105 | 1 | 0.282342718 | 0.092268528 |
| Eri2 | 0.047293397 | -1.529820947 | 0.05 | 0.118 | 1 | 0.123163238 | 1.558490289 | 0.2 | 0.068 | 1 | 0.123163238 | 0.092350128 |
| Prps1 | 0.047309671 | -0.655351829 | 0.252 | 0.353 | 1 | 0.664074918 | -0.248864633 | 0.3 | 0.343 | 1 | 0.664074918 | 0.092381137 |
| P4htm | 0.047386325 | -1 | 0.134 | 0.224 | 1 | 0.160477286 | 1.732519689 | 0.3 | 0.167 | 1 | 0.160477286 | 0.092527186 |
| Daxx | 0.069158326 | -0.644625422 | 0.193 | 0.288 | 1 | 0.04741796 | -2.004445905 | 0 | 0.29 | 1 | 0.069158326 | 0.092587457 |
| Zfp287 | 0.0475103 | -1.432959407 | 0.042 | 0.106 | 1 | 0.815998758 | 0.660369903 | 0.1 | 0.077 | 1 | 0.815998758 | 0.092763371 |
| Plekhg3 | 0.071867809 | -1.020203571 | 0.202 | 0.282 | 1 | 0.047578941 | -1.922244842 | 0.4 | 0.605 | 1 | 0.071867809 | 0.092894126 |
| Ifit3 | 0.047612066 | 2.698997744 | 0.202 | 0.124 | 1 | 0.252969517 | -2.408342847 | 0 | 0.117 | 1 | 0.252969517 | 0.092957224 |
| Shc4 | 0.047628132 | 0.292180751 | 0.361 | 0.247 | 1 | 0.18711205 | 0.832055363 | 0.4 | 0.207 | 1 | 0.18711205 | 0.092987826 |
| Eif1ax | 0.047630252 | -0.30256277 | 0.84 | 0.888 | 1 | 0.18923483 | -0.377255169 | 0.7 | 0.92 | 1 | 0.18923483 | 0.092991863 |
| Klf12 | 0.047642633 | -0.546343932 | 0.874 | 0.929 | 1 | 0.185664896 | 0.426341529 | 0.9 | 0.978 | 1 | 0.185664896 | 0.093015445 |
| Wfdc1 | 0.047680009 | -1.378511623 | 0.042 | 0.106 | 1 | 0.485511734 | -0.441509711 | 0.1 | 0.182 | 1 | 0.485511734 | 0.093086635 |
| Slc6a20a | 0.33261065 | -1.292781749 | 0.008 | 0.024 | 1 | 0.047702188 | 3.280956314 | 0.1 | 0.015 | 1 | 0.33261065 | 0.093128878 |
| Bcl9 | 0.047732473 | -1.311397427 | 0.143 | 0.229 | 1 | 0.462144925 | -0.521236903 | 0.2 | 0.312 | 1 | 0.462144925 | 0.093186558 |
| Fam168b | 0.077083078 | -0.42343994 | 0.58 | 0.612 | 1 | 0.047740367 | -0.693572999 | 0.5 | 0.759 | 1 | 0.077083078 | 0.093201592 |
| Clcn6 | 0.047792583 | -0.622930351 | 0.227 | 0.324 | 1 | 0.61989043 | 0.625604485 | 0.2 | 0.33 | 1 | 0.61989043 | 0.093301035 |
| Rras | 0.047799241 | 0.284143432 | 0.731 | 0.6 | 1 | 0.581924841 | 0.249737583 | 0.6 | 0.531 | 1 | 0.581924841 | 0.093313715 |
| Tbc1d32 | 0.441689596 | -0.150242636 | 0.185 | 0.218 | 1 | 0.047801686 | 1.193493473 | 0.4 | 0.154 | 1 | 0.441689596 | 0.09331837 |
| Mrpl2 | 0.047824424 | -0.602996424 | 0.328 | 0.429 | 1 | 0.780202336 | -0.191531458 | 0.3 | 0.315 | 1 | 0.780202336 | 0.093361673 |
| Slc24a5 | 0.047864325 | 0.893084796 | 0.092 | 0.035 | 1 | 0.168719809 | 1.744903414 | 0.2 | 0.077 | 1 | 0.168719809 | 0.093437656 |
| Tmem67 | 0.047976084 | -0.826463745 | 0.118 | 0.206 | 1 | 0.201730639 | -0.814968106 | 0 | 0.142 | 1 | 0.201730639 | 0.093650464 |
| Zfp959 | 0.04797619 | -1.201633861 | 0.05 | 0.118 | 1 | 0.245373301 | -0.473931188 | 0 | 0.12 | 1 | 0.245373301 | 0.093650666 |
| Prkag1 | 0.048027093 | -0.512699757 | 0.664 | 0.676 | 1 | 0.853075461 | -0.120842162 | 0.8 | 0.66 | 1 | 0.853075461 | 0.093747585 |
| Ndufc1 | 0.048052761 | 0.210239138 | 0.79 | 0.753 | 1 | 0.666410894 | 0.406897111 | 0.4 | 0.673 | 1 | 0.666410894 | 0.093796454 |
| Ampd2 | 0.048077111 | -0.684735635 | 0.176 | 0.276 | 1 | 0.320467377 | 0.494359952 | 0.5 | 0.318 | 1 | 0.320467377 | 0.093842813 |
| Atad1 | 0.048129068 | -0.559851668 | 0.521 | 0.571 | 1 | 0.14450977 | -0.459611517 | 0.4 | 0.62 | 1 | 0.14450977 | 0.093941728 |
| Fbxw17 | 0.048179012 | -1.14839184 | 0.034 | 0.094 | 1 | 0.982938702 | 1.017921908 | 0.1 | 0.111 | 1 | 0.982938702 | 0.094036807 |
| Zfp955b | 0.048337433 | -0.748461233 | 0.21 | 0.306 | 1 | 0.597539426 | 0.708066645 | 0.3 | 0.244 | 1 | 0.597539426 | 0.094338358 |
| Dhrs4 | 0.048374609 | -0.730539325 | 0.202 | 0.3 | 1 | 0.318044366 | 0.684021171 | 0.4 | 0.253 | 1 | 0.318044366 | 0.094409116 |
| Extl1 | 0.126414988 | 0.681223042 | 0.319 | 0.241 | 1 | 0.048381627 | 1.210566986 | 0.5 | 0.222 | 1 | 0.126414988 | 0.094422473 |
| Rigi | 0.284874354 | -0.587306441 | 0.193 | 0.241 | 1 | 0.048399204 | -2.782977992 | 0 | 0.29 | 1 | 0.284874354 | 0.094455925 |
| Tbcd | 0.048411309 | -1.190683562 | 0.16 | 0.247 | 1 | 0.294779005 | -0.959358016 | 0.1 | 0.238 | 1 | 0.294779005 | 0.094478963 |
| Cybc1 | 0.048432389 | -0.790281409 | 0.118 | 0.206 | 1 | 0.190727344 | -0.91281543 | 0 | 0.148 | 1 | 0.190727344 | 0.094519081 |
| Tada3 | 0.048475651 | -0.819845866 | 0.16 | 0.253 | 1 | 1 | 0.558490289 | 0.2 | 0.21 | 1 | 1 | 0.094601412 |
| A930015D03Rik | 0.500134451 | -0.263034406 | 0.504 | 0.494 | 1 | 0.048490404 | 0.501346382 | 1 | 0.568 | 1 | 0.500134451 | 0.094629489 |
| Laptm4b | 0.808302292 | -0.685959815 | 0.345 | 0.318 | 1 | 0.048493177 | -1.750262417 | 0.1 | 0.435 | 1 | 0.808302292 | 0.094634767 |
| Mecp2 | 0.04849455 | -0.590779827 | 0.723 | 0.729 | 1 | 0.882695169 | -0.185018151 | 0.9 | 0.84 | 1 | 0.882695169 | 0.094637378 |
| C530020B09Rik | 0.06000657 | -2.292781749 | 0 | 0.029 | 1 | 0.048521704 | 3.432959407 | 0.1 | 0.015 | 1 | 0.06000657 | 0.094689052 |
| BB218582 | 0.094752821 | -1.807354922 | 0.008 | 0.041 | 1 | 0.048521704 | 3.432959407 | 0.1 | 0.015 | 1 | 0.094752821 | 0.094689052 |
| Gm26826 | 0.147029286 | -1.807354922 | 0 | 0.018 | 1 | 0.048521704 | 3.432959407 | 0.1 | 0.015 | 1 | 0.147029286 | 0.094689052 |
| Gm3604 | 0.147030916 | -1.485426827 | 0 | 0.018 | 1 | 0.048521704 | 3.432959407 | 0.1 | 0.015 | 1 | 0.147030916 | 0.094689052 |
| Il31 | 0.237885589 | -1.070389328 | 0 | 0.012 | 1 | 0.048521704 | 3.432959407 | 0.1 | 0.015 | 1 | 0.237885589 | 0.094689052 |
| Gm26559 | 0.237885589 | -1.070389328 | 0 | 0.012 | 1 | 0.048521704 | 3.432959407 | 0.1 | 0.015 | 1 | 0.237885589 | 0.094689052 |
| 9930004E17Rik | 0.33261065 | -1.070389328 | 0.008 | 0.024 | 1 | 0.048521704 | 3.432959407 | 0.1 | 0.015 | 1 | 0.33261065 | 0.094689052 |
| Gm49893 | 0.33261065 | -1.292781749 | 0.008 | 0.024 | 1 | 0.048521704 | 3.432959407 | 0.1 | 0.015 | 1 | 0.33261065 | 0.094689052 |
| Gm42531 | 0.390871516 | 1.514573173 | 0.025 | 0.012 | 1 | 0.048521704 | 3.432959407 | 0.1 | 0.015 | 1 | 0.390871516 | 0.094689052 |
| Sult2b1 | 0.390871516 | 1 | 0.025 | 0.012 | 1 | 0.048521704 | 3.432959407 | 0.1 | 0.015 | 1 | 0.390871516 | 0.094689052 |
| ENSMUSG00000121346 | 0.508731273 | -0.807354922 | 0.008 | 0.018 | 1 | 0.048521704 | 3.432959407 | 0.1 | 0.015 | 1 | 0.508731273 | 0.094689052 |
| Gm50340 | 0.721162042 | 0.514573173 | 0.017 | 0.012 | 1 | 0.048521704 | 3.432959407 | 0.1 | 0.015 | 1 | 0.721162042 | 0.094689052 |
| Catsperd | 0.048524435 | -1.263034406 | 0.042 | 0.106 | 1 | 0.433721269 | -0.191531458 | 0 | 0.059 | 1 | 0.433721269 | 0.094694249 |
| Ppfibp2 | 0.048524763 | -1.934734229 | 0.076 | 0.147 | 1 | 0.245855205 | -1.425021588 | 0 | 0.12 | 1 | 0.245855205 | 0.094694873 |
| B4galt4 | 0.048540056 | 0.404948682 | 0.303 | 0.194 | 1 | 0.955920248 | 0.223506042 | 0.3 | 0.278 | 1 | 0.955920248 | 0.094723974 |
| Gtf3a | 0.048638753 | -1.03562391 | 0.101 | 0.182 | 1 | 0.478719965 | -0.191531458 | 0.1 | 0.185 | 1 | 0.478719965 | 0.094911778 |
| Zc2hc1a | 0.048694155 | -0.394661363 | 0.202 | 0.312 | 1 | 0.357059944 | 0.466126271 | 0.5 | 0.37 | 1 | 0.357059944 | 0.09501719 |
| Gm11655 | 0.048729099 | 0.80407979 | 0.151 | 0.076 | 1 | 0.062250235 | 2.079322453 | 0.2 | 0.056 | 1 | 0.062250235 | 0.095083674 |
| Copz2 | 0.04874237 | -0.324572816 | 0.866 | 0.824 | 1 | 0.421078047 | -0.206615785 | 0.6 | 0.707 | 1 | 0.421078047 | 0.095108921 |
| Nutf2-ps1 | 0.048767346 | -0.740683882 | 0.244 | 0.335 | 1 | 0.097087847 | -1.304006187 | 0 | 0.219 | 1 | 0.097087847 | 0.095156438 |
| Mettl6 | 0.048790066 | -0.591221491 | 0.277 | 0.382 | 1 | 0.123224287 | -1.236319379 | 0.2 | 0.417 | 1 | 0.123224287 | 0.095199661 |
| Cyb5a | 0.048796362 | 0.63942466 | 0.664 | 0.588 | 1 | 0.067094368 | -0.750262417 | 0.3 | 0.685 | 1 | 0.067094368 | 0.09521164 |
| Best3 | 0.048798367 | 1.929610672 | 0.05 | 0.012 | 1 | 0.199151937 | 1.930459067 | 0.1 | 0.028 | 1 | 0.199151937 | 0.095215454 |
| D5Ertd579e | 0.167346041 | 0.152752351 | 0.756 | 0.665 | 1 | 0.048953163 | -1.116971077 | 0.6 | 0.778 | 1 | 0.167346041 | 0.095509914 |
| Tmem181a | 0.049017937 | -0.718916957 | 0.235 | 0.335 | 1 | 0.913880307 | 0.101445264 | 0.3 | 0.275 | 1 | 0.913880307 | 0.095633115 |
| Desi2 | 0.049053451 | -0.458233864 | 0.866 | 0.912 | 1 | 0.139465786 | -0.562148183 | 0.9 | 0.972 | 1 | 0.139465786 | 0.095700661 |
| Hycc1 | 0.196721134 | -0.364120531 | 0.277 | 0.341 | 1 | 0.049081927 | -1.930445324 | 0.1 | 0.404 | 1 | 0.196721134 | 0.095754818 |
| Ghitm | 0.08299989 | -0.189804495 | 0.798 | 0.841 | 1 | 0.049307616 | -0.650456601 | 0.6 | 0.852 | 1 | 0.08299989 | 0.096183991 |
| Grcc10 | 0.049314414 | -0.344197259 | 0.857 | 0.835 | 1 | 0.595600404 | -0.205228032 | 0.8 | 0.833 | 1 | 0.595600404 | 0.096196917 |
| Pdgfc | 0.049338148 | -0.131789873 | 0.311 | 0.194 | 1 | 0.994117187 | -0.555725279 | 0.5 | 0.435 | 1 | 0.994117187 | 0.096242044 |
| Capn3 | 0.049397229 | 1.736965594 | 0.05 | 0.012 | 1 | 0.049344356 | 3.210566986 | 0.1 | 0.015 | 1 | 0.049397229 | 0.096253846 |
| Timeless | 0.062759149 | -2.070389328 | 0.008 | 0.047 | 1 | 0.049344356 | 3.210566986 | 0.1 | 0.015 | 1 | 0.062759149 | 0.096253846 |
| Zfp658 | 0.063085444 | -1.807354922 | 0.008 | 0.047 | 1 | 0.049344356 | 3.210566986 | 0.1 | 0.015 | 1 | 0.063085444 | 0.096253846 |
| Kntc1 | 0.084054882 | -0.970853654 | 0.017 | 0.059 | 1 | 0.049344356 | 3.210566986 | 0.1 | 0.015 | 1 | 0.084054882 | 0.096253846 |
| Dpysl5 | 0.093244798 | -2.292781749 | 0 | 0.024 | 1 | 0.049344356 | 3.017921908 | 0.1 | 0.015 | 1 | 0.093244798 | 0.096253846 |
| Mlip | 0.147029286 | -2.655351829 | 0 | 0.018 | 1 | 0.049344356 | 3.210566986 | 0.1 | 0.015 | 1 | 0.147029286 | 0.096253846 |
| Tbc1d10c | 0.147029286 | -1.807354922 | 0 | 0.018 | 1 | 0.049344356 | 3.210566986 | 0.1 | 0.015 | 1 | 0.147029286 | 0.096253846 |
| Gm16189 | 0.167062889 | 2.099535674 | 0.025 | 0.006 | 1 | 0.049344356 | 3.017921908 | 0.1 | 0.015 | 1 | 0.167062889 | 0.096253846 |
| Faah | 0.168207387 | -1.807354922 | 0.025 | 0.059 | 1 | 0.049344356 | 3.210566986 | 0.1 | 0.015 | 1 | 0.168207387 | 0.096253846 |
| Nudt12 | 0.201658121 | 1.251538767 | 0.034 | 0.012 | 1 | 0.049344356 | 3.210566986 | 0.1 | 0.015 | 1 | 0.201658121 | 0.096253846 |
| 4930481B07Rik | 0.218485013 | -1.292781749 | 0.008 | 0.029 | 1 | 0.049344356 | 3.017921908 | 0.1 | 0.015 | 1 | 0.218485013 | 0.096253846 |
| Nuak2 | 0.237885589 | -1.485426827 | 0 | 0.012 | 1 | 0.049344356 | 3.210566986 | 0.1 | 0.015 | 1 | 0.237885589 | 0.096253846 |
| Cd101 | 0.33261065 | -1.292781749 | 0.008 | 0.024 | 1 | 0.049344356 | 3.017921908 | 0.1 | 0.015 | 1 | 0.33261065 | 0.096253846 |
| Gm13421 | 0.511005543 | -0.485426827 | 0.008 | 0.018 | 1 | 0.049344356 | 2.695993813 | 0.1 | 0.015 | 1 | 0.511005543 | 0.096253846 |
| 1700028J19Rik | 0.511005543 | -0.485426827 | 0.008 | 0.018 | 1 | 0.049344356 | 3.017921908 | 0.1 | 0.015 | 1 | 0.511005543 | 0.096253846 |
| 4930595D18Rik | 0.659865009 | 0.514573173 | 0.025 | 0.018 | 1 | 0.049344356 | 3.210566986 | 0.1 | 0.015 | 1 | 0.659865009 | 0.096253846 |
| Cenpn | 0.73484547 | -0.359895945 | 0.034 | 0.041 | 1 | 0.049344356 | 3.017921908 | 0.1 | 0.015 | 1 | 0.73484547 | 0.096253846 |
| Gm57069 | 0.781790612 | -0.485426827 | 0.008 | 0.012 | 1 | 0.049344356 | 3.017921908 | 0.1 | 0.015 | 1 | 0.781790612 | 0.096253846 |
| Hk3 | 0.920852644 | 0.514573173 | 0.025 | 0.024 | 1 | 0.049344356 | 3.017921908 | 0.1 | 0.015 | 1 | 0.920852644 | 0.096253846 |
| Rhd | 0.962120409 | 0.192645078 | 0.017 | 0.018 | 1 | 0.049344356 | 2.695993813 | 0.1 | 0.015 | 1 | 0.962120409 | 0.096253846 |
| Zfp212 | 0.267845185 | -0.592342031 | 0.185 | 0.229 | 1 | 0.049348961 | -2.161987182 | 0 | 0.287 | 1 | 0.267845185 | 0.096262602 |
| Ankrd9 | 0.049354807 | 0.929610672 | 0.118 | 0.053 | 1 | 0.632378953 | 1.432959407 | 0.1 | 0.062 | 1 | 0.632378953 | 0.096273716 |
| Plekho1 | 0.049387847 | -1.100136671 | 0.101 | 0.182 | 1 | 0.889715589 | 0.393431043 | 0.1 | 0.12 | 1 | 0.889715589 | 0.096336534 |
| Prxl2c | 0.265980032 | -0.319416876 | 0.454 | 0.482 | 1 | 0.049398082 | -1.11821478 | 0.5 | 0.679 | 1 | 0.265980032 | 0.096355994 |
| Septin2 | 0.04940432 | -1.109917692 | 0.092 | 0.171 | 1 | 0.451476908 | -0.374395515 | 0.1 | 0.191 | 1 | 0.451476908 | 0.096367853 |
| Plcb3 | 0.049405895 | -1.070389328 | 0.092 | 0.171 | 1 | 0.128118413 | -1.457811523 | 0 | 0.191 | 1 | 0.128118413 | 0.096370848 |
| Cavin3 | 0.15474285 | -0.557277112 | 0.639 | 0.665 | 1 | 0.049484734 | -1.220482831 | 0.3 | 0.623 | 1 | 0.15474285 | 0.096520729 |
| Ncapg2 | 0.126769005 | -1.272023189 | 0.084 | 0.141 | 1 | 0.049533052 | 2.14345279 | 0.3 | 0.108 | 1 | 0.126769005 | 0.096612582 |
| Ptpn11 | 0.049616565 | -0.341935234 | 0.748 | 0.771 | 1 | 0.099673291 | -0.627993804 | 0.6 | 0.87 | 1 | 0.099673291 | 0.096771327 |
| Gclc | 0.049669943 | -0.294242706 | 0.387 | 0.529 | 1 | 0.107215016 | -0.432539557 | 0.3 | 0.574 | 1 | 0.107215016 | 0.096872783 |

| | | | | | | | | | | | | |
|---|---|---|---|---|---|---|---|---|---|---|---|---|
| Gid8 | 0.04970294 | -0.568568062 | 0.37 | 0.453 | 1 | 0.590669629 | 0.374065718 | 0.5 | 0.417 | 1 | 0.590669629 | 0.096935498 |
| Zfp410 | 0.049724728 | -0.724892762 | 0.244 | 0.335 | 1 | 0.08164668 | -1.362899876 | 0.2 | 0.454 | 1 | 0.08164668 | 0.096976908 |
| BC065397 | 0.049816865 | -1.122856748 | 0.042 | 0.106 | 1 | 0.292791045 | 0.821524695 | 0.2 | 0.093 | 1 | 0.292791045 | 0.097152009 |
| Oxsr1 | 0.049853087 | -0.762267033 | 0.176 | 0.271 | 1 | 0.360255924 | -0.772155023 | 0.2 | 0.315 | 1 | 0.360255924 | 0.097220843 |
| Slc35b3 | 0.250338676 | 0.131244533 | 0.445 | 0.359 | 1 | 0.049867958 | 0.839118755 | 0.7 | 0.349 | 1 | 0.250338676 | 0.097249102 |
| Elapor1 | 0.049875378 | 0.514573173 | 0.109 | 0.047 | 1 | 0.408705848 | 0.159940913 | 0 | 0.065 | 1 | 0.408705848 | 0.097263203 |
| Lancl2 | 0.194108815 | -0.527247003 | 0.126 | 0.182 | 1 | 0.049933818 | 0.944672926 | 0.5 | 0.21 | 1 | 0.194108815 | 0.097374249 |
| Plekhm1 | 0.075743668 | 0.568012432 | 0.277 | 0.188 | 1 | 0.049935028 | 1.07647409 | 0.6 | 0.293 | 1 | 0.075743668 | 0.09737655 |
| Trpm6 | 0.049978084 | 0.181997834 | 0.227 | 0.129 | 1 | 0.894316574 | -0.391469028 | 0.1 | 0.108 | 1 | 0.894316574 | 0.097458358 |
| Prkra | 0.050006975 | -0.639754974 | 0.21 | 0.312 | 1 | 0.813436767 | 0.558490289 | 0.2 | 0.25 | 1 | 0.813436767 | 0.097513253 |
| Gadd45a | 0.957092435 | -0.549952857 | 0.479 | 0.447 | 1 | 0.050018973 | -1.567040593 | 0.4 | 0.679 | 1 | 0.957092435 | 0.097536047 |
| Mfsd4b4 | 0.338096132 | -0.944858446 | 0.025 | 0.047 | 1 | 0.050020996 | 2.280956314 | 0.2 | 0.052 | 1 | 0.338096132 | 0.097539892 |
| Vps51 | 0.050022556 | -0.932885804 | 0.151 | 0.241 | 1 | 0.926247152 | 0.176619654 | 0.2 | 0.204 | 1 | 0.926247152 | 0.097542856 |
| Srm | 0.050027146 | -0.5318624 | 0.538 | 0.582 | 1 | 0.252488181 | -0.776493958 | 0.3 | 0.438 | 1 | 0.252488181 | 0.097551577 |
| Lhfpl2 | 0.050064829 | -3.109917692 | 0.017 | 0.065 | 1 | 0.597832181 | 0.847996907 | 0 | 0.028 | 1 | 0.597832181 | 0.09762317 |
| Harbi1 | 0.050101152 | 0.762500686 | 0.134 | 0.065 | 1 | 0.744545432 | 1.432959407 | 0.1 | 0.077 | 1 | 0.744545432 | 0.097692178 |
| Slc33a1 | 0.050136345 | -0.63047716 | 0.328 | 0.424 | 1 | 0.117589732 | -0.611434712 | 0.1 | 0.383 | 1 | 0.117589732 | 0.097759037 |
| Gm29684 | 0.075308142 | 1.029146346 | 0.076 | 0.029 | 1 | 0.050189208 | 3.017921908 | 0.1 | 0.015 | 1 | 0.075308142 | 0.09785946 |
| A330015K06Rik | 0.220066125 | 1.592575685 | 0.05 | 0.024 | 1 | 0.050189208 | 2.695993813 | 0.1 | 0.015 | 1 | 0.220066125 | 0.09785946 |
| Anln | 0.312893752 | -2.40599236 | 0.034 | 0.059 | 1 | 0.050189208 | 2.847996907 | 0.1 | 0.015 | 1 | 0.312893752 | 0.09785946 |
| Gm4566 | 0.347966916 | -0.707819249 | 0.017 | 0.035 | 1 | 0.050189208 | 2.558490289 | 0.1 | 0.015 | 1 | 0.347966916 | 0.09785946 |
| Fbxo47 | 0.370241848 | 1.362570079 | 0.042 | 0.024 | 1 | 0.050189208 | 2.432959407 | 0.1 | 0.015 | 1 | 0.370241848 | 0.09785946 |
| Fbxo24 | 0.396130756 | 0.192645078 | 0.025 | 0.012 | 1 | 0.050189208 | 3.017921908 | 0.1 | 0.015 | 1 | 0.396130756 | 0.09785946 |
| Slc17a7 | 0.508731273 | -0.807354922 | 0.008 | 0.018 | 1 | 0.050189208 | 3.017921908 | 0.1 | 0.015 | 1 | 0.508731273 | 0.09785946 |
| A330033J07Rik | 0.627805946 | -0.622930351 | 0.025 | 0.035 | 1 | 0.050189208 | 2.111031312 | 0.1 | 0.015 | 1 | 0.627805946 | 0.09785946 |
| Frs3 | 0.721162042 | 0.514573173 | 0.017 | 0.012 | 1 | 0.050189208 | 3.017921908 | 0.1 | 0.015 | 1 | 0.721162042 | 0.09785946 |
| Ppp1r42 | 0.726456542 | -0.222392421 | 0.017 | 0.012 | 1 | 0.050189208 | 2.558490289 | 0.1 | 0.015 | 1 | 0.726456542 | 0.09785946 |
| Ppp1r14a | 0.050270081 | 1.514573173 | 0.05 | 0.012 | 1 | 0.597824685 | 1.432959407 | 0 | 0.028 | 1 | 0.597824685 | 0.09801308 |
| St3gal2 | 0.050280079 | -0.814049575 | 0.277 | 0.376 | 1 | 0.073111383 | -1.035189428 | 0.2 | 0.512 | 1 | 0.073111383 | 0.098032071 |
| Adck1 | 0.050507383 | -1.032914622 | 0.092 | 0.171 | 1 | 0.810674074 | 0.374065718 | 0.1 | 0.123 | 1 | 0.810674074 | 0.09846377 |
| Fcf1 | 0.050508094 | -0.679437447 | 0.437 | 0.512 | 1 | 0.139496075 | -0.827568143 | 0.2 | 0.454 | 1 | 0.139496075 | 0.09846512 |
| Exosc7 | 0.050679864 | -0.944858446 | 0.185 | 0.271 | 1 | 0.580597587 | -0.210896782 | 0.1 | 0.16 | 1 | 0.580597587 | 0.098791279 |
| Elp2 | 0.05068156 | -0.807354922 | 0.235 | 0.324 | 1 | 0.198807672 | -0.596787936 | 0.1 | 0.312 | 1 | 0.198807672 | 0.0987945 |
| Gm57257 | 0.050739142 | -1.428843299 | 0.076 | 0.147 | 1 | 0.536852613 | -1.069540933 | 0.1 | 0.164 | 1 | 0.536852613 | 0.098903824 |
| Ano8 | 0.112335502 | -1.900464326 | 0.017 | 0.053 | 1 | 0.050745058 | 2.695993813 | 0.2 | 0.056 | 1 | 0.112335502 | 0.098915055 |
| Trmt112 | 0.201199752 | -0.222392421 | 0.782 | 0.765 | 1 | 0.050754078 | -0.699068986 | 0.4 | 0.775 | 1 | 0.201199752 | 0.09893218 |
| Pigl | 0.970435857 | 0.225066556 | 0.101 | 0.1 | 1 | 0.050794982 | 1.558490289 | 0.3 | 0.099 | 1 | 0.970435857 | 0.099009834 |
| Zfp935 | 0.550057298 | -0.378511623 | 0.143 | 0.165 | 1 | 0.050797756 | 1.374065718 | 0.3 | 0.099 | 1 | 0.550057298 | 0.099015099 |
| E430024I08Rik | 0.05089305 | -1.003275132 | 0.16 | 0.247 | 1 | 0.117411796 | -1.473931188 | 0 | 0.201 | 1 | 0.117411796 | 0.099195998 |
| Exosc9 | 0.050922097 | -0.663764068 | 0.269 | 0.371 | 1 | 0.066281338 | 1.017921908 | 0.5 | 0.225 | 1 | 0.066281338 | 0.099251134 |
| Mapk8ip1 | 0.050942901 | -1.418312631 | 0.084 | 0.159 | 1 | 0.374549063 | 1.459926455 | 0.3 | 0.207 | 1 | 0.374549063 | 0.099290623 |
| Tapbpl | 0.050993789 | 0.768329765 | 0.202 | 0.118 | 1 | 0.269196652 | 1.354956895 | 0.2 | 0.093 | 1 | 0.269196652 | 0.099387211 |
| Sh3bgrl2 | 0.050996134 | -2.707819249 | 0.017 | 0.065 | 1 | 0.184723838 | 1.048295557 | 0.2 | 0.077 | 1 | 0.184723838 | 0.099391662 |
| Lrrc27 | 0.358659803 | -0.244418728 | 0.059 | 0.088 | 1 | 0.051013518 | 2.079322453 | 0.2 | 0.052 | 1 | 0.358659803 | 0.099424657 |
| Gm45871 | 0.899314101 | -0.378511623 | 0.05 | 0.053 | 1 | 0.051013518 | 2.079322453 | 0.2 | 0.052 | 1 | 0.899314101 | 0.099424657 |
| Gm44700 | 0.729956195 | -0.622930351 | 0.034 | 0.041 | 1 | 0.051013634 | 2.017921908 | 0.2 | 0.052 | 1 | 0.729956195 | 0.099424877 |
| Gdpd4 | 0.245195939 | 0.292180751 | 0.017 | 0.041 | 1 | 0.051044361 | 2.31748219 | 0.1 | 0.015 | 1 | 0.245195939 | 0.099483195 |
| Gm11290 | 0.070860081 | 1.04508789 | 0.067 | 0.024 | 1 | 0.051044714 | 1.769994395 | 0.1 | 0.015 | 1 | 0.070860081 | 0.099483865 |
| Gpr39 | 0.474027635 | -0.14839184 | 0.025 | 0.041 | 1 | 0.051044714 | 2.515421567 | 0.1 | 0.015 | 1 | 0.474027635 | 0.099483865 |
| Slc35b4 | 0.05109123 | -0.807354922 | 0.151 | 0.241 | 1 | 0.195851971 | -0.982078092 | 0.1 | 0.281 | 1 | 0.195851971 | 0.099572146 |
| BC049762 | 0.051150155 | 0.736965594 | 0.05 | 0.012 | 1 | 0.115118737 | 2.432959407 | 0.1 | 0.022 | 1 | 0.115118737 | 0.099683972 |
| Asgr1 | 0.051150155 | 1 | 0.05 | 0.012 | 1 | 0.274256184 | 2.31748219 | 0.1 | 0.034 | 1 | 0.274256184 | 0.099683972 |
| Gm50295 | 0.051150155 | 1.321928095 | 0.05 | 0.012 | 1 | 0.577155236 | 1.111031312 | 0 | 0.031 | 1 | 0.577155236 | 0.099683972 |
| Vac14 | 0.051154464 | -1.015941544 | 0.059 | 0.129 | 1 | 0.901596368 | 0.176619654 | 0.2 | 0.173 | 1 | 0.901596368 | 0.099692148 |
| Sphk1 | 0.051331231 | 0.836501268 | 0.311 | 0.218 | 1 | 0.871819944 | 0.591657153 | 0.2 | 0.188 | 1 | 0.871819944 | 0.100027567 |
| Arf3 | 0.496620349 | -0.390962143 | 0.471 | 0.459 | 1 | 0.051347296 | -1.135207851 | 0.4 | 0.651 | 1 | 0.496620349 | 0.100058047 |
| Wasf3 | 0.082054654 | 0.598637438 | 0.303 | 0.212 | 1 | 0.051353854 | 1.030860964 | 0.5 | 0.222 | 1 | 0.082054654 | 0.100070489 |
| Tbccd1 | 0.051360112 | -1.008988783 | 0.05 | 0.118 | 1 | 0.542976121 | -0.391469028 | 0.1 | 0.167 | 1 | 0.542976121 | 0.100082363 |
| Zfp711 | 0.051465202 | -2.758445322 | 0.017 | 0.065 | 1 | 0.715089227 | -0.970762779 | 0.1 | 0.133 | 1 | 0.715089227 | 0.100281737 |
| Aurkaip1 | 0.120338584 | -0.287980763 | 0.622 | 0.7 | 1 | 0.051491907 | -0.991066875 | 0.3 | 0.623 | 1 | 0.120338584 | 0.100332397 |
| Gm52988 | 0.051503021 | -1.383547213 | 0.059 | 0.129 | 1 | 0.196834952 | -1.505640048 | 0 | 0.145 | 1 | 0.196834952 | 0.10035348 |
| Ythdc2 | 0.051544994 | -0.771731012 | 0.261 | 0.347 | 1 | 0.772437577 | 0.741797503 | 0.2 | 0.281 | 1 | 0.772437577 | 0.100433101 |
| Cyp39a1 | 0.051671421 | -1.222392421 | 0.059 | 0.129 | 1 | 0.567492749 | -0.321928095 | 0.1 | 0.164 | 1 | 0.567492749 | 0.100672906 |
| Parp9 | 0.051674406 | -0.572889668 | 0.143 | 0.235 | 1 | 0.362064958 | -0.763437806 | 0.1 | 0.213 | 1 | 0.362064958 | 0.100678567 |
| Galk1 | 0.051676509 | -0.642968104 | 0.286 | 0.376 | 1 | 0.446111849 | -0.230005605 | 0.2 | 0.33 | 1 | 0.446111849 | 0.100682556 |
| Mapk14 | 0.328000445 | -0.452259963 | 0.328 | 0.371 | 1 | 0.051680215 | -1.505640048 | 0.3 | 0.562 | 1 | 0.328000445 | 0.100689586 |
| Cx3cr1 | 0.051728157 | -0.422233001 | 0.084 | 0.165 | 1 | 0.667100705 | 0.804142617 | 0.2 | 0.164 | 1 | 0.667100705 | 0.100780512 |
| Tnfrsf1b | 0.051755939 | 0.597035333 | 0.134 | 0.065 | 1 | 0.808436044 | 0.185031894 | 0.1 | 0.123 | 1 | 0.808436044 | 0.100833201 |
| Spred1 | 0.051820844 | 0.200912694 | 0.933 | 0.841 | 1 | 0.947983621 | 0.104772322 | 0.9 | 0.901 | 1 | 0.947983621 | 0.100956288 |
| Upp2 | 0.051871107 | 0.346018903 | 0.605 | 0.494 | 1 | 0.571732281 | -0.68251781 | 0.4 | 0.451 | 1 | 0.571732281 | 0.101051601 |
| Slc31a1 | 0.051887675 | 0.317381306 | 0.765 | 0.665 | 1 | 0.304043972 | -0.492512614 | 0.7 | 0.738 | 1 | 0.304043972 | 0.10108302 |
| Ncaph | 0.063085444 | -1.807354922 | 0.008 | 0.047 | 1 | 0.051911311 | 2.695993813 | 0.1 | 0.015 | 1 | 0.063085444 | 0.101127838 |
| Arsk | 0.05191333 | -0.86393845 | 0.21 | 0.3 | 1 | 0.677511057 | -0.357117523 | 0.3 | 0.327 | 1 | 0.677511057 | 0.101131665 |
| ENSMUSG00000121476 | 0.051935865 | -2.14839184 | 0.017 | 0.065 | 1 | 0.943862173 | 0.432959407 | 0.1 | 0.09 | 1 | 0.943862173 | 0.101174397 |
| Mns1 | 0.051936544 | -2.485426827 | 0.017 | 0.065 | 1 | 0.597838178 | 1.210566986 | 0 | 0.028 | 1 | 0.597838178 | 0.101175683 |
| Taf1d | 0.05197586 | 0.142750454 | 0.79 | 0.859 | 1 | 0.858655281 | 0.290001453 | 0.8 | 0.793 | 1 | 0.858655281 | 0.10125023 |
| Setd7 | 0.05198114 | -0.38714731 | 0.723 | 0.747 | 1 | 0.087413168 | -0.743238191 | 0.8 | 0.883 | 1 | 0.087413168 | 0.101260241 |
| Atp5d | 0.11533154 | -0.214530304 | 0.899 | 0.941 | 1 | 0.052009835 | -0.581207925 | 0.7 | 0.935 | 1 | 0.11533154 | 0.101314646 |
| 1600020E01Rik | 0.052049556 | -0.611289783 | 0.588 | 0.647 | 1 | 0.960376483 | -0.246781318 | 0.7 | 0.639 | 1 | 0.960376483 | 0.101389957 |
| Trappc2 | 0.130827574 | 0.478047297 | 0.294 | 0.218 | 1 | 0.052060393 | 1.255421222 | 0.5 | 0.235 | 1 | 0.130827574 | 0.101410502 |
| Gm16759 | 0.052069554 | -1.807354922 | 0.05 | 0.118 | 1 | 0.682901431 | 1.695993813 | 0.1 | 0.071 | 1 | 0.682901431 | 0.10142787 |
| Ube2d-ps | 0.052071736 | -0.777192952 | 0.21 | 0.306 | 1 | 0.221394238 | 1.197742946 | 0.4 | 0.253 | 1 | 0.221394238 | 0.101432005 |
| Slc37a3 | 0.121606539 | -0.548436625 | 0.353 | 0.418 | 1 | 0.052146056 | -1.030110788 | 0.2 | 0.586 | 1 | 0.121606539 | 0.101572901 |
| Mrps26 | 0.052181659 | -0.678071905 | 0.294 | 0.388 | 1 | 0.38198909 | -0.31299497 | 0.2 | 0.349 | 1 | 0.38198909 | 0.101640393 |
| Tmem216 | 0.096707534 | -0.795766948 | 0.185 | 0.259 | 1 | 0.052189496 | -1.840059087 | 0 | 0.281 | 1 | 0.096707534 | 0.101655249 |
| Ocrl | 0.052253075 | -0.921525942 | 0.345 | 0.4 | 1 | 0.76116593 | 0.22090893 | 0.5 | 0.441 | 1 | 0.76116593 | 0.101775765 |
| Irgq | 0.307569089 | -0.589763487 | 0.126 | 0.165 | 1 | 0.052289048 | 1.75918764 | 0.4 | 0.17 | 1 | 0.307569089 | 0.101843952 |
| Asf1b | 0.052408793 | -2.14839184 | 0.017 | 0.065 | 1 | 0.195226183 | 2.017921908 | 0.1 | 0.028 | 1 | 0.195226183 | 0.102070904 |
| Kank4 | 0.05240902 | -1.987927168 | 0.017 | 0.065 | 1 | 0.410925689 | 2.017921908 | 0.1 | 0.046 | 1 | 0.410925689 | 0.102071335 |
| Fstl3 | 0.05240902 | -1.807354922 | 0.017 | 0.065 | 1 | 0.35375985 | -0.111361109 | 0 | 0.08 | 1 | 0.35375985 | 0.102071335 |
| Vps52 | 0.052424256 | -1.029747343 | 0.092 | 0.171 | 1 | 0.695209309 | 0.744903414 | 0.2 | 0.151 | 1 | 0.695209309 | 0.102100209 |
| Fbxl14 | 0.052424583 | -0.987927168 | 0.092 | 0.171 | 1 | 0.091492948 | -1.374395515 | 0 | 0.225 | 1 | 0.091492948 | 0.102100829 |
| Plcb1 | 0.052438547 | -0.566422559 | 0.832 | 0.835 | 1 | 0.323055234 | -0.553734336 | 0.9 | 0.935 | 1 | 0.323055234 | 0.102127293 |
| Arel1 | 0.052444079 | -1.008988783 | 0.151 | 0.235 | 1 | 0.086640269 | -1.654503434 | 0 | 0.231 | 1 | 0.086640269 | 0.102137776 |
| Nr1d1 | 0.47805518 | -0.112660981 | 0.697 | 0.576 | 1 | 0.052506189 | -0.995540352 | 0.4 | 0.735 | 1 | 0.47805518 | 0.102255477 |
| Adamts3 | 0.052574936 | -0.798017057 | 0.655 | 0.676 | 1 | 0.673582643 | -0.719992516 | 0.7 | 0.679 | 1 | 0.673582643 | 0.102385747 |
| Vapb | 0.05258115 | -0.406438531 | 0.647 | 0.671 | 1 | 0.300640645 | -0.570043081 | 0.6 | 0.71 | 1 | 0.300640645 | 0.102397523 |
| Msantd4 | 0.762605297 | -0.141942589 | 0.345 | 0.347 | 1 | 0.052657418 | 1.395870089 | 0.6 | 0.38 | 1 | 0.762605297 | 0.102542032 |

| | | | | | | | | | | | | |
|---|---|---|---|---|---|---|---|---|---|---|---|---|
| Zswim5 | 0.052686965 | 0.736965594 | 0.05 | 0.012 | 1 | 0.252293596 | 2.930459067 | 0.1 | 0.034 | 1 | 0.252293596 | 0.102598014 |
| Nutf2 | 0.052776339 | -0.970853654 | 0.118 | 0.2 | 1 | 0.670802544 | 0.695993813 | 0.2 | 0.145 | 1 | 0.670802544 | 0.102767336 |
| Stat4 | 0.091361251 | 2.514573173 | 0.017 | 0 | 1 | 0.05278585 | 2.432959407 | 0.1 | 0.015 | 1 | 0.091361251 | 0.102785354 |
| Fam169b | 0.670386669 | -0.163498732 | 0.025 | 0.018 | 1 | 0.052789093 | 1.494359952 | 0.1 | 0.015 | 1 | 0.670386669 | 0.102791498 |
| Vps13c | 0.052795707 | -0.755515991 | 0.336 | 0.418 | 1 | 0.925298926 | 0.466126271 | 0.4 | 0.46 | 1 | 0.925298926 | 0.102804028 |
| Gpam | 0.187968352 | 0.844721775 | 0.176 | 0.124 | 1 | 0.052857387 | 1.732519689 | 0.3 | 0.102 | 1 | 0.187968352 | 0.102920871 |
| Cnih1 | 0.246847265 | -0.255668576 | 0.807 | 0.735 | 1 | 0.052857896 | -0.979257573 | 0.4 | 0.716 | 1 | 0.246847265 | 0.102921835 |
| Clec2l | 0.05288261 | -1.707819249 | 0.017 | 0.065 | 1 | 0.577142568 | 1.432959407 | 0 | 0.031 | 1 | 0.577142568 | 0.10296865 |
| Fxyd7 | 0.05288261 | -1.807354922 | 0.017 | 0.065 | 1 | 0.362872338 | 1.847996907 | 0.1 | 0.04 | 1 | 0.362872338 | 0.10296865 |
| Ccdc85c | 0.05288261 | -1.707819249 | 0.017 | 0.065 | 1 | 0.813207742 | 0.888638891 | 0.1 | 0.077 | 1 | 0.813207742 | 0.10296865 |
| Gm30054 | 0.05288261 | -1.707819249 | 0.017 | 0.065 | 1 | 0.539456618 | 0.695993813 | 0 | 0.037 | 1 | 0.539456618 | 0.10296865 |
| Gm15458 | 0.05288261 | -1.707819249 | 0.017 | 0.065 | 1 | 0.44686405 | 0.374065718 | 0 | 0.056 | 1 | 0.44686405 | 0.10296865 |
| Gnmt | 0.05288261 | -1.707819249 | 0.017 | 0.065 | 1 | 0.154873683 | 2.432959407 | 0.1 | 0.025 | 1 | 0.154873683 | 0.10296865 |
| Scml2 | 0.05288261 | -1.987927168 | 0.017 | 0.065 | 1 | 0.450331524 | 1.625604485 | 0.1 | 0.046 | 1 | 0.450331524 | 0.10296865 |
| Arhgef5 | 0.052887956 | 0.514573173 | 0.395 | 0.288 | 1 | 0.314380406 | -0.596787936 | 0.2 | 0.364 | 1 | 0.314380406 | 0.102978776 |
| Gm28376 | 0.144054748 | -1.485426827 | 0.008 | 0.035 | 1 | 0.053042311 | 1.795529487 | 0.2 | 0.052 | 1 | 0.144054748 | 0.103271136 |
| Cenpt | 0.490319312 | -0.378511623 | 0.109 | 0.135 | 1 | 0.053098955 | 1.385653692 | 0.4 | 0.157 | 1 | 0.490319312 | 0.103378412 |
| Elac2 | 0.053159694 | -0.833350131 | 0.067 | 0.141 | 1 | 0.496793513 | 1.048295557 | 0.2 | 0.123 | 1 | 0.496793513 | 0.103493435 |
| S1pr3 | 0.05317585 | 1.330148602 | 0.168 | 0.094 | 1 | 0.597836679 | 0.930459067 | 0 | 0.028 | 1 | 0.597836679 | 0.103524029 |
| Vps72 | 0.053244718 | -0.690541257 | 0.252 | 0.347 | 1 | 0.830563258 | 0.591657153 | 0.3 | 0.346 | 1 | 0.830563258 | 0.103654437 |
| Farsb | 0.05330928 | -0.748461233 | 0.361 | 0.441 | 1 | 0.791003343 | -0.368659145 | 0.5 | 0.457 | 1 | 0.791003343 | 0.10377668 |
| Zfp623 | 0.053357292 | -1.600904045 | 0.017 | 0.065 | 1 | 0.919812491 | 0.973527789 | 0.1 | 0.09 | 1 | 0.919812491 | 0.103867584 |
| S100a5 | 0.053416159 | -1.86393845 | 0.017 | 0.065 | 1 | 0.362872659 | 1.494359952 | 0.1 | 0.04 | 1 | 0.362872659 | 0.103979032 |
| Clcn5 | 0.387836053 | -0.861126178 | 0.345 | 0.365 | 1 | 0.053422445 | -1.493830746 | 0.1 | 0.441 | 1 | 0.387836053 | 0.103990932 |
| Hsp90aa1 | 0.19684437 | -0.228360145 | 0.975 | 0.988 | 1 | 0.053427045 | -0.624206946 | 1 | 0.997 | 1 | 0.19684437 | 0.103999641 |
| Aff1 | 0.053615546 | 0.234164347 | 0.966 | 0.906 | 1 | 0.972003693 | 0.223506042 | 0.9 | 0.92 | 1 | 0.972003693 | 0.104356465 |
| Itgav | 0.053616877 | 0.249869947 | 0.908 | 0.824 | 1 | 0.201614627 | 0.484635442 | 0.9 | 0.852 | 1 | 0.201614627 | 0.104358985 |
| Tm2d2 | 0.05367379 | 0.208020304 | 0.857 | 0.724 | 1 | 0.257392686 | -0.519825954 | 0.6 | 0.704 | 1 | 0.257392686 | 0.104466704 |
| Ptprm | 0.425561143 | -0.282579427 | 0.882 | 0.888 | 1 | 0.053692726 | 0.543024289 | 1 | 0.963 | 1 | 0.425561143 | 0.104502542 |
| Gm57146 | 0.734009606 | -0.129283017 | 0.403 | 0.353 | 1 | 0.053699912 | 1.238311976 | 0.6 | 0.312 | 1 | 0.734009606 | 0.104516144 |
| Rbm42 | 0.095288458 | -0.521406124 | 0.513 | 0.576 | 1 | 0.053798189 | -1.11821478 | 0.4 | 0.664 | 1 | 0.095288458 | 0.104702132 |
| Gch1 | 0.053844399 | 0.652076697 | 0.244 | 0.147 | 1 | 0.378321355 | -0.668578619 | 0.1 | 0.207 | 1 | 0.378321355 | 0.10478958 |
| Lefty1 | 0.053870521 | 0.617666666 | 0.218 | 0.129 | 1 | 0.23873063 | -0.625934282 | 0 | 0.123 | 1 | 0.23873063 | 0.10483901 |
| Spin1 | 0.053890926 | -0.835369298 | 0.37 | 0.435 | 1 | 0.102236837 | -0.713881981 | 0.4 | 0.654 | 1 | 0.102236837 | 0.104877619 |
| St6galnac3 | 0.053903803 | -2.117695043 | 0.017 | 0.065 | 1 | 0.160108911 | 1.111031312 | 0.1 | 0.025 | 1 | 0.160108911 | 0.104901986 |
| Gm17354 | 0.053916985 | -1.485426827 | 0.017 | 0.065 | 1 | 0.363859515 | 0.210566986 | 0 | 0.077 | 1 | 0.363859515 | 0.104926929 |
| Lman1l | 0.053916985 | -1.485426827 | 0.017 | 0.065 | 1 | 0.878982267 | 0.591657153 | 0.1 | 0.083 | 1 | 0.878982267 | 0.104926929 |
| Nbeal1 | 0.778996826 | -0.116855426 | 0.941 | 0.929 | 1 | 0.053941148 | 0.473939299 | 0.9 | 0.929 | 1 | 0.778996826 | 0.104972649 |
| Aamdc | 0.170064819 | -0.524955191 | 0.37 | 0.429 | 1 | 0.054011201 | -1.852442812 | 0.1 | 0.395 | 1 | 0.170064819 | 0.105105192 |
| Ttf1 | 0.054019278 | -1.182034684 | 0.218 | 0.3 | 1 | 0.526959928 | 0.169925001 | 0.4 | 0.269 | 1 | 0.526959928 | 0.105120474 |
| Trmo | 0.620648735 | -0.102098188 | 0.092 | 0.112 | 1 | 0.054034286 | 1.625604485 | 0.3 | 0.102 | 1 | 0.620648735 | 0.105148868 |
| Lhpp | 0.219157371 | -0.807354922 | 0.05 | 0.088 | 1 | 0.054034956 | 1.695993813 | 0.3 | 0.102 | 1 | 0.219157371 | 0.105150135 |
| Tsc22d2 | 0.05406544 | -0.322041465 | 0.874 | 0.882 | 1 | 0.493115357 | -0.585704437 | 0.9 | 0.892 | 1 | 0.493115357 | 0.105207807 |
| Zcwpw1 | 0.217966241 | -0.944858446 | 0.05 | 0.088 | 1 | 0.054077882 | 1.847996907 | 0.2 | 0.052 | 1 | 0.217966241 | 0.105231346 |
| Midn | 0.054252965 | -0.254101281 | 0.655 | 0.741 | 1 | 0.142301186 | -0.880931369 | 0.7 | 0.806 | 1 | 0.142301186 | 0.105562546 |
| Ppp1r8 | 0.054300532 | -0.84450792 | 0.16 | 0.247 | 1 | 0.126947913 | -1.101019165 | 0.1 | 0.324 | 1 | 0.126947913 | 0.105652516 |
| Slc25a21 | 0.05438491 | -0.250961574 | 0.092 | 0.035 | 1 | 0.474054479 | 1.252387162 | 0.1 | 0.049 | 1 | 0.474054479 | 0.105812101 |
| Mrpl57 | 0.054389973 | 0.309254265 | 0.487 | 0.371 | 1 | 0.08104791 | -1.465893869 | 0.1 | 0.364 | 1 | 0.08104791 | 0.105821677 |
| Gm50333 | 0.054392857 | -2.070389328 | 0.017 | 0.065 | 1 | 0.6104049 | 1.263034406 | 0.1 | 0.059 | 1 | 0.6104049 | 0.105827132 |
| Dclk1 | 0.955309558 | -0.485426827 | 0.134 | 0.129 | 1 | 0.054410705 | 0.542188477 | 0.3 | 0.096 | 1 | 0.955309558 | 0.105860885 |
| Arhgef26 | 0.054447086 | -1.549557165 | 0.067 | 0.135 | 1 | 0.226483722 | -1.141949429 | 0.1 | 0.262 | 1 | 0.226483722 | 0.105929686 |
| Pik3r3 | 0.054534243 | -1.504285854 | 0.076 | 0.147 | 1 | 0.944815301 | -0.242605642 | 0.2 | 0.201 | 1 | 0.944815301 | 0.106094503 |
| Ndufaf7 | 0.054672488 | -0.617877123 | 0.151 | 0.247 | 1 | 0.433438626 | -0.339630097 | 0.1 | 0.198 | 1 | 0.433438626 | 0.106355895 |
| Plekha8 | 0.054691527 | 0.514573173 | 0.303 | 0.206 | 1 | 0.341915916 | 0.642882477 | 0.3 | 0.167 | 1 | 0.341915916 | 0.10639189 |
| Mill2 | 0.054698111 | -1.273922722 | 0.143 | 0.224 | 1 | 0.519355588 | 0.672147071 | 0.2 | 0.123 | 1 | 0.519355588 | 0.106404338 |
| Fam107b | 0.054710237 | -0.626782676 | 0.16 | 0.259 | 1 | 0.311389494 | -0.260062839 | 0.2 | 0.38 | 1 | 0.311389494 | 0.106427265 |
| Fam234a | 0.054735525 | 0.354108501 | 0.471 | 0.353 | 1 | 0.117211651 | -1.083616118 | 0.1 | 0.364 | 1 | 0.117211651 | 0.106475071 |
| Btbd16 | 0.060418894 | -1.758445322 | 0.042 | 0.1 | 1 | 0.054775072 | 3.085036104 | 0.2 | 0.056 | 1 | 0.060418894 | 0.106549835 |
| Rnf149 | 0.054971076 | -0.664750527 | 0.218 | 0.318 | 1 | 0.822235445 | 0.147557188 | 0.3 | 0.247 | 1 | 0.822235445 | 0.106920332 |
| Chdh | 0.054982374 | 0.848992212 | 0.176 | 0.1 | 1 | 0.566286055 | -0.604129911 | 0.3 | 0.343 | 1 | 0.566286055 | 0.106941686 |
| Pon2 | 0.054992273 | 0.171428592 | 0.622 | 0.471 | 1 | 0.341302537 | 0.339850003 | 0.8 | 0.586 | 1 | 0.341302537 | 0.106960396 |
| Mettl16 | 0.111740758 | -0.468553009 | 0.504 | 0.535 | 1 | 0.055024412 | -1.210896782 | 0.3 | 0.59 | 1 | 0.111740758 | 0.107021138 |
| Nubp2 | 0.055044432 | -0.95491211 | 0.092 | 0.171 | 1 | 0.731980265 | 0.31748219 | 0.1 | 0.136 | 1 | 0.731980265 | 0.107058974 |
| 2810004N23Rik | 0.057711713 | -0.67251238 | 0.328 | 0.429 | 1 | 0.055105859 | 1.042725299 | 0.6 | 0.349 | 1 | 0.057711713 | 0.107175063 |
| Znrf2 | 0.055109863 | -0.754887502 | 0.319 | 0.418 | 1 | 0.472888235 | -0.714528205 | 0.5 | 0.525 | 1 | 0.472888235 | 0.10718263 |
| 1700030J22Rik | 0.055140359 | -1.392317423 | 0.017 | 0.065 | 1 | 0.669853054 | 2.210566986 | 0 | 0.019 | 1 | 0.669853054 | 0.107240259 |
| Pcnx | 0.17397162 | -0.268802886 | 0.765 | 0.829 | 1 | 0.055144616 | -0.962969269 | 0.8 | 0.861 | 1 | 0.17397162 | 0.107248304 |
| Atosa | 0.055153951 | -0.890181353 | 0.403 | 0.482 | 1 | 0.939032776 | -0.22708221 | 0.5 | 0.519 | 1 | 0.939032776 | 0.107265943 |
| Fem1b | 0.11342741 | -0.520473774 | 0.361 | 0.441 | 1 | 0.055212328 | -1.521236903 | 0.2 | 0.481 | 1 | 0.11342741 | 0.107376255 |
| Gm20275 | 0.055232949 | -2.276840205 | 0.076 | 0.147 | 1 | 0.519194854 | -0.267480311 | 0.3 | 0.194 | 1 | 0.519194854 | 0.107415219 |
| Rab28 | 0.359086292 | -0.353922044 | 0.706 | 0.641 | 1 | 0.05525716 | -0.864721141 | 0.3 | 0.63 | 1 | 0.359086292 | 0.107460967 |
| Iqcd | 0.144054748 | -1.485426827 | 0.008 | 0.035 | 1 | 0.055268358 | 2.295455884 | 0.2 | 0.056 | 1 | 0.144054748 | 0.107482124 |
| Dtymk | 0.05529603 | -0.602996424 | 0.546 | 0.582 | 1 | 0.807731952 | 0.306427001 | 0.4 | 0.485 | 1 | 0.807731952 | 0.107534408 |
| Txndc16 | 0.055382122 | -1.032914622 | 0.168 | 0.253 | 1 | 1 | 0.942633781 | 0.2 | 0.235 | 1 | 1 | 0.107697065 |
| Cldnd1 | 0.607553416 | -0.171714038 | 0.739 | 0.729 | 1 | 0.055392831 | -0.830460282 | 0.6 | 0.836 | 1 | 0.607553416 | 0.107717296 |
| Spata5l1 | 0.055636933 | -1.292781749 | 0.017 | 0.065 | 1 | 0.298405825 | -0.267480311 | 0 | 0.099 | 1 | 0.298405825 | 0.108178398 |
| Zfp239 | 0.055636933 | -1.292781749 | 0.017 | 0.065 | 1 | 0.48995314 | 0.558490289 | 0 | 0.046 | 1 | 0.48995314 | 0.108178398 |
| Sdhb | 0.290123119 | -0.165026202 | 0.605 | 0.665 | 1 | 0.055697601 | -0.974544419 | 0.5 | 0.694 | 1 | 0.290123119 | 0.108292979 |
| Ism1 | 0.055851367 | -0.690631316 | 0.311 | 0.424 | 1 | 0.325449657 | -1.304006187 | 0.2 | 0.343 | 1 | 0.325449657 | 0.108583358 |
| Arpc5l | 0.097202951 | -0.485426827 | 0.546 | 0.571 | 1 | 0.055874463 | -1.131825212 | 0.4 | 0.623 | 1 | 0.097202951 | 0.10862697 |
| Ly96 | 0.074630997 | -0.925999419 | 0.151 | 0.229 | 1 | 0.055985625 | -1.888968688 | 0 | 0.275 | 1 | 0.074630997 | 0.10883686 |
| Gm44649 | 0.05612203 | 0.625604485 | 0.193 | 0.112 | 1 | 0.930920445 | 0.354956895 | 0.2 | 0.182 | 1 | 0.930920445 | 0.109094377 |
| Trafd1 | 0.056142298 | 0.185265548 | 0.723 | 0.576 | 1 | 0.456806935 | -0.299993124 | 0.5 | 0.608 | 1 | 0.456806935 | 0.109132639 |
| Btbd9 | 0.05627331 | -0.478077371 | 0.857 | 0.912 | 1 | 0.460360318 | -0.424653036 | 0.9 | 0.957 | 1 | 0.460360318 | 0.109379934 |
| Erich1 | 0.05629015 | -1.14839184 | 0.084 | 0.159 | 1 | 0.580893489 | 0.393431043 | 0.2 | 0.13 | 1 | 0.580893489 | 0.109411719 |
| Colq | 0.056321394 | -0.774933444 | 0.042 | 0.106 | 1 | 0.487557793 | 1.625604485 | 0.1 | 0.049 | 1 | 0.487557793 | 0.109470689 |
| Trim24 | 0.987844564 | -0.318607737 | 0.445 | 0.406 | 1 | 0.056359353 | 0.789103218 | 0.8 | 0.469 | 1 | 0.987844564 | 0.10954233 |
| Cabyr | 0.056391829 | -0.86393845 | 0.017 | 0.065 | 1 | 0.518124185 | 2.159940913 | 0.1 | 0.056 | 1 | 0.518124185 | 0.109603619 |
| Ddx19b | 0.479330577 | -0.707819249 | 0.067 | 0.088 | 1 | 0.056444898 | 1.591657153 | 0.3 | 0.102 | 1 | 0.479330577 | 0.109703769 |
| Zc3h12a | 0.180193017 | 0.819427754 | 0.143 | 0.094 | 1 | 0.056446341 | 1.494359952 | 0.3 | 0.102 | 1 | 0.180193017 | 0.109706492 |
| Pde8a | 0.056621112 | -1.026266775 | 0.639 | 0.624 | 1 | 0.943168449 | -0.28665262 | 0.9 | 0.682 | 1 | 0.943168449 | 0.110036274 |
| Emsy | 0.056634065 | -0.715432433 | 0.437 | 0.5 | 1 | 0.061643505 | -0.873861795 | 0.3 | 0.654 | 1 | 0.061643505 | 0.110060712 |
| Phax | 0.082023102 | -0.426264755 | 0.588 | 0.618 | 1 | 0.056675375 | -1.14865965 | 0.5 | 0.67 | 1 | 0.082023102 | 0.110138652 |
| Tsix | 0.05670673 | 0.67211445 | 0.185 | 0.106 | 1 | 0.730750993 | 1.695993813 | 0 | 0.012 | 1 | 0.730750993 | 0.110197806 |
| Flvcr1 | 0.056738987 | -0.934734229 | 0.16 | 0.247 | 1 | 0.242099096 | 0.197742946 | 0.6 | 0.352 | 1 | 0.242099096 | 0.110258661 |
| Trabd | 0.370992869 | -0.422233001 | 0.244 | 0.282 | 1 | 0.056749514 | -1.544320516 | 0.1 | 0.395 | 1 | 0.370992869 | 0.110278521 |
| Spice1 | 0.056749878 | -1.015941544 | 0.143 | 0.224 | 1 | 0.682816447 | 0.432959407 | 0.2 | 0.145 | 1 | 0.682816447 | 0.110279207 |
| Zbtb4 | 0.056785209 | -0.7456626 | 0.277 | 0.365 | 1 | 0.070536944 | -1.591872446 | 0.2 | 0.457 | 1 | 0.070536944 | 0.110345858 |

| | | | | | | | | | | | | |
|---|---|---|---|---|---|---|---|---|---|---|---|---|
| Tes | 0.056878934 | 0.57870351 | 0.134 | 0.065 | 1 | 0.732929433 | 0.847996907 | 0.1 | 0.154 | 1 | 0.732929433 | 0.110522656 |
| Ccdc122 | 0.056901054 | 0.514573173 | 0.235 | 0.147 | 1 | 0.802755022 | 0.542188477 | 0.2 | 0.173 | 1 | 0.802755022 | 0.110564378 |
| Mphosph9 | 0.056919985 | 0.201415288 | 0.218 | 0.129 | 1 | 0.222221227 | 1.14345279 | 0.3 | 0.164 | 1 | 0.222221227 | 0.110600085 |
| Cops3 | 0.056977059 | -0.485426827 | 0.555 | 0.594 | 1 | 0.23660448 | -0.53359411 | 0.5 | 0.676 | 1 | 0.23660448 | 0.110707733 |
| Atf6 | 0.05702706 | -0.443257373 | 0.807 | 0.812 | 1 | 0.892919996 | -0.111361109 | 0.9 | 0.867 | 1 | 0.892919996 | 0.110802035 |
| Spon2 | 0.057165047 | -2.292781749 | 0.025 | 0.076 | 1 | 0.193279696 | 2.432959407 | 0.1 | 0.028 | 1 | 0.193279696 | 0.111062252 |
| Ndufaf1 | 0.057197227 | -1.032914622 | 0.084 | 0.159 | 1 | 0.274994038 | 1.31748219 | 0.2 | 0.093 | 1 | 0.274994038 | 0.111122931 |
| Gm16093 | 0.057279702 | -1.411426246 | 0.109 | 0.182 | 1 | 0.101364941 | 1.185031894 | 0.3 | 0.117 | 1 | 0.101364941 | 0.111278439 |
| Dcaf4 | 0.057288506 | -1.292781749 | 0.067 | 0.135 | 1 | 0.974695342 | 0.558490289 | 0.1 | 0.102 | 1 | 0.974695342 | 0.11129504 |
| Gtf2a2 | 0.649242301 | -0.271124993 | 0.748 | 0.676 | 1 | 0.057347314 | -0.984893108 | 0.6 | 0.747 | 1 | 0.649242301 | 0.111405913 |
| Pde5a | 0.303109214 | -0.511899039 | 0.076 | 0.112 | 1 | 0.057456633 | 0.558490289 | 0.3 | 0.102 | 1 | 0.303109214 | 0.111612002 |
| Abcf1 | 0.086607611 | -0.360620371 | 0.706 | 0.741 | 1 | 0.057498167 | -0.736965594 | 0.5 | 0.796 | 1 | 0.086607611 | 0.111690295 |
| Fbxo25 | 0.333796246 | -0.622930351 | 0.109 | 0.147 | 1 | 0.057562914 | 1.639410285 | 0.3 | 0.108 | 1 | 0.333796246 | 0.111812338 |
| Esco1 | 0.082162134 | -0.485426827 | 0.454 | 0.512 | 1 | 0.057652955 | -1.108413825 | 0.5 | 0.707 | 1 | 0.082162134 | 0.111982048 |
| Efs | 0.0576558 | -0.778208576 | 0.16 | 0.247 | 1 | 0.732158138 | 0.223506042 | 0.3 | 0.231 | 1 | 0.732158138 | 0.111987409 |
| Tmem116 | 0.05770851 | -1.222392421 | 0.067 | 0.135 | 1 | 0.074165461 | 1.667424661 | 0.3 | 0.114 | 1 | 0.074165461 | 0.112086748 |
| Bud23 | 0.057737571 | -0.745293954 | 0.294 | 0.376 | 1 | 0.418467741 | 1.03809979 | 0.4 | 0.318 | 1 | 0.418467741 | 0.112141515 |
| Zfp619 | 0.057865101 | -1.944858446 | 0.025 | 0.076 | 1 | 0.105692031 | 1.515421567 | 0.2 | 0.065 | 1 | 0.105692031 | 0.112381832 |
| Pms1 | 0.057865497 | -2.070389328 | 0.025 | 0.076 | 1 | 0.45800013 | 1.847996907 | 0.1 | 0.049 | 1 | 0.45800013 | 0.112382579 |
| Prx | 0.057865497 | -2.292781749 | 0.025 | 0.076 | 1 | 0.455766715 | 1.210566986 | 0.1 | 0.046 | 1 | 0.455766715 | 0.112382579 |
| Mtf1 | 0.057881071 | -0.790281409 | 0.235 | 0.324 | 1 | 0.626790727 | -0.252607034 | 0.3 | 0.377 | 1 | 0.626790727 | 0.112411924 |
| Rnf141 | 0.83102909 | -0.137503524 | 0.092 | 0.1 | 1 | 0.057991097 | 2.176619654 | 0.3 | 0.111 | 1 | 0.83102909 | 0.112619227 |
| BC031181 | 0.058007341 | 0.22614199 | 0.849 | 0.765 | 1 | 0.433805534 | -0.114106486 | 0.5 | 0.775 | 1 | 0.433805534 | 0.112649831 |
| Clpp | 0.058112763 | -0.410659059 | 0.496 | 0.594 | 1 | 0.662918691 | 0.500073603 | 0.5 | 0.472 | 1 | 0.662918691 | 0.112848432 |
| Anapc7 | 0.058175338 | -0.549557165 | 0.345 | 0.441 | 1 | 0.620766124 | -0.310004928 | 0.4 | 0.454 | 1 | 0.620766124 | 0.112966306 |
| Gpat3 | 0.058201935 | 0.251538767 | 0.277 | 0.171 | 1 | 0.665720356 | -0.374395515 | 0.1 | 0.142 | 1 | 0.665720356 | 0.113016405 |
| Sorbs3 | 0.058302572 | 0.235289415 | 0.538 | 0.394 | 1 | 0.388640884 | -0.709998547 | 0.4 | 0.469 | 1 | 0.388640884 | 0.113205953 |
| Vps29 | 0.058449913 | -0.371094152 | 0.546 | 0.624 | 1 | 0.149355225 | -0.713881981 | 0.5 | 0.667 | 1 | 0.149355225 | 0.113483433 |
| St3gal3 | 0.058474216 | -0.788495895 | 0.353 | 0.453 | 1 | 0.2825462 | -1.489872732 | 0.3 | 0.401 | 1 | 0.2825462 | 0.113529198 |
| Riox1 | 0.058475874 | -0.932885804 | 0.084 | 0.159 | 1 | 0.890218267 | 0.393431043 | 0.2 | 0.176 | 1 | 0.890218267 | 0.113532321 |
| 1700123O20Rik | 0.058481921 | -0.755515991 | 0.244 | 0.329 | 1 | 0.677850277 | -0.171902651 | 0.3 | 0.34 | 1 | 0.677850277 | 0.113543706 |
| Rad9a | 0.058507009 | -0.970853654 | 0.017 | 0.065 | 1 | 0.363906344 | 0.111031312 | 0 | 0.077 | 1 | 0.363906344 | 0.113590948 |
| Dusp1 | 0.058509078 | 0.423726031 | 0.815 | 0.782 | 1 | 0.554671221 | 0.100321669 | 0.8 | 0.867 | 1 | 0.554671221 | 0.113594843 |
| Apbb3 | 0.087607873 | -0.780882711 | 0.067 | 0.129 | 1 | 0.058519675 | 1.695993813 | 0.3 | 0.105 | 1 | 0.087607873 | 0.113614797 |
| Gm42917 | 0.058570531 | -1.733354341 | 0.025 | 0.076 | 1 | 0.979200188 | 0.625604485 | 0.1 | 0.096 | 1 | 0.979200188 | 0.113710555 |
| Usp35 | 0.058570531 | -1.807354922 | 0.025 | 0.076 | 1 | 0.601875777 | 1.432959407 | 0.1 | 0.059 | 1 | 0.601875777 | 0.113710555 |
| Mbnl3 | 0.05857113 | -2.008988783 | 0.025 | 0.076 | 1 | 0.323852509 | 3.111031312 | 0.1 | 0.04 | 1 | 0.323852509 | 0.113711683 |
| 1500004A13Rik | 0.05857133 | -1.655351829 | 0.025 | 0.076 | 1 | 0.219555167 | -0.736965594 | 0 | 0.133 | 1 | 0.219555167 | 0.113712059 |
| Pitpnb | 0.058755778 | -0.222392421 | 0.739 | 0.782 | 1 | 0.16794719 | -0.523451324 | 0.7 | 0.827 | 1 | 0.16794719 | 0.114059315 |
| Zdhhc18 | 0.058783894 | 0.571156701 | 0.479 | 0.382 | 1 | 0.467945027 | 0.22090893 | 0.6 | 0.435 | 1 | 0.467945027 | 0.114112242 |
| Sub1 | 0.254775655 | -0.164724992 | 0.958 | 0.959 | 1 | 0.058831147 | -0.492451276 | 0.9 | 0.981 | 1 | 0.254775655 | 0.11420119 |
| Fbxl20 | 0.222736425 | -0.613750924 | 0.454 | 0.476 | 1 | 0.058946573 | -1.021606456 | 0.3 | 0.667 | 1 | 0.222736425 | 0.114418447 |
| Crip1 | 0.058953937 | 1.041383921 | 0.681 | 0.635 | 1 | 0.062464879 | -1.508772938 | 0.2 | 0.515 | 1 | 0.062464879 | 0.114432306 |
| Edem3 | 0.058995967 | -0.641400383 | 0.571 | 0.624 | 1 | 0.500182003 | -0.290417122 | 0.7 | 0.747 | 1 | 0.500182003 | 0.114511409 |
| Dcun1d4 | 0.059145116 | 0.222807048 | 0.303 | 0.194 | 1 | 0.068601002 | 1.290001453 | 0.4 | 0.185 | 1 | 0.068601002 | 0.114792088 |
| Ttc23 | 0.059223849 | -1.088767857 | 0.168 | 0.247 | 1 | 0.126579685 | 1.06796259 | 0.5 | 0.275 | 1 | 0.126579685 | 0.114940234 |
| Zfp106 | 0.059266191 | -0.243900239 | 0.899 | 0.865 | 1 | 0.085499834 | -0.698040082 | 0.7 | 0.858 | 1 | 0.085499834 | 0.1150199 |
| Slc9a5 | 0.059281554 | -1.87774425 | 0.025 | 0.076 | 1 | 0.577155236 | 1.31748219 | 0 | 0.031 | 1 | 0.577155236 | 0.115048805 |
| Arid3a | 0.059281554 | -1.572889668 | 0.025 | 0.076 | 1 | 0.219917933 | -1.026422211 | 0 | 0.133 | 1 | 0.219917933 | 0.115048805 |
| Ankrd13b | 0.059281554 | -1.572889668 | 0.025 | 0.076 | 1 | 0.413255896 | 0.988174565 | 0.2 | 0.111 | 1 | 0.413255896 | 0.115048805 |
| Gm48045 | 0.059285152 | 1.684498174 | 0.059 | 0.018 | 1 | 0.643850855 | 2.017921908 | 0 | 0.022 | 1 | 0.643850855 | 0.115055575 |
| Gm9922 | 0.059285152 | 1.684498174 | 0.059 | 0.018 | 1 | 0.112266525 | 3.017921908 | 0.1 | 0.022 | 1 | 0.112266525 | 0.115055575 |
| Il11 | 0.059377701 | -2.515174171 | 0.042 | 0.1 | 1 | 0.987182949 | 0.236562194 | 0.1 | 0.099 | 1 | 0.987182949 | 0.115229691 |
| Slc6a6 | 0.059379968 | 0.314864702 | 0.983 | 0.912 | 1 | 0.706682285 | 1.130396637 | 0.9 | 0.923 | 1 | 0.706682285 | 0.115233956 |
| Cyth3 | 0.08185309 | -0.369403031 | 0.908 | 0.876 | 1 | 0.059418003 | -0.52851866 | 0.5 | 0.877 | 1 | 0.08185309 | 0.115305508 |
| Rfc2 | 0.059487412 | -0.959358016 | 0.202 | 0.288 | 1 | 0.455057592 | 1.167065347 | 0.3 | 0.228 | 1 | 0.455057592 | 0.115436071 |
| Nprl2 | 0.05959668 | -1.117695043 | 0.076 | 0.147 | 1 | 0.907531249 | 0.660369903 | 0.1 | 0.111 | 1 | 0.907531249 | 0.115641596 |
| Cubn | 0.420800329 | -0.439623138 | 0.185 | 0.141 | 1 | 0.059622382 | 1.362570079 | 0.3 | 0.105 | 1 | 0.420800329 | 0.115689935 |
| Nop16 | 0.059656196 | -0.645891499 | 0.345 | 0.435 | 1 | 0.400696817 | -0.210896782 | 0.2 | 0.346 | 1 | 0.400696817 | 0.115753531 |
| Med30 | 0.059664202 | 0.477578965 | 0.454 | 0.347 | 1 | 0.141679595 | 0.984498906 | 0.5 | 0.299 | 1 | 0.141679595 | 0.115768588 |
| Chst14 | 0.059796227 | -0.900464326 | 0.084 | 0.159 | 1 | 0.231844057 | -0.473931188 | 0 | 0.127 | 1 | 0.231844057 | 0.116016864 |
| Gm12279 | 0.059996363 | -1.655351829 | 0.025 | 0.076 | 1 | 0.068450024 | 1.930459067 | 0.2 | 0.059 | 1 | 0.068450024 | 0.116393162 |
| 4930444P10Rik | 0.06000657 | -2.485426827 | 0 | 0.029 | 1 | 0.643835681 | 1.847996907 | 0 | 0.022 | 1 | 0.643835681 | 0.116412351 |
| Fam78b | 0.06000657 | -2.292781749 | 0 | 0.029 | 1 | 0.597832556 | 0.769994395 | 0 | 0.028 | 1 | 0.597832556 | 0.116412351 |
| Depdc7 | 0.06000657 | -2.655351829 | 0 | 0.029 | 1 | 0.362872659 | 1.558490289 | 0.1 | 0.04 | 1 | 0.362872659 | 0.116412351 |
| Gm19391 | 0.06000657 | -2.292781749 | 0 | 0.029 | 1 | 0.698529281 | 2.210566986 | 0 | 0.015 | 1 | 0.698529281 | 0.116412351 |
| Vcam1 | 0.06000657 | -2.292781749 | 0 | 0.029 | 1 | 0.669848329 | 1.111031312 | 0 | 0.019 | 1 | 0.669848329 | 0.116412351 |
| Ptch2 | 0.06000657 | -2.292781749 | 0 | 0.029 | 1 | 0.64384652 | 1.111031312 | 0 | 0.022 | 1 | 0.64384652 | 0.116412351 |
| Gm26779 | 0.06000657 | -2.292781749 | 0 | 0.029 | 1 | 0.6199755 | 1.847996907 | 0 | 0.025 | 1 | 0.6199755 | 0.116412351 |
| Gm6598 | 0.06000657 | -2.655351829 | 0 | 0.029 | 1 | 0.73075158 | 2.695993813 | 0 | 0.012 | 1 | 0.73075158 | 0.116412351 |
| Gm29443 | 0.06000657 | -2.655351829 | 0 | 0.029 | 1 | 0.669848329 | 1.432959407 | 0 | 0.019 | 1 | 0.669848329 | 0.116412351 |
| Lmo1 | 0.06000657 | -2.292781749 | 0 | 0.029 | 1 | 0.408673423 | 0.374065718 | 0 | 0.065 | 1 | 0.408673423 | 0.116412351 |
| Pgbd5 | 0.06000657 | -2.485426827 | 0 | 0.029 | 1 | 0.669843604 | 2.017921908 | 0 | 0.019 | 1 | 0.669843604 | 0.116412351 |
| BC050972 | 0.06000657 | -2.292781749 | 0 | 0.029 | 1 | 0.73075158 | 2.695993813 | 0 | 0.012 | 1 | 0.73075158 | 0.116412351 |
| Cacna2d2 | 0.06000657 | -2.292781749 | 0 | 0.029 | 1 | 0.577142568 | 1.432959407 | 0 | 0.031 | 1 | 0.577142568 | 0.116412351 |
| Gm12204 | 0.06000657 | -2.292781749 | 0 | 0.029 | 1 | 0.59781419 | 1.558490289 | 0 | 0.028 | 1 | 0.59781419 | 0.116412351 |
| Abhd15 | 0.06000657 | -3.070389328 | 0 | 0.029 | 1 | 0.684686471 | 0.930459067 | 0.1 | 0.065 | 1 | 0.684686471 | 0.116412351 |
| Gm15893 | 0.06000657 | -2.292781749 | 0 | 0.029 | 1 | 0.698529281 | 2.210566986 | 0 | 0.015 | 1 | 0.698529281 | 0.116412351 |
| Macc1 | 0.06000657 | -2.944858446 | 0 | 0.029 | 1 | 0.643835681 | 1.847996907 | 0 | 0.022 | 1 | 0.643835681 | 0.116412351 |
| Scgn | 0.06000657 | -2.292781749 | 0 | 0.029 | 1 | 0.201133726 | 1.625604485 | 0.1 | 0.028 | 1 | 0.201133726 | 0.116412351 |
| Gm15564 | 0.06000657 | -2.292781749 | 0 | 0.029 | 1 | 0.557756275 | 1.017921908 | 0 | 0.034 | 1 | 0.557756275 | 0.116412351 |
| Tmprss7 | 0.06000657 | -2.485426827 | 0 | 0.029 | 1 | 0.557771538 | 0.625604485 | 0 | 0.034 | 1 | 0.557771538 | 0.116412351 |
| Rsph1 | 0.06000657 | -2.655351829 | 0 | 0.029 | 1 | 0.48995314 | 0.494359952 | 0 | 0.046 | 1 | 0.48995314 | 0.116412351 |
| Pcdhb2 | 0.06000657 | -2.292781749 | 0 | 0.029 | 1 | 0.522066388 | 1.017921908 | 0 | 0.04 | 1 | 0.522066388 | 0.116412351 |
| Gm50462 | 0.06000657 | -2.292781749 | 0 | 0.029 | 1 | 0.669843604 | 2.017921908 | 0 | 0.019 | 1 | 0.669843604 | 0.116412351 |
| Cntf | 0.06000657 | -2.292781749 | 0 | 0.029 | 1 | 0.730749233 | 2.432959407 | 0 | 0.012 | 1 | 0.730749233 | 0.116412351 |
| Nlgn3 | 0.06000657 | -2.292781749 | 0 | 0.029 | 1 | 0.713245375 | 1.017921908 | 0.1 | 0.068 | 1 | 0.713245375 | 0.116412351 |
| Gm16083 | 0.060010312 | -2.944858446 | 0 | 0.029 | 1 | 0.118051503 | 2.111031312 | 0.1 | 0.022 | 1 | 0.118051503 | 0.116419386 |
| Gm26930 | 0.060010312 | -2.485426827 | 0 | 0.029 | 1 | 0.730750993 | 1.695993813 | 0 | 0.012 | 1 | 0.730750993 | 0.116419386 |
| Gm9530 | 0.060010312 | -2.485426827 | 0 | 0.029 | 1 | 0.643835681 | 1.558490289 | 0 | 0.022 | 1 | 0.643835681 | 0.116419386 |
| Rem1 | 0.060010312 | -2.485426827 | 0 | 0.029 | 1 | 0.698529281 | 2.210566986 | 0 | 0.015 | 1 | 0.698529281 | 0.116419386 |
| Gm37459 | 0.060010312 | -3.070389328 | 0 | 0.029 | 1 | 0.643842184 | 1.695993813 | 0 | 0.022 | 1 | 0.643842184 | 0.116419386 |
| Pcdh10 | 0.060010312 | -2.485426827 | 0 | 0.029 | 1 | 0.489955583 | 0.159940913 | 0 | 0.046 | 1 | 0.489955583 | 0.116419386 |
| Gja4 | 0.060010312 | -2.485426827 | 0 | 0.029 | 1 | 0.643842184 | 1.432959407 | 0 | 0.022 | 1 | 0.643842184 | 0.116419386 |
| Hsd17b13 | 0.060010312 | -2.944858446 | 0 | 0.029 | 1 | 0.730749233 | 2.432959407 | 0 | 0.012 | 1 | 0.730749233 | 0.116419386 |
| Gm10399 | 0.060010312 | -2.485426827 | 0 | 0.029 | 1 | 0.080320981 | 2.558490289 | 0.1 | 0.019 | 1 | 0.080320981 | 0.116419386 |
| A930016O22Rik | 0.060010312 | -2.485426827 | 0 | 0.029 | 1 | 0.577155236 | 1.31748219 | 0 | 0.031 | 1 | 0.577155236 | 0.116419386 |
| Gm44837 | 0.060010312 | -2.485426827 | 0 | 0.029 | 1 | 0.669848329 | 1.847996907 | 0 | 0.019 | 1 | 0.669848329 | 0.116419386 |

| | | | | | | | | | | | | |
|---|---|---|---|---|---|---|---|---|---|---|---|---|
| Muc5ac | 0.060010312 | -5.807354922 | 0 | 0.029 | 1 | 0.557756607 | 0.847996907 | 0 | 0.034 | 1 | 0.557756607 | 0.116419386 |
| Gm27201 | 0.060010312 | -2.485426827 | 0 | 0.029 | 1 | 0.73075158 | 2.695993813 | 0 | 0.012 | 1 | 0.73075158 | 0.116419386 |
| Exph5 | 0.060010312 | -2.655351829 | 0 | 0.029 | 1 | 0.343828427 | -0.191531458 | 0 | 0.083 | 1 | 0.343828427 | 0.116419386 |
| Ephb1 | 0.060010312 | -3.655351829 | 0 | 0.029 | 1 | 0.505682638 | -1.069540933 | 0 | 0.043 | 1 | 0.505682638 | 0.116419386 |
| Ano4 | 0.060010312 | -2.485426827 | 0 | 0.029 | 1 | 0.597832556 | 1.31748219 | 0 | 0.028 | 1 | 0.597832556 | 0.116419386 |
| Wnt10b | 0.060010312 | -2.655351829 | 0 | 0.029 | 1 | 0.73075158 | 2.695993813 | 0 | 0.012 | 1 | 0.73075158 | 0.116419386 |
| Ccdc116 | 0.060010312 | -2.655351829 | 0 | 0.029 | 1 | 0.577142568 | 1.432959407 | 0 | 0.031 | 1 | 0.577142568 | 0.116419386 |
| 2810407A14Rik | 0.060010312 | -2.485426827 | 0 | 0.029 | 1 | 0.669850692 | 1.111031312 | 0 | 0.019 | 1 | 0.669850692 | 0.116419386 |
| Hs3st6 | 0.060010312 | -2.655351829 | 0 | 0.029 | 1 | 0.577177049 | 0.432959407 | 0 | 0.031 | 1 | 0.577177049 | 0.116419386 |
| Pik3ap1 | 0.060010312 | -2.655351829 | 0 | 0.029 | 1 | 0.5978438 | -0.152003093 | 0 | 0.028 | 1 | 0.5978438 | 0.116419386 |
| Tsnaxip1 | 0.060011559 | -2.944858446 | 0 | 0.029 | 1 | 0.698529281 | 1.847996907 | 0 | 0.015 | 1 | 0.698529281 | 0.116421731 |
| Gm29677 | 0.060011559 | -2.807354922 | 0 | 0.029 | 1 | 0.557782156 | -0.441509711 | 0 | 0.034 | 1 | 0.557782156 | 0.116421731 |
| Gm28055 | 0.060012183 | -2.655351829 | 0 | 0.029 | 1 | 0.278845643 | 2.017921908 | 0.1 | 0.034 | 1 | 0.278845643 | 0.116422904 |
| Itpripl1 | 0.060012807 | -2.070389328 | 0 | 0.029 | 1 | 0.619961444 | 1.432959407 | 0 | 0.025 | 1 | 0.619961444 | 0.116424076 |
| Gm43768 | 0.060012807 | -2.070389328 | 0 | 0.029 | 1 | 0.193279696 | 2.31748219 | 0.1 | 0.028 | 1 | 0.193279696 | 0.116424076 |
| Cyb561d1 | 0.060012807 | -2.070389328 | 0 | 0.029 | 1 | 0.73075158 | 2.695993813 | 0 | 0.012 | 1 | 0.73075158 | 0.116424076 |
| Selplg | 0.060012807 | -2.070389328 | 0 | 0.029 | 1 | 0.616106286 | 0.930459067 | 0.1 | 0.059 | 1 | 0.616106286 | 0.116424076 |
| Tmem147os | 0.060012807 | -2.070389328 | 0 | 0.029 | 1 | 0.643842184 | 1.558490289 | 0 | 0.022 | 1 | 0.643842184 | 0.116424076 |
| Gdpd3 | 0.060012807 | -2.070389328 | 0 | 0.029 | 1 | 0.112700281 | 2.263034406 | 0.2 | 0.071 | 1 | 0.112700281 | 0.116424076 |
| Pbx4 | 0.060012807 | -2.070389328 | 0 | 0.029 | 1 | 0.398985336 | 2.017921908 | 0.1 | 0.043 | 1 | 0.398985336 | 0.116424076 |
| Unc13a | 0.060012807 | -2.070389328 | 0 | 0.029 | 1 | 0.64384652 | 1.558490289 | 0 | 0.022 | 1 | 0.64384652 | 0.116424076 |
| Gm12111 | 0.060012807 | -2.070389328 | 0 | 0.029 | 1 | 0.643850855 | 2.017921908 | 0 | 0.022 | 1 | 0.643850855 | 0.116424076 |
| N4bp3 | 0.060012807 | -2.070389328 | 0 | 0.029 | 1 | 0.505686333 | 1.111031312 | 0 | 0.043 | 1 | 0.505686333 | 0.116424076 |
| Med9os | 0.060012807 | -2.070389328 | 0 | 0.029 | 1 | 0.643850855 | 2.017921908 | 0 | 0.022 | 1 | 0.643850855 | 0.116424076 |
| Doc2b | 0.060012807 | -2.070389328 | 0 | 0.029 | 1 | 0.64384652 | 1.210566986 | 0 | 0.022 | 1 | 0.64384652 | 0.116424076 |
| 2900041M22Rik | 0.060012807 | -2.070389328 | 0 | 0.029 | 1 | 0.669848329 | 1.847996907 | 0 | 0.019 | 1 | 0.669848329 | 0.116424076 |
| Gpx6 | 0.060012807 | -2.070389328 | 0 | 0.029 | 1 | 0.597824685 | 1.432959407 | 0 | 0.028 | 1 | 0.597824685 | 0.116424076 |
| Gm2237 | 0.060012807 | -2.070389328 | 0 | 0.029 | 1 | 0.698532411 | 2.017921908 | 0 | 0.015 | 1 | 0.698532411 | 0.116424076 |
| Aard | 0.060012807 | -2.070389328 | 0 | 0.029 | 1 | 0.52208609 | 1.017921908 | 0 | 0.04 | 1 | 0.52208609 | 0.116424076 |
| Jrk | 0.060012807 | -2.070389328 | 0 | 0.029 | 1 | 0.505686333 | 1.111031312 | 0 | 0.043 | 1 | 0.505686333 | 0.116424076 |
| Gm39556 | 0.060012807 | -2.070389328 | 0 | 0.029 | 1 | 0.730750993 | 1.558490289 | 0 | 0.012 | 1 | 0.730750993 | 0.116424076 |
| Pdxp | 0.060012807 | -2.070389328 | 0 | 0.029 | 1 | 0.73075158 | 2.695993813 | 0 | 0.012 | 1 | 0.73075158 | 0.116424076 |
| Spink5 | 0.060012807 | -2.070389328 | 0 | 0.029 | 1 | 0.619972287 | 0.930459067 | 0 | 0.025 | 1 | 0.619972287 | 0.116424076 |
| Rps6ka4 | 0.060012807 | -2.070389328 | 0 | 0.029 | 1 | 0.557784478 | 1.432959407 | 0 | 0.034 | 1 | 0.557784478 | 0.116424076 |
| Best1 | 0.060012807 | -2.070389328 | 0 | 0.029 | 1 | 0.505605027 | 0.769994395 | 0 | 0.043 | 1 | 0.505605027 | 0.116424076 |
| Ptpn4 | 0.060042229 | -0.991962681 | 0.294 | 0.376 | 1 | 0.749957338 | 0.287909074 | 0.5 | 0.429 | 1 | 0.749957338 | 0.116479389 |
| Ciao2a | 0.06015167 | -0.529820947 | 0.319 | 0.418 | 1 | 0.342894466 | 0.558490289 | 0.6 | 0.417 | 1 | 0.342894466 | 0.116685118 |
| Ikbkb | 0.060232077 | -0.76106127 | 0.235 | 0.324 | 1 | 0.176132549 | -1.138582578 | 0.2 | 0.389 | 1 | 0.176132549 | 0.116836251 |
| Anxa9 | 0.060242193 | 1.514573173 | 0.059 | 0.018 | 1 | 0.59781419 | 1.432959407 | 0 | 0.028 | 1 | 0.59781419 | 0.116855265 |
| Zbtb6 | 0.060252738 | -0.765534746 | 0.143 | 0.229 | 1 | 0.275441776 | 0.847996907 | 0.4 | 0.244 | 1 | 0.275441776 | 0.116875084 |
| Senp3 | 0.060410885 | -0.617877123 | 0.361 | 0.435 | 1 | 0.880895526 | -0.208146171 | 0.6 | 0.509 | 1 | 0.880895526 | 0.117172295 |
| Map2 | 0.060430487 | 0.262789902 | 0.613 | 0.471 | 1 | 0.785714899 | -0.208757872 | 0.3 | 0.37 | 1 | 0.785714899 | 0.11720913 |
| Serinc1 | 0.060438483 | -0.245508671 | 0.924 | 0.912 | 1 | 0.527448937 | -0.182642366 | 1 | 0.96 | 1 | 0.527448937 | 0.117224156 |
| Tbx2 | 0.060451033 | -1.428843299 | 0.076 | 0.147 | 1 | 0.460671035 | -0.339630097 | 0 | 0.052 | 1 | 0.460671035 | 0.117247738 |
| Plscr4 | 0.430516815 | -0.437598302 | 0.244 | 0.276 | 1 | 0.060527224 | -2.589408406 | 0 | 0.269 | 1 | 0.430516815 | 0.117390904 |
| 0610040B10Rik | 0.060546302 | -1.555816155 | 0.025 | 0.076 | 1 | 0.619953009 | 1.695993813 | 0 | 0.025 | 1 | 0.619953009 | 0.11742675 |
| Smim30 | 0.060551168 | -0.468553009 | 0.487 | 0.547 | 1 | 0.125088852 | -0.959358016 | 0.3 | 0.503 | 1 | 0.125088852 | 0.117435892 |
| Gpc2 | 0.06071555 | -1.392317423 | 0.025 | 0.076 | 1 | 0.669843604 | 2.017921908 | 0 | 0.019 | 1 | 0.669843604 | 0.117744721 |
| Gm56496 | 0.06071555 | -1.392317423 | 0.025 | 0.076 | 1 | 0.670447518 | 1.374065718 | 0.1 | 0.065 | 1 | 0.670447518 | 0.117744721 |
| Pcdhga11 | 0.06071555 | -1.392317423 | 0.025 | 0.076 | 1 | 0.52208609 | 1.017921908 | 0 | 0.04 | 1 | 0.52208609 | 0.117744721 |
| Cstf1 | 0.060726986 | -0.887525271 | 0.109 | 0.188 | 1 | 0.21908128 | -0.505640048 | 0 | 0.133 | 1 | 0.21908128 | 0.117766205 |
| Hsp90b1 | 0.107434942 | -0.174450358 | 1 | 0.982 | 1 | 0.060844801 | -0.614256114 | 1 | 0.994 | 1 | 0.107434942 | 0.117987511 |
| Tfb2m | 0.0608721 | -0.613750924 | 0.303 | 0.394 | 1 | 0.461700497 | -0.604129911 | 0.3 | 0.37 | 1 | 0.461700497 | 0.118038787 |
| Ezh1 | 0.06100028 | -0.568842835 | 0.261 | 0.353 | 1 | 0.375474519 | 0.518076021 | 0.5 | 0.352 | 1 | 0.375474519 | 0.118279526 |
| Efcab8 | 0.061011068 | -1.240314329 | 0.101 | 0.176 | 1 | 0.564091766 | -0.814968106 | 0.1 | 0.164 | 1 | 0.564091766 | 0.118299786 |
| Mlf1 | 0.061025415 | -0.437171659 | 0.319 | 0.2 | 1 | 0.296820298 | 1.31748219 | 0.3 | 0.194 | 1 | 0.296820298 | 0.118326729 |
| Helq | 0.06104448 | -1.544320516 | 0.025 | 0.076 | 1 | 0.274653871 | -0.68251781 | 0 | 0.108 | 1 | 0.274653871 | 0.118362531 |
| Riox2 | 0.061066497 | -1.439623138 | 0.059 | 0.124 | 1 | 0.632378953 | 1.494359952 | 0.1 | 0.062 | 1 | 0.632378953 | 0.118403877 |
| Pced1a | 0.06112219 | -1.117695043 | 0.076 | 0.147 | 1 | 0.252463474 | -0.536666944 | 0 | 0.117 | 1 | 0.252463474 | 0.118508458 |
| Tmem250-ps | 0.061186099 | -1.008988783 | 0.126 | 0.206 | 1 | 0.939142315 | 0.453137289 | 0.2 | 0.185 | 1 | 0.939142315 | 0.11862846 |
| Lcn3 | 0.061210102 | -1.12588444 | 0.109 | 0.188 | 1 | 0.577172123 | -0.888968688 | 0 | 0.031 | 1 | 0.577172123 | 0.118673527 |
| Naf1 | 0.061271062 | -0.648925559 | 0.345 | 0.424 | 1 | 0.141946668 | -1.158666824 | 0.2 | 0.407 | 1 | 0.141946668 | 0.118787981 |
| Wdr36 | 0.123103388 | -0.634804451 | 0.37 | 0.418 | 1 | 0.061315848 | -1.808626579 | 0.1 | 0.383 | 1 | 0.123103388 | 0.118872064 |
| Ccnt1 | 0.200276888 | -0.258574758 | 0.765 | 0.776 | 1 | 0.061419245 | -0.759685671 | 0.8 | 0.873 | 1 | 0.200276888 | 0.119066167 |
| Get3 | 0.061419824 | -0.502914254 | 0.277 | 0.382 | 1 | 0.2401333 | -0.797995028 | 0.2 | 0.361 | 1 | 0.2401333 | 0.119067253 |
| Gm47507 | 0.061446043 | -3.529820947 | 0.008 | 0.047 | 1 | 0.730750993 | 1.111031312 | 0 | 0.012 | 1 | 0.730750993 | 0.11911647 |
| 2500002B13Rik | 0.061459122 | 1.362570079 | 0.059 | 0.018 | 1 | 0.474904369 | 0.769994395 | 0 | 0.049 | 1 | 0.474904369 | 0.11914102 |
| Rfwd3 | 0.829628725 | -0.167250867 | 0.252 | 0.259 | 1 | 0.061460394 | -1.689437224 | 0.1 | 0.386 | 1 | 0.829628725 | 0.119143408 |
| Stmn1 | 0.061465844 | -1.410019425 | 0.118 | 0.194 | 1 | 0.390546309 | -0.596787936 | 0.1 | 0.207 | 1 | 0.390546309 | 0.119153638 |
| Slc39a10 | 0.061490614 | -0.395789615 | 0.303 | 0.382 | 1 | 0.912759247 | -0.125036046 | 0.5 | 0.469 | 1 | 0.912759247 | 0.119200133 |
| Mrnip | 0.115259334 | 0.877143252 | 0.109 | 0.059 | 1 | 0.061516469 | 1.063725598 | 0.3 | 0.102 | 1 | 0.115259334 | 0.119248663 |
| Def6 | 0.061741475 | -0.900464326 | 0.067 | 0.135 | 1 | 0.938517603 | 0.625604485 | 0.1 | 0.09 | 1 | 0.938517603 | 0.11967094 |
| Bend7 | 0.061771615 | -2.485426827 | 0.008 | 0.047 | 1 | 0.433723566 | -0.191531458 | 0 | 0.059 | 1 | 0.433723566 | 0.119727498 |
| Fgd3 | 0.061774832 | -3.070389328 | 0.008 | 0.047 | 1 | 0.597837803 | 0.695993813 | 0 | 0.028 | 1 | 0.597837803 | 0.119733534 |
| Gm29394 | 0.061784696 | -1.333423734 | 0.025 | 0.076 | 1 | 0.577155236 | 1.210566986 | 0 | 0.031 | 1 | 0.577155236 | 0.119752043 |
| 4930417O13Rik | 0.061792708 | 1.707218251 | 0.059 | 0.018 | 1 | 0.730749233 | 1.847996907 | 0 | 0.012 | 1 | 0.730749233 | 0.119767077 |
| Plekhj1 | 0.061913914 | 0.305986551 | 0.672 | 0.565 | 1 | 0.613460812 | 0.582535763 | 0.5 | 0.497 | 1 | 0.613460812 | 0.11999495 |
| Cybrd1 | 0.061948858 | -0.883976204 | 0.134 | 0.218 | 1 | 0.110141944 | 0.894539492 | 0.5 | 0.253 | 1 | 0.110141944 | 0.120060056 |
| Hdac11 | 0.094752821 | -1.807354922 | 0.008 | 0.041 | 1 | 0.062050246 | 2.558490289 | 0.2 | 0.059 | 1 | 0.094752821 | 0.120250259 |
| Agk | 0.06209994 | -0.723586564 | 0.218 | 0.306 | 1 | 0.121091381 | 0.614199722 | 0.5 | 0.235 | 1 | 0.121091381 | 0.120343477 |
| Trim69 | 0.062102212 | -2.292781749 | 0.008 | 0.047 | 1 | 0.619967468 | 1.210566986 | 0 | 0.025 | 1 | 0.619967468 | 0.120347739 |
| Fgf13 | 0.062103288 | -2.807354922 | 0.008 | 0.047 | 1 | 0.460675018 | -0.763437806 | 0 | 0.052 | 1 | 0.460675018 | 0.120349757 |
| Lct | 0.062103646 | -2.185866545 | 0.008 | 0.047 | 1 | 0.15487326 | 2.111031312 | 0.1 | 0.025 | 1 | 0.15487326 | 0.12035043 |
| Rpl41 | 0.06216516 | -0.182462136 | 0.975 | 0.994 | 1 | 0.176561881 | -0.397115591 | 1 | 0.991 | 1 | 0.176561881 | 0.120465812 |
| Hdhd5 | 0.062175299 | -1.622930351 | 0.034 | 0.088 | 1 | 0.248571385 | 1.473601392 | 0.2 | 0.09 | 1 | 0.248571385 | 0.120484831 |
| Wbp4 | 0.107906186 | -0.401362562 | 0.639 | 0.694 | 1 | 0.062255085 | -0.741411499 | 0.5 | 0.793 | 1 | 0.107906186 | 0.120634474 |
| Atp2a2 | 0.197748766 | -0.182073489 | 0.992 | 0.982 | 1 | 0.062290701 | -0.613433498 | 1 | 0.997 | 1 | 0.197748766 | 0.12070127 |
| Hdx | 0.062308079 | -1.678071905 | 0.025 | 0.076 | 1 | 0.522139811 | 0.374065718 | 0 | 0.04 | 1 | 0.522139811 | 0.120733861 |
| Vdac3 | 0.06231006 | -0.380729449 | 0.681 | 0.729 | 1 | 0.12227346 | -0.636388639 | 0.6 | 0.812 | 1 | 0.12227346 | 0.120737576 |
| Cpq | 0.062318675 | 0.149576356 | 0.866 | 0.665 | 1 | 0.193578984 | -0.734640541 | 0.8 | 0.802 | 1 | 0.193578984 | 0.120753734 |
| D230022J07Rik | 0.062394054 | 0.929610672 | 0.059 | 0.018 | 1 | 0.597824685 | 1.210566986 | 0 | 0.028 | 1 | 0.597824685 | 0.12089509 |
| Tulp2 | 0.062431398 | -2.185866545 | 0.008 | 0.047 | 1 | 0.539431168 | 1.111031312 | 0 | 0.037 | 1 | 0.539431168 | 0.120965116 |
| Stk33 | 0.062431757 | -3.185866545 | 0.008 | 0.047 | 1 | 0.698532411 | 0.558490289 | 0 | 0.015 | 1 | 0.698532411 | 0.12096579 |
| Pomc | 0.062431757 | -2.070389328 | 0.008 | 0.047 | 1 | 0.597845674 | 1.695993813 | 0 | 0.028 | 1 | 0.597845674 | 0.12096579 |
| Gsdmd | 0.062431757 | -2.070389328 | 0.008 | 0.047 | 1 | 0.298340756 | -0.111361109 | 0 | 0.099 | 1 | 0.298340756 | 0.12096579 |
| Hltf | 0.062438746 | -0.754887502 | 0.336 | 0.412 | 1 | 0.563970796 | 0.485426827 | 0.4 | 0.324 | 1 | 0.563970796 | 0.120978894 |
| Thsd7a | 0.062475803 | -1.632268215 | 0.034 | 0.088 | 1 | 0.136680051 | -3.0853659 | 0 | 0.185 | 1 | 0.136680051 | 0.12104838 |
| Sertad4 | 0.062519906 | -0.748461233 | 0.277 | 0.371 | 1 | 0.616594055 | -0.171902651 | 0.4 | 0.454 | 1 | 0.616594055 | 0.121131074 |

| | | | | | | | | | | | | |
|---|---|---|---|---|---|---|---|---|---|---|---|---|
| Fbxw8 | 0.062533536 | 0.130232471 | 0.807 | 0.647 | 1 | 0.823190431 | -0.113935053 | 0.7 | 0.701 | 1 | 0.823190431 | 0.121156629 |
| 3110056K07Rik | 0.656151114 | -0.432959407 | 0.286 | 0.288 | 1 | 0.062536634 | -1.625934282 | 0.1 | 0.386 | 1 | 0.656151114 | 0.121162437 |
| Acrbp | 0.062541076 | -1.655351829 | 0.042 | 0.1 | 1 | 0.439498406 | 0.959028219 | 0.2 | 0.114 | 1 | 0.439498406 | 0.121170765 |
| Gm12359 | 0.062541693 | -1.424026283 | 0.042 | 0.1 | 1 | 0.342250455 | 1.463333056 | 0.2 | 0.108 | 1 | 0.342250455 | 0.121171923 |
| Rassf2 | 0.062554654 | -0.817270391 | 0.185 | 0.271 | 1 | 0.342253858 | -0.121629444 | 0.1 | 0.253 | 1 | 0.342253858 | 0.121196223 |
| Mmaa | 0.231501131 | -0.707819249 | 0.101 | 0.147 | 1 | 0.062561107 | 1.591657153 | 0.4 | 0.17 | 1 | 0.231501131 | 0.121208323 |
| Cdk15 | 0.062564366 | -1.544320516 | 0.025 | 0.076 | 1 | 0.698533455 | 1.695993813 | 0 | 0.015 | 1 | 0.698533455 | 0.121214431 |
| Nkiras1 | 0.062573365 | 0.633217669 | 0.261 | 0.171 | 1 | 0.092047986 | -1.505640048 | 0 | 0.225 | 1 | 0.092047986 | 0.121231304 |
| Xrcc4 | 0.062576691 | -0.67305383 | 0.311 | 0.4 | 1 | 0.401256721 | -1.101019165 | 0.2 | 0.309 | 1 | 0.401256721 | 0.121237541 |
| Prorp | 0.771438011 | -0.158616511 | 0.176 | 0.159 | 1 | 0.06259469 | 1.295455884 | 0.3 | 0.105 | 1 | 0.771438011 | 0.121271285 |
| Cyb5rl | 0.563722678 | -0.622930351 | 0.034 | 0.047 | 1 | 0.06262373 | 2.695993813 | 0.2 | 0.059 | 1 | 0.563722678 | 0.121325729 |
| Psmg1 | 0.062665692 | -0.660513534 | 0.227 | 0.324 | 1 | 0.802355431 | -0.339630097 | 0.3 | 0.306 | 1 | 0.802355431 | 0.121404395 |
| Crebzf | 0.235399608 | -0.348924827 | 0.395 | 0.447 | 1 | 0.062698862 | -1.468577954 | 0.2 | 0.475 | 1 | 0.235399608 | 0.121466576 |
| Otud3 | 0.062710206 | 0.691450935 | 0.176 | 0.1 | 1 | 0.407240741 | -0.125036046 | 0.1 | 0.225 | 1 | 0.407240741 | 0.121487841 |
| Lonrf2 | 0.062759149 | -1.944858446 | 0.008 | 0.047 | 1 | 0.59781419 | 1.432959407 | 0 | 0.028 | 1 | 0.59781419 | 0.121579588 |
| Pabpc1l | 0.062759149 | -2.070389328 | 0.008 | 0.047 | 1 | 0.713245375 | 0.888638891 | 0.1 | 0.068 | 1 | 0.713245375 | 0.121579588 |
| Rtn4ip1 | 0.062759149 | -1.944858446 | 0.008 | 0.047 | 1 | 0.295128528 | 2.515421567 | 0.1 | 0.037 | 1 | 0.295128528 | 0.121579588 |
| Rin1 | 0.062759149 | -2.070389328 | 0.008 | 0.047 | 1 | 0.669848329 | 1.558490289 | 0 | 0.019 | 1 | 0.669848329 | 0.121579588 |
| Traf6 | 0.062765713 | -0.549557165 | 0.176 | 0.265 | 1 | 0.164570057 | -0.780819884 | 0.1 | 0.33 | 1 | 0.164570057 | 0.121591891 |
| Ftl1-ps1 | 0.853240938 | -0.265645619 | 0.345 | 0.335 | 1 | 0.062785513 | -1.261301736 | 0.2 | 0.478 | 1 | 0.853240938 | 0.121629005 |
| Stx12 | 0.249689962 | 0.10325876 | 0.815 | 0.694 | 1 | 0.062902909 | -0.83507568 | 0.4 | 0.759 | 1 | 0.249689962 | 0.121849042 |
| Wnt4 | 0.062948806 | -1.600904045 | 0.034 | 0.088 | 1 | 0.100607361 | -1.827568143 | 0 | 0.216 | 1 | 0.100607361 | 0.121935061 |
| Lef1os1 | 0.063085444 | -1.807354922 | 0.008 | 0.047 | 1 | 0.539483006 | 1.31748219 | 0 | 0.037 | 1 | 0.539483006 | 0.122191115 |
| Napepld | 0.063085444 | -1.807354922 | 0.008 | 0.047 | 1 | 0.643842184 | 1.695993813 | 0 | 0.022 | 1 | 0.643842184 | 0.122191115 |
| Gm43378 | 0.063085444 | -1.807354922 | 0.008 | 0.047 | 1 | 0.597824685 | 1.31748219 | 0 | 0.028 | 1 | 0.597824685 | 0.122191115 |
| Il12rb2 | 0.063085444 | -1.944858446 | 0.008 | 0.047 | 1 | 0.489923006 | 0.558490289 | 0 | 0.046 | 1 | 0.489923006 | 0.122191115 |
| Shkbp1 | 0.063085444 | -1.807354922 | 0.008 | 0.047 | 1 | 0.505623509 | 0.769994395 | 0 | 0.043 | 1 | 0.505623509 | 0.122191115 |
| Gm20457 | 0.063085444 | -1.807354922 | 0.008 | 0.047 | 1 | 0.153152701 | 2.558490289 | 0.1 | 0.025 | 1 | 0.153152701 | 0.122191115 |
| Nat2 | 0.063085444 | -1.807354922 | 0.008 | 0.047 | 1 | 0.643842184 | 1.695993813 | 0 | 0.022 | 1 | 0.643842184 | 0.122191115 |
| Pih1d2 | 0.063085444 | -1.807354922 | 0.008 | 0.047 | 1 | 0.539413885 | 1.210566986 | 0 | 0.037 | 1 | 0.539413885 | 0.122191115 |
| Shpk | 0.063085444 | -1.807354922 | 0.008 | 0.047 | 1 | 0.597824685 | 1.432959407 | 0 | 0.028 | 1 | 0.597824685 | 0.122191115 |
| Rapgefl1 | 0.063085444 | -1.807354922 | 0.008 | 0.047 | 1 | 0.317764606 | 1.14345279 | 0.2 | 0.099 | 1 | 0.317764606 | 0.122191115 |
| Zfp229 | 0.063085444 | -1.807354922 | 0.008 | 0.047 | 1 | 0.640912917 | 1.31748219 | 0.1 | 0.062 | 1 | 0.640912917 | 0.122191115 |
| Spdef | 0.063085444 | -2.070389328 | 0.008 | 0.047 | 1 | 0.490317433 | 1.432959407 | 0.1 | 0.049 | 1 | 0.490317433 | 0.122191115 |
| Gm20508 | 0.063085444 | -1.807354922 | 0.008 | 0.047 | 1 | 0.577164735 | 1.111031312 | 0 | 0.031 | 1 | 0.577164735 | 0.122191115 |
| Gm16973 | 0.063094072 | -1.555816155 | 0.034 | 0.088 | 1 | 0.843785996 | 0.769994395 | 0.1 | 0.08 | 1 | 0.843785996 | 0.122207282 |
| 3010001F23Rik | 0.063094072 | -1.411426246 | 0.034 | 0.088 | 1 | 0.239002451 | -0.864721141 | 0 | 0.123 | 1 | 0.239002451 | 0.122207282 |
| Samm50 | 0.06319966 | 0.324748614 | 0.445 | 0.312 | 1 | 0.463309474 | 0.688001022 | 0.5 | 0.37 | 1 | 0.463309474 | 0.122405123 |
| Atrn | 0.063201654 | -0.576125137 | 0.588 | 0.635 | 1 | 0.193024738 | 0.672661309 | 0.9 | 0.852 | 1 | 0.193024738 | 0.122408859 |
| Homer3 | 0.063217873 | -0.665999073 | 0.176 | 0.271 | 1 | 0.492892434 | -0.573039333 | 0.3 | 0.395 | 1 | 0.492892434 | 0.122439246 |
| Tpd52l2 | 0.097496172 | 0.260549142 | 0.866 | 0.829 | 1 | 0.063294596 | 0.677657754 | 0.9 | 0.864 | 1 | 0.097496172 | 0.122582985 |
| Zscan12 | 0.063369665 | -1.021479727 | 0.067 | 0.135 | 1 | 0.298575525 | -0.267480311 | 0 | 0.099 | 1 | 0.298575525 | 0.122723615 |
| Msto1 | 0.063392764 | -1.173482821 | 0.067 | 0.135 | 1 | 0.838483509 | 0.463333056 | 0.1 | 0.12 | 1 | 0.838483509 | 0.122766886 |
| Dus1l | 0.063392764 | -1.070389328 | 0.067 | 0.135 | 1 | 0.810418519 | 0.930459067 | 0.1 | 0.077 | 1 | 0.810418519 | 0.122766886 |
| Ddx23 | 0.265470228 | -0.423142549 | 0.445 | 0.471 | 1 | 0.063393505 | -1.348400306 | 0.3 | 0.543 | 1 | 0.265470228 | 0.122768273 |
| Rab8a | 0.193256318 | -0.313474144 | 0.513 | 0.553 | 1 | 0.063408541 | -0.638247543 | 0.3 | 0.627 | 1 | 0.193256318 | 0.12279644 |
| 1700048O20Rik | 0.083686583 | -1.687060688 | 0.034 | 0.082 | 1 | 0.063410446 | 2.017921908 | 0.2 | 0.056 | 1 | 0.083686583 | 0.122800007 |
| Anxa11os | 0.929389703 | 0.192645078 | 0.025 | 0.024 | 1 | 0.063410824 | 1.902444691 | 0.2 | 0.056 | 1 | 0.929389703 | 0.122800715 |
| Gm11906 | 0.063442928 | -1.610957709 | 0.034 | 0.088 | 1 | 0.364080008 | -0.230005605 | 0 | 0.077 | 1 | 0.364080008 | 0.12286085 |
| Amigo1 | 0.063453834 | -1.655351829 | 0.008 | 0.047 | 1 | 0.522151747 | 1.210566986 | 0 | 0.04 | 1 | 0.522151747 | 0.12288128 |
| Gm36640 | 0.063453834 | -1.655351829 | 0.008 | 0.047 | 1 | 0.408856811 | -0.864721141 | 0 | 0.065 | 1 | 0.408856811 | 0.12288128 |
| Tmem91 | 0.063453834 | -1.655351829 | 0.008 | 0.047 | 1 | 0.420904398 | 0.374065718 | 0 | 0.062 | 1 | 0.420904398 | 0.12288128 |
| Rps29 | 0.063464458 | 0.147848162 | 0.983 | 0.982 | 1 | 0.131917703 | -0.473263119 | 0.9 | 0.994 | 1 | 0.131917703 | 0.122901178 |
| Zfp626 | 0.063541841 | -1.321928095 | 0.05 | 0.112 | 1 | 0.530891827 | 0.821524695 | 0.2 | 0.127 | 1 | 0.530891827 | 0.123046116 |
| Pdk2 | 0.063541841 | -1.321928095 | 0.05 | 0.112 | 1 | 0.353587229 | 0.111031312 | 0 | 0.08 | 1 | 0.353587229 | 0.123046116 |
| Foxred2 | 0.063541841 | -1.321928095 | 0.05 | 0.112 | 1 | 0.857799108 | 0.236562194 | 0.1 | 0.117 | 1 | 0.857799108 | 0.123046116 |
| Slc38a7 | 0.063542395 | -1.263034406 | 0.05 | 0.112 | 1 | 0.654863895 | 0.625604485 | 0.2 | 0.145 | 1 | 0.654863895 | 0.123047154 |
| Rgs20 | 0.063715042 | 2.344648171 | 0.092 | 0.041 | 1 | 0.619967066 | 1.210566986 | 0 | 0.025 | 1 | 0.619967066 | 0.123370478 |
| Tgs1 | 0.063727365 | -0.645504043 | 0.504 | 0.576 | 1 | 0.273187036 | -0.609351398 | 0.4 | 0.543 | 1 | 0.273187036 | 0.123393553 |
| Gm56494 | 0.112335908 | -1.900464326 | 0.017 | 0.053 | 1 | 0.06374617 | 2.374065718 | 0.2 | 0.059 | 1 | 0.112335908 | 0.123428765 |
| Neto2 | 0.063779874 | -0.715724447 | 0.076 | 0.147 | 1 | 0.316165207 | -1.489872732 | 0 | 0.093 | 1 | 0.316165207 | 0.123491875 |
| Bphl | 0.063944732 | -0.637429921 | 0.235 | 0.329 | 1 | 0.641690248 | -0.489872732 | 0.3 | 0.333 | 1 | 0.641690248 | 0.123800536 |
| Il18bp | 0.063947154 | 1.447458977 | 0.118 | 0.059 | 1 | 0.976598201 | 0.660369903 | 0.1 | 0.096 | 1 | 0.976598201 | 0.12380507 |
| Zfp319 | 0.064020597 | -1.411426246 | 0.034 | 0.088 | 1 | 0.815153107 | 0.463333056 | 0.1 | 0.123 | 1 | 0.815153107 | 0.123942556 |
| Igfals | 0.064020597 | -1.485426827 | 0.034 | 0.088 | 1 | 0.364956373 | 1.048295557 | 0.2 | 0.105 | 1 | 0.364956373 | 0.123942556 |
| Ccdc58 | 0.190745061 | 0.318652963 | 0.336 | 0.271 | 1 | 0.06402167 | -2.121629444 | 0 | 0.262 | 1 | 0.190745061 | 0.123944566 |
| Ndufa10 | 0.690188343 | -0.143992418 | 0.605 | 0.588 | 1 | 0.064141516 | -1.022093771 | 0.3 | 0.602 | 1 | 0.690188343 | 0.124168898 |
| Bub3 | 0.064180405 | -0.561777713 | 0.471 | 0.529 | 1 | 0.126647823 | -0.875162888 | 0.6 | 0.654 | 1 | 0.126647823 | 0.124241685 |
| Slc35f6 | 0.064187373 | 0.80407979 | 0.311 | 0.224 | 1 | 0.064261158 | 1.642882477 | 0.4 | 0.185 | 1 | 0.064261158 | 0.124254727 |
| Tmem9b | 0.506184012 | 0.138197294 | 0.513 | 0.494 | 1 | 0.064217497 | -0.789433014 | 0.2 | 0.543 | 1 | 0.506184012 | 0.124311106 |
| Ergic2 | 0.279740345 | -0.226965151 | 0.815 | 0.747 | 1 | 0.064312945 | 0.632059507 | 0.9 | 0.83 | 1 | 0.279740345 | 0.124489734 |
| Phpt1 | 0.064351551 | -0.437332539 | 0.403 | 0.494 | 1 | 0.26498905 | -0.654503434 | 0.3 | 0.448 | 1 | 0.26498905 | 0.12456198 |
| Fsd1l | 0.064370628 | -0.733354341 | 0.092 | 0.171 | 1 | 0.593007977 | 0.135278859 | 0.3 | 0.204 | 1 | 0.593007977 | 0.124597678 |
| Slc19a1 | 0.064378371 | -0.699551633 | 0.185 | 0.276 | 1 | 0.606389335 | 0.526068812 | 0.3 | 0.216 | 1 | 0.606389335 | 0.124612167 |
| Noc2l | 0.064441217 | -0.816632736 | 0.244 | 0.329 | 1 | 0.163195439 | -0.806506527 | 0.1 | 0.33 | 1 | 0.163195439 | 0.124729763 |
| Fhip2b | 0.383374974 | -0.774933444 | 0.067 | 0.094 | 1 | 0.064460505 | 1.695993813 | 0.3 | 0.108 | 1 | 0.383374974 | 0.124765854 |
| Six1 | 0.064573624 | -1.008988783 | 0.134 | 0.212 | 1 | 0.134206245 | -0.989572629 | 0.1 | 0.355 | 1 | 0.134206245 | 0.124977495 |
| Vwce | 0.749417834 | -0.333423734 | 0.05 | 0.059 | 1 | 0.06458666 | 1.602884409 | 0.2 | 0.056 | 1 | 0.749417834 | 0.125001883 |
| Paqr8 | 0.302975942 | 0.321928095 | 0.092 | 0.059 | 1 | 0.064587169 | 1.795529487 | 0.2 | 0.056 | 1 | 0.302975942 | 0.125002835 |
| Mlh3 | 0.44371554 | -0.513995979 | 0.134 | 0.165 | 1 | 0.064591125 | 2.716752373 | 0.3 | 0.12 | 1 | 0.44371554 | 0.125010236 |
| Snrk | 0.094180973 | -0.701155518 | 0.21 | 0.288 | 1 | 0.064624113 | -1.618702713 | 0.1 | 0.383 | 1 | 0.094180973 | 0.125071951 |
| Fkbp3 | 0.064629729 | 0.284668476 | 0.79 | 0.724 | 1 | 0.143905769 | -0.521236903 | 0.4 | 0.719 | 1 | 0.143905769 | 0.125082457 |
| Sike1 | 0.064687476 | -0.485426827 | 0.227 | 0.329 | 1 | 0.196127093 | -0.880931369 | 0.2 | 0.383 | 1 | 0.196127093 | 0.125190483 |
| Dffb | 0.064713836 | -1.359895945 | 0.042 | 0.1 | 1 | 0.374659129 | -0.111361109 | 0 | 0.074 | 1 | 0.374659129 | 0.125239791 |
| Klrg2 | 0.064802212 | -1.5360529 | 0.042 | 0.1 | 1 | 0.534705424 | 0.536795218 | 0.2 | 0.127 | 1 | 0.534705424 | 0.125405097 |
| Nudt4 | 0.064819828 | -0.340553198 | 0.798 | 0.829 | 1 | 0.074557989 | -0.432539557 | 0.5 | 0.87 | 1 | 0.074557989 | 0.125438046 |
| Rnf6 | 0.064870131 | -0.485426827 | 0.513 | 0.571 | 1 | 0.07979684 | -1.008140389 | 0.4 | 0.639 | 1 | 0.07979684 | 0.125532127 |
| BC005624 | 0.064879765 | 0.187870573 | 0.765 | 0.647 | 1 | 0.23123731 | -0.252607034 | 0.4 | 0.654 | 1 | 0.23123731 | 0.125550147 |
| Acy1 | 0.064955424 | -1.250961574 | 0.034 | 0.088 | 1 | 0.374523929 | 0.210566986 | 0 | 0.074 | 1 | 0.374523929 | 0.125691641 |
| Msx2 | 0.065025797 | -1.378511623 | 0.042 | 0.1 | 1 | 0.827409899 | 0.418009066 | 0.2 | 0.176 | 1 | 0.827409899 | 0.125823239 |
| Nupl2 | 0.36119327 | -0.691877705 | 0.092 | 0.124 | 1 | 0.065068365 | 1.403212064 | 0.3 | 0.105 | 1 | 0.36119327 | 0.125902837 |
| Dcbld1 | 0.065068895 | -1.694880193 | 0.042 | 0.1 | 1 | 0.325058394 | -0.473931188 | 0 | 0.09 | 1 | 0.325058394 | 0.125903829 |
| Gm14963 | 0.065079358 | 1.308122295 | 0.134 | 0.071 | 1 | 0.067347313 | 2.532495081 | 0.2 | 0.062 | 1 | 0.067347313 | 0.125923393 |
| Ctsa | 0.065089307 | 0.183122304 | 0.731 | 0.582 | 1 | 0.077300574 | -0.868210127 | 0.3 | 0.639 | 1 | 0.077300574 | 0.125941996 |
| Rbm14 | 0.115589825 | -0.744161096 | 0.21 | 0.282 | 1 | 0.065096407 | -1.457811523 | 0.1 | 0.383 | 1 | 0.115589825 | 0.125955272 |
| Mks1 | 0.065109084 | -1.070389328 | 0.059 | 0.124 | 1 | 0.195226183 | 2.31748219 | 0.1 | 0.028 | 1 | 0.195226183 | 0.125978975 |
| Mterf4 | 0.065111247 | 0.634867407 | 0.185 | 0.106 | 1 | 0.736119637 | 0.345496566 | 0.1 | 0.136 | 1 | 0.736119637 | 0.125983019 |
| Agtr1a | 0.065158879 | -1.485426827 | 0.008 | 0.047 | 1 | 0.643842184 | 1.558490289 | 0 | 0.022 | 1 | 0.643842184 | 0.126072078 |

| | | | | | | | | | | | | |
|---|---|---|---|---|---|---|---|---|---|---|---|---|
| Stx16 | 0.065193873 | -0.597319707 | 0.571 | 0.582 | 1 | 0.802006868 | -0.226550047 | 0.6 | 0.608 | 1 | 0.802006868 | 0.126137505 |
| Ik | 0.065255449 | -0.229246092 | 0.866 | 0.865 | 1 | 0.240035604 | -0.428334322 | 0.9 | 0.883 | 1 | 0.240035604 | 0.126252624 |
| Gpsm1 | 0.065480177 | -0.178999826 | 0.353 | 0.429 | 1 | 0.259150223 | 0.620329543 | 0.7 | 0.481 | 1 | 0.259150223 | 0.1266727 |
| Far1os | 0.065500324 | -1.359895945 | 0.008 | 0.047 | 1 | 0.40687241 | 1.769994395 | 0.1 | 0.043 | 1 | 0.40687241 | 0.126710356 |
| Insyn2a | 0.318674966 | 1.303069068 | 0.067 | 0.041 | 1 | 0.065525828 | 2.539874611 | 0.2 | 0.059 | 1 | 0.318674966 | 0.126758021 |
| G6pdx | 0.065706139 | 0.465663572 | 0.21 | 0.124 | 1 | 0.489468891 | 0.418009066 | 0.1 | 0.21 | 1 | 0.489468891 | 0.127094982 |
| Atp5k | 0.065796855 | -0.456280482 | 0.445 | 0.529 | 1 | 0.478101007 | -0.321928095 | 0.4 | 0.506 | 1 | 0.478101007 | 0.127264484 |
| C920006O11Rik | 0.065819134 | -1.222392421 | 0.042 | 0.1 | 1 | 0.522123697 | 0.625604485 | 0 | 0.04 | 1 | 0.522123697 | 0.127306109 |
| Nup43 | 0.065819134 | -1.222392421 | 0.042 | 0.1 | 1 | 0.324923526 | -0.230005605 | 0 | 0.09 | 1 | 0.324923526 | 0.127306109 |
| Als2cl | 0.065836121 | 0.59946207 | 0.227 | 0.141 | 1 | 0.21381637 | -1.026472211 | 0 | 0.136 | 1 | 0.21381637 | 0.127337847 |
| Hvcn1 | 0.065841031 | -1.222392421 | 0.008 | 0.047 | 1 | 0.619961444 | 1.210566986 | 0 | 0.025 | 1 | 0.619961444 | 0.127347021 |
| Gm26671 | 0.065863243 | 1.707218251 | 0.067 | 0.024 | 1 | 0.730749233 | 2.017921908 | 0 | 0.012 | 1 | 0.730749233 | 0.127388519 |
| Ankrd13c | 0.065899907 | 0.176525413 | 0.706 | 0.588 | 1 | 0.661827105 | -0.272755253 | 0.7 | 0.75 | 1 | 0.661827105 | 0.127457016 |
| Zfp112 | 0.065920977 | -1.432959407 | 0.042 | 0.1 | 1 | 0.878679028 | 0.135278859 | 0.1 | 0.111 | 1 | 0.878679028 | 0.127496379 |
| Ndufs4 | 0.760165526 | -0.192230451 | 0.739 | 0.688 | 1 | 0.065945724 | -0.765820457 | 0.5 | 0.784 | 1 | 0.760165526 | 0.12754261 |
| Mtx1 | 0.280342488 | -0.462706751 | 0.387 | 0.418 | 1 | 0.066009797 | 1.398025617 | 0.7 | 0.41 | 1 | 0.280342488 | 0.127662301 |
| Chmp7 | 0.066028705 | -1.201633861 | 0.05 | 0.112 | 1 | 0.6882351 | -0.125036046 | 0.2 | 0.247 | 1 | 0.6882351 | 0.12769762 |
| Tpd52l1 | 0.066093152 | 0.377069649 | 0.429 | 0.306 | 1 | 0.142408595 | 1.861417422 | 0.4 | 0.231 | 1 | 0.142408595 | 0.127818 |
| Gm46515 | 0.066159764 | 2.230780207 | 0.067 | 0.024 | 1 | 0.238914673 | 1.847996907 | 0.1 | 0.031 | 1 | 0.238914673 | 0.127942413 |
| Fggy | 0.066199372 | -1.304854582 | 0.101 | 0.176 | 1 | 0.264657021 | 0.944672926 | 0.3 | 0.164 | 1 | 0.264657021 | 0.128016388 |
| Cadm2 | 0.066207785 | -1 | 0.008 | 0.047 | 1 | 0.903631786 | -0.668578619 | 0.1 | 0.083 | 1 | 0.903631786 | 0.128032099 |
| Gm47863 | 0.066427141 | 1.514573173 | 0.067 | 0.024 | 1 | 0.318562315 | 2.017921908 | 0.1 | 0.037 | 1 | 0.318562315 | 0.128441717 |
| Gm56741 | 0.06645187 | -1.070389328 | 0.059 | 0.124 | 1 | 0.743727234 | 0.930459067 | 0.1 | 0.071 | 1 | 0.743727234 | 0.128487889 |
| Tmtc3 | 0.06648758 | -0.996388746 | 0.143 | 0.224 | 1 | 0.259374358 | -1.033740212 | 0.2 | 0.349 | 1 | 0.259374358 | 0.128554562 |
| Ormdl1 | 0.066580688 | -0.707819249 | 0.084 | 0.159 | 1 | 0.136365622 | 1.432959407 | 0.3 | 0.133 | 1 | 0.136365622 | 0.128728388 |
| Abraxas2 | 0.06664831 | -0.606442228 | 0.261 | 0.353 | 1 | 0.105444092 | -1.248864633 | 0.2 | 0.432 | 1 | 0.105444092 | 0.128854622 |
| Vcan | 0.389335982 | 1.877143252 | 0.118 | 0.159 | 1 | 0.066681142 | 2.091922489 | 0.2 | 0.059 | 1 | 0.389335982 | 0.128915909 |
| Chka | 0.066708675 | 0.199586687 | 0.857 | 0.818 | 1 | 0.31239378 | 0.163317614 | 1 | 0.914 | 1 | 0.31239378 | 0.128967302 |
| Ccdc191 | 0.066742947 | -0.937939032 | 0.126 | 0.206 | 1 | 0.980741359 | 2.043917117 | 0.1 | 0.114 | 1 | 0.980741359 | 0.129031274 |
| Tspan17 | 0.066779211 | -0.663764068 | 0.118 | 0.2 | 1 | 0.433541956 | 0.625604485 | 0 | 0.059 | 1 | 0.433541956 | 0.129098959 |
| Cpt1a | 0.066820756 | -0.857395605 | 0.109 | 0.188 | 1 | 0.826844939 | -0.171902651 | 0.4 | 0.414 | 1 | 0.826844939 | 0.129176499 |
| Reno1 | 0.066936818 | -1.14839184 | 0.042 | 0.1 | 1 | 0.743726438 | 1.111031312 | 0.1 | 0.071 | 1 | 0.743726438 | 0.129393099 |
| Rnf130 | 0.684906895 | -0.113996771 | 0.874 | 0.865 | 1 | 0.066946711 | -0.829547601 | 0.7 | 0.929 | 1 | 0.684906895 | 0.12941156 |
| Cmtm8 | 0.066963038 | -1.201633861 | 0.126 | 0.2 | 1 | 0.397314192 | -0.603315979 | 0.4 | 0.488 | 1 | 0.397314192 | 0.129442028 |
| Dalrd3 | 0.066982466 | -0.76106127 | 0.134 | 0.218 | 1 | 0.347337489 | 1.503348735 | 0.3 | 0.191 | 1 | 0.347337489 | 0.129478281 |
| Mff | 0.066984057 | -0.485426827 | 0.655 | 0.688 | 1 | 0.482930923 | -0.298446662 | 0.7 | 0.753 | 1 | 0.482930923 | 0.129481251 |
| Ahnak2 | 0.067005841 | -0.347923303 | 0.092 | 0.171 | 1 | 0.878981639 | 0.591657153 | 0.1 | 0.083 | 1 | 0.878981639 | 0.129521899 |
| Gripap1 | 0.067023189 | -0.531523242 | 0.496 | 0.547 | 1 | 0.314864874 | 0.724322536 | 0.7 | 0.605 | 1 | 0.314864874 | 0.12955427 |
| Cby1 | 0.218739523 | -0.671839951 | 0.227 | 0.276 | 1 | 0.067048187 | -1.625934282 | 0 | 0.256 | 1 | 0.218739523 | 0.129600914 |
| Ccne2 | 0.067074406 | -1.269698136 | 0.042 | 0.1 | 1 | 0.44222087 | 1.847996907 | 0.1 | 0.046 | 1 | 0.44222087 | 0.129649835 |
| Apbb2 | 0.067202468 | -0.553870511 | 0.983 | 0.982 | 1 | 0.754514591 | -0.175889651 | 1 | 0.994 | 1 | 0.754514591 | 0.129888765 |
| Tbc1d25 | 0.067297211 | -1.137503524 | 0.05 | 0.112 | 1 | 0.870773711 | 0.660369903 | 0.1 | 0.083 | 1 | 0.870773711 | 0.130065507 |
| Srp54a | 0.067353866 | -0.539874611 | 0.269 | 0.365 | 1 | 0.14197002 | 1.063725598 | 0.5 | 0.281 | 1 | 0.14197002 | 0.130171189 |
| Hsbp1 | 0.067420127 | -0.303096805 | 0.874 | 0.835 | 1 | 0.111396952 | -0.603083343 | 0.5 | 0.836 | 1 | 0.111396952 | 0.13029478 |
| Hnrnpa2b1 | 0.187773699 | -0.209181548 | 1 | 1 | 1 | 0.067534406 | -0.435909904 | 1 | 1 | 1 | 0.187773699 | 0.130507917 |
| Gm15787 | 0.592366805 | -0.770829046 | 0.118 | 0.135 | 1 | 0.067654913 | 1.988174565 | 0.3 | 0.114 | 1 | 0.592366805 | 0.130732638 |
| Rbm12b1 | 0.95111309 | 0.12755005 | 0.084 | 0.082 | 1 | 0.067692047 | 2.245332404 | 0.3 | 0.117 | 1 | 0.95111309 | 0.13080188 |
| Bmi1 | 0.934741647 | -0.281068329 | 0.353 | 0.335 | 1 | 0.067778209 | -1.581990934 | 0.1 | 0.377 | 1 | 0.934741647 | 0.130962533 |
| Senp8 | 0.852932554 | 0.292180751 | 0.118 | 0.112 | 1 | 0.067835469 | 1.263034406 | 0.3 | 0.105 | 1 | 0.852932554 | 0.131069288 |
| 0610030E20Rik | 0.067845239 | -0.867297463 | 0.235 | 0.318 | 1 | 0.533762734 | -0.201246612 | 0.2 | 0.302 | 1 | 0.533762734 | 0.131087502 |
| Mettl26 | 0.067941014 | 0.557008439 | 0.387 | 0.288 | 1 | 0.552161919 | 0.483943336 | 0.2 | 0.312 | 1 | 0.552161919 | 0.131266046 |
| Eef1d | 0.359884592 | -0.122069895 | 0.891 | 0.941 | 1 | 0.068015441 | -0.516056664 | 0.5 | 0.929 | 1 | 0.359884592 | 0.131404781 |
| Gm29418 | 0.068018721 | -0.769219793 | 0.16 | 0.241 | 1 | 0.412907433 | -0.473931188 | 0.2 | 0.306 | 1 | 0.412907433 | 0.131410896 |
| Hspe1 | 0.068110408 | -0.239298132 | 0.95 | 0.918 | 1 | 0.389304166 | -0.428570655 | 0.7 | 0.895 | 1 | 0.389304166 | 0.131581789 |
| Tomm34 | 0.659571935 | -0.244418728 | 0.387 | 0.394 | 1 | 0.068140242 | 0.783866569 | 0.7 | 0.38 | 1 | 0.659571935 | 0.131637391 |
| Csrnp1 | 0.438613907 | 0.186151316 | 0.487 | 0.465 | 1 | 0.06817548 | -1.296774618 | 0.2 | 0.509 | 1 | 0.438613907 | 0.131703064 |
| Itgb6 | 0.912599274 | -0.485426827 | 0.05 | 0.053 | 1 | 0.068211732 | 0.821524695 | 0.2 | 0.056 | 1 | 0.912599274 | 0.131770623 |
| Brf1 | 0.240154123 | -0.542760002 | 0.244 | 0.3 | 1 | 0.068378536 | -1.380109166 | 0.2 | 0.466 | 1 | 0.240154123 | 0.132081447 |
| Rab3gap2 | 0.068409725 | 0.12885124 | 0.723 | 0.588 | 1 | 0.232523393 | 0.725140159 | 0.8 | 0.722 | 1 | 0.232523393 | 0.13213956 |
| Arhgef6 | 0.068496388 | -0.495551044 | 0.412 | 0.494 | 1 | 0.383834207 | -0.754667596 | 0.4 | 0.488 | 1 | 0.383834207 | 0.132301022 |
| Ankrd35 | 0.068498102 | -0.807354922 | 0.067 | 0.135 | 1 | 0.682730486 | 0.135278859 | 0.3 | 0.222 | 1 | 0.682730486 | 0.132304215 |
| Kctd12 | 0.068537819 | -0.719892081 | 0.218 | 0.306 | 1 | 0.238973396 | -1.217294554 | 0.1 | 0.29 | 1 | 0.238973396 | 0.132378204 |
| Senp1 | 0.068637841 | -0.679437447 | 0.403 | 0.465 | 1 | 0.423963555 | -0.208146171 | 0.3 | 0.472 | 1 | 0.423963555 | 0.132564528 |
| Zfp655 | 0.068655092 | 0.201156578 | 0.672 | 0.535 | 1 | 0.845044525 | 0.10782888 | 0.5 | 0.648 | 1 | 0.845044525 | 0.132596662 |
| Ddx49 | 0.068737469 | -0.537265759 | 0.294 | 0.388 | 1 | 0.345122655 | -0.606568957 | 0.2 | 0.327 | 1 | 0.345122655 | 0.132750098 |
| 0610009B22Rik | 0.068808312 | 0.437951891 | 0.37 | 0.271 | 1 | 0.465758965 | 0.20029865 | 0.2 | 0.336 | 1 | 0.465758965 | 0.13288204 |
| Tollip | 0.216243024 | -0.35710273 | 0.294 | 0.359 | 1 | 0.068814427 | -1.176834946 | 0.3 | 0.546 | 1 | 0.216243024 | 0.132893428 |
| Ndufv2 | 0.068870614 | -0.319209875 | 0.739 | 0.771 | 1 | 0.115956247 | -0.637429921 | 0.6 | 0.775 | 1 | 0.115956247 | 0.132998066 |
| Snhg9 | 0.068960179 | -1.008988783 | 0.05 | 0.112 | 1 | 0.635221364 | 1.432959407 | 0.1 | 0.062 | 1 | 0.635221364 | 0.133164852 |
| Gm16268 | 0.069034043 | -1.584962501 | 0.034 | 0.088 | 1 | 0.99241428 | 0.602884409 | 0.1 | 0.102 | 1 | 0.99241428 | 0.133302388 |
| Taf10 | 0.069135002 | 0.279042282 | 0.714 | 0.665 | 1 | 0.885483492 | 0.227844977 | 0.6 | 0.685 | 1 | 0.885483492 | 0.133490355 |
| Map3k14 | 0.069176559 | 0.111217479 | 0.118 | 0.206 | 1 | 0.179394774 | 1.176619654 | 0.4 | 0.228 | 1 | 0.179394774 | 0.133567722 |
| Ecsit | 0.069245123 | -0.987927168 | 0.092 | 0.165 | 1 | 0.802255445 | 0.558490289 | 0.2 | 0.164 | 1 | 0.802255445 | 0.13369536 |
| Dync2i2 | 0.069245518 | -0.944858446 | 0.092 | 0.165 | 1 | 0.926075407 | 0.403212064 | 0.1 | 0.108 | 1 | 0.926075407 | 0.133696094 |
| Ccdc163 | 0.237313032 | -0.642968104 | 0.076 | 0.118 | 1 | 0.069250768 | 0.951832718 | 0.3 | 0.105 | 1 | 0.237313032 | 0.133705866 |
| Mospd2 | 0.069253263 | 0.297543936 | 0.622 | 0.482 | 1 | 0.393474169 | 0.718954962 | 0.7 | 0.512 | 1 | 0.393474169 | 0.133710511 |
| Il6 | 0.069284498 | 0.901596296 | 0.067 | 0.024 | 1 | 0.698534498 | 2.432959407 | 0 | 0.015 | 1 | 0.698534498 | 0.133768655 |
| Hs6st1 | 0.069369634 | -0.399696953 | 0.134 | 0.224 | 1 | 0.144355975 | -1.141949429 | 0.1 | 0.312 | 1 | 0.144355975 | 0.133927123 |
| Stk3 | 0.069427863 | -0.498026864 | 0.756 | 0.765 | 1 | 0.201284334 | -0.687134438 | 0.6 | 0.796 | 1 | 0.201284334 | 0.134035498 |
| Tmem161a | 0.541074615 | -0.352160296 | 0.378 | 0.382 | 1 | 0.069435749 | -1.449683642 | 0.1 | 0.377 | 1 | 0.541074615 | 0.134050175 |
| Tubg1 | 0.069534781 | -0.900464326 | 0.059 | 0.124 | 1 | 0.770508597 | 1.210566986 | 0.1 | 0.074 | 1 | 0.770508597 | 0.134234476 |
| Cenpc1 | 0.069621245 | -0.534336428 | 0.361 | 0.435 | 1 | 0.084585907 | -1.17978625 | 0.3 | 0.568 | 1 | 0.084585907 | 0.134395372 |
| Pawr | 0.06962157 | -1.34869049 | 0.252 | 0.335 | 1 | 0.734503636 | 0.489886475 | 0.3 | 0.383 | 1 | 0.734503636 | 0.134395978 |
| Cdk10 | 0.069627104 | -1.11345805 | 0.084 | 0.153 | 1 | 0.811975214 | 0.828097349 | 0.2 | 0.182 | 1 | 0.811975214 | 0.134406275 |
| Setdb2 | 0.08308731 | -1.306456686 | 0.118 | 0.188 | 1 | 0.069633981 | 1.418009066 | 0.4 | 0.179 | 1 | 0.08308731 | 0.134419072 |
| Ndufa11 | 0.834809474 | -0.125251263 | 0.773 | 0.776 | 1 | 0.069702971 | -0.896961478 | 0.4 | 0.67 | 1 | 0.834809474 | 0.134547437 |
| Zmat5 | 0.069750553 | -0.581642142 | 0.193 | 0.282 | 1 | 0.8090203 | 0.30367639 | 0.3 | 0.25 | 1 | 0.8090203 | 0.134635966 |
| Cd2bp2 | 0.126431169 | -0.669851398 | 0.16 | 0.229 | 1 | 0.0697548 | -1.763437806 | 0 | 0.253 | 1 | 0.126431169 | 0.134643867 |
| B230217C12Rik | 0.069864528 | -0.982926487 | 0.134 | 0.212 | 1 | 0.46716563 | -0.567040593 | 0.1 | 0.185 | 1 | 0.46716563 | 0.134848004 |
| Tbck | 0.069993943 | -0.900464326 | 0.227 | 0.312 | 1 | 0.094177961 | 0.625604485 | 0.7 | 0.41 | 1 | 0.094177961 | 0.135088734 |
| Syn1 | 0.26362158 | 0.120294234 | 0.168 | 0.118 | 1 | 0.070020187 | 2.159940913 | 0.3 | 0.114 | 1 | 0.26362158 | 0.135137548 |
| H2-DMa | 0.174591802 | 0.450442835 | 0.235 | 0.171 | 1 | 0.070034711 | 1.494359952 | 0.3 | 0.108 | 1 | 0.174591802 | 0.135164562 |
| Ptpra | 0.070212956 | 0.213026578 | 0.924 | 0.947 | 1 | 0.164391539 | 0.350606745 | 1 | 0.957 | 1 | 0.164391539 | 0.135496052 |
| Mrs2 | 0.070238299 | -0.959358016 | 0.134 | 0.212 | 1 | 0.388593324 | -0.204470513 | 0.1 | 0.231 | 1 | 0.388593324 | 0.13554318 |
| Bcas3 | 0.070301154 | -0.513995979 | 0.681 | 0.706 | 1 | 0.225925597 | -0.558247474 | 0.5 | 0.735 | 1 | 0.225925597 | 0.135660057 |
| Trerf1 | 0.396943544 | -0.795766948 | 0.134 | 0.165 | 1 | 0.070310085 | -2.141949429 | 0 | 0.253 | 1 | 0.396943544 | 0.135676662 |
| Ap5z1 | 0.070338928 | -0.95491211 | 0.092 | 0.165 | 1 | 0.609526222 | -0.152003093 | 0.1 | 0.154 | 1 | 0.609526222 | 0.135730291 |
| Sgf29 | 0.070338928 | -0.95491211 | 0.092 | 0.165 | 1 | 0.489315033 | 0.71414116 | 0.3 | 0.201 | 1 | 0.489315033 | 0.135730291 |

| | | | | | | | | | | | | |
|---|---|---|---|---|---|---|---|---|---|---|---|---|
| Arfgap2 | 0.070362487 | -0.721494185 | 0.328 | 0.394 | 1 | 0.374082521 | 0.463333056 | 0.5 | 0.315 | 1 | 0.374082521 | 0.135774095 |
| Gria4 | 0.070370714 | -2.106915204 | 0.034 | 0.088 | 1 | 0.353956943 | -1.339630097 | 0 | 0.08 | 1 | 0.353956943 | 0.135789391 |
| Frrs1 | 0.070446221 | 1.514573173 | 0.076 | 0.029 | 1 | 0.099077868 | 0.359710425 | 0.3 | 0.114 | 1 | 0.099077868 | 0.135929772 |
| Dgkd | 0.261844315 | -0.579208973 | 0.378 | 0.406 | 1 | 0.07044703 | -1.5004034 | 0.2 | 0.466 | 1 | 0.261844315 | 0.135931275 |
| Zdhhc23 | 0.070534977 | 1.099535674 | 0.134 | 0.071 | 1 | 0.27460373 | -0.567040593 | 0 | 0.108 | 1 | 0.27460373 | 0.136094772 |
| Hmga2 | 0.070592233 | 1.901596296 | 0.084 | 0.035 | 1 | 0.408802025 | -1.090602549 | 0 | 0.065 | 1 | 0.408802025 | 0.136201202 |
| Kif7 | 0.070721331 | -0.918386234 | 0.059 | 0.124 | 1 | 0.317682216 | 1.982297998 | 0.2 | 0.108 | 1 | 0.317682216 | 0.136441155 |
| Ppp1r13l | 0.070737765 | -0.774933444 | 0.05 | 0.112 | 1 | 0.542084603 | -0.191531458 | 0.1 | 0.17 | 1 | 0.542084603 | 0.136471699 |
| Pcdh17 | 0.101549958 | 0.363706507 | 0.328 | 0.235 | 1 | 0.070740693 | -3.053540455 | 0 | 0.253 | 1 | 0.101549958 | 0.13647714 |
| Mbtps2 | 0.070793822 | -0.726434927 | 0.303 | 0.388 | 1 | 0.949412081 | 0.174225139 | 0.3 | 0.306 | 1 | 0.949412081 | 0.136575878 |
| Polr1c | 0.070814251 | -0.544320516 | 0.462 | 0.518 | 1 | 0.167008542 | -0.820021334 | 0.3 | 0.509 | 1 | 0.167008542 | 0.136613845 |
| Snx8 | 0.109163124 | -0.86393845 | 0.176 | 0.247 | 1 | 0.070910786 | 0.648688098 | 0.7 | 0.349 | 1 | 0.109163124 | 0.136793233 |
| Uba52 | 0.070963833 | 0.175484527 | 0.882 | 0.876 | 1 | 0.078353244 | -0.667500775 | 0.7 | 0.855 | 1 | 0.078353244 | 0.136891801 |
| Abhd1 | 0.070996137 | 1.389042291 | 0.076 | 0.029 | 1 | 0.236765379 | 2.111031312 | 0.1 | 0.031 | 1 | 0.236765379 | 0.136951823 |
| Ppp2r5a | 0.071137095 | -0.442991561 | 0.756 | 0.759 | 1 | 0.073022113 | -0.667969502 | 0.9 | 0.926 | 1 | 0.073022113 | 0.137213703 |
| 4732440D04Rik | 0.206665719 | -0.968208933 | 0.235 | 0.276 | 1 | 0.071191675 | -2.220482831 | 0.1 | 0.364 | 1 | 0.206665719 | 0.137315096 |
| Mir22hg | 0.071239233 | -0.428843299 | 0.462 | 0.535 | 1 | 0.904421396 | 0.207136081 | 0.6 | 0.611 | 1 | 0.904421396 | 0.137403437 |
| Pign | 0.071361834 | -0.831877241 | 0.403 | 0.453 | 1 | 0.404013368 | 0.232197002 | 0.6 | 0.435 | 1 | 0.404013368 | 0.137631156 |
| Alkbh8 | 0.12710407 | -0.748461233 | 0.21 | 0.282 | 1 | 0.071367875 | 1.159940913 | 0.5 | 0.256 | 1 | 0.12710407 | 0.137642377 |
| Dqx1 | 0.374118666 | -1.555816155 | 0.05 | 0.029 | 1 | 0.071481024 | 3.847996907 | 0.1 | 0.019 | 1 | 0.374118666 | 0.13785251 |
| Slbp | 0.178978428 | -0.116501253 | 0.605 | 0.688 | 1 | 0.071492125 | -1.123090402 | 0.4 | 0.679 | 1 | 0.178978428 | 0.137873126 |
| Rmnd1 | 0.071521353 | -1.070389328 | 0.084 | 0.153 | 1 | 0.822433807 | 0.31748219 | 0.2 | 0.164 | 1 | 0.822433807 | 0.137927401 |
| Gsr | 0.071525723 | 1.653134003 | 0.479 | 0.424 | 1 | 0.123398305 | 0.612577805 | 0.4 | 0.645 | 1 | 0.123398305 | 0.137935517 |
| Tufm | 0.08801116 | -0.736965594 | 0.151 | 0.229 | 1 | 0.071561649 | 1.336097868 | 0.4 | 0.179 | 1 | 0.08801116 | 0.138002229 |
| Mpc2 | 0.071577604 | 0.195854634 | 0.916 | 0.906 | 1 | 0.17739443 | 0.312122326 | 0.9 | 0.892 | 1 | 0.17739443 | 0.138031854 |
| Mcm3ap | 0.071585473 | -0.700439718 | 0.193 | 0.282 | 1 | 0.703785053 | -0.101019165 | 0.2 | 0.259 | 1 | 0.703785053 | 0.138046466 |
| Zcchc2 | 0.071599237 | -0.934734229 | 0.185 | 0.265 | 1 | 0.094210868 | 0.878370556 | 0.5 | 0.253 | 1 | 0.094210868 | 0.138072022 |
| Rdh14 | 0.102546732 | 0.298844482 | 0.244 | 0.159 | 1 | 0.07184179 | -1.640289575 | 0 | 0.25 | 1 | 0.102546732 | 0.138522337 |
| Haghl | 0.071900748 | -0.963474124 | 0.109 | 0.182 | 1 | 0.557574822 | 0.847996907 | 0.2 | 0.13 | 1 | 0.557574822 | 0.138631778 |
| Adi1 | 0.071925405 | -1.207892852 | 0.076 | 0.141 | 1 | 0.870361724 | 1.048295557 | 0.1 | 0.13 | 1 | 0.870361724 | 0.138677546 |
| 4930539J05Rik | 0.338096132 | -0.807354922 | 0.025 | 0.047 | 1 | 0.072014428 | 2.210566986 | 0.2 | 0.059 | 1 | 0.338096132 | 0.138842778 |
| Ppip5k1 | 0.07203508 | -0.959358016 | 0.101 | 0.176 | 1 | 0.532309442 | -0.357117523 | 0.1 | 0.17 | 1 | 0.532309442 | 0.138881107 |
| Arl1 | 0.072046225 | -0.349321361 | 0.899 | 0.9 | 1 | 0.592890387 | -0.194128569 | 0.9 | 0.886 | 1 | 0.592890387 | 0.138901792 |
| Ap2s1 | 0.072089914 | -0.464865505 | 0.58 | 0.641 | 1 | 0.709449496 | 0.221650485 | 0.8 | 0.648 | 1 | 0.709449496 | 0.138982872 |
| Gaa | 0.072094242 | 0.343842871 | 0.782 | 0.612 | 1 | 0.14012478 | 0.629043569 | 0.9 | 0.623 | 1 | 0.14012478 | 0.138990904 |
| Tmem242 | 0.717304094 | -0.20269662 | 0.403 | 0.4 | 1 | 0.072268956 | -1.425021588 | 0.1 | 0.373 | 1 | 0.717304094 | 0.139315109 |
| Tmem41b | 0.265199532 | -0.173893904 | 0.555 | 0.6 | 1 | 0.072335036 | -0.706970854 | 0.4 | 0.679 | 1 | 0.265199532 | 0.139437715 |
| Tmem9 | 0.072395291 | -0.6617496 | 0.345 | 0.424 | 1 | 0.319017359 | -0.401616984 | 0.3 | 0.469 | 1 | 0.319017359 | 0.139549503 |
| Champ1 | 0.230199302 | -0.517848305 | 0.16 | 0.212 | 1 | 0.072427479 | -1.947862377 | 0 | 0.25 | 1 | 0.230199302 | 0.139609218 |
| Sh2d4a | 0.0724532 | 0.929610672 | 0.151 | 0.082 | 1 | 0.792962827 | -0.441509711 | 0.1 | 0.12 | 1 | 0.792962827 | 0.139656934 |
| Pin4 | 0.07245445 | -0.577349317 | 0.269 | 0.365 | 1 | 0.215763967 | -0.654503434 | 0.1 | 0.275 | 1 | 0.215763967 | 0.139659253 |
| Mbtps1 | 0.072489772 | -0.478854173 | 0.588 | 0.635 | 1 | 0.666985633 | 0.361298422 | 0.7 | 0.701 | 1 | 0.666985633 | 0.139724777 |
| Lmod1 | 0.236601947 | -1.359895945 | 0.017 | 0.041 | 1 | 0.072537454 | 3.602884409 | 0.1 | 0.019 | 1 | 0.236601947 | 0.139813225 |
| Gm36862 | 0.756080634 | 0.192645078 | 0.034 | 0.041 | 1 | 0.072537454 | 3.602884409 | 0.1 | 0.019 | 1 | 0.756080634 | 0.139813225 |
| Slc25a33 | 0.242082165 | 0.450442835 | 0.252 | 0.194 | 1 | 0.072640201 | -2.026472211 | 0 | 0.25 | 1 | 0.242082165 | 0.140003803 |
| Ptdss2 | 0.072783543 | -0.604725756 | 0.185 | 0.271 | 1 | 0.837260985 | 0.306427001 | 0.3 | 0.269 | 1 | 0.837260985 | 0.140269642 |
| Erbb3 | 0.072821872 | 1.421463768 | 0.084 | 0.035 | 1 | 0.396883838 | 0.210566986 | 0 | 0.068 | 1 | 0.396883838 | 0.140340719 |
| Bloc1s5 | 0.072886769 | -0.900464326 | 0.05 | 0.112 | 1 | 0.921181689 | 0.558490289 | 0.1 | 0.108 | 1 | 0.921181689 | 0.140461058 |
| Efcab2 | 0.749223192 | 0.116023796 | 0.126 | 0.141 | 1 | 0.072949292 | 1.236562194 | 0.3 | 0.108 | 1 | 0.749223192 | 0.140576984 |
| Bcl6 | 0.07300731 | 0.113082205 | 0.756 | 0.682 | 1 | 0.176263483 | 0.322212342 | 1 | 0.858 | 1 | 0.176263483 | 0.140684552 |
| Dohh | 0.372945534 | -0.102098188 | 0.395 | 0.441 | 1 | 0.073070655 | -1.365782384 | 0.1 | 0.373 | 1 | 0.372945534 | 0.14080199 |
| Cpne2 | 0.073263891 | 0.425763906 | 0.319 | 0.218 | 1 | 0.096335569 | -1.489872732 | 0.1 | 0.346 | 1 | 0.096335569 | 0.141160184 |
| D930015M05Rik | 0.124070461 | -1.250961574 | 0.025 | 0.065 | 1 | 0.073299054 | 2.14345279 | 0.2 | 0.059 | 1 | 0.124070461 | 0.141225357 |
| Ciao1 | 0.073354702 | -0.593951284 | 0.218 | 0.306 | 1 | 0.547462655 | 0.515421567 | 0.4 | 0.299 | 1 | 0.547462655 | 0.141328492 |
| Spint2 | 0.073428504 | 1.13498451 | 0.353 | 0.282 | 1 | 0.261069153 | -1.104474723 | 0.2 | 0.343 | 1 | 0.261069153 | 0.141465264 |
| Cldn12 | 0.073540071 | 0.462105753 | 0.21 | 0.129 | 1 | 0.125898525 | 1.450237399 | 0.4 | 0.207 | 1 | 0.125898525 | 0.141671999 |
| Lman2 | 0.073541237 | -0.349479329 | 0.714 | 0.706 | 1 | 0.223429533 | -0.420491929 | 0.5 | 0.741 | 1 | 0.223429533 | 0.14167416 |
| Eftud2 | 0.073568118 | -0.637429921 | 0.319 | 0.4 | 1 | 0.262386433 | -0.473931188 | 0.2 | 0.395 | 1 | 0.262386433 | 0.141723969 |
| Zfp771 | 0.073582868 | -0.652076697 | 0.345 | 0.429 | 1 | 0.084397702 | 0.778455973 | 0.7 | 0.383 | 1 | 0.084397702 | 0.141751298 |
| Heatr5b | 0.073607931 | -1.215819767 | 0.143 | 0.218 | 1 | 0.496997151 | -0.526398608 | 0.2 | 0.281 | 1 | 0.496997151 | 0.141797735 |
| A730036I17Rik | 0.238504649 | -2.343407822 | 0.017 | 0.041 | 1 | 0.073613816 | 3.432959407 | 0.1 | 0.019 | 1 | 0.238504649 | 0.141808638 |
| Syt16 | 0.621077712 | -0.707819249 | 0.025 | 0.035 | 1 | 0.07361683 | 3.31748219 | 0.1 | 0.019 | 1 | 0.621077712 | 0.141814223 |
| Fam89b | 0.151201875 | -0.401903472 | 0.429 | 0.482 | 1 | 0.073630669 | -1.304006187 | 0.2 | 0.491 | 1 | 0.151201875 | 0.141839862 |
| Ddx39a | 0.073662349 | -0.88231698 | 0.202 | 0.276 | 1 | 0.373477845 | -0.484578433 | 0.2 | 0.321 | 1 | 0.373477845 | 0.141898557 |
| Slc25a12 | 0.073667 | -0.835924074 | 0.244 | 0.324 | 1 | 0.996893059 | -0.255096586 | 0.4 | 0.358 | 1 | 0.996893059 | 0.141907174 |
| Kmt5a | 0.412840823 | -0.146289442 | 0.723 | 0.771 | 1 | 0.073768371 | 0.930459067 | 1 | 0.88 | 1 | 0.412840823 | 0.14209497 |
| Steep1 | 0.073811198 | -0.629816737 | 0.345 | 0.435 | 1 | 0.721482949 | -0.536666944 | 0.4 | 0.389 | 1 | 0.721482949 | 0.142174304 |
| Fam13b | 0.138744007 | -0.184136902 | 0.731 | 0.812 | 1 | 0.073842533 | -0.821981708 | 0.6 | 0.852 | 1 | 0.138744007 | 0.142232347 |
| Fgf18 | 0.230628701 | -0.635173947 | 0.218 | 0.282 | 1 | 0.07392008 | -1.518130992 | 0.1 | 0.417 | 1 | 0.230628701 | 0.142375982 |
| Vdac2 | 0.073925591 | -0.299871174 | 0.824 | 0.876 | 1 | 0.321224356 | -0.23908871 | 0.7 | 0.941 | 1 | 0.321224356 | 0.142386188 |
| Nckap5 | 0.595563636 | -0.49848298 | 0.21 | 0.229 | 1 | 0.073977706 | 1.769994395 | 0.5 | 0.29 | 1 | 0.595563636 | 0.14248271 |
| Atg14 | 0.074028406 | -1.163498732 | 0.101 | 0.171 | 1 | 0.542225907 | 0.532495081 | 0.3 | 0.213 | 1 | 0.542225907 | 0.142576608 |
| Fbxo10 | 0.074086135 | -1.129283017 | 0.109 | 0.182 | 1 | 0.185962967 | -1.425021588 | 0 | 0.151 | 1 | 0.185962967 | 0.142683515 |
| Fat2 | 0.074145467 | 4.269460675 | 0.034 | 0.006 | 1 | 0.180146342 | 2.847996907 | 0.1 | 0.028 | 1 | 0.180146342 | 0.142793383 |
| Drc3 | 0.0741692 | 0.652076697 | 0.067 | 0.024 | 1 | 0.452712304 | 2.017921908 | 0.1 | 0.049 | 1 | 0.452712304 | 0.14283733 |
| Armc10 | 0.07421133 | -1.14839184 | 0.084 | 0.153 | 1 | 0.18021444 | -0.888968688 | 0 | 0.154 | 1 | 0.18021444 | 0.142915338 |
| Ap3s2 | 0.074222741 | -1.137503524 | 0.109 | 0.182 | 1 | 0.219556817 | -0.814968106 | 0 | 0.133 | 1 | 0.219556817 | 0.142936466 |
| Vwa1 | 0.074362539 | 0.980236745 | 0.176 | 0.106 | 1 | 0.42087075 | 0.494359952 | 0 | 0.062 | 1 | 0.42087075 | 0.143195291 |
| Ncapd2 | 0.074406369 | -1.778208576 | 0.067 | 0.129 | 1 | 0.306920729 | -0.230005605 | 0 | 0.096 | 1 | 0.306920729 | 0.143276431 |
| Mfsd6 | 0.074417351 | 1.215012891 | 0.076 | 0.029 | 1 | 0.525480919 | -0.131825212 | 0.1 | 0.176 | 1 | 0.525480919 | 0.14329676 |
| Uap1l1 | 0.143482547 | -0.603608254 | 0.252 | 0.324 | 1 | 0.074439117 | -1.567040593 | 0.1 | 0.37 | 1 | 0.143482547 | 0.143337052 |
| Gpcpd1 | 0.074445323 | -0.640496224 | 0.546 | 0.606 | 1 | 0.338304752 | 0.102042529 | 0.8 | 0.58 | 1 | 0.338304752 | 0.143348539 |
| Eif3g | 0.21135177 | -0.29764204 | 0.655 | 0.635 | 1 | 0.074541077 | -0.580337415 | 0.4 | 0.735 | 1 | 0.21135177 | 0.143525782 |
| Washc5 | 0.415162099 | -0.394278939 | 0.277 | 0.312 | 1 | 0.074563582 | -1.526398608 | 0.2 | 0.454 | 1 | 0.415162099 | 0.143567437 |
| Mmab | 0.629925121 | -0.333423734 | 0.084 | 0.1 | 1 | 0.074599625 | 1.959028219 | 0.2 | 0.059 | 1 | 0.629925121 | 0.143634146 |
| 2810432F15Rik | 0.07461065 | 0.901596296 | 0.126 | 0.065 | 1 | 0.790188307 | 0.769994395 | 0.1 | 0.074 | 1 | 0.790188307 | 0.143654552 |
| Sec62 | 0.074629106 | -0.197850237 | 0.975 | 1 | 1 | 0.341835106 | -0.233641562 | 1 | 0.997 | 1 | 0.341835106 | 0.143688708 |
| Gm37982 | 0.074638169 | 1.836501268 | 0.092 | 0.041 | 1 | 0.59781419 | 1.558490289 | 0 | 0.028 | 1 | 0.59781419 | 0.143705482 |
| Taf1 | 0.078659806 | -0.411001986 | 0.748 | 0.753 | 1 | 0.074642157 | 0.556617875 | 0.9 | 0.79 | 1 | 0.078659806 | 0.143712863 |
| Camta2 | 0.074645303 | -0.829381228 | 0.092 | 0.165 | 1 | 0.937522008 | 0.245332404 | 0.2 | 0.204 | 1 | 0.937522008 | 0.143718685 |
| Jdp2 | 0.074677552 | -0.426533138 | 0.412 | 0.524 | 1 | 0.154270847 | -1.242605642 | 0.3 | 0.525 | 1 | 0.154270847 | 0.143778367 |
| Atp6v0d1 | 0.07610615 | -0.503201529 | 0.639 | 0.635 | 1 | 0.074678109 | -0.732505946 | 0.6 | 0.809 | 1 | 0.07610615 | 0.143779398 |
| Pcdhb21 | 0.217412808 | -1.655351829 | 0.008 | 0.029 | 1 | 0.074698996 | 3.14345279 | 0.1 | 0.019 | 1 | 0.217412808 | 0.143818052 |
| Gm30551 | 0.356960217 | 0.652076697 | 0.025 | 0.047 | 1 | 0.074698996 | 3.017921908 | 0.1 | 0.019 | 1 | 0.356960217 | 0.143818052 |
| Bdkrb2 | 0.864020021 | 0.389042291 | 0.042 | 0.047 | 1 | 0.074698996 | 2.795529487 | 0.1 | 0.019 | 1 | 0.864020021 | 0.143818052 |
| Garin2 | 0.76863403 | 1.362570079 | 0.059 | 0.053 | 1 | 0.074705731 | 1.672147071 | 0.3 | 0.114 | 1 | 0.76863403 | 0.143830517 |
| Nlrc3 | 0.074770642 | -0.807354922 | 0.042 | 0.1 | 1 | 0.257956406 | 1.31748219 | 0.2 | 0.096 | 1 | 0.257956406 | 0.143950635 |
| Eif2ak3 | 0.074809777 | -0.658619542 | 0.445 | 0.494 | 1 | 0.202537032 | -1.074247936 | 0.6 | 0.664 | 1 | 0.202537032 | 0.144023051 |

| | | | | | | | | | | | | |
|---|---|---|---|---|---|---|---|---|---|---|---|---|
| Phkg1 | 0.07484372 | -2.222392421 | 0.017 | 0.059 | 1 | 0.535011384 | 0.973527789 | 0.1 | 0.052 | 1 | 0.535011384 | 0.144085857 |
| Lcp1 | 0.07484372 | -2.544320516 | 0.017 | 0.059 | 1 | 0.54547801 | -0.252607034 | 0.1 | 0.191 | 1 | 0.54547801 | 0.144085857 |
| Nhsl2 | 0.074849166 | -3.326729081 | 0.017 | 0.059 | 1 | 0.245950436 | -1.567040593 | 0 | 0.12 | 1 | 0.245950436 | 0.144095934 |
| Ankmy2 | 0.074849463 | -1.237499314 | 0.151 | 0.224 | 1 | 0.599103958 | 0.732519689 | 0.1 | 0.182 | 1 | 0.599103958 | 0.144096484 |
| Gm27253 | 0.074849771 | -1.987927168 | 0.017 | 0.059 | 1 | 0.539456618 | 0.930459067 | 0 | 0.037 | 1 | 0.539456618 | 0.144097054 |
| Gtpbp6 | 0.075160664 | -0.857395605 | 0.134 | 0.212 | 1 | 0.955189183 | 0.732519689 | 0.1 | 0.093 | 1 | 0.955189183 | 0.144672203 |
| Gm40528 | 0.075174255 | 4.269460675 | 0.034 | 0.006 | 1 | 0.07916329 | 2.695993813 | 0.1 | 0.019 | 1 | 0.07916329 | 0.144697341 |
| Gss | 0.075242588 | -0.807354922 | 0.218 | 0.3 | 1 | 0.19329857 | 0.888638891 | 0.5 | 0.327 | 1 | 0.19329857 | 0.144823729 |
| 4930503L19Rik | 0.075292321 | -1.276840205 | 0.101 | 0.171 | 1 | 0.818036108 | 0.795529487 | 0.1 | 0.136 | 1 | 0.818036108 | 0.144915709 |
| Anp32a | 0.216119387 | -0.200939312 | 0.79 | 0.865 | 1 | 0.075298622 | -0.69173634 | 0.5 | 0.818 | 1 | 0.216119387 | 0.144927362 |
| Xrcc6 | 0.108807107 | -0.632268215 | 0.193 | 0.271 | 1 | 0.075304667 | 1.393431043 | 0.4 | 0.179 | 1 | 0.108807107 | 0.144938541 |
| Gm56976 | 0.075308142 | 0.666576266 | 0.076 | 0.029 | 1 | 0.629538555 | 1.558490289 | 0.1 | 0.062 | 1 | 0.629538555 | 0.144944968 |
| Gm45237 | 0.075308142 | 1.029146346 | 0.076 | 0.029 | 1 | 0.557744661 | 1.210566986 | 0 | 0.034 | 1 | 0.557744661 | 0.144944968 |
| Gm4890 | 0.07563317 | 0.877143252 | 0.126 | 0.065 | 1 | 0.318562315 | 2.017921908 | 0.1 | 0.037 | 1 | 0.318562315 | 0.145545964 |
| Cacna1a | 0.07579863 | 0.544320516 | 0.218 | 0.141 | 1 | 0.15226684 | -1.864721141 | 0.1 | 0.296 | 1 | 0.15226684 | 0.145851827 |
| Gm26512 | 0.268222101 | -1.263034406 | 0.05 | 0.082 | 1 | 0.07581449 | 2.31748219 | 0.2 | 0.062 | 1 | 0.268222101 | 0.145881143 |
| Tmem86b | 0.522060045 | -0.485426827 | 0.042 | 0.059 | 1 | 0.075834151 | 2.374065718 | 0.2 | 0.062 | 1 | 0.522060045 | 0.145917484 |
| Mcfd2 | 0.075843212 | 0.192645078 | 0.866 | 0.753 | 1 | 0.76380627 | -0.143965774 | 0.7 | 0.688 | 1 | 0.76380627 | 0.145934232 |
| D030028A08Rik | 0.435188125 | 0.152003093 | 0.076 | 0.053 | 1 | 0.075915843 | 1.959028219 | 0.2 | 0.059 | 1 | 0.435188125 | 0.146068471 |
| Gm35608 | 0.188329318 | 0.444183845 | 0.109 | 0.065 | 1 | 0.075916244 | 1.558490289 | 0.2 | 0.059 | 1 | 0.188329318 | 0.146069211 |
| Gm9993 | 0.360641659 | -0.758445322 | 0.076 | 0.106 | 1 | 0.075916244 | 1.847996907 | 0.2 | 0.059 | 1 | 0.360641659 | 0.146069211 |
| 9130401M01Rik | 0.075977953 | -0.485426827 | 0.218 | 0.312 | 1 | 0.631966711 | -0.138582578 | 0.2 | 0.256 | 1 | 0.631966711 | 0.146183257 |
| Hprt | 0.35289118 | -0.222392421 | 0.655 | 0.676 | 1 | 0.075988233 | -0.811800827 | 0.4 | 0.716 | 1 | 0.35289118 | 0.146202255 |
| Pdcd6ip | 0.076019602 | -0.242263127 | 0.95 | 0.9 | 1 | 0.20213875 | -0.393223776 | 1 | 0.957 | 1 | 0.20213875 | 0.146260224 |
| Sik1 | 0.076039322 | -0.352160296 | 0.176 | 0.271 | 1 | 0.562982491 | 0.341461053 | 0.4 | 0.302 | 1 | 0.562982491 | 0.146296666 |
| Pcdhb20 | 0.076194626 | -1.900464326 | 0.017 | 0.059 | 1 | 0.522102507 | 0.930459067 | 0 | 0.04 | 1 | 0.522102507 | 0.146583631 |
| Gm48512 | 0.076195849 | -1.807354922 | 0.017 | 0.059 | 1 | 0.450330458 | 1.063725598 | 0.1 | 0.046 | 1 | 0.450330458 | 0.14658589 |
| Shroom3 | 0.07620118 | 0.569020957 | 0.143 | 0.076 | 1 | 0.213764069 | 1.150025444 | 0.3 | 0.157 | 1 | 0.213764069 | 0.146595741 |
| Mef2d | 0.076212833 | -0.398146502 | 0.773 | 0.759 | 1 | 0.110979787 | -0.599050988 | 0.9 | 0.938 | 1 | 0.110979787 | 0.146617269 |
| Celsr2 | 0.334866848 | -0.944858446 | 0.025 | 0.047 | 1 | 0.076218018 | 1.785261151 | 0.3 | 0.117 | 1 | 0.334866848 | 0.146626849 |
| Dmrta1 | 0.076220415 | 1.836501268 | 0.034 | 0.006 | 1 | 0.505686333 | 1.111031312 | 0 | 0.043 | 1 | 0.505686333 | 0.146631279 |
| Myc | 0.076231631 | -0.375549209 | 0.462 | 0.553 | 1 | 0.383099905 | -1.304006187 | 0.4 | 0.454 | 1 | 0.383099905 | 0.146652 |
| Chm | 0.200422707 | -0.386648673 | 0.647 | 0.659 | 1 | 0.076272724 | -1.024284652 | 0.7 | 0.759 | 1 | 0.200422707 | 0.146727919 |
| Mettl5 | 0.344856923 | -0.511899039 | 0.202 | 0.241 | 1 | 0.07629084 | -1.611434712 | 0 | 0.244 | 1 | 0.344856923 | 0.146761387 |
| Cap1 | 0.07630704 | -0.270687978 | 0.706 | 0.806 | 1 | 0.308969433 | -0.43658303 | 0.9 | 0.867 | 1 | 0.308969433 | 0.146791315 |
| Fbxl18 | 0.303518965 | -0.527247003 | 0.076 | 0.112 | 1 | 0.076327986 | 2.457206954 | 0.3 | 0.117 | 1 | 0.303518965 | 0.146830011 |
| Osgepl1 | 0.331299826 | -0.584962501 | 0.126 | 0.165 | 1 | 0.076346534 | 1.959028219 | 0.3 | 0.117 | 1 | 0.331299826 | 0.146864275 |
| Ddx59 | 0.455951659 | -0.347923303 | 0.059 | 0.082 | 1 | 0.076505997 | 2.385653692 | 0.2 | 0.062 | 1 | 0.455951659 | 0.147158826 |
| Slc25a14 | 0.610899327 | -0.428843299 | 0.101 | 0.118 | 1 | 0.076537978 | 2.152851488 | 0.3 | 0.127 | 1 | 0.610899327 | 0.147217894 |
| E230016M11Rik | 0.076604997 | -1.135240472 | 0.151 | 0.229 | 1 | 0.41152387 | -0.864721141 | 0.2 | 0.318 | 1 | 0.41152387 | 0.147341668 |
| Alas1 | 0.335295118 | -0.527247003 | 0.252 | 0.3 | 1 | 0.07670987 | 0.990015911 | 0.7 | 0.466 | 1 | 0.335295118 | 0.147535336 |
| Azin2 | 0.55758214 | -0.45845978 | 0.176 | 0.2 | 1 | 0.076728167 | 2.333423734 | 0.3 | 0.12 | 1 | 0.55758214 | 0.147569122 |
| Adcy1 | 0.076872468 | -1.900464326 | 0.017 | 0.059 | 1 | 0.619961444 | 1.558490289 | 0 | 0.025 | 1 | 0.619961444 | 0.147835559 |
| Gm49539 | 0.076872468 | -1.987927168 | 0.017 | 0.059 | 1 | 0.535011813 | 1.111031312 | 0.1 | 0.052 | 1 | 0.535011813 | 0.147835559 |
| Msh5 | 0.076872468 | -1.807354922 | 0.017 | 0.059 | 1 | 0.730749233 | 2.432959407 | 0 | 0.012 | 1 | 0.730749233 | 0.147835559 |
| Foxm1 | 0.076872775 | -1.900464326 | 0.017 | 0.059 | 1 | 0.669848329 | 1.558490289 | 0 | 0.019 | 1 | 0.669848329 | 0.147836126 |
| Wasf1 | 0.076872775 | -1.987927168 | 0.017 | 0.059 | 1 | 0.522066388 | 0.930459067 | 0 | 0.04 | 1 | 0.522066388 | 0.147836126 |
| Timm22 | 0.147064079 | -0.388750808 | 0.361 | 0.429 | 1 | 0.077165569 | -1.399930607 | 0.1 | 0.367 | 1 | 0.147064079 | 0.148376613 |
| Ggnbp1 | 0.480515999 | -0.485426827 | 0.05 | 0.071 | 1 | 0.077249551 | 1.959028219 | 0.2 | 0.059 | 1 | 0.480515999 | 0.148531609 |
| 6720427I07Rik | 0.077302082 | -0.551015169 | 0.227 | 0.324 | 1 | 0.095634188 | -1.776493958 | 0.1 | 0.343 | 1 | 0.095634188 | 0.148628553 |
| Zfp182 | 0.077348833 | 0.362570079 | 0.353 | 0.247 | 1 | 0.479568577 | 0.478763097 | 0.4 | 0.284 | 1 | 0.479568577 | 0.148714825 |
| Pip4k2c | 0.132982601 | 0.205820467 | 0.538 | 0.435 | 1 | 0.077351838 | 0.619890834 | 0.8 | 0.472 | 1 | 0.132982601 | 0.148720369 |
| Gm41790 | 0.077373108 | -0.459431619 | 0.109 | 0.188 | 1 | 0.140879753 | 1.31748219 | 0.4 | 0.21 | 1 | 0.140879753 | 0.148759618 |
| Ucp2 | 0.077439358 | -1.155278225 | 0.084 | 0.153 | 1 | 0.123773987 | -2.31299497 | 0 | 0.194 | 1 | 0.123773987 | 0.148881862 |
| Exosc4 | 0.077471797 | -0.769219793 | 0.185 | 0.265 | 1 | 0.100512397 | -1.425021588 | 0 | 0.216 | 1 | 0.100512397 | 0.148941715 |
| Washc3 | 0.077524201 | 0.24687739 | 0.588 | 0.476 | 1 | 0.66696872 | 0.18865221 | 0.3 | 0.42 | 1 | 0.66696872 | 0.1490384 |
| Cabin1 | 0.077538687 | -0.550521855 | 0.403 | 0.471 | 1 | 0.80148489 | -0.234743524 | 0.5 | 0.506 | 1 | 0.80148489 | 0.149065126 |
| Dnaja4 | 0.265759659 | 0.63005039 | 0.252 | 0.206 | 1 | 0.077539793 | 1.30520386 | 0.5 | 0.265 | 1 | 0.265759659 | 0.149067167 |
| Gpr45 | 0.077551868 | -1.600904045 | 0.017 | 0.059 | 1 | 0.505582845 | 0.930459067 | 0 | 0.043 | 1 | 0.505582845 | 0.149089443 |
| Fbln2 | 0.077551868 | -1.600904045 | 0.017 | 0.059 | 1 | 0.32099467 | 1.930459067 | 0.1 | 0.037 | 1 | 0.32099467 | 0.149089443 |
| Gm56912 | 0.077551868 | -1.600904045 | 0.017 | 0.059 | 1 | 0.505686333 | 1.111031312 | 0 | 0.043 | 1 | 0.505686333 | 0.149089443 |
| Tslp | 0.077551868 | -1.807354922 | 0.017 | 0.059 | 1 | 0.643850855 | 2.017921908 | 0 | 0.022 | 1 | 0.643850855 | 0.149089443 |
| Arfrp1 | 0.077552731 | -0.539874611 | 0.269 | 0.359 | 1 | 0.3811719 | -0.484578433 | 0.2 | 0.318 | 1 | 0.3811719 | 0.149091036 |
| Zfp51 | 0.077609298 | -1.222392421 | 0.067 | 0.129 | 1 | 0.970734692 | 0.354956895 | 0.2 | 0.188 | 1 | 0.970734692 | 0.149195393 |
| Trpv4 | 0.07765194 | -0.121323597 | 0.857 | 0.635 | 1 | 0.15174093 | 0.531655247 | 0.9 | 0.707 | 1 | 0.15174093 | 0.149274056 |
| Cdcp1 | 0.077804613 | 1.514573173 | 0.034 | 0.006 | 1 | 0.274237255 | -0.230005605 | 0 | 0.108 | 1 | 0.274237255 | 0.149555669 |
| Taf1b | 0.077819615 | -0.823296466 | 0.235 | 0.312 | 1 | 0.136531004 | 0.930459067 | 0.5 | 0.284 | 1 | 0.136531004 | 0.149583338 |
| Zcwpw2 | 0.288148609 | -1.807354922 | 0.067 | 0.1 | 1 | 0.077825993 | 2.432959407 | 0.2 | 0.062 | 1 | 0.288148609 | 0.149595102 |
| Rhoq | 0.077991691 | 0.191374541 | 0.941 | 0.835 | 1 | 0.150951276 | -0.711430502 | 0.9 | 0.941 | 1 | 0.150951276 | 0.149900678 |
| Cdca8 | 0.095266307 | -2.343407822 | 0.008 | 0.041 | 1 | 0.078038625 | 3.210566986 | 0.1 | 0.019 | 1 | 0.095266307 | 0.149987223 |
| Lmtk3 | 0.095769202 | -1.485426827 | 0.008 | 0.041 | 1 | 0.078038625 | 3.210566986 | 0.1 | 0.019 | 1 | 0.095769202 | 0.149987223 |
| Espnl | 0.1433296 | -1.655351829 | 0.008 | 0.035 | 1 | 0.078038625 | 3.210566986 | 0.1 | 0.019 | 1 | 0.1433296 | 0.149987223 |
| Notch4 | 0.215141111 | 1.099535674 | 0.059 | 0.029 | 1 | 0.078038625 | 3.210566986 | 0.1 | 0.019 | 1 | 0.215141111 | 0.149987223 |
| Odad4 | 0.329694367 | -0.600904045 | 0.034 | 0.059 | 1 | 0.078038625 | 3.210566986 | 0.1 | 0.019 | 1 | 0.329694367 | 0.149987223 |
| C030010C08Rik | 0.334204811 | -0.807354922 | 0.008 | 0.024 | 1 | 0.078038625 | 3.210566986 | 0.1 | 0.019 | 1 | 0.334204811 | 0.149987223 |
| 4930563E22Rik | 0.347966916 | -0.707819249 | 0.017 | 0.035 | 1 | 0.078038625 | 3.210566986 | 0.1 | 0.019 | 1 | 0.347966916 | 0.149987223 |
| Zfp973 | 0.491791629 | -0.707819249 | 0.017 | 0.029 | 1 | 0.078038625 | 3.210566986 | 0.1 | 0.019 | 1 | 0.491791629 | 0.149987223 |
| Gm2814 | 0.695893386 | -0.222392421 | 0.017 | 0.024 | 1 | 0.078038625 | 3.210566986 | 0.1 | 0.019 | 1 | 0.695893386 | 0.149987223 |
| Gm56859 | 0.929389703 | 0.192645078 | 0.025 | 0.024 | 1 | 0.078038625 | 3.210566986 | 0.1 | 0.019 | 1 | 0.929389703 | 0.149987223 |
| Slc16a10 | 0.71170832 | 0.181589889 | 0.261 | 0.235 | 1 | 0.078063191 | -3.517353469 | 0 | 0.244 | 1 | 0.71170832 | 0.15003252 |
| Gpatch2l | 0.218659204 | -0.600904045 | 0.21 | 0.265 | 1 | 0.078075407 | 0.880418384 | 0.6 | 0.306 | 1 | 0.218659204 | 0.150055045 |
| Pja2 | 0.078104136 | -0.345011708 | 0.824 | 0.818 | 1 | 0.602797965 | -0.117035433 | 0.8 | 0.873 | 1 | 0.602797965 | 0.150108015 |
| Itfg2 | 0.0781916 | -0.900464326 | 0.168 | 0.247 | 1 | 0.640163592 | -0.339630097 | 0.2 | 0.253 | 1 | 0.640163592 | 0.150269273 |
| G3bp1 | 0.078226703 | -0.230385212 | 0.916 | 0.906 | 1 | 0.114493942 | -0.638860456 | 0.8 | 0.917 | 1 | 0.114493942 | 0.150333988 |
| Chmp1b2 | 0.0782286 | -0.755515991 | 0.118 | 0.194 | 1 | 0.196398879 | -1.026472211 | 0 | 0.145 | 1 | 0.196398879 | 0.150337486 |
| Tprn | 0.078232801 | -1.485426827 | 0.017 | 0.059 | 1 | 0.489939838 | 0.625604485 | 0 | 0.046 | 1 | 0.489939838 | 0.15034523 |
| Tmprss5 | 0.078232801 | -1.485426827 | 0.017 | 0.059 | 1 | 0.433636001 | 0.432959407 | 0 | 0.059 | 1 | 0.433636001 | 0.15034523 |
| Zranb2 | 0.262672497 | -0.366187691 | 0.824 | 0.782 | 1 | 0.078235342 | -0.957274701 | 0.6 | 0.781 | 1 | 0.262672497 | 0.150349916 |
| Il6st | 0.078267619 | 0.143063049 | 0.899 | 0.876 | 1 | 0.258338442 | -0.236499368 | 0.9 | 0.923 | 1 | 0.258338442 | 0.150409418 |
| Mier1 | 0.159698354 | -0.281103727 | 0.941 | 0.929 | 1 | 0.078280604 | -0.719531001 | 1 | 0.975 | 1 | 0.159698354 | 0.150433356 |
| Muc4 | 0.078335216 | 0.514573173 | 0.034 | 0.006 | 1 | 0.669853054 | 2.210566986 | 0 | 0.019 | 1 | 0.669853054 | 0.150534026 |
| Sp100 | 0.07834103 | -0.378511623 | 0.034 | 0.006 | 1 | 0.730750993 | 1.111031312 | 0 | 0.012 | 1 | 0.730750993 | 0.150544743 |
| P2ry14 | 0.078402547 | 0.974004791 | 0.084 | 0.035 | 1 | 0.444918466 | 1.695993813 | 0.1 | 0.046 | 1 | 0.444918466 | 0.150658134 |
| Ring1 | 0.078592096 | -0.64689025 | 0.286 | 0.365 | 1 | 0.39086802 | 0.795529487 | 0.5 | 0.395 | 1 | 0.39086802 | 0.151007475 |
| Palmd | 0.927056084 | -1.254101281 | 0.143 | 0.129 | 1 | 0.078656991 | 1.03809979 | 0.4 | 0.176 | 1 | 0.927056084 | 0.15112706 |
| Epcam | 0.078700321 | -0.15680408 | 0.118 | 0.2 | 1 | 0.536774994 | -1.019167411 | 0.2 | 0.265 | 1 | 0.536774994 | 0.151206902 |
| Eml2 | 0.959218734 | -0.168953162 | 0.227 | 0.224 | 1 | 0.078828535 | -1.789433014 | 0 | 0.241 | 1 | 0.959218734 | 0.151443132 |
| Tmem265 | 0.078886097 | -0.854660637 | 0.084 | 0.153 | 1 | 0.749092797 | 0.420986766 | 0.3 | 0.256 | 1 | 0.749092797 | 0.151549177 |

| | | | | | | | | | | | | |
|---|---|---|---|---|---|---|---|---|---|---|---|---|
| Zfp330 | 0.078946923 | -0.454062656 | 0.521 | 0.582 | 1 | 0.418084571 | -0.30914949 | 0.4 | 0.515 | 1 | 0.418084571 | 0.15166123 |
| Fancg | 0.079004402 | -1.359895945 | 0.017 | 0.059 | 1 | 0.248008881 | 3.017921908 | 0.1 | 0.034 | 1 | 0.248008881 | 0.151767108 |
| Trim7 | 0.079004402 | -1.359895945 | 0.017 | 0.059 | 1 | 0.323434771 | 1.847996907 | 0.1 | 0.037 | 1 | 0.323434771 | 0.151767108 |
| Meikin | 0.079004402 | -1.359895945 | 0.017 | 0.059 | 1 | 0.505582845 | 1.017921908 | 0 | 0.043 | 1 | 0.505582845 | 0.151767108 |
| Gm20535 | 0.091361251 | 2.099535674 | 0.017 | 0 | 1 | 0.07916329 | 3.017921908 | 0.1 | 0.019 | 1 | 0.091361251 | 0.152059754 |
| Zfp873 | 0.112335908 | -1.600904045 | 0.017 | 0.053 | 1 | 0.07916329 | 2.695993813 | 0.1 | 0.019 | 1 | 0.112335908 | 0.152059754 |
| Stil | 0.16412139 | -2.292781749 | 0.034 | 0.071 | 1 | 0.07916329 | 2.695993813 | 0.1 | 0.019 | 1 | 0.16412139 | 0.152059754 |
| Gng8 | 0.170630732 | -1.572889668 | 0.025 | 0.059 | 1 | 0.07916329 | 2.111031312 | 0.1 | 0.019 | 1 | 0.170630732 | 0.152059754 |
| Gm30934 | 0.205759512 | 0.836501268 | 0.034 | 0.012 | 1 | 0.07916329 | 2.847996907 | 0.1 | 0.019 | 1 | 0.205759512 | 0.152059754 |
| 1700126G02Rik | 0.218485013 | -1.292781749 | 0.008 | 0.029 | 1 | 0.07916329 | 3.017921908 | 0.1 | 0.019 | 1 | 0.218485013 | 0.152059754 |
| Spc25 | 0.23664108 | -1.600904045 | 0.025 | 0.053 | 1 | 0.07916329 | 3.017921908 | 0.1 | 0.019 | 1 | 0.23664108 | 0.152059754 |
| Kcnc4 | 0.331040183 | -1.292781749 | 0.008 | 0.024 | 1 | 0.07916329 | 3.017921908 | 0.1 | 0.019 | 1 | 0.331040183 | 0.152059754 |
| Gm57243 | 0.334204811 | -0.807354922 | 0.008 | 0.024 | 1 | 0.07916329 | 2.695993813 | 0.1 | 0.019 | 1 | 0.334204811 | 0.152059754 |
| Serpinb6b | 0.337379399 | -0.707819249 | 0.008 | 0.024 | 1 | 0.07916329 | 2.695993813 | 0.1 | 0.019 | 1 | 0.337379399 | 0.152059754 |
| Gm17135 | 0.390876134 | 0.929610672 | 0.025 | 0.012 | 1 | 0.07916329 | 2.847996907 | 0.1 | 0.019 | 1 | 0.390876134 | 0.152059754 |
| Dpf1 | 0.508731273 | -1.070389328 | 0.008 | 0.018 | 1 | 0.07916329 | 3.017921908 | 0.1 | 0.019 | 1 | 0.508731273 | 0.152059754 |
| Jag2 | 0.721161231 | 0.514573173 | 0.017 | 0.012 | 1 | 0.07916329 | 2.847996907 | 0.1 | 0.019 | 1 | 0.721161231 | 0.152059754 |
| Tjp3 | 0.721162042 | 0.514573173 | 0.017 | 0.012 | 1 | 0.07916329 | 2.695993813 | 0.1 | 0.019 | 1 | 0.721162042 | 0.152059754 |
| ENSMUSG00000121456 | 0.926217033 | -0.485426827 | 0.034 | 0.035 | 1 | 0.07916329 | 2.847996907 | 0.1 | 0.019 | 1 | 0.926217033 | 0.152059754 |
| Trpm4 | 0.07932954 | -1.655351829 | 0.059 | 0.118 | 1 | 0.895499827 | 0.403212064 | 0.1 | 0.111 | 1 | 0.895499827 | 0.152365904 |
| Rexo4 | 0.447616633 | -0.469659511 | 0.311 | 0.329 | 1 | 0.079386118 | -1.433289204 | 0.1 | 0.364 | 1 | 0.447616633 | 0.152470081 |
| Cpsf1 | 0.079427914 | -0.970853654 | 0.076 | 0.141 | 1 | 0.744574595 | 0.210566986 | 0.1 | 0.133 | 1 | 0.744574595 | 0.152547034 |
| Ssh3 | 0.079492144 | -0.833350131 | 0.084 | 0.153 | 1 | 0.135281863 | -1.248864633 | 0 | 0.185 | 1 | 0.135281863 | 0.152665287 |
| Irf9 | 0.834988438 | -0.197081333 | 0.378 | 0.371 | 1 | 0.079496682 | -1.833827133 | 0.1 | 0.358 | 1 | 0.834988438 | 0.152673642 |
| Gm54310 | 0.079555112 | 0.80407979 | 0.076 | 0.029 | 1 | 0.149795061 | 2.847996907 | 0.1 | 0.025 | 1 | 0.149795061 | 0.152781207 |
| Nhp2 | 0.25027316 | -0.338200874 | 0.42 | 0.459 | 1 | 0.079680141 | 1.147557188 | 0.5 | 0.269 | 1 | 0.25027316 | 0.153011357 |
| Nexmif | 0.079694088 | -1.687060688 | 0.017 | 0.059 | 1 | 0.201131228 | 1.558490289 | 0.1 | 0.028 | 1 | 0.201131228 | 0.153037029 |
| Trappc10 | 0.079708278 | -0.536773707 | 0.395 | 0.471 | 1 | 0.111264158 | -0.617666666 | 0.3 | 0.577 | 1 | 0.111264158 | 0.153063147 |
| Retreg3 | 0.079729266 | -0.867896464 | 0.21 | 0.288 | 1 | 0.411763664 | -0.428334322 | 0.3 | 0.429 | 1 | 0.411763664 | 0.153101776 |
| Apba3 | 0.199182876 | -0.549557165 | 0.16 | 0.218 | 1 | 0.07976199 | 0.959028219 | 0.5 | 0.231 | 1 | 0.199182876 | 0.153162004 |
| Fam210b | 0.079803961 | 0.988504361 | 0.16 | 0.094 | 1 | 0.710404476 | 0.808468542 | 0.1 | 0.068 | 1 | 0.710404476 | 0.153239249 |
| Tshr | 0.721162042 | 0.514573173 | 0.017 | 0.012 | 1 | 0.079824511 | 2.270687978 | 0.2 | 0.062 | 1 | 0.721162042 | 0.15327707 |
| Bud31 | 0.50624665 | 0.101198723 | 0.782 | 0.741 | 1 | 0.079842974 | -0.837796036 | 0.5 | 0.762 | 1 | 0.50624665 | 0.153311047 |
| Dars | 0.079916548 | -0.553813802 | 0.622 | 0.618 | 1 | 0.307688413 | -0.604129911 | 0.5 | 0.62 | 1 | 0.307688413 | 0.153446441 |
| Exoc3l2 | 0.079991593 | 0.514573173 | 0.084 | 0.035 | 1 | 0.730750993 | 2.210566986 | 0 | 0.012 | 1 | 0.730750993 | 0.153584531 |
| Ccdc68 | 0.157538651 | -0.959358016 | 0.059 | 0.106 | 1 | 0.079998199 | -2.161987182 | 0 | 0.241 | 1 | 0.157538651 | 0.153596686 |
| Ap4s1 | 0.515939596 | 0.262509885 | 0.454 | 0.459 | 1 | 0.080005351 | -0.950168844 | 0.2 | 0.512 | 1 | 0.515939596 | 0.153609846 |
| Agbl2 | 0.080041423 | -1.392317423 | 0.017 | 0.059 | 1 | 0.669852109 | 1.111031312 | 0 | 0.019 | 1 | 0.669852109 | 0.153676216 |
| Smarcd3 | 0.080042107 | -1.03796785 | 0.109 | 0.182 | 1 | 0.843684972 | 0.374065718 | 0.2 | 0.167 | 1 | 0.843684972 | 0.153677475 |
| Gm57402 | 0.080042107 | -0.86393845 | 0.109 | 0.182 | 1 | 0.577022844 | -0.217294554 | 0.2 | 0.269 | 1 | 0.577022844 | 0.153677475 |
| Dnajb9 | 0.080054593 | 0.257360608 | 0.908 | 0.859 | 1 | 0.232137314 | -0.660149997 | 0.8 | 0.907 | 1 | 0.232137314 | 0.153700449 |
| Reg3g | 0.388777089 | 0.462316936 | 0.303 | 0.365 | 1 | 0.080094444 | -1.876895855 | 0.2 | 0.454 | 1 | 0.388777089 | 0.153773769 |
| Itprid2 | 0.080106772 | -0.948398804 | 0.235 | 0.318 | 1 | 0.360987675 | 0.71414116 | 0.5 | 0.454 | 1 | 0.360987675 | 0.15379645 |
| Adh7 | 0.080234283 | 1.035405336 | 0.05 | 0.112 | 1 | 0.081418918 | -2.408342847 | 0 | 0.238 | 1 | 0.081418918 | 0.154031026 |
| Tlnrd1 | 0.080290079 | -0.744813456 | 0.21 | 0.294 | 1 | 0.150781369 | -1.239465935 | 0.1 | 0.306 | 1 | 0.150781369 | 0.154133661 |
| Ercc2 | 0.205718882 | -0.5360529 | 0.092 | 0.141 | 1 | 0.080314922 | 1.231325546 | 0.4 | 0.17 | 1 | 0.205718882 | 0.154179357 |
| Icam2 | 0.147030916 | -2.070389328 | 0 | 0.018 | 1 | 0.080320981 | 2.558490289 | 0.1 | 0.019 | 1 | 0.147030916 | 0.154190502 |
| Zfp660 | 0.218485013 | -1.485426827 | 0.008 | 0.029 | 1 | 0.080320981 | 2.847996907 | 0.1 | 0.019 | 1 | 0.218485013 | 0.154190502 |
| Gm26737 | 0.220673861 | -2.807354922 | 0.008 | 0.029 | 1 | 0.080320981 | 2.695993813 | 0.1 | 0.019 | 1 | 0.220673861 | 0.154190502 |
| Gm57064 | 0.237885589 | -2.292781749 | 0 | 0.012 | 1 | 0.080320981 | 2.695993813 | 0.1 | 0.019 | 1 | 0.237885589 | 0.154190502 |
| Plcxd3 | 0.342819381 | -1.748461233 | 0.017 | 0.035 | 1 | 0.080320981 | 2.31748219 | 0.1 | 0.019 | 1 | 0.342819381 | 0.154190502 |
| Gm32191 | 0.508731273 | -0.807354922 | 0.008 | 0.018 | 1 | 0.080320981 | 2.695993813 | 0.1 | 0.019 | 1 | 0.508731273 | 0.154190502 |
| Ttbk1 | 0.82670638 | -0.485426827 | 0.025 | 0.029 | 1 | 0.080320981 | 2.31748219 | 0.1 | 0.019 | 1 | 0.82670638 | 0.154190502 |
| Wdr18 | 0.08037015 | -0.722466024 | 0.193 | 0.276 | 1 | 0.239853971 | -0.827568143 | 0.1 | 0.259 | 1 | 0.239853971 | 0.154280939 |
| Tjp2 | 0.08040699 | 0.317271732 | 0.387 | 0.282 | 1 | 0.71666024 | 0.348643121 | 0.7 | 0.543 | 1 | 0.71666024 | 0.154348696 |
| Galnt17 | 0.08040858 | -2.185866545 | 0.017 | 0.059 | 1 | 0.539475467 | -0.864721141 | 0 | 0.037 | 1 | 0.539475467 | 0.154351621 |
| Nudt3 | 0.250531115 | -0.443847723 | 0.538 | 0.524 | 1 | 0.080586212 | -1 | 0.4 | 0.608 | 1 | 0.250531115 | 0.154676286 |
| Ranbp10 | 0.080625075 | -0.713232745 | 0.445 | 0.482 | 1 | 0.547438541 | -0.364996817 | 0.6 | 0.62 | 1 | 0.547438541 | 0.154749748 |
| Nucb2 | 0.080632566 | -0.416839742 | 0.622 | 0.659 | 1 | 0.699902862 | 0.35384069 | 0.6 | 0.63 | 1 | 0.699902862 | 0.154763521 |
| Ppp2r3a | 0.080679495 | -0.344180902 | 0.597 | 0.671 | 1 | 0.414334885 | -0.633129783 | 0.7 | 0.747 | 1 | 0.414334885 | 0.154849809 |
| Tmem238 | 0.080737445 | -0.772307975 | 0.176 | 0.259 | 1 | 0.21939837 | -0.970762779 | 0.1 | 0.269 | 1 | 0.21939837 | 0.154956355 |
| Exd2 | 0.080769524 | -1.265645619 | 0.109 | 0.188 | 1 | 0.870507854 | 0.374065718 | 0.1 | 0.114 | 1 | 0.870507854 | 0.155015332 |
| Recql5 | 0.999131864 | -0.211251864 | 0.176 | 0.171 | 1 | 0.080776216 | 1.453137289 | 0.3 | 0.114 | 1 | 0.999131864 | 0.155027635 |
| Id1 | 0.246170632 | 0.25823342 | 0.538 | 0.506 | 1 | 0.080789772 | -1.797141109 | 0.3 | 0.562 | 1 | 0.246170632 | 0.155052557 |
| Usp27x | 0.080924463 | 0.707218251 | 0.126 | 0.065 | 1 | 0.09920356 | 2.079322453 | 0.2 | 0.065 | 1 | 0.09920356 | 0.155300158 |
| Cfap45 | 0.080972186 | -2.070389328 | 0.025 | 0.071 | 1 | 0.175528605 | -1.28585884 | 0 | 0.157 | 1 | 0.175528605 | 0.155387877 |
| Pinx1 | 0.081080661 | -1.041820176 | 0.118 | 0.188 | 1 | 0.489415639 | 0.808468542 | 0.3 | 0.213 | 1 | 0.489415639 | 0.155587249 |
| Wdr5b | 0.081107383 | -1.185866545 | 0.017 | 0.059 | 1 | 0.446953437 | 0.769994395 | 0 | 0.056 | 1 | 0.446953437 | 0.155636359 |
| Slc25a3 | 0.138099214 | -0.135963736 | 0.975 | 0.982 | 1 | 0.081118015 | -0.527610448 | 0.9 | 0.994 | 1 | 0.138099214 | 0.155655897 |
| Ttc1 | 0.680926287 | -0.231243339 | 0.412 | 0.412 | 1 | 0.081183956 | -1.633129783 | 0.1 | 0.361 | 1 | 0.680926287 | 0.155777077 |
| Mfap1a | 0.081199471 | -0.352160296 | 0.286 | 0.376 | 1 | 0.491695226 | 0.190102883 | 0.2 | 0.327 | 1 | 0.491695226 | 0.155805587 |
| Dnajc17 | 0.081235035 | -1.122856748 | 0.067 | 0.129 | 1 | 0.374563499 | 0.111031312 | 0 | 0.074 | 1 | 0.374563499 | 0.155870939 |
| Atp9b | 0.199423939 | -0.19730144 | 0.748 | 0.771 | 1 | 0.081244452 | 0.69951688 | 0.9 | 0.833 | 1 | 0.199423939 | 0.155888244 |
| Gk | 0.982474945 | 0.396736683 | 0.143 | 0.153 | 1 | 0.081248825 | -1.723545078 | 0 | 0.238 | 1 | 0.982474945 | 0.155896278 |
| Pfkfb1 | 0.081454382 | -1.185866545 | 0.017 | 0.059 | 1 | 0.154873683 | 2.432959407 | 0.1 | 0.025 | 1 | 0.154873683 | 0.156273948 |
| Gm36129 | 0.237885589 | -1.070389328 | 0 | 0.012 | 1 | 0.08148995 | 2.558490289 | 0.1 | 0.019 | 1 | 0.237885589 | 0.156339288 |
| 4933406J09Rik | 0.338976313 | -0.485426827 | 0.008 | 0.024 | 1 | 0.081490349 | 1.695993813 | 0.1 | 0.019 | 1 | 0.338976313 | 0.15634002 |
| Serpind1 | 0.609500043 | 0.514573173 | 0.034 | 0.024 | 1 | 0.081490349 | 2.31748219 | 0.1 | 0.019 | 1 | 0.609500043 | 0.15634002 |
| Gm57236 | 0.666161437 | 0.192645078 | 0.025 | 0.018 | 1 | 0.081490349 | 2.695993813 | 0.1 | 0.019 | 1 | 0.666161437 | 0.15634002 |
| Krt8 | 0.114400675 | 0.472752997 | 0.109 | 0.182 | 1 | 0.08149679 | -1.723545078 | 0 | 0.238 | 1 | 0.114400675 | 0.156351853 |
| Ggps1 | 0.933943269 | -0.196430627 | 0.706 | 0.635 | 1 | 0.081621069 | -0.894967428 | 0.4 | 0.688 | 1 | 0.933943269 | 0.156580139 |
| Vti1b | 0.081750052 | 0.182729609 | 0.782 | 0.671 | 1 | 0.391437656 | -0.307608424 | 0.6 | 0.676 | 1 | 0.391437656 | 0.156817033 |
| Stx1b | 0.081816921 | -1.070389328 | 0.017 | 0.059 | 1 | 0.231905241 | 2.017921908 | 0.2 | 0.099 | 1 | 0.231905241 | 0.156939834 |
| Rfx1 | 0.081855076 | -0.868755467 | 0.176 | 0.253 | 1 | 0.500359921 | 0.185031894 | 0.5 | 0.333 | 1 | 0.500359921 | 0.157009898 |
| Atp13a1 | 0.081891989 | -0.59645814 | 0.185 | 0.271 | 1 | 0.56109131 | -0.315978829 | 0.2 | 0.272 | 1 | 0.56109131 | 0.157077681 |
| Gm57325 | 0.081939068 | -1.807354922 | 0.025 | 0.071 | 1 | 0.557729726 | 1.31748219 | 0 | 0.034 | 1 | 0.557729726 | 0.157164126 |
| Zc3h12c | 0.081970809 | -1.180572246 | 0.151 | 0.224 | 1 | 0.091787965 | -1.68251781 | 0.1 | 0.349 | 1 | 0.091787965 | 0.157222405 |
| Miga1 | 0.082105586 | -0.980191519 | 0.16 | 0.235 | 1 | 0.251618696 | -1.033740212 | 0.2 | 0.349 | 1 | 0.251618696 | 0.157469845 |
| Eya4 | 0.142055489 | -0.58345891 | 0.714 | 0.7 | 1 | 0.082261568 | 0.268214645 | 1 | 0.769 | 1 | 0.142055489 | 0.157756171 |
| Nopchap1 | 0.082329783 | -0.59645814 | 0.277 | 0.365 | 1 | 0.113117915 | -2.004445905 | 0.1 | 0.324 | 1 | 0.113117915 | 0.157881374 |
| Emc6 | 0.162950055 | 0.18167396 | 0.647 | 0.541 | 1 | 0.082332983 | -0.898554736 | 0.4 | 0.608 | 1 | 0.162950055 | 0.157887245 |
| Mecom | 0.241128552 | -0.273922722 | 0.193 | 0.271 | 1 | 0.082338878 | -2.080110175 | 0 | 0.238 | 1 | 0.241128552 | 0.157898066 |
| Usb1 | 0.082351155 | -1.070389328 | 0.101 | 0.171 | 1 | 0.232002979 | -0.625934282 | 0 | 0.127 | 1 | 0.232002979 | 0.157920597 |
| Wdr83os | 0.0824709 | 0.227183497 | 0.756 | 0.676 | 1 | 0.704948786 | 0.254156254 | 0.5 | 0.602 | 1 | 0.704948786 | 0.15814035 |
| Gm20257 | 0.082579434 | -0.826463745 | 0.109 | 0.182 | 1 | 0.213477562 | -0.763437866 | 0 | 0.136 | 1 | 0.213477562 | 0.158339505 |
| Mturn | 0.082586516 | -0.86393845 | 0.101 | 0.171 | 1 | 0.978431889 | 0.193493473 | 0.2 | 0.185 | 1 | 0.978431889 | 0.1583525 |
| Ahsa2 | 0.167362535 | -0.721494185 | 0.319 | 0.359 | 1 | 0.082632707 | -1.380109166 | 0.2 | 0.454 | 1 | 0.167362535 | 0.158437251 |
| Cmas | 0.830561328 | -0.212050477 | 0.479 | 0.471 | 1 | 0.082651107 | -0.830074999 | 0.2 | 0.512 | 1 | 0.830561328 | 0.158471008 |

| | | | | | | | | | | | | |
|---|---|---|---|---|---|---|---|---|---|---|---|---|
| Gtf3c6 | 0.082656591 | -0.639754974 | 0.218 | 0.306 | 1 | 0.119734381 | 1.063725598 | 0.5 | 0.296 | 1 | 0.119734381 | 0.15848107 |
| Sox7 | 0.396130756 | 0.514573173 | 0.025 | 0.012 | 1 | 0.082671855 | 1.847996907 | 0.1 | 0.019 | 1 | 0.396130756 | 0.158509074 |
| Edn2 | 0.840043733 | 2.303069068 | 0.05 | 0.047 | 1 | 0.082671855 | -0.567040593 | 0.1 | 0.019 | 1 | 0.840043733 | 0.158509074 |
| Lss | 0.082675926 | 0.347116427 | 0.319 | 0.229 | 1 | 0.125018628 | 1.295455884 | 0.3 | 0.13 | 1 | 0.125018628 | 0.158516544 |
| Mettl7a1 | 0.082683284 | 0.611146433 | 0.504 | 0.453 | 1 | 0.970049567 | 0.3124466 | 0.6 | 0.537 | 1 | 0.970049567 | 0.158530043 |
| Zfp467 | 0.082720225 | -1.292781749 | 0.059 | 0.118 | 1 | 0.202168315 | -1.191531458 | 0 | 0.142 | 1 | 0.202168315 | 0.158597815 |
| Haus6 | 0.082721076 | -1.102098188 | 0.109 | 0.176 | 1 | 0.145862455 | 1.732519689 | 0.3 | 0.139 | 1 | 0.145862455 | 0.158599375 |
| Tmem19 | 0.842811602 | -0.237499314 | 0.244 | 0.224 | 1 | 0.082810663 | 1.159940913 | 0.5 | 0.25 | 1 | 0.842811602 | 0.158763721 |
| Gbe1 | 0.082849493 | -0.788618359 | 0.277 | 0.365 | 1 | 0.676560844 | -0.58364256 | 0.5 | 0.515 | 1 | 0.676560844 | 0.158834948 |
| Cip2a | 0.082917942 | -2.240314329 | 0.025 | 0.071 | 1 | 0.643760887 | 1.111031312 | 0.1 | 0.062 | 1 | 0.643760887 | 0.158960499 |
| 9530077C05Rik | 0.082918201 | -1.655351829 | 0.025 | 0.071 | 1 | 0.307197874 | -0.374395515 | 0 | 0.096 | 1 | 0.307197874 | 0.158960974 |
| Qpct | 0.083055389 | -1.029747343 | 0.076 | 0.141 | 1 | 0.568002172 | 1.409112665 | 0.2 | 0.145 | 1 | 0.568002172 | 0.159212581 |
| Ctu2 | 0.083090633 | -0.936088236 | 0.118 | 0.188 | 1 | 0.576013204 | -0.191531458 | 0.1 | 0.164 | 1 | 0.576013204 | 0.159277212 |
| Cacna1d | 0.083306941 | -1.607417352 | 0.109 | 0.176 | 1 | 0.615352752 | -0.674569057 | 0.3 | 0.361 | 1 | 0.615352752 | 0.159673835 |
| Svopl | 0.088618012 | -2.163498732 | 0.034 | 0.082 | 1 | 0.083307997 | 0.548436625 | 0.6 | 0.306 | 1 | 0.088618012 | 0.159675771 |
| Smc2 | 0.196464919 | -1.42604828 | 0.143 | 0.194 | 1 | 0.083321583 | 1.210566986 | 0.3 | 0.114 | 1 | 0.196464919 | 0.159700679 |
| Polr2j | 0.083363228 | -0.703331858 | 0.319 | 0.388 | 1 | 0.144338084 | -1.152003093 | 0.1 | 0.309 | 1 | 0.144338084 | 0.159777029 |
| Rabgap1l | 0.083373064 | -0.290481411 | 1 | 0.994 | 1 | 0.161747494 | 0.181026515 | 1 | 0.997 | 1 | 0.161747494 | 0.159795061 |
| Nagpa | 0.083390491 | -0.86393845 | 0.076 | 0.141 | 1 | 0.976598179 | 0.732519689 | 0.1 | 0.096 | 1 | 0.976598179 | 0.159827008 |
| Nudt22 | 0.083390491 | -0.86393845 | 0.076 | 0.141 | 1 | 0.984621079 | 0.135278859 | 0.1 | 0.099 | 1 | 0.984621079 | 0.159827008 |
| Proca1 | 0.083515305 | -0.987927168 | 0.076 | 0.141 | 1 | 0.806199775 | 0.159940913 | 0.1 | 0.123 | 1 | 0.806199775 | 0.160055805 |
| Commd9 | 0.12452488 | -0.687060688 | 0.143 | 0.212 | 1 | 0.083539386 | -1.489872732 | 0 | 0.235 | 1 | 0.12452488 | 0.160099942 |
| Mycbpap | 0.083592522 | -1.622930351 | 0.025 | 0.071 | 1 | 0.522115639 | 0.558490289 | 0 | 0.04 | 1 | 0.522115639 | 0.160197334 |
| Cdkl5 | 0.083603127 | -0.867896464 | 0.294 | 0.365 | 1 | 0.342034824 | -1.146985019 | 0.3 | 0.404 | 1 | 0.342034824 | 0.160216771 |
| Zfp971 | 0.083670363 | -1.485426827 | 0.05 | 0.106 | 1 | 0.994871611 | 0.625604485 | 0.1 | 0.099 | 1 | 0.994871611 | 0.160339996 |
| Trmt9b | 0.083686583 | -1.748461233 | 0.034 | 0.082 | 1 | 0.385661066 | -0.473931188 | 0 | 0.071 | 1 | 0.385661066 | 0.160369721 |
| Hspb8 | 0.270883698 | 0.590522026 | 0.193 | 0.147 | 1 | 0.083735155 | -1.750262417 | 0 | 0.235 | 1 | 0.270883698 | 0.160458734 |
| Egflam | 0.083760429 | -1.807354922 | 0.042 | 0.094 | 1 | 0.597836679 | 1.111031312 | 0 | 0.028 | 1 | 0.597836679 | 0.160505049 |
| Pbdc1 | 0.085865044 | -0.366463995 | 0.798 | 0.818 | 1 | 0.083797781 | -0.942466236 | 0.5 | 0.818 | 1 | 0.085865044 | 0.160573494 |
| Nenf | 0.083895017 | -0.294183104 | 0.866 | 0.888 | 1 | 0.100452799 | -0.509555098 | 0.5 | 0.836 | 1 | 0.100452799 | 0.16075166 |
| Gm1123 | 0.083898758 | -1.733354341 | 0.025 | 0.071 | 1 | 0.503837353 | 0.374065718 | 0.2 | 0.12 | 1 | 0.503837353 | 0.160758515 |
| Inca1 | 0.083898758 | -1.733354341 | 0.025 | 0.071 | 1 | 0.522066388 | 0.930459067 | 0 | 0.04 | 1 | 0.522066388 | 0.160758515 |
| Dnajb5 | 0.659299372 | 0.344648171 | 0.059 | 0.047 | 1 | 0.083924373 | 2.210566986 | 0.2 | 0.062 | 1 | 0.659299372 | 0.160805445 |
| Orc5 | 0.083941554 | 0.334000927 | 0.294 | 0.206 | 1 | 0.561366012 | 0.374065718 | 0.1 | 0.188 | 1 | 0.561366012 | 0.160836924 |
| Dgcr6 | 0.083976482 | -0.485426827 | 0.311 | 0.406 | 1 | 0.942140256 | 0.397335498 | 0.3 | 0.299 | 1 | 0.942140256 | 0.160900914 |
| Mrps30 | 0.083984316 | -0.509273569 | 0.227 | 0.318 | 1 | 0.199399955 | -0.840059087 | 0.1 | 0.281 | 1 | 0.199399955 | 0.160915267 |
| Krt5 | 0.567699227 | 0.377069649 | 0.042 | 0.029 | 1 | 0.084001278 | -2.41670632 | 0 | 0.235 | 1 | 0.567699227 | 0.16094634 |
| Rfc5 | 0.084070758 | -1.173482821 | 0.059 | 0.118 | 1 | 0.982277984 | 0.808468542 | 0.1 | 0.102 | 1 | 0.982277984 | 0.161073624 |
| Pigb | 0.08411866 | -0.707819249 | 0.118 | 0.194 | 1 | 0.629951339 | 0.973527789 | 0.2 | 0.151 | 1 | 0.629951339 | 0.16116137 |
| Pnpla2 | 0.964632633 | -0.267835392 | 0.193 | 0.182 | 1 | 0.084155505 | -1.69632361 | 0 | 0.235 | 1 | 0.964632633 | 0.16122886 |
| Unc79 | 0.084157913 | -1.452259963 | 0.059 | 0.118 | 1 | 0.887193786 | 0.210566986 | 0.1 | 0.083 | 1 | 0.887193786 | 0.161233272 |
| Rprd1a | 0.084226015 | -0.707819249 | 0.454 | 0.512 | 1 | 0.212723137 | -1.000741936 | 0.5 | 0.562 | 1 | 0.212723137 | 0.161358008 |
| Avl9 | 0.08423138 | -0.744161096 | 0.454 | 0.494 | 1 | 0.876023451 | -0.181096005 | 0.7 | 0.716 | 1 | 0.876023451 | 0.161367835 |
| Dapk3 | 0.084261231 | -0.520192245 | 0.286 | 0.371 | 1 | 0.130267991 | -1.111361109 | 0.1 | 0.321 | 1 | 0.130267991 | 0.161422507 |
| Wdcp | 0.08431146 | -0.696930932 | 0.126 | 0.2 | 1 | 0.9825718 | 0.127150978 | 0.2 | 0.191 | 1 | 0.9825718 | 0.161514498 |
| Vta1 | 0.266903066 | -0.402010819 | 0.529 | 0.529 | 1 | 0.084327722 | -1.04385429 | 0.4 | 0.599 | 1 | 0.266903066 | 0.16154428 |
| Coq10a | 0.248741027 | -0.559427409 | 0.151 | 0.2 | 1 | 0.084367594 | -1.69632361 | 0 | 0.235 | 1 | 0.248741027 | 0.161617297 |
| Trmt1l | 0.239629343 | -0.376355128 | 0.504 | 0.529 | 1 | 0.084368245 | -1.141949429 | 0.3 | 0.58 | 1 | 0.239629343 | 0.161618489 |
| Smpd3 | 0.084446126 | -3.516645558 | 0.084 | 0.141 | 1 | 0.374732071 | -2.596787936 | 0 | 0.074 | 1 | 0.374732071 | 0.161761103 |
| Zbtb14 | 0.084454263 | -1.185866545 | 0.059 | 0.118 | 1 | 0.600624496 | 1.08718457 | 0.2 | 0.148 | 1 | 0.600624496 | 0.161776003 |
| Casp8ap2 | 0.08447995 | -0.812788808 | 0.294 | 0.365 | 1 | 0.108282931 | -1.489872732 | 0.2 | 0.423 | 1 | 0.108282931 | 0.161823038 |
| Mthfsl | 0.084487652 | -0.461179281 | 0.202 | 0.294 | 1 | 0.763974546 | 0.494359952 | 0.3 | 0.262 | 1 | 0.763974546 | 0.16183714 |
| Meis2 | 0.439514884 | -0.678071905 | 0.067 | 0.094 | 1 | 0.084634596 | -2.465893869 | 0 | 0.235 | 1 | 0.439514884 | 0.162106177 |
| Hsd17b11 | 0.084637386 | 0.132702537 | 0.529 | 0.394 | 1 | 0.084950114 | -1.446964141 | 0.2 | 0.454 | 1 | 0.084950114 | 0.162111286 |
| Gm43445 | 0.08468161 | -0.970853654 | 0.067 | 0.129 | 1 | 0.678986641 | 0.973527789 | 0.1 | 0.065 | 1 | 0.678986641 | 0.162192246 |
| Chchd5 | 0.084730918 | -0.883976204 | 0.076 | 0.141 | 1 | 0.363859515 | 0.159940913 | 0 | 0.077 | 1 | 0.363859515 | 0.162282507 |
| Mmgt1 | 0.084746874 | -0.606442228 | 0.294 | 0.365 | 1 | 0.311658071 | 0.50836681 | 0.6 | 0.414 | 1 | 0.311658071 | 0.162311715 |
| Zbtb17 | 0.084813959 | -0.914270126 | 0.101 | 0.171 | 1 | 0.492003246 | 0.374065718 | 0.4 | 0.272 | 1 | 0.492003246 | 0.162434511 |
| Gm43387 | 0.084886415 | -1.485426827 | 0.025 | 0.071 | 1 | 0.557764571 | 0.930459067 | 0 | 0.034 | 1 | 0.557764571 | 0.162567127 |
| Sms | 0.08505288 | -0.741081703 | 0.311 | 0.394 | 1 | 0.729866563 | 0.614199722 | 0.4 | 0.386 | 1 | 0.729866563 | 0.162871767 |
| Gm38910 | 0.124189855 | -0.839063782 | 0.059 | 0.112 | 1 | 0.085109687 | 2.14345279 | 0.2 | 0.062 | 1 | 0.124189855 | 0.162975715 |
| Mus81 | 0.761526836 | -0.432959407 | 0.109 | 0.118 | 1 | 0.085109687 | 2.14345279 | 0.2 | 0.062 | 1 | 0.761526836 | 0.162975715 |
| Zfp574 | 0.085229871 | -0.857395605 | 0.126 | 0.2 | 1 | 0.347390193 | -0.125036046 | 0.1 | 0.25 | 1 | 0.347390193 | 0.163195612 |
| Mpp2 | 0.085257132 | -1.432959407 | 0.05 | 0.106 | 1 | 0.315846856 | -0.267480311 | 0 | 0.093 | 1 | 0.315846856 | 0.163245485 |
| Vnn1 | 0.085355613 | 1.675037845 | 0.143 | 0.082 | 1 | 0.179618702 | 1.913585248 | 0.2 | 0.083 | 1 | 0.179618702 | 0.163425646 |
| Arrb1 | 0.085371967 | 0.621488377 | 0.101 | 0.047 | 1 | 0.126860282 | 1.432959407 | 0.2 | 0.068 | 1 | 0.126860282 | 0.163455562 |
| Tnfaip6 | 0.085482209 | 1.843696769 | 0.252 | 0.165 | 1 | 0.229657818 | 2.14345279 | 0.2 | 0.096 | 1 | 0.229657818 | 0.163657209 |
| Bpifb1 | 0.23847629 | -1.600904045 | 0.017 | 0.041 | 1 | 0.085507763 | 2.667424661 | 0.2 | 0.068 | 1 | 0.23847629 | 0.163703949 |
| Nagk | 0.125673074 | -0.448900951 | 0.244 | 0.324 | 1 | 0.085516786 | 1.306427001 | 0.5 | 0.293 | 1 | 0.125673074 | 0.16372045 |
| Slc25a19 | 0.085584149 | -1.269698136 | 0.059 | 0.118 | 1 | 0.831639436 | 0.463333056 | 0.1 | 0.12 | 1 | 0.831639436 | 0.163843651 |
| Tmx4 | 0.085612979 | -0.746966986 | 0.613 | 0.647 | 1 | 0.602268418 | -0.630735268 | 0.6 | 0.648 | 1 | 0.602268418 | 0.163896376 |
| Tcirg1 | 0.429235064 | -0.448900951 | 0.143 | 0.176 | 1 | 0.085847841 | 0.667424661 | 0.4 | 0.17 | 1 | 0.429235064 | 0.164325831 |
| Ccnq | 0.678698362 | 0.120294234 | 0.227 | 0.206 | 1 | 0.085865995 | 1.231325546 | 0.4 | 0.176 | 1 | 0.678698362 | 0.164359021 |
| Actr8 | 0.085872507 | -0.900464326 | 0.168 | 0.247 | 1 | 0.642333469 | 0.290001453 | 0.4 | 0.315 | 1 | 0.642333469 | 0.164370926 |
| Or2k2 | 0.085880134 | -1.392317423 | 0.025 | 0.071 | 1 | 0.968372505 | 0.432959407 | 0.1 | 0.093 | 1 | 0.968372505 | 0.16438487 |
| Trim68 | 0.085880134 | -1.655351829 | 0.025 | 0.071 | 1 | 0.290412063 | -0.596787936 | 0 | 0.102 | 1 | 0.290412063 | 0.16438487 |
| Zfp747 | 0.085880134 | -1.392317423 | 0.025 | 0.071 | 1 | 0.191899351 | 1.473601392 | 0.2 | 0.08 | 1 | 0.191899351 | 0.16438487 |
| Tbkbp1 | 0.085880134 | -1.392317423 | 0.025 | 0.071 | 1 | 0.26672884 | -0.374395515 | 0 | 0.111 | 1 | 0.26672884 | 0.16438487 |
| Ccdc18 | 0.085943462 | -2.250961574 | 0.025 | 0.071 | 1 | 0.362872659 | 1.263034406 | 0.1 | 0.04 | 1 | 0.362872659 | 0.164500646 |
| Odad1 | 0.086043597 | -0.698420551 | 0.134 | 0.212 | 1 | 0.439349256 | -0.291933355 | 0.1 | 0.213 | 1 | 0.439349256 | 0.164683694 |
| Rnf113a2 | 0.241295305 | 0.344648171 | 0.269 | 0.206 | 1 | 0.086067798 | 1.503348735 | 0.4 | 0.201 | 1 | 0.241295305 | 0.16472793 |
| Daglb | 0.086104463 | -1.070389328 | 0.059 | 0.118 | 1 | 0.822515869 | 0.290001453 | 0.1 | 0.12 | 1 | 0.822515869 | 0.164794948 |
| Ssr4 | 0.086118421 | 0.236574743 | 0.983 | 0.959 | 1 | 0.921472976 | 0.420264334 | 0.9 | 0.935 | 1 | 0.921472976 | 0.16482046 |
| Flad1 | 0.086130977 | -1.555816155 | 0.034 | 0.082 | 1 | 0.290365769 | -0.505640048 | 0 | 0.102 | 1 | 0.290365769 | 0.164843409 |
| Usp3 | 0.086171987 | -0.498562229 | 0.597 | 0.635 | 1 | 0.147794163 | -0.830074999 | 0.7 | 0.821 | 1 | 0.147794163 | 0.164918362 |
| Nicn1 | 0.086201917 | -0.628384781 | 0.16 | 0.241 | 1 | 0.48571834 | 0.574978412 | 0.3 | 0.198 | 1 | 0.48571834 | 0.164973063 |
| Csnk1g2 | 0.152204214 | -0.31259023 | 0.639 | 0.671 | 1 | 0.086307312 | -0.728032469 | 0.4 | 0.735 | 1 | 0.152204214 | 0.165165673 |
| Dgkz | 0.086398619 | -0.93122658 | 0.202 | 0.276 | 1 | 0.193775056 | 0.804142617 | 0.5 | 0.293 | 1 | 0.193775056 | 0.165332517 |
| Map4k2 | 0.08641939 | -1.070389328 | 0.092 | 0.159 | 1 | 0.124174427 | 2.159940913 | 0.3 | 0.136 | 1 | 0.124174427 | 0.165370469 |
| Mtpap | 0.098420433 | -0.50851044 | 0.378 | 0.441 | 1 | 0.086429015 | 0.739937161 | 0.7 | 0.389 | 1 | 0.098420433 | 0.165388055 |
| Arfip2 | 0.086442456 | -0.56137568 | 0.252 | 0.335 | 1 | 0.306014945 | 0.779517169 | 0.5 | 0.34 | 1 | 0.306014945 | 0.165412614 |
| Eml5 | 0.086493767 | -1.544320516 | 0.059 | 0.118 | 1 | 0.760606608 | 1.057092505 | 0.2 | 0.17 | 1 | 0.760606608 | 0.165506363 |
| Romo1 | 0.12343504 | 0.202629167 | 0.866 | 0.824 | 1 | 0.086519635 | -0.392774044 | 0.5 | 0.784 | 1 | 0.12343504 | 0.165553622 |
| Tctn1 | 0.086600674 | -1.485426827 | 0.042 | 0.094 | 1 | 0.928523052 | 0.591657153 | 0.1 | 0.108 | 1 | 0.928523052 | 0.165701672 |
| Nsmaf | 0.123822609 | 0.255186544 | 0.378 | 0.276 | 1 | 0.086834823 | 0.881383075 | 0.7 | 0.407 | 1 | 0.123822609 | 0.16612936 |
| Pik3r2 | 0.08685501 | -0.691877705 | 0.084 | 0.153 | 1 | 0.701707131 | 0.625604485 | 0.1 | 0.157 | 1 | 0.701707131 | 0.166166227 |
| Ccnj | 0.086879909 | -1.292781749 | 0.025 | 0.071 | 1 | 0.505639431 | 0.769994395 | 0 | 0.043 | 1 | 0.505639431 | 0.1662117 |
| Rab27b | 0.345296183 | 0.270904092 | 0.437 | 0.371 | 1 | 0.08690336 | -0.263776342 | 0.2 | 0.519 | 1 | 0.345296183 | 0.166254526 |

| | | | | | | | | | | | | |
|---|---|---|---|---|---|---|---|---|---|---|---|---|
| Sugct | 0.086989368 | 0.148923701 | 0.395 | 0.288 | 1 | 0.540325152 | -0.129976787 | 0.4 | 0.293 | 1 | 0.540325152 | 0.166411586 |
| Abhd14a | 0.087051696 | -0.548620654 | 0.462 | 0.512 | 1 | 0.913900615 | 0.526068812 | 0.4 | 0.475 | 1 | 0.913900615 | 0.166525395 |
| Ufl1 | 0.087127091 | -0.716752373 | 0.496 | 0.5 | 1 | 0.634854688 | -0.314786026 | 0.5 | 0.574 | 1 | 0.634854688 | 0.166663052 |
| Mid1ip1 | 0.090377066 | 0.736965594 | 0.084 | 0.159 | 1 | 0.087207731 | -1.91281543 | 0 | 0.231 | 1 | 0.090377066 | 0.166810274 |
| Lamtor3 | 0.087287689 | -0.582525515 | 0.487 | 0.541 | 1 | 0.108042503 | -0.615073289 | 0.4 | 0.648 | 1 | 0.108042503 | 0.166956238 |
| Tmc6 | 0.452781322 | -0.551015169 | 0.252 | 0.276 | 1 | 0.087347699 | 1.238311976 | 0.6 | 0.321 | 1 | 0.452781322 | 0.167065777 |
| Gm16845 | 0.087369067 | -1.485426827 | 0.034 | 0.082 | 1 | 0.559081495 | 1.31748219 | 0.1 | 0.056 | 1 | 0.559081495 | 0.16710478 |
| Galnt5 | 0.087369964 | -1.622930351 | 0.034 | 0.082 | 1 | 0.749382172 | 0.973527789 | 0.1 | 0.071 | 1 | 0.749382172 | 0.167106417 |
| N4bp2os | 0.087369964 | -1.411426246 | 0.034 | 0.082 | 1 | 0.36390653 | 0.159940913 | 0 | 0.077 | 1 | 0.36390653 | 0.167106417 |
| Cog8 | 0.087402888 | -0.604071324 | 0.269 | 0.347 | 1 | 0.5935875 | 0.821524695 | 0.4 | 0.352 | 1 | 0.5935875 | 0.167166511 |
| Stambp | 0.087403813 | -0.582288366 | 0.218 | 0.306 | 1 | 0.61694667 | -0.242605642 | 0.2 | 0.256 | 1 | 0.61694667 | 0.1671682 |
| Plb1 | 0.540003619 | -0.436517227 | 0.067 | 0.088 | 1 | 0.087507587 | 2.481869008 | 0.2 | 0.065 | 1 | 0.540003619 | 0.167357596 |
| Ift46 | 0.087509372 | -0.455473605 | 0.42 | 0.494 | 1 | 0.756460164 | 0.187846909 | 0.6 | 0.494 | 1 | 0.756460164 | 0.167360854 |
| Mgme1 | 0.087607873 | -0.780882711 | 0.067 | 0.129 | 1 | 0.186053549 | -1.425021588 | 0 | 0.151 | 1 | 0.186053549 | 0.167540607 |
| Med17 | 0.087744357 | -0.664750527 | 0.202 | 0.288 | 1 | 0.444592813 | -0.28585884 | 0.2 | 0.324 | 1 | 0.444592813 | 0.167789641 |
| Commd6 | 0.087769978 | 0.328706628 | 0.513 | 0.412 | 1 | 0.741071167 | 0.354956895 | 0.4 | 0.349 | 1 | 0.741071167 | 0.167836387 |
| Fam174a | 0.087775459 | 0.155630236 | 0.807 | 0.653 | 1 | 0.129311009 | -0.679740725 | 0.4 | 0.694 | 1 | 0.129311009 | 0.167846386 |
| Cep152 | 0.087861328 | -1.163498732 | 0.042 | 0.094 | 1 | 0.474904369 | 0.695993813 | 0 | 0.049 | 1 | 0.474904369 | 0.168003043 |
| Golm1 | 0.275487408 | -0.485426827 | 0.193 | 0.241 | 1 | 0.08790216 | 1.040641984 | 0.3 | 0.111 | 1 | 0.275487408 | 0.16807753 |
| Sardh | 0.088035112 | -1.185866545 | 0.025 | 0.071 | 1 | 0.364098759 | -0.230005605 | 0 | 0.077 | 1 | 0.364098759 | 0.168320042 |
| Lysmd1 | 0.088035112 | -1.185866545 | 0.025 | 0.071 | 1 | 0.667604536 | 1.432959407 | 0.1 | 0.065 | 1 | 0.667604536 | 0.168320042 |
| Noxo1 | 0.088035112 | -1.185866545 | 0.025 | 0.071 | 1 | 0.619961444 | 1.432959407 | 0 | 0.025 | 1 | 0.619961444 | 0.168320042 |
| 9330179D12Rik | 0.088044807 | -1.359895945 | 0.042 | 0.094 | 1 | 0.446819579 | 0.111031312 | 0 | 0.056 | 1 | 0.446819579 | 0.168337726 |
| Zfp748 | 0.088045392 | -1.424026283 | 0.042 | 0.094 | 1 | 0.815502872 | 0.602884409 | 0.2 | 0.167 | 1 | 0.815502872 | 0.168338794 |
| Atg10 | 0.088045933 | -0.77915803 | 0.16 | 0.241 | 1 | 0.246666496 | -1.258202497 | 0.1 | 0.253 | 1 | 0.246666496 | 0.168339779 |
| Rwdd1 | 0.088054952 | 0.193522363 | 0.857 | 0.812 | 1 | 0.496503993 | 0.253771485 | 0.8 | 0.762 | 1 | 0.496503993 | 0.168356229 |
| Ivd | 0.088129116 | -0.406571406 | 0.252 | 0.347 | 1 | 0.109779089 | -1.230005605 | 0.1 | 0.336 | 1 | 0.109779089 | 0.16849149 |
| Srpr | 0.088146772 | 0.150379351 | 0.992 | 0.965 | 1 | 0.635413721 | 0.120522761 | 1 | 0.966 | 1 | 0.635413721 | 0.16852369 |
| Surf1 | 0.231596016 | -0.457412451 | 0.361 | 0.4 | 1 | 0.088295495 | -1.201246612 | 0.1 | 0.358 | 1 | 0.231596016 | 0.168794896 |
| Cad | 0.088495026 | -1.201633861 | 0.05 | 0.106 | 1 | 0.947574162 | 0.558490289 | 0.1 | 0.105 | 1 | 0.947574162 | 0.169158682 |
| Reep4 | 0.088617786 | -1.411426246 | 0.034 | 0.082 | 1 | 0.886171989 | 1.111031312 | 0.1 | 0.086 | 1 | 0.886171989 | 0.169382459 |
| Ifi30 | 0.162070043 | -0.451318973 | 0.563 | 0.588 | 1 | 0.088640292 | -1.615073289 | 0.3 | 0.522 | 1 | 0.162070043 | 0.169423482 |
| Fam83g | 0.088670088 | -1.250961574 | 0.025 | 0.071 | 1 | 0.29045414 | -0.567040593 | 0 | 0.102 | 1 | 0.29045414 | 0.169477791 |
| Lzic | 0.088704603 | -0.6990223 | 0.303 | 0.371 | 1 | 0.673222623 | 2.063725598 | 0.3 | 0.278 | 1 | 0.673222623 | 0.169540699 |
| Pgm1 | 0.08878902 | 0.59946207 | 0.244 | 0.165 | 1 | 0.635093491 | 1.894539492 | 0.2 | 0.157 | 1 | 0.635093491 | 0.16969455 |
| Cerk | 0.088860175 | -0.929033479 | 0.16 | 0.235 | 1 | 0.121812856 | -1.567040593 | 0.1 | 0.321 | 1 | 0.121812856 | 0.16982422 |
| Nubpl | 0.088956743 | 0.316633795 | 0.244 | 0.159 | 1 | 0.207635132 | -0.959358016 | 0 | 0.139 | 1 | 0.207635132 | 0.170000183 |
| Stk16 | 0.088996998 | -0.470629825 | 0.345 | 0.424 | 1 | 0.370058008 | 0.669193754 | 0.5 | 0.383 | 1 | 0.370058008 | 0.17007353 |
| Ciapin1 | 0.564867992 | -0.227930605 | 0.319 | 0.347 | 1 | 0.089036671 | 1.267150514 | 0.6 | 0.352 | 1 | 0.564867992 | 0.170145813 |
| Gimap6 | 0.089037903 | -1.021479727 | 0.059 | 0.118 | 1 | 0.976598201 | 0.769994395 | 0.1 | 0.096 | 1 | 0.976598201 | 0.170148058 |
| Decr2 | 0.089047327 | 0.477098467 | 0.269 | 0.182 | 1 | 0.799428547 | 0.695993813 | 0.2 | 0.167 | 1 | 0.799428547 | 0.170165227 |
| Cspg5 | 0.089204104 | -1.11345805 | 0.067 | 0.129 | 1 | 0.458661693 | 0.959028219 | 0.2 | 0.117 | 1 | 0.458661693 | 0.170450837 |
| Polg | 0.08928613 | -0.485426827 | 0.244 | 0.329 | 1 | 0.264143231 | 1.085036104 | 0.5 | 0.358 | 1 | 0.264143231 | 0.170600247 |
| Tcp11 | 0.089303409 | -1.321928095 | 0.042 | 0.094 | 1 | 0.577142568 | 1.432959407 | 0 | 0.031 | 1 | 0.577142568 | 0.170631719 |
| Psmd5 | 0.85054582 | -0.485426827 | 0.252 | 0.247 | 1 | 0.089389513 | -1.348400306 | 0.1 | 0.355 | 1 | 0.85054582 | 0.170788541 |
| Gm50232 | 0.644520473 | -0.707819249 | 0.042 | 0.053 | 1 | 0.089481107 | 1.795529487 | 0.2 | 0.062 | 1 | 0.644520473 | 0.170955346 |
| Nme6 | 0.08950598 | -1.359895945 | 0.042 | 0.094 | 1 | 0.673291667 | 1.263034406 | 0.1 | 0.065 | 1 | 0.673291667 | 0.17100064 |
| Rgl2 | 0.089537696 | -0.770829046 | 0.101 | 0.171 | 1 | 0.128235781 | -1.441509711 | 0 | 0.191 | 1 | 0.128235781 | 0.171058392 |
| Ralgps1 | 0.543188037 | -0.555816155 | 0.059 | 0.076 | 1 | 0.089609194 | 0.769994395 | 0.3 | 0.111 | 1 | 0.543188037 | 0.171188581 |
| Tmem134 | 0.08965757 | 0.207518354 | 0.723 | 0.659 | 1 | 0.310791434 | 0.904179742 | 0.6 | 0.59 | 1 | 0.310791434 | 0.171276659 |
| A930037H05Rik | 0.08970361 | -1.250961574 | 0.025 | 0.071 | 1 | 0.597832556 | 1.111031312 | 0 | 0.028 | 1 | 0.597832556 | 0.171360483 |
| Coq7 | 0.089745224 | 0.410737362 | 0.395 | 0.312 | 1 | 0.362650059 | 0.984498906 | 0.4 | 0.293 | 1 | 0.362650059 | 0.171436242 |
| Zfp36 | 0.089784707 | 0.497452903 | 0.782 | 0.788 | 1 | 0.963920036 | 0.521893136 | 0.8 | 0.833 | 1 | 0.963920036 | 0.171508121 |
| Sco1 | 0.089814486 | -1.008988783 | 0.059 | 0.118 | 1 | 0.248571988 | 1.515421567 | 0.2 | 0.09 | 1 | 0.248571988 | 0.17156233 |
| Ubxn11 | 0.089877159 | -1.485426827 | 0.034 | 0.082 | 1 | 0.48480221 | 1.695993813 | 0.1 | 0.049 | 1 | 0.48480221 | 0.171676415 |
| Zbtb48 | 0.089877159 | -1.250961574 | 0.034 | 0.082 | 1 | 0.348486822 | 1.393431043 | 0.2 | 0.105 | 1 | 0.348486822 | 0.171676415 |
| Lrrc73 | 0.089877159 | -1.333423734 | 0.034 | 0.082 | 1 | 0.829966406 | 1.159940913 | 0.1 | 0.08 | 1 | 0.829966406 | 0.171676415 |
| Ccdc127 | 0.414565506 | -0.206710799 | 0.782 | 0.682 | 1 | 0.089949108 | -0.894302132 | 0.4 | 0.698 | 1 | 0.414565506 | 0.171807374 |
| Syce2 | 0.089998769 | -1.185866545 | 0.05 | 0.106 | 1 | 0.792494329 | 1.602884409 | 0.1 | 0.083 | 1 | 0.792494329 | 0.17189776 |
| Alyref2 | 0.090145775 | -1.070389328 | 0.05 | 0.106 | 1 | 0.374600758 | 0.111031312 | 0 | 0.074 | 1 | 0.374600758 | 0.172165289 |
| Wwtr1 | 0.090188671 | -0.218189374 | 0.992 | 1 | 1 | 0.259968565 | -0.35500909 | 1 | 1 | 1 | 0.259968565 | 0.172243345 |
| A330076H08Rik | 0.090193346 | 0.40354186 | 0.286 | 0.194 | 1 | 0.16649103 | 0.515421567 | 0.5 | 0.259 | 1 | 0.16649103 | 0.172251853 |
| Atp11b | 0.17916767 | -0.364959307 | 0.647 | 0.712 | 1 | 0.090213092 | -0.174478126 | 0.6 | 0.83 | 1 | 0.17916767 | 0.172287782 |
| Irs2 | 0.090228581 | -0.582412443 | 0.647 | 0.676 | 1 | 0.425511159 | 0.188337039 | 0.4 | 0.725 | 1 | 0.425511159 | 0.172315966 |
| Jade1 | 0.090230372 | -0.96051171 | 0.336 | 0.388 | 1 | 0.154676431 | -0.750262417 | 0.3 | 0.534 | 1 | 0.154676431 | 0.172319224 |
| Cdh23 | 0.346682404 | -0.694880193 | 0.092 | 0.129 | 1 | 0.090401149 | 2.280956314 | 0.2 | 0.065 | 1 | 0.346682404 | 0.17262993 |
| Ppp1r18 | 0.313817253 | -0.4440069 | 0.345 | 0.388 | 1 | 0.090455277 | -1.321928095 | 0.2 | 0.444 | 1 | 0.313817253 | 0.172728397 |
| Wfdc2 | 0.497139171 | 1.188509303 | 0.269 | 0.318 | 1 | 0.090537183 | -1.385800278 | 0.2 | 0.546 | 1 | 0.497139171 | 0.172877384 |
| Tmem222 | 0.090665504 | -0.524246076 | 0.361 | 0.429 | 1 | 0.382741746 | -0.536666944 | 0.2 | 0.315 | 1 | 0.382741746 | 0.173110775 |
| Creb5 | 0.347497471 | -0.967578522 | 0.218 | 0.247 | 1 | 0.090686626 | -2.574535129 | 0 | 0.228 | 1 | 0.347497471 | 0.173149188 |
| Cox6b2 | 0.090764306 | -0.678071905 | 0.025 | 0.071 | 1 | 0.52208609 | 0.769994395 | 0 | 0.04 | 1 | 0.52208609 | 0.173290453 |
| Pfkfb3 | 0.090807772 | -0.887525271 | 0.277 | 0.341 | 1 | 0.24638552 | 0.568881895 | 0.5 | 0.414 | 1 | 0.24638552 | 0.173369492 |
| Bet1 | 0.090821772 | -0.382333334 | 0.521 | 0.576 | 1 | 0.190614099 | -0.8829449 | 0.3 | 0.469 | 1 | 0.190614099 | 0.173394949 |
| Gpatch4 | 0.090860214 | -1.070389328 | 0.126 | 0.194 | 1 | 0.875033647 | 0.478763097 | 0.2 | 0.185 | 1 | 0.875033647 | 0.17346485 |
| Gipc2 | 0.090869989 | -0.752907138 | 0.042 | 0.094 | 1 | 0.170401245 | -1.840059087 | 0 | 0.16 | 1 | 0.170401245 | 0.173482624 |
| Fam149a | 0.090907914 | 0.163100802 | 0.227 | 0.141 | 1 | 0.8791288 | 1 | 0.2 | 0.194 | 1 | 0.8791288 | 0.173551579 |
| Sec23b | 0.090955502 | -0.547968293 | 0.462 | 0.518 | 1 | 0.167772813 | -0.884955625 | 0.5 | 0.611 | 1 | 0.167772813 | 0.1736381 |
| Gm57455 | 0.091098781 | -0.970853654 | 0.025 | 0.071 | 1 | 0.193279696 | 2.432959407 | 0.1 | 0.028 | 1 | 0.193279696 | 0.173898574 |
| Zfp974 | 0.091147213 | -1.485426827 | 0.034 | 0.082 | 1 | 0.863421649 | 0.210566986 | 0.1 | 0.114 | 1 | 0.863421649 | 0.173986612 |
| Mfhas1 | 0.223490793 | -0.324962155 | 0.143 | 0.2 | 1 | 0.091266966 | 0.565062943 | 0.6 | 0.293 | 1 | 0.223490793 | 0.174204272 |
| Dusp19 | 0.091267511 | -0.765534746 | 0.101 | 0.171 | 1 | 0.928522993 | 0.660369903 | 0.1 | 0.108 | 1 | 0.928522993 | 0.174205263 |
| Maob | 0.091281127 | -0.967295835 | 0.277 | 0.347 | 1 | 0.786993796 | -0.346650524 | 0.3 | 0.33 | 1 | 0.786993796 | 0.17423001 |
| Rgs7 | 0.091361251 | 2.514573173 | 0.017 | 0 | 1 | 0.730749233 | 2.432959407 | 0 | 0.012 | 1 | 0.730749233 | 0.174375624 |
| Ovol2 | 0.091361251 | 2.514573173 | 0.017 | 0 | 1 | 0.619961444 | 1.558490289 | 0 | 0.025 | 1 | 0.619961444 | 0.174375624 |
| Gm56779 | 0.091361251 | 2.099535674 | 0.017 | 0 | 1 | 0.597845674 | 1.695993813 | 0 | 0.028 | 1 | 0.597845674 | 0.174375624 |
| Trim45 | 0.091361251 | 2.099535674 | 0.017 | 0 | 1 | 0.698534498 | 2.432959407 | 0 | 0.015 | 1 | 0.698534498 | 0.174375624 |
| Tek | 0.091361251 | 2.099535674 | 0.017 | 0 | 1 | 0.730749233 | 2.210566986 | 0 | 0.012 | 1 | 0.730749233 | 0.174375624 |
| Pars2 | 0.091361251 | 2.099535674 | 0.017 | 0 | 1 | 0.730749233 | 2.432959407 | 0 | 0.012 | 1 | 0.730749233 | 0.174375624 |
| 4932441J04Rik | 0.091361251 | 2.099535674 | 0.017 | 0 | 1 | 0.698529281 | 2.210566986 | 0 | 0.015 | 1 | 0.698529281 | 0.174375624 |
| Chrna9 | 0.091361251 | 2.099535674 | 0.017 | 0 | 1 | 0.643835681 | 1.558490289 | 0 | 0.022 | 1 | 0.643835681 | 0.174375624 |
| Dao | 0.091361251 | 2.099535674 | 0.017 | 0 | 1 | 0.669853054 | 2.210566986 | 0 | 0.019 | 1 | 0.669853054 | 0.174375624 |
| Gm15406 | 0.091361251 | 2.099535674 | 0.017 | 0 | 1 | 0.113669596 | 2.847996907 | 0.1 | 0.022 | 1 | 0.113669596 | 0.174375624 |
| Cnfn | 0.091361251 | 2.514573173 | 0.017 | 0 | 1 | 0.730749233 | 2.432959407 | 0 | 0.012 | 1 | 0.730749233 | 0.174375624 |
| Scnn1b | 0.091361251 | 2.099535674 | 0.017 | 0 | 1 | 0.698529281 | 2.210566986 | 0 | 0.015 | 1 | 0.698529281 | 0.174375624 |
| Defb1 | 0.091361251 | 2.514573173 | 0.017 | 0 | 1 | 0.460619503 | 0.111031312 | 0 | 0.052 | 1 | 0.460619503 | 0.174375624 |
| Slc12a3 | 0.091361251 | 2.099535674 | 0.017 | 0 | 1 | 0.730750993 | 1.558490289 | 0 | 0.012 | 1 | 0.730750993 | 0.174375624 |
| Zfp810 | 0.091361251 | 2.099535674 | 0.017 | 0 | 1 | 0.730749233 | 2.432959407 | 0 | 0.012 | 1 | 0.730749233 | 0.174375624 |
| Gm28905 | 0.091361251 | 2.514573173 | 0.017 | 0 | 1 | 0.730750993 | 1.847996907 | 0 | 0.012 | 1 | 0.730750993 | 0.174375624 |

| | | | | | | | | | | | | |
|---|---|---|---|---|---|---|---|---|---|---|---|---|
| Gm47922 | 0.091361251 | 2.514573173 | 0.017 | 0 | 1 | 0.698529281 | 2.210566986 | 0 | 0.015 | 1 | 0.698529281 | 0.174375624 |
| Arhgap9 | 0.091361251 | 2.514573173 | 0.017 | 0 | 1 | 0.6199755 | 1.847996907 | 0 | 0.025 | 1 | 0.6199755 | 0.174375624 |
| Asb16 | 0.091361251 | 2.836501268 | 0.017 | 0 | 1 | 0.698534498 | 2.432959407 | 0 | 0.015 | 1 | 0.698534498 | 0.174375624 |
| Rbm24 | 0.091361251 | 3.514573173 | 0.017 | 0 | 1 | 0.669853054 | 2.210566986 | 0 | 0.019 | 1 | 0.669853054 | 0.174375624 |
| Gm15965 | 0.091361251 | 2.099535674 | 0.017 | 0 | 1 | 0.730750993 | 2.017921908 | 0 | 0.012 | 1 | 0.730750993 | 0.174375624 |
| 6330415G19Rik | 0.091361251 | 2.099535674 | 0.017 | 0 | 1 | 0.698534498 | 2.432959407 | 0 | 0.015 | 1 | 0.698534498 | 0.174375624 |
| Rab44 | 0.091361251 | 3.321928095 | 0.017 | 0 | 1 | 0.73075158 | 2.695993813 | 0 | 0.012 | 1 | 0.73075158 | 0.174375624 |
| H2-Q6 | 0.091361251 | 2.099535674 | 0.017 | 0 | 1 | 0.730749233 | 2.432959407 | 0 | 0.012 | 1 | 0.730749233 | 0.174375624 |
| Galnt14 | 0.091361251 | 2.514573173 | 0.017 | 0 | 1 | 0.669852109 | 1.210566986 | 0 | 0.019 | 1 | 0.669852109 | 0.174375624 |
| Tmem17 | 0.091397595 | -1.292781749 | 0.05 | 0.106 | 1 | 0.225422464 | -0.505640048 | 0 | 0.13 | 1 | 0.225422464 | 0.174441669 |
| 4933421O10Rik | 0.782569661 | -0.185866545 | 0.261 | 0.265 | 1 | 0.091622387 | 1.880418384 | 0.4 | 0.207 | 1 | 0.782569661 | 0.174850111 |
| Obp1b | 0.091633164 | -0.719892081 | 0.126 | 0.2 | 1 | 0.148511251 | 0.602884409 | 0.4 | 0.198 | 1 | 0.148511251 | 0.174869692 |
| Septin5 | 0.893287269 | 0.140177658 | 0.185 | 0.182 | 1 | 0.091659188 | 1.190102883 | 0.4 | 0.176 | 1 | 0.893287269 | 0.17491697 |
| Rpl10-ps3 | 0.214658341 | 0.206450877 | 0.756 | 0.665 | 1 | 0.091727325 | -0.926267229 | 0.6 | 0.691 | 1 | 0.214658341 | 0.175040747 |
| Ccpg1 | 0.297611726 | 0.11488992 | 0.874 | 0.824 | 1 | 0.091740381 | -0.731016328 | 0.7 | 0.886 | 1 | 0.297611726 | 0.175064465 |
| Ccdc167 | 0.091774087 | -1.19592021 | 0.084 | 0.147 | 1 | 0.124037212 | -1.339630097 | 0 | 0.194 | 1 | 0.124037212 | 0.17512569 |
| Zscan21 | 0.091775347 | -1.11345805 | 0.084 | 0.147 | 1 | 0.80097483 | 0.263034406 | 0.2 | 0.16 | 1 | 0.80097483 | 0.17512798 |
| Slc31a2 | 0.091776608 | -1.025995209 | 0.084 | 0.147 | 1 | 0.487484474 | 0.769994395 | 0.3 | 0.213 | 1 | 0.487484474 | 0.17513027 |
| Acd | 0.091814832 | -0.534336428 | 0.218 | 0.306 | 1 | 0.365377668 | 0.667424661 | 0.4 | 0.253 | 1 | 0.365377668 | 0.175199701 |
| Ssr2 | 0.09182264 | -0.314795834 | 0.966 | 0.9 | 1 | 0.937208872 | 0.227410496 | 0.9 | 0.892 | 1 | 0.937208872 | 0.175213884 |
| Orc2 | 0.091881536 | -0.916061182 | 0.311 | 0.371 | 1 | 0.286053209 | -0.637429921 | 0.2 | 0.404 | 1 | 0.286053209 | 0.175320855 |
| Gm56517 | 0.091900178 | -0.739183419 | 0.193 | 0.271 | 1 | 0.41655118 | -0.484578433 | 0.2 | 0.302 | 1 | 0.41655118 | 0.175354713 |
| Tuba4a | 0.166965163 | -1.733354341 | 0.025 | 0.059 | 1 | 0.091970626 | -1.5519337 | 0 | 0.225 | 1 | 0.166965163 | 0.175482657 |
| Cln6 | 0.091973719 | -1.008988783 | 0.05 | 0.106 | 1 | 0.305374989 | 1.393431043 | 0.2 | 0.099 | 1 | 0.305374989 | 0.175488273 |
| Fkbp15 | 0.092135397 | -0.642968104 | 0.328 | 0.412 | 1 | 0.776885309 | -0.441509711 | 0.5 | 0.451 | 1 | 0.776885309 | 0.175781862 |
| Otub2 | 0.092147579 | -0.86393845 | 0.025 | 0.071 | 1 | 0.474928012 | 0.494359952 | 0 | 0.049 | 1 | 0.474928012 | 0.175803981 |
| Yod1 | 0.0923349 | -0.783107376 | 0.235 | 0.324 | 1 | 0.386424417 | -0.91281543 | 0.2 | 0.34 | 1 | 0.386424417 | 0.176144067 |
| Snhg16 | 0.127951871 | -0.760049207 | 0.21 | 0.276 | 1 | 0.092504642 | -1.723545078 | 0 | 0.225 | 1 | 0.127951871 | 0.176452176 |
| Arfip1 | 0.751691199 | -0.117039421 | 0.807 | 0.712 | 1 | 0.092551831 | -0.729312022 | 0.6 | 0.833 | 1 | 0.751691199 | 0.17653782 |
| Lmf1 | 0.092567736 | -0.691877705 | 0.193 | 0.276 | 1 | 0.454221257 | -0.191531458 | 0.1 | 0.191 | 1 | 0.454221257 | 0.176566687 |
| Map3k10 | 0.092608149 | -0.765534746 | 0.21 | 0.288 | 1 | 0.462893088 | 0.800691192 | 0.5 | 0.367 | 1 | 0.462893088 | 0.176640029 |
| Pcdhgc5 | 0.092651972 | -1.070389328 | 0.034 | 0.082 | 1 | 0.539413885 | 1.210566986 | 0 | 0.037 | 1 | 0.539413885 | 0.176719556 |
| Ikzf4 | 0.092676246 | -0.900464326 | 0.294 | 0.359 | 1 | 0.479312141 | 0.754887502 | 0.4 | 0.309 | 1 | 0.479312141 | 0.176763605 |
| Rab26os | 0.092693774 | -0.889817082 | 0.134 | 0.206 | 1 | 0.353533258 | 0.210566986 | 0 | 0.08 | 1 | 0.353533258 | 0.176795412 |
| Msi2 | 0.758899424 | -0.104592043 | 1 | 1 | 1 | 0.092703083 | 0.392445874 | 1 | 1 | 1 | 0.758899424 | 0.176812304 |
| Nit1 | 0.092722002 | -0.515174171 | 0.168 | 0.253 | 1 | 0.53908852 | -0.441509711 | 0.2 | 0.272 | 1 | 0.53908852 | 0.176846635 |
| Smg1 | 0.092735266 | 0.252288024 | 0.975 | 0.971 | 1 | 0.820706707 | 0.664455414 | 0.9 | 0.985 | 1 | 0.820706707 | 0.176870703 |
| Mmp9 | 0.092795927 | -3.070389328 | 0.008 | 0.041 | 1 | 0.551967957 | -3.306258639 | 0.1 | 0.052 | 1 | 0.551967957 | 0.17698077 |
| Tcim | 0.092933273 | -0.987927168 | 0.025 | 0.071 | 1 | 0.790244149 | 0.210566986 | 0.1 | 0.127 | 1 | 0.790244149 | 0.177229953 |
| Asrgl1 | 0.129711689 | 0.208764743 | 0.412 | 0.306 | 1 | 0.093101921 | 1.063725598 | 0.5 | 0.262 | 1 | 0.129711689 | 0.177535874 |
| Cdc42bpg | 0.093131512 | -0.839063782 | 0.05 | 0.106 | 1 | 0.522604016 | 1.111031312 | 0.2 | 0.136 | 1 | 0.522604016 | 0.177589546 |
| Tbcel | 0.093132224 | -0.54092194 | 0.294 | 0.382 | 1 | 0.651874453 | 0.277681182 | 0.6 | 0.506 | 1 | 0.651874453 | 0.177590837 |
| Nppc | 0.093241817 | -2.292781749 | 0 | 0.024 | 1 | 0.619961444 | 1.31748219 | 0 | 0.025 | 1 | 0.619961444 | 0.177789598 |
| Mir181a-1hg | 0.093241817 | -2.070389328 | 0 | 0.024 | 1 | 0.730750993 | 2.210566986 | 0 | 0.012 | 1 | 0.730750993 | 0.177789598 |
| Galnt3 | 0.093241817 | -3.392317423 | 0 | 0.024 | 1 | 0.596204587 | 1.374065718 | 0.1 | 0.059 | 1 | 0.596204587 | 0.177789598 |
| Large2 | 0.093241817 | -2.070389328 | 0 | 0.024 | 1 | 0.108064109 | 3.017921908 | 0.1 | 0.022 | 1 | 0.108064109 | 0.177789598 |
| Tnfrsf14 | 0.093241817 | -2.070389328 | 0 | 0.024 | 1 | 0.316137708 | 2.017921908 | 0.1 | 0.037 | 1 | 0.316137708 | 0.177789598 |
| Miat | 0.093241817 | -2.292781749 | 0 | 0.024 | 1 | 0.106693545 | 3.14345279 | 0.1 | 0.022 | 1 | 0.106693545 | 0.177789598 |
| Slc23a4 | 0.093241817 | -2.070389328 | 0 | 0.024 | 1 | 0.669843604 | 2.017921908 | 0 | 0.019 | 1 | 0.669843604 | 0.177789598 |
| Kbtbd8 | 0.093241817 | -2.070389328 | 0 | 0.024 | 1 | 0.619961444 | 1.432959407 | 0 | 0.025 | 1 | 0.619961444 | 0.177789598 |
| Ghrl | 0.093241817 | -2.070389328 | 0 | 0.024 | 1 | 0.577184788 | 1.558490289 | 0 | 0.031 | 1 | 0.577184788 | 0.177789598 |
| Vstm2b | 0.093241817 | -2.070389328 | 0 | 0.024 | 1 | 0.698529281 | 2.210566986 | 0 | 0.015 | 1 | 0.698529281 | 0.177789598 |
| Bean1 | 0.093241817 | -2.070389328 | 0 | 0.024 | 1 | 0.643835681 | 1.847996907 | 0 | 0.022 | 1 | 0.643835681 | 0.177789598 |
| Spata2l | 0.093241817 | -2.070389328 | 0 | 0.024 | 1 | 0.643850855 | 2.017921908 | 0 | 0.022 | 1 | 0.643850855 | 0.177789598 |
| Gm26631 | 0.093241817 | -2.070389328 | 0 | 0.024 | 1 | 0.64384652 | 1.558490289 | 0 | 0.022 | 1 | 0.64384652 | 0.177789598 |
| Adamts7 | 0.093241817 | -2.070389328 | 0 | 0.024 | 1 | 0.643842184 | 1.695993813 | 0 | 0.022 | 1 | 0.643842184 | 0.177789598 |
| Plxnc1 | 0.093241817 | -2.292781749 | 0 | 0.024 | 1 | 0.539456304 | 0.432959407 | 0 | 0.037 | 1 | 0.539456304 | 0.177789598 |
| Lcp2 | 0.093241817 | -3.070389328 | 0 | 0.024 | 1 | 0.669843604 | 1.695993813 | 0 | 0.019 | 1 | 0.669843604 | 0.177789598 |
| 4933427D14Rik | 0.093241817 | -2.070389328 | 0 | 0.024 | 1 | 0.6199755 | 1.847996907 | 0 | 0.025 | 1 | 0.6199755 | 0.177789598 |
| Rundc3a | 0.093241817 | -2.070389328 | 0 | 0.024 | 1 | 0.669853054 | 2.210566986 | 0 | 0.019 | 1 | 0.669853054 | 0.177789598 |
| Adam11 | 0.093241817 | -2.070389328 | 0 | 0.024 | 1 | 0.59781419 | 1.558490289 | 0 | 0.028 | 1 | 0.59781419 | 0.177789598 |
| Milr1 | 0.093241817 | -2.070389328 | 0 | 0.024 | 1 | 0.577165087 | 0.769994395 | 0 | 0.031 | 1 | 0.577165087 | 0.177789598 |
| Fam110c | 0.093241817 | -2.070389328 | 0 | 0.024 | 1 | 0.557756607 | 1.111031312 | 0 | 0.034 | 1 | 0.557756607 | 0.177789598 |
| Gm10457 | 0.093241817 | -2.070389328 | 0 | 0.024 | 1 | 0.698529281 | 1.432959407 | 0 | 0.015 | 1 | 0.698529281 | 0.177789598 |
| Taf9 | 0.093241817 | -2.070389328 | 0 | 0.024 | 1 | 0.577142568 | 1.432959407 | 0 | 0.031 | 1 | 0.577142568 | 0.177789598 |
| Gm15764 | 0.093241817 | -2.070389328 | 0 | 0.024 | 1 | 0.597845674 | 1.695993813 | 0 | 0.028 | 1 | 0.597845674 | 0.177789598 |
| Gm20939 | 0.093241817 | -2.070389328 | 0 | 0.024 | 1 | 0.6199755 | 1.847996907 | 0 | 0.025 | 1 | 0.6199755 | 0.177789598 |
| Cbln2 | 0.093241817 | -2.070389328 | 0 | 0.024 | 1 | 0.698529281 | 2.210566986 | 0 | 0.015 | 1 | 0.698529281 | 0.177789598 |
| Tmeff2 | 0.093244798 | -2.655351829 | 0 | 0.024 | 1 | 0.730749233 | 0.930459067 | 0 | 0.012 | 1 | 0.730749233 | 0.177795004 |
| Dnah7a | 0.093244798 | -2.944858446 | 0 | 0.024 | 1 | 0.669848329 | 1.847996907 | 0 | 0.019 | 1 | 0.669848329 | 0.177795004 |
| Tent5b | 0.093244798 | -2.807354922 | 0 | 0.024 | 1 | 0.522102507 | 0.930459067 | 0 | 0.04 | 1 | 0.522102507 | 0.177795004 |
| Caln1 | 0.093244798 | -2.292781749 | 0 | 0.024 | 1 | 0.730749233 | 0.695993813 | 0 | 0.012 | 1 | 0.730749233 | 0.177795004 |
| Coro1a | 0.093244798 | -4.185866545 | 0 | 0.024 | 1 | 0.439528334 | 1.769994395 | 0.1 | 0.046 | 1 | 0.439528334 | 0.177795004 |
| D7Ertd443e | 0.093244798 | -2.485426827 | 0 | 0.024 | 1 | 0.730749233 | 2.432959407 | 0 | 0.012 | 1 | 0.730749233 | 0.177795004 |
| Slc5a5 | 0.093244798 | -2.292781749 | 0 | 0.024 | 1 | 0.409513437 | 1.374065718 | 0.1 | 0.043 | 1 | 0.409513437 | 0.177795004 |
| Niban3 | 0.093244798 | -2.944858446 | 0 | 0.024 | 1 | 0.619953009 | 1.432959407 | 0 | 0.025 | 1 | 0.619953009 | 0.177795004 |
| Gm57038 | 0.093244798 | -2.292781749 | 0 | 0.024 | 1 | 0.577142568 | 1.432959407 | 0 | 0.031 | 1 | 0.577142568 | 0.177795004 |
| Gask1a | 0.093244798 | -2.655351829 | 0 | 0.024 | 1 | 0.643842184 | 1.432959407 | 0 | 0.022 | 1 | 0.643842184 | 0.177795004 |
| Mmp11 | 0.093244798 | -2.292781749 | 0 | 0.024 | 1 | 0.669843604 | 2.017921908 | 0 | 0.019 | 1 | 0.669843604 | 0.177795004 |
| Gm47561 | 0.093244798 | -2.485426827 | 0 | 0.024 | 1 | 0.669850692 | 1.558490289 | 0 | 0.019 | 1 | 0.669850692 | 0.177795004 |
| Zfp354b | 0.093244798 | -2.292781749 | 0 | 0.024 | 1 | 0.460577419 | 0.494359952 | 0 | 0.052 | 1 | 0.460577419 | 0.177795004 |
| Gm27010 | 0.093244798 | -2.292781749 | 0 | 0.024 | 1 | 0.539413885 | 1.111031312 | 0 | 0.037 | 1 | 0.539413885 | 0.177795004 |
| Blk | 0.093244798 | -3.485426827 | 0 | 0.024 | 1 | 0.698533455 | 0.625604485 | 0 | 0.015 | 1 | 0.698533455 | 0.177795004 |
| Gm6994 | 0.093244798 | -2.292781749 | 0 | 0.024 | 1 | 0.505582845 | 1.017921908 | 0 | 0.043 | 1 | 0.505582845 | 0.177795004 |
| Gm49003 | 0.093244798 | -2.944858446 | 0 | 0.024 | 1 | 0.669851164 | 1.210566986 | 0 | 0.019 | 1 | 0.669851164 | 0.177795004 |
| Ltb | 0.093244798 | -3.392317423 | 0 | 0.024 | 1 | 0.73075158 | 2.695993813 | 0 | 0.012 | 1 | 0.73075158 | 0.177795004 |
| 9430060I03Rik | 0.093245792 | -1.807354922 | 0 | 0.024 | 1 | 0.6199755 | 1.847996907 | 0 | 0.025 | 1 | 0.6199755 | 0.177796806 |
| Myocos | 0.093245792 | -1.807354922 | 0 | 0.024 | 1 | 0.278846016 | 2.111031312 | 0.1 | 0.034 | 1 | 0.278846016 | 0.177796806 |
| Fcgr3 | 0.093245792 | -1.807354922 | 0 | 0.024 | 1 | 0.698529281 | 1.558490289 | 0 | 0.015 | 1 | 0.698529281 | 0.177796806 |
| 8030442B05Rik | 0.093245792 | -2.485426827 | 0 | 0.024 | 1 | 0.619971484 | 1.31748219 | 0 | 0.025 | 1 | 0.619971484 | 0.177796806 |
| Dpp4 | 0.093245792 | -1.807354922 | 0 | 0.024 | 1 | 0.730750993 | 1.210566986 | 0 | 0.012 | 1 | 0.730750993 | 0.177796806 |
| Gm14455 | 0.093245792 | -1.807354922 | 0 | 0.024 | 1 | 0.151443299 | 2.695993813 | 0.1 | 0.025 | 1 | 0.151443299 | 0.177796806 |
| Lor | 0.093245792 | -1.807354922 | 0 | 0.024 | 1 | 0.522066388 | 1.111031312 | 0 | 0.04 | 1 | 0.522066388 | 0.177796806 |
| Gm42702 | 0.093245792 | -1.807354922 | 0 | 0.024 | 1 | 0.113669596 | 2.847996907 | 0.1 | 0.022 | 1 | 0.113669596 | 0.177796806 |
| Ttpa | 0.093245792 | -1.807354922 | 0 | 0.024 | 1 | 0.539413885 | 1.111031312 | 0 | 0.037 | 1 | 0.539413885 | 0.177796806 |
| Tmem275 | 0.093245792 | -2.807354922 | 0 | 0.024 | 1 | 0.730750993 | 2.210566986 | 0 | 0.012 | 1 | 0.730750993 | 0.177796806 |
| Rab42 | 0.093245792 | -1.807354922 | 0 | 0.024 | 1 | 0.73075158 | 2.695993813 | 0 | 0.012 | 1 | 0.73075158 | 0.177796806 |
| Gm42744 | 0.093245792 | -1.807354922 | 0 | 0.024 | 1 | 0.669843604 | 1.847996907 | 0 | 0.019 | 1 | 0.669843604 | 0.177796806 |

| | | | | | | | | | | | | |
|---|---|---|---|---|---|---|---|---|---|---|---|---|
| Zfyve28 | 0.093245792 | -2.944858446 | 0 | 0.024 | 1 | 0.597832556 | 1.31748219 | 0 | 0.028 | 1 | 0.597832556 | 0.177796806 |
| Ncf1 | 0.093245792 | -1.807354922 | 0 | 0.024 | 1 | 0.151443299 | 2.31748219 | 0.1 | 0.025 | 1 | 0.151443299 | 0.177796806 |
| Grid2ip | 0.093245792 | -1.807354922 | 0 | 0.024 | 1 | 0.698534498 | 2.432959407 | 0 | 0.015 | 1 | 0.698534498 | 0.177796806 |
| Pirb | 0.093245792 | -1.807354922 | 0 | 0.024 | 1 | 0.619953009 | 1.558490289 | 0 | 0.025 | 1 | 0.619953009 | 0.177796806 |
| Gm3854 | 0.093245792 | -1.807354922 | 0 | 0.024 | 1 | 0.698529281 | 2.210566986 | 0 | 0.015 | 1 | 0.698529281 | 0.177796806 |
| Zfp108 | 0.093245792 | -1.807354922 | 0 | 0.024 | 1 | 0.669848329 | 1.695993813 | 0 | 0.019 | 1 | 0.669848329 | 0.177796806 |
| Gm38973 | 0.093245792 | -1.807354922 | 0 | 0.024 | 1 | 0.489901827 | 0.695993813 | 0 | 0.046 | 1 | 0.489901827 | 0.177796806 |
| Fbxo27 | 0.093245792 | -1.807354922 | 0 | 0.024 | 1 | 0.698534498 | 2.432959407 | 0 | 0.015 | 1 | 0.698534498 | 0.177796806 |
| Nmb | 0.093245792 | -1.807354922 | 0 | 0.024 | 1 | 0.681837487 | 0.625604485 | 0.1 | 0.065 | 1 | 0.681837487 | 0.177796806 |
| Xrra1 | 0.093245792 | -1.807354922 | 0 | 0.024 | 1 | 0.730750993 | 1.847996907 | 0 | 0.012 | 1 | 0.730750993 | 0.177796806 |
| Gm57240 | 0.093245792 | -1.807354922 | 0 | 0.024 | 1 | 0.698529281 | 2.017921908 | 0 | 0.015 | 1 | 0.698529281 | 0.177796806 |
| Atp2a1 | 0.093245792 | -1.807354922 | 0 | 0.024 | 1 | 0.619961444 | 1.432959407 | 0 | 0.025 | 1 | 0.619961444 | 0.177796806 |
| Gm53374 | 0.093245792 | -1.807354922 | 0 | 0.024 | 1 | 0.698534498 | 2.432959407 | 0 | 0.015 | 1 | 0.698534498 | 0.177796806 |
| Gm56824 | 0.093245792 | -1.807354922 | 0 | 0.024 | 1 | 0.643850855 | 2.017921908 | 0 | 0.022 | 1 | 0.643850855 | 0.177796806 |
| Mettl24 | 0.093245792 | -1.807354922 | 0 | 0.024 | 1 | 0.669843604 | 2.017921908 | 0 | 0.019 | 1 | 0.669843604 | 0.177796806 |
| Ttc41 | 0.093245792 | -1.807354922 | 0 | 0.024 | 1 | 0.557756607 | 1.111031312 | 0 | 0.034 | 1 | 0.557756607 | 0.177796806 |
| 2810021J22Rik | 0.093245792 | -1.807354922 | 0 | 0.024 | 1 | 0.669843604 | 2.017921908 | 0 | 0.019 | 1 | 0.669843604 | 0.177796806 |
| Adora2b | 0.093245792 | -1.807354922 | 0 | 0.024 | 1 | 0.446819579 | 0.432959407 | 0 | 0.056 | 1 | 0.446819579 | 0.177796806 |
| Gm48898 | 0.093245792 | -1.807354922 | 0 | 0.024 | 1 | 0.669850692 | 1.432959407 | 0 | 0.019 | 1 | 0.669850692 | 0.177796806 |
| 2410004P03Rik | 0.093245792 | -1.807354922 | 0 | 0.024 | 1 | 0.669853054 | 2.210566986 | 0 | 0.019 | 1 | 0.669853054 | 0.177796806 |
| Ighm | 0.093245792 | -5.343407822 | 0 | 0.024 | 1 | 0.364080193 | -0.267480311 | 0 | 0.077 | 1 | 0.364080193 | 0.177796806 |
| Gm49267 | 0.093245792 | -2.485426827 | 0 | 0.024 | 1 | 0.619953009 | 1.558490289 | 0 | 0.025 | 1 | 0.619953009 | 0.177796806 |
| Lrrc24 | 0.093245792 | -1.807354922 | 0 | 0.024 | 1 | 0.669853054 | 2.210566986 | 0 | 0.019 | 1 | 0.669853054 | 0.177796806 |
| Apol9b | 0.093245792 | -1.807354922 | 0 | 0.024 | 1 | 0.698534498 | 2.432959407 | 0 | 0.015 | 1 | 0.698534498 | 0.177796806 |
| Panx2 | 0.093245792 | -1.807354922 | 0 | 0.024 | 1 | 0.313847198 | 2.31748219 | 0.1 | 0.037 | 1 | 0.313847198 | 0.177796806 |
| Gm49584 | 0.093245792 | -1.807354922 | 0 | 0.024 | 1 | 0.669853054 | 2.210566986 | 0 | 0.019 | 1 | 0.669853054 | 0.177796806 |
| Prodh | 0.093245792 | -1.807354922 | 0 | 0.024 | 1 | 0.643842184 | 1.695993813 | 0 | 0.022 | 1 | 0.643842184 | 0.177796806 |
| Prr18 | 0.093245792 | -1.807354922 | 0 | 0.024 | 1 | 0.557744661 | 1.111031312 | 0 | 0.034 | 1 | 0.557744661 | 0.177796806 |
| B3galt4 | 0.093245792 | -1.807354922 | 0 | 0.024 | 1 | 0.597845674 | 1.695993813 | 0 | 0.028 | 1 | 0.597845674 | 0.177796806 |
| Guca1a | 0.093245792 | -1.807354922 | 0 | 0.024 | 1 | 0.730750993 | 2.210566986 | 0 | 0.012 | 1 | 0.730750993 | 0.177796806 |
| Gm41604 | 0.093245792 | -1.807354922 | 0 | 0.024 | 1 | 0.490315988 | 1.494359952 | 0.1 | 0.049 | 1 | 0.490315988 | 0.177796806 |
| Gm15956 | 0.093245792 | -1.807354922 | 0 | 0.024 | 1 | 0.433608183 | 0.432959407 | 0 | 0.059 | 1 | 0.433608183 | 0.177796806 |
| Fchsd1 | 0.093245792 | -1.807354922 | 0 | 0.024 | 1 | 0.539413885 | 1.210566986 | 0 | 0.037 | 1 | 0.539413885 | 0.177796806 |
| Kctd16 | 0.093245792 | -3.807354922 | 0 | 0.024 | 1 | 0.597840427 | 0.625604485 | 0 | 0.028 | 1 | 0.597840427 | 0.177796806 |
| Gm39529 | 0.093245792 | -2.485426827 | 0 | 0.024 | 1 | 0.577155236 | 1.210566986 | 0 | 0.031 | 1 | 0.577155236 | 0.177796806 |
| 5430416N02Rik | 0.093286491 | -0.817270391 | 0.21 | 0.282 | 1 | 0.853793526 | -0.111361109 | 0.2 | 0.213 | 1 | 0.853793526 | 0.177870613 |
| Ccdc157 | 0.289731811 | -0.358047521 | 0.437 | 0.465 | 1 | 0.093327388 | -1.171902651 | 0.2 | 0.475 | 1 | 0.289731811 | 0.177944775 |
| Unk | 0.155974209 | -0.665999073 | 0.235 | 0.3 | 1 | 0.093365208 | 0.539874611 | 0.6 | 0.278 | 1 | 0.155974209 | 0.178013355 |
| Ltbp3 | 0.282154947 | -0.327547167 | 0.748 | 0.724 | 1 | 0.093386896 | 0.498847343 | 0.9 | 0.738 | 1 | 0.282154947 | 0.178052679 |
| Adh5 | 0.530517855 | -0.264399948 | 0.555 | 0.541 | 1 | 0.093505015 | -0.888968688 | 0.3 | 0.574 | 1 | 0.530517855 | 0.178266842 |
| 4930579G24Rik | 0.093533261 | -1.070389328 | 0.042 | 0.094 | 1 | 1 | 0.847996907 | 0.1 | 0.099 | 1 | 1 | 0.178318051 |
| Acacb | 0.093709616 | -0.748461233 | 0.059 | 0.118 | 1 | 0.678985981 | 1.063725598 | 0.1 | 0.065 | 1 | 0.678985981 | 0.17863774 |
| Wiz | 0.207435234 | -0.542010356 | 0.202 | 0.259 | 1 | 0.093721106 | -1.28585884 | 0.1 | 0.352 | 1 | 0.207435234 | 0.178658566 |
| Ano1 | 0.093774805 | -2.485426827 | 0.008 | 0.041 | 1 | 0.364055313 | -1.69632361 | 0 | 0.077 | 1 | 0.364055313 | 0.178755895 |
| Klhl3 | 0.093774805 | -2.292781749 | 0.008 | 0.041 | 1 | 0.539456618 | 0.930459067 | 0 | 0.037 | 1 | 0.539456618 | 0.178755895 |
| Zfp12 | 0.093798052 | -0.780882711 | 0.067 | 0.129 | 1 | 0.467730467 | -0.248864633 | 0.1 | 0.188 | 1 | 0.467730467 | 0.178798029 |
| Uhmk1 | 0.365026361 | -0.263034406 | 0.277 | 0.329 | 1 | 0.09387725 | -0.872849022 | 0.1 | 0.395 | 1 | 0.365026361 | 0.178941562 |
| Htt | 0.093879232 | -0.480511337 | 0.655 | 0.659 | 1 | 0.915095213 | -0.12114213 | 0.7 | 0.79 | 1 | 0.915095213 | 0.178945154 |
| 1600002D24Rik | 0.093928832 | -1.201633861 | 0.034 | 0.082 | 1 | 0.698533977 | 1.558490289 | 0 | 0.015 | 1 | 0.698533977 | 0.179035038 |
| Ank1 | 0.355178918 | 1.118644496 | 0.118 | 0.088 | 1 | 0.094005589 | 1.515421567 | 0.2 | 0.062 | 1 | 0.355178918 | 0.179174127 |
| Glce | 0.485819947 | -0.193246076 | 0.412 | 0.441 | 1 | 0.094008943 | 0.60083658 | 0.8 | 0.42 | 1 | 0.485819947 | 0.179180205 |
| Supt20 | 0.346695471 | -0.406571406 | 0.538 | 0.535 | 1 | 0.094028695 | -0.849974556 | 0.2 | 0.534 | 1 | 0.346695471 | 0.179215994 |
| Zfp69 | 0.61215802 | 0.377069649 | 0.067 | 0.053 | 1 | 0.09408953 | 1.432959407 | 0.3 | 0.117 | 1 | 0.61215802 | 0.17932622 |
| Ttll12 | 0.094189415 | -1.222392421 | 0.034 | 0.082 | 1 | 0.735248562 | 1.31748219 | 0.1 | 0.071 | 1 | 0.735248562 | 0.179507185 |
| Kif22 | 0.0942638 | -2.655351829 | 0.008 | 0.041 | 1 | 0.730749233 | 2.432959407 | 0 | 0.012 | 1 | 0.730749233 | 0.179641936 |
| Gm15510 | 0.094264307 | -1.944858446 | 0.008 | 0.041 | 1 | 0.539413885 | 1.210566986 | 0 | 0.037 | 1 | 0.539413885 | 0.179642854 |
| Myadml2os | 0.094264307 | -1.944858446 | 0.008 | 0.041 | 1 | 0.474962821 | 0.494359952 | 0 | 0.049 | 1 | 0.474962821 | 0.179642854 |
| Rab40b | 0.094264307 | -1.944858446 | 0.008 | 0.041 | 1 | 0.52208609 | 0.930459067 | 0 | 0.04 | 1 | 0.52208609 | 0.179642854 |
| Gm15287 | 0.094264307 | -1.944858446 | 0.008 | 0.041 | 1 | 0.698532411 | 0.930459067 | 0 | 0.015 | 1 | 0.698532411 | 0.179642854 |
| Slc8a1 | 0.094264307 | -2.572889668 | 0.008 | 0.041 | 1 | 0.19112519 | -1.567040593 | 0 | 0.148 | 1 | 0.19112519 | 0.179642854 |
| Fam174c | 0.094288918 | 0.344648171 | 0.303 | 0.212 | 1 | 0.750911357 | 0.263034406 | 0.1 | 0.133 | 1 | 0.750911357 | 0.179687436 |
| Abraxas1 | 0.094294783 | -0.987927168 | 0.042 | 0.094 | 1 | 0.539445308 | 1.017921908 | 0 | 0.037 | 1 | 0.539445308 | 0.17969806 |
| Gm20274 | 0.094294783 | -0.987927168 | 0.042 | 0.094 | 1 | 0.37481957 | 0.374065718 | 0 | 0.074 | 1 | 0.37481957 | 0.17969806 |
| Rbmx2 | 0.094442734 | -1 | 0.042 | 0.094 | 1 | 0.5933729 | 1.625604485 | 0.1 | 0.059 | 1 | 0.5933729 | 0.179966039 |
| Smim15 | 0.307610963 | -0.44170545 | 0.412 | 0.441 | 1 | 0.094443932 | -1.019167411 | 0.1 | 0.389 | 1 | 0.307610963 | 0.179968208 |
| Gm45509 | 0.094525946 | -0.987927168 | 0.034 | 0.082 | 1 | 0.832730078 | 1.017921908 | 0.1 | 0.08 | 1 | 0.832730078 | 0.180116738 |
| Wars1 | 0.094536411 | -0.857395605 | 0.126 | 0.2 | 1 | 0.143476623 | -1.191531458 | 0 | 0.179 | 1 | 0.143476623 | 0.180135688 |
| Chuk | 0.094714295 | -0.318776958 | 0.555 | 0.624 | 1 | 0.965691535 | 0.145382817 | 0.7 | 0.75 | 1 | 0.965691535 | 0.180457792 |
| P2ry1 | 0.094752821 | -1.807354922 | 0.008 | 0.041 | 1 | 0.597824685 | 1.210566986 | 0 | 0.028 | 1 | 0.597824685 | 0.180527544 |
| Tmem82 | 0.094752821 | -1.807354922 | 0.008 | 0.041 | 1 | 0.984402643 | 0.558490289 | 0.1 | 0.096 | 1 | 0.984402643 | 0.180527544 |
| Lrrd1 | 0.094752821 | -2.292781749 | 0.008 | 0.041 | 1 | 0.365421266 | 1.625604485 | 0.1 | 0.04 | 1 | 0.365421266 | 0.180527544 |
| A730049H05Rik | 0.094752821 | -1.807354922 | 0.008 | 0.041 | 1 | 0.619967066 | 1.111031312 | 0 | 0.025 | 1 | 0.619967066 | 0.180527544 |
| B130024G19Rik | 0.094752821 | -2.185866545 | 0.008 | 0.041 | 1 | 0.669843604 | 2.017921908 | 0 | 0.019 | 1 | 0.669843604 | 0.180527544 |
| B430319H21Rik | 0.094752821 | -1.807354922 | 0.008 | 0.041 | 1 | 0.539445308 | 1.017921908 | 0 | 0.037 | 1 | 0.539445308 | 0.180527544 |
| Appbp2os | 0.094752821 | -1.944858446 | 0.008 | 0.041 | 1 | 0.19134412 | 2.432959407 | 0.1 | 0.028 | 1 | 0.19134412 | 0.180527544 |
| Ace | 0.094752821 | -1.944858446 | 0.008 | 0.041 | 1 | 0.446790883 | 0.625604485 | 0 | 0.056 | 1 | 0.446790883 | 0.180527544 |
| C230035I16Rik | 0.094752821 | -1.807354922 | 0.008 | 0.041 | 1 | 0.564719271 | 1.494359952 | 0.1 | 0.056 | 1 | 0.564719271 | 0.180527544 |
| Sult1d1 | 0.094757124 | -0.485426827 | 0.025 | 0.071 | 1 | 0.099883994 | -1.625934282 | 0 | 0.216 | 1 | 0.099883994 | 0.180535335 |
| Mfap3 | 0.094845189 | -0.384899951 | 0.319 | 0.406 | 1 | 0.315359415 | 0.526068812 | 0.6 | 0.432 | 1 | 0.315359415 | 0.180694768 |
| Wdr35 | 0.094993565 | -0.79105311 | 0.437 | 0.494 | 1 | 0.544194108 | 0.27308807 | 0.7 | 0.58 | 1 | 0.544194108 | 0.180963352 |
| Gm43848 | 0.095085133 | -0.839063782 | 0.059 | 0.118 | 1 | 0.943858776 | 0.591657153 | 0.1 | 0.09 | 1 | 0.943858776 | 0.181129083 |
| Gyg | 0.095118645 | -0.944858446 | 0.05 | 0.106 | 1 | 0.290452399 | -0.536666944 | 0 | 0.102 | 1 | 0.290452399 | 0.181189733 |
| Zc3h6 | 0.09519821 | 0.807354922 | 0.244 | 0.171 | 1 | 0.422239367 | 0.973527789 | 0.2 | 0.117 | 1 | 0.422239367 | 0.18133372 |
| Vps18 | 0.383646744 | -0.552541023 | 0.168 | 0.2 | 1 | 0.095217024 | -1.640289575 | 0 | 0.222 | 1 | 0.383646744 | 0.181367767 |
| Traf1 | 0.095240327 | -1.807354922 | 0.008 | 0.041 | 1 | 0.669843604 | 1.847996907 | 0 | 0.019 | 1 | 0.669843604 | 0.181409934 |
| 4930417H01Rik | 0.095240327 | -1.655351829 | 0.008 | 0.041 | 1 | 0.669843604 | 1.847996907 | 0 | 0.019 | 1 | 0.669843604 | 0.181409934 |
| Gm14286 | 0.095240327 | -1.807354922 | 0.008 | 0.041 | 1 | 0.446844685 | 0.494359952 | 0 | 0.056 | 1 | 0.446844685 | 0.181409934 |
| Stag3 | 0.095240327 | -1.655351829 | 0.008 | 0.041 | 1 | 0.669853054 | 2.210566986 | 0 | 0.019 | 1 | 0.669853054 | 0.181409934 |
| Fancf | 0.095240327 | -1.655351829 | 0.008 | 0.041 | 1 | 0.597845674 | 1.695993813 | 0 | 0.028 | 1 | 0.597845674 | 0.181409934 |
| Mmp15 | 0.095240327 | -1.655351829 | 0.008 | 0.041 | 1 | 0.642599971 | 1.393431043 | 0.1 | 0.065 | 1 | 0.642599971 | 0.181409934 |
| Gm28229 | 0.095240327 | -1.655351829 | 0.008 | 0.041 | 1 | 0.523811025 | 1.558490289 | 0.1 | 0.052 | 1 | 0.523811025 | 0.181409934 |
| Acvr2b | 0.095240327 | -1.655351829 | 0.008 | 0.041 | 1 | 0.63806535 | 1.31748219 | 0.1 | 0.062 | 1 | 0.63806535 | 0.181409934 |
| Gm30786 | 0.095240327 | -1.807354922 | 0.008 | 0.041 | 1 | 0.669843604 | 2.017921908 | 0 | 0.019 | 1 | 0.669843604 | 0.181409934 |
| Gm26660 | 0.095240327 | -1.807354922 | 0.008 | 0.041 | 1 | 0.669843604 | 2.017921908 | 0 | 0.019 | 1 | 0.669843604 | 0.181409934 |
| Phf11c | 0.095240327 | -1.655351829 | 0.008 | 0.041 | 1 | 0.522066388 | 1.111031312 | 0 | 0.04 | 1 | 0.522066388 | 0.181409934 |
| Cfap91 | 0.095240327 | -1.807354922 | 0.008 | 0.041 | 1 | 0.698529281 | 2.210566986 | 0 | 0.015 | 1 | 0.698529281 | 0.181409934 |
| Gm26724 | 0.095240327 | -1.655351829 | 0.008 | 0.041 | 1 | 0.433577142 | 0.494359952 | 0 | 0.059 | 1 | 0.433577142 | 0.181409934 |
| Magt1 | 0.284967239 | -0.266786541 | 0.815 | 0.712 | 1 | 0.095296532 | -0.734737493 | 0.4 | 0.769 | 1 | 0.284967239 | 0.181511634 |

| | | | | | | | | | | | | |
|---|---|---|---|---|---|---|---|---|---|---|---|---|
| Zfp120 | 0.095386292 | -1.163498732 | 0.076 | 0.135 | 1 | 0.302298355 | 1.210566986 | 0.2 | 0.096 | 1 | 0.302298355 | 0.181674039 |
| Endog | 0.095436041 | 0.563482773 | 0.218 | 0.141 | 1 | 0.610049543 | 0.847996907 | 0.2 | 0.139 | 1 | 0.610049543 | 0.181764045 |
| Rnaset2a | 0.095624705 | 0.247711987 | 0.798 | 0.741 | 1 | 0.606649958 | 0.747669848 | 0.7 | 0.731 | 1 | 0.606649958 | 0.182105326 |
| Gtf2h4 | 0.095641458 | -0.755515991 | 0.109 | 0.182 | 1 | 0.87759965 | 0.558490289 | 0.1 | 0.114 | 1 | 0.87759965 | 0.182135627 |
| 9530026P05Rik | 0.095645338 | -1.03191518 | 0.336 | 0.424 | 1 | 0.350428054 | -0.870353009 | 0.3 | 0.432 | 1 | 0.350428054 | 0.182142645 |
| Cracr2a | 0.095769202 | -1.485426827 | 0.008 | 0.041 | 1 | 0.489922734 | 0.625604485 | 0 | 0.046 | 1 | 0.489922734 | 0.182366664 |
| Lrfn3 | 0.095769202 | -1.485426827 | 0.008 | 0.041 | 1 | 0.490000637 | 1.017921908 | 0 | 0.046 | 1 | 0.490000637 | 0.182366664 |
| Lrrc75b | 0.095769202 | -1.485426827 | 0.008 | 0.041 | 1 | 0.539483006 | 1.31748219 | 0 | 0.037 | 1 | 0.539483006 | 0.182366664 |
| Fzd6 | 0.095818286 | -1.011495639 | 0.16 | 0.229 | 1 | 0.586847726 | 0.802909017 | 0.3 | 0.247 | 1 | 0.586847726 | 0.182455428 |
| Tuft1 | 0.095986787 | 0.436570661 | 0.185 | 0.112 | 1 | 0.509971822 | -0.408342847 | 0.1 | 0.176 | 1 | 0.509971822 | 0.18276011 |
| Snx11 | 0.096022102 | -0.887525271 | 0.109 | 0.176 | 1 | 0.743002921 | 0.744903414 | 0.2 | 0.157 | 1 | 0.743002921 | 0.18282396 |
| Nfe2l3 | 0.096070957 | -1.070389328 | 0.042 | 0.094 | 1 | 0.180629538 | -1.267480311 | 0 | 0.154 | 1 | 0.180629538 | 0.182912286 |
| Ankra2 | 0.096075631 | -0.883976204 | 0.16 | 0.229 | 1 | 0.200319809 | 0.785261151 | 0.4 | 0.207 | 1 | 0.200319809 | 0.182920735 |
| Hspa14 | 0.096141867 | -0.730902862 | 0.42 | 0.482 | 1 | 0.534652612 | 0.44049308 | 0.5 | 0.38 | 1 | 0.534652612 | 0.183040476 |
| Mccc1 | 0.096224764 | -1.070389328 | 0.109 | 0.176 | 1 | 0.508419418 | -0.457811523 | 0.1 | 0.176 | 1 | 0.508419418 | 0.183190323 |
| Tmem63b | 0.096250052 | 0.304355465 | 0.571 | 0.441 | 1 | 0.826661389 | 0.48866884 | 0.5 | 0.478 | 1 | 0.826661389 | 0.183236032 |
| Amd1 | 0.096390508 | 0.3804365 | 0.714 | 0.629 | 1 | 0.610164009 | 0.588305944 | 0.6 | 0.694 | 1 | 0.610164009 | 0.183489886 |
| Yrdc | 0.096393017 | -0.420721138 | 0.597 | 0.629 | 1 | 0.144301131 | -0.847148512 | 0.5 | 0.639 | 1 | 0.144301131 | 0.18349442 |
| Rabepk | 0.096463074 | -0.959358016 | 0.193 | 0.265 | 1 | 0.418869283 | 0.362570079 | 0.4 | 0.256 | 1 | 0.418869283 | 0.183621023 |
| Agfg1 | 0.37720462 | 0.131090407 | 0.874 | 0.871 | 1 | 0.096505932 | -0.638247543 | 0.8 | 0.929 | 1 | 0.37720462 | 0.183698469 |
| Kmt5c | 0.09663232 | -0.980191519 | 0.084 | 0.147 | 1 | 0.207814346 | -0.959358016 | 0 | 0.139 | 1 | 0.207814346 | 0.183926835 |
| Plxnb1 | 0.114018404 | -0.292781749 | 0.126 | 0.206 | 1 | 0.096654541 | 1.083809844 | 0.5 | 0.256 | 1 | 0.114018404 | 0.183966981 |
| Apod | 0.09666749 | 1.281912416 | 0.034 | 0.082 | 1 | 0.364181356 | -2.668578619 | 0 | 0.077 | 1 | 0.364181356 | 0.183990377 |
| Hapln3 | 0.204449765 | 0.344648171 | 0.235 | 0.171 | 1 | 0.096714237 | 1.073063462 | 0.4 | 0.173 | 1 | 0.204449765 | 0.18407483 |
| Ube2u | 0.096770559 | -1.707819249 | 0.008 | 0.041 | 1 | 0.475011905 | -0.505640048 | 0 | 0.049 | 1 | 0.475011905 | 0.184176577 |
| Gm57354 | 0.096848887 | -0.504285854 | 0.235 | 0.324 | 1 | 0.945956787 | 0.245332404 | 0.3 | 0.336 | 1 | 0.945956787 | 0.184318066 |
| Dmac2l | 0.096903212 | 0.514573173 | 0.218 | 0.141 | 1 | 0.274607156 | -0.709998547 | 0 | 0.108 | 1 | 0.274607156 | 0.184416192 |
| Usp39 | 0.096978193 | -0.648925559 | 0.185 | 0.265 | 1 | 0.557424439 | -0.430538593 | 0.2 | 0.269 | 1 | 0.557424439 | 0.184551617 |
| Tnfsf12 | 0.097131471 | -0.827819025 | 0.202 | 0.276 | 1 | 0.172258709 | -0.970762779 | 0.1 | 0.293 | 1 | 0.172258709 | 0.18482842 |
| Gm44899 | 0.09720951 | 0.40354186 | 0.185 | 0.112 | 1 | 0.190933503 | -0.936274402 | 0 | 0.148 | 1 | 0.190933503 | 0.184969331 |
| Stim1 | 0.097219495 | 0.230364744 | 0.571 | 0.441 | 1 | 0.354004306 | 0.937001913 | 0.7 | 0.648 | 1 | 0.354004306 | 0.18498736 |
| Pnpt1 | 0.097245487 | -0.733354341 | 0.244 | 0.318 | 1 | 0.957626514 | 0.263034406 | 0.3 | 0.309 | 1 | 0.957626514 | 0.185034289 |
| Ddc | 0.097272579 | -2.600904045 | 0.008 | 0.041 | 1 | 0.73075158 | 2.695993813 | 0 | 0.012 | 1 | 0.73075158 | 0.185083204 |
| D11Wsu47e | 0.097301124 | -0.707819249 | 0.059 | 0.118 | 1 | 0.642220346 | 0.159940913 | 0.1 | 0.151 | 1 | 0.642220346 | 0.18513474 |
| Foxc1 | 0.149302633 | -0.126081203 | 0.849 | 0.859 | 1 | 0.097340971 | -0.85654721 | 0.9 | 0.858 | 1 | 0.149302633 | 0.185206677 |
| Cfdp1 | 0.097342564 | -0.292127317 | 0.908 | 0.9 | 1 | 0.879599701 | 0.125245172 | 0.9 | 0.932 | 1 | 0.879599701 | 0.185209552 |
| Man2b2 | 0.176991597 | 0.305986551 | 0.319 | 0.247 | 1 | 0.097361758 | -1.152003093 | 0.1 | 0.349 | 1 | 0.176991597 | 0.185244205 |
| Ramp1 | 0.106636566 | 1.330148602 | 0.126 | 0.071 | 1 | 0.097407842 | 1.406487196 | 0.4 | 0.194 | 1 | 0.106636566 | 0.185327397 |
| Has1 | 0.097433372 | 1.514573173 | 0.176 | 0.112 | 1 | 0.181152715 | -2.220482831 | 0 | 0.154 | 1 | 0.181152715 | 0.185373483 |
| Osbpl6 | 0.998822485 | -0.178999826 | 0.655 | 0.612 | 1 | 0.097579907 | -0.91281543 | 0.4 | 0.691 | 1 | 0.998822485 | 0.185637976 |
| E130307A14Rik | 0.097646635 | -1.05177365 | 0.16 | 0.229 | 1 | 0.92174639 | -0.230005605 | 0.2 | 0.198 | 1 | 0.92174639 | 0.185758405 |
| Endov | 0.097656868 | -0.847996907 | 0.092 | 0.159 | 1 | 0.36390653 | 0.159940913 | 0 | 0.077 | 1 | 0.36390653 | 0.185776872 |
| Rccd1 | 0.097668808 | -0.696930932 | 0.092 | 0.159 | 1 | 0.393459672 | 1.017921908 | 0.2 | 0.114 | 1 | 0.393459672 | 0.185798421 |
| Rnf157 | 0.097688321 | -1.025995209 | 0.05 | 0.106 | 1 | 0.841020948 | 0.888638891 | 0.1 | 0.08 | 1 | 0.841020948 | 0.185833633 |
| Sh3glb2 | 0.097734178 | -0.719892081 | 0.126 | 0.2 | 1 | 0.175162596 | -0.959358016 | 0 | 0.157 | 1 | 0.175162596 | 0.185916386 |
| Lmo4 | 0.097975506 | -0.24726709 | 0.882 | 0.959 | 1 | 0.639415506 | -0.237219096 | 0.8 | 0.932 | 1 | 0.639415506 | 0.186351813 |
| Polr2m | 0.108921345 | -0.257797757 | 0.689 | 0.724 | 1 | 0.098069475 | -0.611434712 | 0.4 | 0.735 | 1 | 0.108921345 | 0.186521328 |
| Poc5 | 0.115238092 | -0.86393845 | 0.126 | 0.194 | 1 | 0.09808005 | -1.654503434 | 0 | 0.219 | 1 | 0.115238092 | 0.186540403 |
| Gmppb | 0.978217493 | -0.150242636 | 0.261 | 0.247 | 1 | 0.098203602 | -1.596787936 | 0 | 0.219 | 1 | 0.978217493 | 0.186763256 |
| Gnb5 | 0.266487037 | -0.640705053 | 0.134 | 0.176 | 1 | 0.098276907 | -1.625934282 | 0 | 0.219 | 1 | 0.266487037 | 0.186895464 |
| Cyp2r1 | 0.09828446 | -1.222392421 | 0.008 | 0.041 | 1 | 0.557729726 | 1.31748219 | 0 | 0.034 | 1 | 0.557729726 | 0.186909084 |
| Gm15563 | 0.098292224 | -1.222392421 | 0.008 | 0.041 | 1 | 0.320995017 | 1.769994395 | 0.1 | 0.037 | 1 | 0.320995017 | 0.186923087 |
| Gm32673 | 0.81369306 | -0.519374159 | 0.168 | 0.176 | 1 | 0.098737311 | -2.026472211 | 0 | 0.219 | 1 | 0.81369306 | 0.187725565 |
| Tedc1 | 0.098797453 | -1.070389328 | 0.008 | 0.041 | 1 | 0.234627319 | 2.31748219 | 0.1 | 0.031 | 1 | 0.234627319 | 0.18783397 |
| Krt76 | 0.098797453 | -1.070389328 | 0.008 | 0.041 | 1 | 0.730749233 | 2.210566986 | 0 | 0.012 | 1 | 0.730749233 | 0.18783397 |
| Sephs2 | 0.098846287 | -0.733354341 | 0.118 | 0.188 | 1 | 0.895499855 | 0.432959407 | 0.1 | 0.111 | 1 | 0.895499855 | 0.187921986 |
| Lpcat4 | 0.098902893 | -1.070389328 | 0.076 | 0.135 | 1 | 0.124045071 | -1.230005605 | 0 | 0.194 | 1 | 0.124045071 | 0.188024004 |
| Ptrh1 | 0.09894596 | -0.733354341 | 0.05 | 0.106 | 1 | 0.482051401 | 1.769994395 | 0.1 | 0.049 | 1 | 0.482051401 | 0.188101616 |
| Atp5pb | 0.099031952 | -0.228702743 | 0.924 | 0.924 | 1 | 0.126286703 | -0.511899039 | 0.7 | 0.944 | 1 | 0.126286703 | 0.188256576 |
| Snph | 0.09910821 | -0.774933444 | 0.05 | 0.106 | 1 | 0.248571988 | 1.515421567 | 0.2 | 0.09 | 1 | 0.248571988 | 0.188393982 |
| Hecw2 | 0.225897743 | -1.434800754 | 0.143 | 0.182 | 1 | 0.099271161 | -3.064227133 | 0 | 0.219 | 1 | 0.225897743 | 0.188687558 |
| Pstk | 0.099412774 | -0.485426827 | 0.118 | 0.194 | 1 | 0.190728693 | -0.91281543 | 0 | 0.148 | 1 | 0.190728693 | 0.188942648 |
| Ezr | 0.099429429 | -0.499232627 | 0.756 | 0.735 | 1 | 0.406732462 | 0.131340295 | 0.9 | 0.895 | 1 | 0.406732462 | 0.188972647 |
| Msrb1 | 0.099440697 | 0.32694617 | 0.403 | 0.3 | 1 | 0.421137889 | 1.146436649 | 0.4 | 0.361 | 1 | 0.421137889 | 0.188992941 |
| Etaa1 | 0.099450394 | -0.485426827 | 0.092 | 0.159 | 1 | 0.245614293 | -0.625934282 | 0 | 0.12 | 1 | 0.245614293 | 0.189010407 |
| Aldh16a1 | 0.474205722 | -0.584962501 | 0.05 | 0.071 | 1 | 0.099511414 | 1.236562194 | 0.3 | 0.117 | 1 | 0.474205722 | 0.189120307 |
| Sh3bp5 | 0.099627735 | -0.485426827 | 0.084 | 0.147 | 1 | 0.423822188 | 0.795529487 | 0.2 | 0.111 | 1 | 0.423822188 | 0.189329785 |
| 2810449D17Rik | 0.099681212 | -0.900464326 | 0.109 | 0.176 | 1 | 0.306858329 | -0.111361109 | 0 | 0.096 | 1 | 0.306858329 | 0.18942608 |
| Ttc39aos1 | 0.099733028 | 1.736965594 | 0.042 | 0.012 | 1 | 0.73075158 | 2.695993813 | 0 | 0.012 | 1 | 0.73075158 | 0.18951938 |
| Dcun1d2 | 0.099745326 | -1.292781749 | 0.109 | 0.171 | 1 | 0.409382678 | -0.763437806 | 0.1 | 0.201 | 1 | 0.409382678 | 0.189541523 |
| Ndufaf3 | 0.413407306 | 0.316633795 | 0.227 | 0.188 | 1 | 0.099781901 | 1.558490289 | 0.3 | 0.12 | 1 | 0.413407306 | 0.189607374 |
| Hr | 0.09979922 | 0.666576266 | 0.319 | 0.229 | 1 | 0.169953941 | 0.792955343 | 0.5 | 0.312 | 1 | 0.169953941 | 0.189638556 |
| Itga1 | 0.099890081 | 0.537293249 | 0.277 | 0.194 | 1 | 0.654519418 | -0.654503434 | 0.2 | 0.241 | 1 | 0.654519418 | 0.189802133 |
| Cers6 | 0.099919487 | -1.343407822 | 0.134 | 0.206 | 1 | 0.734561684 | 0.467724825 | 0.3 | 0.256 | 1 | 0.734561684 | 0.18985507 |
| A630072M18Rik | 0.099964469 | -0.86393845 | 0.101 | 0.171 | 1 | 0.448733315 | -0.489872732 | 0.1 | 0.191 | 1 | 0.448733315 | 0.189936043 |
| Acox3 | 0.099981753 | -1.485426827 | 0.084 | 0.147 | 1 | 0.175529674 | -1.391469028 | 0 | 0.157 | 1 | 0.175529674 | 0.189967154 |
| Urb2 | 0.100085448 | -1.070389328 | 0.076 | 0.135 | 1 | 0.499859222 | -0.210896782 | 0.1 | 0.179 | 1 | 0.499859222 | 0.190153799 |
| Ints1 | 0.100086874 | -1.021479727 | 0.076 | 0.135 | 1 | 0.43703994 | 1.210566986 | 0.2 | 0.117 | 1 | 0.43703994 | 0.190156365 |
| Sdhc | 0.301385053 | -0.250961574 | 0.597 | 0.612 | 1 | 0.100166416 | 0.586872091 | 0.9 | 0.574 | 1 | 0.301385053 | 0.190299521 |
| Ntn4 | 0.100261188 | 1.219829907 | 0.185 | 0.118 | 1 | 0.502439396 | 0.625604485 | 0.2 | 0.13 | 1 | 0.502439396 | 0.19047007 |
| Mrpl16 | 0.484699551 | -0.306456686 | 0.202 | 0.229 | 1 | 0.100412283 | -1.339630097 | 0 | 0.216 | 1 | 0.484699551 | 0.19074194 |
| Psmd10 | 0.100493799 | 0.381306642 | 0.235 | 0.153 | 1 | 0.613980018 | 0.847996907 | 0.2 | 0.139 | 1 | 0.613980018 | 0.190888594 |
| 5730522E02Rik | 0.100554912 | -0.929033479 | 0.092 | 0.159 | 1 | 0.105977145 | 0.779517169 | 0.3 | 0.12 | 1 | 0.105977145 | 0.190998533 |
| Tfcp2 | 0.100693338 | -0.570818318 | 0.244 | 0.324 | 1 | 0.578154848 | 0.494359952 | 0.2 | 0.302 | 1 | 0.578154848 | 0.191247528 |
| Cdh1 | 0.1008272 | -0.694880193 | 0.076 | 0.141 | 1 | 0.638668466 | -0.294961047 | 0.2 | 0.269 | 1 | 0.638668466 | 0.191488276 |
| Mup5 | 0.100864308 | 1.514573173 | 0.042 | 0.012 | 1 | 0.318562315 | 2.017921908 | 0.1 | 0.037 | 1 | 0.318562315 | 0.191555007 |
| Adap2 | 0.100864308 | 1.514573173 | 0.042 | 0.012 | 1 | 0.73075158 | 2.695993813 | 0 | 0.012 | 1 | 0.73075158 | 0.191555007 |
| Fam83h | 0.100864308 | 1.514573173 | 0.042 | 0.012 | 1 | 0.151443299 | 2.695993813 | 0.1 | 0.025 | 1 | 0.151443299 | 0.191555007 |
| Scn8a | 0.100864308 | 1.514573173 | 0.042 | 0.012 | 1 | 0.669848329 | 1.695993813 | 0 | 0.019 | 1 | 0.669848329 | 0.191555007 |
| Rasa3 | 0.725049101 | -0.296665184 | 0.58 | 0.535 | 1 | 0.100962072 | 0.608530972 | 0.8 | 0.556 | 1 | 0.725049101 | 0.191730804 |
| Ube2l6 | 0.101006717 | 0.152003093 | 0.092 | 0.165 | 1 | 0.334208749 | -0.408342847 | 0 | 0.086 | 1 | 0.334208749 | 0.191811077 |
| B230217O12Rik | 0.101015466 | -0.86393845 | 0.05 | 0.106 | 1 | 0.245813931 | 1.515421567 | 0.2 | 0.09 | 1 | 0.245813931 | 0.191826807 |
| Aco2 | 0.101150323 | -0.319689623 | 0.613 | 0.694 | 1 | 0.300072806 | -0.39538054 | 0.6 | 0.781 | 1 | 0.300072806 | 0.192069258 |
| Ap1b1 | 0.101244454 | -0.531230517 | 0.202 | 0.288 | 1 | 0.573289833 | -0.327852929 | 0.2 | 0.265 | 1 | 0.573289833 | 0.192238468 |
| Mfsd2a | 0.679845361 | -0.485426827 | 0.076 | 0.088 | 1 | 0.101328442 | 2.017921908 | 0.2 | 0.068 | 1 | 0.679845361 | 0.192389431 |
| Borcs6 | 0.49627785 | -0.333423734 | 0.277 | 0.312 | 1 | 0.101370147 | -1.348400306 | 0.1 | 0.343 | 1 | 0.49627785 | 0.192464387 |
| Mgat5 | 0.101406332 | -0.531720479 | 0.513 | 0.565 | 1 | 0.887400332 | -0.287379868 | 0.8 | 0.772 | 1 | 0.887400332 | 0.192529419 |
| Mindy1 | 0.193581306 | -0.506185387 | 0.412 | 0.465 | 1 | 0.101412191 | -0.768674454 | 0.2 | 0.494 | 1 | 0.193581306 | 0.19253995 |

| | | | | | | | | | | | | |
|---|---|---|---|---|---|---|---|---|---|---|---|---|
| Capg | 0.293662808 | 0.189611018 | 0.429 | 0.371 | 1 | 0.101574177 | -1.258202497 | 0.2 | 0.481 | 1 | 0.293662808 | 0.19283104 |
| Zfp830 | 0.101586605 | -0.56515402 | 0.286 | 0.365 | 1 | 0.143482187 | -1.090602549 | 0.2 | 0.404 | 1 | 0.143482187 | 0.192853372 |
| Fktn | 0.520900123 | -0.456280482 | 0.176 | 0.2 | 1 | 0.101638085 | -1.814968106 | 0 | 0.216 | 1 | 0.520900123 | 0.192945869 |
| Pop1 | 0.101679543 | -1.025995209 | 0.084 | 0.147 | 1 | 0.153840217 | 1.558490289 | 0.3 | 0.142 | 1 | 0.153840217 | 0.193020357 |
| Imp4 | 0.101727508 | -0.796628515 | 0.193 | 0.265 | 1 | 0.184587046 | -0.936274402 | 0.1 | 0.287 | 1 | 0.184587046 | 0.19310653 |
| Lpcat1 | 0.236773792 | -0.424026283 | 0.319 | 0.376 | 1 | 0.101784612 | -1.489872732 | 0.1 | 0.34 | 1 | 0.236773792 | 0.193209117 |
| Babam2 | 0.117313165 | -0.374395515 | 0.639 | 0.682 | 1 | 0.102003171 | 0.492198611 | 0.8 | 0.664 | 1 | 0.117313165 | 0.193601695 |
| Lrrc32 | 0.114195969 | 0.794681092 | 0.118 | 0.065 | 1 | 0.102413136 | 1.695993813 | 0.2 | 0.065 | 1 | 0.114195969 | 0.194337821 |
| Rbks | 0.659565637 | 0.399095955 | 0.084 | 0.071 | 1 | 0.102413136 | 1.847996907 | 0.2 | 0.065 | 1 | 0.659565637 | 0.194337821 |
| C230013L11Rik | 0.908150902 | -0.263034406 | 0.05 | 0.053 | 1 | 0.102413136 | 1.902444691 | 0.2 | 0.065 | 1 | 0.908150902 | 0.194337821 |
| Dffa | 0.246626843 | -0.807354922 | 0.025 | 0.053 | 1 | 0.102413279 | 1.959028219 | 0.2 | 0.065 | 1 | 0.246626843 | 0.194338079 |
| Slc18a1 | 0.10253645 | 1.192645078 | 0.042 | 0.012 | 1 | 0.522144287 | -0.339630097 | 0 | 0.04 | 1 | 0.522144287 | 0.194559176 |
| Spen | 0.392291029 | -0.402136484 | 0.731 | 0.765 | 1 | 0.102557941 | -0.490506745 | 0.7 | 0.895 | 1 | 0.392291029 | 0.194597751 |
| Pla2g12b | 0.483824937 | 0.514573173 | 0.025 | 0.041 | 1 | 0.10265174 | 4.017921908 | 0.1 | 0.022 | 1 | 0.483824937 | 0.1947661 |
| Rabl2 | 0.102747638 | -0.970853654 | 0.109 | 0.176 | 1 | 0.974641528 | 0.193493473 | 0.2 | 0.188 | 1 | 0.974641528 | 0.194938198 |
| Katnal1 | 0.102750954 | -0.780882711 | 0.168 | 0.241 | 1 | 0.776651022 | 0.808468542 | 0.1 | 0.145 | 1 | 0.776651022 | 0.19494415 |
| Abhd10 | 0.102817219 | -0.891419187 | 0.151 | 0.218 | 1 | 0.868833741 | 0.227844977 | 0.2 | 0.216 | 1 | 0.868833741 | 0.195063057 |
| Alad | 0.14317503 | -0.655351829 | 0.168 | 0.235 | 1 | 0.102853901 | 1.769994395 | 0.3 | 0.136 | 1 | 0.14317503 | 0.195128878 |
| Mul1 | 0.103025057 | 0.26459492 | 0.294 | 0.2 | 1 | 0.301155595 | -0.521236903 | 0.1 | 0.238 | 1 | 0.301155595 | 0.195435952 |
| Slc30a4 | 0.103180781 | -0.470319935 | 0.286 | 0.382 | 1 | 0.2683663 | 0.513301516 | 0.6 | 0.395 | 1 | 0.2683663 | 0.195715288 |
| Raf1 | 0.555624422 | 0.12259091 | 0.748 | 0.824 | 1 | 0.103268226 | 0.482712442 | 1 | 0.935 | 1 | 0.555624422 | 0.195872126 |
| Pofut1 | 0.226467723 | -0.565597176 | 0.244 | 0.294 | 1 | 0.103346588 | 1.111031312 | 0.6 | 0.361 | 1 | 0.226467723 | 0.196012658 |
| Atp7a | 0.103517326 | -0.544320516 | 0.496 | 0.518 | 1 | 0.614966188 | -0.345482823 | 0.6 | 0.66 | 1 | 0.614966188 | 0.196318815 |
| Hspb2 | 0.103662468 | 1.099535674 | 0.042 | 0.012 | 1 | 0.116579781 | 2.558490289 | 0.1 | 0.022 | 1 | 0.116579781 | 0.196579028 |
| 9430065F17Rik | 0.103662468 | 1.099535674 | 0.042 | 0.012 | 1 | 0.730750993 | 2.210566986 | 0 | 0.012 | 1 | 0.730750993 | 0.196579028 |
| Srf | 0.20997875 | 0.186998515 | 0.303 | 0.229 | 1 | 0.103886641 | 1.135278859 | 0.5 | 0.265 | 1 | 0.20997875 | 0.196980848 |
| Entpd6 | 0.512030106 | 0.215012891 | 0.092 | 0.071 | 1 | 0.104043387 | 1.795529487 | 0.2 | 0.065 | 1 | 0.512030106 | 0.197261748 |
| Ado | 0.17165204 | -0.694880193 | 0.126 | 0.182 | 1 | 0.104107781 | -1.5519337 | 0 | 0.213 | 1 | 0.17165204 | 0.197377133 |
| Lyrm2 | 0.104265449 | -0.739183419 | 0.193 | 0.265 | 1 | 0.478722715 | -0.230005605 | 0.1 | 0.185 | 1 | 0.478722715 | 0.197659613 |
| Agpat4 | 0.104304383 | -0.510717746 | 0.874 | 0.794 | 1 | 0.582710659 | 0.179202815 | 0.8 | 0.79 | 1 | 0.582710659 | 0.197729362 |
| Tmem109 | 0.104361843 | -0.574236094 | 0.403 | 0.459 | 1 | 0.869271863 | 0.282596672 | 0.5 | 0.522 | 1 | 0.869271863 | 0.197832292 |
| Exosc10 | 0.151116269 | -0.727501614 | 0.37 | 0.406 | 1 | 0.104609935 | -1.217294554 | 0.2 | 0.435 | 1 | 0.151116269 | 0.198276632 |
| Wdr48 | 0.104624404 | -0.584962501 | 0.361 | 0.435 | 1 | 0.636652461 | 0.159940913 | 0.6 | 0.488 | 1 | 0.636652461 | 0.198302542 |
| Tep1 | 0.104761338 | -1.117695043 | 0.067 | 0.124 | 1 | 0.515410835 | 1.362570079 | 0.2 | 0.136 | 1 | 0.515410835 | 0.198547739 |
| Slc16a11 | 0.104825744 | 0.684498174 | 0.042 | 0.012 | 1 | 0.212993607 | -0.625934282 | 0 | 0.136 | 1 | 0.212993607 | 0.198663051 |
| Foxq1 | 0.104825744 | -0.222392421 | 0.042 | 0.012 | 1 | 0.698529281 | 2.017921908 | 0 | 0.015 | 1 | 0.698529281 | 0.198663051 |
| Znhit1 | 0.104831544 | -0.594361199 | 0.345 | 0.412 | 1 | 0.115672755 | -1.171902651 | 0.1 | 0.333 | 1 | 0.115672755 | 0.198673435 |
| Schip1 | 0.104842092 | -0.465389074 | 0.597 | 0.688 | 1 | 0.425628232 | -0.760070954 | 0.8 | 0.852 | 1 | 0.425628232 | 0.19869232 |
| Rnf25 | 0.104947062 | -0.544320516 | 0.387 | 0.453 | 1 | 0.869738141 | -0.175849835 | 0.4 | 0.38 | 1 | 0.869738141 | 0.198880238 |
| Stx17 | 0.10497654 | -0.622930351 | 0.353 | 0.418 | 1 | 0.934928423 | 0.252387162 | 0.4 | 0.429 | 1 | 0.934928423 | 0.198933006 |
| Dbr1 | 0.105121212 | -0.900464326 | 0.168 | 0.235 | 1 | 0.253861541 | -0.776493958 | 0.1 | 0.253 | 1 | 0.253861541 | 0.199191954 |
| Fam32a | 0.105244236 | -0.612183969 | 0.345 | 0.406 | 1 | 0.434840065 | -0.152003093 | 0.3 | 0.472 | 1 | 0.434840065 | 0.199412124 |
| Actr10 | 0.152559499 | -0.352160296 | 0.714 | 0.688 | 1 | 0.105318861 | -0.714528205 | 0.7 | 0.802 | 1 | 0.152559499 | 0.199545659 |
| Ogfrl1 | 0.105421194 | -1.485426827 | 0.092 | 0.153 | 1 | 0.175393422 | -1.131825212 | 0 | 0.157 | 1 | 0.175393422 | 0.19972876 |
| Cul9 | 0.105455767 | -0.780882711 | 0.143 | 0.212 | 1 | 0.442567486 | -0.230005605 | 0.2 | 0.327 | 1 | 0.442567486 | 0.199790615 |
| Pik3c3 | 0.105538972 | -0.485426827 | 0.151 | 0.229 | 1 | 0.292892381 | 1.463333056 | 0.3 | 0.191 | 1 | 0.292892381 | 0.199939469 |
| Tcp11l2 | 0.96377531 | -0.160134498 | 0.622 | 0.606 | 1 | 0.105838343 | -0.609351398 | 0.5 | 0.762 | 1 | 0.96377531 | 0.200474931 |
| Fzr1 | 0.105876501 | -0.563429339 | 0.126 | 0.2 | 1 | 0.540206155 | 0.751135367 | 0.3 | 0.213 | 1 | 0.540206155 | 0.200543169 |
| Rbfa | 0.106065822 | -0.952552838 | 0.134 | 0.2 | 1 | 0.971261163 | 0.245332404 | 0.2 | 0.198 | 1 | 0.971261163 | 0.200881686 |
| Nfkbiz | 0.214764676 | 0.653178411 | 0.588 | 0.512 | 1 | 0.10613842 | 1.471433555 | 0.6 | 0.38 | 1 | 0.214764676 | 0.201011476 |
| Strn4 | 0.106157482 | -0.411426246 | 0.252 | 0.341 | 1 | 0.160442245 | -0.864721141 | 0.2 | 0.435 | 1 | 0.160442245 | 0.201045552 |
| Snrnp25 | 0.198373698 | 0.273565073 | 0.235 | 0.171 | 1 | 0.106482622 | 1.725140159 | 0.3 | 0.13 | 1 | 0.198373698 | 0.201626695 |
| Crlf2 | 0.106518062 | 0.368352432 | 0.445 | 0.335 | 1 | 0.598862105 | 0.351165316 | 0.2 | 0.299 | 1 | 0.598862105 | 0.201690026 |
| Invs | 0.106544476 | -0.707819249 | 0.269 | 0.341 | 1 | 0.386025715 | -0.391469028 | 0.2 | 0.34 | 1 | 0.386025715 | 0.201737227 |
| Emid1 | 0.709935792 | -0.637429921 | 0.067 | 0.076 | 1 | 0.106693545 | 3.14345279 | 0.1 | 0.022 | 1 | 0.709935792 | 0.202003577 |
| 1700003E16Rik | 0.1433296 | -2.070389328 | 0.008 | 0.035 | 1 | 0.106697968 | 3.14345279 | 0.1 | 0.022 | 1 | 0.1433296 | 0.20201148 |
| Prmt2 | 0.106718814 | -0.818002166 | 0.176 | 0.247 | 1 | 0.712329907 | -0.125036046 | 0.2 | 0.241 | 1 | 0.712329907 | 0.202048722 |
| Glis1 | 0.106763131 | -0.421861698 | 0.143 | 0.224 | 1 | 0.151344028 | 0.364479669 | 0.5 | 0.284 | 1 | 0.151344028 | 0.202127896 |
| Dtx3l | 0.106849703 | -0.987927168 | 0.084 | 0.147 | 1 | 0.23908502 | -1.111361109 | 0 | 0.123 | 1 | 0.23908502 | 0.202282547 |
| Plk3 | 0.10688986 | -0.822461814 | 0.126 | 0.194 | 1 | 0.51153589 | -0.441509711 | 0.1 | 0.176 | 1 | 0.51153589 | 0.202354278 |
| Psmg3 | 0.107226828 | -0.600904045 | 0.176 | 0.253 | 1 | 0.725768183 | 0.185031894 | 0.1 | 0.136 | 1 | 0.725768183 | 0.202956063 |
| 4930599N23Rik | 0.107268795 | -0.889817082 | 0.109 | 0.176 | 1 | 0.553860965 | -0.654503434 | 0.1 | 0.164 | 1 | 0.553860965 | 0.203030995 |
| Fgl2 | 0.107556108 | 1.014144182 | 0.126 | 0.071 | 1 | 0.290752693 | -2.481923979 | 0 | 0.102 | 1 | 0.290752693 | 0.2035439 |
| Parp14 | 0.107556816 | -1.770829046 | 0.05 | 0.1 | 1 | 0.298780359 | -0.982078092 | 0 | 0.099 | 1 | 0.298780359 | 0.203545164 |
| Dxo | 0.107557179 | -0.870090677 | 0.143 | 0.212 | 1 | 0.73845289 | 1.040641984 | 0.2 | 0.173 | 1 | 0.73845289 | 0.203545811 |
| Septin10 | 0.107564516 | 0.109731441 | 0.42 | 0.306 | 1 | 0.269045995 | 0.802909017 | 0.4 | 0.244 | 1 | 0.269045995 | 0.203558908 |
| Cnr1 | 0.107566819 | -2.707819249 | 0.017 | 0.053 | 1 | 0.643846086 | -0.374395515 | 0 | 0.022 | 1 | 0.643846086 | 0.203563018 |
| Nostrin | 0.36330301 | -1.185866545 | 0.05 | 0.076 | 1 | 0.107666771 | 2.210566986 | 0.3 | 0.142 | 1 | 0.36330301 | 0.203741408 |
| Lima1 | 0.533708805 | -0.396578481 | 0.739 | 0.712 | 1 | 0.107686858 | -0.447644497 | 0.7 | 0.855 | 1 | 0.533708805 | 0.203777256 |
| Dlg1 | 0.309752268 | -0.226585435 | 0.924 | 0.9 | 1 | 0.107718762 | 0.303260088 | 1 | 0.944 | 1 | 0.309752268 | 0.203834192 |
| Rad17 | 0.107743495 | -0.417255325 | 0.353 | 0.447 | 1 | 0.139708836 | -1.004445905 | 0.2 | 0.438 | 1 | 0.139708836 | 0.203878329 |
| Slc15a4 | 0.107753234 | -0.600904045 | 0.16 | 0.235 | 1 | 0.59822595 | -0.242605642 | 0.2 | 0.262 | 1 | 0.59822595 | 0.203895708 |
| Fcgrt | 0.317628138 | -0.208475641 | 0.908 | 0.841 | 1 | 0.107871871 | -0.54813213 | 0.7 | 0.836 | 1 | 0.317628138 | 0.204107402 |
| Arhgef3 | 0.108091294 | 0.861223697 | 0.336 | 0.265 | 1 | 0.179902277 | 0.968073359 | 0.5 | 0.355 | 1 | 0.179902277 | 0.204498861 |
| Tfg | 0.701517277 | -0.10877042 | 0.866 | 0.829 | 1 | 0.108135159 | -0.55024535 | 0.8 | 0.864 | 1 | 0.701517277 | 0.204577105 |
| Dlat | 0.108143652 | -0.769219793 | 0.328 | 0.382 | 1 | 0.561107418 | -0.134081185 | 0.4 | 0.509 | 1 | 0.561107418 | 0.204592255 |
| Atp13a3 | 0.527424787 | -0.196665596 | 0.731 | 0.753 | 1 | 0.108296371 | -0.663902132 | 0.8 | 0.892 | 1 | 0.527424787 | 0.204864639 |
| Colgalt1 | 0.108364489 | -0.10938346 | 0.874 | 0.782 | 1 | 0.192266532 | 0.307946828 | 1 | 0.784 | 1 | 0.192266532 | 0.204986115 |
| Clstn1 | 0.291863519 | -0.127533235 | 0.647 | 0.659 | 1 | 0.108368575 | -0.761051213 | 0.7 | 0.821 | 1 | 0.291863519 | 0.204993402 |
| Nim1k | 0.108623755 | -1.078124977 | 0.311 | 0.365 | 1 | 0.490056542 | -0.600463594 | 0.4 | 0.46 | 1 | 0.490056542 | 0.20544839 |
| Tug1 | 0.156691955 | -0.270687978 | 0.546 | 0.612 | 1 | 0.108846203 | -0.736965594 | 0.4 | 0.682 | 1 | 0.156691955 | 0.20584491 |
| Oga | 0.109136343 | -0.34552822 | 0.798 | 0.788 | 1 | 0.443275246 | 0.338071459 | 0.9 | 0.892 | 1 | 0.443275246 | 0.206361944 |
| Ccnc | 0.109285384 | -0.456667834 | 0.445 | 0.512 | 1 | 0.715200856 | -0.165299916 | 0.5 | 0.522 | 1 | 0.715200856 | 0.206627474 |
| Tmem165 | 0.109288984 | -0.337493398 | 0.79 | 0.788 | 1 | 0.551988283 | 0.347068986 | 0.9 | 0.84 | 1 | 0.551988283 | 0.206633885 |
| Oxnad1 | 0.142819013 | 0.340543773 | 0.218 | 0.147 | 1 | 0.109615296 | 1.506959989 | 0.3 | 0.13 | 1 | 0.142819013 | 0.207215078 |
| Cmc4 | 0.109630002 | -0.632268215 | 0.202 | 0.282 | 1 | 0.676110446 | 0.526068812 | 0.3 | 0.231 | 1 | 0.676110446 | 0.207241267 |
| Umad1 | 0.109657135 | -1.144389909 | 0.21 | 0.282 | 1 | 0.409954898 | -0.621806396 | 0.3 | 0.417 | 1 | 0.409954898 | 0.207289583 |
| Tysnd1 | 0.109767314 | -0.829381228 | 0.101 | 0.165 | 1 | 0.219337963 | -0.68251781 | 0 | 0.133 | 1 | 0.219337963 | 0.207485765 |
| Akap10 | 0.109779267 | -0.802909017 | 0.345 | 0.394 | 1 | 0.235459462 | -0.929610672 | 0.4 | 0.537 | 1 | 0.235459462 | 0.207507047 |
| Pfkfb2 | 0.110165748 | -0.970853654 | 0.101 | 0.165 | 1 | 0.764433719 | 1.054447784 | 0.2 | 0.302 | 1 | 0.764433719 | 0.208195005 |
| Hyal1 | 0.840272954 | -0.308549065 | 0.101 | 0.106 | 1 | 0.110206181 | 2.374065718 | 0.2 | 0.071 | 1 | 0.840272954 | 0.20826696 |
| Dlg3 | 0.110331586 | -0.696930932 | 0.118 | 0.188 | 1 | 0.725985348 | 0.602884409 | 0.2 | 0.269 | 1 | 0.725985348 | 0.208490114 |
| Pgls | 0.519381327 | -0.197181859 | 0.63 | 0.671 | 1 | 0.110391737 | -0.874469118 | 0.4 | 0.586 | 1 | 0.519381327 | 0.208597138 |
| Akr1b3 | 0.633167243 | 0.514573173 | 0.395 | 0.382 | 1 | 0.110455647 | -0.282201817 | 0.1 | 0.392 | 1 | 0.633167243 | 0.208710843 |
| Elmo2 | 0.35679886 | -0.572889668 | 0.118 | 0.153 | 1 | 0.110604614 | -1.5519337 | 0 | 0.207 | 1 | 0.35679886 | 0.208975848 |
| Tchh | 0.110663996 | 1.140177658 | 0.193 | 0.124 | 1 | 0.539431168 | 0.930459067 | 0 | 0.037 | 1 | 0.539431168 | 0.209081472 |
| Macrod1 | 0.110714356 | -0.485426827 | 0.202 | 0.282 | 1 | 0.599313134 | -0.348400306 | 0.2 | 0.275 | 1 | 0.599313134 | 0.209171043 |
| Nr2f6 | 0.148266269 | -0.783107376 | 0.185 | 0.247 | 1 | 0.110726482 | -1.161987182 | 0.1 | 0.336 | 1 | 0.148266269 | 0.20919261 |

| | | | | | | | | | | | | |
|---|---|---|---|---|---|---|---|---|---|---|---|---|
| Naa40 | 0.153366038 | -0.743224585 | 0.16 | 0.224 | 1 | 0.110726876 | -1.536666944 | 0 | 0.207 | 1 | 0.153366038 | 0.209193312 |
| 9230114K14Rik | 0.380148494 | -0.521050737 | 0.109 | 0.141 | 1 | 0.110735964 | 1.432959407 | 0.2 | 0.065 | 1 | 0.380148494 | 0.209209474 |
| Gm17477 | 0.678847153 | -1.222392421 | 0.017 | 0.024 | 1 | 0.110742227 | 1.354956895 | 0.2 | 0.065 | 1 | 0.678847153 | 0.209220613 |
| Parp2 | 0.110764468 | -0.437598302 | 0.294 | 0.376 | 1 | 0.272099747 | 0.834700084 | 0.4 | 0.247 | 1 | 0.272099747 | 0.209260168 |
| Garin5b | 0.456088296 | -1.163498732 | 0.025 | 0.041 | 1 | 0.110849878 | 2.602884409 | 0.1 | 0.022 | 1 | 0.456088296 | 0.20941206 |
| Mrps33 | 0.144672271 | -0.329307625 | 0.58 | 0.629 | 1 | 0.110945708 | -0.532275175 | 0.3 | 0.58 | 1 | 0.144672271 | 0.209582466 |
| Prdm16os | 0.110971151 | 0.567040593 | 0.202 | 0.129 | 1 | 0.387128844 | 1.299103661 | 0.3 | 0.201 | 1 | 0.387128844 | 0.209627705 |
| Zfp217 | 0.760288773 | -0.418312631 | 0.345 | 0.335 | 1 | 0.111034325 | -1.178475305 | 0.2 | 0.432 | 1 | 0.760288773 | 0.209740029 |
| Slc25a28 | 0.163450863 | -0.267835392 | 0.286 | 0.359 | 1 | 0.111036879 | -1.433289204 | 0.1 | 0.333 | 1 | 0.163450863 | 0.209744569 |
| Plekhg2 | 0.376873362 | -0.669851398 | 0.286 | 0.318 | 1 | 0.111215419 | 2.130396637 | 0.5 | 0.318 | 1 | 0.376873362 | 0.210061969 |
| Ccrl2 | 0.111372969 | -1.987927168 | 0.017 | 0.053 | 1 | 0.643835681 | 1.847996907 | 0 | 0.022 | 1 | 0.643835681 | 0.210342 |
| Tlcd3a | 0.111372969 | -1.900464326 | 0.017 | 0.053 | 1 | 0.420933835 | 0.31748219 | 0 | 0.062 | 1 | 0.420933835 | 0.210342 |
| Cit | 0.111374183 | -2.854660637 | 0.017 | 0.053 | 1 | 0.730749233 | 2.432959407 | 0 | 0.012 | 1 | 0.730749233 | 0.210344157 |
| Dnajc18 | 0.111460244 | -0.702238216 | 0.235 | 0.312 | 1 | 0.379917876 | -0.654503434 | 0.3 | 0.398 | 1 | 0.379917876 | 0.210497101 |
| Exoc3 | 0.120436733 | -0.397963986 | 0.63 | 0.653 | 1 | 0.111618856 | -0.702555563 | 0.5 | 0.765 | 1 | 0.120436733 | 0.210778944 |
| Tmem33 | 0.111860632 | 0.310340121 | 0.807 | 0.7 | 1 | 0.531720996 | -0.272096939 | 0.8 | 0.806 | 1 | 0.531720996 | 0.211208462 |
| Dtnb | 0.867001947 | -0.298341275 | 0.706 | 0.641 | 1 | 0.111922789 | -1.04673105 | 0.7 | 0.778 | 1 | 0.867001947 | 0.211318867 |
| Wwc2 | 0.111966304 | -0.386327739 | 0.899 | 0.882 | 1 | 0.398078104 | -0.305098724 | 1 | 0.898 | 1 | 0.398078104 | 0.211396154 |
| Gprasp1 | 0.112163928 | -0.499101764 | 0.521 | 0.565 | 1 | 0.828692675 | 0.23502003 | 0.7 | 0.605 | 1 | 0.828692675 | 0.211747109 |
| Ap2b1 | 0.112229872 | -0.363325219 | 0.79 | 0.776 | 1 | 0.421658406 | -0.249756562 | 0.7 | 0.84 | 1 | 0.421658406 | 0.2118642 |
| Fgf7 | 0.219589239 | -1.070389328 | 0.008 | 0.029 | 1 | 0.112264464 | 2.017921908 | 0.1 | 0.022 | 1 | 0.219589239 | 0.211925619 |
| Zmynd15 | 0.147029286 | -1.807354922 | 0 | 0.018 | 1 | 0.112266525 | 3.017921908 | 0.1 | 0.022 | 1 | 0.147029286 | 0.211929277 |
| B3gnt6 | 0.147030916 | -1.485426827 | 0 | 0.018 | 1 | 0.112266525 | 3.017921908 | 0.1 | 0.022 | 1 | 0.147030916 | 0.211929277 |
| Birc5 | 0.164849896 | -1.93021167 | 0.017 | 0.047 | 1 | 0.112266525 | 3.017921908 | 0.1 | 0.022 | 1 | 0.164849896 | 0.211929277 |
| Mcrip2 | 0.215728257 | 1.099535674 | 0.042 | 0.018 | 1 | 0.112266525 | 3.017921908 | 0.1 | 0.022 | 1 | 0.215728257 | 0.211929277 |
| Shd | 0.219589239 | -1.070389328 | 0.008 | 0.029 | 1 | 0.112266525 | 3.017921908 | 0.1 | 0.022 | 1 | 0.219589239 | 0.211929277 |
| Gm50146 | 0.225055707 | -0.485426827 | 0.008 | 0.029 | 1 | 0.112266525 | 3.017921908 | 0.1 | 0.022 | 1 | 0.225055707 | 0.211929277 |
| Mrap | 0.384399171 | 0.321928095 | 0.042 | 0.024 | 1 | 0.112266525 | 3.017921908 | 0.1 | 0.022 | 1 | 0.384399171 | 0.211929277 |
| Fam111a | 0.574919344 | -0.244418728 | 0.059 | 0.076 | 1 | 0.112266525 | 3.017921908 | 0.1 | 0.022 | 1 | 0.574919344 | 0.211929277 |
| Gm16861 | 0.687345921 | -0.900464326 | 0.017 | 0.024 | 1 | 0.112266525 | 3.017921908 | 0.1 | 0.022 | 1 | 0.687345921 | 0.211929277 |
| Gm44386 | 0.691605844 | -0.485426827 | 0.017 | 0.024 | 1 | 0.112266525 | 3.017921908 | 0.1 | 0.022 | 1 | 0.691605844 | 0.211929277 |
| AV099323 | 0.112335502 | -1.707819249 | 0.017 | 0.053 | 1 | 0.113669596 | 2.847996907 | 0.1 | 0.022 | 1 | 0.113669596 | 0.212051738 |
| Pik3cd | 0.112335502 | -1.707819249 | 0.017 | 0.053 | 1 | 0.505623509 | 0.695993813 | 0 | 0.043 | 1 | 0.505623509 | 0.212051738 |
| Nrm | 0.112335502 | -1.807354922 | 0.017 | 0.053 | 1 | 0.479305119 | 1.847996907 | 0.1 | 0.049 | 1 | 0.479305119 | 0.212051738 |
| Zbtb45 | 0.112335908 | -1.600904045 | 0.017 | 0.053 | 1 | 0.489901827 | 0.847996907 | 0 | 0.046 | 1 | 0.489901827 | 0.21205246 |
| Gm56844 | 0.112335908 | -1.600904045 | 0.017 | 0.053 | 1 | 0.505623793 | 0.847996907 | 0 | 0.043 | 1 | 0.505623793 | 0.21205246 |
| Efcab5 | 0.112335908 | -1.807354922 | 0.017 | 0.053 | 1 | 0.299732265 | 2.354956895 | 0.1 | 0.037 | 1 | 0.299732265 | 0.21205246 |
| Atg12 | 0.618116644 | -0.222392421 | 0.513 | 0.424 | 1 | 0.112369583 | -1.28128611 | 0.3 | 0.5 | 1 | 0.618116644 | 0.212112242 |
| Zfp57 | 0.112398102 | -0.994440475 | 0.092 | 0.153 | 1 | 0.213665798 | -0.864721141 | 0 | 0.136 | 1 | 0.213665798 | 0.21216287 |
| Gm26911 | 0.445153818 | 0.80407979 | 0.067 | 0.047 | 1 | 0.112461851 | 1.31748219 | 0.2 | 0.065 | 1 | 0.445153818 | 0.212276034 |
| Prepl | 0.112464173 | 0.607682577 | 0.303 | 0.224 | 1 | 0.97528804 | 0.510127268 | 0.2 | 0.201 | 1 | 0.97528804 | 0.212280156 |
| Gm1976 | 0.112666195 | -1.318316841 | 0.126 | 0.188 | 1 | 0.674769437 | 0.732519689 | 0.2 | 0.154 | 1 | 0.674769437 | 0.212638719 |
| Lrrc45 | 0.112833604 | -0.982926487 | 0.126 | 0.188 | 1 | 0.565279233 | 0.720241359 | 0.3 | 0.241 | 1 | 0.565279233 | 0.212935785 |
| Ormdl2 | 0.11287898 | 0.785875195 | 0.21 | 0.141 | 1 | 0.565720925 | 0.542188477 | 0.3 | 0.21 | 1 | 0.565720925 | 0.213016296 |
| Zfp983 | 0.112938994 | -1.070389328 | 0.067 | 0.124 | 1 | 0.810418519 | 0.847996907 | 0.1 | 0.077 | 1 | 0.810418519 | 0.213122771 |
| Ech1 | 0.112994008 | 0.294895136 | 0.655 | 0.582 | 1 | 0.405749443 | -0.255096586 | 0.4 | 0.611 | 1 | 0.405749443 | 0.21322037 |
| Eif2ak4 | 0.61047356 | -0.111031312 | 0.563 | 0.529 | 1 | 0.113018769 | 0.601082166 | 0.8 | 0.571 | 1 | 0.61047356 | 0.213264297 |
| Unc119 | 0.113557594 | 0.334000927 | 0.319 | 0.229 | 1 | 0.113199189 | -1.304006187 | 0 | 0.204 | 1 | 0.113557594 | 0.213584322 |
| Ethe1 | 0.144333789 | -1.025995209 | 0.076 | 0.129 | 1 | 0.113199189 | -1.304006187 | 0 | 0.204 | 1 | 0.144333789 | 0.213584322 |
| Cwf19l2 | 0.948702542 | -0.279842694 | 0.613 | 0.535 | 1 | 0.113216099 | 0.942633781 | 0.8 | 0.528 | 1 | 0.948702542 | 0.213614313 |
| Pde4a | 0.684737648 | -0.597901556 | 0.269 | 0.271 | 1 | 0.113234021 | 0.982297998 | 0.5 | 0.287 | 1 | 0.684737648 | 0.213646098 |
| Bfsp2 | 0.113300556 | -1.600904045 | 0.017 | 0.053 | 1 | 0.643850855 | 2.017921908 | 0 | 0.022 | 1 | 0.643850855 | 0.213764097 |
| Tert | 0.113300556 | -1.485426827 | 0.017 | 0.053 | 1 | 0.557729726 | 1.210566986 | 0 | 0.034 | 1 | 0.557729726 | 0.213764097 |
| Sall2 | 0.113300556 | -1.485426827 | 0.017 | 0.053 | 1 | 0.539444993 | 0.847996907 | 0 | 0.037 | 1 | 0.539444993 | 0.213764097 |
| Stard6 | 0.113300556 | -1.600904045 | 0.017 | 0.053 | 1 | 0.236765379 | 2.111031312 | 0.1 | 0.031 | 1 | 0.236765379 | 0.213764097 |
| Gm56492 | 0.113300556 | -1.600904045 | 0.017 | 0.053 | 1 | 0.365421266 | 1.625604485 | 0.1 | 0.04 | 1 | 0.365421266 | 0.213764097 |
| Zfp865 | 0.113336972 | -1.070389328 | 0.059 | 0.112 | 1 | 0.409771779 | -0.357117523 | 0.1 | 0.204 | 1 | 0.409771779 | 0.213828675 |
| Cptp | 0.1133582 | -0.522901533 | 0.118 | 0.188 | 1 | 0.802052667 | 1.111031312 | 0.1 | 0.077 | 1 | 0.802052667 | 0.213866318 |
| Atmin | 0.148718085 | -0.839063782 | 0.067 | 0.118 | 1 | 0.113408924 | 1.374065718 | 0.3 | 0.123 | 1 | 0.148718085 | 0.213956264 |
| Pitpna | 0.135179418 | -0.312334547 | 0.958 | 0.971 | 1 | 0.113441718 | -0.586281235 | 1 | 0.972 | 1 | 0.135179418 | 0.214014412 |
| Taf7 | 0.113492028 | -0.588304008 | 0.521 | 0.553 | 1 | 0.421429145 | 0.30367639 | 0.7 | 0.559 | 1 | 0.421429145 | 0.214103616 |
| B3gntl1 | 0.291779183 | -0.669851398 | 0.076 | 0.112 | 1 | 0.113639075 | 2.648688098 | 0.2 | 0.071 | 1 | 0.291779183 | 0.214364311 |
| Gm43560 | 0.121112965 | 1.225066556 | 0.092 | 0.047 | 1 | 0.113669596 | 2.695993813 | 0.1 | 0.022 | 1 | 0.121112965 | 0.214418414 |
| Gm7467 | 0.12404457 | 1.192645078 | 0.059 | 0.024 | 1 | 0.113669596 | 2.847996907 | 0.1 | 0.022 | 1 | 0.12404457 | 0.214418414 |
| Gm42788 | 0.13606055 | 0.640104055 | 0.092 | 0.047 | 1 | 0.113669596 | 2.847996907 | 0.1 | 0.022 | 1 | 0.13606055 | 0.214418414 |
| Tk1 | 0.142603094 | -2.572889668 | 0.008 | 0.035 | 1 | 0.113669596 | 2.847996907 | 0.1 | 0.022 | 1 | 0.142603094 | 0.214418414 |
| Gm20548 | 0.222316567 | 0.736965594 | 0.042 | 0.018 | 1 | 0.113669596 | 2.847996907 | 0.1 | 0.022 | 1 | 0.222316567 | 0.214418414 |
| Gm45464 | 0.345346582 | -1.070389328 | 0.017 | 0.035 | 1 | 0.113669596 | 2.847996907 | 0.1 | 0.022 | 1 | 0.345346582 | 0.214418414 |
| Gm43625 | 0.375395897 | 0.777607579 | 0.042 | 0.024 | 1 | 0.113669596 | 2.695993813 | 0.1 | 0.022 | 1 | 0.375395897 | 0.214418414 |
| Gm57301 | 0.630817006 | -0.900464326 | 0.042 | 0.053 | 1 | 0.113669596 | 2.847996907 | 0.1 | 0.022 | 1 | 0.630817006 | 0.214418414 |
| Llgl2 | 0.659871503 | 0.514573173 | 0.025 | 0.018 | 1 | 0.113669596 | 2.847996907 | 0.1 | 0.022 | 1 | 0.659871503 | 0.214418414 |
| Gm21411 | 0.666161437 | 0.192645078 | 0.025 | 0.018 | 1 | 0.113669596 | 2.847996907 | 0.1 | 0.022 | 1 | 0.666161437 | 0.214418414 |
| Tigar | 0.781790612 | -0.807354922 | 0.008 | 0.012 | 1 | 0.113669596 | 2.695993813 | 0.1 | 0.022 | 1 | 0.781790612 | 0.214418414 |
| Syngr3 | 0.781790612 | -0.485426827 | 0.008 | 0.012 | 1 | 0.113669596 | 2.695993813 | 0.1 | 0.022 | 1 | 0.781790612 | 0.214418414 |
| Irag2 | 0.942200939 | -1.070389328 | 0.025 | 0.024 | 1 | 0.113669596 | 2.847996907 | 0.1 | 0.022 | 1 | 0.942200939 | 0.214418414 |
| Krt23 | 0.255815287 | -0.807354922 | 0.067 | 0.106 | 1 | 0.113730316 | -1.654503434 | 0 | 0.204 | 1 | 0.255815287 | 0.214526047 |
| Btla | 0.113839312 | -2.109917692 | 0.017 | 0.053 | 1 | 0.730750993 | 2.210566986 | 0 | 0.012 | 1 | 0.730750993 | 0.214719234 |
| Vdr | 0.113897764 | -0.439623138 | 0.227 | 0.306 | 1 | 0.484025102 | -0.457811523 | 0.1 | 0.182 | 1 | 0.484025102 | 0.214822827 |
| Ctnnbip1 | 0.113905561 | 0.281688378 | 0.655 | 0.529 | 1 | 0.710456239 | 0.350218976 | 0.4 | 0.528 | 1 | 0.710456239 | 0.214836646 |
| Pcdhgb2 | 0.113989317 | 1.684498174 | 0.05 | 0.018 | 1 | 0.19134412 | 2.31748219 | 0.1 | 0.028 | 1 | 0.19134412 | 0.21498507 |
| Uevld | 0.114046817 | -1.070389328 | 0.084 | 0.141 | 1 | 0.21282212 | 1.393431043 | 0.3 | 0.167 | 1 | 0.21282212 | 0.215086957 |
| Bmf | 0.114079836 | -1.944858446 | 0.025 | 0.065 | 1 | 0.930503186 | 0.660369903 | 0.1 | 0.09 | 1 | 0.930503186 | 0.215145463 |
| Krt14 | 0.114100564 | 1.823901231 | 0.05 | 0.106 | 1 | 0.138472048 | -1.026472211 | 0 | 0.182 | 1 | 0.138472048 | 0.215182189 |
| Zfp808 | 0.984061185 | -0.446952679 | 0.109 | 0.106 | 1 | 0.11419596 | 1.473601392 | 0.2 | 0.065 | 1 | 0.984061185 | 0.215351202 |
| Tppp3 | 0.114207321 | 1.292180751 | 0.109 | 0.059 | 1 | 0.927830673 | 0.808468542 | 0.1 | 0.09 | 1 | 0.927830673 | 0.21537133 |
| Ttc16 | 0.447933452 | -0.222392421 | 0.034 | 0.053 | 1 | 0.114234576 | 1.959028219 | 0.2 | 0.068 | 1 | 0.447933452 | 0.215419614 |
| Gm19426 | 0.114268111 | -1.359895945 | 0.017 | 0.053 | 1 | 0.230466554 | 2.558490289 | 0.1 | 0.031 | 1 | 0.230466554 | 0.215479021 |
| Gm12829 | 0.114268111 | -1.359895945 | 0.017 | 0.053 | 1 | 0.698529281 | 2.210566986 | 0 | 0.015 | 1 | 0.698529281 | 0.215479021 |
| Zfp772 | 0.114268111 | -1.485426827 | 0.017 | 0.053 | 1 | 0.151443299 | 2.695993813 | 0.1 | 0.025 | 1 | 0.151443299 | 0.215479021 |
| Slc25a29 | 0.114268111 | -1.359895945 | 0.017 | 0.053 | 1 | 0.559081495 | 1.374065718 | 0.1 | 0.056 | 1 | 0.559081495 | 0.215479021 |
| Dmtn | 0.114268111 | -1.359895945 | 0.017 | 0.053 | 1 | 0.151443299 | 2.558490289 | 0.1 | 0.025 | 1 | 0.151443299 | 0.215479021 |
| Coq5 | 0.320696961 | -0.642968104 | 0.185 | 0.224 | 1 | 0.114287708 | -1.723545078 | 0 | 0.204 | 1 | 0.320696961 | 0.215513736 |
| Ndor1 | 0.114352871 | -0.918386234 | 0.067 | 0.124 | 1 | 0.648368948 | 0.695993813 | 0.2 | 0.142 | 1 | 0.648368948 | 0.215629163 |
| B4galt6 | 0.114364433 | -0.537894247 | 0.168 | 0.247 | 1 | 0.194649818 | -0.947862377 | 0.1 | 0.281 | 1 | 0.194649818 | 0.215649643 |
| Vps13a | 0.742440753 | -0.41091655 | 0.605 | 0.541 | 1 | 0.114385861 | -0.945552216 | 0.3 | 0.667 | 1 | 0.742440753 | 0.215687596 |
| Mtus1 | 0.896834081 | -0.300512792 | 0.874 | 0.924 | 1 | 0.114426095 | -0.904256319 | 0.8 | 0.963 | 1 | 0.896834081 | 0.215758858 |
| Slc25a13 | 0.604462299 | -0.428843299 | 0.218 | 0.235 | 1 | 0.114430345 | 0.538950103 | 0.6 | 0.324 | 1 | 0.604462299 | 0.215766385 |
| Psph | 0.11449896 | 0.451379346 | 0.378 | 0.3 | 1 | 0.432792299 | 1.587934067 | 0.3 | 0.225 | 1 | 0.432792299 | 0.215887908 |

| | | | | | | | | | | | | |
|---|---|---|---|---|---|---|---|---|---|---|---|---|
| G3bp2 | 0.114521497 | -0.361494499 | 0.672 | 0.676 | 1 | 0.979433558 | 0.405891164 | 0.6 | 0.713 | 1 | 0.979433558 | 0.215927821 |
| Gm35546 | 0.802027366 | 0.681223042 | 0.168 | 0.159 | 1 | 0.114664571 | -2.131825212 | 0 | 0.204 | 1 | 0.802027366 | 0.216181179 |
| Ppif | 0.114709411 | 1.369375257 | 0.227 | 0.165 | 1 | 0.315928386 | -0.374395515 | 0 | 0.093 | 1 | 0.315928386 | 0.216260574 |
| Dmd | 0.75352038 | -0.347923303 | 0.05 | 0.059 | 1 | 0.114753686 | -2.959358016 | 0 | 0.204 | 1 | 0.75352038 | 0.216338964 |
| Plekhh3 | 0.11476617 | -1.070389328 | 0.059 | 0.112 | 1 | 0.799489326 | 0.159940913 | 0.1 | 0.123 | 1 | 0.799489326 | 0.216361067 |
| Cd320 | 0.114953397 | 0.443019912 | 0.387 | 0.306 | 1 | 0.913797547 | 0.481869008 | 0.3 | 0.296 | 1 | 0.913797547 | 0.216692511 |
| Pde6d | 0.189747322 | -0.549557165 | 0.303 | 0.347 | 1 | 0.114975361 | -1.310004928 | 0.2 | 0.426 | 1 | 0.189747322 | 0.216731388 |
| Cep192 | 0.11504346 | -1.182864057 | 0.244 | 0.306 | 1 | 0.11694079 | 0.75918764 | 0.5 | 0.25 | 1 | 0.11694079 | 0.216851923 |
| Wfdc12 | 0.115105047 | 1.652076697 | 0.05 | 0.018 | 1 | 0.730749233 | 2.432959407 | 0 | 0.012 | 1 | 0.730749233 | 0.216960922 |
| Lck | 0.219476538 | 0.929610672 | 0.042 | 0.018 | 1 | 0.115118737 | 2.558490289 | 0.1 | 0.022 | 1 | 0.219476538 | 0.21698515 |
| 9330104G04Rik | 0.223965999 | -0.485426827 | 0.008 | 0.029 | 1 | 0.115118737 | 2.31748219 | 0.1 | 0.022 | 1 | 0.223965999 | 0.21698515 |
| Gm56772 | 0.237885589 | -2.485426827 | 0 | 0.012 | 1 | 0.115118737 | 2.432959407 | 0.1 | 0.022 | 1 | 0.237885589 | 0.21698515 |
| Serinc4 | 0.242320046 | -0.900464326 | 0.017 | 0.041 | 1 | 0.115118737 | 2.695993813 | 0.1 | 0.022 | 1 | 0.242320046 | 0.21698515 |
| Aatk | 0.331040183 | -1.292781749 | 0.008 | 0.024 | 1 | 0.115118737 | 2.432959407 | 0.1 | 0.022 | 1 | 0.331040183 | 0.21698515 |
| Gm15345 | 0.629493349 | -0.333423734 | 0.025 | 0.035 | 1 | 0.115118737 | 2.31748219 | 0.1 | 0.022 | 1 | 0.629493349 | 0.21698515 |
| Spata48 | 0.687345921 | -0.707819249 | 0.017 | 0.024 | 1 | 0.115118737 | 1.494359952 | 0.1 | 0.022 | 1 | 0.687345921 | 0.21698515 |
| Cgas | 0.782486655 | 0.514573173 | 0.05 | 0.059 | 1 | 0.115118737 | 2.558490289 | 0.1 | 0.022 | 1 | 0.782486655 | 0.21698515 |
| 4930533B01Rik | 0.817300436 | -0.485426827 | 0.042 | 0.047 | 1 | 0.115118737 | 2.695993813 | 0.1 | 0.022 | 1 | 0.817300436 | 0.21698515 |
| Kif15 | 0.11532534 | -2.424026283 | 0.017 | 0.053 | 1 | 0.156606269 | 1.930459067 | 0.1 | 0.025 | 1 | 0.156606269 | 0.217350746 |
| Gm56867 | 0.11532863 | -1.222392421 | 0.017 | 0.053 | 1 | 0.490000637 | 1.017921908 | 0 | 0.046 | 1 | 0.490000637 | 0.217356567 |
| Hsf4 | 0.11532863 | -1.222392421 | 0.017 | 0.053 | 1 | 0.841021797 | 0.808468542 | 0.1 | 0.08 | 1 | 0.841021797 | 0.217356567 |
| Gzmm | 0.11532863 | -1.222392421 | 0.017 | 0.053 | 1 | 0.324884604 | -0.191531458 | 0 | 0.09 | 1 | 0.324884604 | 0.217356567 |
| L3mbtl2 | 0.11532863 | -1.222392421 | 0.017 | 0.053 | 1 | 0.749384606 | 0.930459067 | 0.1 | 0.071 | 1 | 0.749384606 | 0.217356567 |
| Gm56596 | 0.11532863 | -1.222392421 | 0.017 | 0.053 | 1 | 0.669843604 | 2.017921908 | 0 | 0.019 | 1 | 0.669843604 | 0.217356567 |
| Zfp950 | 0.567736524 | -0.283792966 | 0.462 | 0.465 | 1 | 0.115391579 | 1.19107081 | 0.7 | 0.46 | 1 | 0.567736524 | 0.217467941 |
| Hmgcs1 | 0.115513693 | 0.153778718 | 0.773 | 0.794 | 1 | 0.420205804 | -0.837796036 | 0.8 | 0.883 | 1 | 0.420205804 | 0.217683973 |
| Ppa2 | 0.115545137 | -0.795766948 | 0.176 | 0.247 | 1 | 0.319104705 | -0.640289575 | 0.1 | 0.228 | 1 | 0.319104705 | 0.217739595 |
| Pex10 | 0.115608252 | -1.129283017 | 0.059 | 0.112 | 1 | 0.773317652 | 1.111031312 | 0.1 | 0.074 | 1 | 0.773317652 | 0.217851236 |
| Dnase2a | 0.11561918 | 0.370183263 | 0.134 | 0.076 | 1 | 0.213674789 | -0.789433014 | 0 | 0.136 | 1 | 0.213674789 | 0.217870565 |
| Snhg20 | 0.115622163 | -0.836899198 | 0.227 | 0.3 | 1 | 0.127739955 | -1.248864633 | 0 | 0.191 | 1 | 0.127739955 | 0.217875841 |
| Fem1c | 0.115625612 | -0.463059014 | 0.37 | 0.435 | 1 | 0.469512698 | 0.507864216 | 0.6 | 0.556 | 1 | 0.469512698 | 0.217881942 |
| Rnf185 | 0.478586517 | -0.313946359 | 0.336 | 0.365 | 1 | 0.115653368 | -1.452398027 | 0.2 | 0.42 | 1 | 0.478586517 | 0.217931034 |
| Scpep1 | 0.115710347 | -0.367200746 | 0.782 | 0.747 | 1 | 0.645535547 | -0.261627585 | 0.8 | 0.747 | 1 | 0.645535547 | 0.21803181 |
| Rpl35 | 0.135208272 | -0.18142064 | 0.916 | 0.971 | 1 | 0.115733418 | -0.469918126 | 0.7 | 0.932 | 1 | 0.135208272 | 0.218072611 |
| Lgals8 | 0.115759293 | -0.502105568 | 0.277 | 0.359 | 1 | 0.958372648 | 0.432959407 | 0.3 | 0.33 | 1 | 0.958372648 | 0.218118371 |
| Clspn | 0.115825159 | -1.029747343 | 0.017 | 0.053 | 1 | 0.318562315 | 1.930459067 | 0.1 | 0.037 | 1 | 0.318562315 | 0.218234851 |
| Tma7 | 0.115968065 | -0.201633861 | 0.899 | 0.912 | 1 | 0.202288919 | -0.286932674 | 0.4 | 0.861 | 1 | 0.202288919 | 0.218487537 |
| Dolpp1 | 0.991600982 | -0.185866545 | 0.059 | 0.059 | 1 | 0.115985361 | 1.744903414 | 0.2 | 0.068 | 1 | 0.991600982 | 0.218518117 |
| Pacsin3 | 0.554832416 | 0.110182918 | 0.227 | 0.194 | 1 | 0.115994921 | -1.258202497 | 0.1 | 0.33 | 1 | 0.554832416 | 0.218535021 |
| Gm42047 | 0.116047781 | -0.700932863 | 0.176 | 0.247 | 1 | 0.214175333 | -1.840059087 | 0 | 0.136 | 1 | 0.214175333 | 0.218628475 |
| Ptpn18 | 0.116303552 | -1.292781749 | 0.017 | 0.053 | 1 | 0.505623793 | 0.695993813 | 0 | 0.043 | 1 | 0.505623793 | 0.219080588 |
| Gm56530 | 0.116303552 | -1.392317423 | 0.017 | 0.053 | 1 | 0.698532411 | 1.847996907 | 0 | 0.015 | 1 | 0.698532411 | 0.219080588 |
| Plau | 0.11631305 | -1.333423734 | 0.017 | 0.053 | 1 | 0.577155236 | 1.210566986 | 0 | 0.031 | 1 | 0.577155236 | 0.219097374 |
| Capzb | 0.116350907 | -0.202332198 | 0.933 | 0.959 | 1 | 0.5400876 | -0.228057334 | 0.8 | 0.969 | 1 | 0.5400876 | 0.219164281 |
| Gart | 0.116448042 | -0.688960221 | 0.235 | 0.306 | 1 | 0.466266204 | -0.606568957 | 0.2 | 0.287 | 1 | 0.466266204 | 0.219335937 |
| Lmnb1 | 0.116547739 | -1.544320516 | 0.109 | 0.171 | 1 | 0.482002214 | 0.403212064 | 0.3 | 0.194 | 1 | 0.482002214 | 0.219512103 |
| Nemp1 | 0.116567757 | -0.778208576 | 0.143 | 0.212 | 1 | 0.567494884 | -0.28585884 | 0.1 | 0.164 | 1 | 0.567494884 | 0.219547472 |
| Gm28729 | 0.251035261 | -0.378511623 | 0.017 | 0.041 | 1 | 0.116579362 | 2.432959407 | 0.1 | 0.022 | 1 | 0.251035261 | 0.219567976 |
| Rnf43 | 0.147043407 | -1.222392421 | 0.008 | 0.035 | 1 | 0.116579781 | 2.31748219 | 0.1 | 0.022 | 1 | 0.147043407 | 0.219568717 |
| Epg5 | 0.11665666 | -0.713232745 | 0.252 | 0.324 | 1 | 0.825702775 | -0.167944637 | 0.4 | 0.318 | 1 | 0.825702775 | 0.219704544 |
| Scgb1c1 | 0.246626843 | -0.807354922 | 0.025 | 0.053 | 1 | 0.116727707 | -1.473931188 | 0 | 0.201 | 1 | 0.246626843 | 0.219830057 |
| Tor3a | 0.786213489 | 0.188072348 | 0.303 | 0.347 | 1 | 0.116753065 | -1.41670632 | 0.1 | 0.33 | 1 | 0.786213489 | 0.219874851 |
| Plxna1 | 0.116755587 | -0.753915663 | 0.16 | 0.229 | 1 | 0.500485803 | -0.351311902 | 0.2 | 0.284 | 1 | 0.500485803 | 0.219879306 |
| Dhdh | 0.116795967 | -1.292781749 | 0.017 | 0.053 | 1 | 0.489923006 | 0.494359952 | 0 | 0.046 | 1 | 0.489923006 | 0.219950636 |
| Zfp763 | 0.116807553 | -1.185866545 | 0.017 | 0.053 | 1 | 0.687535637 | 0.973527789 | 0.1 | 0.065 | 1 | 0.687535637 | 0.219971102 |
| Dnajc9 | 0.116882587 | -0.870857864 | 0.378 | 0.435 | 1 | 0.833123345 | -0.104474723 | 0.3 | 0.349 | 1 | 0.833123345 | 0.220103636 |
| Mef2b | 0.464490522 | 0.334000927 | 0.21 | 0.176 | 1 | 0.116905188 | -1.321928095 | 0 | 0.201 | 1 | 0.464490522 | 0.220143553 |
| Metap1 | 0.620539198 | -0.215819767 | 0.269 | 0.288 | 1 | 0.116945069 | -0.572041274 | 0.2 | 0.488 | 1 | 0.620539198 | 0.220213988 |
| Med16 | 0.125459038 | -0.629816737 | 0.126 | 0.194 | 1 | 0.116980597 | -1.321928095 | 0 | 0.201 | 1 | 0.125459038 | 0.220276734 |
| Coq8b | 0.301408447 | -0.497399469 | 0.328 | 0.376 | 1 | 0.117109027 | 1.695993813 | 0.4 | 0.25 | 1 | 0.301408447 | 0.22050353 |
| Gbp3 | 0.117118121 | 1.344648171 | 0.101 | 0.053 | 1 | 0.522102507 | -0.230005605 | 0 | 0.04 | 1 | 0.522102507 | 0.220519589 |
| Sh2b3 | 0.117141176 | -0.774933444 | 0.118 | 0.182 | 1 | 0.390201407 | 0.94753258 | 0.3 | 0.194 | 1 | 0.390201407 | 0.220560297 |
| Mpg | 0.598584123 | -0.31259023 | 0.235 | 0.253 | 1 | 0.117149863 | -1.505640048 | 0 | 0.201 | 1 | 0.598584123 | 0.220575636 |
| 2810013P06Rik | 0.995442303 | -0.269698136 | 0.252 | 0.241 | 1 | 0.117224871 | -1.457811523 | 0 | 0.201 | 1 | 0.995442303 | 0.220708072 |
| Cox11 | 0.117266313 | -0.613750924 | 0.286 | 0.347 | 1 | 0.964785029 | 0.521496082 | 0.3 | 0.33 | 1 | 0.964785029 | 0.220781237 |
| Ei24 | 0.117284078 | -0.605166071 | 0.336 | 0.412 | 1 | 0.660234302 | -0.415037499 | 0.4 | 0.41 | 1 | 0.660234302 | 0.2208126 |
| Spc24 | 0.117336861 | -1.944858446 | 0.034 | 0.076 | 1 | 0.643835681 | 1.695993813 | 0 | 0.022 | 1 | 0.643835681 | 0.220905784 |
| Ppp1r11 | 0.117352664 | -0.387023123 | 0.471 | 0.559 | 1 | 0.836328896 | 0.216303355 | 0.5 | 0.494 | 1 | 0.836328896 | 0.22093368 |
| Slc45a4 | 0.117375736 | 0.464820138 | 0.303 | 0.218 | 1 | 0.734049654 | 0.362570079 | 0.4 | 0.414 | 1 | 0.734049654 | 0.220974408 |
| Alg11 | 0.117438131 | -0.563429339 | 0.118 | 0.188 | 1 | 0.14369286 | -1.230005605 | 0 | 0.179 | 1 | 0.14369286 | 0.221084548 |
| Reps2 | 0.595169987 | -0.632268215 | 0.05 | 0.065 | 1 | 0.117731314 | 1.245332404 | 0.2 | 0.065 | 1 | 0.595169987 | 0.221601965 |
| Gm26944 | 0.533062242 | -0.36994961 | 0.084 | 0.106 | 1 | 0.117753401 | 1.795529487 | 0.2 | 0.068 | 1 | 0.533062242 | 0.221640939 |
| Zdhhc7 | 0.117781876 | -0.854660637 | 0.092 | 0.153 | 1 | 0.169316115 | 1.095089769 | 0.4 | 0.216 | 1 | 0.169316115 | 0.221691181 |
| Arrb2 | 0.117805731 | -1.070389328 | 0.017 | 0.053 | 1 | 0.460577419 | 0.558490289 | 0 | 0.052 | 1 | 0.460577419 | 0.221733271 |
| Naa60 | 0.117865495 | -0.515800476 | 0.303 | 0.376 | 1 | 0.617232049 | 0.778455973 | 0.4 | 0.349 | 1 | 0.617232049 | 0.221838715 |
| Clpb | 0.117877706 | -0.850423644 | 0.218 | 0.288 | 1 | 0.961070131 | -0.289506617 | 0.4 | 0.361 | 1 | 0.961070131 | 0.221860258 |
| G930009F23Rik | 0.1179084 | -0.199122642 | 0.143 | 0.082 | 1 | 0.676915308 | 0.930459067 | 0.1 | 0.068 | 1 | 0.676915308 | 0.22191441 |
| Thap4 | 0.117931424 | -0.807354922 | 0.118 | 0.182 | 1 | 0.304920053 | -0.736965594 | 0.1 | 0.235 | 1 | 0.304920053 | 0.221955027 |
| Bnip1 | 0.118111265 | -0.833350131 | 0.168 | 0.235 | 1 | 0.440445776 | -0.536666944 | 0.1 | 0.191 | 1 | 0.440445776 | 0.22227226 |
| Casp6 | 0.118132246 | -0.669851398 | 0.067 | 0.124 | 1 | 0.202053621 | -0.91281543 | 0 | 0.142 | 1 | 0.202053621 | 0.222309264 |
| Med12l | 0.379191663 | 0.570068285 | 0.16 | 0.124 | 1 | 0.118277953 | -2.618702713 | 0 | 0.201 | 1 | 0.379191663 | 0.222566232 |
| Hs1bp3 | 0.118298188 | -1.185866545 | 0.017 | 0.053 | 1 | 0.489939838 | 0.558490289 | 0 | 0.046 | 1 | 0.489939838 | 0.222601915 |
| Mrps14 | 0.11858451 | 0.219829907 | 0.714 | 0.618 | 1 | 0.15723454 | -0.58694015 | 0.5 | 0.71 | 1 | 0.15723454 | 0.223106734 |
| Chd1l | 0.245953972 | 0.175086707 | 0.185 | 0.241 | 1 | 0.118625738 | 1.367667947 | 0.4 | 0.207 | 1 | 0.245953972 | 0.22317941 |
| Coa8 | 0.118677005 | -0.335679708 | 0.387 | 0.476 | 1 | 0.306312776 | 0.570462931 | 0.6 | 0.451 | 1 | 0.306312776 | 0.223269778 |
| H6pd | 0.118776377 | 0.695145418 | 0.218 | 0.147 | 1 | 0.994182602 | 0.263034406 | 0.2 | 0.191 | 1 | 0.994182602 | 0.223444926 |
| Pp2d1 | 0.118807315 | -0.944858446 | 0.017 | 0.053 | 1 | 0.559081495 | 1.558490289 | 0.1 | 0.056 | 1 | 0.559081495 | 0.223499452 |
| Ndufa13 | 0.118842899 | -0.195425374 | 0.891 | 0.918 | 1 | 0.171384536 | -0.426862935 | 0.8 | 0.889 | 1 | 0.171384536 | 0.223562163 |
| Prpsap1 | 0.527939304 | -0.258357919 | 0.588 | 0.576 | 1 | 0.118868752 | -1 | 0.4 | 0.571 | 1 | 0.527939304 | 0.223607724 |
| Zfp524 | 0.118941427 | -1.292781749 | 0.042 | 0.088 | 1 | 0.433577142 | 0.432959407 | 0 | 0.059 | 1 | 0.433577142 | 0.223735791 |
| Neil2 | 0.118967248 | 1 | 0.05 | 0.018 | 1 | 0.6199755 | 1.847996907 | 0 | 0.025 | 1 | 0.6199755 | 0.223781291 |
| Zfyve26 | 0.319900785 | -0.378511623 | 0.176 | 0.224 | 1 | 0.119055739 | 1.070389328 | 0.4 | 0.188 | 1 | 0.319900785 | 0.22393721 |
| 2500004C02Rik | 0.119072815 | -1.015941544 | 0.059 | 0.112 | 1 | 0.670802544 | 0.695993813 | 0.2 | 0.145 | 1 | 0.670802544 | 0.223967296 |
| Gm44699 | 0.191767884 | -0.662304589 | 0.151 | 0.206 | 1 | 0.119165539 | 1.147557188 | 0.5 | 0.272 | 1 | 0.191767884 | 0.224130653 |
| Maged1 | 0.119262105 | -0.185692317 | 0.958 | 0.935 | 1 | 0.857023489 | 0.128897733 | 0.8 | 0.938 | 1 | 0.857023489 | 0.22430076 |
| Arih2 | 0.191143172 | -0.410976878 | 0.739 | 0.741 | 1 | 0.119300532 | 0.321756845 | 1 | 0.802 | 1 | 0.191143172 | 0.224368447 |
| Sfxn2 | 0.119312254 | -0.528495549 | 0.202 | 0.282 | 1 | 0.340315687 | 1.075407403 | 0.4 | 0.262 | 1 | 0.340315687 | 0.224389094 |
| Ppat | 0.119363129 | -0.567220918 | 0.328 | 0.4 | 1 | 0.882020792 | -0.321928095 | 0.3 | 0.287 | 1 | 0.882020792 | 0.224478702 |

| | | | | | | | | | | | | |
|---|---|---|---|---|---|---|---|---|---|---|---|---|
| Slc5a6 | 0.119425598 | -1.733354341 | 0.025 | 0.065 | 1 | 0.59781419 | 1.558490289 | 0 | 0.028 | 1 | 0.59781419 | 0.224588722 |
| Erich6 | 0.119426926 | -1.485426827 | 0.025 | 0.065 | 1 | 0.673291667 | 1.263034406 | 0.1 | 0.065 | 1 | 0.673291667 | 0.224591061 |
| Egr4 | 0.147029286 | -3.070389328 | 0 | 0.018 | 1 | 0.11953774 | 1.769994395 | 0.1 | 0.022 | 1 | 0.147029286 | 0.224786208 |
| Mmp3 | 0.342923893 | -0.555816155 | 0.126 | 0.088 | 1 | 0.119539013 | 0.31748219 | 0.1 | 0.022 | 1 | 0.342923893 | 0.22478845 |
| Cep41 | 0.597720959 | -0.563429339 | 0.126 | 0.147 | 1 | 0.119564161 | 1.236562194 | 0.3 | 0.123 | 1 | 0.597720959 | 0.224832734 |
| Xpa | 0.119604851 | -0.627445832 | 0.16 | 0.235 | 1 | 0.330264283 | 0.270687978 | 0.5 | 0.312 | 1 | 0.330264283 | 0.224904382 |
| Stap2 | 0.119830721 | 1.514573173 | 0.059 | 0.024 | 1 | 0.698534498 | 2.432959407 | 0 | 0.015 | 1 | 0.698534498 | 0.22530204 |
| Pla2g6 | 0.119866822 | -1.070389328 | 0.05 | 0.1 | 1 | 0.680069523 | 1.648688098 | 0.1 | 0.071 | 1 | 0.680069523 | 0.225365589 |
| Zscan22 | 0.11991697 | -0.980191519 | 0.084 | 0.141 | 1 | 0.266983933 | -0.536666944 | 0 | 0.111 | 1 | 0.266983933 | 0.225453861 |
| Cep112 | 0.634744994 | -0.765534746 | 0.571 | 0.512 | 1 | 0.119949586 | 0.215054377 | 0.8 | 0.481 | 1 | 0.634744994 | 0.225511268 |
| Ints11 | 0.194800781 | -0.467724825 | 0.286 | 0.347 | 1 | 0.119971323 | 1.140177658 | 0.5 | 0.287 | 1 | 0.194800781 | 0.225549528 |
| Grpel1 | 0.120044596 | 0.342512427 | 0.664 | 0.594 | 1 | 0.778063848 | -0.195857383 | 0.7 | 0.623 | 1 | 0.778063848 | 0.225678486 |
| Pot1a | 0.12006251 | -0.936088236 | 0.109 | 0.171 | 1 | 0.219623355 | -0.840059087 | 0 | 0.133 | 1 | 0.219623355 | 0.225710014 |
| Gm48956 | 0.120183375 | 1.868210127 | 0.084 | 0.041 | 1 | 0.643846086 | 1.210566986 | 0 | 0.022 | 1 | 0.643846086 | 0.225922707 |
| Slx1b | 0.120183375 | -1.687060688 | 0.034 | 0.076 | 1 | 0.518235124 | 1.769994395 | 0.1 | 0.052 | 1 | 0.518235124 | 0.225922707 |
| Trp53rkb | 0.120246704 | -0.87774425 | 0.059 | 0.112 | 1 | 0.37374098 | 1.280956314 | 0.2 | 0.108 | 1 | 0.37374098 | 0.226034139 |
| Lrrc8b | 0.530354347 | -0.522901533 | 0.227 | 0.259 | 1 | 0.120270079 | -1.187627003 | 0.1 | 0.367 | 1 | 0.530354347 | 0.226075267 |
| Ifi44 | 0.120377965 | 1.955145764 | 0.067 | 0.029 | 1 | 0.669843604 | -0.111361109 | 0 | 0.019 | 1 | 0.669843604 | 0.226265075 |
| Apool | 0.120648871 | 0.476099025 | 0.227 | 0.153 | 1 | 0.314580102 | 0.295455884 | 0.4 | 0.228 | 1 | 0.314580102 | 0.226741592 |
| Rnls | 0.120682496 | 0.399095955 | 0.134 | 0.076 | 1 | 0.165735208 | 1.08718457 | 0.2 | 0.077 | 1 | 0.165735208 | 0.226800728 |
| Ehhadh | 0.120783297 | -1.485426827 | 0.025 | 0.065 | 1 | 0.643835681 | 1.695993813 | 0 | 0.022 | 1 | 0.643835681 | 0.226977989 |
| Gm21781 | 0.120783631 | -1.392317423 | 0.025 | 0.065 | 1 | 0.420904619 | 0.210566986 | 0 | 0.062 | 1 | 0.420904619 | 0.226978576 |
| Serac1 | 0.120783631 | -1.572889668 | 0.025 | 0.065 | 1 | 0.35382401 | -0.408342847 | 0 | 0.08 | 1 | 0.35382401 | 0.226978576 |
| Gm26672 | 0.120783631 | -1.392317423 | 0.025 | 0.065 | 1 | 0.59781419 | 1.558490289 | 0 | 0.028 | 1 | 0.59781419 | 0.226978576 |
| Dram2 | 0.261208794 | -0.504285854 | 0.218 | 0.271 | 1 | 0.120813529 | -1.145308441 | 0.2 | 0.426 | 1 | 0.261208794 | 0.227031149 |
| Eif3m | 0.120821101 | -0.298056691 | 0.866 | 0.829 | 1 | 0.719578916 | 0.106855636 | 0.6 | 0.883 | 1 | 0.719578916 | 0.227044464 |
| Pigm | 0.120821329 | -0.748461233 | 0.118 | 0.182 | 1 | 0.415749645 | -0.441509711 | 0.1 | 0.201 | 1 | 0.415749645 | 0.227044864 |
| Cln8 | 0.120831255 | -0.426533138 | 0.151 | 0.224 | 1 | 0.824941613 | 0.328622747 | 0.3 | 0.265 | 1 | 0.824941613 | 0.227062317 |
| Nap1l3 | 0.120848405 | -1.222392421 | 0.042 | 0.088 | 1 | 0.743727433 | 1.159940913 | 0.1 | 0.071 | 1 | 0.743727433 | 0.227092474 |
| Gm49692 | 0.120918975 | 0.514573173 | 0.134 | 0.076 | 1 | 0.743726438 | 1.111031312 | 0.1 | 0.071 | 1 | 0.743726438 | 0.227216551 |
| Efcab15 | 0.120942738 | 0.436570661 | 0.126 | 0.071 | 1 | 0.362872659 | 1.930459067 | 0.1 | 0.04 | 1 | 0.362872659 | 0.22725833 |
| Acsf2 | 0.121095302 | 0.208764743 | 0.412 | 0.318 | 1 | 0.290054926 | 1.075407403 | 0.4 | 0.272 | 1 | 0.290054926 | 0.227526532 |
| Nfya | 0.441494463 | -0.411426246 | 0.303 | 0.324 | 1 | 0.121258877 | -1.433289204 | 0.1 | 0.324 | 1 | 0.441494463 | 0.227814039 |
| Slc46a1 | 0.826158434 | -0.163498732 | 0.076 | 0.082 | 1 | 0.12134152 | 1.744903414 | 0.2 | 0.068 | 1 | 0.826158434 | 0.227959276 |
| Gm15411 | 0.890489825 | -0.485426827 | 0.05 | 0.053 | 1 | 0.12134152 | 1.648688098 | 0.2 | 0.068 | 1 | 0.890489825 | 0.227959276 |
| Ndufb7 | 0.121372652 | 0.184424571 | 0.731 | 0.676 | 1 | 0.935678759 | 0.386484977 | 0.6 | 0.645 | 1 | 0.935678759 | 0.228013983 |
| Pop5 | 0.121522419 | 0.263034406 | 0.378 | 0.288 | 1 | 0.418690953 | 0.450237399 | 0.5 | 0.358 | 1 | 0.418690953 | 0.22827714 |
| Nans | 0.1216007 | -0.32788555 | 0.345 | 0.424 | 1 | 0.534659668 | -0.296774618 | 0.3 | 0.395 | 1 | 0.534659668 | 0.22841467 |
| Nexn | 0.468055356 | -0.839063782 | 0.05 | 0.071 | 1 | 0.121668802 | 1.159940913 | 0.3 | 0.123 | 1 | 0.468055356 | 0.228534306 |
| Gm43707 | 0.121786177 | 1.514573173 | 0.067 | 0.029 | 1 | 0.529403237 | 1.31748219 | 0.1 | 0.052 | 1 | 0.529403237 | 0.228740482 |
| Nek11 | 0.121845366 | -1.333423734 | 0.034 | 0.076 | 1 | 0.537822819 | 1.017921908 | 0.1 | 0.052 | 1 | 0.537822819 | 0.22884444 |
| Gas7 | 0.121846047 | -0.36994961 | 0.252 | 0.165 | 1 | 0.929209469 | 0.858050571 | 0.2 | 0.21 | 1 | 0.929209469 | 0.228845636 |
| Gm29358 | 0.121879242 | 1.362570079 | 0.059 | 0.024 | 1 | 0.236765774 | 2.017921908 | 0.1 | 0.031 | 1 | 0.236765774 | 0.228903934 |
| Gm13427 | 0.121879242 | 1.514573173 | 0.059 | 0.024 | 1 | 0.278846016 | 2.111031312 | 0.1 | 0.034 | 1 | 0.278846016 | 0.228903934 |
| Minpp1 | 0.837972321 | -0.226893814 | 0.42 | 0.406 | 1 | 0.122066493 | 0.760534065 | 0.7 | 0.503 | 1 | 0.837972321 | 0.229232758 |
| Rdh11 | 0.122134943 | 0.371833001 | 0.496 | 0.406 | 1 | 0.572437694 | 0.566710796 | 0.4 | 0.327 | 1 | 0.572437694 | 0.229352943 |
| Gnasas1 | 0.122148811 | -1.485426827 | 0.025 | 0.065 | 1 | 0.604715484 | 0.973527789 | 0.1 | 0.059 | 1 | 0.604715484 | 0.22937729 |
| Gm43331 | 0.122148811 | -1.292781749 | 0.025 | 0.065 | 1 | 0.239079302 | -0.888968688 | 0 | 0.123 | 1 | 0.239079302 | 0.22937729 |
| Exoc8 | 0.122148811 | -1.392317423 | 0.025 | 0.065 | 1 | 0.425255145 | 1.111031312 | 0.2 | 0.114 | 1 | 0.425255145 | 0.22937729 |
| Eme1 | 0.122148811 | -1.292781749 | 0.025 | 0.065 | 1 | 0.362872659 | 1.625604485 | 0.1 | 0.04 | 1 | 0.362872659 | 0.22937729 |
| Gm57404 | 0.122148811 | -1.485426827 | 0.025 | 0.065 | 1 | 0.357796814 | 2.111031312 | 0.1 | 0.04 | 1 | 0.357796814 | 0.22937729 |
| Gm2885 | 0.122148811 | -1.292781749 | 0.025 | 0.065 | 1 | 0.539455361 | 0.769994395 | 0 | 0.037 | 1 | 0.539455361 | 0.22937729 |
| B130034C11Rik | 0.122167767 | -1.070389328 | 0.05 | 0.1 | 1 | 0.489972412 | 0.263034406 | 0 | 0.046 | 1 | 0.489972412 | 0.229410571 |
| Eva1c | 0.122377026 | 0.225066556 | 0.168 | 0.106 | 1 | 0.397039386 | -0.230005605 | 0 | 0.068 | 1 | 0.397039386 | 0.229777916 |
| Dennd10 | 0.122479862 | 0.15549208 | 0.538 | 0.406 | 1 | 0.975511211 | 0.290001453 | 0.5 | 0.497 | 1 | 0.975511211 | 0.229958407 |
| Tmem208 | 0.122564255 | -0.302800505 | 0.681 | 0.724 | 1 | 0.612607572 | -0.183711953 | 0.6 | 0.639 | 1 | 0.612607572 | 0.230106513 |
| Usp50 | 0.12259759 | -0.354842717 | 0.328 | 0.229 | 1 | 0.586731859 | 0.675529711 | 0.3 | 0.25 | 1 | 0.586731859 | 0.230165011 |
| Stat5a | 0.122651993 | -0.568842835 | 0.227 | 0.288 | 1 | 0.204134687 | 1.116700062 | 0.5 | 0.346 | 1 | 0.204134687 | 0.230260474 |
| Timm21 | 0.122718501 | -0.552541023 | 0.277 | 0.359 | 1 | 0.981377168 | 0.159940913 | 0.3 | 0.278 | 1 | 0.981377168 | 0.230377171 |
| Prg4 | 0.519601782 | 1.173298671 | 0.345 | 0.329 | 1 | 0.12274394 | -4.976431529 | 0.1 | 0.321 | 1 | 0.519601782 | 0.230421806 |
| 1700113A16Rik | 0.122775433 | -1.222392421 | 0.042 | 0.088 | 1 | 0.324842568 | -0.304006187 | 0 | 0.09 | 1 | 0.324842568 | 0.230477059 |
| Zfp518b | 0.122775433 | -1.222392421 | 0.042 | 0.088 | 1 | 0.27437716 | -0.374395515 | 0 | 0.108 | 1 | 0.27437716 | 0.230477059 |
| Gm16536 | 0.122775674 | -1.14839184 | 0.042 | 0.088 | 1 | 0.902406098 | 0.695993813 | 0.1 | 0.086 | 1 | 0.902406098 | 0.230477481 |
| Ddr1 | 0.122822442 | -0.807354922 | 0.151 | 0.218 | 1 | 0.802067033 | 0.486540447 | 0.4 | 0.392 | 1 | 0.802067033 | 0.230559532 |
| Sema6d | 0.863443455 | -0.364120531 | 0.109 | 0.112 | 1 | 0.122868566 | 0.913585248 | 0.3 | 0.127 | 1 | 0.863443455 | 0.230640448 |
| Prelid3a | 0.122934004 | -1.155278225 | 0.084 | 0.141 | 1 | 0.37172876 | 1.236562194 | 0.2 | 0.111 | 1 | 0.37172876 | 0.230755238 |
| Garin4 | 0.18903364 | -0.807354922 | 0.076 | 0.124 | 1 | 0.123147286 | 1.795529487 | 0.2 | 0.071 | 1 | 0.18903364 | 0.231129319 |
| Rnf146 | 0.131867704 | -0.485426827 | 0.504 | 0.541 | 1 | 0.123203884 | -0.729312022 | 0.4 | 0.63 | 1 | 0.131867704 | 0.231228571 |
| Asb3 | 0.198107202 | -0.436517227 | 0.361 | 0.424 | 1 | 0.123232529 | 1.210566986 | 0.6 | 0.373 | 1 | 0.198107202 | 0.231278802 |
| Gsta3 | 0.823222381 | 0.736965594 | 0.034 | 0.029 | 1 | 0.123362276 | -1.111361109 | 0 | 0.194 | 1 | 0.823222381 | 0.231506302 |
| Gm26879 | 0.123520686 | -1.250961574 | 0.034 | 0.076 | 1 | 0.952551463 | 0.732519689 | 0.1 | 0.093 | 1 | 0.952551463 | 0.231784013 |
| Tmem40 | 0.123522138 | -1.185866545 | 0.025 | 0.065 | 1 | 0.619961444 | 1.432959407 | 0 | 0.025 | 1 | 0.619961444 | 0.231786557 |
| Pus7l | 0.123522138 | -1.185866545 | 0.025 | 0.065 | 1 | 0.433676456 | 0.263034406 | 0 | 0.059 | 1 | 0.433676456 | 0.231786557 |
| Ahcyl | 0.123522138 | -1.185866545 | 0.025 | 0.065 | 1 | 0.557729726 | 1.31748219 | 0 | 0.034 | 1 | 0.557729726 | 0.231786557 |
| Zfp952 | 0.123721889 | -1.029747343 | 0.084 | 0.141 | 1 | 0.749374726 | 0.494359952 | 0.2 | 0.154 | 1 | 0.749374726 | 0.232136672 |
| Iars2 | 0.123877512 | -0.597901556 | 0.227 | 0.306 | 1 | 0.795842597 | 0.131789873 | 0.5 | 0.423 | 1 | 0.795842597 | 0.232409386 |
| Rbm43 | 0.124006159 | -0.718087584 | 0.143 | 0.212 | 1 | 0.827572678 | 0.494359952 | 0.2 | 0.167 | 1 | 0.827572678 | 0.23263479 |
| Gm34655 | 0.12404457 | 1.192645078 | 0.059 | 0.024 | 1 | 0.669843604 | 1.695993813 | 0 | 0.019 | 1 | 0.669843604 | 0.232702084 |
| C330013E15Rik | 0.12404457 | 1.192645078 | 0.059 | 0.024 | 1 | 0.730749233 | 2.432959407 | 0 | 0.012 | 1 | 0.730749233 | 0.232702084 |
| Stbd1 | 0.519851852 | 0.707218251 | 0.05 | 0.035 | 1 | 0.124058736 | 2.385653692 | 0.2 | 0.077 | 1 | 0.519851852 | 0.232726902 |
| A130014A01Rik | 0.124070461 | -1.250961574 | 0.025 | 0.065 | 1 | 0.493083302 | 1.210566986 | 0.1 | 0.049 | 1 | 0.493083302 | 0.232747443 |
| D6Wsu163e | 0.12419098 | -0.925999419 | 0.05 | 0.1 | 1 | 0.293201969 | 1.354956895 | 0.2 | 0.096 | 1 | 0.293201969 | 0.23295856 |
| Pip5kl1 | 0.124193642 | 1.407657969 | 0.067 | 0.029 | 1 | 0.577155236 | 1.31748219 | 0 | 0.031 | 1 | 0.577155236 | 0.232963223 |
| Epb41l5 | 0.124243738 | -1.485426827 | 0.084 | 0.141 | 1 | 0.311522335 | 1.403212064 | 0.3 | 0.194 | 1 | 0.311522335 | 0.233050969 |
| Upf3a | 0.252426978 | -0.393397689 | 0.496 | 0.512 | 1 | 0.124295646 | -0.926936538 | 0.3 | 0.54 | 1 | 0.252426978 | 0.233141884 |
| Pak4 | 0.124420997 | 0.514573173 | 0.185 | 0.118 | 1 | 0.839816919 | 1.31748219 | 0.1 | 0.136 | 1 | 0.839816919 | 0.233361409 |
| Kdsr | 0.124442648 | -0.678071905 | 0.353 | 0.406 | 1 | 0.456464517 | -0.391469028 | 0.4 | 0.5 | 1 | 0.456464517 | 0.233399323 |
| Zfp868 | 0.12452488 | -0.687060688 | 0.143 | 0.212 | 1 | 0.350722492 | 0.769994395 | 0.3 | 0.173 | 1 | 0.350722492 | 0.233543314 |
| Mpv17l | 0.830610802 | -0.718916957 | 0.244 | 0.241 | 1 | 0.124532881 | -1.536666944 | 0 | 0.194 | 1 | 0.830610802 | 0.233557324 |
| Cpt2 | 0.124723089 | -1.070389328 | 0.042 | 0.088 | 1 | 0.770508511 | 1.159940913 | 0.1 | 0.074 | 1 | 0.770508511 | 0.23389033 |
| Ddx1 | 0.124870672 | -0.338801913 | 0.605 | 0.629 | 1 | 0.714050958 | 0.664775082 | 0.5 | 0.586 | 1 | 0.714050958 | 0.23414866 |
| Rnf41 | 0.634636862 | -0.196921734 | 0.454 | 0.453 | 1 | 0.124958485 | -0.74879103 | 0.5 | 0.66 | 1 | 0.634636862 | 0.234302348 |
| Tnfsf9 | 0.125001244 | -1.411426246 | 0.025 | 0.065 | 1 | 0.421020792 | -0.473931188 | 0 | 0.062 | 1 | 0.421020792 | 0.234377176 |
| Mterf2 | 0.240356677 | -1.222392421 | 0.017 | 0.041 | 1 | 0.125004469 | 1.354956895 | 0.2 | 0.068 | 1 | 0.240356677 | 0.234382822 |
| Tsen54 | 0.125057509 | -1.070389328 | 0.025 | 0.065 | 1 | 0.396844932 | 0.374065718 | 0 | 0.068 | 1 | 0.396844932 | 0.234475638 |
| Mrpl23 | 0.125118867 | 0.210566986 | 0.706 | 0.588 | 1 | 0.566657958 | -0.131825212 | 0.5 | 0.571 | 1 | 0.566657958 | 0.234583004 |
| Cxcl1 | 0.125142351 | 0.267933205 | 0.185 | 0.118 | 1 | 0.643842184 | 1.31748219 | 0 | 0.022 | 1 | 0.643842184 | 0.234624094 |
| Wdr81 | 0.12521021 | -1.163498732 | 0.034 | 0.076 | 1 | 0.2691968 | 1.473601392 | 0.2 | 0.093 | 1 | 0.2691968 | 0.234742824 |

| | | | | | | | | | | | | |
|---|---|---|---|---|---|---|---|---|---|---|---|---|
| Tmem41a | 0.12521021 | -1.163498732 | 0.034 | 0.076 | 1 | 0.355033047 | 1.31748219 | 0.2 | 0.105 | 1 | 0.355033047 | 0.234742824 |
| Ptbp3 | 0.837504696 | -0.152630216 | 0.941 | 0.912 | 1 | 0.125282093 | -0.654503434 | 0.9 | 0.954 | 1 | 0.837504696 | 0.234868584 |
| Gorasp1 | 0.125429459 | -0.733354341 | 0.118 | 0.182 | 1 | 0.394127038 | 1.048295557 | 0.2 | 0.108 | 1 | 0.394127038 | 0.235126369 |
| Obp1a | 0.125482321 | -0.59645814 | 0.168 | 0.241 | 1 | 0.892501456 | 0.227844977 | 0.2 | 0.222 | 1 | 0.892501456 | 0.23521883 |
| Gm33302 | 0.125632131 | 0.955145764 | 0.059 | 0.024 | 1 | 0.385755192 | -1.248864633 | 0 | 0.071 | 1 | 0.385755192 | 0.23548083 |
| 2310039H08Rik | 0.125634041 | -0.204526001 | 0.353 | 0.459 | 1 | 0.54298729 | 0.558490289 | 0.4 | 0.327 | 1 | 0.54298729 | 0.235484169 |
| Mre11a | 0.125680818 | -0.870090677 | 0.118 | 0.182 | 1 | 0.428988119 | -0.625934282 | 0.1 | 0.194 | 1 | 0.428988119 | 0.235565968 |
| Mrpl53 | 0.125743227 | 0.278944924 | 0.42 | 0.318 | 1 | 0.702189968 | 0.179978666 | 0.4 | 0.309 | 1 | 0.702189968 | 0.235675096 |
| Gm17484 | 0.125829809 | -1.485426827 | 0.101 | 0.159 | 1 | 0.785821942 | 0.769994395 | 0.2 | 0.182 | 1 | 0.785821942 | 0.235826478 |
| Dpagt1 | 0.125874667 | -0.707819249 | 0.185 | 0.253 | 1 | 0.492898381 | -0.230005605 | 0.1 | 0.182 | 1 | 0.492898381 | 0.235904902 |
| Gm45051 | 0.237677659 | 0.447458977 | 0.16 | 0.112 | 1 | 0.125893763 | 1.403212064 | 0.3 | 0.127 | 1 | 0.237677659 | 0.235938286 |
| Tiam2 | 0.125956359 | -0.288475622 | 0.983 | 0.953 | 1 | 0.410135779 | 0.776691872 | 1 | 0.944 | 1 | 0.410135779 | 0.236047714 |
| Odr4 | 0.12597191 | -0.475966498 | 0.706 | 0.7 | 1 | 0.623813459 | -0.371269414 | 0.7 | 0.627 | 1 | 0.623813459 | 0.236074898 |
| Zfp523 | 0.126080775 | -0.707819249 | 0.16 | 0.229 | 1 | 0.974641329 | 0.193493473 | 0.2 | 0.188 | 1 | 0.974641329 | 0.236265188 |
| Vbp1 | 0.60892712 | -0.239670413 | 0.538 | 0.529 | 1 | 0.126088245 | -0.616284112 | 0.3 | 0.565 | 1 | 0.60892712 | 0.236278245 |
| Mro | 0.126162682 | 0.72102405 | 0.109 | 0.059 | 1 | 0.669853054 | 2.210566986 | 0 | 0.019 | 1 | 0.669853054 | 0.236408342 |
| Preb | 0.126221858 | -0.485426827 | 0.37 | 0.441 | 1 | 0.21307125 | 0.710493383 | 0.6 | 0.395 | 1 | 0.21307125 | 0.236511759 |
| Ttll5 | 0.126422024 | -0.855650479 | 0.479 | 0.512 | 1 | 0.62906621 | 0.179418287 | 0.5 | 0.608 | 1 | 0.62906621 | 0.236861519 |
| Cenpj | 0.126507508 | -1.070389328 | 0.084 | 0.141 | 1 | 0.379926954 | 1.210566986 | 0.2 | 0.111 | 1 | 0.379926954 | 0.237010866 |
| Zfp715 | 0.126805006 | -0.532732542 | 0.092 | 0.153 | 1 | 0.660804483 | 0.210566986 | 0.1 | 0.148 | 1 | 0.660804483 | 0.237530503 |
| Nlrx1 | 0.126832059 | -1 | 0.042 | 0.088 | 1 | 0.564719271 | 1.31748219 | 0.1 | 0.056 | 1 | 0.564719271 | 0.237577747 |
| Mmachc | 0.126913697 | -1.070389328 | 0.034 | 0.076 | 1 | 0.133962084 | 1.744903414 | 0.2 | 0.071 | 1 | 0.133962084 | 0.237720308 |
| Zfp160 | 0.126937917 | -0.659924558 | 0.235 | 0.312 | 1 | 0.150441693 | 0.695993813 | 0.6 | 0.355 | 1 | 0.150441693 | 0.237762599 |
| Pdcd2 | 0.12700269 | -0.722466024 | 0.092 | 0.153 | 1 | 0.500525852 | 0.988174565 | 0.2 | 0.123 | 1 | 0.500525852 | 0.237875696 |
| Zfand4 | 0.127022833 | -1.14839184 | 0.034 | 0.076 | 1 | 0.489922734 | 0.625604485 | 0 | 0.046 | 1 | 0.489922734 | 0.237910866 |
| Ssu72 | 0.127098949 | 0.13606155 | 0.849 | 0.753 | 1 | 0.473522473 | -0.206079766 | 0.6 | 0.762 | 1 | 0.473522473 | 0.238043755 |
| Calr3 | 0.654829089 | -0.622930351 | 0.076 | 0.088 | 1 | 0.127196265 | 2.432959407 | 0.2 | 0.074 | 1 | 0.654829089 | 0.238213641 |
| Lrrc8a | 0.127252801 | -0.494887156 | 0.437 | 0.482 | 1 | 0.920526089 | -0.148355316 | 0.6 | 0.577 | 1 | 0.920526089 | 0.238312326 |
| Trappc2l | 0.127308125 | 0.286905809 | 0.613 | 0.524 | 1 | 0.368092314 | -0.544320516 | 0.3 | 0.41 | 1 | 0.368092314 | 0.238408891 |
| Gm12184 | 0.127403495 | -0.707819249 | 0.092 | 0.153 | 1 | 1 | 0.695993813 | 0.1 | 0.099 | 1 | 1 | 0.238575339 |
| Bysl | 0.414274607 | -0.600904045 | 0.134 | 0.165 | 1 | 0.127572476 | -1.230005605 | 0 | 0.191 | 1 | 0.414274607 | 0.238870215 |
| Ddx52 | 0.148768541 | -0.518593691 | 0.294 | 0.365 | 1 | 0.127580286 | -1.276698841 | 0.1 | 0.321 | 1 | 0.148768541 | 0.238883843 |
| Mllt11 | 0.35761864 | -0.485426827 | 0.261 | 0.294 | 1 | 0.127585657 | -1.31299497 | 0.1 | 0.321 | 1 | 0.35761864 | 0.238893215 |
| Gprin3 | 0.158543148 | -0.222392421 | 0.059 | 0.106 | 1 | 0.127594466 | 0.898980835 | 0.4 | 0.198 | 1 | 0.158543148 | 0.238908584 |
| Gm32655 | 0.127708408 | 0.929610672 | 0.059 | 0.024 | 1 | 0.597845674 | 1.695993813 | 0 | 0.028 | 1 | 0.597845674 | 0.239107378 |
| Aktip | 0.127791485 | 0.192645078 | 0.403 | 0.3 | 1 | 0.953885945 | 0.129178659 | 0.4 | 0.429 | 1 | 0.953885945 | 0.239252307 |
| Hps1 | 0.228146873 | -0.605721061 | 0.16 | 0.212 | 1 | 0.127837547 | 1.494359952 | 0.3 | 0.13 | 1 | 0.228146873 | 0.239332656 |
| Snhg17 | 0.127873213 | -0.567888987 | 0.101 | 0.165 | 1 | 0.862447767 | 0.526068812 | 0.1 | 0.117 | 1 | 0.862447767 | 0.239394867 |
| Mrpl34 | 0.127957948 | 0.18740843 | 0.403 | 0.294 | 1 | 0.516860167 | 0.821524695 | 0.3 | 0.235 | 1 | 0.516860167 | 0.23954266 |
| Hic2 | 0.127959295 | -1.070389328 | 0.034 | 0.076 | 1 | 0.219556817 | -0.91281543 | 0 | 0.133 | 1 | 0.219556817 | 0.239545009 |
| Lrp10 | 0.357665305 | 0.117914203 | 0.462 | 0.388 | 1 | 0.127962697 | 0.837349662 | 0.7 | 0.543 | 1 | 0.357665305 | 0.239550942 |
| Ocel1 | 0.329544702 | -0.537894247 | 0.168 | 0.212 | 1 | 0.127979132 | 1.587934067 | 0.4 | 0.219 | 1 | 0.329544702 | 0.239579606 |
| Paqr6 | 0.128011506 | 0.377069649 | 0.134 | 0.076 | 1 | 0.629177319 | -0.152003093 | 0.1 | 0.151 | 1 | 0.629177319 | 0.239636066 |
| Nr2c2ap | 0.568677868 | -0.273922722 | 0.151 | 0.176 | 1 | 0.128036214 | -1.357117523 | 0 | 0.191 | 1 | 0.568677868 | 0.239679156 |
| Sbsn | 0.128062386 | -1.008988783 | 0.05 | 0.1 | 1 | 0.505582845 | 1.017921908 | 0 | 0.043 | 1 | 0.505582845 | 0.239724797 |
| Atp10d | 0.128175103 | -1.159198595 | 0.101 | 0.159 | 1 | 0.190568424 | -0.982078092 | 0 | 0.148 | 1 | 0.190568424 | 0.239921349 |
| Gm10941 | 0.12818617 | 1.099535674 | 0.067 | 0.029 | 1 | 0.444918734 | 1.769994395 | 0.1 | 0.046 | 1 | 0.444918734 | 0.239940646 |
| Zfp874b | 0.128240814 | -0.663764068 | 0.134 | 0.2 | 1 | 0.529243243 | -0.397115591 | 0.2 | 0.275 | 1 | 0.529243243 | 0.240035922 |
| Nfyb | 0.128288211 | 0.201982943 | 0.597 | 0.476 | 1 | 0.304127484 | -0.785132877 | 0.4 | 0.503 | 1 | 0.304127484 | 0.240118557 |
| Ankrd24 | 0.128327573 | -1.117695043 | 0.076 | 0.129 | 1 | 0.277916787 | 1.280956314 | 0.2 | 0.093 | 1 | 0.277916787 | 0.24018718 |
| Ndufb6 | 0.128341709 | -0.226184 | 0.697 | 0.765 | 1 | 0.168420954 | -0.563278674 | 0.4 | 0.694 | 1 | 0.168420954 | 0.240211824 |
| Srek1ip1 | 0.215101005 | -0.608809243 | 0.185 | 0.241 | 1 | 0.128402189 | -1.611434712 | 0 | 0.191 | 1 | 0.215101005 | 0.240317256 |
| 1700052K11Rik | 0.12842758 | -1.137503524 | 0.042 | 0.088 | 1 | 0.604714968 | 1.063725598 | 0.1 | 0.059 | 1 | 0.604714968 | 0.240361516 |
| Suv39h2 | 0.128441056 | 0.63005039 | 0.101 | 0.053 | 1 | 0.570369307 | 1.159940913 | 0.1 | 0.056 | 1 | 0.570369307 | 0.240385007 |
| Gm16001 | 0.128759008 | -1.070389328 | 0.034 | 0.076 | 1 | 0.891584574 | 0.973527789 | 0.1 | 0.086 | 1 | 0.891584574 | 0.240939134 |
| Stk19 | 0.128890611 | -0.55668551 | 0.504 | 0.529 | 1 | 0.152160681 | -1.058893689 | 0.3 | 0.485 | 1 | 0.152160681 | 0.241168433 |
| Gm56938 | 0.128925668 | 0.929610672 | 0.101 | 0.053 | 1 | 0.276546558 | 2.210566986 | 0.1 | 0.034 | 1 | 0.276546558 | 0.241229508 |
| Ccdc73 | 0.129004627 | -0.900464326 | 0.042 | 0.088 | 1 | 0.539413885 | 1.111031312 | 0 | 0.037 | 1 | 0.539413885 | 0.241367061 |
| Gm14295 | 0.129004627 | -0.900464326 | 0.042 | 0.088 | 1 | 0.770508511 | 1.159940913 | 0.1 | 0.074 | 1 | 0.770508511 | 0.241367061 |
| Dzip1l | 0.647677514 | 0.154677228 | 0.319 | 0.288 | 1 | 0.1290956 | -1.357117523 | 0.1 | 0.318 | 1 | 0.647677514 | 0.241525527 |
| Cyp2a5 | 0.129102985 | -0.807354922 | 0.092 | 0.153 | 1 | 0.176952685 | -0.536666944 | 0.1 | 0.336 | 1 | 0.176952685 | 0.241538389 |
| Ptprk | 0.129105486 | 0.113018903 | 0.992 | 0.971 | 1 | 0.607154853 | -0.368659145 | 1 | 0.963 | 1 | 0.607154853 | 0.241542745 |
| Vps26b | 0.129136944 | -0.883976204 | 0.084 | 0.141 | 1 | 0.909368452 | 0.4480663 | 0.2 | 0.228 | 1 | 0.909368452 | 0.241597537 |
| Det1 | 0.129202249 | -0.987927168 | 0.034 | 0.076 | 1 | 0.334166742 | -0.339630097 | 0 | 0.086 | 1 | 0.334166742 | 0.241711277 |
| Zmat3 | 0.129440022 | -0.944858446 | 0.193 | 0.253 | 1 | 0.737306566 | 0.185031894 | 0.3 | 0.231 | 1 | 0.737306566 | 0.242125324 |
| Kbtbd4 | 0.129469566 | -0.549557165 | 0.067 | 0.124 | 1 | 0.660804483 | 0.210566986 | 0.1 | 0.148 | 1 | 0.660804483 | 0.242176763 |
| Dnpep | 0.129709463 | 0.31955719 | 0.437 | 0.353 | 1 | 0.461463842 | 0.443013072 | 0.2 | 0.34 | 1 | 0.461463842 | 0.242594381 |
| Gatad1 | 0.836844107 | -0.142050313 | 0.697 | 0.641 | 1 | 0.129839051 | 0.671119144 | 0.7 | 0.694 | 1 | 0.836844107 | 0.242819923 |
| Hk2 | 0.129965713 | -0.629392602 | 0.689 | 0.7 | 1 | 0.329457938 | -0.191531458 | 0.4 | 0.676 | 1 | 0.329457938 | 0.24304034 |
| Pigq | 0.130014988 | -0.719892081 | 0.134 | 0.2 | 1 | 0.133885694 | 0.930459067 | 0.4 | 0.188 | 1 | 0.133885694 | 0.243126079 |
| Mtrf1 | 0.130036939 | -1 | 0.042 | 0.088 | 1 | 0.505623793 | 0.847996907 | 0 | 0.043 | 1 | 0.505623793 | 0.243164272 |
| Atp1b1 | 0.130050121 | -0.512907564 | 0.134 | 0.206 | 1 | 0.165107894 | -2.165299916 | 0.1 | 0.321 | 1 | 0.165107894 | 0.243187208 |
| Hsdl1 | 0.130069518 | -0.733354341 | 0.042 | 0.088 | 1 | 0.274556733 | -0.408342847 | 0 | 0.108 | 1 | 0.274556733 | 0.243220956 |
| Fam50a | 0.130075046 | -0.379717386 | 0.479 | 0.535 | 1 | 0.609987545 | -0.204470513 | 0.5 | 0.571 | 1 | 0.609987545 | 0.243230575 |
| Cfh | 0.130178099 | -0.758445322 | 0.042 | 0.088 | 1 | 0.841159024 | -0.993305347 | 0.1 | 0.077 | 1 | 0.841159024 | 0.24340986 |
| Cenpl | 0.13030008 | -0.900464326 | 0.025 | 0.065 | 1 | 0.561898712 | 1.558490289 | 0.1 | 0.056 | 1 | 0.561898712 | 0.24362205 |
| Greb1l | 0.130300275 | 0.57870351 | 0.101 | 0.053 | 1 | 0.449402339 | 1.804928185 | 0.2 | 0.123 | 1 | 0.449402339 | 0.243622388 |
| Thoc5 | 0.130342562 | -0.74176658 | 0.134 | 0.2 | 1 | 0.143218643 | -1.111361109 | 0 | 0.179 | 1 | 0.143218643 | 0.243695941 |
| Epb41l1 | 0.130434219 | -0.63641358 | 0.748 | 0.7 | 1 | 0.133623452 | -0.849700045 | 0.5 | 0.765 | 1 | 0.133623452 | 0.243855352 |
| Zfp932 | 0.130476315 | -0.925999419 | 0.101 | 0.159 | 1 | 0.788045725 | 0.345496566 | 0.1 | 0.127 | 1 | 0.788045725 | 0.243928561 |
| Tox3 | 0.130493492 | -1.698420551 | 0.067 | 0.118 | 1 | 0.538014414 | -1.101019165 | 0.1 | 0.167 | 1 | 0.538014414 | 0.243958432 |
| Angel2 | 0.130564599 | -0.516790998 | 0.571 | 0.582 | 1 | 0.364390598 | -0.700896339 | 0.5 | 0.537 | 1 | 0.364390598 | 0.244082084 |
| Gm13889 | 0.660505326 | 2.41941594 | 0.084 | 0.1 | 1 | 0.13063145 | 0.515421567 | 0.2 | 0.068 | 1 | 0.660505326 | 0.244198324 |
| Mrpl13 | 0.130702308 | -0.383547213 | 0.42 | 0.488 | 1 | 0.164634814 | -0.997028433 | 0.2 | 0.398 | 1 | 0.164634814 | 0.244321523 |
| Mppe1 | 0.13070657 | 0.417275971 | 0.311 | 0.229 | 1 | 0.143304663 | 1.382333334 | 0.4 | 0.213 | 1 | 0.143304663 | 0.244328932 |
| Slc22a18 | 0.13079084 | 0.974004791 | 0.076 | 0.035 | 1 | 0.505605027 | 0.625604485 | 0 | 0.043 | 1 | 0.505605027 | 0.244475437 |
| Fbxl2 | 0.227499514 | -0.252449655 | 0.546 | 0.588 | 1 | 0.130794451 | -0.756865152 | 0.2 | 0.506 | 1 | 0.227499514 | 0.244481713 |
| Gga1 | 0.131108879 | -0.784987109 | 0.202 | 0.265 | 1 | 0.666981268 | 0.172431857 | 0.2 | 0.278 | 1 | 0.666981268 | 0.24502822 |
| Ifi202b | 0.381355455 | 2.514573173 | 0.034 | 0.018 | 1 | 0.131194406 | -2.870821341 | 0 | 0.188 | 1 | 0.381355455 | 0.24517684 |
| Pja1 | 0.131393829 | -0.639460456 | 0.353 | 0.429 | 1 | 0.758024449 | -0.239465935 | 0.4 | 0.451 | 1 | 0.758024449 | 0.245523319 |
| Atp6v0b | 0.263612203 | -0.159656666 | 0.882 | 0.918 | 1 | 0.131580973 | -0.66918748 | 0.8 | 0.944 | 1 | 0.263612203 | 0.245848394 |
| Zfp866 | 0.131609839 | -1.070389328 | 0.118 | 0.176 | 1 | 0.37357326 | -0.5519337 | 0.1 | 0.213 | 1 | 0.37357326 | 0.245898529 |
| Dnajc16 | 0.131616205 | -0.807354922 | 0.109 | 0.171 | 1 | 0.225728289 | -0.654503434 | 0 | 0.13 | 1 | 0.225728289 | 0.245909585 |
| Gpc4 | 0.131627575 | -0.824913293 | 0.252 | 0.329 | 1 | 0.221944423 | 0.695993813 | 0.6 | 0.407 | 1 | 0.221944423 | 0.245929331 |
| Hspg2 | 0.611464292 | -0.131630679 | 0.95 | 0.935 | 1 | 0.131743268 | 0.865063455 | 1 | 0.901 | 1 | 0.611464292 | 0.246130248 |
| Septin1 | 0.302671651 | -0.485426827 | 0.16 | 0.206 | 1 | 0.131766944 | -1.357117523 | 0 | 0.188 | 1 | 0.302671651 | 0.246171361 |
| Klhl42 | 0.251707932 | -0.532732542 | 0.118 | 0.165 | 1 | 0.131771492 | -1.321928095 | 0 | 0.188 | 1 | 0.251707932 | 0.246179259 |
| Prdm5 | 0.131791422 | -0.529016096 | 0.605 | 0.635 | 1 | 0.575000219 | 0.369056355 | 0.7 | 0.623 | 1 | 0.575000219 | 0.246213864 |

| | | | | | | | | | | | | |
|---|---|---|---|---|---|---|---|---|---|---|---|---|
| Dynll2 | 0.131800048 | -0.292781749 | 0.513 | 0.594 | 1 | 0.629726982 | 0.255927519 | 0.7 | 0.673 | 1 | 0.629726982 | 0.246228843 |
| Mtx3 | 0.13180198 | -1.070389328 | 0.076 | 0.129 | 1 | 0.245502526 | -0.567040593 | 0 | 0.12 | 1 | 0.245502526 | 0.246232198 |
| Ints14 | 0.131860803 | -0.826463745 | 0.218 | 0.282 | 1 | 0.593358411 | 0.120681482 | 0.4 | 0.284 | 1 | 0.593358411 | 0.246334334 |
| Zmiz2 | 0.13210128 | -0.887525271 | 0.084 | 0.141 | 1 | 0.672439432 | -0.204470513 | 0.2 | 0.247 | 1 | 0.672439432 | 0.246751811 |
| Ermp1 | 0.157771826 | -0.592342031 | 0.076 | 0.129 | 1 | 0.132229371 | 1.780882711 | 0.3 | 0.145 | 1 | 0.157771826 | 0.246974136 |
| Rab17 | 0.132266052 | 0.929610672 | 0.076 | 0.035 | 1 | 0.505605027 | 0.930459067 | 0 | 0.043 | 1 | 0.505605027 | 0.247037795 |
| Dot1l | 0.132270969 | -0.441572538 | 0.681 | 0.694 | 1 | 0.208045031 | -0.780311624 | 0.9 | 0.824 | 1 | 0.208045031 | 0.247046328 |
| Shroom2 | 0.132328466 | -0.870090677 | 0.185 | 0.247 | 1 | 0.646991726 | 0.336097868 | 0.4 | 0.318 | 1 | 0.646991726 | 0.247146108 |
| Irak2 | 0.304633649 | 0.237732967 | 0.261 | 0.206 | 1 | 0.132470638 | 1.249737583 | 0.5 | 0.306 | 1 | 0.304633649 | 0.247392807 |
| Sft2d3 | 0.1743957 | -0.784987109 | 0.101 | 0.153 | 1 | 0.132527679 | 1.532495081 | 0.3 | 0.136 | 1 | 0.1743957 | 0.247491771 |
| Gm4285 | 0.13264037 | 0.777607579 | 0.16 | 0.1 | 1 | 0.804840839 | 1.017921908 | 0.1 | 0.077 | 1 | 0.804840839 | 0.247687272 |
| Pamr1 | 0.338207799 | -0.431655571 | 0.193 | 0.235 | 1 | 0.132694487 | -2.647414009 | 0 | 0.188 | 1 | 0.338207799 | 0.247781147 |
| Sphk2 | 0.132724789 | -0.807354922 | 0.034 | 0.076 | 1 | 0.976598208 | 0.732519689 | 0.1 | 0.096 | 1 | 0.976598208 | 0.247833708 |
| Nrde2 | 0.132724789 | -0.807354922 | 0.034 | 0.076 | 1 | 0.873509692 | 0.660369903 | 0.1 | 0.083 | 1 | 0.873509692 | 0.247833708 |
| 1110046J04Rik | 0.132842793 | -0.925999419 | 0.084 | 0.141 | 1 | 0.637710823 | 0.558490289 | 0.2 | 0.139 | 1 | 0.637710823 | 0.248038378 |
| Leng9 | 0.132911033 | -0.765534746 | 0.042 | 0.088 | 1 | 0.567542604 | 1.31748219 | 0.1 | 0.056 | 1 | 0.567542604 | 0.248156723 |
| Mgrn1 | 0.132954208 | -0.523561956 | 0.378 | 0.435 | 1 | 0.311795119 | 0.620705686 | 0.7 | 0.552 | 1 | 0.311795119 | 0.248231595 |
| Uspl1 | 0.132975958 | -0.324962155 | 0.395 | 0.471 | 1 | 0.699821923 | -0.24081236 | 0.4 | 0.441 | 1 | 0.699821923 | 0.24826931 |
| Vil1 | 0.13314264 | 0.80407979 | 0.084 | 0.041 | 1 | 0.643842184 | 1.558490289 | 0 | 0.022 | 1 | 0.643842184 | 0.248558317 |
| Plekhb2 | 0.133214804 | 0.514573173 | 0.109 | 0.059 | 1 | 0.911403023 | 0.263034406 | 0.1 | 0.108 | 1 | 0.911403023 | 0.248683424 |
| Dele1 | 0.210201614 | -0.371255807 | 0.311 | 0.382 | 1 | 0.133251292 | -1.201246612 | 0.1 | 0.318 | 1 | 0.210201614 | 0.248746677 |
| Eef1akmt1 | 0.133375175 | -0.640149422 | 0.361 | 0.412 | 1 | 0.611785389 | -0.425021588 | 0.3 | 0.336 | 1 | 0.611785389 | 0.248961412 |
| Dennd6a | 0.265397991 | -0.594203173 | 0.462 | 0.488 | 1 | 0.133624448 | 0.558490289 | 0.8 | 0.568 | 1 | 0.265397991 | 0.249393403 |
| H3c15 | 0.133629804 | 0.377069649 | 0.076 | 0.035 | 1 | 0.597845674 | 1.695993813 | 0 | 0.028 | 1 | 0.597845674 | 0.249402683 |
| Zfp954 | 0.133944892 | -0.921525942 | 0.134 | 0.194 | 1 | 0.93487372 | 0.432959407 | 0.2 | 0.182 | 1 | 0.93487372 | 0.24994855 |
| Spaca1 | 0.522834533 | 0.192645078 | 0.092 | 0.071 | 1 | 0.133962084 | 1.795529487 | 0.2 | 0.071 | 1 | 0.522834533 | 0.249978328 |
| Vangl2 | 0.134031692 | -0.829381228 | 0.084 | 0.141 | 1 | 0.407300575 | -0.339630097 | 0.1 | 0.204 | 1 | 0.407300575 | 0.25009889 |
| Atp8a1 | 0.134365162 | -2.048363022 | 0.067 | 0.118 | 1 | 0.704909746 | -0.419483404 | 0.2 | 0.238 | 1 | 0.704909746 | 0.250676327 |
| Rmdn3 | 0.134411816 | -0.660513534 | 0.235 | 0.3 | 1 | 0.231644818 | -0.91281543 | 0.1 | 0.262 | 1 | 0.231644818 | 0.250757095 |
| Synm | 0.134447918 | -0.987927168 | 0.101 | 0.159 | 1 | 0.24440878 | 0.476548676 | 0.4 | 0.219 | 1 | 0.24440878 | 0.250819594 |
| Elp3 | 0.134455258 | -0.524955191 | 0.345 | 0.418 | 1 | 0.70013015 | 0.119068631 | 0.3 | 0.367 | 1 | 0.70013015 | 0.250832299 |
| Cox20 | 0.134522556 | 0.221791424 | 0.588 | 0.476 | 1 | 0.645522863 | 0.31748219 | 0.3 | 0.426 | 1 | 0.645522863 | 0.250948794 |
| Mrpl32 | 0.134624032 | -0.425720581 | 0.437 | 0.5 | 1 | 0.252026114 | -0.73161236 | 0.4 | 0.519 | 1 | 0.252026114 | 0.251124433 |
| Gas2l1 | 0.21413654 | 0.602036014 | 0.353 | 0.3 | 1 | 0.13480405 | -0.896961478 | 0.1 | 0.352 | 1 | 0.21413654 | 0.251435968 |
| 0610040F04Rik | 0.134847265 | -0.748461233 | 0.092 | 0.047 | 1 | 0.504871086 | 5.054447784 | 0.1 | 0.056 | 1 | 0.504871086 | 0.251510745 |
| Fbxo6 | 0.229225562 | 0.421463768 | 0.21 | 0.153 | 1 | 0.135001619 | -1.111361109 | 0 | 0.185 | 1 | 0.229225562 | 0.2517778 |
| Lpar1 | 0.135087836 | -0.223546116 | 0.899 | 0.812 | 1 | 0.776372104 | 0.249041134 | 0.8 | 0.873 | 1 | 0.776372104 | 0.251926949 |
| Clns1a | 0.135269185 | -0.512907564 | 0.319 | 0.388 | 1 | 0.390796474 | -0.279758641 | 0.3 | 0.444 | 1 | 0.390796474 | 0.252240617 |
| Arl10 | 0.20525694 | -0.519374159 | 0.134 | 0.188 | 1 | 0.135284622 | -1.131825212 | 0 | 0.185 | 1 | 0.20525694 | 0.252267315 |
| Sh3pxd2a | 0.135315978 | -0.157757557 | 0.866 | 0.935 | 1 | 0.370533096 | -0.346809683 | 0.7 | 0.895 | 1 | 0.370533096 | 0.252321542 |
| Tbl2 | 0.42165995 | -0.254101281 | 0.193 | 0.229 | 1 | 0.135358535 | 1.091922489 | 0.4 | 0.194 | 1 | 0.42165995 | 0.252395137 |
| Exosc1 | 0.135366873 | 0.110182918 | 0.471 | 0.353 | 1 | 0.585390211 | -0.449683642 | 0.3 | 0.346 | 1 | 0.585390211 | 0.252409556 |
| Zcchc10 | 0.386926384 | -0.527247003 | 0.126 | 0.159 | 1 | 0.135367037 | -1.28585884 | 0 | 0.185 | 1 | 0.386926384 | 0.25240984 |
| Ddx41 | 0.604453238 | -0.128270244 | 0.328 | 0.353 | 1 | 0.135450484 | 1.299103661 | 0.5 | 0.324 | 1 | 0.604453238 | 0.252554134 |
| Igsf9b | 0.13547406 | 0.192645078 | 0.059 | 0.024 | 1 | 0.474977106 | 0.374065718 | 0 | 0.049 | 1 | 0.474977106 | 0.252594899 |
| 5730480H06Rik | 0.13548629 | -0.770829046 | 0.109 | 0.171 | 1 | 0.584304267 | 0.695993813 | 0.2 | 0.133 | 1 | 0.584304267 | 0.252616046 |
| Repin1 | 0.588567298 | 0.321928095 | 0.101 | 0.082 | 1 | 0.135595114 | -1.267480311 | 0 | 0.185 | 1 | 0.588567298 | 0.252804193 |
| Atf5 | 0.135669312 | -0.282439805 | 0.319 | 0.394 | 1 | 0.149058396 | -1.505640048 | 0.2 | 0.398 | 1 | 0.149058396 | 0.252932461 |
| Tbc1d14 | 0.782696898 | -0.273922722 | 0.193 | 0.206 | 1 | 0.13573279 | 1.197742946 | 0.4 | 0.21 | 1 | 0.782696898 | 0.25304219 |
| Gm14305 | 0.135794752 | -0.748461233 | 0.143 | 0.206 | 1 | 0.213542033 | -0.763437806 | 0 | 0.136 | 1 | 0.213542033 | 0.25314929 |
| Gm56692 | 0.8081821 | -0.359895945 | 0.092 | 0.1 | 1 | 0.135899886 | 1.744903414 | 0.2 | 0.071 | 1 | 0.8081821 | 0.253330993 |
| Pdcd11 | 0.136111371 | -0.485426827 | 0.361 | 0.424 | 1 | 0.393813886 | -0.723545078 | 0.3 | 0.389 | 1 | 0.393813886 | 0.253696437 |
| Imp3 | 0.230879691 | -0.399012075 | 0.555 | 0.594 | 1 | 0.136189453 | -0.767180207 | 0.4 | 0.605 | 1 | 0.230879691 | 0.253831338 |
| Gsdme | 0.596533505 | -0.151007788 | 0.328 | 0.353 | 1 | 0.136350373 | 0.841333176 | 0.6 | 0.361 | 1 | 0.596533505 | 0.254109322 |
| Sorl1 | 0.136411568 | 0.981699183 | 0.202 | 0.141 | 1 | 0.800998995 | 1.617042472 | 0.3 | 0.309 | 1 | 0.800998995 | 0.254215019 |
| Mnt | 0.136433617 | -0.549557165 | 0.227 | 0.3 | 1 | 0.60990746 | -0.516056664 | 0.2 | 0.253 | 1 | 0.60990746 | 0.254253102 |
| 2310033P09Rik | 0.943754746 | -0.103179263 | 0.353 | 0.341 | 1 | 0.136635289 | -1.248864633 | 0.1 | 0.315 | 1 | 0.943754746 | 0.254601376 |
| Inpp5k | 0.19214088 | -0.425305835 | 0.269 | 0.341 | 1 | 0.13664741 | -1.185018151 | 0.2 | 0.414 | 1 | 0.19214088 | 0.254622306 |
| Elof1 | 0.695909237 | -0.281893433 | 0.487 | 0.471 | 1 | 0.13675903 | -0.408342847 | 0.3 | 0.574 | 1 | 0.695909237 | 0.254815028 |
| Gm49077 | 0.563722678 | -0.622930351 | 0.034 | 0.047 | 1 | 0.136823071 | 3.880418384 | 0.1 | 0.025 | 1 | 0.563722678 | 0.25492559 |
| Fam210a | 0.136889921 | -0.378511623 | 0.37 | 0.447 | 1 | 0.251564939 | -0.936274402 | 0.4 | 0.503 | 1 | 0.251564939 | 0.255040992 |
| Mttp | 0.936340801 | -0.442358105 | 0.126 | 0.124 | 1 | 0.136897587 | 1.31748219 | 0.3 | 0.136 | 1 | 0.936340801 | 0.255054225 |
| Emc10 | 0.136942718 | 0.162897735 | 0.798 | 0.706 | 1 | 0.649656714 | 0.370463482 | 0.6 | 0.673 | 1 | 0.649656714 | 0.255132128 |
| 2510039O18Rik | 0.981857919 | -0.103556192 | 0.286 | 0.276 | 1 | 0.13713356 | -0.9669712 | 0.2 | 0.417 | 1 | 0.981857919 | 0.255461507 |
| Ndufb8 | 0.137174174 | 0.218080125 | 0.79 | 0.735 | 1 | 0.282578582 | 0.141404961 | 0.5 | 0.796 | 1 | 0.282578582 | 0.255531595 |
| Utp14a | 0.137228853 | -0.544927839 | 0.513 | 0.535 | 1 | 0.230749645 | -0.84502534 | 0.4 | 0.519 | 1 | 0.230749645 | 0.255625948 |
| Atp6v0a2 | 0.13723333 | -0.347923303 | 0.395 | 0.471 | 1 | 0.553147092 | 0.202661788 | 0.7 | 0.593 | 1 | 0.553147092 | 0.255633673 |
| Sgms1os1 | 0.337681333 | -0.411426246 | 0.067 | 0.1 | 1 | 0.137336027 | 2.31748219 | 0.2 | 0.077 | 1 | 0.337681333 | 0.255810869 |
| Ndrg4 | 0.137382945 | -0.632268215 | 0.084 | 0.141 | 1 | 0.274653734 | -0.654503434 | 0 | 0.108 | 1 | 0.274653734 | 0.255891816 |
| Pex2 | 0.137423904 | 0.344648171 | 0.387 | 0.3 | 1 | 0.590513161 | -0.222392421 | 0.3 | 0.38 | 1 | 0.590513161 | 0.255962479 |
| Dynlt2b | 0.664903344 | -0.209792385 | 0.311 | 0.329 | 1 | 0.137480302 | 0.859492545 | 0.5 | 0.275 | 1 | 0.664903344 | 0.256059771 |
| Saal1 | 0.137565581 | -0.887525271 | 0.101 | 0.159 | 1 | 0.21925749 | -0.567040593 | 0 | 0.133 | 1 | 0.21925749 | 0.256206874 |
| Plin3 | 0.916788519 | -0.101086125 | 0.353 | 0.324 | 1 | 0.137607218 | -0.852442812 | 0.2 | 0.451 | 1 | 0.916788519 | 0.256278689 |
| Paics | 0.137695099 | 0.159439064 | 0.958 | 0.912 | 1 | 0.256033403 | -0.56233666 | 1 | 0.948 | 1 | 0.256033403 | 0.256430257 |
| Smim27 | 0.137755648 | 0.292180751 | 0.319 | 0.229 | 1 | 0.782595554 | 0.888638891 | 0.2 | 0.176 | 1 | 0.782595554 | 0.256534678 |
| Bloc1s3 | 0.137770241 | 0.652076697 | 0.084 | 0.041 | 1 | 0.474877085 | 0.847996907 | 0 | 0.049 | 1 | 0.474877085 | 0.256559843 |
| Atp8a2 | 0.137890762 | -0.881355504 | 0.126 | 0.188 | 1 | 0.90020706 | 0.902444691 | 0.2 | 0.194 | 1 | 0.90020706 | 0.256767662 |
| Gm7628 | 0.138053626 | 0.666576266 | 0.076 | 0.035 | 1 | 0.189486514 | 2.695993813 | 0.1 | 0.028 | 1 | 0.189486514 | 0.257048449 |
| Ubl7 | 0.442122327 | -0.424026283 | 0.261 | 0.288 | 1 | 0.138137626 | 1.111031312 | 0.4 | 0.204 | 1 | 0.442122327 | 0.257193249 |
| Mgst1 | 0.557705469 | -0.241869796 | 0.37 | 0.388 | 1 | 0.138201364 | -0.736965594 | 0.2 | 0.5 | 1 | 0.557705469 | 0.257303112 |
| Rft1 | 0.152202195 | -0.795766948 | 0.176 | 0.241 | 1 | 0.138243722 | 1.744903414 | 0.3 | 0.139 | 1 | 0.152202195 | 0.257376117 |
| Gm525 | 0.138260892 | 0.960416719 | 0.454 | 0.394 | 1 | 0.139214685 | -2.153433629 | 0.3 | 0.478 | 1 | 0.139214685 | 0.257405711 |
| Gm715 | 0.144054748 | -1.655351829 | 0.008 | 0.035 | 1 | 0.138385668 | 3.280956314 | 0.1 | 0.025 | 1 | 0.144054748 | 0.257620742 |
| Tdg | 0.138418722 | -0.497601541 | 0.37 | 0.441 | 1 | 0.672015306 | -0.293664243 | 0.5 | 0.506 | 1 | 0.672015306 | 0.257677701 |
| Clip1 | 0.176789629 | -0.324842958 | 0.916 | 0.882 | 1 | 0.138432021 | -0.582470633 | 0.9 | 0.944 | 1 | 0.176789629 | 0.257700617 |
| Kars | 0.138480136 | -0.584962501 | 0.21 | 0.282 | 1 | 0.37709732 | -0.536666944 | 0.2 | 0.318 | 1 | 0.37709732 | 0.257783524 |
| Plod3 | 0.138491619 | 0.193951275 | 0.748 | 0.647 | 1 | 0.19693327 | 0.538452536 | 0.8 | 0.679 | 1 | 0.19693327 | 0.257803309 |
| Camk2g | 0.138582474 | -0.353119535 | 0.42 | 0.476 | 1 | 0.394838105 | 1.007142069 | 0.5 | 0.571 | 1 | 0.394838105 | 0.257959846 |
| Srp19 | 0.138677465 | -0.190229098 | 0.79 | 0.8 | 1 | 0.231892503 | -0.456730448 | 0.7 | 0.802 | 1 | 0.231892503 | 0.258123491 |
| Wdr11 | 0.231518059 | -0.87108052 | 0.218 | 0.259 | 1 | 0.138707435 | -1.111361109 | 0.1 | 0.315 | 1 | 0.231518059 | 0.258175117 |
| Lyz2 | 0.167499542 | -2.137503524 | 0.042 | 0.082 | 1 | 0.138807831 | -1.91281543 | 0 | 0.182 | 1 | 0.167499542 | 0.258348048 |
| Rps6ka2 | 0.302042502 | -0.276840205 | 0.286 | 0.329 | 1 | 0.138871449 | 0.111031312 | 0.4 | 0.179 | 1 | 0.302042502 | 0.258457619 |
| Hmbs | 0.138882672 | -0.748461233 | 0.109 | 0.171 | 1 | 0.799033398 | 0.526068812 | 0.1 | 0.127 | 1 | 0.799033398 | 0.258476947 |
| Aqp4 | 0.649875936 | 0.707218251 | 0.025 | 0.035 | 1 | 0.13891179 | -1.321928095 | 0 | 0.182 | 1 | 0.649875936 | 0.258527095 |
| Dnajc12 | 0.486156241 | -0.207892852 | 0.151 | 0.118 | 1 | 0.139077324 | 1.290001453 | 0.3 | 0.13 | 1 | 0.486156241 | 0.258812146 |
| Ptpa | 0.139139839 | -0.418999465 | 0.555 | 0.6 | 1 | 0.521725063 | -0.184424571 | 0.5 | 0.642 | 1 | 0.521725063 | 0.258919783 |
| Ebag9 | 0.227478747 | -0.57975421 | 0.269 | 0.324 | 1 | 0.139150004 | -1.321928095 | 0.1 | 0.312 | 1 | 0.227478747 | 0.258937284 |
| Tmx2 | 0.139151485 | -0.503348735 | 0.277 | 0.347 | 1 | 0.537759572 | 1.079322453 | 0.3 | 0.228 | 1 | 0.537759572 | 0.258939834 |

| | | | | | | | | | | | | |
|---|---|---|---|---|---|---|---|---|---|---|---|---|
| Lrrk2 | 0.139259627 | -1.315501826 | 0.126 | 0.182 | 1 | 0.819959586 | -0.191531458 | 0.2 | 0.157 | 1 | 0.819959586 | 0.259126011 |
| Gm41609 | 0.186666079 | -0.23878686 | 0.193 | 0.129 | 1 | 0.139272902 | 1.732519689 | 0.2 | 0.074 | 1 | 0.186666079 | 0.259148863 |
| Tma16 | 0.139471219 | -0.77915803 | 0.21 | 0.276 | 1 | 0.764027856 | 0.532495081 | 0.3 | 0.262 | 1 | 0.764027856 | 0.259490217 |
| Ttc4 | 0.257836224 | -0.528495549 | 0.218 | 0.276 | 1 | 0.139526339 | -1.121629444 | 0.1 | 0.315 | 1 | 0.257836224 | 0.259585079 |
| Dusp10 | 0.700766672 | -0.366782331 | 0.235 | 0.2 | 1 | 0.139672098 | -1.357117523 | 0 | 0.182 | 1 | 0.700766672 | 0.259835902 |
| Mrps24 | 0.438934868 | -0.23388806 | 0.563 | 0.582 | 1 | 0.139673673 | 0.778955262 | 0.7 | 0.596 | 1 | 0.438934868 | 0.259838611 |
| Dhx37 | 0.139692383 | -1.070389328 | 0.067 | 0.118 | 1 | 0.420286532 | 0.847996907 | 0.2 | 0.111 | 1 | 0.420286532 | 0.259870804 |
| Mbd4 | 0.139692383 | -1.173482821 | 0.067 | 0.118 | 1 | 0.334242652 | -0.191531458 | 0 | 0.086 | 1 | 0.334242652 | 0.259870804 |
| Dpy30 | 0.256585628 | -0.402010819 | 0.538 | 0.559 | 1 | 0.13969569 | -0.773892163 | 0.4 | 0.577 | 1 | 0.256585628 | 0.259876494 |
| Mrgprf | 0.590813891 | 0.460125389 | 0.143 | 0.124 | 1 | 0.139730043 | -1.391469028 | 0 | 0.182 | 1 | 0.590813891 | 0.259935602 |
| Gm40437 | 0.147029286 | -2.485426827 | 0 | 0.018 | 1 | 0.13997867 | 3.31748219 | 0.1 | 0.025 | 1 | 0.147029286 | 0.260363313 |
| Atg2a | 0.140034039 | 0.568344429 | 0.37 | 0.288 | 1 | 0.892440113 | 0.299103661 | 0.3 | 0.278 | 1 | 0.892440113 | 0.260458545 |
| Ube2e1 | 0.140035498 | -0.40714207 | 0.731 | 0.682 | 1 | 0.172164651 | -0.291067131 | 0.6 | 0.784 | 1 | 0.172164651 | 0.260461055 |
| Ermard | 0.140099354 | -0.977279923 | 0.118 | 0.176 | 1 | 0.265695397 | 0.909397451 | 0.3 | 0.157 | 1 | 0.265695397 | 0.260570879 |
| Tex10 | 0.140116288 | -0.411426246 | 0.311 | 0.382 | 1 | 0.494543857 | -0.388070452 | 0.3 | 0.41 | 1 | 0.494543857 | 0.260600001 |
| Ctdsp1 | 0.928958725 | -0.213753503 | 0.462 | 0.424 | 1 | 0.140479062 | -0.720845929 | 0.3 | 0.543 | 1 | 0.928958725 | 0.261223757 |
| Il3ra | 0.140551971 | 0.514573173 | 0.067 | 0.029 | 1 | 0.557784478 | 1.432959407 | 0 | 0.034 | 1 | 0.557784478 | 0.261349086 |
| Crcp | 0.472671543 | -0.402964667 | 0.193 | 0.224 | 1 | 0.140632994 | 1 | 0.4 | 0.194 | 1 | 0.472671543 | 0.261488349 |
| Ubald1 | 0.140775626 | 0.289506617 | 0.42 | 0.318 | 1 | 0.145226158 | 0.894539492 | 0.5 | 0.275 | 1 | 0.145226158 | 0.261733474 |
| Lamtor4 | 0.641785186 | -0.130095574 | 0.689 | 0.682 | 1 | 0.14103761 | -0.441509711 | 0.4 | 0.639 | 1 | 0.641785186 | 0.262183613 |
| Fchsd2 | 0.141603305 | -0.173482821 | 0.908 | 0.941 | 1 | 0.430541964 | -0.817584054 | 1 | 0.941 | 1 | 0.430541964 | 0.263155113 |
| Dph6 | 0.217831881 | -0.548162583 | 0.303 | 0.359 | 1 | 0.1416665 | -1.521236903 | 0.1 | 0.309 | 1 | 0.217831881 | 0.263263603 |
| Mcoln1 | 0.141833762 | -0.485426827 | 0.202 | 0.271 | 1 | 0.624372333 | 0.463333056 | 0.3 | 0.219 | 1 | 0.624372333 | 0.263550708 |
| Gstk1 | 0.142220205 | 0.617666666 | 0.193 | 0.129 | 1 | 0.158995328 | 1.432959407 | 0.3 | 0.139 | 1 | 0.158995328 | 0.264213823 |
| Ciao2b | 0.142460623 | -0.448273829 | 0.403 | 0.465 | 1 | 0.376282981 | -0.381249186 | 0.3 | 0.441 | 1 | 0.376282981 | 0.264626217 |
| Usp45 | 0.14250729 | -0.485426827 | 0.328 | 0.4 | 1 | 0.854927824 | 0.197742946 | 0.4 | 0.373 | 1 | 0.854927824 | 0.264706252 |
| Il27 | 0.142603094 | -1.944858446 | 0.008 | 0.035 | 1 | 0.355415506 | 2.210566986 | 0.1 | 0.04 | 1 | 0.355415506 | 0.264870546 |
| Hid1 | 0.142603094 | -1.944858446 | 0.008 | 0.035 | 1 | 0.557756607 | 0.769994395 | 0 | 0.034 | 1 | 0.557756607 | 0.264870546 |
| Lrrc69 | 0.142603794 | -1.807354922 | 0.008 | 0.035 | 1 | 0.154873683 | 2.210566986 | 0.1 | 0.025 | 1 | 0.154873683 | 0.264871746 |
| Homez | 0.142603794 | -1.807354922 | 0.008 | 0.035 | 1 | 0.489940924 | 0.695993813 | 0 | 0.046 | 1 | 0.489940924 | 0.264871746 |
| Ppargc1b | 0.142603794 | -1.807354922 | 0.008 | 0.035 | 1 | 0.597842676 | 0.625604485 | 0 | 0.028 | 1 | 0.597842676 | 0.264871746 |
| Btk | 0.142603794 | -2.392317423 | 0.008 | 0.035 | 1 | 0.643846086 | 0.847996907 | 0 | 0.022 | 1 | 0.643846086 | 0.264871746 |
| Ell | 0.142625249 | 0.105095199 | 0.773 | 0.659 | 1 | 0.694311182 | 0.310562776 | 0.8 | 0.738 | 1 | 0.694311182 | 0.264908536 |
| Lrwd1 | 0.142657723 | -1.222392421 | 0.067 | 0.118 | 1 | 0.211194799 | 1.515421567 | 0.2 | 0.083 | 1 | 0.211194799 | 0.264964221 |
| Zfp687 | 0.142657912 | -0.959358016 | 0.067 | 0.118 | 1 | 0.255499842 | 0.930459067 | 0.3 | 0.157 | 1 | 0.255499842 | 0.264964545 |
| Dmac1 | 0.31504328 | -0.281893433 | 0.361 | 0.406 | 1 | 0.14301867 | -1.026472211 | 0.1 | 0.312 | 1 | 0.31504328 | 0.265583 |
| Etnk1 | 0.143136367 | 0.214210428 | 0.866 | 0.835 | 1 | 0.430287517 | -0.369703819 | 0.9 | 0.867 | 1 | 0.430287517 | 0.265784714 |
| Gnai3 | 0.143185961 | -0.177936116 | 0.916 | 0.935 | 1 | 0.227424671 | -0.37351448 | 1 | 0.975 | 1 | 0.227424671 | 0.265869702 |
| Cyfip2 | 0.603949109 | -0.655351829 | 0.084 | 0.1 | 1 | 0.143218643 | -1.171902651 | 0 | 0.179 | 1 | 0.603949109 | 0.265925707 |
| Prdm14 | 0.1433296 | -1.807354922 | 0.008 | 0.035 | 1 | 0.643835681 | 1.695993813 | 0 | 0.022 | 1 | 0.643835681 | 0.266115826 |
| Gm4673 | 0.1433296 | -1.807354922 | 0.008 | 0.035 | 1 | 0.320995017 | 1.847996907 | 0.1 | 0.037 | 1 | 0.320995017 | 0.266115826 |
| Igf2os | 0.1433296 | -1.655351829 | 0.008 | 0.035 | 1 | 0.698534498 | 2.432959407 | 0 | 0.015 | 1 | 0.698534498 | 0.266115826 |
| Gm45894 | 0.1433296 | -1.655351829 | 0.008 | 0.035 | 1 | 0.401608067 | 1.847996907 | 0.1 | 0.043 | 1 | 0.401608067 | 0.266115826 |
| Kiss1r | 0.1433296 | -1.655351829 | 0.008 | 0.035 | 1 | 0.698529281 | 2.017921908 | 0 | 0.015 | 1 | 0.698529281 | 0.266115826 |
| 4732465J04Rik | 0.1433296 | -1.655351829 | 0.008 | 0.035 | 1 | 0.643842184 | 1.695993813 | 0 | 0.022 | 1 | 0.643842184 | 0.266115826 |
| Tepsin | 0.1433296 | -1.655351829 | 0.008 | 0.035 | 1 | 0.505605027 | 0.847996907 | 0 | 0.043 | 1 | 0.505605027 | 0.266115826 |
| Dlk1 | 0.1433296 | -1.655351829 | 0.008 | 0.035 | 1 | 0.489977569 | -0.888968688 | 0 | 0.046 | 1 | 0.489977569 | 0.266115826 |
| Gm26555 | 0.1433296 | -1.655351829 | 0.008 | 0.035 | 1 | 0.539444993 | 0.930459067 | 0 | 0.037 | 1 | 0.539444993 | 0.266115826 |
| Dusp29 | 0.1433296 | -1.655351829 | 0.008 | 0.035 | 1 | 0.619967066 | 1.111031312 | 0 | 0.025 | 1 | 0.619967066 | 0.266115826 |
| Pced1b | 0.1433296 | -1.655351829 | 0.008 | 0.035 | 1 | 0.597840427 | 0.930459067 | 0 | 0.028 | 1 | 0.597840427 | 0.266115826 |
| A930001C03Rik | 0.1433296 | -1.807354922 | 0.008 | 0.035 | 1 | 0.698529281 | 1.847996907 | 0 | 0.015 | 1 | 0.698529281 | 0.266115826 |
| Rps6ka6 | 0.1433296 | -1.655351829 | 0.008 | 0.035 | 1 | 0.408705208 | 0.263034406 | 0 | 0.065 | 1 | 0.408705208 | 0.266115826 |
| Sprr1a | 0.143404766 | -0.263034406 | 0.059 | 0.112 | 1 | 0.420961055 | -1.111361109 | 0 | 0.062 | 1 | 0.420961055 | 0.266244606 |
| Inpp4a | 0.143459593 | -0.617338503 | 0.294 | 0.371 | 1 | 0.167135275 | 1.115219109 | 0.6 | 0.525 | 1 | 0.167135275 | 0.266338532 |
| Adgre5 | 0.143564897 | -1.070389328 | 0.109 | 0.165 | 1 | 0.369847407 | -1.201246612 | 0.1 | 0.207 | 1 | 0.369847407 | 0.266518914 |
| 3110082I17Rik | 0.143723836 | -0.730539325 | 0.202 | 0.271 | 1 | 0.151936449 | -1.152003093 | 0 | 0.173 | 1 | 0.151936449 | 0.26679113 |
| Ctla4 | 0.143756494 | -0.748461233 | 0.076 | 0.129 | 1 | 0.977222166 | 0.695993813 | 0.1 | 0.102 | 1 | 0.977222166 | 0.266847058 |
| Rrp7a | 0.143795219 | -0.565597176 | 0.261 | 0.329 | 1 | 0.760684162 | -0.201246612 | 0.3 | 0.318 | 1 | 0.760684162 | 0.266913373 |
| Mark2 | 0.143873472 | -0.415962008 | 0.597 | 0.641 | 1 | 0.258636819 | -0.568807519 | 0.8 | 0.852 | 1 | 0.258636819 | 0.267047368 |
| Bzw2 | 0.143937793 | -0.753532223 | 0.756 | 0.747 | 1 | 0.190728779 | 0.889753372 | 0.8 | 0.67 | 1 | 0.190728779 | 0.267157497 |
| 4930590J08Rik | 0.144001366 | -0.315501826 | 0.034 | 0.076 | 1 | 0.723705138 | 0.210566986 | 0.1 | 0.136 | 1 | 0.723705138 | 0.267266338 |
| Gm17056 | 0.821489426 | 0.12755005 | 0.084 | 0.076 | 1 | 0.144015054 | 0.847996907 | 0.3 | 0.127 | 1 | 0.821489426 | 0.267289773 |
| BC055324 | 0.144054748 | -1.485426827 | 0.008 | 0.035 | 1 | 0.730749233 | 2.432959407 | 0 | 0.012 | 1 | 0.730749233 | 0.267357726 |
| Gm10848 | 0.144054748 | -1.485426827 | 0.008 | 0.035 | 1 | 0.460599334 | 0.494359952 | 0 | 0.052 | 1 | 0.460599334 | 0.267357726 |
| Gm15929 | 0.144054748 | -1.485426827 | 0.008 | 0.035 | 1 | 0.698532411 | 2.017921908 | 0 | 0.015 | 1 | 0.698532411 | 0.267357726 |
| Gm16853 | 0.144054748 | -1.485426827 | 0.008 | 0.035 | 1 | 0.315848442 | -0.339630097 | 0 | 0.093 | 1 | 0.315848442 | 0.267357726 |
| Gm19531 | 0.144054748 | -1.485426827 | 0.008 | 0.035 | 1 | 0.474877085 | 0.847996907 | 0 | 0.049 | 1 | 0.474877085 | 0.267357726 |
| Gm3055 | 0.144054748 | -1.485426827 | 0.008 | 0.035 | 1 | 0.669843604 | 2.017921908 | 0 | 0.019 | 1 | 0.669843604 | 0.267357726 |
| Gm57330 | 0.144054748 | -1.485426827 | 0.008 | 0.035 | 1 | 0.73075158 | 2.695993813 | 0 | 0.012 | 1 | 0.73075158 | 0.267357726 |
| Aoc2 | 0.144054748 | -1.485426827 | 0.008 | 0.035 | 1 | 0.320995017 | 1.769994395 | 0.1 | 0.037 | 1 | 0.320995017 | 0.267357726 |
| Acvrl1 | 0.144054748 | -1.485426827 | 0.008 | 0.035 | 1 | 0.539483006 | 1.31748219 | 0 | 0.037 | 1 | 0.539483006 | 0.267357726 |
| Zfp677 | 0.144054748 | -1.485426827 | 0.008 | 0.035 | 1 | 0.557756607 | 0.847996907 | 0 | 0.034 | 1 | 0.557756607 | 0.267357726 |
| Rnf125 | 0.144054748 | -1.485426827 | 0.008 | 0.035 | 1 | 0.398223038 | 3.169925001 | 0.1 | 0.046 | 1 | 0.398223038 | 0.267357726 |
| 2010204K13Rik | 0.144054748 | -1.485426827 | 0.008 | 0.035 | 1 | 0.47676286 | 1.930459067 | 0.1 | 0.049 | 1 | 0.47676286 | 0.267357726 |
| Gm14798 | 0.144054748 | -1.655351829 | 0.008 | 0.035 | 1 | 0.460631204 | 0.374065718 | 0 | 0.052 | 1 | 0.460631204 | 0.267357726 |
| Blnk | 0.144077942 | -2.544320516 | 0.008 | 0.035 | 1 | 0.201481076 | -0.68251781 | 0 | 0.142 | 1 | 0.201481076 | 0.26739743 |
| Trp53bp2 | 0.144081965 | -0.511266832 | 0.403 | 0.471 | 1 | 0.243120555 | -0.388637437 | 0.4 | 0.682 | 1 | 0.243120555 | 0.267404318 |
| Coasy | 0.144116679 | -0.655351829 | 0.109 | 0.171 | 1 | 0.761476353 | 0.494359952 | 0.1 | 0.133 | 1 | 0.761476353 | 0.267463742 |
| Dlgap4 | 0.144131804 | -0.417496973 | 0.866 | 0.853 | 1 | 0.204844688 | -0.495147674 | 1 | 0.938 | 1 | 0.204844688 | 0.267489631 |
| Mboat7 | 0.144144638 | -0.485426827 | 0.218 | 0.288 | 1 | 0.972190936 | 0.169925001 | 0.3 | 0.309 | 1 | 0.972190936 | 0.267511599 |
| Plekha5 | 0.144193182 | -0.264561876 | 0.647 | 0.729 | 1 | 0.50140612 | 0.274530045 | 1 | 0.849 | 1 | 0.50140612 | 0.26759469 |
| Yars | 0.144372942 | -0.590961241 | 0.555 | 0.576 | 1 | 0.30293624 | -0.720845929 | 0.6 | 0.654 | 1 | 0.30293624 | 0.267902337 |
| Tmem18 | 0.14439017 | -0.842978832 | 0.118 | 0.176 | 1 | 0.154389571 | 1.587934067 | 0.4 | 0.213 | 1 | 0.154389571 | 0.267931819 |
| Gm26542 | 0.467461923 | -0.532732542 | 0.109 | 0.135 | 1 | 0.144446262 | -1.814968106 | 0 | 0.179 | 1 | 0.467461923 | 0.268027802 |
| Samd5 | 0.144461794 | -1.235448574 | 0.185 | 0.241 | 1 | 0.816620954 | 0.487407191 | 0.3 | 0.278 | 1 | 0.816620954 | 0.268054378 |
| Cox7a2l | 0.144549444 | 0.171800824 | 0.916 | 0.941 | 1 | 0.171568417 | -0.433289204 | 0.7 | 0.91 | 1 | 0.171568417 | 0.268204346 |
| Eogt | 0.957124317 | -0.366782331 | 0.244 | 0.235 | 1 | 0.144647599 | 0.446379923 | 0.6 | 0.318 | 1 | 0.957124317 | 0.268372271 |
| Neurl2 | 0.144817959 | -1.292781749 | 0.008 | 0.035 | 1 | 0.505686333 | 1.111031312 | 0 | 0.043 | 1 | 0.505686333 | 0.268663676 |
| Myef2l | 0.144817959 | -1.292781749 | 0.008 | 0.035 | 1 | 0.669843604 | 1.847996907 | 0 | 0.019 | 1 | 0.669843604 | 0.268663676 |
| Zfp691 | 0.144817959 | -1.292781749 | 0.008 | 0.035 | 1 | 0.669853054 | 2.210566986 | 0 | 0.019 | 1 | 0.669853054 | 0.268663676 |
| Cmtr2 | 0.144817959 | -1.292781749 | 0.008 | 0.035 | 1 | 0.643850855 | 2.017921908 | 0 | 0.022 | 1 | 0.643850855 | 0.268663676 |
| Gm48163 | 0.144817959 | -1.292781749 | 0.008 | 0.035 | 1 | 0.19522577 | 2.111031312 | 0.1 | 0.028 | 1 | 0.19522577 | 0.268663676 |
| Gm56765 | 0.144817959 | -1.292781749 | 0.008 | 0.035 | 1 | 0.505605027 | 0.847996907 | 0 | 0.043 | 1 | 0.505605027 | 0.268663676 |
| Myo5b | 0.144817959 | -1.292781749 | 0.008 | 0.035 | 1 | 0.761476353 | 0.494359952 | 0.1 | 0.133 | 1 | 0.761476353 | 0.268663676 |
| Tmem151a | 0.144817959 | -1.292781749 | 0.008 | 0.035 | 1 | 0.460521625 | 0.769994395 | 0 | 0.052 | 1 | 0.460521625 | 0.268663676 |
| Bend3 | 0.145034519 | -1.185866545 | 0.059 | 0.106 | 1 | 0.21381637 | -1.131825212 | 0 | 0.136 | 1 | 0.21381637 | 0.269034026 |
| Pdlim7 | 0.338728526 | -0.173042505 | 0.445 | 0.506 | 1 | 0.145138354 | 0.871765072 | 0.6 | 0.475 | 1 | 0.338728526 | 0.269211567 |
| Rab3gap1 | 0.145442909 | -0.246239163 | 0.782 | 0.812 | 1 | 0.559113354 | 0.219797312 | 0.9 | 0.806 | 1 | 0.559113354 | 0.269732178 |

| | | | | | | | | | | | | |
|---|---|---|---|---|---|---|---|---|---|---|---|---|
| Ociad1 | 0.145463856 | -0.202808025 | 0.908 | 0.882 | 1 | 0.320969756 | -0.316680017 | 0.8 | 0.846 | 1 | 0.320969756 | 0.269767979 |
| Mrpl24 | 0.243191528 | -0.287826867 | 0.689 | 0.688 | 1 | 0.145632708 | -0.665072676 | 0.5 | 0.701 | 1 | 0.243191528 | 0.270056531 |
| Gnptg | 0.14573975 | 0.242768557 | 0.496 | 0.406 | 1 | 0.778626993 | 0.326760003 | 0.3 | 0.361 | 1 | 0.778626993 | 0.270239425 |
| Rp1 | 0.146109238 | -1.189970944 | 0.143 | 0.2 | 1 | 0.230005898 | -1.544320516 | 0.1 | 0.259 | 1 | 0.230005898 | 0.270870567 |
| 5031439G07Rik | 0.820387765 | 0.225066556 | 0.395 | 0.388 | 1 | 0.146192135 | -0.789433014 | 0.2 | 0.454 | 1 | 0.820387765 | 0.27101213 |
| Slfn8 | 0.146296808 | -1.807354922 | 0.008 | 0.035 | 1 | 0.730750993 | 2.210566986 | 0 | 0.012 | 1 | 0.730750993 | 0.271190861 |
| Nxn | 0.146506996 | -1.018859027 | 0.555 | 0.553 | 1 | 0.880575438 | 0.505310252 | 0.6 | 0.605 | 1 | 0.880575438 | 0.271549692 |
| Ifih1 | 0.146682355 | -0.687060688 | 0.084 | 0.141 | 1 | 0.213724879 | -1.391469028 | 0 | 0.136 | 1 | 0.213724879 | 0.271848997 |
| Lrch1 | 0.146805059 | 0.195582361 | 0.689 | 0.594 | 1 | 0.324961773 | 0.956016777 | 0.7 | 0.747 | 1 | 0.324961773 | 0.272058393 |
| Gtf3c1 | 0.146975495 | -0.438632616 | 0.445 | 0.518 | 1 | 0.551244009 | -0.162650338 | 0.7 | 0.744 | 1 | 0.551244009 | 0.272349194 |
| Spag16 | 0.147029286 | -1.807354922 | 0 | 0.018 | 1 | 0.730749233 | 2.432959407 | 0 | 0.012 | 1 | 0.730749233 | 0.27244096 |
| Nectin4 | 0.147029286 | -1.807354922 | 0 | 0.018 | 1 | 0.730750993 | 2.210566986 | 0 | 0.012 | 1 | 0.730750993 | 0.27244096 |
| Vash2 | 0.147029286 | -1.807354922 | 0 | 0.018 | 1 | 0.577142568 | 1.432959407 | 0 | 0.031 | 1 | 0.577142568 | 0.27244096 |
| Gm5148 | 0.147029286 | -2.070389328 | 0 | 0.018 | 1 | 0.698534498 | 2.432959407 | 0 | 0.015 | 1 | 0.698534498 | 0.27244096 |
| Gm26594 | 0.147029286 | -1.807354922 | 0 | 0.018 | 1 | 0.73075158 | 2.695993813 | 0 | 0.012 | 1 | 0.73075158 | 0.27244096 |
| 4833413G10Rik | 0.147029286 | -1.807354922 | 0 | 0.018 | 1 | 0.522151747 | 1.210566986 | 0 | 0.04 | 1 | 0.522151747 | 0.27244096 |
| Drc1 | 0.147029286 | -1.807354922 | 0 | 0.018 | 1 | 0.643842184 | 1.31748219 | 0 | 0.022 | 1 | 0.643842184 | 0.27244096 |
| Gm42982 | 0.147029286 | -1.807354922 | 0 | 0.018 | 1 | 0.669853054 | 2.210566986 | 0 | 0.019 | 1 | 0.669853054 | 0.27244096 |
| Gm43661 | 0.147029286 | -1.807354922 | 0 | 0.018 | 1 | 0.643842184 | 1.695993813 | 0 | 0.022 | 1 | 0.643842184 | 0.27244096 |
| Gm34312 | 0.147029286 | -1.807354922 | 0 | 0.018 | 1 | 0.505639431 | 0.769994395 | 0 | 0.043 | 1 | 0.505639431 | 0.27244096 |
| Mfap5 | 0.147029286 | -2.807354922 | 0 | 0.018 | 1 | 0.669843604 | 1.31748219 | 0 | 0.019 | 1 | 0.669843604 | 0.27244096 |
| Gm15704 | 0.147029286 | -2.070389328 | 0 | 0.018 | 1 | 0.698534498 | 2.432959407 | 0 | 0.015 | 1 | 0.698534498 | 0.27244096 |
| Zfp114 | 0.147029286 | -1.807354922 | 0 | 0.018 | 1 | 0.539431168 | 1.111031312 | 0 | 0.037 | 1 | 0.539431168 | 0.27244096 |
| Tyrobp | 0.147029286 | -2.944858446 | 0 | 0.018 | 1 | 0.397024988 | -1.970762779 | 0 | 0.068 | 1 | 0.397024988 | 0.27244096 |
| Il4i1 | 0.147029286 | -1.807354922 | 0 | 0.018 | 1 | 0.539455361 | 0.847996907 | 0 | 0.037 | 1 | 0.539455361 | 0.27244096 |
| Tm6sf1 | 0.147029286 | -1.807354922 | 0 | 0.018 | 1 | 0.698532411 | 1.847996907 | 0 | 0.015 | 1 | 0.698532411 | 0.27244096 |
| Lat | 0.147029286 | -2.070389328 | 0 | 0.018 | 1 | 0.730749233 | 2.432959407 | 0 | 0.012 | 1 | 0.730749233 | 0.27244096 |
| Gm28379 | 0.147029286 | -1.807354922 | 0 | 0.018 | 1 | 0.577142568 | 1.432959407 | 0 | 0.031 | 1 | 0.577142568 | 0.27244096 |
| Hkdc1 | 0.147029286 | -1.807354922 | 0 | 0.018 | 1 | 0.15487326 | 2.017921908 | 0.1 | 0.025 | 1 | 0.15487326 | 0.27244096 |
| Cabcoco1 | 0.147029286 | -1.807354922 | 0 | 0.018 | 1 | 0.557729726 | 1.31748219 | 0 | 0.034 | 1 | 0.557729726 | 0.27244096 |
| Calm4 | 0.147029286 | -1.807354922 | 0 | 0.018 | 1 | 0.730750993 | 1.847996907 | 0 | 0.012 | 1 | 0.730750993 | 0.27244096 |
| Ryr2 | 0.147029286 | -1.807354922 | 0 | 0.018 | 1 | 0.669850692 | 1.432959407 | 0 | 0.019 | 1 | 0.669850692 | 0.27244096 |
| Rem2 | 0.147029286 | -1.807354922 | 0 | 0.018 | 1 | 0.669843604 | 2.017921908 | 0 | 0.019 | 1 | 0.669843604 | 0.27244096 |
| Adam28 | 0.147029286 | -2.292781749 | 0 | 0.018 | 1 | 0.408636939 | 0.374065718 | 0 | 0.065 | 1 | 0.408636939 | 0.27244096 |
| Rxfp3 | 0.147029286 | -1.807354922 | 0 | 0.018 | 1 | 0.42093472 | 0.374065718 | 0 | 0.062 | 1 | 0.42093472 | 0.27244096 |
| Grhl2 | 0.147029286 | -2.292781749 | 0 | 0.018 | 1 | 0.306858329 | -0.152003093 | 0 | 0.096 | 1 | 0.306858329 | 0.27244096 |
| Ly6c2 | 0.147029286 | -1.807354922 | 0 | 0.018 | 1 | 0.669848329 | 1.847996907 | 0 | 0.019 | 1 | 0.669848329 | 0.27244096 |
| Tex49 | 0.147029286 | -4.070389328 | 0 | 0.018 | 1 | 0.6199755 | 1.847996907 | 0 | 0.025 | 1 | 0.6199755 | 0.27244096 |
| B3gnt5 | 0.147029286 | -1.807354922 | 0 | 0.018 | 1 | 0.730750993 | 1.558490289 | 0 | 0.012 | 1 | 0.730750993 | 0.27244096 |
| Hgd | 0.147029286 | -1.807354922 | 0 | 0.018 | 1 | 0.698529281 | 2.210566986 | 0 | 0.015 | 1 | 0.698529281 | 0.27244096 |
| Pcdhb19 | 0.147029286 | -1.807354922 | 0 | 0.018 | 1 | 0.197182351 | 1.930459067 | 0.1 | 0.028 | 1 | 0.197182351 | 0.27244096 |
| Dtx4 | 0.147029286 | -1.807354922 | 0 | 0.018 | 1 | 0.669843604 | 1.847996907 | 0 | 0.019 | 1 | 0.669843604 | 0.27244096 |
| Usp51 | 0.147029286 | -1.807354922 | 0 | 0.018 | 1 | 0.730750993 | 1.695993813 | 0 | 0.012 | 1 | 0.730750993 | 0.27244096 |
| Gm28151 | 0.147030916 | -1.485426827 | 0 | 0.018 | 1 | 0.669853054 | 2.210566986 | 0 | 0.019 | 1 | 0.669853054 | 0.272443742 |
| A030005L19Rik | 0.147030916 | -1.485426827 | 0 | 0.018 | 1 | 0.698532411 | 1.695993813 | 0 | 0.015 | 1 | 0.698532411 | 0.272443742 |
| Gm32250 | 0.147030916 | -1.485426827 | 0 | 0.018 | 1 | 0.232498909 | 2.432959407 | 0.1 | 0.031 | 1 | 0.232498909 | 0.272443742 |
| Fam110a | 0.147030916 | -1.485426827 | 0 | 0.018 | 1 | 0.698529281 | 2.210566986 | 0 | 0.015 | 1 | 0.698529281 | 0.272443742 |
| Fam83c | 0.147030916 | -1.485426827 | 0 | 0.018 | 1 | 0.730749233 | 2.210566986 | 0 | 0.012 | 1 | 0.730749233 | 0.272443742 |
| Ankrd60 | 0.147030916 | -1.485426827 | 0 | 0.018 | 1 | 0.730749233 | 2.432959407 | 0 | 0.012 | 1 | 0.730749233 | 0.272443742 |
| Mccc1os | 0.147030916 | -1.485426827 | 0 | 0.018 | 1 | 0.669843604 | 1.847996907 | 0 | 0.019 | 1 | 0.669843604 | 0.272443742 |
| I830077J02Rik | 0.147030916 | -1.485426827 | 0 | 0.018 | 1 | 0.643835681 | 1.847996907 | 0 | 0.022 | 1 | 0.643835681 | 0.272443742 |
| Cavin4 | 0.147030916 | -1.485426827 | 0 | 0.018 | 1 | 0.698529281 | 2.210566986 | 0 | 0.015 | 1 | 0.698529281 | 0.272443742 |
| Gm57092 | 0.147030916 | -1.485426827 | 0 | 0.018 | 1 | 0.643842184 | 1.558490289 | 0 | 0.022 | 1 | 0.643842184 | 0.272443742 |
| Gm15402 | 0.147030916 | -2.070389328 | 0 | 0.018 | 1 | 0.669853054 | 2.210566986 | 0 | 0.019 | 1 | 0.669853054 | 0.272443742 |
| Zfp109 | 0.147030916 | -1.485426827 | 0 | 0.018 | 1 | 0.698532411 | 2.017921908 | 0 | 0.015 | 1 | 0.698532411 | 0.272443742 |
| Gm45442 | 0.147030916 | -1.485426827 | 0 | 0.018 | 1 | 0.698529281 | 2.210566986 | 0 | 0.015 | 1 | 0.698529281 | 0.272443742 |
| Otud7a | 0.147030916 | -1.485426827 | 0 | 0.018 | 1 | 0.73075158 | 2.695993813 | 0 | 0.012 | 1 | 0.73075158 | 0.272443742 |
| Camsap3 | 0.147030916 | -1.485426827 | 0 | 0.018 | 1 | 0.557744661 | 1.017921908 | 0 | 0.034 | 1 | 0.557744661 | 0.272443742 |
| Zmat4 | 0.147030916 | -2.485426827 | 0 | 0.018 | 1 | 0.518401944 | 1.111031312 | 0.1 | 0.052 | 1 | 0.518401944 | 0.272443742 |
| A230103J11Rik | 0.147030916 | -1.485426827 | 0 | 0.018 | 1 | 0.73075158 | 2.695993813 | 0 | 0.012 | 1 | 0.73075158 | 0.272443742 |
| Odad3 | 0.147030916 | -1.485426827 | 0 | 0.018 | 1 | 0.730749233 | 2.210566986 | 0 | 0.012 | 1 | 0.730749233 | 0.272443742 |
| Tecta | 0.147030916 | -2.807354922 | 0 | 0.018 | 1 | 0.669848329 | 1.847996907 | 0 | 0.019 | 1 | 0.669848329 | 0.272443742 |
| Filip1 | 0.147030916 | -3.807354922 | 0 | 0.018 | 1 | 0.619967066 | 1.111031312 | 0 | 0.025 | 1 | 0.619967066 | 0.272443742 |
| Gm57003 | 0.147030916 | -1.485426827 | 0 | 0.018 | 1 | 0.73075158 | 2.695993813 | 0 | 0.012 | 1 | 0.73075158 | 0.272443742 |
| Matk | 0.147030916 | -2.070389328 | 0 | 0.018 | 1 | 0.560373918 | 1.744903414 | 0.1 | 0.059 | 1 | 0.560373918 | 0.272443742 |
| Ccdc38 | 0.147030916 | -2.070389328 | 0 | 0.018 | 1 | 0.698529281 | 2.017921908 | 0 | 0.015 | 1 | 0.698529281 | 0.272443742 |
| Myl6b | 0.147030916 | -1.485426827 | 0 | 0.018 | 1 | 0.698534498 | 2.432959407 | 0 | 0.015 | 1 | 0.698534498 | 0.272443742 |
| Gm12059 | 0.147030916 | -2.292781749 | 0 | 0.018 | 1 | 0.73075158 | 2.695993813 | 0 | 0.012 | 1 | 0.73075158 | 0.272443742 |
| Dock2 | 0.147030916 | -5.040015679 | 0 | 0.018 | 1 | 0.557729726 | 1.017921908 | 0 | 0.034 | 1 | 0.557729726 | 0.272443742 |
| Zfp454 | 0.147030916 | -1.485426827 | 0 | 0.018 | 1 | 0.698534498 | 2.432959407 | 0 | 0.015 | 1 | 0.698534498 | 0.272443742 |
| Ccdc69 | 0.147030916 | -2.485426827 | 0 | 0.018 | 1 | 0.73075158 | 2.695993813 | 0 | 0.012 | 1 | 0.73075158 | 0.272443742 |
| Vmo1 | 0.147030916 | -1.485426827 | 0 | 0.018 | 1 | 0.522066388 | 0.930459067 | 0 | 0.04 | 1 | 0.522066388 | 0.272443742 |
| Hic1 | 0.147030916 | -1.485426827 | 0 | 0.018 | 1 | 0.73075158 | 2.695993813 | 0 | 0.012 | 1 | 0.73075158 | 0.272443742 |
| Rgs9 | 0.147030916 | -2.944858446 | 0 | 0.018 | 1 | 0.619953009 | 1.695993813 | 0 | 0.025 | 1 | 0.619953009 | 0.272443742 |
| Gm11747 | 0.147030916 | -1.485426827 | 0 | 0.018 | 1 | 0.32099467 | 1.930459067 | 0.1 | 0.037 | 1 | 0.32099467 | 0.272443742 |
| Dtnbos | 0.147030916 | -1.485426827 | 0 | 0.018 | 1 | 0.597824685 | 1.432959407 | 0 | 0.028 | 1 | 0.597824685 | 0.272443742 |
| Mycn | 0.147030916 | -1.485426827 | 0 | 0.018 | 1 | 0.669843604 | 2.017921908 | 0 | 0.019 | 1 | 0.669843604 | 0.272443742 |
| Gm48008 | 0.147030916 | -1.485426827 | 0 | 0.018 | 1 | 0.619961444 | 1.558490289 | 0 | 0.025 | 1 | 0.619961444 | 0.272443742 |
| Gm47882 | 0.147030916 | -1.485426827 | 0 | 0.018 | 1 | 0.73075158 | 2.695993813 | 0 | 0.012 | 1 | 0.73075158 | 0.272443742 |
| Phf2os1 | 0.147030916 | -1.485426827 | 0 | 0.018 | 1 | 0.19134412 | 2.558490289 | 0.1 | 0.028 | 1 | 0.19134412 | 0.272443742 |
| Gm6270 | 0.147030916 | -1.485426827 | 0 | 0.018 | 1 | 0.730749233 | 2.432959407 | 0 | 0.012 | 1 | 0.730749233 | 0.272443742 |
| Gm47814 | 0.147030916 | -1.485426827 | 0 | 0.018 | 1 | 0.149795061 | 2.847996907 | 0.1 | 0.025 | 1 | 0.149795061 | 0.272443742 |
| Itih4 | 0.147030916 | -2.070389328 | 0 | 0.018 | 1 | 0.698534498 | 2.432959407 | 0 | 0.015 | 1 | 0.698534498 | 0.272443742 |
| Stab1 | 0.147030916 | -1.485426827 | 0 | 0.018 | 1 | 0.643835681 | 1.847996907 | 0 | 0.022 | 1 | 0.643835681 | 0.272443742 |
| Gm46447 | 0.147030916 | -1.485426827 | 0 | 0.018 | 1 | 0.489939838 | 0.625604485 | 0 | 0.046 | 1 | 0.489939838 | 0.272443742 |
| Gm49083 | 0.147030916 | -2.070389328 | 0 | 0.018 | 1 | 0.730749233 | 2.017921908 | 0 | 0.012 | 1 | 0.730749233 | 0.272443742 |
| Ltb4r1 | 0.147030916 | -1.485426827 | 0 | 0.018 | 1 | 0.698529281 | 2.017921908 | 0 | 0.015 | 1 | 0.698529281 | 0.272443742 |
| 1700109G14Rik | 0.147030916 | -1.485426827 | 0 | 0.018 | 1 | 0.669851164 | 1.31748219 | 0 | 0.019 | 1 | 0.669851164 | 0.272443742 |
| Gm56752 | 0.147030916 | -1.485426827 | 0 | 0.018 | 1 | 0.730750993 | 2.017921908 | 0 | 0.012 | 1 | 0.730750993 | 0.272443742 |
| 4930592A05Rik | 0.147030916 | -4.392317423 | 0 | 0.018 | 1 | 0.698534498 | 2.432959407 | 0 | 0.015 | 1 | 0.698534498 | 0.272443742 |
| Mal2 | 0.147030916 | -1.485426827 | 0 | 0.018 | 1 | 0.385773258 | 0.432959407 | 0 | 0.071 | 1 | 0.385773258 | 0.272443742 |
| Amigo2 | 0.147030916 | -1.485426827 | 0 | 0.018 | 1 | 0.669843604 | 2.017921908 | 0 | 0.019 | 1 | 0.669843604 | 0.272443742 |
| Gm42477 | 0.147030916 | -1.485426827 | 0 | 0.018 | 1 | 0.669843604 | 1.558490289 | 0 | 0.019 | 1 | 0.669843604 | 0.272443742 |
| Gm49521 | 0.147030916 | -2.070389328 | 0 | 0.018 | 1 | 0.597824685 | 1.432959407 | 0 | 0.028 | 1 | 0.597824685 | 0.272443742 |
| P2rx6 | 0.147030916 | -1.485426827 | 0 | 0.018 | 1 | 0.643835681 | 1.847996907 | 0 | 0.022 | 1 | 0.643835681 | 0.272443742 |
| Stxbp5l | 0.147030916 | -2.070389328 | 0 | 0.018 | 1 | 0.643842184 | 1.558490289 | 0 | 0.022 | 1 | 0.643842184 | 0.272443742 |
| Plcxd2 | 0.147030916 | -1.485426827 | 0 | 0.018 | 1 | 0.730749233 | 2.432959407 | 0 | 0.012 | 1 | 0.730749233 | 0.272443742 |
| Vgll3 | 0.147030916 | -2.070389328 | 0 | 0.018 | 1 | 0.743727035 | 0.374065718 | 0.1 | 0.071 | 1 | 0.743727035 | 0.272443742 |

| | | | | | | | | | | | | |
|---|---|---|---|---|---|---|---|---|---|---|---|---|
| Ptx4 | 0.147030916 | -1.485426827 | 0 | 0.018 | 1 | 0.730750993 | 2.210566986 | 0 | 0.012 | 1 | 0.730750993 | 0.272443742 |
| Cbs | 0.147030916 | -2.070389328 | 0 | 0.018 | 1 | 0.669843604 | 2.017921908 | 0 | 0.019 | 1 | 0.669843604 | 0.272443742 |
| Ly6g6c | 0.147030916 | -1.485426827 | 0 | 0.018 | 1 | 0.698534498 | 2.432959407 | 0 | 0.015 | 1 | 0.698534498 | 0.272443742 |
| 9130008F23Rik | 0.147030916 | -1.485426827 | 0 | 0.018 | 1 | 0.643835681 | 1.695993813 | 0 | 0.022 | 1 | 0.643835681 | 0.272443742 |
| Gm17949 | 0.147030916 | -1.485426827 | 0 | 0.018 | 1 | 0.73075158 | 2.695993813 | 0 | 0.012 | 1 | 0.73075158 | 0.272443742 |
| Prob1 | 0.147030916 | -1.485426827 | 0 | 0.018 | 1 | 0.73075158 | 2.695993813 | 0 | 0.012 | 1 | 0.73075158 | 0.272443742 |
| Gm10053 | 0.147030916 | -1.485426827 | 0 | 0.018 | 1 | 0.669853054 | 2.210566986 | 0 | 0.019 | 1 | 0.669853054 | 0.272443742 |
| Hps6 | 0.147030916 | -1.485426827 | 0 | 0.018 | 1 | 0.489901827 | 0.847996907 | 0 | 0.046 | 1 | 0.489901827 | 0.272443742 |
| Bex1 | 0.147030916 | -1.485426827 | 0 | 0.018 | 1 | 0.669848329 | 1.847996907 | 0 | 0.019 | 1 | 0.669848329 | 0.272443742 |
| Zcchc18 | 0.147030916 | -1.485426827 | 0 | 0.018 | 1 | 0.669848329 | 1.847996907 | 0 | 0.019 | 1 | 0.669848329 | 0.272443742 |
| Abca8a | 0.147047659 | -1.292781749 | 0.008 | 0.035 | 1 | 0.669843604 | 2.017921908 | 0 | 0.019 | 1 | 0.669843604 | 0.272472304 |
| Bag6 | 0.147086905 | -0.40103464 | 0.294 | 0.371 | 1 | 0.166680237 | -0.959358016 | 0.2 | 0.398 | 1 | 0.166680237 | 0.272539252 |
| Tbc1d2b | 0.14711858 | -0.660513534 | 0.21 | 0.276 | 1 | 0.653624842 | -0.291933355 | 0.2 | 0.247 | 1 | 0.653624842 | 0.272593284 |
| Erp27 | 0.147165254 | -0.485426827 | 0.126 | 0.188 | 1 | 0.694884813 | 0.336097868 | 0.2 | 0.145 | 1 | 0.694884813 | 0.272672896 |
| Ndufa3 | 0.147214998 | 0.25766487 | 0.613 | 0.535 | 1 | 0.564217495 | -0.31836148 | 0.4 | 0.448 | 1 | 0.564217495 | 0.27275774 |
| Oxr1 | 0.198743922 | -0.35842155 | 0.714 | 0.724 | 1 | 0.147348925 | -1.002426737 | 0.6 | 0.741 | 1 | 0.198743922 | 0.272986144 |
| Oard1 | 0.982306823 | 0.114035243 | 0.269 | 0.265 | 1 | 0.147437462 | -1.152003093 | 0 | 0.176 | 1 | 0.982306823 | 0.273137118 |
| Mgat2 | 0.586505884 | -0.19592021 | 0.664 | 0.659 | 1 | 0.147437934 | -0.721296138 | 0.6 | 0.728 | 1 | 0.586505884 | 0.273137923 |
| Hacd2 | 0.147710209 | 0.232371356 | 0.655 | 0.559 | 1 | 0.387823916 | 0.405478671 | 0.7 | 0.586 | 1 | 0.387823916 | 0.273602113 |
| Slc27a4 | 0.147766108 | -0.900464326 | 0.092 | 0.147 | 1 | 0.589878624 | 0.526068812 | 0.3 | 0.219 | 1 | 0.589878624 | 0.273697394 |
| Tacstd2 | 0.692072298 | 1.225066556 | 0.042 | 0.053 | 1 | 0.147776142 | -1.736965594 | 0 | 0.176 | 1 | 0.692072298 | 0.273714496 |
| Gm17231 | 0.147787238 | -1.222392421 | 0.008 | 0.035 | 1 | 0.17652086 | 2.94753258 | 0.1 | 0.028 | 1 | 0.17652086 | 0.273733409 |
| Ankrd55 | 0.147787238 | -1.070389328 | 0.008 | 0.035 | 1 | 0.669852109 | 0.558490289 | 0 | 0.019 | 1 | 0.669852109 | 0.273733409 |
| Atf7ip2 | 0.147787238 | -0.944858446 | 0.008 | 0.035 | 1 | 0.669843604 | 2.017921908 | 0 | 0.019 | 1 | 0.669843604 | 0.273733409 |
| Fam53c | 0.147917619 | -0.531230517 | 0.218 | 0.288 | 1 | 0.815164809 | 0.769994395 | 0.3 | 0.336 | 1 | 0.815164809 | 0.273955615 |
| Zdhhc6 | 0.5739218 | -0.396890153 | 0.286 | 0.306 | 1 | 0.147993647 | 0.969558886 | 0.5 | 0.306 | 1 | 0.5739218 | 0.274085174 |
| Lnpk | 0.44010884 | -0.485426827 | 0.387 | 0.394 | 1 | 0.148090566 | 0.804928185 | 0.6 | 0.358 | 1 | 0.44010884 | 0.274250316 |
| Bbs4 | 0.15082099 | -1.04342228 | 0.126 | 0.188 | 1 | 0.148099608 | -1.391469028 | 0 | 0.176 | 1 | 0.15082099 | 0.274265723 |
| Cds2 | 0.613054053 | 0.232802204 | 0.294 | 0.271 | 1 | 0.148274222 | -0.159995884 | 0.1 | 0.361 | 1 | 0.613054053 | 0.2745632 |
| Caml | 0.14838506 | -0.29716019 | 0.487 | 0.571 | 1 | 0.181197692 | -0.830074999 | 0.3 | 0.5 | 1 | 0.181197692 | 0.274751993 |
| Gm39363 | 0.148540368 | -0.900464326 | 0.008 | 0.035 | 1 | 0.522132053 | 0.558490289 | 0 | 0.04 | 1 | 0.522132053 | 0.275016496 |
| Stab2 | 0.148540368 | -0.900464326 | 0.008 | 0.035 | 1 | 0.619953009 | 1.558490289 | 0 | 0.025 | 1 | 0.619953009 | 0.275016496 |
| Dnajc28 | 0.148540368 | -0.900464326 | 0.008 | 0.035 | 1 | 0.522115639 | 0.558490289 | 0 | 0.04 | 1 | 0.522115639 | 0.275016496 |
| Map6 | 0.153790546 | -0.117695043 | 0.067 | 0.118 | 1 | 0.148612307 | -1.763437806 | 0 | 0.176 | 1 | 0.153790546 | 0.275138996 |
| Hibadh | 0.692568911 | -0.158262084 | 0.689 | 0.665 | 1 | 0.148739658 | -0.713019899 | 0.5 | 0.704 | 1 | 0.692568911 | 0.275355831 |
| Las1l | 0.41710424 | 0.624197664 | 0.563 | 0.512 | 1 | 0.148747871 | 0.691192827 | 0.7 | 0.506 | 1 | 0.41710424 | 0.275369813 |
| Ccdc92 | 0.14882213 | 0.389042291 | 0.076 | 0.035 | 1 | 0.362872338 | 1.847996907 | 0.1 | 0.04 | 1 | 0.362872338 | 0.275496234 |
| Spsb1 | 0.148834243 | -0.446952679 | 0.639 | 0.647 | 1 | 0.968866177 | 0.870861015 | 0.6 | 0.679 | 1 | 0.968866177 | 0.275516855 |
| Dynlt2a2 | 0.291023996 | -1.070389328 | 0.042 | 0.071 | 1 | 0.148903646 | 1.744903414 | 0.2 | 0.074 | 1 | 0.291023996 | 0.275634996 |
| Gm3734 | 0.345791776 | -0.970853654 | 0.025 | 0.047 | 1 | 0.148903646 | 1.795529487 | 0.2 | 0.074 | 1 | 0.345791776 | 0.275634996 |
| Inpp1 | 0.148994586 | 0.251538767 | 0.378 | 0.294 | 1 | 0.218995436 | 1.071502948 | 0.5 | 0.358 | 1 | 0.218995436 | 0.275789785 |
| Galnt15 | 0.152147231 | -0.408805546 | 0.168 | 0.106 | 1 | 0.149182272 | 0.979786779 | 0.3 | 0.139 | 1 | 0.152147231 | 0.276109194 |
| Rnf14 | 0.149250013 | -0.460335846 | 0.529 | 0.559 | 1 | 0.510030619 | -0.433289204 | 0.6 | 0.599 | 1 | 0.510030619 | 0.276224459 |
| Gm16152 | 0.149285731 | -0.707819249 | 0.008 | 0.035 | 1 | 0.189486514 | 2.695993813 | 0.1 | 0.028 | 1 | 0.189486514 | 0.276285233 |
| Pgbd1 | 0.149285731 | -0.707819249 | 0.008 | 0.035 | 1 | 0.619953009 | 1.695993813 | 0 | 0.025 | 1 | 0.619953009 | 0.276285233 |
| Zfp174 | 0.149285731 | -0.707819249 | 0.008 | 0.035 | 1 | 0.669843604 | 2.017921908 | 0 | 0.019 | 1 | 0.669843604 | 0.276285233 |
| Cerox1 | 0.149285731 | -0.292781749 | 0.008 | 0.035 | 1 | 0.643850855 | 2.017921908 | 0 | 0.022 | 1 | 0.643850855 | 0.276285233 |
| Mrpl4 | 0.149337856 | 0.189611018 | 0.58 | 0.459 | 1 | 0.562103727 | -0.219117289 | 0.4 | 0.512 | 1 | 0.562103727 | 0.276373916 |
| Nav3 | 0.149359544 | 0.446132117 | 0.622 | 0.524 | 1 | 0.188286031 | -2.540498805 | 0.1 | 0.269 | 1 | 0.188286031 | 0.276410815 |
| Mphosph6 | 0.149429671 | -0.439033922 | 0.42 | 0.482 | 1 | 0.303257897 | -0.260062839 | 0.2 | 0.383 | 1 | 0.303257897 | 0.276530116 |
| Gm20632 | 0.167062889 | 1.836501268 | 0.025 | 0.006 | 1 | 0.149795061 | 2.847996907 | 0.1 | 0.025 | 1 | 0.167062889 | 0.277151562 |
| Tlcd3b | 0.168156185 | 1.514573173 | 0.025 | 0.006 | 1 | 0.149795061 | 2.847996907 | 0.1 | 0.025 | 1 | 0.168156185 | 0.277151562 |
| 2410022M11Rik | 0.30021769 | -0.485426827 | 0.05 | 0.082 | 1 | 0.149795061 | 2.847996907 | 0.1 | 0.025 | 1 | 0.30021769 | 0.277151562 |
| Gm57388 | 0.338041639 | 0.877143252 | 0.059 | 0.035 | 1 | 0.149795061 | 2.847996907 | 0.1 | 0.025 | 1 | 0.338041639 | 0.277151562 |
| Sec31b | 0.345346582 | -0.900464326 | 0.017 | 0.035 | 1 | 0.149795061 | 2.847996907 | 0.1 | 0.025 | 1 | 0.345346582 | 0.277151562 |
| Pcdh1 | 0.375395897 | 0.777607579 | 0.042 | 0.024 | 1 | 0.149795061 | 2.847996907 | 0.1 | 0.025 | 1 | 0.375395897 | 0.277151562 |
| Etaa1os | 0.890081511 | -0.273922722 | 0.067 | 0.071 | 1 | 0.149795061 | 2.847996907 | 0.1 | 0.025 | 1 | 0.890081511 | 0.277151562 |
| Pard3bos1 | 0.929388345 | 0.514573173 | 0.025 | 0.024 | 1 | 0.149795061 | 2.847996907 | 0.1 | 0.025 | 1 | 0.929388345 | 0.277151562 |
| Epn3 | 0.954551343 | -0.222392421 | 0.017 | 0.018 | 1 | 0.149795061 | 2.847996907 | 0.1 | 0.025 | 1 | 0.954551343 | 0.277151562 |
| Ogfr | 0.149805275 | -0.443606651 | 0.429 | 0.494 | 1 | 0.163560027 | -0.614346308 | 0.2 | 0.481 | 1 | 0.163560027 | 0.277168929 |
| Elk1 | 0.149877937 | 0.565199246 | 0.193 | 0.129 | 1 | 0.713159143 | 0.821524695 | 0.2 | 0.154 | 1 | 0.713159143 | 0.277292478 |
| Polr2g | 0.14995111 | -0.346599122 | 0.571 | 0.606 | 1 | 0.33470584 | -0.461049897 | 0.3 | 0.457 | 1 | 0.33470584 | 0.277416885 |
| Lpin1 | 0.149957824 | 0.144204723 | 0.412 | 0.329 | 1 | 0.460096609 | 0.219180116 | 0.4 | 0.265 | 1 | 0.460096609 | 0.277428299 |
| Tmem43 | 0.244429692 | -0.566726929 | 0.437 | 0.441 | 1 | 0.150317189 | -0.950168844 | 0.3 | 0.512 | 1 | 0.244429692 | 0.27803912 |
| Epn2 | 0.150382325 | -0.509539283 | 0.395 | 0.459 | 1 | 0.423894354 | -0.979571236 | 0.6 | 0.599 | 1 | 0.423894354 | 0.278149806 |
| Calm2 | 0.150387132 | 0.108235039 | 0.992 | 1 | 1 | 0.462071513 | -0.203373824 | 1 | 0.997 | 1 | 0.462071513 | 0.278157974 |
| Otulinl | 0.150467206 | -0.883976204 | 0.067 | 0.118 | 1 | 0.446680864 | 0.789103218 | 0.2 | 0.117 | 1 | 0.446680864 | 0.278294032 |
| Rfk | 0.150647751 | 0.160936218 | 0.529 | 0.429 | 1 | 0.779013778 | 0.155425432 | 0.5 | 0.577 | 1 | 0.779013778 | 0.278600757 |
| Prmt3 | 0.150656416 | -0.559427409 | 0.252 | 0.324 | 1 | 0.31032137 | -0.339630097 | 0.1 | 0.259 | 1 | 0.31032137 | 0.278615476 |
| Mon1b | 0.150707399 | -1.129283017 | 0.059 | 0.106 | 1 | 0.433541956 | 0.625604485 | 0 | 0.059 | 1 | 0.433541956 | 0.278702077 |
| Mrps15 | 0.62700499 | -0.139102078 | 0.513 | 0.429 | 1 | 0.15072656 | -0.467504919 | 0.2 | 0.466 | 1 | 0.62700499 | 0.278734624 |
| Col15a1 | 0.644519284 | -0.987927168 | 0.042 | 0.053 | 1 | 0.151071004 | 0.725140159 | 0.3 | 0.133 | 1 | 0.644519284 | 0.27931956 |
| Mif4gd | 0.151150608 | -0.892602209 | 0.176 | 0.235 | 1 | 0.434349365 | -0.457811523 | 0.1 | 0.194 | 1 | 0.434349365 | 0.279454709 |
| Atpsckmt | 0.151313075 | -0.647698256 | 0.16 | 0.224 | 1 | 0.435013519 | 1.210566986 | 0.3 | 0.21 | 1 | 0.435013519 | 0.279730503 |
| Ptcd1 | 0.151384434 | -0.833350131 | 0.067 | 0.118 | 1 | 0.213257061 | -0.68251781 | 0 | 0.136 | 1 | 0.213257061 | 0.279851621 |
| Bco2 | 0.226898421 | 0.514573173 | 0.059 | 0.029 | 1 | 0.151443299 | 2.695993813 | 0.1 | 0.025 | 1 | 0.226898421 | 0.279951526 |
| Hsbp1l1 | 0.23847629 | -1.222392421 | 0.017 | 0.041 | 1 | 0.151443299 | 2.695993813 | 0.1 | 0.025 | 1 | 0.23847629 | 0.279951526 |
| Spata21 | 0.331040183 | -1.292781749 | 0.008 | 0.024 | 1 | 0.151443299 | 2.695993813 | 0.1 | 0.025 | 1 | 0.331040183 | 0.279951526 |
| Gm43113 | 0.334204811 | -0.807354922 | 0.008 | 0.024 | 1 | 0.151443299 | 2.695993813 | 0.1 | 0.025 | 1 | 0.334204811 | 0.279951526 |
| Phlda2 | 0.338096132 | -0.807354922 | 0.025 | 0.047 | 1 | 0.151443299 | 2.695993813 | 0.1 | 0.025 | 1 | 0.338096132 | 0.279951526 |
| Gm5817 | 0.668121723 | -0.678071905 | 0.042 | 0.053 | 1 | 0.151443299 | 2.695993813 | 0.1 | 0.025 | 1 | 0.668121723 | 0.279951526 |
| Gm56589 | 0.929389703 | 0.192645078 | 0.025 | 0.024 | 1 | 0.151443299 | 2.695993813 | 0.1 | 0.025 | 1 | 0.929389703 | 0.279951526 |
| Zfp128 | 0.954551343 | -0.222392421 | 0.017 | 0.018 | 1 | 0.151443299 | 2.695993813 | 0.1 | 0.025 | 1 | 0.954551343 | 0.279951526 |
| Zfp7 | 0.954551343 | -0.222392421 | 0.017 | 0.018 | 1 | 0.151443299 | 2.695993813 | 0.1 | 0.025 | 1 | 0.954551343 | 0.279951526 |
| Gm14634 | 0.954551343 | -0.222392421 | 0.017 | 0.018 | 1 | 0.151443299 | 2.695993813 | 0.1 | 0.025 | 1 | 0.954551343 | 0.279951526 |
| Nkiras2 | 0.15157408 | -0.807354922 | 0.092 | 0.147 | 1 | 0.387282126 | 1.111031312 | 0.2 | 0.108 | 1 | 0.387282126 | 0.280173458 |
| 2900093K20Rik | 0.151661553 | -1.070389328 | 0.059 | 0.106 | 1 | 0.601875777 | 0.973527789 | 0.1 | 0.059 | 1 | 0.601875777 | 0.28032188 |
| Skp2 | 0.15169433 | -0.87244995 | 0.059 | 0.106 | 1 | 0.245667588 | -0.789433014 | 0 | 0.12 | 1 | 0.245667588 | 0.28037749 |
| Gnpda1 | 0.151705514 | -0.555816155 | 0.118 | 0.176 | 1 | 0.613082344 | 0.31748219 | 0.3 | 0.213 | 1 | 0.613082344 | 0.280396465 |
| Grk6 | 0.255425453 | 0.321928095 | 0.37 | 0.3 | 1 | 0.15178164 | 1.19592021 | 0.4 | 0.225 | 1 | 0.255425453 | 0.280525614 |
| Slc16a4 | 0.151806297 | -0.341574575 | 0.714 | 0.724 | 1 | 0.164030395 | -0.515721815 | 0.6 | 0.799 | 1 | 0.164030395 | 0.280567442 |
| Kifbp | 0.151856307 | -0.573864792 | 0.529 | 0.529 | 1 | 0.3171341 | -0.793108672 | 0.5 | 0.562 | 1 | 0.3171341 | 0.280652276 |
| R3hcc1l | 0.152144915 | -0.485426827 | 0.387 | 0.441 | 1 | 0.260706629 | 0.614199722 | 0.8 | 0.531 | 1 | 0.260706629 | 0.281141756 |
| Sgpl1 | 0.404651346 | -0.406816992 | 0.361 | 0.394 | 1 | 0.152192688 | -1.213642159 | 0.6 | 0.66 | 1 | 0.404651346 | 0.281222761 |
| Smg8 | 0.161037717 | -0.637429921 | 0.126 | 0.188 | 1 | 0.152215872 | -1.171902651 | 0 | 0.173 | 1 | 0.161037717 | 0.281262072 |
| Ndufb4 | 0.152294103 | 0.150416843 | 0.597 | 0.494 | 1 | 0.703492149 | 0.710493383 | 0.5 | 0.432 | 1 | 0.703492149 | 0.281394713 |
| Erp29 | 0.152380344 | 0.133668358 | 0.975 | 0.871 | 1 | 0.443898367 | 0.444075439 | 1 | 0.877 | 1 | 0.443898367 | 0.281540919 |
| Susd5 | 0.152434968 | 0.127466606 | 0.983 | 0.888 | 1 | 0.701619029 | -0.152003093 | 1 | 0.87 | 1 | 0.701619029 | 0.281633517 |

| | | | | | | | | | | | | |
|---|---|---|---|---|---|---|---|---|---|---|---|---|
| Tspan32 | 0.225554032 | 0.85160816 | 0.118 | 0.076 | 1 | 0.153060084 | 1.744903414 | 0.2 | 0.074 | 1 | 0.225554032 | 0.282692778 |
| Emc7 | 0.195169425 | 0.137458157 | 0.916 | 0.824 | 1 | 0.153147371 | -0.567040593 | 0.7 | 0.892 | 1 | 0.195169425 | 0.282840625 |
| Gm17359 | 0.334204811 | -0.807354922 | 0.008 | 0.024 | 1 | 0.153152701 | 2.558490289 | 0.1 | 0.025 | 1 | 0.334204811 | 0.282849652 |
| Mm2pr | 0.334204811 | -0.807354922 | 0.008 | 0.024 | 1 | 0.153152701 | 2.31748219 | 0.1 | 0.025 | 1 | 0.334204811 | 0.282849652 |
| 2610029K11Rik | 0.62441508 | -0.485426827 | 0.025 | 0.035 | 1 | 0.153152701 | 2.111031312 | 0.1 | 0.025 | 1 | 0.62441508 | 0.282849652 |
| Platr22 | 0.935790323 | -0.485426827 | 0.025 | 0.024 | 1 | 0.153152701 | 2.558490289 | 0.1 | 0.025 | 1 | 0.935790323 | 0.282849652 |
| Pknox1 | 0.15315803 | -0.635669463 | 0.227 | 0.294 | 1 | 0.736369693 | -0.581990934 | 0.3 | 0.309 | 1 | 0.736369693 | 0.282858677 |
| Dpy19l4 | 0.197511036 | -0.298056691 | 0.378 | 0.453 | 1 | 0.153185236 | -1.080110175 | 0.3 | 0.478 | 1 | 0.197511036 | 0.282904755 |
| Gnl1 | 0.153219578 | -0.560714954 | 0.387 | 0.441 | 1 | 0.569228897 | -0.224571719 | 0.4 | 0.503 | 1 | 0.569228897 | 0.282962917 |
| Eefsec | 0.153336608 | -0.707819249 | 0.16 | 0.224 | 1 | 0.599702147 | 0.506959989 | 0.3 | 0.225 | 1 | 0.599702147 | 0.2831611 |
| Ptpn12 | 0.153575018 | -0.327278994 | 0.639 | 0.7 | 1 | 0.22805107 | -0.747924806 | 0.6 | 0.793 | 1 | 0.22805107 | 0.283564751 |
| Kif3b | 0.153636747 | -0.589763487 | 0.269 | 0.329 | 1 | 0.362647914 | -0.789433014 | 0.3 | 0.398 | 1 | 0.362647914 | 0.283669244 |
| Abr | 0.153756691 | -0.750021747 | 0.361 | 0.429 | 1 | 0.534763129 | -0.297433533 | 0.5 | 0.571 | 1 | 0.534763129 | 0.283872262 |
| Gm34455 | 0.154028519 | -2.089498151 | 0.109 | 0.165 | 1 | 0.872093741 | 0.515421567 | 0.2 | 0.256 | 1 | 0.872093741 | 0.284332253 |
| Pigu | 0.154268479 | -0.587788545 | 0.235 | 0.3 | 1 | 0.901763896 | -0.201246612 | 0.3 | 0.287 | 1 | 0.901763896 | 0.284738195 |
| Tmem65 | 0.895294369 | -0.156289401 | 0.874 | 0.841 | 1 | 0.154450653 | -0.40354186 | 0.8 | 0.951 | 1 | 0.895294369 | 0.285046302 |
| Dag1 | 0.154492133 | -0.283237977 | 0.891 | 0.835 | 1 | 0.842540337 | -0.146266724 | 1 | 0.92 | 1 | 0.842540337 | 0.285116447 |
| Gm14858 | 0.479565188 | -0.572889668 | 0.05 | 0.071 | 1 | 0.154638992 | 2.432959407 | 0.2 | 0.083 | 1 | 0.479565188 | 0.285364767 |
| Tbc1d22b | 0.154753605 | -0.765534746 | 0.126 | 0.188 | 1 | 0.437706179 | -0.709998547 | 0.1 | 0.191 | 1 | 0.437706179 | 0.285558532 |
| Efna3 | 0.243996811 | -0.944858446 | 0.025 | 0.053 | 1 | 0.154873683 | 2.432959407 | 0.1 | 0.025 | 1 | 0.243996811 | 0.285761508 |
| Gm13562 | 0.33261065 | -1.070389328 | 0.008 | 0.024 | 1 | 0.154873683 | 2.432959407 | 0.1 | 0.025 | 1 | 0.33261065 | 0.285761508 |
| Insl6 | 0.382796898 | 1.099535674 | 0.034 | 0.018 | 1 | 0.154873683 | 2.432959407 | 0.1 | 0.025 | 1 | 0.382796898 | 0.285761508 |
| Mrpl38 | 0.154981494 | -0.435386145 | 0.269 | 0.347 | 1 | 0.718016741 | -0.357117523 | 0.3 | 0.324 | 1 | 0.718016741 | 0.285943724 |
| Pip5k1c | 0.155035158 | -0.678071905 | 0.193 | 0.259 | 1 | 0.340167781 | -0.279758641 | 0.1 | 0.25 | 1 | 0.340167781 | 0.286034416 |
| Kctd10 | 0.163711928 | -0.584962501 | 0.277 | 0.341 | 1 | 0.15504225 | -0.840059087 | 0.3 | 0.512 | 1 | 0.163711928 | 0.2860464 |
| Tex2 | 0.155309732 | -0.368613162 | 0.218 | 0.294 | 1 | 0.594651341 | 0.625604485 | 0.5 | 0.525 | 1 | 0.594651341 | 0.286498352 |
| C4b | 0.25496283 | 0.328160049 | 0.109 | 0.071 | 1 | 0.155334127 | 1.210566986 | 0.4 | 0.21 | 1 | 0.25496283 | 0.286539563 |
| Spred3 | 0.155443856 | -1.201633861 | 0.05 | 0.094 | 1 | 0.957826589 | 0.625604485 | 0.1 | 0.093 | 1 | 0.957826589 | 0.28672492 |
| Hyls1 | 0.155445315 | -1.678071905 | 0.05 | 0.094 | 1 | 0.852081709 | 0.494359952 | 0.1 | 0.08 | 1 | 0.852081709 | 0.286727384 |
| ENSMUSG00000121485 | 0.155445315 | -1.263034406 | 0.05 | 0.094 | 1 | 0.23905333 | -1.736965594 | 0 | 0.123 | 1 | 0.23905333 | 0.286727384 |
| Ccdc90b | 0.874471331 | -0.197939378 | 0.597 | 0.559 | 1 | 0.155532874 | -0.926936538 | 0.3 | 0.509 | 1 | 0.874471331 | 0.286875274 |
| Slc35e4 | 0.155704528 | 0.295006796 | 0.714 | 0.582 | 1 | 0.386521321 | -0.513979349 | 0.6 | 0.664 | 1 | 0.386521321 | 0.287165157 |
| Ahr | 0.155714059 | 0.96203215 | 0.16 | 0.106 | 1 | 0.559651619 | -0.248864633 | 0.1 | 0.167 | 1 | 0.559651619 | 0.28718125 |
| Nherf1 | 0.669521295 | -0.485426827 | 0.042 | 0.053 | 1 | 0.155715549 | -1.004445905 | 0 | 0.17 | 1 | 0.669521295 | 0.287183766 |
| Opa1 | 0.46406097 | -0.401213823 | 0.555 | 0.529 | 1 | 0.155771277 | -1.127755547 | 0.3 | 0.503 | 1 | 0.46406097 | 0.287277863 |
| Cadps2 | 0.175676122 | -1.163498732 | 0.025 | 0.059 | 1 | 0.155924918 | -1.026472211 | 0 | 0.17 | 1 | 0.175676122 | 0.287537256 |
| Clk4 | 0.401815517 | -0.279277485 | 0.79 | 0.776 | 1 | 0.156030723 | -0.592575685 | 0.7 | 0.778 | 1 | 0.401815517 | 0.287715859 |
| Mrpl39 | 0.15607466 | -0.608809243 | 0.202 | 0.271 | 1 | 0.489318462 | 0.660369903 | 0.3 | 0.201 | 1 | 0.489318462 | 0.28779002 |
| Vkorc1 | 0.156247416 | -0.177404967 | 0.933 | 0.871 | 1 | 0.728268399 | 0.173397193 | 0.9 | 0.867 | 1 | 0.728268399 | 0.288081577 |
| Hfe | 0.3024216 | -0.555816155 | 0.076 | 0.112 | 1 | 0.156369641 | -1.069540933 | 0 | 0.17 | 1 | 0.3024216 | 0.288287818 |
| Aqp5 | 0.636250625 | 0.321928095 | 0.025 | 0.035 | 1 | 0.156456005 | -1.321928095 | 0 | 0.17 | 1 | 0.636250625 | 0.288433528 |
| Khk | 0.156466554 | 0.285091327 | 0.21 | 0.141 | 1 | 0.175396096 | -0.959358016 | 0 | 0.157 | 1 | 0.175396096 | 0.288451326 |
| Rhno1 | 0.156585639 | -0.847996907 | 0.109 | 0.165 | 1 | 0.190480543 | -0.763437806 | 0 | 0.148 | 1 | 0.190480543 | 0.288652216 |
| Gm15283 | 0.156605262 | 0.860569028 | 0.37 | 0.282 | 1 | 0.417115963 | 0.359710425 | 0.2 | 0.346 | 1 | 0.417115963 | 0.288685315 |
| Dnah2 | 0.216681675 | 2.407657969 | 0.05 | 0.024 | 1 | 0.156606269 | 1.625604485 | 0.1 | 0.025 | 1 | 0.216681675 | 0.288687014 |
| Cfap54 | 0.387377553 | 1.251538767 | 0.025 | 0.012 | 1 | 0.156606269 | 2.017921908 | 0.1 | 0.025 | 1 | 0.387377553 | 0.288687014 |
| Obp2b | 0.913539018 | -0.17055349 | 0.227 | 0.235 | 1 | 0.156606693 | 0.111031312 | 0.1 | 0.025 | 1 | 0.913539018 | 0.288687729 |
| Npc2 | 0.156877752 | 0.257415333 | 0.983 | 0.941 | 1 | 0.799223329 | -0.107491562 | 0.7 | 0.886 | 1 | 0.799223329 | 0.289144875 |
| Cyp4f17 | 0.156966347 | -0.807354922 | 0.185 | 0.247 | 1 | 0.418286619 | -0.339630097 | 0.1 | 0.201 | 1 | 0.418286619 | 0.289294259 |
| Sox8 | 0.156996535 | -0.503348735 | 0.168 | 0.235 | 1 | 0.499781863 | 1.111031312 | 0.2 | 0.136 | 1 | 0.499781863 | 0.289345158 |
| Ifit2 | 0.990433705 | -0.529820947 | 0.109 | 0.106 | 1 | 0.157250184 | 1.660369903 | 0.2 | 0.077 | 1 | 0.990433705 | 0.289772748 |
| Arsg | 0.7852007 | 0.266645659 | 0.084 | 0.094 | 1 | 0.15728729 | 1.515421567 | 0.2 | 0.074 | 1 | 0.7852007 | 0.289835288 |
| Agrn | 0.1573007 | -0.619282574 | 0.252 | 0.312 | 1 | 0.923929162 | 0.558490289 | 0.3 | 0.296 | 1 | 0.923929162 | 0.28985789 |
| Med19 | 0.157498804 | -0.408805546 | 0.336 | 0.406 | 1 | 0.515250554 | 0.874963954 | 0.4 | 0.336 | 1 | 0.515250554 | 0.290191734 |
| Fut8 | 0.157765077 | -0.409573073 | 0.672 | 0.7 | 1 | 0.30906654 | 0.248267695 | 0.8 | 0.91 | 1 | 0.30906654 | 0.290640334 |
| Lfng | 0.157892227 | 0.893084796 | 0.126 | 0.076 | 1 | 0.505639431 | 0.769994395 | 0 | 0.043 | 1 | 0.505639431 | 0.290854499 |
| Gm17251 | 0.707499527 | -0.163498732 | 0.109 | 0.124 | 1 | 0.157901591 | 2.473601392 | 0.2 | 0.08 | 1 | 0.707499527 | 0.290870269 |
| Spout1 | 0.158076521 | -0.244418728 | 0.151 | 0.224 | 1 | 0.336366819 | 1.31748219 | 0.2 | 0.102 | 1 | 0.336366819 | 0.291164855 |
| 4930435F18Rik | 0.158083295 | -2.424026283 | 0.017 | 0.047 | 1 | 0.557771538 | 0.769994395 | 0 | 0.034 | 1 | 0.557771538 | 0.291176261 |
| Rad51d | 0.158141597 | -1.137503524 | 0.05 | 0.094 | 1 | 0.746554885 | 0.973527789 | 0.1 | 0.071 | 1 | 0.746554885 | 0.29127443 |
| Zfand1 | 0.158142332 | -1.378511623 | 0.05 | 0.094 | 1 | 0.216297137 | 1.473601392 | 0.2 | 0.083 | 1 | 0.216297137 | 0.291275666 |
| Aifm1 | 0.158245335 | -0.363436303 | 0.437 | 0.506 | 1 | 0.363084831 | 0.757394358 | 0.5 | 0.438 | 1 | 0.363084831 | 0.291449084 |
| Aasdhppt | 0.158269975 | -0.499502012 | 0.336 | 0.394 | 1 | 0.446803188 | 0.280956314 | 0.2 | 0.34 | 1 | 0.446803188 | 0.291490565 |
| Gm42743 | 0.324971094 | -1.5360529 | 0.034 | 0.059 | 1 | 0.158347926 | 2.111031312 | 0.1 | 0.025 | 1 | 0.324971094 | 0.291621787 |
| Thsd4 | 0.222670078 | -0.158616511 | 0.655 | 0.559 | 1 | 0.158356279 | -1.621806396 | 0.6 | 0.713 | 1 | 0.222670078 | 0.291635846 |
| Zrsr2 | 0.158490174 | -0.480190179 | 0.689 | 0.653 | 1 | 0.804859169 | -0.166245016 | 0.7 | 0.698 | 1 | 0.804859169 | 0.291861213 |
| Scn5a | 0.158535694 | -1.572889668 | 0.042 | 0.082 | 1 | 0.474988533 | 0.210566986 | 0 | 0.049 | 1 | 0.474988533 | 0.291937822 |
| Mipol1 | 0.158723454 | 0.227291221 | 0.378 | 0.282 | 1 | 0.306541227 | -0.831743819 | 0.2 | 0.336 | 1 | 0.306541227 | 0.292253773 |
| Rhoc | 0.243173623 | 0.162400941 | 0.966 | 0.882 | 1 | 0.159021764 | -0.453893435 | 0.7 | 0.901 | 1 | 0.243173623 | 0.292755606 |
| Barx2 | 0.159084391 | 0.173536255 | 0.185 | 0.118 | 1 | 0.743727433 | 0.591657153 | 0.1 | 0.071 | 1 | 0.743727433 | 0.292860938 |
| Ubqln2 | 0.382111096 | -0.35767128 | 0.37 | 0.4 | 1 | 0.159344635 | -0.584962501 | 0.2 | 0.454 | 1 | 0.382111096 | 0.293298558 |
| Snai2 | 0.15943156 | -0.461381353 | 0.42 | 0.465 | 1 | 0.325918603 | 0.582535763 | 0.5 | 0.333 | 1 | 0.325918603 | 0.293444698 |
| Vegfc | 0.159448562 | -1.485426827 | 0.042 | 0.082 | 1 | 0.245813088 | -1.131825212 | 0 | 0.12 | 1 | 0.245813088 | 0.293473281 |
| Pola2 | 0.159448562 | -1.424026283 | 0.042 | 0.082 | 1 | 0.667604536 | 1.31748219 | 0.1 | 0.065 | 1 | 0.667604536 | 0.293473281 |
| 2610020C07Rik | 0.523187922 | -0.141472426 | 0.168 | 0.194 | 1 | 0.159547298 | 0.617042472 | 0.5 | 0.281 | 1 | 0.523187922 | 0.293639256 |
| Eri1 | 0.159585494 | -0.822461814 | 0.261 | 0.318 | 1 | 0.349520121 | -0.362899876 | 0.1 | 0.241 | 1 | 0.349520121 | 0.293703458 |
| Eif2s3y | 0.15980141 | -0.543641441 | 0.328 | 0.4 | 1 | 0.791860818 | 0.151673297 | 0.6 | 0.679 | 1 | 0.791860818 | 0.29406633 |
| Aifm2 | 0.675012197 | -0.207892852 | 0.118 | 0.135 | 1 | 0.159812798 | 1.185031894 | 0.3 | 0.136 | 1 | 0.675012197 | 0.294085466 |
| Limk2 | 0.159962949 | -0.504285854 | 0.202 | 0.265 | 1 | 0.361240817 | 0.835718577 | 0.5 | 0.41 | 1 | 0.361240817 | 0.294337752 |
| C2cd5 | 0.160003605 | -0.727501614 | 0.286 | 0.341 | 1 | 0.896800168 | -0.677886361 | 0.5 | 0.401 | 1 | 0.896800168 | 0.294406057 |
| Esrp1 | 0.355734963 | -0.292781749 | 0.017 | 0.035 | 1 | 0.1602802 | -0.91281543 | 0 | 0.167 | 1 | 0.355734963 | 0.294870657 |
| Tbx3 | 0.16040337 | -0.586964854 | 0.261 | 0.329 | 1 | 0.230735416 | -1.276698841 | 0.1 | 0.259 | 1 | 0.230735416 | 0.295077498 |
| Rala | 0.160551152 | -0.410239331 | 0.613 | 0.641 | 1 | 0.881103116 | 0.105384749 | 0.7 | 0.704 | 1 | 0.881103116 | 0.295325631 |
| Mettl4 | 0.160594548 | -0.847996907 | 0.092 | 0.147 | 1 | 0.731391744 | 0.374065718 | 0.2 | 0.151 | 1 | 0.731391744 | 0.295398486 |
| Prune1 | 0.160621501 | -0.516453723 | 0.143 | 0.206 | 1 | 0.429693212 | 0.249737583 | 0.1 | 0.225 | 1 | 0.429693212 | 0.295443736 |
| Tmed10 | 0.160669994 | 0.119435231 | 1 | 0.988 | 1 | 0.195691494 | -0.354463612 | 1 | 0.981 | 1 | 0.195691494 | 0.295525141 |
| Nanp | 0.160762395 | -0.889817082 | 0.126 | 0.182 | 1 | 0.672105081 | 0.111031312 | 0.1 | 0.145 | 1 | 0.672105081 | 0.295680242 |
| Copz1 | 0.872464905 | -0.163498732 | 0.647 | 0.594 | 1 | 0.160827565 | -0.755923752 | 0.4 | 0.627 | 1 | 0.872464905 | 0.295789624 |
| Sympk | 0.169045485 | -0.744161096 | 0.218 | 0.271 | 1 | 0.160908518 | -1.258202497 | 0.1 | 0.296 | 1 | 0.169045485 | 0.295925485 |
| Cog1 | 0.161037717 | -0.563429339 | 0.126 | 0.188 | 1 | 0.161677071 | 2.079322453 | 0.4 | 0.222 | 1 | 0.161677071 | 0.296142288 |
| Rpusd4 | 0.161040837 | -1.03562391 | 0.109 | 0.165 | 1 | 0.790243401 | 0.31748219 | 0.1 | 0.127 | 1 | 0.790243401 | 0.296147522 |
| Cox5b-ps | 0.161040979 | -0.765534746 | 0.109 | 0.165 | 1 | 0.886171989 | 1.111031312 | 0.1 | 0.086 | 1 | 0.886171989 | 0.296147761 |
| Akip1 | 0.347490014 | -0.485426827 | 0.109 | 0.147 | 1 | 0.161078754 | -1.152003093 | 0 | 0.167 | 1 | 0.347490014 | 0.296211142 |
| Gm35188 | 0.947974969 | 0.621488377 | 0.118 | 0.118 | 1 | 0.161342775 | -1.374395515 | 0 | 0.167 | 1 | 0.947974969 | 0.296654059 |
| Trmt1 | 0.161522022 | -0.502500341 | 0.311 | 0.376 | 1 | 0.598589603 | 0.880418384 | 0.4 | 0.355 | 1 | 0.598589603 | 0.29695468 |
| Gsg1l | 0.443210507 | -0.359895945 | 0.034 | 0.053 | 1 | 0.161590018 | 0.902444691 | 0.2 | 0.074 | 1 | 0.443210507 | 0.297068702 |
| Rpa1 | 0.161608083 | 0.362570079 | 0.261 | 0.188 | 1 | 0.384821966 | 1.017921908 | 0.3 | 0.194 | 1 | 0.384821966 | 0.297098994 |
| Ublcp1 | 0.161920538 | -0.57140005 | 0.311 | 0.376 | 1 | 0.458994564 | 0.486540447 | 0.5 | 0.364 | 1 | 0.458994564 | 0.297622816 |

| | | | | | | | | | | | | |
|---|---|---|---|---|---|---|---|---|---|---|---|---|
| Zfr2 | 0.161955797 | -1.807354922 | 0.034 | 0.071 | 1 | 0.868037657 | 0.888638891 | 0.1 | 0.083 | 1 | 0.868037657 | 0.297681914 |
| Gm42439 | 0.162091423 | 0.440572591 | 0.227 | 0.159 | 1 | 0.518656139 | -0.457811523 | 0.1 | 0.173 | 1 | 0.518656139 | 0.297909217 |
| Cntn3 | 0.162101639 | -1.807354922 | 0.017 | 0.047 | 1 | 0.669848329 | 1.847996907 | 0 | 0.019 | 1 | 0.669848329 | 0.297926336 |
| Ccn3 | 0.1621032 | -2.070389328 | 0.017 | 0.047 | 1 | 0.489984083 | -0.473931188 | 0 | 0.046 | 1 | 0.489984083 | 0.297928953 |
| Rtl8b | 0.23525384 | -0.382333334 | 0.378 | 0.429 | 1 | 0.162235254 | -0.848326703 | 0.2 | 0.404 | 1 | 0.23525384 | 0.29815023 |
| Glrx2 | 0.542566816 | 0.119941884 | 0.487 | 0.465 | 1 | 0.162274907 | -0.730270942 | 0.2 | 0.441 | 1 | 0.542566816 | 0.298216668 |
| Hs2st1 | 0.162437218 | 0.455961013 | 0.723 | 0.688 | 1 | 0.234729614 | 0.782705446 | 0.7 | 0.667 | 1 | 0.234729614 | 0.298488586 |
| Tceal1 | 0.26178811 | -0.637429921 | 0.269 | 0.306 | 1 | 0.162738057 | -1.131825212 | 0.1 | 0.299 | 1 | 0.26178811 | 0.298992439 |
| Tmem218 | 0.162753693 | 0.203371485 | 0.378 | 0.288 | 1 | 0.556174309 | 1.152851488 | 0.3 | 0.231 | 1 | 0.556174309 | 0.299018621 |
| Rogdi | 0.162943225 | 0.695145418 | 0.126 | 0.076 | 1 | 0.767699782 | 1.263034406 | 0.1 | 0.074 | 1 | 0.767699782 | 0.299335956 |
| Lrch4 | 0.163072549 | -0.733354341 | 0.059 | 0.106 | 1 | 0.944635256 | 0.973527789 | 0.1 | 0.093 | 1 | 0.944635256 | 0.299552442 |
| Chmp6 | 0.163122394 | -0.807354922 | 0.143 | 0.2 | 1 | 0.497579023 | 0.982297998 | 0.3 | 0.222 | 1 | 0.497579023 | 0.299635872 |
| Sec61a2 | 0.163137142 | -0.485426827 | 0.487 | 0.524 | 1 | 0.248686806 | -0.922244842 | 0.4 | 0.503 | 1 | 0.248686806 | 0.299660557 |
| Mknk2 | 0.476557833 | -0.302948775 | 0.546 | 0.535 | 1 | 0.163364818 | -0.567040593 | 0.5 | 0.741 | 1 | 0.476557833 | 0.300041572 |
| Trmt2b | 0.163411338 | 0.243271151 | 0.21 | 0.141 | 1 | 0.290445435 | -0.505640048 | 0 | 0.102 | 1 | 0.290445435 | 0.300119411 |
| Katnb1 | 0.163629046 | -0.925999419 | 0.05 | 0.094 | 1 | 0.433608413 | 0.494359952 | 0 | 0.059 | 1 | 0.433608413 | 0.300483627 |
| Mrps25 | 0.163708115 | -0.528928466 | 0.37 | 0.412 | 1 | 0.41017892 | -0.388070452 | 0.3 | 0.435 | 1 | 0.41017892 | 0.300615883 |
| Txndc11 | 0.163722176 | -0.485426827 | 0.529 | 0.547 | 1 | 0.180182293 | -0.885959938 | 0.5 | 0.642 | 1 | 0.180182293 | 0.300639402 |
| Ccl27a | 0.163777662 | -0.854660637 | 0.084 | 0.135 | 1 | 0.907531249 | 0.625604485 | 0.1 | 0.111 | 1 | 0.907531249 | 0.300732202 |
| Snrpb2 | 0.163826313 | -0.214625538 | 0.655 | 0.659 | 1 | 0.291555207 | -0.596787936 | 0.5 | 0.596 | 1 | 0.291555207 | 0.300813565 |
| Gm28050 | 0.164118052 | -1.555816155 | 0.034 | 0.071 | 1 | 0.766391805 | 0.31748219 | 0.1 | 0.071 | 1 | 0.766391805 | 0.301301369 |
| Idh3a | 0.16424322 | -0.608283575 | 0.328 | 0.382 | 1 | 0.320671211 | -0.399930607 | 0.2 | 0.373 | 1 | 0.320671211 | 0.301510605 |
| Tmem205 | 0.336423682 | -0.275503758 | 0.303 | 0.347 | 1 | 0.164317264 | 0.951832718 | 0.5 | 0.309 | 1 | 0.336423682 | 0.301634366 |
| Vrk1 | 0.164527912 | -1.137503524 | 0.101 | 0.153 | 1 | 0.803964451 | 0.695993813 | 0.2 | 0.176 | 1 | 0.803964451 | 0.301986391 |
| 1700085D22Rik | 0.164630923 | -0.669851398 | 0.059 | 0.106 | 1 | 0.94011655 | 0.432959407 | 0.1 | 0.105 | 1 | 0.94011655 | 0.302158505 |
| Abtb3 | 0.604448998 | -0.495843443 | 0.277 | 0.288 | 1 | 0.164671834 | -2.131825212 | 0.1 | 0.287 | 1 | 0.604448998 | 0.302226855 |
| Mipep | 0.164680406 | -0.250961574 | 0.118 | 0.176 | 1 | 0.656340368 | 0.473601392 | 0.2 | 0.142 | 1 | 0.656340368 | 0.302241176 |
| Cmbl | 0.582001671 | -0.263034406 | 0.311 | 0.259 | 1 | 0.164697052 | -0.304006187 | 0.1 | 0.349 | 1 | 0.582001671 | 0.302268986 |
| Sh2b2 | 0.164699155 | -1.070389328 | 0.05 | 0.094 | 1 | 0.965736751 | 0.432959407 | 0.1 | 0.093 | 1 | 0.965736751 | 0.302272499 |
| Dda1 | 0.164761377 | -0.316299556 | 0.597 | 0.629 | 1 | 0.709586045 | 0.125530882 | 0.7 | 0.534 | 1 | 0.709586045 | 0.302376443 |
| Zfp706 | 0.180867833 | -0.211393422 | 0.832 | 0.835 | 1 | 0.164788881 | -0.545666942 | 0.6 | 0.886 | 1 | 0.180867833 | 0.302422386 |
| Scamp3 | 0.164792212 | 0.155170373 | 0.748 | 0.612 | 1 | 0.191694239 | 0.988174565 | 0.6 | 0.503 | 1 | 0.191694239 | 0.302427951 |
| Zfp937 | 0.164808002 | -1.359895945 | 0.017 | 0.047 | 1 | 0.669848329 | 1.847996907 | 0 | 0.019 | 1 | 0.669848329 | 0.302454327 |
| Gm17494 | 0.164808002 | -1.359895945 | 0.017 | 0.047 | 1 | 0.619953009 | 1.558490289 | 0 | 0.025 | 1 | 0.619953009 | 0.302454327 |
| 9530046B11Rik | 0.164808002 | -1.359895945 | 0.017 | 0.047 | 1 | 0.276546558 | 2.210566986 | 0.1 | 0.034 | 1 | 0.276546558 | 0.302454327 |
| Zfp764 | 0.164808002 | -1.359895945 | 0.017 | 0.047 | 1 | 0.370536645 | 1.625604485 | 0.1 | 0.04 | 1 | 0.370536645 | 0.302454327 |
| Gm48604 | 0.164808002 | -1.359895945 | 0.017 | 0.047 | 1 | 0.474928012 | 0.558490289 | 0 | 0.049 | 1 | 0.474928012 | 0.302454327 |
| Acss3 | 0.164808002 | -1.485426827 | 0.017 | 0.047 | 1 | 0.410092532 | -1.037360528 | 0.1 | 0.194 | 1 | 0.410092532 | 0.302454327 |
| 9330136K24Rik | 0.164808002 | -1.707819249 | 0.017 | 0.047 | 1 | 0.643850855 | 2.017921908 | 0 | 0.022 | 1 | 0.643850855 | 0.302454327 |
| Zfp870 | 0.164808002 | -1.359895945 | 0.017 | 0.047 | 1 | 0.539413885 | 1.210566986 | 0 | 0.037 | 1 | 0.539413885 | 0.302454327 |
| Cxcl17 | 0.164841518 | -1.733354341 | 0.017 | 0.047 | 1 | 0.290261107 | -0.267480311 | 0 | 0.102 | 1 | 0.290261107 | 0.302510309 |
| Arhgap27 | 0.164907643 | 2.321928095 | 0.025 | 0.006 | 1 | 0.318562315 | 2.111031312 | 0.1 | 0.037 | 1 | 0.318562315 | 0.302620755 |
| Snrnp40 | 0.164984509 | -0.453246421 | 0.395 | 0.453 | 1 | 0.292778169 | -0.648834684 | 0.3 | 0.463 | 1 | 0.292778169 | 0.30274913 |
| Gcsh | 0.333086021 | -0.336283388 | 0.37 | 0.406 | 1 | 0.165145577 | 0.888638891 | 0.5 | 0.296 | 1 | 0.333086021 | 0.303018092 |
| Prss35 | 0.165148208 | -2.292781749 | 0.025 | 0.059 | 1 | 0.409514316 | 1.31748219 | 0.1 | 0.043 | 1 | 0.409514316 | 0.303022485 |
| Miga2 | 0.481022347 | 0.234465254 | 0.101 | 0.076 | 1 | 0.165155638 | 1.339850003 | 0.3 | 0.145 | 1 | 0.481022347 | 0.303034891 |
| Casz1 | 0.449198523 | -0.263034406 | 0.034 | 0.053 | 1 | 0.165167706 | -1.111361109 | 0 | 0.164 | 1 | 0.449198523 | 0.303055041 |
| Kyat1 | 0.165171017 | -0.944858446 | 0.05 | 0.094 | 1 | 0.837907635 | 1.695993813 | 0.1 | 0.09 | 1 | 0.837907635 | 0.303060569 |
| Gpank1 | 0.165171017 | -1.008988783 | 0.05 | 0.094 | 1 | 0.252267609 | -0.505640048 | 0 | 0.117 | 1 | 0.252267609 | 0.303060569 |
| Unc93b1 | 0.165325111 | -0.581642142 | 0.176 | 0.241 | 1 | 0.990738483 | 0.695993813 | 0.2 | 0.216 | 1 | 0.990738483 | 0.30331783 |
| Tor1aip2 | 0.165388696 | -0.312923282 | 0.899 | 0.888 | 1 | 0.353151504 | -0.355958267 | 0.7 | 0.944 | 1 | 0.353151504 | 0.303423972 |
| Rab15 | 0.165486507 | -0.748461233 | 0.059 | 0.106 | 1 | 0.298735402 | -0.473931188 | 0 | 0.099 | 1 | 0.298735402 | 0.303587229 |
| Abhd14b | 0.165488954 | 0.434402824 | 0.244 | 0.176 | 1 | 0.355007563 | -0.505640048 | 0.1 | 0.219 | 1 | 0.355007563 | 0.303591313 |
| Sh3bp5l | 0.337469985 | -0.610957709 | 0.084 | 0.118 | 1 | 0.165521745 | -1.026472211 | 0 | 0.164 | 1 | 0.337469985 | 0.303646041 |
| Pcyox1 | 0.16555443 | -0.390269594 | 0.294 | 0.365 | 1 | 0.94889839 | 0.236562194 | 0.3 | 0.33 | 1 | 0.94889839 | 0.303700591 |
| Map2k1 | 0.165601072 | -0.633818667 | 0.361 | 0.424 | 1 | 0.198955112 | -0.541045384 | 0.3 | 0.519 | 1 | 0.198955112 | 0.303778429 |
| Ptrhd1 | 0.844170742 | -0.273922722 | 0.252 | 0.253 | 1 | 0.16567287 | -1.069540933 | 0 | 0.164 | 1 | 0.844170742 | 0.30389824 |
| Thap2 | 0.702317195 | -0.253835469 | 0.546 | 0.529 | 1 | 0.165677354 | -0.573704323 | 0.3 | 0.556 | 1 | 0.702317195 | 0.303905722 |
| Pcca | 0.165734474 | -0.977279923 | 0.294 | 0.341 | 1 | 0.936919934 | 0.295455884 | 0.4 | 0.407 | 1 | 0.936919934 | 0.304001032 |
| Nhlrc3 | 0.166459702 | -0.857395605 | 0.126 | 0.182 | 1 | 0.165872673 | -1.171902651 | 0 | 0.164 | 1 | 0.166459702 | 0.304231602 |
| Vav2 | 0.165904919 | -0.881355504 | 0.109 | 0.165 | 1 | 0.383272343 | -0.852442812 | 0.1 | 0.207 | 1 | 0.383272343 | 0.304285396 |
| Atp5c1 | 0.340221848 | -0.116501253 | 0.882 | 0.924 | 1 | 0.165964666 | -0.409867089 | 0.6 | 0.92 | 1 | 0.340221848 | 0.304385062 |
| Gm50250 | 0.165983913 | 2.099535674 | 0.025 | 0.006 | 1 | 0.643850855 | 2.017921908 | 0 | 0.022 | 1 | 0.643850855 | 0.304417167 |
| Lxn | 0.166078598 | 0.302269569 | 0.437 | 0.359 | 1 | 0.417036153 | 0.791853829 | 0.4 | 0.278 | 1 | 0.417036153 | 0.304575095 |
| Gm42939 | 0.1661669 | -1.222392421 | 0.017 | 0.047 | 1 | 0.940117095 | 0.403212064 | 0.1 | 0.105 | 1 | 0.940117095 | 0.304722361 |
| St8sia1 | 0.1661669 | -1.222392421 | 0.017 | 0.047 | 1 | 0.232498909 | 2.432959407 | 0.1 | 0.031 | 1 | 0.232498909 | 0.304722361 |
| Klhl25 | 0.1661669 | -1.222392421 | 0.017 | 0.047 | 1 | 0.539431168 | 1.017921908 | 0 | 0.037 | 1 | 0.539431168 | 0.304722361 |
| Gm44956 | 0.1661669 | -1.222392421 | 0.017 | 0.047 | 1 | 0.59781419 | 1.31748219 | 0 | 0.028 | 1 | 0.59781419 | 0.304722361 |
| Zfp429 | 0.1661669 | -1.222392421 | 0.017 | 0.047 | 1 | 0.396884044 | 0.31748219 | 0 | 0.068 | 1 | 0.396884044 | 0.304722361 |
| Dmxl2 | 0.166347543 | -0.870090677 | 0.126 | 0.182 | 1 | 0.626168585 | 0.127150978 | 0.1 | 0.167 | 1 | 0.626168585 | 0.305023582 |
| Syt2 | 0.166377994 | 0.241554678 | 0.067 | 0.118 | 1 | 0.253088681 | 1.393431043 | 0.2 | 0.093 | 1 | 0.253088681 | 0.305074352 |
| Spryd4 | 0.166419042 | -0.847996907 | 0.05 | 0.094 | 1 | 0.408636939 | 0.432959407 | 0 | 0.065 | 1 | 0.408636939 | 0.305142787 |
| Tmem209 | 0.166459702 | -0.790281409 | 0.126 | 0.182 | 1 | 0.370509459 | 0.542188477 | 0.3 | 0.17 | 1 | 0.370509459 | 0.305210572 |
| Rasal2 | 0.166611276 | -0.333423734 | 0.437 | 0.512 | 1 | 0.71465533 | 0.245332404 | 0.7 | 0.562 | 1 | 0.71465533 | 0.305463235 |
| Tinf2 | 0.166673 | -0.780882711 | 0.084 | 0.135 | 1 | 0.907531249 | 0.660369903 | 0.1 | 0.111 | 1 | 0.907531249 | 0.305566112 |
| D2hgdh | 0.166898464 | -1.070389328 | 0.042 | 0.082 | 1 | 0.385509064 | 0.263034406 | 0 | 0.071 | 1 | 0.385509064 | 0.30594183 |
| Ncald | 0.166898464 | -1.070389328 | 0.042 | 0.082 | 1 | 0.889931078 | 0.210566986 | 0.1 | 0.083 | 1 | 0.889931078 | 0.30594183 |
| Gm27017 | 0.166961482 | -1.655351829 | 0.025 | 0.059 | 1 | 0.505639146 | 0.494359952 | 0 | 0.043 | 1 | 0.505639146 | 0.306046827 |
| Trim17 | 0.166965572 | -1.485426827 | 0.025 | 0.059 | 1 | 0.474927752 | 0.625604485 | 0 | 0.049 | 1 | 0.474927752 | 0.306053642 |
| Gm28818 | 0.167062889 | 1.836501268 | 0.025 | 0.006 | 1 | 0.730749233 | 2.432959407 | 0 | 0.012 | 1 | 0.730749233 | 0.306215769 |
| C030005K06Rik | 0.167062889 | 1.836501268 | 0.025 | 0.006 | 1 | 0.619961444 | 1.558490289 | 0 | 0.025 | 1 | 0.619961444 | 0.306215769 |
| Tcaf2 | 0.167092087 | -0.839063782 | 0.059 | 0.106 | 1 | 0.913226652 | 0.463333056 | 0.1 | 0.086 | 1 | 0.913226652 | 0.306264409 |
| Rnf121 | 0.167257375 | -0.63047716 | 0.345 | 0.4 | 1 | 0.361263771 | 0.331421381 | 0.7 | 0.506 | 1 | 0.361263771 | 0.30653972 |
| Wsb2 | 0.35743316 | 0.123382416 | 0.63 | 0.553 | 1 | 0.167276779 | -0.561016805 | 0.4 | 0.67 | 1 | 0.35743316 | 0.306572037 |
| Fkbp8 | 0.64761884 | -0.129283017 | 0.672 | 0.6 | 1 | 0.167304438 | -0.784271309 | 0.4 | 0.565 | 1 | 0.64761884 | 0.306618101 |
| 6030407O03Rik | 0.341808287 | 0.655435709 | 0.168 | 0.129 | 1 | 0.167431225 | 2.017921908 | 0.2 | 0.08 | 1 | 0.341808287 | 0.306829236 |
| Synrg | 0.167439058 | -0.415037499 | 0.252 | 0.324 | 1 | 0.614704601 | 0.374065718 | 0.4 | 0.336 | 1 | 0.614704601 | 0.306842278 |
| Slc25a40 | 0.20144649 | -0.911691582 | 0.126 | 0.176 | 1 | 0.167514205 | 1.185031894 | 0.3 | 0.136 | 1 | 0.20144649 | 0.306967401 |
| Mars2 | 0.167517146 | -1.008988783 | 0.05 | 0.094 | 1 | 0.364021887 | -0.152003093 | 0 | 0.077 | 1 | 0.364021887 | 0.306972298 |
| Gm56730 | 0.167576301 | 0.287162677 | 0.319 | 0.253 | 1 | 0.622540735 | 0.864116572 | 0.1 | 0.176 | 1 | 0.622540735 | 0.307070786 |
| Ndufb2 | 0.226627065 | -0.182034684 | 0.79 | 0.806 | 1 | 0.167608839 | -0.632332053 | 0.5 | 0.735 | 1 | 0.226627065 | 0.307124955 |
| Ptp4a1.1 | 0.167616515 | -1.070389328 | 0.017 | 0.047 | 1 | 0.577184788 | 1.558490289 | 0 | 0.031 | 1 | 0.577184788 | 0.307137733 |
| Klhl36 | 0.167616515 | -1.070389328 | 0.017 | 0.047 | 1 | 0.577142568 | 1.31748219 | 0 | 0.031 | 1 | 0.577142568 | 0.307137733 |
| Dnah1 | 0.167616515 | -1.070389328 | 0.017 | 0.047 | 1 | 0.460551517 | 0.695993813 | 0 | 0.052 | 1 | 0.460551517 | 0.307137733 |
| Tmem191 | 0.167616515 | -1.070389328 | 0.017 | 0.047 | 1 | 0.827202642 | 1.210566986 | 0.1 | 0.08 | 1 | 0.827202642 | 0.307137733 |
| Gm10184 | 0.167616515 | -1.070389328 | 0.017 | 0.047 | 1 | 0.446758355 | 0.695993813 | 0 | 0.056 | 1 | 0.446758355 | 0.307137733 |
| Thap7 | 0.16777601 | 0.27064759 | 0.261 | 0.188 | 1 | 0.950940311 | 0.299103661 | 0.2 | 0.185 | 1 | 0.950940311 | 0.307403231 |
| Dapk1 | 0.167889364 | -0.634290213 | 0.454 | 0.5 | 1 | 0.241267356 | 0.487407191 | 0.8 | 0.58 | 1 | 0.241267356 | 0.307591889 |

| | | | | | | | | | | | | |
|---|---|---|---|---|---|---|---|---|---|---|---|---|
| Anxa5 | 0.1679963 | 0.243150982 | 0.983 | 0.982 | 1 | 0.278471237 | -0.281469298 | 1 | 1 | 1 | 0.278471237 | 0.307769843 |
| 6330403L08Rik | 0.168010859 | -1.137503524 | 0.042 | 0.082 | 1 | 0.947574312 | 0.660369903 | 0.1 | 0.105 | 1 | 0.947574312 | 0.307794069 |
| Mink1 | 0.168088409 | -0.904139985 | 0.328 | 0.371 | 1 | 0.579308846 | 0.107029382 | 0.4 | 0.534 | 1 | 0.579308846 | 0.307923104 |
| R3hcc1 | 0.168102611 | -0.663764068 | 0.134 | 0.194 | 1 | 0.938824242 | 0.31748219 | 0.2 | 0.182 | 1 | 0.938824242 | 0.307946734 |
| Anapc10 | 0.168145916 | -0.434442898 | 0.429 | 0.482 | 1 | 0.871016035 | 0.40053793 | 0.4 | 0.444 | 1 | 0.871016035 | 0.308018782 |
| Gm43773 | 0.168156185 | 1.514573173 | 0.025 | 0.006 | 1 | 0.505605027 | 0.847996907 | 0 | 0.043 | 1 | 0.505605027 | 0.308035867 |
| Cfap276 | 0.168156185 | 1.514573173 | 0.025 | 0.006 | 1 | 0.730750993 | 1.847996907 | 0 | 0.012 | 1 | 0.730750993 | 0.308035867 |
| Ptprz1 | 0.168156185 | 1.514573173 | 0.025 | 0.006 | 1 | 0.596204587 | 1.558490289 | 0.1 | 0.059 | 1 | 0.596204587 | 0.308035867 |
| 0610038B21Rik | 0.168156185 | 1.514573173 | 0.025 | 0.006 | 1 | 0.730749233 | 2.432959407 | 0 | 0.012 | 1 | 0.730749233 | 0.308035867 |
| Nrg4 | 0.168156185 | 1.514573173 | 0.025 | 0.006 | 1 | 0.597832556 | 0.847996907 | 0 | 0.028 | 1 | 0.597832556 | 0.308035867 |
| Tspan15 | 0.168156185 | 1.514573173 | 0.025 | 0.006 | 1 | 0.643847387 | 1.210566986 | 0 | 0.022 | 1 | 0.643847387 | 0.308035867 |
| 5033403F01Rik | 0.168156185 | 1.514573173 | 0.025 | 0.006 | 1 | 0.698532411 | 1.695993813 | 0 | 0.015 | 1 | 0.698532411 | 0.308035867 |
| Ica1l | 0.16829248 | -1.292781749 | 0.017 | 0.047 | 1 | 0.73075158 | 2.695993813 | 0 | 0.012 | 1 | 0.73075158 | 0.308262602 |
| Dusp4 | 0.168301449 | -1.250961574 | 0.017 | 0.047 | 1 | 0.397058513 | -0.654503434 | 0 | 0.068 | 1 | 0.397058513 | 0.308277521 |
| Pskh1 | 0.168314065 | -0.607417352 | 0.118 | 0.176 | 1 | 0.595315522 | 0.795529487 | 0.2 | 0.136 | 1 | 0.595315522 | 0.308298506 |
| Gm32031 | 0.168409789 | 0.607682577 | 0.126 | 0.076 | 1 | 0.701889643 | 1.374065718 | 0.1 | 0.068 | 1 | 0.701889643 | 0.308457721 |
| Klhl7 | 0.168759686 | -0.408805546 | 0.546 | 0.565 | 1 | 0.229714511 | -0.69173634 | 0.3 | 0.583 | 1 | 0.229714511 | 0.30903954 |
| Gps1 | 0.460256428 | -0.355791547 | 0.42 | 0.435 | 1 | 0.168762489 | 0.637100124 | 0.7 | 0.469 | 1 | 0.460256428 | 0.3090442 |
| Fam81a | 0.168791026 | -1.485426827 | 0.025 | 0.059 | 1 | 0.577172123 | 0.930459067 | 0 | 0.031 | 1 | 0.577172123 | 0.309091641 |
| Mastl | 0.168792258 | -1.87774425 | 0.025 | 0.059 | 1 | 0.433660137 | 0.374065718 | 0 | 0.059 | 1 | 0.433660137 | 0.309093689 |
| Chst8 | 0.168792668 | -2.292781749 | 0.025 | 0.059 | 1 | 0.363828584 | 3.385653692 | 0.1 | 0.043 | 1 | 0.363828584 | 0.309094372 |
| Lurap1 | 0.168993275 | -1.070389328 | 0.017 | 0.047 | 1 | 0.64384652 | 1.558490289 | 0 | 0.022 | 1 | 0.64384652 | 0.309427824 |
| Ripk2 | 0.37676078 | -0.712495736 | 0.227 | 0.265 | 1 | 0.169155111 | -1.647414009 | 0.1 | 0.29 | 1 | 0.37676078 | 0.309696771 |
| Lingo2 | 0.169248346 | 1.514573173 | 0.025 | 0.006 | 1 | 0.730750993 | 1.847996907 | 0 | 0.012 | 1 | 0.730750993 | 0.309851689 |
| Zfp105 | 0.169429949 | -0.987927168 | 0.042 | 0.082 | 1 | 0.950061233 | 0.695993813 | 0.1 | 0.105 | 1 | 0.950061233 | 0.310153391 |
| Lrrc75aos2 | 0.169429949 | -1.222392421 | 0.042 | 0.082 | 1 | 0.334207905 | -0.191531458 | 0 | 0.086 | 1 | 0.334207905 | 0.310153391 |
| Ppwd1 | 0.169877568 | -0.557576613 | 0.261 | 0.324 | 1 | 0.912591666 | 0.415885894 | 0.3 | 0.315 | 1 | 0.912591666 | 0.310896747 |
| Pcdhb7 | 0.17005575 | -1.411426246 | 0.025 | 0.059 | 1 | 0.44693432 | -0.441509711 | 0 | 0.056 | 1 | 0.44693432 | 0.311192542 |
| Lockd | 0.170146453 | -2.273922722 | 0.042 | 0.082 | 1 | 0.522115639 | 0.695993813 | 0 | 0.04 | 1 | 0.522115639 | 0.31134309 |
| Selenbp1 | 0.227966248 | -1.485426827 | 0.034 | 0.065 | 1 | 0.170152187 | -0.91281543 | 0 | 0.16 | 1 | 0.227966248 | 0.311352607 |
| Tmem223 | 0.170275456 | -0.333423734 | 0.521 | 0.559 | 1 | 0.176941852 | 0.583935306 | 0.8 | 0.491 | 1 | 0.176941852 | 0.311557181 |
| Pigx | 0.493981614 | -0.163498732 | 0.412 | 0.429 | 1 | 0.17028477 | -0.654503434 | 0.2 | 0.438 | 1 | 0.493981614 | 0.311572636 |
| Gm10033 | 0.170293958 | -1.008988783 | 0.118 | 0.171 | 1 | 0.185956028 | -1.357117523 | 0 | 0.151 | 1 | 0.185956028 | 0.311587883 |
| Cdipt | 0.170342851 | 0.325539348 | 0.504 | 0.447 | 1 | 0.379130245 | -0.196397213 | 0.4 | 0.531 | 1 | 0.379130245 | 0.311669016 |
| Ghr | 0.918780632 | -0.193388995 | 0.815 | 0.747 | 1 | 0.170460102 | 0.545213861 | 0.9 | 0.722 | 1 | 0.918780632 | 0.311863558 |
| Acot8 | 0.247842471 | -0.67305383 | 0.134 | 0.182 | 1 | 0.170471519 | -1.111361109 | 0 | 0.16 | 1 | 0.247842471 | 0.311882499 |
| Smarcal1 | 0.39344825 | -0.359895945 | 0.076 | 0.106 | 1 | 0.170471519 | -1.111361109 | 0 | 0.16 | 1 | 0.39344825 | 0.311882499 |
| Tbc1d9 | 0.214921791 | 0.666576266 | 0.076 | 0.041 | 1 | 0.170545774 | -1.374395515 | 0 | 0.16 | 1 | 0.214921791 | 0.312005687 |
| H2bc18 | 0.170630732 | -1.572889668 | 0.025 | 0.059 | 1 | 0.276546558 | 1.374065718 | 0.1 | 0.034 | 1 | 0.276546558 | 0.312146617 |
| 4930505N22Rik | 0.170631144 | -1.292781749 | 0.025 | 0.059 | 1 | 0.669848329 | 1.695993813 | 0 | 0.019 | 1 | 0.669848329 | 0.312147301 |
| Ghdc | 0.881055828 | -0.131789873 | 0.151 | 0.141 | 1 | 0.170660906 | -1.111361109 | 0 | 0.16 | 1 | 0.881055828 | 0.312196668 |
| Fam221a | 0.170675822 | -1.021479727 | 0.076 | 0.124 | 1 | 0.460551517 | 0.695993813 | 0 | 0.052 | 1 | 0.460551517 | 0.312221407 |
| Plcd3 | 0.170675822 | -1.021479727 | 0.076 | 0.124 | 1 | 0.803961691 | 0.210566986 | 0.1 | 0.123 | 1 | 0.803961691 | 0.312221407 |
| Iqcb1 | 0.170716151 | -0.583037624 | 0.496 | 0.512 | 1 | 0.753956932 | -0.342334306 | 0.6 | 0.611 | 1 | 0.753956932 | 0.312288298 |
| Zfp446 | 0.17071659 | -1.250961574 | 0.034 | 0.071 | 1 | 0.582686224 | 1.959028219 | 0.1 | 0.062 | 1 | 0.582686224 | 0.312289026 |
| Cdca7 | 0.170716928 | -1.163498732 | 0.034 | 0.071 | 1 | 0.669853054 | 2.210566986 | 0 | 0.019 | 1 | 0.669853054 | 0.312289587 |
| Gm47271 | 0.170716928 | -1.163498732 | 0.034 | 0.071 | 1 | 0.460577419 | 0.374065718 | 0 | 0.052 | 1 | 0.460577419 | 0.312289587 |
| Zfp760 | 0.170716928 | -1.163498732 | 0.034 | 0.071 | 1 | 0.396919237 | 0.263034406 | 0 | 0.068 | 1 | 0.396919237 | 0.312289587 |
| Adam32 | 0.170718266 | -1.572889668 | 0.067 | 0.112 | 1 | 0.949458438 | -0.357117523 | 0.1 | 0.102 | 1 | 0.949458438 | 0.312291805 |
| Hdac7 | 0.292880187 | -0.669851398 | 0.05 | 0.082 | 1 | 0.170814325 | 0.847996907 | 0.3 | 0.133 | 1 | 0.292880187 | 0.312451117 |
| Rbm15 | 0.170921448 | -0.434800754 | 0.336 | 0.406 | 1 | 0.417788989 | -0.360589715 | 0.3 | 0.429 | 1 | 0.417788989 | 0.312628754 |
| Aaas | 0.366498716 | -0.645891499 | 0.134 | 0.171 | 1 | 0.170951447 | 1.515421567 | 0.2 | 0.077 | 1 | 0.366498716 | 0.312678497 |
| Exog | 0.236616883 | -0.970853654 | 0.034 | 0.065 | 1 | 0.170951602 | 1.695993813 | 0.2 | 0.077 | 1 | 0.236616883 | 0.312678753 |
| Unc50 | 0.419349999 | 0.116023796 | 0.748 | 0.659 | 1 | 0.171018842 | -0.533252279 | 0.5 | 0.648 | 1 | 0.419349999 | 0.31279024 |
| Zfpl1 | 0.171044963 | -0.498026864 | 0.37 | 0.429 | 1 | 0.37350452 | -0.526398608 | 0.2 | 0.321 | 1 | 0.37350452 | 0.312833547 |
| Npepl1 | 0.171144099 | -0.900464326 | 0.118 | 0.171 | 1 | 0.470220374 | 0.494359952 | 0.4 | 0.272 | 1 | 0.470220374 | 0.312997896 |
| Ccl19-ps2 | 0.171347213 | -1.292781749 | 0.034 | 0.071 | 1 | 0.474965159 | 0.374065718 | 0 | 0.049 | 1 | 0.474965159 | 0.313334559 |
| Atf7ip | 0.282661503 | -0.327721374 | 0.622 | 0.647 | 1 | 0.171421318 | -0.64385619 | 0.8 | 0.818 | 1 | 0.282661503 | 0.313457368 |
| B930059L03Rik | 0.171447336 | 0.929610672 | 0.025 | 0.006 | 1 | 0.73075158 | 2.695993813 | 0 | 0.012 | 1 | 0.73075158 | 0.313500483 |
| Gm49173 | 0.171447336 | 0.929610672 | 0.025 | 0.006 | 1 | 0.6199755 | 1.847996907 | 0 | 0.025 | 1 | 0.6199755 | 0.313500483 |
| Gm49864 | 0.171447336 | 0.929610672 | 0.025 | 0.006 | 1 | 0.730749233 | 2.210566986 | 0 | 0.012 | 1 | 0.730749233 | 0.313500483 |
| Tmprss11b | 0.171452536 | 1.166649869 | 0.025 | 0.006 | 1 | 0.698534498 | 2.432959407 | 0 | 0.015 | 1 | 0.698534498 | 0.3135091 |
| Zfp957 | 0.171452536 | 0.929610672 | 0.025 | 0.006 | 1 | 0.698532411 | 2.017921908 | 0 | 0.015 | 1 | 0.698532411 | 0.3135091 |
| Gm13391 | 0.334866848 | -1.185866545 | 0.025 | 0.047 | 1 | 0.171659091 | 1.808468542 | 0.2 | 0.08 | 1 | 0.334866848 | 0.313851339 |
| Efnb3 | 0.171783743 | -0.807354922 | 0.017 | 0.047 | 1 | 0.561898712 | 1.374065718 | 0.1 | 0.056 | 1 | 0.561898712 | 0.314057832 |
| 1700109K24Rik | 0.171783743 | -0.807354922 | 0.017 | 0.047 | 1 | 0.355415506 | 2.210566986 | 0.1 | 0.04 | 1 | 0.355415506 | 0.314057832 |
| 4933439C10Rik | 0.883885666 | -0.263034406 | 0.109 | 0.1 | 1 | 0.171859568 | 2.385653692 | 0.2 | 0.083 | 1 | 0.883885666 | 0.314183425 |
| Klf15 | 0.558312301 | -0.678071905 | 0.092 | 0.112 | 1 | 0.171860003 | 2.558490289 | 0.2 | 0.083 | 1 | 0.558312301 | 0.314184145 |
| Gm38551 | 0.171910829 | -1.292781749 | 0.034 | 0.071 | 1 | 0.385705751 | -0.473931188 | 0 | 0.071 | 1 | 0.385705751 | 0.314268325 |
| E130114P18Rik | 0.171926766 | -1.544320516 | 0.025 | 0.059 | 1 | 0.362872659 | 1.930459067 | 0.1 | 0.04 | 1 | 0.362872659 | 0.314294718 |
| Zscan25 | 0.171985674 | -0.900464326 | 0.042 | 0.082 | 1 | 0.944635256 | 0.973527789 | 0.1 | 0.093 | 1 | 0.944635256 | 0.314392277 |
| Nptn | 0.172060765 | -0.221025909 | 1 | 0.994 | 1 | 0.338454136 | -0.233996466 | 1 | 1 | 1 | 0.338454136 | 0.314516623 |
| Dusp7 | 0.260972777 | -0.279842694 | 0.345 | 0.4 | 1 | 0.172137723 | -1.111361109 | 0.4 | 0.525 | 1 | 0.260972777 | 0.31464405 |
| Aldh4a1 | 0.172151625 | -1.255944981 | 0.126 | 0.176 | 1 | 0.656997996 | -0.111361109 | 0.1 | 0.148 | 1 | 0.656997996 | 0.314667068 |
| Rab5b | 0.85144279 | -0.19592021 | 0.487 | 0.435 | 1 | 0.172158609 | -0.579864633 | 0.3 | 0.583 | 1 | 0.85144279 | 0.314678631 |
| Ndufb3 | 0.586405877 | -0.114647248 | 0.798 | 0.835 | 1 | 0.172163702 | -0.111361109 | 0.4 | 0.759 | 1 | 0.586405877 | 0.314687064 |
| Fastkd5 | 0.172480603 | -1.185866545 | 0.025 | 0.059 | 1 | 0.374479548 | 0.210566986 | 0 | 0.074 | 1 | 0.374479548 | 0.315211647 |
| Gm12762 | 0.172480603 | -1.185866545 | 0.025 | 0.059 | 1 | 0.278845643 | 2.017921908 | 0.1 | 0.034 | 1 | 0.278845643 | 0.315211647 |
| Gm49700 | 0.172480603 | -1.292781749 | 0.025 | 0.059 | 1 | 0.320995017 | 2.017921908 | 0.1 | 0.037 | 1 | 0.320995017 | 0.315211647 |
| Gsto2 | 0.172497189 | -0.807354922 | 0.05 | 0.094 | 1 | 0.395903457 | 0.930459067 | 0.2 | 0.111 | 1 | 0.395903457 | 0.315239097 |
| Sod1 | 0.299186766 | 0.117237675 | 0.807 | 0.765 | 1 | 0.17254953 | -0.488430758 | 0.6 | 0.75 | 1 | 0.299186766 | 0.315325719 |
| Gm9929 | 0.241520057 | -1.185866545 | 0.025 | 0.053 | 1 | 0.172749482 | 1.732519689 | 0.2 | 0.08 | 1 | 0.241520057 | 0.315656581 |
| Ccdc15 | 0.361460156 | -0.485426827 | 0.109 | 0.147 | 1 | 0.17278887 | 0.888638891 | 0.3 | 0.136 | 1 | 0.361460156 | 0.315721746 |
| Lonp2 | 0.985495145 | -0.267003308 | 0.597 | 0.553 | 1 | 0.172892508 | 0.599667696 | 0.8 | 0.62 | 1 | 0.985495145 | 0.315893197 |
| Teddm2 | 0.205197403 | 1.142604395 | 0.076 | 0.041 | 1 | 0.172903719 | 3.695993813 | 0.1 | 0.028 | 1 | 0.205197403 | 0.315911743 |
| Fhl1 | 0.501782815 | 0.161901555 | 0.286 | 0.247 | 1 | 0.172949736 | -1.5519337 | 0.2 | 0.38 | 1 | 0.501782815 | 0.315987862 |
| Mterf1a | 0.172951071 | -1.070389328 | 0.034 | 0.071 | 1 | 0.360331175 | 2.017921908 | 0.1 | 0.04 | 1 | 0.360331175 | 0.315990069 |
| Gm17344 | 0.173185841 | -0.655351829 | 0.017 | 0.047 | 1 | 0.44222087 | 1.847996907 | 0.1 | 0.046 | 1 | 0.44222087 | 0.316378347 |
| Fam219b | 0.173451357 | -0.567888987 | 0.134 | 0.194 | 1 | 0.776741883 | 0.536795218 | 0.2 | 0.16 | 1 | 0.776741883 | 0.31681734 |
| Gpr137c | 0.17380357 | -1.432959407 | 0.025 | 0.059 | 1 | 0.47499996 | -0.374395515 | 0 | 0.049 | 1 | 0.47499996 | 0.317399459 |
| D130040H23Rik | 0.173901795 | -0.687060688 | 0.05 | 0.094 | 1 | 0.818788682 | 0.732519689 | 0.1 | 0.077 | 1 | 0.818788682 | 0.317561756 |
| Klhdc4 | 0.173915565 | -0.76106127 | 0.143 | 0.2 | 1 | 0.482708944 | 0.494359952 | 0.3 | 0.191 | 1 | 0.482708944 | 0.317584507 |
| Mrpl14 | 0.173986419 | -0.30560579 | 0.37 | 0.459 | 1 | 0.238976053 | -0.505640048 | 0.2 | 0.407 | 1 | 0.238976053 | 0.317701564 |
| Stk11ip | 0.174205604 | -0.847996907 | 0.042 | 0.082 | 1 | 0.707564904 | 1.263034406 | 0.1 | 0.068 | 1 | 0.707564904 | 0.318063616 |
| Dis3l | 0.17423254 | -0.655351829 | 0.05 | 0.094 | 1 | 0.202249388 | -1.171902651 | 0 | 0.142 | 1 | 0.202249388 | 0.318108102 |
| Gm15862 | 0.174340642 | -1.070389328 | 0.025 | 0.059 | 1 | 0.539431168 | 1.017921908 | 0 | 0.037 | 1 | 0.539431168 | 0.318286625 |
| K230015D01Rik | 0.174340642 | -1.070389328 | 0.025 | 0.059 | 1 | 0.59781419 | 1.558490289 | 0 | 0.028 | 1 | 0.59781419 | 0.318286625 |
| Acot6 | 0.174340642 | -1.070389328 | 0.025 | 0.059 | 1 | 0.482051401 | 1.769994395 | 0.1 | 0.049 | 1 | 0.482051401 | 0.318286625 |

| | | | | | | | | | | | | |
|---|---|---|---|---|---|---|---|---|---|---|---|---|
| Nthl1 | 0.174340642 | -1.070389328 | 0.025 | 0.059 | 1 | 0.577142568 | 1.432959407 | 0 | 0.031 | 1 | 0.577142568 | 0.318286625 |
| Crabp2 | 0.174612932 | 2.514573173 | 0.092 | 0.053 | 1 | 0.397023343 | -0.408342847 | 0 | 0.068 | 1 | 0.397023343 | 0.318736188 |
| 5031415H12Rik | 0.368333876 | 0.514573173 | 0.05 | 0.029 | 1 | 0.174700008 | 3.14345279 | 0.1 | 0.028 | 1 | 0.368333876 | 0.318879922 |
| Rfx5 | 0.174896695 | -0.807354922 | 0.042 | 0.082 | 1 | 0.446819579 | 0.558490289 | 0 | 0.056 | 1 | 0.446819579 | 0.319204536 |
| 1700034P13Rik | 0.17504221 | -2.117695043 | 0.025 | 0.059 | 1 | 0.597838178 | 0.769994395 | 0 | 0.028 | 1 | 0.597838178 | 0.319444644 |
| Gm45025 | 0.175047852 | -0.784987109 | 0.084 | 0.135 | 1 | 0.570895052 | -0.111361109 | 0.1 | 0.164 | 1 | 0.570895052 | 0.319453953 |
| Thumpd2 | 0.175057179 | -1.070389328 | 0.025 | 0.059 | 1 | 0.778938153 | 0.808468542 | 0.1 | 0.074 | 1 | 0.778938153 | 0.319469343 |
| Gm41664 | 0.175057179 | -1.070389328 | 0.025 | 0.059 | 1 | 0.522130859 | 0.494359952 | 0 | 0.04 | 1 | 0.522130859 | 0.319469343 |
| Bcdin3d | 0.175202541 | -0.970853654 | 0.034 | 0.071 | 1 | 0.385546628 | 0.111031312 | 0 | 0.071 | 1 | 0.385546628 | 0.319709151 |
| Slc25a4 | 0.175260217 | 0.110553016 | 0.992 | 0.959 | 1 | 0.188731149 | -0.467778961 | 0.9 | 0.957 | 1 | 0.188731149 | 0.319804291 |
| Ltbp1 | 0.175279165 | 0.529523514 | 0.227 | 0.159 | 1 | 0.37086401 | -0.91281543 | 0.1 | 0.225 | 1 | 0.37086401 | 0.319835545 |
| Shb | 0.175334038 | -1.36994961 | 0.151 | 0.2 | 1 | 1 | 0.336097868 | 0.3 | 0.309 | 1 | 1 | 0.319926052 |
| Daam1 | 0.175440879 | -0.481479643 | 0.882 | 0.835 | 1 | 0.211823557 | 0.650890927 | 0.9 | 0.88 | 1 | 0.211823557 | 0.320102256 |
| Gm57450 | 0.175443909 | -1.359895945 | 0.067 | 0.112 | 1 | 0.231030696 | 1.432959407 | 0.2 | 0.086 | 1 | 0.231030696 | 0.320107253 |
| Prss36 | 0.417750265 | 0.584962501 | 0.134 | 0.106 | 1 | 0.175467868 | 1.515421567 | 0.2 | 0.077 | 1 | 0.417750265 | 0.320146764 |
| Bok | 0.175759461 | -1.137503524 | 0.042 | 0.082 | 1 | 0.238914673 | 2.111031312 | 0.1 | 0.031 | 1 | 0.238914673 | 0.320627533 |
| Pdss1 | 0.244727652 | -0.439623138 | 0.168 | 0.218 | 1 | 0.175774955 | 1.558490289 | 0.3 | 0.145 | 1 | 0.244727652 | 0.320653075 |
| Tbc1d23 | 0.175853057 | -0.387871518 | 0.605 | 0.641 | 1 | 0.202005065 | 0.271071725 | 0.9 | 0.694 | 1 | 0.202005065 | 0.320781817 |
| Dpm1 | 0.336418864 | -0.224636115 | 0.874 | 0.888 | 1 | 0.175924128 | -0.446964141 | 0.7 | 0.87 | 1 | 0.336418864 | 0.320898957 |
| Pccb | 0.182856607 | -0.518593691 | 0.319 | 0.371 | 1 | 0.175945597 | -0.989572629 | 0.2 | 0.389 | 1 | 0.182856607 | 0.320934341 |
| Nek7 | 0.325914759 | -0.276983965 | 0.773 | 0.782 | 1 | 0.175995355 | -0.606125801 | 0.7 | 0.861 | 1 | 0.325914759 | 0.321016345 |
| Cc2d1b | 0.176015217 | -0.622930351 | 0.109 | 0.165 | 1 | 0.312453074 | -0.668578619 | 0.1 | 0.231 | 1 | 0.312453074 | 0.321049078 |
| E130308A19Rik | 0.176095729 | -1.30256277 | 0.151 | 0.2 | 1 | 0.252867638 | 0.905221775 | 0.4 | 0.253 | 1 | 0.252867638 | 0.321181752 |
| Ak2 | 0.266289227 | -0.336563441 | 0.513 | 0.535 | 1 | 0.176231865 | -0.673239997 | 0.3 | 0.562 | 1 | 0.266289227 | 0.32140606 |
| Gm17057 | 0.17632717 | -0.807354922 | 0.025 | 0.059 | 1 | 0.238914673 | 1.847996907 | 0.1 | 0.031 | 1 | 0.238914673 | 0.321563069 |
| Gm13778 | 0.176362599 | -0.944858446 | 0.025 | 0.059 | 1 | 0.474947236 | 0.558490289 | 0 | 0.049 | 1 | 0.474947236 | 0.321621432 |
| Klf7 | 0.176414047 | -0.408051983 | 0.807 | 0.788 | 1 | 0.741330257 | 0.481214576 | 0.8 | 0.833 | 1 | 0.741330257 | 0.321706178 |
| Rbm12b2 | 0.17645202 | -0.854660637 | 0.076 | 0.124 | 1 | 0.701467586 | 0.345496566 | 0.1 | 0.142 | 1 | 0.701467586 | 0.321768725 |
| Arl4c | 0.176501489 | -1.378511623 | 0.034 | 0.071 | 1 | 0.334246869 | -0.374395515 | 0 | 0.086 | 1 | 0.334246869 | 0.321850202 |
| Ergic3 | 0.268830601 | -0.157446062 | 0.84 | 0.806 | 1 | 0.176505856 | -0.443693958 | 0.6 | 0.843 | 1 | 0.268830601 | 0.321857395 |
| Gm31645 | 0.658336322 | -0.707819249 | 0.042 | 0.053 | 1 | 0.176506871 | 2.930459067 | 0.1 | 0.028 | 1 | 0.658336322 | 0.321859067 |
| Adgrv1 | 0.176620785 | 0.573466862 | 0.109 | 0.065 | 1 | 0.730750993 | 1.31748219 | 0 | 0.012 | 1 | 0.730750993 | 0.322046669 |
| Sh2d3c | 0.176824576 | -0.765534746 | 0.042 | 0.082 | 1 | 0.505651086 | 0.374065718 | 0 | 0.043 | 1 | 0.505651086 | 0.322382222 |
| Thyn1 | 0.176838484 | -0.485426827 | 0.286 | 0.347 | 1 | 0.871371724 | 0.345496566 | 0.3 | 0.265 | 1 | 0.871371724 | 0.322405118 |
| Rpp40 | 0.176950337 | -0.970853654 | 0.025 | 0.059 | 1 | 0.357796814 | 2.111031312 | 0.1 | 0.04 | 1 | 0.357796814 | 0.322589252 |
| Fam220a.1 | 0.177025272 | -1.14839184 | 0.034 | 0.071 | 1 | 0.678985981 | 1.063725598 | 0.1 | 0.065 | 1 | 0.678985981 | 0.322712596 |
| Pxn | 0.234249524 | -0.505326385 | 0.42 | 0.435 | 1 | 0.177179942 | -0.928497052 | 0.4 | 0.574 | 1 | 0.234249524 | 0.322967153 |
| Rac1 | 0.17718464 | -0.158906596 | 0.983 | 0.976 | 1 | 0.191250116 | -0.319185328 | 1 | 0.994 | 1 | 0.191250116 | 0.322974884 |
| Tfeb | 0.743038722 | -0.584962501 | 0.252 | 0.253 | 1 | 0.177263701 | -1.198015491 | 0.2 | 0.38 | 1 | 0.743038722 | 0.323104982 |
| Ncdn | 0.37022703 | 0.308122295 | 0.101 | 0.071 | 1 | 0.177281618 | 1.08718457 | 0.3 | 0.139 | 1 | 0.37022703 | 0.323134465 |
| Parp11 | 0.753241924 | -0.292781749 | 0.084 | 0.094 | 1 | 0.17728256 | 1.017921908 | 0.3 | 0.139 | 1 | 0.753241924 | 0.323136014 |
| Nploc4 | 0.177334535 | -0.625151591 | 0.387 | 0.429 | 1 | 0.866841374 | -0.191531458 | 0.7 | 0.636 | 1 | 0.866841374 | 0.323221533 |
| Snx1 | 0.177421093 | 0.242047318 | 0.765 | 0.688 | 1 | 0.946039145 | 0.236562194 | 0.6 | 0.676 | 1 | 0.946039145 | 0.323363942 |
| Txndc2 | 0.177545801 | -0.359895945 | 0.017 | 0.047 | 1 | 0.446819579 | 0.558490289 | 0 | 0.056 | 1 | 0.446819579 | 0.32356909 |
| Nyap1 | 0.177606284 | -0.900464326 | 0.034 | 0.071 | 1 | 0.876246026 | 0.591657153 | 0.1 | 0.083 | 1 | 0.876246026 | 0.323668576 |
| Gm33887 | 0.177704428 | -0.86393845 | 0.034 | 0.071 | 1 | 0.446790883 | 0.625604485 | 0 | 0.056 | 1 | 0.446790883 | 0.323829992 |
| Gm20712 | 0.177704428 | -0.86393845 | 0.034 | 0.071 | 1 | 0.500525852 | 0.988174565 | 0.2 | 0.123 | 1 | 0.500525852 | 0.323829992 |
| Usf2 | 0.928077459 | -0.140515414 | 0.521 | 0.494 | 1 | 0.177718258 | -0.69173634 | 0.4 | 0.58 | 1 | 0.928077459 | 0.323852737 |
| Gm46367 | 0.236604499 | -1.655351829 | 0.025 | 0.053 | 1 | 0.177754291 | 1.245332404 | 0.2 | 0.077 | 1 | 0.236604499 | 0.323911994 |
| Ttbk2 | 0.178258357 | -0.510088881 | 0.218 | 0.282 | 1 | 0.879335223 | -0.111361109 | 0.3 | 0.293 | 1 | 0.879335223 | 0.324740672 |
| Srr | 0.178267352 | -0.578536232 | 0.252 | 0.318 | 1 | 0.232917359 | 0.558490289 | 0.5 | 0.296 | 1 | 0.232917359 | 0.324755455 |
| Gm33696 | 0.498594647 | -0.485426827 | 0.017 | 0.029 | 1 | 0.178315901 | 2.902444691 | 0.1 | 0.028 | 1 | 0.498594647 | 0.324835242 |
| Rap2a | 0.558748664 | 0.326500825 | 0.353 | 0.324 | 1 | 0.178516941 | 1.783456654 | 0.5 | 0.309 | 1 | 0.558748664 | 0.325165584 |
| Peg12 | 0.178742974 | -0.987927168 | 0.034 | 0.071 | 1 | 0.597832181 | 1.210566986 | 0 | 0.028 | 1 | 0.597832181 | 0.325536897 |
| Mrps17 | 0.178767577 | -0.258016331 | 0.546 | 0.571 | 1 | 0.646097602 | -0.198015491 | 0.4 | 0.444 | 1 | 0.646097602 | 0.325577308 |
| AW146154 | 0.178855069 | -0.86393845 | 0.025 | 0.059 | 1 | 0.334207905 | -0.191531458 | 0 | 0.086 | 1 | 0.334207905 | 0.325721002 |
| Atp5g2 | 0.291374243 | 0.129957663 | 0.933 | 0.941 | 1 | 0.178950112 | -0.440294809 | 0.6 | 0.948 | 1 | 0.291374243 | 0.325877081 |
| Smndc1 | 0.49608418 | -0.178947131 | 0.748 | 0.735 | 1 | 0.17956426 | -0.613861449 | 0.6 | 0.787 | 1 | 0.49608418 | 0.326885197 |
| Aox1 | 0.276964127 | -1.321928095 | 0.076 | 0.112 | 1 | 0.179587233 | 1.988174565 | 0.2 | 0.083 | 1 | 0.276964127 | 0.326922892 |
| Map7d3 | 0.295234939 | -0.179323699 | 0.361 | 0.271 | 1 | 0.179728582 | 0.569461407 | 0.4 | 0.198 | 1 | 0.295234939 | 0.3271548 |
| Zfp358 | 0.179729526 | -0.610957709 | 0.084 | 0.135 | 1 | 0.467730467 | -0.248864633 | 0.1 | 0.188 | 1 | 0.467730467 | 0.327156349 |
| Gm43713 | 0.179731198 | 1.099535674 | 0.092 | 0.053 | 1 | 0.241810349 | 1.14345279 | 0.2 | 0.086 | 1 | 0.241810349 | 0.327159093 |
| 4930512B01Rik | 0.179731198 | 1.514573173 | 0.092 | 0.053 | 1 | 0.58430737 | 0.602884409 | 0.2 | 0.133 | 1 | 0.58430737 | 0.327159093 |
| Gda | 0.374548525 | -0.222392421 | 0.017 | 0.006 | 1 | 0.179765786 | -1.248864633 | 0 | 0.154 | 1 | 0.374548525 | 0.327215834 |
| Sestd1 | 0.179772733 | 0.446669108 | 0.723 | 0.671 | 1 | 0.947867954 | 0.199760231 | 0.6 | 0.636 | 1 | 0.947867954 | 0.327227231 |
| Pdzd11 | 0.59612802 | -0.20085926 | 0.462 | 0.482 | 1 | 0.180021382 | -1.111361109 | 0.2 | 0.383 | 1 | 0.59612802 | 0.327635066 |
| Gpr137 | 0.500412712 | 0.292180751 | 0.218 | 0.188 | 1 | 0.180048469 | -0.864721141 | 0 | 0.154 | 1 | 0.500412712 | 0.327679487 |
| Cwf19l1 | 0.180193017 | -0.600904045 | 0.084 | 0.135 | 1 | 0.18713833 | 1.602884409 | 0.2 | 0.08 | 1 | 0.18713833 | 0.327916511 |
| Ptges2 | 0.684132355 | -0.333423734 | 0.076 | 0.088 | 1 | 0.180217735 | -1.004445905 | 0 | 0.154 | 1 | 0.684132355 | 0.327957038 |
| Rnaseh2a | 0.180241497 | -0.911691582 | 0.101 | 0.153 | 1 | 0.306978955 | -0.339630097 | 0 | 0.096 | 1 | 0.306978955 | 0.327995997 |
| Mtif3 | 0.403243536 | -0.485426827 | 0.126 | 0.159 | 1 | 0.180497413 | -1.048167282 | 0 | 0.154 | 1 | 0.403243536 | 0.328415511 |
| Mlf2 | 0.180566083 | 0.204888674 | 0.824 | 0.718 | 1 | 0.236191321 | -0.44905652 | 0.6 | 0.778 | 1 | 0.236191321 | 0.328528055 |
| Lsg1 | 0.180807804 | -0.476915327 | 0.487 | 0.524 | 1 | 0.367964346 | -0.516056664 | 0.4 | 0.509 | 1 | 0.367964346 | 0.328924145 |
| Pitpnm3 | 0.479565188 | -0.572889668 | 0.05 | 0.071 | 1 | 0.180832797 | -1.441509711 | 0 | 0.154 | 1 | 0.479565188 | 0.328965093 |
| Gm16867 | 0.181188472 | 0.862496476 | 0.101 | 0.059 | 1 | 0.52208609 | 1.017921908 | 0 | 0.04 | 1 | 0.52208609 | 0.329547682 |
| Kdm2b | 0.206876142 | -0.541280062 | 0.235 | 0.288 | 1 | 0.181415429 | 1.119801522 | 0.5 | 0.306 | 1 | 0.206876142 | 0.3299193 |
| Gtf3c3 | 0.181478031 | -0.520192245 | 0.286 | 0.347 | 1 | 0.338099058 | -0.68251781 | 0.1 | 0.222 | 1 | 0.338099058 | 0.330021786 |
| Gm17103 | 0.181489765 | -0.807354922 | 0.025 | 0.059 | 1 | 0.489972412 | 0.374065718 | 0 | 0.046 | 1 | 0.489972412 | 0.330040994 |
| Pycard | 0.181534955 | -1.155278225 | 0.084 | 0.129 | 1 | 0.594018878 | -0.279758641 | 0.2 | 0.269 | 1 | 0.594018878 | 0.33011497 |
| Cep95 | 0.181545636 | -0.5360529 | 0.336 | 0.388 | 1 | 0.685971942 | -0.289506617 | 0.3 | 0.355 | 1 | 0.685971942 | 0.330132454 |
| Zfp975 | 0.181733989 | -0.537894247 | 0.185 | 0.247 | 1 | 0.427324778 | 0.973527789 | 0.3 | 0.201 | 1 | 0.427324778 | 0.330440735 |
| Enox2 | 0.181831522 | -0.584962501 | 0.168 | 0.229 | 1 | 0.669017601 | 0.602884409 | 0.3 | 0.238 | 1 | 0.669017601 | 0.330600342 |
| Cstdc2 | 0.840598367 | 0.814133455 | 0.042 | 0.047 | 1 | 0.181995085 | 2.695993813 | 0.1 | 0.028 | 1 | 0.840598367 | 0.33086796 |
| Ptpn6 | 0.18225745 | -0.733354341 | 0.042 | 0.082 | 1 | 0.561898712 | 1.31748219 | 0.1 | 0.056 | 1 | 0.561898712 | 0.331297122 |
| Nol6 | 0.182309612 | -0.485426827 | 0.05 | 0.094 | 1 | 0.997434248 | 0.888638891 | 0.1 | 0.099 | 1 | 0.997434248 | 0.33138243 |
| Fbf1 | 0.182746843 | -0.822461814 | 0.134 | 0.188 | 1 | 0.614403038 | 0.864116572 | 0.1 | 0.179 | 1 | 0.614403038 | 0.332097277 |
| Dgkq | 0.182805385 | -0.833350131 | 0.076 | 0.124 | 1 | 0.368291247 | 1.048295557 | 0.2 | 0.105 | 1 | 0.368291247 | 0.332192962 |
| Cipc | 0.183028293 | 0.675037845 | 0.286 | 0.218 | 1 | 0.643363676 | 0.432959407 | 0.1 | 0.17 | 1 | 0.643363676 | 0.33255723 |
| Pole3 | 0.594964814 | -0.159440516 | 0.504 | 0.488 | 1 | 0.183049011 | 1.574978412 | 0.5 | 0.367 | 1 | 0.594964814 | 0.332591081 |
| Zdhhc9 | 0.183298246 | -0.409477974 | 0.218 | 0.276 | 1 | 0.462739488 | 0.706335757 | 0.4 | 0.287 | 1 | 0.462739488 | 0.332998245 |
| Trp53rka | 0.183954374 | -0.765534746 | 0.101 | 0.153 | 1 | 0.362960976 | 0.789103218 | 0.3 | 0.176 | 1 | 0.362960976 | 0.334069537 |
| Ppil6 | 0.184219916 | 0.798366139 | 0.176 | 0.124 | 1 | 0.50071917 | 1.210566986 | 0.2 | 0.133 | 1 | 0.50071917 | 0.334502855 |
| Hint3 | 0.249813223 | 0.192645078 | 0.378 | 0.306 | 1 | 0.184266299 | 1.403212064 | 0.4 | 0.222 | 1 | 0.249813223 | 0.334578529 |
| Dgat2 | 0.184271132 | 0.853375086 | 0.168 | 0.118 | 1 | 0.247511186 | 3.119801522 | 0.2 | 0.105 | 1 | 0.247511186 | 0.334586414 |
| A830008E24Rik | 0.184506253 | 0.421463768 | 0.101 | 0.059 | 1 | 0.539413885 | 1.210566986 | 0 | 0.037 | 1 | 0.539413885 | 0.334969948 |
| 2310009B15Rik | 0.933371774 | -0.137503524 | 0.336 | 0.312 | 1 | 0.184510843 | 1.611929548 | 0.4 | 0.244 | 1 | 0.933371774 | 0.334977435 |
| Ccdc62 | 0.184628572 | 0.377069649 | 0.252 | 0.182 | 1 | 0.925158688 | 0.684021171 | 0.2 | 0.194 | 1 | 0.925158688 | 0.335169434 |
| Ksr1 | 0.23275291 | -1.977279923 | 0.059 | 0.094 | 1 | 0.184723838 | 1.048295557 | 0.2 | 0.077 | 1 | 0.23275291 | 0.33532478 |

| | | | | | | | | | | | | |
|---|---|---|---|---|---|---|---|---|---|---|---|---|
| Zdhhc24 | 0.19558021 | -0.748461233 | 0.067 | 0.112 | 1 | 0.184784555 | 1.744903414 | 0.2 | 0.08 | 1 | 0.19558021 | 0.335423777 |
| Cat | 0.690203681 | -0.399270183 | 0.429 | 0.412 | 1 | 0.184961373 | -0.201246612 | 0.2 | 0.506 | 1 | 0.690203681 | 0.335712037 |
| Mpzl2 | 0.185092432 | -1.27897595 | 0.109 | 0.159 | 1 | 0.382574577 | -0.536666944 | 0.1 | 0.21 | 1 | 0.382574577 | 0.335925655 |
| Hgsnat | 0.489582889 | 0.165423609 | 0.496 | 0.453 | 1 | 0.18518805 | 0.551947443 | 0.7 | 0.438 | 1 | 0.489582889 | 0.336081486 |
| Gm4631 | 0.217244155 | -0.847996907 | 0.05 | 0.088 | 1 | 0.185322417 | -0.840059087 | 0 | 0.151 | 1 | 0.217244155 | 0.336300435 |
| Cgrrf1 | 0.185323521 | 0.168397532 | 0.588 | 0.488 | 1 | 0.486090148 | 0.210566986 | 0.7 | 0.559 | 1 | 0.486090148 | 0.336302234 |
| Msh2 | 0.185453203 | -0.807354922 | 0.134 | 0.188 | 1 | 0.274766077 | 1.494359952 | 0.3 | 0.176 | 1 | 0.274766077 | 0.336513515 |
| Zfp874a | 0.512772159 | 0.160936218 | 0.126 | 0.1 | 1 | 0.185480222 | -0.959358016 | 0 | 0.151 | 1 | 0.512772159 | 0.336557531 |
| Ttc32 | 0.546497537 | -0.378511623 | 0.143 | 0.165 | 1 | 0.18568104 | -1.026472211 | 0 | 0.151 | 1 | 0.546497537 | 0.336884631 |
| Tmcc3 | 0.185716755 | -1.007847862 | 0.202 | 0.247 | 1 | 0.30460373 | -1.159271094 | 0.3 | 0.519 | 1 | 0.30460373 | 0.336942797 |
| Gfer | 0.263180305 | -0.614709844 | 0.126 | 0.171 | 1 | 0.185734253 | -1.026472211 | 0 | 0.151 | 1 | 0.263180305 | 0.336971293 |
| Ttll1 | 0.251593672 | -0.485426827 | 0.126 | 0.176 | 1 | 0.185747766 | -1.131825212 | 0 | 0.151 | 1 | 0.251593672 | 0.336993299 |
| Rom1 | 0.185786601 | 0.684498174 | 0.202 | 0.147 | 1 | 0.233532583 | 0.973527789 | 0.3 | 0.151 | 1 | 0.233532583 | 0.337056541 |
| Ubald2 | 0.186003153 | -0.299871174 | 0.437 | 0.5 | 1 | 0.460712699 | -0.479264633 | 0.5 | 0.54 | 1 | 0.460712699 | 0.337409134 |
| Cops6 | 0.186010172 | -0.325386415 | 0.563 | 0.6 | 1 | 0.256451659 | -0.571327951 | 0.5 | 0.62 | 1 | 0.256451659 | 0.33742056 |
| Gmpr2 | 0.186020125 | -0.648925559 | 0.193 | 0.253 | 1 | 0.50921148 | 0.245332404 | 0.1 | 0.201 | 1 | 0.50921148 | 0.337436763 |
| Cbfb | 0.186123919 | -0.251291747 | 0.79 | 0.841 | 1 | 0.228110647 | -0.375768856 | 0.8 | 0.87 | 1 | 0.228110647 | 0.337605725 |
| Praf2 | 0.186309061 | -0.342915183 | 0.529 | 0.576 | 1 | 0.79218173 | 0.145093148 | 0.4 | 0.525 | 1 | 0.79218173 | 0.337907055 |
| Il1rap | 0.186497731 | 0.730301864 | 0.168 | 0.112 | 1 | 0.781463531 | -0.210896782 | 0.1 | 0.127 | 1 | 0.781463531 | 0.338214059 |
| Adnp2 | 0.18661353 | -0.839063782 | 0.202 | 0.259 | 1 | 0.330044251 | -0.668578619 | 0.1 | 0.225 | 1 | 0.330044251 | 0.33840245 |
| Qrsl1 | 0.186725655 | 0.984058456 | 0.092 | 0.053 | 1 | 0.194305414 | 1.354956895 | 0.2 | 0.08 | 1 | 0.194305414 | 0.33858484 |
| Manba | 0.186767884 | -0.407424315 | 0.134 | 0.194 | 1 | 0.76989384 | -0.745843746 | 0.2 | 0.219 | 1 | 0.76989384 | 0.338653526 |
| Procr | 0.186773687 | -0.780882711 | 0.076 | 0.124 | 1 | 0.364021887 | -0.152003093 | 0 | 0.077 | 1 | 0.364021887 | 0.338662964 |
| Poldip2 | 0.186835941 | -0.519374159 | 0.303 | 0.353 | 1 | 0.304043136 | -0.589408406 | 0.3 | 0.432 | 1 | 0.304043136 | 0.338764213 |
| Man2a2 | 0.187009772 | -0.782408565 | 0.261 | 0.306 | 1 | 0.931726213 | 0.374065718 | 0.3 | 0.312 | 1 | 0.931726213 | 0.339046889 |
| Fech | 0.563307698 | -0.448900951 | 0.303 | 0.312 | 1 | 0.187010702 | 0.905221775 | 0.5 | 0.315 | 1 | 0.563307698 | 0.339048401 |
| Elp6 | 0.187073615 | -0.584962501 | 0.034 | 0.071 | 1 | 0.596204587 | 1.374065718 | 0.1 | 0.059 | 1 | 0.596204587 | 0.339150693 |
| Dusp12 | 0.187120525 | -0.567888987 | 0.235 | 0.3 | 1 | 0.685741638 | 0.469485283 | 0.3 | 0.244 | 1 | 0.685741638 | 0.339226959 |
| Lrrc28 | 0.848022162 | 0.359294947 | 0.168 | 0.159 | 1 | 0.187184012 | 1.286118019 | 0.3 | 0.148 | 1 | 0.848022162 | 0.339330169 |
| Mterf3 | 0.187291307 | -0.406355256 | 0.479 | 0.518 | 1 | 0.524489576 | -0.304006187 | 0.4 | 0.5 | 1 | 0.524489576 | 0.339504581 |
| Cthrc1 | 0.187410928 | -0.172416797 | 0.689 | 0.753 | 1 | 0.365804126 | -0.413032363 | 0.6 | 0.806 | 1 | 0.365804126 | 0.339698999 |
| Myo1c | 0.187500732 | -0.341416524 | 0.521 | 0.547 | 1 | 0.199184845 | 0.669193754 | 0.7 | 0.543 | 1 | 0.199184845 | 0.339844939 |
| Ccdc71l | 0.187678506 | -0.669851398 | 0.076 | 0.124 | 1 | 0.19086576 | -0.91281543 | 0 | 0.148 | 1 | 0.19086576 | 0.34013379 |
| Rps27rt | 0.238728711 | 0.158429363 | 0.563 | 0.471 | 1 | 0.187799847 | -0.814968106 | 0.2 | 0.392 | 1 | 0.238728711 | 0.340330911 |
| Fam3a | 0.188259513 | -0.537894247 | 0.21 | 0.271 | 1 | 0.522991439 | 0.101445264 | 0.2 | 0.346 | 1 | 0.522991439 | 0.341077381 |
| Pter | 0.189194686 | 0.91190867 | 0.193 | 0.135 | 1 | 0.188300893 | 0.868174788 | 0.3 | 0.139 | 1 | 0.189194686 | 0.34114456 |
| Cry2 | 0.764180143 | -0.177955567 | 0.849 | 0.771 | 1 | 0.189340198 | 0.1754219 | 0.9 | 0.83 | 1 | 0.764180143 | 0.342830685 |
| Tmc4 | 0.189353641 | -0.359895945 | 0.025 | 0.059 | 1 | 0.315892066 | -0.408342847 | 0 | 0.093 | 1 | 0.315892066 | 0.342852481 |
| Atcayos | 0.189353641 | -0.359895945 | 0.025 | 0.059 | 1 | 0.585491729 | 1.930459067 | 0.1 | 0.062 | 1 | 0.585491729 | 0.342852481 |
| Gatd1 | 0.189397743 | 0.477098467 | 0.235 | 0.171 | 1 | 0.824580948 | 0.625604485 | 0.2 | 0.17 | 1 | 0.824580948 | 0.342923982 |
| Gm57027 | 0.343621566 | -0.263034406 | 0.025 | 0.047 | 1 | 0.189486514 | 2.695993813 | 0.1 | 0.028 | 1 | 0.343621566 | 0.343067889 |
| Ppp1r3f | 0.403273565 | -0.600904045 | 0.042 | 0.065 | 1 | 0.189486514 | 2.695993813 | 0.1 | 0.028 | 1 | 0.403273565 | 0.343067889 |
| Galnt12 | 0.933394757 | -0.163498732 | 0.034 | 0.035 | 1 | 0.189486514 | 2.695993813 | 0.1 | 0.028 | 1 | 0.933394757 | 0.343067889 |
| P2ry2 | 0.189963563 | 0.96203215 | 0.092 | 0.053 | 1 | 0.643850855 | 2.017921908 | 0 | 0.022 | 1 | 0.643850855 | 0.343840971 |
| Irak1 | 0.190158704 | -0.50779464 | 0.412 | 0.471 | 1 | 0.470078979 | 0.165479096 | 0.7 | 0.488 | 1 | 0.470078979 | 0.344157075 |
| Aldh3a1 | 0.19022263 | 0.736965594 | 0.168 | 0.118 | 1 | 0.36121732 | 0.306427001 | 0.1 | 0.25 | 1 | 0.36121732 | 0.344260612 |
| Smpd2 | 0.942891649 | 0.514573173 | 0.218 | 0.218 | 1 | 0.190364473 | 0.816288047 | 0.4 | 0.207 | 1 | 0.942891649 | 0.344490313 |
| Pik3c2a | 0.190465958 | -0.565597176 | 0.655 | 0.682 | 1 | 0.618241864 | -0.135514971 | 0.5 | 0.775 | 1 | 0.618241864 | 0.344654634 |
| Ocln | 0.715874401 | 0.929610672 | 0.017 | 0.012 | 1 | 0.190650882 | -0.91281543 | 0 | 0.148 | 1 | 0.715874401 | 0.344954006 |
| Dapl1 | 0.509230724 | 1.602036014 | 0.05 | 0.035 | 1 | 0.190651171 | -1.004445905 | 0 | 0.148 | 1 | 0.509230724 | 0.344954474 |
| Krt7 | 0.383916503 | 1.514573173 | 0.025 | 0.012 | 1 | 0.19072879 | -1.026472211 | 0 | 0.148 | 1 | 0.383916503 | 0.345080108 |
| Bola1 | 0.190790455 | 0.362570079 | 0.269 | 0.2 | 1 | 0.750912484 | 0.345496566 | 0.1 | 0.133 | 1 | 0.750912484 | 0.345179912 |
| Crlf1 | 0.263482981 | 0.432111013 | 0.109 | 0.071 | 1 | 0.19094333 | -1.152003093 | 0 | 0.148 | 1 | 0.263482981 | 0.345427305 |
| 2810403D21Rik | 0.191103387 | 0.268816758 | 0.353 | 0.271 | 1 | 0.516095193 | 0.51967104 | 0.3 | 0.213 | 1 | 0.516095193 | 0.34568627 |
| Dnajc24 | 0.333893753 | -0.735970289 | 0.202 | 0.235 | 1 | 0.191117577 | 0.598383016 | 0.4 | 0.194 | 1 | 0.333893753 | 0.345709226 |
| Hpn | 0.859314287 | 0.362570079 | 0.042 | 0.047 | 1 | 0.191121916 | -1.267480311 | 0 | 0.148 | 1 | 0.859314287 | 0.345716245 |
| Gm49980 | 0.191322589 | -1.292781749 | 0.059 | 0.1 | 1 | 0.252526464 | -0.536666944 | 0 | 0.117 | 1 | 0.252526464 | 0.346040845 |
| Hhipl2 | 0.218485013 | -1.292781749 | 0.008 | 0.029 | 1 | 0.19134412 | 2.558490289 | 0.1 | 0.028 | 1 | 0.218485013 | 0.346075668 |
| Gm43566 | 0.242320046 | -0.900464326 | 0.017 | 0.041 | 1 | 0.19134412 | 2.558490289 | 0.1 | 0.028 | 1 | 0.242320046 | 0.346075668 |
| Mroh5 | 0.343608466 | -0.600904045 | 0.025 | 0.047 | 1 | 0.19134412 | 2.558490289 | 0.1 | 0.028 | 1 | 0.343608466 | 0.346075668 |
| Gm57017 | 0.390876134 | 0.929610672 | 0.025 | 0.012 | 1 | 0.19134412 | 2.432959407 | 0.1 | 0.028 | 1 | 0.390876134 | 0.346075668 |
| Ccdc181 | 0.408579016 | -0.485426827 | 0.042 | 0.065 | 1 | 0.19134412 | 2.558490289 | 0.1 | 0.028 | 1 | 0.408579016 | 0.346075668 |
| Snn | 0.48842591 | -0.900464326 | 0.017 | 0.029 | 1 | 0.19134412 | 2.558490289 | 0.1 | 0.028 | 1 | 0.48842591 | 0.346075668 |
| Tmem79 | 0.55853335 | 2.099535674 | 0.042 | 0.029 | 1 | 0.19134412 | 2.558490289 | 0.1 | 0.028 | 1 | 0.55853335 | 0.346075668 |
| Sting1 | 0.67656539 | -0.185866545 | 0.042 | 0.053 | 1 | 0.19134412 | 2.558490289 | 0.1 | 0.028 | 1 | 0.67656539 | 0.346075668 |
| Tmem169 | 0.695893386 | -0.222392421 | 0.017 | 0.024 | 1 | 0.19134412 | 2.558490289 | 0.1 | 0.028 | 1 | 0.695893386 | 0.346075668 |
| Gm56548 | 0.743002317 | -0.163498732 | 0.034 | 0.041 | 1 | 0.19134412 | 2.558490289 | 0.1 | 0.028 | 1 | 0.743002317 | 0.346075668 |
| Tango2 | 0.881739929 | -0.199122642 | 0.16 | 0.153 | 1 | 0.191344873 | -1.391469028 | 0 | 0.148 | 1 | 0.881739929 | 0.346076885 |
| Elp5 | 0.191368883 | -0.331717412 | 0.563 | 0.582 | 1 | 0.48920537 | -0.495147674 | 0.5 | 0.509 | 1 | 0.48920537 | 0.346115716 |
| Inpp4b | 0.419379487 | 0.478949263 | 0.202 | 0.165 | 1 | 0.191382403 | -1.391469028 | 0 | 0.148 | 1 | 0.419379487 | 0.346137582 |
| Stt3b | 0.30091759 | 0.113669958 | 0.958 | 0.882 | 1 | 0.191395674 | -0.278319543 | 0.9 | 0.966 | 1 | 0.30091759 | 0.346159043 |
| Ntng2 | 0.358965641 | 0.192645078 | 0.151 | 0.112 | 1 | 0.191525558 | 0.385653692 | 0.3 | 0.139 | 1 | 0.358965641 | 0.346369076 |
| Trim12a | 0.191658872 | -0.790281409 | 0.092 | 0.141 | 1 | 0.226262432 | -1.210896782 | 0 | 0.13 | 1 | 0.226262432 | 0.34658462 |
| Faim | 0.594174054 | -0.222392421 | 0.092 | 0.112 | 1 | 0.191899506 | 1.602884409 | 0.2 | 0.08 | 1 | 0.594174054 | 0.346973592 |
| Mrpl19 | 0.192026989 | -0.595051318 | 0.143 | 0.2 | 1 | 0.5726036 | -0.131825212 | 0.1 | 0.164 | 1 | 0.5726036 | 0.347179614 |
| Stam | 0.192098746 | -0.492768803 | 0.546 | 0.565 | 1 | 0.284205562 | -0.41670632 | 0.5 | 0.707 | 1 | 0.284205562 | 0.347295564 |
| Supt3 | 0.192230448 | -0.389502407 | 0.445 | 0.518 | 1 | 0.949583317 | -0.240322896 | 0.5 | 0.475 | 1 | 0.949583317 | 0.347508351 |
| 2410018L13Rik | 0.192313923 | -0.9510904 | 0.168 | 0.218 | 1 | 0.585103807 | -0.505640048 | 0.1 | 0.157 | 1 | 0.585103807 | 0.347643201 |
| Lsm8 | 0.756854279 | -0.218487967 | 0.445 | 0.441 | 1 | 0.19239236 | 0.147557188 | 0.2 | 0.5 | 1 | 0.756854279 | 0.347769901 |
| Akap7 | 0.192445619 | -0.939144795 | 0.185 | 0.229 | 1 | 0.629727104 | 0.245332404 | 0.1 | 0.167 | 1 | 0.629727104 | 0.347855922 |
| Lcmt2 | 0.192622614 | 0.63005039 | 0.101 | 0.059 | 1 | 0.960016185 | 0.847996907 | 0.1 | 0.105 | 1 | 0.960016185 | 0.348141756 |
| Akap1 | 0.192658066 | -0.787989597 | 0.109 | 0.159 | 1 | 0.735268561 | 0.536795218 | 0.1 | 0.148 | 1 | 0.735268561 | 0.348199001 |
| Tmem170b | 0.616285731 | 0.229170954 | 0.193 | 0.171 | 1 | 0.193125573 | 0.795529487 | 0.4 | 0.216 | 1 | 0.616285731 | 0.348953658 |
| ENSMUSG00000095041 | 0.657194237 | -0.349771728 | 0.252 | 0.265 | 1 | 0.193173581 | -0.69173634 | 0.1 | 0.315 | 1 | 0.657194237 | 0.34903113 |
| Gm13703 | 0.201658121 | 1.251538767 | 0.034 | 0.012 | 1 | 0.193279696 | 2.31748219 | 0.1 | 0.028 | 1 | 0.201658121 | 0.349202351 |
| 1700007L15Rik | 0.243996811 | -0.944858446 | 0.025 | 0.053 | 1 | 0.193279696 | 2.31748219 | 0.1 | 0.028 | 1 | 0.243996811 | 0.349202351 |
| Mterf1b | 0.334204811 | -0.807354922 | 0.008 | 0.024 | 1 | 0.193279696 | 2.111031312 | 0.1 | 0.028 | 1 | 0.334204811 | 0.349202351 |
| Mapk12 | 0.334204811 | -0.807354922 | 0.008 | 0.024 | 1 | 0.193279696 | 2.31748219 | 0.1 | 0.028 | 1 | 0.334204811 | 0.349202351 |
| Batf2 | 0.345346582 | -1.070389328 | 0.017 | 0.035 | 1 | 0.193279696 | 2.31748219 | 0.1 | 0.028 | 1 | 0.345346582 | 0.349202351 |
| 9530056K15Rik | 0.432595587 | -0.622930351 | 0.034 | 0.053 | 1 | 0.193279696 | 2.432959407 | 0.1 | 0.028 | 1 | 0.432595587 | 0.349202351 |
| Gm6556 | 0.462887689 | -0.655351829 | 0.025 | 0.041 | 1 | 0.193279696 | 2.432959407 | 0.1 | 0.028 | 1 | 0.462887689 | 0.349202351 |
| Slc37a4 | 0.467093859 | -0.485426827 | 0.025 | 0.041 | 1 | 0.193279696 | 2.31748219 | 0.1 | 0.028 | 1 | 0.467093859 | 0.349202351 |
| Susd1 | 0.474027635 | -0.359895945 | 0.025 | 0.041 | 1 | 0.193279696 | 2.31748219 | 0.1 | 0.028 | 1 | 0.474027635 | 0.349202351 |
| Amz1 | 0.619390946 | -0.944858446 | 0.025 | 0.035 | 1 | 0.193279696 | 2.210566986 | 0.1 | 0.028 | 1 | 0.619390946 | 0.349202351 |
| Rad51c | 0.851095342 | -0.185866545 | 0.059 | 0.053 | 1 | 0.193279696 | 2.432959407 | 0.1 | 0.028 | 1 | 0.851095342 | 0.349202351 |
| Kiz | 0.959580753 | -0.584962501 | 0.294 | 0.276 | 1 | 0.193496835 | 1.40053793 | 0.4 | 0.287 | 1 | 0.959580753 | 0.349552645 |
| Rabac1 | 0.202532912 | 0.10948714 | 0.975 | 0.953 | 1 | 0.193596314 | -0.42815545 | 0.9 | 0.957 | 1 | 0.202532912 | 0.349713096 |
| Trim11 | 0.193644632 | -0.756728849 | 0.21 | 0.265 | 1 | 0.261397263 | 1.185031894 | 0.4 | 0.244 | 1 | 0.261397263 | 0.349791021 |

| | | | | | | | | | | | | |
|---|---|---|---|---|---|---|---|---|---|---|---|---|
| Mier2 | 0.193935163 | -0.774933444 | 0.067 | 0.112 | 1 | 0.385509064 | 0.263034406 | 0 | 0.071 | 1 | 0.385509064 | 0.350259478 |
| 4930430F08Rik | 0.193935163 | -0.839063782 | 0.067 | 0.112 | 1 | 0.743726438 | 1.111031312 | 0.1 | 0.071 | 1 | 0.743726438 | 0.350259478 |
| B930036N10Rik | 0.193935286 | 0.710297644 | 0.294 | 0.229 | 1 | 0.577889808 | 0.982297998 | 0.2 | 0.142 | 1 | 0.577889808 | 0.350259678 |
| Ncstn | 0.194163994 | -0.436517227 | 0.294 | 0.359 | 1 | 0.231983795 | -0.648834684 | 0.3 | 0.494 | 1 | 0.231983795 | 0.350628332 |
| Aldh3a2 | 0.194511313 | -0.428843299 | 0.084 | 0.135 | 1 | 0.860122985 | 0.558490289 | 0.1 | 0.117 | 1 | 0.860122985 | 0.351187976 |
| Mcts2 | 0.194511313 | 0.666576266 | 0.143 | 0.094 | 1 | 1 | 0.808468542 | 0.1 | 0.099 | 1 | 1 | 0.351187976 |
| Acer3 | 0.194544587 | -0.453517579 | 0.613 | 0.659 | 1 | 0.488583806 | 0.419869408 | 0.9 | 0.765 | 1 | 0.488583806 | 0.351241577 |
| Zfp934 | 0.194658022 | -1.250961574 | 0.134 | 0.182 | 1 | 0.629177319 | -0.152003093 | 0.1 | 0.151 | 1 | 0.629177319 | 0.351424298 |
| Hexb | 0.274662605 | -0.387579504 | 0.647 | 0.612 | 1 | 0.19469851 | -1.135883428 | 0.6 | 0.728 | 1 | 0.274662605 | 0.35148951 |
| Gprc5a | 0.194738927 | 0.908852112 | 0.168 | 0.112 | 1 | 0.397023754 | -0.304006187 | 0 | 0.068 | 1 | 0.397023754 | 0.351554604 |
| Fam222b | 0.19495726 | -0.37594835 | 0.571 | 0.606 | 1 | 0.538698042 | -0.390162831 | 0.6 | 0.66 | 1 | 0.538698042 | 0.351906187 |
| Vps41 | 0.194969199 | -0.41072946 | 0.546 | 0.588 | 1 | 0.201398745 | 0.904859244 | 0.8 | 0.599 | 1 | 0.201398745 | 0.351925409 |
| Shq1 | 0.195088983 | -0.807354922 | 0.067 | 0.112 | 1 | 0.308447337 | 1.017921908 | 0.2 | 0.096 | 1 | 0.308447337 | 0.352118254 |
| Cox10 | 0.803234069 | -0.655351829 | 0.126 | 0.129 | 1 | 0.195100754 | 1.41305985 | 0.3 | 0.151 | 1 | 0.803234069 | 0.352137204 |
| Suds3 | 0.982621991 | -0.222392421 | 0.529 | 0.494 | 1 | 0.195207951 | -0.806506527 | 0.5 | 0.602 | 1 | 0.982621991 | 0.352309758 |
| Vps35l | 0.262108033 | -0.628384781 | 0.319 | 0.359 | 1 | 0.195211409 | 0.695993813 | 0.6 | 0.404 | 1 | 0.262108033 | 0.352315324 |
| Gm48768 | 0.498594647 | -0.485426827 | 0.017 | 0.029 | 1 | 0.19522577 | 2.210566986 | 0.1 | 0.028 | 1 | 0.498594647 | 0.352338439 |
| Cdh4 | 0.949508638 | -1.900464326 | 0.017 | 0.018 | 1 | 0.19522577 | 1.695993813 | 0.1 | 0.028 | 1 | 0.949508638 | 0.352338439 |
| Mir670hg | 0.217412808 | -1.655351829 | 0.008 | 0.029 | 1 | 0.195226183 | 2.31748219 | 0.1 | 0.028 | 1 | 0.217412808 | 0.352339103 |
| Ccdc80 | 0.195256755 | 0.286836481 | 0.975 | 0.982 | 1 | 0.263111718 | -0.280010706 | 1 | 0.957 | 1 | 0.263111718 | 0.352388309 |
| Camk2b | 0.769923893 | -0.11345805 | 0.084 | 0.094 | 1 | 0.195403269 | 2.091922489 | 0.2 | 0.086 | 1 | 0.769923893 | 0.352624101 |
| Sf3b5 | 0.195427135 | 0.147153694 | 0.782 | 0.741 | 1 | 0.47763999 | 0.347383676 | 0.4 | 0.679 | 1 | 0.47763999 | 0.352662505 |
| Zfp799 | 0.46299458 | -0.354182294 | 0.143 | 0.176 | 1 | 0.195438999 | 2.091922489 | 0.2 | 0.086 | 1 | 0.46299458 | 0.352681595 |
| Dsc2 | 0.242304531 | -1.572889668 | 0.017 | 0.041 | 1 | 0.195671161 | -0.68251781 | 0 | 0.145 | 1 | 0.242304531 | 0.353055118 |
| Traf7 | 0.19576896 | -0.597901556 | 0.277 | 0.329 | 1 | 0.436567612 | -0.374395515 | 0.3 | 0.401 | 1 | 0.436567612 | 0.353212435 |
| Pex11a | 0.195865014 | 0.158429363 | 0.151 | 0.1 | 1 | 0.474948015 | 0.494359952 | 0 | 0.049 | 1 | 0.474948015 | 0.353366925 |
| Ergic1 | 0.908420759 | -0.436145924 | 0.706 | 0.618 | 1 | 0.196024602 | 0.611362563 | 0.8 | 0.664 | 1 | 0.908420759 | 0.35362356 |
| Gfod2 | 0.238832134 | -1.167250867 | 0.126 | 0.171 | 1 | 0.196334728 | 0.708066645 | 0.4 | 0.207 | 1 | 0.238832134 | 0.354122131 |
| Neat1 | 0.19635655 | -0.327930208 | 1 | 0.976 | 1 | 0.801754352 | 0.121527967 | 1 | 0.997 | 1 | 0.801754352 | 0.354157205 |
| Gm19325 | 0.250480965 | -0.918386234 | 0.067 | 0.106 | 1 | 0.196396505 | -0.982078092 | 0 | 0.145 | 1 | 0.250480965 | 0.354221423 |
| Atg9a | 0.447602814 | -0.347923303 | 0.16 | 0.194 | 1 | 0.196419284 | -0.888968688 | 0.1 | 0.281 | 1 | 0.447602814 | 0.354258032 |
| Ulk4 | 0.19651089 | -0.201633861 | 0.336 | 0.241 | 1 | 0.631885181 | -0.289506617 | 0.3 | 0.21 | 1 | 0.631885181 | 0.35440525 |
| Ncoa7 | 0.196761149 | -0.702238216 | 0.202 | 0.259 | 1 | 0.811756222 | -0.161987182 | 0.2 | 0.228 | 1 | 0.811756222 | 0.354807349 |
| Slc8a2 | 0.394811349 | -0.11345805 | 0.05 | 0.076 | 1 | 0.196842982 | 1.185031894 | 0.2 | 0.083 | 1 | 0.394811349 | 0.354938805 |
| Egln3 | 0.196931943 | 1.152003093 | 0.076 | 0.041 | 1 | 0.318562315 | 2.017921908 | 0.1 | 0.037 | 1 | 0.318562315 | 0.355081697 |
| Slc50a1 | 0.78224408 | -0.255559285 | 0.63 | 0.588 | 1 | 0.197038086 | -0.191531458 | 0.3 | 0.525 | 1 | 0.78224408 | 0.355252165 |
| Raly | 0.540970304 | -0.103055517 | 0.899 | 0.794 | 1 | 0.197057563 | -0.472044079 | 0.7 | 0.861 | 1 | 0.540970304 | 0.355283443 |
| She | 0.219589239 | -1.070389328 | 0.008 | 0.029 | 1 | 0.197183593 | 2.017921908 | 0.1 | 0.028 | 1 | 0.219589239 | 0.355485816 |
| Abhd16a | 0.404414281 | -0.292781749 | 0.395 | 0.418 | 1 | 0.19727163 | -0.880931369 | 0.2 | 0.383 | 1 | 0.404414281 | 0.355627164 |
| Jade2 | 0.331143962 | -0.384497918 | 0.261 | 0.3 | 1 | 0.197306658 | -0.663902132 | 0.1 | 0.312 | 1 | 0.331143962 | 0.355683399 |
| Ccng2 | 0.197368522 | -0.485426827 | 0.118 | 0.176 | 1 | 0.680387269 | 0.336097868 | 0.2 | 0.142 | 1 | 0.680387269 | 0.35578271 |
| Rpn1 | 0.197373344 | -0.256123279 | 0.866 | 0.818 | 1 | 0.441151884 | -0.241514345 | 0.8 | 0.889 | 1 | 0.441151884 | 0.35579045 |
| Cers2 | 0.576066702 | -0.196430627 | 0.723 | 0.682 | 1 | 0.197381398 | -0.604795515 | 0.7 | 0.741 | 1 | 0.576066702 | 0.35580338 |
| Med15 | 0.197448839 | -0.433499035 | 0.58 | 0.576 | 1 | 0.231441766 | -0.345482823 | 0.4 | 0.676 | 1 | 0.231441766 | 0.355911634 |
| Fyb | 0.324311097 | -0.87244995 | 0.084 | 0.053 | 1 | 0.197619546 | 1.874963954 | 0.2 | 0.09 | 1 | 0.324311097 | 0.356185607 |
| Spop | 0.264574864 | -0.16621482 | 0.933 | 0.918 | 1 | 0.197832184 | -0.579864633 | 0.8 | 0.929 | 1 | 0.264574864 | 0.356526795 |
| Tom1 | 0.198179366 | -0.550521855 | 0.395 | 0.435 | 1 | 0.315744914 | -0.68251781 | 0.5 | 0.586 | 1 | 0.315744914 | 0.357083672 |
| Ddx60 | 0.198238628 | 0.514573173 | 0.076 | 0.041 | 1 | 0.669852582 | 0.432959407 | 0 | 0.019 | 1 | 0.669852582 | 0.357178703 |
| Frg1 | 0.198288491 | -0.361168724 | 0.555 | 0.553 | 1 | 0.72308035 | 0.374065718 | 0.6 | 0.537 | 1 | 0.72308035 | 0.357258656 |
| Micos10 | 0.19838789 | -0.247098934 | 0.933 | 0.941 | 1 | 0.419660999 | -0.364053571 | 0.9 | 0.957 | 1 | 0.419660999 | 0.357418025 |
| Tmem94 | 0.198444724 | -1.185866545 | 0.059 | 0.1 | 1 | 0.45030668 | 0.821524695 | 0.2 | 0.114 | 1 | 0.45030668 | 0.35750914 |
| Zbtb9 | 0.19847768 | -0.918386234 | 0.067 | 0.112 | 1 | 0.270488207 | 1.345496566 | 0.2 | 0.096 | 1 | 0.270488207 | 0.35756197 |
| Rtcb | 0.198570254 | -0.320003218 | 0.639 | 0.653 | 1 | 0.213626331 | -0.438227127 | 0.4 | 0.673 | 1 | 0.213626331 | 0.357710362 |
| Kctd9 | 0.198659277 | -0.213347282 | 0.42 | 0.482 | 1 | 0.499519247 | 0.566710796 | 0.6 | 0.457 | 1 | 0.499519247 | 0.357853046 |
| H1f5 | 0.198710494 | -1.682463674 | 0.109 | 0.065 | 1 | 0.643846086 | 1.111031312 | 0 | 0.022 | 1 | 0.643846086 | 0.357935127 |
| Necap2 | 0.198812719 | -0.31444063 | 0.504 | 0.535 | 1 | 0.292856511 | -0.321928095 | 0.4 | 0.602 | 1 | 0.292856511 | 0.35809894 |
| Cdk18 | 0.609500043 | 0.514573173 | 0.034 | 0.024 | 1 | 0.199153598 | 2.111031312 | 0.1 | 0.028 | 1 | 0.609500043 | 0.35864504 |
| Myom1 | 0.199398986 | -0.576574715 | 0.126 | 0.182 | 1 | 0.481035478 | 0.951832718 | 0.2 | 0.123 | 1 | 0.481035478 | 0.359038016 |
| Frmpd3 | 0.199602566 | 1.514573173 | 0.034 | 0.012 | 1 | 0.730750993 | 1.432959407 | 0 | 0.012 | 1 | 0.730750993 | 0.359363947 |
| Abat | 0.548428218 | -0.173482821 | 0.235 | 0.265 | 1 | 0.199857262 | 1.570462931 | 0.4 | 0.25 | 1 | 0.548428218 | 0.359771599 |
| Chpf2 | 0.664701556 | -0.126972856 | 0.319 | 0.329 | 1 | 0.199965107 | -0.542793046 | 0.2 | 0.426 | 1 | 0.664701556 | 0.35994417 |
| 1110059G10Rik | 0.200145855 | -0.702238216 | 0.269 | 0.318 | 1 | 0.853314095 | 0.362570079 | 0.3 | 0.275 | 1 | 0.853314095 | 0.360233347 |
| Rftn2 | 0.271729238 | 0.272912418 | 0.58 | 0.535 | 1 | 0.200339019 | -1.00166882 | 0.4 | 0.562 | 1 | 0.271729238 | 0.360542316 |
| Cox7a2 | 0.20066936 | 0.153003229 | 0.899 | 0.929 | 1 | 0.2320387 | -0.239875849 | 0.6 | 0.917 | 1 | 0.2320387 | 0.361070528 |
| Dhrs11 | 0.200803355 | -1.040015679 | 0.118 | 0.165 | 1 | 0.232144172 | -0.91281543 | 0 | 0.127 | 1 | 0.232144172 | 0.361284722 |
| Agpat1 | 0.20106164 | -0.578536232 | 0.218 | 0.276 | 1 | 0.459488893 | 0.532495081 | 0.4 | 0.272 | 1 | 0.459488893 | 0.361697496 |
| Slc48a1 | 0.201115294 | 0.254929356 | 0.437 | 0.347 | 1 | 0.456034053 | -0.171902651 | 0.3 | 0.426 | 1 | 0.456034053 | 0.361783226 |
| Ckap4 | 0.789749529 | -0.106609516 | 1 | 0.953 | 1 | 0.201132065 | 0.236562194 | 1 | 0.966 | 1 | 0.789749529 | 0.361810023 |
| H1f3 | 0.847726288 | -0.790281409 | 0.076 | 0.082 | 1 | 0.201133309 | 1.625604485 | 0.1 | 0.028 | 1 | 0.847726288 | 0.361812011 |
| Micu2 | 0.201206347 | -0.435550921 | 0.538 | 0.576 | 1 | 0.881806575 | 0.121373257 | 0.5 | 0.59 | 1 | 0.881806575 | 0.3619287 |
| 1700008J07Rik | 0.803106666 | -0.485426827 | 0.059 | 0.065 | 1 | 0.20120644 | 1.744903414 | 0.2 | 0.083 | 1 | 0.803106666 | 0.361928849 |
| Dnlz | 0.247026372 | 0.251538767 | 0.378 | 0.306 | 1 | 0.201239784 | 0.964810572 | 0.4 | 0.216 | 1 | 0.247026372 | 0.361982117 |
| Ppl | 0.490124306 | -0.944858446 | 0.017 | 0.029 | 1 | 0.201293398 | -0.68251781 | 0 | 0.142 | 1 | 0.490124306 | 0.362067764 |
| Nedd1 | 0.201465432 | -0.748461233 | 0.176 | 0.229 | 1 | 0.677634678 | -0.419483404 | 0.2 | 0.235 | 1 | 0.677634678 | 0.362342544 |
| Alx4 | 0.201502846 | -0.485426827 | 0.294 | 0.353 | 1 | 0.262570715 | -0.42170123 | 0.2 | 0.395 | 1 | 0.262570715 | 0.362402295 |
| Bcl11a | 0.333832652 | -1.622930351 | 0.025 | 0.047 | 1 | 0.201568928 | -0.864721141 | 0 | 0.142 | 1 | 0.333832652 | 0.362507823 |
| Serpinb5 | 0.201658121 | 1.251538767 | 0.034 | 0.012 | 1 | 0.619953009 | 1.695993813 | 0 | 0.025 | 1 | 0.619953009 | 0.362650245 |
| Gm13008 | 0.201658121 | 1.251538767 | 0.034 | 0.012 | 1 | 0.52208609 | 1.017921908 | 0 | 0.04 | 1 | 0.52208609 | 0.362650245 |
| Espn | 0.201658121 | 1.251538767 | 0.034 | 0.012 | 1 | 0.412160956 | 1.374065718 | 0.1 | 0.043 | 1 | 0.412160956 | 0.362650245 |
| E230013L22Rik | 0.201658121 | 1.251538767 | 0.034 | 0.012 | 1 | 0.698534498 | 2.432959407 | 0 | 0.015 | 1 | 0.698534498 | 0.362650245 |
| Gm30978 | 0.201658121 | 1.251538767 | 0.034 | 0.012 | 1 | 0.619961444 | 1.558490289 | 0 | 0.025 | 1 | 0.619961444 | 0.362650245 |
| Camp | 0.201658121 | 1.251538767 | 0.034 | 0.012 | 1 | 0.396883838 | 0.159940913 | 0 | 0.068 | 1 | 0.396883838 | 0.362650245 |
| Gm57036 | 0.201658121 | 1.251538767 | 0.034 | 0.012 | 1 | 0.730750993 | 2.210566986 | 0 | 0.012 | 1 | 0.730750993 | 0.362650245 |
| Rph3al | 0.201658121 | 1.251538767 | 0.034 | 0.012 | 1 | 0.489901827 | 0.769994395 | 0 | 0.046 | 1 | 0.489901827 | 0.362650245 |
| Gm48420 | 0.201658121 | 1.251538767 | 0.034 | 0.012 | 1 | 0.73075158 | 2.695993813 | 0 | 0.012 | 1 | 0.73075158 | 0.362650245 |
| Nckap1l | 0.201658121 | 1.251538767 | 0.034 | 0.012 | 1 | 0.698533455 | 1.695993813 | 0 | 0.015 | 1 | 0.698533455 | 0.362650245 |
| Nrg2 | 0.201658121 | 1.251538767 | 0.034 | 0.012 | 1 | 0.669848329 | 1.695993813 | 0 | 0.019 | 1 | 0.669848329 | 0.362650245 |
| Aar2 | 0.201700674 | -0.900464326 | 0.193 | 0.247 | 1 | 0.586922175 | 0.725140159 | 0.3 | 0.231 | 1 | 0.586922175 | 0.362718186 |
| Gm57390 | 0.2253929 | -0.784987109 | 0.101 | 0.147 | 1 | 0.201803389 | -0.814968106 | 0 | 0.142 | 1 | 0.2253929 | 0.362882169 |
| Stat3 | 0.201867399 | -0.197411368 | 0.975 | 0.994 | 1 | 0.231856415 | -0.46096004 | 1 | 0.991 | 1 | 0.231856415 | 0.362984352 |
| Fam122a | 0.201957608 | 0.451837417 | 0.286 | 0.218 | 1 | 0.322562958 | 0.751135367 | 0.3 | 0.167 | 1 | 0.322562958 | 0.363128341 |
| Fundc2 | 0.202240312 | -0.222392421 | 0.731 | 0.706 | 1 | 0.276753153 | -0.377514857 | 0.4 | 0.688 | 1 | 0.276753153 | 0.36357948 |
| Cbr3 | 0.306406703 | -0.687060688 | 0.143 | 0.182 | 1 | 0.2022992 | -1.230005605 | 0 | 0.142 | 1 | 0.306406703 | 0.363673433 |
| Pappa | 0.836533336 | 0.514573173 | 0.025 | 0.029 | 1 | 0.202608854 | -2.191531458 | 0 | 0.142 | 1 | 0.836533336 | 0.36416736 |
| Myrfl | 0.202681485 | 1.614108846 | 0.034 | 0.012 | 1 | 0.643842184 | 1.31748219 | 0 | 0.022 | 1 | 0.643842184 | 0.364283186 |
| Dennd4b | 0.202738265 | -1.444784843 | 0.059 | 0.1 | 1 | 0.353816462 | -0.152003093 | 0 | 0.08 | 1 | 0.353816462 | 0.364373726 |
| Eif3j2 | 0.202754759 | -0.655351829 | 0.092 | 0.141 | 1 | 0.955037198 | 0.695993813 | 0.1 | 0.105 | 1 | 0.955037198 | 0.364400026 |
| Zfp560 | 0.202787265 | -0.296665184 | 0.336 | 0.406 | 1 | 0.240240513 | -0.872849022 | 0.2 | 0.361 | 1 | 0.240240513 | 0.364451854 |

| | | | | | | | | | | | | |
|---|---|---|---|---|---|---|---|---|---|---|---|---|
| Mindy4 | 0.437496032 | -0.557576613 | 0.269 | 0.294 | 1 | 0.203212663 | 0.467724825 | 0.5 | 0.287 | 1 | 0.437496032 | 0.365129939 |
| Ilvbl | 0.606373044 | -0.263034406 | 0.21 | 0.229 | 1 | 0.203331315 | 0.988174565 | 0.4 | 0.228 | 1 | 0.606373044 | 0.365319006 |
| Mast2 | 0.203498777 | -0.444685033 | 0.748 | 0.812 | 1 | 0.881718022 | 0.207993042 | 0.8 | 0.843 | 1 | 0.881718022 | 0.365585801 |
| Cstf2t | 0.425837967 | -0.432959407 | 0.202 | 0.235 | 1 | 0.203519237 | -0.814968106 | 0.1 | 0.278 | 1 | 0.425837967 | 0.365618395 |
| Edem2 | 0.203625155 | 0.514573173 | 0.286 | 0.218 | 1 | 0.498544503 | 0.625604485 | 0.3 | 0.201 | 1 | 0.498544503 | 0.365787107 |
| Clic6 | 0.203706789 | 0.836501268 | 0.034 | 0.012 | 1 | 0.539483006 | 1.31748219 | 0 | 0.037 | 1 | 0.539483006 | 0.365917123 |
| Man2b1 | 0.204031698 | -0.528071165 | 0.361 | 0.4 | 1 | 0.246198373 | -0.327852929 | 0.3 | 0.509 | 1 | 0.246198373 | 0.366434463 |
| Traf5 | 0.204352325 | -0.282609944 | 0.345 | 0.418 | 1 | 0.762563407 | 0.748461233 | 0.3 | 0.293 | 1 | 0.762563407 | 0.366944778 |
| Cmtm4 | 0.59695239 | -0.296981738 | 0.479 | 0.482 | 1 | 0.204365573 | 0.497499659 | 0.7 | 0.559 | 1 | 0.59695239 | 0.366965858 |
| Oat | 0.297882822 | 0.129827031 | 0.941 | 0.924 | 1 | 0.204516417 | -0.492232005 | 0.8 | 0.954 | 1 | 0.297882822 | 0.367205869 |
| Tbx4 | 0.204733108 | 0.777607579 | 0.034 | 0.012 | 1 | 0.643842184 | 1.695993813 | 0 | 0.022 | 1 | 0.643842184 | 0.367550571 |
| Rnf5 | 0.722235126 | 0.251538767 | 0.303 | 0.288 | 1 | 0.204751959 | -0.888968688 | 0.1 | 0.278 | 1 | 0.722235126 | 0.367580554 |
| Asb8 | 0.204931019 | -0.333423734 | 0.151 | 0.212 | 1 | 0.391752957 | -0.536666944 | 0.1 | 0.207 | 1 | 0.391752957 | 0.367865316 |
| Pgs1 | 0.204979282 | 0.408778509 | 0.445 | 0.376 | 1 | 0.264090327 | -0.505640048 | 0.2 | 0.389 | 1 | 0.264090327 | 0.367942057 |
| Fndc4 | 0.802246533 | -0.292781749 | 0.319 | 0.318 | 1 | 0.205081205 | -0.970762779 | 0.1 | 0.275 | 1 | 0.802246533 | 0.36810411 |
| Fxyd6 | 0.205240415 | 0.404630276 | 0.748 | 0.694 | 1 | 0.813712124 | 0.105697868 | 0.6 | 0.593 | 1 | 0.813712124 | 0.368357202 |
| Col23a1 | 0.20527034 | -2.529820947 | 0.05 | 0.088 | 1 | 0.982059529 | -0.171902651 | 0.1 | 0.099 | 1 | 0.982059529 | 0.368404768 |
| Cdc123 | 0.422774622 | -0.109917692 | 0.798 | 0.771 | 1 | 0.205325867 | 0.408575249 | 0.8 | 0.75 | 1 | 0.422774622 | 0.368493023 |
| Gm56773 | 0.205759512 | 0.836501268 | 0.034 | 0.012 | 1 | 0.73075158 | 2.695993813 | 0 | 0.012 | 1 | 0.73075158 | 0.369182047 |
| Aass | 0.205759512 | 0.836501268 | 0.034 | 0.012 | 1 | 0.278845643 | 2.017921908 | 0.1 | 0.034 | 1 | 0.278845643 | 0.369182047 |
| Pkd1l1 | 0.205759512 | 0.836501268 | 0.034 | 0.012 | 1 | 0.730749233 | 2.432959407 | 0 | 0.012 | 1 | 0.730749233 | 0.369182047 |
| Rnf113a1 | 0.205759512 | 0.836501268 | 0.034 | 0.012 | 1 | 0.577142568 | 1.432959407 | 0 | 0.031 | 1 | 0.577142568 | 0.369182047 |
| Myef2 | 0.312514391 | 0.139361654 | 0.504 | 0.435 | 1 | 0.206430381 | 0.829618377 | 0.5 | 0.364 | 1 | 0.312514391 | 0.37024726 |
| Erbb4 | 0.206620145 | 2.192645078 | 0.067 | 0.035 | 1 | 0.535552133 | -2.647414009 | 0.1 | 0.16 | 1 | 0.535552133 | 0.370548405 |
| Sfrp2 | 0.206794694 | -3.765953424 | 0.21 | 0.253 | 1 | 0.353956584 | -2.604129911 | 0 | 0.08 | 1 | 0.353956584 | 0.370825342 |
| Gm16201 | 0.20689069 | -1.137503524 | 0.05 | 0.088 | 1 | 0.41325416 | 1.017921908 | 0.2 | 0.111 | 1 | 0.41325416 | 0.370977623 |
| Serp2 | 0.206968358 | -0.900464326 | 0.059 | 0.1 | 1 | 0.343825642 | -0.111361109 | 0 | 0.083 | 1 | 0.343825642 | 0.371100815 |
| Fam114a2 | 0.411881569 | -0.426533138 | 0.613 | 0.547 | 1 | 0.206998218 | -0.378207129 | 0.4 | 0.639 | 1 | 0.411881569 | 0.371148174 |
| 6230400D17Rik | 0.620760474 | -0.250961574 | 0.134 | 0.153 | 1 | 0.207490875 | -0.736965594 | 0 | 0.139 | 1 | 0.620760474 | 0.371929287 |
| Rsph9 | 0.333027639 | -0.707819249 | 0.118 | 0.153 | 1 | 0.207628562 | -0.814968106 | 0 | 0.139 | 1 | 0.333027639 | 0.372147505 |
| Ppp4c | 0.234270786 | -0.322996362 | 0.555 | 0.588 | 1 | 0.207670508 | -0.315978829 | 0.6 | 0.747 | 1 | 0.234270786 | 0.372213976 |
| Sgsm3 | 0.853331236 | 0.140177658 | 0.286 | 0.288 | 1 | 0.207904204 | 0.642882477 | 0.5 | 0.302 | 1 | 0.853331236 | 0.372584251 |
| Polb | 0.207906361 | -0.410464769 | 0.445 | 0.488 | 1 | 0.244112213 | -0.763437806 | 0.5 | 0.599 | 1 | 0.244112213 | 0.372587668 |
| Noc4l | 0.43825949 | -0.122856748 | 0.118 | 0.153 | 1 | 0.208256419 | 1.457206954 | 0.3 | 0.16 | 1 | 0.43825949 | 0.373142102 |
| Gstp1 | 0.208329163 | -0.833350131 | 0.084 | 0.129 | 1 | 0.644080819 | 0.111031312 | 0.1 | 0.151 | 1 | 0.644080819 | 0.373257285 |
| Thnsl2 | 0.208412017 | -0.600904045 | 0.134 | 0.188 | 1 | 0.213257061 | -0.709998547 | 0 | 0.136 | 1 | 0.213257061 | 0.373388466 |
| Gstm4 | 0.627671129 | -0.418312631 | 0.084 | 0.1 | 1 | 0.208672055 | 1.473601392 | 0.2 | 0.083 | 1 | 0.627671129 | 0.373800083 |
| Rwdd4a | 0.209213194 | -0.448900951 | 0.395 | 0.435 | 1 | 0.657704027 | -0.339630097 | 0.4 | 0.414 | 1 | 0.657704027 | 0.374656228 |
| Sult1e1 | 0.209265315 | 2.251538767 | 0.042 | 0.018 | 1 | 0.266654026 | -0.473931188 | 0 | 0.111 | 1 | 0.266654026 | 0.374738658 |
| Tspyl4 | 0.20936991 | 0.487606125 | 0.303 | 0.241 | 1 | 0.906174647 | 0.299103661 | 0.2 | 0.176 | 1 | 0.906174647 | 0.374904062 |
| Gm57269 | 0.209374352 | -0.137503524 | 0.092 | 0.053 | 1 | 0.479305119 | 1.847996907 | 0.1 | 0.049 | 1 | 0.479305119 | 0.374911084 |
| Tbc1d13 | 0.304957596 | -0.67305383 | 0.118 | 0.159 | 1 | 0.209557815 | 1.190102883 | 0.3 | 0.154 | 1 | 0.304957596 | 0.375201152 |
| Pex6 | 0.20991879 | -0.589763487 | 0.151 | 0.206 | 1 | 0.351299375 | 0.795529487 | 0.4 | 0.269 | 1 | 0.351299375 | 0.375771682 |
| Brme1 | 0.209949382 | 0.514573173 | 0.034 | 0.012 | 1 | 0.698533977 | 1.558490289 | 0 | 0.015 | 1 | 0.698533977 | 0.375820022 |
| Lamtor2 | 0.210077302 | 0.148923701 | 0.815 | 0.759 | 1 | 0.244328005 | -0.290830798 | 0.5 | 0.688 | 1 | 0.244328005 | 0.376022131 |
| Csdc2 | 0.294977787 | -0.544320516 | 0.076 | 0.112 | 1 | 0.210253649 | 2.091922489 | 0.2 | 0.09 | 1 | 0.294977787 | 0.376300701 |
| C430014B12Rik | 0.210665377 | 1.029146346 | 0.067 | 0.035 | 1 | 0.474928012 | 0.695993813 | 0 | 0.049 | 1 | 0.474928012 | 0.376950853 |
| Tbp | 0.210703583 | -0.600904045 | 0.218 | 0.276 | 1 | 0.383786339 | 0.558490289 | 0.4 | 0.256 | 1 | 0.383786339 | 0.377011166 |
| Mrpl48 | 0.210908992 | -0.395131654 | 0.79 | 0.8 | 1 | 0.56272345 | -0.21857071 | 0.7 | 0.753 | 1 | 0.56272345 | 0.377335381 |
| Czib | 0.33641152 | -0.423142549 | 0.479 | 0.488 | 1 | 0.210919855 | -0.846264237 | 0.3 | 0.457 | 1 | 0.33641152 | 0.377352525 |
| Gm56807 | 0.211102229 | -1.14839184 | 0.05 | 0.088 | 1 | 0.655218074 | -0.736965594 | 0.1 | 0.062 | 1 | 0.655218074 | 0.377640307 |
| Prpf38a | 0.211890815 | -0.432959407 | 0.445 | 0.476 | 1 | 0.290039091 | -0.35463226 | 0.3 | 0.525 | 1 | 0.290039091 | 0.378883913 |
| Glrx3 | 0.212030086 | -0.186435544 | 0.748 | 0.741 | 1 | 0.320986375 | -0.478503918 | 0.7 | 0.809 | 1 | 0.320986375 | 0.379103414 |
| Hyi | 0.212353088 | -0.582724029 | 0.303 | 0.347 | 1 | 0.350378717 | -0.58694015 | 0.2 | 0.324 | 1 | 0.350378717 | 0.379612341 |
| Gm1673 | 0.212433903 | -0.807354922 | 0.109 | 0.159 | 1 | 0.225744174 | 1.602884409 | 0.2 | 0.086 | 1 | 0.225744174 | 0.379739643 |
| Ndufaf2 | 0.480596325 | -0.229867542 | 0.378 | 0.435 | 1 | 0.212560653 | 0.930459067 | 0.5 | 0.38 | 1 | 0.480596325 | 0.379939274 |
| Pex13 | 0.212611647 | -0.493332025 | 0.529 | 0.553 | 1 | 0.439638044 | -0.339630097 | 0.5 | 0.568 | 1 | 0.439638044 | 0.380019582 |
| Gm57028 | 0.212902821 | 1.321928095 | 0.042 | 0.018 | 1 | 0.698529281 | 2.210566986 | 0 | 0.015 | 1 | 0.698529281 | 0.380478031 |
| Wnt11 | 0.446795972 | -0.460335846 | 0.294 | 0.324 | 1 | 0.212950357 | 0.898182664 | 0.5 | 0.312 | 1 | 0.446795972 | 0.380552859 |
| Sec13 | 0.3713301 | -0.201633861 | 0.857 | 0.782 | 1 | 0.212975832 | -0.287683882 | 0.6 | 0.793 | 1 | 0.3713301 | 0.38059296 |
| Zgpat | 0.240313282 | -0.839063782 | 0.143 | 0.188 | 1 | 0.213153168 | -0.970762779 | 0.1 | 0.272 | 1 | 0.240313282 | 0.380872062 |
| S100a14 | 0.491791629 | -0.707819249 | 0.017 | 0.029 | 1 | 0.213336472 | -0.789433014 | 0 | 0.136 | 1 | 0.491791629 | 0.381160493 |
| Ndufs7 | 0.519135371 | -0.11345805 | 0.613 | 0.6 | 1 | 0.213340176 | -0.635520331 | 0.4 | 0.577 | 1 | 0.519135371 | 0.381166321 |
| Runx2os1 | 0.213359606 | 0.450442835 | 0.143 | 0.094 | 1 | 0.792440823 | 0.432959407 | 0.1 | 0.127 | 1 | 0.792440823 | 0.381196891 |
| Spring1 | 0.780445949 | 0.234465254 | 0.176 | 0.165 | 1 | 0.21341222 | -0.68251781 | 0 | 0.136 | 1 | 0.780445949 | 0.381279665 |
| Rnd3 | 0.958601157 | 0.520791696 | 0.588 | 0.612 | 1 | 0.213538125 | 0.666592897 | 0.4 | 0.654 | 1 | 0.958601157 | 0.381477719 |
| Gm37640 | 0.832144311 | -0.11345805 | 0.076 | 0.082 | 1 | 0.213605742 | -0.814968106 | 0 | 0.136 | 1 | 0.832144311 | 0.38158407 |
| Tmem14a | 0.213706053 | 0.602036014 | 0.134 | 0.088 | 1 | 0.799264655 | 1.210566986 | 0.1 | 0.077 | 1 | 0.799264655 | 0.381741829 |
| Cyp4v3 | 0.539461866 | 0.140177658 | 0.151 | 0.124 | 1 | 0.213727127 | -1.111361109 | 0 | 0.136 | 1 | 0.539461866 | 0.381774969 |
| Zfp768 | 0.213757656 | -0.925999419 | 0.05 | 0.088 | 1 | 0.748798999 | 0.236562194 | 0.1 | 0.133 | 1 | 0.748798999 | 0.381822976 |
| Lsamp | 0.946477164 | -1.411426246 | 0.025 | 0.024 | 1 | 0.21380157 | -2.159995884 | 0.2 | 0.367 | 1 | 0.946477164 | 0.381892028 |
| Mtmr9 | 0.213844333 | -0.645891499 | 0.126 | 0.176 | 1 | 0.320734601 | 0.710493383 | 0.4 | 0.247 | 1 | 0.320734601 | 0.381959268 |
| Atp6v0d2 | 0.213853217 | 2.099535674 | 0.042 | 0.018 | 1 | 0.643850855 | -2.661558192 | 0 | 0.022 | 1 | 0.643850855 | 0.381973235 |
| Dcbld2 | 0.428969133 | 0.223983044 | 0.546 | 0.471 | 1 | 0.2139888 | -1.121629444 | 0.3 | 0.435 | 1 | 0.428969133 | 0.382186394 |
| Pcdhb22 | 0.628301557 | 0.308122295 | 0.067 | 0.053 | 1 | 0.214223869 | 0.930459067 | 0.2 | 0.08 | 1 | 0.628301557 | 0.382555871 |
| Smim12 | 0.214560014 | 0.381306642 | 0.227 | 0.165 | 1 | 0.366953053 | -0.391469028 | 0.1 | 0.216 | 1 | 0.366953053 | 0.383084028 |
| Agpat2 | 0.214654481 | 0.355374578 | 0.353 | 0.276 | 1 | 0.765744384 | -0.152003093 | 0.2 | 0.228 | 1 | 0.765744384 | 0.383232417 |
| Fam216a | 0.214768765 | -0.678071905 | 0.244 | 0.3 | 1 | 0.405365001 | -0.441509711 | 0.2 | 0.312 | 1 | 0.405365001 | 0.383411908 |
| Trappc13 | 0.341595485 | -0.501728639 | 0.328 | 0.347 | 1 | 0.214973033 | -0.987702641 | 0.3 | 0.444 | 1 | 0.341595485 | 0.383732661 |
| Cep78 | 0.215096734 | -1.424026283 | 0.042 | 0.076 | 1 | 0.365421586 | 1.017921908 | 0.1 | 0.04 | 1 | 0.365421586 | 0.383926863 |
| Fbxo4 | 0.215341403 | 0.21649182 | 0.403 | 0.312 | 1 | 0.808827722 | 0.159940913 | 0.3 | 0.343 | 1 | 0.808827722 | 0.384310886 |
| Ift43 | 0.215354867 | -0.390078951 | 0.496 | 0.535 | 1 | 0.309628617 | -0.408342847 | 0.4 | 0.552 | 1 | 0.309628617 | 0.384332015 |
| Pnrc2 | 0.524108889 | -0.456857675 | 0.353 | 0.359 | 1 | 0.215481036 | -0.888968688 | 0.2 | 0.373 | 1 | 0.524108889 | 0.384529995 |
| Cdc14a | 0.370969596 | -0.507121898 | 0.336 | 0.371 | 1 | 0.215693312 | -0.674569057 | 0.3 | 0.5 | 1 | 0.370969596 | 0.384863019 |
| Gm26788 | 0.215728257 | 1.099535674 | 0.042 | 0.018 | 1 | 0.557729726 | 1.31748219 | 0 | 0.034 | 1 | 0.557729726 | 0.384917832 |
| 1810059H22Rik | 0.215728257 | 1.099535674 | 0.042 | 0.018 | 1 | 0.577142568 | 1.432959407 | 0 | 0.031 | 1 | 0.577142568 | 0.384917832 |
| Cog7 | 0.215788207 | -0.724213687 | 0.193 | 0.241 | 1 | 0.867785371 | -0.165299916 | 0.2 | 0.21 | 1 | 0.867785371 | 0.385011863 |
| Sptssa | 0.215865036 | 0.120794793 | 0.739 | 0.729 | 1 | 0.232864126 | -0.491313267 | 0.7 | 0.753 | 1 | 0.232864126 | 0.385132358 |
| Slf1 | 0.215954229 | -0.876447445 | 0.311 | 0.347 | 1 | 0.408364711 | 0.741081703 | 0.4 | 0.34 | 1 | 0.408364711 | 0.385272229 |
| Mfap1b | 0.216187091 | -0.399012075 | 0.252 | 0.318 | 1 | 0.816647894 | -0.159995884 | 0.4 | 0.392 | 1 | 0.816647894 | 0.385637324 |
| Ramp2 | 0.216236217 | -0.509674373 | 0.294 | 0.347 | 1 | 0.294768651 | -0.763437806 | 0.1 | 0.238 | 1 | 0.294768651 | 0.385714332 |
| Col19a1 | 0.216340691 | -1.655351829 | 0.008 | 0.029 | 1 | 0.487557552 | 1.31748219 | 0.1 | 0.049 | 1 | 0.487557552 | 0.385878088 |
| Tspan1 | 0.216340691 | -1.655351829 | 0.008 | 0.029 | 1 | 0.490000637 | 1.017921908 | 0 | 0.046 | 1 | 0.490000637 | 0.385878088 |
| Pcdhb11 | 0.216340691 | -2.070389328 | 0.008 | 0.029 | 1 | 0.698532411 | 2.017921908 | 0 | 0.015 | 1 | 0.698532411 | 0.385878088 |
| Yif1b | 0.216360182 | -0.227358674 | 0.521 | 0.582 | 1 | 0.687330511 | -0.136896182 | 0.5 | 0.549 | 1 | 0.687330511 | 0.385908636 |
| Tbxas1 | 0.216626974 | 1.362570079 | 0.05 | 0.024 | 1 | 0.643842184 | 1.558490289 | 0 | 0.022 | 1 | 0.643842184 | 0.386326703 |
| Slc4a11 | 0.216724535 | 1.029146346 | 0.059 | 0.029 | 1 | 0.43441669 | 2.930459067 | 0.1 | 0.049 | 1 | 0.43441669 | 0.386479545 |
| Sap18b | 0.216918504 | -0.485426827 | 0.193 | 0.253 | 1 | 0.842884833 | 0.868174788 | 0.2 | 0.188 | 1 | 0.842884833 | 0.38678337 |

| | | | | | | | | | | | | |
|---|---|---|---|---|---|---|---|---|---|---|---|---|
| Nfil3 | 0.21702985 | -0.669851398 | 0.252 | 0.3 | 1 | 0.33897244 | -0.138582578 | 0.2 | 0.37 | 1 | 0.33897244 | 0.386957744 |
| Gm16023 | 0.235608199 | 0.164075926 | 0.252 | 0.188 | 1 | 0.217111813 | 1.089814826 | 0.4 | 0.241 | 1 | 0.235608199 | 0.387086087 |
| Gan | 0.217147268 | 0.275107238 | 0.403 | 0.324 | 1 | 0.430480195 | 0.68163852 | 0.5 | 0.373 | 1 | 0.430480195 | 0.387141601 |
| Akap13 | 0.217346162 | -0.450323026 | 0.824 | 0.853 | 1 | 0.385348913 | 0.316732543 | 1 | 0.877 | 1 | 0.385348913 | 0.38745297 |
| Susd4 | 0.217412808 | -1.485426827 | 0.008 | 0.029 | 1 | 0.522114744 | 0.695993813 | 0 | 0.04 | 1 | 0.522114744 | 0.387557286 |
| 6530402F18Rik | 0.217412808 | -1.485426827 | 0.008 | 0.029 | 1 | 0.398985336 | 2.017921908 | 0.1 | 0.043 | 1 | 0.398985336 | 0.387557286 |
| Nnat | 0.217412808 | -1.807354922 | 0.008 | 0.029 | 1 | 0.619961444 | 1.31748219 | 0 | 0.025 | 1 | 0.619961444 | 0.387557286 |
| 4930537H20Rik | 0.217412808 | -1.485426827 | 0.008 | 0.029 | 1 | 0.597845674 | 1.695993813 | 0 | 0.028 | 1 | 0.597845674 | 0.387557286 |
| Gm43598 | 0.217412808 | -1.655351829 | 0.008 | 0.029 | 1 | 0.669853054 | 2.210566986 | 0 | 0.019 | 1 | 0.669853054 | 0.387557286 |
| Gm20404 | 0.217412808 | -1.485426827 | 0.008 | 0.029 | 1 | 0.557729726 | 1.31748219 | 0 | 0.034 | 1 | 0.557729726 | 0.387557286 |
| Aplp1 | 0.217412808 | -1.655351829 | 0.008 | 0.029 | 1 | 0.669843604 | 2.017921908 | 0 | 0.019 | 1 | 0.669843604 | 0.387557286 |
| Chrna7 | 0.217412808 | -1.807354922 | 0.008 | 0.029 | 1 | 0.557776184 | 0.494359952 | 0 | 0.034 | 1 | 0.557776184 | 0.387557286 |
| Zfp791 | 0.217412808 | -1.655351829 | 0.008 | 0.029 | 1 | 0.450330991 | 1.374065718 | 0.1 | 0.046 | 1 | 0.450330991 | 0.387557286 |
| 4930513N10Rik | 0.217412808 | -1.655351829 | 0.008 | 0.029 | 1 | 0.698534498 | 2.432959407 | 0 | 0.015 | 1 | 0.698534498 | 0.387557286 |
| Slc35f1 | 0.217412808 | -2.572889668 | 0.008 | 0.029 | 1 | 0.669843604 | 2.017921908 | 0 | 0.019 | 1 | 0.669843604 | 0.387557286 |
| Slc47a1 | 0.217412808 | -1.655351829 | 0.008 | 0.029 | 1 | 0.730749233 | 2.017921908 | 0 | 0.012 | 1 | 0.730749233 | 0.387557286 |
| Gm10190 | 0.217412808 | -1.485426827 | 0.008 | 0.029 | 1 | 0.643842184 | 1.695993813 | 0 | 0.022 | 1 | 0.643842184 | 0.387557286 |
| Pcdhb16 | 0.217412808 | -1.485426827 | 0.008 | 0.029 | 1 | 0.364073509 | -0.111361109 | 0 | 0.077 | 1 | 0.364073509 | 0.387557286 |
| Slc9a7 | 0.217412808 | -1.944858446 | 0.008 | 0.029 | 1 | 0.730750993 | 0.847996907 | 0 | 0.012 | 1 | 0.730750993 | 0.387557286 |
| 2700069I18Rik | 0.217575341 | 1 | 0.042 | 0.018 | 1 | 0.491223492 | 1.847996907 | 0.1 | 0.052 | 1 | 0.491223492 | 0.387811653 |
| Styxl1 | 0.217575341 | 1 | 0.042 | 0.018 | 1 | 0.669843604 | 1.432959407 | 0 | 0.019 | 1 | 0.669843604 | 0.387811653 |
| A330035P11Rik | 0.217575341 | 1 | 0.042 | 0.018 | 1 | 0.362872338 | 1.847996907 | 0.1 | 0.04 | 1 | 0.362872338 | 0.387811653 |
| Zfp654 | 0.217639845 | -0.408121101 | 0.471 | 0.506 | 1 | 0.682739469 | 0.185031894 | 0.7 | 0.633 | 1 | 0.682739469 | 0.387912588 |
| Pdcd5 | 0.846808652 | -0.20685157 | 0.714 | 0.676 | 1 | 0.217815071 | 1.136067863 | 0.6 | 0.698 | 1 | 0.846808652 | 0.388186737 |
| Nipal2 | 0.217966241 | -1.343407822 | 0.05 | 0.088 | 1 | 0.424323555 | -0.5519337 | 0.1 | 0.198 | 1 | 0.424323555 | 0.388423199 |
| Dtwd2 | 0.218249139 | -1.424026283 | 0.042 | 0.076 | 1 | 0.236382673 | 1.354956895 | 0.2 | 0.086 | 1 | 0.236382673 | 0.388865591 |
| Rpe | 0.218280802 | -0.584962501 | 0.202 | 0.259 | 1 | 0.323161532 | -0.668578619 | 0.1 | 0.228 | 1 | 0.323161532 | 0.388915095 |
| Serping1 | 0.218308843 | -1.678071905 | 0.042 | 0.076 | 1 | 0.343972692 | -1.982078092 | 0 | 0.083 | 1 | 0.343972692 | 0.388958935 |
| Usp30 | 0.218337102 | -0.707819249 | 0.059 | 0.1 | 1 | 0.737271939 | 0.602884409 | 0.2 | 0.154 | 1 | 0.737271939 | 0.389003114 |
| Platr1 | 0.218485013 | -1.292781749 | 0.008 | 0.029 | 1 | 0.698529281 | 2.210566986 | 0 | 0.015 | 1 | 0.698529281 | 0.389234324 |
| Pla2r1 | 0.218485013 | -2.485426827 | 0.008 | 0.029 | 1 | 0.730749233 | 2.210566986 | 0 | 0.012 | 1 | 0.730749233 | 0.389234324 |
| Gm43260 | 0.218485013 | -1.292781749 | 0.008 | 0.029 | 1 | 0.643850855 | 2.017921908 | 0 | 0.022 | 1 | 0.643850855 | 0.389234324 |
| Fam131c | 0.218485013 | -1.292781749 | 0.008 | 0.029 | 1 | 0.698534498 | 2.432959407 | 0 | 0.015 | 1 | 0.698534498 | 0.389234324 |
| 4933404O12Rik | 0.218485013 | -1.292781749 | 0.008 | 0.029 | 1 | 0.577171068 | 0.847996907 | 0 | 0.031 | 1 | 0.577171068 | 0.389234324 |
| Epo | 0.218485013 | -1.292781749 | 0.008 | 0.029 | 1 | 0.599038642 | 1.432959407 | 0.1 | 0.059 | 1 | 0.599038642 | 0.389234324 |
| Fgd5 | 0.218485013 | -1.655351829 | 0.008 | 0.029 | 1 | 0.557756607 | 0.930459067 | 0 | 0.034 | 1 | 0.557756607 | 0.389234324 |
| Gm32592 | 0.218485013 | -2.292781749 | 0.008 | 0.029 | 1 | 0.489966712 | 0.210566986 | 0 | 0.046 | 1 | 0.489966712 | 0.389234324 |
| Pianp | 0.218485013 | -1.292781749 | 0.008 | 0.029 | 1 | 0.669853054 | 2.210566986 | 0 | 0.019 | 1 | 0.669853054 | 0.389234324 |
| Clec2d | 0.218485013 | -1.292781749 | 0.008 | 0.029 | 1 | 0.446844446 | 0.31748219 | 0 | 0.056 | 1 | 0.446844446 | 0.389234324 |
| Gm44777 | 0.218485013 | -1.292781749 | 0.008 | 0.029 | 1 | 0.44222087 | 1.847996907 | 0.1 | 0.046 | 1 | 0.44222087 | 0.389234324 |
| Lrrc56 | 0.218485013 | -1.292781749 | 0.008 | 0.029 | 1 | 0.698529281 | 2.210566986 | 0 | 0.015 | 1 | 0.698529281 | 0.389234324 |
| Rwdd2a | 0.218485013 | -1.485426827 | 0.008 | 0.029 | 1 | 0.597824685 | 1.432959407 | 0 | 0.028 | 1 | 0.597824685 | 0.389234324 |
| Rab6b | 0.218485013 | -1.944858446 | 0.008 | 0.029 | 1 | 0.557744661 | 1.210566986 | 0 | 0.034 | 1 | 0.557744661 | 0.389234324 |
| Zc3h12d | 0.218485013 | -1.485426827 | 0.008 | 0.029 | 1 | 0.669843604 | 2.017921908 | 0 | 0.019 | 1 | 0.669843604 | 0.389234324 |
| Adora2a | 0.218485013 | -1.292781749 | 0.008 | 0.029 | 1 | 0.597824685 | 1.31748219 | 0 | 0.028 | 1 | 0.597824685 | 0.389234324 |
| Dnah9 | 0.218485013 | -1.292781749 | 0.008 | 0.029 | 1 | 0.669843604 | 1.847996907 | 0 | 0.019 | 1 | 0.669843604 | 0.389234324 |
| Gm11627 | 0.218485013 | -1.292781749 | 0.008 | 0.029 | 1 | 0.6199755 | 1.847996907 | 0 | 0.025 | 1 | 0.6199755 | 0.389234324 |
| Taco1os | 0.218485013 | -1.292781749 | 0.008 | 0.029 | 1 | 0.698529281 | 2.017921908 | 0 | 0.015 | 1 | 0.698529281 | 0.389234324 |
| Zfp184 | 0.218485013 | -1.292781749 | 0.008 | 0.029 | 1 | 0.643835681 | 1.847996907 | 0 | 0.022 | 1 | 0.643835681 | 0.389234324 |
| Ap1g2 | 0.218485013 | -1.292781749 | 0.008 | 0.029 | 1 | 0.698529281 | 2.210566986 | 0 | 0.015 | 1 | 0.698529281 | 0.389234324 |
| Adcy4 | 0.218485013 | -1.292781749 | 0.008 | 0.029 | 1 | 0.505582845 | 1.017921908 | 0 | 0.043 | 1 | 0.505582845 | 0.389234324 |
| Nrros | 0.218485013 | -2.185866545 | 0.008 | 0.029 | 1 | 0.698529281 | 2.017921908 | 0 | 0.015 | 1 | 0.698529281 | 0.389234324 |
| Dnd1 | 0.218485013 | -1.292781749 | 0.008 | 0.029 | 1 | 0.474877085 | 0.847996907 | 0 | 0.049 | 1 | 0.474877085 | 0.389234324 |
| Carns1 | 0.218485013 | -1.292781749 | 0.008 | 0.029 | 1 | 0.460551517 | 0.625604485 | 0 | 0.052 | 1 | 0.460551517 | 0.389234324 |
| ENSMUSG00000121502 | 0.752935768 | -0.333423734 | 0.067 | 0.076 | 1 | 0.218871506 | 1.245332404 | 0.2 | 0.083 | 1 | 0.752935768 | 0.389838276 |
| Tamm41 | 0.218897051 | -0.563429339 | 0.059 | 0.1 | 1 | 0.902716678 | 0.558490289 | 0.1 | 0.111 | 1 | 0.902716678 | 0.389878183 |
| Rnf32 | 0.219074506 | 0.514573173 | 0.076 | 0.041 | 1 | 0.643842184 | 1.558490289 | 0 | 0.022 | 1 | 0.643842184 | 0.390155372 |
| Zfp141 | 0.219092883 | -0.485426827 | 0.059 | 0.1 | 1 | 0.886002664 | 2.14345279 | 0.1 | 0.096 | 1 | 0.886002664 | 0.390184075 |
| A930012L18Rik | 0.21910035 | 0.929610672 | 0.059 | 0.029 | 1 | 0.236765774 | 2.210566986 | 0.1 | 0.031 | 1 | 0.236765774 | 0.390195737 |
| Zfp617 | 0.219161616 | -0.485426827 | 0.176 | 0.235 | 1 | 0.98244984 | 0.263034406 | 0.2 | 0.188 | 1 | 0.98244984 | 0.390291417 |
| Wdr72 | 0.944467697 | -2.485426827 | 0.017 | 0.018 | 1 | 0.21925749 | -1.339630097 | 0 | 0.133 | 1 | 0.944467697 | 0.390441133 |
| Rfng | 0.219304728 | -0.632268215 | 0.109 | 0.159 | 1 | 0.737056954 | 1.185031894 | 0.1 | 0.154 | 1 | 0.737056954 | 0.390514892 |
| Sdhaf1 | 0.865689927 | -0.308549065 | 0.101 | 0.106 | 1 | 0.219415333 | -0.736965594 | 0 | 0.133 | 1 | 0.865689927 | 0.390687577 |
| Asl | 0.376282416 | -0.436517227 | 0.101 | 0.135 | 1 | 0.219556817 | -0.763437806 | 0 | 0.133 | 1 | 0.376282416 | 0.390908437 |
| Bend5 | 0.219589239 | -1.070389328 | 0.008 | 0.029 | 1 | 0.597838178 | 0.930459067 | 0 | 0.028 | 1 | 0.597838178 | 0.390959044 |
| 4930557J02Rik | 0.219589239 | -1.070389328 | 0.008 | 0.029 | 1 | 0.643835681 | 1.432959407 | 0 | 0.022 | 1 | 0.643835681 | 0.390959044 |
| A430102K17Rik | 0.219589239 | -1.070389328 | 0.008 | 0.029 | 1 | 0.59781419 | 1.432959407 | 0 | 0.028 | 1 | 0.59781419 | 0.390959044 |
| Pex26 | 0.219589239 | -1.070389328 | 0.008 | 0.029 | 1 | 0.408673423 | 0.374065718 | 0 | 0.065 | 1 | 0.408673423 | 0.390959044 |
| Gm57192 | 0.219589239 | -1.070389328 | 0.008 | 0.029 | 1 | 0.406872117 | 1.625604485 | 0.1 | 0.043 | 1 | 0.406872117 | 0.390959044 |
| Myo7a | 0.219589239 | -1.070389328 | 0.008 | 0.029 | 1 | 0.619961444 | 1.558490289 | 0 | 0.025 | 1 | 0.619961444 | 0.390959044 |
| Stk32c | 0.219589239 | -1.070389328 | 0.008 | 0.029 | 1 | 0.234627319 | 2.017921908 | 0.1 | 0.031 | 1 | 0.234627319 | 0.390959044 |
| Fam124a | 0.219589239 | -1.070389328 | 0.008 | 0.029 | 1 | 0.730750993 | 2.017921908 | 0 | 0.012 | 1 | 0.730750993 | 0.390959044 |
| Aifm3 | 0.219589239 | -1.070389328 | 0.008 | 0.029 | 1 | 0.232498909 | 2.432959407 | 0.1 | 0.031 | 1 | 0.232498909 | 0.390959044 |
| Scgb2b27 | 0.219592075 | -0.622930351 | 0.277 | 0.329 | 1 | 0.949913265 | 0.732519689 | 0.1 | 0.093 | 1 | 0.949913265 | 0.39096347 |
| Pald1 | 0.219613651 | -0.918386234 | 0.059 | 0.1 | 1 | 0.649373782 | 1.299103661 | 0.2 | 0.173 | 1 | 0.649373782 | 0.390997146 |
| Rtl8a | 0.219851105 | -0.403813062 | 0.555 | 0.553 | 1 | 0.709341436 | 0.124206701 | 0.7 | 0.552 | 1 | 0.709341436 | 0.391367701 |
| Dock7 | 0.37154731 | -0.105630334 | 0.832 | 0.847 | 1 | 0.219921659 | -0.524336142 | 0.8 | 0.873 | 1 | 0.37154731 | 0.391477782 |
| Gm13056 | 0.220074704 | 1 | 0.05 | 0.024 | 1 | 0.669853054 | 2.210566986 | 0 | 0.019 | 1 | 0.669853054 | 0.391716532 |
| Usp20 | 0.394689527 | -0.780882711 | 0.084 | 0.112 | 1 | 0.220238922 | 1.847996907 | 0.2 | 0.09 | 1 | 0.394689527 | 0.391972661 |
| 6430573P05Rik | 0.220389995 | 0.777607579 | 0.042 | 0.018 | 1 | 0.669843604 | 1.847996907 | 0 | 0.019 | 1 | 0.669843604 | 0.39220824 |
| Gm57397 | 0.220389995 | 0.777607579 | 0.042 | 0.018 | 1 | 0.669853054 | 2.210566986 | 0 | 0.019 | 1 | 0.669853054 | 0.39220824 |
| Ism2 | 0.220389995 | 0.777607579 | 0.042 | 0.018 | 1 | 0.365421266 | 1.494359952 | 0.1 | 0.04 | 1 | 0.365421266 | 0.39220824 |
| Them6 | 0.220389995 | 0.777607579 | 0.042 | 0.018 | 1 | 0.669848329 | 1.847996907 | 0 | 0.019 | 1 | 0.669848329 | 0.39220824 |
| Fermt3 | 0.220673861 | -1.359895945 | 0.008 | 0.029 | 1 | 0.971008345 | 0.494359952 | 0.1 | 0.093 | 1 | 0.971008345 | 0.392650769 |
| Emc9 | 0.220877333 | 1.029146346 | 0.05 | 0.024 | 1 | 0.234627319 | 2.210566986 | 0.1 | 0.031 | 1 | 0.234627319 | 0.39296787 |
| Ak1 | 0.221257696 | -0.404808367 | 0.218 | 0.276 | 1 | 0.45738534 | -0.210896782 | 0.2 | 0.324 | 1 | 0.45738534 | 0.393560424 |
| Nxpe3 | 0.221428712 | -1.292781749 | 0.042 | 0.076 | 1 | 0.640912765 | 1.159940913 | 0.1 | 0.062 | 1 | 0.640912765 | 0.393826749 |
| Ly6d | 0.221429694 | -3.090288885 | 0.042 | 0.076 | 1 | 0.577155236 | 1.31748219 | 0 | 0.031 | 1 | 0.577155236 | 0.393828279 |
| Trim13 | 0.294489166 | -0.584962501 | 0.05 | 0.082 | 1 | 0.221457287 | 1.31748219 | 0.2 | 0.083 | 1 | 0.294489166 | 0.393871243 |
| St14 | 0.221503692 | -1.201633861 | 0.042 | 0.076 | 1 | 0.576213673 | 1.409112665 | 0.2 | 0.148 | 1 | 0.576213673 | 0.393943499 |
| Bud13 | 0.221511075 | -0.321928095 | 0.168 | 0.229 | 1 | 0.601142067 | -0.131825212 | 0.1 | 0.157 | 1 | 0.601142067 | 0.393954993 |
| Gm29642 | 0.221730359 | 0.929610672 | 0.05 | 0.024 | 1 | 0.522102507 | 0.930459067 | 0 | 0.04 | 1 | 0.522102507 | 0.394296367 |
| Prrx1 | 0.563786224 | -0.593429477 | 0.672 | 0.647 | 1 | 0.221760209 | -1.000477003 | 0.7 | 0.852 | 1 | 0.563786224 | 0.394342828 |
| Gm15492 | 0.22176705 | -1.070389328 | 0.008 | 0.029 | 1 | 0.619969878 | 1.111031312 | 0 | 0.025 | 1 | 0.619969878 | 0.394353476 |
| Galntl6 | 0.22176705 | -1.070389328 | 0.008 | 0.029 | 1 | 0.698534498 | 2.432959407 | 0 | 0.015 | 1 | 0.698534498 | 0.394353476 |
| Gga2 | 0.221998176 | -0.27897595 | 0.353 | 0.406 | 1 | 0.280473375 | 0.415037499 | 0.6 | 0.423 | 1 | 0.280473375 | 0.394713162 |
| Nfasc | 0.719672462 | -0.167686529 | 0.21 | 0.188 | 1 | 0.222012323 | -1.31299497 | 0.1 | 0.262 | 1 | 0.719672462 | 0.394735175 |
| Map3k13 | 0.222113005 | -0.87774425 | 0.05 | 0.088 | 1 | 0.698532411 | 1.31748219 | 0 | 0.015 | 1 | 0.698532411 | 0.394891823 |

| | | | | | | | | | | | | |
|---|---|---|---|---|---|---|---|---|---|---|---|---|
| Anpep | 0.22220604 | -0.584962501 | 0.084 | 0.129 | 1 | 0.460599334 | 0.432959407 | 0 | 0.052 | 1 | 0.460599334 | 0.395036556 |
| Adamtsl1 | 0.222254178 | -3.240314329 | 0.067 | 0.106 | 1 | 0.910522686 | -0.131825212 | 0.1 | 0.086 | 1 | 0.910522686 | 0.395111437 |
| Bc1 | 0.368025286 | -0.659924558 | 0.101 | 0.135 | 1 | 0.222259489 | 0.868174788 | 0.2 | 0.086 | 1 | 0.368025286 | 0.395119697 |
| 4930458D05Rik | 0.222455988 | 0.514573173 | 0.076 | 0.041 | 1 | 0.597845674 | 1.695993813 | 0 | 0.028 | 1 | 0.597845674 | 0.39542531 |
| Mrpl18 | 0.222536989 | -0.515174171 | 0.437 | 0.459 | 1 | 0.478304418 | -0.351311902 | 0.3 | 0.435 | 1 | 0.478304418 | 0.395551266 |
| Hdac6 | 0.88487267 | -0.319416876 | 0.185 | 0.171 | 1 | 0.222729033 | 0.888638891 | 0.3 | 0.148 | 1 | 0.88487267 | 0.395849843 |
| Lamtor1 | 0.67505855 | -0.146841388 | 0.807 | 0.759 | 1 | 0.22275122 | -0.545410577 | 0.7 | 0.784 | 1 | 0.67505855 | 0.395884335 |
| Ifit3b | 0.222804836 | 2.394278939 | 0.109 | 0.071 | 1 | 0.557777511 | -1.230005605 | 0 | 0.034 | 1 | 0.557777511 | 0.395967677 |
| Pmp22 | 0.222830324 | 0.165972961 | 0.916 | 0.841 | 1 | 0.799085637 | -0.191531458 | 0.8 | 0.781 | 1 | 0.799085637 | 0.396007295 |
| Gm5444 | 0.2228594 | -1.600904045 | 0.008 | 0.029 | 1 | 0.698533977 | 1.210566986 | 0 | 0.015 | 1 | 0.698533977 | 0.396052487 |
| Cdkn3 | 0.222865056 | -1.070389328 | 0.008 | 0.029 | 1 | 0.698529281 | 1.111031312 | 0 | 0.015 | 1 | 0.698529281 | 0.396061279 |
| 1700028E10Rik | 0.222866942 | -1.185866545 | 0.008 | 0.029 | 1 | 0.236765379 | 2.017921908 | 0.1 | 0.031 | 1 | 0.236765379 | 0.39606421 |
| Dph3 | 0.222945546 | 0.136726233 | 0.731 | 0.624 | 1 | 0.322269964 | -0.408342847 | 0.4 | 0.611 | 1 | 0.322269964 | 0.396186376 |
| Apex1 | 0.223114436 | 0.192645078 | 0.504 | 0.429 | 1 | 0.832426343 | 0.483943336 | 0.3 | 0.296 | 1 | 0.832426343 | 0.396448821 |
| Bloc1s6 | 0.223124998 | -0.605721061 | 0.176 | 0.229 | 1 | 0.473131346 | 0.913585248 | 0.3 | 0.216 | 1 | 0.473131346 | 0.396465232 |
| Kbtbd3 | 0.223147667 | -0.739183419 | 0.084 | 0.129 | 1 | 0.44222087 | 1.847996907 | 0.1 | 0.046 | 1 | 0.44222087 | 0.396500453 |
| Larp1b | 0.544186508 | -0.414235779 | 0.521 | 0.524 | 1 | 0.223240733 | 0.103835811 | 0.5 | 0.698 | 1 | 0.544186508 | 0.396645041 |
| Gm47123 | 0.223270748 | 0.321928095 | 0.042 | 0.018 | 1 | 0.619953009 | 1.695993813 | 0 | 0.025 | 1 | 0.619953009 | 0.396691669 |
| Angptl4 | 0.223319364 | -1.26121167 | 0.126 | 0.176 | 1 | 0.478014605 | 1.417017863 | 0.1 | 0.216 | 1 | 0.478014605 | 0.396767189 |
| Itga2 | 0.223471041 | 0.836501268 | 0.05 | 0.024 | 1 | 0.509801007 | 0.210566986 | 0.1 | 0.049 | 1 | 0.509801007 | 0.397002775 |
| Spns2 | 0.223473929 | -0.321928095 | 0.05 | 0.088 | 1 | 0.539431168 | 1.111031312 | 0 | 0.037 | 1 | 0.539431168 | 0.397007262 |
| Mecr | 0.223536488 | -0.537894247 | 0.185 | 0.241 | 1 | 0.54004028 | 0.695993813 | 0.3 | 0.225 | 1 | 0.54004028 | 0.397104415 |
| Ro60 | 0.831616894 | -0.206044427 | 0.378 | 0.376 | 1 | 0.22366812 | -0.413923879 | 0.2 | 0.457 | 1 | 0.831616894 | 0.397308812 |
| Zfp322a | 0.510677086 | -0.605721061 | 0.16 | 0.182 | 1 | 0.223701232 | 1.006694653 | 0.4 | 0.241 | 1 | 0.510677086 | 0.397360223 |
| Cdk5rap1 | 0.223713953 | 0.467267458 | 0.202 | 0.147 | 1 | 0.645676495 | 0.432959407 | 0.2 | 0.139 | 1 | 0.645676495 | 0.397379974 |
| Epyc | 0.223732493 | -0.847996907 | 0.79 | 0.659 | 1 | 0.643933865 | -0.323623389 | 0.5 | 0.577 | 1 | 0.643933865 | 0.397408758 |
| Eqtn | 0.223960336 | -0.707819249 | 0.008 | 0.029 | 1 | 0.539456304 | 0.695993813 | 0 | 0.037 | 1 | 0.539456304 | 0.39776244 |
| Gzf1 | 0.973169026 | -0.100136671 | 0.227 | 0.218 | 1 | 0.223989542 | 0.930459067 | 0.4 | 0.231 | 1 | 0.973169026 | 0.397807769 |
| Clcc1 | 0.644261847 | -0.161154792 | 0.487 | 0.459 | 1 | 0.224086252 | 0.786357841 | 0.6 | 0.46 | 1 | 0.644261847 | 0.397957855 |
| Ntpcr | 0.224127904 | -0.753915663 | 0.16 | 0.212 | 1 | 0.977523303 | 1.176619654 | 0.1 | 0.105 | 1 | 0.977523303 | 0.39802249 |
| Zfp809 | 0.224161768 | -0.707819249 | 0.118 | 0.165 | 1 | 0.251421138 | -0.827568143 | 0.1 | 0.256 | 1 | 0.251421138 | 0.398075038 |
| Mlxipl | 0.224229284 | 0.321928095 | 0.042 | 0.018 | 1 | 0.276546558 | 2.017921908 | 0.1 | 0.034 | 1 | 0.276546558 | 0.398179796 |
| Amdhd2 | 0.224271308 | -0.807354922 | 0.076 | 0.118 | 1 | 0.343644528 | 0.111031312 | 0 | 0.083 | 1 | 0.343644528 | 0.398244997 |
| Ntmt1 | 0.619512312 | -0.318776958 | 0.361 | 0.371 | 1 | 0.224376577 | -0.872849022 | 0.2 | 0.37 | 1 | 0.619512312 | 0.398408306 |
| Gm57266 | 0.224473032 | 0.707218251 | 0.059 | 0.029 | 1 | 0.698534498 | 2.432959407 | 0 | 0.015 | 1 | 0.698534498 | 0.398557922 |
| Zfp563 | 0.224638472 | -0.987927168 | 0.042 | 0.076 | 1 | 0.960463364 | 0.660369903 | 0.1 | 0.093 | 1 | 0.960463364 | 0.3988145 |
| Bmp15 | 0.224638472 | -0.987927168 | 0.042 | 0.076 | 1 | 0.577155236 | 1.210566986 | 0 | 0.031 | 1 | 0.577155236 | 0.3988145 |
| Shld3 | 0.224763609 | -0.678071905 | 0.05 | 0.088 | 1 | 0.42087075 | 0.494359952 | 0 | 0.062 | 1 | 0.42087075 | 0.399008538 |
| Vps39 | 0.224924229 | -0.669851398 | 0.16 | 0.212 | 1 | 0.315359305 | -0.536666944 | 0.1 | 0.231 | 1 | 0.315359305 | 0.39925755 |
| C230034O21Rik | 0.225055707 | -0.485426827 | 0.008 | 0.029 | 1 | 0.276546558 | 2.210566986 | 0.1 | 0.034 | 1 | 0.276546558 | 0.399461342 |
| Frat2 | 0.225055707 | -0.485426827 | 0.008 | 0.029 | 1 | 0.276546558 | 2.210566986 | 0.1 | 0.034 | 1 | 0.276546558 | 0.399461342 |
| Gpr37l1 | 0.225065156 | -0.292781749 | 0.008 | 0.029 | 1 | 0.59781419 | 1.558490289 | 0 | 0.028 | 1 | 0.59781419 | 0.399475987 |
| Gm47438 | 0.225065156 | -0.292781749 | 0.008 | 0.029 | 1 | 0.730749233 | 2.017921908 | 0 | 0.012 | 1 | 0.730749233 | 0.399475987 |
| Nadsyn1 | 0.225145359 | 0.736965594 | 0.05 | 0.024 | 1 | 0.522102209 | 0.625604485 | 0 | 0.04 | 1 | 0.522102209 | 0.399600285 |
| Spdya | 0.225145359 | 0.736965594 | 0.05 | 0.024 | 1 | 0.619961444 | 1.432959407 | 0 | 0.025 | 1 | 0.619961444 | 0.399600285 |
| Gm26633 | 0.225438365 | 0.666576266 | 0.126 | 0.082 | 1 | 0.757881427 | 0.591657153 | 0.1 | 0.071 | 1 | 0.757881427 | 0.400054273 |
| Park7 | 0.225588159 | 0.148225166 | 0.874 | 0.876 | 1 | 0.508809361 | 0.342105977 | 0.8 | 0.849 | 1 | 0.508809361 | 0.4002863 |
| Ckmt1 | 0.551006645 | 0.877143252 | 0.042 | 0.029 | 1 | 0.225659915 | -0.736965594 | 0 | 0.13 | 1 | 0.551006645 | 0.400397433 |
| Gm17491 | 0.330159923 | 0.614108846 | 0.109 | 0.076 | 1 | 0.225744174 | 1.602884409 | 0.2 | 0.086 | 1 | 0.330159923 | 0.400527916 |
| Arl2 | 0.381568046 | -0.437063806 | 0.513 | 0.512 | 1 | 0.225849846 | 1 | 0.5 | 0.33 | 1 | 0.381568046 | 0.400691538 |
| Kbtbd7 | 0.531681693 | -0.36994961 | 0.101 | 0.124 | 1 | 0.225912992 | -0.789433014 | 0 | 0.13 | 1 | 0.531681693 | 0.400789304 |
| Ttpal | 0.226041675 | -0.748461233 | 0.118 | 0.165 | 1 | 0.408647882 | 0.584962501 | 0.4 | 0.256 | 1 | 0.408647882 | 0.400988511 |
| 9430091E24Rik | 0.474205722 | -0.584962501 | 0.05 | 0.071 | 1 | 0.226111238 | -0.959358016 | 0 | 0.13 | 1 | 0.474205722 | 0.401096184 |
| Ndufaf6 | 0.783820105 | -0.292781749 | 0.126 | 0.135 | 1 | 0.226215502 | -0.982078092 | 0 | 0.13 | 1 | 0.783820105 | 0.401257551 |
| Ulk3 | 0.226527747 | 0.225066556 | 0.067 | 0.035 | 1 | 0.46060033 | 0.558490289 | 0 | 0.052 | 1 | 0.46060033 | 0.401740674 |
| Ap1s3 | 0.226595337 | 0.166649869 | 0.067 | 0.035 | 1 | 0.233699361 | 1.111031312 | 0.2 | 0.086 | 1 | 0.233699361 | 0.401845227 |
| Bcr | 0.461123457 | -0.249359469 | 0.689 | 0.594 | 1 | 0.226637587 | -0.68251781 | 0.5 | 0.707 | 1 | 0.461123457 | 0.401910579 |
| Megf9 | 0.226676978 | -0.6412215 | 0.303 | 0.341 | 1 | 0.497435241 | -0.68251781 | 0.3 | 0.367 | 1 | 0.497435241 | 0.401971503 |
| Pdlim3 | 0.226938062 | 1.447458977 | 0.059 | 0.029 | 1 | 0.238914277 | 1.432959407 | 0.1 | 0.031 | 1 | 0.238914277 | 0.402375241 |
| Gm11696 | 0.227253259 | 0.377069649 | 0.076 | 0.041 | 1 | 0.619961444 | 0.625604485 | 0 | 0.025 | 1 | 0.619961444 | 0.402862475 |
| Dnajc2 | 0.227315223 | -0.577873076 | 0.395 | 0.441 | 1 | 0.337270613 | -0.473931188 | 0.3 | 0.457 | 1 | 0.337270613 | 0.402958236 |
| Gm38537 | 0.227695427 | -0.959358016 | 0.059 | 0.029 | 1 | 0.597824685 | 1.432959407 | 0 | 0.028 | 1 | 0.597824685 | 0.403545647 |
| Cdkl3 | 0.227783623 | -0.748461233 | 0.185 | 0.235 | 1 | 0.329592532 | -1.058893689 | 0.1 | 0.222 | 1 | 0.329592532 | 0.403681867 |
| Rwdd2b | 0.227875541 | -0.900464326 | 0.042 | 0.076 | 1 | 0.479305119 | 1.847996907 | 0.1 | 0.049 | 1 | 0.479305119 | 0.403823821 |
| Lama1 | 0.227875541 | -0.900464326 | 0.042 | 0.076 | 1 | 0.365421266 | 1.558490289 | 0.1 | 0.04 | 1 | 0.365421266 | 0.403823821 |
| Ubtd1 | 0.228315828 | -0.446607578 | 0.538 | 0.565 | 1 | 0.35764543 | -0.531075095 | 0.6 | 0.694 | 1 | 0.35764543 | 0.404503538 |
| Hadhb | 0.228833061 | -0.302882339 | 0.639 | 0.635 | 1 | 0.836889448 | 0.10782888 | 0.8 | 0.707 | 1 | 0.836889448 | 0.405301553 |
| Mgat3 | 0.228907949 | 0.13606155 | 0.143 | 0.094 | 1 | 0.251421138 | -0.814968106 | 0.1 | 0.256 | 1 | 0.251421138 | 0.405417048 |
| Dnajc27 | 0.229010426 | -0.891419187 | 0.134 | 0.182 | 1 | 0.622725201 | 0.625604485 | 0.1 | 0.176 | 1 | 0.622725201 | 0.405575077 |
| Aph1c | 0.229139593 | -0.833350131 | 0.076 | 0.118 | 1 | 0.899700754 | 0.847996907 | 0.1 | 0.086 | 1 | 0.899700754 | 0.405774234 |
| Sh3bgr | 0.229539929 | 0.684498174 | 0.05 | 0.024 | 1 | 0.73075158 | 2.695993813 | 0 | 0.012 | 1 | 0.73075158 | 0.406391279 |
| Pax9 | 0.229545074 | -0.163498732 | 0.067 | 0.035 | 1 | 0.557782487 | -0.625934282 | 0 | 0.034 | 1 | 0.557782487 | 0.406399207 |
| Dtna | 0.229986203 | -0.707819249 | 0.042 | 0.076 | 1 | 0.696866015 | -1.058893689 | 0.1 | 0.136 | 1 | 0.696866015 | 0.407078752 |
| Gm32687 | 0.23847629 | -1.222392421 | 0.017 | 0.041 | 1 | 0.230466554 | 2.558490289 | 0.1 | 0.031 | 1 | 0.23847629 | 0.407818275 |
| 6820445E23Rik | 0.327697987 | 0.389042291 | 0.076 | 0.047 | 1 | 0.230466554 | 2.558490289 | 0.1 | 0.031 | 1 | 0.327697987 | 0.407818275 |
| Kics2 | 0.338096132 | -0.807354922 | 0.025 | 0.047 | 1 | 0.230466554 | 2.558490289 | 0.1 | 0.031 | 1 | 0.338096132 | 0.407818275 |
| Gm57238 | 0.616536081 | 0.251538767 | 0.034 | 0.024 | 1 | 0.230466554 | 2.558490289 | 0.1 | 0.031 | 1 | 0.616536081 | 0.407818275 |
| Vsig10l | 0.781790612 | -0.807354922 | 0.008 | 0.012 | 1 | 0.230466554 | 2.558490289 | 0.1 | 0.031 | 1 | 0.781790612 | 0.407818275 |
| Fam207a | 0.29287835 | -0.707819249 | 0.21 | 0.247 | 1 | 0.230602531 | 0.791853829 | 0.5 | 0.321 | 1 | 0.29287835 | 0.408027535 |
| Rab24 | 0.319609142 | 0.117237675 | 0.521 | 0.435 | 1 | 0.230722834 | -0.85654721 | 0.2 | 0.364 | 1 | 0.319609142 | 0.408212643 |
| Rad54b | 0.230827218 | -1.250961574 | 0.034 | 0.065 | 1 | 0.248008881 | 3.754887502 | 0.1 | 0.034 | 1 | 0.248008881 | 0.408373232 |
| Anks6 | 0.230828788 | -1.163498732 | 0.034 | 0.065 | 1 | 0.670447383 | 1.31748219 | 0.1 | 0.065 | 1 | 0.670447383 | 0.408375646 |
| H2-T22 | 0.581745757 | 0.400616984 | 0.328 | 0.3 | 1 | 0.230955316 | -0.896961478 | 0.2 | 0.367 | 1 | 0.581745757 | 0.408570273 |
| Pigw | 0.231140974 | -0.807354922 | 0.042 | 0.076 | 1 | 0.522066388 | 1.111031312 | 0 | 0.04 | 1 | 0.522066388 | 0.408855798 |
| Epb41l4b | 0.231189813 | 0.412693559 | 0.227 | 0.165 | 1 | 0.408755177 | 1.017921908 | 0.5 | 0.41 | 1 | 0.408755177 | 0.408930897 |
| Pkd1 | 0.231341659 | -0.580145484 | 0.765 | 0.676 | 1 | 0.680399448 | -0.128613273 | 0.6 | 0.735 | 1 | 0.680399448 | 0.409164355 |
| Uba3 | 0.231789612 | 0.215012891 | 0.681 | 0.6 | 1 | 0.30149917 | -0.596787936 | 0.6 | 0.645 | 1 | 0.30149917 | 0.4098528 |
| Ccndbp1 | 0.231951426 | -0.301714874 | 0.319 | 0.382 | 1 | 0.404577954 | -0.239465935 | 0.2 | 0.343 | 1 | 0.404577954 | 0.410101388 |
| Trappc11 | 0.23195613 | -0.661276663 | 0.202 | 0.247 | 1 | 0.679976938 | -0.242605642 | 0.2 | 0.241 | 1 | 0.679976938 | 0.410108614 |
| Zfp788 | 0.232006913 | -1.122856748 | 0.067 | 0.106 | 1 | 0.489351369 | 1.111031312 | 0.2 | 0.123 | 1 | 0.489351369 | 0.410186619 |
| Xkr6 | 0.316165555 | 0.315264365 | 0.151 | 0.112 | 1 | 0.232106745 | 1.847996907 | 0.2 | 0.096 | 1 | 0.316165555 | 0.41033995 |
| Nkd1 | 0.518706747 | 0.123382416 | 0.538 | 0.571 | 1 | 0.232118492 | -0.864721141 | 0.4 | 0.552 | 1 | 0.518706747 | 0.41035799 |
| Epop | 0.785106222 | -0.240314329 | 0.126 | 0.112 | 1 | 0.23214127 | -0.625934282 | 0 | 0.127 | 1 | 0.785106222 | 0.410392971 |
| Epb41l4a | 0.653579641 | 0.836501268 | 0.025 | 0.018 | 1 | 0.232208128 | -0.763437806 | 0 | 0.127 | 1 | 0.653579641 | 0.410495642 |
| Gm39469 | 0.503311279 | -0.392317423 | 0.109 | 0.135 | 1 | 0.232270098 | -0.936274402 | 0 | 0.127 | 1 | 0.503311279 | 0.410590797 |
| Pcdhga12 | 0.232440904 | 0.455679484 | 0.168 | 0.118 | 1 | 0.315849236 | -0.441509711 | 0 | 0.093 | 1 | 0.315849236 | 0.410853034 |
| Mgmt | 0.332079647 | -0.527247003 | 0.092 | 0.129 | 1 | 0.232479005 | -1.425021588 | 0 | 0.127 | 1 | 0.332079647 | 0.410911522 |
| AA386476 | 0.251644218 | -0.622930351 | 0.025 | 0.053 | 1 | 0.232498909 | 2.432959407 | 0.1 | 0.031 | 1 | 0.251644218 | 0.410942075 |

| | | | | | | | | | | | | |
|---|---|---|---|---|---|---|---|---|---|---|---|---|
| 6230426M11Rik | 0.327885716 | -0.622930351 | 0.034 | 0.059 | 1 | 0.232498909 | 2.31748219 | 0.1 | 0.031 | 1 | 0.327885716 | 0.410942075 |
| Cyp46a1 | 0.334204811 | -0.807354922 | 0.008 | 0.024 | 1 | 0.232498909 | 2.432959407 | 0.1 | 0.031 | 1 | 0.334204811 | 0.410942075 |
| Kcnh1 | 0.687345921 | -1.707819249 | 0.017 | 0.024 | 1 | 0.232498909 | 2.432959407 | 0.1 | 0.031 | 1 | 0.687345921 | 0.410942075 |
| Zfp672 | 0.648114613 | -0.529820947 | 0.227 | 0.241 | 1 | 0.232571301 | -0.654503434 | 0.1 | 0.29 | 1 | 0.648114613 | 0.411053192 |
| Zfp930 | 0.232592835 | -0.847996907 | 0.042 | 0.076 | 1 | 0.396919237 | 0.263034406 | 0 | 0.068 | 1 | 0.396919237 | 0.411086244 |
| Pex16 | 0.232698994 | 0.450442835 | 0.168 | 0.118 | 1 | 0.245300326 | -0.441509711 | 0 | 0.12 | 1 | 0.245300326 | 0.411249166 |
| Ccdc57 | 0.232787917 | -0.411426246 | 0.092 | 0.141 | 1 | 0.232736427 | -1.304006187 | 0 | 0.127 | 1 | 0.232787917 | 0.41130661 |
| Edem1 | 0.233094693 | -0.525690697 | 0.345 | 0.394 | 1 | 0.976981518 | 0.106230326 | 0.5 | 0.5 | 1 | 0.976981518 | 0.41185625 |
| Gm13830 | 0.233711927 | -1.163498732 | 0.034 | 0.065 | 1 | 0.408636939 | 0.374065718 | 0 | 0.065 | 1 | 0.408636939 | 0.41280259 |
| Gm26903 | 0.233711927 | -1.485426827 | 0.034 | 0.065 | 1 | 0.505623793 | 0.847996907 | 0 | 0.043 | 1 | 0.505623793 | 0.41280259 |
| Tmem176a | 0.233759167 | 0.385586732 | 0.773 | 0.747 | 1 | 0.412110868 | 0.174225139 | 0.5 | 0.704 | 1 | 0.412110868 | 0.412874986 |
| Mrps36 | 0.23399049 | -0.301002256 | 0.311 | 0.376 | 1 | 0.873224509 | 0.432959407 | 0.2 | 0.238 | 1 | 0.873224509 | 0.41322943 |
| Nsun6 | 0.234001758 | -0.665999073 | 0.185 | 0.241 | 1 | 0.481548736 | 0.902444691 | 0.3 | 0.21 | 1 | 0.481548736 | 0.413246693 |
| Lrp12 | 0.733974903 | -0.259150971 | 0.336 | 0.353 | 1 | 0.234022034 | 0.613167382 | 0.6 | 0.423 | 1 | 0.733974903 | 0.413277756 |
| Tmem236 | 0.234168602 | -1.807354922 | 0.025 | 0.053 | 1 | 0.643849988 | 0.111031312 | 0 | 0.022 | 1 | 0.643849988 | 0.41350227 |
| Col5a3 | 0.234168602 | -2.129283017 | 0.025 | 0.053 | 1 | 0.460666555 | -0.152003093 | 0 | 0.052 | 1 | 0.460666555 | 0.41350227 |
| Atp2b1 | 0.234312772 | -0.347402649 | 0.992 | 0.982 | 1 | 0.588399091 | -0.227291221 | 1 | 0.988 | 1 | 0.588399091 | 0.413723068 |
| Wrap53 | 0.234388204 | -0.485426827 | 0.05 | 0.088 | 1 | 0.923628411 | 0.558490289 | 0.1 | 0.108 | 1 | 0.923628411 | 0.413838577 |
| Tmco1 | 0.234388281 | 0.159590797 | 0.79 | 0.747 | 1 | 0.971181568 | 0.497499659 | 0.6 | 0.71 | 1 | 0.971181568 | 0.413838696 |
| Gpr35 | 0.334204811 | -0.807354922 | 0.008 | 0.024 | 1 | 0.234627319 | 2.31748219 | 0.1 | 0.031 | 1 | 0.334204811 | 0.414204659 |
| Lax1 | 0.402360347 | -0.847996907 | 0.042 | 0.065 | 1 | 0.234627319 | 2.017921908 | 0.1 | 0.031 | 1 | 0.402360347 | 0.414204659 |
| Gm47163 | 0.511005543 | -0.485426827 | 0.008 | 0.018 | 1 | 0.234627319 | 2.31748219 | 0.1 | 0.031 | 1 | 0.511005543 | 0.414204659 |
| Racgap1 | 0.585442721 | -1.485426827 | 0.05 | 0.065 | 1 | 0.234627319 | 2.210566986 | 0.1 | 0.031 | 1 | 0.585442721 | 0.414204659 |
| Gm47167 | 0.856663367 | -0.163498732 | 0.034 | 0.029 | 1 | 0.234627319 | 2.210566986 | 0.1 | 0.031 | 1 | 0.856663367 | 0.414204659 |
| Sdcbp2 | 0.234733653 | -1.707819249 | 0.017 | 0.041 | 1 | 0.489901827 | 0.695993813 | 0 | 0.046 | 1 | 0.489901827 | 0.414367419 |
| Galnt16 | 0.234733653 | -2.854660637 | 0.017 | 0.041 | 1 | 0.619971082 | 1.111031312 | 0 | 0.025 | 1 | 0.619971082 | 0.414367419 |
| Ppara | 0.234733653 | -2.14839184 | 0.017 | 0.041 | 1 | 0.539475153 | 0.210566986 | 0 | 0.037 | 1 | 0.539475153 | 0.414367419 |
| Gmeb1 | 0.379822869 | -0.496482016 | 0.336 | 0.371 | 1 | 0.235139192 | -1.236319379 | 0.2 | 0.355 | 1 | 0.379822869 | 0.414987945 |
| Ppil4 | 0.235299141 | -0.194836699 | 0.639 | 0.671 | 1 | 0.462176657 | -0.330806246 | 0.7 | 0.799 | 1 | 0.462176657 | 0.415232597 |
| Fbxl3 | 0.451310156 | -0.175742328 | 0.706 | 0.741 | 1 | 0.235325818 | -0.509815724 | 0.7 | 0.802 | 1 | 0.451310156 | 0.415273395 |
| Chchd4 | 0.235372635 | -0.807354922 | 0.16 | 0.206 | 1 | 0.693795609 | -0.204470513 | 0.2 | 0.244 | 1 | 0.693795609 | 0.415344992 |
| Uimc1 | 0.235512983 | 0.122805453 | 0.824 | 0.682 | 1 | 0.833455431 | 0.291298258 | 0.7 | 0.707 | 1 | 0.833455431 | 0.4155596 |
| Mir99ahg | 0.579429211 | -0.339634576 | 0.933 | 0.929 | 1 | 0.235602203 | -0.422880296 | 0.6 | 0.84 | 1 | 0.579429211 | 0.415696009 |
| Slc9a6 | 0.236049596 | -0.528495549 | 0.227 | 0.282 | 1 | 0.356587212 | 0.383715888 | 0.5 | 0.312 | 1 | 0.356587212 | 0.41637978 |
| Gm16091 | 0.236601303 | -1.987927168 | 0.017 | 0.041 | 1 | 0.577184788 | 1.558490289 | 0 | 0.031 | 1 | 0.577184788 | 0.41722243 |
| Gm10602 | 0.236601947 | -1.359895945 | 0.017 | 0.041 | 1 | 0.398985336 | 2.017921908 | 0.1 | 0.043 | 1 | 0.398985336 | 0.417223412 |
| Gm56874 | 0.236601947 | -1.707819249 | 0.017 | 0.041 | 1 | 0.669843604 | 2.017921908 | 0 | 0.019 | 1 | 0.669843604 | 0.417223412 |
| Lrrn3 | 0.236601947 | -1.900464326 | 0.017 | 0.041 | 1 | 0.505639146 | 0.558490289 | 0 | 0.043 | 1 | 0.505639146 | 0.417223412 |
| Abcd3 | 0.72422253 | -0.324630615 | 0.639 | 0.547 | 1 | 0.236604778 | 0.617699287 | 0.7 | 0.694 | 1 | 0.72422253 | 0.417227735 |
| Megf10 | 0.236605962 | -2.240314329 | 0.025 | 0.053 | 1 | 0.669843604 | 2.017921908 | 0 | 0.019 | 1 | 0.669843604 | 0.417229543 |
| ENSMUSG00000121596 | 0.236616883 | -0.970853654 | 0.034 | 0.065 | 1 | 0.48480221 | 1.695993813 | 0.1 | 0.049 | 1 | 0.48480221 | 0.417246217 |
| Abtb1 | 0.236616883 | -0.970853654 | 0.034 | 0.065 | 1 | 0.604715656 | 1.374065718 | 0.1 | 0.059 | 1 | 0.604715656 | 0.417246217 |
| Garin5a | 0.236616883 | -0.970853654 | 0.034 | 0.065 | 1 | 0.276546558 | 2.210566986 | 0.1 | 0.034 | 1 | 0.276546558 | 0.417246217 |
| Setd4 | 0.236616883 | -1.070389328 | 0.034 | 0.065 | 1 | 0.396883838 | 0.210566986 | 0 | 0.068 | 1 | 0.396883838 | 0.417246217 |
| A230072E10Rik | 0.236713526 | -1 | 0.034 | 0.065 | 1 | 0.374709424 | -0.736965594 | 0 | 0.074 | 1 | 0.374709424 | 0.417393758 |
| Plch2 | 0.236720913 | -1.122856748 | 0.042 | 0.076 | 1 | 0.418559485 | 0.263034406 | 0.2 | 0.108 | 1 | 0.418559485 | 0.417405036 |
| Arg2 | 0.493508373 | -0.655351829 | 0.017 | 0.029 | 1 | 0.236765774 | 1.930459067 | 0.1 | 0.031 | 1 | 0.493508373 | 0.417473516 |
| Prrg4 | 0.561605478 | 0.514573173 | 0.042 | 0.029 | 1 | 0.236765774 | 2.017921908 | 0.1 | 0.031 | 1 | 0.561605478 | 0.417473516 |
| Gm49085 | 0.719955107 | 0.152003093 | 0.05 | 0.041 | 1 | 0.236765774 | 2.017921908 | 0.1 | 0.031 | 1 | 0.719955107 | 0.417473516 |
| Rab5c | 0.237057553 | -0.434201504 | 0.546 | 0.553 | 1 | 0.431963552 | -0.21857071 | 0.5 | 0.688 | 1 | 0.431963552 | 0.417918823 |
| Borcs8 | 0.80794296 | -0.222392421 | 0.328 | 0.329 | 1 | 0.237179365 | 1.097356375 | 0.4 | 0.259 | 1 | 0.80794296 | 0.418104679 |
| Arntl2 | 0.237437188 | -0.584962501 | 0.202 | 0.253 | 1 | 0.954022777 | -0.898554736 | 0.4 | 0.33 | 1 | 0.954022777 | 0.418497957 |
| Gm57175 | 0.237885589 | -1.070389328 | 0 | 0.012 | 1 | 0.73075158 | 2.695993813 | 0 | 0.012 | 1 | 0.73075158 | 0.419181625 |
| Gm10550 | 0.237885589 | -1.070389328 | 0 | 0.012 | 1 | 0.597824685 | 1.432959407 | 0 | 0.028 | 1 | 0.597824685 | 0.419181625 |
| Gm57107 | 0.237885589 | -1.070389328 | 0 | 0.012 | 1 | 0.698532411 | 2.017921908 | 0 | 0.015 | 1 | 0.698532411 | 0.419181625 |
| Kif21b | 0.237885589 | -1.070389328 | 0 | 0.012 | 1 | 0.669853054 | 2.210566986 | 0 | 0.019 | 1 | 0.669853054 | 0.419181625 |
| Ptprc | 0.237885589 | -4.439623138 | 0 | 0.012 | 1 | 0.396978292 | -0.596978936 | 0 | 0.068 | 1 | 0.396978292 | 0.419181625 |
| 4933439K11Rik | 0.237885589 | -1.485426827 | 0 | 0.012 | 1 | 0.730749233 | 2.432959407 | 0 | 0.012 | 1 | 0.730749233 | 0.419181625 |
| Vsig8 | 0.237885589 | -1.485426827 | 0 | 0.012 | 1 | 0.643835681 | 1.558490289 | 0 | 0.022 | 1 | 0.643835681 | 0.419181625 |
| Gm37168 | 0.237885589 | -1.485426827 | 0 | 0.012 | 1 | 0.730749233 | 2.432959407 | 0 | 0.012 | 1 | 0.730749233 | 0.419181625 |
| Irf6 | 0.237885589 | -1.807354922 | 0 | 0.012 | 1 | 0.994867755 | 0.930459067 | 0.1 | 0.099 | 1 | 0.994867755 | 0.419181625 |
| Apbb1ip | 0.237885589 | -3.185866545 | 0 | 0.012 | 1 | 0.557756275 | 0.558490289 | 0 | 0.034 | 1 | 0.557756275 | 0.419181625 |
| Galnt13 | 0.237885589 | -1.070389328 | 0 | 0.012 | 1 | 0.577142568 | 1.111031312 | 0 | 0.031 | 1 | 0.577142568 | 0.419181625 |
| Erich2 | 0.237885589 | -1.485426827 | 0 | 0.012 | 1 | 0.669851164 | 1.432959407 | 0 | 0.019 | 1 | 0.669851164 | 0.419181625 |
| Bpifb4 | 0.237885589 | -1.070389328 | 0 | 0.012 | 1 | 0.539431168 | 0.847996907 | 0 | 0.037 | 1 | 0.539431168 | 0.419181625 |
| Uckl1os | 0.237885589 | -1.485426827 | 0 | 0.012 | 1 | 0.730749233 | 2.017921908 | 0 | 0.012 | 1 | 0.730749233 | 0.419181625 |
| P2ry12 | 0.237885589 | -1.807354922 | 0 | 0.012 | 1 | 0.698529281 | 2.210566986 | 0 | 0.015 | 1 | 0.698529281 | 0.419181625 |
| Lingo4 | 0.237885589 | -1.070389328 | 0 | 0.012 | 1 | 0.669851164 | 1.695993813 | 0 | 0.019 | 1 | 0.669851164 | 0.419181625 |
| Spag17 | 0.237885589 | -1.485426827 | 0 | 0.012 | 1 | 0.730749233 | 2.432959407 | 0 | 0.012 | 1 | 0.730749233 | 0.419181625 |
| Cd53 | 0.237885589 | -3.292781749 | 0 | 0.012 | 1 | 0.577179863 | 0.31748219 | 0 | 0.031 | 1 | 0.577179863 | 0.419181625 |
| Plppr4 | 0.237885589 | -1.807354922 | 0 | 0.012 | 1 | 0.73075158 | 1.558490289 | 0 | 0.012 | 1 | 0.73075158 | 0.419181625 |
| Gm43569 | 0.237885589 | -1.485426827 | 0 | 0.012 | 1 | 0.643835681 | 1.695993813 | 0 | 0.022 | 1 | 0.643835681 | 0.419181625 |
| Gdf6 | 0.237885589 | -1.070389328 | 0 | 0.012 | 1 | 0.698533977 | 1.017921908 | 0 | 0.015 | 1 | 0.698533977 | 0.419181625 |
| Gm6297 | 0.237885589 | -1.070389328 | 0 | 0.012 | 1 | 0.698534498 | 2.432959407 | 0 | 0.015 | 1 | 0.698534498 | 0.419181625 |
| Rasef | 0.237885589 | -1.485426827 | 0 | 0.012 | 1 | 0.698534498 | 2.432959407 | 0 | 0.015 | 1 | 0.698534498 | 0.419181625 |
| Gm42303 | 0.237885589 | -4.614709844 | 0 | 0.012 | 1 | 0.730750993 | 1.847996907 | 0 | 0.012 | 1 | 0.730750993 | 0.419181625 |
| Kcnq4 | 0.237885589 | -1.807354922 | 0 | 0.012 | 1 | 0.557771538 | 0.625604485 | 0 | 0.034 | 1 | 0.557771538 | 0.419181625 |
| Zscan20 | 0.237885589 | -1.070389328 | 0 | 0.012 | 1 | 0.360331175 | 2.017921908 | 0.1 | 0.04 | 1 | 0.360331175 | 0.419181625 |
| Cd52 | 0.237885589 | -3.070389328 | 0 | 0.012 | 1 | 0.396949898 | -0.304006187 | 0 | 0.068 | 1 | 0.396949898 | 0.419181625 |
| Zfp984 | 0.237885589 | -1.485426827 | 0 | 0.012 | 1 | 0.643850855 | 2.017921908 | 0 | 0.022 | 1 | 0.643850855 | 0.419181625 |
| Stap1 | 0.237885589 | -1.070389328 | 0 | 0.012 | 1 | 0.619971484 | 1.31748219 | 0 | 0.025 | 1 | 0.619971484 | 0.419181625 |
| Gm10419 | 0.237885589 | -1.070389328 | 0 | 0.012 | 1 | 0.730749233 | 2.432959407 | 0 | 0.012 | 1 | 0.730749233 | 0.419181625 |
| Gm14508 | 0.237885589 | -1.485426827 | 0 | 0.012 | 1 | 0.73075158 | 2.695993813 | 0 | 0.012 | 1 | 0.73075158 | 0.419181625 |
| Gm16120 | 0.237885589 | -1.070389328 | 0 | 0.012 | 1 | 0.730750993 | 2.017921908 | 0 | 0.012 | 1 | 0.730750993 | 0.419181625 |
| Gpr37 | 0.237885589 | -1.070389328 | 0 | 0.012 | 1 | 0.669843604 | 1.847996907 | 0 | 0.019 | 1 | 0.669843604 | 0.419181625 |
| Fscn3 | 0.237885589 | -1.070389328 | 0 | 0.012 | 1 | 0.669853054 | 2.210566986 | 0 | 0.019 | 1 | 0.669853054 | 0.419181625 |
| Arhgap25 | 0.237885589 | -2.655351829 | 0 | 0.012 | 1 | 0.698529281 | 2.210566986 | 0 | 0.015 | 1 | 0.698529281 | 0.419181625 |
| 4933427D06Rik | 0.237885589 | -1.485426827 | 0 | 0.012 | 1 | 0.730750993 | 0.930459067 | 0 | 0.012 | 1 | 0.730750993 | 0.419181625 |
| Mdfic2 | 0.237885589 | -1.807354922 | 0 | 0.012 | 1 | 0.619961444 | 1.210566986 | 0 | 0.025 | 1 | 0.619961444 | 0.419181625 |
| 4930540M05Rik | 0.237885589 | -1.070389328 | 0 | 0.012 | 1 | 0.698529281 | 2.210566986 | 0 | 0.015 | 1 | 0.698529281 | 0.419181625 |
| Apoc1 | 0.237885589 | -1.485426827 | 0 | 0.012 | 1 | 0.643850855 | 2.017921908 | 0 | 0.022 | 1 | 0.643850855 | 0.419181625 |
| Hipk4 | 0.237885589 | -1.485426827 | 0 | 0.012 | 1 | 0.619961444 | 1.210566986 | 0 | 0.025 | 1 | 0.619961444 | 0.419181625 |
| Dmkn | 0.237885589 | -1.070389328 | 0 | 0.012 | 1 | 0.619961444 | 1.558490289 | 0 | 0.025 | 1 | 0.619961444 | 0.419181625 |
| B230322F03Rik | 0.237885589 | -1.070389328 | 0 | 0.012 | 1 | 0.6199755 | 1.847996907 | 0 | 0.025 | 1 | 0.6199755 | 0.419181625 |
| Gm44997 | 0.237885589 | -1.807354922 | 0 | 0.012 | 1 | 0.643835681 | 1.847996907 | 0 | 0.022 | 1 | 0.643835681 | 0.419181625 |
| Gm57220 | 0.237885589 | -1.485426827 | 0 | 0.012 | 1 | 0.73075158 | 2.695993813 | 0 | 0.012 | 1 | 0.73075158 | 0.419181625 |
| Pgpep1l | 0.237885589 | -1.070389328 | 0 | 0.012 | 1 | 0.619953009 | 1.432959407 | 0 | 0.025 | 1 | 0.619953009 | 0.419181625 |
| Gm21057 | 0.237885589 | -1.070389328 | 0 | 0.012 | 1 | 0.619953009 | 1.558490289 | 0 | 0.025 | 1 | 0.619953009 | 0.419181625 |

| | | | | | | | | | | | | |
|---|---|---|---|---|---|---|---|---|---|---|---|---|
| Rab38 | 0.237885589 | -1.070389328 | 0 | 0.012 | 1 | 0.643848687 | 0.432959407 | 0 | 0.022 | 1 | 0.643848687 | 0.419181625 |
| Ccdc81 | 0.237885589 | -2.292781749 | 0 | 0.012 | 1 | 0.730749233 | 1.847996907 | 0 | 0.012 | 1 | 0.730749233 | 0.419181625 |
| Gm35082 | 0.237885589 | -1.070389328 | 0 | 0.012 | 1 | 0.698532411 | 2.017921908 | 0 | 0.015 | 1 | 0.698532411 | 0.419181625 |
| Muc2 | 0.237885589 | -1.070389328 | 0 | 0.012 | 1 | 0.330809864 | 1.263034406 | 0.1 | 0.037 | 1 | 0.330809864 | 0.419181625 |
| Mrgpre | 0.237885589 | -1.070389328 | 0 | 0.012 | 1 | 0.643850855 | 2.017921908 | 0 | 0.022 | 1 | 0.643850855 | 0.419181625 |
| Gm45370 | 0.237885589 | -1.070389328 | 0 | 0.012 | 1 | 0.577184788 | 1.558490289 | 0 | 0.031 | 1 | 0.577184788 | 0.419181625 |
| Gpm6a | 0.237885589 | -1.070389328 | 0 | 0.012 | 1 | 0.522066388 | 1.111031312 | 0 | 0.04 | 1 | 0.522066388 | 0.419181625 |
| Lpar2 | 0.237885589 | -1.807354922 | 0 | 0.012 | 1 | 0.522151747 | 1.210566986 | 0 | 0.04 | 1 | 0.522151747 | 0.419181625 |
| Lrrc25 | 0.237885589 | -1.070389328 | 0 | 0.012 | 1 | 0.698529281 | 1.847996907 | 0 | 0.015 | 1 | 0.698529281 | 0.419181625 |
| Best2 | 0.237885589 | -1.485426827 | 0 | 0.012 | 1 | 0.577172123 | 0.930459067 | 0 | 0.031 | 1 | 0.577172123 | 0.419181625 |
| Has3 | 0.237885589 | -1.070389328 | 0 | 0.012 | 1 | 0.730749233 | 2.210566986 | 0 | 0.012 | 1 | 0.730749233 | 0.419181625 |
| Calb2 | 0.237885589 | -1.485426827 | 0 | 0.012 | 1 | 0.73075158 | 2.695993813 | 0 | 0.012 | 1 | 0.73075158 | 0.419181625 |
| 4933417D19Rik | 0.237885589 | -1.485426827 | 0 | 0.012 | 1 | 0.643835681 | 1.847996907 | 0 | 0.022 | 1 | 0.643835681 | 0.419181625 |
| Zfp558 | 0.237885589 | -1.070389328 | 0 | 0.012 | 1 | 0.73075158 | 2.695993813 | 0 | 0.012 | 1 | 0.73075158 | 0.419181625 |
| Gm16754 | 0.237885589 | -1.070389328 | 0 | 0.012 | 1 | 0.730750993 | 1.847996907 | 0 | 0.012 | 1 | 0.730750993 | 0.419181625 |
| Gm39326 | 0.237885589 | -1.070389328 | 0 | 0.012 | 1 | 0.52208609 | 0.769994395 | 0 | 0.04 | 1 | 0.52208609 | 0.419181625 |
| Nr2e3 | 0.237885589 | -1.485426827 | 0 | 0.012 | 1 | 0.577164735 | 1.017921908 | 0 | 0.031 | 1 | 0.577164735 | 0.419181625 |
| Col6a5 | 0.237885589 | -1.070389328 | 0 | 0.012 | 1 | 0.669853054 | 2.210566986 | 0 | 0.019 | 1 | 0.669853054 | 0.419181625 |
| Ngp | 0.237885589 | -1.485426827 | 0 | 0.012 | 1 | 0.307031624 | -0.304006187 | 0 | 0.096 | 1 | 0.307031624 | 0.419181625 |
| Gm48249 | 0.237885589 | -1.070389328 | 0 | 0.012 | 1 | 0.698529281 | 2.210566986 | 0 | 0.015 | 1 | 0.698529281 | 0.419181625 |
| Gm48084 | 0.237885589 | -1.485426827 | 0 | 0.012 | 1 | 0.730749233 | 2.210566986 | 0 | 0.012 | 1 | 0.730749233 | 0.419181625 |
| Gm26535 | 0.237885589 | -1.070389328 | 0 | 0.012 | 1 | 0.669843604 | 1.847996907 | 0 | 0.019 | 1 | 0.669843604 | 0.419181625 |
| Mfsd4b1 | 0.237885589 | -2.292781749 | 0 | 0.012 | 1 | 0.619967468 | 1.432959407 | 0 | 0.025 | 1 | 0.619967468 | 0.419181625 |
| Pkib | 0.237885589 | -1.807354922 | 0 | 0.012 | 1 | 0.73075158 | 2.695993813 | 0 | 0.012 | 1 | 0.73075158 | 0.419181625 |
| Gm48086 | 0.237885589 | -1.070389328 | 0 | 0.012 | 1 | 0.73075158 | 2.695993813 | 0 | 0.012 | 1 | 0.73075158 | 0.419181625 |
| Gm30122 | 0.237885589 | -1.070389328 | 0 | 0.012 | 1 | 0.475022551 | 0.930459067 | 0 | 0.049 | 1 | 0.475022551 | 0.419181625 |
| B4galnt1 | 0.237885589 | -1.070389328 | 0 | 0.012 | 1 | 0.52208609 | 0.930459067 | 0 | 0.04 | 1 | 0.52208609 | 0.419181625 |
| Egfros | 0.237885589 | -1.070389328 | 0 | 0.012 | 1 | 0.597832556 | 0.930459067 | 0 | 0.028 | 1 | 0.597832556 | 0.419181625 |
| Gm12166 | 0.237885589 | -1.070389328 | 0 | 0.012 | 1 | 0.669853054 | 2.210566986 | 0 | 0.019 | 1 | 0.669853054 | 0.419181625 |
| Ifi47 | 0.237885589 | -2.070389328 | 0 | 0.012 | 1 | 0.698533977 | 1.210566986 | 0 | 0.015 | 1 | 0.698533977 | 0.419181625 |
| 6720407P12Rik | 0.237885589 | -1.485426827 | 0 | 0.012 | 1 | 0.698534498 | 2.432959407 | 0 | 0.015 | 1 | 0.698534498 | 0.419181625 |
| Mir142hg | 0.237885589 | -3.292781749 | 0 | 0.012 | 1 | 0.669843604 | 1.111031312 | 0 | 0.019 | 1 | 0.669843604 | 0.419181625 |
| Lrrc46 | 0.237885589 | -1.485426827 | 0 | 0.012 | 1 | 0.698529281 | 2.210566986 | 0 | 0.015 | 1 | 0.698529281 | 0.419181625 |
| Card14 | 0.237885589 | -1.070389328 | 0 | 0.012 | 1 | 0.669852109 | 1.31748219 | 0 | 0.019 | 1 | 0.669852109 | 0.419181625 |
| 4930549C15Rik | 0.237885589 | -1.070389328 | 0 | 0.012 | 1 | 0.643842184 | 1.695993813 | 0 | 0.022 | 1 | 0.643842184 | 0.419181625 |
| Gm4756 | 0.237885589 | -1.070389328 | 0 | 0.012 | 1 | 0.669848329 | 1.31748219 | 0 | 0.019 | 1 | 0.669848329 | 0.419181625 |
| 2310002D06Rik | 0.237885589 | -1.070389328 | 0 | 0.012 | 1 | 0.730749233 | 2.432959407 | 0 | 0.012 | 1 | 0.730749233 | 0.419181625 |
| Pnma1 | 0.237885589 | -1.070389328 | 0 | 0.012 | 1 | 0.643835681 | 1.558490289 | 0 | 0.022 | 1 | 0.643835681 | 0.419181625 |
| Ighd | 0.237885589 | -1.807354922 | 0 | 0.012 | 1 | 0.698534498 | 2.432959407 | 0 | 0.015 | 1 | 0.698534498 | 0.419181625 |
| Chrm3 | 0.237885589 | -2.944858446 | 0 | 0.012 | 1 | 0.669853054 | 2.210566986 | 0 | 0.019 | 1 | 0.669853054 | 0.419181625 |
| Gm35161 | 0.237885589 | -1.070389328 | 0 | 0.012 | 1 | 0.669848329 | 1.847996907 | 0 | 0.019 | 1 | 0.669848329 | 0.419181625 |
| Gm34250 | 0.237885589 | -1.070389328 | 0 | 0.012 | 1 | 0.577155236 | 1.31748219 | 0 | 0.031 | 1 | 0.577155236 | 0.419181625 |
| Gm48970 | 0.237885589 | -1.070389328 | 0 | 0.012 | 1 | 0.730749233 | 2.017921908 | 0 | 0.012 | 1 | 0.730749233 | 0.419181625 |
| Fbxo43 | 0.237885589 | -1.485426827 | 0 | 0.012 | 1 | 0.730750993 | 1.695993813 | 0 | 0.012 | 1 | 0.730750993 | 0.419181625 |
| Parp10 | 0.237885589 | -1.485426827 | 0 | 0.012 | 1 | 0.23891309 | 2.017921908 | 0.1 | 0.031 | 1 | 0.23891309 | 0.419181625 |
| Rac2 | 0.237885589 | -1.807354922 | 0 | 0.012 | 1 | 0.557782487 | -0.888968688 | 0 | 0.034 | 1 | 0.557782487 | 0.419181625 |
| Card10 | 0.237885589 | -1.070389328 | 0 | 0.012 | 1 | 0.619961444 | 1.558490289 | 0 | 0.025 | 1 | 0.619961444 | 0.419181625 |
| Kcnj4 | 0.237885589 | -1.485426827 | 0 | 0.012 | 1 | 0.730750993 | 2.210566986 | 0 | 0.012 | 1 | 0.730750993 | 0.419181625 |
| Nfam1 | 0.237885589 | -1.070389328 | 0 | 0.012 | 1 | 0.698532411 | 2.017921908 | 0 | 0.015 | 1 | 0.698532411 | 0.419181625 |
| Arhgap8 | 0.237885589 | -2.070389328 | 0 | 0.012 | 1 | 0.597837803 | 0.930459067 | 0 | 0.028 | 1 | 0.597837803 | 0.419181625 |
| Phf21b | 0.237885589 | -1.070389328 | 0 | 0.012 | 1 | 0.698534498 | 2.432959407 | 0 | 0.015 | 1 | 0.698534498 | 0.419181625 |
| Cntn1 | 0.237885589 | -1.070389328 | 0 | 0.012 | 1 | 0.73075158 | 1.695993813 | 0 | 0.012 | 1 | 0.73075158 | 0.419181625 |
| Fam186b | 0.237885589 | -2.292781749 | 0 | 0.012 | 1 | 0.619961444 | 1.558490289 | 0 | 0.025 | 1 | 0.619961444 | 0.419181625 |
| Fignl2 | 0.237885589 | -1.070389328 | 0 | 0.012 | 1 | 0.364054013 | -0.304054187 | 0 | 0.077 | 1 | 0.364054013 | 0.419181625 |
| A330009N23Rik | 0.237885589 | -1.070389328 | 0 | 0.012 | 1 | 0.643835681 | 1.847996907 | 0 | 0.022 | 1 | 0.643835681 | 0.419181625 |
| Gm49745 | 0.237885589 | -1.070389328 | 0 | 0.012 | 1 | 0.698534498 | 2.432959407 | 0 | 0.015 | 1 | 0.698534498 | 0.419181625 |
| Tm4sf19 | 0.237885589 | -1.485426827 | 0 | 0.012 | 1 | 0.643846086 | 1.111031312 | 0 | 0.022 | 1 | 0.643846086 | 0.419181625 |
| Cfap44 | 0.237885589 | -1.070389328 | 0 | 0.012 | 1 | 0.619967468 | 1.210566986 | 0 | 0.025 | 1 | 0.619967468 | 0.419181625 |
| Cd200r4 | 0.237885589 | -1.807354922 | 0 | 0.012 | 1 | 0.730749233 | 2.432959407 | 0 | 0.012 | 1 | 0.730749233 | 0.419181625 |
| Zpld1 | 0.237885589 | -1.070389328 | 0 | 0.012 | 1 | 0.698532411 | 1.847996907 | 0 | 0.015 | 1 | 0.698532411 | 0.419181625 |
| Baiap3 | 0.237885589 | -1.070389328 | 0 | 0.012 | 1 | 0.698534498 | 2.432959407 | 0 | 0.015 | 1 | 0.698534498 | 0.419181625 |
| Wfikkn1 | 0.237885589 | -1.485426827 | 0 | 0.012 | 1 | 0.669843604 | 2.017921908 | 0 | 0.019 | 1 | 0.669843604 | 0.419181625 |
| Dnah8 | 0.237885589 | -2.070389328 | 0 | 0.012 | 1 | 0.577174234 | 0.847996907 | 0 | 0.031 | 1 | 0.577174234 | 0.419181625 |
| Gm50455 | 0.237885589 | -1.070389328 | 0 | 0.012 | 1 | 0.669843604 | 1.847996907 | 0 | 0.019 | 1 | 0.669843604 | 0.419181625 |
| Gm50107 | 0.237885589 | -1.070389328 | 0 | 0.012 | 1 | 0.730750993 | 2.017921908 | 0 | 0.012 | 1 | 0.730750993 | 0.419181625 |
| Cfb | 0.237885589 | -1.485426827 | 0 | 0.012 | 1 | 0.643842184 | 1.558490289 | 0 | 0.022 | 1 | 0.643842184 | 0.419181625 |
| Acsbg3 | 0.237885589 | -1.070389328 | 0 | 0.012 | 1 | 0.643850855 | 2.017921908 | 0 | 0.022 | 1 | 0.643850855 | 0.419181625 |
| Vav1 | 0.237885589 | -2.292781749 | 0 | 0.012 | 1 | 0.698532411 | 1.847996907 | 0 | 0.015 | 1 | 0.698532411 | 0.419181625 |
| Ttc39d | 0.237885589 | -1.485426827 | 0 | 0.012 | 1 | 0.73075158 | 2.695993813 | 0 | 0.012 | 1 | 0.73075158 | 0.419181625 |
| Gm19689 | 0.237885589 | -1.807354922 | 0 | 0.012 | 1 | 0.730749233 | 2.432959407 | 0 | 0.012 | 1 | 0.730749233 | 0.419181625 |
| Nol4 | 0.237885589 | -1.807354922 | 0 | 0.012 | 1 | 0.669851164 | 0.695993813 | 0 | 0.019 | 1 | 0.669851164 | 0.419181625 |
| Pcdhb3 | 0.237885589 | -1.070389328 | 0 | 0.012 | 1 | 0.577155236 | 1.31748219 | 0 | 0.031 | 1 | 0.577155236 | 0.419181625 |
| Pcdhb15 | 0.237885589 | -2.292781749 | 0 | 0.012 | 1 | 0.698532411 | 2.017921908 | 0 | 0.015 | 1 | 0.698532411 | 0.419181625 |
| Dok6 | 0.237885589 | -2.807354922 | 0 | 0.012 | 1 | 0.619971484 | 0.374065718 | 0 | 0.025 | 1 | 0.619971484 | 0.419181625 |
| Plaat5 | 0.237885589 | -1.070389328 | 0 | 0.012 | 1 | 0.643842184 | 1.31748219 | 0 | 0.022 | 1 | 0.643842184 | 0.419181625 |
| 4930414N06Rik | 0.237885589 | -1.070389328 | 0 | 0.012 | 1 | 0.73075158 | 2.695993813 | 0 | 0.012 | 1 | 0.73075158 | 0.419181625 |
| Gm14703 | 0.237885589 | -1.070389328 | 0 | 0.012 | 1 | 0.643835681 | 1.558490289 | 0 | 0.022 | 1 | 0.643835681 | 0.419181625 |
| Crat | 0.238392794 | -0.854660637 | 0.092 | 0.135 | 1 | 0.87523427 | 0.494359952 | 0.1 | 0.114 | 1 | 0.87523427 | 0.419954464 |
| Nek4 | 0.814408065 | -0.292781749 | 0.076 | 0.082 | 1 | 0.238442079 | -0.536666944 | 0 | 0.123 | 1 | 0.814408065 | 0.420029533 |
| Ypel4 | 0.23847629 | -1.600904045 | 0.017 | 0.041 | 1 | 0.505623793 | 0.695993813 | 0 | 0.043 | 1 | 0.505623793 | 0.420081638 |
| Acad12 | 0.23847629 | -1.222392421 | 0.017 | 0.041 | 1 | 0.698529281 | 2.210566986 | 0 | 0.015 | 1 | 0.698529281 | 0.420081638 |
| Pold1 | 0.23847629 | -1.359895945 | 0.017 | 0.041 | 1 | 0.539431168 | 0.930459067 | 0 | 0.037 | 1 | 0.539431168 | 0.420081638 |
| 4930579K19Rik | 0.23847629 | -1.222392421 | 0.017 | 0.041 | 1 | 0.643842184 | 1.695993813 | 0 | 0.022 | 1 | 0.643842184 | 0.420081638 |
| Ttc21a | 0.23847629 | -1.222392421 | 0.017 | 0.041 | 1 | 0.416075273 | 2.159940913 | 0.1 | 0.046 | 1 | 0.416075273 | 0.420081638 |
| Adam4 | 0.23847629 | -1.222392421 | 0.017 | 0.041 | 1 | 0.730750993 | 2.210566986 | 0 | 0.012 | 1 | 0.730750993 | 0.420081638 |
| Efcab11 | 0.23847629 | -2.944858446 | 0.017 | 0.041 | 1 | 0.577165087 | 0.930459067 | 0 | 0.031 | 1 | 0.577165087 | 0.420081638 |
| 4930506C21Rik | 0.23847629 | -1.222392421 | 0.017 | 0.041 | 1 | 0.643835681 | 1.695993813 | 0 | 0.022 | 1 | 0.643835681 | 0.420081638 |
| Paqr4 | 0.23847629 | -1.222392421 | 0.017 | 0.041 | 1 | 0.577172123 | 0.930459067 | 0 | 0.031 | 1 | 0.577172123 | 0.420081638 |
| Lmx1a | 0.238508516 | -1.411426246 | 0.017 | 0.041 | 1 | 0.40884999 | -0.888968688 | 0 | 0.065 | 1 | 0.40884999 | 0.420130719 |
| Mrpl37 | 0.238782652 | 0.292180751 | 0.261 | 0.2 | 1 | 0.675516286 | 0.31748219 | 0.3 | 0.225 | 1 | 0.675516286 | 0.420548149 |
| Atoh8 | 0.836371502 | 0.219117289 | 0.143 | 0.135 | 1 | 0.238852743 | -0.709998547 | 0 | 0.123 | 1 | 0.836371502 | 0.420654852 |
| Nptxr | 0.267565044 | -0.807354922 | 0.059 | 0.094 | 1 | 0.238909965 | -0.789433014 | 0 | 0.123 | 1 | 0.267565044 | 0.420741959 |
| Plekhn1 | 0.495207728 | -0.485426827 | 0.017 | 0.029 | 1 | 0.23891309 | 2.017921908 | 0.1 | 0.031 | 1 | 0.495207728 | 0.420746715 |
| 2610021A01Rik | 0.285423532 | -1.008988783 | 0.05 | 0.082 | 1 | 0.238949895 | -0.789433014 | 0 | 0.123 | 1 | 0.285423532 | 0.420802738 |
| Ccl19 | 0.239056444 | -1.185866545 | 0.025 | 0.053 | 1 | 0.619953009 | 1.695993813 | 0 | 0.025 | 1 | 0.619953009 | 0.420964905 |
| Gm44710 | 0.239056444 | -1.185866545 | 0.025 | 0.053 | 1 | 0.567543378 | 1.432959407 | 0.1 | 0.056 | 1 | 0.567543378 | 0.420964905 |
| Gm15839 | 0.239056444 | -1.185866545 | 0.025 | 0.053 | 1 | 0.698532411 | 1.695993813 | 0 | 0.015 | 1 | 0.698532411 | 0.420964905 |
| Trnp1 | 0.770562594 | -0.155278225 | 0.126 | 0.112 | 1 | 0.239088237 | -0.959358016 | 0 | 0.123 | 1 | 0.770562594 | 0.421013289 |

| | | | | | | | | | | | | |
|---|---|---|---|---|---|---|---|---|---|---|---|---|
| Larp6 | 0.553219952 | -0.297566812 | 0.437 | 0.447 | 1 | 0.239088354 | 0.484489708 | 0.7 | 0.58 | 1 | 0.553219952 | 0.421013466 |
| Dmc1 | 0.580079944 | -0.292781749 | 0.076 | 0.094 | 1 | 0.239129095 | -1.171902651 | 0 | 0.123 | 1 | 0.580079944 | 0.421075466 |
| Gpr176 | 0.239451286 | -1.655351829 | 0.017 | 0.041 | 1 | 0.557780497 | -0.267480311 | 0 | 0.034 | 1 | 0.557780497 | 0.421565654 |
| Tagap1 | 0.269780997 | -0.707819249 | 0.084 | 0.124 | 1 | 0.239499471 | 1.625604485 | 0.2 | 0.093 | 1 | 0.269780997 | 0.421638945 |
| Slc35c2 | 0.311839696 | -0.607417352 | 0.134 | 0.176 | 1 | 0.239503096 | 1.31748219 | 0.3 | 0.167 | 1 | 0.311839696 | 0.421644459 |
| Zfp286 | 0.239542116 | -0.86393845 | 0.034 | 0.065 | 1 | 0.282346243 | -0.736965594 | 0 | 0.105 | 1 | 0.282346243 | 0.421703806 |
| Cyp2d22 | 0.239542116 | -0.970853654 | 0.034 | 0.065 | 1 | 0.396883838 | 0.210566986 | 0 | 0.068 | 1 | 0.396883838 | 0.421703806 |
| Tesk2 | 0.862632518 | 1.340543773 | 0.134 | 0.129 | 1 | 0.239612216 | 2.097356375 | 0.2 | 0.099 | 1 | 0.862632518 | 0.421810418 |
| Znfx1 | 0.239624046 | -0.726434927 | 0.151 | 0.2 | 1 | 0.333869988 | -0.823380346 | 0.2 | 0.327 | 1 | 0.333869988 | 0.421828408 |
| Klhl29 | 0.773815754 | 0.383013469 | 0.429 | 0.394 | 1 | 0.239753362 | -1.327008012 | 0.3 | 0.463 | 1 | 0.773815754 | 0.422025049 |
| Igfbp2 | 0.239812881 | -1.070389328 | 0.134 | 0.182 | 1 | 0.338139145 | -1.411136062 | 0.1 | 0.238 | 1 | 0.338139145 | 0.422115544 |
| Pcdhb17 | 0.239919666 | -1.250961574 | 0.025 | 0.053 | 1 | 0.244547978 | 1.14345279 | 0.2 | 0.086 | 1 | 0.244547978 | 0.422277886 |
| 9030624G23Rik | 0.239952542 | -0.584962501 | 0.042 | 0.076 | 1 | 0.619967066 | 1.31748219 | 0 | 0.025 | 1 | 0.619967066 | 0.422327862 |
| Prpsap2 | 0.334767633 | -0.28748745 | 0.235 | 0.276 | 1 | 0.240059447 | 0.672147071 | 0.4 | 0.228 | 1 | 0.334767633 | 0.422490356 |
| Jpx | 0.800794122 | -0.333423734 | 0.092 | 0.1 | 1 | 0.240277036 | 0.591657153 | 0.3 | 0.145 | 1 | 0.800794122 | 0.422821018 |
| Etfa | 0.240349638 | -0.283488995 | 0.933 | 0.888 | 1 | 0.249916036 | -0.298300069 | 0.9 | 0.954 | 1 | 0.249916036 | 0.422931327 |
| Gm26982 | 0.240356677 | -1.070389328 | 0.017 | 0.041 | 1 | 0.401608067 | 1.930459067 | 0.1 | 0.043 | 1 | 0.401608067 | 0.422942023 |
| Apobec1 | 0.240356677 | -1.485426827 | 0.017 | 0.041 | 1 | 0.439648252 | 2.159940913 | 0.1 | 0.049 | 1 | 0.439648252 | 0.422942023 |
| Zfp526 | 0.240356677 | -1.070389328 | 0.017 | 0.041 | 1 | 0.401608067 | 1.695993813 | 0.1 | 0.043 | 1 | 0.401608067 | 0.422942023 |
| Kcnn1 | 0.240356677 | -1.070389328 | 0.017 | 0.041 | 1 | 0.669843604 | 2.017921908 | 0 | 0.019 | 1 | 0.669843604 | 0.422942023 |
| Adgrg1 | 0.240356677 | -1.600904045 | 0.017 | 0.041 | 1 | 0.324842732 | -0.111361109 | 0 | 0.09 | 1 | 0.324842732 | 0.422942023 |
| Gm11149 | 0.240356677 | -1.070389328 | 0.017 | 0.041 | 1 | 0.643850855 | 2.017921908 | 0 | 0.022 | 1 | 0.643850855 | 0.422942023 |
| 1700022H01Rik | 0.240356677 | -1.070389328 | 0.017 | 0.041 | 1 | 0.643842184 | 1.695993813 | 0 | 0.022 | 1 | 0.643842184 | 0.422942023 |
| Gm3625 | 0.240356677 | -1.070389328 | 0.017 | 0.041 | 1 | 0.669853054 | 2.210566986 | 0 | 0.019 | 1 | 0.669853054 | 0.422942023 |
| 9530082P21Rik | 0.240356677 | -1.070389328 | 0.017 | 0.041 | 1 | 0.669843604 | 2.017921908 | 0 | 0.019 | 1 | 0.669843604 | 0.422942023 |
| Gm26885 | 0.240356677 | -1.070389328 | 0.017 | 0.041 | 1 | 0.460577419 | 0.374065718 | 0 | 0.052 | 1 | 0.460577419 | 0.422942023 |
| Scd4 | 0.240356677 | -1.070389328 | 0.017 | 0.041 | 1 | 0.482051401 | 1.558490289 | 0.1 | 0.049 | 1 | 0.482051401 | 0.422942023 |
| Arhgef7 | 0.92499886 | -0.572889668 | 0.445 | 0.406 | 1 | 0.240633894 | 1 | 0.5 | 0.33 | 1 | 0.92499886 | 0.423363117 |
| Cpne9 | 0.506471296 | -1.807354922 | 0.008 | 0.018 | 1 | 0.241074418 | 1.695993813 | 0.1 | 0.031 | 1 | 0.506471296 | 0.424031961 |
| Hykk | 0.241123912 | -0.900464326 | 0.034 | 0.065 | 1 | 0.676138615 | 0.973527789 | 0.1 | 0.065 | 1 | 0.676138615 | 0.424107083 |
| Slc39a6 | 0.241322883 | -0.31444063 | 0.496 | 0.553 | 1 | 0.353348579 | 0.327606407 | 0.7 | 0.522 | 1 | 0.353348579 | 0.424409032 |
| Bard1 | 0.241520057 | -1.733354341 | 0.025 | 0.053 | 1 | 0.698534498 | 2.432959407 | 0 | 0.015 | 1 | 0.698534498 | 0.424708177 |
| Gm10353 | 0.241873159 | -0.807354922 | 0.034 | 0.065 | 1 | 0.862565643 | 0.808468542 | 0.1 | 0.083 | 1 | 0.862565643 | 0.425243692 |
| Gm20045 | 0.386945655 | 0.321928095 | 0.092 | 0.065 | 1 | 0.242210977 | 1.374065718 | 0.2 | 0.09 | 1 | 0.386945655 | 0.425755796 |
| 4921511C10Rik | 0.242320046 | -0.900464326 | 0.017 | 0.041 | 1 | 0.460551517 | 0.625604485 | 0 | 0.052 | 1 | 0.460551517 | 0.425921087 |
| A530041M06Rik | 0.242320046 | -0.900464326 | 0.017 | 0.041 | 1 | 0.460577419 | 0.494359952 | 0 | 0.052 | 1 | 0.460577419 | 0.425921087 |
| Clstn3 | 0.242320046 | -0.900464326 | 0.017 | 0.041 | 1 | 0.952551372 | 0.808468542 | 0.1 | 0.093 | 1 | 0.952551372 | 0.425921087 |
| A630052C17Rik | 0.242320046 | -0.900464326 | 0.017 | 0.041 | 1 | 0.318562315 | 2.111031312 | 0.1 | 0.037 | 1 | 0.318562315 | 0.425921087 |
| Gm34552 | 0.242320046 | -0.900464326 | 0.017 | 0.041 | 1 | 0.698529281 | 2.210566986 | 0 | 0.015 | 1 | 0.698529281 | 0.425921087 |
| Il16 | 0.242622913 | -0.337869639 | 0.916 | 0.718 | 1 | 0.669635185 | 0.283331121 | 1 | 0.852 | 1 | 0.669635185 | 0.426379948 |
| A530072M11Rik | 0.242709639 | -0.748461233 | 0.034 | 0.065 | 1 | 0.515666611 | 1.847996907 | 0.1 | 0.052 | 1 | 0.515666611 | 0.426511308 |
| Brca1 | 0.495728344 | -0.572889668 | 0.05 | 0.071 | 1 | 0.243244622 | 1.432959407 | 0.1 | 0.031 | 1 | 0.495728344 | 0.427321298 |
| Phex | 0.243255962 | -3.961160258 | 0.017 | 0.041 | 1 | 0.522126383 | 0.558490289 | 0 | 0.04 | 1 | 0.522126383 | 0.427338461 |
| Gm56796 | 0.915395984 | -0.665999073 | 0.118 | 0.118 | 1 | 0.243506792 | 1.591657153 | 0.2 | 0.093 | 1 | 0.915395984 | 0.427718026 |
| Efhc1 | 0.243996811 | -0.944858446 | 0.025 | 0.053 | 1 | 0.307202183 | -0.408342847 | 0 | 0.096 | 1 | 0.307202183 | 0.428459178 |
| Atxn7l2 | 0.243996811 | -0.944858446 | 0.025 | 0.053 | 1 | 0.827202642 | 1.210566986 | 0.1 | 0.08 | 1 | 0.827202642 | 0.428459178 |
| Gm57314 | 0.243996811 | -1.070389328 | 0.025 | 0.053 | 1 | 0.954546049 | 0.263034406 | 0.1 | 0.09 | 1 | 0.954546049 | 0.428459178 |
| Reep1 | 0.243996811 | -1.070389328 | 0.025 | 0.053 | 1 | 0.643842184 | 1.210566986 | 0 | 0.022 | 1 | 0.643842184 | 0.428459178 |
| 9330160F10Rik | 0.243996811 | -0.944858446 | 0.025 | 0.053 | 1 | 0.48480221 | 1.695993813 | 0.1 | 0.049 | 1 | 0.48480221 | 0.428459178 |
| Vars2 | 0.243996811 | -1.070389328 | 0.025 | 0.053 | 1 | 0.577142568 | 1.432959407 | 0 | 0.031 | 1 | 0.577142568 | 0.428459178 |
| Prelid2 | 0.243996811 | -1.292781749 | 0.025 | 0.053 | 1 | 0.669853054 | 2.210566986 | 0 | 0.019 | 1 | 0.669853054 | 0.428459178 |
| Nqo2 | 0.244069061 | 0.444183845 | 0.134 | 0.088 | 1 | 0.792439835 | 0.374065718 | 0.1 | 0.127 | 1 | 0.792439835 | 0.428568416 |
| Gm15545 | 0.24421652 | -0.944858446 | 0.017 | 0.041 | 1 | 0.698529281 | 2.017921908 | 0 | 0.015 | 1 | 0.698529281 | 0.428791331 |
| Dse | 0.244447506 | 0.165565544 | 0.866 | 0.818 | 1 | 0.855527768 | 0.149377624 | 0.8 | 0.846 | 1 | 0.855527768 | 0.429140429 |
| Ccdc171 | 0.757833833 | 0.421463768 | 0.134 | 0.124 | 1 | 0.244555242 | 0.847996907 | 0.2 | 0.086 | 1 | 0.757833833 | 0.429303218 |
| 2510017J16Rik | 0.244731468 | -0.549557165 | 0.076 | 0.118 | 1 | 0.896995869 | 0.808468542 | 0.1 | 0.086 | 1 | 0.896995869 | 0.429569444 |
| Lrrc14 | 0.24476491 | -0.522901533 | 0.134 | 0.182 | 1 | 0.576516421 | 0.874963954 | 0.2 | 0.133 | 1 | 0.576516421 | 0.429619958 |
| Gm41505 | 0.244921348 | -0.765534746 | 0.025 | 0.053 | 1 | 0.763554956 | 0.185031894 | 0.1 | 0.071 | 1 | 0.763554956 | 0.429856229 |
| Aars | 0.245047823 | -0.397963986 | 0.454 | 0.488 | 1 | 0.988790399 | -0.343144581 | 0.7 | 0.559 | 1 | 0.988790399 | 0.43004721 |
| Riok2 | 0.337211713 | -0.382333334 | 0.513 | 0.512 | 1 | 0.245071172 | -0.859822342 | 0.3 | 0.466 | 1 | 0.337211713 | 0.430082464 |
| Pfkm | 0.40841665 | 0.237039197 | 0.235 | 0.194 | 1 | 0.24508846 | -0.736965594 | 0.1 | 0.259 | 1 | 0.40841665 | 0.430108567 |
| Itgb1bp1 | 0.625671747 | -0.436517227 | 0.336 | 0.341 | 1 | 0.245122676 | 0.732519689 | 0.5 | 0.302 | 1 | 0.625671747 | 0.430160226 |
| Ptpn3 | 0.860855182 | -0.36882907 | 0.563 | 0.529 | 1 | 0.245136216 | -0.269020829 | 0.5 | 0.728 | 1 | 0.860855182 | 0.430180668 |
| Mccc2 | 0.245162314 | -0.605721061 | 0.16 | 0.212 | 1 | 0.843690779 | 0.14345279 | 0.2 | 0.167 | 1 | 0.843690779 | 0.430220068 |
| Mansc4 | 0.245181037 | -0.748461233 | 0.017 | 0.041 | 1 | 0.669848329 | 1.558490289 | 0 | 0.019 | 1 | 0.669848329 | 0.430248332 |
| Tmem30c | 0.245181037 | -0.807354922 | 0.017 | 0.041 | 1 | 0.73075158 | 2.695993813 | 0 | 0.012 | 1 | 0.73075158 | 0.430248332 |
| Kctd8 | 0.729112716 | -0.292781749 | 0.017 | 0.012 | 1 | 0.245427676 | 0.236562194 | 0.1 | 0.031 | 1 | 0.729112716 | 0.430620608 |
| Piezo1 | 0.249868291 | 0.154752779 | 0.958 | 0.924 | 1 | 0.245483323 | 0.371031658 | 1 | 0.843 | 1 | 0.249868291 | 0.430704584 |
| Edc4 | 0.245524865 | -0.441032708 | 0.235 | 0.288 | 1 | 0.809640882 | 0.236562194 | 0.2 | 0.241 | 1 | 0.809640882 | 0.43076727 |
| Oscp1 | 0.245608961 | -0.36994961 | 0.084 | 0.129 | 1 | 0.515566127 | 0.744903414 | 0.2 | 0.123 | 1 | 0.515566127 | 0.430894161 |
| Etfbkmt | 0.35899242 | -0.807354922 | 0.076 | 0.106 | 1 | 0.245620175 | -0.709998547 | 0 | 0.12 | 1 | 0.35899242 | 0.430911079 |
| Trib2 | 0.818884394 | -0.263034406 | 0.345 | 0.347 | 1 | 0.245645492 | -1.279758641 | 0.2 | 0.349 | 1 | 0.818884394 | 0.430949276 |
| Gm48693 | 0.245740898 | -0.970853654 | 0.025 | 0.053 | 1 | 0.539455361 | 0.695993813 | 0 | 0.037 | 1 | 0.539455361 | 0.431093207 |
| Rab1b | 0.279038988 | -0.299871174 | 0.496 | 0.541 | 1 | 0.245768182 | -0.486698484 | 0.3 | 0.503 | 1 | 0.279038988 | 0.431134365 |
| Atp9a | 0.245815262 | -0.510961919 | 0.185 | 0.235 | 1 | 0.716775779 | 0.331421381 | 0.3 | 0.231 | 1 | 0.716775779 | 0.43120538 |
| Tmed5 | 0.256119746 | -0.233239804 | 0.723 | 0.694 | 1 | 0.245981328 | 0.591657153 | 0.7 | 0.688 | 1 | 0.256119746 | 0.431455843 |
| Gm19705 | 0.246100257 | -0.304854582 | 0.134 | 0.088 | 1 | 0.282293785 | -0.654503434 | 0 | 0.105 | 1 | 0.282293785 | 0.431635178 |
| Orm1 | 0.246133955 | -0.807354922 | 0.017 | 0.041 | 1 | 0.355415506 | 2.210566986 | 0.1 | 0.04 | 1 | 0.355415506 | 0.431685986 |
| Gm53018 | 0.246154704 | -0.622930351 | 0.017 | 0.041 | 1 | 0.597832181 | 0.558490289 | 0 | 0.028 | 1 | 0.597832181 | 0.431717269 |
| Nkain1 | 0.246156649 | 0.13606155 | 0.017 | 0.041 | 1 | 0.669848329 | 1.31748219 | 0 | 0.019 | 1 | 0.669848329 | 0.431720202 |
| BC024978 | 0.24639036 | -0.807354922 | 0.067 | 0.106 | 1 | 0.306978493 | -0.230005605 | 0 | 0.096 | 1 | 0.306978493 | 0.43207251 |
| Aknad1 | 0.246626843 | -0.807354922 | 0.025 | 0.053 | 1 | 0.479305119 | 1.847996907 | 0.1 | 0.049 | 1 | 0.479305119 | 0.432428886 |
| Map10 | 0.246626843 | -0.807354922 | 0.025 | 0.053 | 1 | 0.557784478 | 1.432959407 | 0 | 0.034 | 1 | 0.557784478 | 0.432428886 |
| 6330408M09Rik | 0.246626843 | -0.807354922 | 0.025 | 0.053 | 1 | 0.73075158 | 2.695993813 | 0 | 0.012 | 1 | 0.73075158 | 0.432428886 |
| Zfp97 | 0.246626843 | -0.807354922 | 0.025 | 0.053 | 1 | 0.557729726 | 1.31748219 | 0 | 0.034 | 1 | 0.557729726 | 0.432428886 |
| Sel1l | 0.246651386 | -0.23388806 | 0.689 | 0.7 | 1 | 0.571402981 | -0.270120193 | 0.7 | 0.778 | 1 | 0.571402981 | 0.432465865 |
| E130102H24Rik | 0.246800536 | -0.24838763 | 0.185 | 0.241 | 1 | 0.518227108 | 0.418009066 | 0.3 | 0.198 | 1 | 0.518227108 | 0.432690568 |
| Ggta1 | 0.246894713 | 0.192645078 | 0.134 | 0.088 | 1 | 0.316734049 | 1.720241359 | 0.2 | 0.105 | 1 | 0.316734049 | 0.432832427 |
| Zfyve21 | 0.247028107 | -0.341036918 | 0.361 | 0.424 | 1 | 0.906868877 | 0.41305985 | 0.3 | 0.306 | 1 | 0.906868877 | 0.433033328 |
| Strip2 | 0.247103148 | -0.655351829 | 0.017 | 0.041 | 1 | 0.367975074 | 1.263034406 | 0.1 | 0.04 | 1 | 0.367975074 | 0.43314633 |
| Letm1 | 0.247147962 | -0.472251438 | 0.361 | 0.4 | 1 | 0.830554568 | 0.709169202 | 0.4 | 0.463 | 1 | 0.830554568 | 0.433213809 |
| Stxbp1 | 0.434755424 | -0.449626106 | 0.563 | 0.518 | 1 | 0.247212822 | -0.888968688 | 0.5 | 0.63 | 1 | 0.434755424 | 0.433311464 |
| Pth2r | 0.247442064 | -0.402964667 | 0.025 | 0.053 | 1 | 0.505649381 | 0.625604485 | 0 | 0.043 | 1 | 0.505649381 | 0.433656552 |
| 2310022A10Rik | 0.781546994 | -0.102098188 | 0.101 | 0.112 | 1 | 0.247746891 | 1.017921908 | 0.3 | 0.154 | 1 | 0.781546994 | 0.43411526 |
| 1700037C18Rik | 0.387377553 | 1.251538767 | 0.025 | 0.012 | 1 | 0.248008881 | 3.432959407 | 0.1 | 0.034 | 1 | 0.387377553 | 0.434509357 |
| 3930402G23Rik | 0.442576478 | -0.707819249 | 0.059 | 0.082 | 1 | 0.248008881 | 3.017921908 | 0.1 | 0.034 | 1 | 0.442576478 | 0.434509357 |
| Trim72 | 0.24807501 | -0.333423734 | 0.017 | 0.041 | 1 | 0.539445308 | 0.625604485 | 0 | 0.037 | 1 | 0.539445308 | 0.43460881 |
| Tnfsf13b | 0.248258778 | -0.86393845 | 0.025 | 0.053 | 1 | 0.577176345 | 0.769994395 | 0 | 0.031 | 1 | 0.577176345 | 0.434885314 |

| | | | | | | | | | | | | |
|---|---|---|---|---|---|---|---|---|---|---|---|---|
| Ctns | 0.25272315 | -0.175086707 | 0.168 | 0.224 | 1 | 0.248331847 | -0.776493958 | 0.1 | 0.256 | 1 | 0.25272315 | 0.434994987 |
| Lasp1 | 0.596953473 | -0.230497471 | 0.395 | 0.394 | 1 | 0.248984681 | -0.797995028 | 0.4 | 0.549 | 1 | 0.596953473 | 0.43597599 |
| Gm36264 | 0.24903135 | -0.485426827 | 0.017 | 0.041 | 1 | 0.698534498 | 2.432959407 | 0 | 0.015 | 1 | 0.698534498 | 0.436046087 |
| Gm36198 | 0.249108017 | -0.748461233 | 0.025 | 0.053 | 1 | 0.557765898 | 0.494359952 | 0 | 0.034 | 1 | 0.557765898 | 0.43616123 |
| Aven | 0.249287523 | -0.59306755 | 0.403 | 0.435 | 1 | 0.532940671 | -0.556986928 | 0.5 | 0.549 | 1 | 0.532940671 | 0.436430776 |
| Gm39090 | 0.249401202 | 0.514573173 | 0.21 | 0.153 | 1 | 0.294962571 | 0.789103218 | 0.3 | 0.16 | 1 | 0.294962571 | 0.436601445 |
| Get4 | 0.249418195 | -0.390789953 | 0.395 | 0.447 | 1 | 0.277619269 | 0.446379923 | 0.7 | 0.463 | 1 | 0.277619269 | 0.436626954 |
| Etfrf1 | 0.249470792 | 0.402098444 | 0.252 | 0.194 | 1 | 0.529363426 | 0.678071905 | 0.1 | 0.201 | 1 | 0.529363426 | 0.436705907 |
| Tefm | 0.249624576 | -0.707819249 | 0.092 | 0.135 | 1 | 0.293201969 | 1.354956895 | 0.2 | 0.096 | 1 | 0.293201969 | 0.436936723 |
| Zfp825 | 0.249624576 | -0.707819249 | 0.092 | 0.135 | 1 | 0.935975206 | 1.176619654 | 0.1 | 0.099 | 1 | 0.935975206 | 0.436936723 |
| Fpgt | 0.24970484 | -0.485426827 | 0.076 | 0.118 | 1 | 0.767699782 | 1.263034406 | 0.1 | 0.074 | 1 | 0.767699782 | 0.437057172 |
| Hook2 | 0.249766386 | -1.008988783 | 0.059 | 0.094 | 1 | 0.368291381 | 0.821524695 | 0.2 | 0.105 | 1 | 0.368291381 | 0.437149525 |
| Lypd1 | 0.249802223 | 0.910501849 | 0.101 | 0.065 | 1 | 0.309089492 | 1.902444691 | 0.1 | 0.037 | 1 | 0.309089492 | 0.437203295 |
| Lrsam1 | 0.249867367 | -0.691877705 | 0.092 | 0.135 | 1 | 0.438375168 | -0.357117523 | 0.1 | 0.194 | 1 | 0.438375168 | 0.437301033 |
| Hdhd2 | 0.77579956 | -0.166148311 | 0.429 | 0.429 | 1 | 0.249961132 | 0.913585248 | 0.5 | 0.364 | 1 | 0.77579956 | 0.437441696 |
| Fkbp1b | 0.250560853 | -0.448900951 | 0.143 | 0.194 | 1 | 0.920052124 | 1.017921908 | 0.1 | 0.12 | 1 | 0.920052124 | 0.438340966 |
| Ubxn2a | 0.250794448 | 0.131244533 | 0.588 | 0.5 | 1 | 0.368666786 | -0.658538947 | 0.6 | 0.599 | 1 | 0.368666786 | 0.438691041 |
| Ppp1r12c | 0.280331259 | -0.330704232 | 0.387 | 0.435 | 1 | 0.25083739 | -0.75818208 | 0.4 | 0.515 | 1 | 0.280331259 | 0.438755383 |
| Osgep | 0.250938487 | -0.337869639 | 0.479 | 0.482 | 1 | 0.485844255 | 0.663572335 | 0.5 | 0.414 | 1 | 0.485844255 | 0.438906849 |
| F420014N23Rik | 0.556227717 | -0.719892081 | 0.126 | 0.147 | 1 | 0.251063813 | 0.973527789 | 0.3 | 0.154 | 1 | 0.556227717 | 0.439094588 |
| Fam98c | 0.251130077 | -0.563429339 | 0.118 | 0.165 | 1 | 0.877599005 | 0.526068812 | 0.1 | 0.114 | 1 | 0.877599005 | 0.439193839 |
| Gm35019 | 0.251166951 | -1.14839184 | 0.059 | 0.094 | 1 | 0.397068795 | -0.625934282 | 0 | 0.068 | 1 | 0.397068795 | 0.439249065 |
| Nckipsd | 0.809650969 | -0.347923303 | 0.076 | 0.082 | 1 | 0.251346409 | 1.31748219 | 0.2 | 0.09 | 1 | 0.809650969 | 0.439517801 |
| Ankrd33b | 0.251644218 | -0.622930351 | 0.025 | 0.053 | 1 | 0.489967526 | 0.159940913 | 0 | 0.046 | 1 | 0.489967526 | 0.439963623 |
| Pacsin1 | 0.251644218 | -0.622930351 | 0.025 | 0.053 | 1 | 0.87077304 | 0.888638891 | 0.1 | 0.083 | 1 | 0.87077304 | 0.439963623 |
| Fance | 0.251689149 | -0.569491092 | 0.193 | 0.241 | 1 | 0.747730795 | 0.345496566 | 0.3 | 0.238 | 1 | 0.747730795 | 0.44003087 |
| Cntrl | 0.368135205 | -0.584962501 | 0.412 | 0.412 | 1 | 0.251975866 | 1.111031312 | 0.5 | 0.42 | 1 | 0.368135205 | 0.440459895 |
| Rab25 | 0.393128968 | 0.514573173 | 0.034 | 0.018 | 1 | 0.25202959 | -0.304006187 | 0 | 0.117 | 1 | 0.393128968 | 0.440540266 |
| Saxo2 | 0.695893386 | -0.222392421 | 0.017 | 0.024 | 1 | 0.252277061 | 2.795529487 | 0.1 | 0.034 | 1 | 0.695893386 | 0.440910406 |
| Prss53 | 0.767659898 | 0.273565073 | 0.067 | 0.059 | 1 | 0.252277061 | 2.795529487 | 0.1 | 0.034 | 1 | 0.767659898 | 0.440910406 |
| Nol8 | 0.878743232 | -0.50851044 | 0.378 | 0.347 | 1 | 0.252376647 | 1.045229254 | 0.4 | 0.275 | 1 | 0.878743232 | 0.441059322 |
| Msl3l2 | 0.705799754 | 0.351074441 | 0.143 | 0.129 | 1 | 0.252405384 | -0.473931188 | 0 | 0.117 | 1 | 0.705799754 | 0.441102291 |
| Smarcad1 | 0.252588698 | -0.464865505 | 0.538 | 0.541 | 1 | 0.516496642 | 0.239479678 | 0.9 | 0.676 | 1 | 0.516496642 | 0.441376346 |
| Psme3 | 0.694706734 | -0.342062652 | 0.286 | 0.288 | 1 | 0.25261929 | -0.823380346 | 0.2 | 0.355 | 1 | 0.694706734 | 0.441422075 |
| Trim12c | 0.262515828 | -0.683366205 | 0.126 | 0.171 | 1 | 0.252669274 | -0.763437806 | 0 | 0.117 | 1 | 0.262515828 | 0.441496787 |
| Zmiz1os1 | 0.329078398 | 0.377069649 | 0.143 | 0.1 | 1 | 0.252778467 | 1.988174565 | 0.2 | 0.096 | 1 | 0.329078398 | 0.441659981 |
| Ndufs2 | 0.476773107 | -0.225667553 | 0.529 | 0.547 | 1 | 0.253049258 | -0.562740455 | 0.5 | 0.633 | 1 | 0.476773107 | 0.442064589 |
| Nsg1 | 0.254234298 | 0.383679106 | 0.378 | 0.324 | 1 | 0.750427307 | 1.263034406 | 0.2 | 0.173 | 1 | 0.750427307 | 0.443833517 |
| Pdk1 | 0.609582308 | -0.337328188 | 0.269 | 0.282 | 1 | 0.254278455 | 0.808468542 | 0.4 | 0.231 | 1 | 0.609582308 | 0.443899377 |
| Pcgf1 | 0.49663453 | -0.726434927 | 0.067 | 0.088 | 1 | 0.254436177 | 2.602884409 | 0.1 | 0.034 | 1 | 0.49663453 | 0.444134586 |
| Taf12 | 0.901858839 | -0.222392421 | 0.311 | 0.306 | 1 | 0.25446456 | -0.736965594 | 0.2 | 0.361 | 1 | 0.901858839 | 0.444176907 |
| Abi3bp | 0.254774324 | 0.199264527 | 0.824 | 0.776 | 1 | 0.34133969 | -0.149572267 | 1 | 0.889 | 1 | 0.34133969 | 0.444638691 |
| Pole4 | 0.254849799 | -0.393397689 | 0.504 | 0.535 | 1 | 0.27982437 | 0.571665678 | 0.6 | 0.451 | 1 | 0.27982437 | 0.444751179 |
| Alx3 | 0.255060174 | -0.424026283 | 0.185 | 0.235 | 1 | 0.510687509 | -0.654503434 | 0.1 | 0.173 | 1 | 0.510687509 | 0.445064656 |
| Polr2l | 0.25518418 | 0.321928095 | 0.504 | 0.447 | 1 | 0.481520222 | 1.506959989 | 0.3 | 0.259 | 1 | 0.481520222 | 0.445249394 |
| Hspa4l | 0.848651903 | -0.194647431 | 0.244 | 0.218 | 1 | 0.255389019 | -1.510857757 | 0.2 | 0.352 | 1 | 0.848651903 | 0.445554488 |
| Mars1 | 0.682677933 | -0.163498732 | 0.21 | 0.229 | 1 | 0.255476115 | -0.802257054 | 0.1 | 0.253 | 1 | 0.682677933 | 0.445684184 |
| Asah2 | 0.255524051 | -0.459431619 | 0.118 | 0.165 | 1 | 0.698920459 | 0.227844977 | 0.2 | 0.145 | 1 | 0.698920459 | 0.445755561 |
| Ap4b1 | 0.255773081 | -0.333423734 | 0.134 | 0.188 | 1 | 0.3180835 | 1.558490289 | 0.2 | 0.105 | 1 | 0.3180835 | 0.446126292 |
| Gabpb1 | 0.762496467 | -0.138976413 | 0.513 | 0.435 | 1 | 0.255776602 | -0.54813213 | 0.4 | 0.586 | 1 | 0.762496467 | 0.446131534 |
| Bod1 | 0.255888401 | 0.121428279 | 0.689 | 0.618 | 1 | 0.274046888 | -0.722046174 | 0.6 | 0.648 | 1 | 0.274046888 | 0.446297927 |
| Mettl21a | 0.256013726 | -0.485426827 | 0.193 | 0.247 | 1 | 0.541726962 | 0.678071905 | 0.3 | 0.21 | 1 | 0.541726962 | 0.446484424 |
| Pitrm1 | 0.728642681 | -0.707819249 | 0.353 | 0.359 | 1 | 0.256171196 | 0.374065718 | 0.6 | 0.389 | 1 | 0.728642681 | 0.446718711 |
| Rps6ka5 | 0.256396362 | -0.564728947 | 0.706 | 0.718 | 1 | 0.908617205 | 0.398471031 | 0.7 | 0.654 | 1 | 0.908617205 | 0.44705363 |
| Snrnp27 | 0.256450569 | -0.410572832 | 0.546 | 0.547 | 1 | 0.373367279 | -0.665774546 | 0.4 | 0.469 | 1 | 0.373367279 | 0.447134243 |
| Gmnn | 0.897851829 | -1.185866545 | 0.05 | 0.053 | 1 | 0.256603914 | 2.602884409 | 0.1 | 0.034 | 1 | 0.897851829 | 0.447362259 |
| Fam136a | 0.256629333 | 0.148923701 | 0.311 | 0.241 | 1 | 0.70160456 | 0.558490289 | 0.1 | 0.157 | 1 | 0.70160456 | 0.447400051 |
| Trp53cor1 | 0.256795588 | 1.021533162 | 0.101 | 0.065 | 1 | 0.770508597 | 1.210566986 | 0.1 | 0.074 | 1 | 0.770508597 | 0.447647202 |
| Spats2 | 0.257047497 | -0.424885285 | 0.395 | 0.441 | 1 | 0.514724336 | -0.784271309 | 0.4 | 0.438 | 1 | 0.514724336 | 0.448021578 |
| Rnf213 | 0.509227837 | -0.309855263 | 0.151 | 0.182 | 1 | 0.257050541 | -1.769980651 | 0.1 | 0.25 | 1 | 0.509227837 | 0.448026102 |
| Zfp408 | 0.257148134 | -0.807354922 | 0.118 | 0.159 | 1 | 0.536072276 | 0.510127268 | 0.3 | 0.201 | 1 | 0.536072276 | 0.448171105 |
| Sfxn3 | 0.257654604 | -0.566346823 | 0.185 | 0.235 | 1 | 0.514679445 | -0.121629444 | 0.2 | 0.309 | 1 | 0.514679445 | 0.448923313 |
| Klc4 | 0.454784446 | -0.572889668 | 0.109 | 0.135 | 1 | 0.257834691 | -0.668578619 | 0.1 | 0.253 | 1 | 0.454784446 | 0.449190654 |
| Gm43696 | 0.258046661 | -1.070389328 | 0.059 | 0.094 | 1 | 0.932664593 | 0.31748219 | 0.1 | 0.105 | 1 | 0.932664593 | 0.449505242 |
| Il18 | 0.258047729 | 0.176156955 | 0.445 | 0.353 | 1 | 0.2844507 | 1.164470571 | 0.4 | 0.29 | 1 | 0.2844507 | 0.449506828 |
| Galnt4 | 0.258319461 | -0.780882711 | 0.084 | 0.124 | 1 | 0.342815982 | 0.808468542 | 0.3 | 0.173 | 1 | 0.342815982 | 0.449909978 |
| Retsat | 0.258398169 | 0.836501268 | 0.101 | 0.065 | 1 | 0.885920045 | 1.602884409 | 0.1 | 0.096 | 1 | 0.885920045 | 0.450026724 |
| Zfp119a | 0.25856772 | -1.129283017 | 0.059 | 0.094 | 1 | 0.342815159 | 1.14345279 | 0.2 | 0.102 | 1 | 0.342815159 | 0.450278174 |
| Psmd14 | 0.258595 | -0.2348344 | 0.84 | 0.829 | 1 | 0.323809475 | -0.418789634 | 0.6 | 0.796 | 1 | 0.323809475 | 0.450318626 |
| Map2k3os | 0.258712304 | 0.479807755 | 0.227 | 0.176 | 1 | 0.831735067 | 0.625604485 | 0.1 | 0.133 | 1 | 0.831735067 | 0.450492552 |
| Fzd10 | 0.525292084 | 0.929610672 | 0.05 | 0.035 | 1 | 0.258781732 | 1.067309935 | 0.4 | 0.25 | 1 | 0.525292084 | 0.45059548 |
| Reg4 | 0.565402508 | 0.893084796 | 0.042 | 0.059 | 1 | 0.258795395 | 2.017921908 | 0.1 | 0.034 | 1 | 0.565402508 | 0.450615734 |
| Gm28198 | 0.259110098 | -0.485426827 | 0.118 | 0.165 | 1 | 0.732343074 | 1.054447784 | 0.2 | 0.185 | 1 | 0.732343074 | 0.451082153 |
| Tti2 | 0.259126855 | 0.381306642 | 0.218 | 0.165 | 1 | 0.454211821 | -0.248864633 | 0.1 | 0.191 | 1 | 0.454211821 | 0.451106983 |
| Tap1 | 0.475483628 | -0.152851488 | 0.084 | 0.112 | 1 | 0.259355579 | 1.558490289 | 0.2 | 0.096 | 1 | 0.475483628 | 0.451445841 |
| Selenoh | 0.259375837 | -0.4440069 | 0.303 | 0.359 | 1 | 0.680991518 | -0.152003093 | 0.2 | 0.244 | 1 | 0.680991518 | 0.45147585 |
| Hscb | 0.468987371 | 0.192645078 | 0.176 | 0.141 | 1 | 0.259499111 | -0.473931188 | 0 | 0.114 | 1 | 0.468987371 | 0.451658433 |
| Gm57093 | 0.404086076 | -0.739183419 | 0.084 | 0.112 | 1 | 0.259563383 | -0.959358016 | 0 | 0.114 | 1 | 0.404086076 | 0.451753616 |
| Tmem35b | 0.32692511 | -0.584962501 | 0.109 | 0.147 | 1 | 0.259564679 | -0.441509711 | 0 | 0.114 | 1 | 0.32692511 | 0.451755535 |
| Mthfd1l | 0.259653685 | -0.564597041 | 0.546 | 0.571 | 1 | 0.411120761 | 0.162561613 | 0.6 | 0.522 | 1 | 0.411120761 | 0.451887334 |
| Unc13d | 0.712086445 | 0.421463768 | 0.076 | 0.065 | 1 | 0.259731249 | -0.709998547 | 0 | 0.114 | 1 | 0.712086445 | 0.452002176 |
| Wipf2 | 0.842169226 | -0.415037499 | 0.294 | 0.282 | 1 | 0.25991419 | -1.004445905 | 0.3 | 0.435 | 1 | 0.842169226 | 0.452272994 |
| Atp5g3 | 0.260018366 | -0.2003838 | 0.857 | 0.865 | 1 | 0.609657733 | -0.220937725 | 0.7 | 0.861 | 1 | 0.609657733 | 0.452427181 |
| Acaa2 | 0.436832447 | -0.58058406 | 0.151 | 0.182 | 1 | 0.260223896 | -0.936274402 | 0.1 | 0.253 | 1 | 0.436832447 | 0.452731317 |
| Lamb2 | 0.260377491 | -0.184345517 | 0.639 | 0.694 | 1 | 0.340860841 | 0.474538511 | 0.8 | 0.679 | 1 | 0.340860841 | 0.452958545 |
| Phospho2 | 0.260471968 | -0.613182374 | 0.202 | 0.247 | 1 | 0.882826193 | 0.150025444 | 0.3 | 0.324 | 1 | 0.882826193 | 0.45309829 |
| Ppard | 0.260546695 | 0.239713332 | 0.655 | 0.512 | 1 | 0.332108891 | -0.371269414 | 0.5 | 0.63 | 1 | 0.332108891 | 0.453208809 |
| Tprgl | 0.260667433 | -0.294484044 | 0.571 | 0.582 | 1 | 0.342665313 | -0.173698165 | 0.5 | 0.679 | 1 | 0.342665313 | 0.453387356 |
| Cntln | 0.632291472 | -0.64731451 | 0.429 | 0.394 | 1 | 0.260832919 | 0.528537067 | 0.6 | 0.448 | 1 | 0.632291472 | 0.453632026 |
| Gm16751 | 0.331652362 | -1.185866545 | 0.025 | 0.047 | 1 | 0.260963403 | 2.280956314 | 0.1 | 0.034 | 1 | 0.331652362 | 0.453824908 |
| Ccnt2 | 0.261070762 | -0.474022064 | 0.647 | 0.641 | 1 | 0.938175131 | -0.173422077 | 0.9 | 0.704 | 1 | 0.938175131 | 0.453983582 |
| Zfyve16 | 0.261674625 | -0.505604709 | 0.412 | 0.453 | 1 | 0.972861957 | 0.123104145 | 0.4 | 0.401 | 1 | 0.972861957 | 0.454875641 |
| Polr1f | 0.261892765 | -0.403999228 | 0.437 | 0.471 | 1 | 0.90980183 | 0.147557188 | 0.4 | 0.435 | 1 | 0.90980183 | 0.455197709 |
| Rras2 | 0.483815266 | -0.23320394 | 0.824 | 0.759 | 1 | 0.26206658 | -0.781359714 | 0.7 | 0.867 | 1 | 0.483815266 | 0.455454267 |
| Zfp346 | 0.262228346 | -0.724213687 | 0.185 | 0.229 | 1 | 0.448112288 | 0.68163852 | 0.3 | 0.201 | 1 | 0.448112288 | 0.455692987 |
| Pdhx | 0.352649244 | -0.392317423 | 0.319 | 0.359 | 1 | 0.262433422 | -0.944010051 | 0.2 | 0.349 | 1 | 0.352649244 | 0.455995543 |
| Polr3c | 0.262870635 | -0.922832139 | 0.193 | 0.235 | 1 | 0.714023326 | 0.197742946 | 0.3 | 0.228 | 1 | 0.714023326 | 0.4566403 |
| Lyrm4 | 0.496531199 | 0.312939312 | 0.151 | 0.124 | 1 | 0.263492389 | 1.463333056 | 0.2 | 0.096 | 1 | 0.496531199 | 0.457556339 |

| | | | | | | | | | | | | |
|---|---|---|---|---|---|---|---|---|---|---|---|---|
| Casp7 | 0.263502851 | -1.117695043 | 0.076 | 0.112 | 1 | 0.343896307 | -0.339630097 | 0 | 0.083 | 1 | 0.343896307 | 0.457571949 |
| Ccz1 | 0.263573705 | -0.319840761 | 0.479 | 0.512 | 1 | 0.692735759 | 0.18733468 | 0.6 | 0.534 | 1 | 0.692735759 | 0.457676311 |
| Crem | 0.263584719 | -0.453246421 | 0.412 | 0.429 | 1 | 0.339832986 | -0.643143572 | 0.3 | 0.438 | 1 | 0.339832986 | 0.457692533 |
| Mapk9 | 0.263605768 | -0.943010021 | 0.319 | 0.365 | 1 | 0.37844943 | 1.76091129 | 0.4 | 0.318 | 1 | 0.37844943 | 0.457723535 |
| Cdkn2c | 0.872914327 | -0.431655571 | 0.202 | 0.188 | 1 | 0.263643408 | 0.611929548 | 0.4 | 0.213 | 1 | 0.872914327 | 0.45777897 |
| Lbr | 0.26366623 | -0.442358105 | 0.319 | 0.376 | 1 | 0.312436781 | 1.141404961 | 0.6 | 0.46 | 1 | 0.312436781 | 0.457812578 |
| Thumpd3 | 0.313293824 | 0.204233052 | 0.294 | 0.235 | 1 | 0.263677607 | 1.121757719 | 0.4 | 0.256 | 1 | 0.313293824 | 0.457829333 |
| Ankrd29 | 0.263826343 | -0.839063782 | 0.059 | 0.094 | 1 | 0.460652366 | -0.152003093 | 0 | 0.052 | 1 | 0.460652366 | 0.458048347 |
| Ppp1r3b | 0.263865452 | -0.129283017 | 0.059 | 0.094 | 1 | 0.619961444 | 1.432959407 | 0 | 0.025 | 1 | 0.619961444 | 0.458105927 |
| Fabp3 | 0.263973794 | 2.669851398 | 0.143 | 0.106 | 1 | 0.334120172 | -0.111361109 | 0 | 0.086 | 1 | 0.334120172 | 0.458265424 |
| Nudt16l1 | 0.264017067 | -0.456280482 | 0.185 | 0.235 | 1 | 0.662229633 | 0.667424661 | 0.3 | 0.247 | 1 | 0.662229633 | 0.458329122 |
| Pld1 | 0.338367718 | -0.64731451 | 0.378 | 0.406 | 1 | 0.26452266 | 0.882762325 | 0.5 | 0.352 | 1 | 0.338367718 | 0.459073083 |
| Slc10a3 | 0.651751769 | -0.315501826 | 0.193 | 0.212 | 1 | 0.264735571 | 1.31748219 | 0.3 | 0.185 | 1 | 0.651751769 | 0.45938622 |
| Gm12216 | 0.264767457 | -0.839063782 | 0.134 | 0.182 | 1 | 0.438858008 | 0.304733697 | 0.2 | 0.352 | 1 | 0.438858008 | 0.459433108 |
| Fhip1b | 0.913294659 | -0.263034406 | 0.319 | 0.294 | 1 | 0.264934452 | -0.647414009 | 0.2 | 0.38 | 1 | 0.913294659 | 0.459678641 |
| Snapc5 | 0.265018063 | -0.425305835 | 0.336 | 0.382 | 1 | 0.916515774 | 0.111031312 | 0.4 | 0.349 | 1 | 0.916515774 | 0.459801552 |
| Slc4a7 | 0.265086618 | -0.164556039 | 0.765 | 0.812 | 1 | 0.936022596 | 0.242230945 | 0.7 | 0.846 | 1 | 0.936022596 | 0.459902321 |
| Calhm5 | 0.265135626 | -0.86393845 | 0.059 | 0.094 | 1 | 0.328344451 | 1.159940913 | 0.1 | 0.037 | 1 | 0.328344451 | 0.459974352 |
| Gtf2f1 | 0.265327032 | -0.279842694 | 0.605 | 0.629 | 1 | 0.869221818 | 0.219180116 | 0.6 | 0.63 | 1 | 0.869221818 | 0.46025563 |
| Dtd1 | 0.265456059 | -0.731183242 | 0.21 | 0.259 | 1 | 0.363369604 | -0.31836148 | 0.2 | 0.355 | 1 | 0.363369604 | 0.460445199 |
| Fzd3 | 0.265456719 | -0.920363884 | 0.193 | 0.235 | 1 | 0.835789492 | 0.364479669 | 0.3 | 0.269 | 1 | 0.835789492 | 0.460446169 |
| Atxn1l | 0.607327187 | -0.511899039 | 0.193 | 0.212 | 1 | 0.265575339 | -0.864721141 | 0.2 | 0.349 | 1 | 0.607327187 | 0.460620417 |
| Amd2 | 0.428820425 | 0.514573173 | 0.109 | 0.082 | 1 | 0.266321869 | 1.473601392 | 0.2 | 0.093 | 1 | 0.428820425 | 0.461716399 |
| Pde4dip | 0.266600291 | -0.16383863 | 0.983 | 0.918 | 1 | 0.328587078 | 1.013725251 | 1 | 0.969 | 1 | 0.328587078 | 0.462124867 |
| Rrm2b | 0.441895877 | -0.750021747 | 0.479 | 0.471 | 1 | 0.266749603 | 0.688798312 | 0.6 | 0.562 | 1 | 0.441895877 | 0.462343855 |
| Prkar2a | 0.798207694 | -0.382667253 | 0.454 | 0.435 | 1 | 0.266807344 | -0.795859283 | 0.6 | 0.645 | 1 | 0.798207694 | 0.462428529 |
| Emd | 0.266959493 | -0.308080886 | 0.639 | 0.659 | 1 | 0.718469663 | -0.145824519 | 0.7 | 0.719 | 1 | 0.718469663 | 0.462651615 |
| Fitm2 | 0.633616189 | -0.439623138 | 0.126 | 0.141 | 1 | 0.266985265 | -0.441509711 | 0 | 0.111 | 1 | 0.633616189 | 0.462689398 |
| Fam151b | 0.709934092 | -0.774933444 | 0.067 | 0.076 | 1 | 0.267085024 | -0.596787936 | 0 | 0.111 | 1 | 0.709934092 | 0.462835638 |
| 4921531C22Rik | 0.274273472 | -1.021479727 | 0.076 | 0.112 | 1 | 0.267087021 | -0.625934282 | 0 | 0.111 | 1 | 0.274273472 | 0.462838566 |
| R74862 | 0.573917755 | -0.529820947 | 0.109 | 0.129 | 1 | 0.267128965 | -0.625934282 | 0 | 0.111 | 1 | 0.573917755 | 0.462900046 |
| Cftr | 0.506471296 | -1.485426827 | 0.008 | 0.018 | 1 | 0.267176359 | -0.959358016 | 0 | 0.111 | 1 | 0.506471296 | 0.462969512 |
| Shld2 | 0.267311449 | -0.630857267 | 0.193 | 0.235 | 1 | 0.718270774 | 0.140177658 | 0.3 | 0.238 | 1 | 0.718270774 | 0.463167487 |
| Zfp956 | 0.267565044 | -0.807354922 | 0.059 | 0.094 | 1 | 0.269196207 | 1.354956895 | 0.2 | 0.093 | 1 | 0.269196207 | 0.463539035 |
| Abca3 | 0.954284643 | -0.222392421 | 0.109 | 0.106 | 1 | 0.267697626 | 1.403212064 | 0.2 | 0.096 | 1 | 0.954284643 | 0.463733233 |
| Ppp2r5c | 0.267717053 | -0.265645619 | 0.807 | 0.8 | 1 | 0.795417902 | -0.101513323 | 0.9 | 0.83 | 1 | 0.795417902 | 0.463761685 |
| Dync2li1 | 0.368807556 | 0.177538186 | 0.277 | 0.224 | 1 | 0.267975391 | 0.354956895 | 0.5 | 0.29 | 1 | 0.368807556 | 0.464139971 |
| Phldb1 | 0.26817385 | -0.390648602 | 0.908 | 0.829 | 1 | 0.624604852 | -0.140379209 | 1 | 0.895 | 1 | 0.624604852 | 0.464430486 |
| Hpf1 | 0.356673324 | -0.395484702 | 0.555 | 0.529 | 1 | 0.268562108 | 0.908004216 | 0.6 | 0.466 | 1 | 0.356673324 | 0.46499861 |
| Wdr89 | 0.420989806 | -0.182282208 | 0.63 | 0.653 | 1 | 0.2687321 | -0.436818015 | 0.5 | 0.645 | 1 | 0.420989806 | 0.465247259 |
| Dph7 | 0.268823832 | -0.755515991 | 0.126 | 0.171 | 1 | 0.632905675 | -0.131825212 | 0.1 | 0.151 | 1 | 0.632905675 | 0.465381412 |
| N4bp2l1 | 0.269018753 | -0.600904045 | 0.109 | 0.153 | 1 | 0.452221475 | -0.640289575 | 0.1 | 0.188 | 1 | 0.452221475 | 0.465666417 |
| Adra1b | 0.269018753 | -0.707819249 | 0.109 | 0.153 | 1 | 0.973644895 | 0.111031312 | 0.1 | 0.093 | 1 | 0.973644895 | 0.465666417 |
| Bmyc | 0.559113342 | 0.269460675 | 0.319 | 0.288 | 1 | 0.269278288 | -0.611434712 | 0.1 | 0.25 | 1 | 0.559113342 | 0.466045779 |
| Lrriq1 | 0.740152584 | -1.761551232 | 0.345 | 0.312 | 1 | 0.269306806 | -2.0853659 | 0.1 | 0.235 | 1 | 0.740152584 | 0.466087456 |
| Smpd4 | 0.34080683 | -0.450661409 | 0.16 | 0.2 | 1 | 0.270071992 | -0.581990934 | 0.1 | 0.25 | 1 | 0.34080683 | 0.467205103 |
| 1500011B03Rik | 0.270079426 | 0.189611018 | 0.504 | 0.459 | 1 | 0.940763193 | 0.753814467 | 0.4 | 0.488 | 1 | 0.940763193 | 0.467215955 |
| Afap1l2 | 0.270100267 | 0.275786313 | 0.227 | 0.165 | 1 | 0.602533881 | 0.635452271 | 0.1 | 0.179 | 1 | 0.602533881 | 0.467246379 |
| 1600014C10Rik | 0.270363992 | -0.655351829 | 0.084 | 0.124 | 1 | 0.511968279 | 1.048295557 | 0.2 | 0.127 | 1 | 0.511968279 | 0.467631297 |
| Palld | 0.270472898 | 0.244812904 | 0.613 | 0.588 | 1 | 0.590632229 | 1.644878497 | 0.7 | 0.614 | 1 | 0.590632229 | 0.467790207 |
| Dpp3 | 0.270475826 | -0.694880193 | 0.126 | 0.171 | 1 | 0.545343748 | -0.152003093 | 0.1 | 0.17 | 1 | 0.545343748 | 0.46779448 |
| Fbxw5 | 0.270568855 | -0.485426827 | 0.134 | 0.182 | 1 | 0.819633794 | 0.526068812 | 0.1 | 0.123 | 1 | 0.819633794 | 0.467930204 |
| App | 0.513478948 | -0.130723426 | 1 | 0.994 | 1 | 0.27069403 | -0.298952959 | 1 | 1 | 1 | 0.513478948 | 0.468112803 |
| Mgat4b | 0.46063507 | -0.333423734 | 0.328 | 0.359 | 1 | 0.27073141 | -0.452398027 | 0.3 | 0.528 | 1 | 0.46063507 | 0.468167323 |
| Zdhhc12 | 0.271123862 | 0.263034406 | 0.151 | 0.106 | 1 | 0.433635082 | 0.374065718 | 0 | 0.059 | 1 | 0.433635082 | 0.468739576 |
| Tvp23b | 0.881401239 | -0.178449074 | 0.765 | 0.682 | 1 | 0.271128925 | -0.536666944 | 0.5 | 0.651 | 1 | 0.881401239 | 0.468746956 |
| Gm56955 | 0.271714919 | -0.201633861 | 0.16 | 0.112 | 1 | 1 | 0.111031312 | 0.1 | 0.096 | 1 | 1 | 0.469600841 |
| Araf | 0.271768814 | -0.218712075 | 0.723 | 0.682 | 1 | 0.782010753 | -0.173422077 | 0.7 | 0.704 | 1 | 0.782010753 | 0.46967934 |
| S100a7a | 0.338041639 | 0.877143252 | 0.059 | 0.035 | 1 | 0.272080741 | 2.432959407 | 0.1 | 0.034 | 1 | 0.338041639 | 0.470133552 |
| Gm26935 | 0.393128968 | 0.514573173 | 0.034 | 0.018 | 1 | 0.272080741 | 2.432959407 | 0.1 | 0.034 | 1 | 0.393128968 | 0.470133552 |
| Tmem198 | 0.929810917 | -0.222392421 | 0.034 | 0.035 | 1 | 0.272080741 | 2.432959407 | 0.1 | 0.034 | 1 | 0.929810917 | 0.470133552 |
| Rpap1 | 0.398253428 | -0.707819249 | 0.042 | 0.065 | 1 | 0.272087326 | 1.31748219 | 0.2 | 0.093 | 1 | 0.398253428 | 0.470143139 |
| Chtf8 | 0.441422065 | -0.428843299 | 0.101 | 0.129 | 1 | 0.272087919 | 1.432959407 | 0.2 | 0.093 | 1 | 0.441422065 | 0.470144003 |
| D130020L05Rik | 0.272136498 | -0.655351829 | 0.059 | 0.094 | 1 | 0.673291667 | 1.263034406 | 0.1 | 0.065 | 1 | 0.673291667 | 0.470214723 |
| 1190005I06Rik | 0.397925554 | 0.206450877 | 0.134 | 0.1 | 1 | 0.272523112 | 1.695993813 | 0.2 | 0.099 | 1 | 0.397925554 | 0.470777377 |
| Rab20 | 0.624990352 | -0.982926487 | 0.134 | 0.147 | 1 | 0.272617252 | 0.732519689 | 0.3 | 0.157 | 1 | 0.624990352 | 0.470914337 |
| Rbpms | 0.310486888 | -0.640149422 | 0.244 | 0.294 | 1 | 0.272636907 | 1.145382817 | 0.5 | 0.373 | 1 | 0.310486888 | 0.470942931 |
| Il2 | 0.273470513 | 0.514573173 | 0.101 | 0.065 | 1 | 0.557784478 | 1.432959407 | 0 | 0.034 | 1 | 0.557784478 | 0.472154904 |
| 9330020H09Rik | 0.273470513 | 0.514573173 | 0.101 | 0.065 | 1 | 0.333955583 | 0.111031312 | 0 | 0.086 | 1 | 0.333955583 | 0.472154904 |
| Mrps18c | 0.605749836 | -0.336904302 | 0.345 | 0.353 | 1 | 0.273628633 | -0.754667596 | 0.2 | 0.352 | 1 | 0.605749836 | 0.472384637 |
| Gm54621 | 0.334204811 | -0.807354922 | 0.008 | 0.024 | 1 | 0.274256184 | 2.210566986 | 0.1 | 0.034 | 1 | 0.334204811 | 0.473295913 |
| Gm37305 | 0.286155558 | -0.87774425 | 0.05 | 0.082 | 1 | 0.274309548 | -0.304006187 | 0 | 0.108 | 1 | 0.286155558 | 0.473373368 |
| Cd36 | 0.485254257 | -1.240314329 | 0.025 | 0.041 | 1 | 0.274309548 | -0.339630097 | 0 | 0.108 | 1 | 0.485254257 | 0.473373368 |
| Hdac5 | 0.274349441 | -0.180572246 | 0.235 | 0.282 | 1 | 0.674760907 | 0.263034406 | 0.3 | 0.377 | 1 | 0.674760907 | 0.473431266 |
| Tm7sf2 | 0.354971971 | 0.560376862 | 0.185 | 0.147 | 1 | 0.274440092 | -0.374395515 | 0 | 0.108 | 1 | 0.354971971 | 0.473562821 |
| Dipk1b | 0.401173773 | -0.341036918 | 0.134 | 0.171 | 1 | 0.274605786 | -0.763437806 | 0 | 0.108 | 1 | 0.401173773 | 0.473803234 |
| Pcgf5 | 0.314261585 | -0.441572538 | 0.613 | 0.582 | 1 | 0.274703214 | -0.47087463 | 0.6 | 0.765 | 1 | 0.314261585 | 0.473944572 |
| Rfc3 | 0.275030037 | -0.833350131 | 0.076 | 0.112 | 1 | 0.366859967 | 1.290001453 | 0.2 | 0.111 | 1 | 0.366859967 | 0.474418553 |
| Ptpru | 0.275099486 | 0.514573173 | 0.101 | 0.065 | 1 | 0.559081495 | 1.494359952 | 0.1 | 0.056 | 1 | 0.559081495 | 0.474519245 |
| Zdhhc4 | 0.530121033 | -0.418312631 | 0.261 | 0.276 | 1 | 0.275308415 | -0.654503434 | 0.1 | 0.247 | 1 | 0.530121033 | 0.474822107 |
| Idua | 0.275309228 | -0.610957709 | 0.084 | 0.124 | 1 | 0.677102225 | 0.769994395 | 0.2 | 0.148 | 1 | 0.677102225 | 0.474823285 |
| Dennd2c | 0.64800604 | -0.563429339 | 0.286 | 0.306 | 1 | 0.275596523 | 0.69085051 | 0.5 | 0.389 | 1 | 0.64800604 | 0.475239603 |
| Rnaseh2c | 0.275636964 | -0.343407822 | 0.294 | 0.359 | 1 | 0.546179351 | -0.178475305 | 0.2 | 0.278 | 1 | 0.546179351 | 0.475298192 |
| Smim11 | 0.283785201 | -0.321451092 | 0.689 | 0.635 | 1 | 0.275645371 | -0.282201817 | 0.4 | 0.654 | 1 | 0.283785201 | 0.475310371 |
| Stk24 | 0.275708454 | -0.316014055 | 0.689 | 0.729 | 1 | 0.676934903 | 0.172836597 | 0.8 | 0.855 | 1 | 0.676934903 | 0.475401756 |
| Grhl1 | 0.275785942 | -0.442358105 | 0.092 | 0.135 | 1 | 0.707182625 | -0.397115591 | 0.2 | 0.235 | 1 | 0.707182625 | 0.475513998 |
| Zer1 | 0.965249327 | -0.203655859 | 0.227 | 0.218 | 1 | 0.276016232 | 0.754887502 | 0.4 | 0.231 | 1 | 0.965249327 | 0.475847504 |
| Zfhx3 | 0.716147682 | -0.129484075 | 0.908 | 0.912 | 1 | 0.276127584 | 0.423534394 | 0.9 | 0.951 | 1 | 0.716147682 | 0.476008726 |
| Cd200r3 | 0.462887689 | -0.655351829 | 0.025 | 0.041 | 1 | 0.276546558 | 2.210566986 | 0.1 | 0.034 | 1 | 0.462887689 | 0.476615117 |
| Gpr19 | 0.569304225 | -0.485426827 | 0.034 | 0.047 | 1 | 0.276546558 | 2.210566986 | 0.1 | 0.034 | 1 | 0.569304225 | 0.476615117 |
| Tyw3 | 0.752076132 | -0.263034406 | 0.05 | 0.059 | 1 | 0.276546558 | 2.210566986 | 0.1 | 0.034 | 1 | 0.752076132 | 0.476615117 |
| Gm43149 | 0.8327876 | -0.222392421 | 0.042 | 0.047 | 1 | 0.276546558 | 2.111031312 | 0.1 | 0.034 | 1 | 0.8327876 | 0.476615117 |
| Aplf | 0.943948817 | 0.192645078 | 0.084 | 0.082 | 1 | 0.276546558 | 1.930459067 | 0.1 | 0.034 | 1 | 0.943948817 | 0.476615117 |
| Cfp | 0.276680374 | -0.925999419 | 0.05 | 0.082 | 1 | 0.952551463 | 0.769994395 | 0.1 | 0.093 | 1 | 0.952551463 | 0.476808718 |
| Rhobtb2 | 0.276944832 | -0.839063782 | 0.059 | 0.094 | 1 | 0.374523737 | 0.159940913 | 0 | 0.074 | 1 | 0.374523737 | 0.477191224 |
| Pls3 | 0.837239078 | 0.394278939 | 0.588 | 0.594 | 1 | 0.27713814 | -0.100567829 | 0.4 | 0.691 | 1 | 0.837239078 | 0.477470731 |
| Casp8 | 0.277430245 | -0.661276663 | 0.185 | 0.229 | 1 | 0.765706876 | 0.263034406 | 0.3 | 0.241 | 1 | 0.765706876 | 0.477892949 |
| Trps1 | 0.937861701 | -0.152134331 | 1 | 1 | 1 | 0.2775418 | -0.272546552 | 1 | 1 | 1 | 0.937861701 | 0.47805415 |

| | | | | | | | | | | | | |
|---|---|---|---|---|---|---|---|---|---|---|---|---|
| Slc18b1 | 0.277558082 | -0.524955191 | 0.126 | 0.171 | 1 | 0.434361595 | -0.5519337 | 0.1 | 0.194 | 1 | 0.434361595 | 0.478077676 |
| Cmss1 | 0.316231998 | -0.382449818 | 1 | 1 | 1 | 0.277563706 | -0.110058156 | 1 | 1 | 1 | 0.316231998 | 0.478085801 |
| Ankrd49 | 0.277651554 | -0.807354922 | 0.168 | 0.212 | 1 | 0.530768752 | 0.432959407 | 0.2 | 0.123 | 1 | 0.530768752 | 0.478212722 |
| Arf2 | 0.456315681 | -0.104578014 | 0.798 | 0.729 | 1 | 0.277962769 | 0.34807051 | 0.8 | 0.691 | 1 | 0.456315681 | 0.478662238 |
| Nab2 | 0.562608462 | -0.304400832 | 0.504 | 0.506 | 1 | 0.278429023 | -0.574535129 | 0.5 | 0.633 | 1 | 0.562608462 | 0.479335325 |
| Rad52 | 0.278640569 | -0.736965594 | 0.151 | 0.194 | 1 | 0.438375404 | -0.408342847 | 0.1 | 0.194 | 1 | 0.438375404 | 0.479640571 |
| Bbs10 | 0.323940516 | -0.748461233 | 0.034 | 0.059 | 1 | 0.278845643 | 2.017921908 | 0.1 | 0.034 | 1 | 0.323940516 | 0.479936394 |
| B930046K06Rik | 0.563722678 | -0.622930351 | 0.034 | 0.047 | 1 | 0.278845643 | 2.017921908 | 0.1 | 0.034 | 1 | 0.563722678 | 0.479936394 |
| Elfn1 | 0.949508638 | -0.707819249 | 0.017 | 0.018 | 1 | 0.278845643 | 1.847996907 | 0.1 | 0.034 | 1 | 0.949508638 | 0.479936394 |
| Rnd1 | 0.343759729 | 0.616452787 | 0.168 | 0.129 | 1 | 0.278846016 | 1.930459067 | 0.1 | 0.034 | 1 | 0.343759729 | 0.479936931 |
| A430090L17Rik | 0.370184089 | 1.099535674 | 0.017 | 0.006 | 1 | 0.278846016 | 1.930459067 | 0.1 | 0.034 | 1 | 0.370184089 | 0.479936931 |
| Tfb1m | 0.617355833 | -0.485426827 | 0.067 | 0.082 | 1 | 0.278846016 | 2.111031312 | 0.1 | 0.034 | 1 | 0.617355833 | 0.479936931 |
| Nampt | 0.278968112 | -0.325386415 | 0.387 | 0.447 | 1 | 0.336423426 | -0.666576266 | 0.5 | 0.596 | 1 | 0.336423426 | 0.480113016 |
| Igf2bp3 | 0.822688475 | -0.538866086 | 0.244 | 0.218 | 1 | 0.279069881 | 0.385653692 | 0.3 | 0.164 | 1 | 0.822688475 | 0.480259763 |
| Gpsm3 | 0.279097496 | -0.600904045 | 0.143 | 0.188 | 1 | 0.324925979 | -0.339630097 | 0 | 0.09 | 1 | 0.324925979 | 0.48029958 |
| Rufy3 | 0.361342345 | -0.323155398 | 0.403 | 0.441 | 1 | 0.279207653 | -0.668578619 | 0.4 | 0.568 | 1 | 0.361342345 | 0.480458393 |
| Mtch2 | 0.27933336 | -0.161362988 | 0.714 | 0.771 | 1 | 0.432047772 | -0.397115591 | 0.8 | 0.756 | 1 | 0.432047772 | 0.480639594 |
| 2810002D19Rik | 0.279375853 | 0.312939312 | 0.143 | 0.1 | 1 | 0.58821099 | 0.695993813 | 0.2 | 0.133 | 1 | 0.58821099 | 0.480700839 |
| 6330562C20Rik | 0.279761139 | -0.86393845 | 0.076 | 0.112 | 1 | 0.557756607 | 1.111031312 | 0 | 0.034 | 1 | 0.557756607 | 0.481255983 |
| Ric8a | 0.280115846 | -0.513995979 | 0.193 | 0.241 | 1 | 0.57814466 | 0.930459067 | 0.3 | 0.231 | 1 | 0.57814466 | 0.481766805 |
| Tigd2 | 0.347023886 | -0.572889668 | 0.244 | 0.276 | 1 | 0.280351357 | 0.598383016 | 0.4 | 0.228 | 1 | 0.347023886 | 0.48210583 |
| Gamt | 0.280633385 | 0.640104055 | 0.092 | 0.059 | 1 | 0.607556927 | 1.159940913 | 0.1 | 0.059 | 1 | 0.607556927 | 0.482511674 |
| Ccdc130 | 0.356895812 | -0.544320516 | 0.084 | 0.118 | 1 | 0.280855571 | 1.210566986 | 0.2 | 0.093 | 1 | 0.356895812 | 0.482831291 |
| A430033K04Rik | 0.280969056 | -1 | 0.05 | 0.082 | 1 | 0.933175436 | 0.591657153 | 0.1 | 0.09 | 1 | 0.933175436 | 0.482994502 |
| Dhps | 0.367911584 | -0.207892852 | 0.319 | 0.365 | 1 | 0.280988159 | -0.574535129 | 0.2 | 0.383 | 1 | 0.367911584 | 0.483021973 |
| 5330439K02Rik | 0.510175501 | -1.173482821 | 0.042 | 0.059 | 1 | 0.281154556 | 1.769994395 | 0.1 | 0.034 | 1 | 0.510175501 | 0.483261228 |
| Spata32 | 0.702319168 | 0.251538767 | 0.017 | 0.024 | 1 | 0.281154556 | 1.625604485 | 0.1 | 0.034 | 1 | 0.702319168 | 0.483261228 |
| Grin2d | 0.760864275 | 0.726077278 | 0.109 | 0.1 | 1 | 0.281292606 | 1.515421567 | 0.2 | 0.096 | 1 | 0.760864275 | 0.483459681 |
| Bpifb5 | 0.281294119 | 0.731384562 | 0.151 | 0.112 | 1 | 0.892908645 | -0.267480311 | 0.2 | 0.17 | 1 | 0.892908645 | 0.483461857 |
| Rpl9-ps6 | 0.281566392 | -0.316827223 | 0.445 | 0.471 | 1 | 0.429759962 | -0.454565863 | 0.4 | 0.472 | 1 | 0.429759962 | 0.483853151 |
| Cd44 | 0.798609527 | -0.165207075 | 0.378 | 0.388 | 1 | 0.28186263 | -0.117854871 | 0.6 | 0.429 | 1 | 0.798609527 | 0.484278717 |
| Zcrb1 | 0.715798865 | -0.111968432 | 0.857 | 0.888 | 1 | 0.281993156 | -0.207395094 | 0.6 | 0.88 | 1 | 0.715798865 | 0.484466172 |
| Myo18b | 0.281993276 | -0.542010356 | 0.092 | 0.059 | 1 | 0.73075158 | 2.695993813 | 0 | 0.012 | 1 | 0.73075158 | 0.484466344 |
| Tmem69 | 0.469821252 | -0.411426246 | 0.076 | 0.1 | 1 | 0.282040088 | -0.191531458 | 0 | 0.105 | 1 | 0.469821252 | 0.484533565 |
| Cxcl16 | 0.309165704 | -0.485426827 | 0.076 | 0.112 | 1 | 0.282175151 | -0.304006187 | 0 | 0.105 | 1 | 0.309165704 | 0.484727487 |
| Ssbp4 | 0.720402084 | -0.335850471 | 0.429 | 0.365 | 1 | 0.28229169 | 0.505809973 | 0.6 | 0.469 | 1 | 0.720402084 | 0.484894781 |
| Cdr2l | 0.74915534 | 0.213403638 | 0.269 | 0.288 | 1 | 0.282347089 | -0.505640048 | 0 | 0.105 | 1 | 0.74915534 | 0.4849743 |
| Nomo1 | 0.346144151 | -0.501728639 | 0.345 | 0.365 | 1 | 0.282516462 | -0.473931188 | 0.2 | 0.38 | 1 | 0.346144151 | 0.485217373 |
| Gemin7 | 0.529913508 | -0.263034406 | 0.471 | 0.482 | 1 | 0.282539592 | 0.795529487 | 0.6 | 0.466 | 1 | 0.529913508 | 0.485250563 |
| Cemip | 0.775585683 | -0.128270244 | 0.815 | 0.741 | 1 | 0.282574544 | -1.288139781 | 0.8 | 0.784 | 1 | 0.775585683 | 0.485300715 |
| Prr16 | 0.687345921 | -2.707819249 | 0.017 | 0.024 | 1 | 0.28271034 | -1.750262417 | 0 | 0.105 | 1 | 0.687345921 | 0.485495544 |
| Crebrf | 0.282849735 | -0.106915204 | 0.798 | 0.753 | 1 | 0.503335874 | -0.233452907 | 0.6 | 0.799 | 1 | 0.503335874 | 0.485695497 |
| Txlng | 0.283009527 | -0.207324973 | 0.387 | 0.459 | 1 | 0.705467622 | 0.159940913 | 0.5 | 0.426 | 1 | 0.705467622 | 0.485924661 |
| E2f5 | 0.283021396 | -0.655351829 | 0.294 | 0.329 | 1 | 0.316601129 | -0.574535129 | 0.2 | 0.361 | 1 | 0.316601129 | 0.485941681 |
| Gm19303 | 0.283163413 | -1.392317423 | 0.042 | 0.071 | 1 | 0.669848329 | 1.847996907 | 0 | 0.019 | 1 | 0.669848329 | 0.486145307 |
| Sugp2 | 0.283223836 | -0.469996787 | 0.303 | 0.353 | 1 | 0.966960284 | -0.296774618 | 0.4 | 0.343 | 1 | 0.966960284 | 0.48623193 |
| Cacnb4 | 0.283340545 | -1.207892852 | 0.05 | 0.082 | 1 | 0.898150588 | -0.827568143 | 0.1 | 0.083 | 1 | 0.898150588 | 0.486399226 |
| Dok5 | 0.298320054 | 1.277533976 | 0.134 | 0.1 | 1 | 0.283473298 | 1.31748219 | 0.1 | 0.034 | 1 | 0.298320054 | 0.486589486 |
| Axdnd1 | 0.747884097 | 1.099535674 | 0.042 | 0.035 | 1 | 0.283473671 | 1.432959407 | 0.1 | 0.034 | 1 | 0.747884097 | 0.48659002 |
| Man1a | 0.283543762 | -0.232404224 | 0.916 | 0.824 | 1 | 0.911157528 | -0.27959776 | 0.9 | 0.932 | 1 | 0.911157528 | 0.486690459 |
| Sumf1 | 0.291952741 | -0.381591016 | 0.244 | 0.294 | 1 | 0.283871198 | 1.017921908 | 0.4 | 0.275 | 1 | 0.291952741 | 0.487159539 |
| Fbxl12 | 0.284082049 | -0.389849167 | 0.261 | 0.312 | 1 | 0.69841515 | 0.341461053 | 0.4 | 0.349 | 1 | 0.69841515 | 0.487461487 |
| Polr2e | 0.297316026 | -0.389849167 | 0.487 | 0.494 | 1 | 0.284699829 | -0.699068986 | 0.4 | 0.506 | 1 | 0.297316026 | 0.488345666 |
| Ifi27l2a | 0.284813441 | -1.411426246 | 0.05 | 0.082 | 1 | 0.635221364 | 1.017921908 | 0.1 | 0.062 | 1 | 0.635221364 | 0.488508186 |
| Ebf3 | 0.846126662 | -0.659456227 | 0.286 | 0.265 | 1 | 0.285023107 | -0.606568957 | 0.2 | 0.38 | 1 | 0.846126662 | 0.488808043 |
| Dhx40 | 0.806400376 | -0.335233891 | 0.605 | 0.6 | 1 | 0.285187247 | -0.794109501 | 0.6 | 0.688 | 1 | 0.806400376 | 0.489042728 |
| H2ac20 | 0.285298141 | -1.378511623 | 0.05 | 0.082 | 1 | 0.670447518 | 1.374065718 | 0.1 | 0.065 | 1 | 0.670447518 | 0.489201252 |
| Trim32 | 0.285298141 | -0.847996907 | 0.05 | 0.082 | 1 | 0.502048646 | 0.751135367 | 0.3 | 0.204 | 1 | 0.502048646 | 0.489201252 |
| Zfp39 | 0.285298141 | -0.765534746 | 0.05 | 0.082 | 1 | 0.324887875 | -0.152003093 | 0 | 0.09 | 1 | 0.324887875 | 0.489201252 |
| Tpk1 | 0.285323197 | -0.554468472 | 0.269 | 0.312 | 1 | 0.581799183 | 0.644463512 | 0.3 | 0.244 | 1 | 0.581799183 | 0.489237068 |
| Cetn2 | 0.285473041 | 0.255330346 | 0.571 | 0.5 | 1 | 0.618930501 | -0.434937057 | 0.4 | 0.423 | 1 | 0.618930501 | 0.489451225 |
| Add3 | 0.332732474 | -0.738407568 | 0.319 | 0.359 | 1 | 0.285504812 | -1.026472211 | 0.2 | 0.343 | 1 | 0.332732474 | 0.489496626 |
| Qsox2 | 0.285514531 | 0.470179053 | 0.21 | 0.165 | 1 | 0.320198089 | 1.091922489 | 0.3 | 0.182 | 1 | 0.320198089 | 0.489510515 |
| Pdcl3 | 0.721757589 | 0.267306083 | 0.731 | 0.706 | 1 | 0.285719037 | 0.840261258 | 0.8 | 0.79 | 1 | 0.721757589 | 0.489802706 |
| Zfp40 | 0.327741082 | -1.185866545 | 0.059 | 0.088 | 1 | 0.285795154 | 1.695993813 | 0.1 | 0.034 | 1 | 0.327741082 | 0.489911438 |
| Mvb12a | 0.615803617 | -0.248726569 | 0.479 | 0.494 | 1 | 0.285974561 | -0.756865152 | 0.3 | 0.429 | 1 | 0.615803617 | 0.490167672 |
| Rmc1 | 0.285997297 | -0.333423734 | 0.193 | 0.247 | 1 | 0.736727725 | -0.119581616 | 0.3 | 0.346 | 1 | 0.736727725 | 0.49020014 |
| Cox5a | 0.286029582 | -0.217299053 | 0.84 | 0.847 | 1 | 0.536038669 | -0.150180358 | 0.6 | 0.867 | 1 | 0.536038669 | 0.490246242 |
| Evi5l | 0.286031751 | -0.385891154 | 0.252 | 0.312 | 1 | 0.393650386 | -0.876895855 | 0.1 | 0.204 | 1 | 0.393650386 | 0.49024934 |
| Mesd | 0.286144361 | 0.117383675 | 0.714 | 0.629 | 1 | 0.318529925 | 0.338071459 | 0.9 | 0.667 | 1 | 0.318529925 | 0.490410127 |
| Ptgr2 | 0.286632491 | 0.450442835 | 0.168 | 0.124 | 1 | 0.55794495 | 0.526068812 | 0.1 | 0.191 | 1 | 0.55794495 | 0.491106796 |
| Trappc9 | 0.286752098 | -0.517559448 | 0.504 | 0.529 | 1 | 0.910259806 | -0.191531458 | 0.6 | 0.596 | 1 | 0.910259806 | 0.491277431 |
| Rhbdf2 | 0.697511304 | -0.240314329 | 0.227 | 0.194 | 1 | 0.286785417 | 1.14345279 | 0.2 | 0.093 | 1 | 0.697511304 | 0.491324958 |
| Lypla2 | 0.287042025 | -0.173482821 | 0.336 | 0.412 | 1 | 0.363814896 | 0.695993813 | 0.5 | 0.398 | 1 | 0.363814896 | 0.491690926 |
| Sec24c | 0.567010123 | -0.40103464 | 0.311 | 0.324 | 1 | 0.287050972 | -0.772155023 | 0.2 | 0.343 | 1 | 0.567010123 | 0.491703684 |
| Txndc9 | 0.287425772 | 0.125161481 | 0.664 | 0.606 | 1 | 0.617252404 | 0.296822719 | 0.8 | 0.685 | 1 | 0.617252404 | 0.492237969 |
| Pip4p2 | 0.935261602 | -0.331881569 | 0.571 | 0.506 | 1 | 0.287977191 | -0.473931188 | 0.5 | 0.611 | 1 | 0.935261602 | 0.49302352 |
| Itgb8 | 0.672222044 | 0.263034406 | 0.084 | 0.071 | 1 | 0.287991958 | -1.880931369 | 0.2 | 0.321 | 1 | 0.672222044 | 0.493044548 |
| Ccnb1ip1 | 0.288105649 | 0.514573173 | 0.092 | 0.059 | 1 | 0.42083289 | 0.374065718 | 0 | 0.062 | 1 | 0.42083289 | 0.493206433 |
| Gm57315 | 0.745632032 | -0.263034406 | 0.269 | 0.276 | 1 | 0.288171085 | 0.739937161 | 0.4 | 0.235 | 1 | 0.745632032 | 0.493299595 |
| Lonrf3 | 0.288735261 | -0.807354922 | 0.076 | 0.112 | 1 | 0.71280762 | -0.152003093 | 0.1 | 0.145 | 1 | 0.71280762 | 0.494102471 |
| Nle1 | 0.288979814 | -0.87774425 | 0.126 | 0.165 | 1 | 0.29852246 | -0.339630097 | 0 | 0.099 | 1 | 0.29852246 | 0.494450295 |
| Kazn | 0.289213889 | -0.739183419 | 0.185 | 0.224 | 1 | 0.373527882 | -1.239465935 | 0.1 | 0.207 | 1 | 0.373527882 | 0.494783104 |
| Atad5 | 0.289246943 | -0.748461233 | 0.05 | 0.082 | 1 | 0.540641288 | 0.732519689 | 0.1 | 0.052 | 1 | 0.540641288 | 0.494830091 |
| Hps4 | 0.289293037 | -0.436517227 | 0.227 | 0.276 | 1 | 0.420918733 | 0.751135367 | 0.3 | 0.188 | 1 | 0.420918733 | 0.494895613 |
| Adra2a | 0.28974413 | 0.514573173 | 0.092 | 0.059 | 1 | 0.570370849 | 0.930459067 | 0.1 | 0.056 | 1 | 0.570370849 | 0.4955366 |
| Gm30025 | 0.936806234 | 0.215012891 | 0.084 | 0.088 | 1 | 0.290071415 | -0.152003093 | 0 | 0.102 | 1 | 0.936806234 | 0.496001405 |
| Hspa2 | 0.940116403 | -0.201633861 | 0.252 | 0.241 | 1 | 0.290151543 | 0.75918764 | 0.4 | 0.256 | 1 | 0.940116403 | 0.496115168 |
| Mad2l2 | 0.303422377 | -0.527247003 | 0.134 | 0.176 | 1 | 0.290201571 | 1.393431043 | 0.2 | 0.096 | 1 | 0.303422377 | 0.49618619 |
| Bbs2 | 0.476152319 | -0.883976204 | 0.084 | 0.106 | 1 | 0.290201571 | 1.393431043 | 0.2 | 0.096 | 1 | 0.476152319 | 0.49618619 |
| Rfxank | 0.646706704 | 0.312939312 | 0.134 | 0.118 | 1 | 0.290202007 | 1.31748219 | 0.2 | 0.096 | 1 | 0.646706704 | 0.496186809 |
| Gm16159 | 0.609500043 | 0.514573173 | 0.034 | 0.024 | 1 | 0.290259655 | -0.304006187 | 0 | 0.102 | 1 | 0.609500043 | 0.496268643 |
| Tmem127 | 0.290295867 | -0.370784261 | 0.445 | 0.488 | 1 | 0.511666985 | -0.335224918 | 0.5 | 0.577 | 1 | 0.511666985 | 0.496320044 |
| Pheta1 | 0.393886923 | -0.250961574 | 0.05 | 0.076 | 1 | 0.29045414 | -0.567040593 | 0 | 0.102 | 1 | 0.393886923 | 0.496544673 |
| Pafah1b3 | 0.290480518 | -0.58058406 | 0.16 | 0.206 | 1 | 0.67794035 | 0.210566986 | 0.1 | 0.145 | 1 | 0.67794035 | 0.496582104 |
| Cdkl2 | 0.290538245 | -0.617877123 | 0.176 | 0.218 | 1 | 0.737523087 | 0.210566986 | 0.2 | 0.148 | 1 | 0.737523087 | 0.496664018 |
| Acad10 | 0.50416161 | -0.987927168 | 0.042 | 0.059 | 1 | 0.290562315 | 2.695993813 | 0.1 | 0.037 | 1 | 0.50416161 | 0.496698171 |
| Setd6 | 0.290952026 | -0.748461233 | 0.076 | 0.112 | 1 | 0.843784661 | 0.888638891 | 0.1 | 0.08 | 1 | 0.843784661 | 0.49725097 |

| | | | | | | | | | | | | |
|---|---|---|---|---|---|---|---|---|---|---|---|---|
| Crnde | 0.291136805 | -0.833350131 | 0.076 | 0.112 | 1 | 0.343888998 | -0.191531458 | 0 | 0.083 | 1 | 0.343888998 | 0.49751297 |
| Epas1 | 0.886389204 | -0.190298792 | 0.462 | 0.453 | 1 | 0.291241469 | -0.58364256 | 0.4 | 0.608 | 1 | 0.886389204 | 0.497661345 |
| Cnmd | 0.291487836 | 0.120240874 | 1 | 0.941 | 1 | 0.649829294 | -0.161925382 | 1 | 0.92 | 1 | 0.649829294 | 0.498010514 |
| Abca2 | 0.292083791 | 0.726077278 | 0.126 | 0.088 | 1 | 0.664609652 | 0.236562194 | 0.1 | 0.148 | 1 | 0.664609652 | 0.498854641 |
| Zfp639 | 0.292105568 | -0.630857267 | 0.252 | 0.294 | 1 | 0.889131675 | -0.101019165 | 0.3 | 0.293 | 1 | 0.889131675 | 0.498885473 |
| Lrrc40 | 0.292329494 | -0.42129649 | 0.151 | 0.2 | 1 | 0.835629839 | 0.354956895 | 0.2 | 0.167 | 1 | 0.835629839 | 0.499202455 |
| Gm34894 | 0.691605844 | -1.070389328 | 0.017 | 0.024 | 1 | 0.292833419 | 2.602884409 | 0.1 | 0.037 | 1 | 0.691605844 | 0.499915426 |
| Flywch2 | 0.330889747 | -0.807354922 | 0.118 | 0.153 | 1 | 0.292997521 | 1.210566986 | 0.3 | 0.176 | 1 | 0.330889747 | 0.500147494 |
| E130311K13Rik | 0.43258948 | -0.372952098 | 0.118 | 0.147 | 1 | 0.293203422 | 1.393431043 | 0.2 | 0.096 | 1 | 0.43258948 | 0.500438598 |
| Wdr37 | 0.29376691 | -0.496654083 | 0.412 | 0.441 | 1 | 0.844010126 | -0.119581616 | 0.7 | 0.534 | 1 | 0.844010126 | 0.501234823 |
| Slc35f5 | 0.825498472 | -0.147845468 | 0.655 | 0.635 | 1 | 0.293788827 | -0.326777495 | 0.5 | 0.701 | 1 | 0.825498472 | 0.501265779 |
| Ak8 | 0.294281293 | -1.5360529 | 0.042 | 0.071 | 1 | 0.716087628 | 0.847996907 | 0.1 | 0.068 | 1 | 0.716087628 | 0.501961106 |
| Tnfsf13 | 0.331128918 | -0.555816155 | 0.067 | 0.1 | 1 | 0.294315211 | 1.625604485 | 0.2 | 0.102 | 1 | 0.331128918 | 0.502008978 |
| Gpr89 | 0.448466864 | -0.188445089 | 0.269 | 0.312 | 1 | 0.294777006 | -0.802257054 | 0.1 | 0.238 | 1 | 0.448466864 | 0.502660528 |
| Fhl3 | 0.295021169 | -1.14839184 | 0.042 | 0.071 | 1 | 0.77893732 | 1.017921908 | 0.1 | 0.074 | 1 | 0.77893732 | 0.503004849 |
| Rasd1 | 0.608345034 | -0.246239163 | 0.151 | 0.129 | 1 | 0.295080895 | 3.728415291 | 0.2 | 0.114 | 1 | 0.608345034 | 0.503089056 |
| Tut1 | 0.295549156 | -0.628384781 | 0.176 | 0.218 | 1 | 0.749592875 | -0.125036046 | 0.2 | 0.231 | 1 | 0.749592875 | 0.503749008 |
| Mkrn2 | 0.295789315 | -0.567888987 | 0.235 | 0.276 | 1 | 0.736124381 | -0.165299916 | 0.2 | 0.231 | 1 | 0.736124381 | 0.504087312 |
| Gys1 | 0.305531305 | -0.678071905 | 0.076 | 0.112 | 1 | 0.295812998 | 0.847996907 | 0.2 | 0.093 | 1 | 0.305531305 | 0.504120667 |
| Poli | 0.295841525 | -0.637429921 | 0.05 | 0.082 | 1 | 0.446758355 | 0.695993813 | 0 | 0.056 | 1 | 0.446758355 | 0.504160842 |
| Gm56555 | 0.295875308 | 0.156119202 | 0.202 | 0.153 | 1 | 0.908958615 | 0.345496566 | 0.1 | 0.108 | 1 | 0.908958615 | 0.504208418 |
| Map7d1 | 0.435002683 | -0.166197882 | 0.798 | 0.788 | 1 | 0.296001491 | -0.230005605 | 1 | 0.892 | 1 | 0.435002683 | 0.504386099 |
| 2510002D24Rik | 0.29622745 | -0.780882711 | 0.218 | 0.259 | 1 | 0.804338365 | 0.135278859 | 0.3 | 0.244 | 1 | 0.804338365 | 0.504704198 |
| Gm49041 | 0.296361079 | 0.640104055 | 0.084 | 0.053 | 1 | 0.44222087 | 1.769994395 | 0.1 | 0.046 | 1 | 0.44222087 | 0.504892269 |
| Rgs11 | 0.296361079 | 0.640104055 | 0.084 | 0.053 | 1 | 0.44222087 | 1.769994395 | 0.1 | 0.046 | 1 | 0.44222087 | 0.504892269 |
| Fes | 0.296855466 | -0.669851398 | 0.076 | 0.112 | 1 | 0.550047024 | 0.580516596 | 0.2 | 0.127 | 1 | 0.550047024 | 0.505587765 |
| 2010315B03Rik | 0.296859869 | -0.770829046 | 0.126 | 0.165 | 1 | 0.50802954 | 0.795529487 | 0.2 | 0.123 | 1 | 0.50802954 | 0.505593956 |
| Rgs16 | 0.297504972 | -1.201633861 | 0.042 | 0.071 | 1 | 0.669850692 | 1.558490289 | 0 | 0.019 | 1 | 0.669850692 | 0.506500736 |
| Bag5 | 0.360565874 | -0.280956314 | 0.689 | 0.647 | 1 | 0.297532588 | -0.676339235 | 0.6 | 0.623 | 1 | 0.360565874 | 0.506539535 |
| Nudt13 | 0.297546387 | -0.565597176 | 0.218 | 0.259 | 1 | 0.95383112 | 0.930459067 | 0.2 | 0.241 | 1 | 0.95383112 | 0.506558921 |
| Mrpl47 | 0.297554075 | -0.517848305 | 0.143 | 0.188 | 1 | 0.922592259 | 0.354956895 | 0.2 | 0.179 | 1 | 0.922592259 | 0.506569722 |
| Map3k4 | 0.298067721 | 0.753760837 | 0.454 | 0.4 | 1 | 0.633597798 | -0.267480311 | 0.3 | 0.358 | 1 | 0.633597798 | 0.507291075 |
| Kcnj16 | 0.653590196 | 0.777607579 | 0.025 | 0.018 | 1 | 0.298405825 | -0.304006187 | 0 | 0.099 | 1 | 0.653590196 | 0.507765614 |
| Gm57158 | 0.479329584 | -0.839063782 | 0.067 | 0.088 | 1 | 0.298521563 | -0.374395515 | 0 | 0.099 | 1 | 0.479329584 | 0.507928002 |
| Nmnat1 | 0.74780139 | -0.163498732 | 0.084 | 0.094 | 1 | 0.298572536 | -0.304006187 | 0 | 0.099 | 1 | 0.74780139 | 0.507999513 |
| Gm13919 | 0.528185318 | -0.378511623 | 0.042 | 0.059 | 1 | 0.298574031 | -0.339630097 | 0 | 0.099 | 1 | 0.528185318 | 0.50800161 |
| Prps2 | 0.598684756 | -0.584962501 | 0.05 | 0.065 | 1 | 0.298574031 | -0.374395515 | 0 | 0.099 | 1 | 0.598684756 | 0.50800161 |
| Kdm1b | 0.327885716 | -0.622930351 | 0.034 | 0.059 | 1 | 0.298574778 | -0.441509711 | 0 | 0.099 | 1 | 0.327885716 | 0.508002658 |
| Wdr66 | 0.344307415 | -1.137503524 | 0.05 | 0.076 | 1 | 0.298730025 | -0.441509711 | 0 | 0.099 | 1 | 0.344307415 | 0.508220422 |
| Dubr | 0.428171036 | -0.359895945 | 0.067 | 0.094 | 1 | 0.298769755 | -0.567040593 | 0 | 0.099 | 1 | 0.428171036 | 0.508276144 |
| Nsrp1 | 0.386778049 | -0.367163946 | 0.487 | 0.5 | 1 | 0.298892788 | 0.444274721 | 0.7 | 0.562 | 1 | 0.386778049 | 0.508448678 |
| Flrt3 | 0.396447869 | -0.900464326 | 0.101 | 0.129 | 1 | 0.298910851 | -1.004445905 | 0 | 0.099 | 1 | 0.396447869 | 0.508474006 |
| Slc16a3 | 0.299020282 | -0.250961574 | 0.059 | 0.094 | 1 | 0.474948015 | 0.263034406 | 0 | 0.049 | 1 | 0.474948015 | 0.508627435 |
| Shc2 | 0.299051459 | -0.900464326 | 0.042 | 0.071 | 1 | 0.997435548 | 0.732519689 | 0.1 | 0.099 | 1 | 0.997435548 | 0.508671143 |
| Gga3 | 0.304128942 | -0.544320516 | 0.076 | 0.112 | 1 | 0.299253929 | 1.31748219 | 0.2 | 0.096 | 1 | 0.304128942 | 0.508954945 |
| 1600010M07Rik | 0.299430475 | -0.208909192 | 0.235 | 0.288 | 1 | 0.464775341 | -0.616284112 | 0.2 | 0.29 | 1 | 0.464775341 | 0.509202341 |
| Ufsp2 | 0.736696885 | -0.263034406 | 0.454 | 0.447 | 1 | 0.299686647 | 0.296822719 | 0.6 | 0.37 | 1 | 0.736696885 | 0.509561207 |
| 2810001A02Rik | 0.299852363 | -0.103556192 | 0.244 | 0.306 | 1 | 0.734892753 | 0.432959407 | 0.3 | 0.241 | 1 | 0.734892753 | 0.509793287 |
| Lias | 0.765822059 | -0.275503758 | 0.311 | 0.312 | 1 | 0.300209183 | 0.959028219 | 0.4 | 0.287 | 1 | 0.765822059 | 0.510292812 |
| Otud6b | 0.300399033 | -0.45845978 | 0.37 | 0.394 | 1 | 0.758115501 | -0.374395515 | 0.4 | 0.386 | 1 | 0.758115501 | 0.510558487 |
| Slc25a25 | 0.300746547 | -0.485426827 | 0.151 | 0.2 | 1 | 0.386020974 | 1.03809979 | 0.4 | 0.306 | 1 | 0.386020974 | 0.511044608 |
| Slc35a4 | 0.300923283 | -0.385891154 | 0.336 | 0.376 | 1 | 0.530151892 | -0.339630097 | 0.2 | 0.278 | 1 | 0.530151892 | 0.511291744 |
| Fkbp10 | 0.302837188 | -0.139251186 | 0.899 | 0.835 | 1 | 0.301003489 | 0.466747721 | 1 | 0.806 | 1 | 0.302837188 | 0.511403878 |
| Fbxl15 | 0.301007283 | -0.5360529 | 0.101 | 0.141 | 1 | 0.406275623 | 1.14345279 | 0.2 | 0.111 | 1 | 0.406275623 | 0.511409182 |
| Chid1 | 0.30102726 | -0.521952703 | 0.269 | 0.312 | 1 | 0.446099639 | -0.351311902 | 0.2 | 0.302 | 1 | 0.446099639 | 0.511437108 |
| Gli2 | 0.301182873 | -0.983453691 | 0.303 | 0.329 | 1 | 0.991405548 | -0.15815532 | 0.5 | 0.488 | 1 | 0.991405548 | 0.511654623 |
| Ostm1 | 0.301185744 | 0.337695411 | 0.143 | 0.1 | 1 | 0.339737394 | -1.111361109 | 0.1 | 0.219 | 1 | 0.339737394 | 0.511658636 |
| Tti1 | 0.915429626 | 0.453172628 | 0.118 | 0.118 | 1 | 0.301782669 | 2.785261151 | 0.2 | 0.105 | 1 | 0.915429626 | 0.512492558 |
| Rrp15 | 0.301870115 | -0.527247003 | 0.345 | 0.365 | 1 | 0.64413061 | -0.304006187 | 0.3 | 0.34 | 1 | 0.64413061 | 0.512614663 |
| Tatdn2 | 0.324269833 | -0.563429339 | 0.193 | 0.235 | 1 | 0.302013806 | -0.58694015 | 0.2 | 0.343 | 1 | 0.324269833 | 0.512815272 |
| Slk | 0.302360244 | -0.257786328 | 0.95 | 0.924 | 1 | 0.543278466 | 0.152372108 | 0.9 | 0.963 | 1 | 0.543278466 | 0.513298771 |
| Pomp | 0.388887267 | -0.218869355 | 0.866 | 0.847 | 1 | 0.302363923 | -0.18232763 | 0.7 | 0.821 | 1 | 0.388887267 | 0.513303905 |
| Marchf8 | 0.302406565 | -0.519374159 | 0.403 | 0.435 | 1 | 0.657759611 | -0.160881245 | 0.5 | 0.519 | 1 | 0.657759611 | 0.5133634 |
| Trappc14 | 0.3024216 | -0.622930351 | 0.076 | 0.112 | 1 | 0.378682929 | 1.591657153 | 0.2 | 0.117 | 1 | 0.378682929 | 0.513384376 |
| Mrpl27 | 0.302444795 | -0.40053793 | 0.42 | 0.459 | 1 | 0.369589276 | -0.428334322 | 0.3 | 0.441 | 1 | 0.369589276 | 0.513416736 |
| Gm6712 | 0.302643928 | -0.780882711 | 0.076 | 0.112 | 1 | 0.827078451 | 0.236562194 | 0.1 | 0.12 | 1 | 0.827078451 | 0.513694509 |
| Mov10 | 0.302856192 | -1.015941544 | 0.067 | 0.1 | 1 | 0.586883315 | -0.408342847 | 0.1 | 0.157 | 1 | 0.586883315 | 0.513990511 |
| Pcdhga5 | 0.302857204 | -0.839063782 | 0.067 | 0.1 | 1 | 0.96046354 | 0.591657153 | 0.1 | 0.093 | 1 | 0.96046354 | 0.513991922 |
| Rlf | 0.302897256 | -0.193132393 | 0.84 | 0.853 | 1 | 0.414132085 | -0.39025617 | 1 | 0.92 | 1 | 0.414132085 | 0.514047764 |
| Agt | 0.303072733 | -0.413873566 | 0.16 | 0.206 | 1 | 0.669853054 | 2.210566986 | 0 | 0.019 | 1 | 0.669853054 | 0.514292384 |
| Masp2 | 0.303113451 | -0.807354922 | 0.042 | 0.071 | 1 | 0.85982938 | 1.063725598 | 0.1 | 0.083 | 1 | 0.85982938 | 0.514349137 |
| Gm10069 | 0.303113451 | -0.807354922 | 0.042 | 0.071 | 1 | 0.317793426 | 0.672147071 | 0.2 | 0.096 | 1 | 0.317793426 | 0.514349137 |
| Cage1 | 0.303113451 | -0.900464326 | 0.042 | 0.071 | 1 | 0.474928012 | 0.695993813 | 0 | 0.049 | 1 | 0.474928012 | 0.514349137 |
| 4833438C02Rik | 0.303113451 | -0.807354922 | 0.042 | 0.071 | 1 | 0.619953009 | 1.695993813 | 0 | 0.025 | 1 | 0.619953009 | 0.514349137 |
| Rmi1 | 0.303299559 | -0.549557165 | 0.076 | 0.112 | 1 | 0.748799851 | 0.236562194 | 0.1 | 0.133 | 1 | 0.748799851 | 0.514608495 |
| Obp2a | 0.304376588 | -0.358191639 | 0.328 | 0.365 | 1 | 0.518662352 | -1.782977992 | 0.1 | 0.173 | 1 | 0.518662352 | 0.516108068 |
| Rorc | 0.304720429 | 0.80407979 | 0.076 | 0.047 | 1 | 0.7836543 | 0.210566986 | 0.1 | 0.127 | 1 | 0.7836543 | 0.516586318 |
| Nectin2 | 0.304794299 | -0.273922722 | 0.126 | 0.171 | 1 | 0.495411511 | 0.864116572 | 0.3 | 0.216 | 1 | 0.495411511 | 0.516689034 |
| Zfp219 | 0.304881331 | -0.707819249 | 0.092 | 0.129 | 1 | 0.579077979 | 0.695993813 | 0.3 | 0.231 | 1 | 0.579077979 | 0.516810037 |
| Ngef | 0.305024921 | -0.22059914 | 0.639 | 0.471 | 1 | 0.644252115 | 0.31748219 | 0.6 | 0.519 | 1 | 0.644252115 | 0.517009639 |
| Triqk | 0.30512009 | 0.148445274 | 0.277 | 0.218 | 1 | 0.75319459 | 0.494359952 | 0.2 | 0.262 | 1 | 0.75319459 | 0.51714191 |
| Parp3 | 0.50067329 | -0.431655571 | 0.252 | 0.276 | 1 | 0.305127353 | 0.236562194 | 0.5 | 0.287 | 1 | 0.50067329 | 0.517152004 |
| Fam149b | 0.305652076 | -0.644625422 | 0.21 | 0.253 | 1 | 0.547887411 | -0.121629444 | 0.2 | 0.296 | 1 | 0.547887411 | 0.51788096 |
| Cryl1 | 0.933622821 | -0.222392421 | 0.58 | 0.541 | 1 | 0.305858291 | 0.699795383 | 0.6 | 0.583 | 1 | 0.933622821 | 0.518167288 |
| Uqcr10 | 0.306580015 | 0.115302989 | 0.857 | 0.829 | 1 | 0.484285597 | 0.473601392 | 0.7 | 0.756 | 1 | 0.484285597 | 0.519168725 |
| Tlr2 | 0.30677422 | -0.263034406 | 0.16 | 0.212 | 1 | 0.526584835 | -0.516056664 | 0.2 | 0.275 | 1 | 0.526584835 | 0.519438018 |
| Immp1l | 0.306783264 | -0.36746349 | 0.664 | 0.641 | 1 | 0.682421969 | 0.107029382 | 0.7 | 0.608 | 1 | 0.682421969 | 0.519450557 |
| Cgn | 0.508731273 | -1.292781749 | 0.008 | 0.018 | 1 | 0.306858329 | -0.152003093 | 0 | 0.096 | 1 | 0.508731273 | 0.519554623 |
| Stxbp2 | 0.511049521 | 0.308122295 | 0.092 | 0.071 | 1 | 0.306920729 | -0.152003093 | 0 | 0.096 | 1 | 0.511049521 | 0.519641124 |
| Bfsp1 | 0.612842643 | -0.600904045 | 0.084 | 0.1 | 1 | 0.307081665 | -0.567040593 | 0 | 0.096 | 1 | 0.612842643 | 0.519864181 |
| Bloc1s1 | 0.368592076 | 0.164988735 | 0.403 | 0.353 | 1 | 0.307091481 | -0.526398608 | 0.2 | 0.346 | 1 | 0.368592076 | 0.519877785 |
| Paqr7 | 0.994401536 | -0.122856748 | 0.059 | 0.059 | 1 | 0.307126308 | -0.339630097 | 0 | 0.096 | 1 | 0.994401536 | 0.519926047 |
| Msantd1 | 0.30720717 | -0.707819249 | 0.042 | 0.071 | 1 | 0.361269566 | 2.695993813 | 0.1 | 0.043 | 1 | 0.361269566 | 0.520038094 |
| 6330418K02Rik | 0.30720717 | -0.807354922 | 0.042 | 0.071 | 1 | 0.396844932 | 0.374065718 | 0 | 0.068 | 1 | 0.396844932 | 0.520038094 |
| Pex11g | 0.30720717 | -0.707819249 | 0.042 | 0.071 | 1 | 0.521021286 | 1.494359952 | 0.1 | 0.052 | 1 | 0.521021286 | 0.520038094 |
| Car13 | 0.841777167 | 0.251538767 | 0.034 | 0.029 | 1 | 0.307269881 | -0.654503434 | 0 | 0.096 | 1 | 0.841777167 | 0.520124982 |
| Slc37a2 | 0.388295116 | 0.292180751 | 0.042 | 0.024 | 1 | 0.307404299 | -2.111361109 | 0 | 0.096 | 1 | 0.388295116 | 0.520311195 |
| Gosr1 | 0.30743594 | -0.501023682 | 0.319 | 0.359 | 1 | 0.910611498 | 0.203225011 | 0.4 | 0.37 | 1 | 0.910611498 | 0.520355023 |

| | | | | | | | | | | | | |
|---|---|---|---|---|---|---|---|---|---|---|---|---|
| Rgs6 | 0.683093632 | -1.359895945 | 0.017 | 0.024 | 1 | 0.307471634 | -2.833827133 | 0 | 0.096 | 1 | 0.683093632 | 0.520404462 |
| Mpnd | 0.308156617 | -0.576574715 | 0.168 | 0.212 | 1 | 0.81822505 | 0.418009066 | 0.2 | 0.25 | 1 | 0.81822505 | 0.521352733 |
| Gm35417 | 0.851114917 | -0.402964667 | 0.059 | 0.053 | 1 | 0.308449956 | 0.795529487 | 0.2 | 0.096 | 1 | 0.851114917 | 0.521758536 |
| Coq2 | 0.30850483 | -0.303223496 | 0.378 | 0.429 | 1 | 0.985630314 | 0.432959407 | 0.3 | 0.336 | 1 | 0.985630314 | 0.521834429 |
| Mitd1 | 0.308571621 | -0.248986631 | 0.235 | 0.288 | 1 | 0.443206576 | -0.425021588 | 0.1 | 0.191 | 1 | 0.443206576 | 0.521926796 |
| Nlk | 0.308675954 | -0.578938713 | 0.479 | 0.512 | 1 | 0.884573237 | -0.306258639 | 0.7 | 0.67 | 1 | 0.884573237 | 0.522071063 |
| Itgb4 | 0.308986793 | -0.678071905 | 0.042 | 0.071 | 1 | 0.680769929 | 0.494359952 | 0.1 | 0.16 | 1 | 0.680769929 | 0.522500748 |
| Gatc | 0.309129263 | -0.485426827 | 0.118 | 0.159 | 1 | 0.451411735 | 1.111031312 | 0.2 | 0.117 | 1 | 0.451411735 | 0.522697625 |
| Trim39 | 0.827724995 | -0.122856748 | 0.353 | 0.329 | 1 | 0.309162031 | 0.407424315 | 0.6 | 0.367 | 1 | 0.827724995 | 0.522742901 |
| Nfkbie | 0.309180345 | -1.250961574 | 0.034 | 0.059 | 1 | 0.366192339 | 2.515421567 | 0.1 | 0.043 | 1 | 0.366192339 | 0.522768205 |
| Arc | 0.309754039 | 1.736965594 | 0.067 | 0.041 | 1 | 0.331023156 | 2.159940913 | 0.1 | 0.04 | 1 | 0.331023156 | 0.523560514 |
| Stat1 | 0.309796588 | -0.26121167 | 0.227 | 0.276 | 1 | 0.920974242 | -0.11821478 | 0.3 | 0.278 | 1 | 0.920974242 | 0.523619251 |
| Usp24 | 0.477997433 | -0.140488754 | 0.899 | 0.859 | 1 | 0.3100672 | -0.261627585 | 0.9 | 0.941 | 1 | 0.477997433 | 0.523992731 |
| Zdhhc3 | 0.310095856 | 0.150416843 | 0.58 | 0.506 | 1 | 0.875034944 | 0.111031312 | 0.6 | 0.562 | 1 | 0.875034944 | 0.524032272 |
| Mroh1 | 0.310589623 | -0.418312631 | 0.143 | 0.188 | 1 | 0.95440895 | 0.611929548 | 0.2 | 0.201 | 1 | 0.95440895 | 0.524713333 |
| Klhl24 | 0.495032033 | -0.211666015 | 0.639 | 0.635 | 1 | 0.310854611 | 0.443930525 | 0.7 | 0.676 | 1 | 0.495032033 | 0.525078633 |
| Mrm3 | 0.310921415 | 0.432111013 | 0.126 | 0.088 | 1 | 0.835493932 | 1.063725598 | 0.1 | 0.08 | 1 | 0.835493932 | 0.525170703 |
| Srp54c | 0.766788786 | -0.375243909 | 0.269 | 0.265 | 1 | 0.311055042 | 0.808468542 | 0.4 | 0.256 | 1 | 0.766788786 | 0.525354845 |
| Yjefn3 | 0.311066875 | 0.436570661 | 0.126 | 0.088 | 1 | 0.342813781 | 1.14345279 | 0.2 | 0.102 | 1 | 0.342813781 | 0.525371149 |
| Cope | 0.311380642 | -0.139891598 | 0.916 | 0.929 | 1 | 0.859771678 | 0.136807947 | 0.7 | 0.895 | 1 | 0.859771678 | 0.525803379 |
| Pebp1 | 0.311392759 | 0.136871825 | 0.95 | 0.912 | 1 | 0.74458774 | 0.30367639 | 0.8 | 0.914 | 1 | 0.74458774 | 0.525820067 |
| Htra2 | 0.479679704 | 0.175086707 | 0.345 | 0.3 | 1 | 0.311423426 | 0.834700084 | 0.4 | 0.259 | 1 | 0.479679704 | 0.525862302 |
| 1700029J07Rik | 0.402291747 | -0.396159489 | 0.168 | 0.206 | 1 | 0.311539133 | 1.31748219 | 0.2 | 0.099 | 1 | 0.402291747 | 0.526021634 |
| Ralgapb | 0.443626279 | -0.566956713 | 0.496 | 0.465 | 1 | 0.311556009 | -0.367509129 | 0.4 | 0.593 | 1 | 0.443626279 | 0.526044872 |
| Nlrc4 | 0.311627564 | -0.600904045 | 0.042 | 0.071 | 1 | 0.559081495 | 1.494359952 | 0.1 | 0.056 | 1 | 0.559081495 | 0.526143389 |
| Pomgnt1 | 0.47904551 | -0.303588504 | 0.252 | 0.282 | 1 | 0.312071365 | 0.894539492 | 0.4 | 0.275 | 1 | 0.47904551 | 0.526754192 |
| Ndufaf4 | 0.325911355 | -0.323675757 | 0.37 | 0.406 | 1 | 0.3121975 | -0.608517229 | 0.3 | 0.448 | 1 | 0.325911355 | 0.52692772 |
| Tcte2 | 0.312301937 | -0.678071905 | 0.042 | 0.071 | 1 | 0.320995017 | 2.017921908 | 0.1 | 0.037 | 1 | 0.320995017 | 0.527071375 |
| Rpgrip1 | 0.802656338 | -0.347923303 | 0.168 | 0.176 | 1 | 0.312438594 | 1.197742946 | 0.3 | 0.176 | 1 | 0.802656338 | 0.527259313 |
| Psmf1 | 0.504536824 | -0.214124805 | 0.244 | 0.276 | 1 | 0.312445632 | -0.68251781 | 0.1 | 0.231 | 1 | 0.504536824 | 0.527268991 |
| Erlin2 | 0.312604107 | -0.510088881 | 0.218 | 0.265 | 1 | 0.802029995 | 0.494359952 | 0.3 | 0.269 | 1 | 0.802029995 | 0.527486886 |
| Fads2 | 0.998681388 | -0.118369357 | 0.395 | 0.376 | 1 | 0.312879714 | 0.769994395 | 0.4 | 0.247 | 1 | 0.998681388 | 0.527865713 |
| Ppp1r18os | 0.312916234 | 0.292180751 | 0.084 | 0.053 | 1 | 0.698529281 | 2.210566986 | 0 | 0.015 | 1 | 0.698529281 | 0.527915898 |
| Tarbp2 | 0.312982001 | -0.687060688 | 0.16 | 0.2 | 1 | 0.8173934 | 0.526068812 | 0.1 | 0.123 | 1 | 0.8173934 | 0.528006269 |
| Mtg1 | 0.313143129 | -0.584962501 | 0.042 | 0.071 | 1 | 0.947274527 | 0.888638891 | 0.1 | 0.093 | 1 | 0.947274527 | 0.528227639 |
| Pcdhb14 | 0.317038541 | 0.23893873 | 0.126 | 0.088 | 1 | 0.313735272 | 1.695993813 | 0.2 | 0.105 | 1 | 0.317038541 | 0.529040723 |
| Cep128 | 0.498653409 | -1.538538164 | 0.109 | 0.129 | 1 | 0.313763181 | 1.210566986 | 0.2 | 0.102 | 1 | 0.498653409 | 0.529079029 |
| Iws1 | 0.314086512 | -0.358047521 | 0.647 | 0.647 | 1 | 0.617861603 | 0.113862903 | 0.7 | 0.728 | 1 | 0.617861603 | 0.529522687 |
| Bace2 | 0.314353647 | 0.107397791 | 0.252 | 0.2 | 1 | 0.748798999 | 0.236562194 | 0.1 | 0.133 | 1 | 0.748798999 | 0.529889078 |
| Gm16066 | 0.441080022 | -0.180572246 | 0.126 | 0.159 | 1 | 0.314562161 | 1.41305985 | 0.3 | 0.185 | 1 | 0.441080022 | 0.530174969 |
| Sumo3 | 0.314564292 | 0.188697893 | 0.672 | 0.594 | 1 | 0.87875254 | 0.131789873 | 0.6 | 0.676 | 1 | 0.87875254 | 0.53017789 |
| Gm42659 | 0.403100411 | -0.707819249 | 0.067 | 0.094 | 1 | 0.314644262 | 1.280956314 | 0.2 | 0.099 | 1 | 0.403100411 | 0.530287512 |
| Drd3 | 0.314949751 | -0.944858446 | 0.042 | 0.071 | 1 | 0.404313999 | 0.695993813 | 0.1 | 0.043 | 1 | 0.404313999 | 0.530706156 |
| Zfpm1 | 0.9789673 | -0.430979043 | 0.193 | 0.182 | 1 | 0.314961439 | -0.495147674 | 0.1 | 0.253 | 1 | 0.9789673 | 0.530722171 |
| S100pbp | 0.315449319 | -0.6412215 | 0.378 | 0.388 | 1 | 0.491946324 | 0.707134371 | 0.5 | 0.426 | 1 | 0.491946324 | 0.531390366 |
| Gbp6 | 0.315484434 | 0.357031896 | 0.084 | 0.053 | 1 | 0.505677238 | -0.505640048 | 0 | 0.043 | 1 | 0.505677238 | 0.531438441 |
| Tmed1 | 0.315755378 | -0.357419215 | 0.622 | 0.612 | 1 | 0.37463197 | 0.526068812 | 0.7 | 0.531 | 1 | 0.37463197 | 0.531809298 |
| Nelfcd | 0.493487376 | -0.883976204 | 0.084 | 0.106 | 1 | 0.315800844 | -0.191531458 | 0 | 0.093 | 1 | 0.493487376 | 0.531871515 |
| Srrd | 0.444913709 | 0.13606155 | 0.151 | 0.118 | 1 | 0.315848442 | -0.267480311 | 0 | 0.093 | 1 | 0.444913709 | 0.531936646 |
| Ptger1 | 0.962802486 | -0.5360529 | 0.109 | 0.106 | 1 | 0.315930606 | -0.408342847 | 0 | 0.093 | 1 | 0.962802486 | 0.532049065 |
| Dapp1 | 0.315967308 | -0.655351829 | 0.21 | 0.247 | 1 | 0.98674541 | -0.152003093 | 0.3 | 0.293 | 1 | 0.98674541 | 0.532099277 |
| Galm | 0.758918751 | 0.23893873 | 0.109 | 0.1 | 1 | 0.316047777 | -0.536666944 | 0 | 0.093 | 1 | 0.758918751 | 0.532209356 |
| Ccdc149 | 0.319309473 | 0.273565073 | 0.084 | 0.053 | 1 | 0.316066164 | -0.625934282 | 0 | 0.093 | 1 | 0.319309473 | 0.532234507 |
| Esam | 0.341460366 | -0.655351829 | 0.025 | 0.047 | 1 | 0.316137708 | 2.111031312 | 0.1 | 0.037 | 1 | 0.341460366 | 0.532332366 |
| Gm56573 | 0.458788392 | -0.807354922 | 0.025 | 0.041 | 1 | 0.316137708 | 2.210566986 | 0.1 | 0.037 | 1 | 0.458788392 | 0.532332366 |
| Pcdhgc4 | 0.486994077 | -0.392317423 | 0.05 | 0.071 | 1 | 0.316137708 | 2.017921908 | 0.1 | 0.037 | 1 | 0.486994077 | 0.532332366 |
| Gdf9 | 0.729956195 | -0.748461233 | 0.034 | 0.041 | 1 | 0.316137708 | 2.210566986 | 0.1 | 0.037 | 1 | 0.729956195 | 0.532332366 |
| Gm56656 | 0.929389703 | 0.192645078 | 0.025 | 0.024 | 1 | 0.316137708 | 2.210566986 | 0.1 | 0.037 | 1 | 0.929389703 | 0.532332366 |
| Rtp4 | 0.882599202 | -0.394661363 | 0.126 | 0.118 | 1 | 0.316169327 | -1.441509711 | 0 | 0.093 | 1 | 0.882599202 | 0.53237561 |
| Fzd10os | 0.316514938 | -0.970853654 | 0.034 | 0.059 | 1 | 0.818903342 | 0.800691192 | 0.2 | 0.179 | 1 | 0.818903342 | 0.53284817 |
| Ccnf | 0.316569711 | -1.14839184 | 0.034 | 0.059 | 1 | 0.669848329 | 1.847996907 | 0 | 0.019 | 1 | 0.669848329 | 0.532923041 |
| Mettl3 | 0.317276125 | -0.450661409 | 0.151 | 0.194 | 1 | 0.538835794 | -0.321928095 | 0.1 | 0.17 | 1 | 0.538835794 | 0.533888111 |
| Grhpr | 0.317519004 | 0.367731785 | 0.185 | 0.141 | 1 | 0.395948015 | 0.751135367 | 0.3 | 0.182 | 1 | 0.395948015 | 0.534219691 |
| Nedd9 | 0.317550964 | -0.135402001 | 0.916 | 0.859 | 1 | 0.694289182 | -0.412065933 | 0.8 | 0.867 | 1 | 0.694289182 | 0.534263313 |
| Cdk5 | 0.317697121 | -0.368613162 | 0.311 | 0.353 | 1 | 0.563874981 | 0.506959989 | 0.4 | 0.293 | 1 | 0.563874981 | 0.534462782 |
| Atxn7l1 | 0.496630976 | 0.154529705 | 0.454 | 0.418 | 1 | 0.317848446 | 0.266601021 | 0.7 | 0.494 | 1 | 0.496630976 | 0.534669257 |
| Pmm1 | 0.317908567 | 0.362570079 | 0.37 | 0.312 | 1 | 0.365931832 | 0.882762325 | 0.4 | 0.272 | 1 | 0.365931832 | 0.534751277 |
| Anks1b | 0.318308565 | -0.610233283 | 0.252 | 0.206 | 1 | 0.399963158 | 1.210566986 | 0.3 | 0.198 | 1 | 0.399963158 | 0.535296787 |
| Dmap1 | 0.318308698 | -0.544320516 | 0.067 | 0.1 | 1 | 0.960016185 | 0.847996907 | 0.1 | 0.105 | 1 | 0.960016185 | 0.535296968 |
| Gm49463 | 0.318989184 | 0.514573173 | 0.076 | 0.047 | 1 | 0.318562315 | 2.111031312 | 0.1 | 0.037 | 1 | 0.318989184 | 0.535642681 |
| Adm | 0.381101673 | 0.152003093 | 0.05 | 0.029 | 1 | 0.318562315 | 2.111031312 | 0.1 | 0.037 | 1 | 0.381101673 | 0.535642681 |
| Dhrs9 | 0.39315136 | -1.887525271 | 0.034 | 0.018 | 1 | 0.318562315 | 1.847996907 | 0.1 | 0.037 | 1 | 0.39315136 | 0.535642681 |
| ENSMUSG00000121319 | 0.462887689 | -0.655351829 | 0.025 | 0.041 | 1 | 0.318562315 | 2.017921908 | 0.1 | 0.037 | 1 | 0.462887689 | 0.535642681 |
| Sars2 | 0.658336322 | -0.485426827 | 0.042 | 0.053 | 1 | 0.318562315 | 2.017921908 | 0.1 | 0.037 | 1 | 0.658336322 | 0.535642681 |
| Cenps | 0.659008402 | -0.655351829 | 0.059 | 0.071 | 1 | 0.318562315 | 2.111031312 | 0.1 | 0.037 | 1 | 0.659008402 | 0.535642681 |
| Psip1 | 0.716138837 | -0.182557068 | 0.807 | 0.753 | 1 | 0.318784846 | -0.564892126 | 0.6 | 0.781 | 1 | 0.716138837 | 0.535945914 |
| Coq8a | 0.318989184 | 0.514573173 | 0.076 | 0.047 | 1 | 0.420933835 | 0.31748219 | 0 | 0.062 | 1 | 0.420933835 | 0.536224268 |
| Nfatc1 | 0.53282874 | -0.110540429 | 0.983 | 0.906 | 1 | 0.319061266 | -0.430949465 | 1 | 0.981 | 1 | 0.53282874 | 0.536322441 |
| D17H6S53E | 0.319511358 | -0.637429921 | 0.067 | 0.1 | 1 | 0.701889643 | 1.374065718 | 0.1 | 0.068 | 1 | 0.701889643 | 0.536935208 |
| Dctn4 | 0.319521294 | -0.263034406 | 0.798 | 0.735 | 1 | 0.929326207 | -0.106773839 | 0.9 | 0.784 | 1 | 0.929326207 | 0.53694873 |
| Rabl3 | 0.31960216 | 0.370183263 | 0.126 | 0.088 | 1 | 0.353884738 | -0.408342847 | 0 | 0.08 | 1 | 0.353884738 | 0.537058779 |
| Coa7 | 0.320216024 | -0.86393845 | 0.034 | 0.059 | 1 | 0.474904369 | 0.625604485 | 0 | 0.049 | 1 | 0.474904369 | 0.537893746 |
| Zfp688 | 0.320216024 | -0.86393845 | 0.034 | 0.059 | 1 | 0.522066388 | 1.111031312 | 0 | 0.04 | 1 | 0.522066388 | 0.537893746 |
| A330041J22Rik | 0.320216024 | -0.86393845 | 0.034 | 0.059 | 1 | 0.489877116 | 0.930459067 | 0 | 0.046 | 1 | 0.489877116 | 0.537893746 |
| Tmem220 | 0.320216024 | -0.86393845 | 0.034 | 0.059 | 1 | 0.523811244 | 1.625604485 | 0.1 | 0.052 | 1 | 0.523811244 | 0.537893746 |
| Nol7 | 0.853703397 | -0.11345805 | 0.756 | 0.688 | 1 | 0.32024893 | 0.558490289 | 0.4 | 0.753 | 1 | 0.853703397 | 0.537938483 |
| Mllt3 | 0.4950674 | 0.16147706 | 0.899 | 0.918 | 1 | 0.320260734 | 0.267088061 | 0.9 | 0.867 | 1 | 0.4950674 | 0.537954531 |
| 4833422C13Rik | 0.320286451 | -1.070389328 | 0.034 | 0.059 | 1 | 0.669853054 | 2.210566986 | 0 | 0.019 | 1 | 0.669853054 | 0.537989491 |
| Eda2r | 0.320394949 | 1.94753258 | 0.059 | 0.035 | 1 | 0.669851164 | 1.210566986 | 0 | 0.019 | 1 | 0.669851164 | 0.538136974 |
| Coq3 | 0.320608484 | -0.448900951 | 0.134 | 0.176 | 1 | 0.974575842 | -0.152003093 | 0.3 | 0.272 | 1 | 0.974575842 | 0.538427168 |
| Plpp2 | 0.320629713 | 0.344648171 | 0.126 | 0.088 | 1 | 0.493067073 | 0.988174565 | 0.2 | 0.123 | 1 | 0.493067073 | 0.538456014 |
| Mmrn2 | 0.820836204 | -0.807354922 | 0.025 | 0.029 | 1 | 0.32099467 | 1.625604485 | 0.1 | 0.037 | 1 | 0.820836204 | 0.538951762 |
| Gm10382 | 0.844046361 | 0.13606155 | 0.059 | 0.053 | 1 | 0.32099467 | 1.558490289 | 0.1 | 0.037 | 1 | 0.844046361 | 0.538951762 |
| Slc26a6 | 0.399120535 | 0.251538767 | 0.034 | 0.018 | 1 | 0.320995017 | 2.017921908 | 0.1 | 0.037 | 1 | 0.399120535 | 0.538952233 |
| 4931408D14Rik | 0.467093859 | -0.485426827 | 0.025 | 0.041 | 1 | 0.320995017 | 2.017921908 | 0.1 | 0.037 | 1 | 0.467093859 | 0.538952233 |
| Mfsd3 | 0.707123932 | 0.389042291 | 0.076 | 0.065 | 1 | 0.320995017 | 2.017921908 | 0.1 | 0.037 | 1 | 0.707123932 | 0.538952233 |
| B430105G09Rik | 0.321248178 | 0.836501268 | 0.067 | 0.041 | 1 | 0.479305119 | 1.847996907 | 0.1 | 0.049 | 1 | 0.479305119 | 0.539295965 |
| Rcl1 | 0.66646909 | -0.243225925 | 0.286 | 0.312 | 1 | 0.321504664 | -0.823380346 | 0.2 | 0.33 | 1 | 0.66646909 | 0.539644079 |

| | | | | | | | | | | | | |
|---|---|---|---|---|---|---|---|---|---|---|---|---|
| Slc7a6 | 0.321793289 | -0.411426246 | 0.328 | 0.359 | 1 | 0.506007965 | 0.190102883 | 0.4 | 0.265 | 1 | 0.506007965 | 0.540035658 |
| Ptprf | 0.488014746 | -0.178337241 | 0.798 | 0.835 | 1 | 0.322042097 | -0.434662545 | 0.9 | 0.91 | 1 | 0.488014746 | 0.540373081 |
| Gpatch1 | 0.322277564 | -0.822461814 | 0.134 | 0.171 | 1 | 0.889571487 | 0.176619654 | 0.2 | 0.173 | 1 | 0.889571487 | 0.5406923 |
| Ccnh | 0.335601739 | -0.333423734 | 0.412 | 0.447 | 1 | 0.322367304 | -0.5519337 | 0.3 | 0.481 | 1 | 0.335601739 | 0.540813929 |
| Gm38115 | 0.443378777 | -0.572889668 | 0.059 | 0.082 | 1 | 0.322375242 | 2.079322453 | 0.2 | 0.108 | 1 | 0.443378777 | 0.540824688 |
| Gm47644 | 0.322548076 | 0.399095955 | 0.076 | 0.047 | 1 | 0.474948275 | 0.625604485 | 0 | 0.049 | 1 | 0.474948275 | 0.54105889 |
| Cebpzos | 0.824921917 | -0.312095224 | 0.37 | 0.335 | 1 | 0.322746382 | -0.165299916 | 0.1 | 0.259 | 1 | 0.824921917 | 0.541327537 |
| Fam204a | 0.557261607 | -0.24493499 | 0.706 | 0.635 | 1 | 0.323420067 | -0.246781318 | 0.4 | 0.593 | 1 | 0.557261607 | 0.542239594 |
| Enc1 | 0.567218524 | -0.392317423 | 0.076 | 0.094 | 1 | 0.323434771 | 1.695993813 | 0.1 | 0.037 | 1 | 0.567218524 | 0.542259491 |
| Rtl6 | 0.82670638 | -0.292781749 | 0.025 | 0.029 | 1 | 0.323434771 | 1.847996907 | 0.1 | 0.037 | 1 | 0.82670638 | 0.542259491 |
| Gpatch11 | 0.323677932 | -0.485426827 | 0.235 | 0.276 | 1 | 0.353555789 | -0.723545078 | 0.1 | 0.216 | 1 | 0.353555789 | 0.54258846 |
| Fndc10 | 0.323940516 | -0.748461233 | 0.034 | 0.059 | 1 | 0.398985336 | 2.017921908 | 0.1 | 0.043 | 1 | 0.398985336 | 0.542943575 |
| D6Ertd527e | 0.323940516 | -0.748461233 | 0.034 | 0.059 | 1 | 0.629538555 | 1.558490289 | 0.1 | 0.062 | 1 | 0.629538555 | 0.542943575 |
| Lipt2 | 0.323940516 | -0.748461233 | 0.034 | 0.059 | 1 | 0.669848329 | 1.695993813 | 0 | 0.019 | 1 | 0.669848329 | 0.542943575 |
| Fan1 | 0.681126405 | -0.436517227 | 0.109 | 0.124 | 1 | 0.324108708 | 1.374065718 | 0.2 | 0.105 | 1 | 0.681126405 | 0.543170962 |
| Primpol | 0.324161917 | -0.780882711 | 0.084 | 0.118 | 1 | 0.662167716 | 1.558490289 | 0.1 | 0.065 | 1 | 0.662167716 | 0.543242886 |
| Fam177a | 0.462720534 | 0.514573173 | 0.067 | 0.047 | 1 | 0.324792998 | -0.191531458 | 0 | 0.09 | 1 | 0.462720534 | 0.544095505 |
| Gm37768 | 0.328454311 | -1.655351829 | 0.025 | 0.047 | 1 | 0.324842077 | -0.191531458 | 0 | 0.09 | 1 | 0.328454311 | 0.544161779 |
| Tnfrsf18 | 0.398253428 | -0.707819249 | 0.042 | 0.065 | 1 | 0.324842077 | -0.267480311 | 0 | 0.09 | 1 | 0.398253428 | 0.544161779 |
| Gm16552 | 0.834564718 | 0.321928095 | 0.025 | 0.029 | 1 | 0.324842077 | -0.230005605 | 0 | 0.09 | 1 | 0.834564718 | 0.544161779 |
| D730003I15Rik | 0.981839274 | -0.222392421 | 0.042 | 0.041 | 1 | 0.324842732 | -0.111361109 | 0 | 0.09 | 1 | 0.981839274 | 0.544162663 |
| Pdia4 | 0.324890692 | -0.216457141 | 0.882 | 0.882 | 1 | 0.551217609 | -0.235619212 | 0.8 | 0.886 | 1 | 0.551217609 | 0.544227422 |
| Lrp2bp | 0.935195931 | -0.222392421 | 0.034 | 0.035 | 1 | 0.324963586 | -0.441509711 | 0 | 0.09 | 1 | 0.935195931 | 0.544325839 |
| Gm10575 | 0.324969356 | -0.807354922 | 0.034 | 0.059 | 1 | 0.532201836 | 1.31748219 | 0.1 | 0.052 | 1 | 0.532201836 | 0.544333629 |
| Dpyd | 0.715877701 | 1.192645078 | 0.017 | 0.012 | 1 | 0.325056924 | -1.152003093 | 0 | 0.09 | 1 | 0.715877701 | 0.544451844 |
| Gm43328 | 0.325184869 | -0.563429339 | 0.067 | 0.1 | 1 | 0.530179294 | 0.526068812 | 0.3 | 0.204 | 1 | 0.530179294 | 0.544624539 |
| Txnrd1 | 0.325218529 | 0.121982072 | 0.899 | 0.841 | 1 | 0.438059351 | -0.327261539 | 0.9 | 0.969 | 1 | 0.438059351 | 0.544669966 |
| Ccdc30 | 0.325267772 | -1.485426827 | 0.025 | 0.047 | 1 | 0.447621929 | 1.494359952 | 0.1 | 0.046 | 1 | 0.447621929 | 0.54473642 |
| Cyb5d2 | 0.325362796 | 0.124626655 | 0.193 | 0.147 | 1 | 0.783655686 | 0.210566986 | 0.1 | 0.127 | 1 | 0.783655686 | 0.544864643 |
| Zfp612 | 0.651414376 | -0.707819249 | 0.042 | 0.053 | 1 | 0.325884699 | 1.695993813 | 0.1 | 0.037 | 1 | 0.651414376 | 0.545568561 |
| Csmd1 | 0.511005543 | -0.485426827 | 0.008 | 0.018 | 1 | 0.32588574 | -0.191531458 | 0.1 | 0.037 | 1 | 0.511005543 | 0.545569964 |
| Dctn3 | 0.572826855 | 0.166118783 | 0.563 | 0.553 | 1 | 0.326080986 | -0.840059087 | 0.5 | 0.617 | 1 | 0.572826855 | 0.545833163 |
| Gprc5c | 0.32615754 | -0.471212968 | 0.336 | 0.376 | 1 | 0.7502827 | -0.158666824 | 0.4 | 0.423 | 1 | 0.7502827 | 0.545936339 |
| Fmnl2 | 0.976452621 | -0.604193788 | 0.807 | 0.776 | 1 | 0.326409581 | -1.147107619 | 0.8 | 0.83 | 1 | 0.976452621 | 0.546275947 |
| Meaf6 | 0.327039278 | -0.457412451 | 0.345 | 0.371 | 1 | 0.908319932 | 0.494359952 | 0.3 | 0.367 | 1 | 0.908319932 | 0.547123866 |
| Tirap | 0.460231274 | -0.516453723 | 0.176 | 0.206 | 1 | 0.327634038 | 1.054447784 | 0.3 | 0.182 | 1 | 0.460231274 | 0.547924013 |
| Gpatch2 | 0.327648703 | -0.587525015 | 0.445 | 0.453 | 1 | 0.531097816 | 0.403212064 | 0.5 | 0.386 | 1 | 0.531097816 | 0.547943733 |
| Ndfip1 | 0.327825645 | -0.100014497 | 1 | 0.994 | 1 | 0.608188031 | -0.137476979 | 0.9 | 1 | 1 | 0.608188031 | 0.548181637 |
| C130036L24Rik | 0.327829711 | 0.684498174 | 0.067 | 0.041 | 1 | 0.619953009 | 1.695993813 | 0 | 0.025 | 1 | 0.619953009 | 0.548187102 |
| H2ac18 | 0.327829711 | 0.684498174 | 0.067 | 0.041 | 1 | 0.557729726 | 1.31748219 | 0 | 0.034 | 1 | 0.557729726 | 0.548187102 |
| Pdlim5 | 0.327854849 | 0.198137262 | 0.95 | 0.859 | 1 | 0.381164653 | 1.376204629 | 0.8 | 0.88 | 1 | 0.381164653 | 0.548220896 |
| Cfap61 | 0.327877028 | -0.222392421 | 0.034 | 0.059 | 1 | 0.54814124 | 0.769994395 | 0.1 | 0.056 | 1 | 0.54814124 | 0.548250711 |
| Polr1e | 0.327885716 | -0.622930351 | 0.034 | 0.059 | 1 | 0.590778887 | 1.695993813 | 0.1 | 0.059 | 1 | 0.590778887 | 0.548262389 |
| Hdhd3 | 0.327885716 | -0.622930351 | 0.034 | 0.059 | 1 | 0.698534498 | 2.432959407 | 0 | 0.015 | 1 | 0.698534498 | 0.548262389 |
| Pcyox1l | 0.327885716 | -0.622930351 | 0.034 | 0.059 | 1 | 0.556267414 | 1.695993813 | 0.1 | 0.056 | 1 | 0.556267414 | 0.548262389 |
| Hinfp | 0.480296797 | -0.662304589 | 0.168 | 0.194 | 1 | 0.327991521 | -0.709998547 | 0.1 | 0.225 | 1 | 0.480296797 | 0.548404604 |
| Pros1 | 0.328196849 | -0.100997314 | 0.571 | 0.653 | 1 | 0.552477648 | -0.330806246 | 0.6 | 0.627 | 1 | 0.552477648 | 0.548680527 |
| Gm28535 | 0.954551343 | -0.222392421 | 0.017 | 0.018 | 1 | 0.328338207 | 1.695993813 | 0.1 | 0.037 | 1 | 0.954551343 | 0.548870436 |
| Rap1gap | 0.328416316 | -0.485426827 | 0.067 | 0.1 | 1 | 0.490416595 | 1.732519689 | 0.2 | 0.145 | 1 | 0.490416595 | 0.548975355 |
| Rsl1 | 0.32845267 | -1.292781749 | 0.025 | 0.047 | 1 | 0.662167716 | 1.558490289 | 0.1 | 0.065 | 1 | 0.662167716 | 0.549024183 |
| Gm29266 | 0.328480024 | -1.700439718 | 0.025 | 0.047 | 1 | 0.730750993 | 1.017921908 | 0 | 0.012 | 1 | 0.730750993 | 0.549060921 |
| Rasl10a | 0.32860203 | 0.666576266 | 0.067 | 0.041 | 1 | 0.460577668 | 0.625604485 | 0 | 0.052 | 1 | 0.460577668 | 0.549224766 |
| Ctdp1 | 0.32870094 | -0.42129649 | 0.378 | 0.412 | 1 | 0.915995242 | -0.174096864 | 0.6 | 0.522 | 1 | 0.915995242 | 0.549357572 |
| Pcbd2 | 0.532372826 | -0.160804623 | 0.395 | 0.429 | 1 | 0.328802853 | -0.69173634 | 0.2 | 0.33 | 1 | 0.532372826 | 0.549494389 |
| 9430038I01Rik | 0.328863946 | 0.170618772 | 0.168 | 0.124 | 1 | 0.947055009 | 0.769994395 | 0.1 | 0.114 | 1 | 0.947055009 | 0.549576396 |
| Tmub1 | 0.328947692 | -0.534336428 | 0.218 | 0.259 | 1 | 0.580366526 | 0.754887502 | 0.3 | 0.235 | 1 | 0.580366526 | 0.5496888 |
| St8sia4 | 0.329469853 | -1.807354922 | 0.008 | 0.024 | 1 | 0.643849121 | 1.111031312 | 0 | 0.022 | 1 | 0.643849121 | 0.550389322 |
| Gm38825 | 0.329469853 | -1.655351829 | 0.008 | 0.024 | 1 | 0.643850855 | 2.017921908 | 0 | 0.022 | 1 | 0.643850855 | 0.550389322 |
| Mc5r | 0.329694367 | -0.600904045 | 0.034 | 0.059 | 1 | 0.343827383 | -0.536666944 | 0 | 0.083 | 1 | 0.343827383 | 0.550690358 |
| Wdr12 | 0.329739753 | -0.545298283 | 0.353 | 0.382 | 1 | 0.902240388 | 0.299103661 | 0.4 | 0.401 | 1 | 0.902240388 | 0.550751201 |
| 2900089D17Rik | 0.330061685 | -0.388129626 | 0.143 | 0.182 | 1 | 0.907438215 | 0.521496082 | 0.2 | 0.225 | 1 | 0.907438215 | 0.551182654 |
| Tmem120a | 0.330611944 | 0.316633795 | 0.378 | 0.324 | 1 | 0.852653145 | 0.159940913 | 0.3 | 0.253 | 1 | 0.852653145 | 0.55191963 |
| Stard3 | 0.825622469 | 0.175771259 | 0.218 | 0.212 | 1 | 0.330757907 | 0.847996907 | 0.3 | 0.17 | 1 | 0.825622469 | 0.552115021 |
| Slc1a3 | 0.616062867 | -2.250961574 | 0.025 | 0.035 | 1 | 0.330809864 | 1.31748219 | 0.1 | 0.037 | 1 | 0.616062867 | 0.552184561 |
| Tmem37 | 0.667579237 | 0.427110332 | 0.084 | 0.071 | 1 | 0.330969856 | 2.515421567 | 0.1 | 0.04 | 1 | 0.667579237 | 0.552398666 |
| Txndc8 | 0.331040183 | -1.655351829 | 0.008 | 0.024 | 1 | 0.669852582 | 0.769994395 | 0 | 0.019 | 1 | 0.669852582 | 0.552492764 |
| Frem1 | 0.331040183 | -2.070389328 | 0.008 | 0.024 | 1 | 0.669843604 | 1.695993813 | 0 | 0.019 | 1 | 0.669843604 | 0.552492764 |
| Lgals7 | 0.331040183 | -1.807354922 | 0.008 | 0.024 | 1 | 0.577171068 | 0.930459067 | 0 | 0.031 | 1 | 0.577171068 | 0.552492764 |
| Gm26632 | 0.331040183 | -1.292781749 | 0.008 | 0.024 | 1 | 0.557772534 | 0.558490289 | 0 | 0.034 | 1 | 0.557772534 | 0.552492764 |
| Gm26839 | 0.331040183 | -1.292781749 | 0.008 | 0.024 | 1 | 0.730749233 | 2.017921908 | 0 | 0.012 | 1 | 0.730749233 | 0.552492764 |
| Dok2 | 0.331040183 | -1.292781749 | 0.008 | 0.024 | 1 | 0.698534498 | 2.432959407 | 0 | 0.015 | 1 | 0.698534498 | 0.552492764 |
| A430093F15Rik | 0.331040183 | -1.485426827 | 0.008 | 0.024 | 1 | 0.643835681 | 1.847996907 | 0 | 0.022 | 1 | 0.643835681 | 0.552492764 |
| Tenm1 | 0.331040183 | -2.185866545 | 0.008 | 0.024 | 1 | 0.557771538 | -0.68251781 | 0 | 0.034 | 1 | 0.557771538 | 0.552492764 |
| Nkap | 0.470371957 | -0.485426827 | 0.387 | 0.388 | 1 | 0.331465529 | 0.653349476 | 0.5 | 0.377 | 1 | 0.470371957 | 0.553061661 |
| Ptger4 | 0.331652362 | -1.185866545 | 0.025 | 0.047 | 1 | 0.489976754 | 0.111031312 | 0 | 0.046 | 1 | 0.489976754 | 0.553311434 |
| 4931406C07Rik | 0.408226553 | -0.380457268 | 0.345 | 0.371 | 1 | 0.332484268 | -0.663902132 | 0.2 | 0.33 | 1 | 0.408226553 | 0.554422748 |
| G730003C15Rik | 0.33261065 | -1.070389328 | 0.008 | 0.024 | 1 | 0.73075158 | 2.695993813 | 0 | 0.012 | 1 | 0.73075158 | 0.554591456 |
| Gm20753 | 0.33261065 | -1.070389328 | 0.008 | 0.024 | 1 | 0.698534498 | 2.432959407 | 0 | 0.015 | 1 | 0.698534498 | 0.554591456 |
| B3galt2 | 0.33261065 | -1.292781749 | 0.008 | 0.024 | 1 | 0.539455361 | 0.847996907 | 0 | 0.037 | 1 | 0.539455361 | 0.554591456 |
| Ppp1r16b | 0.33261065 | -1.070389328 | 0.008 | 0.024 | 1 | 0.643847387 | 1.210566986 | 0 | 0.022 | 1 | 0.643847387 | 0.554591456 |
| Rpusd3 | 0.33261065 | -1.070389328 | 0.008 | 0.024 | 1 | 0.619953009 | 1.695993813 | 0 | 0.025 | 1 | 0.619953009 | 0.554591456 |
| Gm44448 | 0.33261065 | -1.070389328 | 0.008 | 0.024 | 1 | 0.730749233 | 2.432959407 | 0 | 0.012 | 1 | 0.730749233 | 0.554591456 |
| Smco2 | 0.33261065 | -1.070389328 | 0.008 | 0.024 | 1 | 0.619971082 | 1.111031312 | 0 | 0.025 | 1 | 0.619971082 | 0.554591456 |
| Cd79a | 0.33261065 | -3.614709844 | 0.008 | 0.024 | 1 | 0.781750472 | 0.847996907 | 0.1 | 0.074 | 1 | 0.781750472 | 0.554591456 |
| 1110035H17Rik | 0.33261065 | -1.070389328 | 0.008 | 0.024 | 1 | 0.73075158 | 2.695993813 | 0 | 0.012 | 1 | 0.73075158 | 0.554591456 |
| Svip | 0.33261065 | -1.070389328 | 0.008 | 0.024 | 1 | 0.669851164 | 1.695993813 | 0 | 0.019 | 1 | 0.669851164 | 0.554591456 |
| St8sia2 | 0.33261065 | -1.485426827 | 0.008 | 0.024 | 1 | 0.597845674 | 1.695993813 | 0 | 0.028 | 1 | 0.597845674 | 0.554591456 |
| Tmc7 | 0.33261065 | -1.070389328 | 0.008 | 0.024 | 1 | 0.698534498 | 2.432959407 | 0 | 0.015 | 1 | 0.698534498 | 0.554591456 |
| Vwa3a | 0.33261065 | -1.292781749 | 0.008 | 0.024 | 1 | 0.59781419 | 1.558490289 | 0 | 0.028 | 1 | 0.59781419 | 0.554591456 |
| Gm37612 | 0.33261065 | -1.070389328 | 0.008 | 0.024 | 1 | 0.730749233 | 2.432959407 | 0 | 0.012 | 1 | 0.730749233 | 0.554591456 |
| Arhgap45 | 0.33261065 | -1.070389328 | 0.008 | 0.024 | 1 | 0.59781419 | 1.558490289 | 0 | 0.028 | 1 | 0.59781419 | 0.554591456 |
| Gm47093 | 0.33261065 | -1.070389328 | 0.008 | 0.024 | 1 | 0.577155236 | 1.111031312 | 0 | 0.031 | 1 | 0.577155236 | 0.554591456 |
| Gm57374 | 0.33261065 | -1.485426827 | 0.008 | 0.024 | 1 | 0.73075158 | 2.695993813 | 0 | 0.012 | 1 | 0.73075158 | 0.554591456 |
| Arhgap27os2 | 0.33261065 | -1.070389328 | 0.008 | 0.024 | 1 | 0.643850855 | 2.017921908 | 0 | 0.022 | 1 | 0.643850855 | 0.554591456 |
| Gm47484 | 0.33261065 | -1.292781749 | 0.008 | 0.024 | 1 | 0.73075158 | 2.695993813 | 0 | 0.012 | 1 | 0.73075158 | 0.554591456 |
| Zfp457 | 0.33261065 | -1.070389328 | 0.008 | 0.024 | 1 | 0.730750993 | 2.017921908 | 0 | 0.012 | 1 | 0.730750993 | 0.554591456 |
| Zfp85 | 0.33261065 | -1.070389328 | 0.008 | 0.024 | 1 | 0.619953009 | 1.695993813 | 0 | 0.025 | 1 | 0.619953009 | 0.554591456 |
| 1700023H06Rik | 0.33261065 | -1.070389328 | 0.008 | 0.024 | 1 | 0.669843604 | 2.017921908 | 0 | 0.019 | 1 | 0.669843604 | 0.554591456 |

| | | | | | | | | | | | | |
|---|---|---|---|---|---|---|---|---|---|---|---|---|
| Kcnrg | 0.33261065 | -1.070389328 | 0.008 | 0.024 | 1 | 0.807629212 | 0.973527789 | 0.1 | 0.077 | 1 | 0.807629212 | 0.554591456 |
| Gm9920 | 0.33261065 | -1.070389328 | 0.008 | 0.024 | 1 | 0.643835681 | 1.695993813 | 0 | 0.022 | 1 | 0.643835681 | 0.554591456 |
| Gm53058 | 0.33261065 | -1.485426827 | 0.008 | 0.024 | 1 | 0.421003536 | -0.152003093 | 0 | 0.062 | 1 | 0.421003536 | 0.554591456 |
| 2600002D14Rik | 0.33261065 | -1.070389328 | 0.008 | 0.024 | 1 | 0.698532411 | 2.017921908 | 0 | 0.015 | 1 | 0.698532411 | 0.554591456 |
| D430001F17Rik | 0.33261065 | -1.485426827 | 0.008 | 0.024 | 1 | 0.730750993 | 1.558490289 | 0 | 0.012 | 1 | 0.730750993 | 0.554591456 |
| Prdm9 | 0.33261065 | -1.292781749 | 0.008 | 0.024 | 1 | 0.597824685 | 1.31748219 | 0 | 0.028 | 1 | 0.597824685 | 0.554591456 |
| Tff2 | 0.33261065 | -1.292781749 | 0.008 | 0.024 | 1 | 0.52208609 | 0.930459067 | 0 | 0.04 | 1 | 0.52208609 | 0.554591456 |
| C2 | 0.33261065 | -1.070389328 | 0.008 | 0.024 | 1 | 0.619961444 | 1.432959407 | 0 | 0.025 | 1 | 0.619961444 | 0.554591456 |
| Ablim3 | 0.33261065 | -1.070389328 | 0.008 | 0.024 | 1 | 0.619967468 | 1.432959407 | 0 | 0.025 | 1 | 0.619967468 | 0.554591456 |
| Doc2g | 0.33261065 | -1.292781749 | 0.008 | 0.024 | 1 | 0.669843604 | 1.695993813 | 0 | 0.019 | 1 | 0.669843604 | 0.554591456 |
| Gm39526 | 0.33261065 | -1.070389328 | 0.008 | 0.024 | 1 | 0.669843604 | 2.017921908 | 0 | 0.019 | 1 | 0.669843604 | 0.554591456 |
| 2700054A10Rik | 0.332752594 | -1.070389328 | 0.025 | 0.047 | 1 | 0.698533455 | 1.695993813 | 0 | 0.015 | 1 | 0.698533455 | 0.554780899 |
| Nfib | 0.332799754 | -0.342062652 | 0.798 | 0.688 | 1 | 0.990634489 | -0.173000175 | 0.9 | 0.873 | 1 | 0.990634489 | 0.554843831 |
| Klhdc1 | 0.332808966 | 0.357031896 | 0.16 | 0.118 | 1 | 0.546324534 | -0.391469028 | 0.1 | 0.167 | 1 | 0.546324534 | 0.554856124 |
| Piga | 0.849859613 | -0.359895945 | 0.084 | 0.088 | 1 | 0.333163571 | 1.280956314 | 0.2 | 0.102 | 1 | 0.849859613 | 0.555329176 |
| Mrpl21 | 0.33341356 | -0.327164743 | 0.479 | 0.506 | 1 | 0.79879577 | -0.339630097 | 0.4 | 0.386 | 1 | 0.79879577 | 0.555662518 |
| Lig4 | 0.333495621 | -0.485426827 | 0.034 | 0.059 | 1 | 0.470043448 | 2.558490289 | 0.1 | 0.052 | 1 | 0.470043448 | 0.555771913 |
| Stx4a | 0.601938652 | -0.217493622 | 0.563 | 0.576 | 1 | 0.333548443 | -0.668578619 | 0.6 | 0.599 | 1 | 0.601938652 | 0.555842323 |
| Sult1c1 | 0.342795612 | -1.070389328 | 0.017 | 0.035 | 1 | 0.334014848 | -0.111361109 | 0 | 0.086 | 1 | 0.342795612 | 0.556463777 |
| Dcaf12l1 | 0.77228175 | -0.163498732 | 0.067 | 0.076 | 1 | 0.334165224 | -0.111361109 | 0 | 0.086 | 1 | 0.77228175 | 0.556664051 |
| Dtwd1 | 0.645173409 | -0.748461233 | 0.076 | 0.088 | 1 | 0.334166236 | -0.191531458 | 0 | 0.086 | 1 | 0.645173409 | 0.556665399 |
| Nat14 | 0.665284816 | -0.359895945 | 0.042 | 0.053 | 1 | 0.334166236 | -0.230005605 | 0 | 0.086 | 1 | 0.665284816 | 0.556665399 |
| Prss12 | 0.334203633 | -1.807354922 | 0.008 | 0.024 | 1 | 0.522132053 | 0.374065718 | 0 | 0.04 | 1 | 0.522132053 | 0.556715198 |
| H2ac4 | 0.334203633 | -1.292781749 | 0.008 | 0.024 | 1 | 0.59781419 | 1.558490289 | 0 | 0.028 | 1 | 0.59781419 | 0.556715198 |
| Cpne5 | 0.334203633 | -1.070389328 | 0.008 | 0.024 | 1 | 0.643842184 | 1.558490289 | 0 | 0.022 | 1 | 0.643842184 | 0.556715198 |
| Tox2 | 0.334204811 | -0.807354922 | 0.008 | 0.024 | 1 | 0.643847387 | 0.930459067 | 0 | 0.022 | 1 | 0.643847387 | 0.556716766 |
| Gm11457 | 0.334204811 | -0.807354922 | 0.008 | 0.024 | 1 | 0.52208609 | 1.017921908 | 0 | 0.04 | 1 | 0.52208609 | 0.556716766 |
| Gm42658 | 0.334204811 | -0.807354922 | 0.008 | 0.024 | 1 | 0.460551517 | 0.625604485 | 0 | 0.052 | 1 | 0.460551517 | 0.556716766 |
| Gstm6 | 0.334204811 | -0.807354922 | 0.008 | 0.024 | 1 | 0.619961444 | 1.31748219 | 0 | 0.025 | 1 | 0.619961444 | 0.556716766 |
| Dnali1 | 0.334204811 | -0.807354922 | 0.008 | 0.024 | 1 | 0.730749233 | 2.432959407 | 0 | 0.012 | 1 | 0.730749233 | 0.556716766 |
| Gm43727 | 0.334204811 | -0.807354922 | 0.008 | 0.024 | 1 | 0.522102507 | 0.930459067 | 0 | 0.04 | 1 | 0.522102507 | 0.556716766 |
| Zfp605 | 0.334204811 | -0.807354922 | 0.008 | 0.024 | 1 | 0.698534498 | 2.432959407 | 0 | 0.015 | 1 | 0.698534498 | 0.556716766 |
| Gm43915 | 0.334204811 | -0.807354922 | 0.008 | 0.024 | 1 | 0.669853054 | 2.210566986 | 0 | 0.019 | 1 | 0.669853054 | 0.556716766 |
| C5ar2 | 0.334204811 | -0.807354922 | 0.008 | 0.024 | 1 | 0.460617262 | 0.432959407 | 0 | 0.052 | 1 | 0.460617262 | 0.556716766 |
| Zbtb32 | 0.334204811 | -0.807354922 | 0.008 | 0.024 | 1 | 0.730750993 | 1.847996907 | 0 | 0.012 | 1 | 0.730750993 | 0.556716766 |
| Gm12758 | 0.334204811 | -0.807354922 | 0.008 | 0.024 | 1 | 0.669843604 | 1.847996907 | 0 | 0.019 | 1 | 0.669843604 | 0.556716766 |
| Faap24 | 0.334204811 | -0.807354922 | 0.008 | 0.024 | 1 | 0.577142568 | 1.432959407 | 0 | 0.031 | 1 | 0.577142568 | 0.556716766 |
| Neu3 | 0.334204811 | -0.807354922 | 0.008 | 0.024 | 1 | 0.44222087 | 1.847996907 | 0.1 | 0.046 | 1 | 0.44222087 | 0.556716766 |
| Amt | 0.334204811 | -0.807354922 | 0.008 | 0.024 | 1 | 0.6199755 | 1.847996907 | 0 | 0.025 | 1 | 0.6199755 | 0.556716766 |
| Entpd3 | 0.334204811 | -0.807354922 | 0.008 | 0.024 | 1 | 0.460617262 | 0.432959407 | 0 | 0.052 | 1 | 0.460617262 | 0.556716766 |
| Lgr5 | 0.334204811 | -0.807354922 | 0.008 | 0.024 | 1 | 0.557784146 | -1.750262417 | 0 | 0.034 | 1 | 0.557784146 | 0.556716766 |
| Gm57367 | 0.334204811 | -0.807354922 | 0.008 | 0.024 | 1 | 0.730749233 | 2.432959407 | 0 | 0.012 | 1 | 0.730749233 | 0.556716766 |
| Shmt1 | 0.334204811 | -0.807354922 | 0.008 | 0.024 | 1 | 0.401608067 | 1.769994395 | 0.1 | 0.043 | 1 | 0.401608067 | 0.556716766 |
| Scarf1 | 0.334204811 | -0.807354922 | 0.008 | 0.024 | 1 | 0.421032074 | -0.152003093 | 0 | 0.062 | 1 | 0.421032074 | 0.556716766 |
| Gm56531 | 0.334204811 | -0.807354922 | 0.008 | 0.024 | 1 | 0.577172475 | 0.847996907 | 0 | 0.031 | 1 | 0.577172475 | 0.556716766 |
| Gm56516 | 0.334204811 | -0.807354922 | 0.008 | 0.024 | 1 | 0.698529281 | 2.210566986 | 0 | 0.015 | 1 | 0.698529281 | 0.556716766 |
| Nhlrc1 | 0.334204811 | -0.807354922 | 0.008 | 0.024 | 1 | 0.73075158 | 2.695993813 | 0 | 0.012 | 1 | 0.73075158 | 0.556716766 |
| Gm36445 | 0.334204811 | -0.807354922 | 0.008 | 0.024 | 1 | 0.73075158 | 2.695993813 | 0 | 0.012 | 1 | 0.73075158 | 0.556716766 |
| Gm41291 | 0.334204811 | -0.807354922 | 0.008 | 0.024 | 1 | 0.643850855 | 2.017921908 | 0 | 0.022 | 1 | 0.643850855 | 0.556716766 |
| Gpd1 | 0.334204811 | -0.807354922 | 0.008 | 0.024 | 1 | 0.59781419 | 1.31748219 | 0 | 0.028 | 1 | 0.59781419 | 0.556716766 |
| Gm17058 | 0.334204811 | -0.807354922 | 0.008 | 0.024 | 1 | 0.669848329 | 1.847996907 | 0 | 0.019 | 1 | 0.669848329 | 0.556716766 |
| A230028O05Rik | 0.334204811 | -0.807354922 | 0.008 | 0.024 | 1 | 0.577184788 | 1.558490289 | 0 | 0.031 | 1 | 0.577184788 | 0.556716766 |
| Gp5 | 0.334204811 | -0.807354922 | 0.008 | 0.024 | 1 | 0.73075158 | 2.695993813 | 0 | 0.012 | 1 | 0.73075158 | 0.556716766 |
| Gm16278 | 0.334204811 | -0.807354922 | 0.008 | 0.024 | 1 | 0.619967066 | 0.930459067 | 0 | 0.025 | 1 | 0.619967066 | 0.556716766 |
| Ly6g5b | 0.334204811 | -0.807354922 | 0.008 | 0.024 | 1 | 0.597824685 | 1.31748219 | 0 | 0.028 | 1 | 0.597824685 | 0.556716766 |
| Abcc10 | 0.334204811 | -0.807354922 | 0.008 | 0.024 | 1 | 0.489877116 | 0.930459067 | 0 | 0.046 | 1 | 0.489877116 | 0.556716766 |
| Gm48696 | 0.334204811 | -0.807354922 | 0.008 | 0.024 | 1 | 0.577184788 | 1.558490289 | 0 | 0.031 | 1 | 0.577184788 | 0.556716766 |
| Pnck | 0.334204811 | -0.807354922 | 0.008 | 0.024 | 1 | 0.698534498 | 2.432959407 | 0 | 0.015 | 1 | 0.698534498 | 0.556716766 |
| 1700096K18Rik | 0.815028039 | -0.485426827 | 0.059 | 0.065 | 1 | 0.334207905 | -0.230005605 | 0 | 0.086 | 1 | 0.815028039 | 0.556720887 |
| 1810024B03Rik | 0.381045639 | -0.970853654 | 0.067 | 0.094 | 1 | 0.3342467 | -0.408342847 | 0 | 0.086 | 1 | 0.381045639 | 0.556772544 |
| Ercc6l2 | 0.519172186 | -0.346023771 | 0.387 | 0.418 | 1 | 0.334250463 | -1.062451508 | 0.2 | 0.315 | 1 | 0.519172186 | 0.556777554 |
| Akap17b | 0.962042535 | 0.192645078 | 0.042 | 0.041 | 1 | 0.334309939 | -0.374395515 | 0 | 0.086 | 1 | 0.962042535 | 0.556856742 |
| Mcee | 0.950351423 | -0.175086707 | 0.412 | 0.394 | 1 | 0.334318025 | -0.625934282 | 0.2 | 0.33 | 1 | 0.950351423 | 0.556867508 |
| Usp18 | 0.448121284 | 0.893084796 | 0.101 | 0.076 | 1 | 0.334404174 | -0.959358016 | 0 | 0.086 | 1 | 0.448121284 | 0.556982197 |
| Faap20 | 0.636661913 | -0.329307625 | 0.277 | 0.288 | 1 | 0.334621347 | 1.263034406 | 0.3 | 0.188 | 1 | 0.636661913 | 0.557271248 |
| Irf8 | 0.334756206 | -0.106915204 | 0.076 | 0.047 | 1 | 0.539431168 | 0.847996907 | 0 | 0.037 | 1 | 0.539431168 | 0.557450694 |
| Rslcan18 | 0.334866848 | -0.944858446 | 0.025 | 0.047 | 1 | 0.489923006 | 0.769994395 | 0 | 0.046 | 1 | 0.489923006 | 0.55759789 |
| Gm49741 | 0.334866848 | -0.944858446 | 0.025 | 0.047 | 1 | 0.643846086 | 1.432959407 | 0 | 0.022 | 1 | 0.643846086 | 0.55759789 |
| Gm49915 | 0.334866848 | -0.944858446 | 0.025 | 0.047 | 1 | 0.557765898 | 1.017921908 | 0 | 0.034 | 1 | 0.557765898 | 0.55759789 |
| Fbxo5 | 0.334911132 | -1.687060688 | 0.025 | 0.047 | 1 | 0.597824685 | 1.432959407 | 0 | 0.028 | 1 | 0.597824685 | 0.557656798 |
| Hsd17b4 | 0.335002512 | -0.555816155 | 0.277 | 0.318 | 1 | 0.709780953 | 0.397335498 | 0.5 | 0.454 | 1 | 0.709780953 | 0.55777834 |
| Mrpl36 | 0.335192357 | -0.284514133 | 0.345 | 0.388 | 1 | 0.598414216 | 0.354956895 | 0.4 | 0.309 | 1 | 0.598414216 | 0.558030798 |
| Raver2 | 0.335500324 | -0.392317423 | 0.034 | 0.059 | 1 | 0.610403029 | 0.732519689 | 0.1 | 0.059 | 1 | 0.610403029 | 0.558440181 |
| Kcnq1ot1 | 0.335505861 | -0.410214783 | 0.924 | 0.935 | 1 | 0.43349054 | -0.376757105 | 1 | 0.991 | 1 | 0.43349054 | 0.558447539 |
| Gas2l3 | 0.335588033 | -0.292781749 | 0.076 | 0.047 | 1 | 0.690389141 | 0.930459067 | 0.1 | 0.065 | 1 | 0.690389141 | 0.558556738 |
| Slfn2 | 0.335789682 | -1.600904045 | 0.008 | 0.024 | 1 | 0.643846086 | 1.017921908 | 0 | 0.022 | 1 | 0.643846086 | 0.558824653 |
| Ninj2 | 0.335795703 | -0.300538193 | 0.748 | 0.506 | 1 | 0.851213923 | -0.235689244 | 0.6 | 0.512 | 1 | 0.851213923 | 0.558832651 |
| Epm2a | 0.336298646 | -0.341036918 | 0.092 | 0.129 | 1 | 0.640737411 | 0.290001453 | 0.2 | 0.148 | 1 | 0.640737411 | 0.559500512 |
| Gm35769 | 0.747707899 | -0.347923303 | 0.084 | 0.094 | 1 | 0.336366819 | 1.31748219 | 0.2 | 0.102 | 1 | 0.747707899 | 0.559591001 |
| Cenpe | 0.336457245 | -3.266786541 | 0.017 | 0.035 | 1 | 0.669848329 | 1.847996907 | 0 | 0.019 | 1 | 0.669848329 | 0.559711012 |
| Ercc6 | 0.405370311 | -0.644625422 | 0.193 | 0.224 | 1 | 0.33672005 | 0.748461233 | 0.4 | 0.256 | 1 | 0.405370311 | 0.560059707 |
| Tpgs1 | 0.336855036 | -0.485426827 | 0.235 | 0.276 | 1 | 0.816965012 | 0.159940913 | 0.2 | 0.225 | 1 | 0.816965012 | 0.560238756 |
| Jkamp | 0.33713602 | 0.1602236 | 0.739 | 0.647 | 1 | 0.645505381 | -0.103847055 | 0.5 | 0.611 | 1 | 0.645505381 | 0.560611344 |
| Fcer1g | 0.337379399 | -2.14839184 | 0.008 | 0.024 | 1 | 0.364083721 | -1.230005605 | 0 | 0.077 | 1 | 0.364083721 | 0.560933939 |
| Mapk4 | 0.337379399 | -1.222392421 | 0.008 | 0.024 | 1 | 0.522123697 | 0.695993813 | 0 | 0.04 | 1 | 0.522123697 | 0.560933939 |
| Gm42997 | 0.33738293 | -0.707819249 | 0.008 | 0.024 | 1 | 0.619971082 | 1.111031312 | 0 | 0.025 | 1 | 0.619971082 | 0.560938619 |
| As3mt | 0.33752951 | -0.485426827 | 0.126 | 0.165 | 1 | 0.801502184 | -0.152003093 | 0.2 | 0.219 | 1 | 0.801502184 | 0.56113285 |
| Calhm6 | 0.337717757 | -1.359895945 | 0.017 | 0.035 | 1 | 0.619953009 | 1.558490289 | 0 | 0.025 | 1 | 0.619953009 | 0.561382231 |
| Alg10b | 0.338065559 | -0.430979043 | 0.168 | 0.212 | 1 | 0.358042819 | -0.581990934 | 0.1 | 0.216 | 1 | 0.358042819 | 0.561842795 |
| A430105J06Rik | 0.338096132 | -0.807354922 | 0.025 | 0.047 | 1 | 0.404236993 | 1.847996907 | 0.1 | 0.043 | 1 | 0.404236993 | 0.561883269 |
| Ap3m2 | 0.338096132 | -0.807354922 | 0.025 | 0.047 | 1 | 0.619969878 | 1.111031312 | 0 | 0.025 | 1 | 0.619969878 | 0.561883269 |
| Ybey | 0.338097529 | 0.974004791 | 0.059 | 0.035 | 1 | 0.515666611 | 1.847996907 | 0.1 | 0.052 | 1 | 0.515666611 | 0.561885119 |
| Gm29585 | 0.338155151 | -2.117695043 | 0.025 | 0.047 | 1 | 0.557744661 | 1.210566986 | 0 | 0.034 | 1 | 0.557744661 | 0.561961396 |
| Zfp990 | 0.338530395 | -0.87774425 | 0.059 | 0.088 | 1 | 0.863418778 | 0.263034406 | 0.1 | 0.114 | 1 | 0.863418778 | 0.562457962 |
| Cyc1 | 0.471884218 | -0.121373257 | 0.655 | 0.706 | 1 | 0.338615302 | -0.505640048 | 0.6 | 0.664 | 1 | 0.471884218 | 0.562570281 |
| Atosb | 0.885627576 | -0.23926624 | 0.311 | 0.3 | 1 | 0.33873066 | -0.726239188 | 0.4 | 0.472 | 1 | 0.885627576 | 0.56272286 |
| Zfp646 | 0.342499327 | -0.576574715 | 0.16 | 0.2 | 1 | 0.338936354 | -0.780819884 | 0.2 | 0.324 | 1 | 0.342499327 | 0.562994856 |
| Gm20337 | 0.338976313 | -0.485426827 | 0.008 | 0.024 | 1 | 0.698534498 | 2.432959407 | 0 | 0.015 | 1 | 0.698534498 | 0.563047686 |

| | | | | | | | | | | | | |
|---|---|---|---|---|---|---|---|---|---|---|---|---|
| F2rl1 | 0.338976313 | -0.485426827 | 0.008 | 0.024 | 1 | 0.446819579 | 0.558490289 | 0 | 0.056 | 1 | 0.446819579 | 0.563047686 |
| Dpys | 0.338976313 | -0.485426827 | 0.008 | 0.024 | 1 | 0.698532411 | 2.017921908 | 0 | 0.015 | 1 | 0.698532411 | 0.563047686 |
| Gm15902 | 0.339103263 | 0.929610672 | 0.059 | 0.035 | 1 | 0.730749233 | 2.432959407 | 0 | 0.012 | 1 | 0.730749233 | 0.563215503 |
| Gm17455 | 0.339242628 | -0.970853654 | 0.025 | 0.047 | 1 | 0.577155236 | 1.31748219 | 0 | 0.031 | 1 | 0.577155236 | 0.563399696 |
| Decr1 | 0.339417773 | -0.404808367 | 0.336 | 0.365 | 1 | 0.95174376 | 0.328622747 | 0.3 | 0.29 | 1 | 0.95174376 | 0.563631121 |
| Trmt10a | 0.340049389 | -0.578536232 | 0.227 | 0.265 | 1 | 0.653387069 | 0.515421567 | 0.1 | 0.167 | 1 | 0.653387069 | 0.564465191 |
| Gm17749 | 0.340052008 | -0.163498732 | 0.042 | 0.071 | 1 | 0.460635686 | 0.111031312 | 0 | 0.052 | 1 | 0.460635686 | 0.564468648 |
| Gm57095 | 0.34025168 | -1.359895945 | 0.017 | 0.035 | 1 | 0.643835681 | 1.847996907 | 0 | 0.022 | 1 | 0.643835681 | 0.564732155 |
| Adgb | 0.34025168 | -1.359895945 | 0.017 | 0.035 | 1 | 0.539465415 | 0.847996907 | 0 | 0.037 | 1 | 0.539465415 | 0.564732155 |
| 5430402O13Rik | 0.340571011 | -0.222392421 | 0.008 | 0.024 | 1 | 0.643835681 | 1.847996907 | 0 | 0.022 | 1 | 0.643835681 | 0.565153409 |
| Gm48855 | 0.340571011 | -0.222392421 | 0.008 | 0.024 | 1 | 0.597824685 | 1.432959407 | 0 | 0.028 | 1 | 0.597824685 | 0.565153409 |
| Add2 | 0.34057807 | 0.929610672 | 0.008 | 0.024 | 1 | 0.730749233 | 2.432959407 | 0 | 0.012 | 1 | 0.730749233 | 0.565162719 |
| Gpr15lg | 0.34057807 | 0.251538767 | 0.008 | 0.024 | 1 | 0.698529281 | 2.017921908 | 0 | 0.015 | 1 | 0.698529281 | 0.565162719 |
| Ppfia2 | 0.4698534 | -1.163498732 | 0.025 | 0.041 | 1 | 0.340651062 | 1.754887502 | 0.1 | 0.04 | 1 | 0.4698534 | 0.565258978 |
| Etv1 | 0.485396436 | -0.383547213 | 0.168 | 0.129 | 1 | 0.340669857 | -0.526398608 | 0.1 | 0.241 | 1 | 0.485396436 | 0.565283762 |
| Krcc1 | 0.341044541 | -0.172836597 | 0.63 | 0.688 | 1 | 0.817820561 | -0.168605061 | 0.7 | 0.642 | 1 | 0.817820561 | 0.565777704 |
| Fabp5 | 1 | -0.370867063 | 0.403 | 0.359 | 1 | 0.341442752 | -0.936274402 | 0.2 | 0.34 | 1 | 1 | 0.56630235 |
| Matcap1 | 0.341460366 | -0.655351829 | 0.025 | 0.047 | 1 | 0.522151747 | 1.210566986 | 0 | 0.04 | 1 | 0.522151747 | 0.566325551 |
| A730063M14Rik | 0.341460366 | -0.655351829 | 0.025 | 0.047 | 1 | 0.489939838 | 0.625604485 | 0 | 0.046 | 1 | 0.489939838 | 0.566325551 |
| Hdac10 | 0.341460366 | -0.655351829 | 0.025 | 0.047 | 1 | 0.460577419 | 0.494359952 | 0 | 0.052 | 1 | 0.460577419 | 0.566325551 |
| Mlst8 | 0.341460366 | -0.655351829 | 0.025 | 0.047 | 1 | 0.474904369 | 0.769994395 | 0 | 0.049 | 1 | 0.474904369 | 0.566325551 |
| ENSMUSG00000095742 | 0.341460366 | -0.655351829 | 0.025 | 0.047 | 1 | 0.698534498 | 2.432959407 | 0 | 0.015 | 1 | 0.698534498 | 0.566325551 |
| Dhx36 | 0.341646238 | -0.276331228 | 0.824 | 0.724 | 1 | 0.411107573 | -0.45549394 | 0.8 | 0.765 | 1 | 0.411107573 | 0.566570324 |
| Zfp52 | 0.677126859 | 0.462105753 | 0.269 | 0.259 | 1 | 0.341798979 | 0.753218682 | 0.5 | 0.463 | 1 | 0.677126859 | 0.566771416 |
| Rbp4 | 0.342128371 | -0.381269596 | 0.395 | 0.429 | 1 | 0.470885793 | -0.304006187 | 0.5 | 0.593 | 1 | 0.470885793 | 0.56720492 |
| Cep290 | 0.342342595 | -1.015046294 | 0.328 | 0.335 | 1 | 0.795523314 | 0.140177658 | 0.3 | 0.247 | 1 | 0.795523314 | 0.567486738 |
| Fh1 | 0.342399334 | -0.396890153 | 0.303 | 0.341 | 1 | 0.948643498 | 0.295455884 | 0.3 | 0.296 | 1 | 0.948643498 | 0.567561364 |
| Alg3 | 0.342544168 | -0.354182294 | 0.168 | 0.212 | 1 | 0.7481915 | 0.995554095 | 0.2 | 0.173 | 1 | 0.7481915 | 0.567751829 |
| Fstl5 | 0.342795612 | -1.070389328 | 0.017 | 0.035 | 1 | 0.374633209 | -0.789433014 | 0 | 0.074 | 1 | 0.374633209 | 0.568082393 |
| Tbx15 | 0.342795612 | -1.070389328 | 0.017 | 0.035 | 1 | 0.7522162 | 0.591657153 | 0.1 | 0.071 | 1 | 0.7522162 | 0.568082393 |
| Gm12655 | 0.342795612 | -1.070389328 | 0.017 | 0.035 | 1 | 0.73075158 | 1.432959407 | 0 | 0.012 | 1 | 0.73075158 | 0.568082393 |
| Kctd7 | 0.342795612 | -1.070389328 | 0.017 | 0.035 | 1 | 0.505605027 | 0.930459067 | 0 | 0.043 | 1 | 0.505605027 | 0.568082393 |
| Alox5ap | 0.342795612 | -1.600904045 | 0.017 | 0.035 | 1 | 0.64384652 | 0.494359952 | 0 | 0.022 | 1 | 0.64384652 | 0.568082393 |
| Gm15927 | 0.342795612 | -1.070389328 | 0.017 | 0.035 | 1 | 0.59781419 | 1.558490289 | 0 | 0.028 | 1 | 0.59781419 | 0.568082393 |
| Slc5a11 | 0.342795612 | -1.070389328 | 0.017 | 0.035 | 1 | 0.730749233 | 2.210566986 | 0 | 0.012 | 1 | 0.730749233 | 0.568082393 |
| Gm15910 | 0.342795612 | -1.070389328 | 0.017 | 0.035 | 1 | 0.597845674 | 1.695993813 | 0 | 0.028 | 1 | 0.597845674 | 0.568082393 |
| Evi2b | 0.342795612 | -1.070389328 | 0.017 | 0.035 | 1 | 0.577155236 | 0.847996907 | 0 | 0.031 | 1 | 0.577155236 | 0.568082393 |
| Osr2 | 0.342795612 | -1.222392421 | 0.017 | 0.035 | 1 | 0.397089153 | -0.888968688 | 0 | 0.068 | 1 | 0.397089153 | 0.568082393 |
| Pmaip1 | 0.342795612 | -1.070389328 | 0.017 | 0.035 | 1 | 0.577165087 | 0.930459067 | 0 | 0.031 | 1 | 0.577165087 | 0.568082393 |
| Plp1 | 0.342795612 | -1.070389328 | 0.017 | 0.035 | 1 | 0.669853054 | 2.210566986 | 0 | 0.019 | 1 | 0.669853054 | 0.568082393 |
| Gm53065 | 0.468447396 | -0.485426827 | 0.025 | 0.041 | 1 | 0.3430501 | 2.079322453 | 0.1 | 0.04 | 1 | 0.468447396 | 0.568416829 |
| Necap1 | 0.39170401 | -0.485426827 | 0.244 | 0.271 | 1 | 0.343184543 | -0.577024681 | 0.2 | 0.327 | 1 | 0.39170401 | 0.568593455 |
| Zfp820 | 0.343608466 | -0.622930351 | 0.025 | 0.047 | 1 | 0.64384652 | 1.31748219 | 0 | 0.022 | 1 | 0.64384652 | 0.569150154 |
| Fdxacb1 | 0.443101201 | 0.251538767 | 0.076 | 0.053 | 1 | 0.343787167 | -0.111361109 | 0 | 0.083 | 1 | 0.443101201 | 0.569384718 |
| Tspyl1 | 0.343842772 | -0.250961574 | 0.588 | 0.582 | 1 | 0.66427568 | 0.30520386 | 0.6 | 0.636 | 1 | 0.66427568 | 0.569457692 |
| Spry4 | 0.578808801 | 0.192645078 | 0.101 | 0.082 | 1 | 0.343859585 | -0.152003093 | 0 | 0.083 | 1 | 0.578808801 | 0.569479756 |
| Slc12a7 | 0.474205722 | -0.584962501 | 0.05 | 0.071 | 1 | 0.343863936 | -0.230005605 | 0 | 0.083 | 1 | 0.474205722 | 0.569485466 |
| Gm28791 | 0.870967912 | -0.292781749 | 0.059 | 0.053 | 1 | 0.343865677 | -0.111361109 | 0 | 0.083 | 1 | 0.870967912 | 0.56948775 |
| Igtp | 0.641881888 | -0.648925559 | 0.126 | 0.141 | 1 | 0.343954076 | -1.457811523 | 0 | 0.083 | 1 | 0.641881888 | 0.569603746 |
| Cmtr1 | 0.731617263 | -0.100762977 | 0.319 | 0.282 | 1 | 0.34395853 | -0.797995028 | 0.2 | 0.321 | 1 | 0.731617263 | 0.569609589 |
| Bicd1 | 0.460435947 | 0.713881981 | 0.109 | 0.082 | 1 | 0.344029398 | -0.959358016 | 0 | 0.083 | 1 | 0.460435947 | 0.569702569 |
| D130017N08Rik | 0.344235067 | 0.707218251 | 0.059 | 0.035 | 1 | 0.487557552 | 1.432959407 | 0.1 | 0.049 | 1 | 0.487557552 | 0.569972353 |
| 1500009L16Rik | 0.344235067 | 0.707218251 | 0.059 | 0.035 | 1 | 0.669843604 | 2.017921908 | 0 | 0.019 | 1 | 0.669843604 | 0.569972353 |
| Il1r2 | 0.344246393 | -0.555816155 | 0.034 | 0.059 | 1 | 0.669848329 | 0.625604485 | 0 | 0.019 | 1 | 0.669848329 | 0.569987207 |
| Man2c1os | 0.344317492 | -1.807354922 | 0.05 | 0.076 | 1 | 0.877599489 | 0.210566986 | 0.1 | 0.114 | 1 | 0.877599489 | 0.570080449 |
| Dynlt1c | 0.344507534 | 0.330148602 | 0.151 | 0.112 | 1 | 0.667604536 | 1.432959407 | 0.1 | 0.065 | 1 | 0.667604536 | 0.570329627 |
| Ap1m1 | 0.847105233 | -0.268615438 | 0.311 | 0.306 | 1 | 0.344992087 | 0.494359952 | 0.5 | 0.333 | 1 | 0.847105233 | 0.570964635 |
| Rpgr | 0.345157505 | -0.722466024 | 0.101 | 0.135 | 1 | 0.849313972 | 0.625604485 | 0.1 | 0.08 | 1 | 0.849313972 | 0.571181307 |
| Tmem159 | 0.345287389 | -0.238333965 | 0.319 | 0.376 | 1 | 0.530328566 | 0.219180116 | 0.2 | 0.349 | 1 | 0.530328566 | 0.571351397 |
| Ifi204 | 0.345346582 | -0.900464326 | 0.017 | 0.035 | 1 | 0.619961444 | 1.432959407 | 0 | 0.025 | 1 | 0.619961444 | 0.571428903 |
| Hnmt | 0.345346582 | -0.900464326 | 0.017 | 0.035 | 1 | 0.460577419 | 0.558490289 | 0 | 0.052 | 1 | 0.460577419 | 0.571428903 |
| Rrh | 0.345346582 | -0.900464326 | 0.017 | 0.035 | 1 | 0.669848329 | 1.847996907 | 0 | 0.019 | 1 | 0.669848329 | 0.571428903 |
| 4930534D22Rik | 0.345346582 | -0.900464326 | 0.017 | 0.035 | 1 | 0.597824685 | 1.432959407 | 0 | 0.028 | 1 | 0.597824685 | 0.571428903 |
| Gm36371 | 0.345346582 | -0.900464326 | 0.017 | 0.035 | 1 | 0.669851164 | 1.695993813 | 0 | 0.019 | 1 | 0.669851164 | 0.571428903 |
| Fadd | 0.345346582 | -0.900464326 | 0.017 | 0.035 | 1 | 0.643850855 | 2.017921908 | 0 | 0.022 | 1 | 0.643850855 | 0.571428903 |
| Gm56848 | 0.345346582 | -0.900464326 | 0.017 | 0.035 | 1 | 0.619961444 | 1.558490289 | 0 | 0.025 | 1 | 0.619961444 | 0.571428903 |
| Gm26812 | 0.345346582 | -0.900464326 | 0.017 | 0.035 | 1 | 0.669843604 | 2.017921908 | 0 | 0.019 | 1 | 0.669843604 | 0.571428903 |
| Plppr2 | 0.345346582 | -1.070389328 | 0.017 | 0.035 | 1 | 0.669843604 | 2.017921908 | 0 | 0.019 | 1 | 0.669843604 | 0.571428903 |
| Gm19569 | 0.345346582 | -0.900464326 | 0.017 | 0.035 | 1 | 0.698534498 | 2.432959407 | 0 | 0.015 | 1 | 0.698534498 | 0.571428903 |
| 1700030C12Rik | 0.345346582 | -0.900464326 | 0.017 | 0.035 | 1 | 0.355415506 | 2.210566986 | 0.1 | 0.04 | 1 | 0.355415506 | 0.571428903 |
| Lyrm7 | 0.345346582 | -0.900464326 | 0.017 | 0.035 | 1 | 0.522102209 | 0.847996907 | 0 | 0.04 | 1 | 0.522102209 | 0.571428903 |
| Krt6a | 0.345346582 | -0.900464326 | 0.017 | 0.035 | 1 | 0.824376305 | -0.473931188 | 0.1 | 0.077 | 1 | 0.824376305 | 0.571428903 |
| Hunk | 0.345346582 | -1.485426827 | 0.017 | 0.035 | 1 | 0.489955311 | 0.159940913 | 0 | 0.046 | 1 | 0.489955311 | 0.571428903 |
| Gm49958 | 0.345346582 | -0.900464326 | 0.017 | 0.035 | 1 | 0.52208609 | 0.930459067 | 0 | 0.04 | 1 | 0.52208609 | 0.571428903 |
| Zfp811 | 0.345346582 | -0.900464326 | 0.017 | 0.035 | 1 | 0.557771538 | 0.769994395 | 0 | 0.034 | 1 | 0.557771538 | 0.571428903 |
| Arhgef4 | 0.345379989 | -1.070389328 | 0.017 | 0.035 | 1 | 0.619967468 | 1.432959407 | 0 | 0.025 | 1 | 0.619967468 | 0.571472641 |
| Rxra | 0.582466655 | -0.304854582 | 0.227 | 0.247 | 1 | 0.345406999 | -0.546862711 | 0.2 | 0.33 | 1 | 0.582466655 | 0.571508004 |
| Klhl33 | 0.396430391 | -1.177304532 | 0.084 | 0.112 | 1 | 0.345504381 | 2.017921908 | 0.1 | 0.04 | 1 | 0.396430391 | 0.571635484 |
| Zup1 | 0.345759233 | -0.394914534 | 0.37 | 0.406 | 1 | 0.517026658 | -0.397115591 | 0.3 | 0.407 | 1 | 0.517026658 | 0.571969019 |
| Zfp280b | 0.345791776 | -0.622930351 | 0.025 | 0.047 | 1 | 0.781750802 | 1.017921908 | 0.1 | 0.074 | 1 | 0.781750802 | 0.572011599 |
| Fkbp2 | 0.346217612 | -0.354723136 | 0.613 | 0.6 | 1 | 0.736049231 | -0.210896782 | 0.6 | 0.636 | 1 | 0.736049231 | 0.572568589 |
| Sigmar1 | 0.513620492 | 0.198099508 | 0.336 | 0.3 | 1 | 0.34621888 | 1.280956314 | 0.3 | 0.207 | 1 | 0.513620492 | 0.572570247 |
| Toe1 | 0.346442623 | -0.513995979 | 0.185 | 0.224 | 1 | 0.878099348 | 0.4480663 | 0.2 | 0.235 | 1 | 0.878099348 | 0.572862755 |
| Prdm16 | 0.34645033 | -0.313474144 | 0.798 | 0.8 | 1 | 0.701175554 | 0.105841756 | 0.9 | 0.895 | 1 | 0.701175554 | 0.572872829 |
| Pdcd2l | 0.346481746 | -0.542010356 | 0.143 | 0.182 | 1 | 0.64935734 | 0.345496566 | 0.3 | 0.219 | 1 | 0.64935734 | 0.572913891 |
| Klf3 | 0.373194322 | -0.176921112 | 0.874 | 0.788 | 1 | 0.347458321 | -0.805029869 | 0.9 | 0.935 | 1 | 0.373194322 | 0.574189358 |
| Ndufa5 | 0.347488952 | -0.241190017 | 0.613 | 0.653 | 1 | 0.72907293 | -0.186649236 | 0.6 | 0.574 | 1 | 0.72907293 | 0.574229332 |
| Ap2a1 | 0.34757283 | -0.424026283 | 0.437 | 0.453 | 1 | 0.57120273 | -0.321928095 | 0.5 | 0.528 | 1 | 0.57120273 | 0.574338788 |
| Ascc2 | 0.347643052 | -0.320796125 | 0.353 | 0.394 | 1 | 0.687203849 | -0.105460508 | 0.4 | 0.451 | 1 | 0.687203849 | 0.574430412 |
| Osbpl9 | 0.347674137 | -0.240980424 | 0.908 | 0.906 | 1 | 0.596184969 | -0.300302218 | 1 | 0.948 | 1 | 0.596184969 | 0.574470969 |
| Yipf2 | 0.77736847 | 0.215012891 | 0.227 | 0.218 | 1 | 0.347800863 | 1.642882477 | 0.3 | 0.213 | 1 | 0.77736847 | 0.574636286 |
| Gpr160 | 0.347959498 | -0.807354922 | 0.017 | 0.035 | 1 | 0.643842184 | 1.558490289 | 0 | 0.022 | 1 | 0.643842184 | 0.574843184 |
| Gm57147 | 0.347966916 | -0.707819249 | 0.017 | 0.035 | 1 | 0.518235124 | 1.769994395 | 0.1 | 0.052 | 1 | 0.518235124 | 0.574852858 |
| Naif1 | 0.347966916 | -0.707819249 | 0.017 | 0.035 | 1 | 0.577142568 | 1.432959407 | 0 | 0.031 | 1 | 0.577142568 | 0.574852858 |
| Crybb3 | 0.347966916 | -0.707819249 | 0.017 | 0.035 | 1 | 0.698533455 | 1.558490289 | 0 | 0.015 | 1 | 0.698533455 | 0.574852858 |
| Cldn15 | 0.347966916 | -0.707819249 | 0.017 | 0.035 | 1 | 0.487557552 | 1.494359952 | 0.1 | 0.049 | 1 | 0.487557552 | 0.574852858 |
| A430078I02Rik | 0.347966916 | -0.707819249 | 0.017 | 0.035 | 1 | 0.73075158 | 2.695993813 | 0 | 0.012 | 1 | 0.73075158 | 0.574852858 |
| Trim21 | 0.347966916 | -0.707819249 | 0.017 | 0.035 | 1 | 0.522066388 | 1.111031312 | 0 | 0.04 | 1 | 0.522066388 | 0.574852858 |

| | | | | | | | | | | | | |
|---|---|---|---|---|---|---|---|---|---|---|---|---|
| Ccdc74a | 0.347966916 | -0.707819249 | 0.017 | 0.035 | 1 | 0.597845674 | 1.695993813 | 0 | 0.028 | 1 | 0.597845674 | 0.574852858 |
| Mrpl33 | 0.397846271 | 0.125530882 | 0.807 | 0.706 | 1 | 0.348058324 | -0.355184911 | 0.5 | 0.657 | 1 | 0.397846271 | 0.57497205 |
| Kdm3a | 0.348417068 | -0.367064631 | 0.588 | 0.6 | 1 | 0.591684765 | -0.374395515 | 0.6 | 0.738 | 1 | 0.591684765 | 0.575439682 |
| Zfp1 | 0.38769969 | 0.133144066 | 0.445 | 0.388 | 1 | 0.348896772 | 0.573137065 | 0.5 | 0.352 | 1 | 0.38769969 | 0.576064587 |
| Yap1 | 0.349098056 | -0.431115707 | 0.782 | 0.7 | 1 | 0.497888023 | -0.402229723 | 0.9 | 0.843 | 1 | 0.497888023 | 0.576326659 |
| Phb | 0.349119246 | -0.485426827 | 0.345 | 0.376 | 1 | 0.688615645 | -0.28585884 | 0.3 | 0.333 | 1 | 0.688615645 | 0.576354244 |
| Mrrf | 0.412061134 | 0.106915204 | 0.345 | 0.294 | 1 | 0.349311576 | 0.982297998 | 0.3 | 0.185 | 1 | 0.412061134 | 0.576604575 |
| Arf4os | 0.349688292 | -0.637429921 | 0.059 | 0.088 | 1 | 0.67329207 | 1.159940913 | 0.1 | 0.065 | 1 | 0.67329207 | 0.577094682 |
| Adprm | 0.34989961 | 0.146841388 | 0.21 | 0.159 | 1 | 0.941666367 | 1.450237399 | 0.2 | 0.207 | 1 | 0.941666367 | 0.577369483 |
| Gatd3a | 0.350039259 | -0.175742328 | 0.252 | 0.306 | 1 | 0.450689441 | -0.304006187 | 0.2 | 0.302 | 1 | 0.450689441 | 0.577551035 |
| Clba1 | 0.974388597 | -0.211251864 | 0.151 | 0.153 | 1 | 0.350155284 | -0.750262417 | 0.1 | 0.216 | 1 | 0.974388597 | 0.577701845 |
| Lonrf1 | 0.487131208 | -0.304854582 | 0.353 | 0.382 | 1 | 0.350182501 | -0.676958285 | 0.3 | 0.432 | 1 | 0.487131208 | 0.577737218 |
| Gm29681 | 0.350200104 | 0.152003093 | 0.025 | 0.047 | 1 | 0.460634939 | 0.159940913 | 0 | 0.052 | 1 | 0.460634939 | 0.577760096 |
| Nuggc | 0.350200104 | -0.333423734 | 0.025 | 0.047 | 1 | 0.669850692 | 1.31748219 | 0 | 0.019 | 1 | 0.669850692 | 0.577760096 |
| Ophn1 | 0.458223296 | -0.502774071 | 0.529 | 0.494 | 1 | 0.35031065 | -0.905702703 | 0.4 | 0.525 | 1 | 0.458223296 | 0.577903748 |
| 6820408C15Rik | 0.350543059 | -0.655351829 | 0.017 | 0.035 | 1 | 0.643835681 | 1.847996907 | 0 | 0.022 | 1 | 0.643835681 | 0.578205682 |
| Gm20470 | 0.350543059 | -0.655351829 | 0.017 | 0.035 | 1 | 0.522126383 | 0.558490289 | 0 | 0.04 | 1 | 0.522126383 | 0.578205682 |
| Angptl7 | 0.350596085 | -1.036836768 | 0.076 | 0.106 | 1 | 0.489991953 | -0.654503434 | 0 | 0.046 | 1 | 0.489991953 | 0.578274555 |
| 5033430I15Rik | 0.351259601 | 0.514573173 | 0.059 | 0.035 | 1 | 0.597845674 | 1.695993813 | 0 | 0.028 | 1 | 0.597845674 | 0.579135894 |
| Spata6 | 0.588219713 | -0.36994961 | 0.513 | 0.488 | 1 | 0.351467423 | 0.695993813 | 0.8 | 0.627 | 1 | 0.588219713 | 0.579405496 |
| Arvcf | 0.351832008 | -0.944858446 | 0.017 | 0.035 | 1 | 0.643849121 | 1.432959407 | 0 | 0.022 | 1 | 0.643849121 | 0.579878255 |
| Trim56 | 0.351914373 | -0.547711105 | 0.218 | 0.253 | 1 | 0.369116445 | -0.536666944 | 0.2 | 0.343 | 1 | 0.369116445 | 0.57998502 |
| Nabp1 | 0.352369348 | 0.758498756 | 0.134 | 0.1 | 1 | 0.660248429 | 1.675529711 | 0.2 | 0.16 | 1 | 0.660248429 | 0.580574538 |
| Tex9 | 0.728293952 | 0.152003093 | 0.101 | 0.088 | 1 | 0.352598821 | 0.959028219 | 0.2 | 0.102 | 1 | 0.728293952 | 0.580871714 |
| 9330188P03Rik | 0.352678372 | 1.266645659 | 0.252 | 0.218 | 1 | 0.651560768 | 0.570838682 | 0.3 | 0.269 | 1 | 0.651560768 | 0.58097471 |
| Erp44 | 0.402468869 | -0.175197254 | 0.866 | 0.865 | 1 | 0.353502294 | -0.267480311 | 0.8 | 0.907 | 1 | 0.402468869 | 0.582040717 |
| Rragd | 0.558164171 | -0.970853654 | 0.034 | 0.047 | 1 | 0.353587229 | 0.111031312 | 0 | 0.08 | 1 | 0.558164171 | 0.58215053 |
| Gm17066 | 0.743002317 | -0.163498732 | 0.034 | 0.041 | 1 | 0.353636871 | 0.111031312 | 0 | 0.08 | 1 | 0.743002317 | 0.582214706 |
| Get1 | 0.361745152 | -0.402964667 | 0.059 | 0.088 | 1 | 0.353793999 | -0.191531458 | 0 | 0.08 | 1 | 0.361745152 | 0.582417804 |
| Zfp113 | 0.393887242 | -0.315501826 | 0.05 | 0.076 | 1 | 0.353824728 | -0.267480311 | 0 | 0.08 | 1 | 0.393887242 | 0.582457518 |
| Entrep1 | 0.726318234 | 1.067114196 | 0.067 | 0.059 | 1 | 0.353850423 | -0.304006187 | 0 | 0.08 | 1 | 0.726318234 | 0.582490724 |
| R3hdm4 | 0.64556564 | -0.301002256 | 0.504 | 0.482 | 1 | 0.354376421 | -0.272096939 | 0.4 | 0.59 | 1 | 0.64556564 | 0.583170194 |
| Gm14966 | 0.763303963 | -0.185866545 | 0.101 | 0.112 | 1 | 0.354577235 | 2.625604485 | 0.2 | 0.114 | 1 | 0.763303963 | 0.583429455 |
| Gm34086 | 0.354668871 | 0.929610672 | 0.05 | 0.029 | 1 | 0.535011169 | 1.159940913 | 0.1 | 0.052 | 1 | 0.535011169 | 0.583547734 |
| Polr2a | 0.354691111 | -0.619365205 | 0.706 | 0.724 | 1 | 0.937262719 | -0.139930261 | 0.8 | 0.747 | 1 | 0.937262719 | 0.583576438 |
| Gramd1b | 0.354691115 | 0.147790842 | 0.227 | 0.176 | 1 | 0.809640341 | 0.159940913 | 0.2 | 0.241 | 1 | 0.809640341 | 0.583576443 |
| Wars2 | 0.354716792 | -0.402964667 | 0.294 | 0.341 | 1 | 0.406912746 | 0.695993813 | 0.4 | 0.281 | 1 | 0.406912746 | 0.583609582 |
| Dcaf12 | 0.722125083 | -0.129283017 | 0.487 | 0.418 | 1 | 0.354784919 | -0.408342847 | 0.4 | 0.559 | 1 | 0.722125083 | 0.583697499 |
| Kirrel3 | 0.469479943 | -1.266786541 | 0.092 | 0.118 | 1 | 0.355034862 | 0.861417422 | 0.2 | 0.108 | 1 | 0.469479943 | 0.58401997 |
| Zbtb21 | 0.431673255 | 0.234465254 | 0.412 | 0.365 | 1 | 0.355040632 | -0.689437224 | 0.3 | 0.407 | 1 | 0.431673255 | 0.584027413 |
| Gng10 | 0.954802243 | -0.180813318 | 0.748 | 0.629 | 1 | 0.355158478 | -0.357117523 | 0.6 | 0.654 | 1 | 0.954802243 | 0.584179411 |
| Gm43774 | 0.629493349 | -0.333423734 | 0.025 | 0.035 | 1 | 0.355415506 | 2.210566986 | 0.1 | 0.04 | 1 | 0.629493349 | 0.58451083 |
| C330011M18Rik | 0.629509811 | -0.292781749 | 0.025 | 0.035 | 1 | 0.355415506 | 2.210566986 | 0.1 | 0.04 | 1 | 0.629509811 | 0.58451083 |
| Gm16794 | 0.355734963 | -0.292781749 | 0.017 | 0.035 | 1 | 0.444918466 | 1.494359952 | 0.1 | 0.046 | 1 | 0.444918466 | 0.584922562 |
| Rilp | 0.355734963 | -0.292781749 | 0.017 | 0.035 | 1 | 0.601875604 | 1.263034406 | 0.1 | 0.059 | 1 | 0.601875604 | 0.584922562 |
| Sytl4 | 0.355734963 | -0.292781749 | 0.017 | 0.035 | 1 | 0.539431168 | 1.111031312 | 0 | 0.037 | 1 | 0.539431168 | 0.584922562 |
| Afg1l | 0.355799991 | -0.900464326 | 0.16 | 0.188 | 1 | 0.86159151 | 1.017921908 | 0.1 | 0.13 | 1 | 0.86159151 | 0.585006349 |
| Whamm | 0.379469059 | -0.211251864 | 0.143 | 0.182 | 1 | 0.355805022 | -0.723545078 | 0.1 | 0.216 | 1 | 0.379469059 | 0.58501283 |
| Gm4117 | 0.35618215 | 0.440572591 | 0.034 | 0.059 | 1 | 0.539445308 | 0.558490289 | 0 | 0.037 | 1 | 0.539445308 | 0.585498576 |
| Eef1akmt2 | 0.356570608 | 0.139533741 | 0.412 | 0.335 | 1 | 0.855548392 | 0.20029865 | 0.3 | 0.34 | 1 | 0.855548392 | 0.585998617 |
| Rab4b | 0.676946229 | -0.144389909 | 0.588 | 0.524 | 1 | 0.357033865 | 0.418009066 | 0.7 | 0.522 | 1 | 0.676946229 | 0.58659455 |
| Usp8 | 0.449893537 | -0.155278225 | 0.639 | 0.665 | 1 | 0.357150459 | 0.468924905 | 0.8 | 0.744 | 1 | 0.449893537 | 0.586744467 |
| Tpgs2 | 0.718668392 | -0.381591016 | 0.261 | 0.265 | 1 | 0.357158124 | -0.391469028 | 0.1 | 0.219 | 1 | 0.718668392 | 0.586754323 |
| Gramd4 | 0.540018306 | -0.821029859 | 0.168 | 0.182 | 1 | 0.357210876 | 1.367667947 | 0.3 | 0.204 | 1 | 0.540018306 | 0.586822142 |
| Kxd1 | 0.35723183 | -0.485426827 | 0.252 | 0.282 | 1 | 0.539049278 | 0.31748219 | 0.4 | 0.278 | 1 | 0.539049278 | 0.58684908 |
| Nudt7 | 0.619653652 | -0.273922722 | 0.067 | 0.082 | 1 | 0.357796814 | 2.111031312 | 0.1 | 0.04 | 1 | 0.619653652 | 0.587575068 |
| Ada | 0.687345921 | -0.707819249 | 0.017 | 0.024 | 1 | 0.357796814 | 2.111031312 | 0.1 | 0.04 | 1 | 0.687345921 | 0.587575068 |
| Gm56696 | 0.729956195 | -0.485426827 | 0.034 | 0.041 | 1 | 0.357796814 | 2.017921908 | 0.1 | 0.04 | 1 | 0.729956195 | 0.587575068 |
| Gm19500 | 0.801878508 | -0.807354922 | 0.042 | 0.047 | 1 | 0.357796814 | 2.111031312 | 0.1 | 0.04 | 1 | 0.801878508 | 0.587575068 |
| Rpusd1 | 0.899314017 | -0.485426827 | 0.05 | 0.053 | 1 | 0.357796814 | 1.847996907 | 0.1 | 0.04 | 1 | 0.899314017 | 0.587575068 |
| Lrrc8dos | 0.929389703 | 0.192645078 | 0.025 | 0.024 | 1 | 0.357796814 | 2.111031312 | 0.1 | 0.04 | 1 | 0.929389703 | 0.587575068 |
| Bola2 | 0.960177694 | -0.16002654 | 0.37 | 0.347 | 1 | 0.357922704 | 0.782705446 | 0.4 | 0.265 | 1 | 0.960177694 | 0.587736745 |
| Rsph3b | 0.656859297 | -0.23878686 | 0.218 | 0.235 | 1 | 0.358223933 | 0.913585248 | 0.3 | 0.182 | 1 | 0.656859297 | 0.58812348 |
| Ptpmt1 | 0.358423422 | -0.526068812 | 0.269 | 0.3 | 1 | 0.55464971 | -0.526398608 | 0.2 | 0.269 | 1 | 0.55464971 | 0.588379495 |
| Map4k5 | 0.788421213 | -0.161729639 | 0.908 | 0.924 | 1 | 0.359412888 | -0.101147296 | 1 | 0.972 | 1 | 0.788421213 | 0.589648152 |
| Calhm2 | 0.359884886 | -0.517848305 | 0.118 | 0.153 | 1 | 0.478081765 | -0.441509711 | 0.1 | 0.182 | 1 | 0.478081765 | 0.590252641 |
| Uvssa | 0.359954084 | -0.5360529 | 0.269 | 0.3 | 1 | 0.418168535 | 0.453137289 | 0.4 | 0.269 | 1 | 0.418168535 | 0.590341225 |
| Accs | 0.757769092 | -0.11345805 | 0.067 | 0.076 | 1 | 0.360331175 | 2.017921908 | 0.1 | 0.04 | 1 | 0.757769092 | 0.590823794 |
| Palb2 | 0.933394757 | -0.163498732 | 0.034 | 0.035 | 1 | 0.360331175 | 1.930459067 | 0.1 | 0.04 | 1 | 0.933394757 | 0.590823794 |
| Kcnip1 | 0.954551343 | -0.222392421 | 0.017 | 0.018 | 1 | 0.360331175 | 1.930459067 | 0.1 | 0.04 | 1 | 0.954551343 | 0.590823794 |
| Gm3839 | 0.360394871 | 0.736965594 | 0.05 | 0.029 | 1 | 0.398985336 | 2.017921908 | 0.1 | 0.043 | 1 | 0.398985336 | 0.590905279 |
| Gm53 | 0.360674734 | 0.225066556 | 0.059 | 0.035 | 1 | 0.619961444 | 1.31748219 | 0 | 0.025 | 1 | 0.619961444 | 0.591263205 |
| Pycr2 | 0.360786228 | -0.308549065 | 0.336 | 0.382 | 1 | 0.447077246 | -0.294961047 | 0.2 | 0.324 | 1 | 0.447077246 | 0.591405753 |
| Parva | 0.360940877 | -0.283237977 | 0.798 | 0.771 | 1 | 0.517068351 | -0.506946246 | 0.8 | 0.846 | 1 | 0.517068351 | 0.591603438 |
| Ift57 | 0.436547438 | 0.241554678 | 0.412 | 0.365 | 1 | 0.361310815 | -0.425021588 | 0.2 | 0.352 | 1 | 0.436547438 | 0.592076125 |
| Gm15246 | 0.361463646 | 0.707218251 | 0.05 | 0.029 | 1 | 0.643849121 | 0.625604485 | 0 | 0.022 | 1 | 0.643849121 | 0.592271325 |
| Haus1 | 0.918441518 | 0.251538767 | 0.118 | 0.124 | 1 | 0.361632474 | 1.176619654 | 0.2 | 0.105 | 1 | 0.918441518 | 0.592486902 |
| Yipf1 | 0.496518653 | -0.22762907 | 0.378 | 0.394 | 1 | 0.362019408 | -0.330806246 | 0.2 | 0.355 | 1 | 0.496518653 | 0.592980764 |
| Fhit | 0.362079431 | -0.247927513 | 0.445 | 0.347 | 1 | 0.672920397 | -0.616793628 | 0.7 | 0.515 | 1 | 0.672920397 | 0.593057348 |
| Rbms1 | 0.362131979 | 0.135970063 | 0.983 | 0.971 | 1 | 0.747781164 | 0.265319683 | 1 | 0.988 | 1 | 0.747781164 | 0.593124387 |
| Supt4a | 0.967486082 | -0.204245075 | 0.563 | 0.506 | 1 | 0.362391004 | -0.425021588 | 0.4 | 0.531 | 1 | 0.967486082 | 0.593454769 |
| Ankrd13a | 0.362430683 | -0.40053793 | 0.252 | 0.294 | 1 | 0.639298509 | -0.167944637 | 0.4 | 0.441 | 1 | 0.639298509 | 0.593505366 |
| Slc1a2 | 0.362599853 | 0.514573173 | 0.05 | 0.029 | 1 | 0.39706715 | -0.374395515 | 0 | 0.068 | 1 | 0.39706715 | 0.593721052 |
| Dach1 | 0.362722586 | 1.602036014 | 0.042 | 0.024 | 1 | 0.460685472 | -1.048167282 | 0 | 0.052 | 1 | 0.460685472 | 0.593877498 |
| Npff | 0.403273565 | -0.600904045 | 0.042 | 0.065 | 1 | 0.362872338 | 1.847996907 | 0.1 | 0.04 | 1 | 0.403273565 | 0.594068343 |
| A930001M01Rik | 0.691605844 | -0.485426827 | 0.017 | 0.024 | 1 | 0.362872338 | 1.847996907 | 0.1 | 0.04 | 1 | 0.691605844 | 0.594068343 |
| Galns | 0.744292306 | -0.177304532 | 0.101 | 0.088 | 1 | 0.362872338 | 1.769994395 | 0.1 | 0.04 | 1 | 0.744292306 | 0.594068343 |
| Exoc6 | 0.984059855 | -0.411426246 | 0.067 | 0.065 | 1 | 0.362872659 | 1.930459067 | 0.1 | 0.04 | 1 | 0.984059855 | 0.594068751 |
| Naxd | 0.555200899 | -0.128270244 | 0.63 | 0.553 | 1 | 0.362969706 | -0.492135784 | 0.4 | 0.565 | 1 | 0.555200899 | 0.594192405 |
| Smco4 | 0.363017904 | -0.547711105 | 0.227 | 0.265 | 1 | 0.408679229 | -0.31299497 | 0.2 | 0.336 | 1 | 0.408679229 | 0.59425381 |
| Rasa2 | 0.36331592 | -0.402136484 | 0.714 | 0.659 | 1 | 0.431765469 | 0.36729249 | 1 | 0.793 | 1 | 0.431765469 | 0.594633383 |
| Rrp1b | 0.363348769 | 0.109316694 | 0.244 | 0.188 | 1 | 0.708163291 | -0.165299916 | 0.2 | 0.238 | 1 | 0.708163291 | 0.59467521 |
| AU022252 | 0.417828738 | -0.517848305 | 0.151 | 0.182 | 1 | 0.363637803 | -0.5519337 | 0.1 | 0.216 | 1 | 0.417828738 | 0.595043154 |
| Creld2 | 0.67536163 | -0.186987085 | 0.563 | 0.482 | 1 | 0.363788062 | -0.715432433 | 0.4 | 0.466 | 1 | 0.67536163 | 0.59523437 |
| Ttc14 | 0.363832129 | -0.18339829 | 0.798 | 0.765 | 1 | 0.408633804 | -0.354943152 | 0.7 | 0.772 | 1 | 0.408633804 | 0.59529044 |
| 3110040N11Rik | 0.363977667 | -0.485426827 | 0.185 | 0.224 | 1 | 0.591382622 | 1.339850003 | 0.2 | 0.148 | 1 | 0.591382622 | 0.595475592 |
| Fgfbp3 | 0.793002704 | 0.192645078 | 0.109 | 0.1 | 1 | 0.364021887 | -0.152003093 | 0 | 0.077 | 1 | 0.793002704 | 0.59553184 |
| Gm51425 | 0.5770098 | -0.925999419 | 0.05 | 0.065 | 1 | 0.364023373 | -0.441509711 | 0 | 0.077 | 1 | 0.5770098 | 0.59553373 |

| | | | | | | | | | | | | |
|---|---|---|---|---|---|---|---|---|---|---|---|---|
| Ccdc60 | 0.467093859 | -0.485426827 | 0.025 | 0.041 | 1 | 0.364053642 | -0.152003093 | 0 | 0.077 | 1 | 0.467093859 | 0.59557223 |
| B3galt1 | 0.364123618 | -0.255068096 | 0.513 | 0.429 | 1 | 0.461554258 | 0.607793209 | 0.4 | 0.349 | 1 | 0.461554258 | 0.595661227 |
| Sbk1 | 0.364143092 | -0.428843299 | 0.084 | 0.118 | 1 | 0.42087075 | 0.494359952 | 0 | 0.062 | 1 | 0.42087075 | 0.595685992 |
| H4c16 | 0.364180867 | -1.185866545 | 0.05 | 0.076 | 1 | 0.619972287 | 1.017921908 | 0 | 0.025 | 1 | 0.619972287 | 0.59573403 |
| Cdc20 | 0.364207692 | -1.185866545 | 0.05 | 0.076 | 1 | 0.619961444 | 1.31748219 | 0 | 0.025 | 1 | 0.619961444 | 0.595768141 |
| Rimoc1 | 0.442039218 | -0.415717855 | 0.345 | 0.371 | 1 | 0.364885665 | -0.48032896 | 0.3 | 0.441 | 1 | 0.442039218 | 0.596629782 |
| Wdhd1 | 0.797540729 | -0.485426827 | 0.092 | 0.1 | 1 | 0.364957174 | 0.847996907 | 0.2 | 0.105 | 1 | 0.797540729 | 0.59672061 |
| Tmem192 | 0.733430158 | 0.280107919 | 0.118 | 0.106 | 1 | 0.364957442 | 1.210566986 | 0.2 | 0.105 | 1 | 0.733430158 | 0.596720949 |
| Khnyn | 0.365086091 | -0.376197757 | 0.294 | 0.335 | 1 | 0.635911249 | -0.357117523 | 0.4 | 0.432 | 1 | 0.635911249 | 0.596884328 |
| Tbl3 | 0.365194304 | -0.24726709 | 0.151 | 0.194 | 1 | 0.90419698 | 0.210566986 | 0.2 | 0.207 | 1 | 0.90419698 | 0.597021728 |
| Armc5 | 0.365351964 | -0.359895945 | 0.084 | 0.118 | 1 | 0.799710702 | 0.959028219 | 0.1 | 0.142 | 1 | 0.799710702 | 0.59722187 |
| Tmem30a | 0.365417818 | 0.101254191 | 0.992 | 0.935 | 1 | 0.821565912 | 0.15020191 | 0.9 | 0.963 | 1 | 0.821565912 | 0.597305454 |
| Gm31282 | 0.495207728 | -0.485426827 | 0.017 | 0.029 | 1 | 0.365420305 | 1.769994395 | 0.1 | 0.04 | 1 | 0.495207728 | 0.597308611 |
| 9130230L23Rik | 0.731769749 | -0.222392421 | 0.017 | 0.012 | 1 | 0.365420305 | 1.31748219 | 0.1 | 0.04 | 1 | 0.731769749 | 0.597308611 |
| Rtn1 | 0.948614424 | -0.807354922 | 0.025 | 0.024 | 1 | 0.365421266 | 1.625604485 | 0.1 | 0.04 | 1 | 0.948614424 | 0.59730983 |
| Scgb1b27 | 0.365452675 | -0.544320516 | 0.176 | 0.212 | 1 | 0.420904619 | 0.263034406 | 0 | 0.062 | 1 | 0.420904619 | 0.597349692 |
| Zkscan8 | 0.365652349 | -0.622930351 | 0.076 | 0.106 | 1 | 0.802052667 | 1.111031312 | 0.1 | 0.077 | 1 | 0.802052667 | 0.597603058 |
| Hs3st1 | 0.365849486 | 2.321928095 | 0.017 | 0.006 | 1 | 0.730749233 | 2.432959407 | 0 | 0.012 | 1 | 0.730749233 | 0.597853126 |
| Arfgef3 | 0.365882469 | -0.222392421 | 0.067 | 0.041 | 1 | 0.832755037 | 0.263034406 | 0.1 | 0.077 | 1 | 0.832755037 | 0.597894957 |
| Gm29292 | 0.36591132 | 0.192645078 | 0.059 | 0.035 | 1 | 0.505639146 | 0.625604485 | 0 | 0.043 | 1 | 0.505639146 | 0.597931546 |
| Rbm48 | 0.366052865 | 0.125530882 | 0.294 | 0.235 | 1 | 0.551329394 | -0.374395515 | 0.1 | 0.167 | 1 | 0.551329394 | 0.59811103 |
| Pan2 | 0.366239385 | -0.214779237 | 0.227 | 0.276 | 1 | 0.398635678 | 1.18142064 | 0.3 | 0.213 | 1 | 0.398635678 | 0.598347482 |
| Fam76a | 0.366328374 | 0.280107919 | 0.798 | 0.706 | 1 | 0.559999986 | 0.302028537 | 0.7 | 0.787 | 1 | 0.559999986 | 0.59846027 |
| Gm48027 | 0.3986128 | -0.529820947 | 0.109 | 0.141 | 1 | 0.366816344 | 0.591657153 | 0.3 | 0.173 | 1 | 0.3986128 | 0.599078458 |
| Gm29107 | 0.367142614 | 0.514573173 | 0.05 | 0.029 | 1 | 0.539445308 | 1.017921908 | 0 | 0.037 | 1 | 0.539445308 | 0.599491529 |
| Cyp27a1 | 0.367142614 | 0.514573173 | 0.05 | 0.029 | 1 | 0.577165087 | 0.930459067 | 0 | 0.031 | 1 | 0.577165087 | 0.599491529 |
| Tmem232 | 0.367210365 | 0.514573173 | 0.05 | 0.029 | 1 | 0.619967468 | 0.31748219 | 0 | 0.025 | 1 | 0.619967468 | 0.599577278 |
| 1110065P20Rik | 0.367587905 | -0.226184 | 0.454 | 0.488 | 1 | 0.551296058 | 0.403212064 | 0.5 | 0.417 | 1 | 0.551296058 | 0.600054942 |
| Cenpv | 0.367708716 | -0.485426827 | 0.126 | 0.165 | 1 | 0.639871597 | -0.304006187 | 0.1 | 0.148 | 1 | 0.639871597 | 0.600207732 |
| Ubxn4 | 0.367788314 | -0.194399503 | 0.992 | 0.994 | 1 | 0.656270925 | -0.128390536 | 1 | 0.994 | 1 | 0.656270925 | 0.600308384 |
| 6530409C15Rik | 0.82670638 | -0.292781749 | 0.025 | 0.029 | 1 | 0.367976994 | 1.625604485 | 0.1 | 0.04 | 1 | 0.82670638 | 0.600546919 |
| Gm44577 | 0.918724392 | 1.152003093 | 0.025 | 0.024 | 1 | 0.367977953 | 1.210566986 | 0.1 | 0.04 | 1 | 0.918724392 | 0.600548133 |
| Arhgef38 | 0.368010882 | 1.514573173 | 0.017 | 0.006 | 1 | 0.433541956 | 0.494359952 | 0 | 0.059 | 1 | 0.433541956 | 0.600589755 |
| 9530034E10Rik | 0.368010882 | 1.514573173 | 0.017 | 0.006 | 1 | 0.730749233 | 2.432959407 | 0 | 0.012 | 1 | 0.730749233 | 0.600589755 |
| Hcar2 | 0.368010882 | 1.514573173 | 0.017 | 0.006 | 1 | 0.577184788 | 1.558490289 | 0 | 0.031 | 1 | 0.577184788 | 0.600589755 |
| Gm47772 | 0.368010882 | 1.836501268 | 0.017 | 0.006 | 1 | 0.730749233 | 2.210566986 | 0 | 0.012 | 1 | 0.730749233 | 0.600589755 |
| Dock5 | 0.368684822 | -0.116193018 | 0.143 | 0.106 | 1 | 0.85854053 | -0.135883428 | 0.2 | 0.231 | 1 | 0.85854053 | 0.601441146 |
| Gm10734 | 0.369482921 | 0.344648171 | 0.05 | 0.029 | 1 | 0.539483006 | 1.31748219 | 0 | 0.037 | 1 | 0.539483006 | 0.602448213 |
| Rai2 | 0.36957688 | -1.133125083 | 0.05 | 0.076 | 1 | 0.42105375 | -0.596787936 | 0 | 0.062 | 1 | 0.42105375 | 0.60256669 |
| Ino80c | 0.369732243 | -0.276840205 | 0.16 | 0.2 | 1 | 0.674048896 | 0.159940913 | 0.1 | 0.145 | 1 | 0.674048896 | 0.602762554 |
| Gm14744 | 0.370165731 | -0.347923303 | 0.16 | 0.2 | 1 | 0.696451287 | -0.425021588 | 0.3 | 0.346 | 1 | 0.696451287 | 0.603308794 |
| Kcnj13 | 0.370184089 | 1.099535674 | 0.017 | 0.006 | 1 | 0.687539247 | 0.930459067 | 0.1 | 0.065 | 1 | 0.687539247 | 0.603331918 |
| Cd59a | 0.370184089 | 1.099535674 | 0.017 | 0.006 | 1 | 0.444918466 | 1.625604485 | 0.1 | 0.046 | 1 | 0.444918466 | 0.603331918 |
| Gm14302 | 0.370184089 | 1.099535674 | 0.017 | 0.006 | 1 | 0.73075158 | 2.695993813 | 0 | 0.012 | 1 | 0.73075158 | 0.603331918 |
| Fam209 | 0.370184089 | 1.099535674 | 0.017 | 0.006 | 1 | 0.643850855 | 2.017921908 | 0 | 0.022 | 1 | 0.643850855 | 0.603331918 |
| Lama5 | 0.370184089 | 1.099535674 | 0.017 | 0.006 | 1 | 0.396884044 | 0.210566986 | 0 | 0.068 | 1 | 0.396884044 | 0.603331918 |
| Ppm1j | 0.370184089 | 1.099535674 | 0.017 | 0.006 | 1 | 0.669848329 | 1.847996907 | 0 | 0.019 | 1 | 0.669848329 | 0.603331918 |
| Musk | 0.370184089 | 1.099535674 | 0.017 | 0.006 | 1 | 0.669850692 | 1.432959407 | 0 | 0.019 | 1 | 0.669850692 | 0.603331918 |
| Apobr | 0.370184089 | 1.099535674 | 0.017 | 0.006 | 1 | 0.698532411 | 2.017921908 | 0 | 0.015 | 1 | 0.698532411 | 0.603331918 |
| Ido2 | 0.370184089 | 1.099535674 | 0.017 | 0.006 | 1 | 0.669853054 | 2.210566986 | 0 | 0.019 | 1 | 0.669853054 | 0.603331918 |
| Klf1 | 0.370184089 | 1.099535674 | 0.017 | 0.006 | 1 | 0.730749233 | 2.210566986 | 0 | 0.012 | 1 | 0.730749233 | 0.603331918 |
| Gm10629 | 0.370184089 | 1.099535674 | 0.017 | 0.006 | 1 | 0.73075158 | 2.695993813 | 0 | 0.012 | 1 | 0.73075158 | 0.603331918 |
| Susd2 | 0.370184089 | 1.099535674 | 0.017 | 0.006 | 1 | 0.730749233 | 2.432959407 | 0 | 0.012 | 1 | 0.730749233 | 0.603331918 |
| Tektip1 | 0.370184089 | 1.099535674 | 0.017 | 0.006 | 1 | 0.522102209 | 0.847996907 | 0 | 0.04 | 1 | 0.522102209 | 0.603331918 |
| 1500026H17Rik | 0.370184089 | 1.099535674 | 0.017 | 0.006 | 1 | 0.643850855 | 2.017921908 | 0 | 0.022 | 1 | 0.643850855 | 0.603331918 |
| Kif19a | 0.370184089 | 1.099535674 | 0.017 | 0.006 | 1 | 0.730749233 | 2.432959407 | 0 | 0.012 | 1 | 0.730749233 | 0.603331918 |
| Kcns3 | 0.370184089 | 1.099535674 | 0.017 | 0.006 | 1 | 0.643835681 | 1.847996907 | 0 | 0.022 | 1 | 0.643835681 | 0.603331918 |
| Srl | 0.370184089 | 1.099535674 | 0.017 | 0.006 | 1 | 0.64384652 | 1.31748219 | 0 | 0.022 | 1 | 0.64384652 | 0.603331918 |
| Sidt1 | 0.370184089 | 1.099535674 | 0.017 | 0.006 | 1 | 0.730750993 | 1.31748219 | 0 | 0.012 | 1 | 0.730750993 | 0.603331918 |
| Ripk4 | 0.370184089 | 1.099535674 | 0.017 | 0.006 | 1 | 0.408880042 | 0.558490289 | 0 | 0.065 | 1 | 0.408880042 | 0.603331918 |
| Dynlt1b | 0.370184089 | 1.099535674 | 0.017 | 0.006 | 1 | 0.6199755 | 1.847996907 | 0 | 0.025 | 1 | 0.6199755 | 0.603331918 |
| Gm3435 | 0.370184089 | 1.099535674 | 0.017 | 0.006 | 1 | 0.59781419 | 1.558490289 | 0 | 0.028 | 1 | 0.59781419 | 0.603331918 |
| Crb3 | 0.370184089 | 1.099535674 | 0.017 | 0.006 | 1 | 0.669843604 | 2.017921908 | 0 | 0.019 | 1 | 0.669843604 | 0.603331918 |
| Gm50209 | 0.370184089 | 1.099535674 | 0.017 | 0.006 | 1 | 0.730749233 | 2.017921908 | 0 | 0.012 | 1 | 0.730749233 | 0.603331918 |
| Gm29650 | 0.370184089 | 1.099535674 | 0.017 | 0.006 | 1 | 0.669853054 | 2.210566986 | 0 | 0.019 | 1 | 0.669853054 | 0.603331918 |
| Fads6 | 0.370241848 | 1.192645078 | 0.042 | 0.024 | 1 | 0.698534498 | 2.432959407 | 0 | 0.015 | 1 | 0.698534498 | 0.60340467 |
| 5430414B19Rik | 0.370241848 | 1 | 0.042 | 0.024 | 1 | 0.698534498 | 2.432959407 | 0 | 0.015 | 1 | 0.698534498 | 0.60340467 |
| Lgr4 | 0.722446056 | -0.405373576 | 0.714 | 0.653 | 1 | 0.370256583 | -1.075589978 | 0.7 | 0.741 | 1 | 0.722446056 | 0.603423229 |
| Lif | 0.370273383 | 1.514573173 | 0.042 | 0.024 | 1 | 0.557765566 | -0.267480311 | 0 | 0.034 | 1 | 0.557765566 | 0.603444388 |
| Wdr59 | 0.874866692 | -0.32788555 | 0.361 | 0.318 | 1 | 0.370715121 | 0.751135367 | 0.4 | 0.259 | 1 | 0.874866692 | 0.604000541 |
| Rrp12 | 0.370949067 | -0.584962501 | 0.16 | 0.194 | 1 | 0.740772979 | 0.868174788 | 0.2 | 0.17 | 1 | 0.740772979 | 0.604294923 |
| Stau2 | 0.617609249 | -0.232140398 | 0.328 | 0.282 | 1 | 0.371154072 | 0.909397451 | 0.5 | 0.395 | 1 | 0.617609249 | 0.604552799 |
| Bicral | 0.421143945 | -0.311397427 | 0.538 | 0.529 | 1 | 0.372341989 | -0.636714121 | 0.5 | 0.583 | 1 | 0.421143945 | 0.606045421 |
| 6530401F13Rik | 0.372363304 | 0.929610672 | 0.017 | 0.006 | 1 | 0.73075158 | 2.695993813 | 0 | 0.012 | 1 | 0.73075158 | 0.606072177 |
| Ahrr | 0.372363304 | 0.929610672 | 0.017 | 0.006 | 1 | 0.669850692 | 1.017921908 | 0 | 0.019 | 1 | 0.669850692 | 0.606072177 |
| Spef2 | 0.372363304 | 0.929610672 | 0.017 | 0.006 | 1 | 0.669843604 | 1.695993813 | 0 | 0.019 | 1 | 0.669843604 | 0.606072177 |
| Gm29050 | 0.372363304 | 0.929610672 | 0.017 | 0.006 | 1 | 0.730749233 | 2.432959407 | 0 | 0.012 | 1 | 0.730749233 | 0.606072177 |
| Rxylt1 | 0.44507037 | 0.198931341 | 0.723 | 0.665 | 1 | 0.373154802 | -0.663902132 | 0.7 | 0.633 | 1 | 0.44507037 | 0.607065098 |
| Jmjd8 | 0.477184611 | -0.137503524 | 0.185 | 0.224 | 1 | 0.373582621 | -0.374395515 | 0.1 | 0.213 | 1 | 0.477184611 | 0.607601268 |
| BC049715 | 0.812015907 | -0.485426827 | 0.076 | 0.082 | 1 | 0.373655406 | 2.14345279 | 0.1 | 0.043 | 1 | 0.812015907 | 0.60769245 |
| Cacna2d3 | 0.374088349 | -1.424026283 | 0.084 | 0.112 | 1 | 0.96670128 | 0.295455884 | 0.1 | 0.099 | 1 | 0.96670128 | 0.608234605 |
| Dnmbp | 0.456462022 | 0.402363669 | 0.445 | 0.406 | 1 | 0.374156083 | 0.714935377 | 0.6 | 0.568 | 1 | 0.456462022 | 0.608319391 |
| Atg13 | 0.647590838 | -0.258655965 | 0.319 | 0.276 | 1 | 0.374181546 | 0.383715888 | 0.5 | 0.315 | 1 | 0.647590838 | 0.608351263 |
| Rnf114 | 0.915322807 | -0.199672345 | 0.387 | 0.365 | 1 | 0.374230287 | 0.111031312 | 0.3 | 0.46 | 1 | 0.915322807 | 0.608412266 |
| Thtpa | 0.651414669 | -0.600904045 | 0.042 | 0.053 | 1 | 0.374430929 | 0.263034406 | 0 | 0.074 | 1 | 0.651414669 | 0.608663338 |
| Camk1 | 0.833226962 | -0.111865964 | 0.697 | 0.641 | 1 | 0.374447962 | -0.291933355 | 0.4 | 0.645 | 1 | 0.833226962 | 0.608684648 |
| Zswim1 | 0.736440063 | -0.333423734 | 0.034 | 0.041 | 1 | 0.374479548 | 0.263034406 | 0 | 0.074 | 1 | 0.736440063 | 0.608724164 |
| Gm3235 | 0.964644347 | 0.251538767 | 0.017 | 0.018 | 1 | 0.374523737 | 0.159940913 | 0 | 0.074 | 1 | 0.964644347 | 0.608779444 |
| Gm14320 | 0.374548525 | 0.514573173 | 0.017 | 0.006 | 1 | 0.669848329 | 1.695993813 | 0 | 0.019 | 1 | 0.669848329 | 0.608810452 |
| Pcdh15 | 0.374548525 | 0.514573173 | 0.017 | 0.006 | 1 | 0.73075158 | 2.695993813 | 0 | 0.012 | 1 | 0.73075158 | 0.608810452 |
| Slc26a10 | 0.374548525 | 0.514573173 | 0.017 | 0.006 | 1 | 0.643842184 | 1.695993813 | 0 | 0.022 | 1 | 0.643842184 | 0.608810452 |
| Gm49291 | 0.374548525 | 0.514573173 | 0.017 | 0.006 | 1 | 0.73075158 | 2.695993813 | 0 | 0.012 | 1 | 0.73075158 | 0.608810452 |
| Arhgef33 | 0.374548525 | 0.514573173 | 0.017 | 0.006 | 1 | 0.539483006 | 1.31748219 | 0 | 0.037 | 1 | 0.539483006 | 0.608810452 |
| Flacc1 | 0.374550451 | 0.514573173 | 0.017 | 0.006 | 1 | 0.730749233 | 1.695993813 | 0 | 0.012 | 1 | 0.730749233 | 0.608812862 |
| Cyp4a12b | 0.374550451 | 0.707218251 | 0.017 | 0.006 | 1 | 0.505686333 | 1.111031312 | 0 | 0.043 | 1 | 0.505686333 | 0.608812862 |
| Hap1 | 0.374550451 | 0.514573173 | 0.017 | 0.006 | 1 | 0.698532411 | 2.017921908 | 0 | 0.015 | 1 | 0.698532411 | 0.608812862 |
| Irf7 | 0.903509213 | 0.877143252 | 0.059 | 0.065 | 1 | 0.374563499 | -0.111361109 | 0 | 0.074 | 1 | 0.903509213 | 0.608829183 |
| Gm43330 | 0.926217033 | -0.333423734 | 0.034 | 0.035 | 1 | 0.374600373 | -0.111361109 | 0 | 0.074 | 1 | 0.926217033 | 0.608875307 |

| | | | | | | | | | | | | |
|---|---|---|---|---|---|---|---|---|---|---|---|---|
| Ctc1 | 0.430352465 | -0.807354922 | 0.059 | 0.082 | 1 | 0.374660857 | -0.152003093 | 0 | 0.074 | 1 | 0.430352465 | 0.608950956 |
| Isg15 | 0.797879015 | 0.850176205 | 0.067 | 0.059 | 1 | 0.374785995 | -2.399930607 | 0 | 0.074 | 1 | 0.797879015 | 0.609107447 |
| Dyrk3 | 0.37535919 | 1.514573173 | 0.042 | 0.024 | 1 | 0.698534498 | 2.432959407 | 0 | 0.015 | 1 | 0.698534498 | 0.609823859 |
| Mutyh | 0.37535919 | 1.099535674 | 0.042 | 0.024 | 1 | 0.669853054 | 2.210566986 | 0 | 0.019 | 1 | 0.669853054 | 0.609823859 |
| Zc2hc1c | 0.375395897 | 0.777607579 | 0.042 | 0.024 | 1 | 0.518235124 | 1.769994395 | 0.1 | 0.052 | 1 | 0.518235124 | 0.609869714 |
| Slc7a8 | 0.375395897 | 0.777607579 | 0.042 | 0.024 | 1 | 0.564719077 | 1.263034406 | 0.1 | 0.056 | 1 | 0.564719077 | 0.609869714 |
| Ift81 | 0.486198971 | -0.349365278 | 0.294 | 0.318 | 1 | 0.376052848 | -0.959358016 | 0.2 | 0.306 | 1 | 0.486198971 | 0.610689952 |
| Tada2b | 0.441807137 | -0.276840205 | 0.193 | 0.229 | 1 | 0.376502903 | -0.58694015 | 0.2 | 0.315 | 1 | 0.441807137 | 0.61125137 |
| Gmeb2 | 0.805558089 | -0.214124805 | 0.403 | 0.406 | 1 | 0.377540591 | 0.197742946 | 0.6 | 0.383 | 1 | 0.805558089 | 0.612544284 |
| Bmp2 | 0.406834111 | -0.649251629 | 0.462 | 0.494 | 1 | 0.377861449 | -1.126736335 | 0.5 | 0.623 | 1 | 0.406834111 | 0.612943624 |
| Gm15952 | 0.378629058 | -0.669851398 | 0.076 | 0.106 | 1 | 0.781749978 | 0.769994395 | 0.1 | 0.074 | 1 | 0.781749978 | 0.613898153 |
| Gramd1c | 0.729578475 | -0.762267033 | 0.185 | 0.188 | 1 | 0.378847889 | 0.295455884 | 0.1 | 0.244 | 1 | 0.729578475 | 0.614170055 |
| Hspbap1 | 0.379475942 | -0.519915204 | 0.319 | 0.329 | 1 | 0.888919856 | 0.672147071 | 0.3 | 0.401 | 1 | 0.888919856 | 0.614949894 |
| Ttc17 | 0.381011498 | -0.428201958 | 0.479 | 0.471 | 1 | 0.926961716 | -0.212375711 | 0.7 | 0.617 | 1 | 0.926961716 | 0.616853234 |
| Fcrla | 0.381775243 | -0.600904045 | 0.042 | 0.024 | 1 | 0.669843604 | 2.017921908 | 0 | 0.019 | 1 | 0.669843604 | 0.61779815 |
| Gm3830 | 0.381775243 | 0.514573173 | 0.042 | 0.024 | 1 | 0.669843604 | 1.847996907 | 0 | 0.019 | 1 | 0.669843604 | 0.61779815 |
| Mfsd13a | 0.381775243 | 0.292180751 | 0.042 | 0.024 | 1 | 0.539456304 | 0.625604485 | 0 | 0.037 | 1 | 0.539456304 | 0.61779815 |
| Gm13375 | 0.382348836 | 0.152003093 | 0.21 | 0.165 | 1 | 0.792439835 | 0.403212064 | 0.1 | 0.127 | 1 | 0.792439835 | 0.61850704 |
| Gdi1 | 0.382376402 | -0.168953162 | 0.521 | 0.524 | 1 | 0.663418668 | -0.127755547 | 0.5 | 0.602 | 1 | 0.663418668 | 0.618541091 |
| Glyr1 | 0.384747041 | -0.353829944 | 0.798 | 0.747 | 1 | 0.382387539 | -0.296393003 | 0.8 | 0.852 | 1 | 0.384747041 | 0.618554848 |
| Paxx | 0.864261766 | -0.167686529 | 0.294 | 0.271 | 1 | 0.382567243 | -0.441509711 | 0.1 | 0.21 | 1 | 0.864261766 | 0.618776791 |
| Ankrd26 | 0.399895515 | -0.758445322 | 0.16 | 0.188 | 1 | 0.382675639 | -0.505640048 | 0.1 | 0.225 | 1 | 0.399895515 | 0.618910634 |
| Xrcc2 | 0.382796898 | 1.099535674 | 0.034 | 0.018 | 1 | 0.619961444 | 1.558490289 | 0 | 0.025 | 1 | 0.619961444 | 0.619060331 |
| Mvk | 0.38282696 | -0.642968104 | 0.092 | 0.124 | 1 | 1 | 1.111031312 | 0.1 | 0.108 | 1 | 1 | 0.619097439 |
| Sptbn4 | 0.383102202 | 0.514573173 | 0.042 | 0.024 | 1 | 0.698533455 | 1.695993813 | 0 | 0.015 | 1 | 0.698533455 | 0.619437107 |
| Adipor2 | 0.403799219 | -0.170418146 | 0.58 | 0.588 | 1 | 0.383965178 | -0.191531458 | 0.6 | 0.719 | 1 | 0.403799219 | 0.620501098 |
| Mtmr11 | 0.888318362 | -0.121463512 | 0.387 | 0.376 | 1 | 0.384165588 | -0.759685671 | 0.5 | 0.537 | 1 | 0.888318362 | 0.620747976 |
| Plekho2 | 0.384679248 | -0.485426827 | 0.286 | 0.306 | 1 | 0.814498413 | 0.569461407 | 0.3 | 0.296 | 1 | 0.814498413 | 0.621380372 |
| Mpzl3 | 0.447413382 | -0.424026283 | 0.059 | 0.082 | 1 | 0.385509064 | 0.263034406 | 0 | 0.071 | 1 | 0.447413382 | 0.62240089 |
| Scamp5 | 0.813139637 | 0.192645078 | 0.084 | 0.076 | 1 | 0.385509064 | 0.263034406 | 0 | 0.071 | 1 | 0.813139637 | 0.62240089 |
| Uqcc6 | 0.952043835 | -0.255944981 | 0.118 | 0.118 | 1 | 0.385546031 | 0.159940913 | 0 | 0.071 | 1 | 0.952043835 | 0.622446321 |
| Alg12 | 0.621943472 | -0.36994961 | 0.101 | 0.118 | 1 | 0.385546826 | 0.210566986 | 0 | 0.071 | 1 | 0.621943472 | 0.622447297 |
| Snx21 | 0.385552538 | -0.515174171 | 0.168 | 0.206 | 1 | 0.81682738 | 0.625604485 | 0.3 | 0.275 | 1 | 0.81682738 | 0.622454317 |
| Cd47 | 0.480819506 | -0.278290747 | 0.815 | 0.835 | 1 | 0.385660069 | -0.236319379 | 0.9 | 0.883 | 1 | 0.480819506 | 0.622586449 |
| Gm36447 | 0.825035967 | -0.359895945 | 0.042 | 0.047 | 1 | 0.385675962 | -0.230005605 | 0 | 0.071 | 1 | 0.825035967 | 0.622605976 |
| Ntn1 | 0.957515741 | 0.548520505 | 0.067 | 0.065 | 1 | 0.385681126 | -0.408342847 | 0 | 0.071 | 1 | 0.957515741 | 0.622612321 |
| Slc16a7 | 0.560351215 | 0.152003093 | 0.042 | 0.059 | 1 | 0.385723424 | -0.596787936 | 0 | 0.071 | 1 | 0.560351215 | 0.622664289 |
| Zc3hav1 | 0.926674367 | -0.259745836 | 0.697 | 0.659 | 1 | 0.385950826 | -0.33375353 | 0.5 | 0.645 | 1 | 0.926674367 | 0.622943612 |
| Rpp30 | 0.3860242 | 0.251538767 | 0.218 | 0.176 | 1 | 0.966655593 | 0.263034406 | 0.2 | 0.185 | 1 | 0.966655593 | 0.623033717 |
| Lynx1 | 0.478001056 | 0.212010403 | 0.202 | 0.171 | 1 | 0.386140755 | 1 | 0.3 | 0.194 | 1 | 0.478001056 | 0.623176827 |
| Vasp | 0.386148545 | 0.148445274 | 0.664 | 0.635 | 1 | 0.433205904 | 0.218834602 | 0.8 | 0.809 | 1 | 0.433205904 | 0.623186391 |
| Bfar | 0.386379061 | -0.328701323 | 0.58 | 0.565 | 1 | 0.913143049 | 0.341461053 | 0.5 | 0.565 | 1 | 0.913143049 | 0.623469344 |
| Slc41a1 | 0.386590925 | -0.295885433 | 0.79 | 0.759 | 1 | 0.865966469 | -0.148181382 | 1 | 0.793 | 1 | 0.865966469 | 0.623729306 |
| Gm28501 | 0.387233514 | 0.836501268 | 0.034 | 0.018 | 1 | 0.539445308 | 0.695993813 | 0 | 0.037 | 1 | 0.539445308 | 0.624517234 |
| Cfap126 | 0.387233514 | 0.836501268 | 0.034 | 0.018 | 1 | 0.669853054 | 2.210566986 | 0 | 0.019 | 1 | 0.669853054 | 0.624517234 |
| D630039A03Rik | 0.387233514 | 0.836501268 | 0.034 | 0.018 | 1 | 0.643850855 | 2.017921908 | 0 | 0.022 | 1 | 0.643850855 | 0.624517234 |
| 1700080G18Rik | 0.387233514 | 0.836501268 | 0.034 | 0.018 | 1 | 0.577155236 | 1.210566986 | 0 | 0.031 | 1 | 0.577155236 | 0.624517234 |
| 9130023H24Rik | 0.387233514 | 0.836501268 | 0.034 | 0.018 | 1 | 0.698532411 | 1.847996907 | 0 | 0.015 | 1 | 0.698532411 | 0.624517234 |
| Gpx2 | 0.387233514 | 0.836501268 | 0.034 | 0.018 | 1 | 0.52208609 | 0.930459067 | 0 | 0.04 | 1 | 0.52208609 | 0.624517234 |
| Cdk20 | 0.387233514 | 0.836501268 | 0.034 | 0.018 | 1 | 0.643850855 | 2.017921908 | 0 | 0.022 | 1 | 0.643850855 | 0.624517234 |
| Gm49189 | 0.387233514 | 0.836501268 | 0.034 | 0.018 | 1 | 0.460551517 | 0.695993813 | 0 | 0.052 | 1 | 0.460551517 | 0.624517234 |
| A830018L16Rik | 0.387377553 | 1.251538767 | 0.025 | 0.012 | 1 | 0.619973091 | 0.625604485 | 0 | 0.025 | 1 | 0.619973091 | 0.624693737 |
| Meig1 | 0.387377553 | 1.251538767 | 0.025 | 0.012 | 1 | 0.698532411 | 1.432959407 | 0 | 0.015 | 1 | 0.698532411 | 0.624693737 |
| Gpr87 | 0.387377553 | 1.251538767 | 0.025 | 0.012 | 1 | 0.669853054 | 2.210566986 | 0 | 0.019 | 1 | 0.669853054 | 0.624693737 |
| Krtcap3 | 0.387377553 | 1.251538767 | 0.025 | 0.012 | 1 | 0.619961444 | 1.558490289 | 0 | 0.025 | 1 | 0.619961444 | 0.624693737 |
| 9130230N09Rik | 0.387377553 | 1.251538767 | 0.025 | 0.012 | 1 | 0.597845674 | 1.695993813 | 0 | 0.028 | 1 | 0.597845674 | 0.624693737 |
| Gm56649 | 0.387377553 | 1.251538767 | 0.025 | 0.012 | 1 | 0.643842184 | 1.432959407 | 0 | 0.022 | 1 | 0.643842184 | 0.624693737 |
| Slc35a5 | 0.572648461 | -0.885964757 | 0.193 | 0.206 | 1 | 0.387601025 | 0.591657153 | 0.3 | 0.176 | 1 | 0.572648461 | 0.624967495 |
| Lancl1 | 0.387950415 | 0.138009821 | 0.387 | 0.335 | 1 | 0.908795412 | 0.547601973 | 0.3 | 0.293 | 1 | 0.908795412 | 0.625395306 |
| Mrps18b | 0.495654707 | 0.124626655 | 0.361 | 0.318 | 1 | 0.388066925 | 0.708066645 | 0.4 | 0.278 | 1 | 0.495654707 | 0.625537912 |
| Slc7a14 | 0.388295116 | -0.359895945 | 0.042 | 0.024 | 1 | 0.669848329 | 1.847996907 | 0 | 0.019 | 1 | 0.669848329 | 0.625817135 |
| Npr1 | 0.388295116 | 0.292180751 | 0.042 | 0.024 | 1 | 0.522125488 | 0.558490289 | 0 | 0.04 | 1 | 0.522125488 | 0.625817135 |
| Iigp1 | 0.388330286 | 0.263034406 | 0.042 | 0.024 | 1 | 0.669853054 | -0.814968106 | 0 | 0.019 | 1 | 0.669853054 | 0.625860161 |
| Wdr91 | 0.388372449 | 0.436570661 | 0.126 | 0.094 | 1 | 0.772690975 | -0.425021588 | 0.1 | 0.127 | 1 | 0.772690975 | 0.625911739 |
| Chpf | 0.388540397 | -0.359895945 | 0.571 | 0.541 | 1 | 0.508598302 | 0.535529141 | 0.6 | 0.611 | 1 | 0.508598302 | 0.626117154 |
| Gm11630 | 0.38869288 | 0.777607579 | 0.034 | 0.018 | 1 | 0.698533977 | 1.847996907 | 0 | 0.015 | 1 | 0.698533977 | 0.626303605 |
| Gm48395 | 0.38869288 | 0.777607579 | 0.034 | 0.018 | 1 | 0.698533977 | 1.847996907 | 0 | 0.015 | 1 | 0.698533977 | 0.626303605 |
| Lcn4 | 0.388706181 | 0.836501268 | 0.034 | 0.018 | 1 | 0.619973894 | 0.159940913 | 0 | 0.025 | 1 | 0.619973894 | 0.626319867 |
| Galnt11 | 0.704864694 | -0.188225818 | 0.546 | 0.541 | 1 | 0.388891917 | 0.27838237 | 0.7 | 0.565 | 1 | 0.704864694 | 0.626546911 |
| Lsm6 | 0.388966198 | -0.250961574 | 0.622 | 0.659 | 1 | 0.591992603 | 0.470434113 | 0.5 | 0.583 | 1 | 0.591992603 | 0.626637693 |
| Socs4 | 0.389061237 | -0.263034406 | 0.454 | 0.482 | 1 | 0.634960503 | -0.378682873 | 0.6 | 0.571 | 1 | 0.634960503 | 0.626753828 |
| Gm38134 | 0.389129094 | 1.099535674 | 0.025 | 0.012 | 1 | 0.557729726 | 1.31748219 | 0 | 0.034 | 1 | 0.557729726 | 0.626836736 |
| Gm14636 | 0.389129094 | 1.099535674 | 0.025 | 0.012 | 1 | 0.397022109 | -0.152003093 | 0 | 0.068 | 1 | 0.397022109 | 0.626836736 |
| Odf2l | 0.389305491 | -0.713232745 | 0.261 | 0.276 | 1 | 0.576900743 | 0.249737583 | 0.3 | 0.207 | 1 | 0.576900743 | 0.627052217 |
| Nr2c1 | 0.652561546 | -0.584962501 | 0.235 | 0.241 | 1 | 0.389785408 | 0.695993813 | 0.4 | 0.281 | 1 | 0.652561546 | 0.627638152 |
| Sim1 | 0.390179782 | 0.707218251 | 0.034 | 0.018 | 1 | 0.619972287 | 0.847996907 | 0 | 0.025 | 1 | 0.619972287 | 0.628119302 |
| Prkar2b | 0.390227814 | -0.446952679 | 0.134 | 0.171 | 1 | 0.423008277 | -0.521236903 | 0.1 | 0.198 | 1 | 0.423008277 | 0.628177882 |
| Serhl | 0.390283558 | -0.299871174 | 0.202 | 0.247 | 1 | 0.823550025 | 0.494359952 | 0.2 | 0.167 | 1 | 0.823550025 | 0.628245861 |
| Slc2a13 | 0.390557565 | -0.61667136 | 0.286 | 0.312 | 1 | 0.77183544 | -0.84502534 | 0.3 | 0.306 | 1 | 0.77183544 | 0.628579918 |
| Isoc2b | 0.657824503 | -0.163498732 | 0.151 | 0.129 | 1 | 0.390704404 | 1.14345279 | 0.2 | 0.108 | 1 | 0.657824503 | 0.628758877 |
| 1700066M21Rik | 0.390876134 | 0.929610672 | 0.025 | 0.012 | 1 | 0.460551517 | 0.695993813 | 0 | 0.052 | 1 | 0.460551517 | 0.628968116 |
| 1010001B22Rik | 0.390876134 | 0.929610672 | 0.025 | 0.012 | 1 | 0.669853054 | 2.210566986 | 0 | 0.019 | 1 | 0.669853054 | 0.628968116 |
| Morn3 | 0.390876134 | 0.929610672 | 0.025 | 0.012 | 1 | 0.59781419 | 1.31748219 | 0 | 0.028 | 1 | 0.59781419 | 0.628968116 |
| Limk1 | 0.390876134 | 0.929610672 | 0.025 | 0.012 | 1 | 0.401608067 | 1.930459067 | 0.1 | 0.043 | 1 | 0.401608067 | 0.628968116 |
| Gm34121 | 0.390876134 | 0.929610672 | 0.025 | 0.012 | 1 | 0.730749233 | 2.210566986 | 0 | 0.012 | 1 | 0.730749233 | 0.628968116 |
| Cfap46 | 0.390876134 | 0.929610672 | 0.025 | 0.012 | 1 | 0.701889643 | 1.374065718 | 0.1 | 0.068 | 1 | 0.701889643 | 0.628968116 |
| Gm45570 | 0.390876134 | 0.929610672 | 0.025 | 0.012 | 1 | 0.446790883 | 0.625604485 | 0 | 0.056 | 1 | 0.446790883 | 0.628968116 |
| Tmem25 | 0.390876134 | 0.929610672 | 0.025 | 0.012 | 1 | 0.597824685 | 1.432959407 | 0 | 0.028 | 1 | 0.597824685 | 0.628968116 |
| Gm57399 | 0.390876134 | 0.929610672 | 0.025 | 0.012 | 1 | 0.698534498 | 2.432959407 | 0 | 0.015 | 1 | 0.698534498 | 0.628968116 |
| Dsg2 | 0.390876134 | 0.929610672 | 0.025 | 0.012 | 1 | 0.491419382 | -0.267480311 | 0.1 | 0.182 | 1 | 0.491419382 | 0.628968116 |
| Traf4 | 0.391201026 | 0.107947913 | 0.252 | 0.206 | 1 | 0.595721319 | 0.31748219 | 0.4 | 0.299 | 1 | 0.595721319 | 0.629363809 |
| Dtx2 | 0.721046538 | -0.255944981 | 0.134 | 0.147 | 1 | 0.391734711 | -0.457811523 | 0.1 | 0.207 | 1 | 0.721046538 | 0.630013338 |
| Gstm1 | 0.403908889 | 0.514573173 | 0.345 | 0.312 | 1 | 0.39204498 | -0.204470513 | 0.3 | 0.454 | 1 | 0.403908889 | 0.630390694 |
| Aasdh | 0.392143671 | -0.632268215 | 0.092 | 0.124 | 1 | 0.899701091 | 0.808468542 | 0.1 | 0.086 | 1 | 0.899701091 | 0.630510683 |
| Mydgf | 0.392227037 | -0.231243339 | 0.815 | 0.776 | 1 | 0.567556677 | 0.464949522 | 0.6 | 0.66 | 1 | 0.567556677 | 0.630612025 |
| Zfp758 | 0.392289357 | -0.552541023 | 0.16 | 0.194 | 1 | 0.498363911 | -0.339630097 | 0.1 | 0.179 | 1 | 0.498363911 | 0.630687774 |
| Rpl3l | 0.392622155 | 0.836501268 | 0.025 | 0.012 | 1 | 0.698529281 | 2.017921908 | 0 | 0.015 | 1 | 0.698529281 | 0.631092153 |
| Cd247 | 0.393128968 | -0.163498732 | 0.034 | 0.018 | 1 | 0.643835681 | 1.695993813 | 0 | 0.022 | 1 | 0.643835681 | 0.631707551 |

| | | | | | | | | | | | | |
|---|---|---|---|---|---|---|---|---|---|---|---|---|
| Pam16l | 0.393128968 | 0.514573173 | 0.034 | 0.018 | 1 | 0.577142568 | 1.432959407 | 0 | 0.031 | 1 | 0.577142568 | 0.631707551 |
| Gm28529 | 0.393128968 | 0.514573173 | 0.034 | 0.018 | 1 | 0.557729726 | 1.210566986 | 0 | 0.034 | 1 | 0.557729726 | 0.631707551 |
| Fbxo9 | 0.747670847 | -0.276840205 | 0.513 | 0.494 | 1 | 0.393630147 | -0.536666944 | 0.4 | 0.491 | 1 | 0.747670847 | 0.632315602 |
| 9530062K07Rik | 0.394377538 | 0.514573173 | 0.025 | 0.012 | 1 | 0.643850855 | 2.017921908 | 0 | 0.022 | 1 | 0.643850855 | 0.633221434 |
| Fam3d | 0.394382146 | 0.777607579 | 0.025 | 0.012 | 1 | 0.433541956 | 0.625604485 | 0 | 0.059 | 1 | 0.433541956 | 0.633227015 |
| Psmg2 | 0.650899003 | -0.171318237 | 0.412 | 0.435 | 1 | 0.39492778 | 0.748461233 | 0.4 | 0.296 | 1 | 0.650899003 | 0.633887609 |
| Rraga | 0.395329094 | -0.273922722 | 0.672 | 0.659 | 1 | 0.477673755 | -0.102501132 | 0.5 | 0.682 | 1 | 0.477673755 | 0.634373095 |
| Mal | 0.396130756 | 0.514573173 | 0.025 | 0.012 | 1 | 0.597838178 | 0.930459067 | 0 | 0.028 | 1 | 0.597838178 | 0.635341936 |
| Armh1 | 0.396130756 | 0.514573173 | 0.025 | 0.012 | 1 | 0.73075158 | 2.695993813 | 0 | 0.012 | 1 | 0.73075158 | 0.635341936 |
| Terb1 | 0.396130756 | 0.514573173 | 0.025 | 0.012 | 1 | 0.698534498 | 2.432959407 | 0 | 0.015 | 1 | 0.698534498 | 0.635341936 |
| D630024D03Rik | 0.396130756 | 0.514573173 | 0.025 | 0.012 | 1 | 0.59781419 | 1.432959407 | 0 | 0.028 | 1 | 0.59781419 | 0.635341936 |
| Gm6566 | 0.396130756 | 0.514573173 | 0.025 | 0.012 | 1 | 0.669843604 | 1.847996907 | 0 | 0.019 | 1 | 0.669843604 | 0.635341936 |
| Gm56488 | 0.396130756 | 0.514573173 | 0.025 | 0.012 | 1 | 0.64384652 | 1.111031312 | 0 | 0.022 | 1 | 0.64384652 | 0.635341936 |
| Slc22a21 | 0.396468723 | 0.215012891 | 0.361 | 0.294 | 1 | 0.634716965 | 0.135278859 | 0.3 | 0.21 | 1 | 0.634716965 | 0.635749998 |
| Zbtb7b | 0.396544791 | -0.532732542 | 0.109 | 0.141 | 1 | 0.571244285 | 0.808468542 | 0.3 | 0.247 | 1 | 0.571244285 | 0.635841811 |
| Per1 | 0.396795942 | -0.336563441 | 0.311 | 0.341 | 1 | 0.842859066 | 0.463333056 | 0.4 | 0.37 | 1 | 0.842859066 | 0.636144865 |
| Elac1 | 0.555219608 | -0.485426827 | 0.076 | 0.094 | 1 | 0.396918414 | 0.210566986 | 0 | 0.068 | 1 | 0.555219608 | 0.636292601 |
| Ankrd6 | 0.491812123 | -1.070389328 | 0.017 | 0.029 | 1 | 0.396919031 | 0.159940913 | 0 | 0.068 | 1 | 0.491812123 | 0.636293345 |
| Tmem167b | 0.396967537 | -0.174067974 | 0.521 | 0.535 | 1 | 0.58849542 | -0.195857383 | 0.5 | 0.571 | 1 | 0.58849542 | 0.636351849 |
| Gsap | 0.600299676 | -0.847996907 | 0.042 | 0.029 | 1 | 0.397056868 | -0.505640048 | 0 | 0.068 | 1 | 0.600299676 | 0.636459579 |
| Sntb1 | 0.787038236 | 0.543142325 | 0.067 | 0.076 | 1 | 0.397077021 | -0.864721141 | 0 | 0.068 | 1 | 0.787038236 | 0.636483881 |
| Eola1 | 0.587093033 | -0.244418728 | 0.193 | 0.218 | 1 | 0.397457042 | 1.31748219 | 0.3 | 0.204 | 1 | 0.587093033 | 0.636941984 |
| Zdhhc13 | 0.760148815 | 0.292180751 | 0.151 | 0.141 | 1 | 0.397569712 | 1.048295557 | 0.2 | 0.108 | 1 | 0.760148815 | 0.637077748 |
| Abcd1 | 0.794328197 | 0.432111013 | 0.084 | 0.076 | 1 | 0.397577919 | 0.988174565 | 0.2 | 0.108 | 1 | 0.794328197 | 0.637087636 |
| ENSMUSG00000121445 | 0.397773438 | 0.736965594 | 0.084 | 0.059 | 1 | 0.857092893 | 1.159940913 | 0.1 | 0.083 | 1 | 0.857092893 | 0.637323168 |
| Adam9 | 0.398036777 | -0.398316163 | 0.571 | 0.565 | 1 | 0.977804094 | 0.32277649 | 0.6 | 0.586 | 1 | 0.977804094 | 0.637640278 |
| Spacdr | 0.656590986 | -0.122856748 | 0.622 | 0.612 | 1 | 0.398171961 | 0.185031894 | 0.9 | 0.713 | 1 | 0.656590986 | 0.637803012 |
| Znrf1 | 0.39818483 | 0.156679581 | 0.714 | 0.606 | 1 | 0.558383665 | -0.136333637 | 0.6 | 0.71 | 1 | 0.558383665 | 0.637818501 |
| Brat1 | 0.398253428 | -0.707819249 | 0.042 | 0.065 | 1 | 0.420904619 | 0.432959407 | 0 | 0.062 | 1 | 0.420904619 | 0.637901063 |
| Nudt16 | 0.398253428 | -0.707819249 | 0.042 | 0.065 | 1 | 0.557744661 | 1.210566986 | 0 | 0.034 | 1 | 0.557744661 | 0.637901063 |
| Rapgef5 | 0.398337465 | -0.925999419 | 0.042 | 0.065 | 1 | 0.446923566 | -0.888968688 | 0 | 0.056 | 1 | 0.446923566 | 0.638002194 |
| Lmbrd2 | 0.420020337 | -0.566726929 | 0.429 | 0.435 | 1 | 0.398493551 | 0.147557188 | 0.6 | 0.401 | 1 | 0.420020337 | 0.638189991 |
| Slc12a9 | 0.398791469 | 0.370183263 | 0.126 | 0.094 | 1 | 0.772482357 | 0.263034406 | 0.1 | 0.13 | 1 | 0.772482357 | 0.638548302 |
| Gm19412 | 0.450128493 | -0.402964667 | 0.059 | 0.082 | 1 | 0.398985336 | 1.930459067 | 0.1 | 0.043 | 1 | 0.450128493 | 0.638781374 |
| Rtn2 | 0.958744754 | 0.344648171 | 0.042 | 0.041 | 1 | 0.398985336 | 1.558490289 | 0.1 | 0.043 | 1 | 0.958744754 | 0.638781374 |
| Sdk2 | 0.469834719 | -0.125251263 | 0.992 | 0.929 | 1 | 0.399093609 | 0.237750087 | 1 | 0.966 | 1 | 0.469834719 | 0.63891151 |
| Gm31224 | 0.399120535 | 0.251538767 | 0.034 | 0.018 | 1 | 0.73075158 | 2.695993813 | 0 | 0.012 | 1 | 0.73075158 | 0.638943868 |
| Gm41760 | 0.399120535 | 0.251538767 | 0.034 | 0.018 | 1 | 0.669843604 | 2.017921908 | 0 | 0.019 | 1 | 0.669843604 | 0.638943868 |
| Irx3 | 0.966368454 | -0.142539114 | 0.462 | 0.459 | 1 | 0.3992204 | 1.376204629 | 0.5 | 0.429 | 1 | 0.966368454 | 0.639063872 |
| Ccdc77 | 0.946331675 | -0.352160296 | 0.244 | 0.235 | 1 | 0.39924939 | 1.099058671 | 0.3 | 0.207 | 1 | 0.946331675 | 0.639098704 |
| Kat2b | 0.531063586 | -0.240314329 | 0.277 | 0.306 | 1 | 0.399322154 | -0.579864633 | 0.3 | 0.454 | 1 | 0.531063586 | 0.639186126 |
| Dhcr7 | 0.399382632 | 0.57870351 | 0.21 | 0.171 | 1 | 0.727839674 | 0.159940913 | 0.1 | 0.136 | 1 | 0.727839674 | 0.639258778 |
| Gnptab | 0.399672998 | 0.147790842 | 0.37 | 0.318 | 1 | 0.968103701 | 0.700810829 | 0.4 | 0.389 | 1 | 0.968103701 | 0.63960749 |
| Nprl3 | 0.638237046 | 0.132702537 | 0.193 | 0.171 | 1 | 0.399901406 | 0.769994395 | 0.3 | 0.191 | 1 | 0.638237046 | 0.639881677 |
| Rab8b | 0.40038951 | -0.492694828 | 0.437 | 0.471 | 1 | 0.798143 | 0.345496566 | 0.5 | 0.571 | 1 | 0.798143 | 0.640467261 |
| Bbs9 | 0.401026349 | -0.710393192 | 0.429 | 0.429 | 1 | 0.553050107 | -0.668578616 | 0.6 | 0.639 | 1 | 0.553050107 | 0.641230566 |
| Ankmy1 | 0.405066831 | -0.567888987 | 0.193 | 0.229 | 1 | 0.401106319 | -0.596787936 | 0.1 | 0.204 | 1 | 0.405066831 | 0.641326359 |
| Chkb | 0.401533783 | -0.419838486 | 0.303 | 0.335 | 1 | 0.401111395 | -0.484578433 | 0.2 | 0.312 | 1 | 0.401533783 | 0.641332439 |
| C1ql3 | 0.401446423 | 0.192645078 | 0.025 | 0.012 | 1 | 0.597824685 | 1.31748219 | 0 | 0.028 | 1 | 0.597824685 | 0.641733615 |
| 2810405F17Rik | 0.462720534 | 0.514573173 | 0.067 | 0.047 | 1 | 0.401608067 | 1.847996907 | 0.1 | 0.043 | 1 | 0.462720534 | 0.641927094 |
| Gm45435 | 0.695893386 | -0.222392421 | 0.017 | 0.024 | 1 | 0.401608067 | 1.930459067 | 0.1 | 0.043 | 1 | 0.695893386 | 0.641927094 |
| Crocc | 0.971938394 | -0.263034406 | 0.042 | 0.041 | 1 | 0.401608067 | 1.847996907 | 0.1 | 0.043 | 1 | 0.971938394 | 0.641927094 |
| Mpp1 | 0.624346587 | -0.176098769 | 0.336 | 0.359 | 1 | 0.401810354 | -0.567040593 | 0.3 | 0.42 | 1 | 0.624346587 | 0.642169147 |
| Polr3gl | 0.402544288 | -0.228651412 | 0.311 | 0.347 | 1 | 0.481799279 | -0.49777993 | 0.3 | 0.373 | 1 | 0.481799279 | 0.643046673 |
| Nek8 | 0.40257437 | 0.292180751 | 0.202 | 0.165 | 1 | 0.776742783 | 0.515421567 | 0.2 | 0.16 | 1 | 0.776742783 | 0.643082616 |
| Clu | 0.770804645 | -0.628621463 | 0.563 | 0.524 | 1 | 0.403081586 | -0.407571352 | 0.6 | 0.84 | 1 | 0.770804645 | 0.643688407 |
| Rgp1 | 0.403104811 | 0.450442835 | 0.151 | 0.118 | 1 | 0.90271652 | 0.591657153 | 0.1 | 0.111 | 1 | 0.90271652 | 0.643716133 |
| Cox19 | 0.780261209 | -0.118887846 | 0.471 | 0.429 | 1 | 0.40320162 | -0.339630097 | 0.3 | 0.435 | 1 | 0.780261209 | 0.643831693 |
| Tmem185b | 0.637587543 | 0.185265548 | 0.269 | 0.247 | 1 | 0.403209339 | 1.036069255 | 0.3 | 0.201 | 1 | 0.637587543 | 0.643840906 |
| Gabrr2 | 0.403273565 | -0.807354922 | 0.042 | 0.065 | 1 | 0.849316227 | 0.591657153 | 0.1 | 0.08 | 1 | 0.849316227 | 0.643917561 |
| Gm57395 | 0.719955107 | 0.152003093 | 0.05 | 0.041 | 1 | 0.4042367 | 1.695993813 | 0.1 | 0.043 | 1 | 0.719955107 | 0.64506609 |
| Gpr157 | 0.687345921 | -0.707819249 | 0.017 | 0.024 | 1 | 0.404236993 | 1.847996907 | 0.1 | 0.043 | 1 | 0.687345921 | 0.64506644 |
| Cbarp | 0.954551343 | -0.222392421 | 0.017 | 0.018 | 1 | 0.404236993 | 1.695993813 | 0.1 | 0.043 | 1 | 0.954551343 | 0.64506644 |
| Anxa4 | 0.812505157 | -0.213880921 | 0.748 | 0.653 | 1 | 0.404600571 | 0.474890088 | 0.7 | 0.701 | 1 | 0.812505157 | 0.64549952 |
| Cnot9 | 0.584188737 | -0.173482821 | 0.235 | 0.265 | 1 | 0.404829488 | 0.625604485 | 0.4 | 0.272 | 1 | 0.584188737 | 0.645772062 |
| 6430548M08Rik | 0.404923304 | -1.008988783 | 0.059 | 0.082 | 1 | 0.500552544 | 1.054447784 | 0.2 | 0.13 | 1 | 0.500552544 | 0.645883726 |
| Tor1a | 0.405041054 | -0.299871174 | 0.37 | 0.412 | 1 | 0.689994846 | 0.688798312 | 0.4 | 0.398 | 1 | 0.689994846 | 0.646023853 |
| Nop58 | 0.453540615 | -0.236747714 | 0.966 | 0.924 | 1 | 0.406660205 | -0.505640048 | 0.9 | 0.914 | 1 | 0.453540615 | 0.647947887 |
| Cyld | 0.406733591 | -0.560194596 | 0.303 | 0.318 | 1 | 0.518854295 | 0.362570079 | 0.6 | 0.481 | 1 | 0.518854295 | 0.648034968 |
| Plekha3 | 0.900177896 | -0.208717484 | 0.622 | 0.576 | 1 | 0.406809087 | -0.389042291 | 0.4 | 0.534 | 1 | 0.900177896 | 0.64812454 |
| Gas2 | 0.769211662 | 0.321928095 | 0.042 | 0.035 | 1 | 0.406871237 | 1.558490289 | 0.1 | 0.043 | 1 | 0.769211662 | 0.64819827 |
| Cd248 | 0.508731273 | -0.807354922 | 0.008 | 0.018 | 1 | 0.406872117 | 1.625604485 | 0.1 | 0.043 | 1 | 0.508731273 | 0.648199314 |
| Cyp4f13 | 0.672580325 | -0.392317423 | 0.126 | 0.141 | 1 | 0.406914957 | 1.159940913 | 0.2 | 0.117 | 1 | 0.672580325 | 0.648250132 |
| Fis1 | 0.406978595 | 0.118272236 | 0.891 | 0.812 | 1 | 0.684745783 | 0.267315403 | 0.7 | 0.84 | 1 | 0.684745783 | 0.648325614 |
| D430041D05Rik | 0.407016556 | 0.633217669 | 0.084 | 0.059 | 1 | 0.50699936 | -0.267480311 | 0.1 | 0.049 | 1 | 0.50699936 | 0.648370635 |
| Apex2 | 0.66470915 | -0.110387396 | 0.445 | 0.406 | 1 | 0.407713949 | 0.210566986 | 0.5 | 0.33 | 1 | 0.66470915 | 0.649197233 |
| Erbb2 | 0.452168881 | -0.600904045 | 0.092 | 0.118 | 1 | 0.408420126 | 0.789103218 | 0.3 | 0.185 | 1 | 0.452168881 | 0.650033253 |
| 1500015A07Rik | 0.408443615 | -0.584962501 | 0.042 | 0.065 | 1 | 0.474904369 | 0.695993813 | 0 | 0.049 | 1 | 0.474904369 | 0.650061044 |
| Cyrib | 0.935993062 | -0.539874611 | 0.395 | 0.365 | 1 | 0.408501559 | 0.975147884 | 0.6 | 0.565 | 1 | 0.935993062 | 0.650129594 |
| Gm14327 | 0.408579016 | -0.485426827 | 0.042 | 0.065 | 1 | 0.505623793 | 0.847996907 | 0 | 0.043 | 1 | 0.505623793 | 0.65022122 |
| Slc25a35 | 0.462720534 | 0.514573173 | 0.067 | 0.047 | 1 | 0.408596393 | 0.494359952 | 0 | 0.065 | 1 | 0.462720534 | 0.650241774 |
| 4930509H03Rik | 0.437408619 | -0.485426827 | 0.034 | 0.053 | 1 | 0.408636939 | 0.432959407 | 0 | 0.065 | 1 | 0.437408619 | 0.65028973 |
| Sema3b | 0.82670638 | -0.292781749 | 0.025 | 0.029 | 1 | 0.408636939 | 0.374065718 | 0 | 0.065 | 1 | 0.82670638 | 0.65028973 |
| Pard6a | 0.931091114 | -0.177304532 | 0.092 | 0.094 | 1 | 0.408636939 | 0.432959407 | 0 | 0.065 | 1 | 0.931091114 | 0.65028973 |
| Gm37083 | 0.468235759 | -0.485426827 | 0.076 | 0.1 | 1 | 0.40867321 | 0.31748219 | 0 | 0.065 | 1 | 0.468235759 | 0.650332627 |
| Krt1 | 0.933394757 | -0.163498732 | 0.034 | 0.035 | 1 | 0.40867321 | 0.31748219 | 0 | 0.065 | 1 | 0.933394757 | 0.650332627 |
| Zbtb7c | 0.408711642 | -0.288125387 | 0.63 | 0.541 | 1 | 0.951765977 | 0.534106131 | 0.3 | 0.395 | 1 | 0.951765977 | 0.650378077 |
| Evpl | 0.613389487 | 0.13606155 | 0.067 | 0.053 | 1 | 0.408732936 | 0.210566986 | 0 | 0.065 | 1 | 0.613389487 | 0.650403259 |
| Pcdhgb4 | 0.619563144 | -0.185866545 | 0.05 | 0.065 | 1 | 0.408781983 | -0.111361109 | 0 | 0.065 | 1 | 0.619563144 | 0.650461256 |
| Adssl1 | 0.853671961 | 0.614108846 | 0.059 | 0.065 | 1 | 0.408781983 | -0.596787936 | 0 | 0.065 | 1 | 0.853671961 | 0.650461256 |
| Nudcd2 | 0.409028791 | -0.354182294 | 0.487 | 0.506 | 1 | 0.88539574 | 0.368306449 | 0.4 | 0.441 | 1 | 0.88539574 | 0.65075303 |
| Epdr1 | 0.409093747 | -0.685099172 | 0.361 | 0.371 | 1 | 0.863737736 | 0.290001453 | 0.2 | 0.231 | 1 | 0.863737736 | 0.6508298 |
| Cebpb | 0.409438104 | 0.102871386 | 0.824 | 0.871 | 1 | 0.668418268 | -0.497390161 | 0.9 | 0.92 | 1 | 0.668418268 | 0.651236646 |
| Pithd1 | 0.669167827 | 0.227692025 | 0.328 | 0.312 | 1 | 0.409644168 | -0.408342847 | 0.2 | 0.312 | 1 | 0.669167827 | 0.651479992 |
| 9530052E02Rik | 0.410271646 | 0.455679484 | 0.151 | 0.118 | 1 | 0.974376691 | -0.111361109 | 0.1 | 0.099 | 1 | 0.974376691 | 0.652220468 |
| Cep57 | 0.410381977 | -0.504042505 | 0.42 | 0.435 | 1 | 0.834042909 | -0.158666824 | 0.4 | 0.41 | 1 | 0.834042909 | 0.652350587 |
| Pih1d1 | 0.781793791 | -0.45845978 | 0.21 | 0.212 | 1 | 0.410618379 | -0.425021588 | 0.1 | 0.201 | 1 | 0.781793791 | 0.652629305 |
| Rangrf | 0.411060987 | -0.452259963 | 0.16 | 0.194 | 1 | 0.656993059 | 0.135278859 | 0.1 | 0.148 | 1 | 0.656993059 | 0.653150839 |

| | | | | | | | | | | | | |
|---|---|---|---|---|---|---|---|---|---|---|---|---|
| Mtfr1 | 0.730752997 | -0.100762977 | 0.58 | 0.518 | 1 | 0.411410824 | -0.514573173 | 0.5 | 0.556 | 1 | 0.730752997 | 0.653562782 |
| Trib3 | 0.411860105 | -0.544320516 | 0.067 | 0.094 | 1 | 0.495850843 | 1.31748219 | 0.1 | 0.049 | 1 | 0.495850843 | 0.654091464 |
| Gm33046 | 0.450638771 | -1.292781749 | 0.025 | 0.041 | 1 | 0.412163295 | 1.263034406 | 0.1 | 0.043 | 1 | 0.450638771 | 0.654448008 |
| Xlr3b | 0.450629499 | -1.207892852 | 0.067 | 0.088 | 1 | 0.412163587 | 1.374065718 | 0.1 | 0.043 | 1 | 0.450629499 | 0.654448352 |
| Gm6225 | 0.412706773 | -0.116782233 | 0.277 | 0.224 | 1 | 0.432316013 | 0.95491211 | 0.3 | 0.194 | 1 | 0.432316013 | 0.655086666 |
| Arid1b | 0.412812449 | -0.225257755 | 0.924 | 0.912 | 1 | 0.84249363 | -0.212320963 | 1 | 0.972 | 1 | 0.84249363 | 0.65521078 |
| Nln | 0.413058021 | -0.655351829 | 0.101 | 0.129 | 1 | 0.591309503 | 0.172431857 | 0.1 | 0.176 | 1 | 0.591309503 | 0.655499113 |
| Nipsnap3b | 0.483444595 | -0.359895945 | 0.176 | 0.206 | 1 | 0.41320752 | -0.536666944 | 0.1 | 0.201 | 1 | 0.483444595 | 0.655674586 |
| Pigf | 0.413288344 | 0.133483005 | 0.319 | 0.265 | 1 | 0.531669493 | 0.791853829 | 0.4 | 0.315 | 1 | 0.531669493 | 0.655769433 |
| Dhdds | 0.935280994 | -0.122856748 | 0.403 | 0.371 | 1 | 0.413327249 | 0.453137289 | 0.4 | 0.272 | 1 | 0.935280994 | 0.655815083 |
| Tceanc2 | 0.821317136 | -0.14839184 | 0.227 | 0.206 | 1 | 0.413751642 | 0.409112665 | 0.4 | 0.256 | 1 | 0.821317136 | 0.656312862 |
| Golga7 | 0.413872763 | -0.272578179 | 0.664 | 0.635 | 1 | 0.762093805 | 0.174225139 | 0.7 | 0.63 | 1 | 0.762093805 | 0.656454862 |
| 4930517O19Rik | 0.413944087 | -0.453717967 | 0.134 | 0.165 | 1 | 0.992558317 | 0.667424661 | 0.2 | 0.213 | 1 | 0.992558317 | 0.656538467 |
| Cd99l2 | 0.414395702 | -0.292781749 | 0.294 | 0.324 | 1 | 0.701153283 | 0.391482771 | 0.5 | 0.41 | 1 | 0.701153283 | 0.657067607 |
| Mrpl28 | 0.824125499 | -0.179800545 | 0.403 | 0.388 | 1 | 0.41449057 | 0.190102883 | 0.2 | 0.355 | 1 | 0.824125499 | 0.657178707 |
| Vwa8 | 0.41528737 | -0.277621942 | 0.37 | 0.412 | 1 | 0.925816021 | -0.489872732 | 0.5 | 0.457 | 1 | 0.925816021 | 0.658111141 |
| Glt8d2 | 0.415880845 | -0.453717967 | 0.294 | 0.324 | 1 | 0.437821204 | 0.808468542 | 0.3 | 0.191 | 1 | 0.437821204 | 0.658804813 |
| Tmem42 | 0.416034171 | -0.281068329 | 0.336 | 0.371 | 1 | 0.661524627 | 0.227844977 | 0.4 | 0.309 | 1 | 0.661524627 | 0.65898391 |
| Slc29a3 | 0.416591855 | 0.186998515 | 0.328 | 0.282 | 1 | 0.482538662 | 0.385653692 | 0.4 | 0.269 | 1 | 0.482538662 | 0.659634936 |
| Batf3 | 0.416774963 | 0.321928095 | 0.109 | 0.082 | 1 | 0.607555216 | 1.263034406 | 0.1 | 0.059 | 1 | 0.607555216 | 0.659848556 |
| 5730414N17Rik | 0.417007668 | 0.273565073 | 0.084 | 0.059 | 1 | 0.474877085 | 0.769994395 | 0 | 0.049 | 1 | 0.474877085 | 0.660119941 |
| Ciao3 | 0.417007668 | 0.389042291 | 0.084 | 0.059 | 1 | 0.89158461 | 0.888638891 | 0.1 | 0.086 | 1 | 0.89158461 | 0.660119941 |
| Ccdc71 | 0.675287596 | -0.448900951 | 0.151 | 0.165 | 1 | 0.417018665 | -0.374395515 | 0.1 | 0.201 | 1 | 0.675287596 | 0.660132762 |
| Vps37c | 0.417204061 | -0.76106127 | 0.143 | 0.171 | 1 | 0.917685867 | 0.30367639 | 0.2 | 0.222 | 1 | 0.917685867 | 0.660348894 |
| Rplp1 | 0.417361453 | -0.109862509 | 1 | 1 | 1 | 0.536204331 | -0.162193836 | 1 | 0.997 | 1 | 0.536204331 | 0.660532323 |
| Trmu | 0.417438307 | -0.889817082 | 0.126 | 0.153 | 1 | 0.89790487 | 0.345496566 | 0.1 | 0.111 | 1 | 0.89790487 | 0.660621874 |
| Rnf150 | 0.457512938 | -1.180572246 | 0.126 | 0.094 | 1 | 0.417483039 | 0.847996907 | 0.1 | 0.043 | 1 | 0.457512938 | 0.66067399 |
| Map1a | 0.424612725 | -0.276840205 | 0.294 | 0.318 | 1 | 0.418135765 | -0.195425374 | 0.3 | 0.454 | 1 | 0.424612725 | 0.661434012 |
| Tctn2 | 0.418321265 | 0.12755005 | 0.202 | 0.159 | 1 | 0.613618532 | 0.159940913 | 0.1 | 0.157 | 1 | 0.613618532 | 0.661649849 |
| Lin7a | 0.972021493 | 0.370183263 | 0.059 | 0.059 | 1 | 0.419106384 | 0.708066645 | 0.2 | 0.117 | 1 | 0.972021493 | 0.662562607 |
| Pold2 | 0.419272883 | -0.283792966 | 0.16 | 0.194 | 1 | 0.681830711 | 0.263034406 | 0.1 | 0.145 | 1 | 0.681830711 | 0.662756016 |
| Zfp653 | 0.419669733 | -0.614709844 | 0.118 | 0.147 | 1 | 0.680584459 | 0.290001453 | 0.3 | 0.225 | 1 | 0.680584459 | 0.663216781 |
| Pts | 0.419815789 | 0.18740843 | 0.336 | 0.288 | 1 | 0.732341524 | 0.345496566 | 0.3 | 0.238 | 1 | 0.732341524 | 0.663386282 |
| Prpf31 | 0.419875085 | -0.404808367 | 0.311 | 0.347 | 1 | 0.856687012 | 0.473601392 | 0.3 | 0.309 | 1 | 0.856687012 | 0.663455083 |
| Spata5 | 0.419952957 | 0.333305485 | 0.714 | 0.659 | 1 | 0.868460995 | 0.779517169 | 0.6 | 0.67 | 1 | 0.868460995 | 0.663545428 |
| Cdc42se2 | 0.419953651 | -0.149063642 | 0.496 | 0.482 | 1 | 0.43139803 | 0.50836681 | 0.7 | 0.685 | 1 | 0.43139803 | 0.663546233 |
| Hsd3b7 | 0.420457089 | 0.182729609 | 0.353 | 0.306 | 1 | 0.601087022 | -0.145308441 | 0.3 | 0.38 | 1 | 0.601087022 | 0.664130015 |
| 3110045C21Rik | 0.745529669 | 0.152003093 | 0.143 | 0.129 | 1 | 0.42083289 | 0.558490289 | 0 | 0.062 | 1 | 0.745529669 | 0.664565459 |
| Gm14440 | 0.659871503 | 0.514573173 | 0.025 | 0.018 | 1 | 0.42087075 | 0.432959407 | 0 | 0.062 | 1 | 0.659871503 | 0.664609312 |
| 1700088E04Rik | 0.853677712 | 0.292180751 | 0.076 | 0.071 | 1 | 0.42087075 | 0.374065718 | 0 | 0.062 | 1 | 0.853677712 | 0.664609312 |
| Thg1l | 0.448223156 | -0.563429339 | 0.059 | 0.082 | 1 | 0.420904398 | 0.31748219 | 0 | 0.062 | 1 | 0.448223156 | 0.664648284 |
| Armc6 | 0.563722678 | -0.622930351 | 0.034 | 0.047 | 1 | 0.420904398 | 0.374065718 | 0 | 0.062 | 1 | 0.563722678 | 0.664648284 |
| Cbr4 | 0.565914055 | -0.411426246 | 0.076 | 0.094 | 1 | 0.420904619 | 0.263034406 | 0 | 0.062 | 1 | 0.565914055 | 0.66464854 |
| Gm17024 | 0.8327876 | -0.222392421 | 0.042 | 0.047 | 1 | 0.420904619 | 0.432959407 | 0 | 0.062 | 1 | 0.8327876 | 0.66464854 |
| Zfp709 | 0.735730565 | -0.485426827 | 0.05 | 0.059 | 1 | 0.42093472 | 0.374065718 | 0 | 0.062 | 1 | 0.735730565 | 0.664683402 |
| Dnaaf4 | 0.526968887 | -0.765534746 | 0.042 | 0.059 | 1 | 0.420960391 | 0.210566986 | 0 | 0.062 | 1 | 0.526968887 | 0.664713131 |
| Gm49380 | 0.743002317 | -0.163498732 | 0.034 | 0.041 | 1 | 0.420984509 | 0.263034406 | 0 | 0.062 | 1 | 0.743002317 | 0.664741061 |
| Gm20646 | 0.574974342 | -0.378511623 | 0.034 | 0.047 | 1 | 0.420996899 | 0.159940913 | 0 | 0.062 | 1 | 0.574974342 | 0.664755409 |
| Cntn4 | 0.581554694 | -1.900464326 | 0.042 | 0.029 | 1 | 0.421047336 | -1.026472211 | 0 | 0.062 | 1 | 0.581554694 | 0.664813813 |
| Hexim2 | 0.432595587 | -0.748461233 | 0.034 | 0.053 | 1 | 0.421085155 | 0.625604485 | 0 | 0.062 | 1 | 0.432595587 | 0.664857603 |
| Glb1 | 0.421277025 | -0.55359833 | 0.227 | 0.253 | 1 | 0.896240218 | -0.449683642 | 0.3 | 0.256 | 1 | 0.896240218 | 0.665079718 |
| Cutc | 0.943859233 | -0.333423734 | 0.118 | 0.118 | 1 | 0.421724599 | 1.176619654 | 0.2 | 0.114 | 1 | 0.943859233 | 0.665597561 |
| Wasf2 | 0.422339244 | -0.225410613 | 0.95 | 0.9 | 1 | 0.504751285 | -0.227155367 | 0.8 | 0.954 | 1 | 0.504751285 | 0.66630805 |
| Arap2 | 0.925725731 | -0.304854582 | 0.067 | 0.065 | 1 | 0.423352219 | -0.473931188 | 0.1 | 0.201 | 1 | 0.925725731 | 0.667477337 |
| Zscan29 | 0.849740639 | -0.407424315 | 0.143 | 0.147 | 1 | 0.423665275 | 1.31748219 | 0.2 | 0.12 | 1 | 0.849740639 | 0.667838285 |
| Rchy1 | 0.424030945 | -0.368912818 | 0.361 | 0.388 | 1 | 0.802201382 | 0.478763097 | 0.4 | 0.392 | 1 | 0.802201382 | 0.668259648 |
| Lipo3 | 0.782855568 | -0.552541023 | 0.16 | 0.165 | 1 | 0.425626749 | -0.457811523 | 0.1 | 0.198 | 1 | 0.782855568 | 0.670095368 |
| Ero1a | 0.680654071 | -0.344070978 | 0.437 | 0.418 | 1 | 0.425661463 | -0.448052557 | 0.3 | 0.426 | 1 | 0.680654071 | 0.670135245 |
| Ngrn | 0.527822431 | -0.311397427 | 0.168 | 0.194 | 1 | 0.425871122 | -0.152003093 | 0.1 | 0.201 | 1 | 0.527822431 | 0.670376031 |
| Glrx | 0.425959366 | -0.201633861 | 0.151 | 0.188 | 1 | 0.785609776 | -0.204470513 | 0.2 | 0.225 | 1 | 0.785609776 | 0.670477351 |
| Rela | 0.803783904 | -0.147146527 | 0.639 | 0.6 | 1 | 0.426935162 | 0.43965406 | 0.7 | 0.645 | 1 | 0.803783904 | 0.671596692 |
| Hsd17b10 | 0.540528623 | -0.177304532 | 0.605 | 0.618 | 1 | 0.427238454 | -0.156448998 | 0.4 | 0.571 | 1 | 0.540528623 | 0.671944211 |
| Pdk3 | 0.485158155 | -0.190979469 | 0.387 | 0.406 | 1 | 0.427505757 | 0.453137289 | 0.5 | 0.401 | 1 | 0.485158155 | 0.672250342 |
| H4c3 | 0.427967526 | -0.748461233 | 0.034 | 0.053 | 1 | 0.522151747 | 1.210566986 | 0 | 0.04 | 1 | 0.522151747 | 0.672778848 |
| Gm26621 | 0.427967526 | -0.748461233 | 0.034 | 0.053 | 1 | 0.444833547 | 2.210566986 | 0.1 | 0.049 | 1 | 0.444833547 | 0.672778848 |
| Rdh13 | 0.427993368 | 0.762500686 | 0.109 | 0.082 | 1 | 0.893095748 | 0.263034406 | 0.1 | 0.111 | 1 | 0.893095748 | 0.672808413 |
| Gm41307 | 0.428023634 | -1.222392421 | 0.034 | 0.053 | 1 | 0.522126681 | 0.769994395 | 0 | 0.04 | 1 | 0.522126681 | 0.672843037 |
| Nsun4 | 0.428190141 | -0.442358105 | 0.218 | 0.253 | 1 | 0.56385299 | -0.291933355 | 0.2 | 0.272 | 1 | 0.56385299 | 0.673033485 |
| Lipa | 0.500248059 | -0.669851398 | 0.244 | 0.253 | 1 | 0.428601467 | -0.68251781 | 0.2 | 0.296 | 1 | 0.500248059 | 0.673503716 |
| Bbs7 | 0.755555929 | -0.339575961 | 0.185 | 0.194 | 1 | 0.428970786 | -0.668578619 | 0.1 | 0.194 | 1 | 0.755555929 | 0.673925636 |
| B9d1 | 0.429205831 | 0.212010403 | 0.218 | 0.176 | 1 | 0.631405023 | 0.185031894 | 0.1 | 0.154 | 1 | 0.631405023 | 0.674194017 |
| Ces1d | 0.495189997 | 2.152003093 | 0.025 | 0.041 | 1 | 0.429542466 | -0.654503434 | 0.1 | 0.198 | 1 | 0.495189997 | 0.674578202 |
| Ccdc50 | 0.42964608 | -0.267554603 | 0.807 | 0.759 | 1 | 0.434623358 | -0.374395515 | 1 | 0.849 | 1 | 0.434623358 | 0.674696406 |
| Tle3 | 0.429738649 | -0.292781749 | 0.16 | 0.194 | 1 | 0.653203525 | -0.274859841 | 0.3 | 0.37 | 1 | 0.653203525 | 0.674801992 |
| Vhl | 0.430043086 | -0.569491092 | 0.193 | 0.224 | 1 | 0.466178535 | -0.441509711 | 0.2 | 0.296 | 1 | 0.466178535 | 0.675149116 |
| Gtpbp8 | 0.430175036 | -0.59645814 | 0.193 | 0.218 | 1 | 0.736126198 | -0.242605642 | 0.2 | 0.231 | 1 | 0.736126198 | 0.67529951 |
| Carhsp1 | 0.430256636 | 0.415037499 | 0.336 | 0.306 | 1 | 0.606240231 | -0.465893869 | 0.3 | 0.343 | 1 | 0.606240231 | 0.6753925 |
| Sptlc1 | 0.818729022 | -0.183771127 | 0.378 | 0.371 | 1 | 0.430411773 | -0.55950692 | 0.3 | 0.392 | 1 | 0.818729022 | 0.675569252 |
| Acsl5 | 0.468749824 | -0.58058406 | 0.143 | 0.171 | 1 | 0.431646722 | -0.473931188 | 0.1 | 0.194 | 1 | 0.468749824 | 0.676974551 |
| Jrkl | 0.697081018 | -0.375802336 | 0.269 | 0.276 | 1 | 0.431672802 | -0.581990934 | 0.1 | 0.194 | 1 | 0.697081018 | 0.677004195 |
| Hdac9 | 0.43175308 | 1.157021168 | 0.109 | 0.082 | 1 | 0.808751207 | -0.434937057 | 0.2 | 0.164 | 1 | 0.808751207 | 0.677095438 |
| Sipa1l2 | 0.43203766 | -0.292781749 | 0.277 | 0.224 | 1 | 0.696374135 | -0.433289204 | 0.2 | 0.256 | 1 | 0.696374135 | 0.67741878 |
| Gm48226 | 0.432448405 | -0.292781749 | 0.109 | 0.141 | 1 | 0.981801623 | 0.660369903 | 0.1 | 0.096 | 1 | 0.981801623 | 0.677885187 |
| Gpatch3 | 0.432595587 | -0.622930351 | 0.034 | 0.053 | 1 | 0.505639146 | 0.494359952 | 0 | 0.043 | 1 | 0.505639146 | 0.678052232 |
| Zik1 | 0.432595587 | -0.622930351 | 0.034 | 0.053 | 1 | 0.557765898 | 1.017921908 | 0 | 0.034 | 1 | 0.557765898 | 0.678052232 |
| 9130004C02Rik | 0.432595587 | -0.748461233 | 0.034 | 0.053 | 1 | 0.619953009 | 1.558490289 | 0 | 0.025 | 1 | 0.619953009 | 0.678052232 |
| Apmap | 0.433463805 | -0.380729449 | 0.361 | 0.376 | 1 | 0.554400757 | -0.167944637 | 0.3 | 0.395 | 1 | 0.554400757 | 0.679036739 |
| Mettl23 | 0.680752633 | -0.199122642 | 0.345 | 0.359 | 1 | 0.433523032 | -0.481923979 | 0.3 | 0.392 | 1 | 0.680752633 | 0.679103845 |
| Ccdc120 | 0.658336322 | -0.485426827 | 0.042 | 0.053 | 1 | 0.433608183 | 0.432959407 | 0 | 0.059 | 1 | 0.658336322 | 0.67920031 |
| 9330159M07Rik | 0.783591685 | -0.432959407 | 0.092 | 0.1 | 1 | 0.433636001 | 0.111031312 | 0 | 0.059 | 1 | 0.783591685 | 0.679231821 |
| Rsf1os1 | 0.954551343 | -0.222392421 | 0.017 | 0.018 | 1 | 0.433657839 | 0.31748219 | 0 | 0.059 | 1 | 0.954551343 | 0.679256556 |
| Gm15886 | 0.669421909 | -0.691877705 | 0.092 | 0.106 | 1 | 0.433659218 | 0.263034406 | 0 | 0.059 | 1 | 0.669421909 | 0.679258119 |
| Gm16272 | 0.487658195 | 0.273565073 | 0.059 | 0.041 | 1 | 0.433676456 | 0.263034406 | 0 | 0.059 | 1 | 0.487658195 | 0.679277644 |
| Gm14221 | 0.908150902 | -0.765534746 | 0.05 | 0.053 | 1 | 0.433696681 | -0.152003093 | 0 | 0.059 | 1 | 0.908150902 | 0.67930055 |
| 2900009J06Rik | 0.561550783 | -0.315501826 | 0.059 | 0.076 | 1 | 0.433698059 | -0.152003093 | 0 | 0.059 | 1 | 0.561550783 | 0.679302112 |
| Pthlh | 0.931797332 | 1.131244533 | 0.05 | 0.053 | 1 | 0.43370955 | -0.625934282 | 0 | 0.059 | 1 | 0.931797332 | 0.679315126 |
| 2510046G10Rik | 0.770891247 | 0.292180751 | 0.042 | 0.035 | 1 | 0.433764692 | 0.695993813 | 0 | 0.059 | 1 | 0.770891247 | 0.679377576 |
| Gm36033 | 0.433838095 | -0.600904045 | 0.034 | 0.053 | 1 | 0.858897456 | 1.280956314 | 0.1 | 0.09 | 1 | 0.858897456 | 0.679460697 |

| | | | | | | | | | | | | |
|---|---|---|---|---|---|---|---|---|---|---|---|---|
| Spata7 | 0.434215021 | -0.276840205 | 0.202 | 0.153 | 1 | 0.472143491 | -0.536666944 | 0.1 | 0.182 | 1 | 0.472143491 | 0.679887357 |
| Chmp1b | 0.4344032 | -0.644625422 | 0.21 | 0.235 | 1 | 0.862096937 | 0.31748219 | 0.3 | 0.259 | 1 | 0.862096937 | 0.68010026 |
| Cep76 | 0.435382995 | -0.485426827 | 0.134 | 0.165 | 1 | 0.758020954 | 0.290001453 | 0.3 | 0.238 | 1 | 0.758020954 | 0.681207637 |
| Hif1an | 0.690926911 | -0.448900951 | 0.151 | 0.165 | 1 | 0.435694881 | -0.391469028 | 0.1 | 0.194 | 1 | 0.690926911 | 0.681559733 |
| Mllt6 | 0.756777454 | -0.163498732 | 0.328 | 0.335 | 1 | 0.43630731 | -0.246185533 | 0.3 | 0.457 | 1 | 0.756777454 | 0.682250551 |
| Tab3 | 0.824533127 | 0.570068285 | 0.202 | 0.212 | 1 | 0.436328654 | -0.802257054 | 0.1 | 0.191 | 1 | 0.824533127 | 0.682274614 |
| Zbtb11os1 | 0.436892906 | 0.173536255 | 0.118 | 0.088 | 1 | 0.905123558 | 0.625604485 | 0.1 | 0.111 | 1 | 0.905123558 | 0.6829104 |
| Gm19605 | 0.563722678 | -0.748461233 | 0.034 | 0.047 | 1 | 0.437024249 | 2.017921908 | 0.1 | 0.046 | 1 | 0.563722678 | 0.683058304 |
| Gm56489 | 0.720666617 | 0.251538767 | 0.076 | 0.065 | 1 | 0.437024249 | 2.017921908 | 0.1 | 0.046 | 1 | 0.720666617 | 0.683058304 |
| Shroom1 | 0.843692325 | -0.137503524 | 0.042 | 0.047 | 1 | 0.437024249 | 2.017921908 | 0.1 | 0.046 | 1 | 0.843692325 | 0.683058304 |
| Gm8883 | 0.98180767 | -0.402964667 | 0.059 | 0.059 | 1 | 0.437024249 | 2.017921908 | 0.1 | 0.046 | 1 | 0.98180767 | 0.683058304 |
| Gm57006 | 0.437408619 | -0.485426827 | 0.034 | 0.053 | 1 | 0.643835681 | 1.847996907 | 0 | 0.022 | 1 | 0.643835681 | 0.683490938 |
| Bcat2 | 0.685205143 | -0.313245852 | 0.261 | 0.265 | 1 | 0.438365957 | -0.304006187 | 0.1 | 0.194 | 1 | 0.685205143 | 0.684567202 |
| Ganc | 0.915750808 | -0.258016331 | 0.16 | 0.159 | 1 | 0.438378238 | 0.660369903 | 0.3 | 0.188 | 1 | 0.915750808 | 0.684580997 |
| Ap5m1 | 0.54997316 | -0.333423734 | 0.202 | 0.159 | 1 | 0.438381544 | -0.425021588 | 0.1 | 0.194 | 1 | 0.54997316 | 0.68458471 |
| Smap2 | 0.43862968 | -0.31306278 | 0.597 | 0.588 | 1 | 0.62401536 | 0.165479096 | 0.7 | 0.772 | 1 | 0.62401536 | 0.684863363 |
| Pphln1 | 0.439142949 | -0.203655859 | 0.513 | 0.547 | 1 | 0.492445536 | -0.441509711 | 0.5 | 0.593 | 1 | 0.492445536 | 0.685439368 |
| Wdsub1 | 0.439270793 | -0.667630158 | 0.185 | 0.212 | 1 | 0.800963496 | 0.625604485 | 0.3 | 0.29 | 1 | 0.800963496 | 0.685582757 |
| Gspt2 | 0.884185266 | 0.152003093 | 0.092 | 0.088 | 1 | 0.439498288 | 0.930459067 | 0.2 | 0.114 | 1 | 0.884185266 | 0.68583783 |
| Zfp65 | 0.697642991 | -0.378511623 | 0.092 | 0.106 | 1 | 0.439499712 | 0.902444691 | 0.2 | 0.114 | 1 | 0.697642991 | 0.685839427 |
| 4930473A02Rik | 0.841777167 | 0.251538767 | 0.034 | 0.029 | 1 | 0.439528334 | 1.769994395 | 0.1 | 0.046 | 1 | 0.841777167 | 0.685871511 |
| Ddx54 | 0.440727325 | -0.418312631 | 0.311 | 0.341 | 1 | 0.59737364 | -0.441509711 | 0.3 | 0.346 | 1 | 0.59737364 | 0.687214075 |
| Acsl6 | 0.691605844 | -0.485426827 | 0.017 | 0.024 | 1 | 0.4409452 | 1.111031312 | 0.2 | 0.12 | 1 | 0.691605844 | 0.68745773 |
| Zfand2b | 0.441087651 | -0.292781749 | 0.21 | 0.247 | 1 | 0.73324675 | 0.695993813 | 0.2 | 0.154 | 1 | 0.73324675 | 0.687616986 |
| Junb | 0.441279765 | -0.155148461 | 0.765 | 0.818 | 1 | 0.749012492 | 1.286118019 | 0.7 | 0.799 | 1 | 0.749012492 | 0.687831699 |
| Tmem143 | 0.442027065 | -0.485426827 | 0.034 | 0.053 | 1 | 0.804840839 | 1.017921908 | 0.1 | 0.077 | 1 | 0.804840839 | 0.688666204 |
| Gm9828 | 0.629509811 | -0.292781749 | 0.025 | 0.035 | 1 | 0.44222087 | 1.847996907 | 0.1 | 0.046 | 1 | 0.629509811 | 0.688882442 |
| A330069E16Rik | 0.895003849 | -0.687060688 | 0.076 | 0.071 | 1 | 0.44222087 | 1.494359952 | 0.1 | 0.046 | 1 | 0.895003849 | 0.688882442 |
| Zfp703 | 0.442827429 | 0.176877762 | 0.597 | 0.565 | 1 | 0.773674056 | -0.421067913 | 0.8 | 0.765 | 1 | 0.773674056 | 0.689558726 |
| Zfp790 | 0.443378777 | -0.485426827 | 0.059 | 0.082 | 1 | 0.559646818 | -0.131825212 | 0.1 | 0.167 | 1 | 0.559646818 | 0.690172814 |
| H4c14 | 0.443522758 | -0.485426827 | 0.059 | 0.082 | 1 | 0.643842184 | 1.432959407 | 0 | 0.022 | 1 | 0.643842184 | 0.690333079 |
| Slx4ip | 0.794736253 | -0.347923303 | 0.218 | 0.218 | 1 | 0.443590006 | 0.571665678 | 0.3 | 0.191 | 1 | 0.794736253 | 0.690407919 |
| F3 | 0.443726479 | 0.894578656 | 0.58 | 0.676 | 1 | 0.974019176 | 0.913204037 | 0.5 | 0.698 | 1 | 0.974019176 | 0.69055977 |
| P2rx7 | 0.60313842 | -0.584962501 | 0.193 | 0.212 | 1 | 0.443744263 | -1.026472211 | 0.1 | 0.188 | 1 | 0.60313842 | 0.690579555 |
| Gm14418 | 0.444395758 | -0.137503524 | 0.034 | 0.053 | 1 | 0.640912309 | 1.159940913 | 0.1 | 0.062 | 1 | 0.640912309 | 0.691303926 |
| Irak4 | 0.444905464 | -0.485426827 | 0.176 | 0.206 | 1 | 0.869545745 | 0.41305985 | 0.2 | 0.173 | 1 | 0.869545745 | 0.691870056 |
| Gm11464 | 0.570777422 | 0.514573173 | 0.042 | 0.029 | 1 | 0.444918466 | 1.625604485 | 0.1 | 0.046 | 1 | 0.570777422 | 0.691884491 |
| Nemp2 | 0.580612737 | -0.222392421 | 0.034 | 0.047 | 1 | 0.444918466 | 1.494359952 | 0.1 | 0.046 | 1 | 0.580612737 | 0.691884491 |
| Foxl1 | 0.659871503 | 0.514573173 | 0.025 | 0.018 | 1 | 0.444918466 | 1.494359952 | 0.1 | 0.046 | 1 | 0.659871503 | 0.691884491 |
| Fkbp5 | 0.869647685 | -0.304400832 | 0.429 | 0.406 | 1 | 0.445815989 | -0.685597203 | 0.3 | 0.432 | 1 | 0.869647685 | 0.692880082 |
| Aars2 | 0.446590753 | -1.185866545 | 0.025 | 0.041 | 1 | 0.460577419 | 0.494359952 | 0 | 0.052 | 1 | 0.460577419 | 0.693738205 |
| Maged2 | 0.474332284 | -0.201505055 | 0.866 | 0.818 | 1 | 0.446615413 | -0.173280803 | 0.6 | 0.806 | 1 | 0.474332284 | 0.693765499 |
| Plpp7 | 0.513278666 | -0.485426827 | 0.008 | 0.018 | 1 | 0.446758355 | 0.625604485 | 0 | 0.056 | 1 | 0.513278666 | 0.693923682 |
| Gm52992 | 0.695893386 | -0.222392421 | 0.017 | 0.024 | 1 | 0.446758355 | 0.695993813 | 0 | 0.056 | 1 | 0.695893386 | 0.693923682 |
| Otx1 | 0.715874401 | 0.929610672 | 0.017 | 0.012 | 1 | 0.446758355 | 0.695993813 | 0 | 0.056 | 1 | 0.715874401 | 0.693923682 |
| Amacr | 0.616280491 | 0.407657969 | 0.092 | 0.076 | 1 | 0.446790883 | 0.558490289 | 0 | 0.056 | 1 | 0.616280491 | 0.693959672 |
| Pcdhgb6 | 0.760414269 | -0.137503524 | 0.05 | 0.059 | 1 | 0.446790883 | 0.625604485 | 0 | 0.056 | 1 | 0.760414269 | 0.693959672 |
| Gm15991 | 0.820836204 | -0.485426827 | 0.025 | 0.029 | 1 | 0.446790883 | 0.625604485 | 0 | 0.056 | 1 | 0.820836204 | 0.693959672 |
| Trmt5 | 0.518250144 | -0.315501826 | 0.067 | 0.088 | 1 | 0.44681934 | 0.31748219 | 0 | 0.056 | 1 | 0.518250144 | 0.693991157 |
| F8a | 0.591421701 | -0.678071905 | 0.05 | 0.065 | 1 | 0.446819579 | 0.374065718 | 0 | 0.056 | 1 | 0.591421701 | 0.693991422 |
| Gm12473 | 0.511005543 | -0.485426827 | 0.008 | 0.018 | 1 | 0.446844446 | 0.263034406 | 0 | 0.056 | 1 | 0.511005543 | 0.694018933 |
| Il15 | 0.721162042 | 0.514573173 | 0.017 | 0.012 | 1 | 0.446844446 | 0.31748219 | 0 | 0.056 | 1 | 0.721162042 | 0.694018933 |
| Gm56688 | 0.528185318 | -0.378511623 | 0.042 | 0.059 | 1 | 0.44686405 | 0.374065718 | 0 | 0.056 | 1 | 0.528185318 | 0.69404062 |
| Gm46412 | 0.966646355 | -0.359895945 | 0.092 | 0.088 | 1 | 0.44688413 | 0.210566986 | 0 | 0.056 | 1 | 0.966646355 | 0.694062834 |
| Gm40438 | 0.682146455 | -0.632268215 | 0.059 | 0.071 | 1 | 0.446912573 | -0.267480311 | 0 | 0.056 | 1 | 0.682146455 | 0.694094298 |
| Grid1 | 0.791190296 | 0.514573173 | 0.008 | 0.012 | 1 | 0.446928585 | -0.374395515 | 0 | 0.056 | 1 | 0.791190296 | 0.69411201 |
| Hddc3 | 0.672385695 | -0.222392421 | 0.042 | 0.053 | 1 | 0.446953437 | 0.769994395 | 0 | 0.056 | 1 | 0.672385695 | 0.694139499 |
| Tmem267 | 0.921442135 | -0.122856748 | 0.05 | 0.053 | 1 | 0.447621929 | 1.558490289 | 0.1 | 0.046 | 1 | 0.921442135 | 0.694878467 |
| Pqlc3 | 0.598433805 | -0.271302022 | 0.185 | 0.206 | 1 | 0.449364427 | -0.567040593 | 0.1 | 0.188 | 1 | 0.598433805 | 0.696800465 |
| Tmem184c | 0.517631686 | -0.299013703 | 0.202 | 0.229 | 1 | 0.45023022 | -0.654503434 | 0.2 | 0.29 | 1 | 0.517631686 | 0.697753189 |
| Slfn9 | 0.739199689 | -0.366782331 | 0.084 | 0.094 | 1 | 0.450330458 | 1.31748219 | 0.1 | 0.046 | 1 | 0.739199689 | 0.697863395 |
| Actr1b | 0.450371107 | -0.168236651 | 0.479 | 0.5 | 1 | 0.507622614 | -0.434937057 | 0.4 | 0.441 | 1 | 0.507622614 | 0.69790808 |
| Ddrgk1 | 0.450635776 | -0.309046804 | 0.63 | 0.618 | 1 | 0.455050679 | -0.321928095 | 0.7 | 0.682 | 1 | 0.455050679 | 0.69819895 |
| Sirt3 | 0.450730003 | -0.722466024 | 0.109 | 0.135 | 1 | 0.782668159 | 1.135278859 | 0.1 | 0.145 | 1 | 0.782668159 | 0.698302471 |
| Tbc1d17 | 0.450961676 | -0.578536232 | 0.235 | 0.253 | 1 | 0.727897833 | -0.152003093 | 0.2 | 0.238 | 1 | 0.727897833 | 0.698556918 |
| Hdac8 | 0.684322782 | -0.500300104 | 0.605 | 0.588 | 1 | 0.451511513 | -0.724668328 | 0.8 | 0.735 | 1 | 0.684322782 | 0.69916038 |
| Fbxo17 | 0.451692946 | -0.246239163 | 0.21 | 0.241 | 1 | 0.591575445 | 0.276454922 | 0.3 | 0.207 | 1 | 0.591575445 | 0.699359374 |
| Snx19 | 0.451757874 | -0.389849167 | 0.261 | 0.294 | 1 | 0.735417634 | 0.354956895 | 0.4 | 0.34 | 1 | 0.735417634 | 0.699430571 |
| Ppcdc | 0.451813581 | -0.914270126 | 0.076 | 0.1 | 1 | 0.500075746 | 0.695993813 | 0.2 | 0.12 | 1 | 0.500075746 | 0.69949165 |
| Pgap4 | 0.452099576 | -0.397963986 | 0.118 | 0.147 | 1 | 0.65890161 | 0.185031894 | 0.1 | 0.148 | 1 | 0.65890161 | 0.699805125 |
| Rrn3 | 0.45230909 | -0.425305835 | 0.328 | 0.347 | 1 | 0.548258112 | -0.339630097 | 0.3 | 0.386 | 1 | 0.548258112 | 0.700034667 |
| Ttc39a | 0.452339222 | 0.251538767 | 0.076 | 0.053 | 1 | 0.964594091 | 0.432959407 | 0.1 | 0.102 | 1 | 0.964594091 | 0.700067673 |
| Rpp25l | 0.452404543 | -0.126972856 | 0.168 | 0.206 | 1 | 0.714120306 | 0.249737583 | 0.2 | 0.269 | 1 | 0.714120306 | 0.700139215 |
| Txndc12 | 0.453686247 | -0.19730144 | 0.613 | 0.624 | 1 | 0.770533859 | 0.252387162 | 0.6 | 0.537 | 1 | 0.770533859 | 0.701541283 |
| Chic1 | 0.782273908 | -0.376720568 | 0.319 | 0.282 | 1 | 0.453937983 | 0.336097868 | 0.5 | 0.38 | 1 | 0.782273908 | 0.701816274 |
| Selenos | 0.454649477 | 0.110750461 | 0.975 | 0.935 | 1 | 0.503486784 | -0.241867502 | 0.9 | 0.963 | 1 | 0.503486784 | 0.702592807 |
| Gpr68 | 0.756529862 | -0.177304532 | 0.084 | 0.094 | 1 | 0.455355445 | 1.902444691 | 0.1 | 0.049 | 1 | 0.756529862 | 0.703362309 |
| Hmgxb3 | 0.456396969 | -0.191695624 | 0.328 | 0.341 | 1 | 0.540183596 | 0.526068812 | 0.5 | 0.364 | 1 | 0.540183596 | 0.704495744 |
| Wdr7 | 0.456407949 | -0.575624636 | 0.311 | 0.329 | 1 | 0.80539543 | -0.183711953 | 0.5 | 0.469 | 1 | 0.80539543 | 0.704507683 |
| Tra2a | 0.456607185 | -0.122477839 | 0.966 | 0.959 | 1 | 0.963971388 | -0.111361109 | 1 | 0.957 | 1 | 0.963971388 | 0.704724249 |
| Rcbtb1 | 0.456757926 | -0.189602486 | 0.471 | 0.494 | 1 | 0.897925391 | -0.374395515 | 0.6 | 0.491 | 1 | 0.897925391 | 0.704888049 |
| 2900026A02Rik | 0.457710896 | 0.4639471 | 0.143 | 0.112 | 1 | 0.587504234 | 0.720241359 | 0.2 | 0.136 | 1 | 0.587504234 | 0.705922528 |
| Nat9 | 0.458036779 | 0.260816581 | 0.193 | 0.159 | 1 | 0.463104881 | 1.111031312 | 0.2 | 0.12 | 1 | 0.463104881 | 0.706275867 |
| Inpp5f | 0.458125926 | -0.349105356 | 0.319 | 0.341 | 1 | 0.681214373 | -0.289506617 | 0.3 | 0.352 | 1 | 0.681214373 | 0.706372487 |
| Pygl | 0.458176086 | -0.374395515 | 0.319 | 0.341 | 1 | 0.804509871 | 0.424397394 | 0.3 | 0.284 | 1 | 0.804509871 | 0.706426846 |
| Hs6st2 | 0.614381021 | -0.944858446 | 0.025 | 0.035 | 1 | 0.458496018 | 0.847996907 | 0.1 | 0.046 | 1 | 0.614381021 | 0.706773438 |
| Gdpd1 | 0.458670999 | -1.008988783 | 0.05 | 0.071 | 1 | 0.935846285 | 0.660369903 | 0.1 | 0.09 | 1 | 0.935846285 | 0.706962913 |
| Gm31105 | 0.458788392 | -0.807354922 | 0.025 | 0.041 | 1 | 0.643835681 | 1.847996907 | 0 | 0.022 | 1 | 0.643835681 | 0.707089996 |
| Dynlt1a | 0.458788392 | -0.807354922 | 0.025 | 0.041 | 1 | 0.474927752 | 0.625604485 | 0 | 0.049 | 1 | 0.474927752 | 0.707089996 |
| Haus2 | 0.459701287 | -0.555816155 | 0.227 | 0.253 | 1 | 0.985034612 | 0.359710425 | 0.2 | 0.21 | 1 | 0.985034612 | 0.7080773 |
| Pdzrn4 | 0.459774846 | -4.09423607 | 0.118 | 0.135 | 1 | 0.885340179 | -2.888968688 | 0.1 | 0.08 | 1 | 0.885340179 | 0.708156783 |
| B3gat3 | 0.459923069 | -0.318316841 | 0.58 | 0.565 | 1 | 0.705900236 | 0.115848329 | 0.6 | 0.522 | 1 | 0.705900236 | 0.708316909 |
| Haus7 | 0.459951328 | -0.662304589 | 0.185 | 0.212 | 1 | 0.902186621 | 0.354956895 | 0.2 | 0.176 | 1 | 0.902186621 | 0.708347432 |
| Mrpl42 | 0.460187661 | -0.199672345 | 0.63 | 0.659 | 1 | 0.634406153 | -0.165299916 | 0.6 | 0.599 | 1 | 0.634406153 | 0.708602639 |
| Psmd8 | 0.460275984 | -0.163498732 | 0.79 | 0.765 | 1 | 0.914356379 | 0.182732778 | 0.7 | 0.84 | 1 | 0.914356379 | 0.708697986 |
| Ccdc28a | 0.460309652 | 0.303069068 | 0.109 | 0.082 | 1 | 0.607558467 | 0.973527789 | 0.1 | 0.059 | 1 | 0.607558467 | 0.708734329 |
| Gm11423 | 0.547347962 | -0.485426827 | 0.059 | 0.076 | 1 | 0.460521625 | 0.695993813 | 0 | 0.052 | 1 | 0.547347962 | 0.708963083 |
| Rab3d | 0.929389703 | 0.192645078 | 0.025 | 0.024 | 1 | 0.460551517 | 0.625604485 | 0 | 0.052 | 1 | 0.929389703 | 0.708995334 |

| | | | | | | | | | | | | |
|---|---|---|---|---|---|---|---|---|---|---|---|---|
| Osbp2 | 0.940587367 | -0.263034406 | 0.034 | 0.035 | 1 | 0.460551517 | 0.625604485 | 0 | 0.052 | 1 | 0.940587367 | 0.708995334 |
| Gbp7 | 0.505339217 | -0.900464326 | 0.084 | 0.106 | 1 | 0.460577668 | -0.339630097 | 0 | 0.052 | 1 | 0.505339217 | 0.709023548 |
| Gm14743 | 0.575022075 | -0.333423734 | 0.034 | 0.047 | 1 | 0.460577668 | 0.625604485 | 0 | 0.052 | 1 | 0.575022075 | 0.709023548 |
| Dph1 | 0.82670638 | -0.292781749 | 0.025 | 0.029 | 1 | 0.460577668 | 0.625604485 | 0 | 0.052 | 1 | 0.82670638 | 0.709023548 |
| Mettl22 | 0.831393676 | 0.192645078 | 0.059 | 0.053 | 1 | 0.460577668 | 0.625604485 | 0 | 0.052 | 1 | 0.831393676 | 0.709023548 |
| Bend4 | 0.511005543 | -0.485426827 | 0.008 | 0.018 | 1 | 0.460599334 | 0.432959407 | 0 | 0.052 | 1 | 0.511005543 | 0.709046921 |
| Tor4a | 0.62441508 | -0.485426827 | 0.025 | 0.035 | 1 | 0.460600081 | 0.432959407 | 0 | 0.052 | 1 | 0.62441508 | 0.709047727 |
| Tnfsf13os | 0.959428331 | 1.57870351 | 0.059 | 0.059 | 1 | 0.460600081 | 0.31748219 | 0 | 0.052 | 1 | 0.959428331 | 0.709047727 |
| Pcdhga9 | 0.467093859 | -0.485426827 | 0.025 | 0.041 | 1 | 0.460618756 | 0.374065718 | 0 | 0.052 | 1 | 0.467093859 | 0.709067873 |
| Grin3a | 0.536086759 | 0.152003093 | 0.05 | 0.035 | 1 | 0.460634939 | 0.210566986 | 0 | 0.052 | 1 | 0.536086759 | 0.709085331 |
| Edar | 0.511005543 | -0.485426827 | 0.008 | 0.018 | 1 | 0.460644897 | 0.263034406 | 0 | 0.052 | 1 | 0.511005543 | 0.709096073 |
| Slco1b2 | 0.770881424 | 0.652076697 | 0.034 | 0.041 | 1 | 0.460673275 | -0.374395515 | 0 | 0.052 | 1 | 0.770881424 | 0.709126684 |
| Gm57100 | 0.641768891 | 0.192645078 | 0.092 | 0.076 | 1 | 0.460690947 | 0.847996907 | 0 | 0.052 | 1 | 0.641768891 | 0.709145746 |
| Laptm5 | 0.778665521 | -2.070389328 | 0.008 | 0.012 | 1 | 0.461231174 | -0.763437806 | 0.1 | 0.046 | 1 | 0.778665521 | 0.709728152 |
| Gdf5 | 0.461344327 | 0.514573173 | 0.134 | 0.106 | 1 | 0.905530041 | -1.011825435 | 0.2 | 0.191 | 1 | 0.905530041 | 0.709850066 |
| Rap2b | 0.953845761 | -0.146624914 | 0.235 | 0.235 | 1 | 0.461801569 | -0.11821478 | 0.2 | 0.324 | 1 | 0.953845761 | 0.710342449 |
| Gm56816 | 0.462559356 | 0.514573173 | 0.067 | 0.047 | 1 | 0.678985981 | 1.111031312 | 0.1 | 0.065 | 1 | 0.678985981 | 0.711157555 |
| Copg2 | 0.462798889 | -0.24726709 | 0.277 | 0.312 | 1 | 0.731782648 | 0.415885894 | 0.4 | 0.343 | 1 | 0.731782648 | 0.711414967 |
| AV039307 | 0.462887689 | -0.655351829 | 0.025 | 0.041 | 1 | 0.73075158 | 2.695993813 | 0 | 0.012 | 1 | 0.73075158 | 0.711510366 |
| Gm42797 | 0.462887689 | -0.655351829 | 0.025 | 0.041 | 1 | 0.474974509 | 0.432959407 | 0 | 0.049 | 1 | 0.474974509 | 0.711510366 |
| Tle6 | 0.462887689 | -0.807354922 | 0.025 | 0.041 | 1 | 0.539431168 | 1.017921908 | 0 | 0.037 | 1 | 0.539431168 | 0.711510366 |
| Qrich2 | 0.462887689 | -0.655351829 | 0.025 | 0.041 | 1 | 0.643850855 | 2.017921908 | 0 | 0.022 | 1 | 0.643850855 | 0.711510366 |
| Cep131 | 0.462887689 | -0.807354922 | 0.025 | 0.041 | 1 | 0.52208609 | 0.847996907 | 0 | 0.04 | 1 | 0.52208609 | 0.711510366 |
| Abcc3 | 0.462932621 | -0.748461233 | 0.025 | 0.041 | 1 | 0.669848329 | 1.31748219 | 0 | 0.019 | 1 | 0.669848329 | 0.711558631 |
| Ldlrad3 | 0.463724904 | -0.169838116 | 0.84 | 0.806 | 1 | 0.645435704 | -0.172409304 | 0.9 | 0.87 | 1 | 0.645435704 | 0.712409021 |
| Rarb | 0.464320767 | -1.070389328 | 0.025 | 0.041 | 1 | 0.475011126 | -0.840059087 | 0 | 0.049 | 1 | 0.475011126 | 0.713047759 |
| Rnf145 | 0.464928906 | -0.19592021 | 0.387 | 0.318 | 1 | 0.82647812 | 0.455679484 | 0.4 | 0.398 | 1 | 0.82647812 | 0.713698925 |
| Usp2 | 0.713641087 | 0.248679113 | 0.328 | 0.306 | 1 | 0.464982161 | 1.159940913 | 0.3 | 0.207 | 1 | 0.713641087 | 0.713755912 |
| Mrps23 | 0.521309755 | -0.250961574 | 0.294 | 0.318 | 1 | 0.465528513 | -0.419483404 | 0.2 | 0.293 | 1 | 0.521309755 | 0.71434023 |
| Heatr5a | 0.645613233 | 0.358304616 | 0.824 | 0.818 | 1 | 0.465677796 | 0.276069741 | 0.9 | 0.87 | 1 | 0.645613233 | 0.714499782 |
| Mtus2 | 0.465929249 | -1.185866545 | 0.05 | 0.071 | 1 | 0.718927209 | 0.973527789 | 0.1 | 0.068 | 1 | 0.718927209 | 0.714768433 |
| Tsen15 | 0.46635296 | 0.121428279 | 0.487 | 0.424 | 1 | 0.497072349 | 0.415885894 | 0.5 | 0.367 | 1 | 0.497072349 | 0.715220836 |
| Pfkfb4 | 0.543933546 | -0.807354922 | 0.244 | 0.253 | 1 | 0.466712277 | 1.617042472 | 0.3 | 0.25 | 1 | 0.543933546 | 0.715604204 |
| Gm33508 | 0.467062381 | -0.622930351 | 0.025 | 0.041 | 1 | 0.730750993 | 2.210566986 | 0 | 0.012 | 1 | 0.730750993 | 0.715977494 |
| Gm20544 | 0.467062381 | -0.748461233 | 0.025 | 0.041 | 1 | 0.619961444 | 1.432959407 | 0 | 0.025 | 1 | 0.619961444 | 0.715977494 |
| Zfp324 | 0.467093859 | -0.485426827 | 0.025 | 0.041 | 1 | 0.522151747 | 1.210566986 | 0 | 0.04 | 1 | 0.522151747 | 0.716011044 |
| Gm20036 | 0.467093859 | -0.485426827 | 0.025 | 0.041 | 1 | 0.73075158 | 2.695993813 | 0 | 0.012 | 1 | 0.73075158 | 0.716011044 |
| Gm34961 | 0.467093859 | -0.485426827 | 0.025 | 0.041 | 1 | 0.52208609 | 1.017921908 | 0 | 0.04 | 1 | 0.52208609 | 0.716011044 |
| Gm9776 | 0.467093859 | -0.485426827 | 0.025 | 0.041 | 1 | 0.698534498 | 2.432959407 | 0 | 0.015 | 1 | 0.698534498 | 0.716011044 |
| Klhl26 | 0.46744567 | 0.152003093 | 0.235 | 0.194 | 1 | 0.751609728 | 0.532495081 | 0.3 | 0.259 | 1 | 0.751609728 | 0.716385885 |
| Zdhhc2 | 0.467742685 | -1.539874611 | 0.084 | 0.106 | 1 | 0.969256618 | 0.210566986 | 0.1 | 0.099 | 1 | 0.969256618 | 0.71670215 |
| Fyb2 | 0.468447396 | -0.485426827 | 0.025 | 0.041 | 1 | 0.681612912 | -0.230005605 | 0.1 | 0.142 | 1 | 0.681612912 | 0.717451829 |
| Fuz | 0.582347819 | -0.531230517 | 0.218 | 0.235 | 1 | 0.468604635 | -0.408342847 | 0.1 | 0.185 | 1 | 0.582347819 | 0.717618966 |
| Pde12 | 0.469664964 | -0.375243909 | 0.235 | 0.265 | 1 | 0.920924799 | 0.147557188 | 0.3 | 0.262 | 1 | 0.920924799 | 0.71874475 |
| Mrpl52 | 0.470169363 | -0.179800545 | 0.756 | 0.724 | 1 | 0.911992354 | 0.353601616 | 0.6 | 0.642 | 1 | 0.911992354 | 0.719279497 |
| Smyd3 | 0.834434695 | -0.222392421 | 0.613 | 0.588 | 1 | 0.470732366 | -0.730270942 | 0.7 | 0.688 | 1 | 0.834434695 | 0.719875772 |
| Papln | 0.471242918 | -0.485426827 | 0.025 | 0.041 | 1 | 0.564719271 | 1.494359952 | 0.1 | 0.056 | 1 | 0.564719271 | 0.720415948 |
| Gm28154 | 0.471781614 | -0.137503524 | 0.076 | 0.1 | 1 | 0.906513737 | 0.345496566 | 0.1 | 0.108 | 1 | 0.906513737 | 0.720985337 |
| Ntng1 | 0.959597195 | -0.292781749 | 0.017 | 0.018 | 1 | 0.472246744 | -0.473931188 | 0.1 | 0.046 | 1 | 0.959597195 | 0.721476501 |
| Gm19710 | 0.541457736 | -0.185866545 | 0.084 | 0.106 | 1 | 0.474084773 | 0.959028219 | 0.2 | 0.12 | 1 | 0.541457736 | 0.723413174 |
| Nelfe | 0.817080162 | -0.292781749 | 0.336 | 0.318 | 1 | 0.47441308 | 1.744903414 | 0.3 | 0.269 | 1 | 0.817080162 | 0.72375839 |
| Snapin | 0.474849365 | -0.117275716 | 0.63 | 0.506 | 1 | 0.670361169 | -0.230005605 | 0.5 | 0.531 | 1 | 0.670361169 | 0.724216811 |
| Glra1 | 0.482848516 | 1.225066556 | 0.059 | 0.041 | 1 | 0.474877085 | 0.769994395 | 0 | 0.049 | 1 | 0.482848516 | 0.724245925 |
| Dusp28 | 0.699674204 | -0.263034406 | 0.092 | 0.106 | 1 | 0.474877085 | 0.847996907 | 0 | 0.049 | 1 | 0.699674204 | 0.724245925 |
| Zfp707 | 0.933394757 | -0.163498732 | 0.034 | 0.035 | 1 | 0.474877085 | 0.625604485 | 0 | 0.049 | 1 | 0.933394757 | 0.724245925 |
| Rnf122 | 0.577984721 | -0.292781749 | 0.084 | 0.065 | 1 | 0.474904369 | 0.769994395 | 0 | 0.049 | 1 | 0.577984721 | 0.724274579 |
| Cadm4 | 0.687345921 | -1.359895945 | 0.017 | 0.024 | 1 | 0.474904369 | 0.769994395 | 0 | 0.049 | 1 | 0.687345921 | 0.724274579 |
| Slc6a15 | 0.954551343 | -0.222392421 | 0.017 | 0.018 | 1 | 0.474904369 | 0.625604485 | 0 | 0.049 | 1 | 0.954551343 | 0.724274579 |
| Kctd21 | 0.491791629 | -0.707819249 | 0.017 | 0.029 | 1 | 0.474927752 | 0.625604485 | 0 | 0.049 | 1 | 0.491791629 | 0.724299135 |
| Ric3 | 0.659299372 | 0.344648171 | 0.059 | 0.047 | 1 | 0.474927752 | 0.625604485 | 0 | 0.049 | 1 | 0.659299372 | 0.724299135 |
| 4930570G19Rik | 0.695893386 | -0.222392421 | 0.017 | 0.024 | 1 | 0.474927752 | 0.558490289 | 0 | 0.049 | 1 | 0.695893386 | 0.724299135 |
| Adam8 | 0.954551343 | -0.222392421 | 0.017 | 0.018 | 1 | 0.474927752 | 0.625604485 | 0 | 0.049 | 1 | 0.954551343 | 0.724299135 |
| Duxbl1 | 0.500410556 | -0.555816155 | 0.067 | 0.088 | 1 | 0.474928012 | 0.558490289 | 0 | 0.049 | 1 | 0.500410556 | 0.724299408 |
| 2810457G06Rik | 0.612246903 | 0.421463768 | 0.092 | 0.076 | 1 | 0.474928012 | 0.695993813 | 0 | 0.049 | 1 | 0.612246903 | 0.724299408 |
| Arl4d | 0.745693666 | 0.389042291 | 0.076 | 0.088 | 1 | 0.474928012 | 0.695993813 | 0 | 0.049 | 1 | 0.745693666 | 0.724299408 |
| Xdh | 0.954971566 | 0.652076697 | 0.034 | 0.035 | 1 | 0.474928012 | 0.494359952 | 0 | 0.049 | 1 | 0.954971566 | 0.724299408 |
| Acpp | 0.781790612 | -0.485426827 | 0.008 | 0.012 | 1 | 0.474947236 | 0.31748219 | 0 | 0.049 | 1 | 0.781790612 | 0.724319595 |
| Zfp882 | 0.814974746 | -0.944858446 | 0.025 | 0.029 | 1 | 0.474947236 | 0.558490289 | 0 | 0.049 | 1 | 0.814974746 | 0.724319595 |
| Gm34102 | 0.74698568 | -0.378511623 | 0.05 | 0.041 | 1 | 0.474948275 | 0.625604485 | 0 | 0.049 | 1 | 0.74698568 | 0.724320686 |
| Lama3 | 0.508731273 | -1.944858446 | 0.008 | 0.018 | 1 | 0.474962821 | 0.494359952 | 0 | 0.049 | 1 | 0.508731273 | 0.724335961 |
| Gm8098 | 0.622067527 | 0.251538767 | 0.067 | 0.053 | 1 | 0.474964379 | 0.263034406 | 0 | 0.049 | 1 | 0.622067527 | 0.724337597 |
| ENSMUSG00000121495 | 0.91699934 | -0.378511623 | 0.05 | 0.053 | 1 | 0.474986196 | 0.111031312 | 0 | 0.049 | 1 | 0.91699934 | 0.724360506 |
| Gm34728 | 0.511005543 | -0.485426827 | 0.008 | 0.018 | 1 | 0.47499165 | 0.263034406 | 0 | 0.049 | 1 | 0.511005543 | 0.724366232 |
| Serpinb8 | 0.614381507 | -0.807354922 | 0.025 | 0.035 | 1 | 0.474995545 | 0.111031312 | 0 | 0.049 | 1 | 0.614381507 | 0.724370323 |
| Gm6710 | 0.493791844 | 0.514573173 | 0.059 | 0.041 | 1 | 0.475022551 | 0.930459067 | 0 | 0.049 | 1 | 0.493791844 | 0.724398678 |
| Abca7 | 0.82670638 | -0.292781749 | 0.025 | 0.029 | 1 | 0.475022551 | 0.930459067 | 0 | 0.049 | 1 | 0.82670638 | 0.724398678 |
| Pick1 | 0.476410909 | -0.378511623 | 0.16 | 0.188 | 1 | 0.509966637 | -0.339630097 | 0.1 | 0.176 | 1 | 0.509966637 | 0.725854464 |
| Gtpbp3 | 0.772953872 | 0.225066556 | 0.067 | 0.059 | 1 | 0.47676286 | 1.930459067 | 0.1 | 0.049 | 1 | 0.772953872 | 0.726222896 |
| Coq10b | 0.476879115 | -0.146958462 | 0.916 | 0.818 | 1 | 0.737690713 | 0.206976702 | 1 | 0.898 | 1 | 0.737690713 | 0.72634454 |
| Inhca | 0.477034041 | 0.389042291 | 0.067 | 0.047 | 1 | 0.539413885 | 1.210566986 | 0 | 0.037 | 1 | 0.539413885 | 0.726506605 |
| Smug1 | 0.664542429 | -0.555816155 | 0.076 | 0.088 | 1 | 0.477764505 | 0.959028219 | 0.2 | 0.12 | 1 | 0.664542429 | 0.727270088 |
| Astn2 | 0.478334758 | 1.32901752 | 0.143 | 0.118 | 1 | 0.851630802 | -1.048167282 | 0.1 | 0.114 | 1 | 0.851630802 | 0.727865375 |
| A730081D07Rik | 0.478576589 | -0.669851398 | 0.05 | 0.071 | 1 | 0.959564654 | 1.073063462 | 0.1 | 0.102 | 1 | 0.959564654 | 0.728117627 |
| Unc119b | 0.478642258 | 0.141114777 | 0.311 | 0.271 | 1 | 0.578043218 | 0.403212064 | 0.4 | 0.302 | 1 | 0.578043218 | 0.728186105 |
| Med24 | 0.706644468 | -0.258016331 | 0.185 | 0.2 | 1 | 0.478672634 | 0.864116572 | 0.3 | 0.213 | 1 | 0.706644468 | 0.728217778 |
| Ubxn2b | 0.479019086 | 0.283247627 | 0.16 | 0.129 | 1 | 0.621864716 | 0.580516596 | 0.2 | 0.139 | 1 | 0.621864716 | 0.728578887 |
| Gm9975 | 0.656548497 | 0.377069649 | 0.059 | 0.047 | 1 | 0.479305119 | 1.847996907 | 0.1 | 0.049 | 1 | 0.656548497 | 0.728876841 |
| Spindoc | 0.479308402 | -0.348924827 | 0.395 | 0.4 | 1 | 0.896912019 | -0.182840497 | 0.7 | 0.537 | 1 | 0.896912019 | 0.72888026 |
| Map3k1 | 0.480538623 | -0.326360341 | 0.437 | 0.471 | 1 | 0.522351488 | -0.208757872 | 0.4 | 0.534 | 1 | 0.522351488 | 0.730159878 |
| Tnik | 0.481293537 | 1.799975392 | 0.092 | 0.071 | 1 | 0.502002181 | 1.111031312 | 0.1 | 0.052 | 1 | 0.502002181 | 0.730943605 |
| Samd8 | 0.481497633 | -0.345569164 | 0.504 | 0.5 | 1 | 0.964173353 | -0.198015491 | 0.8 | 0.67 | 1 | 0.964173353 | 0.731155295 |
| Rnf215 | 0.481548679 | -0.272697503 | 0.395 | 0.418 | 1 | 0.571170549 | -0.245112498 | 0.3 | 0.389 | 1 | 0.571170549 | 0.731208228 |
| Kbtbd11 | 0.856663367 | -0.163498732 | 0.034 | 0.029 | 1 | 0.482051401 | 1.695993813 | 0.1 | 0.049 | 1 | 0.856663367 | 0.731729249 |
| B430010I23Rik | 0.791236802 | -0.497203995 | 0.319 | 0.282 | 1 | 0.482718877 | 0.1170551 | 0.5 | 0.327 | 1 | 0.791236802 | 0.73242024 |
| 2810454H06Rik | 0.625689949 | -1.070389328 | 0.059 | 0.071 | 1 | 0.48320251 | 2.017921908 | 0.1 | 0.052 | 1 | 0.625689949 | 0.732920354 |
| Nrbf2 | 0.483411833 | 0.210956956 | 0.504 | 0.465 | 1 | 0.798368456 | 0.283212288 | 0.4 | 0.367 | 1 | 0.798368456 | 0.733166666 |
| Gmppa | 0.499276649 | -0.372952098 | 0.387 | 0.4 | 1 | 0.484388758 | -0.391469028 | 0.3 | 0.377 | 1 | 0.499276649 | 0.734145047 |
| Cdk5rap3 | 0.545957155 | -0.304854582 | 0.345 | 0.359 | 1 | 0.484877664 | 0.20029865 | 0.2 | 0.33 | 1 | 0.545957155 | 0.734648979 |

| | | | | | | | | | | | | |
|---|---|---|---|---|---|---|---|---|---|---|---|---|
| Chrnb1 | 0.485068998 | -1.070389328 | 0.017 | 0.029 | 1 | 0.802052667 | 1.017921908 | 0.1 | 0.077 | 1 | 0.802052667 | 0.734846064 |
| Slc3a1 | 0.485068998 | -1.359895945 | 0.017 | 0.029 | 1 | 0.857614143 | 0.494359952 | 0.1 | 0.08 | 1 | 0.857614143 | 0.734846064 |
| Zkscan1 | 0.523526383 | -0.359895945 | 0.378 | 0.394 | 1 | 0.48513467 | -0.430538593 | 0.4 | 0.475 | 1 | 0.523526383 | 0.734913692 |
| Gm47512 | 0.485250129 | 0.666576266 | 0.059 | 0.041 | 1 | 0.669853054 | 2.210566986 | 0 | 0.019 | 1 | 0.669853054 | 0.73503257 |
| Snhg14 | 0.485640987 | -0.428843299 | 0.092 | 0.118 | 1 | 0.930502981 | 0.625604485 | 0.1 | 0.09 | 1 | 0.930502981 | 0.735434806 |
| Supt7l | 0.485932531 | 0.362570079 | 0.126 | 0.1 | 1 | 0.768686435 | 0.580516596 | 0.2 | 0.16 | 1 | 0.768686435 | 0.735734638 |
| Chst3 | 0.487281696 | 0.514573173 | 0.193 | 0.165 | 1 | 0.785716493 | -0.844198817 | 0.5 | 0.346 | 1 | 0.785716493 | 0.737119941 |
| Bik | 0.721162042 | 0.514573173 | 0.017 | 0.012 | 1 | 0.487556826 | 1.558490289 | 0.1 | 0.049 | 1 | 0.721162042 | 0.737401994 |
| Hmgn5 | 0.48819733 | -0.447952122 | 0.378 | 0.394 | 1 | 0.6556804 | -0.140507455 | 0.3 | 0.401 | 1 | 0.6556804 | 0.738058027 |
| Rsf1os2 | 0.48842591 | -0.900464326 | 0.017 | 0.029 | 1 | 0.698529281 | 2.210566986 | 0 | 0.015 | 1 | 0.698529281 | 0.73829195 |
| Omp | 0.48842591 | -1.359895945 | 0.017 | 0.029 | 1 | 0.698529281 | 2.017921908 | 0 | 0.015 | 1 | 0.698529281 | 0.73829195 |
| Gm15890 | 0.48842591 | -1.070389328 | 0.017 | 0.029 | 1 | 0.597824685 | 1.31748219 | 0 | 0.028 | 1 | 0.597824685 | 0.73829195 |
| Gm1604b | 0.48842591 | -0.900464326 | 0.017 | 0.029 | 1 | 0.53946887 | 0.625604485 | 0 | 0.037 | 1 | 0.53946887 | 0.73829195 |
| Ate1 | 0.488720897 | -0.285128177 | 0.462 | 0.471 | 1 | 0.704083675 | 0.729160677 | 0.4 | 0.417 | 1 | 0.704083675 | 0.738593679 |
| Dhx58 | 0.488764274 | 0.514573173 | 0.092 | 0.071 | 1 | 0.505623793 | 0.432959407 | 0 | 0.043 | 1 | 0.505623793 | 0.738638033 |
| Zbed4 | 0.528392066 | -0.36994961 | 0.084 | 0.106 | 1 | 0.489244633 | -0.457811523 | 0.1 | 0.179 | 1 | 0.528392066 | 0.739128955 |
| C030013C21Rik | 0.826158434 | -0.163498732 | 0.076 | 0.082 | 1 | 0.489351263 | 1.048295557 | 0.2 | 0.123 | 1 | 0.826158434 | 0.739237867 |
| Alg1 | 0.48948765 | 0.114965714 | 0.328 | 0.282 | 1 | 0.546845602 | 0.420986766 | 0.4 | 0.284 | 1 | 0.546845602 | 0.73937714 |
| D130062J10Rik | 0.619390946 | -0.655351829 | 0.025 | 0.035 | 1 | 0.489877116 | 0.930459067 | 0 | 0.046 | 1 | 0.619390946 | 0.739774643 |
| Gm43138 | 0.657855022 | 0.362570079 | 0.059 | 0.047 | 1 | 0.489877116 | 0.930459067 | 0 | 0.046 | 1 | 0.657855022 | 0.739774643 |
| 2310058D17Rik | 0.670922574 | -0.292781749 | 0.042 | 0.053 | 1 | 0.489901827 | 0.847996907 | 0 | 0.046 | 1 | 0.670922574 | 0.739799854 |
| Gp1bb | 0.8327876 | -0.222392421 | 0.042 | 0.047 | 1 | 0.489901827 | 0.847996907 | 0 | 0.046 | 1 | 0.8327876 | 0.739799854 |
| Gm128 | 0.506471296 | -1.070389328 | 0.008 | 0.018 | 1 | 0.489923006 | 0.769994395 | 0 | 0.046 | 1 | 0.506471296 | 0.73982146 |
| Gm56626 | 0.62441508 | -0.485426827 | 0.025 | 0.035 | 1 | 0.489923006 | 0.769994395 | 0 | 0.046 | 1 | 0.62441508 | 0.73982146 |
| Gm7160 | 0.839906751 | 0.292180751 | 0.034 | 0.029 | 1 | 0.489923006 | 0.558490289 | 0 | 0.046 | 1 | 0.839906751 | 0.73982146 |
| Kcnab1 | 0.967819414 | 0.514573173 | 0.059 | 0.059 | 1 | 0.489923006 | 0.494359952 | 0 | 0.046 | 1 | 0.967819414 | 0.73982146 |
| A230087F16Rik | 0.82670638 | -0.485426827 | 0.025 | 0.029 | 1 | 0.489939838 | 0.625604485 | 0 | 0.046 | 1 | 0.82670638 | 0.739838631 |
| Disc1 | 0.948614424 | -0.944858446 | 0.025 | 0.024 | 1 | 0.489940652 | 0.558490289 | 0 | 0.046 | 1 | 0.948614424 | 0.739839462 |
| Gm42829 | 0.659871503 | 0.514573173 | 0.025 | 0.018 | 1 | 0.48995314 | 0.558490289 | 0 | 0.046 | 1 | 0.659871503 | 0.7398522 |
| Dzank1 | 0.542950752 | 0.334000927 | 0.042 | 0.059 | 1 | 0.489954768 | 0.432959407 | 0 | 0.046 | 1 | 0.542950752 | 0.739853862 |
| Zfp341 | 0.558163762 | -1.163498732 | 0.034 | 0.047 | 1 | 0.489954768 | 0.494359952 | 0 | 0.046 | 1 | 0.558163762 | 0.739853862 |
| D530017H19Rik | 0.815893803 | 0.377069649 | 0.059 | 0.053 | 1 | 0.489954768 | 0.374065718 | 0 | 0.046 | 1 | 0.815893803 | 0.739853862 |
| Gm49066 | 0.493401455 | -0.292781749 | 0.05 | 0.071 | 1 | 0.489955583 | 0.159940913 | 0 | 0.046 | 1 | 0.493401455 | 0.739854692 |
| Gm31045 | 0.842422009 | 0.251538767 | 0.025 | 0.029 | 1 | 0.489955854 | 0.625604485 | 0 | 0.046 | 1 | 0.842422009 | 0.739854969 |
| Ngfr | 0.495207728 | -0.485426827 | 0.017 | 0.029 | 1 | 0.489967526 | 0.210566986 | 0 | 0.046 | 1 | 0.495207728 | 0.739866876 |
| Gm49890 | 0.954551343 | -0.222392421 | 0.017 | 0.018 | 1 | 0.489994938 | -1.267480311 | 0 | 0.046 | 1 | 0.954551343 | 0.739894837 |
| Plaat3 | 0.597447999 | -0.351125735 | 0.244 | 0.265 | 1 | 0.49024548 | -0.382957528 | 0.4 | 0.481 | 1 | 0.597447999 | 0.74015033 |
| C330018D20Rik | 0.544321907 | 0.152003093 | 0.05 | 0.035 | 1 | 0.490317433 | 1.374065718 | 0.1 | 0.049 | 1 | 0.544321907 | 0.740223681 |
| 2700097O09Rik | 0.647679843 | -0.464668267 | 0.244 | 0.247 | 1 | 0.490956763 | 0.210566986 | 0.4 | 0.269 | 1 | 0.647679843 | 0.740874983 |
| Tbc1d8 | 0.491342249 | -0.485426827 | 0.05 | 0.071 | 1 | 0.516192301 | -0.408342847 | 0.1 | 0.176 | 1 | 0.516192301 | 0.741267292 |
| Dars2 | 0.491342249 | -0.563429339 | 0.05 | 0.071 | 1 | 0.632378953 | 1.432959407 | 0.1 | 0.062 | 1 | 0.632378953 | 0.741267292 |
| Gm16008 | 0.491791629 | -0.707819249 | 0.017 | 0.029 | 1 | 0.577184788 | 1.558490289 | 0 | 0.031 | 1 | 0.577184788 | 0.741724251 |
| Ephx4 | 0.491791629 | -0.707819249 | 0.017 | 0.029 | 1 | 0.698532411 | 1.847996907 | 0 | 0.015 | 1 | 0.698532411 | 0.741724251 |
| Gm32389 | 0.491791629 | -1.070389328 | 0.017 | 0.029 | 1 | 0.669850692 | 1.558490289 | 0 | 0.019 | 1 | 0.669850692 | 0.741724251 |
| Mfsd12 | 0.491791629 | -0.707819249 | 0.017 | 0.029 | 1 | 0.776128087 | 0.930459067 | 0.1 | 0.074 | 1 | 0.776128087 | 0.741724251 |
| Gm11963 | 0.491791629 | -0.707819249 | 0.017 | 0.029 | 1 | 0.698532411 | 2.017921908 | 0 | 0.015 | 1 | 0.698532411 | 0.741724251 |
| 5830428M24Rik | 0.491791629 | -0.707819249 | 0.017 | 0.029 | 1 | 0.698529281 | 2.210566986 | 0 | 0.015 | 1 | 0.698529281 | 0.741724251 |
| 1700102H20Rik | 0.491791629 | -0.707819249 | 0.017 | 0.029 | 1 | 0.619953009 | 1.695993813 | 0 | 0.025 | 1 | 0.619953009 | 0.741724251 |
| Nme3 | 0.491791629 | -0.707819249 | 0.017 | 0.029 | 1 | 0.597845674 | 1.695993813 | 0 | 0.028 | 1 | 0.597845674 | 0.741724251 |
| Tsx | 0.491791629 | -0.707819249 | 0.017 | 0.029 | 1 | 0.698529281 | 2.210566986 | 0 | 0.015 | 1 | 0.698529281 | 0.741724251 |
| Cdca3 | 0.491818194 | -1.235448574 | 0.017 | 0.029 | 1 | 0.730749233 | 2.432959407 | 0 | 0.012 | 1 | 0.730749233 | 0.741751253 |
| Acad11 | 0.641994176 | -0.244418728 | 0.084 | 0.1 | 1 | 0.492591531 | 0.769994395 | 0.2 | 0.12 | 1 | 0.641994176 | 0.742536646 |
| Dpp7 | 0.492735407 | 0.110182918 | 0.244 | 0.206 | 1 | 0.493918982 | 0.71414116 | 0.3 | 0.201 | 1 | 0.493918982 | 0.742682632 |
| Nhlrc2 | 0.499838719 | -0.301714874 | 0.513 | 0.512 | 1 | 0.49296393 | 0.648688098 | 0.6 | 0.617 | 1 | 0.499838719 | 0.742914424 |
| Ppp1r1c | 0.493503829 | -2.392317423 | 0.017 | 0.029 | 1 | 0.698534498 | 0.930459067 | 0 | 0.015 | 1 | 0.698534498 | 0.743461629 |
| Gm29865 | 0.493503829 | -1.292781749 | 0.017 | 0.029 | 1 | 0.643848687 | 1.017921908 | 0 | 0.022 | 1 | 0.643848687 | 0.743461629 |
| E230001N04Rik | 0.493680852 | 0.514573173 | 0.059 | 0.041 | 1 | 0.698534498 | 2.432959407 | 0 | 0.015 | 1 | 0.698534498 | 0.74364092 |
| Gm56791 | 0.495207728 | -0.485426827 | 0.017 | 0.029 | 1 | 0.59781419 | 1.558490289 | 0 | 0.028 | 1 | 0.59781419 | 0.745184762 |
| Creb3l4 | 0.495207728 | -0.485426827 | 0.017 | 0.029 | 1 | 0.557729726 | 1.31748219 | 0 | 0.034 | 1 | 0.557729726 | 0.745184762 |
| 4933408J17Rik | 0.495207728 | -0.485426827 | 0.017 | 0.029 | 1 | 0.73075158 | 2.695993813 | 0 | 0.012 | 1 | 0.73075158 | 0.745184762 |
| Gm11767 | 0.495207728 | -0.485426827 | 0.017 | 0.029 | 1 | 0.669848329 | 1.847996907 | 0 | 0.019 | 1 | 0.669848329 | 0.745184762 |
| Hhipl1 | 0.495207728 | -0.485426827 | 0.017 | 0.029 | 1 | 0.643850855 | 2.017921908 | 0 | 0.022 | 1 | 0.643850855 | 0.745184762 |
| 5730424H11Rik | 0.495207728 | -0.485426827 | 0.017 | 0.029 | 1 | 0.73075158 | 2.695993813 | 0 | 0.012 | 1 | 0.73075158 | 0.745184762 |
| Gm27042 | 0.495207728 | -0.485426827 | 0.017 | 0.029 | 1 | 0.619961444 | 1.558490289 | 0 | 0.025 | 1 | 0.619961444 | 0.745184762 |
| Pcdhb18 | 0.967167471 | 0.514573173 | 0.017 | 0.018 | 1 | 0.495853236 | 1.263034406 | 0.1 | 0.049 | 1 | 0.967167471 | 0.74583604 |
| Abitram | 0.496410202 | -0.333423734 | 0.143 | 0.171 | 1 | 0.843866241 | 0.185031894 | 0.1 | 0.117 | 1 | 0.843866241 | 0.746397315 |
| Hax1 | 0.496731909 | 0.130786608 | 0.546 | 0.506 | 1 | 0.865078671 | -0.140507455 | 0.5 | 0.469 | 1 | 0.865078671 | 0.746721228 |
| Ints10 | 0.958547113 | -0.237499314 | 0.143 | 0.141 | 1 | 0.496792988 | 0.902444691 | 0.2 | 0.123 | 1 | 0.958547113 | 0.746782703 |
| Kdm5d | 0.497254566 | -0.485426827 | 0.202 | 0.229 | 1 | 0.93457602 | -0.219117289 | 0.5 | 0.42 | 1 | 0.93457602 | 0.747247029 |
| Gnpda2 | 0.498426642 | -0.485426827 | 0.218 | 0.241 | 1 | 0.93224742 | 0.432959407 | 0.2 | 0.222 | 1 | 0.93224742 | 0.748424167 |
| Gm38604 | 0.498594647 | -0.485426827 | 0.017 | 0.029 | 1 | 0.669851164 | 1.695993813 | 0 | 0.019 | 1 | 0.669851164 | 0.748592672 |
| A630014C17Rik | 0.500300313 | -0.292781749 | 0.017 | 0.029 | 1 | 0.643842184 | 1.432959407 | 0 | 0.022 | 1 | 0.643842184 | 0.750300223 |
| Dlk2 | 0.500300313 | -0.292781749 | 0.017 | 0.029 | 1 | 0.643850855 | 2.017921908 | 0 | 0.022 | 1 | 0.643850855 | 0.750300223 |
| Gm50373 | 0.500307072 | 0.836501268 | 0.017 | 0.029 | 1 | 0.730750993 | 1.847996907 | 0 | 0.012 | 1 | 0.730750993 | 0.750306978 |
| Gm37529 | 0.501382816 | -0.485426827 | 0.067 | 0.088 | 1 | 0.675997103 | 0.111031312 | 0.1 | 0.145 | 1 | 0.675997103 | 0.751380904 |
| Klf5 | 0.502252321 | 0.118924659 | 0.597 | 0.553 | 1 | 0.660597177 | 0.123912604 | 0.5 | 0.559 | 1 | 0.660597177 | 0.752247248 |
| Cibar1 | 0.519004038 | -0.242856524 | 0.664 | 0.629 | 1 | 0.502848217 | -0.374395515 | 0.5 | 0.58 | 1 | 0.519004038 | 0.752840105 |
| Gm20186 | 0.503051401 | 0.343092705 | 0.403 | 0.388 | 1 | 0.858278871 | 0.41305985 | 0.3 | 0.417 | 1 | 0.858278871 | 0.75304209 |
| Tmem201 | 0.503394082 | -0.263034406 | 0.092 | 0.118 | 1 | 0.879964264 | 0.558490289 | 0.1 | 0.114 | 1 | 0.879964264 | 0.753382562 |
| Slc38a9 | 0.556541007 | -0.269698136 | 0.252 | 0.206 | 1 | 0.504022096 | 0.172431857 | 0.2 | 0.315 | 1 | 0.556541007 | 0.754005918 |
| Fgd4 | 0.662458041 | -0.418312631 | 0.134 | 0.147 | 1 | 0.504147558 | -0.325485914 | 0.2 | 0.306 | 1 | 0.662458041 | 0.754130355 |
| Gm15337 | 0.692072298 | 0.225066556 | 0.042 | 0.053 | 1 | 0.504201527 | 0.769994395 | 0.1 | 0.049 | 1 | 0.692072298 | 0.754183874 |
| 4631405J19Rik | 0.504214062 | -2.392317423 | 0.008 | 0.018 | 1 | 0.73075158 | 2.695993813 | 0 | 0.012 | 1 | 0.73075158 | 0.754196304 |
| C9 | 0.504214062 | -2.070389328 | 0.008 | 0.018 | 1 | 0.643849121 | 0.625604485 | 0 | 0.022 | 1 | 0.643849121 | 0.754196304 |
| Gcc2 | 0.50422145 | -0.621748298 | 0.529 | 0.506 | 1 | 0.597928483 | -0.175849835 | 0.5 | 0.627 | 1 | 0.597928483 | 0.754203629 |
| Tbcc | 0.818142894 | 0.209718591 | 0.202 | 0.188 | 1 | 0.504275543 | 0.988174565 | 0.2 | 0.123 | 1 | 0.818142894 | 0.754257263 |
| Lrrc1 | 0.504792024 | -0.115713318 | 0.597 | 0.635 | 1 | 0.781054259 | -0.210403469 | 0.9 | 0.821 | 1 | 0.781054259 | 0.75476906 |
| C030034I22Rik | 0.569304225 | -0.485426827 | 0.034 | 0.047 | 1 | 0.504821049 | 2.639410285 | 0.1 | 0.056 | 1 | 0.569304225 | 0.754797807 |
| Fmo5 | 0.651706649 | -0.807354922 | 0.076 | 0.088 | 1 | 0.504836594 | 3.532495081 | 0.1 | 0.056 | 1 | 0.651706649 | 0.754813202 |
| Gm56948 | 0.511005543 | -0.485426827 | 0.008 | 0.018 | 1 | 0.505582845 | 1.017921908 | 0 | 0.043 | 1 | 0.511005543 | 0.755551677 |
| 2310034G01Rik | 0.659299372 | 0.344648171 | 0.059 | 0.047 | 1 | 0.505582845 | 1.017921908 | 0 | 0.043 | 1 | 0.659299372 | 0.755551677 |
| Lrrc75a | 0.929388469 | 0.321928095 | 0.025 | 0.024 | 1 | 0.505582845 | 1.017921908 | 0 | 0.043 | 1 | 0.929388469 | 0.755551677 |
| Nr1h3 | 0.954551343 | -0.222392421 | 0.017 | 0.018 | 1 | 0.505582845 | 0.930459067 | 0 | 0.043 | 1 | 0.954551343 | 0.755551677 |
| Sgsh | 0.933394757 | -0.163498732 | 0.034 | 0.035 | 1 | 0.505605027 | 0.847996907 | 0 | 0.043 | 1 | 0.933394757 | 0.75557361 |
| Pcdhga8 | 0.990094116 | -0.485426827 | 0.042 | 0.041 | 1 | 0.505605027 | 0.847996907 | 0 | 0.043 | 1 | 0.990094116 | 0.75557361 |
| Sox13 | 0.721162042 | 0.514573173 | 0.017 | 0.012 | 1 | 0.505623509 | 0.769994395 | 0 | 0.043 | 1 | 0.721162042 | 0.755591885 |
| 1700085D07Rik | 0.511005543 | -0.485426827 | 0.008 | 0.018 | 1 | 0.505623793 | 0.847996907 | 0 | 0.043 | 1 | 0.511005543 | 0.755592166 |
| Slc4a8 | 0.627805946 | -0.748461233 | 0.025 | 0.035 | 1 | 0.505623793 | 0.695993813 | 0 | 0.043 | 1 | 0.627805946 | 0.755592166 |

| | | | | | | | | | | | | |
|---|---|---|---|---|---|---|---|---|---|---|---|---|
| Pbld2 | 0.629509811 | -0.292781749 | 0.025 | 0.035 | 1 | 0.505638293 | 0.695993813 | 0 | 0.043 | 1 | 0.629509811 | 0.755606503 |
| Coro2b | 0.833998399 | -0.321928095 | 0.101 | 0.106 | 1 | 0.505638293 | 0.695993813 | 0 | 0.043 | 1 | 0.833998399 | 0.755606503 |
| Kif24 | 0.881678885 | -1.137503524 | 0.05 | 0.053 | 1 | 0.505638293 | 0.558490289 | 0 | 0.043 | 1 | 0.881678885 | 0.755606503 |
| Pcdhga1 | 0.629509811 | -0.292781749 | 0.025 | 0.035 | 1 | 0.505639146 | 0.558490289 | 0 | 0.043 | 1 | 0.629509811 | 0.755607346 |
| Snap91 | 0.781790612 | -0.807354922 | 0.008 | 0.012 | 1 | 0.505639146 | 0.31748219 | 0 | 0.043 | 1 | 0.781790612 | 0.755607346 |
| Foxo6 | 0.508731273 | -0.807354922 | 0.008 | 0.018 | 1 | 0.505659614 | 0.374065718 | 0 | 0.043 | 1 | 0.508731273 | 0.755627583 |
| Gm17276 | 0.954551343 | -1.485426827 | 0.017 | 0.018 | 1 | 0.505659614 | 0.31748219 | 0 | 0.043 | 1 | 0.954551343 | 0.755627583 |
| Gm44148 | 0.651414376 | -0.707819249 | 0.042 | 0.053 | 1 | 0.50566132 | 0.159940913 | 0 | 0.043 | 1 | 0.651414376 | 0.75562927 |
| Mapk10 | 0.672494309 | -0.485426827 | 0.025 | 0.018 | 1 | 0.505670416 | 0.210566986 | 0 | 0.043 | 1 | 0.672494309 | 0.755638263 |
| Rnf180 | 0.569374548 | -1.137503524 | 0.034 | 0.047 | 1 | 0.505673259 | -0.709998547 | 0 | 0.043 | 1 | 0.569374548 | 0.755641073 |
| A230056P14Rik | 0.658336322 | -0.807354922 | 0.042 | 0.053 | 1 | 0.505686333 | 1.111031312 | 0 | 0.043 | 1 | 0.658336322 | 0.755653999 |
| Hsdl2 | 0.510644417 | 0.230780207 | 0.269 | 0.235 | 1 | 0.50598674 | 0.494359952 | 0.4 | 0.284 | 1 | 0.510644417 | 0.755950899 |
| Adam12 | 0.988120957 | 0.320749904 | 0.311 | 0.306 | 1 | 0.506209612 | -1.195857383 | 0.3 | 0.38 | 1 | 0.988120957 | 0.756171053 |
| Uqcc1 | 0.506242657 | -0.372045028 | 0.496 | 0.494 | 1 | 0.816144647 | -0.145308441 | 0.5 | 0.549 | 1 | 0.816144647 | 0.756203686 |
| Fmo1 | 0.506471296 | -1.070389328 | 0.008 | 0.018 | 1 | 0.577155236 | 1.017921908 | 0 | 0.031 | 1 | 0.577155236 | 0.756429418 |
| Gm6403 | 0.506471296 | -1.485426827 | 0.008 | 0.018 | 1 | 0.669853054 | 2.210566986 | 0 | 0.019 | 1 | 0.669853054 | 0.756429418 |
| Gcnt7 | 0.506471296 | -1.070389328 | 0.008 | 0.018 | 1 | 0.73075158 | 2.695993813 | 0 | 0.012 | 1 | 0.73075158 | 0.756429418 |
| Oas1b | 0.506471296 | -1.070389328 | 0.008 | 0.018 | 1 | 0.669853054 | 2.210566986 | 0 | 0.019 | 1 | 0.669853054 | 0.756429418 |
| Slc25a18 | 0.506471296 | -1.070389328 | 0.008 | 0.018 | 1 | 0.597824685 | 1.432959407 | 0 | 0.028 | 1 | 0.597824685 | 0.756429418 |
| Gm57242 | 0.506471296 | -1.359895945 | 0.008 | 0.018 | 1 | 0.698532411 | 1.432959407 | 0 | 0.015 | 1 | 0.698532411 | 0.756429418 |
| Aldh1a2 | 0.506471296 | -1.485426827 | 0.008 | 0.018 | 1 | 0.698529281 | 2.017921908 | 0 | 0.015 | 1 | 0.698529281 | 0.756429418 |
| Syk | 0.506471296 | -3.008988783 | 0.008 | 0.018 | 1 | 0.5978438 | 0.31748219 | 0 | 0.028 | 1 | 0.5978438 | 0.756429418 |
| Tigit | 0.506471296 | -1.070389328 | 0.008 | 0.018 | 1 | 0.577172123 | 0.769994395 | 0 | 0.031 | 1 | 0.577172123 | 0.756429418 |
| Gm36279 | 0.506471296 | -1.070389328 | 0.008 | 0.018 | 1 | 0.59781419 | 1.558490289 | 0 | 0.028 | 1 | 0.59781419 | 0.756429418 |
| Tmed8 | 0.668322198 | -0.365132593 | 0.176 | 0.147 | 1 | 0.506848851 | -0.391469028 | 0.1 | 0.176 | 1 | 0.668322198 | 0.756801944 |
| Ccdc152 | 0.733219616 | -1.070389328 | 0.034 | 0.041 | 1 | 0.507439125 | 1.432959407 | 0.1 | 0.052 | 1 | 0.733219616 | 0.757383785 |
| Omd | 0.71636102 | -0.367298122 | 0.983 | 0.947 | 1 | 0.508007008 | -0.454409262 | 1 | 0.91 | 1 | 0.71636102 | 0.757942896 |
| Actr6 | 0.611241902 | -0.222392421 | 0.168 | 0.188 | 1 | 0.508404355 | 1.672147071 | 0.2 | 0.136 | 1 | 0.611241902 | 0.758333722 |
| Mfap3l | 0.508432637 | 0.273565073 | 0.059 | 0.041 | 1 | 0.512834261 | 2.017921908 | 0.1 | 0.056 | 1 | 0.512834261 | 0.758361528 |
| Zfp414 | 0.508511877 | -0.292781749 | 0.143 | 0.171 | 1 | 0.923628199 | 0.558490289 | 0.1 | 0.108 | 1 | 0.923628199 | 0.758439425 |
| BC034090 | 0.508618987 | -0.544320516 | 0.084 | 0.106 | 1 | 0.699481299 | 0.31748219 | 0.1 | 0.142 | 1 | 0.699481299 | 0.7585447 |
| Itpka | 0.508731273 | -0.807354922 | 0.008 | 0.018 | 1 | 0.669848329 | 1.695993813 | 0 | 0.019 | 1 | 0.669848329 | 0.758655038 |
| Slc9b1 | 0.508731273 | -0.807354922 | 0.008 | 0.018 | 1 | 0.557765566 | 0.769994395 | 0 | 0.034 | 1 | 0.557765566 | 0.758655038 |
| Gpr63 | 0.508731273 | -0.807354922 | 0.008 | 0.018 | 1 | 0.669848329 | 1.847996907 | 0 | 0.019 | 1 | 0.669848329 | 0.758655038 |
| Lrrc19 | 0.508731273 | -1.292781749 | 0.008 | 0.018 | 1 | 0.557729726 | 1.31748219 | 0 | 0.034 | 1 | 0.557729726 | 0.758655038 |
| Zmynd12 | 0.508731273 | -0.807354922 | 0.008 | 0.018 | 1 | 0.669851164 | 1.432959407 | 0 | 0.019 | 1 | 0.669851164 | 0.758655038 |
| Tnfrsf25 | 0.508731273 | -1.070389328 | 0.008 | 0.018 | 1 | 0.522066388 | 1.111031312 | 0 | 0.04 | 1 | 0.522066388 | 0.758655038 |
| E130006D01Rik | 0.508731273 | -0.807354922 | 0.008 | 0.018 | 1 | 0.698534498 | 2.432959407 | 0 | 0.015 | 1 | 0.698534498 | 0.758655038 |
| Zfp977 | 0.508731273 | -1.070389328 | 0.008 | 0.018 | 1 | 0.6199755 | 1.847996907 | 0 | 0.025 | 1 | 0.6199755 | 0.758655038 |
| Trim66 | 0.508731273 | -0.807354922 | 0.008 | 0.018 | 1 | 0.64384652 | 1.558490289 | 0 | 0.022 | 1 | 0.64384652 | 0.758655038 |
| Mtnr1a | 0.508731273 | -1.070389328 | 0.008 | 0.018 | 1 | 0.669848329 | 1.847996907 | 0 | 0.019 | 1 | 0.669848329 | 0.758655038 |
| Spata33 | 0.508731273 | -0.807354922 | 0.008 | 0.018 | 1 | 0.698534498 | 2.432959407 | 0 | 0.015 | 1 | 0.698534498 | 0.758655038 |
| 5930403N24Rik | 0.508731273 | -0.807354922 | 0.008 | 0.018 | 1 | 0.669853054 | 2.210566986 | 0 | 0.019 | 1 | 0.669853054 | 0.758655038 |
| Trim16 | 0.508731273 | -1.070389328 | 0.008 | 0.018 | 1 | 0.669843604 | 1.695993813 | 0 | 0.019 | 1 | 0.669843604 | 0.758655038 |
| Evi2a | 0.508731273 | -0.807354922 | 0.008 | 0.018 | 1 | 0.619961444 | 1.31748219 | 0 | 0.025 | 1 | 0.619961444 | 0.758655038 |
| Hoxb9 | 0.508731273 | -0.807354922 | 0.008 | 0.018 | 1 | 0.619971484 | 1.31748219 | 0 | 0.025 | 1 | 0.619971484 | 0.758655038 |
| 6030468B19Rik | 0.508731273 | -0.807354922 | 0.008 | 0.018 | 1 | 0.698529281 | 2.210566986 | 0 | 0.015 | 1 | 0.698529281 | 0.758655038 |
| 9430024F10Rik | 0.508731273 | -0.807354922 | 0.008 | 0.018 | 1 | 0.698532411 | 1.695993813 | 0 | 0.015 | 1 | 0.698532411 | 0.758655038 |
| A330102I10Rik | 0.508731273 | -0.807354922 | 0.008 | 0.018 | 1 | 0.643835681 | 1.847996907 | 0 | 0.022 | 1 | 0.643835681 | 0.758655038 |
| Fam217a | 0.508731273 | -0.807354922 | 0.008 | 0.018 | 1 | 0.730749233 | 2.432959407 | 0 | 0.012 | 1 | 0.730749233 | 0.758655038 |
| Gm41396 | 0.508731273 | -0.807354922 | 0.008 | 0.018 | 1 | 0.643835681 | 1.847996907 | 0 | 0.022 | 1 | 0.643835681 | 0.758655038 |
| Pgap6 | 0.508731273 | -0.807354922 | 0.008 | 0.018 | 1 | 0.669843604 | 2.017921908 | 0 | 0.019 | 1 | 0.669843604 | 0.758655038 |
| Gng5c | 0.508731273 | -0.807354922 | 0.008 | 0.018 | 1 | 0.522066388 | 1.111031312 | 0 | 0.04 | 1 | 0.522066388 | 0.758655038 |
| Gm17315 | 0.508731273 | -0.807354922 | 0.008 | 0.018 | 1 | 0.643835681 | 1.847996907 | 0 | 0.022 | 1 | 0.643835681 | 0.758655038 |
| Pcdhb12 | 0.508731273 | -0.807354922 | 0.008 | 0.018 | 1 | 0.597838178 | 0.930459067 | 0 | 0.028 | 1 | 0.597838178 | 0.758655038 |
| Rnf165 | 0.508731273 | -0.807354922 | 0.008 | 0.018 | 1 | 0.730750993 | 1.558490289 | 0 | 0.012 | 1 | 0.730750993 | 0.758655038 |
| Tent5d | 0.508731273 | -0.807354922 | 0.008 | 0.018 | 1 | 0.730750993 | 2.210566986 | 0 | 0.012 | 1 | 0.730750993 | 0.758655038 |
| Miip | 0.510095062 | -0.807354922 | 0.042 | 0.059 | 1 | 0.73075158 | 2.695993813 | 0 | 0.012 | 1 | 0.73075158 | 0.759993151 |
| Ube2ql1 | 0.510828007 | 0.192645078 | 0.059 | 0.041 | 1 | 0.607556927 | 1.111031312 | 0.1 | 0.059 | 1 | 0.607556927 | 0.760710761 |
| Dlgap5 | 0.51100426 | -1.070389328 | 0.008 | 0.018 | 1 | 0.73075158 | 2.695993813 | 0 | 0.012 | 1 | 0.73075158 | 0.760883167 |
| AI661453 | 0.51100426 | -0.900464326 | 0.008 | 0.018 | 1 | 0.698529281 | 1.695993813 | 0 | 0.015 | 1 | 0.698529281 | 0.760883167 |
| B3gat2 | 0.511005543 | -0.485426827 | 0.008 | 0.018 | 1 | 0.698529281 | 1.847996907 | 0 | 0.015 | 1 | 0.698529281 | 0.760884421 |
| Gm37362 | 0.511005543 | -0.485426827 | 0.008 | 0.018 | 1 | 0.643850855 | 2.017921908 | 0 | 0.022 | 1 | 0.643850855 | 0.760884421 |
| Gm15852 | 0.511005543 | -0.485426827 | 0.008 | 0.018 | 1 | 0.59781419 | 1.558490289 | 0 | 0.028 | 1 | 0.59781419 | 0.760884421 |
| Frmd5 | 0.511005543 | -0.485426827 | 0.008 | 0.018 | 1 | 0.73075158 | 0.625604485 | 0 | 0.012 | 1 | 0.73075158 | 0.760884421 |
| Snap25 | 0.511005543 | -0.485426827 | 0.008 | 0.018 | 1 | 0.698532411 | 1.695993813 | 0 | 0.015 | 1 | 0.698532411 | 0.760884421 |
| Fer1l4 | 0.511005543 | -0.485426827 | 0.008 | 0.018 | 1 | 0.698533977 | 1.847996907 | 0 | 0.015 | 1 | 0.698533977 | 0.760884421 |
| Sptssb | 0.511005543 | -0.485426827 | 0.008 | 0.018 | 1 | 0.64384652 | 1.210566986 | 0 | 0.022 | 1 | 0.64384652 | 0.760884421 |
| Gm42699 | 0.511005543 | -0.485426827 | 0.008 | 0.018 | 1 | 0.730749233 | 2.432959407 | 0 | 0.012 | 1 | 0.730749233 | 0.760884421 |
| Gm12503 | 0.511005543 | -0.485426827 | 0.008 | 0.018 | 1 | 0.698529281 | 2.210566986 | 0 | 0.015 | 1 | 0.698529281 | 0.760884421 |
| Gm13091 | 0.511005543 | -0.485426827 | 0.008 | 0.018 | 1 | 0.698534498 | 2.432959407 | 0 | 0.015 | 1 | 0.698534498 | 0.760884421 |
| Fgfbp1 | 0.511005543 | -0.485426827 | 0.008 | 0.018 | 1 | 0.698534498 | 2.432959407 | 0 | 0.015 | 1 | 0.698534498 | 0.760884421 |
| Gm57291 | 0.511005543 | -0.485426827 | 0.008 | 0.018 | 1 | 0.669843604 | 1.847996907 | 0 | 0.019 | 1 | 0.669843604 | 0.760884421 |
| Gm56980 | 0.511005543 | -0.485426827 | 0.008 | 0.018 | 1 | 0.577172123 | 0.847996907 | 0 | 0.031 | 1 | 0.577172123 | 0.760884421 |
| C3ar1 | 0.511005543 | -0.485426827 | 0.008 | 0.018 | 1 | 0.730749233 | 2.432959407 | 0 | 0.012 | 1 | 0.730749233 | 0.760884421 |
| A230083G16Rik | 0.511005543 | -0.485426827 | 0.008 | 0.018 | 1 | 0.619961444 | 1.558490289 | 0 | 0.025 | 1 | 0.619961444 | 0.760884421 |
| Lrrc4b | 0.511005543 | -0.485426827 | 0.008 | 0.018 | 1 | 0.577164735 | 1.111031312 | 0 | 0.031 | 1 | 0.577164735 | 0.760884421 |
| Ttll13 | 0.511005543 | -0.485426827 | 0.008 | 0.018 | 1 | 0.539431168 | 1.111031312 | 0 | 0.037 | 1 | 0.539431168 | 0.760884421 |
| Fcsk | 0.511005543 | -0.485426827 | 0.008 | 0.018 | 1 | 0.629538555 | 1.558490289 | 0.1 | 0.062 | 1 | 0.629538555 | 0.760884421 |
| 2810408A11Rik | 0.511005543 | -0.485426827 | 0.008 | 0.018 | 1 | 0.698529281 | 2.017921908 | 0 | 0.015 | 1 | 0.698529281 | 0.760884421 |
| Coro6 | 0.511005543 | -0.485426827 | 0.008 | 0.018 | 1 | 0.73075158 | 2.695993813 | 0 | 0.012 | 1 | 0.73075158 | 0.760884421 |
| Gm57341 | 0.511005543 | -0.485426827 | 0.008 | 0.018 | 1 | 0.698534498 | 2.432959407 | 0 | 0.015 | 1 | 0.698534498 | 0.760884421 |
| Gm34868 | 0.511005543 | -0.485426827 | 0.008 | 0.018 | 1 | 0.698534498 | 2.432959407 | 0 | 0.015 | 1 | 0.698534498 | 0.760884421 |
| Slc10a1 | 0.511005543 | -0.485426827 | 0.008 | 0.018 | 1 | 0.619961444 | 1.432959407 | 0 | 0.025 | 1 | 0.619961444 | 0.760884421 |
| Nnt | 0.511005543 | -0.485426827 | 0.008 | 0.018 | 1 | 0.643850855 | 2.017921908 | 0 | 0.022 | 1 | 0.643850855 | 0.760884421 |
| Ccl28 | 0.511005543 | -0.485426827 | 0.008 | 0.018 | 1 | 0.698534498 | 2.432959407 | 0 | 0.015 | 1 | 0.698534498 | 0.760884421 |
| Rec8 | 0.511005543 | -0.485426827 | 0.008 | 0.018 | 1 | 0.669843604 | 1.847996907 | 0 | 0.019 | 1 | 0.669843604 | 0.760884421 |
| Gm49067 | 0.511005543 | -0.485426827 | 0.008 | 0.018 | 1 | 0.539455361 | 0.769994395 | 0 | 0.037 | 1 | 0.539455361 | 0.760884421 |
| Zfp54 | 0.511005543 | -0.485426827 | 0.008 | 0.018 | 1 | 0.539431168 | 1.111031312 | 0 | 0.037 | 1 | 0.539431168 | 0.760884421 |
| C3 | 0.511005543 | -0.485426827 | 0.008 | 0.018 | 1 | 0.619953009 | 1.558490289 | 0 | 0.025 | 1 | 0.619953009 | 0.760884421 |
| Lrrtm2 | 0.511005543 | -0.485426827 | 0.008 | 0.018 | 1 | 0.698534498 | 2.432959407 | 0 | 0.015 | 1 | 0.698534498 | 0.760884421 |
| C130060C02Rik | 0.511005543 | -0.485426827 | 0.008 | 0.018 | 1 | 0.730750993 | 2.210566986 | 0 | 0.012 | 1 | 0.730750993 | 0.760884421 |
| Cyp2c55 | 0.511005543 | -0.485426827 | 0.008 | 0.018 | 1 | 0.73075158 | 2.695993813 | 0 | 0.012 | 1 | 0.73075158 | 0.760884421 |
| Mageh1 | 0.677083422 | -0.124837112 | 0.555 | 0.482 | 1 | 0.511723603 | -0.144698293 | 0.5 | 0.593 | 1 | 0.677083422 | 0.76158616 |
| Fam174b | 0.512326114 | 0.251538767 | 0.017 | 0.029 | 1 | 0.619961444 | 1.558490289 | 0 | 0.025 | 1 | 0.619961444 | 0.762174181 |
| Scn9a | 0.513278666 | -0.485426827 | 0.008 | 0.018 | 1 | 0.698532411 | 1.558490289 | 0 | 0.015 | 1 | 0.698532411 | 0.763102343 |
| Yipf3 | 0.51358867 | -0.261386553 | 0.765 | 0.671 | 1 | 0.603971371 | -0.168605061 | 0.6 | 0.664 | 1 | 0.603971371 | 0.763404018 |
| Tecpr2 | 0.514924542 | -0.396159489 | 0.328 | 0.347 | 1 | 0.948519479 | -0.346650524 | 0.4 | 0.355 | 1 | 0.948519479 | 0.7647018 |

| | | | | | | | | | | | | |
|---|---|---|---|---|---|---|---|---|---|---|---|---|
| Ola1 | 0.515067978 | -0.309046804 | 0.639 | 0.641 | 1 | 0.628913436 | -0.253965504 | 0.5 | 0.648 | 1 | 0.628913436 | 0.764840934 |
| Pafah2 | 0.516061619 | -0.600904045 | 0.042 | 0.059 | 1 | 0.561898712 | 1.558490289 | 0.1 | 0.056 | 1 | 0.561898712 | 0.765803643 |
| Nanos1 | 0.516061619 | -0.600904045 | 0.042 | 0.059 | 1 | 0.964593759 | 0.494359952 | 0.1 | 0.102 | 1 | 0.964593759 | 0.765803643 |
| Hacd4 | 0.861972158 | -0.485426827 | 0.227 | 0.224 | 1 | 0.516144115 | 1.354956895 | 0.3 | 0.219 | 1 | 0.861972158 | 0.765883482 |
| Rnd2 | 0.5968991 | -0.281893433 | 0.143 | 0.165 | 1 | 0.516180601 | -0.152003093 | 0.1 | 0.176 | 1 | 0.5968991 | 0.765918789 |
| Foxj3 | 0.517349946 | -0.153230397 | 0.546 | 0.565 | 1 | 0.521869275 | -0.311201688 | 0.8 | 0.741 | 1 | 0.521869275 | 0.767048925 |
| Pdxk | 0.51851474 | -0.411426246 | 0.134 | 0.159 | 1 | 0.545345363 | -0.210896782 | 0.1 | 0.17 | 1 | 0.545345363 | 0.768171944 |
| Dnai4 | 0.520579061 | 0.116023796 | 0.126 | 0.1 | 1 | 0.91873618 | 0.463333056 | 0.1 | 0.108 | 1 | 0.91873618 | 0.770155563 |
| Zfp784 | 0.52803196 | 0.514573173 | 0.05 | 0.035 | 1 | 0.520825171 | 1.959028219 | 0.1 | 0.056 | 1 | 0.52803196 | 0.770391483 |
| Xrcc3 | 0.704462092 | 0.192645078 | 0.017 | 0.024 | 1 | 0.521021286 | 1.558490289 | 0.1 | 0.052 | 1 | 0.704462092 | 0.770579391 |
| Gm42938 | 0.721162042 | 0.514573173 | 0.017 | 0.012 | 1 | 0.521021286 | 1.695993813 | 0.1 | 0.052 | 1 | 0.721162042 | 0.770579391 |
| Zfp2 | 0.743002317 | -0.163498732 | 0.034 | 0.041 | 1 | 0.521021286 | 1.695993813 | 0.1 | 0.052 | 1 | 0.743002317 | 0.770579391 |
| Acsf3 | 0.837471739 | -0.137503524 | 0.042 | 0.047 | 1 | 0.521021286 | 1.695993813 | 0.1 | 0.052 | 1 | 0.837471739 | 0.770579391 |
| Zfp777 | 0.557333085 | -0.137503524 | 0.176 | 0.206 | 1 | 0.521580462 | 0.847996907 | 0.3 | 0.216 | 1 | 0.557333085 | 0.771114745 |
| 2610306M01Rik | 0.522060045 | -0.485426827 | 0.042 | 0.059 | 1 | 0.590778887 | 1.695993813 | 0.1 | 0.059 | 1 | 0.590778887 | 0.7715734 |
| 5830448L01Rik | 0.522060045 | -0.485426827 | 0.042 | 0.059 | 1 | 0.557729726 | 1.31748219 | 0 | 0.034 | 1 | 0.557729726 | 0.7715734 |
| Gm14403 | 0.609500043 | 0.514573173 | 0.034 | 0.024 | 1 | 0.522066388 | 1.111031312 | 0 | 0.04 | 1 | 0.609500043 | 0.771579463 |
| Gm15494 | 0.657769394 | 0.777607579 | 0.025 | 0.018 | 1 | 0.522066388 | 1.111031312 | 0 | 0.04 | 1 | 0.657769394 | 0.771579463 |
| 9330133O14Rik | 0.832144311 | -0.11345805 | 0.076 | 0.082 | 1 | 0.522066388 | 1.111031312 | 0 | 0.04 | 1 | 0.832144311 | 0.771579463 |
| Smim24 | 0.563722678 | -0.622930351 | 0.034 | 0.047 | 1 | 0.52208609 | 0.847996907 | 0 | 0.04 | 1 | 0.563722678 | 0.771598295 |
| Chek2 | 0.569304225 | -0.485426827 | 0.034 | 0.047 | 1 | 0.52208609 | 1.017921908 | 0 | 0.04 | 1 | 0.569304225 | 0.771598295 |
| A430035B10Rik | 0.619413067 | -0.622930351 | 0.025 | 0.035 | 1 | 0.52208609 | 0.930459067 | 0 | 0.04 | 1 | 0.619413067 | 0.771598295 |
| Taf6l | 0.695893386 | -0.222392421 | 0.017 | 0.024 | 1 | 0.52208609 | 1.017921908 | 0 | 0.04 | 1 | 0.695893386 | 0.771598295 |
| Ttc38 | 0.736440063 | -0.485426827 | 0.034 | 0.041 | 1 | 0.52208609 | 1.017921908 | 0 | 0.04 | 1 | 0.736440063 | 0.771598295 |
| Spns1 | 0.643761544 | 0.247092862 | 0.345 | 0.324 | 1 | 0.522096805 | 0.591657153 | 0.4 | 0.302 | 1 | 0.643761544 | 0.771608536 |
| Gdf15 | 0.606382401 | 2.522747104 | 0.059 | 0.076 | 1 | 0.522102209 | 0.494359952 | 0 | 0.04 | 1 | 0.606382401 | 0.771613701 |
| Hcls1 | 0.666161437 | 0.192645078 | 0.025 | 0.018 | 1 | 0.522102209 | 0.769994395 | 0 | 0.04 | 1 | 0.666161437 | 0.771613701 |
| 2610027K06Rik | 0.61095539 | -0.392317423 | 0.05 | 0.065 | 1 | 0.522102507 | 0.930459067 | 0 | 0.04 | 1 | 0.61095539 | 0.771613986 |
| Ano7 | 0.820836204 | -0.485426827 | 0.025 | 0.029 | 1 | 0.522102507 | 0.769994395 | 0 | 0.04 | 1 | 0.820836204 | 0.771613986 |
| AU040972 | 0.967167471 | 0.514573173 | 0.017 | 0.018 | 1 | 0.522115639 | 0.210566986 | 0 | 0.04 | 1 | 0.967167471 | 0.771626538 |
| Trpv2 | 0.737974447 | -0.707819249 | 0.05 | 0.041 | 1 | 0.522125488 | 0.374065718 | 0 | 0.04 | 1 | 0.737974447 | 0.77163595 |
| Cd274 | 0.780787367 | -0.378511623 | 0.042 | 0.035 | 1 | 0.522126383 | 0.432959407 | 0 | 0.04 | 1 | 0.780787367 | 0.771636806 |
| Kcnn3 | 0.691605844 | -0.485426827 | 0.017 | 0.024 | 1 | 0.52214011 | -0.111361109 | 0 | 0.04 | 1 | 0.691605844 | 0.771649925 |
| Hs3st5 | 0.731769749 | -0.485426827 | 0.017 | 0.012 | 1 | 0.522141303 | -0.789433014 | 0 | 0.04 | 1 | 0.731769749 | 0.771651066 |
| Fam220a | 0.613538034 | -0.263034406 | 0.05 | 0.065 | 1 | 0.522151747 | 1.210566986 | 0 | 0.04 | 1 | 0.613538034 | 0.771661047 |
| Gm43920 | 0.849201806 | -0.163498732 | 0.034 | 0.029 | 1 | 0.522151747 | 1.210566986 | 0 | 0.04 | 1 | 0.849201806 | 0.771661047 |
| Caprin2 | 0.881018668 | -0.27897595 | 0.084 | 0.088 | 1 | 0.522795999 | 0.159940913 | 0.2 | 0.12 | 1 | 0.881018668 | 0.772276341 |
| Fuca2 | 0.522852473 | -0.250961574 | 0.126 | 0.153 | 1 | 0.783642311 | 0.31748219 | 0.3 | 0.244 | 1 | 0.783642311 | 0.772330238 |
| Pigbos1 | 0.52289335 | -0.372952098 | 0.303 | 0.318 | 1 | 0.711898073 | 0.510127268 | 0.3 | 0.238 | 1 | 0.711898073 | 0.772369244 |
| Fbxl4 | 0.523404128 | -0.770829046 | 0.118 | 0.141 | 1 | 0.897905174 | 0.558490289 | 0.1 | 0.111 | 1 | 0.897905174 | 0.772856375 |
| Taf8 | 0.657253221 | -0.326729081 | 0.185 | 0.2 | 1 | 0.5235128 | 0.31748219 | 0.1 | 0.198 | 1 | 0.657253221 | 0.772959948 |
| Kcnj15 | 0.524318795 | -0.647698256 | 0.101 | 0.076 | 1 | 0.670291956 | 0.501615768 | 0.4 | 0.367 | 1 | 0.670291956 | 0.773727391 |
| Mcur1 | 0.525706842 | -0.383547213 | 0.286 | 0.312 | 1 | 0.541922868 | 0.41305985 | 0.2 | 0.312 | 1 | 0.541922868 | 0.775046 |
| Gm43329 | 0.525815167 | -0.11345805 | 0.067 | 0.088 | 1 | 0.611024044 | 0.625604485 | 0.2 | 0.136 | 1 | 0.611024044 | 0.775148744 |
| Rpgrip1l | 0.744304677 | -0.320367581 | 0.261 | 0.229 | 1 | 0.52600992 | 0.75918764 | 0.3 | 0.231 | 1 | 0.744304677 | 0.775333404 |
| Chac1 | 0.526433892 | -0.277318632 | 0.143 | 0.171 | 1 | 0.902664512 | 0.354956895 | 0.1 | 0.117 | 1 | 0.902664512 | 0.775735141 |
| Ipp | 0.527480144 | -0.359895945 | 0.084 | 0.106 | 1 | 0.905123661 | 0.660369903 | 0.1 | 0.111 | 1 | 0.905123661 | 0.776724986 |
| Ttc33 | 0.527557553 | 0.141114777 | 0.286 | 0.247 | 1 | 0.967876186 | 0.210566986 | 0.3 | 0.281 | 1 | 0.967876186 | 0.776798134 |
| Zfp157 | 0.527919153 | -0.453717967 | 0.143 | 0.165 | 1 | 0.851026605 | 0.653349476 | 0.2 | 0.182 | 1 | 0.851026605 | 0.777139674 |
| Mcm4 | 0.800683828 | -0.530750818 | 0.319 | 0.3 | 1 | 0.528311591 | 0.598383016 | 0.3 | 0.213 | 1 | 0.800683828 | 0.777510045 |
| Psat1 | 0.528769511 | -0.144389909 | 0.126 | 0.153 | 1 | 0.670447518 | 1.210566986 | 0.1 | 0.065 | 1 | 0.670447518 | 0.777941826 |
| Fnta | 0.528876744 | -0.174497731 | 0.681 | 0.688 | 1 | 0.787529553 | 0.138338658 | 0.6 | 0.688 | 1 | 0.787529553 | 0.778042878 |
| Zfp976 | 0.529350599 | -0.378511623 | 0.101 | 0.124 | 1 | 0.749513671 | 0.453137289 | 0.1 | 0.145 | 1 | 0.749513671 | 0.778489142 |
| Zfp719 | 0.529775041 | -0.222392421 | 0.143 | 0.171 | 1 | 0.55758256 | 0.672147071 | 0.2 | 0.13 | 1 | 0.55758256 | 0.778888488 |
| Gm29825 | 0.530703346 | 0.225066556 | 0.05 | 0.035 | 1 | 0.577164735 | 1.017921908 | 0 | 0.031 | 1 | 0.577164735 | 0.779760651 |
| Gm56582 | 0.663329737 | -0.137503524 | 0.101 | 0.118 | 1 | 0.530779388 | 1.079322453 | 0.2 | 0.13 | 1 | 0.663329737 | 0.779832017 |
| Nacc2 | 0.884251209 | -0.163498732 | 0.395 | 0.365 | 1 | 0.531016589 | -0.578267848 | 0.6 | 0.552 | 1 | 0.884251209 | 0.78005456 |
| Dcaf11 | 0.531159735 | -0.204245075 | 0.361 | 0.3 | 1 | 0.645176539 | -0.267480311 | 0.3 | 0.349 | 1 | 0.645176539 | 0.780188806 |
| Manea | 0.921063209 | -0.222392421 | 0.16 | 0.159 | 1 | 0.531375843 | -0.171902651 | 0.1 | 0.173 | 1 | 0.921063209 | 0.780391399 |
| Pcbp3 | 0.532093082 | -0.134519665 | 0.193 | 0.218 | 1 | 0.956955183 | -0.161987182 | 0.2 | 0.185 | 1 | 0.956955183 | 0.781063116 |
| Pals1 | 0.610368094 | -0.158049057 | 0.924 | 0.882 | 1 | 0.532196708 | -0.392362061 | 0.9 | 0.91 | 1 | 0.610368094 | 0.78116008 |
| C1rl | 0.532370314 | -0.807354922 | 0.059 | 0.076 | 1 | 0.91873697 | 0.558490289 | 0.1 | 0.108 | 1 | 0.91873697 | 0.781322477 |
| Ccl25 | 0.574729894 | -0.454399932 | 0.261 | 0.276 | 1 | 0.532853498 | -0.152003093 | 0.3 | 0.392 | 1 | 0.574729894 | 0.781774146 |
| Dusp16 | 0.533318292 | -0.537088947 | 0.479 | 0.465 | 1 | 0.98608358 | -0.207687226 | 0.7 | 0.605 | 1 | 0.98608358 | 0.782208184 |
| Aspscr1 | 0.533987992 | -0.288734916 | 0.336 | 0.353 | 1 | 0.659769794 | 0.867207511 | 0.5 | 0.54 | 1 | 0.659769794 | 0.782832809 |
| Rap1a | 0.84804165 | -0.113956189 | 0.975 | 0.935 | 1 | 0.534156839 | -0.249152855 | 1 | 0.978 | 1 | 0.84804165 | 0.78299015 |
| Mfsd14a | 0.534829945 | -0.107478647 | 0.597 | 0.624 | 1 | 0.56942206 | 0.436126686 | 0.6 | 0.688 | 1 | 0.56942206 | 0.78361682 |
| Arid5a | 0.969873156 | -0.154781515 | 0.437 | 0.412 | 1 | 0.535132586 | -0.159995884 | 0.2 | 0.315 | 1 | 0.969873156 | 0.783898287 |
| Exd1 | 0.535426657 | -0.436517227 | 0.101 | 0.124 | 1 | 0.649467485 | 0.888638891 | 0.1 | 0.062 | 1 | 0.649467485 | 0.784171609 |
| Rbck1 | 0.535594791 | -0.329775957 | 0.303 | 0.318 | 1 | 0.887596681 | 0.159940913 | 0.3 | 0.324 | 1 | 0.887596681 | 0.784327802 |
| Mcm8 | 0.536129185 | 0.344648171 | 0.05 | 0.035 | 1 | 0.698534498 | 2.432959407 | 0 | 0.015 | 1 | 0.698534498 | 0.784823867 |
| Paip2b | 0.720642819 | -0.19592021 | 0.345 | 0.353 | 1 | 0.536254627 | 0.443013072 | 0.4 | 0.299 | 1 | 0.720642819 | 0.784940229 |
| Triobp | 0.772864939 | -0.127874823 | 0.261 | 0.276 | 1 | 0.536610172 | 0.280956314 | 0.5 | 0.349 | 1 | 0.772864939 | 0.785269868 |
| Gm31718 | 0.7480211 | -0.392317423 | 0.05 | 0.059 | 1 | 0.537121713 | 1.558490289 | 0.1 | 0.056 | 1 | 0.7480211 | 0.785743692 |
| Dbp | 0.537317465 | -0.165167553 | 0.454 | 0.418 | 1 | 0.99525537 | 0.605140383 | 0.3 | 0.34 | 1 | 0.99525537 | 0.785924872 |
| 9330159F19Rik | 0.537493015 | 0.192645078 | 0.05 | 0.035 | 1 | 0.698532411 | 1.558490289 | 0 | 0.015 | 1 | 0.698532411 | 0.786087289 |
| Sntg2 | 0.556800866 | -0.807354922 | 0.034 | 0.047 | 1 | 0.537821322 | 0.888638891 | 0.1 | 0.052 | 1 | 0.556800866 | 0.78639087 |
| Uros | 0.657164563 | 0.219117289 | 0.16 | 0.141 | 1 | 0.538394642 | 0.988174565 | 0.2 | 0.13 | 1 | 0.657164563 | 0.786920493 |
| Gng4 | 0.616555609 | 0.292180751 | 0.034 | 0.024 | 1 | 0.539413885 | 1.111031312 | 0 | 0.037 | 1 | 0.616555609 | 0.787860431 |
| Vps25 | 0.626109212 | -0.402964667 | 0.067 | 0.082 | 1 | 0.539413885 | 1.210566986 | 0 | 0.037 | 1 | 0.626109212 | 0.787860431 |
| Dppa5a | 0.629509811 | -0.292781749 | 0.025 | 0.035 | 1 | 0.539413885 | 1.210566986 | 0 | 0.037 | 1 | 0.629509811 | 0.787860431 |
| Faim2 | 0.778665521 | -1.070389328 | 0.008 | 0.012 | 1 | 0.539413885 | 1.017921908 | 0 | 0.037 | 1 | 0.778665521 | 0.787860431 |
| Gm13708 | 0.998349055 | -0.359895945 | 0.042 | 0.041 | 1 | 0.539413885 | 1.111031312 | 0 | 0.037 | 1 | 0.998349055 | 0.787860431 |
| Psmb8 | 0.677915106 | 0.745898719 | 0.084 | 0.071 | 1 | 0.539431168 | 0.210566986 | 0 | 0.037 | 1 | 0.677915106 | 0.787876351 |
| Gm52870 | 0.687345921 | -0.707819249 | 0.017 | 0.024 | 1 | 0.539431168 | 1.017921908 | 0 | 0.037 | 1 | 0.687345921 | 0.787876351 |
| Gm43597 | 0.695893386 | -0.222392421 | 0.017 | 0.024 | 1 | 0.539431168 | 1.017921908 | 0 | 0.037 | 1 | 0.695893386 | 0.787876351 |
| Gm26881 | 0.817300263 | -0.600904045 | 0.042 | 0.047 | 1 | 0.539431168 | 1.111031312 | 0 | 0.037 | 1 | 0.817300263 | 0.787876351 |
| 5031434O11Rik | 0.820836204 | -0.655351829 | 0.025 | 0.029 | 1 | 0.539431168 | 0.930459067 | 0 | 0.037 | 1 | 0.820836204 | 0.787876351 |
| Aldh3b1 | 0.856663367 | -0.163498732 | 0.034 | 0.029 | 1 | 0.539431168 | 1.111031312 | 0 | 0.037 | 1 | 0.856663367 | 0.787876351 |
| Tha1 | 0.62441508 | -0.655351829 | 0.025 | 0.035 | 1 | 0.539445308 | 1.017921908 | 0 | 0.037 | 1 | 0.62441508 | 0.787889375 |
| Nudt17 | 0.630794806 | -0.333423734 | 0.034 | 0.024 | 1 | 0.539445308 | 1.017921908 | 0 | 0.037 | 1 | 0.630794806 | 0.787889375 |
| Ncf2 | 0.721161231 | 0.514573173 | 0.017 | 0.012 | 1 | 0.539445308 | 0.210566986 | 0 | 0.037 | 1 | 0.721161231 | 0.787889375 |
| Gpt | 0.721162042 | 0.514573173 | 0.017 | 0.012 | 1 | 0.539445308 | 0.847996907 | 0 | 0.037 | 1 | 0.721162042 | 0.787889375 |
| Mblac2 | 0.742864519 | -0.436517227 | 0.109 | 0.094 | 1 | 0.539445308 | 0.769994395 | 0 | 0.037 | 1 | 0.742864519 | 0.787889375 |
| Muc5b | 0.781790612 | -0.485426827 | 0.008 | 0.012 | 1 | 0.539455361 | 0.847996907 | 0 | 0.037 | 1 | 0.781790612 | 0.787898636 |
| Gm2011 | 0.62441508 | -0.655351829 | 0.025 | 0.035 | 1 | 0.539456618 | 0.930459067 | 0 | 0.037 | 1 | 0.62441508 | 0.787899793 |
| Gm26626 | 0.687345921 | -0.707819249 | 0.017 | 0.024 | 1 | 0.539462273 | 0.695993813 | 0 | 0.037 | 1 | 0.687345921 | 0.787905002 |
| Mtmr7 | 0.935509317 | -0.485426827 | 0.076 | 0.076 | 1 | 0.539470441 | 0.494359952 | 0 | 0.037 | 1 | 0.935509317 | 0.787912525 |

| | | | | | | | | | | | | |
|---|---|---|---|---|---|---|---|---|---|---|---|---|
| Cd72 | 0.954551343 | -1.485426827 | 0.017 | 0.018 | 1 | 0.539475781 | 0.159940913 | 0 | 0.037 | 1 | 0.954551343 | 0.787917444 |
| 1810021B22Rik | 0.575022075 | -0.333423734 | 0.034 | 0.047 | 1 | 0.539483006 | 1.31748219 | 0 | 0.037 | 1 | 0.575022075 | 0.787924098 |
| Gm29491 | 0.653579641 | 0.836501268 | 0.025 | 0.018 | 1 | 0.539483006 | 1.31748219 | 0 | 0.037 | 1 | 0.653579641 | 0.787924098 |
| Gm10135 | 0.695893386 | -0.222392421 | 0.017 | 0.024 | 1 | 0.539483006 | 1.31748219 | 0 | 0.037 | 1 | 0.695893386 | 0.787924098 |
| Zkscan14 | 0.539834531 | -0.655351829 | 0.059 | 0.076 | 1 | 0.710403435 | 1.159940913 | 0.1 | 0.068 | 1 | 0.710403435 | 0.788247741 |
| Lgi4 | 0.540366669 | -0.137503524 | 0.042 | 0.059 | 1 | 0.707564787 | 1.017921908 | 0.1 | 0.068 | 1 | 0.707564787 | 0.788737201 |
| Hemk1 | 0.540366669 | -0.137503524 | 0.042 | 0.059 | 1 | 0.557729726 | 1.210566986 | 0 | 0.034 | 1 | 0.557729726 | 0.788737201 |
| Mcf2l | 0.540446425 | -0.122856748 | 0.042 | 0.059 | 1 | 0.557779501 | 0.159940913 | 0 | 0.034 | 1 | 0.557779501 | 0.788810512 |
| Aldh7a1 | 0.541045664 | -0.210919637 | 0.555 | 0.559 | 1 | 0.672119164 | -0.283247627 | 0.5 | 0.506 | 1 | 0.672119164 | 0.789360917 |
| Stambpl1 | 0.545516192 | -0.533789849 | 0.286 | 0.229 | 1 | 0.541131333 | -0.111361109 | 0.1 | 0.188 | 1 | 0.545516192 | 0.789439546 |
| Surf2 | 0.914049774 | -0.343407822 | 0.235 | 0.229 | 1 | 0.541319942 | 0.678071905 | 0.1 | 0.198 | 1 | 0.914049774 | 0.789612605 |
| Rin3 | 0.542275319 | -0.308549065 | 0.176 | 0.141 | 1 | 0.885121327 | -0.327852929 | 0.2 | 0.204 | 1 | 0.885121327 | 0.790488117 |
| Vps11 | 0.542602386 | -0.347923303 | 0.176 | 0.2 | 1 | 0.849519108 | 0.397335498 | 0.3 | 0.272 | 1 | 0.849519108 | 0.790787423 |
| Tmem50b | 0.543417883 | -0.392317423 | 0.294 | 0.312 | 1 | 0.640289421 | 0.168256181 | 0.3 | 0.389 | 1 | 0.640289421 | 0.79153277 |
| Pdgfrl | 0.600857473 | -0.328701323 | 0.756 | 0.629 | 1 | 0.543805632 | -0.458891789 | 0.6 | 0.633 | 1 | 0.600857473 | 0.791886699 |
| Papolg | 0.543856741 | 0.251538767 | 0.21 | 0.182 | 1 | 0.934159891 | 0.432959407 | 0.2 | 0.222 | 1 | 0.934159891 | 0.791933327 |
| Zfp60 | 0.5440054 | -0.559427409 | 0.143 | 0.165 | 1 | 0.827573814 | 0.473601392 | 0.2 | 0.167 | 1 | 0.827573814 | 0.792068925 |
| Mpv17 | 0.958211849 | -0.105578756 | 0.345 | 0.335 | 1 | 0.544588889 | -0.452398027 | 0.2 | 0.272 | 1 | 0.958211849 | 0.79260072 |
| Mief1 | 0.54525569 | -0.411426246 | 0.202 | 0.224 | 1 | 0.936906035 | 0.558490289 | 0.2 | 0.198 | 1 | 0.936906035 | 0.793207613 |
| Gm7694 | 0.545647726 | 0.273565073 | 0.084 | 0.065 | 1 | 0.554647151 | -0.171902651 | 0.1 | 0.167 | 1 | 0.554647151 | 0.793564011 |
| Napg | 0.682095619 | -0.22650853 | 0.429 | 0.418 | 1 | 0.546300814 | -0.668578619 | 0.5 | 0.494 | 1 | 0.682095619 | 0.794157049 |
| Fbxo21 | 0.605925141 | -0.559427409 | 0.126 | 0.141 | 1 | 0.546968902 | -0.171902651 | 0.1 | 0.17 | 1 | 0.605925141 | 0.794762824 |
| Cndp2 | 0.591318757 | -0.273922722 | 0.16 | 0.182 | 1 | 0.546971357 | -0.171902651 | 0.1 | 0.17 | 1 | 0.591318757 | 0.794765048 |
| Sox5 | 0.906045612 | -0.123199748 | 1 | 0.988 | 1 | 0.549448899 | 0.153440945 | 1 | 0.991 | 1 | 0.906045612 | 0.797003705 |
| Swap70 | 0.549800249 | -0.263034406 | 0.454 | 0.488 | 1 | 0.76801373 | -0.353636955 | 0.5 | 0.552 | 1 | 0.76801373 | 0.797320185 |
| Parm1 | 0.549832743 | -0.30256277 | 0.084 | 0.106 | 1 | 0.681836701 | 0.210566986 | 0.1 | 0.065 | 1 | 0.681836701 | 0.797349441 |
| Plod1 | 0.550649879 | -0.153350777 | 0.622 | 0.529 | 1 | 0.602779605 | -0.200170376 | 0.5 | 0.565 | 1 | 0.602779605 | 0.798084468 |
| Mtg2 | 0.877271662 | -0.442358105 | 0.134 | 0.135 | 1 | 0.553722642 | 0.847996907 | 0.2 | 0.13 | 1 | 0.877271662 | 0.800836519 |
| Abca8b | 0.69920041 | -0.378511623 | 0.202 | 0.212 | 1 | 0.553904104 | 0.602884409 | 0.2 | 0.127 | 1 | 0.69920041 | 0.800998451 |
| Sox2ot | 0.55423075 | -1.284514133 | 0.118 | 0.135 | 1 | 0.881544009 | -1.611434712 | 0.2 | 0.157 | 1 | 0.881544009 | 0.801289776 |
| Arsa | 0.554391976 | 0.192645078 | 0.244 | 0.212 | 1 | 0.954869852 | 0.374065718 | 0.2 | 0.185 | 1 | 0.954869852 | 0.801433489 |
| Wdr55 | 0.554909411 | -0.392317423 | 0.059 | 0.076 | 1 | 0.949913217 | 0.847996907 | 0.1 | 0.093 | 1 | 0.949913217 | 0.801894367 |
| Sdha | 0.759849787 | -0.128000051 | 0.723 | 0.718 | 1 | 0.555016763 | -0.342141316 | 0.9 | 0.762 | 1 | 0.759849787 | 0.801989919 |
| Rep15 | 0.556257834 | 0.370183263 | 0.109 | 0.088 | 1 | 0.749384606 | 0.930459067 | 0.1 | 0.071 | 1 | 0.749384606 | 0.80309289 |
| Zfp960 | 0.566209975 | 0.514573173 | 0.042 | 0.029 | 1 | 0.557729726 | 1.31748219 | 0 | 0.034 | 1 | 0.566209975 | 0.804397005 |
| Catip | 0.602313349 | 0.514573173 | 0.067 | 0.053 | 1 | 0.557729726 | 0.847996907 | 0 | 0.034 | 1 | 0.602313349 | 0.804397005 |
| Mzf1 | 0.695893386 | -0.222392421 | 0.017 | 0.024 | 1 | 0.557729726 | 1.210566986 | 0 | 0.034 | 1 | 0.695893386 | 0.804397005 |
| 9630013D21Rik | 0.726737277 | -0.600904045 | 0.034 | 0.041 | 1 | 0.557729726 | 1.31748219 | 0 | 0.034 | 1 | 0.726737277 | 0.804397005 |
| Gm57182 | 0.74388966 | -0.378511623 | 0.05 | 0.059 | 1 | 0.557729726 | 1.210566986 | 0 | 0.034 | 1 | 0.74388966 | 0.804397005 |
| Snhg10 | 0.825035967 | -0.359895945 | 0.042 | 0.047 | 1 | 0.557729726 | 1.210566986 | 0 | 0.034 | 1 | 0.825035967 | 0.804397005 |
| Dennd6b | 0.902908911 | -0.122856748 | 0.067 | 0.071 | 1 | 0.557729726 | 1.210566986 | 0 | 0.034 | 1 | 0.902908911 | 0.804397005 |
| Amhr2 | 0.573820584 | 0.292180751 | 0.042 | 0.029 | 1 | 0.557744661 | 1.111031312 | 0 | 0.034 | 1 | 0.573820584 | 0.804410215 |
| Pmel | 0.63765316 | -0.987927168 | 0.042 | 0.053 | 1 | 0.557744661 | 1.210566986 | 0 | 0.034 | 1 | 0.63765316 | 0.804410215 |
| Haus5 | 0.91699934 | -0.137503524 | 0.05 | 0.053 | 1 | 0.557744661 | 1.111031312 | 0 | 0.034 | 1 | 0.91699934 | 0.804410215 |
| D730045B01Rik | 0.830633077 | 0.514573173 | 0.034 | 0.029 | 1 | 0.557756275 | 0.930459067 | 0 | 0.034 | 1 | 0.830633077 | 0.804420488 |
| Ccdc189 | 0.648435093 | 0.514573173 | 0.059 | 0.047 | 1 | 0.557756607 | 1.111031312 | 0 | 0.034 | 1 | 0.648435093 | 0.804420781 |
| Armc2 | 0.820836204 | -0.655351829 | 0.025 | 0.029 | 1 | 0.557756607 | 1.111031312 | 0 | 0.034 | 1 | 0.820836204 | 0.804420781 |
| C030014I23Rik | 0.830920729 | -0.185866545 | 0.059 | 0.065 | 1 | 0.557756607 | 0.930459067 | 0 | 0.034 | 1 | 0.830920729 | 0.804420781 |
| 4933424G06Rik | 0.88079732 | 0.514573173 | 0.05 | 0.047 | 1 | 0.557756607 | 0.930459067 | 0 | 0.034 | 1 | 0.88079732 | 0.804420781 |
| C78859 | 0.843628697 | 0.514573173 | 0.034 | 0.029 | 1 | 0.557764571 | 0.695993813 | 0 | 0.034 | 1 | 0.843628697 | 0.804427825 |
| Gm56999 | 0.619390946 | -0.944858446 | 0.025 | 0.035 | 1 | 0.557765566 | 0.847996907 | 0 | 0.034 | 1 | 0.619390946 | 0.804428705 |
| Tbx3os1 | 0.790856346 | -0.544320516 | 0.076 | 0.082 | 1 | 0.557765566 | 0.695993813 | 0 | 0.034 | 1 | 0.790856346 | 0.804428705 |
| Arhgef16 | 0.842422009 | 0.251538767 | 0.025 | 0.029 | 1 | 0.557765898 | 0.558490289 | 0 | 0.034 | 1 | 0.842422009 | 0.804428999 |
| Ptgs2os | 0.968129935 | -0.352160296 | 0.067 | 0.065 | 1 | 0.557772202 | 0.263034406 | 0 | 0.034 | 1 | 0.968129935 | 0.804434575 |
| Ctxnd1 | 0.954551343 | -0.707819249 | 0.017 | 0.018 | 1 | 0.557777511 | -0.596787936 | 0 | 0.034 | 1 | 0.954551343 | 0.80443927 |
| Fkbpl | 0.809580956 | -0.707819249 | 0.042 | 0.047 | 1 | 0.557784478 | 1.432959407 | 0 | 0.034 | 1 | 0.809580956 | 0.804445432 |
| Slc19a3 | 0.942200939 | -0.655351829 | 0.025 | 0.024 | 1 | 0.557784478 | 1.432959407 | 0 | 0.034 | 1 | 0.942200939 | 0.804445432 |
| Gm16192 | 0.558163762 | -1.070389328 | 0.034 | 0.047 | 1 | 0.577155236 | 1.31748219 | 0 | 0.031 | 1 | 0.577155236 | 0.804780738 |
| 4930481A15Rik | 0.558164171 | -0.748461233 | 0.034 | 0.047 | 1 | 0.698532411 | 1.847996907 | 0 | 0.015 | 1 | 0.698532411 | 0.8047811 |
| Gm57453 | 0.55853335 | 0.736965594 | 0.042 | 0.029 | 1 | 0.643850855 | 2.017921908 | 0 | 0.022 | 1 | 0.643850855 | 0.805107197 |
| Slc46a3 | 0.55853335 | 0.736965594 | 0.042 | 0.029 | 1 | 0.643835681 | 1.847996907 | 0 | 0.022 | 1 | 0.643835681 | 0.805107197 |
| Spag4 | 0.752076132 | -0.263034406 | 0.05 | 0.059 | 1 | 0.559081495 | 1.558490289 | 0.1 | 0.056 | 1 | 0.752076132 | 0.805590872 |
| Gm13523 | 0.8327876 | -0.222392421 | 0.042 | 0.047 | 1 | 0.559081495 | 1.625604485 | 0.1 | 0.056 | 1 | 0.8327876 | 0.805590872 |
| Cant1 | 0.660981477 | -0.122856748 | 0.202 | 0.224 | 1 | 0.559652534 | -0.152003093 | 0.1 | 0.167 | 1 | 0.660981477 | 0.806094109 |
| Rufy2 | 0.56097662 | -0.610957709 | 0.286 | 0.3 | 1 | 0.90335801 | 0.299103661 | 0.4 | 0.475 | 1 | 0.90335801 | 0.807258471 |
| 4930556J24Rik | 0.733219616 | -0.600904045 | 0.034 | 0.041 | 1 | 0.561898516 | 1.494359952 | 0.1 | 0.056 | 1 | 0.733219616 | 0.80806709 |
| D3Ertd751e | 0.624727185 | -0.600904045 | 0.227 | 0.188 | 1 | 0.562278809 | 2.091922489 | 0.2 | 0.16 | 1 | 0.624727185 | 0.808400159 |
| Zkscan3 | 0.645717377 | -0.163498732 | 0.647 | 0.612 | 1 | 0.56240357 | 0.114040062 | 0.8 | 0.67 | 1 | 0.645717377 | 0.808509364 |
| Gcdh | 0.562493726 | -0.30256277 | 0.151 | 0.176 | 1 | 0.790243401 | 0.345496566 | 0.1 | 0.127 | 1 | 0.790243401 | 0.80858826 |
| Gdap1 | 0.563722678 | -0.86393845 | 0.034 | 0.047 | 1 | 0.73075158 | 2.695993813 | 0 | 0.012 | 1 | 0.73075158 | 0.809662098 |
| Gm15728 | 0.563722678 | -0.748461233 | 0.034 | 0.047 | 1 | 0.784562821 | 0.808468542 | 0.1 | 0.074 | 1 | 0.784562821 | 0.809662098 |
| H3c6 | 0.563722678 | -0.970853654 | 0.034 | 0.047 | 1 | 0.597845674 | 1.695993813 | 0 | 0.028 | 1 | 0.597845674 | 0.809662098 |
| Ice2 | 0.564038111 | -0.722466024 | 0.193 | 0.206 | 1 | 0.788279795 | -0.230005605 | 0.2 | 0.222 | 1 | 0.788279795 | 0.809937232 |
| Kcnk1 | 0.572985371 | -0.241660755 | 0.479 | 0.406 | 1 | 0.564305233 | -0.563278674 | 0.4 | 0.475 | 1 | 0.572985371 | 0.81017007 |
| Bbs12 | 0.597182466 | 0.215012891 | 0.059 | 0.076 | 1 | 0.564718493 | 1.432959407 | 0.1 | 0.056 | 1 | 0.597182466 | 0.810530009 |
| Itgb3bp | 0.564983782 | -0.102098188 | 0.218 | 0.182 | 1 | 0.631405447 | 0.159940913 | 0.1 | 0.154 | 1 | 0.631405447 | 0.81076089 |
| Fam161b | 0.566169537 | 0.514573173 | 0.042 | 0.029 | 1 | 0.643842184 | 1.558490289 | 0 | 0.022 | 1 | 0.643842184 | 0.81179113 |
| Gm35248 | 0.566169537 | 0.514573173 | 0.042 | 0.029 | 1 | 0.669853054 | 2.210566986 | 0 | 0.019 | 1 | 0.669853054 | 0.81179113 |
| Guca1b | 0.566169537 | 0.514573173 | 0.042 | 0.029 | 1 | 0.577155236 | 1.31748219 | 0 | 0.031 | 1 | 0.577155236 | 0.81179113 |
| Kdm4d | 0.566193127 | 0.514573173 | 0.042 | 0.029 | 1 | 0.64384652 | 1.558490289 | 0 | 0.022 | 1 | 0.64384652 | 0.811811597 |
| Gm17259 | 0.566209975 | 0.514573173 | 0.042 | 0.029 | 1 | 0.669848329 | 1.847996907 | 0 | 0.019 | 1 | 0.669848329 | 0.811826214 |
| Edc3 | 0.566797993 | -0.11345805 | 0.235 | 0.194 | 1 | 0.648412879 | -0.191531458 | 0.2 | 0.25 | 1 | 0.648412879 | 0.812336021 |
| Pbld1 | 0.567953944 | -0.600904045 | 0.034 | 0.047 | 1 | 0.597845674 | 1.695993813 | 0 | 0.028 | 1 | 0.597845674 | 0.813336206 |
| Bhlhe41 | 0.568285456 | -0.500693584 | 0.252 | 0.271 | 1 | 0.792134708 | 0.393431043 | 0.1 | 0.136 | 1 | 0.792134708 | 0.813622553 |
| Mapk1ip1 | 0.970824425 | -0.522901533 | 0.134 | 0.129 | 1 | 0.569190405 | 0.695993813 | 0.2 | 0.13 | 1 | 0.970824425 | 0.814403093 |
| Egfl7 | 0.569304225 | -0.485426827 | 0.034 | 0.047 | 1 | 0.698534498 | 2.432959407 | 0 | 0.015 | 1 | 0.698534498 | 0.814501149 |
| Abcg1 | 0.569358879 | -0.485426827 | 0.034 | 0.047 | 1 | 0.893095315 | 0.345496566 | 0.1 | 0.111 | 1 | 0.893095315 | 0.814548225 |
| Gm47920 | 0.841777167 | 0.251538767 | 0.034 | 0.029 | 1 | 0.570367381 | 1.31748219 | 0.1 | 0.056 | 1 | 0.841777167 | 0.815415812 |
| Gm53610 | 0.93675107 | -0.439623138 | 0.092 | 0.094 | 1 | 0.570371234 | 1.374065718 | 0.1 | 0.056 | 1 | 0.93675107 | 0.815419124 |
| Gt(ROSA)26Sor | 0.571072874 | -0.209217033 | 0.361 | 0.394 | 1 | 0.736038228 | -0.33375353 | 0.5 | 0.472 | 1 | 0.736038228 | 0.81602152 |
| Abcb8 | 0.78506464 | 0.206450877 | 0.151 | 0.141 | 1 | 0.571318392 | 1.185031894 | 0.2 | 0.145 | 1 | 0.78506464 | 0.816232079 |
| 1110002L01Rik | 0.893973372 | -0.116193018 | 0.21 | 0.218 | 1 | 0.572029785 | 0.418009066 | 0.3 | 0.207 | 1 | 0.893973372 | 0.816841495 |
| Ctss | 0.572322988 | -1.129283017 | 0.042 | 0.029 | 1 | 0.681836963 | 0.888638891 | 0.1 | 0.065 | 1 | 0.681836963 | 0.817092374 |
| Catspere2 | 0.938019325 | -0.173482821 | 0.118 | 0.112 | 1 | 0.573092799 | 0.280956314 | 0.2 | 0.13 | 1 | 0.938019325 | 0.817750242 |
| Zfp442 | 0.929660875 | 0.13606155 | 0.067 | 0.065 | 1 | 0.573200928 | 1.111031312 | 0.1 | 0.056 | 1 | 0.929660875 | 0.817842552 |
| Hjurp | 0.573233783 | -0.426533138 | 0.378 | 0.3 | 1 | 0.670262149 | 0.231325546 | 0.4 | 0.302 | 1 | 0.670262149 | 0.817870596 |
| Gm5432 | 0.573820584 | 0.292180751 | 0.042 | 0.029 | 1 | 0.698529281 | 1.847996907 | 0 | 0.015 | 1 | 0.698529281 | 0.818371106 |
| Gm56637 | 0.573820584 | 0.292180751 | 0.042 | 0.029 | 1 | 0.676137417 | 1.159940913 | 0.1 | 0.065 | 1 | 0.676137417 | 0.818371106 |

| | | | | | | | | | | | | |
|---|---|---|---|---|---|---|---|---|---|---|---|---|
| 2810036E18Rik | 0.573850987 | 0.321928095 | 0.042 | 0.029 | 1 | 0.597840427 | 0.930459067 | 0 | 0.028 | 1 | 0.597840427 | 0.818397018 |
| Zwilch | 0.57438583 | -1.222392421 | 0.067 | 0.082 | 1 | 0.59781419 | 1.432959407 | 0 | 0.028 | 1 | 0.59781419 | 0.818852578 |
| Timm10b | 0.574626315 | -0.132411054 | 0.58 | 0.594 | 1 | 0.76203935 | -0.141949429 | 0.5 | 0.549 | 1 | 0.76203935 | 0.819057228 |
| Noxred1 | 0.574974342 | -0.359895945 | 0.034 | 0.047 | 1 | 0.63522121 | 1.31748219 | 0.1 | 0.062 | 1 | 0.63522121 | 0.81935319 |
| 4930545L23Rik | 0.575022075 | -0.333423734 | 0.034 | 0.047 | 1 | 0.896995938 | 0.769994395 | 0.1 | 0.086 | 1 | 0.896995938 | 0.819393763 |
| Sesn3 | 0.694095172 | -0.143507257 | 0.924 | 0.882 | 1 | 0.576206782 | -0.181750437 | 0.9 | 0.895 | 1 | 0.694095172 | 0.820399308 |
| Hmcn2 | 0.576415686 | -0.392317423 | 0.034 | 0.047 | 1 | 0.821582593 | 0.494359952 | 0.1 | 0.077 | 1 | 0.821582593 | 0.820576329 |
| Pmpcb | 0.652479206 | -0.145576824 | 0.487 | 0.488 | 1 | 0.576716222 | 0.154423908 | 0.4 | 0.537 | 1 | 0.652479206 | 0.820830843 |
| Zkscan7 | 0.613868002 | -0.250961574 | 0.076 | 0.059 | 1 | 0.577142568 | 1.432959407 | 0 | 0.031 | 1 | 0.613868002 | 0.821191592 |
| Wdr54 | 0.695893386 | -0.222392421 | 0.017 | 0.024 | 1 | 0.577142568 | 1.432959407 | 0 | 0.031 | 1 | 0.695893386 | 0.821191592 |
| Gm13524 | 0.704462092 | 0.192645078 | 0.017 | 0.024 | 1 | 0.577142568 | 1.210566986 | 0 | 0.031 | 1 | 0.704462092 | 0.821191592 |
| 4930509G22Rik | 0.619390946 | -0.655351829 | 0.025 | 0.035 | 1 | 0.577155236 | 1.31748219 | 0 | 0.031 | 1 | 0.619390946 | 0.821202305 |
| Gm49774 | 0.624459945 | -0.333423734 | 0.076 | 0.059 | 1 | 0.577155236 | 1.017921908 | 0 | 0.031 | 1 | 0.624459945 | 0.821202305 |
| Gm13938 | 0.629509811 | -0.292781749 | 0.025 | 0.035 | 1 | 0.577155236 | 1.31748219 | 0 | 0.031 | 1 | 0.629509811 | 0.821202305 |
| A930033M14Rik | 0.629509811 | -0.292781749 | 0.025 | 0.035 | 1 | 0.577155236 | 1.210566986 | 0 | 0.031 | 1 | 0.629509811 | 0.821202305 |
| Itprid1 | 0.721162042 | 0.514573173 | 0.017 | 0.012 | 1 | 0.577155236 | 1.31748219 | 0 | 0.031 | 1 | 0.721162042 | 0.821202305 |
| Cyria | 0.749411125 | -0.774933444 | 0.05 | 0.059 | 1 | 0.577155236 | 1.31748219 | 0 | 0.031 | 1 | 0.749411125 | 0.821202305 |
| Gm46212 | 0.817300436 | -0.707819249 | 0.042 | 0.047 | 1 | 0.577155236 | 1.31748219 | 0 | 0.031 | 1 | 0.817300436 | 0.821202305 |
| Gm15265 | 0.926217033 | -0.485426827 | 0.034 | 0.035 | 1 | 0.577155236 | 1.111031312 | 0 | 0.031 | 1 | 0.926217033 | 0.821202305 |
| Gm47095 | 0.954551343 | -0.222392421 | 0.017 | 0.018 | 1 | 0.577155236 | 1.210566986 | 0 | 0.031 | 1 | 0.954551343 | 0.821202305 |
| Slc4a5 | 0.611257237 | 0.344648171 | 0.034 | 0.024 | 1 | 0.577164735 | 1.111031312 | 0 | 0.031 | 1 | 0.611257237 | 0.821210339 |
| Gm14471 | 0.721162042 | 0.514573173 | 0.017 | 0.012 | 1 | 0.577164735 | 1.017921908 | 0 | 0.031 | 1 | 0.721162042 | 0.821210339 |
| Hpgds | 0.687345921 | -1.222392421 | 0.017 | 0.024 | 1 | 0.577165087 | 0.930459067 | 0 | 0.031 | 1 | 0.687345921 | 0.821210636 |
| 4921516A02Rik | 0.781790612 | -0.485426827 | 0.008 | 0.012 | 1 | 0.577165087 | 1.210566986 | 0 | 0.031 | 1 | 0.781790612 | 0.821210636 |
| Kdm8 | 0.935790323 | -0.655351829 | 0.025 | 0.024 | 1 | 0.577165087 | 1.017921908 | 0 | 0.031 | 1 | 0.935790323 | 0.821210636 |
| Abhd17a | 0.577172657 | -0.202354668 | 0.521 | 0.535 | 1 | 0.920089177 | 0.153735763 | 0.5 | 0.5 | 1 | 0.920089177 | 0.821217038 |
| Flt1 | 0.589353116 | 0.192645078 | 0.042 | 0.029 | 1 | 0.577177752 | 0.263034406 | 0 | 0.031 | 1 | 0.589353116 | 0.821221347 |
| 9330151L19Rik | 0.616536081 | 0.251538767 | 0.034 | 0.024 | 1 | 0.577184788 | 1.558490289 | 0 | 0.031 | 1 | 0.616536081 | 0.821227297 |
| Gm15441 | 0.695893386 | -0.222392421 | 0.017 | 0.024 | 1 | 0.577184788 | 1.558490289 | 0 | 0.031 | 1 | 0.695893386 | 0.821227297 |
| Ovca2 | 0.695893386 | -0.222392421 | 0.017 | 0.024 | 1 | 0.577184788 | 1.558490289 | 0 | 0.031 | 1 | 0.695893386 | 0.821227297 |
| Hmga1b | 0.951369735 | 0.292180751 | 0.034 | 0.035 | 1 | 0.577184788 | 1.558490289 | 0 | 0.031 | 1 | 0.951369735 | 0.821227297 |
| Ankzf1 | 0.577421493 | 0.116023796 | 0.16 | 0.135 | 1 | 0.97216902 | 0.660369903 | 0.1 | 0.102 | 1 | 0.97216902 | 0.821427406 |
| Ccdc174 | 0.578001324 | -0.290170536 | 0.504 | 0.488 | 1 | 0.629027042 | -0.290417122 | 0.5 | 0.54 | 1 | 0.629027042 | 0.821917117 |
| Me3 | 0.883350772 | -0.333423734 | 0.084 | 0.088 | 1 | 0.57808529 | 0.191373421 | 0.3 | 0.219 | 1 | 0.883350772 | 0.821987977 |
| Lrrc15 | 0.578297036 | -0.510088881 | 0.059 | 0.076 | 1 | 0.73075158 | 2.695993813 | 0 | 0.012 | 1 | 0.73075158 | 0.82216661 |
| Eef1akmt3 | 0.57840222 | -0.432959407 | 0.092 | 0.112 | 1 | 0.905111698 | 0.591657153 | 0.1 | 0.086 | 1 | 0.905111698 | 0.822255312 |
| Trim26 | 0.599237957 | -0.405256478 | 0.269 | 0.282 | 1 | 0.579180318 | -0.339630097 | 0.3 | 0.373 | 1 | 0.599237957 | 0.822910795 |
| Rubcn | 0.730692492 | -0.189970944 | 0.328 | 0.294 | 1 | 0.580455759 | 0.576486719 | 0.5 | 0.373 | 1 | 0.730692492 | 0.82398263 |
| Gm49959 | 0.581576556 | 0.321928095 | 0.134 | 0.112 | 1 | 0.814355306 | 0.41305985 | 0.2 | 0.164 | 1 | 0.814355306 | 0.824921822 |
| Ggact | 0.874760046 | -0.298341275 | 0.252 | 0.229 | 1 | 0.582680506 | -0.315978829 | 0.2 | 0.269 | 1 | 0.874760046 | 0.82584444 |
| Tspan14 | 0.660093998 | -0.419538892 | 0.336 | 0.276 | 1 | 0.582821015 | -0.214738849 | 0.3 | 0.398 | 1 | 0.660093998 | 0.825961695 |
| Snhg7 | 0.915874043 | -0.332226973 | 0.613 | 0.529 | 1 | 0.5833126 | 0.214867123 | 0.6 | 0.5 | 1 | 0.915874043 | 0.826371611 |
| Icmt | 0.857387615 | -0.109917692 | 0.168 | 0.176 | 1 | 0.585381113 | 0.639410285 | 0.3 | 0.228 | 1 | 0.857387615 | 0.828091178 |
| Dcxr | 0.586681172 | 0.877143252 | 0.118 | 0.1 | 1 | 0.770328444 | 0.210566986 | 0.1 | 0.13 | 1 | 0.770328444 | 0.829167547 |
| Snx17 | 0.86985849 | -0.146220741 | 0.571 | 0.553 | 1 | 0.586752861 | 0.322113639 | 0.7 | 0.577 | 1 | 0.86985849 | 0.829226802 |
| Secisbp2 | 0.725581973 | -0.111031312 | 0.487 | 0.418 | 1 | 0.587463484 | 0.18142064 | 0.5 | 0.377 | 1 | 0.725581973 | 0.829813623 |
| Zfp27 | 0.61789105 | -0.539874611 | 0.084 | 0.1 | 1 | 0.588204061 | 0.720241359 | 0.2 | 0.133 | 1 | 0.61789105 | 0.830424105 |
| Tcaim | 0.860740596 | -0.372952098 | 0.151 | 0.153 | 1 | 0.588204061 | 0.720241359 | 0.2 | 0.133 | 1 | 0.860740596 | 0.830424105 |
| Irgm2 | 0.661757414 | -0.870090677 | 0.193 | 0.165 | 1 | 0.589212695 | 0.896906507 | 0.2 | 0.151 | 1 | 0.661757414 | 0.83125379 |
| Thap1 | 0.643915774 | -0.222392421 | 0.134 | 0.153 | 1 | 0.589340857 | 1.210566986 | 0.2 | 0.148 | 1 | 0.643915774 | 0.831359068 |
| Fbxo28 | 0.58937845 | -0.231109638 | 0.555 | 0.535 | 1 | 0.592536727 | -0.465893869 | 0.7 | 0.596 | 1 | 0.592536727 | 0.831389943 |
| Mrpl44 | 0.590491255 | 0.114035243 | 0.185 | 0.159 | 1 | 0.624802901 | 1.135278859 | 0.2 | 0.154 | 1 | 0.624802901 | 0.832302588 |
| Fahd1 | 0.854084332 | -0.222392421 | 0.109 | 0.1 | 1 | 0.590778887 | 1.695993813 | 0.1 | 0.059 | 1 | 0.854084332 | 0.832538081 |
| Tctn3 | 0.981805003 | -0.185866545 | 0.059 | 0.059 | 1 | 0.590778887 | 1.695993813 | 0.1 | 0.059 | 1 | 0.981805003 | 0.832538081 |
| Mtmr4 | 0.591421978 | -0.584962501 | 0.05 | 0.065 | 1 | 0.925765422 | 1.432959407 | 0.1 | 0.099 | 1 | 0.925765422 | 0.833064 |
| Arl3 | 0.875725832 | -0.101333035 | 0.832 | 0.776 | 1 | 0.591511837 | -0.169430165 | 0.6 | 0.738 | 1 | 0.875725832 | 0.83313742 |
| Fbp1 | 0.822639881 | 0.251538767 | 0.084 | 0.076 | 1 | 0.5933729 | 1.625604485 | 0.1 | 0.059 | 1 | 0.822639881 | 0.834654402 |
| Elmo3 | 0.856663367 | -0.163498732 | 0.034 | 0.029 | 1 | 0.5933729 | 1.494359952 | 0.1 | 0.059 | 1 | 0.856663367 | 0.834654402 |
| Ndufs5 | 0.6124823 | 0.105116706 | 0.874 | 0.835 | 1 | 0.594468036 | -0.258918297 | 0.7 | 0.741 | 1 | 0.6124823 | 0.835543827 |
| Fank1 | 0.595456625 | 1 | 0.034 | 0.024 | 1 | 0.597832181 | 1.210566986 | 0 | 0.028 | 1 | 0.597832181 | 0.836344658 |
| Ddit3 | 0.936205458 | 0.340543773 | 0.403 | 0.412 | 1 | 0.596090027 | -0.332575339 | 0.3 | 0.373 | 1 | 0.936205458 | 0.836856734 |
| Prelid3b | 0.596159594 | -0.19592021 | 0.403 | 0.406 | 1 | 0.926527147 | 0.374065718 | 0.4 | 0.423 | 1 | 0.926527147 | 0.836912926 |
| 9030025P20Rik | 0.929389703 | 0.192645078 | 0.025 | 0.024 | 1 | 0.596204587 | 1.558490289 | 0.1 | 0.059 | 1 | 0.929389703 | 0.836949265 |
| Cyb561d2 | 0.596389646 | -0.23878686 | 0.227 | 0.253 | 1 | 0.826469788 | 0.354956895 | 0.2 | 0.164 | 1 | 0.826469788 | 0.837098682 |
| Ift27 | 0.768704427 | -0.222392421 | 0.378 | 0.371 | 1 | 0.597465446 | -0.171902651 | 0.3 | 0.358 | 1 | 0.768704427 | 0.837965933 |
| 3830408C21Rik | 0.597541705 | -0.485426827 | 0.05 | 0.065 | 1 | 0.643835681 | 1.847996907 | 0 | 0.022 | 1 | 0.643835681 | 0.838027321 |
| Serinc2 | 0.634555959 | -0.163498732 | 0.025 | 0.035 | 1 | 0.59781419 | 1.558490289 | 0 | 0.028 | 1 | 0.634555959 | 0.838246574 |
| Lrrc61 | 0.634555959 | -0.163498732 | 0.025 | 0.035 | 1 | 0.59781419 | 1.558490289 | 0 | 0.028 | 1 | 0.634555959 | 0.838246574 |
| Gm56795 | 0.695893386 | -0.222392421 | 0.017 | 0.024 | 1 | 0.59781419 | 1.31748219 | 0 | 0.028 | 1 | 0.695893386 | 0.838246574 |
| Gm43728 | 0.695893386 | -0.222392421 | 0.017 | 0.024 | 1 | 0.59781419 | 1.558490289 | 0 | 0.028 | 1 | 0.695893386 | 0.838246574 |
| Gm56760 | 0.695893386 | -0.222392421 | 0.017 | 0.024 | 1 | 0.59781419 | 1.558490289 | 0 | 0.028 | 1 | 0.695893386 | 0.838246574 |
| Ldhd | 0.718517371 | 0.836501268 | 0.017 | 0.012 | 1 | 0.59781419 | 1.558490289 | 0 | 0.028 | 1 | 0.718517371 | 0.838246574 |
| Mettl18 | 0.791189474 | 0.929610672 | 0.008 | 0.012 | 1 | 0.59781419 | 1.210566986 | 0 | 0.028 | 1 | 0.791189474 | 0.838246574 |
| Sp140 | 0.820836204 | -0.485426827 | 0.025 | 0.029 | 1 | 0.59781419 | 1.558490289 | 0 | 0.028 | 1 | 0.820836204 | 0.838246574 |
| Gm29170 | 0.942200939 | -0.292781749 | 0.025 | 0.024 | 1 | 0.59781419 | 1.558490289 | 0 | 0.028 | 1 | 0.942200939 | 0.838246574 |
| Kcnk7 | 0.954551343 | -0.222392421 | 0.017 | 0.018 | 1 | 0.59781419 | 1.558490289 | 0 | 0.028 | 1 | 0.954551343 | 0.838246574 |
| Il4ra | 0.998349055 | -0.600904045 | 0.042 | 0.041 | 1 | 0.59781419 | 1.558490289 | 0 | 0.028 | 1 | 0.998349055 | 0.838246574 |
| Hecw1 | 0.691615381 | -0.655351829 | 0.017 | 0.024 | 1 | 0.597824685 | 0.769994395 | 0 | 0.028 | 1 | 0.691615381 | 0.838255016 |
| B230208H11Rik | 0.697374687 | -0.292781749 | 0.059 | 0.047 | 1 | 0.597824685 | 1.432959407 | 0 | 0.028 | 1 | 0.697374687 | 0.838255016 |
| Gm56904 | 0.817300436 | -0.485426827 | 0.042 | 0.047 | 1 | 0.597824685 | 1.432959407 | 0 | 0.028 | 1 | 0.817300436 | 0.838255016 |
| Samd13 | 0.933394757 | -0.333423734 | 0.034 | 0.035 | 1 | 0.597824685 | 1.31748219 | 0 | 0.028 | 1 | 0.933394757 | 0.838255016 |
| Zfp473 | 0.949508638 | -1.070389328 | 0.017 | 0.018 | 1 | 0.597824685 | 1.31748219 | 0 | 0.028 | 1 | 0.949508638 | 0.838255016 |
| Cenpk | 0.864460014 | -0.918386234 | 0.067 | 0.071 | 1 | 0.597832181 | 1.210566986 | 0 | 0.028 | 1 | 0.864460014 | 0.838261046 |
| Gpr135 | 0.825035967 | -0.359895945 | 0.042 | 0.047 | 1 | 0.597832556 | 1.31748219 | 0 | 0.028 | 1 | 0.825035967 | 0.838261347 |
| Tbc1d30 | 0.698028986 | -0.655351829 | 0.017 | 0.024 | 1 | 0.597837803 | 1.017921908 | 0 | 0.028 | 1 | 0.698028986 | 0.838265568 |
| Shank2 | 0.864135893 | -0.333423734 | 0.034 | 0.029 | 1 | 0.597837803 | 0.930459067 | 0 | 0.028 | 1 | 0.864135893 | 0.838265568 |
| Sgo1 | 0.919045149 | -0.485426827 | 0.034 | 0.035 | 1 | 0.597838178 | 1.210566986 | 0 | 0.028 | 1 | 0.919045149 | 0.838265869 |
| Rab26 | 0.933394757 | -0.163498732 | 0.034 | 0.035 | 1 | 0.597838178 | 1.210566986 | 0 | 0.028 | 1 | 0.933394757 | 0.838265869 |
| Tnfaip3 | 0.944467697 | -0.707819249 | 0.017 | 0.018 | 1 | 0.597838178 | 1.210566986 | 0 | 0.028 | 1 | 0.944467697 | 0.838265869 |
| 1700010K24Rik | 0.651178013 | 0.862496476 | 0.059 | 0.047 | 1 | 0.597840427 | 0.930459067 | 0 | 0.028 | 1 | 0.651178013 | 0.838267678 |
| Mx2 | 0.954551343 | -0.707819249 | 0.017 | 0.018 | 1 | 0.597843425 | 0.263034406 | 0 | 0.028 | 1 | 0.954551343 | 0.838270089 |
| Oasl2 | 0.63457549 | -0.185866545 | 0.025 | 0.035 | 1 | 0.5978438 | -0.374395515 | 0 | 0.028 | 1 | 0.63457549 | 0.838270391 |
| Gm11175 | 0.62441508 | -0.485426827 | 0.025 | 0.035 | 1 | 0.597845674 | 1.695993813 | 0 | 0.028 | 1 | 0.62441508 | 0.838271898 |
| Thnsl1 | 0.648435093 | 0.514573173 | 0.059 | 0.047 | 1 | 0.597845674 | 1.695993813 | 0 | 0.028 | 1 | 0.648435093 | 0.838271898 |
| Plek2 | 0.653579641 | 1.099535674 | 0.025 | 0.018 | 1 | 0.597845674 | 1.695993813 | 0 | 0.028 | 1 | 0.653579641 | 0.838271898 |
| BC025920 | 0.687345921 | -0.707819249 | 0.017 | 0.024 | 1 | 0.597845674 | 1.695993813 | 0 | 0.028 | 1 | 0.687345921 | 0.838271898 |
| Gm13166 | 0.820836204 | -0.655351829 | 0.025 | 0.029 | 1 | 0.597845674 | 1.695993813 | 0 | 0.028 | 1 | 0.820836204 | 0.838271898 |
| Mroh7 | 0.949508638 | -0.485426827 | 0.017 | 0.018 | 1 | 0.597845674 | 1.695993813 | 0 | 0.028 | 1 | 0.949508638 | 0.838271898 |

| | | | | | | | | | | | | |
|---|---|---|---|---|---|---|---|---|---|---|---|---|
| Sat2 | 0.706962352 | -0.24726709 | 0.168 | 0.182 | 1 | 0.599232526 | 0.648688098 | 0.2 | 0.136 | 1 | 0.706962352 | 0.839385432 |
| Zfp692 | 0.607734737 | -0.235448574 | 0.143 | 0.165 | 1 | 0.599232773 | 0.695993813 | 0.2 | 0.136 | 1 | 0.607734737 | 0.83938563 |
| Arhgef1 | 0.782252743 | -0.180013819 | 0.807 | 0.741 | 1 | 0.599637885 | 0.306427001 | 0.8 | 0.827 | 1 | 0.782252743 | 0.839710177 |
| Nsg2 | 0.602438591 | 1 | 0.034 | 0.024 | 1 | 0.730749233 | 2.210566986 | 0 | 0.012 | 1 | 0.730749233 | 0.841944926 |
| L1cam | 0.602458696 | 0.736965594 | 0.034 | 0.024 | 1 | 0.643835681 | 1.847996907 | 0 | 0.022 | 1 | 0.643835681 | 0.841960912 |
| Myo15a | 0.602963643 | -0.600904045 | 0.084 | 0.1 | 1 | 0.865784955 | 0.111031312 | 0.1 | 0.114 | 1 | 0.865784955 | 0.842362131 |
| Cln3 | 0.603599254 | -0.402964667 | 0.134 | 0.153 | 1 | 0.910678077 | 0.41305985 | 0.2 | 0.179 | 1 | 0.910678077 | 0.842866449 |
| Zbtb18 | 0.603640044 | 0.292180751 | 0.067 | 0.053 | 1 | 0.740899751 | 1.159940913 | 0.1 | 0.071 | 1 | 0.740899751 | 0.842898785 |
| Gm57463 | 0.60429662 | -0.36994961 | 0.092 | 0.112 | 1 | 0.947274527 | 0.808468542 | 0.1 | 0.093 | 1 | 0.947274527 | 0.843418835 |
| Rspo1 | 0.606062622 | -1.250961574 | 0.025 | 0.035 | 1 | 0.669848329 | 1.847996907 | 0 | 0.019 | 1 | 0.669848329 | 0.844813343 |
| Traf2 | 0.618399472 | -0.378511623 | 0.101 | 0.118 | 1 | 0.606437892 | 0.354956895 | 0.1 | 0.176 | 1 | 0.618399472 | 0.845108867 |
| Gm29340 | 0.609500043 | 0.514573173 | 0.034 | 0.024 | 1 | 0.669851164 | 1.695993813 | 0 | 0.019 | 1 | 0.669851164 | 0.847509784 |
| H2bc21 | 0.609500043 | 0.514573173 | 0.034 | 0.024 | 1 | 0.619961444 | 1.558490289 | 0 | 0.025 | 1 | 0.619961444 | 0.847509784 |
| Tril | 0.609500043 | 0.514573173 | 0.034 | 0.024 | 1 | 0.73075158 | 2.695993813 | 0 | 0.012 | 1 | 0.73075158 | 0.847509784 |
| Hhla1 | 0.609500043 | 0.514573173 | 0.034 | 0.024 | 1 | 0.669853054 | 2.210566986 | 0 | 0.019 | 1 | 0.669853054 | 0.847509784 |
| Gm56728 | 0.609500043 | 0.514573173 | 0.034 | 0.024 | 1 | 0.643850855 | 2.017921908 | 0 | 0.022 | 1 | 0.643850855 | 0.847509784 |
| Gdap2 | 0.609690011 | -0.177304532 | 0.311 | 0.276 | 1 | 0.647393638 | -0.385800278 | 0.2 | 0.247 | 1 | 0.647393638 | 0.847658113 |
| Cnep1r1 | 0.776963521 | -0.238116774 | 0.765 | 0.647 | 1 | 0.611360692 | 0.273248219 | 0.6 | 0.741 | 1 | 0.776963521 | 0.848959488 |
| 9030612E09Rik | 0.612185957 | -0.292781749 | 0.05 | 0.065 | 1 | 0.619953009 | 1.695993813 | 0 | 0.025 | 1 | 0.619953009 | 0.849600268 |
| Gm47664 | 0.612185957 | -0.292781749 | 0.05 | 0.065 | 1 | 0.619953009 | 1.695993813 | 0 | 0.025 | 1 | 0.619953009 | 0.849600268 |
| Slco1c1 | 0.612991737 | -0.311397427 | 0.084 | 0.1 | 1 | 0.897033968 | -0.348400306 | 0.2 | 0.17 | 1 | 0.897033968 | 0.850224604 |
| Ggt7 | 0.614381507 | -1.392317423 | 0.025 | 0.035 | 1 | 0.730750993 | 1.432959407 | 0 | 0.012 | 1 | 0.730750993 | 0.851298378 |
| Pcdhb5 | 0.616536081 | 0.251538767 | 0.034 | 0.024 | 1 | 0.669848329 | 1.558490289 | 0 | 0.019 | 1 | 0.669848329 | 0.852955423 |
| A730090N16Rik | 0.616555609 | 0.292180751 | 0.034 | 0.024 | 1 | 0.643850855 | 2.017921908 | 0 | 0.022 | 1 | 0.643850855 | 0.852970399 |
| Mospd3 | 0.652489961 | -0.451479495 | 0.336 | 0.329 | 1 | 0.617568514 | 0.785261151 | 0.3 | 0.244 | 1 | 0.652489961 | 0.853746159 |
| Slc25a36 | 0.729408699 | -0.148031931 | 0.782 | 0.824 | 1 | 0.617658553 | -0.218404319 | 0.8 | 0.852 | 1 | 0.729408699 | 0.853815018 |
| 4930500A05Rik | 0.618327312 | -0.222392421 | 0.034 | 0.024 | 1 | 0.643850855 | 2.017921908 | 0 | 0.022 | 1 | 0.643850855 | 0.85432596 |
| Zfp598 | 0.618558798 | -0.182864057 | 0.261 | 0.282 | 1 | 0.786958765 | 0.326760003 | 0.4 | 0.349 | 1 | 0.786958765 | 0.85450261 |
| Micu1 | 0.95582922 | -0.260632279 | 0.983 | 0.888 | 1 | 0.618795969 | -0.292741361 | 0.9 | 0.92 | 1 | 0.95582922 | 0.854683487 |
| Gm15834 | 0.649543317 | -0.578536232 | 0.076 | 0.088 | 1 | 0.618955758 | 0.973527789 | 0.1 | 0.059 | 1 | 0.649543317 | 0.854805286 |
| Nos1 | 0.619390946 | -0.807354922 | 0.025 | 0.035 | 1 | 0.73075158 | 2.695993813 | 0 | 0.012 | 1 | 0.73075158 | 0.855136748 |
| Gm5608 | 0.619390946 | -0.944858446 | 0.025 | 0.035 | 1 | 0.643850855 | 2.017921908 | 0 | 0.022 | 1 | 0.643850855 | 0.855136748 |
| Gm10644 | 0.619390946 | -0.655351829 | 0.025 | 0.035 | 1 | 0.643842184 | 1.558490289 | 0 | 0.022 | 1 | 0.643842184 | 0.855136748 |
| Zfp953 | 0.619390946 | -0.944858446 | 0.025 | 0.035 | 1 | 0.6199755 | 1.847996907 | 0 | 0.025 | 1 | 0.6199755 | 0.855136748 |
| Acot12 | 0.619390946 | -0.807354922 | 0.025 | 0.035 | 1 | 0.698534498 | 2.432959407 | 0 | 0.015 | 1 | 0.698534498 | 0.855136748 |
| Tigd5 | 0.629509811 | -0.292781749 | 0.025 | 0.035 | 1 | 0.619953009 | 1.695993813 | 0 | 0.025 | 1 | 0.629509811 | 0.855564285 |
| Coa4 | 0.659871503 | 0.514573173 | 0.025 | 0.018 | 1 | 0.619953009 | 1.695993813 | 0 | 0.025 | 1 | 0.659871503 | 0.855564285 |
| 2310040G24Rik | 0.704462092 | 0.192645078 | 0.017 | 0.024 | 1 | 0.619953009 | 1.432959407 | 0 | 0.025 | 1 | 0.704462092 | 0.855564285 |
| Gm13844 | 0.791189474 | 0.514573173 | 0.008 | 0.012 | 1 | 0.619953009 | 1.695993813 | 0 | 0.025 | 1 | 0.791189474 | 0.855564285 |
| Stk35os1 | 0.841777167 | 0.251538767 | 0.034 | 0.029 | 1 | 0.619953009 | 1.695993813 | 0 | 0.025 | 1 | 0.841777167 | 0.855564285 |
| Plac8 | 0.856670374 | -0.765534746 | 0.034 | 0.029 | 1 | 0.619953009 | 1.432959407 | 0 | 0.025 | 1 | 0.856670374 | 0.855564285 |
| Apba2 | 0.949508638 | -1.707819249 | 0.017 | 0.018 | 1 | 0.619953009 | 1.432959407 | 0 | 0.025 | 1 | 0.949508638 | 0.855564285 |
| Cxcr4 | 0.962120409 | 0.251538767 | 0.017 | 0.018 | 1 | 0.619953009 | 1.695993813 | 0 | 0.025 | 1 | 0.962120409 | 0.855564285 |
| Tbc1d2 | 0.659865009 | 0.514573173 | 0.025 | 0.018 | 1 | 0.619961444 | 1.432959407 | 0 | 0.025 | 1 | 0.659865009 | 0.855570696 |
| Myoc | 0.929389703 | 0.192645078 | 0.025 | 0.024 | 1 | 0.619961444 | 1.558490289 | 0 | 0.025 | 1 | 0.929389703 | 0.855570696 |
| Fbp2 | 0.949508638 | -0.485426827 | 0.017 | 0.018 | 1 | 0.619961444 | 1.210566986 | 0 | 0.025 | 1 | 0.949508638 | 0.855570696 |
| Abca6 | 0.981839274 | -0.485426827 | 0.042 | 0.041 | 1 | 0.619961444 | 1.432959407 | 0 | 0.025 | 1 | 0.981839274 | 0.855570696 |
| 6430550D23Rik | 0.830638286 | -0.163498732 | 0.025 | 0.029 | 1 | 0.619967066 | 1.31748219 | 0 | 0.025 | 1 | 0.830638286 | 0.855574969 |
| D830035M03Rik | 0.659871503 | 0.514573173 | 0.025 | 0.018 | 1 | 0.619967468 | 1.432959407 | 0 | 0.025 | 1 | 0.659871503 | 0.855575275 |
| Iqch | 0.682210295 | -1.117695043 | 0.042 | 0.053 | 1 | 0.619971484 | 0.374065718 | 0 | 0.025 | 1 | 0.682210295 | 0.855578327 |
| Gm34471 | 0.780793891 | 0.377069649 | 0.042 | 0.035 | 1 | 0.619972287 | 0.930459067 | 0 | 0.025 | 1 | 0.780793891 | 0.855578938 |
| Acvr1c | 0.954552228 | -0.485426827 | 0.017 | 0.018 | 1 | 0.619973091 | 0.695993813 | 0 | 0.025 | 1 | 0.954552228 | 0.855579548 |
| Trim36 | 0.954551343 | -0.222392421 | 0.017 | 0.018 | 1 | 0.619973492 | 0.930459067 | 0 | 0.025 | 1 | 0.954551343 | 0.855579853 |
| Rskr | 0.919045049 | -0.748461233 | 0.034 | 0.035 | 1 | 0.619973894 | 1.210566986 | 0 | 0.025 | 1 | 0.919045049 | 0.855580159 |
| Spi1 | 0.687345921 | -1.485426827 | 0.017 | 0.024 | 1 | 0.619975098 | -1.267480311 | 0 | 0.025 | 1 | 0.687345921 | 0.855581074 |
| Zbtb42 | 0.704462092 | 0.192645078 | 0.017 | 0.024 | 1 | 0.6199755 | 1.847996907 | 0 | 0.025 | 1 | 0.704462092 | 0.855581379 |
| 1810014B01Rik | 0.770891247 | 0.292180751 | 0.042 | 0.035 | 1 | 0.6199755 | 1.847996907 | 0 | 0.025 | 1 | 0.770891247 | 0.855581379 |
| 1700064N11Rik | 0.856663367 | -0.163498732 | 0.034 | 0.029 | 1 | 0.6199755 | 1.847996907 | 0 | 0.025 | 1 | 0.856663367 | 0.855581379 |
| Aloxe3 | 0.962120594 | 1.251538767 | 0.017 | 0.018 | 1 | 0.6199755 | 1.847996907 | 0 | 0.025 | 1 | 0.962120594 | 0.855581379 |
| Cd4 | 0.620105549 | -0.222392421 | 0.034 | 0.024 | 1 | 0.730749233 | 2.210566986 | 0 | 0.012 | 1 | 0.730749233 | 0.855680206 |
| Tmem106a | 0.622067527 | 0.251538767 | 0.067 | 0.053 | 1 | 0.743727035 | 1.017921908 | 0.1 | 0.071 | 1 | 0.743727035 | 0.857167046 |
| Gm56952 | 0.623307316 | 0.13606155 | 0.067 | 0.053 | 1 | 0.865301582 | 0.973527789 | 0.1 | 0.083 | 1 | 0.865301582 | 0.858102622 |
| Uqcc3 | 0.623641507 | -0.397963986 | 0.37 | 0.365 | 1 | 0.837777619 | 0.210566986 | 0.3 | 0.247 | 1 | 0.837777619 | 0.858354285 |
| Armcx5 | 0.792782687 | 0.246639968 | 0.261 | 0.253 | 1 | 0.62383342 | -0.152003093 | 0.2 | 0.259 | 1 | 0.792782687 | 0.858498704 |
| Ctf1 | 0.623923448 | -0.455053178 | 0.185 | 0.2 | 1 | 0.806699093 | 0.263034406 | 0.3 | 0.247 | 1 | 0.806699093 | 0.858566427 |
| Slc25a37 | 0.624383789 | -0.431655571 | 0.286 | 0.294 | 1 | 0.681009209 | -0.291933355 | 0.2 | 0.244 | 1 | 0.681009209 | 0.858912462 |
| Adam1a | 0.62441508 | -0.485426827 | 0.025 | 0.035 | 1 | 0.669843604 | 2.017921908 | 0 | 0.019 | 1 | 0.669843604 | 0.858935968 |
| Gm57385 | 0.62441508 | -0.485426827 | 0.025 | 0.035 | 1 | 0.64384652 | 1.558490289 | 0 | 0.022 | 1 | 0.64384652 | 0.858935968 |
| Gm5141 | 0.62441508 | -0.485426827 | 0.025 | 0.035 | 1 | 0.643850855 | 2.017921908 | 0 | 0.022 | 1 | 0.643850855 | 0.858935968 |
| Kif6 | 0.62441508 | -0.655351829 | 0.025 | 0.035 | 1 | 0.669848329 | 1.847996907 | 0 | 0.019 | 1 | 0.669848329 | 0.858935968 |
| Slc14a2 | 0.624449806 | -0.315501826 | 0.025 | 0.035 | 1 | 0.669852109 | 1.111031312 | 0 | 0.019 | 1 | 0.669852109 | 0.858962052 |
| Bcl3 | 0.625645141 | -0.35614381 | 0.319 | 0.341 | 1 | 0.667132802 | -0.152003093 | 0.3 | 0.364 | 1 | 0.667132802 | 0.85985844 |
| Gm10785 | 0.626037119 | -1.411426246 | 0.076 | 0.088 | 1 | 0.78456274 | 0.732519689 | 0.1 | 0.074 | 1 | 0.78456274 | 0.860151764 |
| Tmem237 | 0.753084058 | -0.234883366 | 0.429 | 0.424 | 1 | 0.626584376 | -0.175849835 | 0.3 | 0.373 | 1 | 0.753084058 | 0.860560772 |
| Smim26 | 0.78946462 | -0.387246433 | 0.286 | 0.288 | 1 | 0.62710133 | 0.739937161 | 0.3 | 0.244 | 1 | 0.78946462 | 0.860946582 |
| Lcn11 | 0.919683886 | 0.166649869 | 0.067 | 0.071 | 1 | 0.627325888 | -0.425021588 | 0.1 | 0.151 | 1 | 0.919683886 | 0.861114006 |
| Gm47761 | 0.627327552 | -0.19592021 | 0.067 | 0.082 | 1 | 0.669848329 | 1.558490289 | 0 | 0.019 | 1 | 0.669848329 | 0.861115247 |
| Hdc | 0.627805946 | -1.250961574 | 0.025 | 0.035 | 1 | 0.669850692 | 0.769994395 | 0 | 0.019 | 1 | 0.669850692 | 0.861471586 |
| Gm56911 | 0.627805946 | -0.622930351 | 0.025 | 0.035 | 1 | 0.730749233 | 2.432959407 | 0 | 0.012 | 1 | 0.730749233 | 0.861471586 |
| Hibch | 0.928251774 | -0.374395515 | 0.193 | 0.188 | 1 | 0.628268577 | 0.598383016 | 0.3 | 0.235 | 1 | 0.928251774 | 0.861815749 |
| Pgm3 | 0.960932931 | -0.14345279 | 0.689 | 0.635 | 1 | 0.628274852 | 0.17871812 | 0.7 | 0.599 | 1 | 0.960932931 | 0.861820414 |
| Armcx6 | 0.628573237 | -0.378511623 | 0.101 | 0.118 | 1 | 0.935769869 | 0.988174565 | 0.1 | 0.117 | 1 | 0.935769869 | 0.86204216 |
| Gm57475 | 0.629509811 | -0.292781749 | 0.025 | 0.035 | 1 | 0.669853054 | 2.210566986 | 0 | 0.019 | 1 | 0.669853054 | 0.86273702 |
| Pla2g4b | 0.629509811 | -0.292781749 | 0.025 | 0.035 | 1 | 0.63522121 | 1.374065718 | 0.1 | 0.062 | 1 | 0.63522121 | 0.86273702 |
| Hsd17b14 | 0.629509811 | -0.292781749 | 0.025 | 0.035 | 1 | 0.669848329 | -0.267480311 | 0 | 0.019 | 1 | 0.669848329 | 0.86273702 |
| Gm44686 | 0.629509811 | -0.292781749 | 0.025 | 0.035 | 1 | 0.91052109 | 0.625604485 | 0.1 | 0.086 | 1 | 0.91052109 | 0.86273702 |
| Gm19967 | 0.629509811 | -0.292781749 | 0.025 | 0.035 | 1 | 0.640912613 | 1.111031312 | 0.1 | 0.062 | 1 | 0.640912613 | 0.86273702 |
| Mief2 | 0.926311501 | 0.234465254 | 0.084 | 0.088 | 1 | 0.62976973 | 0.695993813 | 0.2 | 0.139 | 1 | 0.926311501 | 0.862929547 |
| Ascc1 | 0.630635104 | -0.158616511 | 0.269 | 0.235 | 1 | 0.696387976 | 0.331421381 | 0.3 | 0.231 | 1 | 0.696387976 | 0.863569574 |
| Rab10os | 0.683860487 | -0.240314329 | 0.328 | 0.335 | 1 | 0.631701795 | 0.808468542 | 0.3 | 0.262 | 1 | 0.683860487 | 0.864356432 |
| Siah1b | 0.828519158 | 0.225066556 | 0.059 | 0.053 | 1 | 0.632378953 | 1.494359952 | 0.1 | 0.062 | 1 | 0.828519158 | 0.864854766 |
| A130010J15Rik | 0.885335088 | 0.344648171 | 0.05 | 0.047 | 1 | 0.632378953 | 1.494359952 | 0.1 | 0.062 | 1 | 0.885335088 | 0.864854766 |
| Mphosph8 | 0.902598638 | -0.197987109 | 0.63 | 0.588 | 1 | 0.632380257 | 0.129178659 | 0.7 | 0.608 | 1 | 0.902598638 | 0.864855724 |
| Prps1l3 | 0.633537455 | -0.333423734 | 0.16 | 0.176 | 1 | 0.836616793 | 0.580516596 | 0.2 | 0.17 | 1 | 0.836616793 | 0.865705203 |
| Trmt61b | 0.634337549 | -0.649813645 | 0.193 | 0.206 | 1 | 1 | 0.374065718 | 0.2 | 0.204 | 1 | 1 | 0.866290972 |
| Bop1 | 0.63445928 | -0.333423734 | 0.185 | 0.2 | 1 | 0.996294322 | 1.14345279 | 0.2 | 0.216 | 1 | 0.996294322 | 0.866379982 |
| Arl5c | 0.719404314 | -0.637429921 | 0.067 | 0.076 | 1 | 0.63522121 | 1.111031312 | 0.1 | 0.062 | 1 | 0.719404314 | 0.866936434 |
| Mtmr1 | 0.635283409 | -0.420331799 | 0.387 | 0.329 | 1 | 0.716242525 | 0.489542936 | 0.4 | 0.472 | 1 | 0.716242525 | 0.866981808 |

| | | | | | | | | | | | | |
|---|---|---|---|---|---|---|---|---|---|---|---|---|
| Gm57349 | 0.636250625 | 0.514573173 | 0.025 | 0.035 | 1 | 0.669843604 | 2.017921908 | 0 | 0.019 | 1 | 0.669843604 | 0.867686393 |
| Ceacam1 | 0.97600602 | -0.551516018 | 0.168 | 0.159 | 1 | 0.63689786 | 0.751135367 | 0.2 | 0.148 | 1 | 0.97600602 | 0.868156836 |
| Snrnp35 | 0.637831555 | -0.642968104 | 0.092 | 0.106 | 1 | 0.952551372 | 0.808468542 | 0.1 | 0.093 | 1 | 0.952551372 | 0.868834017 |
| Nde1 | 0.638317632 | -0.19730144 | 0.37 | 0.371 | 1 | 0.712685563 | 0.450237399 | 0.4 | 0.364 | 1 | 0.712685563 | 0.869185865 |
| Snhg12 | 0.871373168 | -0.237167774 | 0.697 | 0.606 | 1 | 0.638408725 | 0.20029865 | 0.6 | 0.497 | 1 | 0.871373168 | 0.86925175 |
| Dkk3 | 0.980854284 | 0.337332788 | 0.319 | 0.335 | 1 | 0.63852658 | 0.351719559 | 0.6 | 0.605 | 1 | 0.980854284 | 0.869336967 |
| Gm16124 | 0.925605176 | -0.244418728 | 0.16 | 0.159 | 1 | 0.639851846 | -0.171902651 | 0.1 | 0.148 | 1 | 0.925605176 | 0.870293307 |
| Dipk1a | 0.639992957 | -0.461179281 | 0.16 | 0.176 | 1 | 0.678879808 | -0.374395515 | 0.2 | 0.238 | 1 | 0.678879808 | 0.870394929 |
| Cdk5rap2 | 0.948112773 | -0.379298476 | 0.58 | 0.524 | 1 | 0.640534824 | 1.193493473 | 0.5 | 0.469 | 1 | 0.948112773 | 0.870784787 |
| Zfp444 | 0.726528464 | 0.12225575 | 0.126 | 0.112 | 1 | 0.642364859 | 0.494359952 | 0.3 | 0.222 | 1 | 0.726528464 | 0.872097106 |
| Gm16573 | 0.659871503 | 0.514573173 | 0.025 | 0.018 | 1 | 0.643835681 | 1.847996907 | 0 | 0.022 | 1 | 0.659871503 | 0.873146978 |
| Ddo | 0.661963295 | 0.251538767 | 0.025 | 0.018 | 1 | 0.643835681 | 1.558490289 | 0 | 0.022 | 1 | 0.661963295 | 0.873146978 |
| Gm13134 | 0.695893386 | -0.222392421 | 0.017 | 0.024 | 1 | 0.643835681 | 1.847996907 | 0 | 0.022 | 1 | 0.695893386 | 0.873146978 |
| Gm49801 | 0.715874401 | 0.929610672 | 0.017 | 0.012 | 1 | 0.643835681 | 1.558490289 | 0 | 0.022 | 1 | 0.715874401 | 0.873146978 |
| Atp6v0e2 | 0.720666617 | 0.251538767 | 0.076 | 0.065 | 1 | 0.643835681 | 1.847996907 | 0 | 0.022 | 1 | 0.720666617 | 0.873146978 |
| 5730507C01Rik | 0.721162042 | 0.514573173 | 0.017 | 0.012 | 1 | 0.643835681 | 1.847996907 | 0 | 0.022 | 1 | 0.721162042 | 0.873146978 |
| F630206G17Rik | 0.864135893 | -0.333423734 | 0.034 | 0.029 | 1 | 0.643835681 | 1.695993813 | 0 | 0.022 | 1 | 0.864135893 | 0.873146978 |
| Tex26 | 0.903595785 | -0.163498732 | 0.05 | 0.047 | 1 | 0.643835681 | 1.432959407 | 0 | 0.022 | 1 | 0.903595785 | 0.873146978 |
| Gm20492 | 0.929389703 | 0.192645078 | 0.025 | 0.024 | 1 | 0.643835681 | 1.847996907 | 0 | 0.022 | 1 | 0.929389703 | 0.873146978 |
| Psmb9 | 0.97201319 | -0.572889668 | 0.059 | 0.059 | 1 | 0.643835681 | 1.847996907 | 0 | 0.022 | 1 | 0.97201319 | 0.873146978 |
| Gm56897 | 0.687345921 | -0.707819249 | 0.017 | 0.024 | 1 | 0.643842184 | 1.695993813 | 0 | 0.022 | 1 | 0.687345921 | 0.87315161 |
| Mir133a-1hg | 0.721162042 | 0.514573173 | 0.017 | 0.012 | 1 | 0.643842184 | 1.695993813 | 0 | 0.022 | 1 | 0.721162042 | 0.87315161 |
| Angel1 | 0.726456542 | -0.900464326 | 0.017 | 0.012 | 1 | 0.643842184 | 1.210566986 | 0 | 0.022 | 1 | 0.726456542 | 0.87315161 |
| Gm28874 | 0.856663367 | -0.163498732 | 0.034 | 0.029 | 1 | 0.643842184 | 1.558490289 | 0 | 0.022 | 1 | 0.856663367 | 0.87315161 |
| 4931403E22Rik | 0.929389703 | 0.192645078 | 0.025 | 0.024 | 1 | 0.643842184 | 1.695993813 | 0 | 0.022 | 1 | 0.929389703 | 0.87315161 |
| Kif9 | 0.998600264 | -0.122856748 | 0.059 | 0.059 | 1 | 0.643842184 | 1.695993813 | 0 | 0.022 | 1 | 0.998600264 | 0.87315161 |
| Trim30a | 0.920856377 | 0.707218251 | 0.025 | 0.024 | 1 | 0.64384652 | 1.210566986 | 0 | 0.022 | 1 | 0.920856377 | 0.873154698 |
| Slc26a8 | 0.704462092 | 0.192645078 | 0.017 | 0.024 | 1 | 0.643848687 | 1.111031312 | 0 | 0.022 | 1 | 0.704462092 | 0.873156243 |
| Adgrf2 | 0.704462092 | 0.192645078 | 0.017 | 0.024 | 1 | 0.643848687 | 1.111031312 | 0 | 0.022 | 1 | 0.704462092 | 0.873156243 |
| Reln | 0.781790612 | -0.485426827 | 0.008 | 0.012 | 1 | 0.643848687 | 1.017921908 | 0 | 0.022 | 1 | 0.781790612 | 0.873156243 |
| Slc28a1 | 0.781790612 | -0.485426827 | 0.008 | 0.012 | 1 | 0.643848687 | 1.210566986 | 0 | 0.022 | 1 | 0.781790612 | 0.873156243 |
| Mast1 | 0.767588375 | -0.36994961 | 0.042 | 0.035 | 1 | 0.643849121 | 1.111031312 | 0 | 0.022 | 1 | 0.767588375 | 0.873156551 |
| A330032B11Rik | 0.647331896 | 1.684498174 | 0.025 | 0.018 | 1 | 0.643850855 | 2.017921908 | 0 | 0.022 | 1 | 0.647331896 | 0.873157787 |
| Zscan2 | 0.687345921 | -0.900464326 | 0.017 | 0.024 | 1 | 0.643850855 | 2.017921908 | 0 | 0.022 | 1 | 0.687345921 | 0.873157787 |
| Pld4 | 0.687345921 | -1.485426827 | 0.017 | 0.024 | 1 | 0.643850855 | 2.017921908 | 0 | 0.022 | 1 | 0.687345921 | 0.873157787 |
| C230038L03Rik | 0.691605844 | -0.485426827 | 0.017 | 0.024 | 1 | 0.643850855 | 2.017921908 | 0 | 0.022 | 1 | 0.691605844 | 0.873157787 |
| Gm30455 | 0.721161231 | 0.514573173 | 0.017 | 0.012 | 1 | 0.643850855 | 2.017921908 | 0 | 0.022 | 1 | 0.721161231 | 0.873157787 |
| AV356131 | 0.942200939 | -0.655351829 | 0.025 | 0.024 | 1 | 0.643850855 | 2.017921908 | 0 | 0.022 | 1 | 0.942200939 | 0.873157787 |
| Slc26a1 | 0.954551343 | -0.222392421 | 0.017 | 0.018 | 1 | 0.643850855 | 2.017921908 | 0 | 0.022 | 1 | 0.954551343 | 0.873157787 |
| Zfp933 | 0.643902485 | -0.292781749 | 0.084 | 0.1 | 1 | 0.657751416 | 0.828097349 | 0.1 | 0.17 | 1 | 0.657751416 | 0.87319456 |
| Phf7 | 0.643902485 | -0.359895945 | 0.084 | 0.1 | 1 | 0.868037657 | 0.808468542 | 0.1 | 0.083 | 1 | 0.868037657 | 0.87319456 |
| C2cd2l | 0.644919696 | -0.222392421 | 0.084 | 0.1 | 1 | 0.950061233 | 0.695993813 | 0.1 | 0.105 | 1 | 0.950061233 | 0.873917978 |
| Itpripl2 | 0.94919294 | 0.170008881 | 0.613 | 0.635 | 1 | 0.645941583 | -0.427782728 | 0.7 | 0.63 | 1 | 0.94919294 | 0.874642637 |
| Rnasel | 0.646612294 | -0.117695043 | 0.101 | 0.118 | 1 | 0.933175065 | 0.625604485 | 0.1 | 0.09 | 1 | 0.933175065 | 0.875117129 |
| Sema4d | 0.930609841 | -0.552541023 | 0.143 | 0.135 | 1 | 0.646836597 | 0.785261151 | 0.1 | 0.17 | 1 | 0.930609841 | 0.875275611 |
| Mpp7 | 0.722501354 | -0.363254957 | 0.345 | 0.376 | 1 | 0.64684429 | 1.177349209 | 0.6 | 0.571 | 1 | 0.722501354 | 0.875281044 |
| Gcnt1 | 0.646989117 | -0.213054557 | 0.471 | 0.418 | 1 | 0.876159675 | 1.077478753 | 0.3 | 0.315 | 1 | 0.876159675 | 0.875383317 |
| Scrn3 | 0.647663663 | -0.448900951 | 0.151 | 0.165 | 1 | 0.780774352 | 0.393431043 | 0.2 | 0.16 | 1 | 0.780774352 | 0.875859106 |
| Cenpx | 0.907810678 | -0.357876777 | 0.412 | 0.4 | 1 | 0.648175503 | 0.30367639 | 0.3 | 0.426 | 1 | 0.907810678 | 0.876219523 |
| D330023K18Rik | 0.648345132 | -0.222392421 | 0.101 | 0.118 | 1 | 0.994802027 | 0.494359952 | 0.1 | 0.096 | 1 | 0.994802027 | 0.876338854 |
| Mettl27 | 0.813139637 | 0.192645078 | 0.084 | 0.076 | 1 | 0.649466592 | 1.017921908 | 0.1 | 0.062 | 1 | 0.813139637 | 0.87712633 |
| Arhgap18 | 0.651200208 | 0.514573173 | 0.059 | 0.047 | 1 | 0.73566144 | 0.4480663 | 0.2 | 0.16 | 1 | 0.73566144 | 0.878338705 |
| Phox2a | 0.653579641 | 1.321928095 | 0.025 | 0.018 | 1 | 0.669848329 | 1.31748219 | 0 | 0.019 | 1 | 0.669848329 | 0.879992935 |
| Tcea3 | 0.654832922 | -0.23388806 | 0.303 | 0.265 | 1 | 0.776035541 | 0.159940913 | 0.3 | 0.241 | 1 | 0.776035541 | 0.880859688 |
| Polr2f | 0.760425891 | -0.122856748 | 0.639 | 0.624 | 1 | 0.654985639 | 0.40053793 | 0.6 | 0.54 | 1 | 0.760425891 | 0.88096509 |
| Tatdn1 | 0.655210852 | -0.172268942 | 0.269 | 0.288 | 1 | 0.74405179 | 0.123104145 | 0.2 | 0.253 | 1 | 0.74405179 | 0.881120443 |
| Gm30835 | 0.657014966 | -2.185866545 | 0.042 | 0.053 | 1 | 0.730749233 | 2.210566986 | 0 | 0.012 | 1 | 0.730749233 | 0.882361266 |
| Gm16178 | 0.657769394 | 0.736965594 | 0.025 | 0.018 | 1 | 0.730749233 | 2.432959407 | 0 | 0.012 | 1 | 0.730749233 | 0.882878212 |
| Phf11d | 0.657769394 | 0.707218251 | 0.025 | 0.018 | 1 | 0.730750993 | 1.111031312 | 0 | 0.012 | 1 | 0.730750993 | 0.882878212 |
| Pla2g10os | 0.657773312 | 0.321928095 | 0.025 | 0.018 | 1 | 0.669843604 | 2.017921908 | 0 | 0.019 | 1 | 0.669843604 | 0.882880894 |
| Zfp426 | 0.680648329 | -0.222392421 | 0.185 | 0.2 | 1 | 0.659442035 | 0.210566986 | 0.3 | 0.219 | 1 | 0.680648329 | 0.884020272 |
| Gm16098 | 0.659871503 | 0.514573173 | 0.025 | 0.018 | 1 | 0.698529281 | 2.017921908 | 0 | 0.015 | 1 | 0.698529281 | 0.884312606 |
| Gm56971 | 0.659871503 | 0.514573173 | 0.025 | 0.018 | 1 | 0.698534498 | 2.432959407 | 0 | 0.015 | 1 | 0.698534498 | 0.884312606 |
| Fgl1 | 0.659871503 | 0.514573173 | 0.025 | 0.018 | 1 | 0.669853054 | 2.210566986 | 0 | 0.019 | 1 | 0.669853054 | 0.884312606 |
| Pla2g3 | 0.659871503 | 0.514573173 | 0.025 | 0.018 | 1 | 0.698534498 | 2.432959407 | 0 | 0.015 | 1 | 0.698534498 | 0.884312606 |
| Gm12714 | 0.659871503 | 0.514573173 | 0.025 | 0.018 | 1 | 0.698529281 | 2.017921908 | 0 | 0.015 | 1 | 0.698529281 | 0.884312606 |
| Psma8 | 0.659871503 | 0.514573173 | 0.025 | 0.018 | 1 | 0.73075158 | 2.695993813 | 0 | 0.012 | 1 | 0.73075158 | 0.884312606 |
| D330050I16Rik | 0.659871503 | 0.514573173 | 0.025 | 0.018 | 1 | 0.669843604 | 2.017921908 | 0 | 0.019 | 1 | 0.669843604 | 0.884312606 |
| Src | 0.660968347 | -0.365132593 | 0.193 | 0.206 | 1 | 0.939269775 | 0.388565288 | 0.2 | 0.219 | 1 | 0.939269775 | 0.885057538 |
| Scfd2 | 0.684700375 | -0.502914254 | 0.395 | 0.376 | 1 | 0.661852178 | 0.191373421 | 0.2 | 0.306 | 1 | 0.684700375 | 0.885656051 |
| Pkp2 | 0.661963295 | -0.163498732 | 0.025 | 0.018 | 1 | 0.669848329 | 1.695993813 | 0 | 0.019 | 1 | 0.669848329 | 0.885731186 |
| Six3os1 | 0.661963295 | 0.251538767 | 0.025 | 0.018 | 1 | 0.698532411 | 1.695993813 | 0 | 0.015 | 1 | 0.698532411 | 0.885731186 |
| Col17a1 | 0.661963295 | -0.333423734 | 0.025 | 0.018 | 1 | 0.89080433 | 0.930459067 | 0.1 | 0.123 | 1 | 0.89080433 | 0.885731186 |
| Zbtb22 | 0.878456412 | -0.106915204 | 0.101 | 0.106 | 1 | 0.662707401 | 0.236562194 | 0.1 | 0.148 | 1 | 0.878456412 | 0.886233702 |
| Rusc1 | 0.836421481 | -0.24838763 | 0.218 | 0.224 | 1 | 0.662968441 | -0.152003093 | 0.2 | 0.247 | 1 | 0.836421481 | 0.886409728 |
| Mapt | 0.663317101 | 0.292180751 | 0.059 | 0.047 | 1 | 0.727475609 | 0.591657153 | 0.1 | 0.068 | 1 | 0.727475609 | 0.886644626 |
| ENSMUSG00000121487 | 0.663326742 | -0.432959407 | 0.109 | 0.124 | 1 | 0.669853054 | 2.210566986 | 0 | 0.019 | 1 | 0.669853054 | 0.886651117 |
| Trip4 | 0.75049496 | -0.218142948 | 0.403 | 0.394 | 1 | 0.664266739 | -0.246185533 | 0.4 | 0.444 | 1 | 0.75049496 | 0.887283178 |
| Gm26797 | 0.733769298 | -0.610957709 | 0.067 | 0.076 | 1 | 0.664327136 | 0.580516596 | 0.2 | 0.142 | 1 | 0.733769298 | 0.887323729 |
| Iah1 | 0.880224466 | -0.198144875 | 0.235 | 0.235 | 1 | 0.664574935 | 0.625604485 | 0.3 | 0.262 | 1 | 0.880224466 | 0.887490026 |
| Mgll | 0.666074541 | -0.269698136 | 0.597 | 0.606 | 1 | 0.983593256 | -0.177282502 | 0.7 | 0.636 | 1 | 0.983593256 | 0.888493788 |
| Mettl5os | 0.666161437 | 0.192645078 | 0.025 | 0.018 | 1 | 0.669851164 | 1.695993813 | 0 | 0.019 | 1 | 0.669851164 | 0.888551814 |
| Gm57275 | 0.666161437 | 0.192645078 | 0.025 | 0.018 | 1 | 0.669853054 | 2.210566986 | 0 | 0.019 | 1 | 0.669853054 | 0.888551814 |
| Etohd2 | 0.666161437 | 0.192645078 | 0.025 | 0.018 | 1 | 0.669843604 | 2.017921908 | 0 | 0.019 | 1 | 0.669843604 | 0.888551814 |
| Ppm1f | 0.667443205 | -0.485426827 | 0.059 | 0.071 | 1 | 0.709460441 | 0.159940913 | 0.1 | 0.139 | 1 | 0.709460441 | 0.889405978 |
| Donson | 0.667443205 | -0.655351829 | 0.059 | 0.071 | 1 | 0.799264655 | 1.159940913 | 0.1 | 0.077 | 1 | 0.799264655 | 0.889405978 |
| Scap | 0.667786265 | -0.346023771 | 0.277 | 0.288 | 1 | 0.831032868 | -0.31836148 | 0.4 | 0.383 | 1 | 0.831032868 | 0.889634034 |
| Timmdc1 | 0.900959334 | -0.273922722 | 0.294 | 0.282 | 1 | 0.668283293 | 0.403212064 | 0.3 | 0.228 | 1 | 0.900959334 | 0.889964026 |
| Cpsf3 | 0.669153166 | -0.185866545 | 0.277 | 0.288 | 1 | 0.814122732 | 0.87108052 | 0.2 | 0.281 | 1 | 0.814122732 | 0.890540372 |
| Tex14 | 0.687561937 | 0.374065718 | 0.672 | 0.694 | 1 | 0.669364818 | -0.3376739 | 0.9 | 0.667 | 1 | 0.687561937 | 0.890680376 |
| 5430405H02Rik | 0.834936007 | -0.687060688 | 0.118 | 0.124 | 1 | 0.669702134 | -0.230005605 | 0.1 | 0.142 | 1 | 0.834936007 | 0.89090332 |
| Zfp850 | 0.687345921 | -0.707819249 | 0.017 | 0.024 | 1 | 0.669843604 | 2.017921908 | 0 | 0.019 | 1 | 0.687345921 | 0.890996754 |
| Tgm4 | 0.695893386 | -0.222392421 | 0.017 | 0.024 | 1 | 0.669843604 | 2.017921908 | 0 | 0.019 | 1 | 0.695893386 | 0.890996754 |
| Gm43714 | 0.721162042 | 0.514573173 | 0.017 | 0.012 | 1 | 0.669843604 | 2.017921908 | 0 | 0.019 | 1 | 0.721162042 | 0.890996754 |
| Pdf | 0.721162042 | 0.514573173 | 0.017 | 0.012 | 1 | 0.669843604 | 2.017921908 | 0 | 0.019 | 1 | 0.721162042 | 0.890996754 |
| Pde8b | 0.731769749 | -0.485426827 | 0.017 | 0.012 | 1 | 0.669843604 | 1.847996907 | 0 | 0.019 | 1 | 0.731769749 | 0.890996754 |
| 9230116N13Rik | 0.743002317 | -0.163498732 | 0.034 | 0.041 | 1 | 0.669843604 | 2.017921908 | 0 | 0.019 | 1 | 0.743002317 | 0.890996754 |
| Inava | 0.781790612 | -0.807354922 | 0.008 | 0.012 | 1 | 0.669843604 | 1.847996907 | 0 | 0.019 | 1 | 0.781790612 | 0.890996754 |

| | | | | | | | | | | | | |
|---|---|---|---|---|---|---|---|---|---|---|---|---|
| Emx2os | 0.781790612 | -0.807354922 | 0.008 | 0.012 | 1 | 0.669843604 | 2.017921908 | 0 | 0.019 | 1 | 0.781790612 | 0.890996754 |
| Spats1 | 0.962120594 | 0.736965594 | 0.017 | 0.018 | 1 | 0.669843604 | 2.017921908 | 0 | 0.019 | 1 | 0.962120594 | 0.890996754 |
| 9430037G07Rik | 0.717039396 | -0.86393845 | 0.034 | 0.041 | 1 | 0.669848329 | 1.695993813 | 0 | 0.019 | 1 | 0.717039396 | 0.890999874 |
| Fgf12 | 0.778665521 | -1.070389328 | 0.008 | 0.012 | 1 | 0.669848329 | 1.432959407 | 0 | 0.019 | 1 | 0.778665521 | 0.890999874 |
| Myh11 | 0.781790612 | -0.485426827 | 0.008 | 0.012 | 1 | 0.669848329 | 1.847996907 | 0 | 0.019 | 1 | 0.781790612 | 0.890999874 |
| Gm30173 | 0.820836204 | -1.185866545 | 0.025 | 0.029 | 1 | 0.669848329 | 1.695993813 | 0 | 0.019 | 1 | 0.820836204 | 0.890999874 |
| Gm16833 | 0.841777167 | 0.251538767 | 0.034 | 0.029 | 1 | 0.669848329 | 1.847996907 | 0 | 0.019 | 1 | 0.841777167 | 0.890999874 |
| Slc44a4 | 0.954552228 | -0.485426827 | 0.017 | 0.018 | 1 | 0.669848329 | 1.695993813 | 0 | 0.019 | 1 | 0.954552228 | 0.890999874 |
| Gm11520 | 0.998349193 | -0.572889668 | 0.042 | 0.041 | 1 | 0.669848329 | 1.31748219 | 0 | 0.019 | 1 | 0.998349193 | 0.890999874 |
| Irag1 | 0.695893386 | -0.222392421 | 0.017 | 0.024 | 1 | 0.669850692 | 1.432959407 | 0 | 0.019 | 1 | 0.695893386 | 0.891001434 |
| Gm45518 | 0.713085938 | 0.777607579 | 0.017 | 0.024 | 1 | 0.669850692 | 1.31748219 | 0 | 0.019 | 1 | 0.713085938 | 0.891001434 |
| Bnipl | 0.929389703 | 0.192645078 | 0.025 | 0.024 | 1 | 0.669850692 | 1.558490289 | 0 | 0.019 | 1 | 0.929389703 | 0.891001434 |
| Gpr183 | 0.781790612 | -0.485426827 | 0.008 | 0.012 | 1 | 0.669851164 | 0.374065718 | 0 | 0.019 | 1 | 0.781790612 | 0.891001746 |
| 4933412E12Rik | 0.842422009 | 0.251538767 | 0.025 | 0.029 | 1 | 0.669851164 | 1.695993813 | 0 | 0.019 | 1 | 0.842422009 | 0.891001746 |
| Gm54771 | 0.929389703 | 0.192645078 | 0.025 | 0.024 | 1 | 0.669851164 | 1.210566986 | 0 | 0.019 | 1 | 0.929389703 | 0.891001746 |
| Fsip1 | 0.781790612 | -0.485426827 | 0.008 | 0.012 | 1 | 0.669852109 | 0.625604485 | 0 | 0.019 | 1 | 0.781790612 | 0.89100237 |
| H3c7 | 0.687345921 | -1.485426827 | 0.017 | 0.024 | 1 | 0.669853054 | 2.210566986 | 0 | 0.019 | 1 | 0.687345921 | 0.891002994 |
| Mtfp1 | 0.691605844 | -0.485426827 | 0.017 | 0.024 | 1 | 0.669853054 | 2.210566986 | 0 | 0.019 | 1 | 0.691605844 | 0.891002994 |
| 4932443L11Rik | 0.695893386 | -0.222392421 | 0.017 | 0.024 | 1 | 0.669853054 | 2.210566986 | 0 | 0.019 | 1 | 0.695893386 | 0.891002994 |
| Gm30092 | 0.721162042 | 0.514573173 | 0.017 | 0.012 | 1 | 0.669853054 | 2.210566986 | 0 | 0.019 | 1 | 0.721162042 | 0.891002994 |
| Cideb | 0.721162042 | 0.514573173 | 0.017 | 0.012 | 1 | 0.669853054 | 2.210566986 | 0 | 0.019 | 1 | 0.721162042 | 0.891002994 |
| Marveld2 | 0.726456542 | -0.707819249 | 0.017 | 0.012 | 1 | 0.669853054 | 2.210566986 | 0 | 0.019 | 1 | 0.726456542 | 0.891002994 |
| Gm57272 | 0.839906751 | 0.292180751 | 0.034 | 0.029 | 1 | 0.669853054 | 2.210566986 | 0 | 0.019 | 1 | 0.839906751 | 0.891002994 |
| A030001D20Rik | 0.920852644 | 0.514573173 | 0.025 | 0.024 | 1 | 0.669853054 | 2.210566986 | 0 | 0.019 | 1 | 0.920852644 | 0.891002994 |
| Gm43646 | 0.954551343 | -0.222392421 | 0.017 | 0.018 | 1 | 0.669853054 | 2.210566986 | 0 | 0.019 | 1 | 0.954551343 | 0.891002994 |
| Gm34343 | 0.67069931 | 0.436570661 | 0.101 | 0.088 | 1 | 0.876565312 | 1.558490289 | 0.1 | 0.093 | 1 | 0.876565312 | 0.891561055 |
| Skic3 | 0.902584153 | 0.24687739 | 0.655 | 0.576 | 1 | 0.670852198 | 0.584962501 | 0.5 | 0.571 | 1 | 0.902584153 | 0.891661725 |
| Itpk1 | 0.672289116 | -0.516453723 | 0.168 | 0.176 | 1 | 0.755080452 | 0.31748219 | 0.2 | 0.157 | 1 | 0.755080452 | 0.892605577 |
| Aldh9a1 | 0.752244375 | -0.130761946 | 0.252 | 0.265 | 1 | 0.672297518 | 0.135278859 | 0.2 | 0.275 | 1 | 0.752244375 | 0.892611083 |
| Gm6169 | 0.672385695 | -0.222392421 | 0.042 | 0.053 | 1 | 0.770508511 | 1.063725598 | 0.1 | 0.074 | 1 | 0.770508511 | 0.892668867 |
| Exoc6b | 0.673014898 | 0.1667932 | 0.908 | 0.9 | 1 | 0.775117042 | 0.229808782 | 1 | 0.938 | 1 | 0.775117042 | 0.893080743 |
| Gm10501 | 0.753635616 | -0.250961574 | 0.076 | 0.065 | 1 | 0.673292204 | 1.159940913 | 0.1 | 0.065 | 1 | 0.753635616 | 0.893262016 |
| Creld1 | 0.997808749 | -0.201633861 | 0.101 | 0.1 | 1 | 0.673292204 | 1.31748219 | 0.1 | 0.065 | 1 | 0.997808749 | 0.893262016 |
| Fxn | 0.742650324 | -0.205318908 | 0.118 | 0.129 | 1 | 0.674802651 | 0.602884409 | 0.2 | 0.145 | 1 | 0.742650324 | 0.894246684 |
| Oplah | 0.967038687 | -0.222392421 | 0.109 | 0.106 | 1 | 0.674802651 | 0.672147071 | 0.2 | 0.145 | 1 | 0.967038687 | 0.894246684 |
| Scamp2 | 0.74963486 | -0.173148902 | 0.689 | 0.676 | 1 | 0.675421648 | 0.12217187 | 0.5 | 0.651 | 1 | 0.74963486 | 0.894648893 |
| Usp6nl | 0.829810886 | -0.287866259 | 0.697 | 0.641 | 1 | 0.675523019 | -0.293358943 | 0.7 | 0.747 | 1 | 0.829810886 | 0.894714689 |
| Rab29 | 0.675917441 | -0.733354341 | 0.059 | 0.071 | 1 | 0.746553901 | 1.017921908 | 0.1 | 0.071 | 1 | 0.746553901 | 0.894970495 |
| Gprasp2 | 0.67775968 | -0.647698256 | 0.151 | 0.165 | 1 | 0.952657695 | -0.161987182 | 0.3 | 0.275 | 1 | 0.952657695 | 0.896161176 |
| Bcar3 | 0.692043293 | -0.722466024 | 0.076 | 0.088 | 1 | 0.67782495 | -0.463204782 | 0.2 | 0.139 | 1 | 0.692043293 | 0.896203237 |
| Cish | 0.677926639 | -0.137503524 | 0.042 | 0.053 | 1 | 0.698529281 | 2.210566986 | 0 | 0.015 | 1 | 0.698529281 | 0.89626875 |
| Micall2 | 0.830638286 | -0.163498732 | 0.025 | 0.029 | 1 | 0.67898466 | 1.159940913 | 0.1 | 0.065 | 1 | 0.830638286 | 0.896949152 |
| Diablo | 0.870214943 | -0.128270244 | 0.42 | 0.382 | 1 | 0.679273249 | 0.617991301 | 0.5 | 0.429 | 1 | 0.870214943 | 0.897134351 |
| Mrpl58 | 0.962964899 | 0.181149439 | 0.294 | 0.3 | 1 | 0.680570406 | 0.374065718 | 0.3 | 0.225 | 1 | 0.962964899 | 0.897964735 |
| Abcb1a | 0.681646164 | 0.893084796 | 0.05 | 0.041 | 1 | 0.730750993 | 1.695993813 | 0 | 0.012 | 1 | 0.730750993 | 0.898650835 |
| Fanca | 0.683093021 | -1.070389328 | 0.017 | 0.024 | 1 | 0.730749233 | 2.210566986 | 0 | 0.012 | 1 | 0.730749233 | 0.899569967 |
| Ngdn | 0.970716318 | -0.166148311 | 0.487 | 0.435 | 1 | 0.683557866 | 0.293029146 | 0.5 | 0.444 | 1 | 0.970716318 | 0.899864376 |
| Ppp1r12b | 0.890375192 | -0.375802336 | 0.815 | 0.747 | 1 | 0.684442724 | -0.171482101 | 0.7 | 0.824 | 1 | 0.890375192 | 0.900423605 |
| Ptpn23 | 0.902242081 | -0.174225139 | 0.462 | 0.424 | 1 | 0.684480647 | 0.379848071 | 0.5 | 0.441 | 1 | 0.902242081 | 0.900447538 |
| Zfp518a | 0.780477813 | 0.251538767 | 0.168 | 0.159 | 1 | 0.685156579 | -0.230005605 | 0.1 | 0.139 | 1 | 0.780477813 | 0.90087362 |
| Gab2 | 0.686900891 | -0.687735002 | 0.277 | 0.282 | 1 | 0.755903169 | -0.674334606 | 0.5 | 0.54 | 1 | 0.755903169 | 0.901968948 |
| Rab39 | 0.687344614 | -0.359895945 | 0.092 | 0.106 | 1 | 0.696463161 | 1.494359952 | 0.1 | 0.068 | 1 | 0.696463161 | 0.90224661 |
| Chn1os3 | 0.687345921 | -0.707819249 | 0.017 | 0.024 | 1 | 0.730750993 | 2.017921908 | 0 | 0.012 | 1 | 0.730750993 | 0.902247427 |
| Kcnab2 | 0.687345921 | -0.707819249 | 0.017 | 0.024 | 1 | 0.730749233 | 2.432959407 | 0 | 0.012 | 1 | 0.730749233 | 0.902247427 |
| Gm57047 | 0.687345921 | -0.707819249 | 0.017 | 0.024 | 1 | 0.698532411 | 2.017921908 | 0 | 0.015 | 1 | 0.698532411 | 0.902247427 |
| 4933406I18Rik | 0.6875384 | -0.387246433 | 0.286 | 0.253 | 1 | 0.755083981 | 0.299103661 | 0.3 | 0.241 | 1 | 0.755083981 | 0.902367748 |
| C77080 | 0.721162042 | 0.514573173 | 0.017 | 0.012 | 1 | 0.687544921 | 0.695993813 | 0.1 | 0.065 | 1 | 0.721162042 | 0.902371823 |
| Rnf31 | 0.688548539 | -0.485426827 | 0.118 | 0.129 | 1 | 0.721630177 | 0.111031312 | 0.1 | 0.136 | 1 | 0.721630177 | 0.902997988 |
| Ppp1r7 | 0.762713895 | -0.401362562 | 0.353 | 0.341 | 1 | 0.689599391 | 0.349543399 | 0.4 | 0.324 | 1 | 0.762713895 | 0.903651462 |
| Helz2 | 0.911521533 | -0.281893433 | 0.151 | 0.141 | 1 | 0.690166919 | 1.336097868 | 0.2 | 0.179 | 1 | 0.911521533 | 0.904003462 |
| Kif1a | 0.729956195 | -0.485426827 | 0.034 | 0.041 | 1 | 0.690399885 | 0.625604485 | 0.1 | 0.065 | 1 | 0.729956195 | 0.904147769 |
| Gm28875 | 0.69061685 | -0.222392421 | 0.059 | 0.071 | 1 | 0.92362858 | 0.432959407 | 0.1 | 0.108 | 1 | 0.92362858 | 0.904282066 |
| Aarsd1 | 0.691342877 | -0.349771728 | 0.261 | 0.276 | 1 | 0.885127866 | 0.223506042 | 0.2 | 0.222 | 1 | 0.885127866 | 0.90473078 |
| Baat | 0.691605844 | -0.485426827 | 0.017 | 0.024 | 1 | 0.73075158 | 2.695993813 | 0 | 0.012 | 1 | 0.73075158 | 0.904893044 |
| Gm15672 | 0.691605844 | -0.485426827 | 0.017 | 0.024 | 1 | 0.730749233 | 2.210566986 | 0 | 0.012 | 1 | 0.730749233 | 0.904893044 |
| Asb5 | 0.691605844 | -0.707819249 | 0.017 | 0.024 | 1 | 0.698534498 | 2.432959407 | 0 | 0.015 | 1 | 0.698534498 | 0.904893044 |
| 4933431K23Rik | 0.691605844 | -1.070389328 | 0.017 | 0.024 | 1 | 0.73075158 | 1.558490289 | 0 | 0.012 | 1 | 0.73075158 | 0.904893044 |
| 1810041H14Rik | 0.691605844 | -0.707819249 | 0.017 | 0.024 | 1 | 0.698534498 | 2.432959407 | 0 | 0.015 | 1 | 0.698534498 | 0.904893044 |
| Slc25a23 | 0.691605844 | -0.485426827 | 0.017 | 0.024 | 1 | 0.738073905 | 1.263034406 | 0.1 | 0.071 | 1 | 0.738073905 | 0.904893044 |
| Tmem123 | 0.903215166 | -0.134519665 | 0.244 | 0.247 | 1 | 0.691857985 | 0.450237399 | 0.4 | 0.364 | 1 | 0.903215166 | 0.905048499 |
| Rsad1 | 0.692323493 | -0.614709844 | 0.118 | 0.129 | 1 | 0.925746139 | 1.31748219 | 0.1 | 0.099 | 1 | 0.925746139 | 0.905335167 |
| Fam107a | 0.918851277 | -0.485426827 | 0.05 | 0.047 | 1 | 0.694099566 | 0.432959407 | 0.3 | 0.25 | 1 | 0.918851277 | 0.906424925 |
| Aspa | 0.695891031 | -0.292781749 | 0.017 | 0.024 | 1 | 0.698532411 | 1.847996907 | 0 | 0.015 | 1 | 0.698532411 | 0.907517735 |
| 9130604C24Rik | 0.695893386 | -0.222392421 | 0.017 | 0.024 | 1 | 0.698534498 | 2.432959407 | 0 | 0.015 | 1 | 0.698534498 | 0.907519167 |
| Jsrp1 | 0.695893386 | -0.222392421 | 0.017 | 0.024 | 1 | 0.73075158 | 2.695993813 | 0 | 0.012 | 1 | 0.73075158 | 0.907519167 |
| D630036H23Rik | 0.695893386 | -0.222392421 | 0.017 | 0.024 | 1 | 0.730749233 | 2.432959407 | 0 | 0.012 | 1 | 0.730749233 | 0.907519167 |
| Nrxn3 | 0.695893386 | -0.222392421 | 0.017 | 0.024 | 1 | 0.73075158 | 1.558490289 | 0 | 0.012 | 1 | 0.73075158 | 0.907519167 |
| Tprg | 0.695893386 | -0.222392421 | 0.017 | 0.024 | 1 | 0.730750993 | 1.695993813 | 0 | 0.012 | 1 | 0.730750993 | 0.907519167 |
| Prdx3 | 0.943788182 | -0.12609876 | 0.664 | 0.576 | 1 | 0.698254621 | -0.215697769 | 0.5 | 0.512 | 1 | 0.943788182 | 0.908949727 |
| Zfp668 | 0.698446093 | -0.222392421 | 0.092 | 0.106 | 1 | 0.949913217 | 0.847996907 | 0.1 | 0.093 | 1 | 0.949913217 | 0.909065241 |
| Plscr2 | 0.736478911 | 1.04508789 | 0.076 | 0.065 | 1 | 0.698529281 | 2.210566986 | 0 | 0.015 | 1 | 0.736478911 | 0.909115405 |
| 4930444A19Rik | 0.781790612 | -0.485426827 | 0.008 | 0.012 | 1 | 0.698529281 | 2.210566986 | 0 | 0.015 | 1 | 0.781790612 | 0.909115405 |
| Oas1a | 0.784920947 | -0.900464326 | 0.008 | 0.012 | 1 | 0.698529281 | 2.017921908 | 0 | 0.015 | 1 | 0.784920947 | 0.909115405 |
| Gm11266 | 0.791189474 | 0.514573173 | 0.008 | 0.012 | 1 | 0.698529281 | 1.847996907 | 0 | 0.015 | 1 | 0.791189474 | 0.909115405 |
| Tafa4 | 0.830638286 | -0.163498732 | 0.025 | 0.029 | 1 | 0.698529281 | 2.017921908 | 0 | 0.015 | 1 | 0.830638286 | 0.909115405 |
| 2810408I11Rik | 0.926217033 | -0.333423734 | 0.034 | 0.035 | 1 | 0.698529281 | 2.017921908 | 0 | 0.015 | 1 | 0.926217033 | 0.909115405 |
| Nlrp12 | 0.929388469 | 0.321928095 | 0.025 | 0.024 | 1 | 0.698529281 | 2.017921908 | 0 | 0.015 | 1 | 0.929388469 | 0.909115405 |
| Gm39185 | 0.944467697 | -0.707819249 | 0.017 | 0.018 | 1 | 0.698529281 | 2.210566986 | 0 | 0.015 | 1 | 0.944467697 | 0.909115405 |
| Gm37634 | 0.954551343 | -0.222392421 | 0.017 | 0.018 | 1 | 0.698529281 | 2.210566986 | 0 | 0.015 | 1 | 0.954551343 | 0.909115405 |
| Marchf10 | 0.954551343 | -0.222392421 | 0.017 | 0.018 | 1 | 0.698529281 | 1.695993813 | 0 | 0.015 | 1 | 0.954551343 | 0.909115405 |
| Gpr18 | 0.954551343 | -0.707819249 | 0.017 | 0.018 | 1 | 0.698529281 | 2.017921908 | 0 | 0.015 | 1 | 0.954551343 | 0.909115405 |
| Ifi203 | 0.710606097 | 1.251538767 | 0.017 | 0.012 | 1 | 0.698532411 | 1.847996907 | 0 | 0.015 | 1 | 0.710606097 | 0.909117293 |
| Elapor2 | 0.715874401 | 1.251538767 | 0.017 | 0.012 | 1 | 0.698532411 | 1.847996907 | 0 | 0.015 | 1 | 0.715874401 | 0.909117293 |
| Cfap161 | 0.718517371 | 0.836501268 | 0.017 | 0.012 | 1 | 0.698532411 | 1.695993813 | 0 | 0.015 | 1 | 0.718517371 | 0.909117293 |
| Gm29570 | 0.721162042 | 0.514573173 | 0.017 | 0.012 | 1 | 0.698532411 | 1.695993813 | 0 | 0.015 | 1 | 0.721162042 | 0.909117293 |
| Thbs4 | 0.721162042 | 0.514573173 | 0.017 | 0.012 | 1 | 0.698532411 | 1.432959407 | 0 | 0.015 | 1 | 0.721162042 | 0.909117293 |
| 1810034E14Rik | 0.729956195 | -0.748461233 | 0.034 | 0.041 | 1 | 0.698532411 | 1.847996907 | 0 | 0.015 | 1 | 0.729956195 | 0.909117293 |
| Gm12472 | 0.781790612 | -0.485426827 | 0.008 | 0.012 | 1 | 0.698532411 | 1.695993813 | 0 | 0.015 | 1 | 0.781790612 | 0.909117293 |

| | | | | | | | | | | | | |
|---|---|---|---|---|---|---|---|---|---|---|---|---|
| Sema3a | 0.781790612 | -1.070389328 | 0.008 | 0.012 | 1 | 0.698532411 | 1.847996907 | 0 | 0.015 | 1 | 0.781790612 | 0.909117293 |
| ENSMUSG00000121382 | 0.781790612 | -0.485426827 | 0.008 | 0.012 | 1 | 0.698532411 | 1.847996907 | 0 | 0.015 | 1 | 0.781790612 | 0.909117293 |
| Liph | 0.781790612 | -0.807354922 | 0.008 | 0.012 | 1 | 0.698532411 | 1.847996907 | 0 | 0.015 | 1 | 0.781790612 | 0.909117293 |
| Ryr1 | 0.959597195 | -0.292781749 | 0.017 | 0.018 | 1 | 0.698532411 | 1.847996907 | 0 | 0.015 | 1 | 0.959597195 | 0.909117293 |
| Tnfrsf9 | 0.974741666 | 1.321928095 | 0.017 | 0.018 | 1 | 0.698532411 | 1.847996907 | 0 | 0.015 | 1 | 0.974741666 | 0.909117293 |
| Mab21l2 | 0.778665521 | -1.485426827 | 0.008 | 0.012 | 1 | 0.698533977 | 1.210566986 | 0 | 0.015 | 1 | 0.778665521 | 0.909118237 |
| Gpr158 | 0.781790612 | -0.485426827 | 0.008 | 0.012 | 1 | 0.698533977 | 1.210566986 | 0 | 0.015 | 1 | 0.781790612 | 0.909118237 |
| Cldn14 | 0.925119899 | 1.514573173 | 0.025 | 0.024 | 1 | 0.698533977 | 1.847996907 | 0 | 0.015 | 1 | 0.925119899 | 0.909118237 |
| Gm38560 | 0.721161231 | 0.514573173 | 0.017 | 0.012 | 1 | 0.698534498 | 2.432959407 | 0 | 0.015 | 1 | 0.721161231 | 0.909118551 |
| Gm13620 | 0.721162042 | 0.514573173 | 0.017 | 0.012 | 1 | 0.698534498 | 2.432959407 | 0 | 0.015 | 1 | 0.721162042 | 0.909118551 |
| Hoatz | 0.721162042 | 0.514573173 | 0.017 | 0.012 | 1 | 0.698534498 | 2.432959407 | 0 | 0.015 | 1 | 0.721162042 | 0.909118551 |
| Fam228a | 0.721162042 | 0.514573173 | 0.017 | 0.012 | 1 | 0.698534498 | 2.432959407 | 0 | 0.015 | 1 | 0.721162042 | 0.909118551 |
| 1700108F19Rik | 0.721162042 | 0.514573173 | 0.017 | 0.012 | 1 | 0.698534498 | 2.432959407 | 0 | 0.015 | 1 | 0.721162042 | 0.909118551 |
| Mettl21c | 0.781790612 | -0.485426827 | 0.008 | 0.012 | 1 | 0.698534498 | 2.432959407 | 0 | 0.015 | 1 | 0.781790612 | 0.909118551 |
| Dll1 | 0.846810562 | 0.225066556 | 0.042 | 0.047 | 1 | 0.698534498 | 2.432959407 | 0 | 0.015 | 1 | 0.846810562 | 0.909118551 |
| Gm56945 | 0.942200939 | -0.292781749 | 0.025 | 0.024 | 1 | 0.698534498 | 2.432959407 | 0 | 0.015 | 1 | 0.942200939 | 0.909118551 |
| 9330198N18Rik | 0.949508638 | -0.707819249 | 0.017 | 0.018 | 1 | 0.698534498 | 2.432959407 | 0 | 0.015 | 1 | 0.949508638 | 0.909118551 |
| Tmem104 | 0.699203351 | -0.122856748 | 0.059 | 0.071 | 1 | 0.740899751 | 1.111031312 | 0.1 | 0.071 | 1 | 0.740899751 | 0.909521376 |
| Auh | 0.956075636 | -0.32788555 | 0.387 | 0.359 | 1 | 0.700350167 | -0.489872732 | 0.3 | 0.324 | 1 | 0.956075636 | 0.910209978 |
| Cln5 | 0.713512595 | -0.198545679 | 0.395 | 0.394 | 1 | 0.701528 | 0.18142064 | 0.5 | 0.398 | 1 | 0.713512595 | 0.910914465 |
| Chst10 | 0.736440063 | -0.333423734 | 0.034 | 0.041 | 1 | 0.701889643 | 1.210566986 | 0.1 | 0.068 | 1 | 0.736440063 | 0.911130215 |
| Gtpbp2 | 0.98039559 | -0.30560579 | 0.42 | 0.382 | 1 | 0.70206058 | -0.516056664 | 0.5 | 0.46 | 1 | 0.98039559 | 0.911232102 |
| Clasrp | 0.702366872 | -0.292781749 | 0.252 | 0.265 | 1 | 0.750279863 | 0.159940913 | 0.3 | 0.235 | 1 | 0.750279863 | 0.911414521 |
| Rab39b | 0.783113074 | -0.485426827 | 0.193 | 0.194 | 1 | 0.703877869 | -0.327852929 | 0.2 | 0.235 | 1 | 0.783113074 | 0.912311683 |
| Gm28411 | 0.704462092 | 0.192645078 | 0.017 | 0.024 | 1 | 0.73075158 | 2.695993813 | 0 | 0.012 | 1 | 0.73075158 | 0.912657345 |
| Pigg | 0.738417015 | 0.230780207 | 0.16 | 0.147 | 1 | 0.705404156 | 0.111031312 | 0.1 | 0.139 | 1 | 0.738417015 | 0.913213289 |
| 3830406C13Rik | 0.707377823 | -0.253611152 | 0.429 | 0.382 | 1 | 0.801773528 | 0.245332404 | 0.3 | 0.349 | 1 | 0.801773528 | 0.914372262 |
| Gm10549 | 0.742975843 | -0.222392421 | 0.101 | 0.112 | 1 | 0.707564437 | 1.210566986 | 0.1 | 0.068 | 1 | 0.742975843 | 0.914481441 |
| Prxl2b | 0.775208187 | -0.163498732 | 0.151 | 0.135 | 1 | 0.707564437 | 1.210566986 | 0.1 | 0.068 | 1 | 0.775208187 | 0.914481441 |
| Zswim4 | 0.831356128 | -0.24838763 | 0.319 | 0.324 | 1 | 0.708452914 | 0.66624647 | 0.4 | 0.41 | 1 | 0.831356128 | 0.915000297 |
| Spata24 | 0.711008194 | -0.19592021 | 0.168 | 0.147 | 1 | 0.966276887 | 1.14345279 | 0.1 | 0.114 | 1 | 0.966276887 | 0.916483736 |
| Ackr3 | 0.712927937 | 0.211242297 | 0.538 | 0.553 | 1 | 0.792393494 | 1.416657595 | 0.5 | 0.506 | 1 | 0.792393494 | 0.917589631 |
| Kcnd1 | 0.82670638 | -0.292781749 | 0.025 | 0.029 | 1 | 0.713243542 | 1.017921908 | 0.1 | 0.068 | 1 | 0.82670638 | 0.917770734 |
| Trpm3 | 0.802554629 | -1.485426827 | 0.118 | 0.124 | 1 | 0.713848685 | -1.177538186 | 0.2 | 0.241 | 1 | 0.802554629 | 0.918117425 |
| Klhl5 | 0.981892435 | -0.127533235 | 0.37 | 0.353 | 1 | 0.714603108 | 0.571665678 | 0.4 | 0.401 | 1 | 0.981892435 | 0.918548614 |
| Fastk | 0.756623835 | -0.295455884 | 0.479 | 0.453 | 1 | 0.714848205 | -0.11821478 | 0.4 | 0.432 | 1 | 0.756623835 | 0.918688454 |
| Dzip3 | 0.740614169 | -0.423270351 | 0.429 | 0.406 | 1 | 0.717665314 | -0.224571719 | 0.3 | 0.383 | 1 | 0.740614169 | 0.920287125 |
| Rnpep | 0.909797243 | -0.370968103 | 0.361 | 0.329 | 1 | 0.717840041 | -0.262186011 | 0.4 | 0.466 | 1 | 0.909797243 | 0.920385757 |
| Sec14l2 | 0.719564527 | 0.354108501 | 0.118 | 0.106 | 1 | 0.900378755 | 0.388565288 | 0.2 | 0.228 | 1 | 0.900378755 | 0.921355945 |
| Aldh1a1 | 0.918487834 | -0.485426827 | 0.05 | 0.053 | 1 | 0.719601387 | 0.374065718 | 0.1 | 0.139 | 1 | 0.918487834 | 0.921376618 |
| Baz2b | 0.721128943 | -0.189613336 | 0.992 | 0.971 | 1 | 0.835089665 | -0.166893702 | 1 | 0.966 | 1 | 0.835089665 | 0.922230934 |
| Gm16183 | 0.721161231 | 0.514573173 | 0.017 | 0.012 | 1 | 0.730749233 | 2.432959407 | 0 | 0.012 | 1 | 0.730749233 | 0.922248941 |
| P2rx3 | 0.721162042 | 0.514573173 | 0.017 | 0.012 | 1 | 0.73075158 | 2.695993813 | 0 | 0.012 | 1 | 0.73075158 | 0.922249393 |
| Gm43660 | 0.721162042 | 0.514573173 | 0.017 | 0.012 | 1 | 0.730749233 | 2.432959407 | 0 | 0.012 | 1 | 0.730749233 | 0.922249393 |
| Parvaos | 0.721162042 | 0.514573173 | 0.017 | 0.012 | 1 | 0.73075158 | 2.695993813 | 0 | 0.012 | 1 | 0.73075158 | 0.922249393 |
| Icam5 | 0.721162042 | 0.514573173 | 0.017 | 0.012 | 1 | 0.730749233 | 2.210566986 | 0 | 0.012 | 1 | 0.730749233 | 0.922249393 |
| Pgf | 0.721162042 | 0.514573173 | 0.017 | 0.012 | 1 | 0.73075158 | 2.695993813 | 0 | 0.012 | 1 | 0.73075158 | 0.922249393 |
| Acat3 | 0.721162042 | 0.514573173 | 0.017 | 0.012 | 1 | 0.730749233 | 2.432959407 | 0 | 0.012 | 1 | 0.730749233 | 0.922249393 |
| BC037039 | 0.721162042 | 0.514573173 | 0.017 | 0.012 | 1 | 0.73075158 | 2.695993813 | 0 | 0.012 | 1 | 0.73075158 | 0.922249393 |
| Abcc2 | 0.721162042 | 0.514573173 | 0.017 | 0.012 | 1 | 0.73075158 | 2.695993813 | 0 | 0.012 | 1 | 0.73075158 | 0.922249393 |
| Gm16068 | 0.721162042 | 0.514573173 | 0.017 | 0.012 | 1 | 0.730749233 | 2.432959407 | 0 | 0.012 | 1 | 0.730749233 | 0.922249393 |
| Gm21060 | 0.721162042 | 0.514573173 | 0.017 | 0.012 | 1 | 0.73075158 | 2.695993813 | 0 | 0.012 | 1 | 0.73075158 | 0.922249393 |
| 2310061I04Rik | 0.800244188 | -0.201633861 | 0.109 | 0.118 | 1 | 0.721624061 | 0.159940913 | 0.1 | 0.136 | 1 | 0.800244188 | 0.922506836 |
| Olfm2 | 0.723489531 | -1.250961574 | 0.034 | 0.041 | 1 | 0.920257134 | -0.391469028 | 0.1 | 0.105 | 1 | 0.920257134 | 0.923541961 |
| Lypd2 | 0.724131315 | 1.099535674 | 0.118 | 0.135 | 1 | 0.828905415 | 0.785261151 | 0.2 | 0.253 | 1 | 0.828905415 | 0.923896468 |
| Zbtb43 | 0.725662655 | -0.236399279 | 0.252 | 0.259 | 1 | 0.828283226 | 0.359710425 | 0.3 | 0.37 | 1 | 0.828283226 | 0.924739021 |
| Serpinb11 | 0.726456542 | -1.222392421 | 0.017 | 0.012 | 1 | 0.730749233 | 2.432959407 | 0 | 0.012 | 1 | 0.730749233 | 0.925173977 |
| Tgfbr3l | 0.726456542 | -0.222392421 | 0.017 | 0.012 | 1 | 0.73075158 | 2.695993813 | 0 | 0.012 | 1 | 0.73075158 | 0.925173977 |
| Anks1 | 0.728781507 | -0.287098111 | 0.538 | 0.506 | 1 | 0.755996813 | -0.377514857 | 0.7 | 0.642 | 1 | 0.755996813 | 0.926440529 |
| Mlh1 | 0.729331019 | -0.944858446 | 0.084 | 0.094 | 1 | 0.960463364 | 0.660369903 | 0.1 | 0.093 | 1 | 0.960463364 | 0.926738303 |
| Ctu1 | 0.760414269 | -0.137503524 | 0.05 | 0.059 | 1 | 0.729840207 | 1.432959407 | 0.1 | 0.071 | 1 | 0.760414269 | 0.927013686 |
| A530013C23Rik | 0.729956195 | -0.622930351 | 0.034 | 0.041 | 1 | 0.746555377 | 0.847996907 | 0.1 | 0.071 | 1 | 0.746555377 | 0.927076343 |
| Pi15 | 0.731769749 | -1.070389328 | 0.017 | 0.012 | 1 | 0.730749233 | 2.017921908 | 0 | 0.012 | 1 | 0.731769749 | 0.927504024 |
| Cblc | 0.731769749 | -0.222392421 | 0.017 | 0.012 | 1 | 0.730749233 | 2.210566986 | 0 | 0.012 | 1 | 0.731769749 | 0.927504024 |
| E2f7 | 0.778665521 | -1.485426827 | 0.008 | 0.012 | 1 | 0.730749233 | 1.847996907 | 0 | 0.012 | 1 | 0.778665521 | 0.927504024 |
| Gm6345 | 0.781790612 | -0.485426827 | 0.008 | 0.012 | 1 | 0.730749233 | 2.432959407 | 0 | 0.012 | 1 | 0.781790612 | 0.927504024 |
| Prnd | 0.781790612 | -1.070389328 | 0.008 | 0.012 | 1 | 0.730749233 | 2.210566986 | 0 | 0.012 | 1 | 0.781790612 | 0.927504024 |
| Gm52054 | 0.781790612 | -0.485426827 | 0.008 | 0.012 | 1 | 0.730749233 | 2.432959407 | 0 | 0.012 | 1 | 0.781790612 | 0.927504024 |
| Ftcd | 0.791189474 | 0.514573173 | 0.008 | 0.012 | 1 | 0.730749233 | 2.432959407 | 0 | 0.012 | 1 | 0.791189474 | 0.927504024 |
| Fam122b | 0.82670638 | -0.292781749 | 0.025 | 0.029 | 1 | 0.730749233 | 2.432959407 | 0 | 0.012 | 1 | 0.82670638 | 0.927504024 |
| 4930412C18Rik | 0.8327876 | -0.359895945 | 0.042 | 0.047 | 1 | 0.730749233 | 2.432959407 | 0 | 0.012 | 1 | 0.8327876 | 0.927504024 |
| 9930111J21Rik2 | 0.944467697 | -1.485426827 | 0.017 | 0.018 | 1 | 0.730749233 | 2.210566986 | 0 | 0.012 | 1 | 0.944467697 | 0.927504024 |
| Cfap47 | 0.778665521 | -1.070389328 | 0.008 | 0.012 | 1 | 0.730750993 | 2.210566986 | 0 | 0.012 | 1 | 0.778665521 | 0.927504972 |
| Rasgrp1 | 0.781790612 | -1.485426827 | 0.008 | 0.012 | 1 | 0.730750993 | 2.017921908 | 0 | 0.012 | 1 | 0.781790612 | 0.927504972 |
| Gm33732 | 0.842422009 | 0.251538767 | 0.025 | 0.029 | 1 | 0.730750993 | 1.847996907 | 0 | 0.012 | 1 | 0.842422009 | 0.927504972 |
| Gm50186 | 0.954551343 | -0.222392421 | 0.017 | 0.018 | 1 | 0.730750993 | 2.017921908 | 0 | 0.012 | 1 | 0.954551343 | 0.927504972 |
| 1810044D09Rik | 0.770891247 | 0.292180751 | 0.042 | 0.035 | 1 | 0.73075158 | 2.695993813 | 0 | 0.012 | 1 | 0.770891247 | 0.927505289 |
| Klhl41 | 0.781790612 | -0.485426827 | 0.008 | 0.012 | 1 | 0.73075158 | 2.695993813 | 0 | 0.012 | 1 | 0.781790612 | 0.927505289 |
| Mypopos | 0.781790612 | -0.485426827 | 0.008 | 0.012 | 1 | 0.73075158 | 2.695993813 | 0 | 0.012 | 1 | 0.781790612 | 0.927505289 |
| Gm94 | 0.781790612 | -0.485426827 | 0.008 | 0.012 | 1 | 0.73075158 | 2.695993813 | 0 | 0.012 | 1 | 0.781790612 | 0.927505289 |
| Gm21860 | 0.781790612 | -0.485426827 | 0.008 | 0.012 | 1 | 0.73075158 | 2.695993813 | 0 | 0.012 | 1 | 0.781790612 | 0.927505289 |
| Kif14 | 0.809121304 | -1.070389328 | 0.025 | 0.029 | 1 | 0.73075158 | 2.695993813 | 0 | 0.012 | 1 | 0.809121304 | 0.927505289 |
| Gm33694 | 0.839906751 | 0.292180751 | 0.034 | 0.029 | 1 | 0.73075158 | 2.695993813 | 0 | 0.012 | 1 | 0.839906751 | 0.927505289 |
| Swsap1 | 0.841777167 | 0.251538767 | 0.034 | 0.029 | 1 | 0.73075158 | 2.695993813 | 0 | 0.012 | 1 | 0.841777167 | 0.927505289 |
| Gm46210 | 0.841777167 | 0.251538767 | 0.034 | 0.029 | 1 | 0.73075158 | 2.695993813 | 0 | 0.012 | 1 | 0.841777167 | 0.927505289 |
| Myrip | 0.852421231 | -0.11345805 | 0.084 | 0.076 | 1 | 0.73075158 | 0.847996907 | 0 | 0.012 | 1 | 0.852421231 | 0.927505289 |
| Adgrg7 | 0.916593122 | 0.929610672 | 0.025 | 0.024 | 1 | 0.73075158 | 2.695993813 | 0 | 0.012 | 1 | 0.916593122 | 0.927505289 |
| Gm29481 | 0.949508638 | -0.485426827 | 0.017 | 0.018 | 1 | 0.73075158 | 2.695993813 | 0 | 0.012 | 1 | 0.949508638 | 0.927505289 |
| Dmrta2 | 0.954551343 | -0.485426827 | 0.017 | 0.018 | 1 | 0.73075158 | 2.695993813 | 0 | 0.012 | 1 | 0.954551343 | 0.927505289 |
| Tbx21 | 0.954551343 | -0.222392421 | 0.017 | 0.018 | 1 | 0.73075158 | 2.695993813 | 0 | 0.012 | 1 | 0.954551343 | 0.927505289 |
| Oma1 | 0.845455025 | -0.258016331 | 0.16 | 0.147 | 1 | 0.731393023 | 0.354956895 | 0.2 | 0.151 | 1 | 0.845455025 | 0.927850292 |
| Gm17435 | 0.771351101 | -0.222392421 | 0.143 | 0.153 | 1 | 0.731979614 | 0.185031894 | 0.1 | 0.136 | 1 | 0.771351101 | 0.928165073 |
| Cep170b | 0.894995937 | -0.273922722 | 0.227 | 0.229 | 1 | 0.732452172 | -0.357117523 | 0.3 | 0.321 | 1 | 0.894995937 | 0.92841816 |
| Prkaa2 | 0.997075054 | -0.365587057 | 0.277 | 0.265 | 1 | 0.732774205 | -0.649350946 | 0.7 | 0.571 | 1 | 0.997075054 | 0.928590374 |
| Coq9 | 0.782712149 | 0.152003093 | 0.244 | 0.235 | 1 | 0.733243829 | 0.648688098 | 0.2 | 0.154 | 1 | 0.782712149 | 0.928841145 |
| Abcc9 | 0.73484547 | -0.359895945 | 0.034 | 0.041 | 1 | 0.97181614 | 0.210566986 | 0.1 | 0.099 | 1 | 0.97181614 | 0.929693075 |
| Ibtk | 0.791339858 | -0.182415946 | 0.672 | 0.606 | 1 | 0.73635345 | 0.330148602 | 0.6 | 0.685 | 1 | 0.791339858 | 0.930490497 |
| BC004004 | 0.738046229 | -0.209365741 | 0.538 | 0.506 | 1 | 0.993922154 | 0.359710425 | 0.4 | 0.395 | 1 | 0.993922154 | 0.931380222 |

| | | | | | | | | | | | | |
|---|---|---|---|---|---|---|---|---|---|---|---|---|
| A430110L20Rik | 0.902582003 | -0.11345805 | 0.076 | 0.071 | 1 | 0.738073803 | 1.210566986 | 0.1 | 0.071 | 1 | 0.902582003 | 0.931394667 |
| Ficd | 0.798995268 | 0.321928095 | 0.084 | 0.076 | 1 | 0.738190479 | 0.345496566 | 0.1 | 0.136 | 1 | 0.798995268 | 0.931455775 |
| Abcd2 | 0.915783742 | -0.584962501 | 0.05 | 0.047 | 1 | 0.739199376 | 1.591657153 | 0.1 | 0.077 | 1 | 0.915783742 | 0.931983034 |
| Ropn1l | 0.92730367 | 0.308122295 | 0.084 | 0.082 | 1 | 0.740900052 | 1.063725598 | 0.1 | 0.071 | 1 | 0.92730367 | 0.932867217 |
| Zbtb24 | 0.743160593 | -0.182034684 | 0.218 | 0.194 | 1 | 1 | 0.210566986 | 0.2 | 0.194 | 1 | 1 | 0.934033519 |
| Paf1 | 0.84721536 | -0.21041978 | 0.588 | 0.553 | 1 | 0.743239186 | -0.14865965 | 0.6 | 0.651 | 1 | 0.84721536 | 0.934073884 |
| Gpn2 | 0.743347 | -0.173482821 | 0.134 | 0.147 | 1 | 0.77032871 | 0.374065718 | 0.1 | 0.13 | 1 | 0.77032871 | 0.934129238 |
| Wdr44 | 0.743612951 | -0.490292582 | 0.546 | 0.5 | 1 | 0.951281185 | 0.243574092 | 0.7 | 0.685 | 1 | 0.951281185 | 0.934265681 |
| Uqcc5 | 0.74799621 | -0.116193018 | 0.345 | 0.371 | 1 | 0.91035791 | 0.678071905 | 0.3 | 0.327 | 1 | 0.91035791 | 0.93649409 |
| Pop7 | 0.749678593 | 0.109316694 | 0.227 | 0.247 | 1 | 0.757516379 | 1.231325546 | 0.2 | 0.176 | 1 | 0.757516379 | 0.937339193 |
| Slc30a1 | 0.901468271 | 0.185265548 | 0.227 | 0.241 | 1 | 0.750779646 | 0.159940913 | 0.3 | 0.25 | 1 | 0.901468271 | 0.937889215 |
| Gm26728 | 0.779131717 | -0.137503524 | 0.042 | 0.035 | 1 | 0.752214659 | 0.808468542 | 0.1 | 0.071 | 1 | 0.779131717 | 0.938602425 |
| Eci1 | 0.995985306 | -0.267246657 | 0.387 | 0.347 | 1 | 0.752863311 | 0.210566986 | 0.4 | 0.315 | 1 | 0.995985306 | 0.938923457 |
| Zfp58 | 0.753488444 | -0.444784843 | 0.143 | 0.153 | 1 | 0.966655593 | 0.263034406 | 0.2 | 0.185 | 1 | 0.966655593 | 0.939232053 |
| Nin | 0.754189188 | 0.318034179 | 0.445 | 0.429 | 1 | 0.947091324 | 0.629904623 | 0.4 | 0.522 | 1 | 0.947091324 | 0.939577045 |
| Mfsd8 | 0.754337771 | -0.222392421 | 0.084 | 0.094 | 1 | 0.773317994 | 1.159940913 | 0.1 | 0.074 | 1 | 0.773317994 | 0.939650069 |
| Pex7 | 0.931236682 | -0.171318237 | 0.336 | 0.318 | 1 | 0.754770812 | 0.506959989 | 0.2 | 0.262 | 1 | 0.931236682 | 0.939862645 |
| Eif2d | 0.755426191 | 0.114965714 | 0.311 | 0.282 | 1 | 0.901851322 | 0.130396637 | 0.3 | 0.324 | 1 | 0.901851322 | 0.940183652 |
| Sfxn1 | 0.995343462 | -0.279842694 | 0.387 | 0.359 | 1 | 0.756539203 | 0.480487777 | 0.4 | 0.373 | 1 | 0.995343462 | 0.94072684 |
| Ifrd1 | 0.946340668 | 0.159218223 | 0.975 | 0.976 | 1 | 0.760757338 | 0.480823796 | 1 | 0.994 | 1 | 0.946340668 | 0.942762949 |
| Recql | 0.932852538 | -0.304854582 | 0.134 | 0.129 | 1 | 0.763867381 | 0.111031312 | 0.1 | 0.13 | 1 | 0.932852538 | 0.944241386 |
| Smyd5 | 0.765055862 | -0.418312631 | 0.067 | 0.076 | 1 | 0.90271652 | 0.591657153 | 0.1 | 0.111 | 1 | 0.90271652 | 0.944801252 |
| Xab2 | 0.920381825 | -0.201633861 | 0.109 | 0.112 | 1 | 0.772713711 | 0.558490289 | 0.2 | 0.16 | 1 | 0.920381825 | 0.948340943 |
| Coa6 | 0.773386846 | -0.32788555 | 0.193 | 0.2 | 1 | 0.79991673 | 1.135278859 | 0.1 | 0.142 | 1 | 0.79991673 | 0.948646478 |
| Dus2 | 0.774749271 | 0.514573173 | 0.134 | 0.124 | 1 | 0.989745405 | 0.432959407 | 0.1 | 0.099 | 1 | 0.989745405 | 0.949262109 |
| Rasl11a | 0.947172425 | 0.734698987 | 0.168 | 0.165 | 1 | 0.775686745 | 1.190102883 | 0.1 | 0.08 | 1 | 0.947172425 | 0.949683563 |
| Zfp512 | 0.963759138 | -0.432959407 | 0.202 | 0.194 | 1 | 0.776027447 | 0.249737583 | 0.3 | 0.241 | 1 | 0.963759138 | 0.949836296 |
| Shld1 | 0.831393676 | 0.192645078 | 0.059 | 0.053 | 1 | 0.77893732 | 1.017921908 | 0.1 | 0.074 | 1 | 0.831393676 | 0.951131291 |
| Trpc1 | 0.950020442 | -0.163498732 | 0.21 | 0.206 | 1 | 0.781455713 | 0.478763097 | 0.3 | 0.281 | 1 | 0.950020442 | 0.952238395 |
| Pigc | 0.781740974 | -0.563429339 | 0.143 | 0.147 | 1 | 0.89025648 | 0.193493473 | 0.2 | 0.213 | 1 | 0.89025648 | 0.952362998 |
| Fkrp | 0.785007715 | -0.459431619 | 0.21 | 0.212 | 1 | 0.956633159 | 0.463333056 | 0.3 | 0.327 | 1 | 0.956633159 | 0.953778317 |
| Naa80 | 0.7852007 | 0.173536255 | 0.084 | 0.094 | 1 | 0.814156863 | 1.888638891 | 0.1 | 0.086 | 1 | 0.814156863 | 0.953861261 |
| Zhx3 | 0.848050699 | -0.160331453 | 0.681 | 0.641 | 1 | 0.785263559 | -0.281504188 | 0.8 | 0.75 | 1 | 0.848050699 | 0.953888261 |
| Fbxl9 | 0.785768995 | 0.292180751 | 0.109 | 0.1 | 1 | 0.923628623 | 0.625604485 | 0.1 | 0.108 | 1 | 0.923628623 | 0.954105076 |
| Lcat | 0.845214404 | -0.102098188 | 0.092 | 0.1 | 1 | 0.788046783 | 0.263034406 | 0.1 | 0.127 | 1 | 0.845214404 | 0.955075834 |
| Cenpa | 0.791107473 | -0.534336428 | 0.092 | 0.1 | 1 | 1 | 0.463333056 | 0.1 | 0.099 | 1 | 1 | 0.956363912 |
| Sirpa | 0.793742403 | -0.235448574 | 0.168 | 0.176 | 1 | 0.881956326 | -0.165299916 | 0.2 | 0.207 | 1 | 0.881956326 | 0.957457804 |
| Cfap36 | 0.795659659 | -0.173482821 | 0.773 | 0.682 | 1 | 0.946701874 | 0.440996726 | 0.5 | 0.565 | 1 | 0.946701874 | 0.958245025 |
| 2310014F06Rik | 0.81973837 | -0.341036918 | 0.143 | 0.129 | 1 | 0.796928751 | 0.354956895 | 0.2 | 0.16 | 1 | 0.81973837 | 0.958762068 |
| Tbc1d9b | 0.852207552 | -0.13965199 | 0.37 | 0.347 | 1 | 0.799182741 | 0.219180116 | 0.4 | 0.349 | 1 | 0.852207552 | 0.959672429 |
| Dgke | 0.956636546 | -0.592342031 | 0.101 | 0.094 | 1 | 0.799264655 | 1.159940913 | 0.1 | 0.077 | 1 | 0.956636546 | 0.959705321 |
| Mcrs1 | 0.97304451 | -0.172268942 | 0.311 | 0.3 | 1 | 0.803061516 | 0.359710425 | 0.2 | 0.25 | 1 | 0.97304451 | 0.961215233 |
| Adck5 | 0.867089572 | -0.273922722 | 0.084 | 0.088 | 1 | 0.804841125 | 1.111031312 | 0.1 | 0.077 | 1 | 0.867089572 | 0.961913014 |
| Gm56906 | 0.806445602 | 0.688055994 | 0.353 | 0.353 | 1 | 0.825954596 | -0.100075301 | 0.5 | 0.444 | 1 | 0.825954596 | 0.962536695 |
| Pank2 | 0.80707266 | -0.295602268 | 0.445 | 0.429 | 1 | 0.945060187 | 0.227844977 | 0.4 | 0.429 | 1 | 0.945060187 | 0.962779042 |
| Sdhaf2 | 0.936175735 | -0.27897595 | 0.429 | 0.4 | 1 | 0.807802254 | 0.159940913 | 0.4 | 0.343 | 1 | 0.936175735 | 0.963060027 |
| Kcnip2 | 0.809580956 | -0.707819249 | 0.042 | 0.047 | 1 | 0.878982225 | 0.591657153 | 0.1 | 0.083 | 1 | 0.878982225 | 0.963740588 |
| 5430402E10Rik | 0.82670638 | -0.485426827 | 0.025 | 0.029 | 1 | 0.810417824 | 0.847996907 | 0.1 | 0.077 | 1 | 0.82670638 | 0.964058599 |
| Mfsd2b | 0.812352496 | -0.655351829 | 0.059 | 0.065 | 1 | 0.870773487 | 0.808468542 | 0.1 | 0.083 | 1 | 0.870773487 | 0.964788414 |
| Ssx2ip | 0.985443311 | -0.191695624 | 0.244 | 0.235 | 1 | 0.814218228 | -0.127755547 | 0.3 | 0.327 | 1 | 0.985443311 | 0.965485133 |
| Eef1akmt4 | 0.893040156 | 0.152003093 | 0.151 | 0.147 | 1 | 0.815153559 | 0.494359952 | 0.1 | 0.123 | 1 | 0.893040156 | 0.965831793 |
| Zfhx2 | 0.815458199 | -0.222392421 | 0.092 | 0.082 | 1 | 0.997503154 | 0.847996907 | 0.1 | 0.105 | 1 | 0.997503154 | 0.965944324 |
| Ap5s1 | 0.815458483 | -0.292781749 | 0.092 | 0.082 | 1 | 0.896995661 | 0.888638891 | 0.1 | 0.086 | 1 | 0.896995661 | 0.965944429 |
| Gstm2 | 0.949508638 | -1.070389328 | 0.017 | 0.018 | 1 | 0.817393355 | 0.403212064 | 0.1 | 0.123 | 1 | 0.949508638 | 0.966654813 |
| Vps53 | 0.906539718 | -0.114783447 | 0.361 | 0.347 | 1 | 0.817675908 | 0.732519689 | 0.3 | 0.401 | 1 | 0.906539718 | 0.966757926 |
| Gm9968 | 0.895994907 | 0.251538767 | 0.05 | 0.047 | 1 | 0.818789679 | 0.625604485 | 0.1 | 0.077 | 1 | 0.895994907 | 0.96716282 |
| Alkbh6 | 0.824323016 | -0.14839184 | 0.269 | 0.276 | 1 | 0.873815752 | 0.574978412 | 0.2 | 0.188 | 1 | 0.873815752 | 0.969137597 |
| Agtrap | 0.825056477 | 0.116023796 | 0.126 | 0.118 | 1 | 0.853153226 | 0.494359952 | 0.1 | 0.117 | 1 | 0.853153226 | 0.969394764 |
| A930005H10Rik | 0.825868867 | -0.27897595 | 0.118 | 0.124 | 1 | 0.989602979 | 0.591657153 | 0.1 | 0.096 | 1 | 0.989602979 | 0.969678349 |
| Gm43305 | 0.826351242 | -0.655351829 | 0.269 | 0.271 | 1 | 0.943394268 | 0.179978666 | 0.3 | 0.29 | 1 | 0.943394268 | 0.969846109 |
| Ip6k3 | 0.980912009 | -0.222392421 | 0.084 | 0.082 | 1 | 0.832261605 | 0.185031894 | 0.1 | 0.117 | 1 | 0.980912009 | 0.971863831 |
| Lrrc51 | 0.980277813 | -0.244418728 | 0.101 | 0.1 | 1 | 0.832730078 | 1.017921908 | 0.1 | 0.08 | 1 | 0.980277813 | 0.972020773 |
| St6galnac6 | 0.833425941 | 0.198797305 | 0.412 | 0.376 | 1 | 0.874767822 | 0.31748219 | 0.4 | 0.361 | 1 | 0.874767822 | 0.972253083 |
| Gm2a | 0.83407273 | -0.37661334 | 0.58 | 0.518 | 1 | 0.946751866 | -0.105854633 | 0.8 | 0.667 | 1 | 0.946751866 | 0.972468141 |
| Inf2 | 0.840166822 | 0.321928095 | 0.361 | 0.347 | 1 | 0.882323732 | 0.43965406 | 0.4 | 0.398 | 1 | 0.882323732 | 0.974453355 |
| Gm56721 | 0.929389703 | 0.192645078 | 0.025 | 0.024 | 1 | 0.843785495 | 0.769994395 | 0.1 | 0.08 | 1 | 0.929389703 | 0.975597028 |
| Fbxo38 | 0.94619605 | -0.209792385 | 0.639 | 0.547 | 1 | 0.84583938 | 0.334657334 | 0.5 | 0.571 | 1 | 0.94619605 | 0.976234503 |
| Parl | 0.847356646 | -0.202492864 | 0.487 | 0.471 | 1 | 0.99418338 | 0.295455884 | 0.4 | 0.475 | 1 | 0.99418338 | 0.976700007 |
| Rab11fip3 | 0.913970792 | -0.199582538 | 0.58 | 0.541 | 1 | 0.84965706 | -0.224152879 | 0.6 | 0.586 | 1 | 0.913970792 | 0.977397 |
| Gm36975 | 0.851095342 | -0.292781749 | 0.059 | 0.053 | 1 | 0.962069915 | 0.403212064 | 0.1 | 0.102 | 1 | 0.962069915 | 0.977827403 |
| 1700037H04Rik | 0.95840026 | -0.321928095 | 0.101 | 0.1 | 1 | 0.85982938 | 1.111031312 | 0.1 | 0.083 | 1 | 0.95840026 | 0.980352197 |
| Bsdc1 | 0.860459795 | -0.16163839 | 0.555 | 0.512 | 1 | 0.97034554 | 0.159940913 | 0.6 | 0.596 | 1 | 0.97034554 | 0.980528531 |
| Peg13 | 0.962766843 | -0.11345805 | 0.076 | 0.076 | 1 | 0.862565596 | 0.930459067 | 0.1 | 0.083 | 1 | 0.962766843 | 0.981111785 |
| Lage3 | 0.867655974 | -0.112383417 | 0.487 | 0.459 | 1 | 0.94562216 | 0.225363988 | 0.4 | 0.407 | 1 | 0.94562216 | 0.982485059 |
| Gm28403 | 0.868582124 | -0.821029859 | 0.109 | 0.112 | 1 | 0.94945869 | -0.131825212 | 0.1 | 0.102 | 1 | 0.94945869 | 0.982729342 |
| Flii | 0.98613617 | -0.114907995 | 0.639 | 0.594 | 1 | 0.868837106 | 0.133841195 | 0.8 | 0.747 | 1 | 0.98613617 | 0.982796295 |
| Dph5 | 0.968181309 | -0.412670485 | 0.328 | 0.318 | 1 | 0.874024193 | 0.145796731 | 0.2 | 0.188 | 1 | 0.968181309 | 0.984130096 |
| Kantr | 0.876094844 | -0.280607035 | 0.202 | 0.206 | 1 | 0.896304752 | -0.165299916 | 0.2 | 0.204 | 1 | 0.896304752 | 0.984647512 |
| Acot13 | 0.970798276 | -0.14332085 | 0.303 | 0.294 | 1 | 0.879199739 | 0.306427001 | 0.3 | 0.281 | 1 | 0.970798276 | 0.985407297 |
| Gm11944 | 0.879395149 | 0.180154134 | 0.143 | 0.153 | 1 | 0.94617234 | 0.331421381 | 0.2 | 0.194 | 1 | 0.94617234 | 0.98545447 |
| Pramel12 | 0.98448682 | -0.19592021 | 0.084 | 0.082 | 1 | 0.882330095 | 0.591657153 | 0.1 | 0.114 | 1 | 0.98448682 | 0.986153793 |
| Xaf1 | 0.883765714 | 0.221791424 | 0.126 | 0.135 | 1 | 0.939377312 | -0.230005605 | 0.1 | 0.102 | 1 | 0.939377312 | 0.986489591 |
| Gm36220 | 0.946052343 | -0.122856748 | 0.319 | 0.312 | 1 | 0.886838982 | 0.515421567 | 0.2 | 0.179 | 1 | 0.946052343 | 0.987194584 |
| Cep89 | 0.888403756 | -0.854660637 | 0.092 | 0.094 | 1 | 0.918735707 | 0.526068812 | 0.1 | 0.108 | 1 | 0.918735707 | 0.987546278 |
| Unc13b | 0.892670141 | -0.758445322 | 0.067 | 0.071 | 1 | 0.998070471 | 0.159940913 | 0.2 | 0.194 | 1 | 0.998070471 | 0.988480301 |
| Mocs3 | 0.92873938 | -0.14839184 | 0.092 | 0.088 | 1 | 0.902406755 | 0.847996907 | 0.1 | 0.086 | 1 | 0.92873938 | 0.990475558 |
| Ess2 | 0.9038404 | -0.19592021 | 0.076 | 0.071 | 1 | 0.926075407 | 0.591657153 | 0.1 | 0.108 | 1 | 0.926075407 | 0.990753331 |
| Slc2a8 | 0.904640859 | -0.129283017 | 0.126 | 0.129 | 1 | 0.955189183 | 0.732519689 | 0.1 | 0.093 | 1 | 0.955189183 | 0.990906634 |
| Kri1 | 0.908848588 | -0.137503524 | 0.311 | 0.288 | 1 | 0.912253832 | 0.223506042 | 0.2 | 0.222 | 1 | 0.912253832 | 0.99169142 |
| Gm56587 | 0.912587482 | -0.185866545 | 0.05 | 0.053 | 1 | 0.933175065 | 0.625604485 | 0.1 | 0.09 | 1 | 0.933175065 | 0.992359052 |
| Fancl | 0.997578222 | -0.214124805 | 0.21 | 0.206 | 1 | 0.914027159 | 0.832055363 | 0.2 | 0.198 | 1 | 0.997578222 | 0.992608671 |
| Cmc2 | 0.917380054 | -0.209792385 | 0.176 | 0.176 | 1 | 1 | 0.732519689 | 0.1 | 0.099 | 1 | 1 | 0.993173944 |
| Pla2g15 | 0.917588076 | -0.308549065 | 0.092 | 0.094 | 1 | 0.923628199 | 0.591657153 | 0.1 | 0.108 | 1 | 0.923628199 | 0.993208275 |
| Znhit2 | 0.923416138 | 0.225066556 | 0.118 | 0.124 | 1 | 0.952551523 | 0.847996907 | 0.1 | 0.093 | 1 | 0.952551523 | 0.994134912 |
| Cep83os | 0.946247509 | -0.240314329 | 0.118 | 0.112 | 1 | 0.927831027 | 0.591657153 | 0.1 | 0.09 | 1 | 0.946247509 | 0.994791639 |
| Tprkb | 0.92887743 | -0.102098188 | 0.311 | 0.306 | 1 | 0.99013547 | 0.101445264 | 0.3 | 0.302 | 1 | 0.99013547 | 0.99494158 |
| 1700084C06Rik | 0.93550943 | -0.555816155 | 0.076 | 0.076 | 1 | 0.962070425 | 0.135278859 | 0.1 | 0.102 | 1 | 0.962070425 | 0.995840966 |
| Snf8 | 0.941450096 | -0.111611991 | 0.563 | 0.524 | 1 | 0.96005504 | 0.459926455 | 0.4 | 0.451 | 1 | 0.96005504 | 0.996571909 |

| | | | | | | | | | | | | |
|---|---|---|---|---|---|---|---|---|---|---|---|---|
| Dnase1l1 | 0.957582933 | -0.333423734 | 0.185 | 0.182 | 1 | 0.958355597 | 0.574978412 | 0.2 | 0.204 | 1 | 0.958355597 | 0.998200792 |
| Zfp946 | 0.960286596 | -0.347923303 | 0.076 | 0.076 | 1 | 0.9604631 | 0.695993813 | 0.1 | 0.093 | 1 | 0.9604631 | 0.998422846 |
| Drg1 | 0.966543847 | -0.19529876 | 0.689 | 0.588 | 1 | 0.972625027 | 0.399012075 | 0.7 | 0.731 | 1 | 0.972625027 | 0.998880686 |

| scRNA seq_Col2a1/Acan enriched cluster subset: cluster 2 | | | | | | | | | | | | |
|---|---|---|---|---|---|---|---|---|---|---|---|---|
| | cKO cart_p_val | cKO cart_avg_log2FC | cKO cart_pct.1 | cKO cart_pct.2 | cKO cart_p_val_adj | WT cart_p_val | WT cart_avg_log2FC | WT cart_pct.1 | WT cart_pct.2 | WT cart_p_val_adj | max_pval | minimump_p_val |
| Col5a2 | 1.63617E-45 | 5.877465289 | 0.981 | 0.106 | 3.97459E-41 | 1.57861E-12 | 2.592727481 | 0.829 | 0.485 | 3.83475E-08 | 1.57861E-12 | 3.27235E-45 |
| Col3a1 | 1.03302E-43 | 6.23690835 | 0.943 | 0.093 | 2.50941E-39 | 1.06312E-16 | 3.776312529 | 0.805 | 0.29 | 2.58252E-12 | 1.06312E-16 | 2.06604E-43 |
| Lgals1 | 9.63238E-41 | 5.858856796 | 0.943 | 0.11 | 2.3399E-36 | 9.7861E-11 | 2.81659457 | 0.732 | 0.372 | 2.37724E-06 | 9.7861E-11 | 1.92648E-40 |
| Col1a1 | 4.0272E-38 | 6.387210894 | 0.887 | 0.11 | 9.78288E-34 | 2.10465E-17 | 2.995821361 | 0.878 | 0.355 | 5.11262E-13 | 2.10465E-17 | 8.0544E-38 |
| Itgbl1 | 8.83032E-38 | 4.956062101 | 0.83 | 0.059 | 2.14506E-33 | 0.000185614 | 1.996375223 | 0.537 | 0.331 | 1 | 0.000185614 | 1.76606E-37 |
| Cdh11 | 4.1029E-35 | 5.174965815 | 0.943 | 0.191 | 9.96677E-31 | 1.41637E-19 | 4.318239908 | 0.829 | 0.28 | 3.44064E-15 | 1.41637E-19 | 8.2058E-35 |
| Ptprd | 9.66924E-33 | 4.958308227 | 0.981 | 0.275 | 2.34885E-28 | 4.18809E-12 | 2.647645752 | 0.927 | 0.628 | 1.01737E-07 | 4.18809E-12 | 1.93385E-32 |
| Mfap2 | 4.13885E-30 | 3.397993746 | 0.849 | 0.114 | 1.00541E-25 | 2.00856E-05 | 1.476948636 | 0.683 | 0.444 | 0.487918602 | 2.00856E-05 | 8.27769E-30 |
| Robo2 | 1.46426E-29 | 3.637664159 | 0.811 | 0.106 | 3.55698E-25 | 1.15209E-07 | 1.955849346 | 0.659 | 0.283 | 0.002798659 | 1.15209E-07 | 2.92852E-29 |
| Igfbp4 | 1.0842E-28 | 4.172497297 | 0.887 | 0.212 | 2.63374E-24 | 3.38163E-11 | 3.232689552 | 0.756 | 0.372 | 8.21465E-07 | 3.38163E-11 | 2.1684E-28 |
| Gulp1 | 1.54561E-28 | 3.275016829 | 0.943 | 0.225 | 3.7546E-24 | 0.001377962 | 1.206153119 | 0.78 | 0.703 | 1 | 0.001377962 | 3.09123E-28 |
| Tnc | 4.03965E-28 | 9.458503343 | 0.491 | 0.004 | 9.81311E-24 | 6.78534E-05 | 3.403308323 | 0.268 | 0.078 | 1 | 6.78534E-05 | 8.07929E-28 |
| Mmp14 | 7.61897E-28 | 3.57090828 | 1 | 0.555 | 1.8508E-23 | 2.87172E-11 | 2.005762717 | 0.951 | 0.853 | 6.97599E-07 | 2.87172E-11 | 1.52379E-27 |
| Mme | 3.27598E-27 | 7.01270359 | 0.528 | 0.017 | 7.95801E-23 | 1.21866E-09 | 3.540223112 | 0.39 | 0.078 | 2.96036E-05 | 1.21866E-09 | 6.55196E-27 |
| Marcks | 3.71875E-27 | 3.531822809 | 0.962 | 0.347 | 9.03358E-23 | 1.3402E-11 | 1.872361295 | 0.951 | 0.625 | 3.25562E-07 | 1.3402E-11 | 7.4375E-27 |
| Hmcn1 | 8.1047E-27 | 5.304469714 | 0.585 | 0.034 | 1.96879E-22 | 5.17677E-10 | 4.452864148 | 0.341 | 0.055 | 1.25754E-05 | 5.17677E-10 | 1.62094E-26 |
| Mdk | 1.09037E-26 | 4.183166573 | 0.811 | 0.157 | 2.64874E-22 | 4.85389E-08 | 2.413993419 | 0.732 | 0.437 | 0.001179108 | 4.85389E-08 | 2.18075E-26 |
| Mmp13 | 1.04896E-25 | 5.785488785 | 0.604 | 0.047 | 2.54813E-21 | 2.66119E-08 | 6.026495176 | 0.22 | 0.024 | 0.000646455 | 2.66119E-08 | 2.09792E-25 |
| Slc40a1 | 2.72448E-25 | 5.260517259 | 0.604 | 0.051 | 6.6183E-21 | 0.097909772 | 1.164566831 | 0.463 | 0.386 | 1 | 0.097909772 | 5.44896E-25 |
| Nrp2 | 4.76707E-25 | 3.173360136 | 0.981 | 0.445 | 1.15802E-20 | 1.5714E-12 | 1.850692281 | 0.976 | 0.782 | 3.81725E-08 | 1.5714E-12 | 9.53413E-25 |
| Adamts2 | 7.39908E-25 | 3.230894275 | 0.792 | 0.144 | 1.79739E-20 | 0.001798876 | 0.782757066 | 0.78 | 0.573 | 1 | 0.001798876 | 1.47982E-24 |
| Mmp2 | 7.57789E-25 | 2.33606965 | 1 | 0.419 | 1.84082E-20 | 1.89964E-08 | 1.11073612 | 0.976 | 0.809 | 0.000461461 | 1.89964E-08 | 1.51558E-24 |
| Tmsb10 | 8.94615E-25 | 3.165538352 | 1 | 0.648 | 2.1732E-20 | 2.24314E-12 | 1.83720485 | 0.951 | 0.771 | 5.44903E-08 | 2.24314E-12 | 1.78923E-24 |
| Cacna1g | 1.96431E-24 | 4.260517259 | 0.642 | 0.072 | 4.7717E-20 | 5.52476E-08 | 2.648171025 | 0.439 | 0.119 | 0.001342074 | 5.52476E-08 | 3.92862E-24 |
| P3h2 | 4.00182E-24 | -6.853706027 | 0.132 | 0.915 | 9.72122E-20 | 8.46861E-10 | -4.174530665 | 0.171 | 0.689 | 2.0572E-05 | 8.46861E-10 | 8.00364E-24 |
| Twist1 | 4.21697E-24 | 3.707976236 | 0.679 | 0.089 | 1.02439E-19 | 4.53817E-12 | 3.177054853 | 0.683 | 0.259 | 1.10241E-07 | 4.53817E-12 | 8.43393E-24 |
| Snorc | 5.66839E-24 | -4.187958205 | 0.245 | 0.949 | 1.37696E-19 | 1.09454E-15 | -3.639454573 | 0.366 | 0.884 | 2.65885E-11 | 1.09454E-15 | 1.13368E-23 |
| Dapk2 | 5.77366E-24 | 5.402650108 | 0.547 | 0.038 | 1.40254E-19 | 4.35775E-06 | 2.525260843 | 0.463 | 0.188 | 0.105858342 | 4.35775E-06 | 1.15473E-23 |
| Mylk | 1.15187E-23 | 3.459163263 | 0.755 | 0.144 | 2.79811E-19 | 4.54658E-05 | 2.031526506 | 0.561 | 0.331 | 1 | 4.54658E-05 | 2.30373E-23 |
| Thbs2 | 1.25506E-23 | 4.932330173 | 0.585 | 0.051 | 3.0488E-19 | 4.77592E-10 | 4.748668175 | 0.415 | 0.092 | 1.16017E-05 | 4.77592E-10 | 2.51012E-23 |
| Col12a1 | 2.68821E-23 | 2.665398119 | 1 | 0.644 | 6.53021E-19 | 0.047150145 | 0.51703364 | 0.927 | 0.922 | 1 | 0.047150145 | 5.37643E-23 |
| Sema5a | 2.77273E-23 | 3.221393634 | 0.849 | 0.212 | 6.73551E-19 | 4.27111E-05 | 1.374414219 | 0.659 | 0.403 | 1 | 4.27111E-05 | 5.54546E-23 |
| Acan | 2.84754E-23 | -3.211148158 | 0.925 | 0.992 | 6.91725E-19 | 1.28346E-15 | -2.038654183 | 0.829 | 0.983 | 3.11778E-11 | 1.28346E-15 | 5.69508E-23 |
| Kif21a | 3.5701E-23 | -3.767316637 | 0.566 | 0.945 | 8.67248E-19 | 2.45051E-12 | -1.908109326 | 0.634 | 0.973 | 5.95277E-08 | 2.45051E-12 | 7.1402E-23 |
| Col1a2 | 3.92278E-23 | 3.12815863 | 0.962 | 0.733 | 9.52922E-19 | 5.08736E-08 | 0.939496431 | 1 | 0.928 | 0.001235822 | 5.08736E-08 | 7.84556E-23 |
| Gm56768 | 4.60314E-23 | 3.547040018 | 0.906 | 0.343 | 1.1182E-18 | 5.1053E-07 | 1.924140487 | 0.829 | 0.638 | 0.01240179 | 5.1053E-07 | 9.20629E-23 |
| Podxl2 | 4.88294E-23 | 4.842778588 | 0.509 | 0.03 | 1.18616E-18 | 0.003321769 | 1.220533489 | 0.317 | 0.137 | 1 | 0.003321769 | 9.76589E-23 |
| Hapln1 | 5.97663E-23 | -2.989606091 | 0.83 | 0.996 | 1.45184E-18 | 1.6033E-14 | -2.563725098 | 0.707 | 0.959 | 3.89474E-10 | 1.6033E-14 | 1.19533E-22 |
| Fn1 | 8.81734E-23 | -1.976028493 | 1 | 1 | 2.14191E-18 | 1.90672E-14 | -1.663599441 | 1 | 1 | 4.6318E-10 | 1.90672E-14 | 1.76347E-22 |
| Meltf | 1.08185E-22 | -5.620064465 | 0.075 | 0.877 | 2.62802E-18 | 2.01065E-13 | -4.16885145 | 0.098 | 0.758 | 4.88427E-09 | 2.01065E-13 | 2.16369E-22 |
| Tmem119 | 1.09559E-22 | 5.01270359 | 0.491 | 0.025 | 2.66141E-18 | 0.000482673 | 2.100239256 | 0.22 | 0.061 | 1 | 0.000482673 | 2.19118E-22 |
| Fbxl7 | 1.189E-22 | 4.011887817 | 0.755 | 0.165 | 2.88832E-18 | 1.03587E-10 | 2.475548447 | 0.683 | 0.256 | 2.51634E-06 | 1.03587E-10 | 2.378E-22 |
| Gpm6b | 1.26487E-22 | 3.287172891 | 0.887 | 0.28 | 3.07261E-18 | 0.09147324 | 0.763623717 | 0.463 | 0.372 | 1 | 0.09147324 | 2.52973E-22 |
| Dock4 | 1.94234E-22 | 2.558788976 | 0.849 | 0.212 | 4.71834E-18 | 7.19309E-05 | 1.291898549 | 0.756 | 0.522 | 1 | 7.19309E-05 | 3.88469E-22 |
| Eps8l2 | 2.32544E-22 | -4.10985799 | 0.17 | 0.907 | 5.64896E-18 | 7.0107E-13 | -2.073801854 | 0.537 | 0.928 | 1.70304E-08 | 7.0107E-13 | 4.65088E-22 |
| Mtss1 | 2.37487E-22 | 3.812934078 | 0.566 | 0.051 | 5.76903E-18 | 5.59266E-08 | 2.056214632 | 0.488 | 0.147 | 0.001358569 | 5.59266E-08 | 4.74973E-22 |
| Cytl1 | 3.74234E-22 | -4.242827759 | 0.792 | 0.996 | 9.09089E-18 | 1.29302E-11 | -2.390845815 | 0.659 | 0.942 | 3.141E-07 | 1.29302E-11 | 7.48468E-22 |
| Csgalnact1 | 4.48062E-22 | -4.776014743 | 0.283 | 0.936 | 1.08843E-17 | 2.07328E-05 | -1.504390279 | 0.244 | 0.655 | 0.503642094 | 2.07328E-05 | 8.96123E-22 |
| Kalrn | 9.5971E-22 | 2.840086992 | 0.736 | 0.136 | 2.33133E-17 | 7.9171E-08 | 2.83720485 | 0.659 | 0.29 | 0.001923222 | 7.9171E-08 | 1.91942E-21 |
| Cped1 | 1.1737E-21 | 2.387060203 | 1 | 0.458 | 2.85116E-17 | 2.64805E-05 | 1.385993738 | 0.829 | 0.655 | 0.643263122 | 2.64805E-05 | 2.3474E-21 |
| Stk17b | 1.65291E-21 | 3.009292155 | 0.774 | 0.191 | 4.01525E-17 | 1.97599E-08 | 2.51050225 | 0.732 | 0.375 | 0.000480008 | 1.97599E-08 | 3.30582E-21 |
| Xylt1 | 2.48792E-21 | -2.681583701 | 0.755 | 0.983 | 6.04366E-17 | 1.48506E-17 | -2.789066797 | 0.78 | 0.99 | 3.60751E-13 | 1.48506E-17 | 4.97584E-21 |
| Podnl1 | 4.02192E-21 | 4.039245377 | 0.472 | 0.025 | 9.77004E-17 | 0.010038082 | 2.014082612 | 0.171 | 0.061 | 1 | 0.010038082 | 8.04383E-21 |
| Sox4 | 5.62485E-21 | 2.402373968 | 0.943 | 0.39 | 1.36639E-16 | 1.09994E-10 | 2.184278593 | 0.902 | 0.573 | 2.67196E-06 | 1.09994E-10 | 1.12497E-20 |
| Lum | 5.96953E-21 | 3.37422558 | 0.962 | 0.517 | 1.45012E-16 | 0.215059676 | 1.075646383 | 0.732 | 0.761 | 1 | 0.215059676 | 1.19391E-20 |
| Nid1 | 6.19377E-21 | 2.850465526 | 0.736 | 0.14 | 1.50459E-16 | 2.50149E-06 | 2.890316186 | 0.463 | 0.181 | 0.060766295 | 2.50149E-06 | 1.23875E-20 |
| Ncmap | 6.69229E-21 | -3.061214804 | 0.34 | 0.932 | 1.62569E-16 | 7.31077E-10 | -2.061835851 | 0.293 | 0.795 | 1.77593E-05 | 7.31077E-10 | 1.33846E-20 |
| Serpinf1 | 9.40119E-21 | 2.338609749 | 0.925 | 0.331 | 2.28374E-16 | 4.33032E-06 | 1.690363461 | 0.732 | 0.553 | 0.105192165 | 4.33032E-06 | 1.88024E-20 |
| Ssh2 | 1.03103E-20 | 2.670790014 | 0.943 | 0.339 | 2.50457E-16 | 0.00844005 | 0.909745952 | 0.829 | 0.744 | 1 | 0.00844005 | 2.06205E-20 |
| Frmd6 | 1.04298E-20 | 2.488332495 | 0.962 | 0.496 | 2.5336E-16 | 1.05684E-05 | 0.846773559 | 1 | 0.891 | 0.256727616 | 1.05684E-05 | 2.08596E-20 |
| Cdk14 | 1.07442E-20 | 2.553271971 | 0.962 | 0.436 | 2.60999E-16 | 6.46066E-07 | 1.53464208 | 0.902 | 0.737 | 0.01569424 | 6.46066E-07 | 2.14885E-20 |
| Gpc3 | 1.83849E-20 | 3.188405721 | 0.83 | 0.233 | 4.46607E-16 | 0.812917018 | -0.136799942 | 0.488 | 0.522 | 1 | 0.812917018 | 3.67699E-20 |
| Col27a1 | 1.92554E-20 | -2.318527569 | 0.792 | 0.992 | 4.67753E-16 | 3.2353E-15 | -1.973095002 | 0.854 | 0.983 | 7.8592E-11 | 3.2353E-15 | 3.85109E-20 |
| Zfp36l2 | 2.12536E-20 | 2.370451286 | 0.981 | 0.487 | 5.16292E-16 | 1.55304E-09 | 1.69143371 | 0.902 | 0.672 | 3.77264E-05 | 1.55304E-09 | 4.25072E-20 |
| Col5a1 | 2.41996E-20 | 2.529700605 | 0.943 | 0.441 | 5.87857E-16 | 4.16572E-06 | 1.414775216 | 0.756 | 0.584 | 0.101193741 | 4.16572E-06 | 4.83992E-20 |
| Bcorl1 | 2.87926E-20 | 3.413456863 | 0.585 | 0.072 | 6.9943E-16 | 0.000521308 | 1.455059232 | 0.512 | 0.29 | 1 | 0.000521308 | 5.75852E-20 |
| Zfp385b | 3.00591E-20 | -3.343158161 | 0.604 | 0.983 | 7.30197E-16 | 3.25196E-09 | -2.359882483 | 0.585 | 0.908 | 7.89967E-05 | 3.25196E-09 | 6.01183E-20 |
| Dpysl3 | 3.63289E-20 | 4.274021523 | 0.491 | 0.038 | 8.82502E-16 | 2.63613E-14 | 3.727021932 | 0.488 | 0.078 | 6.40369E-10 | 2.63613E-14 | 7.26578E-20 |
| Arhgap28 | 3.82736E-20 | 5.167546635 | 0.472 | 0.034 | 9.29743E-16 | 2.39978E-11 | 3.336775859 | 0.415 | 0.072 | 5.82954E-07 | 2.39978E-11 | 7.65473E-20 |
| Itga11 | 3.91972E-20 | 3.569760094 | 0.811 | 0.229 | 9.52179E-16 | 0.017917476 | 0.868569021 | 0.634 | 0.519 | 1 | 0.017917476 | 7.83945E-20 |
| Amot | 4.61962E-20 | 3.644107436 | 0.585 | 0.076 | 1.1222E-15 | 0.009949082 | 1.64875976 | 0.366 | 0.208 | 1 | 0.009949082 | 9.23924E-20 |
| Axl | 4.91544E-20 | 2.865215978 | 0.755 | 0.174 | 1.19406E-15 | 0.017952331 | 0.52935194 | 0.732 | 0.611 | 1 | 0.017952331 | 9.83087E-20 |
| Comp | 6.56353E-20 | -3.530073402 | 0.604 | 0.962 | 1.59441E-15 | 1.99822E-10 | -1.955634731 | 0.732 | 0.894 | 4.85409E-06 | 1.99822E-10 | 1.31271E-19 |
| Pcolce | 7.23406E-20 | 2.252697754 | 0.943 | 0.373 | 1.7573E-15 | 0.004360179 | 0.624095669 | 0.878 | 0.747 | 1 | 0.004360179 | 1.44681E-19 |
| Papss2 | 8.12539E-20 | -2.331512691 | 0.755 | 0.992 | 1.97382E-15 | 2.11744E-13 | -1.982185798 | 0.756 | 0.99 | 5.14369E-09 | 2.11744E-13 | 1.62508E-19 |
| Cdh13 | 1.49671E-19 | 4.441885272 | 0.472 | 0.034 | 3.6358E-15 | 5.70659E-14 | 5.438108894 | 0.317 | 0.024 | 1.38624E-09 | 5.70659E-14 | 2.99342E-19 |
| Adamts9 | 1.50379E-19 | 2.286428274 | 0.83 | 0.229 | 3.653E-15 | 2.68021E-08 | 2.316372687 | 0.634 | 0.27 | 0.000651076 | 2.68021E-08 | 3.00758E-19 |
| Evl | 1.74219E-19 | 2.731647776 | 0.83 | 0.25 | 4.23213E-15 | 0.028368707 | 1.422167351 | 0.585 | 0.454 | 1 | 0.028368707 | 3.48438E-19 |
| Col9a2 | 2.34251E-19 | -2.191808867 | 0.811 | 0.979 | 5.69043E-15 | 1.73958E-11 | -1.795234768 | 0.707 | 0.952 | 4.22579E-07 | 1.73958E-11 | 4.68502E-19 |
| Lgals3 | 2.39102E-19 | -3.154941734 | 0.566 | 0.953 | 5.80826E-15 | 9.64667E-09 | -1.697527315 | 0.756 | 0.942 | 0.000234337 | 9.64667E-09 | 4.78204E-19 |
| Zfhx4 | 2.88213E-19 | 3.246840797 | 0.887 | 0.339 | 7.00126E-15 | 0.001205507 | 0.751344113 | 0.878 | 0.594 | 1 | 0.001205507 | 5.76425E-19 |
| Sfrp4 | 3.71205E-19 | 8.383541285 | 0.321 | 0 | 9.01732E-15 | 1.26802E-11 | 7.387401932 | 0.22 | 0.01 | 3.08028E-07 | 1.26802E-11 | 7.4241E-19 |
| Scrg1 | 3.92918E-19 | -5.359399665 | 0.113 | 0.814 | 9.54477E-15 | 3.09944E-06 | -4.835220492 | 0.024 | 0.403 | 0.075291579 | 3.09944E-06 | 7.85837E-19 |
| Tfap2b | 4.04138E-19 | 4.054195719 | 0.509 | 0.051 | 9.81731E-15 | 0.715372978 | 0.183529599 | 0.293 | 0.27 | 1 | 0.715372978 | 8.08275E-19 |
| Pcdh18 | 4.77353E-19 | 5.029191713 | 0.396 | 0.017 | 1.15959E-14 | 2.17292E-08 | 2.897325842 | 0.39 | 0.089 | 0.000527846 | 2.17292E-08 | 9.54706E-19 |
| Prrx2 | 5.01231E-19 | 2.527634618 | 0.83 | 0.246 | 1.21759E-14 | 9.94684E-09 | 1.954698413 | 0.805 | 0.42 | 0.000241629 | 9.94684E-09 | 1.00246E-18 |
| Egfr | 6.75155E-19 | 2.983508897 | 0.811 | 0.237 | 1.64009E-14 | 4.41724E-07 | 1.157444595 | 0.976 | 0.778 | 0.010730365 | 4.41724E-07 | 1.35031E-18 |
| Runx1t1 | 7.30374E-19 | 4.167106319 | 0.528 | 0.064 | 1.77423E-14 | 7.06332E-08 | 2.790662264 | 0.415 | 0.109 | 0.001715821 | 7.06332E-08 | 1.46075E-18 |
| Mafb | 8.07872E-19 | 2.514793635 | 0.811 | 0.225 | 1.96248E-14 | 2.84586E-10 | 2.819770256 | 0.756 | 0.341 | 6.91317E-06 | 2.84586E-10 | 1.61574E-18 |
| Myo1b | 8.82357E-19 | 4.088834659 | 0.491 | 0.047 | 2.14342E-14 | 6.09192E-07 | 1.626308067 | 0.439 | 0.14 | 0.014798491 | 6.09192E-07 | 1.76471E-18 |
| Nebl | 1.01821E-18 | -3.027586131 | 0.34 | 0.919 | 2.47345E-14 | 3.49E-10 | -2.687663304 | 0.341 | 0.799 | 8.47792E-06 | 3.49E-10 | 2.03643E-18 |
| Ugdh | 1.43504E-18 | -3.160857773 | 0.887 | 0.983 | 3.48599E-14 | 7.66602E-07 | -1.873065878 | 0.805 | 0.952 | 0.018622289 | 7.66602E-07 | 2.87007E-18 |
| Aebp1 | 1.58288E-18 | 2.234734109 | 0.925 | 0.441 | 3.84512E-14 | 3.00923E-06 | 1.015910737 | 0.854 | 0.778 | 0.073100255 | 3.00923E-06 | 3.16575E-18 |
| Fnbp1l | 1.7207E-18 | 2.558788976 | 0.792 | 0.229 | 4.17991E-14 | 5.16736E-08 | 1.439445342 | 0.829 | 0.536 | 0.001255255 | 5.16736E-08 | 3.44139E-18 |
| Wnt9a | 2.21256E-18 | 3.891688189 | 0.434 | 0.03 | 5.37474E-14 | 7.01095E-06 | 3.19334866 | 0.244 | 0.051 | 0.170310106 | 7.01095E-06 | 4.42511E-18 |
| Pdgfra | 2.73224E-18 | 2.670364035 | 0.83 | 0.275 | 6.63715E-14 | 1.17271E-10 | 2.335807066 | 0.756 | 0.324 | 2.84876E-06 | 1.17271E-10 | 5.46447E-18 |

| | | | | | | | | | | | | |
|---|---|---|---|---|---|---|---|---|---|---|---|---|
| Irf1 | 3.61459E-18 | 2.911451444 | 0.755 | 0.216 | 8.78055E-14 | 3.02039E-09 | 1.897746392 | 0.659 | 0.235 | 7.33714E-05 | 3.02039E-09 | 7.22917E-18 |
| Slc39a14 | 3.82315E-18 | -2.666616557 | 0.528 | 0.962 | 9.28719E-14 | 0.004162721 | -0.781274777 | 0.439 | 0.655 | 1 | 0.004162721 | 7.6463E-18 |
| Slc26a2 | 3.97352E-18 | -2.790789295 | 0.453 | 0.903 | 9.65248E-14 | 1.40041E-05 | -1.445059028 | 0.61 | 0.826 | 0.340188265 | 1.40041E-05 | 7.94704E-18 |
| Bmpr1b | 4.50901E-18 | -3.99594785 | 0.226 | 0.839 | 1.09533E-13 | 1.86535E-05 | -1.779812384 | 0.659 | 0.826 | 0.453131662 | 1.86535E-05 | 9.01802E-18 |
| Pdgfrb | 4.70763E-18 | 2.810768194 | 0.849 | 0.343 | 1.14358E-13 | 2.25036E-06 | 1.337919174 | 0.829 | 0.648 | 0.05466581 | 2.25036E-06 | 9.41527E-18 |
| Tmem204 | 6.7965E-18 | 3.083169334 | 0.566 | 0.081 | 1.65101E-13 | 0.016792081 | 2.029849928 | 0.195 | 0.082 | 1 | 0.016792081 | 1.3593E-17 |
| Bnc2 | 7.10616E-18 | 2.437029931 | 0.906 | 0.403 | 1.72623E-13 | 0.000225959 | 1.26286719 | 0.756 | 0.679 | 1 | 0.000225959 | 1.42123E-17 |
| Prkca | 7.6053E-18 | -2.64419549 | 0.868 | 0.979 | 1.84748E-13 | 2.2082E-11 | -2.027780019 | 0.829 | 0.973 | 5.36416E-07 | 2.2082E-11 | 1.52106E-17 |
| Col8a2 | 7.70004E-18 | 1.979203577 | 0.943 | 0.428 | 1.87049E-13 | 0.000352937 | 0.739357526 | 0.878 | 0.836 | 1 | 0.000352937 | 1.54001E-17 |
| Svil | 8.01167E-18 | 2.184287052 | 0.981 | 0.589 | 1.94619E-13 | 2.07092E-14 | 1.933420165 | 1 | 0.703 | 5.03068E-10 | 2.07092E-14 | 1.60233E-17 |
| Mfap4 | 8.1822E-18 | 3.355419945 | 0.566 | 0.081 | 1.98762E-13 | 2.63498E-07 | 2.721727632 | 0.463 | 0.157 | 0.006400897 | 2.63498E-07 | 1.63644E-17 |
| Mxra7 | 9.94022E-18 | 2.167267848 | 0.925 | 0.458 | 2.41468E-13 | 0.009702568 | 0.994174196 | 0.634 | 0.58 | 1 | 0.009702568 | 1.98804E-17 |
| Chst11 | 1.38868E-17 | -2.217068567 | 0.868 | 1 | 3.37339E-13 | 1.4902E-10 | -1.748343041 | 0.707 | 0.983 | 3.61999E-06 | 1.4902E-10 | 2.77736E-17 |
| Antxr1 | 1.39894E-17 | 2.123561663 | 0.981 | 0.636 | 3.39832E-13 | 1.57783E-12 | 1.920872974 | 0.976 | 0.894 | 3.83286E-08 | 1.57783E-12 | 2.79789E-17 |
| Cacna1c | 1.44206E-17 | 2.900003457 | 0.811 | 0.237 | 3.50306E-13 | 2.76336E-05 | 1.249416305 | 0.659 | 0.413 | 0.671275649 | 2.76336E-05 | 2.88412E-17 |
| Wls | 1.44685E-17 | 1.820876804 | 0.981 | 0.814 | 3.51469E-13 | 0.001041939 | 0.626555914 | 0.976 | 0.952 | 1 | 0.001041939 | 2.8937E-17 |
| Clmp | 1.64054E-17 | 2.245990652 | 0.83 | 0.288 | 3.98521E-13 | 1.49398E-08 | 2.610816041 | 0.683 | 0.321 | 0.000362918 | 1.49398E-08 | 3.28108E-17 |
| Colec12 | 1.84368E-17 | 1.725491214 | 0.981 | 0.886 | 4.47868E-13 | 0.23217992 | 0.296827285 | 0.951 | 0.945 | 1 | 0.23217992 | 3.68737E-17 |
| Sh3pxd2b | 2.04881E-17 | 3.20464782 | 0.698 | 0.174 | 4.97697E-13 | 0.00017961 | 1.578183506 | 0.707 | 0.437 | 1 | 0.00017961 | 4.09762E-17 |
| Tox | 2.17712E-17 | 2.945775705 | 0.849 | 0.326 | 5.28865E-13 | 0.359999238 | 0.887350958 | 0.561 | 0.549 | 1 | 0.359999238 | 4.35423E-17 |
| F13a1 | 2.2503E-17 | -4.144664518 | 0.321 | 0.873 | 5.46644E-13 | 6.91507E-06 | -2.588580525 | 0.341 | 0.696 | 0.167980786 | 6.91507E-06 | 4.50061E-17 |
| Rcan1 | 2.57012E-17 | -4.23435827 | 0.604 | 0.894 | 6.24335E-13 | 1.31466E-05 | -2.026483016 | 0.732 | 0.877 | 0.319356819 | 1.31466E-05 | 5.14025E-17 |
| Prnp | 3.93E-17 | -1.760212214 | 0.792 | 0.979 | 9.54675E-13 | 3.68808E-09 | -1.611068979 | 0.732 | 0.939 | 8.95908E-05 | 3.68808E-09 | 7.86E-17 |
| Epha3 | 4.12898E-17 | 5.643009076 | 0.377 | 0.021 | 1.00301E-12 | 7.41033E-16 | 4.111127572 | 0.415 | 0.041 | 1.80012E-11 | 7.41033E-16 | 8.25796E-17 |
| Col9a3 | 4.44173E-17 | -2.165195284 | 0.679 | 0.97 | 1.07899E-12 | 2.87284E-10 | -1.85312577 | 0.683 | 0.932 | 6.97871E-06 | 2.87284E-10 | 8.88346E-17 |
| Tgm2 | 6.13711E-17 | -4.317855433 | 0.358 | 0.852 | 1.49083E-12 | 0.133073076 | -0.863234868 | 0.463 | 0.56 | 1 | 0.133073076 | 1.22742E-16 |
| Rai14 | 6.23457E-17 | 2.90792883 | 0.83 | 0.305 | 1.5145E-12 | 3.88578E-06 | 1.648061933 | 0.854 | 0.628 | 0.094393316 | 3.88578E-06 | 1.24691E-16 |
| Foxn3 | 6.69547E-17 | 1.344180804 | 1 | 0.958 | 1.62646E-12 | 3.56694E-05 | 0.877607204 | 1 | 0.99 | 0.866482001 | 3.56694E-05 | 1.33909E-16 |
| Tnrc6c | 7.56376E-17 | 1.715379189 | 1 | 0.665 | 1.83739E-12 | 0.000539408 | 0.756089115 | 0.951 | 0.922 | 1 | 0.000539408 | 1.51275E-16 |
| 1700024B18Rik | 8.19281E-17 | 6.564113531 | 0.34 | 0.013 | 1.9902E-12 | 0.010866434 | 1.363273661 | 0.195 | 0.075 | 1 | 0.010866434 | 1.63856E-16 |
| Loxl4 | 8.29973E-17 | -3.68703298 | 0.245 | 0.839 | 2.01617E-12 | 1.93519E-09 | -2.720716632 | 0.317 | 0.799 | 4.70096E-05 | 1.93519E-09 | 1.65995E-16 |
| Ednra | 9.89515E-17 | 5.763531837 | 0.34 | 0.013 | 2.40373E-12 | 9.84915E-15 | 7.281989692 | 0.268 | 0.01 | 2.39255E-10 | 9.84915E-15 | 1.97903E-16 |
| Timp2 | 9.99288E-17 | 1.308019005 | 1 | 0.966 | 2.42747E-12 | 4.11669E-06 | 0.766626799 | 1 | 0.983 | 0.100002567 | 4.11669E-06 | 1.99858E-16 |
| Fstl1 | 1.14541E-16 | 1.868813723 | 0.981 | 0.559 | 2.78243E-12 | 9.63314E-07 | 1.009489501 | 1 | 0.867 | 0.023400826 | 9.63314E-07 | 2.29082E-16 |
| Fat4 | 1.19939E-16 | 2.123612062 | 0.943 | 0.424 | 2.91356E-12 | 0.000147361 | 1.624969044 | 0.756 | 0.567 | 1 | 0.000147361 | 2.39878E-16 |
| Robo1 | 1.30999E-16 | 2.35865348 | 0.792 | 0.237 | 3.18224E-12 | 0.049928402 | 0.582336211 | 0.854 | 0.785 | 1 | 0.049928402 | 2.61999E-16 |
| Anxa8 | 1.51014E-16 | -5.130062956 | 0.17 | 0.775 | 3.66842E-12 | 0.072248696 | -0.975293376 | 0.293 | 0.406 | 1 | 0.072248696 | 3.02027E-16 |
| Sdc4 | 1.62024E-16 | -1.870558447 | 1 | 1 | 3.93589E-12 | 2.12607E-05 | -0.826445883 | 1 | 1 | 0.516465918 | 2.12607E-05 | 3.24048E-16 |
| Atrx | 1.65941E-16 | 1.2925752 | 1 | 0.936 | 4.03105E-12 | 0.051242287 | 0.359077137 | 1 | 0.986 | 1 | 0.051242287 | 3.31883E-16 |
| Ttc3 | 1.94654E-16 | 1.401362562 | 1 | 0.915 | 4.72853E-12 | 0.016012019 | 0.339389675 | 1 | 0.98 | 1 | 0.016012019 | 3.89308E-16 |
| Id3 | 2.52437E-11 | 1.809951959 | 0.887 | 0.538 | 6.13221E-07 | 2.63847E-16 | 2.723588774 | 0.951 | 0.399 | 6.40938E-12 | 2.52437E-11 | 5.27695E-16 |
| Kmt2c | 2.98458E-16 | 1.588210946 | 1 | 0.665 | 7.25014E-12 | 0.003034209 | 0.63266043 | 1 | 0.983 | 1 | 0.003034209 | 5.96916E-16 |
| Tnrc6a | 3.37788E-16 | 1.506101457 | 0.962 | 0.653 | 8.20554E-12 | 0.074642855 | 0.407339933 | 0.976 | 0.962 | 1 | 0.074642855 | 6.75575E-16 |
| Actn1 | 3.77706E-16 | 1.843325167 | 0.943 | 0.483 | 9.17525E-12 | 5.38376E-05 | 0.918087952 | 0.927 | 0.84 | 1 | 5.38376E-05 | 7.55413E-16 |
| Bicc1 | 4.62722E-16 | 2.359307286 | 0.943 | 0.72 | 1.12404E-11 | 4.49731E-06 | 2.698377145 | 0.854 | 0.816 | 0.109248599 | 4.49731E-06 | 9.25444E-16 |
| Sox9 | 4.74448E-16 | -2.359898925 | 0.887 | 0.97 | 1.15253E-11 | 0.000122459 | -1.252331988 | 0.878 | 0.959 | 1 | 0.000122459 | 9.48896E-16 |
| S100b | 5.11425E-16 | -2.929790552 | 0.415 | 0.903 | 1.24235E-11 | 2.87195E-07 | -2.067967229 | 0.22 | 0.676 | 0.006976538 | 2.87195E-07 | 1.02285E-15 |
| Fzd8 | 5.64189E-16 | 3.297680549 | 0.509 | 0.076 | 1.37053E-11 | 1.18357E-05 | 2.062271405 | 0.439 | 0.164 | 0.287513192 | 1.18357E-05 | 1.12838E-15 |
| Rsf1 | 5.74317E-16 | 1.498569046 | 0.981 | 0.636 | 1.39513E-11 | 0.004466819 | 0.768763794 | 0.902 | 0.894 | 1 | 0.004466819 | 1.14863E-15 |
| Hbp1 | 5.90437E-16 | 1.523367189 | 0.981 | 0.674 | 1.43429E-11 | 0.012540859 | 0.495736569 | 0.927 | 0.833 | 1 | 0.012540859 | 1.18087E-15 |
| Clec3a | 6.09367E-16 | -4.07024377 | 0.075 | 0.746 | 1.48028E-11 | 0.002566045 | -1.940402729 | 0.073 | 0.3 | 1 | 0.002566045 | 1.21873E-15 |
| Itgb1 | 6.39854E-16 | 1.097510695 | 1 | 0.992 | 1.55433E-11 | 0.000597445 | 0.482093659 | 0.976 | 0.993 | 1 | 0.000597445 | 1.27971E-15 |
| Ep400 | 7.15879E-16 | 1.845643922 | 0.981 | 0.716 | 1.73901E-11 | 2.20276E-06 | 1.332732267 | 0.951 | 0.884 | 0.053509387 | 2.20276E-06 | 1.43176E-15 |
| Peli2 | 7.58724E-16 | 2.695290976 | 0.604 | 0.11 | 1.84309E-11 | 0.006286158 | 0.83720485 | 0.488 | 0.29 | 1 | 0.006286158 | 1.51745E-15 |
| Cdh2 | 7.70452E-16 | 5.31000082 | 0.396 | 0.034 | 1.87158E-11 | 0.002665574 | 2.625700745 | 0.22 | 0.078 | 1 | 0.002665574 | 1.5409E-15 |
| Hes1 | 8.62729E-16 | 2.168889927 | 0.811 | 0.275 | 2.09574E-11 | 0.000172793 | 1.438108894 | 0.78 | 0.529 | 1 | 0.000172793 | 1.72546E-15 |
| App | 9.07185E-16 | 0.952781931 | 1 | 0.996 | 2.20373E-11 | 0.000135727 | 0.628164353 | 1 | 1 | 1 | 0.000135727 | 1.81437E-15 |
| Tead2 | 9.54379E-16 | 4.232725107 | 0.358 | 0.021 | 2.31838E-11 | 1.09422E-08 | 2.83720485 | 0.366 | 0.075 | 0.000265808 | 1.09422E-08 | 1.90876E-15 |
| 4930523C07Rik | 1.10977E-15 | -2.095360447 | 0.66 | 0.953 | 2.69585E-11 | 1.1671E-12 | -1.717193333 | 0.683 | 0.973 | 2.83512E-08 | 1.1671E-12 | 2.21954E-15 |
| Dennd2a | 1.12253E-15 | 3.019067495 | 0.585 | 0.11 | 2.72684E-11 | 0.051764864 | 0.593279267 | 0.439 | 0.304 | 1 | 0.051764864 | 2.24505E-15 |
| Neo1 | 1.30931E-15 | 1.691104915 | 1 | 0.742 | 3.18058E-11 | 0.000536084 | 0.71345409 | 1 | 0.922 | 1 | 0.000536084 | 2.61863E-15 |
| Gtf2ird1 | 1.42064E-15 | 2.27810501 | 0.792 | 0.284 | 3.45103E-11 | 0.256072108 | 0.57223264 | 0.78 | 0.669 | 1 | 0.256072108 | 2.84129E-15 |
| Nfatc4 | 1.4278E-15 | 2.928446739 | 0.547 | 0.089 | 3.46842E-11 | 0.000411858 | 1.537644568 | 0.463 | 0.239 | 1 | 0.000411858 | 2.85561E-15 |
| Fbn2 | 2.29908E-12 | 5.440124814 | 0.245 | 0.008 | 5.58494E-08 | 1.56106E-15 | 4.893788378 | 0.317 | 0.017 | 3.79214E-11 | 2.29908E-12 | 3.12213E-15 |
| Tle5 | 1.61554E-15 | 1.997482853 | 0.868 | 0.453 | 3.92446E-11 | 0.170148845 | 0.394034212 | 0.659 | 0.648 | 1 | 0.170148845 | 3.23107E-15 |
| Rab3il1 | 1.7492E-15 | 2.884075005 | 0.547 | 0.093 | 4.24916E-11 | 0.064574948 | 1.116358921 | 0.293 | 0.184 | 1 | 0.064574948 | 3.4984E-15 |
| Runx2 | 1.85897E-15 | 2.181497147 | 0.981 | 0.754 | 4.5158E-11 | 1.96205E-05 | 1.620621821 | 0.951 | 0.836 | 0.476620542 | 1.96205E-05 | 3.71793E-15 |
| Crebbp | 1.93271E-15 | 1.105382831 | 1 | 0.97 | 4.69495E-11 | 0.026669729 | 0.26121469 | 1 | 1 | 1 | 0.026669729 | 3.86543E-15 |
| Scube2 | 2.14162E-15 | 3.154722595 | 0.585 | 0.131 | 5.20242E-11 | 0.000133121 | 2.545973552 | 0.366 | 0.154 | 1 | 0.000133121 | 4.28324E-15 |
| Myl9 | 2.1583E-15 | 3.817687608 | 0.491 | 0.076 | 5.24295E-11 | 0.00105299 | 2.60371472 | 0.39 | 0.191 | 1 | 0.00105299 | 4.31661E-15 |
| Ptma | 2.21525E-15 | 1.138117192 | 1 | 0.987 | 5.38128E-11 | 0.000370547 | 0.427762813 | 1 | 0.997 | 1 | 0.000370547 | 4.4305E-15 |
| Slc22a4 | 2.44271E-15 | -5.419465434 | 0.057 | 0.703 | 5.93383E-11 | 4.14156E-05 | -3.810253577 | 0.024 | 0.348 | 1 | 4.14156E-05 | 4.88542E-15 |
| Dock10 | 3.03914E-15 | 3.324647596 | 0.396 | 0.034 | 7.38269E-11 | 0.954860703 | -0.332720152 | 0.195 | 0.191 | 1 | 0.954860703 | 6.07829E-15 |
| Chad | 3.71026E-15 | -2.759601336 | 0.736 | 0.949 | 9.01296E-11 | 1.86046E-08 | -1.491534709 | 0.732 | 0.911 | 0.000451943 | 1.86046E-08 | 7.42052E-15 |
| Kcnma1 | 3.77907E-15 | -1.653227867 | 0.925 | 0.987 | 9.18011E-11 | 5.70487E-06 | -1.415587889 | 0.707 | 0.925 | 0.138582782 | 5.70487E-06 | 7.55813E-15 |
| Fmn1 | 4.15751E-15 | -2.805466227 | 0.642 | 0.928 | 1.00994E-10 | 2.27727E-05 | -1.447995212 | 0.854 | 0.945 | 0.553193637 | 2.27727E-05 | 8.31502E-15 |
| Mex3a | 5.36886E-15 | 3.564113531 | 0.509 | 0.089 | 1.3042E-10 | 0.0007871 | 1.602739596 | 0.439 | 0.239 | 1 | 0.0007871 | 1.07377E-14 |
| Prickle2 | 5.57405E-15 | 2.209689054 | 0.736 | 0.216 | 1.35405E-10 | 0.005238381 | 1.227877801 | 0.561 | 0.379 | 1 | 0.005238381 | 1.11481E-14 |
| Tmem164 | 5.82836E-15 | 1.883238999 | 0.887 | 0.407 | 1.41583E-10 | 0.009973718 | 0.688278039 | 0.902 | 0.788 | 1 | 0.009973718 | 1.16567E-14 |
| P4ha2 | 5.85628E-15 | -3.047623884 | 0.208 | 0.801 | 1.42261E-10 | 2.98288E-05 | -1.649577257 | 0.293 | 0.611 | 0.724601324 | 2.98288E-05 | 1.17126E-14 |
| Fdft1 | 6.07611E-15 | -3.879279919 | 0.189 | 0.754 | 1.47601E-10 | 0.001314155 | -1.607159321 | 0.366 | 0.597 | 1 | 0.001314155 | 1.21522E-14 |
| Tagln | 7.09647E-15 | 5.82374936 | 0.453 | 0.068 | 1.72387E-10 | 0.024372721 | 2.691353983 | 0.195 | 0.089 | 1 | 0.024372721 | 1.41929E-14 |
| Fgd6 | 7.47938E-15 | 2.333059836 | 0.811 | 0.335 | 1.81689E-10 | 2.02186E-07 | 1.349411008 | 0.902 | 0.584 | 0.004911508 | 2.02186E-07 | 1.49588E-14 |
| Adamts12 | 7.56247E-15 | 2.541001101 | 0.604 | 0.127 | 1.83707E-10 | 0.001279746 | 0.735451351 | 0.585 | 0.338 | 1 | 0.001279746 | 1.51249E-14 |
| Ctsk | 8.39328E-15 | 1.854526568 | 0.981 | 0.797 | 2.03889E-10 | 0.001669679 | -0.745380888 | 0.902 | 0.826 | 1 | 0.001669679 | 1.67866E-14 |
| Marcksl1 | 8.47264E-15 | 2.295095251 | 0.774 | 0.25 | 2.05817E-10 | 0.000731949 | 1.08559248 | 0.732 | 0.546 | 1 | 0.000731949 | 1.69453E-14 |
| Gm4258 | 8.87275E-15 | 3.436757963 | 0.358 | 0.025 | 2.15537E-10 | 8.63731E-15 | 2.719635253 | 0.293 | 0.014 | 2.09818E-10 | 8.87275E-15 | 1.72746E-14 |
| Glis3 | 9.22407E-15 | -2.69721569 | 0.396 | 0.843 | 2.24071E-10 | 0.000275856 | -1.184478364 | 0.683 | 0.881 | 1 | 0.000275856 | 1.84481E-14 |
| Hnrnpdl | 1.04258E-14 | 1.129631614 | 1 | 0.941 | 2.53263E-10 | 0.053499223 | 0.333286069 | 0.976 | 0.976 | 1 | 0.053499223 | 2.08515E-14 |
| Gramd2 | 1.05982E-14 | -3.337130502 | 0.132 | 0.75 | 2.57451E-10 | 0.003282801 | -2.574608749 | 0.146 | 0.352 | 1 | 0.003282801 | 2.11963E-14 |
| Aspn | 1.09824E-14 | 2.840614004 | 0.717 | 0.229 | 2.66785E-10 | 0.005269071 | 0.6831338 | 0.756 | 0.56 | 1 | 0.005269071 | 2.19649E-14 |
| Ninj2 | 1.19156E-14 | -3.550752713 | 0.075 | 0.725 | 2.89453E-10 | 4.08354E-08 | -2.942280016 | 0.073 | 0.577 | 0.000991973 | 4.08354E-08 | 2.38312E-14 |
| Lpl | 1.32331E-14 | 4.824573993 | 0.358 | 0.03 | 3.21458E-10 | 0.000307768 | 3.877846834 | 0.195 | 0.048 | 1 | 0.000307768 | 2.64662E-14 |
| Tnfrsf11b | 1.33623E-14 | -3.753047425 | 0.358 | 0.847 | 3.24596E-10 | 0.030975739 | -1.536919456 | 0.317 | 0.505 | 1 | 0.030975739 | 2.67245E-14 |
| Rab11fip4 | 2.37651E-13 | -1.751186242 | 0.604 | 0.932 | 5.77303E-09 | 1.4097E-14 | -2.107185112 | 0.561 | 0.952 | 3.42444E-10 | 2.37651E-13 | 2.8194E-14 |
| Tpm4 | 1.53953E-14 | 1.423394108 | 0.962 | 0.814 | 3.73983E-10 | 0.0051241 | 0.441211769 | 0.927 | 0.956 | 1 | 0.0051241 | 3.07907E-14 |
| Sdk2 | 1.54646E-14 | -1.801347678 | 0.792 | 0.992 | 3.75665E-10 | 4.54454E-13 | -1.730479659 | 0.805 | 0.99 | 1.10396E-08 | 4.54454E-13 | 3.09291E-14 |
| Trp53 | 1.55669E-14 | 1.977844833 | 0.906 | 0.534 | 3.78152E-10 | 0.00042164 | 0.754863402 | 0.927 | 0.782 | 1 | 0.00042164 | 3.11339E-14 |

| | | | | | | | | | | | | |
|---|---|---|---|---|---|---|---|---|---|---|---|---|
| Btg1 | 1.83934E-14 | -1.660391751 | 1 | 0.996 | 4.46811E-10 | 0.032885368 | -0.265045208 | 1 | 1 | 1 | 0.032885368 | 3.67867E-14 |
| Il34 | 1.94598E-14 | 2.928446739 | 0.472 | 0.068 | 4.72717E-10 | 0.265466401 | 1.578470581 | 0.195 | 0.147 | 1 | 0.265466401 | 3.89196E-14 |
| Specc1 | 1.99913E-14 | 2.973357442 | 0.755 | 0.322 | 4.85629E-10 | 0.010194837 | 0.703938319 | 0.854 | 0.758 | 1 | 0.010194837 | 3.99826E-14 |
| Dock8 | 2.09015E-14 | -2.439700235 | 0.283 | 0.86 | 5.0774E-10 | 0.002942769 | -1.063566139 | 0.585 | 0.761 | 1 | 0.002942769 | 4.18031E-14 |
| Cplx2 | 2.13828E-14 | 4.47665069 | 0.283 | 0.008 | 5.19431E-10 | 0.702426751 | 0.614812428 | 0.122 | 0.102 | 1 | 0.702426751 | 4.27656E-14 |
| Abca9 | 2.17772E-14 | 2.290074448 | 0.623 | 0.14 | 5.29012E-10 | 0.809415322 | 1.013235214 | 0.244 | 0.259 | 1 | 0.809415322 | 4.35545E-14 |
| Ppfibp2 | 3.71932E-14 | 4.029191713 | 0.415 | 0.051 | 9.03497E-10 | 2.3711E-14 | 4.013054685 | 0.463 | 0.068 | 5.75988E-10 | 3.71932E-14 | 4.7422E-14 |
| Baiap2l1 | 2.37152E-14 | -2.879224737 | 0.396 | 0.864 | 5.76089E-10 | 5.86281E-06 | -1.39502553 | 0.683 | 0.877 | 0.142419465 | 5.86281E-06 | 4.74304E-14 |
| Rgl1 | 2.55473E-14 | 1.851894599 | 0.925 | 0.686 | 6.20595E-10 | 0.002439447 | 1.051709362 | 0.854 | 0.751 | 1 | 0.002439447 | 5.10946E-14 |
| Maf | 2.64877E-14 | 1.654876708 | 0.962 | 0.521 | 6.4344E-10 | 0.035600502 | 0.465380311 | 0.927 | 0.843 | 1 | 0.035600502 | 5.29755E-14 |
| Trpv4 | 2.81473E-14 | -2.633179965 | 0.283 | 0.826 | 6.83754E-10 | 2.49056E-08 | -2.016624502 | 0.317 | 0.768 | 0.000605008 | 2.49056E-08 | 5.62946E-14 |
| Nasp | 2.84699E-14 | 1.607828135 | 0.943 | 0.623 | 6.91591E-10 | 0.000752451 | 0.876553517 | 0.902 | 0.857 | 1 | 0.000752451 | 5.69398E-14 |
| Cbfa2t3 | 2.85482E-14 | 3.974150349 | 0.415 | 0.051 | 6.93494E-10 | 0.014880547 | 1.351778023 | 0.268 | 0.13 | 1 | 0.014880547 | 5.70965E-14 |
| Ddah2 | 2.86493E-14 | 2.598329246 | 0.623 | 0.161 | 6.95949E-10 | 0.275233748 | 0.589277336 | 0.317 | 0.249 | 1 | 0.275233748 | 5.72986E-14 |
| Chd3 | 2.88035E-14 | 1.951782536 | 0.925 | 0.547 | 6.99696E-10 | 3.38652E-10 | 1.310376525 | 1 | 0.836 | 8.22653E-06 | 3.38652E-10 | 5.76071E-14 |
| Loxl1 | 3.25794E-14 | 2.470107844 | 0.811 | 0.394 | 7.9142E-10 | 4.768E-06 | 1.040810627 | 0.902 | 0.768 | 0.11582426 | 4.768E-06 | 6.51589E-14 |
| ENSMUSG00000121513 | 3.4039E-14 | 1.610023964 | 0.906 | 0.419 | 8.26876E-10 | 0.003955106 | 0.674651837 | 0.854 | 0.795 | 1 | 0.003955106 | 6.8078E-14 |
| Flna | 3.76108E-14 | 2.12037109 | 0.906 | 0.492 | 9.13642E-10 | 0.000142914 | 0.886259074 | 0.878 | 0.823 | 1 | 0.000142914 | 7.52216E-14 |
| Camk2d | 4.093E-14 | -2.003728422 | 0.868 | 1 | 9.9427E-10 | 4.40658E-11 | -2.030407227 | 0.927 | 0.997 | 1.07045E-06 | 4.40658E-11 | 8.18599E-14 |
| Tubb5 | 4.12452E-14 | 1.313103258 | 0.981 | 0.75 | 1.00193E-09 | 6.39905E-06 | 0.974433775 | 0.927 | 0.904 | 0.155445705 | 6.39905E-06 | 8.24904E-14 |
| Tuba1c | 4.14172E-14 | -2.244547589 | 0.717 | 0.958 | 1.00611E-09 | 1.86693E-08 | -1.316806969 | 0.756 | 0.976 | 0.000453516 | 1.86693E-08 | 8.28343E-14 |
| Sema6a | 4.42632E-14 | 2.748097335 | 0.585 | 0.131 | 1.07524E-09 | 0.000668424 | 0.982544213 | 0.683 | 0.461 | 1 | 0.000668424 | 8.85263E-14 |
| Stk26 | 4.58121E-14 | -2.852530789 | 0.264 | 0.818 | 1.11287E-09 | 0.000145475 | -1.619372772 | 0.268 | 0.587 | 1 | 0.000145475 | 9.16242E-14 |
| Nfkbia | 4.94696E-14 | 1.855811153 | 0.925 | 0.551 | 1.20172E-09 | 2.42882E-11 | 1.986609077 | 0.927 | 0.727 | 5.9001E-07 | 2.42882E-11 | 9.89392E-14 |
| Mia | 5.28955E-14 | -1.965712636 | 0.774 | 0.962 | 1.28494E-09 | 3.22781E-09 | -1.808105425 | 0.683 | 0.874 | 7.84099E-05 | 3.22781E-09 | 1.05791E-13 |
| Uap1 | 5.55785E-14 | -2.430900329 | 0.774 | 0.949 | 1.35011E-09 | 0.027304396 | -0.364867587 | 0.902 | 0.901 | 1 | 0.027304396 | 1.11157E-13 |
| Galnt18 | 5.99468E-14 | -3.431575117 | 0.415 | 0.835 | 1.45623E-09 | 3.25953E-06 | -2.151973657 | 0.537 | 0.795 | 0.079180516 | 3.25953E-06 | 1.19894E-13 |
| Insig1 | 6.16075E-14 | -3.732802676 | 0.396 | 0.818 | 1.49657E-09 | 0.092273102 | -0.686357106 | 0.585 | 0.689 | 1 | 0.092273102 | 1.23215E-13 |
| Arid4a | 6.42523E-14 | 1.583665696 | 0.925 | 0.551 | 1.56082E-09 | 2.66563E-07 | 1.300659478 | 0.951 | 0.792 | 0.006475356 | 2.66563E-07 | 1.28505E-13 |
| Chd4 | 7.21605E-14 | 1.223084149 | 0.981 | 0.907 | 1.75292E-09 | 3.5239E-07 | 0.941879844 | 1 | 0.99 | 0.008560247 | 3.5239E-07 | 1.44321E-13 |
| Dst | 7.9582E-14 | -2.093984824 | 0.849 | 0.987 | 1.93321E-09 | 1.56601E-08 | -1.443335007 | 0.902 | 0.983 | 0.000380415 | 1.56601E-08 | 1.59164E-13 |
| Alx1 | 8.4123E-14 | 1.808298369 | 0.962 | 0.78 | 2.04352E-09 | 0.748960683 | 0.340074348 | 0.78 | 0.846 | 1 | 0.748960683 | 1.68246E-13 |
| Fgfr3 | 8.75544E-14 | -2.15710491 | 0.434 | 0.86 | 2.12687E-09 | 6.12697E-07 | -1.475164417 | 0.512 | 0.84 | 0.01488363 | 6.12697E-07 | 1.75109E-13 |
| 2610307P16Rik | 9.1706E-14 | 3.947280515 | 0.434 | 0.064 | 2.22772E-09 | 4.34091E-10 | 4.410390183 | 0.341 | 0.055 | 1.05449E-05 | 4.34091E-10 | 1.83412E-13 |
| Eno3 | 9.21535E-14 | 3.23564259 | 0.472 | 0.076 | 2.23859E-09 | 0.630694111 | -0.292078167 | 0.073 | 0.096 | 1 | 0.630694111 | 1.84307E-13 |
| Fbxo11 | 9.73183E-14 | 1.405530698 | 0.981 | 0.78 | 2.36406E-09 | 0.001554809 | 0.832886693 | 1 | 0.952 | 1 | 0.001554809 | 1.94637E-13 |
| Lamb1 | 9.94423E-14 | 5.427741089 | 0.302 | 0.017 | 2.41565E-09 | 1.50174E-09 | 3.83720485 | 0.244 | 0.024 | 3.64803E-05 | 1.50174E-09 | 1.98885E-13 |
| Mylip | 1.11503E-13 | 2.262975485 | 0.755 | 0.275 | 2.70863E-09 | 4.17349E-09 | 1.738194844 | 0.902 | 0.539 | 0.000101382 | 4.17349E-09 | 2.23006E-13 |
| Cirbp | 1.1322E-13 | 1.697139401 | 0.906 | 0.492 | 2.75034E-09 | 0.015894121 | 0.713822434 | 0.732 | 0.614 | 1 | 0.015894121 | 2.2644E-13 |
| N4bp2 | 1.14419E-13 | 1.834389519 | 0.868 | 0.424 | 2.77946E-09 | 0.107317405 | 0.399624782 | 0.78 | 0.747 | 1 | 0.107317405 | 2.28838E-13 |
| Adamts4 | 1.15956E-13 | 4.017219071 | 0.358 | 0.034 | 2.8168E-09 | 0.00082356 | 2.489281546 | 0.146 | 0.031 | 1 | 0.00082356 | 2.31912E-13 |
| Nr2f1 | 1.33176E-13 | 2.827147937 | 0.472 | 0.076 | 3.23512E-09 | 0.421974442 | 1.33970519 | 0.146 | 0.116 | 1 | 0.421974442 | 2.66353E-13 |
| Etl4 | 1.33469E-13 | 2.223985257 | 0.717 | 0.229 | 3.24224E-09 | 4.6235E-09 | 2.128370691 | 0.756 | 0.358 | 0.000112314 | 4.6235E-09 | 2.66939E-13 |
| Igf1 | 1.35096E-13 | 3.21710138 | 0.604 | 0.157 | 3.28175E-09 | 3.67784E-05 | 2.927659801 | 0.488 | 0.249 | 0.893419716 | 3.67784E-05 | 2.70192E-13 |
| Nfix | 1.62542E-13 | 0.993377318 | 1 | 0.97 | 3.94848E-09 | 0.021563515 | 0.340409209 | 1 | 0.993 | 1 | 0.021563515 | 3.25085E-13 |
| Prrx1 | 1.66225E-13 | 2.221428408 | 0.962 | 0.589 | 4.03793E-09 | 0.001303587 | 0.642991884 | 0.927 | 0.836 | 1 | 0.001303587 | 3.32449E-13 |
| Col6a2 | 1.69059E-13 | -1.416859758 | 0.849 | 0.992 | 4.10677E-09 | 0.024571474 | -0.514061356 | 0.902 | 0.942 | 1 | 0.024571474 | 3.38117E-13 |
| Wwp2 | 1.96383E-13 | -1.876201978 | 0.774 | 0.966 | 4.77054E-09 | 0.00054613 | -0.862713758 | 0.829 | 0.986 | 1 | 0.00054613 | 3.92766E-13 |
| Fhod3 | 2.05637E-13 | 5.551612747 | 0.283 | 0.013 | 4.99534E-09 | 0.000656102 | 3.145327145 | 0.171 | 0.041 | 1 | 0.000656102 | 4.11275E-13 |
| Fbln1 | 2.05945E-13 | 3.209170379 | 0.434 | 0.064 | 5.00281E-09 | 9.20606E-06 | 2.83720485 | 0.22 | 0.041 | 0.223633544 | 9.20606E-06 | 4.1189E-13 |
| Mxra8 | 2.09598E-13 | 1.363309217 | 0.962 | 0.86 | 5.09155E-09 | 0.0148443 | 0.365155255 | 0.951 | 0.857 | 1 | 0.0148443 | 4.19195E-13 |
| Pfkp | 2.11115E-13 | -2.152436041 | 0.434 | 0.86 | 5.1284E-09 | 0.04857525 | -0.994303934 | 0.634 | 0.683 | 1 | 0.04857525 | 4.22229E-13 |
| Mcoln2 | 2.45314E-13 | -3.953801862 | 0.094 | 0.678 | 5.95916E-09 | 0.000722273 | -0.600200463 | 0.146 | 0.454 | 1 | 0.000722273 | 4.90627E-13 |
| Eif4g2 | 2.46748E-13 | 1.018643642 | 1 | 0.975 | 5.994E-09 | 0.006235418 | 0.392498314 | 1 | 0.997 | 1 | 0.006235418 | 4.93496E-13 |
| Chil1 | 2.49402E-13 | -5.801867096 | 0.189 | 0.712 | 6.05847E-09 | 1.995E-05 | -2.600517503 | 0.195 | 0.546 | 0.484625565 | 1.995E-05 | 4.98804E-13 |
| Mxd1 | 2.64816E-13 | 2.373557197 | 0.698 | 0.242 | 6.43291E-09 | 8.51015E-07 | 1.304081281 | 0.756 | 0.41 | 0.02067285 | 8.51015E-07 | 5.29632E-13 |
| Acap2 | 2.72126E-13 | 1.67804603 | 0.906 | 0.487 | 6.6105E-09 | 0.000718343 | 0.552047633 | 0.854 | 0.693 | 1 | 0.000718343 | 5.44253E-13 |
| Cyp51 | 2.82081E-13 | -3.122117611 | 0.434 | 0.814 | 6.85232E-09 | 0.001751304 | -1.480765232 | 0.488 | 0.662 | 1 | 0.001751304 | 5.64162E-13 |
| Dennd2b | 2.87426E-13 | 2.315187267 | 0.566 | 0.123 | 6.98216E-09 | 3.08373E-07 | 2.027307733 | 0.659 | 0.328 | 0.007490987 | 3.08373E-07 | 5.74852E-13 |
| Chadl | 2.99544E-13 | -1.814903756 | 0.491 | 0.903 | 7.27653E-09 | 2.44137E-09 | -2.2931347 | 0.268 | 0.761 | 5.93057E-05 | 2.44137E-09 | 5.99088E-13 |
| Zcchc14 | 3.20884E-13 | 1.551926801 | 0.962 | 0.733 | 7.79493E-09 | 1.68719E-05 | 0.798895493 | 0.976 | 0.932 | 0.409851472 | 1.68719E-05 | 6.41769E-13 |
| Ppp1r1b | 3.50979E-13 | -6.426477987 | 0.038 | 0.619 | 8.52599E-09 | 5.80434E-06 | -3.037264268 | 0.049 | 0.43 | 0.140999071 | 5.80434E-06 | 7.01958E-13 |
| Itsn1 | 3.70133E-13 | 2.248381702 | 0.83 | 0.373 | 8.99128E-09 | 4.3476E-06 | 1.145327145 | 0.854 | 0.7 | 0.105611987 | 4.3476E-06 | 7.40267E-13 |
| Ogn | 3.70188E-13 | 1.507024339 | 0.925 | 0.441 | 8.9926E-09 | 0.01176372 | 1.001624745 | 0.78 | 0.775 | 1 | 0.01176372 | 7.40375E-13 |
| Tmsb4x | 4.17152E-13 | 2.03670062 | 0.962 | 0.767 | 1.01335E-08 | 4.61972E-11 | 2.748384817 | 0.976 | 0.84 | 1.12222E-06 | 4.61972E-11 | 8.34305E-13 |
| Dixdc1 | 4.37206E-13 | -2.337130502 | 0.302 | 0.784 | 1.06206E-08 | 1.58314E-10 | -2.572186086 | 0.244 | 0.792 | 3.84577E-06 | 1.58314E-10 | 8.74413E-13 |
| Tbc1d12 | 5.03856E-13 | 2.220014055 | 0.736 | 0.267 | 1.22397E-08 | 0.056843863 | 0.495098873 | 0.683 | 0.481 | 1 | 0.056843863 | 1.00771E-12 |
| Gja1 | 5.29643E-13 | 1.93418812 | 0.887 | 0.47 | 1.28661E-08 | 1.85011E-08 | 1.633064133 | 0.854 | 0.43 | 0.00044943 | 1.85011E-08 | 1.05929E-12 |
| Aig1 | 5.48295E-13 | -2.309385511 | 0.547 | 0.915 | 1.33192E-08 | 0.032885079 | -0.151588022 | 0.61 | 0.768 | 1 | 0.032885079 | 1.09659E-12 |
| Abca1 | 5.63805E-13 | 2.8327945 | 0.566 | 0.144 | 1.36959E-08 | 3.43135E-08 | 2.208388059 | 0.659 | 0.287 | 0.000833544 | 3.43135E-08 | 1.12761E-12 |
| Smarce1 | 5.67075E-13 | 1.484512807 | 0.906 | 0.708 | 1.37754E-08 | 0.765163678 | 0.135905105 | 0.878 | 0.908 | 1 | 0.765163678 | 1.13415E-12 |
| Cald1 | 6.11849E-13 | 1.601505678 | 0.981 | 0.966 | 1.4863E-08 | 7.45829E-05 | 1.18827929 | 0.976 | 0.925 | 1 | 7.45829E-05 | 1.2237E-12 |
| Pard3b | 6.13536E-13 | 1.696872012 | 0.925 | 0.644 | 1.4904E-08 | 0.005774686 | 0.812026287 | 0.951 | 0.84 | 1 | 0.005774686 | 1.22707E-12 |
| Iqsec1 | 6.18762E-13 | -1.936592572 | 0.604 | 0.958 | 1.5031E-08 | 1.06663E-10 | -1.592682493 | 0.78 | 0.976 | 2.59105E-06 | 1.06663E-10 | 1.23752E-12 |
| Rasl11b | 6.60347E-13 | 3.334046294 | 0.491 | 0.097 | 1.60411E-08 | 2.48231E-09 | 2.510976617 | 0.488 | 0.137 | 6.03003E-05 | 2.48231E-09 | 1.32069E-12 |
| Tmem64 | 6.73289E-13 | 2.256005931 | 0.698 | 0.25 | 1.63555E-08 | 0.271907994 | 0.569982647 | 0.659 | 0.594 | 1 | 0.271907994 | 1.34658E-12 |
| Pard6g | 6.82686E-13 | 3.154722595 | 0.434 | 0.068 | 1.65838E-08 | 6.98242E-05 | 2.232342792 | 0.415 | 0.177 | 1 | 6.98242E-05 | 1.36537E-12 |
| Col6a3 | 6.91928E-13 | -1.82181278 | 0.849 | 0.975 | 1.68083E-08 | 0.000127378 | -1.107813726 | 0.805 | 0.901 | 1 | 0.000127378 | 1.38386E-12 |
| Fscn1 | 7.07209E-13 | 1.547040018 | 0.887 | 0.356 | 1.71795E-08 | 0.007595242 | 0.753483024 | 0.707 | 0.625 | 1 | 0.007595242 | 1.41442E-12 |
| Klf13 | 7.11109E-13 | 1.140448464 | 1 | 0.771 | 1.72743E-08 | 0.076573523 | 0.268276022 | 0.976 | 0.962 | 1 | 0.076573523 | 1.42222E-12 |
| Itga8 | 7.74395E-13 | 4.657222935 | 0.302 | 0.021 | 1.88116E-08 | 1.50965E-10 | 3.974708374 | 0.317 | 0.041 | 3.66723E-06 | 1.50965E-10 | 1.54879E-12 |
| Colgalt2 | 7.86308E-13 | -3.581679337 | 0.17 | 0.716 | 1.9101E-08 | 0.013392919 | -1.431283986 | 0.122 | 0.307 | 1 | 0.013392919 | 1.57262E-12 |
| Nt5e | 8.66293E-13 | -3.993895213 | 0.189 | 0.712 | 2.1044E-08 | 0.003763432 | -1.997630809 | 0.268 | 0.498 | 1 | 0.003763432 | 1.73259E-12 |
| Sdc2 | 8.77235E-13 | 1.896018109 | 0.962 | 0.826 | 2.13098E-08 | 0.138305462 | 1.219124607 | 0.878 | 0.809 | 1 | 0.138305462 | 1.75447E-12 |
| Chsy3 | 9.6296E-13 | -2.450813043 | 0.698 | 0.949 | 2.33922E-08 | 0.000294835 | -1.878610094 | 0.463 | 0.717 | 1 | 0.000294835 | 1.92592E-12 |
| Sesn1 | 9.67014E-13 | 2.308267853 | 0.811 | 0.394 | 2.34907E-08 | 0.05635754 | 0.742770048 | 0.683 | 0.549 | 1 | 0.05635754 | 1.93403E-12 |
| Adam19 | 9.76632E-13 | 3.977844833 | 0.321 | 0.025 | 2.37244E-08 | 0.011563259 | 2.234540347 | 0.195 | 0.078 | 1 | 0.011563259 | 1.95326E-12 |
| Map1b | 1.01849E-12 | 4.228303727 | 0.528 | 0.119 | 2.47413E-08 | 0.000188951 | 2.86719959 | 0.585 | 0.345 | 1 | 0.000188951 | 2.03699E-12 |
| Pax3 | 1.08665E-12 | 7.440124814 | 0.208 | 0 | 2.6397E-08 | 0.00063996 | 2.490554326 | 0.268 | 0.096 | 1 | 0.00063996 | 2.17331E-12 |
| Col2a1 | 1.10416E-12 | -1.913851426 | 0.811 | 0.983 | 2.68222E-08 | 1.97205E-11 | -1.966185628 | 0.634 | 0.973 | 4.7905E-07 | 1.97205E-11 | 2.20831E-12 |
| Gpc6 | 1.25232E-12 | -1.558458932 | 0.981 | 1 | 3.04214E-08 | 7.90453E-10 | -1.232385422 | 0.951 | 0.997 | 1.92017E-05 | 7.90453E-10 | 2.50464E-12 |
| Rbfox2 | 1.28923E-12 | 1.255373213 | 1 | 0.89 | 3.1318E-08 | 4.65167E-06 | 0.644559772 | 1 | 0.969 | 0.112998367 | 4.65167E-06 | 2.57846E-12 |
| Zfp521 | 1.31968E-12 | 1.259883525 | 1 | 0.797 | 3.20577E-08 | 0.00748925 | 0.800773098 | 0.902 | 0.853 | 1 | 0.00748925 | 2.63936E-12 |
| Smarcc1 | 1.37949E-12 | 1.588953659 | 0.943 | 0.623 | 3.35105E-08 | 0.002019847 | 0.747857364 | 0.902 | 0.874 | 1 | 0.002019847 | 2.75897E-12 |
| Cpxm1 | 1.3822E-12 | 1.900676972 | 0.849 | 0.39 | 3.35763E-08 | 0.005209156 | 1.297987939 | 0.732 | 0.638 | 1 | 0.005209156 | 2.76439E-12 |
| Kcne4 | 1.59835E-12 | 5.440124814 | 0.264 | 0.013 | 3.8827E-08 | 0.001085069 | 3.322631677 | 0.073 | 0.007 | 1 | 0.001085069 | 3.19669E-12 |
| Fgf2 | 1.61688E-12 | -2.792613527 | 0.321 | 0.78 | 3.92772E-08 | 6.25967E-08 | -2.474076001 | 0.22 | 0.662 | 0.001520598 | 6.25967E-08 | 3.23376E-12 |
| Mta2 | 1.64482E-12 | 2.043691282 | 0.698 | 0.242 | 3.99559E-08 | 0.037610364 | 0.742047617 | 0.659 | 0.539 | 1 | 0.037610364 | 3.28964E-12 |

| | | | | | | | | | | | | |
|---|---|---|---|---|---|---|---|---|---|---|---|---|
| Clic4 | 1.65649E-12 | -1.544286304 | 1 | 0.996 | 4.02394E-08 | 0.016671504 | -0.444830518 | 0.976 | 0.993 | 1 | 0.016671504 | 3.31298E-12 |
| Phldb2 | 1.72927E-12 | 2.350118284 | 0.717 | 0.233 | 4.20073E-08 | 0.017455138 | 1.229367883 | 0.683 | 0.498 | 1 | 0.017455138 | 3.45853E-12 |
| Slit2 | 1.77854E-12 | 4.176748901 | 0.509 | 0.114 | 4.32044E-08 | 3.05842E-06 | 2.205767919 | 0.659 | 0.314 | 0.074295137 | 3.05842E-06 | 3.55709E-12 |
| Ucma | 3.5041E-10 | -1.526426105 | 0.623 | 0.911 | 8.51217E-06 | 1.86563E-12 | -2.79491291 | 0.293 | 0.836 | 4.532E-08 | 3.5041E-10 | 3.73127E-12 |
| Rbbp6 | 1.95028E-12 | 1.174396014 | 1 | 0.881 | 4.73763E-08 | 0.544020006 | 0.185685573 | 0.927 | 0.98 | 1 | 0.544020006 | 3.90056E-12 |
| Tiam1 | 2.05232E-12 | -1.591093917 | 0.377 | 0.86 | 4.98549E-08 | 5.74396E-06 | -1.528050677 | 0.439 | 0.785 | 0.139532176 | 5.74396E-06 | 4.10464E-12 |
| Cnn2 | 2.12457E-12 | 2.318547397 | 0.736 | 0.292 | 5.161E-08 | 0.001753908 | 0.988466007 | 0.756 | 0.546 | 1 | 0.001753908 | 4.24913E-12 |
| Nbl1 | 2.21959E-12 | 2.062840494 | 0.868 | 0.479 | 5.39184E-08 | 0.137388418 | 0.706957142 | 0.829 | 0.785 | 1 | 0.137388418 | 4.43919E-12 |
| Dusp5 | 2.24181E-12 | 2.139455838 | 0.642 | 0.186 | 5.44581E-08 | 3.60831E-05 | 1.287007767 | 0.561 | 0.259 | 0.876531647 | 3.60831E-05 | 4.48363E-12 |
| Npas3 | 1.91818E-10 | -1.761850554 | 0.472 | 0.907 | 4.65964E-06 | 2.24577E-12 | -2.121743251 | 0.512 | 0.925 | 5.45542E-08 | 1.91818E-10 | 4.49154E-12 |
| Hmgcr | 2.37362E-12 | -3.139898154 | 0.396 | 0.797 | 5.76599E-08 | 0.053865077 | -0.928695645 | 0.585 | 0.676 | 1 | 0.053865077 | 4.74723E-12 |
| Lrrn4cl | 2.54917E-12 | 2.550651271 | 0.491 | 0.097 | 6.19244E-08 | 0.000828451 | 2.614812428 | 0.195 | 0.055 | 1 | 0.000828451 | 5.09834E-12 |
| Parp8 | 2.57828E-12 | 1.488723522 | 0.868 | 0.424 | 6.26316E-08 | 2.88167E-06 | 1.117621334 | 0.829 | 0.57 | 0.070001494 | 2.88167E-06 | 5.15656E-12 |
| Wnt5a | 2.62141E-12 | 3.802420851 | 0.377 | 0.051 | 6.36792E-08 | 0.003123087 | 1.181853021 | 0.317 | 0.137 | 1 | 0.003123087 | 5.24281E-12 |
| Hexim1 | 2.63505E-12 | 1.903761021 | 0.887 | 0.492 | 6.40107E-08 | 0.011064369 | 0.764364043 | 0.878 | 0.782 | 1 | 0.011064369 | 5.27011E-12 |
| Septin11 | 2.65293E-12 | 1.394060908 | 0.962 | 0.729 | 6.44449E-08 | 0.000128056 | 1.044961925 | 0.951 | 0.908 | 1 | 0.000128056 | 5.30585E-12 |
| Sirt1 | 2.70803E-12 | 1.865215978 | 0.868 | 0.504 | 6.57835E-08 | 0.000214857 | 0.990249224 | 0.829 | 0.761 | 1 | 0.000214857 | 5.41606E-12 |
| Fxyd3 | 2.73183E-12 | -4.320335259 | 0.302 | 0.746 | 6.63615E-08 | 0.000943778 | -1.798831836 | 0.341 | 0.601 | 1 | 0.000943778 | 5.46365E-12 |
| Uba7 | 2.78688E-12 | 3.855162313 | 0.321 | 0.03 | 6.76989E-08 | 0.837325339 | 0.452541 | 0.122 | 0.113 | 1 | 0.837325339 | 5.57376E-12 |
| Tpm2 | 2.98969E-12 | 2.329809301 | 0.642 | 0.199 | 7.26255E-08 | 0.008045072 | 1.32379561 | 0.488 | 0.328 | 1 | 0.008045072 | 5.97938E-12 |
| Rbfox1 | 3.03215E-12 | 3.929162894 | 0.434 | 0.076 | 7.36571E-08 | 0.061694361 | 0.799069721 | 0.195 | 0.099 | 1 | 0.061694361 | 6.06431E-12 |
| Dnm3os | 3.243E-12 | 3.265753907 | 0.415 | 0.064 | 7.8779E-08 | 7.67146E-07 | 2.874679555 | 0.341 | 0.085 | 0.018635515 | 7.67146E-07 | 6.486E-12 |
| Col14a1 | 3.28268E-12 | 4.213616284 | 0.321 | 0.03 | 7.97428E-08 | 2.42415E-06 | 5.206438659 | 0.146 | 0.014 | 0.058887485 | 2.42415E-06 | 6.56535E-12 |
| Tacc2 | 3.32065E-12 | -2.033454111 | 0.774 | 0.949 | 8.06651E-08 | 7.14486E-06 | -0.99535642 | 0.854 | 0.969 | 0.173563039 | 7.14486E-06 | 6.64129E-12 |
| Ncoa1 | 3.71183E-12 | 2.049369595 | 0.811 | 0.398 | 9.01679E-08 | 0.003174426 | 0.972529202 | 0.854 | 0.805 | 1 | 0.003174426 | 7.42367E-12 |
| Prrc2b | 4.06861E-12 | 1.205230497 | 0.981 | 0.801 | 9.88348E-08 | 0.015986347 | 0.402066144 | 1 | 0.952 | 1 | 0.015986347 | 8.13723E-12 |
| Bcl2l11 | 4.09611E-12 | 2.314379261 | 0.566 | 0.148 | 9.95026E-08 | 5.7456E-11 | 3.391306387 | 0.659 | 0.222 | 1.39572E-06 | 5.7456E-11 | 8.19221E-12 |
| Hsd17b7 | 4.1025E-12 | -3.083266682 | 0.245 | 0.737 | 9.9658E-08 | 0.000128962 | -1.662261431 | 0.195 | 0.526 | 1 | 0.000128962 | 8.20501E-12 |
| BC006965 | 4.28081E-12 | -6.47554453 | 0.057 | 0.597 | 1.03989E-07 | 0.052834964 | -1.545264787 | 0.098 | 0.229 | 1 | 0.052834964 | 8.56162E-12 |
| Prkg1 | 4.63383E-12 | 1.677212933 | 1 | 0.678 | 1.12565E-07 | 0.037657465 | -0.529617825 | 0.976 | 0.939 | 1 | 0.037657465 | 9.26766E-12 |
| Rfx7 | 4.68444E-12 | 1.698905245 | 0.887 | 0.53 | 1.13794E-07 | 0.00036687 | 0.92707588 | 0.854 | 0.771 | 1 | 0.00036687 | 9.36888E-12 |
| Capn6 | 4.71854E-12 | 1.89866015 | 0.811 | 0.419 | 1.14623E-07 | 0.780158736 | 0.394907578 | 0.61 | 0.73 | 1 | 0.780158736 | 9.43709E-12 |
| Ptprs | 4.79475E-12 | 2.014124265 | 0.792 | 0.377 | 1.16474E-07 | 2.9039E-05 | 1.380690715 | 0.732 | 0.532 | 0.705415633 | 2.9039E-05 | 9.5895E-12 |
| Six4 | 4.93073E-12 | 2.347367673 | 0.396 | 0.055 | 1.19777E-07 | 0.011644022 | 1.088743617 | 0.341 | 0.181 | 1 | 0.011644022 | 9.86146E-12 |
| Pawr | 2.02457E-07 | 2.154722595 | 0.566 | 0.242 | 0.004918075 | 5.28049E-12 | 2.362046358 | 0.78 | 0.324 | 1.28274E-07 | 2.02457E-07 | 1.0561E-11 |
| Tubb2a | 5.37521E-12 | -2.258389294 | 0.774 | 0.936 | 1.30575E-07 | 3.30613E-07 | -1.760035981 | 0.829 | 0.908 | 0.008031254 | 3.30613E-07 | 1.07504E-11 |
| Setd2 | 5.5161E-12 | 0.972345852 | 1 | 0.847 | 1.33997E-07 | 0.188807697 | 0.153409915 | 1 | 0.973 | 1 | 0.188807697 | 1.10322E-11 |
| Ldlr | 5.81027E-12 | -2.69986249 | 0.358 | 0.78 | 1.41143E-07 | 0.002806795 | -1.21365152 | 0.439 | 0.625 | 1 | 0.002806795 | 1.16205E-11 |
| Gjc3 | 5.98905E-12 | -3.629912251 | 0.094 | 0.636 | 1.45486E-07 | 0.139570514 | -2.16279515 | 0 | 0.051 | 1 | 0.139570514 | 1.19781E-11 |
| Pacsin2 | 6.21502E-12 | -1.76565123 | 0.528 | 0.903 | 1.50975E-07 | 0.001129909 | -0.96708594 | 0.732 | 0.881 | 1 | 0.001129909 | 1.243E-11 |
| Wif1 | 6.22565E-12 | -4.079855365 | 0.151 | 0.691 | 1.51234E-07 | 5.56771E-06 | -1.444862468 | 0.415 | 0.778 | 0.135250705 | 5.56771E-06 | 1.24513E-11 |
| Fgd1 | 6.53559E-12 | 2.359081093 | 0.566 | 0.148 | 1.58763E-07 | 0.575040769 | 0.614812428 | 0.268 | 0.235 | 1 | 0.575040769 | 1.30712E-11 |
| Ptgfr | 6.74598E-12 | 4.490325627 | 0.377 | 0.055 | 1.63873E-07 | 4.3131E-08 | 2.325704511 | 0.317 | 0.058 | 0.001047738 | 4.3131E-08 | 1.3492E-11 |
| Thbd | 6.84337E-12 | 3.493524508 | 0.34 | 0.038 | 1.66239E-07 | 1.49441E-08 | 4.744095445 | 0.171 | 0.01 | 0.000363023 | 1.49441E-08 | 1.36867E-11 |
| Mxd4 | 7.31027E-12 | 1.47665069 | 0.925 | 0.627 | 1.77581E-07 | 2.14947E-05 | 0.779653827 | 0.976 | 0.836 | 0.522150261 | 2.14947E-05 | 1.46205E-11 |
| Tgfbr1 | 7.39342E-12 | 1.685659214 | 0.906 | 0.504 | 1.79601E-07 | 6.27876E-06 | 1.319670277 | 0.902 | 0.71 | 0.152523589 | 6.27876E-06 | 1.47868E-11 |
| Chd6 | 7.45924E-12 | 1.477301872 | 0.962 | 0.716 | 1.812E-07 | 0.000103916 | 0.771115659 | 0.927 | 0.87 | 1 | 0.000103916 | 1.49185E-11 |
| Ppp1r14b | 7.71989E-12 | 1.496745297 | 0.943 | 0.653 | 1.87532E-07 | 0.000238661 | 0.821313477 | 0.927 | 0.792 | 1 | 0.000238661 | 1.54398E-11 |
| Serpina3n | 8.06021E-12 | -7.001781891 | 0.019 | 0.564 | 1.95799E-07 | 0.077569551 | -3.777504994 | 0 | 0.072 | 1 | 0.077569551 | 1.61204E-11 |
| Msmo1 | 8.10014E-12 | -3.855347264 | 0.283 | 0.729 | 1.96769E-07 | 0.00025346 | -1.93128725 | 0.293 | 0.584 | 1 | 0.00025346 | 1.62003E-11 |
| Dync1h1 | 8.18881E-12 | 1.185883526 | 0.981 | 0.784 | 1.98923E-07 | 0.01915179 | 0.405938889 | 0.927 | 0.959 | 1 | 0.01915179 | 1.63776E-11 |
| Il16 | 8.58552E-12 | -2.186064527 | 0.434 | 0.881 | 2.08559E-07 | 3.64661E-11 | -2.006153314 | 0.439 | 0.915 | 8.85836E-07 | 3.64661E-11 | 1.7171E-11 |
| Palm | 8.80279E-12 | 2.466666601 | 0.453 | 0.081 | 2.13837E-07 | 0.251959328 | 0.924667691 | 0.244 | 0.167 | 1 | 0.251959328 | 1.76056E-11 |
| Idi1 | 9.45029E-12 | -3.840178251 | 0.208 | 0.708 | 2.29566E-07 | 0.004390389 | -2.107653596 | 0.293 | 0.478 | 1 | 0.004390389 | 1.89006E-11 |
| Smarca2 | 9.64973E-12 | 1.268104394 | 0.962 | 0.75 | 2.34411E-07 | 1.62895E-05 | 1.045522086 | 0.976 | 0.833 | 0.395705534 | 1.62895E-05 | 1.92995E-11 |
| Tamalin | 9.72151E-12 | 2.592843707 | 0.585 | 0.169 | 2.36155E-07 | 0.04545289 | 0.752037467 | 0.415 | 0.276 | 1 | 0.04545289 | 1.9443E-11 |
| Ehmt2 | 1.08753E-11 | 2.154722595 | 0.642 | 0.208 | 2.64183E-07 | 0.187672804 | 0.782757066 | 0.415 | 0.365 | 1 | 0.187672804 | 2.17506E-11 |
| Clip3 | 1.09488E-11 | 2.37231403 | 0.547 | 0.14 | 2.65969E-07 | 0.100662367 | 0.506559538 | 0.463 | 0.331 | 1 | 0.100662367 | 2.18977E-11 |
| Naaladl2 | 1.12098E-11 | 1.843350292 | 0.943 | 0.606 | 2.72308E-07 | 0.957811358 | -0.201065167 | 0.927 | 0.805 | 1 | 0.957811358 | 2.24196E-11 |
| Wipf1 | 1.18382E-11 | 1.697249829 | 0.83 | 0.364 | 2.87574E-07 | 0.005600395 | 1.193943709 | 0.78 | 0.577 | 1 | 0.005600395 | 2.36764E-11 |
| Hs3st3b1 | 1.2458E-11 | -2.09003764 | 0.358 | 0.805 | 3.02631E-07 | 1.34958E-05 | -1.637880033 | 0.268 | 0.659 | 0.327840622 | 1.34958E-05 | 2.49161E-11 |
| Bcat1 | 1.25906E-11 | -3.667582669 | 0.057 | 0.606 | 3.05852E-07 | 0.000235084 | -4.638528581 | 0.024 | 0.294 | 1 | 0.000235084 | 2.51813E-11 |
| Pla2g7 | 1.28995E-11 | -5.616766875 | 0.019 | 0.555 | 3.13355E-07 | 1.58942E-05 | -5.405969134 | 0 | 0.334 | 0.386102701 | 1.58942E-05 | 2.5799E-11 |
| Cspg4 | 1.33304E-11 | -1.748069433 | 0.528 | 0.915 | 3.23822E-07 | 9.076E-10 | -1.607064653 | 0.561 | 0.918 | 2.20474E-05 | 9.076E-10 | 2.66608E-11 |
| Nalcn | 1.36775E-11 | 4.882643049 | 0.264 | 0.017 | 3.32255E-07 | 0.166006825 | 0.410940095 | 0.098 | 0.044 | 1 | 0.166006825 | 2.73551E-11 |
| Alpl | 6.53744E-05 | 1.317104834 | 0.302 | 0.097 | 1 | 1.47359E-11 | 2.659206548 | 0.293 | 0.027 | 3.57964E-07 | 6.53744E-05 | 2.94718E-11 |
| Ctxn1 | 1.58771E-11 | 4.225111923 | 0.264 | 0.017 | 3.85686E-07 | 0.00753319 | 2.007129851 | 0.146 | 0.044 | 1 | 0.00753319 | 3.17541E-11 |
| Hpse2 | 1.59638E-11 | 2.830934158 | 0.509 | 0.119 | 3.87792E-07 | 0.200043229 | -1.037264268 | 0.146 | 0.235 | 1 | 0.200043229 | 3.19276E-11 |
| Tcf12 | 1.59928E-11 | 1.258534293 | 1 | 0.949 | 3.88497E-07 | 0.585696372 | -0.213282255 | 1 | 0.997 | 1 | 0.585696372 | 3.19856E-11 |
| Maml2 | 1.66561E-11 | -1.67478216 | 0.981 | 0.996 | 4.0461E-07 | 2.40192E-06 | -1.117545385 | 1 | 1 | 0.058347419 | 2.40192E-06 | 3.33122E-11 |
| Phip | 1.69947E-11 | 1.844867332 | 0.887 | 0.555 | 4.12834E-07 | 6.46663E-06 | 0.874124313 | 0.927 | 0.727 | 0.15708744 | 6.46663E-06 | 3.39893E-11 |
| Phf23 | 1.70864E-11 | 1.893856026 | 0.736 | 0.347 | 4.15063E-07 | 0.007106313 | 0.860920919 | 0.732 | 0.59 | 1 | 0.007106313 | 3.41728E-11 |
| Irf2 | 1.82911E-11 | 1.71756669 | 0.849 | 0.53 | 4.44327E-07 | 0.005360895 | 0.866033503 | 0.78 | 0.71 | 1 | 0.005360895 | 3.65822E-11 |
| Zfp354c | 1.95518E-11 | 1.517292674 | 0.755 | 0.297 | 4.74952E-07 | 0.6002181 | 0.214571931 | 0.561 | 0.553 | 1 | 0.6002181 | 3.91035E-11 |
| Pik3r3 | 2.00065E-11 | 3.660613525 | 0.377 | 0.059 | 4.85998E-07 | 3.06412E-07 | 2.139233387 | 0.488 | 0.16 | 0.007443367 | 3.06412E-07 | 4.0013E-11 |
| Tcf4 | 2.0191E-11 | 0.838187378 | 1 | 0.983 | 4.90481E-07 | 1.62001E-05 | 0.523803976 | 1 | 1 | 0.393532251 | 1.62001E-05 | 4.03821E-11 |
| 9930021J03Rik | 2.02513E-11 | 1.536029237 | 0.887 | 0.475 | 4.91946E-07 | 6.83558E-05 | 0.950464862 | 0.878 | 0.71 | 1 | 6.83558E-05 | 4.05027E-11 |
| Gna14 | 2.10264E-11 | -4.714693295 | 0.132 | 0.631 | 5.10774E-07 | 0.168505133 | -1.958654433 | 0.366 | 0.433 | 1 | 0.168505133 | 4.20529E-11 |
| Pxdn | 2.10364E-11 | 2.141047658 | 0.623 | 0.203 | 5.11016E-07 | 0.166771228 | 0.502602925 | 0.488 | 0.444 | 1 | 0.166771228 | 4.20728E-11 |
| Podn | 2.16044E-11 | 5.06161319 | 0.208 | 0.004 | 5.24814E-07 | 0.003153551 | 2.904319046 | 0.171 | 0.051 | 1 | 0.003153551 | 4.32088E-11 |
| Acly | 2.19802E-11 | -1.791919158 | 0.642 | 0.932 | 5.33943E-07 | 0.015249843 | -0.827341199 | 0.732 | 0.799 | 1 | 0.015249843 | 4.39604E-11 |
| Sdc1 | 2.23018E-11 | 2.294125651 | 0.604 | 0.195 | 5.41757E-07 | 0.000251437 | 1.204936634 | 0.732 | 0.567 | 1 | 0.000251437 | 4.46037E-11 |
| Ece1 | 2.23056E-11 | 2.204191271 | 0.66 | 0.225 | 5.41849E-07 | 6.83741E-06 | 1.60690723 | 0.683 | 0.372 | 0.166094244 | 6.83741E-06 | 4.46113E-11 |
| Mtch1 | 2.28857E-11 | 1.497947072 | 0.981 | 0.805 | 5.55941E-07 | 0.042923547 | 0.565400234 | 0.902 | 0.86 | 1 | 0.042923547 | 4.57715E-11 |
| Hip1r | 2.3463E-11 | -2.467329225 | 0.226 | 0.716 | 5.69963E-07 | 9.72975E-08 | -1.695290231 | 0.317 | 0.758 | 0.00236355 | 9.72975E-08 | 4.6926E-11 |
| Usp9x | 2.48978E-11 | 1.221993291 | 1 | 0.932 | 6.04817E-07 | 0.000242787 | 0.656732684 | 1 | 0.98 | 1 | 0.000242787 | 4.97956E-11 |
| Ttc7 | 2.52168E-11 | -2.027802443 | 0.321 | 0.809 | 6.12566E-07 | 0.076054162 | -0.630153685 | 0.707 | 0.836 | 1 | 0.076054162 | 5.04335E-11 |
| Man1c1 | 2.72989E-11 | 2.136800687 | 0.623 | 0.195 | 6.63144E-07 | 0.002414365 | 0.790511614 | 0.585 | 0.362 | 1 | 0.002414365 | 5.45977E-11 |
| C030013E06Rik | 2.87737E-11 | 3.417757001 | 0.358 | 0.051 | 6.98972E-07 | 0.007999789 | 2.649577847 | 0.22 | 0.096 | 1 | 0.007999789 | 5.75475E-11 |
| Slit3 | 3.04395E-11 | 5.621848605 | 0.321 | 0.038 | 7.39436E-07 | 6.98332E-09 | 2.558488822 | 0.512 | 0.147 | 0.000169639 | 6.98332E-09 | 6.0879E-11 |
| Lmcd1 | 3.08547E-11 | -3.389597921 | 0.453 | 0.805 | 7.49523E-07 | 8.96335E-07 | -1.891229111 | 0.268 | 0.724 | 0.021773765 | 8.96335E-07 | 6.17095E-11 |
| Topors | 3.09912E-11 | 1.752053653 | 0.849 | 0.458 | 7.52837E-07 | 0.001784898 | 0.793129875 | 0.854 | 0.734 | 1 | 0.001784898 | 6.19823E-11 |
| Tnfsf11 | 3.80946E-06 | 2.19654277 | 0.208 | 0.034 | 0.092539487 | 3.09958E-11 | 6.296636468 | 0.195 | 0.007 | 7.5295E-07 | 3.80946E-06 | 6.19916E-11 |
| Stard4 | 3.13085E-11 | -4.063062045 | 0.113 | 0.614 | 7.60547E-07 | 0.000351915 | -1.841733799 | 0.171 | 0.461 | 1 | 0.000351915 | 6.26171E-11 |
| Sgms2 | 3.14568E-11 | -1.977693312 | 0.698 | 0.936 | 7.64148E-07 | 0.000248515 | -0.989968962 | 0.878 | 0.973 | 1 | 0.000248515 | 6.29136E-11 |
| Nfkbiz | 0.001566098 | 0.523217508 | 0.717 | 0.504 | 1 | 3.27469E-11 | 2.090653207 | 0.756 | 0.334 | 7.95487E-07 | 0.001566098 | 6.54938E-11 |
| Nlgn1 | 3.2837E-11 | 3.689499338 | 0.302 | 0.03 | 7.97676E-07 | 0.179935398 | 2.662118143 | 0.171 | 0.109 | 1 | 0.179935398 | 6.5674E-11 |
| Hmgb1 | 3.35828E-11 | 0.896120435 | 1 | 0.962 | 8.15792E-07 | 0.006708153 | 0.410624603 | 1 | 0.993 | 1 | 0.006708153 | 6.71655E-11 |

| | | | | | | | | | | | | |
|---|---|---|---|---|---|---|---|---|---|---|---|---|
| Maml3 | 3.37836E-11 | 1.839220769 | 0.849 | 0.479 | 8.20671E-07 | 0.142638158 | 0.315017107 | 0.707 | 0.618 | 1 | 0.142638158 | 6.75671E-11 |
| Adam10 | 3.47752E-11 | 1.627272553 | 0.849 | 0.441 | 8.4476E-07 | 2.33812E-05 | 1.307737462 | 0.829 | 0.594 | 0.567975127 | 2.33812E-05 | 6.95505E-11 |
| Scin | 4.08004E-08 | -1.931623177 | 0.377 | 0.788 | 0.000991124 | 3.53727E-11 | -2.980956827 | 0.098 | 0.724 | 8.59274E-07 | 4.08004E-08 | 7.07454E-11 |
| Fgfr1 | 3.80947E-11 | 1.07733442 | 1 | 0.983 | 9.25397E-07 | 0.285718303 | -0.151691871 | 1 | 0.98 | 1 | 0.285718303 | 7.61894E-11 |
| Stk38l | 4.03321E-11 | -2.192296481 | 0.415 | 0.826 | 9.79747E-07 | 0.000122468 | -1.0509849 | 0.537 | 0.785 | 1 | 0.000122468 | 8.06642E-11 |
| Ptpn13 | 4.14114E-11 | 1.993259172 | 0.83 | 0.411 | 1.00597E-06 | 1.283E-07 | 2.250357705 | 0.805 | 0.549 | 0.003116655 | 1.283E-07 | 8.28228E-11 |
| Ankrd11 | 4.20874E-11 | 0.95790908 | 1 | 0.987 | 1.02239E-06 | 0.055243315 | 0.33280783 | 1 | 1 | 1 | 0.055243315 | 8.41747E-11 |
| Mtnr1a | 0.003243872 | 3.739685096 | 0.057 | 0.004 | 1 | 4.28564E-11 | 6.007129851 | 0.146 | 0 | 1.04107E-06 | 0.003243872 | 8.57128E-11 |
| Gm48898 | 2.25643E-05 | 4.47665069 | 0.075 | 0 | 0.548131677 | 4.28709E-11 | 6.422167351 | 0.146 | 0 | 1.04142E-06 | 2.25643E-05 | 8.57418E-11 |
| Mdfi | 4.29421E-11 | -1.775599758 | 0.509 | 0.886 | 1.04315E-06 | 1.20419E-05 | -1.186641892 | 0.463 | 0.785 | 0.292522822 | 1.20419E-05 | 8.58842E-11 |
| Trim33 | 4.35283E-11 | 1.52911811 | 0.887 | 0.564 | 1.05739E-06 | 0.69599101 | 0.237804966 | 0.78 | 0.843 | 1 | 0.69599101 | 8.70567E-11 |
| Ppp1r12a | 4.37391E-11 | 1.034367872 | 0.981 | 0.919 | 1.06251E-06 | 0.002706571 | 0.391780089 | 1 | 0.99 | 1 | 0.002706571 | 8.74782E-11 |
| Adgra2 | 4.40463E-11 | 2.530871081 | 0.528 | 0.144 | 1.06997E-06 | 0.009448981 | 0.989207943 | 0.537 | 0.352 | 1 | 0.009448981 | 8.80927E-11 |
| Ar | 4.49059E-11 | 3.795180208 | 0.302 | 0.03 | 1.09086E-06 | 0.003677438 | 1.828642836 | 0.39 | 0.222 | 1 | 0.003677438 | 8.98119E-11 |
| Has2 | 4.55107E-11 | -3.502736897 | 0.132 | 0.665 | 1.10555E-06 | 0.00303371 | -1.725439271 | 0.171 | 0.427 | 1 | 0.00303371 | 9.10214E-11 |
| Plat | 1.98958E-10 | 2.900149768 | 0.358 | 0.059 | 4.83308E-06 | 4.62476E-11 | 2.907594178 | 0.415 | 0.072 | 1.12345E-06 | 1.98958E-10 | 9.24951E-11 |
| Ssbp3 | 4.67042E-11 | 1.438866027 | 0.925 | 0.597 | 1.13454E-06 | 4.96154E-07 | 1.20438271 | 0.902 | 0.782 | 0.012052562 | 4.96154E-07 | 9.34083E-11 |
| Dnmt3a | 4.69014E-11 | 1.492163689 | 0.943 | 0.581 | 1.13933E-06 | 0.646376021 | 0.17386584 | 0.951 | 0.945 | 1 | 0.646376021 | 9.38028E-11 |
| S100a13 | 4.69968E-11 | -1.822557329 | 0.679 | 0.873 | 1.14165E-06 | 0.002484574 | -0.84522346 | 0.732 | 0.819 | 1 | 0.002484574 | 9.39937E-11 |
| Sqle | 4.76947E-11 | -2.871466929 | 0.264 | 0.72 | 1.1586E-06 | 0.001643746 | -1.454499323 | 0.268 | 0.502 | 1 | 0.001643746 | 9.53893E-11 |
| Cnmd | 1.57938E-10 | -1.06327181 | 0.887 | 0.983 | 3.83662E-06 | 4.89392E-11 | -1.642051853 | 0.78 | 0.942 | 1.18883E-06 | 1.57938E-10 | 9.78785E-11 |
| F5 | 5.00503E-11 | -7.657454711 | 0.019 | 0.534 | 1.21582E-06 | 0.03239195 | -5.766421495 | 0 | 0.102 | 1 | 0.03239195 | 1.00101E-10 |
| Rnf19a | 5.14024E-11 | 1.44606237 | 0.887 | 0.487 | 1.24867E-06 | 6.87827E-06 | 1.027482913 | 0.951 | 0.754 | 0.167087023 | 6.87827E-06 | 1.02805E-10 |
| Safb | 5.27264E-11 | 1.355731146 | 0.887 | 0.682 | 1.28083E-06 | 0.039405276 | 0.615477418 | 0.951 | 0.84 | 1 | 0.039405276 | 1.05453E-10 |
| Adk | 5.3588E-11 | -1.376315501 | 0.906 | 0.996 | 1.30176E-06 | 2.2784E-08 | -1.467699274 | 0.902 | 0.986 | 0.00055347 | 2.2784E-08 | 1.07176E-10 |
| Cd109 | 0.000897329 | -1.141762414 | 0.906 | 0.919 | 1 | 5.36923E-11 | -1.979032977 | 0.732 | 0.962 | 1.30429E-06 | 0.000897329 | 1.07385E-10 |
| Hnrnpl | 5.42369E-11 | 0.92808695 | 1 | 0.932 | 1.31752E-06 | 6.16926E-05 | 0.620755568 | 1 | 0.976 | 1 | 6.16926E-05 | 1.08474E-10 |
| Trpm7 | 5.53571E-11 | 1.088998413 | 1 | 0.877 | 1.34473E-06 | 0.308365254 | 0.325476151 | 1 | 0.976 | 1 | 0.308365254 | 1.10714E-10 |
| H3f3a | 5.61564E-11 | 0.82183853 | 1 | 0.975 | 1.36415E-06 | 0.045593041 | 0.359118614 | 1 | 0.993 | 1 | 0.045593041 | 1.12313E-10 |
| Kcns1 | 5.97215E-11 | 2.71862348 | 0.415 | 0.076 | 1.45076E-06 | 0.061095452 | 1.076392514 | 0.268 | 0.154 | 1 | 0.061095452 | 1.19443E-10 |
| Sos1 | 5.98755E-11 | 1.704061186 | 0.792 | 0.39 | 1.4545E-06 | 0.19399658 | 0.376300342 | 0.732 | 0.666 | 1 | 0.19399658 | 1.19751E-10 |
| Gypc | 6.01935E-11 | 2.602181572 | 0.472 | 0.106 | 1.46222E-06 | 0.020901042 | 1.45387621 | 0.268 | 0.137 | 1 | 0.020901042 | 1.20387E-10 |
| Asap1 | 6.22402E-11 | 1.62237472 | 0.906 | 0.64 | 1.51194E-06 | 0.010496613 | 0.600347067 | 0.976 | 0.87 | 1 | 0.010496613 | 1.2448E-10 |
| Npnt | 6.43592E-11 | 4.21000503 | 0.415 | 0.089 | 1.56341E-06 | 4.0065E-08 | 3.083365437 | 0.634 | 0.283 | 0.000973259 | 4.0065E-08 | 1.28718E-10 |
| Nbea | 6.73235E-11 | 1.80523086 | 0.849 | 0.483 | 1.63542E-06 | 0.115533707 | 0.3762665 | 0.878 | 0.795 | 1 | 0.115533707 | 1.34647E-10 |
| Snx4 | 6.74792E-11 | 1.405188327 | 0.868 | 0.623 | 1.63921E-06 | 0.069336745 | 0.552690717 | 0.878 | 0.799 | 1 | 0.069336745 | 1.34958E-10 |
| Qki | 6.89443E-11 | 0.853706582 | 1 | 0.992 | 1.6748E-06 | 0.002555716 | 0.450884173 | 1 | 1 | 1 | 0.002555716 | 1.37889E-10 |
| Myh9 | 7.29777E-11 | 1.313175463 | 0.943 | 0.733 | 1.77277E-06 | 9.71559E-05 | 0.717905921 | 0.951 | 0.904 | 1 | 9.71559E-05 | 1.45955E-10 |
| Pds5a | 7.58303E-11 | 1.341026146 | 0.906 | 0.653 | 1.84207E-06 | 0.020746876 | 0.519308509 | 0.902 | 0.911 | 1 | 0.020746876 | 1.51661E-10 |
| Morc4 | 7.9006E-11 | 2.502645898 | 0.528 | 0.144 | 1.91921E-06 | 0.00581052 | 0.962735732 | 0.39 | 0.195 | 1 | 0.00581052 | 1.58012E-10 |
| L3mbtl3 | 8.07458E-11 | 2.564113531 | 0.585 | 0.203 | 1.96148E-06 | 0.083945382 | 0.739907648 | 0.439 | 0.324 | 1 | 0.083945382 | 1.61492E-10 |
| Gtf2i | 8.31122E-11 | 1.315187267 | 0.943 | 0.627 | 2.01896E-06 | 0.018253404 | 0.559945804 | 0.951 | 0.863 | 1 | 0.018253404 | 1.66224E-10 |
| Mras | 8.43526E-11 | -1.556124346 | 0.623 | 0.903 | 2.04909E-06 | 8.01992E-06 | -1.220287484 | 0.537 | 0.846 | 0.194819887 | 8.01992E-06 | 1.68705E-10 |
| Plcg2 | 8.63965E-11 | 1.403401708 | 0.509 | 0.114 | 2.09874E-06 | 0.003484755 | 0.965144238 | 0.659 | 0.495 | 1 | 0.003484755 | 1.72793E-10 |
| Pcsk5 | 8.85921E-11 | 1.772149491 | 0.642 | 0.212 | 2.15208E-06 | 2.99487E-06 | 2.204413824 | 0.561 | 0.256 | 0.072751318 | 2.99487E-06 | 1.77184E-10 |
| AW112010 | 9.03633E-11 | -7.237594828 | 0.019 | 0.525 | 2.1951E-06 | 0.188554844 | -1.863234868 | 0 | 0.041 | 1 | 0.188554844 | 1.80727E-10 |
| Rasa1 | 9.17015E-11 | 1.553992778 | 0.849 | 0.551 | 2.22761E-06 | 0.312427483 | 0.429457386 | 0.829 | 0.823 | 1 | 0.312427483 | 1.83403E-10 |
| Apod | 0.020280153 | -0.264816297 | 0.132 | 0.047 | 1 | 9.38889E-11 | 4.565125304 | 0.317 | 0.041 | 2.28075E-06 | 0.020280153 | 1.87778E-10 |
| Atf2 | 9.41571E-11 | 1.247404597 | 0.943 | 0.699 | 2.28726E-06 | 0.001347815 | 0.869069348 | 0.878 | 0.867 | 1 | 0.001347815 | 1.88314E-10 |
| Dbn1 | 9.7141E-11 | 2.685237311 | 0.453 | 0.102 | 2.35975E-06 | 1.01836E-10 | 2.334704509 | 0.512 | 0.119 | 2.47379E-06 | 1.01836E-10 | 1.94282E-10 |
| Tmbim1 | 9.73625E-11 | -1.285849997 | 0.792 | 0.932 | 2.36513E-06 | 4.43905E-05 | -0.816619509 | 0.756 | 0.966 | 1 | 4.43905E-05 | 1.94725E-10 |
| Set | 9.78733E-11 | 0.925289188 | 1 | 0.928 | 2.37754E-06 | 0.215482591 | 0.18026677 | 1 | 0.983 | 1 | 0.215482591 | 1.95747E-10 |
| Pla2g5 | 9.86213E-11 | -3.014023252 | 0.17 | 0.661 | 2.39571E-06 | 2.37637E-05 | -2.118851803 | 0.268 | 0.601 | 0.577268524 | 2.37637E-05 | 1.97243E-10 |
| Rbms3 | 1.04378E-10 | 1.858856796 | 0.887 | 0.534 | 2.53554E-06 | 7.61029E-07 | 1.215509798 | 0.902 | 0.635 | 0.018486924 | 7.61029E-07 | 2.08755E-10 |
| Tiam2 | 1.08112E-10 | -1.973498161 | 0.925 | 0.975 | 2.62626E-06 | 0.262374533 | -0.507827396 | 0.878 | 0.956 | 1 | 0.262374533 | 2.16225E-10 |
| Stk32b | 1.08392E-10 | -2.699106757 | 0.302 | 0.742 | 2.63307E-06 | 4.27898E-06 | -2.339169403 | 0.244 | 0.614 | 0.103944879 | 4.27898E-06 | 2.16785E-10 |
| Basp1 | 1.08878E-10 | 3.920257341 | 0.302 | 0.034 | 2.64486E-06 | 1.01419E-09 | 2.697480086 | 0.463 | 0.109 | 2.46367E-05 | 1.01419E-09 | 2.17756E-10 |
| Fst | 1.10988E-10 | 2.839220769 | 0.34 | 0.047 | 2.69613E-06 | 4.16531E-05 | 6.819057503 | 0.098 | 0.007 | 1 | 4.16531E-05 | 2.21977E-10 |
| Dio2 | 1.11418E-10 | 4.869565378 | 0.415 | 0.081 | 2.70655E-06 | 7.33165E-09 | 2.852311742 | 0.488 | 0.143 | 0.0001781 | 7.33165E-09 | 2.22835E-10 |
| Ngef | 1.13919E-10 | -3.093204919 | 0.132 | 0.631 | 2.76731E-06 | 4.5312E-07 | -2.674907086 | 0.146 | 0.573 | 0.011007188 | 4.5312E-07 | 2.27838E-10 |
| Pde4dip | 1.18633E-10 | -1.484340115 | 0.83 | 0.97 | 2.88184E-06 | 8.85327E-09 | -1.367153649 | 0.902 | 0.98 | 0.000215064 | 8.85327E-09 | 2.37267E-10 |
| Nfatc1 | 2.6781E-10 | -1.442065341 | 0.849 | 0.958 | 6.50564E-06 | 1.1935E-10 | -1.4799231 | 0.927 | 0.99 | 2.89926E-06 | 2.6781E-10 | 2.387E-10 |
| Cacnb2 | 1.23043E-10 | 2.095753669 | 0.849 | 0.428 | 2.98897E-06 | 0.141441071 | 0.719429432 | 0.78 | 0.744 | 1 | 0.141441071 | 2.46087E-10 |
| Ddx6 | 1.26377E-10 | 0.940854246 | 1 | 0.97 | 3.06996E-06 | 0.000787477 | 0.476098921 | 1 | 0.993 | 1 | 0.000787477 | 2.52755E-10 |
| Mib1 | 1.26644E-10 | 1.247388843 | 1 | 0.682 | 3.07644E-06 | 0.000732995 | 0.714753866 | 1 | 0.911 | 1 | 0.000732995 | 2.53288E-10 |
| Tpm1 | 1.27047E-10 | 1.403305844 | 0.981 | 0.97 | 3.08623E-06 | 0.001716149 | 0.979532882 | 0.927 | 0.952 | 1 | 0.001716149 | 2.54095E-10 |
| Gfpt2 | 1.32962E-10 | -1.993551335 | 0.604 | 0.932 | 3.2299E-06 | 1.85247E-08 | -1.759471071 | 0.488 | 0.887 | 0.000450001 | 1.85247E-08 | 2.65923E-10 |
| Plod2 | 1.34361E-10 | -0.966006869 | 1 | 1 | 3.26389E-06 | 0.00032784 | -0.554585368 | 0.976 | 0.942 | 1 | 0.00032784 | 2.68722E-10 |
| Gabre | 1.34864E-10 | 4.690775495 | 0.245 | 0.017 | 3.27613E-06 | 0.931764638 | 0.515276755 | 0.098 | 0.102 | 1 | 0.931764638 | 2.69729E-10 |
| Gnb2 | 1.35876E-10 | 0.902180563 | 1 | 0.924 | 3.30069E-06 | 0.327843398 | 0.177593643 | 0.951 | 0.962 | 1 | 0.327843398 | 2.71752E-10 |
| Smurf2 | 1.4051E-10 | 1.017526977 | 1 | 0.894 | 3.41327E-06 | 0.000699589 | 0.747505114 | 1 | 0.98 | 1 | 0.000699589 | 2.8102E-10 |
| Sparc | 1.40715E-10 | -1.749062769 | 1 | 1 | 3.41825E-06 | 9.95919E-06 | -1.088087114 | 1 | 1 | 0.24192871 | 9.95919E-06 | 2.8143E-10 |
| Sim2 | 1.41001E-10 | -2.494892864 | 0.057 | 0.585 | 3.4252E-06 | 0.000109258 | -1.626319523 | 0.171 | 0.512 | 1 | 0.000109258 | 2.82002E-10 |
| Map3k12 | 1.45227E-10 | 2.720319771 | 0.415 | 0.081 | 3.52787E-06 | 0.780594808 | 0.500169863 | 0.195 | 0.235 | 1 | 0.780594808 | 2.90455E-10 |
| Scd1 | 1.49754E-10 | -3.737205732 | 0.245 | 0.695 | 3.63783E-06 | 0.008443116 | -1.386358115 | 0.268 | 0.481 | 1 | 0.008443116 | 2.99509E-10 |
| Ly6e | 1.7103E-10 | 1.344172334 | 0.962 | 0.547 | 4.15466E-06 | 0.0190681 | 0.447936449 | 0.976 | 0.928 | 1 | 0.0190681 | 3.4206E-10 |
| Usp32 | 1.71573E-10 | 1.482153665 | 0.906 | 0.555 | 4.16786E-06 | 0.024428986 | 0.692026511 | 0.951 | 0.836 | 1 | 0.024428986 | 3.43147E-10 |
| Megf6 | 1.77635E-10 | 3.891688189 | 0.245 | 0.017 | 4.3151E-06 | 8.66397E-05 | 2.289717055 | 0.195 | 0.041 | 1 | 8.66397E-05 | 3.55269E-10 |
| Cyba | 1.8498E-10 | 2.027965453 | 0.679 | 0.263 | 4.49353E-06 | 0.000679266 | 0.971222198 | 0.707 | 0.464 | 1 | 0.000679266 | 3.6996E-10 |
| Akap12 | 1.8646E-10 | 2.249049977 | 0.472 | 0.114 | 4.5295E-06 | 0.016317203 | 1.203568502 | 0.415 | 0.253 | 1 | 0.016317203 | 3.72921E-10 |
| Lsm14a | 1.88183E-10 | 1.436493563 | 0.906 | 0.568 | 4.57135E-06 | 0.111600393 | 0.291616587 | 0.854 | 0.901 | 1 | 0.111600393 | 3.76366E-10 |
| Setbp1 | 1.88501E-10 | 1.049987101 | 1 | 0.941 | 4.57908E-06 | 3.30754E-05 | 0.75480904 | 0.976 | 0.99 | 0.80346701 | 3.30754E-05 | 3.77003E-10 |
| Shisa2 | 1.89367E-10 | 2.406261362 | 0.547 | 0.165 | 4.60011E-06 | 0.264586521 | 0.307256451 | 0.512 | 0.437 | 1 | 0.264586521 | 3.78735E-10 |
| Nxn | 1.8965E-10 | 2.167719935 | 0.83 | 0.462 | 4.60699E-06 | 0.008111482 | 0.708730292 | 0.707 | 0.59 | 1 | 0.008111482 | 3.79301E-10 |
| Ltbp2 | 1.94321E-10 | 2.145733812 | 0.604 | 0.199 | 4.72044E-06 | 1.31941E-07 | 2.539405026 | 0.634 | 0.314 | 0.003205103 | 1.31941E-07 | 3.88642E-10 |
| Fzd1 | 1.95287E-10 | 1.9218378 | 0.698 | 0.297 | 4.74391E-06 | 0.002625542 | 1.609176894 | 0.512 | 0.334 | 1 | 0.002625542 | 3.90574E-10 |
| Lrrc8c | 1.9696E-10 | -1.807429775 | 0.528 | 0.86 | 4.78456E-06 | 1.37543E-08 | -1.943724786 | 0.366 | 0.833 | 0.000334118 | 1.37543E-08 | 3.9392E-10 |
| Stard13 | 1.98149E-10 | -1.981322925 | 0.792 | 0.928 | 4.81342E-06 | 0.003207373 | -0.572872687 | 0.732 | 0.922 | 1 | 0.003207373 | 3.96297E-10 |
| Zeb1 | 1.98641E-10 | 1.104939956 | 0.962 | 0.771 | 4.8254E-06 | 0.048975939 | 0.290965781 | 0.902 | 0.891 | 1 | 0.048975939 | 3.97283E-10 |
| Adcy7 | 2.01723E-10 | 4.657222935 | 0.283 | 0.03 | 4.90026E-06 | 0.080853361 | 1.01123425 | 0.146 | 0.068 | 1 | 0.080853361 | 4.03447E-10 |
| Plxdc2 | 2.04617E-10 | 1.635154512 | 0.925 | 0.763 | 4.97056E-06 | 0.002420703 | 0.906010365 | 0.976 | 0.915 | 1 | 0.002420703 | 4.09234E-10 |
| Cnot6l | 2.09999E-10 | 1.406454352 | 0.906 | 0.551 | 5.10131E-06 | 9.77376E-07 | 0.992112212 | 1 | 0.826 | 0.023742416 | 9.77376E-07 | 4.19999E-10 |
| Atxn1 | 2.13819E-10 | -1.580872702 | 0.774 | 0.975 | 5.1941E-06 | 0.001306285 | -0.953076559 | 0.878 | 0.956 | 1 | 0.001306285 | 4.27638E-10 |
| Steap1 | 2.16749E-10 | -4.764140642 | 0.038 | 0.53 | 5.26528E-06 | 0.057083636 | -1.849295677 | 0.073 | 0.188 | 1 | 0.057083636 | 4.33499E-10 |
| Arid1a | 2.19624E-10 | 1.511719482 | 0.925 | 0.602 | 5.3351E-06 | 0.000944584 | 0.646449353 | 0.902 | 0.874 | 1 | 0.000944584 | 4.39248E-10 |
| Shisal1 | 2.31688E-10 | 5.324647596 | 0.208 | 0.008 | 5.62817E-06 | 4.29016E-09 | 3.974708374 | 0.195 | 0.014 | 0.000104217 | 4.29016E-09 | 4.63376E-10 |
| Fbxo32 | 2.4227E-10 | 2.586438835 | 0.434 | 0.093 | 5.88522E-06 | 1.90146E-05 | 0.854482841 | 0.512 | 0.205 | 0.461902191 | 1.90146E-05 | 4.8454E-10 |
| Hsp90ab1 | 2.46433E-10 | 0.844674032 | 1 | 0.996 | 5.98635E-06 | 0.016876067 | 0.472474752 | 1 | 0.997 | 1 | 0.016876067 | 4.92866E-10 |

| | | | | | | | | | | | | |
|---|---|---|---|---|---|---|---|---|---|---|---|---|
| Stac2 | 2.49674E-10 | 3.039245377 | 0.321 | 0.042 | 6.06507E-06 | 0.333634169 | -0.250257991 | 0.195 | 0.273 | 1 | 0.333634169 | 4.99347E-10 |
| Tnfrsf21 | 2.52919E-10 | -3.440934507 | 0.151 | 0.631 | 6.14391E-06 | 2.04341E-06 | -1.487494994 | 0.488 | 0.816 | 0.049638477 | 2.04341E-06 | 5.05838E-10 |
| Prrc2c | 2.54765E-10 | 0.766955239 | 1 | 0.992 | 6.18874E-06 | 0.087235298 | 0.278252924 | 1 | 1 | 1 | 0.087235298 | 5.09529E-10 |
| Adamts5 | 2.59074E-10 | 3.877188619 | 0.302 | 0.038 | 6.29342E-06 | 8.74255E-09 | 3.136765132 | 0.366 | 0.075 | 0.000212374 | 8.74255E-09 | 5.18147E-10 |
| Ndnf | 2.68563E-10 | 5.242185436 | 0.208 | 0.008 | 6.52393E-06 | 2.45921E-06 | 4.83720485 | 0.146 | 0.014 | 0.059739067 | 2.45921E-06 | 5.37126E-10 |
| Chst5 | 2.74175E-10 | -4.394099314 | 0.094 | 0.572 | 6.66027E-06 | 0.038287497 | -1.16279515 | 0.024 | 0.143 | 1 | 0.038287497 | 5.48351E-10 |
| Plxnd1 | 2.79635E-10 | 3.237184755 | 0.34 | 0.051 | 6.7929E-06 | 5.32614E-07 | 1.744095445 | 0.585 | 0.242 | 0.012938268 | 5.32614E-07 | 5.5927E-10 |
| Satb2 | 2.84769E-10 | 4.010332685 | 0.321 | 0.042 | 6.9176E-06 | 0.000156428 | 3.18867722 | 0.22 | 0.058 | 1 | 0.000156428 | 5.69538E-10 |
| Hspg2 | 2.89104E-10 | -1.541511888 | 0.83 | 0.966 | 7.0229E-06 | 0.000177911 | -1.040060577 | 0.878 | 0.908 | 1 | 0.000177911 | 5.78207E-10 |
| Fzd2 | 2.9088E-10 | 1.8327945 | 0.717 | 0.288 | 7.06605E-06 | 0.014557712 | 1.179409469 | 0.634 | 0.512 | 1 | 0.014557712 | 5.81759E-10 |
| Yy1 | 3.06715E-10 | 1.154722595 | 0.962 | 0.729 | 7.45071E-06 | 0.002624675 | 0.734466744 | 0.951 | 0.922 | 1 | 0.002624675 | 6.13429E-10 |
| Rcor1 | 3.13073E-10 | 2.0734988 | 0.868 | 0.581 | 7.60516E-06 | 0.084542312 | 0.530278918 | 0.829 | 0.85 | 1 | 0.084542312 | 6.26145E-10 |
| Msx1 | 3.14798E-10 | 4.102255175 | 0.34 | 0.055 | 7.64708E-06 | 0.00033493 | 3.181853021 | 0.341 | 0.143 | 1 | 0.00033493 | 6.29597E-10 |
| Ago1 | 3.15219E-10 | 2.041511984 | 0.547 | 0.157 | 7.65731E-06 | 0.211892709 | 0.77191339 | 0.317 | 0.242 | 1 | 0.211892709 | 6.30439E-10 |
| S100a1 | 3.15973E-10 | -1.958078008 | 0.755 | 0.907 | 7.67561E-06 | 0.002173359 | -0.915773062 | 0.683 | 0.826 | 1 | 0.002173359 | 6.31945E-10 |
| Plscr3 | 3.23419E-10 | 2.031340179 | 0.604 | 0.208 | 7.85649E-06 | 0.712248994 | 0.632846351 | 0.244 | 0.3 | 1 | 0.712248994 | 6.46838E-10 |
| Baz2a | 3.23441E-10 | 1.981390992 | 0.774 | 0.369 | 7.85704E-06 | 0.002864673 | 1.006524821 | 0.805 | 0.747 | 1 | 0.002864673 | 6.46883E-10 |
| Farp1 | 3.27619E-10 | -1.210554407 | 0.943 | 1 | 7.95853E-06 | 4.29889E-10 | -1.234744992 | 0.902 | 1 | 1.04429E-05 | 4.29889E-10 | 6.55239E-10 |
| Pten | 3.37518E-10 | 1.159339223 | 0.981 | 0.89 | 8.19898E-06 | 0.000344696 | 0.662707119 | 1 | 0.962 | 1 | 0.000344696 | 6.75036E-10 |
| Il17d | 1.22538E-08 | -1.945300393 | 0.566 | 0.826 | 0.00029767 | 3.40254E-10 | -1.947648074 | 0.39 | 0.857 | 8.26545E-06 | 1.22538E-08 | 6.80508E-10 |
| Eln | 3.55227E-10 | 3.27905073 | 0.585 | 0.195 | 8.62919E-06 | 0.030323552 | 2.479794168 | 0.341 | 0.229 | 1 | 0.030323552 | 7.10455E-10 |
| Il1r1 | 3.5534E-10 | -3.236032624 | 0.189 | 0.644 | 8.63192E-06 | 0.00043472 | -2.256307036 | 0.049 | 0.321 | 1 | 0.00043472 | 7.1068E-10 |
| Tmod3 | 3.58871E-10 | 1.448867533 | 0.906 | 0.703 | 8.7177E-06 | 1.40322E-05 | 0.789615391 | 0.902 | 0.904 | 0.340869797 | 1.40322E-05 | 7.17742E-10 |
| Slmap | 3.75265E-10 | 1.291921785 | 0.943 | 0.737 | 9.11594E-06 | 0.039398529 | 0.540711802 | 0.927 | 0.891 | 1 | 0.039398529 | 7.5053E-10 |
| Ube2e3 | 3.80047E-10 | 1.039803156 | 0.962 | 0.78 | 9.2321E-06 | 0.721617183 | 0.184265555 | 0.854 | 0.85 | 1 | 0.721617183 | 7.60094E-10 |
| Syt11 | 3.8075E-10 | 3.454282877 | 0.302 | 0.038 | 9.24918E-06 | 0.129909575 | 0.944120054 | 0.171 | 0.096 | 1 | 0.129909575 | 7.615E-10 |
| Srsf2 | 3.85031E-10 | 0.847469655 | 1 | 0.945 | 9.35318E-06 | 0.160661995 | 0.194250626 | 1 | 0.983 | 1 | 0.160661995 | 7.70063E-10 |
| Nfib | 4.17016E-10 | 1.650250012 | 0.943 | 0.686 | 1.01302E-05 | 5.84369E-05 | 1.387224026 | 0.976 | 0.86 | 1 | 5.84369E-05 | 8.34033E-10 |
| Lgr5 | 0.209009104 | 1.739685096 | 0.038 | 0.013 | 1 | 4.17906E-10 | 5.866952193 | 0.195 | 0.01 | 1.01518E-05 | 0.209009104 | 8.35812E-10 |
| Pmvk | 4.28913E-10 | -3.03227592 | 0.208 | 0.653 | 1.04192E-05 | 0.000938819 | -2.410722664 | 0.146 | 0.389 | 1 | 0.000938819 | 8.57826E-10 |
| Atxn7l3 | 4.62583E-10 | 2.200526284 | 0.604 | 0.229 | 1.12371E-05 | 0.029600412 | 0.640168003 | 0.585 | 0.413 | 1 | 0.029600412 | 9.25167E-10 |
| Alcam | 4.69678E-10 | 1.817141236 | 0.642 | 0.216 | 1.14094E-05 | 0.000331547 | 0.978902935 | 0.805 | 0.573 | 1 | 0.000331547 | 9.39356E-10 |
| Lrch2 | 4.89828E-10 | 3.669295768 | 0.264 | 0.025 | 1.18989E-05 | 0.005991371 | 1.602739596 | 0.244 | 0.099 | 1 | 0.005991371 | 9.79655E-10 |
| Ccn2 | 5.19545E-10 | -1.646786576 | 0.962 | 0.983 | 1.26208E-05 | 0.643058372 | -0.625957006 | 0.927 | 0.86 | 1 | 0.643058372 | 1.03909E-09 |
| Hspa1b | 5.27098E-10 | 2.075288127 | 0.887 | 0.483 | 1.28043E-05 | 2.41284E-09 | 2.689066983 | 0.902 | 0.577 | 5.86126E-05 | 2.41284E-09 | 1.0542E-09 |
| Ctsc | 5.30039E-10 | 3.704919677 | 0.264 | 0.025 | 1.28757E-05 | 0.009309196 | 1.174239837 | 0.293 | 0.137 | 1 | 0.009309196 | 1.06008E-09 |
| Fgfr2 | 5.71291E-10 | 1.21386608 | 1 | 0.949 | 1.38778E-05 | 0.072166706 | 0.422034168 | 1 | 0.997 | 1 | 0.072166706 | 1.14258E-09 |
| Plekhh2 | 5.85621E-10 | -2.462971108 | 0.434 | 0.801 | 1.42259E-05 | 1.45649E-05 | -1.669532484 | 0.39 | 0.727 | 0.353809895 | 1.45649E-05 | 1.17124E-09 |
| Ltbp4 | 5.99463E-10 | 3.669295768 | 0.321 | 0.047 | 1.45622E-05 | 3.07989E-05 | 1.823908027 | 0.512 | 0.232 | 0.748166344 | 3.07989E-05 | 1.19893E-09 |
| Kdm5b | 6.13932E-10 | 1.07930645 | 0.981 | 0.856 | 1.49136E-05 | 0.835217903 | 0.186471653 | 1 | 0.993 | 1 | 0.835217903 | 1.22786E-09 |
| Cfh | 1.01604E-05 | 2.652222254 | 0.208 | 0.038 | 0.246816252 | 6.19665E-10 | 3.771616508 | 0.317 | 0.044 | 1.50529E-05 | 1.01604E-05 | 1.23933E-09 |
| Enpp2 | 6.2141E-10 | -2.344008679 | 0.698 | 0.949 | 1.50953E-05 | 0.093992135 | -0.884852537 | 0.512 | 0.672 | 1 | 0.093992135 | 1.24282E-09 |
| Phf12 | 6.35804E-10 | 1.479963424 | 0.792 | 0.445 | 1.5445E-05 | 0.07490197 | 0.346006185 | 0.829 | 0.788 | 1 | 0.07490197 | 1.27161E-09 |
| Usp34 | 6.35871E-10 | 0.855624 | 1 | 0.953 | 1.54466E-05 | 0.003585544 | 0.383149607 | 1 | 0.997 | 1 | 0.003585544 | 1.27174E-09 |
| Prkcb | 6.46509E-10 | 3.457285365 | 0.264 | 0.025 | 1.5705E-05 | 0.000220267 | 2.742047617 | 0.244 | 0.068 | 1 | 0.000220267 | 1.29302E-09 |
| Ccnl2 | 6.50719E-10 | 0.905246294 | 0.981 | 0.886 | 1.58073E-05 | 0.262571869 | 0.155578048 | 1 | 0.956 | 1 | 0.262571869 | 1.30144E-09 |
| Macroh2a2 | 6.6096E-10 | 2.174622152 | 0.585 | 0.208 | 1.6056E-05 | 0.002461319 | 0.991714799 | 0.488 | 0.256 | 1 | 0.002461319 | 1.32192E-09 |
| Grik5 | 6.71872E-10 | 2.47665069 | 0.358 | 0.059 | 1.63211E-05 | 0.004764074 | 1.183200705 | 0.415 | 0.232 | 1 | 0.004764074 | 1.34374E-09 |
| 2810001G20Rik | 6.90341E-10 | -1.80412008 | 0.415 | 0.805 | 1.67698E-05 | 0.014578895 | -0.780179129 | 0.366 | 0.563 | 1 | 0.014578895 | 1.38068E-09 |
| Cttnbp2 | 7.02145E-10 | 2.977844833 | 0.321 | 0.047 | 1.70565E-05 | 0.000757007 | 1.377773231 | 0.366 | 0.154 | 1 | 0.000757007 | 1.40429E-09 |
| Tnks2 | 7.08583E-10 | 1.139925593 | 1 | 0.928 | 1.72129E-05 | 0.059775901 | 0.376092748 | 0.976 | 0.983 | 1 | 0.059775901 | 1.41717E-09 |
| Wac | 7.21985E-10 | 0.929328935 | 1 | 0.941 | 1.75384E-05 | 0.354394698 | 0.109678952 | 0.976 | 0.986 | 1 | 0.354394698 | 1.44397E-09 |
| Simc1 | 7.22574E-10 | 2.088488896 | 0.679 | 0.297 | 1.75528E-05 | 0.007507003 | 0.832061547 | 0.634 | 0.427 | 1 | 0.007507003 | 1.44515E-09 |
| Birc6 | 7.24731E-10 | 0.990533963 | 0.981 | 0.949 | 1.76052E-05 | 0.083106764 | 0.214735699 | 1 | 0.993 | 1 | 0.083106764 | 1.44946E-09 |
| Wdfy3 | 7.49807E-10 | 1.307621494 | 0.962 | 0.674 | 1.82143E-05 | 0.013234514 | 0.47463477 | 0.951 | 0.898 | 1 | 0.013234514 | 1.49961E-09 |
| Chd7 | 7.68149E-10 | 1.199116714 | 0.849 | 0.483 | 1.86599E-05 | 0.074768711 | 0.349758855 | 0.78 | 0.768 | 1 | 0.074768711 | 1.5363E-09 |
| Mtus1 | 1.17541E-05 | -1.81114472 | 0.887 | 0.907 | 0.285530801 | 7.91989E-10 | -1.542774238 | 0.829 | 0.976 | 1.9239E-05 | 1.17541E-05 | 1.58398E-09 |
| Enox1 | 8.19513E-10 | 2.524672205 | 0.623 | 0.254 | 1.99076E-05 | 0.002962093 | 1.407060458 | 0.561 | 0.392 | 1 | 0.002962093 | 1.63903E-09 |
| Abi2 | 8.23743E-10 | 1.622708974 | 0.792 | 0.47 | 2.00104E-05 | 0.245277721 | 0.552637282 | 0.707 | 0.724 | 1 | 0.245277721 | 1.64749E-09 |
| Serbp1 | 8.4944E-10 | 0.847942516 | 1 | 0.97 | 2.06346E-05 | 0.556253353 | 0.144182603 | 1 | 0.976 | 1 | 0.556253353 | 1.69888E-09 |
| Wbp1l | 8.51593E-10 | 1.873702443 | 0.679 | 0.267 | 2.06869E-05 | 0.322177752 | 0.259262777 | 0.683 | 0.679 | 1 | 0.322177752 | 1.70319E-09 |
| Brd1 | 8.5584E-10 | 1.157066535 | 0.962 | 0.847 | 2.07901E-05 | 0.338835479 | 0.120830109 | 0.976 | 0.935 | 1 | 0.338835479 | 1.71168E-09 |
| Slc44a2 | 8.64771E-10 | -1.435240587 | 0.679 | 0.886 | 2.1007E-05 | 3.766E-05 | -1.221688839 | 0.61 | 0.785 | 0.914837857 | 3.766E-05 | 1.72954E-09 |
| Shprh | 8.80771E-10 | 2.135093788 | 0.566 | 0.191 | 2.13957E-05 | 0.490119504 | 0.360157688 | 0.39 | 0.358 | 1 | 0.490119504 | 1.76154E-09 |
| Plag1 | 8.88867E-10 | 2.925240749 | 0.321 | 0.047 | 2.15924E-05 | 0.002123045 | 2.08997092 | 0.244 | 0.089 | 1 | 0.002123045 | 1.77773E-09 |
| Pds5b | 8.91435E-10 | 1.275647377 | 0.887 | 0.695 | 2.16547E-05 | 0.658251528 | 0.229367883 | 0.878 | 0.86 | 1 | 0.658251528 | 1.78287E-09 |
| Anxa11 | 4.75466E-07 | -1.391314394 | 0.566 | 0.839 | 0.011550032 | 8.9541E-10 | -1.89476406 | 0.561 | 0.898 | 2.17513E-05 | 4.75466E-07 | 1.79082E-09 |
| Dhx9 | 9.01E-10 | 1.158484514 | 0.962 | 0.763 | 2.18871E-05 | 0.142407174 | 0.336984549 | 0.902 | 0.915 | 1 | 0.142407174 | 1.802E-09 |
| Ankrd10 | 9.19472E-10 | 1.353822695 | 0.83 | 0.492 | 2.23358E-05 | 0.391414924 | 0.191635584 | 0.878 | 0.768 | 1 | 0.391414924 | 1.83894E-09 |
| Ttc23 | 9.29055E-10 | 2.402650108 | 0.509 | 0.148 | 2.25686E-05 | 0.196685789 | 0.776663308 | 0.341 | 0.273 | 1 | 0.196685789 | 1.85811E-09 |
| Rbpj | 9.62597E-10 | 1.612896597 | 0.906 | 0.606 | 2.33834E-05 | 0.000701468 | 1.042879876 | 0.829 | 0.713 | 1 | 0.000701468 | 1.92519E-09 |
| Hnrnpd | 9.9598E-10 | 0.999328912 | 0.981 | 0.822 | 2.41943E-05 | 0.097285659 | 0.347056426 | 0.976 | 0.939 | 1 | 0.097285659 | 1.99196E-09 |
| Flnc | 1.05138E-09 | 3.669295768 | 0.283 | 0.034 | 2.55401E-05 | 1.72326E-09 | 3.87282876 | 0.341 | 0.058 | 4.18615E-05 | 1.72326E-09 | 2.10276E-09 |
| Niban2 | 8.9166E-07 | -1.160919237 | 0.717 | 0.911 | 0.021660209 | 1.07086E-09 | -1.447309283 | 0.78 | 0.959 | 2.60133E-05 | 8.9166E-07 | 2.14171E-09 |
| Olfml2b | 1.41476E-08 | 3.33529484 | 0.302 | 0.051 | 0.000343673 | 1.07535E-09 | 4.266048149 | 0.268 | 0.031 | 2.61223E-05 | 1.41476E-08 | 2.1507E-09 |
| Syne1 | 1.07808E-09 | 1.021581331 | 1 | 0.801 | 2.61887E-05 | 0.3172556 | -0.209199995 | 1 | 0.98 | 1 | 0.3172556 | 2.15616E-09 |
| Nsd1 | 1.09199E-09 | 1.216879071 | 0.887 | 0.75 | 2.65266E-05 | 0.211842329 | 0.232250874 | 0.951 | 0.952 | 1 | 0.211842329 | 2.18397E-09 |
| Gata3 | 0.008804509 | 1.695290976 | 0.094 | 0.021 | 1 | 1.09689E-09 | 6.574170444 | 0.171 | 0.007 | 2.66457E-05 | 0.008804509 | 2.19378E-09 |
| Fat3 | 1.12111E-09 | 3.105812994 | 0.396 | 0.081 | 2.7234E-05 | 0.858846083 | 0.858578501 | 0.171 | 0.164 | 1 | 0.858846083 | 2.24222E-09 |
| Nfic | 1.16973E-09 | 1.038655197 | 0.981 | 0.835 | 2.84151E-05 | 0.121560175 | 0.262013179 | 1 | 0.959 | 1 | 0.121560175 | 2.33946E-09 |
| Eif4e3 | 1.1908E-09 | 2.621848605 | 0.434 | 0.106 | 2.8927E-05 | 0.203220192 | 0.790911198 | 0.268 | 0.191 | 1 | 0.203220192 | 2.38161E-09 |
| Ash1l | 1.2061E-09 | 0.724628521 | 1 | 0.97 | 2.92986E-05 | 0.07859396 | 0.273565454 | 1 | 0.997 | 1 | 0.07859396 | 2.4122E-09 |
| Epb41l2 | 1.21164E-09 | 1.280253477 | 0.981 | 0.746 | 2.94332E-05 | 0.003264373 | 0.742172401 | 0.927 | 0.881 | 1 | 0.003264373 | 2.42328E-09 |
| A830082K12Rik | 1.2301E-09 | 2.762405172 | 0.34 | 0.055 | 2.98815E-05 | 0.113512004 | 1.199774929 | 0.171 | 0.096 | 1 | 0.113512004 | 2.4602E-09 |
| Nrbp1 | 1.24227E-09 | 1.426395919 | 0.849 | 0.525 | 3.01773E-05 | 0.744264052 | 0.195098442 | 0.756 | 0.734 | 1 | 0.744264052 | 2.48455E-09 |
| Adamts1 | 1.27247E-09 | 1.07818104 | 0.585 | 0.174 | 3.09109E-05 | 3.61525E-08 | 2.235754226 | 0.585 | 0.225 | 0.000878216 | 3.61525E-08 | 2.54494E-09 |
| Dlg2 | 1.28628E-09 | -2.201120258 | 0.472 | 0.839 | 3.12463E-05 | 0.000714623 | -1.522337537 | 0.268 | 0.522 | 1 | 0.000714623 | 2.57256E-09 |
| Col11a2 | 1.30212E-09 | -1.040565066 | 0.774 | 0.987 | 3.1631E-05 | 3.88366E-08 | -1.089858403 | 0.659 | 0.973 | 0.00094342 | 3.88366E-08 | 2.60424E-09 |
| Hey1 | 1.32282E-09 | 2.991223863 | 0.34 | 0.055 | 3.2134E-05 | 2.31755E-09 | 2.83720485 | 0.39 | 0.078 | 5.62978E-05 | 2.31755E-09 | 2.64564E-09 |
| Cldn11 | 0.001643111 | -2.211183197 | 0.132 | 0.347 | 1 | 1.33327E-09 | 5.072421311 | 0.317 | 0.048 | 3.23879E-05 | 0.001643111 | 2.66655E-09 |
| Phlpp1 | 1.35314E-09 | 1.220098604 | 0.849 | 0.364 | 3.28705E-05 | 0.000187961 | 1.475278687 | 0.829 | 0.669 | 1 | 0.000187961 | 2.70628E-09 |
| Col10a1 | 1.37787E-09 | -9.758765246 | 0.132 | 0.564 | 3.34712E-05 | 0.10504439 | -6.438454095 | 0.024 | 0.102 | 1 | 0.10504439 | 2.75574E-09 |
| Efnb2 | 1.38643E-09 | 1.83846325 | 0.642 | 0.229 | 3.36792E-05 | 0.000167077 | 1.420119522 | 0.634 | 0.365 | 1 | 0.000167077 | 2.77286E-09 |
| Zfp618 | 1.38679E-09 | 1.991223863 | 0.491 | 0.131 | 3.36879E-05 | 0.000659174 | 1.089012 | 0.561 | 0.324 | 1 | 0.000659174 | 2.77358E-09 |
| Negr1 | 1.39822E-09 | 2.549001534 | 0.34 | 0.055 | 3.39657E-05 | 0.441366869 | 0.295311071 | 0.268 | 0.208 | 1 | 0.441366869 | 2.79645E-09 |
| Susd5 | 0.00117772 | -0.556028481 | 0.83 | 0.949 | 1 | 1.42779E-09 | -1.418202629 | 0.659 | 0.904 | 3.46838E-05 | 0.00117772 | 2.85557E-09 |
| Cert1 | 1.43522E-09 | 1.310640738 | 0.962 | 0.631 | 3.48643E-05 | 0.000114303 | 0.784910342 | 0.878 | 0.775 | 1 | 0.000114303 | 2.87043E-09 |
| Slc43a2 | 1.47446E-09 | 2.365142374 | 0.585 | 0.186 | 3.58176E-05 | 0.107108091 | 0.722872174 | 0.512 | 0.42 | 1 | 0.107108091 | 2.94892E-09 |

| | | | | | | | | | | | | |
|---|---|---|---|---|---|---|---|---|---|---|---|---|
| Atp10a | 1.50343E-09 | -1.718199065 | 0.547 | 0.886 | 3.65212E-05 | 0.005755949 | -0.785512573 | 0.561 | 0.778 | 1 | 0.005755949 | 3.00685E-09 |
| Crabp1 | 1.55175E-09 | 5.739685096 | 0.151 | 0 | 3.76951E-05 | 0.007718533 | 4.83720485 | 0.024 | 0 | 1 | 0.007718533 | 3.1035E-09 |
| 6330403K07Rik | 1.55363E-09 | 5.739685096 | 0.151 | 0 | 3.77407E-05 | 1.11097E-05 | 4.215716473 | 0.122 | 0.01 | 0.269877415 | 1.11097E-05 | 3.10726E-09 |
| Insc | 1.55587E-09 | 7.01270359 | 0.151 | 0 | 3.77951E-05 | 0.262674569 | 4.059597271 | 0.024 | 0.007 | 1 | 0.262674569 | 3.11173E-09 |
| Rnf10 | 1.56298E-09 | 0.864292653 | 1 | 0.881 | 3.7968E-05 | 0.039793356 | 0.328593235 | 0.951 | 0.973 | 1 | 0.039793356 | 3.12596E-09 |
| Hlf | 1.60065E-09 | 3.182736971 | 0.453 | 0.119 | 3.8883E-05 | 3.80211E-06 | 1.353882611 | 0.634 | 0.276 | 0.092360895 | 3.80211E-06 | 3.2013E-09 |
| Usf3 | 1.6173E-09 | 2.017219071 | 0.623 | 0.242 | 3.92875E-05 | 0.01667436 | 0.777222745 | 0.634 | 0.433 | 1 | 0.01667436 | 3.2346E-09 |
| Isg20 | 1.62124E-09 | -4.104549892 | 0.057 | 0.521 | 3.93832E-05 | 0.003218967 | -2.944154864 | 0.024 | 0.218 | 1 | 0.003218967 | 3.24248E-09 |
| Clint1 | 1.73738E-09 | -0.825686677 | 1 | 0.996 | 4.22045E-05 | 1.64036E-07 | -0.620682954 | 1 | 1 | 0.003984762 | 1.64036E-07 | 3.47477E-09 |
| Tgfb3 | 1.73969E-09 | 1.700156731 | 0.717 | 0.309 | 4.22605E-05 | 0.26508492 | 0.512274267 | 0.61 | 0.601 | 1 | 0.26508492 | 3.47938E-09 |
| Eml4 | 1.77677E-09 | 1.73180867 | 0.906 | 0.665 | 4.31613E-05 | 0.03542554 | 0.321657212 | 0.951 | 0.853 | 1 | 0.03542554 | 3.55354E-09 |
| Zbtb7a | 1.79237E-09 | 1.131390091 | 0.925 | 0.627 | 4.35403E-05 | 0.080317829 | 0.264092067 | 0.902 | 0.922 | 1 | 0.080317829 | 3.58475E-09 |
| Sox5 | 2.99608E-05 | -0.741867923 | 0.962 | 1 | 0.727808586 | 1.82493E-09 | -1.182291326 | 0.951 | 0.997 | 4.43312E-05 | 2.99608E-05 | 3.64986E-09 |
| Purg | 1.92154E-09 | 2.10496956 | 0.566 | 0.195 | 4.6678E-05 | 0.415844187 | 0.535085236 | 0.415 | 0.399 | 1 | 0.415844187 | 3.84307E-09 |
| Mgat4a | 1.98129E-09 | -1.621710438 | 0.566 | 0.86 | 4.81294E-05 | 0.000284854 | -0.89177576 | 0.61 | 0.805 | 1 | 0.000284854 | 3.96258E-09 |
| Cdkl4 | 2.01062E-09 | 3.517292674 | 0.226 | 0.017 | 4.88421E-05 | 6.36868E-06 | 3.159132945 | 0.195 | 0.031 | 0.154708072 | 6.36868E-06 | 4.02125E-09 |
| B2m | 2.03029E-09 | 1.131243528 | 1 | 0.911 | 4.93198E-05 | 0.091060405 | 0.423513776 | 0.976 | 0.983 | 1 | 0.091060405 | 4.06058E-09 |
| Arfgef2 | 2.10897E-09 | -1.642689524 | 0.377 | 0.839 | 5.12311E-05 | 4.67272E-05 | -1.123819183 | 0.756 | 0.84 | 1 | 4.67272E-05 | 4.21794E-09 |
| Cnr1 | 0.442404005 | 0.377115016 | 0.057 | 0.034 | 1 | 2.20836E-09 | 7.194756854 | 0.146 | 0.003 | 5.36455E-05 | 0.442404005 | 4.41672E-09 |
| Cd1d1 | 2.91495E-08 | 4.47665069 | 0.151 | 0.004 | 0.0007081 | 2.20847E-09 | 5.422167351 | 0.146 | 0.003 | 5.36481E-05 | 2.91495E-08 | 4.41694E-09 |
| Zfp292 | 2.21092E-09 | 1.710159143 | 0.755 | 0.36 | 5.37078E-05 | 0.005306979 | 0.706808213 | 0.805 | 0.659 | 1 | 0.005306979 | 4.42185E-09 |
| Neto2 | 2.23333E-09 | 2.320732546 | 0.358 | 0.064 | 5.42522E-05 | 1.17517E-05 | 3.088743617 | 0.268 | 0.065 | 0.285473091 | 1.17517E-05 | 4.46667E-09 |
| Cpe | 0.00040576 | -0.797345156 | 0.943 | 0.936 | 1 | 2.25931E-09 | -1.501852406 | 0.902 | 0.962 | 5.48831E-05 | 0.00040576 | 4.51861E-09 |
| Sncaip | 2.31653E-09 | 2.65128998 | 0.472 | 0.127 | 5.62732E-05 | 0.01031514 | 1.949315216 | 0.341 | 0.188 | 1 | 0.01031514 | 4.63307E-09 |
| Lypd6 | 2.40707E-09 | 4.002719501 | 0.208 | 0.013 | 5.84725E-05 | 0.607129262 | 2.489281546 | 0.049 | 0.034 | 1 | 0.607129262 | 4.81414E-09 |
| Anxa3 | 2.45064E-09 | -2.307686724 | 0.208 | 0.653 | 5.9531E-05 | 0.236597911 | -0.490482514 | 0.317 | 0.416 | 1 | 0.236597911 | 4.90129E-09 |
| Syde2 | 2.45918E-09 | 4.06161319 | 0.208 | 0.013 | 5.97383E-05 | 0.008368886 | 1.875678998 | 0.195 | 0.072 | 1 | 0.008368886 | 4.91835E-09 |
| Sfpq | 2.4752E-09 | 0.716884755 | 1 | 0.966 | 6.01275E-05 | 0.703033966 | 0.154061955 | 1 | 0.997 | 1 | 0.703033966 | 4.9504E-09 |
| Eya2 | 2.49517E-09 | 2.117108303 | 0.717 | 0.364 | 6.06126E-05 | 0.000126279 | 1.29753246 | 0.902 | 0.826 | 1 | 0.000126279 | 4.99034E-09 |
| Itpr1 | 2.52032E-09 | 1.54246136 | 0.962 | 0.839 | 6.12237E-05 | 0.002901247 | 0.518216532 | 1 | 0.973 | 1 | 0.002901247 | 5.04064E-09 |
| Adcy5 | 2.56654E-09 | 5.242185436 | 0.17 | 0.004 | 6.23464E-05 | 0.001468631 | 2.952682067 | 0.146 | 0.034 | 1 | 0.001468631 | 5.13308E-09 |
| Nlgn2 | 2.62483E-09 | 1.753360032 | 0.585 | 0.199 | 6.37624E-05 | 0.021216464 | 0.989207943 | 0.512 | 0.355 | 1 | 0.021216464 | 5.24966E-09 |
| Mkx | 2.65219E-09 | 1.8327945 | 0.415 | 0.097 | 6.4427E-05 | 0.000655186 | 2.120997816 | 0.244 | 0.078 | 1 | 0.000655186 | 5.30438E-09 |
| Rps6kb1 | 2.65384E-09 | 1.264173737 | 0.906 | 0.72 | 6.44671E-05 | 0.002804072 | 0.490554326 | 0.976 | 0.898 | 1 | 0.002804072 | 5.30769E-09 |
| Arid2 | 2.66655E-09 | 1.433967228 | 0.83 | 0.513 | 6.47758E-05 | 0.042372443 | 0.304024891 | 0.902 | 0.85 | 1 | 0.042372443 | 5.3331E-09 |
| Elf2 | 2.67792E-09 | 1.394127254 | 0.943 | 0.801 | 6.5052E-05 | 1.4383E-05 | 0.85560943 | 0.976 | 0.945 | 0.349390713 | 1.4383E-05 | 5.35584E-09 |
| Proser2 | 2.74101E-09 | -1.987457416 | 0.283 | 0.746 | 6.65846E-05 | 1.19316E-07 | -1.853110651 | 0.341 | 0.751 | 0.002898431 | 1.19316E-07 | 5.48202E-09 |
| Elovl4 | 2.75768E-09 | 4.47665069 | 0.17 | 0.004 | 6.69896E-05 | 0.121203683 | 1.306690133 | 0.146 | 0.075 | 1 | 0.121203683 | 5.51537E-09 |
| Cbl | 2.75924E-09 | 1.573947891 | 0.774 | 0.394 | 6.70275E-05 | 0.067646184 | 0.418120517 | 0.78 | 0.696 | 1 | 0.067646184 | 5.51848E-09 |
| Rab30 | 2.81742E-09 | 2.569760094 | 0.509 | 0.148 | 6.84408E-05 | 0.056447679 | 0.56635094 | 0.341 | 0.212 | 1 | 0.056447679 | 5.63484E-09 |
| Tent5c | 2.83878E-09 | 2.874614676 | 0.34 | 0.059 | 6.89597E-05 | 1.25138E-08 | 2.410940095 | 0.366 | 0.072 | 0.000303985 | 1.25138E-08 | 5.67756E-09 |
| Gatad2b | 2.85639E-09 | 0.856169008 | 0.981 | 0.822 | 6.93874E-05 | 0.645250064 | 0.102244392 | 0.976 | 0.962 | 1 | 0.645250064 | 5.71278E-09 |
| Clock | 2.89642E-09 | 1.333692736 | 0.943 | 0.657 | 7.03599E-05 | 0.014924714 | 0.677054844 | 0.878 | 0.867 | 1 | 0.014924714 | 5.79285E-09 |
| Fam171b | 3.03097E-09 | 2.285967128 | 0.396 | 0.081 | 7.36284E-05 | 0.052484661 | 1.270164257 | 0.268 | 0.16 | 1 | 0.052484661 | 6.06194E-09 |
| Cdk19 | 3.05266E-09 | 1.425370185 | 0.755 | 0.364 | 7.41552E-05 | 0.001611027 | 1.347620397 | 0.634 | 0.423 | 1 | 0.001611027 | 6.10532E-09 |
| Gm40841 | 3.10942E-09 | 4.402650108 | 0.189 | 0.008 | 7.5534E-05 | 0.000156969 | 4.83720485 | 0.049 | 0 | 1 | 0.000156969 | 6.21883E-09 |
| Hpcal1 | 3.14871E-09 | 2.55911285 | 0.415 | 0.097 | 7.64884E-05 | 0.008948779 | 1.275325962 | 0.293 | 0.137 | 1 | 0.008948779 | 6.29742E-09 |
| Zswim6 | 0.001639283 | 0.700690964 | 0.925 | 0.886 | 1 | 3.18079E-09 | 1.593951069 | 1 | 0.891 | 7.72678E-05 | 0.001639283 | 6.36158E-09 |
| Zfp608 | 3.22544E-09 | -1.492939463 | 0.792 | 0.941 | 7.83524E-05 | 0.000159587 | -1.293560134 | 0.805 | 0.898 | 1 | 0.000159587 | 6.45088E-09 |
| Eeig1 | 3.29169E-09 | -2.880272827 | 0.17 | 0.619 | 7.99617E-05 | 0.773032406 | -0.208309809 | 0.512 | 0.447 | 1 | 0.773032406 | 6.58338E-09 |
| Stx5a | 3.37521E-09 | 1.227889007 | 0.906 | 0.682 | 8.19906E-05 | 0.029527747 | 0.758196975 | 0.902 | 0.826 | 1 | 0.029527747 | 6.75042E-09 |
| Gm11906 | 3.58585E-09 | 3.977844833 | 0.245 | 0.025 | 8.71075E-05 | 0.204687296 | 1.100239256 | 0.122 | 0.068 | 1 | 0.204687296 | 7.1717E-09 |
| Cygb | 3.71476E-09 | 3.209170379 | 0.283 | 0.038 | 9.0239E-05 | 0.139569276 | -2.020776145 | 0 | 0.051 | 1 | 0.139569276 | 7.42953E-09 |
| Pals2 | 3.73221E-09 | -1.586386388 | 0.774 | 0.919 | 9.06628E-05 | 8.41709E-06 | -1.072236931 | 0.756 | 0.932 | 0.204468065 | 8.41709E-06 | 7.46442E-09 |
| Vcp | 3.77543E-09 | 0.803094266 | 1 | 0.966 | 9.17128E-05 | 0.008702047 | 0.455432897 | 1 | 0.993 | 1 | 0.008702047 | 7.55086E-09 |
| Mllt10 | 3.79707E-09 | 1.094107312 | 0.962 | 0.733 | 9.22383E-05 | 0.015762104 | 0.467794098 | 1 | 0.935 | 1 | 0.015762104 | 7.59413E-09 |
| Kif5b | 3.82672E-09 | 0.866707135 | 0.981 | 0.966 | 9.29587E-05 | 0.431114878 | 0.202863982 | 1 | 0.997 | 1 | 0.431114878 | 7.65344E-09 |
| Cask | 4.11089E-09 | 1.333059836 | 0.925 | 0.729 | 9.98618E-05 | 6.85054E-06 | 1.009844236 | 0.927 | 0.857 | 0.166413284 | 6.85054E-06 | 8.22178E-09 |
| Prss23 | 4.2069E-09 | 2.673189684 | 0.472 | 0.144 | 0.000102194 | 2.05277E-05 | 2.43132735 | 0.585 | 0.28 | 0.498658843 | 2.05277E-05 | 8.4138E-09 |
| Frk | 4.4068E-09 | 1.928446739 | 0.434 | 0.106 | 0.00010705 | 0.0202861 | 1.384032222 | 0.366 | 0.212 | 1 | 0.0202861 | 8.81361E-09 |
| Nufip2 | 4.51016E-09 | 0.939399072 | 1 | 0.983 | 0.000109561 | 0.623429478 | 0.100370894 | 1 | 1 | 1 | 0.623429478 | 9.02031E-09 |
| Prkar1a | 4.5455E-09 | 0.950893778 | 0.962 | 0.881 | 0.000110419 | 0.391059719 | 0.153415355 | 0.951 | 0.956 | 1 | 0.391059719 | 9.091E-09 |
| Rbm3 | 4.56213E-09 | 0.774996411 | 1 | 0.945 | 0.000110823 | 0.07803269 | 0.333505511 | 0.976 | 0.993 | 1 | 0.07803269 | 9.12425E-09 |
| Mbtd1 | 4.59432E-09 | 1.408906083 | 0.868 | 0.585 | 0.000111605 | 0.000231436 | 0.74827957 | 0.976 | 0.846 | 1 | 0.000231436 | 9.18864E-09 |
| Zbtb2 | 4.60187E-09 | 1.968856049 | 0.566 | 0.191 | 0.000111789 | 3.78596E-05 | 1.593750813 | 0.585 | 0.317 | 0.919685331 | 3.78596E-05 | 9.20374E-09 |
| Abhd12 | 4.64684E-09 | 1.757651642 | 0.736 | 0.436 | 0.000112881 | 0.245858043 | 0.47463477 | 0.537 | 0.471 | 1 | 0.245858043 | 9.29368E-09 |
| Tgif2 | 4.66567E-09 | 2.363309217 | 0.472 | 0.14 | 0.000113338 | 0.100052398 | 1.074704164 | 0.317 | 0.212 | 1 | 0.100052398 | 9.33133E-09 |
| Tle1 | 4.6969E-09 | 1.43820039 | 0.811 | 0.547 | 0.000114097 | 0.000136133 | 1.081346163 | 0.927 | 0.703 | 1 | 0.000136133 | 9.3938E-09 |
| Fbln5 | 4.7291E-09 | 2.836193076 | 0.472 | 0.144 | 0.000114879 | 6.60542E-05 | 3.309957847 | 0.317 | 0.109 | 1 | 6.60542E-05 | 9.4582E-09 |
| Brwd1 | 4.87368E-09 | 1.174622152 | 0.925 | 0.725 | 0.000118391 | 7.92922E-06 | 1.111379813 | 0.976 | 0.898 | 0.19261657 | 7.92922E-06 | 9.74736E-09 |
| Olfml2a | 0.015350156 | 2.739685096 | 0.057 | 0.008 | 1 | 4.87379E-09 | 4.059597271 | 0.195 | 0.014 | 0.000118394 | 0.015350156 | 9.74759E-09 |
| Cobll1 | 4.90907E-09 | -2.484570507 | 0.453 | 0.754 | 0.000119251 | 0.014441869 | -0.813846841 | 0.463 | 0.669 | 1 | 0.014441869 | 9.81815E-09 |
| Slc43a1 | 4.95103E-09 | 2.820303556 | 0.377 | 0.081 | 0.00012027 | 0.900600623 | -0.123624553 | 0.195 | 0.201 | 1 | 0.900600623 | 9.90206E-09 |
| Atp8a1 | 0.000302781 | 2.53659323 | 0.226 | 0.068 | 1 | 4.98104E-09 | 2.152078187 | 0.585 | 0.188 | 0.000120999 | 0.000302781 | 9.96208E-09 |
| Rev3l | 8.91661E-08 | 1.381111404 | 0.887 | 0.636 | 0.002166023 | 5.36442E-09 | 1.285374589 | 0.976 | 0.843 | 0.000130313 | 8.91661E-08 | 1.07288E-08 |
| Smoc1 | 0.537071855 | -0.317765177 | 0.189 | 0.153 | 1 | 5.5174E-09 | 4.322631677 | 0.293 | 0.044 | 0.000134029 | 0.537071855 | 1.10348E-08 |
| Ywhab | 5.52803E-09 | 0.823692293 | 0.981 | 0.945 | 0.000134287 | 0.331618684 | 0.140959016 | 0.976 | 0.976 | 1 | 0.331618684 | 1.10561E-08 |
| Usp7 | 5.53649E-09 | 1.059199984 | 0.906 | 0.737 | 0.000134492 | 0.583405616 | 0.137520272 | 0.854 | 0.939 | 1 | 0.583405616 | 1.1073E-08 |
| Polg2 | 5.65551E-09 | 2.624207878 | 0.472 | 0.14 | 0.000137384 | 0.004331022 | 0.946829341 | 0.463 | 0.256 | 1 | 0.004331022 | 1.1311E-08 |
| Adamtsl1 | 3.32696E-07 | 5.458503343 | 0.264 | 0.051 | 0.00808186 | 5.81164E-09 | 4.448639562 | 0.317 | 0.055 | 0.000141176 | 3.32696E-07 | 1.16233E-08 |
| Cemip2 | 5.9109E-09 | 1.827035231 | 0.849 | 0.669 | 0.000143588 | 3.82904E-08 | 2.398859977 | 0.854 | 0.594 | 0.000930149 | 3.82904E-08 | 1.18218E-08 |
| Zfp91 | 5.94682E-09 | 1.066064975 | 0.962 | 0.758 | 0.00014446 | 0.340406774 | 0.229305538 | 0.902 | 0.915 | 1 | 0.340406774 | 1.18936E-08 |
| Ndufa4l2 | 5.99308E-09 | -2.528729769 | 0.075 | 0.542 | 0.000145584 | 0.510746787 | -1.278272368 | 0.049 | 0.075 | 1 | 0.510746787 | 1.19862E-08 |
| Pogz | 6.13849E-09 | 1.693588681 | 0.66 | 0.288 | 0.000149116 | 0.287289515 | 0.515276755 | 0.585 | 0.529 | 1 | 0.287289515 | 1.2277E-08 |
| Tbl1x | 6.16791E-09 | 1.027551354 | 0.943 | 0.847 | 0.000149831 | 0.024231662 | 0.438905905 | 0.976 | 0.966 | 1 | 0.024231662 | 1.23358E-08 |
| Jun | 6.1771E-09 | 1.457302564 | 0.981 | 0.941 | 0.000150054 | 0.000116064 | 0.85953241 | 0.976 | 0.986 | 1 | 0.000116064 | 1.23542E-08 |
| Prdx6 | 6.47236E-09 | -1.397818428 | 0.736 | 0.941 | 0.000157227 | 0.061979174 | -0.29011398 | 0.829 | 0.904 | 1 | 0.061979174 | 1.29447E-08 |
| Fdps | 6.51113E-09 | -3.005148742 | 0.226 | 0.631 | 0.000158168 | 0.21095889 | -1.027250479 | 0.512 | 0.543 | 1 | 0.21095889 | 1.30223E-08 |
| Orai1 | 6.52445E-09 | -2.172964769 | 0.226 | 0.678 | 0.000158492 | 0.047685959 | -0.78589278 | 0.366 | 0.509 | 1 | 0.047685959 | 1.30489E-08 |
| Ranbp9 | 6.54627E-09 | 1.221836791 | 0.943 | 0.657 | 0.000159022 | 0.092648657 | 0.384918838 | 0.927 | 0.881 | 1 | 0.092648657 | 1.30925E-08 |
| Shf | 6.59947E-09 | -2.079938664 | 0.34 | 0.699 | 0.000160314 | 0.802927723 | -0.120566915 | 0.244 | 0.256 | 1 | 0.802927723 | 1.31989E-08 |
| Pde4d | 6.70209E-09 | -1.755438493 | 0.849 | 0.928 | 0.000162807 | 2.14033E-05 | -1.562828363 | 0.732 | 0.935 | 0.519928534 | 2.14033E-05 | 1.34042E-08 |
| Pdlim2 | 6.79535E-09 | 2.811834881 | 0.396 | 0.093 | 0.000165073 | 3.89082E-06 | 2.073645046 | 0.463 | 0.174 | 0.094515799 | 3.89082E-06 | 1.35907E-08 |
| Pakap | 6.84078E-09 | 2.332720897 | 0.547 | 0.191 | 0.000166176 | 0.002553405 | 2.591627641 | 0.366 | 0.195 | 1 | 0.002553405 | 1.36816E-08 |
| Abracl | 6.89934E-09 | 1.592843707 | 0.717 | 0.356 | 0.000167599 | 0.004446755 | 0.790911198 | 0.634 | 0.413 | 1 | 0.004446755 | 1.37987E-08 |
| Chd1 | 6.96335E-09 | 0.965987744 | 1 | 0.86 | 0.000169154 | 0.19643785 | 0.282450085 | 0.927 | 0.894 | 1 | 0.19643785 | 1.39267E-08 |
| Rgs2 | 8.45586E-07 | -4.091705495 | 0.377 | 0.636 | 0.020540977 | 7.05374E-09 | 2.51902889 | 0.585 | 0.205 | 0.00017135 | 8.45586E-07 | 1.41075E-08 |
| Dysf | 7.13934E-09 | -3.813368157 | 0.038 | 0.487 | 0.000173429 | 0.017494166 | -1.678494988 | 0.122 | 0.294 | 1 | 0.017494166 | 1.42787E-08 |

| | | | | | | | | | | | | |
|---|---|---|---|---|---|---|---|---|---|---|---|---|
| Phf21a | 7.15152E-09 | 1.294172638 | 0.981 | 0.797 | 0.000173725 | 0.005899488 | 0.50722312 | 0.927 | 0.935 | 1 | 0.005899488 | 1.4303E-08 |
| Tjp1 | 7.16248E-09 | 1.183722467 | 0.925 | 0.644 | 0.000173991 | 0.000181591 | 0.778945452 | 0.951 | 0.805 | 1 | 0.000181591 | 1.4325E-08 |
| Thap3 | 7.18289E-09 | 2.095828906 | 0.509 | 0.148 | 0.000174487 | 0.022566684 | 1.322631677 | 0.317 | 0.184 | 1 | 0.022566684 | 1.43658E-08 |
| Arid4b | 7.32723E-09 | 0.85097116 | 1 | 0.992 | 0.000177993 | 0.011596644 | 0.558833121 | 0.976 | 0.993 | 1 | 0.011596644 | 1.46545E-08 |
| ENSMUSG00000121322 | 7.33844E-09 | -3.678167419 | 0.038 | 0.475 | 0.000178265 | 0.001061073 | -2.196218152 | 0.049 | 0.294 | 1 | 0.001061073 | 1.46769E-08 |
| Olfml3 | 7.33902E-09 | 1.185011217 | 0.906 | 0.648 | 0.000178279 | 0.002810508 | 0.857897245 | 0.732 | 0.573 | 1 | 0.002810508 | 1.4678E-08 |
| Ccnd2 | 7.42653E-09 | 1.062061859 | 0.943 | 0.75 | 0.000180405 | 0.077039123 | 0.624150293 | 0.927 | 0.898 | 1 | 0.077039123 | 1.48531E-08 |
| Phf3 | 7.60617E-09 | 0.944302815 | 0.981 | 0.839 | 0.000184769 | 0.369678582 | 0.216132056 | 0.951 | 0.949 | 1 | 0.369678582 | 1.52123E-08 |
| Syncrip | 7.8425E-09 | 0.882326086 | 1 | 0.907 | 0.00019051 | 0.020167558 | 0.327624161 | 1 | 0.983 | 1 | 0.020167558 | 1.5685E-08 |
| Ppic | 7.92138E-09 | 0.867306282 | 1 | 0.953 | 0.000192426 | 0.299330571 | -0.189084829 | 0.927 | 0.922 | 1 | 0.299330571 | 1.58428E-08 |
| Ier5l | 1.54951E-07 | 2.275016829 | 0.434 | 0.136 | 0.003764078 | 7.92175E-09 | 2.581947795 | 0.488 | 0.137 | 0.000192435 | 1.54951E-07 | 1.58435E-08 |
| Ica1 | 7.94474E-09 | -2.086285505 | 0.283 | 0.742 | 0.000192994 | 0.003734302 | -1.484723245 | 0.244 | 0.481 | 1 | 0.003734302 | 1.58895E-08 |
| Afap1 | 7.94979E-09 | 2.023478062 | 0.604 | 0.25 | 0.000193116 | 0.000163189 | 0.97780318 | 0.756 | 0.471 | 1 | 0.000163189 | 1.58996E-08 |
| Cbx6 | 8.02482E-09 | 1.660613525 | 0.66 | 0.271 | 0.000194939 | 0.000353396 | 0.909460939 | 0.829 | 0.631 | 1 | 0.000353396 | 1.60496E-08 |
| Prkci | 8.13062E-09 | 1.817687608 | 0.604 | 0.229 | 0.000197509 | 0.007353961 | 0.883008539 | 0.683 | 0.539 | 1 | 0.007353961 | 1.62612E-08 |
| Adarb1 | 8.19186E-09 | 2.378845904 | 0.585 | 0.225 | 0.000198997 | 1.58187E-06 | 2.257260424 | 0.634 | 0.334 | 0.038426886 | 1.58187E-06 | 1.63837E-08 |
| Pgm2l1 | 8.27848E-09 | 3.502645898 | 0.264 | 0.034 | 0.000201101 | 0.010188958 | 1.608935862 | 0.341 | 0.177 | 1 | 0.010188958 | 1.6557E-08 |
| Matr3 | 8.49082E-09 | 0.714943238 | 1 | 0.958 | 0.000206259 | 0.027770183 | 0.321283994 | 1 | 0.986 | 1 | 0.027770183 | 1.69816E-08 |
| Cdk4 | 8.56632E-09 | 1.460531024 | 0.792 | 0.597 | 0.000208093 | 0.202555366 | 0.374232873 | 0.732 | 0.696 | 1 | 0.202555366 | 1.71326E-08 |
| Asxl2 | 8.88306E-09 | 1.420012975 | 0.811 | 0.449 | 0.000215787 | 8.15138E-06 | 0.944490023 | 0.927 | 0.655 | 0.198013391 | 8.15138E-06 | 1.77661E-08 |
| Hnrnpll | 8.89738E-09 | 1.326783341 | 0.868 | 0.581 | 0.000216135 | 0.000355764 | 1.169931996 | 0.927 | 0.768 | 1 | 0.000355764 | 1.77948E-08 |
| Kansl2 | 9.01142E-09 | 1.610402079 | 0.679 | 0.331 | 0.000218905 | 0.406939267 | 0.233383322 | 0.537 | 0.481 | 1 | 0.406939267 | 1.80228E-08 |
| Gnao1 | 9.02834E-09 | 2.94321849 | 0.34 | 0.064 | 0.000219316 | 4.96029E-06 | 2.198086579 | 0.439 | 0.164 | 0.120495457 | 4.96029E-06 | 1.80567E-08 |
| Dennd5a | 9.05507E-09 | 1.145952385 | 0.906 | 0.648 | 0.000219966 | 0.016434715 | 0.571743707 | 0.976 | 0.925 | 1 | 0.016434715 | 1.81101E-08 |
| Atpif1 | 9.46082E-09 | 1.052938857 | 0.962 | 0.754 | 0.000229822 | 0.080478895 | 0.401105735 | 0.78 | 0.7 | 1 | 0.080478895 | 1.89216E-08 |
| Ptn | 9.54291E-09 | 2.377115016 | 0.642 | 0.275 | 0.000231816 | 0.20384157 | -0.391475446 | 0.512 | 0.406 | 1 | 0.20384157 | 1.90858E-08 |
| Camk2n1 | 9.76214E-09 | 1.622227514 | 0.698 | 0.347 | 0.000237142 | 0.051674222 | 0.534427855 | 0.756 | 0.686 | 1 | 0.051674222 | 1.95243E-08 |
| Prmt1 | 1.00905E-08 | 1.421368254 | 0.849 | 0.614 | 0.000245117 | 0.365681883 | 0.376231066 | 0.732 | 0.727 | 1 | 0.365681883 | 2.01809E-08 |
| Dcbld1 | 1.02331E-08 | 3.154722595 | 0.264 | 0.034 | 0.000248582 | 0.730207335 | -0.19841906 | 0.073 | 0.089 | 1 | 0.730207335 | 2.04661E-08 |
| Cul7 | 1.0253E-08 | 1.994257923 | 0.547 | 0.191 | 0.000249065 | 0.099695415 | 0.532870815 | 0.488 | 0.362 | 1 | 0.099695415 | 2.05059E-08 |
| Eed | 1.02702E-08 | 1.407109756 | 0.717 | 0.339 | 0.000249484 | 0.027084063 | 0.604544093 | 0.707 | 0.532 | 1 | 0.027084063 | 2.05404E-08 |
| Kcnk2 | 1.07411E-08 | 2.984797593 | 0.472 | 0.144 | 0.000260922 | 0.21529053 | 1.002427232 | 0.39 | 0.345 | 1 | 0.21529053 | 2.14821E-08 |
| Sh3d19 | 0.04496882 | -0.328767029 | 0.981 | 0.966 | 1 | 1.10601E-08 | -1.527367582 | 0.902 | 0.973 | 0.000268672 | 0.04496882 | 2.21202E-08 |
| Dlx5 | 1.13833E-08 | 2.723006354 | 0.415 | 0.102 | 0.000276522 | 0.018964078 | 1.675741427 | 0.268 | 0.143 | 1 | 0.018964078 | 2.27665E-08 |
| Psme4 | 1.14962E-08 | 1.149639695 | 0.962 | 0.886 | 0.000279267 | 0.235934877 | 0.212933056 | 1 | 0.986 | 1 | 0.235934877 | 2.29925E-08 |
| Gapvd1 | 1.17447E-08 | 0.962077517 | 0.962 | 0.729 | 0.000285302 | 0.763130409 | 0.148988387 | 0.878 | 0.898 | 1 | 0.763130409 | 2.34894E-08 |
| Gnai1 | 1.18114E-08 | 2.466666601 | 0.566 | 0.208 | 0.000286922 | 0.022862322 | 0.702045267 | 0.683 | 0.515 | 1 | 0.022862322 | 2.36228E-08 |
| Sp3 | 1.18912E-08 | 1.147149375 | 0.925 | 0.708 | 0.000288862 | 0.220849607 | 0.155256734 | 0.902 | 0.908 | 1 | 0.220849607 | 2.37824E-08 |
| Gm12703 | 1.19787E-08 | 2.880547631 | 0.34 | 0.068 | 0.000290987 | 0.758609712 | 1.808058504 | 0.22 | 0.212 | 1 | 0.758609712 | 2.39574E-08 |
| Kctd4 | 1.21215E-08 | -3.075575025 | 0.132 | 0.576 | 0.000294455 | 0.000391309 | -1.138869311 | 0.146 | 0.447 | 1 | 0.000391309 | 2.4243E-08 |
| Ep300 | 1.21386E-08 | 1.258673559 | 0.868 | 0.657 | 0.00029487 | 0.009838125 | 0.530624719 | 1 | 0.901 | 1 | 0.009838125 | 2.42772E-08 |
| Zkscan17 | 1.2185E-08 | 2.154722595 | 0.434 | 0.114 | 0.000295997 | 0.889131879 | 0.156483366 | 0.268 | 0.3 | 1 | 0.889131879 | 2.43699E-08 |
| Slc7a7 | 1.22425E-08 | 1.927951733 | 0.472 | 0.136 | 0.000297394 | 0.467751963 | 0.331676817 | 0.317 | 0.27 | 1 | 0.467751963 | 2.4485E-08 |
| Kank1 | 1.22484E-08 | -1.323967901 | 0.604 | 0.881 | 0.000297539 | 0.000349324 | -1.299333983 | 0.683 | 0.816 | 1 | 0.000349324 | 2.44969E-08 |
| Itm2a | 1.23349E-08 | 1.651822383 | 0.925 | 0.936 | 0.00029964 | 0.461023218 | -0.208240523 | 0.829 | 0.85 | 1 | 0.461023218 | 2.46698E-08 |
| C1qtnf1 | 1.25759E-08 | 1.845772354 | 0.604 | 0.246 | 0.000305494 | 0.654735669 | 0.142617858 | 0.415 | 0.396 | 1 | 0.654735669 | 2.51518E-08 |
| Gdf11 | 4.13747E-06 | 2.502645898 | 0.208 | 0.034 | 0.10050748 | 1.29361E-08 | 2.997669522 | 0.293 | 0.044 | 0.000314245 | 4.13747E-06 | 2.58723E-08 |
| Elmo1 | 1.33155E-08 | 1.813441545 | 0.811 | 0.521 | 0.00032346 | 0.001413025 | 1.189506594 | 0.561 | 0.317 | 1 | 0.001413025 | 2.6631E-08 |
| Lamc3 | 1.34898E-08 | 2.510866405 | 0.321 | 0.055 | 0.000327694 | 0.029627631 | 1.328191202 | 0.195 | 0.089 | 1 | 0.029627631 | 2.69796E-08 |
| Shank3 | 1.37408E-08 | 3.8327945 | 0.208 | 0.017 | 0.000333791 | 2.83791E-06 | 3.537644568 | 0.171 | 0.02 | 0.068938448 | 2.83791E-06 | 2.74816E-08 |
| Elovl6 | 1.39368E-08 | -1.459332817 | 0.453 | 0.831 | 0.000338552 | 0.471762962 | 0.254830144 | 0.61 | 0.553 | 1 | 0.471762962 | 2.78735E-08 |
| Hells | 1.39717E-08 | 2.556821038 | 0.358 | 0.076 | 0.000339401 | 0.119553235 | 1.050608488 | 0.195 | 0.113 | 1 | 0.119553235 | 2.79434E-08 |
| 1700016P03Rik | 8.52708E-05 | 1.707263618 | 0.17 | 0.03 | 1 | 1.42076E-08 | 5.159132945 | 0.171 | 0.01 | 0.000345131 | 8.52708E-05 | 2.84152E-08 |
| A2m | 1.47012E-08 | -4.490575569 | 0.038 | 0.475 | 0.000357122 | 0.057979938 | -3.717384002 | 0 | 0.082 | 1 | 0.057979938 | 2.94024E-08 |
| Sema4c | 4.24313E-08 | 2.711115943 | 0.302 | 0.055 | 0.00103074 | 1.47092E-08 | 2.515276755 | 0.415 | 0.099 | 0.000357317 | 4.24313E-08 | 2.94185E-08 |
| Chsy1 | 1.4805E-08 | 1.231763631 | 0.83 | 0.525 | 0.000359642 | 0.014107628 | 0.816005684 | 0.756 | 0.72 | 1 | 0.014107628 | 2.96099E-08 |
| Ppfia1 | 1.48115E-08 | 1.325523284 | 0.774 | 0.441 | 0.000359801 | 0.799253345 | 0.255632938 | 0.707 | 0.737 | 1 | 0.799253345 | 2.9623E-08 |
| Itga6 | 1.48772E-08 | -1.918526387 | 0.453 | 0.754 | 0.000361398 | 0.025113473 | -0.785056192 | 0.439 | 0.608 | 1 | 0.025113473 | 2.97545E-08 |
| Timp3 | 1.49358E-08 | -3.459353408 | 1 | 0.941 | 0.000362821 | 0.720366628 | -0.617262794 | 0.902 | 0.935 | 1 | 0.720366628 | 2.98717E-08 |
| Nkd2 | 1.50347E-08 | 3.614154213 | 0.245 | 0.03 | 0.000365224 | 8.33911E-06 | 3.775804305 | 0.171 | 0.024 | 0.202573689 | 8.33911E-06 | 3.00695E-08 |
| Cyp26a1 | 1.52038E-08 | 4.756758609 | 0.208 | 0.017 | 0.00036933 | 0.029009389 | 0.773074512 | 0.341 | 0.198 | 1 | 0.029009389 | 3.04075E-08 |
| Prcp | 1.52362E-08 | 3.06161319 | 0.321 | 0.059 | 0.000370117 | 0.004447756 | 2.007129851 | 0.268 | 0.119 | 1 | 0.004447756 | 3.04723E-08 |
| Camk4 | 1.53614E-08 | 1.681652974 | 0.717 | 0.343 | 0.000373159 | 0.212059577 | 0.491912762 | 0.659 | 0.611 | 1 | 0.212059577 | 3.07228E-08 |
| Cd300a | 1.53682E-08 | 3.817687608 | 0.208 | 0.017 | 0.000373325 | 0.899790943 | 0.292884334 | 0.073 | 0.068 | 1 | 0.899790943 | 3.07365E-08 |
| Hnrnpul1 | 1.53805E-08 | 1.298558367 | 0.849 | 0.602 | 0.000373622 | 0.078043124 | 0.538177157 | 0.878 | 0.867 | 1 | 0.078043124 | 3.07609E-08 |
| Ifitm3 | 1.55431E-08 | 0.612668686 | 0.585 | 0.195 | 0.000377574 | 0.000526792 | 2.416363718 | 0.488 | 0.276 | 1 | 0.000526792 | 3.10863E-08 |
| Bmp1 | 1.56253E-08 | 1.828494362 | 0.717 | 0.449 | 0.000379571 | 0.003924218 | 1.245388678 | 0.561 | 0.454 | 1 | 0.003924218 | 3.12507E-08 |
| Ccn6 | 1.60372E-08 | -3.791696365 | 0.038 | 0.462 | 0.000389575 | 7.48238E-06 | -3.686357106 | 0.024 | 0.382 | 0.181762047 | 7.48238E-06 | 3.20744E-08 |
| Aggf1 | 1.71347E-08 | 1.482608145 | 0.736 | 0.39 | 0.000416235 | 0.028036297 | 0.842851413 | 0.634 | 0.509 | 1 | 0.028036297 | 3.42693E-08 |
| Kcnj2 | 1.71734E-08 | 7.769432439 | 0.132 | 0 | 0.000417177 | 3.96704E-05 | 4.83720485 | 0.098 | 0.007 | 0.963672977 | 3.96704E-05 | 3.43468E-08 |
| Fgf14 | 1.71823E-08 | 5.855162313 | 0.132 | 0 | 0.000417392 | 0.11782599 | 1.962735732 | 0.073 | 0.027 | 1 | 0.11782599 | 3.43646E-08 |
| Nipbl | 1.75808E-08 | 0.878365962 | 1 | 0.975 | 0.000427073 | 0.395706891 | 0.23271224 | 1 | 0.997 | 1 | 0.395706891 | 3.51616E-08 |
| Ehbp1 | 1.80565E-08 | 1.468499849 | 0.868 | 0.678 | 0.000438628 | 2.04308E-06 | 1.035117166 | 0.951 | 0.86 | 0.049630446 | 2.04308E-06 | 3.6113E-08 |
| Bmp6 | 1.81865E-08 | -3.01155463 | 0.453 | 0.814 | 0.000441786 | 0.040606453 | -1.507494553 | 0.22 | 0.389 | 1 | 0.040606453 | 3.63729E-08 |
| Ube2h | 1.82774E-08 | 1.071485714 | 0.962 | 0.89 | 0.000443995 | 3.42218E-05 | 0.740814149 | 0.976 | 0.976 | 0.831314929 | 3.42218E-05 | 3.65549E-08 |
| Tax1bp3 | 1.83475E-08 | 1.860991392 | 0.698 | 0.36 | 0.000445697 | 0.671956733 | 0.259331774 | 0.488 | 0.457 | 1 | 0.671956733 | 3.6695E-08 |
| Zfand5 | 1.84487E-08 | 0.894829606 | 1 | 0.975 | 0.000448157 | 3.46392E-07 | 0.845274866 | 1 | 0.99 | 0.008414563 | 3.46392E-07 | 3.68975E-08 |
| Col24a1 | 1.86326E-08 | 4.999071724 | 0.321 | 0.064 | 0.000452623 | 0.32901212 | 1.377773231 | 0.122 | 0.082 | 1 | 0.32901212 | 3.72652E-08 |
| H2az2 | 1.89584E-08 | 0.91206727 | 0.962 | 0.792 | 0.000460538 | 0.001765626 | 0.54950839 | 0.927 | 0.87 | 1 | 0.001765626 | 3.79168E-08 |
| Psmc3 | 1.96459E-08 | 0.968749652 | 0.943 | 0.712 | 0.000477239 | 0.912659924 | 0.213972332 | 0.78 | 0.829 | 1 | 0.912659924 | 3.92918E-08 |
| Cyp26b1 | 2.01858E-08 | -2.596048799 | 0.132 | 0.581 | 0.000490353 | 0.147851899 | -0.791735348 | 0.341 | 0.457 | 1 | 0.147851899 | 4.03715E-08 |
| Nono | 2.07063E-08 | 1.129890742 | 0.906 | 0.631 | 0.000502997 | 0.603643322 | 0.118386602 | 0.829 | 0.826 | 1 | 0.603643322 | 4.14125E-08 |
| Otor | 2.07066E-08 | -3.279014755 | 0.057 | 0.5 | 0.000503004 | 0.002026602 | -3.314308115 | 0.098 | 0.331 | 1 | 0.002026602 | 4.14132E-08 |
| Sap30l | 2.12678E-08 | 1.650956177 | 0.642 | 0.297 | 0.000516637 | 0.906881226 | 0.165587967 | 0.415 | 0.447 | 1 | 0.906881226 | 4.25355E-08 |
| Pkig | 2.13978E-08 | 1.678881816 | 0.792 | 0.496 | 0.000519794 | 0.025253036 | 0.676930018 | 0.756 | 0.659 | 1 | 0.025253036 | 4.27955E-08 |
| Ell2 | 2.1528E-08 | -1.167968629 | 0.698 | 0.928 | 0.000522958 | 0.012762394 | -0.768931674 | 0.805 | 0.884 | 1 | 0.012762394 | 4.3056E-08 |
| Cdk11b | 2.15669E-08 | 0.957122635 | 0.943 | 0.805 | 0.000523903 | 0.015071193 | 0.492584169 | 0.951 | 0.928 | 1 | 0.015071193 | 4.31338E-08 |
| Lamc1 | 2.19408E-08 | 1.601568832 | 0.717 | 0.326 | 0.000532985 | 0.00013672 | 1.209681887 | 0.805 | 0.631 | 1 | 0.00013672 | 4.38815E-08 |
| Edil3 | 2.31949E-08 | 1.016324099 | 0.925 | 0.682 | 0.000563451 | 0.094810688 | 0.788666852 | 0.829 | 0.823 | 1 | 0.094810688 | 4.63898E-08 |
| Col4a1 | 2.39171E-08 | 2.440124814 | 0.547 | 0.208 | 0.000580994 | 0.011616709 | 2.392902756 | 0.463 | 0.352 | 1 | 0.011616709 | 4.78341E-08 |
| Wnk4 | 2.41885E-08 | -3.076841472 | 0.094 | 0.517 | 0.000587588 | 0.025796506 | -1.432255825 | 0.122 | 0.28 | 1 | 0.025796506 | 4.83771E-08 |
| Ctdspl2 | 2.4364E-08 | 1.326488943 | 0.811 | 0.564 | 0.00059185 | 0.404230681 | 0.350806184 | 0.78 | 0.805 | 1 | 0.404230681 | 4.8728E-08 |
| Kat6b | 2.43859E-08 | 0.965202113 | 0.962 | 0.686 | 0.000592383 | 0.395436758 | -0.31620983 | 0.878 | 0.911 | 1 | 0.395436758 | 4.87719E-08 |
| Hsd11b1 | 2.45824E-08 | -3.965947292 | 0.132 | 0.534 | 0.000597155 | 0.666353766 | -0.555112573 | 0.024 | 0.038 | 1 | 0.666353766 | 4.91648E-08 |
| Dchs1 | 2.48493E-08 | 2.361173472 | 0.377 | 0.085 | 0.00060364 | 0.038293578 | 1.036108918 | 0.415 | 0.294 | 1 | 0.038293578 | 4.96987E-08 |
| Pcdh19 | 2.52199E-08 | 5.614154213 | 0.17 | 0.008 | 0.000612642 | 0.045365236 | 1.654340793 | 0.293 | 0.177 | 1 | 0.045365236 | 5.04398E-08 |
| Hnrnpul2 | 2.57504E-08 | 1.052575708 | 0.943 | 0.725 | 0.00062553 | 0.003341374 | 0.635570989 | 0.927 | 0.904 | 1 | 0.003341374 | 5.15009E-08 |
| Svep1 | 2.66153E-08 | 1.227478937 | 0.66 | 0.246 | 0.000646538 | 0.500294382 | 0.252242349 | 0.634 | 0.59 | 1 | 0.500294382 | 5.32305E-08 |

| | | | | | | | | | | | | |
|---|---|---|---|---|---|---|---|---|---|---|---|---|
| Tanc2 | 2.66616E-08 | 1.27810501 | 0.849 | 0.555 | 0.000647664 | 0.008078151 | 0.404679924 | 0.854 | 0.696 | 1 | 0.008078151 | 5.33232E-08 |
| Rb1 | 2.68643E-08 | 1.560541564 | 0.66 | 0.305 | 0.000652587 | 3.01818E-06 | 1.30412012 | 0.854 | 0.604 | 0.073317673 | 3.01818E-06 | 5.37286E-08 |
| Dync1i2 | 2.69554E-08 | 1.124560624 | 0.962 | 0.758 | 0.000654801 | 0.092978575 | 0.308030127 | 0.902 | 0.87 | 1 | 0.092978575 | 5.39108E-08 |
| Luzp2 | 2.72164E-08 | -5.552636537 | 0.019 | 0.436 | 0.000661142 | 0.002414256 | -3.199884469 | 0.049 | 0.266 | 1 | 0.002414256 | 5.44329E-08 |
| Cxcl12 | 2.76076E-08 | 2.680184084 | 0.396 | 0.106 | 0.000670643 | 8.96564E-08 | 3.465236072 | 0.341 | 0.075 | 0.002177934 | 8.96564E-08 | 5.52151E-08 |
| Syf2 | 2.77759E-08 | 1.361173472 | 0.849 | 0.508 | 0.000674733 | 7.33237E-05 | 1.181650381 | 0.805 | 0.662 | 1 | 7.33237E-05 | 5.55519E-08 |
| Lgr6 | 2.79725E-08 | -3.914950933 | 0.057 | 0.479 | 0.000679509 | 0.004568514 | -2.275495283 | 0.268 | 0.464 | 1 | 0.004568514 | 5.59451E-08 |
| Cavin3 | 2.80534E-08 | 1.251623578 | 0.887 | 0.602 | 0.000681473 | 0.799552598 | 0.103850509 | 0.537 | 0.625 | 1 | 0.799552598 | 5.61068E-08 |
| Fibin | 2.82241E-08 | -1.780442455 | 0.377 | 0.784 | 0.000685621 | 3.94816E-05 | -1.440388446 | 0.195 | 0.58 | 0.959086409 | 3.94816E-05 | 5.64483E-08 |
| Hnrnpab | 2.83758E-08 | 0.776464945 | 1 | 0.928 | 0.000689304 | 0.398777657 | 0.107470825 | 1 | 0.997 | 1 | 0.398777657 | 5.67515E-08 |
| Dlg4 | 2.85559E-08 | 1.350273404 | 0.698 | 0.322 | 0.00069368 | 0.300289536 | 0.334704509 | 0.585 | 0.522 | 1 | 0.300289536 | 5.71118E-08 |
| Pcdhgc3 | 2.87613E-08 | 2.10741688 | 0.415 | 0.106 | 0.000698669 | 0.44787843 | 0.377773231 | 0.244 | 0.191 | 1 | 0.44787843 | 5.75225E-08 |
| Gtpbp1 | 2.91052E-08 | 1.995524 | 0.528 | 0.186 | 0.000707023 | 0.911490812 | 0.452541 | 0.268 | 0.307 | 1 | 0.911490812 | 5.82104E-08 |
| Aplp2 | 2.92024E-08 | 0.800074287 | 1 | 0.987 | 0.000709386 | 0.00034071 | 0.572918648 | 1 | 0.993 | 1 | 0.00034071 | 5.84049E-08 |
| Hmgcs1 | 2.98713E-08 | -2.694842655 | 0.717 | 0.801 | 0.000725634 | 0.016197753 | -1.076030759 | 0.829 | 0.887 | 1 | 0.016197753 | 5.97427E-08 |
| Smarca4 | 3.00783E-08 | 1.108428943 | 0.906 | 0.64 | 0.000730663 | 0.112537694 | 0.386220654 | 0.829 | 0.799 | 1 | 0.112537694 | 6.01567E-08 |
| Pde5a | 0.03480106 | 1.872322864 | 0.17 | 0.081 | 1 | 3.03109E-08 | 4.293583144 | 0.341 | 0.075 | 0.000736312 | 0.03480106 | 6.06217E-08 |
| Pygo1 | 3.08185E-08 | 2.254258268 | 0.453 | 0.14 | 0.000748642 | 0.068406891 | 1.136765132 | 0.39 | 0.28 | 1 | 0.068406891 | 6.16369E-08 |
| Nr2f2 | 3.1036E-08 | 1.86295647 | 0.698 | 0.445 | 0.000753927 | 0.041100142 | 0.679663573 | 0.683 | 0.543 | 1 | 0.041100142 | 6.2072E-08 |
| Lims2 | 0.000424511 | -1.508242418 | 0.226 | 0.487 | 1 | 3.19201E-08 | -3.917682652 | 0.049 | 0.522 | 0.000775402 | 0.000424511 | 6.38401E-08 |
| 4833420G17Rik | 3.23644E-08 | 1.589937976 | 0.717 | 0.411 | 0.000786196 | 0.515440799 | 0.493250449 | 0.512 | 0.529 | 1 | 0.515440799 | 6.47288E-08 |
| Usp53 | 3.27368E-08 | 1.260125275 | 0.849 | 0.551 | 0.000795242 | 0.013006459 | 0.483378403 | 0.976 | 0.812 | 1 | 0.013006459 | 6.54736E-08 |
| Zfp30 | 3.30934E-08 | 3.154722595 | 0.283 | 0.047 | 0.000803904 | 0.393115405 | 0.656632604 | 0.22 | 0.174 | 1 | 0.393115405 | 6.61867E-08 |
| Angpt1 | 3.33642E-08 | 3.582143819 | 0.302 | 0.055 | 0.000810484 | 0.00924986 | 1.782757066 | 0.293 | 0.143 | 1 | 0.00924986 | 6.67285E-08 |
| Rbm26 | 3.33758E-08 | 1.097309352 | 0.981 | 0.814 | 0.000810766 | 0.665451547 | -0.155733486 | 0.902 | 0.915 | 1 | 0.665451547 | 6.67517E-08 |
| Snai1 | 3.40418E-08 | 1.793840866 | 0.585 | 0.233 | 0.000826944 | 0.413566182 | 0.252242349 | 0.39 | 0.338 | 1 | 0.413566182 | 6.80836E-08 |
| Fmnl3 | 3.51616E-08 | 1.439290162 | 0.66 | 0.309 | 0.000854145 | 0.033546871 | 0.705798661 | 0.659 | 0.567 | 1 | 0.033546871 | 7.03232E-08 |
| Fbxl19 | 3.55858E-08 | 2.406261362 | 0.358 | 0.081 | 0.00086445 | 0.449647622 | 1.009385825 | 0.22 | 0.181 | 1 | 0.449647622 | 7.11716E-08 |
| Nhs | 3.56668E-08 | 2.340468114 | 0.566 | 0.216 | 0.000866417 | 0.000508603 | 1.422167351 | 0.732 | 0.478 | 1 | 0.000508603 | 7.13335E-08 |
| Rgs7bp | 4.77436E-05 | 2.444229212 | 0.17 | 0.03 | 1 | 3.57265E-08 | 2.83720485 | 0.244 | 0.031 | 0.000867867 | 4.77436E-05 | 7.14529E-08 |
| Axin1 | 3.58693E-08 | 1.414109224 | 0.679 | 0.305 | 0.000871337 | 0.022978773 | 0.377773231 | 0.805 | 0.614 | 1 | 0.022978773 | 7.17386E-08 |
| Palmd | 3.61069E-08 | 2.719507214 | 0.358 | 0.085 | 0.000877109 | 0.001360781 | 2.196285943 | 0.341 | 0.16 | 1 | 0.001360781 | 7.22138E-08 |
| Cpne8 | 6.29587E-07 | -0.78118758 | 0.925 | 1 | 0.015293928 | 3.61998E-08 | -1.103698863 | 0.927 | 0.993 | 0.000879366 | 6.29587E-07 | 7.23997E-08 |
| Zeb2 | 0.001862144 | 2.634715536 | 0.321 | 0.165 | 1 | 3.72893E-08 | 2.661047895 | 0.439 | 0.116 | 0.00090583 | 0.001862144 | 7.45785E-08 |
| Sbno1 | 3.93807E-08 | 0.93403448 | 0.962 | 0.856 | 0.000956637 | 0.000124331 | 0.788416286 | 1 | 0.952 | 1 | 0.000124331 | 7.87615E-08 |
| Prkn | 3.97641E-08 | -1.854047615 | 0.264 | 0.669 | 0.000965948 | 0.000108575 | -1.400425011 | 0.341 | 0.621 | 1 | 0.000108575 | 7.95281E-08 |
| Itsn2 | 4.06473E-08 | 1.243989933 | 0.868 | 0.623 | 0.000987405 | 0.120837349 | -0.304503909 | 0.78 | 0.898 | 1 | 0.120837349 | 8.12946E-08 |
| Asxl1 | 4.08101E-08 | 1.099038462 | 0.906 | 0.686 | 0.000991358 | 0.122786628 | 0.60080557 | 0.878 | 0.867 | 1 | 0.122786628 | 8.16201E-08 |
| Cys1 | 4.11851E-08 | 3.324647596 | 0.226 | 0.025 | 0.001000468 | 9.50158E-06 | 2.47463477 | 0.293 | 0.075 | 0.230812348 | 9.50158E-06 | 8.23701E-08 |
| Gpbp1l1 | 4.15314E-08 | 1.513176566 | 0.736 | 0.407 | 0.001008881 | 0.003903543 | 0.496167932 | 0.829 | 0.652 | 1 | 0.003903543 | 8.30628E-08 |
| Ehmt1 | 4.2826E-08 | 1.150582864 | 0.887 | 0.644 | 0.001040329 | 0.105367365 | 0.342961206 | 0.902 | 0.87 | 1 | 0.105367365 | 8.5652E-08 |
| Mast4 | 2.6862E-07 | -0.820226642 | 0.887 | 0.945 | 0.006525317 | 4.28605E-08 | -1.743839701 | 0.756 | 0.918 | 0.001041166 | 2.6862E-07 | 8.57209E-08 |
| Dusp4 | 1.89608E-05 | 3.254258268 | 0.132 | 0.013 | 0.460595195 | 4.38463E-08 | 1.795384674 | 0.268 | 0.038 | 0.001065115 | 1.89608E-05 | 8.76927E-08 |
| Rbm25 | 4.40515E-08 | 0.538438483 | 1 | 0.996 | 0.001070098 | 0.496842948 | 0.118676504 | 1 | 0.997 | 1 | 0.496842948 | 8.81029E-08 |
| Git2 | 4.41407E-08 | 1.777487128 | 0.642 | 0.301 | 0.001072266 | 0.461219908 | 0.226886775 | 0.561 | 0.495 | 1 | 0.461219908 | 8.82814E-08 |
| Ccdc138 | 4.45715E-08 | 1.302656024 | 0.679 | 0.284 | 0.001082731 | 0.005989223 | 1.805194455 | 0.561 | 0.42 | 1 | 0.005989223 | 8.9143E-08 |
| Nptn | 4.67518E-08 | 0.521522214 | 1 | 0.996 | 0.001135694 | 5.46073E-05 | 0.500078044 | 1 | 1 | 1 | 5.46073E-05 | 9.35035E-08 |
| Fig4 | 4.72957E-08 | 2.620386167 | 0.34 | 0.072 | 0.001148908 | 0.59577502 | 0.789899135 | 0.171 | 0.15 | 1 | 0.59577502 | 9.45914E-08 |
| Eva1b | 4.78026E-08 | 1.679384585 | 0.755 | 0.424 | 0.00116122 | 0.105209886 | 0.346457691 | 0.659 | 0.512 | 1 | 0.105209886 | 9.56051E-08 |
| Rbbp4 | 4.87929E-08 | 0.681328323 | 0.943 | 0.89 | 0.001185276 | 0.242375588 | 0.21060786 | 0.927 | 0.918 | 1 | 0.242375588 | 9.75857E-08 |
| Pim1 | 4.95528E-08 | 1.393910259 | 0.642 | 0.267 | 0.001203736 | 0.001040172 | 0.862590168 | 0.683 | 0.447 | 1 | 0.001040172 | 9.91055E-08 |
| Map4k3 | 4.99408E-08 | 1.300153034 | 0.811 | 0.47 | 0.001213161 | 0.293216863 | 0.500555249 | 0.756 | 0.758 | 1 | 0.293216863 | 9.98815E-08 |
| Zbtb38 | 5.0577E-08 | -1.512592568 | 0.755 | 0.907 | 0.001228617 | 0.000201965 | -1.154404655 | 0.659 | 0.863 | 1 | 0.000201965 | 1.01154E-07 |
| Tmem117 | 5.2175E-08 | -2.430239906 | 0.132 | 0.568 | 0.001267434 | 0.002289986 | -3.0454382 | 0.073 | 0.29 | 1 | 0.002289986 | 1.0435E-07 |
| Tspan11 | 0.000241473 | 3.614154213 | 0.094 | 0.008 | 1 | 5.44174E-08 | 4.059597271 | 0.244 | 0.034 | 0.001321907 | 0.000241473 | 1.08835E-07 |
| Khdrbs1 | 5.455E-08 | 0.831836217 | 1 | 0.881 | 0.001325128 | 0.135971791 | 0.196628374 | 0.927 | 0.99 | 1 | 0.135971791 | 1.091E-07 |
| Cd276 | 5.46021E-08 | 2.319109413 | 0.491 | 0.178 | 0.001326393 | 0.109302 | 0.795384674 | 0.439 | 0.362 | 1 | 0.109302 | 1.09204E-07 |
| Stk39 | 5.57447E-08 | -1.257020327 | 0.698 | 0.924 | 0.00135415 | 0.004731717 | -0.803339451 | 0.878 | 0.904 | 1 | 0.004731717 | 1.11489E-07 |
| Ndrg2 | 5.59357E-08 | -2.217246183 | 0.208 | 0.623 | 0.00135879 | 0.000240993 | -1.384067032 | 0.317 | 0.604 | 1 | 0.000240993 | 1.11871E-07 |
| Slc12a2 | 0.606337189 | 0.224292442 | 0.66 | 0.644 | 1 | 5.64917E-08 | 2.142419071 | 0.829 | 0.618 | 0.001372295 | 0.606337189 | 1.12983E-07 |
| Rnd3 | 4.75227E-07 | 0.777932565 | 0.906 | 0.534 | 0.011544211 | 5.79714E-08 | 1.3458554 | 0.878 | 0.614 | 0.001408241 | 4.75227E-07 | 1.15943E-07 |
| Atp2b1 | 5.85297E-08 | -0.942204455 | 0.943 | 0.996 | 0.001421804 | 0.014224393 | -0.479323776 | 0.976 | 0.99 | 1 | 0.014224393 | 1.17059E-07 |
| Stard9 | 5.88333E-08 | 2.661682584 | 0.302 | 0.055 | 0.001429178 | 0.434237212 | -0.19841906 | 0.049 | 0.085 | 1 | 0.434237212 | 1.17667E-07 |
| Gm10635 | 5.88473E-08 | 1.46284489 | 0.264 | 0.038 | 0.001429518 | 0.838870453 | -1.105309656 | 0.098 | 0.085 | 1 | 0.838870453 | 1.17695E-07 |
| Mxi1 | 5.94718E-08 | 2.119957177 | 0.528 | 0.203 | 0.001444688 | 2.31096E-07 | 1.597153762 | 0.78 | 0.406 | 0.005613791 | 2.31096E-07 | 1.18944E-07 |
| Fat1 | 5.99508E-08 | 1.199746519 | 0.849 | 0.517 | 0.001456324 | 1.8959E-06 | 1.612678264 | 0.878 | 0.662 | 0.046055169 | 1.8959E-06 | 1.19902E-07 |
| Macf1 | 6.30936E-07 | 0.568835586 | 1 | 0.975 | 0.015326694 | 5.99534E-08 | 0.717728871 | 1 | 0.986 | 0.001456387 | 6.30936E-07 | 1.19907E-07 |
| Sh3rf3 | 6.01923E-08 | 2.308345784 | 0.509 | 0.178 | 0.001462192 | 0.002053174 | 1.681723585 | 0.512 | 0.355 | 1 | 0.002053174 | 1.20385E-07 |
| Smo | 6.07119E-08 | 1.193658257 | 0.868 | 0.551 | 0.001474812 | 0.055719027 | 0.477955119 | 0.805 | 0.778 | 1 | 0.055719027 | 1.21424E-07 |
| Cd9 | 6.15018E-08 | -0.769278456 | 1 | 1 | 0.001494001 | 0.000861521 | -0.442915827 | 1 | 1 | 1 | 0.000861521 | 1.23004E-07 |
| Bmt2 | 6.19731E-08 | 1.353756001 | 0.755 | 0.381 | 0.00150545 | 0.549931825 | 0.359625884 | 0.78 | 0.703 | 1 | 0.549931825 | 1.23946E-07 |
| Evi5 | 6.25202E-08 | 1.422006474 | 0.774 | 0.492 | 0.001518742 | 0.193687247 | 0.321581553 | 0.805 | 0.72 | 1 | 0.193687247 | 1.2504E-07 |
| Rcc2 | 6.35545E-08 | 1.68235192 | 0.717 | 0.394 | 0.001543867 | 0.00016029 | 1.301654543 | 0.78 | 0.59 | 1 | 0.00016029 | 1.27109E-07 |
| Prelp | 6.38651E-08 | -1.167976789 | 0.717 | 0.928 | 0.00155141 | 7.57504E-05 | -0.987048597 | 0.634 | 0.809 | 1 | 7.57504E-05 | 1.2773E-07 |
| Prtn3 | 6.411E-08 | 2.324647596 | 0.226 | 0.025 | 0.001557361 | 0.003075345 | 2.422167351 | 0.171 | 0.051 | 1 | 0.003075345 | 1.2822E-07 |
| Pacs1 | 6.41599E-08 | 1.569760094 | 0.585 | 0.233 | 0.001558572 | 0.015016959 | 0.769849582 | 0.634 | 0.468 | 1 | 0.015016959 | 1.2832E-07 |
| Trp53inp1 | 6.45431E-08 | 1.699640377 | 0.642 | 0.322 | 0.001567881 | 0.041431035 | 0.621887946 | 0.585 | 0.437 | 1 | 0.041431035 | 1.29086E-07 |
| Rdh10 | 6.66451E-08 | 2.329220326 | 0.491 | 0.169 | 0.001618943 | 0.01422676 | 0.808253476 | 0.61 | 0.451 | 1 | 0.01422676 | 1.3329E-07 |
| Arl15 | 6.76191E-08 | 1.189641011 | 0.887 | 0.542 | 0.001642603 | 0.018528643 | 0.543965514 | 0.854 | 0.659 | 1 | 0.018528643 | 1.35238E-07 |
| Mob3b | 6.79435E-08 | 2.868009192 | 0.434 | 0.136 | 0.001650484 | 0.000672644 | 2.325491331 | 0.439 | 0.225 | 1 | 0.000672644 | 1.35887E-07 |
| Tbc1d15 | 6.92382E-08 | 1.126340793 | 0.868 | 0.627 | 0.001681935 | 0.269590476 | 0.507703535 | 0.756 | 0.802 | 1 | 0.269590476 | 1.38476E-07 |
| Dsel | 7.02332E-08 | 1.664671741 | 0.642 | 0.309 | 0.001706104 | 0.392290113 | 0.64128464 | 0.463 | 0.474 | 1 | 0.392290113 | 1.40466E-07 |
| Ago2 | 7.06211E-08 | 1.087772351 | 0.943 | 0.712 | 0.001715527 | 0.888815596 | -0.102757567 | 0.976 | 0.939 | 1 | 0.888815596 | 1.41242E-07 |
| Ptms | 7.08288E-08 | 0.717026544 | 1 | 0.949 | 0.001720574 | 0.117889618 | 0.268564654 | 1 | 0.986 | 1 | 0.117889618 | 1.41658E-07 |
| Mat2a | 7.22684E-08 | 0.960453774 | 1 | 0.864 | 0.001755544 | 0.44989814 | 0.144182603 | 0.927 | 0.922 | 1 | 0.44989814 | 1.44537E-07 |
| Hras | 7.26597E-08 | 1.504665066 | 0.66 | 0.305 | 0.001765049 | 0.131312544 | 0.413993419 | 0.61 | 0.502 | 1 | 0.131312544 | 1.45319E-07 |
| Sh3rf1 | 7.43946E-08 | 1.164415991 | 0.887 | 0.564 | 0.001807193 | 0.118895159 | 0.677127634 | 0.829 | 0.744 | 1 | 0.118895159 | 1.48789E-07 |
| Ddr2 | 7.55893E-08 | -1.441686433 | 0.792 | 0.932 | 0.001836214 | 0.000268435 | -0.948290638 | 0.805 | 0.915 | 1 | 0.000268435 | 1.51179E-07 |
| Cnrip1 | 7.5682E-08 | 2.3688474 | 0.302 | 0.055 | 0.001838467 | 0.347505017 | 0.80158094 | 0.146 | 0.099 | 1 | 0.347505017 | 1.51364E-07 |
| Ccdc117 | 7.61301E-08 | 1.782753817 | 0.585 | 0.258 | 0.001849352 | 0.137455779 | 0.715575405 | 0.561 | 0.474 | 1 | 0.137455779 | 1.5226E-07 |
| F2r | 7.8479E-08 | 3.434830514 | 0.208 | 0.021 | 0.001906411 | 3.43682E-05 | 2.858266465 | 0.415 | 0.171 | 0.834873004 | 3.43682E-05 | 1.56958E-07 |
| Tll1 | 7.90684E-08 | 1.997181318 | 0.491 | 0.165 | 0.001920729 | 0.305431483 | 0.32225207 | 0.561 | 0.522 | 1 | 0.305431483 | 1.58137E-07 |
| Hdac2 | 7.98829E-08 | 1.52115015 | 0.736 | 0.458 | 0.001940515 | 0.978347564 | -0.116991461 | 0.732 | 0.676 | 1 | 0.978347564 | 1.59766E-07 |
| Caln1 | 2.2563E-05 | 4.962077517 | 0.075 | 0 | 0.548100284 | 8.04483E-08 | 7.159132945 | 0.098 | 0 | 0.001954251 | 2.2563E-05 | 1.60897E-07 |
| Acat3 | 0.343006305 | -0.1672055 | 0 | 0.017 | 1 | 8.04483E-08 | 5.422167351 | 0.098 | 0 | 0.001954251 | 0.343006305 | 1.60897E-07 |
| 4933431K23Rik | 0.002005228 | 3.739685096 | 0.075 | 0.008 | 1 | 8.047E-08 | 6.296636468 | 0.098 | 0 | 0.001954778 | 0.002005228 | 1.6094E-07 |
| Cntn1 | 0.249034808 | 2.154722595 | 0.019 | 0.004 | 1 | 8.047E-08 | 6.159132945 | 0.098 | 0 | 0.001954778 | 0.249034808 | 1.6094E-07 |
| Nqo1 | 8.36675E-08 | -3.436238647 | 0.057 | 0.462 | 0.00203245 | 0.707264418 | -0.119726428 | 0.317 | 0.273 | 1 | 0.707264418 | 1.67335E-07 |

| | | | | | | | | | | | | |
|---|---|---|---|---|---|---|---|---|---|---|---|---|
| Rbpms | 8.37068E-08 | 2.070330408 | 0.547 | 0.212 | 0.002033405 | 0.001742549 | 1.619974134 | 0.537 | 0.355 | 1 | 0.001742549 | 1.67414E-07 |
| Tent4b | 8.49204E-08 | 1.807919831 | 0.604 | 0.258 | 0.002062885 | 8.2462E-05 | 1.484533231 | 0.78 | 0.563 | 1 | 8.2462E-05 | 1.69841E-07 |
| Mrps11 | 8.64632E-08 | 1.89946554 | 0.434 | 0.123 | 0.002100365 | 0.292611805 | -0.395455907 | 0.073 | 0.133 | 1 | 0.292611805 | 1.72926E-07 |
| Ccdc12 | 8.64635E-08 | 1.188510908 | 0.83 | 0.517 | 0.002100372 | 0.011510352 | 0.736600909 | 0.634 | 0.451 | 1 | 0.011510352 | 1.72927E-07 |
| Mex3b | 8.66571E-08 | 2.792152515 | 0.302 | 0.059 | 0.002105073 | 0.006236053 | 2.401105735 | 0.171 | 0.058 | 1 | 0.006236053 | 1.73314E-07 |
| Carmil1 | 0.009734838 | -0.926705005 | 0.755 | 0.818 | 1 | 8.77727E-08 | -1.499219261 | 0.756 | 0.959 | 0.002132175 | 0.009734838 | 1.75545E-07 |
| Vcam1 | 0.015178556 | 2.891688189 | 0.057 | 0.008 | 1 | 8.89873E-08 | 5.644559772 | 0.122 | 0.003 | 0.002161678 | 0.015178556 | 1.77975E-07 |
| Eps15 | 8.91119E-08 | 1.30843201 | 0.774 | 0.398 | 0.002164706 | 0.60064311 | 0.136765132 | 0.634 | 0.706 | 1 | 0.60064311 | 1.78224E-07 |
| Ccnyl1 | 8.94137E-08 | 2.074552246 | 0.509 | 0.182 | 0.002172037 | 0.030069432 | 1.177860153 | 0.488 | 0.355 | 1 | 0.030069432 | 1.78827E-07 |
| Prcc | 9.01673E-08 | 1.470224421 | 0.642 | 0.275 | 0.002190343 | 0.602076762 | 0.369056014 | 0.488 | 0.515 | 1 | 0.602076762 | 1.80335E-07 |
| Cbfa2t2 | 9.02382E-08 | 1.248844772 | 0.755 | 0.39 | 0.002192067 | 0.919571174 | -0.128029732 | 0.659 | 0.652 | 1 | 0.919571174 | 1.80476E-07 |
| Pax9 | 0.732136816 | 1.237184755 | 0.057 | 0.047 | 1 | 9.03268E-08 | 5.489281546 | 0.171 | 0.014 | 0.002194219 | 0.732136816 | 1.80654E-07 |
| Hspa1a | 7.02177E-07 | 2.352248922 | 0.755 | 0.449 | 0.017057294 | 9.04615E-08 | 2.337378262 | 0.854 | 0.57 | 0.002197492 | 7.02177E-07 | 1.80923E-07 |
| Ankrd17 | 9.07793E-08 | 0.8327945 | 0.981 | 0.949 | 0.00220521 | 0.004792639 | 0.364871204 | 0.976 | 0.997 | 1 | 0.004792639 | 1.81558E-07 |
| Frem1 | 0.014846242 | 4.029191713 | 0.057 | 0.008 | 1 | 9.08098E-08 | 5.007129851 | 0.122 | 0.003 | 0.002205952 | 0.014846242 | 1.8162E-07 |
| Iigp1 | 0.746406176 | 0.962077517 | 0.038 | 0.03 | 1 | 9.08913E-08 | 6.243197209 | 0.122 | 0.003 | 0.002207933 | 0.746406176 | 1.81783E-07 |
| Rsrp1 | 9.14136E-08 | 1.014422762 | 0.981 | 0.953 | 0.002220619 | 0.01279461 | 0.45676029 | 1 | 0.99 | 1 | 0.01279461 | 1.82827E-07 |
| Slc23a3 | 9.27411E-08 | 2.962077517 | 0.245 | 0.034 | 0.002252866 | 0.000550093 | 1.75474269 | 0.268 | 0.089 | 1 | 0.000550093 | 1.85482E-07 |
| Agbl2 | 0.032198545 | 2.002719501 | 0.094 | 0.03 | 1 | 9.27849E-08 | 4.952682067 | 0.122 | 0.003 | 0.00225393 | 0.032198545 | 1.8557E-07 |
| Rflnb | 9.30242E-08 | 1.347367673 | 0.849 | 0.61 | 0.002259745 | 0.000184105 | 1.015661485 | 0.805 | 0.628 | 1 | 0.000184105 | 1.86048E-07 |
| Tubb2b | 9.33501E-08 | -2.64138757 | 0.189 | 0.585 | 0.002267661 | 0.000507666 | -1.551290862 | 0.146 | 0.44 | 1 | 0.000507666 | 1.867E-07 |
| Fnbp1 | 9.38484E-08 | 1.63859897 | 0.642 | 0.288 | 0.002279764 | 0.001519083 | 1.007129851 | 0.707 | 0.546 | 1 | 0.001519083 | 1.87697E-07 |
| Atad2b | 9.43274E-08 | 1.406430583 | 0.868 | 0.593 | 0.002291401 | 0.019759016 | 0.353882611 | 0.927 | 0.795 | 1 | 0.019759016 | 1.88655E-07 |
| Dhx58os | 9.48707E-08 | -1.600986722 | 0.472 | 0.792 | 0.0023046 | 1.87786E-05 | -1.17201368 | 0.244 | 0.652 | 0.456169446 | 1.87786E-05 | 1.89741E-07 |
| Ldb1 | 9.5443E-08 | 1.334876729 | 0.755 | 0.466 | 0.002318502 | 0.275212455 | 0.347590226 | 0.78 | 0.734 | 1 | 0.275212455 | 1.90886E-07 |
| Nelfb | 9.79633E-08 | 1.386048141 | 0.66 | 0.301 | 0.002379725 | 0.060672567 | 0.784093513 | 0.512 | 0.399 | 1 | 0.060672567 | 1.95927E-07 |
| Cd24a | 9.84468E-08 | -2.668983409 | 0.434 | 0.703 | 0.002391471 | 0.279655853 | -0.337166057 | 0.512 | 0.396 | 1 | 0.279655853 | 1.96894E-07 |
| Slc6a12 | 9.86855E-08 | -6.233294691 | 0 | 0.386 | 0.002397269 | 0.043337147 | -3.466575898 | 0 | 0.092 | 1 | 0.043337147 | 1.97371E-07 |
| Pde7b | 1.00611E-07 | 2.982255399 | 0.434 | 0.144 | 0.002444037 | 4.81089E-05 | 1.747769766 | 0.512 | 0.222 | 1 | 4.81089E-05 | 2.01222E-07 |
| Rspo1 | 0.000123854 | 3.920257341 | 0.113 | 0.013 | 1 | 1.00748E-07 | 4.059597271 | 0.122 | 0.003 | 0.002447377 | 0.000123854 | 2.01497E-07 |
| Pfn1 | 1.03154E-07 | 0.742473893 | 1 | 0.941 | 0.002505815 | 0.365408593 | 0.115716592 | 1 | 0.983 | 1 | 0.365408593 | 2.06308E-07 |
| Arhgap35 | 1.03326E-07 | 1.458590718 | 0.755 | 0.453 | 0.00250999 | 0.538210087 | -0.138547604 | 0.756 | 0.775 | 1 | 0.538210087 | 2.06652E-07 |
| Pum1 | 1.04864E-07 | 0.761807906 | 1 | 0.945 | 0.002547356 | 0.297904049 | 0.201088261 | 1 | 0.986 | 1 | 0.297904049 | 2.09728E-07 |
| Ptgfrn | 1.04921E-07 | 1.954423944 | 0.491 | 0.174 | 0.002548743 | 0.000602264 | 1.83720485 | 0.341 | 0.143 | 1 | 0.000602264 | 2.09842E-07 |
| Cgref1 | 1.07079E-07 | -4.173204241 | 0.038 | 0.428 | 0.002601155 | 0.001202657 | -2.143342788 | 0.073 | 0.321 | 1 | 0.001202657 | 2.14157E-07 |
| Sertad1 | 1.08105E-07 | 1.282932577 | 0.811 | 0.466 | 0.002626098 | 0.033034027 | 1.160449873 | 0.683 | 0.679 | 1 | 0.033034027 | 2.16211E-07 |
| Syn3 | 1.08217E-07 | -3.293259003 | 0.264 | 0.606 | 0.002628819 | 0.029951659 | -2.08562729 | 0.122 | 0.266 | 1 | 0.029951659 | 2.16435E-07 |
| Rad21 | 1.0841E-07 | 1.255354958 | 0.774 | 0.538 | 0.002633488 | 0.002698137 | 0.618113792 | 0.902 | 0.768 | 1 | 0.002698137 | 2.16819E-07 |
| Serinc3 | 1.08813E-07 | 1.130520246 | 0.981 | 0.958 | 0.002643287 | 0.060132894 | 0.280413022 | 1 | 0.993 | 1 | 0.060132894 | 2.17626E-07 |
| Bnip2 | 1.0964E-07 | 0.934275818 | 0.925 | 0.831 | 0.002663373 | 0.582903065 | 0.105280711 | 0.927 | 0.887 | 1 | 0.582903065 | 2.1928E-07 |
| Ppp2r5e | 1.09743E-07 | 1.36726723 | 0.811 | 0.581 | 0.002665885 | 0.111237191 | 0.202722213 | 0.878 | 0.799 | 1 | 0.111237191 | 2.19487E-07 |
| Bmpr2 | 1.09819E-07 | 0.9063847 | 1 | 0.852 | 0.002667731 | 0.029205132 | 0.331771009 | 1 | 0.986 | 1 | 0.029205132 | 2.19639E-07 |
| Arhgap6 | 1.10426E-07 | 2.442524906 | 0.472 | 0.153 | 0.002682465 | 0.52878946 | 0.328655465 | 0.439 | 0.42 | 1 | 0.52878946 | 2.20852E-07 |
| Wscd2 | 1.1051E-07 | -3.126048175 | 0.075 | 0.479 | 0.002684509 | 0.066595255 | -1.155261478 | 0.171 | 0.307 | 1 | 0.066595255 | 2.2102E-07 |
| Sc5d | 1.11647E-07 | -1.814903756 | 0.453 | 0.754 | 0.002712127 | 0.02815574 | -0.852626769 | 0.463 | 0.611 | 1 | 0.02815574 | 2.23294E-07 |
| Lix1l | 1.12141E-07 | 1.099074565 | 0.774 | 0.504 | 0.002724126 | 0.000398054 | 1.096591479 | 0.683 | 0.522 | 1 | 0.000398054 | 2.24282E-07 |
| Fry | 1.12352E-07 | -1.357030059 | 0.717 | 0.941 | 0.002729256 | 0.056542674 | -0.598614212 | 0.683 | 0.863 | 1 | 0.056542674 | 2.24704E-07 |
| Mpped2 | 1.16979E-07 | 1.31362619 | 0.755 | 0.424 | 0.002841664 | 0.003552715 | 0.899668937 | 0.756 | 0.601 | 1 | 0.003552715 | 2.33959E-07 |
| Sdk1 | 1.18913E-07 | 1.824077283 | 0.396 | 0.11 | 0.002888638 | 0.003030085 | 1.547384501 | 0.537 | 0.314 | 1 | 0.003030085 | 2.37826E-07 |
| Tnpo2 | 1.19655E-07 | 1.483345342 | 0.623 | 0.258 | 0.002906649 | 0.318468661 | 0.493250449 | 0.463 | 0.437 | 1 | 0.318468661 | 2.39309E-07 |
| Ybx1 | 1.22845E-07 | 0.628398677 | 1 | 0.966 | 0.002984151 | 0.012219779 | 0.294660444 | 1 | 0.993 | 1 | 0.012219779 | 2.4569E-07 |
| Chchd3 | 1.23846E-07 | 1.380282295 | 0.792 | 0.538 | 0.003008465 | 0.382793429 | 0.353603064 | 0.732 | 0.737 | 1 | 0.382793429 | 2.47692E-07 |
| Alox12 | 1.24937E-07 | -7.399866257 | 0 | 0.381 | 0.003034968 | 0.535786395 | -1.890715605 | 0.024 | 0.044 | 1 | 0.535786395 | 2.49874E-07 |
| Smc3 | 1.27014E-07 | 0.94849233 | 0.981 | 0.771 | 0.003085436 | 0.001561419 | 0.578575623 | 0.927 | 0.918 | 1 | 0.001561419 | 2.54029E-07 |
| Rgmb | 1.29824E-07 | 1.786335189 | 0.528 | 0.199 | 0.003153685 | 0.555675284 | 0.136765132 | 0.439 | 0.423 | 1 | 0.555675284 | 2.59648E-07 |
| Eif3c | 1.31234E-07 | 0.911005886 | 1 | 0.856 | 0.003187933 | 0.324440261 | 0.140925326 | 0.976 | 0.942 | 1 | 0.324440261 | 2.62468E-07 |
| Slc22a23 | 1.31421E-07 | -1.899836504 | 0.151 | 0.568 | 0.003192472 | 0.000392349 | -2.7486966 | 0.293 | 0.526 | 1 | 0.000392349 | 2.62841E-07 |
| Matn1 | 1.32955E-07 | -5.652293308 | 0.151 | 0.534 | 0.003229739 | 1.85384E-06 | -5.465560988 | 0.049 | 0.447 | 0.045033407 | 1.85384E-06 | 2.6591E-07 |
| Fnbp4 | 1.33207E-07 | 1.175186697 | 0.943 | 0.703 | 0.003235862 | 0.002685111 | 0.579188519 | 0.951 | 0.915 | 1 | 0.002685111 | 2.66414E-07 |
| Trim28 | 1.35209E-07 | 1.538051234 | 0.66 | 0.331 | 0.003284509 | 0.156789431 | 0.551153791 | 0.61 | 0.59 | 1 | 0.156789431 | 2.70419E-07 |
| Cobl | 1.80336E-07 | -2.017427072 | 0.264 | 0.648 | 0.004380728 | 1.36032E-07 | -2.034739867 | 0.366 | 0.768 | 0.003304488 | 1.80336E-07 | 2.72064E-07 |
| Fus | 1.3613E-07 | 0.706481175 | 1 | 0.97 | 0.003306881 | 0.100947522 | 0.291972311 | 1 | 1 | 1 | 0.100947522 | 2.72261E-07 |
| Baz1a | 1.36406E-07 | 1.410133759 | 0.736 | 0.386 | 0.003313563 | 0.006603312 | 1.008316766 | 0.707 | 0.563 | 1 | 0.006603312 | 2.72811E-07 |
| Egr1 | 1.36556E-07 | 1.47505567 | 0.83 | 0.547 | 0.003317224 | 0.000277923 | 1.560671177 | 0.805 | 0.666 | 1 | 0.000277923 | 2.73112E-07 |
| Maz | 1.36902E-07 | 1.265753907 | 0.679 | 0.314 | 0.003325621 | 0.989245392 | 0.200580229 | 0.415 | 0.444 | 1 | 0.989245392 | 2.73804E-07 |
| Zfp36l1 | 6.88819E-06 | 0.831489484 | 0.981 | 0.835 | 0.167327878 | 1.38418E-07 | 1.307326607 | 0.976 | 0.819 | 0.003362452 | 6.88819E-06 | 2.76836E-07 |
| Memo1 | 1.3904E-07 | 1.347367673 | 0.83 | 0.538 | 0.003377567 | 0.000193294 | 1.162807266 | 0.805 | 0.713 | 1 | 0.000193294 | 2.78081E-07 |
| Moxd1 | 1.3995E-07 | -4.153616435 | 0.038 | 0.428 | 0.003399662 | 0.037373308 | -1.863234868 | 0.122 | 0.266 | 1 | 0.037373308 | 2.799E-07 |
| Zswim3 | 1.40926E-07 | 3.154722595 | 0.264 | 0.042 | 0.003423365 | 0.03088188 | 1.174239837 | 0.122 | 0.041 | 1 | 0.03088188 | 2.81851E-07 |
| Apaf1 | 1.40985E-07 | 1.814382463 | 0.642 | 0.288 | 0.0034248 | 0.000820409 | 1.077155643 | 0.707 | 0.468 | 1 | 0.000820409 | 2.81969E-07 |
| Gpi1 | 1.44755E-07 | 1.139347369 | 0.925 | 0.644 | 0.003516378 | 0.005255672 | 0.56258247 | 0.927 | 0.761 | 1 | 0.005255672 | 2.89509E-07 |
| Asxl3 | 1.44875E-07 | 3.962077517 | 0.189 | 0.017 | 0.003519307 | 0.193112634 | 0.962735732 | 0.049 | 0.017 | 1 | 0.193112634 | 2.8975E-07 |
| Tsc22d1 | 1.49568E-07 | -1.122821278 | 0.981 | 0.996 | 0.003633297 | 0.442532559 | 0.152557913 | 1 | 0.997 | 1 | 0.442532559 | 2.99135E-07 |
| Entpd3 | 0.209009104 | 1.739685096 | 0.038 | 0.013 | 1 | 1.51143E-07 | 3.185128153 | 0.22 | 0.027 | 0.00367156 | 0.209009104 | 3.02285E-07 |
| Svbp | 1.53452E-07 | 1.091986839 | 0.811 | 0.479 | 0.003727649 | 0.338208729 | -0.190701147 | 0.415 | 0.502 | 1 | 0.338208729 | 3.06903E-07 |
| Fbxo2 | 1.54137E-07 | -4.528271989 | 0.057 | 0.441 | 0.003744291 | 0.015707415 | -1.885261175 | 0.098 | 0.263 | 1 | 0.015707415 | 3.08274E-07 |
| Vps37d | 1.5681E-07 | 2.495759513 | 0.283 | 0.051 | 0.003809229 | 0.10998005 | 1.515276755 | 0.146 | 0.075 | 1 | 0.10998005 | 3.1362E-07 |
| Tgfa | 1.57322E-07 | -3.345123292 | 0.132 | 0.534 | 0.003821655 | 0.002977679 | -1.54256732 | 0.171 | 0.403 | 1 | 0.002977679 | 3.14643E-07 |
| Rock1 | 1.6209E-07 | 0.740021584 | 1 | 0.949 | 0.003937501 | 0.017810919 | 0.284286602 | 1 | 1 | 1 | 0.017810919 | 3.24181E-07 |
| Gbp2 | 0.924182999 | -0.216533212 | 0.113 | 0.11 | 1 | 1.63007E-07 | 3.384692645 | 0.195 | 0.02 | 0.003959766 | 0.924182999 | 3.26014E-07 |
| Jam2 | 1.64302E-07 | -2.098888557 | 0.453 | 0.729 | 0.003991228 | 0.001177811 | -2.101394606 | 0.171 | 0.413 | 1 | 0.001177811 | 3.28604E-07 |
| Foxn2 | 1.64498E-07 | 1.932330173 | 0.585 | 0.267 | 0.003995986 | 0.152647433 | 0.477029286 | 0.61 | 0.539 | 1 | 0.152647433 | 3.28996E-07 |
| Dhx32 | 1.64561E-07 | -1.951476809 | 0.264 | 0.657 | 0.003997514 | 3.23787E-05 | -1.926560804 | 0.122 | 0.488 | 0.786542788 | 3.23787E-05 | 3.29122E-07 |
| Gas1 | 1.65666E-07 | 1.169077888 | 0.717 | 0.356 | 0.004024364 | 0.006422529 | 0.779627574 | 0.659 | 0.464 | 1 | 0.006422529 | 3.31332E-07 |
| Maoa | 1.65742E-07 | 1.739685096 | 0.491 | 0.165 | 0.004026211 | 0.777338118 | 1.439869352 | 0.22 | 0.215 | 1 | 0.777338118 | 3.31485E-07 |
| Aif1l | 1.68612E-07 | -2.545717123 | 0.094 | 0.5 | 0.004095913 | 1.00057E-05 | -3.572186086 | 0.024 | 0.375 | 0.243059061 | 1.00057E-05 | 3.37223E-07 |
| Xist | 1.74807E-07 | -2.660943241 | 0.151 | 0.517 | 0.004246409 | 0.264933923 | 2.422167351 | 0.024 | 0.007 | 1 | 0.264933923 | 3.49614E-07 |
| Ints6 | 1.75653E-07 | 1.269134385 | 0.868 | 0.623 | 0.004266969 | 2.91157E-06 | 1.579839194 | 0.927 | 0.809 | 0.070727952 | 2.91157E-06 | 3.51307E-07 |
| Dennd4c | 1.7708E-07 | 1.728796779 | 0.642 | 0.314 | 0.004301627 | 0.065570828 | 1.203332749 | 0.756 | 0.758 | 1 | 0.065570828 | 3.5416E-07 |
| Creb3l2 | 0.000265732 | -0.616712753 | 1 | 0.987 | 1 | 1.7966E-07 | -0.99903005 | 0.927 | 0.997 | 0.004364293 | 0.000265732 | 3.59319E-07 |
| Insr | 1.82168E-07 | 1.262637934 | 0.792 | 0.462 | 0.004425217 | 0.680572676 | 0.337202683 | 0.659 | 0.703 | 1 | 0.680572676 | 3.64335E-07 |
| Ilf3 | 1.82272E-07 | 1.285306705 | 0.811 | 0.581 | 0.00442776 | 0.005718187 | 0.504388439 | 0.902 | 0.809 | 1 | 0.005718187 | 3.64545E-07 |
| Ttf2 | 1.82402E-07 | 2.984797593 | 0.226 | 0.03 | 0.004430901 | 0.44544556 | -0.747757651 | 0.024 | 0.051 | 1 | 0.44544556 | 3.64803E-07 |
| Lmo4 | 1.24857E-06 | 0.739998759 | 1 | 0.911 | 0.030330143 | 1.85757E-07 | 1.296315239 | 1 | 0.918 | 0.004512415 | 1.24857E-06 | 3.71514E-07 |
| Thy1 | 1.88366E-07 | 5.324647596 | 0.113 | 0 | 0.00457579 | 3.50317E-06 | 7.79140116 | 0.073 | 0 | 0.085098954 | 3.50317E-06 | 3.76732E-07 |
| Lrrn1 | 1.8856E-07 | 6.47665069 | 0.113 | 0 | 0.004580506 | 0.057359562 | 2.515276755 | 0.049 | 0.01 | 1 | 0.057359562 | 3.7712E-07 |
| Ftl1 | 1.89003E-07 | 0.84238245 | 1 | 0.983 | 0.004591252 | 0.022252957 | 0.40749842 | 1 | 1 | 1 | 0.022252957 | 3.78005E-07 |
| Glis2 | 1.89142E-07 | 1.344200394 | 0.604 | 0.25 | 0.004594648 | 0.017929522 | 0.678894477 | 0.683 | 0.543 | 1 | 0.017929522 | 3.78285E-07 |

| | | | | | | | | | | | | |
|---|---|---|---|---|---|---|---|---|---|---|---|---|
| Rad23b | 1.89576E-07 | 0.876665998 | 0.906 | 0.843 | 0.00460517 | 0.445775483 | 0.276361878 | 0.951 | 0.935 | 1 | 0.445775483 | 3.79151E-07 |
| Twist2 | 1.9151E-07 | 3.087608399 | 0.245 | 0.038 | 0.004652149 | 0.305356378 | -0.80665134 | 0.024 | 0.065 | 1 | 0.305356378 | 3.83019E-07 |
| Rarres2 | 1.91573E-07 | 4.242185436 | 0.17 | 0.013 | 0.00465369 | 0.003391254 | 2.252242349 | 0.122 | 0.027 | 1 | 0.003391254 | 3.83146E-07 |
| Lrrc42 | 1.93251E-07 | 1.78699081 | 0.453 | 0.14 | 0.004694458 | 0.79994412 | 0.150144161 | 0.293 | 0.341 | 1 | 0.79994412 | 3.86502E-07 |
| Ssbp2 | 1.97808E-07 | 0.978296027 | 0.868 | 0.551 | 0.004805161 | 0.197097907 | -0.104061325 | 0.927 | 0.857 | 1 | 0.197097907 | 3.95617E-07 |
| Rbm38 | 0.019672433 | 1.569760094 | 0.132 | 0.047 | 1 | 1.98074E-07 | 2.676740178 | 0.268 | 0.044 | 0.004811626 | 0.019672433 | 3.96149E-07 |
| Mmd | 1.9839E-07 | 2.806799291 | 0.189 | 0.017 | 0.004819291 | 0.041522481 | 0.962735732 | 0.098 | 0.031 | 1 | 0.041522481 | 3.9678E-07 |
| Osmr | 2.00412E-07 | -1.375465683 | 0.491 | 0.826 | 0.004868403 | 0.157683905 | 0.165043907 | 0.415 | 0.618 | 1 | 0.157683905 | 4.00823E-07 |
| Has2os | 2.05325E-07 | -2.215735394 | 0.151 | 0.576 | 0.004987748 | 0.001929969 | -2.475678105 | 0.122 | 0.352 | 1 | 0.001929969 | 4.10649E-07 |
| Srek1 | 2.07599E-07 | 0.899811253 | 0.943 | 0.86 | 0.005042997 | 0.003050596 | 0.558413258 | 0.902 | 0.935 | 1 | 0.003050596 | 4.15198E-07 |
| Uty | 2.11002E-07 | 1.47119626 | 0.755 | 0.411 | 0.005125665 | 0.114806276 | 0.566047297 | 0.902 | 0.853 | 1 | 0.114806276 | 4.22004E-07 |
| Ugp2 | 2.13042E-07 | -1.35272484 | 0.925 | 0.932 | 0.005175213 | 0.008323658 | -0.51893896 | 0.732 | 0.86 | 1 | 0.008323658 | 4.26084E-07 |
| Tnrc6b | 2.16589E-07 | 0.80045378 | 1 | 0.966 | 0.005261389 | 0.017829082 | 0.450490133 | 1 | 0.976 | 1 | 0.017829082 | 4.33179E-07 |
| Fam171a2 | 2.16922E-07 | 3.739685096 | 0.17 | 0.013 | 0.005269471 | 3.5574E-07 | 2.730289646 | 0.244 | 0.038 | 0.008641648 | 3.5574E-07 | 4.33844E-07 |
| Tslp | 2.16922E-07 | 3.962077517 | 0.17 | 0.013 | 0.005269471 | 0.872828964 | 1.029849928 | 0.024 | 0.02 | 1 | 0.872828964 | 4.33844E-07 |
| Trps1 | 2.17592E-07 | -1.166089942 | 1 | 1 | 0.005285742 | 0.271616466 | -0.288814667 | 1 | 1 | 1 | 0.271616466 | 4.35184E-07 |
| Dand5 | 2.17997E-07 | 2.10741688 | 0.358 | 0.089 | 0.005295579 | 0.144765332 | 1.413993419 | 0.171 | 0.102 | 1 | 0.144765332 | 4.35994E-07 |
| Ppp4r3a | 2.22237E-07 | 1.035258928 | 0.868 | 0.653 | 0.005398575 | 0.309508297 | -0.153839845 | 0.78 | 0.891 | 1 | 0.309508297 | 4.44473E-07 |
| Dip2c | 2.26001E-07 | 1.285728607 | 0.925 | 0.72 | 0.005490022 | 0.017711965 | 0.29831822 | 0.951 | 0.887 | 1 | 0.017711965 | 4.52002E-07 |
| Snrpe | 2.26971E-07 | 0.831146647 | 0.943 | 0.792 | 0.00551358 | 0.07086526 | 0.369525871 | 0.927 | 0.857 | 1 | 0.07086526 | 4.53942E-07 |
| Syt12 | 2.27784E-07 | 2.70221039 | 0.226 | 0.03 | 0.005533327 | 0.778919402 | 0.544423101 | 0.122 | 0.109 | 1 | 0.778919402 | 4.55568E-07 |
| Fbxw11 | 2.29658E-07 | -1.05853425 | 0.868 | 0.975 | 0.005578844 | 0.002185935 | -0.59019678 | 0.951 | 0.986 | 1 | 0.002185935 | 4.59315E-07 |
| Dusp14 | 2.3006E-07 | -3.389597921 | 0.075 | 0.462 | 0.005588623 | 0.00592124 | -0.970150072 | 0.073 | 0.283 | 1 | 0.00592124 | 4.6012E-07 |
| Nherf2 | 2.31815E-07 | 1.989663349 | 0.434 | 0.131 | 0.005631256 | 0.076818667 | -0.484723245 | 0.073 | 0.195 | 1 | 0.076818667 | 4.6363E-07 |
| Klhl11 | 2.33652E-07 | 1.763932642 | 0.509 | 0.195 | 0.005675871 | 0.126453614 | -0.278272368 | 0.317 | 0.437 | 1 | 0.126453614 | 4.67304E-07 |
| Myh10 | 2.36399E-07 | 1.233332429 | 0.774 | 0.466 | 0.005742605 | 0.1802618 | 0.999700317 | 0.61 | 0.567 | 1 | 0.1802618 | 4.72798E-07 |
| Col6a1 | 2.37438E-07 | -0.861164334 | 0.925 | 0.996 | 0.005767839 | 0.065035666 | -0.389547487 | 0.902 | 0.949 | 1 | 0.065035666 | 4.74876E-07 |
| Heph | 2.39725E-07 | 3.920257341 | 0.17 | 0.013 | 0.005823409 | 0.58792152 | 2.515276755 | 0.024 | 0.014 | 1 | 0.58792152 | 4.79451E-07 |
| Hnrnpa1 | 2.41401E-07 | 0.935861864 | 0.981 | 0.953 | 0.005864109 | 0.000277822 | 0.656715159 | 0.976 | 0.956 | 1 | 0.000277822 | 4.82802E-07 |
| Poln | 2.4352E-07 | 1.721763188 | 0.358 | 0.085 | 0.005915586 | 0.008553045 | 1.747567637 | 0.268 | 0.126 | 1 | 0.008553045 | 4.8704E-07 |
| Gpc4 | 2.44089E-07 | 2.371953311 | 0.547 | 0.242 | 0.005929416 | 0.004339085 | 1.252242349 | 0.537 | 0.396 | 1 | 0.004339085 | 4.88178E-07 |
| Mboat2 | 2.4415E-07 | 2.126153443 | 0.453 | 0.148 | 0.005930887 | 0.287904587 | 0.658401697 | 0.439 | 0.444 | 1 | 0.287904587 | 4.883E-07 |
| Strn3 | 2.44704E-07 | 0.626791039 | 0.981 | 0.966 | 0.005944349 | 0.106619561 | 0.25998119 | 1 | 0.99 | 1 | 0.106619561 | 4.89408E-07 |
| Cdc42ep5 | 2.478E-07 | 1.370451286 | 0.434 | 0.127 | 0.006019567 | 0.814151958 | 0.343665377 | 0.244 | 0.242 | 1 | 0.814151958 | 4.95601E-07 |
| Ccdc50 | 2.53768E-07 | 1.097744654 | 0.868 | 0.758 | 0.006164535 | 0.112706389 | 0.486908242 | 0.854 | 0.853 | 1 | 0.112706389 | 5.07536E-07 |
| Tnfrsf12a | 0.000142403 | -0.972575213 | 0.755 | 0.847 | 1 | 2.54258E-07 | -1.2988567 | 0.854 | 0.942 | 0.00617643 | 0.000142403 | 5.08516E-07 |
| Msi2 | 2.54401E-07 | -0.743154257 | 1 | 1 | 0.006179905 | 0.004370841 | -0.538359158 | 1 | 1 | 1 | 0.004370841 | 5.08802E-07 |
| Synj1 | 2.5615E-07 | 1.836193076 | 0.623 | 0.297 | 0.006222402 | 0.286056891 | 0.673538839 | 0.561 | 0.522 | 1 | 0.286056891 | 5.123E-07 |
| Septin6 | 2.56365E-07 | 2.795180208 | 0.358 | 0.097 | 0.006227624 | 0.000406921 | 1.930314254 | 0.366 | 0.164 | 1 | 0.000406921 | 5.1273E-07 |
| Cnnm4 | 2.59101E-07 | -1.634598404 | 0.226 | 0.657 | 0.006294081 | 0.003677086 | -0.938088863 | 0.244 | 0.526 | 1 | 0.003677086 | 5.18202E-07 |
| Zfp553 | 2.63547E-07 | 2.991223863 | 0.283 | 0.055 | 0.006402083 | 0.421846893 | 0.624211126 | 0.171 | 0.126 | 1 | 0.421846893 | 5.27094E-07 |
| Fzd9 | 2.64233E-07 | -2.108618474 | 0.377 | 0.725 | 0.006418744 | 3.30829E-07 | -1.884420198 | 0.317 | 0.751 | 0.00803649 | 3.30829E-07 | 5.28466E-07 |
| Fbh1 | 2.6457E-07 | 1.877188619 | 0.434 | 0.136 | 0.006426937 | 0.711364825 | 0.334704509 | 0.268 | 0.331 | 1 | 0.711364825 | 5.2914E-07 |
| Irf2bp2 | 2.64749E-07 | 1.138542668 | 0.792 | 0.487 | 0.006431273 | 0.675802938 | 0.323911229 | 0.78 | 0.812 | 1 | 0.675802938 | 5.29497E-07 |
| Prtg | 2.66459E-07 | 2.645708947 | 0.321 | 0.076 | 0.006472813 | 0.546026785 | 0.413993419 | 0.122 | 0.092 | 1 | 0.546026785 | 5.32917E-07 |
| Gstt1 | 2.69975E-07 | 2.395730694 | 0.321 | 0.072 | 0.006558243 | 0.274652187 | 0.811669758 | 0.22 | 0.16 | 1 | 0.274652187 | 5.39951E-07 |
| Gnb1 | 2.71436E-07 | 0.633039882 | 1 | 0.958 | 0.006593733 | 0.019816799 | 0.332016788 | 1 | 0.983 | 1 | 0.019816799 | 5.42873E-07 |
| Zfp609 | 2.73809E-07 | 0.940435223 | 0.887 | 0.581 | 0.006651375 | 0.074241946 | 0.457062141 | 0.78 | 0.761 | 1 | 0.074241946 | 5.47618E-07 |
| Rhod | 2.74204E-07 | 1.553271971 | 0.566 | 0.237 | 0.006660952 | 0.299425451 | 0.776663308 | 0.366 | 0.334 | 1 | 0.299425451 | 5.48407E-07 |
| Ndufv1 | 2.82272E-07 | 1.520372067 | 0.642 | 0.343 | 0.006856948 | 0.009161293 | 0.765414167 | 0.659 | 0.495 | 1 | 0.009161293 | 5.64544E-07 |
| Pou2af1 | 2.82352E-07 | -5.948565214 | 0 | 0.364 | 0.006858894 | 0.001856146 | -4.281736223 | 0 | 0.198 | 1 | 0.001856146 | 5.64704E-07 |
| Mypn | 2.85328E-07 | -5.927426447 | 0 | 0.364 | 0.006931181 | 0.455098275 | -1.16279515 | 0 | 0.014 | 1 | 0.455098275 | 5.70655E-07 |
| Mmp9 | 2.85439E-07 | 5.739685096 | 0.132 | 0.004 | 0.006933874 | 0.00051468 | 0.463503105 | 0.171 | 0.038 | 1 | 0.00051468 | 5.70877E-07 |
| Usp12 | 2.87594E-07 | 1.58098735 | 0.66 | 0.352 | 0.006986229 | 0.054421351 | 0.405031438 | 0.78 | 0.638 | 1 | 0.054421351 | 5.75188E-07 |
| Lrp4 | 2.88162E-07 | 3.569760094 | 0.264 | 0.047 | 0.007000032 | 0.093041697 | 1.795384674 | 0.171 | 0.092 | 1 | 0.093041697 | 5.76324E-07 |
| Lsm7 | 2.88399E-07 | 1.318221327 | 0.755 | 0.47 | 0.007005792 | 0.322379207 | 0.182894302 | 0.61 | 0.536 | 1 | 0.322379207 | 5.76798E-07 |
| Rilpl1 | 2.89251E-07 | 1.525365975 | 0.604 | 0.288 | 0.007026482 | 0.380544315 | 0.562842438 | 0.415 | 0.362 | 1 | 0.380544315 | 5.78502E-07 |
| Mier3 | 2.91363E-07 | 1.491005983 | 0.698 | 0.394 | 0.007077782 | 0.639861901 | 0.196499797 | 0.561 | 0.573 | 1 | 0.639861901 | 5.82725E-07 |
| Igsf10 | 2.97427E-07 | 1.891688189 | 0.585 | 0.246 | 0.0072251 | 0.50866394 | 0.154395026 | 0.415 | 0.375 | 1 | 0.50866394 | 5.94854E-07 |
| Stag1 | 3.00103E-07 | 0.780538263 | 0.981 | 0.915 | 0.007290096 | 0.791290939 | 0.195487571 | 1 | 0.993 | 1 | 0.791290939 | 6.00205E-07 |
| Kifap3 | 3.0265E-07 | 1.389781428 | 0.755 | 0.53 | 0.007351975 | 0.026885284 | 0.784463388 | 0.659 | 0.601 | 1 | 0.026885284 | 6.053E-07 |
| Adamts18 | 3.03626E-07 | 4.8327945 | 0.151 | 0.008 | 0.007375683 | 0.979194004 | 0.83720485 | 0.024 | 0.024 | 1 | 0.979194004 | 6.07252E-07 |
| Kdm6a | 3.04491E-07 | 1.105091827 | 0.906 | 0.669 | 0.007396703 | 0.029461741 | 0.796458507 | 0.829 | 0.829 | 1 | 0.029461741 | 6.08983E-07 |
| Nudt11 | 3.05127E-07 | 4.614154213 | 0.132 | 0.004 | 0.007412139 | 0.252939974 | -0.80665134 | 0.024 | 0.072 | 1 | 0.252939974 | 6.10253E-07 |
| Ctr9 | 3.08803E-07 | 1.425174472 | 0.755 | 0.462 | 0.007501437 | 0.034427943 | 0.708374869 | 0.78 | 0.655 | 1 | 0.034427943 | 6.17605E-07 |
| Porcn | 3.10631E-07 | 2.341135719 | 0.377 | 0.106 | 0.007545839 | 0.942273438 | 0.816143234 | 0.171 | 0.181 | 1 | 0.942273438 | 6.21261E-07 |
| Nsd3 | 3.10727E-07 | 0.858696363 | 0.981 | 0.936 | 0.007548177 | 0.005737094 | 0.379682189 | 1 | 0.99 | 1 | 0.005737094 | 6.21454E-07 |
| Insyn1 | 3.13161E-07 | 3.739685096 | 0.151 | 0.008 | 0.007607315 | 0.842422303 | 0.515276755 | 0.073 | 0.065 | 1 | 0.842422303 | 6.26323E-07 |
| Vgll4 | 3.13605E-07 | 1.267197324 | 0.868 | 0.682 | 0.007618094 | 4.54264E-06 | 1.12364386 | 0.902 | 0.799 | 0.110349705 | 4.54264E-06 | 6.2721E-07 |
| Septin7 | 3.14334E-07 | 0.624954132 | 1 | 0.979 | 0.007635807 | 0.766011858 | 0.113689211 | 1 | 1 | 1 | 0.766011858 | 6.28668E-07 |
| Zfp647 | 3.16115E-07 | 4.154722595 | 0.132 | 0.004 | 0.007679069 | 0.231591308 | -0.747757651 | 0 | 0.034 | 1 | 0.231591308 | 6.3223E-07 |
| Ppp3ca | 3.16369E-07 | -1.458161574 | 1 | 1 | 0.007685246 | 0.642781616 | -0.399335766 | 0.976 | 0.997 | 1 | 0.642781616 | 6.32739E-07 |
| Unc5c | 3.19775E-07 | -1.175089099 | 0.717 | 0.97 | 0.007767976 | 5.20288E-07 | -1.399533504 | 0.561 | 0.928 | 0.012638844 | 5.20288E-07 | 6.3955E-07 |
| Zc3h13 | 3.21119E-07 | 1.028859639 | 0.887 | 0.682 | 0.007800617 | 0.244824554 | 0.351322123 | 0.854 | 0.819 | 1 | 0.244824554 | 6.42237E-07 |
| Anxa2 | 3.26958E-07 | -0.804994662 | 1 | 0.996 | 0.00794246 | 0.000414886 | -0.493674846 | 1 | 0.997 | 1 | 0.000414886 | 6.53916E-07 |
| Xylt2 | 3.27514E-07 | 3.347367673 | 0.208 | 0.025 | 0.00795598 | 0.157045749 | -1.069685746 | 0.024 | 0.089 | 1 | 0.157045749 | 6.55029E-07 |
| Fbxo7 | 3.2922E-07 | -2.079332744 | 0.264 | 0.644 | 0.007997411 | 0.045285907 | -1.049293957 | 0.317 | 0.451 | 1 | 0.045285907 | 6.5844E-07 |
| Chd8 | 3.31544E-07 | 1.331600357 | 0.736 | 0.432 | 0.008053878 | 0.147203718 | 0.416361729 | 0.829 | 0.713 | 1 | 0.147203718 | 6.63089E-07 |
| Rest | 3.3994E-07 | 1.330752959 | 0.792 | 0.53 | 0.008257833 | 0.395860057 | 0.235015935 | 0.78 | 0.775 | 1 | 0.395860057 | 6.79881E-07 |
| Nedd4l | 3.42398E-07 | -1.58574244 | 0.792 | 0.903 | 0.008317539 | 1.60937E-05 | -1.470337077 | 0.78 | 0.942 | 0.390947956 | 1.60937E-05 | 6.84796E-07 |
| Gpc1 | 3.4618E-07 | 1.116179919 | 0.906 | 0.72 | 0.008409414 | 0.064112699 | 0.55894299 | 0.78 | 0.703 | 1 | 0.064112699 | 6.92361E-07 |
| Ehd3 | 3.46633E-07 | -2.086489491 | 0.377 | 0.691 | 0.0084204 | 0.023998769 | -0.970150072 | 0.268 | 0.447 | 1 | 0.023998769 | 6.93265E-07 |
| Msh3 | 3.48509E-07 | 2.811834881 | 0.34 | 0.085 | 0.008465977 | 0.099779582 | 1.451551157 | 0.244 | 0.16 | 1 | 0.099779582 | 6.97018E-07 |
| Map3k3 | 3.48601E-07 | 1.064336358 | 0.925 | 0.682 | 0.008468216 | 0.040822108 | 0.622957517 | 0.951 | 0.863 | 1 | 0.040822108 | 6.97202E-07 |
| Gm32342 | 3.49384E-07 | 3.739685096 | 0.132 | 0.004 | 0.008487229 | 0.446954486 | -0.707115666 | 0.049 | 0.082 | 1 | 0.446954486 | 6.98767E-07 |
| Sarnp | 3.53369E-07 | 0.756363837 | 0.981 | 0.911 | 0.008584033 | 0.262701601 | 0.152154025 | 0.902 | 0.949 | 1 | 0.262701601 | 7.06737E-07 |
| Ppp1r3c | 3.57814E-07 | -2.690767456 | 0.132 | 0.513 | 0.008692027 | 0.014515481 | -1.31254227 | 0.073 | 0.246 | 1 | 0.014515481 | 7.15629E-07 |
| Mideas | 3.60626E-07 | 1.257816088 | 0.717 | 0.381 | 0.008760322 | 1.61899E-05 | 1.377773231 | 0.805 | 0.529 | 0.393285006 | 1.61899E-05 | 7.21251E-07 |
| Il6ra | 3.618E-07 | -3.52334931 | 0.075 | 0.445 | 0.008788845 | 0.017563144 | -1.359192363 | 0.146 | 0.324 | 1 | 0.017563144 | 7.236E-07 |
| Picalm | 3.6278E-07 | 0.848233231 | 0.981 | 0.907 | 0.008812659 | 5.84148E-07 | 0.896225364 | 1 | 0.99 | 0.014190115 | 5.84148E-07 | 7.2556E-07 |
| Fam120a | 3.6347E-07 | 0.874614676 | 0.925 | 0.839 | 0.008829417 | 0.012545201 | 0.447105252 | 0.951 | 0.922 | 1 | 0.012545201 | 7.2694E-07 |
| Zfta | 3.64328E-07 | 1.879088152 | 0.434 | 0.144 | 0.00885025 | 0.001386931 | 1.186654007 | 0.561 | 0.358 | 1 | 0.001386931 | 7.28655E-07 |
| Csrp2 | 3.65084E-07 | 2.629041032 | 0.623 | 0.322 | 0.008868618 | 0.073818332 | 0.888358938 | 0.707 | 0.625 | 1 | 0.073818332 | 7.30168E-07 |
| Spsb2 | 3.65446E-07 | -2.248999591 | 0.302 | 0.627 | 0.008877414 | 0.08751564 | -0.645187917 | 0.39 | 0.495 | 1 | 0.08751564 | 7.30892E-07 |
| Ezh2 | 3.70064E-07 | 1.480492756 | 0.736 | 0.445 | 0.008989602 | 0.010538794 | 1.423646279 | 0.683 | 0.529 | 1 | 0.010538794 | 7.40128E-07 |
| Sf3b2 | 3.70321E-07 | 0.77381778 | 1 | 0.898 | 0.008995848 | 0.002948649 | 0.47463477 | 0.951 | 0.952 | 1 | 0.002948649 | 7.40643E-07 |
| Spast | 3.70371E-07 | 1.446185409 | 0.642 | 0.309 | 0.008997049 | 0.001283445 | 0.956786466 | 0.78 | 0.587 | 1 | 0.001283445 | 7.40742E-07 |
| Dnajb4 | 3.73723E-07 | 1.467961743 | 0.887 | 0.661 | 0.009078486 | 0.001474434 | 0.673358019 | 0.878 | 0.829 | 1 | 0.001474434 | 7.47446E-07 |

| | | | | | | | | | | | | |
|---|---|---|---|---|---|---|---|---|---|---|---|---|
| Dleu2 | 3.76258E-07 | 1.826610039 | 0.792 | 0.496 | 0.009140063 | 0.026998617 | 0.252242349 | 0.878 | 0.771 | 1 | 0.026998617 | 7.52516E-07 |
| Ube2i | 3.79186E-07 | 0.766652143 | 0.962 | 0.907 | 0.009211186 | 0.605971864 | 0.254931437 | 0.951 | 0.949 | 1 | 0.605971864 | 7.58372E-07 |
| Dclk2 | 3.80128E-07 | 3.080722013 | 0.208 | 0.025 | 0.00923407 | 0.768121897 | 0.728680393 | 0.122 | 0.109 | 1 | 0.768121897 | 7.60256E-07 |
| Cks2 | 3.82195E-07 | 1.154722595 | 0.377 | 0.097 | 0.00928429 | 0.000268891 | 2.154395026 | 0.366 | 0.157 | 1 | 0.000268891 | 7.64391E-07 |
| Akt1 | 3.90733E-07 | 1.055186921 | 0.83 | 0.593 | 0.00949169 | 0.801558861 | 0.171430303 | 0.805 | 0.812 | 1 | 0.801558861 | 7.81466E-07 |
| Thrap3 | 3.95667E-07 | 0.681818601 | 0.981 | 0.928 | 0.009611538 | 0.281622639 | 0.262696707 | 0.976 | 0.983 | 1 | 0.281622639 | 7.91333E-07 |
| Mall | 3.95856E-07 | -4.368839361 | 0.019 | 0.381 | 0.009616129 | 0.000685573 | -2.105309656 | 0.098 | 0.369 | 1 | 0.000685573 | 7.91711E-07 |
| Prdx2 | 3.99644E-07 | 1.044463087 | 0.962 | 0.814 | 0.009708158 | 0.045312986 | 0.489281546 | 0.902 | 0.908 | 1 | 0.045312986 | 7.99288E-07 |
| Igsf3 | 3.99816E-07 | 1.270864189 | 0.811 | 0.602 | 0.009712327 | 0.2847316 | 0.271279129 | 0.951 | 0.87 | 1 | 0.2847316 | 7.99632E-07 |
| Pygm | 4.02321E-07 | 2.430357037 | 0.264 | 0.047 | 0.009773181 | 6.16862E-07 | 2.455334214 | 0.366 | 0.096 | 0.014984816 | 6.16862E-07 | 8.04642E-07 |
| Jmy | 4.04675E-07 | 1.227654117 | 0.774 | 0.513 | 0.009830367 | 0.013856455 | 0.495846225 | 0.902 | 0.782 | 1 | 0.013856455 | 8.0935E-07 |
| Pou3f1 | 4.0821E-07 | 2.739685096 | 0.208 | 0.025 | 0.009916245 | 0.007837755 | 1.351778023 | 0.146 | 0.044 | 1 | 0.007837755 | 8.1642E-07 |
| Gxylt2 | 4.0879E-07 | 1.474223952 | 0.83 | 0.648 | 0.009930323 | 0.029004385 | 0.79281073 | 0.732 | 0.689 | 1 | 0.029004385 | 8.1758E-07 |
| Fbxo33 | 4.16713E-07 | 1.960075164 | 0.604 | 0.314 | 0.010122786 | 0.015113198 | 1.042788983 | 0.561 | 0.416 | 1 | 0.015113198 | 8.33425E-07 |
| Fut11 | 4.17981E-07 | 1.644527862 | 0.604 | 0.28 | 0.010153589 | 0.347600645 | 0.380975663 | 0.488 | 0.461 | 1 | 0.347600645 | 8.35961E-07 |
| Gkap1 | 4.18784E-07 | 1.227235095 | 0.736 | 0.424 | 0.01017311 | 0.102781241 | 0.437274243 | 0.659 | 0.539 | 1 | 0.102781241 | 8.37569E-07 |
| Cox4i2 | 4.21138E-07 | -2.254668341 | 0.302 | 0.631 | 0.010230287 | 0.001280194 | -1.096907215 | 0.171 | 0.44 | 1 | 0.001280194 | 8.42276E-07 |
| Ackr3 | 4.29975E-07 | -2.665868154 | 0.264 | 0.61 | 0.010444964 | 0.02074141 | -1.458728037 | 0.341 | 0.529 | 1 | 0.02074141 | 8.59951E-07 |
| Tshz3 | 4.31858E-07 | 2.23878686 | 0.377 | 0.106 | 0.010490696 | 0.018076807 | 1.210422173 | 0.463 | 0.294 | 1 | 0.018076807 | 8.63716E-07 |
| Zfas1 | 4.32038E-07 | 1.121664306 | 0.868 | 0.678 | 0.010495056 | 0.198985182 | 0.564186355 | 0.659 | 0.608 | 1 | 0.198985182 | 8.64075E-07 |
| Yeats2 | 4.32236E-07 | 1.546490315 | 0.623 | 0.309 | 0.010499873 | 0.579733375 | 0.944120054 | 0.439 | 0.464 | 1 | 0.579733375 | 8.64472E-07 |
| Pstpip2 | 3.23185E-05 | 2.102255175 | 0.226 | 0.051 | 0.785080505 | 4.35112E-07 | 2.647380291 | 0.463 | 0.154 | 0.01056975 | 3.23185E-05 | 8.70225E-07 |
| Cav2 | 4.37862E-07 | -2.752168001 | 0.113 | 0.487 | 0.010636554 | 0.088706656 | -0.96369505 | 0.22 | 0.345 | 1 | 0.088706656 | 8.75725E-07 |
| Mocs2 | 4.46554E-07 | 1.517292674 | 0.642 | 0.335 | 0.010847686 | 0.257386873 | 0.403109015 | 0.561 | 0.519 | 1 | 0.257386873 | 8.93107E-07 |
| Rap1gap2 | 4.49254E-07 | 2.438515561 | 0.245 | 0.042 | 0.010913276 | 0.04609121 | 1.363273661 | 0.244 | 0.126 | 1 | 0.04609121 | 8.98508E-07 |
| Stox2 | 4.51516E-07 | 1.280253477 | 0.491 | 0.169 | 0.010968235 | 1.82364E-06 | 1.85900922 | 0.61 | 0.29 | 0.044299855 | 1.82364E-06 | 9.03033E-07 |
| Dek | 4.51984E-07 | 0.623275603 | 0.962 | 0.831 | 0.010979598 | 0.415775518 | 0.211137146 | 0.927 | 0.908 | 1 | 0.415775518 | 9.03968E-07 |
| Ephx2 | 4.58062E-07 | -3.474634025 | 0.019 | 0.39 | 0.011127239 | 0.227738152 | -1.16279515 | 0.024 | 0.075 | 1 | 0.227738152 | 9.16124E-07 |
| Oasl2 | 0.249526738 | 0.47665069 | 0.057 | 0.025 | 1 | 4.60876E-07 | 3.311136038 | 0.146 | 0.01 | 0.011195608 | 0.249526738 | 9.21752E-07 |
| Sema3f | 0.074441876 | 1.47665069 | 0.075 | 0.025 | 1 | 4.67354E-07 | 4.159132945 | 0.146 | 0.01 | 0.011352953 | 0.074441876 | 9.34707E-07 |
| Xpo1 | 0.115306237 | 0.488840099 | 0.604 | 0.5 | 1 | 4.71493E-07 | 1.316540843 | 0.805 | 0.563 | 0.011453508 | 0.115306237 | 9.42986E-07 |
| Arpc3 | 4.7487E-07 | 0.702690888 | 0.981 | 0.894 | 0.01153553 | 0.039646231 | 0.4345968 | 0.902 | 0.918 | 1 | 0.039646231 | 9.49739E-07 |
| Sema3d | 4.76454E-07 | 1.107527427 | 0.736 | 0.436 | 0.011574031 | 0.022645654 | 1.104754737 | 0.634 | 0.529 | 1 | 0.022645654 | 9.52909E-07 |
| Ccn3 | 1.73149E-05 | 3.8327945 | 0.132 | 0.013 | 0.42061249 | 4.81271E-07 | 4.685201756 | 0.195 | 0.024 | 0.011691039 | 1.73149E-05 | 9.62542E-07 |
| Stag2 | 4.88084E-07 | 0.820303556 | 0.962 | 0.831 | 0.011856534 | 0.391131373 | 0.241925058 | 0.976 | 0.969 | 1 | 0.391131373 | 9.76168E-07 |
| Lactb2 | 4.94681E-07 | -2.673942834 | 0.17 | 0.538 | 0.012016785 | 0.012928729 | -1.362467495 | 0.098 | 0.283 | 1 | 0.012928729 | 9.89361E-07 |
| Fap | 4.96889E-07 | 0.79510221 | 0.943 | 0.636 | 0.012070417 | 0.181300583 | -0.369703331 | 0.854 | 0.915 | 1 | 0.181300583 | 9.93777E-07 |
| Ptk2 | 5.01581E-07 | 0.836303286 | 0.943 | 0.775 | 0.012184402 | 0.069289412 | 0.373138062 | 0.976 | 0.942 | 1 | 0.069289412 | 1.00316E-06 |
| Lnx1 | 0.771605102 | -0.430239906 | 0.019 | 0.025 | 1 | 5.01908E-07 | 3.078212949 | 0.22 | 0.031 | 0.012192349 | 0.771605102 | 1.00382E-06 |
| Map2k2 | 5.02114E-07 | 0.853418172 | 0.887 | 0.78 | 0.012197345 | 0.372009245 | 0.210014114 | 0.854 | 0.846 | 1 | 0.372009245 | 1.00423E-06 |
| Rfxap | 5.02769E-07 | 1.631160639 | 0.528 | 0.212 | 0.012213272 | 0.737215785 | 0.448507743 | 0.341 | 0.416 | 1 | 0.737215785 | 1.00554E-06 |
| Kctd12 | 5.0344E-07 | 1.900966003 | 0.528 | 0.212 | 0.01222957 | 2.81631E-06 | 3.178993367 | 0.561 | 0.246 | 0.068413713 | 2.81631E-06 | 1.00688E-06 |
| Srrm2 | 5.07582E-07 | 0.547660835 | 1 | 1 | 0.012330192 | 0.006618892 | -0.210800523 | 1 | 1 | 1 | 0.006618892 | 1.01516E-06 |
| Fam117b | 5.10628E-07 | 1.669295768 | 0.736 | 0.487 | 0.012404176 | 0.000550179 | 1.052264077 | 0.854 | 0.71 | 1 | 0.000550179 | 1.02126E-06 |
| S100a11 | 5.12632E-07 | 0.820027204 | 1 | 0.941 | 0.012452868 | 0.296322117 | 0.31054099 | 0.976 | 0.966 | 1 | 0.296322117 | 1.02526E-06 |
| Ankrd52 | 5.13872E-07 | 1.627974708 | 0.623 | 0.301 | 0.012482984 | 0.623136195 | 0.110020276 | 0.39 | 0.478 | 1 | 0.623136195 | 1.02774E-06 |
| Rev1 | 5.21432E-07 | 1.071982164 | 0.811 | 0.568 | 0.012666622 | 0.997198706 | 0.157914967 | 0.683 | 0.836 | 1 | 0.997198706 | 1.04286E-06 |
| Met | 6.78834E-06 | -2.768109545 | 0.17 | 0.492 | 0.164902307 | 5.24422E-07 | -1.964716186 | 0.415 | 0.765 | 0.012739268 | 6.78834E-06 | 1.04884E-06 |
| Spire1 | 5.29318E-07 | -0.806583692 | 0.774 | 0.962 | 0.012858201 | 0.000430207 | -0.648747645 | 0.829 | 0.908 | 1 | 0.000430207 | 1.05864E-06 |
| Cd248 | 0.003243872 | 3.47665069 | 0.057 | 0.004 | 1 | 5.30651E-07 | 3.744095445 | 0.195 | 0.024 | 0.012890564 | 0.003243872 | 1.0613E-06 |
| Spock1 | 5.30732E-07 | 2.347367673 | 0.283 | 0.055 | 0.012892551 | 0.47105301 | 0.136765132 | 0.122 | 0.085 | 1 | 0.47105301 | 1.06146E-06 |
| Foxp4 | 5.36934E-07 | 1.165693713 | 0.774 | 0.517 | 0.013043208 | 0.003429916 | 0.65571046 | 0.854 | 0.795 | 1 | 0.003429916 | 1.07387E-06 |
| Rnf24 | 5.39694E-07 | 1.398391676 | 0.66 | 0.314 | 0.013110247 | 0.010125707 | 0.847400617 | 0.707 | 0.519 | 1 | 0.010125707 | 1.07939E-06 |
| Adamts17 | 5.43516E-07 | 2.130475049 | 0.415 | 0.14 | 0.013203094 | 3.63507E-06 | 1.858578501 | 0.439 | 0.154 | 0.088303055 | 3.63507E-06 | 1.08703E-06 |
| Efhd2 | 5.47298E-07 | -2.277819306 | 0.377 | 0.695 | 0.013294973 | 0.011408831 | -0.972789954 | 0.537 | 0.686 | 1 | 0.011408831 | 1.0946E-06 |
| Sgcb | 5.53362E-07 | 2.091986839 | 0.453 | 0.169 | 0.013442268 | 0.002662037 | 1.626637864 | 0.39 | 0.215 | 1 | 0.002662037 | 1.10672E-06 |
| Man2a1 | 0.828009926 | -0.461164479 | 0.623 | 0.627 | 1 | 5.55538E-07 | 1.081688859 | 0.829 | 0.471 | 0.013495126 | 0.828009926 | 1.11108E-06 |
| Narf | 5.63465E-07 | 2.184469938 | 0.434 | 0.144 | 0.013687699 | 0.096549946 | 1.377773231 | 0.22 | 0.13 | 1 | 0.096549946 | 1.12693E-06 |
| Mgp | 0.001443697 | -1.491465211 | 0.981 | 0.979 | 1 | 5.65736E-07 | -1.561044674 | 1 | 0.99 | 0.013742863 | 0.001443697 | 1.13147E-06 |
| Taf3 | 5.77525E-07 | 1.459936816 | 0.679 | 0.415 | 0.01402924 | 0.097283969 | 0.434080672 | 0.659 | 0.556 | 1 | 0.097283969 | 1.15505E-06 |
| C630043F03Rik | 5.79058E-07 | 2.034428361 | 0.302 | 0.064 | 0.014066482 | 0.264057548 | 0.471555377 | 0.244 | 0.171 | 1 | 0.264057548 | 1.15812E-06 |
| Vill | 5.79963E-07 | -1.693274312 | 0.208 | 0.602 | 0.014088449 | 1.69012E-05 | -1.546499443 | 0.22 | 0.59 | 0.410562899 | 1.69012E-05 | 1.15992E-06 |
| Papola | 5.85151E-07 | 0.768066897 | 1 | 0.89 | 0.014214489 | 0.734734268 | 0.119604581 | 0.951 | 0.962 | 1 | 0.734734268 | 1.1703E-06 |
| Pgrmc1 | 5.86532E-07 | 0.882835182 | 0.943 | 0.737 | 0.014248036 | 0.000486257 | 0.6255869 | 0.878 | 0.887 | 1 | 0.000486257 | 1.17306E-06 |
| Tbl1xr1 | 5.87892E-07 | 1.565098717 | 0.698 | 0.377 | 0.014281061 | 0.001417855 | 0.594091138 | 0.854 | 0.628 | 1 | 0.001417855 | 1.17578E-06 |
| Fkbp7 | 5.88952E-07 | 1.093938086 | 0.868 | 0.758 | 0.014306834 | 0.305868027 | 0.272610147 | 0.707 | 0.747 | 1 | 0.305868027 | 1.1779E-06 |
| Phgdh | 5.89187E-07 | 2.553271971 | 0.302 | 0.068 | 0.014312542 | 0.140916437 | 0.627751484 | 0.171 | 0.092 | 1 | 0.140916437 | 1.17837E-06 |
| Nol4l | 5.99378E-07 | 2.275016829 | 0.302 | 0.068 | 0.014560098 | 0.131360582 | 0.83720485 | 0.268 | 0.184 | 1 | 0.131360582 | 1.19876E-06 |
| Marf1 | 6.00139E-07 | 1.365401596 | 0.717 | 0.436 | 0.014578566 | 5.64421E-06 | 1.153508395 | 0.805 | 0.543 | 0.137109048 | 5.64421E-06 | 1.20028E-06 |
| Stxbp6 | 6.04536E-07 | -2.365985364 | 0.094 | 0.496 | 0.014685399 | 0.11149326 | -0.529117364 | 0.293 | 0.413 | 1 | 0.11149326 | 1.20907E-06 |
| Btf3l4 | 6.12463E-07 | 1.408009024 | 0.642 | 0.314 | 0.01487794 | 0.75497175 | 0.377773231 | 0.341 | 0.352 | 1 | 0.75497175 | 1.22492E-06 |
| Mapre1 | 6.19166E-07 | 0.72437204 | 0.943 | 0.881 | 0.015040772 | 0.378112817 | 0.289825532 | 0.902 | 0.945 | 1 | 0.378112817 | 1.23833E-06 |
| Grn | 6.20518E-07 | 1.015171242 | 0.83 | 0.644 | 0.015073632 | 0.580648663 | 0.200663562 | 0.732 | 0.775 | 1 | 0.580648663 | 1.24104E-06 |
| Col11a1 | 0.000103415 | -0.707535005 | 1 | 1 | 1 | 6.2101E-07 | -0.96854445 | 0.976 | 0.99 | 0.015085572 | 0.000103415 | 1.24202E-06 |
| Rsbn1 | 6.39705E-07 | 1.111653873 | 0.849 | 0.568 | 0.015539703 | 0.013426609 | 0.724995346 | 0.854 | 0.788 | 1 | 0.013426609 | 1.27941E-06 |
| Agap3 | 6.47208E-07 | 1.849868013 | 0.509 | 0.203 | 0.015721983 | 0.296701681 | 0.422167351 | 0.561 | 0.522 | 1 | 0.296701681 | 1.29442E-06 |
| Cdkn1b | 6.54331E-07 | 1.469924826 | 0.736 | 0.432 | 0.015895 | 0.428523239 | 0.738426995 | 0.683 | 0.765 | 1 | 0.428523239 | 1.30866E-06 |
| Tnfrsf19 | 6.67618E-07 | 5.380142709 | 0.264 | 0.055 | 0.016217774 | 0.000546293 | 4.296636468 | 0.171 | 0.041 | 1 | 0.000546293 | 1.33524E-06 |
| Ifitm2 | 6.7102E-07 | 0.819879409 | 1 | 0.915 | 0.016300426 | 0.008713767 | 0.517550506 | 0.976 | 0.956 | 1 | 0.008713767 | 1.34204E-06 |
| Pdgfa | 6.82065E-07 | 1.347367673 | 0.698 | 0.39 | 0.016568724 | 0.058396941 | 0.24706211 | 0.878 | 0.792 | 1 | 0.058396941 | 1.36413E-06 |
| Atp5d | 6.83169E-07 | 0.763932642 | 1 | 0.907 | 0.016595553 | 0.976199396 | 0.145599789 | 0.854 | 0.939 | 1 | 0.976199396 | 1.36634E-06 |
| Uri1 | 6.83935E-07 | 1.586438835 | 0.642 | 0.326 | 0.016614157 | 0.015706458 | 0.944120054 | 0.634 | 0.505 | 1 | 0.015706458 | 1.36787E-06 |
| Tra2b | 6.85792E-07 | 0.838860501 | 1 | 0.987 | 0.016659248 | 0.062114695 | 0.267455443 | 1 | 0.997 | 1 | 0.062114695 | 1.37158E-06 |
| Trap1 | 6.88372E-07 | 1.267933205 | 0.623 | 0.28 | 0.01672193 | 0.14639642 | 0.853692973 | 0.439 | 0.372 | 1 | 0.14639642 | 1.37674E-06 |
| Eid2 | 6.89657E-07 | 1.614154213 | 0.453 | 0.148 | 0.016753152 | 0.897637051 | 0.156483366 | 0.268 | 0.29 | 1 | 0.897637051 | 1.37931E-06 |
| Zbtb12 | 7.07931E-07 | 2.695290976 | 0.245 | 0.042 | 0.017197064 | 0.224692581 | 1.541748966 | 0.146 | 0.089 | 1 | 0.224692581 | 1.41586E-06 |
| Msl2 | 7.16394E-07 | 1.200526284 | 0.717 | 0.386 | 0.017402652 | 0.287757156 | 0.278120981 | 0.78 | 0.734 | 1 | 0.287757156 | 1.43279E-06 |
| Fam89a | 7.18306E-07 | -4.916739768 | 0.019 | 0.369 | 0.017449094 | 0.114564565 | -2.332720152 | 0 | 0.058 | 1 | 0.114564565 | 1.43661E-06 |
| Taok1 | 7.28022E-07 | 0.8327945 | 0.943 | 0.822 | 0.017685111 | 0.001975535 | 0.527962015 | 0.976 | 0.98 | 1 | 0.001975535 | 1.45604E-06 |
| Six1 | 7.3299E-07 | 2.31000082 | 0.396 | 0.131 | 0.017805798 | 2.03033E-05 | 1.586661388 | 0.61 | 0.311 | 0.493208033 | 2.03033E-05 | 1.46598E-06 |
| Arhgef2 | 7.40563E-07 | 1.515604324 | 0.585 | 0.263 | 0.017989753 | 0.014768224 | 0.79281073 | 0.585 | 0.399 | 1 | 0.014768224 | 1.48113E-06 |
| Apc | 7.44782E-07 | 1.080297754 | 0.868 | 0.644 | 0.018092237 | 0.471896409 | -0.152924906 | 0.805 | 0.853 | 1 | 0.471896409 | 1.48956E-06 |
| Klhl21 | 1.0308E-05 | -1.522571787 | 0.377 | 0.657 | 0.250403011 | 7.49632E-07 | -1.559538021 | 0.293 | 0.717 | 0.018210062 | 1.0308E-05 | 1.49926E-06 |
| Sec63 | 7.52107E-07 | 0.796165578 | 0.962 | 0.924 | 0.018270178 | 0.080692341 | 0.422167351 | 0.976 | 0.959 | 1 | 0.080692341 | 1.50421E-06 |
| Jpt1 | 0.001142339 | 0.900966003 | 0.509 | 0.305 | 1 | 7.55877E-07 | 1.304709769 | 0.756 | 0.382 | 0.018361755 | 0.001142339 | 1.51175E-06 |
| Epyc | 6.01729E-06 | -0.864589097 | 0.377 | 0.788 | 0.146172077 | 7.73535E-07 | -2.859868017 | 0.244 | 0.621 | 0.018790714 | 6.01729E-06 | 1.54707E-06 |
| Ost4 | 7.7678E-07 | -0.854672023 | 0.962 | 0.962 | 0.018869551 | 0.00011068 | -0.736786533 | 0.878 | 0.942 | 1 | 0.00011068 | 1.55356E-06 |
| Lrrc58 | 7.8339E-07 | 0.622891808 | 1 | 0.881 | 0.019030103 | 0.062955087 | 0.246286065 | 0.951 | 0.98 | 1 | 0.062955087 | 1.56678E-06 |

| | | | | | | | | | | | |
|---|---|---|---|---|---|---|---|---|---|---|---|---|
| Dnaja1 | 7.85211E-07 | 1.04118745 | 1 | 0.953 | 0.019074334 | 0.109602991 | 0.367743592 | 1 | 0.986 | 1 | 0.109602991 | 1.57042E-06 |
| Slc12a8 | 0.014067732 | 2.039245377 | 0.151 | 0.055 | 1 | 7.89286E-07 | 3.224227973 | 0.171 | 0.017 | 0.019173332 | 0.014067732 | 1.57857E-06 |
| Eif5 | 7.99968E-07 | 0.63062462 | 1 | 0.975 | 0.019432826 | 0.034345783 | -0.244869209 | 1 | 1 | 1 | 0.034345783 | 1.59994E-06 |
| Tmbim4 | 1.20309E-05 | -0.787281854 | 0.792 | 0.886 | 0.292254254 | 8.05916E-07 | -1.144179472 | 0.585 | 0.867 | 0.019577305 | 1.20309E-05 | 1.61183E-06 |
| Kirrel | 8.23751E-07 | 0.91798915 | 0.868 | 0.602 | 0.02001055 | 0.433156422 | 0.27355858 | 0.707 | 0.754 | 1 | 0.433156422 | 1.6475E-06 |
| Cast | 8.25067E-07 | -1.030632447 | 0.925 | 0.958 | 0.020042538 | 0.004806237 | -0.620287228 | 0.829 | 0.962 | 1 | 0.004806237 | 1.65013E-06 |
| Pcdh10 | 0.000354785 | 3.154722595 | 0.075 | 0.004 | 1 | 8.30389E-07 | 2.83720485 | 0.195 | 0.024 | 0.020171802 | 0.000354785 | 1.66078E-06 |
| Ints6l | 8.31504E-07 | 1.557049042 | 0.623 | 0.258 | 0.020198888 | 0.093307764 | 1.536035315 | 0.61 | 0.56 | 1 | 0.093307764 | 1.66301E-06 |
| Yes1 | 8.3521E-07 | 1.114565468 | 0.792 | 0.475 | 0.020288913 | 0.504853989 | 0.213689108 | 0.707 | 0.662 | 1 | 0.504853989 | 1.67042E-06 |
| Gsto1 | 8.38969E-07 | 1.41149801 | 0.528 | 0.208 | 0.020380225 | 5.50465E-05 | 0.983494292 | 0.707 | 0.399 | 1 | 5.50465E-05 | 1.67794E-06 |
| Setx | 8.41109E-07 | 1.375631525 | 0.66 | 0.343 | 0.020432219 | 0.147303717 | 0.588291552 | 0.659 | 0.577 | 1 | 0.147303717 | 1.68222E-06 |
| Sos2 | 8.42793E-07 | 1.121664306 | 0.774 | 0.547 | 0.020473119 | 0.827730761 | 0.197952194 | 0.756 | 0.812 | 1 | 0.827730761 | 1.68558E-06 |
| Cav1 | 8.44451E-07 | -5.078897082 | 0.038 | 0.39 | 0.020513401 | 0.001229485 | -3.917682652 | 0.049 | 0.28 | 1 | 0.001229485 | 1.6889E-06 |
| Cab39l | 8.46433E-07 | -1.374800919 | 0.358 | 0.72 | 0.020561547 | 0.000507437 | -1.421753701 | 0.561 | 0.744 | 1 | 0.000507437 | 1.69286E-06 |
| Lef1 | 8.47293E-07 | 2.047807391 | 0.509 | 0.195 | 0.020582436 | 0.218282457 | -0.336443238 | 0.366 | 0.433 | 1 | 0.218282457 | 1.69458E-06 |
| Ifitm10 | 0.132300997 | -0.117356951 | 0.528 | 0.686 | 1 | 8.52316E-07 | -1.338237157 | 0.39 | 0.812 | 0.020704464 | 0.132300997 | 1.70463E-06 |
| Smap1 | 8.56237E-07 | 1.041264545 | 0.849 | 0.754 | 0.02079971 | 0.898195922 | 0.178751284 | 0.854 | 0.843 | 1 | 0.898195922 | 1.71247E-06 |
| Nynrin | 8.56752E-07 | 1.950364096 | 0.434 | 0.144 | 0.020812228 | 0.101570317 | 1.013235214 | 0.39 | 0.294 | 1 | 0.101570317 | 1.7135E-06 |
| Slpi | 8.75429E-07 | 0.739685096 | 0.585 | 0.246 | 0.021265914 | 0.998440231 | -0.481834966 | 0.366 | 0.41 | 1 | 0.998440231 | 1.75086E-06 |
| Dbndd2 | 8.76846E-07 | 1.980693195 | 0.434 | 0.153 | 0.021300353 | 0.637811537 | 0.174239837 | 0.293 | 0.263 | 1 | 0.637811537 | 1.75369E-06 |
| Itih5 | 8.9052E-07 | 3.842046813 | 0.34 | 0.097 | 0.021632507 | 1.22304E-05 | 4.602739596 | 0.146 | 0.017 | 0.297100124 | 1.22304E-05 | 1.78104E-06 |
| B4gat1 | 8.99449E-07 | 1.707263618 | 0.472 | 0.169 | 0.021849417 | 0.000326335 | 1.35607816 | 0.415 | 0.177 | 1 | 0.000326335 | 1.7989E-06 |
| Phf6 | 9.00491E-07 | 1.112287328 | 0.604 | 0.263 | 0.021874721 | 0.011565808 | 0.713083538 | 0.634 | 0.457 | 1 | 0.011565808 | 1.80098E-06 |
| Plcl1 | 6.06985E-05 | 1.940128544 | 0.415 | 0.182 | 1 | 9.1364E-07 | 2.547698233 | 0.488 | 0.174 | 0.022194138 | 6.06985E-05 | 1.82728E-06 |
| Mon2 | 9.13793E-07 | 0.856789981 | 0.943 | 0.758 | 0.022197854 | 0.045119513 | 0.509574396 | 0.976 | 0.908 | 1 | 0.045119513 | 1.82758E-06 |
| Pde10a | 9.14362E-07 | 2.099227482 | 0.283 | 0.059 | 0.022211679 | 0.573158132 | 0.441003993 | 0.366 | 0.324 | 1 | 0.573158132 | 1.82872E-06 |
| Osbpl1a | 9.16725E-07 | -1.142052023 | 0.623 | 0.852 | 0.022269084 | 0.000102306 | -1.039084704 | 0.537 | 0.823 | 1 | 0.000102306 | 1.83345E-06 |
| Ext2 | 9.45094E-07 | -1.006065612 | 0.679 | 0.903 | 0.022958224 | 0.750434517 | -0.175038746 | 0.829 | 0.788 | 1 | 0.750434517 | 1.89019E-06 |
| Srsf4 | 9.77732E-07 | 0.961185036 | 0.811 | 0.555 | 0.023751055 | 0.731020362 | 0.168986845 | 0.805 | 0.86 | 1 | 0.731020362 | 1.95546E-06 |
| Rnf111 | 9.82963E-07 | 1.021910766 | 0.868 | 0.631 | 0.023878128 | 0.499929561 | 0.326242931 | 0.854 | 0.84 | 1 | 0.499929561 | 1.96592E-06 |
| Supt16 | 9.85579E-07 | 0.934941387 | 0.906 | 0.682 | 0.023941695 | 0.00685424 | 0.612189346 | 0.902 | 0.853 | 1 | 0.00685424 | 1.97116E-06 |
| Usp46 | 9.88826E-07 | 2.051629102 | 0.321 | 0.076 | 0.024020553 | 0.001213862 | 1.83720485 | 0.268 | 0.099 | 1 | 0.001213862 | 1.97765E-06 |
| Myo9b | 0.001084508 | -0.817756342 | 0.66 | 0.843 | 1 | 9.92198E-07 | -1.173364392 | 0.756 | 0.939 | 0.024102476 | 0.001084508 | 1.9844E-06 |
| Git1 | 9.92446E-07 | 2.417757001 | 0.34 | 0.093 | 0.024108486 | 0.148325082 | 0.854482841 | 0.268 | 0.177 | 1 | 0.148325082 | 1.98489E-06 |
| H13 | 1.00647E-06 | 0.777684864 | 0.962 | 0.877 | 0.024449201 | 0.113835151 | 0.179444515 | 0.976 | 0.935 | 1 | 0.113835151 | 2.01294E-06 |
| Mef2a | 1.01452E-06 | 0.525112873 | 0.981 | 0.915 | 0.024644815 | 0.03716167 | -0.422265878 | 0.976 | 0.99 | 1 | 0.03716167 | 2.02905E-06 |
| 1810055G02Rik | 1.04213E-06 | -1.213428517 | 0.396 | 0.754 | 0.025315342 | 0.002106189 | -0.815734445 | 0.366 | 0.648 | 1 | 0.002106189 | 2.08425E-06 |
| Myl6 | 1.04846E-06 | 0.695095529 | 1 | 0.975 | 0.02546918 | 0.053599516 | 0.471459145 | 1 | 0.993 | 1 | 0.053599516 | 2.09692E-06 |
| Rflna | 0.794383231 | 0.197507461 | 0.566 | 0.691 | 1 | 1.08054E-06 | -1.342830179 | 0.537 | 0.853 | 0.026248374 | 0.794383231 | 2.16107E-06 |
| Trim47 | 1.08478E-06 | -1.512592568 | 0.472 | 0.737 | 0.026351585 | 0.006428471 | -0.65500051 | 0.585 | 0.785 | 1 | 0.006428471 | 2.16957E-06 |
| Il10rb | 1.09274E-06 | 1.984797593 | 0.396 | 0.119 | 0.026544791 | 0.524999404 | 0.618370248 | 0.317 | 0.29 | 1 | 0.524999404 | 2.18547E-06 |
| Rab34 | 1.09773E-06 | 1.051382845 | 0.792 | 0.517 | 0.026666141 | 0.185289411 | 0.613275193 | 0.61 | 0.56 | 1 | 0.185289411 | 2.19547E-06 |
| Gm15559 | 1.09912E-06 | 3.002719501 | 0.208 | 0.03 | 0.026699907 | 0.003474684 | 2.185128153 | 0.122 | 0.027 | 1 | 0.003474684 | 2.19825E-06 |
| Dhx15 | 1.1017E-06 | 0.953088734 | 0.887 | 0.737 | 0.026762414 | 0.002847591 | 0.596381723 | 0.902 | 0.877 | 1 | 0.002847591 | 2.20339E-06 |
| Syt8 | 1.1054E-06 | -3.805279337 | 0.038 | 0.39 | 0.026852258 | 0.054990125 | -0.828131067 | 0.146 | 0.304 | 1 | 0.054990125 | 2.21079E-06 |
| Pura | 1.10611E-06 | 0.673595905 | 1 | 0.911 | 0.026869557 | 0.259053462 | 0.271039945 | 0.951 | 0.956 | 1 | 0.259053462 | 2.21221E-06 |
| Me1 | 1.10612E-06 | -0.977254531 | 0.623 | 0.852 | 0.026869974 | 9.56268E-05 | -0.952872081 | 0.732 | 0.877 | 1 | 9.56268E-05 | 2.21225E-06 |
| Arhgap17 | 1.11045E-06 | -1.541783392 | 0.642 | 0.826 | 0.026975148 | 0.000143983 | -1.324499645 | 0.732 | 0.863 | 1 | 0.000143983 | 2.22091E-06 |
| Srrt | 1.12366E-06 | 1.023478062 | 0.887 | 0.606 | 0.02729596 | 0.682511412 | 0.196747236 | 0.78 | 0.802 | 1 | 0.682511412 | 2.24732E-06 |
| Atp6v0e | 1.13109E-06 | -1.077938162 | 0.981 | 0.97 | 0.027476352 | 0.009920304 | -0.639019711 | 0.902 | 0.956 | 1 | 0.009920304 | 2.26217E-06 |
| Brd3 | 1.13661E-06 | 1.080722013 | 0.755 | 0.466 | 0.02761047 | 0.036179549 | 0.472132263 | 0.927 | 0.778 | 1 | 0.036179549 | 2.27321E-06 |
| Sap30 | 1.14042E-06 | 2.199116714 | 0.34 | 0.089 | 0.02770306 | 0.354341499 | 0.425391251 | 0.415 | 0.341 | 1 | 0.354341499 | 2.28084E-06 |
| Upk1b | 1.14622E-06 | 3.824573993 | 0.17 | 0.017 | 0.027844047 | 0.000228663 | 3.126711467 | 0.122 | 0.017 | 1 | 0.000228663 | 2.29244E-06 |
| Dnm2 | 1.15019E-06 | 0.992892897 | 0.849 | 0.653 | 0.027940462 | 0.000175059 | 0.840947251 | 0.927 | 0.846 | 1 | 0.000175059 | 2.30038E-06 |
| Mtrfr | 1.15467E-06 | 2.226872381 | 0.415 | 0.136 | 0.028049292 | 0.339834243 | -0.213421223 | 0.098 | 0.16 | 1 | 0.339834243 | 2.30934E-06 |
| Pnma8b | 1.16201E-06 | 3.154722595 | 0.208 | 0.03 | 0.028227551 | 0.935120464 | -0.116991461 | 0.073 | 0.075 | 1 | 0.935120464 | 2.32402E-06 |
| Ppm1l | 1.16227E-06 | 2.392882332 | 0.358 | 0.102 | 0.028233782 | 0.077359509 | 0.584224109 | 0.585 | 0.433 | 1 | 0.077359509 | 2.32453E-06 |
| Asphd2 | 1.17025E-06 | 3.417757001 | 0.17 | 0.017 | 0.028427637 | 0.001053884 | 3.574170444 | 0.073 | 0.007 | 1 | 0.001053884 | 2.34049E-06 |
| Ing1 | 1.17476E-06 | 1.435274762 | 0.717 | 0.466 | 0.028537319 | 7.77401E-05 | 1.185841123 | 0.78 | 0.529 | 1 | 7.77401E-05 | 2.34952E-06 |
| Kdm2a | 1.17769E-06 | 1.191716802 | 0.925 | 0.746 | 0.028608565 | 0.000371956 | 0.621135426 | 1 | 0.959 | 1 | 0.000371956 | 2.35539E-06 |
| Ppp1r9b | 1.18575E-06 | 1.932330173 | 0.396 | 0.123 | 0.028804125 | 0.109195736 | 0.847258514 | 0.415 | 0.311 | 1 | 0.109195736 | 2.37149E-06 |
| Mid1 | 1.18664E-06 | 1.587442372 | 0.774 | 0.576 | 0.028825777 | 0.001398012 | 1.618036329 | 0.756 | 0.556 | 1 | 0.001398012 | 2.37327E-06 |
| Ddx50 | 1.18786E-06 | 0.735618709 | 0.981 | 0.932 | 0.028855432 | 0.221296204 | 0.344005328 | 0.976 | 0.959 | 1 | 0.221296204 | 2.37571E-06 |
| Osr2 | 0.617621149 | 0.984797593 | 0.038 | 0.025 | 1 | 1.21157E-06 | 2.504629511 | 0.244 | 0.041 | 0.029431362 | 0.617621149 | 2.42313E-06 |
| Osbpl8 | 1.21525E-06 | 1.105910345 | 0.925 | 0.746 | 0.029520815 | 0.000445104 | 0.966888869 | 0.951 | 0.85 | 1 | 0.000445104 | 2.4305E-06 |
| Ano6 | 1.21706E-06 | -1.117138503 | 0.811 | 0.953 | 0.0295648 | 0.003685928 | -0.689296804 | 0.854 | 0.976 | 1 | 0.003685928 | 2.43412E-06 |
| Srebf2 | 1.22779E-06 | -1.309148089 | 0.547 | 0.792 | 0.029825377 | 0.005471106 | -0.821467337 | 0.634 | 0.771 | 1 | 0.005471106 | 2.45557E-06 |
| Slc1a1 | 1.24851E-06 | -2.349408831 | 0.189 | 0.559 | 0.030328826 | 0.30756876 | 0.252242349 | 0.22 | 0.307 | 1 | 0.30756876 | 2.49702E-06 |
| Zfp131 | 1.25158E-06 | 0.944637069 | 0.906 | 0.712 | 0.030403295 | 0.166780719 | 0.405916196 | 0.854 | 0.887 | 1 | 0.166780719 | 2.50315E-06 |
| Ankhd1 | 1.25528E-06 | 1.038188845 | 0.906 | 0.822 | 0.030493238 | 0.001798639 | 0.828525094 | 0.976 | 0.891 | 1 | 0.001798639 | 2.51056E-06 |
| Rnf217 | 1.25844E-06 | 1.746739853 | 0.717 | 0.436 | 0.030570026 | 0.007101564 | 0.744758601 | 0.805 | 0.683 | 1 | 0.007101564 | 2.51688E-06 |
| Dock11 | 1.26638E-06 | 2.085775241 | 0.585 | 0.258 | 0.030762821 | 0.243629592 | 0.219820871 | 0.561 | 0.444 | 1 | 0.243629592 | 2.53275E-06 |
| Hnrnpa3 | 1.26675E-06 | 0.644118324 | 1 | 0.983 | 0.030771811 | 0.324092381 | 0.128550132 | 1 | 0.993 | 1 | 0.324092381 | 2.53349E-06 |
| Vezf1 | 1.26765E-06 | 0.948878019 | 0.962 | 0.792 | 0.030793827 | 0.16392757 | 0.238039254 | 0.902 | 0.935 | 1 | 0.16392757 | 2.5353E-06 |
| Ptbp1 | 1.27708E-06 | 0.860152616 | 0.925 | 0.801 | 0.031022944 | 0.267154896 | 0.266839725 | 0.805 | 0.925 | 1 | 0.267154896 | 2.55417E-06 |
| Tob2 | 1.2795E-06 | 0.919946449 | 0.925 | 0.742 | 0.031081591 | 0.008963961 | 0.477464401 | 0.951 | 0.918 | 1 | 0.008963961 | 2.559E-06 |
| Bcl2l12 | 1.28227E-06 | 1.962077517 | 0.321 | 0.076 | 0.031149008 | 0.004874322 | 1.363273661 | 0.195 | 0.065 | 1 | 0.004874322 | 2.56455E-06 |
| Slc24a3 | 1.28767E-06 | 3.870929629 | 0.245 | 0.047 | 0.031280149 | 1.57415E-06 | 2.139233387 | 0.293 | 0.061 | 0.038239285 | 1.57415E-06 | 2.57534E-06 |
| Med13 | 1.31048E-06 | 0.728713978 | 0.962 | 0.928 | 0.031834202 | 0.095229545 | 0.391405097 | 1 | 0.983 | 1 | 0.095229545 | 2.62096E-06 |
| 2010109A12Rik | 1.31289E-06 | 2.669295768 | 0.283 | 0.064 | 0.031892715 | 0.167256065 | 1.000703582 | 0.171 | 0.102 | 1 | 0.167256065 | 2.62578E-06 |
| Chd2 | 1.33113E-06 | 1.03721851 | 0.925 | 0.839 | 0.032335926 | 0.002909191 | 0.488697483 | 1 | 0.973 | 1 | 0.002909191 | 2.66227E-06 |
| Slc25a11 | 1.33484E-06 | 1.339147166 | 0.698 | 0.407 | 0.032425932 | 0.197113179 | 0.807457506 | 0.439 | 0.372 | 1 | 0.197113179 | 2.66968E-06 |
| Atm | 1.3463E-06 | 1.776210972 | 0.434 | 0.148 | 0.032704367 | 0.497859501 | 0.568716014 | 0.268 | 0.235 | 1 | 0.497859501 | 2.6926E-06 |
| Dusp3 | 1.34714E-06 | 1.928446739 | 0.472 | 0.182 | 0.032724701 | 0.985139254 | 0.402973785 | 0.39 | 0.403 | 1 | 0.985139254 | 2.69428E-06 |
| 4921524J17Rik | 1.3505E-06 | 1.480950827 | 0.623 | 0.305 | 0.032806309 | 0.709838715 | 0.129385601 | 0.585 | 0.526 | 1 | 0.709838715 | 2.701E-06 |
| Tnfrsf10b | 1.35154E-06 | -2.32424921 | 0.132 | 0.5 | 0.032831595 | 0.009983715 | -1.515311565 | 0.146 | 0.334 | 1 | 0.009983715 | 2.70308E-06 |
| Ubxn1 | 1.36619E-06 | 1.074034749 | 0.849 | 0.585 | 0.033187438 | 0.019377089 | 0.861687535 | 0.78 | 0.71 | 1 | 0.019377089 | 2.73237E-06 |
| Pou2f1 | 1.36957E-06 | 1.376078969 | 0.66 | 0.36 | 0.033269666 | 0.038711478 | 0.52774072 | 0.756 | 0.597 | 1 | 0.038711478 | 2.73914E-06 |
| Bnip3 | 1.37069E-06 | -1.896381746 | 0.283 | 0.644 | 0.033296719 | 5.16205E-05 | -1.274303465 | 0.244 | 0.587 | 1 | 5.16205E-05 | 2.74137E-06 |
| Marchf9 | 1.38131E-06 | 2.932330173 | 0.17 | 0.017 | 0.033554833 | 0.719381525 | 1.059597271 | 0.049 | 0.065 | 1 | 0.719381525 | 2.76262E-06 |
| Ajuba | 1.38131E-06 | 3.029191713 | 0.17 | 0.017 | 0.033554833 | 0.000101236 | 2.351778023 | 0.171 | 0.031 | 1 | 0.000101236 | 2.76262E-06 |
| Prkaca | 1.39442E-06 | 1.709311446 | 0.528 | 0.225 | 0.033873364 | 0.663851423 | 0.887830923 | 0.268 | 0.256 | 1 | 0.663851423 | 2.78885E-06 |
| Ankrd50 | 1.39577E-06 | 1.206437631 | 0.642 | 0.305 | 0.033906101 | 0.346294407 | -0.170614655 | 0.463 | 0.563 | 1 | 0.346294407 | 2.79154E-06 |
| Iqsec2 | 1.39731E-06 | 1.521504925 | 0.453 | 0.161 | 0.033943442 | 0.853648901 | 0.532693808 | 0.293 | 0.321 | 1 | 0.853648901 | 2.79462E-06 |
| Slc25a44 | 1.40176E-06 | 1.569760094 | 0.472 | 0.169 | 0.034051473 | 0.054007949 | -1.090902232 | 0.195 | 0.328 | 1 | 0.054007949 | 2.80351E-06 |
| Atp5h | 1.40889E-06 | 0.620744023 | 1 | 0.936 | 0.03422483 | 0.06274937 | 0.309467218 | 0.927 | 0.918 | 1 | 0.06274937 | 2.81778E-06 |
| A630014C17Rik | 0.094924143 | 1.347367673 | 0.057 | 0.017 | 1 | 1.41423E-06 | 4.574170444 | 0.122 | 0.007 | 0.034354436 | 0.094924143 | 2.82845E-06 |
| Acin1 | 1.41843E-06 | 0.747633849 | 0.981 | 0.903 | 0.034456419 | 0.089125996 | 0.298046039 | 0.976 | 0.966 | 1 | 0.089125996 | 2.83685E-06 |

| | | | | | | | | | | | | |
|---|---|---|---|---|---|---|---|---|---|---|---|---|
| Lyst | 1.43024E-06 | 2.154722595 | 0.472 | 0.178 | 0.034743335 | 0.054213554 | 0.815111079 | 0.585 | 0.478 | 1 | 0.054213554 | 2.86047E-06 |
| Rassf3 | 1.43053E-06 | 1.745331659 | 0.528 | 0.22 | 0.034750437 | 0.068543364 | 0.568171703 | 0.61 | 0.481 | 1 | 0.068543364 | 2.86106E-06 |
| Hivep1 | 1.43613E-06 | 1.306725688 | 0.698 | 0.398 | 0.034886381 | 0.00602828 | 0.900214647 | 0.78 | 0.611 | 1 | 0.00602828 | 2.87225E-06 |
| Boc | 1.44081E-06 | 1.136026501 | 0.717 | 0.381 | 0.035000265 | 0.02771488 | 0.604913967 | 0.756 | 0.645 | 1 | 0.02771488 | 2.88163E-06 |
| Nrk | 0.737817564 | -0.544906848 | 0.623 | 0.64 | 1 | 1.44555E-06 | -1.629986192 | 0.39 | 0.778 | 0.035115219 | 0.737817564 | 2.89109E-06 |
| Pxdc1 | 1.4562E-06 | -1.938482324 | 0.472 | 0.733 | 0.035373899 | 0.055515307 | -0.481111992 | 0.39 | 0.549 | 1 | 0.055515307 | 2.91239E-06 |
| Slc13a5 | 1.4599E-06 | -5.620064465 | 0 | 0.331 | 0.035463845 | 0.01264035 | -3.589059905 | 0.024 | 0.174 | 1 | 0.01264035 | 2.91979E-06 |
| Zfp516 | 1.46148E-06 | 1.033006686 | 0.811 | 0.585 | 0.035502162 | 0.434466963 | 0.266456208 | 0.878 | 0.829 | 1 | 0.434466963 | 2.92295E-06 |
| Rbl2 | 1.46301E-06 | 1.339937467 | 0.679 | 0.369 | 0.035539544 | 0.025071916 | 0.493250449 | 0.707 | 0.505 | 1 | 0.025071916 | 2.92603E-06 |
| Tox2 | 0.287773478 | -0.430239906 | 0 | 0.021 | 1 | 1.46498E-06 | 4.644559772 | 0.122 | 0.007 | 0.035587406 | 0.287773478 | 2.92997E-06 |
| Zfp422 | 1.48451E-06 | 1.891688189 | 0.396 | 0.123 | 0.036061686 | 0.355803348 | 1.029849928 | 0.293 | 0.253 | 1 | 0.355803348 | 2.96902E-06 |
| Slc39a10 | 1.4859E-06 | 1.244359807 | 0.585 | 0.297 | 0.036095474 | 0.007240654 | 1.210765713 | 0.61 | 0.451 | 1 | 0.007240654 | 2.9718E-06 |
| Zfp24 | 1.49626E-06 | 1.312263872 | 0.698 | 0.441 | 0.036347053 | 0.247421796 | -0.305753104 | 0.512 | 0.59 | 1 | 0.247421796 | 2.99251E-06 |
| Zdbf2 | 1.49702E-06 | 1.383663802 | 0.604 | 0.288 | 0.036365507 | 0.007946042 | 1.284303558 | 0.512 | 0.392 | 1 | 0.007946042 | 2.99403E-06 |
| Msl1 | 1.50663E-06 | 1.4226558 | 0.698 | 0.449 | 0.036599146 | 0.587123087 | 0.199986112 | 0.829 | 0.853 | 1 | 0.587123087 | 3.01327E-06 |
| Gstt2 | 1.50711E-06 | 1.956783217 | 0.415 | 0.14 | 0.036610736 | 0.320636885 | -0.126269274 | 0.122 | 0.191 | 1 | 0.320636885 | 3.01422E-06 |
| Fut8 | 1.53055E-06 | 0.96457051 | 0.868 | 0.648 | 0.037180032 | 0.816552363 | -0.112463475 | 0.902 | 0.908 | 1 | 0.816552363 | 3.06109E-06 |
| Nfil3 | 1.54483E-06 | 2.026398498 | 0.509 | 0.229 | 0.037527129 | 1.92459E-05 | 1.455059232 | 0.634 | 0.328 | 0.467520407 | 1.92459E-05 | 3.08967E-06 |
| Smim14 | 1.54486E-06 | -0.841402315 | 1 | 0.975 | 0.037527647 | 0.287821209 | -0.113582176 | 0.951 | 0.983 | 1 | 0.287821209 | 3.08971E-06 |
| Stpg4 | 1.59224E-06 | 2.47665069 | 0.226 | 0.038 | 0.038678776 | 0.033936065 | 1.962735732 | 0.146 | 0.058 | 1 | 0.033936065 | 3.18448E-06 |
| Mtrex | 1.61547E-06 | 1.293885342 | 0.736 | 0.441 | 0.039243052 | 0.696435044 | 0.252242349 | 0.561 | 0.652 | 1 | 0.696435044 | 3.23094E-06 |
| Agbl3 | 1.62048E-06 | 2.687070469 | 0.547 | 0.25 | 0.039364662 | 0.061397473 | 0.898605394 | 0.341 | 0.222 | 1 | 0.061397473 | 3.24095E-06 |
| Tars2 | 1.63495E-06 | 1.845394537 | 0.472 | 0.186 | 0.039716236 | 0.390801152 | 0.222495006 | 0.22 | 0.311 | 1 | 0.390801152 | 3.2699E-06 |
| Yap1 | 1.63541E-06 | 1.198224233 | 0.925 | 0.691 | 0.039727318 | 0.00021442 | 0.885360888 | 0.927 | 0.833 | 1 | 0.00021442 | 3.27081E-06 |
| Gadd45a | 1.63787E-06 | 1.624761483 | 0.717 | 0.403 | 0.039787164 | 0.15044177 | 0.563812358 | 0.732 | 0.662 | 1 | 0.15044177 | 3.27574E-06 |
| Pcbp1 | 1.64023E-06 | 0.598118649 | 1 | 0.941 | 0.039844466 | 0.074760084 | -0.276111674 | 0.951 | 0.983 | 1 | 0.074760084 | 3.28046E-06 |
| Tmem87a | 1.67651E-06 | 1.076720083 | 0.849 | 0.517 | 0.040725752 | 0.009255061 | 0.630511115 | 0.829 | 0.737 | 1 | 0.009255061 | 3.35301E-06 |
| Ninj1 | 1.67799E-06 | -1.709354917 | 0.472 | 0.737 | 0.040761717 | 2.85306E-05 | -1.396667068 | 0.366 | 0.706 | 0.693065012 | 2.85306E-05 | 3.35598E-06 |
| Rab21 | 1.68471E-06 | 1.072884303 | 0.943 | 0.771 | 0.040924987 | 0.352305661 | 0.210219134 | 0.976 | 0.956 | 1 | 0.352305661 | 3.36942E-06 |
| Nradd | 1.6916E-06 | 2.739685096 | 0.245 | 0.047 | 0.041092399 | 0.000631653 | 2.574170444 | 0.268 | 0.096 | 1 | 0.000631653 | 3.3832E-06 |
| Haus4 | 1.71793E-06 | 2.661682584 | 0.264 | 0.055 | 0.04173193 | 0.884674798 | -0.16279515 | 0.073 | 0.078 | 1 | 0.884674798 | 3.43585E-06 |
| Bcl9l | 1.719E-06 | 1.225495669 | 0.717 | 0.415 | 0.041757835 | 0.185715346 | 0.614001242 | 0.78 | 0.737 | 1 | 0.185715346 | 3.43799E-06 |
| Stk4 | 1.72144E-06 | 1.48554709 | 0.547 | 0.263 | 0.041817154 | 0.54133126 | 0.295311071 | 0.439 | 0.403 | 1 | 0.54133126 | 3.44287E-06 |
| Prpf40b | 1.72681E-06 | 1.8327945 | 0.396 | 0.127 | 0.041947727 | 0.44355467 | -0.395455907 | 0.195 | 0.246 | 1 | 0.44355467 | 3.45362E-06 |
| Radil | 1.75386E-06 | -4.054730771 | 0.019 | 0.352 | 0.042604804 | 0.650700755 | -1.0454382 | 0.073 | 0.092 | 1 | 0.650700755 | 3.50772E-06 |
| Cpne1 | 1.76648E-06 | 1.049753035 | 0.792 | 0.564 | 0.042911324 | 0.357303684 | 0.243755883 | 0.878 | 0.836 | 1 | 0.357303684 | 3.53296E-06 |
| Cpped1 | 1.7667E-06 | 1.883420573 | 0.396 | 0.127 | 0.042916566 | 0.789652552 | -0.257952383 | 0.195 | 0.208 | 1 | 0.789652552 | 3.53339E-06 |
| Rapgef6 | 1.77797E-06 | 0.861483259 | 0.962 | 0.881 | 0.043190484 | 0.010992059 | 0.844565555 | 0.976 | 0.945 | 1 | 0.010992059 | 3.55594E-06 |
| Spry1 | 2.38219E-05 | 2.891688189 | 0.189 | 0.034 | 0.57868235 | 1.78078E-06 | 4.190841804 | 0.195 | 0.027 | 0.043258729 | 2.38219E-05 | 3.56156E-06 |
| Nrg1 | 1.80895E-06 | 4.154722595 | 0.151 | 0.013 | 0.043943109 | 1.02519E-05 | 3.775804305 | 0.171 | 0.024 | 0.249039187 | 1.02519E-05 | 3.6179E-06 |
| Apcdd1 | 1.84139E-06 | 2.646575691 | 0.34 | 0.097 | 0.044731105 | 0.273887961 | 0.858578501 | 0.22 | 0.164 | 1 | 0.273887961 | 3.68278E-06 |
| Tcf3 | 1.87367E-06 | 1.855162313 | 0.509 | 0.22 | 0.045515302 | 0.15447136 | 0.609398932 | 0.512 | 0.42 | 1 | 0.15447136 | 3.74735E-06 |
| Aldoc | 1.89881E-06 | -3.464886049 | 0.075 | 0.419 | 0.046125958 | 0.486283127 | 0.287007767 | 0.122 | 0.174 | 1 | 0.486283127 | 3.79762E-06 |
| Crim1 | 0.02867211 | -0.444095877 | 0.962 | 0.979 | 1 | 1.92801E-06 | -0.877965679 | 0.951 | 1 | 0.046835226 | 0.02867211 | 3.85602E-06 |
| Gm56878 | 1.92857E-06 | -1.32971303 | 0.491 | 0.78 | 0.046848931 | 0.516792647 | 0.242046582 | 0.585 | 0.543 | 1 | 0.516792647 | 3.85715E-06 |
| Creb3l1 | 1.93366E-06 | 1.791175411 | 0.66 | 0.377 | 0.046972374 | 0.031346212 | 1.117517247 | 0.585 | 0.515 | 1 | 0.031346212 | 3.86731E-06 |
| Kcna6 | 1.94541E-06 | -1.792106746 | 0.264 | 0.61 | 0.047257847 | 2.32889E-06 | -1.54716382 | 0.244 | 0.679 | 0.056573433 | 2.32889E-06 | 3.89081E-06 |
| Arf5 | 1.9724E-06 | 0.581799344 | 1 | 0.949 | 0.047913477 | 0.208086866 | -0.129628286 | 0.951 | 0.98 | 1 | 0.208086866 | 3.94479E-06 |
| Ncoa2 | 1.9832E-06 | 1.083561566 | 0.868 | 0.691 | 0.048175781 | 0.010186958 | 0.256393992 | 0.976 | 0.857 | 1 | 0.010186958 | 3.96639E-06 |
| Plxna2 | 1.98456E-06 | 1.741201328 | 0.774 | 0.525 | 0.048208872 | 0.004787954 | 1.047086633 | 0.902 | 0.761 | 1 | 0.004787954 | 3.96911E-06 |
| Gm16310 | 2.00051E-06 | 2.495759513 | 0.245 | 0.047 | 0.048596442 | 0.024591692 | 1.450181727 | 0.195 | 0.085 | 1 | 0.024591692 | 4.00102E-06 |
| Pik3r1 | 2.00475E-06 | 0.793349276 | 1 | 0.873 | 0.048699361 | 0.147383032 | 0.261916786 | 0.951 | 0.962 | 1 | 0.147383032 | 4.00949E-06 |
| Tenm4 | 9.11871E-06 | 3.605384004 | 0.302 | 0.089 | 0.22151159 | 2.03504E-06 | 1.99895592 | 0.561 | 0.235 | 0.0494353 | 9.11871E-06 | 4.07008E-06 |
| Camk1g | 2.05987E-06 | 4.962077517 | 0.094 | 0 | 0.050038271 | 0.000156969 | 5.159132945 | 0.049 | 0 | 1 | 0.000156969 | 4.11973E-06 |
| Gm29443 | 2.05987E-06 | 5.324647596 | 0.094 | 0 | 0.050038271 | 0.117592181 | 1.100239256 | 0.049 | 0.014 | 1 | 0.117592181 | 4.11973E-06 |
| Gm28055 | 2.06093E-06 | 5.324647596 | 0.094 | 0 | 0.050064025 | 0.168657861 | 1.83720485 | 0.073 | 0.031 | 1 | 0.168657861 | 4.12185E-06 |
| Rps6ka4 | 2.06104E-06 | 4.739685096 | 0.094 | 0 | 0.050066887 | 0.745976859 | 0.377773231 | 0.024 | 0.034 | 1 | 0.745976859 | 4.12208E-06 |
| Sp4 | 2.06873E-06 | 1.669295768 | 0.528 | 0.233 | 0.050253617 | 0.228832818 | 0.760583568 | 0.439 | 0.375 | 1 | 0.228832818 | 4.13746E-06 |
| Sntb1 | 2.08104E-06 | 2.068992721 | 0.226 | 0.038 | 0.050552662 | 2.34414E-05 | 2.83720485 | 0.22 | 0.044 | 0.569437368 | 2.34414E-05 | 4.16208E-06 |
| Cacnb3 | 2.10067E-06 | 3.417757001 | 0.151 | 0.013 | 0.05102957 | 0.028242983 | 2.252242349 | 0.073 | 0.017 | 1 | 0.028242983 | 4.20134E-06 |
| Gsn | 0.000275801 | -0.743205496 | 0.981 | 0.983 | 1 | 2.132E-06 | -0.713772949 | 0.976 | 1 | 0.051790665 | 0.000275801 | 4.26401E-06 |
| Cep152 | 0.495809695 | 0.47665069 | 0.094 | 0.068 | 1 | 2.13754E-06 | 3.252242349 | 0.195 | 0.027 | 0.051925056 | 0.495809695 | 4.27507E-06 |
| Brd8dc | 2.15953E-06 | 1.757387097 | 0.377 | 0.114 | 0.052459244 | 0.020694296 | -2.060035576 | 0.073 | 0.229 | 1 | 0.020694296 | 4.31905E-06 |
| Gale | 2.16716E-06 | -3.867645218 | 0.019 | 0.347 | 0.052644692 | 0.024060506 | -1.572186086 | 0.049 | 0.195 | 1 | 0.024060506 | 4.33432E-06 |
| Cst3 | 2.17395E-06 | -1.382968487 | 1 | 0.987 | 0.052809608 | 0.005452171 | -0.411466896 | 0.976 | 0.997 | 1 | 0.005452171 | 4.3479E-06 |
| Stc2 | 2.17815E-06 | -2.518750076 | 0.264 | 0.602 | 0.052911668 | 0.177742772 | -0.405651674 | 0.122 | 0.225 | 1 | 0.177742772 | 4.3563E-06 |
| Pwwp2a | 2.17873E-06 | 1.324647596 | 0.642 | 0.331 | 0.052925682 | 0.03628235 | 0.97685684 | 0.634 | 0.539 | 1 | 0.03628235 | 4.35745E-06 |
| Eif4a3 | 2.18412E-06 | 1.055186921 | 0.774 | 0.551 | 0.05305668 | 0.80914718 | 0.30747187 | 0.585 | 0.676 | 1 | 0.80914718 | 4.36824E-06 |
| Fnip1 | 2.18511E-06 | 0.923457769 | 1 | 0.911 | 0.053080709 | 0.018757255 | 0.400060682 | 0.951 | 0.986 | 1 | 0.018757255 | 4.37022E-06 |
| Enho | 2.18555E-06 | -1.951476809 | 0.245 | 0.576 | 0.053091374 | 0.000225215 | -2.298931838 | 0.122 | 0.406 | 1 | 0.000225215 | 4.37109E-06 |
| Zbtb45 | 2.18613E-06 | 3.292226119 | 0.151 | 0.013 | 0.053105542 | 0.360451923 | 0.930314254 | 0.073 | 0.041 | 1 | 0.360451923 | 4.37226E-06 |
| Tead4 | 2.19965E-06 | -3.665456368 | 0.038 | 0.377 | 0.05343381 | 0.000122108 | -4.466575898 | 0.024 | 0.311 | 1 | 0.000122108 | 4.39929E-06 |
| Notch1 | 2.24471E-06 | 1.967444027 | 0.585 | 0.314 | 0.054528499 | 0.002540056 | 1.158174025 | 0.756 | 0.553 | 1 | 0.002540056 | 4.48942E-06 |
| Homer2 | 2.24882E-06 | 2.080722013 | 0.226 | 0.038 | 0.054628401 | 0.895168756 | 0.252242349 | 0.146 | 0.16 | 1 | 0.895168756 | 4.49764E-06 |
| Ankrd28 | 2.26848E-06 | -0.709685704 | 0.943 | 0.987 | 0.055105826 | 0.000541147 | -0.60437688 | 1 | 0.997 | 1 | 0.000541147 | 4.53695E-06 |
| Ncl | 2.326E-06 | 0.485887537 | 1 | 0.987 | 0.056503201 | 0.343332241 | 0.104293325 | 1 | 0.997 | 1 | 0.343332241 | 4.652E-06 |
| Cpm | 2.33146E-06 | -1.067669827 | 0.491 | 0.792 | 0.056635923 | 0.127112085 | -0.863234868 | 0.22 | 0.324 | 1 | 0.127112085 | 4.66292E-06 |
| Slc1a5 | 2.33342E-06 | -1.486382984 | 0.604 | 0.835 | 0.056683401 | 0.001481715 | -0.910988 | 0.537 | 0.788 | 1 | 0.001481715 | 4.66683E-06 |
| Rnf214 | 2.34538E-06 | 1.288578342 | 0.679 | 0.394 | 0.056973898 | 0.607336195 | 0.216618439 | 0.512 | 0.485 | 1 | 0.607336195 | 4.69075E-06 |
| Hs2st1 | 0.009843487 | 0.416262754 | 0.849 | 0.669 | 1 | 2.36289E-06 | 1.231383744 | 0.902 | 0.635 | 0.057399374 | 0.009843487 | 4.72578E-06 |
| Htra4 | 2.36845E-06 | 1.347367673 | 0.264 | 0.055 | 0.057534386 | 0.399993082 | 1.340779024 | 0.171 | 0.13 | 1 | 0.399993082 | 4.73689E-06 |
| Ric1 | 2.36912E-06 | 1.177806208 | 0.792 | 0.551 | 0.057550731 | 0.000110024 | 0.971222198 | 0.829 | 0.679 | 1 | 0.000110024 | 4.73824E-06 |
| Sh3d21 | 2.38085E-06 | 2.974150349 | 0.358 | 0.11 | 0.057835611 | 0.0021045 | 1.494812652 | 0.366 | 0.174 | 1 | 0.0021045 | 4.76169E-06 |
| Kansl1l | 2.40756E-06 | 1.295652636 | 0.849 | 0.525 | 0.058484451 | 0.000517224 | 0.825913689 | 0.927 | 0.857 | 1 | 0.000517224 | 4.81511E-06 |
| Pfdn6 | 2.40985E-06 | 1.291044066 | 0.66 | 0.352 | 0.058540176 | 0.226841065 | -0.387761515 | 0.317 | 0.406 | 1 | 0.226841065 | 4.8197E-06 |
| Col4a2 | 2.41561E-06 | 1.353551544 | 0.717 | 0.534 | 0.058679952 | 0.047082156 | 0.83720485 | 0.634 | 0.601 | 1 | 0.047082156 | 4.83121E-06 |
| Usp3 | 2.45441E-06 | 1.141587193 | 0.792 | 0.581 | 0.059622545 | 0.001539956 | 0.567814949 | 0.927 | 0.802 | 1 | 0.001539956 | 4.90882E-06 |
| Slc35g1 | 2.45554E-06 | -1.865176963 | 0.189 | 0.551 | 0.059649897 | 0.187938135 | -1.136799942 | 0.122 | 0.201 | 1 | 0.187938135 | 4.91107E-06 |
| Psmd8 | 2.46365E-06 | 0.829002921 | 0.906 | 0.746 | 0.059847072 | 0.031636791 | 0.461337948 | 0.854 | 0.833 | 1 | 0.031636791 | 4.9273E-06 |
| Tfap2a | 2.4809E-06 | 1.903183828 | 0.434 | 0.157 | 0.060266021 | 0.544853947 | -0.175619191 | 0.195 | 0.242 | 1 | 0.544853947 | 4.96179E-06 |
| Fam32a | 2.49338E-06 | 1.47665069 | 0.604 | 0.331 | 0.060569277 | 0.562699185 | 0.42657253 | 0.463 | 0.468 | 1 | 0.562699185 | 4.98676E-06 |
| Supt4a | 2.50731E-06 | 1.163234095 | 0.755 | 0.479 | 0.060907625 | 0.09771474 | 0.572864246 | 0.61 | 0.515 | 1 | 0.09771474 | 5.01462E-06 |
| Rarb | 0.008230393 | 2.891688189 | 0.094 | 0.021 | 1 | 2.52432E-06 | 3.381525366 | 0.195 | 0.027 | 0.061320854 | 0.008230393 | 5.04864E-06 |
| Pcdhb9 | 2.53001E-06 | 5.377115016 | 0.132 | 0.008 | 0.061459062 | 0.038776849 | 1.904319046 | 0.098 | 0.031 | 1 | 0.038776849 | 5.06002E-06 |
| Epb41l3 | 2.57153E-06 | 3.533234218 | 0.189 | 0.025 | 0.062467551 | 0.09235202 | 1.59619675 | 0.122 | 0.055 | 1 | 0.09235202 | 5.14305E-06 |
| Bace1 | 2.57995E-06 | 1.617065809 | 0.566 | 0.267 | 0.06267216 | 0.366502206 | 0.413993419 | 0.488 | 0.454 | 1 | 0.366502206 | 5.15989E-06 |
| Arhgap32 | 2.59552E-06 | 0.897122075 | 0.849 | 0.542 | 0.063050469 | 0.12545322 | 0.383948028 | 0.878 | 0.823 | 1 | 0.12545322 | 5.19104E-06 |
| Nt5dc1 | 2.59845E-06 | -2.863604053 | 0.226 | 0.564 | 0.063121614 | 0.519492284 | 0.46797104 | 0.366 | 0.328 | 1 | 0.519492284 | 5.1969E-06 |

| | | | | | | | | | | | | |
|---|---|---|---|---|---|---|---|---|---|---|---|---|
| Trib1 | 2.60974E-06 | -1.34290024 | 0.585 | 0.797 | 0.063395822 | 0.032718464 | -0.745646904 | 0.634 | 0.737 | 1 | 0.032718464 | 5.21947E-06 |
| Vegfc | 2.61837E-06 | 3.377115016 | 0.208 | 0.034 | 0.063605549 | 0.058304253 | 2.185128153 | 0.195 | 0.106 | 1 | 0.058304253 | 5.23674E-06 |
| Esrrg | 2.63165E-06 | -3.254668341 | 0.057 | 0.411 | 0.063928026 | 0.905777747 | -2.077678537 | 0.195 | 0.191 | 1 | 0.905777747 | 5.26329E-06 |
| Pdap1 | 2.64461E-06 | 0.876421432 | 0.925 | 0.771 | 0.064242847 | 0.272444346 | 0.367908611 | 0.756 | 0.812 | 1 | 0.272444346 | 5.28921E-06 |
| Dhcr24 | 2.64801E-06 | -2.786115333 | 0.075 | 0.424 | 0.064325431 | 0.028421816 | -2.605738646 | 0.024 | 0.147 | 1 | 0.028421816 | 5.29601E-06 |
| Nckap5l | 2.65785E-06 | 1.984797593 | 0.453 | 0.165 | 0.064564611 | 0.0646068 | 1.10785244 | 0.488 | 0.396 | 1 | 0.0646068 | 5.3157E-06 |
| Baalc | 2.66268E-06 | -5.496329096 | 0 | 0.318 | 0.064681894 | 0.070383146 | -2.747757651 | 0 | 0.075 | 1 | 0.070383146 | 5.32536E-06 |
| Npm3 | 2.68502E-06 | 1.380998451 | 0.642 | 0.322 | 0.065224619 | 0.627996575 | 0.381753082 | 0.415 | 0.386 | 1 | 0.627996575 | 5.37004E-06 |
| Rnf38 | 2.68795E-06 | 1.370856026 | 0.717 | 0.441 | 0.065295641 | 0.107106491 | 0.510394534 | 0.805 | 0.689 | 1 | 0.107106491 | 5.37589E-06 |
| Ttyh3 | 2.69549E-06 | 1.319353297 | 0.566 | 0.254 | 0.065478857 | 1.71785E-05 | 1.477662463 | 0.561 | 0.276 | 0.417301229 | 1.71785E-05 | 5.39097E-06 |
| Ptprv | 2.72406E-06 | 2.913714495 | 0.189 | 0.025 | 0.066172832 | 0.002303037 | 1.962735732 | 0.171 | 0.048 | 1 | 0.002303037 | 5.44811E-06 |
| Csrnp2 | 2.73682E-06 | 2.849868013 | 0.283 | 0.068 | 0.066482789 | 0.699952729 | 0.359157553 | 0.122 | 0.102 | 1 | 0.699952729 | 5.47363E-06 |
| Ubtf | 2.74188E-06 | 0.884364848 | 0.849 | 0.636 | 0.066605714 | 0.559103718 | -0.172173483 | 0.927 | 0.887 | 1 | 0.559103718 | 5.48375E-06 |
| Rab10 | 2.74872E-06 | 0.690299611 | 0.981 | 0.941 | 0.066771921 | 0.043431893 | 0.352154165 | 1 | 0.983 | 1 | 0.043431893 | 5.49743E-06 |
| Gsdmd | 2.76244E-06 | 4.029191713 | 0.132 | 0.008 | 0.067105179 | 0.273319125 | -0.622226769 | 0.049 | 0.102 | 1 | 0.273319125 | 5.52487E-06 |
| Leo1 | 2.76712E-06 | 1.595295186 | 0.585 | 0.305 | 0.067218959 | 0.646105961 | 0.458693227 | 0.488 | 0.471 | 1 | 0.646105961 | 5.53424E-06 |
| Pald1 | 2.76886E-06 | 1.762405172 | 0.245 | 0.047 | 0.067261149 | 0.617245661 | 0.728680393 | 0.146 | 0.177 | 1 | 0.617245661 | 5.53771E-06 |
| Cdc40 | 2.77493E-06 | 1.154722595 | 0.717 | 0.47 | 0.067408602 | 0.376370926 | 0.13494699 | 0.683 | 0.611 | 1 | 0.376370926 | 5.54985E-06 |
| Iffo1 | 2.78966E-06 | 2.649487287 | 0.302 | 0.081 | 0.06776631 | 0.004166529 | 1.282615998 | 0.268 | 0.106 | 1 | 0.004166529 | 5.5793E-06 |
| Adgre5 | 2.80207E-06 | 2.154722595 | 0.34 | 0.097 | 0.068067867 | 0.000746666 | 1.691353983 | 0.39 | 0.177 | 1 | 0.000746666 | 5.60413E-06 |
| Ak5 | 2.81874E-06 | 3.47665069 | 0.208 | 0.034 | 0.068472837 | 0.035880762 | 0.786578777 | 0.244 | 0.123 | 1 | 0.035880762 | 5.63747E-06 |
| Kat2b | 2.82449E-06 | 0.867841447 | 0.566 | 0.233 | 0.068612397 | 0.006960159 | 0.85796341 | 0.659 | 0.42 | 1 | 0.006960159 | 5.64896E-06 |
| Gnb5 | 2.82863E-06 | 2.195364579 | 0.358 | 0.114 | 0.068713086 | 0.004999113 | 1.382638986 | 0.366 | 0.191 | 1 | 0.004999113 | 5.65725E-06 |
| Plekhm2 | 0.229427834 | -0.132158553 | 0.642 | 0.784 | 1 | 2.84752E-06 | -0.931138933 | 0.683 | 0.898 | 0.069171851 | 0.229427834 | 5.69502E-06 |
| Gm56906 | 2.87845E-06 | 1.092110617 | 0.623 | 0.292 | 0.069923269 | 0.879729493 | 0.277777441 | 0.439 | 0.447 | 1 | 0.879729493 | 5.75689E-06 |
| Sec61a1 | 2.89488E-06 | 1.001537449 | 0.887 | 0.742 | 0.070322377 | 0.514226935 | 0.211296799 | 0.829 | 0.836 | 1 | 0.514226935 | 5.78975E-06 |
| Sec22b | 0.346502668 | -0.186086966 | 0.887 | 0.919 | 1 | 2.92826E-06 | -0.953306562 | 0.732 | 0.911 | 0.071133171 | 0.346502668 | 5.8565E-06 |
| Tcea1 | 2.94304E-06 | 0.751657737 | 0.962 | 0.928 | 0.071492425 | 0.023623685 | 0.364717078 | 0.878 | 0.939 | 1 | 0.023623685 | 5.88608E-06 |
| Cbx4 | 0.000128322 | 1.448453798 | 0.528 | 0.288 | 1 | 2.94915E-06 | 1.413397141 | 0.732 | 0.389 | 0.071640842 | 0.000128322 | 5.8983E-06 |
| Adamts10 | 2.96099E-06 | 1.399141322 | 0.66 | 0.343 | 0.071928377 | 0.153896714 | 0.635570989 | 0.561 | 0.529 | 1 | 0.153896714 | 5.92197E-06 |
| Cds1 | 2.98168E-06 | -2.031143951 | 0.132 | 0.492 | 0.072431037 | 0.878566096 | -0.529577481 | 0.341 | 0.328 | 1 | 0.878566096 | 5.96336E-06 |
| Cnot6 | 2.99919E-06 | 0.76408175 | 0.962 | 0.754 | 0.072856275 | 0.002023774 | 0.533848861 | 0.927 | 0.904 | 1 | 0.002023774 | 5.99837E-06 |
| Slc2a1 | 3.00619E-06 | -2.544760112 | 0.604 | 0.742 | 0.073026324 | 0.743050549 | -0.214883661 | 0.585 | 0.549 | 1 | 0.743050549 | 6.01237E-06 |
| Matn4 | 3.04214E-06 | 1.26688327 | 0.792 | 0.627 | 0.073899653 | 0.087867222 | -0.173343075 | 0.78 | 0.894 | 1 | 0.087867222 | 6.08427E-06 |
| Agtr1a | 3.06399E-06 | 3.614154213 | 0.132 | 0.008 | 0.074430361 | 0.319629309 | -0.622226769 | 0 | 0.024 | 1 | 0.319629309 | 6.12796E-06 |
| Siah1a | 3.08949E-06 | 1.317452095 | 0.679 | 0.39 | 0.075049846 | 0.176215285 | 0.546387287 | 0.683 | 0.604 | 1 | 0.176215285 | 6.17897E-06 |
| Upf2 | 3.09281E-06 | 0.967095592 | 0.925 | 0.623 | 0.075130521 | 0.215346357 | 0.310096735 | 0.829 | 0.799 | 1 | 0.215346357 | 6.18561E-06 |
| Kif1c | 3.11708E-06 | 1.507603618 | 0.566 | 0.275 | 0.07572001 | 0.658044604 | 0.206229951 | 0.366 | 0.457 | 1 | 0.658044604 | 6.23414E-06 |
| Ctdspl | 3.12508E-06 | 0.816726131 | 1 | 0.953 | 0.075914556 | 0.000936918 | -0.566772018 | 1 | 0.997 | 1 | 0.000936918 | 6.25016E-06 |
| Psma7 | 3.15175E-06 | 0.844821584 | 0.962 | 0.835 | 0.076562391 | 0.043488004 | 0.368817926 | 0.927 | 0.911 | 1 | 0.043488004 | 6.3035E-06 |
| Scara3 | 3.16089E-06 | 0.995272387 | 0.906 | 0.805 | 0.076784451 | 0.054725213 | -0.405519292 | 0.976 | 0.945 | 1 | 0.054725213 | 6.32178E-06 |
| Arhgdia | 3.1695E-06 | 0.896120435 | 0.925 | 0.771 | 0.076993375 | 0.318072673 | 0.270332806 | 0.829 | 0.833 | 1 | 0.318072673 | 6.33898E-06 |
| 2900026A02Rik | 0.039219028 | 0.920257341 | 0.208 | 0.106 | 1 | 3.17519E-06 | 2.557096931 | 0.366 | 0.106 | 0.077131802 | 0.039219028 | 6.35038E-06 |
| Hcar2 | 0.507281812 | 1.154722595 | 0.019 | 0.008 | 1 | 3.18068E-06 | 3.322631677 | 0.146 | 0.014 | 0.077265176 | 0.507281812 | 6.36136E-06 |
| Plekhg4 | 3.19027E-06 | 4.324647596 | 0.113 | 0.004 | 0.07749795 | 0.004050657 | 3.422167351 | 0.049 | 0.003 | 1 | 0.004050657 | 6.38052E-06 |
| Gm45894 | 3.19027E-06 | 4.324647596 | 0.113 | 0.004 | 0.07749795 | 0.360451923 | 0.83720485 | 0.073 | 0.041 | 1 | 0.360451923 | 6.38052E-06 |
| Wnt7b | 3.19027E-06 | 4.324647596 | 0.113 | 0.004 | 0.07749795 | 0.004188151 | 2.83720485 | 0.049 | 0.003 | 1 | 0.004188151 | 6.38052E-06 |
| Smchd1 | 3.20605E-06 | 0.695822062 | 0.981 | 0.881 | 0.077881305 | 0.561757452 | 0.167226235 | 0.902 | 0.915 | 1 | 0.561757452 | 6.41208E-06 |
| Rpap2 | 3.21365E-06 | 2.3688474 | 0.321 | 0.089 | 0.078065987 | 0.008884035 | -3.185162963 | 0 | 0.147 | 1 | 0.008884035 | 6.42729E-06 |
| Tenm3 | 3.21762E-06 | 0.664743998 | 1 | 0.996 | 0.078162463 | 0.014184453 | -0.315098339 | 1 | 1 | 1 | 0.014184453 | 6.43523E-06 |
| Plekhg5 | 3.23438E-06 | 2.010332685 | 0.321 | 0.089 | 0.078569441 | 0.612329149 | -0.132421501 | 0.22 | 0.27 | 1 | 0.612329149 | 6.46874E-06 |
| Xrn2 | 3.23783E-06 | 0.681917852 | 0.906 | 0.818 | 0.078653295 | 0.305245026 | 0.346664744 | 0.878 | 0.901 | 1 | 0.305245026 | 6.47564E-06 |
| Gbp5 | 0.03562064 | 3.154722595 | 0.019 | 0 | 1 | 3.24119E-06 | 5.744095445 | 0.098 | 0.003 | 0.078735061 | 0.03562064 | 6.48238E-06 |
| Rexo1 | 3.26414E-06 | 1.502645898 | 0.604 | 0.318 | 0.079292461 | 0.847476227 | 0.238945526 | 0.439 | 0.488 | 1 | 0.847476227 | 6.52827E-06 |
| Prrc2a | 3.29249E-06 | 0.844571668 | 0.906 | 0.703 | 0.079981235 | 0.064795001 | 0.414209829 | 0.854 | 0.877 | 1 | 0.064795001 | 6.58497E-06 |
| Iqgap1 | 0.007895836 | -0.421902459 | 0.962 | 0.987 | 1 | 3.29972E-06 | -0.837016947 | 1 | 0.997 | 0.08015688 | 0.007895836 | 6.59944E-06 |
| Tomm20l | 3.3004E-06 | 3.962077517 | 0.113 | 0.004 | 0.080173304 | 0.435968064 | 1.83720485 | 0.024 | 0.01 | 1 | 0.435968064 | 6.60079E-06 |
| Haus3 | 3.30264E-06 | 1.435904347 | 0.528 | 0.22 | 0.080227659 | 0.325787133 | 0.875006883 | 0.293 | 0.242 | 1 | 0.325787133 | 6.60526E-06 |
| Mab21l2 | 0.029576974 | 4.154722595 | 0.038 | 0.004 | 1 | 3.30639E-06 | 5.537644568 | 0.098 | 0.003 | 0.080318734 | 0.029576974 | 6.61276E-06 |
| Ildr2 | 0.343004841 | -1.304709024 | 0 | 0.017 | 1 | 3.30639E-06 | 6.83720485 | 0.098 | 0.003 | 0.080318734 | 0.343004841 | 6.61276E-06 |
| Sobp | 3.31445E-06 | 0.907550063 | 0.962 | 0.881 | 0.080514547 | 0.019538783 | 0.296384904 | 0.951 | 0.952 | 1 | 0.019538783 | 6.62888E-06 |
| Sfrp1 | 3.33055E-06 | 1.904744342 | 0.358 | 0.11 | 0.080905729 | 0.001658936 | 0.896583397 | 0.463 | 0.225 | 1 | 0.001658936 | 6.66109E-06 |
| Ski | 3.33933E-06 | 0.829146723 | 0.943 | 0.869 | 0.081119037 | 0.007481984 | 0.446706027 | 0.976 | 0.98 | 1 | 0.007481984 | 6.67865E-06 |
| Rbm27 | 3.34905E-06 | 0.924424975 | 0.906 | 0.636 | 0.081355132 | 0.336347959 | 0.291487726 | 0.829 | 0.826 | 1 | 0.336347959 | 6.69809E-06 |
| C1ra | 3.35319E-06 | 1.72408824 | 0.434 | 0.153 | 0.081455625 | 0.439216749 | 0.417665958 | 0.268 | 0.379 | 1 | 0.439216749 | 6.70636E-06 |
| Sox11 | 3.35483E-06 | 4.569760094 | 0.113 | 0.004 | 0.081495632 | 0.058755395 | 1.515276755 | 0.098 | 0.034 | 1 | 0.058755395 | 6.70966E-06 |
| Igfbp6 | 0.000344052 | 3.569760094 | 0.075 | 0.004 | 1 | 3.37242E-06 | 7.263469605 | 0.098 | 0.003 | 0.081922859 | 0.000344052 | 6.74483E-06 |
| Cers6 | 3.37664E-06 | 1.867841447 | 0.396 | 0.127 | 0.082025332 | 0.022202942 | 0.718263777 | 0.415 | 0.235 | 1 | 0.022202942 | 6.75327E-06 |
| Bvht | 0.236091249 | 1.47665069 | 0.057 | 0.025 | 1 | 3.37665E-06 | 2.989207943 | 0.171 | 0.02 | 0.082025529 | 0.236091249 | 6.75328E-06 |
| Anapc5 | 3.38527E-06 | 0.801373402 | 0.925 | 0.856 | 0.082235053 | 0.520680917 | 0.245082558 | 0.878 | 0.894 | 1 | 0.520680917 | 6.77053E-06 |
| Jmjd6 | 3.40532E-06 | 1.234449787 | 0.66 | 0.403 | 0.082722017 | 0.464841362 | 0.406570495 | 0.537 | 0.655 | 1 | 0.464841362 | 6.81063E-06 |
| Sec24a | 3.40748E-06 | 0.920600881 | 0.887 | 0.742 | 0.082774479 | 0.212445021 | 0.402697486 | 0.902 | 0.898 | 1 | 0.212445021 | 6.81495E-06 |
| St6galnac2 | 3.43386E-06 | 2.533234218 | 0.189 | 0.025 | 0.083415443 | 0.854882811 | 1.252242349 | 0.049 | 0.044 | 1 | 0.854882811 | 6.86772E-06 |
| Uqcrc1 | 3.45798E-06 | 1.255416594 | 0.717 | 0.496 | 0.084001286 | 0.462636584 | 0.426682832 | 0.659 | 0.631 | 1 | 0.462636584 | 6.91595E-06 |
| Fam83d | 3.46663E-06 | 3.029191713 | 0.208 | 0.034 | 0.084211415 | 0.025701386 | 1.120997816 | 0.098 | 0.027 | 1 | 0.025701386 | 6.93325E-06 |
| Rspo3 | 3.48112E-06 | -2.305922903 | 0.415 | 0.691 | 0.084563273 | 0.471092629 | -0.508569987 | 0.268 | 0.341 | 1 | 0.471092629 | 6.96222E-06 |
| 6430562O15Rik | 0.000249805 | 4.739685096 | 0.057 | 0 | 1 | 3.50317E-06 | 5.159132945 | 0.073 | 0 | 0.085098954 | 0.000249805 | 7.00632E-06 |
| Necab1 | 0.039851149 | 2.669295768 | 0.075 | 0.021 | 1 | 3.50317E-06 | 6.644559772 | 0.073 | 0 | 0.085098954 | 0.039851149 | 7.00632E-06 |
| Carmn | 0.246443787 | 3.154722595 | 0.019 | 0.004 | 1 | 3.50317E-06 | 5.159132945 | 0.073 | 0 | 0.085098954 | 0.246443787 | 7.00632E-06 |
| 5730419F03Rik | 0.507281812 | 1.154722595 | 0.019 | 0.008 | 1 | 3.50317E-06 | 5.644559772 | 0.073 | 0 | 0.085098954 | 0.507281812 | 7.00632E-06 |
| Gm48089 | 0.015178556 | 2.891688189 | 0.057 | 0.008 | 1 | 3.5034E-06 | 6.537644568 | 0.073 | 0 | 0.085104712 | 0.015178556 | 7.0068E-06 |
| Reep6 | 0.030360955 | 2.739685096 | 0.038 | 0.004 | 1 | 3.5034E-06 | 4.83720485 | 0.073 | 0 | 0.085104712 | 0.030360955 | 7.0068E-06 |
| Gsc | 0.030360955 | 2.739685096 | 0.038 | 0.004 | 1 | 3.5034E-06 | 4.83720485 | 0.073 | 0 | 0.085104712 | 0.030360955 | 7.0068E-06 |
| Vmn2r1 | 0.03562064 | 3.154722595 | 0.019 | 0 | 1 | 3.5034E-06 | 4.83720485 | 0.073 | 0 | 0.085104712 | 0.03562064 | 7.0068E-06 |
| Gm48293 | 0.204653189 | 2.154722595 | 0.038 | 0.013 | 1 | 3.5034E-06 | 5.422167351 | 0.073 | 0 | 0.085104712 | 0.204653189 | 7.0068E-06 |
| Hhip | 0.207551231 | 2.377115016 | 0.038 | 0.013 | 1 | 3.5034E-06 | 7.537644568 | 0.073 | 0 | 0.085104712 | 0.207551231 | 7.0068E-06 |
| 4930544I03Rik | 0.343000446 | -0.430239906 | 0 | 0.017 | 1 | 3.5034E-06 | 6.159132945 | 0.073 | 0 | 0.085104712 | 0.343000446 | 7.0068E-06 |
| F830016B08Rik | 0.722567081 | 1.47665069 | 0.019 | 0.013 | 1 | 3.5034E-06 | 6.644559772 | 0.073 | 0 | 0.085104712 | 0.722567081 | 7.0068E-06 |
| Csnka2ip | 0.736073999 | 0.8327945 | 0.019 | 0.013 | 1 | 3.5034E-06 | 7.537644568 | 0.073 | 0 | 0.085104712 | 0.736073999 | 7.0068E-06 |
| Znrf1 | 0.000190746 | 0.721342515 | 0.792 | 0.619 | 1 | 3.50426E-06 | 1.111599118 | 0.854 | 0.686 | 0.085125525 | 0.000190746 | 7.00851E-06 |
| Ptk7 | 3.50441E-06 | 1.509387476 | 0.585 | 0.301 | 0.085129098 | 0.020156548 | 0.841309248 | 0.732 | 0.56 | 1 | 0.020156548 | 7.00881E-06 |
| Tmem45a | 6.4273E-06 | -0.85499256 | 0.736 | 0.907 | 0.156131939 | 3.52396E-06 | -1.243814501 | 0.585 | 0.843 | 0.085603918 | 6.4273E-06 | 7.0479E-06 |
| Ifi47 | 0.246443787 | 3.47665069 | 0.019 | 0.004 | 1 | 3.57996E-06 | 4.296636468 | 0.098 | 0.003 | 0.086964322 | 0.246443787 | 7.1599E-06 |
| Pxylp1 | 3.61012E-06 | -2.764140642 | 0.132 | 0.458 | 0.087697001 | 0.000356807 | -2.285191782 | 0.073 | 0.355 | 1 | 0.000356807 | 7.22022E-06 |
| Emc3 | 3.63593E-06 | 0.864737392 | 0.887 | 0.686 | 0.088324064 | 0.277330733 | -0.171935804 | 0.707 | 0.761 | 1 | 0.277330733 | 7.27185E-06 |
| Cars | 0.169200042 | -0.304709024 | 0.434 | 0.538 | 1 | 3.6491E-06 | -1.393749585 | 0.293 | 0.713 | 0.088643966 | 0.169200042 | 7.29819E-06 |
| Ppargc1a | 3.65106E-06 | 2.739685096 | 0.132 | 0.008 | 0.088691669 | 0.16031862 | 0.814837037 | 0.293 | 0.201 | 1 | 0.16031862 | 7.30212E-06 |
| Nr6a1 | 3.66406E-06 | 1.619390862 | 0.604 | 0.331 | 0.089007446 | 0.012559188 | 0.678273767 | 0.78 | 0.618 | 1 | 0.012559188 | 7.32811E-06 |

| | | | | | | | | | | | | |
|---|---|---|---|---|---|---|---|---|---|---|---|---|
| Micall1 | 3.67022E-06 | -2.025849651 | 0.226 | 0.555 | 0.089157102 | 0.002025407 | -1.52536523 | 0.171 | 0.43 | 1 | 0.002025407 | 7.34044E-06 |
| Gng11 | 3.67506E-06 | 1.728097121 | 0.566 | 0.318 | 0.089274461 | 0.86222379 | 0.275325962 | 0.439 | 0.457 | 1 | 0.86222379 | 7.3501E-06 |
| Raph1 | 3.67903E-06 | -1.058690595 | 0.868 | 0.949 | 0.089371033 | 9.86502E-06 | -1.003688187 | 0.78 | 0.952 | 0.239641179 | 9.86502E-06 | 7.35805E-06 |
| Rictor | 0.000102077 | 0.922939123 | 0.849 | 0.72 | 1 | 3.68756E-06 | 0.942286896 | 0.902 | 0.833 | 0.089578282 | 0.000102077 | 7.37511E-06 |
| Ank2 | 3.7063E-06 | 1.695290976 | 0.547 | 0.233 | 0.090033379 | 0.002996831 | 1.962735732 | 0.561 | 0.369 | 1 | 0.002996831 | 7.41258E-06 |
| Tmtc1 | 0.005863751 | 1.057861056 | 0.396 | 0.229 | 1 | 3.74006E-06 | 3.604681748 | 0.415 | 0.154 | 0.090853553 | 0.005863751 | 7.48011E-06 |
| Ttc13 | 3.7694E-06 | 2.055186921 | 0.358 | 0.114 | 0.091566333 | 0.714313191 | 0.756284854 | 0.171 | 0.16 | 1 | 0.714313191 | 7.53879E-06 |
| Fdxr | 3.78202E-06 | 2.948271717 | 0.245 | 0.051 | 0.091872878 | 0.015416585 | 2.396632258 | 0.146 | 0.051 | 1 | 0.015416585 | 7.56403E-06 |
| Aspa | 0.04477173 | 1.891688189 | 0.057 | 0.013 | 1 | 3.79878E-06 | 3.422167351 | 0.098 | 0.003 | 0.092279976 | 0.04477173 | 7.59755E-06 |
| Sertad3 | 3.79891E-06 | 1.820303556 | 0.415 | 0.14 | 0.092283242 | 0.172025484 | 0.930314254 | 0.317 | 0.246 | 1 | 0.172025484 | 7.59782E-06 |
| Tlcd2 | 3.81098E-06 | 1.429582436 | 0.528 | 0.229 | 0.092576377 | 0.225208988 | 0.730289646 | 0.341 | 0.259 | 1 | 0.225208988 | 7.62195E-06 |
| Blm | 3.81478E-06 | -3.57319786 | 0.075 | 0.398 | 0.092668699 | 0.333006503 | -1.069685746 | 0.024 | 0.061 | 1 | 0.333006503 | 7.62955E-06 |
| Gm13944 | 3.84534E-06 | -3.617866909 | 0.038 | 0.36 | 0.093410906 | 0.040215224 | -1.863234868 | 0.049 | 0.171 | 1 | 0.040215224 | 7.69066E-06 |
| Csnk1g3 | 3.85659E-06 | 1.017219071 | 0.887 | 0.589 | 0.093684201 | 0.534398109 | 0.197076556 | 0.878 | 0.86 | 1 | 0.534398109 | 7.71316E-06 |
| Eif1a | 0.001206329 | -0.458106699 | 0.962 | 0.949 | 1 | 3.86323E-06 | -0.826335286 | 0.951 | 0.99 | 0.093845649 | 0.001206329 | 7.72645E-06 |
| Lcorl | 3.87824E-06 | 1.630848603 | 0.642 | 0.352 | 0.094210219 | 0.000508279 | 0.456532964 | 0.805 | 0.577 | 1 | 0.000508279 | 7.75647E-06 |
| Ppp2r1a | 3.88774E-06 | 0.971858538 | 0.83 | 0.678 | 0.094440938 | 0.359979416 | 0.388446733 | 0.756 | 0.72 | 1 | 0.359979416 | 7.77546E-06 |
| Otulinl | 3.89662E-06 | 2.59082171 | 0.264 | 0.059 | 0.094656613 | 0.021150363 | 1.891652634 | 0.22 | 0.106 | 1 | 0.021150363 | 7.79322E-06 |
| Sh3bp4 | 4.05295E-06 | 1.86841841 | 0.415 | 0.144 | 0.098454203 | 0.578666175 | -0.372248516 | 0.268 | 0.311 | 1 | 0.578666175 | 8.10588E-06 |
| Hdgfl3 | 4.06915E-06 | 1.543576162 | 0.66 | 0.369 | 0.098847886 | 0.169187803 | 0.495377917 | 0.634 | 0.536 | 1 | 0.169187803 | 8.13829E-06 |
| Oxr1 | 4.07364E-06 | 1.171863728 | 0.868 | 0.686 | 0.098956798 | 0.089886139 | 0.503304113 | 0.756 | 0.734 | 1 | 0.089886139 | 8.14726E-06 |
| Ccnd3 | 4.13539E-06 | 1.314327411 | 0.755 | 0.513 | 0.100456983 | 0.204679307 | 0.345052159 | 0.756 | 0.727 | 1 | 0.204679307 | 8.27077E-06 |
| Imp3 | 4.1529E-06 | 1.189488013 | 0.755 | 0.538 | 0.10088228 | 0.220517485 | 0.392902756 | 0.659 | 0.59 | 1 | 0.220517485 | 8.30579E-06 |
| Tmeff1 | 0.000852474 | 1.614154213 | 0.151 | 0.034 | 1 | 4.15413E-06 | 2.330244861 | 0.293 | 0.068 | 0.100912035 | 0.000852474 | 8.30824E-06 |
| Ubr3 | 4.18427E-06 | 0.773632427 | 0.962 | 0.788 | 0.101644266 | 0.060998257 | 0.342952788 | 0.927 | 0.942 | 1 | 0.060998257 | 8.36852E-06 |
| Rsrc2 | 4.18777E-06 | 0.735603919 | 0.981 | 0.953 | 0.101729387 | 0.387539957 | 0.357324395 | 0.927 | 0.962 | 1 | 0.387539957 | 8.37553E-06 |
| Bmp2k | 4.2504E-06 | 1.594634763 | 0.585 | 0.305 | 0.103250648 | 7.76379E-05 | 1.167353451 | 0.707 | 0.43 | 1 | 7.76379E-05 | 8.50078E-06 |
| Tbx2 | 4.26299E-06 | 2.47665069 | 0.302 | 0.076 | 0.103556616 | 5.30689E-06 | 3.685201756 | 0.195 | 0.031 | 0.128915031 | 5.30689E-06 | 8.52597E-06 |
| Vezt | 4.29622E-06 | 1.246916293 | 0.717 | 0.441 | 0.104363677 | 0.042654062 | 0.950352007 | 0.732 | 0.689 | 1 | 0.042654062 | 8.59241E-06 |
| Itpr3 | 0.050577277 | -0.227747042 | 0.434 | 0.606 | 1 | 4.29837E-06 | -1.219763962 | 0.415 | 0.775 | 0.104416016 | 0.050577277 | 8.59672E-06 |
| Chmp2b | 4.30561E-06 | 1.062464087 | 0.811 | 0.597 | 0.104591907 | 0.19945847 | 0.26666942 | 0.756 | 0.72 | 1 | 0.19945847 | 8.6112E-06 |
| Lats2 | 4.31268E-06 | 1.278812943 | 0.698 | 0.403 | 0.104763628 | 0.023739742 | 0.654726798 | 0.707 | 0.655 | 1 | 0.023739742 | 8.62534E-06 |
| Zfp428 | 4.34682E-06 | 2.782753817 | 0.226 | 0.042 | 0.105592895 | 0.06797635 | 1.667279848 | 0.122 | 0.051 | 1 | 0.06797635 | 8.69362E-06 |
| Tns1 | 4.35157E-06 | -1.330871511 | 0.774 | 0.873 | 0.10570831 | 0.007017494 | -0.688707411 | 0.732 | 0.877 | 1 | 0.007017494 | 8.70312E-06 |
| Tmem39b | 4.36385E-06 | 1.739685096 | 0.434 | 0.161 | 0.106006639 | 0.027998698 | 0.817305292 | 0.439 | 0.287 | 1 | 0.027998698 | 8.72768E-06 |
| Nrp1 | 1.46098E-05 | 1.377115016 | 0.434 | 0.165 | 0.354900847 | 4.37685E-06 | 1.891652634 | 0.561 | 0.27 | 0.106322349 | 1.46098E-05 | 8.75367E-06 |
| Efl1 | 4.39779E-06 | 2.122301117 | 0.509 | 0.25 | 0.106831114 | 0.349756221 | 1.08997092 | 0.488 | 0.457 | 1 | 0.349756221 | 8.79556E-06 |
| Mia2 | 4.47627E-06 | 0.787549484 | 0.962 | 0.881 | 0.108737445 | 0.589536337 | 0.205529604 | 0.951 | 0.932 | 1 | 0.589536337 | 8.95251E-06 |
| Dnajc14 | 4.52157E-06 | 1.910797012 | 0.415 | 0.157 | 0.109837997 | 0.176926551 | 0.937369012 | 0.415 | 0.358 | 1 | 0.176926551 | 9.04312E-06 |
| Ets1 | 4.59405E-06 | -0.684202406 | 0.811 | 0.966 | 0.111598601 | 0.01248036 | -0.27224071 | 0.927 | 0.976 | 1 | 0.01248036 | 9.18807E-06 |
| Tmem88 | 4.64148E-06 | 1.855162313 | 0.283 | 0.068 | 0.112750946 | 0.00298972 | 1.493250449 | 0.195 | 0.061 | 1 | 0.00298972 | 9.28295E-06 |
| B430010I23Rik | 4.73332E-06 | -3.760160792 | 0.038 | 0.356 | 0.114981756 | 0.046364682 | -0.446588116 | 0.171 | 0.355 | 1 | 0.046364682 | 9.46661E-06 |
| Dchs2 | 4.8539E-06 | 2.865215978 | 0.208 | 0.034 | 0.117910954 | 0.01687797 | 1.795384674 | 0.122 | 0.038 | 1 | 0.01687797 | 9.70778E-06 |
| Mrm1 | 4.88173E-06 | 2.47665069 | 0.226 | 0.042 | 0.118586876 | 0.004199752 | 1.635570989 | 0.195 | 0.065 | 1 | 0.004199752 | 9.76343E-06 |
| Rnf13 | 4.89094E-06 | 0.871268648 | 0.868 | 0.699 | 0.118810774 | 0.983805369 | 0.37036845 | 0.707 | 0.823 | 1 | 0.983805369 | 9.78186E-06 |
| Rab22a | 4.90528E-06 | 1.118502407 | 0.774 | 0.593 | 0.119159073 | 0.206305377 | 0.315967947 | 0.878 | 0.775 | 1 | 0.206305377 | 9.81054E-06 |
| Atg4a | 5.04183E-06 | 1.611100889 | 0.358 | 0.11 | 0.122476072 | 0.141075266 | 1.283812428 | 0.317 | 0.229 | 1 | 0.141075266 | 1.00836E-05 |
| Pkp4 | 5.07887E-06 | 1.335601178 | 0.792 | 0.453 | 0.123375973 | 0.000898914 | 1.10301634 | 0.78 | 0.737 | 1 | 0.000898914 | 1.01577E-05 |
| Atn1 | 5.09602E-06 | 1.165369839 | 0.679 | 0.398 | 0.123792599 | 0.009544733 | 0.667279848 | 0.78 | 0.635 | 1 | 0.009544733 | 1.0192E-05 |
| Ccdc3 | 5.24532E-06 | 2.116248447 | 0.245 | 0.051 | 0.127419274 | 0.007020555 | 1.718560353 | 0.341 | 0.184 | 1 | 0.007020555 | 1.04906E-05 |
| Btbd3 | 5.25467E-06 | -1.594984668 | 0.434 | 0.729 | 0.127646326 | 0.009508106 | -0.796256169 | 0.366 | 0.546 | 1 | 0.009508106 | 1.05093E-05 |
| Setd3 | 5.39491E-06 | 0.956078776 | 0.792 | 0.564 | 0.131053145 | 0.056247118 | 0.410940095 | 0.829 | 0.696 | 1 | 0.056247118 | 1.07898E-05 |
| Cpxm2 | 5.42115E-06 | 1.494798036 | 0.547 | 0.28 | 0.131690514 | 0.022971165 | -0.588491131 | 0.439 | 0.628 | 1 | 0.022971165 | 1.08423E-05 |
| Creb1 | 5.49541E-06 | 1.403543141 | 0.66 | 0.377 | 0.133494531 | 0.036852457 | 0.564678995 | 0.732 | 0.587 | 1 | 0.036852457 | 1.09908E-05 |
| Srsf6 | 5.50993E-06 | 0.743028539 | 0.981 | 0.737 | 0.133847189 | 0.082051683 | -0.418556321 | 0.78 | 0.87 | 1 | 0.082051683 | 1.10198E-05 |
| Crkl | 5.51724E-06 | 1.240879239 | 0.642 | 0.347 | 0.134024851 | 0.423045486 | 0.398834847 | 0.61 | 0.625 | 1 | 0.423045486 | 1.10345E-05 |
| Pcolce2 | 5.61391E-06 | -4.384436216 | 0.057 | 0.377 | 0.136372989 | 0.001625709 | -2.516432105 | 0.073 | 0.314 | 1 | 0.001625709 | 1.12278E-05 |
| Nup85 | 5.67224E-06 | 1.635348436 | 0.415 | 0.144 | 0.13778996 | 0.399084792 | -0.332720152 | 0.195 | 0.256 | 1 | 0.399084792 | 1.13444E-05 |
| Olfml1 | 0.007495013 | 2.417757001 | 0.075 | 0.013 | 1 | 5.69655E-06 | 3.602739596 | 0.195 | 0.031 | 0.13838059 | 0.007495013 | 1.13931E-05 |
| Micu1 | 1.93591E-05 | -0.85508058 | 0.755 | 0.966 | 0.470272178 | 5.71372E-06 | -1.07557946 | 0.756 | 0.942 | 0.138797685 | 1.93591E-05 | 1.14274E-05 |
| Gcnt1 | 5.71434E-06 | -2.201298699 | 0.17 | 0.5 | 0.138812832 | 0.000378367 | -2.665295491 | 0.073 | 0.348 | 1 | 0.000378367 | 1.14287E-05 |
| Slc7a2 | 5.77472E-06 | 1.605829349 | 0.623 | 0.343 | 0.14027939 | 0.000599595 | 1.548227177 | 0.61 | 0.41 | 1 | 0.000599595 | 1.15494E-05 |
| Bgn | 5.77653E-06 | 0.909081414 | 0.981 | 0.932 | 0.140323525 | 0.463390537 | 0.1561617 | 0.976 | 0.939 | 1 | 0.463390537 | 1.1553E-05 |
| Npc1 | 5.84652E-06 | -1.260314904 | 0.453 | 0.767 | 0.142023616 | 0.003965947 | -0.812174379 | 0.683 | 0.802 | 1 | 0.003965947 | 1.1693E-05 |
| Tmie | 5.87111E-06 | -3.554935653 | 0.038 | 0.352 | 0.142620975 | 0.077525693 | -2.292078167 | 0 | 0.072 | 1 | 0.077525693 | 1.17422E-05 |
| Klf3 | 5.87214E-06 | -2.006775047 | 0.736 | 0.843 | 0.142645907 | 0.003502612 | -1.155100758 | 0.854 | 0.945 | 1 | 0.003502612 | 1.17442E-05 |
| Caprin1 | 6.05807E-06 | 0.844298472 | 0.925 | 0.869 | 0.147162612 | 0.191632319 | 0.261363739 | 0.951 | 0.959 | 1 | 0.191632319 | 1.21161E-05 |
| Cotl1 | 1.6212E-05 | 1.300573461 | 0.396 | 0.136 | 0.39382232 | 6.07931E-06 | 1.480511337 | 0.537 | 0.229 | 0.147678693 | 1.6212E-05 | 1.21586E-05 |
| Isoc1 | 6.11766E-06 | 1.322450041 | 0.547 | 0.242 | 0.148610238 | 0.934566739 | 0.100239256 | 0.39 | 0.396 | 1 | 0.934566739 | 1.22353E-05 |
| Gnaq | 6.16274E-06 | 0.657603443 | 1 | 0.919 | 0.149705208 | 0.436621792 | -0.169434348 | 0.976 | 0.99 | 1 | 0.436621792 | 1.23254E-05 |
| Gm12940 | 6.16296E-06 | 1.863491297 | 0.509 | 0.242 | 0.149710691 | 0.000294695 | 1.738714986 | 0.561 | 0.334 | 1 | 0.000294695 | 1.23259E-05 |
| Dmxl1 | 6.29783E-06 | 0.932330173 | 0.906 | 0.678 | 0.152986833 | 0.897256599 | 0.119025274 | 0.878 | 0.857 | 1 | 0.897256599 | 1.25956E-05 |
| Brwd3 | 6.31488E-06 | 1.885535962 | 0.491 | 0.233 | 0.15340104 | 9.87284E-06 | 1.815988364 | 0.732 | 0.474 | 0.239830941 | 9.87284E-06 | 1.26297E-05 |
| Xpr1 | 6.3271E-06 | 1.531692053 | 0.736 | 0.564 | 0.153697999 | 0.575086008 | 0.361025339 | 0.854 | 0.853 | 1 | 0.575086008 | 1.26542E-05 |
| Gata2 | 0.001932659 | 4.154722595 | 0.075 | 0.008 | 1 | 6.38015E-06 | 3.169780189 | 0.268 | 0.061 | 0.154986698 | 0.001932659 | 1.27603E-05 |
| Fam180a | 6.41145E-06 | 0.961053239 | 0.868 | 0.602 | 0.155746949 | 0.096426725 | -0.692139459 | 0.805 | 0.805 | 1 | 0.096426725 | 1.28229E-05 |
| Zfpm2 | 0.000182672 | -0.807118597 | 0.755 | 0.936 | 1 | 6.45919E-06 | -0.887216104 | 0.537 | 0.904 | 0.156906653 | 0.000182672 | 1.29183E-05 |
| Cdkn1a | 6.4598E-06 | -1.163217092 | 0.925 | 0.975 | 0.156921497 | 0.124987358 | -0.126269274 | 0.976 | 0.983 | 1 | 0.124987358 | 1.29196E-05 |
| Kdm6b | 6.46523E-06 | 0.787605726 | 0.925 | 0.814 | 0.157053485 | 0.00262809 | 0.529363393 | 0.951 | 0.904 | 1 | 0.00262809 | 1.29304E-05 |
| Cacng7 | 6.51111E-06 | 1.751366901 | 0.377 | 0.123 | 0.158167994 | 0.015977009 | 0.749742009 | 0.415 | 0.246 | 1 | 0.015977009 | 1.30222E-05 |
| Prkrip1 | 6.5637E-06 | 1.391162791 | 0.528 | 0.237 | 0.159445417 | 0.79117163 | 0.186950889 | 0.244 | 0.28 | 1 | 0.79117163 | 1.31274E-05 |
| Akap9 | 6.60883E-06 | 0.762842817 | 0.962 | 0.911 | 0.160541779 | 0.988934408 | 0.15108588 | 0.902 | 0.98 | 1 | 0.988934408 | 1.32176E-05 |
| Ston2 | 0.039810395 | 2.465062715 | 0.302 | 0.203 | 1 | 6.70492E-06 | 3.10399139 | 0.415 | 0.15 | 0.162875933 | 0.039810395 | 1.34098E-05 |
| E130308A19Rik | 6.86434E-06 | 1.941728871 | 0.377 | 0.136 | 0.16674849 | 0.221072116 | 1.385825503 | 0.317 | 0.249 | 1 | 0.221072116 | 1.37286E-05 |
| Nrbp2 | 7.03014E-06 | -1.776014743 | 0.208 | 0.555 | 0.170776086 | 0.10418599 | -0.438919555 | 0.195 | 0.345 | 1 | 0.10418599 | 1.40602E-05 |
| Ralgapa1 | 7.04643E-06 | 0.821498874 | 0.906 | 0.831 | 0.171171808 | 0.155714813 | -0.252958151 | 0.854 | 0.949 | 1 | 0.155714813 | 1.40928E-05 |
| Pacrg | 7.05239E-06 | -5.422706233 | 0 | 0.297 | 0.171316749 | 0.05555427 | -1.747757651 | 0.098 | 0.225 | 1 | 0.05555427 | 1.41047E-05 |
| Kdm4b | 7.05369E-06 | 1.72845784 | 0.396 | 0.136 | 0.171348176 | 0.081897754 | 0.933420165 | 0.341 | 0.242 | 1 | 0.081897754 | 1.41073E-05 |
| Phldb1 | 7.06579E-06 | -0.927071496 | 0.698 | 0.898 | 0.171642232 | 0.005844523 | -0.595754557 | 0.78 | 0.915 | 1 | 0.005844523 | 1.41315E-05 |
| Ahdc1 | 7.15428E-06 | 1.160036395 | 0.774 | 0.547 | 0.173791878 | 0.418644701 | -0.122456221 | 0.805 | 0.86 | 1 | 0.418644701 | 1.43085E-05 |
| 1700025G04Rik | 7.15543E-06 | -0.967142951 | 0.906 | 0.979 | 0.173819803 | 0.047303844 | -0.452086037 | 0.902 | 0.952 | 1 | 0.047303844 | 1.43108E-05 |
| Junb | 0.00298709 | 0.556484863 | 0.868 | 0.78 | 1 | 7.16125E-06 | 1.285032284 | 0.927 | 0.778 | 0.173960983 | 0.00298709 | 1.43224E-05 |
| Pde3a | 0.007056463 | -0.124977265 | 0.623 | 0.86 | 1 | 7.16836E-06 | -1.16279515 | 0.512 | 0.898 | 0.174133788 | 0.007056463 | 1.43367E-05 |
| Acbd6 | 7.1816E-06 | 1.005859209 | 0.868 | 0.64 | 0.17445544 | 0.163076173 | 0.220342265 | 0.732 | 0.645 | 1 | 0.163076173 | 1.43631E-05 |
| Mark4 | 7.21213E-06 | 1.599507437 | 0.509 | 0.237 | 0.175196982 | 0.529692342 | -0.558723827 | 0.39 | 0.403 | 1 | 0.529692342 | 1.44242E-05 |
| Ing4 | 7.21765E-06 | 1.266749213 | 0.679 | 0.432 | 0.175331184 | 0.129108741 | 0.522331512 | 0.561 | 0.454 | 1 | 0.129108741 | 1.44353E-05 |
| Gm8251 | 7.25034E-06 | 2.18574949 | 0.245 | 0.051 | 0.176125353 | 0.481235612 | 0.224227973 | 0.098 | 0.14 | 1 | 0.481235612 | 1.45006E-05 |
| Phf13 | 7.29252E-06 | 1.484871196 | 0.566 | 0.275 | 0.177149859 | 0.479346476 | 0.533424102 | 0.439 | 0.444 | 1 | 0.479346476 | 1.4585E-05 |

| | | | | | | | | | | | | |
|---|---|---|---|---|---|---|---|---|---|---|---|---|
| Usp48 | 7.33654E-06 | 0.880100215 | 0.774 | 0.453 | 0.178219163 | 0.021470097 | 0.538546534 | 0.805 | 0.655 | 1 | 0.021470097 | 1.4673E-05 |
| St13 | 7.36759E-06 | 0.776651422 | 0.962 | 0.924 | 0.178973397 | 0.594699238 | 0.216504568 | 0.927 | 0.966 | 1 | 0.594699238 | 1.47351E-05 |
| Fxyd1 | 7.38722E-06 | 0.82944329 | 0.906 | 0.555 | 0.179450248 | 0.329471705 | -0.208662159 | 0.707 | 0.761 | 1 | 0.329471705 | 1.47744E-05 |
| Mafa | 7.40022E-06 | 3.06161319 | 0.189 | 0.03 | 0.179766025 | 0.340148806 | 0.422167351 | 0.122 | 0.075 | 1 | 0.340148806 | 1.48004E-05 |
| Hacd1 | 7.42692E-06 | 1.509006063 | 0.566 | 0.275 | 0.18041472 | 0.262901774 | 0.691527395 | 0.439 | 0.392 | 1 | 0.262901774 | 1.48538E-05 |
| Syt2 | 7.43913E-06 | 1.659957903 | 0.264 | 0.059 | 0.18071145 | 0.023249884 | 1.557096931 | 0.195 | 0.082 | 1 | 0.023249884 | 1.48782E-05 |
| Ifnar2 | 7.44188E-06 | 1.090016905 | 0.774 | 0.559 | 0.180778053 | 0.339187349 | 0.233383322 | 0.854 | 0.741 | 1 | 0.339187349 | 1.48837E-05 |
| 9530026P05Rik | 7.45425E-06 | 1.491318739 | 0.623 | 0.335 | 0.18107854 | 0.047813203 | 0.898605394 | 0.537 | 0.413 | 1 | 0.047813203 | 1.49084E-05 |
| Hs3st3a1 | 7.52581E-06 | -1.081475626 | 0.321 | 0.682 | 0.18281692 | 0.006665448 | -1.030691614 | 0.22 | 0.464 | 1 | 0.006665448 | 1.50516E-05 |
| Sowahc | 7.53174E-06 | -2.031143951 | 0.245 | 0.564 | 0.182961091 | 0.237252093 | -0.297954733 | 0.244 | 0.341 | 1 | 0.237252093 | 1.50634E-05 |
| Pabpc4l | 7.5755E-06 | 2.984797593 | 0.189 | 0.03 | 0.184024068 | 0.664414892 | -0.250257991 | 0.024 | 0.038 | 1 | 0.664414892 | 1.51509E-05 |
| Fam20a | 7.5838E-06 | 0.953088734 | 0.321 | 0.089 | 0.184225735 | 0.316557688 | 0.568716014 | 0.098 | 0.058 | 1 | 0.316557688 | 1.51675E-05 |
| Sptlc2 | 7.70413E-06 | -1.239762665 | 0.755 | 0.894 | 0.187148778 | 0.000947078 | -1.062072562 | 0.683 | 0.846 | 1 | 0.000947078 | 1.54082E-05 |
| Smim13 | 7.72617E-06 | 1.440124814 | 0.509 | 0.225 | 0.187684175 | 0.268361453 | -0.19841906 | 0.244 | 0.348 | 1 | 0.268361453 | 1.54523E-05 |
| Brip1os | 7.90016E-06 | 1.154722595 | 0.585 | 0.28 | 0.191910747 | 0.726687178 | 0.437274243 | 0.317 | 0.379 | 1 | 0.726687178 | 1.58003E-05 |
| Bach1 | 7.99696E-06 | 1.160325301 | 0.736 | 0.492 | 0.194262141 | 0.079768843 | 0.536809916 | 0.805 | 0.696 | 1 | 0.079768843 | 1.59939E-05 |
| Rcn2 | 8.01835E-06 | -0.66687425 | 0.962 | 0.962 | 0.194781691 | 0.209906065 | -0.283941118 | 0.829 | 0.881 | 1 | 0.209906065 | 1.60366E-05 |
| Parp9 | 8.02363E-06 | 1.521504925 | 0.415 | 0.148 | 0.19490994 | 0.441155805 | 0.545973552 | 0.244 | 0.205 | 1 | 0.441155805 | 1.60472E-05 |
| G3bp2 | 8.03957E-06 | 0.865215978 | 0.811 | 0.644 | 0.195297292 | 0.04704679 | 0.545973552 | 0.756 | 0.703 | 1 | 0.04704679 | 1.60791E-05 |
| Ppfia2 | 0.000532331 | 3.649487287 | 0.113 | 0.017 | 1 | 8.05517E-06 | 4.178241768 | 0.171 | 0.024 | 0.195676297 | 0.000532331 | 1.61103E-05 |
| Csad | 8.10704E-06 | 1.187587229 | 0.736 | 0.479 | 0.196936189 | 0.356183582 | -0.448952114 | 0.659 | 0.669 | 1 | 0.356183582 | 1.6214E-05 |
| Spty2d1 | 8.18837E-06 | 0.998482508 | 0.83 | 0.657 | 0.198911803 | 0.003189307 | 0.791034668 | 0.854 | 0.744 | 1 | 0.003189307 | 1.63767E-05 |
| Tm7sf3 | 8.25792E-06 | 1.316994024 | 0.717 | 0.479 | 0.200601468 | 0.478040682 | 0.208520972 | 0.61 | 0.621 | 1 | 0.478040682 | 1.65158E-05 |
| Cry2 | 8.30355E-06 | -1.035645796 | 0.604 | 0.847 | 0.201709858 | 0.919563011 | 0.113939327 | 0.78 | 0.84 | 1 | 0.919563011 | 1.6607E-05 |
| Gli2 | 8.31666E-06 | 2.26065604 | 0.528 | 0.271 | 0.202028295 | 0.94944779 | 0.227700165 | 0.463 | 0.491 | 1 | 0.94944779 | 1.66332E-05 |
| Dync1li1 | 8.3454E-06 | 0.937365763 | 0.868 | 0.585 | 0.202726488 | 0.258233892 | 0.32089871 | 0.805 | 0.802 | 1 | 0.258233892 | 1.66907E-05 |
| Sem1 | 8.35939E-06 | 0.670337049 | 0.981 | 0.958 | 0.203066312 | 0.391992673 | -0.112467977 | 1 | 0.99 | 1 | 0.391992673 | 1.67187E-05 |
| Tagln2 | 8.40343E-06 | 0.838380935 | 0.925 | 0.695 | 0.204136106 | 0.086862795 | -0.354830853 | 1 | 0.952 | 1 | 0.086862795 | 1.68068E-05 |
| Mrc2 | 8.41528E-06 | 0.789292861 | 0.868 | 0.72 | 0.204423943 | 0.027495548 | 0.642883194 | 0.829 | 0.795 | 1 | 0.027495548 | 1.68305E-05 |
| Aco1 | 8.48838E-06 | 1.502645898 | 0.528 | 0.237 | 0.206199839 | 0.101536149 | -0.721762442 | 0.268 | 0.379 | 1 | 0.101536149 | 1.69767E-05 |
| Mmp23 | 8.5645E-06 | 2.806799291 | 0.226 | 0.047 | 0.208048916 | 0.846926792 | 1.100239256 | 0.024 | 0.031 | 1 | 0.846926792 | 1.71289E-05 |
| Uck1 | 8.56611E-06 | 1.699043111 | 0.434 | 0.169 | 0.20808787 | 0.51819223 | 0.819282942 | 0.268 | 0.246 | 1 | 0.51819223 | 1.71321E-05 |
| Tet1 | 8.57877E-06 | 1.386048141 | 0.528 | 0.237 | 0.208395569 | 0.276543481 | -0.410722664 | 0.171 | 0.246 | 1 | 0.276543481 | 1.71575E-05 |
| Snx2 | 8.58434E-06 | 0.987344275 | 0.868 | 0.665 | 0.20853073 | 0.177318887 | 0.413993419 | 0.829 | 0.846 | 1 | 0.177318887 | 1.71686E-05 |
| Cct5 | 8.58523E-06 | 0.821298861 | 0.906 | 0.703 | 0.208552307 | 0.185560813 | 0.277246355 | 0.805 | 0.84 | 1 | 0.185560813 | 1.71704E-05 |
| Hsf2 | 8.58635E-06 | 1.59400764 | 0.434 | 0.174 | 0.208579711 | 0.011756552 | 1.119792053 | 0.512 | 0.345 | 1 | 0.011756552 | 1.71726E-05 |
| Cdyl | 8.67428E-06 | 1.58002843 | 0.491 | 0.216 | 0.210715638 | 0.18455158 | 0.347399582 | 0.537 | 0.413 | 1 | 0.18455158 | 1.73485E-05 |
| Zcchc8 | 8.67494E-06 | 1.202816883 | 0.623 | 0.314 | 0.210731723 | 0.182464853 | 0.675741427 | 0.585 | 0.522 | 1 | 0.182464853 | 1.73498E-05 |
| Tceal8 | 8.72582E-06 | 0.614529964 | 0.943 | 0.826 | 0.211967555 | 0.667096346 | -0.138547604 | 0.878 | 0.891 | 1 | 0.667096346 | 1.74516E-05 |
| Cdc25a | 8.7913E-06 | 1.974150349 | 0.321 | 0.093 | 0.213558205 | 0.74836577 | 0.441276173 | 0.244 | 0.225 | 1 | 0.74836577 | 1.75825E-05 |
| Sipa1 | 8.82129E-06 | 2.657222935 | 0.226 | 0.047 | 0.214286893 | 0.175359883 | -0.238083278 | 0.073 | 0.16 | 1 | 0.175359883 | 1.76425E-05 |
| Wdr72 | 0.913426097 | 2.261637799 | 0.019 | 0.017 | 1 | 8.84155E-06 | 3.364451853 | 0.341 | 0.099 | 0.214778928 | 0.913426097 | 1.7683E-05 |
| Wsb1 | 8.85479E-06 | 0.765680304 | 1 | 0.932 | 0.215100626 | 0.002757551 | 0.659307978 | 1 | 0.993 | 1 | 0.002757551 | 1.77095E-05 |
| Bnip3l | 8.86353E-06 | 0.664246961 | 0.981 | 0.746 | 0.215312757 | 0.354301163 | 0.380204057 | 0.756 | 0.819 | 1 | 0.354301163 | 1.7727E-05 |
| Ppp1cb | 8.88443E-06 | 0.740567208 | 1 | 0.949 | 0.215820661 | 0.254902585 | -0.104819283 | 0.976 | 0.997 | 1 | 0.254902585 | 1.77688E-05 |
| Hmg20b | 8.90212E-06 | 1.048594244 | 0.811 | 0.619 | 0.216250391 | 0.001024625 | 1.076670784 | 0.732 | 0.546 | 1 | 0.001024625 | 1.78042E-05 |
| Phkg1 | 8.92908E-06 | 3.657222935 | 0.151 | 0.017 | 0.216905156 | 0.039931326 | 1.515276755 | 0.122 | 0.044 | 1 | 0.039931326 | 1.78581E-05 |
| Ube2q2 | 9.06643E-06 | -1.846061266 | 0.736 | 0.86 | 0.220241599 | 0.520084083 | -0.262842327 | 0.732 | 0.805 | 1 | 0.520084083 | 1.81328E-05 |
| Igfbp5 | 9.33423E-06 | 1.660907982 | 0.585 | 0.263 | 0.226747008 | 0.007035199 | 2.232613542 | 0.488 | 0.311 | 1 | 0.007035199 | 1.86684E-05 |
| Dbi | 9.35436E-06 | -1.32201465 | 0.943 | 0.941 | 0.227236167 | 0.50979037 | -0.120566915 | 0.854 | 0.891 | 1 | 0.50979037 | 1.87086E-05 |
| Kif13b | 0.000465597 | -0.961736382 | 1 | 0.975 | 1 | 9.4129E-06 | -1.181744253 | 0.927 | 0.976 | 0.228658214 | 0.000465597 | 1.88257E-05 |
| Lrrk1 | 9.43048E-06 | 0.941588141 | 0.83 | 0.593 | 0.229085203 | 0.609062427 | 0.402627164 | 0.732 | 0.737 | 1 | 0.609062427 | 1.88609E-05 |
| Npat | 9.5467E-06 | 1.005209053 | 0.698 | 0.436 | 0.231908401 | 0.074979257 | 0.726359081 | 0.634 | 0.563 | 1 | 0.074979257 | 1.90933E-05 |
| Gucy1a2 | 9.72896E-06 | 3.595295186 | 0.151 | 0.017 | 0.23633578 | 0.188035462 | 1.08997092 | 0.171 | 0.106 | 1 | 0.188035462 | 1.94578E-05 |
| Cacnb4 | 9.78412E-06 | 3.055186921 | 0.208 | 0.038 | 0.237675893 | 0.626346992 | 2.614812428 | 0.098 | 0.082 | 1 | 0.626346992 | 1.95681E-05 |
| Chst12 | 1.24376E-05 | -0.898388742 | 0.528 | 0.797 | 0.302134953 | 9.83074E-06 | -1.000144813 | 0.268 | 0.686 | 0.23880834 | 1.24376E-05 | 1.96614E-05 |
| Ammecr1 | 9.83623E-06 | 1.6587651 | 0.528 | 0.246 | 0.238941585 | 0.125457081 | 0.531859639 | 0.415 | 0.304 | 1 | 0.125457081 | 1.96724E-05 |
| Tex30 | 9.95224E-06 | 1.78699081 | 0.358 | 0.114 | 0.241759894 | 0.380493813 | 0.212713985 | 0.171 | 0.249 | 1 | 0.380493813 | 1.99044E-05 |
| Gm27241 | 9.95927E-06 | 3.154722595 | 0.151 | 0.017 | 0.241930657 | 0.885202143 | 0.911205431 | 0.049 | 0.044 | 1 | 0.885202143 | 1.99184E-05 |
| Acvr2b | 0.160046644 | 1.347367673 | 0.057 | 0.021 | 1 | 9.96713E-06 | 2.630753972 | 0.22 | 0.041 | 0.242121401 | 0.160046644 | 1.99342E-05 |
| Tmem47 | 1.00402E-05 | 1.8327945 | 0.34 | 0.106 | 0.24389646 | 0.076241907 | 1.225770138 | 0.341 | 0.235 | 1 | 0.076241907 | 2.00803E-05 |
| Atp6v0a4 | 0.188765503 | 0.394393008 | 0.34 | 0.258 | 1 | 1.00833E-05 | -1.100108666 | 0.512 | 0.863 | 0.244942876 | 0.188765503 | 2.01664E-05 |
| Nf2 | 1.02458E-05 | 1.398959405 | 0.547 | 0.267 | 0.248891353 | 0.094929517 | 0.567744175 | 0.634 | 0.539 | 1 | 0.094929517 | 2.04915E-05 |
| Mctp2 | 1.02538E-05 | -2.992118794 | 0.057 | 0.373 | 0.249084961 | 0.554547986 | 0.149148856 | 0.195 | 0.154 | 1 | 0.554547986 | 2.05075E-05 |
| Atox1 | 1.03931E-05 | -1.846319439 | 0.981 | 0.953 | 0.252468071 | 0.087441988 | -0.580203554 | 0.78 | 0.928 | 1 | 0.087441988 | 2.0786E-05 |
| Trim7 | 1.04742E-05 | 3.002719501 | 0.151 | 0.017 | 0.254439295 | 0.631722921 | 0.83720485 | 0.024 | 0.041 | 1 | 0.631722921 | 2.09483E-05 |
| Cebpz | 1.05114E-05 | 1.222537979 | 0.717 | 0.462 | 0.255344019 | 0.002061816 | 0.788445538 | 0.756 | 0.577 | 1 | 0.002061816 | 2.10228E-05 |
| Fam184a | 1.05166E-05 | 1.283046692 | 0.491 | 0.191 | 0.255468421 | 0.823757568 | 0.423807709 | 0.366 | 0.423 | 1 | 0.823757568 | 2.1033E-05 |
| Ahcyl1 | 1.05168E-05 | 0.790234038 | 0.925 | 0.801 | 0.255475046 | 0.000595283 | 0.663944136 | 0.976 | 0.898 | 1 | 0.000595283 | 2.10336E-05 |
| Cep350 | 1.05922E-05 | 0.870455552 | 0.925 | 0.712 | 0.257305564 | 0.59054914 | 0.186034198 | 0.878 | 0.87 | 1 | 0.59054914 | 2.11843E-05 |
| Igfbp3 | 1.0597E-05 | 1.745331659 | 0.245 | 0.051 | 0.257423168 | 0.000458465 | 1.043655727 | 0.439 | 0.205 | 1 | 0.000458465 | 2.1194E-05 |
| Anp32e | 1.06253E-05 | 1.143996188 | 0.811 | 0.585 | 0.258109667 | 0.823794716 | 0.143989458 | 0.756 | 0.768 | 1 | 0.823794716 | 2.12505E-05 |
| Parva | 1.06588E-05 | 0.727588904 | 0.962 | 0.742 | 0.258922677 | 0.19134142 | 0.341208781 | 0.805 | 0.85 | 1 | 0.19134142 | 2.13174E-05 |
| Nexmif | 1.0718E-05 | 3.232725107 | 0.151 | 0.017 | 0.260360743 | 0.077319511 | 2.59619675 | 0.073 | 0.024 | 1 | 0.077319511 | 2.14358E-05 |
| Abhd2 | 0.001185615 | -0.494112089 | 0.887 | 0.949 | 1 | 1.07417E-05 | -0.964075558 | 0.854 | 0.98 | 0.260936578 | 0.001185615 | 2.14832E-05 |
| H2az1 | 1.07479E-05 | 0.599960332 | 0.962 | 0.839 | 0.261086881 | 0.002692767 | 0.611850015 | 0.951 | 0.928 | 1 | 0.002692767 | 2.14956E-05 |
| Slc8a3 | 1.07974E-05 | -2.386374021 | 0.189 | 0.534 | 0.262291093 | 0.351646564 | -1.320885015 | 0.244 | 0.304 | 1 | 0.351646564 | 2.15947E-05 |
| Ncoa3 | 1.0798E-05 | -1.393841145 | 0.679 | 0.864 | 0.262303828 | 0.059039858 | -0.404381137 | 0.902 | 0.925 | 1 | 0.059039858 | 2.15958E-05 |
| Samd1 | 1.08521E-05 | 2.051629102 | 0.34 | 0.11 | 0.263618711 | 0.091749266 | 1.012776414 | 0.293 | 0.198 | 1 | 0.091749266 | 2.1704E-05 |
| Capn2 | 0.098890045 | -0.25203634 | 0.792 | 0.822 | 1 | 1.0881E-05 | -0.82386063 | 0.878 | 0.969 | 0.264321892 | 0.098890045 | 2.17619E-05 |
| Safb2 | 1.09054E-05 | 1.016161764 | 0.83 | 0.593 | 0.264914542 | 0.48186361 | -0.102961901 | 0.805 | 0.823 | 1 | 0.48186361 | 2.18107E-05 |
| Pacsin1 | 1.09568E-05 | 2.891688189 | 0.151 | 0.017 | 0.266161828 | 0.688532549 | 0.979223855 | 0.098 | 0.082 | 1 | 0.688532549 | 2.19134E-05 |
| Rb1cc1 | 1.09637E-05 | 0.694333964 | 0.981 | 0.907 | 0.266330385 | 0.725942502 | 0.188370166 | 0.976 | 0.973 | 1 | 0.725942502 | 2.19273E-05 |
| Nup54 | 1.09754E-05 | 1.053921954 | 0.679 | 0.428 | 0.266614771 | 0.376485548 | -0.337720833 | 0.512 | 0.553 | 1 | 0.376485548 | 2.19507E-05 |
| Nap1l4 | 1.09797E-05 | 1.022694201 | 0.83 | 0.623 | 0.266718401 | 0.436678938 | 0.280966638 | 0.78 | 0.792 | 1 | 0.436678938 | 2.19592E-05 |
| Arhgap12 | 1.10129E-05 | 1.399835093 | 0.528 | 0.237 | 0.267525596 | 0.277356247 | 0.23386382 | 0.537 | 0.43 | 1 | 0.277356247 | 2.20257E-05 |
| Csnk2a2 | 1.10608E-05 | 0.854820806 | 0.887 | 0.665 | 0.268689418 | 0.386602581 | 0.271039945 | 0.854 | 0.816 | 1 | 0.386602581 | 2.21215E-05 |
| Sik3 | 1.10648E-05 | 0.886554982 | 1 | 0.962 | 0.268785721 | 0.000946171 | 0.423187386 | 1 | 0.993 | 1 | 0.000946171 | 2.21294E-05 |
| Sec11a | 1.11843E-05 | 0.767206158 | 0.943 | 0.801 | 0.271689186 | 0.339566696 | 0.147722362 | 0.829 | 0.891 | 1 | 0.339566696 | 2.23685E-05 |
| Cep170 | 1.11986E-05 | 0.959551642 | 0.736 | 0.479 | 0.27203554 | 0.664943527 | 0.176860537 | 0.683 | 0.662 | 1 | 0.664943527 | 2.2397E-05 |
| Mbnl2 | 0.000411375 | 0.36556079 | 1 | 1 | 1 | 1.12415E-05 | 0.607762974 | 1 | 1 | 0.273078321 | 0.000411375 | 2.24829E-05 |
| Erg | 1.13307E-05 | -1.415656583 | 0.66 | 0.907 | 0.275245772 | 0.00053399 | -0.900566494 | 0.732 | 0.939 | 1 | 0.00053399 | 2.26613E-05 |
| Inip | 1.1426E-05 | 2.417757001 | 0.283 | 0.076 | 0.27755983 | 0.29424792 | 0.600165653 | 0.22 | 0.154 | 1 | 0.29424792 | 2.28518E-05 |
| Rftn1 | 2.08414E-05 | 2.312263872 | 0.264 | 0.068 | 0.506278127 | 1.14773E-05 | 2.640168003 | 0.317 | 0.089 | 0.278805836 | 2.08414E-05 | 2.29544E-05 |
| Omd | 0.022215364 | -0.693865325 | 0.943 | 0.966 | 1 | 1.14948E-05 | -1.325863798 | 0.854 | 0.922 | 0.279231953 | 0.022215364 | 2.29895E-05 |
| Magi3 | 1.15498E-05 | 1.264347086 | 0.623 | 0.318 | 0.280567459 | 0.000613055 | 1.248519263 | 0.707 | 0.512 | 1 | 0.000613055 | 2.30994E-05 |
| Stmp1 | 1.16159E-05 | -1.066584685 | 0.66 | 0.847 | 0.282172579 | 0.010486595 | -0.570879889 | 0.537 | 0.747 | 1 | 0.010486595 | 2.32316E-05 |
| BC028528 | 1.16599E-05 | 2.280253477 | 0.151 | 0.017 | 0.283241482 | 0.743829142 | 0.252242349 | 0.024 | 0.034 | 1 | 0.743829142 | 2.33196E-05 |

| | | | | | | | | | | | | |
|---|---|---|---|---|---|---|---|---|---|---|---|---|
| Pi4k2a | 1.16907E-05 | -0.570042546 | 0.547 | 0.843 | 0.283990278 | 0.196468802 | -0.276357709 | 0.659 | 0.727 | 1 | 0.196468802 | 2.33812E-05 |
| Nupr1 | 1.17207E-05 | 0.977347712 | 0.906 | 0.767 | 0.284719982 | 0.038737281 | 0.377773231 | 0.927 | 0.833 | 1 | 0.038737281 | 2.34413E-05 |
| Apbb1 | 1.17268E-05 | 2.237184755 | 0.245 | 0.055 | 0.284868293 | 0.523493397 | 0.493250449 | 0.195 | 0.16 | 1 | 0.523493397 | 2.34535E-05 |
| Rcor3 | 1.1797E-05 | 0.998603393 | 0.66 | 0.39 | 0.286571913 | 0.337843605 | 0.391405097 | 0.537 | 0.546 | 1 | 0.337843605 | 2.35938E-05 |
| Cmtm3 | 1.1817E-05 | 1.61630268 | 0.566 | 0.339 | 0.287058112 | 0.351577822 | 0.584224109 | 0.366 | 0.311 | 1 | 0.351577822 | 2.36338E-05 |
| Ppa1 | 1.18444E-05 | -2.843703842 | 0.736 | 0.826 | 0.287724432 | 0.101901002 | -1.974662729 | 0.634 | 0.734 | 1 | 0.101901002 | 2.36887E-05 |
| Steap2 | 1.18741E-05 | -2.023195197 | 0.302 | 0.589 | 0.288445182 | 0.469245619 | -0.31254227 | 0.293 | 0.362 | 1 | 0.469245619 | 2.3748E-05 |
| Zfp267 | 1.18928E-05 | 2.029191713 | 0.358 | 0.119 | 0.288900462 | 0.671741637 | 0.23386382 | 0.171 | 0.208 | 1 | 0.671741637 | 2.37855E-05 |
| Gcn1 | 1.19349E-05 | 0.892981506 | 0.811 | 0.525 | 0.289921799 | 0.243261531 | 0.555918739 | 0.61 | 0.604 | 1 | 0.243261531 | 2.38696E-05 |
| A730049H05Rik | 0.001025769 | 3.154722595 | 0.094 | 0.013 | 1 | 1.1971E-05 | 3.974708374 | 0.122 | 0.01 | 0.290799649 | 0.001025769 | 2.39419E-05 |
| Brd3os | 1.21754E-05 | 2.270199812 | 0.208 | 0.038 | 0.295765667 | 0.974673471 | 1.334704509 | 0.049 | 0.051 | 1 | 0.974673471 | 2.43507E-05 |
| Pacs2 | 1.22003E-05 | 1.846600299 | 0.415 | 0.161 | 0.296370163 | 0.17818181 | 1.282615998 | 0.317 | 0.253 | 1 | 0.17818181 | 2.44005E-05 |
| Crispld2 | 0.249284487 | 0.451963058 | 0.66 | 0.631 | 1 | 1.22212E-05 | 2.351778023 | 0.488 | 0.218 | 0.296876892 | 0.249284487 | 2.44422E-05 |
| Tia1 | 1.22953E-05 | 1.736009437 | 0.585 | 0.335 | 0.298676263 | 0.00040726 | 1.338953982 | 0.659 | 0.464 | 1 | 0.00040726 | 2.45904E-05 |
| Ulk1 | 1.23042E-05 | 1.32825885 | 0.66 | 0.394 | 0.298893858 | 0.032780593 | 0.757034501 | 0.707 | 0.584 | 1 | 0.032780593 | 2.46083E-05 |
| Nr2c2 | 1.23358E-05 | 1.161675355 | 0.698 | 0.441 | 0.299660082 | 0.009058229 | 0.751089692 | 0.829 | 0.727 | 1 | 0.009058229 | 2.46714E-05 |
| Cap2 | 1.23921E-05 | 2.533234218 | 0.151 | 0.017 | 0.301028664 | 0.60213981 | 1.029849928 | 0.024 | 0.014 | 1 | 0.60213981 | 2.4784E-05 |
| Amer1 | 1.23991E-05 | 1.891688189 | 0.283 | 0.072 | 0.30119787 | 2.01411E-05 | 1.957499084 | 0.341 | 0.102 | 0.489266919 | 2.01411E-05 | 2.4798E-05 |
| Tent4a | 1.24841E-05 | 1.720319771 | 0.415 | 0.157 | 0.303263488 | 0.005111146 | 0.959688856 | 0.537 | 0.331 | 1 | 0.005111146 | 2.4968E-05 |
| Kremen1 | 1.24862E-05 | 1.468476758 | 0.679 | 0.441 | 0.303314604 | 8.9206E-05 | 1.219924629 | 0.756 | 0.526 | 1 | 8.9206E-05 | 2.49722E-05 |
| Stim2 | 1.25577E-05 | 1.169829487 | 0.774 | 0.568 | 0.305052707 | 0.490821707 | 0.203260543 | 0.829 | 0.768 | 1 | 0.490821707 | 2.51153E-05 |
| Knop1 | 1.27006E-05 | 1.47119626 | 0.585 | 0.314 | 0.308523237 | 0.619848399 | 0.486707603 | 0.463 | 0.42 | 1 | 0.619848399 | 2.54011E-05 |
| Kcnq4 | 0.00285365 | 4.47665069 | 0.038 | 0 | 1 | 1.2708E-05 | 3.367719567 | 0.146 | 0.017 | 0.308703334 | 0.00285365 | 2.54159E-05 |
| Dok6 | 0.00285365 | 5.47665069 | 0.038 | 0 | 1 | 1.27111E-05 | 3.515276755 | 0.122 | 0.01 | 0.308777893 | 0.00285365 | 2.5422E-05 |
| Sptan1 | 1.27112E-05 | 1.019533467 | 0.981 | 0.886 | 0.308781175 | 0.00131482 | 0.701952426 | 0.951 | 0.973 | 1 | 0.00131482 | 2.54223E-05 |
| Ly96 | 1.27159E-05 | 1.554330054 | 0.415 | 0.148 | 0.308894861 | 0.06998649 | 0.896098539 | 0.366 | 0.253 | 1 | 0.06998649 | 2.54317E-05 |
| Marchf6 | 1.28898E-05 | 0.853418172 | 0.774 | 0.725 | 0.313120187 | 0.348350192 | -0.148487314 | 0.854 | 0.901 | 1 | 0.348350192 | 2.57795E-05 |
| Tmem106b | 1.28904E-05 | 1.017219071 | 0.774 | 0.559 | 0.313132514 | 0.195548842 | 0.455229371 | 0.683 | 0.662 | 1 | 0.195548842 | 2.57805E-05 |
| Gadd45g | 0.041937072 | 0.788655071 | 0.585 | 0.487 | 1 | 1.29254E-05 | 2.236032122 | 0.561 | 0.266 | 0.313984576 | 0.041937072 | 2.58507E-05 |
| Rbbp7 | 1.30381E-05 | 0.680164284 | 0.981 | 0.877 | 0.316722659 | 0.327510318 | 0.21068267 | 0.976 | 0.959 | 1 | 0.327510318 | 2.60761E-05 |
| Vav3 | 1.30818E-05 | -1.187979793 | 0.491 | 0.788 | 0.317783971 | 0.006052273 | -0.89698366 | 0.634 | 0.778 | 1 | 0.006052273 | 2.61635E-05 |
| Ptpre | 1.3207E-05 | -3.049149739 | 0.038 | 0.347 | 0.320823259 | 0.512556129 | -0.463964685 | 0.171 | 0.208 | 1 | 0.512556129 | 2.64137E-05 |
| Efna1 | 0.000918256 | -0.839910895 | 0.679 | 0.835 | 1 | 1.32207E-05 | -0.987770994 | 0.707 | 0.891 | 0.32115677 | 0.000918256 | 2.64412E-05 |
| Mtx2 | 1.3229E-05 | 1.589125419 | 0.509 | 0.233 | 0.321357916 | 0.821442498 | 0.221545552 | 0.39 | 0.389 | 1 | 0.821442498 | 2.64577E-05 |
| Sbf2 | 1.32564E-05 | 1.274304211 | 0.811 | 0.572 | 0.322024179 | 0.610082082 | 0.270858027 | 0.805 | 0.85 | 1 | 0.610082082 | 2.65126E-05 |
| Mtdh | 1.32686E-05 | 0.675354753 | 1 | 0.97 | 0.322320022 | 0.033094587 | 0.296058581 | 1 | 0.962 | 1 | 0.033094587 | 2.6537E-05 |
| Zfp664 | 1.3302E-05 | 1.222741018 | 0.642 | 0.407 | 0.323132206 | 0.01356307 | 0.780173905 | 0.829 | 0.666 | 1 | 0.01356307 | 2.66038E-05 |
| Cracr2b | 1.35275E-05 | 2.47665069 | 0.264 | 0.068 | 0.328610087 | 0.962913964 | 0.169780189 | 0.244 | 0.235 | 1 | 0.962913964 | 2.70548E-05 |
| Fcho2 | 1.36454E-05 | 0.686364354 | 0.962 | 0.894 | 0.331474051 | 0.029383619 | 0.281867518 | 0.976 | 0.973 | 1 | 0.029383619 | 2.72906E-05 |
| Abraxas2 | 1.36621E-05 | 1.338434548 | 0.547 | 0.263 | 0.331880125 | 0.083437815 | -0.505187348 | 0.293 | 0.444 | 1 | 0.083437815 | 2.7324E-05 |
| Prune2 | 1.36972E-05 | -3.927426447 | 0.075 | 0.381 | 0.332732448 | 0.034440076 | -3.010792057 | 0.073 | 0.208 | 1 | 0.034440076 | 2.73942E-05 |
| Scarf2 | 1.37277E-05 | 1.218852932 | 0.736 | 0.555 | 0.333473484 | 0.018112137 | 0.61396103 | 0.756 | 0.706 | 1 | 0.018112137 | 2.74552E-05 |
| Gxylt1 | 1.37851E-05 | 1.441603743 | 0.566 | 0.297 | 0.334866671 | 0.340103055 | 0.346064371 | 0.488 | 0.447 | 1 | 0.340103055 | 2.75699E-05 |
| Nap1l1 | 1.38235E-05 | 0.684271975 | 0.981 | 0.953 | 0.335800305 | 0.147396206 | -0.222513001 | 1 | 0.976 | 1 | 0.147396206 | 2.76468E-05 |
| Anapc4 | 1.38278E-05 | 1.119957177 | 0.698 | 0.449 | 0.335904252 | 0.458931008 | 0.290044382 | 0.61 | 0.601 | 1 | 0.458931008 | 2.76554E-05 |
| Slc2a3 | 0.002082099 | 1.067259754 | 0.132 | 0.03 | 1 | 1.38978E-05 | 3.377773231 | 0.146 | 0.017 | 0.337606431 | 0.002082099 | 2.77955E-05 |
| Tubgcp3 | 1.39614E-05 | 1.391162791 | 0.472 | 0.195 | 0.339150972 | 0.58837278 | 0.493250449 | 0.317 | 0.297 | 1 | 0.58837278 | 2.79227E-05 |
| Mob1b | 1.40173E-05 | 1.138602929 | 0.698 | 0.415 | 0.340507784 | 0.019007382 | 0.538413254 | 0.854 | 0.71 | 1 | 0.019007382 | 2.80344E-05 |
| Erh | 1.40437E-05 | 0.761911014 | 0.887 | 0.712 | 0.341149367 | 0.568912487 | 0.165942503 | 0.683 | 0.771 | 1 | 0.568912487 | 2.80872E-05 |
| Heyl | 1.41466E-05 | 3.678284551 | 0.17 | 0.025 | 0.34364922 | 0.008422791 | 2.794136128 | 0.195 | 0.075 | 1 | 0.008422791 | 2.8293E-05 |
| Gng5 | 0.011994897 | -0.376359665 | 1 | 0.966 | 1 | 1.41614E-05 | -0.75086892 | 0.951 | 0.976 | 0.344009455 | 0.011994897 | 2.83227E-05 |
| Nectin1 | 1.42766E-05 | 1.84156471 | 0.302 | 0.089 | 0.346806757 | 0.001893832 | 1.210422173 | 0.463 | 0.249 | 1 | 0.001893832 | 2.8553E-05 |
| Ttll4 | 1.43956E-05 | 1.493524508 | 0.434 | 0.178 | 0.3496971 | 0.598691675 | 0.682482255 | 0.39 | 0.44 | 1 | 0.598691675 | 2.87909E-05 |
| Serpinc1 | 1.46126E-05 | 2.685237311 | 0.245 | 0.059 | 0.354968253 | 0.203026202 | 1.876733214 | 0.195 | 0.133 | 1 | 0.203026202 | 2.92249E-05 |
| Enpp1 | 9.6034E-05 | -0.613823317 | 0.679 | 0.924 | 1 | 1.47798E-05 | -1.139456024 | 0.415 | 0.829 | 0.359032105 | 9.6034E-05 | 2.95595E-05 |
| Cry1 | 1.48326E-05 | 1.507832345 | 0.585 | 0.326 | 0.360312793 | 0.326865619 | 0.660882077 | 0.561 | 0.556 | 1 | 0.326865619 | 2.96649E-05 |
| Xxylt1 | 1.49443E-05 | 1.815920681 | 0.415 | 0.165 | 0.363025964 | 0.400638009 | 0.43121249 | 0.317 | 0.266 | 1 | 0.400638009 | 2.98883E-05 |
| Neurl4 | 1.49863E-05 | 1.953088734 | 0.302 | 0.085 | 0.364047353 | 3.36564E-05 | 1.803257518 | 0.366 | 0.123 | 0.817581702 | 3.36564E-05 | 2.99724E-05 |
| Rps6kc1 | 1.50265E-05 | 1.757832442 | 0.453 | 0.195 | 0.365024442 | 0.435151497 | -0.225407127 | 0.366 | 0.427 | 1 | 0.435151497 | 3.00528E-05 |
| Eif3f | 1.50462E-05 | 0.682874174 | 0.981 | 0.89 | 0.365501514 | 0.205110424 | 0.401279984 | 0.951 | 0.922 | 1 | 0.205110424 | 3.00921E-05 |
| Elavl3 | 1.5444E-05 | 2.465062715 | 0.264 | 0.068 | 0.375165767 | 0.851934104 | -0.361575014 | 0.122 | 0.133 | 1 | 0.851934104 | 3.08878E-05 |
| Fbxl2 | 0.084124445 | -0.508242418 | 0.509 | 0.585 | 1 | 1.54695E-05 | -2.11032773 | 0.171 | 0.543 | 0.375785968 | 0.084124445 | 3.09388E-05 |
| Bcl7c | 1.56542E-05 | 0.886233759 | 0.811 | 0.593 | 0.380271067 | 0.773372145 | 0.101462397 | 0.659 | 0.686 | 1 | 0.773372145 | 3.13081E-05 |
| Mafk | 1.56877E-05 | -1.721471204 | 0.245 | 0.564 | 0.381086468 | 0.00897757 | -0.873765536 | 0.585 | 0.73 | 1 | 0.00897757 | 3.13752E-05 |
| Nop56 | 1.57063E-05 | 0.884019823 | 0.83 | 0.695 | 0.381537495 | 0.020408793 | 0.743872273 | 0.756 | 0.683 | 1 | 0.020408793 | 3.14124E-05 |
| Kpna3 | 1.58337E-05 | 1.102526384 | 0.698 | 0.424 | 0.384632445 | 0.53133764 | 0.237292008 | 0.756 | 0.717 | 1 | 0.53133764 | 3.16672E-05 |
| Mamld1 | 1.58578E-05 | -3.304709024 | 0.094 | 0.394 | 0.385218304 | 0.721660257 | 0.377773231 | 0.171 | 0.15 | 1 | 0.721660257 | 3.17154E-05 |
| Gm48024 | 1.59019E-05 | 3.47665069 | 0.17 | 0.025 | 0.38628996 | 0.402372511 | -0.622226769 | 0 | 0.017 | 1 | 0.402372511 | 3.18036E-05 |
| Cxadr | 1.59519E-05 | -2.709099279 | 0.189 | 0.487 | 0.387502796 | 0.795822357 | 0.1105812 | 0.317 | 0.311 | 1 | 0.795822357 | 3.19035E-05 |
| Gm16638 | 0.061202457 | -1.049848549 | 0.283 | 0.403 | 1 | 1.59658E-05 | -1.598614212 | 0.195 | 0.58 | 0.387840648 | 0.061202457 | 3.19313E-05 |
| Mat2b | 1.60159E-05 | 1.163546445 | 0.717 | 0.419 | 0.389058872 | 0.08237752 | 0.571064533 | 0.659 | 0.519 | 1 | 0.08237752 | 3.20316E-05 |
| Tle4 | 1.60984E-05 | 0.883176689 | 0.83 | 0.568 | 0.391061927 | 0.031031923 | 0.919877086 | 0.756 | 0.744 | 1 | 0.031031923 | 3.21965E-05 |
| Fxyd2 | 1.62536E-05 | -6.353072045 | 0.057 | 0.347 | 0.394833258 | 0.124831044 | -3.93758221 | 0.024 | 0.096 | 1 | 0.124831044 | 3.2507E-05 |
| Hectd2 | 1.62712E-05 | -1.130679624 | 0.566 | 0.788 | 0.39526002 | 0.87847762 | 0.140510772 | 0.585 | 0.618 | 1 | 0.87847762 | 3.25421E-05 |
| Clu | 1.63549E-05 | 1.922242767 | 0.717 | 0.5 | 0.397292736 | 0.16393276 | 0.894957617 | 0.805 | 0.836 | 1 | 0.16393276 | 3.27095E-05 |
| Skil | 1.63577E-05 | 0.789002525 | 0.962 | 0.831 | 0.397360267 | 0.042709911 | 0.348025888 | 0.927 | 0.911 | 1 | 0.042709911 | 3.27151E-05 |
| Phaf1 | 1.65028E-05 | 1.621290395 | 0.453 | 0.186 | 0.400885562 | 0.144462548 | 0.679663573 | 0.463 | 0.382 | 1 | 0.144462548 | 3.30053E-05 |
| Col23a1 | 1.66848E-05 | 4.523956404 | 0.208 | 0.042 | 0.405306511 | 0.227742434 | 2.489281546 | 0.146 | 0.092 | 1 | 0.227742434 | 3.33693E-05 |
| Ndrg3 | 1.68436E-05 | 1.272079545 | 0.698 | 0.449 | 0.409163665 | 8.69776E-05 | 1.279209397 | 0.756 | 0.58 | 1 | 8.69776E-05 | 3.36868E-05 |
| Pdzd8 | 1.69393E-05 | 1.069555212 | 0.679 | 0.39 | 0.411489701 | 0.997879807 | 0.16640618 | 0.78 | 0.795 | 1 | 0.997879807 | 3.38783E-05 |
| Srxn1 | 1.71358E-05 | -2.549411607 | 0.17 | 0.483 | 0.41626368 | 0.028840474 | -0.653365281 | 0.22 | 0.403 | 1 | 0.028840474 | 3.42714E-05 |
| Ptpn21 | 1.71399E-05 | 1.695290976 | 0.415 | 0.161 | 0.416361434 | 0.471210206 | 0.439869352 | 0.415 | 0.382 | 1 | 0.471210206 | 3.42794E-05 |
| Serping1 | 0.002506215 | 2.991223863 | 0.151 | 0.042 | 1 | 1.71583E-05 | 5.199774929 | 0.244 | 0.058 | 0.416808374 | 0.002506215 | 3.43162E-05 |
| Washc2 | 1.71881E-05 | 1.266749213 | 0.66 | 0.445 | 0.417534169 | 1.84743E-05 | 1.127460724 | 0.854 | 0.621 | 0.448777963 | 1.84743E-05 | 3.4376E-05 |
| Mrpl43 | 1.72067E-05 | 1.341491106 | 0.604 | 0.377 | 0.417984903 | 0.05435713 | -0.690042153 | 0.293 | 0.447 | 1 | 0.05435713 | 3.44131E-05 |
| Gm15271 | 1.73621E-05 | -4.900559841 | 0 | 0.275 | 0.421758955 | 0.357847088 | -0.332720152 | 0 | 0.02 | 1 | 0.357847088 | 3.47238E-05 |
| Epdr1 | 1.74152E-05 | 1.976724293 | 0.547 | 0.326 | 0.423050436 | 0.000214648 | 2.46797104 | 0.415 | 0.205 | 1 | 0.000214648 | 3.48301E-05 |
| Cnot1 | 1.75608E-05 | 0.826996655 | 0.887 | 0.742 | 0.426586155 | 0.131509208 | 0.428118848 | 0.902 | 0.884 | 1 | 0.131509208 | 3.51212E-05 |
| Clptm1 | 1.75835E-05 | 0.921663248 | 0.755 | 0.53 | 0.427137993 | 0.812929985 | 0.159983841 | 0.659 | 0.683 | 1 | 0.812929985 | 3.51667E-05 |
| Frzb | 1.76316E-05 | -3.237594828 | 0.132 | 0.432 | 0.428305718 | 0.158049817 | -0.727579769 | 0.049 | 0.126 | 1 | 0.158049817 | 3.52628E-05 |
| Cenpc1 | 1.76374E-05 | 1.274683219 | 0.623 | 0.356 | 0.428448325 | 0.521381681 | 0.515276755 | 0.585 | 0.556 | 1 | 0.521381681 | 3.52745E-05 |
| Mmp16 | 1.77101E-05 | 1.268893615 | 0.66 | 0.381 | 0.430214073 | 0.56676964 | -0.152741486 | 0.415 | 0.485 | 1 | 0.56676964 | 3.54199E-05 |
| Gdpd1 | 1.78008E-05 | 3.225111923 | 0.189 | 0.034 | 0.432417217 | 0.009030839 | 1.936740523 | 0.195 | 0.075 | 1 | 0.009030839 | 3.56013E-05 |
| Hmgn5 | 0.618646198 | 0.193611608 | 0.34 | 0.398 | 1 | 1.78413E-05 | 1.447793158 | 0.659 | 0.362 | 0.43340015 | 0.618646198 | 3.56822E-05 |
| Arl5b | 1.79215E-05 | 1.221836791 | 0.66 | 0.445 | 0.435349585 | 0.014158288 | 0.90804509 | 0.756 | 0.645 | 1 | 0.014158288 | 3.58427E-05 |
| Il17rd | 1.79735E-05 | 3.356356456 | 0.189 | 0.034 | 0.436611995 | 0.000494054 | 2.644559772 | 0.195 | 0.051 | 1 | 0.000494054 | 3.59467E-05 |
| Adgra3 | 0.10483554 | -0.407156293 | 0.679 | 0.763 | 1 | 1.7974E-05 | -1.03428041 | 0.561 | 0.881 | 0.436624216 | 0.10483554 | 3.59477E-05 |
| Cpeb1 | 1.80248E-05 | 1.484871196 | 0.396 | 0.14 | 0.437857259 | 0.012189874 | 0.997669522 | 0.439 | 0.27 | 1 | 0.012189874 | 3.60492E-05 |

| | | | | | | | | | | | | |
|---|---|---|---|---|---|---|---|---|---|---|---|---|
| Map4k5 | 0.101076438 | -0.311798449 | 0.925 | 0.915 | 1 | 1.80305E-05 | -0.986779201 | 0.829 | 0.993 | 0.43799652 | 0.101076438 | 3.60606E-05 |
| Dync2h1 | 1.8061E-05 | 1.401734983 | 0.811 | 0.581 | 0.438736845 | 0.090255824 | 0.636107207 | 0.756 | 0.689 | 1 | 0.090255824 | 3.61216E-05 |
| Cachd1 | 1.81805E-05 | 1.811834881 | 0.34 | 0.114 | 0.441640239 | 0.004589967 | 1.199774929 | 0.512 | 0.324 | 1 | 0.004589967 | 3.63606E-05 |
| Nt5c | 1.83747E-05 | 1.44946586 | 0.509 | 0.242 | 0.446357892 | 0.08942642 | 0.887830923 | 0.39 | 0.276 | 1 | 0.08942642 | 3.6749E-05 |
| Zmiz1 | 0.059217012 | -0.549961041 | 0.962 | 0.941 | 1 | 1.84605E-05 | -0.911780327 | 0.951 | 0.983 | 0.448443602 | 0.059217012 | 3.69208E-05 |
| Add3 | 1.85E-05 | 1.534136184 | 0.566 | 0.292 | 0.449402614 | 0.371059459 | 0.436325414 | 0.39 | 0.331 | 1 | 0.371059459 | 3.69997E-05 |
| Zfp651 | 1.87606E-05 | 1.640149422 | 0.396 | 0.148 | 0.455732074 | 0.579046259 | 0.252242349 | 0.463 | 0.437 | 1 | 0.579046259 | 3.75208E-05 |
| Kdm1a | 1.90605E-05 | 0.655648271 | 0.943 | 0.903 | 0.463018581 | 0.870212286 | 0.103142064 | 0.976 | 0.959 | 1 | 0.870212286 | 3.81207E-05 |
| Cercam | 1.93267E-05 | 1.494209061 | 0.377 | 0.131 | 0.46948378 | 0.129166481 | -1.292078167 | 0.073 | 0.16 | 1 | 0.129166481 | 3.8653E-05 |
| Htra1 | 1.9364E-05 | 0.373908702 | 0.906 | 0.61 | 0.470390182 | 0.277884694 | -0.372407657 | 0.951 | 0.901 | 1 | 0.277884694 | 3.87276E-05 |
| Bmper | 1.93748E-05 | 2.048463907 | 0.547 | 0.301 | 0.470652661 | 0.01255397 | 1.803574072 | 0.39 | 0.239 | 1 | 0.01255397 | 3.87492E-05 |
| Atp5o | 1.95632E-05 | 0.791215396 | 0.962 | 0.801 | 0.475230108 | 0.30820035 | -0.201573478 | 0.78 | 0.87 | 1 | 0.30820035 | 3.91261E-05 |
| Meg3 | 1.9693E-05 | -1.451211473 | 0.585 | 0.814 | 0.478383015 | 0.001704989 | -0.85887627 | 0.707 | 0.877 | 1 | 0.001704989 | 3.93857E-05 |
| Serpinb6a | 0.000453978 | -0.698869775 | 0.981 | 0.932 | 1 | 2.01385E-05 | -1.064332511 | 0.683 | 0.925 | 0.489204231 | 0.000453978 | 4.02766E-05 |
| Ndrg1 | 2.01854E-05 | 1.20202831 | 0.547 | 0.258 | 0.490343377 | 0.211475373 | 0.304935243 | 0.659 | 0.587 | 1 | 0.211475373 | 4.03704E-05 |
| Panx1 | 2.02155E-05 | 2.891688189 | 0.132 | 0.013 | 0.491075201 | 0.705656327 | 0.136765132 | 0.049 | 0.065 | 1 | 0.705656327 | 4.04306E-05 |
| Arpc5 | 2.09053E-05 | 0.619800777 | 1 | 0.881 | 0.50783076 | 0.301641966 | 0.247625721 | 0.951 | 0.962 | 1 | 0.301641966 | 4.18101E-05 |
| Ubr5 | 2.09545E-05 | 0.531692053 | 0.981 | 0.992 | 0.509026717 | 0.00088624 | 0.528220858 | 1 | 1 | 1 | 0.00088624 | 4.19086E-05 |
| Psmd1 | 2.09817E-05 | 0.759584653 | 0.962 | 0.847 | 0.509688299 | 0.412555669 | 0.302868422 | 0.854 | 0.925 | 1 | 0.412555669 | 4.1963E-05 |
| Fos | 0.000568595 | 0.68185034 | 0.83 | 0.712 | 1 | 2.12341E-05 | 1.72942989 | 0.951 | 0.85 | 0.515817858 | 0.000568595 | 4.24677E-05 |
| Ugcg | 2.14162E-05 | 1.393910259 | 0.623 | 0.356 | 0.520241458 | 0.082362657 | 0.754063615 | 0.561 | 0.502 | 1 | 0.082362657 | 4.28319E-05 |
| Rabgef1 | 2.15202E-05 | -1.09636526 | 0.491 | 0.758 | 0.522769477 | 0.07055734 | -0.523748402 | 0.707 | 0.747 | 1 | 0.07055734 | 4.304E-05 |
| Luc7l | 2.15478E-05 | 0.713669505 | 0.887 | 0.805 | 0.523439706 | 0.358408621 | 0.189900181 | 0.854 | 0.86 | 1 | 0.358408621 | 4.30952E-05 |
| Cwc25 | 2.1572E-05 | 1.094851139 | 0.642 | 0.364 | 0.52402687 | 0.659679784 | 0.172072 | 0.463 | 0.478 | 1 | 0.659679784 | 4.31435E-05 |
| 2310022B05Rik | 2.16467E-05 | 1.51531231 | 0.528 | 0.292 | 0.525841398 | 0.13285511 | 0.620886943 | 0.463 | 0.375 | 1 | 0.13285511 | 4.32929E-05 |
| Apoe | 2.16571E-05 | 0.887500392 | 0.547 | 0.246 | 0.526094049 | 0.004288403 | -1.314614215 | 0.707 | 0.812 | 1 | 0.004288403 | 4.33137E-05 |
| Nsmce2 | 2.19408E-05 | 0.43203007 | 0.962 | 0.746 | 0.532984835 | 0.20202675 | -0.255989297 | 0.829 | 0.843 | 1 | 0.20202675 | 4.3881E-05 |
| Slc39a8 | 2.21356E-05 | -2.448618435 | 0.245 | 0.542 | 0.537716785 | 0.024131532 | -1.592782991 | 0.098 | 0.259 | 1 | 0.024131532 | 4.42706E-05 |
| Thbs1 | 0.611007622 | 0.204262056 | 0.943 | 0.945 | 1 | 2.22012E-05 | -0.912649427 | 0.927 | 0.973 | 0.539311543 | 0.611007622 | 4.44019E-05 |
| Etfb | 2.2264E-05 | 0.978645366 | 0.83 | 0.597 | 0.540837414 | 0.901373777 | 0.231025863 | 0.585 | 0.587 | 1 | 0.901373777 | 4.45275E-05 |
| Mdh2 | 2.25203E-05 | 0.733879474 | 0.943 | 0.86 | 0.547064003 | 0.656810365 | -0.158826248 | 0.829 | 0.863 | 1 | 0.656810365 | 4.50402E-05 |
| Vstm2b | 2.25591E-05 | 4.739685096 | 0.075 | 0 | 0.548006113 | 0.001133094 | 2.83720485 | 0.073 | 0.007 | 1 | 0.001133094 | 4.51177E-05 |
| Gm266 | 2.25591E-05 | 4.739685096 | 0.075 | 0 | 0.548006113 | 0.007718533 | 3.83720485 | 0.024 | 0 | 1 | 0.007718533 | 4.51177E-05 |
| Dnah7a | 2.2563E-05 | 5.614154213 | 0.075 | 0 | 0.548100284 | 0.107398884 | 2.83720485 | 0.049 | 0.014 | 1 | 0.107398884 | 4.51255E-05 |
| Foxs1 | 2.2563E-05 | 5.154722595 | 0.075 | 0 | 0.548100284 | 0.104856732 | 2.83720485 | 0.024 | 0.003 | 1 | 0.104856732 | 4.51255E-05 |
| Mmp11 | 2.2563E-05 | 4.962077517 | 0.075 | 0 | 0.548100284 | 0.004444719 | 3.159132945 | 0.073 | 0.01 | 1 | 0.004444719 | 4.51255E-05 |
| Krt75 | 2.2563E-05 | 5.154722595 | 0.075 | 0 | 0.548100284 | 0.104856732 | 2.83720485 | 0.024 | 0.003 | 1 | 0.104856732 | 4.51255E-05 |
| Ccser2 | 2.25642E-05 | 0.718151934 | 0.906 | 0.674 | 0.548129706 | 0.299862745 | -0.264490547 | 0.854 | 0.857 | 1 | 0.299862745 | 4.51279E-05 |
| Atp2a1 | 2.25643E-05 | 4.47665069 | 0.075 | 0 | 0.548131677 | 0.027541029 | 2.614812428 | 0.073 | 0.017 | 1 | 0.027541029 | 4.51281E-05 |
| 4930445N18Rik | 2.25643E-05 | 4.47665069 | 0.075 | 0 | 0.548131677 | 0.007718533 | 4.422167351 | 0.024 | 0 | 1 | 0.007718533 | 4.51281E-05 |
| Gm15446 | 2.26235E-05 | 2.817687608 | 0.208 | 0.042 | 0.549569407 | 0.119469353 | 1.547698233 | 0.122 | 0.058 | 1 | 0.119469353 | 4.52464E-05 |
| Pcgf3 | 2.26627E-05 | 1.309050741 | 0.604 | 0.373 | 0.550522167 | 0.130883033 | -0.428139717 | 0.512 | 0.631 | 1 | 0.130883033 | 4.53249E-05 |
| Trappc3 | 2.28929E-05 | 0.922641117 | 0.755 | 0.619 | 0.556115034 | 0.192121541 | 0.288043071 | 0.732 | 0.666 | 1 | 0.192121541 | 4.57853E-05 |
| Grb2 | 2.29303E-05 | 0.595813986 | 0.925 | 0.75 | 0.557023748 | 0.000945196 | 0.74737344 | 0.951 | 0.887 | 1 | 0.000945196 | 4.58601E-05 |
| Zfp81 | 2.3007E-05 | 2.154722595 | 0.264 | 0.068 | 0.55888684 | 0.280370321 | 1.408361551 | 0.146 | 0.099 | 1 | 0.280370321 | 4.60135E-05 |
| Exosc8 | 2.30753E-05 | 1.716601482 | 0.377 | 0.136 | 0.560543996 | 0.782710431 | 0.347820009 | 0.171 | 0.198 | 1 | 0.782710431 | 4.615E-05 |
| Fbxo10 | 2.3123E-05 | 1.44102678 | 0.34 | 0.11 | 0.561704663 | 0.345203971 | 0.554270887 | 0.195 | 0.14 | 1 | 0.345203971 | 4.62455E-05 |
| Eif4enif1 | 2.31321E-05 | 0.988913702 | 0.66 | 0.386 | 0.561925187 | 0.246574216 | -0.143058932 | 0.561 | 0.638 | 1 | 0.246574216 | 4.62637E-05 |
| Hnrnpu | 2.36195E-05 | 0.537356822 | 1 | 0.983 | 0.573764016 | 0.743194043 | 0.150335771 | 1 | 0.997 | 1 | 0.743194043 | 4.72384E-05 |
| Lmo7 | 0.162834464 | -0.263003992 | 0.717 | 0.788 | 1 | 2.39165E-05 | 0.8449822 | 0.878 | 0.628 | 0.580978713 | 0.162834464 | 4.78324E-05 |
| Il11ra1 | 2.40407E-05 | 1.024662054 | 0.849 | 0.686 | 0.583995659 | 0.482048639 | 0.234797215 | 0.659 | 0.792 | 1 | 0.482048639 | 4.80807E-05 |
| Cops4 | 2.40963E-05 | 0.882643049 | 0.811 | 0.606 | 0.585346953 | 0.026219625 | -0.397456409 | 0.585 | 0.734 | 1 | 0.026219625 | 4.8192E-05 |
| Rassf8 | 2.41077E-05 | 1.533234218 | 0.453 | 0.199 | 0.585623869 | 0.007009148 | 1.252242349 | 0.366 | 0.205 | 1 | 0.007009148 | 4.82148E-05 |
| Gli3 | 2.42124E-05 | 0.83118345 | 0.83 | 0.631 | 0.588168076 | 0.008827218 | 0.754215485 | 0.707 | 0.577 | 1 | 0.008827218 | 4.84243E-05 |
| Zdhhc17 | 2.42521E-05 | 1.377115016 | 0.566 | 0.305 | 0.589131167 | 0.528531811 | 0.316948038 | 0.488 | 0.44 | 1 | 0.528531811 | 4.85035E-05 |
| Vps37b | 2.42677E-05 | -1.000555631 | 0.642 | 0.869 | 0.589509979 | 0.003311322 | -0.466826041 | 0.829 | 0.942 | 1 | 0.003311322 | 4.85347E-05 |
| Dusp2 | 2.43561E-05 | 1.187889459 | 0.17 | 0.025 | 0.59165928 | 0.5191986 | 0.252242349 | 0 | 0.01 | 1 | 0.5191986 | 4.87117E-05 |
| Rnf150 | 0.088486065 | 1.031340179 | 0.17 | 0.093 | 1 | 2.4391E-05 | 3.780621321 | 0.171 | 0.027 | 0.592506796 | 0.088486065 | 4.87815E-05 |
| Gm973 | 2.44045E-05 | -1.944813079 | 0.057 | 0.356 | 0.592833423 | 0.280831144 | -0.696773722 | 0.146 | 0.222 | 1 | 0.280831144 | 4.88083E-05 |
| Hif1a | 2.4418E-05 | -0.828283403 | 0.962 | 0.983 | 0.593162153 | 0.032481457 | -0.342298563 | 0.927 | 0.993 | 1 | 0.032481457 | 4.88354E-05 |
| Setd1b | 2.44709E-05 | 1.249359469 | 0.623 | 0.335 | 0.59444749 | 0.963554754 | 0.25943785 | 0.61 | 0.652 | 1 | 0.963554754 | 4.89412E-05 |
| Eif1 | 0.002280908 | -0.303901972 | 1 | 0.996 | 1 | 2.45469E-05 | -0.512418484 | 1 | 0.997 | 0.596294212 | 0.002280908 | 4.90933E-05 |
| Nop53 | 2.48243E-05 | 0.748216729 | 0.849 | 0.669 | 0.603032917 | 0.50263071 | 0.327010117 | 0.732 | 0.703 | 1 | 0.50263071 | 4.96481E-05 |
| Piezo2 | 0.520205973 | 0.177705913 | 0.774 | 0.686 | 1 | 2.50682E-05 | -0.7221557 | 0.61 | 0.935 | 0.608957381 | 0.520205973 | 5.01358E-05 |
| Strip1 | 2.52197E-05 | 1.720319771 | 0.415 | 0.174 | 0.612635894 | 0.058548537 | 0.945581011 | 0.439 | 0.314 | 1 | 0.058548537 | 5.04387E-05 |
| Cetn4 | 2.53731E-05 | -3.015202407 | 0.038 | 0.322 | 0.616363131 | 0.880720437 | 0.19334866 | 0.122 | 0.133 | 1 | 0.880720437 | 5.07455E-05 |
| Notch2 | 2.53989E-05 | 0.594825431 | 1 | 0.89 | 0.616990778 | 0.000459406 | 0.627453868 | 1 | 0.966 | 1 | 0.000459406 | 5.07972E-05 |
| Pitpnm1 | 2.54595E-05 | 2.454282877 | 0.208 | 0.042 | 0.618461306 | 0.116488464 | -1.410722664 | 0.024 | 0.099 | 1 | 0.116488464 | 5.09183E-05 |
| Fgfrl1 | 2.54844E-05 | -0.828938668 | 0.755 | 0.881 | 0.619067988 | 0.010398385 | -0.568208231 | 0.683 | 0.802 | 1 | 0.010398385 | 5.09682E-05 |
| Sdcbp | 0.000304675 | -0.490250088 | 0.943 | 0.975 | 1 | 2.54981E-05 | -0.565868059 | 0.976 | 1 | 0.619400004 | 0.000304675 | 5.09956E-05 |
| Snhg5 | 2.55815E-05 | 1.649799592 | 0.604 | 0.356 | 0.621425577 | 0.963506607 | -0.24525731 | 0.244 | 0.249 | 1 | 0.963506607 | 5.11623E-05 |
| Marchf3 | 2.56182E-05 | 1.804035507 | 0.547 | 0.288 | 0.622317884 | 0.001154856 | 0.955567046 | 0.585 | 0.317 | 1 | 0.001154856 | 5.12358E-05 |
| Cnnm3 | 2.56855E-05 | 1.852159825 | 0.377 | 0.14 | 0.623953121 | 0.352387588 | 0.483567895 | 0.268 | 0.208 | 1 | 0.352387588 | 5.13704E-05 |
| Crot | 2.59198E-05 | 1.900966003 | 0.321 | 0.102 | 0.629643257 | 0.050985426 | 0.497354847 | 0.317 | 0.174 | 1 | 0.050985426 | 5.18389E-05 |
| Ifngr2 | 2.59391E-05 | 1.64423831 | 0.604 | 0.381 | 0.630113084 | 0.067205908 | 0.581947795 | 0.683 | 0.577 | 1 | 0.067205908 | 5.18776E-05 |
| Csf1 | 7.07542E-05 | 1.96568877 | 0.377 | 0.148 | 1 | 2.60789E-05 | 2.057330664 | 0.463 | 0.201 | 0.633507788 | 7.07542E-05 | 5.2157E-05 |
| Tbc1d17 | 2.61014E-05 | 1.595295186 | 0.453 | 0.199 | 0.634054959 | 0.140306596 | -0.849295677 | 0.146 | 0.249 | 1 | 0.140306596 | 5.22021E-05 |
| Phtf1 | 2.61788E-05 | 1.039245377 | 0.679 | 0.394 | 0.635934289 | 0.008061948 | 0.904649371 | 0.707 | 0.58 | 1 | 0.008061948 | 5.23568E-05 |
| Anxa7 | 2.63871E-05 | -0.760221635 | 0.849 | 0.936 | 0.640995306 | 0.004132717 | -0.462968692 | 0.878 | 0.969 | 1 | 0.004132717 | 5.27735E-05 |
| Cdh24 | 2.64431E-05 | 2.526691372 | 0.208 | 0.042 | 0.642355282 | 0.007244896 | 1.580865097 | 0.244 | 0.102 | 1 | 0.007244896 | 5.28855E-05 |
| P2ry1 | 2.64793E-05 | 3.739685096 | 0.113 | 0.008 | 0.643234035 | 6.28796E-05 | 3.422167351 | 0.122 | 0.014 | 1 | 6.28796E-05 | 5.29578E-05 |
| Mitf | 0.098312314 | 0.486867086 | 0.415 | 0.314 | 1 | 2.66409E-05 | 2.324253594 | 0.439 | 0.188 | 0.647161945 | 0.098312314 | 5.32812E-05 |
| Ttc23l | 2.68042E-05 | 3.039245377 | 0.189 | 0.034 | 0.651127007 | 0.000866485 | 2.371541277 | 0.244 | 0.082 | 1 | 0.000866485 | 5.36076E-05 |
| Wasl | 2.68226E-05 | 0.80750048 | 0.849 | 0.716 | 0.651574393 | 0.035429684 | 0.361145079 | 0.902 | 0.918 | 1 | 0.035429684 | 5.36445E-05 |
| Emsy | 2.69949E-05 | 1.074104135 | 0.679 | 0.428 | 0.655759184 | 0.051918717 | 0.650791726 | 0.683 | 0.638 | 1 | 0.051918717 | 5.3989E-05 |
| Sntb2 | 2.70491E-05 | 1.323228319 | 0.849 | 0.703 | 0.657076406 | 0.000683353 | 1.102164133 | 0.878 | 0.741 | 1 | 0.000683353 | 5.40974E-05 |
| Casp12 | 2.70603E-05 | 1.739685096 | 0.396 | 0.157 | 0.657348976 | 0.100352031 | 1.131948115 | 0.366 | 0.27 | 1 | 0.100352031 | 5.41199E-05 |
| Neu2 | 2.71273E-05 | -3.174401001 | 0.038 | 0.326 | 0.658975824 | 0.000151134 | -4.732650759 | 0 | 0.273 | 1 | 0.000151134 | 5.42538E-05 |
| Alkbh5 | 2.72471E-05 | 0.868342079 | 0.849 | 0.712 | 0.661886111 | 0.213039831 | 0.35061409 | 0.805 | 0.939 | 1 | 0.213039831 | 5.44934E-05 |
| Nectin2 | 2.76127E-05 | 1.569760094 | 0.34 | 0.11 | 0.67076652 | 0.000564693 | 1.450181727 | 0.415 | 0.191 | 1 | 0.000564693 | 5.52245E-05 |
| Heca | 2.77508E-05 | 1.47481169 | 0.66 | 0.39 | 0.674123421 | 0.015395837 | 0.882059086 | 0.756 | 0.597 | 1 | 0.015395837 | 5.55009E-05 |
| Pcf11 | 2.78686E-05 | 0.807483411 | 0.962 | 0.784 | 0.676983969 | 0.33093626 | 0.16540827 | 0.927 | 0.925 | 1 | 0.33093626 | 5.57364E-05 |
| Epha4 | 0.000103023 | 0.787665124 | 0.566 | 0.305 | 1 | 2.78914E-05 | 1.99874871 | 0.585 | 0.321 | 0.677537652 | 0.000103023 | 5.5782E-05 |
| Cracr2a | 2.80457E-05 | 3.377115016 | 0.113 | 0.008 | 0.681285853 | 0.346739982 | 1.071670103 | 0.073 | 0.041 | 1 | 0.346739982 | 5.60906E-05 |
| Rnf44 | 2.81962E-05 | 1.75033234 | 0.396 | 0.157 | 0.684942254 | 0.374466547 | -0.15418202 | 0.317 | 0.382 | 1 | 0.374466547 | 5.63916E-05 |
| Ddx3y | 2.84167E-05 | 0.727398722 | 0.849 | 0.487 | 0.690299543 | 4.01702E-05 | -0.686814887 | 0.829 | 0.962 | 0.975813422 | 4.01702E-05 | 5.68327E-05 |
| Snx30 | 2.84711E-05 | 1.540012751 | 0.34 | 0.11 | 0.69161982 | 0.003395276 | 1.268362014 | 0.341 | 0.157 | 1 | 0.003395276 | 5.69414E-05 |
| Cd14 | 2.86063E-05 | -2.25912799 | 0.094 | 0.398 | 0.694905186 | 0.364968557 | -0.707115666 | 0.195 | 0.249 | 1 | 0.364968557 | 5.72119E-05 |

| | | | | | | | | | | | | |
|---|---|---|---|---|---|---|---|---|---|---|---|---|
| 5730471H19Rik | 2.87209E-05 | 1.891688189 | 0.283 | 0.081 | 0.697688978 | 0.779988911 | 0.515276755 | 0.122 | 0.147 | 1 | 0.779988911 | 5.7441E-05 |
| Gm15563 | 2.88309E-05 | 3.324647596 | 0.113 | 0.008 | 0.700359341 | 0.720194014 | 0.749742009 | 0.049 | 0.038 | 1 | 0.720194014 | 5.76609E-05 |
| Foxa3 | 2.88792E-05 | -2.246156841 | 0.132 | 0.428 | 0.701532662 | 0.003165563 | -1.95535307 | 0.098 | 0.321 | 1 | 0.003165563 | 5.77575E-05 |
| Rc3h1 | 2.91137E-05 | 0.880758629 | 0.981 | 0.839 | 0.707228938 | 0.06228214 | 0.488348428 | 0.976 | 0.959 | 1 | 0.06228214 | 5.82265E-05 |
| Lysmd2 | 2.93494E-05 | 1.8327945 | 0.264 | 0.068 | 0.71295604 | 0.166562805 | 1.059597271 | 0.22 | 0.143 | 1 | 0.166562805 | 5.8698E-05 |
| Map7d2 | 2.93546E-05 | -3.425939768 | 0.075 | 0.369 | 0.713081009 | 0.321638138 | -0.448197369 | 0.024 | 0.065 | 1 | 0.321638138 | 5.87083E-05 |
| Scaf1 | 2.94057E-05 | 1.118502407 | 0.66 | 0.369 | 0.714323747 | 0.067904689 | 0.73223529 | 0.561 | 0.464 | 1 | 0.067904689 | 5.88106E-05 |
| Cerk | 0.03435273 | 1.223985257 | 0.302 | 0.182 | 1 | 2.94519E-05 | 1.536035315 | 0.561 | 0.28 | 0.715446719 | 0.03435273 | 5.8903E-05 |
| Adgrl3 | 0.355200887 | -0.634598404 | 0.34 | 0.254 | 1 | 2.95341E-05 | 2.460350544 | 0.39 | 0.14 | 0.717443367 | 0.355200887 | 5.90674E-05 |
| Zfp14 | 2.96411E-05 | 3.154722595 | 0.113 | 0.008 | 0.720042609 | 0.041769421 | 1.458693227 | 0.098 | 0.031 | 1 | 0.041769421 | 5.92814E-05 |
| Sin3a | 2.9642E-05 | 1.242949778 | 0.528 | 0.258 | 0.720063708 | 0.010285027 | 0.736854324 | 0.634 | 0.454 | 1 | 0.010285027 | 5.92831E-05 |
| Ddx11 | 2.96645E-05 | 2.213616284 | 0.264 | 0.072 | 0.720609326 | 0.000101444 | 1.69281494 | 0.244 | 0.065 | 1 | 0.000101444 | 5.93281E-05 |
| Yaf2 | 2.96919E-05 | 0.842285949 | 0.868 | 0.619 | 0.721275025 | 0.043451325 | 0.762911067 | 0.854 | 0.754 | 1 | 0.043451325 | 5.93829E-05 |
| Grsf1 | 3.00435E-05 | 1.011131741 | 0.736 | 0.508 | 0.72981697 | 0.143748914 | 0.619727919 | 0.634 | 0.59 | 1 | 0.143748914 | 6.00861E-05 |
| 1700019D03Rik | 3.00607E-05 | 2.8327945 | 0.113 | 0.008 | 0.730235095 | 0.02065851 | 3.574170444 | 0.049 | 0.007 | 1 | 0.02065851 | 6.01205E-05 |
| Nfxl1 | 3.01037E-05 | 1.327559192 | 0.547 | 0.275 | 0.731279493 | 0.37641628 | 0.478123756 | 0.463 | 0.44 | 1 | 0.37641628 | 6.02065E-05 |
| Slc1a4 | 0.104523157 | -0.7032584 | 0.396 | 0.487 | 1 | 3.02342E-05 | -1.836883541 | 0.293 | 0.608 | 0.734448198 | 0.104523157 | 6.04674E-05 |
| Akap8 | 3.03225E-05 | 1.421116013 | 0.604 | 0.381 | 0.736593404 | 0.565270087 | 0.423646279 | 0.585 | 0.604 | 1 | 0.565270087 | 6.0644E-05 |
| Tbx18 | 3.04679E-05 | 2.640149422 | 0.113 | 0.008 | 0.740126806 | 0.104856732 | 2.83720485 | 0.024 | 0.003 | 1 | 0.104856732 | 6.09349E-05 |
| Zbtb34 | 3.08781E-05 | 2.299112504 | 0.264 | 0.072 | 0.75009098 | 0.989618365 | 0.347820009 | 0.171 | 0.177 | 1 | 0.989618365 | 6.17553E-05 |
| Trp63 | 0.018445361 | 1.806799291 | 0.094 | 0.025 | 1 | 3.10002E-05 | 1.501601818 | 0.366 | 0.123 | 0.753057997 | 0.018445361 | 6.19995E-05 |
| Arl5a | 3.10567E-05 | 1.067259754 | 0.66 | 0.381 | 0.754429946 | 0.184849653 | 0.415378185 | 0.634 | 0.56 | 1 | 0.184849653 | 6.21125E-05 |
| Slc6a8 | 3.11887E-05 | -1.449348729 | 0.302 | 0.589 | 0.75763551 | 0.005926284 | -1.038988949 | 0.244 | 0.502 | 1 | 0.005926284 | 6.23764E-05 |
| Ap2a2 | 3.13589E-05 | 1.168936454 | 0.679 | 0.445 | 0.761770702 | 0.069091296 | 0.313021979 | 0.854 | 0.737 | 1 | 0.069091296 | 6.27168E-05 |
| Lrrk2 | 3.14738E-05 | 1.72941676 | 0.34 | 0.119 | 0.76456213 | 0.542752411 | 0.114738825 | 0.122 | 0.164 | 1 | 0.542752411 | 6.29467E-05 |
| Ccdc83 | 3.17068E-05 | 4.47665069 | 0.094 | 0.004 | 0.77022276 | 0.60213981 | 0.667279848 | 0.024 | 0.014 | 1 | 0.60213981 | 6.34127E-05 |
| Cfl1 | 3.1952E-05 | 0.561018499 | 1 | 0.958 | 0.776178686 | 0.062656318 | 0.305951237 | 1 | 0.986 | 1 | 0.062656318 | 6.3903E-05 |
| Pde6h | 3.22604E-05 | -4.936271937 | 0.038 | 0.314 | 0.783670164 | 0.05261958 | -3.31254227 | 0 | 0.085 | 1 | 0.05261958 | 6.45198E-05 |
| Cdc42se1 | 0.000809378 | -0.632973801 | 0.642 | 0.856 | 1 | 3.26907E-05 | -1.043909111 | 0.732 | 0.874 | 0.794122271 | 0.000809378 | 6.53803E-05 |
| Aplp1 | 3.27343E-05 | 4.324647596 | 0.094 | 0.004 | 0.795181593 | 0.004444719 | 3.159132945 | 0.073 | 0.01 | 1 | 0.004444719 | 6.54675E-05 |
| Tulp4 | 3.28924E-05 | 0.773293488 | 0.925 | 0.877 | 0.799021753 | 0.695737929 | 0.13005491 | 0.976 | 0.959 | 1 | 0.695737929 | 6.57837E-05 |
| Grb10 | 3.29474E-05 | 0.588404642 | 0.981 | 0.886 | 0.800359407 | 0.28873442 | -0.187601224 | 0.927 | 0.962 | 1 | 0.28873442 | 6.58938E-05 |
| Ccs | 3.30322E-05 | 1.311448098 | 0.604 | 0.373 | 0.802417325 | 0.989210882 | 0.274268655 | 0.39 | 0.444 | 1 | 0.989210882 | 6.60632E-05 |
| Chn2 | 0.033719946 | 1.962077517 | 0.094 | 0.03 | 1 | 3.31349E-05 | 2.896098539 | 0.22 | 0.048 | 0.8049131 | 0.033719946 | 6.62687E-05 |
| Pianp | 3.32531E-05 | 3.962077517 | 0.094 | 0.004 | 0.807783504 | 0.004605593 | 2.83720485 | 0.073 | 0.01 | 1 | 0.004605593 | 6.6505E-05 |
| Gm26511 | 3.32531E-05 | 3.962077517 | 0.094 | 0.004 | 0.807783504 | 0.362195044 | 1.100239256 | 0.049 | 0.024 | 1 | 0.362195044 | 6.6505E-05 |
| Gap43 | 3.32531E-05 | 4.614154213 | 0.094 | 0.004 | 0.807783504 | 0.533410994 | 1.83720485 | 0.049 | 0.031 | 1 | 0.533410994 | 6.6505E-05 |
| Zfp207 | 3.33271E-05 | 0.72430987 | 0.962 | 0.86 | 0.809581782 | 0.674241636 | 0.135266228 | 0.829 | 0.935 | 1 | 0.674241636 | 6.66531E-05 |
| Tmem38a | 3.33663E-05 | 2.261637799 | 0.264 | 0.072 | 0.810533194 | 0.483648435 | 0.604544093 | 0.171 | 0.133 | 1 | 0.483648435 | 6.67314E-05 |
| Dcp2 | 3.34111E-05 | 1.261637799 | 0.547 | 0.292 | 0.811621579 | 0.329245499 | -0.218648385 | 0.268 | 0.348 | 1 | 0.329245499 | 6.6821E-05 |
| Heatr5a | 0.000678126 | -0.985851583 | 0.736 | 0.839 | 1 | 3.34674E-05 | -1.153355454 | 0.707 | 0.894 | 0.812989738 | 0.000678126 | 6.69337E-05 |
| Eif2a | 3.37397E-05 | 1.082931912 | 0.774 | 0.538 | 0.819604368 | 0.52581978 | -0.116991461 | 0.634 | 0.676 | 1 | 0.52581978 | 6.74782E-05 |
| Bclaf1 | 3.37573E-05 | 0.538541363 | 0.981 | 0.928 | 0.820031506 | 0.241440026 | 0.303331121 | 1 | 0.98 | 1 | 0.241440026 | 6.75134E-05 |
| Maf1 | 3.37621E-05 | 1.411062348 | 0.491 | 0.233 | 0.820148364 | 0.132714193 | -0.739720332 | 0.293 | 0.392 | 1 | 0.132714193 | 6.7523E-05 |
| Reep2 | 3.38171E-05 | 3.739685096 | 0.094 | 0.004 | 0.821483876 | 0.992196803 | 0.515276755 | 0.024 | 0.024 | 1 | 0.992196803 | 6.7633E-05 |
| Ly6a | 3.40343E-05 | -3.589438501 | 0.245 | 0.496 | 0.82676125 | 0.613423531 | 0.180812491 | 0.561 | 0.655 | 1 | 0.613423531 | 6.80674E-05 |
| 4921531C22Rik | 3.40505E-05 | 2.526691372 | 0.245 | 0.064 | 0.827155555 | 0.008667381 | 1.962735732 | 0.22 | 0.092 | 1 | 0.008667381 | 6.80999E-05 |
| Wipi2 | 3.40599E-05 | 1.207190015 | 0.736 | 0.504 | 0.827382779 | 0.089339167 | 0.619613415 | 0.707 | 0.689 | 1 | 0.089339167 | 6.81186E-05 |
| Fam168a | 3.41693E-05 | 0.627210366 | 0.943 | 0.809 | 0.830039715 | 0.024795351 | 0.427922025 | 0.854 | 0.857 | 1 | 0.024795351 | 6.83374E-05 |
| Pip4k2b | 3.43513E-05 | 1.752624151 | 0.34 | 0.114 | 0.834461293 | 0.604333278 | 0.46797104 | 0.195 | 0.249 | 1 | 0.604333278 | 6.87014E-05 |
| Tmem140 | 3.44905E-05 | -2.193021792 | 0.321 | 0.581 | 0.83784395 | 0.000535514 | -1.944154864 | 0.268 | 0.539 | 1 | 0.000535514 | 6.89799E-05 |
| Kitl | 0.008325974 | 1.649487287 | 0.208 | 0.085 | 1 | 3.44975E-05 | 1.409111198 | 0.512 | 0.229 | 0.838012912 | 0.008325974 | 6.89938E-05 |
| Znrf3 | 3.47185E-05 | 1.125127813 | 0.811 | 0.547 | 0.843382314 | 0.008373896 | 1.10976203 | 0.805 | 0.689 | 1 | 0.008373896 | 6.94358E-05 |
| Cpsf2 | 3.47663E-05 | 1.401815457 | 0.491 | 0.233 | 0.844542088 | 0.674500148 | -0.295820569 | 0.39 | 0.399 | 1 | 0.674500148 | 6.95313E-05 |
| Lrrc1 | 0.261929591 | -0.320788764 | 0.528 | 0.64 | 1 | 3.48682E-05 | -1.27590202 | 0.634 | 0.85 | 0.847018544 | 0.261929591 | 6.97352E-05 |
| Kcnb2 | 3.49183E-05 | 4.690775495 | 0.094 | 0.004 | 0.848235878 | 0.271787759 | 1.029849928 | 0.024 | 0.007 | 1 | 0.271787759 | 6.98354E-05 |
| Gm17167 | 6.95031E-05 | 3.47665069 | 0.132 | 0.017 | 1 | 3.49396E-05 | 2.644559772 | 0.171 | 0.027 | 0.848752856 | 6.95031E-05 | 6.9878E-05 |
| Nr3c1 | 3.51853E-05 | 0.683141893 | 0.981 | 0.89 | 0.854721496 | 0.562424922 | 0.218604752 | 1 | 0.969 | 1 | 0.562424922 | 7.03694E-05 |
| Gsk3b | 3.51855E-05 | 0.313030265 | 1 | 0.932 | 0.854726692 | 0.869981477 | -0.118617766 | 1 | 0.98 | 1 | 0.869981477 | 7.03698E-05 |
| Slc15a2 | 0.839187783 | 0.110328475 | 0.094 | 0.106 | 1 | 3.57597E-05 | 1.456383066 | 0.561 | 0.27 | 0.868673775 | 0.839187783 | 7.15181E-05 |
| Sf3a1 | 3.64235E-05 | 1.403401708 | 0.585 | 0.335 | 0.884799703 | 0.445547603 | 0.457561257 | 0.488 | 0.505 | 1 | 0.445547603 | 7.28457E-05 |
| Ebf4 | 3.65823E-05 | 1.510866405 | 0.151 | 0.021 | 0.888658381 | 0.110420077 | 1.759202338 | 0.171 | 0.096 | 1 | 0.110420077 | 7.31634E-05 |
| Epha5 | 3.66115E-05 | 3.938993904 | 0.151 | 0.021 | 0.889365781 | 0.006416341 | 2.721727632 | 0.098 | 0.02 | 1 | 0.006416341 | 7.32216E-05 |
| S100a16 | 0.026614956 | -0.946815432 | 0.642 | 0.653 | 1 | 3.67343E-05 | -1.138217127 | 0.415 | 0.744 | 0.892350062 | 0.026614956 | 7.34673E-05 |
| Gm43813 | 0.012409259 | 1.055186921 | 0.189 | 0.076 | 1 | 3.68325E-05 | 2.112839292 | 0.293 | 0.082 | 0.894734669 | 0.012409259 | 7.36636E-05 |
| Pnrc2 | 3.69268E-05 | 1.392221909 | 0.566 | 0.309 | 0.897024928 | 0.174071266 | 0.486707043 | 0.463 | 0.355 | 1 | 0.174071266 | 7.38522E-05 |
| Cdc5l | 3.69812E-05 | 0.921866338 | 0.887 | 0.674 | 0.898347052 | 0.034396884 | 0.579223672 | 0.927 | 0.84 | 1 | 0.034396884 | 7.3961E-05 |
| Fzd5 | 3.69884E-05 | 1.652222254 | 0.321 | 0.102 | 0.898521933 | 0.017065874 | 1.30377265 | 0.293 | 0.15 | 1 | 0.017065874 | 7.39754E-05 |
| 1810030O07Rik | 3.71514E-05 | 1.27153626 | 0.547 | 0.263 | 0.902482583 | 0.08958317 | 0.921597037 | 0.463 | 0.365 | 1 | 0.08958317 | 7.43015E-05 |
| Gm16070 | 3.73272E-05 | -1.800974179 | 0.302 | 0.585 | 0.906753152 | 0.001212611 | -1.054812469 | 0.317 | 0.614 | 1 | 0.001212611 | 7.46531E-05 |
| Smg6 | 3.8065E-05 | 0.62893803 | 0.943 | 0.847 | 0.924675425 | 0.105152717 | 0.105779373 | 0.976 | 0.925 | 1 | 0.105152717 | 7.61286E-05 |
| Tmx3 | 3.8206E-05 | 0.992451166 | 0.792 | 0.534 | 0.928101239 | 0.001416979 | 0.753282366 | 0.805 | 0.683 | 1 | 0.001416979 | 7.64106E-05 |
| Erbin | 3.82966E-05 | 0.588845935 | 0.962 | 0.877 | 0.930300672 | 0.166716177 | 0.222068362 | 1 | 0.962 | 1 | 0.166716177 | 7.65917E-05 |
| Lama4 | 3.83768E-05 | 0.81105679 | 0.83 | 0.581 | 0.932248298 | 0.009075209 | 1.559131045 | 0.683 | 0.519 | 1 | 0.009075209 | 7.67521E-05 |
| Adgrl1 | 3.84853E-05 | 1.166218234 | 0.66 | 0.398 | 0.934884812 | 0.21869748 | 0.543021745 | 0.61 | 0.546 | 1 | 0.21869748 | 7.69691E-05 |
| Capns1 | 0.000209853 | -1.065510103 | 0.962 | 0.962 | 1 | 3.8788E-05 | -0.798760554 | 0.878 | 0.949 | 0.942238055 | 0.000209853 | 7.75745E-05 |
| Uvrag | 3.90499E-05 | 0.710715695 | 0.981 | 0.78 | 0.948599487 | 5.64408E-05 | 1.143914988 | 0.951 | 0.874 | 1 | 5.64408E-05 | 7.80982E-05 |
| Tsen34 | 3.90895E-05 | 0.805994441 | 0.849 | 0.669 | 0.949563111 | 0.154205522 | 0.401949071 | 0.805 | 0.744 | 1 | 0.154205522 | 7.81776E-05 |
| Lpgat1 | 3.92257E-05 | 1.03273207 | 0.774 | 0.542 | 0.952870515 | 0.619384356 | -0.137409832 | 0.732 | 0.792 | 1 | 0.619384356 | 7.84498E-05 |
| C2cd2 | 3.9269E-05 | 1.228723176 | 0.377 | 0.131 | 0.953922383 | 0.651860191 | 0.348918368 | 0.171 | 0.215 | 1 | 0.651860191 | 7.85364E-05 |
| Tada1 | 3.93034E-05 | 1.640149422 | 0.377 | 0.14 | 0.954757625 | 0.058322324 | -1.16279515 | 0.098 | 0.222 | 1 | 0.058322324 | 7.86052E-05 |
| Nop2 | 3.93036E-05 | 1.370451286 | 0.453 | 0.195 | 0.954764232 | 0.684373468 | 0.451146417 | 0.244 | 0.29 | 1 | 0.684373468 | 7.86058E-05 |
| Ccdc141 | 3.94949E-05 | 1.395730694 | 0.208 | 0.042 | 0.959409869 | 0.987781007 | 0.269520341 | 0.146 | 0.15 | 1 | 0.987781007 | 7.89882E-05 |
| Loxl3 | 3.95579E-05 | -1.15382647 | 0.528 | 0.771 | 0.960941585 | 0.00224923 | -1.166259005 | 0.366 | 0.611 | 1 | 0.00224923 | 7.91143E-05 |
| Bcl3 | 3.97921E-05 | 1.684271975 | 0.547 | 0.284 | 0.966628934 | 0.078822672 | 0.737669176 | 0.463 | 0.348 | 1 | 0.078822672 | 7.95826E-05 |
| Plekha2 | 4.00637E-05 | 1.598329246 | 0.208 | 0.042 | 0.973228435 | 0.231643049 | -0.622226769 | 0 | 0.034 | 1 | 0.231643049 | 8.01259E-05 |
| Stxbp4 | 4.00792E-05 | 1.551381565 | 0.434 | 0.169 | 0.973604985 | 0.030911844 | 0.934502051 | 0.439 | 0.307 | 1 | 0.030911844 | 8.01569E-05 |
| Ptgds | 4.06639E-05 | -3.46976827 | 0.038 | 0.309 | 0.987806529 | 0.011401301 | -1.970150072 | 0.024 | 0.184 | 1 | 0.011401301 | 8.13261E-05 |
| 1700017B05Rik | 4.07916E-05 | 2.3662267 | 0.245 | 0.064 | 0.990908601 | 0.002279829 | 1.280811501 | 0.415 | 0.218 | 1 | 0.002279829 | 8.15815E-05 |
| Ptov1 | 4.08347E-05 | 0.699927386 | 0.849 | 0.763 | 0.991955691 | 0.420117548 | -0.110631296 | 0.805 | 0.819 | 1 | 0.420117548 | 8.16677E-05 |
| Ccdc186 | 4.08925E-05 | 0.739685096 | 0.849 | 0.737 | 0.993359426 | 0.268650981 | 0.1736019 | 0.854 | 0.833 | 1 | 0.268650981 | 8.17832E-05 |
| Hmgb3 | 4.1148E-05 | 1.857740857 | 0.358 | 0.136 | 0.999566278 | 0.531667115 | 0.343665377 | 0.268 | 0.222 | 1 | 0.531667115 | 8.22942E-05 |
| 5730409E04Rik | 4.11824E-05 | 2.669295768 | 0.151 | 0.021 | 1 | 0.908035876 | 0.334704509 | 0.049 | 0.044 | 1 | 0.908035876 | 8.23632E-05 |
| Gpatch1 | 4.13751E-05 | 1.764776076 | 0.34 | 0.114 | 1 | 0.348552894 | 0.15538081 | 0.122 | 0.181 | 1 | 0.348552894 | 8.27485E-05 |
| Kcnq1ot1 | 0.001708989 | -0.667135287 | 0.925 | 0.932 | 1 | 4.15507E-05 | -0.77663266 | 0.976 | 0.993 | 1 | 0.001708989 | 8.30998E-05 |
| Polr3k | 4.16267E-05 | 1.730914886 | 0.434 | 0.195 | 1 | 0.339287937 | 0.577337723 | 0.293 | 0.235 | 1 | 0.339287937 | 8.32518E-05 |
| Gm50333 | 4.17252E-05 | 1.739685096 | 0.151 | 0.021 | 1 | 0.736872417 | 0.19334866 | 0.073 | 0.058 | 1 | 0.736872417 | 8.34486E-05 |
| Atg101 | 4.19332E-05 | 1.091986839 | 0.566 | 0.301 | 1 | 0.478939218 | 0.476802607 | 0.463 | 0.437 | 1 | 0.478939218 | 8.38646E-05 |

| | | | | | | | | | | | | |
|---|---|---|---|---|---|---|---|---|---|---|---|---|
| Dcaf5 | 4.19956E-05 | 1.744257825 | 0.509 | 0.271 | 1 | 0.010405522 | 0.657655429 | 0.854 | 0.608 | 1 | 0.010405522 | 8.39895E-05 |
| Mfap5 | 0.029966098 | 4.324647596 | 0.038 | 0.004 | 1 | 4.23118E-05 | 4.711673968 | 0.098 | 0.007 | 1 | 0.029966098 | 8.46219E-05 |
| Mccc1os | 0.503971666 | 1.569760094 | 0.019 | 0.008 | 1 | 4.23118E-05 | 4.059597271 | 0.098 | 0.007 | 1 | 0.503971666 | 8.46219E-05 |
| Tgoln1 | 4.23172E-05 | -0.608385414 | 0.943 | 0.962 | 1 | 0.064302237 | -0.191691296 | 0.927 | 0.962 | 1 | 0.064302237 | 8.46327E-05 |
| Khsrp | 4.2537E-05 | 1.246515071 | 0.623 | 0.394 | 1 | 0.045372654 | 0.712049718 | 0.732 | 0.672 | 1 | 0.045372654 | 8.50722E-05 |
| Slc16a6 | 4.26286E-05 | 2.614154213 | 0.151 | 0.021 | 1 | 0.027074279 | 2.43666692 | 0.122 | 0.041 | 1 | 0.027074279 | 8.52554E-05 |
| B930095G15Rik | 0.000759304 | -0.926197401 | 0.283 | 0.538 | 1 | 4.29436E-05 | -1.787286015 | 0.244 | 0.584 | 1 | 0.000759304 | 8.58854E-05 |
| 2310011J03Rik | 4.31169E-05 | 1.528537431 | 0.491 | 0.242 | 1 | 0.771878666 | 1.174239837 | 0.22 | 0.229 | 1 | 0.771878666 | 8.62319E-05 |
| Ube2s | 4.32767E-05 | 1.776210972 | 0.472 | 0.233 | 1 | 0.92865629 | 0.619299819 | 0.366 | 0.406 | 1 | 0.92865629 | 8.65515E-05 |
| AU015336 | 4.3347E-05 | -4.415133014 | 0 | 0.254 | 1 | 0.057944276 | -2.484723245 | 0 | 0.082 | 1 | 0.057944276 | 8.6692E-05 |
| Epb41l1 | 0.000118773 | -0.982780929 | 0.509 | 0.767 | 1 | 4.38153E-05 | -1.260291678 | 0.512 | 0.792 | 1 | 0.000118773 | 8.76287E-05 |
| Itch | 4.39267E-05 | 0.655229436 | 0.981 | 0.839 | 1 | 0.113921261 | -0.24525731 | 0.951 | 0.956 | 1 | 0.113921261 | 8.78515E-05 |
| Gjc1 | 4.40381E-05 | 1.179525986 | 0.623 | 0.398 | 1 | 0.010680795 | 1.051943699 | 0.659 | 0.529 | 1 | 0.010680795 | 8.80744E-05 |
| Tgfb1i1 | 4.40923E-05 | 0.87859819 | 0.66 | 0.36 | 1 | 0.061448372 | 0.799730144 | 0.561 | 0.474 | 1 | 0.061448372 | 8.81827E-05 |
| Bbc3 | 4.41034E-05 | 2.318221327 | 0.283 | 0.085 | 1 | 0.535847356 | 0.589277336 | 0.146 | 0.116 | 1 | 0.535847356 | 8.8205E-05 |
| Cyth1 | 4.4407E-05 | 0.916562858 | 0.736 | 0.466 | 1 | 0.196368249 | 0.130376804 | 0.78 | 0.727 | 1 | 0.196368249 | 8.88119E-05 |
| Dzip1 | 4.4407E-05 | 2.925240749 | 0.226 | 0.055 | 1 | 0.32051672 | 1.136765132 | 0.098 | 0.058 | 1 | 0.32051672 | 8.8812E-05 |
| Dhrs4 | 4.45376E-05 | 1.409295422 | 0.472 | 0.212 | 1 | 0.818608057 | 0.808635698 | 0.244 | 0.259 | 1 | 0.818608057 | 8.90733E-05 |
| Cuedc1 | 4.45548E-05 | 1.154722595 | 0.717 | 0.525 | 1 | 0.398672635 | 0.195755158 | 0.78 | 0.788 | 1 | 0.398672635 | 8.91077E-05 |
| Utrn | 0.028860103 | -0.447394627 | 0.962 | 0.987 | 1 | 4.46488E-05 | -0.959173896 | 0.951 | 0.99 | 1 | 0.028860103 | 8.92956E-05 |
| Baz1b | 4.50185E-05 | 0.9817182 | 0.887 | 0.754 | 1 | 0.43814887 | 0.278182567 | 0.854 | 0.887 | 1 | 0.43814887 | 9.00349E-05 |
| Gm49708 | 4.54236E-05 | 2.395730694 | 0.264 | 0.076 | 1 | 5.05517E-05 | 2.483567895 | 0.244 | 0.061 | 1 | 5.05517E-05 | 9.08452E-05 |
| Htra3 | 0.007510537 | -0.638181171 | 0.34 | 0.555 | 1 | 4.54288E-05 | -2.18959521 | 0.122 | 0.464 | 1 | 0.007510537 | 9.08555E-05 |
| Pcbp2 | 4.58098E-05 | 0.558325199 | 1 | 0.97 | 1 | 0.01167117 | 0.366819962 | 1 | 0.993 | 1 | 0.01167117 | 9.16174E-05 |
| Gm11520 | 0.546805316 | 0.491757582 | 0.057 | 0.038 | 1 | 4.58855E-05 | 2.83720485 | 0.098 | 0.007 | 1 | 0.546805316 | 9.17689E-05 |
| Pnrc1 | 4.61912E-05 | 0.720678046 | 1 | 0.907 | 1 | 4.59568E-05 | 0.883880311 | 0.951 | 0.867 | 1 | 4.61912E-05 | 9.19114E-05 |
| Arpp19 | 4.61245E-05 | 0.869841487 | 0.83 | 0.746 | 1 | 0.008199697 | 0.57010078 | 0.902 | 0.819 | 1 | 0.008199697 | 9.22468E-05 |
| Lonrf3 | 4.61699E-05 | 2.154722595 | 0.245 | 0.064 | 1 | 0.354411168 | 0.737669176 | 0.195 | 0.137 | 1 | 0.354411168 | 9.23377E-05 |
| Rgs16 | 0.014410924 | 1.154722595 | 0.132 | 0.042 | 1 | 4.66375E-05 | 3.322631677 | 0.098 | 0.007 | 1 | 0.014410924 | 9.32728E-05 |
| Eif5b | 4.71088E-05 | 0.515258174 | 0.981 | 0.953 | 1 | 0.182076935 | -0.208029756 | 0.976 | 0.993 | 1 | 0.182076935 | 9.42154E-05 |
| Gon7 | 4.72827E-05 | 1.380998451 | 0.453 | 0.199 | 1 | 0.257828793 | 0.760853964 | 0.317 | 0.253 | 1 | 0.257828793 | 9.45632E-05 |
| Id2 | 0.000396494 | 0.895281454 | 0.811 | 0.614 | 1 | 4.7396E-05 | 2.007129851 | 0.805 | 0.669 | 1 | 0.000396494 | 9.47897E-05 |
| Dstn | 4.75022E-05 | -1.337695679 | 1 | 0.987 | 1 | 0.797976826 | -0.176290993 | 0.976 | 0.993 | 1 | 0.797976826 | 9.50022E-05 |
| Lsp1 | 4.75998E-05 | 1.480122882 | 0.566 | 0.314 | 1 | 0.008281215 | 1.374470608 | 0.634 | 0.505 | 1 | 0.008281215 | 9.51974E-05 |
| Ypel1 | 4.76221E-05 | 2.154722595 | 0.245 | 0.064 | 1 | 0.652439478 | 0.949679579 | 0.122 | 0.102 | 1 | 0.652439478 | 9.5242E-05 |
| Efcab8 | 4.82464E-05 | 1.823516686 | 0.321 | 0.106 | 1 | 0.066105145 | 2.020068907 | 0.244 | 0.15 | 1 | 0.066105145 | 9.64905E-05 |
| Bcl11a | 0.001366773 | 3.669295768 | 0.113 | 0.021 | 1 | 4.83964E-05 | 1.83720485 | 0.341 | 0.109 | 1 | 0.001366773 | 9.67904E-05 |
| Adgrg6 | 0.000151245 | -1.618253535 | 0.415 | 0.669 | 1 | 4.84339E-05 | -1.118541912 | 0.463 | 0.795 | 1 | 0.000151245 | 9.68654E-05 |
| Eif1ad | 4.84484E-05 | 1.331600357 | 0.509 | 0.242 | 1 | 0.496995766 | 0.204936634 | 0.244 | 0.321 | 1 | 0.496995766 | 9.68944E-05 |
| Fkbp9 | 0.004130301 | -0.418616022 | 0.906 | 0.941 | 1 | 4.86754E-05 | -0.774724699 | 0.902 | 0.918 | 1 | 0.004130301 | 9.73484E-05 |
| Hecw2 | 4.87559E-05 | 2.421509135 | 0.34 | 0.127 | 1 | 0.017150309 | 1.195098442 | 0.341 | 0.195 | 1 | 0.017150309 | 9.75093E-05 |
| Gars | 0.000410368 | -1.364006208 | 0.434 | 0.631 | 1 | 4.8963E-05 | -1.557564052 | 0.415 | 0.689 | 1 | 0.000410368 | 9.79236E-05 |
| Eif2ak1 | 4.92172E-05 | 1.127755547 | 0.604 | 0.335 | 1 | 0.475717349 | 0.238167164 | 0.463 | 0.416 | 1 | 0.475717349 | 9.8432E-05 |
| Lasp1 | 4.93185E-05 | 1.074232677 | 0.604 | 0.347 | 1 | 0.001906907 | 0.762657896 | 0.756 | 0.515 | 1 | 0.001906907 | 9.86345E-05 |
| Scaf4 | 4.94266E-05 | 1.040389919 | 0.698 | 0.453 | 1 | 0.430856973 | -0.105805172 | 0.732 | 0.775 | 1 | 0.430856973 | 9.88508E-05 |
| Sap130 | 4.99649E-05 | 1.312602255 | 0.604 | 0.386 | 1 | 5.27753E-05 | 1.373455936 | 0.732 | 0.433 | 1 | 5.27753E-05 | 9.99273E-05 |
| Ero1b | 5.02722E-05 | 2.154722595 | 0.245 | 0.064 | 1 | 0.012737666 | 2.574170444 | 0.171 | 0.065 | 1 | 0.012737666 | 0.000100542 |
| Notch3 | 5.03491E-05 | 2.216123139 | 0.226 | 0.055 | 1 | 0.020136875 | 2.270164257 | 0.171 | 0.068 | 1 | 0.020136875 | 0.000100696 |
| Vstm4 | 5.11055E-05 | 2.70221039 | 0.208 | 0.047 | 1 | 0.459961975 | 0.515276755 | 0.122 | 0.085 | 1 | 0.459961975 | 0.000102208 |
| Trim24 | 0.175659699 | 0.648186741 | 0.472 | 0.411 | 1 | 5.1191E-05 | 0.943292108 | 0.732 | 0.444 | 1 | 0.175659699 | 0.000102379 |
| Pdzd4 | 5.12706E-05 | 1.552058092 | 0.358 | 0.127 | 1 | 0.703663759 | -0.101394606 | 0.22 | 0.249 | 1 | 0.703663759 | 0.000102539 |
| Megf9 | 5.13192E-05 | 1.680042149 | 0.509 | 0.284 | 1 | 0.141803374 | 0.595447104 | 0.439 | 0.355 | 1 | 0.141803374 | 0.000102636 |
| Mcu | 5.15744E-05 | -0.729045443 | 0.792 | 0.958 | 1 | 0.000232846 | -0.801559294 | 0.878 | 0.973 | 1 | 0.000232846 | 0.000103146 |
| Gsk3a | 5.16152E-05 | 0.824801709 | 0.698 | 0.436 | 1 | 0.322113996 | 0.392902756 | 0.659 | 0.625 | 1 | 0.322113996 | 0.000103228 |
| D17H6S53E | 5.16201E-05 | 2.254258268 | 0.226 | 0.055 | 1 | 0.882647938 | 0.699701326 | 0.073 | 0.068 | 1 | 0.882647938 | 0.000103238 |
| Akap7 | 5.16515E-05 | 1.680791406 | 0.396 | 0.169 | 1 | 0.516963191 | 0.232342792 | 0.122 | 0.171 | 1 | 0.516963191 | 0.0001033 |
| Phf20 | 5.17078E-05 | 0.819373984 | 0.755 | 0.525 | 1 | 0.021848126 | 0.470806606 | 0.829 | 0.686 | 1 | 0.021848126 | 0.000103413 |
| Cdc42ep3 | 5.18259E-05 | 0.898761571 | 0.887 | 0.775 | 1 | 0.025472512 | 0.336896223 | 0.927 | 0.898 | 1 | 0.025472512 | 0.000103649 |
| Scara5 | 5.18482E-05 | -2.899354078 | 0.189 | 0.453 | 1 | 0.00199775 | 2.367719567 | 0.244 | 0.092 | 1 | 0.00199775 | 0.000103694 |
| Sufu | 5.18584E-05 | 0.993730718 | 0.585 | 0.309 | 1 | 0.192814334 | 0.975195853 | 0.537 | 0.498 | 1 | 0.192814334 | 0.000103714 |
| Cnot3 | 5.19226E-05 | 1.340278248 | 0.566 | 0.335 | 1 | 0.111161196 | 0.522889939 | 0.537 | 0.382 | 1 | 0.111161196 | 0.000103842 |
| C230034O21Rik | 0.343968402 | 1.154722595 | 0.038 | 0.017 | 1 | 5.19436E-05 | 2.83720485 | 0.146 | 0.02 | 1 | 0.343968402 | 0.000103884 |
| Il17ra | 5.20927E-05 | -1.680976953 | 0.208 | 0.5 | 1 | 0.44462383 | 0.152706676 | 0.268 | 0.205 | 1 | 0.44462383 | 0.000104183 |
| 4833439L19Rik | 5.22669E-05 | 1.145136546 | 0.679 | 0.428 | 1 | 0.653858153 | 0.298338764 | 0.61 | 0.614 | 1 | 0.653858153 | 0.000104531 |
| Tom1l1 | 5.24083E-05 | 2.865215978 | 0.189 | 0.038 | 1 | 0.891353383 | 0.359157553 | 0.098 | 0.106 | 1 | 0.891353383 | 0.000104814 |
| Irx5 | 5.25645E-05 | 1.173338273 | 0.283 | 0.089 | 1 | 0.000227735 | 1.479652845 | 0.195 | 0.044 | 1 | 0.000227735 | 0.000105126 |
| Stxbp5 | 5.25731E-05 | 0.78032708 | 0.849 | 0.576 | 1 | 0.033267254 | 0.554410069 | 0.805 | 0.73 | 1 | 0.033267254 | 0.000105144 |
| Gm16973 | 5.27483E-05 | 2.8327945 | 0.189 | 0.038 | 1 | 0.434130977 | -0.332720152 | 0.049 | 0.085 | 1 | 0.434130977 | 0.000105494 |
| Ncald | 5.27483E-05 | 2.640149422 | 0.189 | 0.038 | 1 | 0.006142124 | 1.317830691 | 0.195 | 0.068 | 1 | 0.006142124 | 0.000105494 |
| Nsdhl | 5.34909E-05 | -2.495531366 | 0.132 | 0.411 | 1 | 0.021338362 | -1.398011612 | 0.073 | 0.232 | 1 | 0.021338362 | 0.000106979 |
| Pa2g4 | 5.37072E-05 | 0.728244155 | 0.868 | 0.784 | 1 | 0.1526214 | 0.378934354 | 0.854 | 0.857 | 1 | 0.1526214 | 0.000107412 |
| Trip12 | 5.40797E-05 | 0.643972199 | 0.962 | 0.958 | 1 | 0.019932242 | 0.309911699 | 1 | 0.986 | 1 | 0.019932242 | 0.000108156 |
| Ptp4a2 | 5.4187E-05 | 0.536100783 | 1 | 0.987 | 1 | 0.275118721 | 0.142446638 | 1 | 0.993 | 1 | 0.275118721 | 0.000108371 |
| Gm10125 | 5.41899E-05 | 2.341135719 | 0.264 | 0.076 | 1 | 0.013060235 | 0.644559772 | 0.244 | 0.102 | 1 | 0.013060235 | 0.000108377 |
| Pip4k2a | 0.005459005 | 1.428287668 | 0.358 | 0.191 | 1 | 5.43419E-05 | 1.592092352 | 0.512 | 0.246 | 1 | 0.005459005 | 0.000108681 |
| Atrnl1 | 0.015304221 | 0.814098406 | 0.547 | 0.428 | 1 | 5.45766E-05 | 1.191789832 | 0.756 | 0.532 | 1 | 0.015304221 | 0.00010915 |
| Ankrd13b | 5.50783E-05 | 2.569760094 | 0.17 | 0.03 | 1 | 0.06635192 | 1.53464208 | 0.195 | 0.102 | 1 | 0.06635192 | 0.000110154 |
| Bend5 | 0.002056378 | 2.891688189 | 0.075 | 0.008 | 1 | 5.52173E-05 | 3.83720485 | 0.122 | 0.014 | 1 | 0.002056378 | 0.000110432 |
| Sppl2a | 5.53285E-05 | -0.669588782 | 0.811 | 0.924 | 1 | 0.1152378 | -0.451276762 | 0.829 | 0.87 | 1 | 0.1152378 | 0.000110654 |
| Ccnl1 | 5.53798E-05 | 0.502997561 | 1 | 0.949 | 1 | 0.229181219 | 0.203721587 | 0.951 | 0.976 | 1 | 0.229181219 | 0.000110757 |
| Akirin2 | 5.56169E-05 | 0.969847252 | 0.83 | 0.631 | 1 | 0.08920217 | 0.325305811 | 0.829 | 0.802 | 1 | 0.08920217 | 0.000111231 |
| Cnksr2 | 0.000549701 | 2.739685096 | 0.113 | 0.017 | 1 | 5.57818E-05 | 2.64875976 | 0.244 | 0.061 | 1 | 0.000549701 | 0.00011156 |
| Shox2 | 0.037573118 | 1.776210972 | 0.113 | 0.042 | 1 | 5.59443E-05 | 3.952682067 | 0.122 | 0.014 | 1 | 0.037573118 | 0.000111886 |
| Antxr2 | 5.60041E-05 | 1.777487128 | 0.509 | 0.275 | 1 | 0.033305733 | 0.943195545 | 0.585 | 0.495 | 1 | 0.033305733 | 0.000112005 |
| Nudt4 | 5.6061E-05 | 0.800579875 | 0.943 | 0.788 | 1 | 0.028501792 | 0.745663396 | 0.878 | 0.857 | 1 | 0.028501792 | 0.000112119 |
| Ube2l6 | 0.004113402 | 0.569760094 | 0.264 | 0.106 | 1 | 5.61996E-05 | 2.169780189 | 0.244 | 0.061 | 1 | 0.004113402 | 0.000112396 |
| Ctif | 5.62688E-05 | 1.329809301 | 0.528 | 0.288 | 1 | 0.005162969 | 1.060056217 | 0.78 | 0.577 | 1 | 0.005162969 | 0.000112534 |
| Col4a5 | 5.63028E-05 | -3.399866257 | 0.075 | 0.352 | 1 | 0.728028858 | -0.444830518 | 0.171 | 0.195 | 1 | 0.728028858 | 0.000112603 |
| H4c16 | 5.63857E-05 | 3.029191713 | 0.189 | 0.038 | 1 | 0.030420195 | 1.574170444 | 0.073 | 0.017 | 1 | 0.030420195 | 0.000112768 |
| Ntng1 | 0.206100734 | 2.417757001 | 0.038 | 0.013 | 1 | 5.75992E-05 | 2.962735732 | 0.171 | 0.031 | 1 | 0.206100734 | 0.000115195 |
| Ralgds | 0.721446283 | -0.397073042 | 0.226 | 0.246 | 1 | 5.76616E-05 | 1.962735732 | 0.512 | 0.249 | 1 | 0.721446283 | 0.00011532 |
| Cfap20 | 5.77172E-05 | 1.030601283 | 0.83 | 0.585 | 1 | 0.416923382 | 0.423228546 | 0.683 | 0.737 | 1 | 0.416923382 | 0.000115431 |
| Taf5 | 5.78029E-05 | 1.47119626 | 0.415 | 0.169 | 1 | 0.061501613 | 0.600165653 | 0.317 | 0.191 | 1 | 0.061501613 | 0.000115602 |
| Cbr3 | 5.8113E-05 | 1.979635888 | 0.34 | 0.127 | 1 | 0.111560721 | 0.789899135 | 0.22 | 0.126 | 1 | 0.111560721 | 0.000116223 |
| Ankrd12 | 5.83148E-05 | 0.725879296 | 0.925 | 0.953 | 1 | 0.130877264 | 0.236375944 | 1 | 0.973 | 1 | 0.130877264 | 0.000116626 |
| Prr5l | 5.85783E-05 | 1.972519264 | 0.377 | 0.153 | 1 | 0.3180689 | -0.221688839 | 0.268 | 0.345 | 1 | 0.3180689 | 0.000117153 |
| Gm41496 | 5.85947E-05 | 0.975398895 | 0.302 | 0.093 | 1 | 0.006654314 | 1.794560512 | 0.293 | 0.147 | 1 | 0.006654314 | 0.000117186 |
| Cep120 | 5.86057E-05 | 1.22061053 | 0.566 | 0.284 | 1 | 0.045242775 | 0.647380291 | 0.585 | 0.502 | 1 | 0.045242775 | 0.000117208 |

| | | | | | | | | | | | | |
|---|---|---|---|---|---|---|---|---|---|---|---|---|
| Dazap2 | 5.87073E-05 | 0.997181318 | 0.774 | 0.597 | 1 | 0.003910091 | 0.697801793 | 0.78 | 0.676 | 1 | 0.003910091 | 0.000117411 |
| Idh2 | 5.87469E-05 | 1.217916421 | 0.698 | 0.466 | 1 | 0.000479421 | 1.023216835 | 0.805 | 0.584 | 1 | 0.000479421 | 0.00011749 |
| Tbc1d1 | 0.544815759 | -0.116263934 | 0.698 | 0.737 | 1 | 5.87963E-05 | -1.008882467 | 0.756 | 0.894 | 1 | 0.544815759 | 0.000117589 |
| Lncppara | 5.88613E-05 | 1.733082361 | 0.528 | 0.297 | 1 | 0.17402062 | 0.66566157 | 0.61 | 0.59 | 1 | 0.17402062 | 0.000117719 |
| Acvr1b | 5.8873E-05 | -1.786725223 | 0.132 | 0.436 | 1 | 0.171013762 | -0.84461919 | 0.195 | 0.297 | 1 | 0.171013762 | 0.000117743 |
| Pcsk7 | 5.90214E-05 | 1.221836791 | 0.434 | 0.174 | 1 | 0.02570682 | 0.83720485 | 0.512 | 0.334 | 1 | 0.02570682 | 0.000118039 |
| Atp2a2 | 0.089112174 | 0.271084087 | 1 | 0.983 | 1 | 5.92882E-05 | 0.701211211 | 1 | 0.997 | 1 | 0.089112174 | 0.000118573 |
| Desi2 | 0.54673921 | -0.233689561 | 0.887 | 0.894 | 1 | 5.95747E-05 | -0.903071958 | 0.927 | 0.976 | 1 | 0.54673921 | 0.000119146 |
| Lonrf2 | 0.042860829 | 1.47665069 | 0.075 | 0.021 | 1 | 5.9657E-05 | 3.515276755 | 0.122 | 0.014 | 1 | 0.042860829 | 0.000119311 |
| Gm12204 | 0.287761733 | -0.652632327 | 0 | 0.021 | 1 | 5.9657E-05 | 3.322631677 | 0.122 | 0.014 | 1 | 0.287761733 | 0.000119311 |
| Sp140 | 0.633487796 | 0.569760094 | 0.038 | 0.025 | 1 | 5.9657E-05 | 3.322631677 | 0.122 | 0.014 | 1 | 0.633487796 | 0.000119311 |
| Lhfpl2 | 0.000606068 | 2.891688189 | 0.132 | 0.025 | 1 | 5.97069E-05 | 3.614812428 | 0.122 | 0.014 | 1 | 0.000606068 | 0.00011941 |
| Kifc3 | 5.99097E-05 | -1.743397791 | 0.132 | 0.436 | 1 | 0.49443761 | 0.478123756 | 0.22 | 0.287 | 1 | 0.49443761 | 0.000119816 |
| Serpini1 | 6.03968E-05 | 2.280253477 | 0.226 | 0.055 | 1 | 0.415156117 | 1.60690723 | 0.146 | 0.109 | 1 | 0.415156117 | 0.00012079 |
| Tut4 | 6.0442E-05 | 0.816674835 | 0.943 | 0.839 | 1 | 0.027877409 | 0.552148394 | 0.902 | 0.87 | 1 | 0.027877409 | 0.00012088 |
| Bccip | 0.26940443 | 0.159022732 | 0.698 | 0.678 | 1 | 6.10537E-05 | -0.857382142 | 0.439 | 0.737 | 1 | 0.26940443 | 0.000122104 |
| Aspscr1 | 0.259164031 | -0.430239906 | 0.283 | 0.36 | 1 | 6.1272E-05 | -1.633753022 | 0.244 | 0.58 | 1 | 0.259164031 | 0.00012254 |
| Usp33 | 6.13939E-05 | 0.890306959 | 0.717 | 0.492 | 1 | 0.871272918 | 0.146155091 | 0.683 | 0.741 | 1 | 0.871272918 | 0.000122784 |
| Ppp2r5c | 6.17839E-05 | 0.777808641 | 0.906 | 0.78 | 1 | 0.000173103 | 0.846790899 | 0.902 | 0.823 | 1 | 0.000173103 | 0.000123564 |
| Metrn | 6.19511E-05 | -2.856504661 | 0.057 | 0.326 | 1 | 0.280264224 | -0.332720152 | 0.098 | 0.167 | 1 | 0.280264224 | 0.000123898 |
| Vit | 0.004933501 | -1.631002311 | 0.17 | 0.364 | 1 | 6.20597E-05 | -2.200930279 | 0.244 | 0.553 | 1 | 0.004933501 | 0.000124116 |
| Metap2 | 6.2545E-05 | 0.803939637 | 0.943 | 0.843 | 1 | 0.714912245 | 0.164315842 | 0.805 | 0.911 | 1 | 0.714912245 | 0.000125086 |
| Rnf227 | 6.29391E-05 | 2.076720083 | 0.226 | 0.055 | 1 | 0.033454942 | 1.730289646 | 0.146 | 0.058 | 1 | 0.033454942 | 0.000125874 |
| Vgll3 | 0.00024982 | 4.739685096 | 0.057 | 0 | 1 | 6.29571E-05 | 3.879025025 | 0.22 | 0.051 | 1 | 0.00024982 | 0.00012591 |
| Gtf2a1 | 6.34649E-05 | 1.575565716 | 0.585 | 0.331 | 1 | 0.124622653 | 0.501601818 | 0.512 | 0.386 | 1 | 0.124622653 | 0.000126926 |
| Fam20b | 0.001867808 | -0.727505947 | 0.547 | 0.754 | 1 | 6.35535E-05 | -1.184562965 | 0.463 | 0.73 | 1 | 0.001867808 | 0.000127103 |
| Rexo2 | 6.36383E-05 | -0.705446908 | 0.962 | 0.97 | 1 | 0.000341993 | -0.602037121 | 0.878 | 0.956 | 1 | 0.000341993 | 0.000127272 |
| Ctnnal1 | 6.37266E-05 | -1.337130502 | 0.415 | 0.661 | 1 | 0.005768306 | -1.062268274 | 0.463 | 0.638 | 1 | 0.005768306 | 0.000127449 |
| Phkg2 | 6.37622E-05 | 1.508359549 | 0.491 | 0.233 | 1 | 0.095474403 | 0.697274589 | 0.415 | 0.297 | 1 | 0.095474403 | 0.00012752 |
| Adgrf5 | 6.37821E-05 | 2.465062715 | 0.283 | 0.089 | 1 | 0.009148179 | 2.883008539 | 0.195 | 0.078 | 1 | 0.009148179 | 0.00012756 |
| Slc7a1 | 6.38513E-05 | -1.388175846 | 0.396 | 0.669 | 1 | 0.000324298 | -1.028920712 | 0.561 | 0.819 | 1 | 0.000324298 | 0.000127699 |
| Immp2l | 6.4211E-05 | -2.041674618 | 0.208 | 0.513 | 1 | 0.10302981 | -1.043213534 | 0.341 | 0.447 | 1 | 0.10302981 | 0.000128418 |
| Etfdh | 6.47341E-05 | 1.545913352 | 0.434 | 0.191 | 1 | 0.029863855 | -0.775238387 | 0.195 | 0.379 | 1 | 0.029863855 | 0.000129464 |
| Tmem101 | 6.47526E-05 | 1.587682002 | 0.396 | 0.161 | 1 | 0.622281961 | 0.711673968 | 0.171 | 0.147 | 1 | 0.622281961 | 0.000129501 |
| Rimoc1 | 0.015747185 | 0.739685096 | 0.491 | 0.331 | 1 | 6.48322E-05 | -1.484723245 | 0.122 | 0.481 | 1 | 0.015747185 | 0.00012966 |
| Sspn | 6.5524E-05 | 0.893676742 | 0.792 | 0.61 | 1 | 0.311873875 | 0.31725234 | 0.707 | 0.662 | 1 | 0.311873875 | 0.000131044 |
| Kcnj15 | 0.003030198 | 2.495759513 | 0.189 | 0.064 | 1 | 6.5824E-05 | 2.739594053 | 0.561 | 0.341 | 1 | 0.003030198 | 0.000131644 |
| Keap1 | 6.61885E-05 | 1.4226558 | 0.528 | 0.301 | 1 | 0.245715306 | 0.660327088 | 0.463 | 0.437 | 1 | 0.245715306 | 0.000132373 |
| Ubn2 | 6.66656E-05 | 0.488963951 | 0.981 | 0.936 | 1 | 0.11553464 | 0.445468105 | 0.927 | 0.956 | 1 | 0.11553464 | 0.000133327 |
| Ppm1n | 0.019420533 | 1.2967416 | 0.094 | 0.025 | 1 | 6.68093E-05 | 1.83720485 | 0.146 | 0.02 | 1 | 0.019420533 | 0.000133614 |
| Intu | 6.69124E-05 | 2.247831999 | 0.189 | 0.038 | 1 | 0.359386644 | 0.962735732 | 0.024 | 0.061 | 1 | 0.359386644 | 0.00013382 |
| Atg3 | 6.70809E-05 | 0.840026069 | 0.83 | 0.708 | 1 | 0.027588333 | 0.395200303 | 0.854 | 0.768 | 1 | 0.027588333 | 0.000134157 |
| Timp1 | 6.75551E-05 | -2.246395276 | 0.717 | 0.818 | 1 | 0.013424935 | -1.80119691 | 0.39 | 0.601 | 1 | 0.013424935 | 0.000135106 |
| Srgn | 6.81565E-05 | -1.024492169 | 0.453 | 0.691 | 1 | 0.006709089 | -0.54220874 | 0.585 | 0.812 | 1 | 0.006709089 | 0.000136308 |
| Cebpd | 6.81959E-05 | -1.971891995 | 0.774 | 0.822 | 1 | 0.004211675 | 0.765769712 | 0.829 | 0.642 | 1 | 0.004211675 | 0.000136387 |
| Sipa1l3 | 6.82093E-05 | 1.538051234 | 0.396 | 0.157 | 1 | 0.138653522 | 0.242461328 | 0.683 | 0.532 | 1 | 0.138653522 | 0.000136414 |
| Cst6 | 6.83315E-05 | 2.039245377 | 0.17 | 0.03 | 1 | 0.441381862 | 1.515276755 | 0.024 | 0.01 | 1 | 0.441381862 | 0.000136658 |
| Clmn | 0.084807514 | -1.093204919 | 0.226 | 0.326 | 1 | 6.85212E-05 | -1.501597064 | 0.512 | 0.706 | 1 | 0.084807514 | 0.000137038 |
| Actr3b | 6.86459E-05 | 3.640149422 | 0.132 | 0.017 | 1 | 0.914200263 | 0.515276755 | 0.024 | 0.027 | 1 | 0.914200263 | 0.000137287 |
| Ackr4 | 6.92169E-05 | 2.237184755 | 0.208 | 0.047 | 1 | 0.000428748 | 2.313642894 | 0.171 | 0.038 | 1 | 0.000428748 | 0.000138429 |
| Fbxl20 | 6.9595E-05 | 1.146817397 | 0.66 | 0.424 | 1 | 0.149124682 | 0.529632048 | 0.756 | 0.642 | 1 | 0.149124682 | 0.000139185 |
| Hdhd5 | 6.975E-05 | 2.047807391 | 0.189 | 0.038 | 1 | 0.322676546 | 0.114738825 | 0.049 | 0.099 | 1 | 0.322676546 | 0.000139495 |
| Btbd7 | 7.00155E-05 | 0.867624484 | 0.906 | 0.771 | 1 | 0.632698393 | 0.207984429 | 0.951 | 0.956 | 1 | 0.632698393 | 0.000140026 |
| Plcg1 | 7.01489E-05 | 1.404700848 | 0.453 | 0.208 | 1 | 0.593430494 | 0.442673006 | 0.341 | 0.341 | 1 | 0.593430494 | 0.000140293 |
| Norad | 7.02144E-05 | 0.581712897 | 0.981 | 0.831 | 1 | 0.096117056 | -0.342402705 | 0.951 | 0.925 | 1 | 0.096117056 | 0.000140424 |
| Isg15 | 0.018918919 | 1.108918905 | 0.132 | 0.047 | 1 | 7.04342E-05 | 2.534135782 | 0.22 | 0.051 | 1 | 0.018918919 | 0.000140863 |
| Tmem150a | 7.0446E-05 | 1.654293604 | 0.453 | 0.216 | 1 | 0.048454917 | 1.401105735 | 0.366 | 0.249 | 1 | 0.048454917 | 0.000140887 |
| Ppp1r15a | 0.096319162 | 0.592747755 | 0.943 | 0.919 | 1 | 7.10103E-05 | 1.392780997 | 0.951 | 0.935 | 1 | 0.096319162 | 0.000142016 |
| Ephb4 | 7.10848E-05 | 1.409295422 | 0.396 | 0.165 | 1 | 0.416182374 | 0.292884334 | 0.488 | 0.42 | 1 | 0.416182374 | 0.000142165 |
| Kctd12b | 7.11295E-05 | 3.154722595 | 0.132 | 0.017 | 1 | 0.478984212 | 0.930314254 | 0.098 | 0.068 | 1 | 0.478984212 | 0.000142254 |
| Nkx6-1 | 7.18383E-05 | 3.154722595 | 0.132 | 0.017 | 1 | 0.513415298 | 2.185128153 | 0.049 | 0.031 | 1 | 0.513415298 | 0.000143671 |
| Nme4 | 7.18383E-05 | 3.154722595 | 0.132 | 0.017 | 1 | 0.003188339 | 2.159132945 | 0.098 | 0.017 | 1 | 0.003188339 | 0.000143671 |
| Gm12349 | 7.18768E-05 | -3.974560422 | 0 | 0.242 | 1 | 0.170373481 | -1.16279515 | 0 | 0.044 | 1 | 0.170373481 | 0.000143748 |
| Senp7 | 7.18815E-05 | 1.080297754 | 0.642 | 0.407 | 1 | 0.587862787 | 0.221047727 | 0.61 | 0.584 | 1 | 0.587862787 | 0.000143758 |
| Fmr1 | 7.19373E-05 | 0.821878722 | 0.849 | 0.691 | 1 | 0.215010854 | 0.458307943 | 0.805 | 0.86 | 1 | 0.215010854 | 0.000143869 |
| Slc35e4 | 0.017195211 | -0.687736128 | 0.509 | 0.665 | 1 | 7.22854E-05 | -1.188947438 | 0.415 | 0.696 | 1 | 0.017195211 | 0.000144566 |
| Ptbp2 | 7.24629E-05 | 0.762802335 | 0.849 | 0.771 | 1 | 0.719886519 | 0.157085116 | 0.951 | 0.887 | 1 | 0.719886519 | 0.00014492 |
| Vars | 7.28704E-05 | 1.510866405 | 0.415 | 0.174 | 1 | 0.201783577 | 0.962735732 | 0.317 | 0.239 | 1 | 0.201783577 | 0.000145735 |
| Tprgl | 7.31784E-05 | 1.015958525 | 0.698 | 0.551 | 1 | 0.578840247 | 0.416629167 | 0.634 | 0.679 | 1 | 0.578840247 | 0.000146351 |
| Smim31 | 7.35386E-05 | -3.639693272 | 0.019 | 0.271 | 1 | 0.032728667 | -1.995685164 | 0.024 | 0.143 | 1 | 0.032728667 | 0.000147072 |
| Tbc1d16 | 7.36117E-05 | 1.411062348 | 0.396 | 0.157 | 1 | 0.388278309 | 0.937731726 | 0.366 | 0.328 | 1 | 0.388278309 | 0.000147218 |
| Klhdc2 | 7.3728E-05 | 0.835142255 | 0.811 | 0.644 | 1 | 0.930354728 | 0.172454323 | 0.732 | 0.782 | 1 | 0.930354728 | 0.000147451 |
| Pecam1 | 7.38215E-05 | 1.988072725 | 0.264 | 0.076 | 1 | 0.000509784 | 0.916639317 | 0.268 | 0.089 | 1 | 0.000509784 | 0.000147638 |
| Tacc1 | 7.41491E-05 | 0.727375954 | 0.906 | 0.869 | 1 | 0.813622908 | 0.13072453 | 0.854 | 0.891 | 1 | 0.813622908 | 0.000148293 |
| Rbpms2 | 7.44504E-05 | 3.029191713 | 0.132 | 0.017 | 1 | 0.641242812 | 0.515276755 | 0.049 | 0.034 | 1 | 0.641242812 | 0.000148895 |
| Foxd1 | 7.44923E-05 | 1.154722595 | 0.377 | 0.136 | 1 | 0.285904916 | 0.59841799 | 0.293 | 0.232 | 1 | 0.285904916 | 0.000148979 |
| Camsap1 | 7.45356E-05 | 1.22106509 | 0.623 | 0.381 | 1 | 0.200881985 | -0.308774456 | 0.659 | 0.703 | 1 | 0.200881985 | 0.000149066 |
| Piezo1 | 0.004019899 | -0.595964387 | 0.887 | 0.949 | 1 | 7.4845E-05 | -1.106784764 | 0.707 | 0.867 | 1 | 0.004019899 | 0.000149684 |
| Nck1 | 7.48587E-05 | 1.062800105 | 0.774 | 0.568 | 1 | 0.990127606 | 0.244160003 | 0.707 | 0.73 | 1 | 0.990127606 | 0.000149712 |
| Kdm3b | 0.000211677 | 1.036725385 | 0.66 | 0.424 | 1 | 7.49289E-05 | 0.917666878 | 0.829 | 0.625 | 1 | 0.000211677 | 0.000149852 |
| Zfp462 | 7.53794E-05 | 1.212825549 | 0.623 | 0.381 | 1 | 0.089219178 | 0.361471419 | 0.512 | 0.341 | 1 | 0.089219178 | 0.000150753 |
| Arhgap22 | 7.6179E-05 | 2.792152515 | 0.132 | 0.017 | 1 | 0.000283397 | 2.111379813 | 0.244 | 0.072 | 1 | 0.000283397 | 0.000152352 |
| Slc25a5 | 7.63391E-05 | 0.92192039 | 0.943 | 0.771 | 1 | 0.010448316 | 0.484108737 | 0.902 | 0.901 | 1 | 0.010448316 | 0.000152672 |
| Zscan18 | 7.63599E-05 | 1.792152515 | 0.208 | 0.047 | 1 | 0.788640814 | 0.486707603 | 0.146 | 0.133 | 1 | 0.788640814 | 0.000152714 |
| Ptger4 | 7.70178E-05 | 3.029191713 | 0.132 | 0.017 | 1 | 0.323872273 | 1.766815522 | 0.073 | 0.041 | 1 | 0.323872273 | 0.00015403 |
| Timm13 | 7.70339E-05 | 0.723110871 | 0.849 | 0.703 | 1 | 0.748403387 | 0.360766806 | 0.634 | 0.672 | 1 | 0.748403387 | 0.000154062 |
| Prc1 | 0.946377329 | -1.932740246 | 0.075 | 0.076 | 1 | 7.72395E-05 | 3.100239256 | 0.171 | 0.031 | 1 | 0.946377329 | 0.000154473 |
| Rp9 | 7.75314E-05 | 0.703686897 | 0.849 | 0.568 | 1 | 0.58472835 | 0.114738825 | 0.61 | 0.693 | 1 | 0.58472835 | 0.000155057 |
| Prr16 | 0.001956492 | 5.377115016 | 0.075 | 0.008 | 1 | 7.77134E-05 | 2.73223529 | 0.268 | 0.078 | 1 | 0.001956492 | 0.000155421 |
| Tekt2 | 7.79076E-05 | 2.891688189 | 0.132 | 0.017 | 1 | 0.546365767 | 1.100239256 | 0.049 | 0.031 | 1 | 0.546365767 | 0.000155809 |
| Arrb2 | 7.79076E-05 | 2.891688189 | 0.132 | 0.017 | 1 | 0.40786703 | -0.555112573 | 0.024 | 0.055 | 1 | 0.40786703 | 0.000155809 |
| Senp6 | 7.79523E-05 | 0.531183062 | 1 | 0.983 | 1 | 0.113147951 | 0.219906367 | 1 | 0.997 | 1 | 0.113147951 | 0.000155899 |
| Fkbp4 | 7.82964E-05 | 1.27829728 | 0.623 | 0.364 | 1 | 0.180047038 | 0.504822397 | 0.659 | 0.573 | 1 | 0.180047038 | 0.000156587 |
| Ophn1 | 7.84681E-05 | 1.084901145 | 0.66 | 0.475 | 1 | 0.322470341 | 0.680103889 | 0.561 | 0.515 | 1 | 0.322470341 | 0.00015693 |
| Pnldc1 | 7.89292E-05 | 2.8327945 | 0.132 | 0.017 | 1 | 0.720202665 | 0.930314254 | 0.049 | 0.038 | 1 | 0.720202665 | 0.000157852 |
| Mfge8 | 7.9053E-05 | -1.609427829 | 0.717 | 0.843 | 1 | 0.232420416 | -0.525094428 | 0.732 | 0.778 | 1 | 0.232420416 | 0.0001581 |
| Tcf7l2 | 7.95186E-05 | 0.61339881 | 1 | 0.894 | 1 | 0.131251405 | 0.290449226 | 0.951 | 0.986 | 1 | 0.131251405 | 0.000159031 |
| Hs6st3 | 7.96204E-05 | -4.067084198 | 0.075 | 0.343 | 1 | 0.555689213 | -0.802205435 | 0.049 | 0.031 | 1 | 0.555689213 | 0.000159235 |

| | | | | | | | | | | | | |
|---|---|---|---|---|---|---|---|---|---|---|---|---|
| Ptp4a3 | 7.96276E-05 | 1.669295768 | 0.226 | 0.055 | 1 | 0.680867383 | -0.16279515 | 0.073 | 0.092 | 1 | 0.680867383 | 0.000159249 |
| Dok3 | 7.97119E-05 | 2.669295768 | 0.132 | 0.017 | 1 | 0.446954486 | -0.970150072 | 0.049 | 0.082 | 1 | 0.446954486 | 0.000159418 |
| Pltp | 8.13664E-05 | 0.964039033 | 0.434 | 0.178 | 1 | 0.004552668 | 1.078212949 | 0.415 | 0.222 | 1 | 0.004552668 | 0.000162726 |
| Pqbp1 | 8.17328E-05 | 0.994257923 | 0.717 | 0.521 | 1 | 0.004385868 | -0.691574816 | 0.244 | 0.532 | 1 | 0.004385868 | 0.000163459 |
| Zfp251 | 0.000280752 | 1.595295186 | 0.283 | 0.097 | 1 | 8.18296E-05 | 1.60690723 | 0.366 | 0.126 | 1 | 0.000280752 | 0.000163652 |
| Gm56530 | 8.25333E-05 | 2.932330173 | 0.132 | 0.017 | 1 | 0.060103117 | 1.614812428 | 0.049 | 0.01 | 1 | 0.060103117 | 0.00016506 |
| Dnah12 | 8.25964E-05 | 2.052842981 | 0.264 | 0.076 | 1 | 0.107929198 | 0.276489895 | 0.098 | 0.041 | 1 | 0.107929198 | 0.000165186 |
| AW011738 | 0.115117141 | 1.614154213 | 0.113 | 0.055 | 1 | 8.26889E-05 | 2.252242349 | 0.268 | 0.075 | 1 | 0.115117141 | 0.000165371 |
| Taf4 | 8.317E-05 | 1.859266711 | 0.283 | 0.089 | 1 | 0.000475667 | 1.342440158 | 0.439 | 0.205 | 1 | 0.000475667 | 0.000166333 |
| Tmcc3 | 8.39808E-05 | -3.223789028 | 0.019 | 0.275 | 1 | 0.021894509 | -1.54205642 | 0.39 | 0.529 | 1 | 0.021894509 | 0.000167955 |
| Cdkn2aip | 0.002378113 | 1.819538403 | 0.377 | 0.212 | 1 | 8.40473E-05 | 1.515276755 | 0.585 | 0.331 | 1 | 0.002378113 | 0.000168088 |
| Trrap | 8.43012E-05 | 1.025439578 | 0.83 | 0.678 | 1 | 0.306320385 | 0.2460372 | 0.854 | 0.853 | 1 | 0.306320385 | 0.000168595 |
| Pou2af3 | 8.45757E-05 | -3.015202407 | 0.038 | 0.301 | 1 | 0.009328784 | -1.987223586 | 0.024 | 0.191 | 1 | 0.009328784 | 0.000169144 |
| Ppp1r2 | 8.56007E-05 | 0.740765361 | 0.906 | 0.822 | 1 | 0.003556115 | 0.69315848 | 0.902 | 0.867 | 1 | 0.003556115 | 0.000171194 |
| Armc10 | 8.56249E-05 | 2.154722595 | 0.283 | 0.089 | 1 | 0.057122107 | 1.220533489 | 0.244 | 0.137 | 1 | 0.057122107 | 0.000171243 |
| Rit1 | 8.57205E-05 | 1.58440687 | 0.547 | 0.318 | 1 | 0.295897672 | 0.296636468 | 0.537 | 0.451 | 1 | 0.295897672 | 0.000171434 |
| Dnajc8 | 8.6083E-05 | 0.636718517 | 0.925 | 0.754 | 1 | 0.506391469 | 0.330244861 | 0.805 | 0.812 | 1 | 0.506391469 | 0.000172159 |
| Hk1 | 0.006517858 | -0.317622343 | 0.642 | 0.809 | 1 | 8.6486E-05 | -1.161239684 | 0.659 | 0.826 | 1 | 0.006517858 | 0.000172965 |
| Atp5l | 8.72505E-05 | 0.458313558 | 1 | 0.945 | 1 | 0.447330358 | 0.149393941 | 0.951 | 0.959 | 1 | 0.447330358 | 0.000174493 |
| Zfp637 | 8.74062E-05 | 1.259059255 | 0.509 | 0.242 | 1 | 0.404954847 | 0.30115195 | 0.415 | 0.358 | 1 | 0.404954847 | 0.000174805 |
| Efcab1 | 8.77647E-05 | -2.396451592 | 0.132 | 0.407 | 1 | 0.000354984 | -1.326025499 | 0.073 | 0.372 | 1 | 0.000354984 | 0.000175522 |
| Spindoc | 8.8188E-05 | 1.165369839 | 0.585 | 0.356 | 1 | 0.431727457 | 0.400218378 | 0.561 | 0.539 | 1 | 0.431727457 | 0.000176368 |
| Fhip2a | 8.84213E-05 | 1.216123139 | 0.604 | 0.364 | 1 | 0.330583302 | -0.132421501 | 0.561 | 0.666 | 1 | 0.330583302 | 0.000176835 |
| Eif2ak2 | 8.87037E-05 | 1.249049977 | 0.528 | 0.28 | 1 | 0.409281188 | 0.640312801 | 0.415 | 0.396 | 1 | 0.409281188 | 0.000177399 |
| Egln2 | 8.886E-05 | 1.43124023 | 0.491 | 0.246 | 1 | 0.696300389 | 0.186440291 | 0.317 | 0.372 | 1 | 0.696300389 | 0.000177712 |
| Gm49454 | 8.94652E-05 | 1.10741688 | 0.132 | 0.017 | 1 | 0.000761857 | 2.252242349 | 0.171 | 0.041 | 1 | 0.000761857 | 0.000178922 |
| Ankrd42 | 8.95869E-05 | 2.312263872 | 0.377 | 0.161 | 1 | 0.027352557 | 1.708620644 | 0.341 | 0.212 | 1 | 0.027352557 | 0.000179166 |
| Mapk1 | 8.99246E-05 | 0.820147124 | 0.906 | 0.801 | 1 | 0.300791043 | 0.176790121 | 0.951 | 0.935 | 1 | 0.300791043 | 0.000179841 |
| Mfsd1 | 9.00364E-05 | 1.465651691 | 0.528 | 0.297 | 1 | 0.579545092 | 0.306690133 | 0.415 | 0.382 | 1 | 0.579545092 | 0.000180065 |
| Atp10d | 0.366907862 | 0.655916738 | 0.17 | 0.127 | 1 | 9.02778E-05 | 1.561570407 | 0.341 | 0.116 | 1 | 0.366907862 | 0.000180547 |
| Klhl22 | 9.05226E-05 | 1.237184755 | 0.453 | 0.199 | 1 | 0.146440186 | 0.974708374 | 0.366 | 0.29 | 1 | 0.146440186 | 0.000181037 |
| Fxr1 | 9.16378E-05 | 0.51531231 | 1 | 0.928 | 1 | 0.236625083 | 0.156880493 | 0.976 | 0.98 | 1 | 0.236625083 | 0.000183267 |
| Slc9a9 | 0.005569608 | 1.669295768 | 0.34 | 0.195 | 1 | 9.18679E-05 | 1.236812309 | 0.659 | 0.369 | 1 | 0.005569608 | 0.000183727 |
| Thsd7a | 0.033178609 | 1.211306123 | 0.132 | 0.051 | 1 | 9.2904E-05 | 1.46736677 | 0.39 | 0.15 | 1 | 0.033178609 | 0.000185799 |
| Armcx4 | 9.30038E-05 | 1.315187267 | 0.472 | 0.225 | 1 | 0.244070701 | -0.126269274 | 0.293 | 0.406 | 1 | 0.244070701 | 0.000185999 |
| Bora | 0.002760321 | 1.762405172 | 0.189 | 0.064 | 1 | 9.34345E-05 | 2.377773231 | 0.171 | 0.031 | 1 | 0.002760321 | 0.00018686 |
| Hira | 9.39543E-05 | 1.454282877 | 0.528 | 0.297 | 1 | 0.531862813 | 0.121710786 | 0.537 | 0.485 | 1 | 0.531862813 | 0.0001879 |
| Cops7b | 9.40253E-05 | 2.47665069 | 0.245 | 0.068 | 1 | 0.01070919 | 1.574170444 | 0.268 | 0.123 | 1 | 0.01070919 | 0.000188042 |
| Col9a1 | 0.142838402 | -0.376411969 | 0.83 | 0.873 | 1 | 9.44576E-05 | -0.745661218 | 0.756 | 0.956 | 1 | 0.142838402 | 0.000188906 |
| Neat1 | 0.000275394 | -0.705861903 | 0.962 | 0.992 | 1 | 9.56273E-05 | -0.960482203 | 1 | 0.997 | 1 | 0.000275394 | 0.000191245 |
| Tfpi | 9.57312E-05 | 2.017219071 | 0.132 | 0.017 | 1 | 0.003983294 | 3.83720485 | 0.049 | 0.003 | 1 | 0.003983294 | 0.000191453 |
| Slc16a2 | 9.58857E-05 | 0.647003093 | 0.868 | 0.636 | 1 | 0.025400253 | 0.675847844 | 0.927 | 0.867 | 1 | 0.025400253 | 0.000191762 |
| Trafd1 | 9.78842E-05 | -1.207333613 | 0.434 | 0.682 | 1 | 0.407469757 | 0.23386382 | 0.61 | 0.604 | 1 | 0.407469757 | 0.000195759 |
| Cdc42bpa | 9.84315E-05 | 0.671232971 | 1 | 0.983 | 1 | 0.509858604 | 0.169464673 | 1 | 0.993 | 1 | 0.509858604 | 0.000196853 |
| Raly | 9.86462E-05 | 0.716697435 | 0.906 | 0.822 | 1 | 0.430848887 | 0.213071752 | 0.829 | 0.86 | 1 | 0.430848887 | 0.000197283 |
| Dgkh | 0.002022071 | 0.725410965 | 0.755 | 0.589 | 1 | 9.88255E-05 | 0.913508217 | 0.829 | 0.594 | 1 | 0.002022071 | 0.000197641 |
| Plcb1 | 0.000678929 | -0.742846443 | 0.66 | 0.873 | 1 | 9.89701E-05 | -0.91444064 | 0.805 | 0.952 | 1 | 0.000678929 | 0.00019793 |
| Synpo2 | 0.000337496 | 1.948271717 | 0.151 | 0.03 | 1 | 9.92209E-05 | 2.252242349 | 0.268 | 0.078 | 1 | 0.000337496 | 0.000198432 |
| Kpnb1 | 9.92518E-05 | 1.029191713 | 0.717 | 0.547 | 1 | 0.285501358 | -0.145236779 | 0.659 | 0.765 | 1 | 0.285501358 | 0.000198494 |
| Cdc42 | 9.92613E-05 | 0.395847384 | 1 | 0.992 | 1 | 0.027359612 | -0.20899095 | 1 | 1 | 1 | 0.027359612 | 0.000198513 |
| Mob2 | 9.93904E-05 | 0.879088152 | 0.623 | 0.352 | 1 | 0.096435054 | 0.616264563 | 0.659 | 0.608 | 1 | 0.096435054 | 0.000198771 |
| Zbtb11 | 9.96993E-05 | 0.916800475 | 0.868 | 0.686 | 1 | 0.04549555 | 0.512020106 | 0.902 | 0.829 | 1 | 0.04549555 | 0.000199389 |
| Sanbr | 0.000100114 | 1.029191713 | 0.585 | 0.314 | 1 | 0.360175344 | 0.679203016 | 0.415 | 0.392 | 1 | 0.360175344 | 0.000200218 |
| 1700123O20Rik | 0.000100209 | 1.444229212 | 0.491 | 0.25 | 1 | 0.903415073 | 0.136765132 | 0.341 | 0.338 | 1 | 0.903415073 | 0.000200408 |
| Ago4 | 0.000100323 | 1.932330173 | 0.245 | 0.068 | 1 | 0.241509658 | 0.568716014 | 0.195 | 0.126 | 1 | 0.241509658 | 0.000200635 |
| Cd82 | 0.000100383 | -0.801334058 | 0.415 | 0.725 | 1 | 0.383690322 | -0.135209319 | 0.439 | 0.502 | 1 | 0.383690322 | 0.000200757 |
| Nos1ap | 0.000100493 | -2.010977821 | 0.321 | 0.64 | 1 | 0.01523289 | -0.922203794 | 0.585 | 0.775 | 1 | 0.01523289 | 0.000200975 |
| Mrps34 | 0.000100508 | 1.201516806 | 0.547 | 0.301 | 1 | 0.791065463 | 0.216618439 | 0.293 | 0.317 | 1 | 0.791065463 | 0.000201006 |
| Ywhah | 0.182302836 | 0.157499679 | 0.83 | 0.725 | 1 | 0.000100662 | -1.03196608 | 0.683 | 0.881 | 1 | 0.182302836 | 0.000201314 |
| Tpst1 | 0.000101099 | 1.082314099 | 0.642 | 0.394 | 1 | 0.039044512 | -0.701534634 | 0.488 | 0.628 | 1 | 0.039044512 | 0.000202187 |
| Map3k1 | 0.011661841 | 0.644527862 | 0.604 | 0.424 | 1 | 0.000101799 | 1.285409213 | 0.732 | 0.502 | 1 | 0.011661841 | 0.000203587 |
| Pank1 | 0.000220898 | -2.187263152 | 0.283 | 0.521 | 1 | 0.000102186 | -2.607809996 | 0.171 | 0.485 | 1 | 0.000220898 | 0.000204362 |
| Ints13 | 0.000103406 | 1.265753907 | 0.566 | 0.297 | 1 | 0.93641099 | 0.323581131 | 0.39 | 0.444 | 1 | 0.93641099 | 0.000206801 |
| Tmco4 | 0.000104528 | -1.652632327 | 0.189 | 0.479 | 1 | 0.050306256 | -0.639608847 | 0.293 | 0.512 | 1 | 0.050306256 | 0.000209045 |
| Cmpk1 | 0.000104625 | 0.542623964 | 0.925 | 0.818 | 1 | 0.028295683 | -0.484723245 | 0.78 | 0.86 | 1 | 0.028295683 | 0.000209238 |
| Gskip | 0.000104743 | 1.471912771 | 0.434 | 0.208 | 1 | 0.266292648 | -0.321224513 | 0.195 | 0.283 | 1 | 0.266292648 | 0.000209475 |
| Lama3 | 0.72256466 | 3.324647596 | 0.019 | 0.013 | 1 | 0.000106212 | 2.100239256 | 0.171 | 0.031 | 1 | 0.72256466 | 0.000212413 |
| Cemip | 0.000106568 | 1.403686105 | 0.792 | 0.767 | 1 | 0.002310703 | 1.225106219 | 0.805 | 0.782 | 1 | 0.002310703 | 0.000213124 |
| Pcmt1 | 0.000106777 | 0.695290976 | 0.906 | 0.847 | 1 | 0.867278277 | 0.12411353 | 0.878 | 0.932 | 1 | 0.867278277 | 0.000213542 |
| Polr2a | 0.000107297 | 1.497867176 | 0.868 | 0.682 | 1 | 0.109897912 | 0.552690717 | 0.829 | 0.737 | 1 | 0.109897912 | 0.000214583 |
| Pcm1 | 0.000107724 | 0.650803814 | 0.943 | 0.89 | 1 | 0.589697462 | 0.157354401 | 0.976 | 0.956 | 1 | 0.589697462 | 0.000215435 |
| B230217C12Rik | 0.00010859 | 1.569760094 | 0.358 | 0.14 | 1 | 0.027979776 | 1.291236481 | 0.293 | 0.167 | 1 | 0.027979776 | 0.000217167 |
| Itgb5 | 0.022018155 | 0.614154213 | 0.868 | 0.894 | 1 | 0.000109167 | 0.616730906 | 0.976 | 0.918 | 1 | 0.022018155 | 0.000218323 |
| Spag9 | 0.000109351 | 0.413965421 | 1 | 0.987 | 1 | 0.395749992 | 0.156822784 | 1 | 0.99 | 1 | 0.395749992 | 0.00021869 |
| Tgds | 0.000109451 | 1.270199812 | 0.396 | 0.161 | 1 | 0.430828873 | -0.448197369 | 0.171 | 0.218 | 1 | 0.430828873 | 0.000218891 |
| Ubl3 | 0.000109709 | 0.64151972 | 0.906 | 0.809 | 1 | 0.068173421 | 0.467154805 | 0.854 | 0.911 | 1 | 0.068173421 | 0.000219407 |
| Prkacb | 0.000109781 | 0.814298156 | 0.83 | 0.669 | 1 | 0.225958059 | 0.221620733 | 0.805 | 0.799 | 1 | 0.225958059 | 0.000219551 |
| Snrpf | 0.000111154 | 0.569760094 | 0.943 | 0.877 | 1 | 0.007798686 | -0.419628217 | 0.732 | 0.884 | 1 | 0.007798686 | 0.000222296 |
| Cspp1 | 0.000111704 | 0.927055231 | 0.717 | 0.449 | 1 | 0.001554018 | 1.159132945 | 0.805 | 0.652 | 1 | 0.001554018 | 0.000223395 |
| Gtf2h1 | 0.000111775 | 1.057111798 | 0.717 | 0.525 | 1 | 0.557585537 | 0.234964358 | 0.756 | 0.724 | 1 | 0.557585537 | 0.000223537 |
| Gm20682 | 0.000112033 | 2.855162313 | 0.151 | 0.025 | 1 | 0.518969492 | 0.589277336 | 0.024 | 0.048 | 1 | 0.518969492 | 0.000224053 |
| Gm42743 | 0.000112039 | 3.739685096 | 0.151 | 0.025 | 1 | 0.37534514 | 0.614812428 | 0.049 | 0.024 | 1 | 0.37534514 | 0.000224066 |
| Gjb2 | 0.00011223 | 4.500497432 | 0.151 | 0.025 | 1 | 0.840070007 | 0.377773231 | 0.024 | 0.031 | 1 | 0.840070007 | 0.000224448 |
| Smad2 | 0.005379676 | 0.888121574 | 0.66 | 0.517 | 1 | 0.00011251 | 0.73286819 | 0.902 | 0.642 | 1 | 0.005379676 | 0.000225008 |
| Rinl | 0.000113264 | 2.792152515 | 0.151 | 0.025 | 1 | 0.323600424 | -0.16279515 | 0.024 | 0.065 | 1 | 0.323600424 | 0.000226515 |
| Ampd1 | 0.000113492 | 1.407488665 | 0.283 | 0.085 | 1 | 0.389400002 | 0.408361551 | 0.195 | 0.143 | 1 | 0.389400002 | 0.000226972 |
| Zdhhc14 | 0.000114072 | 1.792152515 | 0.264 | 0.081 | 1 | 0.042023429 | 0.667279848 | 0.341 | 0.195 | 1 | 0.042023429 | 0.000228131 |
| Foxo1 | 0.000114099 | -0.732118448 | 0.925 | 0.941 | 1 | 0.000451366 | -0.814807301 | 0.878 | 0.98 | 1 | 0.000451366 | 0.000228186 |
| Clip4 | 0.246096337 | 0.874614676 | 0.075 | 0.038 | 1 | 0.000114889 | 2.915207362 | 0.22 | 0.055 | 1 | 0.246096337 | 0.000229765 |
| Obsl1 | 0.000115531 | 1.087608399 | 0.528 | 0.258 | 1 | 0.799400224 | 0.328191202 | 0.341 | 0.365 | 1 | 0.799400224 | 0.000231049 |
| Zfand6 | 0.000115557 | 0.579557721 | 0.962 | 0.86 | 1 | 0.259010477 | 0.188574155 | 0.976 | 0.928 | 1 | 0.259010477 | 0.000231101 |
| C1qtnf2 | 0.000116357 | 1.953088734 | 0.189 | 0.042 | 1 | 0.980939682 | 0.377773231 | 0.146 | 0.147 | 1 | 0.980939682 | 0.0002327 |
| Ptpn9 | 0.000116522 | 0.823365087 | 0.755 | 0.53 | 1 | 0.093116518 | -0.329192778 | 0.634 | 0.785 | 1 | 0.093116518 | 0.00023303 |
| Pus1 | 0.000116821 | 1.46284489 | 0.377 | 0.148 | 1 | 0.072018119 | 1.867578499 | 0.244 | 0.15 | 1 | 0.072018119 | 0.000233629 |
| Ube2a | 0.00011689 | 1.02495472 | 0.698 | 0.521 | 1 | 0.603397296 | 0.166085705 | 0.707 | 0.689 | 1 | 0.603397296 | 0.000233766 |
| Madd | 0.000117451 | 2.669295768 | 0.151 | 0.025 | 1 | 0.043366051 | 1.904319046 | 0.146 | 0.061 | 1 | 0.043366051 | 0.000234889 |
| Ahcy | 0.000117694 | 2.434830514 | 0.208 | 0.051 | 1 | 0.190007084 | -0.917682652 | 0.024 | 0.082 | 1 | 0.190007084 | 0.000235374 |
| Slf2 | 0.000118384 | 1.434339885 | 0.604 | 0.445 | 1 | 0.620055294 | 0.178241768 | 0.756 | 0.672 | 1 | 0.620055294 | 0.000236754 |

| | | | | | | | | | | | | |
|---|---|---|---|---|---|---|---|---|---|---|---|---|
| Vangl1 | 0.000118723 | 1.406261362 | 0.358 | 0.136 | 1 | 0.000652155 | 1.462165418 | 0.439 | 0.215 | 1 | 0.000652155 | 0.000237433 |
| Pik3r2 | 0.000118888 | 1.680791406 | 0.283 | 0.089 | 1 | 0.736798709 | 0.515276755 | 0.171 | 0.154 | 1 | 0.736798709 | 0.000237762 |
| Ppp6r3 | 0.000119846 | 0.795319794 | 0.849 | 0.712 | 1 | 0.000150146 | 0.796312419 | 0.902 | 0.891 | 1 | 0.000150146 | 0.000239677 |
| Gclc | 0.000170448 | 1.199116714 | 0.679 | 0.424 | 1 | 0.000119972 | 1.283352882 | 0.805 | 0.532 | 1 | 0.000170448 | 0.00023993 |
| Rhobtb3 | 0.000119975 | 1.163134835 | 0.623 | 0.403 | 1 | 0.160122406 | 0.689106211 | 0.537 | 0.461 | 1 | 0.160122406 | 0.000239936 |
| Prkd3 | 0.000120345 | 0.886617199 | 0.642 | 0.356 | 1 | 0.688809832 | 0.274784918 | 0.61 | 0.683 | 1 | 0.688809832 | 0.000240676 |
| Tgfbr3 | 0.000120455 | 0.996639206 | 0.887 | 0.784 | 1 | 0.401438254 | -0.199158551 | 0.829 | 0.918 | 1 | 0.401438254 | 0.000240895 |
| Ube2q1 | 0.000120878 | 1.009524679 | 0.679 | 0.508 | 1 | 0.314385053 | -0.17660095 | 0.707 | 0.727 | 1 | 0.314385053 | 0.000241741 |
| Crtc1 | 0.000121122 | 1.508359549 | 0.321 | 0.11 | 1 | 0.107072626 | -0.875513198 | 0.146 | 0.263 | 1 | 0.107072626 | 0.00024223 |
| Cplane1 | 0.00012114 | 1.253732601 | 0.623 | 0.415 | 1 | 0.189832576 | 0.762657896 | 0.585 | 0.611 | 1 | 0.189832576 | 0.000242265 |
| Stoml3 | 0.000121141 | 2.891688189 | 0.17 | 0.034 | 1 | 0.824812458 | 0.422167351 | 0.049 | 0.041 | 1 | 0.824812458 | 0.000242268 |
| Sgip1 | 0.000121144 | 1.874614676 | 0.264 | 0.081 | 1 | 0.0321406 | 2.83720485 | 0.122 | 0.044 | 1 | 0.0321406 | 0.000242274 |
| Rmnd5a | 0.000121491 | 0.870929629 | 0.811 | 0.61 | 1 | 0.036376445 | 0.524737084 | 0.854 | 0.799 | 1 | 0.036376445 | 0.000242967 |
| Jag1 | 0.028486374 | 1.237184755 | 0.302 | 0.182 | 1 | 0.000121918 | 1.360391153 | 0.561 | 0.283 | 1 | 0.028486374 | 0.000243821 |
| Gm11837 | 0.00012249 | 2.640149422 | 0.17 | 0.034 | 1 | 0.151457207 | 0.589277336 | 0.195 | 0.116 | 1 | 0.151457207 | 0.000244964 |
| Polr3b | 0.172003353 | 0.698657827 | 0.358 | 0.271 | 1 | 0.000122696 | 1.321707169 | 0.61 | 0.338 | 1 | 0.172003353 | 0.000245376 |
| Tmeff2 | 0.003437624 | 2.739685096 | 0.057 | 0.004 | 1 | 0.000122962 | 5.83720485 | 0.073 | 0.003 | 1 | 0.003437624 | 0.000245909 |
| Gm42699 | 0.343006305 | -0.1672055 | 0 | 0.017 | 1 | 0.000122962 | 4.159132945 | 0.073 | 0.003 | 1 | 0.343006305 | 0.000245909 |
| Psme1 | 0.000122999 | 0.840614004 | 0.736 | 0.449 | 1 | 0.197504906 | 0.585141562 | 0.634 | 0.556 | 1 | 0.197504906 | 0.000245983 |
| Prss12 | 0.0003388 | 4.47665069 | 0.075 | 0.004 | 1 | 0.000123325 | 3.515276755 | 0.146 | 0.024 | 1 | 0.0003388 | 0.000246635 |
| Casp4 | 0.000123603 | -1.781147068 | 0.132 | 0.428 | 1 | 0.024647735 | -1.201714139 | 0.122 | 0.283 | 1 | 0.024647735 | 0.00024719 |
| Cflar | 0.000123862 | 1.132354782 | 0.792 | 0.568 | 1 | 0.181529383 | 0.497667355 | 0.756 | 0.771 | 1 | 0.181529383 | 0.000247708 |
| Mbd3 | 0.000123993 | 0.945775705 | 0.698 | 0.504 | 1 | 0.786699703 | 0.443638881 | 0.561 | 0.584 | 1 | 0.786699703 | 0.00024797 |
| Psma4 | 0.000124065 | 0.694455842 | 0.906 | 0.75 | 1 | 0.099069915 | 0.352218473 | 0.878 | 0.863 | 1 | 0.099069915 | 0.000248114 |
| Rab27a | 0.000124278 | -2.14644694 | 0.113 | 0.377 | 1 | 0.18916428 | -0.734337135 | 0.146 | 0.235 | 1 | 0.18916428 | 0.00024854 |
| Slx1b | 0.000124729 | 2.782753817 | 0.17 | 0.034 | 1 | 0.177425897 | 1.515276755 | 0.098 | 0.048 | 1 | 0.177425897 | 0.000249443 |
| Tbx21 | 0.204653189 | 2.154722595 | 0.038 | 0.013 | 1 | 0.000125295 | 3.83720485 | 0.073 | 0.003 | 1 | 0.204653189 | 0.000250575 |
| Gm57220 | 0.246443787 | 2.739685096 | 0.019 | 0.004 | 1 | 0.000125295 | 3.83720485 | 0.073 | 0.003 | 1 | 0.246443787 | 0.000250575 |
| Ccdc69 | 0.500671253 | 1.8327945 | 0.019 | 0.008 | 1 | 0.000125295 | 3.83720485 | 0.073 | 0.003 | 1 | 0.500671253 | 0.000250575 |
| Gm56588 | 0.779443586 | 0.154722595 | 0.019 | 0.025 | 1 | 0.000125295 | 3.83720485 | 0.073 | 0.003 | 1 | 0.779443586 | 0.000250575 |
| A4galt | 0.045149504 | -0.748728917 | 0.509 | 0.627 | 1 | 0.000125336 | -1.00104408 | 0.463 | 0.741 | 1 | 0.045149504 | 0.000250657 |
| Plk2 | 0.01003007 | 0.624989615 | 0.453 | 0.271 | 1 | 0.000125535 | 0.863200058 | 0.463 | 0.198 | 1 | 0.01003007 | 0.000251054 |
| Grcc10 | 0.000125552 | 0.684171884 | 0.925 | 0.826 | 1 | 0.021305198 | -0.413169984 | 0.707 | 0.85 | 1 | 0.021305198 | 0.000251089 |
| Btg2 | 0.002257502 | -2.055439538 | 0.736 | 0.809 | 1 | 0.000125618 | 0.646610511 | 0.854 | 0.57 | 1 | 0.002257502 | 0.00025122 |
| Clk1 | 0.000126572 | 0.649524547 | 1 | 0.987 | 1 | 0.024055902 | 0.602299817 | 0.951 | 0.98 | 1 | 0.024055902 | 0.000253128 |
| Zc3hav1l | 0.559783628 | 0.200526284 | 0.132 | 0.102 | 1 | 0.000126916 | 2.271607674 | 0.293 | 0.096 | 1 | 0.559783628 | 0.000253816 |
| Ldhc | 0.000126938 | 4.242185436 | 0.113 | 0.013 | 1 | 0.17794278 | 1.029849928 | 0.073 | 0.031 | 1 | 0.17794278 | 0.00025386 |
| Mapk1ip1l | 0.505627462 | 0.14867355 | 0.83 | 0.797 | 1 | 0.000127005 | -0.768388984 | 0.707 | 0.939 | 1 | 0.505627462 | 0.000253994 |
| Arhgef28 | 0.000127064 | -1.769951832 | 0.302 | 0.597 | 1 | 0.682747218 | -0.354915898 | 0.439 | 0.44 | 1 | 0.682747218 | 0.000254112 |
| 4932422M17Rik | 0.000127473 | 2.541745718 | 0.189 | 0.042 | 1 | 0.006296252 | 2.396632258 | 0.171 | 0.058 | 1 | 0.006296252 | 0.000254929 |
| Chdh | 0.16240763 | -1.077938162 | 0.075 | 0.144 | 1 | 0.000127617 | -2.455576899 | 0.073 | 0.379 | 1 | 0.16240763 | 0.000255218 |
| Chml | 0.00012962 | 2.541745718 | 0.189 | 0.042 | 1 | 0.732726835 | 0.47463477 | 0.098 | 0.082 | 1 | 0.732726835 | 0.000259223 |
| Larp1b | 0.000130003 | 0.956783217 | 0.736 | 0.475 | 1 | 0.112791671 | 0.326515111 | 0.732 | 0.686 | 1 | 0.112791671 | 0.00025999 |
| Btla | 0.001621069 | 2.356356456 | 0.113 | 0.021 | 1 | 0.000130046 | 3.574170444 | 0.073 | 0.003 | 1 | 0.001621069 | 0.000260076 |
| Mir181a-1hg | 0.099079675 | 2.569760094 | 0.038 | 0.008 | 1 | 0.000130046 | 3.574170444 | 0.073 | 0.003 | 1 | 0.099079675 | 0.000260076 |
| Csde1 | 0.000130219 | 0.469712917 | 0.981 | 0.975 | 1 | 0.259786838 | -0.100638674 | 1 | 0.997 | 1 | 0.259786838 | 0.00026042 |
| Tet3 | 0.000130439 | 0.917683397 | 0.755 | 0.436 | 1 | 0.22142908 | 0.138374385 | 0.829 | 0.802 | 1 | 0.22142908 | 0.000260862 |
| Hes6 | 0.000131588 | 2.533234218 | 0.17 | 0.034 | 1 | 0.528139498 | -0.435813645 | 0.049 | 0.075 | 1 | 0.528139498 | 0.000263159 |
| Pgam1 | 0.029920374 | -0.478633831 | 0.906 | 0.907 | 1 | 0.000131885 | -0.74463155 | 0.732 | 0.904 | 1 | 0.029920374 | 0.000263752 |
| Gm44448 | 0.000333511 | 3.739685096 | 0.075 | 0.004 | 1 | 0.000132474 | 3.252242349 | 0.073 | 0.003 | 1 | 0.000333511 | 0.00026493 |
| Gm47507 | 0.00357432 | 3.09332205 | 0.094 | 0.017 | 1 | 0.000132488 | 2.83720485 | 0.073 | 0.003 | 1 | 0.00357432 | 0.000264958 |
| Frmd8 | 0.000132615 | -1.462661384 | 0.245 | 0.542 | 1 | 0.000242902 | -1.584579482 | 0.39 | 0.638 | 1 | 0.000242902 | 0.000265213 |
| 1110038F14Rik | 0.000132919 | 1.395730694 | 0.453 | 0.216 | 1 | 0.271079535 | 1.054796285 | 0.268 | 0.205 | 1 | 0.271079535 | 0.000265821 |
| P4ha1 | 0.000132963 | -0.504750184 | 0.981 | 0.97 | 1 | 0.182217452 | -0.200420222 | 0.976 | 0.993 | 1 | 0.182217452 | 0.000265908 |
| Psd3 | 0.09014489 | -0.396292574 | 0.925 | 0.958 | 1 | 0.000133071 | -0.6815495 | 0.902 | 0.986 | 1 | 0.09014489 | 0.000266124 |
| Qrich1 | 0.000133131 | 0.942418991 | 0.736 | 0.585 | 1 | 0.000473834 | 0.731711721 | 0.951 | 0.809 | 1 | 0.000473834 | 0.000266244 |
| Fam193b | 0.000134275 | 1.494798036 | 0.472 | 0.233 | 1 | 0.608170464 | -0.171783933 | 0.512 | 0.526 | 1 | 0.608170464 | 0.000268531 |
| Cep162 | 0.000134662 | 1.526691372 | 0.415 | 0.182 | 1 | 0.041957134 | 0.814837037 | 0.366 | 0.222 | 1 | 0.041957134 | 0.000269307 |
| Bend7 | 0.000134971 | 3.533234218 | 0.113 | 0.013 | 1 | 0.580382319 | 1.541748966 | 0.073 | 0.055 | 1 | 0.580382319 | 0.000269923 |
| Tsr3 | 0.000135258 | 1.640149422 | 0.415 | 0.182 | 1 | 0.021119624 | 1.093043754 | 0.341 | 0.198 | 1 | 0.021119624 | 0.000270497 |
| Rbm42 | 0.061357102 | 0.531285946 | 0.642 | 0.53 | 1 | 0.000135314 | 0.82453102 | 0.829 | 0.631 | 1 | 0.061357102 | 0.000270611 |
| Pim3 | 0.955393847 | -0.657590047 | 0.679 | 0.568 | 1 | 0.000135974 | 1.146889349 | 0.683 | 0.427 | 1 | 0.955393847 | 0.00027193 |
| Actr2 | 0.000136456 | 0.615788625 | 0.925 | 0.814 | 1 | 0.04467089 | 0.317618358 | 0.927 | 0.928 | 1 | 0.04467089 | 0.000272894 |
| Mif4gd | 0.000136876 | 1.526691372 | 0.396 | 0.169 | 1 | 0.954752992 | 0.118386602 | 0.195 | 0.191 | 1 | 0.954752992 | 0.000273733 |
| Cep85 | 0.0001372 | -1.556772312 | 0.264 | 0.534 | 1 | 0.002875383 | -1.19841906 | 0.146 | 0.416 | 1 | 0.002875383 | 0.000274382 |
| Pir | 0.000138043 | -2.227747042 | 0.283 | 0.517 | 1 | 0.14395291 | -0.599235126 | 0.341 | 0.464 | 1 | 0.14395291 | 0.000276066 |
| Iars | 0.041778002 | -0.594984668 | 0.491 | 0.653 | 1 | 0.000138138 | -1.060710942 | 0.439 | 0.72 | 1 | 0.041778002 | 0.000276257 |
| Tmem97 | 0.000138353 | -1.322483726 | 0.226 | 0.534 | 1 | 0.063421144 | -0.692048218 | 0.146 | 0.287 | 1 | 0.063421144 | 0.000276686 |
| Epb41l5 | 0.000138583 | 2.58098735 | 0.264 | 0.085 | 1 | 0.000634806 | 1.428399304 | 0.39 | 0.171 | 1 | 0.000634806 | 0.000277147 |
| C1d | 0.715388074 | 0.120234218 | 0.755 | 0.742 | 1 | 0.0001389 | -0.93723545 | 0.537 | 0.754 | 1 | 0.715388074 | 0.00027778 |
| ENSMUSG00000121477 | 0.000138982 | 1.428897558 | 0.34 | 0.123 | 1 | 0.812559456 | 0.324754849 | 0.195 | 0.184 | 1 | 0.812559456 | 0.000277945 |
| Stk38 | 0.000139688 | 1.025892614 | 0.736 | 0.513 | 1 | 0.073719765 | 0.449688413 | 0.756 | 0.7 | 1 | 0.073719765 | 0.000279356 |
| Fer | 0.000139921 | 1.32730205 | 0.66 | 0.428 | 1 | 0.304171522 | 0.149727438 | 0.585 | 0.474 | 1 | 0.304171522 | 0.000279822 |
| Gclm | 0.000139921 | 1.080722013 | 0.717 | 0.5 | 1 | 0.015579832 | 1.001918995 | 0.683 | 0.601 | 1 | 0.015579832 | 0.000279822 |
| Paqr7 | 0.000140031 | 2.533234218 | 0.17 | 0.034 | 1 | 0.003000099 | 1.515276755 | 0.22 | 0.075 | 1 | 0.003000099 | 0.000280042 |
| Inka1 | 0.000140031 | 2.739685096 | 0.17 | 0.034 | 1 | 0.324745165 | -0.622226769 | 0.049 | 0.096 | 1 | 0.324745165 | 0.000280042 |
| Prr12 | 0.000140217 | 1.8327945 | 0.283 | 0.093 | 1 | 0.003865383 | 1.163433082 | 0.366 | 0.177 | 1 | 0.003865383 | 0.000280414 |
| Dlgap4 | 0.796102513 | 0.136249935 | 0.906 | 0.847 | 1 | 0.000140353 | -0.911733386 | 0.805 | 0.959 | 1 | 0.796102513 | 0.000280687 |
| Lrpprc | 0.00014075 | 1.142749953 | 0.547 | 0.309 | 1 | 0.758667236 | 0.252242349 | 0.463 | 0.529 | 1 | 0.758667236 | 0.000281479 |
| Bmp5 | 0.000141267 | -3.034203369 | 0.264 | 0.504 | 1 | 0.187658139 | -0.575754907 | 0.659 | 0.71 | 1 | 0.187658139 | 0.000282515 |
| Gm57009 | 0.000141545 | 3.154722595 | 0.113 | 0.013 | 1 | 0.004444719 | 3.159132945 | 0.073 | 0.01 | 1 | 0.004444719 | 0.00028307 |
| Ewsr1 | 0.00014193 | 0.559252436 | 0.943 | 0.886 | 1 | 0.840358951 | 0.114281118 | 0.951 | 0.993 | 1 | 0.840358951 | 0.00028384 |
| Car12 | 0.000141983 | -4.510613322 | 0.019 | 0.254 | 1 | 0.274575496 | -1.31254227 | 0.049 | 0.102 | 1 | 0.274575496 | 0.000283945 |
| Rps6kb2 | 0.000142829 | 1.402650108 | 0.283 | 0.089 | 1 | 0.343461056 | -0.53202896 | 0.098 | 0.154 | 1 | 0.343461056 | 0.000285637 |
| Mknk1 | 0.000143556 | 1.526691372 | 0.377 | 0.153 | 1 | 0.788031076 | -0.200930279 | 0.244 | 0.263 | 1 | 0.788031076 | 0.000287092 |
| Slc25a36 | 0.042266764 | -0.342593784 | 0.736 | 0.822 | 1 | 0.000143621 | -0.604632709 | 0.585 | 0.887 | 1 | 0.042266764 | 0.000287222 |
| Alg10b | 0.000143624 | 1.289652175 | 0.377 | 0.153 | 1 | 0.329443349 | -0.16279515 | 0.146 | 0.222 | 1 | 0.329443349 | 0.000287227 |
| Klc1 | 0.020214862 | -0.525851818 | 0.66 | 0.826 | 1 | 0.000144435 | -0.669802508 | 0.829 | 0.932 | 1 | 0.020214862 | 0.00028885 |
| Enpep | 0.018027274 | 1.962077517 | 0.094 | 0.025 | 1 | 0.000144814 | 3.178241768 | 0.171 | 0.034 | 1 | 0.018027274 | 0.000289607 |
| H2-DMb1 | 0.000145141 | -1.786115333 | 0.189 | 0.47 | 1 | 0.290977274 | 0.326242931 | 0.195 | 0.304 | 1 | 0.290977274 | 0.00029026 |
| Trim27 | 0.00014518 | 1.23878686 | 0.547 | 0.314 | 1 | 0.008718314 | -0.905759485 | 0.293 | 0.502 | 1 | 0.008718314 | 0.000290338 |
| Atp5a1 | 0.000145337 | 0.512664654 | 1 | 0.941 | 1 | 0.19742937 | 0.240160325 | 1 | 0.98 | 1 | 0.19742937 | 0.000290653 |
| Thap11 | 0.000145937 | 1.507024339 | 0.396 | 0.174 | 1 | 0.157801576 | 0.872537923 | 0.39 | 0.307 | 1 | 0.157801576 | 0.000291853 |
| Epc1 | 0.050803077 | 0.352555363 | 0.906 | 0.839 | 1 | 0.00014627 | 0.617824319 | 1 | 0.935 | 1 | 0.050803077 | 0.000292519 |
| Zbtb8a | 0.000147081 | 2.962077517 | 0.113 | 0.013 | 1 | 0.969250601 | 0.667279848 | 0.049 | 0.051 | 1 | 0.969250601 | 0.00029414 |
| Gnai2 | 0.000147185 | 0.412520352 | 1 | 0.97 | 1 | 0.29523817 | 0.147933033 | 1 | 0.983 | 1 | 0.29523817 | 0.000294348 |
| Ing2 | 0.000147624 | 1.119098685 | 0.604 | 0.377 | 1 | 0.026939938 | 0.646262067 | 0.561 | 0.406 | 1 | 0.026939938 | 0.000295227 |
| N4bp1 | 0.000148194 | 1.388313223 | 0.623 | 0.407 | 1 | 0.33339383 | 0.223775694 | 0.805 | 0.724 | 1 | 0.33339383 | 0.000296365 |

| | | | | | | | | | | | | |
|---|---|---|---|---|---|---|---|---|---|---|---|---|
| Ust | 0.000148346 | 0.519218485 | 0.925 | 0.733 | 1 | 0.524278055 | -0.164726038 | 0.951 | 0.949 | 1 | 0.524278055 | 0.000296669 |
| Klf11 | 0.000149085 | 1.417757001 | 0.434 | 0.199 | 1 | 0.852820223 | 0.429120111 | 0.293 | 0.294 | 1 | 0.852820223 | 0.000298149 |
| Tsc22d2 | 0.000149761 | 0.28715015 | 1 | 0.852 | 1 | 0.001199578 | 0.711267861 | 0.854 | 0.898 | 1 | 0.001199578 | 0.000299499 |
| Ubap2l | 0.000632897 | 0.472962503 | 0.962 | 0.945 | 1 | 0.000149815 | 0.503602222 | 1 | 0.973 | 1 | 0.000632897 | 0.000299608 |
| Synj2 | 0.000150408 | 1.763932642 | 0.396 | 0.174 | 1 | 0.146739151 | 0.974708374 | 0.366 | 0.29 | 1 | 0.146739151 | 0.000300792 |
| Ppp1r9a | 0.000150643 | 3.870929629 | 0.113 | 0.013 | 1 | 0.077878427 | -1.440779897 | 0.049 | 0.154 | 1 | 0.077878427 | 0.000301264 |
| Slc13a4 | 0.000150691 | 2.920257341 | 0.113 | 0.013 | 1 | 0.126426307 | -2.20718927 | 0 | 0.055 | 1 | 0.126426307 | 0.000301359 |
| Thoc2 | 0.000151768 | 0.697677569 | 0.925 | 0.932 | 1 | 0.090764109 | -0.278272368 | 0.927 | 0.945 | 1 | 0.090764109 | 0.000303514 |
| Col4a4 | 0.000152284 | -4.776014743 | 0 | 0.225 | 1 | 0.052571469 | -2.589059905 | 0 | 0.085 | 1 | 0.052571469 | 0.000304546 |
| Cd47 | 0.000152629 | 0.738689103 | 0.943 | 0.801 | 1 | 0.00350392 | 0.647497236 | 0.927 | 0.877 | 1 | 0.00350392 | 0.000305234 |
| Epas1 | 0.000152708 | 1.28449047 | 0.66 | 0.411 | 1 | 0.002397833 | 1.008971198 | 0.707 | 0.587 | 1 | 0.002397833 | 0.000305392 |
| Terf1 | 0.000152768 | 1.412520352 | 0.491 | 0.263 | 1 | 0.792769332 | 0.469229387 | 0.366 | 0.379 | 1 | 0.792769332 | 0.000305513 |
| Adam22 | 0.000152891 | 2.254258268 | 0.189 | 0.042 | 1 | 0.456359026 | 0.649577847 | 0.146 | 0.109 | 1 | 0.456359026 | 0.000305758 |
| Ralbp1 | 0.000152895 | 0.815920681 | 0.962 | 0.784 | 1 | 0.335637488 | 0.168826341 | 0.927 | 0.918 | 1 | 0.335637488 | 0.000305767 |
| Alms1 | 0.000153039 | 1.535812762 | 0.321 | 0.119 | 1 | 0.06432467 | 0.895307804 | 0.244 | 0.133 | 1 | 0.06432467 | 0.000306054 |
| Gpha2 | 0.000153478 | 1.092603694 | 0.698 | 0.432 | 1 | 0.057108598 | 0.871440993 | 0.61 | 0.505 | 1 | 0.057108598 | 0.000306933 |
| Kif3a | 0.000153724 | 1.757387097 | 0.377 | 0.165 | 1 | 0.00036702 | 1.749742009 | 0.415 | 0.188 | 1 | 0.00036702 | 0.000307424 |
| Corin | 0.015102354 | -2.56774343 | 0.057 | 0.191 | 1 | 0.000154056 | -5.448197369 | 0 | 0.273 | 1 | 0.015102354 | 0.000308088 |
| Jcad | 0.000154504 | 1.179813576 | 0.415 | 0.174 | 1 | 0.017225916 | 0.972859949 | 0.415 | 0.256 | 1 | 0.017225916 | 0.000308984 |
| Zmiz2 | 0.000154945 | 2.211306123 | 0.264 | 0.085 | 1 | 0.073278263 | 1.132660733 | 0.341 | 0.232 | 1 | 0.073278263 | 0.000309866 |
| St3gal2 | 0.000155009 | 1.513803688 | 0.547 | 0.288 | 1 | 0.406369434 | 0.540811847 | 0.488 | 0.505 | 1 | 0.406369434 | 0.000309994 |
| Mid2 | 0.159195127 | -0.519701533 | 0.585 | 0.665 | 1 | 0.000155152 | -1.225356488 | 0.415 | 0.717 | 1 | 0.159195127 | 0.00031028 |
| Bcl11b | 0.001908644 | 3.739685096 | 0.075 | 0.008 | 1 | 0.000155694 | 3.458693227 | 0.171 | 0.034 | 1 | 0.001908644 | 0.000311363 |
| Sqor | 0.034503322 | -1.093204919 | 0.113 | 0.246 | 1 | 0.000156531 | -2.777504994 | 0.049 | 0.338 | 1 | 0.034503322 | 0.000313037 |
| Map2k5 | 0.000156793 | 1.104096522 | 0.642 | 0.377 | 1 | 0.313559271 | -0.285722672 | 0.61 | 0.672 | 1 | 0.313559271 | 0.000313561 |
| Nox4 | 0.00032444 | 4.595295186 | 0.208 | 0.059 | 1 | 0.000156969 | 8.046658215 | 0.049 | 0 | 1 | 0.00032444 | 0.000313912 |
| Slc5a9 | 0.00285365 | 3.739685096 | 0.038 | 0 | 1 | 0.000156969 | 4.422167351 | 0.049 | 0 | 1 | 0.00285365 | 0.000313912 |
| Fbxo41 | 0.003243872 | 3.47665069 | 0.057 | 0.004 | 1 | 0.000156969 | 4.422167351 | 0.049 | 0 | 1 | 0.003243872 | 0.000313912 |
| Tlr13 | 0.003243872 | 3.47665069 | 0.057 | 0.004 | 1 | 0.000156969 | 4.422167351 | 0.049 | 0 | 1 | 0.003243872 | 0.000313912 |
| Smco3 | 0.00343733 | 2.569760094 | 0.057 | 0.004 | 1 | 0.000156969 | 4.83720485 | 0.049 | 0 | 1 | 0.00343733 | 0.000313912 |
| Hgf | 0.030360955 | 2.739685096 | 0.038 | 0.004 | 1 | 0.000156969 | 4.422167351 | 0.049 | 0 | 1 | 0.030360955 | 0.000313912 |
| Flt4 | 0.030360955 | 2.739685096 | 0.038 | 0.004 | 1 | 0.000156969 | 4.422167351 | 0.049 | 0 | 1 | 0.030360955 | 0.000313912 |
| Tymp | 0.030360955 | 2.739685096 | 0.038 | 0.004 | 1 | 0.000156969 | 4.83720485 | 0.049 | 0 | 1 | 0.030360955 | 0.000313912 |
| Gm57377 | 0.03562064 | 3.154722595 | 0.019 | 0 | 1 | 0.000156969 | 4.422167351 | 0.049 | 0 | 1 | 0.03562064 | 0.000313912 |
| Gm8113 | 0.03562064 | 3.154722595 | 0.019 | 0 | 1 | 0.000156969 | 6.007129851 | 0.049 | 0 | 1 | 0.03562064 | 0.000313912 |
| Gm56759 | 0.03562064 | 3.739685096 | 0.019 | 0 | 1 | 0.000156969 | 4.422167351 | 0.049 | 0 | 1 | 0.03562064 | 0.000313912 |
| I830134H01Rik | 0.03562064 | 3.154722595 | 0.019 | 0 | 1 | 0.000156969 | 4.83720485 | 0.049 | 0 | 1 | 0.03562064 | 0.000313912 |
| Pgm5 | 0.078387299 | 1.984797593 | 0.094 | 0.038 | 1 | 0.000156969 | 4.83720485 | 0.049 | 0 | 1 | 0.078387299 | 0.000313912 |
| Trim5 | 0.085835472 | 2.640149422 | 0.057 | 0.017 | 1 | 0.000156969 | 6.007129851 | 0.049 | 0 | 1 | 0.085835472 | 0.000313912 |
| Cd160 | 0.092905716 | 1.569760094 | 0.057 | 0.017 | 1 | 0.000156969 | 4.83720485 | 0.049 | 0 | 1 | 0.092905716 | 0.000313912 |
| Osr1 | 0.249034808 | 2.154722595 | 0.019 | 0.004 | 1 | 0.000156969 | 7.881598969 | 0.049 | 0 | 1 | 0.249034808 | 0.000313912 |
| Tm4sf4 | 0.251644813 | 0.8327945 | 0.019 | 0.004 | 1 | 0.000156969 | 5.159132945 | 0.049 | 0 | 1 | 0.251644813 | 0.000313912 |
| 2810030D12Rik | 0.336520503 | 1.569760094 | 0.038 | 0.017 | 1 | 0.000156969 | 4.422167351 | 0.049 | 0 | 1 | 0.336520503 | 0.000313912 |
| Armc3 | 0.343006305 | -0.1672055 | 0 | 0.017 | 1 | 0.000156969 | 5.422167351 | 0.049 | 0 | 1 | 0.343006305 | 0.000313912 |
| Gm57357 | 0.413080549 | -0.1672055 | 0 | 0.013 | 1 | 0.000156969 | 5.159132945 | 0.049 | 0 | 1 | 0.413080549 | 0.000313912 |
| C130071C03Rik | 0.413080549 | -0.430239906 | 0 | 0.013 | 1 | 0.000156969 | 4.422167351 | 0.049 | 0 | 1 | 0.413080549 | 0.000313912 |
| Ntrk3 | 0.497380598 | 2.891688189 | 0.019 | 0.008 | 1 | 0.000156969 | 5.422167351 | 0.049 | 0 | 1 | 0.497380598 | 0.000313912 |
| Cdh6 | 0.503971666 | 1.569760094 | 0.019 | 0.008 | 1 | 0.000156969 | 4.422167351 | 0.049 | 0 | 1 | 0.503971666 | 0.000313912 |
| Ifi211 | 0.507281812 | 0.347367673 | 0.019 | 0.008 | 1 | 0.000156969 | 4.83720485 | 0.049 | 0 | 1 | 0.507281812 | 0.000313912 |
| 4930556N13Rik | 0.736073999 | 0.8327945 | 0.019 | 0.013 | 1 | 0.000156969 | 4.422167351 | 0.049 | 0 | 1 | 0.736073999 | 0.000313912 |
| Arpc2 | 0.000157212 | 0.519780245 | 1 | 0.966 | 1 | 0.11783105 | -0.186973688 | 1 | 1 | 1 | 0.11783105 | 0.0003144 |
| Rap1a | 0.670106829 | -0.102958577 | 0.943 | 0.953 | 1 | 0.000157422 | -0.72058745 | 0.976 | 0.98 | 1 | 0.670106829 | 0.000314819 |
| Wdr70 | 0.000157485 | 0.779083913 | 0.811 | 0.653 | 1 | 0.293045075 | 0.512531378 | 0.732 | 0.734 | 1 | 0.293045075 | 0.000314944 |
| Chaserr | 0.000158388 | 0.733355493 | 0.906 | 0.805 | 1 | 0.000303572 | 0.537778753 | 0.951 | 0.891 | 1 | 0.000303572 | 0.000316751 |
| Ppp2r5a | 0.007787643 | 0.629164381 | 0.83 | 0.742 | 1 | 0.000159746 | 0.5733037 | 0.976 | 0.918 | 1 | 0.007787643 | 0.000319466 |
| Cpt1a | 0.000159882 | 1.725879296 | 0.321 | 0.119 | 1 | 0.059661922 | 0.796790581 | 0.537 | 0.396 | 1 | 0.059661922 | 0.000319738 |
| Cct6a | 0.00016011 | 0.542383655 | 0.962 | 0.915 | 1 | 0.007267118 | -0.49132598 | 0.829 | 0.939 | 1 | 0.007267118 | 0.000320194 |
| Pex1 | 0.000160156 | 1.513176566 | 0.321 | 0.114 | 1 | 0.705848648 | 0.608386159 | 0.171 | 0.147 | 1 | 0.705848648 | 0.000320287 |
| Car8 | 0.000305075 | -1.216036263 | 0.283 | 0.606 | 1 | 0.000161342 | -1.0550017 | 0.488 | 0.761 | 1 | 0.000305075 | 0.000322659 |
| Maco1 | 0.000161469 | 1.008173931 | 0.811 | 0.627 | 1 | 0.146837591 | 0.42692088 | 0.78 | 0.754 | 1 | 0.146837591 | 0.000322913 |
| Gipc1 | 0.000161907 | 1.669295768 | 0.415 | 0.208 | 1 | 0.366963225 | 0.660882077 | 0.366 | 0.307 | 1 | 0.366963225 | 0.000323787 |
| Obi1 | 0.000161941 | 1.417757001 | 0.34 | 0.127 | 1 | 0.441003736 | 0.784093513 | 0.122 | 0.191 | 1 | 0.441003736 | 0.000323855 |
| Morf4l1 | 0.000162037 | 0.344844154 | 1 | 0.983 | 1 | 0.001350687 | -0.329977385 | 1 | 0.997 | 1 | 0.001350687 | 0.000324047 |
| Msantd3 | 0.000162053 | 1.844382474 | 0.321 | 0.119 | 1 | 0.194811807 | -0.701953961 | 0.146 | 0.235 | 1 | 0.194811807 | 0.00032408 |
| Phex | 0.00016395 | -1.067669827 | 0.113 | 0.013 | 1 | 0.170414866 | -1.622226769 | 0 | 0.044 | 1 | 0.170414866 | 0.000327874 |
| Dmac2 | 0.000164258 | 1.544669113 | 0.302 | 0.102 | 1 | 0.375356876 | -0.292078167 | 0.049 | 0.092 | 1 | 0.375356876 | 0.000328488 |
| Fbxl22 | 0.00016459 | 2.154722595 | 0.17 | 0.034 | 1 | 0.805509886 | 0.252242349 | 0.073 | 0.085 | 1 | 0.805509886 | 0.000329153 |
| Nmt1 | 0.000165757 | 0.819538403 | 0.698 | 0.453 | 1 | 0.067277607 | 0.83720485 | 0.537 | 0.454 | 1 | 0.067277607 | 0.000331486 |
| Ppp2r2d | 0.647231601 | 0.261637799 | 0.698 | 0.712 | 1 | 0.000166571 | -0.868270458 | 0.732 | 0.887 | 1 | 0.647231601 | 0.000333115 |
| Ppp2r1b | 0.000167087 | 1.434830514 | 0.396 | 0.174 | 1 | 0.150794569 | 0.80158094 | 0.366 | 0.287 | 1 | 0.150794569 | 0.000334146 |
| Api5 | 0.000167244 | 0.924424975 | 0.792 | 0.614 | 1 | 0.766882137 | 0.414233169 | 0.78 | 0.795 | 1 | 0.766882137 | 0.000334461 |
| E2f6 | 0.000167738 | 1.523956404 | 0.377 | 0.153 | 1 | 0.980952361 | 0.278237558 | 0.366 | 0.392 | 1 | 0.980952361 | 0.000335448 |
| Slc22a1 | 0.000168668 | 2.039245377 | 0.189 | 0.042 | 1 | 0.002533587 | 1.88967227 | 0.195 | 0.061 | 1 | 0.002533587 | 0.000337307 |
| Nptxr | 0.105352359 | 1.526691372 | 0.132 | 0.068 | 1 | 0.000170815 | 2.059597271 | 0.293 | 0.096 | 1 | 0.105352359 | 0.000341602 |
| Hectd4 | 0.000170896 | 1.427741089 | 0.528 | 0.305 | 1 | 0.217319699 | 0.781351615 | 0.61 | 0.57 | 1 | 0.217319699 | 0.000341762 |
| Ddit4 | 0.000731678 | 0.962077517 | 0.358 | 0.157 | 1 | 0.000172753 | 2.434658295 | 0.488 | 0.242 | 1 | 0.000731678 | 0.000345475 |
| Bola3 | 0.000173977 | -0.955195097 | 0.321 | 0.631 | 1 | 0.095558811 | -0.484723245 | 0.366 | 0.491 | 1 | 0.095558811 | 0.000347924 |
| Nsd2 | 0.000174231 | 0.888582278 | 0.66 | 0.475 | 1 | 0.199333896 | 0.369765653 | 0.732 | 0.631 | 1 | 0.199333896 | 0.000348432 |
| Cmtm5 | 0.000174436 | -1.924228747 | 0.113 | 0.381 | 1 | 0.059517352 | -1.010792057 | 0.049 | 0.167 | 1 | 0.059517352 | 0.000348841 |
| Mindy2 | 0.003031801 | 0.67104343 | 0.623 | 0.411 | 1 | 0.000175825 | 1.257108104 | 0.756 | 0.549 | 1 | 0.003031801 | 0.00035162 |
| Smoc2 | 0.000176205 | 0.507122411 | 0.887 | 0.669 | 1 | 0.000805433 | -1.347683785 | 0.756 | 0.802 | 1 | 0.000805433 | 0.000352379 |
| Lratd2 | 0.000176263 | 1.758793918 | 0.358 | 0.153 | 1 | 0.068428497 | -0.80665134 | 0.195 | 0.334 | 1 | 0.068428497 | 0.000352495 |
| Cwc27 | 0.000177172 | 0.821878722 | 0.679 | 0.398 | 1 | 0.335446076 | -0.43439157 | 0.415 | 0.526 | 1 | 0.335446076 | 0.000354312 |
| Zfp606 | 0.000178478 | 1.417757001 | 0.34 | 0.123 | 1 | 0.214721655 | -0.863234868 | 0.122 | 0.198 | 1 | 0.214721655 | 0.000356924 |
| Zbp1 | 0.000178786 | 1.948271717 | 0.17 | 0.034 | 1 | 0.008158264 | 1.547698233 | 0.22 | 0.089 | 1 | 0.008158264 | 0.00035754 |
| Tmem245 | 0.000180437 | 1.280253477 | 0.547 | 0.322 | 1 | 0.913862091 | 0.226886775 | 0.512 | 0.536 | 1 | 0.913862091 | 0.000360842 |
| Mcm6 | 0.000180663 | -1.035739338 | 0.264 | 0.564 | 1 | 0.01573178 | -0.913223004 | 0.244 | 0.447 | 1 | 0.01573178 | 0.000361293 |
| Lsmem1 | 0.111857689 | 1.177442671 | 0.226 | 0.144 | 1 | 0.000181069 | 1.553898049 | 0.488 | 0.218 | 1 | 0.111857689 | 0.000362105 |
| Gbp7 | 0.010985144 | 1.427741089 | 0.189 | 0.076 | 1 | 0.000181322 | 1.342440158 | 0.171 | 0.034 | 1 | 0.010985144 | 0.000362612 |
| Bzw1 | 0.33941284 | -0.190992667 | 1 | 0.983 | 1 | 0.00018137 | -0.613840266 | 1 | 0.997 | 1 | 0.33941284 | 0.000362707 |
| Commd8 | 0.000182213 | 1.411062348 | 0.491 | 0.258 | 1 | 0.158352308 | 1.017025887 | 0.366 | 0.3 | 1 | 0.158352308 | 0.000364393 |
| Plekha6 | 0.000182414 | -2.827130058 | 0.038 | 0.284 | 1 | 0.000391553 | 1.578470581 | 0.39 | 0.167 | 1 | 0.000391553 | 0.000364795 |
| Cers4 | 0.000208729 | 1.923397049 | 0.226 | 0.064 | 1 | 0.000182992 | 1.904319046 | 0.317 | 0.109 | 1 | 0.000208729 | 0.00036595 |
| Fmc1 | 0.000183376 | 1.086009845 | 0.547 | 0.297 | 1 | 0.227674133 | 0.261892519 | 0.244 | 0.379 | 1 | 0.227674133 | 0.000366717 |
| Tead3 | 0.000183534 | 1.289652175 | 0.396 | 0.161 | 1 | 0.717381728 | 0.477308905 | 0.171 | 0.205 | 1 | 0.717381728 | 0.000367035 |
| Zfp644 | 0.000185141 | 0.77921346 | 0.868 | 0.695 | 1 | 0.038884839 | 0.491876322 | 0.927 | 0.85 | 1 | 0.038884839 | 0.000370248 |
| Smc6 | 0.00018551 | 0.655035075 | 0.849 | 0.661 | 1 | 0.520465775 | -0.152020679 | 0.756 | 0.775 | 1 | 0.520465775 | 0.000370986 |
| Tshz1 | 0.000186877 | 0.773632427 | 0.717 | 0.453 | 1 | 0.049613377 | 0.791690191 | 0.659 | 0.553 | 1 | 0.049613377 | 0.000373719 |

| | | | | | | | | | | | | |
|---|---|---|---|---|---|---|---|---|---|---|---|---|
| Ino80b | 0.000186906 | 1.806799291 | 0.321 | 0.123 | 1 | 0.542292038 | 0.136765132 | 0.098 | 0.133 | 1 | 0.542292038 | 0.000373777 |
| Plekhb1 | 0.000187188 | -1.944011359 | 0.208 | 0.458 | 1 | 0.010759036 | -2.132421501 | 0.049 | 0.215 | 1 | 0.010759036 | 0.000374341 |
| Ash2l | 0.000187472 | 1.29992051 | 0.509 | 0.263 | 1 | 0.75478588 | 0.368435367 | 0.317 | 0.369 | 1 | 0.75478588 | 0.000374908 |
| Prkcz | 0.003136839 | -1.186314323 | 0.17 | 0.386 | 1 | 0.000188602 | -2.646610927 | 0.049 | 0.341 | 1 | 0.003136839 | 0.000377169 |
| Usf1 | 0.000188719 | 1.549001534 | 0.34 | 0.131 | 1 | 0.03610326 | 0.72678086 | 0.415 | 0.259 | 1 | 0.03610326 | 0.000377403 |
| Tmem213 | 0.246443787 | 2.739685096 | 0.019 | 0.004 | 1 | 0.000189521 | -2.579441902 | 0.073 | 0.372 | 1 | 0.246443787 | 0.000379007 |
| Dyrk1a | 0.000193105 | 0.595849323 | 0.925 | 0.877 | 1 | 0.039072726 | 0.316851431 | 0.927 | 0.966 | 1 | 0.039072726 | 0.000386174 |
| Scml4 | 0.00019316 | 1.557481764 | 0.283 | 0.093 | 1 | 0.625890398 | 0.109284395 | 0.171 | 0.208 | 1 | 0.625890398 | 0.000386283 |
| Mapkapk5 | 0.000194157 | 0.949047568 | 0.623 | 0.381 | 1 | 0.101945539 | 0.597738915 | 0.585 | 0.519 | 1 | 0.101945539 | 0.000388277 |
| Ier5 | 0.596621318 | -0.108123359 | 0.906 | 0.86 | 1 | 0.000196386 | 0.657619737 | 0.951 | 0.778 | 1 | 0.596621318 | 0.000392734 |
| Oaf | 0.316506079 | -0.416735642 | 0.887 | 0.814 | 1 | 0.000196757 | -1.114662655 | 0.634 | 0.809 | 1 | 0.316506079 | 0.000393475 |
| Acat2 | 0.000198531 | -2.185127408 | 0.113 | 0.373 | 1 | 0.572519843 | -0.217242934 | 0.171 | 0.215 | 1 | 0.572519843 | 0.000397022 |
| Tafa2 | 0.00019939 | -4.680275082 | 0.038 | 0.275 | 1 | 0.126451063 | -4.174022406 | 0 | 0.055 | 1 | 0.126451063 | 0.000398741 |
| Arf2 | 0.000200681 | -1.169457952 | 0.642 | 0.784 | 1 | 0.260920844 | -0.574221396 | 0.659 | 0.7 | 1 | 0.260920844 | 0.000401322 |
| Cct2 | 0.000201827 | 0.60001431 | 0.943 | 0.928 | 1 | 0.173182057 | 0.174239837 | 0.976 | 0.952 | 1 | 0.173182057 | 0.000403614 |
| Pde4c | 0.000203141 | 1.932330173 | 0.208 | 0.051 | 1 | 0.123575441 | 0.422167351 | 0.098 | 0.041 | 1 | 0.123575441 | 0.000406241 |
| Cenpb | 0.000203348 | 1.10862618 | 0.66 | 0.419 | 1 | 0.150009954 | 0.437673937 | 0.707 | 0.706 | 1 | 0.150009954 | 0.000406655 |
| Ppp6r1 | 0.000203951 | 1.318221327 | 0.396 | 0.169 | 1 | 0.327813339 | -0.128579435 | 0.22 | 0.307 | 1 | 0.327813339 | 0.00040786 |
| Ier2 | 0.000204265 | 0.543480769 | 0.868 | 0.72 | 1 | 0.178896607 | 0.276128574 | 0.854 | 0.846 | 1 | 0.178896607 | 0.000408487 |
| Malt1 | 0.000205068 | 1.389187848 | 0.283 | 0.093 | 1 | 0.050477573 | 0.763204268 | 0.244 | 0.13 | 1 | 0.050477573 | 0.000410094 |
| Dnajc21 | 0.000205628 | 1.332365509 | 0.547 | 0.36 | 1 | 0.179706634 | 0.494812652 | 0.683 | 0.604 | 1 | 0.179706634 | 0.000411214 |
| Zfp287 | 0.00020676 | 1.46284489 | 0.208 | 0.051 | 1 | 0.479886681 | 0.707921833 | 0.049 | 0.082 | 1 | 0.479886681 | 0.000413476 |
| Mlph | 0.000207763 | -4.237594828 | 0.019 | 0.246 | 1 | 0.026595776 | -3.67058759 | 0 | 0.109 | 1 | 0.026595776 | 0.000415483 |
| Lipa | 0.000207821 | 1.669295768 | 0.415 | 0.212 | 1 | 0.629584356 | 0.61113677 | 0.317 | 0.29 | 1 | 0.629584356 | 0.000415599 |
| 2410004B18Rik | 0.000207952 | 1.017219071 | 0.566 | 0.314 | 1 | 0.553124275 | 0.260855479 | 0.341 | 0.423 | 1 | 0.553124275 | 0.00041586 |
| Gm15337 | 0.08698757 | 1.261637799 | 0.094 | 0.038 | 1 | 0.000208156 | 2.401105735 | 0.171 | 0.034 | 1 | 0.08698757 | 0.000416269 |
| Adcy3 | 0.000208329 | 1.406261362 | 0.358 | 0.14 | 1 | 0.624767614 | -0.101394606 | 0.244 | 0.276 | 1 | 0.624767614 | 0.000416614 |
| Psme3ip1 | 0.000209701 | 1.08568095 | 0.566 | 0.322 | 1 | 0.096859598 | 0.600165653 | 0.512 | 0.41 | 1 | 0.096859598 | 0.000419359 |
| Dennd4a | 0.000376934 | 0.569760094 | 0.623 | 0.398 | 1 | 0.000209799 | 1.116640863 | 0.78 | 0.543 | 1 | 0.000376934 | 0.000419555 |
| Cramp1 | 0.000210654 | 1.397579119 | 0.547 | 0.335 | 1 | 0.000466544 | 1.118042078 | 0.78 | 0.611 | 1 | 0.000466544 | 0.000421263 |
| Fbxl5 | 0.000211104 | -0.954288027 | 0.453 | 0.733 | 1 | 0.001352112 | -0.913095383 | 0.512 | 0.747 | 1 | 0.001352112 | 0.000422163 |
| Cers5 | 0.000211697 | 0.921408887 | 0.717 | 0.534 | 1 | 0.349246371 | 0.338398993 | 0.756 | 0.765 | 1 | 0.349246371 | 0.00042335 |
| Lrba | 0.000211938 | -1.228606045 | 0.679 | 0.835 | 1 | 0.005498831 | -0.701333314 | 0.659 | 0.843 | 1 | 0.005498831 | 0.000423831 |
| Gm527 | 0.000212082 | 2.33529484 | 0.208 | 0.055 | 1 | 0.227183914 | -0.899760744 | 0.049 | 0.109 | 1 | 0.227183914 | 0.00042412 |
| Bpifc | 0.000212459 | -4.474634025 | 0 | 0.216 | 1 | 0.154156177 | -1.747757651 | 0 | 0.048 | 1 | 0.154156177 | 0.000424873 |
| Apof | 0.00021254 | -4.304709024 | 0 | 0.216 | 1 | 0.5191986 | 0.515276755 | 0 | 0.01 | 1 | 0.5191986 | 0.000425036 |
| Man2a2 | 0.000213685 | 1.648831665 | 0.453 | 0.25 | 1 | 0.821479027 | 0.580865097 | 0.293 | 0.314 | 1 | 0.821479027 | 0.000427325 |
| Vrk2 | 0.000214685 | 1.569760094 | 0.302 | 0.106 | 1 | 0.671279014 | -0.188330242 | 0.22 | 0.188 | 1 | 0.671279014 | 0.000429323 |
| Msh6 | 0.000214789 | 1.865215978 | 0.245 | 0.072 | 1 | 0.901160611 | 0.995902596 | 0.122 | 0.119 | 1 | 0.901160611 | 0.000429533 |
| Tssc4 | 0.000217069 | 1.017219071 | 0.528 | 0.292 | 1 | 0.565155538 | 0.165827597 | 0.317 | 0.399 | 1 | 0.565155538 | 0.000434091 |
| Sgk3 | 0.02555531 | -0.671248005 | 0.66 | 0.792 | 1 | 0.000217905 | -1.091613585 | 0.683 | 0.857 | 1 | 0.02555531 | 0.000435763 |
| Vps26b | 0.0002182 | 1.865215978 | 0.264 | 0.085 | 1 | 0.660051918 | 0.567115686 | 0.244 | 0.225 | 1 | 0.660051918 | 0.000436353 |
| Nr4a2 | 0.000218304 | 0.82417201 | 0.868 | 0.636 | 1 | 0.000750705 | 1.200453047 | 0.902 | 0.713 | 1 | 0.000750705 | 0.00043656 |
| Mbd5 | 0.000219497 | 0.603908316 | 0.906 | 0.788 | 1 | 0.629722185 | 0.119836693 | 0.878 | 0.908 | 1 | 0.629722185 | 0.000438945 |
| Gm49980 | 0.000222139 | 2.47665069 | 0.208 | 0.055 | 1 | 0.019562991 | 1.408361551 | 0.22 | 0.099 | 1 | 0.019562991 | 0.000444229 |
| Gfra1 | 0.000223034 | -2.575670345 | 0.189 | 0.449 | 1 | 0.093661861 | -1.080956858 | 0.293 | 0.42 | 1 | 0.093661861 | 0.000446017 |
| Ap2s1 | 0.000223826 | 0.921408887 | 0.774 | 0.581 | 1 | 0.975474311 | 0.145009126 | 0.61 | 0.659 | 1 | 0.975474311 | 0.000447601 |
| Agap1 | 0.000224674 | 0.696517236 | 0.792 | 0.513 | 1 | 0.002271742 | 0.474295193 | 0.927 | 0.747 | 1 | 0.002271742 | 0.000449298 |
| Gorasp2 | 0.071511599 | -0.288541821 | 0.906 | 0.919 | 1 | 0.000226564 | -0.675686236 | 0.878 | 0.939 | 1 | 0.071511599 | 0.000453077 |
| Wdr82 | 0.000227057 | 1.059144935 | 0.642 | 0.411 | 1 | 0.09562235 | 0.816143234 | 0.634 | 0.519 | 1 | 0.09562235 | 0.000454062 |
| Syap1 | 0.000227086 | 0.79075928 | 0.774 | 0.691 | 1 | 0.310322019 | -0.221688839 | 0.732 | 0.823 | 1 | 0.310322019 | 0.000454121 |
| Hpgd | 0.000227199 | -2.308638291 | 0.396 | 0.623 | 1 | 0.619885513 | 0.895449991 | 0.39 | 0.461 | 1 | 0.619885513 | 0.000454346 |
| Ctbp2 | 0.000227348 | 1.154722595 | 0.509 | 0.263 | 1 | 0.993624516 | 0.123509035 | 0.366 | 0.386 | 1 | 0.993624516 | 0.000454645 |
| Elavl1 | 0.000228115 | 0.693188433 | 0.887 | 0.864 | 1 | 0.181632848 | 0.405051192 | 0.951 | 0.956 | 1 | 0.181632848 | 0.000456177 |
| Usp13 | 0.000228851 | 2.242185436 | 0.208 | 0.055 | 1 | 0.368279971 | -0.562725757 | 0.098 | 0.157 | 1 | 0.368279971 | 0.00045765 |
| Ddx21 | 0.000228858 | 0.769732232 | 0.887 | 0.737 | 1 | 0.48968265 | 0.215405581 | 0.829 | 0.843 | 1 | 0.48968265 | 0.000457665 |
| Hid1 | 0.000228867 | 3.891688189 | 0.094 | 0.008 | 1 | 0.000711925 | 1.944120054 | 0.122 | 0.02 | 1 | 0.000711925 | 0.000457682 |
| Ccdc126 | 0.000228875 | 1.544669113 | 0.283 | 0.093 | 1 | 0.188272713 | 0.930314254 | 0.195 | 0.123 | 1 | 0.188272713 | 0.000457698 |
| Itprid1 | 0.732691937 | 0.932330173 | 0.019 | 0.013 | 1 | 0.000231155 | 2.83720485 | 0.122 | 0.017 | 1 | 0.732691937 | 0.000462257 |
| Cracdl | 0.002547209 | -2.814903756 | 0.038 | 0.212 | 1 | 0.000232346 | -2.636726339 | 0.146 | 0.43 | 1 | 0.002547209 | 0.000464638 |
| Slc36a4 | 0.000233337 | 1.154722595 | 0.547 | 0.314 | 1 | 0.080518958 | -0.567549676 | 0.317 | 0.468 | 1 | 0.080518958 | 0.000466619 |
| Ube2e1 | 0.00023335 | 0.876737847 | 0.811 | 0.678 | 1 | 0.528851444 | 0.246317515 | 0.78 | 0.778 | 1 | 0.528851444 | 0.000466645 |
| Fmo1 | 0.725936481 | 1.417757001 | 0.019 | 0.013 | 1 | 0.000233822 | 2.667279848 | 0.122 | 0.017 | 1 | 0.725936481 | 0.00046759 |
| Tufm | 0.000234051 | 1.450178478 | 0.377 | 0.157 | 1 | 0.044877143 | -1.466575898 | 0.073 | 0.201 | 1 | 0.044877143 | 0.000468047 |
| Gm36129 | 0.249034808 | 2.154722595 | 0.019 | 0.004 | 1 | 0.000234983 | 4.007129851 | 0.098 | 0.01 | 1 | 0.249034808 | 0.00046991 |
| Gm4673 | 0.000235029 | 3.739685096 | 0.094 | 0.008 | 1 | 0.7415112 | 0.334704509 | 0.049 | 0.038 | 1 | 0.7415112 | 0.000470002 |
| Pdlim7 | 0.009232395 | 0.627863049 | 0.604 | 0.453 | 1 | 0.0002373 | 0.985727375 | 0.683 | 0.451 | 1 | 0.009232395 | 0.000474543 |
| Flywch1 | 0.000237321 | 1.470224421 | 0.396 | 0.178 | 1 | 0.06696576 | 0.896098539 | 0.366 | 0.246 | 1 | 0.06696576 | 0.000474585 |
| Srsf1 | 0.000238664 | 0.680791406 | 0.868 | 0.669 | 1 | 0.057267626 | -0.197984579 | 0.707 | 0.826 | 1 | 0.057267626 | 0.00047727 |
| Tesc | 0.000238738 | -3.889671525 | 0.057 | 0.288 | 1 | 0.15356791 | -1.970150072 | 0.024 | 0.089 | 1 | 0.15356791 | 0.000477419 |
| Asf1a | 0.00023887 | 1.438515561 | 0.491 | 0.28 | 1 | 0.078302874 | 0.920728205 | 0.439 | 0.328 | 1 | 0.078302874 | 0.000477684 |
| Gpr45 | 0.000239073 | 2.891688189 | 0.132 | 0.021 | 1 | 0.055146445 | 2.185128153 | 0.098 | 0.034 | 1 | 0.055146445 | 0.000478088 |
| Polr2h | 0.00023992 | 1.199116714 | 0.566 | 0.343 | 1 | 0.151662048 | 0.413178567 | 0.488 | 0.375 | 1 | 0.151662048 | 0.000479783 |
| Ablim1 | 0.000240208 | -3.852771942 | 0.038 | 0.267 | 1 | 0.227909888 | 0.311948595 | 0.39 | 0.304 | 1 | 0.227909888 | 0.000480359 |
| Htr6 | 0.000241309 | 3.739685096 | 0.094 | 0.008 | 1 | 0.013354269 | 1.83720485 | 0.073 | 0.014 | 1 | 0.013354269 | 0.00048256 |
| Zfp853 | 0.000241309 | 3.569760094 | 0.094 | 0.008 | 1 | 0.438672228 | 1.83720485 | 0.024 | 0.01 | 1 | 0.438672228 | 0.00048256 |
| Gm15929 | 0.000241309 | 3.377115016 | 0.094 | 0.008 | 1 | 0.605005904 | 1.029849928 | 0.024 | 0.014 | 1 | 0.605005904 | 0.00048256 |
| Igdcc3 | 0.000241309 | 3.377115016 | 0.094 | 0.008 | 1 | 0.51919986 | -0.484723245 | 0 | 0.01 | 1 | 0.51919986 | 0.00048256 |
| Ggt5 | 0.000241309 | 3.377115016 | 0.094 | 0.008 | 1 | 0.433272494 | 2.100239256 | 0.024 | 0.01 | 1 | 0.433272494 | 0.00048256 |
| Psmc4 | 0.000241515 | 0.927807615 | 0.698 | 0.559 | 1 | 0.23803913 | 0.278784137 | 0.732 | 0.696 | 1 | 0.23803913 | 0.000482971 |
| Drosha | 0.00024209 | 1.055186921 | 0.509 | 0.263 | 1 | 0.534716083 | -0.121216046 | 0.293 | 0.348 | 1 | 0.534716083 | 0.000484121 |
| Lgmn | 0.000242741 | 1.012500522 | 0.717 | 0.504 | 1 | 0.072457363 | 0.980162804 | 0.512 | 0.451 | 1 | 0.072457363 | 0.000485422 |
| Rabep1 | 0.000244663 | 0.70005572 | 0.887 | 0.788 | 1 | 0.393890376 | -0.20784163 | 0.78 | 0.884 | 1 | 0.393890376 | 0.000489266 |
| Abca8a | 0.000244739 | 3.533234218 | 0.094 | 0.008 | 1 | 0.115836913 | 1.83720485 | 0.049 | 0.014 | 1 | 0.115836913 | 0.000489418 |
| Tmem63a | 0.181218935 | -0.231694227 | 0.396 | 0.496 | 1 | 0.000245301 | -0.919405274 | 0.366 | 0.676 | 1 | 0.181218935 | 0.000490542 |
| Efnb1 | 0.000246647 | 1.01270359 | 0.547 | 0.301 | 1 | 0.61446334 | 0.439327902 | 0.39 | 0.369 | 1 | 0.61446334 | 0.000493234 |
| Ptpn2 | 0.000247105 | -0.777832884 | 0.698 | 0.86 | 1 | 0.218954379 | -0.289691574 | 0.756 | 0.846 | 1 | 0.218954379 | 0.000494149 |
| Txnl4b | 0.000248058 | 3.154722595 | 0.094 | 0.008 | 1 | 0.231606831 | -0.863234868 | 0 | 0.034 | 1 | 0.231606831 | 0.000496055 |
| Fbxl16 | 0.000248058 | 3.154722595 | 0.094 | 0.008 | 1 | 0.824088343 | 0.377773231 | 0.024 | 0.031 | 1 | 0.824088343 | 0.000496055 |
| G3bp1 | 0.000248669 | 0.615983111 | 0.943 | 0.903 | 1 | 0.062948879 | 0.217596206 | 0.951 | 0.908 | 1 | 0.062948879 | 0.000497276 |
| Vash2 | 0.000249805 | 4.47665069 | 0.057 | 0 | 1 | 0.006468242 | 2.614812428 | 0.098 | 0.02 | 1 | 0.006468242 | 0.000499548 |
| Drc1 | 0.000249805 | 4.47665069 | 0.057 | 0 | 1 | 0.181471053 | 2.422167351 | 0.049 | 0.017 | 1 | 0.181471053 | 0.000499548 |
| Gm15704 | 0.000249805 | 4.739685096 | 0.057 | 0 | 1 | 0.402373368 | 0.252242349 | 0 | 0.017 | 1 | 0.402373368 | 0.000499548 |
| Tm6sf1 | 0.000249805 | 4.47665069 | 0.057 | 0 | 1 | 0.402369942 | -0.332720152 | 0 | 0.017 | 1 | 0.402369942 | 0.000499548 |
| Lat | 0.000249805 | 4.739685096 | 0.057 | 0 | 1 | 0.455094184 | 0.252242349 | 0 | 0.014 | 1 | 0.455094184 | 0.000499548 |
| 4930524B15Rik | 0.000249805 | 4.47665069 | 0.057 | 0 | 1 | 0.455097252 | -0.16279515 | 0 | 0.014 | 1 | 0.455097252 | 0.000499548 |
| Rem2 | 0.000249805 | 4.47665069 | 0.057 | 0 | 1 | 0.115836913 | 1.83720485 | 0.049 | 0.014 | 1 | 0.115836913 | 0.000499548 |
| Gm16253 | 0.00024982 | 4.154722595 | 0.057 | 0 | 1 | 0.007718533 | 3.83720485 | 0.024 | 0 | 1 | 0.007718533 | 0.000499577 |

| | | | | | | | | | | | | |
|---|---|---|---|---|---|---|---|---|---|---|---|---|
| Gm57092 | 0.00024982 | 4.154722595 | 0.057 | 0 | 1 | 0.319629309 | -0.622226769 | 0 | 0.024 | 1 | 0.319629309 | 0.000499577 |
| Otud7a | 0.00024982 | 4.154722595 | 0.057 | 0 | 1 | 0.455098275 | 0.515276755 | 0 | 0.014 | 1 | 0.455098275 | 0.000499577 |
| B3gnt6 | 0.00024982 | 4.154722595 | 0.057 | 0 | 1 | 0.986995563 | 0.83720485 | 0.024 | 0.024 | 1 | 0.986995563 | 0.000499577 |
| Tecta | 0.00024982 | 5.47665069 | 0.057 | 0 | 1 | 0.004604972 | 2.83720485 | 0.073 | 0.01 | 1 | 0.004604972 | 0.000499577 |
| Hic1 | 0.00024982 | 4.154722595 | 0.057 | 0 | 1 | 0.021167579 | 2.83720485 | 0.049 | 0.007 | 1 | 0.021167579 | 0.000499577 |
| Icam2 | 0.00024982 | 4.739685096 | 0.057 | 0 | 1 | 0.878310931 | 0.377773231 | 0.024 | 0.02 | 1 | 0.878310931 | 0.000499577 |
| Mycn | 0.00024982 | 4.154722595 | 0.057 | 0 | 1 | 0.004769158 | 2.515276755 | 0.073 | 0.01 | 1 | 0.004769158 | 0.000499577 |
| Gm47882 | 0.00024982 | 4.154722595 | 0.057 | 0 | 1 | 0.438672228 | 1.83720485 | 0.024 | 0.01 | 1 | 0.438672228 | 0.000499577 |
| 9130008F23Rik | 0.00024982 | 4.154722595 | 0.057 | 0 | 1 | 0.319619815 | -0.484723245 | 0 | 0.024 | 1 | 0.319619815 | 0.000499577 |
| Gm19696 | 0.00024982 | 4.154722595 | 0.057 | 0 | 1 | 0.262674569 | 2.83720485 | 0.024 | 0.007 | 1 | 0.262674569 | 0.000499577 |
| Bex1 | 0.00024982 | 4.154722595 | 0.057 | 0 | 1 | 0.112404454 | 2.252242349 | 0.049 | 0.014 | 1 | 0.112404454 | 0.000499577 |
| Man1a2 | 0.000249871 | 0.700280128 | 0.868 | 0.788 | 1 | 0.007893769 | 0.631344152 | 0.902 | 0.826 | 1 | 0.007893769 | 0.00049968 |
| Pcyt2 | 0.000250178 | -1.331461751 | 0.396 | 0.614 | 1 | 0.721922831 | -0.141421499 | 0.39 | 0.399 | 1 | 0.721922831 | 0.000500294 |
| Sh2b1 | 0.000250609 | 1.489141634 | 0.377 | 0.165 | 1 | 0.062375338 | -0.661046018 | 0.146 | 0.29 | 1 | 0.062375338 | 0.000501155 |
| Ppp1r12b | 0.000254138 | -1.074716476 | 0.585 | 0.818 | 1 | 0.07569702 | -0.663811084 | 0.829 | 0.819 | 1 | 0.07569702 | 0.000508212 |
| Pkia | 0.000254697 | 3.002719501 | 0.094 | 0.008 | 1 | 0.00032942 | 2.730289646 | 0.146 | 0.027 | 1 | 0.00032942 | 0.000509329 |
| Rassf9 | 0.03562064 | 3.739685096 | 0.019 | 0 | 1 | 0.000254855 | 4.296636468 | 0.098 | 0.01 | 1 | 0.03562064 | 0.000509645 |
| Bmf | 0.001339323 | 2.8327945 | 0.132 | 0.03 | 1 | 0.000255721 | 1.786578777 | 0.244 | 0.068 | 1 | 0.001339323 | 0.000511377 |
| Ncor1 | 0.000255798 | 0.407563718 | 1 | 0.983 | 1 | 0.040359146 | 0.295414183 | 1 | 1 | 1 | 0.040359146 | 0.00051153 |
| Atp8a2 | 0.000256146 | 2.029191713 | 0.321 | 0.127 | 1 | 0.222529479 | 0.483567895 | 0.268 | 0.184 | 1 | 0.222529479 | 0.000512226 |
| Dhx36 | 0.000256202 | 0.715242954 | 0.83 | 0.75 | 1 | 0.450931929 | 0.44893426 | 0.732 | 0.771 | 1 | 0.450931929 | 0.000512338 |
| Mfsd11 | 0.000256446 | 1.18386894 | 0.491 | 0.267 | 1 | 0.100524948 | 1.049425466 | 0.463 | 0.396 | 1 | 0.100524948 | 0.000512826 |
| Gyg | 0.000256672 | 2.154722595 | 0.208 | 0.055 | 1 | 0.981361638 | 0.515276755 | 0.098 | 0.099 | 1 | 0.981361638 | 0.000513278 |
| Phtf2 | 0.000256781 | 0.883309771 | 0.774 | 0.576 | 1 | 0.702459023 | -0.18336222 | 0.951 | 0.846 | 1 | 0.702459023 | 0.000513496 |
| Spen | 0.000257526 | 0.843014342 | 0.906 | 0.716 | 1 | 0.199301917 | 0.320548362 | 0.927 | 0.884 | 1 | 0.199301917 | 0.000514987 |
| 5031425E22Rik | 0.000258234 | -0.95121085 | 0.679 | 0.864 | 1 | 0.422460271 | 0.128903014 | 0.732 | 0.857 | 1 | 0.422460271 | 0.000516402 |
| Usp36 | 0.000258501 | 0.974150349 | 0.717 | 0.547 | 1 | 0.925874948 | 0.211019686 | 0.707 | 0.771 | 1 | 0.925874948 | 0.000516935 |
| Mt2 | 0.012728729 | -2.819567286 | 0.358 | 0.458 | 1 | 0.000258773 | 1.439381906 | 0.537 | 0.283 | 1 | 0.012728729 | 0.00051748 |
| Psip1 | 0.000260033 | 0.66802411 | 0.868 | 0.754 | 1 | 0.041871818 | 0.572642974 | 0.805 | 0.771 | 1 | 0.041871818 | 0.000519998 |
| Lsm14b | 0.000260499 | 1.8327945 | 0.283 | 0.097 | 1 | 0.179150634 | -0.555112573 | 0.073 | 0.157 | 1 | 0.179150634 | 0.00052093 |
| Pdcl | 0.000260883 | 1.235341055 | 0.509 | 0.28 | 1 | 0.407783151 | 0.398093216 | 0.366 | 0.314 | 1 | 0.407783151 | 0.000521697 |
| Ip6k2 | 0.000371015 | 0.991223863 | 0.547 | 0.275 | 1 | 0.000261015 | 1.551802631 | 0.61 | 0.386 | 1 | 0.000371015 | 0.000521963 |
| Sbspon | 0.000261541 | -3.551255307 | 0.075 | 0.309 | 1 | 0.103194337 | -1.192542494 | 0.073 | 0.174 | 1 | 0.103194337 | 0.000523014 |
| Clp1 | 0.17760897 | 0.407488665 | 0.321 | 0.237 | 1 | 0.000261846 | 1.696342314 | 0.439 | 0.218 | 1 | 0.17760897 | 0.000523624 |
| Stat2 | 0.176052822 | 0.75033234 | 0.491 | 0.436 | 1 | 0.000262157 | 1.695185845 | 0.488 | 0.273 | 1 | 0.176052822 | 0.000524246 |
| Smap2 | 0.145836737 | 0.415726465 | 0.66 | 0.576 | 1 | 0.00026237 | -1.178061907 | 0.634 | 0.788 | 1 | 0.145836737 | 0.000524671 |
| Uhrf2 | 0.000263959 | 1.051756016 | 0.83 | 0.716 | 1 | 0.00964495 | 0.79281073 | 0.829 | 0.829 | 1 | 0.00964495 | 0.000527848 |
| Slc35a2 | 0.000264276 | -0.966292806 | 0.453 | 0.682 | 1 | 0.103049563 | -0.445729113 | 0.39 | 0.543 | 1 | 0.103049563 | 0.000528482 |
| Carmil2 | 0.000264902 | 2.739685096 | 0.094 | 0.008 | 1 | 0.537088415 | 1.252242349 | 0.049 | 0.031 | 1 | 0.537088415 | 0.000529734 |
| Ptprj | 0.000265043 | -1.412782709 | 0.623 | 0.746 | 1 | 0.526318879 | 0.119891313 | 0.61 | 0.59 | 1 | 0.526318879 | 0.000530016 |
| Zmynd8 | 0.017274034 | 0.805573031 | 0.698 | 0.593 | 1 | 0.000265158 | 0.962735732 | 0.829 | 0.618 | 1 | 0.017274034 | 0.000530245 |
| Gm29683 | 0.003209552 | 3.002719501 | 0.094 | 0.017 | 1 | 0.000265506 | 3.322631677 | 0.098 | 0.01 | 1 | 0.003209552 | 0.000530941 |
| Cxcl1 | 0.813437117 | 0.72941676 | 0.151 | 0.144 | 1 | 0.000265506 | 3.685201756 | 0.098 | 0.01 | 1 | 0.813437117 | 0.000530941 |
| Actn4 | 0.000267116 | -0.668055495 | 1 | 0.983 | 1 | 0.00193746 | -0.479792246 | 0.951 | 1 | 1 | 0.00193746 | 0.000534161 |
| Fancm | 0.000267303 | 1.386896037 | 0.34 | 0.131 | 1 | 0.010001169 | 1.396632258 | 0.366 | 0.208 | 1 | 0.010001169 | 0.000534534 |
| Ipo9 | 0.000268495 | 1.036541168 | 0.491 | 0.237 | 1 | 0.345772481 | 0.515276755 | 0.512 | 0.451 | 1 | 0.345772481 | 0.000536917 |
| Ank | 0.001637005 | -1.126771223 | 0.943 | 0.975 | 1 | 0.000268595 | -1.029233702 | 0.951 | 0.976 | 1 | 0.001637005 | 0.000537119 |
| Etv3 | 0.000268964 | 1.214428841 | 0.453 | 0.229 | 1 | 0.730778266 | 0.446970742 | 0.366 | 0.372 | 1 | 0.730778266 | 0.000537856 |
| Nup155 | 0.000270006 | 1.214428841 | 0.434 | 0.208 | 1 | 0.641385793 | 0.263213467 | 0.244 | 0.304 | 1 | 0.641385793 | 0.000539939 |
| Afmid | 0.000270392 | 1.94321849 | 0.245 | 0.076 | 1 | 0.04640285 | 1.317830691 | 0.22 | 0.113 | 1 | 0.04640285 | 0.000540711 |
| Car5b | 0.00027184 | -2.049149739 | 0.113 | 0.36 | 1 | 0.243915756 | -0.795063366 | 0.073 | 0.14 | 1 | 0.243915756 | 0.000543607 |
| Mb | 0.000273585 | 2.891688189 | 0.151 | 0.03 | 1 | 0.396999105 | 1.635570989 | 0.073 | 0.044 | 1 | 0.396999105 | 0.000547096 |
| Fam72a | 0.000275654 | 2.261637799 | 0.189 | 0.047 | 1 | 0.479272663 | 1.071670103 | 0.073 | 0.048 | 1 | 0.479272663 | 0.000551231 |
| Slfn8 | 0.000275832 | 2.154722595 | 0.094 | 0.008 | 1 | 0.44410834 | 1.252242349 | 0.024 | 0.01 | 1 | 0.44410834 | 0.000551587 |
| Slc39a13 | 0.313725244 | -0.364150715 | 0.642 | 0.682 | 1 | 0.000276579 | -1.16824958 | 0.366 | 0.672 | 1 | 0.313725244 | 0.000553081 |
| Gpd2 | 0.000277078 | -0.889037791 | 0.491 | 0.712 | 1 | 0.004993501 | -0.821006633 | 0.683 | 0.829 | 1 | 0.004993501 | 0.00055408 |
| Agpat4 | 0.000277902 | -1.022007585 | 0.679 | 0.86 | 1 | 0.000371842 | -1.20718927 | 0.61 | 0.816 | 1 | 0.000371842 | 0.000555727 |
| Cdc42ep1 | 0.000277957 | 1.953088734 | 0.264 | 0.089 | 1 | 0.781042988 | 0.486707603 | 0.122 | 0.147 | 1 | 0.781042988 | 0.000555837 |
| Sugt1 | 0.000278316 | 0.666531983 | 0.906 | 0.763 | 1 | 0.886218576 | 0.362365163 | 0.805 | 0.85 | 1 | 0.886218576 | 0.000556555 |
| Galnt17 | 0.000279269 | 0.695290976 | 0.132 | 0.021 | 1 | 0.65086545 | -0.970150072 | 0.049 | 0.034 | 1 | 0.65086545 | 0.000558459 |
| Tex15 | 0.000279565 | 2.395730694 | 0.094 | 0.008 | 1 | 0.752578711 | 0.252242349 | 0.024 | 0.017 | 1 | 0.752578711 | 0.000559053 |
| Mtrf1l | 0.000279734 | 1.598329246 | 0.358 | 0.157 | 1 | 0.214797524 | -0.764831164 | 0.122 | 0.201 | 1 | 0.214797524 | 0.00055939 |
| Pvt1 | 0.000280956 | -1.717402583 | 0.17 | 0.453 | 1 | 0.000892711 | -2.341282165 | 0.146 | 0.41 | 1 | 0.000892711 | 0.000561833 |
| Abtb3 | 0.00028132 | 1.447276088 | 0.472 | 0.242 | 1 | 0.002691146 | 1.546908364 | 0.463 | 0.256 | 1 | 0.002691146 | 0.000562561 |
| Nrd1 | 0.000282923 | 0.771149865 | 0.906 | 0.831 | 1 | 0.03018565 | 0.521300543 | 0.927 | 0.881 | 1 | 0.03018565 | 0.000565766 |
| Ipo11 | 0.000284316 | 1.424811758 | 0.396 | 0.178 | 1 | 0.018134213 | -1.332720152 | 0.146 | 0.321 | 1 | 0.018134213 | 0.000568552 |
| Ube2d3 | 0.138231069 | 0.217439667 | 1 | 0.992 | 1 | 0.000285345 | -0.458921026 | 0.976 | 0.997 | 1 | 0.138231069 | 0.000570608 |
| Slc4a1ap | 0.00028579 | 1.124134275 | 0.585 | 0.386 | 1 | 0.329174583 | 0.758427736 | 0.415 | 0.416 | 1 | 0.329174583 | 0.000571498 |
| Pogk | 0.000287663 | 1.391761792 | 0.415 | 0.195 | 1 | 0.587234882 | -0.246211158 | 0.268 | 0.324 | 1 | 0.587234882 | 0.000575244 |
| Kmt2d | 0.000288548 | 0.89946554 | 0.623 | 0.403 | 1 | 0.800847525 | 0.139688527 | 0.756 | 0.737 | 1 | 0.800847525 | 0.000577012 |
| Ptges3 | 0.000290412 | 0.415313825 | 1 | 0.979 | 1 | 0.762414108 | 0.157914967 | 1 | 0.99 | 1 | 0.762414108 | 0.000580739 |
| Socs3 | 0.000531959 | 1.080974255 | 0.679 | 0.492 | 1 | 0.000291751 | 1.567115686 | 0.707 | 0.495 | 1 | 0.000531959 | 0.000583416 |
| Syt16 | 0.003572946 | 2.154722595 | 0.094 | 0.017 | 1 | 0.000291909 | 2.667279848 | 0.098 | 0.01 | 1 | 0.003572946 | 0.000583733 |
| Pstpip1 | 0.000293747 | -2.600164907 | 0.038 | 0.267 | 1 | 0.103233319 | -1.224571348 | 0.146 | 0.27 | 1 | 0.103233319 | 0.000587408 |
| Gtf3c1 | 0.000293865 | 1.27081966 | 0.66 | 0.449 | 1 | 0.791005164 | 0.124817278 | 0.78 | 0.737 | 1 | 0.791005164 | 0.000587643 |
| Rnf157 | 0.000295303 | 1.817687608 | 0.208 | 0.055 | 1 | 0.823562977 | -0.20718927 | 0.073 | 0.082 | 1 | 0.823562977 | 0.000590518 |
| Tm9sf3 | 0.000295364 | 0.424043263 | 1 | 0.966 | 1 | 0.706529948 | 0.128428021 | 0.976 | 0.997 | 1 | 0.706529948 | 0.00059064 |
| Rnpepl1 | 0.000295755 | 1.517292674 | 0.358 | 0.148 | 1 | 0.196301777 | -0.410722664 | 0.122 | 0.215 | 1 | 0.196301777 | 0.000591422 |
| Zfp180 | 0.000295887 | 2.039245377 | 0.264 | 0.089 | 1 | 1 | 0.220533489 | 0.195 | 0.198 | 1 | 1 | 0.000591686 |
| Slc30a7 | 0.000295995 | 1.154722595 | 0.585 | 0.369 | 1 | 0.460438244 | 0.246908905 | 0.561 | 0.498 | 1 | 0.460438244 | 0.000591903 |
| Afdn | 0.000296741 | 0.875200066 | 0.811 | 0.606 | 1 | 0.021333576 | 0.50593889 | 0.878 | 0.768 | 1 | 0.021333576 | 0.000593394 |
| Peak1 | 0.000297346 | 0.599619383 | 0.792 | 0.564 | 1 | 0.301644549 | 0.682546605 | 0.585 | 0.543 | 1 | 0.301644549 | 0.000594604 |
| Bclaf3 | 0.000297435 | 0.908082627 | 0.585 | 0.322 | 1 | 0.223895483 | 0.712509102 | 0.537 | 0.485 | 1 | 0.223895483 | 0.000594781 |
| Dcaf6 | 0.000298843 | 0.61658595 | 0.868 | 0.665 | 1 | 0.649000259 | -0.317553704 | 0.878 | 0.833 | 1 | 0.649000259 | 0.000597597 |
| Manbal | 0.000298904 | 1.050385935 | 0.604 | 0.411 | 1 | 0.031479616 | -0.622226769 | 0.293 | 0.485 | 1 | 0.031479616 | 0.000597719 |
| Grk5 | 0.000299107 | -1.841351702 | 0.604 | 0.742 | 1 | 0.041644984 | -1.341916645 | 0.683 | 0.741 | 1 | 0.041644984 | 0.000598124 |
| Garem1 | 0.000300613 | -1.949614065 | 0.321 | 0.585 | 1 | 0.007554271 | -1.560672098 | 0.439 | 0.597 | 1 | 0.007554271 | 0.000601135 |
| Ctnnd1 | 0.001141368 | 0.872406356 | 0.868 | 0.801 | 1 | 0.000302298 | 0.67865891 | 0.927 | 0.908 | 1 | 0.001141368 | 0.000604505 |
| Pdrg1 | 0.000302732 | -0.899547068 | 0.623 | 0.805 | 1 | 0.004299589 | -0.754611322 | 0.39 | 0.635 | 1 | 0.004299589 | 0.000605373 |
| Cul3 | 0.000303308 | 0.549407032 | 0.981 | 0.919 | 1 | 0.937127372 | 0.191413455 | 0.976 | 0.983 | 1 | 0.937127372 | 0.000606524 |
| Mical2 | 0.020638591 | 2.361173472 | 0.151 | 0.059 | 1 | 0.000304626 | 1.313642894 | 0.293 | 0.096 | 1 | 0.020638591 | 0.000609159 |
| Mta3 | 0.000304934 | 1.377115016 | 0.491 | 0.263 | 1 | 0.407077195 | -0.121488176 | 0.39 | 0.488 | 1 | 0.407077195 | 0.000609776 |
| Srcap | 0.000309642 | 0.544928241 | 0.83 | 0.585 | 1 | 0.119566991 | 0.317197791 | 0.902 | 0.826 | 1 | 0.119566991 | 0.000619187 |
| Arfgef1 | 0.000310619 | 0.545794509 | 0.943 | 0.826 | 1 | 0.196847538 | -0.183143796 | 0.927 | 0.952 | 1 | 0.196847538 | 0.000621142 |
| Slfn2 | 0.207551231 | 2.891688189 | 0.038 | 0.013 | 1 | 0.000312129 | 1.574170444 | 0.098 | 0.01 | 1 | 0.207551231 | 0.00062416 |
| Gm12258 | 0.000312776 | 2.47665069 | 0.151 | 0.03 | 1 | 0.05398095 | 1.377773231 | 0.073 | 0.02 | 1 | 0.05398095 | 0.000625453 |
| Cyp2d22 | 0.000312776 | 2.614154213 | 0.151 | 0.03 | 1 | 0.003442778 | 1.911205431 | 0.171 | 0.051 | 1 | 0.003442778 | 0.000625453 |
| Tpi1 | 0.015470372 | 0.552166034 | 0.906 | 0.733 | 1 | 0.000313262 | 0.903577872 | 0.829 | 0.805 | 1 | 0.015470372 | 0.000626425 |

| | | | | | | | | | | | | |
|---|---|---|---|---|---|---|---|---|---|---|---|---|
| Eif3b | 0.000313574 | 1.004305752 | 0.66 | 0.508 | 1 | 0.3590287 | -0.14099078 | 0.634 | 0.683 | 1 | 0.3590287 | 0.000627049 |
| B230369F24Rik | 0.00031555 | 1.85638132 | 0.34 | 0.14 | 1 | 0.002476104 | 1.874875268 | 0.463 | 0.263 | 1 | 0.002476104 | 0.000631001 |
| Atosa | 0.000316253 | 1.254718385 | 0.604 | 0.415 | 1 | 0.00120468 | 1.37874507 | 0.707 | 0.491 | 1 | 0.00120468 | 0.000632405 |
| Pcdh17 | 0.142553045 | 0.947678138 | 0.34 | 0.258 | 1 | 0.00031763 | 1.764448507 | 0.463 | 0.215 | 1 | 0.142553045 | 0.000635158 |
| Nr4a3 | 0.00031773 | 0.762405172 | 0.491 | 0.246 | 1 | 0.002826071 | 1.272764673 | 0.61 | 0.451 | 1 | 0.002826071 | 0.000635359 |
| Eya1 | 0.000318682 | 0.485630821 | 0.962 | 0.877 | 1 | 0.021863018 | 0.656673444 | 1 | 0.959 | 1 | 0.021863018 | 0.000637262 |
| Dlat | 0.000319632 | 1.329401126 | 0.528 | 0.322 | 1 | 0.876367444 | 0.252242349 | 0.488 | 0.509 | 1 | 0.876367444 | 0.000639161 |
| Ctsl | 0.000320114 | 0.319859357 | 0.943 | 0.886 | 1 | 0.000545647 | 0.559557316 | 0.976 | 0.959 | 1 | 0.000545647 | 0.000640126 |
| Ilk | 0.000320181 | 0.823313447 | 0.736 | 0.589 | 1 | 0.660306699 | 0.247712691 | 0.561 | 0.58 | 1 | 0.660306699 | 0.00064026 |
| Sec11c | 0.000323413 | -0.927359058 | 0.679 | 0.775 | 1 | 0.033531456 | -0.521481795 | 0.634 | 0.782 | 1 | 0.033531456 | 0.000646722 |
| Cth | 0.000323558 | -2.035101964 | 0.132 | 0.403 | 1 | 0.473646246 | -0.53202896 | 0.073 | 0.109 | 1 | 0.473646246 | 0.000647011 |
| Kcnk5 | 0.000323702 | -1.216725636 | 0.547 | 0.695 | 1 | 0.002696342 | -0.833730874 | 0.39 | 0.669 | 1 | 0.002696342 | 0.000647299 |
| Hipk2 | 0.000324211 | -1.371297105 | 0.755 | 0.826 | 1 | 0.995810349 | -0.177523487 | 0.805 | 0.771 | 1 | 0.995810349 | 0.000648316 |
| Rbmx2 | 0.000324754 | 2.047807391 | 0.189 | 0.047 | 1 | 0.307268086 | -0.555112573 | 0.024 | 0.065 | 1 | 0.307268086 | 0.000649402 |
| Zfat | 0.000325295 | 1.133027524 | 0.377 | 0.157 | 1 | 0.870634236 | 0.1903146 | 0.22 | 0.249 | 1 | 0.870634236 | 0.000650485 |
| Aco2 | 0.000327218 | 0.721212435 | 0.792 | 0.631 | 1 | 0.282974344 | 0.101214379 | 0.805 | 0.771 | 1 | 0.282974344 | 0.00065433 |
| Paxbp1 | 0.000327303 | 0.93417152 | 0.736 | 0.568 | 1 | 0.034163512 | 0.554069238 | 0.829 | 0.72 | 1 | 0.034163512 | 0.000654499 |
| Pigs | 0.000327311 | -1.253362144 | 0.189 | 0.462 | 1 | 0.877462623 | 0.296636468 | 0.317 | 0.307 | 1 | 0.877462623 | 0.000654514 |
| Tcim | 0.000328224 | 2.533234218 | 0.151 | 0.03 | 1 | 0.00035412 | 2.231483789 | 0.293 | 0.102 | 1 | 0.00035412 | 0.000656339 |
| Rem1 | 0.000328465 | 3.962077517 | 0.075 | 0.004 | 1 | 0.60213981 | 1.252242349 | 0.024 | 0.014 | 1 | 0.60213981 | 0.000656823 |
| Chd5 | 0.000328465 | 3.962077517 | 0.075 | 0.004 | 1 | 0.564654905 | 0.721727632 | 0.073 | 0.051 | 1 | 0.564654905 | 0.000656823 |
| Mmp12 | 0.000328465 | 3.962077517 | 0.075 | 0.004 | 1 | 0.5191986 | 0.252242349 | 0 | 0.01 | 1 | 0.5191986 | 0.000656823 |
| Ephb1 | 0.000328465 | 5.242185436 | 0.075 | 0.004 | 1 | 0.005433091 | 2.879025025 | 0.122 | 0.031 | 1 | 0.005433091 | 0.000656823 |
| Wnt10b | 0.000328465 | 4.154722595 | 0.075 | 0.004 | 1 | 0.438672228 | 1.83720485 | 0.024 | 0.01 | 1 | 0.438672228 | 0.000656823 |
| Ints2 | 0.00032938 | 1.790150163 | 0.472 | 0.275 | 1 | 0.536939903 | 1.096292071 | 0.39 | 0.403 | 1 | 0.536939903 | 0.000658652 |
| Gbp6 | 0.747613764 | 0.313420341 | 0.075 | 0.064 | 1 | 0.000329554 | 2.652780279 | 0.146 | 0.027 | 1 | 0.747613764 | 0.000658999 |
| Pias1 | 0.000331085 | 0.619299555 | 0.887 | 0.746 | 1 | 0.696830553 | -0.168744417 | 0.951 | 0.898 | 1 | 0.696830553 | 0.00066206 |
| Rtl3 | 0.000332069 | 1.350447066 | 0.434 | 0.208 | 1 | 0.408243332 | 0.730289646 | 0.268 | 0.235 | 1 | 0.408243332 | 0.000664028 |
| Dtl | 0.023029127 | 0.962077517 | 0.151 | 0.059 | 1 | 0.000332437 | 2.952682067 | 0.146 | 0.027 | 1 | 0.023029127 | 0.000664764 |
| Tmem94 | 0.000332545 | 1.640149422 | 0.208 | 0.055 | 1 | 0.232796044 | 0.898605394 | 0.171 | 0.109 | 1 | 0.232796044 | 0.00066498 |
| St8sia2 | 0.000333511 | 4.154722595 | 0.075 | 0.004 | 1 | 0.359002082 | 1.422167351 | 0.049 | 0.024 | 1 | 0.359002082 | 0.000666911 |
| Col18a1 | 0.000333511 | 3.739685096 | 0.075 | 0.004 | 1 | 0.021167579 | 2.83720485 | 0.049 | 0.007 | 1 | 0.021167579 | 0.000666911 |
| Abhd15 | 0.000333511 | 3.891688189 | 0.075 | 0.004 | 1 | 0.003474311 | 2.351778023 | 0.171 | 0.051 | 1 | 0.003474311 | 0.000666911 |
| Pnp2 | 0.000333511 | 3.739685096 | 0.075 | 0.004 | 1 | 0.87556513 | 0.83720485 | 0.024 | 0.02 | 1 | 0.87556513 | 0.000666911 |
| Gm15564 | 0.000333511 | 3.739685096 | 0.075 | 0.004 | 1 | 0.130804252 | 1.136765132 | 0.073 | 0.027 | 1 | 0.130804252 | 0.000666911 |
| Gm39526 | 0.000333511 | 3.739685096 | 0.075 | 0.004 | 1 | 0.357847088 | -0.16279515 | 0 | 0.02 | 1 | 0.357847088 | 0.000666911 |
| Fbxw2 | 0.000333824 | 0.734390796 | 0.698 | 0.458 | 1 | 0.828933867 | 0.120330674 | 0.585 | 0.594 | 1 | 0.828933867 | 0.000667536 |
| Casp3 | 0.000333922 | 0.671298121 | 0.717 | 0.445 | 1 | 0.036791502 | 0.519526228 | 0.659 | 0.515 | 1 | 0.036791502 | 0.000667732 |
| Wdr74 | 0.000334332 | 1.154722595 | 0.509 | 0.284 | 1 | 0.691279096 | 0.357735478 | 0.39 | 0.382 | 1 | 0.691279096 | 0.000668551 |
| Tbc1d10a | 0.158810494 | -0.243826782 | 0.566 | 0.716 | 1 | 0.000334951 | -0.902013196 | 0.488 | 0.775 | 1 | 0.158810494 | 0.000669789 |
| Sirt4 | 0.000335092 | 2.324647596 | 0.151 | 0.03 | 1 | 0.615241183 | 0.422167351 | 0.122 | 0.096 | 1 | 0.615241183 | 0.000670072 |
| Txnip | 0.000337014 | 0.785128066 | 0.887 | 0.72 | 1 | 0.027864203 | 0.749999978 | 0.902 | 0.833 | 1 | 0.027864203 | 0.000673914 |
| Nsmce4a | 0.00033723 | 0.877882389 | 0.679 | 0.453 | 1 | 0.599482836 | 0.149727438 | 0.537 | 0.587 | 1 | 0.599482836 | 0.000674346 |
| Mnt | 0.000338518 | 1.280253477 | 0.453 | 0.229 | 1 | 0.991833969 | 0.228395607 | 0.244 | 0.253 | 1 | 0.991833969 | 0.000676921 |
| Selplg | 0.000338811 | 3.47665069 | 0.075 | 0.004 | 1 | 0.29958328 | 0.515276755 | 0.098 | 0.055 | 1 | 0.29958328 | 0.000677508 |
| Mlkl | 0.000338811 | 3.47665069 | 0.075 | 0.004 | 1 | 0.267205177 | 2.252242349 | 0.024 | 0.007 | 1 | 0.267205177 | 0.000677508 |
| Abcg4 | 0.000338811 | 3.47665069 | 0.075 | 0.004 | 1 | 0.267205177 | 2.252242349 | 0.024 | 0.007 | 1 | 0.267205177 | 0.000677508 |
| Doc2b | 0.000338811 | 3.47665069 | 0.075 | 0.004 | 1 | 0.881057318 | 0.136765132 | 0.024 | 0.02 | 1 | 0.881057318 | 0.000677508 |
| Fnip2 | 0.06037542 | -0.502996248 | 0.547 | 0.695 | 1 | 0.000339532 | -1.1870917 | 0.683 | 0.843 | 1 | 0.06037542 | 0.000678948 |
| Zbtb44 | 0.00033961 | 1.099440159 | 0.566 | 0.364 | 1 | 0.164815489 | 0.341051616 | 0.707 | 0.608 | 1 | 0.164815489 | 0.000679104 |
| Mlf1 | 0.000340209 | -2.792809985 | 0.057 | 0.292 | 1 | 0.011647677 | -1.787286015 | 0.049 | 0.218 | 1 | 0.011647677 | 0.000680302 |
| Pbx3 | 0.000341852 | 1.308756223 | 0.604 | 0.394 | 1 | 0.004832942 | 0.965144238 | 0.805 | 0.611 | 1 | 0.004832942 | 0.000683587 |
| Tmem243 | 0.013944149 | 1.205348668 | 0.472 | 0.339 | 1 | 0.000342486 | 1.093980265 | 0.707 | 0.505 | 1 | 0.013944149 | 0.000684855 |
| Faf1 | 0.000342655 | 0.582887528 | 0.868 | 0.775 | 1 | 0.122508001 | 0.242527194 | 0.902 | 0.922 | 1 | 0.122508001 | 0.000685192 |
| Fam107a | 0.261818086 | -1.652632327 | 0.019 | 0.055 | 1 | 0.000343463 | -3.79215177 | 0.024 | 0.283 | 1 | 0.261818086 | 0.000686809 |
| Fbxw8 | 0.013655567 | -0.418744267 | 0.547 | 0.75 | 1 | 0.00034649 | -0.871332336 | 0.439 | 0.737 | 1 | 0.013655567 | 0.000692859 |
| Zbtb10 | 0.067576124 | 0.537627473 | 0.566 | 0.462 | 1 | 0.000346997 | 1.074846098 | 0.756 | 0.587 | 1 | 0.067576124 | 0.000693874 |
| Ap1s2 | 0.000347096 | 0.959906418 | 0.604 | 0.352 | 1 | 0.857269698 | 0.393598198 | 0.268 | 0.263 | 1 | 0.857269698 | 0.000694072 |
| Sik1 | 0.068332619 | 0.404700848 | 0.34 | 0.208 | 1 | 0.000347554 | 1.393598198 | 0.512 | 0.276 | 1 | 0.068332619 | 0.000694986 |
| Dmwd | 0.000350852 | 1.333692736 | 0.415 | 0.195 | 1 | 0.956229398 | 0.501601818 | 0.244 | 0.266 | 1 | 0.956229398 | 0.000701581 |
| Pcdhgc5 | 0.000352029 | 2.154722595 | 0.17 | 0.038 | 1 | 0.672172778 | 0.136765132 | 0.024 | 0.038 | 1 | 0.672172778 | 0.000703935 |
| Cyp3a13 | 0.000352286 | -4.237594828 | 0 | 0.203 | 1 | 0.114590127 | -2.410722664 | 0 | 0.058 | 1 | 0.114590127 | 0.000704448 |
| Anxa4 | 0.000352695 | -1.051047606 | 0.509 | 0.733 | 1 | 0.017809917 | -0.743744588 | 0.561 | 0.72 | 1 | 0.017809917 | 0.000705266 |
| Calcoco1 | 0.000355525 | 1.302279783 | 0.415 | 0.203 | 1 | 0.923393301 | 0.308825877 | 0.293 | 0.328 | 1 | 0.923393301 | 0.000710924 |
| Zc3h18 | 0.00035688 | 0.690054328 | 0.755 | 0.593 | 1 | 0.403014941 | 0.340779024 | 0.756 | 0.751 | 1 | 0.403014941 | 0.000713633 |
| Slc35d2 | 0.000358371 | 2.002719501 | 0.151 | 0.03 | 1 | 0.085804408 | 1.149148856 | 0.098 | 0.038 | 1 | 0.085804408 | 0.000716614 |
| Ccnj | 0.000358371 | 2.154722595 | 0.151 | 0.03 | 1 | 0.80551848 | 0.83720485 | 0.049 | 0.041 | 1 | 0.80551848 | 0.000716614 |
| Tradd | 0.000359836 | 1.70221039 | 0.283 | 0.102 | 1 | 0.84750704 | 0.930314254 | 0.098 | 0.092 | 1 | 0.84750704 | 0.000719543 |
| Cacna2d2 | 0.000360208 | 2.891688189 | 0.075 | 0.004 | 1 | 0.007064241 | 2.159132945 | 0.098 | 0.02 | 1 | 0.007064241 | 0.000720286 |
| Gm45847 | 0.00036028 | 2.739685096 | 0.075 | 0.004 | 1 | 0.137917747 | 0.952682067 | 0.171 | 0.096 | 1 | 0.137917747 | 0.00072043 |
| Map2k3 | 0.002373215 | -0.548454686 | 0.925 | 0.907 | 1 | 0.000360458 | -0.773819948 | 0.78 | 0.908 | 1 | 0.002373215 | 0.000720786 |
| Cacul1 | 0.000360579 | 1.033707194 | 0.623 | 0.403 | 1 | 0.038435995 | 0.643600852 | 0.756 | 0.676 | 1 | 0.038435995 | 0.000721027 |
| Ankrd37 | 0.000362322 | 2.154722595 | 0.151 | 0.03 | 1 | 0.404083256 | -0.80665134 | 0.024 | 0.055 | 1 | 0.404083256 | 0.000724513 |
| Get3 | 0.000362515 | 1.137235168 | 0.528 | 0.297 | 1 | 0.201564436 | -0.259656689 | 0.268 | 0.369 | 1 | 0.201564436 | 0.000724899 |
| Kidins220 | 0.000362958 | 0.820083448 | 0.755 | 0.606 | 1 | 0.168893564 | 0.233383322 | 0.878 | 0.809 | 1 | 0.168893564 | 0.000725785 |
| Tgfb1 | 0.000363082 | 0.798101782 | 0.811 | 0.754 | 1 | 0.716096492 | 0.13753069 | 0.805 | 0.853 | 1 | 0.716096492 | 0.000726033 |
| Zfp385c | 0.000426345 | -2.639693272 | 0.038 | 0.263 | 1 | 0.000363445 | -2.762707992 | 0.049 | 0.321 | 1 | 0.000426345 | 0.000726758 |
| Ddx3x | 0.003586918 | 0.438221103 | 1 | 0.936 | 1 | 0.00036386 | 0.525923999 | 1 | 0.962 | 1 | 0.003586918 | 0.000727588 |
| Usp1 | 0.0003665 | 0.569760094 | 0.698 | 0.428 | 1 | 0.109503279 | 0.679663573 | 0.634 | 0.522 | 1 | 0.109503279 | 0.000732866 |
| Ccn1 | 0.042839942 | -1.028179209 | 0.83 | 0.839 | 1 | 0.000368034 | 0.801877913 | 0.805 | 0.611 | 1 | 0.042839942 | 0.000735933 |
| Adcy9 | 0.000368859 | 1.703026625 | 0.415 | 0.203 | 1 | 0.052092842 | 0.983375498 | 0.561 | 0.444 | 1 | 0.052092842 | 0.000737582 |
| Arhgef25 | 0.00037082 | 1.015658557 | 0.679 | 0.508 | 1 | 0.648280161 | 0.437597391 | 0.537 | 0.515 | 1 | 0.648280161 | 0.000741502 |
| Snw1 | 0.000370942 | 0.802269633 | 0.868 | 0.746 | 1 | 0.169859752 | -0.322791034 | 0.756 | 0.85 | 1 | 0.169859752 | 0.000741747 |
| Copb1 | 0.000370962 | 0.474569093 | 0.981 | 0.936 | 1 | 0.046509041 | -0.334540587 | 0.976 | 0.969 | 1 | 0.046509041 | 0.000741787 |
| Pantr1 | 0.00037355 | -2.665456368 | 0.245 | 0.462 | 1 | 0.825421358 | 0.242258261 | 0.268 | 0.294 | 1 | 0.825421358 | 0.000746961 |
| Rhbdf1 | 0.000376475 | 1.399835093 | 0.415 | 0.195 | 1 | 0.582416156 | 0.525260843 | 0.341 | 0.348 | 1 | 0.582416156 | 0.000752809 |
| Mon1a | 0.000377374 | 1.628653783 | 0.321 | 0.131 | 1 | 0.798950398 | 0.386543441 | 0.22 | 0.208 | 1 | 0.798950398 | 0.000754605 |
| Canx | 0.00037996 | 0.39710389 | 1 | 0.992 | 1 | 0.68533808 | 0.104598795 | 1 | 1 | 1 | 0.68533808 | 0.000759776 |
| Tubg1 | 0.000380011 | 1.891688189 | 0.226 | 0.068 | 1 | 0.055796536 | 1.614812428 | 0.146 | 0.065 | 1 | 0.055796536 | 0.000759877 |
| Cidea | 0.000380056 | -2.437734442 | 0.038 | 0.267 | 1 | 0.081425268 | -1.233184478 | 0.049 | 0.15 | 1 | 0.081425268 | 0.000759968 |
| Polm | 0.017364527 | 2.347367673 | 0.094 | 0.025 | 1 | 0.000380126 | 2.47463477 | 0.146 | 0.027 | 1 | 0.017364527 | 0.000760108 |
| Setdb1 | 0.000380386 | 1.167433647 | 0.547 | 0.326 | 1 | 0.603534289 | 0.199131013 | 0.39 | 0.474 | 1 | 0.603534289 | 0.000760628 |
| H1f2 | 0.000380634 | -2.088047826 | 0.434 | 0.631 | 1 | 0.738371497 | 0.484786388 | 0.463 | 0.549 | 1 | 0.738371497 | 0.000761124 |
| Ubqln2 | 0.000381586 | 1.049753035 | 0.585 | 0.343 | 1 | 0.306582258 | 0.543021745 | 0.488 | 0.44 | 1 | 0.306582258 | 0.000763026 |
| Rdx | 0.000382455 | 0.412520352 | 1 | 0.962 | 1 | 0.384769202 | 0.136373042 | 1 | 0.983 | 1 | 0.384769202 | 0.000764763 |
| Epb41 | 0.000382515 | 1.288739943 | 0.509 | 0.292 | 1 | 0.001733636 | 1.040973306 | 0.707 | 0.567 | 1 | 0.001733636 | 0.000764884 |
| Rab11fip1 | 0.000382838 | -2.256210506 | 0.17 | 0.407 | 1 | 0.446011393 | 0.548959881 | 0.341 | 0.29 | 1 | 0.446011393 | 0.000765529 |
| Cisd2 | 0.019575368 | 0.416877577 | 0.868 | 0.831 | 1 | 0.000383127 | -0.76716305 | 0.805 | 0.908 | 1 | 0.019575368 | 0.000766107 |

| | | | | | | | | | | | | |
|---|---|---|---|---|---|---|---|---|---|---|---|---|
| Gm43445 | 0.080589553 | 1.039245377 | 0.17 | 0.089 | 1 | 0.000386305 | 2.252242349 | 0.195 | 0.048 | 1 | 0.080589553 | 0.000772461 |
| Arhgdib | 0.967055092 | -0.108634959 | 0.774 | 0.784 | 1 | 0.000390545 | -0.910242228 | 0.707 | 0.87 | 1 | 0.967055092 | 0.000780938 |
| Usp38 | 0.000393922 | 1.117247889 | 0.509 | 0.284 | 1 | 0.607117273 | 0.289304443 | 0.537 | 0.546 | 1 | 0.607117273 | 0.000787689 |
| Kpna1 | 0.00039439 | 0.891688189 | 0.774 | 0.589 | 1 | 0.499690224 | 0.395436985 | 0.756 | 0.812 | 1 | 0.499690224 | 0.000788625 |
| Itprid2 | 0.000547472 | 1.13912574 | 0.453 | 0.246 | 1 | 0.00039467 | 1.336010707 | 0.634 | 0.43 | 1 | 0.000547472 | 0.000789184 |
| St3gal6 | 0.000396966 | -0.858672999 | 0.491 | 0.767 | 1 | 0.001277007 | -0.701771446 | 0.634 | 0.85 | 1 | 0.001277007 | 0.000793775 |
| Usp14 | 0.000399595 | 0.894599784 | 0.792 | 0.644 | 1 | 0.031236388 | 0.472306716 | 0.805 | 0.7 | 1 | 0.031236388 | 0.00079903 |
| Glis1 | 0.18620908 | -0.240137023 | 0.132 | 0.203 | 1 | 0.000399784 | -1.946537515 | 0.049 | 0.324 | 1 | 0.18620908 | 0.000799408 |
| Raver1 | 0.000401145 | 1.3662267 | 0.472 | 0.254 | 1 | 0.591910269 | 0.271607674 | 0.366 | 0.345 | 1 | 0.591910269 | 0.000802129 |
| Anp32b | 0.000401262 | 0.524452466 | 0.962 | 0.89 | 1 | 0.909878097 | 0.156789427 | 0.902 | 0.952 | 1 | 0.909878097 | 0.000802363 |
| Nuak1 | 0.000402899 | 0.778706984 | 0.566 | 0.356 | 1 | 0.001404759 | 0.925909075 | 0.683 | 0.546 | 1 | 0.001404759 | 0.000805635 |
| Utp14b | 0.000403536 | -1.314391577 | 0.264 | 0.555 | 1 | 0.144990169 | -0.655682066 | 0.415 | 0.512 | 1 | 0.144990169 | 0.00080691 |
| Phka2 | 0.00040357 | -1.888260576 | 0.283 | 0.525 | 1 | 0.153078695 | -0.539171029 | 0.146 | 0.256 | 1 | 0.153078695 | 0.000806978 |
| Dus3l | 0.000405522 | 1.47665069 | 0.358 | 0.157 | 1 | 0.75927208 | 0.694246896 | 0.171 | 0.157 | 1 | 0.75927208 | 0.00081088 |
| Golm2 | 0.000406487 | 0.610574557 | 0.962 | 0.818 | 1 | 0.24105924 | 0.300143894 | 0.854 | 0.857 | 1 | 0.24105924 | 0.000812808 |
| Rps15a | 0.000406507 | 0.360645668 | 1 | 0.966 | 1 | 0.102111906 | -0.173119093 | 0.976 | 0.986 | 1 | 0.102111906 | 0.000812849 |
| 0610009E02Rik | 0.000407229 | 1.906795081 | 0.226 | 0.068 | 1 | 0.012848688 | 1.83720485 | 0.244 | 0.113 | 1 | 0.012848688 | 0.000814292 |
| Kdm4a | 0.000408846 | 1.208493851 | 0.453 | 0.229 | 1 | 0.120447723 | -0.695290231 | 0.244 | 0.372 | 1 | 0.120447723 | 0.000817525 |
| Mamdc2 | 0.000409731 | 1.711115943 | 0.226 | 0.068 | 1 | 0.954638603 | 0.614812428 | 0.098 | 0.106 | 1 | 0.954638603 | 0.000819295 |
| Glipr2 | 0.00041001 | 1.806799291 | 0.17 | 0.038 | 1 | 0.970764619 | 0.515276755 | 0.073 | 0.072 | 1 | 0.970764619 | 0.000819852 |
| Caskin2 | 0.002329586 | 1.569760094 | 0.189 | 0.059 | 1 | 0.000410843 | 1.679663573 | 0.268 | 0.085 | 1 | 0.002329586 | 0.000821517 |
| Tmem158 | 0.000411578 | 2.154722595 | 0.208 | 0.059 | 1 | 0.009560002 | 1.458693227 | 0.244 | 0.106 | 1 | 0.009560002 | 0.000822987 |
| Arid5b | 0.109289155 | 0.291587091 | 0.925 | 0.839 | 1 | 0.00041369 | 1.380425245 | 0.902 | 0.857 | 1 | 0.109289155 | 0.000827209 |
| Ntn3 | 0.000414599 | 1.865215978 | 0.151 | 0.03 | 1 | 0.632655435 | 0.458693227 | 0.098 | 0.075 | 1 | 0.632655435 | 0.000829025 |
| 2600014E21Rik | 0.000416093 | -4.27154216 | 0 | 0.199 | 1 | 0.973993537 | 0.458693227 | 0.024 | 0.024 | 1 | 0.973993537 | 0.000832013 |
| Dhx38 | 0.000419093 | 1.70221039 | 0.377 | 0.182 | 1 | 0.14669727 | 0.604544093 | 0.439 | 0.334 | 1 | 0.14669727 | 0.00083801 |
| Gm11476 | 0.000421043 | 1.948271717 | 0.17 | 0.038 | 1 | 0.304397485 | -0.917682652 | 0.024 | 0.065 | 1 | 0.304397485 | 0.00084191 |
| Chchd10 | 0.083454232 | 1.247831999 | 0.094 | 0.038 | 1 | 0.000423414 | 0.440314697 | 0.341 | 0.123 | 1 | 0.083454232 | 0.000846648 |
| Nup98 | 0.000425908 | 0.4368136 | 0.981 | 0.847 | 1 | 0.748541369 | 0.156483366 | 0.976 | 0.956 | 1 | 0.748541369 | 0.000851636 |
| Rufy3 | 0.000426243 | 1.121555731 | 0.604 | 0.386 | 1 | 0.002077069 | 0.77191339 | 0.78 | 0.532 | 1 | 0.002077069 | 0.000852305 |
| Majin | 0.000427156 | 0.530871081 | 0.66 | 0.428 | 1 | 0.723712948 | -0.231622274 | 0.463 | 0.549 | 1 | 0.723712948 | 0.000854129 |
| Arc | 0.894597189 | -0.719746523 | 0.057 | 0.051 | 1 | 0.000429413 | 1.83720485 | 0.146 | 0.027 | 1 | 0.894597189 | 0.000858641 |
| Ecm2 | 0.000430375 | 0.862413267 | 0.83 | 0.61 | 1 | 0.007499997 | -0.722908689 | 0.683 | 0.829 | 1 | 0.007499997 | 0.000860564 |
| Plod3 | 0.000430914 | -1.005552237 | 0.528 | 0.725 | 1 | 0.000876176 | -1.105626627 | 0.512 | 0.706 | 1 | 0.000876176 | 0.000861642 |
| Tbc1d20 | 0.000432151 | 0.75033234 | 0.698 | 0.462 | 1 | 0.078208175 | -0.167533069 | 0.415 | 0.604 | 1 | 0.078208175 | 0.000864115 |
| Crnkl1 | 0.000434005 | 1.310373465 | 0.453 | 0.242 | 1 | 0.099669596 | 0.990914265 | 0.463 | 0.369 | 1 | 0.099669596 | 0.000867821 |
| Usp15 | 0.000434129 | 0.813106165 | 0.83 | 0.581 | 1 | 0.297719531 | 0.426815854 | 0.854 | 0.823 | 1 | 0.297719531 | 0.00086807 |
| Rragc | 0.000434754 | 0.972030297 | 0.717 | 0.568 | 1 | 0.049285852 | -0.442903069 | 0.659 | 0.812 | 1 | 0.049285852 | 0.000869319 |
| Rab20 | 0.00043478 | 2.377115016 | 0.283 | 0.11 | 1 | 0.84268262 | 0.79281073 | 0.171 | 0.16 | 1 | 0.84268262 | 0.00086937 |
| Ptpn4 | 0.00043614 | 1.321541685 | 0.509 | 0.305 | 1 | 0.008279516 | 0.744758601 | 0.585 | 0.41 | 1 | 0.008279516 | 0.00087209 |
| Tub | 0.000437384 | 1.569760094 | 0.17 | 0.038 | 1 | 0.053472004 | 1.83720485 | 0.146 | 0.065 | 1 | 0.053472004 | 0.000874576 |
| Smc1a | 0.004154573 | 0.370840782 | 0.981 | 0.89 | 1 | 0.000438554 | 0.757351542 | 1 | 0.939 | 1 | 0.004154573 | 0.000876916 |
| Reg4 | 0.000438648 | 1.10741688 | 0.151 | 0.03 | 1 | 0.001521235 | 3.489281546 | 0.122 | 0.024 | 1 | 0.001521235 | 0.000877103 |
| Rasal2 | 0.000439475 | 1.084333267 | 0.679 | 0.436 | 1 | 0.406335742 | 0.334704509 | 0.634 | 0.556 | 1 | 0.406335742 | 0.000878756 |
| Myo18a | 0.230896475 | 0.142195497 | 0.623 | 0.559 | 1 | 0.000439895 | -0.932276513 | 0.756 | 0.901 | 1 | 0.230896475 | 0.000879596 |
| Ints4 | 0.000440529 | 1.100274811 | 0.377 | 0.157 | 1 | 0.442062325 | -0.132421501 | 0.22 | 0.29 | 1 | 0.442062325 | 0.000880864 |
| Lpcat2 | 0.000774984 | 1.690775495 | 0.302 | 0.123 | 1 | 0.000441032 | 1.891652634 | 0.488 | 0.249 | 1 | 0.000774984 | 0.00088187 |
| Slc38a6 | 0.000446905 | 2.105812994 | 0.358 | 0.174 | 1 | 0.511589849 | 0.422167351 | 0.293 | 0.246 | 1 | 0.511589849 | 0.00089361 |
| St3gal3 | 0.000450795 | 0.851653527 | 0.604 | 0.369 | 1 | 0.093313528 | 0.846985871 | 0.463 | 0.389 | 1 | 0.093313528 | 0.000901386 |
| Prpf40a | 0.000453301 | 0.691011249 | 0.925 | 0.928 | 1 | 0.053171675 | 0.344746977 | 0.951 | 0.952 | 1 | 0.053171675 | 0.000906397 |
| Lima1 | 0.000966831 | 0.856641242 | 0.83 | 0.699 | 1 | 0.000455679 | 1.36447955 | 0.902 | 0.843 | 1 | 0.000966831 | 0.00091115 |
| Slbp | 0.06064519 | 0.305983752 | 0.736 | 0.636 | 1 | 0.000456011 | 0.887080756 | 0.78 | 0.655 | 1 | 0.06064519 | 0.000911814 |
| Iqgap2 | 0.000456576 | 1.339147166 | 0.302 | 0.114 | 1 | 0.011438743 | 1.856313673 | 0.366 | 0.205 | 1 | 0.011438743 | 0.000912943 |
| Cnn3 | 0.000457979 | 0.426199797 | 0.981 | 0.97 | 1 | 0.001641822 | 0.394351973 | 0.976 | 0.98 | 1 | 0.001641822 | 0.000915748 |
| Mafg | 0.331334878 | 0.154722595 | 0.792 | 0.801 | 1 | 0.000462077 | -0.695290231 | 0.854 | 0.942 | 1 | 0.331334878 | 0.000923941 |
| Khdrbs3 | 0.000463861 | 1.70221039 | 0.321 | 0.131 | 1 | 0.683396506 | 0.272852629 | 0.537 | 0.488 | 1 | 0.683396506 | 0.000927507 |
| Snrpd2 | 0.000465595 | 0.58556195 | 0.925 | 0.78 | 1 | 0.243088401 | 0.335658357 | 0.756 | 0.775 | 1 | 0.243088401 | 0.000930973 |
| Epc2 | 0.000468039 | 0.548698479 | 0.887 | 0.703 | 1 | 0.198721055 | 0.28609012 | 0.878 | 0.881 | 1 | 0.198721055 | 0.00093586 |
| Tcf7l1 | 0.000469145 | 0.904026524 | 0.717 | 0.487 | 1 | 0.727911672 | -0.178126804 | 0.829 | 0.785 | 1 | 0.727911672 | 0.000938071 |
| Vps72 | 0.000469415 | 1.333059836 | 0.472 | 0.271 | 1 | 0.025357657 | 0.734843132 | 0.488 | 0.324 | 1 | 0.025357657 | 0.000938609 |
| Pwwp3a | 0.000469589 | 1.417757001 | 0.358 | 0.153 | 1 | 0.039507755 | 0.727580359 | 0.341 | 0.195 | 1 | 0.039507755 | 0.000938957 |
| Nr4a1 | 0.000470025 | 0.901815496 | 0.792 | 0.606 | 1 | 0.018929991 | 1.179729849 | 0.78 | 0.737 | 1 | 0.018929991 | 0.000939829 |
| Ngf | 0.000471905 | -4.083682145 | 0.075 | 0.297 | 1 | 0.413137776 | -2.020776145 | 0.049 | 0.085 | 1 | 0.413137776 | 0.000943587 |
| Wdyhv1 | 0.000472597 | 1.541745718 | 0.283 | 0.102 | 1 | 0.728454737 | 0.778311161 | 0.146 | 0.13 | 1 | 0.728454737 | 0.00094497 |
| Arfip2 | 0.000474148 | 1.247831999 | 0.472 | 0.263 | 1 | 0.019764731 | -1.221688839 | 0.195 | 0.365 | 1 | 0.019764731 | 0.000948072 |
| AU020206 | 0.000474495 | 2.010332685 | 0.208 | 0.059 | 1 | 0.001034484 | 1.875678998 | 0.244 | 0.082 | 1 | 0.001034484 | 0.000948766 |
| Pak2 | 0.00047475 | 0.461812181 | 1 | 0.936 | 1 | 0.613273621 | -0.128977365 | 0.927 | 0.993 | 1 | 0.613273621 | 0.000949274 |
| Als2 | 0.000475198 | 1.440124814 | 0.396 | 0.178 | 1 | 0.343533743 | 0.83720485 | 0.293 | 0.239 | 1 | 0.343533743 | 0.000950171 |
| Itprip | 0.000475568 | -1.223789028 | 0.057 | 0.305 | 1 | 0.63600728 | -0.142617268 | 0.146 | 0.177 | 1 | 0.63600728 | 0.000950911 |
| Erg28 | 0.000480947 | -1.551546202 | 0.226 | 0.479 | 1 | 0.982137133 | -0.242522343 | 0.366 | 0.352 | 1 | 0.982137133 | 0.000961663 |
| Tstd2 | 0.000484308 | 1.177442671 | 0.434 | 0.208 | 1 | 0.209440583 | -0.114539982 | 0.22 | 0.348 | 1 | 0.209440583 | 0.000968382 |
| Sema3e | 0.000488197 | -4.758166741 | 0.019 | 0.229 | 1 | 0.298650925 | -2.566517336 | 0.024 | 0.065 | 1 | 0.298650925 | 0.000976155 |
| Elk4 | 0.000489075 | 0.753517369 | 0.83 | 0.695 | 1 | 0.235229209 | -0.188591151 | 0.854 | 0.853 | 1 | 0.235229209 | 0.000977911 |
| Gtf3c2 | 0.000489263 | 1.111909511 | 0.623 | 0.453 | 1 | 0.010869034 | 0.682319248 | 0.732 | 0.655 | 1 | 0.010869034 | 0.000978287 |
| Syngr2 | 0.000490891 | 1.776210972 | 0.264 | 0.093 | 1 | 0.456805468 | 0.328191202 | 0.244 | 0.195 | 1 | 0.456805468 | 0.000981541 |
| Zfp513 | 0.04534964 | 1.2967416 | 0.208 | 0.11 | 1 | 0.000492077 | 1.551802631 | 0.293 | 0.102 | 1 | 0.04534964 | 0.000983912 |
| Pik3ca | 0.000492093 | 0.800004565 | 0.811 | 0.733 | 1 | 0.157099236 | 0.37239338 | 0.854 | 0.863 | 1 | 0.157099236 | 0.000983943 |
| Acad8 | 0.000492329 | 1.497610308 | 0.34 | 0.144 | 1 | 0.432011038 | 0.394261354 | 0.268 | 0.218 | 1 | 0.432011038 | 0.000984417 |
| Irak3 | 0.000492941 | -1.740095169 | 0.208 | 0.449 | 1 | 0.227507614 | -1.332720152 | 0.195 | 0.263 | 1 | 0.227507614 | 0.000985639 |
| Cep97 | 0.000493472 | 1.517292674 | 0.283 | 0.102 | 1 | 0.144684834 | 1.227151368 | 0.22 | 0.137 | 1 | 0.144684834 | 0.0009867 |
| Parp14 | 0.001242996 | 1.970298024 | 0.189 | 0.055 | 1 | 0.000495882 | 2.344164839 | 0.244 | 0.075 | 1 | 0.001242996 | 0.000991517 |
| Abcc5 | 0.000496883 | 1.067961283 | 0.698 | 0.517 | 1 | 0.507709334 | -0.407102348 | 0.707 | 0.734 | 1 | 0.507709334 | 0.000993519 |
| Dedd2 | 0.000501958 | 1.444229212 | 0.283 | 0.102 | 1 | 0.05891639 | 0.885033375 | 0.341 | 0.215 | 1 | 0.05891639 | 0.001003663 |
| Cux2 | 0.000504097 | 3.002719501 | 0.113 | 0.017 | 1 | 0.023468139 | 1.962735732 | 0.098 | 0.027 | 1 | 0.023468139 | 0.00100794 |
| Trac | 0.000504097 | 3.292226119 | 0.113 | 0.017 | 1 | 0.123845731 | 1.721727632 | 0.073 | 0.027 | 1 | 0.123845731 | 0.00100794 |
| Nudcd3 | 0.000504937 | 0.776210972 | 0.868 | 0.648 | 1 | 0.175140307 | 0.235940537 | 0.756 | 0.778 | 1 | 0.175140307 | 0.001009619 |
| Kdm4c | 0.000508215 | 0.734230665 | 0.698 | 0.513 | 1 | 0.541839227 | 0.12478144 | 0.78 | 0.747 | 1 | 0.541839227 | 0.001016171 |
| Zfp667 | 0.000509676 | 2.002719501 | 0.226 | 0.072 | 1 | 0.332012285 | -0.935384654 | 0.122 | 0.174 | 1 | 0.332012285 | 0.001019092 |
| Cntn3 | 0.000509895 | 3.154722595 | 0.113 | 0.017 | 1 | 0.357854401 | -0.332720152 | 0 | 0.02 | 1 | 0.357854401 | 0.001019531 |
| Fzd3 | 0.00050997 | 1.448453798 | 0.377 | 0.182 | 1 | 0.322211272 | 0.569724539 | 0.317 | 0.263 | 1 | 0.322211272 | 0.001019679 |
| Gm14296 | 0.000510502 | 1.891688189 | 0.208 | 0.059 | 1 | 0.893463168 | 0.377773231 | 0.098 | 0.092 | 1 | 0.893463168 | 0.001020744 |
| Cyb5b | 0.000510641 | -0.992920959 | 0.34 | 0.597 | 1 | 0.437992054 | 0.435756415 | 0.585 | 0.577 | 1 | 0.437992054 | 0.001021022 |
| Fubp3 | 0.000510988 | 1.09332205 | 0.528 | 0.301 | 1 | 0.456431935 | 0.321707169 | 0.488 | 0.44 | 1 | 0.456431935 | 0.001021716 |
| Zfp423 | 0.000511695 | 1.865215978 | 0.264 | 0.093 | 1 | 0.001598412 | 1.866952193 | 0.39 | 0.205 | 1 | 0.001598412 | 0.001023128 |
| Trim12a | 0.000513535 | 1.404700848 | 0.264 | 0.089 | 1 | 0.002118536 | 1.977382508 | 0.268 | 0.106 | 1 | 0.002118536 | 0.001026806 |
| Rell1 | 0.000514407 | 1.081141462 | 0.604 | 0.407 | 1 | 0.008288246 | 0.750493217 | 0.854 | 0.679 | 1 | 0.008288246 | 0.00102855 |
| 0610043K17Rik | 0.000516992 | -0.430239906 | 0.17 | 0.445 | 1 | 0.165082955 | 0.641654041 | 0.366 | 0.273 | 1 | 0.165082955 | 0.001033716 |
| Phc2 | 0.000518804 | 0.762405172 | 0.83 | 0.716 | 1 | 0.704713834 | 0.117864926 | 0.878 | 0.908 | 1 | 0.704713834 | 0.001037338 |

| | | | | | | | | | | | | |
|---|---|---|---|---|---|---|---|---|---|---|---|---|
| Thra | 0.000520809 | 0.698972888 | 0.887 | 0.826 | 1 | 0.725928391 | 0.204936634 | 0.878 | 0.901 | 1 | 0.725928391 | 0.001041347 |
| Slc4a3 | 0.000521139 | 1.620386167 | 0.34 | 0.153 | 1 | 0.697464373 | -0.221688839 | 0.171 | 0.195 | 1 | 0.697464373 | 0.001042005 |
| Gpr176 | 0.003471527 | 1.891688189 | 0.094 | 0.017 | 1 | 0.000523333 | 4.235754226 | 0.122 | 0.02 | 1 | 0.003471527 | 0.001046391 |
| Clcn2 | 0.000526901 | 1.602181572 | 0.226 | 0.068 | 1 | 0.615234991 | -0.233184478 | 0.073 | 0.099 | 1 | 0.615234991 | 0.001053524 |
| Otud4 | 0.000528156 | 0.856754399 | 0.698 | 0.5 | 1 | 0.239890042 | 0.372923832 | 0.78 | 0.744 | 1 | 0.239890042 | 0.001056033 |
| Zfp622 | 0.000528829 | 1.223985257 | 0.566 | 0.364 | 1 | 0.532837658 | 0.293890046 | 0.415 | 0.526 | 1 | 0.532837658 | 0.001057379 |
| Nt5c3b | 0.000530609 | 0.814298156 | 0.66 | 0.432 | 1 | 0.023608154 | -0.761054474 | 0.244 | 0.444 | 1 | 0.023608154 | 0.001060936 |
| Map4k4 | 0.000530881 | 0.365644968 | 1 | 0.962 | 1 | 0.090373399 | 0.183696721 | 1 | 0.99 | 1 | 0.090373399 | 0.00106148 |
| Cdo1 | 0.000532288 | -1.46107731 | 0.415 | 0.589 | 1 | 0.931231166 | -0.424559019 | 0.707 | 0.666 | 1 | 0.931231166 | 0.001064293 |
| Rnasel | 0.000533561 | 1.522454379 | 0.245 | 0.081 | 1 | 0.861719075 | 0.174239837 | 0.098 | 0.089 | 1 | 0.861719075 | 0.001066838 |
| Snx6 | 0.000534803 | 0.637931597 | 0.868 | 0.678 | 1 | 0.054055803 | -0.365184547 | 0.78 | 0.891 | 1 | 0.054055803 | 0.001069321 |
| Atf7 | 0.000535111 | 0.777120902 | 0.887 | 0.822 | 1 | 0.243008136 | -0.218329827 | 0.927 | 0.952 | 1 | 0.243008136 | 0.001069937 |
| Efna2 | 0.000535394 | 1.386048141 | 0.321 | 0.127 | 1 | 0.001644336 | 1.857382732 | 0.341 | 0.157 | 1 | 0.001644336 | 0.001070501 |
| Ier3 | 0.929116339 | -0.366343515 | 0.811 | 0.788 | 1 | 0.000535716 | 1.28886094 | 0.78 | 0.666 | 1 | 0.929116339 | 0.001071145 |
| Gm34045 | 0.000537744 | 2.739685096 | 0.113 | 0.017 | 1 | 0.547258706 | -0.332720152 | 0.024 | 0.044 | 1 | 0.547258706 | 0.001075198 |
| Spry4 | 0.22129369 | 1.331600357 | 0.132 | 0.081 | 1 | 0.000537789 | 1.952682067 | 0.22 | 0.061 | 1 | 0.22129369 | 0.001075288 |
| Gm31641 | 0.138460079 | 0.80108564 | 0.151 | 0.089 | 1 | 0.000537866 | 2.747007041 | 0.366 | 0.171 | 1 | 0.138460079 | 0.001075443 |
| Samd14 | 0.000537879 | 2.154722595 | 0.189 | 0.051 | 1 | 0.00446226 | 1.614812428 | 0.293 | 0.13 | 1 | 0.00446226 | 0.001075468 |
| Idh3a | 0.000541138 | 1.169829487 | 0.528 | 0.322 | 1 | 0.141229799 | -0.577832649 | 0.268 | 0.382 | 1 | 0.141229799 | 0.001081983 |
| Commd3 | 0.000541308 | 0.701930174 | 0.774 | 0.576 | 1 | 0.033142591 | -0.602126808 | 0.512 | 0.652 | 1 | 0.033142591 | 0.001082323 |
| Ubxn6 | 0.000542488 | 1.009292155 | 0.698 | 0.564 | 1 | 0.585801299 | 0.257478997 | 0.585 | 0.608 | 1 | 0.585801299 | 0.001084682 |
| Heatr5b | 0.000543397 | 2.002719501 | 0.34 | 0.153 | 1 | 0.569949933 | -0.366667484 | 0.244 | 0.283 | 1 | 0.569949933 | 0.001086498 |
| Unc45a | 0.000547711 | 1.230671448 | 0.453 | 0.237 | 1 | 0.790625728 | 0.493250449 | 0.415 | 0.471 | 1 | 0.790625728 | 0.001095122 |
| Rhpn2 | 0.000549326 | 2.324647596 | 0.113 | 0.017 | 1 | 0.445030093 | 0.635570989 | 0.098 | 0.065 | 1 | 0.445030093 | 0.001098349 |
| 4933427G23Rik | 0.000549823 | 2.640149422 | 0.113 | 0.017 | 1 | 0.170390343 | -1.250257991 | 0 | 0.044 | 1 | 0.170390343 | 0.001099344 |
| Rigi | 0.033762498 | 1.000917259 | 0.321 | 0.199 | 1 | 0.000549965 | 1.483567895 | 0.463 | 0.256 | 1 | 0.033762498 | 0.001099627 |
| Zfp821 | 0.000550079 | 1.669295768 | 0.358 | 0.165 | 1 | 0.706967383 | 0.410940095 | 0.293 | 0.283 | 1 | 0.706967383 | 0.001099855 |
| Scaper | 0.000555011 | 0.804780124 | 0.811 | 0.669 | 1 | 0.856052091 | 0.186839874 | 0.732 | 0.778 | 1 | 0.856052091 | 0.001109713 |
| Cdk9 | 0.000555853 | 1.234892943 | 0.566 | 0.424 | 1 | 0.11586769 | 0.453657637 | 0.683 | 0.608 | 1 | 0.11586769 | 0.001111397 |
| Gadd45b | 0.000557292 | 0.563700887 | 0.981 | 0.822 | 1 | 0.624551182 | 0.116762787 | 0.951 | 0.922 | 1 | 0.624551182 | 0.001114272 |
| Zfp874b | 0.344979884 | -0.600164907 | 0.132 | 0.182 | 1 | 0.000557865 | -2.577832649 | 0.049 | 0.304 | 1 | 0.344979884 | 0.001115418 |
| Cystm1 | 0.000558585 | -2.334662246 | 0.189 | 0.411 | 1 | 0.638635874 | -0.281736223 | 0.317 | 0.256 | 1 | 0.638635874 | 0.001116858 |
| Ap3s2 | 0.000558897 | 1.549001534 | 0.302 | 0.119 | 1 | 0.184443094 | 0.898605394 | 0.195 | 0.119 | 1 | 0.184443094 | 0.001117482 |
| Mapre3 | 0.082821527 | -0.729480406 | 0.472 | 0.597 | 1 | 0.000559233 | -1.192542494 | 0.39 | 0.669 | 1 | 0.082821527 | 0.001118153 |
| Bmp4 | 0.029976142 | 1.46284489 | 0.132 | 0.051 | 1 | 0.000561296 | 2.360766806 | 0.22 | 0.065 | 1 | 0.029976142 | 0.001122277 |
| Actr1a | 0.000561939 | 1.011764641 | 0.623 | 0.436 | 1 | 0.105349876 | 0.338398993 | 0.756 | 0.666 | 1 | 0.105349876 | 0.001123562 |
| Bach2 | 0.000563886 | 0.858737767 | 0.717 | 0.508 | 1 | 0.179998091 | 0.475384028 | 0.805 | 0.717 | 1 | 0.179998091 | 0.001127454 |
| Hsd17b4 | 0.000565776 | 1.491757582 | 0.472 | 0.263 | 1 | 0.928336749 | 0.225770138 | 0.415 | 0.461 | 1 | 0.928336749 | 0.001131232 |
| Ift140 | 0.000567347 | 2.526691372 | 0.208 | 0.064 | 1 | 0.497500298 | 0.904319046 | 0.122 | 0.092 | 1 | 0.497500298 | 0.001134372 |
| Cnksr3 | 0.182171733 | 0.810193071 | 0.453 | 0.403 | 1 | 0.000568113 | 1.388271478 | 0.756 | 0.597 | 1 | 0.182171733 | 0.001135903 |
| 4930435F18Rik | 0.075891445 | 0.984797593 | 0.075 | 0.025 | 1 | 0.000568409 | 3.126711467 | 0.122 | 0.02 | 1 | 0.075891445 | 0.001136495 |
| Gemin7 | 0.000568554 | 0.895988326 | 0.642 | 0.441 | 1 | 0.23274084 | -0.17660095 | 0.366 | 0.485 | 1 | 0.23274084 | 0.001136785 |
| Gm56560 | 0.001102793 | 2.776210972 | 0.17 | 0.047 | 1 | 0.000568997 | 2.334704509 | 0.195 | 0.051 | 1 | 0.001102793 | 0.00113767 |
| Rab5c | 0.000569379 | 0.888442529 | 0.698 | 0.517 | 1 | 0.009534218 | 0.632846351 | 0.78 | 0.669 | 1 | 0.009534218 | 0.001138434 |
| Paip1 | 0.0005718 | 0.816553859 | 0.774 | 0.61 | 1 | 0.289228737 | 0.198944123 | 0.927 | 0.799 | 1 | 0.289228737 | 0.001143274 |
| Scn5a | 0.000572029 | 2.292226119 | 0.17 | 0.042 | 1 | 0.431666094 | 0.574170444 | 0.073 | 0.044 | 1 | 0.431666094 | 0.00114373 |
| Tnpo3 | 0.000572637 | 0.804225348 | 0.774 | 0.564 | 1 | 0.500757012 | -0.10302293 | 0.732 | 0.765 | 1 | 0.500757012 | 0.001144947 |
| Mdga2 | 0.000573652 | 2.234892943 | 0.113 | 0.017 | 1 | 0.853018888 | 1.930314254 | 0.049 | 0.044 | 1 | 0.853018888 | 0.001146974 |
| Zfp36 | 0.550109397 | -0.182270878 | 0.887 | 0.763 | 1 | 0.000575533 | 0.699047882 | 0.976 | 0.812 | 1 | 0.550109397 | 0.001150734 |
| Rtkn | 0.252373652 | 0.480122882 | 0.34 | 0.267 | 1 | 0.000575782 | -1.622226769 | 0.146 | 0.423 | 1 | 0.252373652 | 0.001151232 |
| Pthlh | 0.024739189 | 1.039245377 | 0.113 | 0.038 | 1 | 0.000577803 | 3.83720485 | 0.171 | 0.041 | 1 | 0.024739189 | 0.001155272 |
| P4htm | 0.000578855 | 1.8327945 | 0.34 | 0.153 | 1 | 0.02754041 | 0.91966701 | 0.293 | 0.154 | 1 | 0.02754041 | 0.001157375 |
| Gm36198 | 0.162321801 | 1.640149422 | 0.075 | 0.034 | 1 | 0.000580166 | 3.574170444 | 0.122 | 0.02 | 1 | 0.162321801 | 0.001159996 |
| Vcpkmt | 0.000580209 | 2.361173472 | 0.17 | 0.042 | 1 | 0.193425864 | 1.209173627 | 0.171 | 0.106 | 1 | 0.193425864 | 0.001160081 |
| Cyth3 | 0.003980534 | 0.545071662 | 0.943 | 0.877 | 1 | 0.000580958 | 0.921290331 | 0.951 | 0.853 | 1 | 0.003980534 | 0.001161579 |
| Gm20732 | 0.00058339 | 1.167000925 | 0.491 | 0.28 | 1 | 0.012640992 | 1.117656309 | 0.537 | 0.362 | 1 | 0.012640992 | 0.001166439 |
| Lbh | 0.000584451 | -1.817570476 | 0.208 | 0.445 | 1 | 0.824927279 | -0.640842447 | 0.122 | 0.13 | 1 | 0.824927279 | 0.00116856 |
| Wdr62 | 0.000584928 | 2.377115016 | 0.113 | 0.017 | 1 | 0.004939943 | 1.667279848 | 0.073 | 0.01 | 1 | 0.004939943 | 0.001169514 |
| Hps1 | 0.000585005 | 1.213616284 | 0.358 | 0.153 | 1 | 0.027280655 | -1.747757651 | 0.024 | 0.15 | 1 | 0.027280655 | 0.001169668 |
| Gon4l | 0.000585207 | 1.077646372 | 0.66 | 0.458 | 1 | 0.923758478 | 0.145327145 | 0.537 | 0.539 | 1 | 0.923758478 | 0.001170072 |
| Ccdc84 | 0.000585796 | 1.402650108 | 0.302 | 0.114 | 1 | 0.160741387 | -0.46860358 | 0.122 | 0.222 | 1 | 0.160741387 | 0.001171249 |
| Tube1 | 0.000585813 | 2.154722595 | 0.113 | 0.017 | 1 | 0.175856887 | 2.83720485 | 0.049 | 0.017 | 1 | 0.175856887 | 0.001171282 |
| 1810053B23Rik | 0.000586577 | -5.850901954 | 0 | 0.191 | 1 | 0.208849348 | -2.20718927 | 0 | 0.038 | 1 | 0.208849348 | 0.001172811 |
| Mcm7 | 0.000587259 | 1.624207878 | 0.34 | 0.148 | 1 | 0.450024993 | 0.515276755 | 0.22 | 0.171 | 1 | 0.450024993 | 0.001174173 |
| Lyar | 0.000587524 | 1.039245377 | 0.642 | 0.411 | 1 | 0.527430584 | 0.129845718 | 0.341 | 0.427 | 1 | 0.527430584 | 0.001174704 |
| Trmt13 | 0.000590663 | 1.79482665 | 0.434 | 0.237 | 1 | 0.950481861 | 0.169780189 | 0.195 | 0.188 | 1 | 0.950481861 | 0.001180977 |
| Pcgf2 | 0.000592148 | 1.555260524 | 0.377 | 0.186 | 1 | 0.002100909 | 1.263213467 | 0.512 | 0.311 | 1 | 0.002100909 | 0.001183946 |
| Armc9 | 0.000592773 | 1.657222935 | 0.358 | 0.161 | 1 | 0.372573848 | 0.186440291 | 0.341 | 0.273 | 1 | 0.372573848 | 0.001185194 |
| Fam117a | 0.51711827 | 1.126153443 | 0.17 | 0.14 | 1 | 0.000593529 | 1.721727632 | 0.317 | 0.119 | 1 | 0.51711827 | 0.001186706 |
| Crybg1 | 0.000594946 | -2.503488888 | 0.019 | 0.229 | 1 | 0.879096605 | -0.435813645 | 0.22 | 0.218 | 1 | 0.879096605 | 0.001189539 |
| Zbed6 | 0.000597434 | 0.820083448 | 0.774 | 0.627 | 1 | 0.42096484 | 0.205421492 | 0.878 | 0.799 | 1 | 0.42096484 | 0.001194511 |
| Card19 | 0.178507735 | -0.288220901 | 0.472 | 0.593 | 1 | 0.000597747 | -1.033512133 | 0.463 | 0.693 | 1 | 0.178507735 | 0.001195137 |
| Anpep | 0.000598373 | 1.624207878 | 0.245 | 0.081 | 1 | 0.030881601 | 1.174239837 | 0.122 | 0.041 | 1 | 0.030881601 | 0.001196389 |
| Trub2 | 0.000600137 | 1.727301371 | 0.34 | 0.144 | 1 | 0.270979921 | -0.435813645 | 0.146 | 0.225 | 1 | 0.270979921 | 0.001199914 |
| Vegfa | 0.73354819 | -0.25917308 | 0.585 | 0.504 | 1 | 0.000600619 | 1.190841804 | 0.61 | 0.389 | 1 | 0.73354819 | 0.001200878 |
| Gm31323 | 0.007054001 | 1.8327945 | 0.132 | 0.038 | 1 | 0.000603118 | 2.040738244 | 0.22 | 0.065 | 1 | 0.007054001 | 0.001205872 |
| Phf20l1 | 0.00060578 | 0.692015844 | 0.849 | 0.767 | 1 | 0.044751997 | 0.614812428 | 0.829 | 0.792 | 1 | 0.044751997 | 0.001211193 |
| Pola2 | 0.000607597 | 2.454282877 | 0.17 | 0.042 | 1 | 0.252939974 | -0.863234868 | 0.024 | 0.072 | 1 | 0.252939974 | 0.001214824 |
| Dennd5b | 0.0006091 | 1.334876729 | 0.585 | 0.352 | 1 | 0.235746219 | 0.736275941 | 0.439 | 0.389 | 1 | 0.235746219 | 0.001217828 |
| Myo3b | 0.000611524 | 3.154722595 | 0.132 | 0.025 | 1 | 0.224884123 | 1.30115195 | 0.073 | 0.034 | 1 | 0.224884123 | 0.001222675 |
| Gaa | 0.000613086 | -0.845277405 | 0.472 | 0.729 | 1 | 0.145137511 | -0.441867766 | 0.537 | 0.645 | 1 | 0.145137511 | 0.001225797 |
| Abcf1 | 0.00061376 | 0.740973791 | 0.849 | 0.699 | 1 | 0.688436178 | 0.339374941 | 0.829 | 0.782 | 1 | 0.688436178 | 0.001227144 |
| Xaf1 | 0.457052519 | -0.527101445 | 0.17 | 0.123 | 1 | 0.000615502 | 2.313642894 | 0.244 | 0.082 | 1 | 0.457052519 | 0.001230625 |
| Cbx1 | 0.000616786 | 0.660430453 | 0.868 | 0.826 | 1 | 0.492366469 | 0.369170373 | 0.829 | 0.904 | 1 | 0.492366469 | 0.001233192 |
| Mbd2 | 0.178736688 | 0.289556932 | 1 | 0.962 | 1 | 0.000616829 | -0.607405765 | 0.927 | 0.986 | 1 | 0.178736688 | 0.001233277 |
| Zfp711 | 0.000617004 | 3.739685096 | 0.132 | 0.025 | 1 | 0.160490686 | 0.986348289 | 0.195 | 0.123 | 1 | 0.160490686 | 0.001233628 |
| Ksr1 | 0.000619794 | 3.237184755 | 0.189 | 0.055 | 1 | 0.034053145 | 0.444887427 | 0.171 | 0.068 | 1 | 0.034053145 | 0.001239205 |
| Rarres1 | 0.000622855 | -4.308801778 | 0.057 | 0.267 | 1 | 0.116342058 | -1.513292397 | 0.146 | 0.249 | 1 | 0.116342058 | 0.001245321 |
| Fam110c | 0.343000446 | -0.430239906 | 0 | 0.017 | 1 | 0.000623335 | 2.83720485 | 0.122 | 0.02 | 1 | 0.343000446 | 0.001246281 |
| Magoh | 0.000623676 | 0.494032768 | 0.962 | 0.797 | 1 | 0.109047734 | -0.141208288 | 0.78 | 0.901 | 1 | 0.109047734 | 0.001246963 |
| Ccdc157 | 0.883731495 | 0.234157062 | 0.415 | 0.462 | 1 | 0.000624628 | -1.390330875 | 0.22 | 0.502 | 1 | 0.883731495 | 0.001248866 |
| Pdlim1 | 0.000626072 | 2.045098104 | 0.283 | 0.106 | 1 | 0.195604405 | 0.296636468 | 0.39 | 0.294 | 1 | 0.195604405 | 0.001251752 |
| Mzf1 | 0.917883255 | 0.569760094 | 0.019 | 0.021 | 1 | 0.000629227 | 3.029849928 | 0.122 | 0.02 | 1 | 0.917883255 | 0.001258059 |
| C430049B03Rik | 0.000631443 | 1.734390796 | 0.34 | 0.153 | 1 | 0.00289808 | 1.652780279 | 0.317 | 0.14 | 1 | 0.00289808 | 0.001262488 |
| Timm44 | 0.000632558 | 1.323120126 | 0.528 | 0.335 | 1 | 0.84412766 | 0.291555273 | 0.341 | 0.403 | 1 | 0.84412766 | 0.001264715 |
| Phf2 | 0.000632596 | 0.980693195 | 0.66 | 0.47 | 1 | 0.66257612 | 0.164089718 | 0.707 | 0.795 | 1 | 0.66257612 | 0.001264792 |
| Slc38a3 | 0.000632962 | -1.642290383 | 0.113 | 0.356 | 1 | 0.003920295 | -0.642788091 | 0.171 | 0.437 | 1 | 0.003920295 | 0.001265524 |
| Sumf2 | 0.000633767 | 1.105253919 | 0.491 | 0.271 | 1 | 0.760538085 | 0.450181727 | 0.317 | 0.321 | 1 | 0.760538085 | 0.001267133 |

| | | | | | | | | | | | | |
|---|---|---|---|---|---|---|---|---|---|---|---|---|
| Eif4g3 | 0.000633797 | 0.802584989 | 0.925 | 0.919 | 1 | 0.636973008 | 0.131226948 | 1 | 0.945 | 1 | 0.636973008 | 0.001267192 |
| Cfap141 | 0.000634286 | 2.395730694 | 0.17 | 0.042 | 1 | 0.627074867 | -0.292078167 | 0.073 | 0.096 | 1 | 0.627074867 | 0.001268169 |
| Atxn7l3b | 0.000635426 | 0.459382546 | 0.943 | 0.907 | 1 | 0.005503956 | -0.427848724 | 0.854 | 0.935 | 1 | 0.005503956 | 0.001270448 |
| Suclg1 | 0.000638245 | 0.71862348 | 0.717 | 0.492 | 1 | 0.009166892 | -0.47113418 | 0.439 | 0.652 | 1 | 0.009166892 | 0.001276083 |
| Tcof1 | 0.000640413 | 1.154722595 | 0.453 | 0.237 | 1 | 0.688660118 | 0.20715446 | 0.268 | 0.317 | 1 | 0.688660118 | 0.001280416 |
| St6galnac3 | 0.000640648 | 3.533234218 | 0.132 | 0.025 | 1 | 0.346243518 | 2.83720485 | 0.049 | 0.024 | 1 | 0.346243518 | 0.001280885 |
| Gm44699 | 0.31101939 | -0.385845787 | 0.132 | 0.195 | 1 | 0.000641928 | -2.081658387 | 0.049 | 0.311 | 1 | 0.31101939 | 0.001283445 |
| C78859 | 0.56579457 | -0.652632327 | 0.019 | 0.034 | 1 | 0.00064298 | 3.252242349 | 0.122 | 0.02 | 1 | 0.56579457 | 0.001285547 |
| Osbpl10 | 0.000647164 | 1.513176566 | 0.283 | 0.106 | 1 | 0.383877203 | 0.433205622 | 0.39 | 0.328 | 1 | 0.383877203 | 0.00129391 |
| Pdlim4 | 0.000648209 | 0.969166942 | 0.491 | 0.263 | 1 | 0.416642507 | 0.674933421 | 0.415 | 0.382 | 1 | 0.416642507 | 0.001295998 |
| Cacna1e | 0.000648308 | 1.89946554 | 0.17 | 0.042 | 1 | 0.835815214 | 0.491430013 | 0.146 | 0.167 | 1 | 0.835815214 | 0.001296195 |
| Ttc39c | 0.000968965 | 2.46284489 | 0.226 | 0.081 | 1 | 0.000650932 | 2.322631677 | 0.244 | 0.078 | 1 | 0.000968965 | 0.001301441 |
| Nxt2 | 0.000651202 | 1.813685677 | 0.226 | 0.072 | 1 | 0.439486535 | -0.116991461 | 0.049 | 0.085 | 1 | 0.439486535 | 0.00130198 |
| Sdc3 | 0.197694917 | -0.371687724 | 0.679 | 0.725 | 1 | 0.00065349 | -0.769700204 | 0.634 | 0.833 | 1 | 0.197694917 | 0.001306552 |
| Gm47889 | 0.000654181 | 3.154722595 | 0.151 | 0.034 | 1 | 0.078585422 | 2.100239256 | 0.098 | 0.038 | 1 | 0.078585422 | 0.001307933 |
| mt-Atp6 | 0.000656811 | -0.336998489 | 1 | 1 | 1 | 0.20802093 | -0.214563995 | 1 | 1 | 1 | 0.20802093 | 0.00131319 |
| Pcsk6 | 0.000660375 | -1.494370243 | 0.415 | 0.61 | 1 | 0.018403118 | -2.231274888 | 0.073 | 0.235 | 1 | 0.018403118 | 0.001320314 |
| Zfc3h1 | 0.000660487 | 0.54521791 | 0.925 | 0.877 | 1 | 0.018035225 | 0.43121249 | 0.976 | 0.962 | 1 | 0.018035225 | 0.001320538 |
| Wwc2 | 0.000660677 | 0.712204359 | 0.943 | 0.877 | 1 | 0.755710853 | 0.113265936 | 0.976 | 0.891 | 1 | 0.755710853 | 0.001320918 |
| Msantd4 | 0.000663306 | 1.126153443 | 0.509 | 0.309 | 1 | 0.436210528 | 0.895757032 | 0.39 | 0.386 | 1 | 0.436210528 | 0.001326172 |
| Ccdc25 | 0.000663911 | 0.927807615 | 0.679 | 0.5 | 1 | 0.19481018 | -0.322451816 | 0.512 | 0.648 | 1 | 0.19481018 | 0.001327381 |
| Hnrnph1 | 0.000667109 | 0.487810398 | 0.981 | 0.966 | 1 | 0.018043857 | 0.385359726 | 1 | 0.986 | 1 | 0.018043857 | 0.001333773 |
| Prdm16 | 0.052968909 | 0.322845354 | 0.887 | 0.78 | 1 | 0.000671738 | -1.005438822 | 0.78 | 0.911 | 1 | 0.052968909 | 0.001343025 |
| Sh3bgrl2 | 0.007101514 | 3.017219071 | 0.113 | 0.03 | 1 | 0.000677774 | 1.328191202 | 0.22 | 0.061 | 1 | 0.007101514 | 0.001355089 |
| Rtf1 | 0.000678332 | 0.814770818 | 0.83 | 0.72 | 1 | 0.153726816 | 0.415863608 | 0.902 | 0.891 | 1 | 0.153726816 | 0.001356203 |
| Zfp334 | 0.000680007 | 1.855162313 | 0.189 | 0.051 | 1 | 0.327880046 | 0.83720485 | 0.22 | 0.171 | 1 | 0.327880046 | 0.001359552 |
| Gm2a | 0.000680195 | 1.225425586 | 0.66 | 0.517 | 1 | 0.951777073 | 0.11967323 | 0.61 | 0.679 | 1 | 0.951777073 | 0.001359927 |
| Angptl7 | 0.001570137 | 1.322450041 | 0.208 | 0.068 | 1 | 0.000680295 | 2.040738244 | 0.146 | 0.031 | 1 | 0.001570137 | 0.001360127 |
| Vamp5 | 0.061349391 | -0.910751243 | 0.377 | 0.496 | 1 | 0.000683064 | -2.116991461 | 0.146 | 0.406 | 1 | 0.061349391 | 0.001365661 |
| Acy1 | 0.000684577 | 2.154722595 | 0.17 | 0.042 | 1 | 0.208345579 | -0.917682652 | 0.024 | 0.078 | 1 | 0.208345579 | 0.001368686 |
| Spon2 | 0.000687993 | 2.517292674 | 0.151 | 0.034 | 1 | 0.82181816 | 0.252242349 | 0.024 | 0.031 | 1 | 0.82181816 | 0.001375513 |
| 1110038B12Rik | 0.000689912 | 0.722611582 | 0.755 | 0.64 | 1 | 0.630909061 | 0.377773231 | 0.61 | 0.604 | 1 | 0.630909061 | 0.001379347 |
| Slc20a2 | 0.000690108 | -1.460936703 | 0.34 | 0.581 | 1 | 0.176987171 | 0.933634612 | 0.22 | 0.352 | 1 | 0.176987171 | 0.00137974 |
| Dffb | 0.000693267 | 1.739685096 | 0.189 | 0.051 | 1 | 0.938493157 | 0.515276755 | 0.073 | 0.072 | 1 | 0.938493157 | 0.001386054 |
| Arrdc3 | 0.00163195 | 1.27810501 | 0.491 | 0.292 | 1 | 0.000694855 | 1.774920572 | 0.61 | 0.399 | 1 | 0.00163195 | 0.001389228 |
| Lrp1 | 0.000700035 | 0.324384355 | 1 | 0.983 | 1 | 0.020025361 | 0.365710841 | 1 | 0.986 | 1 | 0.020025361 | 0.00139958 |
| Ube2d2a | 0.000701705 | 0.413809816 | 0.943 | 0.928 | 1 | 0.146606533 | 0.230367849 | 0.976 | 0.966 | 1 | 0.146606533 | 0.001402917 |
| Stat1 | 0.006435572 | 0.866477626 | 0.396 | 0.225 | 1 | 0.000703141 | 1.38230289 | 0.488 | 0.249 | 1 | 0.006435572 | 0.001405788 |
| Rbfa | 0.000703595 | 1.700156731 | 0.321 | 0.14 | 1 | 0.826238249 | 0.614812428 | 0.171 | 0.201 | 1 | 0.826238249 | 0.001406696 |
| Snrpg | 0.000704154 | 0.612294138 | 0.943 | 0.881 | 1 | 0.291970159 | 0.313156729 | 0.902 | 0.898 | 1 | 0.291970159 | 0.001407812 |
| Pcdh9 | 0.006135961 | -1.132733283 | 0.547 | 0.725 | 1 | 0.000706259 | -1.476250198 | 0.463 | 0.72 | 1 | 0.006135961 | 0.001412018 |
| Rab2b | 0.000707036 | 1.098579517 | 0.585 | 0.386 | 1 | 0.405615893 | -0.179096963 | 0.317 | 0.386 | 1 | 0.405615893 | 0.001413571 |
| Sart3 | 0.000709788 | 1.283455909 | 0.434 | 0.233 | 1 | 0.623088515 | 0.476017324 | 0.39 | 0.372 | 1 | 0.623088515 | 0.001419073 |
| Casp6 | 0.000713366 | 1.669295768 | 0.226 | 0.072 | 1 | 0.028132794 | 1.313642894 | 0.244 | 0.123 | 1 | 0.028132794 | 0.001426223 |
| Stx11 | 0.005190065 | 1.782753817 | 0.208 | 0.081 | 1 | 0.00071356 | 2.209173627 | 0.268 | 0.096 | 1 | 0.005190065 | 0.00142661 |
| Tceal9 | 0.000713642 | 0.463393496 | 0.962 | 0.919 | 1 | 0.289678036 | -0.173031106 | 0.951 | 0.959 | 1 | 0.289678036 | 0.001426776 |
| Cfap70 | 0.000713837 | 1.339147166 | 0.245 | 0.085 | 1 | 0.00360508 | -1.923607486 | 0.049 | 0.266 | 1 | 0.00360508 | 0.001427164 |
| Vps33a | 0.000714631 | 1.614154213 | 0.34 | 0.153 | 1 | 0.296524731 | 0.462809335 | 0.366 | 0.304 | 1 | 0.296524731 | 0.001428752 |
| Slc12a1 | 0.000715947 | -3.896939525 | 0.038 | 0.242 | 1 | 0.741726892 | -1.448197369 | 0.024 | 0.034 | 1 | 0.741726892 | 0.001431381 |
| Rtn4rl1 | 0.003170651 | 2.640149422 | 0.132 | 0.034 | 1 | 0.000716574 | 3.660327088 | 0.146 | 0.031 | 1 | 0.003170651 | 0.001432634 |
| Brca1 | 0.321056007 | -0.464187238 | 0.094 | 0.055 | 1 | 0.000719665 | 2.389745873 | 0.122 | 0.02 | 1 | 0.321056007 | 0.001438813 |
| Rapgef5 | 0.000722966 | 2.602181572 | 0.151 | 0.034 | 1 | 0.040586918 | 2.212713985 | 0.122 | 0.044 | 1 | 0.040586918 | 0.00144541 |
| Mrps26 | 0.000723095 | 1.10741688 | 0.528 | 0.309 | 1 | 0.211356184 | -0.104692196 | 0.244 | 0.358 | 1 | 0.211356184 | 0.001445666 |
| U2af1 | 0.201650083 | 0.207556042 | 0.849 | 0.843 | 1 | 0.000726143 | -0.534763928 | 0.78 | 0.942 | 1 | 0.201650083 | 0.001451759 |
| Ptgs1 | 0.000728004 | -1.990327738 | 0.17 | 0.398 | 1 | 0.069203575 | -0.957211017 | 0.073 | 0.195 | 1 | 0.069203575 | 0.001455478 |
| Rab11fip5 | 0.086599295 | -0.677154646 | 0.283 | 0.407 | 1 | 0.000732476 | -1.054519298 | 0.439 | 0.71 | 1 | 0.086599295 | 0.001464415 |
| Pygl | 0.000734075 | -1.545717123 | 0.132 | 0.377 | 1 | 0.951356519 | 0.174239837 | 0.268 | 0.287 | 1 | 0.951356519 | 0.001467611 |
| Gm17354 | 0.00073649 | 2.347367673 | 0.132 | 0.025 | 1 | 0.495369687 | -0.278272368 | 0.049 | 0.078 | 1 | 0.495369687 | 0.001472437 |
| Ndufs7 | 0.000741076 | 0.651060196 | 0.792 | 0.564 | 1 | 0.039155857 | 0.69855131 | 0.659 | 0.56 | 1 | 0.039155857 | 0.001481603 |
| Smpdl3b | 0.000741583 | 2.072260435 | 0.208 | 0.064 | 1 | 0.34180587 | 1.351778023 | 0.073 | 0.041 | 1 | 0.34180587 | 0.001482616 |
| Dpp3 | 0.000741756 | 1.315187267 | 0.302 | 0.119 | 1 | 0.049809518 | 1.091777677 | 0.268 | 0.154 | 1 | 0.049809518 | 0.001482961 |
| Tdrd7 | 0.000744633 | 1.583565894 | 0.377 | 0.174 | 1 | 0.119686106 | -0.517637868 | 0.171 | 0.287 | 1 | 0.119686106 | 0.001488711 |
| Klf6 | 0.985448977 | -0.114522173 | 0.943 | 0.886 | 1 | 0.000744809 | 0.635327886 | 1 | 0.877 | 1 | 0.985448977 | 0.001489063 |
| 2410006H16Rik | 0.000745899 | 0.728663689 | 0.868 | 0.784 | 1 | 0.913855897 | 0.337738569 | 0.634 | 0.717 | 1 | 0.913855897 | 0.001491241 |
| Arl4c | 0.549185868 | 0.953088734 | 0.038 | 0.059 | 1 | 0.000746605 | 1.936740523 | 0.22 | 0.065 | 1 | 0.549185868 | 0.001492653 |
| Sgce | 0.000749378 | 1.055186921 | 0.453 | 0.233 | 1 | 0.988354601 | 0.645584777 | 0.366 | 0.427 | 1 | 0.988354601 | 0.001498194 |
| Ly6c1 | 0.000749552 | -2.982780929 | 0.038 | 0.25 | 1 | 0.05173727 | -2.099433089 | 0.098 | 0.225 | 1 | 0.05173727 | 0.001498543 |
| Ipo8 | 0.000750999 | 1.071306587 | 0.491 | 0.275 | 1 | 0.60688895 | 0.367719567 | 0.463 | 0.433 | 1 | 0.60688895 | 0.001501434 |
| Atg10 | 0.06369536 | 0.727301371 | 0.302 | 0.186 | 1 | 0.000751413 | 1.610433988 | 0.439 | 0.222 | 1 | 0.06369536 | 0.001502261 |
| Nol11 | 0.000755659 | 0.777868289 | 0.642 | 0.428 | 1 | 0.259745089 | 0.279828181 | 0.366 | 0.502 | 1 | 0.259745089 | 0.001510747 |
| Actr5 | 0.000756586 | 2.154722595 | 0.208 | 0.064 | 1 | 0.057552699 | 1.01123425 | 0.195 | 0.096 | 1 | 0.057552699 | 0.001512599 |
| Alpk1 | 0.000756668 | -1.337875888 | 0.302 | 0.551 | 1 | 0.078196943 | -0.671136369 | 0.22 | 0.389 | 1 | 0.078196943 | 0.001512764 |
| Ginm1 | 0.000756767 | 0.634033308 | 0.906 | 0.792 | 1 | 0.917490538 | 0.156483366 | 0.805 | 0.829 | 1 | 0.917490538 | 0.001512962 |
| Trappc9 | 0.000757423 | 0.841746101 | 0.698 | 0.479 | 1 | 0.20264868 | 0.469229387 | 0.634 | 0.59 | 1 | 0.20264868 | 0.001514273 |
| Lrig3 | 0.227265493 | 0.224801031 | 0.755 | 0.691 | 1 | 0.000757866 | 0.968709633 | 0.78 | 0.642 | 1 | 0.227265493 | 0.001515157 |
| Uba5 | 0.00076339 | 0.730785036 | 0.774 | 0.593 | 1 | 0.02779687 | 0.640807637 | 0.78 | 0.686 | 1 | 0.02779687 | 0.001526197 |
| Blvrb | 0.000766037 | 1.493524508 | 0.415 | 0.225 | 1 | 0.019022352 | 0.936740523 | 0.488 | 0.311 | 1 | 0.019022352 | 0.001531488 |
| Ptprk | 0.000770207 | -0.763506513 | 0.981 | 0.979 | 1 | 0.127220868 | -0.147626272 | 0.927 | 0.969 | 1 | 0.127220868 | 0.001539821 |
| Cmss1 | 0.905132392 | -0.138257631 | 1 | 1 | 1 | 0.000771957 | 0.421715826 | 1 | 1 | 1 | 0.905132392 | 0.001543319 |
| Ccne1 | 0.003453733 | 1.920257341 | 0.17 | 0.055 | 1 | 0.000773247 | 2.515276755 | 0.146 | 0.031 | 1 | 0.003453733 | 0.001545895 |
| Slc20a1 | 0.000774876 | -1.822557329 | 0.604 | 0.712 | 1 | 0.193839352 | -0.818146979 | 0.683 | 0.512 | 1 | 0.193839352 | 0.001549151 |
| Lag3 | 0.000777563 | 2.8327945 | 0.151 | 0.034 | 1 | 0.236150728 | 0.589277336 | 0.171 | 0.106 | 1 | 0.236150728 | 0.001554521 |
| Ice1 | 0.000781491 | 1.007380879 | 0.679 | 0.508 | 1 | 0.039684649 | 0.530543512 | 0.756 | 0.648 | 1 | 0.039684649 | 0.001562371 |
| Dab2ip | 0.000781647 | 0.666621633 | 0.698 | 0.462 | 1 | 0.761396767 | -0.180222222 | 0.805 | 0.72 | 1 | 0.761396767 | 0.001562683 |
| Acot6 | 0.000782756 | 2.154722595 | 0.132 | 0.025 | 1 | 0.94215184 | 0.252242349 | 0.049 | 0.051 | 1 | 0.94215184 | 0.001564898 |
| Trip10 | 0.000782813 | 1.201516806 | 0.491 | 0.275 | 1 | 0.51637985 | -0.391613841 | 0.366 | 0.41 | 1 | 0.51637985 | 0.001565014 |
| Kbtbd4 | 0.000783131 | 1.640149422 | 0.226 | 0.072 | 1 | 0.32477683 | -0.56878751 | 0.098 | 0.154 | 1 | 0.32477683 | 0.001565648 |
| Polr2i | 0.000788621 | 1.020866848 | 0.491 | 0.263 | 1 | 0.527459415 | 0.601988388 | 0.317 | 0.297 | 1 | 0.527459415 | 0.00157662 |
| Rchy1 | 0.000792097 | 1.034983351 | 0.547 | 0.339 | 1 | 0.613225429 | 0.328863631 | 0.415 | 0.389 | 1 | 0.613225429 | 0.001583567 |
| Rab40c | 0.000792103 | 1.27153626 | 0.453 | 0.25 | 1 | 0.011766985 | 0.729676385 | 0.512 | 0.321 | 1 | 0.011766985 | 0.00158358 |
| Trim3 | 0.000792261 | 1.036078098 | 0.434 | 0.22 | 1 | 0.553026285 | 0.345351753 | 0.366 | 0.324 | 1 | 0.553026285 | 0.001583894 |
| Dph3 | 0.276147888 | -0.113766241 | 0.585 | 0.686 | 1 | 0.000795378 | -1.054214337 | 0.366 | 0.638 | 1 | 0.276147888 | 0.001590124 |
| Extl2 | 0.000796064 | 1.20255112 | 0.509 | 0.292 | 1 | 0.870651535 | 0.30115195 | 0.268 | 0.294 | 1 | 0.870651535 | 0.001591495 |
| Nmrk1 | 0.000796819 | 1.47665069 | 0.34 | 0.144 | 1 | 0.59431832 | -0.262330824 | 0.122 | 0.154 | 1 | 0.59431832 | 0.001593004 |
| Tmem135 | 0.000797578 | 1.087608399 | 0.528 | 0.331 | 1 | 0.901693128 | 0.420467062 | 0.463 | 0.502 | 1 | 0.901693128 | 0.001594521 |
| Dock3 | 0.000799074 | 2.029191713 | 0.132 | 0.025 | 1 | 0.113990016 | 1.377773231 | 0.122 | 0.058 | 1 | 0.113990016 | 0.001597509 |
| D16Ertd472e | 0.000799163 | 1.351414506 | 0.434 | 0.225 | 1 | 0.451549157 | 0.315967947 | 0.366 | 0.324 | 1 | 0.451549157 | 0.001597688 |
| Ntn1 | 0.118184039 | 0.154722595 | 0.113 | 0.055 | 1 | 0.00080153 | 1.737669176 | 0.195 | 0.051 | 1 | 0.118184039 | 0.001602417 |

| | | | | | | | | | | | | |
|---|---|---|---|---|---|---|---|---|---|---|---|---|
| Unc93b1 | 0.000801922 | 1.341135719 | 0.377 | 0.178 | 1 | 0.872602094 | 0.658867609 | 0.22 | 0.215 | 1 | 0.872602094 | 0.001603201 |
| Orai2 | 0.000803265 | 1.285967128 | 0.302 | 0.119 | 1 | 0.358414592 | 0.551802631 | 0.122 | 0.078 | 1 | 0.358414592 | 0.001605885 |
| Ddx55 | 0.000804828 | 1.18386894 | 0.321 | 0.131 | 1 | 0.304029877 | 0.396632258 | 0.244 | 0.177 | 1 | 0.304029877 | 0.001609007 |
| Lrig1 | 0.002135938 | 1.502645898 | 0.358 | 0.186 | 1 | 0.000805962 | 1.465236072 | 0.537 | 0.3 | 1 | 0.002135938 | 0.001611275 |
| Gmcl1 | 0.000809202 | 1.683416876 | 0.396 | 0.195 | 1 | 0.024674435 | 1.252242349 | 0.512 | 0.348 | 1 | 0.024674435 | 0.001617749 |
| Csnk1d | 0.000812273 | 0.798578785 | 0.792 | 0.564 | 1 | 0.057256824 | 0.459216796 | 0.829 | 0.768 | 1 | 0.057256824 | 0.001623885 |
| Zc3h7b | 0.00082047 | 0.844867332 | 0.585 | 0.331 | 1 | 0.040411013 | 0.484903106 | 0.61 | 0.478 | 1 | 0.040411013 | 0.001640267 |
| Dlg5 | 0.000822237 | 0.922061838 | 0.453 | 0.229 | 1 | 0.636520092 | 0.33109777 | 0.512 | 0.471 | 1 | 0.636520092 | 0.001643798 |
| Mbnl1 | 0.000823254 | -1.034530668 | 1 | 1 | 1 | 0.047074346 | -0.47865603 | 1 | 0.997 | 1 | 0.047074346 | 0.001645831 |
| Ube2v1 | 0.000824826 | 0.910303867 | 0.642 | 0.458 | 1 | 0.91814125 | 0.144182603 | 0.585 | 0.604 | 1 | 0.91814125 | 0.001648972 |
| Pdpr | 0.000827201 | 1.644527862 | 0.34 | 0.153 | 1 | 0.904554548 | 0.412707021 | 0.244 | 0.242 | 1 | 0.904554548 | 0.001653718 |
| Phf1 | 0.000829068 | -1.295310326 | 0.189 | 0.432 | 1 | 0.556052262 | 0.176954222 | 0.341 | 0.44 | 1 | 0.556052262 | 0.00165745 |
| Peg12 | 0.000831604 | 2.029191713 | 0.151 | 0.034 | 1 | 0.355768344 | 1.377773231 | 0.049 | 0.024 | 1 | 0.355768344 | 0.001662516 |
| Ring1 | 0.123558934 | 0.585356949 | 0.415 | 0.314 | 1 | 0.000835981 | -1.563674586 | 0.171 | 0.43 | 1 | 0.123558934 | 0.001671264 |
| Scarb1 | 0.000836891 | -1.922914817 | 0.283 | 0.504 | 1 | 0.482892402 | -0.146307027 | 0.244 | 0.307 | 1 | 0.482892402 | 0.001673081 |
| Usp25 | 0.428721309 | -0.237594828 | 0.811 | 0.669 | 1 | 0.00083984 | 0.774290642 | 0.854 | 0.676 | 1 | 0.428721309 | 0.001678975 |
| Fgd4 | 0.000842622 | 1.513176566 | 0.283 | 0.11 | 1 | 0.003320385 | 1.177280291 | 0.488 | 0.276 | 1 | 0.003320385 | 0.001684534 |
| Pak1ip1 | 0.000843068 | 0.717317282 | 0.792 | 0.585 | 1 | 0.919470269 | 0.167353451 | 0.732 | 0.741 | 1 | 0.919470269 | 0.001685426 |
| Slc35a3 | 0.000843107 | -0.98729641 | 0.34 | 0.606 | 1 | 0.001871647 | -1.013651711 | 0.439 | 0.659 | 1 | 0.001871647 | 0.001685503 |
| Gm56743 | 0.239528771 | 0.434830514 | 0.264 | 0.191 | 1 | 0.000844838 | -1.829551742 | 0.098 | 0.355 | 1 | 0.239528771 | 0.001688961 |
| Rnf41 | 0.633200918 | 0.351759442 | 0.453 | 0.453 | 1 | 0.000845947 | -0.644446934 | 0.39 | 0.693 | 1 | 0.633200918 | 0.001691179 |
| Cldn10 | 0.000846069 | -3.244448499 | 0.057 | 0.267 | 1 | 0.039271642 | -2.484723245 | 0 | 0.096 | 1 | 0.039271642 | 0.001691421 |
| Spata13 | 0.000846344 | -1.183147044 | 0.453 | 0.64 | 1 | 0.181930714 | -0.269294771 | 0.463 | 0.608 | 1 | 0.181930714 | 0.001691973 |
| Pusl1 | 0.00084884 | 1.874614676 | 0.208 | 0.064 | 1 | 0.37346373 | 1.029849928 | 0.098 | 0.061 | 1 | 0.37346373 | 0.00169696 |
| Lat2 | 0.000849855 | 1.776210972 | 0.17 | 0.042 | 1 | 0.376300429 | -0.484723245 | 0.073 | 0.123 | 1 | 0.376300429 | 0.001698988 |
| Rbp4 | 0.004692063 | -1.792809985 | 0.264 | 0.449 | 1 | 0.000849885 | -1.091560245 | 0.317 | 0.628 | 1 | 0.004692063 | 0.001699048 |
| Tns3 | 0.000851221 | 0.609073018 | 0.83 | 0.699 | 1 | 0.001788573 | 1.051523971 | 0.927 | 0.877 | 1 | 0.001788573 | 0.001701717 |
| Fam107b | 0.000852488 | 0.757832442 | 0.396 | 0.178 | 1 | 0.61379894 | 0.343216009 | 0.39 | 0.372 | 1 | 0.61379894 | 0.001704249 |
| Sfxn3 | 0.000853586 | 1.389187848 | 0.377 | 0.178 | 1 | 0.386068179 | 0.574170444 | 0.341 | 0.3 | 1 | 0.386068179 | 0.001706443 |
| Mlxip | 0.000857017 | 0.970298024 | 0.623 | 0.432 | 1 | 0.005202693 | 0.911205431 | 0.78 | 0.642 | 1 | 0.005202693 | 0.001713299 |
| Tgfb2 | 0.000860536 | 0.375984548 | 0.962 | 0.78 | 1 | 0.681920372 | -0.420687357 | 0.878 | 0.901 | 1 | 0.681920372 | 0.001720331 |
| Tmem109 | 0.000860602 | 1.032103308 | 0.604 | 0.398 | 1 | 0.052281053 | 0.75267134 | 0.61 | 0.509 | 1 | 0.052281053 | 0.001720464 |
| Fip1l1 | 0.000861611 | 0.618669695 | 0.792 | 0.669 | 1 | 0.120303305 | -0.194404757 | 0.634 | 0.785 | 1 | 0.120303305 | 0.001722479 |
| Opa3 | 0.000862866 | 1.010332685 | 0.472 | 0.242 | 1 | 0.052903262 | -0.854672855 | 0.195 | 0.341 | 1 | 0.052903262 | 0.001724988 |
| Ogdh | 0.004883766 | 0.569760094 | 0.925 | 0.784 | 1 | 0.000868929 | 0.676339188 | 0.878 | 0.829 | 1 | 0.004883766 | 0.001737103 |
| Golga4 | 0.153642616 | 0.262187292 | 0.906 | 0.877 | 1 | 0.000869589 | -0.542982643 | 0.902 | 0.959 | 1 | 0.153642616 | 0.001738421 |
| Csgalnact2 | 0.000870673 | -0.9442665 | 0.547 | 0.716 | 1 | 0.352203962 | -0.195216628 | 0.415 | 0.529 | 1 | 0.352203962 | 0.001740587 |
| Timm23 | 0.547620537 | -0.13301058 | 0.868 | 0.839 | 1 | 0.000870749 | -0.694176611 | 0.634 | 0.86 | 1 | 0.547620537 | 0.001740739 |
| Whrn | 0.000870992 | 0.953088734 | 0.509 | 0.284 | 1 | 0.729299936 | 0.257260424 | 0.463 | 0.444 | 1 | 0.729299936 | 0.001741225 |
| Ikzf2 | 0.000871907 | 1.412218817 | 0.396 | 0.191 | 1 | 0.071391078 | 0.244974349 | 0.463 | 0.324 | 1 | 0.071391078 | 0.001743054 |
| Ccdc115 | 0.000877543 | 1.121555731 | 0.491 | 0.288 | 1 | 0.536084549 | 0.686645173 | 0.293 | 0.266 | 1 | 0.536084549 | 0.001754315 |
| Myl12a | 0.000877673 | 0.510025429 | 1 | 0.97 | 1 | 0.817345357 | 0.165950052 | 1 | 0.986 | 1 | 0.817345357 | 0.001754575 |
| Yars | 0.000881982 | -1.028877344 | 0.358 | 0.614 | 1 | 0.355738874 | -0.147937191 | 0.537 | 0.669 | 1 | 0.355738874 | 0.001763187 |
| Acbd3 | 0.001363623 | 0.481532911 | 0.962 | 0.907 | 1 | 0.000887024 | -0.607916442 | 0.927 | 0.973 | 1 | 0.001363623 | 0.00177326 |
| Ehd1 | 0.000888762 | -1.004710033 | 0.509 | 0.695 | 1 | 0.018653801 | -0.629494769 | 0.463 | 0.659 | 1 | 0.018653801 | 0.001776734 |
| Adcy2 | 0.000894146 | 0.930792938 | 0.566 | 0.356 | 1 | 0.354701306 | -0.326900478 | 0.756 | 0.778 | 1 | 0.354701306 | 0.001787493 |
| Nisch | 0.000894148 | 0.764671779 | 0.887 | 0.826 | 1 | 0.288764341 | 0.228591081 | 0.902 | 0.901 | 1 | 0.288764341 | 0.001787496 |
| Rhbdf2 | 0.000896509 | -2.20282941 | 0.038 | 0.246 | 1 | 0.970311198 | 0.322631677 | 0.098 | 0.096 | 1 | 0.970311198 | 0.001792213 |
| Mettl8 | 0.000898448 | 1.762405172 | 0.208 | 0.064 | 1 | 0.293411107 | -0.787286015 | 0.049 | 0.099 | 1 | 0.293411107 | 0.00179609 |
| Sgpl1 | 0.000900057 | 1.063092119 | 0.566 | 0.339 | 1 | 0.163209451 | 0.335765705 | 0.756 | 0.645 | 1 | 0.163209451 | 0.001799303 |
| Sco2 | 0.000900309 | 1.544669113 | 0.283 | 0.11 | 1 | 1 | 0.252242349 | 0.122 | 0.123 | 1 | 1 | 0.001799807 |
| Rad54l2 | 0.000904498 | 0.94321849 | 0.604 | 0.364 | 1 | 0.722501839 | 0.183200705 | 0.561 | 0.526 | 1 | 0.722501839 | 0.001808177 |
| Tasor2 | 0.000905085 | 0.868968112 | 0.698 | 0.547 | 1 | 0.257499735 | 0.221093651 | 0.805 | 0.713 | 1 | 0.257499735 | 0.001809351 |
| Ppfibp1 | 0.001292649 | -0.482029208 | 0.943 | 0.949 | 1 | 0.000909118 | -0.840540321 | 0.951 | 0.928 | 1 | 0.001292649 | 0.001817409 |
| Rplp1 | 0.008018922 | -0.352729667 | 1 | 1 | 1 | 0.000910231 | -0.476903741 | 1 | 0.997 | 1 | 0.008018922 | 0.001819633 |
| Kcnk1 | 0.104757762 | -0.736901244 | 0.358 | 0.453 | 1 | 0.000911017 | -1.622226769 | 0.244 | 0.505 | 1 | 0.104757762 | 0.001821204 |
| Dnajb1 | 0.000913819 | -1.329583692 | 0.736 | 0.78 | 1 | 0.159131647 | -0.486460386 | 0.707 | 0.805 | 1 | 0.159131647 | 0.001826804 |
| Nop10 | 0.000915181 | -0.643413797 | 0.906 | 0.886 | 1 | 0.004534335 | -0.597534564 | 0.707 | 0.86 | 1 | 0.004534335 | 0.001829524 |
| Rfc1 | 0.000916892 | 0.588197342 | 0.698 | 0.559 | 1 | 0.697457596 | 0.141211037 | 0.585 | 0.577 | 1 | 0.697457596 | 0.001832943 |
| Pcid2 | 0.000918037 | 1.18386894 | 0.377 | 0.174 | 1 | 0.190671554 | 0.465236072 | 0.317 | 0.222 | 1 | 0.190671554 | 0.00183523 |
| Ebf1 | 0.005298264 | -0.575494664 | 0.887 | 0.987 | 1 | 0.000923448 | -0.685136242 | 1 | 0.962 | 1 | 0.005298264 | 0.001846044 |
| Cd200 | 0.007153149 | 0.758369659 | 0.849 | 0.801 | 1 | 0.000924091 | 1.175340711 | 0.732 | 0.625 | 1 | 0.007153149 | 0.001847327 |
| Utp18 | 0.000929304 | 0.924109667 | 0.642 | 0.445 | 1 | 0.558268721 | 0.743228702 | 0.561 | 0.601 | 1 | 0.558268721 | 0.001857744 |
| Nploc4 | 0.000931479 | 1.050385935 | 0.585 | 0.373 | 1 | 0.88012161 | 0.299667695 | 0.585 | 0.645 | 1 | 0.88012161 | 0.00186209 |
| Sparcl1 | 0.001290158 | -0.565841392 | 0.453 | 0.729 | 1 | 0.000932707 | -1.655753337 | 0.195 | 0.491 | 1 | 0.001290158 | 0.001864543 |
| Gm56599 | 0.00093563 | 1.377115016 | 0.151 | 0.034 | 1 | 0.605005904 | 0.136765152 | 0.024 | 0.014 | 1 | 0.605005904 | 0.001870384 |
| Tfpt | 0.00093648 | 1.181689642 | 0.377 | 0.174 | 1 | 0.255576973 | -0.500664789 | 0.171 | 0.253 | 1 | 0.255576973 | 0.001872083 |
| C2 | 0.016042068 | 2.154722595 | 0.057 | 0.008 | 1 | 0.000936619 | 3.515276755 | 0.098 | 0.014 | 1 | 0.016042068 | 0.00187236 |
| Asb3 | 0.000937395 | 1.174351402 | 0.566 | 0.36 | 1 | 0.25571683 | -0.605738646 | 0.293 | 0.392 | 1 | 0.25571683 | 0.001873912 |
| Cars2 | 0.000937806 | 2.006623956 | 0.321 | 0.144 | 1 | 0.044097841 | -0.863234868 | 0.049 | 0.181 | 1 | 0.044097841 | 0.001874732 |
| Rabgap1l | 0.007356377 | -0.5420601 | 1 | 0.996 | 1 | 0.000938214 | -0.514287703 | 0.976 | 1 | 1 | 0.007356377 | 0.001875548 |
| Tubb6 | 0.50151687 | -0.376800647 | 0.604 | 0.585 | 1 | 0.000940595 | -1.050949183 | 0.366 | 0.625 | 1 | 0.50151687 | 0.001880305 |
| Slc25a34 | 0.200658585 | 1.154722595 | 0.113 | 0.064 | 1 | 0.000942444 | -3.762707992 | 0 | 0.218 | 1 | 0.200658585 | 0.001883999 |
| Barx2 | 0.000943931 | -3.911366596 | 0 | 0.178 | 1 | 0.057889378 | -2.622226769 | 0 | 0.082 | 1 | 0.057889378 | 0.00188697 |
| Ttll7 | 0.000945879 | 1.171596413 | 0.415 | 0.203 | 1 | 0.370866946 | -0.210384609 | 0.366 | 0.474 | 1 | 0.370866946 | 0.001890863 |
| Zfp358 | 0.000948748 | 1.569760094 | 0.245 | 0.085 | 1 | 0.265499271 | 0.622191959 | 0.244 | 0.177 | 1 | 0.265499271 | 0.001896596 |
| Tmem184c | 0.000949047 | 1.409295422 | 0.377 | 0.182 | 1 | 0.21202059 | 0.766815522 | 0.341 | 0.28 | 1 | 0.21202059 | 0.001897194 |
| Slc39a11 | 0.000951871 | -1.508242418 | 0.245 | 0.508 | 1 | 0.423921531 | -0.398011612 | 0.244 | 0.304 | 1 | 0.423921531 | 0.001902836 |
| Atp6v1c2 | 0.000953583 | -4.015202407 | 0 | 0.178 | 1 | 0.537097674 | 1.351778023 | 0.049 | 0.031 | 1 | 0.537097674 | 0.001906256 |
| Dctn1 | 0.000954136 | 0.949902803 | 0.642 | 0.428 | 1 | 0.599426251 | 0.144908305 | 0.488 | 0.567 | 1 | 0.599426251 | 0.001907362 |
| Tmem86a | 0.000956446 | 1.739685096 | 0.151 | 0.034 | 1 | 0.295430251 | 0.515276755 | 0.146 | 0.092 | 1 | 0.295430251 | 0.001911978 |
| Calhm6 | 0.00095849 | 3.47665069 | 0.094 | 0.013 | 1 | 0.27192168 | 1.252242349 | 0.049 | 0.02 | 1 | 0.27192168 | 0.001916062 |
| Bicd2 | 0.000959892 | 1.582143819 | 0.396 | 0.203 | 1 | 0.07597641 | 0.860920919 | 0.488 | 0.399 | 1 | 0.07597641 | 0.001918863 |
| Strn4 | 0.000960614 | 0.86053949 | 0.491 | 0.263 | 1 | 0.015669724 | 0.874485866 | 0.561 | 0.41 | 1 | 0.015669724 | 0.001920305 |
| Rffl | 0.00096222 | 2.225111923 | 0.208 | 0.068 | 1 | 0.001000365 | 1.891652634 | 0.293 | 0.109 | 1 | 0.001000365 | 0.001923514 |
| Fam76b | 0.000967378 | 1.081260433 | 0.566 | 0.377 | 1 | 0.717142863 | 0.194631626 | 0.537 | 0.505 | 1 | 0.717142863 | 0.00193382 |
| Sbno2 | 0.000967574 | -0.770937805 | 0.642 | 0.801 | 1 | 0.2277917 | 0.578470581 | 0.683 | 0.652 | 1 | 0.2277917 | 0.001934212 |
| Sumo1 | 0.000969965 | 0.534952735 | 0.981 | 0.928 | 1 | 0.030908935 | 0.35878582 | 0.927 | 0.956 | 1 | 0.030908935 | 0.001938989 |
| Mcrip1 | 0.000971535 | 0.671732067 | 0.811 | 0.703 | 1 | 0.221059427 | -0.177501649 | 0.61 | 0.727 | 1 | 0.221059427 | 0.001942127 |
| Eng | 0.000978649 | 0.847667776 | 0.623 | 0.369 | 1 | 0.377543602 | -0.110908409 | 0.439 | 0.519 | 1 | 0.377543602 | 0.001956341 |
| Foxc1 | 0.000979378 | 0.637790578 | 0.906 | 0.843 | 1 | 0.0754612 | 0.388630818 | 0.927 | 0.85 | 1 | 0.0754612 | 0.001957797 |
| Sec31a | 0.000980384 | 0.399697739 | 0.981 | 0.97 | 1 | 0.050191188 | -0.305962479 | 1 | 0.986 | 1 | 0.050191188 | 0.001959806 |
| Phactr4 | 0.000980628 | 0.606170734 | 0.774 | 0.542 | 1 | 0.839579906 | 0.10616409 | 0.634 | 0.621 | 1 | 0.839579906 | 0.001960294 |
| Tox4 | 0.000985627 | 0.830049123 | 0.698 | 0.593 | 1 | 0.652449161 | 0.214274499 | 0.732 | 0.747 | 1 | 0.652449161 | 0.001970282 |
| Tlk2 | 0.000988894 | 0.451751388 | 0.925 | 0.877 | 1 | 0.381645297 | -0.106211622 | 0.976 | 0.973 | 1 | 0.381645297 | 0.001976809 |
| Col26a1 | 0.000991627 | 4.242185436 | 0.094 | 0.013 | 1 | 0.776786151 | 2.537644568 | 0.049 | 0.041 | 1 | 0.776786151 | 0.001982271 |
| Higd1a | 0.000994046 | -0.902779874 | 0.774 | 0.839 | 1 | 0.594969841 | -0.170448723 | 0.829 | 0.843 | 1 | 0.594969841 | 0.001987104 |
| Gdf6 | 0.249034808 | 2.154722595 | 0.019 | 0.004 | 1 | 0.000994429 | 5.059597271 | 0.073 | 0.007 | 1 | 0.249034808 | 0.001987869 |

| | | | | | | | | | | | | |
|---|---|---|---|---|---|---|---|---|---|---|---|---|
| Akr1a1 | 0.000994881 | 0.420839213 | 0.981 | 0.945 | 1 | 0.278909328 | 0.181055732 | 1 | 0.983 | 1 | 0.278909328 | 0.001988773 |
| Ankrd29 | 0.00099521 | 1.974150349 | 0.189 | 0.055 | 1 | 0.147416484 | 0.979223855 | 0.098 | 0.044 | 1 | 0.147416484 | 0.00198943 |
| Gm33989 | 0.000996019 | 1.370451286 | 0.453 | 0.258 | 1 | 0.334934177 | 0.456114682 | 0.463 | 0.399 | 1 | 0.334934177 | 0.001991046 |
| Numbl | 0.000997305 | 1.119957177 | 0.453 | 0.237 | 1 | 0.046545077 | -0.229626659 | 0.268 | 0.468 | 1 | 0.046545077 | 0.001993616 |
| Tmem50b | 0.000998928 | 1.302821234 | 0.472 | 0.267 | 1 | 0.283902083 | -0.415182312 | 0.317 | 0.396 | 1 | 0.283902083 | 0.001996858 |
| Ubp1 | 0.00101197 | 0.699982671 | 0.774 | 0.678 | 1 | 0.684938103 | 0.126478492 | 0.878 | 0.792 | 1 | 0.684938103 | 0.002022917 |
| Vps9d1 | 0.001014611 | 1.569760094 | 0.302 | 0.127 | 1 | 0.383128249 | -0.250257991 | 0.122 | 0.181 | 1 | 0.383128249 | 0.002028192 |
| Gab1 | 0.001015663 | 0.861458258 | 0.83 | 0.716 | 1 | 0.170861836 | 0.481573451 | 0.902 | 0.802 | 1 | 0.170861836 | 0.002030294 |
| Large1 | 0.001024217 | 0.394856599 | 0.868 | 0.72 | 1 | 0.199127557 | 0.400506568 | 0.951 | 0.857 | 1 | 0.199127557 | 0.002047384 |
| Ss18l2 | 0.001024262 | -1.382218391 | 0.396 | 0.585 | 1 | 0.251951565 | 0.161094461 | 0.244 | 0.338 | 1 | 0.251951565 | 0.002047474 |
| Tcf24 | 0.001025769 | 3.154722595 | 0.094 | 0.013 | 1 | 0.009536505 | 1.547698233 | 0.146 | 0.044 | 1 | 0.009536505 | 0.002050485 |
| Gm2350 | 0.001025769 | 3.739685096 | 0.094 | 0.013 | 1 | 0.027541029 | 2.614812428 | 0.073 | 0.017 | 1 | 0.027541029 | 0.002050485 |
| Pmaip1 | 0.001025769 | 3.154722595 | 0.094 | 0.013 | 1 | 0.462889204 | 0.515276755 | 0.049 | 0.027 | 1 | 0.462889204 | 0.002050485 |
| Reep3 | 0.020197839 | 0.331600357 | 0.962 | 0.953 | 1 | 0.001026251 | -0.529657777 | 0.976 | 0.993 | 1 | 0.020197839 | 0.002051448 |
| Fam186b | 0.246443787 | 2.891688189 | 0.019 | 0.004 | 1 | 0.001027727 | 2.83720485 | 0.098 | 0.014 | 1 | 0.246443787 | 0.002054398 |
| Tex49 | 0.029966098 | 5.678284551 | 0.038 | 0.004 | 1 | 0.001028208 | 2.83720485 | 0.098 | 0.014 | 1 | 0.029966098 | 0.002055359 |
| Gins1 | 0.100891659 | 2.417757001 | 0.075 | 0.03 | 1 | 0.001028208 | 2.83720485 | 0.098 | 0.014 | 1 | 0.100891659 | 0.002055359 |
| Fzd6 | 0.00102844 | 1.264905513 | 0.358 | 0.165 | 1 | 1 | 0.112312088 | 0.244 | 0.249 | 1 | 1 | 0.002055822 |
| Ibtk | 0.145526272 | -0.466460094 | 0.528 | 0.657 | 1 | 0.001029791 | -0.53113371 | 0.39 | 0.724 | 1 | 0.145526272 | 0.002058522 |
| Pold3 | 0.00103395 | 1.340278248 | 0.358 | 0.165 | 1 | 0.304299616 | -0.343367396 | 0.195 | 0.287 | 1 | 0.304299616 | 0.002066832 |
| Vmp1 | 0.014651178 | 0.391916996 | 0.943 | 0.936 | 1 | 0.001034147 | 0.748171301 | 0.902 | 0.826 | 1 | 0.014651178 | 0.002067225 |
| Atf5 | 0.001038424 | 1.034428361 | 0.528 | 0.326 | 1 | 0.267137652 | -0.16279515 | 0.317 | 0.403 | 1 | 0.267137652 | 0.00207577 |
| Itpripl1 | 0.209009104 | 1.739685096 | 0.038 | 0.013 | 1 | 0.001039676 | 2.614812428 | 0.098 | 0.014 | 1 | 0.209009104 | 0.00207827 |
| Rufy1 | 0.001041158 | 1.05850728 | 0.377 | 0.169 | 1 | 0.017164372 | 0.83720485 | 0.439 | 0.263 | 1 | 0.017164372 | 0.002081232 |
| Hmgn3 | 0.001041641 | 0.946405359 | 0.736 | 0.581 | 1 | 0.255097202 | 0.548528561 | 0.732 | 0.672 | 1 | 0.255097202 | 0.002082196 |
| Prdm10 | 0.001049182 | 1.129187503 | 0.34 | 0.144 | 1 | 0.196584824 | 0.969842618 | 0.293 | 0.208 | 1 | 0.196584824 | 0.002097264 |
| Chac2 | 0.001049784 | 1.377115016 | 0.283 | 0.11 | 1 | 0.528174738 | 0.377773231 | 0.146 | 0.198 | 1 | 0.528174738 | 0.002098465 |
| Pparg | 0.001049835 | -1.338656356 | 0.151 | 0.381 | 1 | 0.074197616 | -1.863234868 | 0.098 | 0.212 | 1 | 0.074197616 | 0.002098568 |
| Cnih1 | 0.001049855 | 0.666882561 | 0.792 | 0.758 | 1 | 0.485812076 | 0.24899669 | 0.756 | 0.7 | 1 | 0.485812076 | 0.002098607 |
| Prox1 | 0.001053176 | 2.232725107 | 0.189 | 0.055 | 1 | 0.543479214 | 1.212713985 | 0.122 | 0.099 | 1 | 0.543479214 | 0.002105243 |
| 1700001K19Rik | 0.001053176 | 2.444229212 | 0.189 | 0.055 | 1 | 0.249741515 | 0.79281073 | 0.122 | 0.072 | 1 | 0.249741515 | 0.002105243 |
| Aldh1a2 | 0.003197361 | 4.154722595 | 0.057 | 0.004 | 1 | 0.001053884 | 3.83720485 | 0.073 | 0.007 | 1 | 0.003197361 | 0.002106657 |
| Cish | 0.016674634 | 1.502645898 | 0.113 | 0.034 | 1 | 0.001053884 | 3.574170444 | 0.073 | 0.007 | 1 | 0.016674634 | 0.002106657 |
| Crybb3 | 0.157246372 | 1.569760094 | 0.057 | 0.021 | 1 | 0.001054062 | 3.83720485 | 0.073 | 0.007 | 1 | 0.157246372 | 0.002107012 |
| Pnpla6 | 0.001054619 | 1.711115943 | 0.283 | 0.114 | 1 | 0.594995998 | 0.647380291 | 0.22 | 0.191 | 1 | 0.594995998 | 0.002108127 |
| Atg12 | 0.189427435 | 0.535202872 | 0.491 | 0.453 | 1 | 0.001057937 | 1.110020276 | 0.659 | 0.471 | 1 | 0.189427435 | 0.002114754 |
| Dnah7b | 0.001058463 | 2.047807391 | 0.226 | 0.076 | 1 | 0.425709359 | 1.667279848 | 0.146 | 0.113 | 1 | 0.425709359 | 0.002115805 |
| Zfp715 | 0.001058708 | 1.384204441 | 0.264 | 0.097 | 1 | 0.359792356 | -0.278272368 | 0.098 | 0.154 | 1 | 0.359792356 | 0.002116295 |
| Gm37233 | 0.001060864 | 3.324647596 | 0.094 | 0.013 | 1 | 0.012973167 | 2.515276755 | 0.073 | 0.014 | 1 | 0.012973167 | 0.002120603 |
| Srsf12 | 0.001060864 | 2.962077517 | 0.094 | 0.013 | 1 | 0.817109602 | 0.721727632 | 0.049 | 0.041 | 1 | 0.817109602 | 0.002120603 |
| Zfp28 | 0.001060864 | 2.962077517 | 0.094 | 0.013 | 1 | 0.672172778 | 0.136765132 | 0.024 | 0.038 | 1 | 0.672172778 | 0.002120603 |
| Mmp15 | 0.001060864 | 2.962077517 | 0.094 | 0.013 | 1 | 0.397816093 | 0.600165653 | 0.098 | 0.061 | 1 | 0.397816093 | 0.002120603 |
| Frmpd4 | 0.001061735 | 3.47665069 | 0.094 | 0.013 | 1 | 0.107398884 | 4.763204268 | 0.049 | 0.014 | 1 | 0.107398884 | 0.002122342 |
| Acox3 | 0.001064447 | 2.8327945 | 0.245 | 0.093 | 1 | 0.977946869 | 0.47463477 | 0.146 | 0.154 | 1 | 0.977946869 | 0.00212776 |
| Sirpa | 0.00106634 | 1.508359549 | 0.321 | 0.14 | 1 | 0.016214435 | 1.252242349 | 0.341 | 0.188 | 1 | 0.016214435 | 0.002131544 |
| Nab2 | 0.001068918 | 1.004305752 | 0.642 | 0.475 | 1 | 0.0681158 | -0.662641037 | 0.537 | 0.642 | 1 | 0.0681158 | 0.002136693 |
| Aox3 | 0.620952062 | 0.216123139 | 0.038 | 0.055 | 1 | 0.001070545 | 2.422167351 | 0.268 | 0.096 | 1 | 0.620952062 | 0.002139943 |
| Zfp248 | 0.001070812 | 1.111653873 | 0.396 | 0.191 | 1 | 0.802755255 | 0.672818032 | 0.22 | 0.253 | 1 | 0.802755255 | 0.002140478 |
| 1700112J16Rik | 0.001073597 | 2.402650108 | 0.17 | 0.047 | 1 | 0.696570733 | 1.489281546 | 0.073 | 0.061 | 1 | 0.696570733 | 0.002146041 |
| Odad2 | 0.00107361 | 3.154722595 | 0.094 | 0.013 | 1 | 0.992196803 | 0.515276755 | 0.024 | 0.024 | 1 | 0.992196803 | 0.002146068 |
| Mcfd2 | 0.226062816 | -0.177353455 | 0.736 | 0.814 | 1 | 0.001075421 | -0.920492186 | 0.537 | 0.71 | 1 | 0.226062816 | 0.002149685 |
| Gm49267 | 0.003339709 | 3.154722595 | 0.057 | 0.004 | 1 | 0.001075996 | 2.351778023 | 0.098 | 0.014 | 1 | 0.003339709 | 0.002150834 |
| Sfrp2 | 0.00107646 | 4.693609225 | 0.377 | 0.203 | 1 | 0.001672306 | 3.613164576 | 0.195 | 0.061 | 1 | 0.001672306 | 0.002151761 |
| Ripk2 | 0.001079431 | 0.927653686 | 0.415 | 0.212 | 1 | 0.033610138 | 0.79281073 | 0.415 | 0.266 | 1 | 0.033610138 | 0.002157696 |
| Spp1 | 0.001080399 | -0.568038026 | 0.491 | 0.263 | 1 | 0.015956725 | 1.165827597 | 0.561 | 0.403 | 1 | 0.015956725 | 0.002159632 |
| Gper1 | 0.001635369 | 2.324647596 | 0.113 | 0.021 | 1 | 0.001085069 | 3.322631677 | 0.073 | 0.007 | 1 | 0.001635369 | 0.002168961 |
| Inhbb | 0.246443787 | 2.739685096 | 0.019 | 0.004 | 1 | 0.00108516 | 3.252242349 | 0.073 | 0.007 | 1 | 0.246443787 | 0.002169143 |
| Gm48913 | 0.413080549 | -1.015202407 | 0 | 0.013 | 1 | 0.00108516 | 3.252242349 | 0.073 | 0.007 | 1 | 0.413080549 | 0.002169143 |
| 1700108F19Rik | 0.732691937 | 1.154722595 | 0.019 | 0.013 | 1 | 0.00108516 | 3.252242349 | 0.073 | 0.007 | 1 | 0.732691937 | 0.002169143 |
| Egf | 0.767066103 | 0.569760094 | 0.038 | 0.03 | 1 | 0.00108516 | 3.252242349 | 0.073 | 0.007 | 1 | 0.767066103 | 0.002169143 |
| Gm15510 | 0.00108584 | 2.8327945 | 0.094 | 0.013 | 1 | 0.672172778 | 0.136765132 | 0.024 | 0.038 | 1 | 0.672172778 | 0.002170501 |
| Malat1 | 0.001088201 | -0.69283202 | 1 | 1 | 1 | 0.068225119 | 0.133831142 | 1 | 1 | 1 | 0.068225119 | 0.002175219 |
| Gosr2 | 0.059598251 | -0.4082136 | 0.264 | 0.407 | 1 | 0.001090843 | -1.524251609 | 0.146 | 0.413 | 1 | 0.059598251 | 0.002180496 |
| Crtac1 | 0.001092394 | 1.454282877 | 0.302 | 0.123 | 1 | 0.666460553 | -1.004097404 | 0.049 | 0.065 | 1 | 0.666460553 | 0.002183594 |
| Tmco3 | 0.001092539 | -0.868609909 | 0.453 | 0.699 | 1 | 0.104723497 | -0.548853583 | 0.317 | 0.468 | 1 | 0.104723497 | 0.002183884 |
| Gm43111 | 0.00109818 | 2.739685096 | 0.094 | 0.013 | 1 | 0.001423246 | 2.59619675 | 0.146 | 0.034 | 1 | 0.001423246 | 0.002195153 |
| Slc25a38 | 0.001098567 | 1.272079545 | 0.415 | 0.216 | 1 | 0.024631304 | -0.89318809 | 0.171 | 0.358 | 1 | 0.024631304 | 0.002195928 |
| Sec24d | 0.001100387 | 0.632856929 | 0.906 | 0.907 | 1 | 0.177479322 | -0.126432947 | 0.902 | 0.956 | 1 | 0.177479322 | 0.002199564 |
| Tmem170 | 0.001104686 | -3.57319786 | 0 | 0.174 | 1 | 0.087679812 | -1.484723245 | 0.024 | 0.109 | 1 | 0.087679812 | 0.002208151 |
| Eapp | 0.001107134 | 0.966061031 | 0.642 | 0.47 | 1 | 0.251184763 | 0.331969542 | 0.488 | 0.403 | 1 | 0.251184763 | 0.002213043 |
| Lsm12 | 0.001109845 | 0.595126814 | 0.887 | 0.788 | 1 | 0.368781146 | 0.168410758 | 0.902 | 0.87 | 1 | 0.368781146 | 0.002218459 |
| Nrip3 | 0.001111113 | -3.822557329 | 0 | 0.174 | 1 | 0.152350708 | 1.479652845 | 0.146 | 0.082 | 1 | 0.152350708 | 0.002220991 |
| Shroom4 | 0.001112581 | -1.752168001 | 0.321 | 0.517 | 1 | 0.00147985 | -1.54256732 | 0.293 | 0.549 | 1 | 0.00147985 | 0.002223924 |
| Ncoa4 | 0.001113009 | 1.806799291 | 0.396 | 0.208 | 1 | 0.805841784 | -0.120090699 | 0.537 | 0.491 | 1 | 0.805841784 | 0.002224779 |
| Ccar1 | 0.001113677 | 0.664500584 | 0.887 | 0.881 | 1 | 0.033999527 | -0.221021543 | 0.951 | 0.956 | 1 | 0.033999527 | 0.002226113 |
| Phyhipl | 0.001549674 | -4.399866257 | 0 | 0.165 | 1 | 0.001116965 | 2.83720485 | 0.073 | 0.007 | 1 | 0.001549674 | 0.002232683 |
| Kifbp | 0.001122137 | 0.866046306 | 0.679 | 0.496 | 1 | 0.226360876 | 0.497894677 | 0.634 | 0.549 | 1 | 0.226360876 | 0.002243015 |
| Opn3 | 0.001122951 | 2.640149422 | 0.094 | 0.013 | 1 | 0.004290705 | 4.159132945 | 0.073 | 0.01 | 1 | 0.004290705 | 0.00224464 |
| Nppc | 0.003243872 | 3.739685096 | 0.057 | 0.004 | 1 | 0.00112698 | 1.989207943 | 0.098 | 0.014 | 1 | 0.003243872 | 0.002252689 |
| 2810408I11Rik | 0.067284646 | 2.154722595 | 0.075 | 0.025 | 1 | 0.001133094 | 2.515276755 | 0.073 | 0.007 | 1 | 0.067284646 | 0.002264905 |
| Zhx2 | 0.001135553 | -0.748510977 | 0.528 | 0.784 | 1 | 0.089069327 | -0.319012732 | 0.756 | 0.836 | 1 | 0.089069327 | 0.002269817 |
| Serpinf2 | 0.00113568 | 2.377115016 | 0.094 | 0.013 | 1 | 0.262674569 | 2.83720485 | 0.024 | 0.007 | 1 | 0.262674569 | 0.002270069 |
| Brip1 | 0.001136333 | 0.757387097 | 0.264 | 0.097 | 1 | 0.894660105 | 1.299239829 | 0.122 | 0.137 | 1 | 0.894660105 | 0.002271375 |
| Gm56501 | 0.488229496 | 0.491757582 | 0.075 | 0.051 | 1 | 0.001141623 | 1.936740523 | 0.22 | 0.068 | 1 | 0.488229496 | 0.002281943 |
| Bmi1 | 0.001141746 | 1.272559085 | 0.491 | 0.309 | 1 | 0.960647675 | 0.425078946 | 0.341 | 0.372 | 1 | 0.960647675 | 0.002282188 |
| Arhgap26 | 0.001144475 | -1.814903756 | 0.302 | 0.517 | 1 | 0.059414239 | -1.185162963 | 0.317 | 0.454 | 1 | 0.059414239 | 0.00228764 |
| Gm13269 | 0.001149047 | 2.569760094 | 0.094 | 0.013 | 1 | 0.281346686 | 1.19334866 | 0.098 | 0.055 | 1 | 0.281346686 | 0.002296774 |
| Nopchap1 | 0.001149207 | 1.043691282 | 0.491 | 0.292 | 1 | 0.766751614 | 0.788531003 | 0.293 | 0.321 | 1 | 0.766751614 | 0.002297093 |
| Palm3 | 0.503971666 | 1.569760094 | 0.019 | 0.008 | 1 | 0.001149798 | 2.83720485 | 0.073 | 0.007 | 1 | 0.503971666 | 0.002298274 |
| Eri2 | 0.001151657 | 1.207190015 | 0.208 | 0.064 | 1 | 0.959995349 | 0.515276755 | 0.073 | 0.072 | 1 | 0.959995349 | 0.002301987 |
| Arfgap3 | 0.543146853 | -0.126321025 | 0.811 | 0.856 | 1 | 0.001156074 | -0.692153398 | 0.902 | 0.904 | 1 | 0.543146853 | 0.002310812 |
| Cryzl2 | 0.076470926 | -0.492167655 | 0.226 | 0.369 | 1 | 0.0011596 | -2.398011612 | 0.049 | 0.283 | 1 | 0.076470926 | 0.002317856 |
| Tfap4 | 0.00116111 | 2.417757001 | 0.094 | 0.013 | 1 | 0.752578711 | 0.667279848 | 0.024 | 0.017 | 1 | 0.752578711 | 0.002320872 |
| Map2k3os | 0.00116117 | -2.582242999 | 0.038 | 0.233 | 1 | 0.221466267 | -1.116991461 | 0.073 | 0.14 | 1 | 0.221466267 | 0.002320992 |
| Dbf4 | 0.00116344 | 2.213616284 | 0.226 | 0.081 | 1 | 0.709278782 | 0.159132945 | 0.098 | 0.078 | 1 | 0.709278782 | 0.002325525 |
| Vps50 | 0.001164205 | 1.47119626 | 0.377 | 0.191 | 1 | 0.072477762 | 1.313642894 | 0.341 | 0.239 | 1 | 0.072477762 | 0.002327054 |
| Aga | 0.001166761 | -0.571595755 | 0.491 | 0.78 | 1 | 0.04132999 | -0.514607525 | 0.488 | 0.672 | 1 | 0.04132999 | 0.002332162 |
| Tlcd4 | 0.00117283 | -1.410062024 | 0.208 | 0.441 | 1 | 0.008369629 | -1.16279515 | 0.268 | 0.474 | 1 | 0.008369629 | 0.002344285 |

| | | | | | | | | | | | | |
|---|---|---|---|---|---|---|---|---|---|---|---|---|
| Rab43 | 0.001173055 | 0.871550543 | 0.585 | 0.394 | 1 | 0.011666272 | 0.770709438 | 0.756 | 0.642 | 1 | 0.011666272 | 0.002344733 |
| Helz | 0.001174497 | 0.605739391 | 0.887 | 0.907 | 1 | 0.432034241 | 0.163980629 | 0.927 | 0.945 | 1 | 0.432034241 | 0.002347614 |
| Qtrt1 | 0.001175701 | 1.245488059 | 0.434 | 0.237 | 1 | 0.494483279 | 0.457062141 | 0.39 | 0.362 | 1 | 0.494483279 | 0.002350019 |
| Txnrd3 | 0.001179418 | 1.792152515 | 0.302 | 0.131 | 1 | 0.613911737 | 1.007129851 | 0.171 | 0.154 | 1 | 0.613911737 | 0.002357445 |
| Prelid3a | 0.001189386 | 1.776210972 | 0.245 | 0.089 | 1 | 0.425043197 | 0.363273661 | 0.073 | 0.119 | 1 | 0.425043197 | 0.002377357 |
| Foxo3 | 0.001192258 | 0.842990267 | 0.906 | 0.792 | 1 | 0.322374139 | 0.483701361 | 0.902 | 0.915 | 1 | 0.322374139 | 0.002383095 |
| Arntl | 0.014019263 | 1.031340179 | 0.415 | 0.267 | 1 | 0.001196661 | 0.937369012 | 0.61 | 0.403 | 1 | 0.014019263 | 0.002391891 |
| Zfp467 | 0.001196729 | 1.994257923 | 0.208 | 0.068 | 1 | 0.264647417 | 1.277777441 | 0.195 | 0.13 | 1 | 0.264647417 | 0.002392027 |
| Micos10 | 0.001198285 | 0.55349504 | 1 | 0.924 | 1 | 0.228860132 | -0.18959521 | 0.976 | 0.952 | 1 | 0.228860132 | 0.002395134 |
| Apba1 | 0.001198572 | 1.948271717 | 0.34 | 0.157 | 1 | 0.726372926 | -0.592782991 | 0.293 | 0.3 | 1 | 0.726372926 | 0.002395706 |
| Gm12536 | 0.012547102 | -2.430239906 | 0.038 | 0.169 | 1 | 0.001199604 | -4.140075074 | 0 | 0.212 | 1 | 0.012547102 | 0.002397769 |
| Tnfsf9 | 0.013444113 | 2.640149422 | 0.113 | 0.034 | 1 | 0.001203423 | 2.711673968 | 0.171 | 0.044 | 1 | 0.013444113 | 0.002405398 |
| St14 | 0.019240161 | 1.767699472 | 0.132 | 0.047 | 1 | 0.001205369 | 1.19334866 | 0.317 | 0.126 | 1 | 0.019240161 | 0.002409285 |
| Abl1 | 0.001206519 | 0.970298024 | 0.547 | 0.331 | 1 | 0.001513111 | 0.787961331 | 0.78 | 0.515 | 1 | 0.001513111 | 0.002411583 |
| Dsn1 | 0.001211015 | 2.261637799 | 0.17 | 0.047 | 1 | 0.075026658 | 1.444887427 | 0.146 | 0.068 | 1 | 0.075026658 | 0.002420564 |
| Paox | 0.001212526 | 1.974150349 | 0.189 | 0.055 | 1 | 0.875793076 | 0.614812428 | 0.098 | 0.092 | 1 | 0.875793076 | 0.002423582 |
| Hectd2os | 0.001221662 | -2.768109545 | 0.038 | 0.242 | 1 | 0.430381562 | -0.80665134 | 0.073 | 0.113 | 1 | 0.430381562 | 0.002441831 |
| Zfp952 | 0.00122292 | 1.721763188 | 0.245 | 0.089 | 1 | 0.555464105 | -0.16279515 | 0.122 | 0.16 | 1 | 0.555464105 | 0.002444345 |
| Uap1l1 | 0.001223069 | 1.087608399 | 0.472 | 0.254 | 1 | 0.729611693 | -0.129628286 | 0.341 | 0.365 | 1 | 0.729611693 | 0.002444642 |
| Zfp740 | 0.001223523 | 1.295585131 | 0.434 | 0.246 | 1 | 0.883425894 | 0.126234464 | 0.512 | 0.526 | 1 | 0.883425894 | 0.002445549 |
| Zc3h10 | 0.001231005 | 2.034428361 | 0.245 | 0.093 | 1 | 0.665139779 | 0.377773231 | 0.122 | 0.157 | 1 | 0.665139779 | 0.002460494 |
| Pam | 0.005924114 | 0.371351732 | 0.981 | 1 | 1 | 0.001231571 | -0.480466396 | 1 | 1 | 1 | 0.005924114 | 0.002461626 |
| Wdr75 | 0.001233967 | 1.067259754 | 0.604 | 0.424 | 1 | 0.650732722 | 0.217596206 | 0.439 | 0.522 | 1 | 0.650732722 | 0.002466412 |
| Ranbp3 | 0.001236408 | 0.781913331 | 0.547 | 0.314 | 1 | 0.951195658 | 0.243680336 | 0.366 | 0.41 | 1 | 0.951195658 | 0.002471287 |
| Sirt7 | 0.001239227 | 1.538051234 | 0.321 | 0.14 | 1 | 0.008139547 | 1.308825877 | 0.341 | 0.167 | 1 | 0.008139547 | 0.002476919 |
| Agrn | 0.00124385 | 1.302821234 | 0.453 | 0.25 | 1 | 0.025434789 | 0.863200058 | 0.439 | 0.276 | 1 | 0.025434789 | 0.002486154 |
| Aff2 | 0.004655978 | 0.974150349 | 0.208 | 0.076 | 1 | 0.001246261 | 1.38550888 | 0.366 | 0.16 | 1 | 0.004655978 | 0.002490969 |
| Srsf3 | 0.001248399 | 0.395247406 | 1 | 0.97 | 1 | 0.112676327 | 0.281976647 | 1 | 0.99 | 1 | 0.112676327 | 0.002495239 |
| Rapgef4 | 0.00125623 | -3.237594828 | 0.038 | 0.233 | 1 | 0.288638437 | -0.813049111 | 0.098 | 0.164 | 1 | 0.288638437 | 0.002510882 |
| Hdhd2 | 0.573232806 | 0.378618607 | 0.453 | 0.424 | 1 | 0.001259288 | -1.605738646 | 0.146 | 0.399 | 1 | 0.573232806 | 0.00251699 |
| Smim3 | 0.001259845 | -0.80412008 | 0.566 | 0.725 | 1 | 0.60565658 | 0.189506594 | 0.512 | 0.642 | 1 | 0.60565658 | 0.002518103 |
| Fbln7 | 0.83931426 | -0.194426969 | 0.434 | 0.39 | 1 | 0.001260146 | -0.758524082 | 0.634 | 0.788 | 1 | 0.83931426 | 0.002518705 |
| Ddr1 | 0.860502843 | -0.282682718 | 0.189 | 0.191 | 1 | 0.001260779 | 0.991117533 | 0.61 | 0.362 | 1 | 0.860502843 | 0.002519969 |
| Pef1 | 0.194917948 | -0.281917159 | 0.358 | 0.475 | 1 | 0.001262173 | -1.06279936 | 0.171 | 0.457 | 1 | 0.194917948 | 0.002522753 |
| Sephs1 | 0.001263922 | 1.228723176 | 0.377 | 0.182 | 1 | 0.219241291 | 0.904319046 | 0.366 | 0.304 | 1 | 0.219241291 | 0.002526247 |
| Cacng4 | 0.001264968 | -2.27154216 | 0.057 | 0.263 | 1 | 0.679848464 | 0.852972166 | 0.146 | 0.126 | 1 | 0.679848464 | 0.002528337 |
| Vcan | 0.001269519 | -0.237594828 | 0.283 | 0.11 | 1 | 0.003204571 | 0.91966701 | 0.171 | 0.048 | 1 | 0.003204571 | 0.002537425 |
| Arl1 | 0.236887361 | 0.253659267 | 0.925 | 0.894 | 1 | 0.001271729 | -0.6416293 | 0.732 | 0.908 | 1 | 0.236887361 | 0.002541841 |
| Fam241a | 0.565960065 | 0.714150003 | 0.094 | 0.072 | 1 | 0.001274483 | 1.766815522 | 0.317 | 0.13 | 1 | 0.565960065 | 0.002547341 |
| Smpd2 | 0.001275127 | 0.962077517 | 0.377 | 0.182 | 1 | 0.153984218 | -0.452301767 | 0.122 | 0.225 | 1 | 0.153984218 | 0.002548629 |
| Meis1 | 0.001277373 | 2.491757582 | 0.17 | 0.047 | 1 | 0.001870301 | 1.985727375 | 0.366 | 0.177 | 1 | 0.001870301 | 0.002553115 |
| Cs | 0.001278814 | 1.312263872 | 0.509 | 0.297 | 1 | 0.048983971 | 0.524465926 | 0.683 | 0.563 | 1 | 0.048983971 | 0.002555993 |
| Gm10762 | 0.001279009 | 1.199116714 | 0.377 | 0.178 | 1 | 0.105845086 | -0.622226769 | 0.098 | 0.205 | 1 | 0.105845086 | 0.002556382 |
| Septin9 | 0.001279644 | -1.033537142 | 0.736 | 0.877 | 1 | 0.007451074 | -0.77274833 | 0.78 | 0.874 | 1 | 0.007451074 | 0.00255765 |
| Nfs1 | 0.001280381 | 1.755452411 | 0.472 | 0.28 | 1 | 0.857320654 | 0.14088124 | 0.341 | 0.369 | 1 | 0.857320654 | 0.002559122 |
| Ncbp3 | 0.001281985 | 0.839520363 | 0.491 | 0.271 | 1 | 0.135936082 | 0.458693327 | 0.537 | 0.44 | 1 | 0.135936082 | 0.002562327 |
| 2900093K20Rik | 0.459891923 | 0.569760094 | 0.113 | 0.081 | 1 | 0.001282957 | 2.589277336 | 0.171 | 0.044 | 1 | 0.459891923 | 0.002564269 |
| Tmem41b | 0.241581032 | 0.120234218 | 0.679 | 0.559 | 1 | 0.001287701 | -0.847293324 | 0.439 | 0.703 | 1 | 0.241581032 | 0.002573744 |
| Hmbs | 0.001289701 | 1.640149422 | 0.283 | 0.114 | 1 | 0.274028343 | -0.589059905 | 0.073 | 0.133 | 1 | 0.274028343 | 0.002577739 |
| Dcaf1 | 0.001289803 | 1.25093791 | 0.566 | 0.386 | 1 | 0.104269212 | 0.679352681 | 0.634 | 0.563 | 1 | 0.104269212 | 0.002577942 |
| Aebp2 | 0.001290917 | 0.496642165 | 0.849 | 0.627 | 1 | 0.009672508 | 0.564507347 | 0.902 | 0.853 | 1 | 0.009672508 | 0.002580168 |
| Psmc2 | 0.001291359 | 0.698905245 | 0.811 | 0.691 | 1 | 0.033281926 | -0.481364233 | 0.659 | 0.744 | 1 | 0.033281926 | 0.002581051 |
| Tial1 | 0.001291836 | 0.60344322 | 0.83 | 0.737 | 1 | 0.067258499 | 0.446745373 | 0.854 | 0.826 | 1 | 0.067258499 | 0.002582003 |
| Lpar1 | 0.001292073 | -1.056529269 | 0.774 | 0.864 | 1 | 0.131740289 | -0.384196369 | 0.78 | 0.884 | 1 | 0.131740289 | 0.002582476 |
| Pla2g15 | 0.001292288 | 1.8327945 | 0.208 | 0.068 | 1 | 0.881271886 | 0.396632258 | 0.098 | 0.109 | 1 | 0.881271886 | 0.002582907 |
| Psmd4 | 0.001302506 | 0.512914234 | 0.887 | 0.818 | 1 | 0.0551986 | -0.345320188 | 0.732 | 0.853 | 1 | 0.0551986 | 0.002603315 |
| Zfp524 | 0.001302911 | 2.047807391 | 0.17 | 0.047 | 1 | 0.611582762 | 1.47463477 | 0.073 | 0.055 | 1 | 0.611582762 | 0.002604125 |
| Lrmda | 0.001302998 | -1.276139872 | 0.302 | 0.542 | 1 | 0.011506353 | -1.849962511 | 0.439 | 0.567 | 1 | 0.011506353 | 0.002604299 |
| Dcun1d4 | 0.807118269 | 0.3688474 | 0.245 | 0.237 | 1 | 0.001304587 | 1.479652845 | 0.366 | 0.167 | 1 | 0.807118269 | 0.002607471 |
| Rims1 | 0.001304903 | 4.108918905 | 0.132 | 0.03 | 1 | 0.618425905 | 1.635570989 | 0.049 | 0.034 | 1 | 0.618425905 | 0.002608104 |
| B4galnt3 | 0.001305538 | -1.818310358 | 0.226 | 0.466 | 1 | 0.851442876 | 0.565400234 | 0.244 | 0.283 | 1 | 0.851442876 | 0.002609372 |
| Ppp1r12c | 0.001307857 | 0.911451444 | 0.585 | 0.377 | 1 | 0.925410885 | 0.299939091 | 0.463 | 0.519 | 1 | 0.925410885 | 0.002614003 |
| Eml5 | 0.001308097 | 2.624207878 | 0.208 | 0.068 | 1 | 0.378409882 | 0.659666664 | 0.22 | 0.164 | 1 | 0.378409882 | 0.002614484 |
| Gm47990 | 0.001310606 | -4.1672055 | 0 | 0.169 | 1 | 0.257308459 | -0.970150072 | 0 | 0.031 | 1 | 0.257308459 | 0.002619493 |
| Slc25a48 | 0.001317227 | -3.974560422 | 0 | 0.169 | 1 | 0.188544916 | -1.250257991 | 0 | 0.041 | 1 | 0.188544916 | 0.002632719 |
| Cir1 | 0.001317274 | 0.935156218 | 0.604 | 0.419 | 1 | 0.152351021 | 0.399799537 | 0.585 | 0.485 | 1 | 0.152351021 | 0.002632814 |
| Pmp22 | 0.001318331 | -0.855626093 | 0.83 | 0.881 | 1 | 0.041417766 | -0.490560185 | 0.61 | 0.805 | 1 | 0.041417766 | 0.002634924 |
| Satb1 | 0.148798744 | 1.891688189 | 0.094 | 0.047 | 1 | 0.001320964 | 2.672145604 | 0.244 | 0.089 | 1 | 0.148798744 | 0.002640183 |
| Irf2bp1 | 0.00132137 | 1.223985257 | 0.396 | 0.203 | 1 | 0.00900217 | -0.755750442 | 0.195 | 0.413 | 1 | 0.00900217 | 0.002640993 |
| Colq | 0.001325427 | 1.685237311 | 0.189 | 0.055 | 1 | 0.114535984 | -1.622226769 | 0 | 0.058 | 1 | 0.114535984 | 0.002649098 |
| Slurp1 | 0.001325785 | -6.68132776 | 0 | 0.169 | 1 | 0.170423449 | -2.372248516 | 0 | 0.044 | 1 | 0.170423449 | 0.002649813 |
| Ruvbl1 | 0.001327283 | 1.009672262 | 0.547 | 0.356 | 1 | 0.875633741 | 0.267839204 | 0.366 | 0.406 | 1 | 0.875633741 | 0.002652805 |
| Tmpo | 0.001329252 | 0.598759966 | 0.623 | 0.398 | 1 | 0.775146199 | 0.241673107 | 0.634 | 0.659 | 1 | 0.775146199 | 0.002656737 |
| Gm3336 | 0.001329537 | 2.051629102 | 0.189 | 0.055 | 1 | 0.051831769 | 1.667279848 | 0.122 | 0.048 | 1 | 0.051831769 | 0.002657307 |
| Mirg | 0.001331153 | -1.466765782 | 0.17 | 0.407 | 1 | 0.082446302 | -0.734337135 | 0.268 | 0.403 | 1 | 0.082446302 | 0.002660534 |
| Cdh19 | 0.001333219 | -3.964218478 | 0.019 | 0.199 | 1 | 0.286427259 | -1.16279515 | 0 | 0.027 | 1 | 0.286427259 | 0.002664661 |
| Herc2 | 0.001334109 | 0.745552103 | 0.792 | 0.716 | 1 | 0.017715927 | 0.413605337 | 0.951 | 0.891 | 1 | 0.017715927 | 0.002666438 |
| Hspe1 | 0.001335646 | 0.51339228 | 0.962 | 0.924 | 1 | 0.604779849 | 0.101124616 | 0.878 | 0.891 | 1 | 0.604779849 | 0.002669508 |
| Abhd5 | 0.001337642 | 0.927312099 | 0.491 | 0.271 | 1 | 0.10923265 | 0.861051592 | 0.561 | 0.491 | 1 | 0.10923265 | 0.002673496 |
| Fcgrt | 0.001338636 | 0.495256569 | 0.925 | 0.856 | 1 | 0.38677724 | -0.100926426 | 0.78 | 0.84 | 1 | 0.38677724 | 0.002675479 |
| Myof | 0.001342113 | 0.854254862 | 0.774 | 0.627 | 1 | 0.202573812 | 0.545438725 | 0.732 | 0.754 | 1 | 0.202573812 | 0.002682424 |
| Gtf3c5 | 0.001348162 | 1.714150003 | 0.302 | 0.131 | 1 | 0.209583224 | 0.624211126 | 0.195 | 0.123 | 1 | 0.209583224 | 0.002694507 |
| Jtb | 0.001353697 | 0.809226029 | 0.679 | 0.559 | 1 | 0.011113875 | -0.668030458 | 0.415 | 0.628 | 1 | 0.011113875 | 0.002705561 |
| Gpr27 | 0.001359018 | 1.33529484 | 0.264 | 0.097 | 1 | 0.035079785 | 1.188112012 | 0.293 | 0.16 | 1 | 0.035079785 | 0.002716189 |
| Gm57146 | 0.001367525 | -0.901545625 | 0.17 | 0.419 | 1 | 0.055365327 | -1.048927186 | 0.195 | 0.338 | 1 | 0.055365327 | 0.002733179 |
| Card6 | 0.001373588 | 2.640149422 | 0.132 | 0.03 | 1 | 0.028943483 | 2.029849928 | 0.073 | 0.017 | 1 | 0.028943483 | 0.00274529 |
| Rnf139 | 0.001373934 | 0.962077517 | 0.434 | 0.233 | 1 | 0.622845855 | 0.183200705 | 0.39 | 0.454 | 1 | 0.622845855 | 0.002745981 |
| Psmd12 | 0.098307577 | 0.364011429 | 0.849 | 0.805 | 1 | 0.001376262 | -0.640842447 | 0.78 | 0.918 | 1 | 0.098307577 | 0.002750629 |
| Tbc1d10b | 0.001377776 | 1.404700848 | 0.283 | 0.11 | 1 | 0.022525153 | -0.795063366 | 0.146 | 0.331 | 1 | 0.022525153 | 0.002753653 |
| Zfp710 | 0.001379632 | 1.553271971 | 0.302 | 0.127 | 1 | 0.296820043 | 0.416873051 | 0.415 | 0.334 | 1 | 0.296820043 | 0.002757361 |
| Mtr | 0.001387069 | 1.221836791 | 0.453 | 0.242 | 1 | 0.699990852 | 0.483567895 | 0.293 | 0.369 | 1 | 0.699990852 | 0.002772214 |
| Crem | 0.599773959 | 0.145733812 | 0.434 | 0.419 | 1 | 0.001393358 | 1.083223279 | 0.634 | 0.406 | 1 | 0.599773959 | 0.002784775 |
| Zfp330 | 0.114281177 | 0.251152357 | 0.66 | 0.534 | 1 | 0.001397852 | -1.069685746 | 0.268 | 0.546 | 1 | 0.114281177 | 0.002793751 |
| Plekhg2 | 0.001398271 | 1.154722595 | 0.472 | 0.267 | 1 | 0.069016064 | 1.120997816 | 0.415 | 0.311 | 1 | 0.069016064 | 0.002794588 |
| Arhgap29 | 0.001400022 | 1.101681632 | 0.755 | 0.716 | 1 | 0.253340902 | 0.529317693 | 0.878 | 0.881 | 1 | 0.253340902 | 0.002798085 |
| C2cd3 | 0.001405309 | 1.61219536 | 0.377 | 0.199 | 1 | 0.832846535 | 0.399083737 | 0.293 | 0.283 | 1 | 0.832846535 | 0.002808643 |
| Ak4 | 0.00141017 | -1.664037091 | 0.132 | 0.352 | 1 | 0.914387501 | 0.578470581 | 0.171 | 0.174 | 1 | 0.914387501 | 0.002818352 |
| Rnf11 | 0.156382948 | 0.277837913 | 0.887 | 0.924 | 1 | 0.001410916 | -0.549149895 | 0.902 | 0.969 | 1 | 0.156382948 | 0.002819841 |

| | | | | | | | | | | | | |
|---|---|---|---|---|---|---|---|---|---|---|---|---|
| Amotl1 | 0.001413807 | 0.387474434 | 0.849 | 0.797 | 1 | 0.187356449 | -0.338608711 | 0.927 | 0.908 | 1 | 0.187356449 | 0.002825615 |
| Bag4 | 0.00142746 | 1.620386167 | 0.302 | 0.131 | 1 | 0.58355004 | 0.644559772 | 0.195 | 0.171 | 1 | 0.58355004 | 0.002852882 |
| Cavin1 | 0.001428948 | 0.507603618 | 0.849 | 0.775 | 1 | 0.717260349 | -0.175804211 | 0.878 | 0.901 | 1 | 0.717260349 | 0.002855853 |
| P3h4 | 0.001430353 | -0.663655839 | 0.811 | 0.898 | 1 | 0.040011899 | -0.540100002 | 0.683 | 0.751 | 1 | 0.040011899 | 0.002858659 |
| Orc1 | 0.001434044 | 2.280253477 | 0.151 | 0.038 | 1 | 0.340924908 | 0.600165653 | 0.098 | 0.058 | 1 | 0.340924908 | 0.002866031 |
| Mier1 | 0.001438243 | 0.419819467 | 0.981 | 0.924 | 1 | 0.365527098 | 0.10463437 | 0.951 | 0.98 | 1 | 0.365527098 | 0.002874417 |
| Mcam | 0.779896917 | -0.982780929 | 0.075 | 0.085 | 1 | 0.001438699 | 1.370078839 | 0.268 | 0.096 | 1 | 0.779896917 | 0.002875329 |
| Hmgxb4 | 0.001439155 | 1.037152998 | 0.509 | 0.271 | 1 | 0.016140714 | -0.935384654 | 0.098 | 0.29 | 1 | 0.016140714 | 0.002876239 |
| Tubgcp4 | 0.001439485 | 0.932330173 | 0.434 | 0.229 | 1 | 0.982006449 | 0.586243276 | 0.244 | 0.259 | 1 | 0.982006449 | 0.002876899 |
| Sorl1 | 0.378742142 | 0.347367673 | 0.208 | 0.157 | 1 | 0.00144632 | 0.885460018 | 0.512 | 0.28 | 1 | 0.378742142 | 0.002890549 |
| Atf6 | 0.001454499 | 0.810119062 | 0.906 | 0.788 | 1 | 0.356120954 | 0.583833988 | 0.878 | 0.867 | 1 | 0.356120954 | 0.002906883 |
| Scmh1 | 0.001456027 | 0.716495468 | 0.792 | 0.623 | 1 | 0.14960802 | 0.361018695 | 0.829 | 0.853 | 1 | 0.14960802 | 0.002909934 |
| Tnik | 0.00145724 | 0.46284489 | 0.189 | 0.055 | 1 | 0.004968499 | 1.970471381 | 0.146 | 0.041 | 1 | 0.004968499 | 0.002912356 |
| Tmem44 | 0.001459208 | 2.739685096 | 0.113 | 0.021 | 1 | 0.452287461 | 1.252242349 | 0.049 | 0.027 | 1 | 0.452287461 | 0.002916287 |
| Flywch2 | 0.575571412 | -0.229941255 | 0.113 | 0.144 | 1 | 0.001461498 | -3.372248516 | 0 | 0.205 | 1 | 0.575571412 | 0.00292086 |
| Nob1 | 0.001463324 | 0.95505025 | 0.604 | 0.419 | 1 | 0.293924173 | -0.190701147 | 0.439 | 0.519 | 1 | 0.293924173 | 0.002924506 |
| Srgap2 | 0.001465646 | 0.349430139 | 0.906 | 0.767 | 1 | 0.993087792 | -0.168067338 | 0.951 | 0.922 | 1 | 0.993087792 | 0.002929144 |
| Acp5 | 0.001467539 | 2.154722595 | 0.151 | 0.038 | 1 | 0.20250665 | -6.647953594 | 0.049 | 0.113 | 1 | 0.20250665 | 0.002932924 |
| Dhrs3 | 0.001468559 | -0.666011357 | 0.302 | 0.597 | 1 | 0.009757117 | -0.147474351 | 0.512 | 0.741 | 1 | 0.009757117 | 0.002934961 |
| Kank4 | 0.007769567 | 2.154722595 | 0.113 | 0.03 | 1 | 0.001481785 | 2.071670103 | 0.146 | 0.034 | 1 | 0.007769567 | 0.002961374 |
| Mir100hg | 0.001483053 | -0.895202631 | 0.679 | 0.801 | 1 | 0.759791994 | -0.126772481 | 0.756 | 0.788 | 1 | 0.759791994 | 0.002963906 |
| Lars2 | 0.330420721 | -0.429506643 | 1 | 1 | 1 | 0.001487202 | 0.496604998 | 1 | 1 | 1 | 0.330420721 | 0.002972193 |
| Itpr2 | 0.001489717 | -0.746015774 | 0.849 | 0.932 | 1 | 0.268333621 | -0.366288326 | 0.78 | 0.86 | 1 | 0.268333621 | 0.002977214 |
| Ano1 | 0.606408808 | 1.502645898 | 0.038 | 0.025 | 1 | 0.001490035 | 0.44893426 | 0.195 | 0.058 | 1 | 0.606408808 | 0.00297785 |
| Vps13d | 0.001499029 | 0.500576529 | 0.736 | 0.559 | 1 | 0.96171068 | -0.139143882 | 0.854 | 0.846 | 1 | 0.96171068 | 0.002995811 |
| Emp1 | 0.012265051 | -0.593633909 | 0.981 | 0.953 | 1 | 0.001499836 | -0.63226669 | 1 | 0.986 | 1 | 0.012265051 | 0.002997423 |
| Arhgef26 | 0.001502097 | 2.104096522 | 0.226 | 0.081 | 1 | 0.011064344 | 0.898605394 | 0.415 | 0.235 | 1 | 0.011064344 | 0.003001938 |
| Ethe1 | 0.001503751 | 2.221836791 | 0.226 | 0.081 | 1 | 0.082722932 | 0.924667691 | 0.293 | 0.184 | 1 | 0.082722932 | 0.00300524 |
| Ssr3 | 0.005031726 | 0.221942813 | 0.981 | 0.966 | 1 | 0.001514294 | -0.422767497 | 0.951 | 0.983 | 1 | 0.005031726 | 0.003026295 |
| Acacb | 0.001517338 | 1.640149422 | 0.208 | 0.068 | 1 | 0.354453741 | 1.635570989 | 0.098 | 0.061 | 1 | 0.354453741 | 0.003032374 |
| Axdnd1 | 0.415175648 | -1.015202407 | 0.019 | 0.042 | 1 | 0.001520836 | 2.614812428 | 0.122 | 0.024 | 1 | 0.415175648 | 0.00303936 |
| Rabif | 0.001521501 | 1.046198138 | 0.415 | 0.216 | 1 | 0.436839324 | -0.432255825 | 0.22 | 0.276 | 1 | 0.436839324 | 0.003040688 |
| Lamp2 | 0.001523804 | 0.32336221 | 1 | 0.987 | 1 | 0.126635275 | 0.259880726 | 1 | 0.997 | 1 | 0.126635275 | 0.003045286 |
| Myl12b | 0.001530989 | 0.640149422 | 0.943 | 0.792 | 1 | 0.047577858 | 0.38122879 | 0.902 | 0.904 | 1 | 0.047577858 | 0.003059634 |
| Dctn2 | 0.001536506 | 0.547817541 | 0.849 | 0.661 | 1 | 0.804907417 | 0.15477654 | 0.707 | 0.72 | 1 | 0.804907417 | 0.003070652 |
| Dlg3 | 0.001540994 | 1.256602209 | 0.302 | 0.127 | 1 | 0.308501786 | 0.712876715 | 0.317 | 0.259 | 1 | 0.308501786 | 0.003079614 |
| Glrx5 | 0.001545138 | 0.798029082 | 0.717 | 0.551 | 1 | 0.162397699 | 0.306135357 | 0.683 | 0.611 | 1 | 0.162397699 | 0.003087889 |
| Vps35l | 0.001546642 | 1.224431567 | 0.509 | 0.305 | 1 | 0.266954314 | -0.282534394 | 0.317 | 0.423 | 1 | 0.266954314 | 0.003090892 |
| Prkar1b | 0.353375086 | 0.370451286 | 0.245 | 0.182 | 1 | 0.001551873 | -1.662641037 | 0.098 | 0.345 | 1 | 0.353375086 | 0.003101338 |
| Mia3 | 0.001563482 | 0.537394568 | 0.943 | 0.869 | 1 | 0.319391398 | -0.135560158 | 0.902 | 0.932 | 1 | 0.319391398 | 0.003124519 |
| Klhl24 | 0.001563741 | 0.751963425 | 0.774 | 0.606 | 1 | 0.249104472 | 0.118386602 | 0.829 | 0.655 | 1 | 0.249104472 | 0.003125036 |
| Trpm4 | 0.001569773 | 1.624207878 | 0.208 | 0.068 | 1 | 0.014889019 | 1.707921833 | 0.22 | 0.096 | 1 | 0.014889019 | 0.003137081 |
| Jade1 | 0.001573516 | 1.389187848 | 0.509 | 0.335 | 1 | 0.004929198 | 1.029849928 | 0.659 | 0.509 | 1 | 0.004929198 | 0.003144556 |
| Srr | 0.001580828 | 1.200526284 | 0.453 | 0.254 | 1 | 0.346724552 | 0.952068807 | 0.341 | 0.297 | 1 | 0.346724552 | 0.003159157 |
| Fign | 0.001582107 | -2.049848549 | 0.151 | 0.369 | 1 | 0.049978494 | -0.606241065 | 0.22 | 0.375 | 1 | 0.049978494 | 0.003161711 |
| Sh3kbp1 | 0.001585334 | -0.90135333 | 0.434 | 0.678 | 1 | 0.880002789 | -0.354415223 | 0.268 | 0.273 | 1 | 0.880002789 | 0.003168156 |
| Tusc3 | 0.001589429 | 0.631687843 | 0.774 | 0.674 | 1 | 0.440921358 | 0.357852537 | 0.683 | 0.71 | 1 | 0.440921358 | 0.003176332 |
| Morn4 | 0.001590232 | 1.8327945 | 0.17 | 0.047 | 1 | 0.370207442 | -0.555112573 | 0.024 | 0.058 | 1 | 0.370207442 | 0.003177935 |
| Glt8d1 | 0.003695212 | 0.550263918 | 0.83 | 0.686 | 1 | 0.001591297 | -0.974437431 | 0.39 | 0.611 | 1 | 0.003695212 | 0.003180062 |
| Myl4 | 0.00159224 | 1.844382474 | 0.226 | 0.081 | 1 | 0.691753484 | -0.352619709 | 0.146 | 0.167 | 1 | 0.691753484 | 0.003181945 |
| Btbd2 | 0.001592402 | 1.110328475 | 0.358 | 0.169 | 1 | 0.757387702 | 0.822254508 | 0.22 | 0.222 | 1 | 0.757387702 | 0.003182267 |
| Hs1bp3 | 0.001603666 | 2.669295768 | 0.113 | 0.021 | 1 | 0.494346678 | -0.555112573 | 0.024 | 0.048 | 1 | 0.494346678 | 0.003204761 |
| Mtarc1 | 0.001605138 | 2.017219071 | 0.113 | 0.021 | 1 | 0.430584486 | 2.83720485 | 0.024 | 0.01 | 1 | 0.430584486 | 0.0032077 |
| 2010007H06Rik | 0.001606748 | -2.564095653 | 0.038 | 0.225 | 1 | 0.048165671 | -1.970150072 | 0.024 | 0.13 | 1 | 0.048165671 | 0.003210914 |
| Dhps | 0.001606989 | 0.945269229 | 0.509 | 0.309 | 1 | 0.602532107 | 0.243680336 | 0.39 | 0.375 | 1 | 0.602532107 | 0.003211395 |
| Ppp4r3b | 0.001614203 | 0.697249829 | 0.849 | 0.695 | 1 | 0.444275237 | 0.109553932 | 0.878 | 0.863 | 1 | 0.444275237 | 0.0032258 |
| Ccne2 | 0.098508297 | 0.569760094 | 0.132 | 0.064 | 1 | 0.00161615 | 2.252242349 | 0.146 | 0.034 | 1 | 0.098508297 | 0.003229688 |
| Plcd1 | 0.002114099 | -0.88591939 | 0.396 | 0.623 | 1 | 0.001616467 | -0.859868017 | 0.366 | 0.631 | 1 | 0.002114099 | 0.003230321 |
| Clasp1 | 0.001618351 | 0.813106165 | 0.755 | 0.644 | 1 | 0.111685433 | -0.236044132 | 0.78 | 0.836 | 1 | 0.111685433 | 0.003234084 |
| Sall2 | 0.001618814 | 2.347367673 | 0.113 | 0.021 | 1 | 0.668293268 | -0.250257991 | 0.024 | 0.038 | 1 | 0.668293268 | 0.003235007 |
| Clic6 | 0.909120594 | -0.845277405 | 0.019 | 0.021 | 1 | 0.001622698 | 2.422167351 | 0.122 | 0.024 | 1 | 0.909120594 | 0.003242762 |
| Klhdc10 | 0.001624016 | 0.895078778 | 0.604 | 0.411 | 1 | 0.029517446 | 0.72975856 | 0.78 | 0.635 | 1 | 0.029517446 | 0.003245395 |
| Nfkbie | 0.00162512 | 2.417757001 | 0.132 | 0.03 | 1 | 0.360451923 | 0.83720485 | 0.073 | 0.041 | 1 | 0.360451923 | 0.003247598 |
| Eya4 | 0.001629928 | 1.03679687 | 0.83 | 0.678 | 1 | 0.160143448 | 0.37036845 | 0.854 | 0.765 | 1 | 0.160143448 | 0.0032572 |
| Col4a3 | 0.001632186 | -3.678167419 | 0.019 | 0.195 | 1 | 0.20883094 | -1.747757651 | 0 | 0.038 | 1 | 0.20883094 | 0.003261707 |
| Zfp300 | 0.001634093 | 2.154722595 | 0.113 | 0.021 | 1 | 0.028242983 | 2.252242349 | 0.073 | 0.017 | 1 | 0.028242983 | 0.003265515 |
| Ppp4r1l-ps | 0.001634303 | 1.173831418 | 0.566 | 0.407 | 1 | 0.44152316 | -0.16279515 | 0.366 | 0.454 | 1 | 0.44152316 | 0.003265935 |
| Pot1b | 0.001635738 | 2.154722595 | 0.151 | 0.038 | 1 | 0.934943007 | -0.385187572 | 0.049 | 0.051 | 1 | 0.934943007 | 0.0032688 |
| Chrdl1 | 0.140022706 | -0.556097893 | 0.528 | 0.602 | 1 | 0.001636355 | -0.920885084 | 0.366 | 0.648 | 1 | 0.140022706 | 0.003270032 |
| Fads3 | 0.001638058 | 0.859979329 | 0.66 | 0.483 | 1 | 0.540279126 | 0.188916414 | 0.537 | 0.481 | 1 | 0.540279126 | 0.003273434 |
| Klhl15 | 0.001639477 | 1.397993746 | 0.453 | 0.275 | 1 | 0.381531151 | 0.184070846 | 0.317 | 0.413 | 1 | 0.381531151 | 0.003276266 |
| Cracd | 0.034521617 | 0.877188619 | 0.302 | 0.178 | 1 | 0.001641998 | 0.876951029 | 0.463 | 0.229 | 1 | 0.034521617 | 0.003281299 |
| Sphk2 | 0.001643815 | 2.017219071 | 0.151 | 0.038 | 1 | 0.021332427 | 1.389745873 | 0.195 | 0.082 | 1 | 0.021332427 | 0.003284928 |
| Rbis | 0.273773012 | -0.208824589 | 0.491 | 0.576 | 1 | 0.001648469 | -1.17660095 | 0.22 | 0.478 | 1 | 0.273773012 | 0.00329422 |
| Gm56901 | 0.001648805 | 2.324647596 | 0.132 | 0.03 | 1 | 0.820932515 | 0.614812428 | 0.049 | 0.041 | 1 | 0.820932515 | 0.00329489 |
| Gm26512 | 0.001650057 | 1.491757582 | 0.17 | 0.047 | 1 | 0.861257699 | 0.136765132 | 0.073 | 0.065 | 1 | 0.861257699 | 0.00329739 |
| Tyw1 | 0.001652055 | 1.221836791 | 0.358 | 0.169 | 1 | 0.721686257 | 0.317830691 | 0.171 | 0.201 | 1 | 0.721686257 | 0.00330138 |
| Arl14ep | 0.281623168 | 0.717858733 | 0.358 | 0.309 | 1 | 0.001654036 | -1.080956858 | 0.195 | 0.471 | 1 | 0.281623168 | 0.003305336 |
| Leng1 | 0.001659764 | 1.026967048 | 0.377 | 0.178 | 1 | 0.031459105 | 1.185128153 | 0.341 | 0.208 | 1 | 0.031459105 | 0.003316772 |
| Tcf20 | 0.00166393 | 0.8327945 | 0.698 | 0.517 | 1 | 0.002817233 | 0.823129665 | 0.902 | 0.713 | 1 | 0.002817233 | 0.003325091 |
| Ulk2 | 0.001664334 | 1.174900477 | 0.736 | 0.585 | 1 | 0.105620569 | 0.734843132 | 0.634 | 0.584 | 1 | 0.105620569 | 0.003325898 |
| Foxj2 | 0.001665532 | 1.223985257 | 0.396 | 0.212 | 1 | 0.467837978 | 0.470422519 | 0.439 | 0.416 | 1 | 0.467837978 | 0.003328291 |
| Efcab7 | 0.001666987 | 1.8327945 | 0.132 | 0.03 | 1 | 0.550075059 | 0.749742009 | 0.049 | 0.031 | 1 | 0.550075059 | 0.003331195 |
| Polr2j | 0.001668636 | 1.025439578 | 0.528 | 0.322 | 1 | 0.111341004 | 0.562197802 | 0.415 | 0.287 | 1 | 0.111341004 | 0.003334488 |
| Myzap | 0.001670419 | 1.20202831 | 0.264 | 0.102 | 1 | 0.972011517 | 0.422167351 | 0.293 | 0.304 | 1 | 0.972011517 | 0.003338048 |
| Ddhd1 | 0.001671833 | 1.271236604 | 0.453 | 0.284 | 1 | 0.011358767 | 0.78473743 | 0.707 | 0.512 | 1 | 0.011358767 | 0.003340872 |
| Egln1 | 0.001677071 | 0.924424975 | 0.642 | 0.521 | 1 | 0.83230401 | 0.188112012 | 0.585 | 0.659 | 1 | 0.83230401 | 0.00335133 |
| Ankrd16 | 0.001678728 | 1.265753907 | 0.321 | 0.14 | 1 | 0.542222544 | 0.493250449 | 0.195 | 0.16 | 1 | 0.542222544 | 0.003354639 |
| Akap11 | 0.035818722 | -0.38312731 | 0.66 | 0.792 | 1 | 0.001679832 | -0.706337724 | 0.463 | 0.765 | 1 | 0.035818722 | 0.003356841 |
| Eif3a | 0.001682248 | 0.413432696 | 1 | 0.992 | 1 | 0.150419985 | -0.175714229 | 1 | 0.99 | 1 | 0.150419985 | 0.003361666 |
| Cdc42bpb | 0.001682737 | 0.715300937 | 0.849 | 0.695 | 1 | 0.124525743 | 0.387449291 | 0.878 | 0.853 | 1 | 0.124525743 | 0.003362642 |
| Tigd2 | 0.001685728 | 1.324647596 | 0.415 | 0.229 | 1 | 0.271242412 | 0.635570989 | 0.146 | 0.246 | 1 | 0.271242412 | 0.003368615 |
| Cfap298 | 0.001686549 | 1.292226119 | 0.377 | 0.191 | 1 | 0.203353957 | 0.773074512 | 0.293 | 0.215 | 1 | 0.203353957 | 0.003370253 |
| Rad9b | 0.001687646 | 1.243989933 | 0.302 | 0.123 | 1 | 0.887046622 | 0.778311161 | 0.195 | 0.198 | 1 | 0.887046622 | 0.003372445 |
| Ddhd2 | 0.001689592 | 0.693672698 | 0.679 | 0.508 | 1 | 0.152077325 | -0.405867824 | 0.561 | 0.669 | 1 | 0.152077325 | 0.00337633 |
| Ccdc34 | 0.001692897 | 0.956496485 | 0.642 | 0.441 | 1 | 0.262653797 | 0.410940095 | 0.415 | 0.324 | 1 | 0.262653797 | 0.003382929 |
| Zfp235 | 0.001695308 | 1.384204441 | 0.264 | 0.102 | 1 | 0.185047927 | 0.574170444 | 0.22 | 0.137 | 1 | 0.185047927 | 0.003387742 |
| Ecm1 | 0.001699037 | 1.695290976 | 0.208 | 0.068 | 1 | 0.026876958 | 1.19334866 | 0.244 | 0.119 | 1 | 0.026876958 | 0.003395188 |

| | | | | | | | | | | | | |
|---|---|---|---|---|---|---|---|---|---|---|---|---|
| Fam227a | 0.001699375 | 1.377115016 | 0.245 | 0.089 | 1 | 0.512923338 | 0.583448258 | 0.146 | 0.116 | 1 | 0.512923338 | 0.003395862 |
| Smurf1 | 0.001700966 | 1.047807391 | 0.491 | 0.28 | 1 | 0.31791083 | 0.607255077 | 0.488 | 0.471 | 1 | 0.31791083 | 0.003399038 |
| Vapb | 0.0017015 | 0.650378554 | 0.755 | 0.64 | 1 | 0.556781168 | 0.319670277 | 0.659 | 0.713 | 1 | 0.556781168 | 0.003400105 |
| Slc35c1 | 0.001702029 | 2.06161319 | 0.189 | 0.059 | 1 | 0.173970007 | -0.717384002 | 0.049 | 0.123 | 1 | 0.173970007 | 0.003401161 |
| Nkrf | 0.00170587 | 1.669295768 | 0.226 | 0.081 | 1 | 0.815649839 | 0.756284854 | 0.122 | 0.143 | 1 | 0.815649839 | 0.003408831 |
| Msi1 | 0.393179448 | 0.377115016 | 0.208 | 0.157 | 1 | 0.001707959 | 1.811669758 | 0.317 | 0.133 | 1 | 0.393179448 | 0.003413001 |
| Foxl1 | 0.917883255 | 0.569760094 | 0.019 | 0.021 | 1 | 0.001709929 | 1.749742009 | 0.146 | 0.034 | 1 | 0.917883255 | 0.003416935 |
| Brox | 0.001710932 | 0.960842861 | 0.623 | 0.424 | 1 | 0.843883153 | 0.104252318 | 0.659 | 0.686 | 1 | 0.843883153 | 0.003418937 |
| Rnf126 | 0.001716739 | 1.094181053 | 0.453 | 0.242 | 1 | 0.306879839 | 0.64319423 | 0.415 | 0.358 | 1 | 0.306879839 | 0.00343053 |
| Atp2c1 | 0.005187188 | 0.754543054 | 0.566 | 0.373 | 1 | 0.001719479 | 0.602299817 | 0.78 | 0.58 | 1 | 0.005187188 | 0.003436002 |
| Igf1r | 0.903568012 | -0.380210793 | 1 | 0.987 | 1 | 0.001721882 | 0.483881559 | 1 | 0.99 | 1 | 0.903568012 | 0.003440799 |
| Kmt5b | 0.001725923 | 0.601124265 | 0.811 | 0.699 | 1 | 0.114777856 | -0.322341006 | 0.78 | 0.877 | 1 | 0.114777856 | 0.003448867 |
| Thbs3 | 0.001729672 | 0.649487287 | 0.509 | 0.288 | 1 | 0.009118172 | 1.127546602 | 0.488 | 0.317 | 1 | 0.009118172 | 0.003456351 |
| Pear1 | 0.001738154 | 0.895988326 | 0.377 | 0.178 | 1 | 0.004384607 | -1.56878751 | 0.171 | 0.386 | 1 | 0.004384607 | 0.003473288 |
| Asb6 | 0.001741653 | 1.228723176 | 0.321 | 0.14 | 1 | 0.451268548 | 0.486707603 | 0.171 | 0.126 | 1 | 0.451268548 | 0.003480273 |
| Tmem87b | 0.001742274 | 0.78829504 | 0.736 | 0.572 | 1 | 0.180059898 | 0.525035721 | 0.732 | 0.717 | 1 | 0.180059898 | 0.003481513 |
| Gm29418 | 0.001742717 | 1.209170379 | 0.358 | 0.174 | 1 | 0.131803076 | -0.555112573 | 0.195 | 0.317 | 1 | 0.131803076 | 0.003482397 |
| Coa7 | 0.001749281 | 2.154722595 | 0.132 | 0.03 | 1 | 0.386576163 | 2.029849928 | 0.073 | 0.044 | 1 | 0.386576163 | 0.003495501 |
| Polr1e | 0.001754074 | 2.154722595 | 0.132 | 0.03 | 1 | 0.279561054 | 1.071670103 | 0.098 | 0.055 | 1 | 0.279561054 | 0.003505071 |
| Map2k1 | 0.001757997 | 0.715610961 | 0.566 | 0.36 | 1 | 0.46000189 | -0.138869311 | 0.463 | 0.519 | 1 | 0.46000189 | 0.003512904 |
| Brms1l | 0.001763551 | 1.070189085 | 0.528 | 0.339 | 1 | 0.519552869 | -0.231957175 | 0.39 | 0.44 | 1 | 0.519552869 | 0.003523992 |
| Slc43a3 | 0.003883431 | -0.772764905 | 0.528 | 0.725 | 1 | 0.001764034 | -1.233184478 | 0.268 | 0.543 | 1 | 0.003883431 | 0.003524956 |
| Lipe | 0.001764517 | 2.154722595 | 0.189 | 0.059 | 1 | 0.188212299 | -1.069685746 | 0.024 | 0.082 | 1 | 0.188212299 | 0.00352592 |
| Gipc2 | 0.01256357 | 1.903183828 | 0.151 | 0.055 | 1 | 0.001770043 | 0.957499084 | 0.317 | 0.133 | 1 | 0.01256357 | 0.003536953 |
| Tmed5 | 0.001772013 | 0.63230156 | 0.792 | 0.686 | 1 | 0.010793887 | 0.664012135 | 0.805 | 0.672 | 1 | 0.010793887 | 0.003540885 |
| Kif28 | 0.001775259 | 1.389187848 | 0.151 | 0.038 | 1 | 0.25154762 | 1.574170444 | 0.098 | 0.055 | 1 | 0.25154762 | 0.003547367 |
| Cdc42ep4 | 0.001775853 | -0.960754623 | 0.377 | 0.619 | 1 | 0.053484808 | -0.505610612 | 0.512 | 0.683 | 1 | 0.053484808 | 0.003548553 |
| Taf13 | 0.001778906 | -0.887284932 | 0.585 | 0.729 | 1 | 0.187080877 | -0.173364392 | 0.61 | 0.7 | 1 | 0.187080877 | 0.003554648 |
| M1ap | 0.001781283 | -2.379255977 | 0.075 | 0.275 | 1 | 0.170402911 | -1.555112573 | 0 | 0.044 | 1 | 0.170402911 | 0.003559393 |
| Cttnbp2nl | 0.001907931 | 0.807919831 | 0.698 | 0.538 | 1 | 0.001790226 | -0.742282521 | 0.61 | 0.812 | 1 | 0.001907931 | 0.003577247 |
| Sv2a | 0.001794349 | 1.962077517 | 0.113 | 0.021 | 1 | 0.716347575 | 1.19334866 | 0.049 | 0.065 | 1 | 0.716347575 | 0.003585479 |
| Ankrd44 | 0.001795061 | 2.08240839 | 0.566 | 0.407 | 1 | 0.002011279 | 0.88967227 | 0.61 | 0.386 | 1 | 0.002011279 | 0.003586899 |
| Peg10 | 0.001811881 | 1.318221327 | 0.132 | 0.03 | 1 | 0.072574324 | 1.515276755 | 0.122 | 0.051 | 1 | 0.072574324 | 0.003620478 |
| Gpsm2 | 0.00181725 | 0.807259408 | 0.66 | 0.47 | 1 | 0.030058642 | -0.642788091 | 0.488 | 0.662 | 1 | 0.030058642 | 0.003631198 |
| Acvr2a | 0.001820217 | 1.007881206 | 0.491 | 0.275 | 1 | 0.099339561 | 0.672633083 | 0.585 | 0.505 | 1 | 0.099339561 | 0.00363712 |
| Plcl2 | 0.001821406 | 1.838695101 | 0.283 | 0.119 | 1 | 0.748078081 | 0.412707021 | 0.146 | 0.181 | 1 | 0.748078081 | 0.003639495 |
| Mpp2 | 0.001829556 | 1.984797593 | 0.189 | 0.059 | 1 | 0.050412123 | 1.252242349 | 0.171 | 0.078 | 1 | 0.050412123 | 0.003655764 |
| Prkx | 0.0103916 | 0.865215978 | 0.396 | 0.233 | 1 | 0.00183122 | 1.100239256 | 0.634 | 0.427 | 1 | 0.0103916 | 0.003659086 |
| Fam53a | 0.001833454 | 1.470224421 | 0.321 | 0.153 | 1 | 0.053329652 | 1.557096931 | 0.317 | 0.218 | 1 | 0.053329652 | 0.003663547 |
| Spaca6 | 0.001834629 | 1.033970435 | 0.509 | 0.335 | 1 | 0.892846382 | 0.127006165 | 0.585 | 0.577 | 1 | 0.892846382 | 0.003665892 |
| Paics | 0.011629355 | -0.439077269 | 0.943 | 0.928 | 1 | 0.001837778 | -0.531016897 | 0.854 | 0.962 | 1 | 0.011629355 | 0.003672179 |
| Ube2r2 | 0.001839041 | 0.506150706 | 0.962 | 0.924 | 1 | 0.109890558 | -0.19841906 | 0.976 | 0.986 | 1 | 0.109890558 | 0.0036747 |
| Gigyf1 | 0.001841748 | 1.465062715 | 0.34 | 0.161 | 1 | 0.150184288 | 0.686262952 | 0.439 | 0.341 | 1 | 0.150184288 | 0.003680105 |
| Depdc5 | 0.001842117 | 1.133027524 | 0.358 | 0.169 | 1 | 0.3572544 | 0.463080544 | 0.39 | 0.331 | 1 | 0.3572544 | 0.003680841 |
| Trappc10 | 0.00184843 | 0.880100215 | 0.604 | 0.403 | 1 | 0.132927291 | 0.420558098 | 0.659 | 0.556 | 1 | 0.132927291 | 0.003693443 |
| Pbx2 | 0.001848485 | 1.247831999 | 0.321 | 0.14 | 1 | 0.204553136 | 1.10681191 | 0.268 | 0.195 | 1 | 0.204553136 | 0.003693553 |
| Sf1 | 0.144703186 | 0.223632304 | 0.849 | 0.839 | 1 | 0.001852319 | -0.525137514 | 0.878 | 0.928 | 1 | 0.144703186 | 0.003701207 |
| Gm49602 | 0.001852603 | 1.984797593 | 0.132 | 0.03 | 1 | 0.457571722 | 1.313642894 | 0.098 | 0.068 | 1 | 0.457571722 | 0.003701774 |
| Sting1 | 0.001856891 | 2.017219071 | 0.132 | 0.03 | 1 | 0.824088343 | 0.377773231 | 0.024 | 0.031 | 1 | 0.824088343 | 0.003710333 |
| Rab11fip2 | 0.001859578 | 1.103375715 | 0.566 | 0.377 | 1 | 0.850848204 | 0.47620719 | 0.39 | 0.433 | 1 | 0.850848204 | 0.003715698 |
| Gm41556 | 0.001859711 | 2.154722595 | 0.208 | 0.072 | 1 | 0.018591525 | 2.408361551 | 0.171 | 0.072 | 1 | 0.018591525 | 0.003715963 |
| Wdr33 | 0.001861364 | 0.518685909 | 0.925 | 0.907 | 1 | 0.169952132 | -0.139435269 | 0.976 | 0.962 | 1 | 0.169952132 | 0.003719263 |
| Vps45 | 0.001873258 | 1.617065809 | 0.321 | 0.148 | 1 | 0.03891567 | 1.377773231 | 0.317 | 0.191 | 1 | 0.03891567 | 0.003743007 |
| Mt1 | 0.001873336 | -2.368013785 | 0.566 | 0.661 | 1 | 0.024423838 | 0.419678657 | 0.585 | 0.433 | 1 | 0.024423838 | 0.003743162 |
| Rsbn1l | 0.001886721 | 1.098139066 | 0.623 | 0.449 | 1 | 0.538866673 | 0.405488609 | 0.707 | 0.703 | 1 | 0.538866673 | 0.003769883 |
| Nr2f6 | 0.001889664 | 1.180717803 | 0.377 | 0.186 | 1 | 0.965840767 | 0.252242349 | 0.317 | 0.331 | 1 | 0.965840767 | 0.003775757 |
| Ninl | 0.001892831 | 1.541745718 | 0.226 | 0.081 | 1 | 0.128491265 | 1.322631677 | 0.22 | 0.14 | 1 | 0.128491265 | 0.003782079 |
| Pou2f3 | 0.00189488 | 1.027610677 | 0.245 | 0.093 | 1 | 0.137193214 | -1.225804948 | 0.073 | 0.16 | 1 | 0.137193214 | 0.003786169 |
| Avpr1a | 0.001896022 | -2.799473716 | 0.038 | 0.22 | 1 | 0.043290284 | -2.520347155 | 0 | 0.092 | 1 | 0.043290284 | 0.00378845 |
| Arhgef12 | 0.001897631 | 0.330392272 | 1 | 0.979 | 1 | 0.013054934 | 0.572810519 | 0.976 | 0.997 | 1 | 0.013054934 | 0.00379166 |
| Itga7 | 0.001898101 | 0.874614676 | 0.113 | 0.021 | 1 | 0.025521447 | 1.422167351 | 0.098 | 0.027 | 1 | 0.025521447 | 0.003792599 |
| Zfp318 | 0.001901002 | 0.948271717 | 0.547 | 0.343 | 1 | 0.4139847 | 0.530543512 | 0.415 | 0.382 | 1 | 0.4139847 | 0.003798391 |
| Ublcp1 | 0.001902867 | 1.178970141 | 0.509 | 0.314 | 1 | 0.840015778 | 0.153940275 | 0.366 | 0.369 | 1 | 0.840015778 | 0.003802113 |
| Grip1 | 0.001904299 | 1.18893831 | 0.321 | 0.153 | 1 | 0.175488001 | 0.186279087 | 0.439 | 0.328 | 1 | 0.175488001 | 0.003804971 |
| Evc2 | 0.001907093 | 1.112902419 | 0.434 | 0.237 | 1 | 0.367881054 | 0.678006255 | 0.39 | 0.352 | 1 | 0.367881054 | 0.00381055 |
| Atl2 | 0.001914443 | 0.609726501 | 0.66 | 0.441 | 1 | 0.58585132 | 0.241974014 | 0.732 | 0.7 | 1 | 0.58585132 | 0.003825221 |
| Rbms2 | 0.001915459 | 0.857042046 | 0.623 | 0.449 | 1 | 0.302054426 | 0.541271963 | 0.488 | 0.481 | 1 | 0.302054426 | 0.003827249 |
| Fbxw7 | 0.001923536 | 0.749968069 | 0.698 | 0.483 | 1 | 0.006405689 | 0.756831433 | 0.78 | 0.614 | 1 | 0.006405689 | 0.003843371 |
| Odc1 | 0.001930445 | -0.696380223 | 0.849 | 0.924 | 1 | 0.211660813 | -0.233418132 | 0.829 | 0.863 | 1 | 0.211660813 | 0.003857164 |
| Tyk2 | 0.001949037 | 2.237184755 | 0.189 | 0.059 | 1 | 0.302185157 | -0.132421501 | 0.073 | 0.137 | 1 | 0.302185157 | 0.003894276 |
| Slc35e3 | 0.001950979 | -1.14644694 | 0.358 | 0.572 | 1 | 0.089671857 | -0.595754557 | 0.293 | 0.427 | 1 | 0.089671857 | 0.003898151 |
| Dner | 0.001956492 | 3.569760094 | 0.075 | 0.008 | 1 | 0.004188151 | 2.83720485 | 0.049 | 0.003 | 1 | 0.004188151 | 0.003909156 |
| Zscan2 | 0.001956492 | 3.569760094 | 0.075 | 0.008 | 1 | 0.872828964 | 1.029849928 | 0.024 | 0.02 | 1 | 0.872828964 | 0.003909156 |
| Zfp791 | 0.001956492 | 3.569760094 | 0.075 | 0.008 | 1 | 0.47519767 | 0.699701326 | 0.024 | 0.051 | 1 | 0.47519767 | 0.003909156 |
| 4930513N10Rik | 0.001956492 | 3.569760094 | 0.075 | 0.008 | 1 | 0.402373368 | 0.252242349 | 0 | 0.017 | 1 | 0.402373368 | 0.003909156 |
| Calml4 | 0.001956492 | 3.377115016 | 0.075 | 0.008 | 1 | 0.916649754 | 0.667279848 | 0.024 | 0.027 | 1 | 0.916649754 | 0.003909156 |
| Fbxl6 | 0.001956685 | 1.340278248 | 0.377 | 0.199 | 1 | 0.269368946 | 0.715214325 | 0.268 | 0.198 | 1 | 0.269368946 | 0.003909541 |
| Rnps1 | 0.001960953 | 0.639469066 | 0.811 | 0.703 | 1 | 0.049220563 | 0.397440265 | 0.878 | 0.785 | 1 | 0.049220563 | 0.003918061 |
| Uba2 | 0.001971549 | 0.837381769 | 0.623 | 0.441 | 1 | 0.675388252 | 0.223992872 | 0.659 | 0.652 | 1 | 0.675388252 | 0.003939211 |
| Kazn | 0.001974388 | 1.758793918 | 0.34 | 0.178 | 1 | 0.008197459 | 0.717905921 | 0.366 | 0.181 | 1 | 0.008197459 | 0.003944878 |
| Mtarc2 | 0.001975222 | 0.598329246 | 0.736 | 0.585 | 1 | 0.608468521 | 0.141818359 | 0.537 | 0.618 | 1 | 0.608468521 | 0.003946543 |
| Zzz3 | 0.00197629 | 0.851231748 | 0.755 | 0.602 | 1 | 0.067970267 | 0.3387791 | 0.951 | 0.867 | 1 | 0.067970267 | 0.003948674 |
| Pik3c3 | 0.004037353 | 1.059565362 | 0.34 | 0.165 | 1 | 0.001988216 | 1.363273661 | 0.366 | 0.171 | 1 | 0.004037353 | 0.00397248 |
| Ensa | 0.001990444 | 0.8327945 | 0.736 | 0.623 | 1 | 0.473525774 | 0.403045197 | 0.707 | 0.788 | 1 | 0.473525774 | 0.003976926 |
| Mrap2 | 0.002005228 | 3.154722595 | 0.075 | 0.008 | 1 | 0.103479965 | 3.422167351 | 0.024 | 0.003 | 1 | 0.103479965 | 0.004006434 |
| Rab6b | 0.002005228 | 3.891688189 | 0.075 | 0.008 | 1 | 0.208817133 | -0.970150072 | 0 | 0.038 | 1 | 0.208817133 | 0.004006434 |
| Engase | 0.002005228 | 3.154722595 | 0.075 | 0.008 | 1 | 0.337213694 | -0.747757651 | 0.024 | 0.061 | 1 | 0.337213694 | 0.004006434 |
| Tes3-ps | 0.002005228 | 3.154722595 | 0.075 | 0.008 | 1 | 0.87556513 | 0.83720485 | 0.024 | 0.02 | 1 | 0.87556513 | 0.004006434 |
| Adcy4 | 0.002005228 | 3.154722595 | 0.075 | 0.008 | 1 | 0.059714165 | 1.574170444 | 0.098 | 0.034 | 1 | 0.059714165 | 0.004006434 |
| Fsd1 | 0.002005228 | 3.377115016 | 0.075 | 0.008 | 1 | 0.257308946 | -0.863234868 | 0 | 0.031 | 1 | 0.257308946 | 0.004006434 |
| Psmc6 | 0.002008492 | 0.472204785 | 0.906 | 0.864 | 1 | 0.1439796 | 0.3384812 | 0.951 | 0.904 | 1 | 0.1439796 | 0.004012949 |
| Egflam | 0.002011015 | 2.315187267 | 0.17 | 0.051 | 1 | 0.35899028 | 1.252242349 | 0.049 | 0.024 | 1 | 0.35899028 | 0.004017986 |
| Rmnd5b | 0.002012567 | 1.337586652 | 0.302 | 0.131 | 1 | 0.550151755 | 0.515276755 | 0.268 | 0.232 | 1 | 0.550151755 | 0.004021084 |
| Eno1 | 0.002016159 | -0.733308974 | 0.698 | 0.805 | 1 | 0.020841248 | -0.505187348 | 0.732 | 0.805 | 1 | 0.020841248 | 0.004028253 |
| Gramd3 | 0.002016597 | 1.106717222 | 0.66 | 0.53 | 1 | 0.224038227 | 0.548528561 | 0.659 | 0.597 | 1 | 0.224038227 | 0.004029128 |
| Zfp469 | 0.714087922 | -0.141518856 | 0.396 | 0.432 | 1 | 0.002026825 | -0.962590081 | 0.439 | 0.693 | 1 | 0.714087922 | 0.004049542 |
| Rnf220 | 0.002027164 | 1.242185436 | 0.396 | 0.203 | 1 | 0.094537168 | 1.048708955 | 0.366 | 0.263 | 1 | 0.094537168 | 0.004050219 |
| Ubn1 | 0.002027719 | 0.539765354 | 0.906 | 0.877 | 1 | 0.234491099 | 0.17882128 | 0.976 | 0.932 | 1 | 0.234491099 | 0.004051327 |

| | | | | | | | | | | | | |
|---|---|---|---|---|---|---|---|---|---|---|---|---|
| Gm15492 | 0.00203087 | 3.154722595 | 0.075 | 0.008 | 1 | 0.260727064 | 1.699701326 | 0.049 | 0.02 | 1 | 0.260727064 | 0.004057616 |
| Adam17 | 0.01781482 | 0.626622431 | 0.547 | 0.403 | 1 | 0.002038738 | 1.228196875 | 0.732 | 0.59 | 1 | 0.01781482 | 0.004073319 |
| Daam2 | 0.00204332 | 1.312904263 | 0.623 | 0.441 | 1 | 0.030258452 | 1.035011633 | 0.512 | 0.389 | 1 | 0.030258452 | 0.004082466 |
| Zfp579 | 0.002046404 | 1.767699472 | 0.189 | 0.059 | 1 | 0.141493889 | -0.665295491 | 0.024 | 0.096 | 1 | 0.141493889 | 0.00408862 |
| Slc25a4 | 0.002048755 | 0.454495899 | 1 | 0.966 | 1 | 0.393532817 | 0.227069278 | 0.976 | 0.952 | 1 | 0.393532817 | 0.004093313 |
| Pick1 | 0.002053783 | 1.261637799 | 0.321 | 0.144 | 1 | 0.70282094 | 0.515276755 | 0.195 | 0.171 | 1 | 0.70282094 | 0.004103347 |
| Fgf7 | 0.002056378 | 2.891688189 | 0.075 | 0.008 | 1 | 0.026641787 | 3.602739596 | 0.073 | 0.017 | 1 | 0.026641787 | 0.004108527 |
| She | 0.002056378 | 2.891688189 | 0.075 | 0.008 | 1 | 0.466469839 | 0.515276755 | 0.049 | 0.027 | 1 | 0.466469839 | 0.004108527 |
| Gm43063 | 0.002056378 | 2.891688189 | 0.075 | 0.008 | 1 | 0.267205177 | 2.252242349 | 0.024 | 0.007 | 1 | 0.267205177 | 0.004108527 |
| Pex26 | 0.002056378 | 2.891688189 | 0.075 | 0.008 | 1 | 0.321379042 | 1.100239256 | 0.098 | 0.058 | 1 | 0.321379042 | 0.004108527 |
| Zfp3 | 0.002056378 | 2.891688189 | 0.075 | 0.008 | 1 | 0.012973167 | 2.515276755 | 0.073 | 0.014 | 1 | 0.012973167 | 0.004108527 |
| Ube2n | 0.002058904 | 0.450797262 | 0.906 | 0.886 | 1 | 0.189667246 | 0.261592317 | 0.878 | 0.908 | 1 | 0.189667246 | 0.004113569 |
| Dnttip2 | 0.002062312 | 0.837043973 | 0.755 | 0.665 | 1 | 0.662667496 | 0.272665067 | 0.756 | 0.724 | 1 | 0.662667496 | 0.00412037 |
| Cnbd2 | 0.002065546 | 1.610402079 | 0.264 | 0.11 | 1 | 0.12960202 | 1.01123425 | 0.268 | 0.181 | 1 | 0.12960202 | 0.004126825 |
| Snx10 | 0.002068753 | -1.346630473 | 0.377 | 0.547 | 1 | 0.365651035 | -1.460475699 | 0.22 | 0.28 | 1 | 0.365651035 | 0.004133226 |
| Kmt2b | 0.002070061 | 1.10741688 | 0.491 | 0.288 | 1 | 0.901506739 | 0.102301722 | 0.488 | 0.485 | 1 | 0.901506739 | 0.004135837 |
| Fosl1 | 0.002070411 | -1.680466535 | 0.094 | 0.309 | 1 | 0.286321756 | -0.84461919 | 0.195 | 0.259 | 1 | 0.286321756 | 0.004136536 |
| Rnf31 | 0.499735438 | 0.526691372 | 0.151 | 0.119 | 1 | 0.002070503 | 1.313642894 | 0.293 | 0.113 | 1 | 0.499735438 | 0.00413672 |
| Appl1 | 0.002071945 | 0.533780797 | 0.83 | 0.725 | 1 | 0.117181077 | 0.424764462 | 0.854 | 0.846 | 1 | 0.117181077 | 0.004139597 |
| Atl1 | 0.002073102 | -1.845277405 | 0.113 | 0.322 | 1 | 0.011237402 | -1.527367582 | 0.049 | 0.222 | 1 | 0.011237402 | 0.004141906 |
| Eif3e | 0.002074844 | 0.441885272 | 0.981 | 0.898 | 1 | 0.407180351 | -0.130227288 | 0.829 | 0.942 | 1 | 0.407180351 | 0.004145382 |
| Cdv3 | 0.00207657 | 0.421155577 | 0.943 | 0.89 | 1 | 0.671147095 | -0.109465648 | 0.951 | 0.966 | 1 | 0.671147095 | 0.004148827 |
| Qars | 0.002076662 | 1.115903346 | 0.528 | 0.36 | 1 | 0.776742985 | 0.185128153 | 0.366 | 0.403 | 1 | 0.776742985 | 0.004149011 |
| Fasn | 0.002077616 | -1.351630071 | 0.34 | 0.53 | 1 | 0.195650328 | -0.500664789 | 0.39 | 0.481 | 1 | 0.195650328 | 0.004150916 |
| Pcdhb21 | 0.002081522 | 2.962077517 | 0.075 | 0.008 | 1 | 0.856409217 | 1.989207943 | 0.024 | 0.02 | 1 | 0.856409217 | 0.004158711 |
| 9330104G04Rik | 0.002081785 | 2.739685096 | 0.075 | 0.008 | 1 | 0.992196803 | 0.136765132 | 0.024 | 0.024 | 1 | 0.992196803 | 0.004159235 |
| Nampt | 0.010672697 | 0.789073122 | 0.566 | 0.39 | 1 | 0.002084871 | 1.278331578 | 0.683 | 0.58 | 1 | 0.010672697 | 0.004165395 |
| Phc1 | 0.00209009 | 1.79717059 | 0.302 | 0.131 | 1 | 0.31957455 | 0.470422519 | 0.317 | 0.253 | 1 | 0.31957455 | 0.004175812 |
| Atp5g1 | 0.002090791 | 0.425271054 | 0.981 | 0.873 | 1 | 0.576113809 | -0.129447893 | 0.854 | 0.857 | 1 | 0.576113809 | 0.004177211 |
| Lypla1 | 0.002100511 | 0.804225348 | 0.509 | 0.292 | 1 | 0.200257143 | 0.600165653 | 0.488 | 0.413 | 1 | 0.200257143 | 0.00419661 |
| Ptges | 0.740832122 | 0.399835093 | 0.132 | 0.157 | 1 | 0.002104598 | 1.289717055 | 0.268 | 0.099 | 1 | 0.740832122 | 0.004204766 |
| Tlr4 | 0.002104832 | -2.489133595 | 0.057 | 0.246 | 1 | 0.691137062 | -0.429581691 | 0.122 | 0.143 | 1 | 0.691137062 | 0.004205234 |
| Phlda1 | 0.002106635 | -1.330359544 | 0.868 | 0.864 | 1 | 0.070150945 | 0.562127253 | 0.78 | 0.741 | 1 | 0.070150945 | 0.004208832 |
| Atg14 | 0.002106882 | 1.925240749 | 0.264 | 0.114 | 1 | 0.226481922 | 0.377773231 | 0.293 | 0.205 | 1 | 0.226481922 | 0.004209326 |
| Pcdhb16 | 0.002107391 | 2.739685096 | 0.075 | 0.008 | 1 | 0.497959937 | 1.252242349 | 0.098 | 0.072 | 1 | 0.497959937 | 0.00421034 |
| Gm15261 | 0.002107391 | 2.739685096 | 0.075 | 0.008 | 1 | 0.889423593 | 0.19334866 | 0.049 | 0.055 | 1 | 0.889423593 | 0.00421034 |
| Rrad | 0.002111144 | 2.877188619 | 0.17 | 0.051 | 1 | 0.023480372 | 2.136765132 | 0.146 | 0.055 | 1 | 0.023480372 | 0.004217831 |
| Idnk | 0.002115472 | 1.569760094 | 0.34 | 0.169 | 1 | 0.195216431 | 1.01123425 | 0.268 | 0.208 | 1 | 0.195216431 | 0.004226468 |
| Swap70 | 0.010149275 | 0.746637856 | 0.642 | 0.436 | 1 | 0.002119371 | 1.270332806 | 0.683 | 0.532 | 1 | 0.010149275 | 0.00423425 |
| Eml1 | 0.0021225 | 1.259692154 | 0.377 | 0.195 | 1 | 0.226676192 | -0.798383724 | 0.341 | 0.444 | 1 | 0.226676192 | 0.004240494 |
| Gm48521 | 0.002130758 | -4.220316837 | 0 | 0.157 | 1 | 0.170422223 | -2.116991461 | 0 | 0.044 | 1 | 0.170422223 | 0.004256977 |
| Cul4b | 0.002130991 | 1.259904831 | 0.528 | 0.347 | 1 | 0.806508653 | 0.207454427 | 0.39 | 0.447 | 1 | 0.806508653 | 0.004257441 |
| Serpina1d | 0.002135421 | -6.349103143 | 0 | 0.157 | 1 | 0.257325016 | -3.982974113 | 0 | 0.031 | 1 | 0.257325016 | 0.004266282 |
| Ikzf5 | 0.002136216 | 0.991784024 | 0.491 | 0.28 | 1 | 0.259197533 | 0.46797104 | 0.415 | 0.348 | 1 | 0.259197533 | 0.004267869 |
| Gmppa | 0.998333974 | 0.247831999 | 0.377 | 0.398 | 1 | 0.002139311 | -1.537834582 | 0.171 | 0.403 | 1 | 0.998333974 | 0.004274046 |
| Senp2 | 0.002155364 | 0.940019439 | 0.943 | 0.814 | 1 | 0.628995706 | 0.438167829 | 0.902 | 0.901 | 1 | 0.628995706 | 0.004306082 |
| Eps8 | 0.463343797 | 0.429620822 | 0.698 | 0.801 | 1 | 0.002156695 | -0.465541832 | 0.707 | 0.918 | 1 | 0.463343797 | 0.004308738 |
| Hip1 | 0.035731324 | 0.537028892 | 0.811 | 0.78 | 1 | 0.00215782 | -0.54104986 | 0.659 | 0.874 | 1 | 0.035731324 | 0.004310984 |
| Adgrb2 | 0.002159563 | 2.47665069 | 0.075 | 0.008 | 1 | 0.462889204 | 0.515276755 | 0.049 | 0.027 | 1 | 0.462889204 | 0.004314463 |
| Rwdd2a | 0.002159563 | 2.154722595 | 0.075 | 0.008 | 1 | 0.365461252 | 1.100239256 | 0.049 | 0.024 | 1 | 0.365461252 | 0.004314463 |
| Cdc42ep2 | 0.002159563 | 2.47665069 | 0.075 | 0.008 | 1 | 0.87556513 | 0.83720485 | 0.024 | 0.02 | 1 | 0.87556513 | 0.004314463 |
| Tmem216 | 0.002159976 | 1.373362881 | 0.377 | 0.195 | 1 | 0.010663763 | -1.136799942 | 0.098 | 0.297 | 1 | 0.010663763 | 0.004315287 |
| Gm56519 | 0.002160241 | 3.002719501 | 0.075 | 0.008 | 1 | 0.593596252 | 1.614812428 | 0.024 | 0.014 | 1 | 0.593596252 | 0.004315815 |
| Pfdn5 | 0.002177019 | 0.340398118 | 1 | 0.953 | 1 | 0.043007514 | 0.290939643 | 0.976 | 0.959 | 1 | 0.043007514 | 0.004349299 |
| Flcn | 0.002181354 | 1.526691372 | 0.358 | 0.178 | 1 | 0.118650951 | -0.421529419 | 0.195 | 0.317 | 1 | 0.118650951 | 0.004357951 |
| Ifngr1 | 0.002185382 | 1.126153443 | 0.566 | 0.428 | 1 | 0.201836744 | 0.675741427 | 0.61 | 0.56 | 1 | 0.201836744 | 0.004365988 |
| Phlpp2 | 0.002191343 | 0.910303867 | 0.566 | 0.347 | 1 | 0.772309482 | 0.204148061 | 0.537 | 0.577 | 1 | 0.772309482 | 0.004377885 |
| Poglut3 | 0.002191739 | 0.803250224 | 0.642 | 0.419 | 1 | 0.649965122 | 0.172515157 | 0.439 | 0.519 | 1 | 0.649965122 | 0.004378675 |
| Bckdk | 0.002192144 | 0.91525666 | 0.491 | 0.292 | 1 | 0.083733202 | -0.561344527 | 0.171 | 0.307 | 1 | 0.083733202 | 0.004379483 |
| Ctsz | 0.002192969 | 0.66139057 | 0.83 | 0.712 | 1 | 0.800405174 | -0.196515768 | 0.707 | 0.744 | 1 | 0.800405174 | 0.004381129 |
| Sh3bp5 | 0.002201039 | 1.039245377 | 0.245 | 0.093 | 1 | 0.005531374 | 1.152706676 | 0.244 | 0.096 | 1 | 0.005531374 | 0.004397233 |
| Calml3 | 0.002204213 | -2.690767456 | 0.057 | 0.242 | 1 | 0.013916965 | -1.435813645 | 0.122 | 0.314 | 1 | 0.013916965 | 0.004403567 |
| Msantd2 | 0.002207209 | 0.953088734 | 0.623 | 0.483 | 1 | 0.045549203 | 0.609719959 | 0.805 | 0.741 | 1 | 0.045549203 | 0.004409546 |
| Gsdme | 0.002212886 | -1.504786859 | 0.17 | 0.381 | 1 | 0.125515441 | -0.257312749 | 0.244 | 0.386 | 1 | 0.125515441 | 0.004420874 |
| Gm56741 | 0.002216608 | 1.8327945 | 0.208 | 0.072 | 1 | 0.045840217 | 1.458693227 | 0.146 | 0.061 | 1 | 0.045840217 | 0.004428303 |
| Pigb | 0.002217469 | 1.261637799 | 0.302 | 0.131 | 1 | 0.161819179 | 1.196747236 | 0.22 | 0.143 | 1 | 0.161819179 | 0.00443002 |
| Eno1b | 0.156268076 | -0.389597921 | 0.17 | 0.271 | 1 | 0.002237492 | -3.185162963 | 0 | 0.191 | 1 | 0.156268076 | 0.004469977 |
| Ylpm1 | 0.002245484 | 0.638147069 | 0.755 | 0.606 | 1 | 0.349204423 | 0.402020777 | 0.829 | 0.778 | 1 | 0.349204423 | 0.004485926 |
| 3110082I17Rik | 0.423096173 | 0.232725107 | 0.283 | 0.233 | 1 | 0.002252228 | -3.332720152 | 0 | 0.191 | 1 | 0.423096173 | 0.004499384 |
| Ccdc71l | 0.026596732 | 1.154722595 | 0.189 | 0.085 | 1 | 0.002252671 | 1.547698233 | 0.293 | 0.123 | 1 | 0.026596732 | 0.004500268 |
| Atp6ap2 | 0.002257163 | 0.633890431 | 0.906 | 0.826 | 1 | 0.728538315 | -0.118359593 | 0.878 | 0.939 | 1 | 0.728538315 | 0.004509232 |
| Dusp10 | 0.019507356 | 0.970298024 | 0.321 | 0.191 | 1 | 0.002258668 | 1.515276755 | 0.341 | 0.154 | 1 | 0.019507356 | 0.004512234 |
| Scand1 | 0.002258801 | 0.821523388 | 0.774 | 0.581 | 1 | 0.008852324 | 0.75768451 | 0.756 | 0.621 | 1 | 0.008852324 | 0.004512499 |
| Gm13832 | 0.002262208 | 1.017219071 | 0.151 | 0.038 | 1 | 0.664414647 | -0.80665134 | 0.024 | 0.038 | 1 | 0.664414647 | 0.004519299 |
| Kin | 0.002265255 | 0.714150003 | 0.604 | 0.403 | 1 | 0.933488134 | 0.543291131 | 0.439 | 0.461 | 1 | 0.933488134 | 0.004525378 |
| Ltbp1 | 0.38434334 | 0.264347086 | 0.226 | 0.178 | 1 | 0.002271428 | 2.050378741 | 0.39 | 0.198 | 1 | 0.38434334 | 0.004537696 |
| Snrpc | 0.002272788 | 0.673595905 | 0.774 | 0.653 | 1 | 0.260317835 | 0.608673291 | 0.659 | 0.638 | 1 | 0.260317835 | 0.004540411 |
| Retreg1 | 0.002280138 | 0.501695819 | 0.943 | 0.877 | 1 | 0.480475223 | 0.216618439 | 0.829 | 0.881 | 1 | 0.480475223 | 0.004555077 |
| Trim41 | 0.002282345 | 1.000917259 | 0.472 | 0.28 | 1 | 0.846000998 | 0.361471419 | 0.39 | 0.41 | 1 | 0.846000998 | 0.004559481 |
| Ggct | 0.002282677 | -1.988235359 | 0.075 | 0.271 | 1 | 0.266251267 | -0.899760744 | 0.049 | 0.102 | 1 | 0.266251267 | 0.004560143 |
| Tubb4b | 0.205986183 | -1.068248641 | 0.868 | 0.797 | 1 | 0.002282828 | -0.924480236 | 0.878 | 0.945 | 1 | 0.205986183 | 0.004560445 |
| Ahctf1 | 0.002283104 | 0.764439143 | 0.83 | 0.729 | 1 | 0.644401309 | 0.327793382 | 0.829 | 0.898 | 1 | 0.644401309 | 0.004560996 |
| Tmem106c | 0.069604582 | -0.589438501 | 0.283 | 0.415 | 1 | 0.002283769 | -1.654648247 | 0.098 | 0.334 | 1 | 0.069604582 | 0.004562321 |
| Fkbp11 | 0.002284813 | 1.045098104 | 0.528 | 0.343 | 1 | 0.442009257 | 0.543021745 | 0.39 | 0.352 | 1 | 0.442009257 | 0.004564406 |
| Arl8a | 0.002287179 | 0.678802853 | 0.698 | 0.572 | 1 | 0.027239687 | -0.450077102 | 0.659 | 0.809 | 1 | 0.027239687 | 0.004569126 |
| Mrpl17 | 0.002289182 | -0.910568865 | 0.491 | 0.708 | 1 | 0.013432739 | -0.686357106 | 0.39 | 0.604 | 1 | 0.013432739 | 0.004573123 |
| Polb | 0.002291613 | 0.85038856 | 0.642 | 0.432 | 1 | 0.817548395 | 0.159983841 | 0.585 | 0.597 | 1 | 0.817548395 | 0.004577975 |
| Hsp90aa1 | 0.002297427 | 0.480251262 | 1 | 0.979 | 1 | 0.00605417 | 0.603516376 | 1 | 0.997 | 1 | 0.00605417 | 0.004589576 |
| Kdm7a | 0.002299254 | 0.669295768 | 0.755 | 0.576 | 1 | 0.0211307 | 0.783993044 | 0.829 | 0.775 | 1 | 0.0211307 | 0.004593221 |
| Slc4a4 | 0.017512007 | -0.153311058 | 0.736 | 0.542 | 1 | 0.002299357 | 0.786229709 | 0.829 | 0.645 | 1 | 0.017512007 | 0.004593427 |
| Camkk2 | 0.002302333 | 0.599203872 | 0.642 | 0.411 | 1 | 0.798609662 | 0.156195662 | 0.537 | 0.532 | 1 | 0.798609662 | 0.004599365 |
| Ptpn3 | 0.246536391 | 0.369547908 | 0.642 | 0.521 | 1 | 0.002304947 | -0.924144903 | 0.585 | 0.741 | 1 | 0.246536391 | 0.004604581 |
| Ncoa7 | 0.002306121 | 1.753360032 | 0.377 | 0.203 | 1 | 0.182773018 | 0.229522273 | 0.317 | 0.215 | 1 | 0.182773018 | 0.004606924 |
| Glmp | 0.002313234 | 0.612464545 | 0.717 | 0.576 | 1 | 0.562005458 | 0.298338764 | 0.61 | 0.604 | 1 | 0.562005458 | 0.004621117 |
| Gstz1 | 0.237116502 | -0.292736382 | 0.396 | 0.475 | 1 | 0.002317574 | -1.341598303 | 0.146 | 0.389 | 1 | 0.237116502 | 0.004629777 |
| Cacna2d1 | 0.002319004 | 0.498099109 | 0.811 | 0.585 | 1 | 0.870570801 | 0.818427643 | 0.634 | 0.737 | 1 | 0.870570801 | 0.00463263 |
| Arih1 | 0.566613112 | -0.312433628 | 0.981 | 0.992 | 1 | 0.002320884 | 0.455010149 | 1 | 0.997 | 1 | 0.566613112 | 0.004636381 |
| Bod1l | 0.002321091 | 0.657816271 | 0.83 | 0.742 | 1 | 0.36608946 | 0.109284395 | 0.829 | 0.765 | 1 | 0.36608946 | 0.004636794 |

| | | | | | | | | | | | | |
|---|---|---|---|---|---|---|---|---|---|---|---|---|
| Ccdc47 | 0.002325351 | 0.589125419 | 0.943 | 0.771 | 1 | 0.303663905 | 0.246293083 | 0.878 | 0.918 | 1 | 0.303663905 | 0.004645294 |
| Lgals9 | 0.002327848 | 1.569760094 | 0.226 | 0.085 | 1 | 0.011795077 | 1.438655473 | 0.195 | 0.075 | 1 | 0.011795077 | 0.004650278 |
| Dpf3 | 0.002332869 | -2.505528033 | 0.075 | 0.267 | 1 | 0.021167286 | 2.83720485 | 0.049 | 0.007 | 1 | 0.021167286 | 0.004660296 |
| Akap13 | 0.00565725 | 0.764999948 | 0.868 | 0.835 | 1 | 0.00233505 | 0.732067459 | 0.976 | 0.867 | 1 | 0.00565725 | 0.004664647 |
| Adra2a | 0.002346127 | 1.948271717 | 0.17 | 0.051 | 1 | 0.634479033 | 0.515276755 | 0.073 | 0.055 | 1 | 0.634479033 | 0.00468675 |
| Aamdc | 0.002347833 | 0.869320376 | 0.585 | 0.364 | 1 | 0.808613721 | 0.363273661 | 0.341 | 0.392 | 1 | 0.808613721 | 0.004690155 |
| Bend3 | 0.002348418 | 2.154722595 | 0.189 | 0.064 | 1 | 0.191081545 | 0.786578777 | 0.195 | 0.123 | 1 | 0.191081545 | 0.004691321 |
| Nup88 | 0.002351013 | 0.889378028 | 0.566 | 0.364 | 1 | 0.603037239 | 0.19334866 | 0.463 | 0.423 | 1 | 0.603037239 | 0.004696498 |
| Adh5 | 0.002352361 | 0.824332921 | 0.698 | 0.513 | 1 | 0.788803336 | 0.306690133 | 0.561 | 0.567 | 1 | 0.788803336 | 0.004699188 |
| Crnde | 0.002354212 | 1.932330173 | 0.208 | 0.072 | 1 | 0.462588913 | 0.114738825 | 0.049 | 0.085 | 1 | 0.462588913 | 0.004702883 |
| Exoc2 | 0.002362504 | 1.362164675 | 0.453 | 0.263 | 1 | 0.330311516 | 0.57131079 | 0.415 | 0.358 | 1 | 0.330311516 | 0.004719426 |
| Phb2 | 0.002370122 | 0.698084191 | 0.736 | 0.636 | 1 | 0.297722832 | -0.104692196 | 0.683 | 0.778 | 1 | 0.297722832 | 0.004734626 |
| Gadd45gip1 | 0.002376763 | 1.423539353 | 0.434 | 0.267 | 1 | 0.91470996 | 0.398093216 | 0.341 | 0.345 | 1 | 0.91470996 | 0.004747877 |
| Slu7 | 0.002385597 | 1.154722595 | 0.547 | 0.39 | 1 | 0.428401067 | 0.522508324 | 0.463 | 0.423 | 1 | 0.428401067 | 0.004765502 |
| Hfe | 0.002393307 | 1.685237311 | 0.208 | 0.072 | 1 | 0.225481555 | -0.555112573 | 0.098 | 0.174 | 1 | 0.225481555 | 0.004780887 |
| Eaf1 | 0.002396649 | 1.068992721 | 0.509 | 0.297 | 1 | 0.467218578 | 0.61149207 | 0.366 | 0.341 | 1 | 0.467218578 | 0.004787553 |
| Fbf1 | 0.413941615 | 0.234157062 | 0.208 | 0.157 | 1 | 0.002411433 | 1.126711467 | 0.341 | 0.154 | 1 | 0.413941615 | 0.004817051 |
| Esyt1 | 0.030672457 | 0.746257749 | 0.509 | 0.381 | 1 | 0.002419547 | 1.231922365 | 0.537 | 0.311 | 1 | 0.030672457 | 0.004833241 |
| 0610040J01Rik | 0.00460403 | -2.102665248 | 0.075 | 0.254 | 1 | 0.002420822 | -0.692048218 | 0.146 | 0.416 | 1 | 0.00460403 | 0.004835785 |
| Mpnd | 0.002421928 | 1.331600357 | 0.34 | 0.161 | 1 | 0.110910381 | -0.701953961 | 0.146 | 0.263 | 1 | 0.110910381 | 0.004837991 |
| Ddx52 | 0.002423242 | 1.020421503 | 0.472 | 0.305 | 1 | 0.341145766 | 0.482362132 | 0.366 | 0.307 | 1 | 0.341145766 | 0.004840612 |
| Riiad1 | 0.002424715 | 1.680791406 | 0.208 | 0.072 | 1 | 0.694862863 | 0.392420007 | 0.146 | 0.126 | 1 | 0.694862863 | 0.004843551 |
| Tpt1 | 0.04659947 | -0.141363742 | 1 | 1 | 1 | 0.002425769 | -0.356886856 | 1 | 1 | 1 | 0.04659947 | 0.004845654 |
| Gm10033 | 0.002432571 | 1.392882332 | 0.283 | 0.119 | 1 | 0.957430737 | 0.387897448 | 0.146 | 0.147 | 1 | 0.957430737 | 0.004859225 |
| Mest | 0.002434467 | 0.505311386 | 0.604 | 0.398 | 1 | 0.433211352 | 0.366413369 | 0.439 | 0.427 | 1 | 0.433211352 | 0.004863007 |
| a | 0.002436263 | 0.772576977 | 0.321 | 0.157 | 1 | 0.005176305 | 1.192121357 | 0.366 | 0.198 | 1 | 0.005176305 | 0.00486659 |
| Serp1 | 0.002442323 | 0.555475868 | 0.906 | 0.801 | 1 | 0.248188258 | -0.274044244 | 0.732 | 0.867 | 1 | 0.248188258 | 0.004878681 |
| Lmo1 | 0.015178556 | 2.891688189 | 0.057 | 0.008 | 1 | 0.002442976 | 1.91966701 | 0.171 | 0.048 | 1 | 0.015178556 | 0.004879983 |
| Gdf10 | 0.002444556 | -1.848279893 | 0.264 | 0.458 | 1 | 0.383008626 | 1.54004466 | 0.341 | 0.334 | 1 | 0.383008626 | 0.004883136 |
| Rpl23 | 0.113057053 | -0.142604315 | 1 | 0.992 | 1 | 0.002444663 | -0.354808581 | 1 | 1 | 1 | 0.113057053 | 0.00488335 |
| Coa8 | 0.002458746 | 0.908082627 | 0.566 | 0.411 | 1 | 0.725112679 | 0.211600365 | 0.463 | 0.454 | 1 | 0.725112679 | 0.004911446 |
| Snrnp200 | 0.002460189 | 1.216123139 | 0.528 | 0.373 | 1 | 0.097667062 | 0.437597391 | 0.634 | 0.546 | 1 | 0.097667062 | 0.004914325 |
| Aars | 0.057128353 | -0.336263758 | 0.34 | 0.504 | 1 | 0.002461589 | -0.919523999 | 0.293 | 0.601 | 1 | 0.057128353 | 0.004917118 |
| Ythdf2 | 0.002462127 | 0.624207878 | 0.774 | 0.682 | 1 | 0.028697378 | 0.642936029 | 0.805 | 0.771 | 1 | 0.028697378 | 0.004918192 |
| Lrp5 | 0.002470885 | 1.370451286 | 0.453 | 0.263 | 1 | 0.768367572 | 0.212713985 | 0.39 | 0.375 | 1 | 0.768367572 | 0.004935665 |
| Rfc3 | 0.002475195 | 1.631160639 | 0.208 | 0.072 | 1 | 0.06622299 | 1.218295017 | 0.195 | 0.102 | 1 | 0.06622299 | 0.004944264 |
| Nt5c3 | 0.765312783 | 0.128250383 | 0.491 | 0.479 | 1 | 0.002476179 | -0.93428462 | 0.439 | 0.659 | 1 | 0.765312783 | 0.004946227 |
| Lmbr1 | 0.002478782 | 1.695290976 | 0.302 | 0.14 | 1 | 0.877402155 | 0.237292008 | 0.171 | 0.167 | 1 | 0.877402155 | 0.00495142 |
| Ppm1h | 0.002479533 | 1.826668397 | 0.302 | 0.144 | 1 | 0.975056766 | 0.351778023 | 0.268 | 0.263 | 1 | 0.975056766 | 0.004952918 |
| Edrf1 | 0.002480592 | 0.975752454 | 0.509 | 0.305 | 1 | 0.88887248 | 0.342440158 | 0.366 | 0.423 | 1 | 0.88887248 | 0.00495503 |
| Leprot | 0.002480606 | 0.499370766 | 0.887 | 0.814 | 1 | 0.124227152 | 0.432509295 | 0.829 | 0.799 | 1 | 0.124227152 | 0.004955059 |
| Zfp280d | 0.002483076 | 0.476160727 | 0.868 | 0.767 | 1 | 0.083932075 | -0.126531606 | 0.854 | 0.932 | 1 | 0.083932075 | 0.004959986 |
| Cd28 | 0.002488627 | -4.430239906 | 0 | 0.153 | 1 | 0.328773791 | -1.555112573 | 0.024 | 0.061 | 1 | 0.328773791 | 0.00497106 |
| Zzef1 | 0.002489452 | 0.59882256 | 0.83 | 0.657 | 1 | 0.232415969 | 0.410390183 | 0.805 | 0.812 | 1 | 0.232415969 | 0.004972707 |
| Atp7a | 0.002497366 | 0.871992388 | 0.604 | 0.487 | 1 | 0.226127122 | 0.424423324 | 0.683 | 0.655 | 1 | 0.226127122 | 0.004988496 |
| Rnf115 | 0.002500117 | 1.053320563 | 0.642 | 0.504 | 1 | 0.090781957 | 0.406342383 | 0.78 | 0.765 | 1 | 0.090781957 | 0.004993983 |
| Acd | 0.002507783 | 1.031340179 | 0.434 | 0.233 | 1 | 0.465708936 | 0.53139642 | 0.293 | 0.253 | 1 | 0.465708936 | 0.005009277 |
| Acp6 | 0.002510428 | 1.404700848 | 0.302 | 0.136 | 1 | 0.27276571 | -0.410722664 | 0.122 | 0.198 | 1 | 0.27276571 | 0.005014553 |
| Slc26a11 | 0.002516545 | 1.243989933 | 0.34 | 0.157 | 1 | 0.771797519 | 0.142617858 | 0.293 | 0.276 | 1 | 0.771797519 | 0.005026757 |
| Mocs1 | 0.002521279 | 1.417757001 | 0.189 | 0.059 | 1 | 0.875461105 | 0.174239837 | 0.049 | 0.055 | 1 | 0.875461105 | 0.005036201 |
| Ube2o | 0.00252604 | 0.914408265 | 0.585 | 0.411 | 1 | 0.098641737 | 0.643896921 | 0.659 | 0.546 | 1 | 0.098641737 | 0.005045699 |
| Slc25a24 | 0.002528117 | 0.748730235 | 0.604 | 0.398 | 1 | 0.142606085 | -0.514470588 | 0.341 | 0.461 | 1 | 0.142606085 | 0.005049843 |
| Capn7 | 0.002530196 | 1.193196743 | 0.472 | 0.309 | 1 | 0.224860878 | 0.366575025 | 0.512 | 0.423 | 1 | 0.224860878 | 0.00505399 |
| Ero1a | 0.046770301 | 0.755315747 | 0.528 | 0.403 | 1 | 0.002536165 | 0.86891371 | 0.61 | 0.396 | 1 | 0.046770301 | 0.005065898 |
| Pin4 | 0.002539903 | 1.062800105 | 0.472 | 0.292 | 1 | 0.020711192 | -1.192542494 | 0.122 | 0.29 | 1 | 0.020711192 | 0.005073355 |
| Scai | 0.00255856 | 1.027343288 | 0.528 | 0.347 | 1 | 0.658098632 | -0.172966955 | 0.561 | 0.594 | 1 | 0.658098632 | 0.005110573 |
| Hdac4 | 0.002563039 | 0.936630311 | 0.717 | 0.619 | 1 | 0.019756035 | 0.565902828 | 0.902 | 0.744 | 1 | 0.019756035 | 0.005119509 |
| Art3 | 0.00256777 | -3.001781891 | 0.038 | 0.212 | 1 | 0.914200263 | 0.252242349 | 0.024 | 0.027 | 1 | 0.914200263 | 0.005128947 |
| Lrfn4 | 0.002572191 | 1.533234218 | 0.208 | 0.072 | 1 | 0.220851649 | 0.883008539 | 0.244 | 0.171 | 1 | 0.220851649 | 0.005137765 |
| Nomo1 | 0.021865118 | 0.724734754 | 0.472 | 0.331 | 1 | 0.002583001 | -1.137991759 | 0.146 | 0.406 | 1 | 0.021865118 | 0.005159329 |
| D030045P18Rik | 0.724777195 | 1.913714495 | 0.057 | 0.047 | 1 | 0.002594567 | -1.829551742 | 0.073 | 0.311 | 1 | 0.724777195 | 0.005182402 |
| Uqcc3 | 0.002600279 | 1.110328475 | 0.509 | 0.335 | 1 | 0.360380731 | 0.742047617 | 0.293 | 0.242 | 1 | 0.360380731 | 0.005193797 |
| Mgst3 | 0.178167434 | 0.8327945 | 0.151 | 0.089 | 1 | 0.002608761 | 0.770090654 | 0.415 | 0.198 | 1 | 0.178167434 | 0.005210716 |
| Capn15 | 0.002609197 | 0.69106639 | 0.623 | 0.424 | 1 | 0.220900357 | 0.361962716 | 0.683 | 0.608 | 1 | 0.220900357 | 0.005211585 |
| Ankrd27 | 0.002619652 | 1.154722595 | 0.34 | 0.161 | 1 | 0.562279315 | 0.887245532 | 0.244 | 0.222 | 1 | 0.562279315 | 0.005232442 |
| Plekhf1 | 0.002620173 | 1.331600357 | 0.434 | 0.263 | 1 | 0.925397645 | 0.590893802 | 0.39 | 0.437 | 1 | 0.925397645 | 0.00523348 |
| Schip1 | 0.580773903 | -0.544751787 | 0.679 | 0.644 | 1 | 0.002628431 | -0.994445659 | 0.756 | 0.863 | 1 | 0.580773903 | 0.005249954 |
| Jade3 | 0.002635359 | 1.695290976 | 0.17 | 0.051 | 1 | 0.578125793 | 0.622191959 | 0.171 | 0.143 | 1 | 0.578125793 | 0.005263772 |
| Lipg | 0.002637282 | -4.399866257 | 0.019 | 0.182 | 1 | 0.357847088 | -0.16279515 | 0 | 0.02 | 1 | 0.357847088 | 0.005267608 |
| Adamts3 | 0.002642474 | -0.477545621 | 0.491 | 0.708 | 1 | 0.580812501 | 0.268470811 | 0.634 | 0.686 | 1 | 0.580812501 | 0.005277965 |
| Polr3h | 0.002644764 | 0.904744342 | 0.509 | 0.305 | 1 | 0.293789847 | 0.463746454 | 0.341 | 0.266 | 1 | 0.293789847 | 0.005282533 |
| Arid1b | 0.002651884 | 0.375528765 | 0.943 | 0.911 | 1 | 0.00895743 | 0.284339224 | 1 | 0.969 | 1 | 0.00895743 | 0.005296735 |
| Pde4b | 0.00265388 | -1.26500355 | 0.642 | 0.763 | 1 | 0.045246971 | -0.643638486 | 0.634 | 0.799 | 1 | 0.045246971 | 0.005300717 |
| Anks3 | 0.002656915 | 1.191248471 | 0.434 | 0.258 | 1 | 0.543202449 | 0.560364644 | 0.39 | 0.403 | 1 | 0.543202449 | 0.005306772 |
| Brpf3 | 0.002670096 | 1.426024617 | 0.358 | 0.182 | 1 | 0.281817952 | 0.695849001 | 0.366 | 0.3 | 1 | 0.281817952 | 0.005333062 |
| Zfp944 | 0.002670973 | 1.517292674 | 0.321 | 0.153 | 1 | 0.550956552 | -0.548853583 | 0.195 | 0.225 | 1 | 0.550956552 | 0.005334812 |
| Renbp | 0.002673403 | 1.391761792 | 0.358 | 0.195 | 1 | 0.757267006 | 0.309957847 | 0.244 | 0.225 | 1 | 0.757267006 | 0.005339658 |
| Ddx47 | 0.002701462 | 1.049753035 | 0.528 | 0.373 | 1 | 0.460545482 | -0.219826095 | 0.463 | 0.509 | 1 | 0.460545482 | 0.005395627 |
| Fhit | 0.332965482 | -0.475327795 | 0.321 | 0.403 | 1 | 0.00270376 | -1.207983924 | 0.317 | 0.549 | 1 | 0.332965482 | 0.005400209 |
| Def8 | 0.002706658 | 1.466666601 | 0.264 | 0.11 | 1 | 0.250949458 | -0.727579769 | 0.098 | 0.167 | 1 | 0.250949458 | 0.005405991 |
| Rhot1 | 0.002709296 | 0.859979329 | 0.623 | 0.453 | 1 | 0.549254117 | 0.120483761 | 0.732 | 0.659 | 1 | 0.549254117 | 0.005411251 |
| Abce1 | 0.045442372 | 0.40402289 | 0.792 | 0.729 | 1 | 0.002709985 | -0.619963717 | 0.61 | 0.823 | 1 | 0.045442372 | 0.005412625 |
| Acat1 | 0.002714559 | 0.538733198 | 0.83 | 0.729 | 1 | 0.924615898 | 0.144351626 | 0.732 | 0.795 | 1 | 0.924615898 | 0.005421749 |
| Tspan3 | 0.002725905 | -0.453251674 | 1 | 0.979 | 1 | 0.548450241 | -0.116157195 | 0.951 | 0.98 | 1 | 0.548450241 | 0.00544438 |
| Cblb | 0.002729315 | 1.019563012 | 0.736 | 0.61 | 1 | 0.607103466 | 0.265605491 | 0.756 | 0.754 | 1 | 0.607103466 | 0.005451181 |
| Ppp2r2a | 0.002735427 | 0.495099089 | 0.868 | 0.873 | 1 | 0.697997667 | -0.130780343 | 0.951 | 0.928 | 1 | 0.697997667 | 0.005463372 |
| Ro60 | 0.002739327 | 1.268379377 | 0.509 | 0.347 | 1 | 0.061084699 | 0.802934948 | 0.537 | 0.437 | 1 | 0.061084699 | 0.005471151 |
| Zfp451 | 0.002744573 | 0.556584705 | 0.717 | 0.559 | 1 | 0.420433938 | -0.267492529 | 0.805 | 0.829 | 1 | 0.420433938 | 0.005481613 |
| Mtcl1 | 0.002747086 | 1.680791406 | 0.189 | 0.064 | 1 | 0.14486162 | 0.930314254 | 0.22 | 0.137 | 1 | 0.14486162 | 0.005486625 |
| Tnrc18 | 0.002749312 | 0.296323609 | 0.962 | 0.894 | 1 | 0.314677525 | -0.171402571 | 0.976 | 0.962 | 1 | 0.314677525 | 0.005491065 |
| Frmd4a | 0.056683969 | -1.411624228 | 0.321 | 0.445 | 1 | 0.002749696 | 0.619974134 | 0.415 | 0.198 | 1 | 0.056683969 | 0.00549183 |
| Ptpn12 | 0.002749823 | 0.633062676 | 0.811 | 0.644 | 1 | 0.220626732 | 0.28239196 | 0.805 | 0.785 | 1 | 0.220626732 | 0.005492085 |
| Krt8 | 0.468340794 | -0.243826782 | 0.189 | 0.144 | 1 | 0.002752142 | 1.715214325 | 0.39 | 0.208 | 1 | 0.468340794 | 0.00549671 |
| Fut4 | 0.002757657 | -3.015202407 | 0.019 | 0.182 | 1 | 0.286433344 | -0.332720152 | 0 | 0.027 | 1 | 0.286433344 | 0.00550771 |
| Gm14325 | 0.002765588 | 1.154722595 | 0.396 | 0.212 | 1 | 0.782188257 | 0.252242349 | 0.244 | 0.229 | 1 | 0.782188257 | 0.005523527 |
| Morf4l2 | 0.00277883 | 0.339501828 | 1 | 0.987 | 1 | 0.075365046 | -0.188330242 | 1 | 0.993 | 1 | 0.075365046 | 0.005549938 |
| Lin54 | 0.002779957 | 1.131638982 | 0.34 | 0.161 | 1 | 0.113970039 | 0.9059176 | 0.366 | 0.287 | 1 | 0.113970039 | 0.005552186 |
| Sqstm1 | 0.002780546 | 0.672005691 | 1 | 0.953 | 1 | 0.152152601 | -0.263505657 | 0.976 | 0.983 | 1 | 0.152152601 | 0.005553361 |

| | | | | | | | | | | | | |
|---|---|---|---|---|---|---|---|---|---|---|---|---|
| Cep85l | 0.002790363 | -1.927639375 | 0.434 | 0.614 | 1 | 0.311944099 | -0.388215264 | 0.366 | 0.478 | 1 | 0.311944099 | 0.005572939 |
| Btbd19 | 0.007950402 | 2.569760094 | 0.094 | 0.021 | 1 | 0.002798243 | 2.83720485 | 0.098 | 0.017 | 1 | 0.007950402 | 0.005588656 |
| Rpl10 | 0.002800827 | 0.348999989 | 1 | 0.987 | 1 | 0.034423466 | -0.261285014 | 1 | 0.993 | 1 | 0.034423466 | 0.00559381 |
| Cfap74 | 0.002807704 | 1.389187848 | 0.302 | 0.144 | 1 | 0.177082512 | -0.395455907 | 0.098 | 0.188 | 1 | 0.177082512 | 0.005607524 |
| Zfand3 | 0.002811199 | 0.457227426 | 1 | 0.979 | 1 | 0.324367776 | 0.137260648 | 1 | 1 | 1 | 0.324367776 | 0.005614495 |
| Togaram1 | 0.277636371 | 0.490239754 | 0.623 | 0.597 | 1 | 0.002815157 | 0.733369039 | 0.878 | 0.758 | 1 | 0.277636371 | 0.005622389 |
| Mex3c | 0.002818044 | 0.645708947 | 0.679 | 0.504 | 1 | 0.708572895 | 0.249543199 | 0.78 | 0.765 | 1 | 0.708572895 | 0.005628147 |
| Ptpn14 | 0.002819801 | 0.618669695 | 0.83 | 0.716 | 1 | 0.833646687 | 0.115607657 | 0.878 | 0.884 | 1 | 0.833646687 | 0.005631651 |
| Pcdh7 | 0.002819939 | 0.710420501 | 0.453 | 0.237 | 1 | 0.16131614 | 0.568481372 | 0.537 | 0.433 | 1 | 0.16131614 | 0.005631926 |
| Entr1 | 0.002821837 | 1.002719501 | 0.547 | 0.386 | 1 | 0.709836048 | 0.285409213 | 0.537 | 0.526 | 1 | 0.709836048 | 0.005635712 |
| Dr1 | 0.002825965 | 1.034428361 | 0.491 | 0.292 | 1 | 0.906144801 | 0.108276575 | 0.39 | 0.41 | 1 | 0.906144801 | 0.005643944 |
| Cnpy4 | 0.002827664 | 1.180257687 | 0.528 | 0.373 | 1 | 0.973831497 | 0.125330237 | 0.415 | 0.447 | 1 | 0.973831497 | 0.005647332 |
| Zbed3 | 0.00283929 | 1.181689642 | 0.358 | 0.182 | 1 | 0.368908918 | -0.651081632 | 0.195 | 0.246 | 1 | 0.368908918 | 0.005670517 |
| 4930558J18Rik | 0.00285365 | 3.739685096 | 0.038 | 0 | 1 | 0.103479965 | 3.83720485 | 0.024 | 0.003 | 1 | 0.103479965 | 0.005699157 |
| Adam23 | 0.00285365 | 4.47665069 | 0.038 | 0 | 1 | 0.007718533 | 4.83720485 | 0.024 | 0 | 1 | 0.007718533 | 0.005699157 |
| Nuak2 | 0.00285365 | 4.154722595 | 0.038 | 0 | 1 | 0.004444719 | 3.159132945 | 0.073 | 0.01 | 1 | 0.004444719 | 0.005699157 |
| Cel | 0.00285365 | 3.739685096 | 0.038 | 0 | 1 | 0.267205177 | 2.252242349 | 0.024 | 0.007 | 1 | 0.267205177 | 0.005699157 |
| Gm56772 | 0.00285365 | 5.154722595 | 0.038 | 0 | 1 | 0.02777124 | 2.422167351 | 0.073 | 0.017 | 1 | 0.02777124 | 0.005699157 |
| Plppr4 | 0.00285365 | 4.47665069 | 0.038 | 0 | 1 | 0.455098275 | -0.622226769 | 0 | 0.014 | 1 | 0.455098275 | 0.005699157 |
| Gm42303 | 0.00285365 | 7.284005612 | 0.038 | 0 | 1 | 0.430587622 | 2.252242349 | 0.024 | 0.01 | 1 | 0.430587622 | 0.005699157 |
| Tex46 | 0.00285365 | 3.739685096 | 0.038 | 0 | 1 | 0.104856732 | 2.83720485 | 0.024 | 0.003 | 1 | 0.104856732 | 0.005699157 |
| Gm57064 | 0.00285365 | 4.962077517 | 0.038 | 0 | 1 | 0.319629309 | -0.622226769 | 0 | 0.024 | 1 | 0.319629309 | 0.005699157 |
| 8030423F21Rik | 0.00285365 | 3.739685096 | 0.038 | 0 | 1 | 0.267205177 | 2.252242349 | 0.024 | 0.007 | 1 | 0.267205177 | 0.005699157 |
| Stap1 | 0.00285365 | 3.739685096 | 0.038 | 0 | 1 | 0.976594067 | 0.962735732 | 0.024 | 0.024 | 1 | 0.976594067 | 0.005699157 |
| Gm14508 | 0.00285365 | 4.154722595 | 0.038 | 0 | 1 | 0.438672228 | 1.83720485 | 0.024 | 0.01 | 1 | 0.438672228 | 0.005699157 |
| Gpr37 | 0.00285365 | 3.739685096 | 0.038 | 0 | 1 | 0.746881137 | 0.83720485 | 0.024 | 0.017 | 1 | 0.746881137 | 0.005699157 |
| Grid2 | 0.00285365 | 3.739685096 | 0.038 | 0 | 1 | 0.5191986 | -0.622226769 | 0 | 0.01 | 1 | 0.5191986 | 0.005699157 |
| Gm38832 | 0.00285365 | 3.739685096 | 0.038 | 0 | 1 | 0.007718533 | 4.83720485 | 0.024 | 0 | 1 | 0.007718533 | 0.005699157 |
| Mdfic2 | 0.00285365 | 4.47665069 | 0.038 | 0 | 1 | 0.286413984 | -0.970150072 | 0 | 0.027 | 1 | 0.286413984 | 0.005699157 |
| 4930540M05Rik | 0.00285365 | 3.739685096 | 0.038 | 0 | 1 | 0.056289003 | 2.83720485 | 0.049 | 0.01 | 1 | 0.056289003 | 0.005699157 |
| Rimklb | 0.00285365 | 3.739685096 | 0.038 | 0 | 1 | 0.007718533 | 3.83720485 | 0.024 | 0 | 1 | 0.007718533 | 0.005699157 |
| Gm10010 | 0.00285365 | 4.47665069 | 0.038 | 0 | 1 | 0.26948834 | 1.83720485 | 0.024 | 0.007 | 1 | 0.26948834 | 0.005699157 |
| Ppfia3 | 0.00285365 | 3.739685096 | 0.038 | 0 | 1 | 0.057901606 | 2.422167351 | 0.049 | 0.01 | 1 | 0.057901606 | 0.005699157 |
| Pgpep1l | 0.00285365 | 3.739685096 | 0.038 | 0 | 1 | 0.286402368 | -0.747757651 | 0 | 0.027 | 1 | 0.286402368 | 0.005699157 |
| Lpar2 | 0.00285365 | 4.47665069 | 0.038 | 0 | 1 | 0.609924905 | 0.136765132 | 0.024 | 0.041 | 1 | 0.609924905 | 0.005699157 |
| Lrrc25 | 0.00285365 | 3.739685096 | 0.038 | 0 | 1 | 0.402364804 | -0.332720152 | 0 | 0.017 | 1 | 0.402364804 | 0.005699157 |
| Gm47326 | 0.00285365 | 3.739685096 | 0.038 | 0 | 1 | 0.455098275 | 0.515276755 | 0 | 0.014 | 1 | 0.455098275 | 0.005699157 |
| Gm16754 | 0.00285365 | 3.739685096 | 0.038 | 0 | 1 | 0.455097252 | -0.332720152 | 0 | 0.014 | 1 | 0.455097252 | 0.005699157 |
| Tnfaip8l3 | 0.00285365 | 4.154722595 | 0.038 | 0 | 1 | 0.103479965 | 3.83720485 | 0.024 | 0.003 | 1 | 0.103479965 | 0.005699157 |
| Nr2e3 | 0.00285365 | 4.154722595 | 0.038 | 0 | 1 | 0.088723073 | 1.136765132 | 0.073 | 0.024 | 1 | 0.088723073 | 0.005699157 |
| Gm47818 | 0.00285365 | 3.739685096 | 0.038 | 0 | 1 | 0.007718533 | 3.83720485 | 0.024 | 0 | 1 | 0.007718533 | 0.005699157 |
| Gm26535 | 0.00285365 | 3.739685096 | 0.038 | 0 | 1 | 0.746881137 | 0.83720485 | 0.024 | 0.017 | 1 | 0.746881137 | 0.005699157 |
| Mfsd4b1 | 0.00285365 | 4.962077517 | 0.038 | 0 | 1 | 0.994797912 | 0.377773231 | 0.024 | 0.024 | 1 | 0.994797912 | 0.005699157 |
| B4galnt1 | 0.00285365 | 3.739685096 | 0.038 | 0 | 1 | 0.208531619 | 2.185128153 | 0.073 | 0.034 | 1 | 0.208531619 | 0.005699157 |
| Egfros | 0.00285365 | 3.739685096 | 0.038 | 0 | 1 | 0.909322542 | -0.16279515 | 0.024 | 0.027 | 1 | 0.909322542 | 0.005699157 |
| 6720407P12Rik | 0.00285365 | 4.154722595 | 0.038 | 0 | 1 | 0.402373368 | 0.252242349 | 0 | 0.017 | 1 | 0.402373368 | 0.005699157 |
| Prcd | 0.00285365 | 3.739685096 | 0.038 | 0 | 1 | 0.103479965 | 4.422167351 | 0.024 | 0.003 | 1 | 0.103479965 | 0.005699157 |
| Jph4 | 0.00285365 | 3.739685096 | 0.038 | 0 | 1 | 0.007718533 | 3.83720485 | 0.024 | 0 | 1 | 0.007718533 | 0.005699157 |
| Rspo2 | 0.00285365 | 4.962077517 | 0.038 | 0 | 1 | 0.007718533 | 3.83720485 | 0.024 | 0 | 1 | 0.007718533 | 0.005699157 |
| Card10 | 0.00285365 | 3.739685096 | 0.038 | 0 | 1 | 0.992196803 | 0.515276755 | 0.024 | 0.024 | 1 | 0.992196803 | 0.005699157 |
| Nfam1 | 0.00285365 | 3.739685096 | 0.038 | 0 | 1 | 0.402369942 | -0.16279515 | 0 | 0.017 | 1 | 0.402369942 | 0.005699157 |
| Arhgap8 | 0.00285365 | 4.739685096 | 0.038 | 0 | 1 | 0.360606599 | 1.029849928 | 0.049 | 0.024 | 1 | 0.360606599 | 0.005699157 |
| Gm15722 | 0.00285365 | 4.154722595 | 0.038 | 0 | 1 | 0.104856732 | 2.83720485 | 0.024 | 0.003 | 1 | 0.104856732 | 0.005699157 |
| Fignl2 | 0.00285365 | 3.739685096 | 0.038 | 0 | 1 | 0.483647624 | -0.825760163 | 0.049 | 0.078 | 1 | 0.483647624 | 0.005699157 |
| Tm4sf19 | 0.00285365 | 4.154722595 | 0.038 | 0 | 1 | 0.856409217 | 2.47463477 | 0.024 | 0.02 | 1 | 0.856409217 | 0.005699157 |
| Dact2 | 0.00285365 | 4.47665069 | 0.038 | 0 | 1 | 0.104856732 | 2.83720485 | 0.024 | 0.003 | 1 | 0.104856732 | 0.005699157 |
| Zcchc12 | 0.00285365 | 3.739685096 | 0.038 | 0 | 1 | 0.267205177 | 2.252242349 | 0.024 | 0.007 | 1 | 0.267205177 | 0.005699157 |
| A630012P03Rik | 0.00285365 | 6.678284551 | 0.038 | 0 | 1 | 0.5191986 | 0.515276755 | 0 | 0.01 | 1 | 0.5191986 | 0.005699157 |
| Tcf7 | 0.002858271 | 1.844382474 | 0.245 | 0.102 | 1 | 0.726565108 | 0.33763384 | 0.293 | 0.273 | 1 | 0.726565108 | 0.005708372 |
| Lin9 | 0.002858789 | 2.039245377 | 0.208 | 0.076 | 1 | 0.706665397 | 0.287007767 | 0.098 | 0.078 | 1 | 0.706665397 | 0.005709405 |
| Osbpl11 | 0.002860913 | 0.752090116 | 0.623 | 0.428 | 1 | 0.334859374 | 0.633845621 | 0.634 | 0.689 | 1 | 0.334859374 | 0.005713641 |
| Gm56862 | 0.00286622 | 2.261637799 | 0.151 | 0.042 | 1 | 0.015454492 | 1.334704509 | 0.22 | 0.092 | 1 | 0.015454492 | 0.005724225 |
| Sox6 | 0.83774729 | -0.125706745 | 0.981 | 0.996 | 1 | 0.002866857 | -0.447232033 | 0.854 | 0.986 | 1 | 0.83774729 | 0.005725496 |
| Tdp2 | 0.003491738 | 1.100274811 | 0.434 | 0.25 | 1 | 0.002885841 | 1.407696868 | 0.537 | 0.328 | 1 | 0.003491738 | 0.005763354 |
| Rngtt | 0.002887311 | 0.959843666 | 0.66 | 0.568 | 1 | 0.007084416 | 0.693800725 | 0.78 | 0.648 | 1 | 0.007084416 | 0.005766285 |
| Fgf13 | 0.003505053 | 2.417757001 | 0.094 | 0.017 | 1 | 0.002892829 | 2.209173627 | 0.146 | 0.038 | 1 | 0.003505053 | 0.005777289 |
| Tbcd | 0.002900162 | 1.918655237 | 0.34 | 0.182 | 1 | 0.394300543 | 0.225770138 | 0.293 | 0.225 | 1 | 0.394300543 | 0.005791913 |
| Dnaaf10 | 0.002901741 | 1.727301371 | 0.283 | 0.127 | 1 | 0.226891521 | 0.873730726 | 0.244 | 0.174 | 1 | 0.226891521 | 0.005795062 |
| Ahsa1 | 0.002902131 | 0.859266711 | 0.642 | 0.475 | 1 | 0.65878874 | -0.244944192 | 0.537 | 0.553 | 1 | 0.65878874 | 0.005795839 |
| Tnn | 0.002910929 | 2.822147256 | 0.151 | 0.042 | 1 | 0.369056111 | -0.80665134 | 0.024 | 0.058 | 1 | 0.369056111 | 0.005813384 |
| Pdia6 | 0.002918114 | -0.42121094 | 0.981 | 0.975 | 1 | 0.057925399 | -0.305753104 | 0.902 | 0.966 | 1 | 0.057925399 | 0.005827714 |
| 5730455P16Rik | 0.002923186 | 1.135863568 | 0.415 | 0.233 | 1 | 0.749992671 | -0.174767792 | 0.268 | 0.287 | 1 | 0.749992671 | 0.005837826 |
| Cpsf7 | 0.002923626 | 0.803500193 | 0.679 | 0.547 | 1 | 0.167660237 | 0.452541 | 0.78 | 0.734 | 1 | 0.167660237 | 0.005838704 |
| Tmcc2 | 0.002924044 | 2.270199812 | 0.151 | 0.042 | 1 | 0.469338706 | 0.80158094 | 0.073 | 0.116 | 1 | 0.469338706 | 0.005839538 |
| Polr2b | 0.002928316 | 0.815412422 | 0.604 | 0.394 | 1 | 0.42739485 | 0.351778023 | 0.561 | 0.556 | 1 | 0.42739485 | 0.005848057 |
| Wnt11 | 0.00293162 | -1.716762665 | 0.151 | 0.347 | 1 | 0.074890632 | -0.96369505 | 0.195 | 0.334 | 1 | 0.074890632 | 0.005854645 |
| Tbcel | 0.002933073 | 1.182203331 | 0.491 | 0.314 | 1 | 0.893808145 | 0.150362735 | 0.488 | 0.512 | 1 | 0.893808145 | 0.005857543 |
| Skap2 | 0.002935174 | -1.778640212 | 0.208 | 0.411 | 1 | 0.163647602 | -0.801395615 | 0.146 | 0.239 | 1 | 0.163647602 | 0.005861732 |
| Plgrkt | 0.00293611 | 0.932330173 | 0.491 | 0.292 | 1 | 0.361015511 | 0.489281546 | 0.366 | 0.3 | 1 | 0.361015511 | 0.0058636 |
| Bltp2 | 0.03083682 | 0.648369929 | 0.679 | 0.589 | 1 | 0.002937793 | 0.725094483 | 0.902 | 0.706 | 1 | 0.03083682 | 0.005866956 |
| Ell | 0.002939324 | -0.674547104 | 0.509 | 0.75 | 1 | 0.24925037 | -0.310693845 | 0.659 | 0.751 | 1 | 0.24925037 | 0.005870009 |
| Mark1 | 0.002939707 | -1.590704578 | 0.189 | 0.398 | 1 | 0.358020071 | 0.198944123 | 0.244 | 0.324 | 1 | 0.358020071 | 0.005870772 |
| Rpl39 | 0.322077345 | -0.115748993 | 1 | 0.966 | 1 | 0.002940751 | -0.433016097 | 1 | 0.986 | 1 | 0.322077345 | 0.005872855 |
| Erlin2 | 0.027168372 | 0.900966003 | 0.358 | 0.22 | 1 | 0.002953638 | -1.686357106 | 0.073 | 0.297 | 1 | 0.027168372 | 0.005898551 |
| Dxo | 0.002954394 | 1.382133091 | 0.321 | 0.153 | 1 | 0.332949689 | -0.508569987 | 0.122 | 0.181 | 1 | 0.332949689 | 0.00590006 |
| Dhfr | 0.002964561 | 1.444229212 | 0.226 | 0.085 | 1 | 0.982921617 | 0.974708374 | 0.098 | 0.102 | 1 | 0.982921617 | 0.005920334 |
| Rfx3 | 0.002965244 | 0.643984679 | 0.642 | 0.453 | 1 | 0.018668069 | 0.812258493 | 0.805 | 0.71 | 1 | 0.018668069 | 0.005921695 |
| Trak2 | 0.002965971 | 0.79717059 | 0.698 | 0.479 | 1 | 0.225212526 | 0.362861522 | 0.756 | 0.676 | 1 | 0.225212526 | 0.005923145 |
| Rpl28 | 0.002967027 | 0.325919253 | 1 | 0.966 | 1 | 0.014268508 | -0.262820454 | 1 | 0.99 | 1 | 0.014268508 | 0.005925251 |
| Calhm5 | 0.005837387 | 2.154722595 | 0.17 | 0.059 | 1 | 0.002968085 | 3.306690133 | 0.122 | 0.027 | 1 | 0.005837387 | 0.005927361 |
| Npas2 | 0.57768449 | 0.772068269 | 0.377 | 0.364 | 1 | 0.00297887 | 1.497354847 | 0.488 | 0.311 | 1 | 0.57768449 | 0.005948866 |
| Mrps35 | 0.002980553 | 1.285967128 | 0.321 | 0.153 | 1 | 0.318430678 | -0.592782991 | 0.146 | 0.212 | 1 | 0.318430678 | 0.005952221 |
| Vps16 | 0.002981066 | 1.182736971 | 0.358 | 0.178 | 1 | 0.807234388 | 0.174239837 | 0.268 | 0.259 | 1 | 0.807234388 | 0.005953245 |
| Zfp974 | 0.002981556 | 2.502645898 | 0.151 | 0.042 | 1 | 0.798360541 | 0.574170444 | 0.122 | 0.113 | 1 | 0.798360541 | 0.005954223 |
| Cat | 0.00298243 | 0.942672118 | 0.566 | 0.386 | 1 | 0.900993258 | 0.224550615 | 0.488 | 0.498 | 1 | 0.900993258 | 0.005955964 |
| Nectin3 | 0.002984821 | 1.30843201 | 0.528 | 0.39 | 1 | 0.453235209 | 0.402802026 | 0.537 | 0.543 | 1 | 0.453235209 | 0.005960733 |
| Egln3 | 0.521955064 | -0.582242999 | 0.038 | 0.059 | 1 | 0.002989727 | 2.83720485 | 0.122 | 0.027 | 1 | 0.521955064 | 0.005970516 |
| Gm3331 | 0.617585569 | -0.1672055 | 0.038 | 0.055 | 1 | 0.002989727 | 2.83720485 | 0.122 | 0.027 | 1 | 0.617585569 | 0.005970516 |

| | | | | | | | | | | | | |
|---|---|---|---|---|---|---|---|---|---|---|---|---|
| Nmt2 | 0.002990373 | 0.939333561 | 0.491 | 0.292 | 1 | 0.007170162 | 0.888086019 | 0.683 | 0.563 | 1 | 0.007170162 | 0.005971804 |
| Dtd1 | 0.002999145 | 1.024325958 | 0.396 | 0.203 | 1 | 0.931750302 | 0.100239256 | 0.341 | 0.352 | 1 | 0.931750302 | 0.005989296 |
| Skp1 | 0.002999965 | 0.304040241 | 0.981 | 0.941 | 1 | 0.053006304 | 0.272514744 | 0.976 | 0.966 | 1 | 0.053006304 | 0.00599093 |
| Taok2 | 0.003000716 | 1.123013735 | 0.472 | 0.292 | 1 | 0.310598649 | -0.355440228 | 0.415 | 0.481 | 1 | 0.310598649 | 0.005992427 |
| BC005537 | 0.003000796 | 0.42380374 | 0.981 | 0.924 | 1 | 0.377039514 | 0.212713985 | 1 | 0.99 | 1 | 0.377039514 | 0.005992588 |
| Mettl14 | 0.003002624 | 1.454282877 | 0.245 | 0.097 | 1 | 0.966485478 | 0.711673968 | 0.122 | 0.126 | 1 | 0.966485478 | 0.005996233 |
| Yod1 | 0.261473089 | -0.797022237 | 0.226 | 0.301 | 1 | 0.003006566 | -1.79700117 | 0.122 | 0.365 | 1 | 0.261473089 | 0.006004093 |
| Sucla2 | 0.003012122 | 0.766992442 | 0.642 | 0.441 | 1 | 0.246204102 | 0.388931021 | 0.61 | 0.608 | 1 | 0.246204102 | 0.006015172 |
| Bmpr1a | 0.003015637 | 0.460014688 | 0.943 | 0.775 | 1 | 0.225920094 | 0.135141998 | 0.951 | 0.935 | 1 | 0.225920094 | 0.00602218 |
| Ebna1bp2 | 0.00302464 | 0.722282049 | 0.774 | 0.648 | 1 | 0.027495287 | -0.501597064 | 0.439 | 0.631 | 1 | 0.027495287 | 0.006040131 |
| Arpc1a | 0.042517875 | -0.38040875 | 0.774 | 0.864 | 1 | 0.003032607 | -0.578609754 | 0.659 | 0.84 | 1 | 0.042517875 | 0.006056017 |
| Zfyve28 | 0.003339709 | 3.154722595 | 0.057 | 0.004 | 1 | 0.003033738 | 2.422167351 | 0.098 | 0.017 | 1 | 0.003339709 | 0.006058272 |
| Ckb | 0.003039787 | -1.246039896 | 0.604 | 0.742 | 1 | 0.873759076 | -1.313157885 | 0.537 | 0.546 | 1 | 0.873759076 | 0.006070333 |
| Cpeb3 | 0.003051126 | 1.04878915 | 0.528 | 0.352 | 1 | 0.010149358 | 0.83720485 | 0.585 | 0.396 | 1 | 0.010149358 | 0.006092942 |
| Tle3 | 0.003053122 | 1.182736971 | 0.321 | 0.148 | 1 | 0.325170338 | 0.640807637 | 0.439 | 0.358 | 1 | 0.325170338 | 0.006096923 |
| Atp6v1g1 | 0.003055176 | 0.557418744 | 0.962 | 0.894 | 1 | 0.68531297 | -0.159859859 | 0.902 | 0.894 | 1 | 0.68531297 | 0.006101019 |
| Zwilch | 0.575545018 | -0.304709024 | 0.057 | 0.081 | 1 | 0.003062748 | 2.159132945 | 0.098 | 0.017 | 1 | 0.575545018 | 0.006116116 |
| Asap2 | 0.003064766 | -0.815893598 | 0.585 | 0.754 | 1 | 0.105031837 | -0.877763642 | 0.415 | 0.515 | 1 | 0.105031837 | 0.006120139 |
| Dhrs11 | 0.003065084 | 1.925240749 | 0.264 | 0.119 | 1 | 0.580773709 | 0.922934724 | 0.146 | 0.119 | 1 | 0.580773709 | 0.006120773 |
| Miip | 0.003073179 | 2.569760094 | 0.132 | 0.034 | 1 | 0.455098275 | 0.515276755 | 0 | 0.014 | 1 | 0.455098275 | 0.006136914 |
| Cct8 | 0.003079228 | 0.515958451 | 0.943 | 0.864 | 1 | 0.003637215 | -0.478028593 | 0.829 | 0.952 | 1 | 0.003637215 | 0.006148975 |
| Tulp2 | 0.003087095 | 3.154722595 | 0.094 | 0.017 | 1 | 0.646976884 | 0.721727632 | 0.049 | 0.034 | 1 | 0.646976884 | 0.006164661 |
| Shank2 | 0.003087136 | 3.002719501 | 0.094 | 0.017 | 1 | 0.352577933 | 1.83720485 | 0.049 | 0.024 | 1 | 0.352577933 | 0.006164742 |
| Vdac2 | 0.003087416 | 0.521261576 | 0.906 | 0.843 | 1 | 0.579184995 | 0.100802918 | 0.927 | 0.935 | 1 | 0.579184995 | 0.0061653 |
| Fgd3 | 0.003088113 | 3.595295186 | 0.094 | 0.017 | 1 | 0.936211935 | 1.83720485 | 0.024 | 0.027 | 1 | 0.936211935 | 0.006166689 |
| Aida | 0.00311143 | 0.753624286 | 0.66 | 0.508 | 1 | 0.117710394 | -0.317730977 | 0.463 | 0.618 | 1 | 0.117710394 | 0.00621318 |
| Naa20 | 0.003115528 | 0.646063461 | 0.736 | 0.585 | 1 | 0.191274399 | 0.272181196 | 0.756 | 0.696 | 1 | 0.191274399 | 0.00622135 |
| Rara | 0.003117835 | 1.470224421 | 0.302 | 0.14 | 1 | 0.003170127 | 1.103850509 | 0.512 | 0.311 | 1 | 0.003170127 | 0.00622595 |
| Pls3 | 0.003131503 | 0.245234888 | 0.755 | 0.555 | 1 | 0.423151733 | 0.411720563 | 0.683 | 0.683 | 1 | 0.423151733 | 0.0062532 |
| Sorbs1 | 0.01911194 | -0.267061737 | 0.792 | 0.907 | 1 | 0.003135241 | -0.861943866 | 0.78 | 0.918 | 1 | 0.01911194 | 0.006260653 |
| Rce1 | 0.003140976 | 1.039245377 | 0.528 | 0.326 | 1 | 0.151203848 | -0.535349318 | 0.268 | 0.382 | 1 | 0.151203848 | 0.006272086 |
| Dis3l2 | 0.003141939 | 0.892752516 | 0.642 | 0.5 | 1 | 0.640956736 | 0.225770138 | 0.585 | 0.584 | 1 | 0.640956736 | 0.006274006 |
| Synpo | 0.003149911 | -0.667899084 | 0.623 | 0.809 | 1 | 0.745386296 | 0.106021608 | 0.634 | 0.73 | 1 | 0.745386296 | 0.006289901 |
| Rrp1 | 0.003150234 | 0.560714954 | 0.792 | 0.678 | 1 | 0.035354071 | -0.453265814 | 0.634 | 0.761 | 1 | 0.035354071 | 0.006290544 |
| Unc13c | 0.003151185 | 5.678284551 | 0.057 | 0.004 | 1 | 0.5191986 | 0.252242349 | 0 | 0.01 | 1 | 0.5191986 | 0.00629244 |
| C9 | 0.003151277 | 4.739685096 | 0.057 | 0.004 | 1 | 0.183764859 | 2.574170444 | 0.049 | 0.017 | 1 | 0.183764859 | 0.006292623 |
| Gm39529 | 0.003151277 | 3.962077517 | 0.057 | 0.004 | 1 | 0.846931821 | 1.377773231 | 0.024 | 0.031 | 1 | 0.846931821 | 0.006292623 |
| Aff3 | 0.003156428 | -0.946028321 | 0.698 | 0.86 | 1 | 0.010441852 | -0.691366469 | 0.732 | 0.846 | 1 | 0.010441852 | 0.006302894 |
| Tlcd1 | 0.003163028 | -0.971809811 | 0.17 | 0.39 | 1 | 0.922696474 | -0.20718927 | 0.244 | 0.242 | 1 | 0.922696474 | 0.006316052 |
| Ppp1cc | 0.003166888 | 0.626343622 | 0.868 | 0.771 | 1 | 0.338588475 | 0.356247719 | 0.829 | 0.853 | 1 | 0.338588475 | 0.006323747 |
| Atp6v1a | 0.003168554 | 0.470818244 | 0.925 | 0.881 | 1 | 0.087224191 | -0.464347109 | 0.951 | 0.959 | 1 | 0.087224191 | 0.006327068 |
| Flvcr1 | 0.003181478 | 1.457285365 | 0.34 | 0.182 | 1 | 0.019713049 | 1.168826341 | 0.512 | 0.338 | 1 | 0.019713049 | 0.006352834 |
| Pcdhga10 | 0.252308722 | -0.783876861 | 0.094 | 0.153 | 1 | 0.003189541 | 2.396632258 | 0.171 | 0.051 | 1 | 0.252308722 | 0.00636891 |
| Dnm1 | 0.003191738 | 0.744257825 | 0.642 | 0.453 | 1 | 0.015599811 | -0.521481795 | 0.537 | 0.737 | 1 | 0.015599811 | 0.006373289 |
| Mdp1 | 0.003192117 | 1.285967128 | 0.321 | 0.153 | 1 | 0.255094321 | 0.737669176 | 0.244 | 0.177 | 1 | 0.255094321 | 0.006374044 |
| Gprin3 | 0.003196565 | 1.792152515 | 0.189 | 0.064 | 1 | 0.716580945 | 0.170448258 | 0.22 | 0.201 | 1 | 0.716580945 | 0.006382911 |
| Timm22 | 0.003196934 | 0.865215978 | 0.547 | 0.369 | 1 | 0.309413448 | -0.360734528 | 0.293 | 0.369 | 1 | 0.309413448 | 0.006383648 |
| Gm6403 | 0.003197361 | 4.154722595 | 0.057 | 0.004 | 1 | 0.114118767 | 2.100239256 | 0.049 | 0.014 | 1 | 0.114118767 | 0.006384499 |
| Hmgcs2 | 0.003197361 | 4.324647596 | 0.057 | 0.004 | 1 | 0.262676008 | 3.644559772 | 0.024 | 0.007 | 1 | 0.262676008 | 0.006384499 |
| Cpne9 | 0.003197361 | 4.47665069 | 0.057 | 0.004 | 1 | 0.760816264 | 0.667279848 | 0.024 | 0.034 | 1 | 0.760816264 | 0.006384499 |
| Lpar5 | 0.003197361 | 3.962077517 | 0.057 | 0.004 | 1 | 0.51919986 | 0.83720485 | 0 | 0.01 | 1 | 0.51919986 | 0.006384499 |
| Gm36660 | 0.003197361 | 3.962077517 | 0.057 | 0.004 | 1 | 0.106248001 | 1.83720485 | 0.024 | 0.003 | 1 | 0.106248001 | 0.006384499 |
| Gm57189 | 0.00319746 | 2.569760094 | 0.132 | 0.034 | 1 | 0.889092853 | 0.809724113 | 0.122 | 0.119 | 1 | 0.889092853 | 0.006384696 |
| Pigv | 0.003200708 | 1.43225657 | 0.283 | 0.127 | 1 | 0.892876108 | 0.413993419 | 0.146 | 0.143 | 1 | 0.892876108 | 0.006391172 |
| Ypel4 | 0.003209552 | 3.292226119 | 0.094 | 0.017 | 1 | 0.265188955 | 2.136765132 | 0.073 | 0.038 | 1 | 0.265188955 | 0.006408803 |
| Plch2 | 0.003213173 | 1.602181572 | 0.151 | 0.042 | 1 | 0.401684222 | -0.863234868 | 0.073 | 0.116 | 1 | 0.401684222 | 0.006416022 |
| Irf9 | 0.18741894 | 0.59082171 | 0.434 | 0.36 | 1 | 0.003220992 | 1.142831132 | 0.512 | 0.328 | 1 | 0.18741894 | 0.006431609 |
| Tuba1b | 0.00322143 | 0.138662744 | 0.943 | 0.826 | 1 | 0.131388451 | 0.464903232 | 0.927 | 0.881 | 1 | 0.131388451 | 0.006432482 |
| Wbp4 | 0.003221608 | 0.830593592 | 0.811 | 0.64 | 1 | 0.109079506 | -0.153014129 | 0.683 | 0.799 | 1 | 0.109079506 | 0.006432838 |
| C1galt1c1 | 0.003225803 | -0.81613106 | 0.302 | 0.551 | 1 | 0.340235872 | -0.236504836 | 0.439 | 0.512 | 1 | 0.340235872 | 0.006441201 |
| Dbr1 | 0.003228831 | 1.427741089 | 0.34 | 0.178 | 1 | 0.448184884 | 0.437274243 | 0.293 | 0.242 | 1 | 0.448184884 | 0.006447238 |
| Rab39b | 0.046505965 | -0.960754623 | 0.094 | 0.216 | 1 | 0.003234136 | -2.081658387 | 0.049 | 0.259 | 1 | 0.046505965 | 0.006457813 |
| Zfp800 | 0.032210393 | 0.631881806 | 0.642 | 0.559 | 1 | 0.0032342 | -0.80665134 | 0.512 | 0.724 | 1 | 0.032210393 | 0.006457939 |
| Spg11 | 0.003235507 | 1.444229212 | 0.283 | 0.127 | 1 | 0.015089037 | 0.974708374 | 0.415 | 0.259 | 1 | 0.015089037 | 0.006460545 |
| Purb | 0.003243708 | 0.424070775 | 0.981 | 0.886 | 1 | 0.205624253 | -0.193462286 | 0.927 | 0.969 | 1 | 0.205624253 | 0.006476895 |
| Slc9b1 | 0.003243872 | 3.47665069 | 0.057 | 0.004 | 1 | 0.538941125 | 1.252242349 | 0.049 | 0.031 | 1 | 0.538941125 | 0.006477221 |
| Gpr63 | 0.003243872 | 3.47665069 | 0.057 | 0.004 | 1 | 0.117592181 | 1.614812428 | 0.049 | 0.014 | 1 | 0.117592181 | 0.006477221 |
| Zmynd12 | 0.003243872 | 3.47665069 | 0.057 | 0.004 | 1 | 0.732718898 | 1.667279848 | 0.024 | 0.017 | 1 | 0.732718898 | 0.006477221 |
| Gm2164 | 0.003243872 | 3.47665069 | 0.057 | 0.004 | 1 | 0.51919986 | 0.83720485 | 0 | 0.01 | 1 | 0.51919986 | 0.006477221 |
| Gm12979 | 0.003243872 | 3.739685096 | 0.057 | 0.004 | 1 | 0.455097252 | -0.16279515 | 0 | 0.014 | 1 | 0.455097252 | 0.006477221 |
| Tnfrsf25 | 0.003243872 | 3.739685096 | 0.057 | 0.004 | 1 | 0.733713289 | 0.721727632 | 0.049 | 0.038 | 1 | 0.733713289 | 0.006477221 |
| Kbtbd8 | 0.003243872 | 3.47665069 | 0.057 | 0.004 | 1 | 0.286413984 | -0.747757651 | 0 | 0.027 | 1 | 0.286413984 | 0.006477221 |
| Ptpro | 0.003243872 | 3.47665069 | 0.057 | 0.004 | 1 | 0.51919986 | 0.83720485 | 0 | 0.01 | 1 | 0.51919986 | 0.006477221 |
| Lcp2 | 0.003243872 | 4.614154213 | 0.057 | 0.004 | 1 | 0.357847088 | -0.484723245 | 0 | 0.02 | 1 | 0.357847088 | 0.006477221 |
| Evi2a | 0.003243872 | 3.47665069 | 0.057 | 0.004 | 1 | 0.973993537 | 0.962735732 | 0.024 | 0.024 | 1 | 0.973993537 | 0.006477221 |
| 6030468B19Rik | 0.003243872 | 3.47665069 | 0.057 | 0.004 | 1 | 0.056289003 | 2.83720485 | 0.049 | 0.01 | 1 | 0.056289003 | 0.006477221 |
| Fam217a | 0.003243872 | 3.47665069 | 0.057 | 0.004 | 1 | 0.455094184 | 0.252242349 | 0 | 0.014 | 1 | 0.455094184 | 0.006477221 |
| Cyp2c23 | 0.003243872 | 3.47665069 | 0.057 | 0.004 | 1 | 0.455094184 | 0.252242349 | 0 | 0.014 | 1 | 0.455094184 | 0.006477221 |
| Rpl22 | 0.139652586 | 0.18880115 | 1 | 0.966 | 1 | 0.003246623 | -0.394829329 | 1 | 0.976 | 1 | 0.139652586 | 0.006482706 |
| Creld1 | 0.00325502 | 1.920257341 | 0.208 | 0.076 | 1 | 0.861257758 | 0.252242349 | 0.073 | 0.065 | 1 | 0.861257758 | 0.006499445 |
| Cxxc1 | 0.003262508 | 1.213616284 | 0.34 | 0.165 | 1 | 0.595871782 | 0.166269126 | 0.268 | 0.328 | 1 | 0.595871782 | 0.006514372 |
| Ilf2 | 0.003270977 | 0.522784478 | 0.849 | 0.767 | 1 | 0.534319973 | 0.64895519 | 0.78 | 0.761 | 1 | 0.534319973 | 0.006531254 |
| Xndc1 | 0.003274596 | 1.840614004 | 0.283 | 0.131 | 1 | 0.843228609 | 0.23386382 | 0.195 | 0.181 | 1 | 0.843228609 | 0.006538468 |
| Stx3 | 0.731363614 | 0.43225657 | 0.302 | 0.292 | 1 | 0.003283267 | -0.601587003 | 0.317 | 0.584 | 1 | 0.731363614 | 0.006555754 |
| Dpp4 | 0.003291662 | 3.154722595 | 0.057 | 0.004 | 1 | 0.455097252 | -0.970150072 | 0 | 0.014 | 1 | 0.455097252 | 0.00657249 |
| Bglap3 | 0.003291662 | 3.154722595 | 0.057 | 0.004 | 1 | 0.357854401 | -0.747757651 | 0 | 0.02 | 1 | 0.357854401 | 0.00657249 |
| Gm42702 | 0.003291662 | 3.154722595 | 0.057 | 0.004 | 1 | 0.260704079 | 2.029849928 | 0.049 | 0.02 | 1 | 0.260704079 | 0.00657249 |
| Ncf1 | 0.003291662 | 3.154722595 | 0.057 | 0.004 | 1 | 0.914200263 | 0.136765132 | 0.024 | 0.027 | 1 | 0.914200263 | 0.00657249 |
| Gm44175 | 0.003291662 | 3.154722595 | 0.057 | 0.004 | 1 | 0.26948834 | 1.029849928 | 0.024 | 0.007 | 1 | 0.26948834 | 0.00657249 |
| Zfp108 | 0.003291662 | 3.154722595 | 0.057 | 0.004 | 1 | 0.729893611 | 2.029849928 | 0.024 | 0.017 | 1 | 0.729893611 | 0.00657249 |
| 1700028J19Rik | 0.003291662 | 3.154722595 | 0.057 | 0.004 | 1 | 0.110710459 | 2.83720485 | 0.049 | 0.014 | 1 | 0.110710459 | 0.00657249 |
| Gm57240 | 0.003291662 | 3.154722595 | 0.057 | 0.004 | 1 | 0.058990144 | 1.83720485 | 0.049 | 0.01 | 1 | 0.058990144 | 0.00657249 |
| 5830408C22Rik | 0.003291662 | 3.154722595 | 0.057 | 0.004 | 1 | 0.262674569 | 3.252242349 | 0.024 | 0.007 | 1 | 0.262674569 | 0.00657249 |
| Gm56824 | 0.003291662 | 3.154722595 | 0.057 | 0.004 | 1 | 0.012973167 | 2.515276755 | 0.073 | 0.014 | 1 | 0.012973167 | 0.00657249 |
| Gm16364.1 | 0.003291662 | 3.154722595 | 0.057 | 0.004 | 1 | 0.828694816 | -0.101394606 | 0.049 | 0.041 | 1 | 0.828694816 | 0.00657249 |
| Gm47163 | 0.003291662 | 3.154722595 | 0.057 | 0.004 | 1 | 0.208817133 | -0.970150072 | 0 | 0.038 | 1 | 0.208817133 | 0.00657249 |
| Adora2b | 0.003291662 | 3.154722595 | 0.057 | 0.004 | 1 | 0.575066969 | 0.444887427 | 0.073 | 0.051 | 1 | 0.575066969 | 0.00657249 |
| 2810408A11Rik | 0.003291662 | 3.154722595 | 0.057 | 0.004 | 1 | 0.60213981 | 1.029849928 | 0.024 | 0.014 | 1 | 0.60213981 | 0.00657249 |

| | | | | | | | | | | | | |
|---|---|---|---|---|---|---|---|---|---|---|---|---|
| Slc6a4 | 0.003291662 | 3.154722595 | 0.057 | 0.004 | 1 | 0.455097252 | -0.332720152 | 0 | 0.014 | 1 | 0.455097252 | 0.00657249 |
| Nnt | 0.003291662 | 3.154722595 | 0.057 | 0.004 | 1 | 0.872828964 | 1.029849928 | 0.024 | 0.02 | 1 | 0.872828964 | 0.00657249 |
| Lrrc24 | 0.003291662 | 3.154722595 | 0.057 | 0.004 | 1 | 0.004605593 | 2.83720485 | 0.073 | 0.01 | 1 | 0.004605593 | 0.00657249 |
| Apol9b | 0.003291662 | 3.154722595 | 0.057 | 0.004 | 1 | 0.057901606 | 2.422167351 | 0.049 | 0.01 | 1 | 0.057901606 | 0.00657249 |
| Sult6b1 | 0.003291662 | 3.154722595 | 0.057 | 0.004 | 1 | 0.587926955 | 2.614812428 | 0.024 | 0.014 | 1 | 0.587926955 | 0.00657249 |
| Hdac8 | 0.003297737 | 0.850072491 | 0.717 | 0.568 | 1 | 0.033976475 | -0.471437554 | 0.61 | 0.754 | 1 | 0.033976475 | 0.006584599 |
| Cnnm2 | 0.107236186 | 0.57654926 | 0.491 | 0.411 | 1 | 0.003297987 | 1.159132945 | 0.488 | 0.297 | 1 | 0.107236186 | 0.006585098 |
| Gpx2 | 0.782077996 | 0.347367673 | 0.019 | 0.025 | 1 | 0.003304785 | 2.515276755 | 0.122 | 0.027 | 1 | 0.782077996 | 0.006598648 |
| Glg1 | 0.003313807 | 0.363309217 | 1 | 0.987 | 1 | 0.21211793 | 0.150411674 | 1 | 0.99 | 1 | 0.21211793 | 0.006616633 |
| Rp2 | 0.003316378 | 1.280253477 | 0.302 | 0.14 | 1 | 0.026670324 | 1.061911137 | 0.341 | 0.201 | 1 | 0.026670324 | 0.006621757 |
| Kif23 | 0.104824324 | 0.446903346 | 0.113 | 0.051 | 1 | 0.003318373 | 1.351778023 | 0.098 | 0.017 | 1 | 0.104824324 | 0.006625735 |
| Sulf1 | 0.003329801 | 0.554078472 | 0.377 | 0.191 | 1 | 0.038731614 | 2.584438779 | 0.341 | 0.222 | 1 | 0.038731614 | 0.006648515 |
| Dexi | 0.003336018 | 1.19654277 | 0.415 | 0.242 | 1 | 0.064097417 | 0.654340793 | 0.512 | 0.372 | 1 | 0.064097417 | 0.006660908 |
| Rapgefl1 | 0.00333608 | 2.640149422 | 0.094 | 0.017 | 1 | 0.500516404 | -0.589059905 | 0.073 | 0.106 | 1 | 0.500516404 | 0.006661031 |
| Gm15545 | 0.00333816 | 2.739685096 | 0.094 | 0.017 | 1 | 0.058990144 | 1.83720485 | 0.049 | 0.01 | 1 | 0.058990144 | 0.006665177 |
| Gm53018 | 0.00333868 | 2.517292674 | 0.094 | 0.017 | 1 | 0.909322449 | -0.555112573 | 0.024 | 0.027 | 1 | 0.909322449 | 0.006666214 |
| Nkain1 | 0.00333881 | 2.029191713 | 0.094 | 0.017 | 1 | 0.749728988 | 0.252242349 | 0.024 | 0.017 | 1 | 0.749728988 | 0.006666473 |
| Ptchd4 | 0.003339613 | 3.377115016 | 0.057 | 0.004 | 1 | 0.003983294 | 5.422167351 | 0.049 | 0.003 | 1 | 0.003983294 | 0.006668073 |
| Col7a1 | 0.003345403 | 3.047807391 | 0.113 | 0.025 | 1 | 0.569723706 | -0.654648247 | 0.073 | 0.099 | 1 | 0.569723706 | 0.006679614 |
| Sumf1 | 0.003348263 | 0.938993904 | 0.434 | 0.237 | 1 | 0.385756636 | -0.190275887 | 0.22 | 0.287 | 1 | 0.385756636 | 0.006685314 |
| Slc25a27 | 0.003350515 | 1.393910259 | 0.321 | 0.157 | 1 | 0.730727078 | 0.73185185 | 0.268 | 0.283 | 1 | 0.730727078 | 0.006689803 |
| Clcc1 | 0.388829505 | 0.370451286 | 0.472 | 0.47 | 1 | 0.003355625 | -0.926094892 | 0.244 | 0.495 | 1 | 0.388829505 | 0.00669999 |
| Rian | 0.006687644 | -0.80479274 | 0.377 | 0.581 | 1 | 0.003359946 | -0.997812511 | 0.439 | 0.696 | 1 | 0.006687644 | 0.006708604 |
| Reep4 | 0.003365997 | 1.669295768 | 0.151 | 0.042 | 1 | 0.400472257 | 0.721727632 | 0.122 | 0.082 | 1 | 0.400472257 | 0.006720664 |
| Hacd3 | 0.003366605 | 1.046662849 | 0.491 | 0.314 | 1 | 0.322304057 | 0.610433988 | 0.488 | 0.457 | 1 | 0.322304057 | 0.006721876 |
| Vldlr | 0.037593965 | 0.602181572 | 0.17 | 0.076 | 1 | 0.003368699 | 0.749742009 | 0.22 | 0.075 | 1 | 0.037593965 | 0.00672605 |
| Cep192 | 0.003369333 | 0.748097335 | 0.434 | 0.246 | 1 | 0.08966749 | 0.623080044 | 0.366 | 0.242 | 1 | 0.08966749 | 0.006727314 |
| Cops5 | 0.003378965 | 0.803648154 | 0.736 | 0.61 | 1 | 0.143244551 | -0.173161866 | 0.61 | 0.72 | 1 | 0.143244551 | 0.006746512 |
| Poc1b | 0.003379067 | 1.627475592 | 0.302 | 0.144 | 1 | 0.6623403 | 0.463746454 | 0.293 | 0.273 | 1 | 0.6623403 | 0.006746716 |
| Gas6 | 0.043444586 | 0.617969187 | 0.377 | 0.233 | 1 | 0.003380785 | 1.664737654 | 0.463 | 0.27 | 1 | 0.043444586 | 0.006750139 |
| Ppargc1b | 0.007807028 | 2.154722595 | 0.075 | 0.013 | 1 | 0.003385722 | 1.737669176 | 0.098 | 0.017 | 1 | 0.007807028 | 0.006759981 |
| Grb14 | 0.003390061 | -1.953801862 | 0.057 | 0.237 | 1 | 0.154173787 | -2.020776145 | 0 | 0.048 | 1 | 0.154173787 | 0.00676863 |
| Cdc42se2 | 0.003390356 | 0.723793723 | 0.604 | 0.462 | 1 | 0.054461827 | 0.641035749 | 0.78 | 0.672 | 1 | 0.054461827 | 0.006769218 |
| Npr1 | 0.225506368 | 1.932330173 | 0.057 | 0.025 | 1 | 0.003391162 | 2.199774929 | 0.122 | 0.027 | 1 | 0.225506368 | 0.006770825 |
| Usp42 | 0.003393132 | 1.316473665 | 0.377 | 0.203 | 1 | 0.010314127 | 0.93278251 | 0.512 | 0.338 | 1 | 0.010314127 | 0.00677475 |
| Pon3 | 0.003394812 | 1.004479959 | 0.434 | 0.254 | 1 | 0.082706639 | -0.122153166 | 0.317 | 0.454 | 1 | 0.082706639 | 0.0067781 |
| D830025C05Rik | 0.003395011 | 1.13912574 | 0.358 | 0.182 | 1 | 0.959034646 | -0.305753104 | 0.415 | 0.382 | 1 | 0.959034646 | 0.006778496 |
| Lgals4 | 0.00340487 | 1.417757001 | 0.245 | 0.097 | 1 | 0.166846671 | 0.930314254 | 0.22 | 0.14 | 1 | 0.166846671 | 0.006798148 |
| Syngap1 | 0.003406485 | 1.549001534 | 0.264 | 0.114 | 1 | 0.021623413 | 1.30377265 | 0.268 | 0.133 | 1 | 0.021623413 | 0.006801366 |
| Dusp15 | 0.003408554 | -3.953801862 | 0 | 0.144 | 1 | 0.231618904 | -1.250257991 | 0 | 0.034 | 1 | 0.231618904 | 0.006805491 |
| Zfp780b | 0.003410674 | 1.640149422 | 0.189 | 0.064 | 1 | 0.7896497 | 0.491430013 | 0.146 | 0.167 | 1 | 0.7896497 | 0.006809716 |
| Otos | 0.003415548 | -5.916739768 | 0 | 0.144 | 1 | 0.103479965 | 3.422167351 | 0.024 | 0.003 | 1 | 0.103479965 | 0.00681943 |
| Cstf1 | 0.003416328 | 1.402650108 | 0.283 | 0.127 | 1 | 0.254443408 | -0.589059905 | 0.073 | 0.137 | 1 | 0.254443408 | 0.006820986 |
| Nxf1 | 0.003418896 | 0.696799283 | 0.717 | 0.555 | 1 | 0.162369015 | 0.451551157 | 0.805 | 0.737 | 1 | 0.162369015 | 0.006826104 |
| Dnaaf4 | 0.00342037 | 2.640149422 | 0.132 | 0.034 | 1 | 0.276640588 | 1.185128153 | 0.098 | 0.055 | 1 | 0.276640588 | 0.006829041 |
| Lama5 | 0.413082433 | 0.154722595 | 0 | 0.013 | 1 | 0.00342063 | 2.126711467 | 0.171 | 0.051 | 1 | 0.413082433 | 0.006829559 |
| Clec16a | 0.003421171 | 1.298458072 | 0.472 | 0.309 | 1 | 0.6530736 | -0.167186919 | 0.488 | 0.522 | 1 | 0.6530736 | 0.006830637 |
| Rgs4 | 0.003437624 | 1.002719501 | 0.057 | 0.004 | 1 | 0.104856732 | 2.83720485 | 0.024 | 0.003 | 1 | 0.104856732 | 0.006863432 |
| Jph2 | 0.003437624 | 1.695290976 | 0.057 | 0.004 | 1 | 0.455098275 | 0.515276755 | 0 | 0.014 | 1 | 0.455098275 | 0.006863432 |
| Blk | 0.003437624 | 0.891688189 | 0.057 | 0.004 | 1 | 0.607875423 | -0.484723245 | 0.024 | 0.014 | 1 | 0.607875423 | 0.006863432 |
| Gpc5 | 0.003437624 | 1.913714495 | 0.057 | 0.004 | 1 | 0.106248001 | -0.16279515 | 0.024 | 0.003 | 1 | 0.106248001 | 0.006863432 |
| Mgrn1 | 0.00344165 | 0.879715547 | 0.566 | 0.377 | 1 | 0.110244827 | -0.131544216 | 0.415 | 0.577 | 1 | 0.110244827 | 0.006871455 |
| mt-Co2 | 0.003453633 | -0.315525502 | 1 | 1 | 1 | 0.358608544 | -0.227141527 | 1 | 1 | 1 | 0.358608544 | 0.006895338 |
| Scarb2 | 0.184058333 | -0.287114964 | 0.566 | 0.657 | 1 | 0.003456327 | -0.606186758 | 0.683 | 0.84 | 1 | 0.184058333 | 0.006900708 |
| Ptx3 | 0.003459002 | -3.586744392 | 0.038 | 0.208 | 1 | 0.032367704 | -3.877040668 | 0 | 0.102 | 1 | 0.032367704 | 0.00690604 |
| Pik3cb | 0.010583889 | -1.221653284 | 0.264 | 0.424 | 1 | 0.003461983 | -1.327181968 | 0.39 | 0.567 | 1 | 0.010583889 | 0.00691198 |
| Tmc8 | 0.003468178 | 2.154722595 | 0.132 | 0.034 | 1 | 0.902330602 | 0.19334866 | 0.049 | 0.044 | 1 | 0.902330602 | 0.006924328 |
| Pabpc1l | 0.003469649 | 2.569760094 | 0.094 | 0.017 | 1 | 0.862535502 | 1.199774929 | 0.073 | 0.068 | 1 | 0.862535502 | 0.006927259 |
| Sytl1 | 0.003472153 | 2.417757001 | 0.094 | 0.017 | 1 | 0.455763738 | 0.962735732 | 0.049 | 0.027 | 1 | 0.455763738 | 0.00693225 |
| Gm44686 | 0.003472153 | 2.417757001 | 0.094 | 0.017 | 1 | 0.801706188 | 0.435106406 | 0.098 | 0.085 | 1 | 0.801706188 | 0.00693225 |
| Dnmt3b | 0.003472811 | 2.154722595 | 0.132 | 0.034 | 1 | 0.547258973 | -0.250257991 | 0.024 | 0.044 | 1 | 0.547258973 | 0.006933561 |
| Ranbp2 | 0.003476245 | 0.445839543 | 0.943 | 0.936 | 1 | 0.397840088 | 0.145027177 | 0.976 | 0.969 | 1 | 0.397840088 | 0.006940406 |
| Mllt6 | 0.003480681 | 1.036541168 | 0.472 | 0.301 | 1 | 0.804209525 | 0.276489895 | 0.39 | 0.461 | 1 | 0.804209525 | 0.006949248 |
| Itpa | 0.00348957 | 1.081473613 | 0.509 | 0.331 | 1 | 0.709281557 | 0.310659035 | 0.293 | 0.331 | 1 | 0.709281557 | 0.006966962 |
| Map1a | 0.630606115 | 0.192524628 | 0.321 | 0.305 | 1 | 0.003494083 | -1.336722082 | 0.244 | 0.478 | 1 | 0.630606115 | 0.006975958 |
| P2rx4 | 0.015289159 | -0.873474297 | 0.208 | 0.394 | 1 | 0.003514635 | -1.25871957 | 0.122 | 0.362 | 1 | 0.015289159 | 0.007016918 |
| A330076H08Rik | 0.003527976 | -1.637835325 | 0.075 | 0.267 | 1 | 0.011390415 | -1.432255825 | 0.098 | 0.29 | 1 | 0.011390415 | 0.007043505 |
| Gm57325 | 0.003530009 | 2.154722595 | 0.132 | 0.034 | 1 | 0.208799874 | -0.863234868 | 0 | 0.038 | 1 | 0.208799874 | 0.007047558 |
| Kcmf1 | 0.003533383 | 0.367085907 | 1 | 0.928 | 1 | 0.383374409 | 0.134082711 | 0.951 | 0.98 | 1 | 0.383374409 | 0.007054281 |
| Gm56999 | 0.003537602 | 1.962077517 | 0.094 | 0.017 | 1 | 0.756560731 | 0.422167351 | 0.024 | 0.034 | 1 | 0.756560731 | 0.007062689 |
| Trim72 | 0.003537602 | 2.154722595 | 0.094 | 0.017 | 1 | 0.180559238 | 0.667279848 | 0.073 | 0.031 | 1 | 0.180559238 | 0.007062689 |
| Anxa6 | 0.003538986 | 0.795394528 | 0.66 | 0.568 | 1 | 0.576000313 | 0.320413852 | 0.537 | 0.577 | 1 | 0.576000313 | 0.007065448 |
| Pde11a | 0.00353944 | -1.332543232 | 0.057 | 0.237 | 1 | 0.253474467 | -0.26713181 | 0.049 | 0.109 | 1 | 0.253474467 | 0.007066353 |
| Slc44a1 | 0.003540102 | -0.701969013 | 0.83 | 0.894 | 1 | 0.78460857 | -0.210827847 | 0.878 | 0.915 | 1 | 0.78460857 | 0.007067672 |
| Gpm6a | 0.249034808 | 2.154722595 | 0.019 | 0.004 | 1 | 0.003559033 | 2.100239256 | 0.122 | 0.027 | 1 | 0.249034808 | 0.0071054 |
| Kat2a | 0.003559936 | 1.618669695 | 0.264 | 0.119 | 1 | 0.156254505 | 0.640807637 | 0.244 | 0.154 | 1 | 0.156254505 | 0.0071072 |
| B830012L14Rik | 0.003565324 | -1.046911266 | 0.17 | 0.381 | 1 | 0.021875306 | -0.717856165 | 0.268 | 0.491 | 1 | 0.021875306 | 0.007117937 |
| Gdpd4 | 0.003574045 | 1.47665069 | 0.094 | 0.017 | 1 | 0.119365389 | 0.83720485 | 0.049 | 0.014 | 1 | 0.119365389 | 0.007135316 |
| Cpeb2 | 0.003577035 | 1.169519597 | 0.415 | 0.225 | 1 | 0.433729713 | 0.237742779 | 0.561 | 0.536 | 1 | 0.433729713 | 0.007141276 |
| Npepps | 0.003589666 | 0.520007059 | 0.906 | 0.839 | 1 | 0.416791943 | 0.187094348 | 0.902 | 0.952 | 1 | 0.416791943 | 0.007166447 |
| Braf | 0.003597049 | 0.625613329 | 0.925 | 0.86 | 1 | 0.604617978 | -0.11126485 | 0.951 | 0.973 | 1 | 0.604617978 | 0.007181159 |
| Casp8ap2 | 0.003602684 | 1.119675648 | 0.491 | 0.301 | 1 | 0.005031384 | 0.92700152 | 0.585 | 0.392 | 1 | 0.005031384 | 0.007192389 |
| Ptgs2 | 0.003604323 | -1.892401317 | 0.151 | 0.352 | 1 | 0.874122727 | -0.686357106 | 0.098 | 0.089 | 1 | 0.874122727 | 0.007195655 |
| Unkl | 0.003610526 | 1.074995402 | 0.472 | 0.301 | 1 | 0.205155276 | -0.292078167 | 0.317 | 0.451 | 1 | 0.205155276 | 0.007208016 |
| Orc2 | 0.003614706 | 1.317275608 | 0.472 | 0.318 | 1 | 0.643005546 | 0.123767791 | 0.415 | 0.396 | 1 | 0.643005546 | 0.007216346 |
| Hsf1 | 0.035006264 | 0.619945851 | 0.491 | 0.343 | 1 | 0.003637942 | -1.380025866 | 0.171 | 0.406 | 1 | 0.035006264 | 0.00726265 |
| Efemp2 | 0.003639016 | -0.625614777 | 0.717 | 0.801 | 1 | 0.206763347 | -0.236044132 | 0.537 | 0.662 | 1 | 0.206763347 | 0.00726479 |
| Lmnb2 | 0.003639371 | 1.640149422 | 0.189 | 0.064 | 1 | 0.596500493 | 0.410940095 | 0.098 | 0.133 | 1 | 0.596500493 | 0.007265497 |
| Shkbp1 | 0.003640503 | 2.154722595 | 0.094 | 0.017 | 1 | 0.269479162 | 1.614812428 | 0.073 | 0.038 | 1 | 0.269479162 | 0.007267753 |
| Esrra | 0.003640503 | 2.154722595 | 0.094 | 0.017 | 1 | 0.049954518 | 1.136765132 | 0.146 | 0.061 | 1 | 0.049954518 | 0.007267753 |
| Adad1 | 0.003643092 | 2.595295186 | 0.113 | 0.025 | 1 | 0.058994294 | 2.185128153 | 0.049 | 0.01 | 1 | 0.058994294 | 0.007272911 |
| A730036I17Rik | 0.00364385 | 0.953088734 | 0.094 | 0.017 | 1 | 0.184911704 | 1.83720485 | 0.049 | 0.017 | 1 | 0.184911704 | 0.007274422 |
| Zfp935 | 0.003644 | 1.468831185 | 0.283 | 0.127 | 1 | 0.491694824 | -0.16279515 | 0.073 | 0.109 | 1 | 0.491694824 | 0.007274721 |
| Agps | 0.003656686 | 0.731223517 | 0.642 | 0.47 | 1 | 0.218012432 | 0.79140116 | 0.707 | 0.703 | 1 | 0.218012432 | 0.007300001 |
| Ccni | 0.004830373 | 0.445312723 | 0.925 | 0.847 | 1 | 0.003658446 | 0.609846176 | 0.878 | 0.891 | 1 | 0.004830373 | 0.007303507 |
| Tafa5 | 0.024538769 | -0.875580165 | 0.396 | 0.564 | 1 | 0.003666596 | -1.127401244 | 0.488 | 0.679 | 1 | 0.024538769 | 0.007319748 |
| Dpy19l1 | 0.00367082 | 0.992451166 | 0.547 | 0.377 | 1 | 0.156872279 | 0.854763221 | 0.512 | 0.471 | 1 | 0.156872279 | 0.007328166 |
| Adcy1 | 0.003670949 | 2.614154213 | 0.113 | 0.025 | 1 | 0.274788535 | 1.252242349 | 0.049 | 0.02 | 1 | 0.274788535 | 0.007328423 |

| | | | | | | | | | | | | |
|---|---|---|---|---|---|---|---|---|---|---|---|---|
| 2610044O15Rik8 | 0.003689302 | 1.78032708 | 0.264 | 0.119 | 1 | 0.164455265 | 1.296636468 | 0.146 | 0.082 | 1 | 0.164455265 | 0.007364994 |
| Pdia4 | 0.003689561 | 0.432878288 | 0.981 | 0.86 | 1 | 0.865694534 | 0.113795579 | 0.829 | 0.891 | 1 | 0.865694534 | 0.00736551 |
| Csf2ra | 0.003689608 | -0.893640427 | 0.113 | 0.322 | 1 | 0.34629187 | 0.334704509 | 0.073 | 0.13 | 1 | 0.34629187 | 0.007365602 |
| Zbtb21 | 0.47625134 | -0.156064942 | 0.34 | 0.394 | 1 | 0.003690058 | -1.106211622 | 0.195 | 0.433 | 1 | 0.47625134 | 0.0073665 |
| Psmc1 | 0.003691533 | 0.553992778 | 0.792 | 0.636 | 1 | 0.155703311 | -0.291200882 | 0.585 | 0.683 | 1 | 0.155703311 | 0.007369439 |
| Sardh | 0.003698097 | 1.984797593 | 0.132 | 0.034 | 1 | 0.209475181 | 1.489281546 | 0.122 | 0.068 | 1 | 0.209475181 | 0.007382518 |
| Noxo1 | 0.003698097 | 1.984797593 | 0.132 | 0.034 | 1 | 0.973993537 | 1.100239256 | 0.024 | 0.024 | 1 | 0.973993537 | 0.007382518 |
| Wnt2b | 0.003698399 | 2.517292674 | 0.113 | 0.025 | 1 | 0.180559238 | 1.136765132 | 0.073 | 0.031 | 1 | 0.180559238 | 0.00738312 |
| C030034I22Rik | 0.003698399 | 2.347367673 | 0.113 | 0.025 | 1 | 0.792969817 | -0.385187572 | 0.049 | 0.058 | 1 | 0.792969817 | 0.00738312 |
| Kctd9 | 0.445070603 | -0.450139463 | 0.415 | 0.466 | 1 | 0.003698599 | -1.275495283 | 0.244 | 0.491 | 1 | 0.445070603 | 0.007383519 |
| Tshz2 | 0.003705789 | -0.871951617 | 0.962 | 0.97 | 1 | 0.037962769 | -0.127073228 | 0.707 | 0.887 | 1 | 0.037962769 | 0.007397845 |
| Myo5c | 0.009815353 | 0.932330173 | 0.094 | 0.021 | 1 | 0.003709094 | 2.907594178 | 0.171 | 0.055 | 1 | 0.009815353 | 0.00740443 |
| Eva1a | 0.003711697 | 0.984797593 | 0.208 | 0.072 | 1 | 0.175071751 | -0.255904555 | 0.122 | 0.218 | 1 | 0.175071751 | 0.007409618 |
| Deaf1 | 0.088181886 | -0.747980204 | 0.283 | 0.398 | 1 | 0.00371306 | -0.962770542 | 0.293 | 0.536 | 1 | 0.088181886 | 0.007412333 |
| Serinc5 | 0.119365269 | -0.731457506 | 0.717 | 0.809 | 1 | 0.003716211 | -0.543467036 | 0.78 | 0.901 | 1 | 0.119365269 | 0.007418611 |
| Micu3 | 0.449987319 | -0.26948655 | 0.415 | 0.513 | 1 | 0.003719908 | -0.663597203 | 0.488 | 0.713 | 1 | 0.449987319 | 0.007425979 |
| Mprip | 0.003726477 | 0.368579432 | 1 | 0.945 | 1 | 0.007461995 | 0.442285993 | 1 | 0.98 | 1 | 0.007461995 | 0.007439067 |
| Dram1 | 0.003728497 | -1.519877118 | 0.151 | 0.352 | 1 | 0.324498573 | -0.577832649 | 0.146 | 0.215 | 1 | 0.324498573 | 0.007443093 |
| Dnajb2 | 0.008110866 | -0.804635421 | 0.34 | 0.542 | 1 | 0.003731716 | -1.452301767 | 0.146 | 0.375 | 1 | 0.008110866 | 0.007449506 |
| Brd2 | 0.003734326 | 0.540568381 | 0.962 | 0.949 | 1 | 0.435985146 | 0.164245868 | 0.976 | 0.986 | 1 | 0.435985146 | 0.007454706 |
| Tbx3 | 0.003742403 | 0.872687227 | 0.453 | 0.267 | 1 | 0.086240215 | 1.405488609 | 0.341 | 0.242 | 1 | 0.086240215 | 0.0074708 |
| Stx18 | 0.003746368 | 0.971500771 | 0.66 | 0.534 | 1 | 0.294955923 | 0.344924352 | 0.61 | 0.621 | 1 | 0.294955923 | 0.007478701 |
| Spop | 0.003751882 | 0.486867086 | 0.962 | 0.915 | 1 | 0.246797685 | 0.19425535 | 0.854 | 0.935 | 1 | 0.246797685 | 0.007489688 |
| Zfp580 | 0.003757708 | 1.59082171 | 0.283 | 0.131 | 1 | 0.342694533 | 1.189506594 | 0.171 | 0.126 | 1 | 0.342694533 | 0.007501296 |
| Zfp341 | 0.003764611 | 1.739685096 | 0.113 | 0.025 | 1 | 0.492842661 | -0.622226769 | 0.024 | 0.048 | 1 | 0.492842661 | 0.00751505 |
| Armcx3 | 0.520090217 | 0.159182243 | 0.623 | 0.678 | 1 | 0.003769208 | -0.803381157 | 0.561 | 0.727 | 1 | 0.520090217 | 0.007524209 |
| Eif2s3y | 0.003774444 | 0.836646825 | 0.528 | 0.335 | 1 | 0.242612473 | -0.226655092 | 0.659 | 0.679 | 1 | 0.242612473 | 0.007534642 |
| Arhgap39 | 0.003774791 | 1.095828906 | 0.321 | 0.153 | 1 | 0.349155548 | -0.40380325 | 0.146 | 0.215 | 1 | 0.349155548 | 0.007535332 |
| Mad2l2 | 0.00378331 | 1.47665069 | 0.283 | 0.131 | 1 | 0.545587835 | -0.332720152 | 0.073 | 0.102 | 1 | 0.545587835 | 0.007552307 |
| Cadm2 | 0.00378724 | 1.614154213 | 0.094 | 0.017 | 1 | 0.663737891 | 1.23386382 | 0.098 | 0.082 | 1 | 0.663737891 | 0.007560138 |
| Arl2bp | 0.003802471 | -0.622000709 | 0.623 | 0.801 | 1 | 0.006083794 | -0.517411864 | 0.561 | 0.765 | 1 | 0.006083794 | 0.007590483 |
| Ptar1 | 0.003804663 | 0.783884899 | 0.604 | 0.407 | 1 | 0.453993268 | 0.298338764 | 0.537 | 0.519 | 1 | 0.453993268 | 0.00759485 |
| Tmod2 | 0.003845844 | 1.984797593 | 0.17 | 0.055 | 1 | 0.903004117 | 0.46797104 | 0.146 | 0.143 | 1 | 0.903004117 | 0.007676897 |
| Ahsa2 | 0.003855722 | 0.964039033 | 0.491 | 0.309 | 1 | 0.93573383 | 0.446564005 | 0.415 | 0.451 | 1 | 0.93573383 | 0.007696578 |
| Ern1 | 0.003860866 | 0.73100534 | 0.736 | 0.534 | 1 | 0.043101973 | 0.636028914 | 0.756 | 0.696 | 1 | 0.043101973 | 0.007706826 |
| Lsm1 | 0.003862808 | 0.896924837 | 0.585 | 0.415 | 1 | 0.692103751 | 0.54962826 | 0.488 | 0.495 | 1 | 0.692103751 | 0.007710695 |
| Tmem265 | 0.003863643 | 1.254258268 | 0.245 | 0.097 | 1 | 0.412667379 | 0.674266279 | 0.293 | 0.253 | 1 | 0.412667379 | 0.007712358 |
| Mecom | 0.648215982 | -1.260314904 | 0.283 | 0.229 | 1 | 0.003870485 | 2.021629421 | 0.39 | 0.208 | 1 | 0.648215982 | 0.00772599 |
| Acrbp | 0.003889286 | 2.242185436 | 0.17 | 0.055 | 1 | 0.540355991 | 0.547698233 | 0.146 | 0.113 | 1 | 0.540355991 | 0.007763445 |
| ENSMUSG00000121485 | 0.003889286 | 2.06161319 | 0.17 | 0.055 | 1 | 0.324449696 | -1.541306773 | 0.073 | 0.126 | 1 | 0.324449696 | 0.007763445 |
| Zfp354a | 0.00389737 | 2.154722595 | 0.113 | 0.025 | 1 | 0.249749409 | -1.020776145 | 0.024 | 0.072 | 1 | 0.249749409 | 0.007779551 |
| Mir99ahg | 0.354385974 | 0.166549702 | 0.906 | 0.936 | 1 | 0.00390851 | 1.001756679 | 0.927 | 0.819 | 1 | 0.354385974 | 0.007801743 |
| Kctd1 | 0.003909135 | 0.921523419 | 0.472 | 0.28 | 1 | 0.218130312 | 0.645253542 | 0.512 | 0.457 | 1 | 0.218130312 | 0.007802988 |
| Arl6ip4 | 0.00391204 | 0.8327945 | 0.528 | 0.356 | 1 | 0.945654128 | 0.373548645 | 0.366 | 0.403 | 1 | 0.945654128 | 0.007808776 |
| Nup107 | 0.003913334 | 1.263951665 | 0.396 | 0.225 | 1 | 0.10461667 | 1.294677615 | 0.39 | 0.283 | 1 | 0.10461667 | 0.007811354 |
| Gm26801 | 0.03562064 | 3.739685096 | 0.019 | 0 | 1 | 0.003917165 | 4.422167351 | 0.049 | 0.003 | 1 | 0.03562064 | 0.007818986 |
| Rasgrf2 | 0.246443787 | 2.739685096 | 0.019 | 0.004 | 1 | 0.003917165 | 4.83720485 | 0.049 | 0.003 | 1 | 0.246443787 | 0.007818986 |
| 4930488L21Rik | 0.413080549 | -0.1672055 | 0 | 0.013 | 1 | 0.003917165 | 4.059597271 | 0.049 | 0.003 | 1 | 0.413080549 | 0.007818986 |
| Shc3 | 0.914958 | 0.347367673 | 0.019 | 0.021 | 1 | 0.003917165 | 4.644559772 | 0.049 | 0.003 | 1 | 0.914958 | 0.007818986 |
| Mgst2 | 0.003919706 | -1.727920455 | 0.132 | 0.322 | 1 | 0.05928562 | -1.787286015 | 0.049 | 0.157 | 1 | 0.05928562 | 0.007824049 |
| Kdm5d | 0.104841625 | 0.8327945 | 0.302 | 0.199 | 1 | 0.003921448 | -0.736262012 | 0.171 | 0.457 | 1 | 0.104841625 | 0.007827518 |
| Cmtm7 | 0.003924208 | 0.766157307 | 0.585 | 0.394 | 1 | 0.177113323 | -0.240038149 | 0.341 | 0.478 | 1 | 0.177113323 | 0.007833016 |
| Eda | 0.003927982 | 1.686573759 | 0.302 | 0.144 | 1 | 0.101915777 | 0.830351179 | 0.463 | 0.334 | 1 | 0.101915777 | 0.007840534 |
| Mkrn2os | 0.003928395 | 2.039245377 | 0.113 | 0.025 | 1 | 0.262676008 | 3.159132945 | 0.024 | 0.007 | 1 | 0.262676008 | 0.007841358 |
| Foxj1 | 0.003935196 | 1.055186921 | 0.094 | 0.017 | 1 | 0.074976089 | 2.071670103 | 0.098 | 0.038 | 1 | 0.074976089 | 0.007854906 |
| Msra | 0.003936932 | 1.034983351 | 0.434 | 0.254 | 1 | 0.429999979 | 0.142059431 | 0.439 | 0.396 | 1 | 0.429999979 | 0.007858365 |
| Dusp6 | 0.365288487 | 1.310373465 | 0.226 | 0.182 | 1 | 0.003938642 | 1.614812428 | 0.268 | 0.113 | 1 | 0.365288487 | 0.007861772 |
| Gramd4 | 0.003943607 | 1.520850494 | 0.302 | 0.148 | 1 | 0.254910264 | -0.995685164 | 0.146 | 0.215 | 1 | 0.254910264 | 0.007871662 |
| Extl1 | 0.003947779 | -1.430239906 | 0.113 | 0.309 | 1 | 0.005548142 | -0.762707992 | 0.049 | 0.256 | 1 | 0.005548142 | 0.007879973 |
| Ubr4 | 0.109487479 | 0.298458072 | 0.943 | 0.847 | 1 | 0.003951024 | -0.40773014 | 0.927 | 0.973 | 1 | 0.109487479 | 0.007886437 |
| Sh3glb2 | 0.003957446 | 1.377115016 | 0.302 | 0.14 | 1 | 0.524561104 | -0.435813645 | 0.122 | 0.157 | 1 | 0.524561104 | 0.00789923 |
| Oxsr1 | 0.003961019 | 1.105812994 | 0.377 | 0.199 | 1 | 0.480304871 | 0.170942247 | 0.366 | 0.304 | 1 | 0.480304871 | 0.007906349 |
| B3galt5 | 0.003961624 | -3.459987249 | 0 | 0.14 | 1 | 0.455097252 | -0.622226769 | 0 | 0.014 | 1 | 0.455097252 | 0.007907554 |
| Cda | 0.003961935 | -3.600164907 | 0 | 0.14 | 1 | 0.606262835 | -0.410722664 | 0.024 | 0.041 | 1 | 0.606262835 | 0.007908172 |
| Pde9a | 0.03562064 | 3.739685096 | 0.019 | 0 | 1 | 0.003983294 | 3.83720485 | 0.049 | 0.003 | 1 | 0.03562064 | 0.007950721 |
| Gm14051 | 0.038976464 | 2.154722595 | 0.075 | 0.021 | 1 | 0.003983294 | 4.159132945 | 0.049 | 0.003 | 1 | 0.038976464 | 0.007950721 |
| Gm57170 | 0.088349056 | 2.154722595 | 0.057 | 0.017 | 1 | 0.003983294 | 4.159132945 | 0.049 | 0.003 | 1 | 0.088349056 | 0.007950721 |
| Gm48202 | 0.413082433 | 0.154722595 | 0 | 0.013 | 1 | 0.003983294 | 4.159132945 | 0.049 | 0.003 | 1 | 0.413082433 | 0.007950721 |
| Slc39a12 | 0.507281812 | 0.8327945 | 0.019 | 0.008 | 1 | 0.003983294 | 4.644559772 | 0.049 | 0.003 | 1 | 0.507281812 | 0.007950721 |
| Plppr1 | 0.507281812 | 0.8327945 | 0.019 | 0.008 | 1 | 0.003983294 | 3.83720485 | 0.049 | 0.003 | 1 | 0.507281812 | 0.007950721 |
| Lypd6b | 0.722567081 | 2.640149422 | 0.019 | 0.013 | 1 | 0.003983294 | 4.422167351 | 0.049 | 0.003 | 1 | 0.722567081 | 0.007950721 |
| Mfsd9 | 0.7394666 | 0.569760094 | 0.019 | 0.013 | 1 | 0.003983294 | 3.83720485 | 0.049 | 0.003 | 1 | 0.7394666 | 0.007950721 |
| Rbm28 | 0.003992358 | 0.79308479 | 0.642 | 0.492 | 1 | 0.161811972 | 0.541406487 | 0.659 | 0.597 | 1 | 0.161811972 | 0.007968777 |
| Ccdc136 | 0.003993249 | 1.2967416 | 0.226 | 0.085 | 1 | 0.717969214 | 0.313642894 | 0.244 | 0.229 | 1 | 0.717969214 | 0.007970552 |
| Rab23 | 0.003994056 | 1.031340179 | 0.34 | 0.169 | 1 | 0.178200026 | -0.970150072 | 0.122 | 0.205 | 1 | 0.178200026 | 0.00797216 |
| Alkbh1 | 0.004004631 | -0.492524184 | 0.509 | 0.75 | 1 | 0.126452923 | -0.316269631 | 0.463 | 0.628 | 1 | 0.126452923 | 0.007993224 |
| Ddx46 | 0.004016238 | 0.698546401 | 0.774 | 0.708 | 1 | 0.065692609 | 0.510223527 | 0.878 | 0.802 | 1 | 0.065692609 | 0.008016345 |
| Mtf2 | 0.004016972 | 0.920257341 | 0.604 | 0.445 | 1 | 0.090229179 | 0.320413852 | 0.78 | 0.625 | 1 | 0.090229179 | 0.008017808 |
| Ercc3 | 0.0040186 | 1.508359549 | 0.264 | 0.119 | 1 | 0.036389802 | 1.372536583 | 0.317 | 0.191 | 1 | 0.036389802 | 0.00802105 |
| Adam9 | 0.137814917 | 0.159604816 | 0.66 | 0.547 | 1 | 0.00402444 | 0.671956336 | 0.78 | 0.56 | 1 | 0.137814917 | 0.008032685 |
| Ythdc1 | 0.004027824 | 0.375749474 | 0.981 | 0.979 | 1 | 0.456414565 | -0.141100079 | 0.976 | 0.983 | 1 | 0.456414565 | 0.008039425 |
| R3hdm4 | 0.157834184 | 0.503958036 | 0.528 | 0.483 | 1 | 0.00404282 | 0.774592873 | 0.732 | 0.563 | 1 | 0.157834184 | 0.008069296 |
| Ddx43 | 0.018669677 | 2.377115016 | 0.075 | 0.017 | 1 | 0.004050657 | 3.422167351 | 0.049 | 0.003 | 1 | 0.018669677 | 0.008084906 |
| Zyg11a | 0.03562064 | 3.154722595 | 0.019 | 0 | 1 | 0.004050657 | 3.422167351 | 0.049 | 0.003 | 1 | 0.03562064 | 0.008084906 |
| 3010003L21Rik | 0.03562064 | 3.154722595 | 0.019 | 0 | 1 | 0.004050657 | 3.422167351 | 0.049 | 0.003 | 1 | 0.03562064 | 0.008084906 |
| 4930500H12Rik | 0.03562064 | 3.154722595 | 0.019 | 0 | 1 | 0.004050657 | 3.422167351 | 0.049 | 0.003 | 1 | 0.03562064 | 0.008084906 |
| Oacyl | 0.207549016 | 1.8327945 | 0.038 | 0.013 | 1 | 0.004050657 | 3.422167351 | 0.049 | 0.003 | 1 | 0.207549016 | 0.008084906 |
| Gm56868 | 0.243102132 | -1.015202407 | 0 | 0.025 | 1 | 0.004050657 | 3.422167351 | 0.049 | 0.003 | 1 | 0.243102132 | 0.008084906 |
| Gm4951 | 0.251644813 | 1.154722595 | 0.019 | 0.004 | 1 | 0.004050657 | 3.644559772 | 0.049 | 0.003 | 1 | 0.251644813 | 0.008084906 |
| Nek2 | 0.287772304 | -1.430239906 | 0 | 0.021 | 1 | 0.004050657 | 3.422167351 | 0.049 | 0.003 | 1 | 0.287772304 | 0.008084906 |
| Gm16230 | 0.324474943 | 1.47665069 | 0.057 | 0.03 | 1 | 0.004050657 | 3.422167351 | 0.049 | 0.003 | 1 | 0.324474943 | 0.008084906 |
| Gm33318 | 0.413082433 | 0.154722595 | 0 | 0.013 | 1 | 0.004050657 | 3.422167351 | 0.049 | 0.003 | 1 | 0.413082433 | 0.008084906 |
| Tmem108 | 0.429421714 | 1.280253477 | 0.057 | 0.034 | 1 | 0.004050657 | 3.422167351 | 0.049 | 0.003 | 1 | 0.429421714 | 0.008084906 |
| Gm36839 | 0.736073999 | -0.304709024 | 0.019 | 0.013 | 1 | 0.004050657 | 3.422167351 | 0.049 | 0.003 | 1 | 0.736073999 | 0.008084906 |
| Gm57180 | 0.926212226 | 0.8327945 | 0.019 | 0.017 | 1 | 0.004050657 | 3.574170444 | 0.049 | 0.003 | 1 | 0.926212226 | 0.008084906 |
| Vdr | 0.004069867 | -1.556772312 | 0.113 | 0.309 | 1 | 0.051582115 | -1.605738646 | 0.073 | 0.195 | 1 | 0.051582115 | 0.008123171 |
| Slc22a15 | 0.004071846 | -1.037922483 | 0.151 | 0.364 | 1 | 0.231020377 | 0.660882077 | 0.317 | 0.239 | 1 | 0.231020377 | 0.008127112 |
| Celf2 | 0.004074734 | 1.984797593 | 0.208 | 0.076 | 1 | 0.067632597 | 0.880273572 | 0.341 | 0.225 | 1 | 0.067632597 | 0.008132864 |

| | | | | | | | | | | | | |
|---|---|---|---|---|---|---|---|---|---|---|---|---|
| Trpm6 | 0.00408354 | -3.155132668 | 0.038 | 0.199 | 1 | 0.017955451 | -3.589059905 | 0 | 0.123 | 1 | 0.017955451 | 0.008150404 |
| Zfp646 | 0.004087546 | 1.331600357 | 0.321 | 0.153 | 1 | 0.902231167 | 0.672460087 | 0.293 | 0.324 | 1 | 0.902231167 | 0.008158384 |
| Sync | 0.004088888 | 1.962077517 | 0.113 | 0.025 | 1 | 0.62803788 | 0.515276755 | 0.024 | 0.041 | 1 | 0.62803788 | 0.008161056 |
| Rgs12 | 0.01101536 | 0.891688189 | 0.226 | 0.097 | 1 | 0.004093405 | 1.30377265 | 0.317 | 0.143 | 1 | 0.01101536 | 0.008170053 |
| Rspry1 | 0.004105234 | 0.594506756 | 0.698 | 0.606 | 1 | 0.01036798 | 0.840188708 | 0.829 | 0.724 | 1 | 0.01036798 | 0.008193615 |
| Nr5a2 | 0.03562064 | 3.739685096 | 0.019 | 0 | 1 | 0.00411894 | 3.252242349 | 0.049 | 0.003 | 1 | 0.03562064 | 0.008220913 |
| Adra2c | 0.069603894 | -1.845277405 | 0.019 | 0.093 | 1 | 0.00411894 | 3.159132945 | 0.049 | 0.003 | 1 | 0.069603894 | 0.008220913 |
| Pik3c2b | 0.101874199 | 1.306725688 | 0.038 | 0.008 | 1 | 0.00411894 | 3.252242349 | 0.049 | 0.003 | 1 | 0.101874199 | 0.008220913 |
| Mfsd10 | 0.013518674 | 0.725879296 | 0.528 | 0.347 | 1 | 0.0041424 | 1.239303293 | 0.463 | 0.294 | 1 | 0.013518674 | 0.008267641 |
| Sned1 | 0.125820171 | -0.443178962 | 0.491 | 0.606 | 1 | 0.004150452 | -1.335075573 | 0.585 | 0.734 | 1 | 0.125820171 | 0.008283678 |
| Upp2 | 0.070887565 | -1.104469745 | 0.472 | 0.555 | 1 | 0.004153211 | -1.296194276 | 0.22 | 0.481 | 1 | 0.070887565 | 0.008289172 |
| Rassf5 | 0.004156565 | 1.669295768 | 0.226 | 0.093 | 1 | 0.202775599 | 0.774920572 | 0.22 | 0.143 | 1 | 0.202775599 | 0.008295853 |
| Gatad1 | 0.004161337 | 0.656603354 | 0.774 | 0.64 | 1 | 0.035249236 | -0.402622165 | 0.585 | 0.71 | 1 | 0.035249236 | 0.008305358 |
| Atg2b | 0.004164675 | 1.07065833 | 0.491 | 0.314 | 1 | 0.197177945 | -0.437802198 | 0.39 | 0.481 | 1 | 0.197177945 | 0.008312006 |
| Hdac5 | 0.004180025 | 1.115194231 | 0.396 | 0.233 | 1 | 0.37466659 | 0.73377526 | 0.39 | 0.372 | 1 | 0.37466659 | 0.008342577 |
| Cyb5r3 | 0.004180884 | 0.586114707 | 0.925 | 0.835 | 1 | 0.075437719 | 0.486448102 | 0.878 | 0.829 | 1 | 0.075437719 | 0.008344289 |
| Irgm2 | 0.923578308 | 1.523367189 | 0.17 | 0.178 | 1 | 0.004187418 | 2.066686696 | 0.293 | 0.133 | 1 | 0.923578308 | 0.008357301 |
| Gm12185 | 0.497382333 | 2.377115016 | 0.019 | 0.008 | 1 | 0.004188265 | 2.515276755 | 0.049 | 0.003 | 1 | 0.497382333 | 0.008358989 |
| Vapa | 0.016422642 | 0.300378581 | 0.981 | 0.962 | 1 | 0.004192528 | -0.335621208 | 0.976 | 0.99 | 1 | 0.016422642 | 0.008367479 |
| Socs4 | 0.084724071 | -0.598184543 | 0.377 | 0.492 | 1 | 0.004203837 | -0.840867055 | 0.341 | 0.604 | 1 | 0.084724071 | 0.008390002 |
| Peg3 | 0.004212683 | 0.327748939 | 1 | 0.962 | 1 | 0.033190579 | -0.326348069 | 1 | 0.993 | 1 | 0.033190579 | 0.008407619 |
| Nav2 | 0.004218637 | -1.447752306 | 0.528 | 0.703 | 1 | 0.081729888 | -0.156703536 | 0.659 | 0.502 | 1 | 0.081729888 | 0.008419476 |
| Flrt3 | 0.004220859 | 1.785488785 | 0.226 | 0.093 | 1 | 0.8736957 | 1.19334866 | 0.098 | 0.096 | 1 | 0.8736957 | 0.008423902 |
| Sfxn5 | 0.004224163 | 0.971858538 | 0.377 | 0.203 | 1 | 0.250554974 | -0.285722672 | 0.415 | 0.519 | 1 | 0.250554974 | 0.008430483 |
| Smg5 | 0.004249026 | 0.925903904 | 0.415 | 0.233 | 1 | 0.491836516 | 0.321505012 | 0.439 | 0.392 | 1 | 0.491836516 | 0.008479998 |
| Arhgef10l | 0.0042567 | 2.154722595 | 0.189 | 0.068 | 1 | 0.997795829 | 0.944120054 | 0.146 | 0.154 | 1 | 0.997795829 | 0.008495281 |
| Jdp2 | 0.0042591 | 1.257292328 | 0.604 | 0.449 | 1 | 0.129556874 | 0.288043071 | 0.61 | 0.505 | 1 | 0.129556874 | 0.008500061 |
| Dhx16 | 0.004274883 | 1.506194965 | 0.377 | 0.208 | 1 | 0.449073363 | 0.774195052 | 0.317 | 0.297 | 1 | 0.449073363 | 0.008531491 |
| Zbtb17 | 0.004287387 | 1.417757001 | 0.264 | 0.114 | 1 | 0.23759352 | -0.213421223 | 0.195 | 0.287 | 1 | 0.23759352 | 0.008556393 |
| Borcs6 | 0.004293149 | 1.154722595 | 0.453 | 0.263 | 1 | 0.769314839 | -0.124321002 | 0.317 | 0.338 | 1 | 0.769314839 | 0.008567867 |
| Abcg1 | 0.004303316 | 1.8327945 | 0.113 | 0.025 | 1 | 0.209595001 | 0.547698233 | 0.171 | 0.102 | 1 | 0.209595001 | 0.008588113 |
| Kif5a | 0.004305963 | 1.906795081 | 0.189 | 0.068 | 1 | 0.352764302 | 0.79281073 | 0.122 | 0.078 | 1 | 0.352764302 | 0.008593385 |
| Tab1 | 0.00430736 | 2.377115016 | 0.151 | 0.047 | 1 | 0.622636773 | -0.577832649 | 0.049 | 0.068 | 1 | 0.622636773 | 0.008596166 |
| Nars2 | 0.004313236 | 1.128250383 | 0.34 | 0.165 | 1 | 0.00724603 | 1.236300805 | 0.341 | 0.167 | 1 | 0.00724603 | 0.008607867 |
| Fam161a | 0.054508986 | 1.607234799 | 0.151 | 0.072 | 1 | 0.004317061 | 1.658867609 | 0.244 | 0.096 | 1 | 0.054508986 | 0.008615485 |
| Itga9 | 0.03434151 | 0.489386678 | 0.509 | 0.364 | 1 | 0.004318353 | 1.003776416 | 0.634 | 0.474 | 1 | 0.03434151 | 0.008618058 |
| Dnal1 | 0.004318967 | 1.154722595 | 0.321 | 0.157 | 1 | 0.194345659 | 0.83720485 | 0.293 | 0.215 | 1 | 0.194345659 | 0.00861928 |
| A630089N07Rik | 0.004321715 | 1.287989126 | 0.321 | 0.161 | 1 | 0.428748 | -0.471547856 | 0.122 | 0.174 | 1 | 0.428748 | 0.008624753 |
| Eif3k | 0.004328483 | 0.371689544 | 0.962 | 0.898 | 1 | 0.305566742 | 0.174239837 | 0.878 | 0.908 | 1 | 0.305566742 | 0.008638229 |
| Mrps31 | 0.004338574 | 1.045098104 | 0.434 | 0.258 | 1 | 0.028021332 | -0.714590787 | 0.146 | 0.321 | 1 | 0.028021332 | 0.008658325 |
| Medag | 0.004342153 | -1.470397032 | 0.264 | 0.458 | 1 | 0.165397712 | -2.328260503 | 0.146 | 0.225 | 1 | 0.165397712 | 0.008665452 |
| Jup | 0.064299403 | 1.059565362 | 0.377 | 0.271 | 1 | 0.004352054 | 1.01123425 | 0.537 | 0.352 | 1 | 0.064299403 | 0.008685169 |
| Chrd | 0.004369896 | 2.154722595 | 0.151 | 0.047 | 1 | 0.740097583 | 0.174239837 | 0.122 | 0.102 | 1 | 0.740097583 | 0.008720695 |
| Rad51d | 0.004376533 | 1.652222254 | 0.17 | 0.055 | 1 | 0.556258075 | -0.116991461 | 0.049 | 0.075 | 1 | 0.556258075 | 0.008733912 |
| Nuggc | 0.986865056 | 0.280253477 | 0.038 | 0.038 | 1 | 0.00439353 | 3.685201756 | 0.073 | 0.01 | 1 | 0.986865056 | 0.008767758 |
| Plekho1 | 0.004400494 | 0.879088152 | 0.283 | 0.119 | 1 | 0.006657446 | 1.326242931 | 0.244 | 0.102 | 1 | 0.006657446 | 0.008781624 |
| Samd8 | 0.247096803 | 0.42498857 | 0.547 | 0.492 | 1 | 0.004403803 | 0.703563966 | 0.854 | 0.648 | 1 | 0.247096803 | 0.008788212 |
| Epn1 | 0.004430558 | 0.8817041 | 0.547 | 0.373 | 1 | 0.399689624 | 0.351778023 | 0.512 | 0.457 | 1 | 0.399689624 | 0.008841486 |
| Cdk2 | 0.004431407 | 1.207190015 | 0.321 | 0.157 | 1 | 0.926045809 | 0.224227973 | 0.268 | 0.266 | 1 | 0.926045809 | 0.008843177 |
| Gm42477 | 0.030360955 | 2.739685096 | 0.038 | 0.004 | 1 | 0.004444719 | 3.83720485 | 0.073 | 0.01 | 1 | 0.030360955 | 0.008869682 |
| Gm50455 | 0.249034808 | 2.154722595 | 0.019 | 0.004 | 1 | 0.004444719 | 3.422167351 | 0.073 | 0.01 | 1 | 0.249034808 | 0.008869682 |
| Amigo2 | 0.503971666 | 1.569760094 | 0.019 | 0.008 | 1 | 0.004444719 | 3.159132945 | 0.073 | 0.01 | 1 | 0.503971666 | 0.008869682 |
| Gm3830 | 0.569866135 | -0.1672055 | 0.019 | 0.034 | 1 | 0.004444719 | 3.422167351 | 0.073 | 0.01 | 1 | 0.569866135 | 0.008869682 |
| Osgin2 | 0.004811135 | -1.484316579 | 0.283 | 0.487 | 1 | 0.004465126 | -1.376142432 | 0.22 | 0.464 | 1 | 0.004811135 | 0.008910314 |
| Jarid2 | 0.010656316 | 0.427260473 | 1 | 0.975 | 1 | 0.004465208 | 0.53452464 | 1 | 0.983 | 1 | 0.010656316 | 0.008910478 |
| Scube1 | 0.004469993 | -1.306725369 | 0.472 | 0.585 | 1 | 0.665587319 | -0.484723245 | 0.293 | 0.311 | 1 | 0.665587319 | 0.008920006 |
| Ankle2 | 0.00447478 | 0.676182153 | 0.698 | 0.547 | 1 | 0.690028362 | 0.132332885 | 0.634 | 0.628 | 1 | 0.690028362 | 0.008929537 |
| Smim27 | 0.004483936 | -1.430239906 | 0.113 | 0.301 | 1 | 0.681533985 | 0.159132945 | 0.146 | 0.181 | 1 | 0.681533985 | 0.008947766 |
| Eid2b | 0.004485665 | 2.541745718 | 0.151 | 0.047 | 1 | 0.451338156 | -0.577832649 | 0.049 | 0.082 | 1 | 0.451338156 | 0.008951208 |
| Tpmt | 0.004508777 | 1.361173472 | 0.226 | 0.089 | 1 | 0.192508176 | -0.787286015 | 0.049 | 0.116 | 1 | 0.192508176 | 0.008997225 |
| Myc | 0.312750251 | -0.24966766 | 0.434 | 0.534 | 1 | 0.004511096 | -1.631564633 | 0.268 | 0.478 | 1 | 0.312750251 | 0.009001842 |
| Ccm2 | 0.004513555 | 0.848619467 | 0.566 | 0.373 | 1 | 0.278187583 | -0.32353098 | 0.39 | 0.505 | 1 | 0.278187583 | 0.009006737 |
| Ccdc134 | 0.004518526 | 0.8327945 | 0.302 | 0.136 | 1 | 0.566034622 | 0.422167351 | 0.171 | 0.137 | 1 | 0.566034622 | 0.009016634 |
| Etnk2 | 0.004530506 | -2.220316837 | 0.019 | 0.174 | 1 | 0.799732736 | 0.930314254 | 0.049 | 0.041 | 1 | 0.799732736 | 0.009040487 |
| Pdpn | 0.004531774 | 1.389187848 | 0.264 | 0.119 | 1 | 0.016610782 | 1.079775153 | 0.463 | 0.307 | 1 | 0.016610782 | 0.009043012 |
| Wdfy2 | 0.004535488 | 1.073192709 | 0.434 | 0.258 | 1 | 0.120211655 | 0.728680393 | 0.463 | 0.375 | 1 | 0.120211655 | 0.009050405 |
| Bag5 | 0.185056067 | 0.409583069 | 0.698 | 0.657 | 1 | 0.004542016 | -0.686357106 | 0.366 | 0.659 | 1 | 0.185056067 | 0.009063402 |
| Trim26 | 0.004552602 | 1.234892943 | 0.415 | 0.246 | 1 | 0.493084007 | 0.413993419 | 0.39 | 0.369 | 1 | 0.493084007 | 0.009084479 |
| Bcl10 | 0.00753905 | 0.507394213 | 0.774 | 0.593 | 1 | 0.004556688 | 0.643896921 | 0.78 | 0.584 | 1 | 0.00753905 | 0.009092613 |
| Riox1 | 0.004558546 | 1.361173472 | 0.245 | 0.102 | 1 | 0.987521713 | 0.252242349 | 0.171 | 0.177 | 1 | 0.987521713 | 0.009096312 |
| Slc25a10 | 0.004558782 | -2.368839361 | 0.019 | 0.169 | 1 | 0.730036921 | -0.20718927 | 0.073 | 0.089 | 1 | 0.730036921 | 0.009096782 |
| Larp1 | 0.004565309 | 0.760349758 | 0.792 | 0.763 | 1 | 0.469694759 | 0.14979508 | 0.902 | 0.942 | 1 | 0.469694759 | 0.009109776 |
| Hs6st1 | 0.004574469 | 1.154722595 | 0.321 | 0.157 | 1 | 0.640310101 | 0.19334866 | 0.268 | 0.311 | 1 | 0.640310101 | 0.009128012 |
| Snhg20 | 0.348750597 | -0.636690783 | 0.226 | 0.28 | 1 | 0.00457493 | -2.410722664 | 0.024 | 0.208 | 1 | 0.348750597 | 0.00912893 |
| Spsb1 | 0.004581892 | 0.769068902 | 0.736 | 0.623 | 1 | 0.013668286 | 0.624211126 | 0.78 | 0.662 | 1 | 0.013668286 | 0.009142789 |
| Trim62 | 0.004603534 | 1.739685096 | 0.208 | 0.081 | 1 | 0.419044044 | 0.855352197 | 0.268 | 0.218 | 1 | 0.419044044 | 0.009185876 |
| Gm26787 | 0.503971666 | 1.569760094 | 0.019 | 0.008 | 1 | 0.004604972 | 2.83720485 | 0.073 | 0.01 | 1 | 0.503971666 | 0.009188739 |
| Txndc8 | 0.014846242 | 3.569760094 | 0.057 | 0.008 | 1 | 0.004605531 | 2.547698233 | 0.073 | 0.01 | 1 | 0.014846242 | 0.009189851 |
| Muc4 | 0.287761733 | -1.932740246 | 0 | 0.021 | 1 | 0.004605593 | 2.83720485 | 0.073 | 0.01 | 1 | 0.287761733 | 0.009189975 |
| Elovl2 | 0.473270726 | 1.669295768 | 0.038 | 0.021 | 1 | 0.004605593 | 2.83720485 | 0.073 | 0.01 | 1 | 0.473270726 | 0.009189975 |
| Srpx | 0.117995623 | 0.347367673 | 0.453 | 0.335 | 1 | 0.0046305 | -1.319299636 | 0.171 | 0.41 | 1 | 0.117995623 | 0.009239559 |
| Dgcr6 | 0.004636876 | 1.108428943 | 0.491 | 0.339 | 1 | 0.052298624 | 0.721727632 | 0.415 | 0.283 | 1 | 0.052298624 | 0.009252252 |
| Peds1 | 0.004647283 | -0.677400232 | 0.358 | 0.589 | 1 | 0.168079436 | -0.372248516 | 0.463 | 0.573 | 1 | 0.168079436 | 0.009272969 |
| Scaf8 | 0.004652763 | 0.442787238 | 0.962 | 0.919 | 1 | 0.836547696 | -0.116202456 | 0.976 | 0.986 | 1 | 0.836547696 | 0.009283877 |
| Nrg2 | 0.917883255 | 0.569760094 | 0.019 | 0.021 | 1 | 0.004659332 | 2.574170444 | 0.073 | 0.01 | 1 | 0.917883255 | 0.009296954 |
| 2810454H06Rik | 0.004673776 | 2.154722595 | 0.151 | 0.047 | 1 | 0.551268628 | 1.322631677 | 0.073 | 0.051 | 1 | 0.551268628 | 0.009325708 |
| Slc10a7 | 0.004675286 | 0.710310493 | 0.585 | 0.373 | 1 | 0.254621544 | 0.209468325 | 0.683 | 0.573 | 1 | 0.254621544 | 0.009328713 |
| Rab3gap2 | 0.004680246 | -0.942138944 | 0.509 | 0.674 | 1 | 0.025585763 | -0.449852785 | 0.61 | 0.741 | 1 | 0.025585763 | 0.009338588 |
| Rps21 | 0.004682775 | -0.278347665 | 1 | 0.992 | 1 | 0.037868444 | -0.265221734 | 1 | 0.99 | 1 | 0.037868444 | 0.009343621 |
| Clec2d | 0.042463172 | 2.47665069 | 0.057 | 0.013 | 1 | 0.004696468 | 2.515276755 | 0.146 | 0.041 | 1 | 0.042463172 | 0.00937088 |
| Tor1b | 0.004723809 | 0.770579103 | 0.604 | 0.424 | 1 | 0.579246292 | 0.119792053 | 0.341 | 0.399 | 1 | 0.579246292 | 0.009425303 |
| Ppp5c | 0.004737658 | 1.023478062 | 0.472 | 0.305 | 1 | 0.233870653 | -0.235203646 | 0.268 | 0.362 | 1 | 0.233870653 | 0.00945287 |
| Nr3c2 | 0.004749341 | -0.771276824 | 0.094 | 0.305 | 1 | 0.518228813 | 0.332584457 | 0.366 | 0.314 | 1 | 0.518228813 | 0.009476125 |
| Tle2 | 0.004754083 | 1.361173472 | 0.226 | 0.089 | 1 | 0.182508323 | 1.212713985 | 0.171 | 0.106 | 1 | 0.182508323 | 0.009485564 |
| Loxl2 | 0.004765601 | -0.783336019 | 0.453 | 0.619 | 1 | 0.862481655 | 0.788295249 | 0.195 | 0.181 | 1 | 0.862481655 | 0.009508491 |
| Hp | 0.03562064 | 4.154722595 | 0.019 | 0 | 1 | 0.004769158 | 1.136765132 | 0.073 | 0.01 | 1 | 0.03562064 | 0.009515572 |
| Dip2b | 0.157811737 | 0.218084656 | 0.604 | 0.483 | 1 | 0.004769905 | 0.668724711 | 0.854 | 0.672 | 1 | 0.157811737 | 0.009517059 |
| Rhbdd1 | 0.004771492 | -1.054730771 | 0.17 | 0.377 | 1 | 0.343596392 | 0.328863631 | 0.22 | 0.311 | 1 | 0.343596392 | 0.009520216 |

| | | | | | | | | | | | | |
|---|---|---|---|---|---|---|---|---|---|---|---|---|
| Erp27 | 0.005723503 | -2.845277405 | 0.038 | 0.191 | 1 | 0.004788837 | -3.466575898 | 0 | 0.167 | 1 | 0.005723503 | 0.009554741 |
| Wsb2 | 0.433360915 | -0.130679624 | 0.509 | 0.602 | 1 | 0.004793611 | -0.856879373 | 0.512 | 0.683 | 1 | 0.433360915 | 0.009564243 |
| Auh | 0.004803582 | 0.991223863 | 0.509 | 0.339 | 1 | 0.005430621 | 1.585141562 | 0.463 | 0.304 | 1 | 0.005430621 | 0.00958409 |
| Samd4b | 0.056920607 | -0.311187234 | 0.736 | 0.826 | 1 | 0.004821069 | -0.661695052 | 0.634 | 0.823 | 1 | 0.056920607 | 0.009618895 |
| Furin | 0.004822861 | -0.696912889 | 0.585 | 0.767 | 1 | 0.112933374 | -0.300795384 | 0.78 | 0.853 | 1 | 0.112933374 | 0.009622462 |
| Mrpl22 | 0.004832049 | 1.340278248 | 0.358 | 0.186 | 1 | 0.402458014 | -0.592782991 | 0.171 | 0.222 | 1 | 0.402458014 | 0.00964075 |
| Lrp8 | 0.094468155 | -0.638826528 | 0.17 | 0.292 | 1 | 0.004836008 | -0.56878751 | 0.122 | 0.372 | 1 | 0.094468155 | 0.00964863 |
| Arglu1 | 0.004839232 | 0.508359549 | 0.943 | 0.89 | 1 | 0.007319592 | -0.445729113 | 0.878 | 0.959 | 1 | 0.007319592 | 0.009655046 |
| Fbl | 0.004844527 | 0.774874524 | 0.66 | 0.521 | 1 | 0.023627217 | -0.285651898 | 0.488 | 0.717 | 1 | 0.023627217 | 0.009665585 |
| Dennd1b | 0.004848575 | 0.983404358 | 0.642 | 0.555 | 1 | 0.093498444 | 0.581190872 | 0.756 | 0.683 | 1 | 0.093498444 | 0.009673642 |
| Rnmt | 0.004872177 | 0.808662837 | 0.642 | 0.517 | 1 | 0.382478282 | -0.217560954 | 0.463 | 0.543 | 1 | 0.382478282 | 0.009720617 |
| Aldh3a1 | 0.147723217 | -0.783876861 | 0.075 | 0.153 | 1 | 0.004875639 | 0.953718859 | 0.415 | 0.222 | 1 | 0.147723217 | 0.009727506 |
| Naalad2 | 0.00489581 | -0.932740246 | 0.113 | 0.318 | 1 | 0.163755924 | -1.144647803 | 0.073 | 0.154 | 1 | 0.163755924 | 0.009767651 |
| Zfp84 | 0.004911436 | 1.185096244 | 0.321 | 0.161 | 1 | 0.148061752 | -0.439635356 | 0.195 | 0.314 | 1 | 0.148061752 | 0.009798749 |
| L2hgdh | 0.004917598 | 1.533234218 | 0.283 | 0.136 | 1 | 0.145753985 | -0.970150072 | 0.073 | 0.16 | 1 | 0.145753985 | 0.009811014 |
| Mapkap1 | 0.004921076 | 0.608440562 | 0.774 | 0.716 | 1 | 0.143976664 | 0.378914153 | 0.854 | 0.792 | 1 | 0.143976664 | 0.009817934 |
| Cox18 | 0.004925017 | 1.134823037 | 0.415 | 0.233 | 1 | 0.045020781 | 0.966487867 | 0.415 | 0.29 | 1 | 0.045020781 | 0.009825778 |
| Adss | 0.015038268 | 0.341925588 | 0.943 | 0.894 | 1 | 0.004926141 | 0.485339471 | 0.927 | 0.809 | 1 | 0.015038268 | 0.009828016 |
| 9430037G07Rik | 0.111557832 | 1.8327945 | 0.075 | 0.03 | 1 | 0.004939943 | 2.029849928 | 0.073 | 0.01 | 1 | 0.111557832 | 0.009855482 |
| Naa16 | 0.004944281 | 0.712500266 | 0.528 | 0.335 | 1 | 0.138888592 | -0.53202896 | 0.39 | 0.505 | 1 | 0.138888592 | 0.009864117 |
| Bicra | 0.004957381 | 1.354031403 | 0.302 | 0.148 | 1 | 0.576699669 | 0.174239837 | 0.39 | 0.341 | 1 | 0.576699669 | 0.009890186 |
| Ndufc2 | 0.004962557 | 0.40955773 | 0.943 | 0.89 | 1 | 0.185430912 | -0.219549928 | 0.829 | 0.874 | 1 | 0.185430912 | 0.009900488 |
| Pcbp3 | 0.004962974 | 1.173831418 | 0.34 | 0.178 | 1 | 0.347711192 | 0.595130063 | 0.244 | 0.177 | 1 | 0.347711192 | 0.009901317 |
| Capza2 | 0.004963166 | 0.451253913 | 0.943 | 0.877 | 1 | 0.364277924 | 0.22461123 | 0.927 | 0.952 | 1 | 0.364277924 | 0.009901698 |
| H2-D1 | 0.004978433 | 0.540012751 | 0.906 | 0.712 | 1 | 0.298056401 | 0.281388695 | 0.878 | 0.812 | 1 | 0.298056401 | 0.009932082 |
| Washc1 | 0.004980798 | 1.228723176 | 0.302 | 0.144 | 1 | 0.834091835 | 0.100239256 | 0.146 | 0.164 | 1 | 0.834091835 | 0.009936787 |
| Ube2f | 0.05248083 | -0.364110317 | 0.717 | 0.809 | 1 | 0.004987485 | -0.344437536 | 0.78 | 0.922 | 1 | 0.05248083 | 0.009950095 |
| Phf8 | 0.373389878 | 0.383541285 | 0.547 | 0.525 | 1 | 0.005020412 | 0.857013331 | 0.805 | 0.635 | 1 | 0.373389878 | 0.010015619 |
| Gmip | 0.005030382 | 0.874614676 | 0.226 | 0.085 | 1 | 0.322447413 | 0.614812428 | 0.195 | 0.137 | 1 | 0.322447413 | 0.01003546 |
| Cope | 0.710357573 | 0.106359573 | 1 | 0.907 | 1 | 0.005033705 | -0.588655619 | 0.829 | 0.898 | 1 | 0.710357573 | 0.010042072 |
| Mgll | 0.141358378 | -0.982780929 | 0.566 | 0.61 | 1 | 0.005042353 | -0.920492186 | 0.463 | 0.662 | 1 | 0.141358378 | 0.01005928 |
| Sox2 | 0.005048979 | -3.517702747 | 0.019 | 0.165 | 1 | 0.017259156 | 1.962735732 | 0.22 | 0.096 | 1 | 0.017259156 | 0.010072466 |
| Lss | 0.005049602 | -1.52334931 | 0.113 | 0.301 | 1 | 0.079005389 | -1.665295491 | 0.049 | 0.147 | 1 | 0.079005389 | 0.010073705 |
| Gpn3 | 0.005077446 | 0.963779812 | 0.547 | 0.364 | 1 | 0.64354584 | -0.172779239 | 0.293 | 0.328 | 1 | 0.64354584 | 0.010129112 |
| Npdc1 | 0.00507846 | 0.402063194 | 0.925 | 0.852 | 1 | 0.223790263 | 0.216514645 | 0.829 | 0.809 | 1 | 0.223790263 | 0.01013113 |
| Nadk | 0.005080039 | 0.739685096 | 0.623 | 0.424 | 1 | 0.357466396 | 0.307585383 | 0.659 | 0.614 | 1 | 0.357466396 | 0.010134271 |
| Rheb | 0.005088995 | -0.478004056 | 0.943 | 0.97 | 1 | 0.027570471 | -0.375788874 | 0.902 | 0.959 | 1 | 0.027570471 | 0.010152093 |
| Magohb | 0.005104639 | 1.052360877 | 0.434 | 0.263 | 1 | 0.102938638 | -0.813049111 | 0.195 | 0.311 | 1 | 0.102938638 | 0.01018322 |
| Dpf2 | 0.005106389 | 0.994257923 | 0.434 | 0.258 | 1 | 0.009945016 | -0.741416124 | 0.268 | 0.498 | 1 | 0.009945016 | 0.010186703 |
| Trpc4ap | 0.005107401 | 0.621848605 | 0.792 | 0.763 | 1 | 0.607267303 | 0.215089351 | 0.78 | 0.867 | 1 | 0.607267303 | 0.010188716 |
| mt-Nd6 | 0.005107817 | -1.089899774 | 0.283 | 0.479 | 1 | 0.005222653 | -0.982974113 | 0.171 | 0.41 | 1 | 0.005222653 | 0.010189544 |
| Tbx3os1 | 0.005127081 | 2.232725107 | 0.17 | 0.059 | 1 | 0.520650529 | 1.83720485 | 0.049 | 0.031 | 1 | 0.520650529 | 0.010227875 |
| Cd63 | 0.005127386 | 0.311743763 | 1 | 0.992 | 1 | 0.028665783 | -0.256825303 | 1 | 0.993 | 1 | 0.028665783 | 0.010228481 |
| Limch1 | 0.005134476 | -1.980436988 | 0.038 | 0.203 | 1 | 0.029635892 | -2.266082959 | 0.098 | 0.246 | 1 | 0.029635892 | 0.010242589 |
| Lpin2 | 0.005140942 | -1.396977452 | 0.623 | 0.771 | 1 | 0.180959155 | -0.513911713 | 0.805 | 0.877 | 1 | 0.180959155 | 0.010255455 |
| Rbak | 0.005160211 | 1.228723176 | 0.245 | 0.102 | 1 | 0.085258157 | -0.695290231 | 0.098 | 0.215 | 1 | 0.085258157 | 0.010293795 |
| 2310010J17Rik | 0.005169177 | -1.337130502 | 0.189 | 0.377 | 1 | 0.551319412 | -0.116991461 | 0.195 | 0.239 | 1 | 0.551319412 | 0.010311634 |
| Mark2 | 0.005179018 | 0.614672516 | 0.774 | 0.589 | 1 | 0.18890882 | -0.283424855 | 0.756 | 0.863 | 1 | 0.18890882 | 0.010331214 |
| Sertad4 | 0.00518074 | 0.891688189 | 0.491 | 0.297 | 1 | 0.949117626 | 0.334704509 | 0.39 | 0.461 | 1 | 0.949117626 | 0.01033464 |
| Rmc1 | 0.738448882 | 0.176096246 | 0.245 | 0.22 | 1 | 0.005180811 | -1.25871957 | 0.146 | 0.372 | 1 | 0.738448882 | 0.010334782 |
| Krt17 | 0.005187746 | 0.997181318 | 0.208 | 0.076 | 1 | 0.093143441 | 1.014082612 | 0.244 | 0.147 | 1 | 0.093143441 | 0.010348579 |
| Gcg | 0.005191326 | 1.906795081 | 0.151 | 0.047 | 1 | 0.962027031 | 0.83720485 | 0.049 | 0.051 | 1 | 0.962027031 | 0.010355702 |
| Serf2 | 0.508275938 | -0.129056925 | 1 | 0.983 | 1 | 0.005217868 | -0.338053596 | 1 | 0.997 | 1 | 0.508275938 | 0.010408511 |
| Edf1 | 0.005218286 | 0.582816247 | 0.887 | 0.805 | 1 | 0.201959505 | -0.155261478 | 0.683 | 0.823 | 1 | 0.201959505 | 0.010409341 |
| D2hgdh | 0.329936506 | 1.154722595 | 0.094 | 0.059 | 1 | 0.00522865 | 1.989207943 | 0.171 | 0.055 | 1 | 0.329936506 | 0.010429962 |
| Gria4 | 0.005231416 | 3.598329246 | 0.151 | 0.047 | 1 | 0.070532845 | 2.016174991 | 0.146 | 0.068 | 1 | 0.070532845 | 0.010435464 |
| Tmx2 | 0.005232255 | 1.082572809 | 0.453 | 0.288 | 1 | 0.349006935 | 0.369056014 | 0.293 | 0.222 | 1 | 0.349006935 | 0.010437134 |
| Ino80c | 0.234285911 | 0.373362881 | 0.245 | 0.169 | 1 | 0.00527153 | -3.0454382 | 0 | 0.164 | 1 | 0.234285911 | 0.01051527 |
| Mlec | 0.005273025 | -0.506932663 | 0.792 | 0.852 | 1 | 0.870648785 | 0.132807118 | 0.805 | 0.826 | 1 | 0.870648785 | 0.010518245 |
| Gm46440 | 0.005315743 | 0.932330173 | 0.17 | 0.055 | 1 | 0.566147642 | 0.667279848 | 0.073 | 0.051 | 1 | 0.566147642 | 0.010603229 |
| Dnajb14 | 0.005330793 | 0.8327945 | 0.604 | 0.504 | 1 | 0.100829109 | 1.022919434 | 0.659 | 0.57 | 1 | 0.100829109 | 0.01063317 |
| Txnl4a | 0.224770723 | 0.389187848 | 0.547 | 0.487 | 1 | 0.005335449 | -0.810767448 | 0.366 | 0.604 | 1 | 0.224770723 | 0.010642431 |
| Bivm | 0.005345678 | 1.739685096 | 0.189 | 0.068 | 1 | 0.603866956 | 0.877846834 | 0.122 | 0.099 | 1 | 0.603866956 | 0.01066278 |
| Ipo4 | 0.179231483 | 0.470224421 | 0.396 | 0.301 | 1 | 0.005353749 | -1.213421223 | 0.098 | 0.321 | 1 | 0.179231483 | 0.010678836 |
| Clca2 | 0.005353775 | -2.974560422 | 0 | 0.131 | 1 | 0.430584486 | 3.159132945 | 0.024 | 0.01 | 1 | 0.430584486 | 0.010678888 |
| Enoph1 | 0.005371021 | 1.444229212 | 0.283 | 0.136 | 1 | 0.456750666 | 1.052933541 | 0.22 | 0.181 | 1 | 0.456750666 | 0.010713194 |
| Trf | 0.212443871 | -2.61890147 | 0.17 | 0.229 | 1 | 0.005376942 | -3.67058979 | 0.024 | 0.201 | 1 | 0.212443871 | 0.010724972 |
| Letmd1 | 0.005388772 | 1.569760094 | 0.189 | 0.068 | 1 | 0.401188856 | -0.484723245 | 0.073 | 0.116 | 1 | 0.401188856 | 0.010748505 |
| Gm11655 | 0.005388824 | -3.130679624 | 0 | 0.131 | 1 | 0.085385065 | -1.80665134 | 0 | 0.068 | 1 | 0.085385065 | 0.010748608 |
| Fpgs | 0.005397267 | 1.680791406 | 0.226 | 0.093 | 1 | 0.25732084 | 1.071670103 | 0.195 | 0.14 | 1 | 0.25732084 | 0.010765403 |
| Gm45187 | 0.005397705 | -3.237594828 | 0 | 0.131 | 1 | 0.441381862 | 1.515276755 | 0.024 | 0.01 | 1 | 0.441381862 | 0.010766275 |
| Oaz2 | 0.005405586 | 0.759862977 | 0.642 | 0.547 | 1 | 0.971933651 | 0.116358921 | 0.634 | 0.676 | 1 | 0.971933651 | 0.010781952 |
| Abhd4 | 0.005409231 | 1.363309217 | 0.302 | 0.148 | 1 | 0.596943695 | 0.377773231 | 0.244 | 0.208 | 1 | 0.596943695 | 0.010789202 |
| St8sia6 | 0.005411193 | -3.678167419 | 0 | 0.131 | 1 | 0.520570945 | 0.898605394 | 0.024 | 0.048 | 1 | 0.520570945 | 0.010793105 |
| Ormdl3 | 0.050288704 | 0.417757001 | 0.717 | 0.695 | 1 | 0.005414884 | 0.493720612 | 0.854 | 0.724 | 1 | 0.050288704 | 0.010800447 |
| Klk10 | 0.005427263 | -5.481902026 | 0 | 0.131 | 1 | 0.286413984 | -0.747757651 | 0 | 0.027 | 1 | 0.286413984 | 0.01082507 |
| Vrk1 | 0.005430487 | 1.337586652 | 0.245 | 0.106 | 1 | 0.058669199 | 1.430029468 | 0.268 | 0.164 | 1 | 0.058669199 | 0.010831484 |
| Spg7 | 0.00544725 | 0.743527162 | 0.566 | 0.373 | 1 | 0.640351757 | 0.191212916 | 0.39 | 0.358 | 1 | 0.640351757 | 0.010864827 |
| Bin1 | 0.005450485 | 0.805573031 | 0.491 | 0.309 | 1 | 0.412040384 | 0.398093216 | 0.341 | 0.287 | 1 | 0.412040384 | 0.010871262 |
| Zc2hc1c | 0.762480493 | 0.739685096 | 0.038 | 0.03 | 1 | 0.005450634 | 1.83720485 | 0.146 | 0.041 | 1 | 0.762480493 | 0.010871559 |
| S100a4 | 0.600739644 | -1.193677711 | 0.566 | 0.517 | 1 | 0.005455507 | 1.017286466 | 0.561 | 0.379 | 1 | 0.600739644 | 0.010881251 |
| Ccdc51 | 0.005461224 | 1.932330173 | 0.151 | 0.047 | 1 | 0.077048992 | 1.679663573 | 0.146 | 0.068 | 1 | 0.077048992 | 0.010892623 |
| Cuedc2 | 0.005469806 | 0.731511164 | 0.66 | 0.508 | 1 | 0.878131954 | 0.143307978 | 0.463 | 0.478 | 1 | 0.878131954 | 0.010909694 |
| Zfp592 | 0.005473614 | 0.814298156 | 0.585 | 0.407 | 1 | 0.148331642 | -0.380318532 | 0.61 | 0.686 | 1 | 0.148331642 | 0.010917267 |
| Mcur1 | 0.005473874 | 0.997181318 | 0.434 | 0.271 | 1 | 0.243611421 | -0.581296505 | 0.244 | 0.317 | 1 | 0.243611421 | 0.010917786 |
| Cluap1 | 0.005478766 | 1.310373465 | 0.415 | 0.25 | 1 | 0.30273245 | -0.265156868 | 0.244 | 0.324 | 1 | 0.30273245 | 0.010927514 |
| Npr3 | 0.00548326 | -1.430239906 | 0.075 | 0.263 | 1 | 0.494347488 | -0.555112573 | 0.024 | 0.048 | 1 | 0.494347488 | 0.010936454 |
| Med21 | 0.005492993 | -0.849279336 | 0.472 | 0.703 | 1 | 0.147932411 | -0.616751639 | 0.39 | 0.498 | 1 | 0.147932411 | 0.010955814 |
| Ola1 | 0.005497086 | 0.71495801 | 0.717 | 0.623 | 1 | 0.460777109 | 0.271452953 | 0.585 | 0.652 | 1 | 0.460777109 | 0.010963955 |
| Ttc21b | 0.005512899 | 1.254258268 | 0.226 | 0.089 | 1 | 0.615676296 | 1.043655727 | 0.146 | 0.126 | 1 | 0.615676296 | 0.010995405 |
| Rad23a | 0.005516323 | 0.770058745 | 0.679 | 0.521 | 1 | 0.201964146 | 0.515276755 | 0.561 | 0.549 | 1 | 0.201964146 | 0.011002217 |
| Tlr2 | 0.838383762 | -0.260314904 | 0.208 | 0.186 | 1 | 0.005523595 | -1.938899138 | 0.098 | 0.297 | 1 | 0.838383762 | 0.011016681 |
| Tor1aip2 | 0.005526988 | 0.574158558 | 0.906 | 0.89 | 1 | 0.131453933 | -0.263595791 | 0.878 | 0.945 | 1 | 0.131453933 | 0.011023428 |
| Ubqln1 | 0.005528529 | 0.661183122 | 0.717 | 0.538 | 1 | 0.661082032 | 0.171079496 | 0.756 | 0.771 | 1 | 0.661082032 | 0.011026493 |
| Tbck | 0.005544446 | 0.888828535 | 0.434 | 0.242 | 1 | 0.539793923 | 0.527349587 | 0.439 | 0.416 | 1 | 0.539793923 | 0.011058151 |
| Znrf2 | 0.005544965 | 0.579813759 | 0.528 | 0.343 | 1 | 0.388508076 | -0.167795831 | 0.439 | 0.536 | 1 | 0.388508076 | 0.011059184 |
| Sass6 | 0.005548004 | 0.8327945 | 0.245 | 0.102 | 1 | 0.280957439 | -1.16279515 | 0.073 | 0.13 | 1 | 0.280957439 | 0.011065228 |
| Zfhx3 | 0.098658709 | 0.464523878 | 0.925 | 0.907 | 1 | 0.005548035 | 0.546749403 | 0.951 | 0.949 | 1 | 0.098658709 | 0.011065269 |

| | | | | | | | | | | | | |
|---|---|---|---|---|---|---|---|---|---|---|---|---|
| Trim2 | 0.005576687 | 0.695290976 | 0.566 | 0.381 | 1 | 0.049927598 | -0.626079094 | 0.39 | 0.567 | 1 | 0.049927598 | 0.011122275 |
| Snrpd3 | 0.005580541 | 0.549001534 | 0.849 | 0.758 | 1 | 0.030890246 | -0.495370489 | 0.683 | 0.857 | 1 | 0.030890246 | 0.011129939 |
| Mvp | 0.005584313 | 1.050385935 | 0.434 | 0.254 | 1 | 0.435219597 | 0.558345477 | 0.366 | 0.324 | 1 | 0.435219597 | 0.011137442 |
| Psme2 | 0.005591695 | 0.68934171 | 0.679 | 0.517 | 1 | 0.561218343 | 0.202773673 | 0.61 | 0.58 | 1 | 0.561218343 | 0.011152123 |
| Hspb1 | 0.02578573 | -1.320254729 | 0.717 | 0.725 | 1 | 0.005593445 | 1.140517267 | 0.805 | 0.761 | 1 | 0.02578573 | 0.011155603 |
| Neb | 0.005593679 | -3.260314904 | 0.038 | 0.191 | 1 | 0.542636497 | 0.930314254 | 0.049 | 0.031 | 1 | 0.542636497 | 0.01115607 |
| Lrrc75aos2 | 0.005597214 | 2.154722595 | 0.151 | 0.047 | 1 | 0.69118247 | 0.930314254 | 0.098 | 0.082 | 1 | 0.69118247 | 0.011163099 |
| Ramp2 | 0.005601557 | 1.166795427 | 0.453 | 0.297 | 1 | 0.366962015 | -0.221688839 | 0.171 | 0.242 | 1 | 0.366962015 | 0.011171737 |
| Casp8 | 0.005602027 | 1.347367673 | 0.34 | 0.182 | 1 | 0.11814576 | -0.863234868 | 0.146 | 0.256 | 1 | 0.11814576 | 0.011172672 |
| Dusp23 | 0.577491074 | -0.268488836 | 0.151 | 0.182 | 1 | 0.005619562 | -2.638528581 | 0.024 | 0.201 | 1 | 0.577491074 | 0.011207544 |
| Rnf135 | 0.005633725 | 1.776210972 | 0.151 | 0.047 | 1 | 0.142325802 | 1.515276755 | 0.098 | 0.044 | 1 | 0.142325802 | 0.011235711 |
| Exoc7 | 0.005647704 | 0.844382474 | 0.396 | 0.212 | 1 | 0.09620671 | 0.661355014 | 0.39 | 0.273 | 1 | 0.09620671 | 0.011263512 |
| Gm16093 | 0.005675991 | 1.502645898 | 0.264 | 0.127 | 1 | 0.612475434 | 0.486707603 | 0.098 | 0.126 | 1 | 0.612475434 | 0.011319764 |
| Akna | 0.005682036 | 1.874614676 | 0.151 | 0.047 | 1 | 0.197390034 | 1.679663573 | 0.122 | 0.068 | 1 | 0.197390034 | 0.011331786 |
| Zfp825 | 0.005685513 | 1.569760094 | 0.226 | 0.093 | 1 | 0.024064336 | -2.654648247 | 0 | 0.113 | 1 | 0.024064336 | 0.011338701 |
| Rnf34 | 0.005723837 | -0.87368582 | 0.472 | 0.644 | 1 | 0.147830481 | -0.34302516 | 0.415 | 0.529 | 1 | 0.147830481 | 0.011414912 |
| Cmya5 | 0.111293723 | 2.154722595 | 0.094 | 0.042 | 1 | 0.00573384 | 1.46797104 | 0.146 | 0.041 | 1 | 0.111293723 | 0.011434802 |
| Tbc1d8 | 0.09620921 | 0.906795081 | 0.113 | 0.051 | 1 | 0.005751791 | 1.661355014 | 0.317 | 0.154 | 1 | 0.09620921 | 0.011470499 |
| Cc2d1a | 0.005771321 | 1.739685096 | 0.151 | 0.047 | 1 | 0.993947333 | 0.152706676 | 0.122 | 0.123 | 1 | 0.993947333 | 0.011509333 |
| Ttc5 | 0.005774097 | 0.970298024 | 0.472 | 0.305 | 1 | 0.707758356 | 0.260855479 | 0.39 | 0.365 | 1 | 0.707758356 | 0.011514854 |
| Slc25a16 | 0.730457285 | 0.458728782 | 0.283 | 0.271 | 1 | 0.00578209 | 1.239642312 | 0.439 | 0.27 | 1 | 0.730457285 | 0.011530748 |
| Itga10 | 0.005782954 | -0.493665286 | 0.66 | 0.809 | 1 | 0.131081196 | -0.373942814 | 0.732 | 0.863 | 1 | 0.131081196 | 0.011532466 |
| Dcaf17 | 0.005784638 | 0.80108564 | 0.34 | 0.169 | 1 | 0.920100572 | 0.146889349 | 0.171 | 0.184 | 1 | 0.920100572 | 0.011535815 |
| Zfp566 | 0.00578946 | 1.502645898 | 0.189 | 0.068 | 1 | 0.685477245 | 0.557096931 | 0.122 | 0.102 | 1 | 0.685477245 | 0.011545402 |
| Sacs | 0.005793557 | 1.252019796 | 0.264 | 0.119 | 1 | 0.021871788 | 1.289717055 | 0.22 | 0.099 | 1 | 0.021871788 | 0.011553548 |
| 2510009E07Rik | 0.005798651 | 0.813031245 | 0.566 | 0.364 | 1 | 0.356281008 | 0.179227552 | 0.659 | 0.597 | 1 | 0.356281008 | 0.011563678 |
| Gm17477 | 0.920805363 | 0.417757001 | 0.019 | 0.021 | 1 | 0.005810809 | 1.564186355 | 0.171 | 0.055 | 1 | 0.920805363 | 0.011587853 |
| Aarsd1 | 0.112947894 | 1.055186921 | 0.34 | 0.254 | 1 | 0.005811801 | 1.399083737 | 0.366 | 0.201 | 1 | 0.112947894 | 0.011589824 |
| Wdr45b | 0.005816248 | 0.587907441 | 0.717 | 0.547 | 1 | 0.570609048 | 0.264626074 | 0.732 | 0.775 | 1 | 0.570609048 | 0.011598667 |
| Isg20l2 | 0.005824689 | 1.067259754 | 0.509 | 0.339 | 1 | 0.163900278 | -0.314104473 | 0.366 | 0.474 | 1 | 0.163900278 | 0.01161545 |
| Rufy4 | 0.005839063 | -2.430239906 | 0.038 | 0.195 | 1 | 0.051682597 | -2.352619709 | 0.024 | 0.126 | 1 | 0.051682597 | 0.011644032 |
| Topbp1 | 0.005841211 | 0.669295768 | 0.396 | 0.22 | 1 | 0.899920228 | 0.564186355 | 0.293 | 0.307 | 1 | 0.899920228 | 0.011648303 |
| Npepl1 | 0.005843397 | 1.569760094 | 0.264 | 0.123 | 1 | 0.551731666 | 0.238167164 | 0.22 | 0.283 | 1 | 0.551731666 | 0.011652648 |
| Orc3 | 0.005843759 | 0.683416876 | 0.509 | 0.326 | 1 | 0.266108896 | 0.980162804 | 0.39 | 0.352 | 1 | 0.266108896 | 0.011653368 |
| Foxp1 | 0.005852068 | 0.313420341 | 1 | 0.97 | 1 | 0.118218742 | 0.197371896 | 1 | 0.993 | 1 | 0.118218742 | 0.011669889 |
| 2700062C07Rik | 0.551664624 | 0.33529484 | 0.245 | 0.212 | 1 | 0.005858506 | -3.093532488 | 0 | 0.16 | 1 | 0.551664624 | 0.011682689 |
| Crk | 0.005864795 | 0.603607865 | 0.83 | 0.678 | 1 | 0.068863355 | -0.268589814 | 0.829 | 0.898 | 1 | 0.068863355 | 0.011695194 |
| Asb7 | 0.005873636 | 1.129187503 | 0.34 | 0.174 | 1 | 0.756975181 | 0.464650682 | 0.293 | 0.304 | 1 | 0.756975181 | 0.011712773 |
| Aldh6a1 | 0.207164383 | -0.54088397 | 0.302 | 0.386 | 1 | 0.005875892 | -1.20718927 | 0.195 | 0.423 | 1 | 0.207164383 | 0.011717258 |
| Celf4 | 0.052258237 | -0.814903756 | 0.132 | 0.254 | 1 | 0.005884057 | -3.271319607 | 0 | 0.16 | 1 | 0.052258237 | 0.011733492 |
| Dicer1 | 0.005886989 | 0.642074299 | 0.66 | 0.475 | 1 | 0.142105791 | -0.378184184 | 0.659 | 0.768 | 1 | 0.142105791 | 0.011739321 |
| Stx6 | 0.005888223 | 1.013168451 | 0.585 | 0.453 | 1 | 0.293142743 | 0.75474269 | 0.561 | 0.515 | 1 | 0.293142743 | 0.011741774 |
| Bckdha | 0.005891316 | 1.361173472 | 0.34 | 0.182 | 1 | 0.047359486 | 1.01123425 | 0.341 | 0.208 | 1 | 0.047359486 | 0.011747924 |
| Rabep2 | 0.005906529 | 1.017219071 | 0.358 | 0.186 | 1 | 0.403704125 | 0.440314697 | 0.244 | 0.188 | 1 | 0.403704125 | 0.011778172 |
| Psmd2 | 0.005912857 | 0.561833178 | 0.811 | 0.712 | 1 | 0.031486898 | -0.385913091 | 0.732 | 0.863 | 1 | 0.031486898 | 0.011790752 |
| 4933407K13Rik | 0.005917709 | 1.454282877 | 0.264 | 0.123 | 1 | 0.327264744 | 0.788841828 | 0.268 | 0.212 | 1 | 0.327264744 | 0.011800399 |
| Tfb2m | 0.0059195 | 0.927312099 | 0.491 | 0.326 | 1 | 0.238015051 | 0.736404209 | 0.415 | 0.362 | 1 | 0.238015051 | 0.011803959 |
| Mrpl40 | 0.005934491 | 0.865215978 | 0.453 | 0.271 | 1 | 0.092035807 | 0.962735732 | 0.293 | 0.184 | 1 | 0.092035807 | 0.011833764 |
| Gm28875 | 0.005938208 | 1.669295768 | 0.151 | 0.047 | 1 | 0.369322099 | 0.83720485 | 0.146 | 0.102 | 1 | 0.369322099 | 0.011841154 |
| Smg7 | 0.005962489 | 0.547305243 | 0.736 | 0.564 | 1 | 0.048670857 | 0.342682567 | 0.829 | 0.751 | 1 | 0.048670857 | 0.011889426 |
| Azi2 | 0.005970154 | 0.68975987 | 0.679 | 0.517 | 1 | 0.759775135 | 0.27208716 | 0.732 | 0.683 | 1 | 0.759775135 | 0.011904664 |
| Tex261 | 0.005975419 | 0.468222068 | 0.792 | 0.653 | 1 | 0.446120578 | -0.123365005 | 0.634 | 0.689 | 1 | 0.446120578 | 0.011915132 |
| Plxna4 | 0.925553644 | -0.639693272 | 0.132 | 0.136 | 1 | 0.005976174 | -1.294823544 | 0.22 | 0.451 | 1 | 0.925553644 | 0.011916634 |
| Dusp8 | 0.00598644 | -2.237594828 | 0.038 | 0.191 | 1 | 0.461502451 | -0.385187572 | 0.122 | 0.167 | 1 | 0.461502451 | 0.011937042 |
| Smad3 | 0.00599446 | 0.647196688 | 0.868 | 0.733 | 1 | 0.338439893 | 0.169380637 | 0.927 | 0.826 | 1 | 0.338439893 | 0.011952986 |
| Vegfd | 0.00599812 | -2.974560422 | 0.019 | 0.161 | 1 | 0.560617452 | -0.221688839 | 0.024 | 0.044 | 1 | 0.560617452 | 0.011960262 |
| Usf2 | 0.882776336 | 0.162759914 | 0.472 | 0.513 | 1 | 0.006003845 | -0.662027777 | 0.341 | 0.608 | 1 | 0.882776336 | 0.011971644 |
| Sgta | 0.006021816 | 0.695290976 | 0.566 | 0.403 | 1 | 0.048397616 | -0.722490892 | 0.463 | 0.567 | 1 | 0.048397616 | 0.01200737 |
| Pbx1 | 0.006055359 | 0.343167684 | 0.962 | 0.945 | 1 | 0.062367647 | 0.213751373 | 0.951 | 0.962 | 1 | 0.062367647 | 0.012074051 |
| Krt5 | 0.901698096 | -0.430239906 | 0.038 | 0.034 | 1 | 0.006067047 | 0.948626132 | 0.39 | 0.205 | 1 | 0.901698096 | 0.012097284 |
| Terf2ip | 0.006070344 | 0.877188619 | 0.415 | 0.237 | 1 | 0.555333085 | 0.334704509 | 0.317 | 0.392 | 1 | 0.555333085 | 0.012103838 |
| Nenf | 0.006079792 | 0.430861988 | 0.962 | 0.86 | 1 | 0.84329699 | 0.160902038 | 0.78 | 0.833 | 1 | 0.84329699 | 0.012122619 |
| Atp8b1 | 0.00648242 | 2.312263872 | 0.189 | 0.076 | 1 | 0.006087363 | 2.215716473 | 0.22 | 0.085 | 1 | 0.00648242 | 0.01213767 |
| Nxt1 | 0.22343671 | 0.154722595 | 0.189 | 0.119 | 1 | 0.006089964 | -1.654648247 | 0.049 | 0.242 | 1 | 0.22343671 | 0.012142841 |
| Arnt2 | 0.006095494 | 1.195364579 | 0.226 | 0.093 | 1 | 0.480474897 | 0.373548645 | 0.22 | 0.283 | 1 | 0.480474897 | 0.012153833 |
| Tm4sf1 | 0.0061163 | -3.212988122 | 0.358 | 0.517 | 1 | 0.062042839 | -0.254371753 | 0.463 | 0.341 | 1 | 0.062042839 | 0.01219519 |
| Hmgn2 | 0.006122347 | 0.355770114 | 0.849 | 0.669 | 1 | 0.3362637 | 0.315807478 | 0.659 | 0.58 | 1 | 0.3362637 | 0.01220721 |
| Ddit4l | 0.00613881 | -2.300179365 | 0.189 | 0.386 | 1 | 0.05008483 | -1.795063366 | 0.098 | 0.242 | 1 | 0.05008483 | 0.012239936 |
| Gtf2h2 | 0.00613943 | 1.3662267 | 0.264 | 0.123 | 1 | 0.241661693 | 0.316948038 | 0.341 | 0.249 | 1 | 0.241661693 | 0.012241167 |
| Arsi | 0.760673286 | -0.119639817 | 0.302 | 0.318 | 1 | 0.006152792 | -1.399834347 | 0.366 | 0.573 | 1 | 0.760673286 | 0.012267728 |
| Hmgb2 | 0.125371146 | -0.266120526 | 0.717 | 0.636 | 1 | 0.006183526 | 0.772161318 | 0.756 | 0.631 | 1 | 0.125371146 | 0.012328815 |
| Srf | 0.006184448 | -1.586744392 | 0.113 | 0.292 | 1 | 0.040484485 | -1.250257991 | 0.146 | 0.29 | 1 | 0.040484485 | 0.012330648 |
| Tmem51 | 0.006207344 | -1.932740246 | 0.113 | 0.284 | 1 | 0.715684974 | 0.640807637 | 0.122 | 0.106 | 1 | 0.715684974 | 0.012376156 |
| Prpsap1 | 0.00621417 | 0.763297653 | 0.698 | 0.555 | 1 | 0.849794884 | 0.28774103 | 0.537 | 0.57 | 1 | 0.849794884 | 0.012389725 |
| Sec16b | 0.179983337 | -0.416026047 | 0.34 | 0.458 | 1 | 0.006216822 | -1.195216628 | 0.122 | 0.345 | 1 | 0.179983337 | 0.012394996 |
| Cdh20 | 0.89708989 | 0.173831418 | 0.075 | 0.072 | 1 | 0.006228711 | 1.000703582 | 0.122 | 0.031 | 1 | 0.89708989 | 0.012418625 |
| Ankrd54 | 0.006242557 | -1.195774652 | 0.151 | 0.339 | 1 | 0.186149432 | -0.255904555 | 0.244 | 0.355 | 1 | 0.186149432 | 0.012446144 |
| Faf2 | 0.006245332 | 0.47665069 | 0.792 | 0.678 | 1 | 0.21891232 | -0.154316996 | 0.683 | 0.785 | 1 | 0.21891232 | 0.01245166 |
| Klhl9 | 0.006261924 | 0.499634008 | 0.755 | 0.572 | 1 | 0.225552979 | -0.320524855 | 0.707 | 0.785 | 1 | 0.225552979 | 0.012484637 |
| Dtnb | 0.654488116 | 0.292226119 | 0.717 | 0.657 | 1 | 0.006267474 | -0.620682954 | 0.61 | 0.799 | 1 | 0.654488116 | 0.012495667 |
| Tuba1a | 0.101017945 | -0.122909922 | 0.981 | 0.818 | 1 | 0.006270612 | 0.587212656 | 1 | 0.881 | 1 | 0.101017945 | 0.012501903 |
| Cep57 | 0.006271871 | 0.69106639 | 0.585 | 0.394 | 1 | 0.028387172 | 0.853324515 | 0.537 | 0.392 | 1 | 0.028387172 | 0.012504406 |
| Rdh5 | 0.006282021 | 2.055186921 | 0.17 | 0.059 | 1 | 0.484769937 | 1.711673968 | 0.098 | 0.072 | 1 | 0.484769937 | 0.012524579 |
| Pik3c2g | 0.006298856 | -3.545717123 | 0 | 0.127 | 1 | 0.365592036 | -1.020776145 | 0.024 | 0.058 | 1 | 0.365592036 | 0.012558037 |
| Soga1 | 0.016787567 | 0.932330173 | 0.528 | 0.386 | 1 | 0.006300595 | 0.99067933 | 0.634 | 0.543 | 1 | 0.016787567 | 0.012561492 |
| Fabp7 | 0.006302336 | -3.489133595 | 0 | 0.127 | 1 | 0.09421266 | -2.250257991 | 0 | 0.065 | 1 | 0.09421266 | 0.012564953 |
| Dnaaf2 | 0.006302853 | 1.285967128 | 0.264 | 0.119 | 1 | 0.415498992 | -0.577832649 | 0.122 | 0.167 | 1 | 0.415498992 | 0.01256598 |
| Gucy2g | 0.006308115 | -3.652632327 | 0 | 0.127 | 1 | 0.357858058 | -0.747757651 | 0 | 0.02 | 1 | 0.357858058 | 0.012576439 |
| Ptp4a1 | 0.583115634 | 0.306725688 | 0.075 | 0.055 | 1 | 0.006315577 | 2.059597271 | 0.122 | 0.031 | 1 | 0.583115634 | 0.012591267 |
| Herc6 | 0.808252115 | -0.617866909 | 0.17 | 0.178 | 1 | 0.006340496 | 1.699701326 | 0.268 | 0.123 | 1 | 0.808252115 | 0.012640791 |
| St3gal1 | 0.006347184 | 0.947280515 | 0.509 | 0.331 | 1 | 0.023932937 | 1.002264096 | 0.39 | 0.229 | 1 | 0.023932937 | 0.012654082 |
| Gpx3 | 0.585248121 | 0.916226865 | 0.509 | 0.64 | 1 | 0.00635252 | 2.366575025 | 0.366 | 0.201 | 1 | 0.585248121 | 0.012664686 |
| Pkn2 | 0.006354251 | 0.591109414 | 0.925 | 0.907 | 1 | 0.229368011 | 0.332185984 | 0.976 | 0.986 | 1 | 0.229368011 | 0.012668125 |
| Wnt4 | 0.742787589 | -1.40599236 | 0.057 | 0.068 | 1 | 0.006354724 | 1.185128153 | 0.366 | 0.188 | 1 | 0.742787589 | 0.012669064 |
| Ets2 | 0.061299805 | 0.222957331 | 0.792 | 0.636 | 1 | 0.006361758 | -0.429581691 | 0.756 | 0.908 | 1 | 0.061299805 | 0.012683044 |
| Pfdn2 | 0.010848888 | 0.404700848 | 0.868 | 0.771 | 1 | 0.006368666 | -0.545923402 | 0.732 | 0.863 | 1 | 0.010848888 | 0.012696772 |
| Gng7 | 0.006369506 | 1.417757001 | 0.151 | 0.047 | 1 | 0.063799075 | -2.16279515 | 0 | 0.078 | 1 | 0.063799075 | 0.012698442 |
| Larp6 | 0.79607843 | 0.177442671 | 0.396 | 0.453 | 1 | 0.006376504 | -1.000738392 | 0.415 | 0.608 | 1 | 0.79607843 | 0.012712348 |

| | | | | | | | | | | | | |
|---|---|---|---|---|---|---|---|---|---|---|---|---|
| D630045J12Rik | 0.006385697 | -1.02488496 | 0.377 | 0.555 | 1 | 0.406416038 | -0.21917762 | 0.488 | 0.625 | 1 | 0.406416038 | 0.012730616 |
| Rnf149 | 0.285489835 | 0.721763188 | 0.321 | 0.267 | 1 | 0.006411765 | 1.0375035 | 0.415 | 0.225 | 1 | 0.285489835 | 0.01278242 |
| Arsb | 0.006415718 | 0.925240749 | 0.396 | 0.22 | 1 | 0.10473158 | -1.754812408 | 0.22 | 0.355 | 1 | 0.10473158 | 0.012790275 |
| Mapre2 | 0.098176156 | -0.520974503 | 0.642 | 0.78 | 1 | 0.006417517 | -0.541967197 | 0.634 | 0.795 | 1 | 0.098176156 | 0.012793849 |
| Fam53b | 0.006424648 | 1.331600357 | 0.208 | 0.081 | 1 | 0.462305128 | 0.667279848 | 0.171 | 0.13 | 1 | 0.462305128 | 0.01280802 |
| Kpna2 | 0.006431603 | 1.280253477 | 0.189 | 0.068 | 1 | 0.767134333 | 0.319356545 | 0.171 | 0.154 | 1 | 0.767134333 | 0.012821841 |
| Cox14 | 0.015897157 | -0.41681939 | 0.83 | 0.869 | 1 | 0.006436234 | -0.504869818 | 0.683 | 0.85 | 1 | 0.015897157 | 0.012831043 |
| 4930578M01Rik | 0.0064438 | 2.154722595 | 0.132 | 0.038 | 1 | 0.494850012 | 0.83720485 | 0.073 | 0.048 | 1 | 0.494850012 | 0.012846077 |
| Cep68 | 0.006463565 | 1.552058092 | 0.283 | 0.14 | 1 | 0.490760601 | -0.16279515 | 0.244 | 0.184 | 1 | 0.490760601 | 0.012885351 |
| Neurl2 | 0.0909501 | 1.8327945 | 0.057 | 0.017 | 1 | 0.006470209 | 2.100239256 | 0.122 | 0.031 | 1 | 0.0909501 | 0.012898554 |
| Gm56912 | 0.178709532 | 1.017219071 | 0.075 | 0.034 | 1 | 0.006470209 | 2.100239256 | 0.122 | 0.031 | 1 | 0.178709532 | 0.012898554 |
| Il27 | 0.494603095 | 0.417757001 | 0.038 | 0.021 | 1 | 0.006470209 | 2.100239256 | 0.122 | 0.031 | 1 | 0.494603095 | 0.012898554 |
| Rapgef2 | 0.006517257 | 0.392011588 | 0.943 | 0.86 | 1 | 0.111518061 | 0.436551269 | 0.976 | 0.983 | 1 | 0.111518061 | 0.01299204 |
| Ubb | 0.006537238 | 0.356721803 | 1 | 0.992 | 1 | 0.068845717 | 0.280527792 | 1 | 0.997 | 1 | 0.068845717 | 0.01303174 |
| Nifk | 0.518138908 | 0.259419973 | 0.491 | 0.479 | 1 | 0.00653975 | -1.221688839 | 0.195 | 0.42 | 1 | 0.518138908 | 0.013036732 |
| Ltf | 0.006540783 | 0.527939918 | 0.472 | 0.275 | 1 | 0.344871914 | 0.710672444 | 0.341 | 0.304 | 1 | 0.344871914 | 0.013038784 |
| Noc3l | 0.006552053 | 1.091986839 | 0.453 | 0.288 | 1 | 0.240932155 | 0.75267134 | 0.366 | 0.294 | 1 | 0.240932155 | 0.013061176 |
| Mex3d | 0.006576371 | 1.191248471 | 0.302 | 0.148 | 1 | 0.49046975 | -0.152741486 | 0.268 | 0.324 | 1 | 0.49046975 | 0.013109494 |
| Zfp142 | 0.006621178 | 1.154722595 | 0.491 | 0.326 | 1 | 0.758517699 | 0.270858027 | 0.439 | 0.454 | 1 | 0.758517699 | 0.013198516 |
| Gtf2f2 | 0.040524369 | -0.422626722 | 0.66 | 0.822 | 1 | 0.006628619 | -0.674201044 | 0.561 | 0.751 | 1 | 0.040524369 | 0.013213299 |
| Cox6c | 0.019642883 | 0.70228063 | 1 | 0.949 | 1 | 0.006643973 | 0.481365695 | 0.951 | 0.949 | 1 | 0.019642883 | 0.013243803 |
| Rgcc | 0.006644254 | -1.826874624 | 0.623 | 0.72 | 1 | 0.138715227 | -0.320256186 | 0.61 | 0.724 | 1 | 0.138715227 | 0.013244361 |
| Ubfd1 | 0.00664728 | 0.89946554 | 0.604 | 0.445 | 1 | 0.325519235 | 0.274268655 | 0.561 | 0.509 | 1 | 0.325519235 | 0.013250375 |
| Tcf25 | 0.006650681 | 0.424562179 | 1 | 0.953 | 1 | 0.220492479 | 0.122575586 | 0.976 | 0.98 | 1 | 0.220492479 | 0.013257131 |
| Mkrn1 | 0.181342376 | 0.275016829 | 0.566 | 0.458 | 1 | 0.006660107 | -0.700760171 | 0.512 | 0.73 | 1 | 0.181342376 | 0.013275857 |
| Tcp1 | 0.00666832 | 0.43982979 | 0.981 | 0.903 | 1 | 0.232950933 | -0.19841906 | 0.829 | 0.939 | 1 | 0.232950933 | 0.013292173 |
| Rnaset2a | 0.006672582 | -0.631873767 | 0.698 | 0.78 | 1 | 0.06414146 | -0.26948435 | 0.659 | 0.741 | 1 | 0.06414146 | 0.013300641 |
| Eps15l1 | 0.006678154 | 0.544992189 | 0.698 | 0.487 | 1 | 0.46848293 | 0.143157234 | 0.756 | 0.747 | 1 | 0.46848293 | 0.013311711 |
| Bcs1l | 0.006695052 | 1.417757001 | 0.226 | 0.093 | 1 | 0.348396049 | -0.707115666 | 0.049 | 0.092 | 1 | 0.348396049 | 0.01334528 |
| Rad51c | 0.556762861 | 0.47665069 | 0.038 | 0.059 | 1 | 0.006702813 | 2.422167351 | 0.098 | 0.02 | 1 | 0.556762861 | 0.013360697 |
| Ckap5 | 0.006709198 | 0.794992467 | 0.679 | 0.568 | 1 | 0.430997726 | 0.274784918 | 0.756 | 0.696 | 1 | 0.430997726 | 0.013373384 |
| Ccdc61 | 0.006709405 | 1.417757001 | 0.151 | 0.047 | 1 | 0.196059041 | 0.898605394 | 0.122 | 0.065 | 1 | 0.196059041 | 0.013373794 |
| Samd9l | 0.006718064 | 2.417757001 | 0.208 | 0.089 | 1 | 0.903372623 | 0.749742009 | 0.244 | 0.263 | 1 | 0.903372623 | 0.013390996 |
| 5730480H06Rik | 0.006754 | 1.232725107 | 0.264 | 0.119 | 1 | 0.506414021 | 0.252242349 | 0.171 | 0.13 | 1 | 0.506414021 | 0.013462383 |
| Mpc1 | 0.03768815 | 0.300800854 | 0.906 | 0.831 | 1 | 0.006764111 | 0.779172728 | 0.902 | 0.819 | 1 | 0.03768815 | 0.013482469 |
| Homez | 0.006766682 | 3.154722595 | 0.075 | 0.013 | 1 | 0.887098934 | 0.749742009 | 0.049 | 0.044 | 1 | 0.887098934 | 0.013487576 |
| Zfp597 | 0.0067737 | 1.339147166 | 0.302 | 0.153 | 1 | 0.52992425 | 0.526864729 | 0.317 | 0.28 | 1 | 0.52992425 | 0.013501518 |
| Ppp1r35 | 0.006774237 | 1.09332205 | 0.34 | 0.178 | 1 | 0.137142061 | 1.055845136 | 0.244 | 0.157 | 1 | 0.137142061 | 0.013502583 |
| Lin37 | 0.00677447 | 1.200526284 | 0.264 | 0.119 | 1 | 0.621307366 | 0.185128153 | 0.098 | 0.13 | 1 | 0.621307366 | 0.013503046 |
| Fto | 0.006781473 | 0.615516918 | 0.962 | 0.936 | 1 | 0.456221078 | 0.131245312 | 0.951 | 0.986 | 1 | 0.456221078 | 0.013516957 |
| Eepd1 | 0.00679374 | 1.940597789 | 0.189 | 0.072 | 1 | 0.007049191 | 1.564186355 | 0.293 | 0.137 | 1 | 0.007049191 | 0.013541325 |
| Arrdc4 | 0.006808871 | 1.389187848 | 0.358 | 0.212 | 1 | 0.068333271 | -0.722222559 | 0.293 | 0.447 | 1 | 0.068333271 | 0.013571382 |
| Matn3 | 0.006812923 | -0.968803151 | 0.509 | 0.699 | 1 | 0.105934636 | -0.442344643 | 0.366 | 0.495 | 1 | 0.105934636 | 0.01357943 |
| Msn | 0.006821583 | -0.458520006 | 0.906 | 0.953 | 1 | 0.026877848 | -0.312643359 | 0.854 | 0.966 | 1 | 0.026877848 | 0.013596632 |
| Pcna | 0.006832017 | 0.556703941 | 0.717 | 0.593 | 1 | 0.418054804 | 0.222495006 | 0.78 | 0.751 | 1 | 0.418054804 | 0.013617358 |
| Ate1 | 0.006832961 | 0.913714495 | 0.604 | 0.436 | 1 | 0.717875292 | 0.437274243 | 0.366 | 0.423 | 1 | 0.717875292 | 0.013619233 |
| Gm10638 | 0.00685564 | 1.891688189 | 0.132 | 0.038 | 1 | 0.231591308 | -0.747757651 | 0 | 0.034 | 1 | 0.231591308 | 0.013664281 |
| Sipa1l1 | 0.006881266 | 0.770457444 | 0.811 | 0.754 | 1 | 0.426306358 | 0.191417156 | 0.854 | 0.85 | 1 | 0.426306358 | 0.013715181 |
| Psmb6 | 0.006888451 | 0.44776957 | 0.962 | 0.839 | 1 | 0.381271438 | -0.153469358 | 0.732 | 0.863 | 1 | 0.381271438 | 0.013729451 |
| Tbc1d8b | 0.006894576 | 0.79778805 | 0.717 | 0.564 | 1 | 0.390174475 | 0.524926925 | 0.585 | 0.648 | 1 | 0.390174475 | 0.013741616 |
| Tmed9 | 0.267088704 | -0.151228648 | 1 | 0.975 | 1 | 0.006905019 | -0.442563572 | 0.976 | 0.98 | 1 | 0.267088704 | 0.013762358 |
| Nup133 | 0.006918853 | 1.339147166 | 0.283 | 0.131 | 1 | 0.904782169 | -0.466575898 | 0.171 | 0.167 | 1 | 0.904782169 | 0.013789836 |
| Tm2d3 | 0.457358604 | -0.278236812 | 0.472 | 0.504 | 1 | 0.006920508 | -0.677368323 | 0.268 | 0.532 | 1 | 0.457358604 | 0.013793124 |
| Trim17 | 0.006940132 | 2.444229212 | 0.113 | 0.03 | 1 | 0.990690153 | 0.174239837 | 0.049 | 0.048 | 1 | 0.990690153 | 0.013832098 |
| Slc27a6 | 0.006977077 | -1.142957954 | 0.132 | 0.326 | 1 | 0.013702032 | -1.10921411 | 0.122 | 0.317 | 1 | 0.013702032 | 0.013905473 |
| Prdm14 | 0.006977505 | 3.154722595 | 0.075 | 0.013 | 1 | 0.87556513 | 0.667279848 | 0.024 | 0.02 | 1 | 0.87556513 | 0.013906324 |
| Rsf1os2 | 0.006977505 | 2.962077517 | 0.075 | 0.013 | 1 | 0.58792152 | 2.100239256 | 0.024 | 0.014 | 1 | 0.58792152 | 0.013906324 |
| Kiss1r | 0.006977505 | 2.962077517 | 0.075 | 0.013 | 1 | 0.056289003 | 3.159132945 | 0.049 | 0.01 | 1 | 0.056289003 | 0.013906324 |
| 4732465J04Rik | 0.006977505 | 2.962077517 | 0.075 | 0.013 | 1 | 0.18261072 | 2.029849928 | 0.049 | 0.017 | 1 | 0.18261072 | 0.013906324 |
| Gm26555 | 0.006977505 | 2.962077517 | 0.075 | 0.013 | 1 | 0.650840959 | 0.515276755 | 0.049 | 0.034 | 1 | 0.650840959 | 0.013906324 |
| Dusp29 | 0.006977505 | 2.962077517 | 0.075 | 0.013 | 1 | 0.02871132 | 1.699701326 | 0.073 | 0.017 | 1 | 0.02871132 | 0.013906324 |
| A930001C03Rik | 0.006977505 | 3.154722595 | 0.075 | 0.013 | 1 | 0.402364804 | -0.332720152 | 0 | 0.017 | 1 | 0.402364804 | 0.013906324 |
| Gm50334 | 0.006977505 | 3.154722595 | 0.075 | 0.013 | 1 | 0.082126335 | 1.711673968 | 0.122 | 0.055 | 1 | 0.082126335 | 0.013906324 |
| Septin8 | 0.006978034 | 0.743789494 | 0.509 | 0.339 | 1 | 0.211665715 | -0.360096591 | 0.39 | 0.526 | 1 | 0.211665715 | 0.013907375 |
| H2bc9 | 0.006988716 | 2.962077517 | 0.113 | 0.03 | 1 | 0.746881137 | 1.029849928 | 0.024 | 0.017 | 1 | 0.746881137 | 0.01392859 |
| BC051537 | 0.006996476 | 3.870929629 | 0.094 | 0.021 | 1 | 0.358996181 | 1.252242349 | 0.049 | 0.024 | 1 | 0.358996181 | 0.013944002 |
| R3hdml | 0.092432065 | 2.014997831 | 0.113 | 0.051 | 1 | 0.007007686 | 0.83720485 | 0.098 | 0.02 | 1 | 0.092432065 | 0.013966264 |
| Cstpp1 | 0.008747855 | 1.079434467 | 0.434 | 0.28 | 1 | 0.00701411 | -0.906956246 | 0.171 | 0.41 | 1 | 0.008747855 | 0.013979023 |
| 6820431F20Rik | 0.572330882 | 0.218609474 | 0.849 | 0.903 | 1 | 0.007017193 | -0.417806733 | 0.854 | 0.976 | 1 | 0.572330882 | 0.013985145 |
| Gprc5a | 0.007021322 | -2.545717123 | 0.019 | 0.161 | 1 | 0.641244846 | -0.372248516 | 0.049 | 0.068 | 1 | 0.641244846 | 0.013993345 |
| Arl6ip5 | 0.957850948 | 0.113573423 | 0.981 | 0.996 | 1 | 0.007029496 | -0.455345655 | 1 | 1 | 1 | 0.957850948 | 0.014009579 |
| B230219D22Rik | 0.007030128 | 0.56356826 | 0.849 | 0.809 | 1 | 0.009530329 | 0.465109968 | 0.951 | 0.853 | 1 | 0.009530329 | 0.014010833 |
| Mpdz | 0.00703059 | 1.109775189 | 0.528 | 0.411 | 1 | 0.186578756 | -0.299457698 | 0.366 | 0.515 | 1 | 0.186578756 | 0.014011751 |
| Zfp934 | 0.007066666 | -1.898388742 | 0.038 | 0.191 | 1 | 0.350585664 | -0.20718927 | 0.098 | 0.157 | 1 | 0.350585664 | 0.014083394 |
| Cstdc2 | 0.237936014 | 0.502645898 | 0.075 | 0.038 | 1 | 0.007070045 | 2.185128153 | 0.098 | 0.02 | 1 | 0.237936014 | 0.014090105 |
| Polr1a | 0.007077356 | 0.992451166 | 0.434 | 0.275 | 1 | 0.210900561 | -0.111811221 | 0.195 | 0.314 | 1 | 0.210900561 | 0.014104623 |
| 5530601H04Rik | 0.007089253 | 0.632769892 | 0.509 | 0.309 | 1 | 0.060064725 | -0.564893594 | 0.244 | 0.406 | 1 | 0.060064725 | 0.014128248 |
| Ccdc88c | 0.19254407 | -0.596048799 | 0.245 | 0.326 | 1 | 0.007097378 | 0.778311161 | 0.463 | 0.266 | 1 | 0.19254407 | 0.014144383 |
| Adar | 0.007099641 | 1.172424597 | 0.415 | 0.25 | 1 | 0.083550272 | 0.591448435 | 0.463 | 0.345 | 1 | 0.083550272 | 0.014148877 |
| Klhl2 | 0.007101791 | 1.57960788 | 0.34 | 0.195 | 1 | 0.963769284 | 0.139613494 | 0.415 | 0.386 | 1 | 0.963769284 | 0.014153147 |
| Rnf213 | 0.037168301 | 1.187144072 | 0.264 | 0.148 | 1 | 0.007107551 | 1.914166832 | 0.39 | 0.225 | 1 | 0.037168301 | 0.014164585 |
| Ap2m1 | 0.007110428 | 0.466666601 | 0.83 | 0.669 | 1 | 0.405484307 | 0.202030903 | 0.878 | 0.788 | 1 | 0.405484307 | 0.014170298 |
| Dmtf1 | 0.007122153 | 1.006623956 | 0.566 | 0.432 | 1 | 0.05588718 | -0.739720332 | 0.512 | 0.601 | 1 | 0.05588718 | 0.014193581 |
| 4933405D12Rik | 0.007125624 | 1.226275856 | 0.208 | 0.085 | 1 | 0.088223603 | 0.694910967 | 0.293 | 0.188 | 1 | 0.088223603 | 0.014200473 |
| Mier2 | 0.007126288 | 1.685237311 | 0.189 | 0.072 | 1 | 0.923177293 | 0.252242349 | 0.073 | 0.068 | 1 | 0.923177293 | 0.014201791 |
| Zc3h12c | 0.007130546 | 1.321832581 | 0.321 | 0.165 | 1 | 0.334529788 | 0.793877418 | 0.366 | 0.338 | 1 | 0.334529788 | 0.014210247 |
| Eif3i | 0.007152896 | 0.34979441 | 0.962 | 0.843 | 1 | 0.162217996 | -0.139542419 | 0.707 | 0.877 | 1 | 0.162217996 | 0.014254628 |
| Plekhh3 | 0.007193708 | 1.154722595 | 0.189 | 0.068 | 1 | 0.319896966 | 0.681926624 | 0.171 | 0.116 | 1 | 0.319896966 | 0.014335667 |
| Ephx4 | 0.007193709 | 2.739685096 | 0.075 | 0.013 | 1 | 0.402369942 | -0.332720152 | 0 | 0.017 | 1 | 0.402369942 | 0.014335668 |
| Gm8066 | 0.007193709 | 2.739685096 | 0.075 | 0.013 | 1 | 0.402373368 | 0.252242349 | 0 | 0.017 | 1 | 0.402373368 | 0.014335668 |
| Aoc2 | 0.007193709 | 2.739685096 | 0.075 | 0.013 | 1 | 0.170390343 | -1.484723245 | 0 | 0.044 | 1 | 0.170390343 | 0.014335668 |
| Samd15 | 0.007193709 | 2.739685096 | 0.075 | 0.013 | 1 | 0.257325989 | -0.484723245 | 0 | 0.031 | 1 | 0.257325989 | 0.014335668 |
| Begain | 0.007193709 | 2.739685096 | 0.075 | 0.013 | 1 | 0.357847088 | -0.16279515 | 0 | 0.02 | 1 | 0.357847088 | 0.014335668 |
| Acvrl1 | 0.007193709 | 2.739685096 | 0.075 | 0.013 | 1 | 0.02398288 | 1.989207943 | 0.098 | 0.027 | 1 | 0.02398288 | 0.014335668 |
| Gm715 | 0.007193709 | 2.962077517 | 0.075 | 0.013 | 1 | 0.25729872 | -0.747757651 | 0 | 0.031 | 1 | 0.25729872 | 0.014335668 |
| Igsf11 | 0.007196753 | 2.669295768 | 0.075 | 0.013 | 1 | 0.402369942 | -0.16279515 | 0 | 0.017 | 1 | 0.402369942 | 0.014341713 |
| Gm826 | 0.007196956 | 2.806799291 | 0.075 | 0.013 | 1 | 0.402373368 | 0.252242349 | 0 | 0.017 | 1 | 0.402373368 | 0.014342117 |
| Snapc5 | 0.007198939 | 0.93713116 | 0.491 | 0.335 | 1 | 0.760319815 | 0.163433082 | 0.317 | 0.355 | 1 | 0.760319815 | 0.014346054 |

| | | | | | | | | | | | | |
|---|---|---|---|---|---|---|---|---|---|---|---|---|
| Tsc22d3 | 0.007201015 | -0.31216055 | 0.792 | 0.936 | 1 | 0.023524384 | -0.617023174 | 0.854 | 0.915 | 1 | 0.023524384 | 0.014350175 |
| Psmd6 | 0.049129424 | 0.618669695 | 0.642 | 0.572 | 1 | 0.007213644 | -0.570298977 | 0.463 | 0.693 | 1 | 0.049129424 | 0.014375252 |
| Arid5a | 0.007216926 | 0.884075005 | 0.547 | 0.394 | 1 | 0.82696479 | 0.624211126 | 0.293 | 0.314 | 1 | 0.82696479 | 0.014381768 |
| Otud3 | 0.007218752 | -2.399866257 | 0.019 | 0.157 | 1 | 0.260888685 | 0.934502051 | 0.268 | 0.215 | 1 | 0.260888685 | 0.014385394 |
| Fam98b | 0.007218787 | 0.623002115 | 0.547 | 0.36 | 1 | 0.537251511 | -0.337881857 | 0.488 | 0.509 | 1 | 0.537251511 | 0.014385463 |
| Pgap1 | 0.007226916 | 2.154722595 | 0.151 | 0.051 | 1 | 0.546786195 | 0.580865097 | 0.146 | 0.116 | 1 | 0.546786195 | 0.014401603 |
| Pprc1 | 0.007228654 | 0.929162894 | 0.566 | 0.411 | 1 | 0.649099286 | 0.509842869 | 0.488 | 0.536 | 1 | 0.649099286 | 0.014405055 |
| Pik3ip1 | 0.408986145 | 0.506194965 | 0.321 | 0.28 | 1 | 0.007237327 | 0.898605394 | 0.561 | 0.355 | 1 | 0.408986145 | 0.014422276 |
| Aen | 0.975532602 | -0.142957954 | 0.34 | 0.331 | 1 | 0.007238505 | -1.917682652 | 0.122 | 0.314 | 1 | 0.975532602 | 0.014424614 |
| Rps11 | 0.143429098 | 0.154409 | 1 | 0.975 | 1 | 0.007250733 | -0.302202962 | 1 | 0.99 | 1 | 0.143429098 | 0.014448894 |
| C030037D09Rik | 0.007266864 | -2.399866257 | 0.019 | 0.157 | 1 | 0.070312671 | -2.069685746 | 0 | 0.075 | 1 | 0.070312671 | 0.014480921 |
| Tspan18 | 0.009141173 | -0.906598853 | 0.642 | 0.742 | 1 | 0.007267041 | -1.208936932 | 0.439 | 0.614 | 1 | 0.009141173 | 0.014481271 |
| Chrnb1 | 0.007270173 | 2.640149422 | 0.075 | 0.013 | 1 | 0.274122622 | 0.614812428 | 0.122 | 0.072 | 1 | 0.274122622 | 0.01448749 |
| Stmn1 | 0.063705638 | -0.430239906 | 0.245 | 0.144 | 1 | 0.00727236 | 1.01123425 | 0.366 | 0.181 | 1 | 0.063705638 | 0.014491833 |
| Trim56 | 0.007278768 | 0.964898036 | 0.377 | 0.208 | 1 | 0.593684557 | 0.346457691 | 0.268 | 0.348 | 1 | 0.593684557 | 0.014504556 |
| Igf2 | 0.007288314 | 2.120775263 | 0.245 | 0.119 | 1 | 0.150500165 | 0.962735732 | 0.195 | 0.119 | 1 | 0.150500165 | 0.014523508 |
| Irak1 | 0.007289505 | 0.925240749 | 0.566 | 0.419 | 1 | 0.120114995 | 0.559866906 | 0.61 | 0.478 | 1 | 0.120114995 | 0.014525872 |
| Rbpjl | 0.00729649 | 1.669295768 | 0.132 | 0.038 | 1 | 0.329079127 | 1.114738825 | 0.122 | 0.078 | 1 | 0.329079127 | 0.01453974 |
| Gm10371 | 0.007302789 | -1.686579659 | 0.075 | 0.242 | 1 | 0.020859607 | -1.470223675 | 0.073 | 0.232 | 1 | 0.020859607 | 0.014552248 |
| Vps4b | 0.007315496 | 0.66461928 | 0.755 | 0.682 | 1 | 0.211429193 | -0.332720152 | 0.683 | 0.768 | 1 | 0.211429193 | 0.014577476 |
| Gm26633 | 0.007325421 | -3.054730771 | 0 | 0.123 | 1 | 0.203695953 | -1.589059905 | 0.024 | 0.078 | 1 | 0.203695953 | 0.01459718 |
| Pcdhga7 | 0.007325421 | -3.054730771 | 0 | 0.123 | 1 | 0.428232129 | 0.515276755 | 0.122 | 0.085 | 1 | 0.428232129 | 0.01459718 |
| Esr1 | 0.176959023 | 0.316641157 | 0.472 | 0.381 | 1 | 0.007338309 | -0.879786045 | 0.439 | 0.642 | 1 | 0.176959023 | 0.014622766 |
| Btk | 0.007344781 | 2.669295768 | 0.075 | 0.013 | 1 | 0.175856887 | 3.614812428 | 0.049 | 0.017 | 1 | 0.175856887 | 0.014635616 |
| Atg5 | 0.007346978 | 0.731865591 | 0.642 | 0.466 | 1 | 0.869855518 | 0.129986599 | 0.561 | 0.584 | 1 | 0.869855518 | 0.014639977 |
| H2bc4 | 0.007361769 | -1.009664223 | 0.792 | 0.831 | 1 | 0.656764373 | -0.156397379 | 0.732 | 0.761 | 1 | 0.656764373 | 0.014669343 |
| Asns | 0.007364063 | -1.26312992 | 0.226 | 0.415 | 1 | 0.022928207 | -1.904262137 | 0.073 | 0.222 | 1 | 0.022928207 | 0.014673896 |
| Cep104 | 0.00736739 | 1.605384004 | 0.34 | 0.178 | 1 | 0.153294809 | -0.659220976 | 0.244 | 0.345 | 1 | 0.153294809 | 0.014680501 |
| Kdm5a | 0.007369728 | 0.510053848 | 0.943 | 0.89 | 1 | 0.192999477 | 0.359908859 | 0.927 | 0.952 | 1 | 0.192999477 | 0.014685143 |
| Adgrl2 | 0.007378012 | 0.655916738 | 0.283 | 0.131 | 1 | 0.107330858 | 0.728680393 | 0.341 | 0.235 | 1 | 0.107330858 | 0.01470159 |
| Pbdc1 | 0.27535378 | -0.176634913 | 0.774 | 0.818 | 1 | 0.007403441 | -0.813231488 | 0.707 | 0.823 | 1 | 0.27535378 | 0.014752071 |
| Akt3 | 0.007413026 | 0.500591655 | 0.943 | 0.835 | 1 | 0.118547752 | -0.260056494 | 0.976 | 0.973 | 1 | 0.118547752 | 0.014771098 |
| Gm48768 | 0.007419035 | 2.640149422 | 0.075 | 0.013 | 1 | 0.083899597 | 1.699701326 | 0.073 | 0.024 | 1 | 0.083899597 | 0.014783029 |
| Gm47400 | 0.007419035 | 2.640149422 | 0.075 | 0.013 | 1 | 0.007718533 | 3.83720485 | 0.024 | 0 | 1 | 0.007718533 | 0.014783029 |
| Atf7ip2 | 0.007419035 | 2.739685096 | 0.075 | 0.013 | 1 | 0.729889663 | 1.83720485 | 0.024 | 0.017 | 1 | 0.729889663 | 0.014783029 |
| Gm4544 | 0.007421114 | 2.47665069 | 0.075 | 0.013 | 1 | 0.51919986 | 0.83720485 | 0 | 0.01 | 1 | 0.51919986 | 0.014787155 |
| H2-Q7 | 0.007421114 | 2.47665069 | 0.075 | 0.013 | 1 | 0.257285084 | -1.069685746 | 0 | 0.031 | 1 | 0.257285084 | 0.014787155 |
| Nemf | 0.007425522 | 0.475432712 | 0.83 | 0.775 | 1 | 0.355104538 | -0.151479837 | 0.78 | 0.826 | 1 | 0.355104538 | 0.014795905 |
| H1f3 | 0.211976407 | -1.626637119 | 0.038 | 0.089 | 1 | 0.007450847 | 1.644559772 | 0.098 | 0.02 | 1 | 0.211976407 | 0.014846179 |
| Plrg1 | 0.325763018 | 0.695290976 | 0.283 | 0.229 | 1 | 0.007454209 | -1.332720152 | 0.122 | 0.328 | 1 | 0.325763018 | 0.014852852 |
| Ncbp2 | 0.098423744 | 0.423539353 | 0.642 | 0.614 | 1 | 0.007464017 | -0.441455791 | 0.439 | 0.7 | 1 | 0.098423744 | 0.014872323 |
| Gm15708 | 0.007484502 | 2.154722595 | 0.113 | 0.03 | 1 | 0.584273352 | -0.16279515 | 0.049 | 0.072 | 1 | 0.584273352 | 0.014912987 |
| Usp39 | 0.007488249 | 1.234157062 | 0.358 | 0.203 | 1 | 0.290053571 | 0.40934331 | 0.341 | 0.256 | 1 | 0.290053571 | 0.014920423 |
| Dlx6 | 0.007495013 | 2.417757001 | 0.075 | 0.013 | 1 | 0.43867119 | 1.422167351 | 0.024 | 0.01 | 1 | 0.43867119 | 0.01493385 |
| 2200002D01Rik | 0.007495013 | 2.417757001 | 0.075 | 0.013 | 1 | 0.9743246 | 0.252242349 | 0.073 | 0.075 | 1 | 0.9743246 | 0.01493385 |
| Tepsin | 0.007495013 | 2.417757001 | 0.075 | 0.013 | 1 | 0.574045084 | 0.930314254 | 0.024 | 0.044 | 1 | 0.574045084 | 0.01493385 |
| Vipas39 | 0.007497675 | 1.128250383 | 0.358 | 0.203 | 1 | 0.631536583 | 0.503304113 | 0.293 | 0.273 | 1 | 0.631536583 | 0.014939135 |
| Nrcam | 0.774217997 | -1.600164907 | 0.019 | 0.025 | 1 | 0.007506182 | 0.422167351 | 0.122 | 0.031 | 1 | 0.774217997 | 0.014956021 |
| Arhgap20 | 0.00750784 | 1.631160639 | 0.132 | 0.038 | 1 | 0.05682354 | 3.322631677 | 0.049 | 0.01 | 1 | 0.05682354 | 0.014959312 |
| Csnk1a1 | 0.007547563 | 0.271795714 | 1 | 0.996 | 1 | 0.104448487 | 0.167277682 | 1 | 1 | 1 | 0.104448487 | 0.015038161 |
| Csk | 0.007550252 | 0.999444369 | 0.321 | 0.161 | 1 | 0.105103261 | -0.520347155 | 0.171 | 0.304 | 1 | 0.105103261 | 0.015043498 |
| Dlx6os1 | 0.007573271 | 2.154722595 | 0.075 | 0.013 | 1 | 0.599290422 | 1.515276755 | 0.024 | 0.014 | 1 | 0.599290422 | 0.015089187 |
| Milr1 | 0.10281534 | 1.739685096 | 0.038 | 0.008 | 1 | 0.00757962 | 1.252242349 | 0.098 | 0.02 | 1 | 0.10281534 | 0.015101789 |
| Gm27003 | 0.007589284 | 1.110328475 | 0.226 | 0.093 | 1 | 0.092096252 | 0.962735732 | 0.195 | 0.106 | 1 | 0.092096252 | 0.015120971 |
| Cep250 | 0.007594854 | 1.120775263 | 0.264 | 0.119 | 1 | 0.693358827 | 0.875678998 | 0.195 | 0.181 | 1 | 0.693358827 | 0.015132026 |
| Ubxn4 | 0.016216438 | -0.318664078 | 0.981 | 0.996 | 1 | 0.007609638 | -0.306612448 | 0.976 | 0.997 | 1 | 0.016216438 | 0.015161369 |
| Relch | 0.007627792 | 0.699718688 | 0.679 | 0.538 | 1 | 0.712796411 | 0.232342792 | 0.659 | 0.679 | 1 | 0.712796411 | 0.015197402 |
| Atg16l1 | 0.0076329 | 0.906795081 | 0.528 | 0.398 | 1 | 0.788590695 | -0.224804812 | 0.512 | 0.498 | 1 | 0.788590695 | 0.015207538 |
| Thnsl2 | 0.007647025 | 1.377115016 | 0.283 | 0.14 | 1 | 0.119624542 | -0.16279515 | 0.049 | 0.143 | 1 | 0.119624542 | 0.015235574 |
| 2700049A03Rik | 0.007652204 | 1.353268274 | 0.34 | 0.186 | 1 | 0.144925032 | 1.074244047 | 0.293 | 0.205 | 1 | 0.144925032 | 0.015245852 |
| Acot11 | 0.118914164 | 1.517292674 | 0.094 | 0.042 | 1 | 0.007659161 | 1.83720485 | 0.22 | 0.089 | 1 | 0.118914164 | 0.01525966 |
| Pcdhb7 | 0.007659872 | 2.154722595 | 0.113 | 0.03 | 1 | 0.522238245 | 1.029849928 | 0.073 | 0.051 | 1 | 0.522238245 | 0.015261071 |
| Clk2 | 0.007675962 | 0.60078699 | 0.547 | 0.356 | 1 | 0.130870176 | -0.417036438 | 0.341 | 0.485 | 1 | 0.130870176 | 0.015293004 |
| Zfp758 | 0.007690834 | 1.382133091 | 0.302 | 0.153 | 1 | 0.218929335 | 0.10681191 | 0.098 | 0.188 | 1 | 0.218929335 | 0.015322518 |
| Resf1 | 0.00769522 | -2.204094599 | 0.698 | 0.729 | 1 | 0.506278521 | -0.575254105 | 0.634 | 0.672 | 1 | 0.506278521 | 0.015331223 |
| Pnpla2 | 0.007695605 | 1.46284489 | 0.302 | 0.161 | 1 | 0.03765005 | 0.547698233 | 0.366 | 0.208 | 1 | 0.03765005 | 0.015331989 |
| Ndufab1 | 0.007698155 | 0.437970222 | 0.83 | 0.746 | 1 | 0.026023702 | -0.237254142 | 0.61 | 0.785 | 1 | 0.026023702 | 0.015337048 |
| Slc22a14 | 0.007704191 | 1.517292674 | 0.132 | 0.038 | 1 | 0.176820381 | 1.515276755 | 0.098 | 0.048 | 1 | 0.176820381 | 0.015349027 |
| Fgf16 | 0.014846242 | 3.154722595 | 0.057 | 0.008 | 1 | 0.007718533 | 3.83720485 | 0.024 | 0 | 1 | 0.014846242 | 0.015377491 |
| Gm16283 | 0.018663137 | 2.417757001 | 0.075 | 0.017 | 1 | 0.007718533 | 3.83720485 | 0.024 | 0 | 1 | 0.018663137 | 0.015377491 |
| Gast | 0.029966098 | 3.154722595 | 0.038 | 0.004 | 1 | 0.007718533 | 3.83720485 | 0.024 | 0 | 1 | 0.029966098 | 0.015377491 |
| Ccdc154 | 0.029966098 | 3.154722595 | 0.038 | 0.004 | 1 | 0.007718533 | 3.83720485 | 0.024 | 0 | 1 | 0.029966098 | 0.015377491 |
| Adam1b | 0.030360955 | 2.739685096 | 0.038 | 0.004 | 1 | 0.007718533 | 3.83720485 | 0.024 | 0 | 1 | 0.030360955 | 0.015377491 |
| Gm15541 | 0.030360955 | 2.739685096 | 0.038 | 0.004 | 1 | 0.007718533 | 3.83720485 | 0.024 | 0 | 1 | 0.030360955 | 0.015377491 |
| Enpp6 | 0.030360955 | 2.739685096 | 0.038 | 0.004 | 1 | 0.007718533 | 3.83720485 | 0.024 | 0 | 1 | 0.030360955 | 0.015377491 |
| Kcnk4 | 0.030360955 | 2.739685096 | 0.038 | 0.004 | 1 | 0.007718533 | 3.83720485 | 0.024 | 0 | 1 | 0.030360955 | 0.015377491 |
| Grem2 | 0.03562064 | 3.154722595 | 0.019 | 0 | 1 | 0.007718533 | 4.422167351 | 0.024 | 0 | 1 | 0.03562064 | 0.015377491 |
| Gm14232 | 0.03562064 | 3.154722595 | 0.019 | 0 | 1 | 0.007718533 | 3.83720485 | 0.024 | 0 | 1 | 0.03562064 | 0.015377491 |
| Gm16337 | 0.03562064 | 3.154722595 | 0.019 | 0 | 1 | 0.007718533 | 4.422167351 | 0.024 | 0 | 1 | 0.03562064 | 0.015377491 |
| Txk | 0.03562064 | 3.739685096 | 0.019 | 0 | 1 | 0.007718533 | 4.83720485 | 0.024 | 0 | 1 | 0.03562064 | 0.015377491 |
| Spata18 | 0.03562064 | 3.154722595 | 0.019 | 0 | 1 | 0.007718533 | 4.83720485 | 0.024 | 0 | 1 | 0.03562064 | 0.015377491 |
| Gm20531 | 0.03562064 | 3.154722595 | 0.019 | 0 | 1 | 0.007718533 | 3.83720485 | 0.024 | 0 | 1 | 0.03562064 | 0.015377491 |
| Gm7457 | 0.03562064 | 3.154722595 | 0.019 | 0 | 1 | 0.007718533 | 3.83720485 | 0.024 | 0 | 1 | 0.03562064 | 0.015377491 |
| Slc17a6 | 0.03562064 | 4.154722595 | 0.019 | 0 | 1 | 0.007718533 | 5.159132945 | 0.024 | 0 | 1 | 0.03562064 | 0.015377491 |
| Fam181b | 0.03562064 | 3.154722595 | 0.019 | 0 | 1 | 0.007718533 | 3.83720485 | 0.024 | 0 | 1 | 0.03562064 | 0.015377491 |
| A630091E08Rik | 0.03562064 | 3.154722595 | 0.019 | 0 | 1 | 0.007718533 | 3.83720485 | 0.024 | 0 | 1 | 0.03562064 | 0.015377491 |
| Art1 | 0.03562064 | 3.739685096 | 0.019 | 0 | 1 | 0.007718533 | 3.83720485 | 0.024 | 0 | 1 | 0.03562064 | 0.015377491 |
| Vipr1 | 0.03562064 | 3.739685096 | 0.019 | 0 | 1 | 0.007718533 | 4.422167351 | 0.024 | 0 | 1 | 0.03562064 | 0.015377491 |
| C1ql1 | 0.03562064 | 3.154722595 | 0.019 | 0 | 1 | 0.007718533 | 3.83720485 | 0.024 | 0 | 1 | 0.03562064 | 0.015377491 |
| Slc7a15 | 0.03562064 | 3.739685096 | 0.019 | 0 | 1 | 0.007718533 | 3.83720485 | 0.024 | 0 | 1 | 0.03562064 | 0.015377491 |
| 5730409N16Rik | 0.03562064 | 3.154722595 | 0.019 | 0 | 1 | 0.007718533 | 3.83720485 | 0.024 | 0 | 1 | 0.03562064 | 0.015377491 |
| H3c4 | 0.03562064 | 3.154722595 | 0.019 | 0 | 1 | 0.007718533 | 3.83720485 | 0.024 | 0 | 1 | 0.03562064 | 0.015377491 |
| AI197445 | 0.03562064 | 3.154722595 | 0.019 | 0 | 1 | 0.007718533 | 3.83720485 | 0.024 | 0 | 1 | 0.03562064 | 0.015377491 |
| Fgf10 | 0.03562064 | 4.962077517 | 0.019 | 0 | 1 | 0.007718533 | 4.83720485 | 0.024 | 0 | 1 | 0.03562064 | 0.015377491 |
| Gm3095 | 0.03562064 | 3.154722595 | 0.019 | 0 | 1 | 0.007718533 | 4.422167351 | 0.024 | 0 | 1 | 0.03562064 | 0.015377491 |
| Ltb4r2 | 0.03562064 | 3.154722595 | 0.019 | 0 | 1 | 0.007718533 | 3.83720485 | 0.024 | 0 | 1 | 0.03562064 | 0.015377491 |
| Cldn1 | 0.03562064 | 3.154722595 | 0.019 | 0 | 1 | 0.007718533 | 3.83720485 | 0.024 | 0 | 1 | 0.03562064 | 0.015377491 |
| Gm28528 | 0.03562064 | 3.154722595 | 0.019 | 0 | 1 | 0.007718533 | 3.83720485 | 0.024 | 0 | 1 | 0.03562064 | 0.015377491 |

| | | | | | | | | | | | | |
|---|---|---|---|---|---|---|---|---|---|---|---|---|
| Gm50011 | 0.03562064 | 3.154722595 | 0.019 | 0 | 1 | 0.007718533 | 4.422167351 | 0.024 | 0 | 1 | 0.03562064 | 0.015377491 |
| Eif2c5 | 0.03562064 | 3.154722595 | 0.019 | 0 | 1 | 0.007718533 | 4.422167351 | 0.024 | 0 | 1 | 0.03562064 | 0.015377491 |
| Dpep1 | 0.039279873 | 2.154722595 | 0.075 | 0.021 | 1 | 0.007718533 | 3.83720485 | 0.024 | 0 | 1 | 0.039279873 | 0.015377491 |
| Brinp1 | 0.044731073 | -3.678167419 | 0 | 0.072 | 1 | 0.007718533 | 5.422167351 | 0.024 | 0 | 1 | 0.044731073 | 0.015377491 |
| Tac1 | 0.04516498 | 1.417757001 | 0.057 | 0.013 | 1 | 0.007718533 | 6.644559772 | 0.024 | 0 | 1 | 0.04516498 | 0.015377491 |
| A930004D18Rik | 0.099079675 | 2.891688189 | 0.038 | 0.008 | 1 | 0.007718533 | 4.422167351 | 0.024 | 0 | 1 | 0.099079675 | 0.015377491 |
| Afap1l1 | 0.100937876 | 2.154722595 | 0.038 | 0.008 | 1 | 0.007718533 | 4.422167351 | 0.024 | 0 | 1 | 0.100937876 | 0.015377491 |
| Ccdc192 | 0.105985328 | -2.799473716 | 0.019 | 0.081 | 1 | 0.007718533 | 3.83720485 | 0.024 | 0 | 1 | 0.105985328 | 0.015377491 |
| Gm42397 | 0.175861997 | -2.368839361 | 0 | 0.034 | 1 | 0.007718533 | 3.83720485 | 0.024 | 0 | 1 | 0.175861997 | 0.015377491 |
| Actg2 | 0.200381013 | 2.47665069 | 0.038 | 0.013 | 1 | 0.007718533 | 4.83720485 | 0.024 | 0 | 1 | 0.200381013 | 0.015377491 |
| 9130015A21Rik | 0.206392421 | -2.1672055 | 0 | 0.03 | 1 | 0.007718533 | 3.83720485 | 0.024 | 0 | 1 | 0.206392421 | 0.015377491 |
| Cnr2 | 0.209009104 | 1.739685096 | 0.038 | 0.013 | 1 | 0.007718533 | 3.83720485 | 0.024 | 0 | 1 | 0.209009104 | 0.015377491 |
| Slco5a1 | 0.243111734 | -0.652632327 | 0 | 0.025 | 1 | 0.007718533 | 4.422167351 | 0.024 | 0 | 1 | 0.243111734 | 0.015377491 |
| Kcng1 | 0.246443787 | 3.154722595 | 0.019 | 0.004 | 1 | 0.007718533 | 3.83720485 | 0.024 | 0 | 1 | 0.246443787 | 0.015377491 |
| Cxcl5 | 0.246443787 | 2.569760094 | 0.019 | 0.004 | 1 | 0.007718533 | 3.83720485 | 0.024 | 0 | 1 | 0.246443787 | 0.015377491 |
| Gm44284 | 0.246443787 | 2.739685096 | 0.019 | 0.004 | 1 | 0.007718533 | 3.83720485 | 0.024 | 0 | 1 | 0.246443787 | 0.015377491 |
| Marchf11 | 0.246443787 | 2.739685096 | 0.019 | 0.004 | 1 | 0.007718533 | 3.83720485 | 0.024 | 0 | 1 | 0.246443787 | 0.015377491 |
| Nell2 | 0.246443787 | 2.739685096 | 0.019 | 0.004 | 1 | 0.007718533 | 4.422167351 | 0.024 | 0 | 1 | 0.246443787 | 0.015377491 |
| C030032O16Rik | 0.249034808 | 2.154722595 | 0.019 | 0.004 | 1 | 0.007718533 | 3.83720485 | 0.024 | 0 | 1 | 0.249034808 | 0.015377491 |
| Atp7b | 0.249034808 | 2.154722595 | 0.019 | 0.004 | 1 | 0.007718533 | 3.83720485 | 0.024 | 0 | 1 | 0.249034808 | 0.015377491 |
| Gm10701 | 0.249034808 | 2.154722595 | 0.019 | 0.004 | 1 | 0.007718533 | 3.83720485 | 0.024 | 0 | 1 | 0.249034808 | 0.015377491 |
| 4930434F21Rik | 0.249034808 | 2.154722595 | 0.019 | 0.004 | 1 | 0.007718533 | 3.83720485 | 0.024 | 0 | 1 | 0.249034808 | 0.015377491 |
| Vstm2a | 0.249034808 | 2.154722595 | 0.019 | 0.004 | 1 | 0.007718533 | 3.83720485 | 0.024 | 0 | 1 | 0.249034808 | 0.015377491 |
| Gm20658 | 0.249034808 | 2.154722595 | 0.019 | 0.004 | 1 | 0.007718533 | 4.422167351 | 0.024 | 0 | 1 | 0.249034808 | 0.015377491 |
| Gm10791 | 0.249034808 | 2.154722595 | 0.019 | 0.004 | 1 | 0.007718533 | 4.83720485 | 0.024 | 0 | 1 | 0.249034808 | 0.015377491 |
| Ly6g6f | 0.249034808 | 2.154722595 | 0.019 | 0.004 | 1 | 0.007718533 | 3.83720485 | 0.024 | 0 | 1 | 0.249034808 | 0.015377491 |
| Gm36914 | 0.249034808 | 2.154722595 | 0.019 | 0.004 | 1 | 0.007718533 | 3.83720485 | 0.024 | 0 | 1 | 0.249034808 | 0.015377491 |
| Tnfrsf13b | 0.251644813 | 1.569760094 | 0.019 | 0.004 | 1 | 0.007718533 | 5.159132945 | 0.024 | 0 | 1 | 0.251644813 | 0.015377491 |
| Tagap | 0.251644813 | 1.569760094 | 0.019 | 0.004 | 1 | 0.007718533 | 3.83720485 | 0.024 | 0 | 1 | 0.251644813 | 0.015377491 |
| Dgat2l6 | 0.287761733 | -0.652632327 | 0 | 0.021 | 1 | 0.007718533 | 5.159132945 | 0.024 | 0 | 1 | 0.287761733 | 0.015377491 |
| Gm57476 | 0.287773478 | -0.430239906 | 0 | 0.021 | 1 | 0.007718533 | 4.422167351 | 0.024 | 0 | 1 | 0.287773478 | 0.015377491 |
| Gm39816 | 0.287773478 | -0.430239906 | 0 | 0.021 | 1 | 0.007718533 | 3.83720485 | 0.024 | 0 | 1 | 0.287773478 | 0.015377491 |
| 4933430I17Rik | 0.343000446 | -0.430239906 | 0 | 0.017 | 1 | 0.007718533 | 3.83720485 | 0.024 | 0 | 1 | 0.343000446 | 0.015377491 |
| Gm34907 | 0.343000446 | -0.845277405 | 0 | 0.017 | 1 | 0.007718533 | 4.83720485 | 0.024 | 0 | 1 | 0.343000446 | 0.015377491 |
| Gm38563 | 0.343000446 | -0.430239906 | 0 | 0.017 | 1 | 0.007718533 | 3.83720485 | 0.024 | 0 | 1 | 0.343000446 | 0.015377491 |
| Gm12796 | 0.413080549 | -0.1672055 | 0 | 0.013 | 1 | 0.007718533 | 3.83720485 | 0.024 | 0 | 1 | 0.413080549 | 0.015377491 |
| Rbp7 | 0.413082433 | 0.154722595 | 0 | 0.013 | 1 | 0.007718533 | 3.83720485 | 0.024 | 0 | 1 | 0.413082433 | 0.015377491 |
| Gm43251 | 0.413082433 | -1.304709024 | 0 | 0.013 | 1 | 0.007718533 | 3.83720485 | 0.024 | 0 | 1 | 0.413082433 | 0.015377491 |
| Il27ra | 0.413082433 | 0.154722595 | 0 | 0.013 | 1 | 0.007718533 | 4.83720485 | 0.024 | 0 | 1 | 0.413082433 | 0.015377491 |
| Ptprb | 0.413082433 | 0.154722595 | 0 | 0.013 | 1 | 0.007718533 | 6.159132945 | 0.024 | 0 | 1 | 0.413082433 | 0.015377491 |
| Gm56535 | 0.413082433 | 0.154722595 | 0 | 0.013 | 1 | 0.007718533 | 3.83720485 | 0.024 | 0 | 1 | 0.413082433 | 0.015377491 |
| Ly6k | 0.413082433 | -0.430239906 | 0 | 0.013 | 1 | 0.007718533 | 3.83720485 | 0.024 | 0 | 1 | 0.413082433 | 0.015377491 |
| Cyp2c65 | 0.413082433 | 0.154722595 | 0 | 0.013 | 1 | 0.007718533 | 4.83720485 | 0.024 | 0 | 1 | 0.413082433 | 0.015377491 |
| Gm41555 | 0.48175124 | 0.417757001 | 0.038 | 0.021 | 1 | 0.007718533 | 4.422167351 | 0.024 | 0 | 1 | 0.48175124 | 0.015377491 |
| Gm57162 | 0.497380598 | 2.154722595 | 0.019 | 0.008 | 1 | 0.007718533 | 3.83720485 | 0.024 | 0 | 1 | 0.497380598 | 0.015377491 |
| Gm43698 | 0.497380598 | 2.569760094 | 0.019 | 0.008 | 1 | 0.007718533 | 3.83720485 | 0.024 | 0 | 1 | 0.497380598 | 0.015377491 |
| 4930517E14Rik | 0.497380598 | 2.154722595 | 0.019 | 0.008 | 1 | 0.007718533 | 3.83720485 | 0.024 | 0 | 1 | 0.497380598 | 0.015377491 |
| Wnt10a | 0.503971666 | 1.569760094 | 0.019 | 0.008 | 1 | 0.007718533 | 3.83720485 | 0.024 | 0 | 1 | 0.503971666 | 0.015377491 |
| Gm15737 | 0.503971666 | 1.569760094 | 0.019 | 0.008 | 1 | 0.007718533 | 3.83720485 | 0.024 | 0 | 1 | 0.503971666 | 0.015377491 |
| Gna15 | 0.503971666 | 1.569760094 | 0.019 | 0.008 | 1 | 0.007718533 | 4.422167351 | 0.024 | 0 | 1 | 0.503971666 | 0.015377491 |
| Slc17a3 | 0.503971666 | 1.154722595 | 0.019 | 0.008 | 1 | 0.007718533 | 5.159132945 | 0.024 | 0 | 1 | 0.503971666 | 0.015377491 |
| Lrfn2 | 0.503971666 | 1.569760094 | 0.019 | 0.008 | 1 | 0.007718533 | 5.159132945 | 0.024 | 0 | 1 | 0.503971666 | 0.015377491 |
| Tmem125 | 0.507281812 | 1.154722595 | 0.019 | 0.008 | 1 | 0.007718533 | 3.83720485 | 0.024 | 0 | 1 | 0.507281812 | 0.015377491 |
| H2bc8 | 0.507281812 | 0.569760094 | 0.019 | 0.008 | 1 | 0.007718533 | 3.83720485 | 0.024 | 0 | 1 | 0.507281812 | 0.015377491 |
| Tff1 | 0.507281812 | 0.154722595 | 0.019 | 0.008 | 1 | 0.007718533 | 4.83720485 | 0.024 | 0 | 1 | 0.507281812 | 0.015377491 |
| H3c3 | 0.663417723 | -2.067669827 | 0.019 | 0.03 | 1 | 0.007718533 | 3.83720485 | 0.024 | 0 | 1 | 0.663417723 | 0.015377491 |
| Nmnat2 | 0.684424342 | 1.154722595 | 0.019 | 0.03 | 1 | 0.007718533 | 3.83720485 | 0.024 | 0 | 1 | 0.684424342 | 0.015377491 |
| ENSMUSG00000121511 | 0.72256466 | 1.739685096 | 0.019 | 0.013 | 1 | 0.007718533 | 3.83720485 | 0.024 | 0 | 1 | 0.72256466 | 0.015377491 |
| Wnt16 | 0.729311193 | 1.154722595 | 0.019 | 0.013 | 1 | 0.007718533 | 3.83720485 | 0.024 | 0 | 1 | 0.729311193 | 0.015377491 |
| Npl | 0.732691937 | 1.154722595 | 0.019 | 0.013 | 1 | 0.007718533 | 3.83720485 | 0.024 | 0 | 1 | 0.732691937 | 0.015377491 |
| Gm4651 | 0.736073999 | 0.8327945 | 0.019 | 0.013 | 1 | 0.007718533 | 4.422167351 | 0.024 | 0 | 1 | 0.736073999 | 0.015377491 |
| Lhfpl3 | 0.912039013 | -0.304709024 | 0.019 | 0.021 | 1 | 0.007718533 | 3.83720485 | 0.024 | 0 | 1 | 0.912039013 | 0.015377491 |
| Aldh3b2 | 0.919816481 | 0.932330173 | 0.019 | 0.017 | 1 | 0.007718533 | 4.83720485 | 0.024 | 0 | 1 | 0.919816481 | 0.015377491 |
| Fmr1nb | 0.932611893 | 0.347367673 | 0.019 | 0.017 | 1 | 0.007718533 | 3.83720485 | 0.024 | 0 | 1 | 0.932611893 | 0.015377491 |
| Reck | 0.007724282 | 0.569760094 | 0.698 | 0.555 | 1 | 0.077851854 | -0.492458894 | 0.659 | 0.771 | 1 | 0.077851854 | 0.015388899 |
| Hspa12b | 0.00772554 | 2.154722595 | 0.075 | 0.013 | 1 | 0.737522908 | -0.250257991 | 0.024 | 0.034 | 1 | 0.737522908 | 0.015391397 |
| Gask1b | 0.00772554 | 1.669295768 | 0.075 | 0.013 | 1 | 0.345194069 | -0.225804948 | 0.171 | 0.116 | 1 | 0.345194069 | 0.015391397 |
| Plppr3 | 0.00772554 | 2.154722595 | 0.075 | 0.013 | 1 | 0.119365389 | 1.252242349 | 0.049 | 0.014 | 1 | 0.119365389 | 0.015391397 |
| Tesmin | 0.00772554 | 2.154722595 | 0.075 | 0.013 | 1 | 0.057743587 | 1.721743582 | 0.098 | 0.034 | 1 | 0.057743587 | 0.015391397 |
| Fhip1a | 0.007728301 | -1.032904408 | 0.264 | 0.475 | 1 | 0.029770326 | -0.574608749 | 0.439 | 0.611 | 1 | 0.029770326 | 0.015396875 |
| Mmgt1 | 0.462485929 | 0.397993746 | 0.358 | 0.331 | 1 | 0.007761503 | -0.892529175 | 0.22 | 0.447 | 1 | 0.462485929 | 0.015462766 |
| Zfp628 | 0.007762402 | 2.614154213 | 0.113 | 0.03 | 1 | 0.014249944 | 2.078212949 | 0.171 | 0.065 | 1 | 0.014249944 | 0.015464548 |
| Mmp28 | 0.007763235 | -1.752168001 | 0.057 | 0.22 | 1 | 0.831038683 | -0.410722664 | 0.122 | 0.106 | 1 | 0.831038683 | 0.015466201 |
| Gm16536 | 0.007776542 | 2.039245377 | 0.151 | 0.051 | 1 | 0.805772606 | 1.159132945 | 0.073 | 0.089 | 1 | 0.805772606 | 0.015492608 |
| Ap1s1 | 0.412848435 | 0.412790748 | 0.358 | 0.318 | 1 | 0.007781044 | -1.10921411 | 0.195 | 0.42 | 1 | 0.412848435 | 0.015501544 |
| D6Wsu163e | 0.007784564 | 1.47665069 | 0.17 | 0.059 | 1 | 0.113140848 | 0.79281073 | 0.171 | 0.089 | 1 | 0.113140848 | 0.015508529 |
| Rtca | 0.00778912 | 0.958325382 | 0.377 | 0.208 | 1 | 0.787547543 | 0.149148856 | 0.293 | 0.276 | 1 | 0.787547543 | 0.01551757 |
| Sesn3 | 0.00779191 | 0.513068088 | 0.962 | 0.886 | 1 | 0.076688061 | 0.498454714 | 0.878 | 0.898 | 1 | 0.076688061 | 0.015523106 |
| Taf1d | 0.007822143 | 0.137149965 | 0.925 | 0.809 | 1 | 0.261043666 | 0.442097453 | 0.805 | 0.792 | 1 | 0.261043666 | 0.0155831 |
| Icam1 | 0.018419898 | 0.330493854 | 0.566 | 0.339 | 1 | 0.007823739 | 0.622761182 | 0.659 | 0.447 | 1 | 0.018419898 | 0.015586267 |
| Dnajc1 | 0.007825868 | 0.3219694 | 1 | 0.979 | 1 | 0.519401991 | 0.253704787 | 0.976 | 0.952 | 1 | 0.519401991 | 0.015590492 |
| Myo1f | 0.007828165 | 1.526691372 | 0.113 | 0.03 | 1 | 0.755974259 | 0.174239837 | 0.049 | 0.061 | 1 | 0.755974259 | 0.01559505 |
| Gfpt1 | 0.007836013 | 0.377274898 | 0.981 | 0.97 | 1 | 0.697416225 | 0.124029819 | 0.976 | 0.983 | 1 | 0.697416225 | 0.015610623 |
| Pcnp | 0.007837189 | 0.470224421 | 0.887 | 0.924 | 1 | 0.362332687 | 0.20784823 | 0.902 | 0.962 | 1 | 0.362332687 | 0.015612956 |
| Tyw5 | 0.007874827 | 1.468831185 | 0.264 | 0.127 | 1 | 0.207265753 | 0.644559772 | 0.244 | 0.164 | 1 | 0.207265753 | 0.01568764 |
| Gm26887 | 0.705883782 | -0.430239906 | 0.132 | 0.148 | 1 | 0.007883031 | 1.561570407 | 0.244 | 0.102 | 1 | 0.705883782 | 0.01570392 |
| Garin5b | 0.007887754 | 2.855162313 | 0.094 | 0.021 | 1 | 0.997398969 | -0.250257991 | 0.024 | 0.024 | 1 | 0.997398969 | 0.015713292 |
| Trim39 | 0.028123684 | -0.676677801 | 0.208 | 0.369 | 1 | 0.007934354 | -1.233184478 | 0.195 | 0.399 | 1 | 0.028123684 | 0.015805755 |
| Snx16 | 0.085074476 | 1.059565362 | 0.226 | 0.136 | 1 | 0.007954072 | -2.221688839 | 0.049 | 0.225 | 1 | 0.085074476 | 0.015844876 |
| Wdfy1 | 0.00795573 | 1.172869942 | 0.585 | 0.415 | 1 | 0.159568497 | -0.149206085 | 0.415 | 0.56 | 1 | 0.159568497 | 0.015848166 |
| Cubn | 0.007956975 | -2.337130502 | 0.038 | 0.186 | 1 | 0.188740461 | -0.286177566 | 0.049 | 0.119 | 1 | 0.188740461 | 0.015850636 |
| Gm10785 | 0.007957528 | 2.517292674 | 0.17 | 0.064 | 1 | 0.571390536 | 0.292884334 | 0.098 | 0.072 | 1 | 0.571390536 | 0.015851734 |
| Sema4a | 0.007958371 | 1.906795081 | 0.208 | 0.085 | 1 | 0.07948816 | 0.83720485 | 0.171 | 0.085 | 1 | 0.07948816 | 0.015853406 |
| Dnajc17 | 0.020215468 | 1.602181572 | 0.189 | 0.085 | 1 | 0.007977035 | 1.974708374 | 0.171 | 0.058 | 1 | 0.020215468 | 0.015890436 |
| Hdx | 0.007979029 | 1.782753817 | 0.132 | 0.038 | 1 | 0.035498737 | 2.159132945 | 0.098 | 0.031 | 1 | 0.035498737 | 0.015894393 |
| Zfp329 | 0.007994577 | 1.276028891 | 0.321 | 0.161 | 1 | 0.504513628 | 0.438655473 | 0.293 | 0.253 | 1 | 0.504513628 | 0.015925241 |
| Cmas | 0.007994719 | -0.705099747 | 0.302 | 0.513 | 1 | 0.088862469 | -0.140768844 | 0.341 | 0.526 | 1 | 0.088862469 | 0.015925522 |
| Slc25a19 | 0.008016111 | 1.71862348 | 0.189 | 0.072 | 1 | 0.130923148 | -1.101394606 | 0.049 | 0.13 | 1 | 0.130923148 | 0.015967963 |

| | | | | | | | | | | | | |
|---|---|---|---|---|---|---|---|---|---|---|---|---|
| Ptch1 | 0.008025442 | 0.840136475 | 0.604 | 0.373 | 1 | 0.440611452 | -0.173211766 | 0.488 | 0.427 | 1 | 0.440611452 | 0.015986476 |
| Dipk1a | 0.0080282 | 1.663736242 | 0.283 | 0.144 | 1 | 0.544083662 | -0.200930279 | 0.195 | 0.242 | 1 | 0.544083662 | 0.015991948 |
| L3mbtl2 | 0.116607058 | 1.47665069 | 0.075 | 0.03 | 1 | 0.008035621 | 1.83720485 | 0.171 | 0.058 | 1 | 0.116607058 | 0.01600667 |
| Ice2 | 0.008039505 | 1.466666601 | 0.321 | 0.174 | 1 | 0.447668824 | 0.559220103 | 0.268 | 0.215 | 1 | 0.447668824 | 0.016014376 |
| Gm17231 | 0.00804668 | 1.47665069 | 0.075 | 0.013 | 1 | 0.815009983 | -0.747757651 | 0.024 | 0.031 | 1 | 0.815009983 | 0.016028612 |
| Lrrc15 | 0.008049706 | 2.48403022 | 0.151 | 0.051 | 1 | 0.021167579 | 2.83720485 | 0.049 | 0.007 | 1 | 0.021167579 | 0.016034614 |
| Cryl1 | 0.00804998 | -1.015202407 | 0.415 | 0.589 | 1 | 0.01053921 | -1.001514243 | 0.439 | 0.604 | 1 | 0.01053921 | 0.016035159 |
| Tada2b | 0.471185117 | 0.487297934 | 0.245 | 0.208 | 1 | 0.008058832 | -0.959808128 | 0.122 | 0.338 | 1 | 0.471185117 | 0.016052719 |
| Gba2 | 0.008063492 | 2.154722595 | 0.113 | 0.03 | 1 | 0.370207442 | -0.555112573 | 0.024 | 0.058 | 1 | 0.370207442 | 0.016061964 |
| Lemd2 | 0.008107695 | 0.768664162 | 0.528 | 0.356 | 1 | 0.31022673 | -0.211704751 | 0.39 | 0.505 | 1 | 0.31022673 | 0.016149656 |
| A430046D13Rik | 0.008124136 | 1.984797593 | 0.113 | 0.03 | 1 | 0.257325989 | -0.484723245 | 0 | 0.031 | 1 | 0.257325989 | 0.01618227 |
| Gm15458 | 0.008124136 | 1.984797593 | 0.113 | 0.03 | 1 | 0.861594903 | -0.101394606 | 0.049 | 0.055 | 1 | 0.861594903 | 0.01618227 |
| Pik3r4 | 0.008129369 | 1.117247889 | 0.245 | 0.11 | 1 | 0.067154911 | 1.515276755 | 0.268 | 0.164 | 1 | 0.067154911 | 0.01619265 |
| Cyp2j6 | 0.008136962 | -1.512033997 | 0.132 | 0.305 | 1 | 0.379295844 | 0.655001519 | 0.195 | 0.147 | 1 | 0.379295844 | 0.016207713 |
| D030056L22Rik | 0.00814229 | 1.211306123 | 0.321 | 0.169 | 1 | 0.109640288 | 0.969308386 | 0.268 | 0.171 | 1 | 0.109640288 | 0.016218282 |
| Fmo5 | 0.040822179 | 1.598329246 | 0.151 | 0.068 | 1 | 0.008145527 | 1.351778023 | 0.146 | 0.044 | 1 | 0.040822179 | 0.016224704 |
| Spink10 | 0.008158003 | 2.932330173 | 0.094 | 0.021 | 1 | 0.622601109 | 0.313642894 | 0.024 | 0.041 | 1 | 0.622601109 | 0.016249452 |
| Baz2b | 0.008161117 | 0.423196741 | 0.962 | 0.983 | 1 | 0.019710923 | 0.488320205 | 0.927 | 0.973 | 1 | 0.019710923 | 0.01625563 |
| C9orf72 | 0.008161464 | -1.304709024 | 0.17 | 0.352 | 1 | 0.57279404 | 0.142617858 | 0.293 | 0.253 | 1 | 0.57279404 | 0.016256318 |
| Irak1bp1 | 0.008164668 | 1.228723176 | 0.396 | 0.242 | 1 | 0.279184257 | -0.395455907 | 0.22 | 0.304 | 1 | 0.279184257 | 0.016262675 |
| Dll1 | 0.008185188 | 1.865215978 | 0.113 | 0.03 | 1 | 0.057901606 | 2.422167351 | 0.049 | 0.01 | 1 | 0.057901606 | 0.016303378 |
| Afg3l2 | 0.008194046 | 0.578693219 | 0.585 | 0.377 | 1 | 0.936403042 | 0.220533489 | 0.512 | 0.526 | 1 | 0.936403042 | 0.016320949 |
| Pycard | 0.008215574 | 1.647762606 | 0.208 | 0.089 | 1 | 0.08681393 | 1.0375035 | 0.366 | 0.253 | 1 | 0.08681393 | 0.016363652 |
| Gm33508 | 0.008227447 | 2.517292674 | 0.094 | 0.021 | 1 | 0.44410834 | 1.252242349 | 0.024 | 0.01 | 1 | 0.44410834 | 0.016387204 |
| Ctbp1 | 0.008272607 | 0.634150681 | 0.736 | 0.597 | 1 | 0.643576904 | 0.252242349 | 0.707 | 0.758 | 1 | 0.643576904 | 0.016476778 |
| Tmem62 | 0.008273228 | 1.948271717 | 0.151 | 0.051 | 1 | 0.97435949 | -0.233184478 | 0.098 | 0.096 | 1 | 0.97435949 | 0.016478009 |
| Iffo2 | 0.083629167 | 0.199391644 | 0.623 | 0.504 | 1 | 0.008282767 | 0.456781106 | 0.829 | 0.597 | 1 | 0.083629167 | 0.016496929 |
| Klhl18 | 0.008285961 | 1.133660979 | 0.358 | 0.199 | 1 | 0.05035072 | -1.093532488 | 0.146 | 0.287 | 1 | 0.05035072 | 0.016503264 |
| Cep131 | 0.008292454 | 2.569760094 | 0.094 | 0.021 | 1 | 0.714406212 | 0.930314254 | 0.049 | 0.038 | 1 | 0.714406212 | 0.016516143 |
| Gm6556 | 0.008292454 | 2.377115016 | 0.094 | 0.021 | 1 | 0.459320496 | 0.962735732 | 0.049 | 0.027 | 1 | 0.459320496 | 0.016516143 |
| 6720489N17Rik | 0.008292454 | 2.377115016 | 0.094 | 0.021 | 1 | 0.319636904 | -0.863234868 | 0 | 0.024 | 1 | 0.319636904 | 0.016516143 |
| Zfp429 | 0.008292454 | 2.377115016 | 0.094 | 0.021 | 1 | 0.37346373 | 1.029849928 | 0.098 | 0.061 | 1 | 0.37346373 | 0.016516143 |
| Gm16156 | 0.008297581 | 2.347367673 | 0.094 | 0.021 | 1 | 0.286422281 | -0.747757651 | 0 | 0.027 | 1 | 0.286422281 | 0.016526312 |
| Gm57387 | 0.008297581 | 2.347367673 | 0.094 | 0.021 | 1 | 0.518949099 | 0.334704509 | 0.024 | 0.048 | 1 | 0.518949099 | 0.016526312 |
| Fbxo8 | 0.00830001 | 1.154722595 | 0.415 | 0.258 | 1 | 0.738058141 | 0.279209397 | 0.244 | 0.232 | 1 | 0.738058141 | 0.01653113 |
| Znfx1 | 0.008300738 | 1.18386894 | 0.302 | 0.153 | 1 | 0.535783281 | 0.621476159 | 0.268 | 0.331 | 1 | 0.535783281 | 0.016532574 |
| Sfr1 | 0.390800133 | 0.126182427 | 1 | 0.949 | 1 | 0.008319644 | -0.475935228 | 0.902 | 0.983 | 1 | 0.390800133 | 0.016570071 |
| Mob3c | 0.008327747 | 1.105812994 | 0.358 | 0.195 | 1 | 0.35955614 | -0.516432105 | 0.146 | 0.208 | 1 | 0.35955614 | 0.016586142 |
| Mrgbp | 0.008348554 | 1.285967128 | 0.283 | 0.14 | 1 | 0.266537396 | 0.858578501 | 0.244 | 0.184 | 1 | 0.266537396 | 0.016627409 |
| Echdc2 | 0.008359873 | 0.909610097 | 0.358 | 0.195 | 1 | 0.341191366 | -0.762707992 | 0.171 | 0.225 | 1 | 0.341191366 | 0.016649859 |
| Trim32 | 0.008362724 | 2.039245377 | 0.151 | 0.051 | 1 | 0.852951204 | 0.167353451 | 0.195 | 0.208 | 1 | 0.852951204 | 0.016655513 |
| Rab3ip | 0.008373614 | -2.678167419 | 0.019 | 0.153 | 1 | 0.226430878 | 0.186950889 | 0.317 | 0.222 | 1 | 0.226430878 | 0.016677111 |
| Ddx39b | 0.008388802 | 0.611929548 | 0.849 | 0.831 | 1 | 0.422020891 | -0.14408289 | 0.829 | 0.887 | 1 | 0.422020891 | 0.016707232 |
| Wdr1 | 0.008391074 | -0.487229885 | 0.698 | 0.839 | 1 | 0.018964022 | -0.430951989 | 0.732 | 0.884 | 1 | 0.018964022 | 0.016711738 |
| Hars2 | 0.008397365 | 1.005859209 | 0.34 | 0.178 | 1 | 0.576206881 | -0.314798244 | 0.171 | 0.205 | 1 | 0.576206881 | 0.016724214 |
| Gm4876 | 0.008398882 | -1.814903756 | 0.038 | 0.186 | 1 | 0.830428412 | -0.134780774 | 0.098 | 0.089 | 1 | 0.830428412 | 0.016727223 |
| Csnk2b | 0.008399925 | 0.484184362 | 0.868 | 0.754 | 1 | 0.291901524 | 0.412282762 | 0.78 | 0.823 | 1 | 0.291901524 | 0.016729291 |
| Fam89b | 0.008403953 | 0.611100889 | 0.585 | 0.432 | 1 | 0.393125103 | 0.383028957 | 0.512 | 0.478 | 1 | 0.393125103 | 0.01673728 |
| Katna1 | 0.008419242 | 0.964619711 | 0.566 | 0.403 | 1 | 0.383028607 | 0.299939091 | 0.561 | 0.509 | 1 | 0.383028607 | 0.016767601 |
| Wnk2 | 0.722666233 | -0.253362144 | 0.132 | 0.148 | 1 | 0.008419352 | -0.930979475 | 0.146 | 0.382 | 1 | 0.722666233 | 0.016767818 |
| Ogg1 | 0.00843264 | 1.71862348 | 0.208 | 0.089 | 1 | 0.408881974 | 1.052933541 | 0.122 | 0.085 | 1 | 0.408881974 | 0.016794171 |
| Map7d1 | 0.008469874 | 0.448168314 | 0.868 | 0.775 | 1 | 0.049075619 | -0.3200091 | 0.854 | 0.901 | 1 | 0.049075619 | 0.016868008 |
| Iah1 | 0.008475559 | 1.244920404 | 0.358 | 0.208 | 1 | 0.232810568 | 1.133189677 | 0.317 | 0.256 | 1 | 0.232810568 | 0.016879283 |
| Rusc2 | 0.008478478 | -1.061933457 | 0.283 | 0.475 | 1 | 0.049081772 | -0.701418936 | 0.341 | 0.519 | 1 | 0.049081772 | 0.016885071 |
| Fstl3 | 0.008500139 | 1.8327945 | 0.113 | 0.03 | 1 | 0.085439336 | 1.029849928 | 0.146 | 0.068 | 1 | 0.085439336 | 0.016928026 |
| Usp28 | 0.008511834 | 0.850569202 | 0.434 | 0.258 | 1 | 0.500810045 | -0.27321914 | 0.268 | 0.338 | 1 | 0.500810045 | 0.016951216 |
| Lpxn | 0.008512345 | 2.324647596 | 0.094 | 0.021 | 1 | 0.979193861 | 0.83720485 | 0.024 | 0.024 | 1 | 0.979193861 | 0.01695223 |
| Birc3 | 0.008516936 | 1.169519597 | 0.377 | 0.22 | 1 | 0.053897507 | 1.20784823 | 0.341 | 0.229 | 1 | 0.053897507 | 0.016961335 |
| Gprasp1 | 0.79057294 | 0.282183504 | 0.528 | 0.551 | 1 | 0.008527912 | -0.599659012 | 0.39 | 0.638 | 1 | 0.79057294 | 0.016983099 |
| Inpp5f | 0.008539098 | 1.179813576 | 0.453 | 0.305 | 1 | 0.328619003 | -0.438429593 | 0.293 | 0.358 | 1 | 0.328619003 | 0.017005279 |
| Gm14963 | 0.008544835 | -3.1672055 | 0 | 0.119 | 1 | 0.619815827 | -0.484723245 | 0.049 | 0.068 | 1 | 0.619815827 | 0.017016655 |
| Cd74 | 0.008546853 | -5.559522923 | 0.038 | 0.178 | 1 | 0.270418241 | -1.410722664 | 0.024 | 0.068 | 1 | 0.270418241 | 0.017020658 |
| Farsb | 0.008547744 | -0.752168001 | 0.245 | 0.445 | 1 | 0.186553708 | -0.509597914 | 0.366 | 0.471 | 1 | 0.186553708 | 0.017022423 |
| Agt | 0.008567092 | 1.502645898 | 0.302 | 0.161 | 1 | 0.744047924 | 1.252242349 | 0.024 | 0.017 | 1 | 0.744047924 | 0.017060789 |
| Camk1 | 0.00856769 | -0.681201479 | 0.528 | 0.695 | 1 | 0.095376353 | -0.326162436 | 0.512 | 0.655 | 1 | 0.095376353 | 0.017061976 |
| Spice1 | 0.0085828 | 0.932330173 | 0.321 | 0.161 | 1 | 0.978697305 | 0.136765132 | 0.146 | 0.147 | 1 | 0.978697305 | 0.017091936 |
| Gm35162 | 0.008587359 | -1.752168001 | 0.019 | 0.157 | 1 | 0.117593704 | -1.20718927 | 0.024 | 0.099 | 1 | 0.117593704 | 0.017100975 |
| Atp11b | 0.012964454 | 0.722916882 | 0.774 | 0.665 | 1 | 0.008603658 | 0.677461225 | 0.854 | 0.819 | 1 | 0.012964454 | 0.017133293 |
| Dnmbp | 0.008605954 | -1.212648471 | 0.264 | 0.458 | 1 | 0.068511184 | -0.652179991 | 0.439 | 0.587 | 1 | 0.068511184 | 0.017137845 |
| Rpl22l1 | 0.047827692 | -0.411669158 | 1 | 0.966 | 1 | 0.008615828 | -0.464175868 | 0.951 | 0.98 | 1 | 0.047827692 | 0.017157424 |
| Gm49797 | 0.008616972 | 0.652222254 | 0.642 | 0.5 | 1 | 0.304636036 | -0.262330824 | 0.756 | 0.775 | 1 | 0.304636036 | 0.017159692 |
| Ctdsp2 | 0.008621285 | 0.426975531 | 0.849 | 0.805 | 1 | 0.180275444 | -0.231052679 | 0.927 | 0.922 | 1 | 0.180275444 | 0.017168243 |
| Lman1 | 0.008621974 | 0.325994022 | 0.962 | 0.928 | 1 | 0.207729714 | -0.178736694 | 0.878 | 0.935 | 1 | 0.207729714 | 0.01716961 |
| Nup214 | 0.00862527 | 0.765310903 | 0.604 | 0.453 | 1 | 0.376870714 | 0.234208426 | 0.634 | 0.57 | 1 | 0.376870714 | 0.017176145 |
| Slco3a1 | 0.010071916 | -1.186745686 | 0.208 | 0.394 | 1 | 0.008632323 | 1.388446733 | 0.366 | 0.188 | 1 | 0.010071916 | 0.01719013 |
| Lrrc8d | 0.008642123 | -0.667707648 | 0.509 | 0.708 | 1 | 0.034411863 | -0.558954639 | 0.707 | 0.874 | 1 | 0.034411863 | 0.017209561 |
| Uso1 | 0.180538191 | 0.344658385 | 0.83 | 0.86 | 1 | 0.008642463 | -0.485966412 | 0.878 | 0.935 | 1 | 0.180538191 | 0.017210233 |
| Osbpl6 | 0.008646616 | 0.791252238 | 0.755 | 0.602 | 1 | 0.644970583 | 0.101049986 | 0.659 | 0.686 | 1 | 0.644970583 | 0.017218468 |
| 9330133O14Rik | 0.008646987 | 1.306725688 | 0.17 | 0.059 | 1 | 0.708632143 | 1.252242349 | 0.049 | 0.038 | 1 | 0.708632143 | 0.017219203 |
| Ubl7 | 0.008649006 | 0.964898036 | 0.415 | 0.246 | 1 | 0.094345676 | 1.170942247 | 0.293 | 0.198 | 1 | 0.094345676 | 0.017223207 |
| Ppp1r26 | 0.008653601 | 2.154722595 | 0.094 | 0.021 | 1 | 0.672172778 | 0.136765132 | 0.024 | 0.038 | 1 | 0.672172778 | 0.017232318 |
| Slc37a4 | 0.008660554 | 2.154722595 | 0.094 | 0.021 | 1 | 0.440032309 | 1.83720485 | 0.049 | 0.027 | 1 | 0.440032309 | 0.017246102 |
| Zxdb | 0.008660554 | 2.154722595 | 0.094 | 0.021 | 1 | 0.040560279 | -1.622226769 | 0.024 | 0.137 | 1 | 0.040560279 | 0.017246102 |
| Sfxn1 | 0.354078102 | -0.430239906 | 0.321 | 0.381 | 1 | 0.008661412 | -1.02786557 | 0.171 | 0.403 | 1 | 0.354078102 | 0.017247804 |
| Med4 | 0.008672867 | 1.065085382 | 0.415 | 0.263 | 1 | 0.25591227 | -0.410722664 | 0.244 | 0.331 | 1 | 0.25591227 | 0.017270516 |
| Smarcad1 | 0.00867979 | 0.575860294 | 0.679 | 0.508 | 1 | 0.25839224 | -0.307845483 | 0.585 | 0.696 | 1 | 0.25839224 | 0.01728424 |
| A430033K04Rik | 0.008680295 | 2.047807391 | 0.151 | 0.051 | 1 | 0.040689782 | 1.83720485 | 0.171 | 0.078 | 1 | 0.040689782 | 0.017285242 |
| Casc1 | 0.008682591 | 1.618669695 | 0.208 | 0.089 | 1 | 0.309072815 | -1.292078167 | 0.122 | 0.177 | 1 | 0.309072815 | 0.017289795 |
| Asrgl1 | 0.360158256 | 0.460135603 | 0.396 | 0.339 | 1 | 0.008688775 | -1.321224513 | 0.098 | 0.294 | 1 | 0.360158256 | 0.017302055 |
| Prss35 | 0.216510852 | 2.517292674 | 0.075 | 0.038 | 1 | 0.008719547 | 3.043655727 | 0.122 | 0.034 | 1 | 0.216510852 | 0.017363064 |
| Phc3 | 0.008725302 | 0.675113094 | 0.623 | 0.475 | 1 | 0.847012217 | 0.149148856 | 0.634 | 0.648 | 1 | 0.847012217 | 0.017374473 |
| Ptcra | 0.008726892 | 1.792152515 | 0.094 | 0.021 | 1 | 0.085488466 | 1.574170444 | 0.073 | 0.024 | 1 | 0.085488466 | 0.017377625 |
| Gm35867 | 0.008726892 | 1.962077517 | 0.094 | 0.021 | 1 | 0.745416242 | 0.334704509 | 0.049 | 0.038 | 1 | 0.745416242 | 0.017377625 |
| Zkscan14 | 0.008733108 | 1.932330173 | 0.151 | 0.051 | 1 | 0.037536938 | 1.547698233 | 0.146 | 0.058 | 1 | 0.037536938 | 0.017389949 |
| Pomk | 0.008733108 | 1.932330173 | 0.151 | 0.051 | 1 | 0.568044235 | 1.19334866 | 0.098 | 0.075 | 1 | 0.568044235 | 0.017389949 |
| Alg14 | 0.008747829 | -0.746499251 | 0.509 | 0.699 | 1 | 0.078021172 | -0.214223389 | 0.317 | 0.505 | 1 | 0.078021172 | 0.017419134 |
| Cep170b | 0.059607551 | -1.015202407 | 0.132 | 0.25 | 1 | 0.008764314 | -1.502645153 | 0.146 | 0.345 | 1 | 0.059607551 | 0.017451814 |
| Saal1 | 0.444883628 | 0.292226119 | 0.17 | 0.127 | 1 | 0.008781941 | -2.747757651 | 0 | 0.147 | 1 | 0.444883628 | 0.017486759 |

| | | | | | | | | | | | | |
|---|---|---|---|---|---|---|---|---|---|---|---|---|
| Fbxw17 | 0.008786295 | 1.806799291 | 0.151 | 0.051 | 1 | 0.447279949 | 0.410940095 | 0.146 | 0.106 | 1 | 0.447279949 | 0.01749539 |
| Tm9sf2 | 0.008811286 | 0.276456909 | 0.962 | 0.949 | 1 | 0.036350033 | -0.313122714 | 0.927 | 0.969 | 1 | 0.036350033 | 0.017544933 |
| Gm15614 | 0.00881436 | 1.225111923 | 0.264 | 0.123 | 1 | 0.801255761 | 0.252242349 | 0.171 | 0.157 | 1 | 0.801255761 | 0.017551028 |
| Telo2 | 0.008814984 | 1.657222935 | 0.208 | 0.085 | 1 | 0.819451157 | 1.450181727 | 0.098 | 0.092 | 1 | 0.819451157 | 0.017552264 |
| Sdhd | 0.008819127 | 0.730914886 | 0.66 | 0.606 | 1 | 0.192533493 | -0.354813922 | 0.561 | 0.638 | 1 | 0.192533493 | 0.017560476 |
| AI506816 | 0.206721143 | -0.247678079 | 0.755 | 0.822 | 1 | 0.008831219 | -0.57921919 | 0.585 | 0.741 | 1 | 0.206721143 | 0.017584447 |
| Slc6a17 | 0.33117623 | -0.667279103 | 0.189 | 0.246 | 1 | 0.008832617 | -1.260827233 | 0.195 | 0.399 | 1 | 0.33117623 | 0.017587218 |
| Morrbid | 0.099340139 | 1.232725107 | 0.132 | 0.064 | 1 | 0.008846545 | 3.199774929 | 0.122 | 0.034 | 1 | 0.099340139 | 0.017614829 |
| Clcn5 | 0.109741892 | 0.373362881 | 0.434 | 0.339 | 1 | 0.008854292 | 1.183655264 | 0.561 | 0.413 | 1 | 0.109741892 | 0.017630186 |
| Ppl | 0.479626322 | 1.154722595 | 0.038 | 0.021 | 1 | 0.00887104 | 1.359157553 | 0.268 | 0.119 | 1 | 0.479626322 | 0.017663384 |
| Mmp25 | 0.008877417 | 2.002719501 | 0.094 | 0.021 | 1 | 0.539566166 | 1.059597271 | 0.073 | 0.051 | 1 | 0.539566166 | 0.017676026 |
| Hltf | 0.00887983 | 0.848619467 | 0.528 | 0.347 | 1 | 0.221898731 | 0.565902828 | 0.39 | 0.317 | 1 | 0.221898731 | 0.017680809 |
| Dhx8 | 0.008887227 | 1.026967048 | 0.509 | 0.347 | 1 | 0.110049444 | 0.876375447 | 0.537 | 0.44 | 1 | 0.110049444 | 0.017695472 |
| Sh3bp1 | 0.008893406 | 1.292226119 | 0.151 | 0.051 | 1 | 0.026172634 | 1.408361551 | 0.22 | 0.102 | 1 | 0.026172634 | 0.017707719 |
| Gm26802 | 0.008907944 | 1.389187848 | 0.283 | 0.144 | 1 | 0.010477546 | 1.087183103 | 0.39 | 0.218 | 1 | 0.010477546 | 0.017736537 |
| Timm29 | 0.008915171 | 1.191248471 | 0.283 | 0.14 | 1 | 0.089585021 | -0.693309867 | 0.146 | 0.276 | 1 | 0.089585021 | 0.017750861 |
| Castor2 | 0.008929419 | 1.133660979 | 0.34 | 0.186 | 1 | 0.214491134 | -0.173364392 | 0.39 | 0.549 | 1 | 0.214491134 | 0.017779103 |
| Zw10 | 0.00894232 | 0.877188619 | 0.415 | 0.246 | 1 | 0.120731042 | -0.305753104 | 0.244 | 0.396 | 1 | 0.120731042 | 0.017804675 |
| Erbb2 | 0.008947583 | 1.377115016 | 0.208 | 0.085 | 1 | 0.741213095 | -0.122153166 | 0.171 | 0.191 | 1 | 0.741213095 | 0.017815107 |
| Rpn2 | 0.008958042 | -0.402556347 | 0.774 | 0.898 | 1 | 0.280748775 | -0.15418202 | 0.805 | 0.86 | 1 | 0.280748775 | 0.017835838 |
| Anp32a | 0.00896421 | 0.548186682 | 0.925 | 0.814 | 1 | 0.02010871 | 0.454927099 | 0.878 | 0.799 | 1 | 0.02010871 | 0.017848063 |
| Tmem80 | 0.871479999 | 0.384204441 | 0.208 | 0.195 | 1 | 0.008987195 | -1.470223675 | 0.073 | 0.263 | 1 | 0.871479999 | 0.017893621 |
| Coro1b | 0.009002911 | 1.3662267 | 0.189 | 0.072 | 1 | 0.045809309 | 1.515276755 | 0.171 | 0.078 | 1 | 0.045809309 | 0.01792477 |
| Gga2 | 0.040138202 | 0.560361396 | 0.491 | 0.36 | 1 | 0.009007497 | -0.957975358 | 0.22 | 0.457 | 1 | 0.040138202 | 0.017933858 |
| 2310047K21Rik | 0.009018564 | 1.962077517 | 0.17 | 0.064 | 1 | 0.971058571 | 0.911205431 | 0.049 | 0.051 | 1 | 0.971058571 | 0.017955793 |
| Pdk1 | 0.309114692 | -0.182312392 | 0.208 | 0.292 | 1 | 0.00901978 | -1.648221977 | 0.073 | 0.259 | 1 | 0.309114692 | 0.017958203 |
| Aoc3 | 0.009022556 | 1.932330173 | 0.094 | 0.021 | 1 | 0.036608095 | 2.100239256 | 0.098 | 0.031 | 1 | 0.036608095 | 0.017963706 |
| Kif27 | 0.009022556 | 1.932330173 | 0.094 | 0.021 | 1 | 0.607145967 | 2.029849928 | 0.049 | 0.034 | 1 | 0.607145967 | 0.017963706 |
| Ppp1r3e | 0.009022556 | 1.739685096 | 0.094 | 0.021 | 1 | 0.529718256 | 1.515276755 | 0.049 | 0.031 | 1 | 0.529718256 | 0.017963706 |
| Polr3g | 0.009023643 | 1.454282877 | 0.113 | 0.03 | 1 | 0.620260362 | 1.377773231 | 0.049 | 0.034 | 1 | 0.620260362 | 0.017965859 |
| Zbtb9 | 0.009033206 | 1.602181572 | 0.189 | 0.072 | 1 | 0.612010787 | 0.667279848 | 0.122 | 0.096 | 1 | 0.612010787 | 0.017984813 |
| Rwdd1 | 0.122691695 | -0.214511215 | 0.792 | 0.839 | 1 | 0.009037218 | 0.562842438 | 0.805 | 0.758 | 1 | 0.122691695 | 0.017992766 |
| Slc29a1 | 0.00903842 | 0.55381855 | 0.849 | 0.89 | 1 | 0.269698881 | -0.149429892 | 0.707 | 0.809 | 1 | 0.269698881 | 0.017995147 |
| Mtres1 | 0.009043382 | 1.081966252 | 0.358 | 0.203 | 1 | 0.327498218 | -0.180717058 | 0.122 | 0.195 | 1 | 0.327498218 | 0.018004981 |
| Sde2 | 0.211200573 | 0.446903346 | 0.66 | 0.72 | 1 | 0.009044411 | 0.689928587 | 0.805 | 0.724 | 1 | 0.211200573 | 0.018007021 |
| Traf3 | 0.009053235 | 0.856064279 | 0.623 | 0.517 | 1 | 0.986690542 | 0.11789312 | 0.707 | 0.751 | 1 | 0.986690542 | 0.01802451 |
| Plekhg3 | 0.01641577 | -1.389597921 | 0.113 | 0.28 | 1 | 0.009067459 | -0.464331456 | 0.463 | 0.618 | 1 | 0.01641577 | 0.018052699 |
| L3mbtl1 | 0.039917687 | 0.859266711 | 0.151 | 0.064 | 1 | 0.009069256 | 1.862295831 | 0.244 | 0.106 | 1 | 0.039917687 | 0.018056261 |
| C87436 | 0.009076503 | 0.945269229 | 0.377 | 0.208 | 1 | 0.021483362 | 1.450181727 | 0.341 | 0.208 | 1 | 0.021483362 | 0.018070622 |
| Stau1 | 0.009078248 | 0.693711671 | 0.83 | 0.78 | 1 | 0.155763059 | 0.449022842 | 0.976 | 0.959 | 1 | 0.155763059 | 0.018074082 |
| Pxk | 0.009128487 | 0.996024849 | 0.415 | 0.242 | 1 | 0.891178824 | 0.116982519 | 0.488 | 0.485 | 1 | 0.891178824 | 0.018173644 |
| Hcfc2 | 0.009157295 | 0.857042046 | 0.528 | 0.347 | 1 | 0.0136447 | -0.833042839 | 0.366 | 0.543 | 1 | 0.0136447 | 0.018230734 |
| Ppif | 0.590054933 | -1.1672055 | 0.17 | 0.195 | 1 | 0.009164965 | 1.786578777 | 0.195 | 0.075 | 1 | 0.590054933 | 0.018245933 |
| Hectd3 | 0.009177121 | 1.347367673 | 0.189 | 0.072 | 1 | 0.789367819 | 0.46797104 | 0.098 | 0.085 | 1 | 0.789367819 | 0.018270023 |
| Nepro | 0.009208188 | 1.154722595 | 0.264 | 0.123 | 1 | 0.655853123 | 0.313642894 | 0.22 | 0.188 | 1 | 0.655853123 | 0.018331585 |
| Ifnar1 | 0.009222364 | 1.127755547 | 0.491 | 0.347 | 1 | 0.434120162 | 0.543845907 | 0.488 | 0.519 | 1 | 0.434120162 | 0.018359676 |
| Ncor2 | 0.009230965 | 0.472557935 | 0.811 | 0.708 | 1 | 0.317600399 | 0.238945526 | 0.854 | 0.87 | 1 | 0.317600399 | 0.018376719 |
| Psmb3 | 0.009237383 | 0.38704098 | 0.906 | 0.839 | 1 | 0.317515382 | 0.19448063 | 0.854 | 0.86 | 1 | 0.317515382 | 0.018389436 |
| Glcci1 | 0.009932543 | 0.782389546 | 0.528 | 0.364 | 1 | 0.009240773 | 0.764225404 | 0.659 | 0.491 | 1 | 0.009932543 | 0.018396154 |
| Dgke | 0.009242964 | 1.94321849 | 0.189 | 0.076 | 1 | 0.069170662 | 1.766815522 | 0.146 | 0.068 | 1 | 0.069170662 | 0.018400495 |
| Gtf3c4 | 0.009268628 | 1.341135719 | 0.358 | 0.216 | 1 | 0.442468369 | 0.515276755 | 0.439 | 0.382 | 1 | 0.442468369 | 0.018451349 |
| Anks1b | 0.009272118 | -3.043547032 | 0.094 | 0.254 | 1 | 0.728800583 | 0.252242349 | 0.22 | 0.198 | 1 | 0.728800583 | 0.018458264 |
| Coro1c | 0.009281591 | 0.414690698 | 0.566 | 0.347 | 1 | 0.065785338 | 0.935385244 | 0.561 | 0.447 | 1 | 0.065785338 | 0.018477034 |
| Sec23a | 0.145042596 | 0.329028082 | 0.774 | 0.775 | 1 | 0.009282083 | -0.386904044 | 0.732 | 0.84 | 1 | 0.145042596 | 0.018478009 |
| Slc35b2 | 0.009305129 | -0.636993203 | 0.528 | 0.699 | 1 | 0.511366845 | -0.263881275 | 0.463 | 0.515 | 1 | 0.511366845 | 0.018523673 |
| Rcan3 | 0.009332925 | 1.331600357 | 0.208 | 0.085 | 1 | 0.028740964 | 1.401105735 | 0.244 | 0.126 | 1 | 0.028740964 | 0.018578746 |
| Parp12 | 0.009364422 | 0.96568877 | 0.302 | 0.148 | 1 | 0.605263547 | 0.686645173 | 0.244 | 0.215 | 1 | 0.605263547 | 0.018641151 |
| Gm49417 | 0.009371301 | -1.878700407 | 0.075 | 0.233 | 1 | 0.01800977 | -1.800868987 | 0.122 | 0.297 | 1 | 0.01800977 | 0.01865478 |
| Eef1a1 | 0.031193485 | 0.224805355 | 1 | 1 | 1 | 0.009394553 | -0.237742857 | 1 | 0.997 | 1 | 0.031193485 | 0.018700849 |
| Rasa2 | 0.009422306 | 0.504022297 | 0.792 | 0.657 | 1 | 0.864053048 | -0.224116598 | 0.805 | 0.799 | 1 | 0.864053048 | 0.018755833 |
| Xkr4 | 0.009433874 | 2.154722595 | 0.17 | 0.064 | 1 | 0.082845297 | 1.252242349 | 0.293 | 0.181 | 1 | 0.082845297 | 0.018778749 |
| Dcun1d2 | 0.009442328 | 1.8327945 | 0.245 | 0.123 | 1 | 0.550788914 | 1.451551157 | 0.22 | 0.195 | 1 | 0.550788914 | 0.018795499 |
| Cenpl | 0.336647531 | 0.389187848 | 0.075 | 0.042 | 1 | 0.009463314 | 1.557096931 | 0.146 | 0.044 | 1 | 0.336647531 | 0.018837073 |
| Srp72 | 0.009472098 | 0.402650108 | 0.906 | 0.864 | 1 | 0.287556553 | -0.128579435 | 0.707 | 0.877 | 1 | 0.287556553 | 0.018854476 |
| Asl | 0.00947539 | 1.347367673 | 0.226 | 0.097 | 1 | 0.114480585 | -0.513292397 | 0.049 | 0.14 | 1 | 0.114480585 | 0.018860997 |
| Itpkb | 0.009493803 | -0.849669174 | 0.377 | 0.551 | 1 | 0.076535545 | 1.181320563 | 0.512 | 0.386 | 1 | 0.076535545 | 0.018897474 |
| Veph1 | 0.232728177 | -0.52334931 | 0.302 | 0.39 | 1 | 0.009499987 | -0.793008732 | 0.22 | 0.437 | 1 | 0.232728177 | 0.018909724 |
| Bax | 0.009508551 | 0.574138535 | 0.698 | 0.657 | 1 | 0.251032078 | -0.264674764 | 0.61 | 0.645 | 1 | 0.251032078 | 0.01892669 |
| Zcchc2 | 0.009519302 | 1.132354782 | 0.358 | 0.203 | 1 | 0.012750109 | 1.488055286 | 0.39 | 0.242 | 1 | 0.012750109 | 0.018947986 |
| Gm7694 | 0.646171646 | 0.228723176 | 0.057 | 0.076 | 1 | 0.009561653 | -2.352619709 | 0.024 | 0.184 | 1 | 0.646171646 | 0.01903188 |
| Sorbs2os | 0.009565085 | 0.984797593 | 0.264 | 0.123 | 1 | 0.018632004 | -1.429581691 | 0.049 | 0.205 | 1 | 0.018632004 | 0.01903868 |
| Tspyl2 | 0.009577073 | 1.036078098 | 0.396 | 0.237 | 1 | 0.631239113 | 0.310659035 | 0.244 | 0.311 | 1 | 0.631239113 | 0.019062427 |
| Pcif1 | 0.009577949 | 0.981390992 | 0.472 | 0.326 | 1 | 0.4273014 | 0.129385601 | 0.366 | 0.427 | 1 | 0.4273014 | 0.019064161 |
| Cyp4f13 | 0.00958258 | 1.10741688 | 0.245 | 0.11 | 1 | 0.279881156 | 0.922934724 | 0.171 | 0.113 | 1 | 0.279881156 | 0.019073333 |
| Cib2 | 0.00958533 | 1.154722595 | 0.245 | 0.11 | 1 | 0.228070883 | -0.648221977 | 0.073 | 0.143 | 1 | 0.228070883 | 0.019078781 |
| Pof1b | 0.009586565 | -2.953801862 | 0.019 | 0.148 | 1 | 0.142447868 | 1.212713985 | 0.195 | 0.116 | 1 | 0.142447868 | 0.019081228 |
| 2900076A07Rik | 0.009606102 | 0.94321849 | 0.283 | 0.136 | 1 | 0.617528011 | 0.377773231 | 0.171 | 0.212 | 1 | 0.617528011 | 0.019119928 |
| Hccs | 0.009625668 | 1.221836791 | 0.358 | 0.195 | 1 | 0.419514646 | 0.581947795 | 0.244 | 0.314 | 1 | 0.419514646 | 0.019158683 |
| 2610035D17Rik | 0.065352095 | -1.031373697 | 0.453 | 0.564 | 1 | 0.009643997 | 0.614812428 | 0.585 | 0.406 | 1 | 0.065352095 | 0.019194987 |
| Ehd2 | 0.905863422 | 0.111095662 | 0.547 | 0.547 | 1 | 0.009645762 | -0.788115389 | 0.585 | 0.72 | 1 | 0.905863422 | 0.019198484 |
| Gm14164 | 0.592586815 | 0.22061053 | 0.396 | 0.36 | 1 | 0.009662771 | -0.932182222 | 0.171 | 0.386 | 1 | 0.592586815 | 0.019232173 |
| Bche | 0.009671214 | -2.1672055 | 0.038 | 0.178 | 1 | 0.019770897 | -2.917682652 | 0 | 0.119 | 1 | 0.019770897 | 0.019248896 |
| Homer3 | 0.009674541 | 0.685237311 | 0.377 | 0.199 | 1 | 0.069797707 | -0.661046018 | 0.268 | 0.41 | 1 | 0.069797707 | 0.019255486 |
| Dyrk2 | 0.009688626 | 0.695290976 | 0.453 | 0.254 | 1 | 0.395242765 | -0.540100002 | 0.39 | 0.44 | 1 | 0.395242765 | 0.019283382 |
| Blmh | 0.009693855 | 0.665836066 | 0.83 | 0.686 | 1 | 0.012585197 | -0.537834582 | 0.707 | 0.819 | 1 | 0.012585197 | 0.019293739 |
| Prkd2 | 0.009706597 | 0.134823037 | 0.453 | 0.246 | 1 | 0.059265286 | -0.590401323 | 0.268 | 0.433 | 1 | 0.059265286 | 0.019318976 |
| Atp6v0e2 | 0.009714931 | 1.624207878 | 0.151 | 0.051 | 1 | 0.188380068 | 1.614812428 | 0.049 | 0.017 | 1 | 0.188380068 | 0.019335482 |
| Vps26c | 0.009734187 | 1.080722013 | 0.283 | 0.136 | 1 | 0.729609522 | 0.370886846 | 0.268 | 0.259 | 1 | 0.729609522 | 0.019373619 |
| N4bp2l2 | 0.009778945 | 0.370182302 | 0.906 | 0.881 | 1 | 0.808879269 | 0.106525518 | 0.854 | 0.901 | 1 | 0.808879269 | 0.019462262 |
| Scx | 0.009788115 | 1.913714495 | 0.17 | 0.064 | 1 | 0.299279682 | 0.648171025 | 0.293 | 0.222 | 1 | 0.299279682 | 0.019480423 |
| Zfp518b | 0.009793526 | 1.707263618 | 0.151 | 0.051 | 1 | 0.376253139 | 0.47463477 | 0.146 | 0.099 | 1 | 0.376253139 | 0.01949114 |
| Pla2g12b | 0.009815262 | 1.154722595 | 0.094 | 0.021 | 1 | 0.27192168 | 1.252242349 | 0.049 | 0.02 | 1 | 0.27192168 | 0.019534184 |
| Eif6 | 0.009879671 | 0.569760094 | 0.755 | 0.631 | 1 | 0.369738071 | 0.322030679 | 0.659 | 0.659 | 1 | 0.369738071 | 0.019661735 |
| Mbip | 0.009906996 | 0.970298024 | 0.415 | 0.25 | 1 | 0.711569926 | 0.515276755 | 0.293 | 0.297 | 1 | 0.711569926 | 0.019715843 |
| Zdhhc20 | 0.009914308 | 0.502645898 | 0.849 | 0.771 | 1 | 0.498300017 | 0.203616995 | 0.927 | 0.901 | 1 | 0.498300017 | 0.019730322 |
| Dgcr2 | 0.009915373 | 0.8327945 | 0.547 | 0.347 | 1 | 0.607893689 | 0.291011919 | 0.463 | 0.549 | 1 | 0.607893689 | 0.019732431 |
| Qsox1 | 0.013062553 | 0.853720339 | 0.698 | 0.585 | 1 | 0.009932092 | 0.830009348 | 0.878 | 0.741 | 1 | 0.013062553 | 0.019765538 |
| Nxpe4 | 0.600639519 | 0.392221909 | 0.264 | 0.242 | 1 | 0.009932468 | -0.758645967 | 0.049 | 0.235 | 1 | 0.600639519 | 0.019766281 |

| | | | | | | | | | | | | |
|---|---|---|---|---|---|---|---|---|---|---|---|---|
| Zfp949 | 0.009939702 | 1.199116714 | 0.34 | 0.195 | 1 | 0.362087408 | 0.597738915 | 0.39 | 0.355 | 1 | 0.362087408 | 0.019780607 |
| Stat6 | 0.009946376 | 1.19695083 | 0.472 | 0.331 | 1 | 0.121180852 | 0.785776611 | 0.585 | 0.509 | 1 | 0.121180852 | 0.019793822 |
| Rbm41 | 0.009956822 | 1.499370766 | 0.34 | 0.195 | 1 | 0.464318865 | -0.221688839 | 0.293 | 0.345 | 1 | 0.464318865 | 0.019814506 |
| Copg1 | 0.009968758 | 0.70524445 | 0.811 | 0.729 | 1 | 0.442686397 | -0.125733056 | 0.707 | 0.761 | 1 | 0.442686397 | 0.019838141 |
| Gm42679 | 0.009974943 | -3.399866257 | 0 | 0.114 | 1 | 0.357854401 | -0.622226769 | 0 | 0.02 | 1 | 0.357854401 | 0.019850386 |
| Gmps | 0.00997714 | 0.757038662 | 0.698 | 0.623 | 1 | 0.547273749 | 0.244068418 | 0.732 | 0.754 | 1 | 0.547273749 | 0.019854736 |
| Bdh2 | 0.00998468 | 2.270199812 | 0.132 | 0.042 | 1 | 0.210202152 | 1.114738825 | 0.122 | 0.068 | 1 | 0.210202152 | 0.019869666 |
| Gm46563 | 0.00999089 | 1.200526284 | 0.245 | 0.11 | 1 | 0.20965574 | -0.80665134 | 0.024 | 0.078 | 1 | 0.20965574 | 0.019881962 |
| Krt16 | 0.009994832 | -5.911366596 | 0 | 0.114 | 1 | 0.188554159 | -4.910988 | 0 | 0.041 | 1 | 0.188554159 | 0.019889768 |
| Mpp1 | 0.74527114 | -0.155617526 | 0.34 | 0.352 | 1 | 0.010040083 | -0.908222323 | 0.22 | 0.444 | 1 | 0.74527114 | 0.019979364 |
| Etv1 | 0.010051945 | 1.925240749 | 0.245 | 0.123 | 1 | 0.731844219 | 0.734111357 | 0.244 | 0.235 | 1 | 0.731844219 | 0.020002848 |
| Kpna4 | 0.01005721 | 0.534427021 | 0.925 | 0.873 | 1 | 0.062325834 | 0.378717094 | 0.976 | 0.959 | 1 | 0.062325834 | 0.020013273 |
| Spats2 | 0.137669874 | 0.524245435 | 0.491 | 0.407 | 1 | 0.01006057 | 0.996985294 | 0.561 | 0.42 | 1 | 0.137669874 | 0.020019925 |
| Letm1 | 0.010065692 | 0.751757928 | 0.528 | 0.352 | 1 | 0.273080459 | 0.317337377 | 0.537 | 0.451 | 1 | 0.273080459 | 0.020030065 |
| R3hdm1 | 0.010073688 | 0.702733959 | 0.774 | 0.733 | 1 | 0.278796137 | -0.125045874 | 0.854 | 0.853 | 1 | 0.278796137 | 0.020045897 |
| Dvl2 | 0.010094858 | 0.995524 | 0.415 | 0.246 | 1 | 0.123219753 | 1.059597271 | 0.341 | 0.273 | 1 | 0.123219753 | 0.020087809 |
| Polk | 0.010098984 | 0.580731212 | 0.453 | 0.258 | 1 | 0.175699412 | 0.535314508 | 0.366 | 0.27 | 1 | 0.175699412 | 0.020095978 |
| Smarcd1 | 0.010100356 | 0.756173218 | 0.434 | 0.254 | 1 | 0.475448174 | 0.618036329 | 0.366 | 0.338 | 1 | 0.475448174 | 0.020098695 |
| Atic | 0.010120951 | 0.776210972 | 0.585 | 0.386 | 1 | 0.123961343 | -0.734337135 | 0.341 | 0.423 | 1 | 0.123961343 | 0.020139469 |
| Rxrb | 0.010150832 | 1.355021245 | 0.321 | 0.174 | 1 | 0.896915463 | 0.551802631 | 0.195 | 0.198 | 1 | 0.896915463 | 0.020198625 |
| Atp9a | 0.010153778 | 1.051629102 | 0.34 | 0.186 | 1 | 0.423476135 | 0.105400961 | 0.171 | 0.242 | 1 | 0.423476135 | 0.020204457 |
| Rps5 | 0.010161521 | 0.293301605 | 1 | 0.97 | 1 | 0.174138355 | -0.129443522 | 1 | 0.986 | 1 | 0.174138355 | 0.020219785 |
| C920021L13Rik | 0.190799923 | 1.100274811 | 0.132 | 0.076 | 1 | 0.010161751 | 1.204936634 | 0.195 | 0.072 | 1 | 0.190799923 | 0.02022024 |
| Snx13 | 0.01016178 | -1.039391872 | 0.434 | 0.593 | 1 | 0.038179262 | -0.639128829 | 0.537 | 0.706 | 1 | 0.038179262 | 0.020220298 |
| Jund | 0.923189342 | -0.116780241 | 1 | 0.983 | 1 | 0.010177862 | -0.443819901 | 1 | 0.993 | 1 | 0.923189342 | 0.020252134 |
| Tdrd3 | 0.010203576 | 1.125378633 | 0.491 | 0.352 | 1 | 0.534883415 | -0.210827847 | 0.512 | 0.58 | 1 | 0.534883415 | 0.020303039 |
| Sertad2 | 0.141648814 | 0.11230419 | 0.943 | 0.907 | 1 | 0.010208851 | 0.504873796 | 0.927 | 0.922 | 1 | 0.141648814 | 0.020313482 |
| Styx | 0.010216983 | 0.690054328 | 0.472 | 0.275 | 1 | 0.393296572 | 0.788295249 | 0.415 | 0.399 | 1 | 0.393296572 | 0.02032958 |
| Gm34455 | 0.010709361 | 0.809894098 | 0.245 | 0.119 | 1 | 0.010254377 | 1.349939023 | 0.415 | 0.232 | 1 | 0.010709361 | 0.020403603 |
| Man1b1 | 0.010270723 | 0.913714495 | 0.415 | 0.258 | 1 | 0.454837044 | -0.212425918 | 0.366 | 0.444 | 1 | 0.454837044 | 0.020435958 |
| Nxph4 | 0.010279983 | 1.526691372 | 0.151 | 0.051 | 1 | 0.015196732 | 1.515276755 | 0.22 | 0.096 | 1 | 0.015196732 | 0.020454287 |
| Mapk8 | 0.010280805 | 0.783628443 | 0.585 | 0.428 | 1 | 0.381283666 | 0.241515942 | 0.683 | 0.645 | 1 | 0.381283666 | 0.020455915 |
| Zfp362 | 0.010292775 | 1.377115016 | 0.321 | 0.186 | 1 | 0.990091622 | 0.150144161 | 0.317 | 0.338 | 1 | 0.990091622 | 0.020479608 |
| Abtb2 | 0.010297011 | -0.484137861 | 0.811 | 0.915 | 1 | 0.030525146 | -0.723962857 | 0.878 | 0.904 | 1 | 0.030525146 | 0.020487993 |
| Homer1 | 0.441877447 | 0.427595193 | 0.528 | 0.487 | 1 | 0.010306343 | 1.098164927 | 0.756 | 0.618 | 1 | 0.441877447 | 0.020506465 |
| Clpb | 0.010306352 | 1.404700848 | 0.377 | 0.233 | 1 | 0.377696931 | 0.252242349 | 0.439 | 0.352 | 1 | 0.377696931 | 0.020506483 |
| Gk5 | 0.010308033 | 1.139455838 | 0.528 | 0.407 | 1 | 0.718997612 | 0.241475918 | 0.61 | 0.594 | 1 | 0.718997612 | 0.020509811 |
| Ap1b1 | 0.01031461 | 0.822147256 | 0.396 | 0.22 | 1 | 0.47530833 | 0.529082554 | 0.293 | 0.259 | 1 | 0.47530833 | 0.020522829 |
| Rnf169 | 0.010337982 | 0.804225348 | 0.547 | 0.403 | 1 | 0.663333243 | 0.135286203 | 0.683 | 0.614 | 1 | 0.663333243 | 0.02056909 |
| Ramac | 0.010344691 | 0.606418565 | 0.736 | 0.669 | 1 | 0.328931931 | 0.124486802 | 0.829 | 0.765 | 1 | 0.328931931 | 0.02058237 |
| Mir17hg | 0.031963796 | 2.47665069 | 0.094 | 0.03 | 1 | 0.010383502 | 2.351778023 | 0.122 | 0.034 | 1 | 0.031963796 | 0.020659187 |
| 5033426O07Rik | 0.024562821 | -2.545717123 | 0 | 0.089 | 1 | 0.010394229 | -1.849295677 | 0.049 | 0.222 | 1 | 0.024562821 | 0.020680419 |
| Ppa2 | 0.010421406 | 1.228723176 | 0.34 | 0.191 | 1 | 0.037822887 | -1.177745492 | 0.098 | 0.242 | 1 | 0.037822887 | 0.020734206 |
| Slitrk6 | 0.010425302 | 1.154722595 | 0.264 | 0.127 | 1 | 0.885198435 | 0.614812428 | 0.049 | 0.044 | 1 | 0.885198435 | 0.020741918 |
| Cnot11 | 0.202248584 | 0.61630268 | 0.302 | 0.229 | 1 | 0.010429341 | -0.747757651 | 0.122 | 0.341 | 1 | 0.202248584 | 0.02074991 |
| Gtf2b | 0.409577369 | 0.100515521 | 0.792 | 0.746 | 1 | 0.010433977 | -0.667139191 | 0.61 | 0.741 | 1 | 0.409577369 | 0.020759087 |
| Kcnmb4 | 0.010440095 | 2.010332685 | 0.17 | 0.064 | 1 | 0.036203725 | 0.915207362 | 0.268 | 0.14 | 1 | 0.036203725 | 0.020771195 |
| Mogs | 0.161996936 | -0.384436216 | 0.208 | 0.309 | 1 | 0.010446781 | -1.188330242 | 0.098 | 0.297 | 1 | 0.161996936 | 0.020784428 |
| Rtn3 | 0.010455817 | 0.211471012 | 0.981 | 0.975 | 1 | 0.070341725 | -0.223233756 | 0.951 | 0.99 | 1 | 0.070341725 | 0.020802309 |
| Ehbp1l1 | 0.064911507 | -0.612300233 | 0.566 | 0.669 | 1 | 0.01046608 | -0.706534988 | 0.561 | 0.747 | 1 | 0.064911507 | 0.020822622 |
| Glyr1 | 0.717401835 | 0.355770114 | 0.698 | 0.784 | 1 | 0.010491203 | 0.345625796 | 0.927 | 0.84 | 1 | 0.717401835 | 0.02087234 |
| Gm15892 | 0.01049128 | 1.607234799 | 0.17 | 0.064 | 1 | 0.243822776 | 0.547698233 | 0.22 | 0.154 | 1 | 0.243822776 | 0.020872492 |
| Chmp5 | 0.038278919 | 0.41149801 | 0.811 | 0.661 | 1 | 0.010505487 | 1.013455489 | 0.756 | 0.737 | 1 | 0.038278919 | 0.020900609 |
| Nufip1 | 0.010507098 | 1.07065833 | 0.472 | 0.339 | 1 | 0.253238721 | 0.446970742 | 0.415 | 0.348 | 1 | 0.253238721 | 0.020903797 |
| Ndc1 | 0.010508483 | 1.017219071 | 0.226 | 0.097 | 1 | 0.922693863 | 1.136765132 | 0.122 | 0.123 | 1 | 0.922693863 | 0.020906538 |
| Cecr2 | 0.338380035 | 1.47665069 | 0.038 | 0.017 | 1 | 0.010517177 | 1.639265472 | 0.244 | 0.106 | 1 | 0.338380035 | 0.020923744 |
| Tbce | 0.049908223 | -0.545717123 | 0.434 | 0.576 | 1 | 0.010528593 | -0.987223586 | 0.268 | 0.481 | 1 | 0.049908223 | 0.020946334 |
| Mrpl49 | 0.010532671 | 1.034428361 | 0.34 | 0.182 | 1 | 0.819631178 | 0.396632258 | 0.22 | 0.212 | 1 | 0.819631178 | 0.020954404 |
| Cinp | 0.010542238 | 1.029191713 | 0.321 | 0.169 | 1 | 0.597443555 | 0.594348326 | 0.22 | 0.191 | 1 | 0.597443555 | 0.020973337 |
| Zfp704 | 0.010547663 | 0.411951379 | 0.717 | 0.576 | 1 | 0.31405455 | -0.180244121 | 0.732 | 0.778 | 1 | 0.31405455 | 0.020984072 |
| Ssr4 | 0.01055202 | -0.518223294 | 0.962 | 0.97 | 1 | 0.148773676 | -0.288941085 | 0.927 | 0.935 | 1 | 0.148773676 | 0.020992694 |
| Tap2 | 0.010563076 | 0.8327945 | 0.302 | 0.153 | 1 | 0.54593315 | 0.72678086 | 0.244 | 0.222 | 1 | 0.54593315 | 0.021014573 |
| Rack1 | 0.155742351 | 0.195326449 | 1 | 0.975 | 1 | 0.01056972 | -0.265258946 | 1 | 0.986 | 1 | 0.155742351 | 0.021027721 |
| Dclre1a | 0.010608389 | 1.517292674 | 0.208 | 0.089 | 1 | 0.24223858 | -0.648221977 | 0.073 | 0.14 | 1 | 0.24223858 | 0.02110424 |
| Eif1ax | 0.081369068 | 0.332476683 | 0.925 | 0.856 | 1 | 0.01061136 | -0.484723245 | 0.805 | 0.928 | 1 | 0.081369068 | 0.021110119 |
| Dnmt1 | 0.010635631 | 0.715838354 | 0.528 | 0.343 | 1 | 0.293673657 | 0.235168836 | 0.512 | 0.44 | 1 | 0.293673657 | 0.021158145 |
| Leng9 | 0.010639181 | 1.417757001 | 0.151 | 0.051 | 1 | 0.216071703 | 1.515276755 | 0.098 | 0.051 | 1 | 0.216071703 | 0.02116517 |
| Agl | 0.010646219 | 1.228723176 | 0.226 | 0.102 | 1 | 0.603884074 | 0.276489895 | 0.171 | 0.14 | 1 | 0.603884074 | 0.021179096 |
| Naa15 | 0.010661148 | 0.349914355 | 0.925 | 0.843 | 1 | 0.542003975 | 0.217907196 | 0.951 | 0.928 | 1 | 0.542003975 | 0.021208636 |
| Traf3ip1 | 0.010665627 | 1.154722595 | 0.302 | 0.157 | 1 | 0.821022231 | 0.494812652 | 0.171 | 0.164 | 1 | 0.821022231 | 0.021217499 |
| Hdac7 | 0.01066931 | 1.331600357 | 0.151 | 0.051 | 1 | 0.149497949 | 1.632846351 | 0.195 | 0.13 | 1 | 0.149497949 | 0.021224785 |
| Zfp706 | 0.010692745 | 0.46700052 | 0.887 | 0.822 | 1 | 0.86533012 | 0.159945562 | 0.805 | 0.887 | 1 | 0.86533012 | 0.021271156 |
| Zcchc9 | 0.010716942 | 1.30324512 | 0.396 | 0.263 | 1 | 0.038446237 | -0.604632709 | 0.195 | 0.358 | 1 | 0.038446237 | 0.02131903 |
| Scrn3 | 0.021994297 | -1.626637119 | 0.057 | 0.182 | 1 | 0.010720919 | -2.391613841 | 0.024 | 0.181 | 1 | 0.021994297 | 0.021326899 |
| Tln2 | 0.010728225 | 0.848736044 | 0.547 | 0.403 | 1 | 0.968605426 | 0.363273661 | 0.195 | 0.205 | 1 | 0.968605426 | 0.021341355 |
| Coro7 | 0.010740769 | 1.261637799 | 0.283 | 0.144 | 1 | 0.755782329 | 0.189506594 | 0.22 | 0.205 | 1 | 0.755782329 | 0.021366174 |
| Urgcp | 0.213068634 | 0.786335189 | 0.415 | 0.356 | 1 | 0.010741203 | -0.72418918 | 0.366 | 0.59 | 1 | 0.213068634 | 0.021367032 |
| Araf | 0.100455892 | -0.366814526 | 0.604 | 0.72 | 1 | 0.010744829 | -0.648221977 | 0.512 | 0.73 | 1 | 0.100455892 | 0.021374207 |
| Snai2 | 0.021799098 | -0.840761924 | 0.321 | 0.475 | 1 | 0.010744963 | -0.603367742 | 0.146 | 0.365 | 1 | 0.021799098 | 0.021374471 |
| Ubc | 0.915228712 | -0.116386893 | 0.981 | 0.987 | 1 | 0.010750403 | 0.306775502 | 1 | 0.99 | 1 | 0.915228712 | 0.021385235 |
| Gpsm3 | 0.010761815 | 1.377115016 | 0.283 | 0.144 | 1 | 0.750173975 | 1.159132945 | 0.098 | 0.085 | 1 | 0.750173975 | 0.021407813 |
| Tusc1 | 0.010770588 | 1.624207878 | 0.208 | 0.089 | 1 | 0.657848822 | 0.43121249 | 0.171 | 0.143 | 1 | 0.657848822 | 0.02142517 |
| Lgi2 | 0.040585941 | 0.169519597 | 0.302 | 0.169 | 1 | 0.01078713 | -1.438919555 | 0.171 | 0.355 | 1 | 0.040585941 | 0.021457897 |
| Slc23a2 | 0.326852159 | -0.31065829 | 0.66 | 0.716 | 1 | 0.010790062 | 0.843178751 | 0.854 | 0.737 | 1 | 0.326852159 | 0.021463699 |
| Marveld1 | 0.010790187 | -0.939253553 | 0.566 | 0.682 | 1 | 0.06139984 | -0.444830518 | 0.439 | 0.625 | 1 | 0.06139984 | 0.021463945 |
| Abhd17b | 0.010798956 | 0.683416876 | 0.585 | 0.407 | 1 | 0.017856502 | 0.862590168 | 0.659 | 0.587 | 1 | 0.017856502 | 0.021481294 |
| Rdh11 | 0.010808724 | -1.096238979 | 0.302 | 0.475 | 1 | 0.778165028 | -0.11032773 | 0.317 | 0.331 | 1 | 0.778165028 | 0.02150062 |
| Bscl2 | 0.010822482 | 0.865215978 | 0.547 | 0.453 | 1 | 0.966510166 | 0.338657101 | 0.488 | 0.526 | 1 | 0.966510166 | 0.021527837 |
| Btrc | 0.010863868 | 0.695290976 | 0.604 | 0.466 | 1 | 0.047098378 | -0.552361962 | 0.659 | 0.775 | 1 | 0.047098378 | 0.021609711 |
| Dzip1l | 0.010864058 | 0.61158027 | 0.453 | 0.267 | 1 | 0.569949641 | 0.10681191 | 0.268 | 0.317 | 1 | 0.569949641 | 0.021610089 |
| Gm16193 | 0.02936866 | -1.218735801 | 0.094 | 0.237 | 1 | 0.010867087 | -1.545264787 | 0.098 | 0.287 | 1 | 0.02936866 | 0.02161608 |
| Acsl6 | 0.932505382 | 1.154722595 | 0.019 | 0.021 | 1 | 0.010874275 | -3.140075074 | 0 | 0.14 | 1 | 0.932505382 | 0.021630301 |
| Rnase4 | 0.010877938 | 0.271875222 | 0.925 | 0.691 | 1 | 0.116525613 | -0.383048026 | 0.902 | 0.891 | 1 | 0.116525613 | 0.021637545 |
| Samhd1 | 0.116388493 | -1.802794074 | 0.66 | 0.623 | 1 | 0.010879885 | 0.731271405 | 0.683 | 0.526 | 1 | 0.116388493 | 0.021641398 |
| Zcchc24 | 0.010886141 | 0.852694057 | 0.566 | 0.424 | 1 | 0.439557247 | 0.559220103 | 0.488 | 0.488 | 1 | 0.439557247 | 0.021653773 |
| Itga5 | 0.288693119 | -0.577993526 | 0.377 | 0.449 | 1 | 0.010888689 | -1.178392005 | 0.268 | 0.481 | 1 | 0.288693119 | 0.021658814 |
| Tmem131 | 0.107194027 | 0.366545774 | 0.755 | 0.737 | 1 | 0.010901057 | 0.500070342 | 0.878 | 0.874 | 1 | 0.107194027 | 0.021683281 |
| Ctsh | 0.010908367 | -1.066276591 | 0.679 | 0.72 | 1 | 0.214421829 | 0.307585383 | 0.707 | 0.58 | 1 | 0.214421829 | 0.021697741 |

| | | | | | | | | | | | | |
|---|---|---|---|---|---|---|---|---|---|---|---|---|
| Ddx19a | 0.861282111 | 0.374503803 | 0.264 | 0.271 | 1 | 0.010913221 | -0.98876575 | 0.171 | 0.389 | 1 | 0.861282111 | 0.021707343 |
| Pycr1 | 0.334316963 | -0.417528854 | 0.226 | 0.297 | 1 | 0.010917891 | -1.661046018 | 0.098 | 0.276 | 1 | 0.334316963 | 0.021716581 |
| Atf4 | 0.316019861 | -0.248692537 | 0.981 | 0.975 | 1 | 0.010943448 | -0.409796375 | 1 | 0.986 | 1 | 0.316019861 | 0.021767136 |
| Smarcb1 | 0.010945155 | 0.679211236 | 0.623 | 0.508 | 1 | 0.336885797 | 0.428740004 | 0.585 | 0.567 | 1 | 0.336885797 | 0.021770514 |
| Zc3h11a | 0.010968051 | 0.86841841 | 0.491 | 0.339 | 1 | 0.429640062 | -0.18959521 | 0.341 | 0.406 | 1 | 0.429640062 | 0.021815805 |
| Mbd6 | 0.010968905 | 1.427741089 | 0.321 | 0.182 | 1 | 0.163820819 | -0.497779398 | 0.195 | 0.297 | 1 | 0.163820819 | 0.021817493 |
| Hunk | 0.628925725 | 0.154722595 | 0.038 | 0.025 | 1 | 0.010976299 | 1.614812428 | 0.122 | 0.034 | 1 | 0.628925725 | 0.021832119 |
| Pla2g6 | 0.956697606 | 0.569760094 | 0.075 | 0.081 | 1 | 0.010994563 | 1.136765132 | 0.171 | 0.058 | 1 | 0.956697606 | 0.021868245 |
| Frg2f1 | 0.934284789 | -0.247375849 | 0.113 | 0.114 | 1 | 0.010994756 | 1.19334866 | 0.171 | 0.058 | 1 | 0.934284789 | 0.021868627 |
| Cyb5a | 0.011018812 | 0.342349598 | 0.792 | 0.581 | 1 | 0.051029571 | 0.861452396 | 0.683 | 0.672 | 1 | 0.051029571 | 0.021916209 |
| Stard7 | 0.011047948 | 0.796268624 | 0.434 | 0.263 | 1 | 0.973940201 | 0.461695715 | 0.366 | 0.396 | 1 | 0.973940201 | 0.021973839 |
| Gm26771 | 0.011100366 | 3.620386167 | 0.17 | 0.068 | 1 | 0.937766637 | 0.435106406 | 0.073 | 0.078 | 1 | 0.937766637 | 0.022077514 |
| Gpsm1 | 0.011123933 | 0.745917049 | 0.509 | 0.373 | 1 | 0.087425151 | 0.484903106 | 0.585 | 0.474 | 1 | 0.087425151 | 0.022124123 |
| Ciz1 | 0.01113614 | 1.052360877 | 0.396 | 0.237 | 1 | 0.267606923 | -0.13108629 | 0.268 | 0.362 | 1 | 0.267606923 | 0.022148266 |
| 4930581F22Rik | 0.011177446 | -1.513655914 | 0.094 | 0.254 | 1 | 0.134666013 | -0.717384002 | 0.049 | 0.133 | 1 | 0.134666013 | 0.022229956 |
| Sirt5 | 0.011190687 | -1.637835325 | 0.075 | 0.233 | 1 | 0.154111598 | -1.484723245 | 0 | 0.048 | 1 | 0.154111598 | 0.022256143 |
| Bcl7a | 0.011200783 | 1.595295186 | 0.208 | 0.089 | 1 | 0.492373798 | -0.648221977 | 0.098 | 0.133 | 1 | 0.492373798 | 0.022276108 |
| Trir | 0.011241466 | 0.483460248 | 0.925 | 0.835 | 1 | 0.285262357 | -0.187179309 | 0.829 | 0.874 | 1 | 0.285262357 | 0.022356561 |
| Tmem131l | 0.011243113 | 0.436698564 | 0.528 | 0.318 | 1 | 0.200467462 | -0.563772706 | 0.634 | 0.676 | 1 | 0.200467462 | 0.022359818 |
| Map9 | 0.011248329 | -1.813096999 | 0.189 | 0.36 | 1 | 0.048466383 | -0.992870149 | 0.268 | 0.413 | 1 | 0.048466383 | 0.022370134 |
| Phf14 | 0.01125204 | 1.000917259 | 0.623 | 0.551 | 1 | 0.082798784 | 0.457271151 | 0.78 | 0.737 | 1 | 0.082798784 | 0.022377473 |
| Ifrd2 | 0.437352348 | 0.610402079 | 0.283 | 0.25 | 1 | 0.011255434 | -1.250257991 | 0.073 | 0.256 | 1 | 0.437352348 | 0.022384182 |
| Zfyve1 | 0.011266382 | 0.445771377 | 0.566 | 0.36 | 1 | 0.474489809 | 0.396632258 | 0.659 | 0.625 | 1 | 0.474489809 | 0.022405834 |
| Colgalt1 | 0.042645308 | -0.527850703 | 0.774 | 0.831 | 1 | 0.011266643 | -0.638279174 | 0.707 | 0.802 | 1 | 0.042645308 | 0.022406348 |
| Cxxc5 | 0.011266767 | 0.769697665 | 0.585 | 0.479 | 1 | 0.429602053 | -0.255904555 | 0.707 | 0.775 | 1 | 0.429602053 | 0.022406593 |
| Zfand4 | 0.914733857 | -0.237594828 | 0.057 | 0.059 | 1 | 0.011276311 | 1.737669176 | 0.122 | 0.034 | 1 | 0.914733857 | 0.022425466 |
| Vopp1 | 0.410698586 | 0.970298024 | 0.208 | 0.174 | 1 | 0.011302978 | -1.004097404 | 0.098 | 0.294 | 1 | 0.410698586 | 0.022478199 |
| Abhd17c | 0.011409383 | 1.242185436 | 0.34 | 0.199 | 1 | 0.285530216 | 0.713822434 | 0.341 | 0.294 | 1 | 0.285530216 | 0.022688591 |
| Mtg1 | 0.011412193 | 1.891688189 | 0.132 | 0.042 | 1 | 0.182124508 | 1.404245443 | 0.146 | 0.085 | 1 | 0.182124508 | 0.022694148 |
| Snx32 | 0.011412193 | 1.891688189 | 0.132 | 0.042 | 1 | 0.434929181 | 0.627751484 | 0.146 | 0.106 | 1 | 0.434929181 | 0.022694148 |
| Ccdc57 | 0.011434991 | 1.002719501 | 0.226 | 0.097 | 1 | 0.345165719 | 0.313642894 | 0.171 | 0.116 | 1 | 0.345165719 | 0.022739223 |
| Zcrb1 | 0.349166938 | 0.179469257 | 0.925 | 0.864 | 1 | 0.011447166 | -0.304486471 | 0.854 | 0.874 | 1 | 0.349166938 | 0.022763294 |
| Cpsf6 | 0.011471528 | 0.403828162 | 0.811 | 0.733 | 1 | 0.392181841 | -0.120893995 | 0.878 | 0.939 | 1 | 0.392181841 | 0.022811461 |
| C030014I23Rik | 0.66868216 | 0.47665069 | 0.075 | 0.059 | 1 | 0.011486559 | 3.159132945 | 0.098 | 0.024 | 1 | 0.66868216 | 0.022841178 |
| Camk1d | 0.011529016 | 0.297599291 | 1 | 1 | 1 | 0.013547988 | 0.663722029 | 1 | 1 | 1 | 0.013547988 | 0.022925114 |
| Gm38910 | 0.011548287 | -2.799473716 | 0 | 0.11 | 1 | 0.070224584 | -1.747757651 | 0 | 0.075 | 1 | 0.070224584 | 0.022963212 |
| Psma6 | 0.011556287 | 0.590108739 | 0.83 | 0.737 | 1 | 0.129311607 | 0.315667729 | 0.805 | 0.823 | 1 | 0.129311607 | 0.022979026 |
| 1700031C06Rik | 0.011558908 | -3.27154216 | 0 | 0.11 | 1 | 0.257325989 | -0.484723245 | 0 | 0.031 | 1 | 0.257325989 | 0.022984207 |
| Stat3 | 0.113011178 | 0.267987429 | 1 | 0.983 | 1 | 0.011564808 | 0.401013854 | 0.976 | 0.993 | 1 | 0.113011178 | 0.022995872 |
| Cited1 | 0.011577389 | -3.20282941 | 0 | 0.11 | 1 | 0.20883094 | -2.589059905 | 0 | 0.038 | 1 | 0.20883094 | 0.023020742 |
| Fam98a | 0.406891022 | 0.244732761 | 0.528 | 0.513 | 1 | 0.01157878 | -0.597617301 | 0.39 | 0.659 | 1 | 0.406891022 | 0.023023492 |
| Gm10602 | 0.038704123 | 2.154722595 | 0.075 | 0.021 | 1 | 0.01158113 | 1.83720485 | 0.122 | 0.034 | 1 | 0.038704123 | 0.023028137 |
| Nipal1 | 0.01160379 | -3.304709024 | 0 | 0.11 | 1 | 0.286430578 | -1.555112573 | 0 | 0.027 | 1 | 0.286430578 | 0.023072933 |
| Wtap | 0.011604324 | 0.73687008 | 0.566 | 0.424 | 1 | 0.656517643 | -0.21906337 | 0.61 | 0.628 | 1 | 0.656517643 | 0.023073987 |
| Akap6 | 0.011608165 | -3.368839361 | 0 | 0.11 | 1 | 0.154227016 | -0.69884805 | 0.024 | 0.092 | 1 | 0.154227016 | 0.023081581 |
| Slco2b1 | 0.011612544 | -1.600164907 | 0.038 | 0.178 | 1 | 0.128315252 | -0.825760163 | 0.024 | 0.099 | 1 | 0.128315252 | 0.023090238 |
| Gm16759 | 0.011615697 | -3.995024525 | 0 | 0.11 | 1 | 0.201684505 | -1.890715605 | 0.024 | 0.078 | 1 | 0.201684505 | 0.02309647 |
| Mtmr3 | 0.011625965 | 0.383138804 | 0.887 | 0.733 | 1 | 0.153494672 | 0.175918854 | 0.951 | 0.904 | 1 | 0.153494672 | 0.023116767 |
| Ap4s1 | 0.423150649 | -0.501323004 | 0.415 | 0.466 | 1 | 0.011632538 | -0.592782991 | 0.268 | 0.536 | 1 | 0.423150649 | 0.02312976 |
| Malsu1 | 0.219545014 | -0.392765201 | 0.453 | 0.568 | 1 | 0.011650785 | -0.957975358 | 0.293 | 0.481 | 1 | 0.219545014 | 0.023165829 |
| D530017H19Rik | 0.182332113 | 0.8327945 | 0.094 | 0.047 | 1 | 0.011675641 | 2.159132945 | 0.122 | 0.034 | 1 | 0.182332113 | 0.023214961 |
| Acaa2 | 0.011680294 | 1.339147166 | 0.283 | 0.144 | 1 | 0.112578076 | 0.674266279 | 0.341 | 0.235 | 1 | 0.112578076 | 0.02322416 |
| Garre1 | 0.019966227 | 0.519428419 | 0.868 | 0.822 | 1 | 0.01168261 | 0.426715368 | 0.854 | 0.843 | 1 | 0.019966227 | 0.023228737 |
| Puf60 | 0.011702778 | 0.490958526 | 0.792 | 0.678 | 1 | 0.203108935 | 0.417490863 | 0.78 | 0.706 | 1 | 0.203108935 | 0.023268601 |
| Tnip2 | 0.177070673 | -0.29778961 | 0.396 | 0.504 | 1 | 0.011704651 | -1.209758629 | 0.244 | 0.437 | 1 | 0.177070673 | 0.023272303 |
| Ipp | 0.011712608 | 1.46284489 | 0.189 | 0.076 | 1 | 0.193633903 | -0.448197369 | 0.049 | 0.119 | 1 | 0.193633903 | 0.023288032 |
| Clstn2 | 0.01172347 | 0.138662744 | 0.415 | 0.229 | 1 | 0.39479124 | 0.438167829 | 0.561 | 0.512 | 1 | 0.39479124 | 0.023309499 |
| Ighm | 0.100937876 | -0.988235359 | 0.038 | 0.008 | 1 | 0.011749432 | 1.782757066 | 0.171 | 0.061 | 1 | 0.100937876 | 0.023360814 |
| Krr1 | 0.01175047 | 0.53659323 | 0.528 | 0.335 | 1 | 0.351888003 | -0.148295581 | 0.341 | 0.44 | 1 | 0.351888003 | 0.023362867 |
| Cibar1 | 0.155338063 | 0.43613553 | 0.642 | 0.644 | 1 | 0.011758355 | -0.538856682 | 0.366 | 0.608 | 1 | 0.155338063 | 0.023378452 |
| B930036N10Rik | 0.011767519 | -2.594215641 | 0.132 | 0.284 | 1 | 0.164184178 | -1.19841906 | 0.073 | 0.154 | 1 | 0.164184178 | 0.023396563 |
| Spata1 | 0.011780693 | -0.749566206 | 0.264 | 0.462 | 1 | 0.377580273 | 0.252242349 | 0.268 | 0.369 | 1 | 0.377580273 | 0.023422601 |
| Pes1 | 0.011797597 | 1.290377694 | 0.377 | 0.242 | 1 | 0.366623776 | -0.204615326 | 0.244 | 0.324 | 1 | 0.366623776 | 0.023456012 |
| Cdhr4 | 0.011810752 | -2.031143951 | 0.075 | 0.22 | 1 | 0.03178054 | 1.252242349 | 0.122 | 0.041 | 1 | 0.03178054 | 0.02348201 |
| Tnip1 | 0.011826167 | 1.649487287 | 0.283 | 0.153 | 1 | 0.314689776 | -0.825760163 | 0.171 | 0.232 | 1 | 0.314689776 | 0.023512476 |
| Dctpp1 | 0.136413197 | 0.550651271 | 0.321 | 0.233 | 1 | 0.011827297 | -2.271319607 | 0.024 | 0.177 | 1 | 0.136413197 | 0.023514708 |
| Pdk4 | 0.011850959 | -2.545717123 | 0.113 | 0.267 | 1 | 0.019869181 | -2.27321914 | 0.098 | 0.266 | 1 | 0.019869181 | 0.023561474 |
| Ttl | 0.110935367 | 0.642823556 | 0.34 | 0.242 | 1 | 0.011862253 | -0.872453398 | 0.146 | 0.348 | 1 | 0.110935367 | 0.023583794 |
| Dennd4b | 0.011886882 | 0.900966003 | 0.17 | 0.064 | 1 | 0.470682201 | -0.20718927 | 0.049 | 0.082 | 1 | 0.470682201 | 0.023632465 |
| Gm39469 | 0.011889567 | -1.237594828 | 0.019 | 0.148 | 1 | 0.63821201 | 0.252242349 | 0.146 | 0.119 | 1 | 0.63821201 | 0.023637773 |
| Mrpl47 | 0.011927139 | 1.313420341 | 0.283 | 0.144 | 1 | 0.174266264 | -0.31254227 | 0.098 | 0.191 | 1 | 0.174266264 | 0.023712022 |
| Ift74 | 0.0906828 | 0.548698479 | 0.623 | 0.5 | 1 | 0.01192754 | -0.653676407 | 0.39 | 0.608 | 1 | 0.0906828 | 0.023712813 |
| Ddx18 | 0.011955454 | 0.795180208 | 0.547 | 0.377 | 1 | 0.704332813 | -0.155261478 | 0.366 | 0.392 | 1 | 0.704332813 | 0.023767975 |
| Lta4h | 0.098357448 | 0.617694571 | 0.472 | 0.386 | 1 | 0.01196752 | -0.802205435 | 0.415 | 0.614 | 1 | 0.098357448 | 0.023791819 |
| Arl5c | 0.011975438 | 2.055186921 | 0.151 | 0.055 | 1 | 0.342867779 | 0.404245443 | 0.098 | 0.058 | 1 | 0.342867779 | 0.023807465 |
| Crocc | 0.872466289 | -0.260314904 | 0.038 | 0.042 | 1 | 0.011988583 | 1.614812428 | 0.122 | 0.034 | 1 | 0.872466289 | 0.023833439 |
| Ppil3 | 0.012014675 | 1.154722595 | 0.358 | 0.216 | 1 | 0.78434725 | 0.582390951 | 0.268 | 0.266 | 1 | 0.78434725 | 0.023884998 |
| Fam25c | 0.051873971 | -2.304709024 | 0 | 0.068 | 1 | 0.012022859 | -3.20718927 | 0 | 0.137 | 1 | 0.051873971 | 0.023901169 |
| Kti12 | 0.012044702 | 1.179813576 | 0.34 | 0.195 | 1 | 0.076212892 | -0.720790603 | 0.146 | 0.276 | 1 | 0.076212892 | 0.023944329 |
| Pde1a | 0.01206625 | -3.067669827 | 0.038 | 0.169 | 1 | 0.103893371 | -3.16279515 | 0 | 0.061 | 1 | 0.103893371 | 0.023986905 |
| Gps2 | 0.012075438 | 1.121807972 | 0.509 | 0.407 | 1 | 0.30600613 | 0.269520341 | 0.634 | 0.529 | 1 | 0.30600613 | 0.02400506 |
| Zfp141 | 0.012080149 | 1.347367673 | 0.17 | 0.064 | 1 | 0.247101343 | 0.83720485 | 0.146 | 0.089 | 1 | 0.247101343 | 0.024014368 |
| Psme3 | 0.012091282 | 0.919094346 | 0.415 | 0.258 | 1 | 0.083591212 | -0.577832649 | 0.22 | 0.369 | 1 | 0.083591212 | 0.024036365 |
| 2410002F23Rik | 0.012097272 | 0.739685096 | 0.528 | 0.415 | 1 | 0.921503473 | 0.284663827 | 0.415 | 0.399 | 1 | 0.921503473 | 0.0240482 |
| Arhgap24 | 0.596247934 | -0.135669927 | 0.208 | 0.169 | 1 | 0.012108459 | 1.236300805 | 0.244 | 0.106 | 1 | 0.596247934 | 0.024070304 |
| Gm56488 | 0.204653189 | 2.154722595 | 0.038 | 0.013 | 1 | 0.012110922 | 3.574170444 | 0.073 | 0.014 | 1 | 0.204653189 | 0.024075169 |
| Mtpn | 0.012114392 | 0.477501586 | 0.811 | 0.712 | 1 | 0.503276008 | 0.20299883 | 0.78 | 0.863 | 1 | 0.503276008 | 0.024082026 |
| Nif3l1 | 0.012116821 | 1.242185436 | 0.245 | 0.114 | 1 | 0.495994869 | -0.508569987 | 0.122 | 0.16 | 1 | 0.495994869 | 0.024086824 |
| 6430590A07Rik | 0.01211793 | 1.200526284 | 0.189 | 0.076 | 1 | 0.171436504 | -1.192542494 | 0.049 | 0.119 | 1 | 0.171436504 | 0.024089016 |
| Spag5 | 0.260667702 | 1.228723176 | 0.094 | 0.055 | 1 | 0.012139487 | 2.644559772 | 0.098 | 0.024 | 1 | 0.260667702 | 0.024131607 |
| Polr3gl | 0.012160333 | 0.707263618 | 0.472 | 0.301 | 1 | 0.866535535 | 0.391405097 | 0.317 | 0.379 | 1 | 0.866535535 | 0.024172792 |
| Atmin | 0.012167918 | 1.417757001 | 0.189 | 0.076 | 1 | 0.543152771 | -0.16279515 | 0.098 | 0.133 | 1 | 0.543152771 | 0.024187777 |
| Zkscan6 | 0.012167918 | 1.417757001 | 0.189 | 0.076 | 1 | 0.210728835 | -0.747757651 | 0.024 | 0.078 | 1 | 0.210728835 | 0.024187777 |
| Irs2 | 0.022907089 | 0.610061459 | 0.717 | 0.653 | 1 | 0.012174002 | 0.68026549 | 0.854 | 0.696 | 1 | 0.022907089 | 0.024199798 |
| Tenm1 | 0.207549016 | 1.962077517 | 0.038 | 0.013 | 1 | 0.012241694 | 4.315252147 | 0.098 | 0.024 | 1 | 0.207549016 | 0.024333528 |
| Naa60 | 0.012247334 | 0.688404591 | 0.491 | 0.314 | 1 | 0.012728264 | -1.095680954 | 0.171 | 0.375 | 1 | 0.012728264 | 0.024344672 |
| Smad7 | 0.016908915 | 0.619087806 | 0.698 | 0.581 | 1 | 0.012252433 | 0.878392896 | 0.78 | 0.625 | 1 | 0.016908915 | 0.024354745 |
| Lgals3bp | 0.012267792 | -1.39371403 | 0.17 | 0.331 | 1 | 0.187033157 | -0.577832649 | 0.195 | 0.294 | 1 | 0.187033157 | 0.024385085 |

| | | | | | | | | | | | | |
|---|---|---|---|---|---|---|---|---|---|---|---|---|
| Map2k6 | 0.012323676 | 2.055186921 | 0.132 | 0.042 | 1 | 0.551133697 | -0.372248516 | 0.049 | 0.075 | 1 | 0.551133697 | 0.024495479 |
| Wdr19 | 0.012326648 | 1.3662267 | 0.17 | 0.064 | 1 | 0.107294683 | -1.385187572 | 0.049 | 0.137 | 1 | 0.107294683 | 0.024501349 |
| Hspa13 | 0.01238477 | 0.806112457 | 0.623 | 0.492 | 1 | 0.177940572 | 0.68745773 | 0.488 | 0.464 | 1 | 0.177940572 | 0.024616157 |
| Uqcrh | 0.012408017 | 0.27919617 | 0.981 | 0.962 | 1 | 0.329464379 | -0.147447186 | 1 | 0.983 | 1 | 0.329464379 | 0.024662074 |
| Eif3d | 0.012411561 | 0.613481472 | 0.623 | 0.504 | 1 | 0.495056771 | -0.245784515 | 0.634 | 0.655 | 1 | 0.495056771 | 0.024669075 |
| Ssbp4 | 0.151140182 | 0.876187095 | 0.434 | 0.381 | 1 | 0.01242236 | -0.923607486 | 0.293 | 0.498 | 1 | 0.151140182 | 0.024690406 |
| Dock6 | 0.01243635 | 1.33529484 | 0.358 | 0.208 | 1 | 0.152331243 | 0.604544093 | 0.463 | 0.372 | 1 | 0.152331243 | 0.024718038 |
| Mboat1 | 0.800113075 | 1.624207878 | 0.057 | 0.051 | 1 | 0.012437929 | 2.489281546 | 0.098 | 0.024 | 1 | 0.800113075 | 0.024721157 |
| Peli3 | 0.03562064 | 3.154722595 | 0.019 | 0 | 1 | 0.012470669 | 2.83720485 | 0.073 | 0.014 | 1 | 0.03562064 | 0.024785821 |
| Kcns3 | 0.503971666 | 1.569760094 | 0.019 | 0.008 | 1 | 0.012470669 | 2.83720485 | 0.073 | 0.014 | 1 | 0.503971666 | 0.024785821 |
| Dpf1 | 0.736073999 | 0.569760094 | 0.019 | 0.013 | 1 | 0.012470669 | 2.83720485 | 0.073 | 0.014 | 1 | 0.736073999 | 0.024785821 |
| Jag2 | 0.7394666 | 0.569760094 | 0.019 | 0.013 | 1 | 0.012470669 | 3.100239256 | 0.073 | 0.014 | 1 | 0.7394666 | 0.024785821 |
| Echs1 | 0.012471997 | 0.735183703 | 0.509 | 0.343 | 1 | 0.850054214 | 0.328863631 | 0.439 | 0.427 | 1 | 0.850054214 | 0.024788443 |
| Srl | 0.413082433 | 0.154722595 | 0 | 0.013 | 1 | 0.012473544 | 2.83720485 | 0.073 | 0.014 | 1 | 0.413082433 | 0.024791499 |
| Ramp1 | 0.112066058 | -2.304709024 | 0.038 | 0.106 | 1 | 0.012476709 | -1.673757069 | 0.049 | 0.222 | 1 | 0.112066058 | 0.02479775 |
| Atp13a3 | 0.012476761 | 0.490641377 | 0.906 | 0.708 | 1 | 0.058931052 | 0.230686206 | 0.976 | 0.877 | 1 | 0.058931052 | 0.024797852 |
| Angel2 | 0.012523384 | 0.672121812 | 0.66 | 0.559 | 1 | 0.520104222 | -0.108159894 | 0.463 | 0.546 | 1 | 0.520104222 | 0.024889933 |
| Zfp712 | 0.0909501 | 1.8327945 | 0.057 | 0.017 | 1 | 0.012531398 | 2.47463477 | 0.098 | 0.024 | 1 | 0.0909501 | 0.024905761 |
| Pbxip1 | 0.071476228 | -0.430239906 | 0.66 | 0.771 | 1 | 0.012531517 | 0.564186355 | 0.878 | 0.734 | 1 | 0.071476228 | 0.024905995 |
| Rgma | 0.012539201 | 1.005859209 | 0.302 | 0.153 | 1 | 0.95629854 | 0.222495006 | 0.22 | 0.222 | 1 | 0.95629854 | 0.024921171 |
| Naxe | 0.331594577 | -0.20282941 | 0.547 | 0.678 | 1 | 0.012546162 | -0.678152183 | 0.341 | 0.57 | 1 | 0.331594577 | 0.024934917 |
| Enpp5 | 0.020848736 | -0.686579659 | 0.189 | 0.369 | 1 | 0.012575492 | -0.992870149 | 0.122 | 0.321 | 1 | 0.020848736 | 0.024992842 |
| Tmcc1 | 0.012590055 | 0.51960789 | 0.83 | 0.767 | 1 | 0.063594379 | -0.406574959 | 0.878 | 0.945 | 1 | 0.063594379 | 0.0250216 |
| Akap1 | 0.208477297 | 0.852159825 | 0.189 | 0.127 | 1 | 0.012598327 | 1.412707021 | 0.268 | 0.13 | 1 | 0.208477297 | 0.025037937 |
| Uba1 | 0.012603713 | 0.66802411 | 0.679 | 0.572 | 1 | 0.403091866 | 0.377773231 | 0.707 | 0.768 | 1 | 0.403091866 | 0.025048573 |
| Cpz | 0.012637954 | 1.292226119 | 0.132 | 0.042 | 1 | 0.028943483 | 1.83720485 | 0.073 | 0.017 | 1 | 0.028943483 | 0.02511619 |
| Prg4 | 0.396946265 | -0.348087437 | 0.283 | 0.347 | 1 | 0.0126409 | -2.311090136 | 0.146 | 0.338 | 1 | 0.396946265 | 0.025122007 |
| Map7 | 0.01265033 | 0.51612213 | 0.585 | 0.436 | 1 | 0.739273662 | -0.359192363 | 0.585 | 0.553 | 1 | 0.739273662 | 0.025140629 |
| Ncam1 | 0.012664624 | 0.877946536 | 0.774 | 0.767 | 1 | 0.967213683 | 0.740775087 | 0.659 | 0.754 | 1 | 0.967213683 | 0.025168856 |
| Rasa3 | 0.10292742 | -0.1672055 | 0.453 | 0.576 | 1 | 0.012669759 | -0.144647803 | 0.341 | 0.594 | 1 | 0.10292742 | 0.025178995 |
| Plxna1 | 0.012695703 | 0.977844833 | 0.321 | 0.174 | 1 | 0.134348447 | 0.735666823 | 0.366 | 0.27 | 1 | 0.134348447 | 0.025230224 |
| Gm26879 | 0.012706454 | 1.624207878 | 0.132 | 0.042 | 1 | 0.191978468 | 1.007129851 | 0.146 | 0.085 | 1 | 0.191978468 | 0.025251454 |
| Oxld1 | 0.012706454 | 1.624207878 | 0.132 | 0.042 | 1 | 0.29852636 | 0.83720485 | 0.073 | 0.038 | 1 | 0.29852636 | 0.025251454 |
| Coa3 | 0.012708842 | 0.80108564 | 0.566 | 0.407 | 1 | 0.668363333 | 0.389059039 | 0.488 | 0.478 | 1 | 0.668363333 | 0.025256169 |
| Gm35550 | 0.503971666 | 1.569760094 | 0.019 | 0.008 | 1 | 0.012719436 | 3.059597271 | 0.073 | 0.014 | 1 | 0.503971666 | 0.025277088 |
| Reln | 0.497380598 | 2.154722595 | 0.019 | 0.008 | 1 | 0.01272186 | 2.667279848 | 0.073 | 0.014 | 1 | 0.497380598 | 0.025280544 |
| Orc4 | 0.454535089 | 0.261637799 | 0.547 | 0.525 | 1 | 0.012724059 | 0.569341912 | 0.829 | 0.662 | 1 | 0.454535089 | 0.025286216 |
| Abhd8 | 0.012747693 | 0.971858538 | 0.453 | 0.301 | 1 | 0.370173856 | 0.370078839 | 0.317 | 0.246 | 1 | 0.370173856 | 0.025332883 |
| Carnmt1 | 0.179807989 | 0.455653608 | 0.642 | 0.581 | 1 | 0.012761274 | -0.525810437 | 0.488 | 0.686 | 1 | 0.179807989 | 0.025359697 |
| Zfp275 | 0.012766339 | 0.751366901 | 0.434 | 0.263 | 1 | 0.830905627 | 0.386543441 | 0.366 | 0.362 | 1 | 0.830905627 | 0.025369699 |
| Bag2 | 0.012769694 | 1.247831999 | 0.283 | 0.144 | 1 | 0.154896638 | -0.679370676 | 0.146 | 0.246 | 1 | 0.154896638 | 0.025376323 |
| Copg2 | 0.012839349 | 0.865215978 | 0.434 | 0.267 | 1 | 0.643351979 | 0.164779508 | 0.39 | 0.338 | 1 | 0.643351979 | 0.02551385 |
| Eif2ak4 | 0.012840321 | -1.308249382 | 0.434 | 0.568 | 1 | 0.066258206 | -0.57501213 | 0.488 | 0.59 | 1 | 0.066258206 | 0.025515768 |
| Adamts7 | 0.10281534 | 1.739685096 | 0.038 | 0.008 | 1 | 0.012844573 | 2.574170444 | 0.073 | 0.014 | 1 | 0.10281534 | 0.025524164 |
| Gm10190 | 0.347746365 | 0.932330173 | 0.038 | 0.017 | 1 | 0.012844573 | 2.574170444 | 0.073 | 0.014 | 1 | 0.347746365 | 0.025524164 |
| Atr | 0.012852362 | 0.978872759 | 0.302 | 0.153 | 1 | 0.290516407 | -0.612102552 | 0.171 | 0.246 | 1 | 0.290516407 | 0.025539541 |
| Fbxl18 | 0.012859824 | 1.112902419 | 0.189 | 0.076 | 1 | 0.928709265 | 0.438655473 | 0.122 | 0.123 | 1 | 0.928709265 | 0.025554273 |
| Ampd3 | 0.012875682 | -2.412961914 | 0.113 | 0.254 | 1 | 0.418671475 | -0.332720152 | 0.073 | 0.116 | 1 | 0.418671475 | 0.025585581 |
| H2-Ab1 | 0.012914866 | -3.922093002 | 0.019 | 0.14 | 1 | 0.345527083 | 1.422167351 | 0.073 | 0.041 | 1 | 0.345527083 | 0.025662939 |
| Arfrp1 | 0.035195524 | 0.756173218 | 0.434 | 0.297 | 1 | 0.012917566 | -1.196218152 | 0.146 | 0.338 | 1 | 0.035195524 | 0.025668268 |
| B3galnt1 | 0.012932669 | 1.569760094 | 0.132 | 0.042 | 1 | 0.494346678 | -0.686357106 | 0.024 | 0.048 | 1 | 0.494346678 | 0.025698084 |
| Tra2a | 0.012942731 | 0.3239087 | 1 | 0.953 | 1 | 0.104964204 | 0.429425379 | 0.951 | 0.959 | 1 | 0.104964204 | 0.025717947 |
| 2810433D01Rik | 0.012944937 | 1.292226119 | 0.17 | 0.064 | 1 | 0.322414763 | 1.252242349 | 0.098 | 0.058 | 1 | 0.322414763 | 0.025722302 |
| Anapc2 | 0.079688534 | 0.569760094 | 0.472 | 0.364 | 1 | 0.012946258 | -0.95069771 | 0.293 | 0.488 | 1 | 0.079688534 | 0.02572491 |
| Nelfa | 0.01296671 | 0.8327945 | 0.415 | 0.258 | 1 | 0.20364861 | -0.16279515 | 0.341 | 0.498 | 1 | 0.20364861 | 0.025765285 |
| Gja4 | 0.015693134 | 2.47665069 | 0.057 | 0.008 | 1 | 0.012970799 | 2.159132945 | 0.073 | 0.014 | 1 | 0.015693134 | 0.025773356 |
| C230038L03Rik | 0.343968402 | 1.154722595 | 0.038 | 0.017 | 1 | 0.012973167 | 2.515276755 | 0.073 | 0.014 | 1 | 0.343968402 | 0.02577803 |
| Med9os | 0.926212226 | 0.8327945 | 0.019 | 0.017 | 1 | 0.012973167 | 2.515276755 | 0.073 | 0.014 | 1 | 0.926212226 | 0.02577803 |
| Socs1 | 0.013012598 | 0.781913331 | 0.358 | 0.208 | 1 | 0.215622414 | 0.735666823 | 0.244 | 0.177 | 1 | 0.215622414 | 0.025855869 |
| Zbtb5 | 0.044854124 | 1.538051234 | 0.17 | 0.085 | 1 | 0.013109296 | 1.930314254 | 0.268 | 0.137 | 1 | 0.044854124 | 0.026046739 |
| Zfp35 | 0.013128203 | 1.306725688 | 0.226 | 0.102 | 1 | 0.463664655 | 0.218295017 | 0.073 | 0.116 | 1 | 0.463664655 | 0.026084056 |
| Lrrc59 | 0.013133345 | 0.429438806 | 0.83 | 0.729 | 1 | 0.020636369 | -0.567636882 | 0.78 | 0.863 | 1 | 0.020636369 | 0.026094205 |
| Rcc1l | 0.01315858 | 1.067259754 | 0.34 | 0.191 | 1 | 0.127056067 | -0.11388555 | 0.146 | 0.276 | 1 | 0.127056067 | 0.026144011 |
| Ass1 | 0.013163603 | -1.193200709 | 0.113 | 0.275 | 1 | 0.872965635 | 0.19334866 | 0.122 | 0.113 | 1 | 0.872965635 | 0.026153925 |
| Lsamp | 0.087097938 | 2.891688189 | 0.057 | 0.017 | 1 | 0.013172112 | -0.615762105 | 0.171 | 0.389 | 1 | 0.087097938 | 0.02617072 |
| Seh1l | 0.323074682 | -0.351911545 | 0.472 | 0.547 | 1 | 0.013184599 | -0.847293324 | 0.317 | 0.515 | 1 | 0.323074682 | 0.026195365 |
| Kdm2b | 0.013199987 | 0.926453607 | 0.396 | 0.237 | 1 | 0.747733267 | 0.505361286 | 0.317 | 0.311 | 1 | 0.747733267 | 0.026225735 |
| Ang | 0.270507009 | -0.600164907 | 0.264 | 0.339 | 1 | 0.013215092 | -1.144647803 | 0.22 | 0.42 | 1 | 0.270507009 | 0.026255545 |
| Med26 | 0.013215121 | 0.789725778 | 0.547 | 0.436 | 1 | 0.089658777 | 0.663067797 | 0.707 | 0.611 | 1 | 0.089658777 | 0.026255603 |
| Pip5k1b | 0.794494793 | 0.228723176 | 0.113 | 0.127 | 1 | 0.013219993 | 1.271607674 | 0.317 | 0.167 | 1 | 0.794494793 | 0.026265217 |
| Gm12743 | 0.013230434 | 1.154722595 | 0.189 | 0.076 | 1 | 0.199711144 | -0.80665134 | 0.049 | 0.116 | 1 | 0.199711144 | 0.026285824 |
| Gm29585 | 0.119161515 | 0.104096522 | 0.075 | 0.03 | 1 | 0.013236535 | 2.252242349 | 0.098 | 0.024 | 1 | 0.119161515 | 0.026297864 |
| Eif2b4 | 0.01324853 | 0.769068902 | 0.377 | 0.225 | 1 | 0.165352281 | 0.502785811 | 0.366 | 0.27 | 1 | 0.165352281 | 0.026321536 |
| Lin52 | 0.013253648 | 0.569760094 | 0.547 | 0.386 | 1 | 0.033470557 | 0.958883406 | 0.512 | 0.389 | 1 | 0.033470557 | 0.026331638 |
| ENSMUSG00000121500 | 0.016034681 | 1.640149422 | 0.17 | 0.068 | 1 | 0.013259526 | 1.483567895 | 0.171 | 0.061 | 1 | 0.016034681 | 0.026343237 |
| Cul9 | 0.013272125 | 1.039245377 | 0.302 | 0.157 | 1 | 0.10632199 | 1.015542091 | 0.39 | 0.314 | 1 | 0.10632199 | 0.026368101 |
| Ssh1 | 0.314826817 | 0.3662267 | 0.415 | 0.364 | 1 | 0.013283062 | -0.671344535 | 0.366 | 0.59 | 1 | 0.314826817 | 0.026389685 |
| Prmt9 | 0.013283188 | 0.846600299 | 0.302 | 0.153 | 1 | 0.652046033 | -0.4233227 | 0.22 | 0.246 | 1 | 0.652046033 | 0.026389933 |
| Arl13b | 0.807052658 | 0.460048722 | 0.642 | 0.712 | 1 | 0.013286842 | -0.644946845 | 0.61 | 0.802 | 1 | 0.807052658 | 0.026397144 |
| Chac1 | 0.02051295 | -2.497354102 | 0.057 | 0.182 | 1 | 0.013307228 | -3.448197369 | 0 | 0.133 | 1 | 0.02051295 | 0.026437373 |
| Pdp1 | 0.013327435 | 1.659957903 | 0.226 | 0.11 | 1 | 0.668831453 | -0.259656689 | 0.146 | 0.174 | 1 | 0.668831453 | 0.026477249 |
| Ipmk | 0.013343896 | 1.394673388 | 0.377 | 0.225 | 1 | 0.015922004 | 1.017777095 | 0.439 | 0.273 | 1 | 0.015922004 | 0.026509732 |
| Cadm1 | 0.013344579 | 0.731511164 | 0.83 | 0.61 | 1 | 0.472573877 | 0.280239735 | 0.976 | 0.952 | 1 | 0.472573877 | 0.026511081 |
| Cldn7 | 0.013345618 | 1.154722595 | 0.17 | 0.064 | 1 | 0.406989874 | 0.120997816 | 0.293 | 0.229 | 1 | 0.406989874 | 0.026513131 |
| Ttc19 | 0.040372285 | 0.60382193 | 0.717 | 0.631 | 1 | 0.013348355 | 0.487938046 | 0.878 | 0.727 | 1 | 0.040372285 | 0.026518531 |
| Psmb9 | 0.224900761 | 0.792152515 | 0.094 | 0.051 | 1 | 0.013354269 | 2.252242349 | 0.073 | 0.014 | 1 | 0.224900761 | 0.026530201 |
| Gm57006 | 0.237970389 | 1.154722595 | 0.075 | 0.038 | 1 | 0.013354269 | 2.252242349 | 0.073 | 0.014 | 1 | 0.237970389 | 0.026530201 |
| Gm15860 | 0.287773478 | -0.430239906 | 0 | 0.021 | 1 | 0.013354269 | 2.252242349 | 0.073 | 0.014 | 1 | 0.287773478 | 0.026530201 |
| I830077J02Rik | 0.503971666 | 1.569760094 | 0.019 | 0.008 | 1 | 0.013354269 | 2.252242349 | 0.073 | 0.014 | 1 | 0.503971666 | 0.026530201 |
| Ttc9c | 0.013388781 | 0.652222254 | 0.528 | 0.339 | 1 | 0.238032455 | -0.115489435 | 0.366 | 0.515 | 1 | 0.238032455 | 0.026598302 |
| Cep164 | 0.013390919 | 0.642823556 | 0.34 | 0.191 | 1 | 0.53132175 | 0.438655473 | 0.293 | 0.253 | 1 | 0.53132175 | 0.026602522 |
| Mcmdc2 | 0.281194124 | 0.762405172 | 0.132 | 0.089 | 1 | 0.013391993 | -2.611225651 | 0.049 | 0.205 | 1 | 0.281194124 | 0.026604641 |
| Nfkb2 | 0.013392491 | 1.027610677 | 0.415 | 0.267 | 1 | 0.110374524 | 0.884202329 | 0.415 | 0.331 | 1 | 0.110374524 | 0.026605624 |
| Arpin | 0.013413147 | 1.280253477 | 0.264 | 0.131 | 1 | 0.194514828 | -0.421529419 | 0.098 | 0.181 | 1 | 0.194514828 | 0.026646382 |
| Plekha4 | 0.013426252 | 1.315187267 | 0.226 | 0.106 | 1 | 0.555925991 | 0.766815522 | 0.122 | 0.096 | 1 | 0.555925991 | 0.02667224 |
| Ap4m1 | 0.013439662 | 1.533234218 | 0.264 | 0.14 | 1 | 0.315756879 | 0.707921833 | 0.22 | 0.16 | 1 | 0.315756879 | 0.026698699 |
| Sf3b6 | 0.013519643 | 0.384842378 | 0.906 | 0.839 | 1 | 0.521235709 | -0.113326474 | 0.854 | 0.891 | 1 | 0.521235709 | 0.026856505 |
| Zfp708 | 0.01351971 | 1.454282877 | 0.132 | 0.042 | 1 | 0.588188971 | 0.204936634 | 0.098 | 0.072 | 1 | 0.588188971 | 0.026856638 |

| | | | | | | | | | | | | |
|---|---|---|---|---|---|---|---|---|---|---|---|---|
| Dgat2 | 0.220681417 | 0.8327945 | 0.189 | 0.127 | 1 | 0.01352237 | 1.252242349 | 0.22 | 0.092 | 1 | 0.220681417 | 0.026861886 |
| Vamp4 | 0.233134443 | -0.240137023 | 0.547 | 0.631 | 1 | 0.013526058 | -0.747757651 | 0.341 | 0.563 | 1 | 0.233134443 | 0.026869162 |
| Slc5a6 | 0.013536077 | 2.533234218 | 0.113 | 0.034 | 1 | 0.362195044 | 1.252242349 | 0.049 | 0.024 | 1 | 0.362195044 | 0.026888929 |
| Fuca1 | 0.013540221 | -0.435310895 | 0.887 | 0.877 | 1 | 0.069692748 | -0.407975544 | 0.829 | 0.833 | 1 | 0.069692748 | 0.026897104 |
| Map3k19 | 0.249034808 | 2.154722595 | 0.019 | 0.004 | 1 | 0.013553388 | 2.306690133 | 0.098 | 0.024 | 1 | 0.249034808 | 0.026923082 |
| Mrpl21 | 0.013579322 | 0.818673392 | 0.585 | 0.475 | 1 | 0.128286932 | 0.564186355 | 0.488 | 0.372 | 1 | 0.128286932 | 0.026974246 |
| Vkorc1 | 0.767496866 | -0.173990105 | 0.906 | 0.894 | 1 | 0.013589686 | -0.548481826 | 0.829 | 0.874 | 1 | 0.767496866 | 0.026994693 |
| Tfcp2 | 0.013615296 | 0.910797012 | 0.415 | 0.263 | 1 | 0.511852658 | 0.387897448 | 0.244 | 0.307 | 1 | 0.511852658 | 0.027045215 |
| Copa | 0.013636581 | 0.404900308 | 1 | 0.979 | 1 | 0.102759248 | -0.209035329 | 1 | 0.99 | 1 | 0.102759248 | 0.027087205 |
| Ccdc173 | 0.949715024 | 0.3662267 | 0.094 | 0.093 | 1 | 0.013636895 | 2.052933541 | 0.171 | 0.065 | 1 | 0.949715024 | 0.027087824 |
| Pole4 | 0.376394923 | 0.478129619 | 0.528 | 0.521 | 1 | 0.013639179 | -0.990614175 | 0.293 | 0.478 | 1 | 0.376394923 | 0.027092331 |
| Twnk | 0.013639551 | 1.80108564 | 0.208 | 0.093 | 1 | 0.30567149 | 1.444887427 | 0.098 | 0.058 | 1 | 0.30567149 | 0.027093065 |
| Txnl1 | 0.018853327 | 0.336995989 | 0.906 | 0.881 | 1 | 0.013642777 | -0.406498078 | 0.902 | 0.952 | 1 | 0.018853327 | 0.027099429 |
| Wdr27 | 0.013660727 | 1.090592257 | 0.17 | 0.064 | 1 | 0.646741372 | 0.377773231 | 0.073 | 0.055 | 1 | 0.646741372 | 0.027134839 |
| Mrpl35 | 0.013684301 | 0.810768194 | 0.396 | 0.229 | 1 | 0.035513157 | -0.930979475 | 0.122 | 0.283 | 1 | 0.035513157 | 0.027181341 |
| Uba3 | 0.013703556 | -0.527537107 | 0.472 | 0.669 | 1 | 0.286023821 | -0.195543733 | 0.537 | 0.659 | 1 | 0.286023821 | 0.027219325 |
| Dact3 | 0.105770772 | 0.555260524 | 0.396 | 0.301 | 1 | 0.013709983 | 1.222495006 | 0.512 | 0.348 | 1 | 0.105770772 | 0.027232002 |
| Stard5 | 0.016417813 | -1.321010836 | 0.094 | 0.246 | 1 | 0.013722108 | -1.659220976 | 0.122 | 0.3 | 1 | 0.016417813 | 0.02725592 |
| Qser1 | 0.013724476 | 0.653324811 | 0.623 | 0.466 | 1 | 0.726181888 | -0.155155088 | 0.78 | 0.747 | 1 | 0.726181888 | 0.027260591 |
| Pip4k2c | 0.071637663 | 0.675253223 | 0.547 | 0.462 | 1 | 0.013725705 | -0.94123738 | 0.317 | 0.505 | 1 | 0.071637663 | 0.027263016 |
| Gm56897 | 0.929579738 | 0.932330173 | 0.019 | 0.021 | 1 | 0.013751449 | 2.029849928 | 0.073 | 0.014 | 1 | 0.929579738 | 0.027313796 |
| Hspb6 | 0.01378562 | -1.799473716 | 0.113 | 0.267 | 1 | 0.246217072 | -0.448197369 | 0.049 | 0.109 | 1 | 0.246217072 | 0.027381196 |
| Tbcb | 0.013829289 | 0.697040758 | 0.698 | 0.581 | 1 | 0.125352989 | 0.357514392 | 0.683 | 0.543 | 1 | 0.125352989 | 0.027467328 |
| Tor2a | 0.013833199 | -0.745741732 | 0.189 | 0.377 | 1 | 0.024861939 | -1.008285201 | 0.146 | 0.317 | 1 | 0.024861939 | 0.02747504 |
| Hsd17b12 | 0.013834925 | -0.576917992 | 0.755 | 0.847 | 1 | 0.868341784 | -0.177954957 | 0.732 | 0.744 | 1 | 0.868341784 | 0.027478445 |
| Postn | 0.173445026 | 1.673423653 | 0.434 | 0.369 | 1 | 0.013836535 | 4.94154151 | 0.195 | 0.085 | 1 | 0.173445026 | 0.027481621 |
| Ahcyl2 | 0.013844338 | 0.62946634 | 0.792 | 0.614 | 1 | 0.243121203 | 0.132660733 | 0.951 | 0.853 | 1 | 0.243121203 | 0.02749701 |
| Gm13986 | 0.039399375 | -0.822557329 | 0.057 | 0.174 | 1 | 0.013852314 | -2.890715605 | 0.049 | 0.205 | 1 | 0.039399375 | 0.027512741 |
| Zbtb41 | 0.013856094 | 0.870269205 | 0.453 | 0.305 | 1 | 0.175602145 | 0.150704323 | 0.293 | 0.471 | 1 | 0.175602145 | 0.027520196 |
| Map3k4 | 0.608068703 | -0.475027828 | 0.396 | 0.428 | 1 | 0.01392192 | -1.265605956 | 0.195 | 0.379 | 1 | 0.608068703 | 0.027650019 |
| Fkbp1a | 0.013923613 | 0.381810304 | 0.925 | 0.894 | 1 | 0.398210555 | -0.100880367 | 0.878 | 0.908 | 1 | 0.398210555 | 0.02765336 |
| Tti2 | 0.552554472 | 0.154722595 | 0.151 | 0.195 | 1 | 0.013929654 | -1.825760163 | 0.049 | 0.208 | 1 | 0.552554472 | 0.027665273 |
| Zfp689 | 0.013943914 | 1.707263618 | 0.151 | 0.055 | 1 | 0.142323729 | -1.16279515 | 0.024 | 0.092 | 1 | 0.142323729 | 0.027693395 |
| Mapk13 | 0.013943914 | 1.707263618 | 0.151 | 0.055 | 1 | 0.531190715 | 0.479652845 | 0.073 | 0.113 | 1 | 0.531190715 | 0.027693395 |
| Immp1l | 0.831722956 | 0.133660979 | 0.623 | 0.657 | 1 | 0.01394438 | -0.677969321 | 0.439 | 0.635 | 1 | 0.831722956 | 0.027694315 |
| Aimp1 | 0.051465954 | 0.601330173 | 0.642 | 0.504 | 1 | 0.013953128 | 0.709449303 | 0.659 | 0.509 | 1 | 0.051465954 | 0.027711566 |
| Ckmt1 | 0.128649744 | -1.752168001 | 0 | 0.042 | 1 | 0.013957216 | 1.145327145 | 0.244 | 0.109 | 1 | 0.128649744 | 0.027719628 |
| Faap20 | 0.013959224 | 0.757832442 | 0.415 | 0.254 | 1 | 0.260764689 | -0.466575898 | 0.122 | 0.201 | 1 | 0.260764689 | 0.027723587 |
| Cstf2 | 0.013972574 | 1.124975251 | 0.302 | 0.161 | 1 | 0.923129958 | 0.500169863 | 0.244 | 0.256 | 1 | 0.923129958 | 0.027749914 |
| Aifm3 | 0.340246113 | 1.417757001 | 0.038 | 0.017 | 1 | 0.013975845 | 1.989207943 | 0.098 | 0.024 | 1 | 0.340246113 | 0.027756365 |
| Ska3 | 0.043642799 | 1.417757001 | 0.113 | 0.042 | 1 | 0.01398455 | 1.83720485 | 0.098 | 0.024 | 1 | 0.043642799 | 0.027773532 |
| Fbxo30 | 0.013992831 | 0.530589497 | 0.755 | 0.623 | 1 | 0.435718148 | -0.164819986 | 0.707 | 0.823 | 1 | 0.435718148 | 0.027789862 |
| Dact1 | 0.014006118 | 0.427155699 | 0.585 | 0.403 | 1 | 0.239429565 | 0.480511337 | 0.537 | 0.485 | 1 | 0.239429565 | 0.027816065 |
| Mok | 0.014008938 | 2.154722595 | 0.113 | 0.034 | 1 | 0.395320196 | -0.410722664 | 0.049 | 0.089 | 1 | 0.395320196 | 0.027821626 |
| Bcl7b | 0.181076378 | 0.44102678 | 0.566 | 0.479 | 1 | 0.014023715 | -0.723510105 | 0.293 | 0.491 | 1 | 0.181076378 | 0.027850765 |
| Lcor | 0.014051396 | 0.40256784 | 0.887 | 0.86 | 1 | 0.547644964 | 0.182935313 | 0.951 | 0.966 | 1 | 0.547644964 | 0.027905351 |
| Tmem26 | 0.016828843 | 3.002719501 | 0.075 | 0.017 | 1 | 0.014093664 | 1.737669176 | 0.098 | 0.024 | 1 | 0.016828843 | 0.027988696 |
| Pcmtd2 | 0.014142151 | 0.795641501 | 0.472 | 0.326 | 1 | 0.452494291 | -0.11032773 | 0.366 | 0.437 | 1 | 0.452494291 | 0.028084302 |
| Rars2 | 0.064181941 | 0.776210972 | 0.34 | 0.225 | 1 | 0.014187516 | -1.006491919 | 0.122 | 0.321 | 1 | 0.064181941 | 0.028173746 |
| Nrgn | 0.213160494 | 0.787940264 | 0.113 | 0.064 | 1 | 0.014203284 | 1.422167351 | 0.098 | 0.024 | 1 | 0.213160494 | 0.028204835 |
| Tnfaip6 | 0.480703359 | -0.313830414 | 0.226 | 0.195 | 1 | 0.014205999 | 2.644559772 | 0.195 | 0.085 | 1 | 0.480703359 | 0.028210187 |
| Mgst1 | 0.845786872 | -0.634857626 | 0.415 | 0.373 | 1 | 0.014240859 | 0.736744668 | 0.634 | 0.471 | 1 | 0.845786872 | 0.028278917 |
| Pdk3 | 0.014261015 | 0.755115136 | 0.528 | 0.369 | 1 | 0.01636584 | 1.128253632 | 0.512 | 0.389 | 1 | 0.01636584 | 0.028318654 |
| Rap1gap | 0.01427215 | 1.782753817 | 0.17 | 0.068 | 1 | 0.235847932 | 1.630753972 | 0.195 | 0.14 | 1 | 0.235847932 | 0.028340606 |
| Spats2l | 0.284543854 | -0.314153056 | 0.887 | 0.898 | 1 | 0.014274932 | -0.443237143 | 0.902 | 0.928 | 1 | 0.284543854 | 0.028346089 |
| Tmem165 | 0.090104425 | -0.38747162 | 0.717 | 0.805 | 1 | 0.01427816 | -0.514955447 | 0.78 | 0.85 | 1 | 0.090104425 | 0.028352454 |
| Arhgef1 | 0.018033372 | -0.416192131 | 0.679 | 0.788 | 1 | 0.014286445 | -0.467306192 | 0.659 | 0.85 | 1 | 0.018033372 | 0.028368787 |
| Lrrc8a | 0.186836915 | 0.387556763 | 0.509 | 0.453 | 1 | 0.014292526 | -0.438429593 | 0.341 | 0.611 | 1 | 0.186836915 | 0.028380775 |
| Gm15246 | 0.018027274 | 1.962077517 | 0.094 | 0.025 | 1 | 0.014296699 | 1.737669176 | 0.073 | 0.014 | 1 | 0.018027274 | 0.028389003 |
| Itgb4 | 0.071478103 | 0.962077517 | 0.113 | 0.047 | 1 | 0.014301015 | 0.911205431 | 0.293 | 0.14 | 1 | 0.071478103 | 0.028397511 |
| Hmbox1 | 0.014320334 | 0.704061186 | 0.585 | 0.462 | 1 | 0.102164156 | 0.509084921 | 0.805 | 0.676 | 1 | 0.102164156 | 0.028435596 |
| Tmem176b | 0.051473881 | 0.329268257 | 0.981 | 0.873 | 1 | 0.014340551 | 0.490579518 | 0.927 | 0.911 | 1 | 0.051473881 | 0.02847545 |
| Ndufb2 | 0.014343634 | 0.481629184 | 0.868 | 0.784 | 1 | 0.07605807 | -0.407267105 | 0.659 | 0.737 | 1 | 0.07605807 | 0.028481528 |
| H2-Aa | 0.014352104 | -3.566376594 | 0.019 | 0.14 | 1 | 0.208817133 | -1.16279515 | 0 | 0.038 | 1 | 0.208817133 | 0.028498225 |
| Ftsj1 | 0.014358339 | 1.084333267 | 0.358 | 0.208 | 1 | 0.488352885 | 0.737669176 | 0.268 | 0.225 | 1 | 0.488352885 | 0.028510516 |
| Thtpa | 0.014386999 | 2.002719501 | 0.113 | 0.034 | 1 | 0.571736654 | 0.635570989 | 0.049 | 0.075 | 1 | 0.571736654 | 0.028567013 |
| R3hdm2 | 0.01440746 | 0.421437347 | 0.887 | 0.809 | 1 | 0.366484965 | 0.179353516 | 0.902 | 0.87 | 1 | 0.366484965 | 0.028607346 |
| Potegl | 0.014420653 | 0.984797593 | 0.132 | 0.042 | 1 | 0.298204415 | 0.970471381 | 0.122 | 0.078 | 1 | 0.298204415 | 0.028633352 |
| Yif1b | 0.557649718 | 0.154722595 | 0.566 | 0.555 | 1 | 0.01450753 | -0.523051364 | 0.341 | 0.577 | 1 | 0.557649718 | 0.028804592 |
| Dnajc4 | 0.384865364 | -0.151938743 | 0.245 | 0.305 | 1 | 0.014514404 | -1.069685746 | 0.073 | 0.256 | 1 | 0.384865364 | 0.02881814 |
| Dctn4 | 0.014516309 | 0.608440562 | 0.811 | 0.75 | 1 | 0.55345315 | 0.153443873 | 0.829 | 0.782 | 1 | 0.55345315 | 0.028821895 |
| Map2 | 0.014526525 | -1.037038208 | 0.377 | 0.564 | 1 | 0.417292945 | -0.559685303 | 0.317 | 0.375 | 1 | 0.417292945 | 0.02884203 |
| Ufl1 | 0.014547691 | 0.923397049 | 0.566 | 0.483 | 1 | 0.16852283 | -0.305753104 | 0.463 | 0.587 | 1 | 0.16852283 | 0.028883747 |
| Khk | 0.227246338 | -0.626637119 | 0.113 | 0.182 | 1 | 0.014565067 | -2.116991461 | 0.024 | 0.171 | 1 | 0.227246338 | 0.028917993 |
| Nog | 0.014572269 | -2.205533619 | 0.113 | 0.267 | 1 | 0.654442505 | -0.863234868 | 0.073 | 0.092 | 1 | 0.654442505 | 0.028932187 |
| Adamts6 | 0.014573944 | 1.934756911 | 0.283 | 0.153 | 1 | 0.153975147 | 1.201168164 | 0.317 | 0.229 | 1 | 0.153975147 | 0.028935488 |
| Gm15270 | 0.014581722 | -1.845277405 | 0.17 | 0.335 | 1 | 0.188689251 | -0.863234868 | 0.073 | 0.15 | 1 | 0.188689251 | 0.028950818 |
| Eml3 | 0.014616546 | 0.909610097 | 0.358 | 0.212 | 1 | 0.418121857 | -0.196218152 | 0.244 | 0.311 | 1 | 0.418121857 | 0.029019449 |
| Erbb4 | 0.686681035 | -1.093204919 | 0.038 | 0.051 | 1 | 0.014646933 | -1.518493982 | 0.024 | 0.177 | 1 | 0.686681035 | 0.029079334 |
| Zfp57 | 0.014650614 | 1.448453798 | 0.226 | 0.106 | 1 | 0.184528319 | 0.83720485 | 0.195 | 0.123 | 1 | 0.184528319 | 0.029086588 |
| Ube2g2 | 0.21023321 | 0.660907982 | 0.453 | 0.428 | 1 | 0.014672623 | -0.381218669 | 0.293 | 0.529 | 1 | 0.21023321 | 0.02912996 |
| Krt18 | 0.014719734 | 0.232725107 | 0.321 | 0.165 | 1 | 0.203584832 | 0.85796341 | 0.341 | 0.259 | 1 | 0.203584832 | 0.029222798 |
| Accs | 0.014758065 | 1.526691372 | 0.151 | 0.055 | 1 | 0.154111598 | -1.250257991 | 0 | 0.048 | 1 | 0.154111598 | 0.029298329 |
| Ndufa11 | 0.014782561 | 0.444618341 | 0.868 | 0.754 | 1 | 0.175010728 | 0.13450208 | 0.512 | 0.683 | 1 | 0.175010728 | 0.029346598 |
| Zc3h12b | 0.014786389 | 1.409651951 | 0.415 | 0.267 | 1 | 0.443748973 | -0.600800211 | 0.366 | 0.403 | 1 | 0.443748973 | 0.029354141 |
| Plaat3 | 0.014839633 | 0.584866986 | 0.396 | 0.225 | 1 | 0.749164604 | -0.408419878 | 0.488 | 0.478 | 1 | 0.749164604 | 0.029459051 |
| Gm9530 | 0.014846242 | 3.154722595 | 0.057 | 0.008 | 1 | 0.188380068 | 1.252242349 | 0.049 | 0.017 | 1 | 0.188380068 | 0.029472073 |
| Cfap206 | 0.014846242 | 3.569760094 | 0.057 | 0.008 | 1 | 0.455098275 | 0.515276755 | 0 | 0.014 | 1 | 0.455098275 | 0.029472073 |
| 2900092C05Rik | 0.014846242 | 3.377115016 | 0.057 | 0.008 | 1 | 0.057901606 | 2.422167351 | 0.049 | 0.01 | 1 | 0.057901606 | 0.029472073 |
| Lgals7 | 0.014846242 | 3.739685096 | 0.057 | 0.008 | 1 | 0.81727826 | -0.16279515 | 0.024 | 0.031 | 1 | 0.81727826 | 0.029472073 |
| Itgal | 0.014846242 | 3.569760094 | 0.057 | 0.008 | 1 | 0.51919986 | 0.83720485 | 0 | 0.01 | 1 | 0.51919986 | 0.029472073 |
| Gm36283 | 0.014846242 | 3.891688189 | 0.057 | 0.008 | 1 | 0.267205177 | 2.252242349 | 0.024 | 0.007 | 1 | 0.267205177 | 0.029472073 |
| Gm26839 | 0.014846242 | 3.154722595 | 0.057 | 0.008 | 1 | 0.430584486 | 3.159132945 | 0.024 | 0.01 | 1 | 0.430584486 | 0.029472073 |
| Gm50321 | 0.014846242 | 3.154722595 | 0.057 | 0.008 | 1 | 0.051733173 | 2.029849928 | 0.073 | 0.02 | 1 | 0.051733173 | 0.029472073 |
| Pcdhga5 | 0.014856536 | 1.607234799 | 0.17 | 0.068 | 1 | 0.917980834 | 0.136765132 | 0.098 | 0.092 | 1 | 0.917980834 | 0.029492355 |
| Clic1 | 0.014874189 | -0.592362319 | 0.962 | 0.945 | 1 | 0.094042673 | -0.190392709 | 0.951 | 0.956 | 1 | 0.094042673 | 0.029527137 |
| Zfp386 | 0.01489967 | 1.022272299 | 0.358 | 0.208 | 1 | 0.288759134 | -0.20718927 | 0.171 | 0.249 | 1 | 0.288759134 | 0.02957734 |
| Sae1 | 0.014899901 | 0.676675298 | 0.679 | 0.568 | 1 | 0.71389469 | 0.208481964 | 0.732 | 0.751 | 1 | 0.71389469 | 0.029577795 |

| | | | | | | | | | | | | |
|---|---|---|---|---|---|---|---|---|---|---|---|---|
| Rnf128 | 0.014907301 | -2.353072045 | 0.057 | 0.191 | 1 | 0.756568101 | 0.898605394 | 0.073 | 0.061 | 1 | 0.756568101 | 0.029592375 |
| Elp3 | 0.164032102 | -0.471462569 | 0.321 | 0.403 | 1 | 0.01490941 | -0.888620187 | 0.195 | 0.389 | 1 | 0.164032102 | 0.029596529 |
| Ccdc30 | 0.98030137 | 1.261637799 | 0.038 | 0.038 | 1 | 0.014936198 | 2.721727632 | 0.122 | 0.038 | 1 | 0.98030137 | 0.029649305 |
| Nod1 | 0.014937919 | -1.353072045 | 0.113 | 0.271 | 1 | 0.10361376 | -0.46860358 | 0.098 | 0.212 | 1 | 0.10361376 | 0.029652696 |
| Ndufa3 | 0.014940335 | 0.635849285 | 0.679 | 0.542 | 1 | 0.508673656 | -0.309636539 | 0.415 | 0.451 | 1 | 0.508673656 | 0.029657457 |
| Wdr3 | 0.01494681 | 1.306725688 | 0.34 | 0.203 | 1 | 0.426593713 | -0.520347155 | 0.244 | 0.287 | 1 | 0.426593713 | 0.029670212 |
| Ripk1 | 0.014952055 | 0.664194896 | 0.774 | 0.678 | 1 | 0.153123994 | 0.515276755 | 0.78 | 0.754 | 1 | 0.153123994 | 0.029680546 |
| Cnpy3 | 0.014961732 | 0.785488785 | 0.547 | 0.424 | 1 | 0.101944308 | -0.484723245 | 0.22 | 0.358 | 1 | 0.101944308 | 0.029699611 |
| Tfip11 | 0.0150053 | 1.280253477 | 0.245 | 0.119 | 1 | 0.322564005 | 0.515276755 | 0.073 | 0.13 | 1 | 0.322564005 | 0.029785441 |
| BC002059 | 0.015027216 | 1.517292674 | 0.189 | 0.081 | 1 | 0.706758 | 0.322631677 | 0.073 | 0.092 | 1 | 0.706758 | 0.029828614 |
| 2510002D24Rik | 0.015066218 | 1.039245377 | 0.358 | 0.216 | 1 | 0.162111157 | 0.210422173 | 0.146 | 0.259 | 1 | 0.162111157 | 0.029905445 |
| Pcnt | 0.015072119 | 0.869320376 | 0.415 | 0.275 | 1 | 0.064880525 | 0.948995675 | 0.512 | 0.43 | 1 | 0.064880525 | 0.029917069 |
| Dyrk1b | 0.015078764 | 2.292226119 | 0.113 | 0.034 | 1 | 0.199752332 | 0.911205431 | 0.098 | 0.048 | 1 | 0.199752332 | 0.029930158 |
| Tmem259 | 0.015094646 | 0.803250224 | 0.585 | 0.453 | 1 | 0.269548661 | 0.40789322 | 0.561 | 0.505 | 1 | 0.269548661 | 0.029961444 |
| Cfap61 | 0.015102942 | 1.948271717 | 0.113 | 0.034 | 1 | 0.062503704 | 0.334704509 | 0.122 | 0.048 | 1 | 0.062503704 | 0.029977786 |
| Pdzrn4 | 0.062908035 | 4.92351767 | 0.189 | 0.114 | 1 | 0.015126219 | 4.320195377 | 0.171 | 0.068 | 1 | 0.062908035 | 0.030023635 |
| Zdhhc1 | 0.028750963 | 0.635348436 | 0.528 | 0.364 | 1 | 0.015138828 | 1.125007161 | 0.488 | 0.345 | 1 | 0.028750963 | 0.030048471 |
| Slc9a8 | 0.015173205 | -0.652632327 | 0.151 | 0.331 | 1 | 0.14332153 | -0.775238387 | 0.22 | 0.331 | 1 | 0.14332153 | 0.030116185 |
| 4930444P10Rik | 0.015178556 | 3.154722595 | 0.057 | 0.008 | 1 | 0.188380068 | 1.614812428 | 0.049 | 0.017 | 1 | 0.188380068 | 0.030126724 |
| Gm20753 | 0.015178556 | 2.891688189 | 0.057 | 0.008 | 1 | 0.599290422 | 1.515276755 | 0.024 | 0.014 | 1 | 0.599290422 | 0.030126724 |
| Depdc7 | 0.015178556 | 3.377115016 | 0.057 | 0.008 | 1 | 0.799741379 | 1.252242349 | 0.049 | 0.041 | 1 | 0.799741379 | 0.030126724 |
| C530020B09Rik | 0.015178556 | 2.891688189 | 0.057 | 0.008 | 1 | 0.114118767 | 2.100239256 | 0.049 | 0.014 | 1 | 0.114118767 | 0.030126724 |
| Fam166b | 0.015178556 | 2.891688189 | 0.057 | 0.008 | 1 | 0.104856732 | 2.83720485 | 0.024 | 0.003 | 1 | 0.104856732 | 0.030126724 |
| Ptch2 | 0.015178556 | 2.891688189 | 0.057 | 0.008 | 1 | 0.182616442 | 2.47463477 | 0.049 | 0.017 | 1 | 0.182616442 | 0.030126724 |
| Gm26779 | 0.015178556 | 2.891688189 | 0.057 | 0.008 | 1 | 0.286433344 | -0.332720152 | 0 | 0.027 | 1 | 0.286433344 | 0.030126724 |
| Tmc7 | 0.015178556 | 2.891688189 | 0.057 | 0.008 | 1 | 0.402373368 | 0.252242349 | 0 | 0.017 | 1 | 0.402373368 | 0.030126724 |
| Arhgap45 | 0.015178556 | 2.891688189 | 0.057 | 0.008 | 1 | 0.936207831 | 1.252242349 | 0.024 | 0.027 | 1 | 0.936207831 | 0.030126724 |
| Slc6a15 | 0.015178556 | 2.891688189 | 0.057 | 0.008 | 1 | 0.450957104 | -0.484723245 | 0.024 | 0.051 | 1 | 0.450957104 | 0.030126724 |
| Gm57374 | 0.015178556 | 3.377115016 | 0.057 | 0.008 | 1 | 0.455098275 | 0.515276755 | 0 | 0.014 | 1 | 0.455098275 | 0.030126724 |
| Gm9920 | 0.015178556 | 2.891688189 | 0.057 | 0.008 | 1 | 0.856404759 | 2.029849928 | 0.024 | 0.02 | 1 | 0.856404759 | 0.030126724 |
| Tmprss7 | 0.015178556 | 3.154722595 | 0.057 | 0.008 | 1 | 0.119148825 | 2.029849928 | 0.073 | 0.027 | 1 | 0.119148825 | 0.030126724 |
| Dscam | 0.015178556 | 2.891688189 | 0.057 | 0.008 | 1 | 0.262676008 | 2.83720485 | 0.024 | 0.007 | 1 | 0.262676008 | 0.030126724 |
| Trim36 | 0.015178556 | 2.891688189 | 0.057 | 0.008 | 1 | 0.973994432 | 1.029849928 | 0.024 | 0.024 | 1 | 0.973994432 | 0.030126724 |
| Cntf | 0.015178556 | 2.891688189 | 0.057 | 0.008 | 1 | 0.02065851 | 3.252242349 | 0.049 | 0.007 | 1 | 0.02065851 | 0.030126724 |
| Nlgn3 | 0.015178556 | 2.891688189 | 0.057 | 0.008 | 1 | 0.834788606 | 1.136765132 | 0.073 | 0.068 | 1 | 0.834788606 | 0.030126724 |
| Iqub | 0.015180537 | 2.962077517 | 0.057 | 0.008 | 1 | 0.5191986 | 0.515276755 | 0 | 0.01 | 1 | 0.5191986 | 0.030130626 |
| Cby1 | 0.015181291 | 1.108918905 | 0.377 | 0.229 | 1 | 0.421262501 | 0.532350268 | 0.293 | 0.242 | 1 | 0.421262501 | 0.030132111 |
| Tmem214 | 0.887549329 | 0.196142522 | 0.509 | 0.547 | 1 | 0.015194702 | -0.429581691 | 0.317 | 0.567 | 1 | 0.887549329 | 0.030158524 |
| Gm4117 | 0.015204103 | 1.47665069 | 0.113 | 0.034 | 1 | 0.171996777 | 1.334704509 | 0.073 | 0.031 | 1 | 0.171996777 | 0.03017704 |
| Zfp9 | 0.15010627 | 0.275404077 | 0.642 | 0.547 | 1 | 0.015224437 | -0.56878751 | 0.439 | 0.631 | 1 | 0.15010627 | 0.03021709 |
| 2810403D21Rik | 0.110679307 | -0.497354102 | 0.208 | 0.326 | 1 | 0.0152265 | -1.995685164 | 0.073 | 0.235 | 1 | 0.110679307 | 0.030221154 |
| Nae1 | 0.015228173 | 0.688404591 | 0.472 | 0.297 | 1 | 0.186474466 | -0.589059905 | 0.439 | 0.495 | 1 | 0.186474466 | 0.030224448 |
| Zfp943 | 0.015243693 | 1.133027524 | 0.358 | 0.216 | 1 | 0.078382848 | -0.725731345 | 0.122 | 0.249 | 1 | 0.078382848 | 0.030255015 |
| Exoc8 | 0.015283463 | 1.695290976 | 0.113 | 0.034 | 1 | 0.144888727 | -1.069685746 | 0.049 | 0.126 | 1 | 0.144888727 | 0.030333342 |
| Rnf185 | 0.015284601 | 0.919506133 | 0.472 | 0.326 | 1 | 0.636005891 | 0.319929157 | 0.463 | 0.406 | 1 | 0.636005891 | 0.030335582 |
| Tnfsf13 | 0.015299135 | 1.685237311 | 0.17 | 0.068 | 1 | 0.13729585 | 0.989207943 | 0.171 | 0.096 | 1 | 0.13729585 | 0.030364206 |
| Gpn2 | 0.015318337 | 1.200526284 | 0.245 | 0.119 | 1 | 0.43516645 | 0.377773231 | 0.171 | 0.123 | 1 | 0.43516645 | 0.030402023 |
| Smagp | 0.015384665 | -1.923279917 | 0.132 | 0.28 | 1 | 0.53893155 | 0.930314254 | 0.049 | 0.031 | 1 | 0.53893155 | 0.030532642 |
| Hddc3 | 0.015413111 | 1.792152515 | 0.113 | 0.034 | 1 | 0.37389475 | -0.332720152 | 0.024 | 0.058 | 1 | 0.37389475 | 0.030588659 |
| Fbxo21 | 0.01543354 | 1.711115943 | 0.226 | 0.114 | 1 | 0.538587779 | 0.583448258 | 0.195 | 0.164 | 1 | 0.538587779 | 0.030628886 |
| Gm48742 | 0.015445441 | -1.686579659 | 0.019 | 0.14 | 1 | 0.452331971 | -0.250257991 | 0.024 | 0.051 | 1 | 0.452331971 | 0.030652321 |
| Tinagl1 | 0.015464757 | -2.517702747 | 0.019 | 0.136 | 1 | 0.084849342 | 1.458693227 | 0.098 | 0.038 | 1 | 0.084849342 | 0.030690356 |
| Eif2s2 | 0.59274146 | -0.123401147 | 0.962 | 0.983 | 1 | 0.01547298 | -0.420886506 | 0.951 | 0.983 | 1 | 0.59274146 | 0.030706546 |
| Gm20274 | 0.01548346 | 1.517292674 | 0.151 | 0.055 | 1 | 0.973847121 | 0.377773231 | 0.073 | 0.072 | 1 | 0.973847121 | 0.030727183 |
| Shld3 | 0.01548346 | 1.517292674 | 0.151 | 0.055 | 1 | 0.306313246 | -0.622226769 | 0.024 | 0.065 | 1 | 0.306313246 | 0.030727183 |
| Optn | 0.015485629 | -0.366109569 | 0.358 | 0.614 | 1 | 0.016396714 | -0.574921054 | 0.293 | 0.536 | 1 | 0.016396714 | 0.030731453 |
| Exph5 | 0.015520433 | 2.739685096 | 0.057 | 0.008 | 1 | 0.709279383 | 0.114738825 | 0.098 | 0.078 | 1 | 0.709279383 | 0.030799983 |
| Tram1 | 0.310290799 | 0.242455967 | 0.962 | 0.941 | 1 | 0.015520895 | -0.362215826 | 0.854 | 0.949 | 1 | 0.310290799 | 0.030800891 |
| Gm43768 | 0.01552144 | 2.569760094 | 0.057 | 0.008 | 1 | 0.44350429 | 1.377773231 | 0.049 | 0.027 | 1 | 0.44350429 | 0.030801965 |
| Gm43727 | 0.01552144 | 2.569760094 | 0.057 | 0.008 | 1 | 0.211360623 | 1.83720485 | 0.073 | 0.034 | 1 | 0.211360623 | 0.030801965 |
| Gdpd3 | 0.01552144 | 2.569760094 | 0.057 | 0.008 | 1 | 0.216244348 | 1.059597271 | 0.122 | 0.068 | 1 | 0.216244348 | 0.030801965 |
| 1700093K21Rik | 0.01552144 | 2.569760094 | 0.057 | 0.008 | 1 | 0.51919986 | 0.83720485 | 0 | 0.01 | 1 | 0.51919986 | 0.030801965 |
| Gpx6 | 0.01552144 | 2.569760094 | 0.057 | 0.008 | 1 | 0.911761241 | 0.377773231 | 0.024 | 0.027 | 1 | 0.911761241 | 0.030801965 |
| Gm54621 | 0.01552144 | 2.569760094 | 0.057 | 0.008 | 1 | 0.620260362 | 1.699701326 | 0.049 | 0.034 | 1 | 0.620260362 | 0.030801965 |
| Gm17058 | 0.01552144 | 2.569760094 | 0.057 | 0.008 | 1 | 0.749728988 | 0.83720485 | 0.024 | 0.017 | 1 | 0.749728988 | 0.030801965 |
| Ly6g5b | 0.01552144 | 2.569760094 | 0.057 | 0.008 | 1 | 0.050256774 | 2.422167351 | 0.073 | 0.02 | 1 | 0.050256774 | 0.030801965 |
| Best1 | 0.01552144 | 2.569760094 | 0.057 | 0.008 | 1 | 0.052077502 | 2.547698233 | 0.098 | 0.034 | 1 | 0.052077502 | 0.030801965 |
| Pnck | 0.01552144 | 2.569760094 | 0.057 | 0.008 | 1 | 0.402373368 | 0.252242349 | 0 | 0.017 | 1 | 0.402373368 | 0.030801965 |
| Angptl4 | 0.015536185 | 1.837240405 | 0.264 | 0.131 | 1 | 0.271610586 | 0.864512196 | 0.268 | 0.205 | 1 | 0.271610586 | 0.030830998 |
| Vdac3 | 0.132449777 | 0.321294161 | 0.792 | 0.691 | 1 | 0.015554682 | -0.465170448 | 0.707 | 0.819 | 1 | 0.132449777 | 0.030867416 |
| Ciao1 | 0.015555121 | 0.842778588 | 0.396 | 0.242 | 1 | 0.246136963 | 0.539524301 | 0.366 | 0.294 | 1 | 0.246136963 | 0.030868279 |
| Prpf3 | 0.169827517 | 0.276713119 | 0.491 | 0.36 | 1 | 0.015573549 | -0.758174267 | 0.317 | 0.519 | 1 | 0.169827517 | 0.030904562 |
| Sft2d1 | 0.055889673 | 0.557481764 | 0.491 | 0.36 | 1 | 0.015580378 | 0.911205431 | 0.585 | 0.451 | 1 | 0.055889673 | 0.030918009 |
| Spsb3 | 0.015614399 | 1.240452469 | 0.321 | 0.178 | 1 | 0.262564073 | -0.332720152 | 0.22 | 0.304 | 1 | 0.262564073 | 0.030984989 |
| Cdkn2b | 0.015633913 | -2.799473716 | 0 | 0.102 | 1 | 0.824812458 | 0.334704509 | 0.049 | 0.041 | 1 | 0.824812458 | 0.031023406 |
| 1700047M11Rik | 0.015645543 | -2.752168001 | 0 | 0.102 | 1 | 0.51919986 | 0.83720485 | 0 | 0.01 | 1 | 0.51919986 | 0.031046303 |
| Ppm1a | 0.015653537 | 0.41175826 | 0.943 | 0.818 | 1 | 0.152495816 | 0.460975371 | 0.878 | 0.911 | 1 | 0.152495816 | 0.031062041 |
| Hdgfl2 | 0.05488268 | 0.721763188 | 0.434 | 0.326 | 1 | 0.015665131 | 1.055205502 | 0.463 | 0.331 | 1 | 0.05488268 | 0.031084866 |
| Gm30489 | 0.015666046 | -3.054730771 | 0 | 0.102 | 1 | 0.231606831 | -0.970150072 | 0 | 0.034 | 1 | 0.231606831 | 0.031086666 |
| Gli1 | 0.015681556 | 1.47665069 | 0.151 | 0.055 | 1 | 0.34180587 | 1.351778023 | 0.073 | 0.041 | 1 | 0.34180587 | 0.031117202 |
| Chmp4c | 0.015688482 | -3.237594828 | 0 | 0.102 | 1 | 0.126389283 | -1.622226769 | 0 | 0.055 | 1 | 0.126389283 | 0.031130835 |
| Gm14009 | 0.015693134 | 2.47665069 | 0.057 | 0.008 | 1 | 0.106248001 | 1.83720485 | 0.024 | 0.003 | 1 | 0.106248001 | 0.031139994 |
| A930016O22Rik | 0.015693134 | 2.47665069 | 0.057 | 0.008 | 1 | 0.82181816 | 0.252242349 | 0.024 | 0.031 | 1 | 0.82181816 | 0.031139994 |
| H2-Eb1 | 0.015699734 | -4.764140642 | 0 | 0.102 | 1 | 0.190537763 | 1.159132945 | 0.098 | 0.048 | 1 | 0.190537763 | 0.031152985 |
| Anapc10 | 0.312819648 | 0.542279358 | 0.491 | 0.453 | 1 | 0.015706315 | -0.948290638 | 0.244 | 0.471 | 1 | 0.312819648 | 0.031165942 |
| Lurap1l | 0.561720372 | -0.752168001 | 0.019 | 0.034 | 1 | 0.015713948 | 2.574170444 | 0.122 | 0.038 | 1 | 0.561720372 | 0.031180969 |
| Atp6v0a1 | 0.015721448 | -0.746814766 | 0.34 | 0.513 | 1 | 0.370442691 | -0.376574441 | 0.488 | 0.522 | 1 | 0.370442691 | 0.031195732 |
| Cpsf4 | 0.015756814 | 0.977844833 | 0.302 | 0.161 | 1 | 0.071279108 | -1.233184478 | 0.098 | 0.215 | 1 | 0.071279108 | 0.031265352 |
| Mettl13 | 0.015812107 | 1.502645898 | 0.17 | 0.068 | 1 | 0.560310843 | -0.484723245 | 0.098 | 0.126 | 1 | 0.560310843 | 0.031374191 |
| A530017D24Rik | 0.015812158 | 1.695290976 | 0.113 | 0.034 | 1 | 0.171658817 | -1.16279515 | 0.024 | 0.085 | 1 | 0.171658817 | 0.031374292 |
| Usp20 | 0.015830038 | 1.538051234 | 0.189 | 0.081 | 1 | 0.189262261 | 1.174239837 | 0.146 | 0.085 | 1 | 0.189262261 | 0.031409486 |
| Ankrd40 | 0.015865586 | 0.674729654 | 0.491 | 0.318 | 1 | 0.67101481 | -0.19841906 | 0.537 | 0.543 | 1 | 0.67101481 | 0.031479455 |
| Add2 | 0.015867524 | 1.306725688 | 0.057 | 0.008 | 1 | 0.430584486 | 2.422167351 | 0.024 | 0.01 | 1 | 0.430584486 | 0.031483269 |
| Atp8b3 | 0.015867524 | 2.154722595 | 0.057 | 0.008 | 1 | 0.079809821 | 1.396632258 | 0.098 | 0.038 | 1 | 0.079809821 | 0.031483269 |
| A430093F15Rik | 0.015867524 | 2.347367673 | 0.057 | 0.008 | 1 | 0.87556513 | 0.83720485 | 0.024 | 0.02 | 1 | 0.87556513 | 0.031483269 |
| Efcab5 | 0.015882612 | 2.806799291 | 0.094 | 0.025 | 1 | 0.599117982 | -0.484723245 | 0.024 | 0.041 | 1 | 0.599117982 | 0.031512967 |
| Penk | 0.073210171 | -1.552636537 | 0.245 | 0.373 | 1 | 0.015904965 | -0.562725757 | 0.317 | 0.515 | 1 | 0.073210171 | 0.031556962 |
| Naprt | 0.01591101 | 1.526691372 | 0.17 | 0.068 | 1 | 0.052525104 | -2.250257991 | 0 | 0.085 | 1 | 0.052525104 | 0.031568859 |

| | | | | | | | | | | | | |
|---|---|---|---|---|---|---|---|---|---|---|---|---|
| Ptprm | 0.198714361 | -0.303313092 | 0.811 | 0.903 | 1 | 0.015912037 | -0.335567109 | 0.951 | 0.966 | 1 | 0.198714361 | 0.031570881 |
| Cdon | 0.015917676 | -1.127312773 | 0.283 | 0.458 | 1 | 0.021120843 | 0.745574374 | 0.561 | 0.423 | 1 | 0.021120843 | 0.031581979 |
| Slc25a39 | 0.015918235 | -0.721345077 | 0.528 | 0.648 | 1 | 0.232141676 | -0.29102086 | 0.561 | 0.645 | 1 | 0.232141676 | 0.031583079 |
| Fez1 | 0.015918457 | 1.569760094 | 0.113 | 0.034 | 1 | 0.387096215 | 0.749742009 | 0.122 | 0.082 | 1 | 0.387096215 | 0.031583516 |
| Gpatch11 | 0.015933974 | 1.090592257 | 0.377 | 0.233 | 1 | 0.462463896 | -0.470223675 | 0.171 | 0.218 | 1 | 0.462463896 | 0.031614056 |
| Pcyt1a | 0.748860808 | 0.158564661 | 0.66 | 0.708 | 1 | 0.015938415 | -0.475445431 | 0.634 | 0.826 | 1 | 0.748860808 | 0.031622797 |
| Nup160 | 0.015957672 | 1.005859209 | 0.434 | 0.297 | 1 | 0.562422081 | 0.644559772 | 0.366 | 0.338 | 1 | 0.562422081 | 0.031660697 |
| Cspg4b | 0.015971221 | 1.444229212 | 0.151 | 0.055 | 1 | 0.224279368 | 0.807457506 | 0.195 | 0.126 | 1 | 0.224279368 | 0.031687362 |
| Crlf1 | 0.015982519 | 1.569760094 | 0.17 | 0.068 | 1 | 0.138037225 | 1.004314836 | 0.22 | 0.133 | 1 | 0.138037225 | 0.031709597 |
| Ide | 0.015984565 | 0.60516543 | 0.792 | 0.661 | 1 | 0.159808521 | 0.301824378 | 0.78 | 0.71 | 1 | 0.159808521 | 0.031713624 |
| Chtop | 0.015986804 | 0.539545481 | 0.66 | 0.517 | 1 | 0.385562917 | 0.247877157 | 0.683 | 0.666 | 1 | 0.385562917 | 0.03171803 |
| Sri | 0.103042352 | 0.353502459 | 0.811 | 0.725 | 1 | 0.016014984 | -0.535537035 | 0.683 | 0.805 | 1 | 0.103042352 | 0.031773489 |
| 9530068E07Rik | 0.016020631 | 0.406485888 | 0.943 | 0.852 | 1 | 0.587192144 | 0.101494319 | 0.976 | 0.87 | 1 | 0.587192144 | 0.031784602 |
| Bop1 | 0.025607321 | 0.932330173 | 0.302 | 0.169 | 1 | 0.016023745 | -1.654648247 | 0.073 | 0.235 | 1 | 0.025607321 | 0.031790729 |
| Adcy6 | 0.016027286 | 1.035423666 | 0.358 | 0.212 | 1 | 0.101084697 | 0.557096931 | 0.512 | 0.375 | 1 | 0.101084697 | 0.031797699 |
| C730034F03Rik | 0.016037543 | 1.669295768 | 0.113 | 0.034 | 1 | 0.718266349 | 0.83720485 | 0.049 | 0.038 | 1 | 0.718266349 | 0.031817883 |
| Fam78b | 0.016042068 | 2.154722595 | 0.057 | 0.008 | 1 | 0.051724793 | 1.944120054 | 0.073 | 0.02 | 1 | 0.051724793 | 0.031826788 |
| Pgbd5 | 0.016042068 | 1.8327945 | 0.057 | 0.008 | 1 | 0.746881137 | 1.029849928 | 0.024 | 0.017 | 1 | 0.746881137 | 0.031826788 |
| Scgn | 0.016042068 | 2.154722595 | 0.057 | 0.008 | 1 | 0.844651838 | 0.989207943 | 0.024 | 0.031 | 1 | 0.844651838 | 0.031826788 |
| Kcnrg | 0.016042068 | 2.154722595 | 0.057 | 0.008 | 1 | 0.643341782 | 0.252242349 | 0.098 | 0.075 | 1 | 0.643341782 | 0.031826788 |
| Popdc2 | 0.016042068 | 2.154722595 | 0.057 | 0.008 | 1 | 0.51919986 | 0.83720485 | 0 | 0.01 | 1 | 0.51919986 | 0.031826788 |
| Parl | 0.154012554 | 0.638707448 | 0.528 | 0.466 | 1 | 0.016044604 | -0.909221074 | 0.317 | 0.495 | 1 | 0.154012554 | 0.031831779 |
| Fzd10 | 0.90991242 | 0.47665069 | 0.038 | 0.042 | 1 | 0.016088384 | -1.236044132 | 0.098 | 0.276 | 1 | 0.90991242 | 0.031917932 |
| Pex11b | 0.016094351 | 0.940597789 | 0.358 | 0.212 | 1 | 0.089856503 | 0.83720485 | 0.293 | 0.188 | 1 | 0.089856503 | 0.031929675 |
| Med14 | 0.016103759 | 0.717658789 | 0.528 | 0.386 | 1 | 0.106604572 | 0.584539417 | 0.683 | 0.57 | 1 | 0.106604572 | 0.031948187 |
| Mon1b | 0.016116859 | 1.533234218 | 0.17 | 0.068 | 1 | 0.637446795 | 0.667279848 | 0.073 | 0.055 | 1 | 0.637446795 | 0.031973965 |
| Slc38a7 | 0.016116915 | 1.491757582 | 0.17 | 0.068 | 1 | 0.604621826 | -0.484723245 | 0.122 | 0.15 | 1 | 0.604621826 | 0.031974075 |
| Gse1 | 0.016150659 | -0.842622952 | 0.415 | 0.606 | 1 | 0.043732035 | -1.090691604 | 0.537 | 0.635 | 1 | 0.043732035 | 0.032040474 |
| Miga2 | 0.811125227 | 0.377115016 | 0.094 | 0.085 | 1 | 0.016172318 | -2.228884341 | 0.024 | 0.167 | 1 | 0.811125227 | 0.032083092 |
| Pgd | 0.016177486 | 0.622501556 | 0.642 | 0.496 | 1 | 0.260736699 | 0.728680393 | 0.634 | 0.58 | 1 | 0.260736699 | 0.03209326 |
| Gm56686 | 0.016194202 | 1.228723176 | 0.264 | 0.136 | 1 | 0.336924056 | 0.614812428 | 0.244 | 0.188 | 1 | 0.336924056 | 0.032126152 |
| 1700096K18Rik | 0.016214441 | 2.261637799 | 0.132 | 0.047 | 1 | 0.324605225 | 1.100239256 | 0.122 | 0.078 | 1 | 0.324605225 | 0.032165974 |
| Slc25a1 | 0.041323376 | -0.759547531 | 0.151 | 0.297 | 1 | 0.016215281 | -2.093532488 | 0.024 | 0.167 | 1 | 0.041323376 | 0.032167626 |
| Gm42997 | 0.016222201 | 2.154722595 | 0.057 | 0.008 | 1 | 0.981794333 | 0.721727632 | 0.024 | 0.024 | 1 | 0.981794333 | 0.032181243 |
| Ccdc91 | 0.016244655 | 1.020866848 | 0.34 | 0.203 | 1 | 0.345885691 | 1.014082612 | 0.244 | 0.195 | 1 | 0.345885691 | 0.032225421 |
| Cacna1d | 0.016287934 | 2.123013735 | 0.245 | 0.127 | 1 | 0.066262841 | 0.817576043 | 0.463 | 0.345 | 1 | 0.066262841 | 0.032310571 |
| Spag1 | 0.016300887 | 1.356356456 | 0.226 | 0.11 | 1 | 0.661698768 | -0.132421501 | 0.146 | 0.177 | 1 | 0.661698768 | 0.032336054 |
| Ranbp1 | 0.016332963 | 0.29467025 | 0.906 | 0.839 | 1 | 0.224625561 | -0.205627066 | 0.829 | 0.911 | 1 | 0.224625561 | 0.032399161 |
| Six3 | 0.779443586 | 0.154722595 | 0.019 | 0.025 | 1 | 0.016339332 | 1.422167351 | 0.195 | 0.078 | 1 | 0.779443586 | 0.032411691 |
| Plscr1 | 0.016348489 | -1.445190247 | 0.321 | 0.47 | 1 | 0.169523729 | 0.603149511 | 0.463 | 0.372 | 1 | 0.169523729 | 0.032429705 |
| Gm17036 | 0.016369897 | 2.47665069 | 0.094 | 0.025 | 1 | 0.699151739 | 0.600165653 | 0.098 | 0.082 | 1 | 0.699151739 | 0.032471821 |
| Fosb | 0.142498592 | 0.458348718 | 0.906 | 0.945 | 1 | 0.016406547 | 0.58949205 | 0.976 | 0.976 | 1 | 0.142498592 | 0.032543919 |
| Heg1 | 0.01641845 | 0.91489558 | 0.358 | 0.212 | 1 | 0.239380574 | 1.370078839 | 0.195 | 0.126 | 1 | 0.239380574 | 0.032567334 |
| Stau2 | 0.023239227 | 0.735497299 | 0.434 | 0.271 | 1 | 0.016440385 | 1.144829407 | 0.561 | 0.375 | 1 | 0.023239227 | 0.032610483 |
| Igsf8 | 0.016440981 | -0.728763396 | 0.358 | 0.538 | 1 | 0.160618934 | -0.410722664 | 0.415 | 0.543 | 1 | 0.160618934 | 0.032611657 |
| Ppp2r3a | 0.016444275 | 0.472462893 | 0.774 | 0.61 | 1 | 0.174751693 | 0.544634935 | 0.805 | 0.737 | 1 | 0.174751693 | 0.032618136 |
| Ranbp17 | 0.016444836 | 0.999444369 | 0.226 | 0.102 | 1 | 0.068152725 | 1.149148856 | 0.293 | 0.191 | 1 | 0.068152725 | 0.03261924 |
| Pkn1 | 0.016461507 | 1.017219071 | 0.302 | 0.165 | 1 | 0.64156796 | -0.254293505 | 0.366 | 0.392 | 1 | 0.64156796 | 0.032652034 |
| Mrpl50 | 0.110245706 | 0.509639102 | 0.528 | 0.445 | 1 | 0.016471757 | -0.976199117 | 0.293 | 0.471 | 1 | 0.110245706 | 0.032672195 |
| Lmln | 0.016517622 | 1.43225657 | 0.17 | 0.068 | 1 | 0.035921968 | 1.111379813 | 0.195 | 0.089 | 1 | 0.035921968 | 0.032762412 |
| Eif2b3 | 0.016519129 | -1.06560736 | 0.358 | 0.492 | 1 | 0.078426023 | -0.560826224 | 0.268 | 0.44 | 1 | 0.078426023 | 0.032765376 |
| Cdip1 | 0.016550792 | 0.614747549 | 0.642 | 0.479 | 1 | 0.616089695 | 0.581947795 | 0.732 | 0.778 | 1 | 0.616089695 | 0.032827655 |
| Clasp2 | 0.016555961 | -0.657010768 | 0.83 | 0.869 | 1 | 0.260313037 | 0.118281046 | 0.878 | 0.874 | 1 | 0.260313037 | 0.032837823 |
| Gpr15lg | 0.016582238 | 1.347367673 | 0.057 | 0.008 | 1 | 0.056289003 | 3.159132945 | 0.049 | 0.01 | 1 | 0.056289003 | 0.032889505 |
| Wwc1 | 0.016584115 | 1.154722595 | 0.151 | 0.055 | 1 | 0.063555886 | 1.042024642 | 0.293 | 0.184 | 1 | 0.063555886 | 0.032893196 |
| Trmt10b | 0.016590296 | 1.714150003 | 0.17 | 0.068 | 1 | 0.62545961 | -0.278272368 | 0.049 | 0.068 | 1 | 0.62545961 | 0.032905353 |
| Gdpgp1 | 0.01660958 | 2.347367673 | 0.094 | 0.025 | 1 | 0.188517184 | -1.069685746 | 0 | 0.041 | 1 | 0.188517184 | 0.032943281 |
| Rnf144a | 0.016612776 | 0.635757653 | 0.566 | 0.436 | 1 | 0.08688351 | 1.271607674 | 0.512 | 0.44 | 1 | 0.08688351 | 0.032949567 |
| Rpp14 | 0.049436965 | 0.771393955 | 0.34 | 0.216 | 1 | 0.016623153 | -1.364429011 | 0.073 | 0.242 | 1 | 0.049436965 | 0.032969977 |
| Zmynd19 | 0.016634224 | 1.181689642 | 0.321 | 0.195 | 1 | 0.025575038 | -0.523542494 | 0.098 | 0.266 | 1 | 0.025575038 | 0.03299175 |
| Cdc37l1 | 0.01664316 | 0.425576505 | 0.792 | 0.691 | 1 | 0.287766197 | -0.227646295 | 0.756 | 0.86 | 1 | 0.287766197 | 0.033009326 |
| Mllt11 | 0.405215324 | -0.214511215 | 0.226 | 0.292 | 1 | 0.016666273 | -0.951291045 | 0.146 | 0.338 | 1 | 0.405215324 | 0.03305478 |
| Bdh1 | 0.016685156 | -1.237594828 | 0.057 | 0.199 | 1 | 0.363969161 | -0.747757651 | 0.073 | 0.119 | 1 | 0.363969161 | 0.033091918 |
| Sh3glb1 | 0.016695042 | 0.287082815 | 1 | 0.979 | 1 | 0.07137117 | -0.200461711 | 1 | 0.99 | 1 | 0.07137117 | 0.033111359 |
| Hsph1 | 0.016713351 | 0.863491297 | 0.811 | 0.72 | 1 | 0.036695168 | 0.408556919 | 0.927 | 0.802 | 1 | 0.036695168 | 0.033147367 |
| Med6 | 0.016731199 | 0.880547631 | 0.509 | 0.369 | 1 | 0.295078801 | 0.118386602 | 0.22 | 0.345 | 1 | 0.295078801 | 0.033182465 |
| Cdr2l | 0.016732885 | 0.615563784 | 0.415 | 0.25 | 1 | 0.223567611 | -1.037264268 | 0.049 | 0.109 | 1 | 0.223567611 | 0.03318578 |
| Zmym6 | 0.016739929 | 0.974150349 | 0.302 | 0.165 | 1 | 0.636789048 | 0.746056962 | 0.195 | 0.167 | 1 | 0.636789048 | 0.033199633 |
| Rps2 | 0.016782217 | 0.285171361 | 1 | 0.987 | 1 | 0.19659314 | -0.107026303 | 1 | 0.997 | 1 | 0.19659314 | 0.033282792 |
| Cldn4 | 0.016838766 | 1.154722595 | 0.151 | 0.055 | 1 | 0.314008923 | -0.174022406 | 0.268 | 0.198 | 1 | 0.314008923 | 0.033393989 |
| Plau | 0.016870117 | 2.533234218 | 0.094 | 0.025 | 1 | 0.82181816 | 0.136765132 | 0.024 | 0.031 | 1 | 0.82181816 | 0.033455633 |
| Wwox | 0.071243011 | -0.701154835 | 0.755 | 0.835 | 1 | 0.016870586 | -1.081658387 | 0.732 | 0.788 | 1 | 0.071243011 | 0.033456554 |
| Arid3b | 0.031574411 | 1.891688189 | 0.094 | 0.03 | 1 | 0.016888528 | 2.363273661 | 0.146 | 0.051 | 1 | 0.031574411 | 0.033491833 |
| Tet2 | 0.016901146 | 0.417757001 | 0.83 | 0.695 | 1 | 0.89198374 | -0.121953055 | 0.878 | 0.853 | 1 | 0.89198374 | 0.033516644 |
| Zfp950 | 0.34605384 | -0.294584807 | 0.396 | 0.479 | 1 | 0.016909758 | -0.430830237 | 0.244 | 0.498 | 1 | 0.34605384 | 0.033533575 |
| Anxa1 | 0.016921357 | -1.238274863 | 0.868 | 0.797 | 1 | 0.035914532 | 1.037131685 | 0.829 | 0.747 | 1 | 0.035914532 | 0.033556381 |
| Abcb10 | 0.016993021 | 1.448453798 | 0.226 | 0.11 | 1 | 0.425526384 | 0.551802631 | 0.244 | 0.198 | 1 | 0.425526384 | 0.03369728 |
| Zbtb20 | 0.264050352 | 0.133395798 | 1 | 1 | 1 | 0.017006124 | -0.272233205 | 1 | 1 | 1 | 0.264050352 | 0.03372304 |
| Asah1 | 0.332301773 | 0.393171362 | 0.585 | 0.572 | 1 | 0.017008395 | 0.591044263 | 0.78 | 0.683 | 1 | 0.332301773 | 0.033727504 |
| Mrpl30 | 0.017023098 | 0.639224914 | 0.679 | 0.602 | 1 | 0.979810214 | 0.220109728 | 0.488 | 0.529 | 1 | 0.979810214 | 0.033756409 |
| Blvra | 0.210912415 | 0.479157545 | 0.415 | 0.343 | 1 | 0.017040961 | -0.79700117 | 0.122 | 0.317 | 1 | 0.210912415 | 0.033791528 |
| Fez2 | 0.017048033 | 0.86841841 | 0.453 | 0.292 | 1 | 0.318966615 | 0.690363461 | 0.366 | 0.311 | 1 | 0.318966615 | 0.03380543 |
| Fkbp15 | 0.017057518 | 0.569760094 | 0.509 | 0.347 | 1 | 0.253059694 | -0.217703491 | 0.341 | 0.468 | 1 | 0.253059694 | 0.033824077 |
| Rps9 | 0.017102478 | 0.30467715 | 1 | 0.983 | 1 | 0.059861651 | -0.141982139 | 1 | 0.986 | 1 | 0.059861651 | 0.033912462 |
| Fdx2 | 0.017105995 | 0.860991392 | 0.415 | 0.271 | 1 | 0.120396894 | 0.430029468 | 0.439 | 0.311 | 1 | 0.120396894 | 0.033919374 |
| Mccc1 | 0.017152571 | 1.569760094 | 0.245 | 0.127 | 1 | 0.606445404 | 0.496167932 | 0.195 | 0.171 | 1 | 0.606445404 | 0.034010931 |
| Tmem115 | 0.086599674 | 0.665975409 | 0.396 | 0.288 | 1 | 0.017177906 | -0.652761137 | 0.244 | 0.454 | 1 | 0.086599674 | 0.034060732 |
| Klhl28 | 0.017235646 | 0.996024849 | 0.453 | 0.314 | 1 | 0.54867199 | 0.749742009 | 0.366 | 0.369 | 1 | 0.54867199 | 0.034174224 |
| Atp6v1h | 0.017267493 | 0.500950936 | 0.849 | 0.72 | 1 | 0.848747739 | 0.121272956 | 0.78 | 0.812 | 1 | 0.848747739 | 0.03423682 |
| Bend6 | 0.017301089 | -1.799473716 | 0.075 | 0.216 | 1 | 0.459505802 | -0.435813645 | 0.024 | 0.051 | 1 | 0.459505802 | 0.03430285 |
| Smc5 | 0.017305251 | 0.550184556 | 0.755 | 0.682 | 1 | 0.813609918 | 0.107192016 | 0.78 | 0.778 | 1 | 0.813609918 | 0.034311031 |
| Stimate | 0.017361488 | 1.002719501 | 0.377 | 0.233 | 1 | 0.289060513 | -0.16279515 | 0.195 | 0.283 | 1 | 0.289060513 | 0.034421556 |
| Gm13563 | 0.017364527 | 2.154722595 | 0.094 | 0.025 | 1 | 0.983241205 | 0.422167351 | 0.049 | 0.048 | 1 | 0.983241205 | 0.034427528 |
| Gfra4 | 0.017364527 | 2.154722595 | 0.094 | 0.025 | 1 | 0.916649754 | 0.667279848 | 0.024 | 0.027 | 1 | 0.916649754 | 0.034427528 |
| Il33 | 0.017397532 | -2.326404095 | 0.019 | 0.136 | 1 | 0.257319172 | -1.484723245 | 0 | 0.031 | 1 | 0.257319172 | 0.03449239 |
| Gm32816 | 0.017422596 | 4.154722595 | 0.075 | 0.017 | 1 | 0.021685149 | 1.100239256 | 0.049 | 0.007 | 1 | 0.021685149 | 0.034541646 |
| Ccdc58 | 0.758553677 | 0.470224421 | 0.302 | 0.297 | 1 | 0.017463636 | -1.228884341 | 0.098 | 0.276 | 1 | 0.758553677 | 0.034622293 |
| Gcc2 | 0.017480215 | 0.784079215 | 0.604 | 0.496 | 1 | 0.822940837 | 0.440314697 | 0.61 | 0.625 | 1 | 0.822940837 | 0.034654873 |

| | | | | | | | | | | | | |
|---|---|---|---|---|---|---|---|---|---|---|---|---|
| Pik3cd | 0.017504397 | 1.984797593 | 0.094 | 0.025 | 1 | 0.26625683 | 2.136765132 | 0.073 | 0.038 | 1 | 0.26625683 | 0.034702391 |
| Cdc25b | 0.017518999 | 1.948271717 | 0.132 | 0.047 | 1 | 0.758237333 | 0.292884334 | 0.073 | 0.089 | 1 | 0.758237333 | 0.034731082 |
| Timm8a1 | 0.04427671 | 0.812330397 | 0.321 | 0.203 | 1 | 0.017544582 | -0.711231775 | 0.098 | 0.28 | 1 | 0.04427671 | 0.034781351 |
| Slc5a3 | 0.017546095 | -0.569376824 | 0.585 | 0.729 | 1 | 0.021803459 | 0.601756276 | 0.756 | 0.628 | 1 | 0.021803459 | 0.034784325 |
| Cnot2 | 0.017568218 | 0.517921163 | 0.811 | 0.86 | 1 | 0.986041015 | 0.113414644 | 0.902 | 0.881 | 1 | 0.986041015 | 0.034827794 |
| Chd9 | 0.270145113 | 0.127801295 | 0.962 | 0.949 | 1 | 0.017597031 | -0.506691722 | 0.976 | 0.962 | 1 | 0.270145113 | 0.034884406 |
| D8Ertd738e | 0.017597057 | 0.301112082 | 0.925 | 0.856 | 1 | 0.022040074 | -0.557192883 | 0.732 | 0.86 | 1 | 0.022040074 | 0.034884457 |
| Fam167a | 0.017609519 | 1.339937467 | 0.396 | 0.267 | 1 | 0.578121111 | -0.16279515 | 0.122 | 0.16 | 1 | 0.578121111 | 0.034908944 |
| Nfe2l3 | 0.017635085 | 1.221836791 | 0.151 | 0.055 | 1 | 0.180559668 | 0.493250449 | 0.22 | 0.14 | 1 | 0.180559668 | 0.034959173 |
| Tlcd3a | 0.017638177 | 2.306725688 | 0.094 | 0.025 | 1 | 0.722625601 | 0.313642894 | 0.073 | 0.058 | 1 | 0.722625601 | 0.034965248 |
| Sppl2b | 0.017657681 | -0.970808287 | 0.151 | 0.318 | 1 | 0.503808841 | -0.296650897 | 0.171 | 0.215 | 1 | 0.503808841 | 0.035003568 |
| Htatip2 | 0.017666534 | -1.032904408 | 0.132 | 0.292 | 1 | 0.296618281 | -0.332720152 | 0.073 | 0.133 | 1 | 0.296618281 | 0.035020962 |
| C4b | 0.393881697 | -1.067669827 | 0.057 | 0.093 | 1 | 0.017669432 | -1.790068456 | 0.073 | 0.235 | 1 | 0.393881697 | 0.035026656 |
| Rpp38 | 0.017686101 | -1.631873767 | 0.075 | 0.212 | 1 | 0.050080034 | -1.278272368 | 0.024 | 0.133 | 1 | 0.050080034 | 0.035059403 |
| Guk1 | 0.108246557 | -0.448161814 | 0.472 | 0.581 | 1 | 0.017768742 | -0.911733386 | 0.317 | 0.488 | 1 | 0.108246557 | 0.035221755 |
| Mga | 0.017773689 | 0.453435976 | 0.792 | 0.665 | 1 | 0.163856947 | 0.24414185 | 0.878 | 0.826 | 1 | 0.163856947 | 0.035231473 |
| Hivep2 | 0.503324566 | -0.283827827 | 1 | 0.97 | 1 | 0.017776516 | 0.384048409 | 1 | 0.976 | 1 | 0.503324566 | 0.035237027 |
| Rp1 | 0.084268268 | -0.946815432 | 0.094 | 0.195 | 1 | 0.017832995 | -0.715336173 | 0.098 | 0.276 | 1 | 0.084268268 | 0.035347974 |
| Dync2i1 | 0.248343296 | 0.657222935 | 0.396 | 0.352 | 1 | 0.017839736 | -0.555112573 | 0.171 | 0.375 | 1 | 0.248343296 | 0.035361216 |
| Slc35g2 | 0.079376758 | -0.600164907 | 0.075 | 0.178 | 1 | 0.017893703 | -2.890175605 | 0 | 0.123 | 1 | 0.079376758 | 0.035467222 |
| Tmtc4 | 0.017899342 | 1.017219071 | 0.302 | 0.161 | 1 | 0.840342415 | 0.201616276 | 0.195 | 0.184 | 1 | 0.840342415 | 0.035478297 |
| Zfp639 | 0.017900839 | 1.195364579 | 0.396 | 0.25 | 1 | 0.397514069 | 0.298784935 | 0.341 | 0.287 | 1 | 0.397514069 | 0.035481238 |
| Tmtc3 | 0.017926577 | 1.098139066 | 0.302 | 0.165 | 1 | 0.185053007 | 0.99285572 | 0.415 | 0.334 | 1 | 0.185053007 | 0.035531792 |
| Tmem183a | 0.017968306 | 0.884633431 | 0.509 | 0.381 | 1 | 0.030422054 | -0.381629752 | 0.341 | 0.546 | 1 | 0.030422054 | 0.035613752 |
| Igbp1 | 0.017969205 | 0.556276865 | 0.679 | 0.564 | 1 | 0.10672622 | 0.536971826 | 0.732 | 0.652 | 1 | 0.10672622 | 0.035615518 |
| Tsix | 0.017993587 | -1.960754623 | 0.038 | 0.161 | 1 | 0.435967024 | 1.422167351 | 0.024 | 0.01 | 1 | 0.435967024 | 0.035663405 |
| Usp49 | 0.018004696 | 0.855162313 | 0.396 | 0.242 | 1 | 0.266048253 | 0.515276755 | 0.415 | 0.348 | 1 | 0.266048253 | 0.035685223 |
| Nfkbil1 | 0.018013047 | 1.444229212 | 0.245 | 0.127 | 1 | 0.789184182 | 0.438655473 | 0.171 | 0.16 | 1 | 0.789184182 | 0.035701624 |
| Flad1 | 0.018031417 | 1.932330173 | 0.132 | 0.047 | 1 | 0.02212569 | 1.883008539 | 0.195 | 0.085 | 1 | 0.02212569 | 0.035737702 |
| Slc5a11 | 0.018034129 | 2.640149422 | 0.075 | 0.017 | 1 | 0.02065851 | 3.574170444 | 0.049 | 0.007 | 1 | 0.02065851 | 0.035743028 |
| Evi2b | 0.018034129 | 2.640149422 | 0.075 | 0.017 | 1 | 0.82181816 | -0.250257991 | 0.024 | 0.031 | 1 | 0.82181816 | 0.035743028 |
| Ace | 0.018034129 | 2.8327945 | 0.075 | 0.017 | 1 | 0.371357099 | -0.484723245 | 0.024 | 0.058 | 1 | 0.371357099 | 0.035743028 |
| Rbmxl1 | 0.036828638 | 0.590224197 | 0.623 | 0.513 | 1 | 0.018046732 | -0.698070527 | 0.415 | 0.611 | 1 | 0.036828638 | 0.035767779 |
| Elovl7 | 0.018078522 | -2.206733864 | 0.132 | 0.267 | 1 | 0.044939522 | -3.147688258 | 0.049 | 0.164 | 1 | 0.044939522 | 0.035830211 |
| Cnst | 0.018094759 | 1.404700848 | 0.226 | 0.11 | 1 | 0.079553011 | 0.79281073 | 0.341 | 0.225 | 1 | 0.079553011 | 0.035862097 |
| Avl9 | 0.018103769 | 0.424329655 | 0.604 | 0.449 | 1 | 0.693922588 | 0.191869731 | 0.732 | 0.713 | 1 | 0.693922588 | 0.035879791 |
| Acap3 | 0.018134831 | 0.945269229 | 0.264 | 0.131 | 1 | 0.028329422 | 0.898605394 | 0.39 | 0.242 | 1 | 0.028329422 | 0.035940791 |
| Meaf6 | 0.157281593 | 0.739685096 | 0.415 | 0.347 | 1 | 0.018136398 | -0.738479837 | 0.171 | 0.392 | 1 | 0.157281593 | 0.035943866 |
| Spred3 | 0.018145157 | 1.962077517 | 0.151 | 0.059 | 1 | 0.196786742 | 1.071670103 | 0.146 | 0.085 | 1 | 0.196786742 | 0.035961068 |
| Kif5c | 0.018161263 | 1.569760094 | 0.094 | 0.025 | 1 | 0.182547274 | 1.082317348 | 0.122 | 0.065 | 1 | 0.182547274 | 0.035992695 |
| Snx31 | 0.018173835 | 1.792152515 | 0.094 | 0.025 | 1 | 0.749728988 | 0.667279848 | 0.024 | 0.017 | 1 | 0.749728988 | 0.036017381 |
| Matcap1 | 0.018176978 | 1.932330173 | 0.094 | 0.025 | 1 | 0.72988856 | 0.83720485 | 0.049 | 0.038 | 1 | 0.72988856 | 0.036023554 |
| Gm49380 | 0.018176978 | 1.932330173 | 0.094 | 0.025 | 1 | 0.732115547 | 0.252242349 | 0.073 | 0.058 | 1 | 0.732115547 | 0.036023554 |
| ENSMUSG00000095742 | 0.018176978 | 1.932330173 | 0.094 | 0.025 | 1 | 0.402373368 | 0.252242349 | 0 | 0.017 | 1 | 0.402373368 | 0.036023554 |
| Pdss2 | 0.018180538 | 0.940943304 | 0.453 | 0.314 | 1 | 0.122085816 | 0.632633706 | 0.512 | 0.389 | 1 | 0.122085816 | 0.036030544 |
| Ube2g1 | 0.018189998 | 0.60768955 | 0.868 | 0.758 | 1 | 0.301984325 | 0.3168859 | 0.927 | 0.922 | 1 | 0.301984325 | 0.03604912 |
| Pspc1 | 0.018190242 | 0.632391082 | 0.66 | 0.568 | 1 | 0.154797209 | 0.703842754 | 0.659 | 0.696 | 1 | 0.154797209 | 0.036049599 |
| Pemt | 0.439203725 | 0.491757582 | 0.094 | 0.064 | 1 | 0.018197174 | 1.91966701 | 0.122 | 0.038 | 1 | 0.439203725 | 0.036063211 |
| Oip5os1 | 0.018198761 | 0.678989164 | 0.755 | 0.716 | 1 | 0.962138612 | 0.111379813 | 0.756 | 0.887 | 1 | 0.962138612 | 0.036066327 |
| Tmem202 | 0.018201144 | -2.20282941 | 0.019 | 0.131 | 1 | 0.570963886 | -0.863234868 | 0.049 | 0.072 | 1 | 0.570963886 | 0.036071006 |
| 4930447C04Rik | 0.018228954 | 1.377115016 | 0.151 | 0.055 | 1 | 0.518178992 | -0.899760744 | 0.049 | 0.075 | 1 | 0.518178992 | 0.036125613 |
| Gm34189 | 0.018239789 | -3.337130502 | 0 | 0.097 | 1 | 0.20883094 | -1.747757651 | 0 | 0.038 | 1 | 0.20883094 | 0.036146888 |
| Rlbp1 | 0.018268947 | -2.27154216 | 0.019 | 0.131 | 1 | 0.077457679 | -1.80665134 | 0 | 0.072 | 1 | 0.077457679 | 0.03620414 |
| Sav1 | 0.026373291 | -0.516600191 | 0.698 | 0.86 | 1 | 0.018303124 | -0.481513689 | 0.78 | 0.887 | 1 | 0.026373291 | 0.036271244 |
| Ctps | 0.018328312 | 1.074552246 | 0.358 | 0.212 | 1 | 0.202280348 | 0.527349587 | 0.317 | 0.239 | 1 | 0.202280348 | 0.036320697 |
| Ephb2 | 0.018337237 | 0.599024689 | 0.491 | 0.339 | 1 | 0.69401199 | -0.467649732 | 0.732 | 0.662 | 1 | 0.69401199 | 0.03633822 |
| Cd4 | 0.018351921 | 2.891688189 | 0.075 | 0.017 | 1 | 0.441381862 | 1.252242349 | 0.024 | 0.01 | 1 | 0.441381862 | 0.036367048 |
| Ppp3r1 | 0.018372861 | 0.454282877 | 0.736 | 0.61 | 1 | 0.227789115 | -0.295689421 | 0.756 | 0.765 | 1 | 0.227789115 | 0.03640816 |
| Foxo4 | 0.018375864 | 1.347367673 | 0.189 | 0.081 | 1 | 0.434461613 | 0.404245443 | 0.195 | 0.147 | 1 | 0.434461613 | 0.036414056 |
| Fmo2 | 0.413080549 | -0.1672055 | 0 | 0.013 | 1 | 0.018378931 | 1.30115195 | 0.22 | 0.096 | 1 | 0.413080549 | 0.036420076 |
| Glod4 | 0.018390078 | 0.708039469 | 0.585 | 0.436 | 1 | 0.16919949 | 0.389059039 | 0.561 | 0.451 | 1 | 0.16919949 | 0.03644196 |
| Ano8 | 0.018436493 | 2.306725688 | 0.094 | 0.025 | 1 | 0.268288619 | 1.252242349 | 0.098 | 0.055 | 1 | 0.268288619 | 0.036533082 |
| Ulk4 | 0.018463238 | -1.455774998 | 0.151 | 0.309 | 1 | 0.70271774 | -0.278272368 | 0.244 | 0.208 | 1 | 0.70271774 | 0.036585584 |
| Ppp2r5d | 0.018493294 | 0.848061257 | 0.302 | 0.161 | 1 | 0.46812586 | 0.494812652 | 0.244 | 0.198 | 1 | 0.46812586 | 0.036644586 |
| Borcs7 | 0.099832837 | 0.483786871 | 0.453 | 0.335 | 1 | 0.018495836 | -1.292078167 | 0.146 | 0.321 | 1 | 0.099832837 | 0.036649575 |
| Atxn7 | 0.018544555 | 0.556276865 | 0.83 | 0.695 | 1 | 0.037989416 | 0.260333761 | 0.902 | 0.853 | 1 | 0.037989416 | 0.03674521 |
| Arhgef17 | 0.018545148 | 0.916858765 | 0.453 | 0.318 | 1 | 0.141666996 | 0.313642894 | 0.512 | 0.403 | 1 | 0.141666996 | 0.036746374 |
| Chmp1a | 0.01859756 | 0.839849257 | 0.472 | 0.331 | 1 | 0.728844457 | -0.105309656 | 0.293 | 0.321 | 1 | 0.728844457 | 0.036849251 |
| Grwd1 | 0.01860041 | 1.389187848 | 0.245 | 0.127 | 1 | 0.292342467 | -0.332720152 | 0.098 | 0.164 | 1 | 0.292342467 | 0.036854845 |
| Mrpl12 | 0.018630165 | 0.783884899 | 0.566 | 0.453 | 1 | 0.127726139 | -0.555112573 | 0.366 | 0.481 | 1 | 0.127726139 | 0.036913246 |
| Eid1 | 0.018634759 | 0.391205476 | 1 | 0.945 | 1 | 0.022569559 | 0.469964339 | 0.927 | 0.939 | 1 | 0.022569559 | 0.036922264 |
| Gm16351 | 0.124227304 | -2.783876861 | 0.094 | 0.174 | 1 | 0.018635658 | -1.751759581 | 0.122 | 0.3 | 1 | 0.124227304 | 0.036924029 |
| Prr14 | 0.018638907 | 0.569760094 | 0.547 | 0.373 | 1 | 0.462663349 | 0.429120111 | 0.463 | 0.447 | 1 | 0.462663349 | 0.036930406 |
| Gm38115 | 0.6159787 | -0.237594828 | 0.057 | 0.076 | 1 | 0.018656393 | 1.252242349 | 0.22 | 0.096 | 1 | 0.6159787 | 0.036964725 |
| Ifi204 | 0.018663137 | 2.417757001 | 0.075 | 0.017 | 1 | 0.027541029 | 2.614812428 | 0.073 | 0.017 | 1 | 0.027541029 | 0.03697796 |
| Fadd | 0.018663137 | 2.417757001 | 0.075 | 0.017 | 1 | 0.872828964 | 1.029849928 | 0.024 | 0.02 | 1 | 0.872828964 | 0.03697796 |
| Gm28229 | 0.018663137 | 2.417757001 | 0.075 | 0.017 | 1 | 0.196649091 | 0.911205431 | 0.098 | 0.048 | 1 | 0.196649091 | 0.03697796 |
| Abca7 | 0.018663137 | 2.417757001 | 0.075 | 0.017 | 1 | 0.420849275 | 1.029849928 | 0.073 | 0.044 | 1 | 0.420849275 | 0.03697796 |
| Dph1 | 0.018663137 | 2.417757001 | 0.075 | 0.017 | 1 | 0.407867291 | -0.484723245 | 0.024 | 0.055 | 1 | 0.407867291 | 0.03697796 |
| Phf11c | 0.018663137 | 2.417757001 | 0.075 | 0.017 | 1 | 0.708632143 | 1.252242349 | 0.049 | 0.038 | 1 | 0.708632143 | 0.03697796 |
| Sema7a | 0.237142171 | 0.347367673 | 0.094 | 0.051 | 1 | 0.018667859 | 1.029849928 | 0.146 | 0.051 | 1 | 0.237142171 | 0.036987229 |
| Igtp | 0.018671318 | 1.614154213 | 0.226 | 0.114 | 1 | 0.081965712 | 2.117312769 | 0.146 | 0.072 | 1 | 0.081965712 | 0.036994018 |
| Slc25a47 | 0.096435945 | 0.8327945 | 0.17 | 0.093 | 1 | 0.018712584 | 1.118386602 | 0.293 | 0.154 | 1 | 0.096435945 | 0.037075007 |
| Ston1 | 0.018713615 | 0.935044558 | 0.566 | 0.432 | 1 | 0.55951015 | 0.321189703 | 0.512 | 0.474 | 1 | 0.55951015 | 0.037077031 |
| Washc4 | 0.018753585 | 0.691155202 | 0.642 | 0.551 | 1 | 0.320666687 | -0.128267467 | 0.585 | 0.741 | 1 | 0.320666687 | 0.037155473 |
| mt-Co1 | 0.018758592 | -0.263981827 | 1 | 1 | 1 | 0.919527305 | -0.280839303 | 1 | 1 | 1 | 0.919527305 | 0.037165299 |
| Cox7b | 0.120748564 | 0.315306464 | 0.906 | 0.86 | 1 | 0.018761915 | -0.440388446 | 0.756 | 0.877 | 1 | 0.120748564 | 0.03717182 |
| Ccdc125 | 0.018765328 | 1.94321849 | 0.151 | 0.059 | 1 | 0.229817256 | 1.174239837 | 0.146 | 0.089 | 1 | 0.229817256 | 0.037178518 |
| Rbm33 | 0.038518863 | 0.307382803 | 0.849 | 0.758 | 1 | 0.018766217 | -0.360332383 | 0.756 | 0.877 | 1 | 0.038518863 | 0.037180263 |
| Ccdc32 | 0.018790106 | 0.977844833 | 0.321 | 0.182 | 1 | 0.773149499 | 0.949679579 | 0.195 | 0.195 | 1 | 0.773149499 | 0.037227143 |
| Med13l | 0.124830353 | 0.168176485 | 1 | 0.958 | 1 | 0.018804579 | -0.466626485 | 1 | 0.997 | 1 | 0.124830353 | 0.037255545 |
| E2f1 | 0.018812445 | 1.948271717 | 0.151 | 0.059 | 1 | 0.523075433 | 0.404245443 | 0.098 | 0.068 | 1 | 0.523075433 | 0.037270982 |
| Zfp990 | 0.018812515 | 2.055186921 | 0.151 | 0.059 | 1 | 0.714971076 | -0.221688839 | 0.098 | 0.116 | 1 | 0.714971076 | 0.03727112 |
| Nid2 | 0.018864289 | -0.66043773 | 0.547 | 0.712 | 1 | 0.053685021 | -0.953432677 | 0.39 | 0.532 | 1 | 0.053685021 | 0.037372717 |
| Sar1a | 0.440334137 | 0.109030504 | 0.887 | 0.894 | 1 | 0.018880572 | -0.41966317 | 0.878 | 0.945 | 1 | 0.440334137 | 0.037404667 |
| Csnk2a1 | 0.018882616 | 0.593663281 | 0.811 | 0.801 | 1 | 0.957295097 | 0.119559866 | 0.829 | 0.891 | 1 | 0.957295097 | 0.037408678 |
| Trnt1 | 0.034063822 | 0.753843159 | 0.547 | 0.445 | 1 | 0.018883476 | -0.548853583 | 0.244 | 0.468 | 1 | 0.034063822 | 0.037410366 |
| Hs6st2 | 0.231357061 | 1.569760094 | 0.057 | 0.025 | 1 | 0.018909686 | 1.541748966 | 0.122 | 0.038 | 1 | 0.231357061 | 0.037461796 |

| | | | | | | | | | | | | |
|---|---|---|---|---|---|---|---|---|---|---|---|---|
| Cyp39a1 | 0.018911859 | 1.358255989 | 0.189 | 0.081 | 1 | 0.271927668 | 0.880273572 | 0.22 | 0.154 | 1 | 0.271927668 | 0.037466059 |
| Zfp511 | 0.018948684 | 0.999444369 | 0.321 | 0.182 | 1 | 0.971560321 | 0.422167351 | 0.22 | 0.225 | 1 | 0.971560321 | 0.037538314 |
| Sec13 | 0.637235222 | 0.134112315 | 0.83 | 0.809 | 1 | 0.018951789 | -0.37471424 | 0.659 | 0.805 | 1 | 0.637235222 | 0.037544408 |
| Nol12 | 0.018954694 | 0.845394537 | 0.491 | 0.369 | 1 | 0.377276642 | -0.262330824 | 0.341 | 0.42 | 1 | 0.377276642 | 0.037550107 |
| Dennd1a | 0.018963684 | 0.268476309 | 0.943 | 0.898 | 1 | 0.742362635 | -0.148855959 | 0.927 | 0.942 | 1 | 0.742362635 | 0.037567747 |
| Slc8a1 | 0.018993345 | 2.002719501 | 0.075 | 0.017 | 1 | 0.040278286 | 1.881598969 | 0.244 | 0.13 | 1 | 0.040278286 | 0.037625943 |
| Ddit3 | 0.019011041 | 0.713017145 | 0.509 | 0.386 | 1 | 0.464561208 | -0.40310948 | 0.317 | 0.379 | 1 | 0.464561208 | 0.037660662 |
| 9330188P03Rik | 0.268348339 | -1.746144213 | 0.189 | 0.242 | 1 | 0.019012466 | -1.921787051 | 0.122 | 0.29 | 1 | 0.268348339 | 0.037663458 |
| Strada | 0.019038639 | 0.968856049 | 0.358 | 0.208 | 1 | 0.262516033 | 0.764448507 | 0.293 | 0.218 | 1 | 0.262516033 | 0.037714808 |
| Lysmd3 | 0.019040359 | 0.445064347 | 0.698 | 0.585 | 1 | 0.200780098 | -0.310592493 | 0.634 | 0.717 | 1 | 0.200780098 | 0.037718182 |
| Mtif3 | 0.019051549 | 1.154722595 | 0.245 | 0.123 | 1 | 0.334093265 | 1.136765132 | 0.195 | 0.143 | 1 | 0.334093265 | 0.037740137 |
| Cbr2 | 0.019054191 | -1.354832503 | 0.396 | 0.551 | 1 | 0.187285291 | 0.954958381 | 0.537 | 0.672 | 1 | 0.187285291 | 0.037745319 |
| Ints7 | 0.01905936 | 0.666784548 | 0.453 | 0.28 | 1 | 0.739840429 | 0.487620412 | 0.317 | 0.317 | 1 | 0.739840429 | 0.037755461 |
| Timm9 | 0.848918404 | -0.158937884 | 0.377 | 0.398 | 1 | 0.019079809 | -1.120566915 | 0.195 | 0.382 | 1 | 0.848918404 | 0.037795579 |
| Ppid | 0.047701096 | 0.617969187 | 0.585 | 0.504 | 1 | 0.019083014 | -0.581925404 | 0.439 | 0.642 | 1 | 0.047701096 | 0.037801867 |
| Tcirg1 | 0.019133535 | 1.404700848 | 0.264 | 0.14 | 1 | 0.192816587 | 0.53139642 | 0.244 | 0.167 | 1 | 0.192816587 | 0.037900978 |
| Mau2 | 0.021405758 | 0.697677569 | 0.547 | 0.398 | 1 | 0.019184805 | 0.667279848 | 0.805 | 0.689 | 1 | 0.021405758 | 0.038001554 |
| Mrpl14 | 0.019220005 | 0.644861549 | 0.547 | 0.394 | 1 | 0.42666866 | 0.903438548 | 0.415 | 0.399 | 1 | 0.42666866 | 0.038070601 |
| Nacc2 | 0.771095397 | 0.417757001 | 0.377 | 0.377 | 1 | 0.019261403 | -0.394647901 | 0.341 | 0.584 | 1 | 0.771095397 | 0.038151805 |
| Hdac3 | 0.019312413 | 0.692095637 | 0.491 | 0.339 | 1 | 0.138812209 | -0.516432105 | 0.293 | 0.413 | 1 | 0.138812209 | 0.038251856 |
| Pdzk1 | 0.019319117 | 2.154722595 | 0.075 | 0.017 | 1 | 0.274788535 | 1.100239256 | 0.049 | 0.02 | 1 | 0.274788535 | 0.038265005 |
| Tbx15 | 0.019319117 | 2.154722595 | 0.075 | 0.017 | 1 | 0.51440413 | 0.136765132 | 0.098 | 0.068 | 1 | 0.51440413 | 0.038265005 |
| Aldh5a1 | 0.019319117 | 2.154722595 | 0.075 | 0.017 | 1 | 0.826384776 | 0.515276755 | 0.024 | 0.031 | 1 | 0.826384776 | 0.038265005 |
| Sema6c | 0.019325634 | 2.154722595 | 0.075 | 0.017 | 1 | 0.989595616 | 0.667279848 | 0.024 | 0.024 | 1 | 0.989595616 | 0.038277788 |
| Lmtk3 | 0.019325634 | 2.154722595 | 0.075 | 0.017 | 1 | 0.872828964 | 1.029849928 | 0.024 | 0.02 | 1 | 0.872828964 | 0.038277788 |
| Trim21 | 0.019325634 | 2.154722595 | 0.075 | 0.017 | 1 | 0.230762368 | 1.252242349 | 0.073 | 0.034 | 1 | 0.230762368 | 0.038277788 |
| 4930533K18Rik | 0.019325634 | 2.154722595 | 0.075 | 0.017 | 1 | 0.15816766 | 2.029849928 | 0.073 | 0.031 | 1 | 0.15816766 | 0.038277788 |
| Spred1 | 0.292904825 | 0.123340469 | 0.868 | 0.881 | 1 | 0.019353549 | -0.438780628 | 0.829 | 0.911 | 1 | 0.292904825 | 0.038332537 |
| Afap1l2 | 0.271002806 | 0.252019796 | 0.245 | 0.178 | 1 | 0.01939665 | 1.707921833 | 0.293 | 0.16 | 1 | 0.271002806 | 0.03841707 |
| Plcxd3 | 0.019491232 | 1.855162313 | 0.075 | 0.017 | 1 | 0.181471053 | 3.029849928 | 0.049 | 0.017 | 1 | 0.181471053 | 0.038602555 |
| Spred2 | 0.019495981 | 0.526691372 | 0.604 | 0.458 | 1 | 0.182096695 | 0.453104332 | 0.756 | 0.768 | 1 | 0.182096695 | 0.038611869 |
| Pcbp4 | 0.025316946 | 0.998603393 | 0.377 | 0.254 | 1 | 0.019547111 | 0.791690191 | 0.512 | 0.365 | 1 | 0.025316946 | 0.038712132 |
| D130043K22Rik | 0.019564485 | 1.154722595 | 0.094 | 0.025 | 1 | 0.674175876 | -0.577832649 | 0.049 | 0.034 | 1 | 0.674175876 | 0.0387462 |
| Csf1r | 0.01965543 | 2.154722595 | 0.075 | 0.017 | 1 | 0.552285888 | -1.508569987 | 0.024 | 0.044 | 1 | 0.552285888 | 0.038924525 |
| Slc8b1 | 0.019657715 | 0.711115943 | 0.321 | 0.174 | 1 | 0.267603643 | -0.662641037 | 0.268 | 0.341 | 1 | 0.267603643 | 0.038929005 |
| Mrtfb | 0.019689109 | 0.607727944 | 0.585 | 0.441 | 1 | 0.093822 | -0.654648247 | 0.61 | 0.706 | 1 | 0.093822 | 0.038990558 |
| Dclre1c | 0.88530492 | 0.203632195 | 0.132 | 0.127 | 1 | 0.019691624 | 1.952682067 | 0.171 | 0.068 | 1 | 0.88530492 | 0.038995487 |
| Txndc16 | 0.019717613 | 1.090592257 | 0.321 | 0.195 | 1 | 0.137361953 | 0.977382508 | 0.317 | 0.222 | 1 | 0.137361953 | 0.039046441 |
| Fbxw5 | 0.019777965 | 1.273367091 | 0.264 | 0.14 | 1 | 0.98425391 | 0.322631677 | 0.122 | 0.123 | 1 | 0.98425391 | 0.039164762 |
| Gm43696 | 0.643144761 | 0.776210972 | 0.094 | 0.076 | 1 | 0.019778898 | -2.890175605 | 0 | 0.119 | 1 | 0.643144761 | 0.039166592 |
| Adat1 | 0.350848329 | -0.373656378 | 0.17 | 0.233 | 1 | 0.019815817 | 1.594348326 | 0.293 | 0.16 | 1 | 0.350848329 | 0.039238968 |
| Rasgrp2 | 0.155354531 | 1.669295768 | 0.057 | 0.021 | 1 | 0.019874433 | 1.635570989 | 0.122 | 0.038 | 1 | 0.155354531 | 0.039353874 |
| Ccnt1 | 0.019886448 | 0.524317123 | 0.811 | 0.763 | 1 | 0.947336889 | 0.148438344 | 0.854 | 0.874 | 1 | 0.947336889 | 0.039377424 |
| Zfp932 | 0.019908678 | 1.47665069 | 0.226 | 0.114 | 1 | 0.726483543 | 0.120997816 | 0.146 | 0.123 | 1 | 0.726483543 | 0.039421 |
| Lca5 | 0.019919809 | 1.399835093 | 0.189 | 0.085 | 1 | 0.066731732 | 1.384692645 | 0.22 | 0.123 | 1 | 0.066731732 | 0.03944282 |
| 2210016F16Rik | 0.019923683 | 0.973115788 | 0.434 | 0.297 | 1 | 0.794759374 | 0.161094461 | 0.317 | 0.348 | 1 | 0.794759374 | 0.039450414 |
| Lmna | 0.019966565 | 0.493695907 | 1 | 0.987 | 1 | 0.975884388 | 0.105759594 | 1 | 0.997 | 1 | 0.975884388 | 0.039534466 |
| Dph2 | 0.0199871 | 1.891688189 | 0.075 | 0.017 | 1 | 0.741726892 | 0.136765132 | 0.024 | 0.034 | 1 | 0.741726892 | 0.039574715 |
| Rab32 | 0.0199871 | 1.891688189 | 0.075 | 0.017 | 1 | 0.023803178 | 2.199774929 | 0.098 | 0.027 | 1 | 0.023803178 | 0.039574715 |
| Gm32036 | 0.0199871 | 1.891688189 | 0.075 | 0.017 | 1 | 0.267205177 | 2.252242349 | 0.024 | 0.007 | 1 | 0.267205177 | 0.039574715 |
| Gm33732 | 0.0199871 | 1.891688189 | 0.075 | 0.017 | 1 | 0.021167286 | 2.83720485 | 0.049 | 0.007 | 1 | 0.021167286 | 0.039574715 |
| Ccdc148 | 0.019997212 | 1.891688189 | 0.075 | 0.017 | 1 | 0.756560731 | -0.435813645 | 0.024 | 0.034 | 1 | 0.756560731 | 0.039594535 |
| Gm47200 | 0.019997402 | 0.8817041 | 0.075 | 0.017 | 1 | 0.267205177 | 2.252242349 | 0.024 | 0.007 | 1 | 0.267205177 | 0.039594909 |
| Txnrd2 | 0.137538287 | -0.652632327 | 0.057 | 0.131 | 1 | 0.020022115 | -1.970150072 | 0.024 | 0.16 | 1 | 0.137538287 | 0.039643344 |
| Sra1 | 0.020033796 | -0.393340323 | 0.755 | 0.826 | 1 | 0.242930033 | -0.179753977 | 0.683 | 0.799 | 1 | 0.242930033 | 0.03966624 |
| Rabl6 | 0.02004669 | 0.532005172 | 0.679 | 0.61 | 1 | 0.380718281 | -0.225235725 | 0.659 | 0.754 | 1 | 0.380718281 | 0.03969151 |
| Zim1 | 0.026944732 | 0.488459992 | 0.415 | 0.258 | 1 | 0.020073465 | -1.083360683 | 0.171 | 0.358 | 1 | 0.026944732 | 0.039743986 |
| Ap3d1 | 0.020098721 | 0.411695248 | 0.811 | 0.746 | 1 | 0.856116475 | 0.105923961 | 0.78 | 0.823 | 1 | 0.856116475 | 0.039793483 |
| G0s2 | 0.020156906 | -0.21991255 | 0.226 | 0.106 | 1 | 0.331197649 | -0.818146979 | 0.049 | 0.096 | 1 | 0.331197649 | 0.039907511 |
| Gm56752 | 0.030360955 | 2.739685096 | 0.038 | 0.004 | 1 | 0.020162119 | 3.83720485 | 0.049 | 0.007 | 1 | 0.030360955 | 0.039917727 |
| D430001F17Rik | 0.204653189 | 2.739685096 | 0.038 | 0.013 | 1 | 0.020162119 | 3.322631677 | 0.049 | 0.007 | 1 | 0.204653189 | 0.039917727 |
| Gm50373 | 0.494603095 | -0.1672055 | 0.038 | 0.021 | 1 | 0.020162119 | 4.059597271 | 0.049 | 0.007 | 1 | 0.494603095 | 0.039917727 |
| Gabrb1 | 0.03562064 | 3.154722595 | 0.019 | 0 | 1 | 0.020162402 | 3.685201756 | 0.049 | 0.007 | 1 | 0.03562064 | 0.039918281 |
| Gm12655 | 0.633487796 | 0.569760094 | 0.038 | 0.025 | 1 | 0.020162402 | 4.007129851 | 0.049 | 0.007 | 1 | 0.633487796 | 0.039918281 |
| Smn1 | 0.02019318 | 0.826396729 | 0.472 | 0.339 | 1 | 0.071286376 | -0.362934765 | 0.244 | 0.423 | 1 | 0.071286376 | 0.039978595 |
| Plpbp | 0.876663998 | 0.111395163 | 0.453 | 0.492 | 1 | 0.02022085 | -0.879786045 | 0.366 | 0.526 | 1 | 0.876663998 | 0.040032818 |
| Maea | 0.02022264 | 0.837107493 | 0.604 | 0.475 | 1 | 0.076334858 | -0.55106574 | 0.585 | 0.662 | 1 | 0.076334858 | 0.040036325 |
| Prkch | 0.020228967 | -1.635354336 | 0.113 | 0.258 | 1 | 0.132564963 | 0.867578499 | 0.244 | 0.15 | 1 | 0.132564963 | 0.040048723 |
| Polr2m | 0.020244917 | 0.406261362 | 0.811 | 0.686 | 1 | 0.400168074 | 0.153178222 | 0.805 | 0.713 | 1 | 0.400168074 | 0.040079978 |
| Katnb1 | 0.020244931 | 1.707263618 | 0.151 | 0.059 | 1 | 0.24180863 | 0.911205431 | 0.098 | 0.051 | 1 | 0.24180863 | 0.040080005 |
| Josd1 | 0.020245727 | 0.592480171 | 0.66 | 0.53 | 1 | 0.652958289 | 0.117082766 | 0.561 | 0.618 | 1 | 0.652958289 | 0.040081565 |
| Nat10 | 0.020265341 | 0.920257341 | 0.358 | 0.22 | 1 | 0.816539257 | 0.538863575 | 0.341 | 0.355 | 1 | 0.816539257 | 0.040119997 |
| Trnp1 | 0.020268475 | 1.484871196 | 0.208 | 0.097 | 1 | 0.412335112 | 0.568716014 | 0.073 | 0.126 | 1 | 0.412335112 | 0.040126138 |
| Kyat1 | 0.020282061 | 1.767699472 | 0.151 | 0.059 | 1 | 0.400227225 | 1.071670103 | 0.122 | 0.085 | 1 | 0.400227225 | 0.04015276 |
| Mtmr14 | 0.020308434 | 1.502645898 | 0.189 | 0.085 | 1 | 0.964436823 | 0.807457506 | 0.122 | 0.13 | 1 | 0.964436823 | 0.040204436 |
| Gle1 | 0.02032825 | 0.636255506 | 0.472 | 0.301 | 1 | 0.577994209 | 0.457062141 | 0.366 | 0.362 | 1 | 0.577994209 | 0.040243263 |
| A130010J15Rik | 0.767318506 | 0.454282877 | 0.057 | 0.047 | 1 | 0.020333392 | 1.47463477 | 0.146 | 0.051 | 1 | 0.767318506 | 0.040253337 |
| Lgalsl | 0.02035235 | -1.053285908 | 0.453 | 0.606 | 1 | 0.085110047 | -0.591032147 | 0.39 | 0.532 | 1 | 0.085110047 | 0.040290482 |
| Mtfr1l | 0.68454332 | 0.11408061 | 0.509 | 0.47 | 1 | 0.020360974 | -0.917682652 | 0.22 | 0.403 | 1 | 0.68454332 | 0.040307378 |
| Tmem212 | 0.020415157 | -2.727920455 | 0.019 | 0.127 | 1 | 0.141940321 | -1.448197369 | 0.073 | 0.157 | 1 | 0.141940321 | 0.040413536 |
| Gm14305 | 0.020460435 | 1.186431455 | 0.283 | 0.157 | 1 | 0.548256432 | 0.252242349 | 0.098 | 0.137 | 1 | 0.548256432 | 0.04050224 |
| Bcor | 0.020494006 | 1.185528642 | 0.453 | 0.343 | 1 | 0.045019734 | 0.421081393 | 0.707 | 0.567 | 1 | 0.045019734 | 0.040568007 |
| Asap3 | 0.139463914 | 0.464407094 | 0.472 | 0.36 | 1 | 0.020516325 | 1.521177356 | 0.341 | 0.205 | 1 | 0.139463914 | 0.040611731 |
| Fdx1 | 0.020551679 | 0.428287668 | 0.679 | 0.475 | 1 | 0.552577717 | 0.265917286 | 0.634 | 0.611 | 1 | 0.552577717 | 0.040680987 |
| Srfbp1 | 0.03004793 | 0.845969889 | 0.453 | 0.326 | 1 | 0.020555407 | -0.80665134 | 0.195 | 0.389 | 1 | 0.03004793 | 0.040688289 |
| Ubash3b | 0.020556037 | -2.226099189 | 0.075 | 0.208 | 1 | 0.057981149 | 0.46797104 | 0.195 | 0.096 | 1 | 0.057981149 | 0.040689524 |
| Ldb2 | 0.020604797 | 1.02113944 | 0.377 | 0.233 | 1 | 0.168579746 | 1.752315952 | 0.22 | 0.16 | 1 | 0.168579746 | 0.040785036 |
| Tm9sf1 | 0.020648226 | 0.614154213 | 0.698 | 0.61 | 1 | 0.94037742 | 0.170942247 | 0.683 | 0.734 | 1 | 0.94037742 | 0.040870102 |
| Wfdc1 | 0.332375765 | 0.510866405 | 0.113 | 0.072 | 1 | 0.020650705 | 1.537644568 | 0.293 | 0.164 | 1 | 0.332375765 | 0.040874958 |
| D430040L24Rik | 0.030360955 | 2.739685096 | 0.038 | 0.004 | 1 | 0.02065851 | 3.574170444 | 0.049 | 0.007 | 1 | 0.030360955 | 0.040890245 |
| Sv2c | 0.081457379 | -3.752168001 | 0 | 0.055 | 1 | 0.02065851 | 4.574170444 | 0.049 | 0.007 | 1 | 0.081457379 | 0.040890245 |
| Uckl1os | 0.246443787 | 2.739685096 | 0.019 | 0.004 | 1 | 0.02065851 | 3.83720485 | 0.049 | 0.007 | 1 | 0.246443787 | 0.040890245 |
| Flacc1 | 0.413082433 | -0.652632327 | 0 | 0.013 | 1 | 0.02065851 | 4.252242349 | 0.049 | 0.007 | 1 | 0.413082433 | 0.040890245 |
| E2f7 | 0.413082433 | -2.093204919 | 0 | 0.013 | 1 | 0.02065851 | 4.059597271 | 0.049 | 0.007 | 1 | 0.413082433 | 0.040890245 |
| Hsd17b13 | 0.932611893 | -0.1672055 | 0.019 | 0.017 | 1 | 0.02065851 | 3.252242349 | 0.049 | 0.007 | 1 | 0.932611893 | 0.040890245 |
| Slc22a17 | 0.05502228 | -0.781147068 | 0.321 | 0.453 | 1 | 0.02067937 | -1.016944284 | 0.244 | 0.423 | 1 | 0.05502228 | 0.040931105 |
| Zfp868 | 0.043547116 | 0.739685096 | 0.283 | 0.161 | 1 | 0.020681475 | -1.84461919 | 0.049 | 0.195 | 1 | 0.043547116 | 0.040935227 |
| Atcayos | 0.059160232 | 1.280253477 | 0.094 | 0.034 | 1 | 0.02071804 | 1.351778023 | 0.146 | 0.051 | 1 | 0.059160232 | 0.041006842 |

| | | | | | | | | | | | | |
|---|---|---|---|---|---|---|---|---|---|---|---|---|
| Rasl11a | 0.20776048 | -0.562690202 | 0.226 | 0.153 | 1 | 0.020728515 | 1.977382508 | 0.171 | 0.068 | 1 | 0.20776048 | 0.041027359 |
| Pml | 0.020737824 | 1.974150349 | 0.132 | 0.047 | 1 | 0.292340691 | 0.279209397 | 0.195 | 0.13 | 1 | 0.292340691 | 0.041045591 |
| Tmed4 | 0.020740099 | 0.596727142 | 0.604 | 0.475 | 1 | 0.695336936 | 0.194756854 | 0.537 | 0.539 | 1 | 0.695336936 | 0.041050046 |
| Trpm3 | 0.020740455 | 3.111653873 | 0.208 | 0.102 | 1 | 0.282073478 | -0.671344535 | 0.171 | 0.249 | 1 | 0.282073478 | 0.041050743 |
| Gdi2 | 0.020751183 | 0.291604401 | 0.943 | 0.932 | 1 | 0.280791641 | -0.143406165 | 0.976 | 0.973 | 1 | 0.280791641 | 0.041071755 |
| Vamp1 | 0.239875555 | 1.084333267 | 0.113 | 0.068 | 1 | 0.0207577 | 1.120997816 | 0.122 | 0.038 | 1 | 0.239875555 | 0.041084518 |
| 4930517O19Rik | 0.384325535 | -0.557995453 | 0.113 | 0.161 | 1 | 0.020794616 | -1.513292397 | 0.073 | 0.232 | 1 | 0.384325535 | 0.041156815 |
| Plxnc1 | 0.099079675 | 2.891688189 | 0.038 | 0.008 | 1 | 0.020826671 | 2.952682067 | 0.098 | 0.027 | 1 | 0.099079675 | 0.041219592 |
| Zfp143 | 0.020826801 | 1.050385935 | 0.245 | 0.119 | 1 | 0.158951279 | 0.94643392 | 0.293 | 0.208 | 1 | 0.158951279 | 0.041219846 |
| Gm17056 | 0.698189964 | 0.154722595 | 0.094 | 0.076 | 1 | 0.020884649 | 1.351778023 | 0.244 | 0.116 | 1 | 0.698189964 | 0.041333129 |
| Plcd3 | 0.020890978 | 1.631160639 | 0.189 | 0.085 | 1 | 0.374150355 | 0.19334866 | 0.171 | 0.116 | 1 | 0.374150355 | 0.041345523 |
| Adprs | 0.020903183 | 0.958325382 | 0.321 | 0.182 | 1 | 0.095815798 | 0.83720485 | 0.244 | 0.143 | 1 | 0.095815798 | 0.041369423 |
| Slc9a7 | 0.043223248 | 2.377115016 | 0.057 | 0.013 | 1 | 0.020911986 | 3.614812428 | 0.049 | 0.007 | 1 | 0.043223248 | 0.04138666 |
| Gm50186 | 0.287761733 | -0.652632327 | 0 | 0.021 | 1 | 0.020911986 | 3.159132945 | 0.049 | 0.007 | 1 | 0.287761733 | 0.04138666 |
| Adgrv1 | 0.362535229 | 1.242185436 | 0.113 | 0.076 | 1 | 0.020911986 | 3.685201756 | 0.049 | 0.007 | 1 | 0.362535229 | 0.04138666 |
| Vamp3 | 0.021800153 | -0.560771469 | 0.491 | 0.657 | 1 | 0.020931398 | -0.644536931 | 0.512 | 0.693 | 1 | 0.021800153 | 0.041424673 |
| Sac3d1 | 0.020931985 | 1.154722595 | 0.245 | 0.123 | 1 | 0.098016708 | 1.048708955 | 0.195 | 0.106 | 1 | 0.098016708 | 0.041425823 |
| Cib1 | 0.906529371 | 0.199116714 | 0.377 | 0.415 | 1 | 0.020941821 | 1.014082612 | 0.488 | 0.352 | 1 | 0.906529371 | 0.041445083 |
| Cyb561a3 | 0.020944152 | 1.090592257 | 0.189 | 0.081 | 1 | 0.20965574 | -0.80665134 | 0.024 | 0.078 | 1 | 0.20965574 | 0.041449647 |
| 0610030E20Rik | 0.02097391 | 1.006623956 | 0.396 | 0.258 | 1 | 0.55629226 | 0.103850509 | 0.244 | 0.307 | 1 | 0.55629226 | 0.041507914 |
| Hdlbp | 0.090107572 | 0.153590625 | 1 | 1 | 1 | 0.021032658 | -0.295915588 | 1 | 1 | 1 | 0.090107572 | 0.041622943 |
| Ccdc85a | 0.02104208 | 1.280253477 | 0.075 | 0.017 | 1 | 0.442708563 | -1.140075074 | 0.049 | 0.082 | 1 | 0.442708563 | 0.041641391 |
| Aff4 | 0.021055029 | 0.26546966 | 0.962 | 0.966 | 1 | 0.335207908 | -0.100307362 | 1 | 1 | 1 | 0.335207908 | 0.041666743 |
| Zmym5 | 0.021066144 | 0.454282877 | 0.868 | 0.784 | 1 | 0.375107137 | -0.114893605 | 0.805 | 0.898 | 1 | 0.375107137 | 0.041688505 |
| Tnfrsf1a | 0.021068339 | -0.457168233 | 0.792 | 0.852 | 1 | 0.444264198 | -0.100015356 | 0.78 | 0.829 | 1 | 0.444264198 | 0.041692803 |
| Kctd3 | 0.226500648 | -0.384436216 | 0.208 | 0.292 | 1 | 0.021103811 | -0.637574733 | 0.195 | 0.399 | 1 | 0.226500648 | 0.04176225 |
| Arhgap10 | 0.02110849 | 0.52453084 | 0.717 | 0.627 | 1 | 0.689158789 | 0.247754958 | 0.732 | 0.652 | 1 | 0.689158789 | 0.041771412 |
| Atp6v0b | 0.021111235 | 0.42652721 | 0.943 | 0.894 | 1 | 0.070737564 | 0.197154789 | 0.951 | 0.939 | 1 | 0.070737564 | 0.041776786 |
| Slfn9 | 0.908073731 | 0.187889459 | 0.094 | 0.089 | 1 | 0.021139139 | 0.962735732 | 0.122 | 0.038 | 1 | 0.908073731 | 0.041831414 |
| Dcun1d5 | 0.021163262 | 0.507759751 | 0.849 | 0.839 | 1 | 0.48022654 | -0.197653907 | 0.878 | 0.877 | 1 | 0.48022654 | 0.04187864 |
| Prima1 | 0.088349056 | 2.154722595 | 0.057 | 0.017 | 1 | 0.021167286 | 2.351778023 | 0.049 | 0.007 | 1 | 0.088349056 | 0.041886518 |
| Sidt1 | 0.413082433 | 0.154722595 | 0 | 0.013 | 1 | 0.021167286 | 2.83720485 | 0.049 | 0.007 | 1 | 0.413082433 | 0.041886518 |
| Ptx4 | 0.503971666 | 1.569760094 | 0.019 | 0.008 | 1 | 0.021167286 | 2.83720485 | 0.049 | 0.007 | 1 | 0.503971666 | 0.041886518 |
| Tent5d | 0.736073999 | 0.8327945 | 0.019 | 0.013 | 1 | 0.021167286 | 2.83720485 | 0.049 | 0.007 | 1 | 0.736073999 | 0.041886518 |
| Gm21860 | 0.029966098 | 3.154722595 | 0.038 | 0.004 | 1 | 0.021167579 | 2.83720485 | 0.049 | 0.007 | 1 | 0.029966098 | 0.041887091 |
| Gm47547 | 0.03562064 | 3.154722595 | 0.019 | 0 | 1 | 0.021167579 | 2.83720485 | 0.049 | 0.007 | 1 | 0.03562064 | 0.041887091 |
| Gm57330 | 0.092905716 | 1.569760094 | 0.057 | 0.017 | 1 | 0.021167579 | 2.83720485 | 0.049 | 0.007 | 1 | 0.092905716 | 0.041887091 |
| Gm9908 | 0.175865986 | -1.015202407 | 0 | 0.034 | 1 | 0.021167579 | 2.83720485 | 0.049 | 0.007 | 1 | 0.175865986 | 0.041887091 |
| Gm6598 | 0.204653189 | 2.739685096 | 0.038 | 0.013 | 1 | 0.021167579 | 2.83720485 | 0.049 | 0.007 | 1 | 0.204653189 | 0.041887091 |
| 1110035H17Rik | 0.211931446 | 1.417757001 | 0.038 | 0.013 | 1 | 0.021167579 | 2.83720485 | 0.049 | 0.007 | 1 | 0.211931446 | 0.041887091 |
| Nos1 | 0.225506368 | 2.154722595 | 0.057 | 0.025 | 1 | 0.021167579 | 2.83720485 | 0.049 | 0.007 | 1 | 0.225506368 | 0.041887091 |
| Ttc39d | 0.246443787 | 2.739685096 | 0.019 | 0.004 | 1 | 0.021167579 | 2.83720485 | 0.049 | 0.007 | 1 | 0.246443787 | 0.041887091 |
| Sult4a1 | 0.413082433 | 0.154722595 | 0 | 0.013 | 1 | 0.021167579 | 2.83720485 | 0.049 | 0.007 | 1 | 0.413082433 | 0.041887091 |
| 5033406O09Rik | 0.485993467 | 0.932330173 | 0.038 | 0.021 | 1 | 0.021167579 | 2.83720485 | 0.049 | 0.007 | 1 | 0.485993467 | 0.041887091 |
| Gm26691 | 0.663401681 | -0.1672055 | 0.019 | 0.03 | 1 | 0.021167579 | 2.83720485 | 0.049 | 0.007 | 1 | 0.663401681 | 0.041887091 |
| Gm38250 | 0.668070194 | 0.154722595 | 0.019 | 0.03 | 1 | 0.021167579 | 2.83720485 | 0.049 | 0.007 | 1 | 0.668070194 | 0.041887091 |
| Parvaos | 0.732691937 | 1.154722595 | 0.019 | 0.013 | 1 | 0.021167579 | 2.83720485 | 0.049 | 0.007 | 1 | 0.732691937 | 0.041887091 |
| Swsap1 | 0.762480493 | 0.739685096 | 0.038 | 0.03 | 1 | 0.021167579 | 2.83720485 | 0.049 | 0.007 | 1 | 0.762480493 | 0.041887091 |
| Adap2 | 0.782077996 | 0.347367673 | 0.019 | 0.025 | 1 | 0.021167579 | 2.83720485 | 0.049 | 0.007 | 1 | 0.782077996 | 0.041887091 |
| 6330408M09Rik | 0.880784741 | 0.280253477 | 0.038 | 0.042 | 1 | 0.021167579 | 2.83720485 | 0.049 | 0.007 | 1 | 0.880784741 | 0.041887091 |
| Gm29481 | 0.916620008 | 1.154722595 | 0.019 | 0.017 | 1 | 0.021167579 | 2.83720485 | 0.049 | 0.007 | 1 | 0.916620008 | 0.041887091 |
| A830008E24Rik | 0.021171539 | -3.776014743 | 0 | 0.093 | 1 | 0.672172778 | 0.136765132 | 0.024 | 0.038 | 1 | 0.672172778 | 0.041894844 |
| Sh2d4a | 0.021197733 | -1.545717123 | 0.019 | 0.131 | 1 | 0.277539989 | -1.589059905 | 0.073 | 0.126 | 1 | 0.277539989 | 0.041946122 |
| Idh1 | 0.025994129 | 0.838050225 | 0.509 | 0.373 | 1 | 0.021212457 | 0.653407674 | 0.634 | 0.457 | 1 | 0.025994129 | 0.041974947 |
| Zfp382 | 0.021214512 | 1.669295768 | 0.189 | 0.085 | 1 | 0.674961871 | 0.557096931 | 0.195 | 0.174 | 1 | 0.674961871 | 0.041978969 |
| Lrrc28 | 0.021220945 | 0.865215978 | 0.264 | 0.14 | 1 | 0.683716842 | 1.093980265 | 0.171 | 0.15 | 1 | 0.683716842 | 0.041991561 |
| Ptpmt1 | 0.021239079 | 1.052360877 | 0.396 | 0.263 | 1 | 0.195579536 | -0.255904555 | 0.171 | 0.28 | 1 | 0.195579536 | 0.04202706 |
| Sgms1 | 0.021259604 | 0.346162917 | 0.868 | 0.784 | 1 | 0.217771615 | 0.308172377 | 0.878 | 0.877 | 1 | 0.217771615 | 0.042067238 |
| Dusp16 | 0.09851991 | 0.415636949 | 0.566 | 0.449 | 1 | 0.021297393 | 0.541406487 | 0.732 | 0.59 | 1 | 0.09851991 | 0.042141206 |
| S100a14 | 0.092905716 | 1.569760094 | 0.057 | 0.017 | 1 | 0.021308648 | 1.351778023 | 0.244 | 0.116 | 1 | 0.092905716 | 0.042163237 |
| Pms2 | 0.021326805 | 0.849868013 | 0.264 | 0.131 | 1 | 0.482549515 | 0.589277336 | 0.171 | 0.137 | 1 | 0.482549515 | 0.042198778 |
| Dnajc22 | 0.021327427 | -2.054730771 | 0.019 | 0.127 | 1 | 0.079643965 | -1.448197369 | 0.024 | 0.113 | 1 | 0.079643965 | 0.042199996 |
| Ogfr | 0.830267412 | 0.145952385 | 0.472 | 0.466 | 1 | 0.021365252 | -0.7050532 | 0.293 | 0.498 | 1 | 0.830267412 | 0.04227403 |
| Atg7 | 0.029206331 | 0.642823556 | 0.358 | 0.212 | 1 | 0.021371003 | 1.042788983 | 0.39 | 0.235 | 1 | 0.029206331 | 0.042285287 |
| Pigp | 0.021382299 | -0.536690848 | 0.66 | 0.742 | 1 | 0.117089536 | -0.236044132 | 0.585 | 0.703 | 1 | 0.117089536 | 0.042307395 |
| Pappa | 0.158183696 | 1.055186921 | 0.057 | 0.021 | 1 | 0.021399939 | 1.53554915 | 0.244 | 0.123 | 1 | 0.158183696 | 0.042341921 |
| Apip | 0.021442463 | 1.19654277 | 0.377 | 0.237 | 1 | 0.46668499 | -0.304150999 | 0.268 | 0.321 | 1 | 0.46668499 | 0.042425148 |
| Gm14167 | 0.116220646 | -0.992118794 | 0.113 | 0.203 | 1 | 0.021450899 | -1.391613841 | 0.024 | 0.164 | 1 | 0.116220646 | 0.042441658 |
| Nhsl1 | 0.054863843 | 1.454282877 | 0.151 | 0.072 | 1 | 0.021520918 | 1.979223855 | 0.195 | 0.085 | 1 | 0.054863843 | 0.042578686 |
| Limd1 | 0.531400441 | -0.1182959 | 0.66 | 0.581 | 1 | 0.021522955 | -0.884870782 | 0.585 | 0.734 | 1 | 0.531400441 | 0.042582673 |
| Hirip3 | 0.021523131 | 1.104096522 | 0.264 | 0.144 | 1 | 0.652219873 | 0.983055716 | 0.098 | 0.123 | 1 | 0.652219873 | 0.042583016 |
| U2af2 | 0.021537562 | 0.717858733 | 0.698 | 0.585 | 1 | 0.574322464 | -0.108504995 | 0.683 | 0.669 | 1 | 0.574322464 | 0.042611258 |
| Catsper2 | 0.258441544 | -0.278236812 | 0.038 | 0.085 | 1 | 0.021538106 | 1.029849928 | 0.171 | 0.065 | 1 | 0.258441544 | 0.042612322 |
| Atp5g3 | 0.021550369 | 0.429987289 | 0.906 | 0.852 | 1 | 0.391006288 | -0.135088221 | 0.829 | 0.86 | 1 | 0.391006288 | 0.042636319 |
| Ndufb6 | 0.021567837 | 0.483172409 | 0.811 | 0.72 | 1 | 0.350896976 | -0.179473891 | 0.659 | 0.689 | 1 | 0.350896976 | 0.042670501 |
| Mars2 | 0.021588089 | 1.874614676 | 0.151 | 0.059 | 1 | 0.983384839 | 0.737669176 | 0.073 | 0.075 | 1 | 0.983384839 | 0.042710132 |
| Junos | 0.021593444 | -1.812110541 | 0.113 | 0.258 | 1 | 0.483913807 | -1.322666487 | 0.122 | 0.16 | 1 | 0.483913807 | 0.042720611 |
| Pwwp3b | 0.021620176 | 0.717317282 | 0.302 | 0.161 | 1 | 0.329757954 | 1.266048149 | 0.268 | 0.239 | 1 | 0.329757954 | 0.042772919 |
| Commd6 | 0.021633348 | -0.324746777 | 0.283 | 0.492 | 1 | 0.16133536 | -0.214883661 | 0.244 | 0.365 | 1 | 0.16133536 | 0.042798694 |
| Fyco1 | 0.119125072 | 0.525365975 | 0.377 | 0.271 | 1 | 0.021653623 | 0.766815522 | 0.439 | 0.273 | 1 | 0.119125072 | 0.042838367 |
| Zbed5 | 0.021683144 | 1.010332685 | 0.245 | 0.123 | 1 | 0.696207799 | 0.578470581 | 0.171 | 0.154 | 1 | 0.696207799 | 0.042896128 |
| Gm6345 | 0.029966098 | 3.154722595 | 0.038 | 0.004 | 1 | 0.021685149 | 2.422167351 | 0.049 | 0.007 | 1 | 0.029966098 | 0.042900053 |
| Padi6 | 0.03562064 | 3.154722595 | 0.019 | 0 | 1 | 0.021685149 | 2.422167351 | 0.049 | 0.007 | 1 | 0.03562064 | 0.042900053 |
| Gm26671 | 0.094748532 | -2.304709024 | 0 | 0.051 | 1 | 0.021685149 | 1.83720485 | 0.049 | 0.007 | 1 | 0.094748532 | 0.042900053 |
| Spag17 | 0.246443787 | 2.739685096 | 0.019 | 0.004 | 1 | 0.021685149 | 2.422167351 | 0.049 | 0.007 | 1 | 0.246443787 | 0.042900053 |
| 1300014J16Rik | 0.413082433 | 0.154722595 | 0 | 0.013 | 1 | 0.021685149 | 2.422167351 | 0.049 | 0.007 | 1 | 0.413082433 | 0.042900053 |
| Mapk11 | 0.503971666 | 1.569760094 | 0.019 | 0.008 | 1 | 0.021685149 | 2.422167351 | 0.049 | 0.007 | 1 | 0.503971666 | 0.042900053 |
| 9930111J21Rik2 | 0.913426097 | 1.962077517 | 0.019 | 0.017 | 1 | 0.021685149 | 2.100239256 | 0.049 | 0.007 | 1 | 0.913426097 | 0.042900053 |
| Adipor2 | 0.021687693 | -0.667279103 | 0.472 | 0.61 | 1 | 0.025490341 | -0.431828297 | 0.585 | 0.734 | 1 | 0.025490341 | 0.042905031 |
| Gm11638 | 0.021697818 | 1.237184755 | 0.132 | 0.047 | 1 | 0.586976522 | 0.136765132 | 0.098 | 0.072 | 1 | 0.586976522 | 0.04292484 |
| Rab1b | 0.021710063 | 0.664396968 | 0.623 | 0.5 | 1 | 0.468650924 | 0.617328026 | 0.512 | 0.495 | 1 | 0.468650924 | 0.042948798 |
| Mrpl46 | 0.039511671 | 1.154722595 | 0.245 | 0.136 | 1 | 0.021727094 | -1.53202896 | 0.049 | 0.195 | 1 | 0.039511671 | 0.042982121 |
| Cxcl16 | 0.615611309 | 0.261637799 | 0.075 | 0.102 | 1 | 0.021731897 | -2.484723245 | 0 | 0.116 | 1 | 0.615611309 | 0.042991518 |
| Ndufaf8 | 0.023245362 | 0.883868685 | 0.509 | 0.364 | 1 | 0.021745727 | -0.79700117 | 0.171 | 0.369 | 1 | 0.023245362 | 0.043018578 |
| Asb4 | 0.405229061 | -1.029701976 | 0.094 | 0.136 | 1 | 0.021767492 | -1.717384002 | 0.073 | 0.232 | 1 | 0.405229061 | 0.043061161 |
| Nek6 | 0.453993424 | -0.442839943 | 0.245 | 0.292 | 1 | 0.021792247 | -1.262330824 | 0.195 | 0.382 | 1 | 0.453993424 | 0.043109592 |
| Wnt5b | 0.220361278 | 0.558355331 | 0.245 | 0.165 | 1 | 0.021802558 | 0.711673968 | 0.171 | 0.065 | 1 | 0.220361278 | 0.043129764 |
| Gsr | 0.416769672 | -0.334991591 | 0.491 | 0.436 | 1 | 0.021816009 | 0.693613996 | 0.683 | 0.631 | 1 | 0.416769672 | 0.04315608 |

| | | | | | | | | | | | | |
|---|---|---|---|---|---|---|---|---|---|---|---|---|
| Stxbp1 | 0.618050093 | 0.118502407 | 0.528 | 0.538 | 1 | 0.021844744 | -0.679942267 | 0.463 | 0.648 | 1 | 0.618050093 | 0.043212295 |
| Slc25a25 | 0.174817513 | 0.906795081 | 0.245 | 0.165 | 1 | 0.021879954 | 0.884510565 | 0.439 | 0.29 | 1 | 0.174817513 | 0.043281176 |
| Rere | 0.021899903 | 0.186948642 | 0.981 | 1 | 1 | 0.751805479 | 0.101264261 | 0.951 | 0.993 | 1 | 0.751805479 | 0.0433202 |
| Pkn3 | 0.022008072 | -1.582242999 | 0.075 | 0.212 | 1 | 0.141427942 | -1.332720152 | 0.024 | 0.092 | 1 | 0.141427942 | 0.043531789 |
| Cd44 | 0.022014656 | -1.721618937 | 0.283 | 0.407 | 1 | 0.856226175 | -0.421198107 | 0.439 | 0.433 | 1 | 0.856226175 | 0.043544668 |
| Ccdc39 | 0.022015552 | 1.569760094 | 0.17 | 0.072 | 1 | 0.518226627 | 1.252242349 | 0.122 | 0.092 | 1 | 0.518226627 | 0.043546419 |
| Armcx1 | 0.022051539 | 1.53659323 | 0.34 | 0.216 | 1 | 0.457517744 | 0.433482664 | 0.341 | 0.29 | 1 | 0.457517744 | 0.043616807 |
| Mgat4b | 0.022055058 | 0.932330173 | 0.453 | 0.322 | 1 | 0.198270329 | -0.344545587 | 0.463 | 0.529 | 1 | 0.198270329 | 0.043623691 |
| Crebl2 | 0.133558004 | -0.492975661 | 0.17 | 0.275 | 1 | 0.022064598 | -1.111811221 | 0.122 | 0.297 | 1 | 0.133558004 | 0.043642349 |
| Zer1 | 0.912762519 | 0.219817623 | 0.226 | 0.22 | 1 | 0.022106739 | -1.425829556 | 0.098 | 0.256 | 1 | 0.912762519 | 0.043724771 |
| Cdkl1 | 0.022143524 | 2.254258268 | 0.113 | 0.038 | 1 | 0.871884828 | 0.624211126 | 0.098 | 0.092 | 1 | 0.871884828 | 0.043796712 |
| Hif1an | 0.022161513 | 0.849868013 | 0.264 | 0.136 | 1 | 0.248539372 | -0.466575898 | 0.122 | 0.201 | 1 | 0.248539372 | 0.043831893 |
| Bphl | 0.305410653 | 0.433876666 | 0.34 | 0.28 | 1 | 0.022195601 | 1.014847764 | 0.463 | 0.314 | 1 | 0.305410653 | 0.043898557 |
| Pdik1l | 0.022195661 | 1.237184755 | 0.208 | 0.097 | 1 | 0.906505461 | 0.185128153 | 0.122 | 0.13 | 1 | 0.906505461 | 0.043898675 |
| Zfp787 | 0.174307745 | 0.685237311 | 0.321 | 0.246 | 1 | 0.022233317 | -0.894599039 | 0.22 | 0.416 | 1 | 0.174307745 | 0.043972314 |
| Cyp2a5 | 0.022237797 | 0.8327945 | 0.226 | 0.106 | 1 | 0.0564259 | 0.515276755 | 0.463 | 0.311 | 1 | 0.0564259 | 0.043981074 |
| Gspt1 | 0.022245589 | 0.369088876 | 0.943 | 0.915 | 1 | 0.251584215 | -0.181422274 | 0.951 | 0.98 | 1 | 0.251584215 | 0.043996312 |
| Slc41a3 | 0.063530508 | 1.377115016 | 0.113 | 0.047 | 1 | 0.022272892 | 2.306690133 | 0.122 | 0.041 | 1 | 0.063530508 | 0.044049703 |
| Eipr1 | 0.812181632 | 0.502645898 | 0.226 | 0.225 | 1 | 0.022290222 | 0.88967227 | 0.341 | 0.184 | 1 | 0.812181632 | 0.044083591 |
| Pim2 | 0.022306187 | 1.491757582 | 0.151 | 0.059 | 1 | 0.7163893 | 0.404245443 | 0.146 | 0.13 | 1 | 0.7163893 | 0.044114809 |
| Spin1 | 0.022322929 | 0.643222256 | 0.509 | 0.386 | 1 | 0.102750839 | 0.397919804 | 0.732 | 0.635 | 1 | 0.102750839 | 0.044147546 |
| Atpaf2 | 0.022337031 | 1.640149422 | 0.226 | 0.119 | 1 | 0.466516809 | -0.292078167 | 0.146 | 0.195 | 1 | 0.466516809 | 0.044175119 |
| Osbpl2 | 0.125712278 | 0.757832442 | 0.321 | 0.233 | 1 | 0.022393164 | 0.913440436 | 0.439 | 0.273 | 1 | 0.125712278 | 0.044284874 |
| Slc44a3 | 0.022497542 | 1.510866405 | 0.189 | 0.085 | 1 | 0.032699804 | 0.564186355 | 0.122 | 0.041 | 1 | 0.032699804 | 0.044488945 |
| Katnal1 | 0.465041772 | 0.359542387 | 0.245 | 0.203 | 1 | 0.022563391 | -1.80665134 | 0.024 | 0.16 | 1 | 0.465041772 | 0.044617675 |
| Gart | 0.223540958 | 0.480122882 | 0.34 | 0.263 | 1 | 0.022605717 | 1.108506872 | 0.415 | 0.266 | 1 | 0.223540958 | 0.044700416 |
| Abca4 | 0.032574092 | 0.781913331 | 0.264 | 0.144 | 1 | 0.022611181 | 1.871970268 | 0.195 | 0.085 | 1 | 0.032574092 | 0.044711096 |
| Snx5 | 0.022625599 | 0.443227688 | 0.887 | 0.758 | 1 | 0.094351787 | -0.341312762 | 0.707 | 0.833 | 1 | 0.094351787 | 0.04473928 |
| Got2 | 0.022642311 | 0.47826174 | 0.642 | 0.487 | 1 | 0.846699658 | 0.133922382 | 0.512 | 0.549 | 1 | 0.846699658 | 0.044771948 |
| Bap1 | 0.022644216 | 1.029191713 | 0.302 | 0.174 | 1 | 0.654748689 | 0.463746454 | 0.341 | 0.324 | 1 | 0.654748689 | 0.044775671 |
| Klhdc8b | 0.969730707 | 0.347367673 | 0.151 | 0.153 | 1 | 0.022690025 | -1.499078538 | 0.073 | 0.229 | 1 | 0.969730707 | 0.044865212 |
| Morc3 | 0.161864725 | 0.632769892 | 0.396 | 0.322 | 1 | 0.022743292 | 0.862008241 | 0.512 | 0.358 | 1 | 0.161864725 | 0.044969326 |
| Arhgap23 | 0.022743575 | 0.54027333 | 0.66 | 0.568 | 1 | 0.109791425 | 0.370078839 | 0.902 | 0.84 | 1 | 0.109791425 | 0.04496988 |
| Sod2 | 0.022791093 | -0.537369781 | 0.774 | 0.814 | 1 | 0.056504868 | 0.730815505 | 0.732 | 0.72 | 1 | 0.056504868 | 0.045062751 |
| Gpr137b | 0.022791725 | 1.112902419 | 0.208 | 0.097 | 1 | 0.249840594 | -1.499078538 | 0.098 | 0.164 | 1 | 0.249840594 | 0.045063987 |
| Smg9 | 0.022803203 | 1.10741688 | 0.226 | 0.11 | 1 | 0.182902503 | -0.372248516 | 0.098 | 0.188 | 1 | 0.182902503 | 0.04508642 |
| Nol10 | 0.430644911 | 0.270199812 | 0.434 | 0.39 | 1 | 0.022811269 | -0.855817397 | 0.22 | 0.406 | 1 | 0.430644911 | 0.045102184 |
| Suco | 0.368767424 | 0.341573709 | 0.774 | 0.775 | 1 | 0.022829238 | 0.405442203 | 0.878 | 0.826 | 1 | 0.368767424 | 0.045137302 |
| Eif4ebp3 | 0.022830917 | 1.389187848 | 0.151 | 0.059 | 1 | 0.377396653 | 0.83720485 | 0.122 | 0.082 | 1 | 0.377396653 | 0.045140584 |
| Crcp | 0.02283403 | 0.984797593 | 0.321 | 0.186 | 1 | 0.405308769 | 0.347820009 | 0.146 | 0.208 | 1 | 0.405308769 | 0.045146667 |
| Atpsckmt | 0.692089444 | 0.569760094 | 0.208 | 0.195 | 1 | 0.022852028 | -1.250257991 | 0.073 | 0.232 | 1 | 0.692089444 | 0.045181841 |
| Shoc2 | 0.022924863 | 0.323322199 | 0.83 | 0.847 | 1 | 0.522764886 | 0.17217796 | 0.902 | 0.942 | 1 | 0.522764886 | 0.045324177 |
| Abhd14a | 0.039171752 | 0.688154795 | 0.604 | 0.466 | 1 | 0.022932108 | -0.511944714 | 0.293 | 0.498 | 1 | 0.039171752 | 0.045338334 |
| Polr3e | 0.977040148 | 0.320732546 | 0.245 | 0.254 | 1 | 0.022936347 | -0.600200463 | 0.22 | 0.423 | 1 | 0.977040148 | 0.045346617 |
| Gm48693 | 0.035512432 | 1.377115016 | 0.094 | 0.03 | 1 | 0.022988526 | 1.83720485 | 0.098 | 0.027 | 1 | 0.035512432 | 0.045448579 |
| Krcc1 | 0.706936174 | -0.141518856 | 0.66 | 0.665 | 1 | 0.023030177 | -0.556873021 | 0.463 | 0.669 | 1 | 0.706936174 | 0.045529965 |
| Ltn1 | 0.023065646 | 0.721763188 | 0.547 | 0.411 | 1 | 0.625566186 | 0.28774103 | 0.561 | 0.543 | 1 | 0.625566186 | 0.045599268 |
| Ndufa8 | 0.225320997 | -0.410062024 | 0.755 | 0.767 | 1 | 0.023080951 | -0.412668308 | 0.537 | 0.727 | 1 | 0.225320997 | 0.045629173 |
| Prdm11 | 0.023113147 | 1.72845784 | 0.264 | 0.148 | 1 | 0.600481727 | 0.100239256 | 0.195 | 0.16 | 1 | 0.600481727 | 0.045692076 |
| Acot2 | 0.023113948 | 1.71862348 | 0.17 | 0.072 | 1 | 0.613065178 | 0.458693227 | 0.049 | 0.072 | 1 | 0.613065178 | 0.045693642 |
| Ttc27 | 0.023122105 | 0.984797593 | 0.321 | 0.182 | 1 | 0.196242506 | 1.388446733 | 0.244 | 0.177 | 1 | 0.196242506 | 0.045709579 |
| Plk3 | 0.023129203 | 1.218852932 | 0.264 | 0.144 | 1 | 0.339798117 | 0.83720485 | 0.22 | 0.167 | 1 | 0.339798117 | 0.045723446 |
| Gm4956 | 0.023133287 | 2.508359549 | 0.113 | 0.038 | 1 | 0.734887054 | 1.604544093 | 0.073 | 0.061 | 1 | 0.734887054 | 0.045731426 |
| Ralb | 0.023135804 | -0.65926542 | 0.377 | 0.555 | 1 | 0.027003787 | -0.576277183 | 0.39 | 0.573 | 1 | 0.027003787 | 0.045736342 |
| Mospd1 | 0.023157036 | 0.879715547 | 0.415 | 0.288 | 1 | 0.731184935 | 0.198944123 | 0.366 | 0.362 | 1 | 0.731184935 | 0.045777823 |
| Zbtb22 | 0.023199141 | 1.347367673 | 0.189 | 0.085 | 1 | 0.32745584 | -0.541306773 | 0.098 | 0.154 | 1 | 0.32745584 | 0.045860083 |
| Rccd1 | 0.580039609 | 0.47665069 | 0.151 | 0.127 | 1 | 0.023199717 | 1.363273661 | 0.22 | 0.102 | 1 | 0.580039609 | 0.045861208 |
| Kat14 | 0.023202078 | 1.306725688 | 0.245 | 0.127 | 1 | 0.677622298 | -0.316600486 | 0.171 | 0.195 | 1 | 0.677622298 | 0.04586582 |
| Rttn | 0.023214565 | 0.454282877 | 0.189 | 0.081 | 1 | 0.688629324 | 0.797676486 | 0.098 | 0.082 | 1 | 0.688629324 | 0.045890214 |
| Psmb10 | 0.023223164 | 0.776210972 | 0.283 | 0.153 | 1 | 0.029600999 | 1.486707603 | 0.244 | 0.126 | 1 | 0.029600999 | 0.045907013 |
| Gpank1 | 0.023229386 | 1.292226119 | 0.151 | 0.059 | 1 | 0.069032735 | 1.600165653 | 0.195 | 0.102 | 1 | 0.069032735 | 0.045919168 |
| Dusp18 | 0.383430045 | -0.501323004 | 0.377 | 0.415 | 1 | 0.023244244 | -0.921494266 | 0.317 | 0.509 | 1 | 0.383430045 | 0.045948192 |
| Anks6 | 0.02325523 | 2.017219071 | 0.113 | 0.038 | 1 | 0.823532891 | 0.635570989 | 0.073 | 0.065 | 1 | 0.823532891 | 0.045969653 |
| Zfp827 | 0.023285369 | 0.680791406 | 0.509 | 0.369 | 1 | 0.045401213 | 0.688341464 | 0.561 | 0.437 | 1 | 0.045401213 | 0.046028529 |
| Klhl29 | 0.023287813 | 0.656120379 | 0.509 | 0.386 | 1 | 0.182725332 | 0.385160319 | 0.537 | 0.447 | 1 | 0.182725332 | 0.046033304 |
| Myo5a | 0.02328909 | -0.63163882 | 0.755 | 0.835 | 1 | 0.565738917 | 0.5568232 | 0.756 | 0.696 | 1 | 0.565738917 | 0.046035798 |
| Ergic1 | 0.777841654 | -0.207221179 | 0.66 | 0.653 | 1 | 0.023293825 | -0.466948543 | 0.512 | 0.689 | 1 | 0.777841654 | 0.046045047 |
| Spns2 | 0.511272335 | 0.417757001 | 0.094 | 0.068 | 1 | 0.023304559 | 2.100239256 | 0.098 | 0.027 | 1 | 0.511272335 | 0.046066015 |
| Ppil4 | 0.023312767 | 0.618669695 | 0.755 | 0.636 | 1 | 0.385526111 | 0.116727378 | 0.829 | 0.792 | 1 | 0.385526111 | 0.046082049 |
| Sema4d | 0.739719753 | 0.154722595 | 0.151 | 0.136 | 1 | 0.023345741 | 0.944120054 | 0.293 | 0.15 | 1 | 0.739719753 | 0.046146458 |
| Odf2 | 0.091708425 | 0.42579432 | 0.509 | 0.424 | 1 | 0.023358237 | 0.985962284 | 0.585 | 0.481 | 1 | 0.091708425 | 0.046170867 |
| Suclg2 | 0.023370712 | 0.683416876 | 0.509 | 0.356 | 1 | 0.031417856 | 0.565400234 | 0.61 | 0.444 | 1 | 0.031417856 | 0.046195233 |
| Rars | 0.023371361 | 0.765680304 | 0.566 | 0.475 | 1 | 0.922606477 | 0.303467672 | 0.537 | 0.553 | 1 | 0.922606477 | 0.046196502 |
| Bahd1 | 0.023405495 | 1.347367673 | 0.17 | 0.072 | 1 | 0.306902365 | -0.262330824 | 0.098 | 0.16 | 1 | 0.306902365 | 0.046263172 |
| Pou6f1 | 0.023419027 | 1.510866405 | 0.264 | 0.153 | 1 | 0.05011143 | 0.88967227 | 0.293 | 0.171 | 1 | 0.05011143 | 0.046289603 |
| Slc25a26 | 0.023445057 | 1.874614676 | 0.151 | 0.059 | 1 | 0.562562137 | -0.26713181 | 0.073 | 0.102 | 1 | 0.562562137 | 0.046340444 |
| Stx2 | 0.023451532 | 1.225111923 | 0.245 | 0.127 | 1 | 0.339076563 | 0.20784823 | 0.146 | 0.229 | 1 | 0.339076563 | 0.04635309 |
| Tnfrsf23 | 0.023603266 | -0.600164907 | 0.283 | 0.466 | 1 | 0.271763663 | -0.477110061 | 0.22 | 0.317 | 1 | 0.271763663 | 0.046649417 |
| Fgl2 | 0.964075615 | 0.280253477 | 0.094 | 0.093 | 1 | 0.023634374 | 1.35607816 | 0.195 | 0.085 | 1 | 0.964075615 | 0.046710165 |
| Otulin | 0.023641335 | -1.153784984 | 0.566 | 0.653 | 1 | 0.196518322 | -0.536253546 | 0.585 | 0.679 | 1 | 0.196518322 | 0.046723757 |
| Or2k2 | 0.023673265 | 2.002719501 | 0.113 | 0.038 | 1 | 0.114805676 | -1.132421501 | 0.024 | 0.102 | 1 | 0.114805676 | 0.046786107 |
| Wwp1 | 0.547991389 | 0.102255175 | 0.623 | 0.593 | 1 | 0.023676849 | 0.90329404 | 0.756 | 0.703 | 1 | 0.547991389 | 0.046793104 |
| Tmem167 | 0.75473688 | -0.12376021 | 0.868 | 0.924 | 1 | 0.023687347 | -0.445134617 | 0.878 | 0.932 | 1 | 0.75473688 | 0.046813603 |
| Pacsin3 | 0.44723462 | -0.326404095 | 0.17 | 0.216 | 1 | 0.023739959 | -0.405651674 | 0.146 | 0.348 | 1 | 0.44723462 | 0.046916332 |
| Grhl1 | 0.023746589 | 1.197791317 | 0.208 | 0.097 | 1 | 0.492893273 | 0.364717078 | 0.268 | 0.229 | 1 | 0.492893273 | 0.046929278 |
| Bak1 | 0.023761517 | 1.211306123 | 0.208 | 0.097 | 1 | 0.275317764 | -0.16279515 | 0.073 | 0.137 | 1 | 0.275317764 | 0.046958424 |
| Macrod2 | 0.023803649 | -0.827789978 | 0.302 | 0.487 | 1 | 0.859368971 | -0.438073152 | 0.293 | 0.3 | 1 | 0.859368971 | 0.047040685 |
| Rcan2 | 0.333491309 | 0.187889459 | 0.113 | 0.072 | 1 | 0.023811975 | 2.007129851 | 0.098 | 0.027 | 1 | 0.333491309 | 0.04705694 |
| Lamb2 | 0.367727702 | -0.162690019 | 0.623 | 0.682 | 1 | 0.023814703 | -0.607579993 | 0.537 | 0.703 | 1 | 0.367727702 | 0.047062267 |
| Gabpa | 0.363566343 | 0.417757001 | 0.321 | 0.271 | 1 | 0.023820807 | 0.785365918 | 0.537 | 0.369 | 1 | 0.363566343 | 0.047074183 |
| Zfp61 | 0.023842054 | 2.029191713 | 0.113 | 0.038 | 1 | 0.165516219 | 1.159132945 | 0.146 | 0.082 | 1 | 0.165516219 | 0.047115665 |
| Mrps5 | 0.023893806 | 0.776210972 | 0.415 | 0.267 | 1 | 0.17372739 | 0.781922414 | 0.366 | 0.283 | 1 | 0.17372739 | 0.047216699 |
| Dguok | 0.02391439 | 1.080722013 | 0.396 | 0.28 | 1 | 0.517778601 | 0.281388695 | 0.39 | 0.358 | 1 | 0.517778601 | 0.047256881 |
| D430041D05Rik | 0.16407236 | -0.675352404 | 0.113 | 0.059 | 1 | 0.023948651 | 1.527876792 | 0.122 | 0.041 | 1 | 0.16407236 | 0.047323764 |
| Chaf1a | 0.227995262 | 1.454282877 | 0.075 | 0.038 | 1 | 0.023969784 | 1.962735732 | 0.098 | 0.027 | 1 | 0.227995262 | 0.047365017 |
| Zfp629 | 0.023972214 | 1.3662267 | 0.151 | 0.059 | 1 | 0.259807619 | -0.53202896 | 0.098 | 0.167 | 1 | 0.259807619 | 0.04736976 |
| Rab25 | 0.206368562 | -1.015202407 | 0 | 0.03 | 1 | 0.024006516 | 1.204936634 | 0.22 | 0.099 | 1 | 0.206368562 | 0.04743672 |
| Gm19951 | 0.550900125 | -0.732802676 | 0.962 | 0.97 | 1 | 0.024006782 | 0.423660978 | 1 | 0.986 | 1 | 0.550900125 | 0.047437228 |

| | | | | | | | | | | | | |
|---|---|---|---|---|---|---|---|---|---|---|---|---|
| Ebag9 | 0.024011721 | 0.824077283 | 0.415 | 0.275 | 1 | 0.244276071 | 0.678006255 | 0.366 | 0.297 | 1 | 0.244276071 | 0.047446879 |
| Nlrc3 | 0.543457596 | -0.430239906 | 0.057 | 0.081 | 1 | 0.024039954 | -2.944154864 | 0 | 0.113 | 1 | 0.543457596 | 0.047501988 |
| Noa1 | 0.02406474 | 1.347367673 | 0.208 | 0.097 | 1 | 0.458485722 | -0.555112573 | 0.073 | 0.109 | 1 | 0.458485722 | 0.047550368 |
| Cyb561d2 | 0.024093958 | 0.767699472 | 0.358 | 0.216 | 1 | 0.84914951 | 0.209173627 | 0.146 | 0.167 | 1 | 0.84914951 | 0.047607396 |
| Rad50 | 0.024145493 | 0.962077517 | 0.491 | 0.347 | 1 | 0.698212033 | -0.292078167 | 0.293 | 0.317 | 1 | 0.698212033 | 0.04770798 |
| 5330439K02Rik | 0.024151136 | 2.002719501 | 0.113 | 0.038 | 1 | 0.654712174 | 0.252242349 | 0.049 | 0.034 | 1 | 0.654712174 | 0.047718994 |
| Gm41077 | 0.02415513 | 1.189488013 | 0.264 | 0.136 | 1 | 0.548896078 | 0.118386602 | 0.195 | 0.154 | 1 | 0.548896078 | 0.047726789 |
| Prpf8 | 0.024168141 | 0.557160057 | 0.679 | 0.61 | 1 | 0.83478564 | 0.411800635 | 0.707 | 0.717 | 1 | 0.83478564 | 0.047752183 |
| Dock9 | 0.02418602 | -1.493565841 | 0.17 | 0.314 | 1 | 0.615805428 | -0.234112555 | 0.39 | 0.41 | 1 | 0.615805428 | 0.047787076 |
| Polh | 0.024207761 | 1.110328475 | 0.208 | 0.097 | 1 | 0.748090517 | 0.904319046 | 0.122 | 0.106 | 1 | 0.748090517 | 0.047829506 |
| Slc17a9 | 0.024232185 | -2.067669827 | 0.038 | 0.153 | 1 | 0.127994637 | -1.020776145 | 0.049 | 0.133 | 1 | 0.127994637 | 0.047877172 |
| Dcaf4 | 0.024234247 | 1.569760094 | 0.189 | 0.089 | 1 | 0.084572163 | 2.066686696 | 0.171 | 0.092 | 1 | 0.084572163 | 0.047881196 |
| Krt15 | 0.771930521 | -1.104011674 | 0.113 | 0.097 | 1 | 0.024248741 | 1.18867722 | 0.415 | 0.27 | 1 | 0.771930521 | 0.04790948 |
| Bpifa1 | 0.609462539 | 0.177442671 | 0.415 | 0.373 | 1 | 0.024281989 | 0.421119261 | 0.805 | 0.713 | 1 | 0.609462539 | 0.047974362 |
| Mgat5 | 0.096869145 | 0.628286171 | 0.566 | 0.538 | 1 | 0.024294369 | 0.668185779 | 0.805 | 0.768 | 1 | 0.096869145 | 0.047998522 |
| Banp | 0.761038257 | 0.333692736 | 0.396 | 0.403 | 1 | 0.024315117 | 0.557712021 | 0.707 | 0.577 | 1 | 0.761038257 | 0.04803901 |
| Tbc1d2b | 0.024337303 | 0.979635888 | 0.358 | 0.225 | 1 | 0.129975536 | -0.497779398 | 0.146 | 0.259 | 1 | 0.129975536 | 0.048082302 |
| Gm42439 | 0.817601149 | 0.311448098 | 0.17 | 0.191 | 1 | 0.024366546 | -2.037264268 | 0.049 | 0.188 | 1 | 0.817601149 | 0.048139364 |
| Zfp120 | 0.024373067 | 1.879088152 | 0.189 | 0.093 | 1 | 0.567039683 | -0.16279515 | 0.073 | 0.102 | 1 | 0.567039683 | 0.048152087 |
| Glb1 | 0.024383185 | 0.874614676 | 0.358 | 0.216 | 1 | 0.777791581 | -0.25471764 | 0.268 | 0.256 | 1 | 0.777791581 | 0.048171829 |
| Endou | 0.024411559 | -1.341703231 | 0.075 | 0.208 | 1 | 0.601776327 | -0.132421501 | 0.122 | 0.157 | 1 | 0.601776327 | 0.048227194 |
| Gm5544 | 0.02445674 | 0.913714495 | 0.245 | 0.119 | 1 | 0.58057174 | 0.500169863 | 0.195 | 0.164 | 1 | 0.58057174 | 0.048315348 |
| Tex10 | 0.024486334 | 0.924424975 | 0.453 | 0.331 | 1 | 0.094274148 | 0.646873638 | 0.512 | 0.392 | 1 | 0.094274148 | 0.048373087 |
| Pofut1 | 0.21081848 | -0.492975661 | 0.208 | 0.288 | 1 | 0.024489668 | -1.069685746 | 0.22 | 0.389 | 1 | 0.21081848 | 0.048379593 |
| Galnt10 | 0.024559643 | 0.911451444 | 0.453 | 0.314 | 1 | 0.599354304 | 0.443126969 | 0.415 | 0.423 | 1 | 0.599354304 | 0.048516111 |
| 9530052E02Rik | 0.024579013 | -1.783876861 | 0.038 | 0.153 | 1 | 0.270926227 | -0.906956246 | 0.049 | 0.106 | 1 | 0.270926227 | 0.048553898 |
| A230057D06Rik | 0.024593349 | -0.637835325 | 0.151 | 0.331 | 1 | 0.140904058 | -0.944154864 | 0.122 | 0.225 | 1 | 0.140904058 | 0.048581864 |
| Crym | 0.024594163 | -2.974560422 | 0 | 0.089 | 1 | 0.368978836 | -0.26713181 | 0.049 | 0.092 | 1 | 0.368978836 | 0.048583453 |
| A630001G21Rik | 0.024597685 | -2.799473716 | 0 | 0.089 | 1 | 0.834695558 | 1.000703582 | 0.049 | 0.058 | 1 | 0.834695558 | 0.048590325 |
| 1600002D24Rik | 0.024608885 | 0.932330173 | 0.132 | 0.047 | 1 | 0.056293038 | 2.83720485 | 0.049 | 0.01 | 1 | 0.056293038 | 0.048612173 |
| Psmc5 | 0.024678017 | 0.412613697 | 0.811 | 0.619 | 1 | 0.167153462 | 0.283048396 | 0.78 | 0.693 | 1 | 0.167153462 | 0.048747029 |
| Tsnax | 0.553446342 | 0.200526284 | 0.509 | 0.462 | 1 | 0.024700891 | -0.80665134 | 0.268 | 0.457 | 1 | 0.553446342 | 0.048791648 |
| Zfand2a | 0.024738005 | -0.796367805 | 0.283 | 0.436 | 1 | 0.03990892 | -1.31118699 | 0.268 | 0.399 | 1 | 0.03990892 | 0.048864042 |
| Mycbpap | 0.024744884 | 1.8327945 | 0.113 | 0.038 | 1 | 0.170403831 | -1.622226769 | 0 | 0.044 | 1 | 0.170403831 | 0.048877459 |
| Nusap1 | 0.024745871 | 0.598329246 | 0.17 | 0.072 | 1 | 0.483789441 | 0.79281073 | 0.098 | 0.068 | 1 | 0.483789441 | 0.048879384 |
| Armcx2 | 0.420800138 | 0.232725107 | 0.283 | 0.225 | 1 | 0.024764764 | -0.522197951 | 0.244 | 0.43 | 1 | 0.420800138 | 0.048916235 |
| Anks1 | 0.827010286 | 0.417757001 | 0.453 | 0.534 | 1 | 0.024775897 | 0.425778604 | 0.805 | 0.621 | 1 | 0.827010286 | 0.048937949 |
| 3110045C21Rik | 0.024785663 | -1.304709024 | 0.038 | 0.157 | 1 | 0.708451968 | 0.589277336 | 0.073 | 0.058 | 1 | 0.708451968 | 0.048956997 |
| Gls | 0.024789457 | 0.221101435 | 1 | 0.975 | 1 | 0.088856196 | 0.400764874 | 1 | 0.993 | 1 | 0.088856196 | 0.048964396 |
| Smarcd2 | 0.142627289 | 0.999444369 | 0.245 | 0.169 | 1 | 0.02479553 | 0.79281073 | 0.439 | 0.276 | 1 | 0.142627289 | 0.048976241 |
| Crtap | 0.065227348 | -0.274350964 | 0.868 | 0.881 | 1 | 0.024839639 | -0.479852014 | 0.707 | 0.816 | 1 | 0.065227348 | 0.04906227 |
| Irf6 | 0.246443787 | 3.154722595 | 0.019 | 0.004 | 1 | 0.024885562 | 1.458693227 | 0.195 | 0.085 | 1 | 0.246443787 | 0.049151832 |
| Csmd3 | 0.497380598 | 2.569760094 | 0.019 | 0.008 | 1 | 0.02490686 | 3.574170444 | 0.073 | 0.017 | 1 | 0.497380598 | 0.049193368 |
| Hycc2 | 0.024935222 | 0.785488785 | 0.472 | 0.322 | 1 | 0.687728814 | 0.239642312 | 0.341 | 0.423 | 1 | 0.687728814 | 0.049248678 |
| Cnppd1 | 0.024976004 | 0.922061838 | 0.415 | 0.288 | 1 | 0.072852271 | 0.53554915 | 0.585 | 0.444 | 1 | 0.072852271 | 0.049328206 |
| Grpel2 | 0.024977994 | 0.984797593 | 0.226 | 0.11 | 1 | 0.648059477 | -0.372248516 | 0.244 | 0.266 | 1 | 0.648059477 | 0.049332088 |
| Pex10 | 0.025002956 | 1.347367673 | 0.17 | 0.072 | 1 | 0.984892814 | 0.404245443 | 0.073 | 0.075 | 1 | 0.984892814 | 0.049380763 |
| Pcyox1 | 0.025050229 | 0.749466116 | 0.453 | 0.309 | 1 | 0.199308856 | 0.718263777 | 0.39 | 0.321 | 1 | 0.199308856 | 0.049472944 |
| Asb13 | 0.025053139 | 1.331600357 | 0.17 | 0.072 | 1 | 0.053576439 | -0.180717058 | 0.073 | 0.201 | 1 | 0.053576439 | 0.049478618 |
| Taf2 | 0.02505668 | 0.960075164 | 0.453 | 0.314 | 1 | 0.045556064 | -0.529577481 | 0.22 | 0.406 | 1 | 0.045556064 | 0.049485522 |
| Nol7 | 0.025063754 | 0.554452841 | 0.792 | 0.699 | 1 | 0.372694215 | 0.291770713 | 0.756 | 0.741 | 1 | 0.372694215 | 0.049499316 |
| H2-Q4 | 0.025117794 | -2.175667079 | 0.17 | 0.318 | 1 | 0.14801619 | 0.458693227 | 0.244 | 0.154 | 1 | 0.14801619 | 0.049604684 |
| Dctn3 | 0.65609025 | -0.686579659 | 0.566 | 0.555 | 1 | 0.025122585 | -0.78044627 | 0.488 | 0.631 | 1 | 0.65609025 | 0.049614025 |
| AU040320 | 0.666417688 | 0.243259269 | 0.245 | 0.22 | 1 | 0.02516483 | -0.759730293 | 0.098 | 0.273 | 1 | 0.666417688 | 0.049696392 |
| Dnah17 | 0.025208004 | 1.962077517 | 0.113 | 0.038 | 1 | 0.630901922 | 1.029849928 | 0.098 | 0.078 | 1 | 0.630901922 | 0.049780564 |
| Epb41l4b | 0.136032543 | -0.595299152 | 0.113 | 0.208 | 1 | 0.025240524 | -0.669147816 | 0.244 | 0.437 | 1 | 0.136032543 | 0.049843963 |
| Yrdc | 0.488474743 | 0.25058261 | 0.604 | 0.619 | 1 | 0.025243458 | -0.47328835 | 0.463 | 0.659 | 1 | 0.488474743 | 0.049849684 |
| Plekhg1 | 0.025258843 | 1.254258268 | 0.189 | 0.085 | 1 | 0.035277012 | 0.567744175 | 0.366 | 0.215 | 1 | 0.035277012 | 0.049879676 |
| Faim | 0.025259104 | 1.261637799 | 0.189 | 0.085 | 1 | 0.039214889 | -2.250257991 | 0 | 0.096 | 1 | 0.039214889 | 0.049880185 |
| Dse | 0.025265961 | -0.683410989 | 0.774 | 0.852 | 1 | 0.455648366 | -0.201055725 | 0.756 | 0.857 | 1 | 0.455648366 | 0.049893553 |
| Rhbdl3 | 0.075950274 | 1.865215978 | 0.132 | 0.064 | 1 | 0.025272128 | 0.313642894 | 0.293 | 0.147 | 1 | 0.075950274 | 0.049905576 |
| Egr4 | 0.507281812 | -0.304709024 | 0.019 | 0.008 | 1 | 0.025332548 | 3.252242349 | 0.073 | 0.017 | 1 | 0.507281812 | 0.050023359 |
| Hmces | 0.02534922 | 0.748730235 | 0.434 | 0.297 | 1 | 0.957812698 | 0.163804384 | 0.341 | 0.386 | 1 | 0.957812698 | 0.050055858 |
| Pcx | 0.494732257 | -0.430239906 | 0.094 | 0.127 | 1 | 0.025372562 | 1.930314254 | 0.122 | 0.041 | 1 | 0.494732257 | 0.050101358 |
| Parm1 | 0.622129999 | 1.117247889 | 0.113 | 0.093 | 1 | 0.025384259 | 3.33970519 | 0.146 | 0.055 | 1 | 0.622129999 | 0.050124157 |
| Snx24 | 0.116370199 | 0.271236604 | 0.717 | 0.691 | 1 | 0.025489684 | -0.320336427 | 0.659 | 0.795 | 1 | 0.116370199 | 0.050329644 |
| Klf5 | 0.53408091 | 0.416446055 | 0.623 | 0.559 | 1 | 0.025495003 | 0.450468459 | 0.732 | 0.532 | 1 | 0.53408091 | 0.050340012 |
| Dennd10 | 0.025521406 | -0.606089741 | 0.321 | 0.492 | 1 | 0.049458887 | -0.549376203 | 0.341 | 0.519 | 1 | 0.049458887 | 0.050391471 |
| Dnal4 | 0.025548736 | 1.228723176 | 0.321 | 0.203 | 1 | 0.930423087 | 0.408361551 | 0.195 | 0.198 | 1 | 0.930423087 | 0.050444734 |
| C1qtnf6 | 0.02558288 | 0.932330173 | 0.509 | 0.407 | 1 | 0.519788542 | 0.443290642 | 0.341 | 0.464 | 1 | 0.519788542 | 0.050511277 |
| Aknad1 | 0.54507254 | 0.8327945 | 0.057 | 0.038 | 1 | 0.025645416 | 1.944120054 | 0.122 | 0.041 | 1 | 0.54507254 | 0.050633145 |
| Iqce | 0.025658747 | 1.844382474 | 0.226 | 0.114 | 1 | 0.81726614 | 1.169780189 | 0.146 | 0.143 | 1 | 0.81726614 | 0.050659122 |
| Rnh1 | 0.723872601 | -0.241060347 | 0.774 | 0.758 | 1 | 0.025689144 | -0.460810625 | 0.659 | 0.843 | 1 | 0.723872601 | 0.050718356 |
| Mthfd2 | 0.12444579 | 0.154722595 | 0.585 | 0.436 | 1 | 0.02571609 | -1.02314316 | 0.268 | 0.437 | 1 | 0.12444579 | 0.050770863 |
| Mospd3 | 0.025730061 | 0.926916677 | 0.434 | 0.309 | 1 | 0.555151305 | 0.667279848 | 0.268 | 0.242 | 1 | 0.555151305 | 0.050798086 |
| Mfn2 | 0.025815304 | 1.05850728 | 0.358 | 0.225 | 1 | 0.930818851 | 0.342440158 | 0.268 | 0.294 | 1 | 0.930818851 | 0.050964179 |
| Nucb2 | 0.051986761 | 0.536483939 | 0.698 | 0.631 | 1 | 0.025821281 | -0.723510105 | 0.537 | 0.642 | 1 | 0.051986761 | 0.050975823 |
| Gapdh | 0.025853674 | -0.453994259 | 0.981 | 0.979 | 1 | 0.263942766 | -0.191746524 | 1 | 0.99 | 1 | 0.263942766 | 0.051038935 |
| Ergic3 | 0.025908142 | 0.355909409 | 0.849 | 0.814 | 1 | 0.707478728 | 0.115721588 | 0.854 | 0.833 | 1 | 0.707478728 | 0.051145051 |
| Krit1 | 0.025945681 | 0.419081182 | 0.736 | 0.678 | 1 | 0.475720061 | 0.211456734 | 0.732 | 0.737 | 1 | 0.475720061 | 0.051218184 |
| Osbpl3 | 0.025947459 | 2.01270359 | 0.17 | 0.076 | 1 | 0.165075562 | 1.15538081 | 0.293 | 0.201 | 1 | 0.165075562 | 0.051221648 |
| Snrpn | 0.025959141 | -1.652632327 | 0.057 | 0.178 | 1 | 0.909652106 | 0.547698233 | 0.122 | 0.119 | 1 | 0.909652106 | 0.051244405 |
| Hvcn1 | 0.042860829 | 1.47665069 | 0.075 | 0.021 | 1 | 0.025975166 | 3.422167351 | 0.073 | 0.017 | 1 | 0.042860829 | 0.051275623 |
| Dtx3l | 0.432287872 | 0.758793918 | 0.151 | 0.114 | 1 | 0.025982289 | 1.585666083 | 0.22 | 0.106 | 1 | 0.432287872 | 0.051289499 |
| Arhgef10 | 0.026012289 | 1.013366746 | 0.34 | 0.208 | 1 | 0.126197922 | -0.825760163 | 0.244 | 0.348 | 1 | 0.126197922 | 0.05134794 |
| Gm56812 | 0.026018725 | 2.039245377 | 0.132 | 0.051 | 1 | 0.879975123 | 0.589277336 | 0.098 | 0.092 | 1 | 0.879975123 | 0.051360476 |
| Acer3 | 0.895802596 | -0.163548476 | 0.679 | 0.631 | 1 | 0.026031434 | -0.581747699 | 0.634 | 0.788 | 1 | 0.895802596 | 0.051385232 |
| Fut10 | 0.026033499 | 1.270199812 | 0.17 | 0.072 | 1 | 0.515329659 | 0.83720485 | 0.171 | 0.137 | 1 | 0.515329659 | 0.051389255 |
| Ppm1g | 0.026066369 | 0.469156153 | 0.623 | 0.479 | 1 | 0.446438902 | 0.185617617 | 0.732 | 0.693 | 1 | 0.446438902 | 0.051453282 |
| Zfp955b | 0.026097869 | 0.891688189 | 0.377 | 0.242 | 1 | 0.962485506 | 0.529082554 | 0.22 | 0.249 | 1 | 0.962485506 | 0.05151464 |
| Nkd1 | 0.09059895 | -0.587479648 | 0.453 | 0.581 | 1 | 0.026113516 | -0.622226769 | 0.39 | 0.57 | 1 | 0.09059895 | 0.051545116 |
| Crbn | 0.026124168 | 1.228723176 | 0.321 | 0.199 | 1 | 0.09123293 | -0.537834582 | 0.098 | 0.218 | 1 | 0.09123293 | 0.051565865 |
| Gm56955 | 0.026162081 | -1.974560422 | 0.038 | 0.153 | 1 | 0.093714829 | -2.069685746 | 0.024 | 0.106 | 1 | 0.093714829 | 0.051639707 |
| Ccdc85b | 0.026193157 | -1.205401824 | 0.453 | 0.581 | 1 | 0.258715694 | -0.327181968 | 0.317 | 0.437 | 1 | 0.258715694 | 0.051700232 |
| Snapin | 0.026205543 | 0.735008608 | 0.585 | 0.551 | 1 | 0.387656672 | 0.447638038 | 0.537 | 0.529 | 1 | 0.387656672 | 0.051724355 |
| Cand2 | 0.02623394 | 1.221836791 | 0.17 | 0.072 | 1 | 0.062499458 | -0.863234868 | 0.024 | 0.126 | 1 | 0.062499458 | 0.05177966 |
| Armc7 | 0.02623394 | 1.221836791 | 0.17 | 0.072 | 1 | 0.614422994 | 0.778311161 | 0.049 | 0.072 | 1 | 0.614422994 | 0.05177966 |
| Rpain | 0.026276468 | 1.104096522 | 0.34 | 0.216 | 1 | 0.221308076 | 0.949679579 | 0.341 | 0.283 | 1 | 0.221308076 | 0.051862483 |

| | | | | | | | | | | | | |
|---|---|---|---|---|---|---|---|---|---|---|---|---|
| Tomm70a | 0.026379783 | 0.394803016 | 0.792 | 0.669 | 1 | 0.109111477 | -0.351456714 | 0.61 | 0.778 | 1 | 0.109111477 | 0.052063673 |
| Tmem120b | 0.026396218 | 1.084333267 | 0.226 | 0.114 | 1 | 0.072674562 | -1.037264268 | 0.073 | 0.188 | 1 | 0.072674562 | 0.052095676 |
| Cdc26 | 0.026404285 | 0.560452352 | 0.736 | 0.695 | 1 | 0.723626249 | 0.183762611 | 0.634 | 0.73 | 1 | 0.723626249 | 0.052111384 |
| Pter | 0.026407371 | -1.736048335 | 0.057 | 0.182 | 1 | 0.937856763 | 0.296636468 | 0.146 | 0.143 | 1 | 0.937856763 | 0.052117394 |
| Cabin1 | 0.026453272 | 0.746257749 | 0.547 | 0.419 | 1 | 0.619108776 | -0.106612315 | 0.439 | 0.515 | 1 | 0.619108776 | 0.052206769 |
| Phyh | 0.04603144 | -0.359850578 | 0.509 | 0.712 | 1 | 0.026477092 | -0.612598068 | 0.537 | 0.679 | 1 | 0.04603144 | 0.052253148 |
| Pde7a | 0.065439592 | 0.622708974 | 0.321 | 0.203 | 1 | 0.026518545 | 0.730289646 | 0.659 | 0.491 | 1 | 0.065439592 | 0.052333857 |
| Rcbtb2 | 0.026527588 | 1.090592257 | 0.358 | 0.242 | 1 | 0.785012724 | -0.224571348 | 0.317 | 0.345 | 1 | 0.785012724 | 0.052351463 |
| Prkab1 | 0.02652857 | 0.932330173 | 0.264 | 0.14 | 1 | 0.589193614 | 0.265662865 | 0.317 | 0.276 | 1 | 0.589193614 | 0.052353374 |
| Setd4 | 0.625902178 | 0.154722595 | 0.038 | 0.055 | 1 | 0.026556239 | 1.444887427 | 0.146 | 0.055 | 1 | 0.625902178 | 0.052407244 |
| Snx25 | 0.412023556 | -0.162222093 | 0.453 | 0.525 | 1 | 0.026559707 | -0.750487444 | 0.415 | 0.608 | 1 | 0.412023556 | 0.052413995 |
| Rab3d | 0.782077996 | 0.347367673 | 0.019 | 0.025 | 1 | 0.026569516 | 1.737669176 | 0.122 | 0.041 | 1 | 0.782077996 | 0.052433094 |
| Samd12 | 0.07373726 | -0.830170513 | 0.038 | 0.123 | 1 | 0.02663639 | -4.20718927 | 0 | 0.109 | 1 | 0.07373726 | 0.052563283 |
| Tysnd1 | 0.026683615 | 1.384204441 | 0.226 | 0.119 | 1 | 0.70787885 | 0.345351753 | 0.146 | 0.126 | 1 | 0.70787885 | 0.052655214 |
| Psma3 | 0.026764323 | 0.275936182 | 1 | 0.97 | 1 | 0.439223191 | 0.162445679 | 0.927 | 0.973 | 1 | 0.439223191 | 0.052812318 |
| Shfl | 0.02676778 | 1.154722595 | 0.226 | 0.114 | 1 | 0.058871847 | -1.140075074 | 0.049 | 0.164 | 1 | 0.058871847 | 0.052819046 |
| Nme7 | 0.026782302 | 1.134823037 | 0.34 | 0.22 | 1 | 0.20205822 | 0.63209042 | 0.366 | 0.28 | 1 | 0.20205822 | 0.052847312 |
| Rbmx | 0.026825654 | 0.507024339 | 0.585 | 0.453 | 1 | 0.959066914 | 0.236645494 | 0.488 | 0.502 | 1 | 0.959066914 | 0.052931691 |
| Psmb8 | 0.026857748 | 0.491757582 | 0.151 | 0.059 | 1 | 0.168643345 | 0.574170444 | 0.073 | 0.031 | 1 | 0.168643345 | 0.052994158 |
| Cenpv | 0.026891458 | 1.039245377 | 0.245 | 0.127 | 1 | 0.66837143 | -0.182694708 | 0.122 | 0.15 | 1 | 0.66837143 | 0.053059765 |
| Ankfy1 | 0.433108754 | 0.201372688 | 0.547 | 0.521 | 1 | 0.026895923 | 0.672460087 | 0.756 | 0.666 | 1 | 0.433108754 | 0.053068455 |
| Gm48678 | 0.026928772 | 1.569760094 | 0.113 | 0.038 | 1 | 0.574764182 | 1.204936634 | 0.122 | 0.096 | 1 | 0.574764182 | 0.053132385 |
| Strbp | 0.364242157 | -0.221338937 | 0.566 | 0.648 | 1 | 0.026962398 | -0.427195098 | 0.707 | 0.863 | 1 | 0.364242157 | 0.053197826 |
| Gm28153 | 0.027061481 | 1.181689642 | 0.283 | 0.161 | 1 | 0.030200254 | -1.890715605 | 0.073 | 0.212 | 1 | 0.030200254 | 0.053390638 |
| Ndufs1 | 0.02707718 | 0.59890644 | 0.566 | 0.419 | 1 | 0.526818572 | 0.265417738 | 0.537 | 0.604 | 1 | 0.526818572 | 0.053421186 |
| Cct6b | 0.473270726 | 1.669295768 | 0.038 | 0.021 | 1 | 0.027081844 | 2.83720485 | 0.073 | 0.017 | 1 | 0.473270726 | 0.053430262 |
| Apba2 | 0.913425467 | 1.984797593 | 0.019 | 0.017 | 1 | 0.027081844 | 3.059597271 | 0.073 | 0.017 | 1 | 0.913425467 | 0.053430262 |
| Thoc7 | 0.027086899 | 0.701636488 | 0.755 | 0.758 | 1 | 0.632812917 | 0.19072796 | 0.805 | 0.788 | 1 | 0.632812917 | 0.053440097 |
| Serpinb9 | 0.027136482 | 1.370451286 | 0.208 | 0.102 | 1 | 0.934996742 | -0.213421223 | 0.146 | 0.147 | 1 | 0.934996742 | 0.053536574 |
| Nol9 | 0.027186827 | 0.805138157 | 0.377 | 0.242 | 1 | 0.287143815 | 0.146042945 | 0.22 | 0.334 | 1 | 0.287143815 | 0.05363453 |
| Polr1d | 0.262889575 | 0.247371883 | 0.925 | 0.941 | 1 | 0.02723227 | -0.293191787 | 0.927 | 0.935 | 1 | 0.262889575 | 0.053722944 |
| Mis12 | 0.027247162 | 1.177442671 | 0.321 | 0.195 | 1 | 0.638194189 | 0.580865097 | 0.244 | 0.222 | 1 | 0.638194189 | 0.053751916 |
| Tspan5 | 0.027247207 | -1.131954653 | 0.547 | 0.661 | 1 | 0.060010356 | 0.814556111 | 0.707 | 0.635 | 1 | 0.060010356 | 0.053752003 |
| Amotl2 | 0.05470271 | 0.514554592 | 0.604 | 0.483 | 1 | 0.027280591 | 0.957499084 | 0.634 | 0.526 | 1 | 0.05470271 | 0.053816951 |
| Vasn | 0.255262067 | 0.157554186 | 0.887 | 0.835 | 1 | 0.027281813 | -0.419332249 | 0.829 | 0.887 | 1 | 0.255262067 | 0.053819328 |
| Dhrs13 | 0.091274665 | -0.982780929 | 0.057 | 0.144 | 1 | 0.027301958 | 1.438655473 | 0.195 | 0.085 | 1 | 0.091274665 | 0.05385852 |
| Eif3m | 0.027360314 | 0.381651148 | 0.868 | 0.839 | 1 | 0.227742269 | -0.151479837 | 0.78 | 0.887 | 1 | 0.227742269 | 0.053972041 |
| Ube3a | 0.027372518 | 0.31740107 | 0.962 | 0.907 | 1 | 0.20035544 | -0.190275887 | 0.951 | 0.969 | 1 | 0.20035544 | 0.053995781 |
| Nme6 | 0.494339628 | 0.8327945 | 0.094 | 0.068 | 1 | 0.027400557 | 1.444887427 | 0.146 | 0.055 | 1 | 0.494339628 | 0.054050323 |
| Sgf29 | 0.027463802 | 1.242185436 | 0.226 | 0.114 | 1 | 0.174295163 | -0.603367742 | 0.122 | 0.215 | 1 | 0.174295163 | 0.054173343 |
| Mto1 | 0.027473981 | 1.154722595 | 0.264 | 0.14 | 1 | 0.250742282 | 0.515276755 | 0.244 | 0.171 | 1 | 0.250742282 | 0.054193143 |
| Ppp3cb | 0.027474235 | 0.662569904 | 0.698 | 0.602 | 1 | 0.714568267 | 0.259331774 | 0.829 | 0.805 | 1 | 0.714568267 | 0.054193637 |
| Prdm6 | 0.725550524 | 0.320866857 | 0.604 | 0.636 | 1 | 0.027536594 | -0.874697886 | 0.512 | 0.689 | 1 | 0.725550524 | 0.054314924 |
| Slc10a1 | 0.100937876 | 2.154722595 | 0.038 | 0.008 | 1 | 0.027541029 | 2.614812428 | 0.073 | 0.017 | 1 | 0.100937876 | 0.054323549 |
| Gm34471 | 0.442381813 | 0.47665069 | 0.057 | 0.034 | 1 | 0.027545835 | 2.515276755 | 0.073 | 0.017 | 1 | 0.442381813 | 0.054332896 |
| Bsg | 0.272049385 | 0.213884667 | 1 | 0.97 | 1 | 0.027608373 | -0.359412773 | 1 | 0.997 | 1 | 0.272049385 | 0.054454525 |
| Epop | 0.027613896 | 0.948271717 | 0.208 | 0.097 | 1 | 0.137455839 | -0.364429011 | 0.049 | 0.133 | 1 | 0.137455839 | 0.054465265 |
| Ercc2 | 0.027626031 | 1.254258268 | 0.208 | 0.102 | 1 | 0.640856999 | 0.159132945 | 0.146 | 0.181 | 1 | 0.640856999 | 0.054488864 |
| Kcnk6 | 0.027626242 | 1.002719501 | 0.151 | 0.059 | 1 | 0.139520591 | -0.572186086 | 0.098 | 0.198 | 1 | 0.139520591 | 0.054489276 |
| Prps1 | 0.137882721 | 0.417757001 | 0.396 | 0.292 | 1 | 0.027690936 | -0.214883661 | 0.171 | 0.365 | 1 | 0.137882721 | 0.054615083 |
| Haghl | 0.02771397 | 1.195364579 | 0.245 | 0.131 | 1 | 0.898145508 | 0.422167351 | 0.122 | 0.133 | 1 | 0.898145508 | 0.054659876 |
| Esyt3 | 0.027734612 | -1.631873767 | 0.057 | 0.178 | 1 | 0.188498353 | -0.970150072 | 0 | 0.041 | 1 | 0.188498353 | 0.054700016 |
| Cox5b | 0.027738006 | 0.336019995 | 1 | 0.89 | 1 | 0.136588475 | 0.34112363 | 0.878 | 0.86 | 1 | 0.136588475 | 0.054706616 |
| Zfp319 | 0.027756034 | 1.913714495 | 0.132 | 0.051 | 1 | 0.033107668 | 1.515276755 | 0.22 | 0.109 | 1 | 0.033107668 | 0.054741671 |
| A230083G16Rik | 0.100937876 | 2.154722595 | 0.038 | 0.008 | 1 | 0.02777124 | 2.351778023 | 0.073 | 0.017 | 1 | 0.100937876 | 0.054771239 |
| Krtcap3 | 0.929410101 | 0.569760094 | 0.019 | 0.017 | 1 | 0.02777124 | 2.351778023 | 0.073 | 0.017 | 1 | 0.929410101 | 0.054771239 |
| Gcfc2 | 0.027788374 | 1.624207878 | 0.189 | 0.093 | 1 | 0.446959877 | 0.100239256 | 0.122 | 0.082 | 1 | 0.446959877 | 0.054804555 |
| Prr5 | 0.027792934 | 1.948271717 | 0.132 | 0.051 | 1 | 0.271903644 | -0.665295491 | 0.049 | 0.102 | 1 | 0.271903644 | 0.054813421 |
| Wee1 | 0.999058719 | 0.841062116 | 0.245 | 0.267 | 1 | 0.027800813 | 1.651338304 | 0.317 | 0.195 | 1 | 0.999058719 | 0.05482874 |
| Zfp236 | 0.027851245 | 0.569760094 | 0.491 | 0.335 | 1 | 0.220776149 | 0.592092352 | 0.537 | 0.495 | 1 | 0.220776149 | 0.054926798 |
| Rbm34 | 0.027853241 | 0.583973953 | 0.434 | 0.271 | 1 | 0.545313984 | -0.506749551 | 0.244 | 0.273 | 1 | 0.545313984 | 0.05493068 |
| Il17rb | 0.027855032 | -1.545717123 | 0.019 | 0.123 | 1 | 0.294063691 | 0.809724113 | 0.171 | 0.116 | 1 | 0.294063691 | 0.054934161 |
| Gm48882 | 0.02787682 | -1.932740246 | 0.038 | 0.148 | 1 | 0.878310931 | 0.515276755 | 0.024 | 0.02 | 1 | 0.878310931 | 0.054976522 |
| Hnrnpf | 0.052360515 | 0.224383473 | 1 | 0.945 | 1 | 0.027881323 | -0.283266754 | 0.976 | 0.99 | 1 | 0.052360515 | 0.054985278 |
| Als2cl | 0.027940752 | -1.500629234 | 0.075 | 0.199 | 1 | 0.987093277 | 1.043655727 | 0.122 | 0.133 | 1 | 0.987093277 | 0.055100818 |
| Zmat1 | 0.027942081 | -0.527101445 | 0.094 | 0.237 | 1 | 0.902455407 | 1.174239837 | 0.122 | 0.119 | 1 | 0.902455407 | 0.055103401 |
| Ubqln4 | 0.02797536 | 0.861940846 | 0.302 | 0.169 | 1 | 0.739101059 | 0.154395026 | 0.244 | 0.287 | 1 | 0.739101059 | 0.055168099 |
| Tmem185a | 0.359592011 | 0.180717803 | 0.434 | 0.356 | 1 | 0.027978239 | -0.784283527 | 0.22 | 0.41 | 1 | 0.359592011 | 0.055173697 |
| Dctn5 | 0.028025897 | 1.017219071 | 0.321 | 0.191 | 1 | 0.802635713 | 0.237292008 | 0.244 | 0.273 | 1 | 0.802635713 | 0.055266343 |
| Zfyve16 | 0.028050387 | 0.751963425 | 0.528 | 0.415 | 1 | 0.473073833 | -0.182694708 | 0.317 | 0.413 | 1 | 0.473073833 | 0.05531395 |
| Ppcs | 0.028095751 | 1.707263618 | 0.132 | 0.051 | 1 | 0.609924905 | 0.136765132 | 0.024 | 0.041 | 1 | 0.609924905 | 0.055402132 |
| 6330411D24Rik | 0.028114769 | 0.953088734 | 0.113 | 0.038 | 1 | 0.122497693 | 1.547698233 | 0.073 | 0.027 | 1 | 0.122497693 | 0.055439098 |
| Ppp6r2 | 0.028115758 | 1.172869942 | 0.377 | 0.258 | 1 | 0.859662472 | 0.597153762 | 0.268 | 0.317 | 1 | 0.859662472 | 0.055441021 |
| Sft2d2 | 0.961124785 | 0.179813576 | 0.321 | 0.318 | 1 | 0.028166869 | 0.811669758 | 0.537 | 0.389 | 1 | 0.961124785 | 0.055540366 |
| 1300002E11Rik | 0.031199332 | 0.804225348 | 0.302 | 0.169 | 1 | 0.028202311 | -1.622226769 | 0.049 | 0.184 | 1 | 0.031199332 | 0.055609252 |
| Pdhx | 0.028202352 | 0.652222254 | 0.453 | 0.318 | 1 | 0.379040428 | 0.690363461 | 0.366 | 0.341 | 1 | 0.379040428 | 0.055609331 |
| Pnkp | 0.945355438 | 0.280253477 | 0.113 | 0.11 | 1 | 0.028217507 | 0.871970268 | 0.341 | 0.201 | 1 | 0.945355438 | 0.055638786 |
| Txn2 | 0.028220306 | -0.459470617 | 0.585 | 0.737 | 1 | 0.636595333 | 0.199390467 | 0.659 | 0.727 | 1 | 0.636595333 | 0.055644226 |
| Ttyh2 | 0.028235037 | 0.932330173 | 0.226 | 0.114 | 1 | 0.384276622 | 0.851144041 | 0.22 | 0.164 | 1 | 0.384276622 | 0.055672856 |
| 1700018L02Rik | 0.209009104 | 1.739685096 | 0.038 | 0.013 | 1 | 0.028242983 | 2.252242349 | 0.073 | 0.017 | 1 | 0.209009104 | 0.0556883 |
| Gm16861 | 0.926653673 | 0.739685096 | 0.019 | 0.021 | 1 | 0.028242983 | 2.252242349 | 0.073 | 0.017 | 1 | 0.926653673 | 0.0556883 |
| Mfsd6 | 0.75990011 | 0.47665069 | 0.057 | 0.047 | 1 | 0.02824574 | 1.004314836 | 0.293 | 0.157 | 1 | 0.75990011 | 0.055693659 |
| N4bp2l1 | 0.028261674 | 1.154722595 | 0.226 | 0.114 | 1 | 0.131885241 | -0.940402729 | 0.098 | 0.198 | 1 | 0.131885241 | 0.055724626 |
| Acta2 | 0.028289472 | 0.410967835 | 0.17 | 0.352 | 1 | 0.505258673 | -1.879002184 | 0.049 | 0.078 | 1 | 0.505258673 | 0.05577865 |
| Pygb | 0.269438213 | 0.595295186 | 0.245 | 0.182 | 1 | 0.028373285 | 0.709449303 | 0.439 | 0.29 | 1 | 0.269438213 | 0.055941527 |
| Cog6 | 0.70587409 | 0.322845354 | 0.283 | 0.271 | 1 | 0.028403017 | -1.152741486 | 0.171 | 0.334 | 1 | 0.70587409 | 0.055999304 |
| Uqcc1 | 0.604137613 | -0.160150743 | 0.472 | 0.5 | 1 | 0.028462361 | -0.615119882 | 0.39 | 0.57 | 1 | 0.604137613 | 0.056114616 |
| Rae1 | 0.028464834 | 0.759421314 | 0.509 | 0.373 | 1 | 0.030386244 | -0.600200463 | 0.22 | 0.423 | 1 | 0.030386244 | 0.05611942 |
| Crppa | 0.036187779 | 0.953088734 | 0.377 | 0.25 | 1 | 0.028499825 | -2.010792057 | 0.098 | 0.239 | 1 | 0.036187779 | 0.056187409 |
| Ldlrad2 | 0.028501494 | -2.430239906 | 0 | 0.085 | 1 | 0.404252812 | -0.20718927 | 0.049 | 0.089 | 1 | 0.404252812 | 0.056190654 |
| Gm47675 | 0.028501494 | -2.368839361 | 0 | 0.085 | 1 | 0.126453962 | -1.250257991 | 0 | 0.055 | 1 | 0.126453962 | 0.056190654 |
| A330048O09Rik | 0.028501494 | -2.368839361 | 0 | 0.085 | 1 | 0.231591308 | -0.863234868 | 0 | 0.034 | 1 | 0.231591308 | 0.056190654 |
| Prkaa2 | 0.186653649 | -0.371000703 | 0.189 | 0.288 | 1 | 0.028504982 | -1.05356608 | 0.463 | 0.59 | 1 | 0.186653649 | 0.056197431 |
| P2ry2 | 0.028517007 | -2.489133595 | 0 | 0.085 | 1 | 0.319641967 | -0.16279515 | 0 | 0.024 | 1 | 0.319641967 | 0.056220795 |
| Pamr1 | 0.028517766 | 1.100274811 | 0.321 | 0.195 | 1 | 0.326024373 | 1.056986058 | 0.22 | 0.177 | 1 | 0.326024373 | 0.056222269 |
| Fhad1 | 0.028542834 | -3.545717123 | 0 | 0.085 | 1 | 0.911761241 | 0.252242349 | 0.024 | 0.027 | 1 | 0.911761241 | 0.056270974 |
| Plce1 | 0.880712889 | -0.218086669 | 0.604 | 0.623 | 1 | 0.028553617 | 1.310575659 | 0.585 | 0.485 | 1 | 0.880712889 | 0.056291924 |
| Zfp830 | 0.028566771 | 0.720319771 | 0.453 | 0.305 | 1 | 0.336165995 | -0.332720152 | 0.341 | 0.406 | 1 | 0.336165995 | 0.056317482 |

| | | | | | | | | | | | | |
|---|---|---|---|---|---|---|---|---|---|---|---|---|
| Pdzk1ip1 | 0.028579494 | -3.054730771 | 0 | 0.085 | 1 | 0.45455539 | 0.515276755 | 0.098 | 0.065 | 1 | 0.45455539 | 0.0563422 |
| Pde1b | 0.028600435 | -2.237594828 | 0 | 0.085 | 1 | 0.402373368 | 0.252242349 | 0 | 0.017 | 1 | 0.402373368 | 0.056382884 |
| Rnf216 | 0.02861846 | 0.434991538 | 0.83 | 0.792 | 1 | 0.763031247 | 0.203815659 | 0.878 | 0.922 | 1 | 0.763031247 | 0.056417904 |
| Uchl5 | 0.028633908 | 0.75033234 | 0.396 | 0.258 | 1 | 0.26277949 | -0.715336173 | 0.293 | 0.358 | 1 | 0.26277949 | 0.056447915 |
| Ppm1d | 0.028638517 | 0.717858733 | 0.415 | 0.28 | 1 | 0.088186037 | 0.849277682 | 0.561 | 0.485 | 1 | 0.088186037 | 0.05645687 |
| Nipsnap3b | 0.461562188 | 0.454282877 | 0.226 | 0.186 | 1 | 0.028646842 | 1.170942247 | 0.317 | 0.181 | 1 | 0.461562188 | 0.056473042 |
| Emid1 | 0.17690986 | 1.444229212 | 0.113 | 0.064 | 1 | 0.02871132 | 1.989207943 | 0.073 | 0.017 | 1 | 0.17690986 | 0.0565983 |
| Rsph3b | 0.02876677 | -1.08474334 | 0.113 | 0.254 | 1 | 0.530212157 | -0.180717058 | 0.146 | 0.191 | 1 | 0.530212157 | 0.056706013 |
| Ppp2r3d | 0.028790691 | 0.857740857 | 0.434 | 0.309 | 1 | 0.804377348 | -0.106211622 | 0.244 | 0.259 | 1 | 0.804377348 | 0.056752479 |
| Dcps | 0.167833773 | 0.569760094 | 0.321 | 0.229 | 1 | 0.02883531 | -0.52536523 | 0.049 | 0.191 | 1 | 0.167833773 | 0.056839144 |
| Zfp583 | 0.028838275 | 1.669295768 | 0.132 | 0.051 | 1 | 0.904354508 | 0.334704509 | 0.098 | 0.092 | 1 | 0.904354508 | 0.056844904 |
| Atg13 | 0.028838865 | 1.169222164 | 0.396 | 0.271 | 1 | 0.344067315 | 0.526417313 | 0.366 | 0.314 | 1 | 0.344067315 | 0.056846049 |
| Inhba | 0.028862275 | -5.53677169 | 0.113 | 0.229 | 1 | 0.051360728 | 1.515276755 | 0.073 | 0.02 | 1 | 0.051360728 | 0.05689152 |
| Dpm1 | 0.028868327 | 0.378278949 | 0.887 | 0.881 | 1 | 0.4420316 | 0.2061727 | 0.902 | 0.86 | 1 | 0.4420316 | 0.056903273 |
| Nab1 | 0.028874429 | 0.472462893 | 0.736 | 0.568 | 1 | 0.074558938 | -0.284424595 | 0.659 | 0.823 | 1 | 0.074558938 | 0.056915125 |
| Galnt4 | 0.028877389 | 1.377115016 | 0.189 | 0.089 | 1 | 0.849087975 | 0.993930353 | 0.171 | 0.177 | 1 | 0.849087975 | 0.056920875 |
| Msrb3 | 0.208431261 | -0.199420125 | 0.755 | 0.831 | 1 | 0.028897107 | 0.210493367 | 0.78 | 0.884 | 1 | 0.208431261 | 0.056959172 |
| Ddi2 | 0.705194074 | -0.217733059 | 0.736 | 0.763 | 1 | 0.028912372 | -0.490623847 | 0.829 | 0.952 | 1 | 0.705194074 | 0.056988819 |
| Zfp941 | 0.030360955 | 2.739685096 | 0.038 | 0.004 | 1 | 0.028943483 | 1.667279848 | 0.073 | 0.017 | 1 | 0.030360955 | 0.05704924 |
| Niban3 | 0.7394666 | -0.1672055 | 0.019 | 0.013 | 1 | 0.028943483 | 1.667279848 | 0.073 | 0.017 | 1 | 0.7394666 | 0.05704924 |
| Rpusd3 | 0.929410101 | 0.569760094 | 0.019 | 0.017 | 1 | 0.028943483 | 2.029849928 | 0.073 | 0.017 | 1 | 0.929410101 | 0.05704924 |
| Zfp397 | 0.081705749 | 0.620744023 | 0.509 | 0.403 | 1 | 0.028971956 | -1.008285201 | 0.268 | 0.44 | 1 | 0.081705749 | 0.057104539 |
| Abcb9 | 0.134153145 | -1.074096096 | 0.132 | 0.216 | 1 | 0.028999171 | -1.31254227 | 0.122 | 0.283 | 1 | 0.134153145 | 0.057157391 |
| Zfp653 | 0.029035976 | 1.154722595 | 0.226 | 0.114 | 1 | 0.430098895 | 0.174239837 | 0.171 | 0.235 | 1 | 0.430098895 | 0.057228865 |
| Alx4 | 0.029085212 | 0.962077517 | 0.434 | 0.305 | 1 | 0.105744111 | -0.421529419 | 0.268 | 0.406 | 1 | 0.105744111 | 0.057324474 |
| 1110019D14Rik | 0.269878427 | -0.430239906 | 0.094 | 0.153 | 1 | 0.029098636 | -2.352619709 | 0.024 | 0.147 | 1 | 0.269878427 | 0.057350541 |
| Ssbp1 | 0.127753816 | 0.389187848 | 0.585 | 0.521 | 1 | 0.029117922 | -0.742021281 | 0.341 | 0.505 | 1 | 0.127753816 | 0.057387991 |
| Adprm | 0.029143359 | -1.282682718 | 0.075 | 0.203 | 1 | 0.069252509 | -0.970150072 | 0.098 | 0.222 | 1 | 0.069252509 | 0.057437382 |
| Rragd | 0.02914791 | 2.614154213 | 0.094 | 0.03 | 1 | 0.455731299 | -0.385187572 | 0.049 | 0.082 | 1 | 0.455731299 | 0.057446219 |
| Wasf1 | 0.02914791 | 2.739685096 | 0.094 | 0.03 | 1 | 0.733713289 | 0.515276755 | 0.049 | 0.038 | 1 | 0.733713289 | 0.057446219 |
| Mocos | 0.02914791 | 2.324647596 | 0.094 | 0.03 | 1 | 0.276574583 | -1.069685746 | 0.024 | 0.068 | 1 | 0.276574583 | 0.057446219 |
| Gm56524 | 0.675133299 | 0.154722595 | 0.075 | 0.059 | 1 | 0.029154486 | -2.20718927 | 0 | 0.106 | 1 | 0.675133299 | 0.057458989 |
| Sergef | 0.029205606 | 1.932330173 | 0.151 | 0.064 | 1 | 0.265737307 | -0.395455907 | 0.049 | 0.106 | 1 | 0.265737307 | 0.057558245 |
| Pecr | 0.564535875 | -0.108311811 | 0.132 | 0.165 | 1 | 0.029216922 | -2.372248516 | 0 | 0.106 | 1 | 0.564535875 | 0.057580215 |
| Gm17178 | 0.772807467 | -0.600164907 | 0.038 | 0.047 | 1 | 0.029250855 | -2.448197369 | 0 | 0.106 | 1 | 0.772807467 | 0.057646097 |
| Obp2a | 0.029318963 | 0.855878113 | 0.453 | 0.326 | 1 | 0.155634518 | -2.622226769 | 0.098 | 0.181 | 1 | 0.155634518 | 0.057778324 |
| Adra1b | 0.3257977 | -0.347777746 | 0.094 | 0.144 | 1 | 0.029332809 | -3.093532488 | 0 | 0.106 | 1 | 0.3257977 | 0.057805204 |
| Trappc8 | 0.02940968 | 0.631160639 | 0.566 | 0.453 | 1 | 0.734549511 | 0.101349448 | 0.61 | 0.683 | 1 | 0.734549511 | 0.057954431 |
| Specc1l | 0.029417071 | 0.619126171 | 0.679 | 0.619 | 1 | 0.64091183 | -0.102188385 | 0.78 | 0.724 | 1 | 0.64091183 | 0.057968778 |
| Ttc1 | 0.029438651 | 0.635348436 | 0.528 | 0.386 | 1 | 0.607646205 | 0.132660733 | 0.293 | 0.362 | 1 | 0.607646205 | 0.058010668 |
| Nav1 | 0.029461338 | -0.447048194 | 0.509 | 0.695 | 1 | 0.451044402 | 0.354422744 | 0.634 | 0.58 | 1 | 0.451044402 | 0.058054706 |
| Pea15a | 0.029469157 | -0.357983817 | 0.472 | 0.657 | 1 | 0.042685444 | -0.562454548 | 0.537 | 0.703 | 1 | 0.042685444 | 0.058069882 |
| Akirin1 | 0.029531963 | 0.478725018 | 0.736 | 0.559 | 1 | 0.383806957 | -0.315079993 | 0.756 | 0.782 | 1 | 0.383806957 | 0.058191788 |
| Coa6 | 0.029540686 | 0.991223863 | 0.302 | 0.174 | 1 | 0.120578778 | 0.893788378 | 0.22 | 0.13 | 1 | 0.120578778 | 0.05820872 |
| Dynlt2b | 0.137277907 | 0.723383283 | 0.396 | 0.305 | 1 | 0.029573302 | -0.126269274 | 0.122 | 0.304 | 1 | 0.137277907 | 0.058272024 |
| Gm42912 | 0.029576974 | 3.47665069 | 0.038 | 0.004 | 1 | 0.51919986 | -0.16279515 | 0 | 0.01 | 1 | 0.51919986 | 0.058279151 |
| Fgf12 | 0.029576974 | 3.739685096 | 0.038 | 0.004 | 1 | 0.749728988 | 0.377773231 | 0.024 | 0.017 | 1 | 0.749728988 | 0.058279151 |
| Degs1 | 0.169938401 | -0.440223994 | 0.83 | 0.835 | 1 | 0.029610715 | 0.520731185 | 0.829 | 0.754 | 1 | 0.169938401 | 0.058344636 |
| Sec22c | 0.831317297 | 0.491757582 | 0.075 | 0.068 | 1 | 0.02963089 | 1.782757066 | 0.171 | 0.072 | 1 | 0.831317297 | 0.0583837 |
| Ech1 | 0.029644777 | 0.540376287 | 0.698 | 0.593 | 1 | 0.246076603 | 0.441276173 | 0.634 | 0.601 | 1 | 0.246076603 | 0.058410742 |
| Gm43466 | 0.241966592 | 0.932330173 | 0.075 | 0.038 | 1 | 0.029673399 | 1.029849928 | 0.073 | 0.017 | 1 | 0.241966592 | 0.058466287 |
| Lck | 0.606409362 | 1.306725688 | 0.038 | 0.025 | 1 | 0.029673399 | 1.667279848 | 0.073 | 0.017 | 1 | 0.606409362 | 0.058466287 |
| Ncbp1 | 0.029712499 | 0.949047568 | 0.509 | 0.377 | 1 | 0.642170558 | 0.231025863 | 0.585 | 0.577 | 1 | 0.642170558 | 0.058542165 |
| Kcnb1 | 0.029810352 | 0.480122882 | 0.283 | 0.161 | 1 | 0.067866097 | -0.728648828 | 0.146 | 0.29 | 1 | 0.067866097 | 0.058732047 |
| Rint1 | 0.478872937 | -0.28137652 | 0.415 | 0.445 | 1 | 0.029823532 | -0.708763519 | 0.341 | 0.529 | 1 | 0.478872937 | 0.058757621 |
| Cntnap2 | 0.029849695 | 1.042022462 | 0.415 | 0.28 | 1 | 0.25274582 | -0.27321914 | 0.195 | 0.3 | 1 | 0.25274582 | 0.058808386 |
| Fra10ac1 | 0.029881445 | 0.939709704 | 0.321 | 0.191 | 1 | 0.139180631 | -0.589059905 | 0.098 | 0.198 | 1 | 0.139180631 | 0.05886999 |
| Ahnak2 | 0.074950654 | 1.359542387 | 0.208 | 0.123 | 1 | 0.029935756 | 1.377773231 | 0.171 | 0.072 | 1 | 0.074950654 | 0.058975362 |
| 4930444A19Rik | 0.029966098 | 3.154722595 | 0.038 | 0.004 | 1 | 0.60213981 | 1.252242349 | 0.024 | 0.014 | 1 | 0.60213981 | 0.05903423 |
| Inava | 0.029966098 | 3.47665069 | 0.038 | 0.004 | 1 | 0.746881137 | 0.83720485 | 0.024 | 0.017 | 1 | 0.746881137 | 0.05903423 |
| Gm13381 | 0.029966098 | 3.154722595 | 0.038 | 0.004 | 1 | 0.51919986 | 0.83720485 | 0 | 0.01 | 1 | 0.51919986 | 0.05903423 |
| Klhl41 | 0.029966098 | 3.154722595 | 0.038 | 0.004 | 1 | 0.455098275 | 0.515276755 | 0 | 0.014 | 1 | 0.455098275 | 0.05903423 |
| 8030451A03Rik | 0.029966098 | 3.154722595 | 0.038 | 0.004 | 1 | 0.267205177 | 2.252242349 | 0.024 | 0.007 | 1 | 0.267205177 | 0.05903423 |
| Frmd3 | 0.029966098 | 3.154722595 | 0.038 | 0.004 | 1 | 0.749728988 | 0.667279848 | 0.024 | 0.017 | 1 | 0.749728988 | 0.05903423 |
| Gm42670 | 0.029966098 | 3.154722595 | 0.038 | 0.004 | 1 | 0.257325989 | -0.484723245 | 0 | 0.031 | 1 | 0.257325989 | 0.05903423 |
| Zfp114 | 0.029966098 | 3.154722595 | 0.038 | 0.004 | 1 | 0.646976884 | 0.721727632 | 0.049 | 0.034 | 1 | 0.646976884 | 0.05903423 |
| Zfp575 | 0.029966098 | 3.154722595 | 0.038 | 0.004 | 1 | 0.267205177 | 2.252242349 | 0.024 | 0.007 | 1 | 0.267205177 | 0.05903423 |
| Tyrobp | 0.029966098 | 4.47665069 | 0.038 | 0.004 | 1 | 0.880265303 | -2.116991461 | 0.073 | 0.065 | 1 | 0.880265303 | 0.05903423 |
| Entrep2 | 0.029966098 | 3.47665069 | 0.038 | 0.004 | 1 | 0.262674569 | 2.83720485 | 0.024 | 0.007 | 1 | 0.262674569 | 0.05903423 |
| Sez6l2 | 0.029966098 | 3.154722595 | 0.038 | 0.004 | 1 | 0.5191986 | 0.515276755 | 0 | 0.01 | 1 | 0.5191986 | 0.05903423 |
| Snap91 | 0.029966098 | 3.47665069 | 0.038 | 0.004 | 1 | 0.284887489 | 1.322631677 | 0.073 | 0.038 | 1 | 0.284887489 | 0.05903423 |
| Hkdc1 | 0.029966098 | 3.154722595 | 0.038 | 0.004 | 1 | 0.928869203 | 1.029849928 | 0.024 | 0.027 | 1 | 0.928869203 | 0.05903423 |
| Ogdhl | 0.029966098 | 3.154722595 | 0.038 | 0.004 | 1 | 0.455098275 | 0.515276755 | 0 | 0.014 | 1 | 0.455098275 | 0.05903423 |
| Gpr183 | 0.029966098 | 3.154722595 | 0.038 | 0.004 | 1 | 0.357858789 | -1.80665134 | 0 | 0.02 | 1 | 0.357858789 | 0.05903423 |
| Smpd5 | 0.029966098 | 3.154722595 | 0.038 | 0.004 | 1 | 0.106248001 | 2.252242349 | 0.024 | 0.003 | 1 | 0.106248001 | 0.05903423 |
| Krt4 | 0.029966098 | 3.154722595 | 0.038 | 0.004 | 1 | 0.104856732 | 2.83720485 | 0.024 | 0.003 | 1 | 0.104856732 | 0.05903423 |
| Syngr3 | 0.029966098 | 3.154722595 | 0.038 | 0.004 | 1 | 0.989595616 | 0.515276755 | 0.024 | 0.024 | 1 | 0.989595616 | 0.05903423 |
| Ms4a6c | 0.029966098 | 4.154722595 | 0.038 | 0.004 | 1 | 0.104856732 | 2.83720485 | 0.024 | 0.003 | 1 | 0.104856732 | 0.05903423 |
| Dtx4 | 0.029966098 | 3.154722595 | 0.038 | 0.004 | 1 | 0.110710459 | 2.83720485 | 0.049 | 0.014 | 1 | 0.110710459 | 0.05903423 |
| Usp51 | 0.029966098 | 3.154722595 | 0.038 | 0.004 | 1 | 0.430587622 | 2.574170444 | 0.024 | 0.01 | 1 | 0.430587622 | 0.05903423 |
| H4c4 | 0.02999467 | -1.392765201 | 0.17 | 0.314 | 1 | 0.451256773 | 0.136765132 | 0.22 | 0.28 | 1 | 0.451256773 | 0.05908966 |
| Ak8 | 0.953755983 | -0.1672055 | 0.057 | 0.059 | 1 | 0.029997125 | 1.721727632 | 0.146 | 0.058 | 1 | 0.953755983 | 0.059094423 |
| Snx14 | 0.030046462 | 0.811238357 | 0.604 | 0.466 | 1 | 0.077928112 | 0.474107566 | 0.683 | 0.58 | 1 | 0.077928112 | 0.059190134 |
| Ralgapb | 0.030098021 | 0.626622431 | 0.566 | 0.458 | 1 | 0.58600061 | 0.32941021 | 0.634 | 0.58 | 1 | 0.58600061 | 0.059290152 |
| Mrpl3 | 0.211104819 | 0.3662267 | 0.472 | 0.39 | 1 | 0.030107244 | -0.638528581 | 0.268 | 0.447 | 1 | 0.211104819 | 0.059308043 |
| Caap1 | 0.030128008 | 0.973115788 | 0.396 | 0.267 | 1 | 0.234179722 | -0.101394606 | 0.195 | 0.29 | 1 | 0.234179722 | 0.059348318 |
| Dcp1a | 0.108299352 | 0.469358197 | 0.453 | 0.343 | 1 | 0.030140653 | -0.404381137 | 0.22 | 0.427 | 1 | 0.108299352 | 0.059372848 |
| Gm41505 | 0.030146987 | 2.154722595 | 0.094 | 0.03 | 1 | 0.189685819 | 0.544423101 | 0.122 | 0.065 | 1 | 0.189685819 | 0.059385133 |
| Zdhhc16 | 0.030177293 | 1.07065833 | 0.321 | 0.199 | 1 | 0.094711181 | -1.020776145 | 0.122 | 0.232 | 1 | 0.094711181 | 0.059443916 |
| Klhl26 | 0.838737051 | -0.210926878 | 0.208 | 0.212 | 1 | 0.030210705 | -1.136799942 | 0.122 | 0.28 | 1 | 0.838737051 | 0.059508723 |
| Upf3b | 0.030230978 | 0.826795759 | 0.623 | 0.538 | 1 | 0.419827339 | 0.312614967 | 0.634 | 0.601 | 1 | 0.419827339 | 0.059548044 |
| Dkk3 | 0.030243725 | 0.62817678 | 0.453 | 0.301 | 1 | 0.076827682 | -0.33568561 | 0.439 | 0.628 | 1 | 0.076827682 | 0.059572767 |
| Lsm6 | 0.030271877 | -0.574380882 | 0.547 | 0.665 | 1 | 0.084137715 | -0.127296469 | 0.439 | 0.601 | 1 | 0.084137715 | 0.059627367 |
| Pde4a | 0.623721263 | 0.68265415 | 0.226 | 0.28 | 1 | 0.030330118 | -0.917682652 | 0.146 | 0.314 | 1 | 0.623721263 | 0.05974032 |
| A030005L19Rik | 0.030360955 | 2.739685096 | 0.038 | 0.004 | 1 | 0.605005904 | 0.667279848 | 0.024 | 0.014 | 1 | 0.605005904 | 0.059800122 |
| Fasl | 0.030360955 | 2.739685096 | 0.038 | 0.004 | 1 | 0.106248001 | 2.252242349 | 0.024 | 0.003 | 1 | 0.106248001 | 0.059800122 |
| Zfp661 | 0.030360955 | 2.739685096 | 0.038 | 0.004 | 1 | 0.267205177 | 2.252242349 | 0.024 | 0.007 | 1 | 0.267205177 | 0.059800122 |
| Myh7b | 0.030360955 | 2.739685096 | 0.038 | 0.004 | 1 | 0.267205177 | 2.252242349 | 0.024 | 0.007 | 1 | 0.267205177 | 0.059800122 |

| | | | | | | | | | | | | |
|---|---|---|---|---|---|---|---|---|---|---|---|---|
| Stmn3 | 0.030360955 | 2.739685096 | 0.038 | 0.004 | 1 | 0.107398884 | 3.100239256 | 0.049 | 0.014 | 1 | 0.107398884 | 0.059800122 |
| 4933429H19Rik | 0.030360955 | 2.739685096 | 0.038 | 0.004 | 1 | 0.51919986 | 0.83720485 | 0 | 0.01 | 1 | 0.51919986 | 0.059800122 |
| Kcnmb3 | 0.030360955 | 2.739685096 | 0.038 | 0.004 | 1 | 0.264933923 | 2.422167351 | 0.024 | 0.007 | 1 | 0.264933923 | 0.059800122 |
| 1110032F04Rik | 0.030360955 | 2.739685096 | 0.038 | 0.004 | 1 | 0.749728988 | 0.83720485 | 0.024 | 0.017 | 1 | 0.749728988 | 0.059800122 |
| Gm42701 | 0.030360955 | 2.739685096 | 0.038 | 0.004 | 1 | 0.51919986 | 0.83720485 | 0 | 0.01 | 1 | 0.51919986 | 0.059800122 |
| Cavin4 | 0.030360955 | 2.739685096 | 0.038 | 0.004 | 1 | 0.056289003 | 2.83720485 | 0.049 | 0.01 | 1 | 0.056289003 | 0.059800122 |
| Abcb4 | 0.030360955 | 2.739685096 | 0.038 | 0.004 | 1 | 0.357854401 | -0.484723245 | 0 | 0.02 | 1 | 0.357854401 | 0.059800122 |
| Gm17216 | 0.030360955 | 2.739685096 | 0.038 | 0.004 | 1 | 0.262674569 | 3.252242349 | 0.024 | 0.007 | 1 | 0.262674569 | 0.059800122 |
| Oscar | 0.030360955 | 2.739685096 | 0.038 | 0.004 | 1 | 0.357860983 | -2.332720152 | 0 | 0.02 | 1 | 0.357860983 | 0.059800122 |
| Gm15873 | 0.030360955 | 2.739685096 | 0.038 | 0.004 | 1 | 0.441381862 | 1.515276755 | 0.024 | 0.01 | 1 | 0.441381862 | 0.059800122 |
| Zfp109 | 0.030360955 | 2.739685096 | 0.038 | 0.004 | 1 | 0.590756788 | 1.83720485 | 0.024 | 0.014 | 1 | 0.590756788 | 0.059800122 |
| Nfkbid | 0.030360955 | 2.739685096 | 0.038 | 0.004 | 1 | 0.51919986 | 0.83720485 | 0 | 0.01 | 1 | 0.51919986 | 0.059800122 |
| Gm15489 | 0.030360955 | 2.739685096 | 0.038 | 0.004 | 1 | 0.319629309 | -0.484723245 | 0 | 0.024 | 1 | 0.319629309 | 0.059800122 |
| Gm20735 | 0.030360955 | 2.739685096 | 0.038 | 0.004 | 1 | 0.666353037 | -0.250257991 | 0.024 | 0.038 | 1 | 0.666353037 | 0.059800122 |
| Trim67 | 0.030360955 | 2.739685096 | 0.038 | 0.004 | 1 | 0.104856732 | 2.83720485 | 0.024 | 0.003 | 1 | 0.104856732 | 0.059800122 |
| Nnmt | 0.030360955 | 2.739685096 | 0.038 | 0.004 | 1 | 0.103479965 | 3.422167351 | 0.024 | 0.003 | 1 | 0.103479965 | 0.059800122 |
| C2cd4b | 0.030360955 | 2.739685096 | 0.038 | 0.004 | 1 | 0.936209655 | 1.515276755 | 0.024 | 0.027 | 1 | 0.936209655 | 0.059800122 |
| Gm17745 | 0.030360955 | 2.739685096 | 0.038 | 0.004 | 1 | 0.87556513 | 0.83720485 | 0.024 | 0.02 | 1 | 0.87556513 | 0.059800122 |
| Myl6b | 0.030360955 | 2.739685096 | 0.038 | 0.004 | 1 | 0.057901606 | 2.422167351 | 0.049 | 0.01 | 1 | 0.057901606 | 0.059800122 |
| Cntnap1 | 0.030360955 | 2.739685096 | 0.038 | 0.004 | 1 | 0.430584486 | 2.83720485 | 0.024 | 0.01 | 1 | 0.430584486 | 0.059800122 |
| Fmnl1 | 0.030360955 | 2.739685096 | 0.038 | 0.004 | 1 | 0.51919986 | 0.83720485 | 0 | 0.01 | 1 | 0.51919986 | 0.059800122 |
| Gm11747 | 0.030360955 | 2.739685096 | 0.038 | 0.004 | 1 | 0.170390036 | -1.332720152 | 0 | 0.044 | 1 | 0.170390036 | 0.059800122 |
| 2210039B01Rik | 0.030360955 | 2.739685096 | 0.038 | 0.004 | 1 | 0.51919986 | 0.83720485 | 0 | 0.01 | 1 | 0.51919986 | 0.059800122 |
| Gm6270 | 0.030360955 | 2.739685096 | 0.038 | 0.004 | 1 | 0.441381862 | 1.515276755 | 0.024 | 0.01 | 1 | 0.441381862 | 0.059800122 |
| Gm47814 | 0.030360955 | 2.739685096 | 0.038 | 0.004 | 1 | 0.916649754 | 0.667279848 | 0.024 | 0.027 | 1 | 0.916649754 | 0.059800122 |
| 1700109G14Rik | 0.030360955 | 2.739685096 | 0.038 | 0.004 | 1 | 0.109050642 | 2.83720485 | 0.049 | 0.014 | 1 | 0.109050642 | 0.059800122 |
| Gm36329 | 0.030360955 | 2.739685096 | 0.038 | 0.004 | 1 | 0.262674569 | 2.83720485 | 0.024 | 0.007 | 1 | 0.262674569 | 0.059800122 |
| Plcxd2 | 0.030360955 | 2.739685096 | 0.038 | 0.004 | 1 | 0.455094184 | 0.252242349 | 0 | 0.014 | 1 | 0.455094184 | 0.059800122 |
| Aif1 | 0.030360955 | 2.739685096 | 0.038 | 0.004 | 1 | 0.319619815 | -0.332720152 | 0 | 0.024 | 1 | 0.319619815 | 0.059800122 |
| Gm17949 | 0.030360955 | 2.739685096 | 0.038 | 0.004 | 1 | 0.455098275 | 0.515276755 | 0 | 0.014 | 1 | 0.455098275 | 0.059800122 |
| Prob1 | 0.030360955 | 2.739685096 | 0.038 | 0.004 | 1 | 0.455098275 | 0.515276755 | 0 | 0.014 | 1 | 0.455098275 | 0.059800122 |
| Hspe1-rs1 | 0.030360955 | 2.739685096 | 0.038 | 0.004 | 1 | 0.51919986 | 0.83720485 | 0 | 0.01 | 1 | 0.51919986 | 0.059800122 |
| Sema4g | 0.030360955 | 2.739685096 | 0.038 | 0.004 | 1 | 0.110710459 | 2.515276755 | 0.049 | 0.014 | 1 | 0.110710459 | 0.059800122 |
| Hps6 | 0.030360955 | 2.739685096 | 0.038 | 0.004 | 1 | 0.080188895 | 1.721727632 | 0.098 | 0.038 | 1 | 0.080188895 | 0.059800122 |
| Arntl2 | 0.030384522 | -1.1672055 | 0.113 | 0.258 | 1 | 0.125718898 | -0.369646721 | 0.22 | 0.348 | 1 | 0.125718898 | 0.059845825 |
| Tmem100 | 0.030404232 | 0.630860546 | 0.774 | 0.597 | 1 | 0.29073548 | -0.301098172 | 0.439 | 0.543 | 1 | 0.29073548 | 0.059884047 |
| Rnf43 | 0.494603095 | 0.417757001 | 0.038 | 0.021 | 1 | 0.030417909 | 1.377773231 | 0.073 | 0.017 | 1 | 0.494603095 | 0.05991057 |
| B230307C23Rik | 0.03048351 | 1.256602209 | 0.245 | 0.136 | 1 | 0.593623769 | -0.122153166 | 0.122 | 0.157 | 1 | 0.593623769 | 0.060037776 |
| Fam162a | 0.03049701 | -0.412752479 | 0.698 | 0.775 | 1 | 0.375695324 | 0.302515556 | 0.707 | 0.727 | 1 | 0.375695324 | 0.060063952 |
| Bpifb5 | 0.383389535 | -1.227747042 | 0.094 | 0.136 | 1 | 0.030509458 | -0.787286015 | 0.049 | 0.188 | 1 | 0.383389535 | 0.060088089 |
| Bcam | 0.287772304 | -1.652632327 | 0 | 0.021 | 1 | 0.030520058 | 1.483567895 | 0.171 | 0.072 | 1 | 0.287772304 | 0.060108641 |
| Vsir | 0.193514436 | 0.588375772 | 0.283 | 0.212 | 1 | 0.030523096 | -1.4233227 | 0.098 | 0.249 | 1 | 0.193514436 | 0.060114533 |
| Gm14547 | 0.030528769 | 2.154722595 | 0.094 | 0.03 | 1 | 0.5191986 | 0.515276755 | 0 | 0.01 | 1 | 0.5191986 | 0.060125533 |
| Gm10353 | 0.030561566 | 1.154722595 | 0.113 | 0.038 | 1 | 0.862957356 | 1.667279848 | 0.073 | 0.085 | 1 | 0.862957356 | 0.060189122 |
| Fabp5 | 0.030568179 | 0.802878922 | 0.472 | 0.356 | 1 | 0.300412345 | 0.712313602 | 0.415 | 0.324 | 1 | 0.300412345 | 0.060201945 |
| Arrdc1 | 0.030572618 | 1.669295768 | 0.132 | 0.051 | 1 | 0.451338156 | -0.577832649 | 0.049 | 0.082 | 1 | 0.451338156 | 0.060210551 |
| Arl6ip1 | 0.030620614 | 0.266655873 | 0.887 | 0.729 | 1 | 0.059350812 | 0.479964274 | 0.878 | 0.785 | 1 | 0.059350812 | 0.060303606 |
| Clcf1 | 0.030640686 | 0.8327945 | 0.189 | 0.089 | 1 | 0.06263815 | 0.658867609 | 0.317 | 0.188 | 1 | 0.06263815 | 0.06034252 |
| Pdxdc1 | 0.030643507 | 0.530871081 | 0.906 | 0.822 | 1 | 0.048626409 | -0.182973032 | 0.854 | 0.932 | 1 | 0.048626409 | 0.06034799 |
| Eef1akmt1 | 0.389350972 | -0.459987249 | 0.358 | 0.398 | 1 | 0.030670498 | -1.087067824 | 0.195 | 0.355 | 1 | 0.389350972 | 0.060400316 |
| Akap8l | 0.031223837 | 0.463475301 | 0.774 | 0.708 | 1 | 0.030711218 | 0.57223264 | 0.878 | 0.7 | 1 | 0.031223837 | 0.060479256 |
| Pcdhb20 | 0.030745068 | 2.002719501 | 0.094 | 0.03 | 1 | 0.716340863 | 1.029849928 | 0.049 | 0.038 | 1 | 0.716340863 | 0.060544876 |
| Mink1 | 0.938356091 | -0.110249615 | 0.34 | 0.356 | 1 | 0.030750037 | -0.626451355 | 0.366 | 0.553 | 1 | 0.938356091 | 0.060554509 |
| Fam114a1 | 0.343349196 | 0.223521592 | 0.849 | 0.903 | 1 | 0.030753457 | -0.428935467 | 0.78 | 0.86 | 1 | 0.343349196 | 0.06056114 |
| Plbd1 | 0.030759595 | 2.569760094 | 0.038 | 0.004 | 1 | 0.106248001 | 1.83720485 | 0.024 | 0.003 | 1 | 0.106248001 | 0.060573038 |
| Itih4 | 0.030759595 | 2.569760094 | 0.038 | 0.004 | 1 | 0.599290422 | 1.515276755 | 0.024 | 0.014 | 1 | 0.599290422 | 0.060573038 |
| Lix1 | 0.030791823 | 1.221836791 | 0.132 | 0.051 | 1 | 0.126502874 | -1.890715605 | 0.024 | 0.096 | 1 | 0.126502874 | 0.060635509 |
| Cep112 | 0.118754643 | -0.375465683 | 0.415 | 0.564 | 1 | 0.030797459 | -0.344813224 | 0.268 | 0.522 | 1 | 0.118754643 | 0.060646435 |
| Sh3bgrl | 0.030814618 | 0.557636535 | 0.774 | 0.742 | 1 | 0.107360202 | 0.279209397 | 0.902 | 0.867 | 1 | 0.107360202 | 0.060679695 |
| Vamp8 | 0.030819029 | -0.494619134 | 0.792 | 0.805 | 1 | 0.656696651 | -0.145236779 | 0.732 | 0.785 | 1 | 0.656696651 | 0.060688246 |
| Hilpda | 0.15586679 | -1.15270593 | 0.321 | 0.407 | 1 | 0.030860576 | -1.928329897 | 0.098 | 0.239 | 1 | 0.15586679 | 0.060768777 |
| H2ax | 0.03088403 | -0.334662246 | 0.472 | 0.314 | 1 | 0.135330698 | -0.747757651 | 0.146 | 0.249 | 1 | 0.135330698 | 0.060814237 |
| Gdi1 | 0.503307076 | 0.168661786 | 0.528 | 0.521 | 1 | 0.030923081 | -0.747757651 | 0.512 | 0.611 | 1 | 0.503307076 | 0.060889925 |
| Zfp326 | 0.030925403 | -0.520357201 | 0.868 | 0.936 | 1 | 0.391524777 | -0.185878763 | 0.878 | 0.928 | 1 | 0.391524777 | 0.060894426 |
| Gm5617 | 0.033296673 | -1.015202407 | 0.151 | 0.288 | 1 | 0.030937735 | -1.767657208 | 0.049 | 0.181 | 1 | 0.033296673 | 0.060918327 |
| Mrps27 | 0.112179473 | 0.538051234 | 0.245 | 0.148 | 1 | 0.03093881 | -1.727579769 | 0.049 | 0.181 | 1 | 0.112179473 | 0.06092041 |
| Hook2 | 0.03094917 | 1.491757582 | 0.151 | 0.064 | 1 | 0.174352866 | 1.045791472 | 0.171 | 0.099 | 1 | 0.174352866 | 0.060940489 |
| Phb | 0.030952021 | 0.974150349 | 0.472 | 0.339 | 1 | 0.596352713 | 0.241825733 | 0.268 | 0.341 | 1 | 0.596352713 | 0.060946015 |
| Ermp1 | 0.03096806 | 1.361173472 | 0.189 | 0.089 | 1 | 0.275815992 | 1.330244861 | 0.195 | 0.143 | 1 | 0.275815992 | 0.060977099 |
| Prpf38a | 0.031004067 | 0.671205358 | 0.547 | 0.445 | 1 | 0.062332893 | -0.544504473 | 0.39 | 0.536 | 1 | 0.062332893 | 0.061046881 |
| Aktip | 0.460480634 | 0.515821287 | 0.377 | 0.335 | 1 | 0.031046187 | 0.90971735 | 0.561 | 0.41 | 1 | 0.460480634 | 0.061128508 |
| Mrto4 | 0.037462812 | 0.760443656 | 0.396 | 0.263 | 1 | 0.031062253 | -0.769452722 | 0.146 | 0.324 | 1 | 0.037462812 | 0.061159643 |
| Edem3 | 0.031068985 | 0.405530698 | 0.698 | 0.581 | 1 | 0.571476994 | 0.199774929 | 0.732 | 0.747 | 1 | 0.571476994 | 0.061172688 |
| Gm46430 | 0.031106123 | 1.47665069 | 0.132 | 0.051 | 1 | 0.074168298 | 1.389745873 | 0.171 | 0.085 | 1 | 0.074168298 | 0.061244655 |
| Ubr7 | 0.0311363 | 0.666436113 | 0.453 | 0.292 | 1 | 0.913902941 | 0.584224109 | 0.317 | 0.348 | 1 | 0.913902941 | 0.061303132 |
| Ppp4c | 0.069819614 | 0.405266056 | 0.642 | 0.559 | 1 | 0.031140455 | 0.480087326 | 0.829 | 0.73 | 1 | 0.069819614 | 0.061311182 |
| Gm57292 | 0.031162042 | 1.154722595 | 0.038 | 0.004 | 1 | 0.231606831 | -0.970150072 | 0 | 0.034 | 1 | 0.231606831 | 0.061353011 |
| Tmed6 | 0.031162042 | 2.154722595 | 0.038 | 0.004 | 1 | 0.104856732 | 2.83720485 | 0.024 | 0.003 | 1 | 0.104856732 | 0.061353011 |
| Mlip | 0.031162042 | 0.932330173 | 0.038 | 0.004 | 1 | 0.110710459 | 2.515276755 | 0.049 | 0.014 | 1 | 0.110710459 | 0.061353011 |
| Dio3os | 0.031162042 | 2.154722595 | 0.038 | 0.004 | 1 | 0.51919986 | 0.83720485 | 0 | 0.01 | 1 | 0.51919986 | 0.061353011 |
| Rxfp3 | 0.031162042 | 2.154722595 | 0.038 | 0.004 | 1 | 0.304397715 | -0.747757651 | 0.024 | 0.065 | 1 | 0.304397715 | 0.061353011 |
| Grhl2 | 0.031162042 | 1.417757001 | 0.038 | 0.004 | 1 | 0.070809483 | 0.979223855 | 0.171 | 0.082 | 1 | 0.070809483 | 0.061353011 |
| Tbc1d10c | 0.031162042 | 2.154722595 | 0.038 | 0.004 | 1 | 0.110710459 | 2.515276755 | 0.049 | 0.014 | 1 | 0.110710459 | 0.061353011 |
| Galc | 0.031168368 | 1.46284489 | 0.17 | 0.076 | 1 | 0.419529726 | 0.807457506 | 0.171 | 0.13 | 1 | 0.419529726 | 0.061365268 |
| Fam50a | 0.031195566 | 0.774691545 | 0.585 | 0.496 | 1 | 0.111753004 | -0.309636539 | 0.463 | 0.584 | 1 | 0.111753004 | 0.061417968 |
| Nubpl | 0.232953386 | -0.340042097 | 0.132 | 0.208 | 1 | 0.031227567 | -1.508569987 | 0.024 | 0.15 | 1 | 0.232953386 | 0.061479972 |
| St6gal1 | 0.588510346 | 0.380282295 | 0.132 | 0.106 | 1 | 0.031230731 | 1.764448507 | 0.195 | 0.092 | 1 | 0.588510346 | 0.061486104 |
| Prkce | 0.031261689 | -0.748464032 | 0.547 | 0.708 | 1 | 0.2022591 | 0.567212893 | 0.805 | 0.71 | 1 | 0.2022591 | 0.061546084 |
| Nln | 0.140585615 | 1.358255989 | 0.17 | 0.106 | 1 | 0.031274177 | 1.068530396 | 0.293 | 0.157 | 1 | 0.140585615 | 0.06157028 |
| Sestd1 | 0.323330102 | -0.576558294 | 0.679 | 0.695 | 1 | 0.031275746 | -0.63075504 | 0.512 | 0.652 | 1 | 0.323330102 | 0.061573319 |
| Oca2 | 0.435798053 | -0.582242999 | 0.057 | 0.089 | 1 | 0.031328467 | -2.292078167 | 0.049 | 0.181 | 1 | 0.435798053 | 0.06167546 |
| Dbt | 0.031344191 | 1.315187267 | 0.226 | 0.114 | 1 | 0.433145568 | -0.448197369 | 0.073 | 0.113 | 1 | 0.433145568 | 0.061705924 |
| Gm39150 | 0.031359353 | -1.771276824 | 0.075 | 0.195 | 1 | 0.157263986 | -0.80665134 | 0.049 | 0.126 | 1 | 0.157263986 | 0.061735298 |
| Parg | 0.031374902 | 0.648088455 | 0.585 | 0.525 | 1 | 0.587553839 | 0.197306674 | 0.756 | 0.744 | 1 | 0.587553839 | 0.061765419 |
| Zfx | 0.031382784 | 0.552058092 | 0.66 | 0.581 | 1 | 0.758746434 | 0.18596913 | 0.805 | 0.843 | 1 | 0.758746434 | 0.061780689 |
| Trp53i11 | 0.708828774 | 0.484871196 | 0.113 | 0.097 | 1 | 0.03140111 | 1.715214325 | 0.146 | 0.058 | 1 | 0.708828774 | 0.06181619 |
| Itga2 | 0.070812095 | 1.8327945 | 0.075 | 0.025 | 1 | 0.031445608 | 0.898605394 | 0.122 | 0.041 | 1 | 0.070812095 | 0.061902389 |
| Zfp866 | 0.93375815 | 0.815920681 | 0.151 | 0.153 | 1 | 0.031492562 | -0.638528581 | 0.073 | 0.229 | 1 | 0.93375815 | 0.061993343 |

| | | | | | | | | | | | | |
|---|---|---|---|---|---|---|---|---|---|---|---|---|
| Nin | 0.031495608 | 0.145620388 | 0.547 | 0.411 | 1 | 0.84711079 | 0.143866188 | 0.488 | 0.522 | 1 | 0.84711079 | 0.061999242 |
| Metap1d | 0.031500309 | -0.376262285 | 0.547 | 0.708 | 1 | 0.062659459 | -0.250257991 | 0.39 | 0.584 | 1 | 0.062659459 | 0.062008348 |
| Gm20646 | 0.031562905 | 2.154722595 | 0.094 | 0.03 | 1 | 0.789697987 | 0.515276755 | 0.049 | 0.061 | 1 | 0.789697987 | 0.062129593 |
| Dnajc27 | 0.031603361 | 0.647762606 | 0.264 | 0.14 | 1 | 0.919635468 | -0.126269274 | 0.171 | 0.174 | 1 | 0.919635468 | 0.062207949 |
| Rfx5 | 0.031658112 | 1.454282877 | 0.132 | 0.051 | 1 | 0.575066969 | 0.589277336 | 0.073 | 0.051 | 1 | 0.575066969 | 0.062313988 |
| Prmt2 | 0.461722868 | 0.589937976 | 0.245 | 0.212 | 1 | 0.031697264 | -0.821006633 | 0.098 | 0.259 | 1 | 0.461722868 | 0.062389811 |
| Isyna1 | 0.031713886 | -1.022007585 | 0.453 | 0.597 | 1 | 0.290763965 | -0.304975161 | 0.463 | 0.567 | 1 | 0.290763965 | 0.062422002 |
| Car2 | 0.031729379 | -3.81106169 | 0.019 | 0.114 | 1 | 0.257323068 | -1.80665134 | 0 | 0.031 | 1 | 0.257323068 | 0.062452005 |
| Cox11 | 0.031736521 | 0.666436113 | 0.434 | 0.297 | 1 | 0.358524078 | 0.667279848 | 0.366 | 0.324 | 1 | 0.358524078 | 0.062465836 |
| Tomm40l | 0.031773209 | 1.47665069 | 0.151 | 0.064 | 1 | 0.184479364 | 0.962735732 | 0.171 | 0.102 | 1 | 0.184479364 | 0.062536881 |
| Trappc6a | 0.031777193 | 0.711779099 | 0.434 | 0.292 | 1 | 0.93867786 | 0.451146417 | 0.268 | 0.297 | 1 | 0.93867786 | 0.062544596 |
| Creb5 | 0.031782476 | -0.38841973 | 0.113 | 0.263 | 1 | 0.202483245 | 0.602739596 | 0.293 | 0.212 | 1 | 0.202483245 | 0.062554827 |
| Flnb | 0.037642699 | -0.884079676 | 0.849 | 0.873 | 1 | 0.031916295 | -0.650417101 | 0.878 | 0.918 | 1 | 0.037642699 | 0.062813941 |
| Sh2d3c | 0.031916899 | 1.417757001 | 0.132 | 0.051 | 1 | 0.836467438 | -0.101394606 | 0.049 | 0.041 | 1 | 0.836467438 | 0.062815109 |
| Prorp | 0.031953713 | 1.086551092 | 0.264 | 0.144 | 1 | 0.421544064 | 0.811669758 | 0.146 | 0.106 | 1 | 0.421544064 | 0.062886386 |
| Tprn | 0.031963796 | 1.962077517 | 0.094 | 0.03 | 1 | 0.328574063 | 1.737669176 | 0.073 | 0.041 | 1 | 0.328574063 | 0.062905908 |
| Atxn7l2 | 0.031963796 | 1.962077517 | 0.094 | 0.03 | 1 | 0.103663347 | 1.120997816 | 0.146 | 0.072 | 1 | 0.103663347 | 0.062905908 |
| Npm2 | 0.031963796 | 1.962077517 | 0.094 | 0.03 | 1 | 0.039195975 | -2.20718927 | 0 | 0.096 | 1 | 0.039195975 | 0.062905908 |
| Hspbap1 | 0.158374335 | 0.640149422 | 0.377 | 0.314 | 1 | 0.031975898 | -0.332720152 | 0.22 | 0.423 | 1 | 0.158374335 | 0.062929337 |
| Gm49539 | 0.031981474 | 1.569760094 | 0.094 | 0.03 | 1 | 0.854674228 | -0.435813645 | 0.049 | 0.055 | 1 | 0.854674228 | 0.062940133 |
| Gm43948 | 0.031989043 | 1.444229212 | 0.151 | 0.064 | 1 | 0.266682704 | -0.435813645 | 0.024 | 0.072 | 1 | 0.266682704 | 0.062954787 |
| Prr3 | 0.032076024 | 0.79482665 | 0.396 | 0.267 | 1 | 0.543112705 | 0.422167351 | 0.293 | 0.263 | 1 | 0.543112705 | 0.063123177 |
| Gm14399 | 0.403474578 | 0.984797593 | 0.094 | 0.064 | 1 | 0.03216596 | -2.16279515 | 0 | 0.102 | 1 | 0.403474578 | 0.063297271 |
| Bicd1 | 0.032169258 | 1.117247889 | 0.17 | 0.076 | 1 | 0.633506765 | 1.593279267 | 0.098 | 0.078 | 1 | 0.633506765 | 0.063303654 |
| Rtl5 | 0.467027607 | -0.430239906 | 0.038 | 0.064 | 1 | 0.032210238 | -2.332720152 | 0 | 0.102 | 1 | 0.467027607 | 0.063382977 |
| Pld1 | 0.032222132 | 0.88695027 | 0.491 | 0.373 | 1 | 0.746575973 | 0.152706676 | 0.366 | 0.355 | 1 | 0.746575973 | 0.063405998 |
| Cdca4 | 0.252719199 | 0.403401708 | 0.396 | 0.318 | 1 | 0.032222777 | -0.693309867 | 0.268 | 0.444 | 1 | 0.252719199 | 0.063407247 |
| Niban1 | 0.032230899 | 0.869320376 | 0.415 | 0.275 | 1 | 0.229644919 | 0.665252167 | 0.561 | 0.491 | 1 | 0.229644919 | 0.063422966 |
| Decr1 | 0.393575733 | 0.436493563 | 0.377 | 0.347 | 1 | 0.032235299 | -0.958654433 | 0.146 | 0.311 | 1 | 0.393575733 | 0.063431484 |
| Cep20 | 0.154455571 | 0.535812762 | 0.377 | 0.275 | 1 | 0.032258069 | 1.002628459 | 0.415 | 0.266 | 1 | 0.154455571 | 0.063475556 |
| Socs7 | 0.032273236 | 0.743296349 | 0.491 | 0.347 | 1 | 0.592780235 | -0.232143197 | 0.488 | 0.512 | 1 | 0.592780235 | 0.063504911 |
| Ccsap | 0.04504193 | -2.015202407 | 0.019 | 0.106 | 1 | 0.032280849 | -2.747757651 | 0 | 0.102 | 1 | 0.04504193 | 0.063519645 |
| Mtm1 | 0.032313707 | -1.29778961 | 0.226 | 0.36 | 1 | 0.096982317 | -0.928329897 | 0.098 | 0.218 | 1 | 0.096982317 | 0.063583239 |
| Reps2 | 0.053161273 | 2.090592257 | 0.113 | 0.047 | 1 | 0.032334624 | 1.583448258 | 0.146 | 0.058 | 1 | 0.053161273 | 0.063623721 |
| P4ha3 | 0.032352373 | 0.964619711 | 0.434 | 0.309 | 1 | 0.640037469 | 0.661355014 | 0.244 | 0.222 | 1 | 0.640037469 | 0.063658069 |
| Gm41609 | 0.032387099 | -1.27154216 | 0.057 | 0.178 | 1 | 0.442583754 | -0.899760744 | 0.049 | 0.082 | 1 | 0.442583754 | 0.063725273 |
| Daglb | 0.032397106 | 1.331600357 | 0.17 | 0.076 | 1 | 0.737111594 | 0.667279848 | 0.098 | 0.123 | 1 | 0.737111594 | 0.06374464 |
| Rnf180 | 0.032418665 | 1.855162313 | 0.094 | 0.03 | 1 | 0.792088554 | 0.313642894 | 0.049 | 0.041 | 1 | 0.792088554 | 0.06378636 |
| Prdx3 | 0.743514181 | 0.154722595 | 0.623 | 0.61 | 1 | 0.032421451 | -0.607013675 | 0.366 | 0.532 | 1 | 0.743514181 | 0.063791751 |
| Zfp574 | 0.032469211 | 1.119098685 | 0.264 | 0.148 | 1 | 0.583796155 | 0.401105735 | 0.195 | 0.253 | 1 | 0.583796155 | 0.063884173 |
| Hps3 | 0.032481078 | -1.389597921 | 0.019 | 0.119 | 1 | 0.844879076 | -0.136799942 | 0.146 | 0.13 | 1 | 0.844879076 | 0.063907136 |
| Ndufv2 | 0.032511679 | 0.43225657 | 0.83 | 0.742 | 1 | 0.143259029 | -0.283497636 | 0.634 | 0.788 | 1 | 0.143259029 | 0.063966349 |
| Mta1 | 0.032574582 | 0.561347854 | 0.585 | 0.504 | 1 | 0.345046469 | -0.28502341 | 0.537 | 0.608 | 1 | 0.345046469 | 0.06408806 |
| Gm19265 | 0.032613361 | 2.06161319 | 0.132 | 0.051 | 1 | 0.090247959 | 1.071670103 | 0.098 | 0.038 | 1 | 0.090247959 | 0.06416309 |
| Wdr66 | 0.03264184 | 1.444229212 | 0.132 | 0.051 | 1 | 0.262016835 | -0.970150072 | 0.049 | 0.102 | 1 | 0.262016835 | 0.06421819 |
| Armh4 | 0.214042707 | -0.27937324 | 0.453 | 0.542 | 1 | 0.03266798 | -0.401934646 | 0.415 | 0.618 | 1 | 0.214042707 | 0.064268763 |
| Ap5z1 | 0.032694421 | 0.974150349 | 0.226 | 0.114 | 1 | 0.933938564 | 0.319356545 | 0.146 | 0.154 | 1 | 0.933938564 | 0.064319918 |
| Abi1 | 0.032697789 | -0.481378755 | 0.868 | 0.945 | 1 | 0.066524304 | -0.470089975 | 0.927 | 0.973 | 1 | 0.066524304 | 0.064326432 |
| Cactin | 0.03790536 | 1.084333267 | 0.17 | 0.076 | 1 | 0.032836495 | 1.370078839 | 0.244 | 0.126 | 1 | 0.03790536 | 0.064594754 |
| Mettl25 | 0.032864035 | 0.569760094 | 0.321 | 0.186 | 1 | 0.181171972 | -1.278272368 | 0.146 | 0.229 | 1 | 0.181171972 | 0.064648025 |
| Aacs | 0.032909939 | -1.036897478 | 0.302 | 0.441 | 1 | 0.522340639 | -0.124727109 | 0.317 | 0.372 | 1 | 0.522340639 | 0.064736814 |
| Lamb3 | 0.330295076 | -0.974560422 | 0.057 | 0.097 | 1 | 0.032912325 | 1.974708374 | 0.146 | 0.058 | 1 | 0.330295076 | 0.064741429 |
| Syde1 | 0.032923467 | 1.010332685 | 0.264 | 0.148 | 1 | 0.138072513 | 0.715214325 | 0.293 | 0.184 | 1 | 0.138072513 | 0.064762979 |
| 2310008N11Rik | 0.032937359 | -2.304709024 | 0.019 | 0.114 | 1 | 0.178315225 | -1.0454382 | 0.049 | 0.119 | 1 | 0.178315225 | 0.064789848 |
| Laptm4a | 0.032938157 | 0.205768889 | 1 | 0.996 | 1 | 0.307438951 | 0.174556252 | 1 | 1 | 1 | 0.307438951 | 0.064791392 |
| Etv5 | 0.032939921 | 1.695290976 | 0.302 | 0.199 | 1 | 0.73264281 | -0.115661175 | 0.341 | 0.386 | 1 | 0.73264281 | 0.064794803 |
| Slc22a5 | 0.239884163 | -0.27244866 | 0.396 | 0.292 | 1 | 0.03300754 | 1.295495547 | 0.488 | 0.355 | 1 | 0.239884163 | 0.064925583 |
| Bbof1 | 0.113798068 | -0.845277405 | 0.038 | 0.11 | 1 | 0.033010901 | 2.252242349 | 0.122 | 0.044 | 1 | 0.113798068 | 0.064932081 |
| Klf9 | 0.033053077 | 0.268893615 | 1 | 0.919 | 1 | 0.143603119 | 0.293202669 | 0.976 | 0.966 | 1 | 0.143603119 | 0.065013648 |
| Gm46367 | 0.033055249 | 1.347367673 | 0.094 | 0.03 | 1 | 0.172798647 | 0.667279848 | 0.024 | 0.089 | 1 | 0.172798647 | 0.065017848 |
| Wdhd1 | 0.033070021 | -2.459987249 | 0.019 | 0.114 | 1 | 0.705906068 | 1.218295017 | 0.122 | 0.106 | 1 | 0.705906068 | 0.065046415 |
| Hexb | 0.03308823 | 0.399835093 | 0.717 | 0.606 | 1 | 0.762282142 | -0.315045259 | 0.634 | 0.737 | 1 | 0.762282142 | 0.065081628 |
| Sympk | 0.033100441 | 1.039245377 | 0.34 | 0.229 | 1 | 0.762766562 | 0.142059431 | 0.317 | 0.287 | 1 | 0.762766562 | 0.065105243 |
| Scube3 | 0.033133834 | -2.489133595 | 0 | 0.081 | 1 | 0.37203631 | 0.614812428 | 0.049 | 0.024 | 1 | 0.37203631 | 0.065169817 |
| Msto1 | 0.033150316 | 1.039245377 | 0.189 | 0.089 | 1 | 0.624558346 | -0.300298674 | 0.098 | 0.123 | 1 | 0.624558346 | 0.065201689 |
| Gm17225 | 0.033179794 | -3.304709024 | 0 | 0.081 | 1 | 0.602692481 | -0.484723245 | 0.024 | 0.041 | 1 | 0.602692481 | 0.06525869 |
| Cpa6 | 0.033193064 | -3.889671525 | 0 | 0.081 | 1 | 0.2088551 | -2.622226769 | 0 | 0.038 | 1 | 0.2088551 | 0.065284349 |
| Brap | 0.033233051 | 0.657222935 | 0.491 | 0.369 | 1 | 0.604738781 | 0.19334866 | 0.512 | 0.625 | 1 | 0.604738781 | 0.065361667 |
| Ddx19b | 0.033268349 | 1.3662267 | 0.151 | 0.064 | 1 | 0.783652209 | -0.233184478 | 0.098 | 0.109 | 1 | 0.783652209 | 0.065429916 |
| Spg20 | 0.03327852 | 0.729115854 | 0.528 | 0.39 | 1 | 0.513362081 | 0.577337723 | 0.39 | 0.379 | 1 | 0.513362081 | 0.065449579 |
| Enc1 | 0.068047114 | 0.810768194 | 0.151 | 0.072 | 1 | 0.033294104 | 2.83720485 | 0.098 | 0.031 | 1 | 0.068047114 | 0.06547971 |
| Fkbp8 | 0.033297933 | 0.68155092 | 0.679 | 0.619 | 1 | 0.345761411 | -0.191592415 | 0.488 | 0.57 | 1 | 0.345761411 | 0.065487113 |
| Smg8 | 0.937735213 | -0.108311811 | 0.17 | 0.161 | 1 | 0.033316779 | -1.292078167 | 0.049 | 0.184 | 1 | 0.937735213 | 0.06552355 |
| Poc5 | 0.03331845 | 1.243989933 | 0.264 | 0.144 | 1 | 0.058722943 | -0.732650759 | 0.098 | 0.229 | 1 | 0.058722943 | 0.065526781 |
| Dipk2a | 0.033326956 | -0.373312838 | 0.868 | 0.903 | 1 | 0.185204165 | -0.187945241 | 0.854 | 0.915 | 1 | 0.185204165 | 0.065543226 |
| Abhd16a | 0.03334173 | 0.650250012 | 0.509 | 0.386 | 1 | 0.391810597 | 0.377773231 | 0.415 | 0.372 | 1 | 0.391810597 | 0.065571789 |
| Adam32 | 0.033359627 | 1.050385935 | 0.17 | 0.076 | 1 | 0.934410222 | -0.240797662 | 0.098 | 0.102 | 1 | 0.934410222 | 0.065606389 |
| Exosc3 | 0.060531027 | -0.479483425 | 0.302 | 0.458 | 1 | 0.03341472 | -0.798383724 | 0.22 | 0.406 | 1 | 0.060531027 | 0.065712896 |
| Ak6 | 0.080294758 | 0.607234799 | 0.434 | 0.331 | 1 | 0.033417215 | -0.760357688 | 0.341 | 0.478 | 1 | 0.080294758 | 0.065717721 |
| Itgav | 0.033429201 | 0.405237531 | 0.906 | 0.847 | 1 | 0.119747374 | 0.23825812 | 0.902 | 0.846 | 1 | 0.119747374 | 0.06574089 |
| Itga1 | 0.063135313 | -1.105804955 | 0.132 | 0.25 | 1 | 0.033480825 | -0.883272622 | 0.098 | 0.259 | 1 | 0.063135313 | 0.065840685 |
| Nt5m | 0.033504812 | 0.655916738 | 0.34 | 0.199 | 1 | 0.137251992 | 0.816143234 | 0.268 | 0.177 | 1 | 0.137251992 | 0.065887051 |
| Fancg | 0.033507074 | 1.739685096 | 0.094 | 0.03 | 1 | 0.643119211 | 0.83720485 | 0.049 | 0.034 | 1 | 0.643119211 | 0.065891425 |
| Poc1a | 0.033507074 | 1.739685096 | 0.094 | 0.03 | 1 | 0.891167418 | 0.377773231 | 0.049 | 0.055 | 1 | 0.891167418 | 0.065891425 |
| 1810021B22Rik | 0.033507074 | 1.739685096 | 0.094 | 0.03 | 1 | 0.674180157 | 0.252242349 | 0.024 | 0.038 | 1 | 0.674180157 | 0.065891425 |
| Avpi1 | 0.033574942 | -1.340972568 | 0.34 | 0.462 | 1 | 0.243340987 | 0.644559772 | 0.39 | 0.307 | 1 | 0.243340987 | 0.066022607 |
| Aldh3a2 | 0.599811324 | -0.260314904 | 0.094 | 0.119 | 1 | 0.033584286 | 0.989207943 | 0.22 | 0.102 | 1 | 0.599811324 | 0.066040668 |
| Ttc17 | 0.033591922 | 0.634855122 | 0.585 | 0.449 | 1 | 0.093717848 | 0.703938319 | 0.634 | 0.618 | 1 | 0.093717848 | 0.066055427 |
| Dhx57 | 0.033609205 | 1.081966252 | 0.453 | 0.322 | 1 | 0.502360363 | 0.151978226 | 0.512 | 0.43 | 1 | 0.502360363 | 0.066088832 |
| Dars | 0.221776485 | -0.140238452 | 0.491 | 0.648 | 1 | 0.033629051 | -0.340712942 | 0.439 | 0.642 | 1 | 0.221776485 | 0.06612719 |
| Inf2 | 0.385287163 | -0.477545621 | 0.321 | 0.36 | 1 | 0.033679379 | -0.863234868 | 0.244 | 0.42 | 1 | 0.385287163 | 0.066224458 |
| Ccng1 | 0.033685462 | -0.932740246 | 0.302 | 0.458 | 1 | 0.403195669 | -0.347219721 | 0.341 | 0.403 | 1 | 0.403195669 | 0.066236214 |
| Pth2r | 0.033715692 | 1.47665069 | 0.094 | 0.03 | 1 | 0.270588322 | 1.737669176 | 0.073 | 0.038 | 1 | 0.270588322 | 0.066294635 |
| Vps13a | 0.033737859 | 0.385728146 | 0.66 | 0.547 | 1 | 0.111450816 | 0.314978104 | 0.756 | 0.642 | 1 | 0.111450816 | 0.066337474 |
| Pdia5 | 0.377547881 | -0.296488517 | 0.34 | 0.403 | 1 | 0.033740098 | -0.567778985 | 0.293 | 0.481 | 1 | 0.377547881 | 0.066341801 |
| Tspan17 | 0.033745484 | 0.976385354 | 0.264 | 0.144 | 1 | 0.234887523 | 1.071670103 | 0.098 | 0.051 | 1 | 0.234887523 | 0.066352211 |
| Tmem29 | 0.033755025 | -0.736901244 | 0.226 | 0.386 | 1 | 0.208336367 | 0.164779508 | 0.146 | 0.256 | 1 | 0.208336367 | 0.066370648 |
| Ippk | 0.033772451 | 0.5310277 | 0.434 | 0.288 | 1 | 0.905459824 | 0.221545552 | 0.341 | 0.372 | 1 | 0.905459824 | 0.066404324 |

| | | | | | | | | | | | | |
|---|---|---|---|---|---|---|---|---|---|---|---|---|
| Shc1 | 0.033780125 | 0.52198437 | 0.698 | 0.657 | 1 | 0.049248551 | 0.41590836 | 0.78 | 0.788 | 1 | 0.049248551 | 0.066419153 |
| Thoc1 | 0.545439209 | 0.129187503 | 0.547 | 0.521 | 1 | 0.033840996 | -0.560989139 | 0.463 | 0.601 | 1 | 0.545439209 | 0.066536778 |
| Mtmr10 | 0.033844843 | -0.493565841 | 0.226 | 0.407 | 1 | 0.135505534 | -0.26713181 | 0.244 | 0.382 | 1 | 0.135505534 | 0.066544212 |
| Gm43707 | 0.761196836 | -0.430239906 | 0.038 | 0.047 | 1 | 0.033937722 | 2.209173627 | 0.122 | 0.044 | 1 | 0.761196836 | 0.066723674 |
| Wdr41 | 0.89637746 | 0.200526284 | 0.264 | 0.288 | 1 | 0.034009581 | -0.374845627 | 0.122 | 0.297 | 1 | 0.89637746 | 0.066862511 |
| Arl8b | 0.038560988 | 0.425619118 | 0.755 | 0.695 | 1 | 0.034061432 | 0.471749379 | 0.854 | 0.785 | 1 | 0.038560988 | 0.066962683 |
| 2900052L18Rik | 0.265220076 | 0.695290976 | 0.113 | 0.068 | 1 | 0.034071634 | 1.898605394 | 0.146 | 0.058 | 1 | 0.265220076 | 0.066982392 |
| Phyhd1 | 0.034091339 | 0.962077517 | 0.264 | 0.148 | 1 | 0.596484337 | -0.17660095 | 0.195 | 0.232 | 1 | 0.596484337 | 0.067020459 |
| Noxred1 | 0.03414753 | 1.640149422 | 0.094 | 0.03 | 1 | 0.319379144 | 1.515276755 | 0.098 | 0.058 | 1 | 0.319379144 | 0.067129007 |
| Wdr5b | 0.03414753 | 1.640149422 | 0.094 | 0.03 | 1 | 0.103916103 | -1.410722664 | 0 | 0.061 | 1 | 0.103916103 | 0.067129007 |
| Slc45a1 | 0.034157605 | 1.569760094 | 0.094 | 0.03 | 1 | 0.059076736 | 1.614812428 | 0.098 | 0.034 | 1 | 0.059076736 | 0.067148468 |
| Kri1 | 0.034167193 | 0.906795081 | 0.396 | 0.275 | 1 | 0.066521703 | 1.644559772 | 0.317 | 0.208 | 1 | 0.066521703 | 0.06716699 |
| Herpud1 | 0.03420043 | -0.414120241 | 0.943 | 0.941 | 1 | 0.050012504 | -0.296218596 | 0.878 | 0.949 | 1 | 0.050012504 | 0.067231191 |
| Efhb | 0.034302114 | 1.104096522 | 0.151 | 0.064 | 1 | 0.178140676 | 1.252242349 | 0.122 | 0.065 | 1 | 0.178140676 | 0.067427593 |
| Ddx42 | 0.034351645 | 0.393562496 | 0.868 | 0.767 | 1 | 0.763940562 | 0.246528698 | 0.829 | 0.867 | 1 | 0.763940562 | 0.067523255 |
| Ltbp3 | 0.03435666 | 0.493138812 | 0.811 | 0.716 | 1 | 0.224522336 | 0.376500452 | 0.732 | 0.744 | 1 | 0.224522336 | 0.06753294 |
| Ube2v2 | 0.068789255 | 0.480950827 | 0.698 | 0.614 | 1 | 0.034399688 | -0.223677392 | 0.561 | 0.747 | 1 | 0.068789255 | 0.067616038 |
| Zscan12 | 0.034399718 | 1.154722595 | 0.189 | 0.089 | 1 | 0.096205992 | -1.410722664 | 0.024 | 0.106 | 1 | 0.096205992 | 0.067616095 |
| Trp53bp2 | 0.034473663 | 0.569760094 | 0.547 | 0.419 | 1 | 0.14243297 | 0.374498099 | 0.78 | 0.659 | 1 | 0.14243297 | 0.067758892 |
| Clk3 | 0.03453035 | 1.002719501 | 0.321 | 0.199 | 1 | 0.08991873 | 1.191488318 | 0.366 | 0.256 | 1 | 0.08991873 | 0.067868356 |
| Ccng2 | 0.037040273 | 0.94321849 | 0.245 | 0.131 | 1 | 0.034531043 | 1.438655473 | 0.244 | 0.13 | 1 | 0.037040273 | 0.067869693 |
| Tnnt3 | 0.034536883 | -3.538764363 | 0.057 | 0.165 | 1 | 0.039948516 | 2.043655727 | 0.146 | 0.061 | 1 | 0.039948516 | 0.06788097 |
| Ncstn | 0.034540256 | 0.794320352 | 0.434 | 0.309 | 1 | 0.292692959 | 0.508582102 | 0.512 | 0.485 | 1 | 0.292692959 | 0.067887483 |
| Trappc5 | 0.171974435 | 0.588375772 | 0.358 | 0.275 | 1 | 0.034555238 | -0.648221977 | 0.146 | 0.317 | 1 | 0.171974435 | 0.067916412 |
| Polr3f | 0.416577372 | -0.359850578 | 0.208 | 0.258 | 1 | 0.034620904 | -0.79700117 | 0.171 | 0.338 | 1 | 0.416577372 | 0.068043202 |
| Slc30a1 | 0.034680272 | 0.821298861 | 0.34 | 0.212 | 1 | 0.256255233 | 0.372082119 | 0.317 | 0.242 | 1 | 0.256255233 | 0.068157823 |
| Phf7 | 0.034717434 | 1.46284489 | 0.17 | 0.076 | 1 | 0.728628616 | 0.377773231 | 0.098 | 0.082 | 1 | 0.728628616 | 0.068229568 |
| Snx33 | 0.034764988 | 1.758793918 | 0.189 | 0.093 | 1 | 0.602541908 | 0.100239256 | 0.171 | 0.205 | 1 | 0.602541908 | 0.068321372 |
| Ptpdc1 | 0.034789034 | 1.337586652 | 0.226 | 0.123 | 1 | 0.155225036 | 1.224227973 | 0.22 | 0.143 | 1 | 0.155225036 | 0.068367791 |
| Il20rb | 0.034809633 | 0.819119563 | 0.264 | 0.148 | 1 | 0.815152438 | 0.252242349 | 0.244 | 0.232 | 1 | 0.815152438 | 0.068407555 |
| Nit2 | 0.034826295 | 0.695290976 | 0.321 | 0.186 | 1 | 0.712772902 | 0.515276755 | 0.195 | 0.181 | 1 | 0.712772902 | 0.068439719 |
| Acsl1 | 0.794361982 | 0.440477077 | 0.491 | 0.475 | 1 | 0.034893345 | 0.535085236 | 0.585 | 0.437 | 1 | 0.794361982 | 0.068569145 |
| Uqcrq | 0.034994713 | 0.365964892 | 0.906 | 0.788 | 1 | 0.055334733 | -0.419134903 | 0.61 | 0.761 | 1 | 0.055334733 | 0.068764797 |
| Zbtb40 | 0.035028282 | 1.221836791 | 0.17 | 0.076 | 1 | 0.776462094 | 0.444887427 | 0.122 | 0.109 | 1 | 0.776462094 | 0.068829583 |
| Mpdu1 | 0.035081395 | 0.790150163 | 0.472 | 0.347 | 1 | 0.77982733 | 0.759202338 | 0.293 | 0.311 | 1 | 0.77982733 | 0.068932086 |
| Zfp46 | 0.096619405 | 0.898382842 | 0.226 | 0.14 | 1 | 0.035089023 | -1.26713181 | 0.073 | 0.215 | 1 | 0.096619405 | 0.068946807 |
| Pkm | 0.035095984 | -0.313426241 | 0.981 | 0.979 | 1 | 0.186962468 | -0.252963891 | 0.976 | 0.983 | 1 | 0.186962468 | 0.068960239 |
| Casp9 | 0.377181002 | 0.417757001 | 0.094 | 0.059 | 1 | 0.035100287 | 1.252242349 | 0.195 | 0.089 | 1 | 0.377181002 | 0.068968544 |
| Gm32618 | 0.035122438 | 1.542793046 | 0.264 | 0.153 | 1 | 0.723816129 | 0.544423101 | 0.22 | 0.198 | 1 | 0.723816129 | 0.06901129 |
| Rad52 | 0.035162072 | 1.450178478 | 0.264 | 0.157 | 1 | 0.620918006 | 0.554270887 | 0.22 | 0.188 | 1 | 0.620918006 | 0.069087772 |
| Brf2 | 0.03518446 | 1.154722595 | 0.17 | 0.076 | 1 | 0.628309298 | 0.377773231 | 0.073 | 0.099 | 1 | 0.628309298 | 0.069130975 |
| Kif3c | 0.035188978 | 1.007881206 | 0.189 | 0.089 | 1 | 0.969981348 | 0.100239256 | 0.146 | 0.15 | 1 | 0.969981348 | 0.069139692 |
| Fastkd1 | 0.035241191 | 1.261637799 | 0.208 | 0.106 | 1 | 0.737064822 | 0.319356545 | 0.171 | 0.15 | 1 | 0.737064822 | 0.06924044 |
| Rsph9 | 0.035250019 | 1.197791317 | 0.226 | 0.119 | 1 | 0.909487215 | 0.742047617 | 0.122 | 0.137 | 1 | 0.909487215 | 0.069257474 |
| Aftph | 0.035250965 | 0.464407094 | 0.755 | 0.636 | 1 | 0.495746346 | -0.145517159 | 0.78 | 0.863 | 1 | 0.495746346 | 0.069259299 |
| Atp13a1 | 0.035365887 | 0.870929629 | 0.34 | 0.212 | 1 | 0.680253533 | -0.136799942 | 0.244 | 0.273 | 1 | 0.680253533 | 0.069481028 |
| Fsd1l | 0.035406006 | 1.154722595 | 0.226 | 0.119 | 1 | 0.396092399 | 0.210422173 | 0.146 | 0.215 | 1 | 0.396092399 | 0.069558427 |
| Dalrd3 | 0.035407363 | 0.984797593 | 0.283 | 0.161 | 1 | 0.52962959 | 0.997669522 | 0.22 | 0.191 | 1 | 0.52962959 | 0.069561045 |
| Atad5 | 0.73304466 | 0.454282877 | 0.057 | 0.072 | 1 | 0.035474967 | 2.33970519 | 0.122 | 0.044 | 1 | 0.73304466 | 0.069691461 |
| 1700008J07Rik | 0.631247702 | 0.874614676 | 0.075 | 0.059 | 1 | 0.035480608 | -2.116991461 | 0 | 0.099 | 1 | 0.631247702 | 0.069702342 |
| 2610027K06Rik | 0.222354209 | 1.247831999 | 0.094 | 0.051 | 1 | 0.035482249 | 2.185128153 | 0.098 | 0.031 | 1 | 0.222354209 | 0.069705508 |
| Idh3b | 0.035563858 | 0.48360345 | 0.679 | 0.551 | 1 | 0.117459547 | -0.255904555 | 0.415 | 0.577 | 1 | 0.117459547 | 0.069862928 |
| Aasdh | 0.284664606 | 1.007881206 | 0.151 | 0.102 | 1 | 0.035581454 | -2.410722664 | 0 | 0.099 | 1 | 0.284664606 | 0.069896869 |
| Lyn | 0.035588353 | 0.762405172 | 0.189 | 0.089 | 1 | 0.071754968 | 0.261175474 | 0.439 | 0.294 | 1 | 0.071754968 | 0.069910176 |
| Gm28323 | 0.03562064 | 3.739685096 | 0.019 | 0 | 1 | 0.104856732 | 2.83720485 | 0.024 | 0.003 | 1 | 0.104856732 | 0.069972449 |
| 1700019A02Rik | 0.03562064 | 3.154722595 | 0.019 | 0 | 1 | 0.262674569 | 2.83720485 | 0.024 | 0.007 | 1 | 0.262674569 | 0.069972449 |
| 5033417F24Rik | 0.03562064 | 3.154722595 | 0.019 | 0 | 1 | 0.455098275 | 0.515276755 | 0 | 0.014 | 1 | 0.455098275 | 0.069972449 |
| Tnfrsf11a | 0.03562064 | 3.154722595 | 0.019 | 0 | 1 | 0.257325016 | -2.890715605 | 0 | 0.031 | 1 | 0.257325016 | 0.069972449 |
| Gm26936 | 0.03562064 | 3.154722595 | 0.019 | 0 | 1 | 0.104856732 | 2.83720485 | 0.024 | 0.003 | 1 | 0.104856732 | 0.069972449 |
| Gm37864 | 0.03562064 | 3.154722595 | 0.019 | 0 | 1 | 0.319629309 | -0.484723245 | 0 | 0.024 | 1 | 0.319629309 | 0.069972449 |
| Cd48 | 0.03562064 | 3.154722595 | 0.019 | 0 | 1 | 0.402369942 | -0.16279515 | 0 | 0.017 | 1 | 0.402369942 | 0.069972449 |
| Slamf1 | 0.03562064 | 3.154722595 | 0.019 | 0 | 1 | 0.438672228 | 1.83720485 | 0.024 | 0.01 | 1 | 0.438672228 | 0.069972449 |
| Slamf9 | 0.03562064 | 3.154722595 | 0.019 | 0 | 1 | 0.455094184 | 0.252242349 | 0 | 0.014 | 1 | 0.455094184 | 0.069972449 |
| Cd34 | 0.03562064 | 3.154722595 | 0.019 | 0 | 1 | 0.262674569 | 3.83720485 | 0.024 | 0.007 | 1 | 0.262674569 | 0.069972449 |
| Lrp1b | 0.03562064 | 3.154722595 | 0.019 | 0 | 1 | 0.186073065 | 1.83720485 | 0.049 | 0.017 | 1 | 0.186073065 | 0.069972449 |
| Mageb3 | 0.03562064 | 3.154722595 | 0.019 | 0 | 1 | 0.106248001 | 2.252242349 | 0.024 | 0.003 | 1 | 0.106248001 | 0.069972449 |
| Tmem74bos | 0.03562064 | 3.154722595 | 0.019 | 0 | 1 | 0.51919986 | 0.83720485 | 0 | 0.01 | 1 | 0.51919986 | 0.069972449 |
| Mmp24 | 0.03562064 | 3.154722595 | 0.019 | 0 | 1 | 0.5191986 | 0.252242349 | 0 | 0.01 | 1 | 0.5191986 | 0.069972449 |
| Edn3 | 0.03562064 | 4.154722595 | 0.019 | 0 | 1 | 0.104856732 | 2.83720485 | 0.024 | 0.003 | 1 | 0.104856732 | 0.069972449 |
| Gm20633 | 0.03562064 | 3.154722595 | 0.019 | 0 | 1 | 0.267205177 | 2.252242349 | 0.024 | 0.007 | 1 | 0.267205177 | 0.069972449 |
| Gm43061 | 0.03562064 | 3.739685096 | 0.019 | 0 | 1 | 0.546365767 | 1.100239256 | 0.049 | 0.031 | 1 | 0.546365767 | 0.069972449 |
| Mybphl | 0.03562064 | 3.154722595 | 0.019 | 0 | 1 | 0.104856732 | 2.83720485 | 0.024 | 0.003 | 1 | 0.104856732 | 0.069972449 |
| Lrrc39 | 0.03562064 | 3.154722595 | 0.019 | 0 | 1 | 0.455094184 | 0.252242349 | 0 | 0.014 | 1 | 0.455094184 | 0.069972449 |
| Spag8 | 0.03562064 | 3.154722595 | 0.019 | 0 | 1 | 0.103479965 | 3.83720485 | 0.024 | 0.003 | 1 | 0.103479965 | 0.069972449 |
| Gm12689 | 0.03562064 | 3.154722595 | 0.019 | 0 | 1 | 0.114118767 | 2.100239256 | 0.049 | 0.014 | 1 | 0.114118767 | 0.069972449 |
| Lactbl1 | 0.03562064 | 3.154722595 | 0.019 | 0 | 1 | 0.26948834 | 1.029849928 | 0.024 | 0.007 | 1 | 0.26948834 | 0.069972449 |
| Pttg1ip2 | 0.03562064 | 3.739685096 | 0.019 | 0 | 1 | 0.51919986 | 0.83720485 | 0 | 0.01 | 1 | 0.51919986 | 0.069972449 |
| 9530036O11Rik | 0.03562064 | 3.154722595 | 0.019 | 0 | 1 | 0.430584486 | 2.422167351 | 0.024 | 0.01 | 1 | 0.430584486 | 0.069972449 |
| 1700126H18Rik | 0.03562064 | 3.154722595 | 0.019 | 0 | 1 | 0.402369942 | -0.622226769 | 0 | 0.017 | 1 | 0.402369942 | 0.069972449 |
| Gm15477 | 0.03562064 | 3.154722595 | 0.019 | 0 | 1 | 0.106248001 | 2.515276755 | 0.024 | 0.003 | 1 | 0.106248001 | 0.069972449 |
| Kit | 0.03562064 | 4.154722595 | 0.019 | 0 | 1 | 0.435967024 | 1.614812428 | 0.024 | 0.01 | 1 | 0.435967024 | 0.069972449 |
| 5430403G16Rik | 0.03562064 | 4.154722595 | 0.019 | 0 | 1 | 0.5191986 | 0.252242349 | 0 | 0.01 | 1 | 0.5191986 | 0.069972449 |
| Lrcol1 | 0.03562064 | 3.154722595 | 0.019 | 0 | 1 | 0.104856732 | 2.83720485 | 0.024 | 0.003 | 1 | 0.104856732 | 0.069972449 |
| Mmp17 | 0.03562064 | 3.739685096 | 0.019 | 0 | 1 | 0.593594236 | 1.614812428 | 0.024 | 0.014 | 1 | 0.593594236 | 0.069972449 |
| Clcn1 | 0.03562064 | 3.154722595 | 0.019 | 0 | 1 | 0.103479965 | 3.422167351 | 0.024 | 0.003 | 1 | 0.103479965 | 0.069972449 |
| Clec4a2 | 0.03562064 | 3.154722595 | 0.019 | 0 | 1 | 0.257323555 | -2.292078167 | 0 | 0.031 | 1 | 0.257323555 | 0.069972449 |
| ENSMUSG00000121470 | 0.03562064 | 3.739685096 | 0.019 | 0 | 1 | 0.186073065 | 1.83720485 | 0.049 | 0.017 | 1 | 0.186073065 | 0.069972449 |
| C5ar1 | 0.03562064 | 4.154722595 | 0.019 | 0 | 1 | 0.362195044 | 0.721727632 | 0.049 | 0.024 | 1 | 0.362195044 | 0.069972449 |
| Gm16175 | 0.03562064 | 3.154722595 | 0.019 | 0 | 1 | 0.872828964 | 1.029849928 | 0.024 | 0.02 | 1 | 0.872828964 | 0.069972449 |
| Gm15883 | 0.03562064 | 4.154722595 | 0.019 | 0 | 1 | 0.455098275 | 0.515276755 | 0 | 0.014 | 1 | 0.455098275 | 0.069972449 |
| Tdrd12 | 0.03562064 | 3.154722595 | 0.019 | 0 | 1 | 0.26948834 | 1.83720485 | 0.024 | 0.007 | 1 | 0.26948834 | 0.069972449 |
| Gm32633 | 0.03562064 | 3.154722595 | 0.019 | 0 | 1 | 0.267205177 | 2.252242349 | 0.024 | 0.007 | 1 | 0.267205177 | 0.069972449 |
| Art5 | 0.03562064 | 3.154722595 | 0.019 | 0 | 1 | 0.110710459 | 2.515276755 | 0.049 | 0.014 | 1 | 0.110710459 | 0.069972449 |
| Gm44987 | 0.03562064 | 3.154722595 | 0.019 | 0 | 1 | 0.51919986 | 0.83720485 | 0 | 0.01 | 1 | 0.51919986 | 0.069972449 |
| Clgn | 0.03562064 | 4.154722595 | 0.019 | 0 | 1 | 0.5191986 | 0.515276755 | 0 | 0.01 | 1 | 0.5191986 | 0.069972449 |
| Cngb1 | 0.03562064 | 4.739685096 | 0.019 | 0 | 1 | 0.51919986 | 0.83720485 | 0 | 0.01 | 1 | 0.51919986 | 0.069972449 |
| Dynlrb2 | 0.03562064 | 3.154722595 | 0.019 | 0 | 1 | 0.856409217 | 1.989207943 | 0.024 | 0.02 | 1 | 0.856409217 | 0.069972449 |
| Hsd17b2 | 0.03562064 | 3.154722595 | 0.019 | 0 | 1 | 0.5191986 | 0.515276755 | 0 | 0.01 | 1 | 0.5191986 | 0.069972449 |

| | | | | | | | | | | | | |
|---|---|---|---|---|---|---|---|---|---|---|---|---|
| Epor | 0.03562064 | 3.154722595 | 0.019 | 0 | 1 | 0.51919986 | 0.83720485 | 0 | 0.01 | 1 | 0.51919986 | 0.069972449 |
| AY074887 | 0.03562064 | 3.154722595 | 0.019 | 0 | 1 | 0.5191986 | 0.515276755 | 0 | 0.01 | 1 | 0.5191986 | 0.069972449 |
| Trcg1 | 0.03562064 | 3.154722595 | 0.019 | 0 | 1 | 0.599290422 | 1.515276755 | 0.024 | 0.014 | 1 | 0.599290422 | 0.069972449 |
| Hcn4 | 0.03562064 | 3.154722595 | 0.019 | 0 | 1 | 0.180328124 | 2.574170444 | 0.049 | 0.017 | 1 | 0.180328124 | 0.069972449 |
| Bcl2a1a | 0.03562064 | 3.739685096 | 0.019 | 0 | 1 | 0.402369942 | -0.622226769 | 0 | 0.017 | 1 | 0.402369942 | 0.069972449 |
| Bcl2a1b | 0.03562064 | 5.154722595 | 0.019 | 0 | 1 | 0.357860251 | -2.020776145 | 0 | 0.02 | 1 | 0.357860251 | 0.069972449 |
| Gadl1 | 0.03562064 | 3.739685096 | 0.019 | 0 | 1 | 0.455094184 | 0.252242349 | 0 | 0.014 | 1 | 0.455094184 | 0.069972449 |
| Lilrb4b | 0.03562064 | 3.739685096 | 0.019 | 0 | 1 | 0.402364804 | -0.747757651 | 0 | 0.017 | 1 | 0.402364804 | 0.069972449 |
| 4930467K11Rik | 0.03562064 | 3.154722595 | 0.019 | 0 | 1 | 0.599290422 | 1.515276755 | 0.024 | 0.014 | 1 | 0.599290422 | 0.069972449 |
| Itgb2 | 0.03562064 | 4.962077517 | 0.019 | 0 | 1 | 0.257320633 | -1.332720152 | 0 | 0.031 | 1 | 0.257320633 | 0.069972449 |
| Osm | 0.03562064 | 3.154722595 | 0.019 | 0 | 1 | 0.267205177 | 2.252242349 | 0.024 | 0.007 | 1 | 0.267205177 | 0.069972449 |
| Gm12122 | 0.03562064 | 3.739685096 | 0.019 | 0 | 1 | 0.106248001 | 1.83720485 | 0.024 | 0.003 | 1 | 0.106248001 | 0.069972449 |
| Myh3 | 0.03562064 | 3.739685096 | 0.019 | 0 | 1 | 0.262674569 | 3.252242349 | 0.024 | 0.007 | 1 | 0.262674569 | 0.069972449 |
| Gm11532 | 0.03562064 | 3.154722595 | 0.019 | 0 | 1 | 0.104856732 | 2.83720485 | 0.024 | 0.003 | 1 | 0.104856732 | 0.069972449 |
| Gm11725 | 0.03562064 | 3.739685096 | 0.019 | 0 | 1 | 0.60213981 | 1.252242349 | 0.024 | 0.014 | 1 | 0.60213981 | 0.069972449 |
| Fscn2 | 0.03562064 | 3.154722595 | 0.019 | 0 | 1 | 0.208841678 | -1.16279515 | 0 | 0.038 | 1 | 0.208841678 | 0.069972449 |
| Gm36752 | 0.03562064 | 3.154722595 | 0.019 | 0 | 1 | 0.103479965 | 3.83720485 | 0.024 | 0.003 | 1 | 0.103479965 | 0.069972449 |
| Gm32219 | 0.03562064 | 3.154722595 | 0.019 | 0 | 1 | 0.104856732 | 2.83720485 | 0.024 | 0.003 | 1 | 0.104856732 | 0.069972449 |
| Gm46378 | 0.03562064 | 3.739685096 | 0.019 | 0 | 1 | 0.104856732 | 2.83720485 | 0.024 | 0.003 | 1 | 0.104856732 | 0.069972449 |
| Dnah11 | 0.03562064 | 3.154722595 | 0.019 | 0 | 1 | 0.51919986 | 0.83720485 | 0 | 0.01 | 1 | 0.51919986 | 0.069972449 |
| Zkscan4 | 0.03562064 | 3.154722595 | 0.019 | 0 | 1 | 0.104856732 | 2.83720485 | 0.024 | 0.003 | 1 | 0.104856732 | 0.069972449 |
| Gm48623 | 0.03562064 | 3.154722595 | 0.019 | 0 | 1 | 0.262674569 | 3.252242349 | 0.024 | 0.007 | 1 | 0.262674569 | 0.069972449 |
| 5430425K12Rik | 0.03562064 | 3.739685096 | 0.019 | 0 | 1 | 0.51919986 | 0.83720485 | 0 | 0.01 | 1 | 0.51919986 | 0.069972449 |
| Ankdd1b | 0.03562064 | 3.739685096 | 0.019 | 0 | 1 | 0.430587622 | 2.515276755 | 0.024 | 0.01 | 1 | 0.430587622 | 0.069972449 |
| Gm30363 | 0.03562064 | 3.154722595 | 0.019 | 0 | 1 | 0.51919986 | -0.16279515 | 0 | 0.01 | 1 | 0.51919986 | 0.069972449 |
| Gm31392 | 0.03562064 | 3.739685096 | 0.019 | 0 | 1 | 0.104856732 | 2.83720485 | 0.024 | 0.003 | 1 | 0.104856732 | 0.069972449 |
| 4933413J09Rik | 0.03562064 | 3.154722595 | 0.019 | 0 | 1 | 0.744047924 | 1.252242349 | 0.024 | 0.017 | 1 | 0.744047924 | 0.069972449 |
| Gm15932 | 0.03562064 | 3.739685096 | 0.019 | 0 | 1 | 0.51919986 | 0.83720485 | 0 | 0.01 | 1 | 0.51919986 | 0.069972449 |
| Erich5 | 0.03562064 | 4.154722595 | 0.019 | 0 | 1 | 0.267205177 | 2.100239256 | 0.024 | 0.007 | 1 | 0.267205177 | 0.069972449 |
| Abra | 0.03562064 | 3.154722595 | 0.019 | 0 | 1 | 0.267205177 | 2.252242349 | 0.024 | 0.007 | 1 | 0.267205177 | 0.069972449 |
| Adamts20 | 0.03562064 | 3.154722595 | 0.019 | 0 | 1 | 0.104856732 | 2.83720485 | 0.024 | 0.003 | 1 | 0.104856732 | 0.069972449 |
| Nckap5los | 0.03562064 | 3.154722595 | 0.019 | 0 | 1 | 0.746881137 | 1.029849928 | 0.024 | 0.017 | 1 | 0.746881137 | 0.069972449 |
| Galnt6 | 0.03562064 | 3.154722595 | 0.019 | 0 | 1 | 0.357860251 | -2.020776145 | 0 | 0.02 | 1 | 0.357860251 | 0.069972449 |
| Smim41 | 0.03562064 | 3.154722595 | 0.019 | 0 | 1 | 0.267205177 | 2.252242349 | 0.024 | 0.007 | 1 | 0.267205177 | 0.069972449 |
| Nfe2 | 0.03562064 | 3.739685096 | 0.019 | 0 | 1 | 0.455097252 | -0.16279515 | 0 | 0.014 | 1 | 0.455097252 | 0.069972449 |
| Gm49482 | 0.03562064 | 3.154722595 | 0.019 | 0 | 1 | 0.267205177 | 2.252242349 | 0.024 | 0.007 | 1 | 0.267205177 | 0.069972449 |
| Pla2g10 | 0.03562064 | 3.154722595 | 0.019 | 0 | 1 | 0.976593771 | 0.83720485 | 0.024 | 0.024 | 1 | 0.976593771 | 0.069972449 |
| Cd86 | 0.03562064 | 3.154722595 | 0.019 | 0 | 1 | 0.455098275 | 0.515276755 | 0 | 0.014 | 1 | 0.455098275 | 0.069972449 |
| BC016579 | 0.03562064 | 3.154722595 | 0.019 | 0 | 1 | 0.402369942 | -0.332720152 | 0 | 0.017 | 1 | 0.402369942 | 0.069972449 |
| Adgrf4 | 0.03562064 | 3.154722595 | 0.019 | 0 | 1 | 0.455097252 | -0.332720152 | 0 | 0.014 | 1 | 0.455097252 | 0.069972449 |
| Gm35097 | 0.03562064 | 3.154722595 | 0.019 | 0 | 1 | 0.455094184 | 0.252242349 | 0 | 0.014 | 1 | 0.455094184 | 0.069972449 |
| ENSMUSG00000121397 | 0.03562064 | 3.154722595 | 0.019 | 0 | 1 | 0.51919986 | 0.83720485 | 0 | 0.01 | 1 | 0.51919986 | 0.069972449 |
| Cyp1b1 | 0.03562064 | 3.739685096 | 0.019 | 0 | 1 | 0.104856732 | 2.83720485 | 0.024 | 0.003 | 1 | 0.104856732 | 0.069972449 |
| Gjd4 | 0.03562064 | 3.154722595 | 0.019 | 0 | 1 | 0.455098275 | 0.515276755 | 0 | 0.014 | 1 | 0.455098275 | 0.069972449 |
| Stk32a | 0.03562064 | 3.739685096 | 0.019 | 0 | 1 | 0.103479965 | 3.159132945 | 0.024 | 0.003 | 1 | 0.103479965 | 0.069972449 |
| Mpeg1 | 0.03562064 | 4.47665069 | 0.019 | 0 | 1 | 0.455098275 | 0.515276755 | 0 | 0.014 | 1 | 0.455098275 | 0.069972449 |
| Entpd1 | 0.03562064 | 3.154722595 | 0.019 | 0 | 1 | 0.269106562 | 0.589277336 | 0.049 | 0.02 | 1 | 0.269106562 | 0.069972449 |
| Gm17203 | 0.03562064 | 3.739685096 | 0.019 | 0 | 1 | 0.587925597 | 2.252242349 | 0.024 | 0.014 | 1 | 0.587925597 | 0.069972449 |
| Lifr | 0.552471758 | -0.430239906 | 0.396 | 0.445 | 1 | 0.035704957 | 1.36186684 | 0.341 | 0.222 | 1 | 0.552471758 | 0.070135071 |
| Arid3a | 0.035705832 | 2.017219071 | 0.113 | 0.042 | 1 | 0.053168571 | 1.043655727 | 0.22 | 0.116 | 1 | 0.053168571 | 0.070136758 |
| Ankrd33b | 0.035728543 | 1.417757001 | 0.094 | 0.03 | 1 | 0.894713448 | 0.515276755 | 0.049 | 0.044 | 1 | 0.894713448 | 0.070180557 |
| Trmu | 0.035729349 | 1.292226119 | 0.226 | 0.123 | 1 | 0.053898198 | 1.75474269 | 0.195 | 0.099 | 1 | 0.053898198 | 0.070182111 |
| 2300009A05Rik | 0.5237135 | 0.20255112 | 0.245 | 0.314 | 1 | 0.035794704 | -0.610254127 | 0.146 | 0.304 | 1 | 0.5237135 | 0.070308146 |
| Gm47237 | 0.109903281 | -1.489133595 | 0.019 | 0.081 | 1 | 0.035815623 | -0.723510105 | 0.024 | 0.147 | 1 | 0.109903281 | 0.070348486 |
| Ccny | 0.035845458 | 0.414985251 | 0.887 | 0.818 | 1 | 0.514116675 | 0.107924004 | 0.902 | 0.932 | 1 | 0.514116675 | 0.070406019 |
| Impa2 | 0.365144349 | -0.525018131 | 0.34 | 0.398 | 1 | 0.035901415 | -0.555112573 | 0.244 | 0.437 | 1 | 0.365144349 | 0.070513918 |
| Gm10382 | 0.961556928 | 0.306725688 | 0.057 | 0.055 | 1 | 0.035935273 | 2.100239256 | 0.098 | 0.031 | 1 | 0.961556928 | 0.070579202 |
| Gm28809 | 0.035952331 | 1.221836791 | 0.151 | 0.064 | 1 | 0.885447308 | 0.604544093 | 0.122 | 0.119 | 1 | 0.885447308 | 0.070612092 |
| C1qtnf4 | 0.730926194 | 0.357215459 | 0.245 | 0.225 | 1 | 0.035967334 | -1.332720152 | 0.122 | 0.266 | 1 | 0.730926194 | 0.070641019 |
| Tmtc2 | 0.035995414 | 0.482297253 | 0.792 | 0.665 | 1 | 0.177917127 | 0.556110472 | 0.732 | 0.683 | 1 | 0.177917127 | 0.070695158 |
| Mast2 | 0.504360392 | -0.506411586 | 0.792 | 0.784 | 1 | 0.036013203 | -0.353997861 | 0.707 | 0.86 | 1 | 0.504360392 | 0.070729456 |
| Pak1 | 0.036023034 | 0.484184362 | 0.755 | 0.627 | 1 | 0.042643558 | -0.410722664 | 0.561 | 0.778 | 1 | 0.042643558 | 0.070748408 |
| Mpzl3 | 0.28312404 | -0.799473716 | 0.038 | 0.081 | 1 | 0.036088838 | 1.515276755 | 0.146 | 0.058 | 1 | 0.28312404 | 0.070875271 |
| Lcat | 0.103896277 | -1.389597921 | 0.038 | 0.11 | 1 | 0.036230829 | -1.917682652 | 0.024 | 0.14 | 1 | 0.103896277 | 0.071148985 |
| Gm6225 | 0.036335822 | -0.895903478 | 0.132 | 0.271 | 1 | 0.040711625 | -1.225804948 | 0.073 | 0.215 | 1 | 0.040711625 | 0.071351353 |
| Gm20712 | 0.171423172 | 1.154722595 | 0.094 | 0.047 | 1 | 0.036337617 | -1.80665361 | 0.024 | 0.14 | 1 | 0.171423172 | 0.071354812 |
| Sco1 | 0.036399086 | 1.084333267 | 0.17 | 0.076 | 1 | 0.204499385 | 0.979223855 | 0.146 | 0.085 | 1 | 0.204499385 | 0.071473278 |
| Sec22a | 0.036418008 | 0.795180208 | 0.472 | 0.377 | 1 | 0.271557694 | -0.113407123 | 0.293 | 0.399 | 1 | 0.271557694 | 0.071509745 |
| Lpcat3 | 0.036445468 | -0.396451592 | 0.302 | 0.508 | 1 | 0.305986186 | -0.445729113 | 0.415 | 0.474 | 1 | 0.305986186 | 0.071562664 |
| Haus8 | 0.094982307 | -0.715216865 | 0.358 | 0.462 | 1 | 0.03645326 | -0.776102277 | 0.22 | 0.396 | 1 | 0.094982307 | 0.07157768 |
| Sdhaf3 | 0.036494725 | 0.948271717 | 0.189 | 0.089 | 1 | 0.98546754 | 0.515276755 | 0.122 | 0.123 | 1 | 0.98546754 | 0.071657584 |
| Prxl2c | 0.03651836 | 0.658569361 | 0.547 | 0.453 | 1 | 0.26987047 | 0.220533489 | 0.78 | 0.659 | 1 | 0.26987047 | 0.071703129 |
| Cep295nl | 0.036552761 | 1.361173472 | 0.189 | 0.093 | 1 | 0.86145439 | 1.055845136 | 0.146 | 0.14 | 1 | 0.86145439 | 0.071769418 |
| Cox20 | 0.835150795 | 0.208161854 | 0.453 | 0.538 | 1 | 0.036571014 | 0.990914265 | 0.512 | 0.41 | 1 | 0.835150795 | 0.071804589 |
| Habp4 | 0.923346281 | 0.117247889 | 0.245 | 0.263 | 1 | 0.036573224 | -0.737703986 | 0.171 | 0.338 | 1 | 0.923346281 | 0.071808848 |
| Zc3hc1 | 0.036619447 | 1.657222935 | 0.189 | 0.093 | 1 | 0.343329314 | 0.88611445 | 0.22 | 0.167 | 1 | 0.343329314 | 0.071897911 |
| Zbtb14 | 0.036642889 | 1.090592257 | 0.17 | 0.076 | 1 | 0.359665845 | -0.213421223 | 0.098 | 0.157 | 1 | 0.359665845 | 0.071943077 |
| Ube4a | 0.03668674 | 0.673595905 | 0.604 | 0.534 | 1 | 0.582634153 | 0.395367291 | 0.561 | 0.645 | 1 | 0.582634153 | 0.072027563 |
| Ap4b1 | 0.036705427 | 0.956783217 | 0.264 | 0.144 | 1 | 0.773848201 | -0.26713181 | 0.098 | 0.109 | 1 | 0.773848201 | 0.072063566 |
| Phactr1 | 0.036719012 | -0.638826528 | 0.094 | 0.216 | 1 | 0.393099334 | 0.800863003 | 0.293 | 0.259 | 1 | 0.393099334 | 0.072089738 |
| Cpeb4 | 0.036755901 | 0.66127875 | 0.887 | 0.797 | 1 | 0.218515404 | 0.20086007 | 0.951 | 0.952 | 1 | 0.218515404 | 0.072160806 |
| Snf8 | 0.036851484 | 0.520007059 | 0.642 | 0.517 | 1 | 0.938569115 | 0.166666617 | 0.439 | 0.451 | 1 | 0.938569115 | 0.072344936 |
| Cdk8 | 0.036883657 | 0.348230271 | 0.962 | 0.945 | 1 | 0.153456304 | 0.383657787 | 0.951 | 0.949 | 1 | 0.153456304 | 0.072406909 |
| Mak16 | 0.036899221 | 0.730696312 | 0.547 | 0.436 | 1 | 0.151513682 | -0.286916462 | 0.39 | 0.529 | 1 | 0.151513682 | 0.07243689 |
| Proser1 | 0.036902391 | 0.887242284 | 0.34 | 0.212 | 1 | 0.326144573 | 0.478123756 | 0.366 | 0.29 | 1 | 0.326144573 | 0.072442996 |
| Cd68 | 0.036902443 | 1.261637799 | 0.189 | 0.089 | 1 | 0.788067132 | -1.188330242 | 0.098 | 0.085 | 1 | 0.788067132 | 0.072443095 |
| Prickle1 | 0.036953342 | 0.789153772 | 0.698 | 0.64 | 1 | 0.420648941 | 0.416629167 | 0.512 | 0.532 | 1 | 0.420648941 | 0.072541135 |
| Ints8 | 0.03695688 | 0.707922533 | 0.547 | 0.466 | 1 | 0.835593082 | 0.139851112 | 0.585 | 0.71 | 1 | 0.835593082 | 0.07254795 |
| Eef2k | 0.044537505 | 1.072260435 | 0.226 | 0.123 | 1 | 0.036991484 | 0.79281073 | 0.341 | 0.201 | 1 | 0.044537505 | 0.072614598 |
| Pafah1b3 | 0.037009071 | 0.962077517 | 0.283 | 0.165 | 1 | 0.637596975 | -0.278272368 | 0.122 | 0.147 | 1 | 0.637596975 | 0.07264847 |
| 1600012H06Rik | 0.543176277 | 0.235341055 | 0.34 | 0.301 | 1 | 0.037029391 | -0.573728251 | 0.146 | 0.324 | 1 | 0.543176277 | 0.072687605 |
| Popdc3 | 0.037031039 | 2.669295768 | 0.075 | 0.021 | 1 | 0.115845783 | 0.83720485 | 0.049 | 0.014 | 1 | 0.115845783 | 0.07269078 |
| Nfkbib | 0.174187695 | 0.450178478 | 0.396 | 0.292 | 1 | 0.03708465 | -0.771604393 | 0.146 | 0.311 | 1 | 0.174187695 | 0.07279403 |
| CJ186046Rik | 0.03709552 | 1.8327945 | 0.208 | 0.106 | 1 | 0.271344616 | -0.513292397 | 0.073 | 0.137 | 1 | 0.271344616 | 0.072814962 |
| 9530059O14Rik | 0.037132527 | -3.27154216 | 0.019 | 0.11 | 1 | 0.551079749 | 1.195658821 | 0.098 | 0.072 | 1 | 0.551079749 | 0.072886229 |
| Llgl1 | 0.037202109 | 1.533234218 | 0.245 | 0.14 | 1 | 0.128766428 | -0.398011612 | 0.171 | 0.297 | 1 | 0.128766428 | 0.073020221 |
| Slain2 | 0.037218668 | 0.601568832 | 0.717 | 0.631 | 1 | 0.691191684 | 0.135688092 | 0.756 | 0.84 | 1 | 0.691191684 | 0.073052107 |
| Acox1 | 0.217705968 | -0.680696246 | 0.698 | 0.75 | 1 | 0.03724105 | 0.660327088 | 0.878 | 0.724 | 1 | 0.217705968 | 0.073095205 |

| | | | | | | | | | | | | |
|---|---|---|---|---|---|---|---|---|---|---|---|---|
| Tubgcp2 | 0.037245985 | 0.895335966 | 0.377 | 0.25 | 1 | 0.919666416 | 0.123509035 | 0.317 | 0.345 | 1 | 0.919666416 | 0.073104707 |
| Epha7 | 0.037281254 | -1.116263934 | 0.358 | 0.5 | 1 | 0.534965473 | 0.896814339 | 0.341 | 0.338 | 1 | 0.534965473 | 0.073172616 |
| Aurka | 0.037307052 | 2.347367673 | 0.075 | 0.021 | 1 | 0.87556513 | 0.83720485 | 0.024 | 0.02 | 1 | 0.87556513 | 0.073222287 |
| Pom121 | 0.037310554 | 0.782304729 | 0.604 | 0.492 | 1 | 0.265378438 | 0.456600848 | 0.585 | 0.563 | 1 | 0.265378438 | 0.07322903 |
| Sars2 | 0.772942585 | 0.347367673 | 0.057 | 0.047 | 1 | 0.037317097 | 2.185128153 | 0.098 | 0.031 | 1 | 0.772942585 | 0.073241628 |
| Wdr35 | 0.609213195 | 0.3292961 | 0.472 | 0.47 | 1 | 0.037321206 | -0.502645153 | 0.39 | 0.611 | 1 | 0.609213195 | 0.07324954 |
| Srp19 | 0.037361151 | 0.291224595 | 0.868 | 0.78 | 1 | 0.070745594 | -0.337427154 | 0.707 | 0.812 | 1 | 0.070745594 | 0.073326446 |
| Med17 | 0.037375112 | 1.036078098 | 0.358 | 0.229 | 1 | 0.863242648 | 0.313642894 | 0.317 | 0.321 | 1 | 0.863242648 | 0.073353325 |
| Stard10 | 0.037421864 | -0.960754623 | 0.17 | 0.309 | 1 | 0.254457579 | -0.305753104 | 0.293 | 0.396 | 1 | 0.254457579 | 0.073443331 |
| Snrpa | 0.03742463 | 0.749083794 | 0.623 | 0.542 | 1 | 0.204147571 | 0.749742009 | 0.61 | 0.536 | 1 | 0.204147571 | 0.073448657 |
| Nudt18 | 0.037480244 | -1.713173869 | 0.075 | 0.191 | 1 | 0.746813312 | 0.359157553 | 0.122 | 0.106 | 1 | 0.746813312 | 0.07355572 |
| Wdr13 | 0.037493481 | 1.53023173 | 0.226 | 0.127 | 1 | 0.04142704 | 1.010041447 | 0.39 | 0.253 | 1 | 0.04142704 | 0.073581202 |
| Tyms | 0.037527004 | 1.154722595 | 0.189 | 0.093 | 1 | 0.072350299 | 1.209173627 | 0.171 | 0.085 | 1 | 0.072350299 | 0.073645732 |
| Adm | 1 | 0.569760094 | 0.038 | 0.038 | 1 | 0.037552986 | 1.962735732 | 0.098 | 0.031 | 1 | 1 | 0.073695745 |
| E230016M11Rik | 0.037624923 | 0.756173218 | 0.302 | 0.174 | 1 | 0.221035095 | -0.686357106 | 0.22 | 0.328 | 1 | 0.221035095 | 0.073834212 |
| Cnnm1 | 0.290483601 | -0.7032584 | 0.094 | 0.148 | 1 | 0.037628064 | 1.759202338 | 0.122 | 0.044 | 1 | 0.290483601 | 0.073840258 |
| Dnttip1 | 0.92510904 | 0.178970141 | 0.472 | 0.513 | 1 | 0.037724706 | -0.597732207 | 0.39 | 0.57 | 1 | 0.92510904 | 0.074026259 |
| Nudt16 | 0.037759341 | 1.865215978 | 0.113 | 0.042 | 1 | 0.208817133 | -0.970150072 | 0 | 0.038 | 1 | 0.208817133 | 0.074092914 |
| Epcam | 0.037793896 | 0.8327945 | 0.264 | 0.144 | 1 | 0.521144445 | 0.444887427 | 0.293 | 0.259 | 1 | 0.521144445 | 0.074159413 |
| Inpp1 | 0.690936036 | -0.228606045 | 0.302 | 0.335 | 1 | 0.037804329 | -0.985917388 | 0.22 | 0.382 | 1 | 0.690936036 | 0.074179491 |
| Rbm20 | 0.037849595 | -1.130679624 | 0.019 | 0.114 | 1 | 0.609087355 | -0.717384002 | 0.073 | 0.096 | 1 | 0.609087355 | 0.074266598 |
| Rbck1 | 0.037864651 | 0.824573993 | 0.415 | 0.288 | 1 | 0.965705687 | 0.136765132 | 0.317 | 0.324 | 1 | 0.965705687 | 0.074295571 |
| A430010J10Rik | 0.037892408 | 0.510866405 | 0.094 | 0.03 | 1 | 0.259050023 | -1.004097404 | 0.024 | 0.072 | 1 | 0.259050023 | 0.074348981 |
| Ppp1r10 | 0.037928775 | 0.485097697 | 0.849 | 0.678 | 1 | 0.298031473 | 0.35349654 | 0.854 | 0.819 | 1 | 0.298031473 | 0.074418958 |
| Chst1 | 0.38910033 | 0.874614676 | 0.075 | 0.047 | 1 | 0.037965503 | 1.265662865 | 0.293 | 0.167 | 1 | 0.38910033 | 0.074489626 |
| Socs5 | 0.037991534 | 0.614154213 | 0.604 | 0.483 | 1 | 0.499291221 | 0.214767644 | 0.585 | 0.553 | 1 | 0.499291221 | 0.074539711 |
| Csdc2 | 0.037991828 | 1.438515561 | 0.17 | 0.081 | 1 | 0.627075744 | -0.448197369 | 0.073 | 0.096 | 1 | 0.627075744 | 0.074540277 |
| Rbm14 | 0.189482114 | 0.424329655 | 0.321 | 0.237 | 1 | 0.037994721 | -0.424060463 | 0.22 | 0.396 | 1 | 0.189482114 | 0.074545843 |
| Axin2 | 0.329401128 | 0.328751995 | 0.34 | 0.28 | 1 | 0.038007462 | 1.113914193 | 0.439 | 0.307 | 1 | 0.329401128 | 0.074570357 |
| Nsmf | 0.038014194 | 0.68265415 | 0.434 | 0.292 | 1 | 0.657701339 | 0.221215453 | 0.366 | 0.328 | 1 | 0.657701339 | 0.074583309 |
| 8030456M14Rik | 0.243102132 | -1.015202407 | 0 | 0.025 | 1 | 0.038036028 | 2.059597271 | 0.098 | 0.031 | 1 | 0.243102132 | 0.074625316 |
| Hebp2 | 0.326693433 | -0.430239906 | 0.208 | 0.271 | 1 | 0.038043019 | -1.622226769 | 0.049 | 0.174 | 1 | 0.326693433 | 0.074638767 |
| Txlna | 0.038062241 | 0.507024339 | 0.736 | 0.665 | 1 | 0.384590591 | 0.174239837 | 0.805 | 0.696 | 1 | 0.384590591 | 0.074675747 |
| Ankib1 | 0.03806558 | 0.637504701 | 0.736 | 0.627 | 1 | 0.521210181 | 0.103484915 | 0.78 | 0.778 | 1 | 0.521210181 | 0.074682172 |
| Urb1 | 0.03809222 | 1.285967128 | 0.245 | 0.14 | 1 | 0.384881434 | 0.10681191 | 0.098 | 0.157 | 1 | 0.384881434 | 0.074733422 |
| Aasdhppt | 0.038149341 | 0.739685096 | 0.472 | 0.347 | 1 | 0.300232682 | 0.126711467 | 0.244 | 0.348 | 1 | 0.300232682 | 0.07484331 |
| Med27 | 0.100715948 | -0.555198176 | 0.358 | 0.504 | 1 | 0.038197974 | -0.684032053 | 0.341 | 0.522 | 1 | 0.100715948 | 0.074936863 |
| Acp2 | 0.038210603 | 0.802737266 | 0.434 | 0.314 | 1 | 0.855477773 | 0.252242349 | 0.317 | 0.362 | 1 | 0.855477773 | 0.074961156 |
| Cept1 | 0.080836665 | 0.578075362 | 0.547 | 0.436 | 1 | 0.038223062 | -0.499830137 | 0.537 | 0.672 | 1 | 0.080836665 | 0.074985122 |
| Spsb4 | 0.943580796 | 0.112902419 | 0.17 | 0.182 | 1 | 0.038234866 | -0.397260404 | 0.122 | 0.297 | 1 | 0.943580796 | 0.075007826 |
| Magt1 | 0.038254657 | 0.48967877 | 0.792 | 0.746 | 1 | 0.505946473 | 0.337278453 | 0.78 | 0.754 | 1 | 0.505946473 | 0.075045895 |
| Klhl7 | 0.038264473 | 0.523589913 | 0.623 | 0.542 | 1 | 0.270227174 | 0.818345823 | 0.61 | 0.57 | 1 | 0.270227174 | 0.075064777 |
| Rpl7 | 0.038285668 | 0.182956276 | 1 | 0.979 | 1 | 0.067196976 | -0.223929125 | 1 | 0.983 | 1 | 0.067196976 | 0.075105544 |
| U2af1l4 | 0.590827978 | 0.292226119 | 0.321 | 0.284 | 1 | 0.0383639 | -1.020776145 | 0.098 | 0.246 | 1 | 0.590827978 | 0.07525601 |
| Batf3 | 0.038376233 | -2.27154216 | 0.019 | 0.11 | 1 | 0.06992677 | 1.685201756 | 0.122 | 0.051 | 1 | 0.06992677 | 0.07527973 |
| Phkb | 0.038392607 | 0.599507437 | 0.472 | 0.318 | 1 | 0.115317073 | 0.356939728 | 0.634 | 0.491 | 1 | 0.115317073 | 0.075311222 |
| Cyth2 | 0.038432396 | 0.569760094 | 0.585 | 0.479 | 1 | 0.455175794 | 0.359157553 | 0.585 | 0.522 | 1 | 0.455175794 | 0.075387743 |
| Prx | 0.038434639 | 1.417757001 | 0.113 | 0.042 | 1 | 0.444067814 | -0.970150072 | 0.024 | 0.051 | 1 | 0.444067814 | 0.075392057 |
| Lmf2 | 0.0384714 | 0.781264199 | 0.34 | 0.208 | 1 | 0.440570945 | -0.410722664 | 0.171 | 0.218 | 1 | 0.440570945 | 0.075462751 |
| AU041133 | 0.038499829 | -2.489133595 | 0 | 0.076 | 1 | 0.278181612 | 1.721727632 | 0.073 | 0.038 | 1 | 0.278181612 | 0.075517422 |
| Gm45397 | 0.038511653 | -2.845277405 | 0 | 0.076 | 1 | 0.319635638 | -0.622226769 | 0 | 0.024 | 1 | 0.319635638 | 0.075540159 |
| Eppk1 | 0.038513716 | -2.974560422 | 0 | 0.076 | 1 | 0.402195522 | 0.252242349 | 0.098 | 0.061 | 1 | 0.402195522 | 0.075544125 |
| Cdk2ap2 | 0.038514343 | 0.569760094 | 0.415 | 0.275 | 1 | 0.052887797 | 0.775428652 | 0.512 | 0.379 | 1 | 0.052887797 | 0.07554533 |
| Tent5b | 0.100006369 | 2.962077517 | 0.038 | 0.008 | 1 | 0.038516927 | 1.83720485 | 0.098 | 0.031 | 1 | 0.100006369 | 0.075550301 |
| Gbp3 | 0.924320687 | 0.154722595 | 0.075 | 0.072 | 1 | 0.038516927 | 0.883008539 | 0.098 | 0.031 | 1 | 0.924320687 | 0.075550301 |
| Zmym1 | 0.038561634 | 0.569760094 | 0.358 | 0.212 | 1 | 0.317192511 | -0.372248516 | 0.171 | 0.246 | 1 | 0.317192511 | 0.075636269 |
| Pomt1 | 0.038575147 | 1.43225657 | 0.226 | 0.123 | 1 | 0.572708492 | 0.678006255 | 0.195 | 0.16 | 1 | 0.572708492 | 0.075662252 |
| Mustn1 | 0.038624971 | -2.126975655 | 0.189 | 0.309 | 1 | 0.923854484 | 0.292884334 | 0.122 | 0.133 | 1 | 0.923854484 | 0.075758054 |
| Tmem168 | 0.038668742 | 0.915935735 | 0.321 | 0.195 | 1 | 0.772491014 | -0.172255479 | 0.293 | 0.317 | 1 | 0.772491014 | 0.075842213 |
| Ret | 0.038686658 | -1.652632327 | 0.038 | 0.14 | 1 | 0.357847088 | -0.16279515 | 0 | 0.02 | 1 | 0.357847088 | 0.075876659 |
| Ckap4 | 0.038783194 | -0.354399964 | 0.962 | 0.975 | 1 | 0.082356867 | -0.350705897 | 0.927 | 0.973 | 1 | 0.082356867 | 0.076062253 |
| Vps13c | 0.038792963 | 0.924424975 | 0.472 | 0.364 | 1 | 0.308325984 | 0.70627398 | 0.488 | 0.454 | 1 | 0.308325984 | 0.076081032 |
| Prkag1 | 0.038819428 | 0.461383933 | 0.717 | 0.661 | 1 | 0.806850503 | 0.142059431 | 0.634 | 0.669 | 1 | 0.806850503 | 0.076131909 |
| Abcc4 | 0.038921514 | -1.027480736 | 0.113 | 0.25 | 1 | 0.226300136 | 0.169780189 | 0.317 | 0.225 | 1 | 0.226300136 | 0.076328144 |
| Znhit6 | 0.038940332 | 0.808546954 | 0.491 | 0.381 | 1 | 0.456723186 | 0.358037013 | 0.39 | 0.341 | 1 | 0.456723186 | 0.076364314 |
| Tspyl3 | 0.038976464 | 2.154722595 | 0.075 | 0.021 | 1 | 0.58792152 | 2.100239256 | 0.024 | 0.014 | 1 | 0.58792152 | 0.076433762 |
| Zfp658 | 0.038976464 | 2.154722595 | 0.075 | 0.021 | 1 | 0.110710459 | 2.515276755 | 0.049 | 0.014 | 1 | 0.110710459 | 0.076433762 |
| Gm20457 | 0.038976464 | 2.154722595 | 0.075 | 0.021 | 1 | 0.25729872 | -0.747757651 | 0 | 0.031 | 1 | 0.25729872 | 0.076433762 |
| Gm11149 | 0.038976464 | 2.154722595 | 0.075 | 0.021 | 1 | 0.319641967 | -0.16279515 | 0 | 0.024 | 1 | 0.319641967 | 0.076433762 |
| Aox1 | 0.588758899 | 1.883420573 | 0.113 | 0.093 | 1 | 0.038984845 | 2.46797104 | 0.171 | 0.075 | 1 | 0.588758899 | 0.076449871 |
| Sil1 | 0.039003475 | -0.347777746 | 0.245 | 0.424 | 1 | 0.298020755 | 0.583448258 | 0.439 | 0.392 | 1 | 0.298020755 | 0.07648568 |
| Ntrk2 | 0.466452234 | 0.84729407 | 0.132 | 0.102 | 1 | 0.039066906 | -1.80665134 | 0.049 | 0.174 | 1 | 0.466452234 | 0.076607589 |
| Sorcs2 | 0.03912188 | 1.506194965 | 0.189 | 0.093 | 1 | 0.436460283 | -0.348661696 | 0.146 | 0.102 | 1 | 0.436460283 | 0.076713239 |
| Soat1 | 0.216784858 | -0.127541283 | 0.434 | 0.589 | 1 | 0.039230968 | -0.159682538 | 0.415 | 0.625 | 1 | 0.216784858 | 0.076922867 |
| Tbc1d25 | 0.831408314 | 0.216123139 | 0.094 | 0.085 | 1 | 0.039231885 | -2.555112573 | 0 | 0.096 | 1 | 0.831408314 | 0.076924629 |
| Gm14964 | 0.471154689 | 1.569760094 | 0.038 | 0.021 | 1 | 0.039251348 | 0.867578499 | 0.146 | 0.058 | 1 | 0.471154689 | 0.076962028 |
| Rnf7 | 0.039400518 | 0.412373804 | 0.792 | 0.746 | 1 | 0.908060936 | 0.17020658 | 0.732 | 0.829 | 1 | 0.908060936 | 0.077248636 |
| Mphosph8 | 0.039461782 | 0.488661669 | 0.698 | 0.585 | 1 | 0.25975292 | 0.457991014 | 0.683 | 0.601 | 1 | 0.25975292 | 0.077366333 |
| Uprt | 0.439203725 | 0.569760094 | 0.094 | 0.064 | 1 | 0.039472155 | 1.47463477 | 0.122 | 0.044 | 1 | 0.439203725 | 0.077386259 |
| Il1rap | 0.376450729 | -0.356239324 | 0.094 | 0.144 | 1 | 0.039494681 | 2.267839204 | 0.22 | 0.113 | 1 | 0.376450729 | 0.077429532 |
| Cse1l | 0.039498013 | 0.650491366 | 0.509 | 0.369 | 1 | 0.990304155 | 0.194756854 | 0.488 | 0.515 | 1 | 0.990304155 | 0.077435934 |
| Nr6a1os | 0.039521374 | 1.569760094 | 0.113 | 0.042 | 1 | 0.487805099 | 0.898605394 | 0.073 | 0.048 | 1 | 0.487805099 | 0.07748081 |
| Bcar3 | 0.039531857 | 2.216123139 | 0.151 | 0.068 | 1 | 0.283258889 | 0.601988388 | 0.195 | 0.133 | 1 | 0.283258889 | 0.077500947 |
| Synm | 0.03955652 | 0.758793918 | 0.226 | 0.114 | 1 | 0.515264586 | -0.756976181 | 0.195 | 0.229 | 1 | 0.515264586 | 0.077548322 |
| A930007I19Rik | 0.613384761 | 0.3662267 | 0.113 | 0.093 | 1 | 0.039591854 | 1.404245443 | 0.146 | 0.058 | 1 | 0.613384761 | 0.077616192 |
| B4galt2 | 0.03965996 | 1.017219071 | 0.226 | 0.119 | 1 | 0.160683958 | 1.422167351 | 0.195 | 0.126 | 1 | 0.160683958 | 0.077747008 |
| Arrdc2 | 0.190671918 | -0.323324702 | 0.057 | 0.123 | 1 | 0.039684072 | 1.120997816 | 0.122 | 0.044 | 1 | 0.190671918 | 0.077793319 |
| Trappc13 | 0.209792335 | 0.496511112 | 0.396 | 0.326 | 1 | 0.03971537 | -0.338872379 | 0.293 | 0.461 | 1 | 0.209792335 | 0.077853429 |
| Gm14403 | 0.175865986 | -1.015202407 | 0 | 0.034 | 1 | 0.039724288 | 1.699701326 | 0.098 | 0.031 | 1 | 0.175865986 | 0.077870558 |
| Gm56656 | 0.48175508 | 1.154722595 | 0.038 | 0.021 | 1 | 0.039724288 | 1.699701326 | 0.098 | 0.031 | 1 | 0.48175508 | 0.077870558 |
| Tgfbi | 0.039728632 | 0.30324512 | 0.208 | 0.102 | 1 | 0.31331221 | 0.540587843 | 0.195 | 0.137 | 1 | 0.31331221 | 0.077878899 |
| Cyria | 0.039749568 | 1.491757582 | 0.113 | 0.042 | 1 | 0.82181816 | 0.252242349 | 0.024 | 0.031 | 1 | 0.82181816 | 0.077919108 |
| Pde8a | 0.474742692 | 0.694533059 | 0.585 | 0.64 | 1 | 0.039818997 | -0.924346383 | 0.61 | 0.7 | 1 | 0.474742692 | 0.078052441 |
| Med30 | 0.569625487 | 0.200526284 | 0.415 | 0.386 | 1 | 0.039831115 | -1.015237962 | 0.171 | 0.324 | 1 | 0.569625487 | 0.078075712 |
| Casp2 | 0.03983138 | 1.707263618 | 0.132 | 0.055 | 1 | 0.873689559 | 0.322631677 | 0.098 | 0.109 | 1 | 0.873689559 | 0.07807622 |
| Gid4 | 0.039832852 | 0.698084191 | 0.528 | 0.394 | 1 | 0.636426037 | 0.545178241 | 0.415 | 0.413 | 1 | 0.636426037 | 0.078079048 |
| Zfp971 | 0.039845057 | 1.569760094 | 0.151 | 0.068 | 1 | 0.095445305 | 1.071670103 | 0.171 | 0.089 | 1 | 0.095445305 | 0.078102486 |
| Fam199x | 0.039880408 | 1.185096244 | 0.264 | 0.157 | 1 | 0.469296536 | 0.10681191 | 0.244 | 0.317 | 1 | 0.469296536 | 0.078170368 |

| | | | | | | | | | | | | |
|---|---|---|---|---|---|---|---|---|---|---|---|---|
| Mcm3ap | 0.039881977 | 0.789073122 | 0.358 | 0.22 | 1 | 0.343229353 | -0.484723245 | 0.195 | 0.266 | 1 | 0.343229353 | 0.078173381 |
| Comtd1 | 0.039910584 | 1.695290976 | 0.113 | 0.042 | 1 | 0.410382697 | -0.332720152 | 0.024 | 0.055 | 1 | 0.410382697 | 0.078228313 |
| Gm56496 | 0.039910584 | 1.695290976 | 0.113 | 0.042 | 1 | 0.631113499 | -0.16279515 | 0.049 | 0.068 | 1 | 0.631113499 | 0.078228313 |
| Pcdhga11 | 0.039910584 | 1.695290976 | 0.113 | 0.042 | 1 | 0.737610035 | 0.614812428 | 0.049 | 0.038 | 1 | 0.737610035 | 0.078228313 |
| Rnf215 | 0.039943486 | 0.692379381 | 0.509 | 0.386 | 1 | 0.431498917 | -0.229909346 | 0.341 | 0.392 | 1 | 0.431498917 | 0.07829149 |
| Pde1c | 0.039945875 | 0.98621687 | 0.547 | 0.445 | 1 | 0.417393871 | -0.801395615 | 0.22 | 0.27 | 1 | 0.417393871 | 0.078296077 |
| Nubp2 | 0.039951424 | 0.974150349 | 0.226 | 0.119 | 1 | 0.233029791 | -0.455576899 | 0.073 | 0.143 | 1 | 0.233029791 | 0.078306731 |
| Usp35 | 0.039953057 | 1.913714495 | 0.113 | 0.042 | 1 | 0.722626107 | 0.377773231 | 0.073 | 0.058 | 1 | 0.722626107 | 0.078309868 |
| Zfp532 | 0.475389844 | 0.361818829 | 0.377 | 0.36 | 1 | 0.040018149 | 0.679203016 | 0.561 | 0.44 | 1 | 0.475389844 | 0.078434846 |
| Stam2 | 0.040034605 | 0.669295768 | 0.528 | 0.377 | 1 | 0.909451125 | 0.164245868 | 0.585 | 0.573 | 1 | 0.909451125 | 0.078466441 |
| Gm33694 | 0.040143819 | 1.932330173 | 0.075 | 0.021 | 1 | 0.438672228 | 1.83720485 | 0.024 | 0.01 | 1 | 0.438672228 | 0.078676111 |
| Ppara | 0.040153397 | 1.624207878 | 0.075 | 0.021 | 1 | 0.165399591 | 1.911205431 | 0.073 | 0.031 | 1 | 0.165399591 | 0.078694499 |
| Trpc1 | 0.040167594 | 0.984797593 | 0.302 | 0.186 | 1 | 0.146595019 | -0.452301767 | 0.171 | 0.297 | 1 | 0.146595019 | 0.078721753 |
| Plec | 0.649033846 | -0.208614998 | 0.981 | 0.932 | 1 | 0.040169616 | -0.453887437 | 0.951 | 0.976 | 1 | 0.649033846 | 0.078725635 |
| Xiap | 0.040175161 | 0.577865144 | 0.755 | 0.682 | 1 | 0.621620461 | 0.168520523 | 0.756 | 0.816 | 1 | 0.621620461 | 0.078736279 |
| Ccz1 | 0.040301951 | 0.611100889 | 0.585 | 0.479 | 1 | 0.845358388 | -0.118248711 | 0.585 | 0.529 | 1 | 0.845358388 | 0.078979654 |
| Gpx8 | 0.04037321 | 0.370962789 | 0.906 | 0.873 | 1 | 0.17488367 | -0.261719383 | 0.732 | 0.84 | 1 | 0.17488367 | 0.079116424 |
| Zfp783 | 0.040375477 | 0.815920681 | 0.264 | 0.144 | 1 | 0.410971732 | -0.541306773 | 0.098 | 0.143 | 1 | 0.410971732 | 0.079120775 |
| Unc13b | 0.04297268 | 2.084333267 | 0.132 | 0.055 | 1 | 0.040399127 | 0.515276755 | 0.317 | 0.177 | 1 | 0.04297268 | 0.079166164 |
| Gm5577 | 0.040399712 | 1.669295768 | 0.113 | 0.042 | 1 | 0.476529618 | 1.000703582 | 0.073 | 0.048 | 1 | 0.476529618 | 0.079167288 |
| Gm26749 | 0.040399871 | 1.624207878 | 0.113 | 0.042 | 1 | 0.616599719 | 0.479652845 | 0.122 | 0.099 | 1 | 0.616599719 | 0.079167592 |
| Igf2bp3 | 0.040420977 | 0.763932642 | 0.321 | 0.208 | 1 | 0.815070516 | 0.39094862 | 0.171 | 0.167 | 1 | 0.815070516 | 0.079208098 |
| Nudt21 | 0.04042421 | 0.431832186 | 0.792 | 0.716 | 1 | 0.078307001 | -0.386963196 | 0.707 | 0.812 | 1 | 0.078307001 | 0.079214304 |
| Oga | 0.172317261 | 0.333277213 | 0.849 | 0.78 | 1 | 0.04043264 | 0.402893338 | 0.927 | 0.887 | 1 | 0.172317261 | 0.079230481 |
| Gm28729 | 0.040450408 | 1.865215978 | 0.075 | 0.021 | 1 | 0.286421728 | -0.863234868 | 0 | 0.027 | 1 | 0.286421728 | 0.07926458 |
| Amigo1 | 0.040453707 | 1.891688189 | 0.075 | 0.021 | 1 | 0.170440921 | -0.970150072 | 0 | 0.044 | 1 | 0.170440921 | 0.079270912 |
| D3Ertd751e | 0.040459729 | -0.502389692 | 0.094 | 0.229 | 1 | 0.802855562 | 0.707921833 | 0.171 | 0.16 | 1 | 0.802855562 | 0.079282469 |
| Nelfcd | 0.040504127 | 1.569760094 | 0.17 | 0.081 | 1 | 0.319799334 | -0.707115666 | 0.049 | 0.096 | 1 | 0.319799334 | 0.07936767 |
| Pdcd5 | 0.040553347 | 0.477501586 | 0.811 | 0.665 | 1 | 0.545147399 | 0.398631836 | 0.732 | 0.689 | 1 | 0.545147399 | 0.079462121 |
| Nek10 | 0.040638111 | 1.47665069 | 0.17 | 0.081 | 1 | 0.206194005 | 0.486707603 | 0.171 | 0.106 | 1 | 0.206194005 | 0.079624766 |
| Gm44649 | 0.040671261 | -1.430239906 | 0.057 | 0.165 | 1 | 0.059687088 | -0.767657208 | 0.073 | 0.198 | 1 | 0.059687088 | 0.079688371 |
| Npr2 | 0.069782804 | 0.627475592 | 0.434 | 0.322 | 1 | 0.040715695 | -0.653120777 | 0.317 | 0.491 | 1 | 0.069782804 | 0.079773621 |
| Taf6l | 0.340246113 | 1.417757001 | 0.038 | 0.017 | 1 | 0.040736816 | 1.574170444 | 0.098 | 0.031 | 1 | 0.340246113 | 0.079814145 |
| Arhgap27 | 0.343006305 | -0.845277405 | 0 | 0.017 | 1 | 0.040736816 | 1.574170444 | 0.098 | 0.031 | 1 | 0.343006305 | 0.079814145 |
| Atl3 | 0.384645326 | 0.305908066 | 0.679 | 0.695 | 1 | 0.04077888 | -0.54927878 | 0.829 | 0.819 | 1 | 0.384645326 | 0.079894844 |
| Eif4a1 | 0.156941248 | 0.199312746 | 0.981 | 0.983 | 1 | 0.040831624 | -0.29988357 | 1 | 0.993 | 1 | 0.156941248 | 0.079996026 |
| 8030462N17Rik | 0.051122099 | 0.507603618 | 0.528 | 0.436 | 1 | 0.040867765 | 0.509278014 | 0.756 | 0.666 | 1 | 0.051122099 | 0.080065355 |
| Zfp408 | 0.040898125 | 1.193196743 | 0.226 | 0.123 | 1 | 0.694500581 | 0.639265472 | 0.22 | 0.201 | 1 | 0.694500581 | 0.080123593 |
| Gm49969 | 0.058295868 | -1.822557329 | 0.057 | 0.153 | 1 | 0.040983002 | -1.050320421 | 0.049 | 0.177 | 1 | 0.058295868 | 0.080286398 |
| Sirt6 | 0.041006383 | 1.669295768 | 0.132 | 0.055 | 1 | 0.796279838 | -0.101394606 | 0.049 | 0.058 | 1 | 0.796279838 | 0.080331242 |
| Pyroxd2 | 0.041006383 | 1.669295768 | 0.132 | 0.055 | 1 | 0.171501658 | -0.116991461 | 0.024 | 0.089 | 1 | 0.171501658 | 0.080331242 |
| Smpd3 | 0.041033851 | -3.941201825 | 0.038 | 0.136 | 1 | 0.057958278 | -4.777504994 | 0 | 0.082 | 1 | 0.057958278 | 0.080383925 |
| Ddx24 | 0.908342044 | 0.192140124 | 0.849 | 0.86 | 1 | 0.041102611 | -0.297280482 | 0.732 | 0.87 | 1 | 0.908342044 | 0.080515797 |
| Dock7 | 0.04112914 | 0.535812762 | 0.868 | 0.835 | 1 | 0.080376608 | 0.413993419 | 0.927 | 0.863 | 1 | 0.080376608 | 0.080566674 |
| Yars2 | 0.584543347 | 0.53023173 | 0.17 | 0.144 | 1 | 0.041137686 | -1.292078167 | 0.073 | 0.205 | 1 | 0.584543347 | 0.080583062 |
| Gpat4 | 0.041276064 | 0.615856509 | 0.585 | 0.441 | 1 | 0.824794794 | 0.107192016 | 0.61 | 0.611 | 1 | 0.824794794 | 0.080848415 |
| Casc3 | 0.041299537 | 0.822147256 | 0.509 | 0.407 | 1 | 0.082416424 | 0.879989716 | 0.707 | 0.58 | 1 | 0.082416424 | 0.080893422 |
| Gng3 | 0.041316766 | 1.714150003 | 0.151 | 0.068 | 1 | 0.415522187 | 0.60690723 | 0.195 | 0.147 | 1 | 0.415522187 | 0.080926457 |
| Phax | 0.041321584 | 0.528283458 | 0.679 | 0.589 | 1 | 0.150620329 | -0.285651898 | 0.561 | 0.679 | 1 | 0.150620329 | 0.080935696 |
| Ahr | 0.181789368 | -1.430239906 | 0.075 | 0.14 | 1 | 0.041322845 | 1.178241768 | 0.268 | 0.15 | 1 | 0.181789368 | 0.080938112 |
| Arhgap11a | 0.041338649 | 1.502645898 | 0.075 | 0.021 | 1 | 0.989595616 | 0.515276755 | 0.024 | 0.024 | 1 | 0.989595616 | 0.080968414 |
| Ccndbp1 | 0.105103246 | 0.461700348 | 0.453 | 0.335 | 1 | 0.041351196 | -0.757741739 | 0.195 | 0.358 | 1 | 0.105103246 | 0.080992471 |
| Mir670hg | 0.041354566 | 2.962077517 | 0.057 | 0.013 | 1 | 0.819548308 | 0.136765132 | 0.024 | 0.031 | 1 | 0.819548308 | 0.080998931 |
| 4930537H20Rik | 0.041354566 | 2.739685096 | 0.057 | 0.013 | 1 | 0.257325989 | -0.484723245 | 0 | 0.031 | 1 | 0.257325989 | 0.080998931 |
| Rad51ap2 | 0.041354566 | 2.739685096 | 0.057 | 0.013 | 1 | 0.262674569 | 2.83720485 | 0.024 | 0.007 | 1 | 0.262674569 | 0.080998931 |
| Dnaaf9 | 0.648827602 | 0.189488013 | 0.66 | 0.623 | 1 | 0.041376744 | -0.173826187 | 0.732 | 0.846 | 1 | 0.648827602 | 0.081041454 |
| Lrrc8b | 0.126996891 | 0.963779812 | 0.321 | 0.229 | 1 | 0.041416593 | -1.646610927 | 0.244 | 0.375 | 1 | 0.126996891 | 0.081117852 |
| Gm47802 | 0.343000446 | -0.430239906 | 0 | 0.017 | 1 | 0.041434534 | 1.47463477 | 0.122 | 0.044 | 1 | 0.343000446 | 0.081152247 |
| Eif2ak3 | 0.041507189 | 0.527235477 | 0.566 | 0.453 | 1 | 0.705012581 | 0.362666339 | 0.659 | 0.662 | 1 | 0.705012581 | 0.081291532 |
| Lin7a | 0.55750758 | 0.756173218 | 0.075 | 0.055 | 1 | 0.041518645 | -3.105309656 | 0.024 | 0.133 | 1 | 0.55750758 | 0.081313491 |
| Sel1l | 0.041527698 | 0.442524906 | 0.755 | 0.682 | 1 | 0.751847746 | 0.290875009 | 0.683 | 0.788 | 1 | 0.751847746 | 0.081330846 |
| Lef1os1 | 0.041626809 | 1.669295768 | 0.075 | 0.021 | 1 | 0.188573672 | -0.863234868 | 0 | 0.041 | 1 | 0.188573672 | 0.081520827 |
| Dpysl4 | 0.041626809 | 1.47665069 | 0.075 | 0.021 | 1 | 0.493019746 | -0.332720152 | 0.049 | 0.078 | 1 | 0.493019746 | 0.081520827 |
| Hsh2d | 0.041626809 | 1.154722595 | 0.075 | 0.021 | 1 | 0.208817133 | -1.16279515 | 0 | 0.038 | 1 | 0.208817133 | 0.081520827 |
| Fads6 | 0.041626809 | 1.47665069 | 0.075 | 0.021 | 1 | 0.057901606 | 2.422167351 | 0.049 | 0.01 | 1 | 0.057901606 | 0.081520827 |
| Gm3625 | 0.041626809 | 1.669295768 | 0.075 | 0.021 | 1 | 0.744047924 | 1.252242349 | 0.024 | 0.017 | 1 | 0.744047924 | 0.081520827 |
| Gm5141 | 0.041626809 | 1.669295768 | 0.075 | 0.021 | 1 | 0.319641967 | -0.16279515 | 0 | 0.024 | 1 | 0.319641967 | 0.081520827 |
| Prrt1 | 0.041626809 | 1.669295768 | 0.075 | 0.021 | 1 | 0.274788535 | 1.100239256 | 0.049 | 0.02 | 1 | 0.274788535 | 0.081520827 |
| Rgl2 | 0.239379778 | 1.115194231 | 0.189 | 0.131 | 1 | 0.041671403 | -1.572186086 | 0.073 | 0.201 | 1 | 0.239379778 | 0.081606299 |
| Rorc | 0.469724974 | -0.347777746 | 0.038 | 0.064 | 1 | 0.041673901 | -1.359192363 | 0.024 | 0.14 | 1 | 0.469724974 | 0.081611088 |
| Krtcap2 | 0.174331783 | 0.135753358 | 0.868 | 0.864 | 1 | 0.041700508 | -0.315747073 | 0.634 | 0.816 | 1 | 0.174331783 | 0.081662085 |
| Emg1 | 0.041935088 | 0.525365975 | 0.66 | 0.551 | 1 | 0.140378514 | 0.396632258 | 0.659 | 0.567 | 1 | 0.140378514 | 0.082111625 |
| Fbxl14 | 0.041967338 | 0.962077517 | 0.226 | 0.119 | 1 | 0.919561519 | 0.367719567 | 0.22 | 0.218 | 1 | 0.919561519 | 0.082173419 |
| Selenop | 0.041973623 | 0.866477626 | 0.453 | 0.343 | 1 | 0.1584555 | 0.798524382 | 0.439 | 0.369 | 1 | 0.1584555 | 0.082185462 |
| Polr1c | 0.285072903 | 0.445771377 | 0.509 | 0.492 | 1 | 0.042015928 | -0.638528581 | 0.366 | 0.522 | 1 | 0.285072903 | 0.082266519 |
| Mad2l1bp | 0.042048634 | 1.280253477 | 0.226 | 0.127 | 1 | 0.051491484 | 1.763204268 | 0.22 | 0.119 | 1 | 0.051491484 | 0.082329181 |
| Mypop | 0.042098806 | 1.640149422 | 0.151 | 0.068 | 1 | 0.929927279 | 0.515276755 | 0.171 | 0.174 | 1 | 0.929927279 | 0.082425302 |
| Nup210l | 0.042107207 | 0.450588393 | 0.566 | 0.428 | 1 | 0.306913096 | -0.378523841 | 0.659 | 0.512 | 1 | 0.306913096 | 0.082441398 |
| Rin2 | 0.042123805 | -1.056694536 | 0.906 | 0.924 | 1 | 0.079709014 | -0.692015099 | 1 | 0.935 | 1 | 0.079709014 | 0.082473195 |
| Stk25 | 0.04216634 | 0.429687211 | 0.679 | 0.661 | 1 | 0.36644821 | -0.209488386 | 0.659 | 0.751 | 1 | 0.36644821 | 0.082554679 |
| Foxred1 | 0.042194999 | -1.060290296 | 0.113 | 0.237 | 1 | 0.272220974 | -0.818146979 | 0.098 | 0.16 | 1 | 0.272220974 | 0.08260958 |
| Snrpd1 | 0.042239347 | 0.234157062 | 0.925 | 0.801 | 1 | 0.641644173 | 0.196747236 | 0.854 | 0.894 | 1 | 0.641644173 | 0.082694532 |
| Trim46 | 0.04226947 | 1.707263618 | 0.132 | 0.055 | 1 | 0.137738323 | 1.564186355 | 0.146 | 0.078 | 1 | 0.137738323 | 0.082752232 |
| Gm33887 | 0.042273726 | 1.502645898 | 0.113 | 0.042 | 1 | 0.903375428 | 0.667279848 | 0.049 | 0.055 | 1 | 0.903375428 | 0.082760384 |
| Gm14327 | 0.042273726 | 1.502645898 | 0.113 | 0.042 | 1 | 0.15412976 | -1.332720152 | 0 | 0.048 | 1 | 0.15412976 | 0.082760384 |
| Pcdhgb6 | 0.042273726 | 1.502645898 | 0.113 | 0.042 | 1 | 0.871991894 | 0.174239837 | 0.049 | 0.055 | 1 | 0.871991894 | 0.082760384 |
| Htatsf1 | 0.042274228 | 0.649194562 | 0.698 | 0.644 | 1 | 0.513512456 | 0.173170778 | 0.756 | 0.713 | 1 | 0.513512456 | 0.082761346 |
| Arsk | 0.643203446 | -0.368839361 | 0.245 | 0.267 | 1 | 0.042322169 | -0.640842447 | 0.171 | 0.348 | 1 | 0.643203446 | 0.082853173 |
| Ptpra | 0.163094229 | -0.239908694 | 0.943 | 0.936 | 1 | 0.042350004 | -0.304092352 | 0.927 | 0.962 | 1 | 0.163094229 | 0.082906486 |
| Zmat5 | 0.042352712 | 0.909610097 | 0.34 | 0.225 | 1 | 0.082841989 | 1.088743617 | 0.341 | 0.239 | 1 | 0.082841989 | 0.082911672 |
| Itgb1bp1 | 0.042387212 | 0.510866405 | 0.453 | 0.314 | 1 | 0.073891282 | -0.825760163 | 0.195 | 0.324 | 1 | 0.073891282 | 0.082977748 |
| Gabbr1 | 0.042424329 | 0.922061838 | 0.509 | 0.441 | 1 | 0.354299708 | 0.539938809 | 0.561 | 0.546 | 1 | 0.354299708 | 0.083048835 |
| Ccdc82 | 0.04243017 | 0.412218817 | 0.585 | 0.403 | 1 | 0.25609411 | -0.16279515 | 0.39 | 0.481 | 1 | 0.25609411 | 0.083060021 |
| Wdr86 | 0.042463172 | 2.47665069 | 0.057 | 0.013 | 1 | 0.599290422 | 1.515276755 | 0.024 | 0.014 | 1 | 0.599290422 | 0.083123223 |
| Epo | 0.042463172 | 2.47665069 | 0.057 | 0.013 | 1 | 0.305356148 | -0.747757651 | 0.024 | 0.065 | 1 | 0.305356148 | 0.083123223 |
| Zfp90 | 0.042463172 | 2.739685096 | 0.057 | 0.013 | 1 | 0.158439491 | 0.989207943 | 0.098 | 0.044 | 1 | 0.158439491 | 0.083123223 |
| Trhde | 0.042463172 | 3.324647596 | 0.057 | 0.013 | 1 | 0.605277123 | 1.711673968 | 0.049 | 0.034 | 1 | 0.605277123 | 0.083123223 |
| Nrros | 0.042463172 | 3.614154213 | 0.057 | 0.013 | 1 | 0.402364804 | -0.16279515 | 0 | 0.017 | 1 | 0.402364804 | 0.083123223 |

| | | | | | | | | | | | | |
|---|---|---|---|---|---|---|---|---|---|---|---|---|
| Slc25a23 | 0.042463172 | 2.47665069 | 0.057 | 0.013 | 1 | 0.225582728 | -0.278272368 | 0.024 | 0.078 | 1 | 0.225582728 | 0.083123223 |
| Prrg3 | 0.042463172 | 2.47665069 | 0.057 | 0.013 | 1 | 0.872828964 | 1.029849928 | 0.024 | 0.02 | 1 | 0.872828964 | 0.083123223 |
| Col19a1 | 0.04246991 | 2.417757001 | 0.057 | 0.013 | 1 | 0.965636861 | 0.635570989 | 0.049 | 0.051 | 1 | 0.965636861 | 0.083136128 |
| Stard8 | 0.084732627 | 1.017219071 | 0.377 | 0.301 | 1 | 0.04247657 | -1.250257991 | 0.171 | 0.321 | 1 | 0.084732627 | 0.083148881 |
| Sec23ip | 0.042532079 | 0.503670904 | 0.642 | 0.496 | 1 | 0.113158513 | -0.463964685 | 0.512 | 0.642 | 1 | 0.113158513 | 0.08325518 |
| Mss51 | 0.042535493 | 1.200526284 | 0.17 | 0.081 | 1 | 0.36392434 | -0.11032773 | 0.073 | 0.123 | 1 | 0.36392434 | 0.083261719 |
| Epb41l4a | 0.914958 | 0.347367673 | 0.019 | 0.021 | 1 | 0.042540147 | 1.252242349 | 0.22 | 0.109 | 1 | 0.914958 | 0.08327063 |
| Atg4b | 0.042638047 | 0.879088152 | 0.491 | 0.39 | 1 | 0.933037291 | 0.156822784 | 0.512 | 0.532 | 1 | 0.933037291 | 0.083458092 |
| H1f4 | 0.042644076 | -1.868361018 | 0.321 | 0.445 | 1 | 0.398824172 | 0.144182603 | 0.415 | 0.362 | 1 | 0.398824172 | 0.083469635 |
| Cd36 | 0.075415174 | 3.09332205 | 0.075 | 0.025 | 1 | 0.042670985 | 1.159132945 | 0.195 | 0.092 | 1 | 0.075415174 | 0.083521157 |
| Fam13a | 0.042737863 | 1.569760094 | 0.132 | 0.055 | 1 | 0.372037778 | 0.174239837 | 0.049 | 0.024 | 1 | 0.372037778 | 0.083649202 |
| Ankrd35 | 0.848124966 | 0.47665069 | 0.113 | 0.106 | 1 | 0.042817732 | 0.807457506 | 0.341 | 0.208 | 1 | 0.848124966 | 0.083802106 |
| Ticam1 | 0.164198307 | 1.055186921 | 0.113 | 0.059 | 1 | 0.042837712 | -0.944154864 | 0.049 | 0.177 | 1 | 0.164198307 | 0.083840354 |
| Nup153 | 0.052518439 | 0.688323291 | 0.792 | 0.78 | 1 | 0.042861756 | 0.329621609 | 0.927 | 0.884 | 1 | 0.052518439 | 0.083886383 |
| Kdelr3 | 0.128023167 | -0.251670013 | 0.698 | 0.822 | 1 | 0.042879958 | -0.383988822 | 0.634 | 0.765 | 1 | 0.128023167 | 0.083921224 |
| Gcnt2 | 0.042886121 | -2.503488888 | 0.189 | 0.297 | 1 | 0.101295954 | 0.515276755 | 0.317 | 0.205 | 1 | 0.101295954 | 0.083933023 |
| Ube2m | 0.042894095 | 0.271463229 | 0.925 | 0.797 | 1 | 0.33758396 | -0.148637087 | 0.707 | 0.809 | 1 | 0.33758396 | 0.083948286 |
| Caprin2 | 0.429872902 | 1.047807391 | 0.113 | 0.081 | 1 | 0.042908994 | 1.029849928 | 0.22 | 0.109 | 1 | 0.429872902 | 0.083976807 |
| Mtmr11 | 0.042925284 | -0.626637119 | 0.264 | 0.407 | 1 | 0.136234482 | -0.33682455 | 0.415 | 0.553 | 1 | 0.136234482 | 0.084007989 |
| Srsf5 | 0.043035981 | 0.291261428 | 0.962 | 0.975 | 1 | 0.124179645 | 0.25328363 | 0.951 | 0.962 | 1 | 0.124179645 | 0.084219866 |
| Eloa | 0.170488092 | 0.3662267 | 0.66 | 0.606 | 1 | 0.043150677 | 0.353477644 | 0.902 | 0.751 | 1 | 0.170488092 | 0.084439374 |
| Hmga1 | 0.978426098 | 0.207190015 | 0.094 | 0.093 | 1 | 0.043213411 | -2.16279515 | 0 | 0.092 | 1 | 0.978426098 | 0.084559423 |
| Cxcl2 | 0.043223248 | 3.533234218 | 0.057 | 0.013 | 1 | 0.51919986 | 0.83720485 | 0 | 0.01 | 1 | 0.51919986 | 0.084578247 |
| Pcdhb4 | 0.043225378 | 1.932330173 | 0.057 | 0.013 | 1 | 0.139550713 | -1.747757651 | 0 | 0.051 | 1 | 0.139550713 | 0.084582322 |
| Fdxacb1 | 0.148562499 | -1.1672055 | 0.019 | 0.072 | 1 | 0.04323174 | -2.292078167 | 0 | 0.092 | 1 | 0.148562499 | 0.084594496 |
| Cct7 | 0.043278713 | 0.39606193 | 0.83 | 0.72 | 1 | 0.264791118 | 0.25514224 | 0.805 | 0.812 | 1 | 0.264791118 | 0.084684379 |
| Ufm1 | 0.043336912 | 0.690319018 | 0.642 | 0.606 | 1 | 0.481624175 | 0.354314258 | 0.634 | 0.645 | 1 | 0.481624175 | 0.084795735 |
| 2210408I21Rik | 0.043342932 | 0.356356456 | 0.679 | 0.551 | 1 | 0.562548636 | 0.172807882 | 0.659 | 0.611 | 1 | 0.562548636 | 0.084807254 |
| Abr | 0.043355133 | -0.317765177 | 0.283 | 0.428 | 1 | 0.164072465 | -0.516824088 | 0.488 | 0.58 | 1 | 0.164072465 | 0.084830599 |
| 1700093J21Rik | 0.043403575 | 1.517292674 | 0.113 | 0.042 | 1 | 0.451338156 | -0.53202896 | 0.049 | 0.082 | 1 | 0.451338156 | 0.08492328 |
| Swt1 | 0.043474164 | 0.604807041 | 0.698 | 0.619 | 1 | 0.120311837 | 0.44749864 | 0.878 | 0.867 | 1 | 0.120311837 | 0.085058325 |
| Anapc15 | 0.043482649 | 0.849868013 | 0.264 | 0.148 | 1 | 0.758820632 | 1.087183103 | 0.122 | 0.116 | 1 | 0.758820632 | 0.085074557 |
| Tbc1d4 | 0.478059247 | 0.59082171 | 0.283 | 0.254 | 1 | 0.043524873 | 0.825977594 | 0.341 | 0.205 | 1 | 0.478059247 | 0.085155332 |
| Jkamp | 0.356353846 | 0.172644503 | 0.679 | 0.686 | 1 | 0.043591924 | -0.476685926 | 0.439 | 0.631 | 1 | 0.356353846 | 0.085283593 |
| Gpc2 | 0.043597347 | 1.377115016 | 0.113 | 0.042 | 1 | 0.746881137 | 1.029849928 | 0.024 | 0.017 | 1 | 0.746881137 | 0.085293965 |
| Fam83e | 0.043597347 | 1.377115016 | 0.113 | 0.042 | 1 | 0.074149471 | 2.422167351 | 0.098 | 0.038 | 1 | 0.074149471 | 0.085293965 |
| Pcdhga6 | 0.043597347 | 1.377115016 | 0.113 | 0.042 | 1 | 0.765010978 | 0.667279848 | 0.098 | 0.085 | 1 | 0.765010978 | 0.085293965 |
| 6530402F18Rik | 0.043605424 | 2.154722595 | 0.057 | 0.013 | 1 | 0.139479918 | -1.250257991 | 0 | 0.051 | 1 | 0.139479918 | 0.085309415 |
| D430018E03Rik | 0.043605424 | 2.154722595 | 0.057 | 0.013 | 1 | 0.746881137 | 1.029849928 | 0.024 | 0.017 | 1 | 0.746881137 | 0.085309415 |
| BC025920 | 0.043605424 | 2.154722595 | 0.057 | 0.013 | 1 | 0.257325989 | -0.484723245 | 0 | 0.031 | 1 | 0.257325989 | 0.085309415 |
| Gm56795 | 0.043609707 | 2.154722595 | 0.057 | 0.013 | 1 | 0.257285084 | -0.863234868 | 0 | 0.031 | 1 | 0.257285084 | 0.085317608 |
| 4930570G19Rik | 0.043609707 | 2.154722595 | 0.057 | 0.013 | 1 | 0.962773283 | 0.589277336 | 0.049 | 0.048 | 1 | 0.962773283 | 0.085317608 |
| Tmod1 | 0.043609707 | 2.154722595 | 0.057 | 0.013 | 1 | 0.391866128 | 0.614812428 | 0.024 | 0.058 | 1 | 0.391866128 | 0.085317608 |
| Gm13134 | 0.043609707 | 2.154722595 | 0.057 | 0.013 | 1 | 0.319619815 | -0.332720152 | 0 | 0.024 | 1 | 0.319619815 | 0.085317608 |
| A430102K17Rik | 0.043609707 | 2.154722595 | 0.057 | 0.013 | 1 | 0.257285084 | -0.747757651 | 0 | 0.031 | 1 | 0.257285084 | 0.085317608 |
| Ssc5d | 0.043609707 | 2.154722595 | 0.057 | 0.013 | 1 | 0.288314827 | 1.252242349 | 0.073 | 0.038 | 1 | 0.288314827 | 0.085317608 |
| Pla2g3 | 0.043609707 | 2.154722595 | 0.057 | 0.013 | 1 | 0.599290422 | 1.515276755 | 0.024 | 0.014 | 1 | 0.599290422 | 0.085317608 |
| Fam124a | 0.043609707 | 2.154722595 | 0.057 | 0.013 | 1 | 0.455097252 | -0.16279515 | 0 | 0.014 | 1 | 0.455097252 | 0.085317608 |
| Map3k8 | 0.043614078 | -0.585757484 | 0.34 | 0.479 | 1 | 0.910829064 | -0.104242968 | 0.317 | 0.3 | 1 | 0.910829064 | 0.085325968 |
| Cldn34c1 | 0.043642799 | 1.417757001 | 0.113 | 0.042 | 1 | 0.170373481 | -1.16279515 | 0 | 0.044 | 1 | 0.170373481 | 0.085380904 |
| Kdelr2 | 0.049097281 | 0.218261677 | 1 | 0.975 | 1 | 0.043682348 | -0.315806769 | 0.976 | 0.973 | 1 | 0.049097281 | 0.085456549 |
| Golm1 | 0.043704048 | 0.946614399 | 0.321 | 0.199 | 1 | 0.505245567 | 0.489281546 | 0.146 | 0.113 | 1 | 0.505245567 | 0.085498052 |
| 5730414N17Rik | 0.043716443 | 1.669295768 | 0.132 | 0.055 | 1 | 0.126329058 | -1.410722664 | 0 | 0.055 | 1 | 0.126329058 | 0.085521759 |
| Snx7 | 0.043764318 | 0.660613525 | 0.547 | 0.407 | 1 | 0.092086558 | 1.100239256 | 0.463 | 0.423 | 1 | 0.092086558 | 0.085613321 |
| Tmem104 | 0.76682112 | 0.306725688 | 0.075 | 0.064 | 1 | 0.043767504 | 1.904319046 | 0.146 | 0.061 | 1 | 0.76682112 | 0.085619414 |
| Galnt2 | 0.043812393 | 0.439228531 | 0.736 | 0.614 | 1 | 0.407821747 | 0.154252558 | 0.78 | 0.775 | 1 | 0.407821747 | 0.08570526 |
| Nudt19 | 0.043838127 | 0.840614004 | 0.415 | 0.309 | 1 | 0.550861784 | 0.658867609 | 0.244 | 0.215 | 1 | 0.550861784 | 0.085754472 |
| Thop1 | 0.043857401 | 1.154722595 | 0.189 | 0.093 | 1 | 0.319407396 | -0.622226769 | 0.073 | 0.126 | 1 | 0.319407396 | 0.08579133 |
| Scn1b | 0.04850846 | 0.684580496 | 0.509 | 0.386 | 1 | 0.043907029 | -0.520347155 | 0.366 | 0.529 | 1 | 0.04850846 | 0.08588623 |
| Fkbp1b | 0.386676888 | -0.319208594 | 0.132 | 0.182 | 1 | 0.043961208 | -1.777504994 | 0.024 | 0.133 | 1 | 0.386676888 | 0.085989828 |
| Edc4 | 0.043972829 | 0.705415193 | 0.377 | 0.242 | 1 | 0.622507527 | -0.389864059 | 0.22 | 0.242 | 1 | 0.622507527 | 0.086012048 |
| Gstm2 | 0.209004666 | 1.669295768 | 0.038 | 0.013 | 1 | 0.043983747 | 0.974708374 | 0.22 | 0.109 | 1 | 0.209004666 | 0.086032924 |
| Togaram2 | 0.04399045 | 1.932330173 | 0.057 | 0.013 | 1 | 0.5191986 | 0.515276755 | 0 | 0.01 | 1 | 0.5191986 | 0.08604574 |
| Acbd5 | 0.043996897 | 0.24997091 | 0.792 | 0.64 | 1 | 0.582807203 | 0.262192009 | 0.732 | 0.792 | 1 | 0.582807203 | 0.086058068 |
| Adamtsl4 | 0.044054763 | 1.337586652 | 0.226 | 0.127 | 1 | 0.834364108 | 0.465236072 | 0.195 | 0.188 | 1 | 0.834364108 | 0.086168704 |
| Dcp1b | 0.044074921 | 1.315187267 | 0.189 | 0.093 | 1 | 0.900266628 | 0.334704509 | 0.098 | 0.092 | 1 | 0.900266628 | 0.086207243 |
| Polq | 0.0441235 | -0.330704232 | 0.075 | 0.021 | 1 | 0.51919986 | 0.83720485 | 0 | 0.01 | 1 | 0.51919986 | 0.086300117 |
| Acot9 | 0.385764122 | -0.132158553 | 0.472 | 0.572 | 1 | 0.044130275 | -0.562326063 | 0.537 | 0.648 | 1 | 0.385764122 | 0.086313069 |
| Pgm3 | 0.095733138 | -0.334991591 | 0.547 | 0.682 | 1 | 0.044176477 | -0.346298558 | 0.439 | 0.625 | 1 | 0.095733138 | 0.086401393 |
| Tatdn2 | 0.619983894 | 0.176096246 | 0.189 | 0.225 | 1 | 0.044178018 | -0.675864732 | 0.195 | 0.358 | 1 | 0.619983894 | 0.086404338 |
| Heatr6 | 0.04418112 | 0.535812762 | 0.377 | 0.242 | 1 | 0.772418745 | -0.136799942 | 0.244 | 0.266 | 1 | 0.772418745 | 0.086410268 |
| Nicn1 | 0.04436375 | 1.011764641 | 0.302 | 0.186 | 1 | 0.71361497 | 0.496167932 | 0.171 | 0.205 | 1 | 0.71361497 | 0.086759358 |
| Ranbp10 | 0.044365225 | 0.760443656 | 0.547 | 0.449 | 1 | 0.288439449 | -0.233184478 | 0.537 | 0.631 | 1 | 0.288439449 | 0.086762178 |
| Tmem68 | 0.044410862 | 0.622227514 | 0.396 | 0.263 | 1 | 0.643594648 | 0.325305811 | 0.366 | 0.328 | 1 | 0.643594648 | 0.0868494 |
| Runx3 | 0.236461932 | -0.417528854 | 0.132 | 0.208 | 1 | 0.044419706 | 0.166850858 | 0.195 | 0.372 | 1 | 0.236461932 | 0.086866303 |
| Fbxl15 | 0.044478001 | 1.104096522 | 0.208 | 0.106 | 1 | 0.400560026 | -0.233184478 | 0.073 | 0.119 | 1 | 0.400560026 | 0.086977709 |
| Zfp771 | 0.044528136 | 0.813685677 | 0.491 | 0.373 | 1 | 0.568272976 | -0.153750011 | 0.341 | 0.399 | 1 | 0.568272976 | 0.087073517 |
| Nherf1 | 0.311426131 | 0.8327945 | 0.075 | 0.042 | 1 | 0.044536585 | 1.635570989 | 0.268 | 0.15 | 1 | 0.311426131 | 0.087089662 |
| Trappc11 | 0.04457215 | 1.055186921 | 0.321 | 0.208 | 1 | 0.390226973 | 0.790911198 | 0.268 | 0.235 | 1 | 0.390226973 | 0.087157623 |
| Tipin | 0.044636299 | 0.518413214 | 0.491 | 0.364 | 1 | 0.144337667 | -0.686357106 | 0.244 | 0.355 | 1 | 0.144337667 | 0.087280198 |
| H2-T23 | 0.213997641 | -1.058271129 | 0.151 | 0.225 | 1 | 0.044647297 | 1.189506594 | 0.22 | 0.109 | 1 | 0.213997641 | 0.087301212 |
| Aldh1l1 | 0.044648689 | -2.304709024 | 0 | 0.072 | 1 | 0.319629309 | -0.622226769 | 0 | 0.024 | 1 | 0.319629309 | 0.087303873 |
| Cage1 | 0.044648689 | -2.237594828 | 0 | 0.072 | 1 | 0.4113861 | 1.159132945 | 0.073 | 0.044 | 1 | 0.4113861 | 0.087303873 |
| Gm8098 | 0.044665451 | -2.237594828 | 0 | 0.072 | 1 | 0.390492216 | 2.209173627 | 0.073 | 0.044 | 1 | 0.390492216 | 0.087335899 |
| Urah | 0.044679819 | -2.600164907 | 0 | 0.072 | 1 | 0.743829142 | 0.252242349 | 0.024 | 0.034 | 1 | 0.743829142 | 0.087363351 |
| Gm8883 | 0.044679978 | -2.489133595 | 0 | 0.072 | 1 | 0.453885966 | -0.16279515 | 0.024 | 0.051 | 1 | 0.453885966 | 0.087363656 |
| Tmem59l | 0.044679978 | -2.304709024 | 0 | 0.072 | 1 | 0.286402368 | -0.484723245 | 0 | 0.027 | 1 | 0.286402368 | 0.087363656 |
| Crlf3 | 0.139767078 | -0.318529518 | 0.585 | 0.699 | 1 | 0.044680541 | -0.544224257 | 0.341 | 0.543 | 1 | 0.139767078 | 0.087364732 |
| Gm47644 | 0.044690835 | -2.430239906 | 0 | 0.072 | 1 | 0.448204037 | -0.484723245 | 0.024 | 0.051 | 1 | 0.448204037 | 0.087384399 |
| Mybph | 0.044691793 | -2.752168001 | 0 | 0.072 | 1 | 0.231632701 | -1.747757651 | 0 | 0.034 | 1 | 0.231632701 | 0.087386229 |
| Polr1has | 0.044693425 | -2.093204919 | 0.019 | 0.106 | 1 | 0.356645148 | 1.029849928 | 0.073 | 0.041 | 1 | 0.356645148 | 0.087389347 |
| Gm19303 | 0.044727719 | -3.054737771 | 0 | 0.072 | 1 | 0.749728988 | 0.83720485 | 0.024 | 0.017 | 1 | 0.749728988 | 0.08745487 |
| Mkks | 0.044738099 | -2.015202407 | 0 | 0.072 | 1 | 0.189411038 | -0.970150072 | 0.024 | 0.082 | 1 | 0.189411038 | 0.087474701 |
| 4930532G15Rik | 0.044764761 | 1.8327945 | 0.057 | 0.013 | 1 | 0.494346678 | -0.555112573 | 0.024 | 0.048 | 1 | 0.494346678 | 0.087525639 |
| Gm57263 | 0.044764761 | 1.154722595 | 0.057 | 0.013 | 1 | 0.057901606 | 2.422167351 | 0.049 | 0.01 | 1 | 0.057901606 | 0.087525639 |
| Alpk3 | 0.044764761 | 0.347367673 | 0.057 | 0.013 | 1 | 0.437109795 | 0.377773231 | 0.073 | 0.044 | 1 | 0.437109795 | 0.087525639 |
| Xylb | 0.044764761 | 1.8327945 | 0.057 | 0.013 | 1 | 0.911761241 | 0.136765132 | 0.024 | 0.027 | 1 | 0.911761241 | 0.087525639 |
| Kif26a | 0.044764761 | 1.8327945 | 0.057 | 0.013 | 1 | 0.205309831 | -0.20718927 | 0.024 | 0.082 | 1 | 0.205309831 | 0.087525639 |

| | | | | | | | | | | | | |
|---|---|---|---|---|---|---|---|---|---|---|---|---|
| Cacna1h | 0.044764761 | 1.8327945 | 0.057 | 0.013 | 1 | 0.51919986 | 0.83720485 | 0 | 0.01 | 1 | 0.51919986 | 0.087525639 |
| Dnd1 | 0.044764761 | 1.8327945 | 0.057 | 0.013 | 1 | 0.126329058 | -1.332720152 | 0 | 0.055 | 1 | 0.126329058 | 0.087525639 |
| Frat2 | 0.044764761 | 1.8327945 | 0.057 | 0.013 | 1 | 0.17794278 | 1.252242349 | 0.073 | 0.031 | 1 | 0.17794278 | 0.087525639 |
| Arhgap4 | 0.044764761 | 1.569760094 | 0.057 | 0.013 | 1 | 0.171157199 | 1.458693227 | 0.073 | 0.031 | 1 | 0.171157199 | 0.087525639 |
| Fermt3 | 0.044771295 | 2.154722595 | 0.057 | 0.013 | 1 | 0.530000087 | 0.287007767 | 0.122 | 0.089 | 1 | 0.530000087 | 0.087538121 |
| Tspan1 | 0.04477173 | 1.891688189 | 0.057 | 0.013 | 1 | 0.083211164 | 1.574170444 | 0.098 | 0.038 | 1 | 0.083211164 | 0.087538953 |
| Rep15 | 0.59720124 | 0.154722595 | 0.075 | 0.102 | 1 | 0.044829216 | 1.458693227 | 0.146 | 0.061 | 1 | 0.59720124 | 0.087648773 |
| 4933406I18Rik | 0.044835436 | -1.075011802 | 0.151 | 0.292 | 1 | 0.16065476 | -0.661046018 | 0.146 | 0.256 | 1 | 0.16065476 | 0.087660656 |
| Laptm4b | 0.072227154 | 0.954189607 | 0.415 | 0.309 | 1 | 0.044867335 | 1.096447676 | 0.512 | 0.413 | 1 | 0.072227154 | 0.087721593 |
| A530046M15Rik | 0.044868025 | -2.015202407 | 0.019 | 0.106 | 1 | 0.590756788 | 1.83720485 | 0.024 | 0.014 | 1 | 0.590756788 | 0.087722909 |
| Nsun4 | 0.572578689 | -0.237594828 | 0.208 | 0.246 | 1 | 0.045020609 | -1.200930279 | 0.146 | 0.287 | 1 | 0.572578689 | 0.088014363 |
| Vcpip1 | 0.914921548 | 0.347367673 | 0.528 | 0.564 | 1 | 0.045097028 | 0.512074323 | 0.805 | 0.724 | 1 | 0.914921548 | 0.088160315 |
| Sgcd | 0.045104179 | 0.215969784 | 0.868 | 0.746 | 1 | 0.801337147 | -0.153867844 | 1 | 0.898 | 1 | 0.801337147 | 0.088173971 |
| Mpp4 | 0.045150456 | 1.762405172 | 0.132 | 0.055 | 1 | 0.822292188 | 0.100239256 | 0.098 | 0.085 | 1 | 0.822292188 | 0.088262348 |
| Paxip1 | 0.453485159 | 0.361173472 | 0.226 | 0.182 | 1 | 0.045151025 | -1.116991461 | 0.146 | 0.29 | 1 | 0.453485159 | 0.088263435 |
| Ephb3 | 0.045165288 | 1.039245377 | 0.321 | 0.212 | 1 | 0.511915877 | -0.195216628 | 0.341 | 0.392 | 1 | 0.511915877 | 0.088290673 |
| Elmod3 | 0.045172181 | 0.869320376 | 0.226 | 0.119 | 1 | 0.61626054 | -0.555112573 | 0.073 | 0.096 | 1 | 0.61626054 | 0.088303836 |
| Cic | 0.045204616 | 0.869320376 | 0.472 | 0.36 | 1 | 0.929914711 | 0.130936053 | 0.488 | 0.505 | 1 | 0.929914711 | 0.088365774 |
| Gpr68 | 0.045213828 | -1.845277405 | 0.019 | 0.106 | 1 | 0.402818967 | -0.917682652 | 0.024 | 0.055 | 1 | 0.402818967 | 0.088383366 |
| Gm4890 | 0.045213937 | -1.752168001 | 0.019 | 0.106 | 1 | 0.710563777 | 1.458693227 | 0.049 | 0.038 | 1 | 0.710563777 | 0.088383573 |
| 2410022M11Rik | 0.045274002 | 1.417757001 | 0.132 | 0.055 | 1 | 0.916649754 | 0.667279848 | 0.024 | 0.027 | 1 | 0.916649754 | 0.08849827 |
| Plac9 | 0.298749209 | -1.30014786 | 0.585 | 0.585 | 1 | 0.045335699 | -0.624746761 | 0.634 | 0.761 | 1 | 0.298749209 | 0.088616073 |
| Utp23 | 0.045338606 | 1.123695699 | 0.245 | 0.144 | 1 | 0.12774759 | -0.622226769 | 0.122 | 0.235 | 1 | 0.12774759 | 0.088621623 |
| Zfp654 | 0.045354374 | 0.540731045 | 0.585 | 0.47 | 1 | 0.047239176 | 0.38418522 | 0.756 | 0.618 | 1 | 0.047239176 | 0.088651728 |
| Esrp1 | 0.160046644 | 1.347367673 | 0.057 | 0.021 | 1 | 0.04539917 | 0.922934724 | 0.268 | 0.147 | 1 | 0.160046644 | 0.088737256 |
| Alx3 | 0.739392541 | 0.154722595 | 0.189 | 0.22 | 1 | 0.045413278 | 1.252242349 | 0.268 | 0.157 | 1 | 0.739392541 | 0.088764189 |
| Hhat | 0.664382361 | 0.190055668 | 0.264 | 0.318 | 1 | 0.045470387 | -0.809396717 | 0.22 | 0.406 | 1 | 0.664382361 | 0.088873218 |
| Prdm4 | 0.045492024 | 0.789073122 | 0.321 | 0.199 | 1 | 0.173378975 | 0.600165653 | 0.39 | 0.294 | 1 | 0.173378975 | 0.088914524 |
| Ift88 | 0.234377538 | -0.356239324 | 0.34 | 0.419 | 1 | 0.045527166 | -0.914536097 | 0.244 | 0.389 | 1 | 0.234377538 | 0.088981609 |
| Cluh | 0.045544961 | 1.09332205 | 0.283 | 0.174 | 1 | 0.687702183 | 0.366884915 | 0.317 | 0.3 | 1 | 0.687702183 | 0.089015579 |
| Ubiad1 | 0.045553653 | 1.389187848 | 0.132 | 0.055 | 1 | 0.397377029 | -0.192542494 | 0.098 | 0.15 | 1 | 0.397377029 | 0.08903217 |
| Pigx | 0.045580064 | 0.548543608 | 0.528 | 0.398 | 1 | 0.093748031 | -0.562725757 | 0.317 | 0.447 | 1 | 0.093748031 | 0.089082586 |
| Reps1 | 0.045671877 | 0.943825812 | 0.434 | 0.309 | 1 | 0.44581349 | 0.477662463 | 0.512 | 0.478 | 1 | 0.44581349 | 0.089257833 |
| Extl3 | 0.045688434 | 0.894855468 | 0.396 | 0.288 | 1 | 0.121186191 | -0.622226769 | 0.244 | 0.369 | 1 | 0.121186191 | 0.089289434 |
| Chst14 | 0.045689563 | 1.203632195 | 0.208 | 0.11 | 1 | 0.126317354 | 0.989207943 | 0.195 | 0.113 | 1 | 0.126317354 | 0.089291589 |
| Mks1 | 0.045718609 | 1.221836791 | 0.17 | 0.081 | 1 | 0.842357354 | 0.83720485 | 0.024 | 0.031 | 1 | 0.842357354 | 0.089347028 |
| Arl4d | 0.045752094 | 1.10741688 | 0.151 | 0.068 | 1 | 0.432970939 | 0.749742009 | 0.073 | 0.044 | 1 | 0.432970939 | 0.089410934 |
| Hmgcl | 0.045752887 | 0.629261106 | 0.434 | 0.301 | 1 | 0.12587948 | -0.385187572 | 0.268 | 0.413 | 1 | 0.12587948 | 0.089412448 |
| Rps6ka2 | 0.045809989 | -0.963274615 | 0.189 | 0.339 | 1 | 0.750948108 | -0.737703986 | 0.171 | 0.188 | 1 | 0.750948108 | 0.089521423 |
| Klhl13 | 0.045837581 | 0.620744023 | 0.415 | 0.275 | 1 | 0.581783311 | 1.019043173 | 0.268 | 0.266 | 1 | 0.581783311 | 0.089574079 |
| Tbc1d30 | 0.045958895 | 0.984797593 | 0.057 | 0.013 | 1 | 0.933764087 | 1.136765132 | 0.024 | 0.027 | 1 | 0.933764087 | 0.08980557 |
| Fes | 0.046002913 | 1.154722595 | 0.17 | 0.081 | 1 | 0.306876945 | -0.286177566 | 0.073 | 0.137 | 1 | 0.306876945 | 0.089889558 |
| Hdac6 | 0.046154435 | 1.221836791 | 0.264 | 0.157 | 1 | 0.593563395 | 1.199774929 | 0.171 | 0.15 | 1 | 0.593563395 | 0.090178637 |
| Dmxl2 | 0.046216243 | 0.984797593 | 0.245 | 0.14 | 1 | 0.287270601 | 0.895307804 | 0.22 | 0.157 | 1 | 0.287270601 | 0.090296546 |
| Kcnip2 | 0.046239828 | 2.444229212 | 0.094 | 0.034 | 1 | 0.402333528 | -0.16279515 | 0.049 | 0.089 | 1 | 0.402333528 | 0.090341534 |
| Nipsnap2 | 0.764011878 | 0.190055668 | 0.34 | 0.381 | 1 | 0.046297526 | -0.832073937 | 0.317 | 0.464 | 1 | 0.764011878 | 0.090451592 |
| Cdk5rap2 | 0.046364307 | 0.377115016 | 0.66 | 0.521 | 1 | 0.786536066 | 0.152706676 | 0.463 | 0.471 | 1 | 0.786536066 | 0.090578966 |
| Plaur | 0.046386747 | -1.553096654 | 0.132 | 0.25 | 1 | 0.613782348 | 0.100239256 | 0.195 | 0.16 | 1 | 0.613782348 | 0.090621763 |
| Vps33b | 0.046395872 | 0.855162313 | 0.226 | 0.119 | 1 | 0.599071773 | 0.392420007 | 0.146 | 0.116 | 1 | 0.599071773 | 0.090639167 |
| Phrf1 | 0.046401786 | 0.598759966 | 0.623 | 0.521 | 1 | 0.449174066 | 0.21693548 | 0.756 | 0.747 | 1 | 0.449174066 | 0.090650446 |
| Mettl5 | 0.859945614 | 0.324647596 | 0.226 | 0.225 | 1 | 0.046405862 | -0.332720152 | 0.098 | 0.256 | 1 | 0.859945614 | 0.090658219 |
| Zswim7 | 0.046410298 | -0.631873767 | 0.094 | 0.225 | 1 | 0.087508341 | 1.438655473 | 0.244 | 0.15 | 1 | 0.087508341 | 0.090666681 |
| Wdr73 | 0.046412746 | 1.090592257 | 0.189 | 0.093 | 1 | 0.089297265 | -0.970150072 | 0.073 | 0.181 | 1 | 0.089297265 | 0.090671348 |
| 4930550C14Rik | 0.536387113 | 0.874614676 | 0.132 | 0.106 | 1 | 0.046412866 | -1.53202896 | 0.049 | 0.167 | 1 | 0.536387113 | 0.090671578 |
| Rps3 | 0.048219292 | 0.224558491 | 1 | 0.97 | 1 | 0.046550076 | -0.201379857 | 1 | 0.99 | 1 | 0.048219292 | 0.090933242 |
| Nars | 0.840871087 | -0.160826588 | 0.887 | 0.818 | 1 | 0.04655468 | -0.305512193 | 0.732 | 0.915 | 1 | 0.840871087 | 0.090942022 |
| Dgkd | 0.046561399 | 0.764776076 | 0.491 | 0.373 | 1 | 0.069366303 | 0.78473743 | 0.561 | 0.444 | 1 | 0.069366303 | 0.090954835 |
| Ssr1 | 0.231018026 | 0.14304079 | 0.943 | 0.945 | 1 | 0.046583531 | -0.308094703 | 0.878 | 0.952 | 1 | 0.231018026 | 0.090997036 |
| Agpat2 | 0.046590955 | -0.639693272 | 0.189 | 0.335 | 1 | 0.735502533 | 0.370078839 | 0.195 | 0.232 | 1 | 0.735502533 | 0.091011193 |
| Nhej1 | 0.046640162 | 1.647762606 | 0.245 | 0.144 | 1 | 0.404492521 | 0.627751484 | 0.195 | 0.16 | 1 | 0.404492521 | 0.09110502 |
| Ric8a | 0.046753158 | 0.859266711 | 0.321 | 0.199 | 1 | 0.918976485 | 0.204936634 | 0.22 | 0.235 | 1 | 0.918976485 | 0.091320458 |
| Ddx10 | 0.222211804 | 0.409651951 | 0.604 | 0.538 | 1 | 0.046763175 | 0.984430803 | 0.585 | 0.474 | 1 | 0.222211804 | 0.091339555 |
| Prr14l | 0.046800614 | 0.457649728 | 0.698 | 0.597 | 1 | 0.05929369 | -0.371248381 | 0.683 | 0.816 | 1 | 0.05929369 | 0.09141093 |
| Fhdc1 | 0.046891983 | -1.112063946 | 0.113 | 0.233 | 1 | 0.969686122 | 0.533424102 | 0.171 | 0.174 | 1 | 0.969686122 | 0.091585109 |
| Tmem248 | 0.145981573 | -0.214511215 | 0.642 | 0.746 | 1 | 0.046933739 | -0.467898622 | 0.683 | 0.84 | 1 | 0.145981573 | 0.091664703 |
| Ccdc93 | 0.047041194 | 0.967095592 | 0.245 | 0.136 | 1 | 0.445814408 | 0.238167164 | 0.317 | 0.253 | 1 | 0.445814408 | 0.091869514 |
| Pdcd2l | 0.047075032 | 1.347367673 | 0.245 | 0.148 | 1 | 0.356003919 | -0.592782991 | 0.171 | 0.229 | 1 | 0.356003919 | 0.091934005 |
| Sh3bp2 | 0.617587111 | -0.330704232 | 0.057 | 0.076 | 1 | 0.047092725 | -0.990614175 | 0.049 | 0.177 | 1 | 0.617587111 | 0.091967726 |
| Lrp10 | 0.047105622 | 0.662869498 | 0.509 | 0.398 | 1 | 0.556842479 | -0.157148587 | 0.512 | 0.553 | 1 | 0.556842479 | 0.091992304 |
| Map2k7 | 0.047108011 | 0.997181318 | 0.302 | 0.195 | 1 | 0.423401842 | -0.114539982 | 0.293 | 0.352 | 1 | 0.423401842 | 0.091996857 |
| Usb1 | 0.257127764 | 1.112902419 | 0.189 | 0.131 | 1 | 0.047135938 | -0.717384002 | 0.024 | 0.137 | 1 | 0.257127764 | 0.092050079 |
| ENSMUSG00000121307 | 0.047222576 | 0.880547631 | 0.415 | 0.292 | 1 | 0.343609318 | -0.376574441 | 0.244 | 0.314 | 1 | 0.343609318 | 0.09221518 |
| Jmjd1c | 0.047368436 | 0.283605984 | 1 | 0.987 | 1 | 0.200224221 | 0.253897006 | 1 | 0.99 | 1 | 0.200224221 | 0.092493103 |
| Slc35e2 | 0.047415966 | 0.776210972 | 0.377 | 0.242 | 1 | 0.458309417 | -0.230539757 | 0.195 | 0.256 | 1 | 0.458309417 | 0.092583659 |
| 2610029K11Rik | 0.243322126 | 1.154722595 | 0.057 | 0.025 | 1 | 0.047434708 | 3.351778023 | 0.073 | 0.02 | 1 | 0.243322126 | 0.092619364 |
| Bglap | 0.047517551 | -2.976432836 | 0.17 | 0.297 | 1 | 0.423536326 | -2.03315987 | 0.073 | 0.044 | 1 | 0.423536326 | 0.092777185 |
| Abt1 | 0.047609619 | 0.8817041 | 0.396 | 0.292 | 1 | 0.060540824 | -0.887687912 | 0.171 | 0.311 | 1 | 0.060540824 | 0.092952563 |
| Kif16b | 0.96987694 | -0.106237483 | 0.453 | 0.47 | 1 | 0.047638937 | -0.474381301 | 0.61 | 0.717 | 1 | 0.96987694 | 0.093008405 |
| Drp2 | 0.219807859 | -1.082316603 | 0.057 | 0.114 | 1 | 0.047672001 | -2.332720152 | 0 | 0.089 | 1 | 0.219807859 | 0.093071383 |
| Lyrm1 | 0.834885587 | -0.142957954 | 0.17 | 0.182 | 1 | 0.047672001 | -2.372248516 | 0 | 0.089 | 1 | 0.834885587 | 0.093071383 |
| Pradc1 | 0.699599866 | -0.134784022 | 0.132 | 0.153 | 1 | 0.047672832 | -2.20718927 | 0 | 0.089 | 1 | 0.699599866 | 0.093072965 |
| Azin1 | 0.159111028 | 0.364940302 | 0.943 | 0.89 | 1 | 0.047729205 | 0.797069738 | 0.902 | 0.925 | 1 | 0.159111028 | 0.093180333 |
| Dnajc7 | 0.047739474 | 0.266912741 | 0.868 | 0.822 | 1 | 0.159412667 | -0.167750032 | 0.854 | 0.904 | 1 | 0.159412667 | 0.09319989 |
| Gtf2ird2 | 0.047743811 | 1.197791317 | 0.189 | 0.093 | 1 | 0.592658325 | 0.422167351 | 0.146 | 0.116 | 1 | 0.592658325 | 0.09320815 |
| Atg2a | 0.047762834 | 0.557787452 | 0.434 | 0.297 | 1 | 0.062757279 | 1.298784935 | 0.366 | 0.266 | 1 | 0.062757279 | 0.093244379 |
| Xrcc6 | 0.047769224 | 0.855162313 | 0.34 | 0.216 | 1 | 0.60130147 | 0.471555377 | 0.146 | 0.191 | 1 | 0.60130147 | 0.093256549 |
| Map3k20 | 0.047774972 | 0.415183057 | 0.491 | 0.343 | 1 | 0.441583213 | 0.109284395 | 0.561 | 0.471 | 1 | 0.441583213 | 0.093267497 |
| Pdcl3 | 0.708070864 | -0.226526591 | 0.736 | 0.712 | 1 | 0.047866251 | -0.338143824 | 0.683 | 0.805 | 1 | 0.708070864 | 0.093441323 |
| Vav2 | 0.048084803 | 1.247831999 | 0.226 | 0.123 | 1 | 0.133005333 | 0.614812428 | 0.293 | 0.191 | 1 | 0.133005333 | 0.093857458 |
| Sugp2 | 0.048110568 | 0.730224766 | 0.434 | 0.309 | 1 | 0.145220179 | -0.622226769 | 0.244 | 0.358 | 1 | 0.145220179 | 0.09390651 |
| Ifit1 | 0.206393216 | -4.531777932 | 0 | 0.03 | 1 | 0.048139335 | 2.515276755 | 0.073 | 0.02 | 1 | 0.206393216 | 0.093961274 |
| Calm3 | 0.602086661 | -0.221293016 | 0.774 | 0.742 | 1 | 0.048149738 | 0.547698233 | 0.805 | 0.788 | 1 | 0.602086661 | 0.09398108 |
| Rel | 0.152894926 | 0.534570666 | 0.453 | 0.339 | 1 | 0.04824958 | 1.136765132 | 0.512 | 0.386 | 1 | 0.152894926 | 0.094171138 |
| Lrrc14 | 0.048301451 | 1.117247889 | 0.245 | 0.144 | 1 | 0.749584433 | -0.221688839 | 0.122 | 0.137 | 1 | 0.749584433 | 0.094269872 |
| Cybc1 | 0.449293242 | 0.331600357 | 0.208 | 0.161 | 1 | 0.048302197 | 1.313642894 | 0.244 | 0.13 | 1 | 0.449293242 | 0.094271292 |
| Pck2 | 0.34089744 | 0.259904831 | 0.34 | 0.275 | 1 | 0.048304766 | -0.811083586 | 0.171 | 0.338 | 1 | 0.34089744 | 0.094276182 |
| Mapk8ip1 | 0.048380693 | 1.547040018 | 0.208 | 0.11 | 1 | 0.793623488 | 0.309957847 | 0.22 | 0.208 | 1 | 0.793623488 | 0.094420695 |
| Ift52 | 0.237817375 | 0.436493563 | 0.396 | 0.318 | 1 | 0.048383892 | -1.037264268 | 0.171 | 0.317 | 1 | 0.237817375 | 0.094426783 |

| | | | | | | | | | | | | |
|---|---|---|---|---|---|---|---|---|---|---|---|---|
| Snhg16 | 0.94915486 | 0.137649081 | 0.245 | 0.25 | 1 | 0.048393539 | -0.622226769 | 0.098 | 0.235 | 1 | 0.94915486 | 0.094445144 |
| Il3ra | 0.048493941 | 2.154722595 | 0.094 | 0.034 | 1 | 0.745976859 | 0.377773231 | 0.024 | 0.034 | 1 | 0.745976859 | 0.09463622 |
| Manba | 0.048499073 | 0.154722595 | 0.264 | 0.148 | 1 | 0.220174515 | 0.200895469 | 0.293 | 0.208 | 1 | 0.220174515 | 0.094645986 |
| Cfp | 0.048520941 | 1.237184755 | 0.132 | 0.055 | 1 | 0.627075525 | -0.332720152 | 0.073 | 0.096 | 1 | 0.627075525 | 0.0946876 |
| Trpv2 | 0.048521896 | 2.860991392 | 0.094 | 0.034 | 1 | 0.243087156 | 0.377773231 | 0.073 | 0.034 | 1 | 0.243087156 | 0.094689418 |
| Golt1b | 0.070447243 | 0.41973465 | 0.679 | 0.606 | 1 | 0.048527514 | -0.648221977 | 0.439 | 0.573 | 1 | 0.070447243 | 0.094700108 |
| Chst3 | 0.317969571 | -0.652632327 | 0.132 | 0.186 | 1 | 0.048533464 | -0.498185505 | 0.195 | 0.372 | 1 | 0.317969571 | 0.094711432 |
| Dsp | 0.048551532 | -0.27154216 | 0.264 | 0.144 | 1 | 0.151544258 | 0.83720485 | 0.341 | 0.249 | 1 | 0.151544258 | 0.094745812 |
| Sipa1l2 | 0.065890933 | -1.410062024 | 0.151 | 0.267 | 1 | 0.048586747 | 0.796562865 | 0.366 | 0.239 | 1 | 0.065890933 | 0.094812821 |
| Syce2 | 0.048642761 | 1.090592257 | 0.151 | 0.068 | 1 | 0.141428497 | -1.20718927 | 0.024 | 0.092 | 1 | 0.141428497 | 0.094919404 |
| Rnd2 | 0.048669329 | 0.852159825 | 0.245 | 0.136 | 1 | 1 | 0.319356545 | 0.171 | 0.174 | 1 | 1 | 0.094969955 |
| Cog5 | 0.14394468 | 0.747812976 | 0.528 | 0.492 | 1 | 0.048788391 | 0.375756886 | 0.732 | 0.536 | 1 | 0.14394468 | 0.095196474 |
| Map3k7 | 0.048790978 | 0.522454379 | 0.604 | 0.487 | 1 | 0.542376765 | 0.237742779 | 0.634 | 0.669 | 1 | 0.542376765 | 0.095201397 |
| Cuta | 0.04895003 | 0.483923925 | 0.736 | 0.653 | 1 | 0.048869801 | 0.633332516 | 0.634 | 0.549 | 1 | 0.04895003 | 0.095351345 |
| Lipo3 | 0.177183041 | 0.154722595 | 0.094 | 0.178 | 1 | 0.048871531 | -0.537834582 | 0.073 | 0.212 | 1 | 0.177183041 | 0.095354635 |
| Bloc1s2 | 0.048881842 | 0.991223863 | 0.302 | 0.186 | 1 | 0.360437673 | -0.385187572 | 0.098 | 0.154 | 1 | 0.360437673 | 0.09537425 |
| Cetn2 | 0.048902385 | 0.435623421 | 0.642 | 0.504 | 1 | 0.614312276 | 0.354812083 | 0.463 | 0.416 | 1 | 0.614312276 | 0.095413326 |
| Atf7ip | 0.048955895 | 0.602181572 | 0.717 | 0.619 | 1 | 0.25821832 | -0.383898876 | 0.732 | 0.829 | 1 | 0.25821832 | 0.095515111 |
| Rnf138 | 0.049014518 | 0.62817678 | 0.491 | 0.377 | 1 | 0.322131523 | 0.527349587 | 0.512 | 0.474 | 1 | 0.322131523 | 0.095626612 |
| Vti1a | 0.049031276 | 1.053277331 | 0.66 | 0.564 | 1 | 0.668478749 | -0.165076091 | 0.756 | 0.686 | 1 | 0.668478749 | 0.095658485 |
| Cox10 | 0.587290004 | 0.256602209 | 0.151 | 0.123 | 1 | 0.04907962 | -1.050320421 | 0.049 | 0.171 | 1 | 0.587290004 | 0.09575043 |
| Aup1 | 0.049088735 | -0.616336198 | 0.321 | 0.462 | 1 | 0.726942854 | 0.319356545 | 0.439 | 0.444 | 1 | 0.726942854 | 0.095767766 |
| Arxes1 | 0.049097572 | 1.438515561 | 0.151 | 0.068 | 1 | 0.669505029 | -0.250257991 | 0.073 | 0.092 | 1 | 0.669505029 | 0.095784573 |
| Magi2 | 0.04910332 | 0.265595812 | 0.849 | 0.712 | 1 | 0.506202661 | -0.232216551 | 0.707 | 0.71 | 1 | 0.506202661 | 0.095795504 |
| Rpl15 | 0.148932899 | 0.153706016 | 1 | 0.966 | 1 | 0.049123606 | -0.235142002 | 0.976 | 1 | 1 | 0.148932899 | 0.095834083 |
| Hace1 | 0.049129285 | 0.8327945 | 0.34 | 0.225 | 1 | 0.063448376 | 0.805355984 | 0.537 | 0.433 | 1 | 0.063448376 | 0.095844883 |
| Tns2 | 0.049146168 | 0.397579119 | 0.679 | 0.525 | 1 | 0.375317792 | -0.210308611 | 0.561 | 0.652 | 1 | 0.375317792 | 0.09587699 |
| Cdk7 | 0.04918461 | 0.628653783 | 0.642 | 0.593 | 1 | 0.942489513 | 0.202317124 | 0.634 | 0.693 | 1 | 0.942489513 | 0.095950094 |
| Ddx20 | 0.049215011 | 1.228723176 | 0.245 | 0.14 | 1 | 0.937729806 | 0.145327145 | 0.22 | 0.229 | 1 | 0.937729806 | 0.096007904 |
| Esco1 | 0.049245138 | 0.459577176 | 0.585 | 0.466 | 1 | 0.901567559 | 0.415247937 | 0.659 | 0.706 | 1 | 0.901567559 | 0.096065192 |
| Gm45159 | 0.049305092 | -1.40599236 | 0.057 | 0.161 | 1 | 0.154111598 | -1.250257991 | 0 | 0.048 | 1 | 0.154111598 | 0.096179191 |
| Hycc1 | 0.338166774 | 0.480950827 | 0.358 | 0.305 | 1 | 0.049311185 | 0.617156369 | 0.512 | 0.379 | 1 | 0.338166774 | 0.096190777 |
| Aldh2 | 0.558465624 | -0.110198921 | 0.962 | 0.869 | 1 | 0.049341959 | -0.338608711 | 0.902 | 0.952 | 1 | 0.558465624 | 0.09624929 |
| Pnp | 0.049346477 | -0.710891725 | 0.906 | 0.911 | 1 | 0.465238624 | -0.177373615 | 0.829 | 0.932 | 1 | 0.465238624 | 0.09625788 |
| Ghr | 0.802460245 | 0.17128644 | 0.717 | 0.788 | 1 | 0.049431281 | -0.752330381 | 0.61 | 0.744 | 1 | 0.802460245 | 0.09641911 |
| Ssna1 | 0.049473201 | 0.640149422 | 0.491 | 0.36 | 1 | 0.736146169 | 0.26892109 | 0.415 | 0.386 | 1 | 0.736146169 | 0.096498805 |
| Eno2 | 0.049488151 | 0.685237311 | 0.208 | 0.106 | 1 | 0.691311839 | 0.410940095 | 0.293 | 0.266 | 1 | 0.691311839 | 0.096527226 |
| Gm44956 | 0.074441876 | 1.47665069 | 0.075 | 0.025 | 1 | 0.049531856 | 2.83720485 | 0.073 | 0.02 | 1 | 0.074441876 | 0.096610308 |
| Kcnk7 | 0.287761733 | -0.652632327 | 0 | 0.021 | 1 | 0.049531856 | 2.351778023 | 0.073 | 0.02 | 1 | 0.287761733 | 0.096610308 |
| Lrrc61 | 0.750969908 | 0.984797593 | 0.038 | 0.03 | 1 | 0.049531856 | 2.351778023 | 0.073 | 0.02 | 1 | 0.750969908 | 0.096610308 |
| Spata21 | 0.932611893 | 0.347367673 | 0.019 | 0.017 | 1 | 0.049531856 | 2.351778023 | 0.073 | 0.02 | 1 | 0.932611893 | 0.096610308 |
| Dtx3 | 0.146596426 | 0.304469714 | 0.698 | 0.627 | 1 | 0.049558057 | -0.316995799 | 0.439 | 0.642 | 1 | 0.146596426 | 0.096660113 |
| Mrpl23 | 0.049564768 | 0.428098945 | 0.736 | 0.614 | 1 | 0.806074125 | 0.181853021 | 0.561 | 0.57 | 1 | 0.806074125 | 0.09667287 |
| C030034L19Rik | 0.049668762 | 2.154722595 | 0.094 | 0.034 | 1 | 0.105459701 | -1.622226769 | 0.049 | 0.137 | 1 | 0.105459701 | 0.096870538 |
| Tnk2 | 0.049712178 | -1.150131987 | 0.245 | 0.377 | 1 | 0.088193922 | -0.819907437 | 0.195 | 0.328 | 1 | 0.088193922 | 0.096953055 |
| Yju2 | 0.684448075 | 0.399835093 | 0.208 | 0.186 | 1 | 0.049712681 | -0.410722664 | 0.073 | 0.205 | 1 | 0.684448075 | 0.096954012 |
| Sap18 | 0.049742933 | 0.380655374 | 0.83 | 0.763 | 1 | 0.520065218 | 0.179905407 | 0.854 | 0.823 | 1 | 0.520065218 | 0.097011507 |
| Ndufaf2 | 0.049816401 | 0.661682584 | 0.528 | 0.386 | 1 | 0.073692113 | -0.455576899 | 0.244 | 0.403 | 1 | 0.073692113 | 0.097151129 |
| Fxyd5 | 0.049904722 | 0.464653912 | 0.811 | 0.712 | 1 | 0.141359018 | -0.320120115 | 0.732 | 0.795 | 1 | 0.141359018 | 0.097318963 |
| Pla2g12a | 0.049910708 | 0.8817041 | 0.396 | 0.288 | 1 | 0.874364712 | 0.142059431 | 0.317 | 0.345 | 1 | 0.874364712 | 0.097330337 |
| Eftud2 | 0.050023764 | 0.609288458 | 0.472 | 0.343 | 1 | 0.282640582 | -0.36790958 | 0.317 | 0.399 | 1 | 0.282640582 | 0.097545152 |
| Krt7 | 0.28776878 | -0.845277405 | 0 | 0.021 | 1 | 0.050102366 | 0.782757066 | 0.244 | 0.13 | 1 | 0.28776878 | 0.097694484 |
| Gm16599 | 0.345721962 | 0.221836791 | 0.17 | 0.119 | 1 | 0.050108295 | 0.422167351 | 0.049 | 0.177 | 1 | 0.345721962 | 0.097705748 |
| Myg1 | 0.050156199 | 0.770058745 | 0.283 | 0.165 | 1 | 0.148756492 | -0.659220976 | 0.122 | 0.222 | 1 | 0.148756492 | 0.097796754 |
| Cyhr1 | 0.05016046 | 1.337586652 | 0.245 | 0.144 | 1 | 0.287459052 | -0.215906487 | 0.171 | 0.256 | 1 | 0.287459052 | 0.097804849 |
| Lpin1 | 0.050169776 | -0.688037663 | 0.245 | 0.39 | 1 | 0.079424645 | 0.909097768 | 0.366 | 0.256 | 1 | 0.079424645 | 0.097822545 |
| Fbxo36 | 0.450947907 | -0.247375849 | 0.17 | 0.22 | 1 | 0.050189863 | -0.16279515 | 0.049 | 0.171 | 1 | 0.450947907 | 0.097860704 |
| Fhl2 | 0.050214868 | -1.239191613 | 0.358 | 0.458 | 1 | 0.111033835 | -1.672852842 | 0.22 | 0.321 | 1 | 0.111033835 | 0.097908203 |
| Zfp263 | 0.050316611 | 1.050385935 | 0.264 | 0.157 | 1 | 0.189390943 | -0.536253546 | 0.146 | 0.246 | 1 | 0.189390943 | 0.098101461 |
| Mturn | 0.05036978 | 0.906795081 | 0.226 | 0.123 | 1 | 0.496308809 | -0.296650897 | 0.146 | 0.191 | 1 | 0.496308809 | 0.098202444 |
| Rps4x | 0.284591068 | 0.104977288 | 1 | 0.987 | 1 | 0.050396242 | -0.267469816 | 1 | 0.986 | 1 | 0.284591068 | 0.098252703 |
| Tmem263 | 0.054610766 | 0.390789953 | 0.792 | 0.784 | 1 | 0.050440559 | -0.40152931 | 0.805 | 0.86 | 1 | 0.054610766 | 0.098336867 |
| Sag | 0.050449319 | 1.459577176 | 0.189 | 0.097 | 1 | 0.748770529 | -0.185162963 | 0.122 | 0.14 | 1 | 0.748770529 | 0.098353505 |
| Wdr48 | 0.050491558 | 0.698084191 | 0.509 | 0.381 | 1 | 0.658653137 | -0.228884341 | 0.488 | 0.491 | 1 | 0.658653137 | 0.098433718 |
| Wdr5 | 0.518832327 | 0.280253477 | 0.302 | 0.258 | 1 | 0.050535081 | -0.16279515 | 0.22 | 0.427 | 1 | 0.518832327 | 0.098516368 |
| G2e3 | 0.050536236 | 0.841062116 | 0.396 | 0.267 | 1 | 0.231634037 | 1.2075733 | 0.39 | 0.355 | 1 | 0.231634037 | 0.098518562 |
| mt-Nd2 | 0.083986484 | -0.152324987 | 1 | 1 | 1 | 0.050592659 | 0.142318475 | 1 | 1 | 1 | 0.083986484 | 0.098625701 |
| Bhlhb9 | 0.05065939 | 0.821298861 | 0.34 | 0.216 | 1 | 0.15621946 | -0.463964685 | 0.22 | 0.334 | 1 | 0.15621946 | 0.098752406 |
| Sult1e1 | 0.656463773 | -2.054730771 | 0.019 | 0.03 | 1 | 0.050702501 | 1.930314254 | 0.195 | 0.096 | 1 | 0.656463773 | 0.098834259 |
| Zfp955a | 0.702779044 | 0.541745718 | 0.189 | 0.174 | 1 | 0.050724439 | -0.795063366 | 0.049 | 0.171 | 1 | 0.702779044 | 0.098875909 |
| Gfod2 | 0.591136794 | 0.842778588 | 0.17 | 0.148 | 1 | 0.050725243 | -1.238083278 | 0.098 | 0.229 | 1 | 0.591136794 | 0.098877436 |
| Dclk1 | 0.050733914 | 1.687596584 | 0.208 | 0.114 | 1 | 0.600733226 | 1.313642894 | 0.122 | 0.099 | 1 | 0.600733226 | 0.098893898 |
| 1700037H04Rik | 0.05076861 | 1.47665069 | 0.17 | 0.085 | 1 | 0.142768149 | -1.069685746 | 0.024 | 0.092 | 1 | 0.142768149 | 0.098959768 |
| Fam185a | 0.554914375 | 0.970298024 | 0.151 | 0.127 | 1 | 0.050775626 | -1.484723245 | 0.049 | 0.167 | 1 | 0.554914375 | 0.098973088 |
| Tmem17 | 0.050836126 | 1.502645898 | 0.151 | 0.068 | 1 | 0.372127461 | 0.551802631 | 0.171 | 0.119 | 1 | 0.372127461 | 0.09908794 |
| Gm12762 | 0.05083669 | 1.984797593 | 0.094 | 0.034 | 1 | 0.6957983 | 1.029849928 | 0.024 | 0.038 | 1 | 0.6957983 | 0.09908901 |
| Fam222b | 0.055104788 | 0.469495971 | 0.679 | 0.572 | 1 | 0.05091071 | -0.559312562 | 0.537 | 0.676 | 1 | 0.055104788 | 0.099229519 |
| Samd13 | 0.338780504 | 0.8327945 | 0.057 | 0.03 | 1 | 0.050983875 | 1.989207943 | 0.073 | 0.02 | 1 | 0.338780504 | 0.099368395 |
| Adh1 | 0.050984403 | -2.237594828 | 0.019 | 0.102 | 1 | 0.383695461 | 0.678006255 | 0.244 | 0.188 | 1 | 0.383695461 | 0.099369397 |
| Pdpk1 | 0.051019285 | 0.392221909 | 0.792 | 0.691 | 1 | 0.516552676 | -0.209337736 | 0.902 | 0.901 | 1 | 0.516552676 | 0.099435603 |
| Gm26827 | 0.997810858 | 0.695290976 | 0.038 | 0.038 | 1 | 0.051041044 | 1.711673968 | 0.122 | 0.048 | 1 | 0.997810858 | 0.099476901 |
| Pcnx3 | 0.932109204 | 0.101611258 | 0.302 | 0.301 | 1 | 0.051082139 | -1.079271795 | 0.22 | 0.358 | 1 | 0.932109204 | 0.099554892 |
| Eif2b1 | 0.462695791 | 0.510866405 | 0.321 | 0.275 | 1 | 0.051107857 | -0.814871847 | 0.098 | 0.242 | 1 | 0.462695791 | 0.099603701 |
| Ppp1r37 | 0.051145713 | 0.670790014 | 0.642 | 0.525 | 1 | 0.879782779 | 0.238167164 | 0.634 | 0.71 | 1 | 0.879782779 | 0.099675543 |
| Prpf19 | 0.051198543 | 0.607979416 | 0.509 | 0.441 | 1 | 0.119512909 | 0.620548798 | 0.61 | 0.526 | 1 | 0.119512909 | 0.099775796 |
| Zfp593 | 0.051207291 | -0.909407743 | 0.17 | 0.297 | 1 | 0.466242739 | -0.484723245 | 0.098 | 0.137 | 1 | 0.466242739 | 0.099792396 |
| Commd10 | 0.227025407 | 0.601468954 | 0.377 | 0.335 | 1 | 0.051279148 | 0.560364644 | 0.537 | 0.372 | 1 | 0.227025407 | 0.099928745 |
| Sptbn1 | 0.051338698 | -0.447613498 | 0.981 | 0.979 | 1 | 0.341619761 | 0.161046431 | 1 | 0.997 | 1 | 0.341619761 | 0.100041733 |
| Papolg | 0.441714378 | 0.544669113 | 0.226 | 0.186 | 1 | 0.051352021 | -0.398011612 | 0.098 | 0.239 | 1 | 0.441714378 | 0.100067012 |
| Nfyb | 0.166164886 | -0.460741994 | 0.453 | 0.542 | 1 | 0.05136215 | -0.76156345 | 0.366 | 0.519 | 1 | 0.166164886 | 0.10008623 |
| Pibf1 | 0.051418765 | -0.357176444 | 0.377 | 0.576 | 1 | 0.349234251 | 0.401105735 | 0.585 | 0.546 | 1 | 0.349234251 | 0.10019364 |
| Tiprl | 0.054140484 | 0.470535715 | 0.736 | 0.712 | 1 | 0.051580536 | -0.352315632 | 0.707 | 0.819 | 1 | 0.054140484 | 0.10050052 |
| Tpd52l1 | 0.051623764 | -0.575670345 | 0.226 | 0.386 | 1 | 0.376009303 | 0.252242349 | 0.293 | 0.229 | 1 | 0.376009303 | 0.100582515 |
| Dipk1b | 0.05164458 | 0.849868013 | 0.245 | 0.136 | 1 | 0.274616159 | 2.209173627 | 0.146 | 0.099 | 1 | 0.274616159 | 0.100621998 |
| Nol3 | 0.051664649 | 1.076720083 | 0.151 | 0.068 | 1 | 0.435914758 | -0.448197369 | 0.073 | 0.113 | 1 | 0.435914758 | 0.100660062 |
| Skic8 | 0.713964504 | 0.124257314 | 0.717 | 0.712 | 1 | 0.051690204 | -0.387289094 | 0.707 | 0.747 | 1 | 0.713964504 | 0.100708531 |
| Pdcd2 | 0.051721569 | 1.06161319 | 0.208 | 0.11 | 1 | 0.139139103 | -0.395455907 | 0.049 | 0.137 | 1 | 0.139139103 | 0.100768018 |
| Amdhd2 | 0.05172571 | 1.395730694 | 0.17 | 0.085 | 1 | 0.106053931 | 1.059597271 | 0.146 | 0.072 | 1 | 0.106053931 | 0.10077587 |

| | | | | | | | | | | | | |
|---|---|---|---|---|---|---|---|---|---|---|---|---|
| Pard3bos1 | 0.089002197 | 1.739685096 | 0.057 | 0.017 | 1 | 0.051733173 | 2.029849928 | 0.073 | 0.02 | 1 | 0.089002197 | 0.100790025 |
| Gm15764 | 0.343000446 | -0.430239906 | 0 | 0.017 | 1 | 0.051733173 | 2.029849928 | 0.073 | 0.02 | 1 | 0.343000446 | 0.100790025 |
| Fkbp14 | 0.241582816 | -0.159385996 | 0.66 | 0.763 | 1 | 0.051769695 | -0.595754557 | 0.439 | 0.56 | 1 | 0.241582816 | 0.100859288 |
| Nfyc | 0.051807117 | 0.479344799 | 0.604 | 0.513 | 1 | 0.60368217 | 0.143942734 | 0.585 | 0.608 | 1 | 0.60368217 | 0.100930257 |
| Pam16 | 0.051826954 | -0.485281269 | 0.377 | 0.542 | 1 | 0.060508493 | -0.872453398 | 0.244 | 0.379 | 1 | 0.060508493 | 0.100967876 |
| Zpr1 | 0.051837562 | 0.552058092 | 0.396 | 0.258 | 1 | 0.919003463 | 0.262296014 | 0.341 | 0.362 | 1 | 0.919003463 | 0.100987991 |
| P2ry14 | 0.051837788 | -2.015202407 | 0 | 0.068 | 1 | 0.470842552 | 0.174239837 | 0.024 | 0.051 | 1 | 0.470842552 | 0.10098842 |
| Bmx | 0.051837788 | -2.304709024 | 0 | 0.068 | 1 | 0.729895585 | 1.83720485 | 0.024 | 0.017 | 1 | 0.729895585 | 0.10098842 |
| 4833419F23Rik | 0.051857271 | -2.1672055 | 0 | 0.068 | 1 | 0.60213981 | 0.667279848 | 0.024 | 0.014 | 1 | 0.60213981 | 0.101025365 |
| Oit3 | 0.051874157 | -2.1672055 | 0 | 0.068 | 1 | 0.170402911 | -1.622226769 | 0 | 0.044 | 1 | 0.170402911 | 0.101057386 |
| Gm11696 | 0.051874157 | -2.1672055 | 0 | 0.068 | 1 | 0.992196803 | -0.484723245 | 0.024 | 0.024 | 1 | 0.992196803 | 0.101057386 |
| Tcap | 0.051888446 | -2.430239906 | 0 | 0.068 | 1 | 0.5191986 | 0.515276755 | 0 | 0.01 | 1 | 0.5191986 | 0.101084481 |
| 4930458D05Rik | 0.051888631 | -2.237594828 | 0 | 0.068 | 1 | 0.257325989 | -0.484723245 | 0 | 0.031 | 1 | 0.257325989 | 0.101084833 |
| Sirt2 | 0.05191569 | 0.666928753 | 0.66 | 0.589 | 1 | 0.761622046 | 0.178993367 | 0.61 | 0.666 | 1 | 0.761622046 | 0.101136141 |
| Ihh | 0.051935214 | -5.288220901 | 0 | 0.068 | 1 | 0.455097252 | -2.069685746 | 0 | 0.014 | 1 | 0.455097252 | 0.101173162 |
| Pfn2 | 0.05194067 | 0.820303556 | 0.453 | 0.36 | 1 | 0.540107504 | 0.826936514 | 0.317 | 0.321 | 1 | 0.540107504 | 0.101183508 |
| 2810405F17Rik | 0.051941711 | -1.932740246 | 0 | 0.068 | 1 | 0.497354299 | -0.332720152 | 0.024 | 0.048 | 1 | 0.497354299 | 0.10118548 |
| Mansc1 | 0.051941711 | -1.932740246 | 0 | 0.068 | 1 | 0.126329058 | -1.484723245 | 0 | 0.055 | 1 | 0.126329058 | 0.10118548 |
| Pja1 | 0.051951147 | 0.162584713 | 0.528 | 0.369 | 1 | 0.195253791 | -0.302346503 | 0.341 | 0.464 | 1 | 0.195253791 | 0.101203373 |
| Bola2 | 0.052057701 | 0.749466116 | 0.453 | 0.335 | 1 | 0.07001368 | -0.60905138 | 0.146 | 0.287 | 1 | 0.07001368 | 0.101405398 |
| Acot8 | 0.052077463 | 1.299112504 | 0.245 | 0.144 | 1 | 0.90501645 | 0.228395607 | 0.146 | 0.157 | 1 | 0.90501645 | 0.101442864 |
| Prep | 0.05226284 | 0.83824893 | 0.491 | 0.377 | 1 | 0.127029106 | 0.726173537 | 0.512 | 0.43 | 1 | 0.127029106 | 0.101794276 |
| Setdb2 | 0.052318205 | 1.46284489 | 0.245 | 0.14 | 1 | 0.281885346 | -0.577832649 | 0.122 | 0.195 | 1 | 0.281885346 | 0.101899215 |
| Nfx1 | 0.052332679 | 0.72062676 | 0.623 | 0.559 | 1 | 0.209596051 | -0.259010465 | 0.707 | 0.758 | 1 | 0.209596051 | 0.101926649 |
| Cdc23 | 0.052360974 | 0.75033234 | 0.415 | 0.301 | 1 | 0.442486821 | -0.20718927 | 0.293 | 0.365 | 1 | 0.442486821 | 0.101980276 |
| Prkag2 | 0.052379298 | -0.857804503 | 0.358 | 0.496 | 1 | 0.368175323 | -0.491918746 | 0.488 | 0.549 | 1 | 0.368175323 | 0.102015006 |
| Bloc1s6 | 0.69493236 | 0.321832581 | 0.226 | 0.203 | 1 | 0.052409937 | -0.350422153 | 0.098 | 0.235 | 1 | 0.69493236 | 0.102073072 |
| Mir155hg | 0.052432397 | -1.7032584 | 0.019 | 0.102 | 1 | 0.188517184 | -1.250257991 | 0 | 0.041 | 1 | 0.188517184 | 0.102115639 |
| Ece2 | 0.265932033 | -1.093204919 | 0.019 | 0.055 | 1 | 0.052462014 | -2.020776145 | 0 | 0.085 | 1 | 0.265932033 | 0.102171765 |
| Zfp512 | 0.50132765 | 0.795641501 | 0.226 | 0.191 | 1 | 0.052474097 | -1.083360683 | 0.122 | 0.259 | 1 | 0.50132765 | 0.102194663 |
| Yipf5 | 0.441093149 | 0.191622178 | 0.811 | 0.89 | 1 | 0.052486095 | -0.447140062 | 0.878 | 0.894 | 1 | 0.441093149 | 0.1022174 |
| Tmx4 | 0.052493899 | -0.761156784 | 0.528 | 0.657 | 1 | 0.068564276 | -0.658490313 | 0.61 | 0.652 | 1 | 0.068564276 | 0.102232189 |
| Bri3bp | 0.052495065 | 0.781913331 | 0.453 | 0.347 | 1 | 0.637457614 | 0.631529823 | 0.366 | 0.372 | 1 | 0.637457614 | 0.102234399 |
| Cyp2f2 | 0.078663301 | -0.876304301 | 0.302 | 0.191 | 1 | 0.052516361 | 0.545178241 | 0.463 | 0.317 | 1 | 0.078663301 | 0.102274755 |
| Arpc1b | 0.08629601 | -0.416863308 | 1 | 0.945 | 1 | 0.052521809 | -0.282120583 | 0.976 | 0.983 | 1 | 0.08629601 | 0.102285078 |
| Fgfbp3 | 0.439885513 | -0.477545621 | 0.075 | 0.11 | 1 | 0.05254193 | -2.332720152 | 0 | 0.085 | 1 | 0.439885513 | 0.102323205 |
| Ccdc60 | 0.89261791 | 0.569760094 | 0.038 | 0.034 | 1 | 0.052557568 | -2.332720152 | 0 | 0.085 | 1 | 0.89261791 | 0.102352838 |
| Prim1 | 0.052615477 | 1.189488013 | 0.208 | 0.114 | 1 | 0.326689845 | -0.410722664 | 0.073 | 0.13 | 1 | 0.326689845 | 0.102462565 |
| Polg | 0.520947675 | 0.320732546 | 0.321 | 0.288 | 1 | 0.052648798 | -0.721762442 | 0.22 | 0.382 | 1 | 0.520947675 | 0.102525701 |
| Prpf6 | 0.226422807 | -0.229420897 | 0.623 | 0.682 | 1 | 0.052673595 | -0.353737933 | 0.585 | 0.73 | 1 | 0.226422807 | 0.102572683 |
| Jmjd8 | 0.070401997 | 0.674729654 | 0.302 | 0.186 | 1 | 0.052698605 | -1.180717058 | 0.098 | 0.225 | 1 | 0.070401997 | 0.102620066 |
| Ttf1 | 0.052810629 | 1.154722595 | 0.358 | 0.246 | 1 | 0.766865812 | 0.307951782 | 0.244 | 0.276 | 1 | 0.766865812 | 0.102832296 |
| Vma21 | 0.107344492 | 0.42680214 | 0.698 | 0.64 | 1 | 0.052821647 | -0.348661696 | 0.512 | 0.662 | 1 | 0.107344492 | 0.102853167 |
| Jak2 | 0.052833049 | -0.601407004 | 0.698 | 0.742 | 1 | 0.162476258 | 0.522055288 | 0.829 | 0.744 | 1 | 0.162476258 | 0.102874767 |
| Serinc2 | 0.236091249 | 1.47665069 | 0.057 | 0.025 | 1 | 0.052833457 | 1.83720485 | 0.073 | 0.02 | 1 | 0.236091249 | 0.102875541 |
| Gm44759 | 0.926212226 | 0.8327945 | 0.019 | 0.017 | 1 | 0.052833457 | 1.83720485 | 0.073 | 0.02 | 1 | 0.926212226 | 0.102875541 |
| Catsperd | 0.115410089 | 1.084333267 | 0.132 | 0.068 | 1 | 0.052870461 | 1.541748966 | 0.122 | 0.048 | 1 | 0.115410089 | 0.102945637 |
| Abhd6 | 0.05295715 | 1.154722595 | 0.189 | 0.097 | 1 | 0.379882221 | -0.53202896 | 0.098 | 0.15 | 1 | 0.379882221 | 0.103109841 |
| Adhfe1 | 0.053005692 | -0.354291053 | 0.094 | 0.22 | 1 | 0.584438431 | 0.707921833 | 0.073 | 0.106 | 1 | 0.584438431 | 0.103201782 |
| Gm43328 | 0.053022653 | -1.545717123 | 0.019 | 0.102 | 1 | 0.067834347 | -1.037264268 | 0.098 | 0.222 | 1 | 0.067834347 | 0.103233904 |
| Kif3b | 0.053138869 | 0.600133743 | 0.415 | 0.28 | 1 | 0.723833036 | 0.169349341 | 0.366 | 0.399 | 1 | 0.723833036 | 0.103453998 |
| Ints12 | 0.053234322 | 0.789073122 | 0.34 | 0.216 | 1 | 0.080008736 | -0.863234868 | 0.146 | 0.273 | 1 | 0.080008736 | 0.10363475 |
| Ganc | 0.543004889 | 0.154722595 | 0.189 | 0.153 | 1 | 0.053264653 | -0.659220976 | 0.073 | 0.208 | 1 | 0.543004889 | 0.103692183 |
| Fah | 0.053271237 | 1.792152515 | 0.094 | 0.034 | 1 | 0.210728835 | -0.747757651 | 0.024 | 0.078 | 1 | 0.210728835 | 0.103704649 |
| Mylpf | 0.053271237 | 1.792152515 | 0.094 | 0.034 | 1 | 0.165335714 | 1.83720485 | 0.073 | 0.031 | 1 | 0.165335714 | 0.103704649 |
| Nthl1 | 0.053271237 | 1.792152515 | 0.094 | 0.034 | 1 | 0.824088343 | 0.377773231 | 0.024 | 0.031 | 1 | 0.824088343 | 0.103704649 |
| Clns1a | 0.053272451 | 1.001248114 | 0.434 | 0.343 | 1 | 0.372737283 | 0.367178116 | 0.488 | 0.433 | 1 | 0.372737283 | 0.103706949 |
| Chst8 | 0.053309369 | 2.33529484 | 0.094 | 0.034 | 1 | 0.073426493 | 1.389745873 | 0.098 | 0.038 | 1 | 0.073426493 | 0.10377685 |
| Lrrd1 | 0.16293479 | 0.569760094 | 0.057 | 0.021 | 1 | 0.053311527 | 2.458693227 | 0.098 | 0.034 | 1 | 0.16293479 | 0.103780936 |
| Hsdl1 | 0.053430609 | 0.906795081 | 0.132 | 0.055 | 1 | 0.882376722 | 0.174239837 | 0.098 | 0.106 | 1 | 0.882376722 | 0.104006388 |
| Rbx1 | 0.053443721 | 0.408625989 | 0.962 | 0.903 | 1 | 0.975589887 | 0.161094461 | 0.951 | 0.904 | 1 | 0.975589887 | 0.104031211 |
| Tmem30b | 0.340246113 | 1.417757001 | 0.038 | 0.017 | 1 | 0.053470525 | 1.773074512 | 0.146 | 0.065 | 1 | 0.340246113 | 0.104081953 |
| Psmd13 | 0.053511845 | 0.490589881 | 0.642 | 0.5 | 1 | 0.237879575 | 0.412022793 | 0.659 | 0.662 | 1 | 0.237879575 | 0.104160172 |
| Erap1 | 0.053522135 | 0.925240749 | 0.34 | 0.233 | 1 | 0.803574343 | 0.347238863 | 0.366 | 0.372 | 1 | 0.803574343 | 0.104179651 |
| Abcc1 | 0.916523617 | -0.144837687 | 0.189 | 0.178 | 1 | 0.053549845 | 0.928352738 | 0.537 | 0.42 | 1 | 0.916523617 | 0.104232104 |
| Rpl6 | 0.053604766 | 0.19795153 | 1 | 0.979 | 1 | 0.111616479 | -0.1504538 | 1 | 0.99 | 1 | 0.111616479 | 0.104336062 |
| Xpo4 | 0.053637331 | 0.588619121 | 0.547 | 0.466 | 1 | 0.199127499 | 0.520093771 | 0.537 | 0.478 | 1 | 0.199127499 | 0.104397698 |
| Tead1 | 0.053755493 | -0.465005324 | 0.981 | 0.962 | 1 | 0.10401155 | 0.370278695 | 1 | 0.969 | 1 | 0.10401155 | 0.104621332 |
| Osbpl5 | 0.053778946 | 0.550651271 | 0.377 | 0.242 | 1 | 0.915405762 | 0.146042945 | 0.366 | 0.348 | 1 | 0.915405762 | 0.104665716 |
| Gm41724 | 0.053818316 | -2.108311811 | 0.057 | 0.157 | 1 | 0.836096179 | 0.515276755 | 0.073 | 0.065 | 1 | 0.836096179 | 0.104740221 |
| Chmp1b | 0.053822171 | 0.85355306 | 0.321 | 0.203 | 1 | 0.645461014 | 0.204936634 | 0.293 | 0.256 | 1 | 0.645461014 | 0.104747516 |
| Tango6 | 0.053842696 | 0.407488665 | 0.208 | 0.106 | 1 | 0.188081245 | 0.383028957 | 0.366 | 0.27 | 1 | 0.188081245 | 0.104786357 |
| Gm56517 | 0.053849932 | 0.821298861 | 0.34 | 0.216 | 1 | 0.090040791 | -0.852094311 | 0.195 | 0.314 | 1 | 0.090040791 | 0.104800048 |
| Cptp | 0.053910356 | 1.002719501 | 0.245 | 0.14 | 1 | 0.453533368 | -0.435813645 | 0.049 | 0.082 | 1 | 0.453533368 | 0.104914386 |
| Ado | 0.05391043 | 0.945269229 | 0.245 | 0.14 | 1 | 0.43446476 | 0.287007767 | 0.146 | 0.215 | 1 | 0.43446476 | 0.104914526 |
| Lox | 0.053929428 | -1.644647343 | 0.679 | 0.716 | 1 | 0.290438564 | 0.47463477 | 0.707 | 0.73 | 1 | 0.290438564 | 0.104950473 |
| Stk16 | 0.67482144 | 0.263428854 | 0.396 | 0.39 | 1 | 0.053964379 | -0.438919555 | 0.22 | 0.41 | 1 | 0.67482144 | 0.105016603 |
| Nhp2 | 0.053965919 | 0.650491366 | 0.528 | 0.424 | 1 | 0.456712465 | 0.357211909 | 0.317 | 0.27 | 1 | 0.456712465 | 0.105019517 |
| Bex3 | 0.130955058 | 0.314105219 | 0.717 | 0.678 | 1 | 0.054109296 | -0.194440777 | 0.512 | 0.73 | 1 | 0.130955058 | 0.105290776 |
| Ldlrad3 | 0.06688358 | -0.247885455 | 0.717 | 0.843 | 1 | 0.054164601 | -0.508750516 | 0.829 | 0.877 | 1 | 0.06688358 | 0.105395399 |
| Mrpl41 | 0.361950017 | 0.429829833 | 0.434 | 0.381 | 1 | 0.054215287 | -0.652179991 | 0.195 | 0.348 | 1 | 0.361950017 | 0.105491277 |
| Mastl | 0.054253096 | 1.569760094 | 0.094 | 0.034 | 1 | 0.656078384 | 0.377773231 | 0.073 | 0.055 | 1 | 0.656078384 | 0.105562794 |
| Chmp2a | 0.054255474 | 0.437253176 | 0.83 | 0.818 | 1 | 0.156034372 | -0.140075074 | 0.756 | 0.833 | 1 | 0.156034372 | 0.105567292 |
| Exosc4 | 0.162929256 | 0.59400764 | 0.302 | 0.216 | 1 | 0.05431659 | -1.215906487 | 0.098 | 0.225 | 1 | 0.162929256 | 0.105682888 |
| Jpx | 0.515277413 | -0.732802676 | 0.075 | 0.102 | 1 | 0.054325321 | -1.589059905 | 0.049 | 0.164 | 1 | 0.515277413 | 0.105699402 |
| Rpl24 | 0.054357622 | 0.168554868 | 1 | 0.979 | 1 | 0.229535511 | -0.112722076 | 1 | 1 | 1 | 0.229535511 | 0.105760493 |
| Arpc4 | 0.069174751 | 0.272494053 | 0.868 | 0.784 | 1 | 0.054481936 | -0.295606979 | 0.756 | 0.84 | 1 | 0.069174751 | 0.105995591 |
| Gal3st1 | 0.054559598 | -1.477545621 | 0.057 | 0.157 | 1 | 0.809543016 | 0.174239837 | 0.049 | 0.058 | 1 | 0.809543016 | 0.106142446 |
| Bcar1 | 0.054588487 | 0.669295768 | 0.491 | 0.364 | 1 | 0.749182092 | 0.565245214 | 0.439 | 0.481 | 1 | 0.749182092 | 0.106197072 |
| Fuz | 0.51266692 | -0.149058153 | 0.189 | 0.237 | 1 | 0.054635643 | -1.215906487 | 0.073 | 0.198 | 1 | 0.51266692 | 0.106286233 |
| Guf1 | 0.054664812 | 1.347367673 | 0.245 | 0.148 | 1 | 0.447044655 | -0.484723245 | 0.098 | 0.14 | 1 | 0.447044655 | 0.106341382 |
| Nhlrc3 | 0.054670602 | 1.010332685 | 0.245 | 0.14 | 1 | 0.848966024 | 0.159132945 | 0.146 | 0.16 | 1 | 0.848966024 | 0.10635233 |
| Tmed2 | 0.054708176 | 0.225391542 | 1 | 0.962 | 1 | 0.056396154 | -0.214160782 | 1 | 0.993 | 1 | 0.056396154 | 0.106423367 |
| Gm31356 | 0.343000446 | -0.430239906 | 0 | 0.017 | 1 | 0.054720963 | 4.159132945 | 0.049 | 0.01 | 1 | 0.343000446 | 0.106447542 |
| Sema3a | 0.507281812 | 0.569760094 | 0.019 | 0.008 | 1 | 0.054720963 | 3.422167351 | 0.049 | 0.01 | 1 | 0.507281812 | 0.106447542 |
| Gins4 | 0.054764614 | 0.932330173 | 0.226 | 0.123 | 1 | 0.673534098 | 0.721727632 | 0.171 | 0.154 | 1 | 0.673534098 | 0.106530066 |
| Fhl3 | 0.054774846 | 2.039245377 | 0.113 | 0.047 | 1 | 0.203282539 | -0.53202896 | 0.024 | 0.082 | 1 | 0.203282539 | 0.106549407 |
| Pheta2 | 0.054866966 | 1.640149422 | 0.094 | 0.034 | 1 | 0.744792403 | 1.100239256 | 0.098 | 0.085 | 1 | 0.744792403 | 0.106723548 |

| | | | | | | | | | | | | |
|---|---|---|---|---|---|---|---|---|---|---|---|---|
| Gnmt | 0.054866966 | 1.640149422 | 0.094 | 0.034 | 1 | 0.909322542 | 0.252242349 | 0.024 | 0.027 | 1 | 0.909322542 | 0.106723548 |
| Nsfl1c | 0.118577276 | 0.386048141 | 0.736 | 0.665 | 1 | 0.054890365 | -0.316918296 | 0.634 | 0.778 | 1 | 0.118577276 | 0.106767778 |
| Nagpa | 0.054908206 | 1.154722595 | 0.189 | 0.097 | 1 | 0.214702526 | 1.564186355 | 0.146 | 0.089 | 1 | 0.214702526 | 0.1068015 |
| S100a5 | 0.054911399 | 1.491757582 | 0.094 | 0.034 | 1 | 0.2673283 | 2.252242349 | 0.073 | 0.038 | 1 | 0.2673283 | 0.106807536 |
| Gpx4 | 0.054967002 | -0.431820942 | 0.943 | 0.907 | 1 | 0.277375925 | 0.244747813 | 0.976 | 0.915 | 1 | 0.277375925 | 0.106912632 |
| Akt2 | 0.116167102 | -0.380800973 | 0.679 | 0.801 | 1 | 0.055047039 | -0.384559083 | 0.707 | 0.86 | 1 | 0.116167102 | 0.107063902 |
| Mcc | 0.055051951 | -0.408738837 | 0.679 | 0.763 | 1 | 0.733151237 | 0.409415853 | 0.659 | 0.73 | 1 | 0.733151237 | 0.107073185 |
| Batf | 0.055058945 | 0.714150003 | 0.34 | 0.225 | 1 | 0.412537726 | 1.47463477 | 0.146 | 0.113 | 1 | 0.412537726 | 0.107086402 |
| Rps6ka5 | 0.624298237 | 0.347367673 | 0.623 | 0.733 | 1 | 0.055083406 | 0.830372816 | 0.756 | 0.642 | 1 | 0.624298237 | 0.107132631 |
| Spon1 | 0.108573991 | -0.655637943 | 0.453 | 0.53 | 1 | 0.055133429 | -0.726224489 | 0.244 | 0.41 | 1 | 0.108573991 | 0.107227163 |
| Nfya | 0.992908173 | 0.154722595 | 0.302 | 0.318 | 1 | 0.055156874 | -0.305753104 | 0.171 | 0.338 | 1 | 0.992908173 | 0.107271467 |
| Hacd4 | 0.055166591 | 1.221836791 | 0.321 | 0.203 | 1 | 0.929738818 | 0.638424986 | 0.195 | 0.225 | 1 | 0.929738818 | 0.10728983 |
| Fxyd7 | 0.055187726 | 1.502645898 | 0.094 | 0.034 | 1 | 0.772963198 | 1.614812428 | 0.049 | 0.041 | 1 | 0.772963198 | 0.107329767 |
| Gm49700 | 0.055187726 | 1.502645898 | 0.094 | 0.034 | 1 | 0.7415112 | 0.515276755 | 0.049 | 0.038 | 1 | 0.7415112 | 0.107329767 |
| Cdc6 | 0.055263206 | 0.695290976 | 0.132 | 0.055 | 1 | 0.899796361 | 0.564186355 | 0.073 | 0.068 | 1 | 0.899796361 | 0.107472389 |
| Ndufaf1 | 0.055275115 | 1.025439578 | 0.208 | 0.11 | 1 | 0.267662921 | -0.863234868 | 0.049 | 0.102 | 1 | 0.267662921 | 0.107494891 |
| Usp5 | 0.453313085 | 0.470224421 | 0.34 | 0.292 | 1 | 0.055295675 | 0.69177441 | 0.463 | 0.355 | 1 | 0.453313085 | 0.107533738 |
| Skic3 | 0.055323657 | -0.704162628 | 0.491 | 0.636 | 1 | 0.237348819 | -0.137828235 | 0.512 | 0.577 | 1 | 0.237348819 | 0.107586607 |
| Gm32282 | 0.055389203 | -1.237594828 | 0.057 | 0.157 | 1 | 0.154111598 | -1.250257991 | 0 | 0.048 | 1 | 0.154111598 | 0.107710442 |
| Rrs1 | 0.055482901 | 0.795180208 | 0.472 | 0.386 | 1 | 0.777498915 | 0.387401932 | 0.366 | 0.444 | 1 | 0.777498915 | 0.107887449 |
| Nabp1 | 0.055604238 | 0.771393955 | 0.189 | 0.097 | 1 | 0.058071154 | 0.644559772 | 0.268 | 0.147 | 1 | 0.058071154 | 0.108116644 |
| Trmt6 | 0.055637601 | 0.569760094 | 0.377 | 0.246 | 1 | 0.612990561 | 1.004314836 | 0.171 | 0.15 | 1 | 0.612990561 | 0.10817966 |
| Akt1s1 | 0.055645577 | 0.991784024 | 0.434 | 0.352 | 1 | 0.291004115 | 0.807457506 | 0.439 | 0.392 | 1 | 0.291004115 | 0.108194723 |
| Tcfl5 | 0.055742304 | 0.809587109 | 0.377 | 0.271 | 1 | 0.239738662 | -0.46860358 | 0.22 | 0.321 | 1 | 0.239738662 | 0.108377404 |
| 9330159F19Rik | 0.17037743 | 1.502645898 | 0.075 | 0.034 | 1 | 0.055763997 | 2.83720485 | 0.049 | 0.01 | 1 | 0.17037743 | 0.108418371 |
| Has1 | 0.855707355 | -0.578631746 | 0.151 | 0.136 | 1 | 0.055850189 | -1.332720152 | 0.049 | 0.164 | 1 | 0.855707355 | 0.108581134 |
| Nutf2 | 0.083744733 | 0.79717059 | 0.245 | 0.148 | 1 | 0.055856485 | -1.484723245 | 0.049 | 0.16 | 1 | 0.083744733 | 0.108593023 |
| Chka | 0.138707408 | 0.389187848 | 0.887 | 0.822 | 1 | 0.055901912 | 0.385963541 | 0.878 | 0.922 | 1 | 0.138707408 | 0.108678801 |
| Tspan13 | 0.05595805 | 0.534465959 | 0.962 | 0.928 | 1 | 0.167093657 | -0.13369992 | 0.902 | 0.956 | 1 | 0.167093657 | 0.108784797 |
| Slc31a1 | 0.055968175 | -0.341132432 | 0.566 | 0.737 | 1 | 0.0973052 | 0.455613834 | 0.829 | 0.724 | 1 | 0.0973052 | 0.108803913 |
| Esf1 | 0.055970124 | 0.264905513 | 0.811 | 0.682 | 1 | 0.523182419 | 0.126965128 | 0.683 | 0.703 | 1 | 0.523182419 | 0.108807592 |
| Gm56765 | 0.0909501 | 1.8327945 | 0.057 | 0.017 | 1 | 0.056112729 | 2.059597271 | 0.098 | 0.034 | 1 | 0.0909501 | 0.10907682 |
| Rangrf | 0.482714097 | -0.482707326 | 0.151 | 0.186 | 1 | 0.056178928 | -1.435813645 | 0.049 | 0.16 | 1 | 0.482714097 | 0.109201783 |
| Rcor2 | 0.056189281 | 1.454282877 | 0.094 | 0.034 | 1 | 0.637411772 | 0.749742009 | 0.049 | 0.034 | 1 | 0.637411772 | 0.109221327 |
| Acadl | 0.056189477 | 0.645167497 | 0.585 | 0.521 | 1 | 0.81968293 | 0.108166171 | 0.756 | 0.758 | 1 | 0.81968293 | 0.109221696 |
| Gm1976 | 0.617505727 | 0.647762606 | 0.132 | 0.169 | 1 | 0.05623828 | -0.825760163 | 0.049 | 0.171 | 1 | 0.617505727 | 0.109313815 |
| Agfg1 | 0.266166207 | -0.268272426 | 0.906 | 0.864 | 1 | 0.056276114 | -0.377808041 | 0.829 | 0.939 | 1 | 0.266166207 | 0.109385228 |
| Gm48302 | 0.056280186 | -1.582242999 | 0.019 | 0.102 | 1 | 0.302474778 | -1.069685746 | 0.024 | 0.065 | 1 | 0.302474778 | 0.109392913 |
| Cdca7l | 0.204653189 | 2.154722595 | 0.038 | 0.013 | 1 | 0.056289003 | 3.83720485 | 0.049 | 0.01 | 1 | 0.204653189 | 0.109409555 |
| Gm6034 | 0.343006305 | -0.845277405 | 0 | 0.017 | 1 | 0.056289003 | 3.159132945 | 0.049 | 0.01 | 1 | 0.343006305 | 0.109409555 |
| Gm16174 | 0.475380264 | 1.347367673 | 0.038 | 0.021 | 1 | 0.056289003 | 3.159132945 | 0.049 | 0.01 | 1 | 0.475380264 | 0.109409555 |
| Zfp786 | 0.497382333 | 2.154722595 | 0.019 | 0.008 | 1 | 0.056289003 | 2.83720485 | 0.049 | 0.01 | 1 | 0.497382333 | 0.109409555 |
| Hgd | 0.507281812 | 1.154722595 | 0.019 | 0.008 | 1 | 0.056289003 | 2.83720485 | 0.049 | 0.01 | 1 | 0.507281812 | 0.109409555 |
| B3gat2 | 0.732691937 | 1.154722595 | 0.019 | 0.013 | 1 | 0.056289003 | 3.422167351 | 0.049 | 0.01 | 1 | 0.732691937 | 0.109409555 |
| Rpl3l | 0.916620008 | 1.154722595 | 0.019 | 0.017 | 1 | 0.056289003 | 3.159132945 | 0.049 | 0.01 | 1 | 0.916620008 | 0.109409555 |
| Ocln | 0.343000446 | -0.430239906 | 0 | 0.017 | 1 | 0.056345411 | 0.778311161 | 0.244 | 0.13 | 1 | 0.343000446 | 0.109516016 |
| Apex2 | 0.056429436 | -0.740580027 | 0.321 | 0.445 | 1 | 0.295092579 | -0.13108629 | 0.244 | 0.348 | 1 | 0.295092579 | 0.109674591 |
| Galnt15 | 0.072194168 | 0.47665069 | 0.208 | 0.114 | 1 | 0.056462892 | -2.472650413 | 0.049 | 0.157 | 1 | 0.072194168 | 0.109737726 |
| Itgb3bp | 0.198796709 | -0.56774343 | 0.132 | 0.212 | 1 | 0.056470894 | -0.622226769 | 0.049 | 0.167 | 1 | 0.198796709 | 0.109752826 |
| Spata5l1 | 0.056499527 | 1.502645898 | 0.094 | 0.034 | 1 | 0.590341795 | -0.332720152 | 0.073 | 0.099 | 1 | 0.590341795 | 0.109806857 |
| Gm56904 | 0.056499527 | 1.502645898 | 0.094 | 0.034 | 1 | 0.25729872 | -0.747757651 | 0 | 0.031 | 1 | 0.25729872 | 0.109806857 |
| 4930505N22Rik | 0.056499527 | 1.502645898 | 0.094 | 0.034 | 1 | 0.357854401 | -0.484723245 | 0 | 0.02 | 1 | 0.357854401 | 0.109806857 |
| Lage3 | 0.111628533 | -0.246156841 | 0.358 | 0.496 | 1 | 0.056667577 | -0.598894265 | 0.268 | 0.427 | 1 | 0.111628533 | 0.11012394 |
| Stoml1 | 0.105293523 | 0.640149422 | 0.208 | 0.119 | 1 | 0.05680209 | 1.185128153 | 0.22 | 0.116 | 1 | 0.105293523 | 0.110377702 |
| Dnase1l1 | 0.056810315 | -1.260314904 | 0.094 | 0.203 | 1 | 0.686893136 | 0.496167932 | 0.171 | 0.208 | 1 | 0.686893136 | 0.110393218 |
| Abcb8 | 0.605478554 | 0.267197324 | 0.17 | 0.14 | 1 | 0.056822455 | -1.435813645 | 0.049 | 0.16 | 1 | 0.605478554 | 0.110416118 |
| Lca5l | 0.056834584 | 1.685237311 | 0.151 | 0.072 | 1 | 0.646741372 | 0.515276755 | 0.073 | 0.055 | 1 | 0.646741372 | 0.110438998 |
| Hspb11 | 0.056861195 | 0.941127122 | 0.434 | 0.339 | 1 | 0.475755558 | 0.264315181 | 0.195 | 0.263 | 1 | 0.475755558 | 0.110489194 |
| Zmat3 | 0.056973273 | 0.695290976 | 0.321 | 0.208 | 1 | 0.519461832 | -0.17660095 | 0.195 | 0.239 | 1 | 0.519461832 | 0.110700592 |
| Flrt1 | 0.056990189 | 0.654293604 | 0.415 | 0.284 | 1 | 0.64000399 | 0.243680336 | 0.317 | 0.3 | 1 | 0.64000399 | 0.110732496 |
| Gpkow | 0.057031379 | 0.960075164 | 0.415 | 0.318 | 1 | 0.367412097 | 0.415247937 | 0.463 | 0.399 | 1 | 0.367412097 | 0.110810179 |
| Bmyc | 0.057096747 | 0.72845784 | 0.396 | 0.28 | 1 | 0.491594495 | 0.819282942 | 0.268 | 0.242 | 1 | 0.491594495 | 0.110933456 |
| Gm15283 | 0.149455156 | -0.845277405 | 0.226 | 0.339 | 1 | 0.057106765 | -0.186127654 | 0.195 | 0.362 | 1 | 0.149455156 | 0.110952347 |
| Gm17103 | 0.057184579 | 1.454282877 | 0.094 | 0.034 | 1 | 0.338230615 | 1.396632258 | 0.073 | 0.041 | 1 | 0.338230615 | 0.111099082 |
| Tnfrsf22 | 0.057186418 | -1.278236812 | 0.057 | 0.157 | 1 | 0.806424891 | 0.778311161 | 0.073 | 0.065 | 1 | 0.806424891 | 0.111102549 |
| Gm15839 | 0.162321801 | 1.640149422 | 0.075 | 0.034 | 1 | 0.057359562 | 2.574170444 | 0.049 | 0.01 | 1 | 0.162321801 | 0.111429005 |
| Gm57107 | 0.249034808 | 2.154722595 | 0.019 | 0.004 | 1 | 0.057359562 | 2.515276755 | 0.049 | 0.01 | 1 | 0.249034808 | 0.111429005 |
| Dpys | 0.916620008 | 1.154722595 | 0.019 | 0.017 | 1 | 0.057359562 | 2.515276755 | 0.049 | 0.01 | 1 | 0.916620008 | 0.111429005 |
| Drd3 | 0.057402143 | 1.70221039 | 0.113 | 0.047 | 1 | 0.495921966 | -1.484723245 | 0.024 | 0.048 | 1 | 0.495921966 | 0.11150928 |
| Slc4a8 | 0.56579457 | -0.752168001 | 0.019 | 0.034 | 1 | 0.05741651 | 1.83720485 | 0.098 | 0.034 | 1 | 0.56579457 | 0.111536364 |
| Cep95 | 0.057444873 | 0.569760094 | 0.472 | 0.343 | 1 | 0.425385796 | 0.523788255 | 0.39 | 0.348 | 1 | 0.425385796 | 0.111589832 |
| Nup93 | 0.057453992 | -1.67309643 | 0.321 | 0.436 | 1 | 0.553542957 | 2.079954984 | 0.293 | 0.283 | 1 | 0.553542957 | 0.111607023 |
| 9130004C02Rik | 0.057461206 | 1.280253477 | 0.094 | 0.034 | 1 | 0.260704079 | 2.351778023 | 0.049 | 0.02 | 1 | 0.260704079 | 0.111620622 |
| Rubcn | 0.830676352 | 0.129187503 | 0.283 | 0.314 | 1 | 0.05747443 | -0.609681516 | 0.22 | 0.399 | 1 | 0.830676352 | 0.11164555 |
| Rpl10a | 0.05748573 | 0.244184222 | 1 | 0.979 | 1 | 0.185179499 | -0.145634599 | 1 | 0.99 | 1 | 0.185179499 | 0.111666851 |
| St3gal5 | 0.057639736 | 1.879088152 | 0.283 | 0.191 | 1 | 0.508553573 | 0.319356545 | 0.195 | 0.154 | 1 | 0.508553573 | 0.111957133 |
| Pag1 | 0.757460333 | 0.154722595 | 0.113 | 0.131 | 1 | 0.0576754 | 0.545178241 | 0.439 | 0.29 | 1 | 0.757460333 | 0.112024348 |
| Src | 0.080508442 | 0.804225348 | 0.283 | 0.182 | 1 | 0.05769473 | -0.732650759 | 0.098 | 0.235 | 1 | 0.080508442 | 0.112060778 |
| Zbtb37 | 0.057716928 | 1.010332685 | 0.226 | 0.127 | 1 | 0.627141184 | 0.458693227 | 0.22 | 0.195 | 1 | 0.627141184 | 0.112102613 |
| Mpzl2 | 0.05772791 | -1.477545621 | 0.057 | 0.157 | 1 | 0.296964633 | -0.622226769 | 0.146 | 0.215 | 1 | 0.296964633 | 0.112123308 |
| Slc16a4 | 0.490364801 | -0.223789028 | 0.698 | 0.725 | 1 | 0.057735088 | -0.453627165 | 0.707 | 0.805 | 1 | 0.490364801 | 0.112136835 |
| Gm26903 | 0.12517633 | 1.067259754 | 0.094 | 0.042 | 1 | 0.057743587 | 1.721727632 | 0.098 | 0.034 | 1 | 0.12517633 | 0.112152851 |
| Gjb6 | 0.057783146 | -1.337130502 | 0.019 | 0.102 | 1 | 0.51919986 | -0.332720152 | 0 | 0.01 | 1 | 0.51919986 | 0.112227399 |
| Maob | 0.05783537 | -1.015202407 | 0.208 | 0.343 | 1 | 0.220360166 | -0.499830137 | 0.244 | 0.341 | 1 | 0.220360166 | 0.11232581 |
| A430110L20Rik | 0.290375571 | -0.430239906 | 0.038 | 0.081 | 1 | 0.057848989 | -2.020776145 | 0 | 0.082 | 1 | 0.290375571 | 0.112351473 |
| Chrna1os | 0.807586016 | 0.417757001 | 0.057 | 0.068 | 1 | 0.05784909 | -1.970150072 | 0 | 0.082 | 1 | 0.807586016 | 0.112351663 |
| Ndufb4 | 0.756452509 | 0.10741688 | 0.566 | 0.53 | 1 | 0.057854847 | -0.242522343 | 0.268 | 0.457 | 1 | 0.756452509 | 0.11236251 |
| Hivep3 | 0.791227759 | -0.302083554 | 0.528 | 0.534 | 1 | 0.0578575 | 0.902299878 | 0.61 | 0.481 | 1 | 0.791227759 | 0.11236751 |
| Fen1 | 0.876757465 | -0.108311811 | 0.075 | 0.068 | 1 | 0.057870241 | -2.020776145 | 0 | 0.082 | 1 | 0.876757465 | 0.112391516 |
| Rdm1 | 0.876981862 | -0.406780933 | 0.358 | 0.335 | 1 | 0.057876759 | -0.970150072 | 0.22 | 0.355 | 1 | 0.876981862 | 0.112403798 |
| Gm43330 | 0.486967902 | -0.430239906 | 0.019 | 0.038 | 1 | 0.057889176 | -2.292078167 | 0 | 0.082 | 1 | 0.486967902 | 0.112427196 |
| Slc27a3 | 0.057893472 | 1.067259754 | 0.283 | 0.169 | 1 | 0.440953085 | 0.854482841 | 0.268 | 0.232 | 1 | 0.440953085 | 0.112435289 |
| Gm57242 | 0.09725594 | 3.002719501 | 0.038 | 0.008 | 1 | 0.057899782 | 1.667279848 | 0.049 | 0.01 | 1 | 0.09725594 | 0.112447179 |
| Thbs4 | 0.732691937 | 1.154722595 | 0.019 | 0.013 | 1 | 0.057899782 | 2.159132945 | 0.049 | 0.01 | 1 | 0.732691937 | 0.112447179 |
| Rorb | 0.413082433 | 0.154722595 | 0 | 0.013 | 1 | 0.05790115 | 2.351778023 | 0.049 | 0.01 | 1 | 0.413082433 | 0.112449756 |
| Acot12 | 0.225506368 | 2.154722595 | 0.057 | 0.025 | 1 | 0.057901606 | 2.422167351 | 0.049 | 0.01 | 1 | 0.225506368 | 0.112450615 |
| Tnnt1 | 0.249034808 | 2.154722595 | 0.019 | 0.004 | 1 | 0.057901606 | 2.422167351 | 0.049 | 0.01 | 1 | 0.249034808 | 0.112450615 |
| Baiap3 | 0.249034808 | 2.154722595 | 0.019 | 0.004 | 1 | 0.057901606 | 2.422167351 | 0.049 | 0.01 | 1 | 0.249034808 | 0.112450615 |

| | | | | | | | | | | | | |
|---|---|---|---|---|---|---|---|---|---|---|---|---|
| Ighd | 0.251644813 | 1.154722595 | 0.019 | 0.004 | 1 | 0.057901606 | 2.422167351 | 0.049 | 0.01 | 1 | 0.251644813 | 0.112450615 |
| 9330198N18Rik | 0.28776878 | -1.015202407 | 0 | 0.021 | 1 | 0.057901606 | 2.422167351 | 0.049 | 0.01 | 1 | 0.28776878 | 0.112450615 |
| Slc17a8 | 0.33449729 | 1.154722595 | 0.057 | 0.03 | 1 | 0.057901606 | 2.422167351 | 0.049 | 0.01 | 1 | 0.33449729 | 0.112450615 |
| Gm9776 | 0.334518863 | 1.154722595 | 0.057 | 0.03 | 1 | 0.057901606 | 2.422167351 | 0.049 | 0.01 | 1 | 0.334518863 | 0.112450615 |
| Gm56971 | 0.340246113 | 1.417757001 | 0.038 | 0.017 | 1 | 0.057901606 | 2.422167351 | 0.049 | 0.01 | 1 | 0.340246113 | 0.112450615 |
| H1f1 | 0.418343931 | 1.3662267 | 0.057 | 0.034 | 1 | 0.057901606 | 2.422167351 | 0.049 | 0.01 | 1 | 0.418343931 | 0.112450615 |
| Fam228a | 0.732691937 | 1.154722595 | 0.019 | 0.013 | 1 | 0.057901606 | 2.422167351 | 0.049 | 0.01 | 1 | 0.732691937 | 0.112450615 |
| Galnt5 | 0.424157953 | -0.368839361 | 0.038 | 0.068 | 1 | 0.057918488 | -2.250257991 | 0 | 0.082 | 1 | 0.424157953 | 0.112482424 |
| Vim | 0.05792259 | -0.318051852 | 1 | 0.992 | 1 | 0.189487678 | 0.4851819 | 0.976 | 0.973 | 1 | 0.189487678 | 0.112490154 |
| Gm5165 | 0.057987413 | 1.254258268 | 0.17 | 0.085 | 1 | 0.094557546 | -0.577832649 | 0.049 | 0.147 | 1 | 0.094557546 | 0.112612285 |
| Gm28376 | 0.779443586 | 0.154722595 | 0.019 | 0.025 | 1 | 0.058044756 | 1.059597271 | 0.122 | 0.048 | 1 | 0.779443586 | 0.112720318 |
| Ap3s1 | 0.086532429 | 0.529980192 | 0.887 | 0.835 | 1 | 0.058116058 | 0.422731013 | 0.878 | 0.887 | 1 | 0.086532429 | 0.11285464 |
| Adrb2 | 0.646760843 | -0.108311811 | 0.094 | 0.119 | 1 | 0.058154182 | 1.007129851 | 0.195 | 0.096 | 1 | 0.646760843 | 0.112926455 |
| Trmt10c | 0.058167684 | 0.721763188 | 0.434 | 0.331 | 1 | 0.065313413 | -0.569970532 | 0.341 | 0.491 | 1 | 0.065313413 | 0.112951889 |
| Rad9a | 0.058169719 | 1.377115016 | 0.094 | 0.034 | 1 | 0.061613439 | 1.547698233 | 0.146 | 0.065 | 1 | 0.061613439 | 0.112955721 |
| Poglut1 | 0.058209673 | 0.664396968 | 0.472 | 0.386 | 1 | 0.390210317 | -0.250257991 | 0.293 | 0.365 | 1 | 0.390210317 | 0.11303098 |
| Isca2 | 0.741602514 | -0.180261653 | 0.34 | 0.356 | 1 | 0.05827144 | -1.128579435 | 0.146 | 0.28 | 1 | 0.741602514 | 0.113147319 |
| Pp2d1 | 0.429421714 | 1.154722595 | 0.057 | 0.034 | 1 | 0.058284872 | 1.252242349 | 0.122 | 0.048 | 1 | 0.429421714 | 0.113172618 |
| Samd4 | 0.058312102 | -0.662223687 | 1 | 0.953 | 1 | 0.995176365 | 0.307514231 | 1 | 0.983 | 1 | 0.995176365 | 0.113223903 |
| Fxr2 | 0.058317139 | 0.621848605 | 0.509 | 0.411 | 1 | 0.232187347 | 0.44609319 | 0.659 | 0.594 | 1 | 0.232187347 | 0.113233389 |
| Gm43154 | 0.058364809 | -3.035101964 | 0.019 | 0.097 | 1 | 0.870087833 | 0.574170444 | 0.024 | 0.02 | 1 | 0.870087833 | 0.113323167 |
| Galm | 0.751722273 | -0.430239906 | 0.094 | 0.106 | 1 | 0.058364999 | 1.087183103 | 0.171 | 0.078 | 1 | 0.751722273 | 0.113323526 |
| Gm11630 | 0.49029049 | 0.739685096 | 0.038 | 0.021 | 1 | 0.058445599 | 2.252242349 | 0.049 | 0.01 | 1 | 0.49029049 | 0.11347531 |
| Mfsd4a | 0.058481756 | 1.306725688 | 0.151 | 0.072 | 1 | 0.504909579 | -0.250257991 | 0.098 | 0.137 | 1 | 0.504909579 | 0.113543397 |
| Mtmr4 | 0.058519528 | 1.624207878 | 0.113 | 0.047 | 1 | 0.115613224 | 0.580865097 | 0.171 | 0.089 | 1 | 0.115613224 | 0.113614521 |
| Tmem121 | 0.058519528 | 1.624207878 | 0.113 | 0.047 | 1 | 0.89988627 | 0.589277336 | 0.049 | 0.055 | 1 | 0.89988627 | 0.113614521 |
| Gnpda2 | 0.058578743 | 0.924424975 | 0.321 | 0.212 | 1 | 0.759086195 | 0.444887427 | 0.195 | 0.225 | 1 | 0.759086195 | 0.113726017 |
| Magi1 | 0.798891683 | -0.74275397 | 0.472 | 0.479 | 1 | 0.058579239 | 0.855052851 | 0.634 | 0.532 | 1 | 0.798891683 | 0.113726951 |
| Tbc1d14 | 0.31851705 | 0.695290976 | 0.245 | 0.191 | 1 | 0.058593323 | -0.635282922 | 0.098 | 0.232 | 1 | 0.31851705 | 0.113753468 |
| Mdm1 | 0.058671809 | 1.395730694 | 0.151 | 0.072 | 1 | 0.877592566 | 1.737669176 | 0.049 | 0.044 | 1 | 0.877592566 | 0.113901238 |
| Rnaseh2a | 0.059349491 | 1.404700848 | 0.208 | 0.114 | 1 | 0.058707408 | 1.83720485 | 0.171 | 0.082 | 1 | 0.059349491 | 0.113968256 |
| Ces1d | 0.352046435 | -0.889671525 | 0.057 | 0.03 | 1 | 0.058830486 | 1.177280291 | 0.293 | 0.181 | 1 | 0.352046435 | 0.114199946 |
| Mbp | 0.058854864 | -2.337130502 | 0.019 | 0.097 | 1 | 0.450179283 | 0.614812428 | 0.22 | 0.171 | 1 | 0.450179283 | 0.114245832 |
| Gorasp1 | 0.560149716 | -0.368839361 | 0.132 | 0.161 | 1 | 0.058867405 | -1.747757651 | 0.024 | 0.123 | 1 | 0.560149716 | 0.114269439 |
| Rgs19 | 0.058952471 | 0.676675298 | 0.245 | 0.136 | 1 | 0.113008642 | -0.648221977 | 0.073 | 0.174 | 1 | 0.113008642 | 0.114429548 |
| Ppm1k | 0.058975385 | 1.10741688 | 0.208 | 0.11 | 1 | 0.506152088 | 0.699701326 | 0.122 | 0.092 | 1 | 0.506152088 | 0.114472673 |
| Gm10457 | 0.099079675 | 2.569760094 | 0.038 | 0.008 | 1 | 0.058990144 | 1.100239256 | 0.049 | 0.01 | 1 | 0.099079675 | 0.11450045 |
| Gm39185 | 0.203229524 | 2.154722595 | 0.038 | 0.013 | 1 | 0.058990144 | 2.100239256 | 0.049 | 0.01 | 1 | 0.203229524 | 0.11450045 |
| Tsnaxip1 | 0.211940354 | 1.154722595 | 0.038 | 0.013 | 1 | 0.058990144 | 1.614812428 | 0.049 | 0.01 | 1 | 0.211940354 | 0.11450045 |
| Gm12714 | 0.340246113 | 1.417757001 | 0.038 | 0.017 | 1 | 0.058990144 | 1.83720485 | 0.049 | 0.01 | 1 | 0.340246113 | 0.11450045 |
| Gm12503 | 0.343006305 | -0.1672055 | 0 | 0.017 | 1 | 0.058990144 | 2.100239256 | 0.049 | 0.01 | 1 | 0.343006305 | 0.11450045 |
| Platr1 | 0.343968402 | 1.154722595 | 0.038 | 0.017 | 1 | 0.058990144 | 2.100239256 | 0.049 | 0.01 | 1 | 0.343968402 | 0.11450045 |
| Gm45442 | 0.503971666 | 1.569760094 | 0.019 | 0.008 | 1 | 0.058990144 | 2.100239256 | 0.049 | 0.01 | 1 | 0.503971666 | 0.11450045 |
| Plscr2 | 0.71903859 | -0.247375849 | 0.057 | 0.072 | 1 | 0.058990144 | 2.100239256 | 0.049 | 0.01 | 1 | 0.71903859 | 0.11450045 |
| Gm5627 | 0.209009104 | 1.739685096 | 0.038 | 0.013 | 1 | 0.058993833 | 2.029849928 | 0.049 | 0.01 | 1 | 0.209009104 | 0.114507393 |
| Arl6 | 0.058996221 | 0.976385354 | 0.264 | 0.161 | 1 | 0.822987837 | 0.574170444 | 0.22 | 0.222 | 1 | 0.822987837 | 0.114511887 |
| Cntrob | 0.418044044 | 0.948271717 | 0.113 | 0.081 | 1 | 0.059057453 | -1.890715605 | 0.024 | 0.123 | 1 | 0.418044044 | 0.114627124 |
| Gm49735 | 0.059116567 | 1.517292674 | 0.113 | 0.047 | 1 | 0.986408398 | 1.450181727 | 0.073 | 0.075 | 1 | 0.986408398 | 0.114738366 |
| Zfp874a | 0.059146035 | -1.508242418 | 0.038 | 0.127 | 1 | 0.146332322 | -1.069685746 | 0.073 | 0.157 | 1 | 0.146332322 | 0.114793817 |
| Ahnak | 0.059149369 | 0.354601508 | 0.981 | 0.975 | 1 | 0.499934284 | -0.151906834 | 1 | 1 | 1 | 0.499934284 | 0.11480009 |
| Fastkd5 | 0.059160232 | 1.280253477 | 0.094 | 0.034 | 1 | 0.538149616 | -0.278272368 | 0.049 | 0.075 | 1 | 0.538149616 | 0.114820531 |
| Cabyr | 0.059160232 | 1.154722595 | 0.094 | 0.034 | 1 | 0.24885456 | 0.351778023 | 0.098 | 0.051 | 1 | 0.24885456 | 0.114820531 |
| Rab33b | 0.460317161 | 0.261637799 | 0.509 | 0.466 | 1 | 0.059216783 | -0.435813645 | 0.39 | 0.584 | 1 | 0.460317161 | 0.114926938 |
| Gm42984 | 0.224148665 | -0.776014743 | 0.113 | 0.182 | 1 | 0.059290042 | -1.0454382 | 0.049 | 0.164 | 1 | 0.224148665 | 0.115064775 |
| Nolc1 | 0.059430952 | 0.282183504 | 0.83 | 0.737 | 1 | 0.378481811 | 0.18753666 | 0.854 | 0.809 | 1 | 0.378481811 | 0.115329866 |
| Slc30a9 | 0.059448939 | 0.693495462 | 0.566 | 0.504 | 1 | 0.198554073 | 0.334704509 | 0.78 | 0.676 | 1 | 0.198554073 | 0.115363701 |
| Chmp3 | 0.24497725 | 0.308635278 | 0.755 | 0.674 | 1 | 0.059520219 | -0.369757531 | 0.61 | 0.72 | 1 | 0.24497725 | 0.115497782 |
| Cfdp1 | 0.059570606 | 0.45646053 | 0.962 | 0.89 | 1 | 0.100871359 | -0.219626373 | 0.878 | 0.939 | 1 | 0.100871359 | 0.115592555 |
| Gm15834 | 0.059589413 | -2.337130502 | 0.019 | 0.097 | 1 | 0.746410507 | -0.116991461 | 0.049 | 0.061 | 1 | 0.746410507 | 0.115627927 |
| Aqp5 | 0.77169508 | 0.154722595 | 0.038 | 0.03 | 1 | 0.059642507 | 0.678006255 | 0.268 | 0.15 | 1 | 0.77169508 | 0.115727786 |
| Rex1bd | 0.059650274 | 0.569760094 | 0.642 | 0.508 | 1 | 0.561913056 | -0.135828103 | 0.439 | 0.481 | 1 | 0.561913056 | 0.115742392 |
| H2ac6 | 0.182555428 | 0.491757582 | 0.226 | 0.153 | 1 | 0.059663647 | 1.189506594 | 0.146 | 0.065 | 1 | 0.182555428 | 0.115767543 |
| Orc6 | 0.06193487 | 0.230671448 | 0.509 | 0.352 | 1 | 0.059724539 | -0.720790603 | 0.244 | 0.41 | 1 | 0.06193487 | 0.115882058 |
| Drg2 | 0.059725109 | 0.754184665 | 0.358 | 0.246 | 1 | 0.132918744 | -0.355440228 | 0.195 | 0.331 | 1 | 0.132918744 | 0.115883129 |
| Efnb3 | 0.343171476 | 0.8327945 | 0.057 | 0.03 | 1 | 0.05974614 | 1.029849928 | 0.122 | 0.048 | 1 | 0.343171476 | 0.115922679 |
| Apbb3 | 0.078483064 | 1.039245377 | 0.17 | 0.089 | 1 | 0.059781374 | -1.555112573 | 0.024 | 0.123 | 1 | 0.078483064 | 0.115988936 |
| Pcdh11x | 0.059790625 | 1.879088152 | 0.113 | 0.047 | 1 | 0.802124775 | 0.866351195 | 0.098 | 0.116 | 1 | 0.802124775 | 0.116006332 |
| Eral1 | 0.05982367 | 0.900966003 | 0.208 | 0.11 | 1 | 0.971303669 | 0.334704509 | 0.146 | 0.154 | 1 | 0.971303669 | 0.116068469 |
| Irgq | 0.206308628 | -0.845277405 | 0.094 | 0.161 | 1 | 0.059905016 | -1.292078167 | 0.073 | 0.191 | 1 | 0.206308628 | 0.116221421 |
| Apbb2 | 0.059930211 | 0.239468526 | 0.981 | 0.983 | 1 | 0.208510589 | -0.293320171 | 0.976 | 0.997 | 1 | 0.208510589 | 0.116268791 |
| Ccl27a | 0.626670448 | 0.614154213 | 0.132 | 0.11 | 1 | 0.059958177 | -1.555112573 | 0.024 | 0.123 | 1 | 0.626670448 | 0.116321371 |
| Akip1 | 0.060025225 | 1.154722595 | 0.208 | 0.114 | 1 | 0.550613491 | 0.181853021 | 0.122 | 0.167 | 1 | 0.550613491 | 0.116447423 |
| Gm35608 | 0.060070909 | -2.130679624 | 0.019 | 0.097 | 1 | 0.276574583 | -1.250257991 | 0.024 | 0.068 | 1 | 0.276574583 | 0.116533304 |
| Gm57047 | 0.243102132 | -1.015202407 | 0 | 0.025 | 1 | 0.060103117 | 1.83720485 | 0.049 | 0.01 | 1 | 0.243102132 | 0.116593849 |
| Cabp1 | 0.776830424 | -1.093204919 | 0.019 | 0.025 | 1 | 0.060103117 | 1.83720485 | 0.049 | 0.01 | 1 | 0.776830424 | 0.116593849 |
| Gps1 | 0.488281321 | 0.290377694 | 0.453 | 0.424 | 1 | 0.060108112 | -0.225804948 | 0.317 | 0.498 | 1 | 0.488281321 | 0.116603238 |
| Pias3 | 0.060193147 | 0.792152515 | 0.226 | 0.123 | 1 | 0.60101237 | 0.553411884 | 0.293 | 0.273 | 1 | 0.60101237 | 0.116763079 |
| H3c15 | 0.060222796 | -2.1672055 | 0 | 0.064 | 1 | 0.359002082 | 1.422167351 | 0.049 | 0.024 | 1 | 0.359002082 | 0.116818807 |
| Rasl10a | 0.060242445 | -2.015202407 | 0 | 0.064 | 1 | 0.938546854 | 0.174239837 | 0.049 | 0.051 | 1 | 0.938546854 | 0.116855738 |
| Car13 | 0.762480493 | 0.739685096 | 0.038 | 0.03 | 1 | 0.060251875 | 1.667279848 | 0.171 | 0.082 | 1 | 0.762480493 | 0.116873462 |
| Clvs1 | 0.060259072 | -2.368839361 | 0 | 0.064 | 1 | 0.357847088 | -0.484723245 | 0 | 0.02 | 1 | 0.357847088 | 0.116886988 |
| Gcnt4 | 0.060259072 | -2.1672055 | 0 | 0.064 | 1 | 0.51919986 | 0.252242349 | 0 | 0.01 | 1 | 0.51919986 | 0.116886988 |
| Gm56974 | 0.060259288 | -2.093204919 | 0 | 0.064 | 1 | 0.257325989 | -0.484723245 | 0 | 0.031 | 1 | 0.257325989 | 0.116887394 |
| Bcan | 0.060272676 | -2.237594828 | 0 | 0.064 | 1 | 0.402369942 | -0.16279515 | 0 | 0.017 | 1 | 0.402369942 | 0.116912556 |
| Gm35330 | 0.060273324 | -2.430239906 | 0 | 0.064 | 1 | 0.435967024 | 1.252242349 | 0.024 | 0.01 | 1 | 0.435967024 | 0.116913774 |
| Gm30085 | 0.060273324 | -2.368839361 | 0 | 0.064 | 1 | 0.864610878 | 1.100239256 | 0.024 | 0.02 | 1 | 0.864610878 | 0.116913774 |
| Rab17 | 0.06027354 | -2.1672055 | 0 | 0.064 | 1 | 0.548898236 | -0.16279515 | 0.024 | 0.044 | 1 | 0.548898236 | 0.11691418 |
| Tank | 0.060281299 | -0.986139941 | 0.434 | 0.525 | 1 | 0.892198424 | 0.252242349 | 0.512 | 0.505 | 1 | 0.892198424 | 0.116928763 |
| Foxa2 | 0.060285201 | -2.430239906 | 0 | 0.064 | 1 | 0.402364804 | -0.16279515 | 0 | 0.017 | 1 | 0.402364804 | 0.116936096 |
| Stpg2 | 0.060292975 | -2.600164907 | 0 | 0.064 | 1 | 0.357854401 | -0.484723245 | 0 | 0.02 | 1 | 0.357854401 | 0.116950706 |
| Gm14168 | 0.06029427 | -2.752168001 | 0 | 0.064 | 1 | 0.966145744 | 0.351778023 | 0.073 | 0.072 | 1 | 0.966145744 | 0.116953142 |
| Cenpw | 0.060308309 | -2.015202407 | 0.019 | 0.097 | 1 | 0.103916103 | -1.410722664 | 0 | 0.061 | 1 | 0.103916103 | 0.116979527 |
| H2ac18 | 0.060321049 | -1.845277405 | 0 | 0.064 | 1 | 0.529718256 | 1.515276755 | 0.049 | 0.031 | 1 | 0.529718256 | 0.117003469 |
| Sort1 | 0.06033687 | -0.552783399 | 0.453 | 0.606 | 1 | 0.189800677 | 0.196499797 | 0.439 | 0.601 | 1 | 0.189800677 | 0.117033202 |
| Spa17 | 0.060373721 | -0.932740246 | 0.057 | 0.157 | 1 | 0.441572754 | -0.116991461 | 0.049 | 0.085 | 1 | 0.441572754 | 0.117102455 |
| Zfp626 | 0.060385646 | 1.417757001 | 0.151 | 0.072 | 1 | 0.89478205 | 0.19334866 | 0.122 | 0.13 | 1 | 0.89478205 | 0.117124865 |
| Plin3 | 0.060429333 | 0.507832345 | 0.434 | 0.314 | 1 | 0.582352198 | -0.253648581 | 0.415 | 0.447 | 1 | 0.582352198 | 0.117206961 |
| Praf2 | 0.771601637 | 0.129187503 | 0.566 | 0.555 | 1 | 0.060449183 | -0.637880033 | 0.39 | 0.539 | 1 | 0.771601637 | 0.117244262 |

| | | | | | | | | | | | | |
|---|---|---|---|---|---|---|---|---|---|---|---|---|
| Hadha | 0.06049504 | 0.400806932 | 0.849 | 0.771 | 1 | 0.951496874 | 0.121335091 | 0.707 | 0.85 | 1 | 0.951496874 | 0.117330431 |
| Mfsd2a | 0.060544708 | -1.799473716 | 0.019 | 0.097 | 1 | 0.561392821 | -0.16279515 | 0.049 | 0.075 | 1 | 0.561392821 | 0.117423755 |
| Il18bp | 0.060544708 | -1.752168001 | 0.019 | 0.097 | 1 | 0.586887905 | -0.448197369 | 0.073 | 0.099 | 1 | 0.586887905 | 0.117423755 |
| Hapln3 | 0.187099207 | -0.557995453 | 0.132 | 0.212 | 1 | 0.060581687 | -1.107653596 | 0.073 | 0.195 | 1 | 0.187099207 | 0.117493234 |
| Pigf | 0.060681896 | -0.845277405 | 0.189 | 0.309 | 1 | 0.100569493 | -0.555112573 | 0.195 | 0.334 | 1 | 0.100569493 | 0.117681499 |
| Gpr137 | 0.888666213 | 0.154722595 | 0.208 | 0.199 | 1 | 0.060765812 | -0.970150072 | 0.049 | 0.164 | 1 | 0.888666213 | 0.11783914 |
| Cdh1 | 0.069588058 | 0.221836791 | 0.189 | 0.097 | 1 | 0.060771474 | 1.356579009 | 0.366 | 0.253 | 1 | 0.069588058 | 0.117849776 |
| Tifa | 0.060792996 | -1.832338349 | 0.113 | 0.216 | 1 | 0.105177328 | -1.26713181 | 0.098 | 0.198 | 1 | 0.105177328 | 0.117890204 |
| Zfp68 | 0.060851167 | 0.691322074 | 0.396 | 0.28 | 1 | 0.993753607 | 0.370078839 | 0.39 | 0.416 | 1 | 0.993753607 | 0.117999469 |
| Asf1b | 0.060897534 | 0.739685096 | 0.094 | 0.034 | 1 | 0.083377301 | 1.83720485 | 0.073 | 0.024 | 1 | 0.083377301 | 0.118086558 |
| Ddx31 | 0.071113388 | 0.932330173 | 0.245 | 0.144 | 1 | 0.060947961 | -1.037264268 | 0.073 | 0.195 | 1 | 0.071113388 | 0.118181268 |
| Coq2 | 0.448974259 | -0.24286977 | 0.358 | 0.419 | 1 | 0.06095356 | 0.649577847 | 0.463 | 0.317 | 1 | 0.448974259 | 0.118191784 |
| Vkorc1l1 | 0.060968872 | 0.12888259 | 0.792 | 0.614 | 1 | 0.261988684 | -0.234456135 | 0.659 | 0.788 | 1 | 0.261988684 | 0.118220541 |
| Lap3 | 0.061046467 | 0.414690698 | 0.528 | 0.398 | 1 | 0.155728242 | 0.521059108 | 0.561 | 0.457 | 1 | 0.155728242 | 0.118366263 |
| Papss1 | 0.061051215 | 0.361488781 | 0.679 | 0.593 | 1 | 0.273982872 | -0.218523298 | 0.634 | 0.809 | 1 | 0.273982872 | 0.118375178 |
| Car11 | 0.601939331 | 1.47665069 | 0.038 | 0.025 | 1 | 0.061088919 | 1.351778023 | 0.098 | 0.034 | 1 | 0.601939331 | 0.118445981 |
| Tada3 | 0.850616316 | 0.242185436 | 0.226 | 0.212 | 1 | 0.061146749 | -1.004097404 | 0.098 | 0.225 | 1 | 0.850616316 | 0.118554573 |
| Foxp2 | 0.386214638 | 0.594254656 | 0.396 | 0.331 | 1 | 0.061191079 | 1.001546278 | 0.537 | 0.403 | 1 | 0.386214638 | 0.118637809 |
| 9330159M07Rik | 0.272433867 | 1.361173472 | 0.132 | 0.089 | 1 | 0.061235389 | 0.778311161 | 0.122 | 0.048 | 1 | 0.272433867 | 0.118721005 |
| Tagap1 | 0.27056741 | 0.569760094 | 0.151 | 0.097 | 1 | 0.061275598 | 1.789899135 | 0.171 | 0.085 | 1 | 0.27056741 | 0.118796497 |
| Bod1 | 0.150081072 | -0.336024564 | 0.566 | 0.665 | 1 | 0.061288329 | -0.268446938 | 0.488 | 0.669 | 1 | 0.150081072 | 0.118820398 |
| Leng8 | 0.696339811 | 0.211947463 | 0.472 | 0.487 | 1 | 0.061429019 | -0.435813645 | 0.317 | 0.519 | 1 | 0.696339811 | 0.119084514 |
| Mvd | 0.061435018 | -0.771276824 | 0.113 | 0.233 | 1 | 0.3432632 | -0.292078167 | 0.049 | 0.096 | 1 | 0.3432632 | 0.119095775 |
| Ptpa | 0.061437896 | 0.523110001 | 0.642 | 0.568 | 1 | 0.066776044 | 0.506937454 | 0.732 | 0.625 | 1 | 0.066776044 | 0.119101177 |
| Pgap2 | 0.061533404 | 0.621848605 | 0.491 | 0.369 | 1 | 0.330237653 | -0.429581691 | 0.39 | 0.454 | 1 | 0.330237653 | 0.119280448 |
| Ndufa6 | 0.115276026 | 0.392882332 | 0.981 | 0.852 | 1 | 0.061625696 | 0.296911398 | 0.829 | 0.802 | 1 | 0.115276026 | 0.119453666 |
| Rps19bp1 | 0.07233062 | 0.964391383 | 0.358 | 0.275 | 1 | 0.061692404 | -0.515311565 | 0.195 | 0.355 | 1 | 0.07233062 | 0.119578855 |
| Mterf4 | 0.0618335 | -0.949614065 | 0.057 | 0.157 | 1 | 0.480257482 | -0.132421501 | 0.098 | 0.14 | 1 | 0.480257482 | 0.119843619 |
| Zfp607b | 0.06184676 | 1.454282877 | 0.113 | 0.047 | 1 | 0.553780906 | 0.83720485 | 0.049 | 0.031 | 1 | 0.553780906 | 0.119868497 |
| Wdr81 | 0.06184676 | 1.454282877 | 0.113 | 0.047 | 1 | 0.568383292 | 0.422167351 | 0.122 | 0.092 | 1 | 0.568383292 | 0.119868497 |
| Nrde2 | 0.06184676 | 1.454282877 | 0.113 | 0.047 | 1 | 0.327299972 | 1.296636468 | 0.122 | 0.078 | 1 | 0.327299972 | 0.119868497 |
| Btbd6 | 0.06184676 | 1.454282877 | 0.113 | 0.047 | 1 | 0.273319125 | -0.622226769 | 0.049 | 0.102 | 1 | 0.273319125 | 0.119868497 |
| Sybu | 0.759606403 | 0.739685096 | 0.094 | 0.085 | 1 | 0.061885891 | -0.800225071 | 0.024 | 0.126 | 1 | 0.759606403 | 0.119941918 |
| Bmp2 | 0.061907783 | -0.246418678 | 0.585 | 0.458 | 1 | 0.538695792 | -0.536629394 | 0.561 | 0.628 | 1 | 0.538695792 | 0.119982992 |
| Prr13 | 0.413600238 | -0.515631397 | 0.491 | 0.5 | 1 | 0.06197922 | -0.114539982 | 0.415 | 0.58 | 1 | 0.413600238 | 0.120117017 |
| Gm52988 | 0.062093891 | 0.984797593 | 0.17 | 0.085 | 1 | 0.275524247 | 0.570418309 | 0.195 | 0.133 | 1 | 0.275524247 | 0.120332131 |
| Nudt22 | 0.062162709 | 0.970298024 | 0.189 | 0.097 | 1 | 0.622587069 | 0.109284395 | 0.122 | 0.096 | 1 | 0.622587069 | 0.120461216 |
| Gm26532 | 0.079360975 | -0.487573081 | 0.283 | 0.432 | 1 | 0.062164614 | -0.511523304 | 0.366 | 0.543 | 1 | 0.079360975 | 0.12046479 |
| Mpp7 | 0.062209198 | 0.116394476 | 0.472 | 0.339 | 1 | 0.955990268 | 0.223387486 | 0.512 | 0.58 | 1 | 0.955990268 | 0.120548412 |
| Cul2 | 0.062296116 | 0.520516575 | 0.491 | 0.364 | 1 | 0.259922051 | 0.385216214 | 0.561 | 0.505 | 1 | 0.259922051 | 0.120711425 |
| Enah | 0.62483508 | -0.166425454 | 0.906 | 0.936 | 1 | 0.06229641 | 0.297661474 | 0.829 | 0.877 | 1 | 0.62483508 | 0.120711978 |
| Calr | 0.062436042 | 0.22638358 | 1 | 0.958 | 1 | 0.126694641 | 0.246720471 | 1 | 0.993 | 1 | 0.126694641 | 0.120973826 |
| Apex1 | 0.062455554 | 0.640149422 | 0.547 | 0.441 | 1 | 0.573137742 | 0.503304113 | 0.317 | 0.294 | 1 | 0.573137742 | 0.121010411 |
| E230029C05Rik | 0.110791028 | 1.962077517 | 0.094 | 0.042 | 1 | 0.062487359 | 0.989207943 | 0.098 | 0.034 | 1 | 0.110791028 | 0.121070048 |
| Dcakd | 0.062643778 | 1.069331104 | 0.377 | 0.271 | 1 | 0.271398337 | 0.458693227 | 0.39 | 0.314 | 1 | 0.271398337 | 0.121363312 |
| Slc11a2 | 0.291961677 | -0.289196024 | 0.302 | 0.411 | 1 | 0.062671087 | 0.566675907 | 0.512 | 0.386 | 1 | 0.291961677 | 0.12141451 |
| Cttn | 0.83439352 | 0.182308426 | 0.566 | 0.657 | 1 | 0.062691133 | -0.283810551 | 0.707 | 0.836 | 1 | 0.83439352 | 0.121452088 |
| Plaa | 0.06269236 | 0.53023173 | 0.566 | 0.496 | 1 | 0.493269905 | -0.16279515 | 0.659 | 0.655 | 1 | 0.493269905 | 0.121454388 |
| Pole3 | 0.704899147 | 0.14690309 | 0.509 | 0.492 | 1 | 0.062701922 | -0.509960537 | 0.22 | 0.392 | 1 | 0.704899147 | 0.121472313 |
| Wiz | 0.067755252 | 0.923397049 | 0.321 | 0.216 | 1 | 0.062810096 | -0.435813645 | 0.195 | 0.365 | 1 | 0.067755252 | 0.121675084 |
| Mipep | 0.062837018 | 1.312263872 | 0.226 | 0.136 | 1 | 0.867140854 | 0.88611445 | 0.146 | 0.143 | 1 | 0.867140854 | 0.121725545 |
| Map3k11 | 0.062869042 | 1.090592257 | 0.17 | 0.085 | 1 | 0.332101251 | 0.60690723 | 0.22 | 0.164 | 1 | 0.332101251 | 0.121785567 |
| Creb3 | 0.454303126 | 0.23564259 | 0.604 | 0.564 | 1 | 0.062890661 | -0.304814155 | 0.366 | 0.577 | 1 | 0.454303126 | 0.121826087 |
| Dnajc16 | 0.062951833 | 0.956783217 | 0.226 | 0.127 | 1 | 0.123422166 | -0.747757651 | 0.049 | 0.137 | 1 | 0.123422166 | 0.121940732 |
| Sec24c | 0.173680661 | 0.816852956 | 0.377 | 0.305 | 1 | 0.063004914 | -0.420182993 | 0.195 | 0.358 | 1 | 0.173680661 | 0.122040209 |
| Bpgm | 0.063024328 | -0.24966766 | 0.396 | 0.555 | 1 | 0.423322585 | -0.170547426 | 0.683 | 0.703 | 1 | 0.423322585 | 0.122076589 |
| Tmem260 | 0.063074372 | 0.772252958 | 0.34 | 0.229 | 1 | 0.98523786 | 0.344164839 | 0.268 | 0.287 | 1 | 0.98523786 | 0.122170368 |
| Rgs10 | 0.063079922 | 0.604525512 | 0.509 | 0.377 | 1 | 0.650410732 | -0.584258919 | 0.244 | 0.266 | 1 | 0.650410732 | 0.122180767 |
| Tcte2 | 0.06312302 | 1.517292674 | 0.113 | 0.047 | 1 | 0.604477838 | -0.16279515 | 0.024 | 0.041 | 1 | 0.604477838 | 0.122261524 |
| Aqp4 | 0.77169508 | 0.154722595 | 0.038 | 0.03 | 1 | 0.063159848 | 1.730289646 | 0.268 | 0.164 | 1 | 0.77169508 | 0.12233053 |
| Zfp213 | 0.063188322 | 1.444229212 | 0.132 | 0.059 | 1 | 0.986965808 | 0.252242349 | 0.049 | 0.048 | 1 | 0.986965808 | 0.12238388 |
| Mrpl38 | 0.063220666 | 0.614154213 | 0.415 | 0.292 | 1 | 0.496534398 | 0.614812428 | 0.341 | 0.321 | 1 | 0.496534398 | 0.122444479 |
| Dhx33 | 0.063229413 | 0.808947758 | 0.321 | 0.208 | 1 | 0.287551084 | -0.257952383 | 0.146 | 0.229 | 1 | 0.287551084 | 0.122460867 |
| Trim8 | 0.06326552 | 0.818673392 | 0.509 | 0.419 | 1 | 0.859749765 | 0.535176312 | 0.61 | 0.621 | 1 | 0.859749765 | 0.122528514 |
| Vps37c | 0.257666935 | 0.657222935 | 0.208 | 0.148 | 1 | 0.063300279 | 1.012776414 | 0.317 | 0.208 | 1 | 0.257666935 | 0.122593633 |
| Plpp5 | 0.747783536 | 0.10137317 | 0.679 | 0.669 | 1 | 0.063419968 | -0.221688839 | 0.415 | 0.638 | 1 | 0.747783536 | 0.122817844 |
| Trabd | 0.063428372 | 0.959171786 | 0.34 | 0.25 | 1 | 0.315645876 | -0.341132391 | 0.317 | 0.396 | 1 | 0.315645876 | 0.122833585 |
| Cbll1 | 0.063442381 | 0.546490315 | 0.491 | 0.369 | 1 | 0.538940489 | 0.644559772 | 0.463 | 0.464 | 1 | 0.538940489 | 0.122859826 |
| Gpd1l | 0.070816288 | 2.029191713 | 0.075 | 0.025 | 1 | 0.063451489 | 0.974708374 | 0.22 | 0.116 | 1 | 0.070816288 | 0.122876886 |
| Zfp212 | 0.06347697 | 0.795641501 | 0.302 | 0.191 | 1 | 0.806431494 | 0.630753972 | 0.268 | 0.28 | 1 | 0.806431494 | 0.122924613 |
| Nlrc4 | 0.063530508 | 1.377115016 | 0.113 | 0.047 | 1 | 0.337213694 | -0.686357106 | 0.024 | 0.061 | 1 | 0.337213694 | 0.12302489 |
| Lonrf1 | 0.063556804 | -0.688037663 | 0.264 | 0.394 | 1 | 0.340119082 | -0.126269274 | 0.366 | 0.437 | 1 | 0.340119082 | 0.12307414 |
| Ptger3 | 0.063616836 | -1.626637119 | 0.075 | 0.178 | 1 | 0.226279267 | -1.332720152 | 0.024 | 0.075 | 1 | 0.226279267 | 0.12318657 |
| Armc1 | 0.063733536 | 0.676675298 | 0.547 | 0.479 | 1 | 0.518328169 | -0.178170376 | 0.463 | 0.567 | 1 | 0.518328169 | 0.123405108 |
| 9030612E09Rik | 0.063759072 | 1.347367673 | 0.113 | 0.047 | 1 | 0.286402368 | -0.484723245 | 0 | 0.027 | 1 | 0.286402368 | 0.123452926 |
| AW046200 | 0.174192683 | -0.214511215 | 0.038 | 0.097 | 1 | 0.063759122 | -2.20718927 | 0 | 0.078 | 1 | 0.174192683 | 0.123453018 |
| Bcdin3d | 0.558272287 | 0.247831999 | 0.038 | 0.059 | 1 | 0.063780097 | -2.069685746 | 0 | 0.078 | 1 | 0.558272287 | 0.123492293 |
| Lgi4 | 0.872144182 | 0.347367673 | 0.057 | 0.051 | 1 | 0.063780097 | -2.20718927 | 0 | 0.078 | 1 | 0.872144182 | 0.123492293 |
| Gfer | 0.421302365 | 0.454282877 | 0.189 | 0.144 | 1 | 0.063780303 | -1.140075074 | 0.049 | 0.16 | 1 | 0.421302365 | 0.12349268 |
| Psmg4 | 0.063791527 | -0.52334931 | 0.264 | 0.407 | 1 | 0.818206188 | 0.242258261 | 0.317 | 0.372 | 1 | 0.818206188 | 0.123513695 |
| Cklf | 0.063819179 | 0.569760094 | 0.245 | 0.14 | 1 | 0.392247349 | -0.132421501 | 0.073 | 0.123 | 1 | 0.392247349 | 0.123565471 |
| 2310057M21Rik | 0.163926926 | 0.589125419 | 0.34 | 0.254 | 1 | 0.063819614 | -0.79215177 | 0.098 | 0.235 | 1 | 0.163926926 | 0.123566285 |
| Dpagt1 | 0.465519954 | 0.370451286 | 0.264 | 0.216 | 1 | 0.063872343 | -1.030691614 | 0.073 | 0.195 | 1 | 0.465519954 | 0.12366501 |
| Spry2 | 0.063880114 | -1.042423875 | 0.396 | 0.487 | 1 | 0.782098723 | -0.1539713 | 0.415 | 0.447 | 1 | 0.782098723 | 0.123679558 |
| Herc3 | 0.424673491 | 1.261637799 | 0.057 | 0.034 | 1 | 0.063896823 | 0.898605394 | 0.146 | 0.065 | 1 | 0.424673491 | 0.123710842 |
| F8a | 0.06407878 | 1.247831999 | 0.113 | 0.047 | 1 | 0.868524543 | -0.101394606 | 0.049 | 0.055 | 1 | 0.868524543 | 0.124051469 |
| Ikbkb | 0.064252795 | 0.714150003 | 0.377 | 0.267 | 1 | 0.764217509 | 0.197200986 | 0.341 | 0.389 | 1 | 0.764217509 | 0.124377168 |
| Apol8 | 0.0642719 | 1.154722595 | 0.132 | 0.059 | 1 | 0.391859748 | 0.711673968 | 0.146 | 0.106 | 1 | 0.391859748 | 0.124412923 |
| Tmem185b | 0.295966246 | -0.508242418 | 0.208 | 0.267 | 1 | 0.064504032 | -1.144647803 | 0.098 | 0.218 | 1 | 0.295966246 | 0.124847293 |
| Ppp2ca | 0.064548127 | 0.233547773 | 0.962 | 0.949 | 1 | 0.217421062 | -0.147513692 | 1 | 0.99 | 1 | 0.217421062 | 0.124929794 |
| Hbegf | 0.123413696 | -1.880492098 | 0.415 | 0.458 | 1 | 0.064652717 | 1.364571049 | 0.463 | 0.358 | 1 | 0.123413696 | 0.12512546 |
| Hlx | 0.064661859 | -1.752168001 | 0.057 | 0.153 | 1 | 0.722131862 | 0.667279848 | 0.049 | 0.038 | 1 | 0.722131862 | 0.125142562 |
| Higd2a | 0.639987508 | 0.126630288 | 0.755 | 0.754 | 1 | 0.064697393 | 0.686645173 | 0.732 | 0.7 | 1 | 0.639987508 | 0.125209034 |
| Gm16062 | 0.103989326 | 0.984797593 | 0.151 | 0.081 | 1 | 0.064757053 | -1.917682652 | 0.049 | 0.154 | 1 | 0.103989326 | 0.12532063 |
| Fis1 | 0.064764397 | 0.281957783 | 0.83 | 0.847 | 1 | 0.157309922 | -0.195638407 | 0.732 | 0.85 | 1 | 0.157309922 | 0.125334367 |
| Zfp26 | 0.064835265 | 0.569760094 | 0.358 | 0.237 | 1 | 0.456669812 | -0.287490898 | 0.268 | 0.331 | 1 | 0.456669812 | 0.125466918 |
| Ypel2 | 0.485939254 | -0.668007553 | 0.453 | 0.415 | 1 | 0.064878325 | 0.396956788 | 0.707 | 0.601 | 1 | 0.485939254 | 0.125547452 |
| Usp22 | 0.125517764 | 0.351261589 | 0.547 | 0.458 | 1 | 0.06492349 | 0.853692973 | 0.585 | 0.526 | 1 | 0.125517764 | 0.125631921 |

| | | | | | | | | | | | | |
|---|---|---|---|---|---|---|---|---|---|---|---|---|
| Trmt11 | 0.064995233 | 0.659957903 | 0.283 | 0.169 | 1 | 0.753055762 | 0.174239837 | 0.293 | 0.273 | 1 | 0.753055762 | 0.125766086 |
| Mosmo | 0.583390686 | 0.374688279 | 0.679 | 0.729 | 1 | 0.065005866 | -0.474908955 | 0.683 | 0.788 | 1 | 0.583390686 | 0.125785969 |
| Spcs3 | 0.065037936 | 0.672329828 | 0.415 | 0.297 | 1 | 0.359378047 | 0.671867118 | 0.439 | 0.362 | 1 | 0.359378047 | 0.125845939 |
| Sgsm3 | 0.799913341 | 0.631160639 | 0.283 | 0.288 | 1 | 0.065151493 | -1.052091686 | 0.195 | 0.324 | 1 | 0.799913341 | 0.126058269 |
| Sprr1a | 0.352620284 | -0.56774343 | 0.057 | 0.097 | 1 | 0.065188715 | 0.228395607 | 0.122 | 0.051 | 1 | 0.352620284 | 0.126127861 |
| BC051226 | 0.065326586 | 1.261637799 | 0.113 | 0.047 | 1 | 0.25729872 | -1.069685746 | 0 | 0.031 | 1 | 0.25729872 | 0.126385609 |
| Myo1c | 0.06536792 | 0.334131846 | 0.66 | 0.508 | 1 | 0.635677201 | 0.422167351 | 0.512 | 0.553 | 1 | 0.635677201 | 0.126462875 |
| Nsun5 | 0.065370231 | 1.154722595 | 0.151 | 0.072 | 1 | 0.129376139 | -1.16279515 | 0.024 | 0.096 | 1 | 0.129376139 | 0.126467194 |
| Taldo1 | 0.65451224 | -0.217552585 | 0.849 | 0.822 | 1 | 0.065403892 | -0.393894267 | 0.634 | 0.805 | 1 | 0.65451224 | 0.126530114 |
| Tram2 | 0.07295422 | 0.569760094 | 0.698 | 0.644 | 1 | 0.065406164 | -0.462918875 | 0.61 | 0.72 | 1 | 0.07295422 | 0.126534362 |
| Caml | 0.065487691 | 0.486867086 | 0.623 | 0.517 | 1 | 0.720364826 | 0.257978719 | 0.488 | 0.495 | 1 | 0.720364826 | 0.126686744 |
| Syn1 | 0.065549638 | -0.845277405 | 0.057 | 0.157 | 1 | 0.900000913 | 0.515276755 | 0.122 | 0.119 | 1 | 0.900000913 | 0.126802522 |
| Coch | 0.065639009 | -0.582242999 | 0.17 | 0.085 | 1 | 0.885193729 | 0.313642894 | 0.049 | 0.044 | 1 | 0.885193729 | 0.126969538 |
| Wrap73 | 0.115294794 | 1.154722595 | 0.132 | 0.068 | 1 | 0.065665849 | -1.555112573 | 0.024 | 0.119 | 1 | 0.115294794 | 0.127019694 |
| Samd5 | 0.066015484 | 1.154722595 | 0.302 | 0.199 | 1 | 0.277452471 | -0.813846841 | 0.22 | 0.287 | 1 | 0.277452471 | 0.127672925 |
| Gid8 | 0.503422595 | 0.260215724 | 0.453 | 0.411 | 1 | 0.066044409 | -0.848361592 | 0.317 | 0.433 | 1 | 0.503422595 | 0.127726954 |
| Mapk8ip3 | 0.066178001 | 0.563362323 | 0.623 | 0.551 | 1 | 0.95174052 | 0.278784137 | 0.585 | 0.652 | 1 | 0.95174052 | 0.127976473 |
| Sdhaf4 | 0.066196848 | -0.477545621 | 0.34 | 0.479 | 1 | 0.156663753 | 0.337972223 | 0.22 | 0.362 | 1 | 0.156663753 | 0.128011674 |
| Slc35e1 | 0.066213794 | -0.674165489 | 0.264 | 0.394 | 1 | 0.763755427 | 0.174239837 | 0.415 | 0.471 | 1 | 0.763755427 | 0.128043322 |
| Mfap1a | 0.332037846 | 0.460825723 | 0.377 | 0.331 | 1 | 0.066524434 | -0.654648247 | 0.195 | 0.341 | 1 | 0.332037846 | 0.128623368 |
| Nup62 | 0.066549506 | 0.822147256 | 0.358 | 0.242 | 1 | 0.681760405 | 0.252242349 | 0.244 | 0.29 | 1 | 0.681760405 | 0.128670176 |
| Hspa14 | 0.066551659 | 0.247831999 | 0.566 | 0.432 | 1 | 0.788272832 | 0.399083737 | 0.317 | 0.392 | 1 | 0.788272832 | 0.128674195 |
| Zbtb7c | 0.066557097 | -0.731008837 | 0.472 | 0.602 | 1 | 0.510078838 | -0.119897152 | 0.463 | 0.382 | 1 | 0.510078838 | 0.128684348 |
| Pknox1 | 0.221389316 | 0.261637799 | 0.34 | 0.25 | 1 | 0.066617572 | 0.944120054 | 0.415 | 0.294 | 1 | 0.221389316 | 0.128797242 |
| Recql | 0.066648584 | 1.466666601 | 0.208 | 0.114 | 1 | 0.678475861 | 0.922934724 | 0.146 | 0.126 | 1 | 0.678475861 | 0.128855135 |
| Igfbp2 | 0.306744973 | 0.41149801 | 0.208 | 0.153 | 1 | 0.066703449 | 0.628170811 | 0.341 | 0.218 | 1 | 0.306744973 | 0.128957547 |
| Ogt | 0.092037654 | -0.281471556 | 0.868 | 0.919 | 1 | 0.066744693 | -0.314798244 | 0.902 | 0.98 | 1 | 0.092037654 | 0.129034532 |
| Zfp383 | 0.161839532 | 0.891688189 | 0.189 | 0.119 | 1 | 0.066760616 | -0.890715605 | 0.049 | 0.16 | 1 | 0.161839532 | 0.129064251 |
| Ik | 0.674087831 | 0.116967673 | 0.906 | 0.856 | 1 | 0.066784253 | -0.248451235 | 0.78 | 0.898 | 1 | 0.674087831 | 0.12910837 |
| Ppp2r3c | 0.066902609 | 0.689059022 | 0.302 | 0.186 | 1 | 0.855418264 | 0.399799537 | 0.171 | 0.16 | 1 | 0.855418264 | 0.129329259 |
| Snph | 0.067076772 | -1.067669827 | 0.019 | 0.097 | 1 | 0.106539077 | -1.292078167 | 0.024 | 0.102 | 1 | 0.106539077 | 0.12965425 |
| Gm19325 | 0.092660621 | 0.721763188 | 0.151 | 0.076 | 1 | 0.067088047 | -1.53202896 | 0.049 | 0.154 | 1 | 0.092660621 | 0.129675287 |
| Stk3 | 0.209533499 | 0.478851598 | 0.774 | 0.758 | 1 | 0.067106499 | 0.43228739 | 0.878 | 0.778 | 1 | 0.209533499 | 0.129709716 |
| Gm20186 | 0.067235525 | 0.47145802 | 0.509 | 0.369 | 1 | 0.18691263 | 0.604544093 | 0.488 | 0.403 | 1 | 0.18691263 | 0.129950435 |
| Tmc6 | 0.715801635 | -0.343528273 | 0.245 | 0.271 | 1 | 0.067300847 | -1.285191782 | 0.22 | 0.345 | 1 | 0.715801635 | 0.130072291 |
| Taco1 | 0.067330003 | 1.154722595 | 0.113 | 0.047 | 1 | 0.838027457 | 0.589277336 | 0.049 | 0.058 | 1 | 0.838027457 | 0.130126676 |
| C1qtnf9 | 0.067330003 | 1.154722595 | 0.113 | 0.047 | 1 | 0.512329427 | 1.19334866 | 0.098 | 0.072 | 1 | 0.512329427 | 0.130126676 |
| Fam151b | 0.067335723 | 1.084333267 | 0.132 | 0.059 | 1 | 0.709546838 | 0.83720485 | 0.122 | 0.106 | 1 | 0.709546838 | 0.130137346 |
| Cbx5 | 0.067363867 | 0.381687746 | 0.811 | 0.792 | 1 | 0.393392689 | -0.106984287 | 0.829 | 0.84 | 1 | 0.393392689 | 0.130189843 |
| Tbca | 0.067420726 | 0.347367673 | 0.906 | 0.903 | 1 | 0.424400275 | 0.242258261 | 0.829 | 0.911 | 1 | 0.424400275 | 0.130295897 |
| Cog2 | 0.151924404 | 0.687596584 | 0.358 | 0.288 | 1 | 0.067485595 | -0.430567475 | 0.244 | 0.406 | 1 | 0.151924404 | 0.130416884 |
| Noc2l | 0.067507655 | 0.887936054 | 0.377 | 0.275 | 1 | 0.83666915 | 0.271607674 | 0.293 | 0.328 | 1 | 0.83666915 | 0.130458027 |
| Gm43462 | 0.067550799 | 0.75033234 | 0.226 | 0.127 | 1 | 0.686621784 | 0.644559772 | 0.171 | 0.15 | 1 | 0.686621784 | 0.130538488 |
| Chpt1 | 0.067582363 | 0.509006063 | 0.66 | 0.661 | 1 | 0.170154099 | -0.152741486 | 0.683 | 0.768 | 1 | 0.170154099 | 0.130597351 |
| Vnn1 | 0.06770409 | -1.960754623 | 0.038 | 0.123 | 1 | 0.127331083 | -1.686357106 | 0.024 | 0.096 | 1 | 0.127331083 | 0.130824337 |
| Rsl1d1 | 0.080210138 | 0.292024639 | 0.925 | 0.843 | 1 | 0.067721709 | -0.367013329 | 0.854 | 0.911 | 1 | 0.080210138 | 0.130857188 |
| Ndufaf6 | 0.067747823 | 1.115194231 | 0.208 | 0.114 | 1 | 0.115268038 | -1.093532488 | 0.049 | 0.137 | 1 | 0.115268038 | 0.130905878 |
| Idh3g | 0.067748413 | 0.47826174 | 0.623 | 0.525 | 1 | 0.961614392 | 0.285106983 | 0.463 | 0.505 | 1 | 0.961614392 | 0.130906979 |
| Helb | 0.067758019 | 1.120775263 | 0.226 | 0.131 | 1 | 0.129757687 | 1.118386602 | 0.268 | 0.184 | 1 | 0.129757687 | 0.130924889 |
| Srp54a | 0.17326529 | 0.50490895 | 0.396 | 0.309 | 1 | 0.067853926 | -0.935384654 | 0.171 | 0.304 | 1 | 0.17326529 | 0.131103696 |
| Gm56829 | 0.776592147 | 0.417757001 | 0.113 | 0.102 | 1 | 0.067961216 | -0.935384654 | 0.024 | 0.123 | 1 | 0.776592147 | 0.131303705 |
| Nup205 | 0.067988134 | 0.727301371 | 0.321 | 0.208 | 1 | 0.075960464 | -0.917682652 | 0.146 | 0.276 | 1 | 0.075960464 | 0.131353882 |
| Unc119 | 0.068020354 | 0.80108564 | 0.358 | 0.246 | 1 | 0.781690394 | 0.578470581 | 0.171 | 0.201 | 1 | 0.781690394 | 0.131413939 |
| Huwe1 | 0.089034228 | 0.274145899 | 1 | 0.987 | 1 | 0.068062266 | -0.128641153 | 1 | 0.997 | 1 | 0.089034228 | 0.13149206 |
| AI987944 | 0.205387602 | -0.652632327 | 0.151 | 0.229 | 1 | 0.06808933 | -0.520347155 | 0.073 | 0.198 | 1 | 0.205387602 | 0.131542502 |
| Six5 | 0.068119757 | 1.154722595 | 0.132 | 0.059 | 1 | 0.116117711 | -1.372248516 | 0.024 | 0.099 | 1 | 0.116117711 | 0.131599213 |
| Prickle3 | 0.608885602 | 0.191248471 | 0.113 | 0.144 | 1 | 0.068253224 | -1.385187572 | 0.049 | 0.154 | 1 | 0.608885602 | 0.131847945 |
| Prrc1 | 0.281251174 | 0.13295478 | 0.849 | 0.852 | 1 | 0.068265421 | -0.322112052 | 0.805 | 0.84 | 1 | 0.281251174 | 0.131870675 |
| Itgam | 0.503281643 | -1.686579659 | 0.038 | 0.021 | 1 | 0.068328784 | -0.56878751 | 0.098 | 0.034 | 1 | 0.503281643 | 0.131988746 |
| 2310002F09Rik | 0.068483021 | -2.459987249 | 0.019 | 0.093 | 1 | 0.114593951 | -2.622226769 | 0 | 0.058 | 1 | 0.114593951 | 0.132276117 |
| Gcat | 0.068655357 | -1.389597921 | 0.038 | 0.123 | 1 | 0.449330947 | 0.667279848 | 0.122 | 0.085 | 1 | 0.449330947 | 0.132597156 |
| Pld3 | 0.068668015 | 0.529402356 | 0.547 | 0.436 | 1 | 0.233104201 | -0.21526257 | 0.341 | 0.485 | 1 | 0.233104201 | 0.132620734 |
| Slc2a10 | 0.635386613 | -0.229941255 | 0.264 | 0.288 | 1 | 0.068853545 | -0.719743275 | 0.195 | 0.341 | 1 | 0.635386613 | 0.132966279 |
| Lims1 | 0.068953929 | -0.220125814 | 0.962 | 0.919 | 1 | 0.099669448 | -0.666095876 | 0.927 | 0.976 | 1 | 0.099669448 | 0.133153213 |
| Sinhcaf | 0.095703663 | 1.055186921 | 0.113 | 0.051 | 1 | 0.068961481 | 2.145327145 | 0.122 | 0.051 | 1 | 0.095703663 | 0.133167276 |
| Tpx2 | 0.069050232 | -3.517702747 | 0.019 | 0.093 | 1 | 0.752578347 | 0.377773231 | 0.024 | 0.017 | 1 | 0.752578347 | 0.133332529 |
| Gm16133 | 0.259729746 | 0.433438623 | 0.34 | 0.267 | 1 | 0.06914951 | -0.859762676 | 0.415 | 0.539 | 1 | 0.259729746 | 0.133517366 |
| Hyal2 | 0.06919367 | 1.039245377 | 0.245 | 0.148 | 1 | 0.204083485 | -0.402261085 | 0.122 | 0.218 | 1 | 0.204083485 | 0.133599577 |
| Rnaseh2c | 0.069244559 | 0.695290976 | 0.434 | 0.309 | 1 | 0.106856606 | -0.693309867 | 0.171 | 0.29 | 1 | 0.106856606 | 0.133694309 |
| Prmt5 | 0.069250493 | 0.812330397 | 0.34 | 0.237 | 1 | 0.106665093 | -0.458251034 | 0.146 | 0.276 | 1 | 0.106665093 | 0.133705356 |
| Espn | 0.243111734 | -0.652632327 | 0 | 0.025 | 1 | 0.069261572 | 2.515276755 | 0.098 | 0.038 | 1 | 0.243111734 | 0.133725978 |
| Mrpl33 | 0.604131832 | 0.245947355 | 0.736 | 0.75 | 1 | 0.069323696 | -0.341132391 | 0.561 | 0.666 | 1 | 0.604131832 | 0.133841618 |
| Il10 | 0.069380208 | 1.055186921 | 0.113 | 0.047 | 1 | 0.522110353 | 1.911205431 | 0.024 | 0.048 | 1 | 0.522110353 | 0.133946803 |
| Lig1 | 0.069398843 | 1.06161319 | 0.226 | 0.131 | 1 | 0.07889756 | -1.863234868 | 0.024 | 0.113 | 1 | 0.07889756 | 0.133981487 |
| Slc25a40 | 0.069465056 | -1.278236812 | 0.075 | 0.174 | 1 | 0.28068511 | 0.652780279 | 0.195 | 0.133 | 1 | 0.28068511 | 0.134104719 |
| Fbxw4 | 0.069484673 | 1.09332205 | 0.264 | 0.161 | 1 | 0.432603894 | 0.97780318 | 0.293 | 0.263 | 1 | 0.432603894 | 0.134141226 |
| Mcee | 0.069517979 | 0.640149422 | 0.472 | 0.386 | 1 | 0.934927918 | 0.338398993 | 0.317 | 0.328 | 1 | 0.934927918 | 0.134203209 |
| Prelid3b | 0.298846618 | -0.108311811 | 0.321 | 0.424 | 1 | 0.069531638 | -0.652761137 | 0.293 | 0.44 | 1 | 0.298846618 | 0.134228627 |
| Pabpc4 | 0.632283613 | 0.154722595 | 0.717 | 0.733 | 1 | 0.069665483 | -0.366484133 | 0.561 | 0.751 | 1 | 0.632283613 | 0.134477687 |
| Srgap1 | 0.069668981 | 0.443036409 | 0.792 | 0.712 | 1 | 0.212829865 | -0.368320927 | 0.927 | 0.935 | 1 | 0.212829865 | 0.134484196 |
| Fkrp | 0.635732636 | -0.274120704 | 0.189 | 0.216 | 1 | 0.069724975 | -0.654648247 | 0.195 | 0.345 | 1 | 0.635732636 | 0.134588377 |
| Pcca | 0.069772578 | 1.046919305 | 0.396 | 0.305 | 1 | 0.565673531 | 0.489854158 | 0.415 | 0.406 | 1 | 0.565673531 | 0.134676943 |
| mt-Nd4l | 0.99195906 | 0.10831213 | 0.887 | 0.941 | 1 | 0.069798025 | 0.186805814 | 0.927 | 0.901 | 1 | 0.99195906 | 0.134724286 |
| Pycrl | 0.069859827 | 0.63687429 | 0.321 | 0.199 | 1 | 0.309278114 | -0.359192363 | 0.098 | 0.16 | 1 | 0.309278114 | 0.134839259 |
| E430018J23Rik | 0.069891378 | 1.932330173 | 0.075 | 0.025 | 1 | 0.278181612 | 2.136765132 | 0.073 | 0.038 | 1 | 0.278181612 | 0.134897951 |
| Dgkz | 0.069985357 | 0.569760094 | 0.34 | 0.225 | 1 | 0.575433448 | 1.208187169 | 0.317 | 0.297 | 1 | 0.575433448 | 0.135072764 |
| Triqk | 0.758451051 | 0.513176566 | 0.245 | 0.242 | 1 | 0.069985951 | -0.970150072 | 0.146 | 0.276 | 1 | 0.758451051 | 0.135073869 |
| Gm13594 | 0.069988491 | -1.845277405 | 0 | 0.059 | 1 | 0.438672228 | 1.83720485 | 0.024 | 0.01 | 1 | 0.438672228 | 0.135078593 |
| Abhd1 | 0.069988491 | -1.845277405 | 0 | 0.059 | 1 | 0.208830556 | -1.16279515 | 0 | 0.038 | 1 | 0.208830556 | 0.135078593 |
| Gm29684 | 0.069988491 | -1.845277405 | 0 | 0.059 | 1 | 0.357854401 | -0.332720152 | 0 | 0.02 | 1 | 0.357854401 | 0.135078593 |
| A330041J22Rik | 0.070008116 | -1.932740246 | 0 | 0.059 | 1 | 0.139479918 | -1.250257991 | 0 | 0.051 | 1 | 0.139479918 | 0.135115096 |
| Pgam5 | 0.537816138 | 0.239114782 | 0.321 | 0.28 | 1 | 0.070013792 | 0.937731726 | 0.366 | 0.249 | 1 | 0.537816138 | 0.135125652 |
| Gm5817 | 0.070024722 | -2.304709024 | 0 | 0.059 | 1 | 0.257285084 | -0.622226769 | 0 | 0.031 | 1 | 0.257285084 | 0.135145982 |
| Nox1 | 0.070038309 | -2.489133595 | 0 | 0.059 | 1 | 0.257308946 | -1.863234868 | 0 | 0.031 | 1 | 0.257308946 | 0.135171253 |
| Gm54310 | 0.07003856 | -2.093204919 | 0 | 0.059 | 1 | 0.257325989 | -0.484723245 | 0 | 0.031 | 1 | 0.257325989 | 0.135171721 |
| A330040F15Rik | 0.070048876 | -2.932740246 | 0 | 0.059 | 1 | 0.357858058 | -1.16279515 | 0 | 0.02 | 1 | 0.357858058 | 0.135190908 |
| Col4a6 | 0.070056425 | -2.545717123 | 0 | 0.059 | 1 | 0.455094184 | -0.484723245 | 0 | 0.014 | 1 | 0.455094184 | 0.135204947 |
| Gm41349 | 0.070080077 | -1.752168001 | 0 | 0.059 | 1 | 0.262674569 | 2.83720485 | 0.024 | 0.007 | 1 | 0.262674569 | 0.135248937 |

| | | | | | | | | | | | | |
|---|---|---|---|---|---|---|---|---|---|---|---|---|
| Rnf19b | 0.070185974 | 0.845394537 | 0.604 | 0.513 | 1 | 0.29490133 | 0.395977827 | 0.61 | 0.57 | 1 | 0.29490133 | 0.135445876 |
| Tmem223 | 0.070247671 | 0.412790748 | 0.642 | 0.521 | 1 | 0.82754912 | 0.406570495 | 0.463 | 0.505 | 1 | 0.82754912 | 0.135560607 |
| Sdhaf1 | 0.070252898 | 1.502645898 | 0.17 | 0.089 | 1 | 0.707925627 | 0.83720485 | 0.146 | 0.126 | 1 | 0.707925627 | 0.135570327 |
| H2ac20 | 0.857199056 | -0.330704232 | 0.075 | 0.068 | 1 | 0.07027352 | -1.863234868 | 0 | 0.075 | 1 | 0.857199056 | 0.135608672 |
| Micall2 | 0.619885227 | 0.984797593 | 0.038 | 0.025 | 1 | 0.070327353 | -2.069685746 | 0 | 0.075 | 1 | 0.619885227 | 0.135708769 |
| Gas2l3 | 0.169426436 | -1.799473716 | 0.019 | 0.068 | 1 | 0.070363324 | -2.292078167 | 0 | 0.075 | 1 | 0.169426436 | 0.135775651 |
| Plek | 0.070365149 | 2.154722595 | 0.075 | 0.025 | 1 | 0.737522269 | -0.332720152 | 0.024 | 0.034 | 1 | 0.737522269 | 0.135779044 |
| Ube2t | 0.070435559 | -1.7032584 | 0.019 | 0.093 | 1 | 0.170373481 | -1.16279515 | 0 | 0.044 | 1 | 0.170373481 | 0.135909951 |
| Tppp3 | 0.070435559 | -2.015202407 | 0.019 | 0.093 | 1 | 0.318153262 | -0.747757651 | 0.049 | 0.096 | 1 | 0.318153262 | 0.135909951 |
| Psmd3 | 0.070491528 | 0.534826304 | 0.623 | 0.564 | 1 | 0.33533569 | 0.261175474 | 0.707 | 0.676 | 1 | 0.33533569 | 0.136014 |
| Rab8b | 0.070642986 | 0.781569177 | 0.528 | 0.441 | 1 | 0.198739663 | 0.318629746 | 0.61 | 0.563 | 1 | 0.198739663 | 0.136295541 |
| Slc25a32 | 0.070698698 | 0.669295768 | 0.321 | 0.203 | 1 | 0.522812207 | 0.526864729 | 0.317 | 0.283 | 1 | 0.522812207 | 0.136399091 |
| Rrm2 | 0.100415114 | -1.093204919 | 0.094 | 0.038 | 1 | 0.07073303 | 2.83720485 | 0.098 | 0.038 | 1 | 0.100415114 | 0.136462899 |
| Zfp384 | 0.07073834 | 0.550651271 | 0.358 | 0.233 | 1 | 0.150674156 | -0.203209419 | 0.268 | 0.427 | 1 | 0.150674156 | 0.136472767 |
| Prrg2 | 0.070739235 | 1.03273207 | 0.208 | 0.119 | 1 | 0.525400465 | 0.811669758 | 0.195 | 0.164 | 1 | 0.525400465 | 0.136474431 |
| Cgn | 0.343000446 | -0.845277405 | 0 | 0.017 | 1 | 0.070809483 | 0.979223855 | 0.171 | 0.082 | 1 | 0.343000446 | 0.136604983 |
| Lurap1 | 0.070812444 | 1.792152515 | 0.075 | 0.025 | 1 | 0.184911704 | 1.83720485 | 0.049 | 0.017 | 1 | 0.184911704 | 0.136610486 |
| Kdm4d | 0.070812444 | 1.792152515 | 0.075 | 0.025 | 1 | 0.881057318 | 0.515276755 | 0.024 | 0.02 | 1 | 0.881057318 | 0.136610486 |
| Atp6v0c | 0.070834506 | 0.244608098 | 0.981 | 0.987 | 1 | 0.425737811 | -0.320459969 | 1 | 0.99 | 1 | 0.425737811 | 0.136651485 |
| Aar2 | 0.490880518 | 0.154722595 | 0.264 | 0.216 | 1 | 0.070841148 | -0.940402729 | 0.122 | 0.249 | 1 | 0.490880518 | 0.136663827 |
| Ttc12 | 0.427821347 | 1.417757001 | 0.057 | 0.034 | 1 | 0.07085378 | 0.737669176 | 0.146 | 0.065 | 1 | 0.427821347 | 0.136687302 |
| Slc16a11 | 0.206392421 | -1.845277405 | 0 | 0.03 | 1 | 0.070882341 | 1.136765132 | 0.22 | 0.119 | 1 | 0.206392421 | 0.136740376 |
| Gprasp2 | 0.511888761 | 0.207190015 | 0.189 | 0.153 | 1 | 0.07093182 | -0.675864732 | 0.146 | 0.294 | 1 | 0.511888761 | 0.136832317 |
| Ammecr1l | 0.070968836 | 0.561767303 | 0.528 | 0.462 | 1 | 0.340454712 | 0.609398932 | 0.78 | 0.696 | 1 | 0.340454712 | 0.136901096 |
| Slc66a1 | 0.070976388 | -1.489133595 | 0.019 | 0.093 | 1 | 0.876550792 | 0.30115195 | 0.171 | 0.16 | 1 | 0.876550792 | 0.136915128 |
| Vps8 | 0.070988175 | 1.13535727 | 0.321 | 0.225 | 1 | 0.668384922 | -0.118595346 | 0.341 | 0.392 | 1 | 0.668384922 | 0.136937029 |
| Gpr89 | 0.129330137 | -0.667279103 | 0.208 | 0.314 | 1 | 0.070997094 | -0.904262137 | 0.122 | 0.249 | 1 | 0.129330137 | 0.1369536 |
| Ttpal | 0.280160115 | 0.752624151 | 0.189 | 0.136 | 1 | 0.071007755 | -0.877040668 | 0.146 | 0.276 | 1 | 0.280160115 | 0.136973409 |
| Ciapin1 | 0.188703799 | 0.527939918 | 0.396 | 0.322 | 1 | 0.07109438 | 0.856313673 | 0.488 | 0.341 | 1 | 0.188703799 | 0.137134349 |
| Pex7 | 0.553901885 | -0.1672055 | 0.283 | 0.335 | 1 | 0.071128075 | -0.957211017 | 0.146 | 0.276 | 1 | 0.553901885 | 0.137196947 |
| Rsrc1 | 0.071128939 | 0.55420528 | 0.717 | 0.678 | 1 | 0.277755665 | 0.507209474 | 0.756 | 0.713 | 1 | 0.277755665 | 0.137198552 |
| Morn2 | 0.071156427 | -0.865176963 | 0.132 | 0.246 | 1 | 0.269066099 | -0.747757651 | 0.122 | 0.188 | 1 | 0.269066099 | 0.137249616 |
| Ccdc68 | 0.725653213 | -0.381330305 | 0.075 | 0.089 | 1 | 0.071170674 | -1.090902232 | 0.122 | 0.249 | 1 | 0.725653213 | 0.137276083 |
| Rnf181 | 0.369214832 | -0.257091004 | 0.34 | 0.411 | 1 | 0.071173068 | -0.790826373 | 0.195 | 0.338 | 1 | 0.369214832 | 0.137280531 |
| Cdadc1 | 0.144560079 | 0.647238847 | 0.472 | 0.419 | 1 | 0.071177545 | -0.349208274 | 0.341 | 0.505 | 1 | 0.144560079 | 0.137288846 |
| Ttll10 | 0.305043961 | -1.237594828 | 0.019 | 0.051 | 1 | 0.071192078 | -1.747757651 | 0.024 | 0.116 | 1 | 0.305043961 | 0.137315845 |
| Il2 | 0.071243125 | -1.430239906 | 0.019 | 0.093 | 1 | 0.745976859 | 0.377773231 | 0.024 | 0.034 | 1 | 0.745976859 | 0.137410668 |
| Gm10687 | 0.071244531 | 1.962077517 | 0.075 | 0.025 | 1 | 0.104856732 | 2.83720485 | 0.024 | 0.003 | 1 | 0.104856732 | 0.13741328 |
| Mapk14 | 0.071253165 | 0.618669695 | 0.453 | 0.331 | 1 | 0.176784095 | 0.515276755 | 0.61 | 0.546 | 1 | 0.176784095 | 0.137429317 |
| Chmp4b | 0.085494143 | 0.21451568 | 0.981 | 0.941 | 1 | 0.071301063 | -0.296218596 | 0.951 | 0.983 | 1 | 0.085494143 | 0.137518283 |
| Ppp2cb | 0.071407946 | 0.55381855 | 0.585 | 0.504 | 1 | 0.160999652 | 0.592582481 | 0.683 | 0.611 | 1 | 0.160999652 | 0.137716797 |
| Mettl9 | 0.356906844 | 0.17471407 | 0.755 | 0.758 | 1 | 0.071553913 | -0.277456133 | 0.683 | 0.819 | 1 | 0.356906844 | 0.137987864 |
| Sdhc | 0.190370508 | 0.3781008 | 0.66 | 0.593 | 1 | 0.071629633 | 0.493250449 | 0.683 | 0.57 | 1 | 0.190370508 | 0.138128462 |
| 6030443J06Rik | 0.416781699 | -0.686579659 | 0.075 | 0.114 | 1 | 0.071652185 | -1.747757651 | 0.024 | 0.116 | 1 | 0.416781699 | 0.138170335 |
| Gm56936 | 0.07166535 | -1.142957954 | 0.094 | 0.195 | 1 | 0.208060389 | -0.825760163 | 0.049 | 0.113 | 1 | 0.208060389 | 0.138194778 |
| Arl9 | 0.071696076 | 1.739685096 | 0.075 | 0.025 | 1 | 0.655636624 | 0.136765132 | 0.049 | 0.068 | 1 | 0.655636624 | 0.138251824 |
| 9530046B11Rik | 0.071696076 | 1.739685096 | 0.075 | 0.025 | 1 | 0.646976884 | 0.721727632 | 0.049 | 0.034 | 1 | 0.646976884 | 0.138251824 |
| Rpp21 | 0.25814464 | 0.438515561 | 0.302 | 0.229 | 1 | 0.071702944 | 0.985727375 | 0.317 | 0.208 | 1 | 0.25814464 | 0.138264577 |
| Trmt9b | 0.071832116 | 2.055186921 | 0.113 | 0.051 | 1 | 0.22554945 | -1.622226769 | 0.024 | 0.075 | 1 | 0.22554945 | 0.13850438 |
| Birc2 | 0.175925191 | 0.569760094 | 0.491 | 0.386 | 1 | 0.071865822 | -0.863234868 | 0.195 | 0.334 | 1 | 0.175925191 | 0.138566948 |
| Dnajc30 | 0.07200839 | 1.005859209 | 0.283 | 0.186 | 1 | 1 | 0.515276755 | 0.146 | 0.15 | 1 | 1 | 0.138831571 |
| Lsm11 | 0.072149962 | 1.569760094 | 0.075 | 0.025 | 1 | 0.488326572 | -0.484723245 | 0.049 | 0.078 | 1 | 0.488326572 | 0.139094307 |
| Gm57383 | 0.07216042 | -0.652632327 | 0.075 | 0.182 | 1 | 0.142502249 | -1.250257991 | 0.049 | 0.126 | 1 | 0.142502249 | 0.139113714 |
| Gm49519 | 0.072171156 | 1.695290976 | 0.075 | 0.025 | 1 | 0.861875503 | 1.252242349 | 0.024 | 0.02 | 1 | 0.861875503 | 0.139133636 |
| Cetn3 | 0.072256639 | 0.439879812 | 0.792 | 0.674 | 1 | 0.267705771 | 0.437082025 | 0.683 | 0.696 | 1 | 0.267705771 | 0.139292256 |
| Traf4 | 0.072393832 | 1.133027524 | 0.302 | 0.208 | 1 | 0.37366461 | -0.16279515 | 0.244 | 0.311 | 1 | 0.37366461 | 0.139546798 |
| Ostm1 | 0.072408138 | 0.974150349 | 0.189 | 0.102 | 1 | 0.558647597 | 0.310659035 | 0.244 | 0.212 | 1 | 0.558647597 | 0.139573338 |
| Retreg3 | 0.072436535 | 0.911866071 | 0.34 | 0.237 | 1 | 0.27145071 | 0.643433106 | 0.463 | 0.42 | 1 | 0.27145071 | 0.139626019 |
| Pmpca | 0.34619388 | -0.265609204 | 0.245 | 0.318 | 1 | 0.072441047 | -0.332720152 | 0.171 | 0.324 | 1 | 0.34619388 | 0.139634389 |
| Zfp324 | 0.072635976 | 1.669295768 | 0.075 | 0.025 | 1 | 0.609924905 | 0.136765132 | 0.024 | 0.041 | 1 | 0.609924905 | 0.139995968 |
| Klhl36 | 0.072635976 | 1.669295768 | 0.075 | 0.025 | 1 | 0.455763738 | 0.962735732 | 0.049 | 0.027 | 1 | 0.455763738 | 0.139995968 |
| Rab36 | 0.072635976 | 1.669295768 | 0.075 | 0.025 | 1 | 0.761280969 | 1.059597271 | 0.073 | 0.061 | 1 | 0.761280969 | 0.139995968 |
| Gm20036 | 0.072635976 | 1.669295768 | 0.075 | 0.025 | 1 | 0.438672228 | 1.83720485 | 0.024 | 0.01 | 1 | 0.438672228 | 0.139995968 |
| Tmem191 | 0.072635976 | 1.669295768 | 0.075 | 0.025 | 1 | 0.682114395 | 0.515276755 | 0.098 | 0.078 | 1 | 0.682114395 | 0.139995968 |
| Tedc2 | 0.072635976 | 1.669295768 | 0.075 | 0.025 | 1 | 0.319619815 | -0.332720152 | 0 | 0.024 | 1 | 0.319619815 | 0.139995968 |
| Rrp15 | 0.248788296 | -0.376997885 | 0.283 | 0.373 | 1 | 0.072639601 | -0.372248516 | 0.195 | 0.358 | 1 | 0.248788296 | 0.140002691 |
| Gpr146 | 0.072657771 | -0.880901315 | 0.151 | 0.267 | 1 | 0.772798826 | 0.749742009 | 0.146 | 0.137 | 1 | 0.772798826 | 0.140036391 |
| Elp4 | 0.072725478 | 0.744090275 | 0.528 | 0.436 | 1 | 0.820006749 | 0.282615998 | 0.415 | 0.399 | 1 | 0.820006749 | 0.140161961 |
| Jade2 | 0.254713732 | -0.44618145 | 0.226 | 0.297 | 1 | 0.072753625 | -1.0405394 | 0.195 | 0.321 | 1 | 0.254713732 | 0.14021416 |
| Zwint | 0.072769513 | -0.286330495 | 0.585 | 0.788 | 1 | 0.214451359 | -0.194273382 | 0.659 | 0.785 | 1 | 0.214451359 | 0.140243623 |
| Crtc3 | 0.072815816 | -0.396687346 | 0.566 | 0.699 | 1 | 0.099658957 | -0.47064806 | 0.756 | 0.833 | 1 | 0.099658957 | 0.140329488 |
| Yipf1 | 0.072822637 | 0.634465783 | 0.472 | 0.369 | 1 | 0.844037599 | 0.116727378 | 0.317 | 0.355 | 1 | 0.844037599 | 0.140342137 |
| Klhdc4 | 0.155547751 | 0.540012751 | 0.245 | 0.161 | 1 | 0.072825086 | 1.000703582 | 0.293 | 0.181 | 1 | 0.155547751 | 0.14034668 |
| Ccp110 | 0.072829102 | 0.859266711 | 0.245 | 0.148 | 1 | 0.155703694 | 0.956786466 | 0.268 | 0.184 | 1 | 0.155703694 | 0.140354125 |
| Gm17024 | 0.659393736 | 0.569760094 | 0.057 | 0.042 | 1 | 0.072849048 | 1.47463477 | 0.122 | 0.051 | 1 | 0.659393736 | 0.140391112 |
| Rps29 | 0.072976546 | -0.173102082 | 1 | 0.979 | 1 | 0.083261602 | -0.252451093 | 1 | 0.99 | 1 | 0.083261602 | 0.140627515 |
| Tbc1d24 | 0.073160222 | 0.8817041 | 0.189 | 0.102 | 1 | 0.137078737 | -0.840867055 | 0.098 | 0.191 | 1 | 0.137078737 | 0.140968026 |
| Gm37529 | 0.506039658 | -0.1672055 | 0.057 | 0.085 | 1 | 0.073181936 | -0.673757069 | 0.049 | 0.157 | 1 | 0.506039658 | 0.141008275 |
| Patl1 | 0.073192541 | 0.617456836 | 0.434 | 0.322 | 1 | 0.416874547 | 0.371230385 | 0.463 | 0.44 | 1 | 0.416874547 | 0.141027934 |
| Lcmt2 | 0.082545374 | -1.368839361 | 0.019 | 0.089 | 1 | 0.073197651 | -1.332720152 | 0.024 | 0.116 | 1 | 0.082545374 | 0.141037407 |
| Prxl2a | 0.202448267 | -0.752168001 | 0.189 | 0.275 | 1 | 0.073222235 | -1.08562729 | 0.098 | 0.218 | 1 | 0.202448267 | 0.141082974 |
| Bbs4 | 0.32801523 | 0.129187503 | 0.208 | 0.153 | 1 | 0.073246879 | 0.707921833 | 0.268 | 0.157 | 1 | 0.32801523 | 0.141128652 |
| Wdr89 | 0.073265953 | 0.20958453 | 0.792 | 0.61 | 1 | 0.397655374 | -0.302346503 | 0.634 | 0.642 | 1 | 0.397655374 | 0.141164006 |
| Pno1 | 0.073326433 | 0.313420341 | 0.528 | 0.369 | 1 | 0.177136593 | -0.16279515 | 0.22 | 0.362 | 1 | 0.177136593 | 0.1412761 |
| Gm11944 | 0.176759835 | 0.714150003 | 0.208 | 0.136 | 1 | 0.073343297 | 1.197794565 | 0.293 | 0.181 | 1 | 0.176759835 | 0.141307355 |
| Atpaf1 | 0.073405778 | 0.569760094 | 0.283 | 0.174 | 1 | 0.637008067 | -0.27321914 | 0.171 | 0.201 | 1 | 0.637008067 | 0.141423148 |
| Hspa2 | 0.073479254 | 0.517292674 | 0.34 | 0.225 | 1 | 0.089184642 | 0.692154517 | 0.366 | 0.246 | 1 | 0.089184642 | 0.141559308 |
| Uxs1 | 0.691016009 | 0.125990876 | 0.698 | 0.703 | 1 | 0.073538448 | -0.339988152 | 0.39 | 0.601 | 1 | 0.691016009 | 0.141668993 |
| Mark3 | 0.349686993 | -0.236653428 | 0.906 | 0.907 | 1 | 0.073539833 | -0.331803864 | 0.927 | 0.983 | 1 | 0.349686993 | 0.141671558 |
| Rilp | 0.628925725 | 0.739685096 | 0.038 | 0.025 | 1 | 0.073540882 | 1.685201756 | 0.122 | 0.051 | 1 | 0.628925725 | 0.141673503 |
| Ptpn1 | 0.073646266 | -0.373848724 | 0.943 | 0.945 | 1 | 0.612194129 | 0.191488318 | 0.951 | 0.973 | 1 | 0.612194129 | 0.141868759 |
| Surf1 | 0.073681814 | 0.493811241 | 0.491 | 0.36 | 1 | 0.210493224 | -0.141421499 | 0.244 | 0.365 | 1 | 0.210493224 | 0.141934619 |
| Ccdc43 | 0.073699345 | 0.874614676 | 0.283 | 0.182 | 1 | 0.405882996 | 0.222495006 | 0.293 | 0.229 | 1 | 0.405882996 | 0.141967097 |
| Nup35 | 0.195511042 | 0.238554186 | 0.453 | 0.343 | 1 | 0.073710684 | -0.775238387 | 0.22 | 0.348 | 1 | 0.195511042 | 0.141988103 |
| Socs2 | 0.119286856 | -0.613244734 | 0.736 | 0.754 | 1 | 0.073803077 | 0.577337723 | 0.659 | 0.512 | 1 | 0.119286856 | 0.142159259 |
| AI480526 | 0.073858312 | 0.946614399 | 0.302 | 0.191 | 1 | 0.253759049 | 0.660882077 | 0.317 | 0.249 | 1 | 0.253759049 | 0.142261573 |
| Mfsd5 | 0.079146188 | 0.614154213 | 0.34 | 0.225 | 1 | 0.07394857 | -0.852094311 | 0.171 | 0.304 | 1 | 0.079146188 | 0.14242875 |
| Hibch | 0.277820865 | 0.442004547 | 0.245 | 0.178 | 1 | 0.07395712 | -0.863234868 | 0.122 | 0.253 | 1 | 0.277820865 | 0.142444584 |

| | | | | | | | | | | | | |
|---|---|---|---|---|---|---|---|---|---|---|---|---|
| Abcc3 | 0.074005219 | 1.640149422 | 0.075 | 0.025 | 1 | 0.117592181 | 0.962735732 | 0.049 | 0.014 | 1 | 0.117592181 | 0.142533665 |
| Sypl | 0.074090192 | -0.261092009 | 0.868 | 0.903 | 1 | 0.122906351 | -0.183312168 | 0.756 | 0.918 | 1 | 0.122906351 | 0.142691027 |
| Procr | 0.074162891 | 1.377115016 | 0.17 | 0.089 | 1 | 0.529642321 | 0.737669176 | 0.098 | 0.072 | 1 | 0.529642321 | 0.142825647 |
| Thumpd3 | 0.074207133 | -1.044057269 | 0.17 | 0.28 | 1 | 0.319292818 | -0.366667484 | 0.195 | 0.27 | 1 | 0.319292818 | 0.142907568 |
| Gigyf2 | 0.210293832 | 0.293550299 | 0.755 | 0.742 | 1 | 0.074270389 | -0.110973954 | 0.634 | 0.881 | 1 | 0.210293832 | 0.143024688 |
| Tomm7 | 0.074297503 | -0.307853293 | 0.906 | 0.89 | 1 | 0.506450192 | -0.109961703 | 0.805 | 0.84 | 1 | 0.506450192 | 0.143074887 |
| Ube2j2 | 0.07436188 | 0.492719058 | 0.698 | 0.669 | 1 | 0.318968284 | 0.436784447 | 0.805 | 0.799 | 1 | 0.318968284 | 0.143194071 |
| Emp2 | 0.074408317 | -0.515407289 | 0.642 | 0.712 | 1 | 0.969845651 | 0.140131983 | 0.707 | 0.782 | 1 | 0.969845651 | 0.143280037 |
| Stum | 0.074441876 | 1.47665069 | 0.075 | 0.025 | 1 | 0.267205177 | 2.252242349 | 0.024 | 0.007 | 1 | 0.267205177 | 0.143342159 |
| Tmem79 | 0.074441876 | 0.228723176 | 0.075 | 0.025 | 1 | 0.080279054 | 2.159132945 | 0.073 | 0.024 | 1 | 0.080279054 | 0.143342159 |
| Gm42939 | 0.074441876 | 1.47665069 | 0.075 | 0.025 | 1 | 0.877271098 | 0.185128153 | 0.098 | 0.106 | 1 | 0.877271098 | 0.143342159 |
| Gm17344 | 0.074441876 | 1.47665069 | 0.075 | 0.025 | 1 | 0.450957104 | -0.332720152 | 0.024 | 0.051 | 1 | 0.450957104 | 0.143342159 |
| Qrich2 | 0.074441876 | 1.47665069 | 0.075 | 0.025 | 1 | 0.872828964 | 1.029849928 | 0.024 | 0.02 | 1 | 0.872828964 | 0.143342159 |
| Rrp12 | 0.074464841 | -0.920565533 | 0.094 | 0.199 | 1 | 0.945208927 | 0.444887427 | 0.171 | 0.171 | 1 | 0.945208927 | 0.14338467 |
| Dnah2 | 0.074476391 | 0.261637799 | 0.075 | 0.025 | 1 | 0.372037778 | 0.100239256 | 0.049 | 0.024 | 1 | 0.372037778 | 0.14340605 |
| Lamc2 | 0.07457944 | -1.867645218 | 0.019 | 0.093 | 1 | 0.222386363 | -1.470223675 | 0.098 | 0.048 | 1 | 0.222386363 | 0.143596786 |
| Slc9a1 | 0.074582873 | 0.398959405 | 0.585 | 0.458 | 1 | 0.563127424 | 0.386259697 | 0.585 | 0.597 | 1 | 0.563127424 | 0.143603141 |
| Cyld | 0.074639472 | 0.543287883 | 0.415 | 0.288 | 1 | 0.7304756 | 0.448162559 | 0.488 | 0.485 | 1 | 0.7304756 | 0.143707893 |
| Gm4419 | 0.074682032 | 2.154722595 | 0.094 | 0.038 | 1 | 0.599290422 | 1.515276755 | 0.024 | 0.014 | 1 | 0.599290422 | 0.143786658 |
| Eif4ebp1 | 0.14946976 | 0.428098945 | 0.396 | 0.305 | 1 | 0.074685338 | -0.753212081 | 0.268 | 0.427 | 1 | 0.14946976 | 0.143792777 |
| Vps18 | 0.074729482 | 0.984797593 | 0.264 | 0.169 | 1 | 0.534826281 | 0.614812428 | 0.244 | 0.212 | 1 | 0.534826281 | 0.143874469 |
| Eefsec | 0.074792799 | 0.845394537 | 0.283 | 0.178 | 1 | 0.965177787 | 0.614812428 | 0.22 | 0.229 | 1 | 0.965177787 | 0.143991635 |
| Ndst2 | 0.074795237 | 0.714150003 | 0.226 | 0.127 | 1 | 0.430854688 | -0.288326032 | 0.195 | 0.253 | 1 | 0.430854688 | 0.143996146 |
| Tpm3-rs7 | 0.144520156 | 1.154722595 | 0.132 | 0.072 | 1 | 0.074829179 | -0.863234868 | 0.024 | 0.119 | 1 | 0.144520156 | 0.144058951 |
| Triobp | 0.074849358 | 0.89946554 | 0.358 | 0.25 | 1 | 0.23822473 | -0.202323514 | 0.244 | 0.369 | 1 | 0.23822473 | 0.14409629 |
| Bms1 | 0.074979641 | 0.798960905 | 0.509 | 0.441 | 1 | 0.170247093 | 0.32941021 | 0.634 | 0.505 | 1 | 0.170247093 | 0.144337336 |
| Kif13a | 0.074981953 | -0.538539521 | 0.472 | 0.614 | 1 | 0.445672536 | -0.195216628 | 0.634 | 0.7 | 1 | 0.445672536 | 0.144341613 |
| Ptgr1 | 0.712757879 | 0.228723176 | 0.057 | 0.072 | 1 | 0.074994234 | 1.204936634 | 0.171 | 0.085 | 1 | 0.712757879 | 0.144364332 |
| Nop58 | 0.075135459 | -0.277528915 | 0.925 | 0.945 | 1 | 0.08459748 | -0.355602392 | 0.805 | 0.928 | 1 | 0.08459748 | 0.14462558 |
| Pfdn1 | 0.950495955 | 0.203632195 | 0.528 | 0.551 | 1 | 0.075144332 | -0.605203621 | 0.512 | 0.614 | 1 | 0.950495955 | 0.144641994 |
| Clip1 | 0.215618321 | 0.212150875 | 0.906 | 0.894 | 1 | 0.075155001 | 0.1231624 | 0.976 | 0.939 | 1 | 0.215618321 | 0.144661728 |
| Net1 | 0.079456977 | 0.781679184 | 0.434 | 0.318 | 1 | 0.075193472 | 0.791562184 | 0.561 | 0.481 | 1 | 0.079456977 | 0.144732887 |
| Vdac1 | 0.075213234 | 0.436048149 | 0.774 | 0.771 | 1 | 0.464368225 | 0.230973732 | 0.805 | 0.833 | 1 | 0.464368225 | 0.144769437 |
| Polr2d | 0.11995471 | 0.258558406 | 0.717 | 0.614 | 1 | 0.07524225 | -0.332720152 | 0.463 | 0.625 | 1 | 0.11995471 | 0.144823103 |
| Lcmt1 | 0.240123938 | 0.661682584 | 0.34 | 0.275 | 1 | 0.075253535 | 1.239964019 | 0.39 | 0.29 | 1 | 0.240123938 | 0.144843975 |
| Crebzf | 0.07537144 | 0.502645898 | 0.509 | 0.407 | 1 | 0.919747465 | 0.384032222 | 0.463 | 0.468 | 1 | 0.919747465 | 0.145062027 |
| Cox15 | 0.075381554 | 1.377115016 | 0.151 | 0.076 | 1 | 0.537652853 | 0.212713985 | 0.073 | 0.109 | 1 | 0.537652853 | 0.145080729 |
| Zfp959 | 0.075476176 | 1.3662267 | 0.151 | 0.076 | 1 | 0.275596727 | 0.589277336 | 0.171 | 0.109 | 1 | 0.275596727 | 0.1452557 |
| Ift172 | 0.075567883 | 0.655916738 | 0.302 | 0.186 | 1 | 0.478647209 | 0.189506594 | 0.171 | 0.229 | 1 | 0.478647209 | 0.145425261 |
| Oard1 | 0.075596188 | 0.699527969 | 0.358 | 0.246 | 1 | 0.22992441 | 0.342440158 | 0.098 | 0.181 | 1 | 0.22992441 | 0.145477593 |
| Trerf1 | 0.344009306 | 0.762405172 | 0.189 | 0.144 | 1 | 0.075822815 | 1.143307978 | 0.341 | 0.232 | 1 | 0.344009306 | 0.14589653 |
| Acyp1 | 0.075847733 | -0.595299152 | 0.132 | 0.25 | 1 | 0.513119262 | 0.209173627 | 0.146 | 0.201 | 1 | 0.513119262 | 0.145942588 |
| Paqr8 | 0.187474181 | 1.3662267 | 0.113 | 0.064 | 1 | 0.075954899 | 1.313642894 | 0.122 | 0.051 | 1 | 0.187474181 | 0.146140651 |
| Ggh | 0.075986358 | 0.467312825 | 0.698 | 0.61 | 1 | 0.286307514 | -0.25471764 | 0.585 | 0.621 | 1 | 0.286307514 | 0.146198789 |
| Otud5 | 0.076046287 | 0.484184362 | 0.585 | 0.508 | 1 | 0.198495558 | -0.402261085 | 0.561 | 0.642 | 1 | 0.198495558 | 0.146309537 |
| Ggps1 | 0.076051147 | 0.560452352 | 0.679 | 0.661 | 1 | 0.398471328 | -0.206364165 | 0.61 | 0.689 | 1 | 0.398471328 | 0.146318517 |
| Fads2 | 0.076083387 | 0.294125651 | 0.491 | 0.36 | 1 | 0.799859718 | 0.252242349 | 0.22 | 0.256 | 1 | 0.799859718 | 0.146378093 |
| Lgals8 | 0.076112268 | 0.540012751 | 0.415 | 0.305 | 1 | 0.763976566 | 0.621476159 | 0.293 | 0.334 | 1 | 0.763976566 | 0.146431458 |
| Smcr8 | 0.076254309 | 0.865215978 | 0.264 | 0.161 | 1 | 0.92787073 | -0.101394606 | 0.171 | 0.174 | 1 | 0.92787073 | 0.146693898 |
| Urm1 | 0.076260773 | 0.687596584 | 0.415 | 0.309 | 1 | 0.877448911 | 0.377773231 | 0.293 | 0.341 | 1 | 0.877448911 | 0.14670584 |
| Nras | 0.608061881 | 0.23878686 | 0.792 | 0.763 | 1 | 0.076321705 | -0.210264724 | 0.732 | 0.826 | 1 | 0.608061881 | 0.146818408 |
| Mical3 | 0.351632763 | -0.322978939 | 0.717 | 0.775 | 1 | 0.076335691 | 0.526137748 | 0.659 | 0.529 | 1 | 0.351632763 | 0.146844245 |
| Ccdc169 | 0.076341719 | 0.776210972 | 0.075 | 0.025 | 1 | 0.319619815 | -0.332720152 | 0 | 0.024 | 1 | 0.319619815 | 0.146855379 |
| Rnf225 | 0.076341719 | 1.306725688 | 0.075 | 0.025 | 1 | 0.319629309 | -0.484723245 | 0 | 0.024 | 1 | 0.319629309 | 0.146855379 |
| Atp2a3 | 0.076341719 | 1.017219071 | 0.075 | 0.025 | 1 | 0.51919986 | 0.83720485 | 0 | 0.01 | 1 | 0.51919986 | 0.146855379 |
| Cep19 | 0.076402614 | 0.79717059 | 0.208 | 0.114 | 1 | 0.50376559 | -0.262330824 | 0.122 | 0.164 | 1 | 0.50376559 | 0.146967868 |
| Mepce | 0.076484992 | 0.955413787 | 0.302 | 0.203 | 1 | 0.466501328 | 0.68395859 | 0.317 | 0.304 | 1 | 0.466501328 | 0.14712003 |
| Vps36 | 0.076500507 | 0.627475592 | 0.509 | 0.407 | 1 | 0.189737214 | 0.573185333 | 0.634 | 0.56 | 1 | 0.189737214 | 0.147148686 |
| Kiz | 0.076513192 | -0.330704232 | 0.17 | 0.309 | 1 | 0.53239969 | 0.491430013 | 0.317 | 0.287 | 1 | 0.53239969 | 0.147172116 |
| Tcaf1 | 0.09829716 | 0.5216186 | 0.792 | 0.78 | 1 | 0.076599684 | -0.277627011 | 0.634 | 0.788 | 1 | 0.09829716 | 0.147331857 |
| Selenbp1 | 0.361072363 | 1.569760094 | 0.075 | 0.047 | 1 | 0.076618737 | 1.059597271 | 0.244 | 0.143 | 1 | 0.361072363 | 0.147367043 |
| Sf3b3 | 0.07672923 | 0.244137516 | 0.774 | 0.733 | 1 | 0.279569259 | 0.44893426 | 0.805 | 0.782 | 1 | 0.279569259 | 0.147571085 |
| Glrx2 | 0.076806111 | 0.435904347 | 0.604 | 0.445 | 1 | 0.809345537 | 0.236812309 | 0.439 | 0.433 | 1 | 0.809345537 | 0.147713044 |
| Mlh1 | 0.076834154 | 1.70221039 | 0.151 | 0.076 | 1 | 0.312064185 | -0.448197369 | 0.049 | 0.099 | 1 | 0.312064185 | 0.14776482 |
| Gm57257 | 0.076870076 | 0.94321849 | 0.189 | 0.102 | 1 | 0.991398301 | 1.388446733 | 0.146 | 0.164 | 1 | 0.991398301 | 0.147831143 |
| Zfp994 | 0.076951995 | 1.280253477 | 0.189 | 0.106 | 1 | 0.884674798 | -0.116991461 | 0.073 | 0.078 | 1 | 0.884674798 | 0.147982381 |
| Amd1 | 0.076953468 | 0.390100658 | 0.717 | 0.653 | 1 | 0.470674508 | 0.511434689 | 0.634 | 0.7 | 1 | 0.470674508 | 0.147985101 |
| Zfp12 | 0.077034778 | 1.211306123 | 0.17 | 0.089 | 1 | 0.111472576 | -1.050320421 | 0.098 | 0.198 | 1 | 0.111472576 | 0.148135199 |
| Palld | 0.092181755 | 0.558871778 | 0.679 | 0.581 | 1 | 0.077047588 | 1.089391873 | 0.683 | 0.608 | 1 | 0.092181755 | 0.148158845 |
| Atp5pb | 0.077140091 | 0.331162801 | 0.962 | 0.915 | 1 | 0.141462651 | -0.201714139 | 0.829 | 0.952 | 1 | 0.141462651 | 0.148329589 |
| 4930539J05Rik | 0.437452221 | 0.280253477 | 0.019 | 0.042 | 1 | 0.077408965 | -1.686357106 | 0 | 0.072 | 1 | 0.437452221 | 0.148825781 |
| Pou5f1 | 0.768911274 | -0.1672055 | 0.038 | 0.047 | 1 | 0.077408965 | -1.686357106 | 0 | 0.072 | 1 | 0.768911274 | 0.148825781 |
| Nfatc2ip | 0.077435556 | 0.80108564 | 0.151 | 0.072 | 1 | 0.139316249 | -1.278272368 | 0.049 | 0.126 | 1 | 0.139316249 | 0.148874847 |
| Zfp503 | 0.077441857 | 0.47665069 | 0.358 | 0.242 | 1 | 0.769528678 | 0.367719567 | 0.341 | 0.365 | 1 | 0.769528678 | 0.148886472 |
| Mmab | 0.623979166 | 0.280253477 | 0.075 | 0.097 | 1 | 0.077457679 | -1.917682652 | 0 | 0.072 | 1 | 0.623979166 | 0.148915667 |
| Pclo | 0.175858672 | -1.845277405 | 0 | 0.034 | 1 | 0.077463951 | 1.124486802 | 0.268 | 0.16 | 1 | 0.175858672 | 0.148927238 |
| Zfp446 | 0.455670339 | 1.055186921 | 0.075 | 0.051 | 1 | 0.077477786 | -1.917682652 | 0 | 0.072 | 1 | 0.455670339 | 0.148952765 |
| Evpl | 0.467027607 | -0.582242999 | 0.038 | 0.064 | 1 | 0.077495329 | -1.970150072 | 0 | 0.072 | 1 | 0.467027607 | 0.148985133 |
| A430018G15Rik | 0.912984262 | 0.154722595 | 0.132 | 0.14 | 1 | 0.07749991 | -1.132421501 | 0.049 | 0.15 | 1 | 0.912984262 | 0.148993583 |
| Mad1l1 | 0.077509524 | 0.817687608 | 0.34 | 0.242 | 1 | 0.105727209 | 0.336143607 | 0.585 | 0.471 | 1 | 0.105727209 | 0.149011321 |
| Smyd4 | 0.084156742 | 0.8327945 | 0.189 | 0.102 | 1 | 0.077524343 | -2.20718927 | 0 | 0.072 | 1 | 0.084156742 | 0.149038663 |
| Adssl1 | 0.687840654 | -0.108311811 | 0.075 | 0.059 | 1 | 0.077526367 | -2.777504994 | 0 | 0.072 | 1 | 0.687840654 | 0.149042397 |
| Ttll1 | 0.734168272 | 0.569760094 | 0.132 | 0.161 | 1 | 0.077543843 | -0.377808041 | 0.049 | 0.16 | 1 | 0.734168272 | 0.149074639 |
| Lmx1a | 0.58643486 | 0.228723176 | 0.019 | 0.034 | 1 | 0.077569416 | -3.069685746 | 0 | 0.072 | 1 | 0.58643486 | 0.149121819 |
| Krt23 | 0.543828205 | 0.154722595 | 0.113 | 0.085 | 1 | 0.07757108 | 0.784093513 | 0.293 | 0.184 | 1 | 0.543828205 | 0.149124887 |
| Dab1 | 0.483884334 | 0.714150003 | 0.038 | 0.021 | 1 | 0.07757387 | -3.636528581 | 0 | 0.072 | 1 | 0.483884334 | 0.149130034 |
| Bhlhe41 | 0.869008941 | 0.542538625 | 0.264 | 0.263 | 1 | 0.077616574 | 1.326242931 | 0.22 | 0.123 | 1 | 0.869008941 | 0.149208816 |
| Impact | 0.077798448 | 0.34058914 | 0.698 | 0.619 | 1 | 0.654706616 | 0.191700807 | 0.585 | 0.686 | 1 | 0.654706616 | 0.149544298 |
| Gm50399 | 0.107309003 | 2.002719501 | 0.075 | 0.03 | 1 | 0.077800883 | 2.47463477 | 0.073 | 0.024 | 1 | 0.107309003 | 0.149548788 |
| Trpt1 | 0.077901906 | 1.100274811 | 0.17 | 0.089 | 1 | 0.423103677 | 1.029849928 | 0.098 | 0.065 | 1 | 0.423103677 | 0.149735104 |
| Tmem203 | 0.178063987 | 0.747547213 | 0.302 | 0.233 | 1 | 0.077914105 | -1.069685746 | 0.098 | 0.212 | 1 | 0.178063987 | 0.149757602 |
| Stxbp2 | 0.077942436 | -0.904171094 | 0.019 | 0.093 | 1 | 0.671462746 | 0.159132945 | 0.073 | 0.096 | 1 | 0.671462746 | 0.149809849 |
| Naa38 | 0.094108972 | -0.338816878 | 0.396 | 0.538 | 1 | 0.077960132 | -0.16279515 | 0.268 | 0.457 | 1 | 0.094108972 | 0.149842482 |
| Dbpht2 | 0.078277786 | 0.669295768 | 0.075 | 0.025 | 1 | 0.5191986 | 0.252242349 | 0 | 0.01 | 1 | 0.5191986 | 0.150428161 |
| Trim38 | 0.078277786 | 0.669295768 | 0.075 | 0.025 | 1 | 0.26948834 | 1.83720485 | 0.024 | 0.007 | 1 | 0.26948834 | 0.150428161 |
| Caskin1 | 0.078277786 | 1.154722595 | 0.075 | 0.025 | 1 | 0.473630309 | 1.644559772 | 0.073 | 0.048 | 1 | 0.473630309 | 0.150428161 |
| Zbtb8os | 0.078311426 | -0.676677801 | 0.245 | 0.364 | 1 | 0.61095925 | 0.19334866 | 0.244 | 0.205 | 1 | 0.61095925 | 0.150490172 |
| Mrps33 | 0.259685118 | 0.455724851 | 0.642 | 0.602 | 1 | 0.078351911 | 0.581443679 | 0.659 | 0.56 | 1 | 0.259685118 | 0.150564801 |

| | | | | | | | | | | | | |
|---|---|---|---|---|---|---|---|---|---|---|---|---|
| Sacm1l | 0.541333647 | -0.210041698 | 0.66 | 0.669 | 1 | 0.07854985 | -0.52606404 | 0.756 | 0.823 | 1 | 0.541333647 | 0.150929621 |
| Zfp809 | 0.303669874 | 0.695290976 | 0.189 | 0.136 | 1 | 0.07868743 | -1.136799942 | 0.146 | 0.266 | 1 | 0.303669874 | 0.151183149 |
| 4933404O12Rik | 0.932505382 | 1.154722595 | 0.019 | 0.021 | 1 | 0.078796479 | 2.685201756 | 0.073 | 0.024 | 1 | 0.932505382 | 0.151384073 |
| Gm56531 | 0.926212226 | 0.8327945 | 0.019 | 0.017 | 1 | 0.078797585 | 2.377773231 | 0.073 | 0.024 | 1 | 0.926212226 | 0.15138611 |
| Arl4a | 0.078802165 | 0.981074507 | 0.755 | 0.72 | 1 | 0.903083209 | -0.106579141 | 0.805 | 0.877 | 1 | 0.903083209 | 0.151394548 |
| Stk11 | 0.078875051 | 0.592843707 | 0.585 | 0.508 | 1 | 0.72320452 | 0.11646557 | 0.732 | 0.775 | 1 | 0.72320452 | 0.151528827 |
| Bud13 | 0.078912085 | 0.896924837 | 0.283 | 0.186 | 1 | 0.270288 | 0.342440158 | 0.22 | 0.147 | 1 | 0.270288 | 0.151597053 |
| Agbl4 | 0.079006321 | -2.727920455 | 0.019 | 0.089 | 1 | 0.272183224 | -1.589059905 | 0.024 | 0.068 | 1 | 0.272183224 | 0.151770644 |
| Tbp | 0.666501055 | 0.669295768 | 0.264 | 0.25 | 1 | 0.079050264 | -0.40380325 | 0.146 | 0.276 | 1 | 0.666501055 | 0.151851584 |
| Creld2 | 0.079129614 | 0.569760094 | 0.604 | 0.496 | 1 | 0.321814061 | -0.168793891 | 0.366 | 0.478 | 1 | 0.321814061 | 0.151997732 |
| Dtnbp1 | 0.079381187 | 0.749880863 | 0.509 | 0.487 | 1 | 0.332277156 | 0.136765132 | 0.732 | 0.625 | 1 | 0.332277156 | 0.152461002 |
| Erp29 | 0.300695212 | -0.15641222 | 0.887 | 0.919 | 1 | 0.079451058 | -0.304050809 | 0.732 | 0.901 | 1 | 0.300695212 | 0.152589645 |
| Tom1 | 0.189164927 | 0.482297253 | 0.472 | 0.407 | 1 | 0.079586596 | 0.710533046 | 0.659 | 0.573 | 1 | 0.189164927 | 0.152839166 |
| Appbp2 | 0.304665503 | 0.278482212 | 0.925 | 0.941 | 1 | 0.079639271 | -0.177981554 | 0.951 | 0.962 | 1 | 0.304665503 | 0.152936128 |
| Gm42659 | 0.344466274 | 1.3662267 | 0.113 | 0.076 | 1 | 0.079643612 | -1.520347155 | 0.024 | 0.113 | 1 | 0.344466274 | 0.15294412 |
| Mief2 | 0.439561627 | 0.631160639 | 0.113 | 0.081 | 1 | 0.079701908 | -0.69884805 | 0.049 | 0.154 | 1 | 0.439561627 | 0.153051422 |
| Gm28375 | 0.361131243 | -0.545717123 | 0.019 | 0.047 | 1 | 0.079774203 | 2.029849928 | 0.098 | 0.038 | 1 | 0.361131243 | 0.153184483 |
| Sugct | 0.925394372 | -0.10977722 | 0.34 | 0.331 | 1 | 0.079830053 | -1.136253363 | 0.171 | 0.314 | 1 | 0.925394372 | 0.153287268 |
| Tarbp1 | 0.357398022 | 1.538051234 | 0.113 | 0.076 | 1 | 0.07988872 | -1.622226769 | 0.024 | 0.113 | 1 | 0.357398022 | 0.153395233 |
| Gm9801 | 0.504721307 | -0.557995453 | 0.151 | 0.182 | 1 | 0.079947358 | -1.84461919 | 0.049 | 0.147 | 1 | 0.504721307 | 0.153503136 |
| Angptl1 | 0.080091902 | 0.207190015 | 0.094 | 0.208 | 1 | 0.190462488 | -0.369246028 | 0.122 | 0.218 | 1 | 0.190462488 | 0.153769092 |
| Ubxn2b | 0.791383167 | 0.328751995 | 0.151 | 0.14 | 1 | 0.080109003 | -0.286177566 | 0.049 | 0.154 | 1 | 0.791383167 | 0.153800554 |
| Tma7 | 0.080233578 | 0.286261586 | 0.943 | 0.898 | 1 | 0.09465374 | -0.239757132 | 0.732 | 0.863 | 1 | 0.09465374 | 0.154029729 |
| Gm56635 | 0.243106933 | -1.304709024 | 0 | 0.025 | 1 | 0.080279054 | 2.159132945 | 0.073 | 0.024 | 1 | 0.243106933 | 0.154113382 |
| Gm41231 | 0.080432773 | 1.695290976 | 0.094 | 0.038 | 1 | 0.622587386 | 0.252242349 | 0.024 | 0.041 | 1 | 0.622587386 | 0.154396115 |
| Phactr2 | 0.080573819 | 0.788228947 | 0.623 | 0.559 | 1 | 0.473164129 | 0.249039917 | 0.732 | 0.71 | 1 | 0.473164129 | 0.154655498 |
| Sdcbp2 | 0.563759422 | -0.752168001 | 0.019 | 0.034 | 1 | 0.080605336 | 1.515276755 | 0.098 | 0.038 | 1 | 0.563759422 | 0.154713452 |
| Snhg3 | 0.080635741 | -0.200372364 | 0.377 | 0.542 | 1 | 0.943223247 | 0.21515303 | 0.439 | 0.457 | 1 | 0.943223247 | 0.154769359 |
| Gm15581 | 0.080641486 | -2.20282941 | 0.019 | 0.089 | 1 | 0.17042253 | -1.863234868 | 0 | 0.044 | 1 | 0.17042253 | 0.154779922 |
| Zfp322a | 0.080747468 | 1.124975251 | 0.245 | 0.157 | 1 | 0.895983101 | 0.371541277 | 0.22 | 0.249 | 1 | 0.895983101 | 0.154974782 |
| Pcgf6 | 0.345577435 | 0.221836791 | 0.283 | 0.216 | 1 | 0.08077722 | -0.654648247 | 0.098 | 0.222 | 1 | 0.345577435 | 0.155029481 |
| Dap | 0.24425283 | -0.179178142 | 0.887 | 0.928 | 1 | 0.080848043 | -0.274920046 | 0.854 | 0.877 | 1 | 0.24425283 | 0.155159681 |
| Rab3a | 0.103822687 | 0.640149422 | 0.113 | 0.051 | 1 | 0.080967876 | 1.000703582 | 0.122 | 0.051 | 1 | 0.103822687 | 0.155379955 |
| Rhof | 0.08098747 | 0.962077517 | 0.17 | 0.089 | 1 | 0.084607555 | 1.537644568 | 0.22 | 0.123 | 1 | 0.084607555 | 0.15541597 |
| Crip1 | 0.853424582 | -0.895202631 | 0.736 | 0.636 | 1 | 0.081009807 | 0.672948598 | 0.585 | 0.495 | 1 | 0.853424582 | 0.155457025 |
| Ifitm1 | 0.081015803 | 1.025439578 | 0.17 | 0.089 | 1 | 0.096706236 | 1.319356545 | 0.293 | 0.191 | 1 | 0.096706236 | 0.155468046 |
| Ier3ip1 | 0.279914494 | 0.178569337 | 0.906 | 0.852 | 1 | 0.081124463 | -0.327854396 | 0.854 | 0.884 | 1 | 0.279914494 | 0.155667747 |
| Pigg | 0.081149034 | -1.046911266 | 0.075 | 0.169 | 1 | 0.099236337 | -0.385187572 | 0.049 | 0.15 | 1 | 0.099236337 | 0.155712903 |
| Dtwd2 | 0.081183316 | 1.855162313 | 0.113 | 0.051 | 1 | 0.853500739 | 0.292884334 | 0.098 | 0.089 | 1 | 0.853500739 | 0.155775902 |
| Pop1 | 0.081201874 | 1.541745718 | 0.189 | 0.106 | 1 | 0.966370685 | 0.515276755 | 0.146 | 0.147 | 1 | 0.966370685 | 0.155810004 |
| Gpat3 | 0.115342143 | -0.7032584 | 0.132 | 0.233 | 1 | 0.08124781 | -1.180717058 | 0.049 | 0.154 | 1 | 0.115342143 | 0.155894413 |
| Runx1 | 0.081304165 | -0.408237883 | 0.981 | 0.975 | 1 | 0.354000667 | -0.457544513 | 0.976 | 0.99 | 1 | 0.354000667 | 0.155997964 |
| Sh3bp5l | 0.081355892 | 1.029191713 | 0.17 | 0.089 | 1 | 0.241130996 | 0.782757066 | 0.22 | 0.15 | 1 | 0.241130996 | 0.156093004 |
| S100a7a | 0.081379558 | -1.752168001 | 0 | 0.055 | 1 | 0.188573672 | -0.863234868 | 0 | 0.041 | 1 | 0.188573672 | 0.156136483 |
| Gm57388 | 0.081379558 | -1.752168001 | 0 | 0.055 | 1 | 0.916649754 | 0.667279848 | 0.024 | 0.027 | 1 | 0.916649754 | 0.156136483 |
| Ranbp3l | 0.081379558 | -2.093204919 | 0 | 0.055 | 1 | 0.622594123 | 0.252242349 | 0.024 | 0.041 | 1 | 0.622594123 | 0.156136483 |
| Gm54798 | 0.08139894 | -1.845277405 | 0 | 0.055 | 1 | 0.787967407 | 0.313642894 | 0.073 | 0.061 | 1 | 0.787967407 | 0.156172092 |
| Scml2 | 0.08139894 | -2.093204919 | 0 | 0.055 | 1 | 0.126404896 | -1.622226769 | 0 | 0.055 | 1 | 0.126404896 | 0.156172092 |
| Mzt2 | 0.081411928 | 0.235859358 | 0.415 | 0.28 | 1 | 0.6186113 | 0.109284395 | 0.22 | 0.27 | 1 | 0.6186113 | 0.156195954 |
| Gm29292 | 0.081415091 | -1.932740246 | 0 | 0.055 | 1 | 0.832579923 | 0.174239837 | 0.049 | 0.041 | 1 | 0.832579923 | 0.156201766 |
| Gm15902 | 0.081415091 | -2.1672055 | 0 | 0.055 | 1 | 0.455094184 | 0.252242349 | 0 | 0.014 | 1 | 0.455094184 | 0.156201766 |
| H2-Q10 | 0.081437704 | -2.489133595 | 0 | 0.055 | 1 | 0.455097252 | -0.16279515 | 0 | 0.014 | 1 | 0.455097252 | 0.156243308 |
| Abcb1a | 0.081444164 | -2.304709024 | 0 | 0.055 | 1 | 0.455097252 | -0.484723245 | 0 | 0.014 | 1 | 0.455097252 | 0.156255177 |
| Lbhd2 | 0.081444164 | -2.545717123 | 0 | 0.055 | 1 | 0.446825814 | -0.80665134 | 0.024 | 0.051 | 1 | 0.446825814 | 0.156255177 |
| Ireb2 | 0.081917256 | 0.523029044 | 0.66 | 0.559 | 1 | 0.081448094 | 0.337738569 | 0.78 | 0.71 | 1 | 0.081917256 | 0.156262396 |
| Cstb | 0.845680559 | -0.446676796 | 0.981 | 0.86 | 1 | 0.081512447 | -0.497779398 | 0.732 | 0.915 | 1 | 0.845680559 | 0.156380615 |
| Gm47761 | 0.081602664 | -1.600164907 | 0.019 | 0.089 | 1 | 0.749728988 | 0.515276755 | 0.024 | 0.017 | 1 | 0.749728988 | 0.156546333 |
| Tmem37 | 0.081603203 | -1.799473716 | 0.019 | 0.089 | 1 | 0.828694816 | 0.334704509 | 0.049 | 0.041 | 1 | 0.828694816 | 0.156547323 |
| Gm48027 | 0.153239186 | 0.8327945 | 0.189 | 0.114 | 1 | 0.081752855 | 1.209173627 | 0.268 | 0.164 | 1 | 0.153239186 | 0.156822181 |
| Ccdc159 | 0.150266341 | -1.1672055 | 0 | 0.038 | 1 | 0.081817456 | 1.989207943 | 0.073 | 0.024 | 1 | 0.150266341 | 0.156940816 |
| Gm34250 | 0.249034808 | 2.154722595 | 0.019 | 0.004 | 1 | 0.081817456 | 1.989207943 | 0.073 | 0.024 | 1 | 0.249034808 | 0.156940816 |
| Coq8b | 0.081821846 | -0.698728742 | 0.245 | 0.381 | 1 | 0.392796981 | -0.255904555 | 0.195 | 0.263 | 1 | 0.392796981 | 0.156948877 |
| Ntn4 | 0.768803235 | -0.458254282 | 0.132 | 0.148 | 1 | 0.081858316 | 0.83720485 | 0.22 | 0.119 | 1 | 0.768803235 | 0.157015849 |
| Gm56626 | 0.569866135 | -0.1672055 | 0.019 | 0.034 | 1 | 0.081880116 | 1.614812428 | 0.098 | 0.038 | 1 | 0.569866135 | 0.157055879 |
| Mrpl20 | 0.081948117 | 0.273181428 | 0.868 | 0.767 | 1 | 0.258431235 | 0.227005058 | 0.805 | 0.792 | 1 | 0.258431235 | 0.15718074 |
| Prpf4 | 0.082012208 | 0.672853587 | 0.358 | 0.25 | 1 | 0.467516082 | 0.46797104 | 0.317 | 0.29 | 1 | 0.467516082 | 0.157298413 |
| Adgrg2 | 0.082055777 | -1.752168001 | 0.151 | 0.246 | 1 | 0.559845024 | -0.356103079 | 0.317 | 0.273 | 1 | 0.559845024 | 0.157378404 |
| Gm56815 | 0.082100671 | 1.640149422 | 0.094 | 0.038 | 1 | 0.997335755 | 0.79281073 | 0.098 | 0.099 | 1 | 0.997335755 | 0.157460822 |
| Bcl9 | 0.082143187 | 1.261637799 | 0.264 | 0.178 | 1 | 0.235227915 | 1.190841804 | 0.366 | 0.3 | 1 | 0.235227915 | 0.15753887 |
| Raver2 | 0.082172169 | 1.517292674 | 0.094 | 0.038 | 1 | 0.281346686 | 1.88967227 | 0.098 | 0.055 | 1 | 0.281346686 | 0.157592073 |
| Nat8f1 | 0.082233882 | -1.489133595 | 0.019 | 0.089 | 1 | 0.672172778 | 0.136765132 | 0.024 | 0.038 | 1 | 0.672172778 | 0.157705353 |
| Gga1 | 0.082265253 | 0.855162313 | 0.321 | 0.22 | 1 | 0.797445264 | 0.348918368 | 0.244 | 0.28 | 1 | 0.797445264 | 0.157762935 |
| Dnajc12 | 0.405743955 | -0.337130502 | 0.094 | 0.14 | 1 | 0.082281329 | -1.332720152 | 0.049 | 0.147 | 1 | 0.405743955 | 0.157792441 |
| Clba1 | 0.37217224 | -0.776014743 | 0.113 | 0.161 | 1 | 0.082289885 | -0.347219721 | 0.098 | 0.229 | 1 | 0.37217224 | 0.157808145 |
| Per3 | 0.434763477 | -0.128211368 | 0.34 | 0.407 | 1 | 0.082312304 | -0.500664789 | 0.22 | 0.369 | 1 | 0.434763477 | 0.157849292 |
| Scyl2 | 0.082317415 | 0.707263618 | 0.415 | 0.331 | 1 | 0.238553475 | -0.257513807 | 0.537 | 0.601 | 1 | 0.238553475 | 0.157858673 |
| Mapk12 | 0.209009104 | 1.739685096 | 0.038 | 0.013 | 1 | 0.082332119 | 1.83720485 | 0.073 | 0.024 | 1 | 0.209009104 | 0.15788566 |
| Mrtfa | 0.082357981 | 0.670337049 | 0.566 | 0.538 | 1 | 0.355739295 | -0.540465797 | 0.707 | 0.693 | 1 | 0.355739295 | 0.157933125 |
| Ankrd13c | 0.083633174 | -0.360451576 | 0.509 | 0.665 | 1 | 0.082378567 | 0.502919548 | 0.732 | 0.751 | 1 | 0.083633174 | 0.157970907 |
| Mthfsl | 0.579648126 | 0.273367091 | 0.283 | 0.25 | 1 | 0.082451618 | -0.622226769 | 0.146 | 0.28 | 1 | 0.579648126 | 0.158104966 |
| Chst15 | 0.082473918 | 1.544669113 | 0.17 | 0.093 | 1 | 0.395031673 | 0.916639317 | 0.195 | 0.15 | 1 | 0.395031673 | 0.158145889 |
| 1110004F10Rik | 0.266064482 | 0.194762955 | 0.868 | 0.818 | 1 | 0.082520972 | -0.312069642 | 0.854 | 0.901 | 1 | 0.266064482 | 0.158232234 |
| Gm42067 | 0.082545374 | -1.368839361 | 0.019 | 0.089 | 1 | 0.231643049 | -0.622226769 | 0 | 0.034 | 1 | 0.231643049 | 0.15827701 |
| Gm50232 | 0.082586873 | 1.569760094 | 0.094 | 0.038 | 1 | 0.355439883 | 1.313642894 | 0.098 | 0.061 | 1 | 0.355439883 | 0.158353155 |
| Tmem35b | 0.082598353 | -0.808751529 | 0.057 | 0.148 | 1 | 0.432197158 | -0.16279515 | 0.073 | 0.116 | 1 | 0.432197158 | 0.158374219 |
| Nbas | 0.08263686 | 0.385335523 | 0.623 | 0.525 | 1 | 0.401675396 | -0.350039628 | 0.512 | 0.56 | 1 | 0.401675396 | 0.15844487 |
| Lrif1 | 0.08264008 | 0.919094346 | 0.321 | 0.225 | 1 | 0.160093648 | 0.728680393 | 0.366 | 0.287 | 1 | 0.160093648 | 0.158450777 |
| Zdhhc9 | 0.082675109 | 0.552686581 | 0.34 | 0.233 | 1 | 0.116438647 | -0.289327556 | 0.171 | 0.307 | 1 | 0.116438647 | 0.158515044 |
| Disc1 | 0.789952486 | 0.154722595 | 0.019 | 0.025 | 1 | 0.082741448 | 1.644559772 | 0.098 | 0.038 | 1 | 0.789952486 | 0.158636748 |
| Pak4 | 0.259291451 | -0.395474488 | 0.094 | 0.157 | 1 | 0.082749639 | -1.305753104 | 0.049 | 0.147 | 1 | 0.259291451 | 0.158651776 |
| Med12l | 0.2772391 | -1.077938162 | 0.094 | 0.148 | 1 | 0.082751468 | 0.600165653 | 0.293 | 0.181 | 1 | 0.2772391 | 0.158655131 |
| Tfeb | 0.133624138 | 1.055186921 | 0.321 | 0.237 | 1 | 0.08287238 | -0.527367582 | 0.244 | 0.392 | 1 | 0.133624138 | 0.158876929 |
| Fjx1 | 0.08292813 | 0.470224421 | 0.302 | 0.203 | 1 | 0.177937456 | 0.515276755 | 0.317 | 0.222 | 1 | 0.177937456 | 0.158979185 |
| Tlnrd1 | 0.33270036 | 0.569760094 | 0.302 | 0.25 | 1 | 0.082935964 | -0.737703986 | 0.171 | 0.317 | 1 | 0.33270036 | 0.158993554 |
| Gm26782 | 0.723856102 | 0.29808677 | 0.396 | 0.373 | 1 | 0.083082431 | -0.309636539 | 0.244 | 0.392 | 1 | 0.723856102 | 0.159262172 |
| Wdr11 | 0.08311295 | 1.193196743 | 0.321 | 0.225 | 1 | 0.567382729 | -0.140075074 | 0.268 | 0.314 | 1 | 0.567382729 | 0.159318138 |
| Rab5if | 0.183763104 | -0.317265196 | 0.83 | 0.809 | 1 | 0.083174818 | -0.358051442 | 0.756 | 0.846 | 1 | 0.183763104 | 0.159431585 |
| Tmem184b | 0.083203094 | 0.733879474 | 0.509 | 0.415 | 1 | 0.092693921 | -0.406720733 | 0.39 | 0.556 | 1 | 0.092693921 | 0.159483432 |

| | | | | | | | | | | | | |
|---|---|---|---|---|---|---|---|---|---|---|---|---|
| 2610507I01Rik | 0.334091523 | 0.874614676 | 0.151 | 0.106 | 1 | 0.08325168 | 1.351778023 | 0.244 | 0.147 | 1 | 0.334091523 | 0.159572519 |
| Kctd18 | 0.083279083 | 0.770058745 | 0.208 | 0.119 | 1 | 0.234852238 | -0.600200463 | 0.098 | 0.171 | 1 | 0.234852238 | 0.15962276 |
| Aak1 | 0.147777844 | -0.297654266 | 0.887 | 0.941 | 1 | 0.083360779 | -0.280673404 | 0.951 | 0.983 | 1 | 0.147777844 | 0.159772538 |
| Rtn1 | 0.083376876 | 3.002719501 | 0.057 | 0.017 | 1 | 0.54561906 | -0.555112573 | 0.024 | 0.044 | 1 | 0.54561906 | 0.159802049 |
| Steap4 | 0.083398474 | -1.304709024 | 0.075 | 0.174 | 1 | 0.128357553 | 2.382638986 | 0.122 | 0.061 | 1 | 0.128357553 | 0.159841642 |
| Gm45509 | 0.083439893 | 1.517292674 | 0.113 | 0.051 | 1 | 0.692944181 | 0.30115195 | 0.098 | 0.078 | 1 | 0.692944181 | 0.15991757 |
| Bmp15 | 0.083439893 | 1.517292674 | 0.113 | 0.051 | 1 | 0.82181816 | 0.136765132 | 0.024 | 0.031 | 1 | 0.82181816 | 0.15991757 |
| Zfp53 | 0.083611268 | 0.776210972 | 0.264 | 0.165 | 1 | 0.898390122 | -0.253397699 | 0.244 | 0.256 | 1 | 0.898390122 | 0.160231691 |
| Gm35188 | 0.760923973 | -0.1672055 | 0.132 | 0.114 | 1 | 0.083626957 | 1.229522273 | 0.244 | 0.15 | 1 | 0.760923973 | 0.160260446 |
| Mtmr6 | 0.306043732 | 0.302279783 | 0.566 | 0.525 | 1 | 0.083737631 | -0.282230381 | 0.488 | 0.638 | 1 | 0.306043732 | 0.160463272 |
| Gcdh | 0.083744733 | 0.762405172 | 0.245 | 0.148 | 1 | 0.624490473 | 0.73286819 | 0.146 | 0.123 | 1 | 0.624490473 | 0.160476286 |
| Hcfc1 | 0.083850764 | 0.751757928 | 0.472 | 0.364 | 1 | 0.172021429 | 0.486130409 | 0.561 | 0.488 | 1 | 0.172021429 | 0.160670578 |
| Gm57404 | 0.083858689 | 1.865215978 | 0.094 | 0.038 | 1 | 0.154090132 | -1.16279515 | 0 | 0.048 | 1 | 0.154090132 | 0.160685098 |
| Gm14471 | 0.343006305 | -0.1672055 | 0 | 0.017 | 1 | 0.083899597 | 1.574170444 | 0.073 | 0.024 | 1 | 0.343006305 | 0.160760051 |
| Ap1s3 | 0.083903744 | 1.806799291 | 0.094 | 0.038 | 1 | 0.167115549 | 0.721727632 | 0.146 | 0.082 | 1 | 0.167115549 | 0.160767651 |
| Gm57027 | 0.424662489 | 1.039245377 | 0.057 | 0.034 | 1 | 0.083915902 | 1.83720485 | 0.073 | 0.024 | 1 | 0.424662489 | 0.160789926 |
| Use1 | 0.387320283 | 0.224431567 | 0.792 | 0.814 | 1 | 0.083923388 | -0.346899087 | 0.585 | 0.717 | 1 | 0.387320283 | 0.160803641 |
| Recql5 | 0.212999922 | 0.962077517 | 0.226 | 0.161 | 1 | 0.08399936 | 1.326242931 | 0.195 | 0.109 | 1 | 0.212999922 | 0.160942828 |
| 6430548M08Rik | 0.292280505 | -0.223789028 | 0.038 | 0.081 | 1 | 0.084145735 | -1.899760744 | 0.049 | 0.143 | 1 | 0.292280505 | 0.161210965 |
| Svopl | 0.084217493 | 1.438515561 | 0.113 | 0.051 | 1 | 0.125807067 | -0.505939731 | 0.195 | 0.331 | 1 | 0.125807067 | 0.1613424 |
| Efr3a | 0.084238841 | 0.347367673 | 0.717 | 0.631 | 1 | 0.282313726 | 0.319134233 | 0.854 | 0.775 | 1 | 0.282313726 | 0.1613815 |
| Ttc30b | 0.084240368 | 1.707263618 | 0.113 | 0.051 | 1 | 0.681377493 | -0.221688839 | 0.049 | 0.065 | 1 | 0.681377493 | 0.161384297 |
| Men1 | 0.084455213 | 0.913714495 | 0.17 | 0.089 | 1 | 0.095494036 | 1.067817778 | 0.317 | 0.218 | 1 | 0.095494036 | 0.161777742 |
| Nptx1 | 0.084560562 | -2.1672055 | 0.019 | 0.089 | 1 | 0.553780906 | 0.422167351 | 0.049 | 0.031 | 1 | 0.553780906 | 0.161970635 |
| Pgghg | 0.453830041 | 0.267197324 | 0.17 | 0.127 | 1 | 0.08460916 | -1.132421501 | 0.049 | 0.147 | 1 | 0.453830041 | 0.162059611 |
| Tspyl1 | 0.084776299 | 0.484290752 | 0.642 | 0.572 | 1 | 0.477586839 | -0.206364165 | 0.61 | 0.638 | 1 | 0.477586839 | 0.162365576 |
| Rrp8 | 0.084786922 | 0.711779099 | 0.377 | 0.271 | 1 | 0.288227486 | -0.284785675 | 0.244 | 0.338 | 1 | 0.288227486 | 0.162385022 |
| Rbm17 | 0.084825322 | 0.411880434 | 0.717 | 0.661 | 1 | 0.189384091 | -0.273716964 | 0.756 | 0.758 | 1 | 0.189384091 | 0.162455308 |
| Rtn2 | 0.560868044 | 0.454282877 | 0.057 | 0.038 | 1 | 0.084849342 | 0.911205431 | 0.098 | 0.038 | 1 | 0.560868044 | 0.162499274 |
| Pbx4 | 0.926212226 | 0.8327945 | 0.019 | 0.017 | 1 | 0.084849342 | 1.458693227 | 0.098 | 0.038 | 1 | 0.926212226 | 0.162499274 |
| Serf1 | 0.084887794 | 0.632953921 | 0.547 | 0.475 | 1 | 0.498729421 | -0.148295581 | 0.366 | 0.42 | 1 | 0.498729421 | 0.16256965 |
| Srebf1 | 0.09053337 | -0.845277405 | 0.283 | 0.394 | 1 | 0.084902864 | -0.823045778 | 0.22 | 0.355 | 1 | 0.09053337 | 0.162597232 |
| Serpine3 | 0.251644813 | 0.8327945 | 0.019 | 0.004 | 1 | 0.08495876 | 1.029849928 | 0.073 | 0.024 | 1 | 0.251644813 | 0.162699529 |
| Gm16287 | 0.62424925 | 0.154722595 | 0.038 | 0.055 | 1 | 0.084959048 | 1.699701326 | 0.073 | 0.024 | 1 | 0.62424925 | 0.162700056 |
| Top1 | 0.084977792 | -0.436672917 | 0.981 | 0.996 | 1 | 0.282941642 | -0.24397919 | 1 | 0.997 | 1 | 0.282941642 | 0.162734359 |
| Dhx30 | 0.085038948 | 0.624207878 | 0.472 | 0.369 | 1 | 0.381751593 | 0.515276755 | 0.463 | 0.44 | 1 | 0.381751593 | 0.162846274 |
| Zfp869 | 0.139877625 | -0.207847485 | 0.358 | 0.462 | 1 | 0.085072548 | -0.435813645 | 0.317 | 0.481 | 1 | 0.139877625 | 0.162907757 |
| Sema4b | 0.446503896 | -1.191052242 | 0.094 | 0.127 | 1 | 0.085122833 | 1.199774929 | 0.146 | 0.068 | 1 | 0.446503896 | 0.162999769 |
| Gm45902 | 0.243111734 | -0.652632327 | 0 | 0.025 | 1 | 0.085336957 | -1.686357106 | 0 | 0.068 | 1 | 0.243111734 | 0.163391517 |
| Fbp1 | 0.48510256 | -0.7032584 | 0.057 | 0.085 | 1 | 0.085336957 | -1.622226769 | 0 | 0.068 | 1 | 0.48510256 | 0.163391517 |
| 9030025P20Rik | 0.206396398 | -0.845277405 | 0 | 0.03 | 1 | 0.085362426 | -1.686357106 | 0 | 0.068 | 1 | 0.206396398 | 0.163438107 |
| Coq8a | 0.172087146 | -1.015202407 | 0.019 | 0.068 | 1 | 0.085404874 | -1.863234868 | 0 | 0.068 | 1 | 0.172087146 | 0.163515755 |
| Upp1 | 0.464650233 | -0.942138944 | 0.094 | 0.131 | 1 | 0.085404874 | -1.970150072 | 0 | 0.068 | 1 | 0.464650233 | 0.163515755 |
| Dok4 | 0.117896765 | -1.368839361 | 0.057 | 0.131 | 1 | 0.085405469 | -1.80665134 | 0 | 0.068 | 1 | 0.117896765 | 0.163516845 |
| Matk | 0.413082433 | -0.430239906 | 0 | 0.013 | 1 | 0.085449555 | -2.116991461 | 0 | 0.068 | 1 | 0.413082433 | 0.163597484 |
| Snn | 0.090287817 | 1.891688189 | 0.057 | 0.017 | 1 | 0.085465061 | 1.667279848 | 0.073 | 0.024 | 1 | 0.090287817 | 0.163625846 |
| Rnf113a1 | 0.24309253 | -0.845277405 | 0 | 0.025 | 1 | 0.085465061 | 1.667279848 | 0.073 | 0.024 | 1 | 0.24309253 | 0.163625846 |
| Gm57038 | 0.725936481 | 1.417757001 | 0.019 | 0.013 | 1 | 0.085465061 | 1.667279848 | 0.073 | 0.024 | 1 | 0.725936481 | 0.163625846 |
| Trmt12 | 0.866605856 | 0.454282877 | 0.057 | 0.051 | 1 | 0.085465061 | 1.667279848 | 0.073 | 0.024 | 1 | 0.866605856 | 0.163625846 |
| Tyro3 | 0.413082433 | 0.154722595 | 0 | 0.013 | 1 | 0.085506748 | -1.555112573 | 0 | 0.068 | 1 | 0.413082433 | 0.163702092 |
| Pigw | 0.085733417 | 1.454282877 | 0.113 | 0.051 | 1 | 0.733713289 | 0.721727632 | 0.049 | 0.038 | 1 | 0.733713289 | 0.164116615 |
| Glmn | 0.085778974 | 0.857042046 | 0.264 | 0.169 | 1 | 0.365662757 | 0.287007767 | 0.122 | 0.191 | 1 | 0.365662757 | 0.164199916 |
| AV356131 | 0.085835472 | 2.8327945 | 0.057 | 0.017 | 1 | 0.872828964 | 1.029849928 | 0.024 | 0.02 | 1 | 0.872828964 | 0.164303215 |
| Sema3g | 0.085835472 | 2.640149422 | 0.057 | 0.017 | 1 | 0.5191986 | 0.515276755 | 0 | 0.01 | 1 | 0.5191986 | 0.164303215 |
| Rps6ka6 | 0.085835472 | 2.417757001 | 0.057 | 0.017 | 1 | 0.681377493 | -0.221688839 | 0.049 | 0.065 | 1 | 0.681377493 | 0.164303215 |
| Lepr | 0.085920688 | 1.299539148 | 0.396 | 0.314 | 1 | 0.165672452 | -0.594968562 | 0.366 | 0.474 | 1 | 0.165672452 | 0.164459011 |
| Erf | 0.458550662 | 0.121299593 | 0.642 | 0.644 | 1 | 0.08610193 | -0.436970114 | 0.707 | 0.833 | 1 | 0.458550662 | 0.164790317 |
| Capg | 0.086102223 | -0.637284363 | 0.302 | 0.415 | 1 | 0.230175827 | -0.353257083 | 0.366 | 0.488 | 1 | 0.230175827 | 0.164790852 |
| Hdhd3 | 0.08615174 | 1.417757001 | 0.094 | 0.038 | 1 | 0.599290422 | 1.515276755 | 0.024 | 0.014 | 1 | 0.599290422 | 0.164881358 |
| Tsen54 | 0.08615174 | 1.417757001 | 0.094 | 0.038 | 1 | 0.369124475 | 1.100239256 | 0.098 | 0.061 | 1 | 0.369124475 | 0.164881358 |
| Gm50069 | 0.08615174 | 1.417757001 | 0.094 | 0.038 | 1 | 0.743829142 | 0.252242349 | 0.024 | 0.034 | 1 | 0.743829142 | 0.164881358 |
| Gla | 0.316197711 | 0.589937976 | 0.283 | 0.22 | 1 | 0.086165719 | 0.966191291 | 0.39 | 0.29 | 1 | 0.316197711 | 0.164906907 |
| Atp5j2 | 0.106149253 | 0.277954361 | 0.981 | 0.86 | 1 | 0.086192234 | 0.336181168 | 0.854 | 0.857 | 1 | 0.106149253 | 0.164955366 |
| Epha2 | 0.273536324 | 0.716601482 | 0.283 | 0.225 | 1 | 0.086216321 | 0.385078032 | 0.512 | 0.369 | 1 | 0.273536324 | 0.164999388 |
| Nexn | 0.086270459 | 1.417757001 | 0.113 | 0.051 | 1 | 0.269608562 | -0.385187572 | 0.073 | 0.137 | 1 | 0.269608562 | 0.165098325 |
| Zfp407 | 0.086345671 | 0.498793573 | 0.679 | 0.61 | 1 | 0.274196895 | 0.106263043 | 0.805 | 0.751 | 1 | 0.274196895 | 0.165235767 |
| Mrps24 | 0.086422575 | 0.569760094 | 0.623 | 0.564 | 1 | 0.145636684 | -0.323259822 | 0.512 | 0.611 | 1 | 0.145636684 | 0.165376289 |
| Slc3a1 | 0.086465956 | 2.569760094 | 0.057 | 0.017 | 1 | 0.827890505 | -0.26713181 | 0.073 | 0.082 | 1 | 0.827890505 | 0.16545555 |
| Zfp975 | 0.646102696 | 0.197791317 | 0.189 | 0.229 | 1 | 0.086498847 | -0.577832649 | 0.098 | 0.218 | 1 | 0.646102696 | 0.165515643 |
| Pwwp2b | 0.676691664 | 0.347367673 | 0.094 | 0.076 | 1 | 0.086530326 | -1.030691614 | 0.073 | 0.181 | 1 | 0.676691664 | 0.165573155 |
| Rmdn2 | 0.575799807 | 1.154722595 | 0.321 | 0.284 | 1 | 0.086565387 | -0.886821689 | 0.171 | 0.307 | 1 | 0.575799807 | 0.165637207 |
| Fam83g | 0.241482942 | -0.508242418 | 0.019 | 0.059 | 1 | 0.086574251 | -1.717384002 | 0.024 | 0.109 | 1 | 0.241482942 | 0.1656534 |
| Lats1 | 0.608540199 | 0.204995801 | 0.585 | 0.623 | 1 | 0.086616145 | -0.240185183 | 0.756 | 0.887 | 1 | 0.608540199 | 0.165729934 |
| Gmpr2 | 0.086717561 | 0.991223863 | 0.302 | 0.212 | 1 | 0.714007889 | 0.232342792 | 0.22 | 0.195 | 1 | 0.714007889 | 0.165915187 |
| Ehhadh | 0.08698757 | 1.261637799 | 0.094 | 0.038 | 1 | 0.87556513 | 0.667279848 | 0.024 | 0.02 | 1 | 0.87556513 | 0.166408303 |
| Klhl12 | 0.087002635 | 0.433438623 | 0.415 | 0.297 | 1 | 0.354225071 | -0.585327186 | 0.293 | 0.355 | 1 | 0.354225071 | 0.166435811 |
| E130202H07Rik | 0.087094062 | 2.377115016 | 0.057 | 0.017 | 1 | 0.104856732 | 2.83720485 | 0.024 | 0.003 | 1 | 0.104856732 | 0.166602749 |
| Ngfr | 0.0909501 | 1.8327945 | 0.057 | 0.017 | 1 | 0.08711311 | 1.185128153 | 0.098 | 0.038 | 1 | 0.0909501 | 0.166637527 |
| Klf15 | 0.087150282 | 1.247831999 | 0.17 | 0.089 | 1 | 0.764914516 | 0.377773231 | 0.073 | 0.089 | 1 | 0.764914516 | 0.166705392 |
| Rpf1 | 0.087153598 | 0.498676996 | 0.434 | 0.318 | 1 | 0.314894255 | -0.123048971 | 0.317 | 0.416 | 1 | 0.314894255 | 0.166711446 |
| Mknk2 | 0.110877183 | -0.185901594 | 0.415 | 0.568 | 1 | 0.087161588 | -0.382473187 | 0.634 | 0.747 | 1 | 0.110877183 | 0.166726034 |
| 2810402E24Rik | 0.087161902 | -0.727920455 | 0.094 | 0.199 | 1 | 0.500914307 | 0.773074512 | 0.073 | 0.109 | 1 | 0.500914307 | 0.166726606 |
| Pfkl | 0.087205356 | -0.468509923 | 0.377 | 0.521 | 1 | 0.721927985 | 0.482362132 | 0.317 | 0.314 | 1 | 0.721927985 | 0.166805937 |
| Ifi27l2a | 0.172452514 | 1.243989933 | 0.113 | 0.059 | 1 | 0.087229089 | 1.773074512 | 0.122 | 0.055 | 1 | 0.172452514 | 0.166849264 |
| Get1 | 0.087236623 | 1.237184755 | 0.132 | 0.064 | 1 | 0.478704236 | 0.114738825 | 0.049 | 0.082 | 1 | 0.478704236 | 0.166863018 |
| Tafazzin | 0.997601494 | 0.232259195 | 0.453 | 0.483 | 1 | 0.087337685 | -0.46860358 | 0.268 | 0.42 | 1 | 0.997601494 | 0.167047498 |
| AU022252 | 0.087380297 | 0.922061838 | 0.245 | 0.153 | 1 | 0.953576626 | 0.703349103 | 0.195 | 0.215 | 1 | 0.953576626 | 0.167125278 |
| Tmem59 | 0.087633396 | 0.203393042 | 1 | 0.979 | 1 | 0.50974974 | 0.174028345 | 0.927 | 0.99 | 1 | 0.50974974 | 0.16758718 |
| Gm16159 | 0.157246372 | 1.569760094 | 0.057 | 0.021 | 1 | 0.087679812 | -1.448197369 | 0.024 | 0.109 | 1 | 0.157246372 | 0.167671874 |
| Mapk1ip1 | 0.575674566 | 0.635348436 | 0.151 | 0.127 | 1 | 0.087713929 | -1.484723245 | 0.049 | 0.143 | 1 | 0.575674566 | 0.167734125 |
| Gm9970 | 0.08772696 | 2.154722595 | 0.057 | 0.017 | 1 | 0.878310931 | 0.667279848 | 0.024 | 0.02 | 1 | 0.878310931 | 0.167757901 |
| Aqp3 | 0.151000758 | -0.845277405 | 0.17 | 0.097 | 1 | 0.087826359 | 1.589277336 | 0.39 | 0.276 | 1 | 0.151000758 | 0.167939249 |
| Foxd2 | 0.57173755 | -0.508242418 | 0.094 | 0.119 | 1 | 0.087870296 | -1.410722664 | 0.049 | 0.147 | 1 | 0.57173755 | 0.168019404 |
| Zxdc | 0.087912681 | 0.370451286 | 0.358 | 0.237 | 1 | 0.390749527 | 0.174239837 | 0.366 | 0.474 | 1 | 0.390749527 | 0.168096722 |
| Ppp2r2b | 0.158435633 | 3.47665069 | 0.075 | 0.034 | 1 | 0.087934666 | 2.145327145 | 0.171 | 0.089 | 1 | 0.158435633 | 0.168136826 |
| Nudt8 | 0.59338774 | 0.377115016 | 0.113 | 0.089 | 1 | 0.087952179 | -1.484723245 | 0.024 | 0.109 | 1 | 0.59338774 | 0.168168773 |
| Atp1a2 | 0.107305355 | -3.112063946 | 0.019 | 0.081 | 1 | 0.087987992 | -2.529117364 | 0.073 | 0.181 | 1 | 0.107305355 | 0.168234097 |
| Sap25 | 0.338780504 | 0.984797593 | 0.057 | 0.03 | 1 | 0.088084868 | 1.635570989 | 0.146 | 0.072 | 1 | 0.338780504 | 0.168410793 |

| | | | | | | | | | | | | |
|---|---|---|---|---|---|---|---|---|---|---|---|---|
| Tst | 0.088122597 | 0.739685096 | 0.208 | 0.119 | 1 | 0.922068358 | -0.264333177 | 0.268 | 0.263 | 1 | 0.922068358 | 0.168479602 |
| Ykt6 | 0.088139784 | 0.545512548 | 0.566 | 0.508 | 1 | 0.157771486 | -0.250257991 | 0.512 | 0.638 | 1 | 0.157771486 | 0.168510947 |
| 9030624G23Rik | 0.088166845 | 1.347367673 | 0.113 | 0.051 | 1 | 0.976593771 | 0.962735732 | 0.024 | 0.024 | 1 | 0.976593771 | 0.168560297 |
| Mtmr2 | 0.088220055 | 0.415183057 | 0.604 | 0.492 | 1 | 0.673987295 | -0.209088802 | 0.537 | 0.594 | 1 | 0.673987295 | 0.168657331 |
| Ttll5 | 0.768374953 | -0.3327234 | 0.491 | 0.5 | 1 | 0.088237605 | 0.722837825 | 0.659 | 0.597 | 1 | 0.768374953 | 0.168689335 |
| Ppp1r18 | 0.088238167 | 0.365142374 | 0.472 | 0.347 | 1 | 0.301907043 | 0.598133687 | 0.463 | 0.433 | 1 | 0.301907043 | 0.168690359 |
| Nat14 | 0.088308085 | 1.280253477 | 0.094 | 0.038 | 1 | 0.157664897 | 0.377773231 | 0.024 | 0.092 | 1 | 0.157664897 | 0.168817852 |
| Gm10848 | 0.088349056 | 2.154722595 | 0.057 | 0.017 | 1 | 0.484946477 | 0.911205431 | 0.073 | 0.048 | 1 | 0.484946477 | 0.168892556 |
| 4930522L14Rik | 0.088349056 | 2.417757001 | 0.057 | 0.017 | 1 | 0.824088343 | 0.377773231 | 0.024 | 0.031 | 1 | 0.824088343 | 0.168892556 |
| Tex45 | 0.088349056 | 2.154722595 | 0.057 | 0.017 | 1 | 0.286421728 | -0.863234868 | 0 | 0.027 | 1 | 0.286421728 | 0.168892556 |
| Gm32389 | 0.088349056 | 2.640149422 | 0.057 | 0.017 | 1 | 0.357858058 | -0.622226769 | 0 | 0.02 | 1 | 0.357858058 | 0.168892556 |
| Zfp964 | 0.088349056 | 2.154722595 | 0.057 | 0.017 | 1 | 0.718283759 | 1.071670103 | 0.049 | 0.038 | 1 | 0.718283759 | 0.168892556 |
| Gm19531 | 0.088349056 | 2.154722595 | 0.057 | 0.017 | 1 | 0.983241205 | 0.422167351 | 0.049 | 0.048 | 1 | 0.983241205 | 0.168892556 |
| B430219N15Rik | 0.088349056 | 2.154722595 | 0.057 | 0.017 | 1 | 0.402373368 | 0.252242349 | 0 | 0.017 | 1 | 0.402373368 | 0.168892556 |
| D430019H16Rik | 0.088349056 | 2.640149422 | 0.057 | 0.017 | 1 | 0.817278758 | -0.555112573 | 0.024 | 0.031 | 1 | 0.817278758 | 0.168892556 |
| 1700102H20Rik | 0.088349056 | 2.154722595 | 0.057 | 0.017 | 1 | 0.968793924 | 1.422167351 | 0.024 | 0.024 | 1 | 0.968793924 | 0.168892556 |
| 2010204K13Rik | 0.088349056 | 2.154722595 | 0.057 | 0.017 | 1 | 0.114609247 | -1.332720152 | 0 | 0.058 | 1 | 0.114609247 | 0.168892556 |
| Gm14798 | 0.088349056 | 2.417757001 | 0.057 | 0.017 | 1 | 0.470914411 | 1.100239256 | 0.073 | 0.048 | 1 | 0.470914411 | 0.168892556 |
| Tmed3 | 0.088441821 | -0.272334487 | 0.736 | 0.831 | 1 | 0.184632293 | -0.248022987 | 0.634 | 0.761 | 1 | 0.184632293 | 0.169061686 |
| Hacd2 | 0.315645812 | 0.257991865 | 0.642 | 0.589 | 1 | 0.088476139 | 0.719938798 | 0.659 | 0.58 | 1 | 0.315645812 | 0.16912425 |
| Gm30025 | 0.83872179 | 0.216123139 | 0.094 | 0.085 | 1 | 0.088489069 | -1.292078167 | 0.024 | 0.109 | 1 | 0.83872179 | 0.169147822 |
| Pnpt1 | 0.088567266 | 0.408479187 | 0.396 | 0.263 | 1 | 0.727764077 | 0.635570989 | 0.293 | 0.311 | 1 | 0.727764077 | 0.169290372 |
| Cep70 | 0.088611768 | -0.752168001 | 0.094 | 0.203 | 1 | 0.749603896 | 0.279209397 | 0.098 | 0.119 | 1 | 0.749603896 | 0.169371491 |
| Cdh4 | 0.28776878 | -2.015202407 | 0 | 0.021 | 1 | 0.088723073 | 0.667279848 | 0.073 | 0.024 | 1 | 0.28776878 | 0.169574363 |
| Dhx34 | 0.602707627 | 0.347367673 | 0.094 | 0.072 | 1 | 0.088779938 | 1.483567895 | 0.122 | 0.055 | 1 | 0.602707627 | 0.169677998 |
| Per1 | 0.3038559 | 0.301563983 | 0.377 | 0.318 | 1 | 0.088806115 | 0.873730726 | 0.463 | 0.358 | 1 | 0.3038559 | 0.169725704 |
| Emilin2 | 0.37533282 | -1.054730771 | 0.057 | 0.093 | 1 | 0.088855034 | 1.730289646 | 0.122 | 0.055 | 1 | 0.37533282 | 0.169814851 |
| C1galt1 | 0.48631404 | 0.325834511 | 0.358 | 0.331 | 1 | 0.088862172 | 0.701068162 | 0.512 | 0.413 | 1 | 0.48631404 | 0.169827858 |
| Pou2f2 | 0.088867931 | 0.865215978 | 0.094 | 0.038 | 1 | 0.089661335 | 0.69281494 | 0.122 | 0.055 | 1 | 0.089661335 | 0.169838354 |
| Il1rapl1 | 0.088881865 | -1.932740246 | 0.057 | 0.14 | 1 | 0.751204029 | -1.348661696 | 0.073 | 0.058 | 1 | 0.751204029 | 0.169863744 |
| Clcn4 | 0.088936799 | 0.589659652 | 0.585 | 0.585 | 1 | 0.210236765 | 0.360948609 | 0.659 | 0.635 | 1 | 0.210236765 | 0.169963844 |
| Bud23 | 0.138502138 | 0.614154213 | 0.415 | 0.326 | 1 | 0.088990912 | -0.570879889 | 0.195 | 0.338 | 1 | 0.138502138 | 0.170062441 |
| Stk35 | 0.089054164 | 0.746637856 | 0.302 | 0.199 | 1 | 0.560868269 | 0.387897448 | 0.317 | 0.29 | 1 | 0.560868269 | 0.170177684 |
| Mfsd8 | 0.089119799 | 1.017219071 | 0.151 | 0.076 | 1 | 0.188812715 | -1.020776165 | 0.024 | 0.082 | 1 | 0.188812715 | 0.17029726 |
| 4930594M22Rik | 0.089292244 | 1.347367673 | 0.094 | 0.038 | 1 | 0.513659043 | 0.159132945 | 0.073 | 0.113 | 1 | 0.513659043 | 0.170611384 |
| N6amt1 | 0.133746829 | 1.034428361 | 0.396 | 0.322 | 1 | 0.089326804 | -1.20718927 | 0.122 | 0.232 | 1 | 0.133746829 | 0.17067433 |
| Stk36 | 0.447253949 | 0.569760094 | 0.057 | 0.034 | 1 | 0.089521939 | 0.944120054 | 0.146 | 0.068 | 1 | 0.447253949 | 0.1710297 |
| Traf6 | 0.384770343 | 0.454282877 | 0.264 | 0.22 | 1 | 0.089543967 | -0.53202896 | 0.195 | 0.341 | 1 | 0.384770343 | 0.171069812 |
| Mrpl10 | 0.288563079 | 0.470224421 | 0.377 | 0.309 | 1 | 0.089630151 | -0.622226769 | 0.171 | 0.311 | 1 | 0.288563079 | 0.171226737 |
| Synj2bp | 0.607687889 | 0.158484514 | 0.736 | 0.725 | 1 | 0.089636838 | -0.358142749 | 0.683 | 0.819 | 1 | 0.607687889 | 0.171238914 |
| Uchl1 | 0.089640617 | 2.569760094 | 0.057 | 0.017 | 1 | 0.930972993 | -0.935384654 | 0.049 | 0.044 | 1 | 0.930972993 | 0.171245793 |
| Gm38604 | 0.089640617 | 2.154722595 | 0.057 | 0.017 | 1 | 0.357858789 | -0.484723245 | 0 | 0.02 | 1 | 0.357858789 | 0.171245793 |
| Sfn | 0.089686182 | -2.156863556 | 0.208 | 0.301 | 1 | 0.624189555 | 0.149148856 | 0.341 | 0.311 | 1 | 0.624189555 | 0.171328753 |
| Acot1 | 0.482570401 | 0.32690357 | 0.245 | 0.199 | 1 | 0.089724691 | -0.863234868 | 0.098 | 0.212 | 1 | 0.482570401 | 0.171398861 |
| Mindy3 | 0.089850239 | 0.561042877 | 0.472 | 0.364 | 1 | 0.857481194 | -0.297637692 | 0.488 | 0.468 | 1 | 0.857481194 | 0.171627412 |
| Nol6 | 0.089864477 | 1.154722595 | 0.132 | 0.064 | 1 | 0.589599113 | 0.159132945 | 0.073 | 0.102 | 1 | 0.589599113 | 0.171653331 |
| Cdr2 | 0.089892246 | 0.616757574 | 0.434 | 0.339 | 1 | 0.525050296 | -0.179096963 | 0.488 | 0.553 | 1 | 0.525050296 | 0.171703876 |
| 1700084C06Rik | 0.089896614 | -0.600164907 | 0.019 | 0.089 | 1 | 0.925228713 | 0.306690133 | 0.098 | 0.102 | 1 | 0.925228713 | 0.171711826 |
| Or8b53 | 0.573775862 | -0.292736382 | 0.17 | 0.131 | 1 | 0.089958222 | -0.80665134 | 0.122 | 0.246 | 1 | 0.573775862 | 0.171823962 |
| Odf2l | 0.089998806 | 0.703026625 | 0.34 | 0.254 | 1 | 0.582967308 | 0.33970519 | 0.171 | 0.215 | 1 | 0.582967308 | 0.171897827 |
| Naa35 | 0.126149431 | 0.837381769 | 0.453 | 0.394 | 1 | 0.090100815 | 0.688527582 | 0.707 | 0.625 | 1 | 0.126149431 | 0.172083473 |
| Nck2 | 0.499760366 | -0.442363465 | 0.623 | 0.669 | 1 | 0.090268963 | -0.465986683 | 0.732 | 0.802 | 1 | 0.499760366 | 0.172389441 |
| Xlr3b | 0.090280906 | 1.817687608 | 0.132 | 0.068 | 1 | 0.139562594 | -1.863234868 | 0 | 0.051 | 1 | 0.139562594 | 0.172411171 |
| Ucp3 | 0.090287817 | 1.891688189 | 0.057 | 0.017 | 1 | 0.262676008 | 2.83720485 | 0.024 | 0.007 | 1 | 0.262676008 | 0.172423744 |
| Grtp1 | 0.090287817 | 1.891688189 | 0.057 | 0.017 | 1 | 0.51919986 | 0.83720485 | 0 | 0.01 | 1 | 0.51919986 | 0.172423744 |
| Dlk1 | 0.090287817 | 1.891688189 | 0.057 | 0.017 | 1 | 0.345569014 | 0.652780279 | 0.073 | 0.041 | 1 | 0.345569014 | 0.172423744 |
| Dlk2 | 0.090287817 | 1.891688189 | 0.057 | 0.017 | 1 | 0.186073065 | 1.83720485 | 0.049 | 0.017 | 1 | 0.186073065 | 0.172423744 |
| Far1 | 0.090359919 | 0.266568562 | 0.849 | 0.788 | 1 | 0.961638546 | 0.111761126 | 0.854 | 0.898 | 1 | 0.961638546 | 0.172554922 |
| Nop16 | 0.090393205 | 0.413965421 | 0.491 | 0.377 | 1 | 0.142509507 | -0.173364392 | 0.22 | 0.358 | 1 | 0.142509507 | 0.172615478 |
| Steep1 | 0.090457174 | 0.860275236 | 0.472 | 0.381 | 1 | 0.527633151 | 0.42692088 | 0.415 | 0.386 | 1 | 0.527633151 | 0.172731847 |
| Dph6 | 0.460070612 | -0.105804955 | 0.283 | 0.347 | 1 | 0.090462857 | -0.913566544 | 0.195 | 0.317 | 1 | 0.460070612 | 0.172742186 |
| Pi4k2b | 0.544693927 | 0.534136184 | 0.358 | 0.335 | 1 | 0.090467814 | -0.360734528 | 0.39 | 0.556 | 1 | 0.544693927 | 0.172751203 |
| Ddx23 | 0.090499919 | 0.579037908 | 0.528 | 0.445 | 1 | 0.941164025 | 0.297096585 | 0.463 | 0.546 | 1 | 0.941164025 | 0.172809602 |
| Celf5 | 0.090605028 | -0.561484439 | 0.057 | 0.144 | 1 | 0.314655312 | -0.305753104 | 0.073 | 0.13 | 1 | 0.314655312 | 0.173000785 |
| A230056P14Rik | 0.090637265 | 0.8327945 | 0.094 | 0.038 | 1 | 0.817109602 | 0.721727632 | 0.049 | 0.041 | 1 | 0.817109602 | 0.173059416 |
| Ccdc120 | 0.090637265 | 1.154722595 | 0.094 | 0.038 | 1 | 0.094139125 | -1.747757651 | 0 | 0.065 | 1 | 0.094139125 | 0.173059416 |
| Il1r2 | 0.090684799 | 0.865215978 | 0.094 | 0.038 | 1 | 0.113256319 | 2.029849928 | 0.049 | 0.014 | 1 | 0.113256319 | 0.173145865 |
| Lamtor4 | 0.090726316 | 0.372723247 | 0.736 | 0.674 | 1 | 0.164468192 | -0.122655759 | 0.537 | 0.645 | 1 | 0.164468192 | 0.173221368 |
| Gtpbp3 | 0.090779589 | 1.261637799 | 0.113 | 0.051 | 1 | 0.490723575 | 0.930314254 | 0.073 | 0.048 | 1 | 0.490723575 | 0.173318245 |
| Sdha | 0.23975009 | 0.339307903 | 0.774 | 0.708 | 1 | 0.090780376 | -0.224434216 | 0.634 | 0.785 | 1 | 0.23975009 | 0.173319675 |
| Slco1c1 | 0.126071338 | -1.330704232 | 0.038 | 0.106 | 1 | 0.090893055 | -1.153750011 | 0.073 | 0.184 | 1 | 0.126071338 | 0.173524562 |
| 2210016L21Rik | 0.528240571 | 0.558871778 | 0.434 | 0.415 | 1 | 0.0908949 | -0.112754468 | 0.244 | 0.399 | 1 | 0.528240571 | 0.173527917 |
| Gm28501 | 0.0909501 | 1.8327945 | 0.057 | 0.017 | 1 | 0.188532591 | -1.484723245 | 0 | 0.041 | 1 | 0.188532591 | 0.173628279 |
| Cmtr2 | 0.0909501 | 1.8327945 | 0.057 | 0.017 | 1 | 0.872828964 | 1.029849928 | 0.024 | 0.02 | 1 | 0.872828964 | 0.173628279 |
| Gm56549 | 0.0909501 | 1.8327945 | 0.057 | 0.017 | 1 | 0.433272494 | 2.100239256 | 0.024 | 0.01 | 1 | 0.433272494 | 0.173628279 |
| Gm50214 | 0.0909501 | 1.8327945 | 0.057 | 0.017 | 1 | 0.909322542 | 0.252242349 | 0.024 | 0.027 | 1 | 0.909322542 | 0.173628279 |
| Flot2 | 0.209582854 | 0.669295768 | 0.34 | 0.267 | 1 | 0.090955882 | 0.676212973 | 0.439 | 0.324 | 1 | 0.209582854 | 0.173638791 |
| Armc8 | 0.090976757 | 0.548065023 | 0.453 | 0.36 | 1 | 0.863657837 | 0.127063044 | 0.561 | 0.587 | 1 | 0.863657837 | 0.173676744 |
| Ncoa5 | 0.318555199 | 0.110328475 | 0.491 | 0.411 | 1 | 0.090992373 | -0.574608749 | 0.415 | 0.553 | 1 | 0.318555199 | 0.173705134 |
| Rasd1 | 0.251262983 | 1.613013292 | 0.189 | 0.127 | 1 | 0.0909973 | 0.570418309 | 0.195 | 0.106 | 1 | 0.251262983 | 0.173714091 |
| Zfp512b | 0.091021184 | 0.869320376 | 0.226 | 0.136 | 1 | 0.901264795 | 0.441276173 | 0.244 | 0.246 | 1 | 0.901264795 | 0.173757512 |
| Mrpl36 | 0.091103982 | 0.555684909 | 0.472 | 0.347 | 1 | 0.747231573 | 0.425778604 | 0.317 | 0.311 | 1 | 0.747231573 | 0.173908028 |
| Ctdsp1 | 0.091120633 | 0.459577176 | 0.509 | 0.424 | 1 | 0.417256004 | -0.121488176 | 0.439 | 0.549 | 1 | 0.417256004 | 0.173938295 |
| Tspyl5 | 0.091152649 | 1.154722595 | 0.094 | 0.038 | 1 | 0.3351125 | -0.970150072 | 0.024 | 0.061 | 1 | 0.3351125 | 0.173996492 |
| Pxn | 0.318626951 | -0.28137652 | 0.358 | 0.445 | 1 | 0.091155379 | -0.447140062 | 0.488 | 0.58 | 1 | 0.318626951 | 0.174001456 |
| Tmem116 | 0.091369799 | 1.154722595 | 0.17 | 0.093 | 1 | 0.255410363 | 0.962735732 | 0.171 | 0.113 | 1 | 0.255410363 | 0.174391159 |
| Tceal5 | 0.091382437 | 1.123695699 | 0.245 | 0.157 | 1 | 0.843077144 | 0.515276755 | 0.171 | 0.171 | 1 | 0.843077144 | 0.174414123 |
| Glrx | 0.259624104 | 0.517292674 | 0.226 | 0.161 | 1 | 0.091395348 | -0.904262137 | 0.122 | 0.239 | 1 | 0.259624104 | 0.174437586 |
| Adora1 | 0.091498451 | -2.130679624 | 0.019 | 0.085 | 1 | 0.414113001 | 0.962735732 | 0.073 | 0.044 | 1 | 0.414113001 | 0.174624935 |
| Fundc2 | 0.476743417 | 0.210432027 | 0.736 | 0.712 | 1 | 0.09152457 | -0.246859415 | 0.512 | 0.703 | 1 | 0.476743417 | 0.174672394 |
| Nsg1 | 0.153772446 | 1.049540358 | 0.434 | 0.326 | 1 | 0.091550662 | 1.685201756 | 0.244 | 0.164 | 1 | 0.153772446 | 0.174719801 |
| Atp5k | 0.131561907 | 0.409295422 | 0.585 | 0.475 | 1 | 0.091668502 | -0.459412157 | 0.39 | 0.519 | 1 | 0.131561907 | 0.174933889 |
| Naa30 | 0.091713781 | 0.569760094 | 0.358 | 0.25 | 1 | 0.217643504 | -0.170289687 | 0.293 | 0.42 | 1 | 0.217643504 | 0.175016145 |
| Gabarapl1 | 0.091794004 | 0.353502459 | 0.736 | 0.636 | 1 | 0.239269009 | -0.160101041 | 0.659 | 0.751 | 1 | 0.239269009 | 0.175161868 |
| Tbpl1 | 0.091812206 | 0.84729407 | 0.396 | 0.314 | 1 | 0.399467383 | 0.416873051 | 0.439 | 0.389 | 1 | 0.399467383 | 0.17519493 |
| Alad | 0.777797691 | 0.311448098 | 0.189 | 0.212 | 1 | 0.091861087 | -0.16279515 | 0.049 | 0.154 | 1 | 0.777797691 | 0.175283716 |
| Znhit3 | 0.091956843 | 0.821298861 | 0.321 | 0.229 | 1 | 0.794776757 | 0.276489895 | 0.146 | 0.167 | 1 | 0.794776757 | 0.175457626 |
| Camta2 | 0.172395619 | 0.962077517 | 0.189 | 0.123 | 1 | 0.092215441 | -0.640842447 | 0.098 | 0.218 | 1 | 0.172395619 | 0.175927194 |

| | | | | | | | | | | | | |
|---|---|---|---|---|---|---|---|---|---|---|---|---|
| Ppp1r1c | 0.092257709 | 0.454282877 | 0.057 | 0.017 | 1 | 0.610750974 | -0.16279515 | 0.024 | 0.014 | 1 | 0.610750974 | 0.176003934 |
| Gm44577 | 0.092257709 | 1.247831999 | 0.057 | 0.017 | 1 | 0.287148249 | 1.377773231 | 0.073 | 0.038 | 1 | 0.287148249 | 0.176003934 |
| Spopl | 0.092531445 | 0.747547213 | 0.453 | 0.356 | 1 | 0.266504557 | -0.300298674 | 0.366 | 0.485 | 1 | 0.266504557 | 0.176500822 |
| Gfus | 0.092637765 | 0.874614676 | 0.283 | 0.191 | 1 | 0.73917707 | 0.678006255 | 0.195 | 0.181 | 1 | 0.73917707 | 0.176693774 |
| Sclt1 | 0.615398608 | -0.285849997 | 0.453 | 0.47 | 1 | 0.092796738 | -0.622226769 | 0.341 | 0.474 | 1 | 0.615398608 | 0.176982241 |
| B4galt3 | 0.651296562 | 0.211306123 | 0.208 | 0.178 | 1 | 0.092812141 | 0.818345823 | 0.317 | 0.212 | 1 | 0.651296562 | 0.177010188 |
| BC055324 | 0.092905716 | 1.569760094 | 0.057 | 0.017 | 1 | 0.441381862 | 1.515276755 | 0.024 | 0.01 | 1 | 0.441381862 | 0.177179959 |
| 1700022I11Rik | 0.092905716 | 1.569760094 | 0.057 | 0.017 | 1 | 0.824088343 | 0.377773231 | 0.024 | 0.031 | 1 | 0.824088343 | 0.177179959 |
| Iqcd | 0.092905716 | 1.569760094 | 0.057 | 0.017 | 1 | 0.732115056 | -0.250257991 | 0.073 | 0.058 | 1 | 0.732115056 | 0.177179959 |
| Abcc6 | 0.092905716 | 1.569760094 | 0.057 | 0.017 | 1 | 0.60213981 | 0.83720485 | 0.024 | 0.014 | 1 | 0.60213981 | 0.177179959 |
| Cerox1 | 0.092905716 | 1.347367673 | 0.057 | 0.017 | 1 | 0.319641967 | -0.16279515 | 0 | 0.024 | 1 | 0.319641967 | 0.177179959 |
| Psmg3 | 0.618519864 | 0.247831999 | 0.245 | 0.216 | 1 | 0.092906736 | -0.944154864 | 0.049 | 0.147 | 1 | 0.618519864 | 0.177181811 |
| Adgrg7 | 0.092919462 | 1.306725688 | 0.057 | 0.017 | 1 | 0.455098275 | 0.515276755 | 0 | 0.014 | 1 | 0.455098275 | 0.177204897 |
| Tacstd2 | 0.093011453 | 0.216123139 | 0.094 | 0.038 | 1 | 0.459414966 | -0.470223675 | 0.22 | 0.164 | 1 | 0.459414966 | 0.177371775 |
| Cdc7 | 0.093011747 | 1.039245377 | 0.094 | 0.038 | 1 | 0.221632028 | 1.557096931 | 0.098 | 0.051 | 1 | 0.221632028 | 0.177372308 |
| Lrp11 | 0.093011747 | 1.039245377 | 0.094 | 0.038 | 1 | 0.124547271 | 1.667279848 | 0.073 | 0.027 | 1 | 0.124547271 | 0.177372308 |
| Capn1 | 0.448671109 | 0.517292674 | 0.113 | 0.081 | 1 | 0.093112976 | 1.159132945 | 0.171 | 0.089 | 1 | 0.448671109 | 0.177555927 |
| Cyfip1 | 0.093195673 | 0.198992862 | 0.755 | 0.631 | 1 | 0.451501474 | 0.22821228 | 0.78 | 0.846 | 1 | 0.451501474 | 0.177705912 |
| Slc31a2 | 0.093316965 | 0.948271717 | 0.189 | 0.106 | 1 | 0.208458799 | -0.894599039 | 0.146 | 0.225 | 1 | 0.208458799 | 0.177925873 |
| Klc2 | 0.093419798 | 1.154722595 | 0.245 | 0.161 | 1 | 0.965867072 | 0.269520341 | 0.22 | 0.225 | 1 | 0.965867072 | 0.178112337 |
| Ifrd1 | 0.727964583 | -0.354591472 | 0.981 | 0.975 | 1 | 0.093669178 | 0.178000696 | 1 | 0.993 | 1 | 0.727964583 | 0.178564441 |
| Slc2a4 | 0.093776296 | -1.7032584 | 0.019 | 0.085 | 1 | 0.989595616 | 0.667279848 | 0.024 | 0.024 | 1 | 0.989595616 | 0.178758598 |
| Fgf2os | 0.093779522 | -2.20282941 | 0.019 | 0.085 | 1 | 0.357847088 | -0.16279515 | 0 | 0.02 | 1 | 0.357847088 | 0.178764445 |
| Cmtm6 | 0.093795827 | 0.569760094 | 0.34 | 0.229 | 1 | 0.235142477 | 0.646063362 | 0.415 | 0.345 | 1 | 0.235142477 | 0.178793996 |
| Sf3a2 | 0.093845458 | 0.676675298 | 0.377 | 0.271 | 1 | 0.190383879 | -0.410722664 | 0.22 | 0.328 | 1 | 0.190383879 | 0.178883945 |
| Smad6 | 0.341058042 | 0.527939918 | 0.321 | 0.258 | 1 | 0.093857074 | 0.883747436 | 0.488 | 0.372 | 1 | 0.341058042 | 0.178904998 |
| Pmpcb | 0.093896459 | 0.523367189 | 0.566 | 0.47 | 1 | 0.902076714 | 0.220946219 | 0.537 | 0.532 | 1 | 0.902076714 | 0.178976373 |
| Ccdc167 | 0.093943545 | 0.767699472 | 0.189 | 0.106 | 1 | 0.872876376 | 0.536035315 | 0.195 | 0.188 | 1 | 0.872876376 | 0.1790617 |
| Cntrl | 0.222662227 | 0.389187848 | 0.453 | 0.403 | 1 | 0.09394564 | 0.967512872 | 0.537 | 0.406 | 1 | 0.222662227 | 0.179065497 |
| Ubald2 | 0.093949525 | 0.664396968 | 0.547 | 0.458 | 1 | 0.345084837 | 0.312113805 | 0.61 | 0.529 | 1 | 0.345084837 | 0.179072536 |
| Fam91a1 | 0.333548355 | 0.324647596 | 0.585 | 0.534 | 1 | 0.093961016 | 0.575818297 | 0.707 | 0.648 | 1 | 0.333548355 | 0.17909336 |
| Atp6v1b2 | 0.149094288 | 0.231890455 | 0.547 | 0.436 | 1 | 0.093965572 | -0.944154864 | 0.634 | 0.689 | 1 | 0.149094288 | 0.179101615 |
| Entrep1 | 0.09409201 | 0.913714495 | 0.113 | 0.051 | 1 | 0.635455346 | 0.292884334 | 0.098 | 0.075 | 1 | 0.635455346 | 0.179330714 |
| Gm16845 | 0.290202443 | 0.792152515 | 0.094 | 0.055 | 1 | 0.094116915 | -1.917682652 | 0 | 0.065 | 1 | 0.290202443 | 0.179375837 |
| C330024C12Rik | 0.324474943 | 1.47665069 | 0.057 | 0.03 | 1 | 0.094116915 | -1.622226769 | 0 | 0.065 | 1 | 0.324474943 | 0.179375837 |
| Gm15956 | 0.732691937 | 1.154722595 | 0.019 | 0.013 | 1 | 0.094139125 | -1.747757651 | 0 | 0.065 | 1 | 0.732691937 | 0.179416075 |
| Prkaa1 | 0.094178886 | 0.739685096 | 0.396 | 0.305 | 1 | 0.329747911 | -0.275857815 | 0.317 | 0.42 | 1 | 0.329747911 | 0.17948811 |
| Itgb3 | 0.253631087 | -0.395474488 | 0.038 | 0.085 | 1 | 0.094203448 | -3.352619709 | 0 | 0.065 | 1 | 0.253631087 | 0.179532606 |
| Hinfp | 0.552867672 | -0.237594828 | 0.151 | 0.191 | 1 | 0.094318123 | -1.013651711 | 0.122 | 0.235 | 1 | 0.552867672 | 0.179740337 |
| 3930402G23Rik | 0.094522549 | -1.652632327 | 0.019 | 0.085 | 1 | 0.672172778 | 0.136765132 | 0.024 | 0.038 | 1 | 0.672172778 | 0.180110586 |
| Cd151 | 0.758838915 | 0.253855787 | 0.528 | 0.597 | 1 | 0.094611278 | -0.640842447 | 0.366 | 0.478 | 1 | 0.758838915 | 0.180271263 |
| Egfl7 | 0.094691859 | -1.652632327 | 0 | 0.051 | 1 | 0.402373368 | 0.252242349 | 0 | 0.017 | 1 | 0.402373368 | 0.180417169 |
| Gm57266 | 0.094691859 | -1.652632327 | 0 | 0.051 | 1 | 0.402373368 | 0.252242349 | 0 | 0.017 | 1 | 0.402373368 | 0.180417169 |
| Gdap1 | 0.09471075 | -1.932740246 | 0 | 0.051 | 1 | 0.455098275 | 0.515276755 | 0 | 0.014 | 1 | 0.455098275 | 0.180451374 |
| Nemp2 | 0.09471075 | -1.752168001 | 0 | 0.051 | 1 | 0.962773283 | 0.83720485 | 0.049 | 0.048 | 1 | 0.962773283 | 0.180451374 |
| Bard1 | 0.09471075 | -2.304709024 | 0 | 0.051 | 1 | 0.599290422 | 1.515276755 | 0.024 | 0.014 | 1 | 0.599290422 | 0.180451374 |
| Spaar | 0.09471075 | -1.932740246 | 0 | 0.051 | 1 | 0.455098275 | 0.515276755 | 0 | 0.014 | 1 | 0.455098275 | 0.180451374 |
| Nupr1l | 0.09471075 | -1.752168001 | 0 | 0.051 | 1 | 0.170390036 | -1.332720152 | 0 | 0.044 | 1 | 0.170390036 | 0.180451374 |
| Gm19605 | 0.09471075 | -1.845277405 | 0 | 0.051 | 1 | 0.453885966 | -0.16279515 | 0.024 | 0.051 | 1 | 0.453885966 | 0.180451374 |
| Pfkfb1 | 0.09471075 | -1.845277405 | 0 | 0.051 | 1 | 0.257308946 | -0.863234868 | 0 | 0.031 | 1 | 0.257308946 | 0.180451374 |
| Bco2 | 0.094725863 | -1.932740246 | 0 | 0.051 | 1 | 0.257285084 | -0.622226769 | 0 | 0.031 | 1 | 0.257285084 | 0.180478737 |
| Gm11290 | 0.094725863 | -2.237594828 | 0 | 0.051 | 1 | 0.357858789 | -1.484723245 | 0 | 0.02 | 1 | 0.357858789 | 0.180478737 |
| Il18r1 | 0.094726206 | -2.237594828 | 0 | 0.051 | 1 | 0.402364804 | -0.16279515 | 0 | 0.017 | 1 | 0.402364804 | 0.180479359 |
| Mcm8 | 0.094726206 | -1.845277405 | 0 | 0.051 | 1 | 0.599290422 | 1.515276755 | 0.024 | 0.014 | 1 | 0.599290422 | 0.180479359 |
| Slc18a2 | 0.094737197 | -2.7032584 | 0 | 0.051 | 1 | 0.257308459 | -0.970150072 | 0 | 0.031 | 1 | 0.257308459 | 0.180499258 |
| Slc45a3 | 0.094738228 | -2.368839361 | 0 | 0.051 | 1 | 0.957192022 | 0.83720485 | 0.049 | 0.048 | 1 | 0.957192022 | 0.180501124 |
| Gm44127 | 0.094738228 | -2.489133595 | 0 | 0.051 | 1 | 0.738375177 | 1.100239256 | 0.024 | 0.017 | 1 | 0.738375177 | 0.180501124 |
| Sntg2 | 0.094748532 | -2.1672055 | 0 | 0.051 | 1 | 0.506557138 | 2.284663827 | 0.073 | 0.051 | 1 | 0.506557138 | 0.180519779 |
| Map10 | 0.094767422 | -1.545717123 | 0 | 0.051 | 1 | 0.745976859 | 0.377773231 | 0.024 | 0.034 | 1 | 0.745976859 | 0.18055398 |
| Prdm16os | 0.804136783 | 0.313420341 | 0.17 | 0.157 | 1 | 0.094781479 | -0.429581691 | 0.098 | 0.218 | 1 | 0.804136783 | 0.18057943 |
| Btg3 | 0.094801342 | 0.286826131 | 0.887 | 0.767 | 1 | 0.629638414 | 0.166512475 | 0.878 | 0.785 | 1 | 0.629638414 | 0.180615389 |
| Mrpl18 | 0.094871031 | 0.613349363 | 0.509 | 0.436 | 1 | 0.154000751 | -0.295245446 | 0.317 | 0.447 | 1 | 0.154000751 | 0.180741549 |
| Gm56938 | 0.094891448 | -1.545717123 | 0.019 | 0.085 | 1 | 0.188517184 | -1.069685746 | 0 | 0.041 | 1 | 0.188517184 | 0.180778509 |
| 4732491K20Rik | 0.094891651 | -1.430239906 | 0.019 | 0.085 | 1 | 0.494850012 | 0.83720485 | 0.073 | 0.048 | 1 | 0.494850012 | 0.180778876 |
| Mef2d | 0.094921562 | 0.285405965 | 0.811 | 0.754 | 1 | 0.556286219 | -0.170289687 | 0.927 | 0.939 | 1 | 0.556286219 | 0.180833021 |
| Gm15853 | 0.094924143 | 1.347367673 | 0.057 | 0.017 | 1 | 0.60213981 | 1.252242349 | 0.024 | 0.014 | 1 | 0.60213981 | 0.180837694 |
| Lrrc43 | 0.094924143 | 0.984797593 | 0.057 | 0.017 | 1 | 0.188380068 | 1.100239256 | 0.049 | 0.017 | 1 | 0.188380068 | 0.180837694 |
| Fam174b | 0.094924143 | 1.347367673 | 0.057 | 0.017 | 1 | 0.263489513 | 1.83720485 | 0.049 | 0.02 | 1 | 0.263489513 | 0.180837694 |
| Gm39363 | 0.094924143 | 1.347367673 | 0.057 | 0.017 | 1 | 0.720213045 | 0.989207943 | 0.049 | 0.038 | 1 | 0.720213045 | 0.180837694 |
| Pced1b | 0.094924143 | 1.347367673 | 0.057 | 0.017 | 1 | 0.355779174 | 1.458693227 | 0.049 | 0.024 | 1 | 0.355779174 | 0.180837694 |
| Ngdn | 0.095011631 | 0.514898159 | 0.528 | 0.441 | 1 | 0.731938168 | 0.238567412 | 0.463 | 0.444 | 1 | 0.731938168 | 0.180996051 |
| Pmm1 | 0.695303526 | 0.113695327 | 0.358 | 0.331 | 1 | 0.095047334 | -0.911733386 | 0.171 | 0.29 | 1 | 0.695303526 | 0.181060672 |
| Tspyl4 | 0.095053615 | -0.970808287 | 0.189 | 0.284 | 1 | 0.732937682 | 0.79281073 | 0.195 | 0.174 | 1 | 0.732937682 | 0.18107204 |
| Oxa1l | 0.095111142 | 0.932330173 | 0.283 | 0.195 | 1 | 0.370106537 | 0.774920572 | 0.366 | 0.331 | 1 | 0.370106537 | 0.181176154 |
| Ube2j1 | 0.095180925 | 0.607603445 | 0.698 | 0.691 | 1 | 0.39672999 | -0.205340845 | 0.829 | 0.785 | 1 | 0.39672999 | 0.181302441 |
| Agpat5 | 0.095265883 | -0.214511215 | 0.283 | 0.436 | 1 | 0.989409962 | 0.145327145 | 0.341 | 0.352 | 1 | 0.989409962 | 0.181456178 |
| Paqr3 | 0.095299816 | -0.693274312 | 0.057 | 0.144 | 1 | 0.934777646 | 0.721727632 | 0.073 | 0.078 | 1 | 0.934777646 | 0.181517577 |
| 5930430L01Rik | 0.113687064 | 1.045098104 | 0.226 | 0.144 | 1 | 0.095326796 | -1.589059905 | 0.073 | 0.171 | 1 | 0.113687064 | 0.181566393 |
| Med25 | 0.095372736 | 0.8327945 | 0.528 | 0.466 | 1 | 0.852377436 | 0.230931843 | 0.585 | 0.58 | 1 | 0.852377436 | 0.181649514 |
| Elmo2 | 0.095559993 | 0.932330173 | 0.208 | 0.123 | 1 | 0.704691347 | 0.287007767 | 0.171 | 0.205 | 1 | 0.704691347 | 0.181988273 |
| Odf3l1 | 0.264906374 | -1.304709024 | 0.019 | 0.055 | 1 | 0.09559395 | -1.747757651 | 0.024 | 0.106 | 1 | 0.264906374 | 0.182049696 |
| Blzf1 | 0.441508122 | 0.126377969 | 0.604 | 0.551 | 1 | 0.095638237 | -0.135683305 | 0.39 | 0.604 | 1 | 0.441508122 | 0.182129802 |
| Rwdd2b | 0.095703663 | 1.055186921 | 0.113 | 0.051 | 1 | 0.94575798 | 0.334704509 | 0.049 | 0.051 | 1 | 0.94575798 | 0.182248134 |
| Nfe2l1 | 0.11670086 | 0.17136286 | 0.962 | 0.97 | 1 | 0.095761882 | 0.268560925 | 0.902 | 0.945 | 1 | 0.11670086 | 0.182353425 |
| Pan2 | 0.095762696 | 0.772252958 | 0.34 | 0.237 | 1 | 0.439443985 | -0.440779897 | 0.171 | 0.222 | 1 | 0.439443985 | 0.182354898 |
| Cdc14b | 0.095830895 | 0.8327945 | 0.189 | 0.11 | 1 | 0.690630713 | 0.478123756 | 0.171 | 0.205 | 1 | 0.690630713 | 0.18247823 |
| Gm16740 | 0.095903694 | 0.974150349 | 0.189 | 0.106 | 1 | 0.213700019 | 0.580865097 | 0.171 | 0.102 | 1 | 0.213700019 | 0.18260987 |
| Crabp2 | 0.689858465 | -1.015202407 | 0.057 | 0.072 | 1 | 0.095912455 | 2.705960317 | 0.122 | 0.058 | 1 | 0.689858465 | 0.182625711 |
| Gm8797 | 0.095925654 | 0.678989164 | 0.415 | 0.309 | 1 | 0.746419207 | 0.170448258 | 0.341 | 0.314 | 1 | 0.746419207 | 0.182649576 |
| Gfm2 | 0.846781124 | 0.377115016 | 0.226 | 0.216 | 1 | 0.095998001 | -0.548853583 | 0.122 | 0.242 | 1 | 0.846781124 | 0.182780386 |
| Hyls1 | 0.096085746 | 0.417757001 | 0.132 | 0.064 | 1 | 0.603831993 | 1.557096931 | 0.098 | 0.078 | 1 | 0.603831993 | 0.182939021 |
| Mnd1 | 0.09620921 | 0.984797593 | 0.113 | 0.051 | 1 | 0.185791946 | -1.654648247 | 0.024 | 0.082 | 1 | 0.185791946 | 0.183162208 |
| Gmds | 0.096258353 | -0.250289186 | 0.887 | 0.949 | 1 | 0.566362819 | -0.113427456 | 0.976 | 0.952 | 1 | 0.566362819 | 0.183251035 |
| Gstcd | 0.096273375 | 0.587682002 | 0.34 | 0.233 | 1 | 0.190896454 | -0.283810551 | 0.122 | 0.215 | 1 | 0.190896454 | 0.183278187 |
| Nubp1 | 0.096300583 | 0.531792244 | 0.358 | 0.246 | 1 | 0.851186073 | 0.642188667 | 0.244 | 0.246 | 1 | 0.851186073 | 0.183327364 |
| Fntb | 0.992159687 | 0.221836791 | 0.226 | 0.237 | 1 | 0.096315332 | -0.473582687 | 0.146 | 0.276 | 1 | 0.992159687 | 0.18335402 |
| Mff | 0.096360006 | 0.283709036 | 0.755 | 0.657 | 1 | 0.16433562 | -0.335741504 | 0.634 | 0.768 | 1 | 0.16433562 | 0.183434762 |
| Spint2 | 0.252436366 | -0.923279917 | 0.264 | 0.322 | 1 | 0.096420407 | 0.642188667 | 0.439 | 0.324 | 1 | 0.252436366 | 0.183543919 |

| | | | | | | | | | | | | |
|---|---|---|---|---|---|---|---|---|---|---|---|---|
| Cenpt | 0.096444158 | -1.015202407 | 0.057 | 0.14 | 1 | 0.200767293 | -0.795063366 | 0.098 | 0.174 | 1 | 0.200767293 | 0.183586841 |
| Chtf8 | 0.1848549 | 0.859266711 | 0.17 | 0.106 | 1 | 0.096508733 | -1.372248516 | 0.024 | 0.106 | 1 | 0.1848549 | 0.183703531 |
| Zfp595 | 0.096645754 | 0.569760094 | 0.113 | 0.051 | 1 | 0.741047133 | 0.88967227 | 0.073 | 0.061 | 1 | 0.741047133 | 0.183951106 |
| Atrn | 0.096668205 | 0.345104621 | 0.698 | 0.597 | 1 | 0.11714462 | -0.285943381 | 0.78 | 0.863 | 1 | 0.11714462 | 0.183991668 |
| Hnrnpr | 0.096727901 | 0.306275479 | 0.887 | 0.89 | 1 | 0.100003821 | 0.290138957 | 0.976 | 0.922 | 1 | 0.100003821 | 0.184099514 |
| Cfap43 | 0.096762976 | -0.464187238 | 0.132 | 0.242 | 1 | 0.338564347 | 0.906467512 | 0.171 | 0.119 | 1 | 0.338564347 | 0.184162878 |
| Arhgap44 | 0.87476158 | -0.120911848 | 0.208 | 0.199 | 1 | 0.096767608 | -0.738479837 | 0.146 | 0.266 | 1 | 0.87476158 | 0.184171247 |
| Elf3 | 0.241893059 | 1.017219071 | 0.075 | 0.038 | 1 | 0.096789097 | 1.043655727 | 0.268 | 0.164 | 1 | 0.241893059 | 0.184210064 |
| Rbbp5 | 0.096843066 | 0.795641501 | 0.283 | 0.191 | 1 | 0.673291719 | 0.122959332 | 0.22 | 0.259 | 1 | 0.673291719 | 0.184307552 |
| Mybl1 | 0.250442599 | 0.524672205 | 0.189 | 0.131 | 1 | 0.096852459 | 2.15131344 | 0.171 | 0.092 | 1 | 0.250442599 | 0.184324519 |
| Dapp1 | 0.97837388 | 0.289652175 | 0.226 | 0.233 | 1 | 0.096965581 | 0.633671456 | 0.39 | 0.28 | 1 | 0.97837388 | 0.184528839 |
| Bcap31 | 0.096989117 | -0.168546918 | 0.962 | 0.936 | 1 | 0.162523605 | -0.116054138 | 0.854 | 0.949 | 1 | 0.162523605 | 0.184571345 |
| Gm28791 | 0.152080548 | 1.624207878 | 0.094 | 0.047 | 1 | 0.097054438 | 1.252242349 | 0.146 | 0.072 | 1 | 0.152080548 | 0.184689312 |
| Smyd3 | 0.392461409 | -0.343682086 | 0.528 | 0.614 | 1 | 0.097085934 | -0.307281545 | 0.585 | 0.703 | 1 | 0.392461409 | 0.18474619 |
| Pi15 | 0.09725594 | 3.739685096 | 0.038 | 0.008 | 1 | 0.441381862 | 1.029849928 | 0.024 | 0.01 | 1 | 0.441381862 | 0.185053162 |
| Dnai1 | 0.09725594 | 3.154722595 | 0.038 | 0.008 | 1 | 0.402373368 | 0.252242349 | 0 | 0.017 | 1 | 0.402373368 | 0.185053162 |
| Asic2 | 0.09725594 | 2.891688189 | 0.038 | 0.008 | 1 | 0.402369942 | -2.250257991 | 0 | 0.017 | 1 | 0.402369942 | 0.185053162 |
| Gm49003 | 0.09725594 | 3.739685096 | 0.038 | 0.008 | 1 | 0.752578711 | 0.136765132 | 0.024 | 0.017 | 1 | 0.752578711 | 0.185053162 |
| Tigit | 0.09725594 | 2.891688189 | 0.038 | 0.008 | 1 | 0.846938525 | 1.252242349 | 0.024 | 0.031 | 1 | 0.846938525 | 0.185053162 |
| 4631405J19Rik | 0.097256955 | 3.855162313 | 0.038 | 0.008 | 1 | 0.438672228 | 1.83720485 | 0.024 | 0.01 | 1 | 0.438672228 | 0.185054995 |
| Spata6 | 0.09726799 | -0.324967863 | 0.377 | 0.525 | 1 | 0.204689144 | -0.235551493 | 0.512 | 0.648 | 1 | 0.204689144 | 0.185074919 |
| Rfc5 | 0.097293064 | 1.211306123 | 0.151 | 0.081 | 1 | 0.6623142 | 0.377773231 | 0.122 | 0.099 | 1 | 0.6623142 | 0.185120188 |
| Fam174a | 0.615623713 | 0.268104394 | 0.623 | 0.737 | 1 | 0.097464122 | -0.270178686 | 0.537 | 0.706 | 1 | 0.615623713 | 0.185428989 |
| 4930430F08Rik | 0.097792432 | 1.417757001 | 0.151 | 0.081 | 1 | 0.96153013 | 0.404245443 | 0.073 | 0.072 | 1 | 0.96153013 | 0.186021504 |
| Synrg | 0.097795422 | 0.473544779 | 0.396 | 0.271 | 1 | 0.757361715 | 0.186440291 | 0.366 | 0.334 | 1 | 0.757361715 | 0.186026899 |
| Klrg2 | 0.998421405 | 0.104096522 | 0.075 | 0.076 | 1 | 0.097883254 | -1.64392184 | 0.049 | 0.14 | 1 | 0.998421405 | 0.186185377 |
| Ccdc90b | 0.32967446 | 0.160458964 | 0.623 | 0.564 | 1 | 0.09789545 | -0.348150192 | 0.366 | 0.522 | 1 | 0.32967446 | 0.18620738 |
| Ppp1r8 | 0.098085353 | 0.96568877 | 0.283 | 0.195 | 1 | 0.512311178 | -0.128579435 | 0.268 | 0.324 | 1 | 0.512311178 | 0.18654997 |
| Dennd3 | 0.846062568 | -0.490612524 | 0.434 | 0.411 | 1 | 0.098242096 | -0.933313304 | 0.268 | 0.386 | 1 | 0.846062568 | 0.186832683 |
| Rwdd4a | 0.098429658 | 0.686943633 | 0.472 | 0.407 | 1 | 0.996874891 | 0.252242349 | 0.39 | 0.416 | 1 | 0.996874891 | 0.187170918 |
| 2010001A14Rik | 0.431872285 | -0.140733289 | 0.151 | 0.203 | 1 | 0.098440597 | -1.305753104 | 0.049 | 0.14 | 1 | 0.431872285 | 0.187190644 |
| Polr1g | 0.098702309 | 0.603312654 | 0.34 | 0.237 | 1 | 0.47123963 | 0.161094461 | 0.22 | 0.297 | 1 | 0.47123963 | 0.187662472 |
| Tceanc | 0.098859881 | 0.906795081 | 0.113 | 0.051 | 1 | 0.203924283 | -0.484723245 | 0.024 | 0.082 | 1 | 0.203924283 | 0.187946487 |
| Acbd7 | 0.099079675 | 2.569760094 | 0.038 | 0.008 | 1 | 0.267205177 | 2.252242349 | 0.024 | 0.007 | 1 | 0.267205177 | 0.188342567 |
| Pm20d2 | 0.099079675 | 2.569760094 | 0.038 | 0.008 | 1 | 0.51919986 | 0.83720485 | 0 | 0.01 | 1 | 0.51919986 | 0.188342567 |
| Ghrl | 0.099079675 | 2.569760094 | 0.038 | 0.008 | 1 | 0.231643049 | -0.622226769 | 0 | 0.034 | 1 | 0.231643049 | 0.188342567 |
| Styk1 | 0.099079675 | 2.891688189 | 0.038 | 0.008 | 1 | 0.103479965 | 4.159132945 | 0.024 | 0.003 | 1 | 0.103479965 | 0.188342567 |
| Gm29571 | 0.099079675 | 2.569760094 | 0.038 | 0.008 | 1 | 0.5191986 | 0.515276755 | 0 | 0.01 | 1 | 0.5191986 | 0.188342567 |
| 5930403N24Rik | 0.099079675 | 2.569760094 | 0.038 | 0.008 | 1 | 0.744047924 | 1.252242349 | 0.024 | 0.017 | 1 | 0.744047924 | 0.188342567 |
| Pctp | 0.099079675 | 2.569760094 | 0.038 | 0.008 | 1 | 0.188380068 | 1.252242349 | 0.049 | 0.017 | 1 | 0.188380068 | 0.188342567 |
| B4galnt2 | 0.099079675 | 2.569760094 | 0.038 | 0.008 | 1 | 0.267205177 | 2.252242349 | 0.024 | 0.007 | 1 | 0.267205177 | 0.188342567 |
| Hoxb9 | 0.099079675 | 2.569760094 | 0.038 | 0.008 | 1 | 0.269112057 | 1.515276755 | 0.049 | 0.02 | 1 | 0.269112057 | 0.188342567 |
| Adam11 | 0.099079675 | 2.569760094 | 0.038 | 0.008 | 1 | 0.257285084 | -0.622226769 | 0 | 0.031 | 1 | 0.257285084 | 0.188342567 |
| Gm57347 | 0.099079675 | 3.154722595 | 0.038 | 0.008 | 1 | 0.104856732 | 2.83720485 | 0.024 | 0.003 | 1 | 0.104856732 | 0.188342567 |
| A330102I10Rik | 0.099079675 | 2.569760094 | 0.038 | 0.008 | 1 | 0.87556513 | 0.83720485 | 0.024 | 0.02 | 1 | 0.87556513 | 0.188342567 |
| Pgap6 | 0.099079675 | 2.569760094 | 0.038 | 0.008 | 1 | 0.746881137 | 1.029849928 | 0.024 | 0.017 | 1 | 0.746881137 | 0.188342567 |
| Tex43 | 0.099079675 | 2.569760094 | 0.038 | 0.008 | 1 | 0.357854401 | -0.332720152 | 0 | 0.02 | 1 | 0.357854401 | 0.188342552 |
| Was | 0.099079675 | 2.569760094 | 0.038 | 0.008 | 1 | 0.51919986 | 0.83720485 | 0 | 0.01 | 1 | 0.51919986 | 0.188342567 |
| Gtf2a2 | 0.099148527 | 0.463826649 | 0.698 | 0.708 | 1 | 0.183589522 | -0.104527495 | 0.634 | 0.758 | 1 | 0.183589522 | 0.188466623 |
| Ralgapa2 | 0.09919394 | 0.406261362 | 0.434 | 0.326 | 1 | 0.507517081 | 0.350327349 | 0.512 | 0.498 | 1 | 0.507517081 | 0.188548443 |
| Rab11fip3 | 0.099411198 | -0.805531226 | 0.491 | 0.572 | 1 | 0.163452373 | -0.255053658 | 0.463 | 0.604 | 1 | 0.163452373 | 0.188939809 |
| Dsg2 | 0.287773478 | -0.430239906 | 0 | 0.021 | 1 | 0.099594597 | 0.83720485 | 0.268 | 0.167 | 1 | 0.287773478 | 0.18927011 |
| Dtx2 | 0.09966771 | 1.10741688 | 0.208 | 0.127 | 1 | 0.967566863 | 0.252242349 | 0.195 | 0.205 | 1 | 0.967566863 | 0.189401767 |
| Eif3g | 0.574072228 | -0.1182959 | 0.623 | 0.648 | 1 | 0.099819646 | -0.387582087 | 0.659 | 0.734 | 1 | 0.574072228 | 0.18967533 |
| Tef | 0.099873823 | 0.591455165 | 0.585 | 0.492 | 1 | 0.457958897 | 0.277972624 | 0.659 | 0.631 | 1 | 0.457958897 | 0.189772866 |
| Fance | 0.099901991 | 0.714150003 | 0.302 | 0.203 | 1 | 0.30027283 | 0.642188867 | 0.293 | 0.232 | 1 | 0.30027283 | 0.189823574 |
| Rras2 | 0.099959101 | 0.719966861 | 0.774 | 0.788 | 1 | 0.464325167 | 0.109970794 | 0.854 | 0.863 | 1 | 0.464325167 | 0.189926379 |
| Gm47561 | 0.100006369 | 2.47665069 | 0.038 | 0.008 | 1 | 0.752578347 | 0.515276755 | 0.024 | 0.017 | 1 | 0.752578347 | 0.190011465 |
| Tmem275 | 0.100007399 | 2.417757001 | 0.038 | 0.008 | 1 | 0.44410834 | 1.252242349 | 0.024 | 0.01 | 1 | 0.44410834 | 0.190013319 |
| 1600010M07Rik | 0.100121715 | 0.466666601 | 0.358 | 0.246 | 1 | 0.898726184 | 0.626308067 | 0.268 | 0.29 | 1 | 0.898726184 | 0.190219072 |
| Zcchc3 | 0.100237272 | 0.776210972 | 0.226 | 0.136 | 1 | 0.961292008 | 0.35607816 | 0.171 | 0.174 | 1 | 0.961292008 | 0.190427032 |
| Sirt3 | 0.100261215 | 1.33529484 | 0.189 | 0.11 | 1 | 0.673569058 | -0.16279515 | 0.122 | 0.147 | 1 | 0.673569058 | 0.190470119 |
| Desi1 | 0.100287296 | -0.441128222 | 0.453 | 0.597 | 1 | 0.150230487 | -0.540465797 | 0.512 | 0.604 | 1 | 0.150230487 | 0.190517051 |
| Sdr39u1 | 0.460365572 | 0.434830514 | 0.245 | 0.203 | 1 | 0.100331481 | -1.19841906 | 0.098 | 0.201 | 1 | 0.460365572 | 0.190596555 |
| Ftx | 0.152902355 | -0.759547531 | 0.189 | 0.275 | 1 | 0.100395052 | -0.597423378 | 0.171 | 0.3 | 1 | 0.152902355 | 0.190710938 |
| H4c9 | 0.100416345 | -1.108311811 | 0.057 | 0.14 | 1 | 0.894652652 | 0.444887427 | 0.049 | 0.055 | 1 | 0.894652652 | 0.190749248 |
| Zfp933 | 0.100427108 | 1.347367673 | 0.151 | 0.081 | 1 | 0.155119504 | 0.906467512 | 0.244 | 0.157 | 1 | 0.155119504 | 0.190768612 |
| Ttbk2 | 0.100471725 | 0.655916738 | 0.34 | 0.237 | 1 | 0.494196325 | -0.26713181 | 0.244 | 0.3 | 1 | 0.494196325 | 0.190848882 |
| Spata6l | 0.100516628 | -0.464187238 | 0.038 | 0.114 | 1 | 0.608968477 | 0.515276755 | 0.049 | 0.072 | 1 | 0.608968477 | 0.190929664 |
| Fam131a | 0.180522518 | 0.645708947 | 0.302 | 0.229 | 1 | 0.10055769 | -0.665295491 | 0.122 | 0.249 | 1 | 0.180522518 | 0.19100353 |
| Hspb8 | 0.100579489 | -1.389597921 | 0.094 | 0.182 | 1 | 0.124839939 | 1.145327145 | 0.317 | 0.215 | 1 | 0.124839939 | 0.191042744 |
| Mob1a | 0.100601219 | 0.632156631 | 0.547 | 0.47 | 1 | 0.892177075 | 0.147905689 | 0.707 | 0.706 | 1 | 0.892177075 | 0.191081833 |
| Diablo | 0.100734354 | -0.418744267 | 0.283 | 0.424 | 1 | 0.786137539 | 0.10301634 | 0.39 | 0.437 | 1 | 0.786137539 | 0.191321299 |
| Hars | 0.441532822 | 0.441285536 | 0.566 | 0.568 | 1 | 0.10076409 | -0.104061325 | 0.415 | 0.618 | 1 | 0.441532822 | 0.191374778 |
| Cdan1 | 0.100801058 | 1.221836791 | 0.151 | 0.081 | 1 | 0.943127655 | 0.282615998 | 0.122 | 0.13 | 1 | 0.943127655 | 0.191441263 |
| Selenoh | 0.10082977 | 0.569760094 | 0.415 | 0.318 | 1 | 0.388636256 | -0.499078538 | 0.195 | 0.249 | 1 | 0.388636256 | 0.191492897 |
| Fem1b | 0.710976309 | 0.193196743 | 0.434 | 0.403 | 1 | 0.100893859 | -0.52456351 | 0.366 | 0.488 | 1 | 0.710976309 | 0.191608147 |
| Ncf2 | 0.100936842 | 2.154722595 | 0.038 | 0.008 | 1 | 0.180559238 | 0.19334866 | 0.073 | 0.031 | 1 | 0.180559238 | 0.191685438 |
| Cxxc4 | 0.100936842 | 2.154722595 | 0.038 | 0.008 | 1 | 0.587925597 | 2.252242349 | 0.024 | 0.014 | 1 | 0.587925597 | 0.191685438 |
| Dpysl5 | 0.100936842 | 2.154722595 | 0.038 | 0.008 | 1 | 0.115836913 | 1.614812428 | 0.049 | 0.014 | 1 | 0.115836913 | 0.191685438 |
| Oas1b | 0.100936842 | 2.154722595 | 0.038 | 0.008 | 1 | 0.114118767 | 2.100239256 | 0.049 | 0.014 | 1 | 0.114118767 | 0.191685438 |
| Art4 | 0.100936842 | 2.154722595 | 0.038 | 0.008 | 1 | 0.267205177 | 2.252242349 | 0.024 | 0.007 | 1 | 0.267205177 | 0.191685438 |
| Gm16183 | 0.100936842 | 2.154722595 | 0.038 | 0.008 | 1 | 0.455094184 | 0.252242349 | 0 | 0.014 | 1 | 0.455094184 | 0.191685438 |
| Slc35f3 | 0.100936842 | 2.154722595 | 0.038 | 0.008 | 1 | 0.106248001 | 1.83720485 | 0.024 | 0.003 | 1 | 0.106248001 | 0.191685438 |
| Gask1a | 0.100936842 | 2.154722595 | 0.038 | 0.008 | 1 | 0.190737323 | 1.100239256 | 0.049 | 0.017 | 1 | 0.190737323 | 0.191685438 |
| Gm27010 | 0.100936842 | 2.154722595 | 0.038 | 0.008 | 1 | 0.188498353 | -1.069685746 | 0 | 0.041 | 1 | 0.188498353 | 0.191685438 |
| Gm6994 | 0.100936842 | 2.154722595 | 0.038 | 0.008 | 1 | 0.793962817 | 1.136765132 | 0.049 | 0.041 | 1 | 0.793962817 | 0.191685438 |
| Gm37362 | 0.100937876 | 2.154722595 | 0.038 | 0.008 | 1 | 0.872828964 | 1.029849928 | 0.024 | 0.02 | 1 | 0.872828964 | 0.191687298 |
| Gm57465 | 0.100937876 | 2.154722595 | 0.038 | 0.008 | 1 | 0.60213981 | 1.252242349 | 0.024 | 0.014 | 1 | 0.60213981 | 0.191687298 |
| Gm14024 | 0.100937876 | 2.154722595 | 0.038 | 0.008 | 1 | 0.104856732 | 2.83720485 | 0.024 | 0.003 | 1 | 0.104856732 | 0.191687298 |
| Fer1l4 | 0.100937876 | 2.154722595 | 0.038 | 0.008 | 1 | 0.607876076 | 0.83720485 | 0.024 | 0.014 | 1 | 0.607876076 | 0.191687298 |
| Gm14455 | 0.100937876 | 2.154722595 | 0.038 | 0.008 | 1 | 0.257285084 | -0.622226769 | 0 | 0.031 | 1 | 0.257285084 | 0.191687298 |
| Rab42 | 0.100937876 | 2.154722595 | 0.038 | 0.008 | 1 | 0.455098275 | 0.515276755 | 0 | 0.014 | 1 | 0.455098275 | 0.191687298 |
| Atad3aos | 0.100937876 | 2.154722595 | 0.038 | 0.008 | 1 | 0.51919986 | 0.83720485 | 0 | 0.01 | 1 | 0.51919986 | 0.191687298 |
| Gm42744 | 0.100937876 | 2.154722595 | 0.038 | 0.008 | 1 | 0.357847088 | -0.332720152 | 0 | 0.02 | 1 | 0.357847088 | 0.191687298 |
| Bend4 | 0.100937876 | 2.154722595 | 0.038 | 0.008 | 1 | 0.462553972 | 1.174239837 | 0.073 | 0.048 | 1 | 0.462553972 | 0.191687298 |
| Pirb | 0.100937876 | 2.154722595 | 0.038 | 0.008 | 1 | 0.989595616 | 0.515276755 | 0.024 | 0.024 | 1 | 0.989595616 | 0.191687298 |
| Gm38973 | 0.100937876 | 2.154722595 | 0.038 | 0.008 | 1 | 0.908035876 | 0.252242349 | 0.049 | 0.044 | 1 | 0.908035876 | 0.191687298 |

| | | | | | | | | | | | | |
|---|---|---|---|---|---|---|---|---|---|---|---|---|
| Gm57214 | 0.100937876 | 2.154722595 | 0.038 | 0.008 | 1 | 0.5191986 | 0.515276755 | 0 | 0.01 | 1 | 0.5191986 | 0.191687298 |
| Fbxo27 | 0.100937876 | 2.154722595 | 0.038 | 0.008 | 1 | 0.402373368 | 0.252242349 | 0 | 0.017 | 1 | 0.402373368 | 0.191687298 |
| Lrrc4b | 0.100937876 | 2.154722595 | 0.038 | 0.008 | 1 | 0.441777736 | 1.699701326 | 0.049 | 0.027 | 1 | 0.441777736 | 0.191687298 |
| Xrra1 | 0.100937876 | 2.154722595 | 0.038 | 0.008 | 1 | 0.455097252 | -0.332720152 | 0 | 0.014 | 1 | 0.455097252 | 0.191687298 |
| Gm53374 | 0.100937876 | 2.154722595 | 0.038 | 0.008 | 1 | 0.599290422 | 1.515276755 | 0.024 | 0.014 | 1 | 0.599290422 | 0.191687298 |
| Csmd1 | 0.100937876 | 2.154722595 | 0.038 | 0.008 | 1 | 0.74932726 | -1.767657208 | 0.049 | 0.038 | 1 | 0.74932726 | 0.191687298 |
| Gm28731 | 0.100937876 | 2.154722595 | 0.038 | 0.008 | 1 | 0.5191986 | 0.252242349 | 0 | 0.01 | 1 | 0.5191986 | 0.191687298 |
| Gm17163 | 0.100937876 | 2.154722595 | 0.038 | 0.008 | 1 | 0.5191986 | 0.515276755 | 0 | 0.01 | 1 | 0.5191986 | 0.191687298 |
| Mettl24 | 0.100937876 | 2.154722595 | 0.038 | 0.008 | 1 | 0.729889663 | 1.83720485 | 0.024 | 0.017 | 1 | 0.729889663 | 0.191687298 |
| Hcn2 | 0.100937876 | 2.154722595 | 0.038 | 0.008 | 1 | 0.402364804 | -0.16279515 | 0 | 0.017 | 1 | 0.402364804 | 0.191687298 |
| 2810021J22Rik | 0.100937876 | 2.154722595 | 0.038 | 0.008 | 1 | 0.746881137 | 1.029849928 | 0.024 | 0.017 | 1 | 0.746881137 | 0.191687298 |
| Coro6 | 0.100937876 | 2.154722595 | 0.038 | 0.008 | 1 | 0.455098275 | 0.515276755 | 0 | 0.014 | 1 | 0.455098275 | 0.191687298 |
| Gm57341 | 0.100937876 | 2.154722595 | 0.038 | 0.008 | 1 | 0.599290422 | 1.515276755 | 0.024 | 0.014 | 1 | 0.599290422 | 0.191687298 |
| 2410004P03Rik | 0.100937876 | 2.154722595 | 0.038 | 0.008 | 1 | 0.114118767 | 2.100239256 | 0.049 | 0.014 | 1 | 0.114118767 | 0.191687298 |
| Gm34868 | 0.100937876 | 2.154722595 | 0.038 | 0.008 | 1 | 0.599290422 | 1.515276755 | 0.024 | 0.014 | 1 | 0.599290422 | 0.191687298 |
| Pgf | 0.100937876 | 2.154722595 | 0.038 | 0.008 | 1 | 0.438672228 | 1.83720485 | 0.024 | 0.01 | 1 | 0.438672228 | 0.191687298 |
| Tshr | 0.100937876 | 2.154722595 | 0.038 | 0.008 | 1 | 0.648487669 | -0.455576899 | 0.049 | 0.068 | 1 | 0.648487669 | 0.191687298 |
| Gm48765 | 0.100937876 | 2.154722595 | 0.038 | 0.008 | 1 | 0.267205177 | 2.252242349 | 0.024 | 0.007 | 1 | 0.267205177 | 0.191687298 |
| Gm49067 | 0.100937876 | 2.154722595 | 0.038 | 0.008 | 1 | 0.633586667 | 0.83720485 | 0.049 | 0.034 | 1 | 0.633586667 | 0.191687298 |
| Gm49584 | 0.100937876 | 2.154722595 | 0.038 | 0.008 | 1 | 0.744047924 | 1.252242349 | 0.024 | 0.017 | 1 | 0.744047924 | 0.191687298 |
| Prodh | 0.100937876 | 2.154722595 | 0.038 | 0.008 | 1 | 0.319629309 | -0.484723245 | 0 | 0.024 | 1 | 0.319629309 | 0.191687298 |
| B3galt4 | 0.100937876 | 2.154722595 | 0.038 | 0.008 | 1 | 0.257325989 | -0.484723245 | 0 | 0.031 | 1 | 0.257325989 | 0.191687298 |
| C3 | 0.100937876 | 2.154722595 | 0.038 | 0.008 | 1 | 0.27192168 | 1.252242349 | 0.049 | 0.02 | 1 | 0.27192168 | 0.191687298 |
| Cyp2c55 | 0.100937876 | 2.154722595 | 0.038 | 0.008 | 1 | 0.438672228 | 1.83720485 | 0.024 | 0.01 | 1 | 0.438672228 | 0.191687298 |
| Gm56646 | 0.100937876 | 2.154722595 | 0.038 | 0.008 | 1 | 0.44410834 | 0.515276755 | 0.024 | 0.01 | 1 | 0.44410834 | 0.191687298 |
| Dlc1 | 0.100974616 | 0.325361131 | 0.906 | 0.89 | 1 | 0.559863265 | -0.192007071 | 0.854 | 0.87 | 1 | 0.559863265 | 0.191753359 |
| Atp11c | 0.219920267 | 0.305139437 | 0.585 | 0.581 | 1 | 0.100984109 | 0.478310429 | 0.707 | 0.689 | 1 | 0.219920267 | 0.191770427 |
| Nedd9 | 0.433005882 | 0.520291418 | 0.962 | 0.864 | 1 | 0.101024853 | 0.497762791 | 0.902 | 0.86 | 1 | 0.433005882 | 0.191843685 |
| Rtp4 | 0.101235162 | 0.602181572 | 0.189 | 0.106 | 1 | 0.137523656 | 1.861452396 | 0.146 | 0.082 | 1 | 0.137523656 | 0.192221766 |
| Fnta | 0.169912215 | 0.361818829 | 0.736 | 0.674 | 1 | 0.101288202 | -0.322872366 | 0.585 | 0.7 | 1 | 0.169912215 | 0.192317104 |
| S100pbp | 0.101303067 | 0.849220048 | 0.453 | 0.369 | 1 | 0.566671111 | -0.115489435 | 0.415 | 0.43 | 1 | 0.566671111 | 0.192343823 |
| Zfp51 | 0.849955492 | 0.417757001 | 0.094 | 0.106 | 1 | 0.101310185 | -1.050320421 | 0.098 | 0.201 | 1 | 0.849955492 | 0.192356617 |
| Kctd15 | 0.101393603 | 0.748730235 | 0.283 | 0.186 | 1 | 0.860844316 | 0.499335211 | 0.195 | 0.225 | 1 | 0.860844316 | 0.192506544 |
| Aamp | 0.101461669 | 0.293818204 | 0.849 | 0.822 | 1 | 0.694567338 | -0.111395998 | 0.878 | 0.85 | 1 | 0.694567338 | 0.192628868 |
| Calhm2 | 0.101538915 | 0.817687608 | 0.208 | 0.123 | 1 | 0.107534542 | 0.83720485 | 0.268 | 0.167 | 1 | 0.107534542 | 0.192767678 |
| Rtel1 | 0.101648743 | 1.8327945 | 0.113 | 0.055 | 1 | 0.103792432 | -1.622226769 | 0 | 0.061 | 1 | 0.103792432 | 0.192965019 |
| Lfng | 0.101754481 | -1.686579659 | 0.038 | 0.11 | 1 | 0.154144895 | -1.410722664 | 0 | 0.048 | 1 | 0.154144895 | 0.193154988 |
| Clk4 | 0.101775752 | 0.334428617 | 0.868 | 0.763 | 1 | 0.157298057 | 0.355671939 | 0.805 | 0.771 | 1 | 0.157298057 | 0.1931932 |
| Atf1 | 0.101780249 | 0.389187848 | 0.547 | 0.436 | 1 | 0.903340254 | 0.355001924 | 0.634 | 0.693 | 1 | 0.903340254 | 0.193201279 |
| Lamtor1 | 0.101860776 | 0.421460969 | 0.774 | 0.78 | 1 | 0.888486052 | 0.189011588 | 0.78 | 0.782 | 1 | 0.888486052 | 0.193345933 |
| Syk | 0.101874199 | 1.067259754 | 0.038 | 0.008 | 1 | 0.257323555 | -1.863234868 | 0 | 0.031 | 1 | 0.257323555 | 0.193370045 |
| Gm1673 | 0.1021931 | 0.8327945 | 0.208 | 0.123 | 1 | 0.680867482 | -0.116991461 | 0.073 | 0.092 | 1 | 0.680867482 | 0.19394277 |
| Lcn11 | 0.110329043 | -1.368839361 | 0.019 | 0.081 | 1 | 0.102260336 | 1.713822434 | 0.22 | 0.14 | 1 | 0.110329043 | 0.194063496 |
| Slc35a5 | 0.102423044 | 1.543287883 | 0.264 | 0.186 | 1 | 0.706708186 | 0.270858027 | 0.195 | 0.177 | 1 | 0.706708186 | 0.194355607 |
| Greb1l | 0.102462777 | -1.20282941 | 0.019 | 0.085 | 1 | 0.989233534 | 1.021629421 | 0.122 | 0.126 | 1 | 0.989233534 | 0.194426933 |
| Ormdl2 | 0.102571459 | 0.687596584 | 0.245 | 0.153 | 1 | 0.324825356 | 0.570418309 | 0.268 | 0.205 | 1 | 0.324825356 | 0.194622014 |
| Irak4 | 0.102584222 | 0.984797593 | 0.264 | 0.178 | 1 | 0.359710668 | -0.250257991 | 0.122 | 0.181 | 1 | 0.359710668 | 0.194644921 |
| Fem1c | 0.662621043 | 0.173338273 | 0.415 | 0.407 | 1 | 0.102618596 | 0.370078839 | 0.683 | 0.539 | 1 | 0.662621043 | 0.194706617 |
| Gltp | 0.102635149 | -0.669026766 | 0.302 | 0.424 | 1 | 0.272974755 | -0.475073075 | 0.293 | 0.365 | 1 | 0.272974755 | 0.194736325 |
| Large2 | 0.10281534 | 1.739685096 | 0.038 | 0.008 | 1 | 0.994797912 | 0.136765132 | 0.024 | 0.024 | 1 | 0.994797912 | 0.195059685 |
| Itpka | 0.10281534 | 1.739685096 | 0.038 | 0.008 | 1 | 0.729893611 | 2.029849928 | 0.024 | 0.017 | 1 | 0.729893611 | 0.195059685 |
| Pex5l | 0.10281534 | 1.739685096 | 0.038 | 0.008 | 1 | 0.5191986 | 0.252242349 | 0 | 0.01 | 1 | 0.5191986 | 0.195059685 |
| Miat | 0.10281534 | 1.417757001 | 0.038 | 0.008 | 1 | 0.994797906 | 0.252242349 | 0.024 | 0.024 | 1 | 0.994797906 | 0.195059685 |
| Spata2l | 0.10281534 | 1.739685096 | 0.038 | 0.008 | 1 | 0.319641967 | -0.16279515 | 0 | 0.024 | 1 | 0.319641967 | 0.195059685 |
| Gm6980 | 0.10281534 | 1.739685096 | 0.038 | 0.008 | 1 | 0.262676008 | 3.029849928 | 0.024 | 0.007 | 1 | 0.262676008 | 0.195059685 |
| Apc2 | 0.10281534 | 1.739685096 | 0.038 | 0.008 | 1 | 0.402373368 | 0.252242349 | 0 | 0.017 | 1 | 0.402373368 | 0.195059685 |
| Gm49226 | 0.10281534 | 1.739685096 | 0.038 | 0.008 | 1 | 0.599290422 | 1.515276755 | 0.024 | 0.014 | 1 | 0.599290422 | 0.195059685 |
| Gm17315 | 0.10281534 | 1.739685096 | 0.038 | 0.008 | 1 | 0.180328124 | 2.252242349 | 0.049 | 0.017 | 1 | 0.180328124 | 0.195059685 |
| Znhit2 | 0.102858545 | 0.756173218 | 0.189 | 0.106 | 1 | 0.208249271 | 0.930314254 | 0.146 | 0.085 | 1 | 0.208249271 | 0.195137209 |
| Rab4a | 0.321478357 | -0.108311811 | 0.113 | 0.178 | 1 | 0.102885258 | -0.278272368 | 0.098 | 0.208 | 1 | 0.321478357 | 0.19518514 |
| Ercc6l2 | 0.103038033 | -0.461948766 | 0.302 | 0.428 | 1 | 0.294004875 | -0.380025866 | 0.244 | 0.321 | 1 | 0.294004875 | 0.19545923 |
| Zdhhc23 | 0.103180216 | -1.46976827 | 0.038 | 0.11 | 1 | 0.134736465 | 1.212713985 | 0.171 | 0.096 | 1 | 0.134736465 | 0.195714276 |
| Commd7 | 0.103215102 | 0.755115136 | 0.472 | 0.398 | 1 | 0.144728716 | -0.308774456 | 0.317 | 0.471 | 1 | 0.144728716 | 0.195776847 |
| Rab8a | 0.103293247 | 0.482776792 | 0.623 | 0.517 | 1 | 0.383246632 | -0.288732139 | 0.585 | 0.621 | 1 | 0.383246632 | 0.195917 |
| Slc35a1 | 0.115987857 | 0.647762606 | 0.434 | 0.343 | 1 | 0.10340396 | -0.21445727 | 0.293 | 0.437 | 1 | 0.115987857 | 0.196115542 |
| Rpl27 | 0.103424904 | 0.222752061 | 1 | 0.962 | 1 | 0.419668895 | 0.144152678 | 1 | 0.986 | 1 | 0.419668895 | 0.196153097 |
| Mttp | 0.103445697 | -0.719746523 | 0.057 | 0.14 | 1 | 0.817848127 | 0.962735732 | 0.146 | 0.14 | 1 | 0.817848127 | 0.196190381 |
| Gm26698 | 0.128609464 | -1.430239906 | 0 | 0.042 | 1 | 0.103479965 | 3.83720485 | 0.024 | 0.003 | 1 | 0.128609464 | 0.196251827 |
| Gm49171 | 0.209009104 | 1.739685096 | 0.038 | 0.013 | 1 | 0.103479965 | 3.422167351 | 0.024 | 0.003 | 1 | 0.209009104 | 0.196251827 |
| Hcn1 | 0.214901224 | 0.932330173 | 0.038 | 0.013 | 1 | 0.103479965 | 3.159132945 | 0.024 | 0.003 | 1 | 0.214901224 | 0.196251827 |
| C2cd4a | 0.246443787 | 2.891688189 | 0.019 | 0.004 | 1 | 0.103479965 | 3.422167351 | 0.024 | 0.003 | 1 | 0.246443787 | 0.196251827 |
| Fbxo40 | 0.246443787 | 2.739685096 | 0.019 | 0.004 | 1 | 0.103479965 | 3.83720485 | 0.024 | 0.003 | 1 | 0.246443787 | 0.196251827 |
| Naip2 | 0.249034808 | 2.154722595 | 0.019 | 0.004 | 1 | 0.103479965 | 3.422167351 | 0.024 | 0.003 | 1 | 0.249034808 | 0.196251827 |
| Ascl3 | 0.251644813 | 1.569760094 | 0.019 | 0.004 | 1 | 0.103479965 | 3.83720485 | 0.024 | 0.003 | 1 | 0.251644813 | 0.196251827 |
| 4931428L18Rik | 0.287761733 | -0.652632327 | 0 | 0.021 | 1 | 0.103479965 | 3.422167351 | 0.024 | 0.003 | 1 | 0.287761733 | 0.196251827 |
| Mycl | 0.287761733 | -0.652632327 | 0 | 0.021 | 1 | 0.103479965 | 3.422167351 | 0.024 | 0.003 | 1 | 0.287761733 | 0.196251827 |
| Gm11592 | 0.287773478 | -0.430239906 | 0 | 0.021 | 1 | 0.103479965 | 3.422167351 | 0.024 | 0.003 | 1 | 0.287773478 | 0.196251827 |
| En2 | 0.343000446 | -0.430239906 | 0 | 0.017 | 1 | 0.103479965 | 3.422167351 | 0.024 | 0.003 | 1 | 0.343000446 | 0.196251827 |
| Alkal2 | 0.343000446 | -0.430239906 | 0 | 0.017 | 1 | 0.103479965 | 4.159132945 | 0.024 | 0.003 | 1 | 0.343000446 | 0.196251827 |
| Jhy | 0.343004841 | -1.015202407 | 0 | 0.017 | 1 | 0.103479965 | 3.252242349 | 0.024 | 0.003 | 1 | 0.343004841 | 0.196251827 |
| Gm38287 | 0.343968402 | 1.154722595 | 0.038 | 0.017 | 1 | 0.103479965 | 3.422167351 | 0.024 | 0.003 | 1 | 0.343968402 | 0.196251827 |
| Tnfrsf26 | 0.351545549 | 0.569760094 | 0.038 | 0.017 | 1 | 0.103479965 | 3.83720485 | 0.024 | 0.003 | 1 | 0.351545549 | 0.196251827 |
| Zfp296 | 0.49029049 | 0.739685096 | 0.038 | 0.021 | 1 | 0.103479965 | 3.83720485 | 0.024 | 0.003 | 1 | 0.49029049 | 0.196251827 |
| Npas1 | 0.503971666 | 1.569760094 | 0.019 | 0.008 | 1 | 0.103479965 | 3.422167351 | 0.024 | 0.003 | 1 | 0.503971666 | 0.196251827 |
| Pcsk2 | 0.510606779 | 0.569760094 | 0.019 | 0.008 | 1 | 0.103479965 | 4.644559772 | 0.024 | 0.003 | 1 | 0.510606779 | 0.196251827 |
| Cdh17 | 0.510606779 | -1.093204919 | 0.019 | 0.008 | 1 | 0.103479965 | 3.422167351 | 0.024 | 0.003 | 1 | 0.510606779 | 0.196251827 |
| Tfap2c | 0.628925725 | 0.739685096 | 0.038 | 0.025 | 1 | 0.103479965 | 3.422167351 | 0.024 | 0.003 | 1 | 0.628925725 | 0.196251827 |
| Lvrn | 0.725936481 | 1.417757001 | 0.019 | 0.013 | 1 | 0.103479965 | 3.574170444 | 0.024 | 0.003 | 1 | 0.725936481 | 0.196251827 |
| A930038B10Rik | 0.7394666 | 0.569760094 | 0.019 | 0.013 | 1 | 0.103479965 | 4.159132945 | 0.024 | 0.003 | 1 | 0.7394666 | 0.196251827 |
| Cenpi | 0.89261791 | 0.569760094 | 0.038 | 0.034 | 1 | 0.103479965 | 3.422167351 | 0.024 | 0.003 | 1 | 0.89261791 | 0.196251827 |
| 2010002M09Rik | 0.926212226 | 0.8327945 | 0.019 | 0.017 | 1 | 0.103479965 | 3.422167351 | 0.024 | 0.003 | 1 | 0.926212226 | 0.196251827 |
| Mppe1 | 0.103529675 | -0.527101445 | 0.17 | 0.284 | 1 | 0.42456901 | -0.233184478 | 0.171 | 0.225 | 1 | 0.42456901 | 0.196340956 |
| Aip | 0.103747594 | 0.782753817 | 0.396 | 0.305 | 1 | 0.681794956 | 0.338398993 | 0.341 | 0.328 | 1 | 0.681794956 | 0.196731624 |
| Scrn2 | 0.324442569 | 0.714150003 | 0.113 | 0.072 | 1 | 0.103767696 | -1.622226769 | 0 | 0.061 | 1 | 0.324442569 | 0.196767658 |
| Gm29157 | 0.547552679 | -0.330704232 | 0.075 | 0.102 | 1 | 0.103767696 | -1.484723245 | 0 | 0.061 | 1 | 0.547552679 | 0.196767658 |
| Gm57147 | 0.624418548 | 0.932330173 | 0.038 | 0.025 | 1 | 0.103767696 | -1.484723245 | 0 | 0.061 | 1 | 0.624418548 | 0.196767658 |
| Gm56915 | 0.905020105 | 0.569760094 | 0.075 | 0.072 | 1 | 0.103792432 | -1.555112573 | 0 | 0.061 | 1 | 0.905020105 | 0.196811996 |
| A130051J06Rik | 0.196612515 | -0.7032584 | 0.038 | 0.093 | 1 | 0.103814258 | -1.622226769 | 0 | 0.061 | 1 | 0.196612515 | 0.196851115 |

| | | | | | | | | | | | | |
|---|---|---|---|---|---|---|---|---|---|---|---|---|
| Ppp1r14c | 0.464333985 | -0.783876861 | 0.038 | 0.064 | 1 | 0.103833173 | -1.80665134 | 0 | 0.061 | 1 | 0.464333985 | 0.196885018 |
| Gm46412 | 0.35921294 | -0.381330305 | 0.057 | 0.097 | 1 | 0.103863363 | -1.970150072 | 0 | 0.061 | 1 | 0.35921294 | 0.196939128 |
| 2700069I18Rik | 0.175842714 | -1.304709024 | 0 | 0.034 | 1 | 0.103872457 | -2.020776145 | 0 | 0.061 | 1 | 0.175842714 | 0.196955426 |
| Zmat4 | 0.413082433 | -0.845277405 | 0 | 0.013 | 1 | 0.103905556 | -2.717384002 | 0 | 0.061 | 1 | 0.413082433 | 0.197014747 |
| Usp37 | 0.103930676 | 0.417757001 | 0.642 | 0.568 | 1 | 0.821684448 | 0.202611581 | 0.756 | 0.829 | 1 | 0.821684448 | 0.197059767 |
| Wdtc1 | 0.103976384 | 0.5917864 | 0.321 | 0.216 | 1 | 0.524496593 | 0.383028957 | 0.341 | 0.314 | 1 | 0.524496593 | 0.197141679 |
| Cyfip2 | 0.103989326 | 0.984797593 | 0.151 | 0.081 | 1 | 0.112999113 | 0.46797104 | 0.268 | 0.16 | 1 | 0.112999113 | 0.197164872 |
| mt-Nd1 | 0.682314285 | 0.241383262 | 1 | 1 | 1 | 0.104071493 | 0.206193337 | 1 | 1 | 1 | 0.682314285 | 0.19731211 |
| Plbd2 | 0.785502438 | 0.209864149 | 0.453 | 0.47 | 1 | 0.104076111 | -0.484723245 | 0.341 | 0.491 | 1 | 0.785502438 | 0.197320384 |
| Bsdc1 | 0.46748804 | 0.380282295 | 0.566 | 0.521 | 1 | 0.104077786 | -0.238083278 | 0.463 | 0.614 | 1 | 0.46748804 | 0.197323387 |
| Afg1l | 0.132785844 | 0.739685096 | 0.245 | 0.161 | 1 | 0.104085923 | -1.440779897 | 0.049 | 0.14 | 1 | 0.132785844 | 0.197337967 |
| Ism1 | 0.477497852 | -1.600164907 | 0.358 | 0.381 | 1 | 0.10409203 | 0.412071473 | 0.439 | 0.324 | 1 | 0.477497852 | 0.197348909 |
| Gm56830 | 0.177057772 | 0.430932389 | 0.321 | 0.237 | 1 | 0.104143385 | 0.847258514 | 0.366 | 0.266 | 1 | 0.177057772 | 0.197440925 |
| Cwc22 | 0.24871908 | 0.499370766 | 0.415 | 0.347 | 1 | 0.104177951 | -0.478150591 | 0.293 | 0.451 | 1 | 0.24871908 | 0.197502856 |
| Prex1 | 0.756716266 | 0.527939918 | 0.226 | 0.258 | 1 | 0.104306174 | -0.172779239 | 0.293 | 0.44 | 1 | 0.756716266 | 0.19773257 |
| Hmgxb3 | 0.604709083 | 0.199391644 | 0.34 | 0.335 | 1 | 0.104446708 | -0.233184478 | 0.22 | 0.389 | 1 | 0.604709083 | 0.197984301 |
| Mettl6 | 0.6437637 | 0.276028891 | 0.358 | 0.335 | 1 | 0.10449057 | -0.56878751 | 0.293 | 0.427 | 1 | 0.6437637 | 0.198062862 |
| Ndufaf7 | 0.104571594 | 0.940597789 | 0.283 | 0.191 | 1 | 0.624365324 | 0.515276755 | 0.22 | 0.191 | 1 | 0.624365324 | 0.19820797 |
| P2ry10b | 0.104701374 | 0.739685096 | 0.302 | 0.199 | 1 | 0.835956367 | 0.136765132 | 0.341 | 0.331 | 1 | 0.835956367 | 0.19844037 |
| Cdh15 | 0.104727895 | 1.417757001 | 0.038 | 0.008 | 1 | 0.106248001 | 2.252242349 | 0.024 | 0.003 | 1 | 0.106248001 | 0.198487858 |
| St8sia5 | 0.104727895 | 0.739685096 | 0.038 | 0.008 | 1 | 0.26948834 | 1.83720485 | 0.024 | 0.007 | 1 | 0.26948834 | 0.198487858 |
| Cmbl | 0.104744075 | 1.154722595 | 0.34 | 0.267 | 1 | 0.720173229 | 0.402576622 | 0.366 | 0.338 | 1 | 0.720173229 | 0.19851683 |
| Gm57030 | 0.204653189 | 2.154722595 | 0.038 | 0.013 | 1 | 0.104856732 | 2.83720485 | 0.024 | 0.003 | 1 | 0.204653189 | 0.19871853 |
| Alk | 0.204653189 | 2.154722595 | 0.038 | 0.013 | 1 | 0.104856732 | 2.83720485 | 0.024 | 0.003 | 1 | 0.204653189 | 0.19871853 |
| Gls2 | 0.206368562 | -1.015202407 | 0 | 0.03 | 1 | 0.104856732 | 2.83720485 | 0.024 | 0.003 | 1 | 0.206368562 | 0.19871853 |
| Apold1 | 0.206380492 | -2.368839361 | 0 | 0.03 | 1 | 0.104856732 | 2.83720485 | 0.024 | 0.003 | 1 | 0.206380492 | 0.19871853 |
| Gm5468.1 | 0.206380492 | -1.1672055 | 0 | 0.03 | 1 | 0.104856732 | 2.83720485 | 0.024 | 0.003 | 1 | 0.206380492 | 0.19871853 |
| Gm56593 | 0.209009104 | 1.739685096 | 0.038 | 0.013 | 1 | 0.104856732 | 2.83720485 | 0.024 | 0.003 | 1 | 0.209009104 | 0.19871853 |
| Melk | 0.243102132 | -1.1672055 | 0 | 0.025 | 1 | 0.104856732 | 2.83720485 | 0.024 | 0.003 | 1 | 0.243102132 | 0.19871853 |
| Gm12043 | 0.243111734 | -0.652632327 | 0 | 0.025 | 1 | 0.104856732 | 2.83720485 | 0.024 | 0.003 | 1 | 0.243111734 | 0.19871853 |
| 4930412B13Rik | 0.246443787 | 3.47665069 | 0.019 | 0.004 | 1 | 0.104856732 | 2.83720485 | 0.024 | 0.003 | 1 | 0.246443787 | 0.19871853 |
| Dlgap2 | 0.246443787 | 3.154722595 | 0.019 | 0.004 | 1 | 0.104856732 | 2.83720485 | 0.024 | 0.003 | 1 | 0.246443787 | 0.19871853 |
| Zfp456 | 0.246443787 | 2.739685096 | 0.019 | 0.004 | 1 | 0.104856732 | 2.83720485 | 0.024 | 0.003 | 1 | 0.246443787 | 0.19871853 |
| Adgre1 | 0.246443787 | 2.739685096 | 0.019 | 0.004 | 1 | 0.104856732 | 2.83720485 | 0.024 | 0.003 | 1 | 0.246443787 | 0.19871853 |
| Ell3 | 0.249034808 | 2.154722595 | 0.019 | 0.004 | 1 | 0.104856732 | 2.83720485 | 0.024 | 0.003 | 1 | 0.249034808 | 0.19871853 |
| Chrna10 | 0.249034808 | 2.154722595 | 0.019 | 0.004 | 1 | 0.104856732 | 2.83720485 | 0.024 | 0.003 | 1 | 0.249034808 | 0.19871853 |
| Ap1m2 | 0.249034808 | 2.154722595 | 0.019 | 0.004 | 1 | 0.104856732 | 2.83720485 | 0.024 | 0.003 | 1 | 0.249034808 | 0.19871853 |
| Rasgrp3 | 0.249034808 | 2.154722595 | 0.019 | 0.004 | 1 | 0.104856732 | 2.83720485 | 0.024 | 0.003 | 1 | 0.249034808 | 0.19871853 |
| Plac8l1 | 0.249034808 | 2.154722595 | 0.019 | 0.004 | 1 | 0.104856732 | 2.83720485 | 0.024 | 0.003 | 1 | 0.249034808 | 0.19871853 |
| Onecut2 | 0.249034808 | 2.154722595 | 0.019 | 0.004 | 1 | 0.104856732 | 2.83720485 | 0.024 | 0.003 | 1 | 0.249034808 | 0.19871853 |
| Slc9a2 | 0.251644813 | 1.569760094 | 0.019 | 0.004 | 1 | 0.104856732 | 2.83720485 | 0.024 | 0.003 | 1 | 0.251644813 | 0.19871853 |
| Platr12 | 0.251644813 | 1.569760094 | 0.019 | 0.004 | 1 | 0.104856732 | 2.83720485 | 0.024 | 0.003 | 1 | 0.251644813 | 0.19871853 |
| Gfi1 | 0.251644813 | 1.154722595 | 0.019 | 0.004 | 1 | 0.104856732 | 2.83720485 | 0.024 | 0.003 | 1 | 0.251644813 | 0.19871853 |
| Gm48236 | 0.251644813 | 1.569760094 | 0.019 | 0.004 | 1 | 0.104856732 | 2.83720485 | 0.024 | 0.003 | 1 | 0.251644813 | 0.19871853 |
| Pi16 | 0.251644813 | 1.569760094 | 0.019 | 0.004 | 1 | 0.104856732 | 2.83720485 | 0.024 | 0.003 | 1 | 0.251644813 | 0.19871853 |
| F12 | 0.287761733 | -1.430239906 | 0 | 0.021 | 1 | 0.104856732 | 2.83720485 | 0.024 | 0.003 | 1 | 0.287761733 | 0.19871853 |
| S100a2 | 0.28776878 | -1.015202407 | 0 | 0.021 | 1 | 0.104856732 | 2.83720485 | 0.024 | 0.003 | 1 | 0.28776878 | 0.19871853 |
| Gm45447 | 0.287773478 | -0.430239906 | 0 | 0.021 | 1 | 0.104856732 | 2.83720485 | 0.024 | 0.003 | 1 | 0.287773478 | 0.19871853 |
| Ddias | 0.287773478 | -0.430239906 | 0 | 0.021 | 1 | 0.104856732 | 2.83720485 | 0.024 | 0.003 | 1 | 0.287773478 | 0.19871853 |
| Gm15943 | 0.287773478 | -0.430239906 | 0 | 0.021 | 1 | 0.104856732 | 2.83720485 | 0.024 | 0.003 | 1 | 0.287773478 | 0.19871853 |
| Gm33373 | 0.338375114 | 1.347367673 | 0.038 | 0.017 | 1 | 0.104856732 | 2.83720485 | 0.024 | 0.003 | 1 | 0.338375114 | 0.19871853 |
| Angpt2 | 0.340246113 | 1.417757001 | 0.038 | 0.017 | 1 | 0.104856732 | 2.83720485 | 0.024 | 0.003 | 1 | 0.340246113 | 0.19871853 |
| Tfr2 | 0.343000446 | -0.430239906 | 0 | 0.017 | 1 | 0.104856732 | 2.83720485 | 0.024 | 0.003 | 1 | 0.343000446 | 0.19871853 |
| Tas1r1 | 0.343004841 | -1.304709024 | 0 | 0.017 | 1 | 0.104856732 | 2.83720485 | 0.024 | 0.003 | 1 | 0.343004841 | 0.19871853 |
| Gm36988 | 0.343006305 | -0.1672055 | 0 | 0.017 | 1 | 0.104856732 | 2.83720485 | 0.024 | 0.003 | 1 | 0.343006305 | 0.19871853 |
| Clcnka | 0.413080549 | -0.430239906 | 0 | 0.013 | 1 | 0.104856732 | 2.83720485 | 0.024 | 0.003 | 1 | 0.413080549 | 0.19871853 |
| AI847159 | 0.413082433 | 0.154722595 | 0 | 0.013 | 1 | 0.104856732 | 2.83720485 | 0.024 | 0.003 | 1 | 0.413082433 | 0.19871853 |
| Dmrt3 | 0.413082433 | 0.154722595 | 0 | 0.013 | 1 | 0.104856732 | 2.83720485 | 0.024 | 0.003 | 1 | 0.413082433 | 0.19871853 |
| Ednrb | 0.469074933 | 1.569760094 | 0.038 | 0.021 | 1 | 0.104856732 | 2.83720485 | 0.024 | 0.003 | 1 | 0.469074933 | 0.19871853 |
| Ufsp1 | 0.48175508 | 1.154722595 | 0.038 | 0.021 | 1 | 0.104856732 | 2.83720485 | 0.024 | 0.003 | 1 | 0.48175508 | 0.19871853 |
| Gm57328 | 0.497380598 | 2.154722595 | 0.019 | 0.008 | 1 | 0.104856732 | 2.83720485 | 0.024 | 0.003 | 1 | 0.497380598 | 0.19871853 |
| 0610039K10Rik | 0.503971666 | 1.569760094 | 0.019 | 0.008 | 1 | 0.104856732 | 2.83720485 | 0.024 | 0.003 | 1 | 0.503971666 | 0.19871853 |
| Gm48860 | 0.503971666 | 1.569760094 | 0.019 | 0.008 | 1 | 0.104856732 | 2.83720485 | 0.024 | 0.003 | 1 | 0.503971666 | 0.19871853 |
| Bdkrb1 | 0.507281812 | 1.154722595 | 0.019 | 0.008 | 1 | 0.104856732 | 2.83720485 | 0.024 | 0.003 | 1 | 0.507281812 | 0.19871853 |
| Acer1 | 0.507281812 | 1.154722595 | 0.019 | 0.008 | 1 | 0.104856732 | 2.83720485 | 0.024 | 0.003 | 1 | 0.507281812 | 0.19871853 |
| 2010308F09Rik | 0.507281812 | 0.569760094 | 0.019 | 0.008 | 1 | 0.104856732 | 2.83720485 | 0.024 | 0.003 | 1 | 0.507281812 | 0.19871853 |
| Nup62cl | 0.507281812 | 1.154722595 | 0.019 | 0.008 | 1 | 0.104856732 | 2.83720485 | 0.024 | 0.003 | 1 | 0.507281812 | 0.19871853 |
| Spdl1 | 0.580201714 | -0.1672055 | 0.019 | 0.034 | 1 | 0.104856732 | 2.83720485 | 0.024 | 0.003 | 1 | 0.580201714 | 0.19871853 |
| 1700057G04Rik | 0.61989169 | 0.669295768 | 0.038 | 0.025 | 1 | 0.104856732 | 2.83720485 | 0.024 | 0.003 | 1 | 0.61989169 | 0.19871853 |
| Lzts1 | 0.628925725 | 0.739685096 | 0.038 | 0.025 | 1 | 0.104856732 | 2.83720485 | 0.024 | 0.003 | 1 | 0.628925725 | 0.19871853 |
| Nkapl | 0.72256466 | 1.739685096 | 0.019 | 0.013 | 1 | 0.104856732 | 2.83720485 | 0.024 | 0.003 | 1 | 0.72256466 | 0.19871853 |
| 4932412D23Rik | 0.72256466 | 1.739685096 | 0.019 | 0.013 | 1 | 0.104856732 | 2.83720485 | 0.024 | 0.003 | 1 | 0.72256466 | 0.19871853 |
| Fkbp6 | 0.725937279 | 1.347367673 | 0.019 | 0.013 | 1 | 0.104856732 | 2.83720485 | 0.024 | 0.003 | 1 | 0.725937279 | 0.19871853 |
| Ttk | 0.729311982 | 0.389187848 | 0.019 | 0.013 | 1 | 0.104856732 | 2.83720485 | 0.024 | 0.003 | 1 | 0.729311982 | 0.19871853 |
| Ccdc146 | 0.732691937 | 1.154722595 | 0.019 | 0.013 | 1 | 0.104856732 | 2.83720485 | 0.024 | 0.003 | 1 | 0.732691937 | 0.19871853 |
| Gm29675 | 0.732691937 | 1.154722595 | 0.019 | 0.013 | 1 | 0.104856732 | 2.83720485 | 0.024 | 0.003 | 1 | 0.732691937 | 0.19871853 |
| 1110013H19Rik | 0.732691937 | 1.154722595 | 0.019 | 0.013 | 1 | 0.104856732 | 2.83720485 | 0.024 | 0.003 | 1 | 0.732691937 | 0.19871853 |
| A830052D11Rik | 0.732691937 | 1.154722595 | 0.019 | 0.013 | 1 | 0.104856732 | 2.83720485 | 0.024 | 0.003 | 1 | 0.732691937 | 0.19871853 |
| Them7 | 0.736073999 | 0.569760094 | 0.019 | 0.013 | 1 | 0.104856732 | 2.83720485 | 0.024 | 0.003 | 1 | 0.736073999 | 0.19871853 |
| Gm20621 | 0.736073999 | 0.8327945 | 0.019 | 0.013 | 1 | 0.104856732 | 2.83720485 | 0.024 | 0.003 | 1 | 0.736073999 | 0.19871853 |
| Chrna4 | 0.787324952 | 0.280253477 | 0.019 | 0.025 | 1 | 0.104856732 | 2.83720485 | 0.024 | 0.003 | 1 | 0.787324952 | 0.19871853 |
| Gm49518 | 0.797844282 | 0.932330173 | 0.019 | 0.025 | 1 | 0.104856732 | 2.83720485 | 0.024 | 0.003 | 1 | 0.797844282 | 0.19871853 |
| Tex29 | 0.916620008 | 1.154722595 | 0.019 | 0.017 | 1 | 0.104856732 | 2.83720485 | 0.024 | 0.003 | 1 | 0.916620008 | 0.19871853 |
| Lin7b | 0.926212226 | 0.8327945 | 0.019 | 0.017 | 1 | 0.104856732 | 2.83720485 | 0.024 | 0.003 | 1 | 0.926212226 | 0.19871853 |
| Slc39a4 | 0.926212226 | 0.8327945 | 0.019 | 0.017 | 1 | 0.104856732 | 2.83720485 | 0.024 | 0.003 | 1 | 0.926212226 | 0.19871853 |
| Dym | 0.104858812 | 1.113499932 | 0.453 | 0.369 | 1 | 0.378949204 | 0.391894339 | 0.561 | 0.536 | 1 | 0.378949204 | 0.198722254 |
| Tmem256 | 0.106435559 | 0.429229785 | 0.736 | 0.657 | 1 | 0.104860572 | 0.688974919 | 0.634 | 0.59 | 1 | 0.106435559 | 0.198725404 |
| Zfp738 | 0.104907359 | 0.417757001 | 0.245 | 0.148 | 1 | 0.669399154 | -0.250257991 | 0.146 | 0.171 | 1 | 0.669399154 | 0.198809164 |
| Ddx39a | 0.138274252 | 0.406261362 | 0.321 | 0.229 | 1 | 0.104934718 | -0.439635356 | 0.195 | 0.334 | 1 | 0.138274252 | 0.198858141 |
| Tm2d2 | 0.104936468 | 0.174583017 | 0.83 | 0.767 | 1 | 0.441470533 | -0.107653596 | 0.634 | 0.71 | 1 | 0.441470533 | 0.198861274 |
| Tpgs1 | 0.6305605 | 0.286826131 | 0.283 | 0.254 | 1 | 0.105035902 | 0.83720485 | 0.317 | 0.212 | 1 | 0.6305605 | 0.199039263 |
| Acss3 | 0.506983348 | 0.695290976 | 0.019 | 0.038 | 1 | 0.10510891 | -0.80665134 | 0.098 | 0.205 | 1 | 0.506983348 | 0.199169937 |
| Nek8 | 0.263131017 | 0.879088152 | 0.226 | 0.169 | 1 | 0.105249088 | 1.169780189 | 0.244 | 0.15 | 1 | 0.263131017 | 0.199420806 |
| Tspan14 | 0.727196251 | 0.176748901 | 0.302 | 0.301 | 1 | 0.105283043 | 0.33763384 | 0.512 | 0.379 | 1 | 0.727196251 | 0.199481567 |
| Rpia | 0.10529036 | 0.678284551 | 0.283 | 0.191 | 1 | 0.707999217 | 0.422167351 | 0.268 | 0.263 | 1 | 0.707999217 | 0.199494661 |
| Serp2 | 0.14523548 | 1.347367673 | 0.132 | 0.072 | 1 | 0.105396607 | 1.029849928 | 0.146 | 0.072 | 1 | 0.14523548 | 0.19968477 |
| Eif4e | 0.10541805 | -0.468152322 | 0.943 | 0.915 | 1 | 0.535540225 | -0.129210924 | 0.902 | 0.966 | 1 | 0.535540225 | 0.199723135 |
| Fzd4 | 0.334518863 | 1.154722595 | 0.057 | 0.03 | 1 | 0.105452763 | 2.351778023 | 0.122 | 0.058 | 1 | 0.334518863 | 0.19978524 |

| | | | | | | | | | | | | |
|---|---|---|---|---|---|---|---|---|---|---|---|---|
| Ocel1 | 0.71718895 | 0.402650108 | 0.17 | 0.199 | 1 | 0.105456232 | -0.777504994 | 0.122 | 0.239 | 1 | 0.71718895 | 0.199791447 |
| 4930556J24Rik | 0.105519369 | 2.306725688 | 0.075 | 0.03 | 1 | 0.226583045 | 1.174239837 | 0.098 | 0.051 | 1 | 0.226583045 | 0.199904401 |
| Uqcc2 | 0.322096991 | -0.282682718 | 0.623 | 0.695 | 1 | 0.105522596 | -0.391613841 | 0.415 | 0.56 | 1 | 0.322096991 | 0.199910174 |
| Dnaaf5 | 0.35437551 | 1.084333267 | 0.113 | 0.076 | 1 | 0.105533372 | -1.555112573 | 0.024 | 0.102 | 1 | 0.35437551 | 0.199929451 |
| Zfp37 | 0.105541015 | 0.8327945 | 0.264 | 0.174 | 1 | 0.989428516 | 0.174239837 | 0.268 | 0.276 | 1 | 0.989428516 | 0.199943124 |
| Snrk | 0.105617574 | 0.733258826 | 0.34 | 0.237 | 1 | 0.814880351 | 0.382010224 | 0.366 | 0.375 | 1 | 0.814880351 | 0.200080077 |
| Tsc1 | 0.105644292 | 0.63230156 | 0.491 | 0.394 | 1 | 0.433299387 | -0.17660095 | 0.463 | 0.529 | 1 | 0.433299387 | 0.200127867 |
| Scnm1 | 0.105686522 | 0.739685096 | 0.245 | 0.153 | 1 | 0.122515773 | 0.995902596 | 0.22 | 0.133 | 1 | 0.122515773 | 0.200203402 |
| Gatd3a | 0.105746195 | 0.519134021 | 0.377 | 0.263 | 1 | 0.115390854 | -0.661046018 | 0.195 | 0.314 | 1 | 0.115390854 | 0.200310131 |
| Wbp2 | 0.339228291 | -0.263590037 | 0.623 | 0.644 | 1 | 0.105772871 | -0.25871957 | 0.537 | 0.659 | 1 | 0.339228291 | 0.200357842 |
| Prpf18 | 0.105792154 | 0.68265415 | 0.377 | 0.284 | 1 | 0.995929284 | 0.506559538 | 0.317 | 0.355 | 1 | 0.995929284 | 0.200392329 |
| Zbtb33 | 0.107869624 | 0.395730694 | 0.358 | 0.25 | 1 | 0.105811235 | -0.515311565 | 0.195 | 0.317 | 1 | 0.107869624 | 0.200426453 |
| Riok2 | 0.105840497 | -0.457461375 | 0.415 | 0.534 | 1 | 0.600316515 | 0.655454413 | 0.463 | 0.461 | 1 | 0.600316515 | 0.200478784 |
| Rasgef1b | 0.105843586 | -1.7032584 | 0.113 | 0.199 | 1 | 0.735265116 | -0.385187572 | 0.049 | 0.061 | 1 | 0.735265116 | 0.200484308 |
| Erc1 | 0.105992629 | -0.527952844 | 0.962 | 0.953 | 1 | 0.98411648 | -0.124418478 | 1 | 0.98 | 1 | 0.98411648 | 0.200750821 |
| Tmem160 | 0.10602458 | 0.569760094 | 0.547 | 0.487 | 1 | 0.995963465 | 0.200895469 | 0.366 | 0.372 | 1 | 0.995963465 | 0.200807949 |
| Usp29 | 0.882815226 | 0.173831418 | 0.17 | 0.161 | 1 | 0.10616816 | 1.504629511 | 0.22 | 0.13 | 1 | 0.882815226 | 0.201064642 |
| Pank4 | 0.10618355 | 1.203632195 | 0.189 | 0.11 | 1 | 0.154659913 | 0.906467512 | 0.244 | 0.16 | 1 | 0.154659913 | 0.201092153 |
| Actr10 | 0.106205257 | 0.446903346 | 0.698 | 0.699 | 1 | 0.293180907 | -0.123266786 | 0.732 | 0.809 | 1 | 0.293180907 | 0.201130957 |
| Carm1 | 0.106220249 | 0.609561102 | 0.491 | 0.381 | 1 | 0.488670355 | 0.115669296 | 0.512 | 0.58 | 1 | 0.488670355 | 0.201157757 |
| Gdnf | 0.172851729 | 0.318221327 | 0.075 | 0.034 | 1 | 0.106248001 | 2.422167351 | 0.024 | 0.003 | 1 | 0.172851729 | 0.201207365 |
| 5730420D15Rik | 0.227851115 | 1.984797593 | 0.057 | 0.025 | 1 | 0.106248001 | 1.83720485 | 0.024 | 0.003 | 1 | 0.227851115 | 0.201207365 |
| 4930565N06Rik | 0.243106933 | -1.304709024 | 0 | 0.025 | 1 | 0.106248001 | 2.252242349 | 0.024 | 0.003 | 1 | 0.243106933 | 0.201207365 |
| 4930547M16Rik | 0.249034808 | 2.154722595 | 0.019 | 0.004 | 1 | 0.106248001 | 2.252242349 | 0.024 | 0.003 | 1 | 0.249034808 | 0.201207365 |
| Gm5533 | 0.251644813 | 1.569760094 | 0.019 | 0.004 | 1 | 0.106248001 | 1.83720485 | 0.024 | 0.003 | 1 | 0.251644813 | 0.201207365 |
| Atp8b4 | 0.251644813 | 1.739685096 | 0.019 | 0.004 | 1 | 0.106248001 | 1.252242349 | 0.024 | 0.003 | 1 | 0.251644813 | 0.201207365 |
| Gm41821 | 0.343004841 | -1.1672055 | 0 | 0.017 | 1 | 0.106248001 | 1.029849928 | 0.024 | 0.003 | 1 | 0.343004841 | 0.201207365 |
| Gm50215 | 0.413080549 | -0.430239906 | 0 | 0.013 | 1 | 0.106248001 | 2.252242349 | 0.024 | 0.003 | 1 | 0.413080549 | 0.201207365 |
| D830013O20Rik | 0.413082433 | 0.154722595 | 0 | 0.013 | 1 | 0.106248001 | 2.252242349 | 0.024 | 0.003 | 1 | 0.413082433 | 0.201207365 |
| Gm48968 | 0.413082433 | 0.154722595 | 0 | 0.013 | 1 | 0.106248001 | 2.252242349 | 0.024 | 0.003 | 1 | 0.413082433 | 0.201207365 |
| Rhoh | 0.507281812 | 0.569760094 | 0.019 | 0.008 | 1 | 0.106248001 | 2.252242349 | 0.024 | 0.003 | 1 | 0.507281812 | 0.201207365 |
| Rundc3b | 0.736077083 | -0.260314904 | 0.019 | 0.013 | 1 | 0.106248001 | 0.83720485 | 0.024 | 0.003 | 1 | 0.736077083 | 0.201207365 |
| Phtf1os | 0.779443586 | -0.1672055 | 0.019 | 0.025 | 1 | 0.106248001 | 2.252242349 | 0.024 | 0.003 | 1 | 0.779443586 | 0.201207365 |
| Tgtp1 | 0.912039013 | -0.304709024 | 0.019 | 0.021 | 1 | 0.106248001 | 1.515276755 | 0.024 | 0.003 | 1 | 0.912039013 | 0.201207365 |
| Zdhhc12 | 0.106527683 | -0.582242999 | 0.057 | 0.14 | 1 | 0.354892874 | -0.101394606 | 0.024 | 0.061 | 1 | 0.354892874 | 0.201707218 |
| Edc3 | 0.106735703 | 0.915935735 | 0.283 | 0.195 | 1 | 0.231023557 | -0.389864059 | 0.171 | 0.259 | 1 | 0.231023557 | 0.202078895 |
| Glra1 | 0.701077609 | 0.216123139 | 0.038 | 0.051 | 1 | 0.106796283 | 1.944120054 | 0.098 | 0.041 | 1 | 0.701077609 | 0.202187121 |
| Rpusd4 | 0.106823257 | 1.232725107 | 0.208 | 0.127 | 1 | 0.26651401 | -0.80665134 | 0.073 | 0.133 | 1 | 0.26651401 | 0.202235306 |
| Cdca2 | 0.106866362 | -3.384436216 | 0.019 | 0.081 | 1 | 0.992196803 | 0.515276755 | 0.024 | 0.024 | 1 | 0.992196803 | 0.202312304 |
| Gm14326 | 0.106941238 | 0.631160639 | 0.264 | 0.165 | 1 | 0.199589714 | 1.170942247 | 0.244 | 0.177 | 1 | 0.199589714 | 0.202446048 |
| Slc25a14 | 0.107007396 | 1.211306123 | 0.17 | 0.097 | 1 | 0.224689994 | -0.917682652 | 0.073 | 0.14 | 1 | 0.224689994 | 0.202564209 |
| Gm10138 | 0.107044187 | -1.341703231 | 0.094 | 0.182 | 1 | 0.414637695 | -1.030691614 | 0.098 | 0.14 | 1 | 0.414637695 | 0.202629915 |
| Fam120c | 0.107097254 | 0.771393955 | 0.396 | 0.309 | 1 | 0.61410139 | -0.156563196 | 0.317 | 0.372 | 1 | 0.61410139 | 0.202724687 |
| 2810449D17Rik | 0.184615916 | 0.739685096 | 0.208 | 0.136 | 1 | 0.107198349 | -1.250257991 | 0.024 | 0.102 | 1 | 0.184615916 | 0.202905212 |
| Med7 | 0.120591814 | -0.319208594 | 0.302 | 0.436 | 1 | 0.107280133 | -0.127691349 | 0.268 | 0.44 | 1 | 0.120591814 | 0.203051239 |
| Gnb1l | 0.16898621 | 1.007881206 | 0.17 | 0.106 | 1 | 0.107293059 | -1.460475699 | 0.049 | 0.137 | 1 | 0.16898621 | 0.203074317 |
| Lgr4 | 0.107321736 | -0.695212116 | 0.623 | 0.691 | 1 | 0.577911967 | 0.340528882 | 0.732 | 0.741 | 1 | 0.577911967 | 0.203125518 |
| Nol4 | 0.249034808 | 2.154722595 | 0.019 | 0.004 | 1 | 0.107402092 | 4.159132945 | 0.049 | 0.014 | 1 | 0.249034808 | 0.203268975 |
| Nmnat3 | 0.273196595 | -0.678167419 | 0.057 | 0.11 | 1 | 0.107461096 | -1.08562729 | 0.049 | 0.143 | 1 | 0.273196595 | 0.203374304 |
| Slc25a22 | 0.107477907 | 0.984797593 | 0.208 | 0.127 | 1 | 0.277111276 | 0.804783372 | 0.171 | 0.113 | 1 | 0.277111276 | 0.203404313 |
| Nek4 | 0.533762035 | 0.261637799 | 0.057 | 0.085 | 1 | 0.107481494 | 1.408361551 | 0.195 | 0.109 | 1 | 0.533762035 | 0.203410716 |
| Nub1 | 0.68366932 | 0.254621165 | 0.604 | 0.623 | 1 | 0.107707856 | 0.590290109 | 0.683 | 0.628 | 1 | 0.68366932 | 0.203814731 |
| Akap17b | 0.182154182 | 0.776210972 | 0.075 | 0.034 | 1 | 0.107739591 | 1.630753972 | 0.146 | 0.075 | 1 | 0.182154182 | 0.203871362 |
| A530013C23Rik | 0.10787897 | 2.154722595 | 0.075 | 0.03 | 1 | 0.163071656 | 1.351778023 | 0.122 | 0.065 | 1 | 0.163071656 | 0.204120068 |
| Gm56492 | 0.10787897 | 2.154722595 | 0.075 | 0.03 | 1 | 0.832579923 | 0.100239256 | 0.049 | 0.041 | 1 | 0.832579923 | 0.204120068 |
| Hus1 | 0.668873349 | 0.806799291 | 0.113 | 0.097 | 1 | 0.107900337 | -1.250257991 | 0.049 | 0.137 | 1 | 0.668873349 | 0.204158192 |
| Phospho2 | 0.107916012 | 0.867841447 | 0.302 | 0.212 | 1 | 0.262782221 | -0.239610747 | 0.244 | 0.334 | 1 | 0.262782221 | 0.204186159 |
| Kat5 | 0.434418497 | 0.347367673 | 0.264 | 0.22 | 1 | 0.107996721 | -0.610254127 | 0.195 | 0.321 | 1 | 0.434418497 | 0.20433015 |
| Fam3a | 0.656728499 | 0.302279783 | 0.264 | 0.242 | 1 | 0.108069757 | -0.292078167 | 0.22 | 0.358 | 1 | 0.656728499 | 0.204460442 |
| Nrtn | 0.873681762 | -0.1672055 | 0.075 | 0.081 | 1 | 0.108151276 | -0.747757651 | 0.024 | 0.106 | 1 | 0.873681762 | 0.204605854 |
| Trnau1ap | 0.108160987 | -0.418744267 | 0.264 | 0.407 | 1 | 0.464232613 | 0.269520341 | 0.268 | 0.375 | 1 | 0.464232613 | 0.204623175 |
| Opn1mw | 0.108170944 | -1.652632327 | 0.019 | 0.081 | 1 | 0.15412976 | -1.622226769 | 0 | 0.048 | 1 | 0.15412976 | 0.204640935 |
| Slc2a12 | 0.108384928 | -0.727920455 | 0.094 | 0.191 | 1 | 0.372505086 | -0.410722664 | 0.024 | 0.058 | 1 | 0.372505086 | 0.205022564 |
| Smpd1 | 0.108556184 | 0.555260524 | 0.434 | 0.326 | 1 | 0.61602313 | 0.324135268 | 0.39 | 0.375 | 1 | 0.61602313 | 0.205327923 |
| Ttc4 | 0.108867077 | 0.772252958 | 0.34 | 0.233 | 1 | 0.201680637 | -0.151479837 | 0.22 | 0.321 | 1 | 0.201680637 | 0.205882113 |
| Nfat5 | 0.182427013 | 0.155222229 | 1 | 0.992 | 1 | 0.108951709 | 0.287747021 | 1 | 1 | 1 | 0.182427013 | 0.206032942 |
| Psen1 | 0.108961091 | 0.437122325 | 0.642 | 0.585 | 1 | 0.892551407 | 0.413065143 | 0.659 | 0.696 | 1 | 0.892551407 | 0.206049663 |
| Jpt2 | 0.108971771 | 0.617065809 | 0.377 | 0.267 | 1 | 0.310012832 | -0.196218152 | 0.22 | 0.304 | 1 | 0.310012832 | 0.206068695 |
| Arhgap1 | 0.109005772 | 0.440477077 | 0.434 | 0.318 | 1 | 0.627520641 | 0.402267793 | 0.366 | 0.352 | 1 | 0.627520641 | 0.206129286 |
| Gm6277 | 0.109042924 | -1.799473716 | 0.019 | 0.081 | 1 | 0.208799874 | -1.069685746 | 0 | 0.038 | 1 | 0.208799874 | 0.206195489 |
| Fbxo45 | 0.109126863 | 1.110328475 | 0.302 | 0.216 | 1 | 0.397451288 | -0.30718506 | 0.317 | 0.386 | 1 | 0.397451288 | 0.206345055 |
| Alg6 | 0.218983836 | -0.356239324 | 0.132 | 0.212 | 1 | 0.109129932 | -1.16279515 | 0.049 | 0.137 | 1 | 0.218983836 | 0.206350521 |
| Psmb4 | 0.109140453 | 0.277870826 | 0.943 | 0.869 | 1 | 0.741590199 | 0.176691316 | 0.732 | 0.846 | 1 | 0.741590199 | 0.206369268 |
| Snx27 | 0.109158632 | 0.435174054 | 0.604 | 0.564 | 1 | 0.452143 | -0.246211158 | 0.683 | 0.717 | 1 | 0.452143 | 0.206401658 |
| Rnaseh1 | 0.109193211 | -0.988235359 | 0.094 | 0.182 | 1 | 0.532538836 | -0.192542494 | 0.098 | 0.133 | 1 | 0.532538836 | 0.206463265 |
| Gramd1a | 0.295794076 | 0.545913352 | 0.415 | 0.364 | 1 | 0.109209654 | 0.683081704 | 0.561 | 0.491 | 1 | 0.295794076 | 0.20649256 |
| Brf1 | 0.109210562 | 0.733879474 | 0.358 | 0.258 | 1 | 0.411290737 | -0.307845483 | 0.39 | 0.468 | 1 | 0.411290737 | 0.206494178 |
| Clip2 | 0.109315321 | 0.739685096 | 0.208 | 0.123 | 1 | 0.773446479 | 0.410940095 | 0.195 | 0.225 | 1 | 0.773446479 | 0.206680802 |
| Camp | 0.243111734 | -0.652632327 | 0 | 0.025 | 1 | 0.109331674 | 2.048708955 | 0.122 | 0.058 | 1 | 0.243111734 | 0.206709933 |
| 1110017D15Rik | 0.10937249 | 1.154722595 | 0.113 | 0.055 | 1 | 0.676908292 | 0.136765132 | 0.098 | 0.078 | 1 | 0.676908292 | 0.206782639 |
| 8430426J06Rik | 0.429454595 | -0.508242418 | 0.019 | 0.042 | 1 | 0.109378456 | 1.396632258 | 0.098 | 0.041 | 1 | 0.429454595 | 0.206793266 |
| Gm57269 | 0.109475251 | -1.932740246 | 0.019 | 0.081 | 1 | 0.114472375 | -1.410722664 | 0 | 0.058 | 1 | 0.114472375 | 0.206965672 |
| Nudc | 0.109510236 | 0.320554842 | 0.717 | 0.691 | 1 | 0.129697137 | -0.389566012 | 0.634 | 0.703 | 1 | 0.129697137 | 0.207027981 |
| Zfp157 | 0.109675342 | 0.787940264 | 0.226 | 0.14 | 1 | 0.260373918 | -0.622226769 | 0.122 | 0.191 | 1 | 0.260373918 | 0.207322003 |
| Gm16751 | 0.109708284 | 1.984797593 | 0.075 | 0.03 | 1 | 0.69185719 | 0.589277336 | 0.024 | 0.038 | 1 | 0.69185719 | 0.207380661 |
| Chkb | 0.207079549 | 0.340278248 | 0.396 | 0.305 | 1 | 0.109772887 | -0.321224513 | 0.195 | 0.324 | 1 | 0.207079549 | 0.207495687 |
| Tmem201 | 0.109908306 | 0.859266711 | 0.17 | 0.093 | 1 | 0.74898193 | 0.100239256 | 0.098 | 0.116 | 1 | 0.74898193 | 0.207736777 |
| Gsta4 | 0.109966959 | 0.404700848 | 0.189 | 0.106 | 1 | 0.157606248 | 0.492376353 | 0.366 | 0.266 | 1 | 0.157606248 | 0.207841187 |
| Rnf170 | 0.495951482 | 0.382133091 | 0.302 | 0.263 | 1 | 0.109974961 | 0.721727632 | 0.341 | 0.229 | 1 | 0.495951482 | 0.20785543 |
| Entrep3 | 0.109988498 | 0.770058745 | 0.245 | 0.157 | 1 | 0.73395792 | 0.667279848 | 0.244 | 0.239 | 1 | 0.73395792 | 0.207879526 |
| Hsd17b11 | 0.122440176 | 0.535812762 | 0.528 | 0.432 | 1 | 0.110033143 | 0.712876715 | 0.512 | 0.437 | 1 | 0.122440176 | 0.207958993 |
| Grk3 | 0.120650931 | 1.011764641 | 0.245 | 0.169 | 1 | 0.110050678 | -0.715767537 | 0.22 | 0.365 | 1 | 0.120650931 | 0.207990205 |
| Fbn1 | 0.110207868 | 0.248778339 | 0.849 | 0.839 | 1 | 0.49326962 | 0.585286476 | 0.878 | 0.901 | 1 | 0.49326962 | 0.208269961 |
| Cnksr1 | 0.11028652 | -1.545717123 | 0 | 0.047 | 1 | 0.674180157 | 0.252242349 | 0.024 | 0.038 | 1 | 0.674180157 | 0.208409924 |
| Gm33424 | 0.11028652 | -1.545717123 | 0 | 0.047 | 1 | 0.455097252 | -0.16279515 | 0 | 0.014 | 1 | 0.455097252 | 0.208409924 |
| Tsbp1 | 0.11028652 | -1.545717123 | 0 | 0.047 | 1 | 0.154144895 | -1.410722664 | 0 | 0.048 | 1 | 0.154144895 | 0.208409924 |
| Arsj | 0.110304646 | -1.752168001 | 0 | 0.047 | 1 | 0.486825794 | -1.332720152 | 0.024 | 0.048 | 1 | 0.486825794 | 0.208442177 |
| B230303O12Rik | 0.110304646 | -1.652632327 | 0 | 0.047 | 1 | 0.733713289 | 0.614812428 | 0.049 | 0.038 | 1 | 0.733713289 | 0.208442177 |

| | | | | | | | | | | | | |
|---|---|---|---|---|---|---|---|---|---|---|---|---|
| Gm44700 | 0.110304646 | -1.752168001 | 0 | 0.047 | 1 | 0.802906569 | -0.16279515 | 0.049 | 0.058 | 1 | 0.802906569 | 0.208442177 |
| Tert | 0.110304646 | -1.652632327 | 0 | 0.047 | 1 | 0.743829142 | 0.136765132 | 0.024 | 0.034 | 1 | 0.743829142 | 0.208442177 |
| Sh3tc2 | 0.110304646 | -1.752168001 | 0 | 0.047 | 1 | 0.51919986 | 0.252242349 | 0 | 0.01 | 1 | 0.51919986 | 0.208442177 |
| Stard6 | 0.110304646 | -1.752168001 | 0 | 0.047 | 1 | 0.208830556 | -1.16279515 | 0 | 0.038 | 1 | 0.208830556 | 0.208442177 |
| Gm26728 | 0.110318743 | -1.932740246 | 0 | 0.047 | 1 | 0.461198188 | 0.883008539 | 0.098 | 0.068 | 1 | 0.461198188 | 0.208467261 |
| Gm10734 | 0.110318743 | -1.845277405 | 0 | 0.047 | 1 | 0.188573672 | -0.863234868 | 0 | 0.041 | 1 | 0.188573672 | 0.208467261 |
| Gm56933 | 0.110319146 | -1.752168001 | 0 | 0.047 | 1 | 0.746881137 | 0.83720485 | 0.024 | 0.017 | 1 | 0.746881137 | 0.208467978 |
| Pllp | 0.110319146 | -1.752168001 | 0 | 0.047 | 1 | 0.319641967 | -0.16279515 | 0 | 0.024 | 1 | 0.319641967 | 0.208467978 |
| Amdhd1 | 0.110319146 | -2.015202407 | 0 | 0.047 | 1 | 0.824088343 | 0.377773231 | 0.024 | 0.031 | 1 | 0.824088343 | 0.208467978 |
| 5031415H12Rik | 0.110319146 | -1.752168001 | 0 | 0.047 | 1 | 0.844641919 | 0.962735732 | 0.024 | 0.031 | 1 | 0.844641919 | 0.208467978 |
| B430212C06Rik | 0.110319146 | -1.752168001 | 0 | 0.047 | 1 | 0.188380068 | 1.614812428 | 0.049 | 0.017 | 1 | 0.188380068 | 0.208467978 |
| Zfp820 | 0.110319146 | 1.792152515 | 0.075 | 0.03 | 1 | 0.319635638 | -0.863234868 | 0 | 0.024 | 1 | 0.319635638 | 0.208467978 |
| Cfap53 | 0.110319146 | 1.792152515 | 0.075 | 0.03 | 1 | 0.155817442 | 0.962735732 | 0.098 | 0.044 | 1 | 0.155817442 | 0.208467979 |
| Efhd1 | 0.110328812 | -2.093204919 | 0 | 0.047 | 1 | 0.739624823 | -1.020776145 | 0.024 | 0.034 | 1 | 0.739624823 | 0.208485178 |
| Gm37245 | 0.110337271 | -2.1672055 | 0 | 0.047 | 1 | 0.26948834 | 1.515276755 | 0.024 | 0.007 | 1 | 0.26948834 | 0.208500228 |
| Gm3455 | 0.110337271 | -2.600164907 | 0 | 0.047 | 1 | 0.357854401 | -0.622226769 | 0 | 0.02 | 1 | 0.357854401 | 0.208500228 |
| Aipl1 | 0.110338882 | -2.932740246 | 0 | 0.047 | 1 | 0.267205177 | 2.252242349 | 0.024 | 0.007 | 1 | 0.267205177 | 0.208503094 |
| Iqgap3 | 0.11034452 | -2.845277405 | 0 | 0.047 | 1 | 0.455098275 | 0.515276755 | 0 | 0.014 | 1 | 0.455098275 | 0.208513128 |
| Azin2 | 0.479452994 | 0.154722595 | 0.226 | 0.182 | 1 | 0.110351423 | -1.192542494 | 0.049 | 0.137 | 1 | 0.479452994 | 0.208525409 |
| C330013E15Rik | 0.110352978 | -1.430239906 | 0 | 0.047 | 1 | 0.441381862 | 1.515276755 | 0.024 | 0.01 | 1 | 0.441381862 | 0.208528177 |
| 3830406C13Rik | 0.22106236 | 0.276713119 | 0.472 | 0.386 | 1 | 0.11040259 | -0.756976181 | 0.244 | 0.362 | 1 | 0.22106236 | 0.208616448 |
| Sars | 0.110407479 | 0.243773783 | 0.642 | 0.538 | 1 | 0.266190091 | -0.398308493 | 0.585 | 0.652 | 1 | 0.266190091 | 0.208625146 |
| Cyp4f16 | 0.241699822 | 0.884633431 | 0.208 | 0.144 | 1 | 0.110430567 | -0.771604393 | 0.073 | 0.174 | 1 | 0.241699822 | 0.208666223 |
| Uba6 | 0.110511831 | 0.954025245 | 0.472 | 0.407 | 1 | 0.19293543 | 0.270164257 | 0.683 | 0.563 | 1 | 0.19293543 | 0.208810797 |
| Isy1 | 0.110595603 | 0.385335523 | 0.547 | 0.453 | 1 | 0.191031293 | 0.316372687 | 0.683 | 0.642 | 1 | 0.191031293 | 0.208959819 |
| Gm30786 | 0.152568749 | 2.154722595 | 0.057 | 0.021 | 1 | 0.110710459 | 2.515276755 | 0.049 | 0.014 | 1 | 0.152568749 | 0.209164112 |
| Slfn5 | 0.251644813 | 1.154722595 | 0.019 | 0.004 | 1 | 0.110710459 | 2.515276755 | 0.049 | 0.014 | 1 | 0.251644813 | 0.209164112 |
| Gm50462 | 0.287761733 | -0.652632327 | 0 | 0.021 | 1 | 0.110710459 | 2.515276755 | 0.049 | 0.014 | 1 | 0.287761733 | 0.209164112 |
| Zc3h12d | 0.332821087 | 2.154722595 | 0.038 | 0.017 | 1 | 0.110710459 | 2.515276755 | 0.049 | 0.014 | 1 | 0.332821087 | 0.209164112 |
| D330050I16Rik | 0.340246113 | 1.417757001 | 0.038 | 0.017 | 1 | 0.110710459 | 2.515276755 | 0.049 | 0.014 | 1 | 0.340246113 | 0.209164112 |
| Gm57069 | 0.507281812 | 1.154722595 | 0.019 | 0.008 | 1 | 0.110710459 | 2.83720485 | 0.049 | 0.014 | 1 | 0.507281812 | 0.209164112 |
| Cbs | 0.510606779 | 0.8327945 | 0.019 | 0.008 | 1 | 0.110710459 | 2.515276755 | 0.049 | 0.014 | 1 | 0.510606779 | 0.209164112 |
| Pde8b | 0.722567081 | 1.8327945 | 0.019 | 0.013 | 1 | 0.110710459 | 2.83720485 | 0.049 | 0.014 | 1 | 0.722567081 | 0.209164112 |
| Rec8 | 0.732691937 | 1.154722595 | 0.019 | 0.013 | 1 | 0.110710459 | 2.83720485 | 0.049 | 0.014 | 1 | 0.732691937 | 0.209164112 |
| Kcnn1 | 0.767066103 | 0.569760094 | 0.038 | 0.03 | 1 | 0.110710459 | 2.515276755 | 0.049 | 0.014 | 1 | 0.767066103 | 0.209164112 |
| Mfsd7a | 0.779443586 | 0.154722595 | 0.019 | 0.025 | 1 | 0.110710459 | 2.515276755 | 0.049 | 0.014 | 1 | 0.779443586 | 0.209164112 |
| Megf10 | 0.922440134 | 2.624207878 | 0.038 | 0.042 | 1 | 0.110710459 | 2.515276755 | 0.049 | 0.014 | 1 | 0.922440134 | 0.209164112 |
| Myorg | 0.926212226 | 0.8327945 | 0.019 | 0.017 | 1 | 0.110710459 | 2.515276755 | 0.049 | 0.014 | 1 | 0.926212226 | 0.209164112 |
| Ncaph | 0.243322126 | 1.154722595 | 0.057 | 0.025 | 1 | 0.110720768 | 2.351778023 | 0.049 | 0.014 | 1 | 0.243322126 | 0.209182448 |
| Fam169b | 0.338381019 | 1.47665069 | 0.038 | 0.017 | 1 | 0.110720768 | 1.749742009 | 0.049 | 0.014 | 1 | 0.338381019 | 0.209182448 |
| Ddx60 | 0.987183403 | -1.015202407 | 0.057 | 0.055 | 1 | 0.110720768 | 2.489281546 | 0.049 | 0.014 | 1 | 0.987183403 | 0.209182448 |
| Sox12 | 0.110748107 | 1.739685096 | 0.094 | 0.042 | 1 | 0.567913408 | 0.721727632 | 0.049 | 0.075 | 1 | 0.567913408 | 0.20923107 |
| Eny2 | 0.110754085 | 0.253427462 | 0.887 | 0.839 | 1 | 0.157304082 | 0.303226278 | 0.878 | 0.887 | 1 | 0.157304082 | 0.209241702 |
| Pigq | 0.416596902 | 0.669295768 | 0.208 | 0.165 | 1 | 0.11081923 | -0.730479659 | 0.098 | 0.208 | 1 | 0.416596902 | 0.209357557 |
| Akap10 | 0.110939766 | 0.403951201 | 0.453 | 0.356 | 1 | 0.913715483 | 0.312678203 | 0.488 | 0.539 | 1 | 0.913715483 | 0.2095719 |
| Aaas | 0.880426804 | -0.19891436 | 0.151 | 0.157 | 1 | 0.110954678 | 0.737669176 | 0.146 | 0.072 | 1 | 0.880426804 | 0.209598416 |
| C030013C21Rik | 0.213025208 | -0.783876861 | 0.038 | 0.089 | 1 | 0.110962133 | -1.132421501 | 0.049 | 0.137 | 1 | 0.213025208 | 0.209611671 |
| Zfp105 | 0.111200679 | 1.517292674 | 0.113 | 0.055 | 1 | 0.467255202 | -0.410722664 | 0.073 | 0.109 | 1 | 0.467255202 | 0.210035768 |
| Golga1 | 0.111201032 | 0.73289593 | 0.453 | 0.369 | 1 | 0.849035995 | -0.107085718 | 0.439 | 0.437 | 1 | 0.849035995 | 0.210036394 |
| Frg1 | 0.11126552 | 0.407030594 | 0.623 | 0.538 | 1 | 0.862709186 | 0.47390632 | 0.463 | 0.549 | 1 | 0.862709186 | 0.210151023 |
| Abca8b | 0.930293511 | 0.154722595 | 0.208 | 0.208 | 1 | 0.111391937 | -0.795063366 | 0.049 | 0.14 | 1 | 0.930293511 | 0.21037571 |
| Nmd3 | 0.531566644 | 0.270199812 | 0.509 | 0.496 | 1 | 0.111439973 | -0.188330242 | 0.439 | 0.608 | 1 | 0.531566644 | 0.210461079 |
| Taok3 | 0.111443318 | 0.406261362 | 0.566 | 0.504 | 1 | 0.220718712 | -0.11208433 | 0.585 | 0.71 | 1 | 0.220718712 | 0.210467023 |
| 1700120C14Rik | 0.394320115 | 0.454282877 | 0.113 | 0.076 | 1 | 0.111480555 | 1.334704509 | 0.098 | 0.041 | 1 | 0.394320115 | 0.210533196 |
| C1qtnf12 | 0.111501294 | 1.100274811 | 0.17 | 0.097 | 1 | 0.673079312 | 0.410940095 | 0.146 | 0.123 | 1 | 0.673079312 | 0.210570049 |
| Ccdc152 | 0.111557832 | 2.640149422 | 0.075 | 0.03 | 1 | 0.548375203 | 0.883008539 | 0.073 | 0.051 | 1 | 0.548375203 | 0.210670514 |
| Setd1a | 0.111590708 | 0.739685096 | 0.34 | 0.254 | 1 | 0.253904528 | 0.501918372 | 0.488 | 0.433 | 1 | 0.253904528 | 0.210728931 |
| Zfp445 | 0.111771936 | 0.351119808 | 0.604 | 0.508 | 1 | 0.355770678 | -0.188330242 | 0.488 | 0.601 | 1 | 0.355770678 | 0.211050906 |
| Taf5l | 0.111790568 | 1.034428361 | 0.264 | 0.182 | 1 | 0.489530489 | 0.653983026 | 0.317 | 0.28 | 1 | 0.489530489 | 0.211084005 |
| Supt7l | 0.279353546 | 0.859266711 | 0.151 | 0.102 | 1 | 0.111947073 | -0.53202896 | 0.073 | 0.174 | 1 | 0.279353546 | 0.211361998 |
| Tbc1d22a | 0.112089434 | 0.709937752 | 0.358 | 0.275 | 1 | 0.444779624 | 0.539524301 | 0.39 | 0.358 | 1 | 0.444779624 | 0.211614826 |
| A430105J06Rik | 0.11213375 | 1.739685096 | 0.075 | 0.03 | 1 | 0.139521753 | -1.410722664 | 0 | 0.051 | 1 | 0.139521753 | 0.211693522 |
| Zfp772 | 0.11213375 | 1.962077517 | 0.075 | 0.03 | 1 | 0.362195044 | 1.252242349 | 0.049 | 0.024 | 1 | 0.362195044 | 0.211693522 |
| Ttc38 | 0.11213375 | 1.962077517 | 0.075 | 0.03 | 1 | 0.220910219 | 1.574170444 | 0.073 | 0.034 | 1 | 0.220910219 | 0.211693522 |
| 1700056N10Rik | 0.11213375 | 2.324647596 | 0.075 | 0.03 | 1 | 0.94215184 | 0.174239837 | 0.049 | 0.051 | 1 | 0.94215184 | 0.211693522 |
| 6030458C11Rik | 0.288368048 | 0.705415193 | 0.264 | 0.208 | 1 | 0.11216332 | -0.584258919 | 0.098 | 0.205 | 1 | 0.288368048 | 0.21174603 |
| Sat1 | 0.112276251 | -0.710664449 | 0.906 | 0.852 | 1 | 0.182302042 | -0.177601085 | 0.902 | 0.915 | 1 | 0.182302042 | 0.211946545 |
| Flvcr2 | 0.112392044 | -0.904171094 | 0.075 | 0.161 | 1 | 0.293427309 | 0.742047617 | 0.049 | 0.102 | 1 | 0.293427309 | 0.212152116 |
| Cenpe | 0.175864657 | -3.7032584 | 0 | 0.034 | 1 | 0.112404454 | 2.252242349 | 0.049 | 0.014 | 1 | 0.175864657 | 0.212174147 |
| Zcchc18 | 0.413082433 | 0.154722595 | 0 | 0.013 | 1 | 0.112404454 | 2.252242349 | 0.049 | 0.014 | 1 | 0.413082433 | 0.212174147 |
| Gm16833 | 0.762480493 | 0.739685096 | 0.038 | 0.03 | 1 | 0.112404454 | 2.252242349 | 0.049 | 0.014 | 1 | 0.762480493 | 0.212174147 |
| Kcnip3 | 0.209009104 | 1.739685096 | 0.038 | 0.013 | 1 | 0.112407187 | 2.614812428 | 0.049 | 0.014 | 1 | 0.209009104 | 0.212178998 |
| Tut1 | 0.591575238 | 0.223985257 | 0.226 | 0.195 | 1 | 0.112500192 | -0.177745492 | 0.122 | 0.246 | 1 | 0.591575238 | 0.212344091 |
| Zfp507 | 0.416250083 | -0.243826782 | 0.283 | 0.347 | 1 | 0.112544895 | -0.475678105 | 0.195 | 0.321 | 1 | 0.416250083 | 0.212423437 |
| 1700123M08Rik | 0.631876444 | 0.510866405 | 0.057 | 0.076 | 1 | 0.11266859 | 2.059597271 | 0.073 | 0.027 | 1 | 0.631876444 | 0.212642969 |
| Sec61b | 0.112711133 | 0.164905278 | 1 | 0.919 | 1 | 0.209677223 | -0.132748014 | 0.902 | 0.956 | 1 | 0.209677223 | 0.212718467 |
| Eef1d | 0.258286206 | 0.17440008 | 0.962 | 0.911 | 1 | 0.112746162 | -0.307100033 | 0.878 | 0.922 | 1 | 0.258286206 | 0.212780627 |
| Wfs1 | 0.115733083 | 0.865215978 | 0.151 | 0.081 | 1 | 0.112842528 | 1.404245443 | 0.146 | 0.075 | 1 | 0.115733083 | 0.21295162 |
| Mrps6 | 0.211721379 | -0.14585618 | 0.792 | 0.818 | 1 | 0.112876837 | 0.493581684 | 0.854 | 0.792 | 1 | 0.211721379 | 0.213012494 |
| Prdx5 | 0.435181175 | 0.191018724 | 0.943 | 0.864 | 1 | 0.113133613 | -0.264597431 | 0.78 | 0.911 | 1 | 0.435181175 | 0.213468012 |
| Bok | 0.113147793 | 2.055186921 | 0.113 | 0.055 | 1 | 0.52065088 | 1.574170444 | 0.049 | 0.031 | 1 | 0.52065088 | 0.213493163 |
| Tgif1 | 0.150443715 | -0.350512713 | 0.642 | 0.716 | 1 | 0.11324053 | -0.26713181 | 0.61 | 0.747 | 1 | 0.150443715 | 0.213657642 |
| Dgkg | 0.209009104 | 1.739685096 | 0.038 | 0.013 | 1 | 0.113256319 | 2.029849928 | 0.049 | 0.014 | 1 | 0.209009104 | 0.213685645 |
| Abtb1 | 0.113375819 | 1.695290976 | 0.094 | 0.042 | 1 | 0.696067731 | 0.699701326 | 0.073 | 0.058 | 1 | 0.696067731 | 0.213897562 |
| Pqlc3 | 0.277440276 | 0.781264199 | 0.245 | 0.186 | 1 | 0.113436814 | -1.101394606 | 0.098 | 0.198 | 1 | 0.277440276 | 0.214005717 |
| Cks1b | 0.11349598 | 0.387168681 | 0.434 | 0.343 | 1 | 0.551133349 | 0.304709769 | 0.366 | 0.345 | 1 | 0.551133349 | 0.214110623 |
| Sp3os | 0.955079974 | 0.43225657 | 0.208 | 0.212 | 1 | 0.11350284 | -0.659220976 | 0.098 | 0.208 | 1 | 0.955079974 | 0.214122785 |
| Srp9 | 0.194990823 | 0.229471462 | 0.887 | 0.898 | 1 | 0.113693594 | -0.268529093 | 0.805 | 0.925 | 1 | 0.194990823 | 0.214460954 |
| Zfp280b | 0.114023709 | 1.792152515 | 0.075 | 0.03 | 1 | 0.972816755 | 0.30115195 | 0.073 | 0.075 | 1 | 0.972816755 | 0.215046012 |
| Smim36 | 0.114023709 | 1.792152515 | 0.075 | 0.03 | 1 | 0.279285383 | 1.458693227 | 0.073 | 0.038 | 1 | 0.279285383 | 0.215046012 |
| St7 | 0.114080626 | 0.884270717 | 0.585 | 0.534 | 1 | 0.117997342 | -0.410722664 | 0.463 | 0.584 | 1 | 0.117997342 | 0.215146862 |
| Mtmr1 | 0.506518398 | -0.330704232 | 0.321 | 0.36 | 1 | 0.114105386 | -0.688863962 | 0.39 | 0.481 | 1 | 0.506518398 | 0.215190733 |
| Mettl5os | 0.932505382 | 1.154722595 | 0.019 | 0.021 | 1 | 0.114116566 | 2.029849928 | 0.049 | 0.014 | 1 | 0.932505382 | 0.21521054 |
| Gm57275 | 0.24309253 | -0.845277405 | 0 | 0.025 | 1 | 0.114118767 | 2.100239256 | 0.049 | 0.014 | 1 | 0.24309253 | 0.215214442 |
| Gm47512 | 0.265932462 | -0.932740246 | 0.019 | 0.055 | 1 | 0.114118767 | 2.100239256 | 0.049 | 0.014 | 1 | 0.265932462 | 0.215214442 |
| Gm28151 | 0.413082433 | 0.154722595 | 0 | 0.013 | 1 | 0.114118767 | 2.100239256 | 0.049 | 0.014 | 1 | 0.413082433 | 0.215214442 |
| Capza1 | 0.114164594 | 0.388545079 | 0.849 | 0.818 | 1 | 0.445725573 | -0.161092855 | 0.829 | 0.904 | 1 | 0.445725573 | 0.215295634 |
| Stk40 | 0.800403072 | 0.201516806 | 0.679 | 0.691 | 1 | 0.114363765 | -0.482535686 | 0.78 | 0.809 | 1 | 0.800403072 | 0.21564846 |

| | | | | | | | | | | | | |
|---|---|---|---|---|---|---|---|---|---|---|---|---|
| Gm35019 | 0.317289525 | 0.645708947 | 0.113 | 0.072 | 1 | 0.114453919 | 1.574170444 | 0.122 | 0.058 | 1 | 0.317289525 | 0.215808139 |
| Fbxo44 | 0.374932142 | -0.304709024 | 0.038 | 0.072 | 1 | 0.114472375 | -1.410722664 | 0 | 0.058 | 1 | 0.374932142 | 0.215840826 |
| Traf3ip2 | 0.114479402 | -0.889671525 | 0.038 | 0.11 | 1 | 0.477243365 | -0.309636539 | 0.098 | 0.14 | 1 | 0.477243365 | 0.215853271 |
| Gm49189 | 0.782077996 | 0.347367673 | 0.019 | 0.025 | 1 | 0.114496531 | -1.484723245 | 0 | 0.058 | 1 | 0.782077996 | 0.215883607 |
| Hdac10 | 0.991244785 | 0.417757001 | 0.038 | 0.038 | 1 | 0.114517465 | -1.686357106 | 0 | 0.058 | 1 | 0.991244785 | 0.215920681 |
| Gm14743 | 0.362626638 | -0.430239906 | 0.019 | 0.047 | 1 | 0.114517667 | -1.555112573 | 0 | 0.058 | 1 | 0.362626638 | 0.215921037 |
| Bik | 0.732691937 | 1.154722595 | 0.019 | 0.013 | 1 | 0.114535179 | -1.686357106 | 0 | 0.058 | 1 | 0.732691937 | 0.21595205 |
| Apobec1 | 0.767066103 | 0.154722595 | 0.038 | 0.03 | 1 | 0.114535782 | -2.16279515 | 0 | 0.058 | 1 | 0.767066103 | 0.215953119 |
| Trib3 | 0.41260394 | -0.889671525 | 0.057 | 0.089 | 1 | 0.114572012 | -1.917682652 | 0 | 0.058 | 1 | 0.41260394 | 0.216017279 |
| Pms1 | 0.156080464 | 1.526691372 | 0.094 | 0.047 | 1 | 0.114575434 | -2.020776145 | 0 | 0.058 | 1 | 0.156080464 | 0.216023338 |
| Sytl2 | 0.198145345 | -2.326404095 | 0.038 | 0.089 | 1 | 0.114582881 | -2.250257991 | 0 | 0.058 | 1 | 0.198145345 | 0.216036525 |
| Rsph10b | 0.114598312 | 1.039245377 | 0.132 | 0.068 | 1 | 0.407867291 | -0.622226769 | 0.024 | 0.055 | 1 | 0.407867291 | 0.216063851 |
| Cpq | 0.166009116 | 0.325780811 | 0.774 | 0.742 | 1 | 0.114631315 | -0.13220683 | 0.707 | 0.816 | 1 | 0.166009116 | 0.216122291 |
| U2surp | 0.114709755 | 0.262436412 | 0.887 | 0.797 | 1 | 0.591060159 | 0.236167586 | 0.854 | 0.918 | 1 | 0.591060159 | 0.216261182 |
| Itm2b | 0.114731574 | 0.124165128 | 1 | 0.996 | 1 | 0.26448423 | 0.329264769 | 1 | 0.997 | 1 | 0.26448423 | 0.216299814 |
| Gpr155 | 0.552726057 | 0.762405172 | 0.094 | 0.072 | 1 | 0.114758964 | 1.029849928 | 0.22 | 0.13 | 1 | 0.552726057 | 0.216348308 |
| Rnf167 | 0.550280002 | 0.166596695 | 0.623 | 0.61 | 1 | 0.114865855 | 0.935667489 | 0.585 | 0.601 | 1 | 0.550280002 | 0.216537545 |
| Oxsm | 0.115087754 | 1.324647596 | 0.113 | 0.055 | 1 | 0.94215184 | -0.101394606 | 0.049 | 0.051 | 1 | 0.94215184 | 0.216930317 |
| Heatr1 | 0.115138584 | -0.592511335 | 0.189 | 0.301 | 1 | 0.918513766 | 0.422167351 | 0.293 | 0.304 | 1 | 0.918513766 | 0.217020274 |
| Ngrn | 0.579734439 | 0.407488665 | 0.208 | 0.178 | 1 | 0.115158184 | -0.421529419 | 0.098 | 0.212 | 1 | 0.579734439 | 0.21705496 |
| Coro1a | 0.729311193 | 1.154722595 | 0.019 | 0.013 | 1 | 0.115159882 | 1.159132945 | 0.098 | 0.041 | 1 | 0.729311193 | 0.217057966 |
| Mus81 | 0.121444117 | 1.361173472 | 0.17 | 0.102 | 1 | 0.115173176 | 1.47463477 | 0.122 | 0.058 | 1 | 0.121444117 | 0.217081491 |
| Tardbp | 0.144316584 | 0.211306123 | 0.83 | 0.792 | 1 | 0.115192494 | -0.366272711 | 0.854 | 0.884 | 1 | 0.144316584 | 0.217115677 |
| Farp2 | 0.115318957 | 0.344099959 | 0.509 | 0.432 | 1 | 0.696941645 | 0.202489314 | 0.585 | 0.628 | 1 | 0.696941645 | 0.217339452 |
| Nt5dc3 | 0.115370192 | -0.40860989 | 0.491 | 0.64 | 1 | 0.758162372 | 0.173530373 | 0.78 | 0.713 | 1 | 0.758162372 | 0.217430103 |
| Wdr83 | 0.115388501 | 0.891688189 | 0.208 | 0.127 | 1 | 0.272324688 | -0.188330242 | 0.098 | 0.167 | 1 | 0.272324688 | 0.217462495 |
| Ddx27 | 0.629860647 | 0.179813576 | 0.585 | 0.559 | 1 | 0.115436553 | 0.731163212 | 0.537 | 0.505 | 1 | 0.629860647 | 0.217547509 |
| Snu13 | 0.535825954 | -0.174446143 | 0.925 | 0.911 | 1 | 0.115442773 | -0.298266095 | 0.829 | 0.904 | 1 | 0.535825954 | 0.217558512 |
| Rpl7l1 | 0.285250456 | 0.172511908 | 0.774 | 0.75 | 1 | 0.115615597 | -0.248910308 | 0.634 | 0.768 | 1 | 0.285250456 | 0.217864228 |
| Wrnip1 | 0.115720921 | 0.8327945 | 0.283 | 0.195 | 1 | 0.174213273 | 0.962735732 | 0.341 | 0.27 | 1 | 0.174213273 | 0.218050511 |
| Cyc1 | 0.115801638 | 0.278528796 | 0.792 | 0.661 | 1 | 0.682352103 | 0.261175474 | 0.683 | 0.659 | 1 | 0.682352103 | 0.218193257 |
| Gm56596 | 0.116607058 | 1.47665069 | 0.075 | 0.03 | 1 | 0.115836913 | 1.83720485 | 0.049 | 0.014 | 1 | 0.116607058 | 0.218255636 |
| 1700023H06Rik | 0.204653189 | 2.154722595 | 0.038 | 0.013 | 1 | 0.115836913 | 1.83720485 | 0.049 | 0.014 | 1 | 0.204653189 | 0.218255636 |
| Ifi44 | 0.312288875 | -1.875024749 | 0.019 | 0.051 | 1 | 0.115836913 | -0.622226769 | 0.049 | 0.014 | 1 | 0.312288875 | 0.218255636 |
| 4930481B07Rik | 0.343968402 | 1.154722595 | 0.038 | 0.017 | 1 | 0.115836913 | 1.614812428 | 0.049 | 0.014 | 1 | 0.343968402 | 0.218255636 |
| Gm3055 | 0.485993467 | 0.932330173 | 0.038 | 0.021 | 1 | 0.115836913 | 1.83720485 | 0.049 | 0.014 | 1 | 0.485993467 | 0.218255636 |
| Kntc1 | 0.860061344 | -0.260314904 | 0.038 | 0.042 | 1 | 0.115836913 | 1.83720485 | 0.049 | 0.014 | 1 | 0.860061344 | 0.218255636 |
| Doc2g | 0.929410101 | 0.347367673 | 0.019 | 0.017 | 1 | 0.115836913 | 1.422167351 | 0.049 | 0.014 | 1 | 0.929410101 | 0.218255636 |
| Zfp101 | 0.163005014 | 0.695290976 | 0.264 | 0.182 | 1 | 0.115886157 | -0.652179991 | 0.171 | 0.3 | 1 | 0.163005014 | 0.218342712 |
| Lrrc47 | 0.115949881 | 0.647762606 | 0.302 | 0.203 | 1 | 0.506582884 | 0.406570495 | 0.244 | 0.317 | 1 | 0.506582884 | 0.218455386 |
| Wdpcp | 0.116019511 | 0.569760094 | 0.321 | 0.225 | 1 | 0.96468578 | 0.374232873 | 0.244 | 0.253 | 1 | 0.96468578 | 0.218578496 |
| Vmn2r29 | 0.116121209 | 1.569760094 | 0.094 | 0.042 | 1 | 0.44682528 | -0.747757651 | 0.024 | 0.051 | 1 | 0.44682528 | 0.218758283 |
| Gm17092 | 0.116140444 | 1.115194231 | 0.208 | 0.131 | 1 | 0.616258969 | -0.448197369 | 0.073 | 0.096 | 1 | 0.616258969 | 0.218792286 |
| Mbd4 | 0.135791612 | 0.859266711 | 0.151 | 0.085 | 1 | 0.116202298 | 1.351778023 | 0.146 | 0.075 | 1 | 0.135791612 | 0.218901621 |
| Herc4 | 0.186476794 | -0.460613555 | 0.604 | 0.729 | 1 | 0.116221834 | 0.384573063 | 0.756 | 0.747 | 1 | 0.186476794 | 0.218936153 |
| Gm10648 | 0.11640703 | -0.904171094 | 0.094 | 0.186 | 1 | 0.372670599 | -0.825760163 | 0.146 | 0.201 | 1 | 0.372670599 | 0.219263463 |
| Gstm1 | 0.197389403 | 0.394673388 | 0.396 | 0.309 | 1 | 0.116473099 | 0.83720485 | 0.561 | 0.433 | 1 | 0.197389403 | 0.219380215 |
| Parp10 | 0.251644813 | 1.569760094 | 0.019 | 0.004 | 1 | 0.116518671 | 2.185128153 | 0.073 | 0.027 | 1 | 0.251644813 | 0.219460741 |
| Dhdh | 0.116572331 | 1.502645898 | 0.075 | 0.03 | 1 | 0.522093435 | 1.252242349 | 0.024 | 0.048 | 1 | 0.522093435 | 0.219555553 |
| Cenpk | 0.116592449 | -1.347777746 | 0.019 | 0.081 | 1 | 0.257308459 | -0.970150072 | 0 | 0.031 | 1 | 0.257308459 | 0.219591099 |
| Hsf4 | 0.116607058 | 1.47665069 | 0.075 | 0.03 | 1 | 0.468117655 | 0.377773231 | 0.049 | 0.085 | 1 | 0.468117655 | 0.219616909 |
| Gzmm | 0.116607058 | 1.47665069 | 0.075 | 0.03 | 1 | 0.718177571 | -0.250257991 | 0.073 | 0.089 | 1 | 0.718177571 | 0.219616909 |
| Mlst8 | 0.116607058 | 1.47665069 | 0.075 | 0.03 | 1 | 0.986965808 | 0.334704509 | 0.049 | 0.048 | 1 | 0.986965808 | 0.219616909 |
| Ahrr | 0.413082433 | -0.430239906 | 0 | 0.013 | 1 | 0.116717525 | 1.136765132 | 0.049 | 0.014 | 1 | 0.413082433 | 0.21981207 |
| Bloc1s4 | 0.116726277 | 0.599203872 | 0.434 | 0.339 | 1 | 0.167210778 | 0.131948115 | 0.244 | 0.399 | 1 | 0.167210778 | 0.21982753 |
| Zcwpw2 | 0.762452419 | 1.468831185 | 0.094 | 0.085 | 1 | 0.11675642 | 1.212713985 | 0.122 | 0.058 | 1 | 0.762452419 | 0.219880778 |
| Kif1a | 0.121757426 | 1.154722595 | 0.075 | 0.03 | 1 | 0.116764546 | 1.252242349 | 0.122 | 0.058 | 1 | 0.121757426 | 0.219895133 |
| Coq4 | 0.559398622 | 0.695290976 | 0.094 | 0.072 | 1 | 0.116859493 | -1.448197369 | 0.024 | 0.099 | 1 | 0.559398622 | 0.220062844 |
| Cdkn2d | 0.116889517 | -0.27154216 | 0.434 | 0.542 | 1 | 0.341355439 | 0.195098442 | 0.439 | 0.348 | 1 | 0.341355439 | 0.220115875 |
| Smim5 | 0.116916791 | -1.686579659 | 0.038 | 0.106 | 1 | 0.15414462 | -2.020776145 | 0 | 0.048 | 1 | 0.15414462 | 0.220164046 |
| Rrp9 | 0.116976796 | 0.569760094 | 0.396 | 0.292 | 1 | 0.623976962 | 0.145327145 | 0.268 | 0.229 | 1 | 0.623976962 | 0.220270022 |
| Plagl1 | 0.351994284 | -0.524789639 | 0.83 | 0.852 | 1 | 0.116988867 | -0.334150687 | 0.902 | 0.911 | 1 | 0.351994284 | 0.22029134 |
| Zfp677 | 0.485993467 | 0.932330173 | 0.038 | 0.021 | 1 | 0.117162876 | 2.059597271 | 0.073 | 0.027 | 1 | 0.485993467 | 0.220598612 |
| Zfp672 | 0.657994237 | 0.191248471 | 0.264 | 0.229 | 1 | 0.117163668 | -0.395455907 | 0.171 | 0.3 | 1 | 0.657994237 | 0.22060001 |
| Rims2 | 0.117237117 | 1.417757001 | 0.075 | 0.03 | 1 | 0.868757657 | 0.786578777 | 0.073 | 0.068 | 1 | 0.868757657 | 0.220729693 |
| Cadm4 | 0.243102132 | -1.545717123 | 0 | 0.025 | 1 | 0.117351913 | 1.252242349 | 0.098 | 0.041 | 1 | 0.243102132 | 0.220932354 |
| Zfp365 | 0.767066103 | 0.569760094 | 0.038 | 0.03 | 1 | 0.117351913 | 1.252242349 | 0.098 | 0.041 | 1 | 0.767066103 | 0.220932354 |
| Gm4221 | 0.190794962 | -0.942138944 | 0.113 | 0.186 | 1 | 0.117574766 | -0.495778434 | 0.098 | 0.205 | 1 | 0.190794962 | 0.221325706 |
| Gm24474 | 0.249034808 | 2.154722595 | 0.019 | 0.004 | 1 | 0.117592181 | 1.614812428 | 0.049 | 0.014 | 1 | 0.249034808 | 0.221356441 |
| Dscaml1 | 0.343000446 | -2.1672055 | 0 | 0.017 | 1 | 0.117592181 | 0.721727632 | 0.049 | 0.014 | 1 | 0.343000446 | 0.221356441 |
| Igf1os | 0.347593463 | 0.695290976 | 0.057 | 0.03 | 1 | 0.117592181 | 1.614812428 | 0.049 | 0.014 | 1 | 0.347593463 | 0.221356441 |
| Gnb3 | 0.503971666 | 1.569760094 | 0.019 | 0.008 | 1 | 0.117592181 | 1.100239256 | 0.049 | 0.014 | 1 | 0.503971666 | 0.221356441 |
| Ddx54 | 0.223374229 | 0.744846801 | 0.377 | 0.318 | 1 | 0.117664095 | 0.886786879 | 0.415 | 0.334 | 1 | 0.223374229 | 0.221483351 |
| Foxa1 | 0.343006305 | -0.1672055 | 0 | 0.017 | 1 | 0.117708875 | 1.19334866 | 0.146 | 0.075 | 1 | 0.343006305 | 0.221562371 |
| Rsl1 | 0.117865727 | 1.502645898 | 0.075 | 0.03 | 1 | 0.842422303 | 0.515276755 | 0.073 | 0.065 | 1 | 0.842422303 | 0.221839125 |
| Slc10a3 | 0.585469773 | 0.491757582 | 0.226 | 0.199 | 1 | 0.117916969 | -0.863234868 | 0.098 | 0.201 | 1 | 0.585469773 | 0.221929527 |
| Clcn6 | 0.346178337 | 0.20255112 | 0.34 | 0.271 | 1 | 0.117968714 | -0.601906784 | 0.22 | 0.341 | 1 | 0.346178337 | 0.222020811 |
| Gm37305 | 0.11800028 | -0.960754623 | 0.019 | 0.081 | 1 | 0.473132449 | -0.372248516 | 0.073 | 0.109 | 1 | 0.473132449 | 0.222076495 |
| Arfgap1 | 0.549344235 | 0.303866034 | 0.698 | 0.746 | 1 | 0.11806608 | -0.387896859 | 0.561 | 0.72 | 1 | 0.549344235 | 0.22219256 |
| H1f10 | 0.118136322 | -0.652632327 | 0.245 | 0.144 | 1 | 0.529627549 | 0.196747236 | 0.171 | 0.222 | 1 | 0.529627549 | 0.222316454 |
| Ppil1 | 0.11814633 | 0.739685096 | 0.245 | 0.157 | 1 | 0.990561755 | 0.225770138 | 0.171 | 0.174 | 1 | 0.990561755 | 0.222334105 |
| Mapt | 0.118391686 | 1.47665069 | 0.094 | 0.042 | 1 | 0.136446906 | 1.690363461 | 0.122 | 0.061 | 1 | 0.136446906 | 0.22276678 |
| Acad9 | 0.153355359 | 0.491757582 | 0.208 | 0.127 | 1 | 0.118473845 | -0.747757651 | 0.073 | 0.171 | 1 | 0.153355359 | 0.222911638 |
| Olfm2 | 0.11850161 | 0.739685096 | 0.075 | 0.03 | 1 | 0.193682915 | -1.970150072 | 0.049 | 0.113 | 1 | 0.193682915 | 0.222960589 |
| Bcas2 | 0.118543908 | 0.412054641 | 0.792 | 0.712 | 1 | 0.397992732 | -0.20859884 | 0.707 | 0.788 | 1 | 0.397992732 | 0.223035158 |
| Manea | 0.125086002 | 0.8327945 | 0.226 | 0.144 | 1 | 0.118635238 | 1.277777441 | 0.244 | 0.16 | 1 | 0.125086002 | 0.223196156 |
| Fam210b | 0.118947929 | -0.771276824 | 0.057 | 0.136 | 1 | 0.648024459 | 1.667279848 | 0.049 | 0.072 | 1 | 0.648024459 | 0.223747248 |
| Pgap3 | 0.118964647 | 0.932330173 | 0.132 | 0.068 | 1 | 0.424903183 | -0.300298674 | 0.073 | 0.116 | 1 | 0.424903183 | 0.223776707 |
| Zfp119b | 0.784690809 | 0.347367673 | 0.057 | 0.068 | 1 | 0.119109634 | 1.989207943 | 0.073 | 0.027 | 1 | 0.784690809 | 0.224032164 |
| Gm19426 | 0.119116409 | 1.306725688 | 0.075 | 0.03 | 1 | 0.745976859 | 0.377773231 | 0.024 | 0.034 | 1 | 0.745976859 | 0.224044098 |
| Slc25a29 | 0.119116409 | 1.306725688 | 0.075 | 0.03 | 1 | 0.611582762 | 1.589277336 | 0.073 | 0.055 | 1 | 0.611582762 | 0.224044098 |
| Dmtn | 0.119116409 | 1.306725688 | 0.075 | 0.03 | 1 | 0.914200263 | 0.377773231 | 0.024 | 0.027 | 1 | 0.914200263 | 0.224044098 |
| Spin2c | 0.119116409 | 1.306725688 | 0.075 | 0.03 | 1 | 0.450957104 | -0.80665134 | 0.024 | 0.051 | 1 | 0.450957104 | 0.224044098 |
| Gm17057 | 0.780566168 | -0.430239906 | 0.038 | 0.047 | 1 | 0.119143537 | 1.944120054 | 0.073 | 0.027 | 1 | 0.780566168 | 0.224091892 |
| Cyp4f17 | 0.119379369 | 0.59082171 | 0.302 | 0.203 | 1 | 0.524688878 | 0.252242349 | 0.146 | 0.205 | 1 | 0.524688878 | 0.224507305 |
| Rrnad1 | 0.968244127 | 0.258558406 | 0.226 | 0.229 | 1 | 0.119479267 | -0.798383724 | 0.122 | 0.229 | 1 | 0.968244127 | 0.224683238 |
| Gnl3 | 0.207644378 | 0.265606701 | 0.925 | 0.843 | 1 | 0.1195229 | 0.372465947 | 0.854 | 0.823 | 1 | 0.207644378 | 0.224760076 |
| Fam13c | 0.119532729 | 1.039245377 | 0.208 | 0.123 | 1 | 0.975971463 | 0.252242349 | 0.171 | 0.174 | 1 | 0.975971463 | 0.224777385 |

| | | | | | | | | | | | | |
|---|---|---|---|---|---|---|---|---|---|---|---|---|
| Gpr161 | 0.347746365 | 0.932330173 | 0.038 | 0.017 | 1 | 0.119573021 | 1.071670103 | 0.098 | 0.041 | 1 | 0.347746365 | 0.224848334 |
| Zdhhc3 | 0.300741011 | 0.33529484 | 0.566 | 0.53 | 1 | 0.119603021 | 0.592092352 | 0.634 | 0.553 | 1 | 0.300741011 | 0.224901159 |
| Trmt5 | 0.119702178 | 1.067259754 | 0.132 | 0.068 | 1 | 0.868524474 | -0.16279515 | 0.049 | 0.055 | 1 | 0.868524474 | 0.225075745 |
| Minpp1 | 0.861815789 | 0.247831999 | 0.415 | 0.411 | 1 | 0.119765466 | -0.152741486 | 0.341 | 0.532 | 1 | 0.861815789 | 0.225187165 |
| Rbm44 | 0.883533598 | 0.776210972 | 0.038 | 0.034 | 1 | 0.119812863 | 1.930314254 | 0.073 | 0.027 | 1 | 0.883533598 | 0.225270604 |
| Zfp763 | 0.119813369 | 1.280253477 | 0.075 | 0.03 | 1 | 0.644124708 | -0.116991461 | 0.049 | 0.068 | 1 | 0.644124708 | 0.225271494 |
| Dok5 | 0.119962068 | -2.254668341 | 0.057 | 0.127 | 1 | 0.664414647 | -0.863234868 | 0.024 | 0.038 | 1 | 0.664414647 | 0.225533239 |
| Lcn3 | 0.119972815 | 1.029191713 | 0.226 | 0.14 | 1 | 0.231627527 | -3.069685746 | 0 | 0.034 | 1 | 0.231627527 | 0.225552154 |
| Chuk | 0.169610313 | 0.401362562 | 0.642 | 0.585 | 1 | 0.120013674 | 0.477978213 | 0.829 | 0.737 | 1 | 0.169610313 | 0.225624066 |
| Kcng2 | 0.120109229 | 1.237184755 | 0.113 | 0.055 | 1 | 0.677889541 | 0.944120054 | 0.098 | 0.082 | 1 | 0.677889541 | 0.225792231 |
| Zfp69 | 0.960852989 | 0.47665069 | 0.057 | 0.059 | 1 | 0.120226446 | -1.192542494 | 0.049 | 0.133 | 1 | 0.960852989 | 0.225998494 |
| Sh2b3 | 0.120249488 | 0.75033234 | 0.226 | 0.14 | 1 | 0.405747493 | -0.314798244 | 0.146 | 0.205 | 1 | 0.405747493 | 0.226039037 |
| Lrrfip1 | 0.120266677 | -0.164737244 | 0.792 | 0.648 | 1 | 0.448724297 | -0.219990552 | 0.854 | 0.85 | 1 | 0.448724297 | 0.226069281 |
| Mcm3 | 0.120288956 | 0.920257341 | 0.189 | 0.11 | 1 | 0.497720212 | -0.19841906 | 0.073 | 0.109 | 1 | 0.497720212 | 0.226108478 |
| Thoc6 | 0.120397949 | 0.60628597 | 0.415 | 0.322 | 1 | 0.290814973 | -0.238083278 | 0.293 | 0.396 | 1 | 0.290814973 | 0.226300232 |
| Trib2 | 0.120398369 | 0.337586652 | 0.434 | 0.326 | 1 | 0.807514533 | 0.729145104 | 0.317 | 0.348 | 1 | 0.807514533 | 0.226300971 |
| Slc26a7 | 0.377182288 | -0.237594828 | 0.038 | 0.072 | 1 | 0.120452595 | 2.515276755 | 0.073 | 0.027 | 1 | 0.377182288 | 0.226396363 |
| Mast1 | 0.120481862 | 0.2967416 | 0.075 | 0.03 | 1 | 0.195508378 | 0.721727632 | 0.049 | 0.017 | 1 | 0.195508378 | 0.226447845 |
| Lrrcc1 | 0.120498215 | 1.009292155 | 0.283 | 0.203 | 1 | 0.545668406 | 0.941902228 | 0.22 | 0.188 | 1 | 0.545668406 | 0.226476609 |
| Rgs11 | 0.120524599 | 1.247831999 | 0.113 | 0.055 | 1 | 0.450957104 | -0.410722664 | 0.024 | 0.051 | 1 | 0.450957104 | 0.226523018 |
| B230354K17Rik | 0.120636574 | 0.747547213 | 0.283 | 0.195 | 1 | 0.882841115 | 0.235168836 | 0.195 | 0.212 | 1 | 0.882841115 | 0.226719964 |
| Magee1 | 0.120651803 | 0.8327945 | 0.189 | 0.11 | 1 | 0.615362039 | -0.332720152 | 0.146 | 0.174 | 1 | 0.615362039 | 0.226746749 |
| Nfatc2 | 0.120669084 | -1.432060342 | 0.547 | 0.619 | 1 | 0.780075743 | -0.607097244 | 0.634 | 0.621 | 1 | 0.780075743 | 0.22677714 |
| Wipi1 | 0.135966807 | -0.120457241 | 0.547 | 0.678 | 1 | 0.120952844 | -0.484723245 | 0.39 | 0.532 | 1 | 0.135966807 | 0.227276097 |
| Kbtbd7 | 0.322108007 | -0.52334931 | 0.075 | 0.123 | 1 | 0.121030988 | -0.890715605 | 0.049 | 0.137 | 1 | 0.322108007 | 0.227413477 |
| Tedc1 | 0.663401681 | -0.1672055 | 0.019 | 0.03 | 1 | 0.121121415 | 1.83720485 | 0.073 | 0.027 | 1 | 0.663401681 | 0.227572434 |
| Ank3 | 0.162614425 | 0.75033234 | 0.189 | 0.114 | 1 | 0.121126941 | 1.437274243 | 0.293 | 0.198 | 1 | 0.162614425 | 0.227582146 |
| Gpr107 | 0.134509076 | -0.400796128 | 0.396 | 0.53 | 1 | 0.12113373 | -0.443889528 | 0.585 | 0.676 | 1 | 0.134509076 | 0.22759408 |
| Tmem129 | 0.322747099 | 0.533234218 | 0.189 | 0.136 | 1 | 0.121133967 | 1.164779508 | 0.22 | 0.133 | 1 | 0.322747099 | 0.227594495 |
| Tarbp2 | 0.323101243 | 0.627475592 | 0.226 | 0.174 | 1 | 0.121561696 | -1.037264268 | 0.049 | 0.133 | 1 | 0.323101243 | 0.228346146 |
| Adam30 | 0.659862295 | 0.306725688 | 0.132 | 0.11 | 1 | 0.121571703 | 0.656632604 | 0.244 | 0.143 | 1 | 0.659862295 | 0.228363727 |
| Tmem127 | 0.121614177 | 0.634715536 | 0.528 | 0.458 | 1 | 0.341154864 | -0.35680577 | 0.537 | 0.58 | 1 | 0.341154864 | 0.228438346 |
| Atxn2l | 0.1216684 | 0.384204441 | 0.698 | 0.631 | 1 | 0.359963606 | 0.132556976 | 0.829 | 0.782 | 1 | 0.359963606 | 0.228533601 |
| Ccdc9 | 0.178788961 | 0.980224864 | 0.34 | 0.275 | 1 | 0.121690275 | 0.626637864 | 0.463 | 0.355 | 1 | 0.178788961 | 0.228572027 |
| ENSMUSG00000121596 | 0.121693932 | 1.377115016 | 0.094 | 0.042 | 1 | 0.479272663 | 1.071670103 | 0.073 | 0.048 | 1 | 0.479272663 | 0.228578452 |
| Garin5a | 0.121693932 | 1.377115016 | 0.094 | 0.042 | 1 | 0.188517184 | -1.069685746 | 0 | 0.041 | 1 | 0.188517184 | 0.228578452 |
| Zfp747 | 0.121693932 | 1.377115016 | 0.094 | 0.042 | 1 | 0.37936884 | -0.707115666 | 0.049 | 0.089 | 1 | 0.37936884 | 0.228578452 |
| Gm10658 | 0.121693932 | 1.377115016 | 0.094 | 0.042 | 1 | 0.957194286 | 1.071670103 | 0.049 | 0.048 | 1 | 0.957194286 | 0.228578452 |
| C230013L11Rik | 0.121693932 | 1.377115016 | 0.094 | 0.042 | 1 | 0.229188679 | -0.917682652 | 0.024 | 0.075 | 1 | 0.229188679 | 0.228578452 |
| Nanos1 | 0.121693932 | 1.377115016 | 0.094 | 0.042 | 1 | 0.33375249 | 0.515276755 | 0.146 | 0.096 | 1 | 0.33375249 | 0.228578452 |
| Pgp | 0.121763301 | -0.582242999 | 0.34 | 0.449 | 1 | 0.727045216 | 0.206438659 | 0.439 | 0.406 | 1 | 0.727045216 | 0.228700301 |
| Acpp | 0.413080549 | -0.1672055 | 0 | 0.013 | 1 | 0.121823396 | 0.699701326 | 0.098 | 0.041 | 1 | 0.413080549 | 0.228805853 |
| Rhebl1 | 0.121958554 | 0.725879296 | 0.132 | 0.068 | 1 | 0.654447162 | -0.890715605 | 0.073 | 0.092 | 1 | 0.654447162 | 0.229043219 |
| Ooep | 0.859433366 | 0.166795427 | 0.283 | 0.275 | 1 | 0.122137832 | -0.814871847 | 0.317 | 0.427 | 1 | 0.859433366 | 0.229358014 |
| 2700097O09Rik | 0.681210884 | 0.338434548 | 0.208 | 0.254 | 1 | 0.122185566 | -0.677368323 | 0.171 | 0.287 | 1 | 0.681210884 | 0.22944182 |
| Bglap2 | 0.92348286 | 0.662326213 | 0.132 | 0.131 | 1 | 0.12242413 | -0.484723245 | 0.098 | 0.041 | 1 | 0.92348286 | 0.229860593 |
| Gm49085 | 0.241893059 | 1.017219071 | 0.075 | 0.038 | 1 | 0.122479108 | 1.83720485 | 0.073 | 0.027 | 1 | 0.241893059 | 0.229957084 |
| Pip4p1 | 0.543672408 | 0.32730205 | 0.528 | 0.525 | 1 | 0.122613152 | -0.697384128 | 0.293 | 0.399 | 1 | 0.543672408 | 0.230192318 |
| Ccnt2 | 0.836082248 | 0.111510528 | 0.604 | 0.653 | 1 | 0.122992861 | 0.529420937 | 0.732 | 0.706 | 1 | 0.836082248 | 0.230858477 |
| Pgk1 | 0.12301025 | -0.341510987 | 0.774 | 0.818 | 1 | 0.337558928 | -0.172642936 | 0.707 | 0.802 | 1 | 0.337558928 | 0.230888979 |
| 9530018F02Rik | 0.334518863 | 1.154722595 | 0.057 | 0.03 | 1 | 0.123156684 | 1.699701326 | 0.073 | 0.027 | 1 | 0.334518863 | 0.231145799 |
| Ppip5k2 | 0.123159182 | 0.8327945 | 0.358 | 0.275 | 1 | 0.607235861 | 0.125330237 | 0.366 | 0.334 | 1 | 0.607235861 | 0.23115018 |
| Gm45767 | 0.12319145 | 1.154722595 | 0.113 | 0.055 | 1 | 0.130736995 | 1.635570989 | 0.122 | 0.061 | 1 | 0.130736995 | 0.231206767 |
| Duxbl1 | 0.123225666 | 1.084333267 | 0.132 | 0.068 | 1 | 0.990690158 | 0.100239256 | 0.049 | 0.048 | 1 | 0.990690158 | 0.231266767 |
| Aldoa | 0.589878459 | -0.278901802 | 0.981 | 0.941 | 1 | 0.123395445 | -0.282987061 | 0.878 | 0.959 | 1 | 0.589878459 | 0.231564454 |
| Gnrh1 | 0.123445956 | -1.015202407 | 0.038 | 0.106 | 1 | 0.133147141 | -0.863234868 | 0.049 | 0.133 | 1 | 0.133147141 | 0.231653008 |
| 4933406C10Rik | 0.123466297 | -1.237594828 | 0.057 | 0.131 | 1 | 0.201398 | -1.305753104 | 0.049 | 0.113 | 1 | 0.201398 | 0.231688668 |
| Pepd | 0.302468607 | 0.552058092 | 0.453 | 0.398 | 1 | 0.123548516 | -0.26622474 | 0.39 | 0.529 | 1 | 0.302468607 | 0.231832795 |
| Tmem219 | 0.123575184 | 0.683416876 | 0.302 | 0.208 | 1 | 0.772413065 | 0.364717078 | 0.268 | 0.253 | 1 | 0.772413065 | 0.231879541 |
| D730045B01Rik | 0.584341866 | 0.280253477 | 0.019 | 0.034 | 1 | 0.123836867 | 1.458693227 | 0.073 | 0.027 | 1 | 0.584341866 | 0.232338164 |
| Dkk2 | 0.123939671 | -1.870812497 | 0.038 | 0.106 | 1 | 0.451651068 | 2.685201756 | 0.073 | 0.048 | 1 | 0.451651068 | 0.232518299 |
| Mgat2 | 0.486750697 | 0.194371671 | 0.698 | 0.653 | 1 | 0.124006553 | -0.441511178 | 0.683 | 0.73 | 1 | 0.486750697 | 0.232635481 |
| Coq3 | 0.124028112 | 0.967095592 | 0.226 | 0.144 | 1 | 0.564629923 | -0.467649732 | 0.244 | 0.276 | 1 | 0.564629923 | 0.232673251 |
| Agk | 0.124029785 | 0.340278248 | 0.358 | 0.25 | 1 | 0.657458079 | 0.338953982 | 0.195 | 0.249 | 1 | 0.657458079 | 0.232676182 |
| Tspan7 | 0.124055172 | -1.20517335 | 0.377 | 0.436 | 1 | 0.871478306 | -0.150722318 | 0.366 | 0.396 | 1 | 0.871478306 | 0.232720659 |
| Cmc2 | 0.124109487 | 0.898382842 | 0.245 | 0.161 | 1 | 0.994672181 | 0.292884334 | 0.098 | 0.099 | 1 | 0.994672181 | 0.23281581 |
| Lsm2 | 0.134019479 | 0.403401708 | 0.415 | 0.301 | 1 | 0.124125121 | -0.484723245 | 0.268 | 0.392 | 1 | 0.134019479 | 0.232843196 |
| Klhl8 | 0.124270023 | 1.095828906 | 0.17 | 0.097 | 1 | 0.837677854 | -0.364429011 | 0.098 | 0.106 | 1 | 0.837677854 | 0.233097007 |
| Gm40438 | 0.124272867 | -2.27154216 | 0.019 | 0.076 | 1 | 0.364441285 | -1.410722664 | 0.024 | 0.058 | 1 | 0.364441285 | 0.233101988 |
| Adrm1b | 0.124316978 | 1.154722595 | 0.113 | 0.055 | 1 | 0.475031087 | 1.159132945 | 0.073 | 0.048 | 1 | 0.475031087 | 0.233179246 |
| Gm30551 | 0.12443947 | 0.47665069 | 0.075 | 0.03 | 1 | 0.319635005 | -0.970150072 | 0 | 0.024 | 1 | 0.319635005 | 0.233393759 |
| Mroh5 | 0.124439896 | 0.669295768 | 0.075 | 0.03 | 1 | 0.231591308 | -0.747757651 | 0 | 0.034 | 1 | 0.231591308 | 0.233394505 |
| Fabp4 | 0.756189816 | 1.377115016 | 0.057 | 0.047 | 1 | 0.12454029 | 1.000703582 | 0.073 | 0.027 | 1 | 0.756189816 | 0.233570296 |
| Lysmd1 | 0.124685691 | 1.280253477 | 0.094 | 0.042 | 1 | 0.665756646 | 0.377773231 | 0.049 | 0.068 | 1 | 0.665756646 | 0.23382486 |
| Gm19744 | 0.124685691 | 1.280253477 | 0.094 | 0.042 | 1 | 0.369056111 | -0.863234868 | 0.024 | 0.058 | 1 | 0.369056111 | 0.23382486 |
| Ghitm | 0.124732935 | 0.246949887 | 0.906 | 0.805 | 1 | 0.74380693 | -0.116430646 | 0.829 | 0.846 | 1 | 0.74380693 | 0.233907565 |
| Mfap3 | 0.124741777 | 0.396695385 | 0.453 | 0.352 | 1 | 0.239448593 | -0.233184478 | 0.341 | 0.451 | 1 | 0.239448593 | 0.233923044 |
| Ciao2b | 0.124792878 | 0.678284551 | 0.491 | 0.428 | 1 | 0.506328041 | -0.16279515 | 0.39 | 0.444 | 1 | 0.506328041 | 0.234012493 |
| Ctbs | 0.124872118 | -0.7032584 | 0.189 | 0.288 | 1 | 0.145821061 | -0.385187572 | 0.122 | 0.229 | 1 | 0.145821061 | 0.23415119 |
| Pcbd2 | 0.124903355 | -0.290062248 | 0.321 | 0.436 | 1 | 0.672699568 | 0.262734723 | 0.341 | 0.324 | 1 | 0.672699568 | 0.234205863 |
| Klc3 | 0.330103143 | -0.368839361 | 0.038 | 0.076 | 1 | 0.124928916 | -1.020776145 | 0.073 | 0.164 | 1 | 0.330103143 | 0.234250598 |
| Gtf2h3 | 0.124945893 | 0.633890431 | 0.321 | 0.22 | 1 | 0.86719888 | 0.146889349 | 0.195 | 0.212 | 1 | 0.86719888 | 0.234280311 |
| Opa1 | 0.331042804 | 0.186712709 | 0.434 | 0.564 | 1 | 0.125018666 | -0.328260503 | 0.39 | 0.512 | 1 | 0.331042804 | 0.234407664 |
| Gm13830 | 0.12517633 | 1.154722595 | 0.094 | 0.042 | 1 | 0.751853856 | 0.766815522 | 0.073 | 0.061 | 1 | 0.751853856 | 0.234683546 |
| Tulp3 | 0.125261407 | -0.352014739 | 0.472 | 0.648 | 1 | 0.671012671 | 0.278237558 | 0.439 | 0.539 | 1 | 0.671012671 | 0.234832393 |
| 6330403L08Rik | 0.125262144 | 1.002719501 | 0.113 | 0.055 | 1 | 0.464327149 | -0.448197369 | 0.073 | 0.109 | 1 | 0.464327149 | 0.234833683 |
| Dtymk | 0.125370892 | 0.380282295 | 0.623 | 0.555 | 1 | 0.249493466 | -0.121488176 | 0.366 | 0.498 | 1 | 0.249493466 | 0.235023924 |
| Wfdc18 | 0.393142571 | -0.1672055 | 0.377 | 0.297 | 1 | 0.125400188 | 0.86891371 | 0.268 | 0.174 | 1 | 0.393142571 | 0.235075169 |
| Exo5 | 0.12560134 | 0.865215978 | 0.132 | 0.068 | 1 | 0.322799813 | 1.029849928 | 0.122 | 0.078 | 1 | 0.322799813 | 0.235426984 |
| Zfp286 | 0.127470223 | 1.154722595 | 0.094 | 0.042 | 1 | 0.125604778 | 0.667279848 | 0.171 | 0.092 | 1 | 0.127470223 | 0.235432997 |
| Sike1 | 0.139097264 | 0.395730694 | 0.377 | 0.267 | 1 | 0.125694613 | -0.180717058 | 0.244 | 0.396 | 1 | 0.139097264 | 0.235590091 |
| Gm15983 | 0.125780136 | 2.154722595 | 0.094 | 0.042 | 1 | 0.934943043 | 0.100239256 | 0.049 | 0.051 | 1 | 0.934943043 | 0.23573963 |
| 2810032G03Rik | 0.125784675 | -1.652632327 | 0.019 | 0.076 | 1 | 0.257325989 | -0.484723245 | 0 | 0.031 | 1 | 0.257325989 | 0.235747565 |
| Mydgf | 0.312155989 | -0.108311811 | 0.736 | 0.805 | 1 | 0.125795363 | -0.385187572 | 0.585 | 0.669 | 1 | 0.312155989 | 0.235766253 |
| Tmsb15b1 | 0.786228092 | 0.635348436 | 0.17 | 0.161 | 1 | 0.125821607 | -0.622226769 | 0.049 | 0.137 | 1 | 0.786228092 | 0.235812137 |
| Pi4kb | 0.125933135 | 0.625794129 | 0.396 | 0.292 | 1 | 0.752510322 | 0.130612905 | 0.439 | 0.43 | 1 | 0.752510322 | 0.236007115 |
| Fam118b | 0.125951071 | -1.015202407 | 0.208 | 0.297 | 1 | 0.713813523 | 0.126711467 | 0.268 | 0.311 | 1 | 0.713813523 | 0.23603847 |
| 0610040F04Rik | 0.1262877 | -2.459987249 | 0.019 | 0.076 | 1 | 0.811288594 | -1.050320421 | 0.049 | 0.058 | 1 | 0.811288594 | 0.236626816 |

| | | | | | | | | | | | | |
|---|---|---|---|---|---|---|---|---|---|---|---|---|
| Slc52a2 | 0.1975444 | 0.8327945 | 0.132 | 0.076 | 1 | 0.126329058 | -1.332720152 | 0 | 0.055 | 1 | 0.1975444 | 0.236699085 |
| Gm57071 | 0.451976832 | -0.330704232 | 0.075 | 0.11 | 1 | 0.126329058 | -1.410722664 | 0 | 0.055 | 1 | 0.451976832 | 0.236699085 |
| Neu3 | 0.287773478 | -0.430239906 | 0 | 0.021 | 1 | 0.126352479 | -1.410722664 | 0 | 0.055 | 1 | 0.287773478 | 0.23674001 |
| Tincr | 0.437537757 | 0.984797593 | 0.057 | 0.034 | 1 | 0.126352479 | -1.410722664 | 0 | 0.055 | 1 | 0.437537757 | 0.23674001 |
| Rftn2 | 0.126362499 | 0.554163239 | 0.642 | 0.534 | 1 | 0.961594568 | 0.200480921 | 0.463 | 0.57 | 1 | 0.961594568 | 0.236757516 |
| Mcf2l | 0.126367001 | 1.247831999 | 0.094 | 0.042 | 1 | 0.208857401 | -2.020776145 | 0 | 0.038 | 1 | 0.208857401 | 0.236765384 |
| C430014B12Rik | 0.317243945 | 0.891688189 | 0.075 | 0.042 | 1 | 0.126372777 | -1.484723245 | 0 | 0.055 | 1 | 0.317243945 | 0.236775476 |
| Otub2 | 0.605896064 | -0.1672055 | 0.038 | 0.055 | 1 | 0.126372777 | -1.686357106 | 0 | 0.055 | 1 | 0.605896064 | 0.236775476 |
| Tmem267 | 0.392962724 | 0.652222254 | 0.075 | 0.047 | 1 | 0.126389952 | -1.686357106 | 0 | 0.055 | 1 | 0.392962724 | 0.236805484 |
| Myadml2os | 0.601939331 | 1.47665069 | 0.038 | 0.025 | 1 | 0.126402665 | -1.686357106 | 0 | 0.055 | 1 | 0.601939331 | 0.236827697 |
| Igsf9b | 0.431044319 | -0.1672055 | 0.019 | 0.042 | 1 | 0.126414932 | -1.80665134 | 0 | 0.055 | 1 | 0.431044319 | 0.236849129 |
| Gm48321 | 0.569866135 | -0.1672055 | 0.019 | 0.034 | 1 | 0.126422738 | -1.863234868 | 0 | 0.055 | 1 | 0.569866135 | 0.236862768 |
| Gm56783 | 0.150219314 | -1.304709024 | 0 | 0.038 | 1 | 0.126453962 | -1.250257991 | 0 | 0.055 | 1 | 0.150219314 | 0.236917319 |
| Shroom1 | 0.246023494 | 0.776210972 | 0.075 | 0.038 | 1 | 0.126453962 | -1.250257991 | 0 | 0.055 | 1 | 0.246023494 | 0.236917319 |
| Gm6710 | 0.312692399 | 1.017219071 | 0.075 | 0.042 | 1 | 0.126453962 | -1.250257991 | 0 | 0.055 | 1 | 0.312692399 | 0.236917319 |
| Gm56489 | 0.844324661 | 0.389187848 | 0.075 | 0.068 | 1 | 0.126453962 | -1.250257991 | 0 | 0.055 | 1 | 0.844324661 | 0.236917319 |
| Tbc1d9b | 0.906123384 | 0.299670931 | 0.358 | 0.356 | 1 | 0.126488035 | -0.268994554 | 0.22 | 0.369 | 1 | 0.906123384 | 0.236976848 |
| Dock2 | 0.503971666 | 0.569760094 | 0.019 | 0.008 | 1 | 0.126562667 | 1.136765132 | 0.073 | 0.027 | 1 | 0.503971666 | 0.237107226 |
| Ints3 | 0.126627945 | 0.765680304 | 0.264 | 0.178 | 1 | 0.24861262 | -0.230539757 | 0.171 | 0.263 | 1 | 0.24861262 | 0.237221253 |
| Mtcp1 | 0.126731036 | -0.477545621 | 0.038 | 0.11 | 1 | 0.267565071 | -0.840867055 | 0.098 | 0.16 | 1 | 0.267565071 | 0.237401317 |
| E130317F20Rik | 0.126786203 | -1.430239906 | 0.019 | 0.076 | 1 | 0.622166556 | 1.83720485 | 0.049 | 0.034 | 1 | 0.622166556 | 0.237497664 |
| Irf3 | 0.126859191 | 0.874614676 | 0.321 | 0.233 | 1 | 0.990364497 | 0.653983026 | 0.268 | 0.283 | 1 | 0.990364497 | 0.237625128 |
| Smyd2 | 0.711493384 | 0.22310957 | 0.358 | 0.335 | 1 | 0.126880526 | -0.242522343 | 0.146 | 0.256 | 1 | 0.711493384 | 0.237662383 |
| Trp53inp2 | 0.297980775 | -0.366514308 | 0.34 | 0.415 | 1 | 0.126947961 | -0.440133094 | 0.317 | 0.44 | 1 | 0.297980775 | 0.237780138 |
| Iqank1 | 0.126955034 | 1.154722595 | 0.094 | 0.042 | 1 | 0.650841212 | 0.174239837 | 0.049 | 0.034 | 1 | 0.650841212 | 0.237792488 |
| Cdk2ap1 | 0.127031621 | 0.280253477 | 0.906 | 0.847 | 1 | 0.544667081 | 0.306690133 | 0.756 | 0.792 | 1 | 0.544667081 | 0.237926208 |
| Gm36862 | 0.12717834 | 0.640149422 | 0.075 | 0.03 | 1 | 0.190737323 | 1.422167351 | 0.049 | 0.017 | 1 | 0.190737323 | 0.238182351 |
| F11r | 0.127209887 | 1.055186921 | 0.113 | 0.055 | 1 | 0.603144629 | 0.10681191 | 0.22 | 0.184 | 1 | 0.603144629 | 0.238237419 |
| 3010001F23Rik | 0.753937509 | -0.1672055 | 0.057 | 0.068 | 1 | 0.127274814 | -1.410722664 | 0.049 | 0.13 | 1 | 0.753937509 | 0.238350749 |
| Ccdc149 | 0.127283999 | -1.304709024 | 0.019 | 0.076 | 1 | 0.32861307 | -0.717384002 | 0.049 | 0.096 | 1 | 0.32861307 | 0.238366781 |
| Tcea2 | 0.127470223 | 1.154722595 | 0.094 | 0.042 | 1 | 0.362195044 | 1.252242349 | 0.049 | 0.024 | 1 | 0.362195044 | 0.238691789 |
| Chpf | 0.186505929 | -0.187669603 | 0.453 | 0.576 | 1 | 0.127627606 | -0.320647319 | 0.488 | 0.628 | 1 | 0.186505929 | 0.238966406 |
| Gm42788 | 0.127778252 | -1.237594828 | 0.019 | 0.076 | 1 | 0.286402368 | -0.484723245 | 0 | 0.027 | 1 | 0.286402368 | 0.239229222 |
| Ubl5 | 0.43029829 | -0.122536451 | 0.906 | 0.915 | 1 | 0.127815886 | 0.309169417 | 0.854 | 0.857 | 1 | 0.43029829 | 0.239294872 |
| Prmt3 | 0.127861602 | 0.536970159 | 0.377 | 0.275 | 1 | 0.76829584 | 0.252242349 | 0.22 | 0.259 | 1 | 0.76829584 | 0.239374614 |
| Fggy | 0.127912061 | 0.8327945 | 0.208 | 0.131 | 1 | 0.183087564 | -1.236044132 | 0.098 | 0.177 | 1 | 0.183087564 | 0.239462626 |
| Ccn5 | 0.551438809 | 0.358255989 | 0.377 | 0.335 | 1 | 0.127967124 | -0.11526678 | 0.098 | 0.208 | 1 | 0.551438809 | 0.239558664 |
| Napa | 0.232333703 | 0.198224233 | 0.811 | 0.797 | 1 | 0.128221322 | -0.229536701 | 0.683 | 0.809 | 1 | 0.232333703 | 0.240001937 |
| Ubox5 | 0.128229798 | -1.054730771 | 0.038 | 0.106 | 1 | 0.66647708 | -0.228884341 | 0.122 | 0.147 | 1 | 0.66647708 | 0.240016714 |
| Asb8 | 0.128300854 | 0.514264981 | 0.264 | 0.169 | 1 | 0.670788134 | 0.287007767 | 0.171 | 0.208 | 1 | 0.670788134 | 0.240140599 |
| Uimc1 | 0.130911891 | -0.441921711 | 0.66 | 0.758 | 1 | 0.128302786 | -0.296960389 | 0.585 | 0.724 | 1 | 0.130911891 | 0.240143968 |
| Kctd2 | 0.145139141 | 0.550651271 | 0.358 | 0.263 | 1 | 0.128324093 | -0.577832649 | 0.146 | 0.263 | 1 | 0.145139141 | 0.240181114 |
| Mitd1 | 0.128373491 | 0.502645898 | 0.358 | 0.246 | 1 | 0.740641096 | -0.144647803 | 0.171 | 0.191 | 1 | 0.740641096 | 0.240267228 |
| Siae | 0.128466301 | 0.842778588 | 0.358 | 0.284 | 1 | 0.849413356 | 0.343665377 | 0.244 | 0.287 | 1 | 0.849413356 | 0.240429011 |
| Hyi | 0.128500564 | 0.408479187 | 0.415 | 0.309 | 1 | 0.210220449 | -0.119726428 | 0.22 | 0.334 | 1 | 0.210220449 | 0.240488734 |
| Hmga1b | 0.128609464 | -1.430239906 | 0 | 0.042 | 1 | 0.231643049 | -0.622226769 | 0 | 0.034 | 1 | 0.231643049 | 0.240678534 |
| Amhr2 | 0.128609464 | -1.430239906 | 0 | 0.042 | 1 | 0.553780906 | 0.721727632 | 0.049 | 0.031 | 1 | 0.553780906 | 0.240678534 |
| Apoa1 | 0.128626524 | -1.652632327 | 0 | 0.042 | 1 | 0.530820926 | 1.174239837 | 0.073 | 0.051 | 1 | 0.530820926 | 0.240708266 |
| Fam161b | 0.128626524 | -1.545717123 | 0 | 0.042 | 1 | 0.319629309 | -0.622226769 | 0 | 0.024 | 1 | 0.319629309 | 0.240708266 |
| Emc9 | 0.128626524 | -1.845277405 | 0 | 0.042 | 1 | 0.531564619 | 1.699701326 | 0.049 | 0.031 | 1 | 0.531564619 | 0.240708266 |
| Guca1b | 0.128626524 | -1.545717123 | 0 | 0.042 | 1 | 0.231606831 | -0.863234868 | 0 | 0.034 | 1 | 0.231606831 | 0.240708266 |
| Spata17 | 0.128639319 | -1.845277405 | 0 | 0.042 | 1 | 0.989595616 | 0.667279848 | 0.024 | 0.024 | 1 | 0.989595616 | 0.240730563 |
| Prrg4 | 0.128639319 | -1.752168001 | 0 | 0.042 | 1 | 0.739624823 | -0.16279515 | 0.024 | 0.034 | 1 | 0.739624823 | 0.240730563 |
| S100a3 | 0.128639793 | -1.652632327 | 0 | 0.042 | 1 | 0.60213981 | 1.252242349 | 0.024 | 0.014 | 1 | 0.60213981 | 0.240731389 |
| Myl3 | 0.128639793 | -1.652632327 | 0 | 0.042 | 1 | 0.605005904 | 1.029849928 | 0.024 | 0.014 | 1 | 0.605005904 | 0.240731389 |
| Sgo1 | 0.128639793 | -1.652632327 | 0 | 0.042 | 1 | 0.257316251 | -0.970150072 | 0 | 0.031 | 1 | 0.257316251 | 0.240731389 |
| Lypd3 | 0.12864927 | -2.1672055 | 0 | 0.042 | 1 | 0.402369942 | -0.16279515 | 0 | 0.017 | 1 | 0.402369942 | 0.240747905 |
| Neurl1b | 0.12864927 | -1.932740246 | 0 | 0.042 | 1 | 0.599290422 | 1.515276755 | 0.024 | 0.014 | 1 | 0.599290422 | 0.240747905 |
| Nek5 | 0.128649744 | -1.752168001 | 0 | 0.042 | 1 | 0.455097252 | -0.16279515 | 0 | 0.014 | 1 | 0.455097252 | 0.240748731 |
| Psrc1 | 0.128656378 | -2.237594828 | 0 | 0.042 | 1 | 0.402373368 | 0.252242349 | 0 | 0.017 | 1 | 0.402373368 | 0.240760292 |
| Gpr39 | 0.128656852 | -2.752168001 | 0 | 0.042 | 1 | 0.752578711 | 0.334704509 | 0.024 | 0.017 | 1 | 0.752578711 | 0.240761118 |
| Gpr3 | 0.128666329 | -1.304709024 | 0 | 0.042 | 1 | 0.826384776 | 0.515276755 | 0.024 | 0.031 | 1 | 0.826384776 | 0.240777634 |
| Gm13056 | 0.128666329 | -1.304709024 | 0 | 0.042 | 1 | 0.744047924 | 1.252242349 | 0.024 | 0.017 | 1 | 0.744047924 | 0.240777634 |
| Gm17259 | 0.128666329 | -1.304709024 | 0 | 0.042 | 1 | 0.357854401 | -0.332720152 | 0 | 0.02 | 1 | 0.357854401 | 0.240777634 |
| 4930452G13Rik | 0.128666329 | -1.304709024 | 0 | 0.042 | 1 | 0.208799874 | -0.863234868 | 0 | 0.038 | 1 | 0.208799874 | 0.240777634 |
| Gm31814 | 0.128666329 | -1.304709024 | 0 | 0.042 | 1 | 0.5191986 | 0.515276755 | 0 | 0.01 | 1 | 0.5191986 | 0.240777634 |
| Rps6ka3 | 0.128722355 | -0.340636445 | 0.868 | 0.877 | 1 | 0.306416811 | -0.105714055 | 0.854 | 0.918 | 1 | 0.306416811 | 0.240875266 |
| 4833418N02Rik | 0.128948588 | -0.713173869 | 0.113 | 0.208 | 1 | 0.550678023 | 0.429120111 | 0.146 | 0.191 | 1 | 0.550678023 | 0.241269438 |
| Llph | 0.128950274 | 0.183635287 | 0.698 | 0.589 | 1 | 0.313193508 | 0.279828181 | 0.61 | 0.526 | 1 | 0.313193508 | 0.241272374 |
| Cmklr1 | 0.128960873 | 0.83846325 | 0.396 | 0.309 | 1 | 0.153547245 | -0.481714495 | 0.488 | 0.614 | 1 | 0.153547245 | 0.241290839 |
| Cog7 | 0.192281463 | 0.659957903 | 0.283 | 0.208 | 1 | 0.129211519 | 0.635570989 | 0.293 | 0.198 | 1 | 0.192281463 | 0.241727421 |
| Shroom2 | 0.328542491 | 0.540012751 | 0.264 | 0.212 | 1 | 0.129251238 | 1.408361551 | 0.415 | 0.307 | 1 | 0.328542491 | 0.241796593 |
| Tbc1d13 | 0.824162189 | -0.138059154 | 0.132 | 0.144 | 1 | 0.129294105 | -0.707115666 | 0.073 | 0.171 | 1 | 0.824162189 | 0.241871244 |
| Wdr90 | 0.527892651 | -0.783876861 | 0.038 | 0.059 | 1 | 0.12937601 | -1.20718927 | 0.024 | 0.096 | 1 | 0.527892651 | 0.242013867 |
| Gm15787 | 0.450459274 | -0.19891436 | 0.094 | 0.136 | 1 | 0.129464027 | -1.278272368 | 0.049 | 0.13 | 1 | 0.450459274 | 0.242167119 |
| Dop1b | 0.129471691 | 0.165369839 | 0.321 | 0.22 | 1 | 0.474654789 | -0.343367396 | 0.22 | 0.276 | 1 | 0.474654789 | 0.242180462 |
| Rnf113a2 | 0.129479493 | -0.595299152 | 0.151 | 0.25 | 1 | 0.268713947 | -0.677368323 | 0.146 | 0.215 | 1 | 0.268713947 | 0.242194047 |
| Pcmtd1 | 0.13003175 | 0.547392281 | 0.717 | 0.72 | 1 | 0.346512866 | 0.382010224 | 0.854 | 0.816 | 1 | 0.346512866 | 0.243155243 |
| Psph | 0.130052083 | -0.231335838 | 0.245 | 0.352 | 1 | 0.137112112 | -0.425829556 | 0.122 | 0.242 | 1 | 0.137112112 | 0.243190622 |
| Gnl3l | 0.609585373 | 0.146359122 | 0.509 | 0.5 | 1 | 0.130064088 | -0.443751464 | 0.366 | 0.495 | 1 | 0.609585373 | 0.24321151 |
| Mtfmt | 0.130417615 | 1.017219071 | 0.189 | 0.114 | 1 | 0.597902727 | 0.893788378 | 0.171 | 0.143 | 1 | 0.597902727 | 0.243826476 |
| Gm57455 | 0.130481779 | 1.039245377 | 0.094 | 0.042 | 1 | 0.82181816 | 0.252242349 | 0.024 | 0.031 | 1 | 0.82181816 | 0.243938063 |
| Smug1 | 0.130568442 | 1.491757582 | 0.132 | 0.072 | 1 | 0.288910755 | -0.777504994 | 0.073 | 0.13 | 1 | 0.288910755 | 0.244088766 |
| Trim25 | 0.130699742 | 0.792152515 | 0.283 | 0.195 | 1 | 0.765219002 | 0.235168836 | 0.341 | 0.324 | 1 | 0.765219002 | 0.244317062 |
| Rfwd3 | 0.346637765 | 0.294900253 | 0.302 | 0.246 | 1 | 0.130805654 | -0.675245151 | 0.268 | 0.392 | 1 | 0.346637765 | 0.24450119 |
| H2-T22 | 0.130857875 | 0.304298951 | 0.396 | 0.292 | 1 | 0.590661675 | 0.695849001 | 0.366 | 0.362 | 1 | 0.590661675 | 0.244591967 |
| Rnf121 | 0.130966601 | 0.440124814 | 0.453 | 0.36 | 1 | 0.816936108 | 0.182894302 | 0.439 | 0.522 | 1 | 0.816936108 | 0.244780951 |
| Prkar2a | 0.672226259 | 0.162584713 | 0.453 | 0.441 | 1 | 0.130989411 | -0.127691349 | 0.512 | 0.662 | 1 | 0.672226259 | 0.244820596 |
| Gm41604 | 0.732691937 | 1.154722595 | 0.019 | 0.013 | 1 | 0.131030883 | 1.749742009 | 0.098 | 0.044 | 1 | 0.732691937 | 0.244892673 |
| Sat2 | 0.131076497 | 0.75033234 | 0.245 | 0.161 | 1 | 0.440656127 | 1.33970519 | 0.171 | 0.133 | 1 | 0.440656127 | 0.244971947 |
| Ilrun | 0.238150394 | 0.265341767 | 0.774 | 0.75 | 1 | 0.131277291 | -0.268099304 | 0.878 | 0.894 | 1 | 0.238150394 | 0.245320856 |
| Gm16153 | 0.163351484 | 0.984797593 | 0.113 | 0.059 | 1 | 0.131599703 | 0.97213443 | 0.195 | 0.116 | 1 | 0.163351484 | 0.245880924 |
| Gtf3a | 0.13160904 | 1.115194231 | 0.208 | 0.136 | 1 | 0.786079022 | 0.422167351 | 0.195 | 0.181 | 1 | 0.786079022 | 0.245897141 |
| Sec24b | 0.18975773 | 0.113553406 | 0.736 | 0.657 | 1 | 0.131619225 | -0.148581291 | 0.756 | 0.867 | 1 | 0.18975773 | 0.24591483 |
| Nfasc | 0.131809901 | -0.538299652 | 0.113 | 0.216 | 1 | 0.510625529 | 0.966487867 | 0.268 | 0.256 | 1 | 0.510625529 | 0.246245952 |
| Pex14 | 0.131882513 | 0.526691372 | 0.509 | 0.445 | 1 | 0.69631382 | 0.125709943 | 0.561 | 0.621 | 1 | 0.69631382 | 0.246372029 |
| Gm12905 | 0.607941221 | 0.404700848 | 0.132 | 0.106 | 1 | 0.132198155 | -1.185162963 | 0.073 | 0.16 | 1 | 0.607941221 | 0.246919957 |
| Cip2a | 0.132361498 | 0.377115016 | 0.094 | 0.042 | 1 | 0.269443216 | 2.100239256 | 0.098 | 0.058 | 1 | 0.269443216 | 0.247203429 |

| | | | | | | | | | | | | |
|---|---|---|---|---|---|---|---|---|---|---|---|---|
| Prkcd | 0.132426707 | 0.992451166 | 0.189 | 0.114 | 1 | 0.242641133 | 0.494060269 | 0.415 | 0.321 | 1 | 0.242641133 | 0.247316581 |
| Arap2 | 0.338601192 | -1.719746523 | 0.038 | 0.072 | 1 | 0.1324319 | 0.23386382 | 0.293 | 0.184 | 1 | 0.338601192 | 0.247325592 |
| Mgat3 | 0.665739254 | 0.484871196 | 0.132 | 0.11 | 1 | 0.13260718 | 0.659666664 | 0.341 | 0.239 | 1 | 0.665739254 | 0.247629696 |
| Platr25 | 0.385979878 | 0.247831999 | 0.226 | 0.174 | 1 | 0.133109463 | -1.004097404 | 0.049 | 0.13 | 1 | 0.385979878 | 0.248500796 |
| Gm10561 | 0.133161347 | -0.577081294 | 0.189 | 0.292 | 1 | 0.26763843 | -0.701953961 | 0.146 | 0.218 | 1 | 0.26763843 | 0.24859075 |
| Zfp940 | 0.133543194 | 0.8327945 | 0.094 | 0.042 | 1 | 0.337213694 | -0.686357106 | 0.024 | 0.061 | 1 | 0.337213694 | 0.249252603 |
| Mbtps2 | 0.886922802 | 0.2967416 | 0.321 | 0.36 | 1 | 0.133687007 | -0.170614655 | 0.195 | 0.321 | 1 | 0.886922802 | 0.249501798 |
| Scfd2 | 0.412221485 | 0.205096117 | 0.434 | 0.373 | 1 | 0.13379545 | 0.551802631 | 0.39 | 0.29 | 1 | 0.412221485 | 0.249689678 |
| Armh3 | 0.366359404 | 0.96034455 | 0.34 | 0.309 | 1 | 0.134040569 | -0.867851496 | 0.366 | 0.468 | 1 | 0.366359404 | 0.250114264 |
| Nol8 | 0.305607987 | 0.401078259 | 0.415 | 0.347 | 1 | 0.134098749 | -0.503832068 | 0.171 | 0.294 | 1 | 0.305607987 | 0.250215024 |
| 1810024B03Rik | 0.13412369 | 1.377115016 | 0.132 | 0.072 | 1 | 0.303059962 | 1.252242349 | 0.122 | 0.078 | 1 | 0.303059962 | 0.250258216 |
| Ap5m1 | 0.13434541 | 0.8327945 | 0.245 | 0.161 | 1 | 0.92662174 | 0.289717055 | 0.195 | 0.191 | 1 | 0.92662174 | 0.25064213 |
| 4732440D04Rik | 0.134351769 | -0.214511215 | 0.17 | 0.28 | 1 | 0.206773643 | 0.386169152 | 0.439 | 0.345 | 1 | 0.206773643 | 0.250653141 |
| Gmeb1 | 0.751727879 | 0.408170952 | 0.377 | 0.352 | 1 | 0.134371614 | -0.649577257 | 0.244 | 0.365 | 1 | 0.751727879 | 0.250687497 |
| Mettl22 | 0.967052971 | 0.347367673 | 0.057 | 0.055 | 1 | 0.134503296 | 1.737669176 | 0.098 | 0.044 | 1 | 0.967052971 | 0.250915456 |
| Krt19 | 0.307371127 | 0.154722595 | 0.094 | 0.055 | 1 | 0.134518149 | 1.911205431 | 0.268 | 0.177 | 1 | 0.307371127 | 0.250941166 |
| Ptcd1 | 0.134520812 | 0.984797593 | 0.151 | 0.085 | 1 | 0.870172807 | 0.345351753 | 0.122 | 0.133 | 1 | 0.870172807 | 0.250945775 |
| Tbcc | 0.134522328 | 0.669295768 | 0.264 | 0.178 | 1 | 0.292611805 | -0.395455907 | 0.073 | 0.133 | 1 | 0.292611805 | 0.250948399 |
| Gm56764 | 0.763636454 | 0.47665069 | 0.057 | 0.047 | 1 | 0.134522772 | 2.007129851 | 0.098 | 0.044 | 1 | 0.763636454 | 0.250949168 |
| Cln6 | 0.134525339 | 1.292226119 | 0.132 | 0.072 | 1 | 0.229598851 | -0.787286015 | 0.049 | 0.109 | 1 | 0.229598851 | 0.250953612 |
| Abcd4 | 0.134603353 | 0.932330173 | 0.17 | 0.097 | 1 | 0.939515018 | 0.365899131 | 0.146 | 0.147 | 1 | 0.939515018 | 0.251088643 |
| Taf12 | 0.260734113 | 0.687596584 | 0.358 | 0.297 | 1 | 0.134812765 | -0.284785675 | 0.244 | 0.372 | 1 | 0.260734113 | 0.251451049 |
| 2310033P09Rik | 0.134902868 | 0.582816247 | 0.415 | 0.331 | 1 | 0.932212467 | 0.230547278 | 0.293 | 0.311 | 1 | 0.932212467 | 0.251606952 |
| Zscan29 | 0.134997996 | 1.232725107 | 0.208 | 0.131 | 1 | 0.348672295 | 0.452541 | 0.171 | 0.116 | 1 | 0.348672295 | 0.251771533 |
| Cln3 | 0.134997996 | 0.984797593 | 0.208 | 0.131 | 1 | 0.548881623 | -0.250257991 | 0.146 | 0.184 | 1 | 0.548881623 | 0.251771533 |
| Coasy | 0.134997996 | 0.984797593 | 0.208 | 0.131 | 1 | 0.782930807 | 0.287007767 | 0.146 | 0.13 | 1 | 0.782930807 | 0.251771533 |
| Dtwd1 | 0.135055342 | 1.46284489 | 0.132 | 0.072 | 1 | 0.379368341 | -0.707115666 | 0.049 | 0.089 | 1 | 0.379368341 | 0.251870739 |
| Sharpin | 0.377800503 | 0.2967416 | 0.245 | 0.191 | 1 | 0.13513732 | 1.351778023 | 0.268 | 0.174 | 1 | 0.377800503 | 0.252012545 |
| Acsl4 | 0.135205255 | -0.925004598 | 0.528 | 0.631 | 1 | 0.355593876 | -0.352137605 | 0.488 | 0.567 | 1 | 0.355593876 | 0.252130049 |
| Gm32673 | 0.701009961 | 0.509006063 | 0.189 | 0.169 | 1 | 0.135296493 | 0.796177582 | 0.293 | 0.201 | 1 | 0.701009961 | 0.252287844 |
| Bcl2l2 | 0.906990039 | 0.280253477 | 0.189 | 0.208 | 1 | 0.135351234 | -0.711231775 | 0.195 | 0.3 | 1 | 0.906990039 | 0.252382511 |
| Tmed10 | 0.305457144 | -0.116579427 | 1 | 0.992 | 1 | 0.135462887 | -0.157614911 | 0.951 | 0.986 | 1 | 0.305457144 | 0.25257558 |
| Sgpp1 | 0.976731231 | -0.502996248 | 0.34 | 0.347 | 1 | 0.135489485 | -0.681523953 | 0.244 | 0.352 | 1 | 0.976731231 | 0.25262157 |
| St6galnac4 | 0.135554013 | -0.49928155 | 0.189 | 0.301 | 1 | 0.534128619 | 0.19334866 | 0.244 | 0.198 | 1 | 0.534128619 | 0.252733136 |
| Gm12216 | 0.13571738 | 0.946614399 | 0.226 | 0.148 | 1 | 0.159548801 | -0.819907437 | 0.268 | 0.358 | 1 | 0.159548801 | 0.253015553 |
| Pmm2 | 0.175419524 | 0.459577176 | 0.321 | 0.229 | 1 | 0.135751375 | -0.410722664 | 0.171 | 0.294 | 1 | 0.175419524 | 0.253074313 |
| Fam234a | 0.433110563 | -0.285849997 | 0.358 | 0.411 | 1 | 0.135946428 | -0.178392005 | 0.244 | 0.372 | 1 | 0.433110563 | 0.253411425 |
| Swi5 | 0.136208562 | -0.25404876 | 0.962 | 0.932 | 1 | 0.246085247 | -0.172718197 | 0.854 | 0.945 | 1 | 0.246085247 | 0.253864351 |
| Zdhhc15 | 0.308773971 | 1.318221327 | 0.113 | 0.072 | 1 | 0.136300661 | 1.30115195 | 0.146 | 0.078 | 1 | 0.308773971 | 0.254023452 |
| Ints1 | 0.136355778 | 0.806799291 | 0.17 | 0.097 | 1 | 0.151739708 | -0.16279515 | 0.049 | 0.13 | 1 | 0.151739708 | 0.254118658 |
| Dusp19 | 0.136382767 | 0.725879296 | 0.208 | 0.127 | 1 | 0.435914758 | -0.448197369 | 0.073 | 0.113 | 1 | 0.435914758 | 0.254165274 |
| Tomm34 | 0.136388537 | 0.751000409 | 0.453 | 0.377 | 1 | 0.485884971 | 0.593279267 | 0.39 | 0.389 | 1 | 0.485884971 | 0.254175241 |
| Amer2 | 0.136574337 | -0.783876861 | 0.019 | 0.076 | 1 | 0.274788535 | 1.252242349 | 0.049 | 0.02 | 1 | 0.274788535 | 0.254496124 |
| Sult1d1 | 0.136655855 | 0.569760094 | 0.094 | 0.042 | 1 | 0.314951298 | 1.922093747 | 0.268 | 0.201 | 1 | 0.314951298 | 0.254636886 |
| Arhgef5 | 0.352204228 | -0.120284453 | 0.264 | 0.347 | 1 | 0.136663929 | -0.179096963 | 0.244 | 0.375 | 1 | 0.352204228 | 0.254650828 |
| Slc25a28 | 0.136681144 | 0.394673388 | 0.415 | 0.309 | 1 | 0.913570767 | 0.425391251 | 0.293 | 0.331 | 1 | 0.913570767 | 0.254680552 |
| Chpf2 | 0.768039097 | 0.112078257 | 0.34 | 0.322 | 1 | 0.136842864 | -0.273826463 | 0.293 | 0.437 | 1 | 0.768039097 | 0.254959758 |
| Afg3l1 | 0.136916954 | 0.739685096 | 0.377 | 0.305 | 1 | 0.158737636 | -0.340333336 | 0.244 | 0.382 | 1 | 0.158737636 | 0.255087657 |
| Mtor | 0.136996654 | 0.825774072 | 0.377 | 0.314 | 1 | 0.39639869 | -0.16279515 | 0.293 | 0.386 | 1 | 0.39639869 | 0.255225224 |
| Fbrsl1 | 0.137073271 | 0.665684514 | 0.528 | 0.449 | 1 | 0.923348455 | 0.124085998 | 0.634 | 0.652 | 1 | 0.923348455 | 0.255357461 |
| Ptpn6 | 0.13710505 | 0.640149422 | 0.113 | 0.055 | 1 | 0.646741372 | 0.252242349 | 0.073 | 0.055 | 1 | 0.646741372 | 0.255412305 |
| Ifit3 | 0.137484316 | -3.257091004 | 0.094 | 0.169 | 1 | 0.815449804 | -0.457415899 | 0.098 | 0.116 | 1 | 0.815449804 | 0.256066694 |
| Ralgps1 | 0.385213857 | 1.084333267 | 0.094 | 0.064 | 1 | 0.137748752 | -1.84461919 | 0.049 | 0.126 | 1 | 0.385213857 | 0.256522786 |
| Ciart | 0.137945745 | -1.347777746 | 0.038 | 0.102 | 1 | 0.452331971 | -0.250257991 | 0.024 | 0.051 | 1 | 0.452331971 | 0.256862461 |
| Ogfod2 | 0.315370244 | 0.891688189 | 0.132 | 0.089 | 1 | 0.138054254 | 0.83720485 | 0.146 | 0.075 | 1 | 0.315370244 | 0.257049531 |
| Trp53i13 | 0.138258822 | 0.587354154 | 0.566 | 0.53 | 1 | 0.503931751 | -0.101394606 | 0.439 | 0.495 | 1 | 0.503931751 | 0.257402142 |
| Zdhhc21 | 0.138335046 | 0.555260524 | 0.358 | 0.263 | 1 | 0.262611825 | 0.774195052 | 0.317 | 0.263 | 1 | 0.262611825 | 0.257533508 |
| Ubr1 | 0.266770448 | 0.173054418 | 0.736 | 0.699 | 1 | 0.138528137 | 0.594030866 | 0.78 | 0.744 | 1 | 0.266770448 | 0.25786623 |
| Adprh | 0.146997599 | 0.550651271 | 0.528 | 0.47 | 1 | 0.138695889 | 0.752315952 | 0.439 | 0.352 | 1 | 0.146997599 | 0.258155228 |
| Zfp954 | 0.162917867 | 1.050385935 | 0.226 | 0.157 | 1 | 0.138831642 | -0.686357106 | 0.098 | 0.195 | 1 | 0.162917867 | 0.25838906 |
| Maff | 0.191632563 | -0.649917943 | 0.509 | 0.572 | 1 | 0.138864424 | 0.268362014 | 0.683 | 0.546 | 1 | 0.191632563 | 0.25844552 |
| Stk11ip | 0.355831567 | -0.719746523 | 0.038 | 0.072 | 1 | 0.138920764 | 1.444887427 | 0.122 | 0.061 | 1 | 0.355831567 | 0.258542549 |
| Pwp1 | 0.138927844 | 0.939709704 | 0.302 | 0.225 | 1 | 0.536779703 | 0.129385601 | 0.195 | 0.249 | 1 | 0.536779703 | 0.258554742 |
| Ino80e | 0.138999556 | 0.769068902 | 0.358 | 0.271 | 1 | 0.14044692 | -0.508569987 | 0.22 | 0.338 | 1 | 0.14044692 | 0.258678235 |
| Cep290 | 0.445033767 | 0.257292328 | 0.264 | 0.347 | 1 | 0.139067958 | 0.813358108 | 0.341 | 0.235 | 1 | 0.445033767 | 0.258796018 |
| Coq6 | 0.499735438 | 0.484871196 | 0.151 | 0.119 | 1 | 0.139083439 | 0.635570989 | 0.22 | 0.13 | 1 | 0.499735438 | 0.258822675 |
| Pold4 | 0.311085597 | -0.344083262 | 0.321 | 0.39 | 1 | 0.139200659 | -0.622226769 | 0.171 | 0.283 | 1 | 0.311085597 | 0.259024494 |
| Lrp2bp | 0.858654164 | 1.417757001 | 0.038 | 0.034 | 1 | 0.139214502 | -0.970150072 | 0.024 | 0.096 | 1 | 0.858654164 | 0.259048326 |
| Gm15879 | 0.139226325 | 1.569760094 | 0.113 | 0.059 | 1 | 0.856647753 | 0.109284395 | 0.146 | 0.133 | 1 | 0.856647753 | 0.259068681 |
| Gm3839 | 0.421517727 | -0.304709024 | 0.019 | 0.042 | 1 | 0.139479918 | -1.250257991 | 0 | 0.051 | 1 | 0.421517727 | 0.259505189 |
| Msantd1 | 0.469724974 | -0.347777746 | 0.038 | 0.064 | 1 | 0.139479918 | -1.250257991 | 0 | 0.051 | 1 | 0.469724974 | 0.259505189 |
| Itgb1bp2 | 0.247902855 | 0.491757582 | 0.094 | 0.051 | 1 | 0.139502445 | -1.332720152 | 0 | 0.051 | 1 | 0.247902855 | 0.259543958 |
| Gm26688 | 0.295239083 | 0.8327945 | 0.094 | 0.055 | 1 | 0.139502445 | -1.332720152 | 0 | 0.051 | 1 | 0.295239083 | 0.259543958 |
| Rph3al | 0.340246113 | 1.417757001 | 0.038 | 0.017 | 1 | 0.139502445 | -1.410722664 | 0 | 0.051 | 1 | 0.340246113 | 0.259543958 |
| Tnfaip2 | 0.74913526 | -0.430239906 | 0.057 | 0.068 | 1 | 0.139502445 | -1.410722664 | 0 | 0.051 | 1 | 0.74913526 | 0.259543958 |
| Il17re | 0.913425467 | 1.569760094 | 0.019 | 0.017 | 1 | 0.139521505 | -1.484723245 | 0 | 0.051 | 1 | 0.913425467 | 0.25957676 |
| Gpr157 | 0.347746365 | 0.932330173 | 0.038 | 0.017 | 1 | 0.139521753 | -1.410722664 | 0 | 0.051 | 1 | 0.347746365 | 0.259577186 |
| Rslcan18 | 0.982488248 | 0.154722595 | 0.038 | 0.038 | 1 | 0.139521753 | -1.410722664 | 0 | 0.051 | 1 | 0.982488248 | 0.259577186 |
| Zfp239 | 0.670470453 | 0.347367673 | 0.057 | 0.042 | 1 | 0.139549228 | -1.622226769 | 0 | 0.051 | 1 | 0.670470453 | 0.259624469 |
| Slc5a5 | 0.7394666 | 0.569760094 | 0.019 | 0.013 | 1 | 0.139550713 | -1.863234868 | 0 | 0.051 | 1 | 0.7394666 | 0.259627025 |
| Gm32592 | 0.914958 | -0.652632327 | 0.019 | 0.021 | 1 | 0.139561604 | -1.970150072 | 0 | 0.051 | 1 | 0.914958 | 0.259645766 |
| 4833411C07Rik | 0.139807644 | -1.214511215 | 0.038 | 0.102 | 1 | 0.188498353 | -0.970150072 | 0 | 0.041 | 1 | 0.188498353 | 0.260069111 |
| Col25a1 | 0.139833724 | -1.419268788 | 0.151 | 0.237 | 1 | 0.995994343 | -0.494101577 | 0.366 | 0.365 | 1 | 0.995994343 | 0.260113978 |
| Dglucy | 0.143356305 | -1.077938162 | 0.057 | 0.127 | 1 | 0.140065855 | -1.410722664 | 0.024 | 0.092 | 1 | 0.143356305 | 0.260513267 |
| Nyap1 | 0.52492136 | -0.508242418 | 0.038 | 0.059 | 1 | 0.140069174 | -1.589059905 | 0.024 | 0.092 | 1 | 0.52492136 | 0.260518975 |
| Ggcx | 0.140172513 | -0.255909725 | 0.472 | 0.581 | 1 | 0.225604296 | -0.425829556 | 0.488 | 0.553 | 1 | 0.225604296 | 0.260696692 |
| Otud1 | 0.140192359 | 0.855162313 | 0.151 | 0.085 | 1 | 0.209621103 | 0.811669758 | 0.22 | 0.147 | 1 | 0.209621103 | 0.260730821 |
| Unc119b | 0.215913325 | -0.604269306 | 0.226 | 0.301 | 1 | 0.14041902 | -0.474381301 | 0.195 | 0.321 | 1 | 0.215913325 | 0.261120539 |
| Tomm20 | 0.576805964 | -0.131930025 | 0.981 | 0.936 | 1 | 0.140442741 | -0.171474906 | 0.902 | 0.959 | 1 | 0.576805964 | 0.261161318 |
| Ccdc6 | 0.140483568 | 0.882469659 | 0.491 | 0.432 | 1 | 0.481171384 | 0.295055433 | 0.659 | 0.618 | 1 | 0.481171384 | 0.261231503 |
| Irf2bpl | 0.599695943 | -0.20282941 | 0.604 | 0.614 | 1 | 0.140600329 | 0.261892519 | 0.902 | 0.758 | 1 | 0.599695943 | 0.261432205 |
| Gm9967 | 0.246023494 | 0.891688189 | 0.075 | 0.038 | 1 | 0.140611281 | 1.30115195 | 0.122 | 0.061 | 1 | 0.246023494 | 0.261451029 |
| Luzp1 | 0.140828516 | 0.237785903 | 0.981 | 0.953 | 1 | 0.288730229 | 0.305919668 | 0.951 | 0.99 | 1 | 0.288730229 | 0.261824361 |
| Ppp1r16a | 0.140877301 | 0.891688189 | 0.151 | 0.085 | 1 | 0.15507078 | -1.20718927 | 0.024 | 0.089 | 1 | 0.15507078 | 0.261908189 |
| AW146154 | 0.324505834 | -0.1672055 | 0.019 | 0.051 | 1 | 0.140976325 | -1.332720152 | 0.024 | 0.092 | 1 | 0.324505834 | 0.262078326 |
| Bbs1 | 0.90155955 | 0.695290976 | 0.038 | 0.042 | 1 | 0.14106199 | 1.483567895 | 0.122 | 0.061 | 1 | 0.90155955 | 0.262225494 |
| G930009F23Rik | 0.523104508 | 0.510866405 | 0.132 | 0.102 | 1 | 0.141161399 | 1.589277336 | 0.122 | 0.061 | 1 | 0.523104508 | 0.262396258 |
| Rnd1 | 0.14121303 | 0.63687429 | 0.208 | 0.131 | 1 | 0.631672177 | 0.930314254 | 0.049 | 0.034 | 1 | 0.631672177 | 0.26248494 |

| | | | | | | | | | | | | |
|---|---|---|---|---|---|---|---|---|---|---|---|---|
| Epm2aip1 | 0.14134441 | 0.717858733 | 0.415 | 0.347 | 1 | 0.712260602 | 0.124486802 | 0.341 | 0.392 | 1 | 0.712260602 | 0.262710577 |
| Zfr2 | 0.16923322 | 1.154722595 | 0.094 | 0.047 | 1 | 0.141427942 | -1.292078167 | 0.024 | 0.092 | 1 | 0.16923322 | 0.262854021 |
| Cct4 | 0.141447806 | 0.318648856 | 0.868 | 0.788 | 1 | 0.266328061 | -0.290015701 | 0.829 | 0.86 | 1 | 0.266328061 | 0.26288813 |
| Atp5md | 0.141511728 | -0.245815335 | 0.736 | 0.843 | 1 | 0.14567676 | -0.300941443 | 0.659 | 0.754 | 1 | 0.14567676 | 0.262997887 |
| Lins1 | 0.141556265 | 1.270199812 | 0.151 | 0.089 | 1 | 0.50899571 | 0.483567895 | 0.146 | 0.113 | 1 | 0.50899571 | 0.263074353 |
| Shroom3 | 0.233609367 | -0.134784022 | 0.057 | 0.114 | 1 | 0.141653485 | 0.883008539 | 0.244 | 0.15 | 1 | 0.233609367 | 0.263241261 |
| Trp53rkb | 0.141670377 | -0.960754623 | 0.038 | 0.102 | 1 | 0.216233519 | 0.212713985 | 0.049 | 0.119 | 1 | 0.216233519 | 0.263270258 |
| Mob4 | 0.141821083 | 0.558747101 | 0.679 | 0.657 | 1 | 0.390390437 | 0.294567759 | 0.878 | 0.863 | 1 | 0.390390437 | 0.263528946 |
| Tinf2 | 0.141989837 | 1.039245377 | 0.17 | 0.102 | 1 | 0.204671491 | 0.707921833 | 0.171 | 0.102 | 1 | 0.204671491 | 0.26381856 |
| Cpsf1 | 0.141989837 | 1.039245377 | 0.17 | 0.102 | 1 | 0.907467977 | 0.574170444 | 0.122 | 0.133 | 1 | 0.907467977 | 0.26381856 |
| Asah2 | 0.142016525 | 0.402650108 | 0.208 | 0.131 | 1 | 0.182984402 | 0.816143234 | 0.22 | 0.137 | 1 | 0.182984402 | 0.263864357 |
| Fam3c | 0.340558174 | 0.312130493 | 0.736 | 0.716 | 1 | 0.142121328 | -0.35994162 | 0.512 | 0.608 | 1 | 0.340558174 | 0.264044185 |
| Dhx29 | 0.142219579 | 0.72941676 | 0.358 | 0.28 | 1 | 0.151432564 | 1.019581592 | 0.39 | 0.3 | 1 | 0.151432564 | 0.264212749 |
| Rtl8a | 0.142229207 | 0.340941404 | 0.604 | 0.542 | 1 | 0.145404956 | -0.276686117 | 0.463 | 0.57 | 1 | 0.145404956 | 0.264229267 |
| Timm8b | 0.277355838 | -0.149875622 | 0.604 | 0.708 | 1 | 0.14237894 | -0.183856766 | 0.439 | 0.577 | 1 | 0.277355838 | 0.264486117 |
| Trim37 | 0.142448294 | 0.399835093 | 0.453 | 0.343 | 1 | 0.944659347 | 0.129385601 | 0.366 | 0.389 | 1 | 0.944659347 | 0.264605071 |
| Trappc12 | 0.451661097 | 0.466666601 | 0.226 | 0.182 | 1 | 0.142449211 | -0.572186086 | 0.122 | 0.222 | 1 | 0.451661097 | 0.264606645 |
| Tspan2 | 0.231980555 | -0.981054184 | 0.226 | 0.288 | 1 | 0.142453301 | 1.0375035 | 0.268 | 0.181 | 1 | 0.231980555 | 0.264613658 |
| Dnajc19 | 0.142506766 | 0.470750088 | 0.585 | 0.508 | 1 | 0.884437862 | 0.391218762 | 0.512 | 0.546 | 1 | 0.884437862 | 0.264705354 |
| Mrps36 | 0.142647773 | 0.42680214 | 0.434 | 0.331 | 1 | 0.901539369 | 0.444887427 | 0.22 | 0.239 | 1 | 0.901539369 | 0.264947159 |
| Zmiz1os1 | 0.142733275 | -0.845277405 | 0.057 | 0.131 | 1 | 0.59343528 | 0.483567895 | 0.122 | 0.096 | 1 | 0.59343528 | 0.265093763 |
| Mrpl42 | 0.272410359 | 0.344622296 | 0.66 | 0.644 | 1 | 0.142838294 | 0.452541 | 0.659 | 0.59 | 1 | 0.272410359 | 0.26527381 |
| Terf2 | 0.142957993 | 0.410561499 | 0.472 | 0.364 | 1 | 0.359940018 | 0.456383066 | 0.537 | 0.488 | 1 | 0.359940018 | 0.265478998 |
| Tmem14c | 0.143047853 | 0.325094044 | 0.774 | 0.754 | 1 | 0.594886827 | 0.156719738 | 0.585 | 0.696 | 1 | 0.594886827 | 0.265633019 |
| Slc3a2 | 0.143087258 | 0.10644258 | 0.962 | 0.835 | 1 | 0.507273509 | -0.223824584 | 0.902 | 0.945 | 1 | 0.507273509 | 0.265700553 |
| Plscr4 | 0.353243091 | 0.408479187 | 0.302 | 0.254 | 1 | 0.143244348 | -0.863234868 | 0.171 | 0.273 | 1 | 0.353243091 | 0.265969752 |
| Mrpl52 | 0.143318836 | 0.347367673 | 0.755 | 0.733 | 1 | 0.175475479 | 0.354540925 | 0.707 | 0.631 | 1 | 0.175475479 | 0.266097384 |
| Gpr173 | 0.143406786 | 0.906795081 | 0.189 | 0.114 | 1 | 0.180671984 | 0.989207943 | 0.195 | 0.123 | 1 | 0.180671984 | 0.266248066 |
| Fxyd6 | 0.447661659 | -0.124303844 | 0.642 | 0.733 | 1 | 0.143683423 | -0.420716633 | 0.512 | 0.604 | 1 | 0.447661659 | 0.26672192 |
| C1rl | 0.143739358 | 1.526691372 | 0.113 | 0.059 | 1 | 0.459188435 | -0.19841906 | 0.073 | 0.113 | 1 | 0.459188435 | 0.266817714 |
| Lrwd1 | 0.143829008 | 0.806799291 | 0.151 | 0.085 | 1 | 0.759542931 | 0.159132945 | 0.073 | 0.089 | 1 | 0.759542931 | 0.266971232 |
| Vwa5a | 0.964913454 | 0.262478775 | 0.453 | 0.521 | 1 | 0.143958399 | -0.230539757 | 0.463 | 0.631 | 1 | 0.964913454 | 0.267192778 |
| Tmem231 | 0.14686216 | 0.695290976 | 0.189 | 0.114 | 1 | 0.144160273 | -0.970150072 | 0.024 | 0.092 | 1 | 0.14686216 | 0.267538362 |
| Yipf2 | 0.744695251 | -0.227747042 | 0.208 | 0.225 | 1 | 0.144197377 | -0.537834582 | 0.122 | 0.229 | 1 | 0.744695251 | 0.26760187 |
| Gm14636 | 0.919816481 | 0.932330173 | 0.019 | 0.017 | 1 | 0.14425054 | 0.46797104 | 0.122 | 0.058 | 1 | 0.919816481 | 0.267692862 |
| 2010320M18Rik | 0.787948437 | 0.906795081 | 0.057 | 0.068 | 1 | 0.144306924 | 1.136765132 | 0.146 | 0.078 | 1 | 0.787948437 | 0.26778936 |
| 1500015L24Rik | 0.144532988 | -1.489133595 | 0.038 | 0.102 | 1 | 0.866453105 | 0.204936634 | 0.073 | 0.082 | 1 | 0.866453105 | 0.268176191 |
| Unc5b | 0.176818004 | 0.697374291 | 0.377 | 0.314 | 1 | 0.144538703 | 0.473800119 | 0.463 | 0.355 | 1 | 0.176818004 | 0.268185969 |
| Irf7 | 0.145100504 | -1.752168001 | 0.019 | 0.072 | 1 | 0.199454674 | 0.515276755 | 0.122 | 0.065 | 1 | 0.199454674 | 0.269146852 |
| Tnfrsf1b | 0.145156542 | -0.430239906 | 0.038 | 0.106 | 1 | 0.314531573 | -0.595754557 | 0.073 | 0.13 | 1 | 0.314531573 | 0.269242662 |
| Dnai4 | 0.389521236 | -0.27154216 | 0.075 | 0.119 | 1 | 0.145310299 | 1.048708955 | 0.171 | 0.099 | 1 | 0.389521236 | 0.269505514 |
| Ppme1 | 0.145451017 | -0.307620619 | 0.679 | 0.771 | 1 | 0.381749139 | -0.146554509 | 0.707 | 0.73 | 1 | 0.381749139 | 0.269746035 |
| Qsox2 | 0.145487319 | -0.678167419 | 0.113 | 0.199 | 1 | 0.763955425 | 0.814837037 | 0.195 | 0.184 | 1 | 0.763955425 | 0.269808079 |
| Helz2 | 0.864170297 | 0.569760094 | 0.132 | 0.148 | 1 | 0.145497978 | -0.84461919 | 0.098 | 0.191 | 1 | 0.864170297 | 0.269826294 |
| Rhog | 0.145542533 | 0.739685096 | 0.321 | 0.242 | 1 | 0.550576125 | -0.192542494 | 0.22 | 0.266 | 1 | 0.550576125 | 0.269902437 |
| Anapc13 | 0.475562902 | -0.160150743 | 0.509 | 0.568 | 1 | 0.145595379 | -0.19841906 | 0.244 | 0.375 | 1 | 0.475562902 | 0.269992744 |
| Mtus2 | 0.145682257 | -1.845277405 | 0.019 | 0.072 | 1 | 0.225539557 | -1.20718927 | 0.024 | 0.075 | 1 | 0.225539557 | 0.270141194 |
| Nacc1 | 0.14575383 | 0.792152515 | 0.358 | 0.288 | 1 | 0.148523419 | -0.446588116 | 0.317 | 0.454 | 1 | 0.148523419 | 0.270263481 |
| Grk4 | 0.145767583 | 0.874614676 | 0.283 | 0.199 | 1 | 0.318099093 | 0.118386602 | 0.171 | 0.253 | 1 | 0.318099093 | 0.270286978 |
| Tomm40 | 0.146054615 | 0.245488059 | 0.528 | 0.411 | 1 | 0.930279827 | 0.175891463 | 0.561 | 0.577 | 1 | 0.930279827 | 0.27077728 |
| Sdhb | 0.146119159 | 0.31392119 | 0.698 | 0.627 | 1 | 0.672010218 | 0.328863631 | 0.61 | 0.7 | 1 | 0.672010218 | 0.270887509 |
| Golph3l | 0.14612178 | 0.732489594 | 0.283 | 0.203 | 1 | 0.491242538 | 0.626308067 | 0.268 | 0.229 | 1 | 0.491242538 | 0.270891985 |
| Fhl1 | 0.380059062 | 0.819119563 | 0.302 | 0.254 | 1 | 0.146385604 | 0.9413937 | 0.439 | 0.365 | 1 | 0.380059062 | 0.271342462 |
| Ndufa2 | 0.14655449 | 0.2967416 | 0.962 | 0.869 | 1 | 0.174387498 | 0.240560544 | 0.829 | 0.768 | 1 | 0.174387498 | 0.271630761 |
| Qpct | 0.567267181 | 0.917683397 | 0.132 | 0.11 | 1 | 0.146565489 | 0.944120054 | 0.22 | 0.137 | 1 | 0.567267181 | 0.271649535 |
| Gpn1 | 0.189932025 | 0.859266711 | 0.264 | 0.199 | 1 | 0.146566185 | -0.80665134 | 0.098 | 0.191 | 1 | 0.189932025 | 0.271650723 |
| Dapl1 | 0.548538879 | 0.154722595 | 0.057 | 0.038 | 1 | 0.146645254 | 0.667279848 | 0.22 | 0.133 | 1 | 0.548538879 | 0.271785678 |
| Hyou1 | 0.146827523 | 0.600133743 | 0.396 | 0.305 | 1 | 0.227301578 | 0.53464208 | 0.415 | 0.338 | 1 | 0.227301578 | 0.272096724 |
| Zbed4 | 0.339010327 | 0.721763188 | 0.132 | 0.089 | 1 | 0.146831898 | -0.987223586 | 0.098 | 0.188 | 1 | 0.339010327 | 0.272104189 |
| D330050G23Rik | 0.14684297 | -1.304709024 | 0.019 | 0.072 | 1 | 0.286433344 | -0.332720152 | 0 | 0.027 | 1 | 0.286433344 | 0.272123081 |
| Bcl6 | 0.867344906 | -0.114404084 | 0.755 | 0.703 | 1 | 0.146844601 | 0.120246803 | 0.902 | 0.857 | 1 | 0.867344906 | 0.272125865 |
| Ogfrl1 | 0.146879465 | 0.770058745 | 0.189 | 0.114 | 1 | 0.994484319 | 0.655001519 | 0.146 | 0.154 | 1 | 0.994484319 | 0.272185353 |
| Hmgcll1 | 0.959797248 | 0.200526284 | 0.113 | 0.119 | 1 | 0.147016766 | -0.622226769 | 0.122 | 0.222 | 1 | 0.959797248 | 0.272419603 |
| Exosc10 | 0.147023912 | 0.709311446 | 0.434 | 0.381 | 1 | 0.957458124 | 0.21515303 | 0.39 | 0.433 | 1 | 0.957458124 | 0.272431793 |
| Mrpl1 | 0.147143535 | 0.586438835 | 0.34 | 0.254 | 1 | 0.302492643 | -0.194273382 | 0.195 | 0.294 | 1 | 0.302492643 | 0.27263585 |
| Brpf1 | 0.147218926 | 1.046198138 | 0.321 | 0.246 | 1 | 0.353910341 | 0.313642894 | 0.463 | 0.403 | 1 | 0.353910341 | 0.272764441 |
| Zfp59 | 0.340246113 | 1.417757001 | 0.038 | 0.017 | 1 | 0.147369519 | 1.334704509 | 0.098 | 0.044 | 1 | 0.340246113 | 0.273021263 |
| Nudt1 | 0.147756771 | 0.948271717 | 0.17 | 0.102 | 1 | 0.913896025 | 0.292884334 | 0.098 | 0.092 | 1 | 0.913896025 | 0.273681478 |
| Rps6ka1 | 0.147863734 | 1.337586652 | 0.208 | 0.14 | 1 | 0.720663204 | 0.690363461 | 0.195 | 0.177 | 1 | 0.720663204 | 0.273863784 |
| Slc35b1 | 0.309038088 | 0.472462893 | 0.698 | 0.648 | 1 | 0.147863781 | 0.37036845 | 0.683 | 0.57 | 1 | 0.309038088 | 0.273863865 |
| Ubxn11 | 0.14799203 | -1.368839361 | 0.019 | 0.072 | 1 | 0.969250601 | 0.667279848 | 0.049 | 0.051 | 1 | 0.969250601 | 0.27408242 |
| 9430091E24Rik | 0.14799203 | -1.1672055 | 0.019 | 0.072 | 1 | 0.311923187 | -0.188330242 | 0.073 | 0.133 | 1 | 0.311923187 | 0.27408242 |
| 5031434O11Rik | 0.148045042 | 2.377115016 | 0.057 | 0.021 | 1 | 0.691833411 | 0.515276755 | 0.024 | 0.038 | 1 | 0.691833411 | 0.27417275 |
| Mmrn2 | 0.148045042 | 2.569760094 | 0.057 | 0.021 | 1 | 0.170390036 | -1.622226769 | 0 | 0.044 | 1 | 0.170390036 | 0.27417275 |
| Wdr59 | 0.830214892 | 0.295320925 | 0.302 | 0.343 | 1 | 0.148169935 | -0.80665134 | 0.171 | 0.276 | 1 | 0.830214892 | 0.274385541 |
| Helq | 0.148217519 | 2.06161319 | 0.094 | 0.047 | 1 | 0.720623372 | 0.120997816 | 0.122 | 0.102 | 1 | 0.720623372 | 0.274466605 |
| Gpalpp1 | 0.1482856 | 0.678989164 | 0.377 | 0.288 | 1 | 0.882273598 | 0.211019686 | 0.317 | 0.321 | 1 | 0.882273598 | 0.274582581 |
| Fech | 0.148384328 | 0.602181572 | 0.377 | 0.292 | 1 | 0.56595899 | 0.486707603 | 0.341 | 0.317 | 1 | 0.56595899 | 0.274750747 |
| Nudt3 | 0.187953036 | 0.456103312 | 0.547 | 0.525 | 1 | 0.148430434 | 0.423902402 | 0.659 | 0.594 | 1 | 0.187953036 | 0.274829274 |
| Gm16268 | 0.792882556 | -0.52334931 | 0.075 | 0.064 | 1 | 0.148443437 | 0.326242931 | 0.171 | 0.092 | 1 | 0.792882556 | 0.274851419 |
| Lyrm2 | 0.148468085 | 0.739685096 | 0.302 | 0.22 | 1 | 0.729336747 | 0.644559772 | 0.195 | 0.181 | 1 | 0.729336747 | 0.274893398 |
| Scrib | 0.605478554 | 0.191248471 | 0.17 | 0.14 | 1 | 0.148523479 | 0.970471381 | 0.244 | 0.16 | 1 | 0.605478554 | 0.274987734 |
| Cdcp1 | 0.28776878 | -0.845277405 | 0 | 0.021 | 1 | 0.148524906 | 0.883008539 | 0.171 | 0.096 | 1 | 0.28776878 | 0.274990164 |
| Fastkd2 | 0.941565696 | 0.247831999 | 0.113 | 0.119 | 1 | 0.148547926 | -0.917682652 | 0.073 | 0.157 | 1 | 0.941565696 | 0.275029366 |
| Hrct1 | 0.148562499 | -1.093204919 | 0.019 | 0.072 | 1 | 0.548898236 | -0.332720152 | 0.024 | 0.044 | 1 | 0.548898236 | 0.275054182 |
| Ggnbp1 | 0.148562499 | -1.093204919 | 0.019 | 0.072 | 1 | 0.764400411 | 0.635570989 | 0.073 | 0.061 | 1 | 0.764400411 | 0.275054182 |
| Igip | 0.148614146 | 0.33529484 | 0.283 | 0.186 | 1 | 0.801590201 | 0.136765132 | 0.317 | 0.304 | 1 | 0.801590201 | 0.275142128 |
| Oat | 0.503052449 | -0.189463509 | 0.962 | 0.924 | 1 | 0.148907516 | -0.218492956 | 0.878 | 0.959 | 1 | 0.503052449 | 0.275641584 |
| Secisbp2l | 0.149755324 | 0.171210718 | 0.887 | 0.847 | 1 | 0.360510897 | 0.246436727 | 0.829 | 0.891 | 1 | 0.360510897 | 0.27708399 |
| Fam133b | 0.30365191 | 0.232556735 | 0.792 | 0.746 | 1 | 0.14977469 | 0.577337723 | 0.829 | 0.754 | 1 | 0.30365191 | 0.277116922 |
| Stim1 | 0.149788684 | 0.433623406 | 0.547 | 0.483 | 1 | 0.914816333 | 0.361243126 | 0.61 | 0.655 | 1 | 0.914816333 | 0.277140718 |
| Gm57095 | 0.149856081 | 2.154722595 | 0.057 | 0.021 | 1 | 0.87556513 | 0.83720485 | 0.024 | 0.02 | 1 | 0.87556513 | 0.277255317 |
| Umad1 | 0.539974703 | 0.495759513 | 0.283 | 0.246 | 1 | 0.149874371 | -0.453791386 | 0.317 | 0.427 | 1 | 0.539974703 | 0.277286414 |
| Pigu | 0.149977382 | 0.79482665 | 0.34 | 0.258 | 1 | 0.465314232 | 0.410940095 | 0.317 | 0.283 | 1 | 0.465314232 | 0.27746155 |
| Rrm2b | 0.19418009 | 0.782304729 | 0.491 | 0.47 | 1 | 0.149977413 | 0.954018515 | 0.634 | 0.553 | 1 | 0.19418009 | 0.277461602 |
| Usp11 | 0.181634716 | 0.617065809 | 0.358 | 0.28 | 1 | 0.1499906 | -0.265156868 | 0.22 | 0.334 | 1 | 0.181634716 | 0.27748402 |
| Eef1akmt2 | 0.150034387 | 0.47665069 | 0.434 | 0.352 | 1 | 0.227746762 | -0.185162963 | 0.244 | 0.352 | 1 | 0.227746762 | 0.277558457 |
| Tor3a | 0.15012805 | -0.822557329 | 0.264 | 0.343 | 1 | 0.362820112 | 0.785116339 | 0.366 | 0.317 | 1 | 0.362820112 | 0.277717668 |

| | | | | | | | | | | | | |
|---|---|---|---|---|---|---|---|---|---|---|---|---|
| Klhl23 | 0.150215458 | 0.526691372 | 0.17 | 0.097 | 1 | 0.634215795 | 0.422167351 | 0.122 | 0.099 | 1 | 0.634215795 | 0.277866232 |
| Gm43920 | 0.150219314 | -1.430239906 | 0 | 0.038 | 1 | 0.609924905 | 0.136765132 | 0.024 | 0.041 | 1 | 0.609924905 | 0.277872786 |
| Adgrg1 | 0.150219314 | -1.752168001 | 0 | 0.038 | 1 | 0.420814977 | 0.515276755 | 0.122 | 0.082 | 1 | 0.420814977 | 0.277872786 |
| Gm12002 | 0.150219314 | -1.304709024 | 0 | 0.038 | 1 | 0.154190296 | -1.069685746 | 0 | 0.048 | 1 | 0.154190296 | 0.277872786 |
| Pcdhgb2 | 0.150219314 | -1.430239906 | 0 | 0.038 | 1 | 0.846926792 | 1.83720485 | 0.024 | 0.031 | 1 | 0.846926792 | 0.277872786 |
| Slc7a14 | 0.15023499 | -1.845277405 | 0 | 0.038 | 1 | 0.749728988 | 0.83720485 | 0.024 | 0.017 | 1 | 0.749728988 | 0.277899428 |
| Mutyh | 0.15023499 | -1.652632327 | 0 | 0.038 | 1 | 0.744047924 | 1.252242349 | 0.024 | 0.017 | 1 | 0.744047924 | 0.277899428 |
| Lsr | 0.15023499 | -1.430239906 | 0 | 0.038 | 1 | 0.572537732 | 0.351778023 | 0.098 | 0.072 | 1 | 0.572537732 | 0.277899428 |
| Gm5608 | 0.15023499 | -1.652632327 | 0 | 0.038 | 1 | 0.319641967 | -0.16279515 | 0 | 0.024 | 1 | 0.319641967 | 0.277899428 |
| Gm10644 | 0.15023499 | -1.430239906 | 0 | 0.038 | 1 | 0.878310931 | 0.515276755 | 0.024 | 0.02 | 1 | 0.878310931 | 0.277899428 |
| Gm26664 | 0.15023499 | -1.545717123 | 0 | 0.038 | 1 | 0.208830556 | -1.16279515 | 0 | 0.038 | 1 | 0.208830556 | 0.277899428 |
| 1700019J19Rik | 0.15023499 | -1.545717123 | 0 | 0.038 | 1 | 0.438672228 | 1.83720485 | 0.024 | 0.01 | 1 | 0.438672228 | 0.277899428 |
| Zfp953 | 0.15023499 | -1.652632327 | 0 | 0.038 | 1 | 0.286433344 | -0.332720152 | 0 | 0.027 | 1 | 0.286433344 | 0.277899428 |
| 4930509G22Rik | 0.15023499 | -1.430239906 | 0 | 0.038 | 1 | 0.231606831 | -0.863234868 | 0 | 0.034 | 1 | 0.231606831 | 0.277899428 |
| Fiz1 | 0.291274566 | 0.8327945 | 0.208 | 0.153 | 1 | 0.150240587 | -0.555112573 | 0.122 | 0.225 | 1 | 0.291274566 | 0.27790894 |
| A330033J07Rik | 0.150246187 | -1.752168001 | 0 | 0.038 | 1 | 0.357854401 | -1.16279515 | 0 | 0.02 | 1 | 0.357854401 | 0.277918458 |
| Mas1 | 0.150246187 | -1.652632327 | 0 | 0.038 | 1 | 0.170353247 | -1.069685746 | 0 | 0.044 | 1 | 0.170353247 | 0.277918458 |
| Mansc4 | 0.150246747 | -1.845277405 | 0 | 0.038 | 1 | 0.357854401 | -0.622226769 | 0 | 0.02 | 1 | 0.357854401 | 0.277919409 |
| F630206G17Rik | 0.150246747 | -1.545717123 | 0 | 0.038 | 1 | 0.319619815 | -0.484723245 | 0 | 0.024 | 1 | 0.319619815 | 0.277919409 |
| Pomc | 0.150246747 | -1.545717123 | 0 | 0.038 | 1 | 0.916649754 | 0.667279848 | 0.024 | 0.027 | 1 | 0.916649754 | 0.277919409 |
| Dsc2 | 0.150252905 | -2.1672055 | 0 | 0.038 | 1 | 0.296893019 | 0.547698233 | 0.195 | 0.133 | 1 | 0.296893019 | 0.277929875 |
| Sfrp5 | 0.150254585 | -3.093204919 | 0 | 0.038 | 1 | 0.286402368 | -1.410722664 | 0 | 0.027 | 1 | 0.286402368 | 0.277932729 |
| Lct | 0.150255145 | -1.652632327 | 0 | 0.038 | 1 | 0.928869203 | 0.614812428 | 0.024 | 0.027 | 1 | 0.928869203 | 0.277933681 |
| A230108P19Rik | 0.150255145 | -1.652632327 | 0 | 0.038 | 1 | 0.319629309 | -0.484723245 | 0 | 0.024 | 1 | 0.319629309 | 0.277933681 |
| Ablim2 | 0.150255145 | -2.093204919 | 0 | 0.038 | 1 | 0.51919986 | 0.83720485 | 0 | 0.01 | 1 | 0.51919986 | 0.277933681 |
| 4921511C10Rik | 0.150266341 | -1.1672055 | 0 | 0.038 | 1 | 0.430024119 | 0.174239837 | 0.024 | 0.055 | 1 | 0.430024119 | 0.277952708 |
| Gm42809 | 0.150266341 | -1.1672055 | 0 | 0.038 | 1 | 0.5191986 | 0.252242349 | 0 | 0.01 | 1 | 0.5191986 | 0.277952708 |
| Gm57313 | 0.150266341 | -1.1672055 | 0 | 0.038 | 1 | 0.51919986 | 0.83720485 | 0 | 0.01 | 1 | 0.51919986 | 0.277952708 |
| Gm39167 | 0.150266341 | -1.1672055 | 0 | 0.038 | 1 | 0.357854401 | -0.484723245 | 0 | 0.02 | 1 | 0.357854401 | 0.277952708 |
| A930033M14Rik | 0.150266341 | -1.1672055 | 0 | 0.038 | 1 | 0.231606831 | -0.970150072 | 0 | 0.034 | 1 | 0.231606831 | 0.277952708 |
| Gm47920 | 0.150266341 | -1.1672055 | 0 | 0.038 | 1 | 0.8229637 | 0.252242349 | 0.049 | 0.058 | 1 | 0.8229637 | 0.277952708 |
| Pcdh1 | 0.150266341 | -1.1672055 | 0 | 0.038 | 1 | 0.916649754 | 0.667279848 | 0.024 | 0.027 | 1 | 0.916649754 | 0.277952708 |
| Fmnl2 | 0.186146396 | 0.214148873 | 0.887 | 0.767 | 1 | 0.150487093 | 0.251678025 | 0.902 | 0.819 | 1 | 0.186146396 | 0.278327821 |
| Noct | 0.150499794 | -0.745442138 | 0.434 | 0.504 | 1 | 0.291480836 | -0.692318664 | 0.585 | 0.604 | 1 | 0.291480836 | 0.2783494 |
| Thsd4 | 0.150512775 | -0.124350021 | 0.472 | 0.627 | 1 | 0.611226588 | 0.559034972 | 0.61 | 0.724 | 1 | 0.611226588 | 0.278371454 |
| Rab40b | 0.150764114 | 1.932330173 | 0.057 | 0.021 | 1 | 0.606262835 | -0.16279515 | 0.024 | 0.041 | 1 | 0.606262835 | 0.278798409 |
| Riok1 | 0.150831937 | 0.439363457 | 0.604 | 0.538 | 1 | 0.528982751 | 0.208388059 | 0.61 | 0.577 | 1 | 0.528982751 | 0.278913601 |
| Dcaf13 | 0.150844151 | 0.460135603 | 0.453 | 0.373 | 1 | 0.538248361 | 0.235754226 | 0.341 | 0.416 | 1 | 0.538248361 | 0.278934345 |
| Lockd | 0.151170084 | -1.149058153 | 0.113 | 0.055 | 1 | 0.22485586 | 1.515276755 | 0.073 | 0.034 | 1 | 0.22485586 | 0.279487774 |
| Snx19 | 0.151317571 | 1.017219071 | 0.34 | 0.267 | 1 | 0.866370078 | 0.13149019 | 0.317 | 0.345 | 1 | 0.866370078 | 0.279738135 |
| Nipa1 | 0.336724592 | 0.874614676 | 0.17 | 0.123 | 1 | 0.151321402 | 0.721727632 | 0.195 | 0.116 | 1 | 0.336724592 | 0.279744638 |
| Pstk | 0.151323358 | 0.776210972 | 0.226 | 0.148 | 1 | 0.632601175 | -0.385187572 | 0.122 | 0.147 | 1 | 0.632601175 | 0.279747957 |
| Cep83os | 0.15142322 | -0.771276824 | 0.057 | 0.127 | 1 | 0.714475009 | 0.83720485 | 0.073 | 0.092 | 1 | 0.714475009 | 0.279917448 |
| Tmem230 | 0.151514091 | -0.216836268 | 0.396 | 0.555 | 1 | 0.523890983 | 0.356772933 | 0.317 | 0.399 | 1 | 0.523890983 | 0.280071662 |
| Gm37083 | 0.151531956 | -0.7032584 | 0.038 | 0.102 | 1 | 0.27744837 | -0.80665134 | 0.024 | 0.068 | 1 | 0.27744837 | 0.280101978 |
| Nudt10 | 0.151679481 | 1.962077517 | 0.057 | 0.021 | 1 | 0.441381862 | 1.515276755 | 0.024 | 0.01 | 1 | 0.441381862 | 0.280352297 |
| Slc35f5 | 0.951493427 | 0.15974067 | 0.642 | 0.644 | 1 | 0.151763271 | -0.291200882 | 0.585 | 0.71 | 1 | 0.951493427 | 0.280494451 |
| Hint3 | 0.151862731 | -0.477545621 | 0.245 | 0.356 | 1 | 0.844611327 | 0.120997816 | 0.244 | 0.225 | 1 | 0.844611327 | 0.280663173 |
| Psmd5 | 0.151873479 | 0.47665069 | 0.321 | 0.233 | 1 | 0.198530961 | 0.646063362 | 0.415 | 0.338 | 1 | 0.198530961 | 0.280681404 |
| Zfp385a | 0.151885074 | 0.435904347 | 0.321 | 0.225 | 1 | 0.83068402 | -0.286177566 | 0.146 | 0.126 | 1 | 0.83068402 | 0.280701073 |
| Rnls | 0.707521342 | 0.645708947 | 0.113 | 0.097 | 1 | 0.152050579 | -2.140075074 | 0.024 | 0.089 | 1 | 0.707521342 | 0.28098178 |
| Tmub1 | 0.520815516 | -0.147840175 | 0.208 | 0.25 | 1 | 0.152124891 | -0.516432105 | 0.146 | 0.249 | 1 | 0.520815516 | 0.2811078 |
| Ergic2 | 0.560873291 | 0.242398113 | 0.717 | 0.788 | 1 | 0.152170006 | 0.457711467 | 0.854 | 0.829 | 1 | 0.560873291 | 0.281184301 |
| Spryd7 | 0.152275603 | 0.701004627 | 0.283 | 0.203 | 1 | 0.903048731 | 0.328191202 | 0.195 | 0.188 | 1 | 0.903048731 | 0.281363347 |
| Ncapg2 | 0.152361651 | -0.827130058 | 0.057 | 0.131 | 1 | 0.928839407 | -0.442903069 | 0.122 | 0.113 | 1 | 0.928839407 | 0.28150923 |
| Dpp9 | 0.268772074 | 0.652222254 | 0.226 | 0.165 | 1 | 0.152466977 | -0.238083278 | 0.195 | 0.321 | 1 | 0.268772074 | 0.281687775 |
| 1810062O18Rik | 0.232775423 | -0.545717123 | 0.057 | 0.114 | 1 | 0.152564924 | -0.899760744 | 0.024 | 0.092 | 1 | 0.232775423 | 0.281853792 |
| Traf1 | 0.152568749 | 2.154722595 | 0.057 | 0.021 | 1 | 0.729889663 | 2.252242349 | 0.024 | 0.017 | 1 | 0.729889663 | 0.281860276 |
| Rrh | 0.152568749 | 1.891688189 | 0.057 | 0.021 | 1 | 0.732716552 | 1.614812428 | 0.024 | 0.017 | 1 | 0.732716552 | 0.281860276 |
| Stag3 | 0.152568749 | 1.891688189 | 0.057 | 0.021 | 1 | 0.744047924 | 1.252242349 | 0.024 | 0.017 | 1 | 0.744047924 | 0.281860276 |
| 6530437J22Rik | 0.152568749 | 1.891688189 | 0.057 | 0.021 | 1 | 0.878310931 | 0.667279848 | 0.024 | 0.02 | 1 | 0.878310931 | 0.281860276 |
| 1700030C12Rik | 0.152568749 | 1.891688189 | 0.057 | 0.021 | 1 | 0.288314827 | 1.252242349 | 0.073 | 0.038 | 1 | 0.288314827 | 0.281860276 |
| Ttbk1 | 0.152568749 | 2.154722595 | 0.057 | 0.021 | 1 | 0.183755686 | 1.83720485 | 0.049 | 0.017 | 1 | 0.183755686 | 0.281860276 |
| Sec31b | 0.152568749 | 1.891688189 | 0.057 | 0.021 | 1 | 0.916649754 | 0.667279848 | 0.024 | 0.027 | 1 | 0.916649754 | 0.281860276 |
| Fam122b | 0.152568749 | 1.891688189 | 0.057 | 0.021 | 1 | 0.441381862 | 1.515276755 | 0.024 | 0.01 | 1 | 0.441381862 | 0.281860276 |
| Srprb | 0.152609154 | 0.354492107 | 0.604 | 0.547 | 1 | 0.277876306 | -0.150722318 | 0.39 | 0.505 | 1 | 0.277876306 | 0.281928754 |
| Dynlt1f | 0.15269861 | 0.990335777 | 0.302 | 0.225 | 1 | 0.490636399 | 0.252242349 | 0.122 | 0.171 | 1 | 0.490636399 | 0.282080355 |
| Scamp1 | 0.152788727 | 0.760190751 | 0.472 | 0.432 | 1 | 0.870882053 | 0.194631626 | 0.512 | 0.536 | 1 | 0.870882053 | 0.282233059 |
| ENSMUSG00000121476 | 0.226045728 | 1.891688189 | 0.075 | 0.038 | 1 | 0.152803103 | 1.328191202 | 0.146 | 0.082 | 1 | 0.226045728 | 0.282257417 |
| Psmf1 | 0.153074648 | 0.549001534 | 0.34 | 0.246 | 1 | 0.805875963 | -0.27321914 | 0.22 | 0.229 | 1 | 0.805875963 | 0.282717448 |
| Adsl | 0.153086206 | 1.180717803 | 0.264 | 0.199 | 1 | 0.78374919 | 0.646521288 | 0.22 | 0.27 | 1 | 0.78374919 | 0.282737025 |
| Actr8 | 0.153099533 | 0.480950827 | 0.283 | 0.199 | 1 | 0.647585408 | 0.483567895 | 0.268 | 0.324 | 1 | 0.647585408 | 0.2827596 |
| Pkp1 | 0.343000446 | -0.430239906 | 0 | 0.017 | 1 | 0.153109368 | 1.071670103 | 0.098 | 0.044 | 1 | 0.343000446 | 0.282776257 |
| Scd4 | 0.767066103 | 0.569760094 | 0.038 | 0.03 | 1 | 0.153109368 | 0.911205431 | 0.098 | 0.044 | 1 | 0.767066103 | 0.282776257 |
| Snx29 | 0.153193753 | 0.45589213 | 0.509 | 0.436 | 1 | 0.30717345 | 0.202854322 | 0.634 | 0.58 | 1 | 0.30717345 | 0.28291918 |
| Ankmy1 | 0.167049705 | 0.533234218 | 0.283 | 0.199 | 1 | 0.153195373 | 1.328191202 | 0.268 | 0.191 | 1 | 0.167049705 | 0.282921923 |
| Bysl | 0.600624856 | 0.535812762 | 0.17 | 0.148 | 1 | 0.153298184 | -0.271319607 | 0.098 | 0.198 | 1 | 0.600624856 | 0.283096034 |
| Pfkfb4 | 0.856896162 | 0.140367302 | 0.226 | 0.254 | 1 | 0.153301168 | -1.146307027 | 0.171 | 0.263 | 1 | 0.856896162 | 0.283101087 |
| Zfp938 | 0.606521232 | 0.178970141 | 0.208 | 0.178 | 1 | 0.153320325 | -0.648221977 | 0.073 | 0.16 | 1 | 0.606521232 | 0.283133528 |
| Yipf6 | 0.48458902 | -0.207847485 | 0.283 | 0.335 | 1 | 0.15340195 | -0.529117364 | 0.244 | 0.362 | 1 | 0.48458902 | 0.283271742 |
| Klhl3 | 0.153517158 | 1.792152515 | 0.057 | 0.021 | 1 | 0.609002981 | 1.458693227 | 0.049 | 0.034 | 1 | 0.609002981 | 0.283466798 |
| Gpr160 | 0.153517791 | 1.739685096 | 0.057 | 0.021 | 1 | 0.878310931 | 0.515276755 | 0.024 | 0.02 | 1 | 0.878310931 | 0.283467869 |
| Faap100 | 0.542187926 | 0.80108564 | 0.113 | 0.089 | 1 | 0.153601789 | -0.185162963 | 0.073 | 0.171 | 1 | 0.542187926 | 0.283610068 |
| Ercc1 | 0.153651309 | 0.810768194 | 0.321 | 0.246 | 1 | 0.260062112 | -0.221688839 | 0.22 | 0.324 | 1 | 0.260062112 | 0.283693894 |
| Impdh1 | 0.153796157 | 0.752624151 | 0.208 | 0.131 | 1 | 0.618074001 | 0.359157553 | 0.244 | 0.212 | 1 | 0.618074001 | 0.283939055 |
| Abhd17a | 0.153890004 | 0.450178478 | 0.566 | 0.521 | 1 | 0.951657214 | 0.181159251 | 0.488 | 0.502 | 1 | 0.951657214 | 0.284097874 |
| 4930453N24Rik | 0.655198284 | 0.241137347 | 0.472 | 0.466 | 1 | 0.15397901 | 0.667279848 | 0.537 | 0.444 | 1 | 0.655198284 | 0.284248496 |
| Tspan8 | 0.153991622 | -1.845277405 | 0.038 | 0.097 | 1 | 0.829279853 | -0.38886323 | 0.244 | 0.218 | 1 | 0.829279853 | 0.284269825 |
| Cdin1 | 0.230028758 | 1.104096522 | 0.189 | 0.131 | 1 | 0.154051294 | -1.050320421 | 0.049 | 0.126 | 1 | 0.230028758 | 0.284370787 |
| Usp27x | 0.154056102 | -0.545717123 | 0.038 | 0.102 | 1 | 0.230630589 | -0.747757651 | 0.024 | 0.075 | 1 | 0.230630589 | 0.284378922 |
| Pdcd10 | 0.154075736 | 0.418818196 | 0.83 | 0.826 | 1 | 0.75749083 | 0.15834221 | 0.756 | 0.881 | 1 | 0.75749083 | 0.28441214 |
| Gm45238 | 0.157246372 | 1.569760094 | 0.057 | 0.021 | 1 | 0.154090132 | -1.16279515 | 0 | 0.048 | 1 | 0.157246372 | 0.284436495 |
| 2310034G01Rik | 0.232194313 | -0.752168001 | 0.019 | 0.059 | 1 | 0.154090132 | -1.16279515 | 0 | 0.048 | 1 | 0.232194313 | 0.284436495 |
| Nudt7 | 0.251917817 | 0.932330173 | 0.113 | 0.068 | 1 | 0.154090132 | -1.16279515 | 0 | 0.048 | 1 | 0.251917817 | 0.284436495 |
| Gm4013 | 0.312692399 | 1.017219071 | 0.075 | 0.042 | 1 | 0.154090132 | -1.250257991 | 0 | 0.048 | 1 | 0.312692399 | 0.284436495 |
| Gm56696 | 0.418292339 | -0.545717123 | 0.019 | 0.042 | 1 | 0.154090132 | -1.250257991 | 0 | 0.048 | 1 | 0.418292339 | 0.284436495 |
| Gm19500 | 0.628582732 | 1.154722595 | 0.057 | 0.042 | 1 | 0.154090132 | -1.16279515 | 0 | 0.048 | 1 | 0.628582732 | 0.284436495 |
| Hgh1 | 0.321169157 | 0.865215978 | 0.132 | 0.089 | 1 | 0.154111598 | -1.250257991 | 0 | 0.048 | 1 | 0.321169157 | 0.284472811 |

| | | | | | | | | | | | | |
|---|---|---|---|---|---|---|---|---|---|---|---|---|
| Mbnl3 | 0.989012386 | -0.368839361 | 0.057 | 0.055 | 1 | 0.154111598 | -1.332720152 | 0 | 0.048 | 1 | 0.989012386 | 0.284472811 |
| Acot13 | 0.405229203 | 0.433876666 | 0.34 | 0.288 | 1 | 0.154119299 | -0.818146979 | 0.195 | 0.294 | 1 | 0.405229203 | 0.28448584 |
| E130114P18Rik | 0.225110966 | 1.874614676 | 0.075 | 0.038 | 1 | 0.15412976 | -1.332720152 | 0 | 0.048 | 1 | 0.225110966 | 0.284503537 |
| 9130230L23Rik | 0.725936481 | 1.417757001 | 0.019 | 0.013 | 1 | 0.154143794 | -1.917682652 | 0 | 0.048 | 1 | 0.725936481 | 0.284527279 |
| Gm34312 | 0.507281812 | 1.154722595 | 0.019 | 0.008 | 1 | 0.154144895 | -1.410722664 | 0 | 0.048 | 1 | 0.507281812 | 0.284529141 |
| Klhl40 | 0.776830424 | -0.652632327 | 0.019 | 0.025 | 1 | 0.154154526 | -1.555112573 | 0 | 0.048 | 1 | 0.776830424 | 0.284545434 |
| Gm53065 | 0.503327126 | 0.154722595 | 0.019 | 0.038 | 1 | 0.154156177 | -1.80665134 | 0 | 0.048 | 1 | 0.503327126 | 0.284548227 |
| Gm17276 | 0.929410101 | -0.545717123 | 0.019 | 0.017 | 1 | 0.154164432 | -1.863234868 | 0 | 0.048 | 1 | 0.929410101 | 0.284562191 |
| Ptk2b | 0.50699757 | 0.669295768 | 0.019 | 0.038 | 1 | 0.154172136 | -2.116991461 | 0 | 0.048 | 1 | 0.50699757 | 0.284575224 |
| Gm13199 | 0.343968402 | 1.154722595 | 0.038 | 0.017 | 1 | 0.154190296 | -1.069685746 | 0 | 0.048 | 1 | 0.343968402 | 0.284605944 |
| Tmem95 | 0.550306015 | 0.669295768 | 0.057 | 0.038 | 1 | 0.154190296 | -1.069685746 | 0 | 0.048 | 1 | 0.550306015 | 0.284605944 |
| Orm1 | 0.567830665 | -0.430239906 | 0.019 | 0.034 | 1 | 0.154190296 | -1.069685746 | 0 | 0.048 | 1 | 0.567830665 | 0.284605944 |
| Rexo4 | 0.326476951 | 0.773293488 | 0.34 | 0.318 | 1 | 0.154422556 | -0.305753104 | 0.244 | 0.372 | 1 | 0.326476951 | 0.284998786 |
| Cplx1 | 0.154426139 | 1.932330173 | 0.057 | 0.021 | 1 | 0.286413984 | -0.863234868 | 0 | 0.027 | 1 | 0.286413984 | 0.285004846 |
| Arvcf | 0.154426139 | 2.347367673 | 0.057 | 0.021 | 1 | 0.319639436 | -0.747757651 | 0 | 0.024 | 1 | 0.319639436 | 0.285004846 |
| Zfp867 | 0.46991905 | 0.569760094 | 0.17 | 0.136 | 1 | 0.154566836 | -1.292078167 | 0.024 | 0.089 | 1 | 0.46991905 | 0.285242766 |
| Gm28050 | 0.154760744 | 1.707263618 | 0.094 | 0.047 | 1 | 0.213504175 | -1.250257991 | 0.024 | 0.078 | 1 | 0.213504175 | 0.2855706 |
| Dimt1 | 0.154787362 | 0.633185474 | 0.358 | 0.284 | 1 | 0.558275552 | 0.114738825 | 0.244 | 0.304 | 1 | 0.558275552 | 0.285615596 |
| Glce | 0.15482752 | 0.59400764 | 0.509 | 0.411 | 1 | 0.820553723 | 0.146889349 | 0.366 | 0.44 | 1 | 0.820553723 | 0.28568348 |
| Kif17 | 0.154930625 | 1.287989126 | 0.151 | 0.089 | 1 | 0.432606101 | 0.88967227 | 0.171 | 0.126 | 1 | 0.432606101 | 0.285857751 |
| Zfp775 | 0.155037799 | -1.430239906 | 0.038 | 0.097 | 1 | 0.69297937 | 0.979223855 | 0.073 | 0.058 | 1 | 0.69297937 | 0.286038879 |
| Zfyve19 | 0.155297009 | 1.417757001 | 0.113 | 0.059 | 1 | 0.477010767 | 0.667279848 | 0.049 | 0.085 | 1 | 0.477010767 | 0.286476856 |
| Cnep1r1 | 0.155521491 | -0.14961144 | 0.585 | 0.72 | 1 | 0.995760678 | 0.139613494 | 0.634 | 0.751 | 1 | 0.995760678 | 0.286856047 |
| Stbd1 | 0.361131243 | -0.545717123 | 0.019 | 0.047 | 1 | 0.155567084 | -1.250257991 | 0.024 | 0.089 | 1 | 0.361131243 | 0.28693305 |
| Ubald1 | 0.155584851 | -0.351168335 | 0.264 | 0.381 | 1 | 0.936212114 | 0.39094862 | 0.268 | 0.283 | 1 | 0.936212114 | 0.286963056 |
| Plekha8 | 0.502983574 | -0.357727406 | 0.208 | 0.254 | 1 | 0.155608312 | -1.19841906 | 0.098 | 0.181 | 1 | 0.502983574 | 0.287002678 |
| Lekr1 | 0.155632647 | -0.776014743 | 0.057 | 0.131 | 1 | 0.240183821 | -0.787286015 | 0.024 | 0.075 | 1 | 0.240183821 | 0.287043774 |
| Tmem258 | 0.155746853 | -0.355033562 | 0.736 | 0.831 | 1 | 0.796995057 | 0.127006165 | 0.683 | 0.724 | 1 | 0.796995057 | 0.287236624 |
| Aph1c | 0.155783418 | 1.154722595 | 0.151 | 0.089 | 1 | 0.829120528 | 0.114738825 | 0.098 | 0.085 | 1 | 0.829120528 | 0.287298362 |
| Uqcc5 | 0.859911639 | 0.341491106 | 0.358 | 0.36 | 1 | 0.155913833 | -0.332720152 | 0.22 | 0.341 | 1 | 0.859911639 | 0.287518542 |
| Fam217b | 0.156004119 | 1.569760094 | 0.094 | 0.047 | 1 | 0.957192022 | 0.83720485 | 0.049 | 0.048 | 1 | 0.957192022 | 0.287670954 |
| Fam234b | 0.156004119 | 1.569760094 | 0.094 | 0.047 | 1 | 0.170353247 | -1.16279515 | 0 | 0.044 | 1 | 0.170353247 | 0.287670954 |
| Gm12279 | 0.156004119 | 1.891688189 | 0.094 | 0.047 | 1 | 0.672416858 | -0.707115666 | 0.049 | 0.065 | 1 | 0.672416858 | 0.287670954 |
| Exog | 0.385114122 | 1.039245377 | 0.075 | 0.047 | 1 | 0.156062583 | -1.116991461 | 0.024 | 0.089 | 1 | 0.385114122 | 0.287769637 |
| Gm48512 | 0.156102092 | 1.8327945 | 0.075 | 0.034 | 1 | 0.466583276 | -0.484723245 | 0.024 | 0.051 | 1 | 0.466583276 | 0.287836321 |
| As3mt | 0.755941787 | 0.119098685 | 0.132 | 0.153 | 1 | 0.156126892 | -0.20718927 | 0.122 | 0.232 | 1 | 0.755941787 | 0.287878178 |
| Usp2 | 0.489105964 | -0.102665248 | 0.264 | 0.326 | 1 | 0.156157983 | 0.778311161 | 0.293 | 0.198 | 1 | 0.489105964 | 0.287930651 |
| Gm37982 | 0.156177221 | -1.260314904 | 0.019 | 0.072 | 1 | 0.257285084 | -0.622226769 | 0 | 0.031 | 1 | 0.257285084 | 0.287963118 |
| Gm42047 | 0.70650009 | -0.905973337 | 0.208 | 0.22 | 1 | 0.156178415 | 0.883008539 | 0.195 | 0.123 | 1 | 0.70650009 | 0.287965133 |
| Armc2 | 0.156287154 | 1.47665069 | 0.057 | 0.021 | 1 | 0.20883094 | -1.069685746 | 0 | 0.038 | 1 | 0.20883094 | 0.288148633 |
| F930017D23Rik | 0.156296076 | 1.739685096 | 0.057 | 0.021 | 1 | 0.5191986 | -0.332720152 | 0 | 0.01 | 1 | 0.5191986 | 0.288163688 |
| Tph2 | 0.15630436 | 0.739685096 | 0.057 | 0.021 | 1 | 0.492843202 | -0.747757651 | 0.024 | 0.048 | 1 | 0.492843202 | 0.288177667 |
| Sprtn | 0.156350042 | 0.739685096 | 0.226 | 0.148 | 1 | 0.424433582 | 0.458693227 | 0.146 | 0.212 | 1 | 0.424433582 | 0.288254749 |
| Gm12089 | 0.338780504 | 0.984797593 | 0.057 | 0.03 | 1 | 0.156555327 | -1.069685746 | 0.024 | 0.089 | 1 | 0.338780504 | 0.288601083 |
| Crat | 0.717783041 | 0.43225657 | 0.132 | 0.114 | 1 | 0.15677102 | -1.069685746 | 0.049 | 0.123 | 1 | 0.717783041 | 0.288964888 |
| Mtfr2 | 0.156804309 | 1.055186921 | 0.151 | 0.089 | 1 | 0.280833036 | 1.109284395 | 0.171 | 0.119 | 1 | 0.280833036 | 0.289021027 |
| Shisa5 | 0.168620333 | -0.287023715 | 0.83 | 0.835 | 1 | 0.15690387 | -0.312898711 | 0.659 | 0.737 | 1 | 0.168620333 | 0.289188915 |
| Atp6v0d1 | 0.205252398 | 0.293596416 | 0.66 | 0.631 | 1 | 0.156926185 | -0.256394517 | 0.707 | 0.816 | 1 | 0.205252398 | 0.289226542 |
| Nqo2 | 0.777125669 | 0.110328475 | 0.094 | 0.11 | 1 | 0.157170927 | 0.766815522 | 0.195 | 0.116 | 1 | 0.777125669 | 0.289639153 |
| Wdr91 | 0.869366983 | 0.2967416 | 0.113 | 0.106 | 1 | 0.157178758 | 0.289717055 | 0.195 | 0.116 | 1 | 0.869366983 | 0.289652354 |
| Cldn15 | 0.157246372 | 1.569760094 | 0.057 | 0.021 | 1 | 0.969252137 | 0.83720485 | 0.049 | 0.051 | 1 | 0.969252137 | 0.289766323 |
| Tril | 0.157246372 | 1.569760094 | 0.057 | 0.021 | 1 | 0.438672228 | 1.83720485 | 0.024 | 0.01 | 1 | 0.438672228 | 0.289766323 |
| Kifc2 | 0.157246372 | 1.569760094 | 0.057 | 0.021 | 1 | 0.561713804 | 0.83720485 | 0.073 | 0.051 | 1 | 0.561713804 | 0.289766323 |
| Eme2 | 0.157246372 | 1.569760094 | 0.057 | 0.021 | 1 | 0.305356148 | -0.747757651 | 0.024 | 0.065 | 1 | 0.305356148 | 0.289766323 |
| Gm4566 | 0.157246372 | 1.569760094 | 0.057 | 0.021 | 1 | 0.357854401 | -0.747757651 | 0 | 0.02 | 1 | 0.357854401 | 0.289766323 |
| Rbms1 | 0.923059628 | -0.246848837 | 1 | 0.97 | 1 | 0.157397436 | 0.192709026 | 0.951 | 0.993 | 1 | 0.923059628 | 0.290020919 |
| Atg4c | 0.352747923 | 0.610402079 | 0.17 | 0.123 | 1 | 0.157407194 | -0.885261175 | 0.098 | 0.184 | 1 | 0.352747923 | 0.290037364 |
| Gm56698 | 0.157431992 | -0.719746523 | 0.019 | 0.072 | 1 | 0.643119211 | 0.721727632 | 0.049 | 0.034 | 1 | 0.643119211 | 0.290079151 |
| Rabggta | 0.157507282 | 0.906795081 | 0.132 | 0.072 | 1 | 0.250552398 | -1.116991461 | 0.024 | 0.072 | 1 | 0.250552398 | 0.29020602 |
| Ccn4 | 0.157522514 | 0.932330173 | 0.472 | 0.381 | 1 | 0.325222716 | 0.57223264 | 0.317 | 0.273 | 1 | 0.325222716 | 0.290231686 |
| Otub1 | 0.157600173 | 0.441435997 | 0.547 | 0.453 | 1 | 0.311680773 | -0.290284885 | 0.39 | 0.474 | 1 | 0.311680773 | 0.290362532 |
| Msrb1 | 0.375741961 | 0.544669113 | 0.377 | 0.335 | 1 | 0.157678747 | -0.550525303 | 0.268 | 0.375 | 1 | 0.375741961 | 0.290494907 |
| Rbm47 | 0.306679482 | -0.796367805 | 0.208 | 0.263 | 1 | 0.157728771 | 0.387897448 | 0.366 | 0.263 | 1 | 0.306679482 | 0.290579177 |
| Clpp | 0.157832771 | 0.467398376 | 0.604 | 0.542 | 1 | 0.389256003 | 0.358233045 | 0.488 | 0.471 | 1 | 0.389256003 | 0.290754359 |
| Klf8 | 0.295306833 | 0.484871196 | 0.094 | 0.055 | 1 | 0.157848015 | 0.580865097 | 0.024 | 0.092 | 1 | 0.295306833 | 0.290780034 |
| Lmtk2 | 0.158078436 | 0.442847982 | 0.321 | 0.225 | 1 | 0.181426857 | 0.115920878 | 0.561 | 0.427 | 1 | 0.181426857 | 0.291168079 |
| Setd7 | 0.158118154 | 0.294784341 | 0.792 | 0.725 | 1 | 0.471788277 | 0.147373231 | 0.878 | 0.881 | 1 | 0.471788277 | 0.291234957 |
| Gm20470 | 0.158174734 | 1.47665069 | 0.057 | 0.021 | 1 | 0.74932726 | 0.100239256 | 0.049 | 0.038 | 1 | 0.74932726 | 0.291330221 |
| Ube2u | 0.158180495 | 1.640149422 | 0.057 | 0.021 | 1 | 0.416810083 | 0.949679579 | 0.073 | 0.044 | 1 | 0.416810083 | 0.291339921 |
| Whamm | 0.188662726 | 0.541745718 | 0.226 | 0.153 | 1 | 0.158307139 | 1.162605137 | 0.293 | 0.201 | 1 | 0.188662726 | 0.291553128 |
| Rhot2 | 0.520550951 | 0.618669695 | 0.132 | 0.102 | 1 | 0.158351678 | 1.159132945 | 0.171 | 0.099 | 1 | 0.520550951 | 0.291628101 |
| Ptcd2 | 0.158371357 | 1.324647596 | 0.113 | 0.059 | 1 | 0.586888021 | -0.448197369 | 0.073 | 0.099 | 1 | 0.586888021 | 0.291661227 |
| Mdga1 | 0.158470515 | -0.494370243 | 0.208 | 0.309 | 1 | 0.365125634 | 0.969842618 | 0.293 | 0.249 | 1 | 0.365125634 | 0.291828126 |
| Mindy1 | 0.158519954 | 0.485695829 | 0.509 | 0.428 | 1 | 0.390770429 | -0.122797082 | 0.415 | 0.495 | 1 | 0.390770429 | 0.291911332 |
| Cap1 | 0.158557968 | 0.200083128 | 0.83 | 0.75 | 1 | 0.21096109 | -0.206738498 | 0.756 | 0.884 | 1 | 0.21096109 | 0.291975308 |
| Hal | 0.158593074 | -3.021866137 | 0.075 | 0.144 | 1 | 0.515698409 | -1.069685746 | 0.049 | 0.075 | 1 | 0.515698409 | 0.292034384 |
| 2310014F06Rik | 0.158781663 | -0.988235359 | 0.075 | 0.148 | 1 | 0.545350841 | 0.209173627 | 0.122 | 0.167 | 1 | 0.545350841 | 0.292351709 |
| Med23 | 0.158788762 | 0.855162313 | 0.283 | 0.212 | 1 | 0.912861235 | 0.313642894 | 0.293 | 0.304 | 1 | 0.912861235 | 0.292363652 |
| Tbkbp1 | 0.385114122 | 1.039245377 | 0.075 | 0.047 | 1 | 0.158826311 | 1.114738825 | 0.171 | 0.099 | 1 | 0.385114122 | 0.292426825 |
| Ndufb7 | 0.159066125 | 0.393910259 | 0.736 | 0.691 | 1 | 0.440822165 | 0.26618154 | 0.634 | 0.645 | 1 | 0.440822165 | 0.292830218 |
| H2ac25 | 0.159095446 | 0.8327945 | 0.17 | 0.102 | 1 | 0.60296698 | 0.604544093 | 0.146 | 0.119 | 1 | 0.60296698 | 0.292879531 |
| Mipol1 | 0.307829713 | 0.209170379 | 0.377 | 0.309 | 1 | 0.15910362 | 0.721727632 | 0.415 | 0.321 | 1 | 0.307829713 | 0.292893279 |
| Gdap2 | 0.159118017 | -0.968659821 | 0.226 | 0.305 | 1 | 0.266162083 | -0.213421223 | 0.171 | 0.256 | 1 | 0.266162083 | 0.29291749 |
| Nedd8 | 0.159279281 | 0.198992862 | 0.981 | 0.877 | 1 | 0.212815732 | 0.232980447 | 0.854 | 0.881 | 1 | 0.212815732 | 0.293188673 |
| Gas7 | 0.544857305 | -0.214511215 | 0.17 | 0.208 | 1 | 0.159371107 | -0.832646549 | 0.122 | 0.222 | 1 | 0.544857305 | 0.293343064 |
| Gm26910 | 0.159409442 | 1.454282877 | 0.094 | 0.047 | 1 | 0.959997791 | 0.930314254 | 0.073 | 0.072 | 1 | 0.959997791 | 0.293407514 |
| Ift20 | 0.159501581 | 0.130228033 | 0.925 | 0.864 | 1 | 0.545899872 | -0.11032773 | 0.951 | 0.894 | 1 | 0.545899872 | 0.293562408 |
| Fbrs | 0.159534282 | 0.631160639 | 0.396 | 0.309 | 1 | 0.608671431 | -0.124993117 | 0.439 | 0.488 | 1 | 0.608671431 | 0.293617376 |
| Nme2 | 0.159549924 | 0.188094053 | 1 | 0.958 | 1 | 0.382299357 | -0.208936932 | 0.951 | 0.983 | 1 | 0.382299357 | 0.29364367 |
| St7l | 0.30861784 | 0.385020214 | 0.528 | 0.47 | 1 | 0.159794803 | -0.335987865 | 0.512 | 0.642 | 1 | 0.30861784 | 0.294055226 |
| Gnl1 | 0.245472958 | 0.410561499 | 0.472 | 0.407 | 1 | 0.159802898 | -0.174767792 | 0.366 | 0.519 | 1 | 0.245472958 | 0.29406883 |
| Ufd1 | 0.159894251 | 0.417757001 | 0.642 | 0.602 | 1 | 0.529218588 | 0.136765132 | 0.537 | 0.679 | 1 | 0.529218588 | 0.29422233 |
| Nova1 | 0.159972144 | 0.466666601 | 0.019 | 0.072 | 1 | 0.554222599 | 0.962735732 | 0.073 | 0.051 | 1 | 0.554222599 | 0.294353202 |
| Arhgef16 | 0.160046644 | 1.347367673 | 0.057 | 0.021 | 1 | 0.535257741 | 1.644559772 | 0.049 | 0.031 | 1 | 0.535257741 | 0.294478359 |
| Gm36371 | 0.160046644 | 1.347367673 | 0.057 | 0.021 | 1 | 0.735545437 | 1.422167351 | 0.024 | 0.017 | 1 | 0.735545437 | 0.294478359 |
| Fancf | 0.160046644 | 1.347367673 | 0.057 | 0.021 | 1 | 0.359002082 | 1.422167351 | 0.049 | 0.024 | 1 | 0.359002082 | 0.294478359 |
| F730043M19Rik | 0.160046644 | 1.347367673 | 0.057 | 0.021 | 1 | 0.267205177 | 2.252242349 | 0.024 | 0.007 | 1 | 0.267205177 | 0.294478359 |
| R3hcc1 | 0.16032151 | 0.976385354 | 0.226 | 0.157 | 1 | 0.874342486 | 0.313642894 | 0.171 | 0.184 | 1 | 0.874342486 | 0.294940033 |

| | | | | | | | | | | | | |
|---|---|---|---|---|---|---|---|---|---|---|---|---|
| S1pr2 | 0.160390529 | 0.522454379 | 0.283 | 0.195 | 1 | 0.254885525 | -0.366078749 | 0.244 | 0.338 | 1 | 0.254885525 | 0.295055936 |
| Mrgprf | 0.194368 | 0.533234218 | 0.189 | 0.119 | 1 | 0.160411329 | -0.659220976 | 0.098 | 0.188 | 1 | 0.194368 | 0.295090863 |
| Utp4 | 0.160609388 | 0.560541564 | 0.472 | 0.403 | 1 | 0.746727209 | -0.151824032 | 0.512 | 0.522 | 1 | 0.746727209 | 0.295423401 |
| Pomt2 | 0.160626991 | 0.758793918 | 0.245 | 0.169 | 1 | 0.205209552 | -0.332720152 | 0.195 | 0.297 | 1 | 0.205209552 | 0.295452951 |
| Cdkn2c | 0.160659694 | 0.300573461 | 0.264 | 0.178 | 1 | 0.281364896 | -0.347219721 | 0.146 | 0.229 | 1 | 0.281364896 | 0.295507851 |
| Epb41l4aos | 0.160759095 | -0.515128804 | 0.17 | 0.263 | 1 | 0.683510392 | -0.332720152 | 0.146 | 0.167 | 1 | 0.683510392 | 0.295674703 |
| Vwce | 0.160849228 | 1.526691372 | 0.094 | 0.047 | 1 | 0.692964516 | 0.564186355 | 0.073 | 0.058 | 1 | 0.692964516 | 0.295825981 |
| Zmym3 | 0.199061375 | 1.154722595 | 0.113 | 0.064 | 1 | 0.160997312 | 1.064615346 | 0.195 | 0.119 | 1 | 0.199061375 | 0.29607449 |
| Tll2 | 0.161033404 | 1.154722595 | 0.057 | 0.021 | 1 | 0.722142177 | 0.635570989 | 0.049 | 0.038 | 1 | 0.722142177 | 0.296135051 |
| Inpp5e | 0.161053743 | 0.491757582 | 0.17 | 0.102 | 1 | 0.878349089 | 0.109284395 | 0.146 | 0.137 | 1 | 0.878349089 | 0.296169178 |
| Srd5a3 | 0.161147197 | 0.8327945 | 0.226 | 0.153 | 1 | 0.412911906 | -0.144647803 | 0.146 | 0.205 | 1 | 0.412911906 | 0.296325975 |
| Ube2d-ps | 0.39609062 | 0.154722595 | 0.321 | 0.254 | 1 | 0.16115641 | -0.346016974 | 0.171 | 0.27 | 1 | 0.39609062 | 0.296341431 |
| Zfp367 | 0.271278143 | 0.154722595 | 0.264 | 0.203 | 1 | 0.161259286 | 1.082317348 | 0.293 | 0.212 | 1 | 0.271278143 | 0.296514015 |
| Thrb | 0.161409902 | -0.345123292 | 0.717 | 0.835 | 1 | 0.219995732 | -0.524530095 | 0.805 | 0.836 | 1 | 0.219995732 | 0.296766648 |
| Rac3 | 0.979468923 | 0.154722595 | 0.075 | 0.076 | 1 | 0.161506624 | -0.707115666 | 0.073 | 0.16 | 1 | 0.979468923 | 0.296928859 |
| Dhx35 | 0.161514697 | 0.762405172 | 0.132 | 0.072 | 1 | 0.403242332 | -0.250257991 | 0.049 | 0.089 | 1 | 0.403242332 | 0.296942397 |
| Septin5 | 0.262825766 | -0.600164907 | 0.132 | 0.195 | 1 | 0.161666974 | -0.463964685 | 0.098 | 0.195 | 1 | 0.262825766 | 0.297197737 |
| Ripor1 | 0.683020254 | 0.318221327 | 0.264 | 0.246 | 1 | 0.161688126 | -0.509597914 | 0.415 | 0.502 | 1 | 0.683020254 | 0.297233202 |
| Ifit3b | 0.161728777 | -2.330704232 | 0.038 | 0.097 | 1 | 0.555685551 | -1.352619709 | 0.049 | 0.031 | 1 | 0.555685551 | 0.297301356 |
| Mrpl54 | 0.277919044 | 0.371820716 | 0.585 | 0.572 | 1 | 0.161814148 | -0.208807548 | 0.341 | 0.464 | 1 | 0.277919044 | 0.297444478 |
| Lbr | 0.161844152 | 0.438515561 | 0.434 | 0.335 | 1 | 0.321789438 | 0.258937002 | 0.512 | 0.457 | 1 | 0.321789438 | 0.297494775 |
| Commd9 | 0.161864748 | 0.739685096 | 0.245 | 0.169 | 1 | 0.3619072 | -0.19841906 | 0.171 | 0.235 | 1 | 0.3619072 | 0.297529299 |
| Gm50012 | 0.820242667 | 0.569760094 | 0.113 | 0.106 | 1 | 0.161945624 | 0.589277336 | 0.268 | 0.181 | 1 | 0.820242667 | 0.297664863 |
| Zfp64 | 0.535844724 | -0.506007414 | 0.396 | 0.419 | 1 | 0.161952207 | -0.263059274 | 0.293 | 0.447 | 1 | 0.535844724 | 0.297675896 |
| Rfk | 0.810576619 | 0.110328475 | 0.491 | 0.466 | 1 | 0.162039305 | 0.632385058 | 0.659 | 0.563 | 1 | 0.810576619 | 0.297821874 |
| Serpine1 | 0.162071419 | -0.326131503 | 0.472 | 0.331 | 1 | 0.771572255 | -0.47412813 | 0.244 | 0.215 | 1 | 0.771572255 | 0.297875693 |
| Pole | 0.162174986 | 1.569760094 | 0.094 | 0.047 | 1 | 0.878310931 | 0.515276755 | 0.024 | 0.02 | 1 | 0.878310931 | 0.298049245 |
| Zfp219 | 0.162344807 | 0.756173218 | 0.17 | 0.102 | 1 | 0.563700678 | -0.116991461 | 0.195 | 0.239 | 1 | 0.563700678 | 0.298333777 |
| Zbtb6 | 0.162514495 | 0.674729654 | 0.264 | 0.178 | 1 | 0.903462579 | 0.222495006 | 0.244 | 0.249 | 1 | 0.903462579 | 0.29861803 |
| Sgtb | 0.162820175 | 1.002719501 | 0.113 | 0.059 | 1 | 0.190244126 | 1.082317348 | 0.122 | 0.065 | 1 | 0.190244126 | 0.299129941 |
| Elp6 | 0.162865867 | 1.347367673 | 0.094 | 0.047 | 1 | 0.713168757 | 0.313642894 | 0.073 | 0.058 | 1 | 0.713168757 | 0.299206443 |
| 2810029C07Rik | 0.162916707 | -0.430239906 | 0.038 | 0.102 | 1 | 0.82181816 | 0.136765132 | 0.024 | 0.031 | 1 | 0.82181816 | 0.29929156 |
| Rad54l | 0.16293479 | 0.984797593 | 0.057 | 0.021 | 1 | 0.548898236 | -0.16279515 | 0.024 | 0.044 | 1 | 0.548898236 | 0.299321834 |
| Gm13166 | 0.16293479 | 0.984797593 | 0.057 | 0.021 | 1 | 0.916649754 | 0.667279848 | 0.024 | 0.027 | 1 | 0.916649754 | 0.299321834 |
| Hopx | 0.16293479 | 1.154722595 | 0.057 | 0.021 | 1 | 0.231618904 | -1.250257991 | 0 | 0.034 | 1 | 0.231618904 | 0.299321834 |
| Appbp2os | 0.16293479 | 0.984797593 | 0.057 | 0.021 | 1 | 0.824088343 | 0.252242349 | 0.024 | 0.031 | 1 | 0.824088343 | 0.299321834 |
| Gm20559 | 0.939016638 | -0.1672055 | 0.019 | 0.017 | 1 | 0.162944451 | 1.944120054 | 0.073 | 0.031 | 1 | 0.939016638 | 0.299338008 |
| Ccdc112 | 0.162955454 | 0.904179133 | 0.283 | 0.22 | 1 | 0.44165826 | 0.504221566 | 0.293 | 0.253 | 1 | 0.44165826 | 0.299356428 |
| Lrrc20 | 0.16301986 | -0.997280499 | 0.113 | 0.195 | 1 | 0.887779406 | -0.16279515 | 0.098 | 0.089 | 1 | 0.887779406 | 0.299464245 |
| Zfp62 | 0.163208935 | 0.669295768 | 0.34 | 0.263 | 1 | 0.817007142 | 0.266456208 | 0.39 | 0.427 | 1 | 0.817007142 | 0.299780714 |
| Snx9 | 0.266893026 | 0.105006835 | 0.887 | 0.801 | 1 | 0.163249673 | 0.428716136 | 0.927 | 0.901 | 1 | 0.266893026 | 0.299848891 |
| Stt3b | 0.163314613 | 0.169829487 | 0.981 | 0.898 | 1 | 0.48703123 | 0.451146417 | 0.951 | 0.966 | 1 | 0.48703123 | 0.299957563 |
| Cdk5rap3 | 0.384168212 | 0.267197324 | 0.396 | 0.343 | 1 | 0.163524836 | -0.185162963 | 0.22 | 0.341 | 1 | 0.384168212 | 0.3003093 |
| Zkscan5 | 0.163553442 | -1.077938162 | 0.057 | 0.123 | 1 | 0.765425678 | -0.520347155 | 0.073 | 0.085 | 1 | 0.765425678 | 0.300357155 |
| Proca1 | 0.383132106 | 0.306725688 | 0.151 | 0.106 | 1 | 0.163801853 | 0.334704509 | 0.195 | 0.113 | 1 | 0.383132106 | 0.300772659 |
| Gmeb2 | 0.163871946 | -0.341703231 | 0.321 | 0.424 | 1 | 0.641782421 | 0.178993367 | 0.415 | 0.386 | 1 | 0.641782421 | 0.300889878 |
| Bambi | 0.302958321 | -0.395474488 | 0.623 | 0.674 | 1 | 0.164108228 | 1.120997816 | 0.561 | 0.481 | 1 | 0.302958321 | 0.301284945 |
| Gm14295 | 0.164198307 | 1.055186921 | 0.113 | 0.059 | 1 | 0.572537583 | 0.458693227 | 0.098 | 0.072 | 1 | 0.572537583 | 0.301435529 |
| Taf1c | 0.164198307 | 1.055186921 | 0.113 | 0.059 | 1 | 0.337213694 | -0.555112573 | 0.024 | 0.061 | 1 | 0.337213694 | 0.301435529 |
| Nmral1 | 0.164198307 | 1.055186921 | 0.113 | 0.059 | 1 | 0.711767985 | 0.404245443 | 0.049 | 0.065 | 1 | 0.711767985 | 0.301435529 |
| Stard3nl | 0.164222765 | 0.389187848 | 0.491 | 0.394 | 1 | 0.69761947 | 0.280811501 | 0.463 | 0.43 | 1 | 0.69761947 | 0.301476413 |
| Pwp2 | 0.174195296 | 0.8327945 | 0.151 | 0.089 | 1 | 0.164263955 | -0.364429011 | 0.049 | 0.13 | 1 | 0.174195296 | 0.301545264 |
| Crtc2 | 0.164338109 | 0.548065023 | 0.321 | 0.229 | 1 | 0.670820167 | 0.680700364 | 0.293 | 0.29 | 1 | 0.670820167 | 0.301669203 |
| Ocrl | 0.164390205 | 0.71862348 | 0.415 | 0.369 | 1 | 0.474833196 | -0.321622443 | 0.415 | 0.447 | 1 | 0.474833196 | 0.301756271 |
| Dip2a | 0.164417704 | 0.669295768 | 0.226 | 0.153 | 1 | 0.454009506 | 0.261892519 | 0.22 | 0.3 | 1 | 0.454009506 | 0.301802226 |
| E130307A14Rik | 0.164527595 | 0.803250224 | 0.264 | 0.186 | 1 | 0.412219522 | -0.292078167 | 0.146 | 0.205 | 1 | 0.412219522 | 0.301985861 |
| B3galt2 | 0.929410101 | 0.347367673 | 0.019 | 0.017 | 1 | 0.164564102 | 1.721727632 | 0.073 | 0.031 | 1 | 0.929410101 | 0.302046861 |
| Zfp668 | 0.164832152 | 1.017219071 | 0.151 | 0.089 | 1 | 0.205111975 | 0.930314254 | 0.146 | 0.085 | 1 | 0.205111975 | 0.302494665 |
| Atxn7l1 | 0.427335558 | -0.489133595 | 0.396 | 0.441 | 1 | 0.164945368 | -0.574070731 | 0.415 | 0.512 | 1 | 0.427335558 | 0.302683762 |
| Washc5 | 0.16510041 | 0.767699472 | 0.358 | 0.284 | 1 | 0.796382885 | -0.185878763 | 0.439 | 0.447 | 1 | 0.796382885 | 0.302942674 |
| Utp15 | 0.556142788 | 0.285967128 | 0.321 | 0.288 | 1 | 0.165113606 | -0.693309867 | 0.146 | 0.246 | 1 | 0.556142788 | 0.302964709 |
| Tspan9 | 0.277927848 | 0.627475592 | 0.34 | 0.267 | 1 | 0.165129579 | 0.557096931 | 0.488 | 0.427 | 1 | 0.277927848 | 0.30299138 |
| Mrps7 | 0.165165786 | 0.442004547 | 0.472 | 0.398 | 1 | 0.900886904 | 0.30115195 | 0.366 | 0.399 | 1 | 0.900886904 | 0.303051835 |
| Inhca | 0.524920047 | -0.582242999 | 0.038 | 0.059 | 1 | 0.165335714 | 1.83720485 | 0.073 | 0.031 | 1 | 0.524920047 | 0.30333553 |
| Mfsd13a | 0.569866135 | -0.304709024 | 0.019 | 0.034 | 1 | 0.165379938 | 1.737669176 | 0.073 | 0.031 | 1 | 0.569866135 | 0.303409352 |
| Dnaaf3 | 0.243102132 | -1.1672055 | 0 | 0.025 | 1 | 0.165389437 | 1.721727632 | 0.073 | 0.031 | 1 | 0.243102132 | 0.303425208 |
| Gm57314 | 0.165465021 | 1.792152515 | 0.075 | 0.034 | 1 | 0.33733901 | -0.221688839 | 0.049 | 0.096 | 1 | 0.33733901 | 0.303551368 |
| Gpr19 | 0.165465021 | 1.569760094 | 0.075 | 0.034 | 1 | 0.188517184 | -1.069685746 | 0 | 0.041 | 1 | 0.188517184 | 0.303551368 |
| Lst1 | 0.165465021 | 1.569760094 | 0.075 | 0.034 | 1 | 0.992196803 | 0.515276755 | 0.024 | 0.024 | 1 | 0.992196803 | 0.303551368 |
| Prelid2 | 0.165465021 | 2.154722595 | 0.075 | 0.034 | 1 | 0.744047924 | 1.252242349 | 0.024 | 0.017 | 1 | 0.744047924 | 0.303551368 |
| Efcab2 | 0.16549671 | -0.752168001 | 0.075 | 0.148 | 1 | 0.806790342 | 1.532350268 | 0.122 | 0.113 | 1 | 0.806790342 | 0.303604258 |
| H3f3b | 0.367847745 | -0.336315358 | 1 | 0.996 | 1 | 0.165680655 | 0.111744545 | 1 | 1 | 1 | 0.367847745 | 0.30391123 |
| Tenm2 | 0.165712582 | -1.369698364 | 0.226 | 0.309 | 1 | 0.213889961 | -1.995685164 | 0.341 | 0.396 | 1 | 0.213889961 | 0.303964505 |
| Usp45 | 0.165733165 | 0.58098735 | 0.434 | 0.356 | 1 | 0.349515671 | -0.360332383 | 0.293 | 0.386 | 1 | 0.349515671 | 0.303998849 |
| Spns1 | 0.165781793 | 0.585879759 | 0.396 | 0.318 | 1 | 0.330162461 | 0.553411884 | 0.366 | 0.297 | 1 | 0.330162461 | 0.304079983 |
| Tmem123 | 0.165814422 | 0.620386167 | 0.321 | 0.229 | 1 | 0.370079432 | 0.20299883 | 0.439 | 0.355 | 1 | 0.370079432 | 0.304134421 |
| Gm31812 | 0.165865786 | 0.776210972 | 0.057 | 0.021 | 1 | 0.402369942 | -1.16279515 | 0 | 0.017 | 1 | 0.402369942 | 0.304220114 |
| Rps27rt | 0.76925626 | 0.247831999 | 0.472 | 0.517 | 1 | 0.16603063 | -0.466575898 | 0.293 | 0.399 | 1 | 0.76925626 | 0.30449509 |
| Trp53bp1 | 0.166095795 | 0.414690698 | 0.491 | 0.407 | 1 | 0.313198436 | 0.649577847 | 0.512 | 0.488 | 1 | 0.313198436 | 0.304603777 |
| Klhl42 | 0.166131468 | 0.652222254 | 0.208 | 0.131 | 1 | 0.522137983 | 0.429120111 | 0.22 | 0.177 | 1 | 0.522137983 | 0.304663272 |
| Prkab2 | 0.166149885 | 0.808272181 | 0.302 | 0.22 | 1 | 0.388122674 | 0.707921833 | 0.39 | 0.369 | 1 | 0.388122674 | 0.304693986 |
| Gpam | 0.258208217 | -0.473308628 | 0.094 | 0.157 | 1 | 0.166194694 | 0.91966701 | 0.171 | 0.099 | 1 | 0.258208217 | 0.304768712 |
| Rabepk | 0.16625429 | 0.891688189 | 0.302 | 0.22 | 1 | 0.927359353 | 0.528452144 | 0.244 | 0.263 | 1 | 0.927359353 | 0.304868091 |
| Timmdc1 | 0.166271161 | 0.50490895 | 0.358 | 0.271 | 1 | 0.170149055 | -0.717384002 | 0.146 | 0.242 | 1 | 0.170149055 | 0.304896223 |
| Zfp976 | 0.166419782 | 0.756173218 | 0.17 | 0.102 | 1 | 0.887300553 | 0.46797104 | 0.146 | 0.143 | 1 | 0.887300553 | 0.30514402 |
| Rela | 0.166489105 | -0.321010836 | 0.528 | 0.636 | 1 | 0.20791359 | -0.325578082 | 0.561 | 0.659 | 1 | 0.20791359 | 0.305259588 |
| Mccc2 | 0.166546665 | 1.124975251 | 0.245 | 0.178 | 1 | 0.562035408 | 0.817305292 | 0.195 | 0.164 | 1 | 0.562035408 | 0.305355539 |
| Tmc4 | 0.664924942 | 0.454282877 | 0.057 | 0.042 | 1 | 0.166619142 | 1.422167351 | 0.146 | 0.082 | 1 | 0.664924942 | 0.305476345 |
| Vegfb | 0.359812059 | 0.41149801 | 0.283 | 0.229 | 1 | 0.166765878 | 0.716189449 | 0.293 | 0.205 | 1 | 0.359812059 | 0.305720897 |
| Vwa1 | 0.166835882 | -0.719746523 | 0.075 | 0.148 | 1 | 0.682087988 | 0.911205431 | 0.073 | 0.058 | 1 | 0.682087988 | 0.305837552 |
| Uhrf1bp1 | 0.664496109 | 0.280253477 | 0.132 | 0.11 | 1 | 0.166870026 | 1.326242931 | 0.244 | 0.164 | 1 | 0.664496109 | 0.305894446 |
| Eola1 | 0.814252285 | 0.154722595 | 0.189 | 0.212 | 1 | 0.166920393 | -0.221688839 | 0.122 | 0.218 | 1 | 0.814252285 | 0.305978368 |
| Saxo2 | 0.340246113 | 1.417757001 | 0.038 | 0.017 | 1 | 0.166992451 | 1.962735732 | 0.073 | 0.031 | 1 | 0.340246113 | 0.306098423 |
| Mterf1a | 0.166999014 | 1.261637799 | 0.094 | 0.047 | 1 | 0.797811701 | 1.029849928 | 0.049 | 0.041 | 1 | 0.797811701 | 0.306109358 |
| Col5a3 | 0.167146717 | 2.454282877 | 0.075 | 0.034 | 1 | 0.496423701 | 0.737669176 | 0.073 | 0.048 | 1 | 0.496423701 | 0.306355409 |
| Zc3h7a | 0.167166448 | 0.366668229 | 0.906 | 0.898 | 1 | 0.435196692 | -0.128751687 | 0.902 | 0.918 | 1 | 0.435196692 | 0.306388274 |
| Ubap1 | 0.167257779 | -0.518833829 | 0.491 | 0.589 | 1 | 0.716656075 | -0.175619191 | 0.634 | 0.676 | 1 | 0.716656075 | 0.306540393 |
| Naaa | 0.167557732 | -0.819282197 | 0.113 | 0.191 | 1 | 0.977665808 | 0.721727632 | 0.098 | 0.102 | 1 | 0.977665808 | 0.30703987 |

| | | | | | | | | | | | | |
|---|---|---|---|---|---|---|---|---|---|---|---|---|
| Wdr25 | 0.167679743 | 0.962077517 | 0.113 | 0.059 | 1 | 0.77193712 | 0.614812428 | 0.049 | 0.061 | 1 | 0.77193712 | 0.307242989 |
| Wdr55 | 0.167679743 | 0.962077517 | 0.113 | 0.059 | 1 | 0.516329798 | 0.422167351 | 0.122 | 0.089 | 1 | 0.516329798 | 0.307242989 |
| Gm26881 | 0.226026934 | 1.377115016 | 0.075 | 0.038 | 1 | 0.167816374 | 1.699701326 | 0.073 | 0.031 | 1 | 0.226026934 | 0.307470412 |
| 2610021A01Rik | 0.167837111 | 1.292226119 | 0.113 | 0.059 | 1 | 0.313183952 | 0.422167351 | 0.171 | 0.113 | 1 | 0.313183952 | 0.307504926 |
| Lrrc4 | 0.167853121 | -0.6337733 | 0.132 | 0.216 | 1 | 0.584698736 | -0.316600486 | 0.171 | 0.205 | 1 | 0.584698736 | 0.307531572 |
| Gm15728 | 0.167910296 | 1.640149422 | 0.075 | 0.034 | 1 | 0.561140381 | 0.644559772 | 0.098 | 0.072 | 1 | 0.561140381 | 0.307626725 |
| Pcdhga8 | 0.167910296 | 1.8327945 | 0.075 | 0.034 | 1 | 0.548898236 | -0.250257991 | 0.024 | 0.044 | 1 | 0.548898236 | 0.307626725 |
| Cfap68 | 0.168020641 | -0.418167074 | 0.302 | 0.398 | 1 | 0.967156933 | 0.237292008 | 0.244 | 0.249 | 1 | 0.967156933 | 0.307810347 |
| E2f8 | 0.168082993 | -2.27154216 | 0.019 | 0.068 | 1 | 0.51919986 | 0.83720485 | 0 | 0.01 | 1 | 0.51919986 | 0.307914093 |
| Zdhhc13 | 0.586138248 | 0.417757001 | 0.17 | 0.14 | 1 | 0.168602902 | -1.132421501 | 0.049 | 0.119 | 1 | 0.586138248 | 0.308778865 |
| Klhl25 | 0.338780504 | 0.984797593 | 0.057 | 0.03 | 1 | 0.168643345 | 1.574170444 | 0.073 | 0.031 | 1 | 0.338780504 | 0.308846113 |
| Gm43597 | 0.917883255 | 0.569760094 | 0.019 | 0.021 | 1 | 0.168643345 | 1.574170444 | 0.073 | 0.031 | 1 | 0.917883255 | 0.308846113 |
| Galnt1 | 0.253605532 | 0.101510789 | 0.906 | 0.864 | 1 | 0.168684532 | 0.429341383 | 0.902 | 0.898 | 1 | 0.253605532 | 0.308914593 |
| Vps53 | 0.644157985 | 0.534136184 | 0.34 | 0.356 | 1 | 0.16871634 | -0.629921161 | 0.317 | 0.41 | 1 | 0.644157985 | 0.308967476 |
| Cyb5rl | 0.1687232 | 1.417757001 | 0.075 | 0.034 | 1 | 0.302745174 | 1.614812428 | 0.098 | 0.058 | 1 | 0.302745174 | 0.308978881 |
| Med18 | 0.1687232 | 1.417757001 | 0.075 | 0.034 | 1 | 0.205693707 | -1.16279515 | 0.024 | 0.078 | 1 | 0.205693707 | 0.308978881 |
| AA386476 | 0.1687232 | 1.417757001 | 0.075 | 0.034 | 1 | 0.743829142 | 0.252242349 | 0.024 | 0.034 | 1 | 0.743829142 | 0.308978881 |
| Cpt1b | 0.1687232 | 1.417757001 | 0.075 | 0.034 | 1 | 0.894661632 | 0.574170444 | 0.049 | 0.055 | 1 | 0.894661632 | 0.308978881 |
| Ankrd2 | 0.1687232 | 1.417757001 | 0.075 | 0.034 | 1 | 0.331950705 | -1.069685746 | 0.024 | 0.061 | 1 | 0.331950705 | 0.308978881 |
| Eps8l1 | 0.168755115 | -1.545717123 | 0.019 | 0.068 | 1 | 0.834677311 | 0.515276755 | 0.049 | 0.058 | 1 | 0.834677311 | 0.309031942 |
| Ift57 | 0.697398569 | -0.200120123 | 0.377 | 0.386 | 1 | 0.168797933 | -0.502645153 | 0.244 | 0.362 | 1 | 0.697398569 | 0.309103123 |
| Mov10 | 0.168928241 | 1.395730694 | 0.132 | 0.076 | 1 | 0.414261829 | 0.667279848 | 0.195 | 0.15 | 1 | 0.414261829 | 0.309319732 |
| Usp40 | 0.585507038 | 0.454282877 | 0.34 | 0.322 | 1 | 0.169043029 | -0.484723245 | 0.268 | 0.372 | 1 | 0.585507038 | 0.309510512 |
| 1500015A07Rik | 0.169154244 | 1.347367673 | 0.094 | 0.047 | 1 | 0.450957104 | -0.410722664 | 0.024 | 0.051 | 1 | 0.450957104 | 0.30969533 |
| Chaf1b | 0.169154582 | 1.417757001 | 0.094 | 0.047 | 1 | 0.666353766 | -0.410722664 | 0.024 | 0.038 | 1 | 0.666353766 | 0.309695891 |
| Rpl9-ps6 | 0.973374567 | 0.164245369 | 0.434 | 0.466 | 1 | 0.169274612 | -0.534354013 | 0.39 | 0.481 | 1 | 0.973374567 | 0.30989533 |
| Rps24 | 0.261651981 | -0.142732568 | 1 | 0.992 | 1 | 0.169316255 | -0.128210479 | 1 | 0.997 | 1 | 0.261651981 | 0.309964516 |
| Smpd4 | 0.169482724 | 0.75033234 | 0.245 | 0.169 | 1 | 0.275106554 | -0.316600486 | 0.171 | 0.256 | 1 | 0.275106554 | 0.310241054 |
| Them4 | 0.264753267 | -0.268488836 | 0.113 | 0.182 | 1 | 0.169553483 | -1.250257991 | 0.049 | 0.119 | 1 | 0.264753267 | 0.310358583 |
| Cbfb | 0.169561057 | 0.157319706 | 0.868 | 0.809 | 1 | 0.347790703 | -0.129903269 | 0.78 | 0.881 | 1 | 0.347790703 | 0.310371162 |
| Golga5 | 0.422107436 | 0.353626663 | 0.547 | 0.555 | 1 | 0.169920849 | 0.612029988 | 0.61 | 0.59 | 1 | 0.422107436 | 0.310968603 |
| Egr2 | 0.169941344 | -0.113766241 | 0.132 | 0.072 | 1 | 0.670233172 | -1.332720152 | 0.024 | 0.038 | 1 | 0.670233172 | 0.311002627 |
| Zbtb39 | 0.169995419 | 0.948271717 | 0.17 | 0.106 | 1 | 0.207212079 | -0.46860358 | 0.146 | 0.239 | 1 | 0.207212079 | 0.311092395 |
| Tmem241 | 0.170003961 | 0.464407094 | 0.245 | 0.165 | 1 | 0.19696121 | -0.460475699 | 0.122 | 0.212 | 1 | 0.19696121 | 0.311106576 |
| Tnfrsf14 | 0.72256466 | 1.739685096 | 0.019 | 0.013 | 1 | 0.170353247 | -1.250257991 | 0 | 0.044 | 1 | 0.72256466 | 0.311686265 |
| Gm39326 | 0.249034808 | 2.154722595 | 0.019 | 0.004 | 1 | 0.170373481 | -1.410722664 | 0 | 0.044 | 1 | 0.249034808 | 0.31171984 |
| Gm13008 | 0.340246113 | 1.417757001 | 0.038 | 0.017 | 1 | 0.170373481 | -1.16279515 | 0 | 0.044 | 1 | 0.340246113 | 0.31171984 |
| Catsperg1 | 0.503971666 | 1.569760094 | 0.019 | 0.008 | 1 | 0.170373481 | -1.16279515 | 0 | 0.044 | 1 | 0.503971666 | 0.31171984 |
| Gm34961 | 0.89261791 | 0.569760094 | 0.038 | 0.034 | 1 | 0.170373481 | -1.16279515 | 0 | 0.044 | 1 | 0.89261791 | 0.31171984 |
| Gm49463 | 0.966184735 | 0.569760094 | 0.057 | 0.059 | 1 | 0.170373481 | -1.16279515 | 0 | 0.044 | 1 | 0.966184735 | 0.31171984 |
| 2900005J15Rik | 0.199211269 | -0.489133595 | 0.038 | 0.093 | 1 | 0.170390343 | -1.484723245 | 0 | 0.044 | 1 | 0.199211269 | 0.311747816 |
| Mfsd3 | 0.3177504 | -0.582242999 | 0.038 | 0.076 | 1 | 0.170390343 | -1.250257991 | 0 | 0.044 | 1 | 0.3177504 | 0.311747816 |
| Isoc2a | 0.306896082 | -0.1672055 | 0.057 | 0.106 | 1 | 0.170402911 | -1.410722664 | 0 | 0.044 | 1 | 0.306896082 | 0.31176867 |
| Acad10 | 0.228430171 | -1.237594828 | 0.019 | 0.059 | 1 | 0.170403831 | -1.686357106 | 0 | 0.044 | 1 | 0.228430171 | 0.311770196 |
| Dnajc28 | 0.49029049 | 0.739685096 | 0.038 | 0.021 | 1 | 0.170403831 | -1.622226769 | 0 | 0.044 | 1 | 0.49029049 | 0.311770196 |
| Rhbdl2 | 0.195531196 | -1.545717123 | 0.019 | 0.064 | 1 | 0.170404137 | -1.332720152 | 0 | 0.044 | 1 | 0.195531196 | 0.311770705 |
| Arhgef37 | 0.982488248 | -0.1672055 | 0.038 | 0.038 | 1 | 0.170404137 | -1.686357106 | 0 | 0.044 | 1 | 0.982488248 | 0.311770705 |
| Pcdhb17 | 0.170411229 | 1.347367673 | 0.075 | 0.034 | 1 | 0.405126994 | 0.949679579 | 0.122 | 0.085 | 1 | 0.405126994 | 0.311782471 |
| Gm42567 | 0.725936481 | 1.569760094 | 0.019 | 0.013 | 1 | 0.170413947 | -1.622226769 | 0 | 0.044 | 1 | 0.725936481 | 0.31178698 |
| Lypd1 | 0.204874398 | -1.617866909 | 0.038 | 0.089 | 1 | 0.17042866 | -1.970150072 | 0 | 0.044 | 1 | 0.204874398 | 0.311811393 |
| Gm29966 | 0.485184337 | -0.752168001 | 0.019 | 0.038 | 1 | 0.170433258 | -2.250257991 | 0 | 0.044 | 1 | 0.485184337 | 0.311819021 |
| Slc18a1 | 0.795213727 | 0.984797593 | 0.019 | 0.025 | 1 | 0.170433258 | -2.520347155 | 0 | 0.044 | 1 | 0.795213727 | 0.311819021 |
| 4833413G10Rik | 0.413080549 | -0.1672055 | 0 | 0.013 | 1 | 0.170440921 | -0.970150072 | 0 | 0.044 | 1 | 0.413080549 | 0.311831735 |
| Fbxl12 | 0.417255393 | 0.20255112 | 0.34 | 0.28 | 1 | 0.170450445 | 0.728083127 | 0.415 | 0.341 | 1 | 0.417255393 | 0.311847536 |
| Jazf1 | 0.293777376 | 0.219817623 | 0.264 | 0.216 | 1 | 0.170549135 | -0.830747016 | 0.293 | 0.399 | 1 | 0.293777376 | 0.312011262 |
| Gm12992 | 0.170582465 | 1.154722595 | 0.094 | 0.047 | 1 | 0.186194355 | 1.614812428 | 0.098 | 0.048 | 1 | 0.186194355 | 0.312066553 |
| Zfp760 | 0.170582465 | 1.154722595 | 0.094 | 0.047 | 1 | 0.628284978 | -0.221688839 | 0.049 | 0.068 | 1 | 0.628284978 | 0.312066553 |
| Snx17 | 0.935889253 | 0.114458725 | 0.528 | 0.568 | 1 | 0.170675444 | -0.157777075 | 0.463 | 0.597 | 1 | 0.935889253 | 0.312220781 |
| Rnaset2b | 0.224660053 | -0.21229339 | 0.585 | 0.695 | 1 | 0.170744008 | -0.242859539 | 0.585 | 0.679 | 1 | 0.224660053 | 0.312334499 |
| Siah2 | 0.193400493 | 0.212543249 | 0.604 | 0.551 | 1 | 0.170767084 | -0.438238343 | 0.488 | 0.587 | 1 | 0.193400493 | 0.312372772 |
| Cgrrf1 | 0.170860706 | -0.356239324 | 0.434 | 0.551 | 1 | 0.579249504 | 0.256701997 | 0.439 | 0.58 | 1 | 0.579249504 | 0.312528031 |
| Zfp189 | 0.190396754 | 0.756173218 | 0.17 | 0.106 | 1 | 0.170948386 | -0.767657208 | 0.098 | 0.184 | 1 | 0.190396754 | 0.312673421 |
| Adrm1 | 0.255871632 | -0.28137652 | 0.453 | 0.534 | 1 | 0.171327362 | -0.257952383 | 0.463 | 0.584 | 1 | 0.255871632 | 0.31330166 |
| Gpcpd1 | 0.249143111 | -0.620291607 | 0.547 | 0.589 | 1 | 0.171397646 | 0.42674008 | 0.585 | 0.587 | 1 | 0.249143111 | 0.31341814 |
| 4921504A21Rik | 0.171582584 | 0.874614676 | 0.113 | 0.059 | 1 | 0.432070036 | -0.395455907 | 0.049 | 0.085 | 1 | 0.432070036 | 0.313724585 |
| Rpgrip1 | 0.854181209 | -0.466765782 | 0.189 | 0.169 | 1 | 0.171615297 | -0.800225071 | 0.098 | 0.191 | 1 | 0.854181209 | 0.313778783 |
| Tspan32 | 0.294953953 | -1.130679624 | 0.057 | 0.102 | 1 | 0.171658977 | -1.069685746 | 0.024 | 0.085 | 1 | 0.294953953 | 0.313851149 |
| Alg2 | 0.171961755 | -0.590704578 | 0.264 | 0.352 | 1 | 0.741274069 | 0.334704509 | 0.341 | 0.406 | 1 | 0.741274069 | 0.314352665 |
| Gm44710 | 0.172024441 | 1.280253477 | 0.075 | 0.034 | 1 | 0.827983455 | 0.377773231 | 0.049 | 0.058 | 1 | 0.827983455 | 0.314456474 |
| Zfand1 | 0.973162777 | 0.953088734 | 0.075 | 0.076 | 1 | 0.172059598 | 0.690363461 | 0.146 | 0.078 | 1 | 0.973162777 | 0.31451469 |
| Gm27253 | 0.172062369 | 1.261637799 | 0.075 | 0.034 | 1 | 0.666354009 | -0.16279515 | 0.024 | 0.038 | 1 | 0.666354009 | 0.31451928 |
| Dffa | 0.172077986 | 1.306725688 | 0.075 | 0.034 | 1 | 0.401822916 | 1.252242349 | 0.098 | 0.065 | 1 | 0.401822916 | 0.314545139 |
| Meikin | 0.172077986 | 1.306725688 | 0.075 | 0.034 | 1 | 0.820932515 | 0.614812428 | 0.049 | 0.041 | 1 | 0.820932515 | 0.314545139 |
| Rras | 0.172094906 | 0.46396543 | 0.679 | 0.648 | 1 | 0.638724655 | 0.166009951 | 0.463 | 0.543 | 1 | 0.638724655 | 0.314573156 |
| Zbed5-ps | 0.172503738 | -0.52334931 | 0.038 | 0.097 | 1 | 0.431656591 | 1.614812428 | 0.122 | 0.092 | 1 | 0.431656591 | 0.315249937 |
| Appl2 | 0.454789236 | 0.253500749 | 0.623 | 0.564 | 1 | 0.172649034 | -0.406066301 | 0.61 | 0.662 | 1 | 0.454789236 | 0.31549038 |
| Nes | 0.172676421 | -1.785720561 | 0.17 | 0.246 | 1 | 0.4113861 | 1.159132945 | 0.073 | 0.044 | 1 | 0.4113861 | 0.315535696 |
| Gm3734 | 0.982488248 | -0.260314904 | 0.038 | 0.038 | 1 | 0.172743061 | -1.020776145 | 0.024 | 0.085 | 1 | 0.982488248 | 0.315645957 |
| Nip7 | 0.653462554 | 0.210217707 | 0.491 | 0.504 | 1 | 0.172836763 | -0.219826095 | 0.415 | 0.512 | 1 | 0.653462554 | 0.315800979 |
| Cnot8 | 0.172851109 | 0.269365161 | 0.509 | 0.39 | 1 | 0.700316712 | 0.143717892 | 0.439 | 0.44 | 1 | 0.700316712 | 0.315824713 |
| Ptdss2 | 0.172910827 | 0.569760094 | 0.302 | 0.22 | 1 | 0.608483287 | 0.214767644 | 0.293 | 0.266 | 1 | 0.608483287 | 0.3159235 |
| Mtss2 | 0.976490022 | 0.311448098 | 0.547 | 0.631 | 1 | 0.172995702 | -0.320796984 | 0.488 | 0.618 | 1 | 0.976490022 | 0.316063891 |
| D17H6S56E-5 | 0.172996346 | -1.15816036 | 0.132 | 0.072 | 1 | 0.199046585 | 0.47463477 | 0.098 | 0.048 | 1 | 0.199046585 | 0.316064957 |
| Sub1 | 0.173032405 | 0.231771259 | 1 | 0.949 | 1 | 0.296237383 | -0.116381656 | 0.976 | 0.98 | 1 | 0.296237383 | 0.316124597 |
| Hmgn2-ps | 0.542187926 | 0.80108564 | 0.113 | 0.089 | 1 | 0.17328147 | -0.917682652 | 0.024 | 0.085 | 1 | 0.542187926 | 0.316536472 |
| Gm16279 | 0.173491509 | 1.154722595 | 0.094 | 0.047 | 1 | 0.871991894 | 0.174239837 | 0.049 | 0.055 | 1 | 0.871991894 | 0.316883715 |
| Map4k2 | 0.180256037 | 0.714150003 | 0.189 | 0.119 | 1 | 0.173567142 | -1.116991461 | 0.073 | 0.15 | 1 | 0.180256037 | 0.317008731 |
| Acer2 | 0.475388 | 1.569760094 | 0.038 | 0.021 | 1 | 0.173695166 | 1.252242349 | 0.073 | 0.031 | 1 | 0.475388 | 0.317220322 |
| 4931414P19Rik | 0.173711986 | 0.962077517 | 0.075 | 0.034 | 1 | 0.831329289 | 0.444887427 | 0.049 | 0.058 | 1 | 0.831329289 | 0.317248118 |
| Mrpl45 | 0.173895343 | 0.53023173 | 0.34 | 0.254 | 1 | 0.414642739 | 0.492376353 | 0.341 | 0.29 | 1 | 0.414642739 | 0.317551096 |
| 3110009E18Rik | 0.173961536 | 0.2967416 | 0.132 | 0.072 | 1 | 0.334510406 | 1.007129851 | 0.122 | 0.078 | 1 | 0.334510406 | 0.317660456 |
| Vac14 | 0.174195296 | 0.8327945 | 0.151 | 0.089 | 1 | 0.304228017 | 0.75474269 | 0.22 | 0.167 | 1 | 0.304228017 | 0.31804659 |
| Map3k6 | 0.624515373 | -0.189231806 | 0.151 | 0.178 | 1 | 0.174263831 | -0.970150072 | 0.049 | 0.119 | 1 | 0.624515373 | 0.318159779 |
| Pced1a | 0.940269558 | 0.324647596 | 0.113 | 0.119 | 1 | 0.174465349 | -0.622226769 | 0.049 | 0.123 | 1 | 0.940269558 | 0.318492539 |
| Gm1604b | 0.460722276 | 1.891688189 | 0.038 | 0.021 | 1 | 0.174558027 | 1.422167351 | 0.073 | 0.031 | 1 | 0.460722276 | 0.318645549 |
| Paxx | 0.174574985 | 0.8327945 | 0.34 | 0.267 | 1 | 0.281895127 | 0.270858027 | 0.122 | 0.218 | 1 | 0.281895127 | 0.318673545 |
| Ehf | 0.583452454 | -0.875024749 | 0.17 | 0.195 | 1 | 0.174598819 | 1.252242349 | 0.317 | 0.246 | 1 | 0.583452454 | 0.31871289 |
| Mir133a-1hg | 0.732691937 | 1.154722595 | 0.019 | 0.013 | 1 | 0.174743691 | 2.574170444 | 0.049 | 0.017 | 1 | 0.732691937 | 0.318952025 |

| | | | | | | | | | | | | |
|---|---|---|---|---|---|---|---|---|---|---|---|---|
| Acads | 0.985704031 | 0.466666601 | 0.113 | 0.119 | 1 | 0.174854015 | -0.944154864 | 0.073 | 0.15 | 1 | 0.985704031 | 0.319134104 |
| Vwa8 | 0.1748541 | 0.80108564 | 0.434 | 0.386 | 1 | 0.571365685 | 0.310225533 | 0.488 | 0.454 | 1 | 0.571365685 | 0.319134243 |
| Tmem154 | 0.3621258 | -0.697720217 | 0.132 | 0.178 | 1 | 0.174854715 | -0.840867055 | 0.073 | 0.154 | 1 | 0.3621258 | 0.319135259 |
| Gm31718 | 0.174945555 | 0.962077517 | 0.094 | 0.047 | 1 | 0.237614931 | 1.029849928 | 0.098 | 0.051 | 1 | 0.237614931 | 0.319285163 |
| Smim7 | 0.745237739 | 0.165920801 | 0.736 | 0.771 | 1 | 0.174947658 | -0.200269856 | 0.561 | 0.754 | 1 | 0.745237739 | 0.319288632 |
| Zfp619 | 0.174968091 | 1.237184755 | 0.094 | 0.047 | 1 | 0.89822728 | 0.46797104 | 0.073 | 0.068 | 1 | 0.89822728 | 0.319322349 |
| Rrp7a | 0.175178175 | 0.480122882 | 0.377 | 0.284 | 1 | 0.719149301 | 0.410940095 | 0.317 | 0.317 | 1 | 0.719149301 | 0.319668957 |
| Dnajb12 | 0.420902216 | 0.403602906 | 0.585 | 0.627 | 1 | 0.175183449 | -0.229909346 | 0.488 | 0.631 | 1 | 0.420902216 | 0.319677657 |
| Arhgap18 | 0.407599676 | 0.569760094 | 0.075 | 0.047 | 1 | 0.175280313 | 0.111379813 | 0.244 | 0.15 | 1 | 0.407599676 | 0.319837438 |
| Reep1 | 0.175308325 | 1.017219071 | 0.075 | 0.034 | 1 | 0.856407545 | 2.252242349 | 0.024 | 0.02 | 1 | 0.856407545 | 0.319883641 |
| Enkd1 | 0.175308325 | 1.154722595 | 0.075 | 0.034 | 1 | 0.208863153 | -0.747757651 | 0 | 0.038 | 1 | 0.208863153 | 0.319883641 |
| Abca6 | 0.175308325 | 0.776210972 | 0.075 | 0.034 | 1 | 0.992196803 | 0.377773231 | 0.024 | 0.024 | 1 | 0.992196803 | 0.319883641 |
| Nprl3 | 0.175344316 | -0.741441594 | 0.113 | 0.195 | 1 | 0.661985174 | 0.218295017 | 0.22 | 0.191 | 1 | 0.661985174 | 0.319943003 |
| Pcdhga2 | 0.300962247 | 1.280253477 | 0.075 | 0.042 | 1 | 0.175351127 | 1.252242349 | 0.073 | 0.031 | 1 | 0.300962247 | 0.319954236 |
| Zfp870 | 0.343171476 | 0.8327945 | 0.057 | 0.03 | 1 | 0.175351127 | 1.377773231 | 0.073 | 0.031 | 1 | 0.343171476 | 0.319954236 |
| Isl1 | 0.413082433 | -0.430239906 | 0 | 0.013 | 1 | 0.175351127 | 1.377773231 | 0.073 | 0.031 | 1 | 0.413082433 | 0.319954236 |
| Trim65 | 0.794190052 | 0.347367673 | 0.038 | 0.047 | 1 | 0.175351127 | 1.252242349 | 0.073 | 0.031 | 1 | 0.794190052 | 0.319954236 |
| 1700037C18Rik | 0.929410101 | 0.569760094 | 0.019 | 0.017 | 1 | 0.175351127 | 1.252242349 | 0.073 | 0.031 | 1 | 0.929410101 | 0.319954236 |
| Lsm10 | 0.175377512 | 0.739685096 | 0.189 | 0.119 | 1 | 0.209660448 | 0.334704509 | 0.073 | 0.154 | 1 | 0.209660448 | 0.319997752 |
| Megf11 | 0.175390527 | 0.399835093 | 0.075 | 0.034 | 1 | 0.720214199 | 1.185128153 | 0.049 | 0.038 | 1 | 0.720214199 | 0.320019217 |
| Scap | 0.322690802 | 0.285967128 | 0.34 | 0.271 | 1 | 0.175504963 | -0.16279515 | 0.268 | 0.399 | 1 | 0.322690802 | 0.320207935 |
| Rnf122 | 0.175611841 | 1.891688189 | 0.113 | 0.064 | 1 | 0.428849875 | 0.83720485 | 0.073 | 0.044 | 1 | 0.428849875 | 0.320384163 |
| 2810001A02Rik | 0.1756155 | -0.607778091 | 0.208 | 0.297 | 1 | 0.898729492 | -0.179096963 | 0.244 | 0.242 | 1 | 0.898729492 | 0.320390197 |
| Tsg101 | 0.175655248 | 0.294125651 | 0.774 | 0.708 | 1 | 0.949970946 | 0.101879614 | 0.78 | 0.836 | 1 | 0.949970946 | 0.32045573 |
| Cgas | 0.175755477 | 0.659957903 | 0.094 | 0.047 | 1 | 0.968794676 | 1.667279848 | 0.024 | 0.024 | 1 | 0.968794676 | 0.320620966 |
| Cutal | 0.17582875 | -1.1672055 | 0 | 0.034 | 1 | 0.51919986 | 0.83720485 | 0 | 0.01 | 1 | 0.51919986 | 0.32074175 |
| BB557941 | 0.17582875 | -1.1672055 | 0 | 0.034 | 1 | 0.5191986 | 0.515276755 | 0 | 0.01 | 1 | 0.5191986 | 0.32074175 |
| 4930417H01Rik | 0.17582875 | -1.1672055 | 0 | 0.034 | 1 | 0.357847088 | -0.332720152 | 0 | 0.02 | 1 | 0.357847088 | 0.32074175 |
| 4930534D22Rik | 0.17582875 | -1.1672055 | 0 | 0.034 | 1 | 0.365461252 | 1.100239256 | 0.049 | 0.024 | 1 | 0.365461252 | 0.32074175 |
| Gm12724 | 0.17582875 | -1.1672055 | 0 | 0.034 | 1 | 0.455098275 | 0.515276755 | 0 | 0.014 | 1 | 0.455098275 | 0.32074175 |
| Hpse | 0.17582875 | -1.304709024 | 0 | 0.034 | 1 | 0.402369942 | -0.16279515 | 0 | 0.017 | 1 | 0.402369942 | 0.32074175 |
| Asgr1 | 0.17582875 | -1.304709024 | 0 | 0.034 | 1 | 0.188498353 | -0.970150072 | 0 | 0.041 | 1 | 0.188498353 | 0.32074175 |
| Krt222 | 0.17582875 | -1.1672055 | 0 | 0.034 | 1 | 0.51919986 | 0.83720485 | 0 | 0.01 | 1 | 0.51919986 | 0.32074175 |
| Gm49958 | 0.17582875 | -1.1672055 | 0 | 0.034 | 1 | 0.606262835 | -0.16279515 | 0.024 | 0.041 | 1 | 0.606262835 | 0.32074175 |
| Gm50295 | 0.17582875 | -1.1672055 | 0 | 0.034 | 1 | 0.231606831 | -1.069685746 | 0 | 0.034 | 1 | 0.231606831 | 0.32074175 |
| 5430402E10Rik | 0.17582875 | -1.304709024 | 0 | 0.034 | 1 | 0.648706972 | 0.114738825 | 0.098 | 0.075 | 1 | 0.648706972 | 0.32074175 |
| Ano7 | 0.175842714 | -1.304709024 | 0 | 0.034 | 1 | 0.604477838 | -0.332720152 | 0.024 | 0.041 | 1 | 0.604477838 | 0.320764768 |
| Gm30173 | 0.175842714 | -1.845277405 | 0 | 0.034 | 1 | 0.357854401 | -0.484723245 | 0 | 0.02 | 1 | 0.357854401 | 0.320764768 |
| Tmem82 | 0.175842714 | -1.304709024 | 0 | 0.034 | 1 | 0.576568947 | -0.555112573 | 0.073 | 0.099 | 1 | 0.576568947 | 0.320764768 |
| A330035P11Rik | 0.175842714 | -1.304709024 | 0 | 0.034 | 1 | 0.547258706 | -0.332720152 | 0.024 | 0.044 | 1 | 0.547258706 | 0.320764768 |
| Gdpd2 | 0.175842714 | -1.304709024 | 0 | 0.034 | 1 | 0.51919986 | 0.83720485 | 0 | 0.01 | 1 | 0.51919986 | 0.320764768 |
| Ly75 | 0.175852023 | -1.545717123 | 0 | 0.034 | 1 | 0.430584486 | 2.422167351 | 0.024 | 0.01 | 1 | 0.430584486 | 0.320780112 |
| Necab3 | 0.175852023 | -1.932740246 | 0 | 0.034 | 1 | 0.357847088 | -0.16279515 | 0 | 0.02 | 1 | 0.357847088 | 0.320780112 |
| Atp6v0d2 | 0.175852023 | -2.093204919 | 0 | 0.034 | 1 | 0.319641967 | -4.84227525 | 0 | 0.024 | 1 | 0.319641967 | 0.320780112 |
| Plet1os | 0.175852023 | -1.545717123 | 0 | 0.034 | 1 | 0.51919986 | 0.83720485 | 0 | 0.01 | 1 | 0.51919986 | 0.320780112 |
| 4930500A05Rik | 0.175852023 | -1.752168001 | 0 | 0.034 | 1 | 0.319641967 | -0.16279515 | 0 | 0.024 | 1 | 0.319641967 | 0.320780112 |
| Rassf6 | 0.175852688 | -1.430239906 | 0 | 0.034 | 1 | 0.402373368 | 0.252242349 | 0 | 0.017 | 1 | 0.402373368 | 0.320781208 |
| Gm20548 | 0.175852688 | -1.430239906 | 0 | 0.034 | 1 | 0.286402368 | -0.484723245 | 0 | 0.027 | 1 | 0.286402368 | 0.320781208 |
| Gm17655 | 0.175852688 | -1.430239906 | 0 | 0.034 | 1 | 0.5191986 | 0.515276755 | 0 | 0.01 | 1 | 0.5191986 | 0.320781208 |
| Ppp1r14a | 0.175852688 | -1.652632327 | 0 | 0.034 | 1 | 0.911761241 | 0.377773231 | 0.024 | 0.027 | 1 | 0.911761241 | 0.320781208 |
| Cpn1 | 0.175852688 | -1.430239906 | 0 | 0.034 | 1 | 0.51919986 | 0.83720485 | 0 | 0.01 | 1 | 0.51919986 | 0.320781208 |
| Kif14 | 0.175856678 | -1.752168001 | 0 | 0.034 | 1 | 0.438672228 | 1.83720485 | 0.024 | 0.01 | 1 | 0.438672228 | 0.320787784 |
| Optc | 0.175860667 | -2.015202407 | 0 | 0.034 | 1 | 0.257308459 | -1.069685746 | 0 | 0.031 | 1 | 0.257308459 | 0.32079436 |
| Lcn2 | 0.175863327 | -3.368839361 | 0 | 0.034 | 1 | 0.621195238 | -0.572186086 | 0.171 | 0.14 | 1 | 0.621195238 | 0.320798744 |
| Esco2 | 0.175863327 | -2.7032584 | 0 | 0.034 | 1 | 0.51919986 | 0.83720485 | 0 | 0.01 | 1 | 0.51919986 | 0.320798744 |
| Zswim5 | 0.175863992 | -2.489133595 | 0 | 0.034 | 1 | 0.624072438 | 1.83720485 | 0.049 | 0.034 | 1 | 0.624072438 | 0.32079984 |
| Cdk18 | 0.175865986 | -1.015202407 | 0 | 0.034 | 1 | 0.837789621 | 0.614812428 | 0.024 | 0.031 | 1 | 0.837789621 | 0.320803128 |
| Gm29340 | 0.175865986 | -1.015202407 | 0 | 0.034 | 1 | 0.357858789 | -0.484723245 | 0 | 0.02 | 1 | 0.357858789 | 0.320803128 |
| Dok7 | 0.175865986 | -1.015202407 | 0 | 0.034 | 1 | 0.565083593 | 0.334704509 | 0.049 | 0.031 | 1 | 0.565083593 | 0.320803128 |
| A430078I02Rik | 0.175865986 | -1.015202407 | 0 | 0.034 | 1 | 0.438672228 | 1.83720485 | 0.024 | 0.01 | 1 | 0.438672228 | 0.320803128 |
| Hyal3 | 0.175865986 | -1.015202407 | 0 | 0.034 | 1 | 0.295050944 | 1.029849928 | 0.073 | 0.038 | 1 | 0.295050944 | 0.320803128 |
| Gm56728 | 0.175865986 | -1.015202407 | 0 | 0.034 | 1 | 0.319641967 | -0.16279515 | 0 | 0.024 | 1 | 0.319641967 | 0.320803128 |
| Ccdc74a | 0.175865986 | -1.015202407 | 0 | 0.034 | 1 | 0.916649754 | 0.667279848 | 0.024 | 0.027 | 1 | 0.916649754 | 0.320803128 |
| Mcrip2 | 0.175865986 | -1.015202407 | 0 | 0.034 | 1 | 0.986995563 | 0.83720485 | 0.024 | 0.024 | 1 | 0.986995563 | 0.320803128 |
| Bcr | 0.221502678 | 0.116587466 | 0.698 | 0.619 | 1 | 0.175979989 | 0.409697999 | 0.756 | 0.693 | 1 | 0.221502678 | 0.320991021 |
| Gm56555 | 0.49832147 | 0.717317282 | 0.208 | 0.165 | 1 | 0.176036426 | 0.73286819 | 0.171 | 0.099 | 1 | 0.49832147 | 0.321084029 |
| Usp54 | 0.176148751 | 0.819738347 | 0.377 | 0.314 | 1 | 0.511213736 | -0.541306773 | 0.463 | 0.488 | 1 | 0.511213736 | 0.321269119 |
| Rrp36 | 0.176571379 | 0.569760094 | 0.358 | 0.271 | 1 | 0.328374623 | -0.138547604 | 0.195 | 0.283 | 1 | 0.328374623 | 0.321965307 |
| Il18 | 0.375209464 | -0.211599619 | 0.321 | 0.407 | 1 | 0.176720274 | -0.429581691 | 0.195 | 0.307 | 1 | 0.375209464 | 0.322210493 |
| Pglyrp1 | 0.176734272 | 0.471912771 | 0.226 | 0.148 | 1 | 0.261370689 | 0.235168836 | 0.268 | 0.195 | 1 | 0.261370689 | 0.32223354 |
| Uqcc6 | 0.176800064 | 0.962077517 | 0.17 | 0.106 | 1 | 0.887328571 | 0.83720485 | 0.073 | 0.068 | 1 | 0.887328571 | 0.322341866 |
| Arl10 | 0.353834133 | 0.446903346 | 0.208 | 0.157 | 1 | 0.176816728 | -0.377808041 | 0.098 | 0.191 | 1 | 0.353834133 | 0.322369301 |
| Sema3c | 0.176852152 | -0.247375849 | 0.377 | 0.496 | 1 | 0.39038036 | -0.174767792 | 0.488 | 0.526 | 1 | 0.39038036 | 0.32242762 |
| Gm57236 | 0.914958 | 0.347367673 | 0.019 | 0.021 | 1 | 0.176970806 | 2.351778023 | 0.049 | 0.017 | 1 | 0.914958 | 0.322622946 |
| Gm10131 | 0.176999872 | 0.865215978 | 0.151 | 0.089 | 1 | 0.938546301 | 1.47463477 | 0.098 | 0.109 | 1 | 0.938546301 | 0.322670789 |
| Ankrd39 | 0.177413817 | 0.861940846 | 0.245 | 0.169 | 1 | 0.713802607 | 0.228395607 | 0.122 | 0.15 | 1 | 0.713802607 | 0.323351971 |
| Plekha3 | 0.177430086 | 0.479344799 | 0.604 | 0.593 | 1 | 0.565071443 | 0.422167351 | 0.512 | 0.532 | 1 | 0.565071443 | 0.323378736 |
| Suox | 0.22221683 | -0.494370243 | 0.151 | 0.229 | 1 | 0.177449258 | -0.863234868 | 0.073 | 0.15 | 1 | 0.22221683 | 0.323410278 |
| Mrps10 | 0.177518334 | 0.669295768 | 0.358 | 0.28 | 1 | 0.202881918 | 0.481724195 | 0.293 | 0.208 | 1 | 0.202881918 | 0.323523909 |
| Gm56688 | 0.855418007 | 0.569760094 | 0.057 | 0.051 | 1 | 0.177526694 | 1.396632258 | 0.098 | 0.048 | 1 | 0.855418007 | 0.323537661 |
| Plekha7 | 0.177709701 | -2.067669827 | 0.038 | 0.093 | 1 | 0.454986006 | 0.119348079 | 0.171 | 0.225 | 1 | 0.454986006 | 0.323838664 |
| Prpsap2 | 0.177759217 | 0.587682002 | 0.321 | 0.246 | 1 | 0.310877465 | -0.307845483 | 0.171 | 0.242 | 1 | 0.310877465 | 0.323920094 |
| Angptl2 | 0.258752717 | 0.536970159 | 0.415 | 0.347 | 1 | 0.177764039 | 0.580865097 | 0.488 | 0.369 | 1 | 0.258752717 | 0.323928025 |
| Gm38882 | 0.177781126 | -0.347777746 | 0.057 | 0.123 | 1 | 0.637446795 | 0.667279848 | 0.073 | 0.055 | 1 | 0.637446795 | 0.323956123 |
| Cgnl1 | 0.45464203 | 0.211947463 | 0.302 | 0.381 | 1 | 0.177843888 | -1.037264268 | 0.293 | 0.386 | 1 | 0.45464203 | 0.324059327 |
| Kcnq1 | 0.177909201 | 0.714150003 | 0.075 | 0.034 | 1 | 0.737522269 | -0.332720152 | 0.024 | 0.034 | 1 | 0.737522269 | 0.324166719 |
| Pold1 | 0.567830665 | -0.430239906 | 0.019 | 0.034 | 1 | 0.17794278 | 1.029849928 | 0.073 | 0.031 | 1 | 0.567830665 | 0.324221927 |
| 1700001P01Rik | 0.628925725 | 0.739685096 | 0.038 | 0.025 | 1 | 0.17794278 | 1.029849928 | 0.073 | 0.031 | 1 | 0.628925725 | 0.324221927 |
| Sox7 | 0.913424206 | 1.417757001 | 0.019 | 0.017 | 1 | 0.178092808 | 2.547698233 | 0.049 | 0.017 | 1 | 0.913424206 | 0.324468568 |
| Gm15417 | 0.821165746 | 0.417757001 | 0.189 | 0.182 | 1 | 0.178189713 | -0.935384654 | 0.122 | 0.205 | 1 | 0.821165746 | 0.324627852 |
| Zfp260 | 0.226965316 | 0.41637046 | 0.66 | 0.678 | 1 | 0.178316517 | -0.206621391 | 0.61 | 0.741 | 1 | 0.226965316 | 0.324836254 |
| Nanp | 0.440362837 | 0.75033234 | 0.189 | 0.153 | 1 | 0.178358349 | -0.673757069 | 0.073 | 0.154 | 1 | 0.440362837 | 0.324904998 |
| Rsu1 | 0.178391658 | 0.339973013 | 0.774 | 0.797 | 1 | 0.243068461 | -0.117491788 | 0.683 | 0.826 | 1 | 0.243068461 | 0.324959732 |
| Ift27 | 0.178396604 | 0.512616187 | 0.434 | 0.36 | 1 | 0.589708781 | 0.296636468 | 0.39 | 0.352 | 1 | 0.589708781 | 0.324967859 |
| Mdfic | 0.22921056 | 0.417757001 | 0.434 | 0.331 | 1 | 0.178501664 | -0.318073376 | 0.366 | 0.519 | 1 | 0.22921056 | 0.325140484 |
| Adal | 0.178545164 | 0.569760094 | 0.226 | 0.148 | 1 | 0.922670096 | 0.351778023 | 0.122 | 0.133 | 1 | 0.922670096 | 0.325211952 |
| Brat1 | 0.1785663 | 0.932330173 | 0.094 | 0.047 | 1 | 0.254061981 | 1.47463477 | 0.098 | 0.055 | 1 | 0.254061981 | 0.325246677 |
| Stx1b | 0.178709532 | 1.017219071 | 0.075 | 0.034 | 1 | 0.327974058 | 0.627751484 | 0.146 | 0.096 | 1 | 0.327974058 | 0.325481967 |

| | | | | | | | | | | | | |
|---|---|---|---|---|---|---|---|---|---|---|---|---|
| Ints15 | 0.179068732 | 0.707263618 | 0.189 | 0.119 | 1 | 0.517317396 | -0.16279515 | 0.146 | 0.191 | 1 | 0.517317396 | 0.326071854 |
| Agpat3 | 0.179160122 | 0.576746524 | 0.547 | 0.525 | 1 | 0.843592636 | 0.174239837 | 0.707 | 0.693 | 1 | 0.843592636 | 0.326221894 |
| Rnf26 | 0.179173694 | -0.732802676 | 0.075 | 0.144 | 1 | 0.355625518 | -0.20718927 | 0.122 | 0.188 | 1 | 0.355625518 | 0.326244175 |
| Rab4b | 0.179179603 | 0.593219067 | 0.585 | 0.542 | 1 | 0.301329855 | 0.391793701 | 0.585 | 0.519 | 1 | 0.301329855 | 0.326253875 |
| Rnf152 | 0.17922613 | -1.223789028 | 0.057 | 0.119 | 1 | 0.527279176 | 0.877846834 | 0.098 | 0.072 | 1 | 0.527279176 | 0.326330254 |
| S100a10 | 0.179778797 | -0.31559734 | 0.981 | 0.97 | 1 | 0.537310618 | -0.167705064 | 0.976 | 0.976 | 1 | 0.537310618 | 0.327237178 |
| Ints11 | 0.511830059 | 0.270199812 | 0.358 | 0.314 | 1 | 0.179981844 | -0.515311565 | 0.195 | 0.307 | 1 | 0.511830059 | 0.327570224 |
| Glt28d2 | 0.979446593 | 0.552058092 | 0.226 | 0.237 | 1 | 0.180030298 | -0.26713181 | 0.244 | 0.372 | 1 | 0.979446593 | 0.327649688 |
| Fzd7 | 0.180117125 | 0.494209061 | 0.34 | 0.254 | 1 | 0.19946149 | 1.082317348 | 0.293 | 0.232 | 1 | 0.19946149 | 0.327792072 |
| Hmcn2 | 0.368179284 | -0.582242999 | 0.019 | 0.047 | 1 | 0.180142662 | -1.069685746 | 0.024 | 0.085 | 1 | 0.368179284 | 0.327833945 |
| Gm45570 | 0.287773478 | -0.430239906 | 0 | 0.021 | 1 | 0.180311132 | 1.422167351 | 0.098 | 0.048 | 1 | 0.287773478 | 0.328110159 |
| Cd247 | 0.471154689 | 2.377115016 | 0.038 | 0.021 | 1 | 0.180328124 | 2.574170444 | 0.049 | 0.017 | 1 | 0.471154689 | 0.328138016 |
| Gm20492 | 0.48175508 | 1.154722595 | 0.038 | 0.021 | 1 | 0.180328124 | 2.252242349 | 0.049 | 0.017 | 1 | 0.48175508 | 0.328138016 |
| Gm57065 | 0.563757816 | -1.015202407 | 0.019 | 0.034 | 1 | 0.180328124 | 2.252242349 | 0.049 | 0.017 | 1 | 0.563757816 | 0.328138016 |
| Gm57243 | 0.926212226 | 0.8327945 | 0.019 | 0.017 | 1 | 0.180328124 | 2.83720485 | 0.049 | 0.017 | 1 | 0.926212226 | 0.328138016 |
| Stil | 0.97619858 | -0.347777746 | 0.057 | 0.055 | 1 | 0.180328124 | 2.83720485 | 0.049 | 0.017 | 1 | 0.97619858 | 0.328138016 |
| Acox2 | 0.997810945 | 0.891688189 | 0.038 | 0.038 | 1 | 0.180328124 | 2.252242349 | 0.049 | 0.017 | 1 | 0.997810945 | 0.328138016 |
| Zfp112 | 0.55809939 | -0.381330305 | 0.057 | 0.081 | 1 | 0.180337238 | -0.995685164 | 0.049 | 0.119 | 1 | 0.55809939 | 0.328152957 |
| Slc26a8 | 0.24309253 | -0.845277405 | 0 | 0.025 | 1 | 0.18034522 | 2.100239256 | 0.049 | 0.017 | 1 | 0.24309253 | 0.328166042 |
| Aldh4a1 | 0.180429057 | 0.695290976 | 0.208 | 0.144 | 1 | 0.570919757 | 0.515276755 | 0.171 | 0.143 | 1 | 0.570919757 | 0.328303469 |
| Bbs10 | 0.767318506 | 0.454282877 | 0.057 | 0.047 | 1 | 0.1805589 | 1.029849928 | 0.073 | 0.031 | 1 | 0.767318506 | 0.328516284 |
| Mien1 | 0.232171936 | -0.492975661 | 0.585 | 0.61 | 1 | 0.180640638 | -0.122153166 | 0.366 | 0.488 | 1 | 0.232171936 | 0.328650236 |
| Chchd4 | 0.436935547 | -0.430239906 | 0.151 | 0.195 | 1 | 0.180995151 | 0.309957847 | 0.146 | 0.256 | 1 | 0.436935547 | 0.329231058 |
| Mettl15 | 0.181243067 | -0.795236723 | 0.151 | 0.233 | 1 | 0.523136554 | 0.15538081 | 0.122 | 0.171 | 1 | 0.523136554 | 0.329637085 |
| Haus7 | 0.399950268 | 0.272079545 | 0.151 | 0.212 | 1 | 0.181406539 | -0.52536523 | 0.098 | 0.188 | 1 | 0.399950268 | 0.329904745 |
| Myrfl | 0.243110774 | -2.237594828 | 0 | 0.025 | 1 | 0.181471053 | 2.422167351 | 0.049 | 0.017 | 1 | 0.243110774 | 0.330010363 |
| Plaat5 | 0.249034808 | 2.154722595 | 0.019 | 0.004 | 1 | 0.181471053 | 2.422167351 | 0.049 | 0.017 | 1 | 0.249034808 | 0.330010363 |
| Fhl4 | 0.343000446 | -0.430239906 | 0 | 0.017 | 1 | 0.181471053 | 2.614812428 | 0.049 | 0.017 | 1 | 0.343000446 | 0.330010363 |
| St3gal4 | 0.181855763 | -0.480280588 | 0.642 | 0.712 | 1 | 0.685518239 | 0.330061882 | 0.61 | 0.625 | 1 | 0.685518239 | 0.330640008 |
| Bloc1s5 | 0.182008745 | 1.154722595 | 0.132 | 0.076 | 1 | 0.403708081 | 0.589277336 | 0.146 | 0.102 | 1 | 0.403708081 | 0.330890307 |
| Snhg15 | 0.45763248 | 0.46284489 | 0.189 | 0.148 | 1 | 0.18204078 | 0.883008539 | 0.146 | 0.082 | 1 | 0.45763248 | 0.330942715 |
| Pcdhga3 | 0.182081857 | 1.526691372 | 0.113 | 0.064 | 1 | 0.328505351 | 1.000703582 | 0.098 | 0.058 | 1 | 0.328505351 | 0.331009911 |
| Rgl3 | 0.18218924 | 1.533234218 | 0.113 | 0.064 | 1 | 0.987649386 | 0.19334866 | 0.146 | 0.147 | 1 | 0.987649386 | 0.33118556 |
| Ubxn8 | 0.949664717 | 0.154722595 | 0.34 | 0.347 | 1 | 0.18230016 | -0.353936638 | 0.244 | 0.358 | 1 | 0.949664717 | 0.331366971 |
| Slc9a5 | 0.182332113 | 0.491757582 | 0.094 | 0.047 | 1 | 0.82181816 | 0.252242349 | 0.024 | 0.031 | 1 | 0.82181816 | 0.331419227 |
| Rbm12b1 | 0.182499022 | -1.015202407 | 0.038 | 0.093 | 1 | 0.296959488 | -0.589059905 | 0.073 | 0.13 | 1 | 0.296959488 | 0.331692151 |
| Osgep | 0.182511055 | 0.392882332 | 0.547 | 0.466 | 1 | 0.250790978 | 0.164779508 | 0.317 | 0.43 | 1 | 0.250790978 | 0.331711824 |
| 2810004N23Rik | 0.99167413 | 0.171596413 | 0.377 | 0.39 | 1 | 0.18256905 | -0.555112573 | 0.268 | 0.369 | 1 | 0.99167413 | 0.331806641 |
| Napepld | 0.243322126 | 1.154722595 | 0.057 | 0.025 | 1 | 0.18261072 | 2.029849928 | 0.049 | 0.017 | 1 | 0.243322126 | 0.331874766 |
| Zfp660 | 0.914958 | 0.154722595 | 0.019 | 0.021 | 1 | 0.18261072 | 2.029849928 | 0.049 | 0.017 | 1 | 0.914958 | 0.331874766 |
| Tbx4 | 0.932506754 | 1.347367673 | 0.019 | 0.021 | 1 | 0.18261072 | 2.029849928 | 0.049 | 0.017 | 1 | 0.932506754 | 0.331874766 |
| Hsf5 | 0.610863297 | 1.347367673 | 0.038 | 0.025 | 1 | 0.18261587 | 2.422167351 | 0.049 | 0.017 | 1 | 0.610863297 | 0.331883184 |
| Ntng2 | 0.674311707 | -0.368839361 | 0.113 | 0.131 | 1 | 0.182692679 | -0.899760744 | 0.073 | 0.154 | 1 | 0.674311707 | 0.332008743 |
| Gm16124 | 0.182855385 | -0.430239906 | 0.094 | 0.174 | 1 | 0.744521103 | 0.296636468 | 0.122 | 0.15 | 1 | 0.744521103 | 0.332274679 |
| Ociad2 | 0.182969399 | -0.368839361 | 0.019 | 0.068 | 1 | 0.7193606 | 0.444887427 | 0.049 | 0.065 | 1 | 0.7193606 | 0.332460996 |
| Snx3 | 0.183165796 | 0.16750875 | 0.943 | 0.898 | 1 | 0.26397042 | -0.118056835 | 0.927 | 0.932 | 1 | 0.26397042 | 0.332781882 |
| F2rl1 | 0.932611893 | 0.347367673 | 0.019 | 0.017 | 1 | 0.183226966 | 1.334704509 | 0.098 | 0.048 | 1 | 0.932611893 | 0.332881811 |
| Rad51b | 0.194234407 | -1.123262153 | 0.132 | 0.208 | 1 | 0.183257726 | 1.690363461 | 0.122 | 0.065 | 1 | 0.194234407 | 0.332932059 |
| Aox2 | 0.518992757 | -1.960754623 | 0.038 | 0.059 | 1 | 0.183315445 | 1.635570989 | 0.098 | 0.048 | 1 | 0.518992757 | 0.333026338 |
| Fktn | 0.183352477 | 0.913714495 | 0.245 | 0.178 | 1 | 0.881085068 | 0.100239256 | 0.195 | 0.212 | 1 | 0.881085068 | 0.333086824 |
| Parp11 | 0.509113633 | 0.739685096 | 0.113 | 0.085 | 1 | 0.18337955 | -0.286177566 | 0.073 | 0.154 | 1 | 0.509113633 | 0.333131041 |
| Etv4 | 0.183540215 | -1.138059154 | 0.075 | 0.14 | 1 | 0.32762653 | 0.91966701 | 0.146 | 0.099 | 1 | 0.32762653 | 0.333393419 |
| Cpsf3 | 0.183582909 | 0.539063297 | 0.34 | 0.271 | 1 | 0.305783933 | 0.753140585 | 0.317 | 0.273 | 1 | 0.305783933 | 0.333463134 |
| Suz12 | 0.183619171 | 0.45911603 | 0.679 | 0.597 | 1 | 0.555790234 | 0.249348276 | 0.732 | 0.7 | 1 | 0.555790234 | 0.333522342 |
| Zbtb24 | 0.183735236 | 0.859266711 | 0.264 | 0.191 | 1 | 0.800755181 | 0.19334866 | 0.171 | 0.198 | 1 | 0.800755181 | 0.333711836 |
| Thada | 0.485188119 | 0.638147069 | 0.434 | 0.449 | 1 | 0.184089387 | 0.550803189 | 0.585 | 0.519 | 1 | 0.485188119 | 0.334289871 |
| Gng10 | 0.955209533 | 0.17031945 | 0.623 | 0.691 | 1 | 0.184191362 | -0.390205646 | 0.561 | 0.666 | 1 | 0.955209533 | 0.334456267 |
| Gm12236 | 0.828131282 | 0.261637799 | 0.075 | 0.068 | 1 | 0.184380802 | 1.252242349 | 0.122 | 0.065 | 1 | 0.828131282 | 0.334765324 |
| Gm29488 | 0.767066103 | 0.569760094 | 0.038 | 0.03 | 1 | 0.184405761 | -0.80665134 | 0.049 | 0.119 | 1 | 0.767066103 | 0.334806038 |
| Zfp33b | 0.184442051 | 0.813685677 | 0.189 | 0.123 | 1 | 0.914201521 | 0.377773231 | 0.195 | 0.208 | 1 | 0.914201521 | 0.334865232 |
| Lias | 0.343942195 | 0.340278248 | 0.377 | 0.297 | 1 | 0.18444465 | 0.848520163 | 0.366 | 0.28 | 1 | 0.343942195 | 0.334869472 |
| Yae1d1 | 0.184508773 | 0.336560918 | 0.509 | 0.411 | 1 | 0.241782792 | -0.254293505 | 0.317 | 0.41 | 1 | 0.241782792 | 0.334974059 |
| Gm56507 | 0.877748789 | 0.247831999 | 0.057 | 0.051 | 1 | 0.184683918 | 1.059597271 | 0.098 | 0.048 | 1 | 0.877748789 | 0.335259686 |
| 4930599N23Rik | 0.184909048 | 0.631160639 | 0.208 | 0.136 | 1 | 0.488257334 | -0.395455907 | 0.122 | 0.167 | 1 | 0.488257334 | 0.335626739 |
| Zfp266 | 0.205386937 | 0.448453798 | 0.604 | 0.589 | 1 | 0.184911602 | -0.160539176 | 0.707 | 0.836 | 1 | 0.205386937 | 0.335630904 |
| Unc13a | 0.926212226 | 0.8327945 | 0.019 | 0.017 | 1 | 0.184911704 | 1.83720485 | 0.049 | 0.017 | 1 | 0.926212226 | 0.33563107 |
| Elof1 | 0.272579508 | 0.417757001 | 0.528 | 0.466 | 1 | 0.184954018 | -0.357173195 | 0.488 | 0.577 | 1 | 0.272579508 | 0.335700047 |
| Mrps18a | 0.835239977 | 0.209170379 | 0.377 | 0.377 | 1 | 0.184971665 | 0.650791726 | 0.415 | 0.345 | 1 | 0.835239977 | 0.335728813 |
| Vps11 | 0.421557469 | 0.507024339 | 0.226 | 0.182 | 1 | 0.185094533 | -0.188330242 | 0.171 | 0.287 | 1 | 0.421557469 | 0.335929079 |
| Gstt3 | 0.263607288 | -0.274120704 | 0.17 | 0.242 | 1 | 0.185277563 | -0.863234868 | 0.098 | 0.177 | 1 | 0.263607288 | 0.336227351 |
| Rdh13 | 0.965479387 | 0.154722595 | 0.094 | 0.093 | 1 | 0.185347991 | 1.111379813 | 0.171 | 0.102 | 1 | 0.965479387 | 0.336342104 |
| Fbxo46 | 0.185460869 | 0.739685096 | 0.151 | 0.089 | 1 | 0.270542168 | 0.100239256 | 0.049 | 0.109 | 1 | 0.270542168 | 0.336526004 |
| Zbtb11os1 | 0.185460869 | 0.680791406 | 0.151 | 0.089 | 1 | 0.463387713 | 0.435106406 | 0.146 | 0.106 | 1 | 0.463387713 | 0.336526004 |
| Tcaf2 | 0.185546097 | 1.417757001 | 0.132 | 0.076 | 1 | 0.344979528 | 1.048708955 | 0.122 | 0.082 | 1 | 0.344979528 | 0.33666484 |
| Eef2kmt | 0.593176225 | 0.377115016 | 0.226 | 0.208 | 1 | 0.185734655 | -0.727579769 | 0.098 | 0.181 | 1 | 0.593176225 | 0.336971949 |
| Rab15 | 0.185745115 | 1.090592257 | 0.132 | 0.076 | 1 | 0.557180722 | 0.589277336 | 0.122 | 0.092 | 1 | 0.557180722 | 0.336988983 |
| Stat5a | 0.185839257 | 0.58440687 | 0.321 | 0.25 | 1 | 0.282217201 | 0.69315848 | 0.415 | 0.341 | 1 | 0.282217201 | 0.337142285 |
| Cherp | 0.185951906 | 0.338434548 | 0.491 | 0.403 | 1 | 0.249024531 | 0.707246256 | 0.512 | 0.478 | 1 | 0.249024531 | 0.3373257 |
| Tpbg | 0.18601126 | 0.111395163 | 0.396 | 0.301 | 1 | 0.658513286 | -0.195216628 | 0.293 | 0.328 | 1 | 0.658513286 | 0.337422331 |
| Gm10974 | 0.413082433 | 0.154722595 | 0 | 0.013 | 1 | 0.186066733 | 1.667279848 | 0.049 | 0.017 | 1 | 0.413082433 | 0.337512637 |
| Sytl5 | 0.28143885 | 1.029191713 | 0.094 | 0.055 | 1 | 0.186070187 | 1.944120054 | 0.049 | 0.017 | 1 | 0.28143885 | 0.337518259 |
| Cxcr5 | 0.209009104 | 1.739685096 | 0.038 | 0.013 | 1 | 0.186073065 | 1.83720485 | 0.049 | 0.017 | 1 | 0.209009104 | 0.337522944 |
| Fgf20 | 0.287773478 | -0.430239906 | 0 | 0.021 | 1 | 0.186073065 | 1.83720485 | 0.049 | 0.017 | 1 | 0.287773478 | 0.337522944 |
| Gm12781 | 0.471154689 | 1.569760094 | 0.038 | 0.021 | 1 | 0.186073065 | 1.83720485 | 0.049 | 0.017 | 1 | 0.471154689 | 0.337522944 |
| Hhipl1 | 0.48175508 | 1.154722595 | 0.038 | 0.021 | 1 | 0.186073065 | 1.83720485 | 0.049 | 0.017 | 1 | 0.48175508 | 0.337522944 |
| 1500026H17Rik | 0.503971666 | 1.569760094 | 0.019 | 0.008 | 1 | 0.186073065 | 1.83720485 | 0.049 | 0.017 | 1 | 0.503971666 | 0.337522944 |
| 4930563E22Rik | 0.668070194 | 0.154722595 | 0.019 | 0.03 | 1 | 0.186073065 | 1.83720485 | 0.049 | 0.017 | 1 | 0.668070194 | 0.337522944 |
| A730090N16Rik | 0.679745214 | 0.417757001 | 0.019 | 0.03 | 1 | 0.186073065 | 1.83720485 | 0.049 | 0.017 | 1 | 0.679745214 | 0.337522944 |
| D630039A03Rik | 0.782077996 | 0.347367673 | 0.019 | 0.025 | 1 | 0.186073065 | 1.83720485 | 0.049 | 0.017 | 1 | 0.782077996 | 0.337522944 |
| Rbl1 | 0.259305799 | 0.510866405 | 0.17 | 0.114 | 1 | 0.186147378 | 1.199774929 | 0.146 | 0.085 | 1 | 0.259305799 | 0.337643909 |
| Gatb | 0.186167104 | 1.002719501 | 0.132 | 0.076 | 1 | 0.953187697 | 0.136765132 | 0.073 | 0.075 | 1 | 0.953187697 | 0.337676017 |
| Rab28 | 0.186176213 | -0.257255105 | 0.566 | 0.691 | 1 | 0.581129047 | 0.136765132 | 0.561 | 0.628 | 1 | 0.581129047 | 0.337690844 |
| Gprc5b | 0.697421791 | -0.375792122 | 0.132 | 0.153 | 1 | 0.186184714 | -0.410722664 | 0.024 | 0.085 | 1 | 0.697421791 | 0.337704681 |
| Zscan22 | 0.186252471 | 0.8327945 | 0.17 | 0.106 | 1 | 0.205741141 | -0.622226769 | 0.049 | 0.116 | 1 | 0.205741141 | 0.337814959 |
| Me3 | 0.186468671 | 0.8327945 | 0.132 | 0.076 | 1 | 0.645507194 | 0.223387486 | 0.244 | 0.218 | 1 | 0.645507194 | 0.338166776 |
| Ctso | 0.186730891 | 0.855162313 | 0.189 | 0.123 | 1 | 0.613142284 | 0.60690723 | 0.195 | 0.174 | 1 | 0.613142284 | 0.338593357 |
| Hnrnph3 | 0.187057405 | 0.211947463 | 0.811 | 0.852 | 1 | 0.583704028 | 0.162605137 | 0.805 | 0.881 | 1 | 0.583704028 | 0.339124337 |
| 4933406J09Rik | 0.207549016 | 1.8327945 | 0.038 | 0.013 | 1 | 0.187228242 | 0.667279848 | 0.049 | 0.017 | 1 | 0.207549016 | 0.33940207 |

| | | | | | | | | | | | | |
|---|---|---|---|---|---|---|---|---|---|---|---|---|
| Fitm2 | 0.197370522 | 0.767699472 | 0.189 | 0.123 | 1 | 0.187386282 | -0.970150072 | 0.049 | 0.116 | 1 | 0.197370522 | 0.339658945 |
| Copz2 | 0.493507562 | -0.178625309 | 0.849 | 0.839 | 1 | 0.187555953 | -0.335041464 | 0.634 | 0.713 | 1 | 0.493507562 | 0.339934671 |
| Zpbp | 0.273166284 | -0.845277405 | 0.038 | 0.081 | 1 | 0.187609799 | -1.484723245 | 0.024 | 0.082 | 1 | 0.273166284 | 0.340022162 |
| Lacc1 | 0.187923286 | 1.211306123 | 0.245 | 0.178 | 1 | 0.753022409 | 0.568716014 | 0.146 | 0.133 | 1 | 0.753022409 | 0.340531411 |
| Mast3 | 0.188037008 | 1.090592257 | 0.151 | 0.093 | 1 | 0.977366645 | 0.136765132 | 0.098 | 0.099 | 1 | 0.977366645 | 0.3407161 |
| H2-K1 | 0.400256664 | -0.112757716 | 0.792 | 0.669 | 1 | 0.188153804 | 0.208008378 | 0.78 | 0.727 | 1 | 0.400256664 | 0.340905754 |
| Gm17135 | 0.209009104 | 1.739685096 | 0.038 | 0.013 | 1 | 0.188380068 | 1.422167351 | 0.049 | 0.017 | 1 | 0.209009104 | 0.341273086 |
| 4930506C21Rik | 0.247022813 | 0.984797593 | 0.057 | 0.025 | 1 | 0.188380068 | 1.422167351 | 0.049 | 0.017 | 1 | 0.247022813 | 0.341273086 |
| Vsig8 | 0.251644813 | 1.569760094 | 0.019 | 0.004 | 1 | 0.188380068 | 1.252242349 | 0.049 | 0.017 | 1 | 0.251644813 | 0.341273086 |
| Gpr21 | 0.325530479 | 2.154722595 | 0.038 | 0.017 | 1 | 0.188380068 | 1.614812428 | 0.049 | 0.017 | 1 | 0.325530479 | 0.341273086 |
| Zfp184 | 0.343968402 | 1.154722595 | 0.038 | 0.017 | 1 | 0.188380068 | 1.614812428 | 0.049 | 0.017 | 1 | 0.343968402 | 0.341273086 |
| Spc25 | 0.359852535 | -1.752168001 | 0.019 | 0.047 | 1 | 0.188380068 | 1.614812428 | 0.049 | 0.017 | 1 | 0.359852535 | 0.341273086 |
| P2rx6 | 0.503971666 | 1.569760094 | 0.019 | 0.008 | 1 | 0.188380068 | 1.614812428 | 0.049 | 0.017 | 1 | 0.503971666 | 0.341273086 |
| Hormad2 | 0.188408576 | 0.417757001 | 0.208 | 0.14 | 1 | 0.215108214 | 0.863200058 | 0.122 | 0.068 | 1 | 0.215108214 | 0.34131936 |
| Vps25 | 0.559233506 | 0.984797593 | 0.094 | 0.072 | 1 | 0.188498353 | -0.970150072 | 0 | 0.041 | 1 | 0.559233506 | 0.341465076 |
| Pih1d2 | 0.569866135 | -0.1672055 | 0.019 | 0.034 | 1 | 0.188498353 | -0.970150072 | 0 | 0.041 | 1 | 0.569866135 | 0.341465076 |
| Fchsd1 | 0.732691937 | 1.154722595 | 0.019 | 0.013 | 1 | 0.188498353 | -0.970150072 | 0 | 0.041 | 1 | 0.732691937 | 0.341465076 |
| Rad54b | 0.855425466 | 0.739685096 | 0.057 | 0.051 | 1 | 0.188498353 | -1.16279515 | 0 | 0.041 | 1 | 0.855425466 | 0.341465076 |
| Tyw3 | 0.199383551 | -0.932740246 | 0.019 | 0.064 | 1 | 0.188517184 | -1.069685746 | 0 | 0.041 | 1 | 0.199383551 | 0.341495639 |
| Bpifb4 | 0.249034808 | 2.154722595 | 0.019 | 0.004 | 1 | 0.188517184 | -1.332720152 | 0 | 0.041 | 1 | 0.249034808 | 0.341495639 |
| Zfp202 | 0.418326033 | 1.569760094 | 0.057 | 0.034 | 1 | 0.188517184 | -1.069685746 | 0 | 0.041 | 1 | 0.418326033 | 0.341495639 |
| Prss53 | 0.451077244 | 0.228723176 | 0.038 | 0.068 | 1 | 0.188517184 | -1.16279515 | 0 | 0.041 | 1 | 0.451077244 | 0.341495639 |
| Cd200r3 | 0.488750271 | -0.304709024 | 0.019 | 0.038 | 1 | 0.188517184 | -1.069685746 | 0 | 0.041 | 1 | 0.488750271 | 0.341495639 |
| Mlxipl | 0.68209151 | 0.695290976 | 0.019 | 0.03 | 1 | 0.188517184 | -1.250257991 | 0 | 0.041 | 1 | 0.68209151 | 0.341495639 |
| Gm52870 | 0.929579738 | 0.932330173 | 0.019 | 0.021 | 1 | 0.188517184 | -1.16279515 | 0 | 0.041 | 1 | 0.929579738 | 0.341495639 |
| Aass | 0.24309253 | -0.845277405 | 0 | 0.025 | 1 | 0.188532249 | -1.250257991 | 0 | 0.041 | 1 | 0.24309253 | 0.341520088 |
| B930046K06Rik | 0.359776551 | -0.652632327 | 0.019 | 0.047 | 1 | 0.188532249 | -1.250257991 | 0 | 0.041 | 1 | 0.359776551 | 0.341520088 |
| Hao1 | 0.48151808 | -0.545717123 | 0.057 | 0.085 | 1 | 0.188532249 | -1.410722664 | 0 | 0.041 | 1 | 0.48151808 | 0.341520088 |
| Nudt17 | 0.677401546 | 0.280253477 | 0.019 | 0.03 | 1 | 0.188532591 | -1.16279515 | 0 | 0.041 | 1 | 0.677401546 | 0.341520644 |
| Rnft2 | 0.80199058 | 0.891688189 | 0.038 | 0.047 | 1 | 0.188532591 | -1.16279515 | 0 | 0.041 | 1 | 0.80199058 | 0.341520644 |
| Gm13427 | 0.986865056 | 0.154722595 | 0.038 | 0.038 | 1 | 0.188532591 | -1.16279515 | 0 | 0.041 | 1 | 0.986865056 | 0.341520644 |
| Gmnn | 0.227539261 | -1.845277405 | 0.019 | 0.059 | 1 | 0.188543546 | -1.332720152 | 0 | 0.041 | 1 | 0.227539261 | 0.341538424 |
| Gm2885 | 0.281986656 | -0.1672055 | 0.019 | 0.055 | 1 | 0.188543546 | -1.410722664 | 0 | 0.041 | 1 | 0.281986656 | 0.341538424 |
| Il4i1 | 0.413080549 | -0.1672055 | 0 | 0.013 | 1 | 0.188543546 | -1.332720152 | 0 | 0.041 | 1 | 0.413080549 | 0.341538424 |
| Muc5b | 0.507281812 | 1.154722595 | 0.019 | 0.008 | 1 | 0.188543546 | -1.332720152 | 0 | 0.041 | 1 | 0.507281812 | 0.341538424 |
| Gm47460 | 0.206368562 | -1.015202407 | 0 | 0.03 | 1 | 0.188544573 | -1.410722664 | 0 | 0.041 | 1 | 0.206368562 | 0.341540091 |
| Gm30054 | 0.246023494 | 0.891688189 | 0.075 | 0.038 | 1 | 0.188544916 | -1.484723245 | 0 | 0.041 | 1 | 0.246023494 | 0.341540646 |
| Gm26626 | 0.912039013 | 0.154722595 | 0.019 | 0.021 | 1 | 0.188551078 | -1.484723245 | 0 | 0.041 | 1 | 0.912039013 | 0.341550647 |
| Gm12811 | 0.327325725 | 2.002719501 | 0.057 | 0.03 | 1 | 0.188553817 | -1.917682652 | 0 | 0.041 | 1 | 0.327325725 | 0.341555092 |
| Adgb | 0.684440425 | 0.8327945 | 0.019 | 0.03 | 1 | 0.188554502 | -1.332720152 | 0 | 0.041 | 1 | 0.684440425 | 0.341556203 |
| Mtmr7 | 0.567381671 | -0.1672055 | 0.057 | 0.081 | 1 | 0.188559979 | -1.686357106 | 0 | 0.041 | 1 | 0.567381671 | 0.341565092 |
| Gm29491 | 0.24309253 | -0.845277405 | 0 | 0.025 | 1 | 0.188573672 | -0.863234868 | 0 | 0.041 | 1 | 0.24309253 | 0.341587315 |
| Tstd3 | 0.312692399 | 1.017219071 | 0.075 | 0.042 | 1 | 0.188573672 | -0.863234868 | 0 | 0.041 | 1 | 0.312692399 | 0.341587315 |
| 9530077C14Rik | 0.469724974 | -0.347777746 | 0.038 | 0.064 | 1 | 0.188573672 | -0.863234868 | 0 | 0.041 | 1 | 0.469724974 | 0.341587315 |
| Lrrc75b | 0.668070194 | 0.154722595 | 0.019 | 0.03 | 1 | 0.188573672 | -0.863234868 | 0 | 0.041 | 1 | 0.668070194 | 0.341587315 |
| Gm10135 | 0.917883255 | 0.569760094 | 0.019 | 0.021 | 1 | 0.188573672 | -0.863234868 | 0 | 0.041 | 1 | 0.917883255 | 0.341587315 |
| Moap1 | 0.917883255 | 0.569760094 | 0.019 | 0.021 | 1 | 0.188573672 | -0.863234868 | 0 | 0.041 | 1 | 0.917883255 | 0.341587315 |
| Ctns | 0.395006511 | 0.242185436 | 0.245 | 0.191 | 1 | 0.188789908 | -0.527367582 | 0.171 | 0.263 | 1 | 0.395006511 | 0.341938187 |
| Lpin3 | 0.752453414 | 0.739685096 | 0.057 | 0.047 | 1 | 0.18881203 | -1.116991461 | 0.024 | 0.082 | 1 | 0.752453414 | 0.341974078 |
| Aopep | 0.188856151 | 0.198091558 | 1 | 1 | 1 | 0.877140794 | -0.116107037 | 1 | 1 | 1 | 0.877140794 | 0.342045656 |
| Nckipsd | 0.629258964 | 0.865215978 | 0.094 | 0.076 | 1 | 0.189071225 | 1.342440158 | 0.146 | 0.085 | 1 | 0.629258964 | 0.342394521 |
| Hspa12a | 0.189127774 | -1.046911266 | 0.057 | 0.119 | 1 | 0.288957625 | 0.515276755 | 0.049 | 0.102 | 1 | 0.288957625 | 0.342486234 |
| Rab5b | 0.189192133 | -0.210274222 | 0.358 | 0.479 | 1 | 0.533015875 | 0.43368922 | 0.61 | 0.57 | 1 | 0.533015875 | 0.342590602 |
| Pofut2 | 0.555529534 | 0.165583587 | 0.717 | 0.754 | 1 | 0.189296644 | 0.351157375 | 0.756 | 0.727 | 1 | 0.555529534 | 0.342760068 |
| Pals1 | 0.445159251 | 0.275870048 | 0.925 | 0.894 | 1 | 0.189396473 | 0.292543714 | 0.927 | 0.908 | 1 | 0.445159251 | 0.342921923 |
| Gm31763 | 0.359307617 | -0.214511215 | 0.057 | 0.097 | 1 | 0.189408523 | -0.695290231 | 0.098 | 0.184 | 1 | 0.359307617 | 0.342941457 |
| Spaca1 | 0.644375912 | 0.640149422 | 0.094 | 0.076 | 1 | 0.189410866 | -1.020776145 | 0.024 | 0.082 | 1 | 0.644375912 | 0.342945256 |
| Zfp931 | 0.189460616 | 1.039245377 | 0.151 | 0.093 | 1 | 0.985544352 | 0.911205431 | 0.098 | 0.102 | 1 | 0.985544352 | 0.343025908 |
| Hyal1 | 0.455315314 | 0.569760094 | 0.132 | 0.097 | 1 | 0.190007084 | -0.917682652 | 0.024 | 0.082 | 1 | 0.455315314 | 0.343911477 |
| Fpgt | 0.747889121 | 0.261637799 | 0.113 | 0.097 | 1 | 0.190007084 | -0.917682652 | 0.024 | 0.082 | 1 | 0.747889121 | 0.343911477 |
| Kmt5c | 0.190022159 | 1.154722595 | 0.17 | 0.11 | 1 | 0.453069113 | -0.435813645 | 0.098 | 0.14 | 1 | 0.453069113 | 0.343935897 |
| Dnah10 | 0.190023427 | 0.417757001 | 0.094 | 0.047 | 1 | 0.410112287 | 0.614812428 | 0.073 | 0.044 | 1 | 0.410112287 | 0.343937951 |
| Vps26a | 0.829963551 | 0.143667406 | 0.528 | 0.568 | 1 | 0.190250321 | -0.222337096 | 0.78 | 0.87 | 1 | 0.829963551 | 0.344305457 |
| Slc9a6 | 0.190262013 | 0.310373465 | 0.34 | 0.242 | 1 | 0.685515152 | 0.592092352 | 0.317 | 0.317 | 1 | 0.685515152 | 0.344324392 |
| Tmem41a | 0.960852989 | 0.47665069 | 0.057 | 0.059 | 1 | 0.190458117 | -0.863234868 | 0.049 | 0.116 | 1 | 0.960852989 | 0.34464194 |
| Galnt7 | 0.19050519 | 0.100274811 | 0.509 | 0.411 | 1 | 0.289153373 | 0.206918359 | 0.512 | 0.437 | 1 | 0.289153373 | 0.344718153 |
| Otx1 | 0.736073999 | 0.8327945 | 0.019 | 0.013 | 1 | 0.190537763 | 1.159132945 | 0.098 | 0.048 | 1 | 0.736073999 | 0.344770887 |
| Gm52992 | 0.917883255 | 0.569760094 | 0.019 | 0.021 | 1 | 0.190537763 | 1.159132945 | 0.098 | 0.048 | 1 | 0.917883255 | 0.344770887 |
| Gm49207 | 0.391848483 | 1.154722595 | 0.094 | 0.064 | 1 | 0.190600673 | -0.917682652 | 0.024 | 0.082 | 1 | 0.391848483 | 0.344872729 |
| Kcnj6 | 0.206396398 | -0.845277405 | 0 | 0.03 | 1 | 0.190737323 | 0.721727632 | 0.049 | 0.017 | 1 | 0.206396398 | 0.34509392 |
| 1700022H01Rik | 0.232506641 | 1.669295768 | 0.057 | 0.025 | 1 | 0.190737323 | 1.422167351 | 0.049 | 0.017 | 1 | 0.232506641 | 0.34509392 |
| Gm26737 | 0.920805363 | -1.1672055 | 0.019 | 0.021 | 1 | 0.190737323 | 1.252242349 | 0.049 | 0.017 | 1 | 0.920805363 | 0.34509392 |
| Tmem147os | 0.926212226 | 0.8327945 | 0.019 | 0.017 | 1 | 0.190737323 | 1.252242349 | 0.049 | 0.017 | 1 | 0.926212226 | 0.34509392 |
| Uqcrfs1 | 0.190737949 | 0.393171362 | 0.66 | 0.589 | 1 | 0.265526329 | -0.374299255 | 0.659 | 0.696 | 1 | 0.265526329 | 0.345094933 |
| Tmem159 | 0.190910428 | 0.334543632 | 0.434 | 0.335 | 1 | 0.528576605 | -0.181654177 | 0.293 | 0.352 | 1 | 0.528576605 | 0.345374065 |
| Rnf123 | 0.190948898 | 0.614154213 | 0.17 | 0.106 | 1 | 0.955837617 | 0.742047617 | 0.122 | 0.137 | 1 | 0.955837617 | 0.345436314 |
| Map3k5 | 0.628155526 | -0.197620463 | 0.868 | 0.907 | 1 | 0.191107821 | -0.493752211 | 0.927 | 0.898 | 1 | 0.628155526 | 0.345693443 |
| Lrrc71 | 0.606005219 | -0.489133595 | 0.038 | 0.055 | 1 | 0.191169615 | -0.332720152 | 0.049 | 0.123 | 1 | 0.606005219 | 0.345793408 |
| Tex14 | 0.191228333 | -1.058758773 | 0.698 | 0.682 | 1 | 0.282320371 | -0.546761204 | 0.61 | 0.683 | 1 | 0.282320371 | 0.34588839 |
| Pum3 | 0.191481296 | 0.660907982 | 0.434 | 0.39 | 1 | 0.806580828 | 0.465645987 | 0.439 | 0.42 | 1 | 0.806580828 | 0.346297505 |
| Zfp335 | 0.191683002 | -0.347777746 | 0.057 | 0.123 | 1 | 0.388257291 | -0.240797662 | 0.146 | 0.208 | 1 | 0.388257291 | 0.346623631 |
| Cep295 | 0.191762308 | 1.004975475 | 0.245 | 0.186 | 1 | 0.728841038 | 0.435106406 | 0.244 | 0.218 | 1 | 0.728841038 | 0.346751833 |
| E2f3 | 0.191779059 | 0.672853587 | 0.264 | 0.195 | 1 | 0.240744258 | -0.308472605 | 0.268 | 0.355 | 1 | 0.240744258 | 0.34677891 |
| Apoo | 0.192000289 | 0.810768194 | 0.189 | 0.123 | 1 | 0.994764071 | 0.252242349 | 0.171 | 0.174 | 1 | 0.994764071 | 0.347136466 |
| Smyd5 | 0.218263436 | 0.569760094 | 0.113 | 0.064 | 1 | 0.192069421 | -0.520347155 | 0.049 | 0.119 | 1 | 0.218263436 | 0.34724818 |
| Ccdc137 | 0.192079465 | 0.631160639 | 0.302 | 0.225 | 1 | 0.993372271 | 0.268362014 | 0.22 | 0.225 | 1 | 0.993372271 | 0.34726441 |
| Trim12c | 0.238028207 | 0.704061186 | 0.208 | 0.14 | 1 | 0.19273005 | -0.221688839 | 0.049 | 0.123 | 1 | 0.238028207 | 0.348315228 |
| Ddx51 | 0.192867466 | 1.237184755 | 0.113 | 0.064 | 1 | 0.302618984 | 0.88967227 | 0.122 | 0.075 | 1 | 0.302618984 | 0.348537073 |
| Zfp768 | 0.192867466 | 1.237184755 | 0.113 | 0.064 | 1 | 0.4371226 | 0.773074512 | 0.171 | 0.126 | 1 | 0.4371226 | 0.348537073 |
| Krt10 | 0.193046318 | 0.598329246 | 0.264 | 0.182 | 1 | 0.504265055 | 0.557096931 | 0.195 | 0.154 | 1 | 0.504265055 | 0.348825756 |
| Tmem145 | 0.336520503 | 1.569760094 | 0.038 | 0.017 | 1 | 0.193113216 | 1.252242349 | 0.049 | 0.017 | 1 | 0.336520503 | 0.348933718 |
| Gm35546 | 0.193275797 | -0.101617158 | 0.226 | 0.148 | 1 | 0.423341791 | -0.366667484 | 0.146 | 0.205 | 1 | 0.423341791 | 0.34919606 |
| Gm11508 | 0.20258154 | 0.940597789 | 0.226 | 0.161 | 1 | 0.193330741 | -0.233184478 | 0.146 | 0.249 | 1 | 0.20258154 | 0.349284707 |
| Sesn2 | 0.424235692 | -0.737294724 | 0.283 | 0.322 | 1 | 0.193564847 | -0.405651674 | 0.171 | 0.28 | 1 | 0.424235692 | 0.349662343 |
| Lrch4 | 0.19357165 | 0.906795081 | 0.132 | 0.076 | 1 | 0.297996186 | -0.577832649 | 0.049 | 0.099 | 1 | 0.297996186 | 0.349673316 |
| Tesk1 | 0.193794928 | 0.600133743 | 0.245 | 0.169 | 1 | 0.342942497 | 0.422167351 | 0.146 | 0.225 | 1 | 0.342942497 | 0.350033382 |
| Zfp568 | 0.193822549 | 0.347367673 | 0.415 | 0.322 | 1 | 0.994691674 | 0.184265555 | 0.439 | 0.468 | 1 | 0.994691674 | 0.350077918 |
| Reg3g | 0.618898762 | -1.85766113 | 0.377 | 0.331 | 1 | 0.19404302 | 0.515276755 | 0.488 | 0.44 | 1 | 0.618898762 | 0.350433346 |

| | | | | | | | | | | | | |
|---|---|---|---|---|---|---|---|---|---|---|---|---|
| Ror2 | 0.925102122 | -0.136141488 | 0.698 | 0.767 | 1 | 0.194353556 | -0.188330242 | 0.512 | 0.683 | 1 | 0.925102122 | 0.350933808 |
| Trappc2l | 0.702611832 | 0.19695083 | 0.566 | 0.559 | 1 | 0.194372108 | -0.121216046 | 0.317 | 0.42 | 1 | 0.702611832 | 0.3509637 |
| Megf8 | 0.194373307 | -0.392765201 | 0.17 | 0.25 | 1 | 0.845866289 | 0.168826341 | 0.22 | 0.235 | 1 | 0.845866289 | 0.350965632 |
| C2cd5 | 0.703839187 | -0.364006208 | 0.302 | 0.322 | 1 | 0.194489447 | -0.26814815 | 0.293 | 0.42 | 1 | 0.703839187 | 0.351152749 |
| Rpn1 | 0.194709719 | -0.206543648 | 0.811 | 0.843 | 1 | 0.707322592 | 0.197509616 | 0.854 | 0.891 | 1 | 0.707322592 | 0.351507564 |
| Rad51 | 0.194755936 | -2.054730771 | 0.019 | 0.064 | 1 | 0.599290422 | 1.515276755 | 0.024 | 0.014 | 1 | 0.599290422 | 0.351581998 |
| Alkbh7 | 0.194928405 | 0.756173218 | 0.17 | 0.106 | 1 | 0.196788309 | 1.052933541 | 0.146 | 0.085 | 1 | 0.196788309 | 0.351859728 |
| Prkar2b | 0.723252451 | 0.428897558 | 0.17 | 0.153 | 1 | 0.195003992 | 0.83720485 | 0.268 | 0.184 | 1 | 0.723252451 | 0.351981427 |
| Plpp6 | 0.195136308 | 0.865215978 | 0.132 | 0.076 | 1 | 0.719596325 | 0.252242349 | 0.073 | 0.092 | 1 | 0.719596325 | 0.352194437 |
| Dele1 | 0.322038443 | -0.317765177 | 0.302 | 0.364 | 1 | 0.19521462 | 0.571860283 | 0.39 | 0.3 | 1 | 0.322038443 | 0.352320492 |
| Wipf2 | 0.54898844 | 0.124975251 | 0.245 | 0.297 | 1 | 0.195351478 | 0.630242469 | 0.488 | 0.423 | 1 | 0.54898844 | 0.352540756 |
| Myadm | 0.244053711 | -0.823805875 | 0.755 | 0.737 | 1 | 0.195438882 | -0.661046018 | 0.707 | 0.71 | 1 | 0.244053711 | 0.352681407 |
| Marchf5 | 0.19573266 | -0.208682136 | 0.736 | 0.801 | 1 | 0.786647827 | 0.170942247 | 0.756 | 0.86 | 1 | 0.786647827 | 0.353154047 |
| Pdk2 | 0.195779678 | 0.80108564 | 0.132 | 0.076 | 1 | 0.627334271 | 0.458693227 | 0.098 | 0.075 | 1 | 0.627334271 | 0.353229674 |
| Slc35b4 | 0.195818116 | 0.643760676 | 0.264 | 0.191 | 1 | 0.686754269 | 0.461337948 | 0.293 | 0.273 | 1 | 0.686754269 | 0.353291497 |
| Nhlrc2 | 0.248913437 | -0.264816297 | 0.434 | 0.53 | 1 | 0.195830561 | -0.460475699 | 0.537 | 0.628 | 1 | 0.248913437 | 0.353311513 |
| Parp6 | 0.371728624 | 0.385335523 | 0.283 | 0.225 | 1 | 0.196165742 | -0.416551742 | 0.171 | 0.27 | 1 | 0.371728624 | 0.353850486 |
| Gulo | 0.196305582 | -1.845277405 | 0.019 | 0.064 | 1 | 0.357854401 | -0.747757651 | 0 | 0.02 | 1 | 0.357854401 | 0.354075283 |
| Teddm2 | 0.196306645 | -1.545717123 | 0.019 | 0.064 | 1 | 0.824088343 | 0.252242349 | 0.024 | 0.031 | 1 | 0.824088343 | 0.354076991 |
| Gm57352 | 0.382167301 | 0.124975251 | 0.113 | 0.165 | 1 | 0.19636614 | 0.377773231 | 0.049 | 0.123 | 1 | 0.382167301 | 0.354172619 |
| Top2a | 0.196371804 | -2.290062248 | 0.094 | 0.047 | 1 | 0.815008472 | -0.622226769 | 0.024 | 0.031 | 1 | 0.815008472 | 0.354181722 |
| Sh3yl1 | 0.19638739 | 1.3662267 | 0.113 | 0.064 | 1 | 0.609086074 | 0.557096931 | 0.098 | 0.075 | 1 | 0.609086074 | 0.354206773 |
| Dpy19l3 | 0.27483013 | 0.306725688 | 0.226 | 0.161 | 1 | 0.19645285 | -0.835220492 | 0.146 | 0.232 | 1 | 0.27483013 | 0.354311978 |
| Plin2 | 0.662360793 | -0.321426419 | 0.377 | 0.326 | 1 | 0.196920597 | -1.069685746 | 0.22 | 0.3 | 1 | 0.662360793 | 0.355063473 |
| Gm10549 | 0.197024584 | -0.652632327 | 0.057 | 0.119 | 1 | 0.245628692 | -0.332720152 | 0.024 | 0.075 | 1 | 0.245628692 | 0.355230481 |
| Rnf144b | 0.390776147 | -0.515128804 | 0.132 | 0.182 | 1 | 0.19703965 | -0.242522343 | 0.122 | 0.218 | 1 | 0.390776147 | 0.355254676 |
| Banf2 | 0.197849573 | -1.237594828 | 0.019 | 0.064 | 1 | 0.319641967 | -0.16279515 | 0 | 0.024 | 1 | 0.319641967 | 0.356554693 |
| Fut2 | 0.197849573 | -1.1672055 | 0.019 | 0.064 | 1 | 0.438672228 | 1.83720485 | 0.024 | 0.01 | 1 | 0.438672228 | 0.356554693 |
| Scp2 | 0.331709035 | 0.24131028 | 0.792 | 0.771 | 1 | 0.198004895 | -0.139658651 | 0.707 | 0.758 | 1 | 0.331709035 | 0.356803851 |
| Zadh2 | 0.445936408 | 0.569760094 | 0.245 | 0.208 | 1 | 0.198038811 | 0.824821125 | 0.341 | 0.263 | 1 | 0.445936408 | 0.356858251 |
| Copb2 | 0.198199834 | 0.107801547 | 0.981 | 0.903 | 1 | 0.274076542 | -0.232824749 | 0.927 | 0.956 | 1 | 0.274076542 | 0.357116494 |
| Ppip5k1 | 0.831767916 | 0.333692736 | 0.151 | 0.144 | 1 | 0.19822298 | -0.640842447 | 0.098 | 0.177 | 1 | 0.831767916 | 0.357153611 |
| Gm26911 | 0.198618088 | -1.015202407 | 0.019 | 0.064 | 1 | 0.565661834 | -0.863234868 | 0.049 | 0.072 | 1 | 0.565661834 | 0.35778703 |
| Siah1b | 0.198618088 | -1.015202407 | 0.019 | 0.064 | 1 | 0.278320434 | -0.686357106 | 0.024 | 0.068 | 1 | 0.278320434 | 0.35778703 |
| Pou3f3 | 0.198790086 | 1.276028891 | 0.264 | 0.199 | 1 | 0.822353017 | 0.133189677 | 0.317 | 0.352 | 1 | 0.822353017 | 0.358062674 |
| Get4 | 0.198834257 | 0.55911285 | 0.472 | 0.415 | 1 | 0.478115272 | 0.311743361 | 0.488 | 0.468 | 1 | 0.478115272 | 0.358133452 |
| Sec14l1 | 0.198869766 | -0.538155245 | 0.811 | 0.792 | 1 | 0.800100791 | 0.499940445 | 0.854 | 0.84 | 1 | 0.800100791 | 0.358190349 |
| Ess2 | 0.199238042 | 1.228723176 | 0.113 | 0.064 | 1 | 0.433145056 | -0.520347155 | 0.073 | 0.113 | 1 | 0.433145056 | 0.358780286 |
| Gabrr2 | 0.199383551 | -1.093204919 | 0.019 | 0.064 | 1 | 0.261904818 | 1.690363461 | 0.122 | 0.075 | 1 | 0.261904818 | 0.359013301 |
| Syt17 | 0.496240311 | -0.719746523 | 0.057 | 0.085 | 1 | 0.199428918 | -0.970150072 | 0.024 | 0.082 | 1 | 0.496240311 | 0.359085943 |
| Nlrx1 | 0.45789401 | 0.347367673 | 0.094 | 0.064 | 1 | 0.199742661 | 1.989207943 | 0.098 | 0.051 | 1 | 0.45789401 | 0.359588191 |
| Ppp1r7 | 0.815275961 | 0.119098685 | 0.321 | 0.352 | 1 | 0.199824512 | 0.633671456 | 0.39 | 0.317 | 1 | 0.815275961 | 0.359719188 |
| Ints10 | 0.200026403 | 1.154722595 | 0.189 | 0.131 | 1 | 0.586694479 | 0.282615998 | 0.098 | 0.13 | 1 | 0.586694479 | 0.360042244 |
| Gm56807 | 0.200124145 | 1.067259754 | 0.113 | 0.064 | 1 | 0.699638409 | -0.192542494 | 0.049 | 0.065 | 1 | 0.699638409 | 0.360198616 |
| Por | 0.200329473 | -0.463254963 | 0.491 | 0.551 | 1 | 0.606304581 | 0.112690997 | 0.659 | 0.635 | 1 | 0.606304581 | 0.360527048 |
| Zfp473 | 0.200381013 | 3.154722595 | 0.038 | 0.013 | 1 | 0.25729872 | -0.863234868 | 0 | 0.031 | 1 | 0.25729872 | 0.360609475 |
| Pdzd2 | 0.200408504 | -1.044057269 | 0.434 | 0.504 | 1 | 0.884004311 | -0.343733144 | 0.61 | 0.532 | 1 | 0.884004311 | 0.360653439 |
| H6pd | 0.346657195 | -0.456712117 | 0.132 | 0.186 | 1 | 0.201045379 | -0.863234868 | 0.122 | 0.201 | 1 | 0.346657195 | 0.361671514 |
| Zfp87 | 0.201211632 | 0.792152515 | 0.189 | 0.123 | 1 | 0.438545867 | -0.410722664 | 0.122 | 0.167 | 1 | 0.438545867 | 0.361937144 |
| Amy1 | 0.448111812 | 0.267197324 | 0.245 | 0.195 | 1 | 0.201260063 | -0.917682652 | 0.146 | 0.235 | 1 | 0.448111812 | 0.362014514 |
| Gm15952 | 0.992807017 | -0.1672055 | 0.094 | 0.093 | 1 | 0.201304031 | -0.787286015 | 0.024 | 0.082 | 1 | 0.992807017 | 0.362084749 |
| Dck | 0.201473505 | 0.526691372 | 0.151 | 0.089 | 1 | 0.603576896 | 0.773074512 | 0.146 | 0.123 | 1 | 0.603576896 | 0.362355438 |
| Surf6 | 0.201513004 | 0.683416876 | 0.283 | 0.212 | 1 | 0.203827189 | 0.370886846 | 0.171 | 0.276 | 1 | 0.203827189 | 0.362418518 |
| Nup50 | 0.296234949 | 0.452522419 | 0.585 | 0.551 | 1 | 0.201537542 | 0.599137359 | 0.683 | 0.703 | 1 | 0.296234949 | 0.362457704 |
| Slc7a6 | 0.496418275 | -0.290062248 | 0.302 | 0.356 | 1 | 0.201541637 | 0.170448258 | 0.171 | 0.283 | 1 | 0.496418275 | 0.362464242 |
| Aifm2 | 0.476460592 | 0.945269229 | 0.151 | 0.123 | 1 | 0.201660759 | 0.306690133 | 0.073 | 0.15 | 1 | 0.476460592 | 0.362654456 |
| Dapk1 | 0.201887286 | 0.569760094 | 0.547 | 0.466 | 1 | 0.461107832 | -0.239146036 | 0.512 | 0.597 | 1 | 0.461107832 | 0.363016096 |
| Tmem138 | 0.456708741 | 0.254258268 | 0.226 | 0.182 | 1 | 0.202034098 | -0.747757651 | 0.098 | 0.177 | 1 | 0.456708741 | 0.363250419 |
| Cryzl1 | 0.202151414 | 0.503958036 | 0.528 | 0.432 | 1 | 0.431370197 | 0.276489895 | 0.585 | 0.56 | 1 | 0.431370197 | 0.363437634 |
| 5430405H02Rik | 0.202229816 | 0.510866405 | 0.17 | 0.11 | 1 | 0.490004305 | 0.766815522 | 0.171 | 0.137 | 1 | 0.490004305 | 0.363562734 |
| Ndufaf5 | 0.202361489 | 0.491757582 | 0.189 | 0.119 | 1 | 0.866794156 | 0.483567895 | 0.122 | 0.113 | 1 | 0.866794156 | 0.363772805 |
| Tmem43 | 0.202442266 | 0.237184755 | 0.509 | 0.424 | 1 | 0.44112058 | -0.322666487 | 0.463 | 0.512 | 1 | 0.44112058 | 0.363901661 |
| 1110065P20Rik | 0.202602014 | 0.383663802 | 0.528 | 0.462 | 1 | 0.58748774 | 0.662953923 | 0.39 | 0.423 | 1 | 0.58748774 | 0.364156452 |
| Dcaf12l1 | 0.202674222 | 1.237184755 | 0.113 | 0.064 | 1 | 0.406241801 | -0.16279515 | 0.049 | 0.089 | 1 | 0.406241801 | 0.364271604 |
| Dmac2l | 0.202793104 | -0.600164907 | 0.113 | 0.186 | 1 | 0.203614517 | -1.250257991 | 0.049 | 0.113 | 1 | 0.203614517 | 0.364461165 |
| Lig4 | 0.305590208 | 1.154722595 | 0.075 | 0.042 | 1 | 0.202884897 | 0.766815522 | 0.098 | 0.048 | 1 | 0.305590208 | 0.364607512 |
| Lyplal1 | 0.20291565 | 0.382133091 | 0.189 | 0.119 | 1 | 1 | 0.898605394 | 0.073 | 0.075 | 1 | 1 | 0.364656539 |
| Rbm12b2 | 0.229496088 | 0.614154213 | 0.151 | 0.093 | 1 | 0.202924838 | -0.425829556 | 0.073 | 0.15 | 1 | 0.229496088 | 0.364671186 |
| Fbxl9 | 0.203148209 | -0.974560422 | 0.057 | 0.114 | 1 | 0.464949359 | -0.126269274 | 0.073 | 0.113 | 1 | 0.464949359 | 0.365027222 |
| Rabl3 | 0.203150217 | -1.015202407 | 0.057 | 0.114 | 1 | 0.892056604 | -0.126269274 | 0.073 | 0.078 | 1 | 0.892056604 | 0.365030423 |
| Arhgef6 | 0.374096367 | 0.163134835 | 0.509 | 0.449 | 1 | 0.203251004 | 0.616560091 | 0.561 | 0.474 | 1 | 0.374096367 | 0.365191038 |
| Hax1 | 0.203370555 | 0.355021245 | 0.585 | 0.508 | 1 | 0.861405435 | 0.397581712 | 0.463 | 0.471 | 1 | 0.861405435 | 0.365381528 |
| Dclre1b | 0.203391832 | 1.067259754 | 0.113 | 0.064 | 1 | 0.828239242 | 0.351778023 | 0.073 | 0.065 | 1 | 0.828239242 | 0.365415427 |
| Ndst1 | 0.20394636 | -0.540739447 | 0.585 | 0.665 | 1 | 0.891647904 | -0.194273382 | 0.829 | 0.771 | 1 | 0.891647904 | 0.366298603 |
| Inpp4b | 0.204051626 | 0.242185436 | 0.245 | 0.165 | 1 | 0.898555555 | 0.47463477 | 0.146 | 0.143 | 1 | 0.898555555 | 0.366466186 |
| Ankrd49 | 0.204208353 | -0.554228623 | 0.132 | 0.208 | 1 | 0.597459184 | 1.47463477 | 0.146 | 0.123 | 1 | 0.597459184 | 0.366715654 |
| Tcn2 | 0.545816683 | 0.385335523 | 0.283 | 0.254 | 1 | 0.204209974 | 1.083844817 | 0.39 | 0.307 | 1 | 0.545816683 | 0.366718234 |
| Sh3pxd2a | 0.204348968 | -0.359850578 | 0.925 | 0.903 | 1 | 0.425245581 | -0.245921664 | 0.976 | 0.877 | 1 | 0.425245581 | 0.366939435 |
| Abhd3 | 0.432162471 | -0.799473716 | 0.057 | 0.089 | 1 | 0.204633319 | -0.448197369 | 0.024 | 0.082 | 1 | 0.432162471 | 0.367391843 |
| Tsga10 | 0.204650267 | 1.047807391 | 0.132 | 0.081 | 1 | 0.659673142 | -0.286177566 | 0.122 | 0.096 | 1 | 0.659673142 | 0.367418802 |
| Gm13562 | 0.204653189 | 2.154722595 | 0.038 | 0.013 | 1 | 0.354167655 | 1.515276755 | 0.049 | 0.024 | 1 | 0.354167655 | 0.36742345 |
| Nr1h3 | 0.204653189 | 2.154722595 | 0.038 | 0.013 | 1 | 0.574025125 | 1.029849928 | 0.024 | 0.044 | 1 | 0.574025125 | 0.36742345 |
| Tdrkh | 0.204653189 | 2.47665069 | 0.038 | 0.013 | 1 | 0.402373368 | 0.252242349 | 0 | 0.017 | 1 | 0.402373368 | 0.36742345 |
| Cd101 | 0.204653189 | 2.47665069 | 0.038 | 0.013 | 1 | 0.746881137 | 0.83720485 | 0.024 | 0.017 | 1 | 0.746881137 | 0.36742345 |
| Fbxo24 | 0.204653189 | 2.47665069 | 0.038 | 0.013 | 1 | 0.732716552 | 1.614812428 | 0.024 | 0.017 | 1 | 0.732716552 | 0.36742345 |
| Vwa3a | 0.204653189 | 2.47665069 | 0.038 | 0.013 | 1 | 0.257285084 | -0.622226769 | 0 | 0.031 | 1 | 0.257285084 | 0.36742345 |
| Adam8 | 0.204653189 | 2.154722595 | 0.038 | 0.013 | 1 | 0.990690153 | 0.174239837 | 0.049 | 0.048 | 1 | 0.990690153 | 0.36742345 |
| BC050972 | 0.204653189 | 2.154722595 | 0.038 | 0.013 | 1 | 0.438672228 | 1.83720485 | 0.024 | 0.01 | 1 | 0.438672228 | 0.36742345 |
| Cbarp | 0.204653189 | 2.154722595 | 0.038 | 0.013 | 1 | 0.881394782 | 1.071670103 | 0.049 | 0.044 | 1 | 0.881394782 | 0.36742345 |
| D630024D03Rik | 0.204653189 | 2.154722595 | 0.038 | 0.013 | 1 | 0.257285084 | -0.747757651 | 0 | 0.031 | 1 | 0.257285084 | 0.36742345 |
| Zfp457 | 0.204653189 | 2.154722595 | 0.038 | 0.013 | 1 | 0.430587622 | 2.515276755 | 0.024 | 0.01 | 1 | 0.430587622 | 0.36742345 |
| Zfp7 | 0.204653189 | 2.154722595 | 0.038 | 0.013 | 1 | 0.257285084 | -0.622226769 | 0 | 0.031 | 1 | 0.257285084 | 0.36742345 |
| Mx2 | 0.204653189 | 2.739685096 | 0.038 | 0.013 | 1 | 0.902008896 | -0.863234868 | 0.024 | 0.027 | 1 | 0.902008896 | 0.36742345 |
| Ablim3 | 0.204653189 | 2.154722595 | 0.038 | 0.013 | 1 | 0.286422281 | -0.747757651 | 0 | 0.027 | 1 | 0.286422281 | 0.36742345 |
| Garin4 | 0.204708128 | 0.932330173 | 0.151 | 0.093 | 1 | 0.921594231 | -1.020776145 | 0.073 | 0.075 | 1 | 0.921594231 | 0.367510839 |
| Dvl1 | 0.204815735 | -0.395474488 | 0.321 | 0.398 | 1 | 0.217205692 | -0.360096591 | 0.415 | 0.519 | 1 | 0.217205692 | 0.367681985 |
| Abca2 | 0.204928697 | -1.054730771 | 0.057 | 0.114 | 1 | 0.631316319 | 0.422167351 | 0.171 | 0.143 | 1 | 0.631316319 | 0.367861623 |
| Prorsd1 | 0.477798802 | -0.26312992 | 0.377 | 0.428 | 1 | 0.20509225 | -0.545264787 | 0.244 | 0.345 | 1 | 0.477798802 | 0.368121169 |

| | | | | | | | | | | | | |
|---|---|---|---|---|---|---|---|---|---|---|---|---|
| Klhl5 | 0.205218654 | 0.778574109 | 0.415 | 0.347 | 1 | 0.419172323 | -0.318713294 | 0.341 | 0.41 | 1 | 0.419172323 | 0.368322611 |
| Lysmd4 | 0.441027632 | 0.569760094 | 0.132 | 0.097 | 1 | 0.205394181 | -0.747757651 | 0.122 | 0.205 | 1 | 0.441027632 | 0.368601592 |
| Nsun7 | 0.394300775 | 0.714150003 | 0.075 | 0.047 | 1 | 0.205699173 | -1.448197369 | 0.024 | 0.078 | 1 | 0.394300775 | 0.369086197 |
| Mterf2 | 0.588476383 | 0.984797593 | 0.019 | 0.034 | 1 | 0.20569972 | -1.448197369 | 0.024 | 0.078 | 1 | 0.588476383 | 0.369087065 |
| Pex5 | 0.518137771 | -0.216836268 | 0.453 | 0.504 | 1 | 0.205768384 | -0.324476158 | 0.488 | 0.567 | 1 | 0.518137771 | 0.36919614 |
| Ddost | 0.205789283 | 0.198006693 | 0.981 | 0.919 | 1 | 0.442140884 | 0.164995464 | 0.854 | 0.911 | 1 | 0.442140884 | 0.369229336 |
| Mpg | 0.211573879 | 0.701004627 | 0.302 | 0.233 | 1 | 0.205839997 | -0.781704983 | 0.122 | 0.205 | 1 | 0.211573879 | 0.369309889 |
| Pcdhb14 | 0.265162361 | 0.280253477 | 0.057 | 0.114 | 1 | 0.206003556 | -0.484723245 | 0.049 | 0.116 | 1 | 0.265162361 | 0.369569647 |
| Thoc5 | 0.328831804 | 1.010332685 | 0.208 | 0.165 | 1 | 0.206061146 | -0.355440228 | 0.098 | 0.184 | 1 | 0.328831804 | 0.369661096 |
| Fcer1g | 0.206100734 | 3.920257341 | 0.038 | 0.013 | 1 | 0.598714593 | -1.010792057 | 0.098 | 0.072 | 1 | 0.598714593 | 0.369723956 |
| Ccdc116 | 0.206100734 | 2.154722595 | 0.038 | 0.013 | 1 | 0.824088343 | 0.377773231 | 0.024 | 0.031 | 1 | 0.824088343 | 0.369723956 |
| Gtf2a1l | 0.206100734 | 2.154722595 | 0.038 | 0.013 | 1 | 0.729895585 | 1.989207943 | 0.024 | 0.017 | 1 | 0.729895585 | 0.369723956 |
| Pik3ap1 | 0.206100734 | 2.154722595 | 0.038 | 0.013 | 1 | 0.257323555 | -2.332720152 | 0 | 0.031 | 1 | 0.257323555 | 0.369723956 |
| St8sia4 | 0.206102945 | 1.669295768 | 0.038 | 0.013 | 1 | 0.861876576 | 1.252242349 | 0.024 | 0.02 | 1 | 0.861876576 | 0.369727466 |
| Noc4l | 0.534440846 | -0.323324702 | 0.113 | 0.144 | 1 | 0.206197146 | -0.723510105 | 0.098 | 0.174 | 1 | 0.534440846 | 0.369877028 |
| Plb1 | 0.206227354 | -1.617866909 | 0.038 | 0.089 | 1 | 0.227008706 | -1.16279515 | 0.024 | 0.075 | 1 | 0.227008706 | 0.369924986 |
| Perp | 0.774982626 | -0.356239324 | 0.17 | 0.148 | 1 | 0.206233984 | 0.826635608 | 0.366 | 0.276 | 1 | 0.774982626 | 0.369935512 |
| Btbd10 | 0.206321376 | 0.681969598 | 0.415 | 0.331 | 1 | 0.554298817 | -0.106211622 | 0.512 | 0.56 | 1 | 0.554298817 | 0.370074242 |
| Gm16152 | 0.206368562 | -1.015202407 | 0 | 0.03 | 1 | 0.231643049 | -0.622226769 | 0 | 0.034 | 1 | 0.231643049 | 0.37014914 |
| 4930556G22Rik | 0.206368562 | -1.015202407 | 0 | 0.03 | 1 | 0.455094184 | 0.252242349 | 0 | 0.014 | 1 | 0.455094184 | 0.37014914 |
| Gca | 0.206368562 | -1.015202407 | 0 | 0.03 | 1 | 0.267205177 | 2.252242349 | 0.024 | 0.007 | 1 | 0.267205177 | 0.37014914 |
| Zfp973 | 0.206368562 | -1.015202407 | 0 | 0.03 | 1 | 0.319641967 | -0.16279515 | 0 | 0.024 | 1 | 0.319641967 | 0.37014914 |
| Gm16008 | 0.206368562 | -1.015202407 | 0 | 0.03 | 1 | 0.231643049 | -0.622226769 | 0 | 0.034 | 1 | 0.231643049 | 0.37014914 |
| Cd27 | 0.206368562 | -1.015202407 | 0 | 0.03 | 1 | 0.5191986 | 0.252242349 | 0 | 0.01 | 1 | 0.5191986 | 0.37014914 |
| Kctd21 | 0.206368562 | -1.015202407 | 0 | 0.03 | 1 | 0.934901982 | 1.071670103 | 0.049 | 0.048 | 1 | 0.934901982 | 0.37014914 |
| Kdm8 | 0.206368562 | -1.430239906 | 0 | 0.03 | 1 | 0.429722366 | 1.962735732 | 0.049 | 0.027 | 1 | 0.429722366 | 0.37014914 |
| Gm45250 | 0.206368562 | -1.015202407 | 0 | 0.03 | 1 | 0.93376172 | 1.100239256 | 0.024 | 0.027 | 1 | 0.93376172 | 0.37014914 |
| Hspb2 | 0.206368562 | -1.015202407 | 0 | 0.03 | 1 | 0.973993984 | 1.100239256 | 0.024 | 0.024 | 1 | 0.973993984 | 0.37014914 |
| Gm57050 | 0.206368562 | -1.015202407 | 0 | 0.03 | 1 | 0.231591308 | -0.747757651 | 0 | 0.034 | 1 | 0.231591308 | 0.37014914 |
| Pam16l | 0.206368562 | -1.015202407 | 0 | 0.03 | 1 | 0.455763738 | 1.100239256 | 0.049 | 0.027 | 1 | 0.455763738 | 0.37014914 |
| Gm10778 | 0.206368562 | -1.015202407 | 0 | 0.03 | 1 | 0.51919986 | 0.83720485 | 0 | 0.01 | 1 | 0.51919986 | 0.37014914 |
| Gm11963 | 0.206368562 | -1.015202407 | 0 | 0.03 | 1 | 0.605005904 | 1.029849928 | 0.024 | 0.014 | 1 | 0.605005904 | 0.37014914 |
| 9330111N05Rik | 0.206368562 | -1.304709024 | 0 | 0.03 | 1 | 0.208863153 | -0.747757651 | 0 | 0.038 | 1 | 0.208863153 | 0.37014914 |
| Slc14a1 | 0.206368562 | -1.015202407 | 0 | 0.03 | 1 | 0.5191986 | 0.252242349 | 0 | 0.01 | 1 | 0.5191986 | 0.37014914 |
| Insl6 | 0.206368562 | -1.015202407 | 0 | 0.03 | 1 | 0.257308946 | -0.863234868 | 0 | 0.031 | 1 | 0.257308946 | 0.37014914 |
| Slc19a3 | 0.206380492 | -1.430239906 | 0 | 0.03 | 1 | 0.745976859 | 0.377773231 | 0.024 | 0.034 | 1 | 0.745976859 | 0.370168076 |
| Gm29170 | 0.206380492 | -1.1672055 | 0 | 0.03 | 1 | 0.257285084 | -0.622226769 | 0 | 0.031 | 1 | 0.257285084 | 0.370168076 |
| Syt14 | 0.206380492 | -1.652632327 | 0 | 0.03 | 1 | 0.26948834 | 1.83720485 | 0.024 | 0.007 | 1 | 0.26948834 | 0.370168076 |
| Kcnk3 | 0.206380492 | -1.845277405 | 0 | 0.03 | 1 | 0.231634426 | -2.16279515 | 0 | 0.034 | 1 | 0.231634426 | 0.370168076 |
| Igf2os | 0.206380492 | -1.1672055 | 0 | 0.03 | 1 | 0.599290422 | 1.515276755 | 0.024 | 0.014 | 1 | 0.599290422 | 0.370168076 |
| Gm31224 | 0.206380492 | -1.1672055 | 0 | 0.03 | 1 | 0.455098275 | 0.515276755 | 0 | 0.014 | 1 | 0.455098275 | 0.370168076 |
| 4933400C23Rik | 0.206380492 | -1.430239906 | 0 | 0.03 | 1 | 0.455097252 | -0.747757651 | 0 | 0.014 | 1 | 0.455097252 | 0.370168076 |
| Gm56945 | 0.206380492 | -1.1672055 | 0 | 0.03 | 1 | 0.402373368 | 0.252242349 | 0 | 0.017 | 1 | 0.402373368 | 0.370168076 |
| Gm48395 | 0.206380492 | -1.1672055 | 0 | 0.03 | 1 | 0.402372511 | -0.332720152 | 0 | 0.017 | 1 | 0.402372511 | 0.370168076 |
| Gm33696 | 0.206380492 | -1.304709024 | 0 | 0.03 | 1 | 0.231618473 | -1.069685746 | 0 | 0.034 | 1 | 0.231618473 | 0.370168076 |
| Fancd2 | 0.206387649 | -1.545717123 | 0 | 0.03 | 1 | 0.968795213 | 1.699701326 | 0.024 | 0.024 | 1 | 0.968795213 | 0.370179437 |
| Tk1 | 0.206387649 | -2.015202407 | 0 | 0.03 | 1 | 0.989595616 | 0.667279848 | 0.024 | 0.024 | 1 | 0.989595616 | 0.370179437 |
| Nlrp12 | 0.206388445 | -1.652632327 | 0 | 0.03 | 1 | 0.402364804 | -0.16279515 | 0 | 0.017 | 1 | 0.402364804 | 0.370180699 |
| Gm48877 | 0.206388445 | -1.845277405 | 0 | 0.03 | 1 | 0.5191986 | 0.515276755 | 0 | 0.01 | 1 | 0.5191986 | 0.370180699 |
| Gm33906 | 0.206388445 | -2.237594828 | 0 | 0.03 | 1 | 0.402369942 | -0.332720152 | 0 | 0.017 | 1 | 0.402369942 | 0.370180699 |
| Shcbp1 | 0.206392421 | -1.845277405 | 0 | 0.03 | 1 | 0.455098275 | 0.515276755 | 0 | 0.014 | 1 | 0.455098275 | 0.370187011 |
| Foxq1 | 0.206392421 | -2.368839361 | 0 | 0.03 | 1 | 0.60213981 | 1.029849928 | 0.024 | 0.014 | 1 | 0.60213981 | 0.370187011 |
| Cdr1os | 0.206392421 | -2.845277405 | 0 | 0.03 | 1 | 0.51919986 | 0.83720485 | 0 | 0.01 | 1 | 0.51919986 | 0.370187011 |
| Lcn4 | 0.206393216 | -1.932740246 | 0 | 0.03 | 1 | 1 | -0.970150072 | 0.024 | 0.024 | 1 | 1 | 0.370188273 |
| Cep55 | 0.206394807 | -2.093204919 | 0 | 0.03 | 1 | 0.267205177 | 2.252242349 | 0.024 | 0.007 | 1 | 0.267205177 | 0.370190798 |
| Blnk | 0.206394807 | -2.600164907 | 0 | 0.03 | 1 | 0.276475896 | 0.547698233 | 0.195 | 0.13 | 1 | 0.276475896 | 0.370190798 |
| Cenpf | 0.206395602 | -4.237594828 | 0 | 0.03 | 1 | 0.51919986 | 0.83720485 | 0 | 0.01 | 1 | 0.51919986 | 0.37019206 |
| Cfap126 | 0.206396398 | -0.845277405 | 0 | 0.03 | 1 | 0.744047924 | 1.252242349 | 0.024 | 0.017 | 1 | 0.744047924 | 0.370193322 |
| Creb3l4 | 0.206396398 | -0.845277405 | 0 | 0.03 | 1 | 0.208799874 | -0.863234868 | 0 | 0.038 | 1 | 0.208799874 | 0.370193322 |
| Mup5 | 0.206396398 | -0.845277405 | 0 | 0.03 | 1 | 0.606262835 | -0.16279515 | 0.024 | 0.041 | 1 | 0.606262835 | 0.370193322 |
| Cdca3 | 0.206396398 | -3.399866257 | 0 | 0.03 | 1 | 0.430584486 | 2.422167351 | 0.024 | 0.01 | 1 | 0.430584486 | 0.370193322 |
| Gm54771 | 0.206396398 | -0.845277405 | 0 | 0.03 | 1 | 0.357858789 | -0.970150072 | 0 | 0.02 | 1 | 0.357858789 | 0.370193322 |
| Cdk20 | 0.206396398 | -0.845277405 | 0 | 0.03 | 1 | 0.872828964 | 1.029849928 | 0.024 | 0.02 | 1 | 0.872828964 | 0.370193322 |
| Fam83h | 0.206396398 | -0.845277405 | 0 | 0.03 | 1 | 0.257285084 | -0.622226769 | 0 | 0.031 | 1 | 0.257285084 | 0.370193322 |
| Myo5b | 0.206396398 | -0.845277405 | 0 | 0.03 | 1 | 0.209931505 | 0.721727632 | 0.195 | 0.123 | 1 | 0.209931505 | 0.370193322 |
| Tmem151a | 0.206396398 | -0.845277405 | 0 | 0.03 | 1 | 0.494850012 | 0.83720485 | 0.073 | 0.048 | 1 | 0.494850012 | 0.370193322 |
| Ercc6l | 0.206396398 | -0.845277405 | 0 | 0.03 | 1 | 0.51919986 | 0.83720485 | 0 | 0.01 | 1 | 0.51919986 | 0.370193322 |
| Trpc3 | 0.87372059 | -0.430239906 | 0.075 | 0.081 | 1 | 0.206719391 | -0.20718927 | 0.024 | 0.082 | 1 | 0.87372059 | 0.370705875 |
| Lmbrd2 | 0.207022563 | 0.578693219 | 0.472 | 0.424 | 1 | 0.96713678 | 0.500921462 | 0.39 | 0.41 | 1 | 0.96713678 | 0.371186785 |
| Gm49692 | 0.670181184 | 1.017219071 | 0.113 | 0.097 | 1 | 0.20702469 | -1.069685746 | 0.024 | 0.078 | 1 | 0.670181184 | 0.371190158 |
| Nap1l3 | 0.837789196 | 0.417757001 | 0.075 | 0.068 | 1 | 0.207026516 | -1.020776145 | 0.024 | 0.078 | 1 | 0.837789196 | 0.371193053 |
| Rapgef3 | 0.207072232 | 0.953088734 | 0.208 | 0.144 | 1 | 0.22792473 | -0.20718927 | 0.195 | 0.283 | 1 | 0.22792473 | 0.371265555 |
| Eif4e2 | 0.254530408 | 0.312706665 | 0.698 | 0.712 | 1 | 0.207237644 | -0.240154033 | 0.659 | 0.741 | 1 | 0.254530408 | 0.371527847 |
| Gm6169 | 0.312692399 | 1.017219071 | 0.075 | 0.042 | 1 | 0.207238811 | 0.667279848 | 0.024 | 0.082 | 1 | 0.312692399 | 0.371529698 |
| Mrps17 | 0.673808915 | 0.119788805 | 0.566 | 0.559 | 1 | 0.207439328 | -0.325733721 | 0.341 | 0.457 | 1 | 0.673808915 | 0.371847581 |
| Rhbdd2 | 0.556009061 | -0.253362144 | 0.208 | 0.242 | 1 | 0.207485379 | -0.64392184 | 0.098 | 0.177 | 1 | 0.556009061 | 0.371920575 |
| Spag6 | 0.207549016 | 1.8327945 | 0.038 | 0.013 | 1 | 0.51919986 | 0.83720485 | 0 | 0.01 | 1 | 0.51919986 | 0.372021438 |
| Kcnc4 | 0.207549016 | 1.8327945 | 0.038 | 0.013 | 1 | 0.87556513 | 0.83720485 | 0.024 | 0.02 | 1 | 0.87556513 | 0.372021438 |
| Gm44837 | 0.207549016 | 1.8327945 | 0.038 | 0.013 | 1 | 0.749728988 | 0.83720485 | 0.024 | 0.017 | 1 | 0.749728988 | 0.372021438 |
| Gm26632 | 0.207549016 | 1.8327945 | 0.038 | 0.013 | 1 | 0.529762985 | 1.334704509 | 0.049 | 0.031 | 1 | 0.529762985 | 0.372021438 |
| Ano4 | 0.207549016 | 1.8327945 | 0.038 | 0.013 | 1 | 0.909322542 | 0.252242349 | 0.024 | 0.027 | 1 | 0.909322542 | 0.372021438 |
| Aatk | 0.207549016 | 1.8327945 | 0.038 | 0.013 | 1 | 0.286413984 | -0.863234868 | 0 | 0.027 | 1 | 0.286413984 | 0.372021438 |
| Gm34917 | 0.207549016 | 1.8327945 | 0.038 | 0.013 | 1 | 0.402369942 | -0.332720152 | 0 | 0.017 | 1 | 0.402369942 | 0.372021438 |
| Slc44a4 | 0.207551231 | 2.154722595 | 0.038 | 0.013 | 1 | 0.749728988 | 0.667279848 | 0.024 | 0.017 | 1 | 0.749728988 | 0.372024948 |
| Gnptg | 0.207708817 | -0.295310326 | 0.358 | 0.462 | 1 | 0.523431195 | 0.100239256 | 0.293 | 0.369 | 1 | 0.523431195 | 0.372274681 |
| Mad2l1 | 0.20774093 | 0.154722595 | 0.132 | 0.076 | 1 | 0.893463168 | 0.422167351 | 0.098 | 0.092 | 1 | 0.893463168 | 0.372325565 |
| Gm19412 | 0.207784127 | 0.984797593 | 0.113 | 0.064 | 1 | 0.498856313 | -0.250257991 | 0.024 | 0.048 | 1 | 0.498856313 | 0.372394011 |
| Gm57184 | 0.2079853 | -0.696133966 | 0.151 | 0.229 | 1 | 0.267112265 | 0.222495006 | 0.049 | 0.109 | 1 | 0.267112265 | 0.372712715 |
| Gtf2e2 | 0.20815375 | 0.351954942 | 0.509 | 0.436 | 1 | 0.73433563 | 0.174239837 | 0.439 | 0.526 | 1 | 0.73433563 | 0.372979516 |
| Plcb3 | 0.208154464 | 0.874614676 | 0.189 | 0.127 | 1 | 0.706478631 | 0.627751484 | 0.195 | 0.184 | 1 | 0.706478631 | 0.372980647 |
| Mrm3 | 0.208503957 | -0.799473716 | 0.057 | 0.114 | 1 | 0.870768967 | 0.351778023 | 0.073 | 0.082 | 1 | 0.870768967 | 0.373534015 |
| Ntmt1 | 0.208523502 | 0.454282877 | 0.434 | 0.352 | 1 | 0.347798112 | 1.019581592 | 0.39 | 0.362 | 1 | 0.347798112 | 0.373564953 |
| Smim24 | 0.534564335 | 1.017219071 | 0.057 | 0.038 | 1 | 0.208531619 | 2.377773231 | 0.073 | 0.034 | 1 | 0.534564335 | 0.373577801 |
| Gm15726 | 0.208626283 | 0.154722595 | 0.113 | 0.064 | 1 | 0.267704969 | 1.19334866 | 0.049 | 0.02 | 1 | 0.267704969 | 0.37372764 |
| Gm56546 | 0.243111734 | -0.652632327 | 0 | 0.025 | 1 | 0.208799874 | -0.863234868 | 0 | 0.038 | 1 | 0.243111734 | 0.374002361 |
| Cyp46a1 | 0.287773478 | -0.430239906 | 0 | 0.021 | 1 | 0.208799874 | -0.863234868 | 0 | 0.038 | 1 | 0.287773478 | 0.374002321 |
| Gm20404 | 0.336520503 | 1.569760094 | 0.038 | 0.017 | 1 | 0.208799874 | -0.863234868 | 0 | 0.038 | 1 | 0.336520503 | 0.374002361 |
| Zfp11 | 0.503971666 | 1.569760094 | 0.019 | 0.008 | 1 | 0.208799874 | -0.970150072 | 0 | 0.038 | 1 | 0.503971666 | 0.374002361 |

| | | | | | | | | | | | | |
|---|---|---|---|---|---|---|---|---|---|---|---|---|
| Gm50163 | 0.503971666 | 1.569760094 | 0.019 | 0.008 | 1 | 0.208799874 | -0.863234868 | 0 | 0.038 | 1 | 0.503971666 | 0.374002361 |
| 5830448L01Rik | 0.605896064 | -0.1672055 | 0.038 | 0.055 | 1 | 0.208799874 | -0.863234868 | 0 | 0.038 | 1 | 0.605896064 | 0.374002361 |
| Lrrc19 | 0.736073999 | 0.347367673 | 0.019 | 0.013 | 1 | 0.208799874 | -0.863234868 | 0 | 0.038 | 1 | 0.736073999 | 0.374002361 |
| St8sia1 | 0.874462081 | 0.984797593 | 0.038 | 0.034 | 1 | 0.208799874 | -0.863234868 | 0 | 0.038 | 1 | 0.874462081 | 0.374002361 |
| Cdnf | 0.991244785 | 0.417757001 | 0.038 | 0.038 | 1 | 0.208799874 | -0.863234868 | 0 | 0.038 | 1 | 0.991244785 | 0.374002361 |
| Gm45237 | 0.267977656 | -0.752168001 | 0.019 | 0.055 | 1 | 0.208817133 | -0.970150072 | 0 | 0.038 | 1 | 0.267977656 | 0.374029671 |
| Pla1a | 0.308041209 | -0.960754623 | 0.038 | 0.076 | 1 | 0.208817133 | -1.332720152 | 0 | 0.038 | 1 | 0.308041209 | 0.374029671 |
| Racgap1 | 0.456281772 | -1.652632327 | 0.038 | 0.064 | 1 | 0.208817133 | -1.069685746 | 0 | 0.038 | 1 | 0.456281772 | 0.374029671 |
| Camsap3 | 0.503971666 | 1.569760094 | 0.019 | 0.008 | 1 | 0.208817133 | -1.16279515 | 0 | 0.038 | 1 | 0.503971666 | 0.374029671 |
| Ankrd23 | 0.520823689 | 1.377115016 | 0.057 | 0.038 | 1 | 0.208817133 | -0.970150072 | 0 | 0.038 | 1 | 0.520823689 | 0.374029671 |
| Gm47167 | 0.567830665 | -0.304709024 | 0.019 | 0.034 | 1 | 0.208817133 | -1.069685746 | 0 | 0.038 | 1 | 0.567830665 | 0.374029671 |
| Gm39929 | 0.675049917 | 0.389187848 | 0.075 | 0.059 | 1 | 0.208830556 | -1.16279515 | 0 | 0.038 | 1 | 0.675049917 | 0.374050911 |
| Gm14443 | 0.239732556 | 1.347367673 | 0.057 | 0.025 | 1 | 0.20883094 | -1.332720152 | 0 | 0.038 | 1 | 0.239732556 | 0.374051518 |
| Ttc41 | 0.343006305 | -0.1672055 | 0 | 0.017 | 1 | 0.20883094 | -1.069685746 | 0 | 0.038 | 1 | 0.343006305 | 0.374051518 |
| A930012L18Rik | 0.362626638 | -0.430239906 | 0.019 | 0.047 | 1 | 0.20883094 | -1.069685746 | 0 | 0.038 | 1 | 0.362626638 | 0.374051518 |
| Ccdc189 | 0.605896064 | -0.1672055 | 0.038 | 0.055 | 1 | 0.20883094 | -1.069685746 | 0 | 0.038 | 1 | 0.605896064 | 0.374051518 |
| Pak6 | 0.243111734 | -0.652632327 | 0 | 0.025 | 1 | 0.208840144 | -1.410722664 | 0 | 0.038 | 1 | 0.243111734 | 0.374066082 |
| Plekhn1 | 0.48175508 | 1.154722595 | 0.038 | 0.021 | 1 | 0.208840144 | -1.250257991 | 0 | 0.038 | 1 | 0.48175508 | 0.374066082 |
| Gm56606 | 0.530868628 | -0.347777746 | 0.038 | 0.059 | 1 | 0.208840144 | -1.250257991 | 0 | 0.038 | 1 | 0.530868628 | 0.374066082 |
| Pdlim3 | 0.882856291 | -0.278236812 | 0.038 | 0.042 | 1 | 0.208841294 | -1.80665134 | 0 | 0.038 | 1 | 0.882856291 | 0.374067902 |
| Zik1 | 0.228899781 | 1.417757001 | 0.075 | 0.038 | 1 | 0.208841678 | -1.16279515 | 0 | 0.038 | 1 | 0.228899781 | 0.374068509 |
| Gm49915 | 0.429421714 | 1.154722595 | 0.057 | 0.034 | 1 | 0.208841678 | -1.16279515 | 0 | 0.038 | 1 | 0.429421714 | 0.374068509 |
| Gm46515 | 0.864193501 | -1.015202407 | 0.038 | 0.042 | 1 | 0.208841678 | -1.410722664 | 0 | 0.038 | 1 | 0.864193501 | 0.374068509 |
| 9630014M24Rik | 0.226045728 | 2.270199812 | 0.075 | 0.038 | 1 | 0.208848197 | -1.686357106 | 0 | 0.038 | 1 | 0.226045728 | 0.374078825 |
| Gm42951 | 0.855931477 | -0.652632327 | 0.038 | 0.042 | 1 | 0.208858934 | -2.717384002 | 0 | 0.038 | 1 | 0.855931477 | 0.374095814 |
| 2210406O10Rik | 0.410365984 | -0.719746523 | 0.038 | 0.068 | 1 | 0.208863153 | -0.747757651 | 0 | 0.038 | 1 | 0.410365984 | 0.374102489 |
| AI597479 | 0.265279113 | -0.799473716 | 0.057 | 0.106 | 1 | 0.208869059 | 0.83720485 | 0.146 | 0.085 | 1 | 0.265279113 | 0.374111834 |
| Emc1 | 0.208891511 | -0.341703231 | 0.189 | 0.275 | 1 | 0.2857598 | -0.516432105 | 0.195 | 0.266 | 1 | 0.2857598 | 0.374147358 |
| Hs3st6 | 0.209004666 | 1.569760094 | 0.038 | 0.013 | 1 | 0.457573452 | 0.444887427 | 0.049 | 0.027 | 1 | 0.457573452 | 0.374326382 |
| Emx2 | 0.209004666 | 1.154722595 | 0.038 | 0.013 | 1 | 0.605005904 | 1.029849928 | 0.024 | 0.014 | 1 | 0.605005904 | 0.374326382 |
| Cpne5 | 0.209007994 | 1.669295768 | 0.038 | 0.013 | 1 | 0.856407545 | 1.83720485 | 0.024 | 0.02 | 1 | 0.856407545 | 0.374331647 |
| 1700066M21Rik | 0.209009104 | 1.739685096 | 0.038 | 0.013 | 1 | 0.409125717 | -0.410722664 | 0.024 | 0.055 | 1 | 0.409125717 | 0.374333402 |
| Gpr35 | 0.209009104 | 1.739685096 | 0.038 | 0.013 | 1 | 0.553780906 | 0.83720485 | 0.049 | 0.031 | 1 | 0.553780906 | 0.374333402 |
| Gm9821 | 0.209009104 | 1.739685096 | 0.038 | 0.013 | 1 | 0.267205177 | 2.252242349 | 0.024 | 0.007 | 1 | 0.267205177 | 0.374333402 |
| Gstm6 | 0.209009104 | 1.739685096 | 0.038 | 0.013 | 1 | 0.274788535 | 0.962735732 | 0.049 | 0.02 | 1 | 0.274788535 | 0.374333402 |
| Dnali1 | 0.209009104 | 1.739685096 | 0.038 | 0.013 | 1 | 0.455094184 | 0.252242349 | 0 | 0.014 | 1 | 0.455094184 | 0.374333402 |
| Gm12963 | 0.209009104 | 1.739685096 | 0.038 | 0.013 | 1 | 0.989595616 | 0.667279848 | 0.024 | 0.024 | 1 | 0.989595616 | 0.374333402 |
| Gm43113 | 0.209009104 | 1.739685096 | 0.038 | 0.013 | 1 | 0.257285084 | -0.622226769 | 0 | 0.031 | 1 | 0.257285084 | 0.374333402 |
| 1010001B22Rik | 0.209009104 | 1.739685096 | 0.038 | 0.013 | 1 | 0.744047924 | 1.252242349 | 0.024 | 0.017 | 1 | 0.744047924 | 0.374333402 |
| Zfp605 | 0.209009104 | 1.739685096 | 0.038 | 0.013 | 1 | 0.402373368 | 0.252242349 | 0 | 0.017 | 1 | 0.402373368 | 0.374333402 |
| Gm40323 | 0.209009104 | 1.739685096 | 0.038 | 0.013 | 1 | 0.267205177 | 2.252242349 | 0.024 | 0.007 | 1 | 0.267205177 | 0.374333402 |
| C5ar2 | 0.209009104 | 1.739685096 | 0.038 | 0.013 | 1 | 0.960216817 | 0.444887427 | 0.049 | 0.051 | 1 | 0.960216817 | 0.374333402 |
| Faap24 | 0.209009104 | 1.739685096 | 0.038 | 0.013 | 1 | 0.824088343 | 0.377773231 | 0.024 | 0.031 | 1 | 0.824088343 | 0.374333402 |
| Meak7 | 0.209009104 | 1.739685096 | 0.038 | 0.013 | 1 | 0.402373368 | 0.252242349 | 0 | 0.017 | 1 | 0.402373368 | 0.374333402 |
| Iho1 | 0.209009104 | 1.739685096 | 0.038 | 0.013 | 1 | 0.856409403 | 1.83720485 | 0.024 | 0.02 | 1 | 0.856409403 | 0.374333402 |
| Gm12111 | 0.209009104 | 1.739685096 | 0.038 | 0.013 | 1 | 0.872828964 | 1.029849928 | 0.024 | 0.02 | 1 | 0.872828964 | 0.374333402 |
| Shmt1 | 0.209009104 | 1.739685096 | 0.038 | 0.013 | 1 | 0.908035876 | 0.252242349 | 0.049 | 0.044 | 1 | 0.908035876 | 0.374333402 |
| Gm51903 | 0.209009104 | 1.739685096 | 0.038 | 0.013 | 1 | 0.51919986 | 0.83720485 | 0 | 0.01 | 1 | 0.51919986 | 0.374333402 |
| Scarf1 | 0.209009104 | 1.739685096 | 0.038 | 0.013 | 1 | 0.722681838 | -0.665295491 | 0.049 | 0.061 | 1 | 0.722681838 | 0.374333402 |
| Gm36445 | 0.209009104 | 1.739685096 | 0.038 | 0.013 | 1 | 0.455098275 | 0.515276755 | 0 | 0.014 | 1 | 0.455098275 | 0.374333402 |
| Pdxp | 0.209009104 | 1.739685096 | 0.038 | 0.013 | 1 | 0.455098275 | 0.515276755 | 0 | 0.014 | 1 | 0.455098275 | 0.374333402 |
| A230028O05Rik | 0.209009104 | 1.739685096 | 0.038 | 0.013 | 1 | 0.231643049 | -0.622226769 | 0 | 0.034 | 1 | 0.231643049 | 0.374333402 |
| Abcc10 | 0.209009104 | 1.739685096 | 0.038 | 0.013 | 1 | 0.33812456 | 1.458693227 | 0.073 | 0.041 | 1 | 0.33812456 | 0.374333402 |
| Gm56952 | 0.209121018 | 1.517292674 | 0.094 | 0.051 | 1 | 0.342487713 | 0.83720485 | 0.122 | 0.078 | 1 | 0.342487713 | 0.374510436 |
| Eif3j2 | 0.220283019 | 0.806799291 | 0.17 | 0.11 | 1 | 0.209173343 | -0.863234868 | 0.049 | 0.113 | 1 | 0.220283019 | 0.374593198 |
| Nt5c2 | 0.345321393 | 0.561198081 | 0.434 | 0.424 | 1 | 0.209192104 | 0.388744349 | 0.537 | 0.461 | 1 | 0.345321393 | 0.374622872 |
| Nprl2 | 0.209238715 | 0.522454379 | 0.17 | 0.106 | 1 | 0.827836237 | 0.212713985 | 0.122 | 0.109 | 1 | 0.827836237 | 0.37469659 |
| Ndufaf4 | 0.209255922 | 0.397292898 | 0.472 | 0.373 | 1 | 0.315087725 | -0.345487448 | 0.366 | 0.454 | 1 | 0.315087725 | 0.374723803 |
| Nudt9 | 0.943280499 | 0.204110622 | 0.528 | 0.568 | 1 | 0.209260309 | -0.42868921 | 0.341 | 0.447 | 1 | 0.943280499 | 0.374730741 |
| Ccdc15 | 0.383847999 | -0.316583124 | 0.094 | 0.14 | 1 | 0.209701214 | 1.199774929 | 0.195 | 0.133 | 1 | 0.383847999 | 0.375427829 |
| Hic2 | 0.600201153 | 0.280253477 | 0.075 | 0.055 | 1 | 0.209706349 | 0.279209397 | 0.195 | 0.119 | 1 | 0.600201153 | 0.375435945 |
| Krt14 | 0.432375074 | -1.227747042 | 0.113 | 0.076 | 1 | 0.209833299 | 1.164779508 | 0.244 | 0.167 | 1 | 0.432375074 | 0.375636584 |
| Gm5914 | 0.210023327 | 1.076720083 | 0.113 | 0.064 | 1 | 0.980363019 | 0.351778023 | 0.073 | 0.075 | 1 | 0.980363019 | 0.375936855 |
| Anapc7 | 0.210030135 | 0.547040018 | 0.453 | 0.39 | 1 | 0.368993667 | -0.250257991 | 0.39 | 0.461 | 1 | 0.368993667 | 0.375947613 |
| Decr2 | 0.210034837 | -0.430239906 | 0.151 | 0.233 | 1 | 0.431630977 | -0.16279515 | 0.122 | 0.174 | 1 | 0.431630977 | 0.375955041 |
| Urod | 0.210322415 | 0.425370185 | 0.302 | 0.216 | 1 | 0.840113855 | 0.129385601 | 0.244 | 0.273 | 1 | 0.840113855 | 0.376409312 |
| Slc25a53 | 0.210347764 | 0.533234218 | 0.17 | 0.106 | 1 | 0.871301821 | 0.136765132 | 0.146 | 0.16 | 1 | 0.871301821 | 0.376449346 |
| Nedd1 | 0.210385145 | 0.63687429 | 0.264 | 0.195 | 1 | 0.723652038 | 0.338953982 | 0.244 | 0.232 | 1 | 0.723652038 | 0.37650838 |
| Morn1 | 0.400690149 | 0.104096522 | 0.226 | 0.169 | 1 | 0.210407577 | -0.217242934 | 0.122 | 0.218 | 1 | 0.400690149 | 0.376543805 |
| Ppm1f | 0.329936506 | 1.154722595 | 0.094 | 0.059 | 1 | 0.210593767 | -0.622226769 | 0.073 | 0.147 | 1 | 0.329936506 | 0.376837799 |
| Slc25a3 | 0.211091337 | 0.135431256 | 0.981 | 0.979 | 1 | 0.265787466 | -0.164797503 | 0.976 | 0.993 | 1 | 0.265787466 | 0.377623121 |
| Cspg5 | 0.211254675 | 0.695290976 | 0.151 | 0.093 | 1 | 0.988989914 | -0.132421501 | 0.122 | 0.119 | 1 | 0.988989914 | 0.377880812 |
| 4931408D14Rik | 0.334518863 | 1.154722595 | 0.057 | 0.03 | 1 | 0.211360623 | 1.83720485 | 0.073 | 0.034 | 1 | 0.334518863 | 0.378047933 |
| Zfp78 | 0.343000446 | -0.430239906 | 0 | 0.017 | 1 | 0.211360623 | 1.83720485 | 0.073 | 0.034 | 1 | 0.343000446 | 0.378047933 |
| E2f4 | 0.211576473 | 0.510866405 | 0.321 | 0.246 | 1 | 0.401139138 | -0.348661696 | 0.341 | 0.399 | 1 | 0.401139138 | 0.378388342 |
| Alg1 | 0.364851018 | -0.28137652 | 0.245 | 0.314 | 1 | 0.211917168 | -0.282534394 | 0.195 | 0.3 | 1 | 0.364851018 | 0.378925449 |
| Cdc34 | 0.211926331 | 0.450178478 | 0.377 | 0.284 | 1 | 0.905613854 | -0.146307027 | 0.415 | 0.399 | 1 | 0.905613854 | 0.378939893 |
| Ppp1r16b | 0.211931446 | 1.417757001 | 0.038 | 0.013 | 1 | 0.319636904 | -0.970150072 | 0 | 0.024 | 1 | 0.319636904 | 0.378947954 |
| Smco2 | 0.211931446 | 1.417757001 | 0.038 | 0.013 | 1 | 0.286427259 | -1.069685746 | 0 | 0.027 | 1 | 0.286427259 | 0.378947954 |
| Svip | 0.211931446 | 1.417757001 | 0.038 | 0.013 | 1 | 0.752578711 | 0.667279848 | 0.024 | 0.017 | 1 | 0.752578711 | 0.378947954 |
| Gm48855 | 0.211931446 | 1.417757001 | 0.038 | 0.013 | 1 | 0.93376172 | 1.100239256 | 0.024 | 0.027 | 1 | 0.93376172 | 0.378947954 |
| Slc6a20a | 0.211931446 | 1.154722595 | 0.038 | 0.013 | 1 | 0.357858058 | -0.747757651 | 0 | 0.02 | 1 | 0.357858058 | 0.378947954 |
| 9930004E17Rik | 0.211931446 | 1.417757001 | 0.038 | 0.013 | 1 | 0.744047924 | 1.252242349 | 0.024 | 0.017 | 1 | 0.744047924 | 0.378947954 |
| Gm9195 | 0.211931446 | 1.417757001 | 0.038 | 0.013 | 1 | 0.267205177 | 1.614812428 | 0.024 | 0.007 | 1 | 0.267205177 | 0.378947954 |
| Gm56767 | 0.211931446 | 1.417757001 | 0.038 | 0.013 | 1 | 0.455098275 | 0.515276755 | 0 | 0.014 | 1 | 0.455098275 | 0.378947954 |
| Pcdhb18 | 0.211931446 | 1.417757001 | 0.038 | 0.013 | 1 | 0.402821579 | -0.917682652 | 0.024 | 0.055 | 1 | 0.402821579 | 0.378947954 |
| Mrpl48 | 0.211933003 | 0.249823297 | 0.868 | 0.78 | 1 | 0.419246845 | 0.433205622 | 0.78 | 0.747 | 1 | 0.419246845 | 0.378950408 |
| Rapgef1 | 0.327708077 | -0.326404095 | 0.642 | 0.661 | 1 | 0.212303887 | -0.19841906 | 0.634 | 0.734 | 1 | 0.327708077 | 0.379534833 |
| Tspan12 | 0.212404654 | 1.8327945 | 0.094 | 0.051 | 1 | 0.265387061 | 1.252242349 | 0.146 | 0.092 | 1 | 0.265387061 | 0.37969357 |
| Ppp2r5b | 0.212446561 | 0.40997965 | 0.434 | 0.356 | 1 | 0.689791597 | 0.43121249 | 0.39 | 0.386 | 1 | 0.689791597 | 0.37975958 |
| Ccdc66 | 0.212604294 | 0.322845354 | 0.321 | 0.237 | 1 | 0.780849275 | 0.439612485 | 0.293 | 0.297 | 1 | 0.780849275 | 0.380008002 |
| Cd79a | 0.287761733 | -3.015202407 | 0 | 0.021 | 1 | 0.212889431 | 1.149148856 | 0.122 | 0.068 | 1 | 0.287761733 | 0.380456953 |
| Cdk16 | 0.325867934 | 0.399835093 | 0.34 | 0.271 | 1 | 0.212899033 | -0.440779897 | 0.293 | 0.403 | 1 | 0.325867934 | 0.380472068 |
| D130020L05Rik | 0.213025208 | -0.783876861 | 0.038 | 0.089 | 1 | 0.352310473 | 1.444887427 | 0.098 | 0.061 | 1 | 0.352310473 | 0.380670677 |
| Fancc | 0.236943219 | 0.247109025 | 0.623 | 0.508 | 1 | 0.213196923 | -0.498617687 | 0.512 | 0.662 | 1 | 0.236943219 | 0.380940919 |
| Mef2b | 0.270857012 | -0.278236812 | 0.132 | 0.203 | 1 | 0.213277286 | -0.584258919 | 0.122 | 0.205 | 1 | 0.270857012 | 0.381067372 |
| Slc1a3 | 0.229023186 | 3.690775495 | 0.057 | 0.025 | 1 | 0.21328888 | 1.759202338 | 0.073 | 0.034 | 1 | 0.229023186 | 0.381085613 |
| Gas2l1 | 0.213349498 | -0.417528854 | 0.245 | 0.339 | 1 | 0.778346201 | 0.168178084 | 0.366 | 0.341 | 1 | 0.778346201 | 0.381180988 |

| | | | | | | | | | | | | |
|---|---|---|---|---|---|---|---|---|---|---|---|---|
| Naglu | 0.213539209 | 0.417757001 | 0.34 | 0.254 | 1 | 0.250941023 | -0.537834582 | 0.146 | 0.225 | 1 | 0.250941023 | 0.381479425 |
| Cyb5r4 | 0.610916846 | 0.589937976 | 0.245 | 0.237 | 1 | 0.213543476 | -0.513292397 | 0.244 | 0.338 | 1 | 0.610916846 | 0.381486135 |
| 2900089D17Rik | 0.677283246 | -0.330704232 | 0.151 | 0.169 | 1 | 0.213800271 | -0.381963671 | 0.146 | 0.235 | 1 | 0.677283246 | 0.381889986 |
| Aph1a | 0.21421339 | 0.329401126 | 0.585 | 0.538 | 1 | 0.838653543 | 0.167792027 | 0.585 | 0.631 | 1 | 0.838653543 | 0.382539404 |
| Rrp1b | 0.214308838 | 0.614154213 | 0.264 | 0.199 | 1 | 0.291055624 | 0.804783372 | 0.293 | 0.229 | 1 | 0.291055624 | 0.382689398 |
| Alg3 | 0.214406861 | -0.482707326 | 0.132 | 0.208 | 1 | 0.683079103 | 0.100239256 | 0.146 | 0.177 | 1 | 0.683079103 | 0.382843419 |
| Eeig2 | 0.270049647 | -0.346501024 | 0.642 | 0.784 | 1 | 0.214445569 | -0.416757651 | 0.707 | 0.765 | 1 | 0.270049647 | 0.382904236 |
| Dnm3 | 0.214608684 | -0.486382984 | 0.17 | 0.25 | 1 | 0.488800797 | 0.506559538 | 0.244 | 0.205 | 1 | 0.488800797 | 0.383160481 |
| Gpx1 | 0.214637362 | -0.59537342 | 1 | 0.932 | 1 | 0.788540733 | 0.22061074 | 0.976 | 0.959 | 1 | 0.788540733 | 0.383205527 |
| Med15 | 0.214708868 | 0.475344492 | 0.604 | 0.572 | 1 | 0.515065211 | -0.140476771 | 0.61 | 0.676 | 1 | 0.515065211 | 0.383317838 |
| Gbe1 | 0.301136859 | 0.785488785 | 0.358 | 0.322 | 1 | 0.214745789 | 0.750740759 | 0.585 | 0.505 | 1 | 0.301136859 | 0.383375824 |
| Gm26930 | 0.214901224 | 1.154722595 | 0.038 | 0.013 | 1 | 0.44410834 | 0.667279848 | 0.024 | 0.01 | 1 | 0.44410834 | 0.383619913 |
| Gm20337 | 0.214901224 | 1.154722595 | 0.038 | 0.013 | 1 | 0.402373368 | 0.252242349 | 0 | 0.017 | 1 | 0.402373368 | 0.383619913 |
| Cfap418 | 0.546686593 | 0.189488013 | 0.264 | 0.229 | 1 | 0.215324608 | 0.627751484 | 0.341 | 0.263 | 1 | 0.546686593 | 0.384284529 |
| Ndufb11 | 0.321952571 | 0.152972813 | 0.981 | 0.898 | 1 | 0.215402449 | -0.180244121 | 0.829 | 0.945 | 1 | 0.321952571 | 0.384406684 |
| B3glct | 0.307190892 | -0.171706892 | 0.528 | 0.61 | 1 | 0.215483455 | 0.70627398 | 0.463 | 0.427 | 1 | 0.307190892 | 0.384533791 |
| Rangap1 | 0.356211817 | 0.223435345 | 0.472 | 0.411 | 1 | 0.215488353 | 0.781922414 | 0.61 | 0.539 | 1 | 0.356211817 | 0.384541476 |
| Snx12 | 0.215519576 | -0.490993937 | 0.358 | 0.441 | 1 | 0.990310026 | 0.169186609 | 0.463 | 0.519 | 1 | 0.990310026 | 0.384590464 |
| Trp53cor1 | 0.313470077 | 0.810768194 | 0.113 | 0.072 | 1 | 0.215541947 | 1.120997816 | 0.122 | 0.068 | 1 | 0.313470077 | 0.384625563 |
| Pex11g | 0.215587606 | 1.261637799 | 0.094 | 0.051 | 1 | 0.542392144 | 1.749742009 | 0.073 | 0.051 | 1 | 0.542392144 | 0.384697196 |
| Retreg2 | 0.215844967 | 0.386896037 | 0.491 | 0.407 | 1 | 0.362562339 | -0.380025866 | 0.39 | 0.437 | 1 | 0.362562339 | 0.385100884 |
| Cdt1 | 0.215984756 | 0.361173472 | 0.208 | 0.136 | 1 | 0.771970126 | 0.544423101 | 0.098 | 0.119 | 1 | 0.771970126 | 0.385320097 |
| Srcin1 | 0.532479883 | -0.7032584 | 0.038 | 0.059 | 1 | 0.216369067 | -1.484723245 | 0.049 | 0.109 | 1 | 0.532479883 | 0.385922561 |
| Fam122a | 0.46014523 | 0.397292898 | 0.283 | 0.237 | 1 | 0.216415454 | -0.16279515 | 0.098 | 0.181 | 1 | 0.46014523 | 0.38599526 |
| Gpr137c | 0.216511295 | 2.324647596 | 0.075 | 0.038 | 1 | 0.975801516 | 0.47463477 | 0.049 | 0.048 | 1 | 0.975801516 | 0.386145449 |
| Ogfod1 | 0.216560762 | 1.082931912 | 0.377 | 0.314 | 1 | 0.754928221 | 0.649577847 | 0.366 | 0.375 | 1 | 0.754928221 | 0.386222961 |
| Lhpp | 0.216756417 | 0.8327945 | 0.113 | 0.064 | 1 | 0.883807496 | 0.435106406 | 0.098 | 0.109 | 1 | 0.883807496 | 0.38652949 |
| Ripk3 | 0.216825422 | 0.645708947 | 0.208 | 0.14 | 1 | 0.359371268 | 1.082317348 | 0.122 | 0.082 | 1 | 0.359371268 | 0.386637581 |
| Bsn | 0.888969647 | -0.237594828 | 0.057 | 0.051 | 1 | 0.216890963 | -1.777504994 | 0.073 | 0.137 | 1 | 0.888969647 | 0.386740237 |
| Icmt | 0.217168859 | 0.898382842 | 0.226 | 0.161 | 1 | 0.790482307 | 0.298046039 | 0.244 | 0.229 | 1 | 0.790482307 | 0.387175404 |
| Ctnnd2 | 0.933062077 | -0.362676622 | 0.283 | 0.292 | 1 | 0.217785003 | 0.272420231 | 0.195 | 0.119 | 1 | 0.933062077 | 0.388139699 |
| Pcdhgc4 | 0.302217547 | 0.652222254 | 0.094 | 0.055 | 1 | 0.217986098 | 2.185128153 | 0.073 | 0.034 | 1 | 0.302217547 | 0.388454257 |
| Bicdl1 | 0.267977656 | -0.752168001 | 0.019 | 0.055 | 1 | 0.218040571 | 1.911205431 | 0.073 | 0.034 | 1 | 0.267977656 | 0.388539451 |
| Gm12353 | 0.328915185 | 0.657222935 | 0.17 | 0.123 | 1 | 0.218055331 | -0.484723245 | 0.073 | 0.15 | 1 | 0.328915185 | 0.388562535 |
| Sbds | 0.218103746 | -0.206613885 | 0.736 | 0.839 | 1 | 0.625675585 | -0.118672597 | 0.78 | 0.836 | 1 | 0.625675585 | 0.388638248 |
| 3300002I08Rik | 0.21820182 | 0.739685096 | 0.208 | 0.144 | 1 | 0.36252357 | 0.83720485 | 0.195 | 0.143 | 1 | 0.36252357 | 0.388791606 |
| Kctd6 | 0.218402412 | 0.767699472 | 0.17 | 0.11 | 1 | 0.860858733 | 0.168826341 | 0.22 | 0.208 | 1 | 0.860858733 | 0.38910521 |
| Tpk1 | 0.218422811 | 0.445312723 | 0.358 | 0.28 | 1 | 0.722833605 | 0.195098442 | 0.268 | 0.242 | 1 | 0.722833605 | 0.389137097 |
| Mrps28 | 0.218504062 | 0.475823693 | 0.66 | 0.691 | 1 | 0.790112173 | 0.210422173 | 0.732 | 0.771 | 1 | 0.790112173 | 0.389264098 |
| Snx1 | 0.544225401 | -0.128731352 | 0.679 | 0.729 | 1 | 0.218612338 | -0.21697029 | 0.537 | 0.693 | 1 | 0.544225401 | 0.389433322 |
| Nostrin | 0.338517886 | 1.640149422 | 0.094 | 0.059 | 1 | 0.21867314 | 0.721727632 | 0.073 | 0.157 | 1 | 0.338517886 | 0.389528337 |
| Rps8 | 0.219146879 | 0.154306012 | 1 | 0.992 | 1 | 0.301236779 | -0.119115279 | 1 | 0.997 | 1 | 0.301236779 | 0.390268404 |
| Gm28154 | 0.728881193 | 0.104096522 | 0.075 | 0.093 | 1 | 0.219519873 | -0.425829556 | 0.049 | 0.116 | 1 | 0.728881193 | 0.390850771 |
| Mapkbp1 | 0.219528789 | 0.487297934 | 0.415 | 0.347 | 1 | 0.628728967 | 0.633030286 | 0.463 | 0.488 | 1 | 0.628728967 | 0.390864689 |
| Gtf3c3 | 0.849657842 | 0.225805693 | 0.321 | 0.322 | 1 | 0.219624104 | -0.592782991 | 0.146 | 0.229 | 1 | 0.849657842 | 0.391013461 |
| Oxct1 | 0.243712715 | 0.297757225 | 0.925 | 0.873 | 1 | 0.219652149 | 0.37705756 | 0.829 | 0.891 | 1 | 0.243712715 | 0.391057231 |
| ENSMUSG00000121319 | 0.488750271 | -0.304709024 | 0.019 | 0.038 | 1 | 0.219939713 | 1.83720485 | 0.073 | 0.034 | 1 | 0.488750271 | 0.391505949 |
| Gng2 | 0.22042268 | 0.71862348 | 0.151 | 0.093 | 1 | 0.58836895 | -0.505187348 | 0.098 | 0.126 | 1 | 0.58836895 | 0.392259202 |
| Ntan1 | 0.220760788 | -0.258759438 | 0.547 | 0.644 | 1 | 0.329340444 | -0.126269274 | 0.585 | 0.655 | 1 | 0.329340444 | 0.392786251 |
| A730081D07Rik | 0.631302606 | 0.776210972 | 0.075 | 0.059 | 1 | 0.220818886 | -1.448197369 | 0.049 | 0.109 | 1 | 0.631302606 | 0.392876792 |
| Mreg | 0.343000446 | -0.430239906 | 0 | 0.017 | 1 | 0.220910219 | 1.721727632 | 0.073 | 0.034 | 1 | 0.343000446 | 0.393019113 |
| Stx12 | 0.580537121 | 0.166662208 | 0.755 | 0.742 | 1 | 0.221129904 | -0.121845522 | 0.659 | 0.761 | 1 | 0.580537121 | 0.393361374 |
| Mib2 | 0.618410763 | 0.522454379 | 0.132 | 0.11 | 1 | 0.221696848 | 1.042024642 | 0.22 | 0.154 | 1 | 0.618410763 | 0.394244204 |
| Etfrf1 | 0.368366014 | 0.442004547 | 0.264 | 0.208 | 1 | 0.222247843 | -0.429581691 | 0.122 | 0.208 | 1 | 0.368366014 | 0.395101582 |
| H2bc6 | 0.882410767 | 0.650250012 | 0.226 | 0.233 | 1 | 0.222358972 | -0.835220492 | 0.146 | 0.085 | 1 | 0.882410767 | 0.395274431 |
| Def6 | 0.222422401 | 0.900966003 | 0.151 | 0.097 | 1 | 0.438320217 | 0.667279848 | 0.122 | 0.085 | 1 | 0.438320217 | 0.395373078 |
| Nipa2 | 0.577760757 | -0.1182959 | 0.623 | 0.708 | 1 | 0.222639448 | -0.278272368 | 0.683 | 0.744 | 1 | 0.577760757 | 0.395710572 |
| Zfp169 | 0.586206676 | -0.207847485 | 0.075 | 0.102 | 1 | 0.222679923 | -0.723510105 | 0.073 | 0.143 | 1 | 0.586206676 | 0.395773497 |
| BC005624 | 0.608949857 | 0.119098685 | 0.623 | 0.712 | 1 | 0.222817366 | 0.294306409 | 0.707 | 0.638 | 1 | 0.608949857 | 0.395987154 |
| Prps2 | 0.555862324 | 1.055186921 | 0.075 | 0.055 | 1 | 0.223048999 | 1.114738825 | 0.146 | 0.089 | 1 | 0.555862324 | 0.396347143 |
| 4930512B01Rik | 0.677640974 | -0.545717123 | 0.057 | 0.072 | 1 | 0.223180347 | -0.795063366 | 0.073 | 0.143 | 1 | 0.677640974 | 0.396551226 |
| Med31 | 0.846697766 | 0.170489911 | 0.509 | 0.513 | 1 | 0.223189356 | -0.151113345 | 0.39 | 0.543 | 1 | 0.846697766 | 0.396565224 |
| Cables1 | 0.223317284 | -0.877698883 | 0.113 | 0.182 | 1 | 0.941164401 | 0.404245443 | 0.146 | 0.15 | 1 | 0.941164401 | 0.396763959 |
| Gm17484 | 0.420959194 | 0.272079545 | 0.17 | 0.127 | 1 | 0.223408818 | 0.983055716 | 0.244 | 0.174 | 1 | 0.420959194 | 0.396906135 |
| 1190005I06Rik | 0.343921463 | 0.767699472 | 0.151 | 0.106 | 1 | 0.223563406 | -0.935384654 | 0.049 | 0.109 | 1 | 0.343921463 | 0.397146216 |
| Snhg17 | 0.223597247 | 0.813685677 | 0.189 | 0.127 | 1 | 0.381589372 | 0.551802631 | 0.073 | 0.123 | 1 | 0.381589372 | 0.397198766 |
| Phf10 | 0.223751249 | 0.347367673 | 0.566 | 0.521 | 1 | 0.546788311 | 0.155669089 | 0.561 | 0.683 | 1 | 0.546788311 | 0.397437877 |
| BC024063 | 0.752453414 | 0.669295768 | 0.057 | 0.047 | 1 | 0.223858419 | 1.458693227 | 0.073 | 0.034 | 1 | 0.752453414 | 0.397604246 |
| Tes | 0.507271194 | 1.007881206 | 0.113 | 0.089 | 1 | 0.224148503 | 0.604544093 | 0.22 | 0.143 | 1 | 0.507271194 | 0.398054454 |
| Znrd2 | 0.224480109 | -0.454086648 | 0.132 | 0.208 | 1 | 0.843961831 | 0.188112012 | 0.171 | 0.188 | 1 | 0.843961831 | 0.398568899 |
| Yjefn3 | 0.435888471 | -0.52334931 | 0.075 | 0.11 | 1 | 0.224535913 | -0.589059905 | 0.049 | 0.113 | 1 | 0.435888471 | 0.398655449 |
| Armt1 | 0.365513344 | 0.855162313 | 0.151 | 0.11 | 1 | 0.224685005 | -0.863234868 | 0.073 | 0.14 | 1 | 0.365513344 | 0.398886659 |
| Gm20257 | 0.394952512 | 0.533234218 | 0.189 | 0.144 | 1 | 0.224685005 | -0.863234868 | 0.073 | 0.14 | 1 | 0.394952512 | 0.398886659 |
| Ftsj3 | 0.224745868 | 0.58098735 | 0.434 | 0.394 | 1 | 0.33971411 | 0.822558074 | 0.463 | 0.433 | 1 | 0.33971411 | 0.398981031 |
| Cdh23 | 0.608182413 | -0.337130502 | 0.094 | 0.119 | 1 | 0.224816156 | -1.622226769 | 0.024 | 0.075 | 1 | 0.608182413 | 0.399090009 |
| Ube2cbp | 0.327566396 | 0.913714495 | 0.113 | 0.072 | 1 | 0.22481635 | -1.589059905 | 0.024 | 0.075 | 1 | 0.327566396 | 0.399090309 |
| Polr1b | 0.234619368 | 0.739685096 | 0.132 | 0.081 | 1 | 0.224839276 | -0.69884805 | 0.073 | 0.143 | 1 | 0.234619368 | 0.399125853 |
| Gm3235 | 0.923013829 | 0.739685096 | 0.019 | 0.017 | 1 | 0.224844589 | -0.332720152 | 0.024 | 0.078 | 1 | 0.923013829 | 0.399134088 |
| Rasl12 | 0.224853418 | -1.652632327 | 0.019 | 0.059 | 1 | 0.666778333 | -0.429581691 | 0.073 | 0.096 | 1 | 0.666778333 | 0.399147777 |
| Gm51425 | 0.519157514 | 1.47665069 | 0.075 | 0.055 | 1 | 0.224917245 | 0.83720485 | 0.122 | 0.068 | 1 | 0.519157514 | 0.399246723 |
| Srbd1 | 0.61957356 | 0.213274777 | 0.642 | 0.725 | 1 | 0.225025918 | -0.296194276 | 0.561 | 0.686 | 1 | 0.61957356 | 0.399415173 |
| Mrpl32 | 0.401231896 | 0.228723176 | 0.509 | 0.466 | 1 | 0.225070975 | -0.297954733 | 0.439 | 0.526 | 1 | 0.401231896 | 0.399485007 |
| Mrpl51 | 0.854774626 | 0.10741688 | 0.491 | 0.496 | 1 | 0.225258714 | -0.449957827 | 0.366 | 0.457 | 1 | 0.854774626 | 0.39977594 |
| Slc16a10 | 0.225381063 | 0.798237278 | 0.302 | 0.233 | 1 | 0.501146899 | -0.622226769 | 0.293 | 0.229 | 1 | 0.501146899 | 0.399965503 |
| Cops3 | 0.856380652 | 0.238380524 | 0.528 | 0.589 | 1 | 0.225413919 | -0.363093801 | 0.634 | 0.676 | 1 | 0.856380652 | 0.400016403 |
| Slc14a2 | 0.225535028 | 2.039245377 | 0.057 | 0.025 | 1 | 0.357860251 | -1.069685746 | 0 | 0.02 | 1 | 0.357860251 | 0.400204008 |
| A430035B10Rik | 0.225535665 | 1.792152515 | 0.057 | 0.025 | 1 | 0.710563777 | 1.458693227 | 0.049 | 0.038 | 1 | 0.710563777 | 0.400204994 |
| Abcd1 | 0.225601086 | -0.1182959 | 0.038 | 0.089 | 1 | 0.727142728 | -0.395455907 | 0.098 | 0.113 | 1 | 0.727142728 | 0.400306321 |
| Mdc1 | 0.225728458 | 0.689059022 | 0.245 | 0.178 | 1 | 0.806915048 | -0.17660095 | 0.195 | 0.215 | 1 | 0.806915048 | 0.40050358 |
| Vps41 | 0.225768716 | 0.341491106 | 0.623 | 0.559 | 1 | 0.429302492 | 0.357424586 | 0.585 | 0.608 | 1 | 0.429302492 | 0.400565919 |
| Pgs1 | 0.2257859 | 0.283455909 | 0.472 | 0.39 | 1 | 0.529418824 | 0.322631677 | 0.415 | 0.379 | 1 | 0.529418824 | 0.400592527 |
| Fam220a | 0.225912448 | 1.039245377 | 0.094 | 0.051 | 1 | 0.609924905 | 0.136765132 | 0.024 | 0.041 | 1 | 0.609924905 | 0.400788462 |
| Rnf114 | 0.899957989 | 0.111653873 | 0.358 | 0.377 | 1 | 0.226218326 | 0.415247937 | 0.512 | 0.447 | 1 | 0.899957989 | 0.401261921 |
| Top1mt | 0.271969145 | 0.714150003 | 0.113 | 0.068 | 1 | 0.226277326 | -1.16279515 | 0.024 | 0.075 | 1 | 0.271969145 | 0.401353224 |
| Ttll12 | 0.870598372 | 0.984797593 | 0.057 | 0.064 | 1 | 0.22627927 | -0.221688839 | 0.024 | 0.078 | 1 | 0.870598372 | 0.401356231 |
| Capn11 | 0.359776551 | -0.652632327 | 0.019 | 0.047 | 1 | 0.226281208 | -1.069685746 | 0.024 | 0.075 | 1 | 0.359776551 | 0.401359231 |
| Ap2b1 | 0.241770425 | -0.232056644 | 0.717 | 0.797 | 1 | 0.226337657 | -0.257070055 | 0.78 | 0.843 | 1 | 0.241770425 | 0.40144658 |
| Nsf | 0.226368423 | 0.327738866 | 0.585 | 0.538 | 1 | 0.435452031 | -0.28965143 | 0.707 | 0.703 | 1 | 0.435452031 | 0.401494183 |

| | | | | | | | | | | | | |
|---|---|---|---|---|---|---|---|---|---|---|---|---|
| Sstr2 | 0.226423846 | -1.368839361 | 0.057 | 0.11 | 1 | 0.535256365 | 1.458693227 | 0.049 | 0.031 | 1 | 0.535256365 | 0.401579934 |
| Lrrn3 | 0.226678839 | 2.614154213 | 0.057 | 0.025 | 1 | 0.832579923 | 0.100239256 | 0.049 | 0.041 | 1 | 0.832579923 | 0.401974382 |
| Septin10 | 0.468354921 | -0.330704232 | 0.321 | 0.36 | 1 | 0.226862054 | -0.297954733 | 0.171 | 0.259 | 1 | 0.468354921 | 0.402257717 |
| Ovgp1 | 0.373358787 | 0.695290976 | 0.132 | 0.093 | 1 | 0.227158876 | -0.300298674 | 0.049 | 0.113 | 1 | 0.373358787 | 0.402716597 |
| Ctla4 | 0.886688014 | 0.207190015 | 0.113 | 0.106 | 1 | 0.227183506 | -0.863234868 | 0.049 | 0.109 | 1 | 0.886688014 | 0.402754666 |
| 1110059E24Rik | 0.227504627 | 0.25093791 | 0.358 | 0.275 | 1 | 0.740895406 | 0.927402659 | 0.268 | 0.273 | 1 | 0.740895406 | 0.403250899 |
| Gm49575 | 0.227535349 | -1.368839361 | 0.019 | 0.059 | 1 | 0.231630114 | -1.484723245 | 0 | 0.034 | 1 | 0.231630114 | 0.403298363 |
| Gm35853 | 0.227588869 | -2.237594828 | 0.019 | 0.059 | 1 | 0.402369942 | -0.16279515 | 0 | 0.017 | 1 | 0.402369942 | 0.403303802 |
| Mettl16 | 0.227540791 | 0.478129619 | 0.528 | 0.521 | 1 | 0.33293056 | 0.450529574 | 0.61 | 0.577 | 1 | 0.33293056 | 0.40330677 |
| Aph1b | 0.227667644 | 0.79717059 | 0.208 | 0.144 | 1 | 0.984304534 | 0.276489895 | 0.146 | 0.147 | 1 | 0.984304534 | 0.403502732 |
| Muc1 | 0.416734858 | -0.752168001 | 0.019 | 0.042 | 1 | 0.227736597 | -1.116991461 | 0.024 | 0.075 | 1 | 0.416734858 | 0.403609236 |
| Ficd | 0.227742276 | -0.1672055 | 0.038 | 0.089 | 1 | 0.777183196 | -0.132421501 | 0.122 | 0.137 | 1 | 0.777183196 | 0.403618007 |
| Gm5535 | 0.22784154 | 1.962077517 | 0.057 | 0.025 | 1 | 0.605005904 | 0.667279848 | 0.024 | 0.014 | 1 | 0.605005904 | 0.403771313 |
| 2610020C07Rik | 0.537863531 | 0.454282877 | 0.208 | 0.178 | 1 | 0.227852022 | -0.397260404 | 0.195 | 0.3 | 1 | 0.537863531 | 0.4037875 |
| H4c3 | 0.641178868 | 0.891688189 | 0.057 | 0.042 | 1 | 0.227863168 | 1.377773231 | 0.073 | 0.034 | 1 | 0.641178868 | 0.403804713 |
| Rragb | 0.227995262 | 1.454282877 | 0.075 | 0.038 | 1 | 0.3351125 | -0.747757651 | 0.024 | 0.061 | 1 | 0.3351125 | 0.404008685 |
| 1700030J22Rik | 0.227995711 | 1.377115016 | 0.075 | 0.038 | 1 | 0.744047924 | 1.252242349 | 0.024 | 0.017 | 1 | 0.744047924 | 0.404009377 |
| Gm43329 | 0.488696298 | -0.430239906 | 0.057 | 0.085 | 1 | 0.228115562 | -0.484723245 | 0.073 | 0.147 | 1 | 0.488696298 | 0.404194414 |
| Pigyl | 0.228118679 | 0.8327945 | 0.283 | 0.22 | 1 | 0.966195215 | 0.136765132 | 0.195 | 0.198 | 1 | 0.966195215 | 0.404199227 |
| Pitpnm2 | 0.228186469 | 0.47665069 | 0.34 | 0.267 | 1 | 0.938922786 | 0.331676817 | 0.293 | 0.307 | 1 | 0.938922786 | 0.404303873 |
| Rassf1 | 0.228355795 | 0.262398112 | 0.623 | 0.547 | 1 | 0.431017429 | 0.30361074 | 0.732 | 0.71 | 1 | 0.431017429 | 0.404565221 |
| Nek11 | 0.228360488 | 0.962077517 | 0.094 | 0.051 | 1 | 0.887691052 | 0.30115195 | 0.049 | 0.055 | 1 | 0.887691052 | 0.404572463 |
| Entpd6 | 0.485102313 | -0.489133595 | 0.057 | 0.085 | 1 | 0.228463905 | -1.020776145 | 0.024 | 0.075 | 1 | 0.485102313 | 0.404732054 |
| Olfm1 | 0.228548298 | 0.412790748 | 0.358 | 0.28 | 1 | 0.331057302 | 0.870371714 | 0.244 | 0.191 | 1 | 0.331057302 | 0.404862271 |
| Gm33302 | 0.407395106 | -2.237594828 | 0.019 | 0.042 | 1 | 0.228668766 | -1.80665134 | 0.024 | 0.075 | 1 | 0.407395106 | 0.405048127 |
| Tnfaip1 | 0.354544433 | 0.358863312 | 0.434 | 0.39 | 1 | 0.228823168 | 0.46797104 | 0.585 | 0.529 | 1 | 0.354544433 | 0.405286294 |
| Muc2 | 0.249034808 | 2.154722595 | 0.019 | 0.004 | 1 | 0.228843009 | 0.898605394 | 0.073 | 0.034 | 1 | 0.249034808 | 0.405316895 |
| Zfp623 | 0.228899781 | 1.417757001 | 0.075 | 0.038 | 1 | 0.719596325 | 0.252242349 | 0.073 | 0.092 | 1 | 0.719596325 | 0.405404452 |
| Stx8 | 0.22896267 | 0.47145802 | 0.453 | 0.411 | 1 | 0.23080549 | 0.414795362 | 0.585 | 0.546 | 1 | 0.23080549 | 0.405501436 |
| Gm36037 | 0.229007934 | 1.154722595 | 0.075 | 0.038 | 1 | 0.538950358 | 0.944120054 | 0.049 | 0.031 | 1 | 0.538950358 | 0.405571234 |
| Tmprss5 | 0.529359278 | 1.154722595 | 0.057 | 0.038 | 1 | 0.229100126 | 1.174239837 | 0.098 | 0.051 | 1 | 0.529359278 | 0.405713384 |
| Rfesd | 0.505581759 | 0.2967416 | 0.075 | 0.11 | 1 | 0.229188679 | -0.917682652 | 0.024 | 0.075 | 1 | 0.505581759 | 0.405849907 |
| Jmjd4 | 0.919472562 | 0.347367673 | 0.075 | 0.072 | 1 | 0.229188679 | -0.917682652 | 0.024 | 0.075 | 1 | 0.919472562 | 0.405849907 |
| Gm16001 | 0.229221835 | 0.874614676 | 0.094 | 0.051 | 1 | 0.382976968 | 0.83720485 | 0.122 | 0.082 | 1 | 0.382976968 | 0.40590102 |
| Rpusd1 | 0.229319728 | -1.093204919 | 0.019 | 0.059 | 1 | 0.574025125 | 1.614812428 | 0.024 | 0.044 | 1 | 0.574025125 | 0.406051919 |
| Gm13919 | 0.22932012 | -1.015202407 | 0.019 | 0.059 | 1 | 0.95276834 | 0.515276755 | 0.098 | 0.096 | 1 | 0.95276834 | 0.406052523 |
| 1500011B03Rik | 0.668963086 | 0.285967128 | 0.472 | 0.479 | 1 | 0.229363021 | -0.238898129 | 0.39 | 0.498 | 1 | 0.668963086 | 0.406118647 |
| Tmem69 | 0.346600384 | -0.489133595 | 0.057 | 0.097 | 1 | 0.229598851 | -0.707115666 | 0.049 | 0.109 | 1 | 0.346600384 | 0.406482069 |
| Clic5 | 0.779362808 | -0.27154216 | 0.094 | 0.106 | 1 | 0.229828583 | 1.252242349 | 0.073 | 0.034 | 1 | 0.779362808 | 0.406835989 |
| Kcnn3 | 0.343968402 | 1.154722595 | 0.038 | 0.017 | 1 | 0.229843619 | 1.652780279 | 0.073 | 0.034 | 1 | 0.343968402 | 0.406859149 |
| Trim11 | 0.229901942 | 0.665337754 | 0.302 | 0.229 | 1 | 0.51521652 | 0.210422173 | 0.195 | 0.256 | 1 | 0.51521652 | 0.406948981 |
| Tonsl | 0.22995404 | 1.261637799 | 0.075 | 0.038 | 1 | 0.441381862 | 1.515276755 | 0.024 | 0.01 | 1 | 0.441381862 | 0.407029219 |
| Gnal | 0.22995404 | 1.154722595 | 0.075 | 0.038 | 1 | 0.249107223 | 0.939084464 | 0.122 | 0.072 | 1 | 0.249107223 | 0.407029219 |
| Zbtb26 | 0.230207531 | -0.932740246 | 0.019 | 0.059 | 1 | 0.365228117 | 0.962735732 | 0.122 | 0.082 | 1 | 0.365228117 | 0.407419554 |
| Gm50240 | 0.230437825 | 1.292226119 | 0.113 | 0.068 | 1 | 0.936982722 | 0.136765132 | 0.098 | 0.102 | 1 | 0.936982722 | 0.407774058 |
| Ubxn2a | 0.230442524 | -0.213428517 | 0.453 | 0.555 | 1 | 0.911248828 | 0.148133946 | 0.585 | 0.601 | 1 | 0.911248828 | 0.407781291 |
| Myoz3 | 0.430735738 | -0.134784022 | 0.208 | 0.157 | 1 | 0.230524075 | -0.700760171 | 0.146 | 0.229 | 1 | 0.430735738 | 0.407906801 |
| Ppcdc | 0.230594569 | 0.569760094 | 0.132 | 0.081 | 1 | 0.909468515 | 0.59619675 | 0.122 | 0.123 | 1 | 0.909468515 | 0.408015282 |
| H3c14 | 0.282558974 | -0.389597921 | 0.057 | 0.106 | 1 | 0.230762368 | 1.252242349 | 0.073 | 0.034 | 1 | 0.282558974 | 0.408273466 |
| Zfp688 | 0.305590208 | 1.154722595 | 0.075 | 0.042 | 1 | 0.230762368 | 1.252242349 | 0.073 | 0.034 | 1 | 0.305590208 | 0.408273466 |
| Supt6 | 0.231006189 | 0.390535532 | 0.811 | 0.784 | 1 | 0.400785068 | 0.235033059 | 0.878 | 0.881 | 1 | 0.400785068 | 0.408648518 |
| Gm2415 | 0.559233506 | 0.792152515 | 0.094 | 0.072 | 1 | 0.231067302 | -0.863234868 | 0.098 | 0.167 | 1 | 0.559233506 | 0.408742507 |
| Tmem86b | 0.23109239 | -0.845277405 | 0.019 | 0.059 | 1 | 0.269239723 | -0.278272368 | 0.024 | 0.072 | 1 | 0.269239723 | 0.408781088 |
| Gm47096 | 0.23109239 | -0.845277405 | 0.019 | 0.059 | 1 | 0.938546854 | 0.174239837 | 0.049 | 0.051 | 1 | 0.938546854 | 0.408781088 |
| Nfu1 | 0.231183544 | 0.424329655 | 0.491 | 0.411 | 1 | 0.409545338 | 0.907594178 | 0.512 | 0.498 | 1 | 0.409545338 | 0.408921256 |
| Pde2a | 0.231366669 | 1.569760094 | 0.057 | 0.025 | 1 | 0.880719545 | 0.136765132 | 0.049 | 0.055 | 1 | 0.880719545 | 0.409202802 |
| Spout1 | 0.231391309 | -0.430239906 | 0.132 | 0.208 | 1 | 0.467255202 | -0.410722664 | 0.073 | 0.109 | 1 | 0.467255202 | 0.40924068 |
| Mrps2 | 0.231562242 | 0.707263618 | 0.17 | 0.11 | 1 | 0.32373352 | -0.520347155 | 0.073 | 0.126 | 1 | 0.32373352 | 0.409503412 |
| Gm34788 | 0.249034808 | 2.154722595 | 0.019 | 0.004 | 1 | 0.231591308 | -1.16279515 | 0 | 0.034 | 1 | 0.249034808 | 0.409548082 |
| Tcp11 | 0.267991969 | -1.1182959 | 0.038 | 0.081 | 1 | 0.231591308 | -0.747757651 | 0 | 0.034 | 1 | 0.267991969 | 0.409548082 |
| Gm41293 | 0.340246113 | 1.417757001 | 0.038 | 0.017 | 1 | 0.231591308 | -0.863234868 | 0 | 0.034 | 1 | 0.340246113 | 0.409548082 |
| Upb1 | 0.413082433 | 0.154722595 | 0 | 0.013 | 1 | 0.231591308 | -0.747757651 | 0 | 0.034 | 1 | 0.413082433 | 0.409548082 |
| Vstm2l | 0.732691937 | 1.154722595 | 0.019 | 0.013 | 1 | 0.231591308 | -0.747757651 | 0 | 0.034 | 1 | 0.732691937 | 0.409548082 |
| Gm43566 | 0.762480493 | 0.739685096 | 0.038 | 0.03 | 1 | 0.231591308 | -0.747757651 | 0 | 0.034 | 1 | 0.762480493 | 0.409548082 |
| Pcdhgb1 | 0.776830424 | -0.545717123 | 0.019 | 0.025 | 1 | 0.231591308 | -0.747757651 | 0 | 0.034 | 1 | 0.776830424 | 0.409548082 |
| Wdr54 | 0.917883255 | 0.569760094 | 0.019 | 0.021 | 1 | 0.231591308 | -0.747757651 | 0 | 0.034 | 1 | 0.917883255 | 0.409548082 |
| Gm57017 | 0.926212226 | 0.8327945 | 0.019 | 0.017 | 1 | 0.231591308 | -0.863234868 | 0 | 0.034 | 1 | 0.926212226 | 0.409548082 |
| Gm56548 | 0.991244785 | 0.417757001 | 0.038 | 0.038 | 1 | 0.231591308 | -0.747757651 | 0 | 0.034 | 1 | 0.991244785 | 0.409548082 |
| Pcdhb3 | 0.249034808 | 2.154722595 | 0.019 | 0.004 | 1 | 0.231606831 | -0.863234868 | 0 | 0.034 | 1 | 0.249034808 | 0.409571938 |
| Inha | 0.340246113 | 1.417757001 | 0.038 | 0.017 | 1 | 0.231606831 | -0.970150072 | 0 | 0.034 | 1 | 0.340246113 | 0.409571938 |
| Sptb | 0.413082433 | 0.154722595 | 0 | 0.013 | 1 | 0.231606831 | -1.069685746 | 0 | 0.034 | 1 | 0.413082433 | 0.409571938 |
| Gm17455 | 0.418323927 | 1.695290976 | 0.057 | 0.034 | 1 | 0.231606831 | -0.863234868 | 0 | 0.034 | 1 | 0.418323927 | 0.409571938 |
| Gm15265 | 0.506983348 | 0.695290976 | 0.019 | 0.038 | 1 | 0.231606831 | -1.069685746 | 0 | 0.034 | 1 | 0.506983348 | 0.409571938 |
| 4930434J08Rik | 0.653990165 | 0.695290976 | 0.057 | 0.042 | 1 | 0.231606831 | -0.863234868 | 0 | 0.034 | 1 | 0.653990165 | 0.409571938 |
| Batf2 | 0.684424342 | 1.154722595 | 0.019 | 0.03 | 1 | 0.231606831 | -0.970150072 | 0 | 0.034 | 1 | 0.684424342 | 0.409571938 |
| Gm13938 | 0.762480493 | 0.739685096 | 0.038 | 0.03 | 1 | 0.231606831 | -0.863234868 | 0 | 0.034 | 1 | 0.762480493 | 0.409571938 |
| E430024P14Rik | 0.897146191 | 0.417757001 | 0.038 | 0.034 | 1 | 0.231606831 | -0.863234868 | 0 | 0.034 | 1 | 0.897146191 | 0.409571938 |
| 4930404H11Rik | 0.279875847 | -0.304709024 | 0.019 | 0.055 | 1 | 0.231618473 | -1.250257991 | 0 | 0.034 | 1 | 0.279875847 | 0.409589829 |
| Gm29825 | 0.566212 | 0.154722595 | 0.057 | 0.038 | 1 | 0.231618473 | -1.16279515 | 0 | 0.034 | 1 | 0.566212 | 0.409589829 |
| Slc4a5 | 0.658774723 | -0.752168001 | 0.019 | 0.03 | 1 | 0.231618473 | -1.069685746 | 0 | 0.034 | 1 | 0.658774723 | 0.409589829 |
| Cyp27a1 | 0.419848106 | -0.430239906 | 0.019 | 0.042 | 1 | 0.231618904 | -1.250257991 | 0 | 0.034 | 1 | 0.419848106 | 0.409590491 |
| D030016M11Rik | 0.641178868 | 0.891688189 | 0.057 | 0.042 | 1 | 0.231618904 | -1.16279515 | 0 | 0.034 | 1 | 0.641178868 | 0.409590491 |
| 4930593A02Rik | 0.430509759 | 0.438515561 | 0.038 | 0.068 | 1 | 0.231627527 | -1.250257991 | 0 | 0.034 | 1 | 0.430509759 | 0.409603743 |
| Paqr4 | 0.567830665 | -0.304709024 | 0.019 | 0.034 | 1 | 0.231627527 | -1.250257991 | 0 | 0.034 | 1 | 0.567830665 | 0.409603743 |
| Gm56980 | 0.732691937 | 1.154722595 | 0.019 | 0.013 | 1 | 0.231627527 | -1.332720152 | 0 | 0.034 | 1 | 0.732691937 | 0.409603743 |
| 2610206C17Rik | 0.765052878 | -0.582242999 | 0.038 | 0.047 | 1 | 0.231633995 | -1.622226769 | 0 | 0.034 | 1 | 0.765052878 | 0.409613682 |
| Cd53 | 0.251644813 | 1.154722595 | 0.019 | 0.004 | 1 | 0.231637013 | -1.863234868 | 0 | 0.034 | 1 | 0.251644813 | 0.40961832 |
| Exoc3l4 | 0.242083189 | -0.904171094 | 0.038 | 0.085 | 1 | 0.231643049 | -0.622226769 | 0 | 0.034 | 1 | 0.242083189 | 0.409627595 |
| Gm7628 | 0.24429165 | -0.260314904 | 0.019 | 0.059 | 1 | 0.231643049 | -0.622226769 | 0 | 0.034 | 1 | 0.24429165 | 0.409627595 |
| Gm42869 | 0.305590208 | 1.154722595 | 0.075 | 0.042 | 1 | 0.231643049 | -0.622226769 | 0 | 0.034 | 1 | 0.305590208 | 0.409627595 |
| Galnt12 | 0.324474943 | 1.47665069 | 0.057 | 0.03 | 1 | 0.231643049 | -0.622226769 | 0 | 0.034 | 1 | 0.324474943 | 0.409627595 |
| Ptp4a1.1 | 0.490620203 | -0.1672055 | 0.019 | 0.038 | 1 | 0.231643049 | -0.622226769 | 0 | 0.034 | 1 | 0.490620203 | 0.409627595 |
| Gm16091 | 0.77632945 | -0.347777746 | 0.038 | 0.03 | 1 | 0.231643049 | -0.622226769 | 0 | 0.034 | 1 | 0.77632945 | 0.409627595 |
| Ovca2 | 0.917883255 | 0.569760094 | 0.019 | 0.021 | 1 | 0.231643049 | -0.622226769 | 0 | 0.034 | 1 | 0.917883255 | 0.409627595 |
| Cenpo | 0.824909603 | 0.438515561 | 0.075 | 0.068 | 1 | 0.231649566 | 1.252242349 | 0.098 | 0.051 | 1 | 0.824909603 | 0.409637611 |
| Rab39 | 0.38147512 | 0.891688189 | 0.132 | 0.093 | 1 | 0.231742716 | -0.686357106 | 0.024 | 0.075 | 1 | 0.38147512 | 0.409780746 |
| Fuca2 | 0.231848504 | 0.906795081 | 0.189 | 0.131 | 1 | 0.601609755 | 0.609398932 | 0.268 | 0.242 | 1 | 0.601609755 | 0.40994328 |
| Exosc9 | 0.510967468 | 0.442847982 | 0.358 | 0.322 | 1 | 0.232033915 | 0.80158094 | 0.293 | 0.225 | 1 | 0.510967468 | 0.410228092 |

| | | | | | | | | | | | | |
|---|---|---|---|---|---|---|---|---|---|---|---|---|
| Slc27a4 | 0.232324381 | 0.900966003 | 0.17 | 0.114 | 1 | 0.414582341 | -0.233184478 | 0.171 | 0.229 | 1 | 0.414582341 | 0.410674144 |
| Alkbh8 | 0.23234097 | 0.680791406 | 0.302 | 0.242 | 1 | 0.322689156 | -0.281736223 | 0.195 | 0.273 | 1 | 0.322689156 | 0.410699613 |
| Tor4a | 0.232506641 | 1.669295768 | 0.057 | 0.025 | 1 | 0.969252137 | 0.83720485 | 0.049 | 0.051 | 1 | 0.969252137 | 0.410953943 |
| Adam1a | 0.232506641 | 1.669295768 | 0.057 | 0.025 | 1 | 0.729889663 | 1.83720485 | 0.024 | 0.017 | 1 | 0.729889663 | 0.410953943 |
| Shpk | 0.232506641 | 1.669295768 | 0.057 | 0.025 | 1 | 0.25729872 | -0.747757651 | 0 | 0.031 | 1 | 0.25729872 | 0.410953943 |
| Gm49179 | 0.232506641 | 1.669295768 | 0.057 | 0.025 | 1 | 0.746881137 | 1.029849928 | 0.024 | 0.017 | 1 | 0.746881137 | 0.410953943 |
| Tmem161b | 0.232522895 | 0.379428882 | 0.34 | 0.263 | 1 | 0.517758726 | 0.505998941 | 0.317 | 0.3 | 1 | 0.517758726 | 0.410978894 |
| Zfp426 | 0.232644658 | 0.756173218 | 0.245 | 0.182 | 1 | 0.589216825 | 0.659666664 | 0.244 | 0.218 | 1 | 0.589216825 | 0.411165779 |
| Zcchc17 | 0.451129449 | 0.265979346 | 0.566 | 0.581 | 1 | 0.23277061 | 0.538546534 | 0.537 | 0.461 | 1 | 0.451129449 | 0.411359063 |
| Slc16a1 | 0.232786632 | 0.732031523 | 0.396 | 0.326 | 1 | 0.352868244 | 0.909555694 | 0.39 | 0.355 | 1 | 0.352868244 | 0.411383647 |
| Slc45a4 | 0.952284303 | 0.13912574 | 0.245 | 0.254 | 1 | 0.232804394 | -0.610254127 | 0.341 | 0.423 | 1 | 0.952284303 | 0.411410902 |
| Gm56582 | 0.316405629 | 0.569760094 | 0.151 | 0.102 | 1 | 0.232826734 | -0.654648247 | 0.073 | 0.14 | 1 | 0.316405629 | 0.41144518 |
| Gm36447 | 0.232873406 | 1.280253477 | 0.075 | 0.038 | 1 | 0.416504493 | 1.30115195 | 0.098 | 0.065 | 1 | 0.416504493 | 0.411516789 |
| Clec2l | 0.232873406 | 1.280253477 | 0.075 | 0.038 | 1 | 0.455763738 | 1.100239256 | 0.049 | 0.027 | 1 | 0.455763738 | 0.411516789 |
| Gm12248 | 0.232873406 | 1.280253477 | 0.075 | 0.038 | 1 | 0.921481726 | -0.385187572 | 0.122 | 0.123 | 1 | 0.921481726 | 0.411516789 |
| Ccdc85c | 0.232873406 | 1.280253477 | 0.075 | 0.038 | 1 | 0.471804215 | -0.20718927 | 0.049 | 0.082 | 1 | 0.471804215 | 0.411516789 |
| Snrnp25 | 0.657678744 | 0.104096522 | 0.17 | 0.203 | 1 | 0.233029791 | -0.455576899 | 0.073 | 0.143 | 1 | 0.657678744 | 0.411756698 |
| Ilvbl | 0.23321828 | 0.466666601 | 0.283 | 0.208 | 1 | 0.868815299 | 0.152706676 | 0.244 | 0.232 | 1 | 0.868815299 | 0.412045793 |
| Irak2 | 0.435485307 | 0.450178478 | 0.264 | 0.22 | 1 | 0.233452798 | -0.226298093 | 0.415 | 0.297 | 1 | 0.435485307 | 0.412405388 |
| Cmtm4 | 0.233648733 | 0.468222068 | 0.509 | 0.475 | 1 | 0.899159205 | 0.338244867 | 0.512 | 0.57 | 1 | 0.899159205 | 0.412705736 |
| Gm57446 | 0.340246113 | 1.417757001 | 0.038 | 0.017 | 1 | 0.233806971 | 1.136765132 | 0.073 | 0.034 | 1 | 0.340246113 | 0.412948242 |
| Dhrs9 | 0.771605102 | -2.237594828 | 0.019 | 0.025 | 1 | 0.233806971 | 0.83720485 | 0.073 | 0.034 | 1 | 0.771605102 | 0.412948242 |
| Nrxn1 | 0.615989618 | 3.899261409 | 0.113 | 0.093 | 1 | 0.233918176 | 1.416873051 | 0.244 | 0.171 | 1 | 0.615989618 | 0.413118639 |
| Tlr3 | 0.233940218 | -0.845277405 | 0.094 | 0.157 | 1 | 0.506717129 | -0.863234868 | 0.073 | 0.106 | 1 | 0.506717129 | 0.41315241 |
| Metap1 | 0.719375878 | 0.417757001 | 0.283 | 0.28 | 1 | 0.233982206 | -0.473996838 | 0.39 | 0.491 | 1 | 0.719375878 | 0.413216739 |
| Leprotl1 | 0.234270645 | 0.318221327 | 0.585 | 0.542 | 1 | 0.86011403 | 0.241595105 | 0.61 | 0.57 | 1 | 0.86011403 | 0.413658555 |
| Zfp58 | 0.234315353 | 0.221836791 | 0.094 | 0.161 | 1 | 0.256251415 | -0.622226769 | 0.122 | 0.195 | 1 | 0.256251415 | 0.413727021 |
| Gsto2 | 0.240745263 | 1.029191713 | 0.113 | 0.068 | 1 | 0.23431566 | 0.313642894 | 0.171 | 0.106 | 1 | 0.240745263 | 0.413727491 |
| Tlcd5 | 0.234341522 | -0.799473716 | 0.057 | 0.11 | 1 | 0.522167156 | 0.252242349 | 0.049 | 0.078 | 1 | 0.522167156 | 0.413767095 |
| Prkra | 0.317850719 | 0.41149801 | 0.321 | 0.258 | 1 | 0.234460379 | -0.555112573 | 0.171 | 0.259 | 1 | 0.317850719 | 0.413949088 |
| Arhgef38 | 0.413080549 | -0.1672055 | 0 | 0.013 | 1 | 0.234887523 | 0.911205431 | 0.098 | 0.051 | 1 | 0.413080549 | 0.414602897 |
| Fam3d | 0.923013829 | 0.739685096 | 0.019 | 0.017 | 1 | 0.234887523 | 1.071670103 | 0.098 | 0.051 | 1 | 0.923013829 | 0.414602897 |
| Gm30371 | 0.234928933 | 0.752624151 | 0.057 | 0.025 | 1 | 0.402373368 | -1.410722664 | 0 | 0.017 | 1 | 0.402373368 | 0.414666263 |
| Xrcc5 | 0.235023162 | 0.402650108 | 0.358 | 0.288 | 1 | 0.339501258 | 0.604544093 | 0.39 | 0.352 | 1 | 0.339501258 | 0.414810438 |
| Ypel5 | 0.510678088 | -0.17477872 | 0.642 | 0.691 | 1 | 0.235102181 | 0.252242349 | 0.659 | 0.567 | 1 | 0.510678088 | 0.414931326 |
| Mrpl44 | 0.235164323 | -0.489133595 | 0.113 | 0.182 | 1 | 0.291084876 | -0.410722664 | 0.098 | 0.164 | 1 | 0.291084876 | 0.415026387 |
| Ube2e2 | 0.235284537 | 0.20274411 | 0.849 | 0.822 | 1 | 0.430634922 | -0.249039327 | 0.829 | 0.857 | 1 | 0.430634922 | 0.415210261 |
| Abca3 | 0.235289341 | 1.100274811 | 0.151 | 0.097 | 1 | 0.580769483 | 0.547698233 | 0.122 | 0.096 | 1 | 0.580769483 | 0.415217608 |
| Gm45025 | 0.675354755 | 0.154722595 | 0.132 | 0.11 | 1 | 0.235860429 | -0.622226769 | 0.098 | 0.171 | 1 | 0.675354755 | 0.416090717 |
| Fzd10os | 0.679455468 | -0.260314904 | 0.038 | 0.051 | 1 | 0.23595121 | 0.479652845 | 0.098 | 0.191 | 1 | 0.679455468 | 0.416229446 |
| Kbtbd11 | 0.236091249 | 1.47665069 | 0.057 | 0.025 | 1 | 0.476434136 | 1.071670103 | 0.073 | 0.048 | 1 | 0.476434136 | 0.416443421 |
| Elmo3 | 0.236091249 | 1.47665069 | 0.057 | 0.025 | 1 | 0.708451968 | 0.444887427 | 0.073 | 0.058 | 1 | 0.708451968 | 0.416443421 |
| Rtn4ip1 | 0.236091249 | 1.47665069 | 0.057 | 0.025 | 1 | 0.624419251 | 0.334704509 | 0.024 | 0.041 | 1 | 0.624419251 | 0.416443421 |
| Polrmt | 0.236091249 | 1.47665069 | 0.057 | 0.025 | 1 | 0.409125717 | -0.410722664 | 0.024 | 0.055 | 1 | 0.409125717 | 0.416443421 |
| Xkr6 | 0.555484956 | 0.884633431 | 0.151 | 0.123 | 1 | 0.236240362 | -1.410722664 | 0.049 | 0.106 | 1 | 0.555484956 | 0.416671216 |
| Ncdn | 0.716332981 | 0.8327945 | 0.094 | 0.081 | 1 | 0.236281389 | 1.452541 | 0.195 | 0.137 | 1 | 0.716332981 | 0.416733884 |
| Nsmce1 | 0.779410121 | 0.392221909 | 0.302 | 0.301 | 1 | 0.236407155 | -0.520347155 | 0.22 | 0.3 | 1 | 0.779410121 | 0.416925967 |
| Tbc1d19 | 0.236614571 | 0.695290976 | 0.283 | 0.22 | 1 | 0.535445643 | 0.34953955 | 0.317 | 0.27 | 1 | 0.535445643 | 0.417242687 |
| Hcls1 | 0.332821087 | 1.8327945 | 0.038 | 0.017 | 1 | 0.236875607 | 0.83720485 | 0.073 | 0.034 | 1 | 0.332821087 | 0.417641161 |
| Acsf3 | 0.236890229 | 1.154722595 | 0.075 | 0.038 | 1 | 0.371357099 | -0.484723245 | 0.024 | 0.058 | 1 | 0.371357099 | 0.417663478 |
| Gal3st4 | 0.732315363 | -0.757814564 | 0.226 | 0.237 | 1 | 0.236898584 | 0.625154373 | 0.317 | 0.246 | 1 | 0.732315363 | 0.417676228 |
| Gm41664 | 0.681605053 | 0.154722595 | 0.057 | 0.042 | 1 | 0.236907891 | 0.911205431 | 0.073 | 0.034 | 1 | 0.681605053 | 0.417690433 |
| Tmem107 | 0.236912726 | 0.672329828 | 0.34 | 0.284 | 1 | 0.582836027 | -0.318073376 | 0.195 | 0.232 | 1 | 0.582836027 | 0.417697813 |
| Dot1l | 0.237018208 | -0.101222386 | 0.547 | 0.72 | 1 | 0.528550708 | -0.288326032 | 0.805 | 0.829 | 1 | 0.528550708 | 0.417858786 |
| Pinx1 | 0.576948666 | -0.193200709 | 0.132 | 0.165 | 1 | 0.237302913 | 0.280811501 | 0.146 | 0.225 | 1 | 0.576948666 | 0.418293153 |
| Ctps2 | 0.67938589 | 0.268379377 | 0.321 | 0.314 | 1 | 0.237361435 | -0.402261085 | 0.341 | 0.447 | 1 | 0.67938589 | 0.41838242 |
| Dcaf11 | 0.237460847 | 0.584866986 | 0.377 | 0.314 | 1 | 0.353565227 | 0.600165653 | 0.39 | 0.341 | 1 | 0.353565227 | 0.41853404 |
| Trub1 | 0.37341076 | 0.865215978 | 0.113 | 0.076 | 1 | 0.237489964 | 1.01123425 | 0.146 | 0.092 | 1 | 0.37341076 | 0.418578444 |
| Nrep | 0.329552482 | -0.173483947 | 0.868 | 0.784 | 1 | 0.237497473 | 0.427595395 | 0.805 | 0.805 | 1 | 0.329552482 | 0.418589896 |
| Ptgis | 0.861317891 | -0.187755337 | 0.943 | 0.822 | 1 | 0.237641935 | -0.309367504 | 0.756 | 0.846 | 1 | 0.861317891 | 0.418810181 |
| Ybey | 0.237895302 | 1.039245377 | 0.075 | 0.038 | 1 | 0.37389475 | -0.332720152 | 0.024 | 0.058 | 1 | 0.37389475 | 0.419196429 |
| Igf2bp1 | 0.237936014 | 1.055186921 | 0.075 | 0.038 | 1 | 0.940361588 | -0.116991461 | 0.049 | 0.051 | 1 | 0.940361588 | 0.419258482 |
| Brms1 | 0.820836852 | 0.198665942 | 0.302 | 0.305 | 1 | 0.237951161 | -0.536253546 | 0.195 | 0.283 | 1 | 0.820836852 | 0.419281567 |
| Gm13778 | 0.237970389 | 1.154722595 | 0.075 | 0.038 | 1 | 0.448202969 | -0.555112573 | 0.024 | 0.051 | 1 | 0.448202969 | 0.419310872 |
| Lman1l | 0.237970389 | 1.154722595 | 0.075 | 0.038 | 1 | 0.712566948 | 0.627751484 | 0.098 | 0.082 | 1 | 0.712566948 | 0.419310872 |
| Camk2b | 0.238000476 | 0.8327945 | 0.132 | 0.081 | 1 | 0.424965286 | 0.707921833 | 0.122 | 0.085 | 1 | 0.424965286 | 0.419356726 |
| Rbbp8 | 0.238169379 | 0.20839126 | 0.585 | 0.525 | 1 | 0.334703834 | 0.308095584 | 0.659 | 0.59 | 1 | 0.334703834 | 0.419614105 |
| Gm11210 | 0.238299361 | 1.216123139 | 0.113 | 0.068 | 1 | 0.54534378 | 0.915207362 | 0.195 | 0.164 | 1 | 0.54534378 | 0.419812137 |
| Mfn1 | 0.325030249 | 0.490142926 | 0.434 | 0.39 | 1 | 0.238465782 | -0.339672912 | 0.39 | 0.485 | 1 | 0.325030249 | 0.420065634 |
| Trim35 | 0.793958706 | 0.10741688 | 0.358 | 0.364 | 1 | 0.238505813 | 0.711096081 | 0.463 | 0.423 | 1 | 0.793958706 | 0.420126603 |
| Dusp22 | 0.238511706 | -0.55223043 | 0.245 | 0.314 | 1 | 0.269196127 | -0.764831164 | 0.122 | 0.191 | 1 | 0.269196127 | 0.420135579 |
| Kcnq3 | 0.510488781 | -0.697720217 | 0.094 | 0.127 | 1 | 0.238593511 | 0.614812428 | 0.146 | 0.089 | 1 | 0.510488781 | 0.420260159 |
| Jmjd7 | 0.238834387 | 0.652222254 | 0.094 | 0.051 | 1 | 0.89988627 | 0.589277336 | 0.049 | 0.055 | 1 | 0.89988627 | 0.420626909 |
| Gm47664 | 0.238834738 | 0.739685096 | 0.094 | 0.051 | 1 | 0.260704079 | 2.029849928 | 0.049 | 0.02 | 1 | 0.260704079 | 0.420627444 |
| Stub1 | 0.23884972 | 0.361173472 | 0.774 | 0.703 | 1 | 0.651354108 | 0.15834221 | 0.683 | 0.666 | 1 | 0.651354108 | 0.420650252 |
| Mtch2 | 0.238880181 | 0.259253178 | 0.755 | 0.746 | 1 | 0.685876348 | 0.255451906 | 0.732 | 0.761 | 1 | 0.685876348 | 0.420696622 |
| Mrrf | 0.238937653 | -0.357727406 | 0.245 | 0.331 | 1 | 0.493152368 | -0.107653596 | 0.146 | 0.195 | 1 | 0.493152368 | 0.420784104 |
| Dnajc2 | 0.609984853 | -0.142957954 | 0.377 | 0.432 | 1 | 0.238979641 | 0.545178241 | 0.512 | 0.444 | 1 | 0.609984853 | 0.420848013 |
| Entpd5 | 0.239123632 | 1.110328475 | 0.189 | 0.136 | 1 | 0.41932075 | 0.186950889 | 0.341 | 0.28 | 1 | 0.41932075 | 0.421067153 |
| Bmp7 | 0.258516252 | 0.848914165 | 0.264 | 0.199 | 1 | 0.239294276 | 0.401105735 | 0.268 | 0.188 | 1 | 0.258516252 | 0.421326801 |
| Rock2 | 0.674774168 | -0.142052023 | 1 | 0.992 | 1 | 0.239397128 | 0.165381509 | 1 | 1 | 1 | 0.674774168 | 0.421483272 |
| Ggt7 | 0.239709976 | 1.039245377 | 0.057 | 0.025 | 1 | 0.455097252 | -0.747757651 | 0 | 0.014 | 1 | 0.455097252 | 0.421959079 |
| Gm15345 | 0.239709976 | 1.306725688 | 0.057 | 0.025 | 1 | 0.286413984 | -0.970150072 | 0 | 0.027 | 1 | 0.286413984 | 0.421959079 |
| Pla2g4b | 0.239732556 | 1.347367673 | 0.057 | 0.025 | 1 | 0.27744837 | -0.80665134 | 0.024 | 0.068 | 1 | 0.27744837 | 0.421993414 |
| Tmem91 | 0.239732556 | 1.347367673 | 0.057 | 0.025 | 1 | 0.741574404 | -0.101394606 | 0.049 | 0.061 | 1 | 0.741574404 | 0.421993414 |
| C330011M18Rik | 0.239732556 | 1.347367673 | 0.057 | 0.025 | 1 | 0.288314827 | 1.252242349 | 0.073 | 0.038 | 1 | 0.288314827 | 0.421993414 |
| Mc1r | 0.239732556 | 1.347367673 | 0.057 | 0.025 | 1 | 0.402364804 | -0.16279515 | 0 | 0.017 | 1 | 0.402364804 | 0.421993414 |
| BC065397 | 0.332470232 | 0.517292674 | 0.113 | 0.072 | 1 | 0.239762478 | 1.045791472 | 0.146 | 0.089 | 1 | 0.332470232 | 0.422038911 |
| Gtf3c6 | 0.263968929 | 0.645708947 | 0.321 | 0.258 | 1 | 0.239792192 | -0.238083278 | 0.22 | 0.314 | 1 | 0.263968929 | 0.422084088 |
| Hsdl2 | 0.536096253 | 0.292226119 | 0.283 | 0.242 | 1 | 0.239891048 | 0.586243276 | 0.366 | 0.276 | 1 | 0.536096253 | 0.422234381 |
| Fendrr | 0.240125386 | -0.312403416 | 0.17 | 0.263 | 1 | 0.487558021 | -0.577832649 | 0.22 | 0.259 | 1 | 0.487558021 | 0.422590571 |
| G6pdx | 0.240449503 | 0.849868013 | 0.208 | 0.148 | 1 | 0.555694186 | -0.348661696 | 0.171 | 0.212 | 1 | 0.555694186 | 0.423083043 |
| Tbl2 | 0.24050192 | 0.748730235 | 0.264 | 0.203 | 1 | 0.643485232 | 0.578470581 | 0.22 | 0.198 | 1 | 0.643485232 | 0.423162667 |
| Ctf1 | 0.406504033 | -0.292736382 | 0.151 | 0.203 | 1 | 0.240622679 | -0.221688839 | 0.171 | 0.259 | 1 | 0.406504033 | 0.423346084 |
| Gm9968 | 0.305657989 | 1.055186921 | 0.075 | 0.042 | 1 | 0.240766646 | 1.071670103 | 0.122 | 0.072 | 1 | 0.305657989 | 0.423564714 |
| Nfe2l2 | 0.241042055 | 0.168721902 | 0.906 | 0.86 | 1 | 0.391114321 | 0.224101869 | 0.951 | 0.925 | 1 | 0.391114321 | 0.423982837 |
| Fam135a | 0.296157924 | -0.198648548 | 0.642 | 0.725 | 1 | 0.241272473 | -0.324258573 | 0.561 | 0.669 | 1 | 0.296157924 | 0.42433254 |
| Tubb3 | 0.241367687 | 1.154722595 | 0.132 | 0.085 | 1 | 0.806223616 | 0.100239256 | 0.049 | 0.058 | 1 | 0.806223616 | 0.424477014 |

| | | | | | | | | | | | | |
|---|---|---|---|---|---|---|---|---|---|---|---|---|
| Ttc36 | 0.241482942 | -0.652632327 | 0.019 | 0.059 | 1 | 0.402373368 | 0.252242349 | 0 | 0.017 | 1 | 0.402373368 | 0.424651873 |
| Gm48226 | 0.294375565 | 0.739685096 | 0.17 | 0.119 | 1 | 0.241568232 | 1.114738825 | 0.146 | 0.089 | 1 | 0.294375565 | 0.424781254 |
| Tmem181a | 0.385899991 | 0.569760094 | 0.321 | 0.288 | 1 | 0.241749157 | 0.580865097 | 0.341 | 0.266 | 1 | 0.385899991 | 0.42505566 |
| K230015D01Rik | 0.241893059 | 1.017219071 | 0.075 | 0.038 | 1 | 0.257285084 | -0.622226769 | 0 | 0.031 | 1 | 0.257285084 | 0.425273866 |
| Vps28 | 0.242347997 | -0.268488836 | 0.642 | 0.686 | 1 | 0.978689271 | 0.391218762 | 0.537 | 0.59 | 1 | 0.978689271 | 0.425963443 |
| Ftl1-ps1 | 0.266729528 | 0.734390796 | 0.377 | 0.331 | 1 | 0.242741111 | 0.544423101 | 0.512 | 0.464 | 1 | 0.266729528 | 0.426558975 |
| Tmem126b | 0.242784489 | 0.652222254 | 0.189 | 0.127 | 1 | 0.381576586 | -0.188330242 | 0.098 | 0.15 | 1 | 0.381576586 | 0.42662467 |
| Cadps2 | 0.242988571 | 0.8327945 | 0.075 | 0.038 | 1 | 0.344006811 | 0.326242931 | 0.22 | 0.157 | 1 | 0.344006811 | 0.426933696 |
| Cd274 | 0.997810947 | 1.261637799 | 0.038 | 0.038 | 1 | 0.24308817 | 0.444887427 | 0.073 | 0.034 | 1 | 0.997810947 | 0.427084482 |
| Gm41307 | 0.761196836 | -0.783876861 | 0.038 | 0.047 | 1 | 0.243088508 | 0.83720485 | 0.073 | 0.034 | 1 | 0.761196836 | 0.427084993 |
| Vwa3b | 0.24309253 | -1.015202407 | 0 | 0.025 | 1 | 0.27192168 | 1.422167351 | 0.049 | 0.02 | 1 | 0.27192168 | 0.427091082 |
| Gm28411 | 0.24309253 | -0.845277405 | 0 | 0.025 | 1 | 0.455098275 | 0.515276755 | 0 | 0.014 | 1 | 0.455098275 | 0.427091082 |
| Hhipl2 | 0.24309253 | -0.845277405 | 0 | 0.025 | 1 | 0.824088343 | 0.377773231 | 0.024 | 0.031 | 1 | 0.824088343 | 0.427091082 |
| Tgm5 | 0.24309253 | -0.845277405 | 0 | 0.025 | 1 | 0.402373368 | 0.252242349 | 0 | 0.017 | 1 | 0.402373368 | 0.427091082 |
| Fabp12 | 0.24309253 | -0.845277405 | 0 | 0.025 | 1 | 0.435967024 | 1.614812428 | 0.024 | 0.01 | 1 | 0.435967024 | 0.427091082 |
| Atp8b5 | 0.24309253 | -0.845277405 | 0 | 0.025 | 1 | 0.402373368 | 0.252242349 | 0 | 0.017 | 1 | 0.402373368 | 0.427091082 |
| Gm57068 | 0.24309253 | -0.845277405 | 0 | 0.025 | 1 | 0.441381862 | 1.252242349 | 0.024 | 0.01 | 1 | 0.441381862 | 0.427091082 |
| C530005A16Rik | 0.24309253 | -0.845277405 | 0 | 0.025 | 1 | 0.51919986 | 0.83720485 | 0 | 0.01 | 1 | 0.51919986 | 0.427091082 |
| 1700019G24Rik | 0.24309253 | -2.652632327 | 0 | 0.025 | 1 | 0.257325989 | -0.484723245 | 0 | 0.031 | 1 | 0.257325989 | 0.427091082 |
| 2310040G24Rik | 0.24309253 | -0.845277405 | 0 | 0.025 | 1 | 0.968793924 | 2.159132945 | 0.024 | 0.024 | 1 | 0.968793924 | 0.427091082 |
| Nrn1l | 0.24309253 | -0.845277405 | 0 | 0.025 | 1 | 0.51919986 | 0.83720485 | 0 | 0.01 | 1 | 0.51919986 | 0.427091082 |
| Pkd1l1 | 0.24309253 | -0.845277405 | 0 | 0.025 | 1 | 0.430584486 | 2.422167351 | 0.024 | 0.01 | 1 | 0.430584486 | 0.427091082 |
| Plek2 | 0.24309253 | -1.015202407 | 0 | 0.025 | 1 | 0.916649754 | 0.667279848 | 0.024 | 0.027 | 1 | 0.916649754 | 0.427091082 |
| A930001M01Rik | 0.24309253 | -0.845277405 | 0 | 0.025 | 1 | 0.803586789 | 0.83720485 | 0.049 | 0.041 | 1 | 0.803586789 | 0.427091082 |
| Rmi2 | 0.24309253 | -0.845277405 | 0 | 0.025 | 1 | 0.319635005 | -0.747757651 | 0 | 0.024 | 1 | 0.319635005 | 0.427091082 |
| Gm20522 | 0.24309253 | -0.845277405 | 0 | 0.025 | 1 | 0.51919986 | 0.83720485 | 0 | 0.01 | 1 | 0.51919986 | 0.427091082 |
| Frmpd3 | 0.24309253 | -0.845277405 | 0 | 0.025 | 1 | 0.430587622 | 2.159132945 | 0.024 | 0.01 | 1 | 0.430587622 | 0.427091082 |
| Gpr37l1 | 0.243102132 | -1.1672055 | 0 | 0.025 | 1 | 0.914200263 | 0.515276755 | 0.024 | 0.027 | 1 | 0.914200263 | 0.427105618 |
| Bbox1 | 0.243102132 | -1.015202407 | 0 | 0.025 | 1 | 0.5191986 | 0.515276755 | 0 | 0.01 | 1 | 0.5191986 | 0.427105618 |
| Tbc1d2 | 0.243102132 | -1.015202407 | 0 | 0.025 | 1 | 0.286413984 | -0.747757651 | 0 | 0.027 | 1 | 0.286413984 | 0.427105618 |
| Ccdc158 | 0.243102132 | -1.015202407 | 0 | 0.025 | 1 | 0.455094184 | 0.252242349 | 0 | 0.014 | 1 | 0.455094184 | 0.427105618 |
| Mapk10 | 0.243102132 | -1.304709024 | 0 | 0.025 | 1 | 0.293979622 | 0.898605394 | 0.073 | 0.038 | 1 | 0.293979622 | 0.427105618 |
| Gm45518 | 0.243102132 | -1.1672055 | 0 | 0.025 | 1 | 0.752578347 | 0.252242349 | 0.024 | 0.017 | 1 | 0.752578347 | 0.427105618 |
| Ccdc113 | 0.243102132 | -1.015202407 | 0 | 0.025 | 1 | 0.362195044 | 1.252242349 | 0.049 | 0.024 | 1 | 0.362195044 | 0.427105618 |
| Klhl31 | 0.243102132 | -1.015202407 | 0 | 0.025 | 1 | 0.319638803 | -1.332720152 | 0 | 0.024 | 1 | 0.319638803 | 0.427105618 |
| Spata32 | 0.243102132 | -1.1672055 | 0 | 0.025 | 1 | 0.666353766 | -0.555112573 | 0.024 | 0.038 | 1 | 0.666353766 | 0.427105618 |
| Gm5444 | 0.243102132 | -1.752168001 | 0 | 0.025 | 1 | 0.402372511 | -0.970150072 | 0 | 0.017 | 1 | 0.402372511 | 0.427105618 |
| Gm41541 | 0.243102132 | -1.1672055 | 0 | 0.025 | 1 | 0.267205177 | 2.252242349 | 0.024 | 0.007 | 1 | 0.267205177 | 0.427105618 |
| A330032B11Rik | 0.243102132 | -1.430239906 | 0 | 0.025 | 1 | 0.319641967 | -0.16279515 | 0 | 0.024 | 1 | 0.319641967 | 0.427105618 |
| Galntl6 | 0.243106933 | -1.304709024 | 0 | 0.025 | 1 | 0.402373368 | 0.252242349 | 0 | 0.017 | 1 | 0.402373368 | 0.427112886 |
| Ngb | 0.243106933 | -1.304709024 | 0 | 0.025 | 1 | 0.455098275 | 0.515276755 | 0 | 0.014 | 1 | 0.455098275 | 0.427112886 |
| Gm41031 | 0.243106933 | -1.845277405 | 0 | 0.025 | 1 | 0.51919986 | 0.252242349 | 0 | 0.01 | 1 | 0.51919986 | 0.427112886 |
| Vwc2 | 0.243107894 | -1.752168001 | 0 | 0.025 | 1 | 0.267205177 | 2.252242349 | 0.024 | 0.007 | 1 | 0.267205177 | 0.427114339 |
| Kcnj16 | 0.243107894 | -1.1672055 | 0 | 0.025 | 1 | 0.254570244 | 0.557096931 | 0.146 | 0.089 | 1 | 0.254570244 | 0.427114339 |
| Rgs6 | 0.243107894 | -1.545717123 | 0 | 0.025 | 1 | 0.611660813 | -1.359192363 | 0.073 | 0.096 | 1 | 0.611660813 | 0.427114339 |
| Cdkn3 | 0.243107894 | -1.752168001 | 0 | 0.025 | 1 | 0.58792152 | 3.974708374 | 0.024 | 0.014 | 1 | 0.58792152 | 0.427114339 |
| T2 | 0.243107894 | -1.1672055 | 0 | 0.025 | 1 | 0.319629309 | -0.747757651 | 0 | 0.024 | 1 | 0.319629309 | 0.427114339 |
| H2ac24 | 0.243109814 | -1.932740246 | 0 | 0.025 | 1 | 0.51919986 | 0.83720485 | 0 | 0.01 | 1 | 0.51919986 | 0.427117246 |
| Tmem169 | 0.243111734 | -0.652632327 | 0 | 0.025 | 1 | 0.455763738 | 1.100239256 | 0.049 | 0.027 | 1 | 0.455763738 | 0.427120153 |
| Serpinb5 | 0.243111734 | -0.652632327 | 0 | 0.025 | 1 | 0.27192168 | 1.422167351 | 0.049 | 0.02 | 1 | 0.27192168 | 0.427120153 |
| Gm16098 | 0.243111734 | -0.652632327 | 0 | 0.025 | 1 | 0.402364804 | -0.16279515 | 0 | 0.017 | 1 | 0.402364804 | 0.427120153 |
| 9130604C24Rik | 0.243111734 | -0.652632327 | 0 | 0.025 | 1 | 0.402373368 | 0.252242349 | 0 | 0.017 | 1 | 0.402373368 | 0.427120153 |
| Ano2 | 0.243111734 | -0.652632327 | 0 | 0.025 | 1 | 0.402372511 | -0.332720152 | 0 | 0.017 | 1 | 0.402372511 | 0.427120153 |
| Gm15635 | 0.243111734 | -0.652632327 | 0 | 0.025 | 1 | 0.51919986 | 0.83720485 | 0 | 0.01 | 1 | 0.51919986 | 0.427120153 |
| Irag1 | 0.243111734 | -0.652632327 | 0 | 0.025 | 1 | 0.357858058 | -0.747757651 | 0 | 0.02 | 1 | 0.357858058 | 0.427120153 |
| Gm2814 | 0.243111734 | -0.652632327 | 0 | 0.025 | 1 | 0.319641967 | -0.16279515 | 0 | 0.024 | 1 | 0.319641967 | 0.427120153 |
| Gm30978 | 0.243111734 | -0.652632327 | 0 | 0.025 | 1 | 0.286413984 | -0.622226769 | 0 | 0.027 | 1 | 0.286413984 | 0.427120153 |
| D830035M03Rik | 0.243111734 | -0.652632327 | 0 | 0.025 | 1 | 0.25523664 | 2.159132945 | 0.049 | 0.02 | 1 | 0.25523664 | 0.427120153 |
| Gm57036 | 0.243111734 | -0.652632327 | 0 | 0.025 | 1 | 0.433272494 | 2.100239256 | 0.024 | 0.01 | 1 | 0.433272494 | 0.427120153 |
| 4933433G19Rik | 0.243111734 | -0.652632327 | 0 | 0.025 | 1 | 0.875461105 | 0.252242349 | 0.049 | 0.055 | 1 | 0.875461105 | 0.427120153 |
| Tprg | 0.243111734 | -0.652632327 | 0 | 0.025 | 1 | 0.430587622 | 2.574170444 | 0.024 | 0.01 | 1 | 0.430587622 | 0.427120153 |
| Zfp474 | 0.243111734 | -0.652632327 | 0 | 0.025 | 1 | 0.51919986 | 0.83720485 | 0 | 0.01 | 1 | 0.51919986 | 0.427120153 |
| A730017L22Rik | 0.243322126 | 1.154722595 | 0.057 | 0.025 | 1 | 0.990690153 | 0.174239837 | 0.049 | 0.048 | 1 | 0.990690153 | 0.427438595 |
| Il12rb2 | 0.243322126 | 0.984797593 | 0.057 | 0.025 | 1 | 0.91185603 | 0.100239256 | 0.049 | 0.044 | 1 | 0.91185603 | 0.427438595 |
| Zfp526 | 0.243322126 | 1.154722595 | 0.057 | 0.025 | 1 | 0.323788749 | 2.306690133 | 0.073 | 0.041 | 1 | 0.323788749 | 0.427438595 |
| Nat2 | 0.243322126 | 1.154722595 | 0.057 | 0.025 | 1 | 0.878310931 | 0.667279848 | 0.024 | 0.02 | 1 | 0.878310931 | 0.427438595 |
| Zfp229 | 0.243322126 | 1.154722595 | 0.057 | 0.025 | 1 | 0.329522958 | 0.962735732 | 0.098 | 0.058 | 1 | 0.329522958 | 0.427438595 |
| 9530082P21Rik | 0.243322126 | 1.154722595 | 0.057 | 0.025 | 1 | 0.746881137 | 1.029849928 | 0.024 | 0.017 | 1 | 0.746881137 | 0.427438595 |
| Spdef | 0.243322126 | 0.8327945 | 0.057 | 0.025 | 1 | 0.958410232 | 0.377773231 | 0.049 | 0.051 | 1 | 0.958410232 | 0.427438595 |
| Gm20508 | 0.243322126 | 1.154722595 | 0.057 | 0.025 | 1 | 0.462888791 | 0.721727632 | 0.049 | 0.027 | 1 | 0.462888791 | 0.427438595 |
| Mettl21a | 0.243352133 | 0.640149422 | 0.283 | 0.212 | 1 | 0.724755376 | 0.614812428 | 0.22 | 0.212 | 1 | 0.724755376 | 0.427484006 |
| Tmem53 | 0.243779426 | 1.491757582 | 0.113 | 0.068 | 1 | 0.622636015 | -0.484723245 | 0.049 | 0.068 | 1 | 0.622636015 | 0.428130444 |
| Atxn3 | 0.24388179 | 0.444229212 | 0.302 | 0.225 | 1 | 0.290446363 | 0.539524301 | 0.317 | 0.249 | 1 | 0.290446363 | 0.428285252 |
| Gga3 | 0.258662883 | -0.889671525 | 0.057 | 0.106 | 1 | 0.244034643 | -0.863234868 | 0.049 | 0.106 | 1 | 0.258662883 | 0.428516379 |
| Zfp598 | 0.794460213 | 0.209864149 | 0.283 | 0.271 | 1 | 0.24407808 | 0.526003162 | 0.415 | 0.341 | 1 | 0.794460213 | 0.428582051 |
| Arfgef3 | 0.244369049 | -0.1672055 | 0.019 | 0.059 | 1 | 0.964417771 | 1.111379813 | 0.073 | 0.078 | 1 | 0.964417771 | 0.429021865 |
| Cc2d1b | 0.521036556 | -0.1672055 | 0.113 | 0.148 | 1 | 0.244525779 | 0.749742009 | 0.293 | 0.218 | 1 | 0.521036556 | 0.429258702 |
| Mfap3l | 0.750625248 | 0.569760094 | 0.057 | 0.047 | 1 | 0.244535097 | 1.029849928 | 0.098 | 0.051 | 1 | 0.750625248 | 0.42927278 |
| Trim69 | 0.244584181 | 1.017219071 | 0.057 | 0.025 | 1 | 0.971394446 | 1.377773231 | 0.024 | 0.024 | 1 | 0.971394446 | 0.42934694 |
| Fxyd4 | 0.244584181 | 1.017219071 | 0.057 | 0.025 | 1 | 0.40660741 | -0.686357106 | 0.024 | 0.055 | 1 | 0.40660741 | 0.42934694 |
| Nelfe | 0.244659574 | -0.247375849 | 0.245 | 0.343 | 1 | 0.900863183 | 0.404245443 | 0.244 | 0.273 | 1 | 0.900863183 | 0.42946084 |
| Sephs2 | 0.2446654 | 0.776210972 | 0.208 | 0.148 | 1 | 0.487163827 | 0.218295017 | 0.146 | 0.106 | 1 | 0.487163827 | 0.429469642 |
| Pomgnt1 | 0.270151491 | 0.551381565 | 0.321 | 0.258 | 1 | 0.244693142 | -0.320336427 | 0.195 | 0.29 | 1 | 0.270151491 | 0.429511551 |
| Hbs1l | 0.37060411 | -0.307278383 | 0.585 | 0.623 | 1 | 0.244767554 | 0.273303965 | 0.732 | 0.703 | 1 | 0.37060411 | 0.429623952 |
| Hspa4l | 0.308746343 | 0.197791317 | 0.283 | 0.216 | 1 | 0.244865981 | 0.252242349 | 0.415 | 0.338 | 1 | 0.308746343 | 0.429772613 |
| 3110056K07Rik | 0.828493607 | 0.18386894 | 0.283 | 0.288 | 1 | 0.244869561 | 0.819282942 | 0.439 | 0.369 | 1 | 0.828493607 | 0.42977802 |
| Pde6d | 0.244966986 | 0.625794129 | 0.358 | 0.322 | 1 | 0.974376017 | 0.398631836 | 0.39 | 0.423 | 1 | 0.974376017 | 0.429925147 |
| Gys1 | 0.588686021 | 0.115194231 | 0.075 | 0.102 | 1 | 0.245238242 | 0.773074512 | 0.146 | 0.089 | 1 | 0.588686021 | 0.430334689 |
| Ivns1abp | 0.245267871 | 0.133804688 | 0.943 | 0.932 | 1 | 0.81702871 | 0.118975818 | 0.927 | 0.969 | 1 | 0.81702871 | 0.430379414 |
| 2010004M13Rik | 0.58732968 | -0.20282941 | 0.132 | 0.161 | 1 | 0.245338642 | -0.825760163 | 0.049 | 0.106 | 1 | 0.58732968 | 0.430486235 |
| Aspm | 0.473741189 | 0.122301117 | 0.113 | 0.081 | 1 | 0.245391209 | 0.667279848 | 0.098 | 0.051 | 1 | 0.473741189 | 0.430565573 |
| Zfp40 | 0.245427429 | 1.46284489 | 0.113 | 0.068 | 1 | 0.609011024 | 1.737669176 | 0.049 | 0.034 | 1 | 0.609011024 | 0.430620235 |
| Thoc2l | 0.887723438 | 0.137988579 | 0.736 | 0.831 | 1 | 0.245579349 | 0.434858072 | 0.854 | 0.829 | 1 | 0.887723438 | 0.430849481 |
| Plekhm3 | 0.245682787 | 0.932330173 | 0.245 | 0.186 | 1 | 0.449049784 | 0.35607816 | 0.171 | 0.126 | 1 | 0.449049784 | 0.431005542 |
| Dus1l | 0.246004563 | 0.970298024 | 0.151 | 0.097 | 1 | 0.446954486 | -0.622226769 | 0.049 | 0.082 | 1 | 0.446954486 | 0.431490882 |
| Tmem143 | 0.246023494 | 0.776210972 | 0.075 | 0.038 | 1 | 0.451337649 | -0.53202896 | 0.049 | 0.082 | 1 | 0.451337649 | 0.431519428 |
| Spata5 | 0.907381116 | 0.140828153 | 0.623 | 0.695 | 1 | 0.246134777 | 0.136765132 | 0.78 | 0.652 | 1 | 0.907381116 | 0.431687225 |
| Dpm3 | 0.246315484 | 0.185973529 | 0.755 | 0.631 | 1 | 0.857181321 | 0.322030679 | 0.683 | 0.652 | 1 | 0.857181321 | 0.43195965 |

| | | | | | | | | | | | | |
|---|---|---|---|---|---|---|---|---|---|---|---|---|
| Gm11713 | 0.566042372 | -0.278236812 | 0.057 | 0.081 | 1 | 0.246318788 | -0.300298674 | 0.049 | 0.109 | 1 | 0.566042372 | 0.431964631 |
| Gtf2h5 | 0.511837519 | -0.102232643 | 0.755 | 0.814 | 1 | 0.246340298 | 0.350580268 | 0.854 | 0.744 | 1 | 0.511837519 | 0.431997054 |
| Slc11a1 | 0.246443787 | 2.739685096 | 0.019 | 0.004 | 1 | 0.438672228 | 1.83720485 | 0.024 | 0.01 | 1 | 0.438672228 | 0.432153034 |
| Apbb1ip | 0.246443787 | 3.002719501 | 0.019 | 0.004 | 1 | 0.739624612 | -0.555112573 | 0.024 | 0.034 | 1 | 0.739624612 | 0.432153034 |
| Cytip | 0.246443787 | 3.324647596 | 0.019 | 0.004 | 1 | 0.5191986 | 0.252242349 | 0 | 0.01 | 1 | 0.5191986 | 0.432153034 |
| P2ry12 | 0.246443787 | 3.154722595 | 0.019 | 0.004 | 1 | 0.58792152 | 2.100239256 | 0.024 | 0.014 | 1 | 0.58792152 | 0.432153034 |
| Zfp984 | 0.246443787 | 2.739685096 | 0.019 | 0.004 | 1 | 0.872828964 | 1.029849928 | 0.024 | 0.02 | 1 | 0.872828964 | 0.432153034 |
| Tmem150c | 0.246443787 | 2.739685096 | 0.019 | 0.004 | 1 | 0.840075108 | 0.749742009 | 0.024 | 0.031 | 1 | 0.840075108 | 0.432153034 |
| Nptx2 | 0.246443787 | 2.739685096 | 0.019 | 0.004 | 1 | 0.402373368 | 0.252242349 | 0 | 0.017 | 1 | 0.402373368 | 0.432153034 |
| Stk31 | 0.246443787 | 3.739685096 | 0.019 | 0.004 | 1 | 0.455098275 | 0.515276755 | 0 | 0.014 | 1 | 0.455098275 | 0.432153034 |
| 4930479D17Rik | 0.246443787 | 2.739685096 | 0.019 | 0.004 | 1 | 0.262676008 | 2.83720485 | 0.024 | 0.007 | 1 | 0.262676008 | 0.432153034 |
| Apoc1 | 0.246443787 | 2.739685096 | 0.019 | 0.004 | 1 | 0.319641967 | -0.16279515 | 0 | 0.024 | 1 | 0.319641967 | 0.432153034 |
| Ccdc81 | 0.246443787 | 3.739685096 | 0.019 | 0.004 | 1 | 0.430584486 | 3.422167351 | 0.024 | 0.01 | 1 | 0.430584486 | 0.432153034 |
| Tomt | 0.246443787 | 2.739685096 | 0.019 | 0.004 | 1 | 0.5191986 | -0.16279515 | 0 | 0.01 | 1 | 0.5191986 | 0.432153034 |
| Adam5 | 0.246443787 | 3.739685096 | 0.019 | 0.004 | 1 | 0.271787759 | 1.515276755 | 0.024 | 0.007 | 1 | 0.271787759 | 0.432153034 |
| B230317F23Rik | 0.246443787 | 3.154722595 | 0.019 | 0.004 | 1 | 0.26948834 | 1.83720485 | 0.024 | 0.007 | 1 | 0.26948834 | 0.432153034 |
| Lilrb4a | 0.246443787 | 3.47665069 | 0.019 | 0.004 | 1 | 0.26948834 | 0.515276755 | 0.024 | 0.007 | 1 | 0.26948834 | 0.432153034 |
| Ascl4 | 0.246443787 | 2.739685096 | 0.019 | 0.004 | 1 | 0.269117553 | 1.614812428 | 0.049 | 0.02 | 1 | 0.269117553 | 0.432153034 |
| Abca13 | 0.246443787 | 3.154722595 | 0.019 | 0.004 | 1 | 0.396115178 | 0.778311161 | 0.122 | 0.082 | 1 | 0.396115178 | 0.432153034 |
| Tc2n | 0.246443787 | 2.739685096 | 0.019 | 0.004 | 1 | 0.51919986 | 0.83720485 | 0 | 0.01 | 1 | 0.51919986 | 0.432153034 |
| Samsn1 | 0.246443787 | 2.891688189 | 0.019 | 0.004 | 1 | 0.5191986 | -1.332720152 | 0 | 0.01 | 1 | 0.5191986 | 0.432153034 |
| Fpr3 | 0.246443787 | 3.154722595 | 0.019 | 0.004 | 1 | 0.262674569 | 3.252242349 | 0.024 | 0.007 | 1 | 0.262674569 | 0.432153034 |
| Dnah8 | 0.246443787 | 2.569760094 | 0.019 | 0.004 | 1 | 0.457570128 | 0.930314254 | 0.049 | 0.027 | 1 | 0.457570128 | 0.432153034 |
| Mesd | 0.279258448 | 0.420704285 | 0.66 | 0.665 | 1 | 0.246502951 | 0.402893338 | 0.659 | 0.676 | 1 | 0.279258448 | 0.432242197 |
| Gemin5 | 0.246616621 | 0.500219161 | 0.321 | 0.25 | 1 | 0.456332382 | 0.91966701 | 0.22 | 0.276 | 1 | 0.456332382 | 0.432413484 |
| Gemin6 | 0.292065803 | 0.8327945 | 0.094 | 0.055 | 1 | 0.246642531 | -0.825760163 | 0.049 | 0.106 | 1 | 0.292065803 | 0.432452523 |
| Patz1 | 0.246673171 | 0.42680214 | 0.226 | 0.157 | 1 | 0.376227582 | -0.132421501 | 0.171 | 0.242 | 1 | 0.376227582 | 0.432498689 |
| Zfp113 | 0.779828588 | 0.154722595 | 0.075 | 0.064 | 1 | 0.246757312 | 1.252242349 | 0.122 | 0.072 | 1 | 0.779828588 | 0.432625453 |
| Gm7160 | 0.247022813 | 0.984797593 | 0.057 | 0.025 | 1 | 0.91185603 | 0.100239256 | 0.049 | 0.044 | 1 | 0.91185603 | 0.433025356 |
| Gm57272 | 0.247022813 | 0.984797593 | 0.057 | 0.025 | 1 | 0.744047924 | 1.252242349 | 0.024 | 0.017 | 1 | 0.744047924 | 0.433025356 |
| Ccdc106 | 0.247022813 | 0.8327945 | 0.057 | 0.025 | 1 | 0.672172778 | 0.136765132 | 0.024 | 0.038 | 1 | 0.672172778 | 0.433025356 |
| Gm32687 | 0.247022813 | 0.984797593 | 0.057 | 0.025 | 1 | 0.546365767 | 1.100239256 | 0.049 | 0.031 | 1 | 0.546365767 | 0.433025356 |
| Gm57349 | 0.247022813 | 0.569760094 | 0.057 | 0.025 | 1 | 0.746881137 | 1.029849928 | 0.024 | 0.017 | 1 | 0.746881137 | 0.433025356 |
| Hsbp1l1 | 0.247022813 | 0.984797593 | 0.057 | 0.025 | 1 | 0.347784376 | 1.83720485 | 0.049 | 0.024 | 1 | 0.347784376 | 0.433025356 |
| Rin1 | 0.247022813 | 0.8327945 | 0.057 | 0.025 | 1 | 0.357854401 | -0.622226769 | 0 | 0.02 | 1 | 0.357854401 | 0.433025356 |
| Miga1 | 0.5458686 | 0.176096246 | 0.17 | 0.212 | 1 | 0.247296896 | 1.034437197 | 0.39 | 0.338 | 1 | 0.5458686 | 0.433438037 |
| Kcnj13 | 0.503971666 | 1.569760094 | 0.019 | 0.008 | 1 | 0.247336483 | -1.250257991 | 0.024 | 0.072 | 1 | 0.503971666 | 0.43349763 |
| Mettl1 | 0.247346978 | 0.618669695 | 0.547 | 0.517 | 1 | 0.731889374 | 0.150362735 | 0.39 | 0.471 | 1 | 0.731889374 | 0.433513429 |
| Nipsnap1 | 0.247808368 | 0.631160639 | 0.132 | 0.081 | 1 | 0.9339635 | 0.396632258 | 0.122 | 0.119 | 1 | 0.9339635 | 0.434207748 |
| Zbtb18 | 0.247902855 | 0.417757001 | 0.094 | 0.051 | 1 | 0.489719381 | 0.778311161 | 0.098 | 0.068 | 1 | 0.489719381 | 0.434349884 |
| Dubr | 0.760979569 | 0.207190015 | 0.094 | 0.081 | 1 | 0.247936015 | 0.685201756 | 0.146 | 0.089 | 1 | 0.760979569 | 0.434399762 |
| Zdhhc2 | 0.991514454 | 1.122301117 | 0.094 | 0.097 | 1 | 0.248407689 | -1.537834582 | 0.049 | 0.106 | 1 | 0.991514454 | 0.435108999 |
| Cox19 | 0.338603974 | 0.338029808 | 0.491 | 0.436 | 1 | 0.248594935 | 0.361471419 | 0.488 | 0.423 | 1 | 0.338603974 | 0.435390428 |
| Rsf1os1 | 0.929410101 | 0.569760094 | 0.019 | 0.017 | 1 | 0.248852431 | 0.699701326 | 0.098 | 0.051 | 1 | 0.929410101 | 0.435777329 |
| Gpr153 | 0.248879265 | 0.51531231 | 0.34 | 0.271 | 1 | 0.457666294 | -0.332720152 | 0.439 | 0.474 | 1 | 0.457666294 | 0.435817641 |
| Gm57175 | 0.249034808 | 2.154722595 | 0.019 | 0.004 | 1 | 0.455098275 | 0.515276755 | 0 | 0.014 | 1 | 0.455098275 | 0.43605128 |
| Card9 | 0.249034808 | 2.154722595 | 0.019 | 0.004 | 1 | 0.267205177 | 2.252242349 | 0.024 | 0.007 | 1 | 0.267205177 | 0.43605128 |
| Galnt13 | 0.249034808 | 2.154722595 | 0.019 | 0.004 | 1 | 0.43828695 | 2.47463477 | 0.049 | 0.027 | 1 | 0.43828695 | 0.43605128 |
| Pard6b | 0.249034808 | 2.154722595 | 0.019 | 0.004 | 1 | 0.319619815 | -0.332720152 | 0 | 0.024 | 1 | 0.319619815 | 0.43605128 |
| Oaz3 | 0.249034808 | 2.154722595 | 0.019 | 0.004 | 1 | 0.60213981 | 0.83720485 | 0.024 | 0.014 | 1 | 0.60213981 | 0.43605128 |
| Mcoln3 | 0.249034808 | 2.154722595 | 0.019 | 0.004 | 1 | 0.5191986 | 0.515276755 | 0 | 0.01 | 1 | 0.5191986 | 0.43605128 |
| Wdr31 | 0.249034808 | 2.154722595 | 0.019 | 0.004 | 1 | 0.51919986 | 0.83720485 | 0 | 0.01 | 1 | 0.51919986 | 0.43605128 |
| Bspry | 0.249034808 | 2.154722595 | 0.019 | 0.004 | 1 | 0.51919986 | 0.83720485 | 0 | 0.01 | 1 | 0.51919986 | 0.43605128 |
| Zscan20 | 0.249034808 | 2.154722595 | 0.019 | 0.004 | 1 | 0.797811701 | 1.029849928 | 0.049 | 0.041 | 1 | 0.797811701 | 0.43605128 |
| Tmem156 | 0.249034808 | 2.154722595 | 0.019 | 0.004 | 1 | 0.357854401 | -0.484723245 | 0 | 0.02 | 1 | 0.357854401 | 0.43605128 |
| Gm10419 | 0.249034808 | 2.154722595 | 0.019 | 0.004 | 1 | 0.441381862 | 1.515276755 | 0.024 | 0.01 | 1 | 0.441381862 | 0.43605128 |
| Gm16120 | 0.249034808 | 2.154722595 | 0.019 | 0.004 | 1 | 0.455097252 | -0.16279515 | 0 | 0.014 | 1 | 0.455097252 | 0.43605128 |
| Akr1b8 | 0.249034808 | 2.154722595 | 0.019 | 0.004 | 1 | 0.357847088 | -0.16279515 | 0 | 0.02 | 1 | 0.357847088 | 0.43605128 |
| Ephb6 | 0.249034808 | 2.154722595 | 0.019 | 0.004 | 1 | 0.498856313 | -0.16279515 | 0.024 | 0.048 | 1 | 0.498856313 | 0.43605128 |
| 4930597O21Rik | 0.249034808 | 2.154722595 | 0.019 | 0.004 | 1 | 0.51919986 | 0.83720485 | 0 | 0.01 | 1 | 0.51919986 | 0.43605128 |
| Meiosin | 0.249034808 | 2.154722595 | 0.019 | 0.004 | 1 | 0.267205177 | 2.100239256 | 0.024 | 0.007 | 1 | 0.267205177 | 0.43605128 |
| Gm26852 | 0.249034808 | 2.154722595 | 0.019 | 0.004 | 1 | 0.357847088 | -0.332720152 | 0 | 0.02 | 1 | 0.357847088 | 0.43605128 |
| Dmkn | 0.249034808 | 2.154722595 | 0.019 | 0.004 | 1 | 0.286413984 | -0.622226769 | 0 | 0.027 | 1 | 0.286413984 | 0.43605128 |
| B230322F03Rik | 0.249034808 | 2.154722595 | 0.019 | 0.004 | 1 | 0.286433344 | -0.332720152 | 0 | 0.027 | 1 | 0.286433344 | 0.43605128 |
| Gm57226 | 0.249034808 | 2.154722595 | 0.019 | 0.004 | 1 | 0.267205177 | 2.252242349 | 0.024 | 0.007 | 1 | 0.267205177 | 0.43605128 |
| Rab38 | 0.249034808 | 2.154722595 | 0.019 | 0.004 | 1 | 0.889300905 | -0.686357106 | 0.024 | 0.02 | 1 | 0.889300905 | 0.43605128 |
| Gm35082 | 0.249034808 | 2.154722595 | 0.019 | 0.004 | 1 | 0.605005904 | 1.029849928 | 0.024 | 0.014 | 1 | 0.605005904 | 0.43605128 |
| Tmc5 | 0.249034808 | 2.154722595 | 0.019 | 0.004 | 1 | 0.402373368 | 0.252242349 | 0 | 0.017 | 1 | 0.402373368 | 0.43605128 |
| Cend1 | 0.249034808 | 2.154722595 | 0.019 | 0.004 | 1 | 0.267205177 | 2.252242349 | 0.024 | 0.007 | 1 | 0.267205177 | 0.43605128 |
| Mrgpre | 0.249034808 | 2.154722595 | 0.019 | 0.004 | 1 | 0.872828964 | 1.029849928 | 0.024 | 0.02 | 1 | 0.872828964 | 0.43605128 |
| Gm45370 | 0.249034808 | 2.154722595 | 0.019 | 0.004 | 1 | 0.452287461 | 1.252242349 | 0.049 | 0.027 | 1 | 0.452287461 | 0.43605128 |
| Gm56852 | 0.249034808 | 2.154722595 | 0.019 | 0.004 | 1 | 0.51919986 | 0.83720485 | 0 | 0.01 | 1 | 0.51919986 | 0.43605128 |
| Marchf1 | 0.249034808 | 2.154722595 | 0.019 | 0.004 | 1 | 0.51919986 | 0.83720485 | 0 | 0.01 | 1 | 0.51919986 | 0.43605128 |
| G430095P16Rik | 0.249034808 | 2.154722595 | 0.019 | 0.004 | 1 | 0.51919986 | 0.83720485 | 0 | 0.01 | 1 | 0.51919986 | 0.43605128 |
| Cdh3 | 0.249034808 | 2.154722595 | 0.019 | 0.004 | 1 | 0.430584486 | 2.83720485 | 0.024 | 0.01 | 1 | 0.430584486 | 0.43605128 |
| Gm47350 | 0.249034808 | 2.154722595 | 0.019 | 0.004 | 1 | 0.402371655 | -0.484723245 | 0 | 0.017 | 1 | 0.402371655 | 0.43605128 |
| Ccr9 | 0.249034808 | 2.154722595 | 0.019 | 0.004 | 1 | 0.402369942 | -0.332720152 | 0 | 0.017 | 1 | 0.402369942 | 0.43605128 |
| Gm30122 | 0.249034808 | 2.154722595 | 0.019 | 0.004 | 1 | 0.979524515 | 0.515276755 | 0.049 | 0.048 | 1 | 0.979524515 | 0.43605128 |
| Gm47101 | 0.249034808 | 2.154722595 | 0.019 | 0.004 | 1 | 0.455098275 | 0.515276755 | 0 | 0.014 | 1 | 0.455098275 | 0.43605128 |
| Gpr75 | 0.249034808 | 2.154722595 | 0.019 | 0.004 | 1 | 0.986995563 | 0.83720485 | 0.024 | 0.024 | 1 | 0.986995563 | 0.43605128 |
| Gm46280 | 0.249034808 | 2.154722595 | 0.019 | 0.004 | 1 | 0.402369942 | -0.484723245 | 0 | 0.017 | 1 | 0.402369942 | 0.43605128 |
| Card14 | 0.249034808 | 2.154722595 | 0.019 | 0.004 | 1 | 0.357860251 | -0.863234868 | 0 | 0.02 | 1 | 0.357860251 | 0.43605128 |
| Fn3krp | 0.249034808 | 2.154722595 | 0.019 | 0.004 | 1 | 0.402373368 | 0.252242349 | 0 | 0.017 | 1 | 0.402373368 | 0.43605128 |
| Gm29968 | 0.249034808 | 2.154722595 | 0.019 | 0.004 | 1 | 0.51919986 | 0.83720485 | 0 | 0.01 | 1 | 0.51919986 | 0.43605128 |
| Lrfn5 | 0.249034808 | 2.154722595 | 0.019 | 0.004 | 1 | 0.5191986 | -0.16279515 | 0 | 0.01 | 1 | 0.5191986 | 0.43605128 |
| Gm4756 | 0.249034808 | 2.154722595 | 0.019 | 0.004 | 1 | 0.744044166 | 0.962735732 | 0.024 | 0.017 | 1 | 0.744044166 | 0.43605128 |
| Tmem229b | 0.249034808 | 2.154722595 | 0.019 | 0.004 | 1 | 0.992196803 | 0.136765132 | 0.024 | 0.024 | 1 | 0.992196803 | 0.43605128 |
| Gm48807 | 0.249034808 | 2.154722595 | 0.019 | 0.004 | 1 | 0.51919986 | 0.83720485 | 0 | 0.01 | 1 | 0.51919986 | 0.43605128 |
| Gm35161 | 0.249034808 | 2.154722595 | 0.019 | 0.004 | 1 | 0.749728988 | 0.83720485 | 0.024 | 0.017 | 1 | 0.749728988 | 0.43605128 |
| Tmem253 | 0.249034808 | 2.154722595 | 0.019 | 0.004 | 1 | 0.286422281 | -1.069685746 | 0 | 0.027 | 1 | 0.286422281 | 0.43605128 |
| Trdv4 | 0.249034808 | 2.154722595 | 0.019 | 0.004 | 1 | 0.267205177 | 2.252242349 | 0.024 | 0.007 | 1 | 0.267205177 | 0.43605128 |
| Stmn4 | 0.249034808 | 2.154722595 | 0.019 | 0.004 | 1 | 0.5191986 | 0.515276755 | 0 | 0.01 | 1 | 0.5191986 | 0.43605128 |
| Gm17753 | 0.249034808 | 2.154722595 | 0.019 | 0.004 | 1 | 0.357858789 | -0.484723245 | 0 | 0.02 | 1 | 0.357858789 | 0.43605128 |
| Gm17546 | 0.249034808 | 2.154722595 | 0.019 | 0.004 | 1 | 0.51919986 | 0.83720485 | 0 | 0.01 | 1 | 0.51919986 | 0.43605128 |
| A330009N23Rik | 0.249034808 | 2.154722595 | 0.019 | 0.004 | 1 | 0.319619815 | -0.332720152 | 0 | 0.024 | 1 | 0.319619815 | 0.43605128 |
| Krt6b | 0.249034808 | 2.154722595 | 0.019 | 0.004 | 1 | 0.7415112 | 0.515276755 | 0.049 | 0.038 | 1 | 0.7415112 | 0.43605128 |
| Zpld1 | 0.249034808 | 2.154722595 | 0.019 | 0.004 | 1 | 0.402369942 | -0.332720152 | 0 | 0.017 | 1 | 0.402369942 | 0.43605128 |
| Gm50107 | 0.249034808 | 2.154722595 | 0.019 | 0.004 | 1 | 0.455097252 | -0.16279515 | 0 | 0.014 | 1 | 0.455097252 | 0.43605128 |

| | | | | | | | | | | | | |
|---|---|---|---|---|---|---|---|---|---|---|---|---|
| Tnxb | 0.249034808 | 2.154722595 | 0.019 | 0.004 | 1 | 0.264933923 | 2.422167351 | 0.024 | 0.007 | 1 | 0.264933923 | 0.43605128 |
| Tmem151b | 0.249034808 | 2.154722595 | 0.019 | 0.004 | 1 | 0.441381862 | 1.515276755 | 0.024 | 0.01 | 1 | 0.441381862 | 0.43605128 |
| Vav1 | 0.249034808 | 2.154722595 | 0.019 | 0.004 | 1 | 0.587925597 | 2.252242349 | 0.024 | 0.014 | 1 | 0.587925597 | 0.43605128 |
| Gm19689 | 0.249034808 | 2.154722595 | 0.019 | 0.004 | 1 | 0.430584486 | 2.422167351 | 0.024 | 0.01 | 1 | 0.430584486 | 0.43605128 |
| Pcdhb15 | 0.249034808 | 2.154722595 | 0.019 | 0.004 | 1 | 0.590756788 | 1.83720485 | 0.024 | 0.014 | 1 | 0.590756788 | 0.43605128 |
| Sptbn2 | 0.249034808 | 2.154722595 | 0.019 | 0.004 | 1 | 0.357854401 | -0.484723245 | 0 | 0.02 | 1 | 0.357854401 | 0.43605128 |
| 4930414N06Rik | 0.249034808 | 2.154722595 | 0.019 | 0.004 | 1 | 0.438672228 | 1.83720485 | 0.024 | 0.01 | 1 | 0.438672228 | 0.43605128 |
| Gm4219 | 0.249034808 | 2.154722595 | 0.019 | 0.004 | 1 | 0.51919986 | 0.83720485 | 0 | 0.01 | 1 | 0.51919986 | 0.43605128 |
| B3galnt2 | 0.644792964 | -0.130679624 | 0.283 | 0.314 | 1 | 0.249333739 | -0.21906337 | 0.268 | 0.369 | 1 | 0.644792964 | 0.436500164 |
| Plcd4 | 0.249526738 | 0.569760094 | 0.057 | 0.025 | 1 | 0.796279439 | -0.435813645 | 0.049 | 0.058 | 1 | 0.796279439 | 0.436789883 |
| Fam20c | 0.729919591 | -0.106542718 | 0.528 | 0.538 | 1 | 0.249595285 | -0.236996092 | 0.585 | 0.734 | 1 | 0.729919591 | 0.436892763 |
| Gm9856 | 0.410635679 | 0.347367673 | 0.132 | 0.093 | 1 | 0.249752505 | -1.292078167 | 0.024 | 0.072 | 1 | 0.410635679 | 0.437128696 |
| Dusp28 | 0.249922726 | -0.56774343 | 0.057 | 0.11 | 1 | 0.452331971 | -0.250257991 | 0.024 | 0.051 | 1 | 0.452331971 | 0.437384083 |
| Lman2l | 0.438272076 | 0.870929629 | 0.245 | 0.212 | 1 | 0.250356727 | -0.225804948 | 0.195 | 0.28 | 1 | 0.438272076 | 0.438034964 |
| Necap1 | 0.647427129 | 0.228723176 | 0.283 | 0.254 | 1 | 0.250417681 | 0.538546534 | 0.39 | 0.314 | 1 | 0.647427129 | 0.438126347 |
| Endod1 | 0.250433282 | 0.422418378 | 0.358 | 0.284 | 1 | 0.268707715 | 0.156027034 | 0.512 | 0.406 | 1 | 0.268707715 | 0.438149736 |
| Cacna2d3 | 0.824670081 | -0.790135851 | 0.094 | 0.102 | 1 | 0.250538294 | -1.174022406 | 0.049 | 0.106 | 1 | 0.824670081 | 0.438307151 |
| Snapc3 | 0.250569763 | 0.470224421 | 0.377 | 0.326 | 1 | 0.56031897 | 0.577337723 | 0.488 | 0.488 | 1 | 0.56031897 | 0.43835432 |
| Wdr6 | 0.250649163 | 0.598329246 | 0.245 | 0.178 | 1 | 0.299379612 | 0.739594053 | 0.341 | 0.294 | 1 | 0.299379612 | 0.438473324 |
| Ndp | 0.25074741 | 0.407488665 | 0.17 | 0.11 | 1 | 0.568512908 | 0.404245443 | 0.146 | 0.119 | 1 | 0.568512908 | 0.438620557 |
| Lmod1 | 0.250757901 | 0.8327945 | 0.057 | 0.025 | 1 | 0.878310931 | 0.667279848 | 0.024 | 0.02 | 1 | 0.878310931 | 0.438636277 |
| Kif20a | 0.250757901 | 0.154722595 | 0.057 | 0.025 | 1 | 0.402364804 | -0.332720152 | 0 | 0.017 | 1 | 0.402364804 | 0.438636277 |
| Supt3 | 0.250798278 | -0.316898433 | 0.415 | 0.504 | 1 | 0.829883355 | -0.377808041 | 0.463 | 0.478 | 1 | 0.829883355 | 0.43869678 |
| A930005H10Rik | 0.250925395 | -0.693274312 | 0.075 | 0.131 | 1 | 0.516992978 | 1.029849928 | 0.122 | 0.092 | 1 | 0.516992978 | 0.438887236 |
| Tiparp | 0.850250739 | -0.276031699 | 0.849 | 0.818 | 1 | 0.251010568 | -0.188499533 | 0.878 | 0.898 | 1 | 0.850250739 | 0.439014831 |
| Cmtr1 | 0.251106824 | 0.585879759 | 0.34 | 0.288 | 1 | 0.706345313 | 0.212713985 | 0.268 | 0.324 | 1 | 0.706345313 | 0.439159011 |
| Spring1 | 0.251300016 | -0.403272858 | 0.113 | 0.182 | 1 | 0.855175746 | 0.120997816 | 0.122 | 0.133 | 1 | 0.855175746 | 0.439448334 |
| Erich6 | 0.314969793 | 0.932330173 | 0.075 | 0.042 | 1 | 0.251349623 | -0.917682652 | 0.024 | 0.072 | 1 | 0.314969793 | 0.439522614 |
| Zfp930 | 0.860358653 | 0.154722595 | 0.057 | 0.064 | 1 | 0.25135045 | -0.863234868 | 0.024 | 0.072 | 1 | 0.860358653 | 0.439523851 |
| Cxcr2 | 0.251644813 | 0.932330173 | 0.019 | 0.004 | 1 | 0.455094184 | -0.484723245 | 0 | 0.014 | 1 | 0.455094184 | 0.439964514 |
| Tmem163 | 0.251644813 | 1.417757001 | 0.019 | 0.004 | 1 | 0.455098275 | 0.515276755 | 0 | 0.014 | 1 | 0.455098275 | 0.439964514 |
| Ptprc | 0.251644813 | 1.607234799 | 0.019 | 0.004 | 1 | 0.870754939 | -0.686357106 | 0.073 | 0.065 | 1 | 0.870754939 | 0.439964514 |
| 4933439K11Rik | 0.251644813 | 1.569760094 | 0.019 | 0.004 | 1 | 0.430584486 | 2.422167351 | 0.024 | 0.01 | 1 | 0.430584486 | 0.439964514 |
| Mndal | 0.251644813 | 1.306725688 | 0.019 | 0.004 | 1 | 0.267205177 | 2.252242349 | 0.024 | 0.007 | 1 | 0.267205177 | 0.439964514 |
| 4930578G10Rik | 0.251644813 | 1.154722595 | 0.019 | 0.004 | 1 | 0.5191986 | 0.515276755 | 0 | 0.01 | 1 | 0.5191986 | 0.439964514 |
| Gm830 | 0.251644813 | 1.569760094 | 0.019 | 0.004 | 1 | 0.51919986 | 0.83720485 | 0 | 0.01 | 1 | 0.51919986 | 0.439964514 |
| Cd52 | 0.251644813 | 1.932330173 | 0.019 | 0.004 | 1 | 0.866004546 | -0.372248516 | 0.073 | 0.065 | 1 | 0.866004546 | 0.439964514 |
| Trp73 | 0.251644813 | 0.8327945 | 0.019 | 0.004 | 1 | 0.267205177 | 2.252242349 | 0.024 | 0.007 | 1 | 0.267205177 | 0.439964514 |
| Gbx1 | 0.251644813 | 1.154722595 | 0.019 | 0.004 | 1 | 0.989595616 | 0.667279848 | 0.024 | 0.024 | 1 | 0.989595616 | 0.439964514 |
| Gm15756 | 0.251644813 | 0.569760094 | 0.019 | 0.004 | 1 | 0.271787759 | 1.029849928 | 0.024 | 0.007 | 1 | 0.271787759 | 0.439964514 |
| Nup210 | 0.251644813 | 1.569760094 | 0.019 | 0.004 | 1 | 0.5191986 | -0.484723245 | 0 | 0.01 | 1 | 0.5191986 | 0.439964514 |
| Hipk4 | 0.251644813 | 1.569760094 | 0.019 | 0.004 | 1 | 0.264884094 | 1.83720485 | 0.049 | 0.02 | 1 | 0.264884094 | 0.439964514 |
| Selenov | 0.251644813 | 0.569760094 | 0.019 | 0.004 | 1 | 0.51919986 | 0.83720485 | 0 | 0.01 | 1 | 0.51919986 | 0.439964514 |
| Gm44997 | 0.251644813 | 1.154722595 | 0.019 | 0.004 | 1 | 0.319619815 | -0.332720152 | 0 | 0.024 | 1 | 0.319619815 | 0.439964514 |
| Ngp | 0.251644813 | 1.569760094 | 0.019 | 0.004 | 1 | 0.489024353 | 0.557096931 | 0.122 | 0.089 | 1 | 0.489024353 | 0.439964514 |
| Vpreb3 | 0.251644813 | -0.430239906 | 0.019 | 0.004 | 1 | 0.264933923 | 2.422167351 | 0.024 | 0.007 | 1 | 0.264933923 | 0.439964514 |
| Havcr2 | 0.251644813 | 1.417757001 | 0.019 | 0.004 | 1 | 0.51919986 | 0.83720485 | 0 | 0.01 | 1 | 0.51919986 | 0.439964514 |
| Mir142hg | 0.251644813 | 0.695290976 | 0.019 | 0.004 | 1 | 0.357847088 | -1.069685746 | 0 | 0.02 | 1 | 0.357847088 | 0.439964514 |
| Akap5 | 0.251644813 | 1.569760094 | 0.019 | 0.004 | 1 | 0.51919986 | 0.83720485 | 0 | 0.01 | 1 | 0.51919986 | 0.439964514 |
| Cdh18 | 0.251644813 | 1.154722595 | 0.019 | 0.004 | 1 | 0.51919986 | -0.16279515 | 0 | 0.01 | 1 | 0.51919986 | 0.439964514 |
| Fbxo43 | 0.251644813 | 1.569760094 | 0.019 | 0.004 | 1 | 0.455097252 | -0.484723245 | 0 | 0.014 | 1 | 0.455097252 | 0.439964514 |
| Csf2rb2 | 0.251644813 | 1.569760094 | 0.019 | 0.004 | 1 | 0.455094184 | 0.252242349 | 0 | 0.014 | 1 | 0.455094184 | 0.439964514 |
| Rac2 | 0.251644813 | 1.154722595 | 0.019 | 0.004 | 1 | 0.576479046 | -1.435813645 | 0.049 | 0.031 | 1 | 0.576479046 | 0.439964514 |
| Parvb | 0.251644813 | 1.569760094 | 0.019 | 0.004 | 1 | 0.355779174 | 1.458693227 | 0.049 | 0.024 | 1 | 0.355779174 | 0.439964514 |
| Kcnh3 | 0.251644813 | 1.569760094 | 0.019 | 0.004 | 1 | 0.262674569 | 3.252242349 | 0.024 | 0.007 | 1 | 0.262674569 | 0.439964514 |
| Cd200r4 | 0.251644813 | 1.154722595 | 0.019 | 0.004 | 1 | 0.455094184 | 0.252242349 | 0 | 0.014 | 1 | 0.455094184 | 0.439964514 |
| Wfikkn1 | 0.251644813 | 1.569760094 | 0.019 | 0.004 | 1 | 0.357847088 | -0.16279515 | 0 | 0.02 | 1 | 0.357847088 | 0.439964514 |
| Cfb | 0.251644813 | 1.569760094 | 0.019 | 0.004 | 1 | 0.861873715 | 1.252242349 | 0.024 | 0.02 | 1 | 0.861873715 | 0.439964514 |
| Kcng3 | 0.251644813 | 1.569760094 | 0.019 | 0.004 | 1 | 0.26948834 | 1.83720485 | 0.024 | 0.007 | 1 | 0.26948834 | 0.439964514 |
| Fndc1 | 0.482376966 | -0.213999712 | 0.396 | 0.458 | 1 | 0.251764631 | 1.063480706 | 0.39 | 0.331 | 1 | 0.482376966 | 0.440143832 |
| Dnajb11 | 0.251772798 | 0.353383088 | 0.717 | 0.716 | 1 | 0.405116986 | 0.273303965 | 0.78 | 0.799 | 1 | 0.405116986 | 0.440156055 |
| Tmem161a | 0.436367225 | 0.347367673 | 0.415 | 0.373 | 1 | 0.251876952 | -0.415182312 | 0.293 | 0.379 | 1 | 0.436367225 | 0.440311905 |
| Mtrr | 0.956697606 | 0.569760094 | 0.075 | 0.081 | 1 | 0.251921424 | 1.252242349 | 0.122 | 0.072 | 1 | 0.956697606 | 0.440378444 |
| Casz1 | 0.676030438 | 0.247831999 | 0.057 | 0.042 | 1 | 0.252402634 | 0.679663573 | 0.22 | 0.15 | 1 | 0.676030438 | 0.441098179 |
| Fam204a | 0.793721033 | 0.144153353 | 0.642 | 0.669 | 1 | 0.252427959 | 0.326720565 | 0.659 | 0.577 | 1 | 0.793721033 | 0.441136043 |
| Zkscan1 | 0.666617597 | 0.308893525 | 0.396 | 0.386 | 1 | 0.252464153 | -0.320796984 | 0.415 | 0.481 | 1 | 0.666617597 | 0.441190157 |
| Cd46 | 0.294550482 | -1.356239324 | 0.075 | 0.123 | 1 | 0.252653899 | -0.425829556 | 0.098 | 0.174 | 1 | 0.294550482 | 0.441473805 |
| Hmox2 | 0.643438315 | 0.389187848 | 0.491 | 0.475 | 1 | 0.252868563 | 0.275549723 | 0.707 | 0.631 | 1 | 0.643438315 | 0.441794616 |
| Gspt2 | 0.253167671 | 0.569760094 | 0.132 | 0.081 | 1 | 0.459676233 | 1.180092563 | 0.146 | 0.113 | 1 | 0.459676233 | 0.442241473 |
| Nudt6 | 0.253167911 | 0.510866405 | 0.132 | 0.081 | 1 | 0.464673518 | 1.19334866 | 0.098 | 0.068 | 1 | 0.464673518 | 0.442241831 |
| Ckap2 | 0.253390183 | -0.582242999 | 0.075 | 0.038 | 1 | 0.729895585 | 1.83720485 | 0.024 | 0.017 | 1 | 0.729895585 | 0.442573781 |
| Uspl1 | 0.32198035 | 0.137649081 | 0.509 | 0.424 | 1 | 0.253450112 | 0.521563018 | 0.463 | 0.437 | 1 | 0.32198035 | 0.442663265 |
| Zfp449 | 0.453009872 | 0.237184755 | 0.151 | 0.11 | 1 | 0.253552426 | -0.995685164 | 0.073 | 0.133 | 1 | 0.453009872 | 0.442816019 |
| Bcl2l13 | 0.253842941 | 0.772252958 | 0.302 | 0.246 | 1 | 0.270515507 | 0.829030918 | 0.415 | 0.379 | 1 | 0.270515507 | 0.443249643 |
| A830018L16Rik | 0.929410101 | 0.569760094 | 0.019 | 0.017 | 1 | 0.253885121 | 2.574170444 | 0.049 | 0.02 | 1 | 0.929410101 | 0.443312587 |
| Slc25a13 | 0.717159697 | 0.535812762 | 0.208 | 0.233 | 1 | 0.253993881 | -0.498398182 | 0.244 | 0.345 | 1 | 0.717159697 | 0.443474871 |
| Mettl2 | 0.254138076 | 0.544669113 | 0.264 | 0.199 | 1 | 0.985089113 | 0.304709769 | 0.195 | 0.198 | 1 | 0.985089113 | 0.44368999 |
| Pla2g4a | 0.254421736 | -0.42039212 | 0.377 | 0.475 | 1 | 0.302026562 | 0.332860809 | 0.366 | 0.29 | 1 | 0.302026562 | 0.444113053 |
| Aff1 | 0.273641087 | 0.316742398 | 0.925 | 0.932 | 1 | 0.254826307 | 0.139348293 | 0.927 | 0.918 | 1 | 0.273641087 | 0.444716168 |
| Scly | 0.254919974 | 0.544669113 | 0.208 | 0.144 | 1 | 0.782665388 | 0.201616276 | 0.195 | 0.174 | 1 | 0.782665388 | 0.444855754 |
| Manf | 0.254944185 | 0.130520246 | 0.962 | 0.903 | 1 | 0.650502449 | 0.2917334 | 0.854 | 0.901 | 1 | 0.650502449 | 0.444891833 |
| Champ1 | 0.825806199 | 0.361173472 | 0.17 | 0.195 | 1 | 0.255513133 | -0.410722664 | 0.171 | 0.253 | 1 | 0.825806199 | 0.445739305 |
| Gm15867 | 0.255513922 | -0.556772312 | 0.226 | 0.305 | 1 | 0.529844413 | 0.371541277 | 0.293 | 0.253 | 1 | 0.529844413 | 0.445740479 |
| Tmsb15b2 | 0.822395308 | 0.347367673 | 0.17 | 0.161 | 1 | 0.255818106 | 0.222495006 | 0.073 | 0.14 | 1 | 0.822395308 | 0.446193309 |
| Map3k14 | 0.255947819 | 0.228723176 | 0.226 | 0.157 | 1 | 0.531299549 | -0.309636539 | 0.195 | 0.239 | 1 | 0.531299549 | 0.446386352 |
| Zfp280c | 0.256162859 | 0.624207878 | 0.434 | 0.386 | 1 | 0.662374583 | 0.510475769 | 0.488 | 0.485 | 1 | 0.662374583 | 0.446706308 |
| Rpp30 | 0.533440568 | 0.261637799 | 0.226 | 0.186 | 1 | 0.256251415 | -0.622226769 | 0.122 | 0.195 | 1 | 0.533440568 | 0.446838043 |
| Snhg1 | 0.455198538 | 0.12817544 | 0.925 | 0.847 | 1 | 0.256305774 | -0.151113345 | 0.78 | 0.85 | 1 | 0.455198538 | 0.446918898 |
| Morn3 | 0.287773478 | -0.430239906 | 0 | 0.021 | 1 | 0.257285084 | -0.863234868 | 0 | 0.031 | 1 | 0.287773478 | 0.448374553 |
| Ssc4d | 0.292183883 | -0.368839361 | 0.038 | 0.081 | 1 | 0.257285084 | -0.622226769 | 0 | 0.031 | 1 | 0.292183883 | 0.448374553 |
| Gm26672 | 0.310259246 | 1.039245377 | 0.075 | 0.042 | 1 | 0.257285084 | -0.622226769 | 0 | 0.031 | 1 | 0.310259246 | 0.448374553 |
| Gm15852 | 0.343006305 | -0.1672055 | 0 | 0.017 | 1 | 0.257285084 | -0.622226769 | 0 | 0.031 | 1 | 0.343006305 | 0.448374553 |
| Phlda2 | 0.419848106 | -0.430239906 | 0.019 | 0.042 | 1 | 0.257285084 | -0.622226769 | 0 | 0.031 | 1 | 0.419848106 | 0.448374553 |
| Gm56791 | 0.48175508 | 1.154722595 | 0.038 | 0.021 | 1 | 0.257285084 | -0.622226769 | 0 | 0.031 | 1 | 0.48175508 | 0.448374553 |
| Gm56589 | 0.48175508 | 1.154722595 | 0.038 | 0.021 | 1 | 0.257285084 | -0.622226769 | 0 | 0.031 | 1 | 0.48175508 | 0.448374553 |
| Mettl18 | 0.507281812 | 0.8327945 | 0.019 | 0.008 | 1 | 0.257285084 | -0.970150072 | 0 | 0.031 | 1 | 0.507281812 | 0.448374553 |
| Zfp128 | 0.913424206 | 1.417757001 | 0.019 | 0.017 | 1 | 0.257285084 | -0.622226769 | 0 | 0.031 | 1 | 0.913424206 | 0.448374553 |
| Gm29994 | 0.307417922 | -0.932740246 | 0.019 | 0.051 | 1 | 0.25729872 | -0.970150072 | 0 | 0.031 | 1 | 0.307417922 | 0.448394808 |

| | | | | | | | | | | | | |
|---|---|---|---|---|---|---|---|---|---|---|---|---|
| Gm49521 | 0.413082433 | -0.430239906 | 0 | 0.013 | 1 | 0.25729872 | -0.747757651 | 0 | 0.031 | 1 | 0.413082433 | 0.448394808 |
| Gins3 | 0.679455468 | -0.260314904 | 0.038 | 0.051 | 1 | 0.25729872 | -0.863234868 | 0 | 0.031 | 1 | 0.679455468 | 0.448394808 |
| Gm43378 | 0.767066103 | 0.569760094 | 0.038 | 0.03 | 1 | 0.25729872 | -0.863234868 | 0 | 0.031 | 1 | 0.767066103 | 0.448394808 |
| Platr22 | 0.779443586 | -0.1672055 | 0.019 | 0.025 | 1 | 0.25729872 | -0.747757651 | 0 | 0.031 | 1 | 0.779443586 | 0.448394808 |
| Prss46 | 0.779443586 | 0.154722595 | 0.019 | 0.025 | 1 | 0.25729872 | -0.863234868 | 0 | 0.031 | 1 | 0.779443586 | 0.448394808 |
| Adora2a | 0.914958 | 0.347367673 | 0.019 | 0.021 | 1 | 0.25729872 | -0.863234868 | 0 | 0.031 | 1 | 0.914958 | 0.448394808 |
| Gm17359 | 0.926212226 | 0.8327945 | 0.019 | 0.017 | 1 | 0.25729872 | -0.747757651 | 0 | 0.031 | 1 | 0.926212226 | 0.448394808 |
| Fank1 | 0.663401681 | -0.1672055 | 0.019 | 0.03 | 1 | 0.257308459 | -0.970150072 | 0 | 0.031 | 1 | 0.663401681 | 0.448409275 |
| Mal | 0.287761733 | -0.652632327 | 0 | 0.021 | 1 | 0.257316251 | -1.250257991 | 0 | 0.031 | 1 | 0.287761733 | 0.448420848 |
| Obp2b | 0.339336339 | 0.436493563 | 0.283 | 0.22 | 1 | 0.257316251 | -3.093532488 | 0 | 0.031 | 1 | 0.339336339 | 0.448420848 |
| Gm47283 | 0.989078441 | -0.28621268 | 0.943 | 0.915 | 1 | 0.257317903 | 0.100635655 | 0.927 | 0.922 | 1 | 0.989078441 | 0.448423303 |
| 2810036E18Rik | 0.906251902 | 0.154722595 | 0.038 | 0.034 | 1 | 0.257319172 | -1.250257991 | 0 | 0.031 | 1 | 0.906251902 | 0.448425188 |
| Snhg11 | 0.503971666 | 1.569760094 | 0.019 | 0.008 | 1 | 0.257320146 | -1.484723245 | 0 | 0.031 | 1 | 0.503971666 | 0.448426635 |
| Mst1r | 0.503971666 | 1.569760094 | 0.019 | 0.008 | 1 | 0.257323068 | -2.116991461 | 0 | 0.031 | 1 | 0.503971666 | 0.448430974 |
| Pbld1 | 0.357060703 | -0.932740246 | 0.019 | 0.047 | 1 | 0.257325989 | -0.484723245 | 0 | 0.031 | 1 | 0.357060703 | 0.448435314 |
| Gm32655 | 0.419848106 | -0.430239906 | 0.019 | 0.042 | 1 | 0.257325989 | -0.484723245 | 0 | 0.031 | 1 | 0.419848106 | 0.448435314 |
| Thnsl1 | 0.605896064 | -0.1672055 | 0.038 | 0.055 | 1 | 0.257325989 | -0.484723245 | 0 | 0.031 | 1 | 0.605896064 | 0.448435314 |
| Etaa1os | 0.821680601 | 0.438515561 | 0.075 | 0.068 | 1 | 0.257325989 | -0.484723245 | 0 | 0.031 | 1 | 0.821680601 | 0.448435314 |
| Dnase1l2 | 0.917883255 | 0.569760094 | 0.019 | 0.021 | 1 | 0.257325989 | -0.484723245 | 0 | 0.031 | 1 | 0.917883255 | 0.448435314 |
| 1700052K11Rik | 0.423548797 | 1.228723176 | 0.094 | 0.064 | 1 | 0.25763405 | 1.83720485 | 0.098 | 0.055 | 1 | 0.423548797 | 0.448892796 |
| Ilkap | 0.49578161 | 0.265426059 | 0.491 | 0.479 | 1 | 0.257745144 | -0.123624553 | 0.512 | 0.611 | 1 | 0.49578161 | 0.449057729 |
| Prss36 | 0.535073776 | -0.389597921 | 0.094 | 0.123 | 1 | 0.257942453 | 1.59619675 | 0.122 | 0.075 | 1 | 0.535073776 | 0.449350596 |
| 6430550D23Rik | 0.661088008 | -0.304709024 | 0.019 | 0.03 | 1 | 0.257969546 | 1.989207943 | 0.049 | 0.02 | 1 | 0.661088008 | 0.449390806 |
| Jrkl | 0.258037873 | -0.401670754 | 0.208 | 0.288 | 1 | 0.393498596 | -0.67058979 | 0.146 | 0.198 | 1 | 0.393498596 | 0.449492202 |
| Wfdc2 | 0.528445339 | -1.439458436 | 0.358 | 0.284 | 1 | 0.258133367 | -0.137704169 | 0.61 | 0.526 | 1 | 0.528445339 | 0.449633898 |
| Cog3 | 0.727117945 | 0.209864149 | 0.453 | 0.436 | 1 | 0.258389813 | 0.417879992 | 0.61 | 0.567 | 1 | 0.727117945 | 0.45001433 |
| Grik3 | 0.258390936 | -0.752168001 | 0.094 | 0.153 | 1 | 0.549376113 | 0.10681191 | 0.171 | 0.225 | 1 | 0.549376113 | 0.450015997 |
| Hsd17b8 | 0.258424527 | -0.147840175 | 0.17 | 0.25 | 1 | 0.80564835 | 0.181853021 | 0.195 | 0.177 | 1 | 0.80564835 | 0.450065818 |
| Pop4 | 0.258651958 | 0.601124265 | 0.377 | 0.326 | 1 | 0.272608877 | 0.83720485 | 0.293 | 0.232 | 1 | 0.272608877 | 0.45040308 |
| Mis18a | 0.258813978 | 0.669295768 | 0.151 | 0.097 | 1 | 0.371357099 | -0.484723245 | 0.024 | 0.058 | 1 | 0.371357099 | 0.45064328 |
| Cilk1 | 0.258932449 | 0.154722595 | 0.528 | 0.436 | 1 | 0.478646977 | 0.186950889 | 0.634 | 0.573 | 1 | 0.478646977 | 0.450818885 |
| Saraf | 0.915149522 | 0.102961167 | 0.962 | 0.949 | 1 | 0.259072987 | 0.324033032 | 0.927 | 0.942 | 1 | 0.915149522 | 0.451027161 |
| Cables2 | 0.259168526 | 0.756173218 | 0.17 | 0.114 | 1 | 0.402172901 | 0.491430013 | 0.22 | 0.164 | 1 | 0.402172901 | 0.451168727 |
| Ptrhd1 | 0.762913908 | 0.347367673 | 0.264 | 0.25 | 1 | 0.259236849 | -0.555112573 | 0.098 | 0.167 | 1 | 0.762913908 | 0.451269954 |
| Fam43a | 0.259408968 | -1.738362201 | 0.113 | 0.165 | 1 | 0.575233611 | 0.252242349 | 0.171 | 0.137 | 1 | 0.575233611 | 0.451524924 |
| Scamp3 | 0.259419847 | -0.192296481 | 0.604 | 0.682 | 1 | 0.559769244 | 0.245758316 | 0.537 | 0.502 | 1 | 0.559769244 | 0.451541037 |
| Folr2 | 0.259444172 | -1.652632327 | 0.038 | 0.081 | 1 | 0.968795535 | 2.252242349 | 0.024 | 0.024 | 1 | 0.968795535 | 0.451577066 |
| Mcm9 | 0.259554598 | 0.412790748 | 0.264 | 0.195 | 1 | 0.612110499 | 0.394982521 | 0.244 | 0.218 | 1 | 0.612110499 | 0.451740607 |
| Med9 | 0.582587824 | -0.260314904 | 0.226 | 0.254 | 1 | 0.259881074 | -0.364429011 | 0.171 | 0.249 | 1 | 0.582587824 | 0.452223976 |
| Coq5 | 0.259983827 | 0.47665069 | 0.264 | 0.195 | 1 | 0.7509763 | 0.416873051 | 0.171 | 0.201 | 1 | 0.7509763 | 0.452376064 |
| Gm7467 | 0.421517727 | -0.304709024 | 0.019 | 0.042 | 1 | 0.260704079 | 2.029849928 | 0.049 | 0.02 | 1 | 0.421517727 | 0.453441541 |
| Tigar | 0.497380598 | 2.569760094 | 0.019 | 0.008 | 1 | 0.260704079 | 2.351778023 | 0.049 | 0.02 | 1 | 0.497380598 | 0.453441541 |
| Gm45464 | 0.665715777 | -0.1672055 | 0.019 | 0.03 | 1 | 0.260704079 | 2.029849928 | 0.049 | 0.02 | 1 | 0.665715777 | 0.453441541 |
| Plac8 | 0.739544426 | 1.669295768 | 0.038 | 0.03 | 1 | 0.260704079 | 2.614812428 | 0.049 | 0.02 | 1 | 0.739544426 | 0.453441541 |
| B630019K06Rik | 0.26085577 | -0.889671525 | 0.057 | 0.106 | 1 | 0.476434136 | 1.071670103 | 0.073 | 0.048 | 1 | 0.476434136 | 0.453665808 |
| Bola1 | 0.260956128 | -0.4082136 | 0.17 | 0.242 | 1 | 0.906262622 | 0.345351753 | 0.122 | 0.133 | 1 | 0.906262622 | 0.453814156 |
| Pkdcc | 0.260969458 | -0.988235359 | 0.226 | 0.301 | 1 | 0.797071653 | -0.370688002 | 0.341 | 0.355 | 1 | 0.797071653 | 0.453833858 |
| Slc7a6os | 0.261364072 | 0.784610966 | 0.396 | 0.36 | 1 | 0.56785742 | 0.582783574 | 0.366 | 0.348 | 1 | 0.56785742 | 0.454416966 |
| Ttll3 | 0.969605466 | -0.932740246 | 0.226 | 0.212 | 1 | 0.261441545 | -0.372248516 | 0.22 | 0.317 | 1 | 0.969605466 | 0.454531408 |
| Pex19 | 0.26149509 | 0.444229212 | 0.434 | 0.364 | 1 | 0.428990697 | 0.715741337 | 0.439 | 0.413 | 1 | 0.428990697 | 0.454610497 |
| Aldh1l2 | 0.261871082 | -1.195774652 | 0.057 | 0.106 | 1 | 0.392436731 | -0.600200463 | 0.049 | 0.089 | 1 | 0.392436731 | 0.4551657 |
| Lama1 | 0.262357648 | 1.347367673 | 0.094 | 0.055 | 1 | 0.282653135 | 1.83720485 | 0.073 | 0.038 | 1 | 0.282653135 | 0.45588376 |
| Muc16 | 0.343000446 | -0.430239906 | 0 | 0.017 | 1 | 0.262428917 | 0.962735732 | 0.122 | 0.072 | 1 | 0.343000446 | 0.455988898 |
| Cd99l2 | 0.262467519 | 0.627475592 | 0.358 | 0.301 | 1 | 0.410175278 | 0.172072 | 0.317 | 0.427 | 1 | 0.410175278 | 0.456045839 |
| Zfp369 | 0.262551048 | 0.287989126 | 0.226 | 0.161 | 1 | 0.801236796 | 0.438655473 | 0.22 | 0.266 | 1 | 0.801236796 | 0.456169044 |
| B4galt5 | 0.262551902 | -0.553622321 | 0.34 | 0.403 | 1 | 0.745639554 | 1.061911137 | 0.415 | 0.423 | 1 | 0.745639554 | 0.456170303 |
| Ccdc86 | 0.262657279 | 0.538051234 | 0.415 | 0.386 | 1 | 0.450232886 | -0.421529419 | 0.268 | 0.314 | 1 | 0.450232886 | 0.456325711 |
| Hey2 | 0.343006305 | -0.1672055 | 0 | 0.017 | 1 | 0.262674569 | 3.574170444 | 0.024 | 0.007 | 1 | 0.343006305 | 0.456351209 |
| 4833445I07Rik | 0.412009681 | 1.792152515 | 0.057 | 0.034 | 1 | 0.262674569 | 2.83720485 | 0.024 | 0.007 | 1 | 0.412009681 | 0.456351209 |
| Adap2os | 0.413080549 | -0.1672055 | 0 | 0.013 | 1 | 0.262674569 | 3.252242349 | 0.024 | 0.007 | 1 | 0.413080549 | 0.456351209 |
| Ccdc96 | 0.413082433 | 0.154722595 | 0 | 0.013 | 1 | 0.262674569 | 3.252242349 | 0.024 | 0.007 | 1 | 0.413082433 | 0.456351209 |
| D430040D24Rik | 0.503971666 | 1.569760094 | 0.019 | 0.008 | 1 | 0.262674569 | 2.83720485 | 0.024 | 0.007 | 1 | 0.503971666 | 0.456351209 |
| Enpp3 | 0.929410101 | 0.569760094 | 0.019 | 0.017 | 1 | 0.262674569 | 2.83720485 | 0.024 | 0.007 | 1 | 0.929410101 | 0.456351209 |
| Gm15608 | 0.929410101 | 0.569760094 | 0.019 | 0.017 | 1 | 0.262674569 | 2.83720485 | 0.024 | 0.007 | 1 | 0.929410101 | 0.456351209 |
| Trim71 | 0.926654567 | 0.739685096 | 0.019 | 0.021 | 1 | 0.262676008 | 2.83720485 | 0.024 | 0.007 | 1 | 0.926654567 | 0.456353331 |
| Cpsf4l | 0.263063806 | -0.974560422 | 0.057 | 0.106 | 1 | 0.993251604 | -0.16279515 | 0.098 | 0.096 | 1 | 0.993251604 | 0.456925046 |
| Wdr47 | 0.263168667 | -0.686579659 | 0.226 | 0.309 | 1 | 0.413018446 | 0.363273661 | 0.415 | 0.375 | 1 | 0.413018446 | 0.457079586 |
| Tecpr1 | 0.26524591 | -0.600164907 | 0.17 | 0.237 | 1 | 0.263234974 | 1.051329655 | 0.293 | 0.239 | 1 | 0.26524591 | 0.457177297 |
| Tmem9b | 0.263272276 | 0.308267853 | 0.566 | 0.487 | 1 | 0.65157583 | -0.145788725 | 0.512 | 0.536 | 1 | 0.65157583 | 0.45723226 |
| Myo7a | 0.917883255 | 0.569760094 | 0.019 | 0.021 | 1 | 0.263489513 | 1.83720485 | 0.049 | 0.02 | 1 | 0.917883255 | 0.457552303 |
| Fgf1 | 0.263793882 | -0.411624228 | 0.094 | 0.161 | 1 | 0.284887811 | 1.334704509 | 0.073 | 0.038 | 1 | 0.284887811 | 0.458000553 |
| Rft1 | 0.263898788 | 0.844382474 | 0.264 | 0.203 | 1 | 0.335719997 | -0.723510105 | 0.098 | 0.15 | 1 | 0.335719997 | 0.458155005 |
| Palb2 | 0.897146191 | 0.417757001 | 0.038 | 0.034 | 1 | 0.264102656 | 2.059597271 | 0.073 | 0.038 | 1 | 0.897146191 | 0.458455099 |
| Tsacc | 0.264265675 | -0.247375849 | 0.075 | 0.136 | 1 | 0.277675489 | -0.665295491 | 0.073 | 0.137 | 1 | 0.277675489 | 0.458695002 |
| Naf1 | 0.264527356 | 0.448453798 | 0.434 | 0.381 | 1 | 0.272089004 | -0.407102348 | 0.317 | 0.413 | 1 | 0.272089004 | 0.459079989 |
| Greb1 | 0.287761733 | -0.652632327 | 0 | 0.021 | 1 | 0.264933923 | 2.422167351 | 0.024 | 0.007 | 1 | 0.287761733 | 0.459677863 |
| Gm19466 | 0.638059929 | 0.280253477 | 0.038 | 0.025 | 1 | 0.264933923 | 2.422167351 | 0.024 | 0.007 | 1 | 0.638059929 | 0.459677863 |
| Vamp2 | 0.424610113 | 0.162256267 | 0.528 | 0.492 | 1 | 0.265121409 | -0.152147906 | 0.415 | 0.526 | 1 | 0.424610113 | 0.459953457 |
| 1700048O20Rik | 0.26528608 | 1.306725688 | 0.094 | 0.055 | 1 | 0.738418148 | -0.16279515 | 0.049 | 0.061 | 1 | 0.738418148 | 0.460195456 |
| Xrcc1 | 0.265469569 | 0.970298024 | 0.226 | 0.169 | 1 | 0.949617711 | 0.757477657 | 0.22 | 0.235 | 1 | 0.949617711 | 0.460465045 |
| Mphosph10 | 0.265737593 | 0.502645898 | 0.453 | 0.428 | 1 | 0.419285735 | 0.326242931 | 0.317 | 0.447 | 1 | 0.419285735 | 0.460858718 |
| Cd2bp2 | 0.265750234 | 0.874614676 | 0.245 | 0.191 | 1 | 0.668809296 | -0.20718927 | 0.22 | 0.249 | 1 | 0.668809296 | 0.460877281 |
| Gm44148 | 0.265932033 | -1.015202407 | 0.019 | 0.055 | 1 | 0.274951624 | 1.83720485 | 0.073 | 0.038 | 1 | 0.274951624 | 0.46114422 |
| Tex26 | 0.265932462 | -1.430239906 | 0.019 | 0.055 | 1 | 0.87556513 | 0.377773231 | 0.024 | 0.02 | 1 | 0.87556513 | 0.46114485 |
| Gm16272 | 0.265932462 | -1.304709024 | 0.019 | 0.055 | 1 | 0.352702675 | -0.221688839 | 0.024 | 0.061 | 1 | 0.352702675 | 0.46114485 |
| 4933412O06Rik | 0.265932462 | -1.093204919 | 0.019 | 0.055 | 1 | 0.5191986 | 0.515276755 | 0 | 0.01 | 1 | 0.5191986 | 0.46114485 |
| Gm16196 | 0.265932462 | -0.932740246 | 0.019 | 0.055 | 1 | 0.599290422 | 1.515276755 | 0.024 | 0.014 | 1 | 0.599290422 | 0.46114485 |
| Mfsd2b | 0.26611511 | 1.707263618 | 0.094 | 0.055 | 1 | 0.815235757 | 0.644559772 | 0.073 | 0.085 | 1 | 0.815235757 | 0.461412969 |
| Kptn | 0.266564008 | 0.472898555 | 0.264 | 0.212 | 1 | 0.682340672 | -0.359192363 | 0.268 | 0.294 | 1 | 0.682340672 | 0.462071645 |
| Gm56816 | 0.530868628 | -0.347777746 | 0.038 | 0.059 | 1 | 0.266662906 | -0.435813645 | 0.024 | 0.072 | 1 | 0.530868628 | 0.462216706 |
| Mthfr | 0.442855436 | 0.552058092 | 0.283 | 0.242 | 1 | 0.266817886 | 0.812542796 | 0.341 | 0.287 | 1 | 0.442855436 | 0.462443988 |
| Gm45669 | 0.432054479 | 0.510866405 | 0.189 | 0.144 | 1 | 0.266825161 | 0.515276755 | 0.244 | 0.171 | 1 | 0.432054479 | 0.462454655 |
| E230001N04Rik | 0.266956226 | -0.845277405 | 0.019 | 0.055 | 1 | 0.599290422 | 1.515276755 | 0.024 | 0.014 | 1 | 0.599290422 | 0.462646825 |
| Inpp4a | 0.739698319 | 0.438515561 | 0.34 | 0.339 | 1 | 0.267018929 | -0.363557556 | 0.463 | 0.536 | 1 | 0.739698319 | 0.462738749 |
| Rsad1 | 0.47205486 | 0.891688189 | 0.151 | 0.119 | 1 | 0.267048857 | -0.410722664 | 0.049 | 0.106 | 1 | 0.47205486 | 0.462782622 |
| Rfc2 | 0.267174803 | 0.292226119 | 0.321 | 0.237 | 1 | 0.336230515 | -0.455576899 | 0.171 | 0.239 | 1 | 0.336230515 | 0.46296723 |
| Espl1 | 0.287761733 | -1.1672055 | 0 | 0.021 | 1 | 0.267205177 | 2.252242349 | 0.024 | 0.007 | 1 | 0.287761733 | 0.463011747 |
| Pcyt1b | 0.28776878 | -0.845277405 | 0 | 0.021 | 1 | 0.267205177 | 2.252242349 | 0.024 | 0.007 | 1 | 0.28776878 | 0.463011747 |

| | | | | | | | | | | | | |
|---|---|---|---|---|---|---|---|---|---|---|---|---|
| Ubxn10 | 0.287773478 | -0.430239906 | 0 | 0.021 | 1 | 0.267205177 | 2.252242349 | 0.024 | 0.007 | 1 | 0.287773478 | 0.463011747 |
| Acy3 | 0.343000446 | -0.430239906 | 0 | 0.017 | 1 | 0.267205177 | 2.252242349 | 0.024 | 0.007 | 1 | 0.343000446 | 0.463011747 |
| Vtcn1 | 0.343006305 | -0.1672055 | 0 | 0.017 | 1 | 0.267205177 | 2.252242349 | 0.024 | 0.007 | 1 | 0.343006305 | 0.463011747 |
| Lratd1 | 0.343006305 | -0.1672055 | 0 | 0.017 | 1 | 0.267205177 | 2.252242349 | 0.024 | 0.007 | 1 | 0.343006305 | 0.463011747 |
| Ciita | 0.343006305 | -0.1672055 | 0 | 0.017 | 1 | 0.267205177 | 2.252242349 | 0.024 | 0.007 | 1 | 0.343006305 | 0.463011747 |
| 1700064H15Rik | 0.413080549 | -0.1672055 | 0 | 0.013 | 1 | 0.267205177 | 2.252242349 | 0.024 | 0.007 | 1 | 0.413080549 | 0.463011747 |
| Gm29128 | 0.413082433 | 0.154722595 | 0 | 0.013 | 1 | 0.267205177 | 2.252242349 | 0.024 | 0.007 | 1 | 0.413082433 | 0.463011747 |
| Gm57150 | 0.413082433 | 0.154722595 | 0 | 0.013 | 1 | 0.267205177 | 2.252242349 | 0.024 | 0.007 | 1 | 0.413082433 | 0.463011747 |
| Gm43017 | 0.413082433 | 0.154722595 | 0 | 0.013 | 1 | 0.267205177 | 2.252242349 | 0.024 | 0.007 | 1 | 0.413082433 | 0.463011747 |
| 2310016G11Rik | 0.413082433 | 0.154722595 | 0 | 0.013 | 1 | 0.267205177 | 2.252242349 | 0.024 | 0.007 | 1 | 0.413082433 | 0.463011747 |
| Sec1 | 0.413082433 | 0.154722595 | 0 | 0.013 | 1 | 0.267205177 | 2.252242349 | 0.024 | 0.007 | 1 | 0.413082433 | 0.463011747 |
| Tpbgl | 0.413082433 | 0.154722595 | 0 | 0.013 | 1 | 0.267205177 | 2.252242349 | 0.024 | 0.007 | 1 | 0.413082433 | 0.463011747 |
| Stra6 | 0.413082433 | 0.154722595 | 0 | 0.013 | 1 | 0.267205177 | 2.252242349 | 0.024 | 0.007 | 1 | 0.413082433 | 0.463011747 |
| Gm50203 | 0.413082433 | 0.154722595 | 0 | 0.013 | 1 | 0.267205177 | 2.252242349 | 0.024 | 0.007 | 1 | 0.413082433 | 0.463011747 |
| Gm39302 | 0.435857866 | 0.154722595 | 0.019 | 0.042 | 1 | 0.267205177 | 2.252242349 | 0.024 | 0.007 | 1 | 0.435857866 | 0.463011747 |
| Skap1 | 0.440752287 | 0.776210972 | 0.057 | 0.034 | 1 | 0.267205177 | 2.252242349 | 0.024 | 0.007 | 1 | 0.440752287 | 0.463011747 |
| Lmo3 | 0.497380598 | 2.154722595 | 0.019 | 0.008 | 1 | 0.267205177 | 2.252242349 | 0.024 | 0.007 | 1 | 0.497380598 | 0.463011747 |
| Knstrn | 0.503280895 | -0.904171094 | 0.038 | 0.021 | 1 | 0.267205177 | 2.252242349 | 0.024 | 0.007 | 1 | 0.503280895 | 0.463011747 |
| Skida1 | 0.503971666 | 1.569760094 | 0.019 | 0.008 | 1 | 0.267205177 | 2.252242349 | 0.024 | 0.007 | 1 | 0.503971666 | 0.463011747 |
| Gm50019 | 0.503971666 | 1.569760094 | 0.019 | 0.008 | 1 | 0.267205177 | 2.252242349 | 0.024 | 0.007 | 1 | 0.503971666 | 0.463011747 |
| Gm48796 | 0.507281812 | 1.154722595 | 0.019 | 0.008 | 1 | 0.267205177 | 2.252242349 | 0.024 | 0.007 | 1 | 0.507281812 | 0.463011747 |
| Gm10857 | 0.677401546 | 0.280253477 | 0.019 | 0.03 | 1 | 0.267205177 | 2.252242349 | 0.024 | 0.007 | 1 | 0.677401546 | 0.463011747 |
| Ptgir | 0.7394666 | 0.154722595 | 0.019 | 0.013 | 1 | 0.267205177 | 2.252242349 | 0.024 | 0.007 | 1 | 0.7394666 | 0.463011747 |
| Gm32856 | 0.782077996 | 0.347367673 | 0.019 | 0.025 | 1 | 0.267205177 | 2.252242349 | 0.024 | 0.007 | 1 | 0.782077996 | 0.463011747 |
| Cenpu | 0.879000798 | 1.377115016 | 0.038 | 0.034 | 1 | 0.267205177 | 2.252242349 | 0.024 | 0.007 | 1 | 0.879000798 | 0.463011747 |
| Cep72 | 0.912039013 | 0.154722595 | 0.019 | 0.021 | 1 | 0.267205177 | 2.252242349 | 0.024 | 0.007 | 1 | 0.912039013 | 0.463011747 |
| Gm56847 | 0.914958 | 0.347367673 | 0.019 | 0.021 | 1 | 0.267205177 | 2.252242349 | 0.024 | 0.007 | 1 | 0.914958 | 0.463011747 |
| A330074K22Rik | 0.914958 | 0.347367673 | 0.019 | 0.021 | 1 | 0.267205177 | 2.252242349 | 0.024 | 0.007 | 1 | 0.914958 | 0.463011747 |
| Gm10244 | 0.926212226 | 0.8327945 | 0.019 | 0.017 | 1 | 0.267205177 | 2.252242349 | 0.024 | 0.007 | 1 | 0.926212226 | 0.463011747 |
| Gm48952 | 0.926212226 | 0.8327945 | 0.019 | 0.017 | 1 | 0.267205177 | 2.252242349 | 0.024 | 0.007 | 1 | 0.926212226 | 0.463011747 |
| Gm16618 | 0.926212226 | 0.8327945 | 0.019 | 0.017 | 1 | 0.267205177 | 2.252242349 | 0.024 | 0.007 | 1 | 0.926212226 | 0.463011747 |
| Rgs22 | 0.929410101 | 0.347367673 | 0.019 | 0.017 | 1 | 0.267205177 | 2.100239256 | 0.024 | 0.007 | 1 | 0.929410101 | 0.463011747 |
| Arl2 | 0.267485385 | 0.489386678 | 0.509 | 0.513 | 1 | 0.766168575 | 0.412183262 | 0.341 | 0.334 | 1 | 0.766168575 | 0.463422338 |
| Rfc4 | 0.26748703 | 0.891688189 | 0.132 | 0.085 | 1 | 0.409125717 | -0.410722664 | 0.024 | 0.055 | 1 | 0.409125717 | 0.463424749 |
| Ebp | 0.267630402 | 0.400632823 | 0.585 | 0.568 | 1 | 0.53463874 | 0.243680336 | 0.415 | 0.386 | 1 | 0.53463874 | 0.463634772 |
| Rufy2 | 0.267639526 | 0.218287724 | 0.358 | 0.28 | 1 | 0.702378812 | 0.292884334 | 0.463 | 0.474 | 1 | 0.702378812 | 0.463648137 |
| Serac1 | 0.706522032 | 0.47665069 | 0.038 | 0.051 | 1 | 0.267833449 | 1.252242349 | 0.122 | 0.072 | 1 | 0.706522032 | 0.463932141 |
| Fam210a | 0.26784368 | 0.300573461 | 0.472 | 0.403 | 1 | 0.757722658 | 0.214487427 | 0.415 | 0.512 | 1 | 0.757722658 | 0.463947124 |
| Rc3h2 | 0.268092569 | 0.222135516 | 0.792 | 0.801 | 1 | 0.645668895 | 0.219399092 | 0.902 | 0.939 | 1 | 0.645668895 | 0.464311513 |
| Ptprz1 | 0.343006305 | -0.1672055 | 0 | 0.017 | 1 | 0.268288619 | 1.252242349 | 0.098 | 0.055 | 1 | 0.343006305 | 0.464598454 |
| Gamt | 0.93236326 | 0.306725688 | 0.075 | 0.072 | 1 | 0.268403557 | 1.059597271 | 0.098 | 0.055 | 1 | 0.93236326 | 0.464766645 |
| Ywhag | 0.693993317 | -0.120756577 | 0.792 | 0.775 | 1 | 0.268452456 | -0.113344708 | 0.854 | 0.908 | 1 | 0.693993317 | 0.464838191 |
| Eva1c | 0.650695347 | 0.695290976 | 0.113 | 0.136 | 1 | 0.268459991 | -0.292078167 | 0.024 | 0.072 | 1 | 0.650695347 | 0.464849215 |
| Cldn12 | 0.532612823 | 0.154722595 | 0.132 | 0.169 | 1 | 0.268951695 | -0.520347155 | 0.146 | 0.222 | 1 | 0.532612823 | 0.465568376 |
| Dpp7 | 0.432887055 | 0.287989126 | 0.264 | 0.212 | 1 | 0.269035449 | 0.536035315 | 0.268 | 0.195 | 1 | 0.432887055 | 0.465690825 |
| Gm11627 | 0.332821087 | 1.8327945 | 0.038 | 0.017 | 1 | 0.269117553 | 1.614812428 | 0.049 | 0.02 | 1 | 0.332821087 | 0.465810849 |
| Dynlt1b | 0.413082433 | 0.154722595 | 0 | 0.013 | 1 | 0.269117553 | 1.614812428 | 0.049 | 0.02 | 1 | 0.413082433 | 0.465810849 |
| Birc5 | 0.478056755 | -2.652632327 | 0.019 | 0.038 | 1 | 0.269117553 | 1.614812428 | 0.049 | 0.02 | 1 | 0.478056755 | 0.465810849 |
| Zfp977 | 0.72256466 | 2.154722595 | 0.019 | 0.013 | 1 | 0.269117553 | 1.614812428 | 0.049 | 0.02 | 1 | 0.72256466 | 0.465810849 |
| Rtl4 | 0.269179131 | -2.889671525 | 0.019 | 0.055 | 1 | 0.435968064 | 1.252242349 | 0.024 | 0.01 | 1 | 0.435968064 | 0.465900857 |
| Dhh | 0.269192031 | -0.652632327 | 0.019 | 0.055 | 1 | 0.826384776 | 0.515276755 | 0.024 | 0.031 | 1 | 0.826384776 | 0.465919712 |
| Cyp7b1 | 0.269292203 | 1.025439578 | 0.132 | 0.085 | 1 | 0.450631276 | -0.355440228 | 0.098 | 0.14 | 1 | 0.450631276 | 0.466066116 |
| Stn1 | 0.385086159 | -0.130679624 | 0.264 | 0.347 | 1 | 0.269340427 | -0.582334042 | 0.171 | 0.249 | 1 | 0.385086159 | 0.466136588 |
| Fam186a | 0.287773478 | -0.430239906 | 0 | 0.021 | 1 | 0.26948834 | 1.83720485 | 0.024 | 0.007 | 1 | 0.287773478 | 0.466352715 |
| Gm30541 | 0.287773478 | -0.430239906 | 0 | 0.021 | 1 | 0.26948834 | 1.252242349 | 0.024 | 0.007 | 1 | 0.287773478 | 0.466352715 |
| Pcp4l1 | 0.327350685 | 2.154722595 | 0.038 | 0.017 | 1 | 0.26948834 | 1.515276755 | 0.024 | 0.007 | 1 | 0.327350685 | 0.466352715 |
| Gm17227 | 0.343000446 | -0.430239906 | 0 | 0.017 | 1 | 0.26948834 | 1.83720485 | 0.024 | 0.007 | 1 | 0.343000446 | 0.466352715 |
| Sntg1 | 0.343004841 | -0.652632327 | 0 | 0.017 | 1 | 0.26948834 | 1.83720485 | 0.024 | 0.007 | 1 | 0.343004841 | 0.466352715 |
| Abcg5 | 0.343006305 | -0.1672055 | 0 | 0.017 | 1 | 0.26948834 | 1.83720485 | 0.024 | 0.007 | 1 | 0.343006305 | 0.466352715 |
| Dennd2d | 0.413080549 | -0.1672055 | 0 | 0.013 | 1 | 0.26948834 | 1.83720485 | 0.024 | 0.007 | 1 | 0.413080549 | 0.466352715 |
| Necab2 | 0.413080549 | -0.1672055 | 0 | 0.013 | 1 | 0.26948834 | 1.515276755 | 0.024 | 0.007 | 1 | 0.413080549 | 0.466352715 |
| Dusp13 | 0.413082433 | 0.154722595 | 0 | 0.013 | 1 | 0.26948834 | 1.83720485 | 0.024 | 0.007 | 1 | 0.413082433 | 0.466352715 |
| Klhl32 | 0.418311293 | 1.502645898 | 0.057 | 0.034 | 1 | 0.26948834 | 1.515276755 | 0.024 | 0.007 | 1 | 0.418311293 | 0.466352715 |
| H4c1 | 0.486967902 | -0.430239906 | 0.019 | 0.038 | 1 | 0.26948834 | 1.83720485 | 0.024 | 0.007 | 1 | 0.486967902 | 0.466352715 |
| Bdnf | 0.503971666 | 1.569760094 | 0.019 | 0.008 | 1 | 0.26948834 | 1.515276755 | 0.024 | 0.007 | 1 | 0.503971666 | 0.466352715 |
| Syne4 | 0.507281812 | 1.154722595 | 0.019 | 0.008 | 1 | 0.26948834 | 1.83720485 | 0.024 | 0.007 | 1 | 0.507281812 | 0.466352715 |
| Soat2 | 0.532834334 | 1.280253477 | 0.057 | 0.038 | 1 | 0.26948834 | 1.515276755 | 0.024 | 0.007 | 1 | 0.532834334 | 0.466352715 |
| Gm15675 | 0.586427152 | 0.454282877 | 0.019 | 0.034 | 1 | 0.26948834 | 0.83720485 | 0.024 | 0.007 | 1 | 0.586427152 | 0.466352715 |
| Fbxl13 | 0.682084573 | 0.8327945 | 0.019 | 0.03 | 1 | 0.26948834 | 1.83720485 | 0.024 | 0.007 | 1 | 0.682084573 | 0.466352715 |
| Gm31816 | 0.732691937 | 1.154722595 | 0.019 | 0.013 | 1 | 0.26948834 | 1.515276755 | 0.024 | 0.007 | 1 | 0.732691937 | 0.466352715 |
| Gm36246 | 0.732691937 | 1.154722595 | 0.019 | 0.013 | 1 | 0.26948834 | 1.83720485 | 0.024 | 0.007 | 1 | 0.732691937 | 0.466352715 |
| Slc30a10 | 0.7394666 | 0.347367673 | 0.019 | 0.013 | 1 | 0.26948834 | 1.252242349 | 0.024 | 0.007 | 1 | 0.7394666 | 0.466352715 |
| Slc5a10 | 0.909120409 | -0.1672055 | 0.019 | 0.021 | 1 | 0.26948834 | 1.515276755 | 0.024 | 0.007 | 1 | 0.909120409 | 0.466352715 |
| Gm5089 | 0.914958 | 0.347367673 | 0.019 | 0.021 | 1 | 0.26948834 | 1.83720485 | 0.024 | 0.007 | 1 | 0.914958 | 0.466352715 |
| Gm56633 | 0.917883255 | 0.569760094 | 0.019 | 0.021 | 1 | 0.26948834 | 1.83720485 | 0.024 | 0.007 | 1 | 0.917883255 | 0.466352715 |
| Gm6260 | 0.932505382 | 1.154722595 | 0.019 | 0.021 | 1 | 0.26948834 | 0.962735732 | 0.024 | 0.007 | 1 | 0.932505382 | 0.466352715 |
| A2ml1 | 1 | 0.47665069 | 0.038 | 0.038 | 1 | 0.26948834 | 0.252242349 | 0.024 | 0.007 | 1 | 1 | 0.466352715 |
| D130040H23Rik | 0.269566579 | 0.953088734 | 0.113 | 0.068 | 1 | 0.440399914 | -0.825760163 | 0.049 | 0.082 | 1 | 0.440399914 | 0.466467017 |
| Dohh | 0.269597783 | 0.359542387 | 0.472 | 0.411 | 1 | 0.568081031 | -0.323259822 | 0.341 | 0.369 | 1 | 0.568081031 | 0.466512602 |
| Zfp458 | 0.269711516 | -1.015202407 | 0.038 | 0.081 | 1 | 0.472947647 | -0.16279515 | 0.049 | 0.082 | 1 | 0.472947647 | 0.46667873 |
| Donson | 0.741101491 | 0.714150003 | 0.075 | 0.064 | 1 | 0.270013856 | 0.778311161 | 0.122 | 0.072 | 1 | 0.741101491 | 0.467120229 |
| Rcc1 | 0.270062231 | 0.466666601 | 0.226 | 0.161 | 1 | 0.881888777 | 0.252242349 | 0.195 | 0.188 | 1 | 0.881888777 | 0.467190853 |
| Plpp2 | 0.471960761 | 0.454282877 | 0.132 | 0.097 | 1 | 0.270261388 | -0.747757651 | 0.073 | 0.133 | 1 | 0.471960761 | 0.467481558 |
| Prkg2 | 0.295397457 | -2.872569097 | 0.151 | 0.208 | 1 | 0.270420203 | -3.484723245 | 0.024 | 0.068 | 1 | 0.295397457 | 0.467713319 |
| Chmp1b2 | 0.270736622 | 0.762405172 | 0.208 | 0.153 | 1 | 0.761682862 | 0.916639317 | 0.146 | 0.14 | 1 | 0.761682862 | 0.468174925 |
| C79798 | 0.524261996 | 1.154722595 | 0.057 | 0.038 | 1 | 0.27115967 | 1.007129851 | 0.098 | 0.055 | 1 | 0.524261996 | 0.468791774 |
| 4930579G24Rik | 0.271439132 | -1.1182959 | 0.038 | 0.081 | 1 | 0.615241183 | 0.422167351 | 0.122 | 0.096 | 1 | 0.615241183 | 0.469199062 |
| Lingo1 | 0.271439132 | -1.067669827 | 0.038 | 0.081 | 1 | 0.627883358 | 1.422167351 | 0.049 | 0.034 | 1 | 0.627883358 | 0.469199062 |
| 4933424G05Rik | 0.271439132 | -1.214511215 | 0.038 | 0.081 | 1 | 0.485983635 | -0.53202896 | 0.049 | 0.078 | 1 | 0.485983635 | 0.469199062 |
| Trappc4 | 0.271477703 | 0.251790013 | 0.679 | 0.602 | 1 | 0.446026964 | 0.43435676 | 0.585 | 0.584 | 1 | 0.446026964 | 0.469255263 |
| Snx11 | 0.382379275 | 0.454282877 | 0.189 | 0.14 | 1 | 0.271628329 | -0.359192363 | 0.098 | 0.167 | 1 | 0.382379275 | 0.469474709 |
| Mrps18c | 0.374799563 | 0.43225657 | 0.396 | 0.339 | 1 | 0.271752195 | 0.869385256 | 0.39 | 0.341 | 1 | 0.374799563 | 0.469655135 |
| Syvn1 | 0.271779294 | 0.458114738 | 0.358 | 0.288 | 1 | 0.350112316 | 0.505008419 | 0.415 | 0.362 | 1 | 0.350112316 | 0.469694604 |
| C330002G04Rik | 0.287771129 | -1.430239906 | 0 | 0.021 | 1 | 0.271787759 | 1.029849928 | 0.024 | 0.007 | 1 | 0.287771129 | 0.469706932 |
| Prph | 0.336525426 | 1.669295768 | 0.038 | 0.017 | 1 | 0.271787759 | 1.252242349 | 0.024 | 0.007 | 1 | 0.336525426 | 0.469706932 |
| Amph | 0.343004841 | -0.652632327 | 0 | 0.017 | 1 | 0.271787759 | 1.100239256 | 0.024 | 0.007 | 1 | 0.343004841 | 0.469706932 |
| 1700061H18Rik | 0.736073999 | 0.154722595 | 0.019 | 0.013 | 1 | 0.271787759 | 0.83720485 | 0.024 | 0.007 | 1 | 0.736073999 | 0.469706932 |
| Cartpt | 0.913424206 | 1.417757001 | 0.019 | 0.017 | 1 | 0.271787759 | 1.029849928 | 0.024 | 0.007 | 1 | 0.913424206 | 0.469706932 |

| | | | | | | | | | | | | |
|---|---|---|---|---|---|---|---|---|---|---|---|---|
| Gm17494 | 0.50514607 | 0.280253477 | 0.019 | 0.038 | 1 | 0.27192168 | 1.252242349 | 0.049 | 0.02 | 1 | 0.50514607 | 0.46990196 |
| Ccl19 | 0.539790273 | 0.891688189 | 0.057 | 0.038 | 1 | 0.27192168 | 1.422167351 | 0.049 | 0.02 | 1 | 0.539790273 | 0.46990196 |
| Rad1 | 0.340804648 | 0.624207878 | 0.132 | 0.089 | 1 | 0.272063045 | 0.435106406 | 0.049 | 0.106 | 1 | 0.340804648 | 0.470107789 |
| Msh2 | 0.483911497 | 0.8327945 | 0.189 | 0.161 | 1 | 0.272149896 | 0.713822434 | 0.244 | 0.171 | 1 | 0.483911497 | 0.470234226 |
| Ddt | 0.526370656 | -0.160319115 | 0.472 | 0.538 | 1 | 0.272297522 | 0.109284395 | 0.317 | 0.427 | 1 | 0.526370656 | 0.470449103 |
| Pde3b | 0.567939333 | 0.598329246 | 0.434 | 0.386 | 1 | 0.272411007 | -0.171357164 | 0.39 | 0.509 | 1 | 0.567939333 | 0.470614258 |
| Nr2c1 | 0.272424988 | 0.869320376 | 0.283 | 0.229 | 1 | 0.625531628 | 0.693369077 | 0.293 | 0.283 | 1 | 0.625531628 | 0.470634601 |
| Limd2 | 0.272569556 | 0.8327945 | 0.189 | 0.136 | 1 | 0.755248421 | 0.551802631 | 0.098 | 0.119 | 1 | 0.755248421 | 0.47084495 |
| Sin3b | 0.633399116 | 0.130180411 | 0.83 | 0.788 | 1 | 0.272779147 | 0.354968651 | 0.878 | 0.833 | 1 | 0.633399116 | 0.471149831 |
| Rab9 | 0.272914269 | 0.325341366 | 0.623 | 0.576 | 1 | 0.820160445 | 0.107125641 | 0.634 | 0.672 | 1 | 0.820160445 | 0.471346339 |
| Shq1 | 0.27302858 | 0.8327945 | 0.132 | 0.085 | 1 | 0.526045905 | -0.717384002 | 0.073 | 0.102 | 1 | 0.526045905 | 0.471512555 |
| Exd1 | 0.622002663 | 0.195364579 | 0.094 | 0.119 | 1 | 0.273061637 | -1.292078167 | 0.024 | 0.068 | 1 | 0.622002663 | 0.471560616 |
| Zswim9 | 0.683023708 | -0.347777746 | 0.038 | 0.051 | 1 | 0.27346637 | 0.287007767 | 0.049 | 0.109 | 1 | 0.683023708 | 0.472148885 |
| Agtpbp1 | 0.441068838 | 0.381493457 | 0.491 | 0.424 | 1 | 0.273731996 | 0.274560729 | 0.683 | 0.645 | 1 | 0.441068838 | 0.472534786 |
| Sorbs3 | 0.407373141 | -0.308451426 | 0.415 | 0.462 | 1 | 0.274334684 | -0.39309277 | 0.39 | 0.478 | 1 | 0.407373141 | 0.473409849 |
| Gnl2 | 0.274354355 | 0.195785251 | 0.755 | 0.729 | 1 | 0.393213665 | -0.177745492 | 0.659 | 0.758 | 1 | 0.393213665 | 0.473438397 |
| Casp7 | 0.656287548 | 0.324647596 | 0.113 | 0.093 | 1 | 0.274511252 | 1.159132945 | 0.122 | 0.075 | 1 | 0.656287548 | 0.473666077 |
| Ly6d | 0.274646579 | -0.804635421 | 0.094 | 0.055 | 1 | 0.844641919 | 0.962735732 | 0.024 | 0.031 | 1 | 0.844641919 | 0.473862414 |
| Mcph1 | 0.274691448 | -0.527101445 | 0.113 | 0.182 | 1 | 0.790430655 | 0.415741081 | 0.171 | 0.16 | 1 | 0.790430655 | 0.473927505 |
| Acbd4 | 0.295945425 | 1.100274811 | 0.151 | 0.102 | 1 | 0.27473555 | -0.622226769 | 0.049 | 0.102 | 1 | 0.295945425 | 0.473991478 |
| Ildr1 | 0.343006305 | -0.1672055 | 0 | 0.017 | 1 | 0.274788535 | 1.252242349 | 0.049 | 0.02 | 1 | 0.343006305 | 0.474068331 |
| Arhgap40 | 0.347746365 | 0.739685096 | 0.038 | 0.017 | 1 | 0.274788535 | 1.252242349 | 0.049 | 0.02 | 1 | 0.347746365 | 0.474068331 |
| Spdya | 0.874462081 | 0.984797593 | 0.038 | 0.034 | 1 | 0.274788535 | 1.100239256 | 0.049 | 0.02 | 1 | 0.874462081 | 0.474068331 |
| Sim1 | 0.797852601 | 0.695290976 | 0.019 | 0.025 | 1 | 0.27480341 | 0.930314254 | 0.049 | 0.02 | 1 | 0.797852601 | 0.474089906 |
| Slc2a8 | 0.27499384 | 0.8817041 | 0.17 | 0.119 | 1 | 0.858309577 | 0.79281073 | 0.098 | 0.092 | 1 | 0.858309577 | 0.474366067 |
| Acss2 | 0.410545609 | -0.430239906 | 0.057 | 0.093 | 1 | 0.275081053 | 0.930314254 | 0.171 | 0.113 | 1 | 0.410545609 | 0.47449252 |
| Ptges3l | 0.539565946 | -0.489133595 | 0.057 | 0.081 | 1 | 0.275698423 | -0.970150072 | 0.024 | 0.068 | 1 | 0.539565946 | 0.475387225 |
| Rexo5 | 0.275916427 | 1.154722595 | 0.151 | 0.102 | 1 | 0.920631033 | 0.544423101 | 0.122 | 0.119 | 1 | 0.920631033 | 0.475702979 |
| Gar1 | 0.276028665 | 0.513803688 | 0.509 | 0.453 | 1 | 0.459533461 | -0.181988715 | 0.39 | 0.461 | 1 | 0.459533461 | 0.475865506 |
| Zfp13 | 0.563400654 | 0.739685096 | 0.075 | 0.055 | 1 | 0.276033035 | 1.252242349 | 0.073 | 0.038 | 1 | 0.563400654 | 0.475871833 |
| Alg11 | 0.276307974 | 0.704061186 | 0.208 | 0.148 | 1 | 0.98744342 | 0.252242349 | 0.171 | 0.174 | 1 | 0.98744342 | 0.476269851 |
| Aldh1a1 | 0.410227083 | 0.228723176 | 0.075 | 0.047 | 1 | 0.276476329 | 0.580865097 | 0.195 | 0.13 | 1 | 0.410227083 | 0.476513498 |
| D030028A08Rik | 0.400685124 | -1.1182959 | 0.038 | 0.068 | 1 | 0.276573927 | -0.863234868 | 0.024 | 0.068 | 1 | 0.400685124 | 0.476654717 |
| 1700029H14Rik | 0.318933047 | 0.739685096 | 0.113 | 0.072 | 1 | 0.276574583 | -0.970150072 | 0.024 | 0.068 | 1 | 0.318933047 | 0.476655666 |
| A330069E16Rik | 0.276626224 | -1.1672055 | 0.038 | 0.081 | 1 | 0.450957104 | -0.686357106 | 0.024 | 0.051 | 1 | 0.450957104 | 0.476730381 |
| Akr1b3 | 0.836385841 | -0.281917159 | 0.415 | 0.381 | 1 | 0.277074944 | 0.773878914 | 0.439 | 0.375 | 1 | 0.836385841 | 0.477379363 |
| Cbwd1 | 0.374835604 | -0.242612903 | 0.415 | 0.475 | 1 | 0.277128703 | -0.528444623 | 0.195 | 0.276 | 1 | 0.374835604 | 0.477457088 |
| Tpcn2 | 0.599114706 | -0.738362201 | 0.113 | 0.136 | 1 | 0.277143318 | -1.098254898 | 0.171 | 0.242 | 1 | 0.599114706 | 0.477478217 |
| Agrp | 0.286299986 | 1.221836791 | 0.113 | 0.072 | 1 | 0.277582718 | 1.136765132 | 0.122 | 0.075 | 1 | 0.286299986 | 0.478113271 |
| D830036C21Rik | 0.997810858 | 0.695290976 | 0.038 | 0.038 | 1 | 0.277587118 | 0.515276755 | 0.122 | 0.072 | 1 | 0.997810858 | 0.478119627 |
| Emc10 | 0.277691736 | 0.247831999 | 0.736 | 0.746 | 1 | 0.847780946 | 0.148704745 | 0.659 | 0.672 | 1 | 0.847780946 | 0.478270772 |
| Fam216a | 0.277878788 | 0.84156471 | 0.321 | 0.267 | 1 | 0.79871701 | 0.396632258 | 0.268 | 0.314 | 1 | 0.79871701 | 0.478540956 |
| Nom1 | 0.277982518 | 0.352403263 | 0.415 | 0.364 | 1 | 0.728074836 | 0.252242349 | 0.366 | 0.433 | 1 | 0.728074836 | 0.478690755 |
| Strn | 0.278093686 | 0.208615603 | 0.585 | 0.538 | 1 | 0.462727094 | 0.269116168 | 0.683 | 0.696 | 1 | 0.462727094 | 0.478851274 |
| B9d2 | 0.278180766 | 0.792152515 | 0.189 | 0.136 | 1 | 0.97385589 | 0.35607816 | 0.171 | 0.174 | 1 | 0.97385589 | 0.478976994 |
| Tada2a | 0.278229277 | 0.261637799 | 0.377 | 0.309 | 1 | 0.91019747 | 0.262296014 | 0.268 | 0.3 | 1 | 0.91019747 | 0.479047024 |
| Riox2 | 0.348109245 | 0.417757001 | 0.132 | 0.089 | 1 | 0.278320434 | -0.686357106 | 0.024 | 0.068 | 1 | 0.348109245 | 0.479178604 |
| Sema3b | 0.628925725 | 0.739685096 | 0.038 | 0.025 | 1 | 0.278320434 | -0.747757651 | 0.024 | 0.068 | 1 | 0.628925725 | 0.479178604 |
| Dars2 | 0.27850928 | 0.8327945 | 0.094 | 0.055 | 1 | 0.338667992 | 0.766815522 | 0.098 | 0.058 | 1 | 0.338667992 | 0.479451141 |
| Mrps12 | 0.278656079 | 0.53023173 | 0.472 | 0.441 | 1 | 0.394258108 | 0.359157553 | 0.268 | 0.386 | 1 | 0.394258108 | 0.479662947 |
| 2810002D19Rik | 0.278967432 | -0.771276824 | 0.075 | 0.127 | 1 | 0.463665412 | -0.16279515 | 0.098 | 0.14 | 1 | 0.463665412 | 0.480112035 |
| Slc35c2 | 0.278985709 | 0.906795081 | 0.208 | 0.148 | 1 | 0.437671635 | 0.252242349 | 0.122 | 0.177 | 1 | 0.437671635 | 0.480138392 |
| Vasp | 0.429488791 | 0.232259195 | 0.698 | 0.636 | 1 | 0.279065906 | -0.269366814 | 0.732 | 0.819 | 1 | 0.429488791 | 0.480254032 |
| Pigc | 0.417804239 | 1.076720083 | 0.17 | 0.14 | 1 | 0.279185752 | -0.484723245 | 0.146 | 0.222 | 1 | 0.417804239 | 0.480426821 |
| Gm56976 | 0.686596267 | -0.260314904 | 0.038 | 0.051 | 1 | 0.279190116 | -0.622226769 | 0.024 | 0.068 | 1 | 0.686596267 | 0.480433111 |
| Brix1 | 0.705091666 | 0.17408792 | 0.642 | 0.661 | 1 | 0.279363128 | -0.195810207 | 0.61 | 0.696 | 1 | 0.705091666 | 0.480682498 |
| Spx | 0.279369347 | 1.055186921 | 0.094 | 0.055 | 1 | 0.522228536 | 0.30115195 | 0.049 | 0.078 | 1 | 0.522228536 | 0.480691462 |
| Parp16 | 0.279369347 | 1.055186921 | 0.094 | 0.055 | 1 | 0.329784344 | 1.749742009 | 0.073 | 0.041 | 1 | 0.329784344 | 0.480691462 |
| Cybrd1 | 0.41163426 | 0.370451286 | 0.226 | 0.174 | 1 | 0.279480408 | 0.212713985 | 0.171 | 0.273 | 1 | 0.41163426 | 0.480851517 |
| Chchd7 | 0.372460598 | 0.251398614 | 0.566 | 0.538 | 1 | 0.279835739 | -0.197065052 | 0.366 | 0.457 | 1 | 0.372460598 | 0.481363438 |
| Kcnip1 | 0.287761733 | -0.652632327 | 0 | 0.021 | 1 | 0.280392128 | 1.351778023 | 0.073 | 0.038 | 1 | 0.287761733 | 0.482164511 |
| Slc22a18 | 0.849840916 | 0.669295768 | 0.057 | 0.051 | 1 | 0.280392128 | 1.071670103 | 0.073 | 0.038 | 1 | 0.849840916 | 0.482164511 |
| Zhx3 | 0.28686419 | 0.218852932 | 0.717 | 0.644 | 1 | 0.28085723 | 0.461531183 | 0.732 | 0.754 | 1 | 0.28686419 | 0.482833677 |
| Chd1l | 0.822232137 | -0.312403416 | 0.208 | 0.22 | 1 | 0.281129491 | -0.648221977 | 0.146 | 0.222 | 1 | 0.822232137 | 0.483225192 |
| Psmg2 | 0.281428853 | 0.438515561 | 0.491 | 0.411 | 1 | 0.991423292 | 0.228395607 | 0.293 | 0.3 | 1 | 0.991423292 | 0.483655506 |
| Rfng | 0.298600205 | -0.545717123 | 0.094 | 0.148 | 1 | 0.28163501 | -0.648221977 | 0.098 | 0.16 | 1 | 0.298600205 | 0.483951742 |
| Zfp617 | 0.281734851 | 0.569760094 | 0.264 | 0.199 | 1 | 0.306864116 | -0.221688839 | 0.122 | 0.198 | 1 | 0.306864116 | 0.484095176 |
| Slc2a13 | 0.281840062 | 0.190346505 | 0.358 | 0.288 | 1 | 0.571177873 | -0.10875531 | 0.268 | 0.311 | 1 | 0.571177873 | 0.484246304 |
| Ulk3 | 0.281986656 | 0.154722595 | 0.019 | 0.055 | 1 | 0.483530927 | 0.989207943 | 0.073 | 0.048 | 1 | 0.483530927 | 0.484456838 |
| Upf3a | 0.282093715 | 0.539545481 | 0.528 | 0.5 | 1 | 0.682657085 | 0.147373231 | 0.439 | 0.546 | 1 | 0.682657085 | 0.484610567 |
| Tsc2 | 0.282231438 | 0.514264981 | 0.264 | 0.195 | 1 | 0.660740151 | 0.471555377 | 0.146 | 0.184 | 1 | 0.660740151 | 0.484808292 |
| Polr2k | 0.282498317 | 0.2967416 | 0.717 | 0.653 | 1 | 0.957695422 | 0.275893617 | 0.537 | 0.556 | 1 | 0.957695422 | 0.485191335 |
| Efcab15 | 0.975550414 | 0.2967416 | 0.094 | 0.093 | 1 | 0.28263577 | 1.351778023 | 0.073 | 0.038 | 1 | 0.975550414 | 0.485388562 |
| Lsm3 | 0.816814967 | 0.154722595 | 0.472 | 0.504 | 1 | 0.282834306 | 0.856313673 | 0.439 | 0.389 | 1 | 0.816814967 | 0.485673367 |
| Med10 | 0.283043857 | 0.385335523 | 0.491 | 0.424 | 1 | 0.329101496 | -0.405651674 | 0.293 | 0.355 | 1 | 0.329101496 | 0.485973889 |
| Eif2b2 | 0.283051173 | 0.8327945 | 0.264 | 0.208 | 1 | 0.81461822 | 0.316948038 | 0.293 | 0.287 | 1 | 0.81461822 | 0.485984379 |
| 2310058D17Rik | 0.283110345 | 0.389187848 | 0.019 | 0.055 | 1 | 0.360451923 | 0.930314254 | 0.073 | 0.041 | 1 | 0.360451923 | 0.486069223 |
| Gngt2 | 0.283133698 | 0.491757582 | 0.019 | 0.055 | 1 | 0.66053601 | -0.747757651 | 0.024 | 0.038 | 1 | 0.66053601 | 0.486102706 |
| Ibsp | 0.95405956 | -6.864868134 | 0.038 | 0.038 | 1 | 0.283494073 | -1.952872081 | 0.049 | 0.02 | 1 | 0.95405956 | 0.486619257 |
| Itga2b | 0.394083223 | 0.997181318 | 0.151 | 0.114 | 1 | 0.283563212 | 0.493250449 | 0.195 | 0.133 | 1 | 0.394083223 | 0.486718329 |
| Vrk3 | 0.710383136 | 0.240452469 | 0.34 | 0.326 | 1 | 0.283879492 | -0.320336427 | 0.195 | 0.287 | 1 | 0.710383136 | 0.487171418 |
| Ucp2 | 0.284103786 | 0.306725688 | 0.17 | 0.114 | 1 | 0.613910785 | -0.941768271 | 0.22 | 0.184 | 1 | 0.613910785 | 0.487492611 |
| Zglp1 | 0.284266016 | 0.962077517 | 0.094 | 0.055 | 1 | 0.30567149 | 1.766815522 | 0.098 | 0.058 | 1 | 0.30567149 | 0.487724864 |
| Polr2f | 0.284402898 | 0.278205648 | 0.679 | 0.619 | 1 | 0.862377823 | 0.287866259 | 0.537 | 0.543 | 1 | 0.862377823 | 0.487920788 |
| Pxmp4 | 0.608350988 | -0.181560793 | 0.302 | 0.331 | 1 | 0.284792093 | -0.136799942 | 0.098 | 0.171 | 1 | 0.608350988 | 0.48847765 |
| Gm28198 | 0.2849274 | 0.752624151 | 0.189 | 0.136 | 1 | 0.538796679 | 0.408361551 | 0.22 | 0.181 | 1 | 0.538796679 | 0.488671177 |
| Plekha5 | 0.285021387 | 0.315338819 | 0.755 | 0.682 | 1 | 0.689974699 | 0.265181405 | 0.756 | 0.867 | 1 | 0.689974699 | 0.488805583 |
| Utp11 | 0.285081398 | 0.166218234 | 0.623 | 0.593 | 1 | 0.49219801 | 0.202254213 | 0.463 | 0.601 | 1 | 0.49219801 | 0.488891392 |
| Coa5 | 0.285189715 | 0.526691372 | 0.17 | 0.114 | 1 | 0.59528192 | -0.213421223 | 0.122 | 0.154 | 1 | 0.59528192 | 0.489046256 |
| Dgki | 0.28536692 | -1.027480736 | 0.094 | 0.153 | 1 | 0.656824467 | -0.210100865 | 0.122 | 0.099 | 1 | 0.656824467 | 0.48929956 |
| Rpap3 | 0.285789516 | 0.690054328 | 0.302 | 0.237 | 1 | 0.87407844 | 0.601988388 | 0.22 | 0.249 | 1 | 0.87407844 | 0.489903385 |
| Lynx1 | 0.285798574 | -0.356239324 | 0.132 | 0.195 | 1 | 0.476532722 | 0.212713985 | 0.146 | 0.205 | 1 | 0.476532722 | 0.489916324 |
| Gm15738 | 0.972529586 | 0.247831999 | 0.057 | 0.055 | 1 | 0.285802944 | -1.16279515 | 0.049 | 0.099 | 1 | 0.972529586 | 0.489922566 |
| Coprs | 0.920350007 | 0.129187503 | 0.208 | 0.203 | 1 | 0.285850284 | -0.332720152 | 0.098 | 0.164 | 1 | 0.920350007 | 0.489990183 |
| Mthfsd | 0.461591142 | 0.454282877 | 0.189 | 0.148 | 1 | 0.286298195 | -0.440779897 | 0.171 | 0.249 | 1 | 0.461591142 | 0.490629733 |
| Efcab9 | 0.287773478 | -0.430239906 | 0 | 0.021 | 1 | 0.286402368 | -0.622226769 | 0 | 0.027 | 1 | 0.287773478 | 0.49077842 |
| Irag2 | 0.47749495 | 0.569760094 | 0.038 | 0.021 | 1 | 0.286402368 | -0.484723245 | 0 | 0.027 | 1 | 0.47749495 | 0.49077842 |
| Gm56817 | 0.668070194 | 0.154722595 | 0.019 | 0.03 | 1 | 0.286402368 | -0.484723245 | 0 | 0.027 | 1 | 0.668070194 | 0.49077842 |

| | | | | | | | | | | | | |
|---|---|---|---|---|---|---|---|---|---|---|---|---|
| Ube2c | 0.668779201 | -3.001781891 | 0.038 | 0.051 | 1 | 0.286402368 | -0.747757651 | 0 | 0.027 | 1 | 0.668779201 | 0.49077842 |
| Gm43560 | 0.765272199 | 0.216123139 | 0.075 | 0.064 | 1 | 0.286402368 | -0.622226769 | 0 | 0.027 | 1 | 0.765272199 | 0.49077842 |
| Cxcr4 | 0.916620008 | 1.347367673 | 0.019 | 0.017 | 1 | 0.286402368 | -0.484723245 | 0 | 0.027 | 1 | 0.916620008 | 0.49077842 |
| Llgl2 | 0.917883255 | 0.569760094 | 0.019 | 0.021 | 1 | 0.286402368 | -0.484723245 | 0 | 0.027 | 1 | 0.917883255 | 0.49077842 |
| Gm48529 | 0.926212226 | 0.8327945 | 0.019 | 0.017 | 1 | 0.286402368 | -0.484723245 | 0 | 0.027 | 1 | 0.926212226 | 0.49077842 |
| Gm21411 | 0.932505382 | 1.154722595 | 0.019 | 0.021 | 1 | 0.286402368 | -0.484723245 | 0 | 0.027 | 1 | 0.932505382 | 0.49077842 |
| Fbp2 | 0.28776878 | -0.845277405 | 0 | 0.021 | 1 | 0.286413984 | -0.970150072 | 0 | 0.027 | 1 | 0.28776878 | 0.490794998 |
| Gm43150 | 0.370966835 | -0.430239906 | 0.019 | 0.047 | 1 | 0.286413984 | -0.863234868 | 0 | 0.027 | 1 | 0.370966835 | 0.490794998 |
| Sycp3 | 0.503971666 | 1.569760094 | 0.019 | 0.008 | 1 | 0.286413984 | -0.622226769 | 0 | 0.027 | 1 | 0.503971666 | 0.490794998 |
| Gm48008 | 0.503971666 | 1.569760094 | 0.019 | 0.008 | 1 | 0.286413984 | -0.622226769 | 0 | 0.027 | 1 | 0.503971666 | 0.490794998 |
| BC049762 | 0.684424342 | 1.47665069 | 0.019 | 0.03 | 1 | 0.286413984 | -0.863234868 | 0 | 0.027 | 1 | 0.684424342 | 0.490794998 |
| Xrcc2 | 0.779443586 | 0.154722595 | 0.019 | 0.025 | 1 | 0.286413984 | -0.622226769 | 0 | 0.027 | 1 | 0.779443586 | 0.490794998 |
| Ppp1r3b | 0.861498586 | -0.693274312 | 0.075 | 0.081 | 1 | 0.286413984 | -0.747757651 | 0 | 0.027 | 1 | 0.861498586 | 0.490794998 |
| Spata48 | 0.912039013 | 0.154722595 | 0.019 | 0.021 | 1 | 0.286413984 | -1.747757651 | 0 | 0.027 | 1 | 0.912039013 | 0.490794998 |
| Gm16278 | 0.926212226 | 0.8327945 | 0.019 | 0.017 | 1 | 0.286421728 | -1.250257991 | 0 | 0.027 | 1 | 0.926212226 | 0.49080605 |
| Odf3l2 | 0.343000446 | -0.430239906 | 0 | 0.017 | 1 | 0.286422281 | -0.747757651 | 0 | 0.027 | 1 | 0.343000446 | 0.490806839 |
| Arhgef4 | 0.658774723 | -1.015202407 | 0.019 | 0.03 | 1 | 0.286422281 | -0.747757651 | 0 | 0.027 | 1 | 0.658774723 | 0.490806839 |
| Cfap299 | 0.923013642 | 0.228723176 | 0.019 | 0.017 | 1 | 0.286422281 | -1.555112573 | 0 | 0.027 | 1 | 0.923013642 | 0.490806839 |
| 2010009K17Rik | 0.661097298 | -0.845277405 | 0.019 | 0.03 | 1 | 0.2864256 | -1.332720152 | 0 | 0.027 | 1 | 0.661097298 | 0.490811576 |
| Galnt16 | 0.764778716 | -0.752168001 | 0.038 | 0.03 | 1 | 0.286427259 | -1.069685746 | 0 | 0.027 | 1 | 0.764778716 | 0.490813944 |
| 8030442B05Rik | 0.343006305 | -0.845277405 | 0 | 0.017 | 1 | 0.286427813 | -0.863234868 | 0 | 0.027 | 1 | 0.343006305 | 0.490814733 |
| 1700003E16Rik | 0.776830424 | -0.430239906 | 0.019 | 0.025 | 1 | 0.286427813 | -0.863234868 | 0 | 0.027 | 1 | 0.776830424 | 0.490814733 |
| Iqch | 0.77860339 | -0.693274312 | 0.057 | 0.047 | 1 | 0.286427813 | -1.80665134 | 0 | 0.027 | 1 | 0.77860339 | 0.490814733 |
| Pnpla3 | 0.302504397 | -0.508242418 | 0.075 | 0.127 | 1 | 0.286428919 | -1.332720152 | 0 | 0.027 | 1 | 0.302504397 | 0.490816312 |
| Spink5 | 0.926212226 | 0.8327945 | 0.019 | 0.017 | 1 | 0.286428919 | -1.250257991 | 0 | 0.027 | 1 | 0.926212226 | 0.490816312 |
| Mmp3 | 0.471584468 | 0.492891331 | 0.132 | 0.097 | 1 | 0.286430578 | -2.890715605 | 0 | 0.027 | 1 | 0.471584468 | 0.49081868 |
| Spi1 | 0.932506754 | 1.962077517 | 0.019 | 0.021 | 1 | 0.28643279 | -3.448197369 | 0 | 0.027 | 1 | 0.932506754 | 0.490821837 |
| Gm50146 | 0.343968402 | 1.154722595 | 0.038 | 0.017 | 1 | 0.286433344 | -0.332720152 | 0 | 0.027 | 1 | 0.343968402 | 0.490822627 |
| Zmynd15 | 0.413080549 | -0.1672055 | 0 | 0.013 | 1 | 0.286433344 | -0.332720152 | 0 | 0.027 | 1 | 0.413080549 | 0.490822627 |
| Ect2 | 0.450583858 | -0.477545621 | 0.057 | 0.034 | 1 | 0.286433344 | -0.332720152 | 0 | 0.027 | 1 | 0.450583858 | 0.490822627 |
| Poll | 0.529359278 | 1.154722595 | 0.057 | 0.038 | 1 | 0.286433344 | -0.332720152 | 0 | 0.027 | 1 | 0.529359278 | 0.490822627 |
| Mrap | 0.588500957 | 0.695290976 | 0.019 | 0.034 | 1 | 0.286433344 | -0.332720152 | 0 | 0.027 | 1 | 0.588500957 | 0.490822627 |
| Gm9922 | 0.897146191 | 0.417757001 | 0.038 | 0.034 | 1 | 0.286433344 | -0.332720152 | 0 | 0.027 | 1 | 0.897146191 | 0.490822627 |
| Gm42595 | 0.926212226 | 0.8327945 | 0.019 | 0.017 | 1 | 0.286433344 | -0.332720152 | 0 | 0.027 | 1 | 0.926212226 | 0.490822627 |
| Cep128 | 0.286436373 | 0.101611258 | 0.075 | 0.131 | 1 | 0.497057746 | -0.442903069 | 0.073 | 0.109 | 1 | 0.497057746 | 0.49082695 |
| Ran | 0.286471342 | 0.193444593 | 0.925 | 0.894 | 1 | 0.392582032 | -0.119013458 | 0.854 | 0.925 | 1 | 0.392582032 | 0.490876854 |
| Kars | 0.46109942 | 0.655916738 | 0.283 | 0.246 | 1 | 0.286549209 | 0.119348079 | 0.22 | 0.328 | 1 | 0.46109942 | 0.490987969 |
| Ctu2 | 0.286660234 | 0.569760094 | 0.208 | 0.148 | 1 | 0.508908435 | 0.690363461 | 0.195 | 0.157 | 1 | 0.508908435 | 0.491146378 |
| Ptprn2 | 0.311343629 | -2.441128222 | 0.094 | 0.144 | 1 | 0.286788914 | -1.949391512 | 0.049 | 0.099 | 1 | 0.311343629 | 0.491329947 |
| Zdhhc18 | 0.286832249 | -0.438507522 | 0.358 | 0.436 | 1 | 0.552061511 | 0.246389623 | 0.341 | 0.454 | 1 | 0.552061511 | 0.491391759 |
| Alkbh6 | 0.493901834 | 0.377115016 | 0.302 | 0.267 | 1 | 0.287060824 | 0.715214325 | 0.122 | 0.198 | 1 | 0.493901834 | 0.491717731 |
| Usp30 | 0.815010891 | -0.1672055 | 0.075 | 0.085 | 1 | 0.287090666 | -0.262330824 | 0.098 | 0.164 | 1 | 0.815010891 | 0.491760281 |
| Tmem240 | 0.28739801 | 0.932330173 | 0.094 | 0.055 | 1 | 0.542016385 | 0.721727632 | 0.098 | 0.072 | 1 | 0.542016385 | 0.492198405 |
| 2010310C07Rik | 0.572347278 | 0.491757582 | 0.038 | 0.059 | 1 | 0.287488435 | 0.707921833 | 0.122 | 0.075 | 1 | 0.572347278 | 0.492327269 |
| Slco2a1 | 0.287540064 | -0.681778673 | 0.057 | 0.106 | 1 | 0.828937904 | -1.088794569 | 0.098 | 0.106 | 1 | 0.828937904 | 0.49240084 |
| 4930549G23Rik | 0.287647022 | -0.545717123 | 0.038 | 0.081 | 1 | 0.979754959 | -0.233184478 | 0.098 | 0.096 | 1 | 0.979754959 | 0.492553234 |
| G730003C15Rik | 0.287761733 | -0.652632327 | 0 | 0.021 | 1 | 0.455098275 | 0.515276755 | 0 | 0.014 | 1 | 0.455098275 | 0.49271665 |
| Meig1 | 0.287761733 | -0.652632327 | 0 | 0.021 | 1 | 0.590756788 | 2.159132945 | 0.024 | 0.014 | 1 | 0.590756788 | 0.49271665 |
| Gpr87 | 0.287761733 | -0.652632327 | 0 | 0.021 | 1 | 0.744047924 | 1.252242349 | 0.024 | 0.017 | 1 | 0.744047924 | 0.49271665 |
| Cd72 | 0.287761733 | -1.652632327 | 0 | 0.021 | 1 | 0.66636882 | -0.332720152 | 0.049 | 0.034 | 1 | 0.66636882 | 0.49271665 |
| Gm49890 | 0.287761733 | -0.652632327 | 0 | 0.021 | 1 | 0.938628024 | -1.825760163 | 0.049 | 0.044 | 1 | 0.938628024 | 0.49271665 |
| Armh1 | 0.287761733 | -0.652632327 | 0 | 0.021 | 1 | 0.438672228 | 1.83720485 | 0.024 | 0.01 | 1 | 0.438672228 | 0.49271665 |
| Slc26a1 | 0.287761733 | -0.652632327 | 0 | 0.021 | 1 | 0.319641967 | -0.16279515 | 0 | 0.024 | 1 | 0.319641967 | 0.49271665 |
| 5430402O13Rik | 0.287761733 | -0.652632327 | 0 | 0.021 | 1 | 0.319619815 | -0.332720152 | 0 | 0.024 | 1 | 0.319619815 | 0.49271665 |
| Sox5os4 | 0.287761733 | -0.652632327 | 0 | 0.021 | 1 | 0.5191986 | 0.515276755 | 0 | 0.01 | 1 | 0.5191986 | 0.49271665 |
| Gm26708 | 0.287761733 | -0.652632327 | 0 | 0.021 | 1 | 0.51919986 | 0.252242349 | 0 | 0.01 | 1 | 0.51919986 | 0.49271665 |
| Ctxnd1 | 0.287761733 | -1.015202407 | 0 | 0.021 | 1 | 0.733321373 | -1.747757651 | 0.024 | 0.034 | 1 | 0.733321373 | 0.49271665 |
| Tex12 | 0.287761733 | -0.652632327 | 0 | 0.021 | 1 | 0.402369942 | -0.16279515 | 0 | 0.017 | 1 | 0.402369942 | 0.49271665 |
| Sox30 | 0.287761733 | -0.652632327 | 0 | 0.021 | 1 | 0.455098275 | 0.515276755 | 0 | 0.014 | 1 | 0.455098275 | 0.49271665 |
| Gm57409 | 0.287761733 | -0.652632327 | 0 | 0.021 | 1 | 0.5191986 | 0.515276755 | 0 | 0.01 | 1 | 0.5191986 | 0.49271665 |
| Zfp85 | 0.287761733 | -0.652632327 | 0 | 0.021 | 1 | 0.989595616 | 0.667279848 | 0.024 | 0.024 | 1 | 0.989595616 | 0.49271665 |
| Gm10037 | 0.287761733 | -0.652632327 | 0 | 0.021 | 1 | 0.51919986 | 0.83720485 | 0 | 0.01 | 1 | 0.51919986 | 0.49271665 |
| Gpr18 | 0.287761733 | -1.015202407 | 0 | 0.021 | 1 | 0.58792152 | 2.515276755 | 0.024 | 0.014 | 1 | 0.58792152 | 0.49271665 |
| Gm53058 | 0.287761733 | -1.015202407 | 0 | 0.021 | 1 | 0.728967958 | -0.665295491 | 0.049 | 0.061 | 1 | 0.728967958 | 0.49271665 |
| A930007A09Rik | 0.287761733 | -0.652632327 | 0 | 0.021 | 1 | 0.438672228 | 1.83720485 | 0.024 | 0.01 | 1 | 0.438672228 | 0.49271665 |
| Tff2 | 0.287761733 | -0.845277405 | 0 | 0.021 | 1 | 0.606262835 | -0.16279515 | 0.024 | 0.041 | 1 | 0.606262835 | 0.49271665 |
| Rsph1 | 0.287761733 | -1.015202407 | 0 | 0.021 | 1 | 0.492841038 | -0.622226769 | 0.024 | 0.048 | 1 | 0.492841038 | 0.49271665 |
| Gm56649 | 0.287761733 | -0.652632327 | 0 | 0.021 | 1 | 0.856407545 | 2.159132945 | 0.024 | 0.02 | 1 | 0.856407545 | 0.49271665 |
| Gm32261 | 0.287761733 | -0.652632327 | 0 | 0.021 | 1 | 0.438672228 | 1.83720485 | 0.024 | 0.01 | 1 | 0.438672228 | 0.49271665 |
| Cdkn2a | 0.28776878 | -0.845277405 | 0 | 0.021 | 1 | 0.5191986 | 0.515276755 | 0 | 0.01 | 1 | 0.5191986 | 0.49272669 |
| Mroh7 | 0.28776878 | -0.845277405 | 0 | 0.021 | 1 | 0.916649754 | 0.667279848 | 0.024 | 0.027 | 1 | 0.916649754 | 0.49272669 |
| Rhd | 0.28776878 | -0.845277405 | 0 | 0.021 | 1 | 0.357847088 | -0.622226769 | 0 | 0.02 | 1 | 0.357847088 | 0.49272669 |
| Elfn1 | 0.28776878 | -1.015202407 | 0 | 0.021 | 1 | 0.629768283 | 0.83720485 | 0.049 | 0.034 | 1 | 0.629768283 | 0.49272669 |
| 9530062K07Rik | 0.28776878 | -1.015202407 | 0 | 0.021 | 1 | 0.872828964 | 1.029849928 | 0.024 | 0.02 | 1 | 0.872828964 | 0.49272669 |
| Sult2b1 | 0.28776878 | -1.304709024 | 0 | 0.021 | 1 | 0.744047924 | 1.252242349 | 0.024 | 0.017 | 1 | 0.744047924 | 0.49272669 |
| Gm27201 | 0.28776878 | -0.845277405 | 0 | 0.021 | 1 | 0.455098275 | 0.515276755 | 0 | 0.014 | 1 | 0.455098275 | 0.49272669 |
| Gm40528 | 0.28776878 | -2.652632327 | 0 | 0.021 | 1 | 0.87556513 | 0.515276755 | 0.024 | 0.02 | 1 | 0.87556513 | 0.49272669 |
| Slc39a2 | 0.28776878 | -1.304709024 | 0 | 0.021 | 1 | 0.5191986 | 0.515276755 | 0 | 0.01 | 1 | 0.5191986 | 0.49272669 |
| Sp100 | 0.287771129 | -2.093204919 | 0 | 0.021 | 1 | 0.435967024 | 1.252242349 | 0.024 | 0.01 | 1 | 0.435967024 | 0.492730036 |
| Acvr1c | 0.287771129 | -1.304709024 | 0 | 0.021 | 1 | 0.992197063 | 0.252242349 | 0.024 | 0.024 | 1 | 0.992197063 | 0.492730036 |
| Ksr2 | 0.287771129 | -2.015202407 | 0 | 0.021 | 1 | 0.522145483 | 1.564186355 | 0.024 | 0.048 | 1 | 0.522145483 | 0.492730036 |
| Gm29677 | 0.287771129 | -1.1672055 | 0 | 0.021 | 1 | 0.527945596 | 0.83720485 | 0.049 | 0.031 | 1 | 0.527945596 | 0.492730036 |
| Gm2447 | 0.287772304 | -2.1672055 | 0 | 0.021 | 1 | 0.51919986 | 0.83720485 | 0 | 0.01 | 1 | 0.51919986 | 0.492731709 |
| Vmn1r4 | 0.287772304 | -1.015202407 | 0 | 0.021 | 1 | 0.441381862 | 1.515276755 | 0.024 | 0.01 | 1 | 0.441381862 | 0.492731709 |
| Rad51ap1 | 0.287772304 | -1.845277405 | 0 | 0.021 | 1 | 0.455094184 | 0.252242349 | 0 | 0.014 | 1 | 0.455094184 | 0.492731709 |
| H2ac4 | 0.287772304 | -2.974560422 | 0 | 0.021 | 1 | 0.362195044 | 1.252242349 | 0.049 | 0.024 | 1 | 0.362195044 | 0.492731709 |
| Cyb561d1 | 0.287773478 | -0.430239906 | 0 | 0.021 | 1 | 0.455098275 | 0.515276755 | 0 | 0.014 | 1 | 0.455098275 | 0.492733382 |
| Dmrta1 | 0.287773478 | -0.430239906 | 0 | 0.021 | 1 | 0.817109602 | 0.721727632 | 0.049 | 0.041 | 1 | 0.817109602 | 0.492733382 |
| Gm12758 | 0.287773478 | -0.430239906 | 0 | 0.021 | 1 | 0.357847088 | -0.332720152 | 0 | 0.02 | 1 | 0.357847088 | 0.492733382 |
| Cfap46 | 0.287773478 | -0.430239906 | 0 | 0.021 | 1 | 0.615748201 | 0.313642894 | 0.049 | 0.072 | 1 | 0.615748201 | 0.492733382 |
| Osgin1 | 0.287773478 | -1.752168001 | 0 | 0.021 | 1 | 0.735264387 | -0.278272368 | 0.049 | 0.061 | 1 | 0.735264387 | 0.492733382 |
| Gm57013 | 0.287773478 | -0.430239906 | 0 | 0.021 | 1 | 0.872828964 | 1.029849928 | 0.024 | 0.02 | 1 | 0.872828964 | 0.492733382 |
| Gm15915 | 0.287773478 | -0.430239906 | 0 | 0.021 | 1 | 0.51919986 | 0.83720485 | 0 | 0.01 | 1 | 0.51919986 | 0.492733382 |
| Gm56994 | 0.287773478 | -0.430239906 | 0 | 0.021 | 1 | 0.51919986 | 0.83720485 | 0 | 0.01 | 1 | 0.51919986 | 0.492733382 |
| Fat2 | 0.287773478 | -2.652632327 | 0 | 0.021 | 1 | 0.81727826 | -0.484723245 | 0.024 | 0.031 | 1 | 0.81727826 | 0.492733382 |
| Gm57337 | 0.287773478 | -0.430239906 | 0 | 0.021 | 1 | 0.599290422 | 1.515276755 | 0.024 | 0.014 | 1 | 0.599290422 | 0.492733382 |
| Gm32934 | 0.287773478 | -0.430239906 | 0 | 0.021 | 1 | 0.51919986 | -0.16279515 | 0 | 0.01 | 1 | 0.51919986 | 0.492733382 |
| Nhlrc1 | 0.287773478 | -0.430239906 | 0 | 0.021 | 1 | 0.455098275 | 0.515276755 | 0 | 0.014 | 1 | 0.455098275 | 0.492733382 |

| | | | | | | | | | | | | |
|---|---|---|---|---|---|---|---|---|---|---|---|---|
| Gm41291 | 0.287773478 | -0.430239906 | 0 | 0.021 | 1 | 0.872828964 | 1.029849928 | 0.024 | 0.02 | 1 | 0.872828964 | 0.492733382 |
| Gm39556 | 0.287773478 | -0.430239906 | 0 | 0.021 | 1 | 0.455097252 | -0.622226769 | 0 | 0.014 | 1 | 0.455097252 | 0.492733382 |
| Taf9b | 0.287773478 | -0.430239906 | 0 | 0.021 | 1 | 0.674180157 | 0.252242349 | 0.024 | 0.038 | 1 | 0.674180157 | 0.492733382 |
| Atxn10 | 0.399785234 | 0.322349213 | 0.887 | 0.894 | 1 | 0.287861168 | -0.103586152 | 0.829 | 0.942 | 1 | 0.399785234 | 0.492858284 |
| Mill2 | 0.522318616 | 0.3662267 | 0.226 | 0.182 | 1 | 0.288052399 | -0.435813645 | 0.073 | 0.133 | 1 | 0.522318616 | 0.493130614 |
| Cyp4a12b | 0.413082433 | -1.652632327 | 0 | 0.013 | 1 | 0.288314827 | 1.252242349 | 0.073 | 0.038 | 1 | 0.413082433 | 0.493504215 |
| Gm43774 | 0.565795103 | -0.430239906 | 0.019 | 0.034 | 1 | 0.288314827 | 1.252242349 | 0.073 | 0.038 | 1 | 0.565795103 | 0.493504215 |
| Gm56730 | 0.288646204 | -0.147840175 | 0.358 | 0.263 | 1 | 0.347226956 | 0.856313673 | 0.22 | 0.167 | 1 | 0.347226956 | 0.493975778 |
| Arap1 | 0.288710476 | 0.347367673 | 0.264 | 0.203 | 1 | 0.317276024 | 0.562197802 | 0.317 | 0.253 | 1 | 0.317276024 | 0.494067213 |
| Cyp2u1 | 0.41630927 | -1.015202407 | 0.075 | 0.11 | 1 | 0.288749523 | -1.088794569 | 0.073 | 0.13 | 1 | 0.41630927 | 0.494122759 |
| Gm34343 | 0.314526532 | -0.52334931 | 0.057 | 0.102 | 1 | 0.288842413 | -1.069685746 | 0.049 | 0.099 | 1 | 0.314526532 | 0.494254887 |
| 4933431E20Rik | 0.289451254 | -0.545717123 | 0.038 | 0.081 | 1 | 0.350310865 | 0.159132945 | 0.049 | 0.096 | 1 | 0.350310865 | 0.495120479 |
| Klf4 | 0.289587966 | 0.120129201 | 1 | 0.958 | 1 | 0.334264322 | 0.262835578 | 0.951 | 0.99 | 1 | 0.334264322 | 0.495314741 |
| Eme1 | 0.289713138 | 1.502645898 | 0.075 | 0.042 | 1 | 0.771060519 | 2.199774929 | 0.049 | 0.041 | 1 | 0.771060519 | 0.495492574 |
| Borcs8 | 0.441575067 | 0.50490895 | 0.358 | 0.322 | 1 | 0.289914028 | 0.788841828 | 0.317 | 0.256 | 1 | 0.441575067 | 0.495777912 |
| Coq9 | 0.289914239 | 0.488459992 | 0.302 | 0.225 | 1 | 0.730554617 | 0.541748966 | 0.171 | 0.154 | 1 | 0.730554617 | 0.495778212 |
| Speg | 0.290202443 | 0.792152515 | 0.094 | 0.055 | 1 | 0.448281635 | 0.334704509 | 0.122 | 0.174 | 1 | 0.448281635 | 0.496187428 |
| Carf | 0.585232464 | 0.122301117 | 0.189 | 0.153 | 1 | 0.290292802 | 0.564186355 | 0.171 | 0.113 | 1 | 0.585232464 | 0.496315693 |
| Sys1 | 0.849089917 | -0.195376173 | 0.698 | 0.644 | 1 | 0.290292853 | -0.287950282 | 0.585 | 0.628 | 1 | 0.849089917 | 0.496315766 |
| Slc25a17 | 0.290481216 | 0.369735486 | 0.642 | 0.589 | 1 | 0.35548188 | 0.562025014 | 0.634 | 0.662 | 1 | 0.35548188 | 0.496583095 |
| Stam | 0.290606203 | 0.470224421 | 0.566 | 0.555 | 1 | 0.597128058 | 0.255409628 | 0.756 | 0.693 | 1 | 0.597128058 | 0.496760441 |
| Slk | 0.290834269 | -0.236558781 | 0.906 | 0.941 | 1 | 0.319435667 | 0.329291013 | 0.902 | 0.969 | 1 | 0.319435667 | 0.497083966 |
| Smim20 | 0.291027329 | -0.278236812 | 0.132 | 0.199 | 1 | 0.924157653 | 0.287007767 | 0.122 | 0.13 | 1 | 0.924157653 | 0.497357752 |
| Sik2 | 0.291101334 | 0.692095637 | 0.491 | 0.411 | 1 | 0.928064332 | 0.19448063 | 0.707 | 0.7 | 1 | 0.928064332 | 0.497462682 |
| Zfp961 | 0.291195087 | 0.714150003 | 0.094 | 0.055 | 1 | 0.328782493 | -1.250257991 | 0.024 | 0.061 | 1 | 0.328782493 | 0.497595595 |
| Slc37a2 | 0.750969908 | 0.984797593 | 0.038 | 0.03 | 1 | 0.291422946 | -2.260827233 | 0.049 | 0.099 | 1 | 0.750969908 | 0.497918558 |
| Rhobtb2 | 0.291454641 | 1.607234799 | 0.113 | 0.072 | 1 | 0.535642424 | -0.332720152 | 0.049 | 0.075 | 1 | 0.535642424 | 0.497963475 |
| Evc | 0.957893518 | -0.207847485 | 0.113 | 0.114 | 1 | 0.291497355 | 0.983055716 | 0.195 | 0.14 | 1 | 0.957893518 | 0.498024003 |
| Zcchc10 | 0.291775321 | 0.714150003 | 0.189 | 0.136 | 1 | 0.921618955 | 0.188112012 | 0.171 | 0.181 | 1 | 0.921618955 | 0.498417804 |
| Rogdi | 0.291884007 | -0.278236812 | 0.057 | 0.106 | 1 | 0.995466755 | 0.574170444 | 0.073 | 0.075 | 1 | 0.995466755 | 0.49857174 |
| Bfsp1 | 0.599888051 | 0.247831999 | 0.113 | 0.089 | 1 | 0.291886353 | -1.101394606 | 0.049 | 0.099 | 1 | 0.599888051 | 0.498575063 |
| Tnks1bp1 | 0.29192533 | -0.128211368 | 0.302 | 0.394 | 1 | 0.468215141 | 0.262510685 | 0.585 | 0.549 | 1 | 0.468215141 | 0.498630261 |
| Haus6 | 0.292202355 | 0.782753817 | 0.189 | 0.14 | 1 | 0.894035711 | 0.594348326 | 0.146 | 0.143 | 1 | 0.894035711 | 0.499022494 |
| Insyn2a | 0.414345737 | 0.154722595 | 0.075 | 0.047 | 1 | 0.292387884 | -0.555112573 | 0.024 | 0.068 | 1 | 0.414345737 | 0.499285093 |
| Cdk10 | 0.876328301 | 0.526691372 | 0.113 | 0.127 | 1 | 0.292501087 | -0.484723245 | 0.122 | 0.191 | 1 | 0.876328301 | 0.499445288 |
| Rpgr | 0.293212617 | -0.130679624 | 0.075 | 0.131 | 1 | 0.292578142 | 1.252242349 | 0.122 | 0.075 | 1 | 0.293212617 | 0.499554314 |
| 2010315B03Rik | 0.418025187 | -0.399866257 | 0.113 | 0.157 | 1 | 0.292675244 | -0.305753104 | 0.073 | 0.133 | 1 | 0.418025187 | 0.499691689 |
| Eef1akmt4 | 0.407411643 | -0.395474488 | 0.113 | 0.157 | 1 | 0.292925206 | -0.622226769 | 0.073 | 0.13 | 1 | 0.407411643 | 0.500045235 |
| Klf12 | 0.996361914 | -0.127387677 | 0.943 | 0.898 | 1 | 0.293104488 | -0.373236701 | 0.976 | 0.976 | 1 | 0.996361914 | 0.500298735 |
| Clasrp | 0.884742316 | 0.172869942 | 0.245 | 0.263 | 1 | 0.293214513 | 0.681926624 | 0.293 | 0.229 | 1 | 0.884742316 | 0.500454276 |
| Gm14418 | 0.642992906 | 0.776210972 | 0.057 | 0.042 | 1 | 0.293244655 | -0.385187572 | 0.024 | 0.068 | 1 | 0.642992906 | 0.500496882 |
| Cwf19l2 | 0.769071755 | 0.180348402 | 0.528 | 0.576 | 1 | 0.29325142 | 0.357211909 | 0.585 | 0.529 | 1 | 0.769071755 | 0.500506445 |
| Zfp420 | 0.758747385 | -0.1672055 | 0.057 | 0.068 | 1 | 0.293411107 | -0.747757651 | 0.049 | 0.099 | 1 | 0.758747385 | 0.500732136 |
| Pggt1b | 0.29365239 | 0.490471184 | 0.509 | 0.504 | 1 | 0.385921148 | -0.107653596 | 0.537 | 0.628 | 1 | 0.385921148 | 0.501073053 |
| Pnpo | 0.293768758 | -0.40599236 | 0.132 | 0.195 | 1 | 0.453533537 | -0.385187572 | 0.049 | 0.082 | 1 | 0.453533537 | 0.501237433 |
| Plxnb1 | 0.293769077 | 0.389187848 | 0.226 | 0.161 | 1 | 0.548350344 | -0.120566915 | 0.22 | 0.27 | 1 | 0.548350344 | 0.501237884 |
| Zfp764 | 0.328746305 | 1.306725688 | 0.057 | 0.03 | 1 | 0.293964142 | 0.989207943 | 0.073 | 0.038 | 1 | 0.328746305 | 0.501513368 |
| Nckap1 | 0.35799845 | -0.175562052 | 0.962 | 0.966 | 1 | 0.294445761 | -0.13585267 | 0.951 | 0.983 | 1 | 0.35799845 | 0.502193215 |
| Gm12184 | 0.294684842 | 0.756173218 | 0.17 | 0.119 | 1 | 0.583953617 | 0.252242349 | 0.073 | 0.102 | 1 | 0.583953617 | 0.502530528 |
| Kbtbd3 | 0.294694844 | 0.900966003 | 0.151 | 0.102 | 1 | 0.986965808 | 0.334704509 | 0.049 | 0.048 | 1 | 0.986965808 | 0.502544637 |
| Tapt1 | 0.326873182 | 0.459936816 | 0.491 | 0.47 | 1 | 0.295046104 | -0.224891271 | 0.659 | 0.72 | 1 | 0.326873182 | 0.503040005 |
| Pex16 | 0.295199779 | -0.545717123 | 0.094 | 0.148 | 1 | 0.55557743 | 0.396632258 | 0.146 | 0.113 | 1 | 0.55557743 | 0.503256648 |
| Eprs | 0.369040543 | 0.147202013 | 0.981 | 0.953 | 1 | 0.295219233 | -0.213707104 | 0.951 | 0.956 | 1 | 0.369040543 | 0.50328407 |
| Alas1 | 0.29532952 | 0.984797593 | 0.321 | 0.271 | 1 | 0.791253745 | 0.227789305 | 0.488 | 0.471 | 1 | 0.791253745 | 0.503439515 |
| Gne | 0.314483564 | -0.147840175 | 0.151 | 0.216 | 1 | 0.295493846 | -0.116991461 | 0.146 | 0.235 | 1 | 0.314483564 | 0.503671079 |
| Polr1f | 0.381386935 | -0.189231806 | 0.415 | 0.466 | 1 | 0.29578302 | -0.414020254 | 0.39 | 0.44 | 1 | 0.381386935 | 0.504078445 |
| Krt1 | 0.488750271 | -0.304709024 | 0.019 | 0.038 | 1 | 0.296019272 | -0.16279515 | 0.024 | 0.068 | 1 | 0.488750271 | 0.504411135 |
| B230217O12Rik | 0.829895379 | 0.347367673 | 0.075 | 0.085 | 1 | 0.296466794 | -0.665295491 | 0.049 | 0.099 | 1 | 0.829895379 | 0.505041027 |
| Itgb6 | 0.377514817 | 0.776210972 | 0.075 | 0.047 | 1 | 0.296644421 | -0.278272368 | 0.098 | 0.055 | 1 | 0.377514817 | 0.50529093 |
| Kansl3 | 0.499695902 | 0.282878946 | 0.547 | 0.504 | 1 | 0.296759583 | -0.219231096 | 0.61 | 0.696 | 1 | 0.499695902 | 0.505452916 |
| D930015M05Rik | 0.728468844 | 0.739685096 | 0.038 | 0.051 | 1 | 0.296931774 | 1.47463477 | 0.098 | 0.058 | 1 | 0.728468844 | 0.505695069 |
| Hdac11 | 0.61536875 | 1.154722595 | 0.038 | 0.025 | 1 | 0.296969383 | -0.101394606 | 0.024 | 0.068 | 1 | 0.61536875 | 0.505747951 |
| Zbtb48 | 0.297115888 | 0.739685096 | 0.094 | 0.055 | 1 | 0.858435718 | 0.334704509 | 0.098 | 0.109 | 1 | 0.858435718 | 0.505953926 |
| Ccnq | 0.737877148 | 0.347367673 | 0.226 | 0.212 | 1 | 0.297124851 | -0.421529419 | 0.122 | 0.191 | 1 | 0.737877148 | 0.505966524 |
| Rnpc3 | 0.488815121 | 0.303480029 | 0.585 | 0.627 | 1 | 0.297457985 | -0.354813922 | 0.488 | 0.543 | 1 | 0.488815121 | 0.506434717 |
| Slc41a2 | 0.371875831 | 0.209864149 | 0.566 | 0.538 | 1 | 0.297474525 | -0.203822418 | 0.585 | 0.696 | 1 | 0.371875831 | 0.506457957 |
| Ccnb1ip1 | 0.297624472 | -0.1672055 | 0.038 | 0.081 | 1 | 0.307268086 | -0.747757651 | 0.024 | 0.065 | 1 | 0.307268086 | 0.506668618 |
| Fahd2a | 0.78442244 | -0.1182959 | 0.094 | 0.106 | 1 | 0.297749795 | -0.16279515 | 0.049 | 0.102 | 1 | 0.78442244 | 0.50684465 |
| Gas5 | 0.536133832 | 0.116117408 | 1 | 0.975 | 1 | 0.297820587 | -0.158389971 | 0.976 | 0.98 | 1 | 0.536133832 | 0.506944071 |
| Wnk3 | 0.781328153 | 0.906795081 | 0.132 | 0.127 | 1 | 0.297921283 | 0.174239837 | 0.146 | 0.239 | 1 | 0.781328153 | 0.507085476 |
| Sox2ot | 0.298114494 | 0.769068902 | 0.17 | 0.119 | 1 | 0.752777536 | -1.049293957 | 0.146 | 0.16 | 1 | 0.752777536 | 0.507356737 |
| Ptpn18 | 0.424662489 | 1.154722595 | 0.057 | 0.034 | 1 | 0.298526682 | 0.749742009 | 0.073 | 0.038 | 1 | 0.424662489 | 0.507935185 |
| Dhx58 | 0.683834454 | 0.280253477 | 0.094 | 0.076 | 1 | 0.298526682 | 0.444887427 | 0.073 | 0.038 | 1 | 0.683834454 | 0.507935185 |
| Hsp90b1 | 0.298746769 | 0.122571234 | 1 | 0.987 | 1 | 0.47337878 | 0.117028212 | 0.976 | 0.997 | 1 | 0.47337878 | 0.508243905 |
| Kctd14 | 0.347166718 | -1.015202407 | 0.038 | 0.072 | 1 | 0.298937964 | 1.59619675 | 0.024 | 0.068 | 1 | 0.347166718 | 0.508512021 |
| Wbp11 | 0.299160906 | -0.240137023 | 0.679 | 0.775 | 1 | 0.659133062 | 0.160181796 | 0.829 | 0.782 | 1 | 0.659133062 | 0.508824564 |
| Nosip | 0.299213346 | 0.502645898 | 0.226 | 0.165 | 1 | 0.393751222 | 1.093043754 | 0.22 | 0.184 | 1 | 0.393751222 | 0.508898065 |
| Cacfd1 | 0.910612475 | -0.101617158 | 0.302 | 0.305 | 1 | 0.29936779 | -0.549818273 | 0.244 | 0.314 | 1 | 0.910612475 | 0.509114506 |
| Tbl3 | 0.350064104 | 0.240452469 | 0.226 | 0.165 | 1 | 0.299786026 | -0.466575898 | 0.146 | 0.215 | 1 | 0.350064104 | 0.509700391 |
| Prag1 | 0.299871187 | -0.218735801 | 0.094 | 0.157 | 1 | 0.650840959 | 0.515276755 | 0.049 | 0.034 | 1 | 0.650840959 | 0.509819646 |
| Lmbrd1 | 0.727496083 | -0.143526504 | 0.774 | 0.754 | 1 | 0.300123113 | 0.100975512 | 0.732 | 0.833 | 1 | 0.727496083 | 0.510172343 |
| Scyl3 | 0.300334455 | 0.701004627 | 0.264 | 0.208 | 1 | 0.314063111 | 0.674266279 | 0.317 | 0.246 | 1 | 0.314063111 | 0.510468126 |
| Atrip | 0.574469885 | 0.106359573 | 0.321 | 0.28 | 1 | 0.300701108 | -0.673757069 | 0.195 | 0.263 | 1 | 0.574469885 | 0.51098106 |
| D6Ertd527e | 0.300962247 | 1.280253477 | 0.075 | 0.042 | 1 | 0.334496323 | 0.911205431 | 0.098 | 0.058 | 1 | 0.334496323 | 0.51134622 |
| Lipt2 | 0.300962247 | 1.280253477 | 0.075 | 0.042 | 1 | 0.357854401 | -0.484723245 | 0 | 0.02 | 1 | 0.357854401 | 0.51134622 |
| Eid3 | 0.300962247 | 1.280253477 | 0.075 | 0.042 | 1 | 0.305356378 | -0.686357106 | 0.024 | 0.065 | 1 | 0.305356378 | 0.51134622 |
| Srd5a1 | 0.300962247 | 1.280253477 | 0.075 | 0.042 | 1 | 0.606262835 | -0.250257991 | 0.024 | 0.041 | 1 | 0.606262835 | 0.51134622 |
| Ahcyl | 0.300962247 | 1.280253477 | 0.075 | 0.042 | 1 | 0.743829142 | 0.252242349 | 0.024 | 0.034 | 1 | 0.743829142 | 0.51134622 |
| Tma16 | 0.501984813 | 0.587682002 | 0.264 | 0.246 | 1 | 0.301083996 | -0.292078167 | 0.195 | 0.273 | 1 | 0.501984813 | 0.511516419 |
| Mllt3 | 0.301095004 | -0.418592396 | 0.887 | 0.915 | 1 | 0.35156765 | -0.283810551 | 0.829 | 0.874 | 1 | 0.35156765 | 0.511531806 |
| Vhl | 0.301582892 | 0.520007059 | 0.264 | 0.199 | 1 | 0.724523665 | 0.227151368 | 0.268 | 0.297 | 1 | 0.724523665 | 0.512213543 |
| A230072C01Rik | 0.30179302 | -0.260314904 | 0.151 | 0.22 | 1 | 0.302335378 | -0.505187348 | 0.122 | 0.188 | 1 | 0.302335378 | 0.512507012 |
| Coro2b | 0.837864131 | 0.200526284 | 0.094 | 0.106 | 1 | 0.302024498 | 0.749742009 | 0.073 | 0.038 | 1 | 0.837864131 | 0.512830199 |
| Myo19 | 0.665829106 | 0.355021245 | 0.283 | 0.258 | 1 | 0.302206402 | -0.600200463 | 0.22 | 0.29 | 1 | 0.665829106 | 0.513084094 |
| Tapbp | 0.302257943 | -0.264685387 | 0.83 | 0.805 | 1 | 0.81191812 | 0.35013455 | 0.683 | 0.771 | 1 | 0.81191812 | 0.513156021 |
| Cfap157 | 0.336520503 | 1.569760094 | 0.038 | 0.017 | 1 | 0.302474778 | -1.069685746 | 0.024 | 0.065 | 1 | 0.336520503 | 0.513458565 |
| Chst2 | 0.357059697 | -0.681778673 | 0.245 | 0.297 | 1 | 0.302722381 | 1.043655727 | 0.171 | 0.119 | 1 | 0.357059697 | 0.513803922 |
| Fan1 | 0.302754342 | -0.508242418 | 0.075 | 0.127 | 1 | 0.729990277 | 0.515276755 | 0.122 | 0.106 | 1 | 0.729990277 | 0.513848492 |

| | | | | | | | | | | | | |
|---|---|---|---|---|---|---|---|---|---|---|---|---|
| Srpx2 | 0.303309165 | -0.899272295 | 0.472 | 0.483 | 1 | 0.371056553 | -0.344545587 | 0.244 | 0.304 | 1 | 0.371056553 | 0.51462188 |
| Gm1123 | 0.398253154 | 0.652222254 | 0.075 | 0.047 | 1 | 0.303319559 | -0.747757651 | 0.073 | 0.13 | 1 | 0.398253154 | 0.514636364 |
| Kif19b | 0.499041755 | 0.247831999 | 0.17 | 0.131 | 1 | 0.303388986 | -0.185162963 | 0.146 | 0.225 | 1 | 0.499041755 | 0.514733095 |
| Gnasas1 | 0.74502569 | 1.261637799 | 0.057 | 0.047 | 1 | 0.303437258 | -1.20718927 | 0.024 | 0.065 | 1 | 0.74502569 | 0.514800346 |
| Fastk | 0.613035538 | 0.342989232 | 0.472 | 0.462 | 1 | 0.303456505 | -0.170289687 | 0.341 | 0.444 | 1 | 0.613035538 | 0.51482716 |
| Rida | 0.31476168 | -0.112063946 | 0.189 | 0.267 | 1 | 0.303736151 | -0.116991461 | 0.171 | 0.242 | 1 | 0.31476168 | 0.515216653 |
| Pex2 | 0.567926241 | 0.207190015 | 0.358 | 0.331 | 1 | 0.304112636 | -0.452301767 | 0.317 | 0.386 | 1 | 0.567926241 | 0.515740777 |
| Anxa11os | 0.48175508 | 1.154722595 | 0.038 | 0.021 | 1 | 0.304397485 | -0.917682652 | 0.024 | 0.065 | 1 | 0.48175508 | 0.51613714 |
| Epm2a | 0.30456639 | 0.849868013 | 0.151 | 0.106 | 1 | 0.402533445 | 0.282615998 | 0.195 | 0.143 | 1 | 0.402533445 | 0.516372095 |
| Limk2 | 0.415155884 | -0.182312392 | 0.189 | 0.25 | 1 | 0.304571255 | -0.529912019 | 0.366 | 0.42 | 1 | 0.415155884 | 0.516378861 |
| Gstm4 | 0.304790107 | -0.652632327 | 0.057 | 0.102 | 1 | 0.768967689 | 0.930314254 | 0.073 | 0.089 | 1 | 0.768967689 | 0.516683204 |
| Mri1 | 0.30489787 | 0.397292898 | 0.321 | 0.254 | 1 | 0.396338655 | -0.300298674 | 0.146 | 0.205 | 1 | 0.396338655 | 0.516833029 |
| Evi5l | 0.304939782 | 0.554809753 | 0.34 | 0.275 | 1 | 0.671245829 | 0.746056962 | 0.22 | 0.198 | 1 | 0.671245829 | 0.516891293 |
| Ghdc | 0.304983951 | 0.470224421 | 0.189 | 0.136 | 1 | 0.552761046 | -0.140075074 | 0.122 | 0.16 | 1 | 0.552761046 | 0.516952691 |
| Armc6 | 0.534564335 | 1.017219071 | 0.057 | 0.038 | 1 | 0.305356148 | -0.747757651 | 0.024 | 0.065 | 1 | 0.534564335 | 0.517469919 |
| Zfp94 | 0.404836717 | 0.669295768 | 0.075 | 0.047 | 1 | 0.305356378 | -0.686357106 | 0.024 | 0.065 | 1 | 0.404836717 | 0.517470239 |
| St6galnac6 | 0.497071498 | -0.382934191 | 0.34 | 0.403 | 1 | 0.305533939 | -0.16279515 | 0.293 | 0.372 | 1 | 0.497071498 | 0.51771689 |
| Tubg2 | 0.305590208 | 1.154722595 | 0.075 | 0.042 | 1 | 0.914200263 | 0.515276755 | 0.024 | 0.027 | 1 | 0.914200263 | 0.517795041 |
| Selenof | 0.41511732 | 0.114458725 | 1 | 0.983 | 1 | 0.305939073 | 0.163720982 | 0.976 | 0.986 | 1 | 0.41511732 | 0.518279429 |
| Setd6 | 0.331821369 | 0.891688189 | 0.132 | 0.089 | 1 | 0.30613173 | 0.979223855 | 0.122 | 0.075 | 1 | 0.331821369 | 0.518546824 |
| Gm27017 | 0.306231257 | -1.1672055 | 0.019 | 0.051 | 1 | 0.54561906 | -0.622226769 | 0.024 | 0.044 | 1 | 0.54561906 | 0.518684931 |
| Foxd2os | 0.306719687 | 0.29992051 | 0.396 | 0.326 | 1 | 0.486701709 | -0.970150072 | 0.171 | 0.205 | 1 | 0.486701709 | 0.519362408 |
| Snrnp40 | 0.30690641 | 0.442524906 | 0.472 | 0.419 | 1 | 0.996189393 | 0.252242349 | 0.415 | 0.464 | 1 | 0.996189393 | 0.519621276 |
| Pip5kl1 | 0.307416057 | -1.015202407 | 0.019 | 0.051 | 1 | 0.459320496 | 0.962735732 | 0.049 | 0.027 | 1 | 0.459320496 | 0.520327482 |
| Map3k21 | 0.307416057 | -1.015202407 | 0.019 | 0.051 | 1 | 0.796279439 | -0.278272368 | 0.049 | 0.058 | 1 | 0.796279439 | 0.520327482 |
| Gm5820 | 0.307417922 | -0.932740246 | 0.019 | 0.051 | 1 | 0.319636904 | -0.970150072 | 0 | 0.024 | 1 | 0.319636904 | 0.520330066 |
| Drg1 | 0.481482222 | 0.18109801 | 0.623 | 0.631 | 1 | 0.307593729 | 0.385695794 | 0.756 | 0.727 | 1 | 0.481482222 | 0.520573556 |
| Rtn4 | 0.307642909 | 0.108180009 | 1 | 0.975 | 1 | 0.524706939 | 0.107992722 | 1 | 0.997 | 1 | 0.524706939 | 0.520641659 |
| Ptpn23 | 0.484959979 | -0.177120969 | 0.415 | 0.445 | 1 | 0.307879467 | -0.265888643 | 0.341 | 0.457 | 1 | 0.484959979 | 0.520969167 |
| Fbxo25 | 0.307902333 | 1.154722595 | 0.17 | 0.123 | 1 | 0.864299847 | 0.222495006 | 0.122 | 0.113 | 1 | 0.864299847 | 0.52100082 |
| H2ac8 | 0.307975 | -0.992118794 | 0.075 | 0.042 | 1 | 0.438672228 | 1.83720485 | 0.024 | 0.01 | 1 | 0.438672228 | 0.5211014 |
| Gm57301 | 0.30799238 | 1.055186921 | 0.075 | 0.042 | 1 | 0.989595616 | 0.667279848 | 0.024 | 0.024 | 1 | 0.989595616 | 0.521125454 |
| Unc13d | 0.308044316 | -1.067669827 | 0.038 | 0.076 | 1 | 0.390822065 | -0.80665134 | 0.073 | 0.116 | 1 | 0.390822065 | 0.521197332 |
| 1700036A12Rik | 0.308230949 | 0.852159825 | 0.17 | 0.123 | 1 | 0.329453331 | -0.813049111 | 0.171 | 0.232 | 1 | 0.329453331 | 0.52145558 |
| Nagk | 0.496258467 | 0.331600357 | 0.321 | 0.284 | 1 | 0.308324145 | -0.233184478 | 0.22 | 0.311 | 1 | 0.496258467 | 0.521584512 |
| 2610306M01Rik | 0.37991914 | 1.154722595 | 0.075 | 0.047 | 1 | 0.308527224 | -0.484723245 | 0.024 | 0.065 | 1 | 0.37991914 | 0.5218654 |
| 4933417E11Rik | 0.30859836 | -1.093204919 | 0.019 | 0.051 | 1 | 0.51919986 | 0.83720485 | 0 | 0.01 | 1 | 0.51919986 | 0.521963772 |
| Hpn | 0.30859836 | -0.932740246 | 0.019 | 0.051 | 1 | 0.886170233 | 0.744095445 | 0.146 | 0.143 | 1 | 0.886170233 | 0.521963772 |
| Gabbr2 | 0.308598826 | -1.093204919 | 0.019 | 0.051 | 1 | 0.402372511 | -1.410722664 | 0 | 0.017 | 1 | 0.402372511 | 0.521964417 |
| Cyp2j9 | 0.308598826 | -0.845277405 | 0.019 | 0.051 | 1 | 0.455098275 | 0.515276755 | 0 | 0.014 | 1 | 0.455098275 | 0.521964417 |
| Gm46212 | 0.308598826 | -1.015202407 | 0.019 | 0.051 | 1 | 0.82181816 | 0.252242349 | 0.024 | 0.031 | 1 | 0.82181816 | 0.521964417 |
| Trmt2a | 0.361310564 | -0.545717123 | 0.17 | 0.22 | 1 | 0.308682504 | -0.466575898 | 0.146 | 0.215 | 1 | 0.361310564 | 0.52208012 |
| Garnl3 | 0.308974054 | 0.637664159 | 0.226 | 0.174 | 1 | 0.360495803 | 0.984762038 | 0.171 | 0.123 | 1 | 0.360495803 | 0.522483142 |
| Slc30a6 | 0.309224627 | 0.3662267 | 0.321 | 0.254 | 1 | 0.361039083 | 0.721727632 | 0.317 | 0.273 | 1 | 0.361039083 | 0.522829383 |
| Pnpla8 | 0.309301524 | 0.250070471 | 0.736 | 0.627 | 1 | 0.854380961 | 0.279209397 | 0.805 | 0.792 | 1 | 0.854380961 | 0.522935615 |
| Dram2 | 0.736503706 | 0.379428882 | 0.264 | 0.246 | 1 | 0.309334553 | 0.2445067 | 0.488 | 0.41 | 1 | 0.736503706 | 0.52298124 |
| Ext1 | 0.798122371 | -0.341087528 | 1 | 0.958 | 1 | 0.30940161 | 0.339770721 | 0.976 | 0.952 | 1 | 0.798122371 | 0.523073864 |
| Slirp | 0.475779897 | -0.288220901 | 0.453 | 0.483 | 1 | 0.309470488 | -0.152741486 | 0.268 | 0.375 | 1 | 0.475779897 | 0.523168993 |
| Hint2 | 0.629960687 | 0.163767734 | 0.491 | 0.483 | 1 | 0.309769711 | -0.428139717 | 0.268 | 0.338 | 1 | 0.629960687 | 0.523582148 |
| Igflr1 | 0.309778164 | -0.752168001 | 0.019 | 0.051 | 1 | 0.881389038 | 0.749742009 | 0.049 | 0.044 | 1 | 0.881389038 | 0.523593817 |
| Gsg1l | 0.309778164 | -0.752168001 | 0.019 | 0.051 | 1 | 0.946637396 | 0.667279848 | 0.073 | 0.078 | 1 | 0.946637396 | 0.523593817 |
| Mrpl13 | 0.657479607 | 0.211306123 | 0.472 | 0.458 | 1 | 0.30985209 | -0.476685926 | 0.341 | 0.399 | 1 | 0.657479607 | 0.523695862 |
| Kif4 | 0.30987377 | -1.652632327 | 0.019 | 0.051 | 1 | 0.5191986 | 0.252242349 | 0 | 0.01 | 1 | 0.5191986 | 0.523725786 |
| Wdr12 | 0.30996655 | 0.505629757 | 0.415 | 0.36 | 1 | 0.825237376 | 0.384692645 | 0.39 | 0.403 | 1 | 0.825237376 | 0.523853837 |
| Rcbtb1 | 0.996823786 | 0.200526284 | 0.472 | 0.487 | 1 | 0.309975319 | 0.291011919 | 0.366 | 0.512 | 1 | 0.996823786 | 0.523865939 |
| Nit1 | 0.524625918 | 0.434830514 | 0.245 | 0.212 | 1 | 0.31024654 | -0.39309277 | 0.195 | 0.28 | 1 | 0.524625918 | 0.524240165 |
| Gm26759 | 0.310259246 | 1.039245377 | 0.075 | 0.042 | 1 | 0.518021606 | 0.627751484 | 0.122 | 0.092 | 1 | 0.518021606 | 0.524257692 |
| Odad4 | 0.310259246 | 1.039245377 | 0.075 | 0.042 | 1 | 0.319641967 | -0.16279515 | 0 | 0.024 | 1 | 0.319641967 | 0.524257692 |
| Rala | 0.416032791 | 0.198078964 | 0.642 | 0.627 | 1 | 0.310454422 | 0.431860747 | 0.683 | 0.706 | 1 | 0.416032791 | 0.524526895 |
| Paqr6 | 0.685534188 | 0.659957903 | 0.113 | 0.097 | 1 | 0.310458537 | -0.665295491 | 0.098 | 0.157 | 1 | 0.685534188 | 0.524532571 |
| Poldip2 | 0.997373689 | 0.182736971 | 0.321 | 0.335 | 1 | 0.310642528 | -0.246211158 | 0.341 | 0.44 | 1 | 0.997373689 | 0.524786275 |
| Ak3 | 0.310837931 | 0.294900253 | 0.358 | 0.284 | 1 | 0.314485164 | 0.576080034 | 0.585 | 0.549 | 1 | 0.314485164 | 0.525055643 |
| A330023F24Rik | 0.926169972 | 0.510866405 | 0.151 | 0.148 | 1 | 0.310874517 | -0.323787027 | 0.171 | 0.259 | 1 | 0.926169972 | 0.525106068 |
| Il1rl2 | 0.310955466 | -0.652632327 | 0.019 | 0.051 | 1 | 0.60213981 | 1.252242349 | 0.024 | 0.014 | 1 | 0.60213981 | 0.52521763 |
| 4930412C18Rik | 0.310955466 | -0.752168001 | 0.019 | 0.051 | 1 | 0.455094184 | 0.252242349 | 0 | 0.014 | 1 | 0.455094184 | 0.52521763 |
| Taf4b | 0.607190864 | 0.218685654 | 0.585 | 0.555 | 1 | 0.310995378 | 0.424709545 | 0.585 | 0.502 | 1 | 0.607190864 | 0.525272631 |
| 2310015A10Rik | 0.311029212 | 0.47665069 | 0.151 | 0.102 | 1 | 1 | 0.759202338 | 0.098 | 0.099 | 1 | 1 | 0.525319253 |
| Macrod1 | 0.887990693 | -0.1672055 | 0.245 | 0.25 | 1 | 0.311070989 | -0.143686327 | 0.195 | 0.283 | 1 | 0.887990693 | 0.525376818 |
| Mios | 0.311168991 | 0.645708947 | 0.283 | 0.229 | 1 | 0.709381607 | 0.437274243 | 0.195 | 0.229 | 1 | 0.709381607 | 0.525511841 |
| Cryz | 0.311353341 | 0.607234799 | 0.208 | 0.153 | 1 | 0.38239465 | -0.455576899 | 0.098 | 0.147 | 1 | 0.38239465 | 0.525765779 |
| Qrsl1 | 0.311919061 | -1.067669827 | 0.038 | 0.076 | 1 | 0.403242332 | -0.372248516 | 0.049 | 0.089 | 1 | 0.403242332 | 0.526544622 |
| Ten1 | 0.312092574 | 0.624207878 | 0.226 | 0.169 | 1 | 0.947540988 | -0.116991461 | 0.146 | 0.14 | 1 | 0.947540988 | 0.526783374 |
| Eda2r | 0.31228701 | -1.347777746 | 0.019 | 0.051 | 1 | 0.758283818 | 0.136765132 | 0.024 | 0.017 | 1 | 0.758283818 | 0.527050843 |
| Dhtkd1 | 0.312300068 | -0.545717123 | 0.019 | 0.051 | 1 | 0.529718256 | 1.83720485 | 0.049 | 0.031 | 1 | 0.529718256 | 0.527068803 |
| Derl3 | 0.312300068 | -0.545717123 | 0.019 | 0.051 | 1 | 0.599290422 | 1.515276755 | 0.024 | 0.014 | 1 | 0.599290422 | 0.527068803 |
| Gm10941 | 0.312300068 | -0.545717123 | 0.019 | 0.051 | 1 | 0.990690158 | 0.252242349 | 0.049 | 0.048 | 1 | 0.990690158 | 0.527068803 |
| Gmpr | 0.312300068 | -0.545717123 | 0.019 | 0.051 | 1 | 0.329039295 | 0.883008539 | 0.122 | 0.078 | 1 | 0.329039295 | 0.527068803 |
| Utp25 | 0.312571694 | 0.46284489 | 0.283 | 0.225 | 1 | 0.436450405 | 0.57131079 | 0.293 | 0.253 | 1 | 0.436450405 | 0.527442324 |
| Xpc | 0.312580883 | 0.874614676 | 0.226 | 0.174 | 1 | 0.424212674 | -0.53202896 | 0.171 | 0.218 | 1 | 0.424212674 | 0.527454957 |
| Sepsecs | 0.774442191 | 0.731511164 | 0.208 | 0.203 | 1 | 0.312581302 | -0.395455907 | 0.171 | 0.246 | 1 | 0.774442191 | 0.527455534 |
| Gm36640 | 0.762480493 | 0.739685096 | 0.038 | 0.03 | 1 | 0.312634129 | 1.120997816 | 0.098 | 0.058 | 1 | 0.762480493 | 0.527528159 |
| Fam222a | 0.312692399 | 1.017219071 | 0.075 | 0.042 | 1 | 0.38441392 | 0.721727632 | 0.098 | 0.061 | 1 | 0.38441392 | 0.527608261 |
| Rnf4 | 0.631793887 | 0.195364579 | 0.698 | 0.665 | 1 | 0.313188122 | -0.182385879 | 0.78 | 0.805 | 1 | 0.631793887 | 0.528289444 |
| Slc25a35 | 0.967052971 | 0.347367673 | 0.057 | 0.055 | 1 | 0.313349742 | 1.252242349 | 0.098 | 0.058 | 1 | 0.967052971 | 0.528511423 |
| Aars2 | 0.313377909 | 1.640149422 | 0.057 | 0.03 | 1 | 0.40786703 | -0.622226769 | 0.024 | 0.055 | 1 | 0.40786703 | 0.528550104 |
| Anapc16 | 0.62904345 | 0.11177045 | 0.755 | 0.729 | 1 | 0.3137357 | -0.148855959 | 0.732 | 0.761 | 1 | 0.62904345 | 0.52904131 |
| Atf6b | 0.313762497 | 0.438515561 | 0.321 | 0.258 | 1 | 0.769679 | 0.649577847 | 0.22 | 0.205 | 1 | 0.769679 | 0.529078089 |
| ENSMUSG00000121445 | 0.313859655 | -0.783876861 | 0.038 | 0.076 | 1 | 0.387007894 | -0.385187572 | 0.049 | 0.089 | 1 | 0.387007894 | 0.529211428 |
| Rdh12 | 0.313859966 | -0.719746523 | 0.038 | 0.076 | 1 | 0.32154479 | -0.435813645 | 0.024 | 0.065 | 1 | 0.32154479 | 0.529211854 |
| Elmod2 | 0.313920823 | 1.017219071 | 0.17 | 0.123 | 1 | 0.980459991 | 0.679663573 | 0.146 | 0.16 | 1 | 0.980459991 | 0.529295363 |
| Ankrd9 | 0.31419656 | 0.8327945 | 0.113 | 0.072 | 1 | 0.318344907 | 1.100239256 | 0.098 | 0.058 | 1 | 0.318344907 | 0.529673642 |
| Bace2 | 0.314507041 | 0.51531231 | 0.264 | 0.212 | 1 | 0.683521218 | 0.604544093 | 0.146 | 0.13 | 1 | 0.683521218 | 0.530099403 |
| Tmem276 | 0.314608352 | 0.589937976 | 0.302 | 0.25 | 1 | 0.773681056 | 1.064615346 | 0.244 | 0.229 | 1 | 0.773681056 | 0.530238289 |
| Acad11 | 0.597385401 | 0.399835093 | 0.113 | 0.089 | 1 | 0.314704597 | -0.332720152 | 0.073 | 0.13 | 1 | 0.597385401 | 0.530370211 |
| Cntn4 | 0.314720163 | 4.06161319 | 0.057 | 0.03 | 1 | 0.68528621 | 2.088743617 | 0.073 | 0.058 | 1 | 0.68528621 | 0.530391545 |
| 9330136K24Rik | 0.314720163 | 2.324647596 | 0.057 | 0.03 | 1 | 0.319641967 | -0.16279515 | 0 | 0.024 | 1 | 0.319641967 | 0.530391545 |
| Smarcal1 | 0.939643391 | 0.454282877 | 0.094 | 0.093 | 1 | 0.314722284 | -0.140075074 | 0.098 | 0.164 | 1 | 0.939643391 | 0.530394451 |
| Cdk6 | 0.314745234 | -0.454569235 | 0.623 | 0.699 | 1 | 0.431389093 | 0.33607394 | 0.488 | 0.457 | 1 | 0.431389093 | 0.530425905 |

| | | | | | | | | | | | | |
|---|---|---|---|---|---|---|---|---|---|---|---|---|
| Astn2 | 0.314825296 | 0.154722595 | 0.17 | 0.119 | 1 | 0.762368087 | 1.136765132 | 0.122 | 0.113 | 1 | 0.762368087 | 0.530535625 |
| Mvb12b | 0.44964864 | 0.776210972 | 0.321 | 0.292 | 1 | 0.314898155 | 0.856833657 | 0.366 | 0.314 | 1 | 0.44964864 | 0.530635463 |
| Cbx8 | 0.314969793 | 0.932330173 | 0.075 | 0.042 | 1 | 0.964635009 | 0.667279848 | 0.049 | 0.048 | 1 | 0.964635009 | 0.530733616 |
| Pik3c2a | 0.32107774 | 0.580329336 | 0.698 | 0.665 | 1 | 0.315127416 | 0.266825672 | 0.78 | 0.765 | 1 | 0.32107774 | 0.530949544 |
| Gm9750 | 0.315926645 | -0.783876861 | 0.019 | 0.051 | 1 | 0.441381862 | 1.515276755 | 0.024 | 0.01 | 1 | 0.441381862 | 0.532043645 |
| Pkd2 | 0.622518272 | -0.125659055 | 0.83 | 0.839 | 1 | 0.316375637 | 0.175700794 | 0.878 | 0.891 | 1 | 0.622518272 | 0.532657731 |
| Dolk | 0.316708502 | 0.669295768 | 0.264 | 0.208 | 1 | 0.488685403 | -0.314798244 | 0.171 | 0.215 | 1 | 0.488685403 | 0.533112728 |
| Tfdp2 | 0.316848024 | 0.728958689 | 0.472 | 0.436 | 1 | 0.461128898 | 0.587728549 | 0.61 | 0.618 | 1 | 0.461128898 | 0.533303378 |
| Fndc10 | 0.317243945 | 0.891688189 | 0.075 | 0.042 | 1 | 0.9042167 | 0.515276755 | 0.049 | 0.044 | 1 | 0.9042167 | 0.533844169 |
| Zfp433 | 0.317243945 | 0.891688189 | 0.075 | 0.042 | 1 | 0.459320496 | 0.962735732 | 0.049 | 0.027 | 1 | 0.459320496 | 0.533844169 |
| 1600020E01Rik | 0.402554088 | 0.515821287 | 0.623 | 0.623 | 1 | 0.317303086 | -0.401761796 | 0.585 | 0.648 | 1 | 0.402554088 | 0.533924923 |
| Cenph | 0.317369156 | 0.640149422 | 0.075 | 0.042 | 1 | 0.402369942 | -0.622226769 | 0 | 0.017 | 1 | 0.402369942 | 0.534015131 |
| Klk8 | 0.363535861 | -2.074096096 | 0.057 | 0.093 | 1 | 0.317506163 | -0.825760163 | 0.024 | 0.065 | 1 | 0.363535861 | 0.534202162 |
| Traf7 | 0.318302722 | 0.553271971 | 0.358 | 0.297 | 1 | 0.91103323 | 0.47463477 | 0.366 | 0.403 | 1 | 0.91103323 | 0.535288821 |
| Vipr2 | 0.610886353 | 1.154722595 | 0.038 | 0.025 | 1 | 0.318344907 | 1.377773231 | 0.098 | 0.058 | 1 | 0.610886353 | 0.535346334 |
| Zc3h12a | 0.318984663 | -0.56774343 | 0.075 | 0.123 | 1 | 0.457238139 | -0.332720152 | 0.073 | 0.113 | 1 | 0.457238139 | 0.536218111 |
| Szt2 | 0.35245935 | 0.614154213 | 0.151 | 0.106 | 1 | 0.319407027 | -0.589059905 | 0.073 | 0.126 | 1 | 0.35245935 | 0.536793205 |
| Chic1 | 0.319438908 | 0.134258492 | 0.34 | 0.288 | 1 | 0.984998786 | 0.149727438 | 0.366 | 0.386 | 1 | 0.984998786 | 0.5368366 |
| Catspere2 | 0.319490745 | -0.617866909 | 0.075 | 0.123 | 1 | 0.589401251 | 1.275325962 | 0.098 | 0.137 | 1 | 0.589401251 | 0.536907154 |
| Gm56858 | 0.347746365 | 0.932330173 | 0.038 | 0.017 | 1 | 0.319619815 | -0.332720152 | 0 | 0.024 | 1 | 0.347746365 | 0.537082804 |
| 3830408C21Rik | 0.458961249 | -0.652632327 | 0.038 | 0.064 | 1 | 0.319619815 | -0.332720152 | 0 | 0.024 | 1 | 0.458961249 | 0.537082804 |
| Gm16045 | 0.503971666 | 1.569760094 | 0.019 | 0.008 | 1 | 0.319619815 | -0.332720152 | 0 | 0.024 | 1 | 0.503971666 | 0.537082804 |
| L1cam | 0.661088008 | -0.304709024 | 0.019 | 0.03 | 1 | 0.319619815 | -0.332720152 | 0 | 0.024 | 1 | 0.661088008 | 0.537082804 |
| Tjp3 | 0.732691937 | 1.154722595 | 0.019 | 0.013 | 1 | 0.319619815 | -0.622226769 | 0 | 0.024 | 1 | 0.732691937 | 0.537082804 |
| Gm41396 | 0.736073999 | 0.8327945 | 0.019 | 0.013 | 1 | 0.319619815 | -0.332720152 | 0 | 0.024 | 1 | 0.736073999 | 0.537082804 |
| Slc24a4 | 0.80004514 | 0.776210972 | 0.038 | 0.047 | 1 | 0.319619815 | -0.747757651 | 0 | 0.024 | 1 | 0.80004514 | 0.537082804 |
| C1qtnf3 | 0.88363765 | 0.522454379 | 0.075 | 0.085 | 1 | 0.319619815 | -0.332720152 | 0 | 0.024 | 1 | 0.88363765 | 0.537082804 |
| Gm31105 | 0.901698096 | 0.280253477 | 0.038 | 0.034 | 1 | 0.319619815 | -0.332720152 | 0 | 0.024 | 1 | 0.901698096 | 0.537082804 |
| 1700126G02Rik | 0.914958 | 0.347367673 | 0.019 | 0.021 | 1 | 0.319619815 | -0.332720152 | 0 | 0.024 | 1 | 0.914958 | 0.537082804 |
| Gm16573 | 0.917883255 | 0.569760094 | 0.019 | 0.021 | 1 | 0.319619815 | -0.332720152 | 0 | 0.024 | 1 | 0.917883255 | 0.537082804 |
| Macc1 | 0.929410101 | -0.1672055 | 0.019 | 0.017 | 1 | 0.319619815 | -0.332720152 | 0 | 0.024 | 1 | 0.929410101 | 0.537082804 |
| Gm30934 | 0.932505382 | 1.154722595 | 0.019 | 0.021 | 1 | 0.319619815 | -0.484723245 | 0 | 0.024 | 1 | 0.932505382 | 0.537082804 |
| Ccrl2 | 0.978112783 | 1.569760094 | 0.038 | 0.038 | 1 | 0.319619815 | -0.332720152 | 0 | 0.024 | 1 | 0.978112783 | 0.537082804 |
| Ankrd61 | 0.340246113 | 1.417757001 | 0.038 | 0.017 | 1 | 0.319629309 | -0.484723245 | 0 | 0.024 | 1 | 0.340246113 | 0.537095723 |
| Angel1 | 0.343000446 | -1.1672055 | 0 | 0.017 | 1 | 0.319629309 | -0.970150072 | 0 | 0.024 | 1 | 0.343000446 | 0.537095723 |
| Gm43661 | 0.413080549 | -0.1672055 | 0 | 0.013 | 1 | 0.319629309 | -0.484723245 | 0 | 0.024 | 1 | 0.413080549 | 0.537095723 |
| 4931403E22Rik | 0.48175508 | 1.154722595 | 0.038 | 0.021 | 1 | 0.319629309 | -0.484723245 | 0 | 0.024 | 1 | 0.48175508 | 0.537095723 |
| Tbxas1 | 0.488750271 | -0.430239906 | 0.019 | 0.038 | 1 | 0.319629309 | -0.622226769 | 0 | 0.024 | 1 | 0.488750271 | 0.537095723 |
| Rnf32 | 0.52195177 | -0.652632327 | 0.038 | 0.059 | 1 | 0.319629309 | -0.622226769 | 0 | 0.024 | 1 | 0.52195177 | 0.537095723 |
| Kif9 | 0.583115634 | 0.306725688 | 0.075 | 0.055 | 1 | 0.319629309 | -0.484723245 | 0 | 0.024 | 1 | 0.583115634 | 0.537095723 |
| 4930579K19Rik | 0.586409164 | 0.417757001 | 0.019 | 0.034 | 1 | 0.319629309 | -0.484723245 | 0 | 0.024 | 1 | 0.586409164 | 0.537095723 |
| Pkmyt1 | 0.596092797 | -0.52334931 | 0.075 | 0.097 | 1 | 0.319629309 | -0.622226769 | 0 | 0.024 | 1 | 0.596092797 | 0.537095723 |
| Gm10399 | 0.916620008 | 1.154722595 | 0.019 | 0.017 | 1 | 0.319629309 | -0.747757651 | 0 | 0.024 | 1 | 0.916620008 | 0.537095723 |
| Gm38505 | 0.413082433 | 0.154722595 | 0 | 0.013 | 1 | 0.319635005 | -0.747757651 | 0 | 0.024 | 1 | 0.413082433 | 0.537103474 |
| H1f5 | 0.421426794 | -3.061214804 | 0.057 | 0.089 | 1 | 0.319635005 | -1.069685746 | 0 | 0.024 | 1 | 0.421426794 | 0.537103474 |
| Gm49741 | 0.429421714 | 1.154722595 | 0.057 | 0.034 | 1 | 0.319635005 | -0.747757651 | 0 | 0.024 | 1 | 0.429421714 | 0.537103474 |
| Gm48956 | 0.458959906 | -1.1182959 | 0.038 | 0.064 | 1 | 0.319635005 | -0.970150072 | 0 | 0.024 | 1 | 0.458959906 | 0.537103474 |
| Gm2449 | 0.969362106 | -0.430239906 | 0.038 | 0.038 | 1 | 0.319635005 | -0.747757651 | 0 | 0.024 | 1 | 0.969362106 | 0.537103474 |
| Trim30a | 0.498931247 | 0.154722595 | 0.038 | 0.021 | 1 | 0.319635638 | -0.970150072 | 0 | 0.024 | 1 | 0.498931247 | 0.537104335 |
| Pcdhb10 | 0.503971666 | 1.569760094 | 0.019 | 0.008 | 1 | 0.319635638 | -1.069685746 | 0 | 0.024 | 1 | 0.503971666 | 0.537104335 |
| Gm57385 | 0.588476383 | 0.569760094 | 0.019 | 0.034 | 1 | 0.319635638 | -0.622226769 | 0 | 0.024 | 1 | 0.588476383 | 0.537104335 |
| Slc2a9 | 0.589572453 | -2.389597921 | 0.038 | 0.055 | 1 | 0.319635638 | -2.372248516 | 0 | 0.024 | 1 | 0.589572453 | 0.537104335 |
| Serpind1 | 0.624418548 | 0.932330173 | 0.038 | 0.025 | 1 | 0.319635638 | -0.970150072 | 0 | 0.024 | 1 | 0.624418548 | 0.537104335 |
| Dlec1 | 0.725936481 | 1.417757001 | 0.019 | 0.013 | 1 | 0.319635638 | -0.622226769 | 0 | 0.024 | 1 | 0.725936481 | 0.537104335 |
| Sptssb | 0.732691937 | 1.154722595 | 0.019 | 0.013 | 1 | 0.319635638 | -0.970150072 | 0 | 0.024 | 1 | 0.732691937 | 0.537104335 |
| Trim66 | 0.736073999 | 0.8327945 | 0.019 | 0.013 | 1 | 0.319635638 | -0.622226769 | 0 | 0.024 | 1 | 0.736073999 | 0.537104335 |
| Gm26631 | 0.736073999 | 0.8327945 | 0.019 | 0.013 | 1 | 0.319635638 | -0.622226769 | 0 | 0.024 | 1 | 0.736073999 | 0.537104335 |
| Slc28a1 | 0.413080549 | -0.1672055 | 0 | 0.013 | 1 | 0.319638803 | -0.970150072 | 0 | 0.024 | 1 | 0.413080549 | 0.537108641 |
| Gm29865 | 0.477502668 | 1.502645898 | 0.038 | 0.021 | 1 | 0.319638803 | -1.16279515 | 0 | 0.024 | 1 | 0.477502668 | 0.537108641 |
| Edn2 | 0.335451174 | -0.686579659 | 0.075 | 0.042 | 1 | 0.319639436 | -3.762707992 | 0 | 0.024 | 1 | 0.335451174 | 0.537109502 |
| Gm57453 | 0.338780504 | 0.984797593 | 0.057 | 0.03 | 1 | 0.319641967 | -0.16279515 | 0 | 0.024 | 1 | 0.338780504 | 0.537112947 |
| Gm30455 | 0.343004841 | -0.652632327 | 0 | 0.017 | 1 | 0.319641967 | -0.16279515 | 0 | 0.024 | 1 | 0.343004841 | 0.537112947 |
| A730011C13Rik | 0.413082433 | 0.154722595 | 0 | 0.013 | 1 | 0.319641967 | -0.16279515 | 0 | 0.024 | 1 | 0.413082433 | 0.537112947 |
| 9130221H12Rik | 0.413082433 | 0.154722595 | 0 | 0.013 | 1 | 0.319641967 | -0.16279515 | 0 | 0.024 | 1 | 0.413082433 | 0.537112947 |
| Fam209 | 0.503971666 | 1.569760094 | 0.019 | 0.008 | 1 | 0.319641967 | -0.16279515 | 0 | 0.024 | 1 | 0.503971666 | 0.537112947 |
| Cdca8 | 0.654152175 | -2.845277405 | 0.019 | 0.03 | 1 | 0.319641967 | -0.16279515 | 0 | 0.024 | 1 | 0.654152175 | 0.537112947 |
| Gm56859 | 0.782077996 | 0.347367673 | 0.019 | 0.025 | 1 | 0.319641967 | -0.16279515 | 0 | 0.024 | 1 | 0.782077996 | 0.537112947 |
| Notch4 | 0.876606655 | 0.154722595 | 0.038 | 0.042 | 1 | 0.319641967 | -0.16279515 | 0 | 0.024 | 1 | 0.876606655 | 0.537112947 |
| Gm43260 | 0.914958 | 0.347367673 | 0.019 | 0.021 | 1 | 0.319641967 | -0.16279515 | 0 | 0.024 | 1 | 0.914958 | 0.537112947 |
| C030010C08Rik | 0.926212226 | 0.8327945 | 0.019 | 0.017 | 1 | 0.319641967 | -0.16279515 | 0 | 0.024 | 1 | 0.926212226 | 0.537112947 |
| Arhgap27os2 | 0.929410101 | 0.569760094 | 0.019 | 0.017 | 1 | 0.319641967 | -0.16279515 | 0 | 0.024 | 1 | 0.929410101 | 0.537112947 |
| Zfpl1 | 0.319703391 | 0.569760094 | 0.434 | 0.398 | 1 | 0.340107671 | -0.151479837 | 0.244 | 0.328 | 1 | 0.340107671 | 0.537196524 |
| Iqcc | 0.31987574 | 0.762405172 | 0.113 | 0.072 | 1 | 1 | 0.557096931 | 0.098 | 0.099 | 1 | 1 | 0.537430991 |
| Ccser1 | 0.965440754 | 0.21498312 | 0.321 | 0.309 | 1 | 0.319884694 | -1.051538399 | 0.293 | 0.345 | 1 | 0.965440754 | 0.537443171 |
| Rhoj | 0.319951541 | -0.393976362 | 0.774 | 0.847 | 1 | 0.755361591 | 0.415158292 | 0.585 | 0.771 | 1 | 0.755361591 | 0.537534093 |
| Ralgps2 | 0.320248407 | 0.547040018 | 0.358 | 0.305 | 1 | 0.906184959 | 0.304709769 | 0.171 | 0.164 | 1 | 0.906184959 | 0.537937771 |
| Sugp1 | 0.320361795 | 0.648831665 | 0.264 | 0.208 | 1 | 0.72086259 | 0.5766773 | 0.268 | 0.259 | 1 | 0.72086259 | 0.53809191 |
| Polr3c | 0.321051386 | 0.459577176 | 0.264 | 0.208 | 1 | 0.466391605 | 0.470422519 | 0.268 | 0.225 | 1 | 0.466391605 | 0.539028779 |
| Romo1 | 0.368597137 | 0.168554868 | 0.849 | 0.839 | 1 | 0.321086852 | -0.134338704 | 0.683 | 0.788 | 1 | 0.368597137 | 0.539076938 |
| Septin4 | 0.579510289 | 0.569760094 | 0.094 | 0.072 | 1 | 0.321447309 | -0.622226769 | 0.049 | 0.096 | 1 | 0.579510289 | 0.539566246 |
| 3222401L13Rik | 0.354382417 | -0.134784022 | 0.057 | 0.102 | 1 | 0.32154479 | -0.278272368 | 0.024 | 0.065 | 1 | 0.354382417 | 0.539698528 |
| Pgap4 | 0.662895684 | 0.685237311 | 0.151 | 0.131 | 1 | 0.322108791 | -0.595754557 | 0.098 | 0.154 | 1 | 0.662895684 | 0.540463509 |
| Invs | 0.409744403 | 0.547040018 | 0.34 | 0.305 | 1 | 0.322178472 | 0.284232463 | 0.244 | 0.348 | 1 | 0.409744403 | 0.540557976 |
| Ormdl1 | 0.322286411 | 0.569760094 | 0.17 | 0.119 | 1 | 0.420625129 | -0.395455907 | 0.098 | 0.143 | 1 | 0.420625129 | 0.540704292 |
| AB124611 | 0.322352926 | 0.200526284 | 0.132 | 0.085 | 1 | 0.664753994 | 0.429120111 | 0.146 | 0.123 | 1 | 0.664753994 | 0.540794443 |
| Mkln1os | 0.621365635 | -0.430239906 | 0.075 | 0.097 | 1 | 0.322502854 | 1.000703582 | 0.098 | 0.058 | 1 | 0.621365635 | 0.540997617 |
| Crlf2 | 0.884324237 | 0.177442671 | 0.34 | 0.39 | 1 | 0.322616136 | 0.351778023 | 0.366 | 0.287 | 1 | 0.884324237 | 0.541151101 |
| Zfyve26 | 0.322710085 | 0.154722595 | 0.264 | 0.191 | 1 | 0.385906043 | 0.407217009 | 0.244 | 0.188 | 1 | 0.385906043 | 0.541278371 |
| H1f0 | 0.322951835 | -1.334583819 | 0.755 | 0.682 | 1 | 0.485491466 | -0.195412102 | 0.732 | 0.648 | 1 | 0.485491466 | 0.541605782 |
| Tmem192 | 0.323000726 | 0.517292674 | 0.151 | 0.102 | 1 | 0.669277944 | 0.83720485 | 0.122 | 0.106 | 1 | 0.669277944 | 0.541671984 |
| Gm17491 | 0.910889192 | 0.216123139 | 0.094 | 0.089 | 1 | 0.323095443 | -0.577832649 | 0.049 | 0.096 | 1 | 0.910889192 | 0.54180022 |
| Rpp40 | 0.323284582 | -0.260314904 | 0.019 | 0.051 | 1 | 0.820932515 | 0.614812428 | 0.049 | 0.041 | 1 | 0.820932515 | 0.542056244 |
| Scgb1b27 | 0.356344543 | -0.36312571 | 0.151 | 0.208 | 1 | 0.323541405 | -0.221688839 | 0.024 | 0.065 | 1 | 0.356344543 | 0.542403769 |
| Tjap1 | 0.502152011 | 0.458114738 | 0.34 | 0.314 | 1 | 0.323698086 | 0.743545743 | 0.439 | 0.399 | 1 | 0.502152011 | 0.542615721 |
| Pola1 | 0.324019533 | -0.253362144 | 0.283 | 0.216 | 1 | 0.596153662 | -0.101394606 | 0.341 | 0.287 | 1 | 0.596153662 | 0.543050408 |
| Ppwd1 | 0.3240565 | 0.55381855 | 0.34 | 0.288 | 1 | 0.706261247 | 0.164779508 | 0.341 | 0.311 | 1 | 0.706261247 | 0.543100385 |
| D11Wsu47e | 0.59338774 | 0.438515561 | 0.113 | 0.089 | 1 | 0.324111702 | 0.83720485 | 0.195 | 0.143 | 1 | 0.59338774 | 0.543175009 |
| Bcl2 | 0.657849108 | -0.556247791 | 0.377 | 0.398 | 1 | 0.324222939 | -0.465781681 | 0.366 | 0.454 | 1 | 0.657849108 | 0.543325364 |

| | | | | | | | | | | | | |
|---|---|---|---|---|---|---|---|---|---|---|---|---|
| Spef1 | 0.324474943 | 1.47665069 | 0.057 | 0.03 | 1 | 0.807969784 | 0.422167351 | 0.073 | 0.065 | 1 | 0.807969784 | 0.543665897 |
| Tle6 | 0.324474943 | 1.739685096 | 0.057 | 0.03 | 1 | 0.646976884 | 0.614812428 | 0.049 | 0.034 | 1 | 0.646976884 | 0.543665897 |
| Sgsh | 0.324474943 | 1.47665069 | 0.057 | 0.03 | 1 | 0.574045084 | 0.930314254 | 0.024 | 0.044 | 1 | 0.574045084 | 0.543665897 |
| Zfp160 | 0.484744555 | 0.247831999 | 0.321 | 0.271 | 1 | 0.324488746 | 0.658867609 | 0.415 | 0.355 | 1 | 0.484744555 | 0.543684546 |
| Gm20342 | 0.324552512 | 0.417757001 | 0.17 | 0.119 | 1 | 0.670504853 | -0.210100865 | 0.244 | 0.273 | 1 | 0.670504853 | 0.543770692 |
| Mboat7 | 0.324577477 | 0.510866405 | 0.302 | 0.25 | 1 | 0.955837507 | 0.229522273 | 0.293 | 0.311 | 1 | 0.955837507 | 0.543804415 |
| Btbd16 | 0.971591433 | -0.1182959 | 0.075 | 0.076 | 1 | 0.324691595 | 0.73286819 | 0.024 | 0.065 | 1 | 0.971591433 | 0.543958558 |
| Kcnab1 | 0.595477844 | 0.228723176 | 0.075 | 0.055 | 1 | 0.324993891 | 2.351778023 | 0.073 | 0.041 | 1 | 0.595477844 | 0.544366753 |
| Osgepl1 | 0.325117608 | 0.739685096 | 0.189 | 0.14 | 1 | 0.33637573 | -0.221688839 | 0.073 | 0.13 | 1 | 0.33637573 | 0.544533758 |
| Mrpl16 | 0.325378778 | 0.499370766 | 0.264 | 0.208 | 1 | 0.463208204 | -0.391613841 | 0.171 | 0.215 | 1 | 0.463208204 | 0.544886207 |
| Gsta3 | 0.567830665 | -0.304709024 | 0.019 | 0.034 | 1 | 0.32541576 | 0.679663573 | 0.244 | 0.181 | 1 | 0.567830665 | 0.544936103 |
| Zfp850 | 0.325530479 | 2.154722595 | 0.038 | 0.017 | 1 | 0.746881137 | 1.029849928 | 0.024 | 0.017 | 1 | 0.746881137 | 0.545090866 |
| Slc35f1 | 0.325530479 | 3.640149422 | 0.038 | 0.017 | 1 | 0.357847088 | -0.16279515 | 0 | 0.02 | 1 | 0.357847088 | 0.545090866 |
| AW822252 | 0.325530479 | 2.154722595 | 0.038 | 0.017 | 1 | 0.402369942 | -0.747757651 | 0 | 0.017 | 1 | 0.402369942 | 0.545090866 |
| Hecw1 | 0.325536394 | 2.377115016 | 0.038 | 0.017 | 1 | 0.911761241 | -0.332720152 | 0.024 | 0.027 | 1 | 0.911761241 | 0.545098844 |
| Med19 | 0.549258084 | 0.380282295 | 0.396 | 0.373 | 1 | 0.325723206 | 0.576244772 | 0.39 | 0.331 | 1 | 0.549258084 | 0.545350805 |
| Ica1l | 0.325936131 | 1.154722595 | 0.057 | 0.03 | 1 | 0.455098275 | 0.515276755 | 0 | 0.014 | 1 | 0.455098275 | 0.5456379 |
| Slc25a15 | 0.566288393 | 1.090592257 | 0.113 | 0.089 | 1 | 0.326167917 | 0.88967227 | 0.195 | 0.143 | 1 | 0.566288393 | 0.545950324 |
| Ctnnbip1 | 0.666499205 | 0.136498836 | 0.585 | 0.581 | 1 | 0.326292936 | 0.385825503 | 0.537 | 0.522 | 1 | 0.666499205 | 0.546118792 |
| Scgb1c1 | 0.362626638 | -0.430239906 | 0.019 | 0.047 | 1 | 0.326294192 | 1.592092352 | 0.244 | 0.188 | 1 | 0.362626638 | 0.546120484 |
| Gm56692 | 0.32647098 | 1.090592257 | 0.132 | 0.089 | 1 | 0.984892814 | 0.404245443 | 0.073 | 0.075 | 1 | 0.984892814 | 0.54635866 |
| Slc29a3 | 0.326564286 | -0.372524408 | 0.245 | 0.314 | 1 | 0.675212518 | -0.200930279 | 0.244 | 0.276 | 1 | 0.675212518 | 0.546484339 |
| Ccnf | 0.32691973 | 0.491757582 | 0.075 | 0.042 | 1 | 0.357854401 | -0.332720152 | 0 | 0.02 | 1 | 0.357854401 | 0.54696295 |
| Grem1 | 0.327036932 | 0.865215978 | 0.019 | 0.051 | 1 | 0.54731036 | -0.899760744 | 0.024 | 0.044 | 1 | 0.54731036 | 0.547120709 |
| Tmem144 | 0.327350685 | 2.154722595 | 0.038 | 0.017 | 1 | 0.523148091 | -0.622226769 | 0.049 | 0.075 | 1 | 0.523148091 | 0.547542899 |
| Susd1 | 0.327364111 | 1.280253477 | 0.057 | 0.03 | 1 | 0.440032309 | 1.83720485 | 0.049 | 0.027 | 1 | 0.440032309 | 0.54756096 |
| Psmc3ip | 0.328052493 | -0.489133595 | 0.038 | 0.076 | 1 | 0.626818216 | 0.766815522 | 0.073 | 0.055 | 1 | 0.626818216 | 0.548486548 |
| Cox6b1 | 0.384456565 | 0.100016703 | 0.962 | 0.903 | 1 | 0.328269088 | 0.126002352 | 0.878 | 0.918 | 1 | 0.384456565 | 0.548777582 |
| Cdc73 | 0.32850138 | 0.201265181 | 0.774 | 0.818 | 1 | 0.366722863 | -0.154648911 | 0.805 | 0.894 | 1 | 0.366722863 | 0.549089603 |
| Iqcn | 0.32862862 | 0.454282877 | 0.132 | 0.089 | 1 | 0.671540682 | 0.564186355 | 0.195 | 0.174 | 1 | 0.671540682 | 0.54926047 |
| Gm35248 | 0.328746305 | 1.306725688 | 0.057 | 0.03 | 1 | 0.744047924 | 1.252242349 | 0.024 | 0.017 | 1 | 0.744047924 | 0.549418476 |
| Dynlt1a | 0.328746305 | 1.306725688 | 0.057 | 0.03 | 1 | 0.449580774 | -0.484723245 | 0.024 | 0.051 | 1 | 0.449580774 | 0.549418476 |
| Dok1 | 0.328750173 | -0.349897798 | 0.321 | 0.403 | 1 | 0.524566271 | -0.344124915 | 0.195 | 0.239 | 1 | 0.524566271 | 0.54942367 |
| Paip2 | 0.424694227 | 0.100547455 | 0.981 | 0.975 | 1 | 0.328898848 | 0.236687535 | 0.976 | 0.986 | 1 | 0.424694227 | 0.549623244 |
| Pitpnm3 | 0.426679191 | -0.237594828 | 0.038 | 0.068 | 1 | 0.328915336 | -0.712992233 | 0.098 | 0.157 | 1 | 0.426679191 | 0.549645374 |
| Pcdhb11 | 0.329173506 | 2.154722595 | 0.038 | 0.017 | 1 | 0.605005904 | 1.029849928 | 0.024 | 0.014 | 1 | 0.605005904 | 0.549991815 |
| Cript | 0.329506091 | 0.226615513 | 0.472 | 0.398 | 1 | 0.92167143 | 0.353171258 | 0.415 | 0.447 | 1 | 0.92167143 | 0.550437918 |
| Btbd9 | 0.496733313 | -0.105706171 | 0.943 | 0.877 | 1 | 0.329569387 | -0.326325925 | 0.902 | 0.962 | 1 | 0.496733313 | 0.550522793 |
| 5730522E02Rik | 0.574905166 | 1.084333267 | 0.151 | 0.127 | 1 | 0.329833629 | 0.904319046 | 0.171 | 0.119 | 1 | 0.574905166 | 0.550877035 |
| Ggact | 0.329992458 | 0.531792244 | 0.283 | 0.229 | 1 | 0.977680893 | 0.702555323 | 0.244 | 0.27 | 1 | 0.977680893 | 0.551089893 |
| Gm57200 | 0.3301007 | -1.195774652 | 0.075 | 0.119 | 1 | 0.648957759 | -0.470223675 | 0.146 | 0.174 | 1 | 0.648957759 | 0.551234928 |
| Zfp111 | 0.330253102 | 0.640149422 | 0.113 | 0.072 | 1 | 0.540093871 | 0.174239837 | 0.073 | 0.106 | 1 | 0.540093871 | 0.551439092 |
| Ripk4 | 0.413082433 | 0.154722595 | 0 | 0.013 | 1 | 0.330691785 | 0.989207943 | 0.098 | 0.058 | 1 | 0.413082433 | 0.552026513 |
| Ift22 | 0.331018419 | 0.417757001 | 0.226 | 0.169 | 1 | 0.952969365 | 0.326242931 | 0.146 | 0.157 | 1 | 0.952969365 | 0.552463645 |
| Dph5 | 0.332357405 | -0.196749776 | 0.245 | 0.339 | 1 | 0.461735542 | -1.354415223 | 0.146 | 0.195 | 1 | 0.461735542 | 0.554253365 |
| Gm11827 | 0.332821087 | 1.8327945 | 0.038 | 0.017 | 1 | 0.51919986 | 0.83720485 | 0 | 0.01 | 1 | 0.51919986 | 0.554872298 |
| Fam131c | 0.332821087 | 1.8327945 | 0.038 | 0.017 | 1 | 0.402373368 | 0.252242349 | 0 | 0.017 | 1 | 0.402373368 | 0.554872298 |
| Gm44777 | 0.332821087 | 1.8327945 | 0.038 | 0.017 | 1 | 0.450957104 | -0.332720152 | 0.024 | 0.051 | 1 | 0.450957104 | 0.554872298 |
| Mtfp1 | 0.332821087 | 1.8327945 | 0.038 | 0.017 | 1 | 0.744047924 | 1.252242349 | 0.024 | 0.017 | 1 | 0.744047924 | 0.554872298 |
| Taco1os | 0.332821087 | 1.8327945 | 0.038 | 0.017 | 1 | 0.402364804 | -0.16279515 | 0 | 0.017 | 1 | 0.402364804 | 0.554872298 |
| Etohd2 | 0.332821087 | 1.8327945 | 0.038 | 0.017 | 1 | 0.746881137 | 1.029849928 | 0.024 | 0.017 | 1 | 0.746881137 | 0.554872298 |
| Zfpm1 | 0.332898426 | 1.043691282 | 0.226 | 0.178 | 1 | 0.844288026 | 0.635570989 | 0.244 | 0.249 | 1 | 0.844288026 | 0.55497549 |
| Tmem132b | 0.332999856 | -0.214511215 | 0.075 | 0.042 | 1 | 0.485324194 | -1.995685164 | 0.024 | 0.048 | 1 | 0.485324194 | 0.555110808 |
| Zfp784 | 0.362626638 | -0.430239906 | 0.019 | 0.047 | 1 | 0.333002878 | -0.863234868 | 0.024 | 0.061 | 1 | 0.362626638 | 0.55511484 |
| Fyb2 | 0.333048769 | 1.154722595 | 0.057 | 0.03 | 1 | 0.415029055 | -0.505187348 | 0.098 | 0.147 | 1 | 0.415029055 | 0.555176056 |
| Clcn3 | 0.333063913 | 0.521780065 | 0.528 | 0.483 | 1 | 0.668609581 | 0.101600931 | 0.537 | 0.621 | 1 | 0.668609581 | 0.555196256 |
| Armcx6 | 0.689493841 | 0.154722595 | 0.094 | 0.114 | 1 | 0.334197171 | -0.747757651 | 0.073 | 0.123 | 1 | 0.689493841 | 0.556706593 |
| Rskr | 0.334484705 | 0.891688189 | 0.057 | 0.03 | 1 | 0.979194243 | 0.83720485 | 0.024 | 0.024 | 1 | 0.979194243 | 0.557089393 |
| Abcb6 | 0.334518863 | 1.154722595 | 0.057 | 0.03 | 1 | 0.894652652 | 0.377773231 | 0.049 | 0.055 | 1 | 0.894652652 | 0.557134856 |
| Gm31728 | 0.334518863 | 1.154722595 | 0.057 | 0.03 | 1 | 0.859141609 | 1.699701326 | 0.024 | 0.02 | 1 | 0.859141609 | 0.557134856 |
| Tpgs2 | 0.334640417 | 0.938993904 | 0.302 | 0.254 | 1 | 0.860053644 | 0.328191202 | 0.195 | 0.218 | 1 | 0.860053644 | 0.557296626 |
| Nnat | 0.334672444 | 2.154722595 | 0.038 | 0.017 | 1 | 0.973993537 | 0.962735732 | 0.024 | 0.024 | 1 | 0.973993537 | 0.557339243 |
| Six3os1 | 0.334672444 | 1.891688189 | 0.038 | 0.017 | 1 | 0.593594236 | 1.422167351 | 0.024 | 0.014 | 1 | 0.593594236 | 0.557339243 |
| Atp6v1f | 0.334762544 | 0.214940118 | 0.925 | 0.873 | 1 | 0.723102143 | -0.131723704 | 0.829 | 0.867 | 1 | 0.723102143 | 0.557459128 |
| Abhd18 | 0.334817767 | -0.20282941 | 0.075 | 0.123 | 1 | 0.954825159 | 0.23386382 | 0.146 | 0.143 | 1 | 0.954825159 | 0.557532597 |
| Snhg7 | 0.334865629 | -0.209160817 | 0.491 | 0.581 | 1 | 0.616461981 | -0.273640919 | 0.488 | 0.505 | 1 | 0.616461981 | 0.557596269 |
| Rgs3 | 0.334958071 | 0.352572832 | 0.604 | 0.559 | 1 | 0.723003826 | 0.617824319 | 0.634 | 0.706 | 1 | 0.723003826 | 0.557719232 |
| Papln | 0.337391903 | 0.8327945 | 0.057 | 0.03 | 1 | 0.335113466 | -0.686357106 | 0.024 | 0.061 | 1 | 0.337391903 | 0.557925898 |
| H2-DMa | 0.335192889 | -0.689626535 | 0.151 | 0.208 | 1 | 0.404146736 | 0.218295017 | 0.073 | 0.119 | 1 | 0.404146736 | 0.558031505 |
| Hlcs | 0.918803784 | 0.256602209 | 0.132 | 0.127 | 1 | 0.335682043 | 0.326242931 | 0.171 | 0.116 | 1 | 0.918803784 | 0.558681652 |
| 2500004C02Rik | 0.335694972 | -0.799473716 | 0.057 | 0.097 | 1 | 0.585391668 | -0.410722664 | 0.122 | 0.15 | 1 | 0.585391668 | 0.55869883 |
| Fbxl3 | 0.335740731 | 0.212825549 | 0.755 | 0.72 | 1 | 0.622097507 | 0.218633956 | 0.829 | 0.795 | 1 | 0.622097507 | 0.558759623 |
| Pitrm1 | 0.971808752 | -0.296973375 | 0.358 | 0.356 | 1 | 0.336006649 | -0.226298093 | 0.293 | 0.41 | 1 | 0.971808752 | 0.559112829 |
| Gm5113 | 0.52492136 | -0.508242418 | 0.038 | 0.059 | 1 | 0.33616415 | -0.686357106 | 0.024 | 0.061 | 1 | 0.52492136 | 0.559321964 |
| Lrrc27 | 0.531815432 | -0.799473716 | 0.057 | 0.081 | 1 | 0.33616415 | -0.747757651 | 0.024 | 0.061 | 1 | 0.531815432 | 0.559321964 |
| Pafah2 | 0.877748789 | 0.247831999 | 0.057 | 0.051 | 1 | 0.336164391 | -0.622226769 | 0.024 | 0.061 | 1 | 0.877748789 | 0.559322285 |
| Nipal3 | 0.938741618 | 0.228723176 | 0.075 | 0.072 | 1 | 0.336164391 | -0.622226769 | 0.024 | 0.061 | 1 | 0.938741618 | 0.559322285 |
| Ebf3 | 0.357738988 | 0.816852956 | 0.302 | 0.267 | 1 | 0.336224431 | 0.825232208 | 0.415 | 0.369 | 1 | 0.357738988 | 0.559401994 |
| Susd4 | 0.336520503 | 1.569760094 | 0.038 | 0.017 | 1 | 0.602691701 | -0.410722664 | 0.024 | 0.041 | 1 | 0.602691701 | 0.559794957 |
| Chn1os3 | 0.336520503 | 1.569760094 | 0.038 | 0.017 | 1 | 0.455097252 | -0.16279515 | 0 | 0.014 | 1 | 0.455097252 | 0.559794957 |
| Eqtn | 0.336520503 | 1.569760094 | 0.038 | 0.017 | 1 | 0.691851529 | 0.749742009 | 0.024 | 0.038 | 1 | 0.691851529 | 0.559794957 |
| Kcnab2 | 0.336520503 | 1.569760094 | 0.038 | 0.017 | 1 | 0.430584486 | 2.422167351 | 0.024 | 0.01 | 1 | 0.430584486 | 0.559794957 |
| Gm20324 | 0.336520503 | 1.569760094 | 0.038 | 0.017 | 1 | 0.5191986 | 0.515276755 | 0 | 0.01 | 1 | 0.5191986 | 0.559794957 |
| Ydjc | 0.336520503 | 1.569760094 | 0.038 | 0.017 | 1 | 0.455094184 | 0.252242349 | 0 | 0.014 | 1 | 0.455094184 | 0.559794957 |
| Ikbke | 0.336825881 | 0.409295422 | 0.245 | 0.186 | 1 | 0.380549595 | -0.138547604 | 0.195 | 0.259 | 1 | 0.380549595 | 0.560200088 |
| Pisd | 0.337046272 | 0.384204441 | 0.226 | 0.169 | 1 | 0.602085017 | -0.250257991 | 0.244 | 0.283 | 1 | 0.602085017 | 0.560492354 |
| Mfsd4b4 | 0.442326061 | 0.695290976 | 0.057 | 0.034 | 1 | 0.337213694 | -0.555112573 | 0.024 | 0.061 | 1 | 0.442326061 | 0.560714312 |
| Ptpru | 0.958257527 | 0.454282877 | 0.075 | 0.081 | 1 | 0.337213694 | -0.686357106 | 0.024 | 0.061 | 1 | 0.958257527 | 0.560714312 |
| Clhc1 | 0.337396694 | 0.906795081 | 0.057 | 0.03 | 1 | 0.89280767 | 0.458693227 | 0.049 | 0.044 | 1 | 0.89280767 | 0.560956859 |
| Slc8a2 | 0.377181966 | 0.280253477 | 0.094 | 0.059 | 1 | 0.337804198 | -1.435813645 | 0.049 | 0.092 | 1 | 0.377181966 | 0.56149672 |
| Gnpnat1 | 0.468259388 | 0.374035623 | 0.434 | 0.39 | 1 | 0.337889405 | 0.231483789 | 0.244 | 0.355 | 1 | 0.468259388 | 0.56160956 |
| Arl6ip6 | 0.337932322 | 0.369461444 | 0.472 | 0.419 | 1 | 0.833209222 | 0.589277336 | 0.366 | 0.416 | 1 | 0.833209222 | 0.56166639 |
| Gm56587 | 0.395615987 | 0.8327945 | 0.075 | 0.047 | 1 | 0.33805041 | -0.484723245 | 0.049 | 0.096 | 1 | 0.395615987 | 0.561822741 |
| Zfp799 | 0.547543999 | 0.417757001 | 0.189 | 0.157 | 1 | 0.338050569 | -0.448197369 | 0.049 | 0.096 | 1 | 0.547543999 | 0.561822951 |
| Emc4 | 0.338072533 | -0.141894412 | 0.509 | 0.597 | 1 | 0.379268537 | -0.140075074 | 0.463 | 0.539 | 1 | 0.379268537 | 0.561852029 |
| Pcyox1l | 0.761797609 | 0.569760094 | 0.057 | 0.047 | 1 | 0.338261133 | -0.484723245 | 0.024 | 0.061 | 1 | 0.761797609 | 0.562101671 |
| Fibp | 0.33830904 | 0.491757582 | 0.453 | 0.462 | 1 | 0.470656563 | 0.242258261 | 0.268 | 0.362 | 1 | 0.470656563 | 0.562165074 |
| Pigl | 0.39233877 | 0.569760094 | 0.132 | 0.093 | 1 | 0.338329262 | 0.911205431 | 0.146 | 0.099 | 1 | 0.39233877 | 0.562191834 |

| | | | | | | | | | | | | |
|---|---|---|---|---|---|---|---|---|---|---|---|---|
| Sec14l2 | 0.663657426 | -0.1672055 | 0.094 | 0.114 | 1 | 0.338342398 | -0.380025866 | 0.171 | 0.235 | 1 | 0.663657426 | 0.562209217 |
| Kcnh1 | 0.338375114 | 0.891688189 | 0.038 | 0.017 | 1 | 0.767181773 | 0.962735732 | 0.024 | 0.034 | 1 | 0.767181773 | 0.56225251 |
| Gm16178 | 0.338375114 | 1.47665069 | 0.038 | 0.017 | 1 | 0.455094184 | 0.252242349 | 0 | 0.014 | 1 | 0.455094184 | 0.56225251 |
| Impg2 | 0.338381019 | 1.569760094 | 0.038 | 0.017 | 1 | 0.883302434 | 1.174239837 | 0.049 | 0.044 | 1 | 0.883302434 | 0.562260324 |
| Dnajc24 | 0.581593219 | 0.280253477 | 0.245 | 0.216 | 1 | 0.338487043 | -0.513292397 | 0.146 | 0.208 | 1 | 0.581593219 | 0.562400608 |
| Adam28 | 0.413080549 | -0.652632327 | 0 | 0.013 | 1 | 0.338667992 | 0.766815522 | 0.098 | 0.058 | 1 | 0.413080549 | 0.562639976 |
| Gm15832 | 0.338780504 | 0.8327945 | 0.057 | 0.03 | 1 | 0.455098275 | 0.515276755 | 0 | 0.014 | 1 | 0.455098275 | 0.562788778 |
| Ryr3 | 0.338780504 | 0.984797593 | 0.057 | 0.03 | 1 | 0.815220306 | 0.422167351 | 0.073 | 0.085 | 1 | 0.815220306 | 0.562788778 |
| AV039307 | 0.338780504 | 0.984797593 | 0.057 | 0.03 | 1 | 0.455098275 | 0.515276755 | 0 | 0.014 | 1 | 0.455098275 | 0.562788778 |
| Zfp707 | 0.338780504 | 0.984797593 | 0.057 | 0.03 | 1 | 0.953470771 | 1.737669176 | 0.049 | 0.048 | 1 | 0.953470771 | 0.562788778 |
| Rab26 | 0.338780504 | 0.984797593 | 0.057 | 0.03 | 1 | 0.357381956 | 1.377773231 | 0.049 | 0.024 | 1 | 0.357381956 | 0.562788778 |
| Mysm1 | 0.413362138 | 0.123695699 | 0.491 | 0.602 | 1 | 0.338972728 | 0.433626466 | 0.683 | 0.672 | 1 | 0.413362138 | 0.563042946 |
| 2310061I04Rik | 0.339334745 | 0.721763188 | 0.151 | 0.106 | 1 | 0.845211542 | 0.334704509 | 0.122 | 0.137 | 1 | 0.845211542 | 0.56352142 |
| 2510046G10Rik | 0.421517727 | -0.304709024 | 0.019 | 0.042 | 1 | 0.339581736 | -0.410722664 | 0.024 | 0.061 | 1 | 0.421517727 | 0.563847717 |
| Secisbp2 | 0.340111441 | 0.414242516 | 0.472 | 0.441 | 1 | 0.758292843 | 0.540223112 | 0.317 | 0.389 | 1 | 0.758292843 | 0.56454709 |
| Rgs5 | 0.340246113 | 1.417757001 | 0.038 | 0.017 | 1 | 0.711735899 | 0.19334866 | 0.049 | 0.065 | 1 | 0.711735899 | 0.564724809 |
| Gm14440 | 0.340246113 | 1.417757001 | 0.038 | 0.017 | 1 | 0.713168757 | 0.444887427 | 0.073 | 0.058 | 1 | 0.713168757 | 0.564724809 |
| Gm15441 | 0.340246113 | 1.417757001 | 0.038 | 0.017 | 1 | 0.826384776 | 0.515276755 | 0.024 | 0.031 | 1 | 0.826384776 | 0.564724809 |
| Lrriq3 | 0.340246113 | 1.417757001 | 0.038 | 0.017 | 1 | 0.92332264 | -0.16279515 | 0.049 | 0.044 | 1 | 0.92332264 | 0.564724809 |
| Gm43728 | 0.340246113 | 1.417757001 | 0.038 | 0.017 | 1 | 0.936207831 | 1.252242349 | 0.024 | 0.027 | 1 | 0.936207831 | 0.564724809 |
| 4930557J02Rik | 0.340246113 | 1.417757001 | 0.038 | 0.017 | 1 | 0.87556513 | 0.377773231 | 0.024 | 0.02 | 1 | 0.87556513 | 0.564724809 |
| Gm57192 | 0.340246113 | 1.417757001 | 0.038 | 0.017 | 1 | 0.883298666 | 0.989207943 | 0.049 | 0.044 | 1 | 0.883298666 | 0.564724809 |
| Vstm5 | 0.340246113 | 1.417757001 | 0.038 | 0.017 | 1 | 0.51919986 | 0.83720485 | 0 | 0.01 | 1 | 0.51919986 | 0.564724809 |
| Jsrp1 | 0.340246113 | 1.417757001 | 0.038 | 0.017 | 1 | 0.438672228 | 1.83720485 | 0.024 | 0.01 | 1 | 0.438672228 | 0.564724809 |
| D630036H23Rik | 0.340246113 | 1.417757001 | 0.038 | 0.017 | 1 | 0.430584486 | 2.422167351 | 0.024 | 0.01 | 1 | 0.430584486 | 0.564724809 |
| Nrxn3 | 0.340246113 | 1.417757001 | 0.038 | 0.017 | 1 | 0.435968064 | 1.83720485 | 0.024 | 0.01 | 1 | 0.435968064 | 0.564724809 |
| Gm48420 | 0.340246113 | 1.417757001 | 0.038 | 0.017 | 1 | 0.455098275 | 0.515276755 | 0 | 0.014 | 1 | 0.455098275 | 0.564724809 |
| Gm56760 | 0.340246113 | 1.417757001 | 0.038 | 0.017 | 1 | 0.362195044 | 1.252242349 | 0.049 | 0.024 | 1 | 0.362195044 | 0.564724809 |
| 4921515G04Rik | 0.340246113 | 1.417757001 | 0.038 | 0.017 | 1 | 0.856404759 | 1.614812428 | 0.024 | 0.02 | 1 | 0.856404759 | 0.564724809 |
| Gm49785 | 0.340246113 | 1.417757001 | 0.038 | 0.017 | 1 | 0.5191986 | 0.252242349 | 0 | 0.01 | 1 | 0.5191986 | 0.564724809 |
| Gm33046 | 0.340283291 | 0.776210972 | 0.057 | 0.03 | 1 | 0.894713448 | 0.574170444 | 0.049 | 0.044 | 1 | 0.894713448 | 0.564773864 |
| Zfp317 | 0.478505112 | 0.489141634 | 0.302 | 0.267 | 1 | 0.34088299 | -0.385187572 | 0.22 | 0.283 | 1 | 0.478505112 | 0.565564767 |
| Kctd10 | 0.340925897 | 0.450707422 | 0.358 | 0.305 | 1 | 0.361825795 | -0.252992959 | 0.439 | 0.515 | 1 | 0.361825795 | 0.565621326 |
| Rad18 | 0.341152009 | -0.752168001 | 0.132 | 0.182 | 1 | 0.36154551 | -0.970150072 | 0.073 | 0.119 | 1 | 0.36154551 | 0.565919325 |
| Lsg1 | 0.341321595 | -0.100091304 | 0.434 | 0.525 | 1 | 0.78481415 | -0.140075074 | 0.512 | 0.505 | 1 | 0.78481415 | 0.566142759 |
| A930037H05Rik | 0.387740096 | 1.347367673 | 0.075 | 0.047 | 1 | 0.341522536 | 2.100239256 | 0.049 | 0.024 | 1 | 0.387740096 | 0.56640743 |
| Cxcl17 | 0.34174933 | 0.652222254 | 0.057 | 0.03 | 1 | 0.627119409 | 0.334704509 | 0.122 | 0.096 | 1 | 0.627119409 | 0.566706055 |
| Limk1 | 0.926212226 | 0.8327945 | 0.019 | 0.017 | 1 | 0.34180587 | 1.351778023 | 0.073 | 0.041 | 1 | 0.926212226 | 0.566780487 |
| Nrxn2 | 0.342053322 | 0.318221327 | 0.113 | 0.072 | 1 | 0.411629078 | 0.160743995 | 0.317 | 0.249 | 1 | 0.411629078 | 0.567106168 |
| Isca1 | 0.766723903 | 0.217653851 | 0.453 | 0.466 | 1 | 0.342183507 | 0.323581131 | 0.366 | 0.474 | 1 | 0.766723903 | 0.567277461 |
| Tent5a | 0.342501803 | -0.965295687 | 0.604 | 0.631 | 1 | 0.772471584 | -0.213421223 | 0.415 | 0.41 | 1 | 0.772471584 | 0.567696121 |
| Mecr | 0.342874284 | -0.410062024 | 0.17 | 0.229 | 1 | 0.861015361 | 0.136765132 | 0.22 | 0.229 | 1 | 0.861015361 | 0.568185794 |
| Gm28818 | 0.343000446 | -0.430239906 | 0 | 0.017 | 1 | 0.441381862 | 1.515276755 | 0.024 | 0.01 | 1 | 0.441381862 | 0.568351586 |
| Serpinb11 | 0.343000446 | -1.430239906 | 0 | 0.017 | 1 | 0.455094184 | 0.252242349 | 0 | 0.014 | 1 | 0.455094184 | 0.568351586 |
| Oip5 | 0.343000446 | -0.430239906 | 0 | 0.017 | 1 | 0.5191986 | 0.515276755 | 0 | 0.01 | 1 | 0.5191986 | 0.568351586 |
| Gm42705 | 0.343000446 | -0.430239906 | 0 | 0.017 | 1 | 0.5191986 | 0.515276755 | 0 | 0.01 | 1 | 0.5191986 | 0.568351586 |
| Gm52698 | 0.343000446 | -0.430239906 | 0 | 0.017 | 1 | 0.749728988 | 0.83720485 | 0.024 | 0.017 | 1 | 0.749728988 | 0.568351586 |
| E130006D01Rik | 0.343000446 | -0.430239906 | 0 | 0.017 | 1 | 0.402373368 | 0.252242349 | 0 | 0.017 | 1 | 0.402373368 | 0.568351586 |
| Kcna1 | 0.343000446 | -0.430239906 | 0 | 0.017 | 1 | 0.455763738 | 1.100239256 | 0.049 | 0.027 | 1 | 0.455763738 | 0.568351586 |
| Syt3 | 0.343000446 | -0.430239906 | 0 | 0.017 | 1 | 0.402369942 | -0.16279515 | 0 | 0.017 | 1 | 0.402369942 | 0.568351586 |
| Tgfbr3l | 0.343000446 | -0.652632327 | 0 | 0.017 | 1 | 0.438672228 | 1.83720485 | 0.024 | 0.01 | 1 | 0.438672228 | 0.568351586 |
| Gm45820 | 0.343000446 | -0.652632327 | 0 | 0.017 | 1 | 0.51919986 | 0.83720485 | 0 | 0.01 | 1 | 0.51919986 | 0.568351586 |
| Gm35721 | 0.343000446 | -0.430239906 | 0 | 0.017 | 1 | 0.5191986 | 0.515276755 | 0 | 0.01 | 1 | 0.5191986 | 0.568351586 |
| Trpm2 | 0.343000446 | -0.430239906 | 0 | 0.017 | 1 | 0.5191986 | 0.252242349 | 0 | 0.01 | 1 | 0.5191986 | 0.568351586 |
| Tcam1 | 0.343000446 | -0.652632327 | 0 | 0.017 | 1 | 0.60213981 | 1.252242349 | 0.024 | 0.014 | 1 | 0.60213981 | 0.568351586 |
| 9430024F10Rik | 0.343000446 | -0.430239906 | 0 | 0.017 | 1 | 0.605005904 | 0.667279848 | 0.024 | 0.014 | 1 | 0.605005904 | 0.568351586 |
| H3c1 | 0.343000446 | -0.430239906 | 0 | 0.017 | 1 | 0.51919986 | 0.83720485 | 0 | 0.01 | 1 | 0.51919986 | 0.568351586 |
| Gm48250 | 0.343000446 | -0.430239906 | 0 | 0.017 | 1 | 0.51919986 | -0.16279515 | 0 | 0.01 | 1 | 0.51919986 | 0.568351586 |
| Marveld2 | 0.343000446 | -1.015202407 | 0 | 0.017 | 1 | 0.744047924 | 1.252242349 | 0.024 | 0.017 | 1 | 0.744047924 | 0.568351586 |
| Depdc1b | 0.343000446 | -1.015202407 | 0 | 0.017 | 1 | 0.51919986 | 0.83720485 | 0 | 0.01 | 1 | 0.51919986 | 0.568351586 |
| 9930012K11Rik | 0.343000446 | -0.430239906 | 0 | 0.017 | 1 | 0.402373368 | 0.252242349 | 0 | 0.017 | 1 | 0.402373368 | 0.568351586 |
| Cyyr1 | 0.343000446 | -0.845277405 | 0 | 0.017 | 1 | 0.5191986 | 0.252242349 | 0 | 0.01 | 1 | 0.5191986 | 0.568351586 |
| Gm49801 | 0.343000446 | -0.430239906 | 0 | 0.017 | 1 | 0.87556513 | 0.515276755 | 0.024 | 0.02 | 1 | 0.87556513 | 0.568351586 |
| Gng5c | 0.343000446 | -0.430239906 | 0 | 0.017 | 1 | 0.733713289 | 0.721727632 | 0.049 | 0.038 | 1 | 0.733713289 | 0.568351586 |
| C030005K06Rik | 0.343000446 | -0.430239906 | 0 | 0.017 | 1 | 0.992196803 | 0.515276755 | 0.024 | 0.024 | 1 | 0.992196803 | 0.568351586 |
| Gm9895 | 0.343000446 | -0.430239906 | 0 | 0.017 | 1 | 0.5191986 | 0.515276755 | 0 | 0.01 | 1 | 0.5191986 | 0.568351586 |
| Gm16189 | 0.343000446 | -0.652632327 | 0 | 0.017 | 1 | 0.729889663 | 2.252242349 | 0.024 | 0.017 | 1 | 0.729889663 | 0.568351586 |
| Ppef1 | 0.343000446 | -0.430239906 | 0 | 0.017 | 1 | 0.51919986 | -0.332720152 | 0 | 0.01 | 1 | 0.51919986 | 0.568351586 |
| Iqca | 0.343004841 | -1.015202407 | 0 | 0.017 | 1 | 0.51919986 | 0.83720485 | 0 | 0.01 | 1 | 0.51919986 | 0.568357361 |
| Ifi203 | 0.343004841 | -0.652632327 | 0 | 0.017 | 1 | 0.587925597 | 2.252242349 | 0.024 | 0.014 | 1 | 0.587925597 | 0.568357361 |
| Gcnt7 | 0.343004841 | -0.652632327 | 0 | 0.017 | 1 | 0.455098275 | 0.515276755 | 0 | 0.014 | 1 | 0.455098275 | 0.568357361 |
| Chrnb2 | 0.343004841 | -0.652632327 | 0 | 0.017 | 1 | 0.455098275 | 0.515276755 | 0 | 0.014 | 1 | 0.455098275 | 0.568357361 |
| Gm128 | 0.343004841 | -0.652632327 | 0 | 0.017 | 1 | 0.495851362 | -0.332720152 | 0.024 | 0.048 | 1 | 0.495851362 | 0.568357361 |
| Cfap161 | 0.343004841 | -0.845277405 | 0 | 0.017 | 1 | 0.402369942 | -0.484723245 | 0 | 0.017 | 1 | 0.402369942 | 0.568357361 |
| D7Ertd443e | 0.343004841 | -0.845277405 | 0 | 0.017 | 1 | 0.455094184 | 0.252242349 | 0 | 0.014 | 1 | 0.455094184 | 0.568357361 |
| Gm56614 | 0.343004841 | -1.1672055 | 0 | 0.017 | 1 | 0.689891827 | 0.404245443 | 0.024 | 0.038 | 1 | 0.689891827 | 0.568357361 |
| Dlgap5 | 0.343004841 | -1.304709024 | 0 | 0.017 | 1 | 0.455098275 | 0.515276755 | 0 | 0.014 | 1 | 0.455098275 | 0.568357361 |
| Gm50250 | 0.343004841 | -0.652632327 | 0 | 0.017 | 1 | 0.872828964 | 1.029849928 | 0.024 | 0.02 | 1 | 0.872828964 | 0.568357361 |
| AI661453 | 0.343004841 | -1.1672055 | 0 | 0.017 | 1 | 0.402364804 | -0.484723245 | 0 | 0.017 | 1 | 0.402364804 | 0.568357361 |
| Tnfaip8l1 | 0.343004841 | -1.304709024 | 0 | 0.017 | 1 | 0.455097252 | -0.332720152 | 0 | 0.014 | 1 | 0.455097252 | 0.568357361 |
| Cdc25c | 0.343004841 | -1.845277405 | 0 | 0.017 | 1 | 0.455094184 | 0.252242349 | 0 | 0.014 | 1 | 0.455094184 | 0.568357361 |
| 1700016C15Rik | 0.343006305 | -0.1672055 | 0 | 0.017 | 1 | 0.5191986 | 0.515276755 | 0 | 0.01 | 1 | 0.5191986 | 0.568359285 |
| Gm13421 | 0.343006305 | -0.1672055 | 0 | 0.017 | 1 | 0.729889663 | 2.83720485 | 0.024 | 0.017 | 1 | 0.729889663 | 0.568359285 |
| P2rx3 | 0.343006305 | -0.1672055 | 0 | 0.017 | 1 | 0.438672228 | 1.83720485 | 0.024 | 0.01 | 1 | 0.438672228 | 0.568359285 |
| Frmd5 | 0.343006305 | -0.1672055 | 0 | 0.017 | 1 | 0.430588668 | 3.644559772 | 0.024 | 0.01 | 1 | 0.430588668 | 0.568359285 |
| Gm43714 | 0.343006305 | -0.1672055 | 0 | 0.017 | 1 | 0.357847088 | -0.16279515 | 0 | 0.02 | 1 | 0.357847088 | 0.568359285 |
| Gm43773 | 0.343006305 | -0.1672055 | 0 | 0.017 | 1 | 0.548898236 | -0.250257991 | 0.024 | 0.044 | 1 | 0.548898236 | 0.568359285 |
| Gm57103 | 0.343006305 | -0.1672055 | 0 | 0.017 | 1 | 0.51919986 | 0.83720485 | 0 | 0.01 | 1 | 0.51919986 | 0.568359285 |
| Gm12473 | 0.343006305 | -0.1672055 | 0 | 0.017 | 1 | 0.369056111 | -0.863234868 | 0.024 | 0.058 | 1 | 0.369056111 | 0.568359285 |
| C77080 | 0.343006305 | -0.1672055 | 0 | 0.017 | 1 | 0.885024635 | -0.410722664 | 0.073 | 0.065 | 1 | 0.885024635 | 0.568359285 |
| Gm13091 | 0.343006305 | -0.1672055 | 0 | 0.017 | 1 | 0.402373368 | 0.252242349 | 0 | 0.017 | 1 | 0.402373368 | 0.568359285 |
| Gm43660 | 0.343006305 | -0.1672055 | 0 | 0.017 | 1 | 0.430584486 | 2.422167351 | 0.024 | 0.01 | 1 | 0.430584486 | 0.568359285 |
| Tmprss11b | 0.343006305 | -1.932740246 | 0 | 0.017 | 1 | 0.402373368 | 0.252242349 | 0 | 0.017 | 1 | 0.402373368 | 0.568359285 |
| Gm57291 | 0.343006305 | -0.1672055 | 0 | 0.017 | 1 | 0.729889663 | 2.252242349 | 0.024 | 0.017 | 1 | 0.729889663 | 0.568359285 |
| Grid2ip | 0.343006305 | -0.1672055 | 0 | 0.017 | 1 | 0.599290422 | 1.515276755 | 0.024 | 0.014 | 1 | 0.599290422 | 0.568359285 |
| C3ar1 | 0.343006305 | -0.1672055 | 0 | 0.017 | 1 | 0.455094184 | 0.252242349 | 0 | 0.014 | 1 | 0.455094184 | 0.568359285 |
| Ttll13 | 0.343006305 | -0.1672055 | 0 | 0.017 | 1 | 0.693809887 | 0.721727632 | 0.024 | 0.038 | 1 | 0.693809887 | 0.568359285 |
| Tlcd3b | 0.343006305 | -0.1672055 | 0 | 0.017 | 1 | 0.916649754 | 0.667279848 | 0.024 | 0.027 | 1 | 0.916649754 | 0.568359285 |

| | | | | | | | | | | | | |
|---|---|---|---|---|---|---|---|---|---|---|---|---|
| Spaca7b | 0.343006305 | -0.1672055 | 0 | 0.017 | 1 | 0.51919986 | 0.83720485 | 0 | 0.01 | 1 | 0.51919986 | 0.568359285 |
| 0610038B21Rik | 0.343006305 | -0.1672055 | 0 | 0.017 | 1 | 0.455094184 | 0.252242349 | 0 | 0.014 | 1 | 0.455094184 | 0.568359285 |
| Gm26537 | 0.343006305 | -0.1672055 | 0 | 0.017 | 1 | 0.51919986 | 0.83720485 | 0 | 0.01 | 1 | 0.51919986 | 0.568359285 |
| Hoatz | 0.343006305 | -0.1672055 | 0 | 0.017 | 1 | 0.402373368 | 0.252242349 | 0 | 0.017 | 1 | 0.402373368 | 0.568359285 |
| Gm32742 | 0.343006305 | -0.1672055 | 0 | 0.017 | 1 | 0.51919986 | 0.83720485 | 0 | 0.01 | 1 | 0.51919986 | 0.568359285 |
| Nrg4 | 0.343006305 | -0.1672055 | 0 | 0.017 | 1 | 0.931315883 | 1.351778023 | 0.024 | 0.027 | 1 | 0.931315883 | 0.568359285 |
| Fam241b | 0.343006305 | -0.1672055 | 0 | 0.017 | 1 | 0.359002082 | 1.422167351 | 0.049 | 0.024 | 1 | 0.359002082 | 0.568359285 |
| Tspan15 | 0.343006305 | -0.1672055 | 0 | 0.017 | 1 | 0.883803856 | 0.136765132 | 0.024 | 0.02 | 1 | 0.883803856 | 0.568359285 |
| Gm56948 | 0.343006305 | -0.1672055 | 0 | 0.017 | 1 | 0.793962817 | 1.136765132 | 0.049 | 0.041 | 1 | 0.793962817 | 0.568359285 |
| 5033403F01Rik | 0.343006305 | -0.1672055 | 0 | 0.017 | 1 | 0.402369942 | -0.484723245 | 0 | 0.017 | 1 | 0.402369942 | 0.568359285 |
| Gm30655 | 0.343006305 | -0.1672055 | 0 | 0.017 | 1 | 0.5191986 | 0.252242349 | 0 | 0.01 | 1 | 0.5191986 | 0.568359285 |
| Zfp957 | 0.343006305 | -1.545717123 | 0 | 0.017 | 1 | 0.402369942 | -0.16279515 | 0 | 0.017 | 1 | 0.402369942 | 0.568359285 |
| 1700085D07Rik | 0.343006305 | -0.1672055 | 0 | 0.017 | 1 | 0.828694929 | 0.422167351 | 0.049 | 0.041 | 1 | 0.828694929 | 0.568359285 |
| Gm36738 | 0.343006305 | -0.1672055 | 0 | 0.017 | 1 | 0.51919986 | 0.83720485 | 0 | 0.01 | 1 | 0.51919986 | 0.568359285 |
| 4930483P17Rik | 0.343006305 | -0.1672055 | 0 | 0.017 | 1 | 0.51919986 | 0.83720485 | 0 | 0.01 | 1 | 0.51919986 | 0.568359285 |
| Pdxk-ps | 0.343006305 | -0.1672055 | 0 | 0.017 | 1 | 0.441381862 | 1.515276755 | 0.024 | 0.01 | 1 | 0.441381862 | 0.568359285 |
| Lrrtm2 | 0.343006305 | -0.1672055 | 0 | 0.017 | 1 | 0.402373368 | 0.252242349 | 0 | 0.017 | 1 | 0.402373368 | 0.568359285 |
| Gm36419 | 0.343006305 | -0.1672055 | 0 | 0.017 | 1 | 0.51919986 | 0.83720485 | 0 | 0.01 | 1 | 0.51919986 | 0.568359285 |
| Abcc2 | 0.343006305 | -0.1672055 | 0 | 0.017 | 1 | 0.455098275 | 0.515276755 | 0 | 0.014 | 1 | 0.455098275 | 0.568359285 |
| Gm16068 | 0.343006305 | -0.1672055 | 0 | 0.017 | 1 | 0.441381862 | 1.515276755 | 0.024 | 0.01 | 1 | 0.441381862 | 0.568359285 |
| Gm21060 | 0.343006305 | -0.1672055 | 0 | 0.017 | 1 | 0.455098275 | 0.515276755 | 0 | 0.014 | 1 | 0.455098275 | 0.568359285 |
| Smim12 | 0.872795249 | 0.417757001 | 0.189 | 0.191 | 1 | 0.343126354 | 0.454735213 | 0.268 | 0.205 | 1 | 0.872795249 | 0.568517013 |
| Gm16675 | 0.343171476 | 0.8327945 | 0.057 | 0.03 | 1 | 0.455094184 | -0.16279515 | 0 | 0.014 | 1 | 0.455094184 | 0.568576291 |
| 1700109K24Rik | 0.343171476 | 0.8327945 | 0.057 | 0.03 | 1 | 0.817109602 | 0.721727632 | 0.049 | 0.041 | 1 | 0.817109602 | 0.568576291 |
| Tep1 | 0.343261237 | 1.047807391 | 0.132 | 0.093 | 1 | 0.811735567 | 0.614812428 | 0.122 | 0.14 | 1 | 0.811735567 | 0.568694197 |
| Cog1 | 0.343464192 | 0.428897558 | 0.208 | 0.153 | 1 | 0.722510137 | 0.377773231 | 0.195 | 0.232 | 1 | 0.722510137 | 0.568960733 |
| Spacdr | 0.771945601 | -0.145179194 | 0.604 | 0.619 | 1 | 0.343647735 | -0.227174378 | 0.659 | 0.727 | 1 | 0.771945601 | 0.569201705 |
| Tmub2 | 0.343765693 | 0.445771377 | 0.34 | 0.284 | 1 | 0.435258111 | 0.42692088 | 0.22 | 0.304 | 1 | 0.435258111 | 0.569356534 |
| Gm19710 | 0.343766385 | 0.659957903 | 0.132 | 0.089 | 1 | 0.576567467 | -0.425829556 | 0.098 | 0.126 | 1 | 0.576567467 | 0.569357443 |
| Phox2a | 0.343968402 | 0.739685096 | 0.038 | 0.017 | 1 | 0.357854401 | -0.863234868 | 0 | 0.02 | 1 | 0.357854401 | 0.569622543 |
| Dnah9 | 0.343968402 | 1.154722595 | 0.038 | 0.017 | 1 | 0.746881137 | 0.83720485 | 0.024 | 0.017 | 1 | 0.746881137 | 0.569622543 |
| Gm26583 | 0.343968402 | 1.154722595 | 0.038 | 0.017 | 1 | 0.44410834 | 1.029849928 | 0.024 | 0.01 | 1 | 0.44410834 | 0.569622543 |
| Zdhhc19 | 0.343984134 | 0.280253477 | 0.038 | 0.017 | 1 | 0.906883853 | -0.16279515 | 0.024 | 0.027 | 1 | 0.906883853 | 0.569643183 |
| Ptger1 | 0.478732386 | 0.785488785 | 0.132 | 0.102 | 1 | 0.344234562 | -0.126269274 | 0.049 | 0.096 | 1 | 0.478732386 | 0.569971691 |
| Tsen2 | 0.344466274 | 1.228723176 | 0.113 | 0.076 | 1 | 0.969378959 | 0.667279848 | 0.098 | 0.099 | 1 | 0.969378959 | 0.570275534 |
| Flt3l | 0.880784741 | 0.280253477 | 0.038 | 0.042 | 1 | 0.344856459 | -0.825760163 | 0.049 | 0.092 | 1 | 0.880784741 | 0.57078694 |
| Spryd3 | 0.344939701 | 0.444229212 | 0.396 | 0.335 | 1 | 0.508322401 | 0.637532505 | 0.366 | 0.362 | 1 | 0.508322401 | 0.570896005 |
| Plet1 | 0.345023796 | -1.508242418 | 0.038 | 0.072 | 1 | 0.658201928 | -1.020776145 | 0.073 | 0.092 | 1 | 0.658201928 | 0.571006172 |
| Ltv1 | 0.520221512 | -0.279905633 | 0.415 | 0.453 | 1 | 0.345281916 | -0.268589814 | 0.317 | 0.416 | 1 | 0.520221512 | 0.571344231 |
| A730063M14Rik | 0.437537757 | 0.984797593 | 0.057 | 0.034 | 1 | 0.345527083 | 1.422167351 | 0.073 | 0.041 | 1 | 0.437537757 | 0.571665201 |
| Ankzf1 | 0.346052786 | 0.292226119 | 0.189 | 0.136 | 1 | 0.6084686 | 0.707921833 | 0.122 | 0.099 | 1 | 0.6084686 | 0.572353041 |
| Tdp1 | 0.393245768 | 0.347367673 | 0.17 | 0.123 | 1 | 0.346203962 | 0.83720485 | 0.122 | 0.078 | 1 | 0.393245768 | 0.57255074 |
| Nup37 | 0.346290655 | 0.624207878 | 0.132 | 0.089 | 1 | 0.514808015 | 1.19334866 | 0.098 | 0.072 | 1 | 0.514808015 | 0.572664092 |
| Tfcp2l1 | 0.809876017 | 0.569760094 | 0.038 | 0.047 | 1 | 0.346609471 | 0.489281546 | 0.171 | 0.119 | 1 | 0.809876017 | 0.573080817 |
| Oscp1 | 0.932034295 | 0.2967416 | 0.113 | 0.11 | 1 | 0.346720471 | 0.807457506 | 0.171 | 0.119 | 1 | 0.932034295 | 0.573225857 |
| B130034C11Rik | 0.474382806 | -0.845277405 | 0.057 | 0.085 | 1 | 0.346795173 | 0.962735732 | 0.073 | 0.041 | 1 | 0.474382806 | 0.573323454 |
| Rybp | 0.522912994 | -0.144167964 | 0.792 | 0.809 | 1 | 0.347000948 | 0.22079271 | 0.854 | 0.836 | 1 | 0.522912994 | 0.573592237 |
| Elapor1 | 0.898683122 | 0.776210972 | 0.075 | 0.072 | 1 | 0.347096313 | 0.515276755 | 0.098 | 0.058 | 1 | 0.898683122 | 0.573716775 |
| Gm16541 | 0.347184584 | -3.326404095 | 0.038 | 0.072 | 1 | 0.828239242 | 0.252242349 | 0.073 | 0.065 | 1 | 0.828239242 | 0.573832033 |
| Harbi1 | 0.347592474 | 0.154722595 | 0.057 | 0.102 | 1 | 0.568078062 | 1.667279848 | 0.098 | 0.075 | 1 | 0.568078062 | 0.57436442 |
| Gm34934 | 0.347593463 | 0.695290976 | 0.057 | 0.03 | 1 | 0.824812458 | 0.515276755 | 0.049 | 0.041 | 1 | 0.824812458 | 0.574365711 |
| Tacc3 | 0.347593766 | 0.154722595 | 0.151 | 0.106 | 1 | 0.408803696 | -0.825760163 | 0.049 | 0.085 | 1 | 0.408803696 | 0.574366106 |
| Phf19 | 0.347746365 | 0.739685096 | 0.038 | 0.017 | 1 | 0.744047924 | 1.252242349 | 0.024 | 0.017 | 1 | 0.744047924 | 0.574565195 |
| Ckm | 0.347746365 | 0.932330173 | 0.038 | 0.017 | 1 | 0.5191986 | 0.515276755 | 0 | 0.01 | 1 | 0.5191986 | 0.574565195 |
| Gm36279 | 0.7394666 | 0.569760094 | 0.019 | 0.013 | 1 | 0.347784376 | 1.83720485 | 0.049 | 0.024 | 1 | 0.7394666 | 0.574614779 |
| Gpd1 | 0.926212226 | 0.8327945 | 0.019 | 0.017 | 1 | 0.347784376 | 2.422167351 | 0.049 | 0.024 | 1 | 0.926212226 | 0.574614779 |
| Fbxl12os | 0.932506754 | 1.669295768 | 0.019 | 0.021 | 1 | 0.347784376 | 1.83720485 | 0.049 | 0.024 | 1 | 0.932506754 | 0.574614779 |
| Tmem242 | 0.347902762 | -0.376997885 | 0.358 | 0.411 | 1 | 0.986268026 | 0.435106406 | 0.341 | 0.369 | 1 | 0.986268026 | 0.574769192 |
| Hacl1 | 0.348160563 | 0.318221327 | 0.132 | 0.195 | 1 | 0.444785902 | -0.136799942 | 0.098 | 0.147 | 1 | 0.444785902 | 0.575105348 |
| Nsrp1 | 0.706395689 | 0.395262363 | 0.472 | 0.5 | 1 | 0.348384254 | -0.204135946 | 0.488 | 0.577 | 1 | 0.706395689 | 0.575396919 |
| Cfap410 | 0.348979222 | -0.142957954 | 0.151 | 0.212 | 1 | 0.976959824 | 0.491430013 | 0.171 | 0.174 | 1 | 0.976959824 | 0.576171947 |
| Cdkl2 | 0.349223337 | 0.402650108 | 0.245 | 0.191 | 1 | 0.987779961 | 0.19334866 | 0.146 | 0.15 | 1 | 0.987779961 | 0.576489735 |
| D230022J07Rik | 0.488750271 | -0.430239906 | 0.019 | 0.038 | 1 | 0.349381124 | 2.252242349 | 0.049 | 0.024 | 1 | 0.488750271 | 0.576695078 |
| C1ql3 | 0.916620008 | 1.154722595 | 0.019 | 0.017 | 1 | 0.349381124 | 1.989207943 | 0.049 | 0.024 | 1 | 0.916620008 | 0.576695078 |
| Dera | 0.349393024 | -0.534576566 | 0.283 | 0.343 | 1 | 0.75500134 | 0.504629511 | 0.293 | 0.294 | 1 | 0.75500134 | 0.576710562 |
| Mlycd | 0.61103347 | -0.278236812 | 0.075 | 0.097 | 1 | 0.349463147 | 0.114738825 | 0.049 | 0.096 | 1 | 0.61103347 | 0.576801803 |
| Gm11973 | 0.411970653 | -0.57319786 | 0.132 | 0.174 | 1 | 0.349916226 | -0.366667484 | 0.195 | 0.263 | 1 | 0.411970653 | 0.577391087 |
| Prune1 | 0.543543162 | 0.399835093 | 0.208 | 0.174 | 1 | 0.349939056 | -0.693309867 | 0.171 | 0.229 | 1 | 0.543543162 | 0.577420769 |
| Mfap1b | 0.746509809 | -0.13946051 | 0.321 | 0.284 | 1 | 0.350078472 | -0.136799942 | 0.293 | 0.406 | 1 | 0.746509809 | 0.577602008 |
| Ak2 | 0.350100129 | 0.271678741 | 0.547 | 0.521 | 1 | 0.569785624 | -0.122153166 | 0.512 | 0.56 | 1 | 0.569785624 | 0.577630157 |
| Ociad1 | 0.350279172 | 0.202774724 | 0.906 | 0.89 | 1 | 0.957840441 | 0.105738236 | 0.829 | 0.846 | 1 | 0.957840441 | 0.577862846 |
| Kazald1 | 0.403997363 | 0.219262847 | 0.528 | 0.458 | 1 | 0.35049295 | 0.494443251 | 0.488 | 0.44 | 1 | 0.403997363 | 0.578140592 |
| Gnpat | 0.350877833 | 0.569760094 | 0.208 | 0.161 | 1 | 0.777902877 | 0.308825877 | 0.171 | 0.198 | 1 | 0.777902877 | 0.578640412 |
| Ank1 | 0.812451721 | -0.430239906 | 0.113 | 0.097 | 1 | 0.351357076 | 1.199774929 | 0.098 | 0.061 | 1 | 0.812451721 | 0.579262357 |
| Tmco6 | 0.35149249 | -0.323324702 | 0.094 | 0.144 | 1 | 0.354892874 | -0.16279515 | 0.024 | 0.061 | 1 | 0.354892874 | 0.579438009 |
| Nlrp5-ps | 0.351545549 | 0.154722595 | 0.038 | 0.017 | 1 | 0.428023103 | 1.962735732 | 0.049 | 0.027 | 1 | 0.428023103 | 0.579506824 |
| Pla2g10os | 0.35154653 | 0.154722595 | 0.038 | 0.017 | 1 | 0.357847088 | -0.16279515 | 0 | 0.02 | 1 | 0.357847088 | 0.579508097 |
| Gstp1 | 0.542787842 | 0.806799291 | 0.132 | 0.106 | 1 | 0.352145075 | 0.778311161 | 0.195 | 0.143 | 1 | 0.542787842 | 0.580283996 |
| Lrrc69 | 0.479622472 | 1.154722595 | 0.038 | 0.021 | 1 | 0.352571034 | 1.699701326 | 0.049 | 0.024 | 1 | 0.479622472 | 0.580835734 |
| Apobec2 | 0.35268327 | -2.81106169 | 0.038 | 0.072 | 1 | 0.51919986 | 0.83720485 | 0 | 0.01 | 1 | 0.51919986 | 0.58098105 |
| Fkbp3 | 0.373331391 | 0.115803606 | 0.792 | 0.742 | 1 | 0.352688819 | 0.756766234 | 0.707 | 0.71 | 1 | 0.373331391 | 0.580988235 |
| Arsa | 0.942157775 | 0.132354782 | 0.226 | 0.225 | 1 | 0.352893421 | 0.230547278 | 0.122 | 0.195 | 1 | 0.942157775 | 0.581253076 |
| Edem2 | 0.353121367 | -0.247375849 | 0.189 | 0.258 | 1 | 0.369947653 | -0.296650897 | 0.146 | 0.212 | 1 | 0.369947653 | 0.581548034 |
| Dmc1 | 0.831509183 | 0.261637799 | 0.094 | 0.085 | 1 | 0.353145557 | -0.421529419 | 0.073 | 0.126 | 1 | 0.831509183 | 0.581579329 |
| Kif7 | 0.353658063 | 0.517292674 | 0.132 | 0.089 | 1 | 0.750946477 | -0.19841906 | 0.098 | 0.113 | 1 | 0.750946477 | 0.5822421 |
| Exoc6 | 0.35370685 | 0.762405172 | 0.094 | 0.059 | 1 | 0.547258973 | -0.250257991 | 0.024 | 0.044 | 1 | 0.547258973 | 0.582305164 |
| Pcdhb12 | 0.736073999 | 0.8327945 | 0.019 | 0.013 | 1 | 0.354175043 | 1.83720485 | 0.049 | 0.024 | 1 | 0.736073999 | 0.582910125 |
| Il6 | 0.354345216 | -1.652632327 | 0.019 | 0.047 | 1 | 0.599290422 | 1.515276755 | 0.024 | 0.014 | 1 | 0.599290422 | 0.583129899 |
| Dynlt3 | 0.354625066 | 0.340437179 | 0.736 | 0.691 | 1 | 0.923800192 | 0.237103752 | 0.707 | 0.802 | 1 | 0.923800192 | 0.583491194 |
| Tmem199 | 0.35537264 | -0.285849997 | 0.094 | 0.144 | 1 | 0.847034704 | 0.649577847 | 0.122 | 0.113 | 1 | 0.847034704 | 0.584455567 |
| Tmem8b | 0.355437423 | 0.953088734 | 0.132 | 0.093 | 1 | 0.879508038 | -0.332720152 | 0.122 | 0.126 | 1 | 0.879508038 | 0.584539084 |
| Pold2 | 0.547458019 | -0.251269765 | 0.151 | 0.186 | 1 | 0.355472334 | -0.513292397 | 0.098 | 0.15 | 1 | 0.547458019 | 0.584584088 |
| Vps39 | 0.355696714 | 0.776210972 | 0.226 | 0.182 | 1 | 0.985849631 | 0.287007767 | 0.22 | 0.229 | 1 | 0.985849631 | 0.584873276 |
| 1700027I24Rik | 0.355699537 | -1.015202407 | 0.019 | 0.047 | 1 | 0.822976752 | 0.159132945 | 0.049 | 0.058 | 1 | 0.822976752 | 0.584876914 |
| Ngly1 | 0.374181636 | 0.796268624 | 0.358 | 0.326 | 1 | 0.355729874 | -0.269710354 | 0.39 | 0.468 | 1 | 0.374181636 | 0.584916005 |
| Cfap54 | 0.929410101 | 0.569760094 | 0.019 | 0.017 | 1 | 0.355775728 | 1.458693227 | 0.049 | 0.024 | 1 | 0.929410101 | 0.584975088 |
| Stard3 | 0.355818665 | 0.33529484 | 0.264 | 0.203 | 1 | 0.559344887 | -0.463964685 | 0.146 | 0.177 | 1 | 0.559344887 | 0.585030408 |
| Zkscan7 | 0.355831243 | -0.845277405 | 0.038 | 0.072 | 1 | 0.846926792 | 1.100239256 | 0.024 | 0.031 | 1 | 0.846926792 | 0.585046612 |

| | | | | | | | | | | | | |
|---|---|---|---|---|---|---|---|---|---|---|---|---|
| Nemp1 | 0.356429675 | 0.42680214 | 0.226 | 0.174 | 1 | 0.488699569 | -0.16279515 | 0.122 | 0.167 | 1 | 0.488699569 | 0.585817237 |
| E2f5 | 0.356675291 | 0.354492107 | 0.358 | 0.305 | 1 | 0.722950382 | 0.302868422 | 0.366 | 0.355 | 1 | 0.722950382 | 0.586133318 |
| Myo15a | 0.567746321 | 0.669295768 | 0.113 | 0.089 | 1 | 0.356716792 | -1.020776145 | 0.073 | 0.119 | 1 | 0.567746321 | 0.586186714 |
| Prdm2 | 0.493915239 | 0.258949964 | 0.906 | 0.86 | 1 | 0.35676242 | 0.17081475 | 0.976 | 0.918 | 1 | 0.493915239 | 0.586245416 |
| Gm4107 | 0.357062705 | -1.974560422 | 0.019 | 0.047 | 1 | 0.605005904 | 0.377773231 | 0.024 | 0.014 | 1 | 0.605005904 | 0.586631634 |
| Tnfaip3 | 0.919816481 | 0.932330173 | 0.019 | 0.017 | 1 | 0.357381956 | 1.377773231 | 0.049 | 0.024 | 1 | 0.919816481 | 0.587042049 |
| Gm13483 | 0.357540258 | 0.328751995 | 0.226 | 0.174 | 1 | 0.980391336 | -0.543616934 | 0.22 | 0.201 | 1 | 0.980391336 | 0.58724548 |
| Aimp2 | 0.357614546 | 0.491757582 | 0.264 | 0.212 | 1 | 0.634469889 | 0.280811501 | 0.22 | 0.273 | 1 | 0.634469889 | 0.587340928 |
| Yif1a | 0.588362164 | -0.103293736 | 0.566 | 0.653 | 1 | 0.357759556 | 0.338398993 | 0.415 | 0.556 | 1 | 0.588362164 | 0.587527213 |
| Gm57270 | 0.413082433 | 0.154722595 | 0 | 0.013 | 1 | 0.357847088 | -0.332720152 | 0 | 0.02 | 1 | 0.413082433 | 0.587639638 |
| Gm34655 | 0.421517727 | -0.304709024 | 0.019 | 0.042 | 1 | 0.357847088 | -0.484723245 | 0 | 0.02 | 1 | 0.421517727 | 0.587639638 |
| Spef2 | 0.500671253 | 1.739685096 | 0.019 | 0.008 | 1 | 0.357847088 | -0.484723245 | 0 | 0.02 | 1 | 0.500671253 | 0.587639638 |
| Cd83 | 0.503971666 | 1.569760094 | 0.019 | 0.008 | 1 | 0.357847088 | -0.16279515 | 0 | 0.02 | 1 | 0.503971666 | 0.587639638 |
| Car15 | 0.503971666 | 1.569760094 | 0.019 | 0.008 | 1 | 0.357847088 | -0.332720152 | 0 | 0.02 | 1 | 0.503971666 | 0.587639638 |
| Fcrla | 0.569866135 | -0.932740246 | 0.019 | 0.034 | 1 | 0.357847088 | -0.16279515 | 0 | 0.02 | 1 | 0.569866135 | 0.587639638 |
| B130024G19Rik | 0.633487796 | 0.154722595 | 0.038 | 0.025 | 1 | 0.357847088 | -0.16279515 | 0 | 0.02 | 1 | 0.633487796 | 0.587639638 |
| 1500009L16Rik | 0.653990165 | 0.695290976 | 0.057 | 0.042 | 1 | 0.357847088 | -0.16279515 | 0 | 0.02 | 1 | 0.653990165 | 0.587639638 |
| 6430573P05Rik | 0.684424342 | 0.739685096 | 0.019 | 0.03 | 1 | 0.357847088 | -0.332720152 | 0 | 0.02 | 1 | 0.684424342 | 0.587639638 |
| Pdf | 0.732691937 | 1.154722595 | 0.019 | 0.013 | 1 | 0.357847088 | -0.16279515 | 0 | 0.02 | 1 | 0.732691937 | 0.587639638 |
| Timeless | 0.753264159 | 0.8327945 | 0.038 | 0.03 | 1 | 0.357847088 | -0.16279515 | 0 | 0.02 | 1 | 0.753264159 | 0.587639638 |
| Myef2l | 0.782077996 | 0.347367673 | 0.019 | 0.025 | 1 | 0.357847088 | -0.332720152 | 0 | 0.02 | 1 | 0.782077996 | 0.587639638 |
| Gm41760 | 0.795213727 | 0.739685096 | 0.019 | 0.025 | 1 | 0.357847088 | -0.16279515 | 0 | 0.02 | 1 | 0.795213727 | 0.587639638 |
| Nudt12 | 0.917883255 | 0.569760094 | 0.019 | 0.021 | 1 | 0.357847088 | -0.16279515 | 0 | 0.02 | 1 | 0.917883255 | 0.587639638 |
| Spats1 | 0.923013455 | 0.417757001 | 0.019 | 0.017 | 1 | 0.357847088 | -0.16279515 | 0 | 0.02 | 1 | 0.923013455 | 0.587639638 |
| Gm6566 | 0.929410101 | 0.569760094 | 0.019 | 0.017 | 1 | 0.357847088 | -0.332720152 | 0 | 0.02 | 1 | 0.929410101 | 0.587639638 |
| 4930595D18Rik | 0.929579738 | 0.932330173 | 0.019 | 0.021 | 1 | 0.357847088 | -0.16279515 | 0 | 0.02 | 1 | 0.929579738 | 0.587639638 |
| 9230116N13Rik | 0.991244785 | 0.417757001 | 0.038 | 0.038 | 1 | 0.357847088 | -0.16279515 | 0 | 0.02 | 1 | 0.991244785 | 0.587639638 |
| Cenpn | 0.997811016 | 0.454282877 | 0.038 | 0.038 | 1 | 0.357847088 | -0.332720152 | 0 | 0.02 | 1 | 0.997811016 | 0.587639638 |
| Gm14320 | 0.413080549 | -0.1672055 | 0 | 0.013 | 1 | 0.357854401 | -0.484723245 | 0 | 0.02 | 1 | 0.413080549 | 0.58764903 |
| Ly6c2 | 0.413080549 | -0.1672055 | 0 | 0.013 | 1 | 0.357854401 | -0.332720152 | 0 | 0.02 | 1 | 0.413080549 | 0.58764903 |
| Myh11 | 0.413080549 | -0.1672055 | 0 | 0.013 | 1 | 0.357854401 | -0.332720152 | 0 | 0.02 | 1 | 0.413080549 | 0.58764903 |
| Ppm1j | 0.413082433 | 0.154722595 | 0 | 0.013 | 1 | 0.357854401 | -0.332720152 | 0 | 0.02 | 1 | 0.413082433 | 0.58764903 |
| Gm56660 | 0.442326061 | 0.8327945 | 0.057 | 0.034 | 1 | 0.357854401 | -0.332720152 | 0 | 0.02 | 1 | 0.442326061 | 0.58764903 |
| Gm11767 | 0.48175508 | 1.154722595 | 0.038 | 0.021 | 1 | 0.357854401 | -0.332720152 | 0 | 0.02 | 1 | 0.48175508 | 0.58764903 |
| Foxm1 | 0.538051855 | 1.039245377 | 0.057 | 0.038 | 1 | 0.357854401 | -0.622226769 | 0 | 0.02 | 1 | 0.538051855 | 0.58764903 |
| Fbxo47 | 0.569866135 | -0.430239906 | 0.019 | 0.034 | 1 | 0.357854401 | -0.863234868 | 0 | 0.02 | 1 | 0.569866135 | 0.58764903 |
| Pate2 | 0.631221588 | 0.347367673 | 0.038 | 0.025 | 1 | 0.357854401 | -0.747757651 | 0 | 0.02 | 1 | 0.631221588 | 0.58764903 |
| Anln | 0.661687831 | -2.093204919 | 0.038 | 0.051 | 1 | 0.357854401 | -0.484723245 | 0 | 0.02 | 1 | 0.661687831 | 0.58764903 |
| Pnpla1 | 0.732691937 | 1.154722595 | 0.019 | 0.013 | 1 | 0.357854401 | -0.332720152 | 0 | 0.02 | 1 | 0.732691937 | 0.58764903 |
| Kif6 | 0.767066103 | 0.417757001 | 0.038 | 0.03 | 1 | 0.357854401 | -0.332720152 | 0 | 0.02 | 1 | 0.767066103 | 0.58764903 |
| Scn8a | 0.782077996 | 0.347367673 | 0.019 | 0.025 | 1 | 0.357854401 | -0.484723245 | 0 | 0.02 | 1 | 0.782077996 | 0.58764903 |
| Cntd1 | 0.869949949 | 0.891688189 | 0.038 | 0.034 | 1 | 0.357854401 | -0.747757651 | 0 | 0.02 | 1 | 0.869949949 | 0.58764903 |
| Pkp2 | 0.912039013 | -0.304709024 | 0.019 | 0.021 | 1 | 0.357854401 | -0.484723245 | 0 | 0.02 | 1 | 0.912039013 | 0.58764903 |
| 2900041M22Rik | 0.926212226 | 0.8327945 | 0.019 | 0.017 | 1 | 0.357854401 | -0.332720152 | 0 | 0.02 | 1 | 0.926212226 | 0.58764903 |
| Musk | 0.413082433 | 0.154722595 | 0 | 0.013 | 1 | 0.357858058 | -0.747757651 | 0 | 0.02 | 1 | 0.413082433 | 0.587653726 |
| Ccdc14 | 0.47749495 | 1.154722595 | 0.038 | 0.021 | 1 | 0.357858058 | -0.747757651 | 0 | 0.02 | 1 | 0.47749495 | 0.587653726 |
| Knl1 | 0.479836037 | -1.389597921 | 0.019 | 0.038 | 1 | 0.357858058 | -0.622226769 | 0 | 0.02 | 1 | 0.479836037 | 0.587653726 |
| Bnipl | 0.48175508 | 1.154722595 | 0.038 | 0.021 | 1 | 0.357858058 | -0.622226769 | 0 | 0.02 | 1 | 0.48175508 | 0.587653726 |
| Ripor2 | 0.764778716 | -1.489133595 | 0.038 | 0.03 | 1 | 0.357858058 | -1.332720152 | 0 | 0.02 | 1 | 0.764778716 | 0.587653726 |
| Ankdd1a | 0.906251902 | 0.154722595 | 0.038 | 0.034 | 1 | 0.357858058 | -0.622226769 | 0 | 0.02 | 1 | 0.906251902 | 0.587653726 |
| 2810407A14Rik | 0.916620008 | 1.154722595 | 0.019 | 0.017 | 1 | 0.357858058 | -1.069685746 | 0 | 0.02 | 1 | 0.916620008 | 0.587653726 |
| Mfsd13b | 0.628925725 | 0.280253477 | 0.038 | 0.025 | 1 | 0.357858789 | -1.069685746 | 0 | 0.02 | 1 | 0.628925725 | 0.587654665 |
| Gm56786 | 0.358003231 | -0.783876861 | 0.038 | 0.072 | 1 | 0.455098275 | 0.515276755 | 0 | 0.014 | 1 | 0.455098275 | 0.587840148 |
| Cd164l2 | 0.358003231 | -0.719746523 | 0.038 | 0.072 | 1 | 0.761330249 | -0.372248516 | 0.073 | 0.085 | 1 | 0.761330249 | 0.587840148 |
| Glp2r | 0.358003231 | -0.845277405 | 0.038 | 0.072 | 1 | 0.604477578 | -0.250257991 | 0.024 | 0.041 | 1 | 0.604477578 | 0.587840148 |
| Zdhhc24 | 0.596093515 | -0.381330305 | 0.075 | 0.097 | 1 | 0.358149327 | 0.721727632 | 0.122 | 0.078 | 1 | 0.596093515 | 0.588027713 |
| Sult1c1 | 0.682084573 | 0.569760094 | 0.019 | 0.03 | 1 | 0.358149327 | 0.515276755 | 0.122 | 0.078 | 1 | 0.682084573 | 0.588027713 |
| Las1l | 0.831922848 | -0.291263493 | 0.547 | 0.53 | 1 | 0.358350162 | -0.167938453 | 0.439 | 0.522 | 1 | 0.831922848 | 0.588285485 |
| Gm16192 | 0.358419706 | -1.015202407 | 0.019 | 0.047 | 1 | 0.82181816 | 0.252242349 | 0.024 | 0.031 | 1 | 0.82181816 | 0.588374726 |
| Slc41a1 | 0.466125652 | 0.14482979 | 0.83 | 0.758 | 1 | 0.358749128 | -0.146116409 | 0.707 | 0.812 | 1 | 0.466125652 | 0.588797319 |
| H3c6 | 0.872466289 | -0.260314904 | 0.038 | 0.042 | 1 | 0.359002082 | 1.422167351 | 0.049 | 0.024 | 1 | 0.872466289 | 0.589121669 |
| Tcf19 | 0.888969925 | -0.489133595 | 0.057 | 0.051 | 1 | 0.359002082 | 1.422167351 | 0.049 | 0.024 | 1 | 0.888969925 | 0.589121669 |
| A130023E24Rik | 0.876606655 | 0.154722595 | 0.038 | 0.042 | 1 | 0.3593261 | 1.100239256 | 0.024 | 0.061 | 1 | 0.876606655 | 0.589536954 |
| Sptlc1 | 0.475375132 | -0.256210506 | 0.34 | 0.381 | 1 | 0.359343903 | 0.199774929 | 0.463 | 0.379 | 1 | 0.475375132 | 0.589559766 |
| Bbs12 | 0.451620383 | 0.216123139 | 0.094 | 0.064 | 1 | 0.359386644 | 0.377773231 | 0.024 | 0.061 | 1 | 0.451620383 | 0.589614529 |
| Ano10 | 0.787608878 | 0.526691372 | 0.302 | 0.301 | 1 | 0.359389045 | -0.455576809 | 0.293 | 0.352 | 1 | 0.787608878 | 0.589617604 |
| Rab29 | 0.360179034 | -0.845277405 | 0.038 | 0.072 | 1 | 0.528136181 | -0.53202896 | 0.049 | 0.075 | 1 | 0.528136181 | 0.590629132 |
| Gatd1 | 0.493541608 | 0.442004547 | 0.226 | 0.191 | 1 | 0.360561236 | -0.484723245 | 0.122 | 0.177 | 1 | 0.493541608 | 0.591118067 |
| Cx3cr1 | 0.36070424 | -1.214511215 | 0.094 | 0.14 | 1 | 0.532190708 | 0.2426563 | 0.195 | 0.16 | 1 | 0.532190708 | 0.591300931 |
| D730003I15Rik | 0.361131243 | -0.652632327 | 0.019 | 0.047 | 1 | 0.78390515 | 0.515276755 | 0.098 | 0.085 | 1 | 0.78390515 | 0.591846711 |
| 9330160F10Rik | 0.361131243 | -0.545717123 | 0.019 | 0.047 | 1 | 0.430051013 | 0.174239837 | 0.024 | 0.055 | 1 | 0.430051013 | 0.591846711 |
| Gm47863 | 0.361131243 | -0.545717123 | 0.019 | 0.047 | 1 | 0.714406212 | 1.029849928 | 0.049 | 0.038 | 1 | 0.714406212 | 0.591846711 |
| P2rx7 | 0.36166808 | 1.174351402 | 0.245 | 0.195 | 1 | 0.459659253 | 0.782757066 | 0.22 | 0.181 | 1 | 0.459659253 | 0.59253236 |
| Babam2 | 0.361914704 | -0.178476612 | 0.585 | 0.682 | 1 | 0.424620015 | 0.358110984 | 0.707 | 0.662 | 1 | 0.424620015 | 0.592847155 |
| 2510017J16Rik | 0.361916615 | 0.891688189 | 0.132 | 0.093 | 1 | 0.722125564 | -0.292078167 | 0.073 | 0.089 | 1 | 0.722125564 | 0.592849594 |
| Gm15927 | 0.597481259 | 1.669295768 | 0.038 | 0.025 | 1 | 0.362195044 | 1.252242349 | 0.049 | 0.024 | 1 | 0.597481259 | 0.593204838 |
| 1700034P13Rik | 0.765053098 | -1.304709024 | 0.038 | 0.047 | 1 | 0.362226486 | 0.83720485 | 0.049 | 0.024 | 1 | 0.765053098 | 0.593244944 |
| Septin1 | 0.362293181 | 0.674729654 | 0.226 | 0.178 | 1 | 0.937509062 | 0.749742009 | 0.171 | 0.184 | 1 | 0.937509062 | 0.593330013 |
| Gm20300 | 0.362357689 | -0.508242418 | 0.038 | 0.072 | 1 | 1 | 0.185128153 | 0.122 | 0.123 | 1 | 1 | 0.593412283 |
| Fosl2 | 0.636428381 | 0.204380481 | 0.755 | 0.746 | 1 | 0.362455063 | -0.233521202 | 0.854 | 0.812 | 1 | 0.636428381 | 0.593536453 |
| Echdc3 | 0.362626638 | -0.430239906 | 0.019 | 0.047 | 1 | 0.465362733 | 1.322631677 | 0.073 | 0.048 | 1 | 0.465362733 | 0.593755197 |
| 4930545L23Rik | 0.362626638 | -0.430239906 | 0.019 | 0.047 | 1 | 0.372552715 | -0.332720152 | 0.049 | 0.092 | 1 | 0.372552715 | 0.593755197 |
| Lrfn1 | 0.362626638 | -0.430239906 | 0.019 | 0.047 | 1 | 0.826384776 | 0.515276755 | 0.024 | 0.031 | 1 | 0.826384776 | 0.593755197 |
| Zmpste24 | 0.363259732 | 0.20958453 | 0.642 | 0.606 | 1 | 0.691684044 | 0.493250449 | 0.707 | 0.706 | 1 | 0.691684044 | 0.594561831 |
| Adck1 | 0.363375239 | 0.891688189 | 0.17 | 0.131 | 1 | 1 | 0.580865097 | 0.122 | 0.123 | 1 | 1 | 0.594708914 |
| Capn5 | 0.664141793 | -0.489133595 | 0.057 | 0.072 | 1 | 0.364013539 | 0.649577847 | 0.098 | 0.061 | 1 | 0.664141793 | 0.595521222 |
| Zfp777 | 0.364034267 | 0.228723176 | 0.245 | 0.182 | 1 | 0.913239382 | 0.422167351 | 0.22 | 0.218 | 1 | 0.913239382 | 0.595547587 |
| AA986860 | 0.855419041 | -0.732802676 | 0.075 | 0.081 | 1 | 0.364440781 | -1.250257991 | 0.024 | 0.058 | 1 | 0.855419041 | 0.596064479 |
| Atp5mpl | 0.364556624 | 0.122301117 | 0.736 | 0.716 | 1 | 0.857327158 | -0.16279515 | 0.585 | 0.59 | 1 | 0.857327158 | 0.596211716 |
| Pip5k1c | 0.364906152 | 0.212825549 | 0.283 | 0.22 | 1 | 0.498436785 | 0.822254508 | 0.268 | 0.242 | 1 | 0.498436785 | 0.596655804 |
| Fbxw10 | 0.485993467 | 0.932330173 | 0.038 | 0.021 | 1 | 0.365461252 | 0.962735732 | 0.049 | 0.024 | 1 | 0.485993467 | 0.597360578 |
| Rbm4 | 0.365551874 | 0.374503803 | 0.472 | 0.424 | 1 | 0.998525302 | 0.552475373 | 0.488 | 0.556 | 1 | 0.998525302 | 0.597475576 |
| Ywhaq | 0.691904132 | -0.315185357 | 1 | 0.97 | 1 | 0.365664143 | -0.221864232 | 0.976 | 0.997 | 1 | 0.691904132 | 0.59761802 |
| Tmem250-ps | 0.365760172 | 0.865215978 | 0.208 | 0.165 | 1 | 0.501906513 | -0.20718927 | 0.146 | 0.191 | 1 | 0.501906513 | 0.59773984 |
| G530011O06Rik | 0.365765614 | 0.652222254 | 0.094 | 0.059 | 1 | 0.60213981 | 1.029849928 | 0.024 | 0.014 | 1 | 0.60213981 | 0.597746744 |
| Uros | 0.366309736 | 0.53023173 | 0.189 | 0.14 | 1 | 0.473330185 | -0.395455907 | 0.098 | 0.137 | 1 | 0.473330185 | 0.598436649 |
| Fam229b | 0.383191341 | -0.395474488 | 0.094 | 0.14 | 1 | 0.36640097 | -0.654648247 | 0.073 | 0.119 | 1 | 0.383191341 | 0.598552269 |
| Tmprss9 | 0.906251902 | 0.154722595 | 0.038 | 0.034 | 1 | 0.366745891 | -0.917682652 | 0.024 | 0.058 | 1 | 0.906251902 | 0.598989234 |

| | | | | | | | | | | | | |
|---|---|---|---|---|---|---|---|---|---|---|---|---|
| Adam2 | 0.366790997 | -1.368839361 | 0.019 | 0.047 | 1 | 0.402372511 | -0.970150072 | 0 | 0.017 | 1 | 0.402372511 | 0.599046358 |
| Hddc2 | 0.898160429 | 0.47665069 | 0.208 | 0.212 | 1 | 0.36685874 | -0.11032773 | 0.098 | 0.157 | 1 | 0.898160429 | 0.599132145 |
| Zfp963 | 0.564907667 | 0.569760094 | 0.075 | 0.055 | 1 | 0.366923935 | -0.332720152 | 0.073 | 0.123 | 1 | 0.564907667 | 0.599214696 |
| Trim13 | 0.427354314 | 0.739685096 | 0.094 | 0.064 | 1 | 0.367032083 | -0.410722664 | 0.049 | 0.092 | 1 | 0.427354314 | 0.599351617 |
| Cops8 | 0.367102507 | 0.261637799 | 0.623 | 0.619 | 1 | 0.714071435 | 0.140596993 | 0.659 | 0.659 | 1 | 0.714071435 | 0.599440764 |
| Poldip3 | 0.440022336 | 0.211712573 | 0.604 | 0.648 | 1 | 0.367340395 | 0.302868422 | 0.78 | 0.785 | 1 | 0.440022336 | 0.599741824 |
| Mocs3 | 0.367745886 | -0.223789028 | 0.057 | 0.097 | 1 | 0.371690742 | -0.250257991 | 0.049 | 0.092 | 1 | 0.371690742 | 0.600254735 |
| Scn2a | 0.885889171 | 0.569760094 | 0.075 | 0.072 | 1 | 0.36790411 | -1.069685746 | 0.024 | 0.058 | 1 | 0.885889171 | 0.600454786 |
| Far1os | 0.755567413 | 0.8327945 | 0.038 | 0.03 | 1 | 0.368128933 | 0.749742009 | 0.073 | 0.041 | 1 | 0.755567413 | 0.600738954 |
| Ptgs2os | 0.368233176 | -0.652632327 | 0.038 | 0.072 | 1 | 0.767206154 | 1.252242349 | 0.024 | 0.034 | 1 | 0.767206154 | 0.60087068 |
| 2310039H08Rik | 0.624656586 | 0.185973529 | 0.434 | 0.411 | 1 | 0.368239832 | 0.252242349 | 0.244 | 0.341 | 1 | 0.624656586 | 0.600879091 |
| Kif26b | 0.641087737 | -0.216115101 | 0.453 | 0.458 | 1 | 0.368341391 | -0.418095337 | 0.415 | 0.488 | 1 | 0.641087737 | 0.601007402 |
| Efna3 | 0.550237964 | 0.695290976 | 0.057 | 0.038 | 1 | 0.368742168 | 0.962735732 | 0.049 | 0.024 | 1 | 0.550237964 | 0.601513549 |
| Zfp93 | 0.550237964 | 0.695290976 | 0.057 | 0.038 | 1 | 0.369055355 | -0.686357106 | 0.024 | 0.058 | 1 | 0.550237964 | 0.601908855 |
| Gm16023 | 0.849702961 | 0.134823037 | 0.208 | 0.216 | 1 | 0.369271993 | -0.64392184 | 0.195 | 0.253 | 1 | 0.849702961 | 0.602182181 |
| Dap3 | 0.769354928 | 0.228723176 | 0.604 | 0.606 | 1 | 0.369481679 | 0.320413852 | 0.707 | 0.676 | 1 | 0.769354928 | 0.602446646 |
| Tmem236 | 0.369573887 | -0.652632327 | 0.019 | 0.047 | 1 | 0.886552366 | -1.020776145 | 0.024 | 0.02 | 1 | 0.886552366 | 0.602562916 |
| Dkc1 | 0.53710578 | 0.309232544 | 0.358 | 0.322 | 1 | 0.369740968 | -0.152147906 | 0.195 | 0.273 | 1 | 0.53710578 | 0.602773553 |
| Agtrap | 0.872378326 | 0.292226119 | 0.113 | 0.123 | 1 | 0.370171802 | -0.26713181 | 0.073 | 0.123 | 1 | 0.872378326 | 0.603316441 |
| Tmem220 | 0.746876889 | 0.776210972 | 0.057 | 0.047 | 1 | 0.370207442 | -0.555112573 | 0.024 | 0.058 | 1 | 0.746876889 | 0.603361334 |
| 9330179D12Rik | 0.925902203 | 0.280253477 | 0.075 | 0.072 | 1 | 0.370207442 | -1.020776145 | 0.024 | 0.058 | 1 | 0.925902203 | 0.603361334 |
| Isoc2b | 0.370364969 | -0.101617158 | 0.094 | 0.148 | 1 | 0.826714817 | 0.322631677 | 0.098 | 0.113 | 1 | 0.826714817 | 0.603559728 |
| Pak3 | 0.370856644 | -0.531777932 | 0.189 | 0.25 | 1 | 0.375652506 | -1.584258919 | 0.122 | 0.171 | 1 | 0.375652506 | 0.604178638 |
| Prkcg | 0.371050443 | -0.921121163 | 0.302 | 0.339 | 1 | 0.432759313 | -0.666620888 | 0.244 | 0.324 | 1 | 0.432759313 | 0.604422455 |
| Gm43387 | 0.37106248 | 1.261637799 | 0.075 | 0.047 | 1 | 0.561305882 | 0.515276755 | 0.049 | 0.031 | 1 | 0.561305882 | 0.604437596 |
| Selenom | 0.371227195 | -0.192399582 | 0.887 | 0.852 | 1 | 0.755482965 | 0.184399703 | 0.732 | 0.799 | 1 | 0.755482965 | 0.604644759 |
| Garin1a | 0.421693172 | -0.368839361 | 0.038 | 0.068 | 1 | 0.371357099 | -0.686357106 | 0.024 | 0.058 | 1 | 0.421693172 | 0.604808103 |
| Rab35 | 0.371721523 | 0.225111923 | 0.491 | 0.415 | 1 | 0.613048114 | -0.242152942 | 0.585 | 0.597 | 1 | 0.613048114 | 0.605266156 |
| Gm9929 | 0.89531717 | 0.454282877 | 0.038 | 0.042 | 1 | 0.371762646 | -0.899760744 | 0.049 | 0.089 | 1 | 0.89531717 | 0.605317827 |
| Irx3 | 0.926421362 | 0.325523284 | 0.434 | 0.466 | 1 | 0.371913993 | 0.118386602 | 0.341 | 0.444 | 1 | 0.926421362 | 0.605507968 |
| Stoml2 | 0.372371709 | 0.61630268 | 0.396 | 0.381 | 1 | 0.916526548 | 0.225770138 | 0.366 | 0.386 | 1 | 0.916526548 | 0.606082729 |
| Gm12592 | 0.372389973 | -0.260314904 | 0.019 | 0.047 | 1 | 0.410846513 | -0.577832649 | 0.049 | 0.085 | 1 | 0.410846513 | 0.606105654 |
| Poli | 0.440573353 | 0.502645898 | 0.094 | 0.064 | 1 | 0.372505086 | -0.410722664 | 0.024 | 0.058 | 1 | 0.440573353 | 0.606250133 |
| Plpp7 | 0.7394666 | 0.347367673 | 0.019 | 0.013 | 1 | 0.372505086 | -0.484723245 | 0.024 | 0.058 | 1 | 0.7394666 | 0.606250133 |
| Gm20536 | 0.799206618 | 0.792152515 | 0.075 | 0.068 | 1 | 0.372505086 | -0.410722664 | 0.024 | 0.058 | 1 | 0.799206618 | 0.606250133 |
| Gm10184 | 0.89261791 | 0.569760094 | 0.038 | 0.034 | 1 | 0.372505086 | -0.410722664 | 0.024 | 0.058 | 1 | 0.89261791 | 0.606250133 |
| Dnhd1 | 0.372550101 | 0.211306123 | 0.189 | 0.136 | 1 | 0.799027953 | -0.305753104 | 0.122 | 0.133 | 1 | 0.799027953 | 0.606306624 |
| Naa40 | 0.811011986 | 0.200526284 | 0.208 | 0.195 | 1 | 0.372688886 | -0.250257991 | 0.146 | 0.208 | 1 | 0.811011986 | 0.606480766 |
| Uevld | 0.747297884 | 0.195364579 | 0.132 | 0.114 | 1 | 0.37277508 | -0.332720152 | 0.122 | 0.177 | 1 | 0.747297884 | 0.6065889 |
| Primpol | 0.372951631 | 1.211306123 | 0.132 | 0.097 | 1 | 0.842422303 | 0.515276755 | 0.073 | 0.065 | 1 | 0.842422303 | 0.606810343 |
| Abhd14b | 0.575917445 | 0.104096522 | 0.17 | 0.212 | 1 | 0.373098127 | 0.65788115 | 0.268 | 0.208 | 1 | 0.575917445 | 0.606994042 |
| Mcm5 | 0.435690605 | 0.510866405 | 0.094 | 0.064 | 1 | 0.37346373 | 1.029849928 | 0.098 | 0.061 | 1 | 0.435690605 | 0.607452302 |
| Fastkd3 | 0.58073812 | 0.154722595 | 0.057 | 0.081 | 1 | 0.373501708 | -0.292078167 | 0.049 | 0.092 | 1 | 0.58073812 | 0.60749989 |
| Gm9887 | 0.373623877 | 1.347367673 | 0.075 | 0.047 | 1 | 0.672172778 | 0.136765132 | 0.024 | 0.038 | 1 | 0.672172778 | 0.607652952 |
| Pcdhb22 | 0.548462127 | 0.8327945 | 0.075 | 0.055 | 1 | 0.373794727 | 0.313642894 | 0.122 | 0.078 | 1 | 0.548462127 | 0.607866956 |
| Cdpf1 | 0.374296037 | 0.491757582 | 0.189 | 0.14 | 1 | 0.893594919 | 0.786578777 | 0.098 | 0.096 | 1 | 0.893594919 | 0.608494551 |
| Gm56860 | 0.375072044 | 0.776210972 | 0.132 | 0.093 | 1 | 0.911204201 | 0.185128153 | 0.122 | 0.13 | 1 | 0.911204201 | 0.609465049 |
| Tmem141 | 0.375294434 | -0.430239906 | 0.113 | 0.161 | 1 | 0.622356447 | 0.212713985 | 0.073 | 0.099 | 1 | 0.622356447 | 0.609742955 |
| Arv1 | 0.375328821 | -0.799473716 | 0.057 | 0.093 | 1 | 0.586944228 | -0.101394606 | 0.049 | 0.072 | 1 | 0.586944228 | 0.609785918 |
| Acsf2 | 0.375368156 | -0.288884057 | 0.283 | 0.373 | 1 | 0.793716138 | 0.186950889 | 0.244 | 0.28 | 1 | 0.793716138 | 0.60983506 |
| Dlst | 0.375440529 | 0.205187662 | 0.66 | 0.657 | 1 | 0.963956177 | -0.101394606 | 0.756 | 0.761 | 1 | 0.963956177 | 0.609925467 |
| Ubac1 | 0.771127604 | 0.2967416 | 0.264 | 0.258 | 1 | 0.375623137 | 1.140131983 | 0.317 | 0.273 | 1 | 0.771127604 | 0.610153533 |
| Mbd1 | 0.375782034 | 0.397292898 | 0.302 | 0.25 | 1 | 0.566161174 | 0.435106406 | 0.317 | 0.28 | 1 | 0.566161174 | 0.61035193 |
| Cdkn2aipnl | 0.377292453 | 0.444229212 | 0.17 | 0.123 | 1 | 0.804496433 | 0.252242349 | 0.122 | 0.109 | 1 | 0.804496433 | 0.61223531 |
| Ahi1 | 0.377684008 | 0.446024726 | 0.509 | 0.475 | 1 | 0.682651084 | 0.231778247 | 0.317 | 0.392 | 1 | 0.682651084 | 0.612722807 |
| Vta1 | 0.37779342 | 0.376078969 | 0.547 | 0.525 | 1 | 0.457171288 | -0.106211622 | 0.537 | 0.601 | 1 | 0.457171288 | 0.612858972 |
| Grin3a | 0.377976475 | 0.569760094 | 0.019 | 0.047 | 1 | 0.484981146 | 1.185128153 | 0.073 | 0.048 | 1 | 0.484981146 | 0.613086734 |
| 1700007L15Rik | 0.377976475 | 0.154722595 | 0.019 | 0.047 | 1 | 0.440032309 | 1.83720485 | 0.049 | 0.027 | 1 | 0.440032309 | 0.613086734 |
| ENSMUSG00000121502 | 0.477925251 | 0.8327945 | 0.094 | 0.068 | 1 | 0.37839063 | 0.877846834 | 0.122 | 0.082 | 1 | 0.477925251 | 0.613601791 |
| Gm1604a | 0.378755743 | 0.526691372 | 0.113 | 0.076 | 1 | 0.946193611 | 0.43121249 | 0.098 | 0.102 | 1 | 0.946193611 | 0.614055573 |
| Srgap3 | 0.378941909 | 0.640149422 | 0.226 | 0.174 | 1 | 0.73435373 | -0.426347672 | 0.537 | 0.481 | 1 | 0.73435373 | 0.614286848 |
| Pth1r | 0.379697285 | -1.257091004 | 0.547 | 0.5 | 1 | 0.389502983 | -0.55696813 | 0.39 | 0.485 | 1 | 0.389502983 | 0.615224542 |
| Gm15886 | 0.379849917 | 0.948271717 | 0.132 | 0.093 | 1 | 0.629871081 | 0.635570989 | 0.073 | 0.055 | 1 | 0.629871081 | 0.615413874 |
| Fam177a | 0.537052636 | -0.1672055 | 0.038 | 0.059 | 1 | 0.380140443 | 0.690363461 | 0.049 | 0.092 | 1 | 0.537052636 | 0.615774129 |
| Mplkip | 0.96842187 | 0.222409402 | 0.547 | 0.572 | 1 | 0.380254903 | 0.300070874 | 0.415 | 0.529 | 1 | 0.96842187 | 0.615916015 |
| Mgme1 | 0.463730159 | 0.466666601 | 0.132 | 0.097 | 1 | 0.38085138 | -0.296650897 | 0.098 | 0.154 | 1 | 0.463730159 | 0.616654987 |
| Kcnc3 | 0.380880107 | 0.154722595 | 0.057 | 0.097 | 1 | 0.452677794 | -0.686357106 | 0.073 | 0.109 | 1 | 0.452677794 | 0.616690558 |
| Ankmy2 | 0.403246511 | 1.102255175 | 0.226 | 0.186 | 1 | 0.381053961 | 0.902299878 | 0.22 | 0.174 | 1 | 0.403246511 | 0.6169058 |
| 1600014C10Rik | 0.885441525 | 0.200526284 | 0.113 | 0.106 | 1 | 0.381863234 | 0.649577847 | 0.171 | 0.123 | 1 | 0.885441525 | 0.617906938 |
| Dph7 | 0.38301283 | -0.489133595 | 0.113 | 0.161 | 1 | 0.536084624 | -0.64392184 | 0.122 | 0.154 | 1 | 0.536084624 | 0.619326832 |
| Rassf2 | 0.383346953 | 0.207190015 | 0.283 | 0.225 | 1 | 0.50419385 | 0.354121963 | 0.293 | 0.242 | 1 | 0.50419385 | 0.619739019 |
| Timm21 | 0.678703932 | 0.289652175 | 0.34 | 0.322 | 1 | 0.383964256 | -0.203822418 | 0.22 | 0.287 | 1 | 0.678703932 | 0.620499962 |
| Me2 | 0.449605028 | 0.491757582 | 0.245 | 0.212 | 1 | 0.383982031 | -0.502645153 | 0.122 | 0.174 | 1 | 0.449605028 | 0.620521861 |
| Tango2 | 0.384620557 | 1.252019796 | 0.189 | 0.148 | 1 | 0.636102147 | -0.259656689 | 0.122 | 0.147 | 1 | 0.636102147 | 0.621308142 |
| Wars1 | 0.384868679 | 0.407488665 | 0.208 | 0.161 | 1 | 0.402486701 | 0.444887427 | 0.22 | 0.167 | 1 | 0.402486701 | 0.621613458 |
| Nudt16l1 | 0.385625162 | 0.2967416 | 0.264 | 0.203 | 1 | 0.897889994 | 0.432814595 | 0.244 | 0.249 | 1 | 0.897889994 | 0.622543559 |
| Rabl2 | 0.385697757 | 0.669295768 | 0.189 | 0.14 | 1 | 0.874879183 | 0.308825877 | 0.195 | 0.188 | 1 | 0.874879183 | 0.622632754 |
| Zfp746 | 0.385764201 | -0.359850578 | 0.208 | 0.263 | 1 | 0.803175819 | 0.324135268 | 0.268 | 0.3 | 1 | 0.803175819 | 0.622714383 |
| Stambpl1 | 0.38604819 | 0.749466116 | 0.283 | 0.246 | 1 | 0.574059435 | 0.119348079 | 0.146 | 0.191 | 1 | 0.574059435 | 0.623063175 |
| Cdipt | 0.512777875 | -0.353072045 | 0.453 | 0.475 | 1 | 0.386401314 | 0.416542801 | 0.561 | 0.522 | 1 | 0.512777875 | 0.623496653 |
| Nr1h2 | 0.386548367 | 0.569760094 | 0.415 | 0.39 | 1 | 0.768579536 | 0.168826341 | 0.366 | 0.403 | 1 | 0.768579536 | 0.623677094 |
| Itih2 | 0.386654903 | 0.610402079 | 0.17 | 0.127 | 1 | 0.580911574 | 0.225770138 | 0.122 | 0.157 | 1 | 0.580911574 | 0.623807792 |
| Aplf | 0.386706609 | 0.640149422 | 0.113 | 0.076 | 1 | 0.62595663 | 1.351778023 | 0.049 | 0.034 | 1 | 0.62595663 | 0.623871217 |
| Fhip2b | 0.386934795 | 0.631160639 | 0.113 | 0.076 | 1 | 0.802994279 | 0.435106406 | 0.098 | 0.116 | 1 | 0.802994279 | 0.624151055 |
| Sidt2 | 0.387242722 | -0.105513178 | 0.66 | 0.75 | 1 | 0.572226725 | -0.115661175 | 0.732 | 0.765 | 1 | 0.572226725 | 0.624528519 |
| Cfap45 | 0.387813168 | 1.076720083 | 0.075 | 0.047 | 1 | 0.948204846 | 0.83720485 | 0.146 | 0.154 | 1 | 0.948204846 | 0.625227283 |
| 4933439C10Rik | 0.844569019 | 0.247831999 | 0.094 | 0.106 | 1 | 0.387872452 | 0.786578777 | 0.122 | 0.082 | 1 | 0.844569019 | 0.625299865 |
| Taf8 | 0.642921392 | 0.598329246 | 0.208 | 0.191 | 1 | 0.387918669 | 0.450181727 | 0.244 | 0.188 | 1 | 0.642921392 | 0.625356444 |
| A430073D23Rik | 0.960557568 | 0.932330173 | 0.075 | 0.076 | 1 | 0.38819584 | -0.16279515 | 0.024 | 0.058 | 1 | 0.960557568 | 0.62569567 |
| Abcd3 | 0.388238774 | -0.123993433 | 0.509 | 0.602 | 1 | 0.796730792 | 0.143989458 | 0.732 | 0.689 | 1 | 0.796730792 | 0.625748203 |
| 2310009B15Rik | 0.38826496 | 0.47665069 | 0.358 | 0.314 | 1 | 0.850181064 | 0.295311071 | 0.22 | 0.253 | 1 | 0.850181064 | 0.625780241 |
| Pcdhga12 | 0.842312554 | -0.20282941 | 0.132 | 0.14 | 1 | 0.388485579 | 1.029849928 | 0.122 | 0.085 | 1 | 0.842312554 | 0.626050112 |
| Zkscan3 | 0.389324727 | 0.395335706 | 0.604 | 0.631 | 1 | 0.422246297 | -0.16279515 | 0.61 | 0.683 | 1 | 0.422246297 | 0.627075712 |
| E430024I08Rik | 0.389748052 | 0.154722595 | 0.264 | 0.199 | 1 | 0.746505823 | 0.118386602 | 0.171 | 0.198 | 1 | 0.746505823 | 0.62759256 |
| Carhsp1 | 0.999114647 | -0.407156293 | 0.34 | 0.314 | 1 | 0.389957115 | 0.486707603 | 0.39 | 0.334 | 1 | 0.999114647 | 0.627847678 |
| Cox6b2 | 0.390346302 | 0.962077517 | 0.075 | 0.047 | 1 | 0.710563777 | 1.614812428 | 0.049 | 0.038 | 1 | 0.710563777 | 0.628322369 |
| Gm10076 | 0.390545542 | -0.151307873 | 0.943 | 0.89 | 1 | 0.391663743 | -0.213421223 | 0.732 | 0.812 | 1 | 0.391663743 | 0.628565263 |
| Zdhhc6 | 0.39083298 | 0.285306705 | 0.34 | 0.288 | 1 | 0.707571099 | -0.196742482 | 0.293 | 0.314 | 1 | 0.707571099 | 0.628915542 |

| | | | | | | | | | | | | |
|---|---|---|---|---|---|---|---|---|---|---|---|---|
| Il31ra | 0.390924926 | 0.859266711 | 0.094 | 0.064 | 1 | 0.617676868 | 1.100239256 | 0.073 | 0.055 | 1 | 0.617676868 | 0.629027555 |
| Cryab | 0.53039856 | -2.058872878 | 0.453 | 0.432 | 1 | 0.391749455 | -0.12402558 | 0.268 | 0.222 | 1 | 0.53039856 | 0.630031275 |
| Sp7 | 0.905470136 | -0.237594828 | 0.17 | 0.174 | 1 | 0.391902179 | 0.944120054 | 0.146 | 0.106 | 1 | 0.905470136 | 0.63021704 |
| Erlin1 | 0.39202077 | 0.242185436 | 0.245 | 0.191 | 1 | 0.966290768 | 0.180092563 | 0.293 | 0.314 | 1 | 0.966290768 | 0.630361256 |
| Gm9725 | 0.392292247 | -0.108311811 | 0.038 | 0.072 | 1 | 0.828694929 | 0.422167351 | 0.049 | 0.041 | 1 | 0.828694929 | 0.630691287 |
| Cdk5rap1 | 0.98775452 | 0.124975251 | 0.17 | 0.169 | 1 | 0.392537695 | -0.442903069 | 0.098 | 0.147 | 1 | 0.98775452 | 0.630989549 |
| Itgb7 | 0.392552036 | 0.399835093 | 0.189 | 0.144 | 1 | 0.62415002 | -0.466575898 | 0.317 | 0.345 | 1 | 0.62415002 | 0.631006971 |
| Alyref2 | 0.393064389 | 0.569760094 | 0.113 | 0.076 | 1 | 0.923149946 | 0.778311161 | 0.073 | 0.072 | 1 | 0.923149946 | 0.631629164 |
| Gdf15 | 0.393268394 | -0.407519829 | 0.094 | 0.064 | 1 | 0.604477578 | -0.622226769 | 0.024 | 0.041 | 1 | 0.604477578 | 0.631876759 |
| Mmgt2 | 0.526289851 | 0.377115016 | 0.321 | 0.284 | 1 | 0.393381016 | -0.221688839 | 0.244 | 0.314 | 1 | 0.526289851 | 0.632013408 |
| Lemd3 | 0.394003897 | 0.111220956 | 0.566 | 0.487 | 1 | 0.791315325 | 0.304983811 | 0.61 | 0.669 | 1 | 0.791315325 | 0.632768724 |
| Lamtor2 | 0.394119828 | 0.225658502 | 0.792 | 0.78 | 1 | 0.666484887 | 0.282143835 | 0.683 | 0.683 | 1 | 0.666484887 | 0.632909218 |
| Dcaf15 | 0.394320115 | 0.510866405 | 0.113 | 0.076 | 1 | 0.410579345 | 0.614812428 | 0.122 | 0.082 | 1 | 0.410579345 | 0.633151877 |
| Grid1 | 0.510606779 | 0.569760094 | 0.019 | 0.008 | 1 | 0.39435929 | 0.91966701 | 0.024 | 0.058 | 1 | 0.510606779 | 0.63319933 |
| C130036L24Rik | 0.394388685 | 0.891688189 | 0.075 | 0.047 | 1 | 0.989595616 | 0.667279848 | 0.024 | 0.024 | 1 | 0.989595616 | 0.633234936 |
| Gm17435 | 0.670446368 | 0.122301117 | 0.17 | 0.144 | 1 | 0.394527229 | 1.284663827 | 0.171 | 0.13 | 1 | 0.670446368 | 0.633402724 |
| Cdc20 | 0.394669687 | -0.752168001 | 0.094 | 0.059 | 1 | 0.992196803 | 0.252242349 | 0.024 | 0.024 | 1 | 0.992196803 | 0.633575213 |
| Gm15411 | 0.395615987 | 0.8327945 | 0.075 | 0.047 | 1 | 0.488539607 | 1.363273661 | 0.098 | 0.068 | 1 | 0.488539607 | 0.634719965 |
| Gm9975 | 0.395615987 | 0.8327945 | 0.075 | 0.047 | 1 | 0.494850012 | 0.83720485 | 0.073 | 0.048 | 1 | 0.494850012 | 0.634719965 |
| Tmem170b | 0.813754918 | -0.142957954 | 0.17 | 0.182 | 1 | 0.396052161 | 1.284663827 | 0.244 | 0.218 | 1 | 0.813754918 | 0.635247008 |
| Yipf3 | 0.421720128 | 0.239743179 | 0.717 | 0.708 | 1 | 0.396212519 | -0.126994429 | 0.61 | 0.669 | 1 | 0.421720128 | 0.635440677 |
| Tasp1 | 0.778696382 | -0.324174846 | 0.604 | 0.589 | 1 | 0.396414221 | 0.101640611 | 0.512 | 0.642 | 1 | 0.778696382 | 0.635684207 |
| Slc25a21 | 0.998220543 | 1.502645898 | 0.057 | 0.059 | 1 | 0.396514443 | -2.31254227 | 0.024 | 0.055 | 1 | 0.998220543 | 0.635805182 |
| Rbsn | 0.539509392 | 0.470224421 | 0.34 | 0.314 | 1 | 0.396721205 | -0.238083278 | 0.317 | 0.389 | 1 | 0.539509392 | 0.636054695 |
| Ceacam1 | 0.396901946 | 1.021456064 | 0.189 | 0.157 | 1 | 0.518087774 | -0.80665134 | 0.122 | 0.154 | 1 | 0.518087774 | 0.636272737 |
| Ifit2 | 0.396927521 | -0.686579659 | 0.075 | 0.114 | 1 | 0.60891903 | 1.377773231 | 0.098 | 0.078 | 1 | 0.60891903 | 0.636303585 |
| Mrps25 | 0.7744007 | 0.272079545 | 0.396 | 0.394 | 1 | 0.397717759 | 0.358037013 | 0.341 | 0.444 | 1 | 0.7744007 | 0.637256102 |
| Uggt1 | 0.397764668 | 0.260952921 | 0.698 | 0.678 | 1 | 0.480073346 | 0.245289588 | 0.732 | 0.693 | 1 | 0.480073346 | 0.637312605 |
| Cyrib | 0.492404917 | 0.370451286 | 0.396 | 0.373 | 1 | 0.398209354 | -0.373942814 | 0.488 | 0.577 | 1 | 0.492404917 | 0.637848018 |
| Inca1 | 0.398253154 | 0.652222254 | 0.075 | 0.047 | 1 | 0.608047495 | -0.16279515 | 0.024 | 0.041 | 1 | 0.608047495 | 0.637900733 |
| Otoa | 0.909701062 | -0.282682718 | 0.132 | 0.136 | 1 | 0.398953908 | 0.667279848 | 0.098 | 0.061 | 1 | 0.909701062 | 0.638743596 |
| Cwc15 | 0.949626717 | 0.10496956 | 0.849 | 0.792 | 1 | 0.399508167 | 0.235992997 | 0.854 | 0.857 | 1 | 0.949626717 | 0.639409558 |
| Efemp1 | 0.906913463 | -0.672128503 | 0.321 | 0.318 | 1 | 0.39990763 | -0.538433832 | 0.293 | 0.365 | 1 | 0.906913463 | 0.639889148 |
| Pramel12 | 0.689557822 | 1.002719501 | 0.094 | 0.081 | 1 | 0.400425697 | 1.209173627 | 0.146 | 0.109 | 1 | 0.689557822 | 0.640510655 |
| 9830144P21Rik | 0.400685124 | -0.904171094 | 0.038 | 0.068 | 1 | 0.997398972 | -0.250257991 | 0.024 | 0.024 | 1 | 0.997398972 | 0.64082168 |
| Gm17251 | 0.429599096 | 0.417757001 | 0.151 | 0.11 | 1 | 0.401223474 | -0.372248516 | 0.049 | 0.089 | 1 | 0.429599096 | 0.641466672 |
| Hjurp | 0.401332204 | -0.418744267 | 0.283 | 0.343 | 1 | 0.568378077 | 0.527349587 | 0.341 | 0.3 | 1 | 0.568378077 | 0.64159687 |
| Pdgfd | 0.813368953 | 0.556085157 | 0.151 | 0.178 | 1 | 0.401695125 | -0.770477727 | 0.146 | 0.205 | 1 | 0.813368953 | 0.642031276 |
| Zfp1 | 0.401966746 | 0.446185409 | 0.434 | 0.407 | 1 | 0.910242605 | 0.402576622 | 0.317 | 0.362 | 1 | 0.910242605 | 0.642356227 |
| Etaa1 | 0.402323126 | 0.187889459 | 0.17 | 0.123 | 1 | 0.547552316 | 0.410940095 | 0.146 | 0.113 | 1 | 0.547552316 | 0.642782354 |
| Cenpm | 0.479836037 | -1.489133595 | 0.019 | 0.038 | 1 | 0.402364804 | -0.16279515 | 0 | 0.017 | 1 | 0.479836037 | 0.642832172 |
| Ccdc38 | 0.500671253 | 1.739685096 | 0.019 | 0.008 | 1 | 0.402364804 | -0.16279515 | 0 | 0.017 | 1 | 0.500671253 | 0.642832172 |
| Gm17090 | 0.503971666 | 1.569760094 | 0.019 | 0.008 | 1 | 0.402364804 | -0.16279515 | 0 | 0.017 | 1 | 0.503971666 | 0.642832172 |
| Tafa4 | 0.619885227 | 0.984797593 | 0.038 | 0.025 | 1 | 0.402364804 | -0.16279515 | 0 | 0.017 | 1 | 0.619885227 | 0.642832172 |
| Fcgr3 | 0.732691937 | 1.154722595 | 0.019 | 0.013 | 1 | 0.402364804 | -0.622226769 | 0 | 0.017 | 1 | 0.732691937 | 0.642832172 |
| Omp | 0.792580307 | 0.8327945 | 0.019 | 0.025 | 1 | 0.402364804 | -0.16279515 | 0 | 0.017 | 1 | 0.792580307 | 0.642832172 |
| Rnf138rt1 | 0.929410101 | 0.569760094 | 0.019 | 0.017 | 1 | 0.402364804 | -0.16279515 | 0 | 0.017 | 1 | 0.929410101 | 0.642832172 |
| Gm12472 | 0.413080549 | -0.1672055 | 0 | 0.013 | 1 | 0.402369942 | -0.484723245 | 0 | 0.017 | 1 | 0.413080549 | 0.642838314 |
| ENSMUSG00000121382 | 0.413080549 | -0.1672055 | 0 | 0.013 | 1 | 0.402369942 | -0.332720152 | 0 | 0.017 | 1 | 0.413080549 | 0.642838314 |
| Apobr | 0.413082433 | 0.154722595 | 0 | 0.013 | 1 | 0.402369942 | -0.16279515 | 0 | 0.017 | 1 | 0.413082433 | 0.642838314 |
| Cep112it | 0.416735385 | -0.652632327 | 0.019 | 0.042 | 1 | 0.402369942 | -0.332720152 | 0 | 0.017 | 1 | 0.416735385 | 0.642838314 |
| Map3k13 | 0.503362835 | 0.502645898 | 0.094 | 0.068 | 1 | 0.402369942 | -0.863234868 | 0 | 0.017 | 1 | 0.503362835 | 0.642838314 |
| Liph | 0.507281812 | 0.8327945 | 0.019 | 0.008 | 1 | 0.402369942 | -0.332720152 | 0 | 0.017 | 1 | 0.507281812 | 0.642838314 |
| Stk33 | 0.565795103 | -1.545717123 | 0.019 | 0.034 | 1 | 0.402369942 | -1.622226769 | 0 | 0.017 | 1 | 0.565795103 | 0.642838314 |
| Gm15287 | 0.661088008 | -0.304709024 | 0.019 | 0.03 | 1 | 0.402369942 | -1.250257991 | 0 | 0.017 | 1 | 0.661088008 | 0.642838314 |
| Fer1l5 | 0.909120409 | -1.652632327 | 0.019 | 0.021 | 1 | 0.402369942 | -0.332720152 | 0 | 0.017 | 1 | 0.909120409 | 0.642838314 |
| Gm2237 | 0.926212226 | 0.8327945 | 0.019 | 0.017 | 1 | 0.402369942 | -0.16279515 | 0 | 0.017 | 1 | 0.926212226 | 0.642838314 |
| 2600002D14Rik | 0.929410101 | 0.569760094 | 0.019 | 0.017 | 1 | 0.402369942 | -0.16279515 | 0 | 0.017 | 1 | 0.929410101 | 0.642838314 |
| Dnajb3 | 0.503971666 | 1.569760094 | 0.019 | 0.008 | 1 | 0.402371655 | -0.484723245 | 0 | 0.017 | 1 | 0.503971666 | 0.642840361 |
| Sptbn4 | 0.776329749 | 0.280253477 | 0.038 | 0.03 | 1 | 0.402371655 | -0.484723245 | 0 | 0.017 | 1 | 0.776329749 | 0.642840361 |
| Nckap1l | 0.917883255 | 0.569760094 | 0.019 | 0.021 | 1 | 0.402371655 | -0.484723245 | 0 | 0.017 | 1 | 0.917883255 | 0.642840361 |
| Cldn14 | 0.776830424 | -1.093204919 | 0.019 | 0.025 | 1 | 0.402372511 | -0.332720152 | 0 | 0.017 | 1 | 0.776830424 | 0.642841385 |
| 4930447K03Rik | 0.912039013 | -0.304709024 | 0.019 | 0.021 | 1 | 0.402372511 | -1.410722664 | 0 | 0.017 | 1 | 0.912039013 | 0.642841385 |
| Brme1 | 0.929579738 | 0.932330173 | 0.019 | 0.021 | 1 | 0.402372511 | -0.622226769 | 0 | 0.017 | 1 | 0.929579738 | 0.642841385 |
| 4930592A05Rik | 0.413082433 | -2.752168001 | 0 | 0.013 | 1 | 0.402373368 | 0.252242349 | 0 | 0.017 | 1 | 0.413082433 | 0.642842408 |
| Ly6g6c | 0.413082433 | 0.154722595 | 0 | 0.013 | 1 | 0.402373368 | 0.252242349 | 0 | 0.017 | 1 | 0.413082433 | 0.642842408 |
| 1810010H24Rik | 0.464892121 | 1.669295768 | 0.038 | 0.021 | 1 | 0.402373368 | 0.252242349 | 0 | 0.017 | 1 | 0.464892121 | 0.642842408 |
| Gm43672 | 0.471154689 | 1.891688189 | 0.038 | 0.021 | 1 | 0.402373368 | 0.252242349 | 0 | 0.017 | 1 | 0.471154689 | 0.642842408 |
| Mettl21c | 0.497380598 | 2.154722595 | 0.019 | 0.008 | 1 | 0.402373368 | 0.252242349 | 0 | 0.017 | 1 | 0.497380598 | 0.642842408 |
| Treh | 0.503971666 | 1.569760094 | 0.019 | 0.008 | 1 | 0.402373368 | 0.252242349 | 0 | 0.017 | 1 | 0.503971666 | 0.642842408 |
| Gm5148 | 0.507281812 | 0.8327945 | 0.019 | 0.008 | 1 | 0.402373368 | 0.252242349 | 0 | 0.017 | 1 | 0.507281812 | 0.642842408 |
| A630066F11Rik | 0.550237964 | 0.695290976 | 0.057 | 0.038 | 1 | 0.402373368 | 0.252242349 | 0 | 0.017 | 1 | 0.550237964 | 0.642842408 |
| Gm19569 | 0.610863297 | 1.347367673 | 0.038 | 0.025 | 1 | 0.402373368 | 0.252242349 | 0 | 0.017 | 1 | 0.610863297 | 0.642842408 |
| Gm56977 | 0.624418548 | 0.932330173 | 0.038 | 0.025 | 1 | 0.402373368 | 0.252242349 | 0 | 0.017 | 1 | 0.624418548 | 0.642842408 |
| Gpld1 | 0.684433992 | 0.984797593 | 0.019 | 0.03 | 1 | 0.402373368 | 0.252242349 | 0 | 0.017 | 1 | 0.684433992 | 0.642842408 |
| Gm56735 | 0.72256466 | 1.739685096 | 0.019 | 0.013 | 1 | 0.402373368 | 0.252242349 | 0 | 0.017 | 1 | 0.72256466 | 0.642842408 |
| Gm38560 | 0.725936481 | 1.417757001 | 0.019 | 0.013 | 1 | 0.402373368 | 0.252242349 | 0 | 0.017 | 1 | 0.725936481 | 0.642842408 |
| Fgfbp1 | 0.732691937 | 1.154722595 | 0.019 | 0.013 | 1 | 0.402373368 | 0.252242349 | 0 | 0.017 | 1 | 0.732691937 | 0.642842408 |
| Spata33 | 0.736073999 | 0.8327945 | 0.019 | 0.013 | 1 | 0.402373368 | 0.252242349 | 0 | 0.017 | 1 | 0.736073999 | 0.642842408 |
| Dyrk3 | 0.753264159 | 0.347367673 | 0.038 | 0.03 | 1 | 0.402373368 | 0.252242349 | 0 | 0.017 | 1 | 0.753264159 | 0.642842408 |
| Gm36264 | 0.767066103 | 0.569760094 | 0.038 | 0.03 | 1 | 0.402373368 | 0.252242349 | 0 | 0.017 | 1 | 0.767066103 | 0.642842408 |
| Arhgdig | 0.767066103 | 0.569760094 | 0.038 | 0.03 | 1 | 0.402373368 | 0.252242349 | 0 | 0.017 | 1 | 0.767066103 | 0.642842408 |
| E230013L22Rik | 0.917883255 | 0.569760094 | 0.019 | 0.021 | 1 | 0.402373368 | 0.252242349 | 0 | 0.017 | 1 | 0.917883255 | 0.642842408 |
| Gm57399 | 0.926212226 | 0.8327945 | 0.019 | 0.017 | 1 | 0.402373368 | 0.252242349 | 0 | 0.017 | 1 | 0.926212226 | 0.642842408 |
| Lrrc40 | 0.855147106 | 0.234157062 | 0.189 | 0.178 | 1 | 0.402725772 | -0.31254227 | 0.122 | 0.174 | 1 | 0.855147106 | 0.643263496 |
| Edar | 0.732691937 | 1.154722595 | 0.019 | 0.013 | 1 | 0.402818967 | -0.863234868 | 0.024 | 0.055 | 1 | 0.732691937 | 0.643374814 |
| Spcs2 | 0.418509377 | 0.142459175 | 0.943 | 0.903 | 1 | 0.402931495 | -0.187309299 | 0.829 | 0.942 | 1 | 0.418509377 | 0.6435092 |
| Bex2 | 0.403100898 | -1.067669827 | 0.038 | 0.068 | 1 | 0.810504515 | 0.515276755 | 0.098 | 0.116 | 1 | 0.810504515 | 0.643711462 |
| Col20a1 | 0.403167988 | -0.609043059 | 0.264 | 0.318 | 1 | 0.503591066 | -0.43741753 | 0.39 | 0.427 | 1 | 0.503591066 | 0.64379155 |
| Asb1 | 0.677640974 | -0.237594828 | 0.057 | 0.072 | 1 | 0.403405135 | 1.351778023 | 0.073 | 0.044 | 1 | 0.677640974 | 0.644074567 |
| Ndrg4 | 0.851548464 | -0.285849997 | 0.113 | 0.119 | 1 | 0.403411405 | 0.152706676 | 0.146 | 0.099 | 1 | 0.851548464 | 0.644082048 |
| Il17rc | 0.403474578 | 0.984797593 | 0.094 | 0.064 | 1 | 0.796511241 | 0.930314254 | 0.098 | 0.089 | 1 | 0.796511241 | 0.644157421 |
| Arel1 | 0.403600667 | 0.286826131 | 0.245 | 0.191 | 1 | 0.999052101 | 0.168826341 | 0.22 | 0.225 | 1 | 0.999052101 | 0.644307836 |
| Rab3gap1 | 0.664927981 | 0.101457445 | 0.811 | 0.797 | 1 | 0.403877698 | -0.138816287 | 0.78 | 0.812 | 1 | 0.664927981 | 0.644638201 |
| Mettl7a1 | 0.403902767 | 0.227367897 | 0.528 | 0.462 | 1 | 0.684233512 | 0.456383066 | 0.512 | 0.543 | 1 | 0.684233512 | 0.644668088 |
| Rom1 | 0.40406473 | -0.557995453 | 0.132 | 0.178 | 1 | 0.853076459 | -0.116991461 | 0.171 | 0.154 | 1 | 0.853076459 | 0.644861154 |
| Reep5 | 0.578472814 | -0.145479313 | 0.981 | 0.962 | 1 | 0.404312613 | -0.13630726 | 0.927 | 0.993 | 1 | 0.578472814 | 0.645156537 |
| Gda | 0.413080549 | -0.652632327 | 0 | 0.013 | 1 | 0.404564934 | -0.101394606 | 0.195 | 0.143 | 1 | 0.413080549 | 0.645457082 |

| | | | | | | | | | | | | |
|---|---|---|---|---|---|---|---|---|---|---|---|---|
| Ttc32 | 0.404661166 | 0.695290976 | 0.189 | 0.148 | 1 | 0.537207104 | 0.782757066 | 0.171 | 0.143 | 1 | 0.537207104 | 0.645571672 |
| Dbnl | 0.703636419 | 0.374503803 | 0.283 | 0.263 | 1 | 0.404769745 | 0.707921833 | 0.39 | 0.348 | 1 | 0.703636419 | 0.645700944 |
| Zfp316 | 0.404836717 | 0.669295768 | 0.075 | 0.047 | 1 | 0.672172778 | 0.136765132 | 0.024 | 0.038 | 1 | 0.672172778 | 0.645780667 |
| Acot7 | 0.997652223 | 0.200526284 | 0.472 | 0.517 | 1 | 0.404889106 | 0.346218497 | 0.488 | 0.447 | 1 | 0.997652223 | 0.645843023 |
| Tkfc | 0.404964703 | 1.002719501 | 0.113 | 0.081 | 1 | 0.424747458 | 0.930314254 | 0.073 | 0.044 | 1 | 0.424747458 | 0.645932996 |
| Mettl26 | 0.982419472 | 0.141425772 | 0.321 | 0.331 | 1 | 0.405083055 | -0.278272368 | 0.244 | 0.317 | 1 | 0.982419472 | 0.646073829 |
| Gm57093 | 0.405147636 | 0.569760094 | 0.132 | 0.093 | 1 | 0.746167976 | -0.723510105 | 0.098 | 0.113 | 1 | 0.746167976 | 0.646150665 |
| Taf1 | 0.40533634 | -0.197437702 | 0.736 | 0.754 | 1 | 0.834033644 | 0.160374329 | 0.78 | 0.795 | 1 | 0.834033644 | 0.646375132 |
| Ift81 | 0.738966912 | 0.205348668 | 0.321 | 0.305 | 1 | 0.405432226 | 0.460484282 | 0.22 | 0.314 | 1 | 0.738966912 | 0.646489162 |
| Rbm48 | 0.932806215 | 0.385335523 | 0.245 | 0.263 | 1 | 0.40553747 | 1.114738825 | 0.195 | 0.16 | 1 | 0.932806215 | 0.6466143 |
| Ascc1 | 0.521880357 | 0.648831665 | 0.283 | 0.242 | 1 | 0.405599267 | 0.32941021 | 0.171 | 0.242 | 1 | 0.521880357 | 0.646687769 |
| Mlf2 | 0.405766647 | 0.203378401 | 0.774 | 0.758 | 1 | 0.947070039 | 0.128253632 | 0.756 | 0.775 | 1 | 0.947070039 | 0.646886722 |
| Nceh1 | 0.40602292 | -0.138059154 | 0.094 | 0.144 | 1 | 0.577839073 | 1.383486882 | 0.146 | 0.123 | 1 | 0.577839073 | 0.647191228 |
| Emb | 0.578784212 | -0.595299152 | 0.755 | 0.72 | 1 | 0.406203807 | -0.224891271 | 0.78 | 0.672 | 1 | 0.578784212 | 0.647406082 |
| Ttc33 | 0.406203893 | 0.392882332 | 0.302 | 0.254 | 1 | 0.940332045 | 0.659666664 | 0.268 | 0.283 | 1 | 0.940332045 | 0.647406184 |
| Anapc1 | 0.406317163 | 0.287172891 | 0.547 | 0.508 | 1 | 0.557197954 | 0.252242349 | 0.659 | 0.683 | 1 | 0.557197954 | 0.647540689 |
| D130017N08Rik | 0.653990165 | 0.695290976 | 0.057 | 0.042 | 1 | 0.40660741 | -0.747757651 | 0.024 | 0.055 | 1 | 0.653990165 | 0.647885234 |
| Fcf1 | 0.407844295 | 0.488840099 | 0.509 | 0.475 | 1 | 0.524480174 | -0.16279515 | 0.415 | 0.451 | 1 | 0.524480174 | 0.649351621 |
| Zfp354b | 0.725936481 | 1.417757001 | 0.019 | 0.013 | 1 | 0.40786703 | -0.622226769 | 0.024 | 0.055 | 1 | 0.725936481 | 0.649378546 |
| Gm26885 | 0.767066103 | 0.569760094 | 0.038 | 0.03 | 1 | 0.40786703 | -0.747757651 | 0.024 | 0.055 | 1 | 0.767066103 | 0.649378546 |
| Cd38 | 0.407939982 | -0.783876861 | 0.038 | 0.068 | 1 | 0.51919986 | 0.83720485 | 0 | 0.01 | 1 | 0.51919986 | 0.649464935 |
| Arhgap21 | 0.670368908 | -0.1672055 | 0.906 | 0.86 | 1 | 0.408646741 | -0.161865279 | 0.951 | 0.952 | 1 | 0.670368908 | 0.650301323 |
| Gm56721 | 0.782077996 | 0.347367673 | 0.019 | 0.025 | 1 | 0.408802174 | -0.787286015 | 0.049 | 0.085 | 1 | 0.782077996 | 0.650485131 |
| Snhg14 | 0.481663295 | 0.859266711 | 0.132 | 0.102 | 1 | 0.408967986 | 1.583448258 | 0.122 | 0.085 | 1 | 0.481663295 | 0.650681158 |
| Fam221a | 0.764727501 | 0.618669695 | 0.113 | 0.102 | 1 | 0.409125717 | -0.410722664 | 0.024 | 0.055 | 1 | 0.764727501 | 0.650867582 |
| 9430015G10Rik | 0.40918063 | 0.962077517 | 0.094 | 0.064 | 1 | 0.898226899 | 0.404245443 | 0.073 | 0.068 | 1 | 0.898226899 | 0.650932472 |
| B9d1 | 0.409235236 | 0.598329246 | 0.226 | 0.186 | 1 | 0.770240849 | 0.164779508 | 0.171 | 0.15 | 1 | 0.770240849 | 0.650996994 |
| Dis3l | 0.595789058 | 0.417757001 | 0.094 | 0.072 | 1 | 0.410035756 | -0.665295491 | 0.098 | 0.143 | 1 | 0.595789058 | 0.651942191 |
| Gm48604 | 0.87899663 | 0.8327945 | 0.038 | 0.034 | 1 | 0.410058594 | 1.334704509 | 0.073 | 0.044 | 1 | 0.87899663 | 0.651969138 |
| Gm29266 | 0.410506039 | -3.112063946 | 0.019 | 0.042 | 1 | 0.455097252 | -1.16279515 | 0 | 0.014 | 1 | 0.455097252 | 0.65249687 |
| Dync2i2 | 0.41056936 | 0.526691372 | 0.17 | 0.127 | 1 | 0.848418482 | 1.289717055 | 0.098 | 0.109 | 1 | 0.848418482 | 0.652571521 |
| Aldh1a3 | 0.410638789 | -0.175922717 | 0.226 | 0.288 | 1 | 0.616177071 | 0.422167351 | 0.171 | 0.147 | 1 | 0.616177071 | 0.652653363 |
| Bc1 | 0.803460347 | -0.357727406 | 0.132 | 0.119 | 1 | 0.411111063 | 1.174239837 | 0.122 | 0.085 | 1 | 0.803460347 | 0.65320982 |
| Det1 | 0.461645427 | -0.582242999 | 0.038 | 0.064 | 1 | 0.411341917 | 0.515276755 | 0.049 | 0.089 | 1 | 0.461645427 | 0.653481662 |
| Adck5 | 0.411375766 | 0.932330173 | 0.113 | 0.081 | 1 | 0.570568845 | 1.000703582 | 0.098 | 0.075 | 1 | 0.570568845 | 0.653521511 |
| Gm57100 | 0.438760049 | -0.368839361 | 0.057 | 0.089 | 1 | 0.411849875 | -0.250257991 | 0.024 | 0.055 | 1 | 0.438760049 | 0.65407943 |
| Upk1a | 0.412009681 | 1.792152515 | 0.057 | 0.034 | 1 | 0.523398916 | 0.351778023 | 0.049 | 0.078 | 1 | 0.523398916 | 0.654267385 |
| 2700054A10Rik | 0.412051988 | -1.015202407 | 0.019 | 0.042 | 1 | 0.596439243 | 1.422167351 | 0.024 | 0.014 | 1 | 0.596439243 | 0.654317136 |
| Orc5 | 0.412239942 | -0.652632327 | 0.208 | 0.25 | 1 | 0.859994978 | 0.515276755 | 0.171 | 0.188 | 1 | 0.859994978 | 0.654538114 |
| Klf10 | 0.804713469 | -0.430239906 | 0.604 | 0.547 | 1 | 0.412501635 | -0.515861757 | 0.561 | 0.597 | 1 | 0.804713469 | 0.654845671 |
| Cpt1c | 0.412542809 | 0.8327945 | 0.208 | 0.169 | 1 | 0.448733233 | 0.450181727 | 0.195 | 0.157 | 1 | 0.448733233 | 0.654894049 |
| Thap2 | 0.412709978 | 0.297462767 | 0.566 | 0.53 | 1 | 0.733600376 | 0.271150812 | 0.561 | 0.546 | 1 | 0.733600376 | 0.655090431 |
| Spag16 | 0.413080549 | -0.1672055 | 0 | 0.013 | 1 | 0.455094184 | 0.252242349 | 0 | 0.014 | 1 | 0.455094184 | 0.655525558 |
| Rgsl1 | 0.413080549 | -0.430239906 | 0 | 0.013 | 1 | 0.51919986 | 0.83720485 | 0 | 0.01 | 1 | 0.51919986 | 0.655525558 |
| Gpr158 | 0.413080549 | -0.1672055 | 0 | 0.013 | 1 | 0.587927635 | 1.83720485 | 0.024 | 0.014 | 1 | 0.587927635 | 0.655525558 |
| Fsip1 | 0.413080549 | -0.1672055 | 0 | 0.013 | 1 | 0.755430318 | -0.484723245 | 0.024 | 0.017 | 1 | 0.755430318 | 0.655525558 |
| Gm12500 | 0.413080549 | -0.1672055 | 0 | 0.013 | 1 | 0.441381862 | 1.515276755 | 0.024 | 0.01 | 1 | 0.441381862 | 0.655525558 |
| Gm11266 | 0.413080549 | -0.1672055 | 0 | 0.013 | 1 | 0.60213981 | 0.83720485 | 0.024 | 0.014 | 1 | 0.60213981 | 0.655525558 |
| Lrp8os3 | 0.413080549 | -0.1672055 | 0 | 0.013 | 1 | 0.605005904 | 0.136765132 | 0.024 | 0.014 | 1 | 0.605005904 | 0.655525558 |
| 9530034E10Rik | 0.413080549 | -0.1672055 | 0 | 0.013 | 1 | 0.441381862 | 1.515276755 | 0.024 | 0.01 | 1 | 0.441381862 | 0.655525558 |
| Spink2 | 0.413080549 | -0.1672055 | 0 | 0.013 | 1 | 0.51919986 | 0.83720485 | 0 | 0.01 | 1 | 0.51919986 | 0.655525558 |
| Oas1c | 0.413080549 | -0.1672055 | 0 | 0.013 | 1 | 0.51919986 | 0.252242349 | 0 | 0.01 | 1 | 0.51919986 | 0.655525558 |
| Gm13844 | 0.413080549 | -0.1672055 | 0 | 0.013 | 1 | 0.989595616 | 0.667279848 | 0.024 | 0.024 | 1 | 0.989595616 | 0.655525558 |
| Gm10610 | 0.413080549 | -0.845277405 | 0 | 0.013 | 1 | 0.455094184 | 0.252242349 | 0 | 0.014 | 1 | 0.455094184 | 0.655525558 |
| Gm47772 | 0.413080549 | -0.430239906 | 0 | 0.013 | 1 | 0.441381862 | 1.252242349 | 0.024 | 0.01 | 1 | 0.441381862 | 0.655525558 |
| Gsta2 | 0.413080549 | -0.430239906 | 0 | 0.013 | 1 | 0.51919986 | 0.83720485 | 0 | 0.01 | 1 | 0.51919986 | 0.655525558 |
| Gm57061 | 0.413080549 | -0.845277405 | 0 | 0.013 | 1 | 0.51919986 | 0.83720485 | 0 | 0.01 | 1 | 0.51919986 | 0.655525558 |
| 4930432B10Rik | 0.413080549 | -0.1672055 | 0 | 0.013 | 1 | 0.51919986 | 0.252242349 | 0 | 0.01 | 1 | 0.51919986 | 0.655525558 |
| Txlnb | 0.413080549 | -0.1672055 | 0 | 0.013 | 1 | 0.51919986 | -0.16279515 | 0 | 0.01 | 1 | 0.51919986 | 0.655525558 |
| Pcdh15 | 0.413080549 | -0.1672055 | 0 | 0.013 | 1 | 0.455098275 | 0.515276755 | 0 | 0.014 | 1 | 0.455098275 | 0.655525558 |
| Klhl10 | 0.413080549 | -0.1672055 | 0 | 0.013 | 1 | 0.60213981 | 1.252242349 | 0.024 | 0.014 | 1 | 0.60213981 | 0.655525558 |
| Gm49291 | 0.413080549 | -0.1672055 | 0 | 0.013 | 1 | 0.455098275 | 0.515276755 | 0 | 0.014 | 1 | 0.455098275 | 0.655525558 |
| Gm15327 | 0.413080549 | -0.1672055 | 0 | 0.013 | 1 | 0.51919986 | 0.83720485 | 0 | 0.01 | 1 | 0.51919986 | 0.655525558 |
| B3gnt5 | 0.413080549 | -0.1672055 | 0 | 0.013 | 1 | 0.455097252 | -0.622226769 | 0 | 0.014 | 1 | 0.455097252 | 0.655525558 |
| Dynlt2a3 | 0.413080549 | -0.1672055 | 0 | 0.013 | 1 | 0.438672228 | 1.83720485 | 0.024 | 0.01 | 1 | 0.438672228 | 0.655525558 |
| Gm30984 | 0.413080549 | -0.1672055 | 0 | 0.013 | 1 | 0.51919986 | 0.83720485 | 0 | 0.01 | 1 | 0.51919986 | 0.655525558 |
| Gm94 | 0.413080549 | -0.1672055 | 0 | 0.013 | 1 | 0.455098275 | 0.515276755 | 0 | 0.014 | 1 | 0.455098275 | 0.655525558 |
| Slc16a12 | 0.413080549 | -0.1672055 | 0 | 0.013 | 1 | 0.51919986 | 0.252242349 | 0 | 0.01 | 1 | 0.51919986 | 0.655525558 |
| Emx2os | 0.413080549 | -0.430239906 | 0 | 0.013 | 1 | 0.746881137 | 1.029849928 | 0.024 | 0.017 | 1 | 0.746881137 | 0.655525558 |
| Gm26901 | 0.413082433 | 0.154722595 | 0 | 0.013 | 1 | 0.430584486 | 2.422167351 | 0.024 | 0.01 | 1 | 0.430584486 | 0.65552777 |
| Gm19497 | 0.413082433 | 0.154722595 | 0 | 0.013 | 1 | 0.438672228 | 1.83720485 | 0.024 | 0.01 | 1 | 0.438672228 | 0.65552777 |
| Gm37748 | 0.413082433 | 0.154722595 | 0 | 0.013 | 1 | 0.5191986 | 0.515276755 | 0 | 0.01 | 1 | 0.5191986 | 0.65552777 |
| Gm32250 | 0.413082433 | 0.154722595 | 0 | 0.013 | 1 | 0.529718256 | 1.515276755 | 0.049 | 0.031 | 1 | 0.529718256 | 0.65552777 |
| C8g | 0.413082433 | 0.154722595 | 0 | 0.013 | 1 | 0.744047924 | 1.252242349 | 0.024 | 0.017 | 1 | 0.744047924 | 0.65552777 |
| 1700084E18Rik | 0.413082433 | 0.154722595 | 0 | 0.013 | 1 | 0.455094184 | 0.252242349 | 0 | 0.014 | 1 | 0.455094184 | 0.65552777 |
| Ttc30a1 | 0.413082433 | 0.154722595 | 0 | 0.013 | 1 | 0.51919986 | 0.83720485 | 0 | 0.01 | 1 | 0.51919986 | 0.65552777 |
| Gm14302 | 0.413082433 | 0.154722595 | 0 | 0.013 | 1 | 0.455098275 | 0.515276755 | 0 | 0.014 | 1 | 0.455098275 | 0.65552777 |
| Tmbim7 | 0.413082433 | 0.154722595 | 0 | 0.013 | 1 | 0.51919986 | -0.622226769 | 0 | 0.01 | 1 | 0.51919986 | 0.65552777 |
| Oas1a | 0.413082433 | -1.1672055 | 0 | 0.013 | 1 | 0.60213981 | 1.029849928 | 0.024 | 0.014 | 1 | 0.60213981 | 0.65552777 |
| Gm15747 | 0.413082433 | 0.154722595 | 0 | 0.013 | 1 | 0.744047924 | 1.252242349 | 0.024 | 0.017 | 1 | 0.744047924 | 0.65552777 |
| Gm15997 | 0.413082433 | -0.430239906 | 0 | 0.013 | 1 | 0.51919986 | 0.83720485 | 0 | 0.01 | 1 | 0.51919986 | 0.65552777 |
| Epha1 | 0.413082433 | 0.154722595 | 0 | 0.013 | 1 | 0.550031708 | 0.721727632 | 0.049 | 0.031 | 1 | 0.550031708 | 0.65552777 |
| Gm15402 | 0.413082433 | -0.430239906 | 0 | 0.013 | 1 | 0.744047924 | 1.252242349 | 0.024 | 0.017 | 1 | 0.744047924 | 0.65552777 |
| Slco1a6 | 0.413082433 | 0.154722595 | 0 | 0.013 | 1 | 0.51919986 | 0.252242349 | 0 | 0.01 | 1 | 0.51919986 | 0.65552777 |
| Klk14 | 0.413082433 | 0.154722595 | 0 | 0.013 | 1 | 0.51919986 | 0.83720485 | 0 | 0.01 | 1 | 0.51919986 | 0.65552777 |
| 1500002F19Rik | 0.413082433 | 0.154722595 | 0 | 0.013 | 1 | 0.599290422 | 1.515276755 | 0.024 | 0.014 | 1 | 0.599290422 | 0.65552777 |
| Rassf7 | 0.413082433 | 0.154722595 | 0 | 0.013 | 1 | 0.60213981 | 1.252242349 | 0.024 | 0.014 | 1 | 0.60213981 | 0.65552777 |
| Proscos | 0.413082433 | 0.154722595 | 0 | 0.013 | 1 | 0.51919986 | 0.83720485 | 0 | 0.01 | 1 | 0.51919986 | 0.65552777 |
| Snx20 | 0.413082433 | 0.154722595 | 0 | 0.013 | 1 | 0.5191986 | 0.515276755 | 0 | 0.01 | 1 | 0.5191986 | 0.65552777 |
| Adgrg3 | 0.413082433 | 0.154722595 | 0 | 0.013 | 1 | 0.815009983 | -1.116991461 | 0.024 | 0.031 | 1 | 0.815009983 | 0.65552777 |
| Gm10629 | 0.413082433 | 0.154722595 | 0 | 0.013 | 1 | 0.438672228 | 1.83720485 | 0.024 | 0.01 | 1 | 0.438672228 | 0.65552777 |
| Angptl6 | 0.413082433 | 0.154722595 | 0 | 0.013 | 1 | 0.51919986 | 0.83720485 | 0 | 0.01 | 1 | 0.51919986 | 0.65552777 |
| Gm47204 | 0.413082433 | 0.154722595 | 0 | 0.013 | 1 | 0.914200263 | 0.515276755 | 0.024 | 0.027 | 1 | 0.914200263 | 0.65552777 |
| Nbeal2 | 0.413082433 | 0.154722595 | 0 | 0.013 | 1 | 0.60213981 | 1.252242349 | 0.024 | 0.014 | 1 | 0.60213981 | 0.65552777 |
| Gm20597 | 0.413082433 | 0.154722595 | 0 | 0.013 | 1 | 0.455097252 | -0.332720152 | 0 | 0.014 | 1 | 0.455097252 | 0.65552777 |
| Susd2 | 0.413082433 | 0.154722595 | 0 | 0.013 | 1 | 0.455094184 | 0.252242349 | 0 | 0.014 | 1 | 0.455094184 | 0.65552777 |
| Mmp19 | 0.413082433 | 0.154722595 | 0 | 0.013 | 1 | 0.455098275 | 0.515276755 | 0 | 0.014 | 1 | 0.455098275 | 0.65552777 |
| Gm12059 | 0.413082433 | -0.652632327 | 0 | 0.013 | 1 | 0.438672228 | 1.83720485 | 0.024 | 0.01 | 1 | 0.438672228 | 0.65552777 |

| | | | | | | | | | | | | |
|---|---|---|---|---|---|---|---|---|---|---|---|---|
| 1700003D09Rik | 0.413082433 | 0.154722595 | 0 | 0.013 | 1 | 0.455098275 | 0.515276755 | 0 | 0.014 | 1 | 0.455098275 | 0.65552777 |
| Kif19a | 0.413082433 | 0.154722595 | 0 | 0.013 | 1 | 0.441381862 | 1.515276755 | 0.024 | 0.01 | 1 | 0.441381862 | 0.65552777 |
| Zfp750 | 0.413082433 | 0.154722595 | 0 | 0.013 | 1 | 0.455098275 | 0.515276755 | 0 | 0.014 | 1 | 0.455098275 | 0.65552777 |
| Myt1l | 0.413082433 | 0.154722595 | 0 | 0.013 | 1 | 0.452283702 | 0.88967227 | 0.049 | 0.027 | 1 | 0.452283702 | 0.65552777 |
| Tmem63c | 0.413082433 | 0.154722595 | 0 | 0.013 | 1 | 0.51919986 | 0.83720485 | 0 | 0.01 | 1 | 0.51919986 | 0.65552777 |
| 3300002A11Rik | 0.413082433 | -0.430239906 | 0 | 0.013 | 1 | 0.746881137 | 1.029849928 | 0.024 | 0.017 | 1 | 0.746881137 | 0.65552777 |
| Serpinb6c | 0.413082433 | 0.154722595 | 0 | 0.013 | 1 | 0.60213981 | 1.029849928 | 0.024 | 0.014 | 1 | 0.60213981 | 0.65552777 |
| Gm5083 | 0.413082433 | 0.154722595 | 0 | 0.013 | 1 | 0.455098275 | 0.515276755 | 0 | 0.014 | 1 | 0.455098275 | 0.65552777 |
| Gm48342 | 0.413082433 | 0.154722595 | 0 | 0.013 | 1 | 0.455098275 | 0.515276755 | 0 | 0.014 | 1 | 0.455098275 | 0.65552777 |
| Scel | 0.413082433 | -0.652632327 | 0 | 0.013 | 1 | 0.51919986 | 0.83720485 | 0 | 0.01 | 1 | 0.51919986 | 0.65552777 |
| Mal2 | 0.413082433 | 0.154722595 | 0 | 0.013 | 1 | 0.440689737 | 0.83720485 | 0.098 | 0.065 | 1 | 0.440689737 | 0.65552777 |
| Zfp41 | 0.413082433 | 0.154722595 | 0 | 0.013 | 1 | 0.856404759 | 2.029849928 | 0.024 | 0.02 | 1 | 0.856404759 | 0.65552777 |
| Muc13 | 0.413082433 | 0.154722595 | 0 | 0.013 | 1 | 0.51919986 | 0.83720485 | 0 | 0.01 | 1 | 0.51919986 | 0.65552777 |
| Gm3435 | 0.413082433 | 0.154722595 | 0 | 0.013 | 1 | 0.914200263 | 0.515276755 | 0.024 | 0.027 | 1 | 0.914200263 | 0.65552777 |
| Gm20468 | 0.413082433 | 0.154722595 | 0 | 0.013 | 1 | 0.746881137 | 0.136765132 | 0.024 | 0.017 | 1 | 0.746881137 | 0.65552777 |
| Plin4 | 0.413082433 | 0.154722595 | 0 | 0.013 | 1 | 0.455098275 | 0.515276755 | 0 | 0.014 | 1 | 0.455098275 | 0.65552777 |
| Crb3 | 0.413082433 | 0.154722595 | 0 | 0.013 | 1 | 0.746881137 | 1.029849928 | 0.024 | 0.017 | 1 | 0.746881137 | 0.65552777 |
| Gm50209 | 0.413082433 | 0.154722595 | 0 | 0.013 | 1 | 0.455094184 | -0.16279515 | 0 | 0.014 | 1 | 0.455094184 | 0.65552777 |
| Syt7 | 0.413082433 | 0.154722595 | 0 | 0.013 | 1 | 0.605005904 | 0.83720485 | 0.024 | 0.014 | 1 | 0.605005904 | 0.65552777 |
| Gm34235 | 0.413082433 | 0.154722595 | 0 | 0.013 | 1 | 0.430584486 | 2.422167351 | 0.024 | 0.01 | 1 | 0.430584486 | 0.65552777 |
| Gm1141 | 0.413082433 | 0.154722595 | 0 | 0.013 | 1 | 0.51919986 | 0.83720485 | 0 | 0.01 | 1 | 0.51919986 | 0.65552777 |
| Peg13 | 0.579510289 | 0.569760094 | 0.094 | 0.072 | 1 | 0.413298036 | 0.786578777 | 0.049 | 0.089 | 1 | 0.579510289 | 0.655780806 |
| Ppp1r13l | 0.413504844 | 0.865215978 | 0.113 | 0.081 | 1 | 0.449074365 | 0.274268655 | 0.122 | 0.174 | 1 | 0.449074365 | 0.656023432 |
| Sumo3 | 0.546693408 | 0.267197324 | 0.623 | 0.627 | 1 | 0.414697236 | 0.275701322 | 0.707 | 0.669 | 1 | 0.546693408 | 0.657420675 |
| Dqx1 | 0.415177231 | -2.489133595 | 0.019 | 0.042 | 1 | 0.864610878 | 1.100239256 | 0.024 | 0.02 | 1 | 0.864610878 | 0.657982328 |
| Ube4bos1 | 0.415177758 | -0.752168001 | 0.019 | 0.042 | 1 | 0.990690158 | 0.252242349 | 0.049 | 0.048 | 1 | 0.990690158 | 0.657982946 |
| Clspn | 0.415204138 | -1.719746523 | 0.019 | 0.042 | 1 | 0.606262835 | -0.250257991 | 0.024 | 0.041 | 1 | 0.606262835 | 0.6580138 |
| Bbln | 0.792947192 | -0.1672055 | 0.472 | 0.466 | 1 | 0.415333892 | 0.328191202 | 0.39 | 0.345 | 1 | 0.792947192 | 0.658165543 |
| Gm10643 | 0.415548353 | -0.347777746 | 0.038 | 0.068 | 1 | 0.476459603 | 1.749742009 | 0.073 | 0.048 | 1 | 0.476459603 | 0.658416272 |
| Lclat1 | 0.491583401 | -0.353072045 | 0.396 | 0.453 | 1 | 0.415826067 | 0.686645173 | 0.293 | 0.256 | 1 | 0.491583401 | 0.658740816 |
| Dtna | 0.645249533 | 0.695290976 | 0.075 | 0.059 | 1 | 0.415873019 | -1.377808041 | 0.098 | 0.14 | 1 | 0.645249533 | 0.65879567 |
| Nmb | 0.732691937 | 1.154722595 | 0.019 | 0.013 | 1 | 0.415956482 | -0.126269274 | 0.098 | 0.061 | 1 | 0.732691937 | 0.658893169 |
| Scn2b | 0.416734858 | -0.752168001 | 0.019 | 0.042 | 1 | 0.983241205 | 0.422167351 | 0.049 | 0.048 | 1 | 0.983241205 | 0.659801774 |
| Mycbp | 0.796578244 | -0.135954566 | 0.358 | 0.373 | 1 | 0.417444699 | 0.45387621 | 0.317 | 0.263 | 1 | 0.796578244 | 0.660629321 |
| Gm13684 | 0.768911057 | -0.582242999 | 0.038 | 0.047 | 1 | 0.417566431 | 0.944120054 | 0.098 | 0.065 | 1 | 0.768911057 | 0.660771138 |
| Tk2 | 0.417734289 | 0.8327945 | 0.132 | 0.097 | 1 | 0.888866644 | 0.47463477 | 0.098 | 0.109 | 1 | 0.888866644 | 0.660966642 |
| Snhg9 | 0.417839588 | 0.865215978 | 0.113 | 0.081 | 1 | 0.787967501 | 0.377773231 | 0.073 | 0.061 | 1 | 0.787967501 | 0.661089254 |
| Gm45494 | 0.418292339 | -0.545717123 | 0.019 | 0.042 | 1 | 0.878310931 | 0.515276755 | 0.024 | 0.02 | 1 | 0.878310931 | 0.661616197 |
| Gm11613 | 0.418292339 | -0.932740246 | 0.019 | 0.042 | 1 | 0.665727056 | -0.332720152 | 0.073 | 0.092 | 1 | 0.665727056 | 0.661616197 |
| Map1s | 0.418785003 | 0.491757582 | 0.245 | 0.195 | 1 | 0.637533643 | 0.294677615 | 0.244 | 0.311 | 1 | 0.637533643 | 0.662189127 |
| Hspbp1 | 0.675513661 | 0.311448098 | 0.245 | 0.225 | 1 | 0.419113584 | -0.380025866 | 0.195 | 0.249 | 1 | 0.675513661 | 0.662570972 |
| 4933433G15Rik | 0.419848106 | -0.430239906 | 0.019 | 0.042 | 1 | 0.972078811 | 0.515276755 | 0.049 | 0.048 | 1 | 0.972078811 | 0.66342378 |
| 1700110C19Rik | 0.419848106 | -0.545717123 | 0.019 | 0.042 | 1 | 0.481233368 | 1.120997816 | 0.098 | 0.068 | 1 | 0.481233368 | 0.66342378 |
| Bag6 | 0.419870661 | 0.332720897 | 0.377 | 0.331 | 1 | 0.987318908 | 0.346457691 | 0.366 | 0.396 | 1 | 0.987318908 | 0.66344995 |
| Abcf3 | 0.419871317 | 0.377115016 | 0.358 | 0.309 | 1 | 0.57318004 | -0.143686327 | 0.317 | 0.369 | 1 | 0.57318004 | 0.663450711 |
| Enpp4 | 0.419980609 | 0.631160639 | 0.208 | 0.169 | 1 | 0.660969823 | 0.313642894 | 0.098 | 0.123 | 1 | 0.660969823 | 0.663577506 |
| Gm48677 | 0.420442892 | -0.430239906 | 0.038 | 0.068 | 1 | 0.441381862 | 1.252242349 | 0.024 | 0.01 | 1 | 0.441381862 | 0.664113558 |
| Ttc21a | 0.77169508 | 0.417757001 | 0.038 | 0.03 | 1 | 0.420803657 | 0.944120054 | 0.073 | 0.044 | 1 | 0.77169508 | 0.664531596 |
| Tatdn1 | 0.859257966 | 0.44102678 | 0.283 | 0.28 | 1 | 0.421079369 | -0.123624553 | 0.195 | 0.259 | 1 | 0.859257966 | 0.664850903 |
| Mpv17l | 0.583031535 | -0.311595409 | 0.208 | 0.25 | 1 | 0.421368229 | -0.605738646 | 0.146 | 0.195 | 1 | 0.583031535 | 0.665185274 |
| 1810014B01Rik | 0.421517727 | -0.304709024 | 0.019 | 0.042 | 1 | 0.986995563 | 0.83720485 | 0.024 | 0.024 | 1 | 0.986995563 | 0.66535826 |
| Psmg1 | 0.422181824 | 0.472898555 | 0.321 | 0.275 | 1 | 0.57954288 | 0.515276755 | 0.317 | 0.304 | 1 | 0.57954288 | 0.666126155 |
| Adam12 | 0.9351967 | -0.354688873 | 0.321 | 0.305 | 1 | 0.422231561 | -0.27906484 | 0.317 | 0.386 | 1 | 0.9351967 | 0.666183631 |
| Pus7 | 0.423025638 | -0.240578686 | 0.264 | 0.331 | 1 | 0.540882157 | -0.254293505 | 0.268 | 0.334 | 1 | 0.540882157 | 0.667100586 |
| Klhdc1 | 0.423331006 | 0.313420341 | 0.17 | 0.127 | 1 | 0.893489698 | 0.454735213 | 0.146 | 0.167 | 1 | 0.893489698 | 0.667452871 |
| Mcrs1 | 0.606872461 | 0.44102678 | 0.321 | 0.301 | 1 | 0.423702901 | -0.257952383 | 0.195 | 0.256 | 1 | 0.606872461 | 0.667881653 |
| Cldn3 | 0.424607473 | 1.306725688 | 0.057 | 0.034 | 1 | 0.749278309 | 0.444887427 | 0.122 | 0.106 | 1 | 0.749278309 | 0.66892344 |
| Bloc1s3 | 0.960852989 | 0.47665069 | 0.057 | 0.059 | 1 | 0.424747458 | 0.930314254 | 0.073 | 0.044 | 1 | 0.960852989 | 0.669084513 |
| Slc30a4 | 0.42480443 | 0.360306728 | 0.377 | 0.335 | 1 | 0.837289657 | -0.183454621 | 0.39 | 0.403 | 1 | 0.837289657 | 0.669150056 |
| Lrrc32 | 0.775146659 | 0.377115016 | 0.075 | 0.089 | 1 | 0.425426486 | 1.422167351 | 0.098 | 0.065 | 1 | 0.775146659 | 0.669865277 |
| Adamtsl3 | 0.570917816 | -1.057464443 | 0.66 | 0.708 | 1 | 0.42581839 | 0.461143318 | 0.683 | 0.792 | 1 | 0.570917816 | 0.670315478 |
| Ppp3cc | 0.425943242 | -0.783876861 | 0.415 | 0.445 | 1 | 0.510401281 | 0.658401697 | 0.268 | 0.249 | 1 | 0.510401281 | 0.670458839 |
| D330041H03Rik | 0.425949632 | 0.80108564 | 0.094 | 0.064 | 1 | 0.503613225 | -0.484723255 | 0.073 | 0.106 | 1 | 0.503613225 | 0.670466175 |
| Atp6v1c1 | 0.426948578 | 0.319781841 | 0.415 | 0.36 | 1 | 0.987633703 | -0.16279515 | 0.415 | 0.437 | 1 | 0.987633703 | 0.671612068 |
| Ssx2ip | 0.428024008 | -0.132853995 | 0.189 | 0.25 | 1 | 0.462760468 | 0.118386602 | 0.268 | 0.334 | 1 | 0.462760468 | 0.672843465 |
| Bst2 | 0.605137469 | -1.446395674 | 0.34 | 0.284 | 1 | 0.428329954 | -0.190275887 | 0.341 | 0.287 | 1 | 0.605137469 | 0.673193358 |
| Ttll11 | 0.428356002 | 0.640149422 | 0.094 | 0.064 | 1 | 0.858365504 | -0.555112573 | 0.073 | 0.078 | 1 | 0.858365504 | 0.67322314 |
| Lmbr1l | 0.428737498 | 0.446903346 | 0.453 | 0.403 | 1 | 0.479960461 | 0.189958071 | 0.463 | 0.433 | 1 | 0.479960461 | 0.673659154 |
| Sec23b | 0.507438488 | 0.339754489 | 0.509 | 0.492 | 1 | 0.428772915 | 0.433091 | 0.659 | 0.601 | 1 | 0.507438488 | 0.673699617 |
| 2510039O18Rik | 0.428807855 | 0.406261362 | 0.321 | 0.271 | 1 | 0.967052485 | 0.217052921 | 0.39 | 0.413 | 1 | 0.967052485 | 0.673739534 |
| Adck2 | 0.428877595 | -0.600164907 | 0.057 | 0.089 | 1 | 0.695253371 | 1.029849928 | 0.146 | 0.13 | 1 | 0.695253371 | 0.673819199 |
| Rundc1 | 0.470381763 | 0.389187848 | 0.151 | 0.114 | 1 | 0.429196713 | 0.656632604 | 0.22 | 0.177 | 1 | 0.470381763 | 0.674183608 |
| Celsr2 | 0.429421714 | 1.154722595 | 0.057 | 0.034 | 1 | 0.987896638 | 0.152706676 | 0.122 | 0.123 | 1 | 0.987896638 | 0.674440419 |
| Wrap53 | 0.529499066 | 0.154722595 | 0.094 | 0.068 | 1 | 0.430381562 | 0.359157553 | 0.146 | 0.102 | 1 | 0.529499066 | 0.675534835 |
| Ftcd | 0.507281812 | 1.154722595 | 0.019 | 0.008 | 1 | 0.430584486 | 2.422167351 | 0.024 | 0.01 | 1 | 0.507281812 | 0.675765972 |
| Hs3st1 | 0.510606779 | 0.347367673 | 0.019 | 0.008 | 1 | 0.430584486 | 2.422167351 | 0.024 | 0.01 | 1 | 0.510606779 | 0.675765972 |
| Kif22 | 0.638059929 | -0.347777746 | 0.038 | 0.025 | 1 | 0.430584486 | 2.422167351 | 0.024 | 0.01 | 1 | 0.638059929 | 0.675765972 |
| Clec4a1 | 0.732691937 | 1.154722595 | 0.019 | 0.013 | 1 | 0.430584486 | 2.422167351 | 0.024 | 0.01 | 1 | 0.732691937 | 0.675765972 |
| 4833408A19Rik | 0.926212226 | 0.8327945 | 0.019 | 0.017 | 1 | 0.430584486 | 2.83720485 | 0.024 | 0.01 | 1 | 0.926212226 | 0.675765972 |
| Lingo2 | 0.729311193 | 1.154722595 | 0.019 | 0.013 | 1 | 0.430587622 | 2.83720485 | 0.024 | 0.01 | 1 | 0.729311193 | 0.675769544 |
| Elac1 | 0.430925503 | 0.739685096 | 0.113 | 0.081 | 1 | 0.628284378 | -0.278272368 | 0.049 | 0.068 | 1 | 0.628284378 | 0.676154217 |
| Gm13391 | 0.431025553 | 0.891688189 | 0.057 | 0.034 | 1 | 0.703206616 | 0.79281073 | 0.098 | 0.082 | 1 | 0.703206616 | 0.676268079 |
| Rnf5 | 0.682763936 | -0.105144532 | 0.264 | 0.301 | 1 | 0.431041102 | 1.156027034 | 0.293 | 0.27 | 1 | 0.682763936 | 0.676285773 |
| Gm26549 | 0.669818222 | 0.197791317 | 0.189 | 0.22 | 1 | 0.431090434 | -0.303276374 | 0.146 | 0.201 | 1 | 0.669818222 | 0.676341905 |
| Plekhm1 | 0.431316667 | -0.477545621 | 0.189 | 0.233 | 1 | 0.784685473 | 0.136765132 | 0.317 | 0.3 | 1 | 0.784685473 | 0.676599267 |
| Otud6b | 0.431377991 | 0.318221327 | 0.434 | 0.373 | 1 | 0.984149184 | 0.156027034 | 0.366 | 0.389 | 1 | 0.984149184 | 0.676669011 |
| Dmap1 | 0.432017134 | 0.721763188 | 0.113 | 0.081 | 1 | 0.479037536 | -0.250257991 | 0.073 | 0.109 | 1 | 0.479037536 | 0.677395464 |
| 5330438D12Rik | 0.984224548 | 0.137649081 | 0.264 | 0.28 | 1 | 0.432046803 | 0.183762611 | 0.268 | 0.345 | 1 | 0.984224548 | 0.677429166 |
| Mtbp | 0.432172505 | -0.932740246 | 0.075 | 0.11 | 1 | 0.81082337 | -0.448197369 | 0.073 | 0.082 | 1 | 0.81082337 | 0.677571936 |
| Dhx37 | 0.911029143 | -0.1672055 | 0.094 | 0.097 | 1 | 0.432243933 | 0.222495006 | 0.073 | 0.119 | 1 | 0.911029143 | 0.677653048 |
| Zfp52 | 0.432512063 | -1.064445926 | 0.245 | 0.267 | 1 | 0.618872442 | 0.273706858 | 0.488 | 0.461 | 1 | 0.618872442 | 0.677957442 |
| Wdr76 | 0.659664406 | -0.7032584 | 0.057 | 0.072 | 1 | 0.432679313 | 0.667279848 | 0.024 | 0.055 | 1 | 0.659664406 | 0.678142738 |
| Kctd11 | 0.794581651 | -0.323324702 | 0.113 | 0.123 | 1 | 0.432720149 | 0.43121249 | 0.098 | 0.154 | 1 | 0.794581651 | 0.678193571 |
| Mpv17l2 | 0.432782941 | -0.343528273 | 0.283 | 0.331 | 1 | 0.544652729 | -0.228884341 | 0.146 | 0.184 | 1 | 0.544652729 | 0.678264808 |
| Cd59a | 0.503971666 | 1.569760094 | 0.019 | 0.008 | 1 | 0.432970668 | 0.589277336 | 0.073 | 0.044 | 1 | 0.503971666 | 0.678477737 |
| Xdh | 0.485184878 | -1.093204919 | 0.019 | 0.038 | 1 | 0.432970939 | 0.515276755 | 0.073 | 0.044 | 1 | 0.485184878 | 0.678478044 |
| Sord | 0.433002157 | 0.657222935 | 0.189 | 0.153 | 1 | 0.582112694 | 0.667279848 | 0.22 | 0.184 | 1 | 0.582112694 | 0.678513446 |
| Vamp9 | 0.668070194 | 0.154722595 | 0.019 | 0.03 | 1 | 0.433272494 | 2.100239256 | 0.024 | 0.01 | 1 | 0.668070194 | 0.678819934 |

| | | | | | | | | | | | | |
|---|---|---|---|---|---|---|---|---|---|---|---|---|
| Adam4 | 0.77169508 | 0.417757001 | 0.038 | 0.03 | 1 | 0.433272494 | 2.100239256 | 0.024 | 0.01 | 1 | 0.77169508 | 0.678819934 |
| Lrguk | 0.797844282 | 1.932330173 | 0.019 | 0.025 | 1 | 0.433272494 | 2.100239256 | 0.024 | 0.01 | 1 | 0.797844282 | 0.678819934 |
| Cox7a2 | 0.433656042 | 0.106655936 | 0.981 | 0.903 | 1 | 0.765656537 | 0.14504329 | 0.854 | 0.915 | 1 | 0.765656537 | 0.679254521 |
| Itpripl2 | 0.53276272 | -0.21745369 | 0.736 | 0.602 | 1 | 0.433758488 | -0.409233045 | 0.634 | 0.631 | 1 | 0.53276272 | 0.67937055 |
| Polr2e | 0.433901785 | 0.428897558 | 0.491 | 0.492 | 1 | 0.7378719 | 0.56635094 | 0.463 | 0.509 | 1 | 0.7378719 | 0.679532811 |
| Mybbp1a | 0.434219922 | 0.326609113 | 0.604 | 0.538 | 1 | 0.500891728 | -0.167082509 | 0.585 | 0.631 | 1 | 0.500891728 | 0.679892903 |
| Zfp873 | 0.434262349 | 1.017219071 | 0.057 | 0.034 | 1 | 0.87556513 | 0.515276755 | 0.024 | 0.02 | 1 | 0.87556513 | 0.67994091 |
| Gm37768 | 0.434276384 | 1.347367673 | 0.057 | 0.034 | 1 | 0.722125564 | -0.250257991 | 0.073 | 0.089 | 1 | 0.722125564 | 0.67995679 |
| Gm34728 | 0.732691937 | 1.154722595 | 0.019 | 0.013 | 1 | 0.434407618 | 0.635570989 | 0.073 | 0.044 | 1 | 0.732691937 | 0.680105257 |
| Parp2 | 0.434912832 | 0.347367673 | 0.377 | 0.335 | 1 | 0.81421993 | -0.264333177 | 0.244 | 0.253 | 1 | 0.81421993 | 0.680676493 |
| Atp13a2 | 0.901698096 | 0.280253477 | 0.038 | 0.034 | 1 | 0.435238881 | -0.20718927 | 0.049 | 0.085 | 1 | 0.901698096 | 0.681044878 |
| Ruvbl2 | 0.704818705 | 0.182736971 | 0.208 | 0.182 | 1 | 0.435381781 | -0.16279515 | 0.146 | 0.198 | 1 | 0.704818705 | 0.681206267 |
| Tmem198b | 0.435456367 | -0.430239906 | 0.057 | 0.089 | 1 | 0.897875248 | 0.136765132 | 0.073 | 0.078 | 1 | 0.897875248 | 0.681290487 |
| 4930527F14Rik | 0.435857866 | 0.154722595 | 0.019 | 0.042 | 1 | 0.450520938 | 0.911205431 | 0.049 | 0.027 | 1 | 0.450520938 | 0.681743652 |
| Pcdhgb7 | 0.435857866 | 0.154722595 | 0.019 | 0.042 | 1 | 0.674180157 | 0.252242349 | 0.024 | 0.038 | 1 | 0.674180157 | 0.681743652 |
| Tctn1 | 0.933942263 | 0.154722595 | 0.075 | 0.072 | 1 | 0.435915141 | -0.520347155 | 0.073 | 0.113 | 1 | 0.933942263 | 0.681808272 |
| Gm47923 | 0.864193501 | -0.845277405 | 0.038 | 0.042 | 1 | 0.435968064 | 1.83720485 | 0.024 | 0.01 | 1 | 0.864193501 | 0.681867976 |
| Hcfc1r1 | 0.70022973 | -0.109550708 | 0.774 | 0.754 | 1 | 0.436517985 | 0.158405963 | 0.732 | 0.669 | 1 | 0.70022973 | 0.682488019 |
| Gm41611 | 0.437452221 | 0.280253477 | 0.019 | 0.042 | 1 | 0.51919986 | 0.83720485 | 0 | 0.01 | 1 | 0.51919986 | 0.683539996 |
| Frem2 | 0.437537757 | 0.984797593 | 0.057 | 0.034 | 1 | 0.438672228 | 1.83720485 | 0.024 | 0.01 | 1 | 0.438672228 | 0.683636225 |
| Gm56867 | 0.437537757 | 0.984797593 | 0.057 | 0.034 | 1 | 0.900433136 | 0.614812428 | 0.049 | 0.044 | 1 | 0.900433136 | 0.683636225 |
| Zfp2 | 0.437537757 | 0.984797593 | 0.057 | 0.034 | 1 | 0.903375428 | 0.667279848 | 0.049 | 0.055 | 1 | 0.903375428 | 0.683636225 |
| Bnip1 | 0.437807982 | 0.417757001 | 0.245 | 0.199 | 1 | 0.4967839 | 0.515276755 | 0.22 | 0.184 | 1 | 0.4967839 | 0.683940135 |
| Fstl5 | 0.61536875 | 1.154722595 | 0.038 | 0.025 | 1 | 0.438021043 | 1.83720485 | 0.098 | 0.068 | 1 | 0.61536875 | 0.684179652 |
| Pfkfb2 | 0.438160896 | 0.669295768 | 0.17 | 0.131 | 1 | 0.915380293 | -0.294500829 | 0.317 | 0.297 | 1 | 0.915380293 | 0.684336821 |
| 9130410C08Rik | 0.438209469 | -0.247375849 | 0.038 | 0.068 | 1 | 0.523152099 | -0.825760163 | 0.049 | 0.075 | 1 | 0.523152099 | 0.684391399 |
| Gna12 | 0.438226793 | -0.176056418 | 0.604 | 0.691 | 1 | 0.719044933 | 0.198855136 | 0.683 | 0.683 | 1 | 0.719044933 | 0.684410864 |
| Ccdc181 | 0.48113705 | 0.776210972 | 0.075 | 0.051 | 1 | 0.43828695 | 1.667279848 | 0.049 | 0.027 | 1 | 0.48113705 | 0.684478449 |
| Nudt5 | 0.973756676 | -0.123812905 | 0.377 | 0.373 | 1 | 0.438635043 | -0.101394606 | 0.244 | 0.304 | 1 | 0.973756676 | 0.684869385 |
| Gm36411 | 0.447253949 | 0.695290976 | 0.057 | 0.034 | 1 | 0.438672228 | 1.83720485 | 0.024 | 0.01 | 1 | 0.447253949 | 0.684911133 |
| Gm49492 | 0.471154689 | 1.569760094 | 0.038 | 0.021 | 1 | 0.438672228 | 1.83720485 | 0.024 | 0.01 | 1 | 0.471154689 | 0.684911133 |
| 4933408J17Rik | 0.48175508 | 1.154722595 | 0.038 | 0.021 | 1 | 0.438672228 | 1.83720485 | 0.024 | 0.01 | 1 | 0.48175508 | 0.684911133 |
| 5730424H11Rik | 0.48175508 | 1.154722595 | 0.038 | 0.021 | 1 | 0.438672228 | 1.83720485 | 0.024 | 0.01 | 1 | 0.48175508 | 0.684911133 |
| Eldr | 0.497857761 | -0.1672055 | 0.019 | 0.038 | 1 | 0.438672228 | 1.83720485 | 0.024 | 0.01 | 1 | 0.497857761 | 0.684911133 |
| A230103J11Rik | 0.503971666 | 1.569760094 | 0.019 | 0.008 | 1 | 0.438672228 | 1.83720485 | 0.024 | 0.01 | 1 | 0.503971666 | 0.684911133 |
| Gm57003 | 0.503971666 | 1.569760094 | 0.019 | 0.008 | 1 | 0.438672228 | 1.83720485 | 0.024 | 0.01 | 1 | 0.503971666 | 0.684911133 |
| Nacad | 0.503971666 | 1.569760094 | 0.019 | 0.008 | 1 | 0.438672228 | 1.83720485 | 0.024 | 0.01 | 1 | 0.503971666 | 0.684911133 |
| Gm26531 | 0.503971666 | 1.569760094 | 0.019 | 0.008 | 1 | 0.438672228 | 1.83720485 | 0.024 | 0.01 | 1 | 0.503971666 | 0.684911133 |
| 1110006O24Rik | 0.571087469 | 0.669295768 | 0.075 | 0.055 | 1 | 0.438672228 | 1.83720485 | 0.024 | 0.01 | 1 | 0.571087469 | 0.684911133 |
| Ddc | 0.679745214 | -0.108311811 | 0.019 | 0.03 | 1 | 0.438672228 | 1.83720485 | 0.024 | 0.01 | 1 | 0.679745214 | 0.684911133 |
| BC037039 | 0.732691937 | 1.154722595 | 0.019 | 0.013 | 1 | 0.438672228 | 1.83720485 | 0.024 | 0.01 | 1 | 0.732691937 | 0.684911133 |
| Tmem30c | 0.776329749 | 0.280253477 | 0.038 | 0.03 | 1 | 0.438672228 | 1.83720485 | 0.024 | 0.01 | 1 | 0.776329749 | 0.684911133 |
| Ttc39aos1 | 0.779443586 | 0.154722595 | 0.019 | 0.025 | 1 | 0.438672228 | 1.83720485 | 0.024 | 0.01 | 1 | 0.779443586 | 0.684911133 |
| Sh3bgr | 0.869949949 | 0.891688189 | 0.038 | 0.034 | 1 | 0.438672228 | 1.83720485 | 0.024 | 0.01 | 1 | 0.869949949 | 0.684911133 |
| Gm56773 | 0.914958 | 0.347367673 | 0.019 | 0.021 | 1 | 0.438672228 | 1.83720485 | 0.024 | 0.01 | 1 | 0.914958 | 0.684911133 |
| 2810459M11Rik | 0.926212226 | 0.8327945 | 0.019 | 0.017 | 1 | 0.438672228 | 1.83720485 | 0.024 | 0.01 | 1 | 0.926212226 | 0.684911133 |
| Fam178b | 0.929410101 | -0.545717123 | 0.019 | 0.017 | 1 | 0.438672228 | 1.83720485 | 0.024 | 0.01 | 1 | 0.929410101 | 0.684911133 |
| Dmrta2 | 0.929410101 | 0.347367673 | 0.019 | 0.017 | 1 | 0.438672228 | 1.83720485 | 0.024 | 0.01 | 1 | 0.929410101 | 0.684911133 |
| Gm47484 | 0.929410101 | 0.347367673 | 0.019 | 0.017 | 1 | 0.438672228 | 1.83720485 | 0.024 | 0.01 | 1 | 0.929410101 | 0.684911133 |
| Slc29a4 | 0.935814214 | 0.154722595 | 0.019 | 0.017 | 1 | 0.438672228 | 1.83720485 | 0.024 | 0.01 | 1 | 0.935814214 | 0.684911133 |
| 1810044D09Rik | 0.991244785 | 0.417757001 | 0.038 | 0.038 | 1 | 0.438672228 | 1.83720485 | 0.024 | 0.01 | 1 | 0.991244785 | 0.684911133 |
| Tchh | 0.438996384 | -0.147840175 | 0.113 | 0.161 | 1 | 0.646976884 | 0.515276755 | 0.049 | 0.034 | 1 | 0.646976884 | 0.685274943 |
| Smad5 | 0.43950174 | 0.212825549 | 0.604 | 0.602 | 1 | 0.555990322 | -0.177265633 | 0.634 | 0.693 | 1 | 0.555990322 | 0.6858417 |
| Grina | 0.687977597 | -0.27154216 | 0.377 | 0.419 | 1 | 0.439679385 | 0.42692088 | 0.561 | 0.519 | 1 | 0.687977597 | 0.686040808 |
| Acsl5 | 0.51228372 | 0.306725688 | 0.189 | 0.153 | 1 | 0.439718844 | -0.36790958 | 0.146 | 0.198 | 1 | 0.51228372 | 0.686085026 |
| Mrpl4 | 0.56783844 | 0.120775263 | 0.547 | 0.5 | 1 | 0.440730971 | 0.264626074 | 0.39 | 0.526 | 1 | 0.56783844 | 0.687218154 |
| Gm29724 | 0.440752287 | 0.776210972 | 0.057 | 0.034 | 1 | 0.58792152 | 2.100239256 | 0.024 | 0.014 | 1 | 0.58792152 | 0.687241996 |
| Bloc1s1 | 0.441225922 | 0.327559192 | 0.415 | 0.364 | 1 | 0.503831799 | 0.130936053 | 0.268 | 0.352 | 1 | 0.503831799 | 0.68777153 |
| Aldh3b3 | 0.497380598 | 2.154722595 | 0.019 | 0.008 | 1 | 0.441381862 | 1.252242349 | 0.024 | 0.01 | 1 | 0.497380598 | 0.687945776 |
| Gm49083 | 0.500671253 | 1.739685096 | 0.019 | 0.008 | 1 | 0.441381862 | 1.029849928 | 0.024 | 0.01 | 1 | 0.500671253 | 0.687945776 |
| Gm29050 | 0.500671253 | 1.739685096 | 0.019 | 0.008 | 1 | 0.441381862 | 1.515276755 | 0.024 | 0.01 | 1 | 0.500671253 | 0.687945776 |
| Asic4 | 0.503971666 | 1.569760094 | 0.019 | 0.008 | 1 | 0.441381862 | 1.515276755 | 0.024 | 0.01 | 1 | 0.503971666 | 0.687945776 |
| Fam83c | 0.503971666 | 1.569760094 | 0.019 | 0.008 | 1 | 0.441381862 | 1.252242349 | 0.024 | 0.01 | 1 | 0.503971666 | 0.687945776 |
| Gm49864 | 0.72256466 | 1.739685096 | 0.019 | 0.013 | 1 | 0.441381862 | 1.252242349 | 0.024 | 0.01 | 1 | 0.72256466 | 0.687945776 |
| Aknaos | 0.74286271 | -0.304709024 | 0.019 | 0.013 | 1 | 0.441381862 | 1.515276755 | 0.024 | 0.01 | 1 | 0.74286271 | 0.687945776 |
| Gm56911 | 0.750976519 | 1.417757001 | 0.038 | 0.03 | 1 | 0.441381862 | 1.515276755 | 0.024 | 0.01 | 1 | 0.750976519 | 0.687945776 |
| Pla2r1 | 0.914958 | -0.845277405 | 0.019 | 0.021 | 1 | 0.441381862 | 1.252242349 | 0.024 | 0.01 | 1 | 0.914958 | 0.687945776 |
| Stmn2 | 0.926211509 | 0.569760094 | 0.019 | 0.017 | 1 | 0.441381862 | 1.515276755 | 0.024 | 0.01 | 1 | 0.926211509 | 0.687945776 |
| Gm47438 | 0.926653673 | 0.739685096 | 0.019 | 0.021 | 1 | 0.441381862 | 1.029849928 | 0.024 | 0.01 | 1 | 0.926653673 | 0.687945776 |
| Tap1 | 0.441516061 | 0.195364579 | 0.132 | 0.093 | 1 | 0.564782929 | 0.911205431 | 0.122 | 0.096 | 1 | 0.564782929 | 0.68809569 |
| Zswim1 | 0.442326061 | 0.8327945 | 0.057 | 0.034 | 1 | 0.94614879 | 0.635570989 | 0.073 | 0.072 | 1 | 0.94614879 | 0.688999778 |
| Zfp82 | 0.442326061 | 0.8327945 | 0.057 | 0.034 | 1 | 0.738418148 | -0.221688839 | 0.049 | 0.061 | 1 | 0.738418148 | 0.688999778 |
| Kics2 | 0.442326061 | 0.8327945 | 0.057 | 0.034 | 1 | 0.745976859 | 0.377773231 | 0.024 | 0.034 | 1 | 0.745976859 | 0.688999778 |
| Ankrd13d | 0.442326061 | 0.569760094 | 0.057 | 0.034 | 1 | 0.95551802 | 0.690363461 | 0.073 | 0.078 | 1 | 0.95551802 | 0.688999778 |
| Baiap2 | 0.457616956 | -0.1672055 | 0.792 | 0.631 | 1 | 0.442376234 | 0.148542299 | 0.854 | 0.744 | 1 | 0.457616956 | 0.689055736 |
| Ctsf | 0.690542121 | -0.151380533 | 0.623 | 0.593 | 1 | 0.442396659 | 0.385993738 | 0.61 | 0.614 | 1 | 0.690542121 | 0.689078514 |
| Ppp1r36 | 0.635884378 | 0.997181318 | 0.113 | 0.093 | 1 | 0.442769278 | 0.114738825 | 0.049 | 0.085 | 1 | 0.635884378 | 0.689493922 |
| Fahd1 | 0.443071943 | 0.569760094 | 0.132 | 0.097 | 1 | 0.751210902 | 0.174239837 | 0.049 | 0.061 | 1 | 0.751210902 | 0.68983114 |
| Mindy4 | 0.443552963 | 0.139772253 | 0.321 | 0.275 | 1 | 0.907163874 | -0.250257991 | 0.293 | 0.294 | 1 | 0.907163874 | 0.690366695 |
| Trmt61a | 0.443899187 | -0.477545621 | 0.075 | 0.11 | 1 | 0.87507634 | 0.404245443 | 0.073 | 0.082 | 1 | 0.87507634 | 0.690751886 |
| Rnf165 | 0.736073999 | 0.8327945 | 0.019 | 0.013 | 1 | 0.44410834 | 1.252242349 | 0.024 | 0.01 | 1 | 0.736073999 | 0.690984462 |
| 1700030K09Rik | 0.445137732 | 0.324647596 | 0.151 | 0.11 | 1 | 0.893549725 | 0.614812428 | 0.073 | 0.068 | 1 | 0.893549725 | 0.692127863 |
| Gm15336 | 0.792583859 | 0.569760094 | 0.019 | 0.025 | 1 | 0.445266316 | 1.614812428 | 0.049 | 0.027 | 1 | 0.792583859 | 0.69227054 |
| Zfp992 | 0.798247423 | -0.130679624 | 0.132 | 0.144 | 1 | 0.445824819 | -0.116991461 | 0.122 | 0.174 | 1 | 0.798247423 | 0.692889869 |
| Sh2b2 | 0.992106036 | 0.318221327 | 0.075 | 0.076 | 1 | 0.446108759 | 1.01123425 | 0.122 | 0.089 | 1 | 0.992106036 | 0.693204493 |
| Zfp438 | 0.782887603 | 0.104096522 | 0.151 | 0.169 | 1 | 0.446688327 | 0.995902596 | 0.122 | 0.085 | 1 | 0.782887603 | 0.693846193 |
| Camk2a | 0.779443586 | 0.154722595 | 0.019 | 0.025 | 1 | 0.446825814 | -0.80665134 | 0.024 | 0.051 | 1 | 0.779443586 | 0.69399832 |
| Adcy10 | 0.756188968 | 1.237184755 | 0.057 | 0.047 | 1 | 0.446826082 | -0.917682652 | 0.024 | 0.051 | 1 | 0.756188968 | 0.693998616 |
| Msl3l2 | 0.592435282 | -0.337130502 | 0.113 | 0.14 | 1 | 0.446876348 | 0.797676486 | 0.146 | 0.109 | 1 | 0.592435282 | 0.694054226 |
| Calr3 | 0.706543665 | 0.80108564 | 0.094 | 0.081 | 1 | 0.446952789 | -0.577832649 | 0.049 | 0.082 | 1 | 0.706543665 | 0.694138782 |
| Clcn7 | 0.446978679 | 0.652222254 | 0.226 | 0.191 | 1 | 0.829583675 | 0.357735478 | 0.268 | 0.311 | 1 | 0.829583675 | 0.694167418 |
| Gm48163 | 0.48175508 | 1.154722595 | 0.038 | 0.021 | 1 | 0.447003204 | 1.136765132 | 0.049 | 0.027 | 1 | 0.48175508 | 0.694194544 |
| Cndp2 | 0.447013717 | 0.438515561 | 0.208 | 0.165 | 1 | 0.748374379 | 0.136765132 | 0.146 | 0.171 | 1 | 0.748374379 | 0.694206171 |
| Gemin2 | 0.769642794 | 0.635348436 | 0.151 | 0.14 | 1 | 0.447044759 | -0.435813645 | 0.098 | 0.14 | 1 | 0.769642794 | 0.694240502 |
| Stap2 | 0.447253949 | 0.695290976 | 0.057 | 0.034 | 1 | 0.599290422 | 1.515276755 | 0.024 | 0.014 | 1 | 0.599290422 | 0.694471803 |
| Nphp4 | 0.447295572 | 0.569760094 | 0.057 | 0.034 | 1 | 0.87556513 | 0.83720485 | 0.024 | 0.02 | 1 | 0.87556513 | 0.694517815 |
| Tex2 | 0.447331744 | 0.251398614 | 0.302 | 0.254 | 1 | 0.487328912 | -0.145517159 | 0.463 | 0.532 | 1 | 0.487328912 | 0.694557799 |
| Vps37a | 0.447865522 | 0.230357033 | 0.642 | 0.695 | 1 | 0.642901046 | -0.21251091 | 0.854 | 0.819 | 1 | 0.642901046 | 0.695147518 |

| | | | | | | | | | | | | |
|---|---|---|---|---|---|---|---|---|---|---|---|---|
| Kctd13 | 0.447929896 | -0.381330305 | 0.075 | 0.11 | 1 | 0.643341782 | 0.30115195 | 0.098 | 0.075 | 1 | 0.643341782 | 0.695218601 |
| Gm34102 | 0.819693881 | 0.932330173 | 0.038 | 0.047 | 1 | 0.448204037 | -0.484723245 | 0.024 | 0.051 | 1 | 0.819693881 | 0.695521215 |
| Cyp4v3 | 0.955983779 | 0.240452469 | 0.132 | 0.136 | 1 | 0.44873847 | -0.885261175 | 0.098 | 0.137 | 1 | 0.955983779 | 0.696110725 |
| Gm31645 | 0.748729549 | 0.569760094 | 0.057 | 0.047 | 1 | 0.448753228 | 0.749742009 | 0.049 | 0.027 | 1 | 0.748729549 | 0.696126996 |
| Spata24 | 0.448827587 | 0.502645898 | 0.189 | 0.148 | 1 | 0.800654532 | 0.759202338 | 0.122 | 0.113 | 1 | 0.800654532 | 0.696208971 |
| Pigh | 0.449036123 | 0.569760094 | 0.132 | 0.097 | 1 | 0.938668711 | 0.941040661 | 0.146 | 0.164 | 1 | 0.938668711 | 0.696438806 |
| Ercc8 | 0.823773582 | 0.46284489 | 0.113 | 0.131 | 1 | 0.449065762 | 0.109284395 | 0.098 | 0.147 | 1 | 0.823773582 | 0.696471465 |
| Gan | 0.449181815 | -0.540422824 | 0.321 | 0.364 | 1 | 0.964953381 | 0.236300805 | 0.341 | 0.382 | 1 | 0.964953381 | 0.696599327 |
| Rnf125 | 0.779443586 | 0.154722595 | 0.019 | 0.025 | 1 | 0.449580774 | -0.555112573 | 0.024 | 0.051 | 1 | 0.779443586 | 0.697038676 |
| Emc2 | 0.449764105 | 0.383809669 | 0.528 | 0.555 | 1 | 0.681709336 | 0.323433397 | 0.512 | 0.526 | 1 | 0.681709336 | 0.69724046 |
| Srp68 | 0.449843165 | 0.32730205 | 0.547 | 0.517 | 1 | 0.813580334 | 0.265417738 | 0.537 | 0.611 | 1 | 0.813580334 | 0.697327457 |
| Scgb2b27 | 0.691130569 | 0.112078257 | 0.283 | 0.314 | 1 | 0.449970348 | 1.204936634 | 0.122 | 0.089 | 1 | 0.691130569 | 0.697467382 |
| Pcdhb19 | 0.497380598 | 2.154722595 | 0.019 | 0.008 | 1 | 0.450513825 | 0.930314254 | 0.049 | 0.027 | 1 | 0.497380598 | 0.698064944 |
| Tmem232 | 0.45058181 | 0.569760094 | 0.057 | 0.034 | 1 | 0.994797912 | -0.80665134 | 0.024 | 0.024 | 1 | 0.994797912 | 0.698139652 |
| Mkrn2 | 0.580450338 | 0.458728782 | 0.283 | 0.254 | 1 | 0.450806174 | 0.144182603 | 0.171 | 0.239 | 1 | 0.580450338 | 0.698386141 |
| Dpy30 | 0.621794543 | 0.24427418 | 0.566 | 0.547 | 1 | 0.450848361 | 0.478123756 | 0.585 | 0.57 | 1 | 0.621794543 | 0.698432477 |
| Brd7 | 0.451097422 | 0.178569337 | 0.585 | 0.559 | 1 | 0.889777195 | 0.156483366 | 0.707 | 0.683 | 1 | 0.889777195 | 0.69870596 |
| Gm11714 | 0.490620203 | -0.1672055 | 0.019 | 0.038 | 1 | 0.451338156 | -0.53202896 | 0.049 | 0.082 | 1 | 0.490620203 | 0.698970181 |
| Osbp2 | 0.885804251 | 0.669295768 | 0.038 | 0.034 | 1 | 0.451529477 | 1.737669176 | 0.073 | 0.048 | 1 | 0.885804251 | 0.699180085 |
| Dnah1 | 0.89261791 | 0.569760094 | 0.038 | 0.034 | 1 | 0.451529477 | 1.515276755 | 0.073 | 0.048 | 1 | 0.89261791 | 0.699180085 |
| Sdccag8 | 0.58924865 | 0.20316295 | 0.453 | 0.432 | 1 | 0.451738076 | 0.265181405 | 0.537 | 0.495 | 1 | 0.58924865 | 0.699408863 |
| Gm48696 | 0.926212226 | 0.8327945 | 0.019 | 0.017 | 1 | 0.452287461 | 1.252242349 | 0.049 | 0.027 | 1 | 0.926212226 | 0.700010975 |
| Gm10509 | 0.729878071 | 0.739685096 | 0.075 | 0.064 | 1 | 0.452331971 | -0.410722664 | 0.024 | 0.051 | 1 | 0.729878071 | 0.70005973 |
| Sult5a1 | 0.878177639 | -0.223789028 | 0.17 | 0.174 | 1 | 0.453508074 | 0.515276755 | 0.098 | 0.065 | 1 | 0.878177639 | 0.701346575 |
| Napb | 0.752839678 | -0.292736382 | 0.151 | 0.165 | 1 | 0.453548653 | -0.192542494 | 0.073 | 0.113 | 1 | 0.752839678 | 0.701390925 |
| Rcsd1 | 0.453968807 | -0.1672055 | 0.132 | 0.093 | 1 | 0.933316456 | 0.30115195 | 0.22 | 0.239 | 1 | 0.933316456 | 0.701849937 |
| Mlh3 | 0.454220368 | 0.247831999 | 0.189 | 0.144 | 1 | 0.912826771 | 0.880273572 | 0.122 | 0.126 | 1 | 0.912826771 | 0.702124593 |
| Ndufa9 | 0.454804758 | 0.465423434 | 0.453 | 0.432 | 1 | 0.541626797 | 0.217052921 | 0.463 | 0.413 | 1 | 0.541626797 | 0.702762148 |
| 4930417O13Rik | 0.481616787 | -1.237594828 | 0.019 | 0.038 | 1 | 0.455094184 | -0.332720152 | 0 | 0.014 | 1 | 0.481616787 | 0.703077652 |
| Gm52054 | 0.497380598 | 2.154722595 | 0.019 | 0.008 | 1 | 0.455094184 | 0.252242349 | 0 | 0.014 | 1 | 0.497380598 | 0.703077652 |
| Ankrd60 | 0.503971666 | 1.569760094 | 0.019 | 0.008 | 1 | 0.455094184 | 0.252242349 | 0 | 0.014 | 1 | 0.503971666 | 0.703077652 |
| Acot4 | 0.503971666 | 1.569760094 | 0.019 | 0.008 | 1 | 0.455094184 | 0.252242349 | 0 | 0.014 | 1 | 0.503971666 | 0.703077652 |
| Msh5 | 0.517412033 | 1.502645898 | 0.057 | 0.038 | 1 | 0.455094184 | 0.252242349 | 0 | 0.014 | 1 | 0.517412033 | 0.703077652 |
| Wfdc12 | 0.56579457 | -0.652632327 | 0.019 | 0.034 | 1 | 0.455094184 | 0.252242349 | 0 | 0.014 | 1 | 0.56579457 | 0.703077652 |
| Rbm46 | 0.917883255 | 0.569760094 | 0.019 | 0.021 | 1 | 0.455094184 | 0.252242349 | 0 | 0.014 | 1 | 0.917883255 | 0.703077652 |
| Fndc5 | 0.917883255 | 0.569760094 | 0.019 | 0.021 | 1 | 0.455094184 | 0.252242349 | 0 | 0.014 | 1 | 0.917883255 | 0.703077652 |
| Gm47270 | 0.917883255 | 0.569760094 | 0.019 | 0.021 | 1 | 0.455094184 | -0.16279515 | 0 | 0.014 | 1 | 0.917883255 | 0.703077652 |
| Gm57367 | 0.926212226 | 0.8327945 | 0.019 | 0.017 | 1 | 0.455094184 | 0.252242349 | 0 | 0.014 | 1 | 0.926212226 | 0.703077652 |
| P2ry6 | 0.929410101 | 0.569760094 | 0.019 | 0.017 | 1 | 0.455094184 | 0.252242349 | 0 | 0.014 | 1 | 0.929410101 | 0.703077652 |
| Gm37612 | 0.929410101 | 0.569760094 | 0.019 | 0.017 | 1 | 0.455094184 | 0.252242349 | 0 | 0.014 | 1 | 0.929410101 | 0.703077652 |
| Cit | 0.973736879 | -1.214511215 | 0.038 | 0.038 | 1 | 0.455094184 | 0.252242349 | 0 | 0.014 | 1 | 0.973736879 | 0.703077652 |
| Agap2 | 0.486967902 | -0.430239906 | 0.019 | 0.038 | 1 | 0.455097252 | -0.332720152 | 0 | 0.014 | 1 | 0.486967902 | 0.703080995 |
| Pclaf | 0.494238239 | -0.904171094 | 0.019 | 0.038 | 1 | 0.455097252 | -0.747757651 | 0 | 0.014 | 1 | 0.494238239 | 0.703080995 |
| Gm57096 | 0.503971666 | 1.569760094 | 0.019 | 0.008 | 1 | 0.455097252 | -0.16279515 | 0 | 0.014 | 1 | 0.503971666 | 0.703080995 |
| Rasgrp1 | 0.507281812 | 0.154722595 | 0.019 | 0.008 | 1 | 0.455097252 | -0.16279515 | 0 | 0.014 | 1 | 0.507281812 | 0.703080995 |
| Calm4 | 0.507281812 | 1.154722595 | 0.019 | 0.008 | 1 | 0.455097252 | -0.332720152 | 0 | 0.014 | 1 | 0.507281812 | 0.703080995 |
| Cfap276 | 0.732691937 | 1.154722595 | 0.019 | 0.013 | 1 | 0.455097252 | -0.332720152 | 0 | 0.014 | 1 | 0.732691937 | 0.703080995 |
| Ankrd45 | 0.736073999 | 0.8327945 | 0.019 | 0.013 | 1 | 0.455097252 | -0.332720152 | 0 | 0.014 | 1 | 0.736073999 | 0.703080995 |
| Kcnh4 | 0.912039013 | 0.154722595 | 0.019 | 0.021 | 1 | 0.455097252 | -0.332720152 | 0 | 0.014 | 1 | 0.912039013 | 0.703080995 |
| Zbtb32 | 0.926212226 | 0.8327945 | 0.019 | 0.017 | 1 | 0.455097252 | -0.332720152 | 0 | 0.014 | 1 | 0.926212226 | 0.703080995 |
| Elp2 | 0.45509736 | 0.428897558 | 0.321 | 0.28 | 1 | 0.904119585 | 0.136765132 | 0.293 | 0.307 | 1 | 0.904119585 | 0.703081113 |
| Ccna2 | 0.479836582 | -2.015202407 | 0.019 | 0.038 | 1 | 0.455098275 | 0.515276755 | 0 | 0.014 | 1 | 0.479836582 | 0.70308211 |
| Grin1 | 0.481613526 | -0.845277405 | 0.019 | 0.038 | 1 | 0.455098275 | 0.515276755 | 0 | 0.014 | 1 | 0.481613526 | 0.70308211 |
| C230012O17Rik | 0.48175508 | 1.154722595 | 0.038 | 0.021 | 1 | 0.455098275 | 0.515276755 | 0 | 0.014 | 1 | 0.48175508 | 0.70308211 |
| Mypopos | 0.497380598 | 2.154722595 | 0.019 | 0.008 | 1 | 0.455098275 | 0.515276755 | 0 | 0.014 | 1 | 0.497380598 | 0.70308211 |
| 6530401F13Rik | 0.500671253 | 1.739685096 | 0.019 | 0.008 | 1 | 0.455098275 | 0.515276755 | 0 | 0.014 | 1 | 0.500671253 | 0.70308211 |
| Gm26594 | 0.507281812 | 1.154722595 | 0.019 | 0.008 | 1 | 0.455098275 | 0.515276755 | 0 | 0.014 | 1 | 0.507281812 | 0.70308211 |
| Cdc45 | 0.550237964 | 0.454282877 | 0.057 | 0.038 | 1 | 0.455098275 | 0.515276755 | 0 | 0.014 | 1 | 0.550237964 | 0.70308211 |
| Myo18b | 0.596455809 | -1.693274312 | 0.057 | 0.076 | 1 | 0.455098275 | 0.515276755 | 0 | 0.014 | 1 | 0.596455809 | 0.70308211 |
| B930059L03Rik | 0.72256466 | 1.739685096 | 0.019 | 0.013 | 1 | 0.455098275 | 0.515276755 | 0 | 0.014 | 1 | 0.72256466 | 0.70308211 |
| 5830454E08Rik | 0.732691937 | 1.154722595 | 0.019 | 0.013 | 1 | 0.455098275 | 0.515276755 | 0 | 0.014 | 1 | 0.732691937 | 0.70308211 |
| Gm30881 | 0.736073999 | 0.8327945 | 0.019 | 0.013 | 1 | 0.455098275 | 0.515276755 | 0 | 0.014 | 1 | 0.736073999 | 0.70308211 |
| Bub1 | 0.906202711 | -1.430239906 | 0.019 | 0.021 | 1 | 0.455098275 | 0.515276755 | 0 | 0.014 | 1 | 0.906202711 | 0.70308211 |
| Baat | 0.914958 | 0.347367673 | 0.019 | 0.021 | 1 | 0.455098275 | 0.515276755 | 0 | 0.014 | 1 | 0.914958 | 0.70308211 |
| Psma8 | 0.917883255 | 0.569760094 | 0.019 | 0.021 | 1 | 0.455098275 | 0.515276755 | 0 | 0.014 | 1 | 0.917883255 | 0.70308211 |
| Myrip | 0.921217478 | 0.280253477 | 0.075 | 0.081 | 1 | 0.455098275 | -1.332720152 | 0 | 0.014 | 1 | 0.921217478 | 0.70308211 |
| Gp5 | 0.926212226 | 0.8327945 | 0.019 | 0.017 | 1 | 0.455098275 | 0.515276755 | 0 | 0.014 | 1 | 0.926212226 | 0.70308211 |
| Gm15908 | 0.926653673 | 0.154722595 | 0.019 | 0.021 | 1 | 0.455098275 | 0.515276755 | 0 | 0.014 | 1 | 0.926653673 | 0.70308211 |
| Dcaf7 | 0.455630743 | 0.470224421 | 0.396 | 0.364 | 1 | 0.796153701 | 0.282615998 | 0.366 | 0.396 | 1 | 0.796153701 | 0.703662112 |
| 9130019O22Rik | 0.991244785 | 0.417757001 | 0.038 | 0.038 | 1 | 0.455731299 | -0.435813645 | 0.049 | 0.082 | 1 | 0.991244785 | 0.703771581 |
| Aspg | 0.553736606 | 0.792152515 | 0.038 | 0.059 | 1 | 0.455763738 | 1.100239256 | 0.049 | 0.027 | 1 | 0.553736606 | 0.703806891 |
| Naif1 | 0.624418548 | 0.932330173 | 0.038 | 0.025 | 1 | 0.455763738 | 1.100239256 | 0.049 | 0.027 | 1 | 0.624418548 | 0.703806891 |
| 1810059H22Rik | 0.624418548 | 0.932330173 | 0.038 | 0.025 | 1 | 0.455763738 | 1.100239256 | 0.049 | 0.027 | 1 | 0.624418548 | 0.703806891 |
| Taf9 | 0.736073999 | 0.8327945 | 0.019 | 0.013 | 1 | 0.455763738 | 1.100239256 | 0.049 | 0.027 | 1 | 0.736073999 | 0.703806891 |
| Ankra2 | 0.470831129 | 0.664396968 | 0.226 | 0.195 | 1 | 0.456250304 | 0.763204268 | 0.244 | 0.208 | 1 | 0.470831129 | 0.704336268 |
| Itpkc | 0.456310413 | 0.100274811 | 0.302 | 0.25 | 1 | 0.458427315 | 0.726173537 | 0.293 | 0.27 | 1 | 0.458427315 | 0.704401633 |
| Eef1akmt3 | 0.456531042 | -0.389597921 | 0.075 | 0.11 | 1 | 0.708986965 | 1.377773231 | 0.098 | 0.085 | 1 | 0.708986965 | 0.704641491 |
| Pdgfrl | 0.912319977 | 0.494531463 | 0.566 | 0.708 | 1 | 0.456550626 | 0.775690461 | 0.537 | 0.645 | 1 | 0.912319977 | 0.704662778 |
| Clstn3 | 0.762480493 | 0.739685096 | 0.038 | 0.03 | 1 | 0.457260871 | 0.883008539 | 0.122 | 0.089 | 1 | 0.762480493 | 0.705434238 |
| Plcb4 | 0.699965951 | -0.1182959 | 0.283 | 0.242 | 1 | 0.457304697 | 0.209173627 | 0.512 | 0.423 | 1 | 0.699965951 | 0.705481809 |
| Tcta | 0.553126861 | 0.63687429 | 0.245 | 0.225 | 1 | 0.457411355 | -0.101394606 | 0.146 | 0.201 | 1 | 0.553126861 | 0.705597562 |
| Lztfl1 | 0.457715971 | -0.1672055 | 0.717 | 0.742 | 1 | 0.799510042 | -0.10285261 | 0.756 | 0.771 | 1 | 0.799510042 | 0.705928032 |
| Exoc6b | 0.457833953 | -0.182089737 | 0.925 | 0.898 | 1 | 0.900570196 | -0.20627819 | 0.951 | 0.939 | 1 | 0.900570196 | 0.706055978 |
| Laptm5 | 0.497382333 | 2.154722595 | 0.019 | 0.008 | 1 | 0.458099588 | -1.917682652 | 0.073 | 0.044 | 1 | 0.497382333 | 0.706343943 |
| Btd | 0.458211857 | -0.106542718 | 0.415 | 0.483 | 1 | 0.809828488 | 0.295311071 | 0.439 | 0.457 | 1 | 0.809828488 | 0.706465608 |
| Emd | 0.458427814 | 0.245367959 | 0.66 | 0.648 | 1 | 0.700055085 | 0.241673107 | 0.585 | 0.737 | 1 | 0.700055085 | 0.706699568 |
| Bmp8a | 0.936447367 | -1.088133929 | 0.208 | 0.186 | 1 | 0.458535036 | -0.814871847 | 0.073 | 0.113 | 1 | 0.936447367 | 0.706815693 |
| Bbs9 | 0.66539089 | 0.459577176 | 0.434 | 0.428 | 1 | 0.458823915 | 0.268209097 | 0.683 | 0.631 | 1 | 0.66539089 | 0.707128445 |
| Dynlt2a2 | 0.458961249 | -0.652632327 | 0.038 | 0.064 | 1 | 0.627334271 | 0.404245443 | 0.098 | 0.075 | 1 | 0.627334271 | 0.70727707 |
| Thap4 | 0.459376534 | 0.428897558 | 0.189 | 0.148 | 1 | 0.703144399 | 0.174239837 | 0.195 | 0.235 | 1 | 0.703144399 | 0.707726268 |
| Shc4 | 0.601055679 | -0.38369732 | 0.264 | 0.301 | 1 | 0.459509407 | -0.435813645 | 0.171 | 0.218 | 1 | 0.601055679 | 0.707869919 |
| Dcbld2 | 0.47627248 | 0.227367897 | 0.491 | 0.504 | 1 | 0.459532275 | 0.298784935 | 0.463 | 0.427 | 1 | 0.47627248 | 0.707894638 |
| Fyb | 0.830659588 | 1.721763188 | 0.057 | 0.068 | 1 | 0.459645218 | 1.152706676 | 0.122 | 0.089 | 1 | 0.830659588 | 0.708016709 |
| Sgms1os1 | 0.45964867 | 0.502645898 | 0.113 | 0.081 | 1 | 0.844886174 | 0.136765132 | 0.073 | 0.082 | 1 | 0.844886174 | 0.70802044 |
| Dnajc13 | 0.45996575 | 0.333383443 | 0.642 | 0.631 | 1 | 0.561885238 | -0.125800943 | 0.732 | 0.775 | 1 | 0.561885238 | 0.708363009 |
| Chchd1 | 0.460127386 | 0.182736971 | 0.698 | 0.653 | 1 | 0.837658161 | 0.174239837 | 0.61 | 0.652 | 1 | 0.837658161 | 0.708537561 |
| Dda1 | 0.460621468 | 0.18893831 | 0.604 | 0.619 | 1 | 0.974900123 | 0.332860809 | 0.512 | 0.543 | 1 | 0.974900123 | 0.709070799 |
| Fbxo6 | 0.460659108 | 0.507024339 | 0.208 | 0.169 | 1 | 0.569430494 | -0.140075074 | 0.146 | 0.184 | 1 | 0.569430494 | 0.709111402 |

| | | | | | | | | | | | | |
|---|---|---|---|---|---|---|---|---|---|---|---|---|
| Grik4 | 0.587408118 | -0.417528854 | 0.151 | 0.178 | 1 | 0.461048972 | -0.26713181 | 0.146 | 0.106 | 1 | 0.587408118 | 0.709531789 |
| Fam220a.1 | 0.461262272 | 0.792152515 | 0.075 | 0.051 | 1 | 0.622636015 | -0.484723245 | 0.049 | 0.068 | 1 | 0.622636015 | 0.709761661 |
| Itpk1 | 0.621511234 | -0.268488836 | 0.151 | 0.178 | 1 | 0.461420511 | 1.029849928 | 0.195 | 0.154 | 1 | 0.621511234 | 0.709932134 |
| Pigm | 0.461591142 | 0.454282877 | 0.189 | 0.148 | 1 | 0.635671706 | 0.515276755 | 0.22 | 0.195 | 1 | 0.635671706 | 0.710115902 |
| Slc36a1 | 0.461702194 | 0.154722595 | 0.057 | 0.089 | 1 | 0.611746269 | -0.364429011 | 0.098 | 0.123 | 1 | 0.611746269 | 0.710235472 |
| Pappa2 | 0.760992231 | 0.333692736 | 0.113 | 0.102 | 1 | 0.462817568 | -1.041940755 | 0.146 | 0.184 | 1 | 0.760992231 | 0.711435035 |
| Thegl | 0.661289389 | -0.223789028 | 0.057 | 0.042 | 1 | 0.462889204 | 0.422167351 | 0.049 | 0.027 | 1 | 0.661289389 | 0.711511993 |
| Hpgds | 0.912039013 | -0.304709024 | 0.019 | 0.021 | 1 | 0.462889204 | 0.515276755 | 0.049 | 0.027 | 1 | 0.912039013 | 0.711511993 |
| Utp3 | 0.538613967 | 0.199810484 | 0.547 | 0.555 | 1 | 0.463092914 | -0.19841906 | 0.659 | 0.679 | 1 | 0.538613967 | 0.71173078 |
| Spc24 | 0.463094925 | -0.974560422 | 0.038 | 0.064 | 1 | 0.87556513 | 0.667279848 | 0.024 | 0.02 | 1 | 0.87556513 | 0.711732941 |
| Ppp1r3f | 0.463963499 | 1.039245377 | 0.075 | 0.051 | 1 | 0.826384776 | 0.515276755 | 0.024 | 0.031 | 1 | 0.826384776 | 0.71266487 |
| ENSMUSG00000121479 | 0.756168608 | 0.47665069 | 0.057 | 0.047 | 1 | 0.463970634 | 1.515276755 | 0.073 | 0.048 | 1 | 0.756168608 | 0.712672518 |
| Nipal2 | 0.599153699 | -0.889671525 | 0.057 | 0.076 | 1 | 0.464131275 | 0.145327145 | 0.244 | 0.188 | 1 | 0.599153699 | 0.71284471 |
| Tnfsf13os | 0.464334319 | -1.214511215 | 0.038 | 0.064 | 1 | 0.483530684 | 1.029849928 | 0.073 | 0.048 | 1 | 0.483530684 | 0.713062279 |
| Zfp27 | 0.553213024 | 0.569760094 | 0.113 | 0.089 | 1 | 0.464546297 | -0.136799942 | 0.098 | 0.14 | 1 | 0.553213024 | 0.713289331 |
| Gtpbp6 | 0.487472041 | 0.538051234 | 0.208 | 0.174 | 1 | 0.464613233 | 0.79281073 | 0.122 | 0.089 | 1 | 0.487472041 | 0.71336101 |
| Usp16 | 0.464933797 | 0.146846169 | 0.868 | 0.847 | 1 | 0.54537359 | 0.178139015 | 0.878 | 0.867 | 1 | 0.54537359 | 0.713704159 |
| Rnf20 | 0.465495121 | 0.353104042 | 0.604 | 0.606 | 1 | 0.858963429 | 0.307800326 | 0.683 | 0.706 | 1 | 0.858963429 | 0.714304534 |
| Gm17529 | 0.465569538 | 1.280253477 | 0.094 | 0.068 | 1 | 0.664521036 | 0.809724113 | 0.146 | 0.126 | 1 | 0.664521036 | 0.714384081 |
| S1pr1 | 0.647136736 | 0.485928503 | 0.321 | 0.297 | 1 | 0.465818968 | 0.810732638 | 0.22 | 0.191 | 1 | 0.647136736 | 0.714650626 |
| Glb1l | 0.584378896 | 0.270199812 | 0.17 | 0.14 | 1 | 0.465967797 | -0.144647803 | 0.146 | 0.198 | 1 | 0.584378896 | 0.714809606 |
| Gm38190 | 0.473473819 | 0.318221327 | 0.113 | 0.081 | 1 | 0.466034165 | 0.515276755 | 0.073 | 0.113 | 1 | 0.473473819 | 0.714880487 |
| Kirrel3 | 0.466331343 | 2.11408061 | 0.132 | 0.102 | 1 | 0.472698783 | -0.236044132 | 0.073 | 0.116 | 1 | 0.472698783 | 0.715197765 |
| Gm2000 | 0.533658184 | -0.127541283 | 0.585 | 0.665 | 1 | 0.466447094 | -0.211158172 | 0.585 | 0.601 | 1 | 0.533658184 | 0.715321297 |
| Tmem60 | 0.976569112 | 0.199116714 | 0.377 | 0.394 | 1 | 0.466789735 | -0.275004654 | 0.268 | 0.328 | 1 | 0.976569112 | 0.715686814 |
| 9230114K14Rik | 0.467005003 | -0.195774652 | 0.094 | 0.136 | 1 | 0.608981053 | 1.30115195 | 0.049 | 0.072 | 1 | 0.608981053 | 0.715916333 |
| AW209491 | 0.467006111 | 0.321832581 | 0.245 | 0.199 | 1 | 0.713163944 | 0.252242349 | 0.195 | 0.174 | 1 | 0.713163944 | 0.715917514 |
| Gm10069 | 0.467027607 | -0.430239906 | 0.038 | 0.064 | 1 | 0.997337913 | 0.225770138 | 0.098 | 0.099 | 1 | 0.997337913 | 0.715940428 |
| Pyroxd1 | 0.812592853 | -0.187669603 | 0.226 | 0.203 | 1 | 0.467031524 | 0.620886943 | 0.244 | 0.195 | 1 | 0.812592853 | 0.715944604 |
| C2cd2l | 0.574920744 | 0.569760094 | 0.113 | 0.089 | 1 | 0.467255202 | -0.410722664 | 0.073 | 0.109 | 1 | 0.574920744 | 0.71618298 |
| Ptrh2 | 0.905852067 | 0.176096246 | 0.226 | 0.246 | 1 | 0.468094679 | -0.221688839 | 0.146 | 0.198 | 1 | 0.905852067 | 0.71707673 |
| Gins2 | 0.469056418 | -0.130679624 | 0.094 | 0.131 | 1 | 0.79248022 | 1.100239256 | 0.098 | 0.089 | 1 | 0.79248022 | 0.718098912 |
| Zfp882 | 0.68209151 | 0.695290976 | 0.019 | 0.03 | 1 | 0.469427653 | 0.100239256 | 0.024 | 0.051 | 1 | 0.68209151 | 0.718492985 |
| Vil1 | 0.469724974 | -0.347777746 | 0.038 | 0.064 | 1 | 0.861873715 | 1.252242349 | 0.024 | 0.02 | 1 | 0.861873715 | 0.718808397 |
| Ndfip2 | 0.870884474 | 0.164966626 | 0.83 | 0.822 | 1 | 0.470107277 | -0.186763798 | 0.854 | 0.877 | 1 | 0.870884474 | 0.719213702 |
| Smim15 | 0.486928507 | 0.226872381 | 0.434 | 0.428 | 1 | 0.470634474 | 0.390521724 | 0.415 | 0.375 | 1 | 0.486928507 | 0.71977214 |
| Tsku | 0.471154689 | 1.569760094 | 0.038 | 0.021 | 1 | 0.608540312 | 1.749742009 | 0.073 | 0.055 | 1 | 0.608540312 | 0.720322637 |
| Nme3 | 0.471154689 | 1.569760094 | 0.038 | 0.021 | 1 | 0.916649754 | 0.667279848 | 0.024 | 0.027 | 1 | 0.916649754 | 0.720322637 |
| Lig3 | 0.970084624 | 0.154722595 | 0.264 | 0.271 | 1 | 0.471546275 | -0.278272368 | 0.268 | 0.324 | 1 | 0.970084624 | 0.72073666 |
| Abhd11 | 0.471597554 | 0.333692736 | 0.226 | 0.182 | 1 | 0.903141963 | 0.222495006 | 0.146 | 0.14 | 1 | 0.903141963 | 0.720790856 |
| Kat8 | 0.53943143 | 0.446903346 | 0.208 | 0.174 | 1 | 0.471783161 | 0.317830691 | 0.293 | 0.239 | 1 | 0.53943143 | 0.720986971 |
| Flot1 | 0.983816362 | 0.329556397 | 0.434 | 0.445 | 1 | 0.472443101 | 0.736404209 | 0.39 | 0.372 | 1 | 0.983816362 | 0.721683718 |
| 4632404H12Rik | 0.473270726 | 1.669295768 | 0.038 | 0.021 | 1 | 0.714416985 | 1.252242349 | 0.049 | 0.038 | 1 | 0.714416985 | 0.722556272 |
| Ankrd6 | 0.473280032 | 1.017219071 | 0.038 | 0.021 | 1 | 0.861257699 | 0.136765132 | 0.073 | 0.065 | 1 | 0.861257699 | 0.722566075 |
| Cebpa | 0.473473819 | 0.318221327 | 0.113 | 0.081 | 1 | 0.79927949 | 0.308825877 | 0.195 | 0.181 | 1 | 0.79927949 | 0.722770181 |
| 4930556M19Rik | 0.675049917 | 0.306725688 | 0.075 | 0.059 | 1 | 0.475031087 | 1.159132945 | 0.073 | 0.048 | 1 | 0.675049917 | 0.72440764 |
| Ptrh1 | 0.750708181 | 0.347367673 | 0.094 | 0.081 | 1 | 0.475031087 | 1.159132945 | 0.073 | 0.048 | 1 | 0.750708181 | 0.72440764 |
| Gm42797 | 0.897146191 | 0.417757001 | 0.038 | 0.034 | 1 | 0.475207277 | 0.444887427 | 0.024 | 0.051 | 1 | 0.897146191 | 0.724592598 |
| Got1 | 0.475267829 | -0.1672055 | 0.151 | 0.195 | 1 | 0.876333542 | 0.795384674 | 0.171 | 0.167 | 1 | 0.876333542 | 0.724656148 |
| Gm26620 | 0.475380264 | 1.347367673 | 0.038 | 0.021 | 1 | 0.51919986 | 0.83720485 | 0 | 0.01 | 1 | 0.51919986 | 0.724774132 |
| Syt9 | 0.475380264 | 1.347367673 | 0.038 | 0.021 | 1 | 0.607875423 | 0.667279848 | 0.024 | 0.014 | 1 | 0.607875423 | 0.724774132 |
| Slc26a6 | 0.475380264 | 1.347367673 | 0.038 | 0.021 | 1 | 0.716340863 | 1.029849928 | 0.049 | 0.038 | 1 | 0.716340863 | 0.724774132 |
| Zfp748 | 0.475421542 | 1.002719501 | 0.094 | 0.068 | 1 | 0.678775307 | 0.149148856 | 0.195 | 0.164 | 1 | 0.678775307 | 0.724817441 |
| Coq10a | 0.475446993 | 0.602181572 | 0.208 | 0.174 | 1 | 0.918169843 | 0.44893426 | 0.22 | 0.229 | 1 | 0.918169843 | 0.724844143 |
| Tbccd1 | 0.475541021 | 0.865215978 | 0.113 | 0.085 | 1 | 0.87554304 | 0.328191202 | 0.171 | 0.164 | 1 | 0.87554304 | 0.724942779 |
| Cep135 | 0.475749133 | 0.635348436 | 0.132 | 0.102 | 1 | 0.549185102 | 0.389745873 | 0.171 | 0.137 | 1 | 0.549185102 | 0.725161028 |
| Gm16853 | 0.485993467 | 0.932330173 | 0.038 | 0.021 | 1 | 0.47612871 | 0.252242349 | 0.122 | 0.085 | 1 | 0.485993467 | 0.725558871 |
| Tnfaip8 | 0.476195174 | 0.18481466 | 0.698 | 0.597 | 1 | 0.681471799 | -0.458251034 | 0.829 | 0.768 | 1 | 0.681471799 | 0.725628504 |
| Exoc1 | 0.842888914 | 0.216123139 | 0.132 | 0.144 | 1 | 0.476453384 | 0.416873051 | 0.268 | 0.218 | 1 | 0.842888914 | 0.725898941 |
| 2810432F15Rik | 0.70960347 | 0.154722595 | 0.075 | 0.093 | 1 | 0.476646183 | -0.787286015 | 0.049 | 0.078 | 1 | 0.70960347 | 0.726100782 |
| Ddx59 | 0.476709722 | 1.084333267 | 0.094 | 0.068 | 1 | 0.831400855 | 1.029849928 | 0.073 | 0.065 | 1 | 0.831400855 | 0.726167285 |
| Mapk3 | 0.477077131 | 0.242949778 | 0.566 | 0.572 | 1 | 0.612732577 | 0.206072168 | 0.61 | 0.536 | 1 | 0.612732577 | 0.726551673 |
| BC049715 | 0.477945573 | -0.845277405 | 0.057 | 0.085 | 1 | 0.494347488 | -0.686357106 | 0.024 | 0.048 | 1 | 0.494347488 | 0.727459175 |
| Gm35417 | 0.478184586 | 0.984797593 | 0.075 | 0.051 | 1 | 0.881947238 | 1.763204268 | 0.098 | 0.099 | 1 | 0.881947238 | 0.727708674 |
| Ccdc191 | 0.478904133 | 0.254258268 | 0.208 | 0.165 | 1 | 0.740909756 | -0.136799942 | 0.098 | 0.116 | 1 | 0.740909756 | 0.728459098 |
| Mphosph9 | 0.923781609 | 0.2967416 | 0.17 | 0.165 | 1 | 0.479557107 | 0.551802631 | 0.122 | 0.174 | 1 | 0.923781609 | 0.729139196 |
| Cplane2 | 0.479622472 | 1.154722595 | 0.038 | 0.021 | 1 | 0.498856313 | -0.250257991 | 0.024 | 0.048 | 1 | 0.498856313 | 0.729207228 |
| Myo1g | 0.479622472 | 1.154722595 | 0.038 | 0.021 | 1 | 0.60213981 | 1.252242349 | 0.024 | 0.014 | 1 | 0.60213981 | 0.729207228 |
| Gm53028 | 0.479954615 | -0.1672055 | 0.189 | 0.237 | 1 | 0.795712643 | -0.179096963 | 0.22 | 0.232 | 1 | 0.795712643 | 0.729552797 |
| Gm47271 | 0.954239151 | 0.47665069 | 0.057 | 0.055 | 1 | 0.480682218 | 1.100239256 | 0.073 | 0.048 | 1 | 0.954239151 | 0.730309041 |
| 4933423P22Rik | 0.48175508 | 1.154722595 | 0.038 | 0.021 | 1 | 0.5191986 | 0.515276755 | 0 | 0.01 | 1 | 0.5191986 | 0.731422203 |
| Myo1h | 0.48175508 | 1.154722595 | 0.038 | 0.021 | 1 | 0.94215184 | 0.252242349 | 0.049 | 0.051 | 1 | 0.94215184 | 0.731422203 |
| 9130023H24Rik | 0.48175508 | 1.154722595 | 0.038 | 0.021 | 1 | 0.587925597 | 2.252242349 | 0.024 | 0.014 | 1 | 0.587925597 | 0.731422203 |
| Gm27042 | 0.48175508 | 1.154722595 | 0.038 | 0.021 | 1 | 0.992196803 | 0.515276755 | 0.024 | 0.024 | 1 | 0.992196803 | 0.731422203 |
| Mb21d2 | 0.973742649 | -0.178701139 | 0.34 | 0.326 | 1 | 0.481811833 | 0.166666617 | 0.341 | 0.28 | 1 | 0.973742649 | 0.731481024 |
| Zfp865 | 0.482431187 | 0.80108564 | 0.113 | 0.085 | 1 | 0.763017656 | 0.367719567 | 0.22 | 0.198 | 1 | 0.763017656 | 0.732122524 |
| Coq7 | 0.742046584 | 0.167661651 | 0.321 | 0.352 | 1 | 0.482588257 | 0.264315181 | 0.341 | 0.29 | 1 | 0.742046584 | 0.732285088 |
| Txndc2 | 0.483399552 | -0.752168001 | 0.019 | 0.038 | 1 | 0.868524543 | 0.100239256 | 0.049 | 0.055 | 1 | 0.868524543 | 0.733123978 |
| Akr1b10 | 0.858271359 | 0.154722595 | 0.17 | 0.186 | 1 | 0.483650504 | 0.19334866 | 0.098 | 0.143 | 1 | 0.858271359 | 0.733383198 |
| Eif1b | 0.591051482 | 0.167569474 | 0.774 | 0.797 | 1 | 0.483685784 | -0.18959521 | 0.854 | 0.84 | 1 | 0.591051482 | 0.73341963 |
| Zfp808 | 0.484219558 | 0.704061186 | 0.132 | 0.102 | 1 | 0.617298283 | 0.114738825 | 0.049 | 0.072 | 1 | 0.617298283 | 0.733970536 |
| Ddx56 | 0.484413265 | 0.232725107 | 0.321 | 0.267 | 1 | 0.783898152 | 0.498402936 | 0.195 | 0.229 | 1 | 0.783898152 | 0.734170318 |
| Adam15 | 0.484621523 | 0.417757001 | 0.434 | 0.411 | 1 | 0.854521677 | 0.377773231 | 0.268 | 0.266 | 1 | 0.854521677 | 0.734385025 |
| Arsg | 0.484720015 | 0.624207878 | 0.113 | 0.085 | 1 | 0.880281089 | -0.20718927 | 0.073 | 0.078 | 1 | 0.880281089 | 0.734486537 |
| Gm21859 | 0.485102394 | 0.154722595 | 0.038 | 0.064 | 1 | 0.807443647 | 0.252242349 | 0.049 | 0.041 | 1 | 0.807443647 | 0.734880455 |
| 4930405A21Rik | 0.485184337 | -0.752168001 | 0.019 | 0.038 | 1 | 0.872828964 | 1.029849928 | 0.024 | 0.02 | 1 | 0.872828964 | 0.734964833 |
| Mybl2 | 0.485184878 | -0.752168001 | 0.019 | 0.038 | 1 | 0.765070435 | 0.930314254 | 0.024 | 0.034 | 1 | 0.765070435 | 0.73496539 |
| Dmp1 | 0.485184878 | -3.517702747 | 0.019 | 0.038 | 1 | 0.5191986 | -0.16279515 | 0 | 0.01 | 1 | 0.5191986 | 0.73496539 |
| Dennd11 | 0.485369227 | 0.792152515 | 0.075 | 0.051 | 1 | 0.695788449 | 1.515276755 | 0.024 | 0.038 | 1 | 0.695788449 | 0.735155168 |
| Repin1 | 0.498467127 | 0.695290976 | 0.113 | 0.085 | 1 | 0.48554133 | -0.52536523 | 0.146 | 0.184 | 1 | 0.498467127 | 0.735332277 |
| A230083N12Rik | 0.485993467 | 0.932330173 | 0.038 | 0.021 | 1 | 0.51919986 | 0.83720485 | 0 | 0.01 | 1 | 0.51919986 | 0.735797284 |
| Gm26935 | 0.485993467 | 0.932330173 | 0.038 | 0.021 | 1 | 0.674180157 | 0.252242349 | 0.024 | 0.038 | 1 | 0.674180157 | 0.735797284 |
| BC037032 | 0.485993467 | 0.932330173 | 0.038 | 0.021 | 1 | 0.51919986 | 0.83720485 | 0 | 0.01 | 1 | 0.51919986 | 0.735797284 |
| Zfp174 | 0.485993467 | 0.932330173 | 0.038 | 0.021 | 1 | 0.746881137 | 1.029849928 | 0.024 | 0.017 | 1 | 0.746881137 | 0.735797284 |
| C130050O18Rik | 0.486676589 | 0.669295768 | 0.075 | 0.051 | 1 | 0.607249896 | 0.322631677 | 0.073 | 0.102 | 1 | 0.607249896 | 0.736499076 |
| Gm29642 | 0.486967902 | -0.430239906 | 0.019 | 0.038 | 1 | 0.716340863 | 1.029849928 | 0.049 | 0.038 | 1 | 0.716340863 | 0.736798066 |
| Gm33037 | 0.486967902 | -0.752168001 | 0.019 | 0.038 | 1 | 0.58792152 | 2.100239256 | 0.024 | 0.014 | 1 | 0.58792152 | 0.736798066 |
| Trip13 | 0.486967902 | -0.932740246 | 0.019 | 0.038 | 1 | 0.5191986 | 0.515276755 | 0 | 0.01 | 1 | 0.5191986 | 0.736798066 |

| | | | | | | | | | | | | |
|---|---|---|---|---|---|---|---|---|---|---|---|---|
| Wdr24 | 0.486967902 | -0.430239906 | 0.019 | 0.038 | 1 | 0.672172778 | 0.136765132 | 0.024 | 0.038 | 1 | 0.672172778 | 0.736798066 |
| Grin2d | 0.487069644 | 0.110328475 | 0.132 | 0.097 | 1 | 0.555454952 | -0.20718927 | 0.073 | 0.102 | 1 | 0.555454952 | 0.73690245 |
| Klf7 | 0.487372862 | -0.281013897 | 0.755 | 0.805 | 1 | 0.760032626 | 0.127697251 | 0.878 | 0.826 | 1 | 0.760032626 | 0.737213417 |
| Zfp839 | 0.594700767 | 0.347367673 | 0.151 | 0.123 | 1 | 0.487894095 | 0.989207943 | 0.122 | 0.092 | 1 | 0.594700767 | 0.737747542 |
| Mfsd12 | 0.779443586 | 0.154722595 | 0.019 | 0.025 | 1 | 0.488328255 | -0.622226769 | 0.049 | 0.078 | 1 | 0.779443586 | 0.738192026 |
| Smc2 | 0.962533826 | -0.310400136 | 0.17 | 0.174 | 1 | 0.488615785 | 1.174239837 | 0.146 | 0.116 | 1 | 0.962533826 | 0.738486185 |
| 1700085D22Rik | 0.808308185 | 0.347367673 | 0.094 | 0.085 | 1 | 0.488622947 | -0.332720152 | 0.073 | 0.109 | 1 | 0.808308185 | 0.738493509 |
| Brca2 | 0.721577188 | -1.232300528 | 0.151 | 0.127 | 1 | 0.488724449 | 0.537644568 | 0.098 | 0.068 | 1 | 0.721577188 | 0.738597311 |
| Setmar | 0.488750271 | -0.304709024 | 0.019 | 0.038 | 1 | 0.684370942 | -0.101394606 | 0.049 | 0.065 | 1 | 0.684370942 | 0.738623714 |
| Nadsyn1 | 0.488750271 | -0.304709024 | 0.019 | 0.038 | 1 | 0.741511021 | 0.174239837 | 0.049 | 0.038 | 1 | 0.741511021 | 0.738623714 |
| Gm48045 | 0.488750271 | -0.304709024 | 0.019 | 0.038 | 1 | 0.872828964 | 1.029849928 | 0.024 | 0.02 | 1 | 0.872828964 | 0.738623714 |
| Pole2 | 0.488960784 | 0.676675298 | 0.151 | 0.119 | 1 | 0.959813146 | 0.773074512 | 0.098 | 0.106 | 1 | 0.959813146 | 0.73883892 |
| Ddx28 | 0.489198959 | 0.792152515 | 0.094 | 0.068 | 1 | 0.566018719 | 0.749742009 | 0.073 | 0.051 | 1 | 0.566018719 | 0.739082297 |
| Tapbpl | 0.489445117 | 0.247831999 | 0.113 | 0.161 | 1 | 0.662319999 | 0.557096931 | 0.073 | 0.099 | 1 | 0.662319999 | 0.739333712 |
| Gm12227 | 0.489599381 | 0.610402079 | 0.113 | 0.085 | 1 | 0.700250611 | -0.233184478 | 0.073 | 0.092 | 1 | 0.700250611 | 0.739491208 |
| Dmd | 0.489691464 | 0.631160639 | 0.075 | 0.051 | 1 | 0.774024661 | -0.785232356 | 0.22 | 0.195 | 1 | 0.774024661 | 0.739585198 |
| Mapk6 | 0.734789323 | 0.141744741 | 0.83 | 0.814 | 1 | 0.48976613 | 0.435811249 | 0.854 | 0.853 | 1 | 0.734789323 | 0.739661398 |
| Med20 | 0.490069471 | 0.235341055 | 0.321 | 0.275 | 1 | 0.682649246 | 0.628805701 | 0.366 | 0.379 | 1 | 0.682649246 | 0.739970856 |
| Gm13274 | 0.49029049 | 0.739685096 | 0.038 | 0.021 | 1 | 0.846935592 | 1.574170444 | 0.024 | 0.031 | 1 | 0.846935592 | 0.740196215 |
| Ankrd55 | 0.49029049 | 0.569760094 | 0.038 | 0.021 | 1 | 0.72989677 | 3.215716473 | 0.024 | 0.017 | 1 | 0.72989677 | 0.740196215 |
| Zfp72 | 0.490620203 | -0.1672055 | 0.019 | 0.038 | 1 | 0.549777959 | 1.100239256 | 0.073 | 0.051 | 1 | 0.549777959 | 0.740532222 |
| Zfp960 | 0.490620203 | -0.1672055 | 0.019 | 0.038 | 1 | 0.743829142 | 0.252242349 | 0.024 | 0.034 | 1 | 0.743829142 | 0.740532222 |
| Abcb7 | 0.490821386 | 0.852159825 | 0.226 | 0.195 | 1 | 0.648957387 | 0.404245443 | 0.293 | 0.263 | 1 | 0.648957387 | 0.740737139 |
| Gpaa1 | 0.987529791 | 0.216123139 | 0.302 | 0.322 | 1 | 0.491567067 | 0.100239256 | 0.22 | 0.266 | 1 | 0.987529791 | 0.741495952 |
| Tjp2 | 0.492026859 | -0.65926542 | 0.302 | 0.331 | 1 | 0.958407446 | 0.20211887 | 0.537 | 0.549 | 1 | 0.958407446 | 0.741963288 |
| Gm20275 | 0.651164202 | 1.958325382 | 0.132 | 0.114 | 1 | 0.492124218 | -0.235203646 | 0.146 | 0.205 | 1 | 0.651164202 | 0.74206219 |
| Cpt2 | 0.824853803 | 0.569760094 | 0.075 | 0.068 | 1 | 0.493019579 | -0.385187572 | 0.049 | 0.078 | 1 | 0.824853803 | 0.742970853 |
| N4bp2os | 0.831453313 | -0.1672055 | 0.057 | 0.064 | 1 | 0.493019746 | -0.332720152 | 0.049 | 0.078 | 1 | 0.831453313 | 0.742971022 |
| Endov | 0.970303881 | 0.154722595 | 0.132 | 0.131 | 1 | 0.493019746 | -0.332720152 | 0.049 | 0.078 | 1 | 0.970303881 | 0.742971022 |
| Ssh3 | 0.493270688 | 0.618669695 | 0.151 | 0.119 | 1 | 0.613537904 | 0.377773231 | 0.146 | 0.184 | 1 | 0.613537904 | 0.743225405 |
| Dolpp1 | 0.493346902 | 0.47665069 | 0.038 | 0.064 | 1 | 0.909349126 | 1.699701326 | 0.073 | 0.072 | 1 | 0.909349126 | 0.743302638 |
| Ppat | 0.495621735 | 0.322450041 | 0.396 | 0.364 | 1 | 0.493359348 | 0.272420231 | 0.22 | 0.297 | 1 | 0.495621735 | 0.74331525 |
| Ift122 | 0.988980239 | 0.154722595 | 0.132 | 0.136 | 1 | 0.493708433 | -0.140075074 | 0.122 | 0.167 | 1 | 0.988980239 | 0.743668849 |
| Mrpl24 | 0.494259103 | 0.165214969 | 0.755 | 0.674 | 1 | 0.960442406 | 0.28412641 | 0.683 | 0.696 | 1 | 0.960442406 | 0.744226145 |
| Gm46447 | 0.503971666 | 1.569760094 | 0.019 | 0.008 | 1 | 0.494346678 | -0.484723245 | 0.024 | 0.048 | 1 | 0.503971666 | 0.744314718 |
| Dcdc2b | 0.982488248 | -0.1672055 | 0.038 | 0.038 | 1 | 0.494347488 | -0.686357106 | 0.024 | 0.048 | 1 | 0.982488248 | 0.744315537 |
| Gtpbp2 | 0.494384901 | 0.558871778 | 0.396 | 0.398 | 1 | 0.660547826 | 0.269944351 | 0.39 | 0.471 | 1 | 0.660547826 | 0.744353372 |
| Tmed7 | 0.494602782 | 0.210533458 | 0.774 | 0.831 | 1 | 0.683005083 | 0.114738825 | 0.829 | 0.85 | 1 | 0.683005083 | 0.744573652 |
| Gm56935 | 0.89116038 | 0.154722595 | 0.038 | 0.042 | 1 | 0.495851362 | -0.555112573 | 0.024 | 0.048 | 1 | 0.89116038 | 0.745834151 |
| Igfals | 0.787929599 | 0.154722595 | 0.057 | 0.068 | 1 | 0.49596326 | 0.218295017 | 0.073 | 0.113 | 1 | 0.787929599 | 0.745946964 |
| Dhcr7 | 0.49646301 | -0.285849997 | 0.151 | 0.195 | 1 | 0.749582434 | -0.332720152 | 0.122 | 0.137 | 1 | 0.749582434 | 0.7464505 |
| Chmp6 | 0.496466524 | 0.600133743 | 0.208 | 0.169 | 1 | 0.992403687 | 0.19334866 | 0.22 | 0.225 | 1 | 0.992403687 | 0.746454039 |
| Sft2d3 | 0.647078441 | 0.347367673 | 0.151 | 0.127 | 1 | 0.496755377 | 0.652780279 | 0.171 | 0.137 | 1 | 0.647078441 | 0.746744849 |
| Enox2 | 0.496854967 | 0.672853587 | 0.226 | 0.199 | 1 | 0.88233563 | -0.598894265 | 0.244 | 0.239 | 1 | 0.88233563 | 0.746845076 |
| Gm45435 | 0.917883255 | 0.569760094 | 0.019 | 0.021 | 1 | 0.497354299 | -0.250257991 | 0.024 | 0.048 | 1 | 0.917883255 | 0.747347299 |
| Pla2g4c | 0.497380598 | 2.154722595 | 0.019 | 0.008 | 1 | 0.5191986 | 0.515276755 | 0 | 0.01 | 1 | 0.5191986 | 0.747373737 |
| Vsig10l | 0.497380598 | 2.569760094 | 0.019 | 0.008 | 1 | 0.546365767 | 1.100239256 | 0.049 | 0.031 | 1 | 0.546365767 | 0.747373737 |
| Gm28379 | 0.497380598 | 2.154722595 | 0.019 | 0.008 | 1 | 0.846926792 | 1.100239256 | 0.024 | 0.031 | 1 | 0.846926792 | 0.747373737 |
| Cabcoco1 | 0.497380598 | 2.154722595 | 0.019 | 0.008 | 1 | 0.743829142 | 0.252242349 | 0.024 | 0.034 | 1 | 0.743829142 | 0.747373737 |
| 4921516A02Rik | 0.497380598 | 2.154722595 | 0.019 | 0.008 | 1 | 0.819548308 | 0.136765132 | 0.024 | 0.031 | 1 | 0.819548308 | 0.747373737 |
| Slc26a10 | 0.497380598 | 2.154722595 | 0.019 | 0.008 | 1 | 0.878310931 | 0.667279848 | 0.024 | 0.02 | 1 | 0.878310931 | 0.747373737 |
| Ryr2 | 0.497380598 | 2.154722595 | 0.019 | 0.008 | 1 | 0.738375177 | 1.100239256 | 0.024 | 0.017 | 1 | 0.738375177 | 0.747373737 |
| Gm56602 | 0.497380598 | 2.569760094 | 0.019 | 0.008 | 1 | 0.51919986 | 0.83720485 | 0 | 0.01 | 1 | 0.51919986 | 0.747373737 |
| Arhgef33 | 0.497380598 | 2.154722595 | 0.019 | 0.008 | 1 | 0.674180157 | 0.252242349 | 0.024 | 0.038 | 1 | 0.674180157 | 0.747373737 |
| Hap1 | 0.497382333 | 2.154722595 | 0.019 | 0.008 | 1 | 0.605005904 | 1.029849928 | 0.024 | 0.014 | 1 | 0.605005904 | 0.74737548 |
| Farsa | 0.498286645 | 0.483786871 | 0.377 | 0.352 | 1 | 0.805027359 | 0.203332749 | 0.39 | 0.416 | 1 | 0.805027359 | 0.74828371 |
| Gem | 0.908238381 | -0.368839361 | 0.66 | 0.657 | 1 | 0.49857029 | 0.363621341 | 0.756 | 0.771 | 1 | 0.908238381 | 0.748568246 |
| D130062J10Rik | 0.77169508 | 0.417757001 | 0.038 | 0.03 | 1 | 0.498856313 | -0.16279515 | 0.024 | 0.048 | 1 | 0.77169508 | 0.748855005 |
| Gm43138 | 0.877748789 | 0.247831999 | 0.057 | 0.051 | 1 | 0.498856313 | -0.16279515 | 0.024 | 0.048 | 1 | 0.877748789 | 0.748855005 |
| Micos13 | 0.499983308 | -0.137728299 | 0.792 | 0.771 | 1 | 0.659416902 | 0.232007865 | 0.659 | 0.734 | 1 | 0.659416902 | 0.749983308 |
| Vps52 | 0.500054035 | 0.267197324 | 0.17 | 0.131 | 1 | 0.98012274 | 0.306690133 | 0.146 | 0.154 | 1 | 0.98012274 | 0.750054032 |
| Trmt10a | 0.64758919 | -0.242167558 | 0.226 | 0.254 | 1 | 0.500201067 | 0.811669758 | 0.195 | 0.16 | 1 | 0.64758919 | 0.750201026 |
| Suv39h1 | 0.573265034 | 0.652222254 | 0.151 | 0.127 | 1 | 0.500329547 | 1.422167351 | 0.122 | 0.096 | 1 | 0.573265034 | 0.750329439 |
| Stxbp5l | 0.500671253 | 1.739685096 | 0.019 | 0.008 | 1 | 0.878310931 | 0.515276755 | 0.024 | 0.02 | 1 | 0.878310931 | 0.750670802 |
| Gm26542 | 0.500813115 | 0.652222254 | 0.151 | 0.119 | 1 | 0.98328985 | 0.100239256 | 0.171 | 0.174 | 1 | 0.98328985 | 0.750812454 |
| Rnf168 | 0.500996092 | 0.486344086 | 0.453 | 0.445 | 1 | 0.801088361 | 0.180451666 | 0.439 | 0.526 | 1 | 0.801088361 | 0.7509951 |
| Fbxo31 | 0.821691149 | -0.278236812 | 0.17 | 0.178 | 1 | 0.501220823 | -0.305753104 | 0.098 | 0.137 | 1 | 0.821691149 | 0.751219332 |
| Tbc1d31 | 0.502167138 | 0.154722595 | 0.094 | 0.068 | 1 | 0.980364501 | 0.46797104 | 0.073 | 0.075 | 1 | 0.980364501 | 0.752162441 |
| Spin4 | 0.502892343 | 0.569760094 | 0.132 | 0.102 | 1 | 0.932212486 | 0.865219268 | 0.146 | 0.15 | 1 | 0.932212486 | 0.752883977 |
| ENSMUSG00000121456 | 0.503312306 | 0.154722595 | 0.019 | 0.038 | 1 | 0.87556513 | 0.667279848 | 0.024 | 0.02 | 1 | 0.87556513 | 0.753301335 |
| A830082N09Rik | 0.812595221 | 0.569760094 | 0.094 | 0.085 | 1 | 0.503543529 | 0.515276755 | 0.073 | 0.048 | 1 | 0.812595221 | 0.753530973 |
| Ccdc187 | 0.503971666 | 1.569760094 | 0.019 | 0.008 | 1 | 0.856407545 | 2.159132945 | 0.024 | 0.02 | 1 | 0.856407545 | 0.753955892 |
| Phactr3 | 0.503971666 | 1.569760094 | 0.019 | 0.008 | 1 | 0.51919986 | 0.252242349 | 0 | 0.01 | 1 | 0.51919986 | 0.753955892 |
| 4930589L23Rik | 0.503971666 | 1.569760094 | 0.019 | 0.008 | 1 | 0.51919986 | 0.83720485 | 0 | 0.01 | 1 | 0.51919986 | 0.753955892 |
| Slc9b2 | 0.503971666 | 1.569760094 | 0.019 | 0.008 | 1 | 0.648942779 | -1.057612914 | 0.049 | 0.034 | 1 | 0.648942779 | 0.753955892 |
| Hook1 | 0.503971666 | 1.569760094 | 0.019 | 0.008 | 1 | 0.567356669 | 0.404245443 | 0.098 | 0.072 | 1 | 0.567356669 | 0.753955892 |
| ENSMUSG00000121439 | 0.503971666 | 1.569760094 | 0.019 | 0.008 | 1 | 0.989595616 | 0.515276755 | 0.024 | 0.024 | 1 | 0.989595616 | 0.753955892 |
| Fam184b | 0.503971666 | 1.417757001 | 0.019 | 0.008 | 1 | 0.51919986 | 0.83720485 | 0 | 0.01 | 1 | 0.51919986 | 0.753955892 |
| Ccdc142os | 0.503971666 | 1.569760094 | 0.019 | 0.008 | 1 | 0.51919986 | 0.83720485 | 0 | 0.01 | 1 | 0.51919986 | 0.753955892 |
| D6Ertd474e | 0.503971666 | 1.569760094 | 0.019 | 0.008 | 1 | 0.587925597 | 2.351778023 | 0.024 | 0.014 | 1 | 0.587925597 | 0.753955892 |
| Gm45502 | 0.503971666 | 1.569760094 | 0.019 | 0.008 | 1 | 0.51919986 | 0.83720485 | 0 | 0.01 | 1 | 0.51919986 | 0.753955892 |
| Gm40630 | 0.503971666 | 1.154722595 | 0.019 | 0.008 | 1 | 0.5191986 | -1.069685746 | 0 | 0.01 | 1 | 0.5191986 | 0.753955892 |
| Tbxa2r | 0.503971666 | 1.569760094 | 0.019 | 0.008 | 1 | 0.599290422 | 1.515276755 | 0.024 | 0.014 | 1 | 0.599290422 | 0.753955892 |
| Tektip1 | 0.503971666 | 1.569760094 | 0.019 | 0.008 | 1 | 0.718266349 | 0.930314254 | 0.049 | 0.038 | 1 | 0.718266349 | 0.753955892 |
| Syt1 | 0.503971666 | 1.569760094 | 0.019 | 0.008 | 1 | 0.756560731 | 0.174239837 | 0.024 | 0.034 | 1 | 0.756560731 | 0.753955892 |
| Zfp454 | 0.503971666 | 1.569760094 | 0.019 | 0.008 | 1 | 0.599290422 | 1.515276755 | 0.024 | 0.014 | 1 | 0.599290422 | 0.753955892 |
| Vmo1 | 0.503971666 | 1.569760094 | 0.019 | 0.008 | 1 | 0.733713289 | 0.515276755 | 0.049 | 0.038 | 1 | 0.733713289 | 0.753955892 |
| 4933428G20Rik | 0.503971666 | 1.569760094 | 0.019 | 0.008 | 1 | 0.732716552 | 1.614812428 | 0.024 | 0.017 | 1 | 0.732716552 | 0.753955892 |
| Rgs9 | 0.503971666 | 1.669295768 | 0.019 | 0.008 | 1 | 0.989595616 | 0.667279848 | 0.024 | 0.024 | 1 | 0.989595616 | 0.753955892 |
| Dtnbos | 0.503971666 | 1.569760094 | 0.019 | 0.008 | 1 | 0.911761241 | 0.377773231 | 0.024 | 0.027 | 1 | 0.911761241 | 0.753955892 |
| Gm47819 | 0.503971666 | 1.569760094 | 0.019 | 0.008 | 1 | 0.60213981 | 1.252242349 | 0.024 | 0.014 | 1 | 0.60213981 | 0.753955892 |
| Phf2os1 | 0.503971666 | 1.569760094 | 0.019 | 0.008 | 1 | 0.824088343 | 0.377773231 | 0.024 | 0.031 | 1 | 0.824088343 | 0.753955892 |
| A430090L17Rik | 0.503971666 | 1.569760094 | 0.019 | 0.008 | 1 | 0.650841212 | 0.422167351 | 0.049 | 0.034 | 1 | 0.650841212 | 0.753955892 |
| D030051J21Rik | 0.503971666 | 1.569760094 | 0.019 | 0.008 | 1 | 0.51919986 | 0.83720485 | 0 | 0.01 | 1 | 0.51919986 | 0.753955892 |
| Stab1 | 0.503971666 | 1.569760094 | 0.019 | 0.008 | 1 | 0.87556513 | 0.83720485 | 0.024 | 0.02 | 1 | 0.87556513 | 0.753955892 |
| Gm10101 | 0.503971666 | 1.569760094 | 0.019 | 0.008 | 1 | 0.5191986 | 0.515276755 | 0 | 0.01 | 1 | 0.5191986 | 0.753955892 |
| Ltb4r1 | 0.503971666 | 1.569760094 | 0.019 | 0.008 | 1 | 0.58792152 | 2.515276755 | 0.024 | 0.014 | 1 | 0.58792152 | 0.753955892 |
| Mapk8ip2 | 0.503971666 | 1.569760094 | 0.019 | 0.008 | 1 | 0.872828964 | 1.029849928 | 0.024 | 0.02 | 1 | 0.872828964 | 0.753955892 |

| | | | | | | | | | | | | |
|---|---|---|---|---|---|---|---|---|---|---|---|---|
| Gm50071 | 0.503971666 | 1.569760094 | 0.019 | 0.008 | 1 | 0.983241205 | 0.422167351 | 0.049 | 0.048 | 1 | 0.983241205 | 0.753955892 |
| Egfl8 | 0.503971666 | 1.569760094 | 0.019 | 0.008 | 1 | 0.826384776 | 0.515276755 | 0.024 | 0.031 | 1 | 0.826384776 | 0.753955892 |
| Gm50020 | 0.503971666 | 1.569760094 | 0.019 | 0.008 | 1 | 0.5191986 | 0.252242349 | 0 | 0.01 | 1 | 0.5191986 | 0.753955892 |
| Gm2629 | 0.503971666 | 1.569760094 | 0.019 | 0.008 | 1 | 0.60213981 | 1.029849928 | 0.024 | 0.014 | 1 | 0.60213981 | 0.753955892 |
| Gm29650 | 0.503971666 | 1.569760094 | 0.019 | 0.008 | 1 | 0.744047924 | 1.252242349 | 0.024 | 0.017 | 1 | 0.744047924 | 0.753955892 |
| Agr2 | 0.504094136 | -2.557995453 | 0.075 | 0.102 | 1 | 0.975749067 | 0.432814595 | 0.146 | 0.154 | 1 | 0.975749067 | 0.754077374 |
| Tdg | 0.504529103 | 0.459577176 | 0.434 | 0.407 | 1 | 0.587352508 | -0.145788725 | 0.463 | 0.512 | 1 | 0.587352508 | 0.75450859 |
| 2500002B13Rik | 0.50514607 | 0.280253477 | 0.019 | 0.038 | 1 | 0.986965808 | 0.334704509 | 0.049 | 0.048 | 1 | 0.986965808 | 0.755119588 |
| Zfp946 | 0.59998885 | 0.631160639 | 0.057 | 0.081 | 1 | 0.505456779 | 0.515276755 | 0.122 | 0.089 | 1 | 0.59998885 | 0.755427002 |
| Ccdc71 | 0.505457708 | 0.470224421 | 0.189 | 0.153 | 1 | 0.80401249 | 0.816143234 | 0.195 | 0.198 | 1 | 0.80401249 | 0.755427921 |
| Neu1 | 0.50624816 | 0.63687429 | 0.264 | 0.233 | 1 | 0.820196974 | 0.178993367 | 0.22 | 0.246 | 1 | 0.820196974 | 0.75620912 |
| Gm42941 | 0.639630093 | 0.404700848 | 0.132 | 0.11 | 1 | 0.506717502 | -0.555112573 | 0.073 | 0.106 | 1 | 0.639630093 | 0.756672378 |
| Gsap | 0.507017057 | 1.324647596 | 0.019 | 0.038 | 1 | 0.815790012 | 1.126711467 | 0.073 | 0.065 | 1 | 0.815790012 | 0.756967818 |
| Fem1a | 0.850692736 | 0.306725688 | 0.075 | 0.068 | 1 | 0.507047055 | 0.493250449 | 0.22 | 0.177 | 1 | 0.850692736 | 0.756997394 |
| Nudcd1 | 0.587984097 | 0.324647596 | 0.34 | 0.301 | 1 | 0.507210123 | 0.796177582 | 0.268 | 0.229 | 1 | 0.587984097 | 0.757158137 |
| Kmo | 0.507281812 | 1.154722595 | 0.019 | 0.008 | 1 | 0.51919986 | 0.83720485 | 0 | 0.01 | 1 | 0.51919986 | 0.757228787 |
| Cacna1b | 0.507281812 | 1.154722595 | 0.019 | 0.008 | 1 | 0.605005904 | 0.83720485 | 0.024 | 0.014 | 1 | 0.605005904 | 0.757228787 |
| Coro2a | 0.507281812 | 0.739685096 | 0.019 | 0.008 | 1 | 0.5191986 | 0.515276755 | 0 | 0.01 | 1 | 0.5191986 | 0.757228787 |
| Prom1 | 0.507281812 | 0.8327945 | 0.019 | 0.008 | 1 | 0.656078223 | -0.116991461 | 0.073 | 0.055 | 1 | 0.656078223 | 0.757228787 |
| Tfpi2 | 0.507281812 | 1.154722595 | 0.019 | 0.008 | 1 | 0.51919986 | 0.83720485 | 0 | 0.01 | 1 | 0.51919986 | 0.757228787 |
| B4galnt4 | 0.507281812 | 1.154722595 | 0.019 | 0.008 | 1 | 0.51919986 | 0.83720485 | 0 | 0.01 | 1 | 0.51919986 | 0.757228787 |
| Gm16104 | 0.507281812 | 1.154722595 | 0.019 | 0.008 | 1 | 0.5191986 | 0.515276755 | 0 | 0.01 | 1 | 0.5191986 | 0.757228787 |
| 4930505A04Rik | 0.507281812 | 1.154722595 | 0.019 | 0.008 | 1 | 0.58792152 | 2.515276755 | 0.024 | 0.014 | 1 | 0.58792152 | 0.757228787 |
| Abi3 | 0.507281812 | 1.154722595 | 0.019 | 0.008 | 1 | 0.51919986 | 0.83720485 | 0 | 0.01 | 1 | 0.51919986 | 0.757228787 |
| Zfp85os | 0.507281812 | 1.154722595 | 0.019 | 0.008 | 1 | 0.51919986 | 0.83720485 | 0 | 0.01 | 1 | 0.51919986 | 0.757228787 |
| Gm47467 | 0.507281812 | 1.154722595 | 0.019 | 0.008 | 1 | 0.51919986 | 0.83720485 | 0 | 0.01 | 1 | 0.51919986 | 0.757228787 |
| Gm49164 | 0.507281812 | 1.154722595 | 0.019 | 0.008 | 1 | 0.5191986 | 0.252242349 | 0 | 0.01 | 1 | 0.5191986 | 0.757228787 |
| Trem3 | 0.507281812 | 0.154722595 | 0.019 | 0.008 | 1 | 0.51919986 | 0.83720485 | 0 | 0.01 | 1 | 0.51919986 | 0.757228787 |
| Zfp719 | 0.5072929 | 0.428897558 | 0.189 | 0.153 | 1 | 0.953070443 | 1.000703582 | 0.122 | 0.133 | 1 | 0.953070443 | 0.757239713 |
| Rab24 | 0.862966179 | 0.106788118 | 0.472 | 0.47 | 1 | 0.507957981 | 0.466367154 | 0.39 | 0.355 | 1 | 0.862966179 | 0.757894652 |
| Fkbp5 | 0.515264682 | 0.10496956 | 0.377 | 0.424 | 1 | 0.508490789 | 0.294062525 | 0.463 | 0.423 | 1 | 0.515264682 | 0.758418696 |
| Elk1 | 0.553448095 | -0.459987249 | 0.132 | 0.161 | 1 | 0.508836108 | -0.278272368 | 0.122 | 0.16 | 1 | 0.553448095 | 0.758758031 |
| Gm16201 | 0.509865945 | 0.502645898 | 0.094 | 0.068 | 1 | 0.822590838 | 0.410940095 | 0.122 | 0.113 | 1 | 0.822590838 | 0.759768608 |
| Gm19409 | 0.510090659 | -0.1672055 | 0.075 | 0.051 | 1 | 0.701437198 | -0.262330824 | 0.122 | 0.143 | 1 | 0.701437198 | 0.759988838 |
| Toe1 | 0.958882966 | 0.130875853 | 0.208 | 0.208 | 1 | 0.510329389 | -0.16279515 | 0.195 | 0.239 | 1 | 0.958882966 | 0.760222693 |
| Sgk2 | 0.510606779 | 0.8327945 | 0.019 | 0.008 | 1 | 0.51919986 | 0.83720485 | 0 | 0.01 | 1 | 0.51919986 | 0.760494275 |
| Cdc20b | 0.510606779 | 0.8327945 | 0.019 | 0.008 | 1 | 0.916649754 | 0.667279848 | 0.024 | 0.027 | 1 | 0.916649754 | 0.760494275 |
| Sod1 | 0.510658419 | 0.192393013 | 0.792 | 0.78 | 1 | 0.592431444 | -0.128181395 | 0.683 | 0.754 | 1 | 0.592431444 | 0.760544817 |
| Alkbh4 | 0.662550611 | 0.569760094 | 0.075 | 0.059 | 1 | 0.510746073 | 0.515276755 | 0.122 | 0.089 | 1 | 0.662550611 | 0.760630595 |
| Zfp395 | 0.511429866 | 0.2967416 | 0.302 | 0.258 | 1 | 0.537543742 | 0.412707021 | 0.366 | 0.341 | 1 | 0.537543742 | 0.761299224 |
| Gm39090 | 0.755540713 | 0.219817623 | 0.189 | 0.174 | 1 | 0.511549151 | 0.209173627 | 0.122 | 0.171 | 1 | 0.755540713 | 0.761415768 |
| Cfap97 | 0.512182989 | 0.231343876 | 0.358 | 0.305 | 1 | 0.959096583 | 0.345351753 | 0.22 | 0.222 | 1 | 0.959096583 | 0.762034564 |
| Lancl2 | 0.512394956 | 0.417757001 | 0.189 | 0.153 | 1 | 0.859123358 | -0.440779897 | 0.22 | 0.218 | 1 | 0.859123358 | 0.762241321 |
| Dnaja4 | 0.819337513 | -0.464187238 | 0.226 | 0.225 | 1 | 0.512402451 | 0.330244861 | 0.317 | 0.266 | 1 | 0.819337513 | 0.76224863 |
| Ccdc22 | 0.516262239 | 0.491757582 | 0.094 | 0.068 | 1 | 0.51246171 | 0.392420007 | 0.098 | 0.137 | 1 | 0.516262239 | 0.762306415 |
| Hmga2 | 0.513088673 | -1.719746523 | 0.038 | 0.059 | 1 | 0.789604246 | -0.863234868 | 0.073 | 0.061 | 1 | 0.789604246 | 0.76291736 |
| Nupl2 | 0.513564211 | 0.785488785 | 0.132 | 0.106 | 1 | 0.823027036 | 0.120997816 | 0.122 | 0.109 | 1 | 0.823027036 | 0.763380223 |
| Taf7 | 0.966135242 | 0.194250959 | 0.509 | 0.547 | 1 | 0.513687158 | -0.144275062 | 0.512 | 0.57 | 1 | 0.966135242 | 0.76349982 |
| Cradd | 0.514348467 | -0.500629234 | 0.189 | 0.233 | 1 | 0.524115085 | 0.149880631 | 0.341 | 0.29 | 1 | 0.524115085 | 0.764142588 |
| Pdss1 | 0.514520133 | 0.267197324 | 0.226 | 0.191 | 1 | 0.731968828 | 0.721727632 | 0.122 | 0.154 | 1 | 0.731968828 | 0.764309299 |
| Aste1 | 0.514984487 | 0.242185436 | 0.264 | 0.22 | 1 | 0.803325588 | 0.788841828 | 0.22 | 0.225 | 1 | 0.803325588 | 0.764759952 |
| 1700028E10Rik | 0.932507852 | 1.640149422 | 0.019 | 0.021 | 1 | 0.515216473 | 2.185128153 | 0.049 | 0.031 | 1 | 0.932507852 | 0.764984932 |
| Tex9 | 0.964042 | 0.318221327 | 0.094 | 0.093 | 1 | 0.515581526 | -0.132421501 | 0.073 | 0.109 | 1 | 0.964042 | 0.765338742 |
| Gm31305 | 0.563757816 | -0.652632327 | 0.019 | 0.034 | 1 | 0.515886997 | 0.174239837 | 0.024 | 0.048 | 1 | 0.563757816 | 0.7656346 |
| Etnk1 | 0.515910175 | -0.171814753 | 0.83 | 0.852 | 1 | 0.587987007 | 0.255808964 | 0.829 | 0.874 | 1 | 0.587987007 | 0.765657041 |
| Gm57457 | 0.516262239 | 0.491757582 | 0.094 | 0.068 | 1 | 0.878993361 | 0.334704509 | 0.049 | 0.055 | 1 | 0.878993361 | 0.765997779 |
| Adat2 | 0.89261791 | 0.569760094 | 0.038 | 0.034 | 1 | 0.517811386 | 1.422167351 | 0.073 | 0.051 | 1 | 0.89261791 | 0.767494141 |
| Zfp182 | 0.93450284 | -0.180261653 | 0.302 | 0.288 | 1 | 0.51788793 | -0.283810551 | 0.244 | 0.294 | 1 | 0.93450284 | 0.767567952 |
| Bpifb1 | 0.567830665 | -0.652632327 | 0.019 | 0.034 | 1 | 0.518181178 | -1.435813645 | 0.049 | 0.075 | 1 | 0.567830665 | 0.767850623 |
| Rnf182 | 0.753377891 | 0.195364579 | 0.113 | 0.097 | 1 | 0.518616834 | 1.126711467 | 0.049 | 0.078 | 1 | 0.753377891 | 0.768270248 |
| Ccl19-ps2 | 0.518992757 | -0.845277405 | 0.038 | 0.059 | 1 | 0.96463636 | 0.766815522 | 0.049 | 0.048 | 1 | 0.96463636 | 0.768632032 |
| Sema4f | 0.569866135 | -0.1672055 | 0.019 | 0.034 | 1 | 0.5191986 | 0.515276755 | 0 | 0.01 | 1 | 0.569866135 | 0.768830014 |
| Gm35315 | 0.610863297 | 1.347367673 | 0.038 | 0.025 | 1 | 0.5191986 | 0.252242349 | 0 | 0.01 | 1 | 0.610863297 | 0.768830014 |
| AA465934 | 0.628925725 | 0.739685096 | 0.038 | 0.025 | 1 | 0.5191986 | 0.515276755 | 0 | 0.01 | 1 | 0.628925725 | 0.768830014 |
| Fancb | 0.628925725 | 0.739685096 | 0.038 | 0.025 | 1 | 0.5191986 | 0.515276755 | 0 | 0.01 | 1 | 0.628925725 | 0.768830014 |
| Map3k15 | 0.638060446 | 0.417757001 | 0.038 | 0.025 | 1 | 0.5191986 | 0.252242349 | 0 | 0.01 | 1 | 0.638060446 | 0.768830014 |
| Dnajc6 | 0.645965914 | 0.280253477 | 0.094 | 0.076 | 1 | 0.5191986 | 0.515276755 | 0 | 0.01 | 1 | 0.645965914 | 0.768830014 |
| Gm36201 | 0.722567081 | 1.8327945 | 0.019 | 0.013 | 1 | 0.5191986 | 0.515276755 | 0 | 0.01 | 1 | 0.722567081 | 0.768830014 |
| Nsl1 | 0.732691937 | 1.154722595 | 0.019 | 0.013 | 1 | 0.5191986 | 0.515276755 | 0 | 0.01 | 1 | 0.732691937 | 0.768830014 |
| Srrm3 | 0.736073999 | 0.8327945 | 0.019 | 0.013 | 1 | 0.5191986 | 0.515276755 | 0 | 0.01 | 1 | 0.736073999 | 0.768830014 |
| Tmem130 | 0.736073999 | 0.154722595 | 0.019 | 0.013 | 1 | 0.5191986 | 0.515276755 | 0 | 0.01 | 1 | 0.736073999 | 0.768830014 |
| Bcl2l14 | 0.7394666 | 0.569760094 | 0.019 | 0.013 | 1 | 0.5191986 | 0.252242349 | 0 | 0.01 | 1 | 0.7394666 | 0.768830014 |
| Plxna4os1 | 0.762480493 | 0.739685096 | 0.038 | 0.03 | 1 | 0.5191986 | 0.515276755 | 0 | 0.01 | 1 | 0.762480493 | 0.768830014 |
| Col13a1 | 0.797852948 | 2.010332685 | 0.019 | 0.025 | 1 | 0.5191986 | -0.484723245 | 0 | 0.01 | 1 | 0.797852948 | 0.768830014 |
| Cacng5 | 0.912039013 | -0.304709024 | 0.019 | 0.021 | 1 | 0.5191986 | 0.515276755 | 0 | 0.01 | 1 | 0.912039013 | 0.768830014 |
| Ankfn1 | 0.916620008 | 1.154722595 | 0.019 | 0.017 | 1 | 0.5191986 | 0.515276755 | 0 | 0.01 | 1 | 0.916620008 | 0.768830014 |
| ENSMUSG00000121482 | 0.917883255 | 0.569760094 | 0.019 | 0.021 | 1 | 0.5191986 | 0.515276755 | 0 | 0.01 | 1 | 0.917883255 | 0.768830014 |
| Wdr38 | 0.926212226 | 0.8327945 | 0.019 | 0.017 | 1 | 0.5191986 | 0.515276755 | 0 | 0.01 | 1 | 0.926212226 | 0.768830014 |
| Sgsm1 | 0.929410101 | 0.569760094 | 0.019 | 0.017 | 1 | 0.5191986 | 0.515276755 | 0 | 0.01 | 1 | 0.929410101 | 0.768830014 |
| Ankle1 | 0.932611893 | 0.154722595 | 0.019 | 0.017 | 1 | 0.5191986 | 0.252242349 | 0 | 0.01 | 1 | 0.932611893 | 0.768830014 |
| Gm14023 | 0.569866135 | -0.1672055 | 0.019 | 0.034 | 1 | 0.51919986 | 0.83720485 | 0 | 0.01 | 1 | 0.569866135 | 0.768831226 |
| E130304I02Rik | 0.569866135 | -0.1672055 | 0.019 | 0.034 | 1 | 0.51919986 | 0.83720485 | 0 | 0.01 | 1 | 0.569866135 | 0.768831226 |
| Duoxa1 | 0.582274458 | 0.389187848 | 0.019 | 0.034 | 1 | 0.51919986 | -0.16279515 | 0 | 0.01 | 1 | 0.582274458 | 0.768831226 |
| Rassf4 | 0.589569635 | -0.783876861 | 0.038 | 0.055 | 1 | 0.51919986 | 0.83720485 | 0 | 0.01 | 1 | 0.589569635 | 0.768831226 |
| Rab3b | 0.610863297 | 1.669295768 | 0.038 | 0.025 | 1 | 0.51919986 | 0.83720485 | 0 | 0.01 | 1 | 0.610863297 | 0.768831226 |
| Gm32585 | 0.626679848 | 0.47665069 | 0.038 | 0.025 | 1 | 0.51919986 | 0.83720485 | 0 | 0.01 | 1 | 0.626679848 | 0.768831226 |
| Gm56727 | 0.668070194 | 0.154722595 | 0.019 | 0.03 | 1 | 0.51919986 | 0.83720485 | 0 | 0.01 | 1 | 0.668070194 | 0.768831226 |
| 4930471E19Rik | 0.679455468 | -1.015202407 | 0.038 | 0.051 | 1 | 0.51919986 | -1.410722664 | 0 | 0.01 | 1 | 0.679455468 | 0.768831226 |
| Gm49313 | 0.72256466 | 1.739685096 | 0.019 | 0.013 | 1 | 0.51919986 | 0.83720485 | 0 | 0.01 | 1 | 0.72256466 | 0.768831226 |
| Sphkap | 0.725936481 | 1.417757001 | 0.019 | 0.013 | 1 | 0.51919986 | -0.332720152 | 0 | 0.01 | 1 | 0.725936481 | 0.768831226 |
| Gm44957 | 0.732691937 | 1.154722595 | 0.019 | 0.013 | 1 | 0.51919986 | 0.83720485 | 0 | 0.01 | 1 | 0.732691937 | 0.768831226 |
| Sdr42e1 | 0.732691937 | 1.154722595 | 0.019 | 0.013 | 1 | 0.51919986 | 0.83720485 | 0 | 0.01 | 1 | 0.732691937 | 0.768831226 |
| Acaa1b | 0.732691937 | 1.154722595 | 0.019 | 0.013 | 1 | 0.51919986 | 0.83720485 | 0 | 0.01 | 1 | 0.732691937 | 0.768831226 |
| Ccdc170 | 0.732691937 | 1.154722595 | 0.019 | 0.013 | 1 | 0.51919986 | -0.622226769 | 0 | 0.01 | 1 | 0.732691937 | 0.768831226 |
| Dsg3 | 0.732691937 | 1.154722595 | 0.019 | 0.013 | 1 | 0.51919986 | 0.83720485 | 0 | 0.01 | 1 | 0.732691937 | 0.768831226 |
| Creg2 | 0.736073999 | 0.569760094 | 0.019 | 0.013 | 1 | 0.51919986 | 0.83720485 | 0 | 0.01 | 1 | 0.736073999 | 0.768831226 |
| Gm11739 | 0.736073999 | 0.8327945 | 0.019 | 0.013 | 1 | 0.51919986 | 0.83720485 | 0 | 0.01 | 1 | 0.736073999 | 0.768831226 |
| Gm41335 | 0.736073999 | 0.8327945 | 0.019 | 0.013 | 1 | 0.51919986 | -1.16279515 | 0 | 0.01 | 1 | 0.736073999 | 0.768831226 |

| | | | | | | | | | | | | |
|---|---|---|---|---|---|---|---|---|---|---|---|---|
| Lrrc3 | 0.776830424 | -0.304709024 | 0.019 | 0.025 | 1 | 0.51919986 | 0.83720485 | 0 | 0.01 | 1 | 0.776830424 | 0.768831226 |
| Gm42918 | 0.782077996 | 0.347367673 | 0.019 | 0.025 | 1 | 0.51919986 | 0.83720485 | 0 | 0.01 | 1 | 0.782077996 | 0.768831226 |
| Irf5 | 0.782077996 | 0.347367673 | 0.019 | 0.025 | 1 | 0.51919986 | 0.83720485 | 0 | 0.01 | 1 | 0.782077996 | 0.768831226 |
| Gm13270 | 0.797844282 | 1.347367673 | 0.019 | 0.025 | 1 | 0.51919986 | 0.83720485 | 0 | 0.01 | 1 | 0.797844282 | 0.768831226 |
| Tnr | 0.837801119 | 1.216123139 | 0.075 | 0.068 | 1 | 0.51919986 | 0.252242349 | 0 | 0.01 | 1 | 0.837801119 | 0.768831226 |
| Xk | 0.89261791 | 0.569760094 | 0.038 | 0.034 | 1 | 0.51919986 | 0.83720485 | 0 | 0.01 | 1 | 0.89261791 | 0.768831226 |
| Gm36457 | 0.90155955 | 0.695290976 | 0.038 | 0.042 | 1 | 0.51919986 | 0.83720485 | 0 | 0.01 | 1 | 0.90155955 | 0.768831226 |
| Tg | 0.90620214 | -1.1672055 | 0.019 | 0.021 | 1 | 0.51919986 | 0.252242349 | 0 | 0.01 | 1 | 0.90620214 | 0.768831226 |
| Awat2 | 0.913424206 | 2.640149422 | 0.019 | 0.017 | 1 | 0.51919986 | 0.83720485 | 0 | 0.01 | 1 | 0.913424206 | 0.768831226 |
| Gm49719 | 0.914958 | 0.347367673 | 0.019 | 0.021 | 1 | 0.51919986 | 0.83720485 | 0 | 0.01 | 1 | 0.914958 | 0.768831226 |
| Igsf6 | 0.929410101 | 0.569760094 | 0.019 | 0.017 | 1 | 0.51919986 | 0.83720485 | 0 | 0.01 | 1 | 0.929410101 | 0.768831226 |
| Gm56533 | 0.929410101 | 0.569760094 | 0.019 | 0.017 | 1 | 0.51919986 | 0.83720485 | 0 | 0.01 | 1 | 0.929410101 | 0.768831226 |
| 1700105P06Rik | 0.932611893 | 0.154722595 | 0.019 | 0.017 | 1 | 0.51919986 | 0.83720485 | 0 | 0.01 | 1 | 0.932611893 | 0.768831226 |
| Unc80 | 0.96498869 | -0.582242999 | 0.038 | 0.038 | 1 | 0.51919986 | 0.83720485 | 0 | 0.01 | 1 | 0.96498869 | 0.768831226 |
| Il4ra | 0.520823689 | 1.377115016 | 0.057 | 0.038 | 1 | 0.914200263 | 0.515276755 | 0.024 | 0.027 | 1 | 0.914200263 | 0.770390063 |
| Trim59 | 0.727252586 | 0.180717803 | 0.151 | 0.136 | 1 | 0.520922423 | 0.630753972 | 0.171 | 0.137 | 1 | 0.727252586 | 0.770484675 |
| Garin2 | 0.521954735 | -0.904171094 | 0.038 | 0.059 | 1 | 0.945021736 | 0.326242931 | 0.122 | 0.119 | 1 | 0.945021736 | 0.771472724 |
| Tmem268 | 0.991244785 | 0.417757001 | 0.038 | 0.038 | 1 | 0.522109572 | 0.989207943 | 0.024 | 0.048 | 1 | 0.991244785 | 0.771620739 |
| Cpox | 0.786407409 | -0.237594828 | 0.302 | 0.305 | 1 | 0.522115289 | 0.442673006 | 0.366 | 0.352 | 1 | 0.786407409 | 0.771626203 |
| Dzank1 | 0.607656166 | -0.652632327 | 0.038 | 0.055 | 1 | 0.522123885 | 0.83720485 | 0.024 | 0.048 | 1 | 0.607656166 | 0.771634419 |
| Thg1l | 0.522862101 | 0.280253477 | 0.094 | 0.068 | 1 | 0.683651486 | 1.322631677 | 0.073 | 0.058 | 1 | 0.683651486 | 0.772339426 |
| Arrb1 | 0.772193541 | 1.154722595 | 0.075 | 0.068 | 1 | 0.523150596 | -0.747757651 | 0.049 | 0.075 | 1 | 0.772193541 | 0.772614646 |
| Nmnat1 | 0.523385329 | 0.200526284 | 0.113 | 0.085 | 1 | 0.5799992 | -0.372248516 | 0.073 | 0.099 | 1 | 0.5799992 | 0.772838456 |
| Msrb2 | 0.850604434 | -0.130679624 | 0.132 | 0.14 | 1 | 0.524023309 | -0.112754468 | 0.171 | 0.212 | 1 | 0.850604434 | 0.77344619 |
| Cyb561 | 0.524096498 | -0.7032584 | 0.057 | 0.081 | 1 | 0.550536867 | -0.250257991 | 0.024 | 0.044 | 1 | 0.550536867 | 0.773515857 |
| Tpd52 | 0.524247543 | 0.115194231 | 0.245 | 0.199 | 1 | 0.82931337 | 0.107470825 | 0.341 | 0.375 | 1 | 0.82931337 | 0.773659599 |
| Mcm2 | 0.524921031 | -0.652632327 | 0.038 | 0.059 | 1 | 0.902331879 | 0.252242349 | 0.049 | 0.044 | 1 | 0.902331879 | 0.774299974 |
| Zc3h3 | 0.526200401 | 0.137649081 | 0.302 | 0.254 | 1 | 0.634854308 | 0.439612485 | 0.317 | 0.304 | 1 | 0.634854308 | 0.77551394 |
| Rfxank | 0.526260575 | 0.43225657 | 0.151 | 0.119 | 1 | 0.583953617 | 0.252242349 | 0.073 | 0.102 | 1 | 0.583953617 | 0.775570957 |
| Slc38a1 | 0.527324855 | 0.131638982 | 0.66 | 0.691 | 1 | 0.603855069 | 0.252242349 | 0.244 | 0.205 | 1 | 0.603855069 | 0.776578207 |
| Gm57198 | 0.527414315 | 0.46284489 | 0.17 | 0.14 | 1 | 1 | 0.493250449 | 0.171 | 0.174 | 1 | 1 | 0.776662771 |
| Jam3 | 0.52748583 | 0.277733485 | 0.528 | 0.504 | 1 | 0.712074853 | 0.142059431 | 0.463 | 0.532 | 1 | 0.712074853 | 0.77673036 |
| Amd2 | 0.608559479 | 0.318221327 | 0.113 | 0.089 | 1 | 0.527746174 | 0.690363461 | 0.122 | 0.092 | 1 | 0.608559479 | 0.776976323 |
| Dnajb6 | 0.528108679 | 0.174825498 | 0.925 | 0.962 | 1 | 0.692481431 | 0.128583881 | 1 | 0.983 | 1 | 0.692481431 | 0.777318582 |
| Denr | 0.543750672 | 0.12144644 | 0.698 | 0.657 | 1 | 0.528197786 | 0.162894863 | 0.561 | 0.666 | 1 | 0.543750672 | 0.777402671 |
| Brcc3 | 0.968882535 | 0.186431455 | 0.283 | 0.288 | 1 | 0.52842785 | 0.593279267 | 0.39 | 0.382 | 1 | 0.968882535 | 0.777619707 |
| Vars2 | 0.529359278 | 1.417757001 | 0.057 | 0.038 | 1 | 0.824088343 | 0.377773231 | 0.024 | 0.031 | 1 | 0.824088343 | 0.778497311 |
| Slc22a21 | 0.814502235 | -0.354291053 | 0.321 | 0.322 | 1 | 0.529393901 | 0.486707603 | 0.244 | 0.208 | 1 | 0.814502235 | 0.778529899 |
| Gm38134 | 0.919816481 | 0.932330173 | 0.019 | 0.017 | 1 | 0.529718256 | 1.515276755 | 0.049 | 0.031 | 1 | 0.919816481 | 0.778835081 |
| Rmdn1 | 1 | 0.260215724 | 0.358 | 0.386 | 1 | 0.530175068 | -0.130227288 | 0.317 | 0.379 | 1 | 1 | 0.779264533 |
| Mcat | 0.941333459 | 0.247831999 | 0.057 | 0.059 | 1 | 0.530635728 | -0.484723245 | 0.049 | 0.075 | 1 | 0.941333459 | 0.77969718 |
| Nkap | 0.531540593 | 0.165064539 | 0.321 | 0.403 | 1 | 0.630207939 | 0.416873051 | 0.39 | 0.379 | 1 | 0.630207939 | 0.780545784 |
| Stk32c | 0.917883255 | 0.569760094 | 0.019 | 0.021 | 1 | 0.531564619 | 2.185128153 | 0.049 | 0.031 | 1 | 0.917883255 | 0.780568294 |
| Il15 | 0.732691937 | 1.154722595 | 0.019 | 0.013 | 1 | 0.532248915 | 1.029849928 | 0.073 | 0.051 | 1 | 0.732691937 | 0.781208923 |
| Epg5 | 0.636235201 | -0.180880437 | 0.264 | 0.301 | 1 | 0.532437022 | -0.224571348 | 0.268 | 0.328 | 1 | 0.636235201 | 0.781384862 |
| Mrps18b | 0.534520072 | -0.193200709 | 0.302 | 0.343 | 1 | 0.532603324 | 0.83720485 | 0.317 | 0.276 | 1 | 0.534520072 | 0.781540348 |
| Gm19705 | 0.748628929 | -0.359850578 | 0.094 | 0.11 | 1 | 0.533536804 | -0.395455907 | 0.073 | 0.106 | 1 | 0.748628929 | 0.782412087 |
| Hk1os | 0.533848988 | -0.260314904 | 0.038 | 0.059 | 1 | 0.744047924 | 1.252242349 | 0.024 | 0.017 | 1 | 0.744047924 | 0.782703234 |
| 2810013P06Rik | 0.534518442 | 0.3662267 | 0.264 | 0.242 | 1 | 0.900242788 | 0.515276755 | 0.195 | 0.195 | 1 | 0.900242788 | 0.783326919 |
| Grk2 | 0.535170394 | 0.549001534 | 0.283 | 0.258 | 1 | 0.897775672 | 0.747567637 | 0.366 | 0.406 | 1 | 0.897775672 | 0.783933437 |
| Prr18 | 0.732691937 | 1.154722595 | 0.019 | 0.013 | 1 | 0.535235731 | 1.252242349 | 0.049 | 0.031 | 1 | 0.732691937 | 0.783994175 |
| Plekhb2 | 0.536286207 | 0.318221327 | 0.057 | 0.085 | 1 | 0.768953216 | -0.513292397 | 0.098 | 0.109 | 1 | 0.768953216 | 0.784969518 |
| Prxl2b | 0.537369199 | -0.304709024 | 0.113 | 0.148 | 1 | 0.57893969 | -0.332720152 | 0.049 | 0.072 | 1 | 0.57893969 | 0.785972742 |
| Gm42917 | 0.961556928 | 0.306725688 | 0.057 | 0.055 | 1 | 0.537851449 | 0.667279848 | 0.122 | 0.092 | 1 | 0.961556928 | 0.786418717 |
| Ptgr2 | 0.538464569 | 0.347367673 | 0.17 | 0.136 | 1 | 0.872083355 | 0.322631677 | 0.171 | 0.191 | 1 | 0.872083355 | 0.786985046 |
| B3gntl1 | 0.904012755 | -0.1672055 | 0.094 | 0.097 | 1 | 0.539633009 | 0.159132945 | 0.049 | 0.078 | 1 | 0.904012755 | 0.788062234 |
| Nudt14 | 0.873614749 | 0.228723176 | 0.113 | 0.123 | 1 | 0.540659399 | -0.16279515 | 0.049 | 0.075 | 1 | 0.873614749 | 0.789006212 |
| Arg2 | 0.789955003 | 0.417757001 | 0.019 | 0.025 | 1 | 0.540778536 | 0.930314254 | 0.049 | 0.031 | 1 | 0.789955003 | 0.789115647 |
| Fabp3 | 0.540811687 | -1.412961914 | 0.151 | 0.114 | 1 | 0.683192894 | 1.029849928 | 0.098 | 0.082 | 1 | 0.683192894 | 0.789146093 |
| Zfp560 | 0.541467546 | 0.14233887 | 0.415 | 0.369 | 1 | 0.729202835 | 0.325305811 | 0.293 | 0.365 | 1 | 0.729202835 | 0.789747989 |
| Zbtb43 | 0.790719585 | 0.154722595 | 0.226 | 0.263 | 1 | 0.541531426 | -0.186127654 | 0.317 | 0.375 | 1 | 0.790719585 | 0.789806567 |
| Aagab | 0.541552582 | 0.819119563 | 0.208 | 0.182 | 1 | 0.706338447 | 0.268362014 | 0.244 | 0.218 | 1 | 0.706338447 | 0.789825965 |
| Gna11 | 0.541665869 | 0.158262953 | 0.774 | 0.771 | 1 | 0.571889259 | 0.207029772 | 0.805 | 0.775 | 1 | 0.571889259 | 0.789929824 |
| Vangl2 | 0.553620571 | -0.323324702 | 0.094 | 0.123 | 1 | 0.541745061 | -0.410722664 | 0.171 | 0.205 | 1 | 0.553620571 | 0.790002411 |
| Gm17382 | 0.796143932 | 0.739685096 | 0.038 | 0.047 | 1 | 0.542339903 | -0.484723245 | 0.024 | 0.044 | 1 | 0.796143932 | 0.790547235 |
| Ctu1 | 0.967052971 | 0.347367673 | 0.057 | 0.055 | 1 | 0.543544585 | -0.101394606 | 0.049 | 0.075 | 1 | 0.967052971 | 0.791648454 |
| Abitram | 0.543713421 | -0.229941255 | 0.132 | 0.165 | 1 | 0.80536539 | 0.544423101 | 0.122 | 0.116 | 1 | 0.80536539 | 0.791802558 |
| Naa80 | 0.543828205 | 0.207190015 | 0.113 | 0.085 | 1 | 0.722125834 | -0.292078167 | 0.073 | 0.089 | 1 | 0.722125834 | 0.791907294 |
| Gm15991 | 0.633487796 | 0.569760094 | 0.038 | 0.025 | 1 | 0.543855984 | 1.071670103 | 0.073 | 0.051 | 1 | 0.633487796 | 0.791932637 |
| Strip2 | 0.586409164 | 0.417757001 | 0.019 | 0.034 | 1 | 0.543976862 | -0.917682652 | 0.024 | 0.044 | 1 | 0.586409164 | 0.792042898 |
| 6530409C15Rik | 0.610863297 | 1.347367673 | 0.038 | 0.025 | 1 | 0.543978477 | -0.555112573 | 0.024 | 0.044 | 1 | 0.610863297 | 0.79204437 |
| Zfp97 | 0.54507254 | 0.8327945 | 0.057 | 0.038 | 1 | 0.550031708 | 0.962735732 | 0.049 | 0.031 | 1 | 0.550031708 | 0.793041006 |
| Ighmbp2 | 0.54507254 | 0.8327945 | 0.057 | 0.038 | 1 | 0.949388172 | 0.422167351 | 0.049 | 0.051 | 1 | 0.949388172 | 0.793041006 |
| Ism2 | 0.628925725 | 0.739685096 | 0.038 | 0.025 | 1 | 0.54561906 | -0.686357106 | 0.024 | 0.044 | 1 | 0.628925725 | 0.793537961 |
| Mrps15 | 0.786918109 | 0.429582436 | 0.434 | 0.47 | 1 | 0.54576309 | 0.16204454 | 0.512 | 0.451 | 1 | 0.786918109 | 0.79366883 |
| Fkbpl | 0.796151811 | 0.569760094 | 0.038 | 0.047 | 1 | 0.546365767 | 1.100239256 | 0.049 | 0.031 | 1 | 0.796151811 | 0.794215983 |
| Mlx | 0.830137754 | 0.240452469 | 0.321 | 0.318 | 1 | 0.546953918 | -0.286177566 | 0.268 | 0.307 | 1 | 0.830137754 | 0.794749248 |
| Sfi1 | 0.54712417 | 0.792152515 | 0.245 | 0.225 | 1 | 0.922883531 | 0.440314697 | 0.268 | 0.287 | 1 | 0.922883531 | 0.794903483 |
| Mrpl19 | 0.547124514 | -0.347777746 | 0.151 | 0.182 | 1 | 0.892586639 | 0.181853021 | 0.171 | 0.16 | 1 | 0.892586639 | 0.794903794 |
| Galns | 0.958287491 | 0.207190015 | 0.094 | 0.093 | 1 | 0.547258706 | -0.410722664 | 0.024 | 0.044 | 1 | 0.958287491 | 0.79502532 |
| Oxnad1 | 0.548086447 | -0.555770788 | 0.151 | 0.182 | 1 | 0.775099199 | 0.279209397 | 0.146 | 0.133 | 1 | 0.775099199 | 0.795774141 |
| Zbtb49 | 0.709332007 | 1.652222254 | 0.075 | 0.064 | 1 | 0.548870133 | -0.292078167 | 0.073 | 0.102 | 1 | 0.709332007 | 0.796481843 |
| Kctd7 | 0.597481259 | 1.669295768 | 0.038 | 0.025 | 1 | 0.548898236 | -0.16279515 | 0.024 | 0.044 | 1 | 0.597481259 | 0.796507199 |
| Tspan32os | 0.901698096 | 0.280253477 | 0.038 | 0.034 | 1 | 0.548898236 | -0.16279515 | 0.024 | 0.044 | 1 | 0.901698096 | 0.796507199 |
| Enkur | 0.549185868 | 0.438515561 | 0.038 | 0.059 | 1 | 0.858133882 | -0.435813645 | 0.049 | 0.055 | 1 | 0.858133882 | 0.796766618 |
| Eaf2 | 0.549905985 | 0.336560918 | 0.151 | 0.195 | 1 | 0.573491877 | 0.717465606 | 0.195 | 0.167 | 1 | 0.573491877 | 0.797415378 |
| Hemk1 | 0.605896064 | -0.1672055 | 0.038 | 0.055 | 1 | 0.550031708 | 0.83720485 | 0.049 | 0.031 | 1 | 0.605896064 | 0.797528536 |
| Cyp2r1 | 0.663401681 | -0.304709024 | 0.019 | 0.03 | 1 | 0.550031708 | 0.962735732 | 0.049 | 0.031 | 1 | 0.663401681 | 0.797528536 |
| Ccno | 0.992810097 | 0.228723176 | 0.094 | 0.093 | 1 | 0.550031708 | 0.962735732 | 0.049 | 0.031 | 1 | 0.992810097 | 0.797528536 |
| Actr6 | 0.55021179 | 0.347367673 | 0.208 | 0.174 | 1 | 0.803974818 | 0.489281546 | 0.146 | 0.137 | 1 | 0.803974818 | 0.797690566 |
| Chek2 | 0.550237964 | 0.695290976 | 0.057 | 0.038 | 1 | 0.737610035 | 0.614812428 | 0.049 | 0.038 | 1 | 0.737610035 | 0.797714111 |
| Tecr | 0.680508456 | -0.275805986 | 0.83 | 0.814 | 1 | 0.550290188 | 0.167074966 | 0.756 | 0.771 | 1 | 0.680508456 | 0.797761085 |
| Snx18 | 0.886040997 | -0.264229954 | 0.698 | 0.746 | 1 | 0.550974859 | -0.11526678 | 0.902 | 0.843 | 1 | 0.886040997 | 0.798376422 |
| Rbks | 0.551270122 | -0.237594828 | 0.057 | 0.081 | 1 | 0.615779071 | 0.83720485 | 0.049 | 0.072 | 1 | 0.615779071 | 0.798641496 |
| Gm16867 | 0.551270122 | -0.237594828 | 0.057 | 0.081 | 1 | 0.737610035 | 0.614812428 | 0.049 | 0.038 | 1 | 0.737610035 | 0.798641496 |
| Ikzf4 | 0.612694334 | 0.195364579 | 0.358 | 0.326 | 1 | 0.552273986 | 0.427813914 | 0.341 | 0.307 | 1 | 0.612694334 | 0.799541417 |
| Smim1 | 0.969050772 | -0.1672055 | 0.132 | 0.131 | 1 | 0.553369223 | 0.392420007 | 0.171 | 0.137 | 1 | 0.969050772 | 0.800520949 |
| Rasa4 | 0.554142306 | -0.430239906 | 0.057 | 0.081 | 1 | 0.878043813 | 0.690363461 | 0.073 | 0.068 | 1 | 0.878043813 | 0.801210917 |

| | | | | | | | | | | | | |
|---|---|---|---|---|---|---|---|---|---|---|---|---|
| Urb2 | 0.555111138 | 0.569760094 | 0.132 | 0.106 | 1 | 0.816644007 | -0.271319607 | 0.171 | 0.177 | 1 | 0.816644007 | 0.8020739 |
| Dmac1 | 0.555215033 | 0.311448098 | 0.396 | 0.386 | 1 | 0.749403986 | -0.150722318 | 0.293 | 0.307 | 1 | 0.749403986 | 0.802166333 |
| Ppih | 0.55628892 | 0.8327945 | 0.132 | 0.106 | 1 | 0.807144138 | 0.574170444 | 0.146 | 0.137 | 1 | 0.807144138 | 0.803120477 |
| Gm12359 | 0.556741021 | 0.762405172 | 0.094 | 0.072 | 1 | 0.80950686 | 0.185128153 | 0.098 | 0.113 | 1 | 0.80950686 | 0.803521478 |
| Stradb | 0.674095102 | 0.318221327 | 0.113 | 0.093 | 1 | 0.557067995 | 0.865219226 | 0.171 | 0.14 | 1 | 0.674095102 | 0.803811239 |
| Pcdhgb4 | 0.557359948 | 0.8327945 | 0.075 | 0.055 | 1 | 0.780128581 | 0.204936634 | 0.073 | 0.061 | 1 | 0.780128581 | 0.804069784 |
| Tctn3 | 0.557359948 | 0.8327945 | 0.075 | 0.055 | 1 | 0.703921817 | 0.667279848 | 0.073 | 0.058 | 1 | 0.703921817 | 0.804069784 |
| 4933424G06Rik | 0.679455197 | -0.347777746 | 0.038 | 0.051 | 1 | 0.557539301 | 0.515276755 | 0.049 | 0.031 | 1 | 0.679455197 | 0.80422853 |
| Mki67 | 0.557662211 | -3.498719644 | 0.019 | 0.034 | 1 | 0.60213981 | 0.252242349 | 0.024 | 0.014 | 1 | 0.60213981 | 0.80433728 |
| Tusc2 | 0.59577707 | 0.373362881 | 0.226 | 0.195 | 1 | 0.558651643 | 0.568716014 | 0.195 | 0.164 | 1 | 0.59577707 | 0.805211628 |
| Thap1 | 0.732132973 | -0.1672055 | 0.132 | 0.148 | 1 | 0.558968845 | -0.332720152 | 0.122 | 0.154 | 1 | 0.732132973 | 0.80549152 |
| Txlng | 0.844785173 | 0.119098685 | 0.434 | 0.428 | 1 | 0.559211854 | 0.769460243 | 0.439 | 0.427 | 1 | 0.844785173 | 0.80570581 |
| Mfsd14b | 0.560106612 | 0.642272594 | 0.472 | 0.483 | 1 | 0.752702838 | 0.176816079 | 0.683 | 0.672 | 1 | 0.752702838 | 0.806493807 |
| Gm57084 | 0.560391874 | 1.389187848 | 0.075 | 0.055 | 1 | 0.976480397 | 0.83720485 | 0.049 | 0.051 | 1 | 0.976480397 | 0.806744696 |
| Chid1 | 0.608334933 | -0.241206082 | 0.264 | 0.301 | 1 | 0.560414473 | 0.735666823 | 0.317 | 0.297 | 1 | 0.608334933 | 0.806764565 |
| Ankrd26 | 0.858565395 | -0.125385324 | 0.17 | 0.178 | 1 | 0.560647514 | 0.692154517 | 0.244 | 0.218 | 1 | 0.858565395 | 0.806969393 |
| Zfp790 | 0.938741618 | 0.228723176 | 0.075 | 0.072 | 1 | 0.560705384 | 0.342440158 | 0.195 | 0.16 | 1 | 0.938741618 | 0.807020241 |
| Tnfsf13b | 0.560867636 | 0.347367673 | 0.057 | 0.038 | 1 | 0.815009479 | -0.332720152 | 0.024 | 0.031 | 1 | 0.815009479 | 0.807162767 |
| Rasgef1c | 0.560907182 | -0.46976827 | 0.057 | 0.038 | 1 | 0.997398969 | -0.622226769 | 0.024 | 0.024 | 1 | 0.997398969 | 0.807197497 |
| Gm56962 | 0.633501905 | 0.247831999 | 0.038 | 0.025 | 1 | 0.561305882 | 0.515276755 | 0.049 | 0.031 | 1 | 0.633501905 | 0.80754747 |
| Lif | 0.56579457 | -0.845277405 | 0.019 | 0.034 | 1 | 0.561306871 | -0.787286015 | 0.049 | 0.031 | 1 | 0.56579457 | 0.807548339 |
| Rbm43 | 0.56167106 | 0.482297253 | 0.208 | 0.178 | 1 | 0.599116661 | 0.365899131 | 0.195 | 0.164 | 1 | 0.599116661 | 0.80786774 |
| 9330154J02Rik | 0.561723594 | -0.932740246 | 0.019 | 0.034 | 1 | 0.973994253 | 1.136765132 | 0.024 | 0.024 | 1 | 0.973994253 | 0.807913792 |
| Atoh8 | 0.561922689 | -0.20282941 | 0.113 | 0.144 | 1 | 0.982876127 | -0.16279515 | 0.122 | 0.119 | 1 | 0.982876127 | 0.80808827 |
| Vps51 | 0.609593732 | 0.324647596 | 0.226 | 0.199 | 1 | 0.562355183 | -0.144647803 | 0.171 | 0.208 | 1 | 0.609593732 | 0.808467015 |
| Srrd | 0.562671052 | 0.810768194 | 0.151 | 0.127 | 1 | 0.66950401 | -0.332720152 | 0.073 | 0.092 | 1 | 0.66950401 | 0.808743391 |
| Scrn1 | 0.874462081 | 0.984797593 | 0.038 | 0.034 | 1 | 0.563653543 | 0.83720485 | 0.122 | 0.096 | 1 | 0.874462081 | 0.809601769 |
| Wdr4 | 0.67166853 | 0.237184755 | 0.151 | 0.127 | 1 | 0.564713918 | 0.667279848 | 0.171 | 0.14 | 1 | 0.67166853 | 0.810526026 |
| Zfp942 | 0.566793567 | 0.499370766 | 0.245 | 0.22 | 1 | 0.744559114 | 0.541748966 | 0.268 | 0.266 | 1 | 0.744559114 | 0.812332187 |
| Gm31282 | 0.782077996 | 0.347367673 | 0.019 | 0.025 | 1 | 0.567316456 | 0.252242349 | 0.024 | 0.044 | 1 | 0.782077996 | 0.81278495 |
| Bub1b | 0.567830665 | -0.652632327 | 0.019 | 0.034 | 1 | 0.87556513 | 0.83720485 | 0.024 | 0.02 | 1 | 0.87556513 | 0.813229666 |
| Gm28874 | 0.567830665 | -0.304709024 | 0.019 | 0.034 | 1 | 0.878310931 | 0.515276755 | 0.024 | 0.02 | 1 | 0.878310931 | 0.813229666 |
| 1700064N11Rik | 0.567830665 | -0.304709024 | 0.019 | 0.034 | 1 | 0.986995563 | 0.83720485 | 0.024 | 0.024 | 1 | 0.986995563 | 0.813229666 |
| Efcab11 | 0.567830665 | -1.932740246 | 0.019 | 0.034 | 1 | 0.819548308 | -0.16279515 | 0.024 | 0.031 | 1 | 0.819548308 | 0.813229666 |
| Panx3 | 0.568858791 | -0.811768143 | 0.113 | 0.14 | 1 | 0.664427337 | -2.755252187 | 0.049 | 0.034 | 1 | 0.664427337 | 0.814117258 |
| Gpatch3 | 0.659393736 | 0.569760094 | 0.057 | 0.042 | 1 | 0.569003293 | 0.515276755 | 0.024 | 0.044 | 1 | 0.659393736 | 0.814241839 |
| Mrpl53 | 0.777466206 | 0.284783135 | 0.358 | 0.36 | 1 | 0.569234444 | 0.159132945 | 0.268 | 0.317 | 1 | 0.777466206 | 0.814441036 |
| C130013H08Rik | 0.569866135 | -0.1672055 | 0.019 | 0.034 | 1 | 0.799741379 | 1.159132945 | 0.049 | 0.041 | 1 | 0.799741379 | 0.814984858 |
| Tha1 | 0.569866135 | -0.304709024 | 0.019 | 0.034 | 1 | 0.650841212 | 0.614812428 | 0.049 | 0.034 | 1 | 0.650841212 | 0.814984858 |
| 5430414B19Rik | 0.569866135 | -0.1672055 | 0.019 | 0.034 | 1 | 0.599290422 | 1.515276755 | 0.024 | 0.014 | 1 | 0.599290422 | 0.814984858 |
| Cep78 | 0.649930681 | 0.502645898 | 0.075 | 0.059 | 1 | 0.570692001 | 1.136765132 | 0.024 | 0.044 | 1 | 0.649930681 | 0.815694642 |
| 4921536K21Rik | 0.836526844 | -0.237594828 | 0.057 | 0.064 | 1 | 0.570959056 | -0.484723245 | 0.049 | 0.072 | 1 | 0.836526844 | 0.815923869 |
| Snapc2 | 0.586701325 | 0.306725688 | 0.17 | 0.14 | 1 | 0.571251052 | 0.351778023 | 0.122 | 0.167 | 1 | 0.586701325 | 0.81617434 |
| Ipo13 | 0.571372899 | 0.762405172 | 0.094 | 0.072 | 1 | 0.617591246 | 0.279209397 | 0.171 | 0.14 | 1 | 0.617591246 | 0.816278808 |
| Clybl | 0.571636314 | 0.268893615 | 0.245 | 0.212 | 1 | 0.618282394 | 0.746602301 | 0.22 | 0.198 | 1 | 0.618282394 | 0.816504553 |
| Sphk1 | 0.64893557 | 0.154722595 | 0.283 | 0.25 | 1 | 0.572095134 | 0.627751484 | 0.22 | 0.184 | 1 | 0.64893557 | 0.816897426 |
| Slc39a6 | 0.572115301 | 0.147069022 | 0.566 | 0.521 | 1 | 0.772997544 | -0.103238306 | 0.512 | 0.529 | 1 | 0.772997544 | 0.816914684 |
| Fndc3a | 0.572169471 | 0.104806117 | 1 | 0.983 | 1 | 0.625172909 | 0.123387283 | 0.829 | 0.952 | 1 | 0.625172909 | 0.816961038 |
| Mrpl57 | 0.572199932 | -0.195774652 | 0.377 | 0.428 | 1 | 0.747905153 | 0.724995346 | 0.341 | 0.358 | 1 | 0.747905153 | 0.816987102 |
| Gm17690 | 0.572756752 | 0.216123139 | 0.057 | 0.081 | 1 | 0.604477578 | -0.250257991 | 0.024 | 0.041 | 1 | 0.604477578 | 0.817463207 |
| Rfx1 | 0.573026345 | 0.402650108 | 0.245 | 0.216 | 1 | 0.975291318 | 0.233133526 | 0.317 | 0.341 | 1 | 0.975291318 | 0.817693498 |
| Ccdc130 | 0.78442244 | -0.214511215 | 0.094 | 0.106 | 1 | 0.573139763 | -0.520347155 | 0.073 | 0.099 | 1 | 0.78442244 | 0.817790338 |
| Ccdc92 | 0.639445448 | 0.306725688 | 0.038 | 0.055 | 1 | 0.574061715 | 0.83720485 | 0.024 | 0.044 | 1 | 0.639445448 | 0.818576578 |
| Sox13 | 0.732691937 | 1.154722595 | 0.019 | 0.013 | 1 | 0.574061715 | 0.83720485 | 0.024 | 0.044 | 1 | 0.732691937 | 0.818576578 |
| Abat | 0.575205204 | 0.357215459 | 0.283 | 0.246 | 1 | 0.778497932 | 0.448162559 | 0.22 | 0.259 | 1 | 0.778497932 | 0.819549381 |
| Zfp346 | 0.913367358 | 0.176096246 | 0.208 | 0.212 | 1 | 0.575379635 | -0.27321914 | 0.171 | 0.208 | 1 | 0.913367358 | 0.819697546 |
| Zfp523 | 0.857844918 | 0.154722595 | 0.208 | 0.199 | 1 | 0.576334002 | 0.554270887 | 0.146 | 0.195 | 1 | 0.857844918 | 0.820507123 |
| 5430416N02Rik | 0.576738787 | 0.531792244 | 0.264 | 0.25 | 1 | 0.763845441 | -0.16279515 | 0.195 | 0.215 | 1 | 0.763845441 | 0.820849946 |
| Catip | 0.576971578 | 0.569760094 | 0.075 | 0.055 | 1 | 0.743829142 | -0.250257991 | 0.024 | 0.034 | 1 | 0.743829142 | 0.821046954 |
| Ccdc28a | 0.577593818 | 0.522454379 | 0.113 | 0.089 | 1 | 0.732115424 | -0.116991461 | 0.073 | 0.058 | 1 | 0.732115424 | 0.821573017 |
| Ifi202b | 0.774217997 | -1.368839361 | 0.019 | 0.025 | 1 | 0.578171261 | -1.673757069 | 0.22 | 0.177 | 1 | 0.774217997 | 0.822060515 |
| Zc2hc1a | 0.578438508 | -0.226706512 | 0.321 | 0.254 | 1 | 0.703618118 | -0.103416603 | 0.341 | 0.379 | 1 | 0.703618118 | 0.822285909 |
| Tpra1 | 0.980890598 | 0.154722595 | 0.283 | 0.297 | 1 | 0.57844229 | 0.478123756 | 0.317 | 0.283 | 1 | 0.980890598 | 0.822289097 |
| Sfxn2 | 0.76534461 | 0.310373465 | 0.264 | 0.246 | 1 | 0.578597526 | -0.11388555 | 0.22 | 0.273 | 1 | 0.76534461 | 0.822419955 |
| Dhrs1 | 0.578634288 | -0.293737906 | 0.377 | 0.403 | 1 | 0.66837425 | 0.430029468 | 0.317 | 0.311 | 1 | 0.66837425 | 0.822450937 |
| Pcdhgb5 | 0.588900999 | 0.491757582 | 0.094 | 0.072 | 1 | 0.578940167 | -0.484723245 | 0.049 | 0.072 | 1 | 0.588900999 | 0.822708617 |
| Tmed8 | 0.578964783 | -0.142957954 | 0.132 | 0.165 | 1 | 0.639846675 | -0.107653596 | 0.146 | 0.177 | 1 | 0.639846675 | 0.822729346 |
| 1110046J04Rik | 0.579084907 | -0.101617158 | 0.094 | 0.123 | 1 | 0.792437066 | 0.276489895 | 0.122 | 0.143 | 1 | 0.792437066 | 0.822830484 |
| Il11 | 0.579622494 | -0.19891436 | 0.094 | 0.072 | 1 | 0.925554239 | -0.541306773 | 0.098 | 0.099 | 1 | 0.925554239 | 0.823282752 |
| Cep41 | 0.987886313 | 1.076720083 | 0.132 | 0.14 | 1 | 0.58039919 | 0.164779508 | 0.098 | 0.133 | 1 | 0.987886313 | 0.82393516 |
| Rdh14 | 0.580582579 | 0.817687608 | 0.208 | 0.191 | 1 | 0.667286353 | 0.30115195 | 0.195 | 0.249 | 1 | 0.667286353 | 0.824089027 |
| Fxn | 0.783562778 | 0.526691372 | 0.132 | 0.123 | 1 | 0.580627224 | -0.508569987 | 0.122 | 0.15 | 1 | 0.783562778 | 0.824126474 |
| Snap29 | 0.754476317 | 0.569760094 | 0.302 | 0.322 | 1 | 0.580709256 | 0.291236481 | 0.317 | 0.283 | 1 | 0.754476317 | 0.824195272 |
| Aven | 0.581128761 | 0.464790534 | 0.415 | 0.424 | 1 | 0.763453801 | 0.236812309 | 0.439 | 0.563 | 1 | 0.763453801 | 0.824546886 |
| Smad9 | 0.581275657 | -0.134784022 | 0.094 | 0.123 | 1 | 0.766359018 | 0.319356545 | 0.146 | 0.133 | 1 | 0.766359018 | 0.824669925 |
| Hdc | 0.582266689 | -0.508242418 | 0.019 | 0.034 | 1 | 0.741208854 | 0.334704509 | 0.024 | 0.017 | 1 | 0.741208854 | 0.825498881 |
| Fbxo28 | 0.582380247 | 0.154722595 | 0.547 | 0.542 | 1 | 0.687788893 | 0.109284395 | 0.561 | 0.604 | 1 | 0.687788893 | 0.825593742 |
| Tefm | 0.975314454 | 0.370451286 | 0.113 | 0.119 | 1 | 0.583998563 | 0.292884334 | 0.073 | 0.102 | 1 | 0.975314454 | 0.826942805 |
| Zgpat | 0.590090321 | 0.446903346 | 0.189 | 0.165 | 1 | 0.584269707 | 0.30377265 | 0.293 | 0.263 | 1 | 0.590090321 | 0.827168323 |
| Mef2c | 0.584863215 | -0.78254165 | 0.283 | 0.301 | 1 | 0.718999979 | -1.055879946 | 0.146 | 0.171 | 1 | 0.718999979 | 0.82766145 |
| Septin2 | 0.737215654 | 0.3662267 | 0.151 | 0.136 | 1 | 0.585116982 | 0.347820009 | 0.22 | 0.184 | 1 | 0.737215654 | 0.827872082 |
| B4galt4 | 0.585612597 | 0.393910259 | 0.264 | 0.233 | 1 | 0.942789681 | 0.309957847 | 0.268 | 0.28 | 1 | 0.942789681 | 0.82828308 |
| Agbl5 | 0.642974908 | 0.569760094 | 0.132 | 0.11 | 1 | 0.586248643 | 0.334704509 | 0.073 | 0.102 | 1 | 0.642974908 | 0.828809814 |
| Arhgef7 | 0.667362798 | 0.632044372 | 0.434 | 0.419 | 1 | 0.586298517 | 0.535314508 | 0.341 | 0.334 | 1 | 0.667362798 | 0.828851083 |
| Kcnu1 | 0.586409164 | 0.417757001 | 0.019 | 0.034 | 1 | 0.746881137 | 1.029849928 | 0.024 | 0.017 | 1 | 0.746881137 | 0.828942621 |
| Ccnb2 | 0.586528827 | -0.944813079 | 0.094 | 0.119 | 1 | 0.65317831 | -0.292078167 | 0.146 | 0.174 | 1 | 0.65317831 | 0.829041589 |
| Slc25a37 | 0.58731547 | -0.255561375 | 0.264 | 0.297 | 1 | 0.679508303 | 0.398631836 | 0.268 | 0.239 | 1 | 0.679508303 | 0.829691479 |
| Lnx2 | 0.58769384 | 0.438515561 | 0.094 | 0.072 | 1 | 0.691070576 | -0.348661696 | 0.146 | 0.171 | 1 | 0.691070576 | 0.830003631 |
| Gm11110 | 0.732691937 | 1.154722595 | 0.019 | 0.013 | 1 | 0.58792152 | 2.515276755 | 0.024 | 0.014 | 1 | 0.732691937 | 0.830191326 |
| Acad12 | 0.750969908 | 0.984797593 | 0.038 | 0.03 | 1 | 0.58792152 | 2.100239256 | 0.024 | 0.014 | 1 | 0.750969908 | 0.830191326 |
| Snap25 | 0.732691937 | 1.154722595 | 0.019 | 0.013 | 1 | 0.587925597 | 2.574170444 | 0.024 | 0.014 | 1 | 0.732691937 | 0.830194686 |
| BC051019 | 0.732691937 | 1.154722595 | 0.019 | 0.013 | 1 | 0.587925597 | 2.83720485 | 0.024 | 0.014 | 1 | 0.732691937 | 0.830194686 |
| Elapor2 | 0.736073999 | 0.569760094 | 0.019 | 0.013 | 1 | 0.587925597 | 2.252242349 | 0.024 | 0.014 | 1 | 0.736073999 | 0.830194686 |
| Mis18bp1 | 0.913088291 | -0.799473716 | 0.038 | 0.034 | 1 | 0.587925597 | 2.351778023 | 0.024 | 0.014 | 1 | 0.913088291 | 0.830194686 |
| 1810034E14Rik | 0.995621732 | 1.280253477 | 0.038 | 0.038 | 1 | 0.587925597 | 2.252242349 | 0.024 | 0.014 | 1 | 0.995621732 | 0.830194686 |
| Gm2011 | 0.588476383 | 0.984797593 | 0.019 | 0.034 | 1 | 0.68987748 | 0.515276755 | 0.024 | 0.038 | 1 | 0.68987748 | 0.830648313 |
| Gm31834 | 0.588476383 | 0.569760094 | 0.019 | 0.034 | 1 | 0.911761241 | 0.252242349 | 0.024 | 0.027 | 1 | 0.911761241 | 0.830648313 |
| Neil2 | 0.588476383 | 0.569760094 | 0.019 | 0.034 | 1 | 0.986995563 | 0.83720485 | 0.024 | 0.024 | 1 | 0.986995563 | 0.830648313 |

| | | | | | | | | | | | | |
|---|---|---|---|---|---|---|---|---|---|---|---|---|
| Gimap6 | 0.588746185 | 0.417757001 | 0.113 | 0.089 | 1 | 0.627108882 | 0.334704509 | 0.073 | 0.099 | 1 | 0.627108882 | 0.8308703 |
| D5Ertd579e | 0.596836664 | -0.199306343 | 0.66 | 0.712 | 1 | 0.589900266 | -0.202552859 | 0.756 | 0.775 | 1 | 0.596836664 | 0.831818208 |
| Gm26608 | 0.732691937 | 1.154722595 | 0.019 | 0.013 | 1 | 0.590756788 | 1.83720485 | 0.024 | 0.014 | 1 | 0.732691937 | 0.832519993 |
| Gm4631 | 0.945185506 | 0.154722595 | 0.075 | 0.072 | 1 | 0.591016961 | 0.711673968 | 0.171 | 0.143 | 1 | 0.945185506 | 0.832732874 |
| Zmat2 | 0.918400894 | 0.270641882 | 0.528 | 0.631 | 1 | 0.59102933 | -0.199453621 | 0.561 | 0.577 | 1 | 0.918400894 | 0.832742991 |
| Ctc1 | 0.885848516 | 0.8327945 | 0.075 | 0.072 | 1 | 0.591760082 | 0.737669176 | 0.049 | 0.075 | 1 | 0.885848516 | 0.833340169 |
| Qpctl | 0.642743588 | 0.122301117 | 0.132 | 0.161 | 1 | 0.593935932 | 0.652780279 | 0.171 | 0.143 | 1 | 0.642743588 | 0.835111973 |
| Abca5 | 0.931623121 | 0.131638982 | 0.321 | 0.326 | 1 | 0.59462434 | 0.141598285 | 0.244 | 0.314 | 1 | 0.931623121 | 0.835670574 |
| 1700010K24Rik | 0.596086858 | -0.719746523 | 0.038 | 0.055 | 1 | 0.921535259 | 0.515276755 | 0.024 | 0.027 | 1 | 0.921535259 | 0.836854174 |
| Ccar2 | 0.765243115 | 0.216123139 | 0.094 | 0.081 | 1 | 0.596623371 | 0.319356545 | 0.146 | 0.181 | 1 | 0.765243115 | 0.837287295 |
| Bbs2 | 0.59720124 | 0.154722595 | 0.075 | 0.102 | 1 | 1 | 0.334704509 | 0.098 | 0.099 | 1 | 1 | 0.837753159 |
| Retsat | 0.66957246 | 0.347367673 | 0.094 | 0.076 | 1 | 0.597269879 | -0.116991461 | 0.073 | 0.099 | 1 | 0.66957246 | 0.83780845 |
| Ppt1 | 0.846465797 | 0.154722595 | 0.604 | 0.653 | 1 | 0.597955977 | 0.205767919 | 0.634 | 0.618 | 1 | 0.846465797 | 0.838360604 |
| D830050J10Rik | 0.598026031 | 0.404700848 | 0.151 | 0.123 | 1 | 0.94575798 | 0.334704509 | 0.049 | 0.051 | 1 | 0.94575798 | 0.838416929 |
| Sgo2a | 0.878702404 | -0.304709024 | 0.038 | 0.042 | 1 | 0.599289758 | 0.83720485 | 0.024 | 0.014 | 1 | 0.878702404 | 0.839431302 |
| Gm13620 | 0.732691937 | 1.154722595 | 0.019 | 0.013 | 1 | 0.599290422 | 1.515276755 | 0.024 | 0.014 | 1 | 0.732691937 | 0.839431834 |
| Ccl28 | 0.732691937 | 1.154722595 | 0.019 | 0.013 | 1 | 0.599290422 | 1.515276755 | 0.024 | 0.014 | 1 | 0.732691937 | 0.839431834 |
| Asb5 | 0.914958 | 0.154722595 | 0.019 | 0.021 | 1 | 0.599290422 | 1.515276755 | 0.024 | 0.014 | 1 | 0.914958 | 0.839431834 |
| 1810041H14Rik | 0.914958 | 0.154722595 | 0.019 | 0.021 | 1 | 0.599290422 | 1.515276755 | 0.024 | 0.014 | 1 | 0.914958 | 0.839431834 |
| Dok2 | 0.916620008 | 1.154722595 | 0.019 | 0.017 | 1 | 0.599290422 | 1.515276755 | 0.024 | 0.014 | 1 | 0.916620008 | 0.839431834 |
| Sptbn5 | 0.920805363 | 0.417757001 | 0.019 | 0.021 | 1 | 0.599290422 | 1.515276755 | 0.024 | 0.014 | 1 | 0.920805363 | 0.839431834 |
| Terb1 | 0.929410101 | 0.569760094 | 0.019 | 0.017 | 1 | 0.599290422 | 1.515276755 | 0.024 | 0.014 | 1 | 0.929410101 | 0.839431834 |
| AW554918 | 0.59954431 | 0.146548663 | 0.396 | 0.36 | 1 | 0.962269496 | 0.301995384 | 0.439 | 0.461 | 1 | 0.962269496 | 0.839635241 |
| 2900009J06Rik | 0.677640974 | -0.237594828 | 0.057 | 0.072 | 1 | 0.599658621 | 1.404245443 | 0.073 | 0.055 | 1 | 0.677640974 | 0.83972678 |
| Actb | 0.5997314 | 0.102847193 | 1 | 0.996 | 1 | 0.879208921 | 0.139132302 | 1 | 0.997 | 1 | 0.879208921 | 0.839785048 |
| Trak1 | 0.664508266 | 0.102255175 | 0.66 | 0.644 | 1 | 0.60042065 | 0.143866188 | 0.829 | 0.765 | 1 | 0.664508266 | 0.840336343 |
| C920006O11Rik | 0.99526215 | 0.417757001 | 0.075 | 0.076 | 1 | 0.600905177 | -0.484723245 | 0.024 | 0.041 | 1 | 0.99526215 | 0.840723322 |
| Cep76 | 0.939532196 | 0.347367673 | 0.151 | 0.153 | 1 | 0.601052205 | 0.174239837 | 0.195 | 0.246 | 1 | 0.939532196 | 0.840840657 |
| Gm50013 | 0.60105781 | -0.330704232 | 0.075 | 0.097 | 1 | 0.833360163 | -0.11032773 | 0.122 | 0.133 | 1 | 0.833360163 | 0.840845129 |
| Txn1 | 0.80618165 | -0.146205201 | 1 | 0.953 | 1 | 0.601730494 | 0.287470837 | 0.951 | 0.952 | 1 | 0.80618165 | 0.8413814 |
| Apeh | 0.601779682 | 0.347367673 | 0.132 | 0.106 | 1 | 0.847601063 | 0.345351753 | 0.146 | 0.137 | 1 | 0.847601063 | 0.841420579 |
| Pigt | 0.601842935 | -0.244684253 | 0.453 | 0.458 | 1 | 0.765349238 | 0.132660733 | 0.39 | 0.437 | 1 | 0.765349238 | 0.841470951 |
| 5830428M24Rik | 0.779443586 | 0.154722595 | 0.019 | 0.025 | 1 | 0.60213981 | 1.252242349 | 0.024 | 0.014 | 1 | 0.779443586 | 0.841707269 |
| 1700030N03Rik | 0.868328394 | -0.260314904 | 0.038 | 0.042 | 1 | 0.60213981 | 1.252242349 | 0.024 | 0.014 | 1 | 0.868328394 | 0.841707269 |
| Gm5432 | 0.874462081 | 0.984797593 | 0.038 | 0.034 | 1 | 0.60213981 | 0.83720485 | 0.024 | 0.014 | 1 | 0.874462081 | 0.841707269 |
| Gm37634 | 0.913424206 | 1.417757001 | 0.019 | 0.017 | 1 | 0.60213981 | 1.252242349 | 0.024 | 0.014 | 1 | 0.913424206 | 0.841707269 |
| Marchf10 | 0.913424206 | 1.417757001 | 0.019 | 0.017 | 1 | 0.60213981 | 0.667279848 | 0.024 | 0.014 | 1 | 0.913424206 | 0.841707269 |
| Gm14221 | 0.602621844 | -0.652632327 | 0.038 | 0.055 | 1 | 0.816281931 | -0.20718927 | 0.049 | 0.058 | 1 | 0.816281931 | 0.842090601 |
| Kyat3 | 0.602621844 | -0.260314904 | 0.038 | 0.055 | 1 | 0.976480397 | 1.159132945 | 0.049 | 0.051 | 1 | 0.976480397 | 0.842090601 |
| Pkd2l2 | 0.602621844 | -0.347777746 | 0.038 | 0.055 | 1 | 0.689850603 | 0.944120054 | 0.073 | 0.058 | 1 | 0.689850603 | 0.842090601 |
| Ptpn22 | 0.672332674 | -1.1672055 | 0.038 | 0.051 | 1 | 0.602692741 | -0.484723245 | 0.024 | 0.041 | 1 | 0.672332674 | 0.842146942 |
| Slc35f6 | 0.602756661 | -0.464187238 | 0.245 | 0.263 | 1 | 0.825332128 | 0.608386159 | 0.195 | 0.191 | 1 | 0.825332128 | 0.842197729 |
| Zdhhc4 | 0.840868153 | 0.318221327 | 0.264 | 0.271 | 1 | 0.602852843 | 0.675741427 | 0.268 | 0.239 | 1 | 0.840868153 | 0.842274135 |
| Mcub | 0.602935833 | -0.207847485 | 0.132 | 0.169 | 1 | 0.748407724 | -0.385187572 | 0.146 | 0.16 | 1 | 0.748407724 | 0.842340047 |
| Daam1 | 0.799582273 | -0.117237041 | 0.811 | 0.864 | 1 | 0.603095761 | -0.125659384 | 0.902 | 0.877 | 1 | 0.799582273 | 0.842467025 |
| Smad1 | 0.603893447 | -0.148589822 | 0.585 | 0.678 | 1 | 0.63853302 | -0.16889535 | 0.683 | 0.713 | 1 | 0.63853302 | 0.843099598 |
| Fetub | 0.683806282 | -0.617866909 | 0.057 | 0.072 | 1 | 0.604854178 | -0.16279515 | 0.122 | 0.154 | 1 | 0.683806282 | 0.84385978 |
| Scn9a | 0.722567081 | 1.8327945 | 0.019 | 0.013 | 1 | 0.605005904 | 0.515276755 | 0.024 | 0.014 | 1 | 0.722567081 | 0.843979664 |
| Gm29570 | 0.732691937 | 1.154722595 | 0.019 | 0.013 | 1 | 0.605005904 | 0.667279848 | 0.024 | 0.014 | 1 | 0.732691937 | 0.843979664 |
| 2700099C18Rik | 0.778599904 | -0.330704232 | 0.057 | 0.047 | 1 | 0.605005904 | 0.667279848 | 0.024 | 0.014 | 1 | 0.778599904 | 0.843979664 |
| Kif11 | 0.914746621 | -1.40599236 | 0.057 | 0.059 | 1 | 0.605005904 | 0.377773231 | 0.024 | 0.014 | 1 | 0.914746621 | 0.843979664 |
| Cdk5r1 | 0.88484015 | 0.2967416 | 0.094 | 0.089 | 1 | 0.605084784 | 0.624211126 | 0.146 | 0.119 | 1 | 0.88484015 | 0.844041972 |
| Slc18b1 | 0.633485883 | -0.19891436 | 0.132 | 0.157 | 1 | 0.60519603 | 0.269520341 | 0.22 | 0.188 | 1 | 0.633485883 | 0.844129826 |
| Mtg2 | 0.606176398 | 0.645708947 | 0.151 | 0.131 | 1 | 0.782053996 | 0.222495006 | 0.146 | 0.13 | 1 | 0.782053996 | 0.844902971 |
| Zbtb25 | 0.606202373 | 0.110328475 | 0.113 | 0.089 | 1 | 0.695010541 | 0.795384674 | 0.195 | 0.171 | 1 | 0.695010541 | 0.844923429 |
| Zfp846 | 0.606768997 | 1.237184755 | 0.075 | 0.059 | 1 | 0.772272944 | 0.564186355 | 0.073 | 0.061 | 1 | 0.772272944 | 0.845369379 |
| Dnpep | 0.768847938 | 0.176748901 | 0.396 | 0.386 | 1 | 0.60693948 | -0.119726428 | 0.293 | 0.341 | 1 | 0.768847938 | 0.845503428 |
| Zfp697 | 0.93671561 | -0.304709024 | 0.151 | 0.148 | 1 | 0.607818019 | 0.276489895 | 0.098 | 0.13 | 1 | 0.93671561 | 0.846193294 |
| Hgsnat | 0.643647061 | 0.23824595 | 0.491 | 0.466 | 1 | 0.607821731 | 0.399799537 | 0.463 | 0.444 | 1 | 0.643647061 | 0.846196205 |
| Cdk15 | 0.946937754 | 0.631160639 | 0.057 | 0.055 | 1 | 0.607875423 | 0.667279848 | 0.024 | 0.014 | 1 | 0.946937754 | 0.846238316 |
| Gm42209 | 0.778655755 | -1.545717123 | 0.038 | 0.03 | 1 | 0.607876076 | -0.250257991 | 0.024 | 0.014 | 1 | 0.778655755 | 0.846238829 |
| Ccdc28b | 0.608063366 | -0.1672055 | 0.302 | 0.339 | 1 | 0.806360447 | 0.199774929 | 0.293 | 0.276 | 1 | 0.806360447 | 0.846385675 |
| Fndc4 | 0.964593376 | 0.127241858 | 0.302 | 0.322 | 1 | 0.608317837 | 0.47463477 | 0.293 | 0.266 | 1 | 0.964593376 | 0.846585083 |
| Bbx | 0.608756674 | 0.251584134 | 0.868 | 0.894 | 1 | 0.720043817 | 0.111046011 | 0.902 | 0.928 | 1 | 0.720043817 | 0.84692866 |
| Pcgf1 | 0.60883182 | 1.270199812 | 0.094 | 0.076 | 1 | 0.629768283 | 0.930314254 | 0.049 | 0.034 | 1 | 0.629768283 | 0.846987455 |
| Atad1 | 0.609193466 | 0.127241858 | 0.547 | 0.551 | 1 | 0.978330726 | 0.176954252 | 0.585 | 0.618 | 1 | 0.978330726 | 0.847270253 |
| Gosr1 | 0.609563179 | 0.102255175 | 0.302 | 0.352 | 1 | 0.928710936 | 0.614812428 | 0.341 | 0.375 | 1 | 0.928710936 | 0.847559089 |
| Panx2 | 0.732691937 | 1.154722595 | 0.019 | 0.013 | 1 | 0.609924905 | 0.136765132 | 0.024 | 0.041 | 1 | 0.732691937 | 0.84784142 |
| Aldh7a1 | 0.610407512 | 0.168143111 | 0.566 | 0.555 | 1 | 0.959717518 | 0.10414371 | 0.488 | 0.509 | 1 | 0.959717518 | 0.848217693 |
| Gm57238 | 0.610863297 | 1.347367673 | 0.038 | 0.025 | 1 | 0.745976859 | 0.377773231 | 0.024 | 0.034 | 1 | 0.745976859 | 0.848572626 |
| A230087F16Rik | 0.610863297 | 1.669295768 | 0.038 | 0.025 | 1 | 0.888999736 | 0.667279848 | 0.049 | 0.044 | 1 | 0.888999736 | 0.848572626 |
| Pcdhb5 | 0.610863297 | 1.347367673 | 0.038 | 0.025 | 1 | 0.729893611 | 2.351778023 | 0.024 | 0.017 | 1 | 0.729893611 | 0.848572626 |
| Cdkl5 | 0.726855916 | 0.550651271 | 0.283 | 0.347 | 1 | 0.612008946 | 0.22897257 | 0.415 | 0.399 | 1 | 0.726855916 | 0.849462942 |
| Scamp5 | 0.676691664 | 0.347367673 | 0.094 | 0.076 | 1 | 0.613013048 | 0.252242349 | 0.049 | 0.072 | 1 | 0.676691664 | 0.850241099 |
| Natd1 | 0.613803461 | 0.280253477 | 0.132 | 0.106 | 1 | 0.727655634 | 0.893788378 | 0.146 | 0.133 | 1 | 0.727655634 | 0.850852233 |
| Incenp | 0.614285535 | -0.330704232 | 0.038 | 0.055 | 1 | 0.966495121 | 0.589277336 | 0.049 | 0.048 | 1 | 0.966495121 | 0.851224352 |
| BB218582 | 0.61536875 | 1.154722595 | 0.038 | 0.025 | 1 | 0.744047924 | 1.252242349 | 0.024 | 0.017 | 1 | 0.744047924 | 0.852058801 |
| Gm36266 | 0.61536875 | 1.154722595 | 0.038 | 0.025 | 1 | 0.674180157 | 0.252242349 | 0.024 | 0.038 | 1 | 0.674180157 | 0.852058801 |
| B430319H21Rik | 0.61536875 | 1.154722595 | 0.038 | 0.025 | 1 | 0.691843603 | 0.614812428 | 0.024 | 0.038 | 1 | 0.691843603 | 0.852058801 |
| Gpr156 | 0.61536875 | 1.154722595 | 0.038 | 0.025 | 1 | 0.979193861 | -0.101394606 | 0.024 | 0.024 | 1 | 0.979193861 | 0.852058801 |
| 2410080I02Rik | 0.61536875 | 1.154722595 | 0.038 | 0.025 | 1 | 0.6898875 | 0.515276755 | 0.024 | 0.038 | 1 | 0.6898875 | 0.852058801 |
| Gm50431 | 0.6159787 | -0.1672055 | 0.057 | 0.076 | 1 | 0.934943043 | 0.100239256 | 0.049 | 0.051 | 1 | 0.934943043 | 0.852527641 |
| Etfbkmt | 0.616199059 | 0.154722595 | 0.113 | 0.089 | 1 | 0.684656986 | -0.16279515 | 0.098 | 0.119 | 1 | 0.684656986 | 0.852696837 |
| H2aj | 0.617881009 | -0.105340245 | 0.698 | 0.716 | 1 | 0.629692555 | -0.101394606 | 0.585 | 0.645 | 1 | 0.629692555 | 0.853985077 |
| Ppm1b | 0.727872069 | 0.131638982 | 0.623 | 0.614 | 1 | 0.618046036 | -0.127691349 | 0.707 | 0.809 | 1 | 0.727872069 | 0.854111169 |
| Irgm1 | 0.618046825 | 0.474001111 | 0.34 | 0.305 | 1 | 0.825887159 | 0.795384674 | 0.22 | 0.249 | 1 | 0.825887159 | 0.854111772 |
| Senp3 | 0.641075733 | 0.197791317 | 0.415 | 0.403 | 1 | 0.618609752 | 0.18867722 | 0.512 | 0.512 | 1 | 0.641075733 | 0.854541479 |
| Dna2 | 0.641178868 | 0.891688189 | 0.057 | 0.042 | 1 | 0.619172754 | 0.766815522 | 0.073 | 0.055 | 1 | 0.641178868 | 0.854970609 |
| 6820408C15Rik | 0.619885227 | 0.984797593 | 0.038 | 0.025 | 1 | 0.87556513 | 0.83720485 | 0.024 | 0.02 | 1 | 0.87556513 | 0.85551276 |
| Bcat2 | 0.976471627 | 0.218852932 | 0.245 | 0.267 | 1 | 0.62002032 | 0.429120111 | 0.22 | 0.188 | 1 | 0.976471627 | 0.855615443 |
| Mms19 | 0.84168773 | -0.189231806 | 0.208 | 0.186 | 1 | 0.621148028 | 0.174239837 | 0.195 | 0.239 | 1 | 0.84168773 | 0.856471183 |
| Odr4 | 0.621965813 | 0.318798521 | 0.66 | 0.712 | 1 | 0.768627389 | 0.266456208 | 0.634 | 0.628 | 1 | 0.768627389 | 0.857090154 |
| Dpcd | 0.622057077 | -0.183147044 | 0.264 | 0.288 | 1 | 0.743466335 | 0.212713985 | 0.268 | 0.239 | 1 | 0.743466335 | 0.857159147 |
| Psd | 0.767066103 | 0.569760094 | 0.038 | 0.03 | 1 | 0.622166556 | 1.574170444 | 0.049 | 0.034 | 1 | 0.767066103 | 0.857241889 |
| Tmem186 | 0.837800081 | 0.104096522 | 0.113 | 0.102 | 1 | 0.622544362 | 0.185128153 | 0.146 | 0.116 | 1 | 0.837800081 | 0.857527241 |
| Gm10130 | 0.641859861 | 0.154722595 | 0.057 | 0.076 | 1 | 0.622634499 | -0.435813645 | 0.049 | 0.068 | 1 | 0.641859861 | 0.857595279 |
| Suv39h2 | 0.938741618 | 0.154722595 | 0.075 | 0.072 | 1 | 0.623785099 | 1.313642894 | 0.073 | 0.055 | 1 | 0.938741618 | 0.858462348 |
| Zfp54 | 0.732691937 | 1.154722595 | 0.019 | 0.013 | 1 | 0.624060349 | 1.252242349 | 0.049 | 0.034 | 1 | 0.732691937 | 0.858669379 |

| | | | | | | | | | | | | |
|---|---|---|---|---|---|---|---|---|---|---|---|---|
| Mblac2 | 0.713924126 | 0.389187848 | 0.113 | 0.097 | 1 | 0.624072438 | 1.614812428 | 0.049 | 0.034 | 1 | 0.713924126 | 0.858678469 |
| Ecd | 0.835968556 | 0.355635289 | 0.264 | 0.246 | 1 | 0.624241456 | 0.149148856 | 0.293 | 0.263 | 1 | 0.835968556 | 0.858805517 |
| Gm26788 | 0.624418548 | 0.932330173 | 0.038 | 0.025 | 1 | 0.743829142 | 0.252242349 | 0.024 | 0.034 | 1 | 0.743829142 | 0.858938573 |
| Capn3 | 0.624418548 | 0.932330173 | 0.038 | 0.025 | 1 | 0.746881137 | 1.029849928 | 0.024 | 0.017 | 1 | 0.746881137 | 0.858938573 |
| H2bc21 | 0.624418548 | 0.932330173 | 0.038 | 0.025 | 1 | 0.992196803 | 0.515276755 | 0.024 | 0.024 | 1 | 0.992196803 | 0.858938573 |
| Lrfn3 | 0.624418548 | 0.932330173 | 0.038 | 0.025 | 1 | 0.900433136 | 0.614812428 | 0.049 | 0.044 | 1 | 0.900433136 | 0.858938573 |
| Cenpa | 0.913878478 | -1.395474488 | 0.094 | 0.097 | 1 | 0.624753797 | 1.367719567 | 0.073 | 0.102 | 1 | 0.913878478 | 0.859190287 |
| Cmtm8 | 0.680193691 | 0.154722595 | 0.189 | 0.165 | 1 | 0.624811281 | -0.137828235 | 0.439 | 0.491 | 1 | 0.680193691 | 0.859233425 |
| Zfp563 | 0.625057589 | 0.962077517 | 0.075 | 0.059 | 1 | 0.708290322 | 0.707921833 | 0.073 | 0.096 | 1 | 0.708290322 | 0.859418188 |
| Gm53610 | 0.625283795 | -0.130679624 | 0.113 | 0.089 | 1 | 0.660763023 | 0.313642894 | 0.073 | 0.055 | 1 | 0.660763023 | 0.859587765 |
| Gm56637 | 0.874462081 | 0.984797593 | 0.038 | 0.034 | 1 | 0.625458103 | -0.385187572 | 0.049 | 0.068 | 1 | 0.874462081 | 0.859718368 |
| Bcap29 | 0.99298492 | 0.464407094 | 0.208 | 0.225 | 1 | 0.626143435 | 0.197794565 | 0.195 | 0.16 | 1 | 0.99298492 | 0.860231269 |
| Gm17106 | 0.626632759 | 0.110328475 | 0.132 | 0.106 | 1 | 0.836704118 | 0.515276755 | 0.146 | 0.171 | 1 | 0.836704118 | 0.860596903 |
| H2bc18 | 0.626762308 | 0.8327945 | 0.057 | 0.042 | 1 | 0.646976884 | -0.16279515 | 0.049 | 0.034 | 1 | 0.646976884 | 0.860693625 |
| Sh2d4b | 0.805982999 | -0.330704232 | 0.075 | 0.064 | 1 | 0.628284378 | -0.332720152 | 0.049 | 0.068 | 1 | 0.805982999 | 0.861827497 |
| Hnmt | 0.628925725 | 0.739685096 | 0.038 | 0.025 | 1 | 0.967442771 | 0.589277336 | 0.049 | 0.051 | 1 | 0.967442771 | 0.862303882 |
| Plppr2 | 0.628925725 | 0.569760094 | 0.038 | 0.025 | 1 | 0.729889663 | 1.83720485 | 0.024 | 0.017 | 1 | 0.729889663 | 0.862303882 |
| 4933412E12Rik | 0.628925725 | 0.739685096 | 0.038 | 0.025 | 1 | 0.735545437 | 1.422167351 | 0.024 | 0.017 | 1 | 0.735545437 | 0.862303882 |
| Psmd10 | 0.630369351 | 0.417757001 | 0.208 | 0.182 | 1 | 0.775755056 | 0.222495006 | 0.122 | 0.143 | 1 | 0.775755056 | 0.863373183 |
| Raf1 | 0.63053335 | -0.233974512 | 0.849 | 0.78 | 1 | 0.634934708 | -0.148067864 | 1 | 0.928 | 1 | 0.634934708 | 0.863494395 |
| BC024978 | 0.947371719 | -0.1182959 | 0.094 | 0.089 | 1 | 0.630693784 | -0.292078167 | 0.073 | 0.096 | 1 | 0.947371719 | 0.863612919 |
| Gm43062 | 0.89261791 | 0.569760094 | 0.038 | 0.034 | 1 | 0.63111335 | -0.221688839 | 0.049 | 0.068 | 1 | 0.89261791 | 0.863922639 |
| Psat1 | 0.782134192 | -0.20282941 | 0.132 | 0.144 | 1 | 0.631113499 | -0.332720152 | 0.049 | 0.068 | 1 | 0.782134192 | 0.863922749 |
| AU040972 | 0.929410101 | 0.569760094 | 0.019 | 0.017 | 1 | 0.631723655 | 2.334704509 | 0.024 | 0.041 | 1 | 0.929410101 | 0.864372533 |
| Gm53 | 0.632172478 | 0.932330173 | 0.057 | 0.042 | 1 | 0.992196803 | 0.252242349 | 0.024 | 0.024 | 1 | 0.992196803 | 0.864702914 |
| Gm42726 | 0.632188235 | 1.247831999 | 0.057 | 0.042 | 1 | 0.942334653 | 1.120997816 | 0.049 | 0.048 | 1 | 0.942334653 | 0.864714506 |
| Alg13 | 0.886557546 | -0.274120704 | 0.17 | 0.178 | 1 | 0.632926916 | -0.332720152 | 0.195 | 0.222 | 1 | 0.886557546 | 0.865257351 |
| Ttc39a | 0.646717977 | 0.739685096 | 0.075 | 0.059 | 1 | 0.632989178 | 0.444887427 | 0.122 | 0.099 | 1 | 0.646717977 | 0.865303057 |
| Klf16 | 0.633122404 | 0.589937976 | 0.264 | 0.242 | 1 | 0.903559005 | 0.448162559 | 0.244 | 0.273 | 1 | 0.903559005 | 0.86540083 |
| Pask | 0.633487796 | 0.417757001 | 0.038 | 0.025 | 1 | 0.976594201 | 1.029849928 | 0.024 | 0.024 | 1 | 0.976594201 | 0.865668804 |
| Mmaa | 0.856836395 | 0.645708947 | 0.132 | 0.127 | 1 | 0.633776585 | -0.233184478 | 0.146 | 0.181 | 1 | 0.856836395 | 0.86588041 |
| Dynlt1c | 0.924977223 | 0.267197324 | 0.132 | 0.127 | 1 | 0.633945044 | -0.101394606 | 0.049 | 0.068 | 1 | 0.924977223 | 0.866003769 |
| Gm14858 | 0.634338601 | 0.792152515 | 0.075 | 0.059 | 1 | 0.771804212 | 0.252242349 | 0.073 | 0.089 | 1 | 0.771804212 | 0.866291742 |
| Rpp25l | 0.910735322 | -0.113766241 | 0.189 | 0.191 | 1 | 0.634347596 | 0.33970519 | 0.293 | 0.263 | 1 | 0.910735322 | 0.866298319 |
| Gtf2h4 | 0.635490099 | 0.363309217 | 0.17 | 0.148 | 1 | 0.700802611 | -0.19841906 | 0.098 | 0.116 | 1 | 0.700802611 | 0.867132532 |
| Ccdc162 | 0.702904456 | 0.228723176 | 0.038 | 0.051 | 1 | 0.635502471 | 0.911205431 | 0.049 | 0.034 | 1 | 0.702904456 | 0.867141552 |
| Mtln | 0.797783427 | 0.192524628 | 0.358 | 0.347 | 1 | 0.635521749 | 0.222495006 | 0.195 | 0.242 | 1 | 0.797783427 | 0.867155604 |
| Hexim2 | 0.635728131 | 1.280253477 | 0.057 | 0.042 | 1 | 0.751210902 | 0.174239837 | 0.049 | 0.061 | 1 | 0.751210902 | 0.867306005 |
| 4732471J01Rik | 0.881934828 | -0.20517335 | 0.151 | 0.144 | 1 | 0.638277175 | -0.410722664 | 0.073 | 0.096 | 1 | 0.881934828 | 0.869156598 |
| Tmem198 | 0.888073109 | 0.569760094 | 0.038 | 0.034 | 1 | 0.639342184 | 0.962735732 | 0.049 | 0.034 | 1 | 0.888073109 | 0.86992594 |
| Zfp995 | 0.917867709 | 0.417757001 | 0.075 | 0.072 | 1 | 0.639909796 | 0.883008539 | 0.098 | 0.078 | 1 | 0.917867709 | 0.870335045 |
| Gpr135 | 0.641178868 | 0.891688189 | 0.057 | 0.042 | 1 | 0.931315883 | 0.962735732 | 0.024 | 0.027 | 1 | 0.931315883 | 0.871247395 |
| Snhg10 | 0.641178868 | 0.891688189 | 0.057 | 0.042 | 1 | 0.743829142 | 0.136765132 | 0.024 | 0.034 | 1 | 0.743829142 | 0.871247395 |
| Slc12a7 | 0.872298139 | 0.389187848 | 0.057 | 0.064 | 1 | 0.641218506 | 0.883008539 | 0.098 | 0.078 | 1 | 0.872298139 | 0.87127584 |
| Ccdc8 | 0.642624038 | 0.569760094 | 0.151 | 0.127 | 1 | 0.761649039 | 0.922934724 | 0.146 | 0.14 | 1 | 0.761649039 | 0.872282422 |
| B230208H11Rik | 0.642803358 | 0.389187848 | 0.038 | 0.055 | 1 | 0.93376172 | 1.100239256 | 0.024 | 0.027 | 1 | 0.93376172 | 0.872410559 |
| Kif18a | 0.643039013 | 0.739685096 | 0.057 | 0.042 | 1 | 0.82181816 | 0.252242349 | 0.024 | 0.031 | 1 | 0.82181816 | 0.872578854 |
| Ccdc73 | 0.844324661 | 0.389187848 | 0.075 | 0.068 | 1 | 0.643119211 | 0.721727632 | 0.049 | 0.034 | 1 | 0.844324661 | 0.872636102 |
| Fbxo22 | 0.899412243 | -0.219401711 | 0.396 | 0.415 | 1 | 0.644214776 | 0.280256725 | 0.293 | 0.352 | 1 | 0.899412243 | 0.873416874 |
| Scnn1a | 0.792583859 | 0.569760094 | 0.019 | 0.025 | 1 | 0.644425228 | 0.359157553 | 0.146 | 0.119 | 1 | 0.792583859 | 0.873566582 |
| Selenot | 0.645110843 | 0.192970064 | 0.792 | 0.699 | 1 | 0.749677901 | 0.160993287 | 0.829 | 0.881 | 1 | 0.749677901 | 0.874053686 |
| Ccl25 | 0.808971154 | -0.179696444 | 0.264 | 0.271 | 1 | 0.6454051 | 0.1108865 | 0.341 | 0.396 | 1 | 0.808971154 | 0.874262457 |
| Man2c1 | 0.646687354 | 0.454282877 | 0.226 | 0.208 | 1 | 0.90669681 | 0.363273661 | 0.146 | 0.16 | 1 | 0.90669681 | 0.875170174 |
| Gm15862 | 0.659393736 | 0.569760094 | 0.057 | 0.042 | 1 | 0.646976884 | 0.614812428 | 0.049 | 0.034 | 1 | 0.659393736 | 0.87537468 |
| Mrpl39 | 0.70374314 | 0.174900477 | 0.264 | 0.237 | 1 | 0.647113542 | 0.347820009 | 0.171 | 0.208 | 1 | 0.70374314 | 0.875471148 |
| Chst10 | 0.986865056 | 0.280253477 | 0.038 | 0.038 | 1 | 0.647946199 | 1.377773231 | 0.049 | 0.072 | 1 | 0.986865056 | 0.876058121 |
| Efhc1 | 0.90155955 | 0.695290976 | 0.038 | 0.042 | 1 | 0.648029111 | -0.126629274 | 0.073 | 0.096 | 1 | 0.90155955 | 0.876116493 |
| Supv3l1 | 0.648949222 | 0.377115016 | 0.226 | 0.208 | 1 | 0.952202181 | 0.252242349 | 0.268 | 0.287 | 1 | 0.952202181 | 0.876763351 |
| D330023K18Rik | 0.758424309 | 0.370451286 | 0.094 | 0.114 | 1 | 0.649068014 | 0.721727632 | 0.073 | 0.099 | 1 | 0.758424309 | 0.876846741 |
| Dut | 0.64952768 | 0.736934485 | 0.585 | 0.585 | 1 | 0.798800878 | 0.580865097 | 0.61 | 0.652 | 1 | 0.798800878 | 0.877169153 |
| Mif | 0.649709903 | -0.241794817 | 0.755 | 0.75 | 1 | 0.900827112 | 0.304983811 | 0.61 | 0.648 | 1 | 0.900827112 | 0.877296848 |
| Gm35769 | 0.71096955 | 0.377115016 | 0.075 | 0.093 | 1 | 0.650019355 | 0.749742009 | 0.122 | 0.102 | 1 | 0.71096955 | 0.877513548 |
| Ptges2 | 0.689557822 | 0.8327945 | 0.094 | 0.081 | 1 | 0.650350628 | 0.351778023 | 0.171 | 0.147 | 1 | 0.689557822 | 0.877745317 |
| Mettl17 | 0.651385437 | 0.569760094 | 0.094 | 0.076 | 1 | 0.93978668 | 0.114738825 | 0.098 | 0.092 | 1 | 0.93978668 | 0.878467886 |
| C230035I16Rik | 0.663401681 | -0.1672055 | 0.019 | 0.03 | 1 | 0.651404528 | 0.444887427 | 0.073 | 0.055 | 1 | 0.663401681 | 0.878481197 |
| Hykk | 0.855425466 | 0.739685096 | 0.057 | 0.051 | 1 | 0.652738383 | -0.116991461 | 0.049 | 0.068 | 1 | 0.855425466 | 0.879409369 |
| Nr2c2ap | 0.653419886 | 0.300573461 | 0.189 | 0.161 | 1 | 0.802384281 | 0.515276755 | 0.195 | 0.184 | 1 | 0.802384281 | 0.879882224 |
| Fam81a | 0.653951506 | 0.569760094 | 0.057 | 0.042 | 1 | 0.835502813 | 0.515276755 | 0.024 | 0.031 | 1 | 0.835502813 | 0.88025044 |
| 4930558N01Rik | 0.654566199 | 0.417757001 | 0.075 | 0.059 | 1 | 0.787967407 | 0.313642894 | 0.073 | 0.061 | 1 | 0.787967407 | 0.880675489 |
| Glul | 0.721512262 | -0.657667917 | 0.943 | 0.792 | 1 | 0.65506106 | 0.104900633 | 0.805 | 0.812 | 1 | 0.721512262 | 0.881017128 |
| Qtrt2 | 0.656280575 | 0.172000586 | 0.283 | 0.25 | 1 | 0.763834449 | 0.252242349 | 0.317 | 0.3 | 1 | 0.763834449 | 0.881856957 |
| Ankrd24 | 0.888118588 | 0.110328475 | 0.113 | 0.106 | 1 | 0.656295755 | 0.47463477 | 0.073 | 0.099 | 1 | 0.888118588 | 0.881867392 |
| Lcp1 | 0.891167808 | 1.739685096 | 0.038 | 0.042 | 1 | 0.656401123 | -1.395455907 | 0.22 | 0.184 | 1 | 0.891167808 | 0.881939812 |
| Gm36899 | 0.656496624 | 0.154722595 | 0.057 | 0.076 | 1 | 0.846931821 | 1.377773231 | 0.024 | 0.031 | 1 | 0.846931821 | 0.88200543 |
| Spr | 0.696139061 | 0.166795427 | 0.358 | 0.356 | 1 | 0.657235595 | 0.285409213 | 0.341 | 0.314 | 1 | 0.696139061 | 0.882512563 |
| Dpysl2 | 0.892444655 | -0.276993646 | 0.434 | 0.462 | 1 | 0.65801748 | 0.515276755 | 0.366 | 0.379 | 1 | 0.892444655 | 0.883047956 |
| Cfap300 | 0.658384643 | -0.20282941 | 0.113 | 0.136 | 1 | 0.788022388 | 0.600165653 | 0.098 | 0.085 | 1 | 0.788022388 | 0.883298948 |
| BC004004 | 0.658441453 | 0.295766476 | 0.509 | 0.521 | 1 | 0.96607825 | 0.195339958 | 0.39 | 0.396 | 1 | 0.96607825 | 0.883337759 |
| Rbm15 | 0.905737945 | -0.126847762 | 0.377 | 0.377 | 1 | 0.65903014 | 0.228395607 | 0.341 | 0.437 | 1 | 0.905737945 | 0.883739555 |
| 1110059G10Rik | 0.659173645 | 0.255249471 | 0.321 | 0.292 | 1 | 0.741398908 | 0.277777441 | 0.244 | 0.28 | 1 | 0.741398908 | 0.883837396 |
| Zfp970 | 0.659393736 | 0.569760094 | 0.057 | 0.042 | 1 | 0.96563688 | 0.589277336 | 0.049 | 0.051 | 1 | 0.96563688 | 0.883987373 |
| E530011L22Rik | 0.659393736 | 0.569760094 | 0.057 | 0.042 | 1 | 0.901627466 | 0.515276755 | 0.049 | 0.055 | 1 | 0.901627466 | 0.883987373 |
| Taf6 | 0.659576228 | 0.333692736 | 0.208 | 0.182 | 1 | 0.82424513 | 0.458693227 | 0.171 | 0.195 | 1 | 0.82424513 | 0.884111655 |
| Dus4l | 0.785055709 | 0.228723176 | 0.094 | 0.11 | 1 | 0.660027706 | 0.667279848 | 0.049 | 0.068 | 1 | 0.785055709 | 0.88441884 |
| 2410018L13Rik | 0.744550492 | 0.495759513 | 0.208 | 0.195 | 1 | 0.661166084 | 0.65788115 | 0.171 | 0.154 | 1 | 0.744550492 | 0.885191577 |
| Arf4os | 0.993684223 | 0.216123139 | 0.075 | 0.076 | 1 | 0.661431287 | 0.458693227 | 0.049 | 0.068 | 1 | 0.993684223 | 0.885371227 |
| Cmc1 | 0.802535307 | 0.110328475 | 0.208 | 0.233 | 1 | 0.66152156 | 0.574170444 | 0.244 | 0.242 | 1 | 0.802535307 | 0.885432346 |
| Mpi | 0.662550611 | 0.569760094 | 0.075 | 0.059 | 1 | 0.684370942 | -0.101394606 | 0.049 | 0.065 | 1 | 0.684370942 | 0.88612791 |
| Gm29055 | 0.663401681 | -0.304709024 | 0.019 | 0.03 | 1 | 0.87556513 | 0.83720485 | 0.024 | 0.02 | 1 | 0.87556513 | 0.886701572 |
| Slc7a11 | 0.896371449 | -0.52334931 | 0.075 | 0.068 | 1 | 0.663487562 | -1.686357106 | 0.049 | 0.065 | 1 | 0.896371449 | 0.886759379 |
| Zranb3 | 0.663820972 | -0.330704232 | 0.075 | 0.093 | 1 | 0.785812954 | 1.126711467 | 0.073 | 0.092 | 1 | 0.785812954 | 0.886983661 |
| Slc38a10 | 0.664225544 | -0.162690019 | 0.623 | 0.661 | 1 | 0.748459782 | 0.329997507 | 0.561 | 0.58 | 1 | 0.748459782 | 0.887255515 |
| Lmf1 | 0.664877119 | 0.43225657 | 0.264 | 0.237 | 1 | 0.804381767 | 0.114738825 | 0.171 | 0.191 | 1 | 0.804381767 | 0.887692655 |
| Gm13293 | 0.664924942 | 0.454282877 | 0.057 | 0.042 | 1 | 0.74789451 | 0.100239256 | 0.049 | 0.061 | 1 | 0.74789451 | 0.887724706 |
| Fbxl4 | 0.665078483 | -0.399866257 | 0.113 | 0.136 | 1 | 0.761867419 | 0.589277336 | 0.122 | 0.109 | 1 | 0.761867419 | 0.887827578 |
| Ltbr | 0.931158111 | 0.192790636 | 0.491 | 0.551 | 1 | 0.665286165 | 0.336939514 | 0.488 | 0.481 | 1 | 0.931158111 | 0.887966649 |
| Pigo | 0.665481273 | 0.438515561 | 0.094 | 0.076 | 1 | 0.969378959 | 0.667279848 | 0.098 | 0.099 | 1 | 0.969378959 | 0.888097221 |
| Nkapd1 | 0.665594839 | 0.445064347 | 0.377 | 0.394 | 1 | 0.941921627 | 0.354422744 | 0.39 | 0.423 | 1 | 0.941921627 | 0.888173188 |

| | | | | | | | | | | | | |
|---|---|---|---|---|---|---|---|---|---|---|---|---|
| Gm14286 | 0.665715777 | -0.1672055 | 0.019 | 0.03 | 1 | 0.896398673 | 0.515276755 | 0.049 | 0.055 | 1 | 0.896398673 | 0.888254058 |
| Gm26660 | 0.665715777 | -0.1672055 | 0.019 | 0.03 | 1 | 0.729889663 | 1.83720485 | 0.024 | 0.017 | 1 | 0.729889663 | 0.888254058 |
| Smco4 | 0.667379298 | 0.239114782 | 0.264 | 0.246 | 1 | 0.863888893 | 0.282615998 | 0.293 | 0.338 | 1 | 0.863888893 | 0.889363469 |
| Ccdc17 | 0.668070194 | 0.154722595 | 0.019 | 0.03 | 1 | 0.826384776 | 0.515276755 | 0.024 | 0.031 | 1 | 0.826384776 | 0.889822604 |
| Irx3os | 0.668070194 | 0.154722595 | 0.019 | 0.03 | 1 | 0.824088343 | 0.377773231 | 0.024 | 0.031 | 1 | 0.824088343 | 0.889822604 |
| Gm35612 | 0.668070194 | 0.154722595 | 0.019 | 0.03 | 1 | 0.684370806 | -0.16279515 | 0.049 | 0.065 | 1 | 0.684370806 | 0.889822604 |
| Hdac9 | 0.672032977 | -1.260314904 | 0.113 | 0.089 | 1 | 0.809788604 | 1.483567895 | 0.171 | 0.164 | 1 | 0.809788604 | 0.892437632 |
| Dppa5a | 0.762480493 | 0.739685096 | 0.038 | 0.03 | 1 | 0.672172778 | 0.136765132 | 0.024 | 0.038 | 1 | 0.762480493 | 0.892529313 |
| Gab3 | 0.672332674 | -0.783876861 | 0.038 | 0.051 | 1 | 0.735421977 | -0.484723245 | 0.024 | 0.034 | 1 | 0.735421977 | 0.892634124 |
| Fads1 | 0.945899417 | -0.210926878 | 0.396 | 0.369 | 1 | 0.673131387 | 0.161094461 | 0.366 | 0.334 | 1 | 0.945899417 | 0.89315691 |
| Fam111a | 0.673218486 | -0.430239906 | 0.057 | 0.072 | 1 | 0.986995563 | 0.83720485 | 0.024 | 0.024 | 1 | 0.986995563 | 0.893213842 |
| Eml6 | 0.67322644 | 0.528537431 | 0.17 | 0.157 | 1 | 0.92932982 | -0.475073075 | 0.244 | 0.242 | 1 | 0.92932982 | 0.893219041 |
| Wdr53 | 0.6734413 | 0.216123139 | 0.264 | 0.237 | 1 | 0.788426862 | 0.545973552 | 0.195 | 0.229 | 1 | 0.788426862 | 0.893359415 |
| Man2b2 | 0.673907532 | -0.295938814 | 0.264 | 0.28 | 1 | 0.780738016 | 0.263737988 | 0.293 | 0.348 | 1 | 0.780738016 | 0.893663702 |
| Aldh16a1 | 0.675049917 | 0.389187848 | 0.075 | 0.059 | 1 | 0.96733343 | 0.544423101 | 0.122 | 0.123 | 1 | 0.96733343 | 0.894407443 |
| Tceal1 | 0.926259258 | 0.216123139 | 0.264 | 0.297 | 1 | 0.675406471 | 0.441903569 | 0.244 | 0.3 | 1 | 0.926259258 | 0.894639041 |
| Gnpda1 | 0.67658588 | -0.223789028 | 0.132 | 0.157 | 1 | 0.786353588 | 0.416873051 | 0.22 | 0.215 | 1 | 0.786353588 | 0.895403307 |
| Slc16a3 | 0.676691664 | 0.347367673 | 0.094 | 0.076 | 1 | 0.994414267 | -0.221688839 | 0.049 | 0.048 | 1 | 0.994414267 | 0.89547172 |
| Snhg12 | 0.741230324 | 0.251584134 | 0.585 | 0.657 | 1 | 0.67685484 | 0.181853021 | 0.439 | 0.509 | 1 | 0.741230324 | 0.895577206 |
| Ears2 | 0.677263881 | -0.125385324 | 0.094 | 0.114 | 1 | 0.885367882 | 0.252242349 | 0.122 | 0.133 | 1 | 0.885367882 | 0.895841398 |
| Gm47123 | 0.677401546 | 0.154722595 | 0.019 | 0.03 | 1 | 0.968793924 | 1.422167351 | 0.024 | 0.024 | 1 | 0.968793924 | 0.895930238 |
| Zfp612 | 0.679455197 | -0.347777746 | 0.038 | 0.051 | 1 | 0.727936015 | 0.667279848 | 0.049 | 0.038 | 1 | 0.727936015 | 0.897251029 |
| Abi3bp | 0.679591434 | 0.112494359 | 0.83 | 0.788 | 1 | 0.813230194 | 1.046534514 | 0.829 | 0.901 | 1 | 0.813230194 | 0.897338351 |
| Tpp1 | 0.918824429 | -0.1182959 | 0.528 | 0.576 | 1 | 0.681020315 | -0.214075655 | 0.561 | 0.553 | 1 | 0.918824429 | 0.89825196 |
| Mre11a | 0.746223686 | 0.289652175 | 0.17 | 0.153 | 1 | 0.681252975 | 0.185128153 | 0.22 | 0.188 | 1 | 0.746223686 | 0.898400334 |
| Trim23 | 0.862131696 | 0.776210972 | 0.226 | 0.22 | 1 | 0.681534138 | 0.240269707 | 0.268 | 0.253 | 1 | 0.862131696 | 0.898579494 |
| Styxl1 | 0.682084573 | 0.569760094 | 0.019 | 0.03 | 1 | 0.746881137 | 0.377773231 | 0.024 | 0.017 | 1 | 0.746881137 | 0.898929781 |
| Gm15910 | 0.682084573 | 0.569760094 | 0.019 | 0.03 | 1 | 0.916649754 | 0.667279848 | 0.024 | 0.027 | 1 | 0.916649754 | 0.898929781 |
| C130046K22Rik | 0.682084573 | 0.569760094 | 0.019 | 0.03 | 1 | 0.824812458 | 0.334704509 | 0.049 | 0.041 | 1 | 0.824812458 | 0.898929781 |
| Zfp39 | 0.682157794 | -0.1672055 | 0.057 | 0.072 | 1 | 0.789367819 | 0.46797104 | 0.098 | 0.085 | 1 | 0.789367819 | 0.898976332 |
| Gm11423 | 0.682157794 | -0.1672055 | 0.057 | 0.072 | 1 | 0.94575798 | 0.252242349 | 0.049 | 0.051 | 1 | 0.94575798 | 0.898976332 |
| Serpinb1a | 0.779443586 | 0.154722595 | 0.019 | 0.025 | 1 | 0.68364805 | 0.410940095 | 0.098 | 0.078 | 1 | 0.779443586 | 0.899921444 |
| Gm37640 | 0.683834454 | 0.280253477 | 0.094 | 0.076 | 1 | 0.8958181 | 0.392420007 | 0.122 | 0.133 | 1 | 0.8958181 | 0.900039347 |
| Rusf1 | 0.684048386 | 0.614154213 | 0.132 | 0.114 | 1 | 0.708212712 | 0.644559772 | 0.073 | 0.096 | 1 | 0.708212712 | 0.900174578 |
| Them6 | 0.684424342 | 0.739685096 | 0.019 | 0.03 | 1 | 0.749728988 | 0.83720485 | 0.024 | 0.017 | 1 | 0.749728988 | 0.900412004 |
| Rtl6 | 0.684424342 | 0.739685096 | 0.019 | 0.03 | 1 | 0.722131097 | 0.83720485 | 0.049 | 0.038 | 1 | 0.722131097 | 0.900412004 |
| Rrn3 | 0.770729821 | 0.241434228 | 0.302 | 0.347 | 1 | 0.684428388 | 0.465236072 | 0.39 | 0.382 | 1 | 0.770729821 | 0.900414558 |
| Zscan26 | 0.685222483 | 0.382133091 | 0.415 | 0.445 | 1 | 0.798111219 | 0.162605137 | 0.488 | 0.502 | 1 | 0.798111219 | 0.900915115 |
| Cp | 0.903754264 | -0.382519013 | 0.66 | 0.606 | 1 | 0.686428612 | 1.848886655 | 0.195 | 0.184 | 1 | 0.903754264 | 0.901672985 |
| Hs3st5 | 0.722567081 | 1.8327945 | 0.019 | 0.013 | 1 | 0.687705661 | 2.167353451 | 0.049 | 0.038 | 1 | 0.722567081 | 0.902472246 |
| Ctss | 0.881269634 | 2.595295186 | 0.038 | 0.034 | 1 | 0.689378239 | 0.737669176 | 0.049 | 0.068 | 1 | 0.881269634 | 0.903514122 |
| Fbln2 | 0.89739913 | 0.569760094 | 0.038 | 0.042 | 1 | 0.689575365 | 1.721727632 | 0.049 | 0.038 | 1 | 0.89739913 | 0.903636546 |
| Rai2 | 0.750817599 | -1.067669827 | 0.057 | 0.068 | 1 | 0.689926087 | 0.218295017 | 0.073 | 0.058 | 1 | 0.750817599 | 0.903854168 |
| Slc38a9 | 0.690459553 | -0.313426241 | 0.208 | 0.229 | 1 | 0.99410919 | 0.348918368 | 0.293 | 0.314 | 1 | 0.99410919 | 0.904184712 |
| Med24 | 0.985190142 | 0.289652175 | 0.189 | 0.195 | 1 | 0.690575935 | -0.233184478 | 0.195 | 0.218 | 1 | 0.985190142 | 0.904256748 |
| Tfb1m | 0.996841423 | 0.347367673 | 0.075 | 0.076 | 1 | 0.691843603 | 0.614812428 | 0.024 | 0.038 | 1 | 0.996841423 | 0.905039635 |
| Tmem209 | 0.777892928 | -0.319208594 | 0.151 | 0.161 | 1 | 0.69237049 | 0.676740178 | 0.195 | 0.171 | 1 | 0.777892928 | 0.905364085 |
| Lrrc51 | 0.934575394 | 0.624207878 | 0.094 | 0.102 | 1 | 0.692944181 | 0.30115195 | 0.098 | 0.078 | 1 | 0.934575394 | 0.905716724 |
| Gpt | 0.732691937 | 1.154722595 | 0.019 | 0.013 | 1 | 0.693820021 | 0.930314254 | 0.024 | 0.038 | 1 | 0.732691937 | 0.906253821 |
| Hscb | 0.695216286 | 0.187889459 | 0.132 | 0.161 | 1 | 0.789769931 | 0.797676486 | 0.122 | 0.109 | 1 | 0.789769931 | 0.907106887 |
| Rfx2 | 0.875747477 | 0.154722595 | 0.17 | 0.182 | 1 | 0.695279745 | 0.748078959 | 0.317 | 0.328 | 1 | 0.875747477 | 0.907145566 |
| Cntfr | 0.792025927 | 0.267197324 | 0.283 | 0.267 | 1 | 0.695538608 | -0.372248516 | 0.195 | 0.218 | 1 | 0.792025927 | 0.907303261 |
| Amigo3 | 0.880784741 | 0.280253477 | 0.038 | 0.042 | 1 | 0.695776133 | 1.574170444 | 0.024 | 0.038 | 1 | 0.880784741 | 0.907447839 |
| Gm43112 | 0.732691937 | 1.154722595 | 0.019 | 0.013 | 1 | 0.695788449 | 1.458693227 | 0.024 | 0.038 | 1 | 0.732691937 | 0.907455332 |
| Tubgcp6 | 0.873180917 | 0.502645898 | 0.094 | 0.089 | 1 | 0.696271586 | 0.707921833 | 0.122 | 0.106 | 1 | 0.873180917 | 0.90774905 |
| Mrpl37 | 0.942081523 | 0.311448098 | 0.226 | 0.225 | 1 | 0.696625228 | 0.416873051 | 0.195 | 0.232 | 1 | 0.942081523 | 0.907963748 |
| Rgs20 | 0.699130418 | -0.389597921 | 0.075 | 0.059 | 1 | 0.976593771 | 0.83720485 | 0.024 | 0.024 | 1 | 0.976593771 | 0.909477494 |
| Cad | 0.765242991 | 0.154722595 | 0.094 | 0.081 | 1 | 0.699952729 | 0.359157553 | 0.122 | 0.102 | 1 | 0.765242991 | 0.909971635 |
| Gm43331 | 0.746876889 | 0.776210972 | 0.057 | 0.047 | 1 | 0.700519221 | 0.109284395 | 0.098 | 0.123 | 1 | 0.746876889 | 0.910311263 |
| Sun2 | 0.701585244 | 0.335748589 | 0.491 | 0.496 | 1 | 0.710058972 | 0.430029468 | 0.463 | 0.468 | 1 | 0.710058972 | 0.910948633 |
| Trappc1 | 0.836733231 | 0.267197324 | 0.245 | 0.28 | 1 | 0.702449347 | 0.304709769 | 0.195 | 0.232 | 1 | 0.836733231 | 0.911463609 |
| Mapk4 | 0.932611893 | -0.304709024 | 0.019 | 0.017 | 1 | 0.702926594 | 1.159132945 | 0.049 | 0.038 | 1 | 0.932611893 | 0.911747392 |
| Mavs | 0.827127062 | 0.207190015 | 0.094 | 0.085 | 1 | 0.704336089 | 1.363273661 | 0.098 | 0.082 | 1 | 0.827127062 | 0.912582852 |
| Zfp518a | 0.832697916 | 0.261637799 | 0.17 | 0.161 | 1 | 0.705542819 | -0.292078167 | 0.122 | 0.14 | 1 | 0.832697916 | 0.913294969 |
| Cenpp | 0.707153229 | -0.53715511 | 0.245 | 0.225 | 1 | 0.959540073 | 0.295311071 | 0.22 | 0.229 | 1 | 0.959540073 | 0.914240768 |
| Zfp119a | 0.948980228 | 0.510866405 | 0.075 | 0.081 | 1 | 0.707226064 | 0.322631677 | 0.122 | 0.102 | 1 | 0.948980228 | 0.914283422 |
| Comt | 0.708284479 | 0.456378295 | 0.434 | 0.445 | 1 | 0.94510486 | 0.210663245 | 0.561 | 0.608 | 1 | 0.94510486 | 0.914902055 |
| Phykpl | 0.993975553 | -0.104011674 | 0.226 | 0.22 | 1 | 0.708836184 | 0.298046039 | 0.268 | 0.246 | 1 | 0.993975553 | 0.915223633 |
| Samm50 | 0.708943726 | 0.320146204 | 0.358 | 0.369 | 1 | 0.805513646 | -0.171357164 | 0.366 | 0.375 | 1 | 0.805513646 | 0.915286245 |
| Lypd2 | 0.97621125 | -0.678167419 | 0.132 | 0.127 | 1 | 0.711453858 | 0.667279848 | 0.268 | 0.249 | 1 | 0.97621125 | 0.916741124 |
| Pop5 | 0.831577533 | -0.1672055 | 0.321 | 0.326 | 1 | 0.712025488 | 0.584224109 | 0.366 | 0.362 | 1 | 0.831577533 | 0.917070681 |
| Gm11457 | 0.926212226 | 0.8327945 | 0.019 | 0.017 | 1 | 0.712484083 | 1.136765132 | 0.049 | 0.038 | 1 | 0.926212226 | 0.917334597 |
| 4930511M06Rik | 0.89532551 | 0.669295768 | 0.038 | 0.042 | 1 | 0.714779374 | 0.252242349 | 0.049 | 0.065 | 1 | 0.89532551 | 0.918649195 |
| Smim19 | 0.714788175 | -0.888051429 | 0.566 | 0.547 | 1 | 0.935521361 | 0.129845718 | 0.463 | 0.481 | 1 | 0.935521361 | 0.918654215 |
| Msl3 | 0.715627067 | 0.308635278 | 0.396 | 0.428 | 1 | 0.922907077 | 0.296636468 | 0.366 | 0.42 | 1 | 0.922907077 | 0.919132035 |
| Rnf40 | 0.864536461 | 0.286826131 | 0.264 | 0.263 | 1 | 0.717185194 | 0.282615998 | 0.341 | 0.331 | 1 | 0.864536461 | 0.920015786 |
| Pard6a | 0.939643391 | 0.454282877 | 0.094 | 0.093 | 1 | 0.7193606 | 0.444887427 | 0.049 | 0.065 | 1 | 0.939643391 | 0.921241527 |
| Ift43 | 0.71952858 | 0.176526965 | 0.509 | 0.521 | 1 | 0.947597484 | 0.252242349 | 0.512 | 0.553 | 1 | 0.947597484 | 0.921335783 |
| Gm34894 | 0.914958 | -0.1672055 | 0.019 | 0.021 | 1 | 0.720202665 | 0.930314254 | 0.049 | 0.038 | 1 | 0.914958 | 0.921713451 |
| Mrpl34 | 0.978198602 | 0.240452469 | 0.321 | 0.343 | 1 | 0.721518322 | 0.659666664 | 0.244 | 0.235 | 1 | 0.978198602 | 0.922447955 |
| Tab3 | 0.722318617 | -0.101617158 | 0.226 | 0.203 | 1 | 0.9767475 | -0.134780774 | 0.195 | 0.188 | 1 | 0.9767475 | 0.922893049 |
| Mettl4 | 0.790920555 | 0.491757582 | 0.132 | 0.123 | 1 | 0.722464802 | 0.766815522 | 0.171 | 0.15 | 1 | 0.790920555 | 0.922974214 |
| Ppp1r42 | 0.72256466 | 2.154722595 | 0.019 | 0.013 | 1 | 0.749728988 | 0.377773231 | 0.024 | 0.017 | 1 | 0.749728988 | 0.923029632 |
| Gm32191 | 0.72256466 | 1.739685096 | 0.019 | 0.013 | 1 | 0.878310931 | 0.515276755 | 0.024 | 0.02 | 1 | 0.878310931 | 0.923029632 |
| Slc17a7 | 0.72256466 | 1.739685096 | 0.019 | 0.013 | 1 | 0.732716552 | 1.614812428 | 0.024 | 0.017 | 1 | 0.732716552 | 0.923029632 |
| Gm49173 | 0.72256466 | 1.739685096 | 0.019 | 0.013 | 1 | 0.986995563 | 0.83720485 | 0.024 | 0.024 | 1 | 0.986995563 | 0.923029632 |
| Kctd8 | 0.722567081 | 1.891688189 | 0.019 | 0.013 | 1 | 0.758694497 | 0.773074512 | 0.024 | 0.034 | 1 | 0.758694497 | 0.923030976 |
| Tamm41 | 0.723486387 | 0.569760094 | 0.094 | 0.081 | 1 | 0.750946477 | -0.19841906 | 0.098 | 0.113 | 1 | 0.750946477 | 0.923540222 |
| Slx4ip | 0.723639959 | 0.193196743 | 0.189 | 0.225 | 1 | 0.938166052 | 0.222495006 | 0.195 | 0.195 | 1 | 0.938166052 | 0.923625128 |
| Mertk | 0.724455941 | 0.115194231 | 0.528 | 0.521 | 1 | 0.794369083 | -0.334203643 | 0.659 | 0.669 | 1 | 0.794369083 | 0.924075471 |
| 2310022A10Rik | 0.865364543 | 0.466666601 | 0.113 | 0.106 | 1 | 0.725456001 | 0.756284854 | 0.171 | 0.157 | 1 | 0.865364543 | 0.924625593 |
| Gm9993 | 0.971243602 | -0.389597921 | 0.094 | 0.093 | 1 | 0.725551616 | 1.377773231 | 0.073 | 0.061 | 1 | 0.971243602 | 0.924678084 |
| Thap12 | 0.725572239 | 0.177929986 | 0.434 | 0.513 | 1 | 0.861306192 | 0.232746173 | 0.561 | 0.577 | 1 | 0.861306192 | 0.924689404 |
| Cdc123 | 0.72578928 | 0.124975251 | 0.755 | 0.788 | 1 | 0.880214661 | 0.212713985 | 0.659 | 0.765 | 1 | 0.880214661 | 0.924808481 |
| Cftr | 0.725936481 | 1.569760094 | 0.019 | 0.013 | 1 | 0.824695335 | -0.435813645 | 0.122 | 0.106 | 1 | 0.824695335 | 0.924889188 |
| Gm21781 | 0.726611811 | 1.039245377 | 0.057 | 0.047 | 1 | 0.741574522 | -0.278272368 | 0.049 | 0.061 | 1 | 0.741574522 | 0.925258898 |
| Dnlz | 0.726769769 | 0.139772253 | 0.358 | 0.331 | 1 | 0.78787049 | 0.174239837 | 0.195 | 0.225 | 1 | 0.78787049 | 0.925345241 |
| Eogt | 0.727863197 | 0.423539353 | 0.245 | 0.237 | 1 | 0.968457801 | 0.686962214 | 0.293 | 0.331 | 1 | 0.968457801 | 0.92594156 |

| | | | | | | | | | | | | |
|---|---|---|---|---|---|---|---|---|---|---|---|---|
| Srp54c | 0.728059401 | 0.385335523 | 0.283 | 0.263 | 1 | 0.774771365 | 0.470422519 | 0.268 | 0.259 | 1 | 0.774771365 | 0.926048311 |
| Ak7 | 0.728477277 | 0.874614676 | 0.057 | 0.047 | 1 | 0.925762561 | 0.870371714 | 0.073 | 0.082 | 1 | 0.925762561 | 0.926275411 |
| Gm28535 | 0.929410101 | 0.569760094 | 0.019 | 0.017 | 1 | 0.729874414 | 0.667279848 | 0.049 | 0.038 | 1 | 0.929410101 | 0.927032168 |
| Ciao3 | 0.850692736 | 0.306725688 | 0.075 | 0.068 | 1 | 0.730036921 | -0.20718927 | 0.073 | 0.089 | 1 | 0.850692736 | 0.927119936 |
| Cacnb1 | 0.932769215 | -0.1182959 | 0.094 | 0.089 | 1 | 0.731642967 | 0.152706676 | 0.122 | 0.102 | 1 | 0.932769215 | 0.927984503 |
| 0610009L18Rik | 0.732089493 | 0.438515561 | 0.094 | 0.081 | 1 | 0.974673471 | 1.071670103 | 0.049 | 0.051 | 1 | 0.974673471 | 0.92822396 |
| Cenpj | 0.946797688 | 0.228723176 | 0.113 | 0.119 | 1 | 0.732650632 | 1.120997816 | 0.122 | 0.113 | 1 | 0.946797688 | 0.928524315 |
| 9430060I03Rik | 0.732691937 | 1.154722595 | 0.019 | 0.013 | 1 | 0.986995563 | 0.83720485 | 0.024 | 0.024 | 1 | 0.986995563 | 0.928546399 |
| Lor | 0.732691937 | 1.154722595 | 0.019 | 0.013 | 1 | 0.733713289 | 0.721727632 | 0.049 | 0.038 | 1 | 0.733713289 | 0.928546399 |
| Gm42938 | 0.732691937 | 1.154722595 | 0.019 | 0.013 | 1 | 0.903375428 | 0.667279848 | 0.049 | 0.055 | 1 | 0.903375428 | 0.928546399 |
| Fcsk | 0.732691937 | 1.154722595 | 0.019 | 0.013 | 1 | 0.745607362 | 0.911205431 | 0.073 | 0.061 | 1 | 0.745607362 | 0.928546399 |
| 5730507C01Rik | 0.732691937 | 1.154722595 | 0.019 | 0.013 | 1 | 0.87556513 | 0.83720485 | 0.024 | 0.02 | 1 | 0.87556513 | 0.928546399 |
| 5033425G24Rik | 0.732691937 | 1.154722595 | 0.019 | 0.013 | 1 | 0.872828964 | 1.029849928 | 0.024 | 0.02 | 1 | 0.872828964 | 0.928546399 |
| Frs3 | 0.732691937 | 1.154722595 | 0.019 | 0.013 | 1 | 0.732716552 | 1.614812428 | 0.024 | 0.017 | 1 | 0.732716552 | 0.928546399 |
| Gm50340 | 0.732691937 | 1.154722595 | 0.019 | 0.013 | 1 | 0.744047924 | 1.252242349 | 0.024 | 0.017 | 1 | 0.744047924 | 0.928546399 |
| B230319C09Rik | 0.736073999 | 0.8327945 | 0.019 | 0.013 | 1 | 0.732716552 | 1.614812428 | 0.024 | 0.017 | 1 | 0.736073999 | 0.928559558 |
| Cyren | 0.733039588 | 0.417757001 | 0.094 | 0.114 | 1 | 0.792136996 | 0.707921833 | 0.073 | 0.089 | 1 | 0.792136996 | 0.928732139 |
| Arl16 | 0.733492164 | 0.631160639 | 0.17 | 0.165 | 1 | 0.850265609 | -0.132421501 | 0.122 | 0.13 | 1 | 0.850265609 | 0.928973573 |
| Crispld1 | 0.927315 | 0.346796874 | 0.792 | 0.881 | 1 | 0.733995858 | -0.210048337 | 0.805 | 0.816 | 1 | 0.927315 | 0.929241796 |
| Ntpcr | 0.735978719 | 0.154722595 | 0.208 | 0.186 | 1 | 0.832615981 | -0.19841906 | 0.098 | 0.106 | 1 | 0.832615981 | 0.930292763 |
| Gm20632 | 0.736073999 | 0.8327945 | 0.019 | 0.013 | 1 | 0.916649754 | 0.667279848 | 0.024 | 0.027 | 1 | 0.916649754 | 0.930343066 |
| Foxo6 | 0.736073999 | 0.8327945 | 0.019 | 0.013 | 1 | 0.784426307 | 1.100239256 | 0.049 | 0.041 | 1 | 0.784426307 | 0.930343066 |
| Slc23a4 | 0.736073999 | 0.8327945 | 0.019 | 0.013 | 1 | 0.746881137 | 1.029849928 | 0.024 | 0.017 | 1 | 0.746881137 | 0.930343066 |
| Bean1 | 0.736073999 | 0.8327945 | 0.019 | 0.013 | 1 | 0.87556513 | 0.83720485 | 0.024 | 0.02 | 1 | 0.87556513 | 0.930343066 |
| Trim16 | 0.736073999 | 0.569760094 | 0.019 | 0.013 | 1 | 0.746881137 | 0.667279848 | 0.024 | 0.017 | 1 | 0.746881137 | 0.930343066 |
| 4933427D14Rik | 0.736073999 | 0.8327945 | 0.019 | 0.013 | 1 | 0.986995563 | 0.83720485 | 0.024 | 0.024 | 1 | 0.986995563 | 0.930343066 |
| Gm20939 | 0.736073999 | 0.8327945 | 0.019 | 0.013 | 1 | 0.986995563 | 0.83720485 | 0.024 | 0.024 | 1 | 0.986995563 | 0.930343066 |
| Tmem255a | 0.736073999 | 0.8327945 | 0.019 | 0.013 | 1 | 0.878310931 | 0.667279848 | 0.024 | 0.02 | 1 | 0.878310931 | 0.930343066 |
| Gm49041 | 0.736283004 | 0.652222254 | 0.075 | 0.064 | 1 | 0.95533271 | 1.159132945 | 0.049 | 0.048 | 1 | 0.95533271 | 0.930453346 |
| Cnot10 | 0.895759863 | 0.116042127 | 0.491 | 0.483 | 1 | 0.737338122 | -0.139079081 | 0.537 | 0.573 | 1 | 0.895759863 | 0.931008738 |
| Man2c1os | 0.737969899 | 0.997181318 | 0.075 | 0.064 | 1 | 0.848263359 | 1.911205431 | 0.098 | 0.116 | 1 | 0.848263359 | 0.931340226 |
| Slc25a18 | 0.7394666 | 0.569760094 | 0.019 | 0.013 | 1 | 0.93376172 | 1.100239256 | 0.024 | 0.027 | 1 | 0.93376172 | 0.932122347 |
| Ldhd | 0.7394666 | 0.347367673 | 0.019 | 0.013 | 1 | 0.914200263 | 0.515276755 | 0.024 | 0.027 | 1 | 0.914200263 | 0.932122347 |
| Fgd5 | 0.932505382 | 1.891688189 | 0.019 | 0.021 | 1 | 0.739624823 | -0.16279515 | 0.024 | 0.034 | 1 | 0.932505382 | 0.932204767 |
| Muc5ac | 0.932611893 | -3.149058153 | 0.019 | 0.017 | 1 | 0.739624823 | -0.250257991 | 0.024 | 0.034 | 1 | 0.932611893 | 0.932204767 |
| Hmg20a | 0.741197993 | 0.329665671 | 0.585 | 0.593 | 1 | 0.885090898 | 0.283048396 | 0.585 | 0.635 | 1 | 0.885090898 | 0.933021521 |
| Pih1d1 | 0.928553721 | 0.569760094 | 0.208 | 0.212 | 1 | 0.74129546 | 0.15538081 | 0.22 | 0.195 | 1 | 0.928553721 | 0.933071961 |
| Gm4681 | 0.919816481 | 0.932330173 | 0.019 | 0.017 | 1 | 0.741726892 | 0.136765132 | 0.024 | 0.034 | 1 | 0.919816481 | 0.933295002 |
| Selenoo | 0.741864846 | 0.502645898 | 0.17 | 0.157 | 1 | 0.907988607 | -0.16279515 | 0.171 | 0.174 | 1 | 0.907988607 | 0.933366242 |
| Cbr4 | 0.742432686 | -0.108311811 | 0.075 | 0.089 | 1 | 0.773530306 | 0.635570989 | 0.049 | 0.061 | 1 | 0.773530306 | 0.933659079 |
| Ap5s1 | 0.742495533 | 1.154722595 | 0.094 | 0.085 | 1 | 0.766287268 | 0.46797104 | 0.073 | 0.089 | 1 | 0.766287268 | 0.93369145 |
| Armc5 | 0.821649301 | 0.207190015 | 0.113 | 0.102 | 1 | 0.742924285 | 0.120997816 | 0.122 | 0.143 | 1 | 0.821649301 | 0.933912077 |
| H2-K2 | 0.756622516 | 0.104096522 | 0.094 | 0.081 | 1 | 0.743829142 | 0.252242349 | 0.024 | 0.034 | 1 | 0.756622516 | 0.934376492 |
| 6230426M11Rik | 0.761797609 | 0.569760094 | 0.057 | 0.047 | 1 | 0.743829142 | 0.136765132 | 0.024 | 0.034 | 1 | 0.761797609 | 0.934376492 |
| Gm28529 | 0.797844282 | 0.932330173 | 0.019 | 0.025 | 1 | 0.743829142 | 0.136765132 | 0.024 | 0.034 | 1 | 0.797844282 | 0.934376492 |
| H3c7 | 0.912039013 | -0.545717123 | 0.019 | 0.021 | 1 | 0.744047924 | 1.252242349 | 0.024 | 0.017 | 1 | 0.912039013 | 0.934488535 |
| Gm43915 | 0.926212226 | 0.8327945 | 0.019 | 0.017 | 1 | 0.744047924 | 1.252242349 | 0.024 | 0.017 | 1 | 0.926212226 | 0.934488535 |
| Gm43646 | 0.929410101 | 0.569760094 | 0.019 | 0.017 | 1 | 0.744047924 | 1.252242349 | 0.024 | 0.017 | 1 | 0.929410101 | 0.934488535 |
| Gm49893 | 0.929410101 | 0.347367673 | 0.019 | 0.017 | 1 | 0.744047924 | 1.252242349 | 0.024 | 0.017 | 1 | 0.929410101 | 0.934488535 |
| Gm42531 | 0.932611893 | -0.1672055 | 0.019 | 0.017 | 1 | 0.744047924 | 1.252242349 | 0.024 | 0.017 | 1 | 0.932611893 | 0.934488535 |
| L3hypdh | 0.852965394 | 0.272079545 | 0.208 | 0.199 | 1 | 0.744092825 | 0.630753972 | 0.171 | 0.154 | 1 | 0.852965394 | 0.934511518 |
| Slc50a1 | 0.744584698 | 0.285509202 | 0.547 | 0.619 | 1 | 0.775439471 | 0.155837865 | 0.488 | 0.522 | 1 | 0.775439471 | 0.934763024 |
| Mamstr | 0.798088617 | 0.454282877 | 0.038 | 0.047 | 1 | 0.745976859 | 0.377773231 | 0.024 | 0.034 | 1 | 0.798088617 | 0.935472244 |
| 6820445E23Rik | 0.992880642 | 0.962077517 | 0.057 | 0.059 | 1 | 0.745976859 | 0.377773231 | 0.024 | 0.034 | 1 | 0.992880642 | 0.935472244 |
| Slc25a45 | 0.941268385 | 0.377115016 | 0.075 | 0.081 | 1 | 0.746273068 | 0.136765132 | 0.122 | 0.102 | 1 | 0.941268385 | 0.935622644 |
| Ubap1l | 0.746876889 | 0.776210972 | 0.057 | 0.047 | 1 | 0.794802136 | 0.377773231 | 0.073 | 0.089 | 1 | 0.794802136 | 0.935928691 |
| Gm56874 | 0.760166624 | 0.454282877 | 0.038 | 0.03 | 1 | 0.746881137 | 1.029849928 | 0.024 | 0.017 | 1 | 0.760166624 | 0.935930841 |
| Hk3 | 0.779443586 | 0.154722595 | 0.019 | 0.025 | 1 | 0.746881137 | 0.83720485 | 0.024 | 0.017 | 1 | 0.779443586 | 0.935930841 |
| Faah | 0.780574205 | -0.1672055 | 0.038 | 0.047 | 1 | 0.746881137 | 1.029849928 | 0.024 | 0.017 | 1 | 0.780574205 | 0.935930841 |
| Tgm4 | 0.917883255 | 0.569760094 | 0.019 | 0.021 | 1 | 0.746881137 | 1.029849928 | 0.024 | 0.017 | 1 | 0.917883255 | 0.935930841 |
| Tnnc1 | 0.935978094 | 0.906795081 | 0.057 | 0.055 | 1 | 0.746881137 | 1.029849928 | 0.024 | 0.017 | 1 | 0.935978094 | 0.935930841 |
| Gcc1 | 0.747597654 | 0.272079545 | 0.226 | 0.208 | 1 | 0.920734619 | 0.156483366 | 0.268 | 0.287 | 1 | 0.920734619 | 0.936293056 |
| Zfp983 | 0.747889121 | 0.104096522 | 0.113 | 0.097 | 1 | 0.922972772 | 0.422167351 | 0.073 | 0.078 | 1 | 0.922972772 | 0.936440105 |
| 1700113A16Rik | 0.828092466 | 0.714150003 | 0.075 | 0.068 | 1 | 0.748790861 | 0.79281073 | 0.098 | 0.085 | 1 | 0.828092466 | 0.936893968 |
| Camkk1 | 0.897146191 | 0.417757001 | 0.038 | 0.034 | 1 | 0.749728988 | 0.83720485 | 0.024 | 0.017 | 1 | 0.897146191 | 0.93736442 |
| Zfp937 | 0.901698096 | 0.280253477 | 0.038 | 0.034 | 1 | 0.749728988 | 0.83720485 | 0.024 | 0.017 | 1 | 0.901698096 | 0.93736442 |
| Camk2n2 | 0.905020105 | 0.569760094 | 0.075 | 0.072 | 1 | 0.751853856 | 1.100239256 | 0.049 | 0.065 | 1 | 0.905020105 | 0.938423491 |
| 2700038G22Rik | 0.752333887 | 0.10741688 | 0.189 | 0.174 | 1 | 0.883849994 | 0.188112012 | 0.171 | 0.164 | 1 | 0.883849994 | 0.938661496 |
| Ccdc122 | 0.752874909 | -0.214511215 | 0.17 | 0.186 | 1 | 0.936293691 | 0.340779024 | 0.171 | 0.174 | 1 | 0.936293691 | 0.938929189 |
| Rpa3 | 0.753853733 | -0.444315091 | 0.283 | 0.288 | 1 | 0.989570328 | 0.162605137 | 0.22 | 0.225 | 1 | 0.989570328 | 0.939412015 |
| Selenok-ps1 | 0.755567413 | 0.8327945 | 0.038 | 0.03 | 1 | 0.916649754 | 0.667279848 | 0.024 | 0.027 | 1 | 0.916649754 | 0.940252711 |
| Rnf145 | 0.767195817 | 0.661183122 | 0.34 | 0.347 | 1 | 0.756662356 | 0.119792053 | 0.415 | 0.396 | 1 | 0.767195817 | 0.940786791 |
| Ropn1l | 0.757967314 | 0.280253477 | 0.094 | 0.081 | 1 | 0.912424882 | 1.422167351 | 0.073 | 0.072 | 1 | 0.912424882 | 0.941420179 |
| Gtdc1 | 0.915511617 | 0.232401959 | 0.358 | 0.394 | 1 | 0.759116879 | 0.610854415 | 0.463 | 0.44 | 1 | 0.915511617 | 0.941975322 |
| Apba3 | 0.760689874 | 0.333692736 | 0.208 | 0.191 | 1 | 0.849953713 | 0.44893426 | 0.22 | 0.242 | 1 | 0.849953713 | 0.942730664 |
| Uckl1 | 0.761640591 | 0.185096244 | 0.189 | 0.169 | 1 | 0.802461614 | 0.109284395 | 0.146 | 0.164 | 1 | 0.802461614 | 0.943184792 |
| Kdm1b | 0.761797609 | 0.569760094 | 0.057 | 0.047 | 1 | 0.956812693 | 0.396632258 | 0.098 | 0.096 | 1 | 0.956812693 | 0.943259621 |
| Serinc4 | 0.762480493 | 0.739685096 | 0.038 | 0.03 | 1 | 0.992196803 | 0.515276755 | 0.024 | 0.024 | 1 | 0.992196803 | 0.943584484 |
| Lax1 | 0.989012386 | -0.237594828 | 0.057 | 0.055 | 1 | 0.762928098 | 0.515276755 | 0.024 | 0.034 | 1 | 0.989012386 | 0.943796913 |
| Rmnd1 | 0.764034925 | 0.739685096 | 0.132 | 0.123 | 1 | 0.891314564 | 0.699701326 | 0.146 | 0.167 | 1 | 0.891314564 | 0.944320483 |
| Napg | 0.76416501 | 0.367130928 | 0.396 | 0.428 | 1 | 0.952872925 | 0.214487427 | 0.463 | 0.498 | 1 | 0.952872925 | 0.944381858 |
| Tctn2 | 0.764500005 | 0.300573461 | 0.189 | 0.174 | 1 | 0.767865096 | 0.334704509 | 0.171 | 0.154 | 1 | 0.767865096 | 0.944539752 |
| Tcp11l1 | 0.967329516 | 0.347367673 | 0.208 | 0.212 | 1 | 0.76483593 | 0.169780189 | 0.268 | 0.253 | 1 | 0.967329516 | 0.94469786 |
| Plekho2 | 0.764908477 | 0.324647596 | 0.302 | 0.297 | 1 | 0.84441179 | -0.186641892 | 0.293 | 0.297 | 1 | 0.84441179 | 0.944731976 |
| Gm29358 | 0.986865056 | 0.280253477 | 0.038 | 0.038 | 1 | 0.765065249 | 1.029849928 | 0.024 | 0.034 | 1 | 0.986865056 | 0.944805663 |
| Dock5 | 0.765204508 | 0.495759513 | 0.132 | 0.119 | 1 | 0.794777975 | 0.486707603 | 0.244 | 0.229 | 1 | 0.794777975 | 0.944871077 |
| Gm26982 | 0.767066103 | 0.569760094 | 0.038 | 0.03 | 1 | 0.879490456 | 0.930314254 | 0.049 | 0.044 | 1 | 0.879490456 | 0.9457418 |
| Gm11175 | 0.767066103 | 0.569760094 | 0.038 | 0.03 | 1 | 0.916649754 | 0.667279848 | 0.024 | 0.027 | 1 | 0.916649754 | 0.9457418 |
| Gm57182 | 0.948751313 | 0.569760094 | 0.057 | 0.055 | 1 | 0.767181773 | 1.377773231 | 0.024 | 0.034 | 1 | 0.948751313 | 0.945795673 |
| 9630013D21Rik | 1 | 0.669295768 | 0.038 | 0.038 | 1 | 0.767181773 | 0.962735732 | 0.024 | 0.034 | 1 | 1 | 0.945795673 |
| Tmem40 | 0.767318506 | 0.454282877 | 0.057 | 0.047 | 1 | 0.992196803 | 0.377773231 | 0.024 | 0.024 | 1 | 0.992196803 | 0.945859323 |
| Pus7l | 0.767318506 | 0.454282877 | 0.057 | 0.047 | 1 | 0.7929683 | -0.221688839 | 0.049 | 0.058 | 1 | 0.7929683 | 0.945859323 |
| Dis3 | 0.768982354 | -0.112757716 | 0.396 | 0.347 | 1 | 0.940698805 | 0.278237558 | 0.317 | 0.358 | 1 | 0.940698805 | 0.946630847 |
| Dnajc15 | 0.949307838 | 0.201720074 | 0.415 | 0.479 | 1 | 0.769517152 | 0.755169081 | 0.415 | 0.437 | 1 | 0.949307838 | 0.946877657 |
| Rtl8b | 0.771444339 | 0.197791317 | 0.415 | 0.407 | 1 | 0.800887012 | 0.223673197 | 0.366 | 0.403 | 1 | 0.800887012 | 0.94776231 |
| Plekhg6 | 0.77169508 | 0.154722595 | 0.038 | 0.03 | 1 | 0.824088343 | 0.377773231 | 0.024 | 0.031 | 1 | 0.824088343 | 0.947876863 |
| Npff | 0.943264487 | 0.669295768 | 0.057 | 0.055 | 1 | 0.772963198 | 1.614812428 | 0.049 | 0.041 | 1 | 0.943264487 | 0.948454291 |
| Susd6 | 0.772994868 | 0.21090543 | 0.774 | 0.801 | 1 | 0.932439637 | 0.209173627 | 0.78 | 0.87 | 1 | 0.932439637 | 0.94846867 |

| | | | | | | | | | | | | |
|---|---|---|---|---|---|---|---|---|---|---|---|---|
| Cyb5r1 | 0.864446111 | 0.195364579 | 0.321 | 0.36 | 1 | 0.775393796 | 0.162605137 | 0.171 | 0.201 | 1 | 0.864446111 | 0.949552053 |
| Sbk1 | 0.807045346 | 0.247831999 | 0.113 | 0.102 | 1 | 0.776732269 | 0.515276755 | 0.049 | 0.061 | 1 | 0.807045346 | 0.95015152 |
| Gm15890 | 0.776830424 | -0.1672055 | 0.019 | 0.025 | 1 | 0.936209655 | 1.515276755 | 0.024 | 0.027 | 1 | 0.936209655 | 0.95019534 |
| Sbf1 | 0.776951782 | 0.154722595 | 0.17 | 0.153 | 1 | 0.984651805 | 0.252242349 | 0.244 | 0.259 | 1 | 0.984651805 | 0.950249492 |
| Atp1b2 | 0.924297469 | 1.280253477 | 0.075 | 0.081 | 1 | 0.778450341 | 0.564186355 | 0.098 | 0.085 | 1 | 0.924297469 | 0.950915749 |
| Pgbd1 | 0.779443586 | 0.154722595 | 0.019 | 0.025 | 1 | 0.989595616 | 0.667279848 | 0.024 | 0.024 | 1 | 0.989595616 | 0.951354868 |
| Gpnmb | 0.779455635 | -0.260314904 | 0.019 | 0.025 | 1 | 0.881057163 | 0.377773231 | 0.024 | 0.02 | 1 | 0.881057163 | 0.951360183 |
| Prkdc | 0.879385603 | -0.234319696 | 0.226 | 0.233 | 1 | 0.780235959 | 0.252242349 | 0.195 | 0.232 | 1 | 0.879385603 | 0.951703766 |
| Syne3 | 0.841941318 | 0.954785024 | 0.245 | 0.246 | 1 | 0.78045967 | 0.795864054 | 0.293 | 0.341 | 1 | 0.841941318 | 0.951802043 |
| Myoc | 0.782077996 | 0.347367673 | 0.019 | 0.025 | 1 | 0.992196803 | 0.515276755 | 0.024 | 0.024 | 1 | 0.992196803 | 0.95251 |
| 1700080G18Rik | 0.782077996 | 0.347367673 | 0.019 | 0.025 | 1 | 0.846931821 | 1.377773231 | 0.024 | 0.031 | 1 | 0.846931821 | 0.95251 |
| Ei24 | 0.78233687 | 0.208762435 | 0.377 | 0.381 | 1 | 0.92194801 | 0.327010117 | 0.39 | 0.413 | 1 | 0.92194801 | 0.952622762 |
| Acss1 | 0.926212226 | 0.8327945 | 0.019 | 0.017 | 1 | 0.783215246 | 0.313642894 | 0.073 | 0.061 | 1 | 0.926212226 | 0.95300437 |
| Gpatch4 | 0.784031888 | 0.228723176 | 0.151 | 0.169 | 1 | 0.911595387 | -0.129628286 | 0.195 | 0.184 | 1 | 0.911595387 | 0.953357775 |
| Rap2c | 0.784538777 | -0.139008608 | 0.377 | 0.39 | 1 | 0.871865685 | 0.195339958 | 0.39 | 0.375 | 1 | 0.871865685 | 0.953576461 |
| Ggta1 | 0.784984695 | 0.695290976 | 0.113 | 0.106 | 1 | 0.901656478 | 0.83720485 | 0.098 | 0.109 | 1 | 0.901656478 | 0.953768418 |
| Cxcl14 | 0.876609362 | -0.1672055 | 0.094 | 0.102 | 1 | 0.785197302 | 0.797676486 | 0.098 | 0.116 | 1 | 0.876609362 | 0.953859801 |
| Nphp1 | 0.785407239 | 0.207190015 | 0.283 | 0.275 | 1 | 0.864322047 | 0.136765132 | 0.244 | 0.253 | 1 | 0.864322047 | 0.953949947 |
| Fras1 | 0.805628541 | -0.783876861 | 0.094 | 0.102 | 1 | 0.785426384 | 0.440314697 | 0.146 | 0.133 | 1 | 0.805628541 | 0.953958163 |
| Lsm8 | 0.785641675 | 0.270915612 | 0.434 | 0.445 | 1 | 0.935012692 | 0.174239837 | 0.463 | 0.495 | 1 | 0.935012692 | 0.954050508 |
| Mns1 | 0.786396395 | -0.368839361 | 0.038 | 0.047 | 1 | 0.928870364 | 0.83720485 | 0.024 | 0.027 | 1 | 0.928870364 | 0.9543735 |
| Zfp185 | 0.826446784 | 0.221836791 | 0.151 | 0.14 | 1 | 0.788134566 | 0.727580359 | 0.195 | 0.188 | 1 | 0.826446784 | 0.955113038 |
| Ntm | 0.907425869 | 1.154722595 | 0.075 | 0.081 | 1 | 0.788255483 | 1.252242349 | 0.049 | 0.041 | 1 | 0.907425869 | 0.95516426 |
| Cnot9 | 0.86941578 | 0.267197324 | 0.264 | 0.25 | 1 | 0.788560922 | 0.264953401 | 0.244 | 0.28 | 1 | 0.86941578 | 0.955293516 |
| Zscan21 | 0.934663314 | 0.526691372 | 0.113 | 0.123 | 1 | 0.79240953 | 0.635570989 | 0.171 | 0.16 | 1 | 0.934663314 | 0.956906197 |
| Gm56494 | 0.997810947 | 1.039245377 | 0.038 | 0.038 | 1 | 0.792724891 | 0.19334866 | 0.073 | 0.061 | 1 | 0.997810947 | 0.957037029 |
| Pdgfc | 0.930476029 | 0.283709036 | 0.245 | 0.242 | 1 | 0.793095758 | 0.465756619 | 0.439 | 0.437 | 1 | 0.930476029 | 0.957190635 |
| Zfp414 | 0.83546256 | 0.154722595 | 0.17 | 0.157 | 1 | 0.79349386 | -0.19841906 | 0.098 | 0.109 | 1 | 0.83546256 | 0.957355214 |
| Chek1 | 0.79385604 | -0.260314904 | 0.132 | 0.144 | 1 | 0.904100276 | 0.204936634 | 0.195 | 0.212 | 1 | 0.904100276 | 0.957504668 |
| 4930533B01Rik | 0.794190052 | 0.347367673 | 0.038 | 0.047 | 1 | 0.992196803 | 0.515276755 | 0.024 | 0.024 | 1 | 0.992196803 | 0.957642265 |
| Inafm1 | 0.854351088 | 0.275016829 | 0.264 | 0.292 | 1 | 0.794886042 | 0.547698233 | 0.244 | 0.239 | 1 | 0.854351088 | 0.957928264 |
| Stab2 | 0.795213727 | 0.739685096 | 0.019 | 0.025 | 1 | 0.989595616 | 0.515276755 | 0.024 | 0.024 | 1 | 0.989595616 | 0.958062582 |
| Bbs5 | 0.824091747 | 0.444229212 | 0.132 | 0.123 | 1 | 0.795696851 | 0.164779508 | 0.146 | 0.13 | 1 | 0.824091747 | 0.958260223 |
| Fam110b | 0.797505304 | -0.361527156 | 0.528 | 0.487 | 1 | 0.870795434 | -0.10875531 | 0.415 | 0.413 | 1 | 0.870795434 | 0.958995898 |
| Kif24 | 0.866587402 | 0.216123139 | 0.057 | 0.051 | 1 | 0.799748057 | 1.644559772 | 0.049 | 0.041 | 1 | 0.866587402 | 0.959899159 |
| 5430427G11Rik | 0.80004514 | 0.776210972 | 0.038 | 0.047 | 1 | 0.888999736 | 0.589277336 | 0.049 | 0.044 | 1 | 0.888999736 | 0.960018054 |
| Gm45871 | 0.855418007 | 0.569760094 | 0.057 | 0.051 | 1 | 0.802906472 | -0.101394606 | 0.049 | 0.058 | 1 | 0.855418007 | 0.961154141 |
| Tnfrsf18 | 0.978023932 | 0.154722595 | 0.057 | 0.055 | 1 | 0.803001574 | 1.052933541 | 0.073 | 0.089 | 1 | 0.978023932 | 0.96119162 |
| Ivd | 0.803696851 | 0.141425772 | 0.321 | 0.305 | 1 | 0.975029725 | 0.327343804 | 0.317 | 0.331 | 1 | 0.975029725 | 0.961465074 |
| Foxred2 | 0.805362391 | 0.569760094 | 0.075 | 0.089 | 1 | 0.896350094 | 0.422167351 | 0.122 | 0.116 | 1 | 0.896350094 | 0.962116201 |
| Grhpr | 0.805507962 | 0.331600357 | 0.17 | 0.157 | 1 | 0.854007242 | 0.578470581 | 0.171 | 0.188 | 1 | 0.854007242 | 0.962172847 |
| Pskh1 | 0.970907506 | 0.187889459 | 0.151 | 0.153 | 1 | 0.8071757 | 0.164779508 | 0.122 | 0.14 | 1 | 0.970907506 | 0.962818789 |
| Atxn1l | 0.807398827 | 0.219817623 | 0.208 | 0.203 | 1 | 0.955775216 | 0.334704509 | 0.317 | 0.348 | 1 | 0.955775216 | 0.962904788 |
| Nek9 | 0.807421717 | 0.105091827 | 0.491 | 0.555 | 1 | 0.872152354 | -0.170448723 | 0.634 | 0.625 | 1 | 0.872152354 | 0.962913605 |
| Nsun3 | 0.953749455 | 0.491757582 | 0.057 | 0.059 | 1 | 0.811810001 | -0.116991461 | 0.073 | 0.061 | 1 | 0.953749455 | 0.964584524 |
| Gm20033 | 0.921038405 | 0.522454379 | 0.094 | 0.093 | 1 | 0.81243016 | 0.252242349 | 0.073 | 0.085 | 1 | 0.921038405 | 0.964817555 |
| Atp6v1g2 | 0.877748789 | 0.247831999 | 0.057 | 0.051 | 1 | 0.820790572 | 0.786578777 | 0.073 | 0.085 | 1 | 0.877748789 | 0.967883981 |
| Zcwpw1 | 0.933942263 | 0.154722595 | 0.075 | 0.072 | 1 | 0.821290924 | 0.136765132 | 0.049 | 0.058 | 1 | 0.933942263 | 0.968063066 |
| Gm13703 | 0.917883255 | 0.569760094 | 0.019 | 0.021 | 1 | 0.82181816 | 0.136765132 | 0.024 | 0.031 | 1 | 0.917883255 | 0.968251232 |
| Gm47095 | 0.929410101 | 0.569760094 | 0.019 | 0.017 | 1 | 0.82181816 | 0.136765132 | 0.024 | 0.031 | 1 | 0.929410101 | 0.968251232 |
| Gm13524 | 0.914958 | 0.347367673 | 0.019 | 0.021 | 1 | 0.824088343 | 0.136765132 | 0.024 | 0.031 | 1 | 0.914958 | 0.969055089 |
| 2310068J16Rik | 0.897146191 | 0.417757001 | 0.038 | 0.034 | 1 | 0.826384776 | 0.515276755 | 0.024 | 0.031 | 1 | 0.897146191 | 0.969857754 |
| Twf1 | 0.983011751 | 0.154722595 | 0.509 | 0.576 | 1 | 0.828472361 | 0.19880309 | 0.634 | 0.642 | 1 | 0.983011751 | 0.970578269 |
| Itga3 | 0.951630882 | -0.108311811 | 0.075 | 0.072 | 1 | 0.832579593 | 0.252242349 | 0.049 | 0.041 | 1 | 0.951630882 | 0.971970407 |
| Arhgef3 | 0.83940729 | -0.20517335 | 0.321 | 0.288 | 1 | 0.976614303 | 0.246243608 | 0.366 | 0.358 | 1 | 0.976614303 | 0.974209981 |
| Fam120aos | 0.84028294 | 0.259419973 | 0.321 | 0.326 | 1 | 0.868311713 | 0.219820871 | 0.317 | 0.352 | 1 | 0.868311713 | 0.974490461 |
| Mvk | 0.998666354 | -0.1672055 | 0.113 | 0.11 | 1 | 0.84091406 | 0.515276755 | 0.098 | 0.109 | 1 | 0.998666354 | 0.974691664 |
| 4930403P22Rik | 0.917883255 | 0.569760094 | 0.019 | 0.021 | 1 | 0.842357354 | 0.83720485 | 0.024 | 0.031 | 1 | 0.917883255 | 0.975148796 |
| Lsm4 | 0.984593797 | -0.131434369 | 0.604 | 0.585 | 1 | 0.845686588 | 0.278473891 | 0.488 | 0.505 | 1 | 0.984593797 | 0.976187371 |
| Lzic | 0.851455512 | 0.256602209 | 0.34 | 0.343 | 1 | 0.861948786 | -0.116991461 | 0.293 | 0.276 | 1 | 0.861948786 | 0.977934535 |
| Slc46a3 | 0.897146191 | 0.417757001 | 0.038 | 0.034 | 1 | 0.856404759 | 1.614812428 | 0.024 | 0.02 | 1 | 0.897146191 | 0.979380407 |
| Hr | 0.856696849 | -0.260314904 | 0.264 | 0.267 | 1 | 1 | 0.524046965 | 0.293 | 0.321 | 1 | 1 | 0.979464207 |
| Efcab14 | 0.917462451 | 0.184779829 | 0.66 | 0.758 | 1 | 0.857781452 | 0.238330042 | 0.707 | 0.802 | 1 | 0.917462451 | 0.979773885 |
| Gm56813 | 0.897146191 | 0.417757001 | 0.038 | 0.034 | 1 | 0.859139786 | 1.422167351 | 0.024 | 0.02 | 1 | 0.897146191 | 0.9801584 |
| Cntln | 0.884141336 | 0.393509454 | 0.377 | 0.415 | 1 | 0.859362231 | 0.109284395 | 0.439 | 0.454 | 1 | 0.884141336 | 0.980221018 |
| Tmem38b | 0.861433281 | 0.487297934 | 0.189 | 0.191 | 1 | 0.974004867 | -0.203822418 | 0.268 | 0.263 | 1 | 0.974004867 | 0.980799264 |
| Cd59b | 0.905123087 | 0.104096522 | 0.094 | 0.089 | 1 | 0.861892873 | -0.26713181 | 0.146 | 0.154 | 1 | 0.905123087 | 0.980926421 |
| Ric3 | 0.866605856 | 0.454282877 | 0.057 | 0.051 | 1 | 0.990690153 | 0.174239837 | 0.049 | 0.048 | 1 | 0.990690153 | 0.982206002 |
| Gm38537 | 0.86832853 | -1.214511215 | 0.038 | 0.042 | 1 | 0.911761241 | 0.377773231 | 0.024 | 0.027 | 1 | 0.911761241 | 0.982662624 |
| Lzts3 | 0.90155955 | 1.280253477 | 0.038 | 0.042 | 1 | 0.868524543 | 0.100239256 | 0.049 | 0.055 | 1 | 0.90155955 | 0.982714204 |
| B430105G09Rik | 0.872144182 | 0.347367673 | 0.057 | 0.051 | 1 | 0.94575798 | 0.334704509 | 0.049 | 0.051 | 1 | 0.94575798 | 0.98365289 |
| Pld4 | 0.912039013 | -0.545717123 | 0.019 | 0.021 | 1 | 0.872828964 | 1.029849928 | 0.024 | 0.02 | 1 | 0.912039013 | 0.983827528 |
| Serpinb6b | 0.932611893 | 0.154722595 | 0.019 | 0.017 | 1 | 0.87556513 | 0.515276755 | 0.024 | 0.02 | 1 | 0.932611893 | 0.984515963 |
| Krba1 | 0.967451273 | 0.454282877 | 0.094 | 0.097 | 1 | 0.877499685 | 0.479652845 | 0.098 | 0.109 | 1 | 0.967451273 | 0.984993673 |
| Slc16a9 | 0.877748789 | 0.247831999 | 0.057 | 0.051 | 1 | 0.987914946 | 0.667279848 | 0.073 | 0.075 | 1 | 0.987914946 | 0.985054641 |
| Gm37459 | 0.932611893 | -0.304709024 | 0.019 | 0.017 | 1 | 0.878310931 | 0.667279848 | 0.024 | 0.02 | 1 | 0.932611893 | 0.985191771 |
| Il12rb1 | 0.885801298 | 0.389187848 | 0.038 | 0.034 | 1 | 0.881057163 | 0.136765132 | 0.024 | 0.02 | 1 | 0.885801298 | 0.985852602 |
| Fstl4 | 0.881273141 | 0.569760094 | 0.038 | 0.034 | 1 | 0.981881939 | 1.26892109 | 0.073 | 0.075 | 1 | 0.981881939 | 0.985903933 |
| 1700102P08Rik | 0.943259908 | -0.285849997 | 0.113 | 0.106 | 1 | 0.888929525 | 0.334704509 | 0.171 | 0.167 | 1 | 0.943259908 | 0.98766335 |
| Fgfr4 | 0.88895204 | 0.110328475 | 0.094 | 0.102 | 1 | 0.970217098 | 0.515276755 | 0.049 | 0.048 | 1 | 0.970217098 | 0.987668351 |
| Kras | 0.888957519 | -0.223454723 | 0.906 | 0.852 | 1 | 0.945301898 | -0.258046539 | 0.976 | 0.918 | 1 | 0.945301898 | 0.987669567 |
| Anxa9 | 0.89261791 | 0.569760094 | 0.038 | 0.034 | 1 | 0.914200263 | 0.377773231 | 0.024 | 0.027 | 1 | 0.914200263 | 0.988469087 |
| Pcdhga9 | 0.89261791 | 0.569760094 | 0.038 | 0.034 | 1 | 0.958410232 | 0.377773231 | 0.049 | 0.051 | 1 | 0.958410232 | 0.988469087 |
| Pgm2 | 0.917668245 | 0.128250383 | 0.302 | 0.305 | 1 | 0.894206051 | 0.10681191 | 0.39 | 0.427 | 1 | 0.917668245 | 0.98880764 |
| Rab26os | 0.98903571 | 0.377115016 | 0.17 | 0.178 | 1 | 0.902281785 | 0.19334866 | 0.073 | 0.078 | 1 | 0.98903571 | 0.99045115 |
| Gm17749 | 0.904133521 | -0.304709024 | 0.057 | 0.059 | 1 | 0.958411801 | 0.458693227 | 0.049 | 0.051 | 1 | 0.958411801 | 0.990809618 |
| Nek3 | 0.911442241 | 0.640149422 | 0.075 | 0.072 | 1 | 0.94866804 | -0.16279515 | 0.073 | 0.075 | 1 | 0.94866804 | 0.992157523 |
| Tmem25 | 0.926212226 | 0.8327945 | 0.019 | 0.017 | 1 | 0.911761241 | 0.377773231 | 0.024 | 0.027 | 1 | 0.926212226 | 0.992213921 |
| Prdm9 | 0.929410101 | 0.347367673 | 0.019 | 0.017 | 1 | 0.911761241 | 0.252242349 | 0.024 | 0.027 | 1 | 0.929410101 | 0.992213921 |
| Epn3 | 0.913424206 | 1.417757001 | 0.019 | 0.017 | 1 | 0.916649754 | 0.667279848 | 0.024 | 0.027 | 1 | 0.916649754 | 0.992504632 |
| Gm16083 | 0.913425467 | 1.669295768 | 0.019 | 0.017 | 1 | 0.979194004 | 0.614812428 | 0.024 | 0.024 | 1 | 0.979194004 | 0.99250485 |
| Gm14634 | 0.929410101 | 0.569760094 | 0.019 | 0.017 | 1 | 0.914200263 | 0.515276755 | 0.024 | 0.027 | 1 | 0.929410101 | 0.992638405 |
| Gpatch2l | 0.968948381 | 0.154722595 | 0.245 | 0.242 | 1 | 0.914467756 | 0.458693227 | 0.293 | 0.317 | 1 | 0.968948381 | 0.992684235 |
| Gm26797 | 0.9146894 | 0.399835093 | 0.075 | 0.072 | 1 | 0.9243431 | 0.463746454 | 0.146 | 0.143 | 1 | 0.9243431 | 0.992722102 |
| Gm44386 | 0.914958 | 0.347367673 | 0.019 | 0.021 | 1 | 0.986995563 | 0.83720485 | 0.024 | 0.024 | 1 | 0.986995563 | 0.992767858 |
| Zbtb42 | 0.914958 | 0.347367673 | 0.019 | 0.021 | 1 | 0.986995563 | 0.83720485 | 0.024 | 0.024 | 1 | 0.986995563 | 0.992767858 |
| Carns1 | 0.914958 | 0.347367673 | 0.019 | 0.021 | 1 | 0.94215184 | 0.174239837 | 0.049 | 0.051 | 1 | 0.94215184 | 0.992767858 |
| Fanci | 0.916038253 | -1.713173869 | 0.075 | 0.068 | 1 | 0.941978923 | -0.885261175 | 0.073 | 0.068 | 1 | 0.941978923 | 0.992950425 |

| | | | | | | | | | | | | |
|---|---|---|---|---|---|---|---|---|---|---|---|---|
| 9130230N09Rik | 0.929410101 | 0.569760094 | 0.019 | 0.017 | 1 | 0.916649754 | 0.667279848 | 0.024 | 0.027 | 1 | 0.929410101 | 0.993052737 |
| Klhdc8a | 0.950579237 | 0.389187848 | 0.057 | 0.055 | 1 | 0.916649754 | 0.667279848 | 0.024 | 0.027 | 1 | 0.950579237 | 0.993052737 |
| Glt8d2 | 0.917325305 | 0.271236604 | 0.302 | 0.314 | 1 | 0.92798605 | -0.101394606 | 0.195 | 0.195 | 1 | 0.92798605 | 0.993164895 |
| Gm42829 | 0.917883255 | 0.569760094 | 0.019 | 0.021 | 1 | 0.919498987 | 0.100239256 | 0.049 | 0.044 | 1 | 0.919498987 | 0.99325684 |
| Coa4 | 0.917883255 | 0.569760094 | 0.019 | 0.021 | 1 | 0.989595616 | 0.667279848 | 0.024 | 0.024 | 1 | 0.989595616 | 0.99325684 |
| Shd | 0.917883255 | 0.569760094 | 0.019 | 0.021 | 1 | 0.986995563 | 0.83720485 | 0.024 | 0.024 | 1 | 0.986995563 | 0.99325684 |
| Kif15 | 0.969363387 | -1.430239906 | 0.038 | 0.038 | 1 | 0.923979186 | 0.422167351 | 0.024 | 0.027 | 1 | 0.969363387 | 0.994220836 |
| Zfp956 | 0.925829179 | 0.280253477 | 0.075 | 0.081 | 1 | 0.943323924 | 1.007129851 | 0.098 | 0.096 | 1 | 0.943323924 | 0.994498689 |
| Gm10516 | 0.9960514 | 0.324647596 | 0.113 | 0.114 | 1 | 0.925848222 | 0.477308905 | 0.171 | 0.174 | 1 | 0.9960514 | 0.994501514 |
| Gm42658 | 0.926212226 | 0.8327945 | 0.019 | 0.017 | 1 | 0.974673471 | 1.071670103 | 0.049 | 0.051 | 1 | 0.974673471 | 0.994555364 |
| Mm2pr | 0.926212226 | 0.8327945 | 0.019 | 0.017 | 1 | 0.936209655 | 1.83720485 | 0.024 | 0.027 | 1 | 0.936209655 | 0.994555364 |
| 4930557K07Rik | 0.926212226 | 0.8327945 | 0.019 | 0.017 | 1 | 0.986995563 | 0.83720485 | 0.024 | 0.024 | 1 | 0.986995563 | 0.994555364 |
| Amt | 0.926212226 | 0.8327945 | 0.019 | 0.017 | 1 | 0.986995563 | 0.83720485 | 0.024 | 0.024 | 1 | 0.986995563 | 0.994555364 |
| Gm47071 | 0.926212226 | 0.8327945 | 0.019 | 0.017 | 1 | 0.926424166 | 0.721727632 | 0.024 | 0.027 | 1 | 0.926424166 | 0.994555364 |
| Cd80 | 0.970708181 | 0.228723176 | 0.057 | 0.055 | 1 | 0.928869203 | 0.614812428 | 0.024 | 0.027 | 1 | 0.970708181 | 0.99494041 |
| Obp1a | 0.983657297 | 0.177442671 | 0.208 | 0.212 | 1 | 0.936476728 | 0.47463477 | 0.22 | 0.222 | 1 | 0.983657297 | 0.995964794 |
| Rmi1 | 0.957559435 | 0.207190015 | 0.094 | 0.097 | 1 | 0.960092216 | 0.773074512 | 0.122 | 0.133 | 1 | 0.960092216 | 0.998198798 |
| Cisd3 | 0.958692287 | 0.132354782 | 0.245 | 0.254 | 1 | 1 | 0.46797104 | 0.171 | 0.181 | 1 | 1 | 0.998293673 |
| Dcaf12 | 0.968524345 | 0.198311863 | 0.415 | 0.453 | 1 | 0.964887511 | 0.139128269 | 0.537 | 0.556 | 1 | 0.968524345 | 0.998767113 |
| Gm4924 | 0.991510144 | 0.707263618 | 0.094 | 0.097 | 1 | 0.96649646 | 0.699701326 | 0.049 | 0.048 | 1 | 0.991510144 | 0.998877513 |
| Gm13889 | 0.968379475 | -2.792809985 | 0.094 | 0.093 | 1 | 0.972303856 | -0.355440228 | 0.073 | 0.072 | 1 | 0.972303856 | 0.999000142 |
| Borcs5 | 0.998666354 | -0.323324702 | 0.113 | 0.11 | 1 | 0.968545992 | 0.483567895 | 0.122 | 0.123 | 1 | 0.998666354 | 0.999010645 |
| Nrm | 0.978112164 | -0.1672055 | 0.038 | 0.038 | 1 | 0.976480397 | 0.83720485 | 0.049 | 0.051 | 1 | 0.978112164 | 0.999446828 |
| Tirap | 0.982034447 | 0.514264981 | 0.189 | 0.195 | 1 | 0.982675819 | -0.202323514 | 0.195 | 0.184 | 1 | 0.982675819 | 0.999677239 |
| AV099323 | 1 | 0.454282877 | 0.038 | 0.038 | 1 | 0.989595616 | 0.667279848 | 0.024 | 0.024 | 1 | 1 | 0.999891749 |

**scRNA seq_Col2a1/Acan enriched cluster subset: cluster 3**

| | cKO cart_p_val | cKO cart_avg_log2FC | cKO cart_pct.1 | cKO cart_pct.2 | cKO cart_p_val_adj | WT cart_p_val | WT cart_avg_log2FC | WT cart_pct.1 | WT cart_pct.2 | WT cart_p_val_adj | max_pval | minimump_p_val |
|---|---|---|---|---|---|---|---|---|---|---|---|---|
| Ank3 | 4.77854E-10 | 2.150209812 | 0.571 | 0.093 | 1.1608E-05 | 4.51895E-24 | 2.351229022 | 0.778 | 0.121 | 1.09774E-19 | 4.77854E-10 | 9.03791E-24 |
| Pclo | 0.000757052 | 3.843696769 | 0.143 | 0.019 | 1 | 2.38576E-23 | 2.268035087 | 0.711 | 0.09 | 5.79548E-19 | 0.000757052 | 4.77152E-23 |
| Esrp1 | 0.052165518 | 2.258734268 | 0.095 | 0.022 | 1 | 8.91111E-23 | 3.057468101 | 0.667 | 0.083 | 2.16469E-18 | 0.052165518 | 1.78222E-22 |
| Sparc | 2.34756E-11 | -3.221662987 | 1 | 1 | 5.70269E-07 | 2.21615E-22 | -3.119304958 | 1 | 1 | 5.38348E-18 | 2.34756E-11 | 4.4323E-22 |
| Mecom | 9.8862E-07 | 2.214340149 | 0.714 | 0.201 | 0.024015548 | 1.37298E-21 | 2.132875504 | 0.778 | 0.145 | 3.33523E-17 | 9.8862E-07 | 2.74595E-21 |
| Serpinh1 | 2.14532E-09 | -2.048406609 | 0.762 | 0.996 | 5.2114E-05 | 1.62777E-21 | -2.58774631 | 0.778 | 0.993 | 3.95417E-17 | 2.14532E-09 | 3.25554E-21 |
| Celf2 | 0.000334491 | 1.247507013 | 0.333 | 0.082 | 1 | 2.21956E-21 | 2.997181177 | 0.8 | 0.152 | 5.39174E-17 | 0.000334491 | 4.43911E-21 |
| Cst3 | 6.35532E-11 | -2.435100297 | 0.857 | 1 | 1.54383E-06 | 3.43038E-21 | -2.459423927 | 0.978 | 0.997 | 8.33308E-17 | 6.35532E-11 | 6.86076E-21 |
| Mia | 2.19185E-10 | -3.464642261 | 0.571 | 0.955 | 5.32444E-06 | 1.38094E-20 | -3.686789986 | 0.244 | 0.945 | 3.35458E-16 | 2.19185E-10 | 2.76188E-20 |
| Fmod | 3.01011E-10 | -3.590490372 | 0.381 | 0.925 | 7.31215E-06 | 2.44539E-20 | -4.147178512 | 0.378 | 0.927 | 5.94034E-16 | 3.01011E-10 | 4.89078E-20 |
| Fmo2 | 0.63207476 | 1.351843673 | 0 | 0.011 | 1 | 5.08499E-20 | 3.361144491 | 0.511 | 0.048 | 1.23525E-15 | 0.63207476 | 1.017E-19 |
| Rps19 | 1.41373E-10 | -2.371701406 | 0.905 | 0.993 | 3.43424E-06 | 7.80342E-20 | -1.866868749 | 0.956 | 0.993 | 1.89561E-15 | 1.41373E-10 | 1.56068E-19 |
| Elf3 | 0.000766448 | 2.936806174 | 0.19 | 0.034 | 1 | 9.96285E-20 | 2.467343895 | 0.667 | 0.1 | 2.42018E-15 | 0.000766448 | 1.99257E-19 |
| Gsta3 | 0.662059486 | 1.214340149 | 0.048 | 0.03 | 1 | 1.10102E-19 | 2.479539192 | 0.689 | 0.111 | 2.67461E-15 | 0.662059486 | 2.20205E-19 |
| Rps3a1 | 1.8582E-09 | -2.084530399 | 0.952 | 0.996 | 4.51394E-05 | 1.15596E-19 | -1.95279234 | 0.956 | 1 | 2.80807E-15 | 1.8582E-09 | 2.31193E-19 |
| Itm2a | 1.95031E-10 | -3.354275954 | 0.476 | 0.97 | 4.73769E-06 | 1.15603E-19 | -3.212426111 | 0.2 | 0.948 | 2.80824E-15 | 1.95031E-10 | 2.31207E-19 |
| Casz1 | 0.263742123 | 1.258734268 | 0.095 | 0.041 | 1 | 2.04256E-19 | 2.479539192 | 0.622 | 0.087 | 4.96179E-15 | 0.263742123 | 4.08512E-19 |
| Aqp4 | 0.083583389 | 1.673771768 | 0.095 | 0.026 | 1 | 2.07508E-19 | 1.964843555 | 0.667 | 0.1 | 5.04077E-15 | 0.083583389 | 4.15015E-19 |
| Bgn | 1.15454E-10 | -2.936096075 | 0.571 | 0.97 | 2.80461E-06 | 2.19124E-19 | -2.242787246 | 0.6 | 0.997 | 5.32295E-15 | 1.15454E-10 | 4.38247E-19 |
| Meis2 | 0.000504688 | 2.903253614 | 0.286 | 0.067 | 1 | 2.43818E-19 | 1.223640968 | 0.778 | 0.142 | 5.92284E-15 | 0.000504688 | 4.87637E-19 |
| Rplp1 | 1.59653E-10 | -2.455185364 | 1 | 1 | 3.87829E-06 | 3.14273E-19 | -2.117157901 | 0.978 | 1 | 7.63432E-15 | 1.59653E-10 | 6.28546E-19 |
| Cd63 | 4.53247E-10 | -1.915745629 | 0.905 | 1 | 1.10103E-05 | 3.48277E-19 | -1.988699805 | 0.956 | 1 | 8.46034E-15 | 4.53247E-10 | 6.96553E-19 |
| Myo5b | 0.472861597 | 1.866416846 | 0.048 | 0.022 | 1 | 3.6363E-19 | 2.924080686 | 0.556 | 0.066 | 8.8333E-15 | 0.472861597 | 7.2726E-19 |
| Ucp2 | 0.022236787 | 1.195724471 | 0.286 | 0.112 | 1 | 5.11467E-19 | 0.71623945 | 0.689 | 0.111 | 1.24245E-14 | 0.022236787 | 1.02293E-18 |
| Rpl23a | 1.33906E-09 | -2.047327421 | 0.81 | 0.996 | 3.25285E-05 | 7.31406E-19 | -1.834034348 | 0.978 | 1 | 1.77673E-14 | 1.33906E-09 | 1.46281E-18 |
| Ecrg4 | 8.70128E-07 | -5.021166447 | 0.286 | 0.765 | 0.02113716 | 8.6381E-19 | -4.897136385 | 0.089 | 0.865 | 2.09837E-14 | 8.70128E-07 | 1.72762E-18 |
| Rps21 | 1.71579E-11 | -2.505189428 | 0.905 | 1 | 4.16799E-07 | 9.91123E-19 | -1.749060299 | 0.956 | 0.997 | 2.40764E-14 | 1.71579E-11 | 1.98225E-18 |
| Rps4x | 5.07314E-10 | -2.387284343 | 0.905 | 0.996 | 1.23237E-05 | 1.14032E-18 | -1.956852552 | 0.933 | 0.997 | 2.77007E-14 | 5.07314E-10 | 2.28065E-18 |
| Rpl6 | 1.8856E-10 | -2.150085352 | 0.857 | 0.993 | 4.58049E-06 | 1.35884E-18 | -1.730429314 | 0.956 | 0.997 | 3.3009E-14 | 1.8856E-10 | 2.71768E-18 |
| Ostc | 1.24714E-08 | -2.07486836 | 0.524 | 0.951 | 0.000302956 | 1.38907E-18 | -2.306542932 | 0.711 | 0.955 | 3.37432E-14 | 1.24714E-08 | 2.77814E-18 |
| Gsta4 | 2.5184E-07 | 2.536268244 | 0.476 | 0.093 | 0.006117686 | 1.76642E-18 | 1.905465008 | 0.822 | 0.194 | 4.29099E-14 | 2.5184E-07 | 3.53284E-18 |
| Nherf1 | 0.312655833 | 1.088809267 | 0.095 | 0.045 | 1 | 1.84588E-18 | 2.331600216 | 0.622 | 0.093 | 4.48402E-14 | 0.312655833 | 3.69177E-18 |
| Maged2 | 3.65178E-10 | -3.647028781 | 0.19 | 0.888 | 8.87091E-06 | 1.91488E-18 | -3.429083578 | 0.178 | 0.896 | 4.65162E-14 | 3.65178E-10 | 3.82975E-18 |
| Krt19 | 1.23719E-05 | 2.673771768 | 0.286 | 0.045 | 0.300537838 | 2.89512E-18 | 2.042614973 | 0.667 | 0.114 | 7.03284E-14 | 1.23719E-05 | 5.79025E-18 |
| Rps12 | 1.27294E-10 | -2.61354729 | 0.905 | 0.993 | 3.09223E-06 | 2.95567E-18 | -1.785155981 | 0.978 | 1 | 7.17992E-14 | 1.27294E-10 | 5.91135E-18 |
| Rpl8 | 6.99346E-10 | -2.342119605 | 0.762 | 0.989 | 1.69885E-05 | 3.29534E-18 | -1.647655232 | 1 | 1 | 8.00505E-14 | 6.99346E-10 | 6.59069E-18 |
| Snorc | 8.42421E-07 | -2.415311776 | 0.524 | 0.843 | 0.020464083 | 3.66508E-18 | -3.970523745 | 0.356 | 0.893 | 8.90321E-14 | 8.42421E-07 | 7.33016E-18 |
| Rps9 | 4.48508E-10 | -2.320680922 | 0.905 | 0.993 | 1.08952E-05 | 3.95076E-18 | -1.645974921 | 0.933 | 0.997 | 9.59719E-14 | 4.48508E-10 | 7.90153E-18 |
| Rpl14 | 5.30362E-10 | -2.320581669 | 0.857 | 0.993 | 1.28836E-05 | 3.95809E-18 | -1.676023417 | 0.956 | 1 | 9.615E-14 | 5.30362E-10 | 7.91619E-18 |
| Cnmd | 1.48223E-11 | -3.021590386 | 0.81 | 0.978 | 3.60063E-07 | 3.95957E-18 | -2.46638599 | 0.622 | 0.969 | 9.61859E-14 | 1.48223E-11 | 7.91914E-18 |
| Ikbip | 1.09092E-06 | -1.612862997 | 0.476 | 0.933 | 0.026500647 | 4.04989E-18 | -2.66806743 | 0.244 | 0.934 | 9.838E-14 | 1.09092E-06 | 8.09979E-18 |
| Rps16 | 8.668E-10 | -2.208871282 | 0.905 | 0.993 | 2.10563E-05 | 4.1183E-18 | -1.716419362 | 0.956 | 1 | 1.00042E-13 | 8.668E-10 | 8.23661E-18 |
| Rpsa | 7.54524E-10 | -2.089993886 | 0.905 | 0.993 | 1.83289E-05 | 4.41232E-18 | -1.770904602 | 0.978 | 1 | 1.07184E-13 | 7.54524E-10 | 8.82463E-18 |
| Xkr4 | 0.894071418 | -0.798716004 | 0.095 | 0.082 | 1 | 4.4487E-18 | 1.513147585 | 0.689 | 0.118 | 1.08068E-13 | 0.894071418 | 8.8974E-18 |
| Hspa8 | 5.31428E-10 | -2.299718071 | 0.857 | 0.996 | 1.29094E-05 | 4.71066E-18 | -1.867750922 | 0.978 | 0.997 | 1.14431E-13 | 5.31428E-10 | 9.42132E-18 |
| Rpl27a | 2.20541E-10 | -2.374528672 | 0.81 | 0.989 | 5.35737E-06 | 4.74831E-18 | -1.754632632 | 0.978 | 0.997 | 1.15346E-13 | 2.20541E-10 | 9.49662E-18 |
| Blnk | 0.029715341 | 1.472137907 | 0.095 | 0.019 | 1 | 5.10986E-18 | 2.846571318 | 0.556 | 0.073 | 1.24129E-13 | 0.029715341 | 1.02197E-17 |
| F11r | 2.10164E-05 | 2.866416846 | 0.286 | 0.049 | 0.510530891 | 5.1213E-18 | 2.545569062 | 0.667 | 0.114 | 1.24407E-13 | 2.10164E-05 | 1.02426E-17 |
| Arap2 | 0.01575824 | 2.286748645 | 0.19 | 0.056 | 1 | 5.26151E-18 | 2.111166238 | 0.689 | 0.121 | 1.27813E-13 | 0.01575824 | 1.0523E-17 |
| Islr | 0.000611506 | -2.188528677 | 0.286 | 0.642 | 1 | 5.37188E-18 | -3.508469467 | 0.133 | 0.882 | 1.30494E-13 | 0.000611506 | 1.07438E-17 |
| Rpl15 | 2.53583E-10 | -2.271637641 | 0.762 | 0.989 | 6.16003E-06 | 5.58573E-18 | -1.724031728 | 1 | 0.997 | 1.35689E-13 | 2.53583E-10 | 1.11715E-17 |
| Cyp2f2 | 5.03482E-09 | 0.711839809 | 0.714 | 0.172 | 0.000122306 | 6.41872E-18 | 0.520801157 | 0.889 | 0.249 | 1.55924E-13 | 5.03482E-09 | 1.28374E-17 |
| Meis1 | 0.01916162 | 2.133203386 | 0.19 | 0.06 | 1 | 6.65584E-18 | 1.975854335 | 0.689 | 0.125 | 1.61684E-13 | 0.01916162 | 1.33117E-17 |
| Ppib | 1.37658E-10 | -2.26891779 | 0.857 | 0.989 | 3.34398E-06 | 6.71684E-18 | -1.815621007 | 0.933 | 0.997 | 1.63165E-13 | 1.37658E-10 | 1.34337E-17 |
| Col9a1 | 6.2195E-09 | -3.790649463 | 0.524 | 0.892 | 0.000151084 | 6.90416E-18 | -2.51350882 | 0.756 | 0.958 | 1.67716E-13 | 6.2195E-09 | 1.38083E-17 |
| Rplp0 | 5.83801E-10 | -2.258033273 | 0.952 | 1 | 1.41817E-05 | 7.14412E-18 | -1.86579871 | 0.978 | 1 | 1.73545E-13 | 5.83801E-10 | 1.42882E-17 |
| Tpt1 | 7.82514E-11 | -2.436852027 | 1 | 1 | 1.90088E-06 | 8.45081E-18 | -1.94127278 | 1 | 1 | 2.05287E-13 | 7.82514E-11 | 1.69016E-17 |
| Rpl13a | 1.55495E-10 | -2.405077943 | 0.952 | 0.996 | 3.77729E-06 | 9.71626E-18 | -1.779020643 | 1 | 1 | 2.36027E-13 | 1.55495E-10 | 1.94325E-17 |
| Pglyrp1 | 0.000482225 | 1.55829455 | 0.429 | 0.142 | 1 | 1.06244E-17 | 2.182643596 | 0.689 | 0.128 | 2.58087E-13 | 0.000482225 | 2.12487E-17 |
| Rps2 | 4.58866E-10 | -2.350841922 | 0.905 | 0.996 | 1.11468E-05 | 1.09064E-17 | -1.677211624 | 0.978 | 1 | 2.64938E-13 | 4.58866E-10 | 2.18128E-17 |
| Col9a2 | 1.14918E-09 | -2.71335659 | 0.762 | 0.963 | 2.79158E-05 | 1.09776E-17 | -2.586674848 | 0.689 | 0.958 | 2.66668E-13 | 1.14918E-09 | 2.19552E-17 |
| Gda | 0.07959214 | 4.410737362 | 0.048 | 0.007 | 1 | 1.49705E-17 | 1.765534746 | 0.578 | 0.083 | 3.63663E-13 | 0.07959214 | 2.99409E-17 |
| Unc13b | 0.670601477 | -0.611630451 | 0.048 | 0.071 | 1 | 1.62018E-17 | 2.490427508 | 0.667 | 0.121 | 3.93574E-13 | 0.670601477 | 3.24036E-17 |
| Ucma | 8.0315E-11 | -5.322309354 | 0.238 | 0.907 | 1.95101E-06 | 1.65795E-17 | -4.468160571 | 0.2 | 0.858 | 4.02748E-13 | 8.0315E-11 | 3.31589E-17 |
| Slc25a4 | 1.18912E-11 | -2.561664802 | 0.714 | 0.993 | 2.88861E-07 | 1.76276E-17 | -1.975487247 | 0.822 | 0.976 | 4.2821E-13 | 1.18912E-11 | 3.52552E-17 |
| Serf2 | 1.18853E-10 | -2.204442237 | 0.905 | 0.993 | 2.88717E-06 | 1.87034E-17 | -1.483693458 | 0.978 | 1 | 4.54344E-13 | 1.18853E-10 | 3.74069E-17 |
| Ddx5 | 3.05843E-09 | -1.411405917 | 0.952 | 1 | 7.42954E-05 | 1.89038E-17 | -1.265973657 | 1 | 1 | 4.5921E-13 | 3.05843E-09 | 3.78075E-17 |
| Rps3 | 6.85718E-10 | -2.471070723 | 0.81 | 0.989 | 1.66575E-05 | 1.8976E-17 | -1.740405768 | 0.956 | 0.997 | 4.60964E-13 | 6.85718E-10 | 3.79519E-17 |
| Ifitm1 | 0.000438003 | 1.638147858 | 0.333 | 0.086 | 1 | 2.19401E-17 | 2.168499413 | 0.689 | 0.128 | 5.3297E-13 | 0.000438003 | 4.38803E-17 |
| Rplp2 | 2.91136E-09 | -1.964263183 | 0.857 | 0.989 | 7.07226E-05 | 2.21576E-17 | -1.739022881 | 0.978 | 0.997 | 5.38253E-13 | 2.91136E-09 | 4.43153E-17 |
| Rpl4 | 1.22725E-08 | -1.546095322 | 0.952 | 0.996 | 0.000298123 | 2.26292E-17 | -1.572523756 | 0.956 | 0.997 | 5.49709E-13 | 1.22725E-08 | 4.52585E-17 |
| Selenof | 3.00324E-10 | -1.841112297 | 0.952 | 0.989 | 7.29546E-06 | 2.6801E-17 | -1.609489424 | 0.933 | 0.993 | 6.51049E-13 | 3.00324E-10 | 5.36019E-17 |
| Npm1 | 5.92559E-07 | -1.284732936 | 0.952 | 0.985 | 0.014394431 | 2.81178E-17 | -1.61241184 | 0.978 | 0.997 | 6.83038E-13 | 5.92559E-07 | 5.62356E-17 |
| Actg1 | 2.6867E-08 | -1.986414426 | 0.952 | 1 | 0.000652653 | 2.94161E-17 | -1.830194926 | 0.956 | 1 | 7.14576E-13 | 2.6867E-08 | 5.88322E-17 |
| Rpl35a | 1.64969E-10 | -2.469544308 | 0.81 | 0.981 | 4.00742E-06 | 3.07301E-17 | -1.517431235 | 0.956 | 0.997 | 7.46495E-13 | 1.64969E-10 | 6.14601E-17 |
| Rpl41 | 2.48095E-09 | -1.805984004 | 0.952 | 0.989 | 6.02672E-05 | 3.28708E-17 | -1.54214465 | 0.956 | 0.997 | 7.98497E-13 | 2.48095E-09 | 6.57416E-17 |
| Eef1a1 | 3.38331E-11 | -2.440541517 | 1 | 1 | 8.21873E-07 | 3.35761E-17 | -1.817778263 | 0.978 | 1 | 8.1563E-13 | 3.38331E-11 | 6.71522E-17 |
| Omd | 8.00507E-09 | -3.110479481 | 0.762 | 0.978 | 0.000194459 | 4.06318E-17 | -2.90368903 | 0.6 | 0.962 | 9.87027E-13 | 8.00507E-09 | 8.12635E-17 |
| Rpl19 | 1.56391E-10 | -2.284014513 | 0.905 | 0.996 | 3.79905E-06 | 4.19905E-17 | -1.55707102 | 0.978 | 1 | 1.02003E-12 | 1.56391E-10 | 8.39809E-17 |
| Rpl7a | 2.32543E-10 | -2.395576279 | 0.857 | 0.978 | 5.64894E-06 | 4.46665E-17 | -1.557730493 | 0.956 | 0.997 | 1.08504E-12 | 2.32543E-10 | 8.9333E-17 |
| Krt23 | 0.001162117 | 1.918884266 | 0.286 | 0.075 | 1 | 5.63975E-17 | 1.724892762 | 0.689 | 0.121 | 1.37001E-12 | 0.001162117 | 1.12795E-16 |
| Ppia | 2.38071E-10 | -2.310186498 | 0.81 | 1 | 5.78323E-06 | 6.59534E-17 | -1.682669964 | 0.978 | 1 | 1.60214E-12 | 2.38071E-10 | 1.31907E-16 |
| Gcnt2 | 0.000577045 | 2.418514712 | 0.571 | 0.254 | 1 | 9.48187E-17 | 2.009882902 | 0.711 | 0.142 | 2.30334E-12 | 0.000577045 | 1.89637E-16 |
| Tmed9 | 4.18202E-11 | -1.967905124 | 0.762 | 0.996 | 1.0159E-06 | 9.5261E-17 | -1.657060104 | 0.889 | 0.993 | 2.31408E-12 | 4.18202E-11 | 1.90522E-16 |
| Vkorc1 | 1.8863E-09 | -2.653882938 | 0.333 | 0.94 | 4.5822E-05 | 1.00775E-16 | -2.344112243 | 0.467 | 0.931 | 2.44802E-12 | 1.8863E-09 | 2.0155E-16 |
| Rps27a | 5.31078E-11 | -2.524188821 | 0.905 | 0.993 | 1.29009E-06 | 1.02607E-16 | -1.639309115 | 0.933 | 1 | 2.49254E-12 | 5.31078E-11 | 2.05215E-16 |
| Atp6v0c | 6.0239E-10 | -1.825727526 | 0.905 | 0.993 | 1.46332E-05 | 1.21304E-16 | -1.791529467 | 0.978 | 0.993 | 2.94671E-12 | 6.0239E-10 | 2.42607E-16 |
| Dsp | 5.13358E-07 | 1.811275291 | 0.571 | 0.134 | 0.012470484 | 1.3528E-16 | 1.496659462 | 0.778 | 0.18 | 3.28623E-12 | 5.13358E-07 | 2.70561E-16 |
| Itm2c | 8.90437E-10 | -2.295854583 | 0.619 | 0.963 | 2.16305E-05 | 1.36767E-16 | -1.95830829 | 0.756 | 0.948 | 3.32233E-12 | 8.90437E-10 | 2.73533E-16 |
| Ppl | 0.457304706 | 2.214340149 | 0.048 | 0.022 | 1 | 1.45861E-16 | 2.846571318 | 0.533 | 0.076 | 3.54326E-12 | 0.457304706 | 2.91723E-16 |
| Cdh1 | 1.14464E-07 | 1.740885964 | 0.476 | 0.086 | 0.002780563 | 1.49876E-16 | 1.709544798 | 0.778 | 0.187 | 3.64078E-12 | 1.14464E-07 | 2.99751E-16 |
| Bsg | 1.49828E-11 | -2.350759818 | 0.762 | 0.993 | 3.63961E-07 | 1.51943E-16 | -1.682 | 0.978 | 1 | 3.69099E-12 | 1.49828E-11 | 3.03885E-16 |
| Nme2 | 2.89396E-10 | -2.499791811 | 0.81 | 0.978 | 7.03E-06 | 1.61012E-16 | -1.798777612 | 0.933 | 0.986 | 3.9113E-12 | 2.89396E-10 | 3.22024E-16 |
| Wwc1 | 0.000101027 | 2.451379346 | 0.286 | 0.056 | 1 | 1.61092E-16 | 2.1454158 | 0.667 | 0.125 | 3.91325E-12 | 0.000101027 | 3.22184E-16 |
| Rpl26 | 1.78254E-09 | -2.208098892 | 0.81 | 0.989 | 4.33014E-05 | 1.6225E-16 | -1.579293751 | 0.956 | 0.993 | 3.94137E-12 | 1.78254E-09 | 3.24499E-16 |
| Rps15 | 3.03161E-11 | -2.32755363 | 0.81 | 0.993 | 7.36439E-07 | 1.7052E-16 | -1.503606745 | 0.956 | 1 | 4.14226E-12 | 3.03161E-11 | 3.41039E-16 |

| | | | | | | | | | | | | |
|---|---|---|---|---|---|---|---|---|---|---|---|---|
| Oat | 5.27826E-10 | -2.531437126 | 0.571 | 0.959 | 1.28219E-05 | 1.74133E-16 | -1.946455467 | 0.844 | 0.965 | 4.23005E-12 | 5.27826E-10 | 3.48267E-16 |
| Rabac1 | 3.12531E-11 | -2.368799616 | 0.714 | 0.981 | 7.59201E-07 | 1.74204E-16 | -1.729525868 | 0.889 | 0.965 | 4.23175E-12 | 3.12531E-11 | 3.48407E-16 |
| Krt18 | 3.86519E-07 | 1.75177428 | 0.619 | 0.16 | 0.009389317 | 1.76609E-16 | 1.914888261 | 0.778 | 0.19 | 4.29018E-12 | 3.86519E-07 | 3.53218E-16 |
| Pfn1 | 2.7314E-06 | -1.182492755 | 0.81 | 0.963 | 0.066351244 | 1.80191E-16 | -1.6384502 | 0.956 | 0.99 | 4.37719E-12 | 2.7314E-06 | 3.60382E-16 |
| Selenos | 2.86657E-07 | -1.654507159 | 0.714 | 0.97 | 0.006963462 | 1.91253E-16 | -1.832251795 | 0.867 | 0.976 | 4.64591E-12 | 2.86657E-07 | 3.82505E-16 |
| Selenok | 3.5157E-11 | -2.272347224 | 0.857 | 0.993 | 8.54034E-07 | 2.14946E-16 | -1.661548095 | 0.933 | 0.986 | 5.22148E-12 | 3.5157E-11 | 4.29893E-16 |
| Ces1d | 0.005240437 | 0.995699863 | 0.143 | 0.026 | 1 | 2.27128E-16 | 1.765534746 | 0.667 | 0.121 | 5.5174E-12 | 0.005240437 | 4.54257E-16 |
| Rps8 | 6.8481E-10 | -2.107135619 | 0.952 | 0.996 | 1.66354E-05 | 2.50329E-16 | -1.644173829 | 0.978 | 1 | 6.08098E-12 | 6.8481E-10 | 5.00657E-16 |
| Epcam | 1.79873E-10 | 2.143257051 | 0.667 | 0.127 | 4.36946E-06 | 2.51667E-16 | 1.211320696 | 0.778 | 0.183 | 6.11349E-12 | 1.79873E-10 | 5.03333E-16 |
| Rps10 | 1.59026E-10 | -2.257767067 | 1 | 0.993 | 3.86306E-06 | 2.54209E-16 | -1.631548156 | 0.978 | 1 | 6.17524E-12 | 1.59026E-10 | 5.08418E-16 |
| Rpl30 | 3.9178E-10 | -2.136734292 | 0.81 | 0.996 | 9.51712E-06 | 2.71474E-16 | -1.572099702 | 0.978 | 0.997 | 6.59464E-12 | 3.9178E-10 | 5.42947E-16 |
| Rpl10 | 2.94435E-10 | -2.283007704 | 0.952 | 0.993 | 7.15241E-06 | 3.03508E-16 | -1.627420614 | 0.978 | 0.997 | 7.3728E-12 | 2.94435E-10 | 6.07015E-16 |
| Clec11a | 7.48074E-08 | -3.183431704 | 0.238 | 0.821 | 0.001817221 | 3.14462E-16 | -2.656777417 | 0.2 | 0.907 | 7.6389E-12 | 7.48074E-08 | 6.28923E-16 |
| Vasn | 3.54679E-08 | -2.621997167 | 0.286 | 0.888 | 0.000861586 | 3.1856E-16 | -2.121509521 | 0.378 | 0.958 | 7.73845E-12 | 3.54679E-08 | 6.37119E-16 |
| Rpl39 | 3.74595E-10 | -2.327837485 | 0.81 | 0.985 | 9.09966E-06 | 3.22488E-16 | -1.51385985 | 0.933 | 0.997 | 7.83387E-12 | 3.74595E-10 | 6.44975E-16 |
| Rps23 | 1.17538E-10 | -2.380646462 | 0.952 | 0.996 | 2.85524E-06 | 3.3415E-16 | -1.585641403 | 0.933 | 0.997 | 8.11717E-12 | 1.17538E-10 | 6.683E-16 |
| Rpl24 | 1.79517E-08 | -1.717693754 | 0.857 | 0.993 | 0.000436083 | 3.73409E-16 | -1.42896025 | 1 | 1 | 9.07086E-12 | 1.79517E-08 | 7.46818E-16 |
| Rpl10a | 2.95928E-09 | -1.971864429 | 0.905 | 0.989 | 7.18869E-05 | 3.81299E-16 | -1.464363633 | 0.978 | 0.993 | 9.26252E-12 | 2.95928E-09 | 7.62599E-16 |
| Rps24 | 1.33567E-09 | -2.07113362 | 0.952 | 0.996 | 3.24462E-05 | 4.01125E-16 | -1.592241158 | 0.978 | 1 | 9.74413E-12 | 1.33567E-09 | 8.0225E-16 |
| Rpl28 | 3.46827E-08 | -1.835593114 | 0.857 | 0.981 | 0.000842513 | 4.49052E-16 | -1.514295811 | 0.956 | 0.997 | 1.09084E-11 | 3.46827E-08 | 8.98104E-16 |
| H3f3a | 2.58762E-10 | -2.219890915 | 0.81 | 0.993 | 6.28586E-06 | 4.57474E-16 | -1.494186925 | 0.956 | 1 | 1.1113E-11 | 2.58762E-10 | 9.14949E-16 |
| Sox2 | 0.197781623 | -1.998653574 | 0.048 | 0.146 | 1 | 4.793E-16 | 2.39927962 | 0.467 | 0.055 | 1.16432E-11 | 0.197781623 | 9.586E-16 |
| Mbp | 0.787787966 | 0.901182264 | 0.095 | 0.082 | 1 | 4.96684E-16 | 1.733698659 | 0.622 | 0.107 | 1.20655E-11 | 0.787787966 | 9.93368E-16 |
| Actb | 1.9591E-09 | -2.050565052 | 0.952 | 1 | 4.75904E-05 | 5.03956E-16 | -1.729502634 | 0.978 | 1 | 1.22421E-11 | 1.9591E-09 | 1.00791E-15 |
| Rps14 | 4.99306E-10 | -2.175430592 | 0.952 | 1 | 1.21291E-05 | 5.12167E-16 | -1.567059479 | 1 | 1 | 1.24416E-11 | 4.99306E-10 | 1.02433E-15 |
| Ptms | 1.40614E-05 | -1.292158605 | 0.905 | 0.963 | 0.341579315 | 5.6928E-16 | -1.550136203 | 0.933 | 0.997 | 1.3829E-11 | 1.40614E-05 | 1.13856E-15 |
| Lcp1 | 0.872226982 | 0.614878079 | 0.048 | 0.041 | 1 | 6.21702E-16 | 0.197645759 | 0.644 | 0.118 | 1.51024E-11 | 0.872226982 | 1.2434E-15 |
| Rps6 | 1.53388E-10 | -2.330182908 | 0.857 | 0.996 | 3.72611E-06 | 7.33755E-16 | -1.451475391 | 0.978 | 1 | 1.78244E-11 | 1.53388E-10 | 1.46751E-15 |
| Dsc2 | 0.64258579 | 1.088809267 | 0.048 | 0.03 | 1 | 7.69638E-16 | 2.519573854 | 0.533 | 0.08 | 1.8696E-11 | 0.64258579 | 1.53928E-15 |
| Krt8 | 3.41551E-05 | 1.275222391 | 0.476 | 0.127 | 0.829694682 | 7.71392E-16 | 1.9745354 | 0.711 | 0.156 | 1.87387E-11 | 3.41551E-05 | 1.54278E-15 |
| Oaz1 | 1.71471E-10 | -2.197990605 | 0.905 | 0.996 | 4.16538E-06 | 7.74204E-16 | -1.58753395 | 0.956 | 0.997 | 1.8807E-11 | 1.71471E-10 | 1.54841E-15 |
| Ctsk | 3.37932E-06 | -2.229414859 | 0.333 | 0.869 | 0.082090536 | 7.94111E-16 | -4.078755635 | 0.244 | 0.927 | 1.92906E-11 | 3.37932E-06 | 1.58822E-15 |
| Rpl37a | 1.31203E-09 | -2.092669961 | 0.857 | 0.985 | 3.18718E-05 | 8.30093E-16 | -1.513324627 | 0.956 | 0.99 | 2.01646E-11 | 1.31203E-09 | 1.66019E-15 |
| Eef1b2 | 2.53246E-10 | -2.231178417 | 0.857 | 0.989 | 6.15184E-06 | 8.32581E-16 | -1.512799915 | 0.956 | 0.993 | 2.02251E-11 | 2.53246E-10 | 1.66516E-15 |
| Rpl9 | 4.90266E-10 | -2.307981632 | 0.952 | 0.989 | 1.19095E-05 | 8.45308E-16 | -1.58022283 | 1 | 1 | 2.05342E-11 | 4.90266E-10 | 1.69062E-15 |
| Ubb | 1.189E-10 | -2.610529073 | 0.952 | 0.996 | 2.88832E-06 | 8.48457E-16 | -1.73491823 | 0.978 | 1 | 2.06107E-11 | 1.189E-10 | 1.69691E-15 |
| Myl12a | 1.74012E-09 | -2.010652192 | 0.857 | 0.985 | 4.22711E-05 | 9.02772E-16 | -1.49087948 | 0.956 | 0.993 | 2.19301E-11 | 1.74012E-09 | 1.80554E-15 |
| Psap | 4.55179E-11 | -1.962322288 | 0.857 | 0.996 | 1.10572E-06 | 9.89753E-16 | -1.483204638 | 0.956 | 0.997 | 2.40431E-11 | 4.55179E-11 | 1.97951E-15 |
| Rpl32 | 6.32768E-10 | -2.36067533 | 0.81 | 0.996 | 1.53712E-05 | 1.01753E-15 | -1.536853498 | 0.956 | 0.993 | 2.47178E-11 | 6.32768E-10 | 2.03506E-15 |
| Hspa5 | 4.28552E-09 | -1.684192534 | 0.905 | 0.996 | 0.000104104 | 1.02873E-15 | -1.681684481 | 0.978 | 1 | 2.49898E-11 | 4.28552E-09 | 2.05745E-15 |
| Rpl11 | 4.75448E-11 | -2.510395156 | 0.81 | 0.996 | 1.15496E-06 | 1.05865E-15 | -1.410132332 | 0.956 | 0.997 | 2.57166E-11 | 4.75448E-11 | 2.11729E-15 |
| Rbm47 | 0.065804434 | 1.201283996 | 0.429 | 0.239 | 1 | 1.05905E-15 | 2.162240423 | 0.778 | 0.197 | 2.57265E-11 | 0.065804434 | 2.1181E-15 |
| Rps7 | 1.27091E-09 | -2.078724467 | 0.905 | 0.993 | 3.08729E-05 | 1.14525E-15 | -1.584174071 | 0.956 | 0.997 | 2.78205E-11 | 1.27091E-09 | 2.2905E-15 |
| Tceal9 | 1.10307E-07 | -2.02597418 | 0.714 | 0.944 | 0.002679574 | 1.16466E-15 | -1.882062329 | 0.867 | 0.972 | 2.82919E-11 | 1.10307E-07 | 2.32932E-15 |
| Mmp2 | 0.000175263 | -3.600282773 | 0.143 | 0.556 | 1 | 1.20638E-15 | -2.372393691 | 0.267 | 0.917 | 2.93055E-11 | 0.000175263 | 2.41277E-15 |
| Tmed10 | 5.86823E-07 | -1.168663523 | 0.905 | 1 | 0.014255112 | 1.27189E-15 | -1.472316322 | 0.956 | 0.986 | 3.08968E-11 | 5.86823E-07 | 2.54379E-15 |
| Rps5 | 3.25107E-10 | -2.410353256 | 0.81 | 0.989 | 7.89749E-06 | 1.35071E-15 | -1.52107117 | 0.933 | 0.997 | 3.28114E-11 | 3.25107E-10 | 2.70141E-15 |
| Malat1 | 3.42445E-05 | 1.018561183 | 1 | 1 | 0.83186681 | 1.35427E-15 | 1.142149093 | 1 | 1 | 3.2898E-11 | 3.42445E-05 | 2.70855E-15 |
| Sult1d1 | 0.050253997 | 2.088809267 | 0.143 | 0.045 | 1 | 1.47497E-15 | 1.704446237 | 0.667 | 0.138 | 3.583E-11 | 0.050253997 | 2.94994E-15 |
| Gabarap | 9.18183E-10 | -2.049679368 | 0.857 | 0.981 | 2.23045E-05 | 1.53555E-15 | -1.4734319 | 0.933 | 0.983 | 3.73016E-11 | 9.18183E-10 | 3.0711E-15 |
| Mrfap1 | 4.03386E-09 | -2.011650915 | 0.619 | 0.966 | 9.79905E-05 | 1.72231E-15 | -1.663377828 | 0.8 | 0.969 | 4.18385E-11 | 4.03386E-09 | 3.44463E-15 |
| Maged1 | 4.38098E-09 | -1.960434252 | 0.571 | 0.974 | 0.000106423 | 1.78929E-15 | -1.803868882 | 0.711 | 0.969 | 4.34654E-11 | 4.38098E-09 | 3.57857E-15 |
| Cldn4 | 1.66564E-06 | 3.133203386 | 0.333 | 0.052 | 0.040461637 | 1.87767E-15 | 0.976119561 | 0.667 | 0.135 | 4.56124E-11 | 1.66564E-06 | 3.75535E-15 |
| Gnai2 | 1.0863E-09 | -1.901697447 | 0.857 | 0.985 | 2.63885E-05 | 1.88077E-15 | -1.499899485 | 0.933 | 0.993 | 4.56877E-11 | 1.0863E-09 | 3.76154E-15 |
| Cd36 | 0.754598222 | -0.370622352 | 0.048 | 0.034 | 1 | 1.95814E-15 | 3.098110085 | 0.444 | 0.052 | 4.7567E-11 | 0.754598222 | 3.91627E-15 |
| Crtap | 1.50966E-12 | -4.479780264 | 0.095 | 0.94 | 3.66726E-08 | 1.95913E-15 | -2.452304189 | 0.244 | 0.889 | 4.75912E-11 | 1.50966E-12 | 3.91826E-15 |
| Rps25 | 3.48933E-10 | -2.180734248 | 0.905 | 0.993 | 8.47628E-06 | 2.04661E-15 | -1.459598806 | 0.956 | 0.99 | 4.97163E-11 | 3.48933E-10 | 4.09322E-15 |
| Tspan3 | 1.13786E-07 | -1.505523017 | 0.81 | 0.996 | 0.002764093 | 2.08377E-15 | -1.527939607 | 0.911 | 0.986 | 5.06189E-11 | 1.13786E-07 | 4.16754E-15 |
| Prelp | 2.73086E-07 | -2.097628473 | 0.571 | 0.914 | 0.006633811 | 2.33182E-15 | -2.515340972 | 0.2 | 0.879 | 5.66445E-11 | 2.73086E-07 | 4.66363E-15 |
| Spcs1 | 1.95862E-09 | -1.83445129 | 0.667 | 0.974 | 4.75788E-05 | 2.33907E-15 | -1.752556008 | 0.844 | 0.962 | 5.68207E-11 | 1.95862E-09 | 4.67814E-15 |
| Col11a1 | 3.56121E-06 | -1.378068165 | 1 | 1 | 0.086508997 | 2.35875E-15 | -1.709542024 | 0.933 | 0.997 | 5.72987E-11 | 3.56121E-06 | 4.71749E-15 |
| Hnrnpa0 | 0.000266564 | -1.033734226 | 0.714 | 0.937 | 1 | 2.36274E-15 | -1.498629583 | 0.911 | 0.979 | 5.73957E-11 | 0.000266564 | 4.72548E-15 |
| Cdcp1 | 0.532432377 | 0.673771768 | 0 | 0.019 | 1 | 2.40234E-15 | 2.905465008 | 0.444 | 0.052 | 5.83576E-11 | 0.532432377 | 4.80468E-15 |
| Rcn3 | 3.29028E-09 | -2.3838668858 | 0.286 | 0.959 | 7.99275E-05 | 2.44832E-15 | -2.19696679 | 0.289 | 0.931 | 5.94745E-11 | 3.29028E-09 | 4.89664E-15 |
| Map1lc3b | 5.88243E-09 | -2.012944232 | 0.762 | 0.985 | 0.000142896 | 2.51011E-15 | -1.537058203 | 0.956 | 1 | 6.09757E-11 | 5.88243E-09 | 5.02023E-15 |
| Tacstd2 | 0.312655833 | 0.55829455 | 0.095 | 0.045 | 1 | 2.54233E-15 | 1.416286045 | 0.6 | 0.104 | 6.17582E-11 | 0.312655833 | 5.08466E-15 |
| Kdelr2 | 1.74032E-08 | -1.667306061 | 0.81 | 0.993 | 0.000422758 | 2.554E-15 | -1.591913017 | 0.889 | 0.986 | 6.20416E-11 | 1.74032E-08 | 5.10799E-15 |
| Col2a1 | 0.000243494 | -1.741215142 | 0.905 | 0.955 | 1 | 2.77942E-15 | -2.840803823 | 0.844 | 0.945 | 6.75176E-11 | 0.000243494 | 5.55884E-15 |
| Laptm4a | 2.97317E-13 | -2.432944639 | 0.952 | 1 | 7.22243E-09 | 2.81595E-15 | -1.2804597 | 1 | 1 | 6.84051E-11 | 2.97317E-13 | 5.6319E-15 |
| Chad | 2.0352E-07 | -3.122694838 | 0.762 | 0.922 | 0.004943896 | 2.85158E-15 | -2.359003485 | 0.444 | 0.958 | 6.92706E-11 | 2.0352E-07 | 5.70316E-15 |
| Calm1 | 2.26782E-08 | -1.841186689 | 0.905 | 0.993 | 0.0005509 | 3.12226E-15 | -1.475668677 | 1 | 1 | 7.58459E-11 | 2.26782E-08 | 6.24452E-15 |
| Rpl34 | 2.54311E-10 | -2.443991813 | 0.857 | 0.989 | 6.17773E-06 | 3.15421E-15 | -1.626822953 | 0.933 | 0.993 | 7.66222E-11 | 2.54311E-10 | 6.30843E-15 |
| Col9a3 | 6.63441E-09 | -2.834727313 | 0.667 | 0.937 | 0.000161163 | 3.28345E-15 | -2.617346668 | 0.578 | 0.952 | 7.97616E-11 | 6.63441E-09 | 6.5669E-15 |
| Chchd2 | 1.70058E-09 | -2.107947227 | 0.81 | 0.989 | 4.13104E-05 | 3.47173E-15 | -1.571442587 | 0.933 | 0.986 | 8.43354E-11 | 1.70058E-09 | 6.94347E-15 |
| Ybx1 | 3.41657E-08 | -1.720784127 | 0.857 | 0.981 | 0.000829954 | 3.49161E-15 | -1.364900535 | 0.978 | 0.997 | 8.48182E-11 | 3.41657E-08 | 6.98322E-15 |
| Rpl13 | 2.92702E-10 | -2.241744665 | 0.905 | 1 | 7.11032E-06 | 3.70412E-15 | -1.574103992 | 0.978 | 1 | 8.99804E-11 | 2.92702E-10 | 7.40824E-15 |
| Cd81 | 1.679E-11 | -2.498091038 | 0.81 | 0.974 | 4.07862E-07 | 3.89365E-15 | -1.686353519 | 0.844 | 0.972 | 9.45845E-11 | 1.679E-11 | 7.7873E-15 |
| Aqp3 | 2.93776E-08 | 1.351843673 | 0.476 | 0.082 | 0.00071364 | 4.01665E-15 | 1.717288302 | 0.778 | 0.215 | 9.75726E-11 | 2.93776E-08 | 8.03331E-15 |
| Rpl27 | 1.53663E-08 | -2.006471236 | 0.857 | 0.978 | 0.000373277 | 4.02441E-15 | -1.450737505 | 0.978 | 0.99 | 9.77609E-11 | 1.53663E-08 | 8.04881E-15 |
| Rbm3 | 2.67144E-05 | -1.27289873 | 0.857 | 0.963 | 0.648945766 | 4.33256E-15 | -1.432135246 | 0.956 | 0.997 | 1.05247E-10 | 2.67144E-05 | 8.66513E-15 |
| Plaur | 0.584877441 | -0.99156415 | 0.19 | 0.231 | 1 | 4.47642E-15 | 2.045642666 | 0.578 | 0.1 | 1.08741E-10 | 0.584877441 | 8.95283E-15 |
| Cadps2 | 0.024430871 | 2.351843673 | 0.143 | 0.037 | 1 | 4.47652E-15 | 2.116031993 | 0.578 | 0.1 | 1.08744E-10 | 0.024430871 | 8.95305E-15 |
| Smoc2 | 1.05098E-05 | -3.848295942 | 0.286 | 0.743 | 0.255303427 | 4.8882E-15 | -3.56403823 | 0.356 | 0.865 | 1.18744E-10 | 1.05098E-05 | 9.77641E-15 |
| Hsp90ab1 | 6.27752E-08 | -1.593176284 | 0.952 | 1 | 0.001524936 | 4.95504E-15 | -1.49162089 | 1 | 0.997 | 1.20368E-10 | 6.27752E-08 | 9.91008E-15 |
| Ccdc3 | 0.000960363 | 2.673771768 | 0.286 | 0.071 | 1 | 5.21449E-15 | 1.315340802 | 0.667 | 0.131 | 1.2667E-10 | 0.000960363 | 1.0429E-14 |
| Ptma | 4.11351E-10 | -2.292075584 | 0.905 | 0.996 | 9.99254E-06 | 5.77925E-15 | -1.529751624 | 0.978 | 1 | 1.40389E-10 | 4.11351E-10 | 1.15585E-14 |
| Rack1 | 2.12622E-09 | -2.161063891 | 0.905 | 0.985 | 5.16502E-05 | 5.82771E-15 | -1.463721888 | 0.956 | 0.993 | 1.41567E-10 | 2.12622E-09 | 1.16554E-14 |
| Ifitm3 | 0.00181539 | -0.39661756 | 0.619 | 0.239 | 1 | 6.10221E-15 | 1.757073168 | 0.822 | 0.221 | 1.48235E-10 | 0.00181539 | 1.22044E-14 |
| Rps20 | 8.7831E-10 | -2.285334885 | 0.905 | 0.993 | 2.13359E-05 | 6.36675E-15 | -1.508280802 | 0.978 | 0.993 | 1.54661E-10 | 8.7831E-10 | 1.27335E-14 |
| Ssr4 | 1.62591E-08 | -1.883883387 | 0.857 | 0.978 | 0.000394967 | 6.39531E-15 | -1.914584987 | 0.8 | 0.955 | 1.55355E-10 | 1.62591E-08 | 1.27906E-14 |
| Hmgb1 | 3.06612E-08 | -1.911190733 | 0.857 | 0.978 | 0.000744823 | 6.43845E-15 | -1.411163484 | 0.978 | 0.997 | 1.56403E-10 | 3.06612E-08 | 1.28769E-14 |
| Mgp | 3.16462E-06 | -2.658957124 | 1 | 0.978 | 0.076874963 | 6.85965E-15 | -2.102769076 | 0.978 | 0.993 | 1.66635E-10 | 3.16462E-06 | 1.37193E-14 |
| Eif1 | 1.10122E-10 | -2.390008495 | 0.952 | 1 | 2.67508E-06 | 7.58582E-15 | -1.465551264 | 0.978 | 1 | 1.84275E-10 | 1.10122E-10 | 1.51716E-14 |
| Nrxn1 | 0.424932388 | -0.283523781 | 0.143 | 0.093 | 1 | 7.59493E-15 | 1.433094333 | 0.6 | 0.114 | 1.84496E-10 | 0.424932388 | 1.51899E-14 |
| Dcn | 0.000153629 | -2.532292107 | 0.619 | 0.843 | 1 | 7.80173E-15 | -3.419775164 | 0.756 | 0.92 | 1.8952E-10 | 0.000153629 | 1.56035E-14 |
| Dynll1 | 9.4311E-09 | -1.918329268 | 0.857 | 0.985 | 0.0002291 | 7.88022E-15 | -1.532302139 | 0.956 | 0.997 | 1.91426E-10 | 9.4311E-09 | 1.57604E-14 |
| Krt15 | 0.003522588 | 0.995699863 | 0.286 | 0.086 | 1 | 8.21912E-15 | 1.849722456 | 0.778 | 0.211 | 1.99659E-10 | 0.003522588 | 1.64382E-14 |
| Tceal8 | 1.19832E-09 | -3.015838017 | 0.238 | 0.896 | 2.91096E-05 | 8.47006E-15 | -2.087644017 | 0.622 | 0.931 | 2.05755E-10 | 1.19832E-09 | 1.69401E-14 |

| | | | | | | | | | | | | |
|---|---|---|---|---|---|---|---|---|---|---|---|---|
| Gk | 0.000182867 | 2.423228306 | 0.429 | 0.127 | 1 | 9.56096E-15 | 1.561082062 | 0.711 | 0.156 | 2.32255E-10 | 0.000182867 | 1.91219E-14 |
| Pmp22 | 8.7868E-08 | -2.102502059 | 0.381 | 0.91 | 0.00213449 | 1.01627E-14 | -2.748622646 | 0.267 | 0.862 | 2.46871E-10 | 8.7868E-08 | 2.03253E-14 |
| P4hb | 9.6499E-09 | -1.593582428 | 0.905 | 0.989 | 0.000234415 | 1.05409E-14 | -1.40332906 | 0.956 | 0.986 | 2.56058E-10 | 9.6499E-09 | 2.10817E-14 |
| Aldh1a1 | 0.267985206 | -0.849790188 | 0 | 0.056 | 1 | 1.07583E-14 | 2.572041274 | 0.511 | 0.08 | 2.61341E-10 | 0.267985206 | 2.15167E-14 |
| Rpl7 | 5.28514E-09 | -2.152198833 | 0.857 | 0.993 | 0.000128387 | 1.12413E-14 | -1.618723288 | 0.956 | 0.99 | 2.73075E-10 | 5.28514E-09 | 2.24827E-14 |
| Pdia3 | 6.13231E-11 | -2.016440058 | 0.905 | 1 | 1.48966E-06 | 1.12881E-14 | -1.363099659 | 1 | 1 | 2.74212E-10 | 6.13231E-11 | 2.25763E-14 |
| Aox3 | 0.268051241 | -1.133583154 | 0 | 0.056 | 1 | 1.22597E-14 | 2.055041364 | 0.467 | 0.062 | 2.97813E-10 | 0.268051241 | 2.45194E-14 |
| Fth1 | 6.95259E-11 | -2.708007295 | 0.905 | 1 | 1.68892E-06 | 1.23279E-14 | -1.584594554 | 1 | 1 | 2.9947E-10 | 6.95259E-11 | 2.46559E-14 |
| Rpl23 | 1.60209E-10 | -2.194593852 | 0.905 | 1 | 3.8918E-06 | 1.31003E-14 | -1.511741389 | 1 | 1 | 3.18233E-10 | 1.60209E-10 | 2.62006E-14 |
| Emp3 | 1.24491E-08 | -2.556969235 | 0.286 | 0.914 | 0.000302413 | 1.31245E-14 | -1.855465577 | 0.289 | 0.945 | 3.18822E-10 | 1.24491E-08 | 2.62491E-14 |
| Rpl12 | 1.22387E-09 | -2.045293242 | 0.857 | 0.985 | 2.97302E-05 | 1.37008E-14 | -1.475213692 | 0.956 | 0.997 | 3.32819E-10 | 1.22387E-09 | 2.74015E-14 |
| Cdkn1c | 1.46016E-06 | -3.889742314 | 0.143 | 0.731 | 0.035470137 | 1.51698E-14 | -2.718391434 | 0.333 | 0.92 | 3.68506E-10 | 1.46016E-06 | 3.03397E-14 |
| Cfl1 | 3.81204E-09 | -1.923060586 | 0.81 | 0.978 | 9.26021E-05 | 1.64879E-14 | -1.479505766 | 0.933 | 0.997 | 4.00525E-10 | 3.81204E-09 | 3.29758E-14 |
| Susd5 | 1.45939E-07 | -1.578578287 | 0.571 | 0.955 | 0.003545158 | 1.80816E-14 | -1.963550922 | 0.489 | 0.934 | 4.39238E-10 | 1.45939E-07 | 3.61632E-14 |
| Filip1l | 0.015444346 | -0.747692001 | 0.857 | 0.933 | 1 | 1.83875E-14 | -1.328812383 | 0.956 | 1 | 4.4667E-10 | 0.015444346 | 3.67751E-14 |
| Mgat2 | 7.54846E-07 | -2.977279923 | 0.095 | 0.705 | 0.018336709 | 1.87771E-14 | -2.938415791 | 0.178 | 0.81 | 4.56132E-10 | 7.54846E-07 | 3.75541E-14 |
| Myl6 | 3.29016E-09 | -1.950621615 | 0.905 | 0.985 | 7.99245E-05 | 2.10337E-14 | -1.419435879 | 0.978 | 0.997 | 5.1095E-10 | 3.29016E-09 | 4.20674E-14 |
| Pebp1 | 8.25393E-10 | -2.280424543 | 0.571 | 0.955 | 2.00504E-05 | 2.12765E-14 | -1.665482011 | 0.822 | 0.924 | 5.1685E-10 | 8.25393E-10 | 4.25531E-14 |
| Tmed3 | 1.6816E-09 | -3.168129488 | 0.19 | 0.862 | 4.08494E-05 | 2.30375E-14 | -2.778406861 | 0.244 | 0.824 | 5.59628E-10 | 1.6816E-09 | 4.60751E-14 |
| Six3 | 0.026779265 | 3.088809267 | 0.095 | 0.019 | 1 | 2.75572E-14 | 3.123645178 | 0.4 | 0.045 | 6.69419E-10 | 0.026779265 | 5.51144E-14 |
| Iqgap2 | 0.104672229 | 0.526930379 | 0.286 | 0.138 | 1 | 2.7741E-14 | 1.586211047 | 0.689 | 0.152 | 6.73883E-10 | 0.104672229 | 5.54819E-14 |
| Morf4l2 | 1.9159E-05 | -0.934042475 | 0.952 | 0.993 | 0.465410777 | 2.79033E-14 | -1.190981427 | 0.978 | 0.997 | 6.77827E-10 | 1.9159E-05 | 5.58066E-14 |
| Rcn1 | 2.0186E-07 | -2.03588648 | 0.381 | 0.903 | 0.004903589 | 2.85462E-14 | -2.592017258 | 0.133 | 0.837 | 6.93445E-10 | 2.0186E-07 | 5.70924E-14 |
| Rpl21 | 2.75398E-10 | -2.333846319 | 0.905 | 0.993 | 6.68998E-06 | 2.94855E-14 | -1.384690403 | 0.956 | 0.997 | 7.16262E-10 | 2.75398E-10 | 5.8971E-14 |
| Ckap4 | 2.06165E-09 | -1.820545369 | 0.762 | 0.989 | 5.00816E-05 | 3.13403E-14 | -1.606568404 | 0.889 | 0.979 | 7.61318E-10 | 2.06165E-09 | 6.26805E-14 |
| Eif4a1 | 0.000582868 | -0.772807249 | 1 | 0.981 | 1 | 3.29724E-14 | -1.301160098 | 1 | 0.993 | 8.00965E-10 | 0.000582868 | 6.59448E-14 |
| Rpl5 | 7.7753E-10 | -2.041041405 | 0.952 | 0.989 | 1.88878E-05 | 3.63842E-14 | -1.422989933 | 0.978 | 0.997 | 8.83845E-10 | 7.7753E-10 | 7.27684E-14 |
| Atp1b1 | 5.45905E-08 | 1.807038299 | 0.619 | 0.142 | 0.001326112 | 3.79891E-14 | 0.184434885 | 0.8 | 0.239 | 9.22832E-10 | 5.45905E-08 | 7.59783E-14 |
| Grhl2 | 0.07959214 | 4.410737362 | 0.048 | 0.007 | 1 | 4.03938E-14 | 2.761075098 | 0.4 | 0.045 | 9.81247E-10 | 0.07959214 | 8.07877E-14 |
| Ftl1 | 2.45305E-08 | -1.791323656 | 0.81 | 1 | 0.000595895 | 4.18892E-14 | -1.373342708 | 1 | 1 | 1.01757E-09 | 2.45305E-08 | 8.37783E-14 |
| Rpl18a | 1.56514E-08 | -1.847765353 | 0.81 | 0.978 | 0.000380204 | 4.48186E-14 | -1.263685393 | 0.933 | 0.997 | 1.08873E-09 | 1.56514E-08 | 8.96372E-14 |
| Tmem59 | 3.95681E-11 | -2.019134278 | 0.857 | 0.993 | 9.61189E-07 | 4.79937E-14 | -1.296102264 | 0.956 | 0.986 | 1.16586E-09 | 3.95681E-11 | 9.59873E-14 |
| Itm2b | 1.30759E-12 | -2.735911775 | 0.952 | 1 | 3.1764E-08 | 4.99407E-14 | -1.401250168 | 0.978 | 1 | 1.21316E-09 | 1.30759E-12 | 9.98815E-14 |
| Fkbp7 | 1.59271E-07 | -2.695462042 | 0.19 | 0.825 | 0.003869009 | 5.37082E-14 | -2.56285535 | 0.178 | 0.83 | 1.30468E-09 | 1.59271E-07 | 1.07416E-13 |
| Rpl3 | 6.66972E-10 | -2.202579247 | 0.905 | 0.985 | 1.62021E-05 | 5.43258E-14 | -1.307220103 | 0.978 | 0.997 | 1.31968E-09 | 6.66972E-10 | 1.08652E-13 |
| Rpl17 | 1.09583E-09 | -2.150953366 | 0.905 | 0.996 | 2.66199E-05 | 5.48417E-14 | -1.389005092 | 1 | 1 | 1.33221E-09 | 1.09583E-09 | 1.09683E-13 |
| Rps18 | 2.18092E-09 | -2.017921908 | 0.81 | 0.996 | 5.29789E-05 | 5.632E-14 | -1.3532028 | 0.978 | 0.993 | 1.36813E-09 | 2.18092E-09 | 1.1264E-13 |
| Ifitm2 | 9.74194E-10 | -2.816156452 | 0.619 | 0.955 | 2.36651E-05 | 5.73108E-14 | -1.750151317 | 0.933 | 0.962 | 1.39219E-09 | 9.74194E-10 | 1.14622E-13 |
| Pofut2 | 0.000538126 | -1.236121316 | 0.381 | 0.776 | 1 | 5.95112E-14 | -2.372209849 | 0.133 | 0.824 | 1.44564E-09 | 0.000538126 | 1.19022E-13 |
| Rps13 | 7.12722E-09 | -1.926471551 | 0.857 | 0.981 | 0.000173134 | 6.2299E-14 | -1.351552201 | 0.978 | 0.986 | 1.51337E-09 | 7.12722E-09 | 1.24598E-13 |
| mt-Cytb | 0.00240725 | 1.085978294 | 1 | 1 | 1 | 6.23819E-14 | 1.605326121 | 1 | 1 | 1.51538E-09 | 0.00240725 | 1.24764E-13 |
| Shroom3 | 0.484228121 | 1.38426515 | 0.143 | 0.101 | 1 | 6.38174E-14 | 2.005000681 | 0.556 | 0.1 | 1.55025E-09 | 0.484228121 | 1.27635E-13 |
| Rps29 | 2.50903E-09 | -1.911190733 | 0.857 | 0.993 | 6.09495E-05 | 6.38752E-14 | -1.268342624 | 0.978 | 0.993 | 1.55166E-09 | 2.50903E-09 | 1.2775E-13 |
| mt-Nd1 | 0.007052929 | 1.087164099 | 1 | 1 | 1 | 6.43465E-14 | 1.520957531 | 1 | 1 | 1.5631E-09 | 0.007052929 | 1.28693E-13 |
| Fkbp9 | 5.86212E-09 | -1.847436049 | 0.524 | 0.966 | 0.000142403 | 6.76564E-14 | -1.596211171 | 0.556 | 0.972 | 1.64351E-09 | 5.86212E-09 | 1.35313E-13 |
| Eif4g2 | 3.78941E-07 | -1.370061755 | 0.905 | 0.985 | 0.009205245 | 8.17208E-14 | -1.19298733 | 0.978 | 1 | 1.98516E-09 | 3.78941E-07 | 1.63442E-13 |
| Lamp1 | 2.99417E-10 | -1.768599798 | 0.857 | 1 | 7.27345E-06 | 8.70738E-14 | -1.208522349 | 0.956 | 0.997 | 2.1152E-09 | 2.99417E-10 | 1.74148E-13 |
| Ccdc80 | 5.32957E-08 | -1.864602133 | 0.857 | 0.989 | 0.00129466 | 8.782E-14 | -1.449917405 | 0.733 | 0.993 | 2.13332E-09 | 5.32957E-08 | 1.7564E-13 |
| Hsp90b1 | 1.04957E-10 | -1.867705999 | 0.857 | 1 | 2.54963E-06 | 9.46647E-14 | -1.398549849 | 0.978 | 0.997 | 2.2996E-09 | 1.04957E-10 | 1.89329E-13 |
| Sub1 | 5.68517E-08 | -1.668399547 | 0.762 | 0.974 | 0.00138104 | 1.18076E-13 | -1.3016929 | 0.933 | 0.986 | 2.86829E-09 | 5.68517E-08 | 2.36151E-13 |
| Fbxo32 | 0.127921069 | 0.320134813 | 0.286 | 0.146 | 1 | 1.21234E-13 | 1.277557861 | 0.711 | 0.17 | 2.94501E-09 | 0.127921069 | 2.42467E-13 |
| Pdia6 | 2.98858E-09 | -1.971780713 | 0.81 | 0.989 | 7.25986E-05 | 1.46954E-13 | -1.539319835 | 0.822 | 0.979 | 3.56981E-09 | 2.98858E-09 | 2.93908E-13 |
| Fcgrt | 2.25017E-11 | -3.689632963 | 0.143 | 0.925 | 5.46611E-07 | 1.54309E-13 | -1.949195629 | 0.267 | 0.92 | 3.74848E-09 | 2.25017E-11 | 3.08619E-13 |
| Plac9 | 0.003145312 | -2.411992786 | 0.286 | 0.608 | 1 | 1.59677E-13 | -2.912135811 | 0.222 | 0.827 | 3.87887E-09 | 0.003145312 | 3.19354E-13 |
| Calr | 9.55081E-09 | -1.720803748 | 0.81 | 0.978 | 0.000232008 | 1.60253E-13 | -1.386777517 | 1 | 0.993 | 3.89287E-09 | 9.55081E-09 | 3.20507E-13 |
| Rps26 | 6.57873E-10 | -2.204832238 | 0.81 | 0.981 | 1.59811E-05 | 1.6889E-13 | -1.335348611 | 0.978 | 0.986 | 4.10267E-09 | 6.57873E-10 | 3.3778E-13 |
| Surf4 | 0.000141522 | -1.300961474 | 0.524 | 0.873 | 1 | 1.74331E-13 | -1.834902935 | 0.689 | 0.9 | 4.23485E-09 | 0.000141522 | 3.48662E-13 |
| Cthrc1 | 3.44316E-07 | -3.243207646 | 0.143 | 0.772 | 0.008364134 | 1.89704E-13 | -2.336602375 | 0.267 | 0.882 | 4.60829E-09 | 3.44316E-07 | 3.79408E-13 |
| Ddost | 2.65538E-08 | -1.855751747 | 0.619 | 0.955 | 0.000645045 | 2.05965E-13 | -1.606259613 | 0.8 | 0.92 | 5.00329E-09 | 2.65538E-08 | 4.11929E-13 |
| Fkbp10 | 2.33504E-08 | -2.216492509 | 0.238 | 0.91 | 0.000567228 | 2.34478E-13 | -2.044847868 | 0.289 | 0.893 | 5.69594E-09 | 2.33504E-08 | 4.68956E-13 |
| Sec61b | 2.0043E-08 | -2.138122314 | 0.667 | 0.955 | 0.000486886 | 2.60904E-13 | -1.565769472 | 0.956 | 0.948 | 6.33787E-09 | 2.0043E-08 | 5.21807E-13 |
| Dapl1 | 0.213671712 | 0.79930265 | 0.095 | 0.037 | 1 | 2.6867E-13 | 1.875717664 | 0.511 | 0.087 | 6.52654E-09 | 0.213671712 | 5.37341E-13 |
| Rps11 | 3.28657E-10 | -2.221939029 | 0.81 | 0.993 | 7.98374E-06 | 2.96528E-13 | -1.295774973 | 0.956 | 0.997 | 7.20327E-09 | 3.28657E-10 | 5.93057E-13 |
| Krt5 | 0.109251422 | 2.97333205 | 0.095 | 0.03 | 1 | 3.21782E-13 | 1.03099589 | 0.667 | 0.159 | 7.81673E-09 | 0.109251422 | 6.43564E-13 |
| Dnaja1 | 4.42076E-09 | -2.129898108 | 0.762 | 0.978 | 0.000107389 | 3.39478E-13 | -1.557353174 | 1 | 0.986 | 8.24659E-09 | 4.42076E-09 | 6.78955E-13 |
| Krt7 | 0.532432377 | 0.673771768 | 0 | 0.019 | 1 | 3.42349E-13 | 1.841770332 | 0.511 | 0.087 | 8.31634E-09 | 0.532432377 | 6.84698E-13 |
| Igfbp7 | 0.001416983 | -2.042507864 | 0.571 | 0.765 | 1 | 3.58241E-13 | -1.67365318 | 0.822 | 0.972 | 8.70239E-09 | 0.001416983 | 7.16482E-13 |
| mt-Co3 | 0.005649634 | 0.934196873 | 1 | 1 | 1 | 3.62769E-13 | 1.43170306 | 1 | 1 | 8.81237E-09 | 0.005649634 | 7.25537E-13 |
| Gas5 | 3.09983E-05 | -1.0570416 | 0.905 | 0.985 | 0.753009549 | 3.66409E-13 | -1.339295227 | 0.933 | 0.986 | 8.9008E-09 | 3.09983E-05 | 7.32817E-13 |
| Sar1a | 6.30585E-07 | -1.605928092 | 0.524 | 0.922 | 0.015318175 | 3.74061E-13 | -1.499392712 | 0.778 | 0.962 | 9.0867E-09 | 6.30585E-07 | 7.48123E-13 |
| Norad | 1.76765E-05 | -1.475975352 | 0.524 | 0.884 | 0.429397809 | 4.15018E-13 | -1.528672591 | 0.756 | 0.955 | 1.00816E-08 | 1.76765E-05 | 8.30036E-13 |
| Rpl37 | 1.72906E-08 | -1.92814956 | 0.905 | 0.978 | 0.000420024 | 4.28496E-13 | -1.116462831 | 0.956 | 0.997 | 1.0409E-08 | 1.72906E-08 | 8.56993E-13 |
| Rps17 | 2.35897E-09 | -2.072770086 | 0.857 | 0.978 | 5.7304E-05 | 4.40183E-13 | -1.365346653 | 0.933 | 0.983 | 1.06929E-08 | 2.35897E-09 | 8.80366E-13 |
| Neat1 | 1.10202E-08 | 2.310706767 | 1 | 0.985 | 0.000267703 | 4.47313E-13 | 1.777514633 | 1 | 0.997 | 1.08661E-08 | 1.10202E-08 | 8.94626E-13 |
| Btf3 | 2.82963E-08 | -1.657454072 | 0.857 | 0.974 | 0.000687375 | 4.6328E-13 | -1.159033062 | 0.956 | 0.983 | 1.1254E-08 | 2.82963E-08 | 9.26561E-13 |
| Fgfrl1 | 1.08502E-07 | -2.163577895 | 0.238 | 0.907 | 0.002635732 | 4.96997E-13 | -2.188661564 | 0.244 | 0.872 | 1.2073E-08 | 1.08502E-07 | 9.93994E-13 |
| Prdx4 | 2.70009E-05 | -2.168578576 | 0.143 | 0.69 | 0.655905572 | 5.1821E-13 | -2.642130641 | 0.089 | 0.72 | 1.25884E-08 | 2.70009E-05 | 1.03642E-12 |
| Tsc22d1 | 1.25066E-07 | -1.888243907 | 0.952 | 0.996 | 0.003038095 | 5.47945E-13 | -1.623788061 | 1 | 0.997 | 1.33107E-08 | 1.25066E-07 | 1.09589E-12 |
| Snhg18 | 1.25421E-07 | -2.359651234 | 0.238 | 0.854 | 0.003046722 | 5.66212E-13 | -2.579544313 | 0.178 | 0.799 | 1.37544E-08 | 1.25421E-07 | 1.13242E-12 |
| Cnpy2 | 8.87409E-08 | -1.842534372 | 0.476 | 0.948 | 0.002155693 | 5.696E-13 | -1.665960739 | 0.689 | 0.927 | 1.38367E-08 | 8.87409E-08 | 1.1392E-12 |
| Cnbp | 0.003595464 | -0.638055737 | 0.905 | 0.985 | 1 | 5.72093E-13 | -0.975852925 | 0.978 | 0.997 | 1.38973E-08 | 0.003595464 | 1.14419E-12 |
| Adh7 | 0.01337844 | 0.614878079 | 0.238 | 0.075 | 1 | 5.77594E-13 | 0.754965505 | 0.667 | 0.163 | 1.40309E-08 | 0.01337844 | 1.15519E-12 |
| Chadl | 1.4722E-08 | -2.955584852 | 0.333 | 0.866 | 0.000357626 | 5.79677E-13 | -3.103523776 | 0.2 | 0.779 | 1.40815E-08 | 1.4722E-08 | 1.15935E-12 |
| mt-Nd4 | 0.001559139 | 1.075050685 | 1 | 1 | 1 | 5.98931E-13 | 1.589782856 | 1 | 1 | 1.45492E-08 | 0.001559139 | 1.19786E-12 |
| Wdr72 | 0.27197965 | 1.150209812 | 0.048 | 0.015 | 1 | 6.23142E-13 | 1.55062229 | 0.467 | 0.076 | 1.51374E-08 | 0.27197965 | 1.24628E-12 |
| Cox4i1 | 2.71119E-09 | -1.925810522 | 0.762 | 0.985 | 6.58603E-05 | 6.45542E-13 | -1.218589495 | 0.956 | 0.993 | 1.56815E-08 | 2.71119E-09 | 1.29108E-12 |
| Ppic | 8.68848E-11 | -2.5554031 | 0.619 | 0.989 | 2.11061E-06 | 6.71254E-13 | -1.432183546 | 0.533 | 0.983 | 1.63061E-08 | 8.68848E-11 | 1.34251E-12 |
| Spint2 | 0.000467595 | 0.683294542 | 0.714 | 0.28 | 1 | 6.98621E-13 | 1.784001495 | 0.8 | 0.266 | 1.69709E-08 | 0.000467595 | 1.39724E-12 |
| Bcl11a | 0.346873332 | -1.02666795 | 0 | 0.041 | 1 | 7.19223E-13 | 1.997181177 | 0.489 | 0.083 | 1.74714E-08 | 0.346873332 | 1.43845E-12 |
| Ubc | 9.80495E-08 | -2.050680073 | 0.905 | 0.993 | 0.002381819 | 7.95705E-13 | -1.430020202 | 0.956 | 0.997 | 1.93293E-08 | 9.80495E-08 | 1.59141E-12 |
| Calu | 7.09944E-05 | -0.679244005 | 0.952 | 0.996 | 1 | 8.3592E-13 | -1.141782241 | 0.911 | 1 | 2.03062E-08 | 7.09944E-05 | 1.67184E-12 |
| Jund | 6.11884E-07 | -1.70242599 | 0.857 | 0.996 | 0.014863885 | 9.02643E-13 | -1.343933293 | 0.956 | 1 | 2.1927E-08 | 6.11884E-07 | 1.80529E-12 |
| mt-Co2 | 0.00154488 | 1.031863049 | 1 | 1 | 1 | 9.62999E-13 | 1.492072214 | 1 | 1 | 2.33932E-08 | 0.00154488 | 1.926E-12 |
| Tmem30a | 5.10809E-09 | -1.725600413 | 0.667 | 0.981 | 0.000124086 | 1.00155E-12 | -1.339748121 | 0.867 | 0.976 | 2.43295E-08 | 5.10809E-09 | 2.00309E-12 |
| Atf4 | 1.34517E-05 | -1.28523086 | 0.952 | 0.978 | 0.326767783 | 1.01213E-12 | -1.3357396 | 0.933 | 0.997 | 2.45866E-08 | 1.34517E-05 | 2.02425E-12 |
| Chchd10 | 0.036753894 | 1.825774861 | 0.143 | 0.041 | 1 | 1.03485E-12 | 1.371870898 | 0.511 | 0.093 | 2.51386E-08 | 0.036753894 | 2.0697E-12 |
| Naca | 1.48634E-08 | -1.670389617 | 1 | 0.978 | 0.000361062 | 1.08052E-12 | -1.128734339 | 0.956 | 0.99 | 2.6248E-08 | 1.48634E-08 | 2.16104E-12 |
| Ctsf | 9.58007E-06 | -2.661618587 | 0.095 | 0.638 | 0.232719113 | 1.09829E-12 | -3.404390255 | 0.089 | 0.696 | 2.66798E-08 | 9.58007E-06 | 2.19659E-12 |

| | | | | | | | | | | | | |
|---|---|---|---|---|---|---|---|---|---|---|---|---|
| Fxyd1 | 1.1009E-05 | -2.962852853 | 0.143 | 0.657 | 0.267431234 | 1.30495E-12 | -2.093757469 | 0.178 | 0.844 | 3.16998E-08 | 1.1009E-05 | 2.6099E-12 |
| Trp63 | 0.815669668 | 0.79930265 | 0.048 | 0.037 | 1 | 1.36358E-12 | 2.761075098 | 0.511 | 0.097 | 3.31242E-08 | 0.815669668 | 2.72717E-12 |
| Krt14 | 2.07399E-05 | 0.866416846 | 0.333 | 0.063 | 0.503812964 | 1.3819E-12 | 2.0259603 | 0.556 | 0.118 | 3.35691E-08 | 2.07399E-05 | 2.7638E-12 |
| Rps15a | 1.12535E-08 | -2.136597223 | 0.905 | 0.978 | 0.000273371 | 1.39905E-12 | -1.251754886 | 0.956 | 0.99 | 3.39856E-08 | 1.12535E-08 | 2.79809E-12 |
| Srsf7 | 0.028313054 | -0.562894785 | 0.857 | 0.944 | 1 | 1.43086E-12 | -1.443936679 | 0.911 | 0.976 | 3.47584E-08 | 0.028313054 | 2.86171E-12 |
| Tmed2 | 3.31297E-06 | -1.161125614 | 0.857 | 0.978 | 0.080478557 | 1.48362E-12 | -1.139194045 | 0.978 | 0.997 | 3.60402E-08 | 3.31297E-06 | 2.96725E-12 |
| Sumo2 | 1.00715E-06 | -1.360091684 | 0.81 | 0.966 | 0.024465729 | 1.5371E-12 | -1.216017722 | 0.933 | 0.986 | 3.73393E-08 | 1.00715E-06 | 3.0742E-12 |
| Gapdh | 2.69205E-09 | -2.061038942 | 0.905 | 0.985 | 6.53954E-05 | 1.55025E-12 | -1.148394792 | 0.956 | 0.997 | 3.76587E-08 | 2.69205E-09 | 3.1005E-12 |
| Slc16a11 | 0.45422894 | 2.79930265 | 0.048 | 0.022 | 1 | 1.58212E-12 | 2.28452321 | 0.467 | 0.08 | 3.8433E-08 | 0.45422894 | 3.16425E-12 |
| Osbpl3 | 0.378300217 | 1.789248985 | 0.143 | 0.09 | 1 | 1.5995E-12 | 1.953720176 | 0.622 | 0.149 | 3.8855E-08 | 0.378300217 | 3.19899E-12 |
| Cggbp1 | 3.53493E-05 | -1.415922621 | 0.524 | 0.862 | 0.858705623 | 1.73748E-12 | -1.413788953 | 0.822 | 0.952 | 4.22068E-08 | 3.53493E-05 | 3.47495E-12 |
| Col1a2 | 0.000290442 | -3.530431228 | 0.429 | 0.802 | 1 | 1.75978E-12 | -1.910668528 | 0.667 | 0.979 | 4.27487E-08 | 0.000290442 | 3.51957E-12 |
| Tspan13 | 1.10419E-10 | -3.548262798 | 0.619 | 0.959 | 2.68229E-06 | 1.87141E-12 | -1.765437299 | 0.933 | 0.952 | 4.54603E-08 | 1.10419E-10 | 3.74282E-12 |
| Aqp5 | 0.662059486 | 0.97333205 | 0.048 | 0.03 | 1 | 1.9089E-12 | 2.612683258 | 0.533 | 0.107 | 4.6371E-08 | 0.662059486 | 3.8178E-12 |
| Fau | 1.74958E-10 | -2.348108154 | 0.905 | 0.993 | 4.25009E-06 | 1.92475E-12 | -1.270825369 | 0.978 | 1 | 4.67559E-08 | 1.74958E-10 | 3.84949E-12 |
| mt-Co1 | 0.001104586 | 0.946344118 | 1 | 1 | 1 | 1.97858E-12 | 1.449837515 | 1 | 1 | 4.80638E-08 | 0.001104586 | 3.95717E-12 |
| Aga | 1.10384E-08 | -3.749694354 | 0.095 | 0.776 | 0.000268145 | 2.05968E-12 | -2.60848186 | 0.111 | 0.734 | 5.00337E-08 | 1.10384E-08 | 4.11935E-12 |
| Dipk2a | 1.08654E-05 | -1.531777143 | 0.429 | 0.933 | 0.263942979 | 2.1068E-12 | -1.63193911 | 0.6 | 0.955 | 5.11783E-08 | 1.08654E-05 | 4.21359E-12 |
| Tubb2a | 5.31143E-09 | -3.509317693 | 0.619 | 0.929 | 0.000129025 | 2.16772E-12 | -2.090700633 | 0.711 | 0.927 | 5.26581E-08 | 5.31143E-09 | 4.33543E-12 |
| mt-Atp6 | 0.006611028 | 0.921576421 | 1 | 1 | 1 | 2.40267E-12 | 1.41910614 | 1 | 1 | 5.83657E-08 | 0.006611028 | 4.80535E-12 |
| Bcam | 0.277253676 | 1.673771768 | 0.048 | 0.015 | 1 | 2.53594E-12 | 2.863644832 | 0.356 | 0.042 | 6.1603E-08 | 0.277253676 | 5.07187E-12 |
| Thra | 1.62308E-06 | -1.472276132 | 0.286 | 0.881 | 0.039427971 | 2.62308E-12 | -1.545125457 | 0.622 | 0.941 | 6.372E-08 | 1.62308E-06 | 5.24617E-12 |
| Lypd2 | 0.025610415 | 0.840881754 | 0.286 | 0.116 | 1 | 2.63634E-12 | 1.616983396 | 0.689 | 0.183 | 6.40419E-08 | 0.025610415 | 5.27267E-12 |
| Sfr1 | 2.80041E-07 | -1.518446313 | 0.857 | 0.966 | 0.006802749 | 2.71901E-12 | -1.395369858 | 0.978 | 0.972 | 6.60503E-08 | 2.80041E-07 | 5.43803E-12 |
| Gng5 | 1.8422E-10 | -2.149733933 | 0.81 | 0.985 | 4.47508E-06 | 2.767E-12 | -1.305383362 | 0.933 | 0.979 | 6.7216E-08 | 1.8422E-10 | 5.53401E-12 |
| Auts2 | 3.14486E-07 | 1.74984635 | 0.952 | 1 | 0.007639501 | 2.79201E-12 | 1.323991493 | 1 | 1 | 6.78235E-08 | 3.14486E-07 | 5.58402E-12 |
| Rpl22 | 6.05313E-10 | -2.096447388 | 0.81 | 0.985 | 1.47043E-05 | 2.81806E-12 | -1.140749547 | 0.933 | 0.986 | 6.84564E-08 | 6.05313E-10 | 5.63612E-12 |
| Epb41l5 | 0.703449108 | 0.79930265 | 0.143 | 0.116 | 1 | 2.93107E-12 | 2.018256778 | 0.6 | 0.135 | 7.12016E-08 | 0.703449108 | 5.86215E-12 |
| Rab25 | 0.4759797 | 1.673771768 | 0.048 | 0.022 | 1 | 3.32441E-12 | 2.612683258 | 0.422 | 0.066 | 8.07566E-08 | 0.4759797 | 6.64882E-12 |
| Gpx8 | 3.51315E-10 | -2.854607205 | 0.19 | 0.933 | 8.53415E-06 | 3.33521E-12 | -1.874182985 | 0.444 | 0.886 | 8.1019E-08 | 3.51315E-10 | 6.67042E-12 |
| Tgoln1 | 1.5954E-06 | -1.266285692 | 0.714 | 0.978 | 0.038755507 | 3.79531E-12 | -1.337363908 | 0.889 | 0.969 | 9.21957E-08 | 1.5954E-06 | 7.59062E-12 |
| S100a14 | 0.454205211 | 2.451379346 | 0.048 | 0.022 | 1 | 3.83999E-12 | 2.013221188 | 0.467 | 0.08 | 9.32809E-08 | 0.454205211 | 7.67997E-12 |
| Ehf | 6.42258E-06 | 1.754390228 | 0.571 | 0.16 | 0.156017233 | 3.96532E-12 | 1.393565969 | 0.711 | 0.183 | 9.63255E-08 | 6.42258E-06 | 7.93063E-12 |
| Hmgn1 | 6.95067E-06 | -1.190263349 | 0.714 | 0.978 | 0.168845582 | 3.98596E-12 | -1.230591238 | 0.911 | 0.948 | 9.6827E-08 | 6.95067E-06 | 7.97193E-12 |
| Col4a2 | 0.000106809 | -1.966874866 | 0.095 | 0.604 | 1 | 4.1456E-12 | -3.104829973 | 0.067 | 0.689 | 1.00705E-07 | 0.000106809 | 8.29121E-12 |
| Selenom | 7.00567E-12 | -4.519753593 | 0.095 | 0.918 | 1.70182E-07 | 4.25471E-12 | -1.84048937 | 0.244 | 0.875 | 1.03356E-07 | 7.00567E-12 | 8.50943E-12 |
| Anxa5 | 1.3556E-06 | -1.27140154 | 0.857 | 0.993 | 0.032930202 | 4.33269E-12 | -1.046866363 | 1 | 1 | 1.0525E-07 | 1.3556E-06 | 8.66539E-12 |
| Aplp2 | 3.31891E-05 | -1.037270481 | 0.905 | 0.996 | 0.806230497 | 4.54932E-12 | -1.166993124 | 0.956 | 1 | 1.10512E-07 | 3.31891E-05 | 9.09864E-12 |
| Sdc4 | 1.21826E-06 | -1.60399424 | 1 | 1 | 0.029593996 | 4.78325E-12 | -1.347230174 | 1 | 1 | 1.16195E-07 | 1.21826E-06 | 9.5665E-12 |
| Nenf | 2.79274E-09 | -2.379701646 | 0.333 | 0.922 | 6.78412E-05 | 4.79944E-12 | -1.474779583 | 0.511 | 0.875 | 1.16588E-07 | 2.79274E-09 | 9.59887E-12 |
| Col11a2 | 1.51007E-06 | -1.45535686 | 0.905 | 0.951 | 0.036682564 | 4.87596E-12 | -1.399483542 | 0.844 | 0.948 | 1.18447E-07 | 1.51007E-06 | 9.75193E-12 |
| Ssr2 | 2.39356E-08 | -1.810823775 | 0.476 | 0.963 | 0.000581444 | 4.91718E-12 | -1.547734657 | 0.667 | 0.927 | 1.19448E-07 | 2.39356E-08 | 9.83435E-12 |
| Rpl18 | 1.3531E-09 | -2.092879184 | 0.905 | 0.985 | 3.28695E-05 | 4.92299E-12 | -1.175550177 | 0.956 | 0.997 | 1.19589E-07 | 1.3531E-09 | 9.84598E-12 |
| Arpc1b | 3.83478E-05 | -0.988490006 | 0.81 | 0.966 | 0.931544321 | 5.24061E-12 | -1.209907115 | 0.933 | 0.99 | 1.27305E-07 | 3.83478E-05 | 1.04812E-11 |
| Ift20 | 2.78779E-06 | -1.76585137 | 0.524 | 0.903 | 0.067721029 | 5.45575E-12 | -1.615436621 | 0.667 | 0.938 | 1.32531E-07 | 2.78779E-06 | 1.09115E-11 |
| Copz2 | 4.35862E-09 | -2.682179909 | 0.238 | 0.888 | 0.00010588 | 5.53076E-12 | -2.243358196 | 0.156 | 0.789 | 1.34353E-07 | 4.35862E-09 | 1.10615E-11 |
| Eid1 | 4.22516E-10 | -2.388724158 | 0.762 | 0.97 | 1.02638E-05 | 5.56931E-12 | -1.420878378 | 0.822 | 0.955 | 1.3529E-07 | 4.22516E-10 | 1.11386E-11 |
| Cldn7 | 0.000467252 | 2.214340149 | 0.286 | 0.067 | 1 | 5.77065E-12 | 1.408450206 | 0.667 | 0.17 | 1.40181E-07 | 0.000467252 | 1.15413E-11 |
| Rpl29 | 1.63732E-09 | -1.928311922 | 0.857 | 1 | 3.97739E-05 | 6.132E-12 | -1.087568458 | 0.956 | 0.997 | 1.48959E-07 | 1.63732E-09 | 1.2264E-11 |
| Polr2d | 2.67412E-06 | -2.718545655 | 0.095 | 0.675 | 0.064959675 | 6.51788E-12 | -3.090773074 | 0.089 | 0.685 | 1.58332E-07 | 2.67412E-06 | 1.30358E-11 |
| Arf4 | 1.59958E-05 | -0.870654981 | 1 | 0.996 | 0.388569029 | 6.65081E-12 | -0.916561937 | 1 | 0.993 | 1.61562E-07 | 1.59958E-05 | 1.33016E-11 |
| Cnn3 | 7.2607E-06 | -1.348596045 | 0.857 | 0.981 | 0.176376997 | 7.01208E-12 | -1.290467045 | 0.933 | 0.986 | 1.70337E-07 | 7.2607E-06 | 1.40242E-11 |
| H3f3b | 7.0895E-08 | -1.622893416 | 0.952 | 1 | 0.001722181 | 7.0126E-12 | -1.235009635 | 1 | 1 | 1.7035E-07 | 7.0895E-08 | 1.40252E-11 |
| Muc16 | 0.578156445 | 1.088809267 | 0 | 0.015 | 1 | 7.02095E-12 | 3.098110085 | 0.333 | 0.038 | 1.70553E-07 | 0.578156445 | 1.40419E-11 |
| Ecm2 | 0.002812832 | -1.53338614 | 0.333 | 0.675 | 1 | 7.28574E-12 | -2.038626251 | 0.333 | 0.886 | 1.76985E-07 | 0.002812832 | 1.45715E-11 |
| Cox7c | 2.0286E-07 | -1.460352512 | 0.905 | 0.981 | 0.004927863 | 8.18897E-12 | -1.121417002 | 0.933 | 0.979 | 1.98926E-07 | 2.0286E-07 | 1.63779E-11 |
| Cdkn1a | 2.95607E-06 | -1.87137291 | 0.952 | 0.966 | 0.071808867 | 8.76982E-12 | -1.619123885 | 0.956 | 0.986 | 2.13036E-07 | 2.95607E-06 | 1.75396E-11 |
| Arf2 | 0.000159453 | -1.15277672 | 0.238 | 0.799 | 1 | 9.67649E-12 | -2.524684489 | 0.178 | 0.775 | 2.35061E-07 | 0.000159453 | 1.9353E-11 |
| Serinc1 | 2.02068E-06 | -1.432136741 | 0.524 | 0.948 | 0.049086323 | 9.73139E-12 | -1.221967214 | 0.867 | 0.976 | 2.36395E-07 | 2.02068E-06 | 1.94628E-11 |
| Mxra8 | 6.95552E-09 | -2.843640875 | 0.286 | 0.925 | 0.000168964 | 1.00156E-11 | -1.490514744 | 0.289 | 0.958 | 2.43298E-07 | 6.95552E-09 | 2.00311E-11 |
| Ston2 | 0.050766442 | -2.529168291 | 0.048 | 0.235 | 1 | 1.02733E-11 | 0.795547315 | 0.578 | 0.121 | 2.49559E-07 | 0.050766442 | 2.05466E-11 |
| Perp | 0.000487416 | 1.162809848 | 0.429 | 0.131 | 1 | 1.1226E-11 | 1.325520582 | 0.733 | 0.218 | 2.72701E-07 | 0.000487416 | 2.24519E-11 |
| Sec61g | 3.97644E-08 | -1.779087197 | 0.762 | 0.955 | 0.000965956 | 1.16513E-11 | -1.421877682 | 0.822 | 0.941 | 2.83033E-07 | 3.97644E-08 | 2.33026E-11 |
| Pla2g5 | 0.000175143 | -3.288545887 | 0.19 | 0.601 | 1 | 1.17867E-11 | -4.366775963 | 0.044 | 0.64 | 2.86323E-07 | 0.000175143 | 2.35734E-11 |
| Tsn | 3.82757E-06 | -1.485725862 | 0.429 | 0.922 | 0.092979264 | 1.22648E-11 | -1.385264544 | 0.844 | 0.927 | 2.97936E-07 | 3.82757E-06 | 2.45296E-11 |
| Cldn3 | 0.81881728 | 1.088809267 | 0.048 | 0.037 | 1 | 1.28891E-11 | 2.567595369 | 0.4 | 0.062 | 3.13103E-07 | 0.81881728 | 2.57783E-11 |
| Gnb2 | 0.000237013 | -0.999938826 | 0.762 | 0.951 | 1 | 1.3033E-11 | -1.212518809 | 0.933 | 0.965 | 3.16597E-07 | 0.000237013 | 2.60659E-11 |
| Ifi27 | 5.10558E-07 | -1.842199893 | 0.571 | 0.94 | 0.012402485 | 1.38385E-11 | -1.45378365 | 0.733 | 0.924 | 3.36165E-07 | 5.10558E-07 | 2.7677E-11 |
| Copb2 | 0.000902347 | -0.931896716 | 0.714 | 0.933 | 1 | 1.39036E-11 | -1.35494074 | 0.889 | 0.962 | 3.37746E-07 | 0.000902347 | 2.78072E-11 |
| Comp | 0.000329782 | -1.806911532 | 0.714 | 0.91 | 1 | 1.50795E-11 | -1.963953057 | 0.467 | 0.938 | 3.66312E-07 | 0.000329782 | 3.01591E-11 |
| Ckmt1 | 0.11409323 | 2.088809267 | 0.095 | 0.03 | 1 | 1.59544E-11 | 1.987927168 | 0.444 | 0.076 | 3.87565E-07 | 0.11409323 | 3.19089E-11 |
| Sult1e1 | 0.425636763 | -1.574155746 | 0 | 0.03 | 1 | 1.74568E-11 | 2.176112597 | 0.4 | 0.062 | 4.24061E-07 | 0.425636763 | 3.49136E-11 |
| Cct2 | 0.000520698 | -0.892003713 | 0.857 | 0.937 | 1 | 1.83452E-11 | -1.019804117 | 0.889 | 0.965 | 4.45641E-07 | 0.000520698 | 3.66903E-11 |
| Igfbp3 | 0.845641408 | -0.66862043 | 0.095 | 0.086 | 1 | 1.86608E-11 | 0.950768369 | 0.667 | 0.166 | 4.53307E-07 | 0.845641408 | 3.73215E-11 |
| Cytl1 | 5.1829E-07 | -3.822946332 | 1 | 0.955 | 0.012590308 | 1.9417E-11 | -1.899143758 | 0.711 | 0.938 | 4.71677E-07 | 5.1829E-07 | 3.8834E-11 |
| Npdc1 | 4.40334E-06 | -1.540988882 | 0.429 | 0.899 | 0.106966029 | 2.02872E-11 | -1.618364983 | 0.333 | 0.886 | 4.92817E-07 | 4.40334E-06 | 4.05745E-11 |
| Rpl31 | 7.87354E-09 | -1.665441986 | 0.857 | 0.981 | 0.000191264 | 2.12388E-11 | -1.067321172 | 0.956 | 0.993 | 5.15933E-07 | 7.87354E-09 | 4.24776E-11 |
| Cilp2 | 0.01733795 | -1.519999976 | 0.381 | 0.604 | 1 | 2.35029E-11 | -2.099996819 | 0.333 | 0.844 | 5.70933E-07 | 0.01733795 | 4.70058E-11 |
| Ldha | 4.07907E-09 | -1.854060626 | 0.857 | 0.978 | 9.90888E-05 | 2.53114E-11 | -1.181887329 | 0.911 | 0.979 | 6.14865E-07 | 4.07907E-09 | 5.06229E-11 |
| Tspyl1 | 0.006389113 | -1.169207064 | 0.286 | 0.608 | 1 | 2.55696E-11 | -2.301820521 | 0.089 | 0.72 | 6.21136E-07 | 0.006389113 | 5.11392E-11 |
| Fscn1 | 0.004604435 | -2.02666795 | 0.143 | 0.478 | 1 | 2.70456E-11 | -2.737884885 | 0.133 | 0.713 | 6.56991E-07 | 0.004604435 | 5.40912E-11 |
| Elob | 5.55261E-09 | -1.79713439 | 0.81 | 0.974 | 0.000134884 | 2.71377E-11 | -1.204033357 | 0.933 | 0.972 | 6.5923E-07 | 5.55261E-09 | 5.42755E-11 |
| Aldh3a1 | 0.584468473 | 0.33137957 | 0.095 | 0.142 | 1 | 2.84084E-11 | 1.209141398 | 0.667 | 0.18 | 6.90097E-07 | 0.584468473 | 5.68168E-11 |
| Msi2 | 3.20402E-07 | 1.221075758 | 1 | 1 | 0.007783204 | 2.92095E-11 | 1.158387246 | 1 | 1 | 7.09558E-07 | 3.20402E-07 | 5.84191E-11 |
| Gpx1 | 8.71443E-08 | -2.650678837 | 0.81 | 0.955 | 0.002116909 | 3.10671E-11 | -1.365818798 | 0.933 | 0.965 | 7.54681E-07 | 8.71443E-08 | 6.21341E-11 |
| Dbi | 9.84165E-10 | -3.090170893 | 0.667 | 0.963 | 2.39073E-05 | 3.12154E-11 | -1.790858602 | 0.822 | 0.896 | 7.58284E-07 | 9.84165E-10 | 6.24308E-11 |
| Lum | 0.000254215 | -2.901409712 | 0.19 | 0.631 | 1 | 3.39425E-11 | -2.447314112 | 0.333 | 0.824 | 8.2453E-07 | 0.000254215 | 6.78849E-11 |
| Cd302 | 4.68202E-09 | -2.896083841 | 0.19 | 0.881 | 0.000113736 | 3.53372E-11 | -1.798054104 | 0.244 | 0.848 | 8.58412E-07 | 4.68202E-09 | 7.06744E-11 |
| Emc7 | 5.30545E-08 | -2.004702619 | 0.381 | 0.899 | 0.001288799 | 3.6746E-11 | -1.46797095 | 0.733 | 0.91 | 8.92634E-07 | 5.30545E-08 | 7.3492E-11 |
| Tnfrsf12a | 0.000213085 | -1.596269586 | 0.571 | 0.851 | 1 | 3.734E-11 | -1.756416685 | 0.867 | 0.941 | 9.07064E-07 | 0.000213085 | 7.468E-11 |
| C1d | 1.20305E-08 | -3.202745179 | 0.095 | 0.795 | 0.000292244 | 3.92301E-11 | -2.066461681 | 0.289 | 0.796 | 9.52978E-07 | 1.20305E-08 | 7.84602E-11 |
| Tst | 0.217829703 | -1.054148687 | 0.048 | 0.142 | 1 | 3.99283E-11 | 2.045642666 | 0.667 | 0.201 | 9.69938E-07 | 0.217829703 | 7.98566E-11 |
| S100a10 | 1.18841E-07 | -1.894143865 | 0.857 | 0.981 | 0.002886888 | 3.9988E-11 | -1.544089079 | 0.933 | 0.983 | 9.71389E-07 | 1.18841E-07 | 7.99761E-11 |
| Slc25a3 | 6.28404E-06 | -1.00627883 | 0.952 | 0.981 | 0.152651907 | 4.03459E-11 | -1.06070622 | 0.978 | 0.993 | 9.80084E-07 | 6.28404E-06 | 8.06919E-11 |
| Morf4l1 | 9.33656E-08 | -1.331903354 | 0.905 | 0.993 | 0.002268038 | 4.10533E-11 | -0.806260518 | 1 | 0.997 | 9.97266E-07 | 9.33656E-08 | 8.21066E-11 |
| Ikzf2 | 0.061931621 | 1.339352729 | 0.381 | 0.216 | 1 | 4.14228E-11 | 2.154896112 | 0.778 | 0.273 | 1.00624E-06 | 0.061931621 | 8.28457E-11 |
| Wnt4 | 0.572090711 | 0.400753273 | 0.095 | 0.063 | 1 | 4.42439E-11 | 1.352427274 | 0.6 | 0.149 | 1.07477E-06 | 0.572090711 | 8.84877E-11 |
| Atp5g2 | 6.35656E-08 | -1.677735416 | 0.81 | 0.948 | 0.001544136 | 4.49104E-11 | -1.201157412 | 0.911 | 0.941 | 1.09096E-06 | 6.35656E-08 | 8.98208E-11 |

| | | | | | | | | | | | | |
|---|---|---|---|---|---|---|---|---|---|---|---|---|
| Zbtb20 | 3.5639E-05 | 1.003614616 | 1 | 1 | 0.865741777 | 5.23005E-11 | 0.785075735 | 1 | 1 | 1.27048E-06 | 3.5639E-05 | 1.04601E-10 |
| Timp2 | 5.32982E-11 | -2.697065928 | 0.762 | 0.989 | 1.29472E-06 | 2.85369E-10 | -1.040090462 | 0.911 | 0.997 | 6.93219E-06 | 2.85369E-10 | 1.06596E-10 |
| Mcl1 | 6.39883E-05 | -0.966764064 | 0.857 | 0.951 | 1 | 5.33948E-11 | -1.202036034 | 0.978 | 0.983 | 1.29707E-06 | 6.39883E-05 | 1.0679E-10 |
| Ran | 7.64333E-06 | -1.506458242 | 0.667 | 0.918 | 0.18567175 | 5.37583E-11 | -1.298018951 | 0.822 | 0.931 | 1.3059E-06 | 7.64333E-06 | 1.07517E-10 |
| Pcbp1 | 1.16766E-07 | -1.63000898 | 0.762 | 0.966 | 0.002836487 | 5.50505E-11 | -1.244414914 | 0.911 | 0.99 | 1.33729E-06 | 1.16766E-07 | 1.10101E-10 |
| Il11ra1 | 3.97359E-06 | -2.537846182 | 0.19 | 0.757 | 0.096526511 | 5.65006E-11 | -1.817356404 | 0.289 | 0.851 | 1.37251E-06 | 3.97359E-06 | 1.13001E-10 |
| Emilin1 | 0.003416536 | -1.208871282 | 0.19 | 0.571 | 1 | 6.16887E-11 | -1.865749322 | 0.267 | 0.834 | 1.49854E-06 | 0.003416536 | 1.23377E-10 |
| mt-Nd2 | 0.002791126 | 0.982790607 | 1 | 1 | 1 | 6.41552E-11 | 1.426344524 | 1 | 1 | 1.55846E-06 | 0.002791126 | 1.2831E-10 |
| Col12a1 | 0.001494427 | -3.001270472 | 0.524 | 0.724 | 1 | 6.46157E-11 | -1.396654606 | 0.733 | 0.952 | 1.56965E-06 | 0.001494427 | 1.29231E-10 |
| Metrnl | 0.000660744 | -1.815163845 | 0.19 | 0.623 | 1 | 6.94148E-11 | -3.564854927 | 0.089 | 0.64 | 1.68622E-06 | 0.000660744 | 1.3883E-10 |
| Atp10d | 0.70122657 | 0.866416846 | 0.095 | 0.138 | 1 | 7.18242E-11 | 2.005000681 | 0.467 | 0.093 | 1.74475E-06 | 0.70122657 | 1.43648E-10 |
| Gstm2 | 0.261635242 | 3.188344941 | 0.048 | 0.015 | 1 | 7.18688E-11 | 2.39927962 | 0.422 | 0.076 | 1.74584E-06 | 0.261635242 | 1.43738E-10 |
| Polr1h | 8.44538E-05 | -2.10233222 | 0.095 | 0.586 | 1 | 7.24621E-11 | -2.357341682 | 0.111 | 0.699 | 1.76025E-06 | 8.44538E-05 | 1.44924E-10 |
| Tuba1b | 7.69122E-07 | -2.383678505 | 0.524 | 0.873 | 0.018683522 | 7.50491E-11 | -1.586209967 | 0.8 | 0.9 | 1.82309E-06 | 7.69122E-07 | 1.50098E-10 |
| Rpl36a | 3.1072E-10 | -2.226452729 | 0.762 | 0.989 | 7.548E-06 | 7.65384E-11 | -1.203270632 | 0.933 | 0.986 | 1.85927E-06 | 3.1072E-10 | 1.53077E-10 |
| P3h4 | 6.28397E-10 | -2.838892734 | 0.286 | 0.929 | 1.5265E-05 | 7.75482E-11 | -1.829039349 | 0.244 | 0.82 | 1.8838E-06 | 6.28397E-10 | 1.55096E-10 |
| Ccn2 | 1.96798E-05 | -2.086552092 | 0.952 | 0.981 | 0.478061397 | 8.03879E-11 | -2.743092311 | 0.556 | 0.917 | 1.95278E-06 | 1.96798E-05 | 1.60776E-10 |
| Rpl36 | 2.0363E-08 | -1.691512696 | 0.905 | 0.985 | 0.000494659 | 8.0881E-11 | -1.04118672 | 0.956 | 0.983 | 1.96476E-06 | 2.0363E-08 | 1.61762E-10 |
| Ocln | 0.578156445 | 1.088809267 | 0 | 0.015 | 1 | 8.83879E-11 | 2.020107573 | 0.467 | 0.093 | 2.14712E-06 | 0.578156445 | 1.76776E-10 |
| Leprot | 5.70734E-08 | -2.217735795 | 0.286 | 0.869 | 0.001386426 | 8.97628E-11 | -1.567154043 | 0.444 | 0.858 | 2.18052E-06 | 5.70734E-08 | 1.79526E-10 |
| Il17d | 2.60315E-06 | -2.448988126 | 0.286 | 0.817 | 0.063235623 | 9.01989E-11 | -1.981935284 | 0.356 | 0.869 | 2.19111E-06 | 2.60315E-06 | 1.80398E-10 |
| Gm2a | 3.39951E-05 | -2.098817736 | 0.048 | 0.582 | 0.825809088 | 9.07672E-11 | -2.088590629 | 0.156 | 0.751 | 2.20492E-06 | 3.39951E-05 | 1.81534E-10 |
| Colgalt1 | 3.82882E-08 | -2.519999976 | 0.238 | 0.866 | 0.000930098 | 9.56267E-11 | -1.744150452 | 0.4 | 0.851 | 2.32296E-06 | 3.82882E-08 | 1.91253E-10 |
| Hsp90aa1 | 1.65028E-08 | -1.787220365 | 0.905 | 0.989 | 0.000400887 | 9.84076E-11 | -1.280556093 | 0.978 | 1 | 2.39052E-06 | 1.65028E-08 | 1.96815E-10 |
| Hnrnpk | 1.17171E-06 | -1.252694366 | 0.905 | 0.985 | 0.028463238 | 1.01783E-10 | -0.984383729 | 0.978 | 0.993 | 2.4725E-06 | 1.17171E-06 | 2.03565E-10 |
| Fam180a | 0.000323329 | -2.973686659 | 0.333 | 0.675 | 1 | 1.05687E-10 | -1.962705916 | 0.378 | 0.872 | 2.56736E-06 | 0.000323329 | 2.11375E-10 |
| Rps27l | 3.33298E-07 | -1.825349329 | 0.619 | 0.918 | 0.008096485 | 1.05854E-10 | -1.318536667 | 0.8 | 0.907 | 2.57141E-06 | 3.33298E-07 | 2.11708E-10 |
| Col8a2 | 0.001488947 | -2.23071233 | 0.19 | 0.549 | 1 | 1.09629E-10 | -1.604010116 | 0.422 | 0.907 | 2.6631E-06 | 0.001488947 | 2.19257E-10 |
| Snrpg | 0.000310834 | -1.055026506 | 0.667 | 0.91 | 1 | 1.12719E-10 | -1.294207337 | 0.8 | 0.913 | 2.73817E-06 | 0.000310834 | 2.25438E-10 |
| Gpx7 | 9.04091E-07 | -2.756680784 | 0.095 | 0.705 | 0.021962184 | 1.2119E-10 | -2.794974711 | 0.067 | 0.626 | 2.94394E-06 | 9.04091E-07 | 2.42379E-10 |
| Pdia4 | 1.65606E-08 | -1.590568836 | 0.381 | 0.922 | 0.000402291 | 1.23453E-10 | -1.39718037 | 0.689 | 0.913 | 2.99893E-06 | 1.65606E-08 | 2.46906E-10 |
| Pfdn5 | 2.31615E-08 | -1.686555541 | 0.762 | 0.978 | 0.000562638 | 1.23513E-10 | -1.099258312 | 0.911 | 0.969 | 3.00037E-06 | 2.31615E-08 | 2.47025E-10 |
| B2m | 5.65852E-07 | -2.077772291 | 0.81 | 0.937 | 0.013745686 | 1.29223E-10 | -0.686658876 | 0.933 | 0.99 | 3.13909E-06 | 5.65852E-07 | 2.58446E-10 |
| Sec13 | 2.00198E-06 | -1.845602391 | 0.238 | 0.858 | 0.048632113 | 1.29741E-10 | -1.831235361 | 0.511 | 0.83 | 3.15166E-06 | 2.00198E-06 | 2.59481E-10 |
| Vamp3 | 9.00782E-07 | -3.107587946 | 0.048 | 0.672 | 0.021881786 | 1.36129E-10 | -1.737884885 | 0.156 | 0.751 | 3.30683E-06 | 9.00782E-07 | 2.72257E-10 |
| Vim | 4.06049E-09 | -2.3703758 | 1 | 0.993 | 9.86375E-05 | 1.44027E-10 | -1.664584709 | 0.956 | 0.976 | 3.49871E-06 | 4.06049E-09 | 2.88054E-10 |
| Eef1g | 6.15635E-06 | -1.173111519 | 0.81 | 0.97 | 0.14955016 | 1.4413E-10 | -1.003160271 | 1 | 0.979 | 3.5012E-06 | 6.15635E-06 | 2.8826E-10 |
| Cpxm1 | 0.000990871 | -3.192476843 | 0.143 | 0.5 | 1 | 1.49671E-10 | -2.66459307 | 0.156 | 0.727 | 3.63582E-06 | 0.000990871 | 2.99343E-10 |
| Adh1 | 0.144977306 | -1.752492987 | 0 | 0.093 | 1 | 1.53542E-10 | 1.576157382 | 0.556 | 0.138 | 3.72985E-06 | 0.144977306 | 3.07085E-10 |
| Tram1 | 2.07646E-05 | -1.249477517 | 0.667 | 0.966 | 0.504414733 | 1.55713E-10 | -1.244282112 | 0.933 | 0.938 | 3.78258E-06 | 2.07646E-05 | 3.11426E-10 |
| Thbs1 | 0.000212916 | -2.139782031 | 0.81 | 0.955 | 1 | 1.58846E-10 | -1.70216932 | 0.933 | 0.972 | 3.85869E-06 | 0.000212916 | 3.17693E-10 |
| Npc2 | 4.86533E-06 | -1.31387984 | 0.714 | 0.978 | 0.118188626 | 1.5979E-10 | -1.805505378 | 0.8 | 0.893 | 3.88162E-06 | 4.86533E-06 | 3.1958E-10 |
| Fto | 0.007226582 | 0.96946596 | 1 | 0.937 | 1 | 1.6234E-10 | 0.993439623 | 1 | 0.979 | 3.94355E-06 | 0.007226582 | 3.24679E-10 |
| Kdelr1 | 2.07577E-09 | -1.914654588 | 0.619 | 0.959 | 5.04246E-05 | 1.72435E-10 | -1.157212611 | 0.911 | 0.955 | 4.18879E-06 | 2.07577E-09 | 3.4487E-10 |
| Pof1b | 0.624476205 | -0.828728573 | 0.095 | 0.127 | 1 | 1.77438E-10 | 2.147019686 | 0.422 | 0.08 | 4.31032E-06 | 0.624476205 | 3.54875E-10 |
| Dnajc3 | 5.1108E-05 | -0.974321415 | 0.81 | 0.981 | 1 | 1.8737E-10 | -1.271123724 | 0.844 | 0.979 | 4.5516E-06 | 5.1108E-05 | 3.74741E-10 |
| Fkbp14 | 2.08579E-06 | -2.087779465 | 0.19 | 0.787 | 0.050667981 | 1.88057E-10 | -3.442082545 | 0.089 | 0.616 | 4.56828E-06 | 2.08579E-06 | 3.76114E-10 |
| Pdrg1 | 0.001964868 | -0.906890596 | 0.333 | 0.806 | 1 | 2.0336E-10 | -2.529626439 | 0.111 | 0.682 | 4.94001E-06 | 0.001964868 | 4.06719E-10 |
| Snx17 | 0.005944345 | -1.189726232 | 0.238 | 0.586 | 1 | 2.06235E-10 | -2.81335324 | 0.133 | 0.651 | 5.00987E-06 | 0.005944345 | 4.12471E-10 |
| Dap | 4.62864E-06 | -1.429132124 | 0.571 | 0.948 | 0.112438807 | 2.10906E-10 | -1.433313988 | 0.622 | 0.913 | 5.12332E-06 | 4.62864E-06 | 4.21811E-10 |
| Srp9 | 8.40779E-08 | -1.902145593 | 0.571 | 0.922 | 0.00204242 | 2.22457E-10 | -1.335646611 | 0.822 | 0.924 | 5.40394E-06 | 8.40779E-08 | 4.44915E-10 |
| Tubb5 | 0.000131654 | -1.649157851 | 0.524 | 0.813 | 1 | 2.22997E-10 | -1.489354922 | 0.844 | 0.917 | 5.41704E-06 | 0.000131654 | 4.45994E-10 |
| Cd200 | 1.05029E-05 | -2.191157205 | 0.476 | 0.836 | 0.255136694 | 2.2425E-10 | -2.442771347 | 0.156 | 0.713 | 5.44749E-06 | 1.05029E-05 | 4.485E-10 |
| Selenow | 2.10787E-07 | -1.639644827 | 0.762 | 0.948 | 0.005120426 | 2.35821E-10 | -0.99113559 | 0.911 | 0.976 | 5.72857E-06 | 2.10787E-07 | 4.71643E-10 |
| Ndufc2 | 3.46185E-06 | -1.413691074 | 0.619 | 0.922 | 0.084095267 | 2.37704E-10 | -1.439155673 | 0.778 | 0.882 | 5.77431E-06 | 3.46185E-06 | 4.75408E-10 |
| Ctsb | 4.00204E-06 | -1.200472754 | 0.81 | 0.993 | 0.097217467 | 2.42391E-10 | -1.158420286 | 0.956 | 0.983 | 5.88816E-06 | 4.00204E-06 | 4.84782E-10 |
| Erp29 | 2.60273E-10 | -2.51300913 | 0.381 | 0.955 | 6.32256E-06 | 2.51498E-10 | -1.329983567 | 0.689 | 0.91 | 6.10939E-06 | 2.60273E-10 | 5.02996E-10 |
| Bag5 | 2.86217E-05 | -1.876975018 | 0.143 | 0.705 | 0.695279486 | 2.56109E-10 | -1.751158478 | 0.133 | 0.699 | 6.22141E-06 | 2.86217E-05 | 5.12219E-10 |
| Nab2 | 0.000746284 | -2.174225139 | 0.143 | 0.534 | 1 | 2.58008E-10 | -2.48434556 | 0.156 | 0.702 | 6.26754E-06 | 0.000746284 | 5.16016E-10 |
| St14 | 0.758436286 | -0.133583154 | 0.048 | 0.063 | 1 | 2.6169E-10 | 2.42003818 | 0.467 | 0.1 | 6.35697E-06 | 0.758436286 | 5.2338E-10 |
| Hspd1 | 2.37654E-06 | -1.741668213 | 0.762 | 0.922 | 0.057731021 | 2.68001E-10 | -1.360897024 | 0.933 | 0.965 | 6.51027E-06 | 2.37654E-06 | 5.36002E-10 |
| Rpl38 | 3.07233E-09 | -2.138726458 | 0.81 | 0.963 | 7.46331E-05 | 2.68533E-10 | -1.02251962 | 0.933 | 0.972 | 6.52321E-06 | 3.07233E-09 | 5.37067E-10 |
| Skp1 | 0.000621227 | -0.647154108 | 0.857 | 0.955 | 1 | 2.72819E-10 | -1.050073137 | 0.911 | 0.976 | 6.62732E-06 | 0.000621227 | 5.45638E-10 |
| Clic1 | 0.000102169 | -1.183270278 | 0.762 | 0.963 | 1 | 2.79224E-10 | -1.104829973 | 0.911 | 0.962 | 6.78291E-06 | 0.000102169 | 5.58448E-10 |
| Mfsd6 | 0.97673807 | 0.425844254 | 0.048 | 0.049 | 1 | 2.82849E-10 | 2.081621963 | 0.511 | 0.121 | 6.87098E-06 | 0.97673807 | 5.65699E-10 |
| Ndn | 7.05819E-06 | -2.531777143 | 0.095 | 0.672 | 0.171457532 | 2.92333E-10 | -3.149817428 | 0.067 | 0.609 | 7.10136E-06 | 7.05819E-06 | 5.84667E-10 |
| Armcx3 | 0.036177392 | -0.667015354 | 0.381 | 0.69 | 1 | 3.03044E-10 | -1.878906597 | 0.2 | 0.785 | 7.36155E-06 | 0.036177392 | 6.06089E-10 |
| Tm9sf3 | 4.87723E-05 | -1.004702619 | 0.857 | 0.981 | 1 | 3.13472E-10 | -0.923327085 | 1 | 0.993 | 7.61486E-06 | 4.87723E-05 | 6.26944E-10 |
| Ralgapa2 | 0.007407916 | 1.383379246 | 0.571 | 0.328 | 1 | 3.19672E-10 | 1.664576243 | 0.956 | 0.429 | 7.76547E-06 | 0.007407916 | 6.39344E-10 |
| Dad1 | 3.67036E-10 | -2.150906247 | 0.667 | 0.981 | 8.91604E-06 | 3.21222E-10 | -1.165120752 | 0.867 | 0.962 | 7.80313E-06 | 3.67036E-10 | 6.42445E-10 |
| Serpine2 | 0.11169552 | -0.12290352 | 0.952 | 0.989 | 1 | 3.41751E-10 | -2.036891697 | 0.756 | 0.965 | 8.30181E-06 | 0.11169552 | 6.83501E-10 |
| Ost4 | 7.38046E-10 | -2.027234046 | 0.762 | 0.978 | 1.79286E-05 | 3.46331E-10 | -1.275184479 | 0.844 | 0.948 | 8.41306E-06 | 7.38046E-10 | 6.92661E-10 |
| Kcnj16 | 0.492604708 | 0.351843673 | 0 | 0.022 | 1 | 3.5581E-10 | 2.185572927 | 0.356 | 0.055 | 8.64333E-06 | 0.492604708 | 7.1162E-10 |
| Canx | 4.82619E-06 | -0.898479681 | 0.905 | 1 | 0.117237686 | 3.61088E-10 | -0.953771706 | 1 | 1 | 8.77154E-06 | 4.82619E-06 | 7.22176E-10 |
| Rpl22l1 | 2.54375E-09 | -2.039455617 | 0.857 | 0.981 | 6.17928E-05 | 3.63323E-10 | -1.085201775 | 0.933 | 0.983 | 8.82583E-06 | 2.54375E-09 | 7.26645E-10 |
| Kcnma1 | 2.20837E-05 | 1.658440114 | 0.952 | 0.978 | 0.536457182 | 3.63627E-10 | 1.864124987 | 0.978 | 0.886 | 8.83322E-06 | 2.20837E-05 | 7.27253E-10 |
| Tmem98 | 2.23227E-05 | -4.262866171 | 0 | 0.507 | 0.542262271 | 3.65772E-10 | -2.975138897 | 0.089 | 0.619 | 8.88533E-06 | 2.23227E-05 | 7.31544E-10 |
| Sdf2 | 3.68929E-10 | -2.356895369 | 0.524 | 0.959 | 8.96203E-06 | 6.2543E-06 | -0.845432933 | 0.778 | 0.907 | 0.151929443 | 6.2543E-06 | 7.37858E-10 |
| Bod1 | 9.0825E-06 | -1.880817084 | 0.095 | 0.69 | 0.220632101 | 3.76772E-10 | -2.301820521 | 0.2 | 0.716 | 9.15255E-06 | 9.0825E-06 | 7.53545E-10 |
| Eif1b | 2.04676E-05 | -1.654507159 | 0.333 | 0.828 | 0.497199028 | 3.76779E-10 | -1.556577825 | 0.578 | 0.882 | 9.15271E-06 | 2.04676E-05 | 7.53557E-10 |
| Rab11fip1 | 0.710157178 | 0.332303486 | 0.429 | 0.358 | 1 | 3.81341E-10 | 1.37431988 | 0.711 | 0.232 | 9.26353E-06 | 0.710157178 | 7.62682E-10 |
| Ppargc1a | 0.639355656 | 1.673771768 | 0.048 | 0.03 | 1 | 3.8426E-10 | 0.660704773 | 0.6 | 0.152 | 9.33445E-06 | 0.639355656 | 7.68521E-10 |
| P3h3 | 0.000446786 | -1.481769316 | 0.286 | 0.72 | 1 | 4.03224E-10 | -1.758362603 | 0.244 | 0.813 | 9.79511E-06 | 0.000446786 | 8.06448E-10 |
| Marveld1 | 3.81662E-05 | -0.68836265 | 0.143 | 0.701 | 0.927132488 | 4.03506E-10 | -2.706118736 | 0.156 | 0.671 | 9.80198E-06 | 3.81662E-05 | 8.07013E-10 |
| Mal2 | 0.632076134 | 1.673771768 | 0 | 0.011 | 1 | 4.11284E-10 | 3.03099589 | 0.289 | 0.035 | 9.99091E-06 | 0.632076134 | 8.22568E-10 |
| Camk1 | 0.000124996 | -1.637295335 | 0.19 | 0.701 | 1 | 4.11815E-10 | -2.109047822 | 0.133 | 0.716 | 1.00038E-05 | 0.000124996 | 8.2363E-10 |
| Jkamp | 2.65444E-07 | -2.959223429 | 0.095 | 0.731 | 0.006448164 | 4.17647E-10 | -2.309393741 | 0.133 | 0.682 | 1.01455E-05 | 2.65444E-07 | 8.35294E-10 |
| Eef2 | 4.62971E-09 | -1.791230109 | 1 | 0.974 | 0.000112465 | 4.24616E-10 | -0.956595807 | 0.956 | 1 | 1.03148E-05 | 4.62971E-09 | 8.49232E-10 |
| Lyz2 | 0.754649812 | -0.326228232 | 0.048 | 0.067 | 1 | 4.46408E-10 | 0.852997588 | 0.511 | 0.125 | 1.08441E-05 | 0.754649812 | 8.92816E-10 |
| Cyfip2 | 0.952859852 | 0.544488751 | 0.095 | 0.093 | 1 | 4.4686E-10 | 1.712218932 | 0.511 | 0.121 | 1.08551E-05 | 0.952859852 | 8.93721E-10 |
| Fibin | 1.02173E-06 | -3.341643285 | 0.19 | 0.75 | 0.024819763 | 4.52335E-10 | -3.728202994 | 0.067 | 0.606 | 1.09881E-05 | 1.02173E-06 | 9.04669E-10 |
| 2900026A02Rik | 0.26101443 | -1.107587946 | 0.048 | 0.131 | 1 | 4.5759E-10 | 1.717838004 | 0.444 | 0.09 | 1.11158E-05 | 0.26101443 | 9.15179E-10 |
| Rps27 | 3.5263E-08 | -1.849322009 | 0.81 | 0.985 | 0.000856608 | 4.59684E-10 | -1.000142622 | 0.956 | 0.99 | 1.11666E-05 | 3.5263E-08 | 9.19367E-10 |
| Six2 | 0.000929789 | -1.544651751 | 0.286 | 0.679 | 1 | 4.63098E-10 | -1.681924231 | 0.2 | 0.796 | 1.12496E-05 | 0.000929789 | 9.26196E-10 |
| Herpud1 | 0.000699061 | -0.997109759 | 0.905 | 0.944 | 1 | 4.72748E-10 | -1.335591258 | 0.844 | 0.955 | 1.1484E-05 | 0.000699061 | 9.45496E-10 |
| Plod2 | 0.014619189 | -0.334971865 | 1 | 1 | 1 | 4.76964E-10 | -1.262264445 | 0.689 | 0.986 | 1.15864E-05 | 0.014619189 | 9.53929E-10 |
| Tpm4 | 0.014931795 | -0.880029802 | 0.81 | 0.843 | 1 | 4.80203E-10 | -1.166684778 | 0.867 | 0.965 | 1.16651E-05 | 0.014931795 | 9.60405E-10 |
| Celsr1 | 0.019692488 | 4.258734268 | 0.048 | 0.004 | 1 | 4.95391E-10 | 2.761075098 | 0.356 | 0.059 | 1.2034E-05 | 0.019692488 | 9.90782E-10 |

| | | | | | | | | | | | | |
|---|---|---|---|---|---|---|---|---|---|---|---|---|
| Cyp2a5 | 0.026418672 | 1.316219763 | 0.286 | 0.116 | 1 | 4.98488E-10 | 1.881618266 | 0.756 | 0.263 | 1.21093E-05 | 0.026418672 | 9.96976E-10 |
| Pcolce | 0.000911002 | -3.676220246 | 0.143 | 0.504 | 1 | 5.10474E-10 | -2.175989484 | 0.467 | 0.81 | 1.24004E-05 | 0.000911002 | 1.02095E-09 |
| Pstpip2 | 0.761764764 | 1.119182916 | 0.095 | 0.082 | 1 | 5.33886E-10 | 1.30164348 | 0.556 | 0.135 | 1.29692E-05 | 0.761764764 | 1.06777E-09 |
| Ptges3 | 8.57326E-07 | -1.307250558 | 0.857 | 0.993 | 0.020826164 | 5.35988E-10 | -0.907603566 | 1 | 0.99 | 1.30202E-05 | 8.57326E-07 | 1.07198E-09 |
| Sox9 | 0.000197214 | -1.374803837 | 0.762 | 0.97 | 1 | 5.37609E-10 | -1.218490559 | 0.822 | 0.969 | 1.30596E-05 | 0.000197214 | 1.07522E-09 |
| Mfap2 | 0.027446066 | -2.674956387 | 0.048 | 0.265 | 1 | 5.544E-10 | -5.848308874 | 0.022 | 0.543 | 1.34675E-05 | 0.027446066 | 1.1088E-09 |
| Prdx1 | 1.78967E-07 | -1.555516016 | 0.81 | 0.981 | 0.004347465 | 5.61477E-10 | -1.021819135 | 0.978 | 0.997 | 1.36394E-05 | 1.78967E-07 | 1.12295E-09 |
| Paics | 6.20668E-09 | -2.068657978 | 0.667 | 0.951 | 0.000150773 | 5.6666E-10 | -1.218902519 | 0.889 | 0.958 | 1.37653E-05 | 6.20668E-09 | 1.13332E-09 |
| Pigk | 5.65713E-05 | -1.960434252 | 0.143 | 0.657 | 1 | 6.18017E-10 | -1.434924623 | 0.111 | 0.702 | 1.50129E-05 | 5.65713E-05 | 1.23603E-09 |
| Col16a1 | 0.041201963 | -0.709932525 | 0.429 | 0.784 | 1 | 6.28193E-10 | -1.448249576 | 0.489 | 0.92 | 1.52601E-05 | 0.041201963 | 1.25639E-09 |
| Tuba1a | 1.12339E-05 | -1.813953443 | 0.524 | 0.873 | 0.272893867 | 6.45072E-10 | -1.5581395 | 0.867 | 0.9 | 1.56701E-05 | 1.12339E-05 | 1.29014E-09 |
| Tmem50a | 6.60704E-10 | -2.540547353 | 0.524 | 0.918 | 1.60498E-05 | 0.001171891 | -0.578774905 | 0.8 | 0.858 | 1 | 0.001171891 | 1.32141E-09 |
| Ssr3 | 4.87158E-06 | -1.128804432 | 0.857 | 0.978 | 0.118340508 | 6.70709E-10 | -1.052164194 | 0.933 | 0.986 | 1.62929E-05 | 4.87158E-06 | 1.34142E-09 |
| mt-Nd5 | 0.003915075 | 0.946396668 | 1 | 1 | 1 | 6.71647E-10 | 1.205215493 | 1 | 1 | 1.63157E-05 | 0.003915075 | 1.34329E-09 |
| Irf6 | 0.020707684 | 2.673771768 | 0.048 | 0.004 | 1 | 6.93419E-10 | 2.522607914 | 0.356 | 0.059 | 1.68445E-05 | 0.020707684 | 1.38684E-09 |
| Klf4 | 0.00021782 | -1.437672443 | 0.905 | 0.97 | 1 | 6.98199E-10 | -0.994000272 | 0.956 | 0.99 | 1.69606E-05 | 0.00021782 | 1.3964E-09 |
| Dleu2 | 0.00014941 | 1.138076812 | 0.857 | 0.526 | 1 | 7.21391E-10 | 1.714893352 | 1 | 0.751 | 1.7524E-05 | 0.00014941 | 1.44278E-09 |
| Slc41a1 | 0.031604246 | -0.596480292 | 0.429 | 0.799 | 1 | 7.2331E-10 | -1.555087151 | 0.289 | 0.879 | 1.75706E-05 | 0.031604246 | 1.44662E-09 |
| Ubl5 | 8.07189E-08 | -1.738701851 | 0.619 | 0.937 | 0.001960824 | 7.62927E-10 | -1.269098889 | 0.756 | 0.872 | 1.8533E-05 | 8.07189E-08 | 1.52585E-09 |
| Prnp | 1.29513E-05 | -1.356969921 | 0.81 | 0.955 | 0.314612783 | 7.80296E-10 | -1.690283748 | 0.8 | 0.931 | 1.8955E-05 | 1.29513E-05 | 1.56059E-09 |
| Tubb4b | 7.69532E-06 | -2.694933098 | 0.571 | 0.828 | 0.186934752 | 7.87437E-10 | -1.43709938 | 0.889 | 0.945 | 1.91284E-05 | 7.69532E-06 | 1.57487E-09 |
| Atp6v0e | 8.18932E-10 | -2.37541271 | 0.762 | 0.989 | 1.98935E-05 | 1.40826E-06 | -0.916596413 | 0.867 | 0.962 | 0.034209496 | 1.40826E-06 | 1.63786E-09 |
| Hnrnpab | 5.67569E-06 | -1.059392256 | 0.762 | 0.955 | 0.137873928 | 8.30354E-10 | -0.92110783 | 1 | 0.997 | 2.0171E-05 | 5.67569E-06 | 1.66071E-09 |
| Arl1 | 5.78702E-08 | -1.967196143 | 0.524 | 0.929 | 0.001405782 | 8.32048E-10 | -1.298242851 | 0.733 | 0.91 | 2.02121E-05 | 5.78702E-08 | 1.6641E-09 |
| Akr1a1 | 4.46424E-07 | -1.3970364 | 0.905 | 0.955 | 0.010844526 | 8.32827E-10 | -0.985911779 | 0.956 | 0.99 | 2.0231E-05 | 4.46424E-07 | 1.66565E-09 |
| Fam107b | 0.013907221 | 1.519966432 | 0.429 | 0.201 | 1 | 8.42895E-10 | 1.258050998 | 0.822 | 0.304 | 2.04756E-05 | 0.013907221 | 1.68579E-09 |
| Tango6 | 0.002786881 | 1.489347197 | 0.333 | 0.108 | 1 | 8.45682E-10 | 1.477603468 | 0.711 | 0.215 | 2.05433E-05 | 0.002786881 | 1.69136E-09 |
| Calm2 | 2.05741E-09 | -1.758554133 | 1 | 0.996 | 4.99786E-05 | 8.69887E-10 | -0.953299534 | 0.978 | 1 | 2.11313E-05 | 2.05741E-09 | 1.73977E-09 |
| Gsto1 | 4.62081E-05 | 1.221259563 | 0.667 | 0.235 | 1 | 8.79869E-10 | 1.288441298 | 0.822 | 0.377 | 2.13738E-05 | 4.62081E-05 | 1.75974E-09 |
| Bzw1 | 5.92616E-05 | -0.904980241 | 1 | 0.985 | 1 | 8.80415E-10 | -0.888322836 | 1 | 0.997 | 2.1387E-05 | 5.92616E-05 | 1.76083E-09 |
| Atp5l | 8.72195E-09 | -1.721254942 | 0.857 | 0.963 | 0.000211874 | 9.55821E-10 | -0.987808941 | 0.911 | 0.965 | 2.32188E-05 | 8.72195E-09 | 1.91164E-09 |
| Kdelr3 | 1.43103E-07 | -2.13701406 | 0.143 | 0.851 | 0.003476266 | 9.59461E-10 | -1.649123844 | 0.244 | 0.827 | 2.33072E-05 | 1.43103E-07 | 1.91892E-09 |
| Swi5 | 2.25989E-08 | -1.801961663 | 0.714 | 0.955 | 0.000548972 | 9.77659E-10 | -1.162417465 | 0.867 | 0.945 | 2.37493E-05 | 2.25989E-08 | 1.95532E-09 |
| Gnl1 | 0.003106887 | -1.735619168 | 0.095 | 0.444 | 1 | 9.9366E-10 | -3.055695251 | 0.044 | 0.571 | 2.4138E-05 | 0.003106887 | 1.98732E-09 |
| Gipc2 | 0.673171208 | 0.281454345 | 0.095 | 0.071 | 1 | 9.98463E-10 | 2.70773464 | 0.467 | 0.107 | 2.42547E-05 | 0.673171208 | 1.99693E-09 |
| Eif1a | 0.002388317 | -0.722315438 | 0.952 | 0.951 | 1 | 9.98782E-10 | -1.044561307 | 0.956 | 0.99 | 2.42624E-05 | 0.002388317 | 1.99756E-09 |
| Ndufb11 | 1.06488E-09 | -2.224607328 | 0.619 | 0.937 | 2.58682E-05 | 3.8807E-08 | -0.883742568 | 0.889 | 0.938 | 0.000942699 | 3.8807E-08 | 2.12977E-09 |
| Slc25a25 | 0.734738016 | 0.845952743 | 0.19 | 0.179 | 1 | 1.09212E-09 | 1.58153456 | 0.711 | 0.246 | 2.65297E-05 | 0.734738016 | 2.18423E-09 |
| Lmna | 0.000552108 | -0.972231518 | 1 | 0.989 | 1 | 1.1559E-09 | -1.05518541 | 1 | 0.997 | 2.80792E-05 | 0.000552108 | 2.31181E-09 |
| Nop10 | 1.15945E-09 | -2.594185317 | 0.381 | 0.929 | 2.81654E-05 | 1.49986E-07 | -1.152433376 | 0.711 | 0.862 | 0.00364346 | 1.49986E-07 | 2.3189E-09 |
| Lcn2 | 0.583271652 | -0.818081329 | 0.048 | 0.026 | 1 | 1.16153E-09 | 0.905465008 | 0.444 | 0.097 | 2.82159E-05 | 0.583271652 | 2.32306E-09 |
| Rps28 | 3.04124E-08 | -1.657476955 | 0.857 | 0.981 | 0.000738777 | 1.18038E-09 | -0.97029442 | 0.956 | 0.969 | 2.86739E-05 | 3.04124E-08 | 2.36077E-09 |
| Cotl1 | 0.059844218 | 1.59848364 | 0.333 | 0.172 | 1 | 1.19967E-09 | 1.176112597 | 0.667 | 0.204 | 2.91424E-05 | 0.059844218 | 2.39934E-09 |
| Ccnd2 | 2.58234E-05 | -2.329334143 | 0.571 | 0.802 | 0.627301762 | 1.25591E-09 | -1.77578368 | 0.867 | 0.907 | 3.05087E-05 | 2.58234E-05 | 2.51183E-09 |
| Crispld1 | 2.36842E-07 | -2.978176843 | 0.333 | 0.907 | 0.005753377 | 1.26287E-09 | -1.128052119 | 0.378 | 0.882 | 3.06776E-05 | 2.36842E-07 | 2.52574E-09 |
| Shb | 0.333747872 | 1.407877708 | 0.238 | 0.175 | 1 | 1.30853E-09 | 1.545569062 | 0.711 | 0.246 | 3.17869E-05 | 0.333747872 | 2.61706E-09 |
| Eif5a | 1.3625E-08 | -1.798854266 | 0.81 | 0.985 | 0.000330979 | 1.32066E-09 | -0.906536279 | 0.911 | 0.979 | 3.20814E-05 | 1.3625E-08 | 2.64131E-09 |
| Peg3 | 0.05509231 | -0.702997289 | 1 | 0.966 | 1 | 1.35886E-09 | -1.009324603 | 0.956 | 1 | 3.30094E-05 | 0.05509231 | 2.71771E-09 |
| Ddx3x | 1.25631E-05 | -1.205607264 | 0.81 | 0.959 | 0.305183344 | 1.412E-09 | -1.079155267 | 0.933 | 0.972 | 3.43003E-05 | 1.25631E-05 | 2.824E-09 |
| Gm56768 | 0.022882332 | -1.679865187 | 0.19 | 0.466 | 1 | 1.42688E-09 | -1.80725304 | 0.156 | 0.74 | 3.46618E-05 | 0.022882332 | 2.85376E-09 |
| Sec62 | 1.98055E-07 | -1.11692581 | 1 | 0.989 | 0.004811155 | 1.42927E-09 | -0.849579412 | 1 | 0.997 | 3.47198E-05 | 1.98055E-07 | 2.85854E-09 |
| Pmepa1 | 0.001304449 | -0.831302517 | 0.952 | 0.963 | 1 | 1.48434E-09 | -0.939088533 | 1 | 0.993 | 3.60575E-05 | 0.001304449 | 2.96867E-09 |
| Srsf3 | 7.43298E-06 | -1.24822572 | 0.952 | 0.978 | 0.180561935 | 1.48591E-09 | -0.850016074 | 0.978 | 0.993 | 3.60958E-05 | 7.43298E-06 | 2.97183E-09 |
| Mesd | 5.11122E-06 | -2.448624864 | 0.143 | 0.705 | 0.12416184 | 1.4891E-09 | -1.846450928 | 0.2 | 0.747 | 3.61733E-05 | 5.11122E-06 | 2.9782E-09 |
| Ncl | 0.000111954 | -0.884411402 | 0.952 | 0.993 | 1 | 1.53011E-09 | -0.897031182 | 1 | 0.997 | 3.71695E-05 | 0.000111954 | 3.06022E-09 |
| Yif1b | 1.22622E-06 | -4.59769126 | 0 | 0.601 | 0.029787337 | 1.54883E-09 | -2.296749532 | 0.089 | 0.619 | 3.76243E-05 | 1.22622E-06 | 3.09767E-09 |
| 9530068E07Rik | 1.58741E-09 | -2.518247005 | 0.238 | 0.918 | 3.85615E-05 | 4.76957E-09 | -1.143695039 | 0.644 | 0.92 | 0.000115862 | 4.76957E-09 | 3.17483E-09 |
| Tgfb3 | 0.001706666 | -2.724259306 | 0.048 | 0.41 | 1 | 1.58862E-09 | -2.043551064 | 0.133 | 0.675 | 3.85907E-05 | 0.001706666 | 3.17724E-09 |
| Cope | 2.06867E-08 | -1.852401402 | 0.571 | 0.951 | 0.000502521 | 1.59956E-09 | -1.222240431 | 0.733 | 0.913 | 3.88565E-05 | 2.06867E-08 | 3.19912E-09 |
| Tm2d2 | 1.62824E-09 | -3.623144439 | 0.095 | 0.832 | 3.95532E-05 | 1.73175E-06 | -1.115422359 | 0.356 | 0.754 | 0.04206764 | 1.73175E-06 | 3.25648E-09 |
| Set | 0.00253991 | -0.686881696 | 0.857 | 0.948 | 1 | 1.65981E-09 | -0.941887982 | 0.978 | 0.986 | 4.03202E-05 | 0.00253991 | 3.31963E-09 |
| Aldoa | 7.15547E-06 | -1.287195501 | 0.81 | 0.959 | 0.173820593 | 1.74423E-09 | -1.043713228 | 0.933 | 0.952 | 4.23709E-05 | 7.15547E-06 | 3.48847E-09 |
| Mageh1 | 0.000198339 | -2.359651234 | 0.095 | 0.545 | 1 | 1.921E-09 | -2.350350415 | 0.133 | 0.661 | 4.66648E-05 | 0.000198339 | 3.84199E-09 |
| Ctsd | 3.00964E-09 | -2.207807739 | 0.667 | 0.97 | 7.31102E-05 | 1.99134E-09 | -1.102770115 | 0.911 | 0.979 | 4.83736E-05 | 3.00964E-09 | 3.98268E-09 |
| Unc50 | 1.37445E-08 | -3.99156415 | 0.048 | 0.746 | 0.000333882 | 2.00119E-09 | -1.92295153 | 0.178 | 0.716 | 4.86129E-05 | 1.37445E-08 | 4.00238E-09 |
| Nap1l1 | 0.033242386 | -0.605182641 | 0.952 | 0.959 | 1 | 2.01737E-09 | -1.097695333 | 0.956 | 0.983 | 4.9006E-05 | 0.033242386 | 4.03475E-09 |
| Yipf3 | 2.37047E-08 | -3.359651234 | 0.048 | 0.761 | 0.000575834 | 2.09889E-09 | -1.666076977 | 0.178 | 0.737 | 5.09861E-05 | 2.37047E-08 | 4.19777E-09 |
| Fuca1 | 2.26533E-09 | -2.56830302 | 0.381 | 0.918 | 5.50293E-05 | 4.77707E-08 | -1.206223949 | 0.622 | 0.865 | 0.001160445 | 4.77707E-08 | 4.53066E-09 |
| Tpd52 | 0.055369212 | 0.793353383 | 0.381 | 0.194 | 1 | 2.27406E-09 | 1.645104736 | 0.8 | 0.304 | 5.52414E-05 | 0.055369212 | 4.54812E-09 |
| Ncmap | 7.2796E-07 | -2.19339255 | 0.381 | 0.858 | 0.017683599 | 2.32403E-09 | -1.847316015 | 0.356 | 0.792 | 5.64554E-05 | 7.2796E-07 | 4.64806E-09 |
| Tmx1 | 2.3468E-09 | -3.063193826 | 0.19 | 0.858 | 5.70085E-05 | 1.83601E-07 | -1.088852846 | 0.689 | 0.879 | 0.004460046 | 1.83601E-07 | 4.69361E-09 |
| Hnrnpa1 | 0.012566982 | -0.643710422 | 0.952 | 0.959 | 1 | 2.35242E-09 | -1.05716814 | 0.956 | 0.958 | 5.71449E-05 | 0.012566982 | 4.70483E-09 |
| Praf2 | 0.000494683 | -1.632615476 | 0.143 | 0.59 | 1 | 2.36113E-09 | -2.521498558 | 0.067 | 0.592 | 5.73566E-05 | 0.000494683 | 4.72226E-09 |
| Cd55 | 0.158039963 | 0.866416846 | 0.286 | 0.164 | 1 | 2.48314E-09 | 1.098110085 | 0.711 | 0.228 | 6.03206E-05 | 0.158039963 | 4.96629E-09 |
| Mydgf | 0.000757266 | -1.052715309 | 0.333 | 0.828 | 1 | 2.49339E-09 | -1.951133434 | 0.244 | 0.723 | 6.05693E-05 | 0.000757266 | 4.98677E-09 |
| Ninj1 | 9.93578E-05 | -2.187521961 | 0.286 | 0.72 | 1 | 2.49514E-09 | -1.907888655 | 0.2 | 0.737 | 6.0612E-05 | 9.93578E-05 | 4.99029E-09 |
| Sf3b4 | 0.000114316 | -2.455511249 | 0.048 | 0.522 | 1 | 2.55892E-09 | -2.167234481 | 0.222 | 0.689 | 6.21614E-05 | 0.000114316 | 5.11785E-09 |
| Tm9sf1 | 7.88788E-05 | -1.915478091 | 0.143 | 0.664 | 1 | 2.62107E-09 | -1.58596056 | 0.289 | 0.796 | 6.36711E-05 | 7.88788E-05 | 5.24215E-09 |
| Copb1 | 0.001443043 | -0.939759885 | 0.714 | 0.963 | 1 | 2.64031E-09 | -1.040450745 | 0.933 | 0.976 | 6.41384E-05 | 0.001443043 | 5.28062E-09 |
| Krt17 | 0.000197728 | 2.351843673 | 0.333 | 0.082 | 1 | 2.64944E-09 | 1.933050839 | 0.467 | 0.111 | 6.43602E-05 | 0.000197728 | 5.29888E-09 |
| Lyn | 0.337380639 | -1.735619168 | 0.048 | 0.112 | 1 | 2.77268E-09 | 1.178841358 | 0.733 | 0.246 | 6.73539E-05 | 0.337380639 | 5.54535E-09 |
| Lman1 | 1.35553E-07 | -1.738041831 | 0.619 | 0.959 | 0.003292843 | 2.78663E-09 | -1.120507292 | 0.733 | 0.958 | 6.76928E-05 | 1.35553E-07 | 5.57326E-09 |
| Emc10 | 6.04647E-06 | -1.897770217 | 0.238 | 0.784 | 0.146880757 | 2.84026E-09 | -1.873320762 | 0.311 | 0.727 | 6.89957E-05 | 6.04647E-06 | 5.68053E-09 |
| Nrep | 8.85108E-07 | -2.702023199 | 0.286 | 0.84 | 0.021501056 | 2.8546E-09 | -1.436906023 | 0.333 | 0.879 | 6.9344E-05 | 8.85108E-07 | 5.7092E-09 |
| Ctnnb1 | 0.000721258 | -0.926787588 | 0.81 | 0.959 | 1 | 2.86167E-09 | -0.835955644 | 1 | 0.997 | 6.95157E-05 | 0.000721258 | 5.72334E-09 |
| Ext1 | 0.011693233 | 1.884596158 | 0.905 | 0.97 | 1 | 2.88382E-09 | 1.224501251 | 1 | 0.948 | 7.00536E-05 | 0.011693233 | 5.76763E-09 |
| Mcfd2 | 3.544E-05 | -1.379339569 | 0.238 | 0.843 | 0.860908591 | 2.91601E-09 | -1.542995493 | 0.222 | 0.761 | 7.08357E-05 | 3.544E-05 | 5.83202E-09 |
| Epb41l4a | 0.492590603 | 0.673771768 | 0 | 0.022 | 1 | 2.98376E-09 | 2.055041364 | 0.4 | 0.08 | 7.24814E-05 | 0.492590603 | 5.96751E-09 |
| U2af1 | 0.002083102 | -0.877638173 | 0.524 | 0.869 | 1 | 3.01712E-09 | -1.015757879 | 0.889 | 0.927 | 7.3292E-05 | 0.002083102 | 6.03425E-09 |
| Golm2 | 2.64041E-05 | -0.849790188 | 0.381 | 0.881 | 0.641409396 | 3.03563E-09 | -1.297257434 | 0.6 | 0.896 | 7.37416E-05 | 2.64041E-05 | 6.07126E-09 |
| Stt3a | 0.085724371 | -0.310798192 | 0.714 | 0.884 | 1 | 3.0991E-09 | -0.9655282 | 0.778 | 0.941 | 7.52833E-05 | 0.085724371 | 6.1982E-09 |
| Hexa | 3.18191E-07 | -2.321210158 | 0.238 | 0.828 | 0.007729492 | 3.17057E-09 | -2.242926832 | 0.222 | 0.73 | 7.70195E-05 | 3.18191E-07 | 6.34114E-09 |
| Slc16a4 | 1.82716E-05 | -1.929112641 | 0.19 | 0.761 | 0.443853005 | 3.17635E-09 | -1.572117701 | 0.378 | 0.858 | 7.71599E-05 | 1.82716E-05 | 6.3527E-09 |
| Rcn2 | 1.42575E-08 | -1.775535634 | 0.667 | 0.985 | 0.000346344 | 3.23908E-09 | -1.326911502 | 0.578 | 0.92 | 7.86837E-05 | 1.42575E-08 | 6.47816E-09 |
| Btg1 | 0.000520483 | -1.065898896 | 1 | 0.996 | 1 | 3.30416E-09 | -0.99147424 | 1 | 1 | 8.02647E-05 | 0.000520483 | 6.60832E-09 |
| Magoh | 8.93654E-05 | -0.900416261 | 0.381 | 0.862 | 1 | 3.35529E-09 | -1.145090798 | 0.822 | 0.896 | 8.15067E-05 | 8.93654E-05 | 6.71058E-09 |
| Hnrnph1 | 0.051461102 | -0.354636648 | 0.952 | 0.97 | 1 | 3.46027E-09 | -0.915100788 | 0.978 | 0.99 | 8.40568E-05 | 0.051461102 | 6.92054E-09 |
| Mctp2 | 0.305227842 | 0.766881172 | 0.381 | 0.31 | 1 | 3.61522E-09 | 1.747202924 | 0.467 | 0.111 | 8.7821E-05 | 0.305227842 | 7.23044E-09 |

| | | | | | | | | | | | | |
|---|---|---|---|---|---|---|---|---|---|---|---|---|
| Chst12 | 3.62621E-09 | -3.857609693 | 0.095 | 0.799 | 8.80879E-05 | 3.7309E-07 | -1.564854927 | 0.222 | 0.699 | 0.009063095 | 3.7309E-07 | 7.25242E-09 |
| Fkbp1a | 0.000102808 | -1.142921019 | 0.667 | 0.918 | 1 | 3.74433E-09 | -1.156770189 | 0.756 | 0.927 | 9.09573E-05 | 0.000102808 | 7.48866E-09 |
| Rpl36al | 2.05043E-07 | -1.429132124 | 0.81 | 0.989 | 0.004980913 | 3.89058E-09 | -0.989644357 | 0.911 | 0.972 | 9.451E-05 | 2.05043E-07 | 7.78116E-09 |
| Dtymk | 0.000423991 | -1.880817084 | 0.143 | 0.601 | 1 | 3.94319E-09 | -2.787247349 | 0.044 | 0.55 | 9.57879E-05 | 0.000423991 | 7.88637E-09 |
| Fis1 | 4.00873E-09 | -2.471905688 | 0.381 | 0.881 | 9.73801E-05 | 8.94331E-06 | -0.836001979 | 0.8 | 0.841 | 0.217250982 | 8.94331E-06 | 8.01746E-09 |
| Ltf | 0.078272979 | 0.731257262 | 0.476 | 0.299 | 1 | 4.02735E-09 | 1.61532798 | 0.689 | 0.249 | 9.78324E-05 | 0.078272979 | 8.0547E-09 |
| Srpr | 3.11307E-08 | -1.339470809 | 0.857 | 0.985 | 0.000756228 | 4.21778E-09 | -0.985424239 | 0.933 | 0.972 | 0.000102458 | 3.11307E-08 | 8.43555E-09 |
| Gng10 | 2.2205E-06 | -2.570988467 | 0.143 | 0.72 | 0.053940348 | 4.22843E-09 | -1.815887397 | 0.178 | 0.727 | 0.000102717 | 2.2205E-06 | 8.45686E-09 |
| Pde4d | 6.80024E-06 | 2.10712945 | 0.952 | 0.91 | 0.165191321 | 4.25493E-09 | 1.978774361 | 1 | 0.896 | 0.000103361 | 6.80024E-06 | 8.50986E-09 |
| Strap | 0.000181188 | -1.19464468 | 0.429 | 0.825 | 1 | 4.25768E-09 | -1.309547883 | 0.733 | 0.872 | 0.000103428 | 0.000181188 | 8.51536E-09 |
| Col17a1 | 0.359280535 | 3.673771768 | 0.048 | 0.019 | 1 | 4.52015E-09 | 1.203079645 | 0.4 | 0.08 | 0.000109803 | 0.359280535 | 9.0403E-09 |
| Ctsl | 1.74145E-06 | -2.508125875 | 0.81 | 0.903 | 0.042303191 | 4.55778E-09 | -1.152479228 | 0.956 | 0.962 | 0.000110718 | 1.74145E-06 | 9.11556E-09 |
| Slc39a1 | 0.000181721 | -1.593014773 | 0.143 | 0.631 | 1 | 4.6058E-09 | -1.583067731 | 0.2 | 0.765 | 0.000111884 | 0.000181721 | 9.21161E-09 |
| Ddah1 | 0.359158144 | 1.000656636 | 0.381 | 0.597 | 1 | 4.63094E-09 | -1.396154105 | 0.089 | 0.644 | 0.000112495 | 0.359158144 | 9.26187E-09 |
| Hint1 | 3.36008E-07 | -1.620236242 | 0.762 | 0.933 | 0.008162301 | 4.63485E-09 | -1.070329898 | 0.867 | 0.92 | 0.00011259 | 3.36008E-07 | 9.2697E-09 |
| Atp5j | 8.63052E-08 | -1.723275807 | 0.714 | 0.948 | 0.002096526 | 4.77857E-09 | -1.035745661 | 0.911 | 0.958 | 0.000116081 | 8.63052E-08 | 9.55713E-09 |
| Eif3c | 0.000665929 | -0.718545655 | 0.81 | 0.888 | 1 | 5.0318E-09 | -1.063528305 | 0.933 | 0.948 | 0.000122233 | 0.000665929 | 1.00636E-08 |
| Fam3d | 0.532436372 | 0.351843673 | 0 | 0.019 | 1 | 5.07168E-09 | 3.213587303 | 0.244 | 0.028 | 0.000123201 | 0.532436372 | 1.01434E-08 |
| Ogn | 0.000347196 | -3.430564892 | 0.143 | 0.56 | 1 | 5.12001E-09 | -1.723975361 | 0.333 | 0.844 | 0.000124375 | 0.000347196 | 1.024E-08 |
| Loxl1 | 0.000794852 | -2.677903671 | 0.095 | 0.5 | 1 | 5.14746E-09 | -1.646723752 | 0.333 | 0.855 | 0.000125042 | 0.000794852 | 1.02949E-08 |
| Srsf9 | 2.90979E-05 | -1.535681598 | 0.286 | 0.81 | 0.706847231 | 5.15064E-09 | -1.33219417 | 0.667 | 0.882 | 0.000125119 | 2.90979E-05 | 1.03013E-08 |
| Pgam1 | 5.16025E-09 | -2.247474121 | 0.524 | 0.937 | 0.000125353 | 5.4842E-09 | -1.259634189 | 0.8 | 0.896 | 0.000133222 | 5.4842E-09 | 1.03205E-08 |
| Tfcp2l1 | 0.921804813 | 1.351843673 | 0.048 | 0.045 | 1 | 5.17248E-09 | 2.0259603 | 0.4 | 0.083 | 0.00012565 | 0.921804813 | 1.0345E-08 |
| Snrpb | 4.74635E-05 | -1.22480356 | 0.667 | 0.918 | 1 | 5.34632E-09 | -1.068341052 | 0.822 | 0.962 | 0.000129873 | 4.74635E-05 | 1.06926E-08 |
| Tceal3 | 0.002009912 | -3.184209227 | 0.048 | 0.384 | 1 | 5.54753E-09 | -3.37942334 | 0.089 | 0.571 | 0.00013476 | 0.002009912 | 1.10951E-08 |
| Lurap1l | 0.39695844 | -0.326228232 | 0 | 0.034 | 1 | 5.55451E-09 | 2.683072586 | 0.222 | 0.021 | 0.00013493 | 0.39695844 | 1.1109E-08 |
| Ndufa4 | 5.55929E-09 | -1.925397226 | 0.81 | 0.97 | 0.000135046 | 6.00611E-08 | -1.001559165 | 0.889 | 0.955 | 0.001459004 | 6.00611E-08 | 1.11186E-08 |
| Tm7sf3 | 0.000153804 | -2.337455488 | 0.095 | 0.556 | 1 | 5.66242E-09 | -1.79913034 | 0.133 | 0.696 | 0.000137551 | 0.000153804 | 1.13248E-08 |
| Matn4 | 0.000133334 | -2.676136023 | 0.238 | 0.69 | 1 | 5.90544E-09 | -1.563159035 | 0.578 | 0.927 | 0.000143455 | 0.000133334 | 1.18109E-08 |
| Nfix | 1.77742E-05 | -1.031508311 | 0.905 | 0.981 | 0.431770812 | 5.94084E-09 | -0.825320186 | 1 | 0.993 | 0.000144315 | 1.77742E-05 | 1.18817E-08 |
| Sdf4 | 1.0074E-07 | -1.678973919 | 0.476 | 0.955 | 0.002447166 | 5.95029E-09 | -1.061466228 | 0.756 | 0.907 | 0.000144544 | 1.0074E-07 | 1.19006E-08 |
| P3h1 | 2.90433E-07 | -2.252227651 | 0.19 | 0.828 | 0.007055191 | 6.17563E-09 | -1.401650093 | 0.244 | 0.775 | 0.000150018 | 2.90433E-07 | 1.23513E-08 |
| Smim19 | 0.000665396 | -2.461387816 | 0.143 | 0.582 | 1 | 6.29812E-09 | -3.205670663 | 0.067 | 0.543 | 0.000152994 | 0.000665396 | 1.25962E-08 |
| Etl4 | 0.228335692 | -1.443685814 | 0.524 | 0.302 | 1 | 6.31595E-09 | 1.187377424 | 0.844 | 0.339 | 0.000153427 | 0.228335692 | 1.26319E-08 |
| Atp5d | 6.37582E-09 | -2.063193826 | 0.667 | 0.944 | 0.000154881 | 1.54364E-07 | -0.974931866 | 0.911 | 0.931 | 0.003749803 | 1.54364E-07 | 1.27516E-08 |
| Arhgef38 | 0.63207476 | 1.351843673 | 0 | 0.011 | 1 | 6.41717E-09 | 2.683072586 | 0.244 | 0.028 | 0.000155886 | 0.63207476 | 1.28343E-08 |
| Ccdc127 | 0.046788239 | -0.543458949 | 0.381 | 0.75 | 1 | 6.50302E-09 | -1.652661308 | 0.222 | 0.761 | 0.000157971 | 0.046788239 | 1.3006E-08 |
| Cpxm2 | 0.004067437 | -3.326228232 | 0.048 | 0.351 | 1 | 6.81047E-09 | -1.73341851 | 0.156 | 0.675 | 0.00016544 | 0.004067437 | 1.36209E-08 |
| Bcl7b | 0.000377739 | -2.04047375 | 0.095 | 0.526 | 1 | 6.82801E-09 | -3.294207337 | 0.044 | 0.533 | 0.000165866 | 0.000377739 | 1.3656E-08 |
| Hspe1 | 6.94794E-09 | -2.121163895 | 0.667 | 0.951 | 0.000168779 | 2.53307E-08 | -1.30591909 | 0.844 | 0.896 | 0.000615333 | 2.53307E-08 | 1.38959E-08 |
| Hspb8 | 0.70287302 | -0.496153234 | 0.143 | 0.168 | 1 | 7.23246E-09 | 1.522607914 | 0.578 | 0.173 | 0.000175691 | 0.70287302 | 1.44649E-08 |
| Ergic3 | 2.68573E-08 | -2.214294954 | 0.286 | 0.862 | 0.000652418 | 7.63501E-09 | -1.269988191 | 0.622 | 0.869 | 0.00018547 | 2.68573E-08 | 1.527E-08 |
| Uqcrh | 4.70261E-07 | -1.260830509 | 0.857 | 0.974 | 0.011423576 | 7.78248E-09 | -0.849821194 | 0.978 | 0.986 | 0.000189052 | 4.70261E-07 | 1.5565E-08 |
| Lrpap1 | 9.20349E-09 | -2.455511249 | 0.238 | 0.896 | 0.000223571 | 7.83903E-09 | -1.310148881 | 0.467 | 0.872 | 0.000190426 | 9.20349E-09 | 1.56781E-08 |
| 2610035D17Rik | 0.048169339 | 1.27343078 | 0.762 | 0.526 | 1 | 7.89883E-09 | 0.902853794 | 0.844 | 0.363 | 0.000191878 | 0.048169339 | 1.57977E-08 |
| Drap1 | 1.49813E-06 | -1.687914036 | 0.429 | 0.884 | 0.036392572 | 7.95626E-09 | -1.264175747 | 0.733 | 0.862 | 0.000193274 | 1.49813E-06 | 1.59125E-08 |
| 1810037I17Rik | 7.98036E-09 | -2.16543202 | 0.571 | 0.925 | 0.000193859 | 5.4397E-05 | -0.637116274 | 0.8 | 0.886 | 1 | 5.4397E-05 | 1.59607E-08 |
| Ece1 | 0.014589035 | 0.875815544 | 0.571 | 0.284 | 1 | 8.33665E-09 | 1.345631492 | 0.8 | 0.349 | 0.000202514 | 0.014589035 | 1.66733E-08 |
| Tmem263 | 0.004427438 | -0.805200037 | 0.381 | 0.817 | 1 | 8.41863E-09 | -1.279387697 | 0.422 | 0.92 | 0.000204505 | 0.004427438 | 1.68373E-08 |
| Cd164 | 0.002706223 | -0.722156909 | 0.905 | 0.955 | 1 | 8.50289E-09 | -1.154555347 | 0.844 | 0.917 | 0.000206552 | 0.002706223 | 1.70058E-08 |
| Farp1 | 1.21066E-06 | 1.466757448 | 1 | 0.989 | 0.029409337 | 8.74941E-09 | 1.251121704 | 1 | 0.986 | 0.000212541 | 1.21066E-06 | 1.74988E-08 |
| Rfxap | 0.016705453 | -2.171718283 | 0.048 | 0.287 | 1 | 8.76032E-09 | -5.199570463 | 0 | 0.471 | 0.000212806 | 0.016705453 | 1.75206E-08 |
| Tenm4 | 0.613898991 | -1.353028292 | 0.095 | 0.131 | 1 | 8.84715E-09 | 1.180572246 | 0.667 | 0.215 | 0.000214915 | 0.613898991 | 1.76943E-08 |
| Atp5b | 4.42402E-08 | -1.530255886 | 0.857 | 0.963 | 0.001074683 | 8.87572E-09 | -0.829996996 | 1 | 0.983 | 0.000215609 | 4.42402E-08 | 1.77514E-08 |
| S100a11 | 1.73071E-08 | -2.063451428 | 0.762 | 0.966 | 0.000420425 | 8.92375E-09 | -0.94828282 | 0.933 | 0.972 | 0.000216776 | 1.73071E-08 | 1.78475E-08 |
| Fndc3b | 2.43486E-07 | 1.160577625 | 1 | 1 | 0.005914753 | 8.98406E-09 | 0.933534518 | 1 | 1 | 0.000218241 | 2.43486E-07 | 1.79681E-08 |
| Tmem251 | 0.020565989 | -1.126203624 | 0.19 | 0.47 | 1 | 9.04882E-09 | -1.455991452 | 0.111 | 0.626 | 0.000219814 | 0.020565989 | 1.80976E-08 |
| Tomm20 | 6.65576E-07 | -1.281648722 | 0.714 | 0.963 | 0.016168171 | 9.2347E-09 | -0.854892435 | 0.889 | 0.962 | 0.000224329 | 6.65576E-07 | 1.84694E-08 |
| Ykt6 | 0.001311688 | -1.605451876 | 0.143 | 0.549 | 1 | 9.23803E-09 | -1.486852415 | 0.156 | 0.696 | 0.00022441 | 0.001311688 | 1.84761E-08 |
| Sem1 | 9.42845E-09 | -1.743099389 | 0.81 | 0.974 | 0.000229036 | 6.02971E-06 | -0.613711521 | 1 | 0.99 | 0.146473597 | 6.02971E-06 | 1.88569E-08 |
| AI506816 | 4.13783E-05 | -1.631834022 | 0.381 | 0.843 | 1 | 9.44295E-09 | -1.644854249 | 0.267 | 0.792 | 0.000229388 | 4.13783E-05 | 1.88859E-08 |
| Srp19 | 0.00025218 | -1.202186968 | 0.381 | 0.828 | 1 | 9.93472E-09 | -1.399916779 | 0.556 | 0.837 | 0.000241334 | 0.00025218 | 1.98694E-08 |
| Tes | 0.493350967 | 0.316219763 | 0.048 | 0.097 | 1 | 1.02211E-08 | 1.905465008 | 0.444 | 0.107 | 0.000248292 | 0.493350967 | 2.04423E-08 |
| Srp14 | 5.53697E-08 | -1.675637064 | 0.619 | 0.948 | 0.001345041 | 1.04858E-08 | -1.009530638 | 0.844 | 0.927 | 0.000254722 | 5.53697E-08 | 2.09717E-08 |
| Nedd8 | 3.98011E-08 | -1.921174822 | 0.571 | 0.922 | 0.000966848 | 1.0795E-08 | -1.192567075 | 0.822 | 0.886 | 0.000262232 | 3.98011E-08 | 2.159E-08 |
| Atp5j2 | 1.12226E-08 | -2.244174016 | 0.571 | 0.907 | 0.000272619 | 2.05762E-05 | -0.847669524 | 0.822 | 0.862 | 0.499837655 | 2.05762E-05 | 2.24452E-08 |
| 1500011B03Rik | 7.44693E-05 | -3.054148687 | 0.048 | 0.511 | 1 | 1.13442E-08 | -2.845707079 | 0.067 | 0.55 | 0.000275574 | 7.44693E-05 | 2.26885E-08 |
| Krtcap2 | 6.89418E-08 | -2.045617053 | 0.381 | 0.903 | 0.001674734 | 1.15798E-08 | -1.509283461 | 0.578 | 0.827 | 0.000281296 | 6.89418E-08 | 2.31596E-08 |
| Aopep | 1.9347E-06 | 1.181096857 | 1 | 1 | 0.046997704 | 1.16316E-08 | 1.092215705 | 1 | 1 | 0.000282555 | 1.9347E-06 | 2.32632E-08 |
| Psmd9 | 0.004166904 | -0.99156415 | 0.095 | 0.463 | 1 | 1.1665E-08 | -3.253565353 | 0.067 | 0.533 | 0.000283366 | 0.004166904 | 2.333E-08 |
| Hnrnpa2b1 | 0.000157344 | -0.765742376 | 1 | 1 | 1 | 1.18779E-08 | -0.702548793 | 1 | 1 | 0.000288537 | 0.000157344 | 2.37557E-08 |
| Serpinf1 | 0.000887431 | -3.897348808 | 0.095 | 0.466 | 1 | 1.21898E-08 | -3.055019673 | 0.156 | 0.64 | 0.000296115 | 0.000887431 | 2.43796E-08 |
| Aldh7a1 | 0.012831295 | -0.402331211 | 0.19 | 0.586 | 1 | 1.22168E-08 | -2.420215222 | 0.067 | 0.574 | 0.00029677 | 0.012831295 | 2.44336E-08 |
| Sspn | 9.9009E-06 | -2.403043829 | 0.143 | 0.683 | 0.240512572 | 1.25686E-08 | -1.528977891 | 0.2 | 0.74 | 0.000305317 | 9.9009E-06 | 2.51373E-08 |
| Atraid | 9.46011E-07 | -1.906559284 | 0.238 | 0.847 | 0.02298049 | 1.30319E-08 | -1.442672967 | 0.511 | 0.81 | 0.00031657 | 9.46011E-07 | 2.60637E-08 |
| Nbdy | 2.28722E-05 | -1.763633545 | 0.19 | 0.743 | 0.555610809 | 1.35168E-08 | -2.10265232 | 0.111 | 0.612 | 0.000328351 | 2.28722E-05 | 2.70336E-08 |
| Eef1d | 9.6356E-06 | -1.298327917 | 0.714 | 0.937 | 0.234068004 | 1.38441E-08 | -1.084332956 | 0.822 | 0.931 | 0.000336302 | 9.6356E-06 | 2.76883E-08 |
| Foxc1 | 1.46005E-05 | -1.700572221 | 0.476 | 0.884 | 0.354675 | 1.40992E-08 | -1.115669206 | 0.4 | 0.931 | 0.000342497 | 1.46005E-05 | 2.81983E-08 |
| Fam241a | 0.173729397 | -0.970084422 | 0 | 0.082 | 1 | 1.41691E-08 | 1.717838004 | 0.444 | 0.107 | 0.000344197 | 0.173729397 | 2.83383E-08 |
| Uqcrb | 2.43178E-06 | -1.429919535 | 0.619 | 0.933 | 0.059072896 | 1.60811E-08 | -1.149817428 | 0.8 | 0.917 | 0.000390642 | 2.43178E-06 | 3.21622E-08 |
| Lox | 0.063189449 | -0.622524015 | 0.476 | 0.728 | 1 | 1.6168E-08 | -0.627896297 | 0.289 | 0.796 | 0.000392753 | 0.063189449 | 3.2336E-08 |
| Taf7 | 0.055358748 | -0.207807739 | 0.238 | 0.563 | 1 | 1.61888E-08 | -2.459885368 | 0.156 | 0.626 | 0.000393258 | 0.055358748 | 3.23776E-08 |
| Tmem176b | 1.63675E-08 | -2.234620853 | 0.571 | 0.918 | 0.000397598 | 8.07502E-07 | -0.833090918 | 0.889 | 0.917 | 0.019615843 | 8.07502E-07 | 3.27349E-08 |
| Tmem167b | 0.000540458 | -1.76585137 | 0.143 | 0.56 | 1 | 1.70859E-08 | -2.10265232 | 0.133 | 0.637 | 0.000415051 | 0.000540458 | 3.41719E-08 |
| Tsc22d3 | 0.00717559 | -1.136483045 | 0.81 | 0.918 | 1 | 1.75223E-08 | -1.318196833 | 0.778 | 0.927 | 0.000425652 | 0.00717559 | 3.50447E-08 |
| Med28 | 1.79149E-08 | -3.535681598 | 0.095 | 0.765 | 0.00043519 | 1.16489E-07 | -1.004983408 | 0.422 | 0.775 | 0.002829763 | 1.16489E-07 | 3.58299E-08 |
| Cystm1 | 0.892568042 | -0.599246727 | 0.381 | 0.369 | 1 | 1.81393E-08 | 1.65505821 | 0.622 | 0.208 | 0.000440641 | 0.892568042 | 3.62787E-08 |
| Tagln2 | 0.006522237 | -1.388210103 | 0.667 | 0.743 | 1 | 1.81688E-08 | -1.215634142 | 0.978 | 0.955 | 0.000441356 | 0.006522237 | 3.63376E-08 |
| Cox6c | 3.352E-08 | -1.607625743 | 0.81 | 0.97 | 0.000814267 | 1.85702E-08 | -0.85021388 | 0.933 | 0.952 | 0.000451108 | 3.352E-08 | 3.71404E-08 |
| Leo1 | 0.183278147 | -0.480033568 | 0.19 | 0.369 | 1 | 1.9022E-08 | -3.093031402 | 0.067 | 0.536 | 0.000462081 | 0.183278147 | 3.80439E-08 |
| Lncpint | 0.000485452 | 1.523418822 | 0.952 | 0.933 | 1 | 1.93675E-08 | 1.438174977 | 0.978 | 0.955 | 0.000470475 | 0.000485452 | 3.87349E-08 |
| Rpl35 | 4.68103E-08 | -1.700797807 | 0.762 | 0.963 | 0.001137115 | 1.94828E-08 | -1.132727404 | 0.889 | 0.931 | 0.000473277 | 4.68103E-08 | 3.89656E-08 |
| H19 | 0.186098499 | -1.33828985 | 0.381 | 0.504 | 1 | 2.00697E-08 | -2.768138526 | 0.178 | 0.657 | 0.000487534 | 0.186098499 | 4.01395E-08 |
| Bet1l | 0.000871302 | -1.180888869 | 0.238 | 0.724 | 1 | 2.08633E-08 | -1.66806743 | 0.156 | 0.692 | 0.000506811 | 0.000871302 | 4.17266E-08 |
| Lrrc58 | 5.44718E-05 | -1.081115734 | 0.571 | 0.929 | 1 | 2.09458E-08 | -0.916150608 | 0.933 | 0.983 | 0.000508815 | 5.44718E-05 | 4.18916E-08 |
| Snu13 | 3.25445E-05 | -1.228457451 | 0.667 | 0.933 | 0.790569863 | 2.09868E-08 | -1.139727394 | 0.8 | 0.91 | 0.000509811 | 3.25445E-05 | 4.19735E-08 |
| Dynlrb1 | 1.28487E-06 | -1.521688326 | 0.667 | 0.918 | 0.031212001 | 2.18695E-08 | -1.100586243 | 0.8 | 0.896 | 0.000531255 | 1.28487E-06 | 4.37391E-08 |
| Mfge8 | 1.8192E-06 | -2.699093292 | 0.333 | 0.858 | 0.044191959 | 2.20843E-08 | -1.747718984 | 0.444 | 0.824 | 0.000536471 | 1.8192E-06 | 4.41685E-08 |

| | | | | | | | | | | | | |
|---|---|---|---|---|---|---|---|---|---|---|---|---|
| Ebp | 7.34063E-06 | -2.921174822 | 0.048 | 0.612 | 0.178318622 | 2.21977E-08 | -4.931637258 | 0 | 0.45 | 0.000539226 | 7.34063E-06 | 4.43954E-08 |
| Arhgap15 | 0.492599419 | -1.752492987 | 0 | 0.022 | 1 | 2.28992E-08 | 1.335149283 | 0.222 | 0.024 | 0.000556267 | 0.492599419 | 4.57984E-08 |
| Ap1s3 | 0.285569273 | -0.911190733 | 0 | 0.052 | 1 | 2.30415E-08 | 3.168499413 | 0.311 | 0.055 | 0.000559724 | 0.285569273 | 4.6083E-08 |
| Xbp1 | 2.39124E-05 | -1.359003268 | 0.524 | 0.925 | 0.580878972 | 2.31066E-08 | -1.339967655 | 0.756 | 0.879 | 0.000561307 | 2.39124E-05 | 4.62133E-08 |
| Grcc10 | 3.23308E-07 | -1.959742478 | 0.381 | 0.881 | 0.007853797 | 2.3223E-08 | -1.371916648 | 0.778 | 0.841 | 0.000564132 | 3.23308E-07 | 4.64459E-08 |
| Mrpl4 | 0.046589244 | -0.399477214 | 0.238 | 0.53 | 1 | 2.34594E-08 | -2.180425414 | 0.089 | 0.574 | 0.000569876 | 0.046589244 | 4.69188E-08 |
| Fzd9 | 8.86158E-07 | -3.683780237 | 0.095 | 0.705 | 0.021526553 | 2.3598E-08 | -1.412994824 | 0.244 | 0.768 | 0.000573244 | 8.86158E-07 | 4.71961E-08 |
| Klhdc2 | 0.007495421 | -0.928763362 | 0.333 | 0.701 | 1 | 2.36443E-08 | -1.263756755 | 0.356 | 0.841 | 0.000574367 | 0.007495421 | 4.72886E-08 |
| Capn6 | 8.19583E-05 | -3.785659851 | 0.048 | 0.526 | 1 | 2.39039E-08 | -1.644981612 | 0.267 | 0.785 | 0.000580673 | 8.19583E-05 | 4.78078E-08 |
| Ablim1 | 0.028258969 | 1.927220125 | 0.381 | 0.213 | 1 | 2.39994E-08 | 1.460680165 | 0.711 | 0.253 | 0.000582993 | 0.028258969 | 4.79987E-08 |
| Gps1 | 0.002354197 | -1.793833782 | 0.095 | 0.455 | 1 | 2.40734E-08 | -2.503925929 | 0.067 | 0.54 | 0.000584791 | 0.002354197 | 4.81468E-08 |
| Rnase4 | 0.004310051 | -1.723820597 | 0.524 | 0.75 | 1 | 2.42201E-08 | -0.682136376 | 0.644 | 0.931 | 0.000588354 | 0.004310051 | 4.84401E-08 |
| Fbln7 | 0.001154097 | -2.990142074 | 0.048 | 0.425 | 1 | 2.49411E-08 | -1.374508965 | 0.356 | 0.834 | 0.000605869 | 0.001154097 | 4.98822E-08 |
| Gaa | 5.91403E-05 | -1.849790188 | 0.238 | 0.716 | 1 | 2.4982E-08 | -1.705805753 | 0.2 | 0.699 | 0.000606862 | 5.91403E-05 | 4.99639E-08 |
| Os9 | 2.53432E-08 | -1.942777076 | 0.429 | 0.925 | 0.000615636 | 0.000533512 | -0.427351404 | 0.889 | 0.917 | 1 | 0.000533512 | 5.06863E-08 |
| Reck | 0.000254372 | -1.409828434 | 0.143 | 0.616 | 1 | 2.60422E-08 | -1.434384995 | 0.311 | 0.827 | 0.000632618 | 0.000254372 | 5.20845E-08 |
| Cox8a | 2.60629E-08 | -1.615067487 | 0.81 | 0.978 | 0.000633121 | 7.15609E-08 | -0.885094672 | 0.933 | 0.958 | 0.001738358 | 7.15609E-08 | 5.21259E-08 |
| Trp53i13 | 8.86152E-06 | -3.081115734 | 0.048 | 0.575 | 0.21526414 | 2.70502E-08 | -2.294207337 | 0.067 | 0.554 | 0.000657104 | 8.86152E-06 | 5.41004E-08 |
| Txndc5 | 0.000367205 | -1.215377246 | 0.524 | 0.918 | 1 | 2.72634E-08 | -1.166593141 | 0.422 | 0.92 | 0.000662282 | 0.000367205 | 5.45268E-08 |
| Rer1 | 5.57874E-08 | -1.826074119 | 0.476 | 0.922 | 0.001355187 | 2.75631E-08 | -0.989816421 | 0.644 | 0.893 | 0.000669562 | 5.57874E-08 | 5.51261E-08 |
| Clcn4 | 7.14513E-05 | -2.154047257 | 0.143 | 0.619 | 1 | 2.83448E-08 | -1.668912743 | 0.244 | 0.699 | 0.000688552 | 7.14513E-05 | 5.66896E-08 |
| Cmip | 2.84649E-08 | 1.537177219 | 1 | 0.929 | 0.00069147 | 3.85075E-07 | 1.092881029 | 0.978 | 0.83 | 0.009354231 | 3.85075E-07 | 5.69299E-08 |
| Spcs2 | 1.72526E-07 | -1.658220011 | 0.571 | 0.937 | 0.004190996 | 2.88525E-08 | -1.03528904 | 0.867 | 0.938 | 0.000700884 | 1.72526E-07 | 5.77049E-08 |
| Sik1 | 0.086860043 | 0.866416846 | 0.381 | 0.22 | 1 | 2.9082E-08 | 1.355710605 | 0.711 | 0.242 | 0.00070646 | 0.086860043 | 5.8164E-08 |
| Ldlrad4 | 7.15379E-07 | 2.30961982 | 1 | 0.925 | 0.017377993 | 2.97825E-08 | 1.175500791 | 1 | 0.972 | 0.000723475 | 7.15379E-07 | 5.95649E-08 |
| Eloc | 7.0073E-05 | -1.31387984 | 0.667 | 0.873 | 1 | 3.0382E-08 | -0.968751869 | 0.844 | 0.927 | 0.00073804 | 7.0073E-05 | 6.07641E-08 |
| Col6a1 | 0.00136547 | -0.732024276 | 1 | 0.981 | 1 | 3.05439E-08 | -1.06549007 | 0.689 | 0.983 | 0.000741973 | 0.00136547 | 6.10878E-08 |
| Anapc11 | 3.06778E-08 | -2.780184721 | 0.19 | 0.81 | 0.000745226 | 8.56377E-06 | -1.121335052 | 0.444 | 0.765 | 0.208030987 | 8.56377E-06 | 6.13557E-08 |
| Atp5g1 | 1.19617E-07 | -2.011991374 | 0.571 | 0.918 | 0.002905744 | 3.08921E-08 | -1.356663166 | 0.8 | 0.865 | 0.00075043 | 1.19617E-07 | 6.17841E-08 |
| Cd9 | 0.001099806 | -0.665092833 | 1 | 1 | 1 | 3.20793E-08 | -0.720900705 | 1 | 1 | 0.000779271 | 0.001099806 | 6.41587E-08 |
| Park7 | 2.43713E-07 | -1.722357269 | 0.619 | 0.896 | 0.005920276 | 3.22392E-08 | -1.205380761 | 0.778 | 0.858 | 0.000783154 | 2.43713E-07 | 6.44784E-08 |
| Dnajb11 | 0.00028008 | -1.436360149 | 0.286 | 0.75 | 1 | 3.22655E-08 | -1.333886241 | 0.556 | 0.834 | 0.000783794 | 0.00028008 | 6.4531E-08 |
| Med13l | 3.2421E-08 | 1.138562302 | 1 | 0.963 | 0.00078757 | 2.7376E-05 | 1.130268753 | 1 | 0.997 | 0.665017346 | 2.7376E-05 | 6.48419E-08 |
| Nsmce3 | 0.002099281 | -2.107587946 | 0.095 | 0.437 | 1 | 3.2588E-08 | -3.071814916 | 0.067 | 0.516 | 0.000791629 | 0.002099281 | 6.51761E-08 |
| Tmem106b | 2.40969E-05 | -2.303508156 | 0.095 | 0.638 | 0.585362384 | 3.30108E-08 | -1.555714273 | 0.222 | 0.734 | 0.000801898 | 2.40969E-05 | 6.60216E-08 |
| Cuedc2 | 0.000215806 | -1.899117901 | 0.095 | 0.571 | 1 | 3.33507E-08 | -2.686161223 | 0.067 | 0.54 | 0.000810156 | 0.000215806 | 6.67015E-08 |
| Med31 | 0.056297951 | -0.260340297 | 0.238 | 0.534 | 1 | 3.39797E-08 | -2.055695251 | 0.111 | 0.588 | 0.000825435 | 0.056297951 | 6.79594E-08 |
| Smim11 | 3.10493E-06 | -2.379339569 | 0.095 | 0.701 | 0.075424847 | 3.42012E-08 | -1.505810573 | 0.222 | 0.713 | 0.000830816 | 3.10493E-06 | 6.84024E-08 |
| Fbn1 | 6.27161E-06 | -1.888979454 | 0.429 | 0.873 | 0.152349925 | 3.4612E-08 | -1.462207213 | 0.6 | 0.945 | 0.000840794 | 6.27161E-06 | 6.9224E-08 |
| Cd151 | 1.68684E-06 | -3.559847909 | 0.048 | 0.627 | 0.040976645 | 3.55429E-08 | -2.960783604 | 0.067 | 0.526 | 0.000863408 | 1.68684E-06 | 7.10858E-08 |
| Ndufa13 | 7.35851E-08 | -1.783933106 | 0.571 | 0.933 | 0.001787529 | 3.6105E-08 | -1.025134015 | 0.822 | 0.896 | 0.000877061 | 7.35851E-08 | 7.22099E-08 |
| Bnip3 | 6.1515E-05 | -2.8244791 | 0.143 | 0.612 | 1 | 3.79091E-08 | -2.040953952 | 0.111 | 0.612 | 0.000920888 | 6.1515E-05 | 7.58182E-08 |
| 1110065P20Rik | 0.000688696 | -1.962852853 | 0.095 | 0.504 | 1 | 3.80004E-08 | -2.960783604 | 0.022 | 0.481 | 0.000923105 | 0.000688696 | 7.60008E-08 |
| Rbbp7 | 0.000633417 | -0.937343021 | 0.667 | 0.914 | 1 | 3.81132E-08 | -0.976810841 | 0.867 | 0.976 | 0.000925847 | 0.000633417 | 7.62265E-08 |
| Gar1 | 0.164530468 | -0.471278565 | 0.286 | 0.478 | 1 | 3.978E-08 | -3.288470968 | 0.067 | 0.512 | 0.000966335 | 0.164530468 | 7.956E-08 |
| Ranbp1 | 7.09853E-06 | -1.695462042 | 0.571 | 0.873 | 0.172437599 | 3.9923E-08 | -1.146836652 | 0.844 | 0.91 | 0.000969808 | 7.09853E-06 | 7.98459E-08 |
| Acan | 0.06698873 | -0.996110483 | 0.905 | 0.985 | 1 | 4.01955E-08 | -1.031130127 | 0.911 | 0.972 | 0.000976429 | 0.06698873 | 8.0391E-08 |
| Nbl1 | 0.00219788 | -1.958819596 | 0.19 | 0.578 | 1 | 4.02641E-08 | -1.553561758 | 0.378 | 0.855 | 0.000978096 | 0.00219788 | 8.05282E-08 |
| Mxra7 | 0.002103178 | -2.146407195 | 0.238 | 0.567 | 1 | 4.18851E-08 | -1.885094672 | 0.156 | 0.654 | 0.001017473 | 0.002103178 | 8.37702E-08 |
| Atp5h | 4.26779E-08 | -1.562324923 | 0.857 | 0.955 | 0.001036731 | 4.90306E-08 | -0.96462567 | 0.867 | 0.927 | 0.00119105 | 4.90306E-08 | 8.53558E-08 |
| Tmed1 | 2.41179E-05 | -2.208871282 | 0.143 | 0.653 | 0.585872236 | 4.27184E-08 | -1.80878051 | 0.111 | 0.602 | 0.001037715 | 2.41179E-05 | 8.54367E-08 |
| Rnasek | 4.2905E-08 | -2.40115672 | 0.571 | 0.925 | 0.001042248 | 0.0020742 | -0.789160315 | 0.8 | 0.782 | 1 | 0.0020742 | 8.581E-08 |
| Eif3k | 0.002341447 | -0.798354924 | 0.762 | 0.922 | 1 | 4.29204E-08 | -1.000831764 | 0.822 | 0.917 | 0.001042623 | 0.002341447 | 8.58409E-08 |
| Inppl1 | 0.04535297 | -1.096302138 | 0.143 | 0.377 | 1 | 4.36447E-08 | -1.9862062 | 0.156 | 0.654 | 0.001060217 | 0.04535297 | 8.72894E-08 |
| Tbc1d8 | 0.460417827 | 2.159198595 | 0.095 | 0.06 | 1 | 4.37954E-08 | 1.875717664 | 0.467 | 0.128 | 0.001063877 | 0.460417827 | 8.75908E-08 |
| S100b | 6.03487E-07 | -2.96430207 | 0.381 | 0.847 | 0.014659916 | 4.50083E-08 | -2.119120631 | 0.2 | 0.685 | 0.00109334 | 6.03487E-07 | 9.00165E-08 |
| AI837181 | 0.026499083 | -1.159118246 | 0.19 | 0.44 | 1 | 4.57314E-08 | -2.776359032 | 0.067 | 0.522 | 0.001110908 | 0.026499083 | 9.14629E-08 |
| Thumpd1 | 0.007388781 | -1.260340297 | 0.143 | 0.496 | 1 | 4.79162E-08 | -3.017367132 | 0.067 | 0.519 | 0.00116398 | 0.007388781 | 9.58323E-08 |
| Tspan31 | 6.48496E-07 | -1.950719097 | 0.286 | 0.851 | 0.015753263 | 4.80791E-08 | -1.323674246 | 0.578 | 0.848 | 0.001167937 | 6.48496E-07 | 9.61581E-08 |
| Creb3 | 1.21201E-05 | -2.763633545 | 0.095 | 0.608 | 0.294421924 | 4.85555E-08 | -1.824722054 | 0.133 | 0.616 | 0.001179511 | 1.21201E-05 | 9.7111E-08 |
| Serinc3 | 4.31393E-07 | -1.661962127 | 0.762 | 0.978 | 0.010479387 | 4.9418E-08 | -0.796687965 | 0.978 | 0.997 | 0.001200463 | 4.31393E-07 | 9.88361E-08 |
| Sra1 | 5.00485E-08 | -2.38800443 | 0.238 | 0.858 | 0.001215778 | 2.05146E-06 | -1.019677293 | 0.578 | 0.817 | 0.049834168 | 2.05146E-06 | 1.00097E-07 |
| Adamts2 | 0.154368031 | -1.718545655 | 0.143 | 0.272 | 1 | 5.05332E-08 | -1.783126965 | 0.2 | 0.661 | 0.001227552 | 0.154368031 | 1.01066E-07 |
| Rab3ip | 0.289489561 | 0.780686972 | 0.048 | 0.134 | 1 | 5.20976E-08 | 2.171173548 | 0.556 | 0.183 | 0.001265554 | 0.289489561 | 1.04195E-07 |
| Cct8 | 0.000379383 | -1.047563175 | 0.571 | 0.903 | 1 | 5.33088E-08 | -0.99001047 | 0.911 | 0.941 | 0.001294978 | 0.000379383 | 1.06618E-07 |
| Slc35b2 | 7.59643E-07 | -3.02666795 | 0.143 | 0.709 | 0.018453249 | 5.33125E-08 | -2.491853096 | 0.111 | 0.571 | 0.001295066 | 7.59643E-07 | 1.06625E-07 |
| Epas1 | 0.074194913 | 0.604609743 | 0.667 | 0.44 | 1 | 5.35503E-08 | 1.417287569 | 0.911 | 0.554 | 0.001300843 | 0.074194913 | 1.07101E-07 |
| Slc50a1 | 0.000372513 | -1.739856161 | 0.19 | 0.638 | 1 | 5.36475E-08 | -2.351573557 | 0.111 | 0.581 | 0.001303206 | 0.000372513 | 1.07295E-07 |
| Stmp1 | 5.37892E-08 | -2.661618587 | 0.238 | 0.858 | 0.001306648 | 5.17316E-07 | -1.254678973 | 0.422 | 0.768 | 0.012566643 | 5.17316E-07 | 1.07578E-07 |
| Psmd7 | 0.004753543 | -0.840350492 | 0.714 | 0.888 | 1 | 5.40168E-08 | -1.008512334 | 0.844 | 0.927 | 0.001312175 | 0.004753543 | 1.08034E-07 |
| Snrpe | 2.50883E-05 | -1.547815354 | 0.571 | 0.84 | 0.609444873 | 5.44063E-08 | -1.134976937 | 0.822 | 0.872 | 0.001321638 | 2.50883E-05 | 1.08813E-07 |
| Cox14 | 5.45252E-08 | -2.167530486 | 0.429 | 0.896 | 0.001324527 | 1.69697E-07 | -1.243358196 | 0.711 | 0.848 | 0.00412228 | 1.69697E-07 | 1.0905E-07 |
| Txnrd1 | 0.014647209 | -0.656004189 | 0.714 | 0.877 | 1 | 5.46107E-08 | -0.797268022 | 0.933 | 0.972 | 0.001326603 | 0.014647209 | 1.09221E-07 |
| Ormdl3 | 8.7873E-05 | -1.233118828 | 0.19 | 0.739 | 1 | 5.56401E-08 | -1.4734319 | 0.422 | 0.789 | 0.001351609 | 8.7873E-05 | 1.1128E-07 |
| Pigp | 5.58509E-08 | -3.09969716 | 0.095 | 0.776 | 0.00135673 | 0.000169759 | -1.031172931 | 0.556 | 0.709 | 1 | 0.000169759 | 1.11702E-07 |
| Rpn2 | 5.59399E-08 | -1.91306802 | 0.429 | 0.91 | 0.001358892 | 2.54106E-07 | -1.097772706 | 0.733 | 0.872 | 0.006172748 | 2.54106E-07 | 1.1188E-07 |
| Gna11 | 1.21153E-07 | -2.490416864 | 0.19 | 0.817 | 0.002943061 | 5.65258E-08 | -1.262746376 | 0.444 | 0.83 | 0.001373125 | 1.21153E-07 | 1.13052E-07 |
| Bmp2 | 0.316104789 | 1.345070444 | 0.333 | 0.493 | 1 | 5.68829E-08 | -1.835252722 | 0.2 | 0.685 | 0.001381799 | 0.316104789 | 1.13766E-07 |
| Lamb2 | 0.001108961 | -1.276541108 | 0.286 | 0.701 | 1 | 5.76959E-08 | -1.340011027 | 0.244 | 0.751 | 0.001401548 | 0.001108961 | 1.15392E-07 |
| Saraf | 4.68344E-05 | -0.982112519 | 0.762 | 0.966 | 1 | 5.79186E-08 | -0.936159817 | 0.844 | 0.955 | 0.001406959 | 4.68344E-05 | 1.15837E-07 |
| Prkar1a | 2.62149E-05 | -1.310537135 | 0.667 | 0.914 | 0.636812857 | 5.79317E-08 | -0.953841994 | 0.956 | 0.955 | 0.001407277 | 2.62149E-05 | 1.15863E-07 |
| Scarf2 | 5.45729E-05 | -2.370622352 | 0.143 | 0.623 | 1 | 5.8698E-08 | -1.255526869 | 0.244 | 0.785 | 0.001425893 | 5.45729E-05 | 1.17396E-07 |
| Bnip2 | 5.08523E-06 | -1.582567986 | 0.333 | 0.888 | 0.123530313 | 6.19806E-08 | -1.004808227 | 0.711 | 0.92 | 0.001505632 | 5.08523E-06 | 1.23961E-07 |
| Fstl1 | 0.032592042 | -1.321995383 | 0.429 | 0.653 | 1 | 6.21789E-08 | -1.196011314 | 0.533 | 0.938 | 0.001510449 | 0.032592042 | 1.24358E-07 |
| Hsd17b10 | 1.55348E-06 | -2.984439715 | 0.048 | 0.657 | 0.037737166 | 6.23445E-08 | -2.062881791 | 0.178 | 0.626 | 0.001514474 | 1.55348E-06 | 1.24689E-07 |
| Trir | 6.24321E-08 | -1.967677924 | 0.381 | 0.888 | 0.0015166 | 2.98637E-06 | -0.872977223 | 0.778 | 0.882 | 0.07254496 | 2.98637E-06 | 1.24864E-07 |
| Gpx4 | 6.33657E-08 | -1.827805839 | 0.81 | 0.922 | 0.00153928 | 1.56204E-07 | -0.875695974 | 0.911 | 0.924 | 0.003794498 | 1.56204E-07 | 1.26731E-07 |
| Ptov1 | 3.91537E-06 | -1.908876867 | 0.286 | 0.817 | 0.095112102 | 6.37349E-08 | -1.190057604 | 0.6 | 0.851 | 0.001548247 | 3.91537E-06 | 1.2747E-07 |
| Efemp2 | 5.42697E-06 | -1.626351957 | 0.286 | 0.825 | 0.131831877 | 6.4262E-08 | -1.605688645 | 0.222 | 0.713 | 0.001561052 | 5.42697E-06 | 1.28524E-07 |
| Manf | 1.65695E-06 | -1.320719653 | 0.667 | 0.933 | 0.040250526 | 6.46843E-08 | -1.193916524 | 0.778 | 0.913 | 0.001571312 | 1.65695E-06 | 1.29369E-07 |
| Cavin3 | 1.10801E-07 | -4.832039786 | 0.048 | 0.701 | 0.002691571 | 6.47426E-08 | -1.646298492 | 0.2 | 0.678 | 0.001572727 | 1.10801E-07 | 1.29485E-07 |
| Ggcx | 0.002598375 | -1.297771786 | 0.19 | 0.59 | 1 | 6.47676E-08 | -1.869153128 | 0.111 | 0.612 | 0.001573335 | 0.002598375 | 1.29535E-07 |
| Itgb4 | 0.446328101 | 1.425844254 | 0.095 | 0.056 | 1 | 6.49886E-08 | 1.591924698 | 0.444 | 0.114 | 0.001578703 | 0.446328101 | 1.29977E-07 |
| Tubb6 | 0.002514936 | -1.117641611 | 0.238 | 0.616 | 1 | 6.53385E-08 | -1.307569359 | 0.156 | 0.661 | 0.001587203 | 0.002514936 | 1.30677E-07 |
| Snx18 | 0.028904989 | -0.595095083 | 0.476 | 0.757 | 1 | 6.58061E-08 | -1.348405645 | 0.6 | 0.889 | 0.001598563 | 0.028904989 | 1.31612E-07 |
| Ftl1-ps1 | 0.000792314 | -3.280424543 | 0 | 0.366 | 1 | 6.78502E-08 | -2.34877628 | 0.067 | 0.533 | 0.001648218 | 0.000792314 | 1.357E-07 |
| Matn3 | 0.00032221 | -2.958314645 | 0.333 | 0.69 | 1 | 6.80191E-08 | -3.506751973 | 0.089 | 0.54 | 0.001652321 | 0.00032221 | 1.36038E-07 |
| Rsl24d1 | 0.00716057 | -1.447243633 | 0.095 | 0.41 | 1 | 6.9039E-08 | -2.416463087 | 0.089 | 0.55 | 0.001677095 | 0.00716057 | 1.38078E-07 |

| | | | | | | | | | | | | |
|---|---|---|---|---|---|---|---|---|---|---|---|---|
| Cct6a | 9.06417E-05 | -1.196945215 | 0.762 | 0.937 | 1 | 6.94773E-08 | -0.964021978 | 0.867 | 0.934 | 0.001687743 | 9.06417E-05 | 1.38955E-07 |
| Slc8a1 | 0.425625368 | -0.496153234 | 0 | 0.03 | 1 | 7.03589E-08 | 0.827462496 | 0.422 | 0.1 | 0.001709158 | 0.425625368 | 1.40718E-07 |
| Trak1 | 7.10854E-08 | 1.791953194 | 0.952 | 0.623 | 0.001726807 | 3.66942E-06 | 1.324984828 | 0.889 | 0.754 | 0.089137652 | 3.66942E-06 | 1.42171E-07 |
| Ndrg2 | 0.016605567 | -0.886943187 | 0.238 | 0.571 | 1 | 7.18127E-08 | -2.085111739 | 0.156 | 0.633 | 0.001744474 | 0.016605567 | 1.43625E-07 |
| Phpt1 | 2.76123E-05 | -3.801961663 | 0 | 0.493 | 0.670758178 | 7.29028E-08 | -2.989352756 | 0.067 | 0.502 | 0.001770956 | 2.76123E-05 | 1.45806E-07 |
| Smg1 | 0.003910178 | 0.892704621 | 0.952 | 0.974 | 1 | 7.31453E-08 | 0.896633441 | 1 | 0.979 | 0.001776845 | 0.003910178 | 1.46291E-07 |
| Cox6a1 | 7.39471E-08 | -1.912496931 | 0.667 | 0.929 | 0.001796323 | 1.22066E-07 | -1.07504966 | 0.933 | 0.917 | 0.00296522 | 1.22066E-07 | 1.47894E-07 |
| Ripk4 | 0.082280431 | 3.088809267 | 0.048 | 0.007 | 1 | 7.42669E-08 | 2.808603468 | 0.244 | 0.035 | 0.001804091 | 0.082280431 | 1.48534E-07 |
| Gnas | 7.43747E-08 | -1.021293063 | 0.952 | 0.996 | 0.001806711 | 2.45829E-07 | -0.661195047 | 1 | 1 | 0.005971682 | 2.45829E-07 | 1.48749E-07 |
| Ldhb | 0.000389028 | -2.492140171 | 0.143 | 0.545 | 1 | 7.48872E-08 | -1.612725777 | 0.578 | 0.81 | 0.001819159 | 0.000389028 | 1.49774E-07 |
| Rtl8a | 4.99977E-05 | -2.481046341 | 0.095 | 0.59 | 1 | 7.54239E-08 | -1.745427173 | 0.133 | 0.623 | 0.001832197 | 4.99977E-05 | 1.50848E-07 |
| Stox2 | 0.218804152 | 1.811275291 | 0.333 | 0.22 | 1 | 7.56826E-08 | 0.609072005 | 0.756 | 0.263 | 0.001838481 | 0.218804152 | 1.51365E-07 |
| Ntan1 | 7.91415E-08 | -4.841928071 | 0 | 0.675 | 0.001922505 | 0.005529433 | -0.486852415 | 0.422 | 0.682 | 1 | 0.005529433 | 1.58283E-07 |
| Psmb4 | 1.84386E-06 | -1.484657595 | 0.619 | 0.903 | 0.044791005 | 8.29617E-08 | -1.208075273 | 0.778 | 0.841 | 0.002015305 | 1.84386E-06 | 1.65923E-07 |
| Rraga | 3.99875E-05 | -1.073462162 | 0.143 | 0.705 | 0.971375421 | 8.37074E-08 | -1.629810369 | 0.333 | 0.73 | 0.00203342 | 3.99875E-05 | 1.67415E-07 |
| Armh4 | 0.012153431 | -0.579208973 | 0.19 | 0.552 | 1 | 8.39857E-08 | -1.815733271 | 0.156 | 0.661 | 0.002040181 | 0.012153431 | 1.67971E-07 |
| Lrrc59 | 0.010775258 | -0.673893889 | 0.476 | 0.769 | 1 | 8.43177E-08 | -1.177721595 | 0.689 | 0.879 | 0.002048246 | 0.010775258 | 1.68635E-07 |
| Rhoc | 8.58196E-08 | -1.644661707 | 0.429 | 0.955 | 0.00208473 | 4.07919E-06 | -0.984630346 | 0.756 | 0.917 | 0.099091739 | 4.07919E-06 | 1.71639E-07 |
| Gabarapl2 | 3.52937E-05 | -1.167983808 | 0.667 | 0.937 | 0.857354718 | 8.72583E-08 | -0.919949886 | 0.844 | 0.962 | 0.002119679 | 3.52937E-05 | 1.74517E-07 |
| Ccdc90b | 0.004248928 | -0.747692001 | 0.19 | 0.604 | 1 | 8.77848E-08 | -2.394929926 | 0.111 | 0.564 | 0.002132467 | 0.004248928 | 1.7557E-07 |
| Slc39a7 | 1.60261E-06 | -1.604677691 | 0.19 | 0.843 | 0.038930526 | 8.86783E-08 | -1.050119845 | 0.622 | 0.913 | 0.002154173 | 1.60261E-06 | 1.77357E-07 |
| B3gnt9 | 0.000879825 | -3.639111188 | 0 | 0.366 | 1 | 9.08032E-08 | -2.913862556 | 0.067 | 0.509 | 0.002205792 | 0.000879825 | 1.81606E-07 |
| Mrps25 | 0.032640431 | -0.678744647 | 0.143 | 0.414 | 1 | 9.13914E-08 | -2.223818009 | 0.044 | 0.491 | 0.002220079 | 0.032640431 | 1.82783E-07 |
| Slc6a8 | 0.113215216 | -0.354425124 | 0.286 | 0.556 | 1 | 9.4047E-08 | -2.022402722 | 0.067 | 0.533 | 0.002284591 | 0.113215216 | 1.88094E-07 |
| Rxylt1 | 0.000961612 | -1.388724158 | 0.333 | 0.716 | 1 | 9.45968E-08 | -1.356455778 | 0.178 | 0.706 | 0.002297945 | 0.000961612 | 1.89194E-07 |
| Atp6ap1 | 3.9838E-05 | -1.109970597 | 0.476 | 0.91 | 0.967745327 | 9.60625E-08 | -1.14894439 | 0.756 | 0.889 | 0.002333551 | 3.9838E-05 | 1.92125E-07 |
| P4ha1 | 9.65919E-08 | -1.495437434 | 0.714 | 0.993 | 0.00234641 | 0.000359745 | -0.512501003 | 0.956 | 0.997 | 1 | 0.000359745 | 1.93184E-07 |
| Kazald1 | 0.000153499 | -3.322466314 | 0.048 | 0.504 | 1 | 9.68228E-08 | -2.49129832 | 0.044 | 0.509 | 0.002352019 | 0.000153499 | 1.93646E-07 |
| Zfp9 | 0.122906022 | -0.492140171 | 0.333 | 0.582 | 1 | 9.71272E-08 | -1.570684006 | 0.178 | 0.675 | 0.002359414 | 0.122906022 | 1.94254E-07 |
| Chpf | 9.82E-05 | -2.297771786 | 0.143 | 0.586 | 1 | 9.89514E-08 | -1.713056451 | 0.2 | 0.675 | 0.002403727 | 9.82E-05 | 1.97903E-07 |
| Cavin1 | 0.040776706 | -0.613940612 | 0.476 | 0.813 | 1 | 1.0041E-07 | -1.096125323 | 0.8 | 0.913 | 0.002439153 | 0.040776706 | 2.00819E-07 |
| Pdgfrl | 3.18912E-05 | -1.993501087 | 0.238 | 0.716 | 0.774701666 | 1.00532E-07 | -1.5274944 | 0.2 | 0.699 | 0.002442121 | 3.18912E-05 | 2.01064E-07 |
| Fzd2 | 0.011095316 | -1.345421798 | 0.095 | 0.388 | 1 | 1.00859E-07 | -2.10003479 | 0.111 | 0.592 | 0.002450057 | 0.011095316 | 2.01717E-07 |
| Cox7a2 | 1.90469E-06 | -1.432727854 | 0.667 | 0.937 | 0.046268749 | 1.02405E-07 | -0.981617107 | 0.867 | 0.913 | 0.002487623 | 1.90469E-06 | 2.0481E-07 |
| Dph3 | 2.2961E-06 | -2.570354176 | 0.143 | 0.709 | 0.055776959 | 1.03073E-07 | -1.886783022 | 0.244 | 0.661 | 0.002503841 | 2.2961E-06 | 2.06145E-07 |
| Scamp3 | 0.093816717 | -0.282638964 | 0.381 | 0.69 | 1 | 1.03884E-07 | -2.118545966 | 0.111 | 0.567 | 0.002523554 | 0.093816717 | 2.07768E-07 |
| Vma21 | 1.04734E-07 | -3.748292998 | 0.048 | 0.698 | 0.002544197 | 3.14173E-06 | -1.367012645 | 0.333 | 0.692 | 0.076319015 | 3.14173E-06 | 2.09468E-07 |
| Rtraf | 1.05504E-07 | -1.677368248 | 0.667 | 0.944 | 0.002562899 | 6.44564E-06 | -0.810665155 | 0.844 | 0.882 | 0.156577483 | 6.44564E-06 | 2.11008E-07 |
| Igf1r | 1.05685E-07 | 1.937182414 | 0.952 | 0.993 | 0.002567297 | 0.010009269 | 0.680119269 | 1 | 0.99 | 1 | 0.010009269 | 2.1137E-07 |
| Rexo2 | 1.07382E-07 | -1.649845996 | 0.81 | 0.981 | 0.002608526 | 1.51447E-07 | -0.89183625 | 0.889 | 0.955 | 0.00367895 | 1.51447E-07 | 2.14764E-07 |
| Bcap31 | 1.08711E-07 | -1.44977488 | 0.714 | 0.959 | 0.00264081 | 1.41581E-05 | -0.658450989 | 0.933 | 0.938 | 0.343929683 | 1.41581E-05 | 2.17422E-07 |
| Tm2d3 | 0.000626131 | -2.107587946 | 0.143 | 0.526 | 1 | 1.09918E-07 | -2.299920989 | 0.111 | 0.561 | 0.002670118 | 0.000626131 | 2.19835E-07 |
| Ndufa7 | 1.11371E-07 | -1.761295303 | 0.667 | 0.922 | 0.002705423 | 0.0001843 | -0.815419152 | 0.8 | 0.834 | 1 | 0.0001843 | 2.22742E-07 |
| Ccdc88c | 0.280233984 | -0.436652222 | 0.19 | 0.321 | 1 | 1.13967E-07 | 1.317423114 | 0.667 | 0.232 | 0.002768483 | 0.280233984 | 2.27934E-07 |
| Wif1 | 0.010943216 | -3.894184308 | 0.429 | 0.604 | 1 | 1.16367E-07 | -1.211031854 | 0.378 | 0.789 | 0.002826783 | 0.010943216 | 2.32734E-07 |
| Arpc5 | 0.006560506 | -0.810776528 | 0.857 | 0.907 | 1 | 1.19806E-07 | -0.878500094 | 0.911 | 0.969 | 0.002910322 | 0.006560506 | 2.39612E-07 |
| Stmn1 | 0.836047971 | -0.820992924 | 0.143 | 0.164 | 1 | 1.22554E-07 | 1.661377515 | 0.511 | 0.156 | 0.002977089 | 0.836047971 | 2.45109E-07 |
| Maml2 | 0.002045686 | 1.06557789 | 1 | 0.993 | 1 | 1.23076E-07 | 1.31604596 | 1 | 1 | 0.002989766 | 0.002045686 | 2.46152E-07 |
| Purb | 0.001023131 | -0.894564414 | 0.81 | 0.91 | 1 | 1.23285E-07 | -0.817051969 | 0.933 | 0.969 | 0.002994844 | 0.001023131 | 2.4657E-07 |
| Nipal2 | 0.649049492 | 2.614878079 | 0.095 | 0.071 | 1 | 1.2465E-07 | 1.078210528 | 0.511 | 0.145 | 0.003028008 | 0.649049492 | 2.49301E-07 |
| Trim47 | 0.00313865 | -1.287566612 | 0.381 | 0.713 | 1 | 1.29092E-07 | -1.453064102 | 0.356 | 0.824 | 0.003135904 | 0.00313865 | 2.58184E-07 |
| Fam98a | 0.018739719 | -0.65329963 | 0.19 | 0.541 | 1 | 1.29184E-07 | -1.529377032 | 0.2 | 0.692 | 0.003138142 | 0.018739719 | 2.58368E-07 |
| Foxa1 | 0.57816106 | 1.351843673 | 0 | 0.015 | 1 | 1.29236E-07 | 2.242499995 | 0.289 | 0.052 | 0.003139406 | 0.57816106 | 2.58472E-07 |
| S100a13 | 1.30959E-07 | -2.517092571 | 0.381 | 0.873 | 0.003181261 | 0.001970875 | -0.618626184 | 0.689 | 0.827 | 1 | 0.001970875 | 2.61918E-07 |
| Penk | 0.02582449 | -1.087779465 | 0.095 | 0.369 | 1 | 1.31943E-07 | -2.762237924 | 0.089 | 0.554 | 0.003205166 | 0.02582449 | 2.63887E-07 |
| Atp6v1g1 | 1.33651E-07 | -1.731842478 | 0.667 | 0.925 | 0.003246654 | 5.14157E-07 | -1.159626951 | 0.844 | 0.903 | 0.012489913 | 5.14157E-07 | 2.67302E-07 |
| Tbc1d8b | 0.276161256 | -0.31869456 | 0.381 | 0.608 | 1 | 1.36811E-07 | -1.673071224 | 0.222 | 0.706 | 0.003323423 | 0.276161256 | 2.73623E-07 |
| mt-Nd4l | 0.036642255 | 1.136196769 | 1 | 0.925 | 1 | 1.38312E-07 | 1.282969418 | 0.956 | 0.896 | 0.003359885 | 0.036642255 | 2.76625E-07 |
| Romo1 | 1.39157E-07 | -2.058678345 | 0.381 | 0.877 | 0.003380407 | 4.23958E-06 | -1.11339402 | 0.6 | 0.803 | 0.102987929 | 4.23958E-06 | 2.78314E-07 |
| Atp5o | 9.41371E-05 | -1.368191525 | 0.619 | 0.847 | 1 | 1.39165E-07 | -1.122133869 | 0.778 | 0.872 | 0.003380598 | 9.41371E-05 | 2.7833E-07 |
| Ginm1 | 1.40509E-07 | -2.18917548 | 0.238 | 0.858 | 0.003413246 | 6.35516E-06 | -0.751555641 | 0.578 | 0.865 | 0.154379541 | 6.35516E-06 | 2.81018E-07 |
| Ptpn13 | 0.30976473 | 0.204536974 | 0.619 | 0.478 | 1 | 1.40538E-07 | 0.702777996 | 0.956 | 0.522 | 0.003413958 | 0.30976473 | 2.81077E-07 |
| Lgr6 | 0.146865698 | 1.764646168 | 0.476 | 0.396 | 1 | 1.41974E-07 | 1.8992283 | 0.8 | 0.384 | 0.003448832 | 0.146865698 | 2.83948E-07 |
| Tmco1 | 1.43662E-07 | -2.552296312 | 0.238 | 0.806 | 0.003489843 | 0.000534212 | -0.895376608 | 0.556 | 0.73 | 1 | 0.000534212 | 2.87324E-07 |
| Rbp1 | 2.63118E-06 | -4.527739577 | 0.048 | 0.634 | 0.063916711 | 1.44019E-07 | -1.230183211 | 0.378 | 0.855 | 0.003498503 | 2.63118E-06 | 2.88037E-07 |
| Myadm | 0.003899777 | -1.076528465 | 0.381 | 0.769 | 1 | 1.4569E-07 | -1.749341482 | 0.311 | 0.772 | 0.003539102 | 0.003899777 | 2.9138E-07 |
| Ly6e | 0.119584821 | -1.235521318 | 0.714 | 0.616 | 1 | 1.47054E-07 | -0.916763191 | 0.933 | 0.934 | 0.00357224 | 0.119584821 | 2.94108E-07 |
| Prdx2 | 6.12718E-05 | -1.24509147 | 0.667 | 0.854 | 1 | 1.47925E-07 | -0.970111406 | 0.867 | 0.913 | 0.003593382 | 6.12718E-05 | 2.95849E-07 |
| Oxct1 | 1.49277E-07 | -1.006949716 | 0.333 | 0.925 | 0.003626247 | 0.00045119 | -0.555499762 | 0.844 | 0.889 | 1 | 0.00045119 | 2.98555E-07 |
| Pgrmc2 | 1.51675E-07 | -3.614709844 | 0.095 | 0.72 | 0.003684499 | 5.08377E-05 | -0.791130783 | 0.556 | 0.827 | 1 | 5.08377E-05 | 3.03351E-07 |
| Mrpl20 | 1.75635E-05 | -1.501866885 | 0.333 | 0.821 | 0.42665187 | 1.54447E-07 | -1.134550671 | 0.6 | 0.824 | 0.003751815 | 1.75635E-05 | 3.08893E-07 |
| Chchd1 | 0.448444911 | -0.109604291 | 0.476 | 0.675 | 1 | 1.54471E-07 | -1.540675115 | 0.289 | 0.702 | 0.003752412 | 0.448444911 | 3.08942E-07 |
| Crybg1 | 0.787515686 | 1.400753273 | 0.19 | 0.19 | 1 | 1.55556E-07 | 1.492741374 | 0.533 | 0.17 | 0.003778775 | 0.787515686 | 3.11113E-07 |
| Josd1 | 0.084300369 | -0.419740118 | 0.286 | 0.575 | 1 | 1.58495E-07 | -1.512131824 | 0.178 | 0.678 | 0.003850168 | 0.084300369 | 3.16991E-07 |
| Chchd7 | 1.97688E-05 | -2.724259306 | 0.048 | 0.582 | 0.480223419 | 1.60266E-07 | -2.367553487 | 0.067 | 0.505 | 0.003893176 | 1.97688E-05 | 3.20532E-07 |
| Mmp14 | 0.000332616 | -2.008903671 | 0.238 | 0.668 | 1 | 1.60995E-07 | -1.28569489 | 0.556 | 0.913 | 0.003910888 | 0.000332616 | 3.2199E-07 |
| Tnfrsf1a | 1.61867E-07 | -2.169925001 | 0.381 | 0.877 | 0.003932065 | 1.03842E-05 | -0.970699072 | 0.667 | 0.848 | 0.252254172 | 1.03842E-05 | 3.23733E-07 |
| Cetn2 | 1.30615E-05 | -3.439970398 | 0.048 | 0.567 | 0.317290884 | 1.68499E-07 | -1.592205415 | 0.044 | 0.481 | 0.004093167 | 1.30615E-05 | 3.36997E-07 |
| Cwc15 | 0.004107735 | -0.875692052 | 0.524 | 0.825 | 1 | 1.68539E-07 | -1.109485334 | 0.756 | 0.872 | 0.004094156 | 0.004107735 | 3.37079E-07 |
| Reg3g | 0.033805315 | -1.434752689 | 0.619 | 0.317 | 1 | 1.7095E-07 | 1.033258941 | 0.778 | 0.394 | 0.004152708 | 0.033805315 | 3.41899E-07 |
| Nucb1 | 1.73071E-07 | -1.597936972 | 0.429 | 0.918 | 0.00420425 | 1.78942E-06 | -0.683889262 | 0.8 | 0.907 | 0.043468476 | 1.78942E-06 | 3.46143E-07 |
| Tspan4 | 3.63752E-05 | -1.714723944 | 0.333 | 0.806 | 0.883625786 | 1.76073E-07 | -1.74765872 | 0.267 | 0.716 | 0.004277155 | 3.63752E-05 | 3.52145E-07 |
| Sdc3 | 0.000306594 | -1.32873074 | 0.286 | 0.75 | 1 | 1.77599E-07 | -1.154197914 | 0.4 | 0.872 | 0.004314236 | 0.000306594 | 3.55198E-07 |
| Yipf4 | 9.39267E-05 | -1.282749596 | 0.381 | 0.847 | 1 | 1.78136E-07 | -1.005897605 | 0.667 | 0.896 | 0.004327271 | 9.39267E-05 | 3.56271E-07 |
| Psenen | 1.82457E-07 | -2.12324121 | 0.333 | 0.854 | 0.004432255 | 3.06769E-05 | -1.006226574 | 0.689 | 0.779 | 0.745203077 | 3.06769E-05 | 3.64915E-07 |
| Alyref | 0.002444901 | -1.049190967 | 0.476 | 0.81 | 1 | 1.86235E-07 | -1.067092506 | 0.8 | 0.907 | 0.004524032 | 0.002444901 | 3.72471E-07 |
| Fkbp11 | 0.012650767 | -2.208871282 | 0.143 | 0.396 | 1 | 1.86589E-07 | -5.265294645 | 0 | 0.412 | 0.004532613 | 0.012650767 | 3.73177E-07 |
| Zfand3 | 0.000416478 | 0.902590458 | 0.952 | 0.985 | 1 | 1.89732E-07 | 0.967425713 | 1 | 1 | 0.004608974 | 0.000416478 | 3.79464E-07 |
| Eva1b | 1.49031E-05 | -4.520985087 | 0 | 0.522 | 0.362026676 | 1.90279E-07 | -1.580752554 | 0.133 | 0.592 | 0.004622256 | 1.49031E-05 | 3.80558E-07 |
| Phf13 | 0.072698202 | -1.107587946 | 0.143 | 0.343 | 1 | 1.90749E-07 | -2.856086225 | 0.067 | 0.502 | 0.004633678 | 0.072698202 | 3.81498E-07 |
| Pja1 | 0.000782562 | -3.15277672 | 0.048 | 0.425 | 1 | 1.90797E-07 | -2.574315257 | 0.067 | 0.509 | 0.004634838 | 0.000782562 | 3.81594E-07 |
| Arf5 | 1.20813E-06 | -1.342906973 | 0.81 | 0.97 | 0.029347879 | 1.92274E-07 | -0.781775916 | 0.956 | 0.979 | 0.004670721 | 1.20813E-06 | 3.84548E-07 |
| Cxadr | 0.776355689 | 0.497574169 | 0.381 | 0.437 | 1 | 1.94828E-07 | 0.793255504 | 0.711 | 0.249 | 0.004732757 | 0.776355689 | 3.89656E-07 |
| Fundc2 | 2.18799E-06 | -2.208871282 | 0.19 | 0.757 | 0.053148543 | 1.94945E-07 | -1.545746104 | 0.378 | 0.727 | 0.004735596 | 2.18799E-06 | 3.89889E-07 |
| Ssna1 | 0.04065103 | -0.593014773 | 0.143 | 0.403 | 1 | 1.96419E-07 | -3.361321533 | 0.022 | 0.446 | 0.004771414 | 0.04065103 | 3.92838E-07 |
| Fut11 | 0.067280238 | -1.008052272 | 0.143 | 0.354 | 1 | 1.97341E-07 | -2.328154669 | 0.089 | 0.522 | 0.004793815 | 0.067280238 | 3.94683E-07 |
| Pdgfrb | 0.027545407 | -1.526990638 | 0.19 | 0.455 | 1 | 2.00539E-07 | -1.26787319 | 0.244 | 0.737 | 0.004871503 | 0.027545407 | 4.01079E-07 |
| Arhgap26 | 0.001982471 | 2.311201688 | 0.667 | 0.463 | 1 | 2.01648E-07 | 1.410546731 | 0.8 | 0.381 | 0.004898437 | 0.001982471 | 4.03296E-07 |

| | | | | | | | | | | | | |
|---|---|---|---|---|---|---|---|---|---|---|---|---|
| Psmb1 | 2.78669E-07 | -1.502816964 | 0.714 | 0.955 | 0.006769424 | 2.03716E-07 | -0.690261733 | 0.933 | 0.952 | 0.00494866 | 2.78669E-07 | 4.07431E-07 |
| Thoc7 | 2.05411E-07 | -2.582268392 | 0.238 | 0.799 | 0.004989847 | 0.000149749 | -0.875195048 | 0.667 | 0.81 | 1 | 0.000149749 | 4.10822E-07 |
| Dnajb9 | 1.66581E-05 | -1.588009755 | 0.667 | 0.896 | 0.404658682 | 2.05968E-07 | -1.190740612 | 0.822 | 0.917 | 0.005003385 | 1.66581E-05 | 4.11937E-07 |
| Fam89b | 0.001601696 | -1.8244791 | 0.095 | 0.489 | 1 | 2.07645E-07 | -2.620708162 | 0.111 | 0.54 | 0.005044109 | 0.001601696 | 4.1529E-07 |
| Ctnnbip1 | 0.001076606 | -1.506137322 | 0.19 | 0.612 | 1 | 2.09077E-07 | -1.550547091 | 0.133 | 0.585 | 0.005078902 | 0.001076606 | 4.18154E-07 |
| Smim7 | 2.12832E-07 | -2.214971481 | 0.143 | 0.813 | 0.005170124 | 2.15102E-05 | -0.996777862 | 0.489 | 0.768 | 0.522526985 | 2.15102E-05 | 4.25665E-07 |
| Adrb2 | 0.311307311 | -1.133583154 | 0.048 | 0.119 | 1 | 2.12985E-07 | 2.039216396 | 0.333 | 0.073 | 0.005173839 | 0.311307311 | 4.25971E-07 |
| Adrm1 | 0.082192172 | -0.305268614 | 0.238 | 0.541 | 1 | 2.20623E-07 | -1.854892435 | 0.2 | 0.626 | 0.005359372 | 0.082192172 | 4.41246E-07 |
| Eif1ax | 0.001093933 | -0.451759114 | 0.667 | 0.884 | 1 | 2.21645E-07 | -0.952515988 | 0.8 | 0.931 | 0.00538421 | 0.001093933 | 4.43291E-07 |
| Abhd14a | 0.000204814 | -2.413691074 | 0.095 | 0.522 | 1 | 2.32073E-07 | -2.143475901 | 0.089 | 0.533 | 0.005637526 | 0.000204814 | 4.64147E-07 |
| Yif1a | 2.38606E-06 | -2.748292998 | 0.095 | 0.679 | 0.057962155 | 2.40272E-07 | -1.699397051 | 0.156 | 0.599 | 0.005836679 | 2.38606E-06 | 4.80543E-07 |
| Fam193a | 0.004139518 | 1.189143996 | 0.952 | 0.799 | 1 | 2.42159E-07 | 0.995217204 | 1 | 0.983 | 0.005882535 | 0.004139518 | 4.84319E-07 |
| Reep5 | 2.47981E-07 | -1.431635273 | 0.714 | 0.985 | 0.006023959 | 0.000170598 | -0.500503091 | 0.978 | 0.986 | 1 | 0.000170598 | 4.95962E-07 |
| Cftr | 0.167544534 | 3.088809267 | 0.048 | 0.011 | 1 | 2.48499E-07 | 2.041526557 | 0.333 | 0.073 | 0.006036545 | 0.167544534 | 4.96999E-07 |
| Dusp18 | 0.023250632 | -1.190414377 | 0.143 | 0.429 | 1 | 2.52663E-07 | -2.127957994 | 0.089 | 0.547 | 0.006137683 | 0.023250632 | 5.05325E-07 |
| Polr1d | 0.000806757 | -0.757060891 | 0.857 | 0.944 | 1 | 2.55971E-07 | -0.780724225 | 0.867 | 0.945 | 0.006218058 | 0.000806757 | 5.11943E-07 |
| D8Ertd738e | 2.57973E-07 | -1.831222428 | 0.429 | 0.903 | 0.006266677 | 0.000148889 | -0.640944857 | 0.756 | 0.858 | 1 | 0.000148889 | 5.15946E-07 |
| Timm8b | 2.58714E-07 | -2.948280052 | 0.143 | 0.731 | 0.006284675 | 9.01485E-05 | -1.336826971 | 0.311 | 0.599 | 1 | 9.01485E-05 | 5.17428E-07 |
| Myo1e | 0.000119948 | 1.125962265 | 1 | 0.993 | 1 | 2.59697E-07 | 0.818998382 | 1 | 1 | 0.006308552 | 0.000119948 | 5.19393E-07 |
| Bax | 2.60002E-07 | -3.509450056 | 0.095 | 0.709 | 0.006315959 | 0.027233744 | -0.499130745 | 0.556 | 0.654 | 1 | 0.027233744 | 5.20003E-07 |
| Pawr | 0.358454845 | 1.383379246 | 0.381 | 0.295 | 1 | 2.64457E-07 | 0.725507852 | 0.8 | 0.315 | 0.006424186 | 0.358454845 | 5.28914E-07 |
| Odc1 | 0.000166044 | -1.398957699 | 0.714 | 0.925 | 1 | 2.65228E-07 | -1.502353509 | 0.778 | 0.872 | 0.00644291 | 0.000166044 | 5.30455E-07 |
| Psmb6 | 7.18674E-06 | -1.541825092 | 0.619 | 0.881 | 0.174580216 | 2.65362E-07 | -1.068248301 | 0.756 | 0.862 | 0.006446167 | 7.18674E-06 | 5.30723E-07 |
| Rrs1 | 0.013097331 | -1.527862093 | 0.143 | 0.422 | 1 | 2.70912E-07 | -2.748773201 | 0.067 | 0.491 | 0.006581 | 0.013097331 | 5.41824E-07 |
| Gstm5 | 2.51864E-06 | -2.271086678 | 0.19 | 0.769 | 0.06118279 | 2.81576E-07 | -1.558237049 | 0.156 | 0.651 | 0.006840033 | 2.51864E-06 | 5.63151E-07 |
| Dhrs7 | 2.85925E-07 | -4.511103575 | 0 | 0.634 | 0.006945691 | 0.000166847 | -1.399916779 | 0.378 | 0.612 | 1 | 0.000166847 | 5.7185E-07 |
| Cibar1 | 0.003866451 | -0.75918764 | 0.286 | 0.672 | 1 | 2.89859E-07 | -1.301160098 | 0.156 | 0.644 | 0.007041246 | 0.003866451 | 5.79717E-07 |
| Zfp706 | 2.70042E-05 | -1.411331801 | 0.476 | 0.862 | 0.655986868 | 2.90125E-07 | -1.007931552 | 0.822 | 0.886 | 0.007047721 | 2.70042E-05 | 5.8025E-07 |
| Abi1 | 2.91267E-07 | 1.258326324 | 1 | 0.925 | 0.007075462 | 2.48182E-06 | 1.237272833 | 1 | 0.962 | 0.060288401 | 2.48182E-06 | 5.82534E-07 |
| Cd44 | 0.002577696 | 1.789248985 | 0.619 | 0.366 | 1 | 2.92569E-07 | 1.840952246 | 0.822 | 0.374 | 0.007107094 | 0.002577696 | 5.85139E-07 |
| Trps1 | 2.98724E-07 | 1.638321362 | 1 | 1 | 0.007256614 | 4.16107E-06 | 1.044870241 | 1 | 1 | 0.101080767 | 4.16107E-06 | 5.97449E-07 |
| Rgs3 | 0.184678073 | -0.936725825 | 0.429 | 0.578 | 1 | 3.01317E-07 | -1.174908409 | 0.267 | 0.765 | 0.007319589 | 0.184678073 | 6.02634E-07 |
| Lamtor4 | 3.03601E-07 | -2.818081329 | 0.095 | 0.731 | 0.007375082 | 0.041162452 | -0.674479418 | 0.578 | 0.64 | 1 | 0.041162452 | 6.07202E-07 |
| Wnk1 | 0.008473108 | 1.092362705 | 1 | 0.985 | 1 | 3.13743E-07 | 0.767697793 | 1 | 0.993 | 0.007621445 | 0.008473108 | 6.27486E-07 |
| Snx15 | 0.036439166 | -0.672003069 | 0.19 | 0.478 | 1 | 3.13787E-07 | -3.863821874 | 0.022 | 0.426 | 0.007622505 | 0.036439166 | 6.27573E-07 |
| Selenbp1 | 0.268033526 | -0.911190733 | 0 | 0.056 | 1 | 3.25621E-07 | 1.612683258 | 0.422 | 0.114 | 0.007909984 | 0.268033526 | 6.51242E-07 |
| Ptprm | 3.32771E-07 | 1.479892034 | 1 | 0.877 | 0.008083677 | 0.000300667 | 0.959939265 | 0.956 | 0.965 | 1 | 0.000300667 | 6.65542E-07 |
| Snhg3 | 0.013287961 | -0.987293712 | 0.19 | 0.537 | 1 | 3.42289E-07 | -2.303988358 | 0.089 | 0.512 | 0.008314886 | 0.013287961 | 6.84578E-07 |
| Isyna1 | 0.000200357 | -2.379960216 | 0.143 | 0.604 | 1 | 3.54651E-07 | -1.674479418 | 0.156 | 0.616 | 0.008615171 | 0.000200357 | 7.09301E-07 |
| Ltbr | 0.000314459 | -1.351763324 | 0.095 | 0.575 | 1 | 3.55255E-07 | -2.253565353 | 0.111 | 0.54 | 0.008629859 | 0.000314459 | 7.1051E-07 |
| Rnf144a | 0.001659805 | -2.022036502 | 0.095 | 0.489 | 1 | 3.58194E-07 | -3.113198837 | 0.089 | 0.505 | 0.008701252 | 0.001659805 | 7.16388E-07 |
| Esrrg | 0.898241106 | 0.728913322 | 0.286 | 0.351 | 1 | 3.59893E-07 | 2.528626317 | 0.489 | 0.145 | 0.008742528 | 0.898241106 | 7.19786E-07 |
| Prpsap1 | 0.000432956 | -1.233118828 | 0.143 | 0.616 | 1 | 3.67804E-07 | -1.606946261 | 0.156 | 0.63 | 0.008934694 | 0.000432956 | 7.35608E-07 |
| Cldnd1 | 0.028365933 | -0.742245491 | 0.476 | 0.754 | 1 | 3.72049E-07 | -1.001134105 | 0.644 | 0.858 | 0.009037803 | 0.028365933 | 7.44097E-07 |
| Sdk1 | 0.169309442 | 2.075870211 | 0.048 | 0.172 | 1 | 3.75372E-07 | 0.989175714 | 0.711 | 0.284 | 0.009118527 | 0.169309442 | 7.50743E-07 |
| Eif3h | 0.000490563 | -0.907177671 | 0.81 | 0.944 | 1 | 3.79182E-07 | -0.64110796 | 0.956 | 0.976 | 0.009211087 | 0.000490563 | 7.58364E-07 |
| Tob1 | 1.98693E-05 | -1.542932993 | 0.81 | 0.937 | 0.482664531 | 3.90977E-07 | -1.021188843 | 0.978 | 0.976 | 0.009497614 | 1.98693E-05 | 7.81954E-07 |
| Fnta | 0.000687084 | -0.849790188 | 0.238 | 0.72 | 1 | 3.94194E-07 | -1.465110636 | 0.356 | 0.737 | 0.009575759 | 0.000687084 | 7.88388E-07 |
| Nav3 | 0.048751771 | 1.206744779 | 0.714 | 0.552 | 1 | 3.97055E-07 | 2.446535983 | 0.511 | 0.225 | 0.009645269 | 0.048751771 | 7.94111E-07 |
| Sik3 | 0.081009756 | 0.743387469 | 0.952 | 0.97 | 1 | 3.98872E-07 | 0.95820647 | 1 | 0.993 | 0.009689401 | 0.081009756 | 7.97744E-07 |
| Chil1 | 0.080782108 | -1.683083283 | 0.429 | 0.631 | 1 | 4.02E-07 | -3.813487513 | 0.133 | 0.561 | 0.009765392 | 0.080782108 | 8.04E-07 |
| Bambi | 4.04918E-07 | -3.90365706 | 0.095 | 0.709 | 0.009836271 | 1.54735E-06 | -1.466674533 | 0.133 | 0.547 | 0.037588147 | 1.54735E-06 | 8.09836E-07 |
| Slc15a2 | 0.516567901 | 1.393663848 | 0.143 | 0.101 | 1 | 4.06043E-07 | 0.651594355 | 0.689 | 0.246 | 0.009863599 | 0.516567901 | 8.12086E-07 |
| Zcchc17 | 0.057683836 | -0.388512511 | 0.286 | 0.601 | 1 | 4.06175E-07 | -2.486852415 | 0.111 | 0.526 | 0.009866801 | 0.057683836 | 8.1235E-07 |
| Foxf1 | 0.000354514 | -2.543458949 | 0.095 | 0.526 | 1 | 4.09063E-07 | -2.005637839 | 0.133 | 0.595 | 0.00993696 | 0.000354514 | 8.18126E-07 |
| Rbm18 | 0.00044839 | -1.752492987 | 0.19 | 0.619 | 1 | 4.12757E-07 | -1.606024116 | 0.267 | 0.685 | 0.010026705 | 0.00044839 | 8.25515E-07 |
| Cox6b1 | 4.15335E-07 | -1.593708543 | 0.714 | 0.929 | 0.010089327 | 5.62276E-05 | -0.734433169 | 0.911 | 0.913 | 1 | 5.62276E-05 | 8.30671E-07 |
| C1qtnf1 | 0.078725236 | -1.184209227 | 0.143 | 0.325 | 1 | 4.16077E-07 | -3.446210431 | 0.044 | 0.453 | 0.010107343 | 0.078725236 | 8.32154E-07 |
| Emc3 | 1.1389E-05 | -1.586755783 | 0.19 | 0.765 | 0.276662106 | 4.16565E-07 | -1.337145693 | 0.533 | 0.789 | 0.010119198 | 1.1389E-05 | 8.3313E-07 |
| Pttg1ip | 1.33262E-06 | -1.6498728 | 0.571 | 0.903 | 0.032372018 | 4.2067E-07 | -0.829470369 | 0.689 | 0.903 | 0.010218923 | 1.33262E-06 | 8.4134E-07 |
| Cxcl17 | 0.00458782 | 2.48112669 | 0.143 | 0.026 | 1 | 4.21945E-07 | 2.098110085 | 0.311 | 0.066 | 0.01024988 | 0.00458782 | 8.43889E-07 |
| Impdh2 | 6.38595E-07 | -1.896083841 | 0.333 | 0.84 | 0.015512741 | 4.25968E-07 | -1.283993524 | 0.622 | 0.806 | 0.010347618 | 6.38595E-07 | 8.51936E-07 |
| Atp5e | 4.36282E-07 | -1.661772598 | 0.714 | 0.929 | 0.010598174 | 1.94529E-06 | -0.912778231 | 0.933 | 0.91 | 0.047255028 | 1.94529E-06 | 8.72565E-07 |
| Snrpd2 | 1.82496E-05 | -1.533242552 | 0.381 | 0.84 | 0.443319155 | 4.40257E-07 | -1.356169045 | 0.689 | 0.785 | 0.010694727 | 1.82496E-05 | 8.80514E-07 |
| Gnaq | 0.040247463 | 0.621998118 | 0.952 | 0.933 | 1 | 4.42799E-07 | 1.122357632 | 1 | 0.986 | 0.01075647 | 0.040247463 | 8.85598E-07 |
| Eif3f | 4.44165E-07 | -1.70576705 | 0.762 | 0.918 | 0.010789664 | 1.12084E-05 | -0.714730376 | 0.933 | 0.924 | 0.272274387 | 1.12084E-05 | 8.8833E-07 |
| Exoc6b | 0.017382775 | 1.317670172 | 0.905 | 0.903 | 1 | 4.4742E-07 | 1.193009349 | 1 | 0.931 | 0.010868728 | 0.017382775 | 8.9484E-07 |
| Sec22b | 0.021049771 | -0.59376403 | 0.619 | 0.937 | 1 | 4.54535E-07 | -0.959471466 | 0.622 | 0.931 | 0.01104156 | 0.021049771 | 9.09069E-07 |
| Cnih1 | 2.74631E-05 | -1.404569684 | 0.238 | 0.806 | 0.667133093 | 4.54536E-07 | -1.449151706 | 0.489 | 0.74 | 0.01104159 | 2.74631E-05 | 9.09072E-07 |
| Dennd1a | 0.000982858 | 1.122311011 | 0.905 | 0.907 | 1 | 4.61401E-07 | 1.298071461 | 0.956 | 0.938 | 0.011208351 | 0.000982858 | 9.22802E-07 |
| Ncbp2 | 0.000103623 | -1.82721234 | 0.143 | 0.657 | 1 | 4.68731E-07 | -1.536095934 | 0.333 | 0.72 | 0.011386417 | 0.000103623 | 9.37462E-07 |
| Tmem223 | 0.000859831 | -1.455511249 | 0.143 | 0.575 | 1 | 4.8505E-07 | -1.871516266 | 0.133 | 0.557 | 0.011782826 | 0.000859831 | 9.70099E-07 |
| Rsl1d1 | 0.02736162 | -0.45092398 | 0.619 | 0.877 | 1 | 4.87099E-07 | -0.969453198 | 0.8 | 0.92 | 0.011832615 | 0.02736162 | 9.74198E-07 |
| Scrg1 | 4.88645E-07 | -4.163856165 | 0.143 | 0.728 | 0.011870162 | 6.66646E-06 | -3.16420852 | 0.044 | 0.405 | 0.161941733 | 6.66646E-06 | 9.7729E-07 |
| Amotl2 | 0.01186464 | -1.172682974 | 0.238 | 0.526 | 1 | 4.98542E-07 | -2.137106376 | 0.156 | 0.599 | 0.012110583 | 0.01186464 | 9.97084E-07 |
| Ctsa | 5.12648E-06 | -2.39661756 | 0.095 | 0.687 | 0.12453242 | 5.01266E-07 | -1.497740731 | 0.267 | 0.685 | 0.012176755 | 5.12648E-06 | 1.00253E-06 |
| Gsto2 | 0.173959863 | -1.413691074 | 0 | 0.082 | 1 | 5.04428E-07 | 2.897197392 | 0.333 | 0.08 | 0.012253554 | 0.173959863 | 1.00885E-06 |
| Pfdn4 | 0.00566941 | -1.159118246 | 0.095 | 0.448 | 1 | 5.0558E-07 | -2.782901878 | 0.089 | 0.488 | 0.012281554 | 0.00566941 | 1.01116E-06 |
| Cops8 | 0.00945016 | -0.718545655 | 0.333 | 0.642 | 1 | 5.07548E-07 | -0.919991721 | 0.2 | 0.73 | 0.012329347 | 0.00945016 | 1.0151E-06 |
| Clta | 5.89538E-05 | -1.051779729 | 0.762 | 0.959 | 1 | 5.12624E-07 | -0.696571007 | 0.956 | 0.972 | 0.012452664 | 5.89538E-05 | 1.02525E-06 |
| Tvp23b | 0.002045267 | -1.155950967 | 0.333 | 0.746 | 1 | 5.31307E-07 | -1.421842424 | 0.244 | 0.709 | 0.01290651 | 0.002045267 | 1.06261E-06 |
| C1galt1c1 | 0.000128309 | -2.509450056 | 0.095 | 0.537 | 1 | 5.33256E-07 | -1.86124793 | 0.133 | 0.561 | 0.012953847 | 0.000128309 | 1.06651E-06 |
| Bri3bp | 0.003020341 | -1.839297815 | 0.048 | 0.392 | 1 | 5.46144E-07 | -3.149817428 | 0.022 | 0.426 | 0.013266928 | 0.003020341 | 1.09229E-06 |
| Samd4 | 5.47048E-07 | 1.583722163 | 1 | 0.959 | 0.01328889 | 0.000120224 | 0.951951615 | 1 | 0.983 | 1 | 0.000120224 | 1.0941E-06 |
| Mlec | 1.17623E-06 | -1.777005386 | 0.333 | 0.881 | 0.028573014 | 5.57159E-07 | -1.047152862 | 0.6 | 0.858 | 0.013534498 | 1.17623E-06 | 1.11432E-06 |
| Ctsc | 0.196608492 | -2.081115734 | 0 | 0.075 | 1 | 5.61677E-07 | 1.223640968 | 0.422 | 0.114 | 0.013644246 | 0.196608492 | 1.12335E-06 |
| Kitl | 0.194385609 | 1.133203386 | 0.19 | 0.101 | 1 | 5.65292E-07 | 1.361144491 | 0.6 | 0.211 | 0.013732061 | 0.194385609 | 1.13058E-06 |
| Rwdd1 | 6.13834E-05 | -1.354636648 | 0.524 | 0.854 | 1 | 5.69656E-07 | -1.404390255 | 0.578 | 0.792 | 0.013838089 | 6.13834E-05 | 1.13931E-06 |
| Med7 | 0.011511845 | -1.430564892 | 0.143 | 0.433 | 1 | 5.7043E-07 | -2.802354241 | 0.067 | 0.474 | 0.013856891 | 0.011511845 | 1.14086E-06 |
| Map9 | 0.026408165 | -1.855007898 | 0.095 | 0.347 | 1 | 5.73668E-07 | -3.660335236 | 0.044 | 0.45 | 0.013935548 | 0.026408165 | 1.14734E-06 |
| Tmed4 | 0.000713979 | -1.704739856 | 0.095 | 0.53 | 1 | 5.78314E-07 | -1.654230735 | 0.178 | 0.595 | 0.014048411 | 0.000713979 | 1.15663E-06 |
| Cnpy4 | 0.042130824 | -1.089993886 | 0.19 | 0.418 | 1 | 5.7925E-07 | -2.466674533 | 0.089 | 0.498 | 0.014071144 | 0.042130824 | 1.1585E-06 |
| Atp1b3 | 5.87122E-07 | -1.525620143 | 0.714 | 0.963 | 0.014262378 | 0.023859152 | -0.157660894 | 0.978 | 0.958 | 1 | 0.023859152 | 1.17424E-06 |
| Tfrc | 0.260858733 | -0.765020085 | 0.381 | 0.567 | 1 | 5.88501E-07 | -1.447539491 | 0.6 | 0.83 | 0.014295861 | 0.260858733 | 1.177E-06 |
| App | 1.21211E-06 | -1.256334746 | 0.952 | 1 | 0.029444507 | 6.0006E-07 | -0.627484988 | 1 | 1 | 0.014576659 | 1.21211E-06 | 1.20012E-06 |
| Atox1 | 6.0572E-07 | -2.507628994 | 0.857 | 0.966 | 0.01471415 | 0.000127048 | -0.854675276 | 0.889 | 0.913 | 1 | 0.000127048 | 1.21144E-06 |
| Vat1 | 0.079979614 | -0.960434252 | 0.143 | 0.343 | 1 | 6.17468E-07 | -2.100303473 | 0.133 | 0.574 | 0.014999539 | 0.079979614 | 1.23494E-06 |
| Add3 | 0.722572136 | 0.705480627 | 0.286 | 0.347 | 1 | 6.18149E-07 | 1.650651108 | 0.689 | 0.284 | 0.015016066 | 0.722572136 | 1.2363E-06 |

| | | | | | | | | | | | | |
|---|---|---|---|---|---|---|---|---|---|---|---|---|
| Eci2 | 0.00207061 | -1.109970597 | 0.238 | 0.653 | 1 | 6.29708E-07 | -2.017367132 | 0.133 | 0.561 | 0.015296873 | 0.00207061 | 1.25942E-06 |
| Hspa4l | 0.005312326 | 1.921699281 | 0.429 | 0.213 | 1 | 6.31451E-07 | 1.258574758 | 0.733 | 0.287 | 0.015339199 | 0.005312326 | 1.2629E-06 |
| Glmp | 3.72175E-05 | -2.003360577 | 0.095 | 0.642 | 0.904088018 | 6.38768E-07 | -1.194244894 | 0.2 | 0.668 | 0.015516956 | 3.72175E-05 | 1.27754E-06 |
| Tspan6 | 9.01369E-07 | -3.193506972 | 0.143 | 0.701 | 0.021896057 | 6.54083E-07 | -1.674479418 | 0.178 | 0.633 | 0.01588899 | 9.01369E-07 | 1.30817E-06 |
| Zcchc7 | 0.019711472 | 0.900939611 | 0.81 | 0.896 | 1 | 6.56676E-07 | 1.334706127 | 0.978 | 0.858 | 0.015951964 | 0.019711472 | 1.31335E-06 |
| Ptdss1 | 0.736882902 | 0.311201688 | 0.286 | 0.388 | 1 | 6.7008E-07 | -1.375821103 | 0.111 | 0.567 | 0.016277586 | 0.736882902 | 1.34016E-06 |
| Rnf187 | 6.74662E-07 | -2.370622352 | 0.095 | 0.735 | 0.01638889 | 1.16526E-05 | -1.07019482 | 0.467 | 0.751 | 0.283064274 | 1.16526E-05 | 1.34932E-06 |
| Gpsm2 | 0.905139283 | 0.475358209 | 0.429 | 0.511 | 1 | 6.75443E-07 | -1.593599228 | 0.244 | 0.702 | 0.016407859 | 0.905139283 | 1.35089E-06 |
| Ctsz | 6.76418E-07 | -2.452563965 | 0.19 | 0.776 | 0.016431556 | 0.00014076 | -1.101704488 | 0.511 | 0.775 | 1 | 0.00014076 | 1.35284E-06 |
| Tomm7 | 6.79106E-07 | -1.565087865 | 0.571 | 0.918 | 0.016496835 | 5.42176E-05 | -0.836794886 | 0.778 | 0.844 | 1 | 5.42176E-05 | 1.35821E-06 |
| Uqcc2 | 1.32868E-06 | -2.574155746 | 0.143 | 0.724 | 0.032276191 | 6.90452E-07 | -1.718375848 | 0.178 | 0.599 | 0.016772464 | 1.32868E-06 | 1.3809E-06 |
| Pigt | 0.012754705 | -1.20069735 | 0.19 | 0.478 | 1 | 6.9158E-07 | -2.624355939 | 0.089 | 0.484 | 0.01679986 | 0.012754705 | 1.38316E-06 |
| Tob2 | 0.004294559 | -1.299686445 | 0.571 | 0.791 | 1 | 6.92529E-07 | -1.070766827 | 0.867 | 0.931 | 0.016822904 | 0.004294559 | 1.38506E-06 |
| Erlec1 | 3.06906E-06 | -1.588546838 | 0.381 | 0.914 | 0.074553641 | 6.96665E-07 | -1.11808607 | 0.533 | 0.841 | 0.016923376 | 3.06906E-06 | 1.39333E-06 |
| Arf1 | 2.32761E-06 | -1.068877465 | 0.952 | 0.989 | 0.056542231 | 7.03317E-07 | -0.637467635 | 1 | 0.993 | 0.017084971 | 2.32761E-06 | 1.40663E-06 |
| Rbfox2 | 0.006204858 | 0.961697349 | 0.952 | 0.907 | 1 | 7.22494E-07 | 0.998295657 | 0.978 | 0.972 | 0.017550825 | 0.006204858 | 1.44499E-06 |
| Psma7 | 0.00023015 | -1.270763602 | 0.714 | 0.869 | 1 | 7.22839E-07 | -0.953449922 | 0.889 | 0.917 | 0.017559214 | 0.00023015 | 1.44568E-06 |
| H2az2 | 8.94383E-06 | -1.701267664 | 0.381 | 0.858 | 0.21726346 | 7.31333E-07 | -1.041213874 | 0.822 | 0.886 | 0.017765543 | 8.94383E-06 | 1.46267E-06 |
| Sema3d | 0.594836257 | -0.116904413 | 0.381 | 0.5 | 1 | 7.34192E-07 | -1.105762248 | 0.133 | 0.606 | 0.017834988 | 0.594836257 | 1.46838E-06 |
| Hax1 | 0.006076186 | -1.196592952 | 0.19 | 0.549 | 1 | 7.35416E-07 | -1.328900055 | 0.089 | 0.529 | 0.017864719 | 0.006076186 | 1.47083E-06 |
| Ociad1 | 7.42409E-07 | -1.502467008 | 0.476 | 0.925 | 0.018034604 | 0.00037943 | -0.540077354 | 0.822 | 0.848 | 1 | 0.00037943 | 1.48482E-06 |
| Pcdh7 | 0.235633305 | 0.136337637 | 0.429 | 0.265 | 1 | 7.5853E-07 | 1.168499413 | 0.822 | 0.388 | 0.018426201 | 0.235633305 | 1.51706E-06 |
| Pura | 0.053788898 | -0.405319512 | 0.857 | 0.933 | 1 | 7.8205E-07 | -0.827889333 | 0.867 | 0.969 | 0.018997568 | 0.053788898 | 1.5641E-06 |
| Ccs | 0.083328164 | -0.216999163 | 0.19 | 0.433 | 1 | 7.90173E-07 | -2.526380779 | 0.089 | 0.491 | 0.019194893 | 0.083328164 | 1.58035E-06 |
| Rel | 0.85905623 | 0.351843673 | 0.333 | 0.362 | 1 | 7.90251E-07 | 1.194572925 | 0.778 | 0.343 | 0.019196773 | 0.85905623 | 1.5805E-06 |
| Svbp | 0.001097253 | -1.403043829 | 0.143 | 0.571 | 1 | 7.94287E-07 | -1.997047148 | 0.133 | 0.547 | 0.019294823 | 0.001097253 | 1.58857E-06 |
| Mmgt1 | 0.031174928 | -0.935037475 | 0.095 | 0.354 | 1 | 8.10298E-07 | -2.203059449 | 0.067 | 0.474 | 0.019683761 | 0.031174928 | 1.6206E-06 |
| Etv6 | 0.003824084 | 1.142820202 | 1 | 0.951 | 1 | 8.10588E-07 | 1.494154869 | 1 | 0.889 | 0.019690808 | 0.003824084 | 1.62118E-06 |
| Cdv3 | 0.448326514 | -0.144832296 | 0.81 | 0.907 | 1 | 8.18368E-07 | -0.79983391 | 0.933 | 0.969 | 0.019879804 | 0.448326514 | 1.63674E-06 |
| Fam174a | 8.26728E-07 | -2.377890352 | 0.143 | 0.761 | 0.020082883 | 4.05667E-05 | -0.815475163 | 0.333 | 0.74 | 0.985445354 | 4.05667E-05 | 1.65346E-06 |
| Arhgef3 | 0.978456248 | -0.167530486 | 0.286 | 0.295 | 1 | 8.32999E-07 | 1.199038994 | 0.733 | 0.301 | 0.020235213 | 0.978456248 | 1.666E-06 |
| Pabpn1 | 0.419110455 | -0.181493139 | 0.857 | 0.918 | 1 | 8.38504E-07 | -0.846596025 | 0.867 | 0.907 | 0.020368928 | 0.419110455 | 1.67701E-06 |
| Lamtor2 | 8.41691E-07 | -2.023581971 | 0.333 | 0.817 | 0.020446351 | 0.001985867 | -0.966020252 | 0.622 | 0.692 | 1 | 0.001985867 | 1.68338E-06 |
| Txndc12 | 0.020107614 | -0.708697869 | 0.286 | 0.646 | 1 | 8.47862E-07 | -1.80878051 | 0.178 | 0.595 | 0.020596271 | 0.020107614 | 1.69572E-06 |
| Cbx3 | 8.57821E-07 | -1.92222177 | 0.524 | 0.899 | 0.020838188 | 1.33526E-05 | -0.662458408 | 0.778 | 0.934 | 0.324360631 | 1.33526E-05 | 1.71564E-06 |
| Lpp | 0.000849258 | 0.817533977 | 1 | 0.974 | 1 | 8.59666E-07 | 1.001429294 | 1 | 0.99 | 0.020883008 | 0.000849258 | 1.71933E-06 |
| Prdm16 | 0.016312695 | 1.520443277 | 0.81 | 0.799 | 1 | 8.65681E-07 | 1.494543617 | 0.889 | 0.896 | 0.021029112 | 0.016312695 | 1.73136E-06 |
| Scoc | 8.67492E-07 | -2.725399326 | 0.095 | 0.724 | 0.021073108 | 7.59047E-06 | -0.825654329 | 0.2 | 0.654 | 0.184387692 | 7.59047E-06 | 1.73498E-06 |
| Snapin | 4.85426E-06 | -3.460654553 | 0.048 | 0.597 | 0.117919652 | 8.68023E-07 | -1.434035717 | 0.133 | 0.592 | 0.021086024 | 4.85426E-06 | 1.73605E-06 |
| Golt1b | 0.190211803 | -0.462767065 | 0.476 | 0.631 | 1 | 8.70546E-07 | -1.676547799 | 0.178 | 0.616 | 0.021147305 | 0.190211803 | 1.74109E-06 |
| Lmnb1 | 0.438413506 | 1.896164189 | 0.19 | 0.142 | 1 | 8.80277E-07 | 1.42003818 | 0.489 | 0.152 | 0.021383697 | 0.438413506 | 1.76055E-06 |
| Sec11c | 8.81617E-07 | -2.531777143 | 0.286 | 0.795 | 0.021416235 | 0.004694061 | -0.675614059 | 0.711 | 0.772 | 1 | 0.004694061 | 1.76323E-06 |
| Spryd3 | 0.00567656 | -1.091762979 | 0.048 | 0.369 | 1 | 9.00419E-07 | -3.191396532 | 0.022 | 0.415 | 0.021872974 | 0.00567656 | 1.80084E-06 |
| Chrdl1 | 0.000259174 | -3.26011556 | 0.238 | 0.616 | 1 | 9.04962E-07 | -1.516468896 | 0.222 | 0.675 | 0.021983341 | 0.000259174 | 1.80992E-06 |
| Anapc13 | 9.05061E-07 | -4.25696557 | 0 | 0.601 | 0.021985734 | 1.06666E-05 | -2.80878051 | 0.067 | 0.405 | 0.259112811 | 1.06666E-05 | 1.81012E-06 |
| Fbxw11 | 9.21974E-07 | 1.346350674 | 0.952 | 0.955 | 0.022396585 | 1.01311E-05 | 1.000083329 | 1 | 0.979 | 0.246104744 | 1.01311E-05 | 1.84395E-06 |
| Tpst1 | 0.005308454 | -1.690800665 | 0.143 | 0.463 | 1 | 9.44437E-07 | -1.401122541 | 0.2 | 0.675 | 0.02294226 | 0.005308454 | 1.88887E-06 |
| Zfp260 | 0.025253176 | -0.611630451 | 0.333 | 0.701 | 1 | 9.44617E-07 | -1.206617269 | 0.311 | 0.789 | 0.022946648 | 0.025253176 | 1.88923E-06 |
| Calm3 | 9.68311E-07 | -2.238452272 | 0.238 | 0.787 | 0.023522199 | 1.46808E-05 | -1.041524451 | 0.711 | 0.803 | 0.356625949 | 1.46808E-05 | 1.93662E-06 |
| Atp5a1 | 9.70643E-07 | -1.329980367 | 0.857 | 0.959 | 0.023578871 | 3.67927E-05 | -0.515234217 | 1 | 0.979 | 0.893768622 | 3.67927E-05 | 1.94129E-06 |
| Snf8 | 0.001004555 | -1.345421798 | 0.143 | 0.571 | 1 | 9.73081E-07 | -2.051637034 | 0.089 | 0.505 | 0.023638077 | 0.001004555 | 1.94616E-06 |
| Cbr2 | 0.128369301 | 0.369545675 | 0.714 | 0.507 | 1 | 9.75529E-07 | 0.528349991 | 0.844 | 0.626 | 0.023697553 | 0.128369301 | 1.95106E-06 |
| Slc29a1 | 0.000335414 | -0.111662536 | 0.524 | 0.91 | 1 | 9.76217E-07 | -0.916302572 | 0.378 | 0.862 | 0.023714265 | 0.000335414 | 1.95243E-06 |
| Creg1 | 1.01742E-06 | -2.600489894 | 0.19 | 0.731 | 0.024715249 | 0.000940789 | -0.805213895 | 0.578 | 0.709 | 1 | 0.000940789 | 2.03485E-06 |
| Btf3l4 | 0.67173385 | 0.802758208 | 0.333 | 0.377 | 1 | 1.03026E-06 | -3.700631706 | 0.022 | 0.401 | 0.025027174 | 0.67173385 | 2.06053E-06 |
| Smim30 | 0.001354073 | -1.478875242 | 0.143 | 0.552 | 1 | 1.03117E-06 | -1.937513824 | 0.133 | 0.554 | 0.025049076 | 0.001354073 | 2.06233E-06 |
| Cdipt | 0.005401073 | -1.212360268 | 0.143 | 0.496 | 1 | 1.03853E-06 | -0.93291681 | 0.156 | 0.585 | 0.025228 | 0.005401073 | 2.07706E-06 |
| Ndufa1 | 1.03868E-06 | -1.855007898 | 0.429 | 0.858 | 0.025231647 | 0.001336087 | -0.782446362 | 0.733 | 0.768 | 1 | 0.001336087 | 2.07736E-06 |
| Gnai3 | 0.094744962 | -0.37129478 | 0.81 | 0.937 | 1 | 1.04855E-06 | -0.653116202 | 0.978 | 0.976 | 0.025471489 | 0.094744962 | 2.09711E-06 |
| Map7 | 0.456850892 | 1.688670644 | 0.571 | 0.455 | 1 | 1.04917E-06 | 1.494310943 | 0.867 | 0.509 | 0.025486398 | 0.456850892 | 2.09834E-06 |
| Mcrip1 | 1.06816E-06 | -2.410657013 | 0.19 | 0.765 | 0.025947623 | 1.94139E-05 | -1.039953817 | 0.489 | 0.747 | 0.471602987 | 1.94139E-05 | 2.13631E-06 |
| Edf1 | 1.08555E-06 | -1.79320666 | 0.381 | 0.854 | 0.026370079 | 0.000291137 | -0.798460325 | 0.8 | 0.806 | 1 | 0.000291137 | 2.17109E-06 |
| Sc5d | 2.6878E-05 | -2.167530486 | 0.238 | 0.735 | 0.652919644 | 1.091E-06 | -1.378441803 | 0.2 | 0.654 | 0.026502625 | 2.6878E-05 | 2.182E-06 |
| Eif3i | 1.64599E-05 | -1.272647192 | 0.571 | 0.888 | 0.399844985 | 1.12767E-06 | -0.975138897 | 0.822 | 0.862 | 0.027393326 | 1.64599E-05 | 2.25534E-06 |
| Nectin1 | 0.729413059 | 0.995699863 | 0.095 | 0.131 | 1 | 1.144E-06 | 1.572648596 | 0.578 | 0.228 | 0.027790008 | 0.729413059 | 2.288E-06 |
| Taf11 | 0.026388475 | -0.726158839 | 0.19 | 0.496 | 1 | 1.16121E-06 | -1.885401792 | 0.111 | 0.533 | 0.02820806 | 0.026388475 | 2.32241E-06 |
| Snrpd1 | 1.4944E-05 | -1.619009982 | 0.476 | 0.851 | 0.363020716 | 1.1838E-06 | -0.841552129 | 0.8 | 0.903 | 0.028756768 | 1.4944E-05 | 2.36759E-06 |
| Irf2bp1 | 0.032119498 | -1.834022873 | 0.048 | 0.254 | 1 | 1.18657E-06 | -2.572428147 | 0.044 | 0.439 | 0.028824047 | 0.032119498 | 2.37313E-06 |
| Rere | 1.38876E-05 | 1.060738501 | 1 | 0.996 | 0.337357731 | 1.19363E-06 | 0.931346943 | 1 | 0.986 | 0.028995696 | 1.38876E-05 | 2.38726E-06 |
| Tln1 | 0.073185443 | -0.452431856 | 0.905 | 0.955 | 1 | 1.19549E-06 | -0.71942099 | 0.911 | 0.976 | 0.02904092 | 0.073185443 | 2.39098E-06 |
| Jagn1 | 0.00739755 | -1.447243633 | 0.143 | 0.455 | 1 | 1.19558E-06 | -2.901889915 | 0.067 | 0.45 | 0.029043027 | 0.00739755 | 2.39116E-06 |
| Glt8d1 | 0.012027946 | -0.718545655 | 0.381 | 0.739 | 1 | 1.20617E-06 | -1.128569694 | 0.178 | 0.647 | 0.029300371 | 0.012027946 | 2.41235E-06 |
| Rtl8b | 0.000585721 | -2.683780237 | 0.048 | 0.437 | 1 | 1.20705E-06 | -2.75987091 | 0.067 | 0.45 | 0.029321585 | 0.000585721 | 2.41409E-06 |
| Tex261 | 1.21626E-06 | -2.648156327 | 0.143 | 0.72 | 0.029545394 | 6.74122E-05 | -1.017367132 | 0.467 | 0.716 | 1 | 6.74122E-05 | 2.43252E-06 |
| Txndc15 | 1.21635E-06 | -2.355975576 | 0.19 | 0.75 | 0.029547621 | 1.93375E-06 | -0.845933509 | 0.378 | 0.796 | 0.046974673 | 1.93375E-06 | 2.4327E-06 |
| Erp44 | 3.89799E-05 | -1.302417271 | 0.476 | 0.896 | 0.946900139 | 1.21913E-06 | -0.831987901 | 0.778 | 0.924 | 0.029615159 | 3.89799E-05 | 2.43826E-06 |
| Hs3st3b1 | 0.011950935 | -0.737591411 | 0.333 | 0.754 | 1 | 1.2327E-06 | -1.675782078 | 0.244 | 0.668 | 0.029944644 | 0.011950935 | 2.46539E-06 |
| Higd2a | 1.2338E-06 | -2.120644099 | 0.286 | 0.791 | 0.029971521 | 0.03219687 | -0.686161223 | 0.711 | 0.702 | 1 | 0.03219687 | 2.4676E-06 |
| Snd1 | 1.27072E-06 | 1.376855561 | 1 | 0.959 | 0.030868288 | 1.1226E-05 | 1.235478532 | 1 | 0.99 | 0.272702844 | 1.1226E-05 | 2.54143E-06 |
| Tmem134 | 1.27921E-06 | -2.430564892 | 0.143 | 0.728 | 0.031074613 | 0.175121587 | -0.503340538 | 0.556 | 0.595 | 1 | 0.175121587 | 2.55842E-06 |
| Polr2g | 1.28897E-06 | -3.578893665 | 0.048 | 0.634 | 0.031311662 | 2.40347E-06 | -2.098287127 | 0.111 | 0.505 | 0.058385064 | 2.40347E-06 | 2.57794E-06 |
| Dnajc25 | 0.00057332 | -0.996631577 | 0.429 | 0.881 | 1 | 1.29986E-06 | -1.104917011 | 0.444 | 0.872 | 0.03157625 | 0.00057332 | 2.59972E-06 |
| Gpha2 | 0.013963266 | -1.36695417 | 0.19 | 0.504 | 1 | 1.30933E-06 | -2.943366551 | 0.156 | 0.574 | 0.031806224 | 0.013963266 | 2.61866E-06 |
| Srpx | 0.002177432 | -3.005708332 | 0.048 | 0.381 | 1 | 1.31105E-06 | -3.11134328 | 0.044 | 0.433 | 0.03184813 | 0.002177432 | 2.62211E-06 |
| Tpm3 | 0.000430582 | -0.719187566 | 0.952 | 0.981 | 1 | 1.31172E-06 | -0.725844925 | 1 | 1 | 0.031864418 | 0.000430582 | 2.62345E-06 |
| Stk26 | 0.297888354 | 1.317896341 | 0.524 | 0.731 | 1 | 1.33415E-06 | -1.530706705 | 0.156 | 0.609 | 0.032409239 | 0.297888354 | 2.6683E-06 |
| Psma4 | 2.65749E-05 | -1.584244563 | 0.381 | 0.81 | 0.645558161 | 1.34097E-06 | -0.859895826 | 0.822 | 0.872 | 0.032574763 | 2.65749E-05 | 2.68193E-06 |
| Mrpl54 | 2.56006E-06 | -3.555046923 | 0.048 | 0.616 | 0.062188901 | 1.35284E-06 | -2.434384995 | 0.111 | 0.502 | 0.03286315 | 2.56006E-06 | 2.70567E-06 |
| Gm2000 | 1.36938E-06 | -2.744080747 | 0.095 | 0.694 | 0.0332649 | 0.061148469 | -0.518561275 | 0.533 | 0.609 | 1 | 0.061148469 | 2.73875E-06 |
| Rdx | 0.741135878 | 0.335164932 | 0.952 | 0.97 | 1 | 1.37361E-06 | -0.723068513 | 0.978 | 0.986 | 0.03336782 | 0.741135878 | 2.74723E-06 |
| Glis3 | 1.30367E-05 | 1.523212091 | 0.905 | 0.75 | 0.316686943 | 1.38151E-06 | 1.541921557 | 0.978 | 0.837 | 0.03355953 | 1.30367E-05 | 2.76301E-06 |
| Sox6 | 0.010665208 | 1.050829025 | 1 | 0.993 | 1 | 1.38943E-06 | 1.083281063 | 0.956 | 0.972 | 0.033752106 | 0.010665208 | 2.77886E-06 |
| Mlf2 | 0.000404475 | -1.174851173 | 0.333 | 0.795 | 1 | 1.39185E-06 | -1.255526869 | 0.644 | 0.792 | 0.033810729 | 0.000404475 | 2.78369E-06 |
| Ppp1r1b | 2.93944E-05 | -4.907428814 | 0.048 | 0.549 | 0.714048252 | 1.39255E-06 | -2.964899712 | 0.044 | 0.436 | 0.033827881 | 2.93944E-05 | 2.7851E-06 |
| Ndufc1 | 1.40806E-06 | -2.184209227 | 0.381 | 0.799 | 0.03420469 | 0.212554942 | -0.361321533 | 0.667 | 0.664 | 1 | 0.212554942 | 2.81613E-06 |
| Pnp | 0.016952559 | -0.950022648 | 0.81 | 0.918 | 1 | 1.41054E-06 | -0.790497883 | 0.867 | 0.927 | 0.034264747 | 0.016952559 | 2.82107E-06 |
| Igsf10 | 0.001922201 | -4.239117569 | 0 | 0.332 | 1 | 1.41529E-06 | -3.108741485 | 0.044 | 0.433 | 0.034380107 | 0.001922201 | 2.83057E-06 |
| Mrps21 | 1.41676E-06 | -2.144582153 | 0.143 | 0.739 | 0.034416013 | 8.52979E-05 | -1.162417465 | 0.422 | 0.661 | 1 | 8.52979E-05 | 2.83352E-06 |

| | | | | | | | | | | | | |
|---|---|---|---|---|---|---|---|---|---|---|---|---|
| Mtfr1l | 0.208378404 | -0.259114036 | 0.286 | 0.493 | 1 | 1.45805E-06 | -2.72631835 | 0.044 | 0.433 | 0.035418841 | 0.208378404 | 2.91609E-06 |
| Gid8 | 0.055628627 | -0.785659851 | 0.19 | 0.437 | 1 | 1.46821E-06 | -1.960783604 | 0.067 | 0.474 | 0.035665824 | 0.055628627 | 2.93642E-06 |
| Dctn2 | 1.47384E-06 | -2.175893959 | 0.143 | 0.739 | 0.035802424 | 0.021514305 | -0.557935513 | 0.667 | 0.727 | 1 | 0.021514305 | 2.94767E-06 |
| Armc1 | 0.019019161 | -0.66251162 | 0.19 | 0.515 | 1 | 1.49192E-06 | -1.555332153 | 0.178 | 0.612 | 0.036241764 | 0.019019161 | 2.98384E-06 |
| Fundc1 | 0.000147089 | -2.46236492 | 0.095 | 0.534 | 1 | 1.50175E-06 | -1.871516266 | 0.089 | 0.519 | 0.036480407 | 0.000147089 | 3.00349E-06 |
| Slc40a1 | 0.045258302 | -4.107587946 | 0 | 0.164 | 1 | 1.51822E-06 | -3.837873422 | 0.067 | 0.446 | 0.036880538 | 0.045258302 | 3.03643E-06 |
| Dda1 | 0.001148203 | -1.252227651 | 0.19 | 0.649 | 1 | 1.53073E-06 | -1.409373663 | 0.156 | 0.599 | 0.037184487 | 0.001148203 | 3.06146E-06 |
| Txn1 | 1.53427E-06 | -1.619587175 | 0.857 | 0.97 | 0.037270535 | 9.41969E-06 | -0.866266005 | 0.956 | 0.952 | 0.228823211 | 9.41969E-06 | 3.06854E-06 |
| 1810055G02Rik | 1.54174E-06 | -2.651158815 | 0.19 | 0.728 | 0.037451868 | 5.02501E-06 | -1.142353909 | 0.244 | 0.671 | 0.122067487 | 5.02501E-06 | 3.08347E-06 |
| Prdx3 | 1.54451E-06 | -3.07218261 | 0.048 | 0.657 | 0.037519235 | 0.006003949 | -0.332752381 | 0.267 | 0.55 | 1 | 0.006003949 | 3.08902E-06 |
| Cat | 0.062652111 | -0.726158839 | 0.19 | 0.437 | 1 | 1.56703E-06 | -1.594912161 | 0.133 | 0.554 | 0.038066375 | 0.062652111 | 3.13406E-06 |
| Rheb | 0.036726369 | -0.535161614 | 0.952 | 0.966 | 1 | 1.57398E-06 | -0.735070824 | 0.867 | 0.965 | 0.038235179 | 0.036726369 | 3.14796E-06 |
| Pdzd11 | 0.014336316 | -1.01272876 | 0.19 | 0.496 | 1 | 1.57546E-06 | -3.024286546 | 0.044 | 0.429 | 0.038271083 | 0.014336316 | 3.15092E-06 |
| Nucb2 | 1.58562E-06 | -2.980864261 | 0.095 | 0.687 | 0.038517844 | 0.008900773 | -0.515707278 | 0.444 | 0.657 | 1 | 0.008900773 | 3.17123E-06 |
| Sparcl1 | 3.7256E-05 | -2.431403425 | 0.19 | 0.716 | 0.905021632 | 1.61086E-06 | -2.696305781 | 0.111 | 0.509 | 0.039130934 | 3.7256E-05 | 3.22171E-06 |
| Zfpm2 | 9.50397E-06 | 1.786369118 | 0.905 | 0.903 | 0.230870523 | 1.61896E-06 | 0.98095434 | 0.911 | 0.851 | 0.039327779 | 9.50397E-06 | 3.23792E-06 |
| Cox7a2l | 1.00333E-05 | -1.158434018 | 0.81 | 0.94 | 0.243728418 | 1.63717E-06 | -0.944325181 | 0.933 | 0.9 | 0.039770121 | 1.00333E-05 | 3.27434E-06 |
| Vps13b | 0.001365793 | 1.125118831 | 0.905 | 0.84 | 1 | 1.66974E-06 | 1.088802343 | 0.978 | 0.851 | 0.040561302 | 0.001365793 | 3.33948E-06 |
| Ssbp1 | 0.000731666 | -1.752492987 | 0.143 | 0.563 | 1 | 1.67906E-06 | -1.633295983 | 0.111 | 0.543 | 0.040787815 | 0.000731666 | 3.35812E-06 |
| Emc6 | 1.68196E-06 | -3.525900577 | 0.048 | 0.627 | 0.040858099 | 0.008861306 | -0.750914016 | 0.489 | 0.619 | 1 | 0.008861306 | 3.36391E-06 |
| Rftn2 | 0.00049226 | -1.656149118 | 0.143 | 0.586 | 1 | 1.70118E-06 | -1.418948351 | 0.178 | 0.616 | 0.041324946 | 0.00049226 | 3.40235E-06 |
| Washc3 | 0.003170773 | -1.101015292 | 0.143 | 0.552 | 1 | 1.70541E-06 | -1.946284034 | 0.067 | 0.471 | 0.04142774 | 0.003170773 | 3.41081E-06 |
| Arpc4 | 1.70856E-06 | -1.948945655 | 0.381 | 0.832 | 0.041504312 | 0.06618809 | -0.295693579 | 0.822 | 0.83 | 1 | 0.06618809 | 3.41711E-06 |
| Tfe3 | 0.176290435 | -0.34612779 | 0.19 | 0.373 | 1 | 1.71724E-06 | -1.756039048 | 0.133 | 0.561 | 0.04171516 | 0.176290435 | 3.43447E-06 |
| B3gat3 | 0.001252757 | -0.866796614 | 0.143 | 0.604 | 1 | 1.72678E-06 | -1.609709163 | 0.156 | 0.581 | 0.041946936 | 0.001252757 | 3.45356E-06 |
| Lsm5 | 0.005584769 | -0.83578333 | 0.143 | 0.5 | 1 | 1.72897E-06 | -2.064161343 | 0.067 | 0.471 | 0.042000208 | 0.005584769 | 3.45794E-06 |
| Bex3 | 1.74267E-06 | -2.53242862 | 0.143 | 0.728 | 0.042332963 | 3.53201E-05 | -0.893669408 | 0.422 | 0.747 | 0.857995075 | 3.53201E-05 | 3.48534E-06 |
| Hmgcs1 | 0.394118424 | -0.942454947 | 0.857 | 0.78 | 1 | 1.74363E-06 | -1.335164527 | 0.867 | 0.882 | 0.042356276 | 0.394118424 | 3.48726E-06 |
| Rbp4 | 0.000431222 | -4.488619561 | 0.048 | 0.444 | 1 | 1.75633E-06 | -1.815565115 | 0.222 | 0.647 | 0.042664805 | 0.000431222 | 3.51266E-06 |
| Tmem33 | 1.76939E-06 | -1.995507019 | 0.19 | 0.787 | 0.04298208 | 0.001281219 | -0.676330214 | 0.711 | 0.82 | 1 | 0.001281219 | 3.53878E-06 |
| Nudcd2 | 0.004563405 | -1.127846785 | 0.143 | 0.526 | 1 | 1.83973E-06 | -1.966542873 | 0.089 | 0.495 | 0.044690789 | 0.004563405 | 3.67946E-06 |
| Ptprk | 1.84436E-06 | 1.395804457 | 0.952 | 0.981 | 0.04480311 | 0.000857222 | 0.780569943 | 1 | 0.958 | 1 | 0.000857222 | 3.68871E-06 |
| Cers2 | 0.063775327 | -0.552296312 | 0.429 | 0.72 | 1 | 1.85245E-06 | -1.204452685 | 0.467 | 0.782 | 0.044999742 | 0.063775327 | 3.7049E-06 |
| Ppp1ca | 0.000931294 | -1.181442793 | 0.524 | 0.795 | 1 | 1.85713E-06 | -0.988220786 | 0.756 | 0.855 | 0.045113336 | 0.000931294 | 3.71425E-06 |
| Srprb | 0.01198078 | -0.826798546 | 0.238 | 0.582 | 1 | 1.86859E-06 | -1.393030393 | 0.111 | 0.55 | 0.045391823 | 0.01198078 | 3.73718E-06 |
| Ubtf | 0.068377682 | -0.229679744 | 0.429 | 0.694 | 1 | 1.90239E-06 | -1.024846128 | 0.8 | 0.907 | 0.046212883 | 0.068377682 | 3.80478E-06 |
| Mrps17 | 0.006354871 | -0.970084422 | 0.19 | 0.59 | 1 | 1.90507E-06 | -1.493516146 | 0.089 | 0.498 | 0.046277983 | 0.006354871 | 3.81014E-06 |
| Slc35a2 | 0.000730007 | -1.425217327 | 0.286 | 0.668 | 1 | 1.909E-06 | -1.158229668 | 0.156 | 0.581 | 0.046373429 | 0.000730007 | 3.818E-06 |
| Pqbp1 | 0.00445753 | -1.211585666 | 0.19 | 0.586 | 1 | 1.90939E-06 | -1.786412697 | 0.156 | 0.55 | 0.046382989 | 0.00445753 | 3.81878E-06 |
| Mdk | 0.164144765 | -1.275244304 | 0.143 | 0.287 | 1 | 1.912E-06 | -2.219000993 | 0.133 | 0.526 | 0.04644639 | 0.164144765 | 3.824E-06 |
| Mef2a | 0.096282803 | 1.054861935 | 0.952 | 0.925 | 1 | 1.92339E-06 | 0.896175246 | 1 | 0.986 | 0.046722926 | 0.096282803 | 3.84677E-06 |
| Raly | 1.93161E-06 | -1.761412305 | 0.333 | 0.877 | 0.046922783 | 3.58391E-05 | -0.754332726 | 0.756 | 0.872 | 0.870602606 | 3.58391E-05 | 3.86323E-06 |
| Bcas2 | 1.9461E-06 | -2.273760812 | 0.286 | 0.761 | 0.047274748 | 0.000417239 | -0.770883902 | 0.689 | 0.792 | 1 | 0.000417239 | 3.8922E-06 |
| Capns1 | 6.38512E-06 | -1.650408779 | 0.81 | 0.974 | 0.155107378 | 1.96149E-06 | -0.784366611 | 0.933 | 0.941 | 0.047648461 | 6.38512E-06 | 3.92297E-06 |
| Tmem258 | 1.9649E-06 | -1.845602391 | 0.333 | 0.851 | 0.047731406 | 0.00068805 | -0.920268444 | 0.622 | 0.734 | 1 | 0.00068805 | 3.9298E-06 |
| Txndc17 | 1.96745E-06 | -1.801961663 | 0.381 | 0.862 | 0.047793399 | 4.97276E-05 | -0.689022295 | 0.8 | 0.858 | 1 | 4.97276E-05 | 3.9349E-06 |
| Chst11 | 1.99343E-06 | 1.718773892 | 1 | 0.974 | 0.048424515 | 0.000376047 | 1.121241456 | 0.956 | 0.948 | 1 | 0.000376047 | 3.98687E-06 |
| Rras | 2.0161E-06 | -2.589262638 | 0.095 | 0.698 | 0.048975071 | 3.08127E-05 | -0.967691973 | 0.178 | 0.588 | 0.748502043 | 3.08127E-05 | 4.03219E-06 |
| Serinc5 | 2.02923E-06 | -1.536899576 | 0.19 | 0.84 | 0.049294062 | 0.000517385 | -0.319699163 | 0.644 | 0.924 | 1 | 0.000517385 | 4.05846E-06 |
| S100a1 | 2.02936E-06 | -2.349560736 | 0.667 | 0.896 | 0.049297306 | 6.45111E-05 | -0.861891847 | 0.733 | 0.82 | 1 | 6.45111E-05 | 4.05872E-06 |
| Akt3 | 2.03768E-06 | 1.347914406 | 0.952 | 0.847 | 0.049499251 | 1.49553E-05 | 0.871276321 | 1 | 0.969 | 0.363293367 | 1.49553E-05 | 4.07535E-06 |
| Hsbp1 | 5.80945E-06 | -1.438437736 | 0.381 | 0.888 | 0.141123229 | 2.07057E-06 | -0.993230226 | 0.711 | 0.844 | 0.050298401 | 5.80945E-06 | 4.14115E-06 |
| Snrpf | 2.64315E-06 | -1.552908012 | 0.619 | 0.91 | 0.064207478 | 2.12624E-06 | -0.937079343 | 0.844 | 0.869 | 0.051650514 | 2.64315E-06 | 4.25247E-06 |
| Eif3m | 2.13227E-06 | -1.623144439 | 0.524 | 0.869 | 0.051797081 | 2.66777E-06 | -0.873707147 | 0.844 | 0.879 | 0.064805526 | 2.66777E-06 | 4.26453E-06 |
| Nfic | 2.1363E-06 | -2.057855828 | 0.476 | 0.892 | 0.051894929 | 3.36463E-06 | -0.864523062 | 0.889 | 0.976 | 0.081733628 | 3.36463E-06 | 4.27259E-06 |
| Ap2m1 | 0.002510412 | -1.073973665 | 0.333 | 0.728 | 1 | 2.14863E-06 | -1.143141363 | 0.622 | 0.827 | 0.052194638 | 0.002510412 | 4.29727E-06 |
| Irf2bpl | 2.17081E-06 | -3.669043694 | 0.095 | 0.653 | 0.052733374 | 1.23137E-05 | -1.206223949 | 0.578 | 0.806 | 0.299125565 | 1.23137E-05 | 4.34162E-06 |
| Eno1 | 2.18409E-06 | -2.051878513 | 0.381 | 0.817 | 0.053055797 | 0.001870666 | -0.603604662 | 0.733 | 0.806 | 1 | 0.001870666 | 4.36817E-06 |
| Naf1 | 0.009678828 | -1.298920886 | 0.095 | 0.414 | 1 | 2.19241E-06 | -2.244282112 | 0.067 | 0.453 | 0.053258144 | 0.009678828 | 4.38483E-06 |
| Commd2 | 0.006187695 | -2.337455488 | 0.048 | 0.336 | 1 | 2.22732E-06 | -2.439324045 | 0.089 | 0.471 | 0.054106019 | 0.006187695 | 4.45463E-06 |
| Trnt1 | 0.002796717 | -1.837190152 | 0.143 | 0.489 | 1 | 2.23034E-06 | -1.517354504 | 0.089 | 0.495 | 0.054179354 | 0.002796717 | 4.46067E-06 |
| Magt1 | 2.23425E-06 | -2.090798288 | 0.19 | 0.799 | 0.05427435 | 0.015134887 | -0.508878722 | 0.667 | 0.772 | 1 | 0.015134887 | 4.46849E-06 |
| Fam98b | 0.193683205 | 0.512779891 | 0.238 | 0.407 | 1 | 2.241E-06 | -1.521498558 | 0.133 | 0.564 | 0.05443848 | 0.193683205 | 4.482E-06 |
| Slirp | 0.029595773 | -0.474126927 | 0.19 | 0.5 | 1 | 2.25218E-06 | -3.035745661 | 0.044 | 0.412 | 0.054709877 | 0.029595773 | 4.50435E-06 |
| Peli1 | 0.528680752 | 0.202466049 | 0.714 | 0.884 | 1 | 2.25777E-06 | 1.774975219 | 0.956 | 0.862 | 0.0548458 | 0.528680752 | 4.51554E-06 |
| Smim10l1 | 0.000133842 | -1.268742738 | 0.524 | 0.854 | 1 | 2.29188E-06 | -1.069834562 | 0.667 | 0.851 | 0.055674312 | 0.000133842 | 4.58375E-06 |
| S100a6 | 2.31797E-06 | -2.44571547 | 0.857 | 0.944 | 0.056308173 | 0.006983496 | -0.735924168 | 0.956 | 0.931 | 1 | 0.006983496 | 4.63594E-06 |
| Ptbp1 | 0.010670369 | -0.687415758 | 0.524 | 0.847 | 1 | 2.32694E-06 | -0.873409815 | 0.867 | 0.917 | 0.056525916 | 0.010670369 | 4.65387E-06 |
| Arl2 | 0.00062866 | -1.34303652 | 0.095 | 0.545 | 1 | 2.34044E-06 | -3.72631835 | 0.022 | 0.384 | 0.056854066 | 0.00062866 | 4.68088E-06 |
| Pabpc1 | 0.000965114 | -0.594174962 | 0.952 | 0.996 | 1 | 2.3814E-06 | -0.606551128 | 1 | 0.993 | 0.057848942 | 0.000965114 | 4.76279E-06 |
| Pmpcb | 0.000468076 | -2.005708332 | 0.095 | 0.519 | 1 | 2.38682E-06 | -1.515197041 | 0.178 | 0.588 | 0.057980662 | 0.000468076 | 4.77364E-06 |
| Ahr | 0.857229584 | 0.33137957 | 0.143 | 0.127 | 1 | 2.3962E-06 | 1.122357632 | 0.422 | 0.125 | 0.058208527 | 0.857229584 | 4.7924E-06 |
| Apbb2 | 0.006360804 | 1.296831423 | 1 | 0.981 | 1 | 2.44124E-06 | 1.090607485 | 1 | 0.993 | 0.059302628 | 0.006360804 | 4.88248E-06 |
| Tmem106c | 0.000938978 | -2.574155746 | 0.048 | 0.418 | 1 | 2.44463E-06 | -4.446210431 | 0 | 0.353 | 0.059385047 | 0.000938978 | 4.88926E-06 |
| Atp6v1f | 3.88938E-06 | -1.480805269 | 0.571 | 0.907 | 0.094480845 | 2.44789E-06 | -0.975890496 | 0.8 | 0.872 | 0.059464176 | 3.88938E-06 | 4.89578E-06 |
| F2r | 0.251704191 | -0.849790188 | 0 | 0.06 | 1 | 2.4494E-06 | 0.373744528 | 0.489 | 0.156 | 0.059500783 | 0.251704191 | 4.89879E-06 |
| Ctdsp1 | 0.047572627 | -0.735619168 | 0.19 | 0.459 | 1 | 2.45432E-06 | -1.633295983 | 0.178 | 0.592 | 0.059620266 | 0.047572627 | 4.90863E-06 |
| Fxyd6 | 0.000121282 | -1.89208191 | 0.333 | 0.746 | 1 | 2.48983E-06 | -0.765153578 | 0.2 | 0.654 | 0.060482861 | 0.000121282 | 4.97965E-06 |
| Rex1bd | 2.49964E-06 | -4.274595464 | 0 | 0.575 | 0.060721142 | 1.05652E-05 | -1.5585134 | 0.156 | 0.526 | 0.256650689 | 1.05652E-05 | 4.99926E-06 |
| Tmbim6 | 2.50971E-06 | -1.100802771 | 0.905 | 0.989 | 0.06096577 | 3.36232E-06 | -0.586732778 | 0.978 | 0.997 | 0.081677381 | 3.36232E-06 | 5.01941E-06 |
| Ext2 | 0.102456896 | 0.566856564 | 0.81 | 0.866 | 1 | 2.52808E-06 | -0.812228989 | 0.356 | 0.862 | 0.061412233 | 0.102456896 | 5.05616E-06 |
| Tmem167 | 5.31624E-05 | -1.250279379 | 0.667 | 0.933 | 1 | 2.5369E-06 | -0.815398389 | 0.778 | 0.948 | 0.06162648 | 5.31624E-05 | 5.0738E-06 |
| Lsm3 | 0.000342567 | -2.081115734 | 0.095 | 0.53 | 1 | 2.55577E-06 | -2.510699157 | 0.067 | 0.446 | 0.062084856 | 0.000342567 | 5.11154E-06 |
| Ccpg1 | 2.56702E-06 | -1.769171728 | 0.429 | 0.877 | 0.062357997 | 1.36947E-05 | -0.741593614 | 0.822 | 0.889 | 0.332671124 | 1.36947E-05 | 5.13403E-06 |
| Anxa3 | 0.01356061 | 0.575143735 | 0.81 | 0.552 | 1 | 2.56944E-06 | 0.83507568 | 0.778 | 0.346 | 0.062416723 | 0.01356061 | 5.13886E-06 |
| Castor2 | 0.244302408 | 0.289107917 | 0.095 | 0.224 | 1 | 2.57181E-06 | -1.638855509 | 0.156 | 0.588 | 0.062474384 | 0.244302408 | 5.14361E-06 |
| Nfe2l1 | 2.59214E-06 | -1.064580367 | 0.857 | 0.978 | 0.062968259 | 0.000727558 | -0.430981153 | 0.911 | 0.945 | 1 | 0.000727558 | 5.18427E-06 |
| Tma7 | 0.000409353 | -0.93725303 | 0.714 | 0.922 | 1 | 2.59895E-06 | -1.024145665 | 0.822 | 0.851 | 0.063133776 | 0.000409353 | 5.1979E-06 |
| Txnl4a | 0.00297428 | -1.25358293 | 0.143 | 0.526 | 1 | 2.60771E-06 | -0.613909152 | 0.178 | 0.637 | 0.063346407 | 0.00297428 | 5.21541E-06 |
| Atp5pb | 2.60921E-06 | -1.355467853 | 0.667 | 0.944 | 0.063382989 | 8.79906E-06 | -0.765721601 | 0.911 | 0.941 | 0.213746792 | 8.79906E-06 | 5.21842E-06 |
| Serf1 | 0.000695143 | -2.059582573 | 0.095 | 0.519 | 1 | 2.61751E-06 | -1.729708939 | 0.067 | 0.467 | 0.0635846 | 0.000695143 | 5.23502E-06 |
| Xylt1 | 2.65031E-06 | 1.417928103 | 1 | 0.937 | 0.064381349 | 5.3595E-05 | 0.949217686 | 1 | 0.958 | 1 | 5.3595E-05 | 5.30061E-06 |
| Frmd6 | 0.003413344 | -1.828728573 | 0.238 | 0.608 | 1 | 2.65116E-06 | -0.903401381 | 0.778 | 0.924 | 0.064401901 | 0.003413344 | 5.30231E-06 |
| Nuak1 | 0.251129249 | 1.171271427 | 0.238 | 0.407 | 1 | 2.66112E-06 | -0.679061831 | 0.2 | 0.619 | 0.064643961 | 0.251129249 | 5.32224E-06 |
| Gpr108 | 2.58857E-05 | -3.91868527 | 0 | 0.496 | 0.628815258 | 2.68767E-06 | -1.517354504 | 0.133 | 0.547 | 0.065288781 | 2.58857E-05 | 5.37532E-06 |
| Polr1c | 0.045960866 | -0.587895802 | 0.238 | 0.515 | 1 | 2.69766E-06 | -1.420215222 | 0.133 | 0.561 | 0.06553154 | 0.045960866 | 5.39531E-06 |
| Gpc1 | 0.000106264 | -1.487414897 | 0.286 | 0.791 | 1 | 2.70412E-06 | -1.095004543 | 0.289 | 0.779 | 0.0656886 | 0.000106264 | 5.40824E-06 |
| Mest | 0.059958097 | -1.566906007 | 0.238 | 0.451 | 1 | 2.73323E-06 | -2.179874662 | 0.089 | 0.481 | 0.066395741 | 0.059958097 | 5.46646E-06 |

| | | | | | | | | | | | | |
|---|---|---|---|---|---|---|---|---|---|---|---|---|
| Aif1l | 0.002913627 | -1.63000898 | 0.095 | 0.451 | 1 | 2.75301E-06 | -3.72631835 | 0.022 | 0.381 | 0.066876015 | 0.002913627 | 5.506E-06 |
| Foxp1 | 4.15158E-06 | 1.340942366 | 1 | 0.974 | 0.100850238 | 2.76982E-06 | 1.000938033 | 1 | 0.993 | 0.06728445 | 4.15158E-06 | 5.53963E-06 |
| Usp34 | 0.001717745 | 0.634952519 | 1 | 0.959 | 1 | 2.78796E-06 | 0.746463372 | 1 | 0.997 | 0.067725134 | 0.001717745 | 5.57591E-06 |
| Lgalsl | 7.93726E-05 | -2.836662754 | 0.143 | 0.612 | 1 | 2.79468E-06 | -1.901889915 | 0.178 | 0.567 | 0.06788826 | 7.93726E-05 | 5.58934E-06 |
| Adam28 | 0.07959214 | 4.410737362 | 0.048 | 0.007 | 1 | 2.80297E-06 | 2.235613609 | 0.222 | 0.038 | 0.06808984 | 0.07959214 | 5.60594E-06 |
| Cox5b | 2.84183E-06 | -1.398927721 | 0.714 | 0.925 | 0.069033835 | 0.001087703 | -0.714865004 | 0.911 | 0.855 | 1 | 0.001087703 | 5.68366E-06 |
| Tmem109 | 0.000266242 | -2.933558546 | 0.048 | 0.466 | 1 | 2.84627E-06 | -1.921789472 | 0.2 | 0.571 | 0.069141471 | 0.000266242 | 5.69252E-06 |
| Arhgdia | 2.19645E-05 | -1.593014773 | 0.381 | 0.832 | 0.533560579 | 2.86114E-06 | -1.044847868 | 0.733 | 0.848 | 0.069502723 | 2.19645E-05 | 5.72226E-06 |
| Gclc | 0.419082693 | -0.249060372 | 0.619 | 0.459 | 1 | 2.87611E-06 | 1.066660447 | 0.889 | 0.516 | 0.069866493 | 0.419082693 | 5.75221E-06 |
| Tbrg1 | 0.001593926 | -0.738041831 | 0.714 | 0.944 | 1 | 2.88992E-06 | -0.90633582 | 0.8 | 0.952 | 0.070202042 | 0.001593926 | 5.77984E-06 |
| Scp2 | 2.89749E-06 | -1.914193221 | 0.381 | 0.806 | 0.070385863 | 0.1453022 | -0.235581459 | 0.822 | 0.74 | 1 | 0.1453022 | 5.79497E-06 |
| Mettl9 | 5.1351E-05 | -1.664707503 | 0.333 | 0.791 | 1 | 2.89855E-06 | -1.048394028 | 0.667 | 0.824 | 0.070411492 | 5.1351E-05 | 5.79708E-06 |
| Dsg2 | 0.532436372 | 1.088809267 | 0 | 0.019 | 1 | 2.90566E-06 | 1.19274696 | 0.444 | 0.138 | 0.070584304 | 0.532436372 | 5.81131E-06 |
| Wdr75 | 0.194568285 | -0.295854583 | 0.286 | 0.47 | 1 | 2.9297E-06 | -1.115293553 | 0.133 | 0.571 | 0.071168187 | 0.194568285 | 5.85938E-06 |
| Cisd1 | 2.95789E-06 | -3.625436251 | 0.048 | 0.612 | 0.071853035 | 0.50971476 | -0.316927414 | 0.533 | 0.519 | 1 | 0.50971476 | 5.91577E-06 |
| Gng11 | 0.001868171 | -3.081115734 | 0.048 | 0.388 | 1 | 2.9686E-06 | -0.722919774 | 0.089 | 0.512 | 0.072113235 | 0.001868171 | 5.93719E-06 |
| Ier3ip1 | 2.99997E-06 | -1.34612779 | 0.476 | 0.892 | 0.072875216 | 5.3648E-06 | -0.839695153 | 0.711 | 0.907 | 0.130321707 | 5.3648E-06 | 5.99993E-06 |
| Cacybp | 0.002209141 | -1.358916851 | 0.429 | 0.72 | 1 | 3.01884E-06 | -0.979892427 | 0.689 | 0.848 | 0.073333575 | 0.002209141 | 6.03766E-06 |
| Cct3 | 0.067856829 | -0.583330308 | 0.667 | 0.84 | 1 | 3.05107E-06 | -0.930744777 | 0.822 | 0.886 | 0.074116642 | 0.067856829 | 6.10213E-06 |
| Tuba1c | 8.84765E-06 | -2.040714782 | 0.762 | 0.925 | 0.214927113 | 3.05865E-06 | -0.795659588 | 0.933 | 0.952 | 0.074300747 | 8.84765E-06 | 6.11729E-06 |
| Slc35e4 | 0.0103092 | -0.908023454 | 0.333 | 0.66 | 1 | 3.07188E-06 | -1.234706326 | 0.244 | 0.727 | 0.074622192 | 0.0103092 | 6.14376E-06 |
| Cops7a | 0.003854387 | -0.889164427 | 0.19 | 0.631 | 1 | 3.1153E-06 | -1.321928095 | 0.222 | 0.619 | 0.075676942 | 0.003854387 | 6.2306E-06 |
| Cdk2ap1 | 0.001756636 | -0.970084422 | 0.524 | 0.884 | 1 | 3.12015E-06 | -1.115093164 | 0.6 | 0.817 | 0.075794686 | 0.001756636 | 6.24029E-06 |
| Gla | 0.00958426 | -2.880817084 | 0 | 0.25 | 1 | 3.15401E-06 | -4.776359032 | 0 | 0.349 | 0.076617119 | 0.00958426 | 6.308E-06 |
| Eif3g | 3.18173E-06 | -2.434752689 | 0.095 | 0.687 | 0.077290598 | 0.009249749 | -0.36496011 | 0.511 | 0.758 | 1 | 0.009249749 | 6.36345E-06 |
| Ndufa8 | 3.20861E-06 | -2.161647073 | 0.333 | 0.799 | 0.077943548 | 0.010732192 | -0.526380779 | 0.622 | 0.716 | 1 | 0.010732192 | 6.41721E-06 |
| Nrp2 | 0.00946789 | -2.303508156 | 0.286 | 0.563 | 1 | 3.23165E-06 | -1.410121561 | 0.467 | 0.858 | 0.078503241 | 0.00946789 | 6.46329E-06 |
| Mien1 | 3.28298E-06 | -3.081115734 | 0.095 | 0.646 | 0.07975017 | 0.008671354 | -1.104829973 | 0.333 | 0.495 | 1 | 0.008671354 | 6.56595E-06 |
| Dpy19l1 | 0.010781358 | -1.640924758 | 0.143 | 0.429 | 1 | 3.2852E-06 | -1.238924902 | 0.111 | 0.533 | 0.079803998 | 0.010781358 | 6.57038E-06 |
| Uba52 | 3.32274E-06 | -1.395123314 | 0.619 | 0.899 | 0.080715992 | 0.000133505 | -0.539319835 | 0.822 | 0.855 | 1 | 0.000133505 | 6.64547E-06 |
| Tmed7 | 0.012705352 | -0.582567986 | 0.524 | 0.843 | 1 | 3.35541E-06 | -0.920268444 | 0.756 | 0.862 | 0.081509559 | 0.012705352 | 6.7108E-06 |
| Dnajc8 | 3.42271E-06 | -1.860206804 | 0.333 | 0.821 | 0.083144504 | 0.299182601 | -0.195704373 | 0.867 | 0.803 | 1 | 0.299182601 | 6.84541E-06 |
| Cdk19 | 0.013813272 | 1.468657338 | 0.619 | 0.422 | 1 | 3.43094E-06 | 1.290755163 | 0.8 | 0.394 | 0.083344275 | 0.013813272 | 6.86186E-06 |
| Ppp1r11 | 5.64572E-05 | -2.331852782 | 0.048 | 0.56 | 1 | 3.44938E-06 | -1.960783604 | 0.178 | 0.543 | 0.0837923 | 5.64572E-05 | 6.89874E-06 |
| Snhg1 | 7.84037E-05 | -1.321210158 | 0.524 | 0.888 | 1 | 3.50966E-06 | -1.026741402 | 0.778 | 0.851 | 0.085256566 | 7.84037E-05 | 7.0193E-06 |
| Mbd3 | 0.002314627 | -1.098817736 | 0.143 | 0.571 | 1 | 3.51583E-06 | -1.119794945 | 0.2 | 0.64 | 0.085406432 | 0.002314627 | 7.03164E-06 |
| Trmt10c | 0.014892176 | -1.496153234 | 0.095 | 0.369 | 1 | 3.54463E-06 | -1.195248857 | 0.111 | 0.529 | 0.086106141 | 0.014892176 | 7.08925E-06 |
| Atp5g3 | 9.47119E-05 | -1.124594371 | 0.619 | 0.881 | 1 | 3.5514E-06 | -1.021188843 | 0.822 | 0.862 | 0.086270497 | 9.47119E-05 | 7.10278E-06 |
| Anapc16 | 3.55591E-06 | -1.992353586 | 0.238 | 0.772 | 0.086380057 | 0.001823409 | -0.486852415 | 0.578 | 0.785 | 1 | 0.001823409 | 7.1118E-06 |
| Gm525 | 0.000483332 | -4.680844946 | 0.048 | 0.448 | 1 | 3.56499E-06 | -1.529377032 | 0.111 | 0.529 | 0.086600781 | 0.000483332 | 7.12997E-06 |
| Commd7 | 0.003663473 | -1.709932525 | 0.095 | 0.437 | 1 | 3.5696E-06 | -1.994999319 | 0.111 | 0.505 | 0.086712655 | 0.003663473 | 7.13918E-06 |
| Odr4 | 0.00150556 | -0.775789607 | 0.286 | 0.735 | 1 | 3.58676E-06 | -1.197939378 | 0.289 | 0.682 | 0.087129565 | 0.00150556 | 7.17351E-06 |
| 1110038B12Rik | 3.61099E-06 | -2.586755783 | 0.143 | 0.701 | 0.087718252 | 0.578175857 | -0.196073019 | 0.622 | 0.602 | 1 | 0.578175857 | 7.22197E-06 |
| Gm10076 | 3.62628E-06 | -1.401760863 | 0.571 | 0.925 | 0.08808961 | 3.66622E-05 | -0.9728117 | 0.756 | 0.81 | 0.890598621 | 3.66622E-05 | 7.25255E-06 |
| Uqcrq | 3.64386E-06 | -1.822654058 | 0.524 | 0.832 | 0.08851659 | 0.001625107 | -0.826788459 | 0.756 | 0.74 | 1 | 0.001625107 | 7.2877E-06 |
| Ncbp2as2 | 1.19503E-05 | -3.998653574 | 0 | 0.522 | 0.290296241 | 3.70121E-06 | -2.924257728 | 0.067 | 0.426 | 0.089909762 | 1.19503E-05 | 7.4024E-06 |
| Mcee | 0.000155399 | -3.65714511 | 0 | 0.433 | 1 | 3.71443E-06 | -3.555332153 | 0.022 | 0.374 | 0.090230988 | 0.000155399 | 7.42885E-06 |
| Cyb5r3 | 4.32585E-06 | -1.725759145 | 0.476 | 0.881 | 0.105083647 | 3.76235E-06 | -1.03380159 | 0.778 | 0.844 | 0.091394938 | 4.32585E-06 | 7.52468E-06 |
| Mgat1 | 0.260774506 | 0.392485657 | 0.238 | 0.433 | 1 | 3.77127E-06 | -1.061088509 | 0.178 | 0.619 | 0.091611574 | 0.260774506 | 7.54252E-06 |
| Gusb | 0.001272446 | -3.268742738 | 0 | 0.347 | 1 | 3.78515E-06 | -2.946284034 | 0.022 | 0.388 | 0.091948937 | 0.001272446 | 7.57029E-06 |
| G6pdx | 0.4987111 | 0.24081236 | 0.095 | 0.164 | 1 | 3.79186E-06 | 1.953720176 | 0.467 | 0.166 | 0.092111844 | 0.4987111 | 7.5837E-06 |
| Abi3bp | 0.018781182 | -1.4932879 | 0.81 | 0.795 | 1 | 3.81384E-06 | -1.622781894 | 0.889 | 0.893 | 0.092645706 | 0.018781182 | 7.62766E-06 |
| Gm19303 | 3.87165E-06 | 3.057100407 | 0.286 | 0.041 | 0.094050141 | 0.809040217 | 1.460680165 | 0.022 | 0.017 | 1 | 0.809040217 | 7.74329E-06 |
| Polr2e | 0.000411823 | -2.107587946 | 0.095 | 0.522 | 1 | 3.87596E-06 | -2.006226574 | 0.2 | 0.55 | 0.094154776 | 0.000411823 | 7.7519E-06 |
| Spag9 | 3.8811E-06 | 1.023142349 | 1 | 0.989 | 0.094279605 | 1.89278E-05 | 0.746086523 | 1 | 0.99 | 0.459794425 | 1.89278E-05 | 7.76218E-06 |
| Rhod | 0.146962561 | -0.637814383 | 0.143 | 0.31 | 1 | 3.89311E-06 | -2.454430938 | 0.022 | 0.388 | 0.094571534 | 0.146962561 | 7.78621E-06 |
| Cdc42 | 0.000299777 | -0.614094491 | 1 | 0.993 | 1 | 3.89446E-06 | -0.519719101 | 1 | 1 | 0.094604139 | 0.000299777 | 7.7889E-06 |
| Sdk2 | 3.9211E-06 | 1.567462866 | 0.952 | 0.955 | 0.09525145 | 7.96011E-06 | 0.956746251 | 0.978 | 0.965 | 0.193367003 | 7.96011E-06 | 7.84219E-06 |
| Derl1 | 3.93708E-06 | -1.697250877 | 0.238 | 0.843 | 0.095639443 | 3.56718E-05 | -0.856660661 | 0.733 | 0.848 | 0.86654009 | 3.56718E-05 | 7.87414E-06 |
| Efna1 | 3.95479E-06 | -1.950719097 | 0.333 | 0.843 | 0.096069808 | 9.10766E-06 | -0.880972947 | 0.622 | 0.907 | 0.22124321 | 9.10766E-06 | 7.90957E-06 |
| Rabgap1l | 3.95599E-06 | 1.263365717 | 1 | 0.996 | 0.096098855 | 4.50476E-05 | 0.851099754 | 1 | 0.997 | 1 | 4.50476E-05 | 7.91196E-06 |
| Tasor | 3.97243E-06 | 1.28058499 | 0.905 | 0.672 | 0.096498251 | 0.519136912 | 0.361144491 | 0.889 | 0.803 | 1 | 0.519136912 | 7.94484E-06 |
| Ndufa6 | 3.98077E-06 | -1.440889215 | 0.667 | 0.892 | 0.096700886 | 0.002619144 | -0.572000534 | 0.8 | 0.806 | 1 | 0.002619144 | 7.96153E-06 |
| Aco1 | 0.214542189 | -0.208871282 | 0.143 | 0.302 | 1 | 3.99883E-06 | -2.166593141 | 0.044 | 0.415 | 0.097139591 | 0.214542189 | 7.99765E-06 |
| Rnaset2b | 4.017E-06 | -2.092520498 | 0.143 | 0.716 | 0.097580912 | 0.000832231 | -0.787247349 | 0.489 | 0.696 | 1 | 0.000832231 | 8.03398E-06 |
| Arf6 | 0.001198185 | -1.214971481 | 0.381 | 0.754 | 1 | 4.03564E-06 | -0.944059369 | 0.756 | 0.865 | 0.098033759 | 0.001198185 | 8.07126E-06 |
| Ppt1 | 5.09628E-05 | -1.722833014 | 0.143 | 0.683 | 1 | 4.04201E-06 | -1.363221066 | 0.311 | 0.668 | 0.098188516 | 5.09628E-05 | 8.084E-06 |
| Ndufb10 | 4.05449E-06 | -1.797533951 | 0.381 | 0.817 | 0.098491576 | 2.39913E-05 | -0.939514987 | 0.778 | 0.817 | 0.582797292 | 2.39913E-05 | 8.10896E-06 |
| Tspan8 | 0.144945336 | -1.970084422 | 0 | 0.093 | 1 | 4.05627E-06 | 1.388889482 | 0.511 | 0.176 | 0.098534898 | 0.144945336 | 8.11252E-06 |
| Snhg6 | 0.010638558 | -0.871760588 | 0.429 | 0.731 | 1 | 4.08399E-06 | -1.138266565 | 0.156 | 0.602 | 0.099208271 | 0.010638558 | 8.16796E-06 |
| Nucks1 | 4.09305E-06 | -1.478875242 | 0.667 | 0.933 | 0.099428317 | 9.02661E-06 | -0.779515916 | 0.844 | 0.924 | 0.219274302 | 9.02661E-06 | 8.18608E-06 |
| Rflna | 3.64265E-05 | -2.693521631 | 0.238 | 0.701 | 0.88487197 | 4.09438E-06 | -1.182580976 | 0.533 | 0.858 | 0.099460763 | 3.64265E-05 | 8.18875E-06 |
| Med12 | 0.040190897 | -0.794997716 | 0.143 | 0.414 | 1 | 4.15479E-06 | -1.526380779 | 0.133 | 0.536 | 0.100928083 | 0.040190897 | 8.30956E-06 |
| Ik | 0.004074414 | -0.854263665 | 0.619 | 0.884 | 1 | 4.16482E-06 | -0.9159651 | 0.844 | 0.889 | 0.10117182 | 0.004074414 | 8.32962E-06 |
| Bad | 6.57862E-05 | -3.865387043 | 0 | 0.466 | 1 | 4.20248E-06 | -2.32442195 | 0.067 | 0.446 | 0.102086646 | 6.57862E-05 | 8.40494E-06 |
| Krr1 | 0.812634041 | 0.719575457 | 0.286 | 0.377 | 1 | 4.29083E-06 | -2.065120263 | 0.089 | 0.481 | 0.104232812 | 0.812634041 | 8.58164E-06 |
| Armcx2 | 0.034166139 | -1.801961663 | 0.048 | 0.25 | 1 | 4.31864E-06 | -2.432404631 | 0.089 | 0.457 | 0.104908329 | 0.034166139 | 8.63726E-06 |
| Gm27201 | 4.38626E-06 | 4.673771768 | 0.143 | 0.007 | 0.106550985 | 0.430487676 | 0.361144491 | 0 | 0.014 | 1 | 0.430487676 | 8.7725E-06 |
| Larp7 | 0.007372247 | -0.64213254 | 0.143 | 0.493 | 1 | 4.41543E-06 | -1.638855509 | 0.111 | 0.498 | 0.107259587 | 0.007372247 | 8.83084E-06 |
| Ppdpf | 2.17551E-05 | -2.413691074 | 0.143 | 0.649 | 0.528474261 | 4.43187E-06 | -1.622112245 | 0.178 | 0.557 | 0.107658963 | 2.17551E-05 | 8.86372E-06 |
| Cox4i2 | 4.43635E-06 | -4.381510668 | 0.048 | 0.612 | 0.107767919 | 8.75201E-06 | -2.407650584 | 0.089 | 0.457 | 0.212603927 | 8.75201E-06 | 8.87269E-06 |
| Zfp830 | 0.457753695 | -0.156303231 | 0.238 | 0.34 | 1 | 4.44101E-06 | -1.975138897 | 0.067 | 0.45 | 0.107881034 | 0.457753695 | 8.882E-06 |
| Mtch1 | 4.5217E-06 | -1.978304929 | 0.429 | 0.869 | 0.109841093 | 0.000185204 | -0.829860228 | 0.822 | 0.872 | 1 | 0.000185204 | 9.04338E-06 |
| Col4a1 | 0.077524622 | -1.704739856 | 0.095 | 0.284 | 1 | 4.55129E-06 | -3.139657562 | 0.044 | 0.415 | 0.110560023 | 0.077524622 | 9.10257E-06 |
| Kcnip1 | 4.64425E-06 | 4.410737362 | 0.143 | 0.007 | 0.112818158 | 0.500244121 | 0.776181991 | 0.022 | 0.045 | 1 | 0.500244121 | 9.28848E-06 |
| Cers6 | 0.381959846 | 0.629377648 | 0.095 | 0.183 | 1 | 4.66761E-06 | 0.776181991 | 0.578 | 0.208 | 0.113385551 | 0.381959846 | 9.3352E-06 |
| Ssr1 | 0.002217564 | -0.780794096 | 0.619 | 0.97 | 1 | 4.72373E-06 | -0.664657002 | 0.889 | 0.952 | 0.114748791 | 0.002217564 | 9.44743E-06 |
| P2ry10b | 0.045333197 | -2.348596045 | 0.048 | 0.231 | 1 | 4.73188E-06 | -3.104829973 | 0.022 | 0.381 | 0.114946846 | 0.045333197 | 9.46374E-06 |
| Mical2 | 0.732276218 | 0.323274521 | 0.095 | 0.075 | 1 | 4.76749E-06 | 1.15951063 | 0.333 | 0.087 | 0.115811887 | 0.732276218 | 9.53496E-06 |
| Nans | 0.018387432 | -1.298920886 | 0.143 | 0.41 | 1 | 4.79831E-06 | -2.361321533 | 0.067 | 0.443 | 0.116560559 | 0.018387432 | 9.5966E-06 |
| Atxn7l3b | 2.89933E-05 | -1.273991109 | 0.762 | 0.925 | 0.704304775 | 4.86098E-06 | -0.664166597 | 0.867 | 0.934 | 0.118082921 | 2.89933E-05 | 9.72194E-06 |
| Pard3 | 6.74151E-05 | 1.269528584 | 1 | 0.914 | 1 | 4.86229E-06 | 0.931609475 | 1 | 0.997 | 0.118114806 | 6.74151E-05 | 9.72456E-06 |
| Snrpc | 4.8839E-06 | -2.225081509 | 0.143 | 0.716 | 0.118639673 | 0.097889542 | -0.291542095 | 0.556 | 0.654 | 1 | 0.097889542 | 9.76777E-06 |
| Rab13 | 0.43034024 | 0.422233001 | 0.286 | 0.459 | 1 | 4.98784E-06 | -2.038026603 | 0.067 | 0.446 | 0.121164496 | 0.43034024 | 9.97565E-06 |
| Alg14 | 0.00298231 | -1.12324121 | 0.286 | 0.694 | 1 | 4.99913E-06 | -1.404390255 | 0.133 | 0.536 | 0.121438927 | 0.00298231 | 9.99824E-06 |
| Bag1 | 7.48544E-05 | -1.471905688 | 0.571 | 0.91 | 1 | 5.02857E-06 | -0.94461079 | 0.733 | 0.869 | 0.122154026 | 7.48544E-05 | 1.00571E-05 |
| Mdh1 | 5.0457E-06 | -1.38052421 | 0.667 | 0.922 | 0.122570234 | 0.008262326 | -0.4734319 | 0.889 | 0.837 | 1 | 0.008262326 | 1.00914E-05 |
| Dusp7 | 0.011957419 | -1.413691074 | 0.095 | 0.399 | 1 | 5.05255E-06 | -1.543607194 | 0.156 | 0.578 | 0.122736474 | 0.011957419 | 1.01051E-05 |

| | | | | | | | | | | | | |
|---|---|---|---|---|---|---|---|---|---|---|---|---|
| Hyou1 | 0.005378409 | -2.280424543 | 0.048 | 0.343 | 1 | 5.07345E-06 | -3.183176025 | 0.044 | 0.394 | 0.123244297 | 0.005378409 | 1.01469E-05 |
| Etfa | 0.008563193 | -0.39035857 | 0.714 | 0.922 | 1 | 5.07434E-06 | -0.76826535 | 0.933 | 0.955 | 0.123265894 | 0.008563193 | 1.01487E-05 |
| Stk16 | 0.056228725 | -0.870548749 | 0.19 | 0.407 | 1 | 5.12279E-06 | -1.867124496 | 0.067 | 0.436 | 0.124442735 | 0.056228725 | 1.02455E-05 |
| Polr2m | 5.15886E-06 | -2.067695219 | 0.19 | 0.75 | 0.12531892 | 0.000539607 | -0.779478374 | 0.578 | 0.747 | 1 | 0.000539607 | 1.03177E-05 |
| Rflnb | 0.003614325 | -1.041071016 | 0.286 | 0.683 | 1 | 5.17137E-06 | -0.972711252 | 0.4 | 0.689 | 0.125622922 | 0.003614325 | 1.03427E-05 |
| Vezf1 | 0.289478144 | -0.238995394 | 0.667 | 0.836 | 1 | 5.18072E-06 | -0.861923507 | 0.911 | 0.934 | 0.125849988 | 0.289478144 | 1.03614E-05 |
| Dpyd | 0.178098064 | 1.214340149 | 0.048 | 0.011 | 1 | 5.19284E-06 | 3.202446745 | 0.267 | 0.059 | 0.126144491 | 0.178098064 | 1.03857E-05 |
| Mt2 | 0.608064605 | -1.366642501 | 0.476 | 0.437 | 1 | 5.21879E-06 | -0.998751454 | 0.689 | 0.256 | 0.126774851 | 0.608064605 | 1.04376E-05 |
| Grn | 5.23402E-06 | -2.399477214 | 0.19 | 0.716 | 0.127144894 | 1.78392E-05 | -1.273571045 | 0.6 | 0.796 | 0.433348775 | 1.78392E-05 | 1.0468E-05 |
| ENSMUSG00000095041 | 0.255710987 | -0.44170545 | 0.143 | 0.269 | 1 | 5.27252E-06 | 1.049611568 | 0.622 | 0.26 | 0.128080035 | 0.255710987 | 1.0545E-05 |
| Extl2 | 0.007251431 | -1.359651234 | 0.048 | 0.354 | 1 | 5.28774E-06 | -4.393743011 | 0 | 0.336 | 0.12844979 | 0.007251431 | 1.05755E-05 |
| Smg6 | 0.074791256 | 0.881429416 | 0.81 | 0.869 | 1 | 5.29002E-06 | 1.231033102 | 0.978 | 0.924 | 0.128505279 | 0.074791256 | 1.058E-05 |
| Car8 | 0.021883451 | -0.600709165 | 0.286 | 0.567 | 1 | 5.291E-06 | -0.923244955 | 0.356 | 0.785 | 0.128528878 | 0.021883451 | 1.0582E-05 |
| Map4 | 0.017467347 | 1.142681561 | 0.905 | 0.914 | 1 | 5.31804E-06 | 0.862682028 | 1 | 0.958 | 0.129185719 | 0.017467347 | 1.0636E-05 |
| Scnn1a | 0.460388202 | 2.088809267 | 0.048 | 0.022 | 1 | 5.34009E-06 | 1.776181991 | 0.333 | 0.09 | 0.129721375 | 0.460388202 | 1.06801E-05 |
| Bscl2 | 0.001317171 | -1.732220592 | 0.095 | 0.5 | 1 | 5.34241E-06 | -1.050281754 | 0.156 | 0.578 | 0.129777763 | 0.001317171 | 1.06848E-05 |
| Degs1 | 5.39333E-06 | -1.620411337 | 0.333 | 0.873 | 0.131014877 | 1.08818E-05 | -1.171263704 | 0.622 | 0.785 | 0.264341189 | 1.08818E-05 | 1.07866E-05 |
| Foxo1 | 0.077232956 | 1.487206501 | 0.952 | 0.937 | 1 | 5.41921E-06 | 1.340867967 | 0.978 | 0.965 | 0.131643417 | 0.077232956 | 1.08384E-05 |
| Plekha6 | 0.771311837 | 1.955914996 | 0.238 | 0.239 | 1 | 5.45337E-06 | 0.952679646 | 0.467 | 0.152 | 0.132473284 | 0.771311837 | 1.09067E-05 |
| Htatsf1 | 0.000888851 | -1.448624864 | 0.286 | 0.683 | 1 | 5.46432E-06 | -1.157242535 | 0.489 | 0.754 | 0.132739225 | 0.000888851 | 1.09286E-05 |
| Slc43a3 | 0.017624061 | -0.503105994 | 0.333 | 0.716 | 1 | 5.46772E-06 | -1.634606036 | 0.156 | 0.564 | 0.132821838 | 0.017624061 | 1.09354E-05 |
| Slc22a23 | 0.447240056 | 0.333734292 | 0.571 | 0.485 | 1 | 5.48258E-06 | 1.548510338 | 0.822 | 0.446 | 0.133182765 | 0.447240056 | 1.09651E-05 |
| Lrig3 | 0.307325197 | 0.197333724 | 0.524 | 0.716 | 1 | 5.5128E-06 | -0.844324627 | 0.244 | 0.723 | 0.133916991 | 0.307325197 | 1.10256E-05 |
| Metap2 | 0.043572459 | -0.496153234 | 0.81 | 0.866 | 1 | 5.53096E-06 | -0.818846374 | 0.844 | 0.907 | 0.134358186 | 0.043572459 | 1.10619E-05 |
| Snrpa1 | 0.178148068 | -0.246793765 | 0.286 | 0.511 | 1 | 5.53732E-06 | -1.350350415 | 0.133 | 0.547 | 0.134512457 | 0.178148068 | 1.10746E-05 |
| Rab24 | 0.002345613 | -0.834022873 | 0.095 | 0.5 | 1 | 5.57956E-06 | -1.935837246 | 0.044 | 0.408 | 0.135538791 | 0.002345613 | 1.11591E-05 |
| Mcu | 0.028171071 | 1.0038446 | 0.905 | 0.929 | 1 | 5.58859E-06 | 1.158949015 | 0.978 | 0.958 | 0.135758038 | 0.028171071 | 1.11771E-05 |
| Gpc6 | 5.64102E-06 | 1.520020812 | 1 | 0.996 | 0.13703171 | 0.000371909 | 0.698966786 | 0.978 | 0.993 | 1 | 0.000371909 | 1.1282E-05 |
| Pdcd5 | 5.71887E-06 | -2.097717702 | 0.19 | 0.731 | 0.138922824 | 0.037084292 | -0.581268017 | 0.622 | 0.706 | 1 | 0.037084292 | 1.14377E-05 |
| Snrpa | 0.071779764 | -0.376076782 | 0.286 | 0.578 | 1 | 5.7654E-06 | -1.638855509 | 0.244 | 0.592 | 0.140053053 | 0.071779764 | 1.15308E-05 |
| Pdcl3 | 0.000107606 | -1.562426453 | 0.238 | 0.754 | 1 | 5.80062E-06 | -0.991324998 | 0.6 | 0.82 | 0.140908559 | 0.000107606 | 1.16012E-05 |
| Cops9 | 5.8017E-06 | -1.992984824 | 0.286 | 0.787 | 0.14093494 | 0.000762555 | -0.979892427 | 0.556 | 0.699 | 1 | 0.000762555 | 1.16034E-05 |
| Fam20c | 0.263258882 | 0.754390228 | 0.286 | 0.556 | 1 | 5.84632E-06 | -0.355607882 | 0.267 | 0.785 | 0.142018832 | 0.263258882 | 1.16926E-05 |
| Ywhah | 0.015610458 | -0.333329289 | 0.524 | 0.761 | 1 | 5.85611E-06 | -1.089516918 | 0.822 | 0.862 | 0.142256562 | 0.015610458 | 1.17122E-05 |
| Jun | 0.00124768 | -1.597565068 | 0.857 | 0.955 | 1 | 5.8683E-06 | -0.963264335 | 1 | 0.983 | 0.142552743 | 0.00124768 | 1.17366E-05 |
| Cnih4 | 0.000185358 | -1.211924606 | 0.286 | 0.795 | 1 | 5.87072E-06 | -1.04756737 | 0.689 | 0.81 | 0.142611626 | 0.000185358 | 1.17414E-05 |
| Ccnh | 0.000293705 | -2.926141075 | 0.048 | 0.463 | 1 | 5.95059E-06 | -1.518561275 | 0.133 | 0.529 | 0.144551631 | 0.000293705 | 1.19011E-05 |
| Gas1 | 0.002504329 | -3.326228232 | 0.095 | 0.448 | 1 | 6.00136E-06 | -2.447146942 | 0.156 | 0.54 | 0.145785102 | 0.002504329 | 1.20027E-05 |
| Gadd45gip1 | 0.041209171 | -1.337455488 | 0.095 | 0.313 | 1 | 6.03576E-06 | -3.089516918 | 0.044 | 0.391 | 0.146620786 | 0.041209171 | 1.20715E-05 |
| Lysmd3 | 0.145654909 | -0.537537155 | 0.381 | 0.623 | 1 | 6.03683E-06 | -1.258375231 | 0.422 | 0.751 | 0.146646622 | 0.145654909 | 1.20736E-05 |
| Slc25a39 | 6.05718E-06 | -2.400904919 | 0.095 | 0.668 | 0.147140945 | 0.011648716 | -0.711325147 | 0.533 | 0.651 | 1 | 0.011648716 | 1.21143E-05 |
| Rab33b | 0.000627567 | -2.15277672 | 0.095 | 0.504 | 1 | 6.0734E-06 | -1.223818009 | 0.178 | 0.619 | 0.147535126 | 0.000627567 | 1.21468E-05 |
| Sgta | 0.002613134 | -1.455511249 | 0.095 | 0.459 | 1 | 6.08708E-06 | -1.409373663 | 0.2 | 0.609 | 0.147867244 | 0.002613134 | 1.21741E-05 |
| Snrpb2 | 0.000233386 | -1.206989096 | 0.19 | 0.694 | 1 | 6.09581E-06 | -1.439687307 | 0.289 | 0.64 | 0.148079398 | 0.000233386 | 1.21916E-05 |
| Hypk | 0.000163481 | -1.233118828 | 0.333 | 0.802 | 1 | 6.09632E-06 | -1.124282336 | 0.533 | 0.768 | 0.148091732 | 0.000163481 | 1.21926E-05 |
| Marchf5 | 0.090051435 | -0.186825176 | 0.524 | 0.81 | 1 | 6.18054E-06 | -0.901889915 | 0.733 | 0.865 | 0.15013758 | 0.090051435 | 1.2361E-05 |
| Nkap | 0.01886382 | -1.405179574 | 0.143 | 0.407 | 1 | 6.21816E-06 | -2.446210431 | 0.067 | 0.429 | 0.151051485 | 0.01886382 | 1.24363E-05 |
| Bola3 | 6.24023E-06 | -2.901137068 | 0.048 | 0.616 | 0.151587689 | 1.46294E-05 | -1.577454964 | 0.156 | 0.526 | 0.355377549 | 1.46294E-05 | 1.24804E-05 |
| Gid4 | 0.11450883 | -0.540007523 | 0.238 | 0.433 | 1 | 6.24818E-06 | -2.11134328 | 0.089 | 0.464 | 0.15178079 | 0.11450883 | 1.24963E-05 |
| 1110004F10Rik | 0.005793871 | -0.596983675 | 0.619 | 0.843 | 1 | 6.4548E-06 | -0.815733271 | 0.8 | 0.91 | 0.156799989 | 0.005793871 | 1.29096E-05 |
| Mier1 | 6.46858E-06 | -1.355975576 | 0.762 | 0.948 | 0.157134718 | 0.000340913 | -0.527971403 | 1 | 0.972 | 1 | 0.000340913 | 1.29371E-05 |
| Ublcp1 | 0.037258496 | -1.438928365 | 0.143 | 0.366 | 1 | 6.48199E-06 | -2.879169838 | 0.067 | 0.415 | 0.157460415 | 0.037258496 | 1.29639E-05 |
| Pik3r3 | 0.299602397 | -1.280424543 | 0.048 | 0.123 | 1 | 6.54036E-06 | 1.495445583 | 0.467 | 0.159 | 0.158878517 | 0.299602397 | 1.30807E-05 |
| Sowahc | 0.075481953 | -0.776261153 | 0.286 | 0.522 | 1 | 6.58073E-06 | -2.798054104 | 0.022 | 0.377 | 0.159859158 | 0.075481953 | 1.31614E-05 |
| Igsf8 | 6.6835E-06 | -4.24509147 | 0 | 0.545 | 0.162355489 | 4.80582E-05 | -1.361321533 | 0.222 | 0.574 | 1 | 4.80582E-05 | 1.33669E-05 |
| Mettl1 | 0.028707677 | -0.238765391 | 0.19 | 0.549 | 1 | 6.69068E-06 | -1.589945908 | 0.111 | 0.516 | 0.162529976 | 0.028707677 | 1.33813E-05 |
| Tmem147 | 5.98273E-05 | -1.863169218 | 0.143 | 0.668 | 1 | 6.70182E-06 | -1.432404631 | 0.244 | 0.616 | 0.162800504 | 5.98273E-05 | 1.34036E-05 |
| Arl3 | 6.70928E-06 | -1.481939375 | 0.286 | 0.84 | 0.162981875 | 0.002372607 | -0.511099962 | 0.578 | 0.758 | 1 | 0.002372607 | 1.34185E-05 |
| Bhlhe40 | 0.000142498 | -1.686709568 | 0.524 | 0.84 | 1 | 6.87956E-06 | -0.998487538 | 0.889 | 0.889 | 0.167118243 | 0.000142498 | 1.37591E-05 |
| Pkdcc | 0.010922189 | -3.196592952 | 0.048 | 0.306 | 1 | 6.88183E-06 | -2.697749198 | 0.044 | 0.401 | 0.1671735 | 0.010922189 | 1.37636E-05 |
| Casd1 | 0.001407053 | -2.355975576 | 0.048 | 0.422 | 1 | 6.96336E-06 | -1.195248857 | 0.089 | 0.481 | 0.169154058 | 0.001407053 | 1.39267E-05 |
| Hdgf | 0.000524885 | -0.849790188 | 0.857 | 0.951 | 1 | 7.00175E-06 | -0.675151638 | 0.933 | 0.976 | 0.170086394 | 0.000524885 | 1.40034E-05 |
| Tpm2 | 0.615673259 | -0.484657595 | 0.238 | 0.284 | 1 | 7.05627E-06 | -3.19344436 | 0.044 | 0.394 | 0.171410903 | 0.615673259 | 1.41125E-05 |
| Gemin7 | 0.000155917 | -2.288160191 | 0.048 | 0.511 | 1 | 7.07093E-06 | -1.539319835 | 0.133 | 0.522 | 0.171767103 | 0.000155917 | 1.41418E-05 |
| Ndufa2 | 7.07692E-06 | -1.496153234 | 0.571 | 0.91 | 0.171912483 | 0.000279057 | -0.88306604 | 0.778 | 0.775 | 1 | 0.000279057 | 1.41538E-05 |
| Adk | 7.08187E-06 | 1.199130067 | 1 | 0.978 | 0.172032843 | 1.80041E-05 | 0.944717112 | 0.978 | 0.976 | 0.437356311 | 1.80041E-05 | 1.41637E-05 |
| Ankrd54 | 0.007978811 | -2.107587946 | 0.048 | 0.325 | 1 | 7.10101E-06 | -2.86124793 | 0.044 | 0.388 | 0.172497641 | 0.007978811 | 1.4202E-05 |
| Fam3a | 0.092268318 | -1.063193826 | 0.095 | 0.257 | 1 | 7.11359E-06 | -2.989352756 | 0.044 | 0.388 | 0.172803403 | 0.092268318 | 1.42271E-05 |
| Polr1f | 0.034674148 | -0.555046923 | 0.19 | 0.478 | 1 | 7.205E-06 | -1.852980314 | 0.111 | 0.484 | 0.175023821 | 0.034674148 | 1.44099E-05 |
| Lpcat2 | 0.149395917 | -1.392317423 | 0.048 | 0.164 | 1 | 7.20533E-06 | 0.761682421 | 0.6 | 0.228 | 0.175031984 | 0.149395917 | 1.44106E-05 |
| Abcc4 | 0.626425122 | 1.659965968 | 0.238 | 0.224 | 1 | 7.22313E-06 | 1.488056604 | 0.511 | 0.194 | 0.175464334 | 0.626425122 | 1.44462E-05 |
| Baz2b | 7.26231E-06 | 1.062104943 | 1 | 0.978 | 0.176416122 | 0.001561583 | 0.693514333 | 1 | 0.962 | 1 | 0.001561583 | 1.45246E-05 |
| Jam3 | 0.000233431 | -2.22709504 | 0.095 | 0.541 | 1 | 7.26959E-06 | -1.356455778 | 0.178 | 0.578 | 0.176592957 | 0.000233431 | 1.45391E-05 |
| Hnrnpu | 0.012735731 | -0.492331523 | 1 | 0.985 | 1 | 7.26977E-06 | -0.509319755 | 1 | 0.997 | 0.176597177 | 0.012735731 | 1.45395E-05 |
| Bccip | 0.008768995 | -0.644545074 | 0.286 | 0.713 | 1 | 7.33703E-06 | -0.782308299 | 0.4 | 0.747 | 0.178231051 | 0.008768995 | 1.4674E-05 |
| Srm | 6.20853E-05 | -2.666078235 | 0.143 | 0.597 | 1 | 7.35981E-06 | -1.908553645 | 0.111 | 0.484 | 0.178784493 | 6.20853E-05 | 1.47196E-05 |
| Mrpl28 | 0.000848798 | -2.475975352 | 0.048 | 0.422 | 1 | 7.45311E-06 | -2.476798751 | 0.044 | 0.398 | 0.181050883 | 0.000848798 | 1.49062E-05 |
| Zfand5 | 0.000544363 | -0.776986308 | 0.905 | 0.985 | 1 | 7.45326E-06 | -0.618281969 | 1 | 0.99 | 0.181054583 | 0.000544363 | 1.49065E-05 |
| Txnl1 | 2.88593E-05 | -1.334553097 | 0.619 | 0.907 | 0.701050262 | 7.46686E-06 | -0.710817256 | 0.889 | 0.955 | 0.181385016 | 2.88593E-05 | 1.49337E-05 |
| Snai2 | 0.000312076 | -3.216999163 | 0.048 | 0.478 | 1 | 7.52585E-06 | -3.674479418 | 0.044 | 0.384 | 0.182817938 | 0.000312076 | 1.50516E-05 |
| Smarca1 | 0.135761307 | 0.357403198 | 0.238 | 0.433 | 1 | 7.54857E-06 | -1.169752852 | 0.178 | 0.606 | 0.183369862 | 0.135761307 | 1.50971E-05 |
| Dohh | 0.001681931 | -1.960434252 | 0.095 | 0.448 | 1 | 7.84704E-06 | -2.683249628 | 0.067 | 0.412 | 0.190620238 | 0.001681931 | 1.5694E-05 |
| Vdac2 | 4.70823E-05 | -1.356750177 | 0.667 | 0.869 | 1 | 7.87135E-06 | -0.752459617 | 0.956 | 0.931 | 0.191210873 | 4.70823E-05 | 1.57426E-05 |
| Enho | 2.17983E-05 | -3.674956387 | 0.048 | 0.552 | 0.529525029 | 7.89783E-06 | -2.722919774 | 0.067 | 0.419 | 0.191854103 | 2.17983E-05 | 1.57956E-05 |
| Gpat4 | 0.097574578 | -0.481506458 | 0.238 | 0.485 | 1 | 7.9201E-06 | -1.212230035 | 0.244 | 0.668 | 0.192395027 | 0.097574578 | 1.58401E-05 |
| Eif4b | 0.022637405 | -0.742392397 | 0.476 | 0.72 | 1 | 7.96523E-06 | -0.84707703 | 0.822 | 0.92 | 0.193491336 | 0.022637405 | 1.59304E-05 |
| Pgp | 0.012112262 | -1.31977321 | 0.143 | 0.451 | 1 | 7.98544E-06 | -1.888469399 | 0.089 | 0.46 | 0.193982277 | 0.012112262 | 1.59708E-05 |
| Uqcr10 | 8.02116E-06 | -1.521853744 | 0.476 | 0.869 | 0.19484996 | 0.006154817 | -0.798054104 | 0.8 | 0.747 | 1 | 0.006154817 | 1.60423E-05 |
| Tspan7 | 0.059988076 | -1.361852142 | 0.238 | 0.44 | 1 | 8.09303E-06 | -1.112786697 | 0.067 | 0.443 | 0.196596003 | 0.059988076 | 1.6186E-05 |
| Bcl7c | 0.037307842 | -0.636083495 | 0.381 | 0.653 | 1 | 8.16084E-06 | -1.031172931 | 0.333 | 0.737 | 0.198243053 | 0.037307842 | 1.63216E-05 |
| Sgpp1 | 0.002965558 | -3.04047375 | 0.048 | 0.369 | 1 | 8.22624E-06 | -3.044847868 | 0.044 | 0.384 | 0.199831704 | 0.002965558 | 1.64524E-05 |
| Atp8a1 | 0.938908218 | 1.766881172 | 0.095 | 0.097 | 1 | 8.26307E-06 | 1.141178807 | 0.533 | 0.19 | 0.200726463 | 0.938908218 | 1.65261E-05 |
| Ankrd6 | 0.454226567 | 2.936806174 | 0.048 | 0.022 | 1 | 8.26864E-06 | 1.894576691 | 0.222 | 0.042 | 0.200861763 | 0.454226567 | 1.65372E-05 |
| Eef1akmt2 | 0.001778842 | -2.506137322 | 0.048 | 0.392 | 1 | 8.37587E-06 | -2.89183625 | 0.044 | 0.384 | 0.203466526 | 0.001778842 | 1.67517E-05 |
| Ebpl | 8.39811E-06 | -4.067695219 | 0 | 0.534 | 0.204006995 | 5.41341E-05 | -1.030364427 | 0.133 | 0.502 | 1 | 5.41341E-05 | 1.67962E-05 |
| Dennd2c | 0.517865401 | 0.866416846 | 0.19 | 0.306 | 1 | 8.42842E-06 | 1.35429082 | 0.711 | 0.343 | 0.204743227 | 0.517865401 | 1.68568E-05 |
| Prkca | 8.48376E-06 | 1.585362793 | 1 | 0.955 | 0.206087476 | 0.00041189 | 1.384973431 | 0.978 | 0.952 | 1 | 0.00041189 | 1.69674E-05 |
| Nolc1 | 0.002657238 | -1.057694846 | 0.381 | 0.784 | 1 | 8.53094E-06 | -0.980546858 | 0.711 | 0.83 | 0.207233686 | 0.002657238 | 1.70618E-05 |

| | | | | | | | | | | | | |
|---|---|---|---|---|---|---|---|---|---|---|---|---|
| Bpnt2 | 0.005161029 | -0.691300819 | 0.333 | 0.836 | 1 | 8.59234E-06 | -0.873875538 | 0.667 | 0.879 | 0.208725126 | 0.005161029 | 1.71846E-05 |
| Pole3 | 0.00012539 | -2.15277672 | 0.048 | 0.53 | 1 | 8.62409E-06 | -2.526380779 | 0.067 | 0.419 | 0.20949641 | 0.00012539 | 1.72481E-05 |
| Ndufb4 | 3.26881E-05 | -2.15277672 | 0.048 | 0.575 | 0.794059289 | 8.6275E-06 | -2.160624183 | 0.133 | 0.481 | 0.20957914 | 3.26881E-05 | 1.72549E-05 |
| Bpgm | 0.002039118 | -1.44170545 | 0.143 | 0.556 | 1 | 8.6924E-06 | -1.285091327 | 0.333 | 0.758 | 0.211155771 | 0.002039118 | 1.73847E-05 |
| Akt1s1 | 0.021559057 | -1.506137322 | 0.143 | 0.384 | 1 | 8.70726E-06 | -2.199570463 | 0.089 | 0.446 | 0.211516866 | 0.021559057 | 1.74145E-05 |
| Dsel | 0.105947362 | -0.83578333 | 0.19 | 0.384 | 1 | 8.74742E-06 | -1.409373663 | 0.133 | 0.526 | 0.212492238 | 0.105947362 | 1.74948E-05 |
| Ndufs7 | 8.83816E-06 | -2.434752689 | 0.095 | 0.646 | 0.214696583 | 0.266207339 | -0.234465254 | 0.533 | 0.578 | 1 | 0.266207339 | 1.76762E-05 |
| Acadvl | 0.001029057 | -1.911190733 | 0.095 | 0.489 | 1 | 8.9085E-06 | -1.638855509 | 0.089 | 0.46 | 0.216405174 | 0.001029057 | 1.78169E-05 |
| Tcp1 | 0.000229782 | -1.092128727 | 0.762 | 0.929 | 1 | 8.91806E-06 | -0.638855509 | 0.889 | 0.931 | 0.216637402 | 0.000229782 | 1.7836E-05 |
| Prkcsh | 1.43699E-05 | -2.030625964 | 0.19 | 0.713 | 0.349073888 | 8.98015E-06 | -1.136595597 | 0.244 | 0.654 | 0.218145877 | 1.43699E-05 | 1.79602E-05 |
| Brd2 | 8.99373E-06 | -1.450349544 | 0.905 | 0.955 | 0.218475714 | 1.55747E-05 | -0.65434659 | 0.956 | 0.99 | 0.378339831 | 1.55747E-05 | 1.79874E-05 |
| Tprgl | 9.02146E-06 | -3.107587946 | 0.095 | 0.616 | 0.219149371 | 4.47632E-05 | -1.212677452 | 0.489 | 0.702 | 1 | 4.47632E-05 | 1.80428E-05 |
| Nono | 0.010272015 | -0.867109665 | 0.381 | 0.705 | 1 | 9.04475E-06 | -1.000379778 | 0.733 | 0.841 | 0.219715109 | 0.010272015 | 1.80894E-05 |
| 2810001G20Rik | 9.06048E-06 | -2.213310347 | 0.286 | 0.769 | 0.220097157 | 2.24361E-05 | -1.189052591 | 0.2 | 0.592 | 0.545017905 | 2.24361E-05 | 1.81209E-05 |
| Septin7 | 0.129451077 | -0.260625123 | 0.952 | 0.985 | 1 | 9.10209E-06 | -0.587662078 | 1 | 1 | 0.221107963 | 0.129451077 | 1.82041E-05 |
| Mllt10 | 0.077294722 | 0.490227157 | 0.857 | 0.769 | 1 | 9.13978E-06 | 0.784336803 | 1 | 0.934 | 0.222023588 | 0.077294722 | 1.82795E-05 |
| Ccdc34 | 0.012966157 | -1.560283571 | 0.19 | 0.5 | 1 | 9.15306E-06 | -3.383016604 | 0.044 | 0.381 | 0.222346025 | 0.012966157 | 1.8306E-05 |
| Rab35 | 0.211458079 | -0.31112134 | 0.238 | 0.444 | 1 | 9.2611E-06 | -1.212230035 | 0.222 | 0.654 | 0.224970573 | 0.211458079 | 1.85221E-05 |
| Dlg1 | 0.002700793 | 0.880479558 | 1 | 0.903 | 1 | 9.31491E-06 | 0.913591224 | 1 | 0.938 | 0.22627778 | 0.002700793 | 1.86297E-05 |
| Cwc25 | 0.161934083 | -0.150656668 | 0.238 | 0.429 | 1 | 9.31632E-06 | -1.547225033 | 0.156 | 0.526 | 0.226312138 | 0.161934083 | 1.86326E-05 |
| Map1b | 0.854923676 | -2.529168291 | 0.238 | 0.19 | 1 | 9.31876E-06 | -0.183460272 | 0.756 | 0.315 | 0.226371405 | 0.854923676 | 1.86374E-05 |
| Ift27 | 0.010790567 | -0.763633545 | 0.095 | 0.396 | 1 | 9.35929E-06 | -2.174908409 | 0.044 | 0.405 | 0.227355828 | 0.010790567 | 1.87185E-05 |
| Higd1a | 0.000194498 | -1.421385465 | 0.524 | 0.851 | 1 | 9.38279E-06 | -1.035745661 | 0.711 | 0.862 | 0.227926747 | 0.000194498 | 1.87655E-05 |
| Rbbp9 | 0.050668883 | -1.164171474 | 0.143 | 0.354 | 1 | 9.41306E-06 | -2.387316742 | 0.067 | 0.419 | 0.228662068 | 0.050668883 | 1.8826E-05 |
| Trim41 | 0.944460625 | 0.516230491 | 0.286 | 0.317 | 1 | 9.4195E-06 | -1.975138897 | 0.089 | 0.457 | 0.228818596 | 0.944460625 | 1.88389E-05 |
| Hnrnpdl | 0.129630787 | -0.411485117 | 1 | 0.948 | 1 | 9.48921E-06 | -0.556482209 | 1 | 0.972 | 0.230511954 | 0.129630787 | 1.89783E-05 |
| Rspo3 | 0.000444173 | -1.236721065 | 0.19 | 0.675 | 1 | 9.51421E-06 | -4.107004344 | 0.044 | 0.377 | 0.231119121 | 0.000444173 | 1.90283E-05 |
| Pdcl | 0.026361485 | -1.337455488 | 0.095 | 0.34 | 1 | 9.52216E-06 | -2.665655568 | 0.022 | 0.367 | 0.231312351 | 0.026361485 | 1.90442E-05 |
| Septin8 | 0.011426561 | -0.785659851 | 0.095 | 0.392 | 1 | 9.62307E-06 | -1.098287127 | 0.156 | 0.564 | 0.233763656 | 0.011426561 | 1.92461E-05 |
| Slc25a38 | 0.024540228 | -1.801961663 | 0.048 | 0.269 | 1 | 9.68175E-06 | -2.998751454 | 0.044 | 0.381 | 0.235189033 | 0.024540228 | 1.93634E-05 |
| Map2k7 | 0.051280692 | -1.683780237 | 0.048 | 0.228 | 1 | 9.69884E-06 | -2.656777417 | 0.044 | 0.391 | 0.235604273 | 0.051280692 | 1.93976E-05 |
| Rps19bp1 | 0.052433842 | -0.946814643 | 0.095 | 0.306 | 1 | 9.74685E-06 | -2.998751454 | 0.044 | 0.381 | 0.236770558 | 0.052433842 | 1.94936E-05 |
| Pkm | 4.07917E-05 | -1.042732985 | 0.905 | 0.985 | 0.99091233 | 9.80198E-06 | -0.65489552 | 0.956 | 0.986 | 0.238109814 | 4.07917E-05 | 1.96039E-05 |
| Gm2296 | 0.003557166 | -2.280424543 | 0.048 | 0.362 | 1 | 9.81845E-06 | -4.328154669 | 0 | 0.322 | 0.238509833 | 0.003557166 | 1.96368E-05 |
| Isg20l2 | 0.882695972 | 0.393663848 | 0.333 | 0.373 | 1 | 9.85272E-06 | -1.547225033 | 0.133 | 0.512 | 0.239342257 | 0.882695972 | 1.97053E-05 |
| Ifi30 | 0.00017616 | -2.017390137 | 0.143 | 0.612 | 1 | 9.87711E-06 | -2.002493855 | 0.2 | 0.564 | 0.23993476 | 0.00017616 | 1.97541E-05 |
| Map7d1 | 9.88837E-06 | -1.733495997 | 0.333 | 0.828 | 0.240208246 | 0.001307745 | -0.54592257 | 0.756 | 0.917 | 1 | 0.001307745 | 1.97766E-05 |
| Lrrc8a | 0.195326283 | -0.502816964 | 0.286 | 0.478 | 1 | 9.915E-06 | -1.246718412 | 0.2 | 0.637 | 0.240855144 | 0.195326283 | 1.98299E-05 |
| Csnk2b | 0.000211204 | -1.251040736 | 0.381 | 0.806 | 1 | 1.00457E-05 | -0.966909706 | 0.756 | 0.827 | 0.244030871 | 0.000211204 | 2.00914E-05 |
| Arhgdib | 1.008E-05 | -2.128149268 | 0.333 | 0.817 | 0.244862797 | 0.000122425 | -0.680586073 | 0.844 | 0.851 | 1 | 0.000122425 | 2.01599E-05 |
| Phyh | 7.24682E-05 | -1.849790188 | 0.238 | 0.709 | 1 | 1.01274E-05 | -1.358195432 | 0.444 | 0.696 | 0.246014758 | 7.24682E-05 | 2.02547E-05 |
| Lzts2 | 0.052782602 | -0.779087197 | 0.238 | 0.489 | 1 | 1.01743E-05 | -0.991015805 | 0.222 | 0.678 | 0.247154374 | 0.052782602 | 2.03485E-05 |
| Aup1 | 0.006734314 | -1.455511249 | 0.143 | 0.459 | 1 | 1.03251E-05 | -1.578594984 | 0.111 | 0.495 | 0.250817486 | 0.006734314 | 2.06501E-05 |
| Ssb | 0.001155068 | -0.800159421 | 0.952 | 0.978 | 1 | 1.04135E-05 | -0.693011279 | 0.956 | 0.965 | 0.252963549 | 0.001155068 | 2.08268E-05 |
| Nme1 | 1.05482E-05 | -1.709932525 | 0.381 | 0.821 | 0.256238 | 0.00018833 | -0.753478808 | 0.667 | 0.792 | 1 | 0.00018833 | 2.10964E-05 |
| Ccnc | 0.520923789 | 0.467867469 | 0.476 | 0.485 | 1 | 1.05911E-05 | -1.223818009 | 0.156 | 0.578 | 0.257278897 | 0.520923789 | 2.11821E-05 |
| Poldip2 | 0.120205688 | -0.437736548 | 0.143 | 0.347 | 1 | 1.06173E-05 | -1.894356242 | 0.111 | 0.478 | 0.257915743 | 0.120205688 | 2.12345E-05 |
| Dctn6 | 0.002521435 | -1.286230164 | 0.19 | 0.593 | 1 | 1.07181E-05 | -1.451353734 | 0.178 | 0.557 | 0.260365197 | 0.002521435 | 2.14362E-05 |
| Mrps14 | 1.07393E-05 | -2.098817736 | 0.143 | 0.698 | 0.260877963 | 0.000135357 | -0.966020252 | 0.556 | 0.727 | 1 | 0.000135357 | 2.14784E-05 |
| Npepl1 | 0.203736014 | -0.44170545 | 0.048 | 0.157 | 1 | 1.07897E-05 | -4.211745177 | 0 | 0.318 | 0.262103988 | 0.203736014 | 2.15793E-05 |
| Aldh2 | 1.09086E-05 | -1.545062834 | 0.667 | 0.903 | 0.264991039 | 6.9129E-05 | -0.625327124 | 0.956 | 0.945 | 1 | 6.9129E-05 | 2.1817E-05 |
| Hdlbp | 0.00343339 | -0.567973097 | 1 | 1 | 1 | 1.09298E-05 | -0.498356387 | 1 | 1 | 0.265507348 | 0.00343339 | 2.18595E-05 |
| Igip | 0.282054762 | 0.565247311 | 0.095 | 0.213 | 1 | 1.09435E-05 | -3.372209849 | 0.022 | 0.349 | 0.265839572 | 0.282054762 | 2.18869E-05 |
| Dmac1 | 0.056396056 | -0.970084422 | 0.19 | 0.403 | 1 | 1.09968E-05 | -3.361321533 | 0.022 | 0.349 | 0.267133229 | 0.056396056 | 2.19934E-05 |
| Tomm34 | 0.026607251 | -1.142145168 | 0.143 | 0.41 | 1 | 1.10125E-05 | -1.854361544 | 0.067 | 0.439 | 0.267514521 | 0.026607251 | 2.20248E-05 |
| Rpl10-ps3 | 1.10225E-05 | -1.849790188 | 0.19 | 0.743 | 0.267757968 | 0.003623011 | -0.842280425 | 0.6 | 0.702 | 1 | 0.003623011 | 2.20448E-05 |
| Stk33 | 0.629723166 | 2.825774861 | 0.048 | 0.03 | 1 | 1.10785E-05 | 6.075390009 | 0.089 | 0.003 | 0.269118609 | 0.629723166 | 2.21569E-05 |
| Gprc5c | 0.020679451 | -0.93109029 | 0.095 | 0.381 | 1 | 1.10899E-05 | -2.065120263 | 0.111 | 0.471 | 0.2693952 | 0.020679451 | 2.21796E-05 |
| Polr2k | 0.000222388 | -1.339690492 | 0.19 | 0.701 | 1 | 1.1203E-05 | -1.581034854 | 0.267 | 0.599 | 0.272142168 | 0.000222388 | 2.24058E-05 |
| Gm11808 | 1.1283E-05 | -2.397690595 | 0.095 | 0.638 | 0.274086612 | 0.383333638 | -0.156887001 | 0.4 | 0.457 | 1 | 0.383333638 | 2.25659E-05 |
| Pno1 | 0.029537818 | -0.620848981 | 0.143 | 0.418 | 1 | 1.13004E-05 | -2.37003875 | 0.044 | 0.391 | 0.274510371 | 0.029537818 | 2.26008E-05 |
| Micos10 | 1.13756E-05 | -1.243207646 | 0.762 | 0.951 | 0.276335522 | 0.00142377 | -0.438696377 | 0.911 | 0.962 | 1 | 0.00142377 | 2.2751E-05 |
| Rpl9-ps6 | 2.55661E-05 | -3.880817084 | 0 | 0.496 | 0.621051771 | 1.14094E-05 | -1.756550551 | 0.156 | 0.519 | 0.277158236 | 2.55661E-05 | 2.28188E-05 |
| Cald1 | 0.030215126 | -0.591166349 | 0.857 | 0.978 | 1 | 1.14762E-05 | -1.000423369 | 0.844 | 0.945 | 0.278780148 | 0.030215126 | 2.29523E-05 |
| Etfb | 1.14802E-05 | -2.38800443 | 0.19 | 0.675 | 0.278876834 | 0.077494114 | -0.584962501 | 0.578 | 0.588 | 1 | 0.077494114 | 2.29603E-05 |
| G6pc3 | 0.001176918 | -2.02666795 | 0.048 | 0.433 | 1 | 1.14972E-05 | -2.86124793 | 0.044 | 0.377 | 0.279290011 | 0.001176918 | 2.29943E-05 |
| Pnrc1 | 1.15579E-05 | -1.541112075 | 0.762 | 0.937 | 0.280764842 | 0.25783867 | -0.289682578 | 0.911 | 0.872 | 1 | 0.25783867 | 2.31157E-05 |
| Hikeshi | 0.000910799 | -1.837190152 | 0.095 | 0.515 | 1 | 1.1562E-05 | -1.329751454 | 0.156 | 0.526 | 0.280864475 | 0.000910799 | 2.31239E-05 |
| Elof1 | 0.011793231 | -1.169925001 | 0.19 | 0.5 | 1 | 1.15909E-05 | -1.475356776 | 0.289 | 0.609 | 0.281564995 | 0.011793231 | 2.31816E-05 |
| Bloc1s1 | 0.045283538 | -0.818081329 | 0.143 | 0.392 | 1 | 1.16747E-05 | -2.435868487 | 0.044 | 0.388 | 0.283601743 | 0.045283538 | 2.33493E-05 |
| Ooep | 0.142918078 | -1.174225139 | 0.143 | 0.287 | 1 | 1.16874E-05 | -1.486852415 | 0.089 | 0.464 | 0.283910304 | 0.142918078 | 2.33747E-05 |
| Timm17a | 1.1695E-05 | -1.91868527 | 0.19 | 0.743 | 0.284095285 | 0.000123138 | -0.84674836 | 0.6 | 0.792 | 1 | 0.000123138 | 2.33899E-05 |
| Rab5if | 0.002255071 | -1.039924047 | 0.571 | 0.832 | 1 | 1.17274E-05 | -0.842727842 | 0.711 | 0.855 | 0.284881708 | 0.002255071 | 2.34546E-05 |
| Czib | 0.180960896 | -0.225701356 | 0.286 | 0.5 | 1 | 1.17535E-05 | -1.310472392 | 0.111 | 0.505 | 0.285516832 | 0.180960896 | 2.35069E-05 |
| Nfu1 | 0.145631282 | -0.350612391 | 0.238 | 0.44 | 1 | 1.17845E-05 | -1.301820521 | 0.156 | 0.554 | 0.286270056 | 0.145631282 | 2.35689E-05 |
| 4921524J17Rik | 0.096650846 | -1.04047375 | 0.19 | 0.377 | 1 | 1.18109E-05 | -1.4734319 | 0.178 | 0.588 | 0.286910952 | 0.096650846 | 2.36217E-05 |
| Brk1 | 1.18788E-05 | -1.457933912 | 0.429 | 0.873 | 0.288560264 | 6.30588E-05 | -0.805213895 | 0.756 | 0.837 | 1 | 6.30588E-05 | 2.37575E-05 |
| Tiprl | 0.006284987 | -0.856742949 | 0.381 | 0.743 | 1 | 1.1984E-05 | -0.743192169 | 0.578 | 0.841 | 0.291114186 | 0.006284987 | 2.39678E-05 |
| Tssc4 | 0.202195879 | -0.233118828 | 0.19 | 0.347 | 1 | 1.2203E-05 | -2.32442195 | 0.089 | 0.436 | 0.296435474 | 0.202195879 | 2.44059E-05 |
| Sf3b1 | 0.000753018 | -0.64572959 | 1 | 0.993 | 1 | 1.22531E-05 | -0.450401659 | 1 | 1 | 0.297651833 | 0.000753018 | 2.4506E-05 |
| Eif4g1 | 0.237110295 | -0.267776478 | 0.905 | 0.981 | 1 | 1.22965E-05 | -0.587522506 | 1 | 0.993 | 0.298706736 | 0.237110295 | 2.45929E-05 |
| Diaph1 | 0.038021114 | 0.985469517 | 0.81 | 0.784 | 1 | 1.23866E-05 | 1.064426959 | 0.978 | 0.785 | 0.30089528 | 0.038021114 | 2.4773E-05 |
| Lgals1 | 0.020995493 | -4.037034666 | 0.048 | 0.28 | 1 | 1.23877E-05 | -1.798054104 | 0.089 | 0.467 | 0.300922341 | 0.020995493 | 2.47753E-05 |
| Diaph2 | 0.010410079 | 1.168916006 | 0.81 | 0.679 | 1 | 1.24293E-05 | 1.132447493 | 0.978 | 0.792 | 0.301932881 | 0.010410079 | 2.48585E-05 |
| Ubr3 | 1.24965E-05 | 1.062577725 | 1 | 0.806 | 0.303566071 | 0.001993369 | 0.758524815 | 0.978 | 0.934 | 1 | 0.001993369 | 2.49929E-05 |
| Vbp1 | 0.001026413 | -1.502816964 | 0.143 | 0.563 | 1 | 1.25327E-05 | -1.144746438 | 0.222 | 0.609 | 0.304445047 | 0.001026413 | 2.50653E-05 |
| Nsun2 | 0.98493128 | 0.425844254 | 0.333 | 0.377 | 1 | 1.25452E-05 | -1.188412673 | 0.178 | 0.585 | 0.304747211 | 0.98493128 | 2.50902E-05 |
| Nrk | 0.208124792 | 0.240198317 | 0.381 | 0.657 | 1 | 1.27698E-05 | -0.704039677 | 0.4 | 0.782 | 0.310204957 | 0.208124792 | 2.55395E-05 |
| Eif4ebp1 | 0.268196568 | -0.27618755 | 0.19 | 0.332 | 1 | 1.2809E-05 | -2.108741485 | 0.089 | 0.457 | 0.311156161 | 0.268196568 | 2.56178E-05 |
| Xxylt1 | 0.055624058 | -1.555046923 | 0.048 | 0.224 | 1 | 1.28335E-05 | -4.282711698 | 0 | 0.315 | 0.311750845 | 0.055624058 | 2.56668E-05 |
| Naa38 | 5.2654E-05 | -2.701267664 | 0.048 | 0.549 | 1 | 1.29225E-05 | -2.017367132 | 0.133 | 0.481 | 0.313913461 | 5.2654E-05 | 2.58448E-05 |
| Nop16 | 0.005178678 | -1.65714511 | 0.095 | 0.422 | 1 | 1.30886E-05 | -2.214167839 | 0.044 | 0.388 | 0.317948675 | 0.005178678 | 2.61771E-05 |
| Plk2 | 0.529770274 | 1.026881518 | 0.333 | 0.302 | 1 | 1.30948E-05 | 1.776181991 | 0.489 | 0.19 | 0.318099594 | 0.529770274 | 2.61895E-05 |
| Pabpc4 | 0.008503252 | -0.666078235 | 0.381 | 0.757 | 1 | 1.31032E-05 | -1.107004344 | 0.511 | 0.761 | 0.318303682 | 0.008503252 | 2.62063E-05 |
| Clptm1l | 1.65037E-05 | -1.58868452 | 0.286 | 0.817 | 0.400907852 | 1.31926E-05 | -0.935837246 | 0.578 | 0.792 | 0.32047386 | 1.65037E-05 | 2.6385E-05 |
| Mmut | 0.050754307 | -1.509450056 | 0.095 | 0.287 | 1 | 1.32109E-05 | -2.620708162 | 0.044 | 0.384 | 0.320918259 | 0.050754307 | 2.64215E-05 |
| Tmem126a | 0.100893581 | -0.118042303 | 0.286 | 0.593 | 1 | 1.33123E-05 | -0.629810369 | 0.178 | 0.606 | 0.323383545 | 0.100893581 | 2.66245E-05 |
| Tmem9 | 0.000940843 | -2.65714511 | 0.048 | 0.418 | 1 | 1.34447E-05 | -1.302988223 | 0.133 | 0.516 | 0.326598214 | 0.000940843 | 2.68892E-05 |

| | | | | | | | | | | | | |
|---|---|---|---|---|---|---|---|---|---|---|---|---|
| Tcim | 0.918519165 | 0.281454345 | 0.048 | 0.052 | 1 | 1.3642E-05 | 1.290755163 | 0.333 | 0.093 | 0.331390921 | 0.918519165 | 2.72838E-05 |
| Arpc2 | 0.003195776 | -0.653641866 | 0.857 | 0.981 | 1 | 1.36833E-05 | -0.52805567 | 1 | 1 | 0.332394881 | 0.003195776 | 2.73664E-05 |
| Ccdc124 | 0.008363934 | -0.933558546 | 0.143 | 0.507 | 1 | 1.3789E-05 | -1.701591264 | 0.067 | 0.439 | 0.334961983 | 0.008363934 | 2.75778E-05 |
| Rbis | 0.019181459 | -0.308080886 | 0.238 | 0.586 | 1 | 1.39936E-05 | -1.330733213 | 0.111 | 0.498 | 0.339933718 | 0.019181459 | 2.79871E-05 |
| Zfp598 | 0.566644938 | 0.198038337 | 0.19 | 0.28 | 1 | 1.42266E-05 | -2.008089318 | 0.044 | 0.398 | 0.34559268 | 0.566644938 | 2.8453E-05 |
| Marchf3 | 0.032337364 | 1.108572522 | 0.571 | 0.317 | 1 | 1.42688E-05 | 1.66404776 | 0.667 | 0.301 | 0.346616781 | 0.032337364 | 2.85373E-05 |
| Tmco3 | 0.004774521 | -1.294673498 | 0.333 | 0.679 | 1 | 1.43082E-05 | -1.404390255 | 0.111 | 0.502 | 0.347574196 | 0.004774521 | 2.86161E-05 |
| Fam76a | 1.43918E-05 | -1.585971496 | 0.238 | 0.784 | 0.349604914 | 0.002029303 | -0.588390442 | 0.622 | 0.81 | 1 | 0.002029303 | 2.87833E-05 |
| Hp1bp3 | 0.007397618 | -0.594000557 | 0.762 | 0.91 | 1 | 1.45333E-05 | -0.640283213 | 0.889 | 0.958 | 0.353043204 | 0.007397618 | 2.90664E-05 |
| Psma2 | 1.45664E-05 | -1.263815625 | 0.667 | 0.937 | 0.353847315 | 0.000260353 | -0.557935513 | 0.933 | 0.955 | 1 | 0.000260353 | 2.91326E-05 |
| Pold4 | 0.009570196 | -1.213753503 | 0.095 | 0.399 | 1 | 1.46868E-05 | -4.211745177 | 0 | 0.311 | 0.356772163 | 0.009570196 | 2.93734E-05 |
| Rbx1 | 1.46945E-05 | -1.264224428 | 0.714 | 0.929 | 0.356958039 | 0.000102938 | -0.651346453 | 0.822 | 0.924 | 1 | 0.000102938 | 2.93887E-05 |
| Zfp637 | 0.002699733 | -3.25696557 | 0 | 0.313 | 1 | 1.48039E-05 | -2.089516918 | 0.067 | 0.412 | 0.359616976 | 0.002699733 | 2.96076E-05 |
| Gmds | 1.49035E-05 | 1.032490834 | 1 | 0.933 | 0.362035938 | 0.002439829 | 0.770988431 | 0.978 | 0.952 | 1 | 0.002439829 | 2.98068E-05 |
| Mrps18a | 0.009157322 | -1.314912919 | 0.095 | 0.399 | 1 | 1.49931E-05 | -2.611548163 | 0.067 | 0.398 | 0.364211915 | 0.009157322 | 2.99859E-05 |
| Ndufs6 | 1.49997E-05 | -1.989193245 | 0.19 | 0.701 | 0.364373722 | 0.199334483 | -0.409097258 | 0.6 | 0.612 | 1 | 0.199334483 | 2.99993E-05 |
| Gtf2a2 | 2.39127E-05 | -1.921174822 | 0.238 | 0.743 | 0.580886556 | 1.50297E-05 | -0.866266005 | 0.511 | 0.779 | 0.365102046 | 2.39127E-05 | 3.00592E-05 |
| Nudt4 | 0.003778364 | -1.057547263 | 0.476 | 0.843 | 1 | 1.50388E-05 | -1.186704836 | 0.867 | 0.858 | 0.365323364 | 0.003778364 | 3.00774E-05 |
| Magi1 | 0.016409472 | 1.122529885 | 0.714 | 0.459 | 1 | 1.50733E-05 | 0.994320983 | 0.889 | 0.491 | 0.366160756 | 0.016409472 | 3.01464E-05 |
| Xpot | 0.685953207 | 0.522284014 | 0.429 | 0.627 | 1 | 1.51576E-05 | -1.106507634 | 0.4 | 0.73 | 0.368209279 | 0.685953207 | 3.0315E-05 |
| Plekha3 | 0.14586457 | -0.28315951 | 0.333 | 0.616 | 1 | 1.52094E-05 | -1.306873749 | 0.178 | 0.585 | 0.369465722 | 0.14586457 | 3.04185E-05 |
| Nop58 | 0.000141107 | -1.170366632 | 0.857 | 0.948 | 1 | 1.52222E-05 | -0.823887403 | 0.911 | 0.913 | 0.369777575 | 0.000141107 | 3.04442E-05 |
| Nf1 | 0.062885395 | 0.881429416 | 0.905 | 0.888 | 1 | 1.52396E-05 | 0.788851183 | 1 | 0.958 | 0.370201001 | 0.062885395 | 3.0479E-05 |
| Ing3 | 0.92723038 | 0.608816326 | 0.333 | 0.377 | 1 | 1.54714E-05 | -1.166102512 | 0.089 | 0.478 | 0.375831588 | 0.92723038 | 3.09426E-05 |
| Tecr | 1.55685E-05 | -1.750394521 | 0.476 | 0.843 | 0.378190627 | 0.001063442 | -0.624355939 | 0.733 | 0.775 | 1 | 0.001063442 | 3.11368E-05 |
| Klhl9 | 2.56648E-05 | -2.480033568 | 0.143 | 0.642 | 0.623449032 | 1.56071E-05 | -0.977517619 | 0.622 | 0.799 | 0.379126661 | 2.56648E-05 | 3.12139E-05 |
| Prkce | 1.56492E-05 | 1.36358888 | 0.905 | 0.66 | 0.380150016 | 0.000549595 | 0.990229545 | 0.978 | 0.682 | 1 | 0.000549595 | 3.12981E-05 |
| Hnrnpa3 | 1.58114E-05 | -0.912416993 | 0.905 | 0.993 | 0.384091237 | 2.85203E-05 | -0.525339081 | 1 | 0.993 | 0.69281561 | 2.85203E-05 | 3.16226E-05 |
| Maz | 0.002527588 | -1.555046923 | 0.048 | 0.407 | 1 | 1.58973E-05 | -1.638855509 | 0.111 | 0.491 | 0.386176435 | 0.002527588 | 3.17943E-05 |
| Ndufa5 | 1.59071E-05 | -2.07218261 | 0.143 | 0.675 | 0.386415584 | 0.000916904 | -1.064161343 | 0.4 | 0.602 | 1 | 0.000916904 | 3.1814E-05 |
| Tomm6 | 1.59142E-05 | -1.420375387 | 0.524 | 0.866 | 0.386586921 | 0.000104561 | -0.904683129 | 0.733 | 0.806 | 1 | 0.000104561 | 3.18281E-05 |
| Tns2 | 0.08919733 | -0.438437736 | 0.333 | 0.571 | 1 | 1.5994E-05 | -1.293468441 | 0.311 | 0.692 | 0.388525666 | 0.08919733 | 3.19877E-05 |
| Rnaset2a | 1.60228E-05 | -1.702604111 | 0.286 | 0.802 | 0.389226544 | 0.006844074 | -0.367975019 | 0.556 | 0.758 | 1 | 0.006844074 | 3.20454E-05 |
| Ilk | 8.1249E-05 | -1.808892157 | 0.143 | 0.653 | 1 | 1.61444E-05 | -0.776359032 | 0.222 | 0.633 | 0.392179675 | 8.1249E-05 | 3.22885E-05 |
| Mea1 | 0.0009737 | -1.834022873 | 0.095 | 0.493 | 1 | 1.61967E-05 | -1.316927414 | 0.067 | 0.426 | 0.393451078 | 0.0009737 | 3.23932E-05 |
| Med19 | 0.004899665 | -2.107587946 | 0.095 | 0.399 | 1 | 1.62143E-05 | -1.84048937 | 0.044 | 0.384 | 0.39387835 | 0.004899665 | 3.24284E-05 |
| Lamtor5 | 1.6264E-05 | -1.957665163 | 0.19 | 0.716 | 0.395085912 | 4.13001E-05 | -1.067698808 | 0.378 | 0.678 | 1 | 4.13001E-05 | 3.25278E-05 |
| Prkch | 0.358498393 | 1.483947209 | 0.286 | 0.228 | 1 | 1.64615E-05 | 0.475477167 | 0.4 | 0.125 | 0.399882797 | 0.358498393 | 3.29227E-05 |
| Ankrd40 | 0.017110954 | -1.12324121 | 0.095 | 0.369 | 1 | 1.64915E-05 | -0.825941061 | 0.178 | 0.599 | 0.400611397 | 0.017110954 | 3.29827E-05 |
| Yipf5 | 0.01967495 | -0.410544518 | 0.571 | 0.899 | 1 | 1.65026E-05 | -0.571227578 | 0.711 | 0.92 | 0.400881544 | 0.01967495 | 3.3005E-05 |
| Psd3 | 1.67518E-05 | 1.161659832 | 0.952 | 0.951 | 0.406934599 | 0.000104229 | 0.871197262 | 1 | 0.972 | 1 | 0.000104229 | 3.35033E-05 |
| Plekhf1 | 0.00963212 | -2.475975352 | 0.048 | 0.313 | 1 | 1.67759E-05 | -1.509572492 | 0.111 | 0.481 | 0.407520357 | 0.00963212 | 3.35515E-05 |
| Bcas3 | 0.012425046 | 1.144631177 | 0.81 | 0.687 | 1 | 1.68929E-05 | 1.193623372 | 0.956 | 0.692 | 0.410361568 | 0.012425046 | 3.37855E-05 |
| Snrnp27 | 1.70677E-05 | -2.735619168 | 0.048 | 0.586 | 0.414609257 | 0.000108319 | -0.913862556 | 0.156 | 0.516 | 1 | 0.000108319 | 3.41352E-05 |
| H2bc4 | 1.70914E-05 | -2.213956571 | 0.524 | 0.847 | 0.415183489 | 0.004447822 | -0.523621149 | 0.756 | 0.758 | 1 | 0.004447822 | 3.41824E-05 |
| Tiam1 | 0.002119413 | 0.933852856 | 0.905 | 0.761 | 1 | 1.72714E-05 | 1.254572827 | 0.978 | 0.706 | 0.419555988 | 0.002119413 | 3.45424E-05 |
| Fmc1 | 0.000728445 | -3.303508156 | 0 | 0.369 | 1 | 1.72758E-05 | -2.124282336 | 0.067 | 0.408 | 0.419663249 | 0.000728445 | 3.45513E-05 |
| Trip11 | 0.064557128 | -0.348596045 | 0.81 | 0.955 | 1 | 1.73424E-05 | -0.750875985 | 0.978 | 0.969 | 0.421282431 | 0.064557128 | 3.46846E-05 |
| Psmd6 | 1.7351E-05 | -2.593014773 | 0.095 | 0.623 | 0.421490264 | 0.003888976 | -0.814427073 | 0.533 | 0.685 | 1 | 0.003888976 | 3.47017E-05 |
| Smo | 0.009720183 | -1.063193826 | 0.286 | 0.634 | 1 | 1.7356E-05 | -0.577454964 | 0.378 | 0.844 | 0.421611949 | 0.009720183 | 3.47117E-05 |
| Msantd4 | 0.017954186 | -1.203972482 | 0.095 | 0.366 | 1 | 1.7396E-05 | -1.996407513 | 0.089 | 0.433 | 0.422584181 | 0.017954186 | 3.47917E-05 |
| Maf1 | 0.078867623 | -0.785659851 | 0.095 | 0.295 | 1 | 1.74202E-05 | -1.856086225 | 0.067 | 0.429 | 0.423172242 | 0.078867623 | 3.48402E-05 |
| Lmtk2 | 0.691912175 | 1.295260144 | 0.238 | 0.243 | 1 | 1.74318E-05 | 1.25152 | 0.778 | 0.391 | 0.423454341 | 0.691912175 | 3.48634E-05 |
| Zpr1 | 0.057371779 | -1.04047375 | 0.095 | 0.299 | 1 | 1.74377E-05 | -2.334849322 | 0.067 | 0.405 | 0.423597098 | 0.057371779 | 3.48751E-05 |
| Mrpl42 | 1.75122E-05 | -2.116305163 | 0.143 | 0.687 | 0.425406747 | 0.000748071 | -0.953841994 | 0.422 | 0.626 | 1 | 0.000748071 | 3.50241E-05 |
| Gkap1 | 1.75244E-05 | -4.530799377 | 0 | 0.519 | 0.425703197 | 0.006459113 | -0.529921137 | 0.289 | 0.595 | 1 | 0.006459113 | 3.50485E-05 |
| Tmem208 | 0.005038143 | -0.921174822 | 0.381 | 0.731 | 1 | 1.75523E-05 | -1.017367132 | 0.333 | 0.685 | 0.426380505 | 0.005038143 | 3.51043E-05 |
| Alg5 | 0.001351456 | -1.351763324 | 0.095 | 0.522 | 1 | 1.76151E-05 | -1.638855509 | 0.156 | 0.519 | 0.427906281 | 0.001351456 | 3.52299E-05 |
| Ube2m | 0.001677776 | -0.866061358 | 0.476 | 0.847 | 1 | 1.78269E-05 | -0.935056778 | 0.733 | 0.806 | 0.433050289 | 0.001677776 | 3.56534E-05 |
| Tsr2 | 0.014047691 | -1.303508156 | 0.095 | 0.377 | 1 | 1.79397E-05 | -2.316927414 | 0.067 | 0.405 | 0.435790106 | 0.014047691 | 3.5879E-05 |
| Fam120b | 0.208685561 | -0.301424841 | 0.238 | 0.388 | 1 | 1.81007E-05 | -1.457939723 | 0.111 | 0.491 | 0.439702938 | 0.208685561 | 3.62011E-05 |
| Eif5 | 0.004254507 | -0.745434169 | 0.905 | 0.985 | 1 | 1.81135E-05 | -0.615987683 | 1 | 1 | 0.440013083 | 0.004254507 | 3.62267E-05 |
| Elp5 | 0.006496125 | -0.987293712 | 0.238 | 0.601 | 1 | 1.81851E-05 | -1.017367132 | 0.156 | 0.564 | 0.441753524 | 0.006496125 | 3.637E-05 |
| Mrps7 | 0.014737022 | -0.950719097 | 0.143 | 0.433 | 1 | 1.84813E-05 | -1.594912161 | 0.089 | 0.443 | 0.448947376 | 0.014737022 | 3.69622E-05 |
| Bud31 | 4.37675E-05 | -1.571232351 | 0.286 | 0.795 | 1 | 1.84839E-05 | -0.928629763 | 0.578 | 0.782 | 0.449011549 | 4.37675E-05 | 3.69675E-05 |
| Klhl13 | 0.002207829 | -3.865387043 | 0 | 0.325 | 1 | 1.85284E-05 | -4.734779929 | 0 | 0.308 | 0.450092543 | 0.002207829 | 3.70565E-05 |
| Dhx38 | 0.142646763 | -1.184209227 | 0.095 | 0.228 | 1 | 1.86815E-05 | -1.867124496 | 0.044 | 0.394 | 0.453809979 | 0.142646763 | 3.73626E-05 |
| Utrn | 0.000252267 | 0.837191015 | 1 | 0.981 | 1 | 1.87939E-05 | 0.975904153 | 1 | 0.983 | 0.456541922 | 0.000252267 | 3.75875E-05 |
| Zfta | 0.087720671 | -1.081115734 | 0.048 | 0.209 | 1 | 1.88137E-05 | -2.158229668 | 0.089 | 0.429 | 0.457022656 | 0.087720671 | 3.76271E-05 |
| Siah2 | 0.000430819 | -1.696915639 | 0.143 | 0.593 | 1 | 1.88699E-05 | -0.934311392 | 0.267 | 0.623 | 0.458386894 | 0.000430819 | 3.77394E-05 |
| Nupr1 | 0.224885553 | -1.104305362 | 0.857 | 0.787 | 1 | 1.90491E-05 | -1.172894074 | 0.889 | 0.837 | 0.462741672 | 0.224885553 | 3.80979E-05 |
| Cdc14a | 0.240669704 | 0.421853394 | 0.476 | 0.347 | 1 | 1.90784E-05 | 1.235613609 | 0.8 | 0.446 | 0.463452096 | 0.240669704 | 3.81564E-05 |
| Aebp1 | 2.44223E-05 | -3.977279923 | 0.048 | 0.567 | 0.593266702 | 1.92373E-05 | -0.942531899 | 0.467 | 0.837 | 0.467312322 | 2.44223E-05 | 3.84742E-05 |
| Atxn1 | 4.6821E-05 | 1.346736782 | 1 | 0.933 | 1 | 1.93695E-05 | 1.291545032 | 0.978 | 0.941 | 0.470522832 | 4.6821E-05 | 3.87385E-05 |
| Dact3 | 0.02891702 | -1.337455488 | 0.095 | 0.336 | 1 | 1.93905E-05 | -2.064161343 | 0.067 | 0.415 | 0.47103493 | 0.02891702 | 3.87807E-05 |
| Actn4 | 1.94517E-05 | 1.138506651 | 1 | 0.985 | 0.472521361 | 0.253333498 | 0.357826996 | 1 | 0.993 | 1 | 0.253333498 | 3.89031E-05 |
| Qdpr | 3.84451E-05 | -2.948280052 | 0.048 | 0.53 | 0.933908996 | 1.96186E-05 | -2.307882274 | 0.089 | 0.419 | 0.476574355 | 3.84451E-05 | 3.92368E-05 |
| Zfyve21 | 0.00774934 | -1.430564892 | 0.095 | 0.422 | 1 | 1.96487E-05 | -2.526380779 | 0.022 | 0.349 | 0.477305607 | 0.00774934 | 3.9297E-05 |
| Ndufb2 | 1.9835E-05 | -1.535681598 | 0.476 | 0.825 | 0.481832863 | 8.52939E-05 | -1.061761251 | 0.578 | 0.751 | 1 | 8.52939E-05 | 3.96697E-05 |
| Zhx1 | 0.41801848 | 0.226312791 | 0.19 | 0.302 | 1 | 1.99068E-05 | -1.960783604 | 0.067 | 0.415 | 0.483574848 | 0.41801848 | 3.98131E-05 |
| Ccser1 | 0.642837231 | -0.292012517 | 0.381 | 0.306 | 1 | 2.00424E-05 | 0.697286445 | 0.689 | 0.284 | 0.486869071 | 0.642837231 | 4.00843E-05 |
| Itpripl2 | 0.039190715 | -0.730982758 | 0.333 | 0.649 | 1 | 2.01518E-05 | -1.06747598 | 0.267 | 0.689 | 0.489528355 | 0.039190715 | 4.03033E-05 |
| Sf3b5 | 2.01683E-05 | -1.63701577 | 0.238 | 0.799 | 0.489927635 | 0.001380771 | -0.321020168 | 0.511 | 0.696 | 1 | 0.001380771 | 4.03361E-05 |
| Commd10 | 0.668068645 | 1.809426867 | 0.333 | 0.343 | 1 | 2.0185E-05 | -1.921789472 | 0.089 | 0.439 | 0.490333646 | 0.668068645 | 4.03696E-05 |
| Mrps18c | 0.016235056 | -1.292012517 | 0.095 | 0.369 | 1 | 2.05573E-05 | -2.665655568 | 0.067 | 0.391 | 0.499377385 | 0.016235056 | 4.11141E-05 |
| Commd3 | 2.06462E-05 | -2.516052791 | 0.143 | 0.649 | 0.501537527 | 0.00226232 | -0.663377828 | 0.422 | 0.668 | 1 | 0.00226232 | 4.1292E-05 |
| Mrpl9 | 0.311657463 | -0.376854305 | 0.19 | 0.306 | 1 | 2.07603E-05 | -1.924257728 | 0.089 | 0.429 | 0.504309841 | 0.311657463 | 4.15202E-05 |
| Rbm17 | 2.07878E-05 | -2.033587364 | 0.19 | 0.709 | 0.50497764 | 9.3656E-05 | -0.841823346 | 0.533 | 0.792 | 1 | 9.3656E-05 | 4.15752E-05 |
| Minpp1 | 0.000114133 | -3.735619168 | 0 | 0.444 | 1 | 2.07947E-05 | -1.394929926 | 0.178 | 0.561 | 0.505144212 | 0.000114133 | 4.15889E-05 |
| Hexb | 2.08367E-05 | -2.167530486 | 0.143 | 0.664 | 0.506165899 | 0.002681144 | -1.161082254 | 0.6 | 0.744 | 1 | 0.002681144 | 4.1673E-05 |
| Sox5 | 0.000150309 | 1.26252409 | 1 | 0.993 | 1 | 2.09001E-05 | 0.845716487 | 1 | 0.99 | 0.507705409 | 0.000150309 | 4.17998E-05 |
| 0610009B22Rik | 0.006863504 | -2.120644099 | 0.048 | 0.332 | 1 | 2.10488E-05 | -1.86124793 | 0.044 | 0.377 | 0.511317679 | 0.006863504 | 4.20972E-05 |
| Iftap | 0.0043295 | -1.405179574 | 0.095 | 0.437 | 1 | 2.11683E-05 | -1.792660845 | 0.089 | 0.45 | 0.514219225 | 0.0043295 | 4.23361E-05 |
| Cox7b | 0.00012927 | -1.19339255 | 0.667 | 0.884 | 1 | 2.12825E-05 | -0.702455671 | 0.8 | 0.872 | 0.516995001 | 0.00012927 | 4.25646E-05 |
| Hivep2 | 7.71947E-05 | 1.727086547 | 1 | 0.974 | 1 | 2.1411E-05 | 1.025523433 | 0.978 | 0.979 | 0.520116349 | 7.71947E-05 | 4.28216E-05 |
| Sgce | 0.024766919 | -0.718545655 | 0.048 | 0.291 | 1 | 2.14159E-05 | -1.156131202 | 0.089 | 0.471 | 0.520235478 | 0.024766919 | 4.28314E-05 |
| Tfam | 0.016553054 | -1.555046923 | 0.095 | 0.362 | 1 | 2.1463E-05 | -2.191396532 | 0.044 | 0.377 | 0.521378208 | 0.016553054 | 4.29255E-05 |
| Cmss1 | 0.271946933 | 1.537508623 | 1 | 1 | 1 | 2.15142E-05 | 0.309586702 | 1 | 1 | 0.522622357 | 0.271946933 | 4.30279E-05 |

| | | | | | | | | | | | | |
|---|---|---|---|---|---|---|---|---|---|---|---|---|
| Ephx1 | 2.16106E-05 | -3.752492987 | 0.095 | 0.586 | 0.524964749 | 0.578305049 | -0.713532195 | 0.578 | 0.495 | 1 | 0.578305049 | 4.32207E-05 |
| Fastk | 0.000904616 | -2.02666795 | 0.095 | 0.493 | 1 | 2.1635E-05 | -1.688486276 | 0.111 | 0.481 | 0.52555658 | 0.000904616 | 4.32695E-05 |
| Lage3 | 0.000977043 | -1.619009982 | 0.095 | 0.5 | 1 | 2.16708E-05 | -1.572428147 | 0.111 | 0.453 | 0.526425876 | 0.000977043 | 4.3341E-05 |
| Ccdc85b | 3.62873E-05 | -3.16543202 | 0.095 | 0.593 | 0.881491435 | 2.21063E-05 | -1.845707079 | 0.111 | 0.471 | 0.537006558 | 3.62873E-05 | 4.42121E-05 |
| Blzf1 | 0.4185685 | -0.233118828 | 0.429 | 0.571 | 1 | 2.22096E-05 | -1.223818009 | 0.222 | 0.633 | 0.539515442 | 0.4185685 | 4.44187E-05 |
| Pfkp | 2.23273E-05 | 1.440955204 | 0.905 | 0.772 | 0.542374415 | 0.114543675 | 0.961727973 | 0.778 | 0.661 | 1 | 0.114543675 | 4.46541E-05 |
| Nr6a1 | 0.014593132 | 1.113555929 | 0.667 | 0.358 | 1 | 2.23305E-05 | 1.437182984 | 0.956 | 0.588 | 0.54245335 | 0.014593132 | 4.46606E-05 |
| Mvb12a | 5.07854E-05 | -2.99156415 | 0.048 | 0.522 | 1 | 2.2349E-05 | -1.58541625 | 0.111 | 0.474 | 0.542901783 | 5.07854E-05 | 4.46975E-05 |
| Rpn1 | 0.00016575 | -1.254993328 | 0.524 | 0.862 | 1 | 2.26905E-05 | -0.804441574 | 0.8 | 0.9 | 0.551197397 | 0.00016575 | 4.53805E-05 |
| Ufc1 | 2.2787E-05 | -1.636083495 | 0.476 | 0.847 | 0.553541804 | 8.65408E-05 | -0.829377415 | 0.667 | 0.806 | 1 | 8.65408E-05 | 4.55735E-05 |
| Hars | 0.097243698 | -0.527862093 | 0.333 | 0.586 | 1 | 2.28246E-05 | -1.0794281 | 0.244 | 0.647 | 0.554454816 | 0.097243698 | 4.56486E-05 |
| Rnf7 | 2.29093E-05 | -1.693345101 | 0.381 | 0.784 | 0.556513555 | 6.13376E-05 | -0.848450656 | 0.733 | 0.83 | 1 | 6.13376E-05 | 4.58181E-05 |
| Tmem43 | 0.199699659 | -0.283999997 | 0.238 | 0.455 | 1 | 2.30122E-05 | -1.214167839 | 0.156 | 0.561 | 0.559012421 | 0.199699659 | 4.60239E-05 |
| Bckdk | 0.004563151 | -2.221045996 | 0.048 | 0.351 | 1 | 2.32025E-05 | -3.282711698 | 0.022 | 0.332 | 0.563633917 | 0.004563151 | 4.64044E-05 |
| Tm9sf2 | 2.32163E-05 | -0.9003608 | 0.762 | 0.966 | 0.563971196 | 0.000383151 | -0.444467694 | 0.911 | 0.972 | 1 | 0.000383151 | 4.64321E-05 |
| Fkbp8 | 2.33349E-05 | -2.14385149 | 0.19 | 0.664 | 0.566850995 | 0.174207721 | -0.294207337 | 0.489 | 0.571 | 1 | 0.174207721 | 4.66692E-05 |
| Mfap1b | 0.193280862 | 0.451379346 | 0.143 | 0.302 | 1 | 2.33372E-05 | -1.614607962 | 0.089 | 0.439 | 0.566906491 | 0.193280862 | 4.66738E-05 |
| Hltf | 0.546088323 | -0.101521945 | 0.286 | 0.388 | 1 | 2.34365E-05 | -2.683249628 | 0.044 | 0.37 | 0.569318703 | 0.546088323 | 4.68724E-05 |
| Cybrd1 | 0.090450526 | -1.611630451 | 0.048 | 0.194 | 1 | 2.34732E-05 | -4.316927414 | 0 | 0.301 | 0.570210642 | 0.090450526 | 4.69458E-05 |
| Aldh6a1 | 0.008972458 | -1.718545655 | 0.095 | 0.392 | 1 | 2.34989E-05 | -1.361321533 | 0.089 | 0.443 | 0.570836124 | 0.008972458 | 4.69973E-05 |
| Dbp | 7.49936E-05 | -4.455511249 | 0 | 0.466 | 1 | 2.35162E-05 | -2.065120263 | 0.044 | 0.384 | 0.571255895 | 7.49936E-05 | 4.70319E-05 |
| Med25 | 0.244828949 | -0.220237537 | 0.286 | 0.493 | 1 | 2.35997E-05 | -0.780327935 | 0.222 | 0.637 | 0.573284666 | 0.244828949 | 4.71989E-05 |
| Clasp1 | 0.730205783 | 0.422396953 | 0.571 | 0.672 | 1 | 2.3739E-05 | 0.885956849 | 0.978 | 0.806 | 0.576668801 | 0.730205783 | 4.74775E-05 |
| Ss18l2 | 0.029923419 | -0.924014773 | 0.286 | 0.571 | 1 | 2.38488E-05 | -2.691966845 | 0.044 | 0.37 | 0.579334362 | 0.029923419 | 4.7697E-05 |
| Mgst3 | 0.981306717 | 0.719575457 | 0.095 | 0.101 | 1 | 2.39273E-05 | 0.934611353 | 0.489 | 0.183 | 0.58124206 | 0.981306717 | 4.7854E-05 |
| Prelid1 | 0.000255885 | -1.238765391 | 0.429 | 0.787 | 1 | 2.39579E-05 | -0.917831458 | 0.778 | 0.83 | 0.581984268 | 0.000255885 | 4.79151E-05 |
| C1qbp | 0.001940891 | -1.050254771 | 0.286 | 0.69 | 1 | 2.39808E-05 | -1.088416883 | 0.556 | 0.765 | 0.582542379 | 0.001940891 | 4.79611E-05 |
| Tmem70 | 0.002673948 | -1.527862093 | 0.143 | 0.5 | 1 | 2.40338E-05 | -1.104829973 | 0.111 | 0.495 | 0.583830184 | 0.002673948 | 4.80671E-05 |
| Sdcbp | 0.002284051 | -0.707162066 | 0.905 | 0.974 | 1 | 2.40388E-05 | -0.532926426 | 1 | 0.997 | 0.5839511 | 0.002284051 | 4.80771E-05 |
| Pex19 | 0.145062103 | -0.381510668 | 0.19 | 0.392 | 1 | 2.40749E-05 | -1.404390255 | 0.111 | 0.464 | 0.584826966 | 0.145062103 | 4.81492E-05 |
| Lsm12 | 0.020259506 | -0.3935835 | 0.571 | 0.825 | 1 | 2.41043E-05 | -0.749154674 | 0.8 | 0.886 | 0.585540785 | 0.020259506 | 4.82079E-05 |
| Atp5c1 | 2.41803E-05 | -1.145655987 | 0.81 | 0.914 | 0.587388015 | 0.000122636 | -0.676932287 | 0.911 | 0.91 | 1 | 0.000122636 | 4.836E-05 |
| Hnrnph2 | 0.00239838 | -0.800487269 | 0.381 | 0.817 | 1 | 2.43702E-05 | -0.918100142 | 0.756 | 0.851 | 0.592001503 | 0.00239838 | 4.87399E-05 |
| ENSMUSG00000121513 | 0.032304692 | 1.513694552 | 0.714 | 0.493 | 1 | 2.44004E-05 | 1.120577426 | 0.978 | 0.775 | 0.592735713 | 0.032304692 | 4.88003E-05 |
| Fam162a | 2.44408E-05 | -1.452932705 | 0.238 | 0.802 | 0.593715436 | 0.000288857 | -0.789098144 | 0.511 | 0.758 | 1 | 0.000288857 | 4.8881E-05 |
| Shisa5 | 2.49355E-05 | -1.574155746 | 0.524 | 0.858 | 0.605732551 | 0.002360508 | -0.644502072 | 0.6 | 0.747 | 1 | 0.002360508 | 4.98703E-05 |
| Cpox | 0.452680658 | 0.150209812 | 0.19 | 0.313 | 1 | 2.49355E-05 | -2.286553765 | 0.067 | 0.398 | 0.605733971 | 0.452680658 | 4.98704E-05 |
| Zfpl1 | 0.174951781 | -0.505031385 | 0.238 | 0.418 | 1 | 2.49486E-05 | -2.86124793 | 0.044 | 0.36 | 0.606050678 | 0.174951781 | 4.98965E-05 |
| Uty | 0.287339178 | 0.576910228 | 0.571 | 0.466 | 1 | 2.49494E-05 | 0.941088917 | 1 | 0.837 | 0.606071524 | 0.287339178 | 4.98982E-05 |
| Cebpd | 2.49697E-05 | -3.265611548 | 0.667 | 0.825 | 0.606564919 | 0.267462686 | -1.376318498 | 0.822 | 0.64 | 1 | 0.267462686 | 4.99389E-05 |
| Myzap | 0.222680203 | -1.801961663 | 0.048 | 0.138 | 1 | 2.51926E-05 | -2.526380779 | 0.022 | 0.346 | 0.611979702 | 0.222680203 | 5.03847E-05 |
| Ppp1r15a | 0.001323782 | -0.904931743 | 0.762 | 0.937 | 1 | 2.54413E-05 | -1.046126125 | 0.956 | 0.934 | 0.618018958 | 0.001323782 | 5.08819E-05 |
| C1qtnf6 | 0.007847288 | -1.535681598 | 0.143 | 0.448 | 1 | 2.55337E-05 | -1.398721505 | 0.133 | 0.498 | 0.620264284 | 0.007847288 | 5.10667E-05 |
| Ube2g2 | 0.011436799 | -1.241111619 | 0.143 | 0.455 | 1 | 2.55741E-05 | -1.393030393 | 0.178 | 0.55 | 0.621244991 | 0.011436799 | 5.11475E-05 |
| Bag3 | 0.00013332 | -1.242298255 | 0.762 | 0.918 | 1 | 2.5654E-05 | -0.675297505 | 0.911 | 0.962 | 0.623187721 | 0.00013332 | 5.13074E-05 |
| Ndufb7 | 2.57004E-05 | -1.922417988 | 0.333 | 0.728 | 0.624314824 | 0.216613448 | -0.321428806 | 0.667 | 0.64 | 1 | 0.216613448 | 5.14002E-05 |
| Luc7l2 | 2.57436E-05 | 0.979120507 | 1 | 0.993 | 0.625364225 | 0.002379714 | 0.493877558 | 1 | 1 | 1 | 0.002379714 | 5.14866E-05 |
| Gtf2h5 | 7.07405E-05 | -1.333329289 | 0.429 | 0.832 | 1 | 2.57597E-05 | -1.001034561 | 0.6 | 0.782 | 0.625753684 | 7.07405E-05 | 5.15187E-05 |
| Stat6 | 0.130821596 | -0.708697869 | 0.19 | 0.369 | 1 | 2.59978E-05 | -1.467869312 | 0.2 | 0.567 | 0.631539554 | 0.130821596 | 5.1995E-05 |
| Psen2 | 0.000723915 | -3.506137322 | 0 | 0.373 | 1 | 2.60824E-05 | -2.429627547 | 0.067 | 0.394 | 0.633592551 | 0.000723915 | 5.2164E-05 |
| Pttg1 | 0.000277405 | -3.91868527 | 0 | 0.414 | 1 | 2.60901E-05 | -2.462604869 | 0.067 | 0.394 | 0.633780252 | 0.000277405 | 5.21795E-05 |
| Lrrc8c | 0.489188993 | 0.759826415 | 0.667 | 0.81 | 1 | 2.61747E-05 | 1.375061272 | 0.822 | 0.768 | 0.635835029 | 0.489188993 | 5.23487E-05 |
| Klf12 | 0.002028142 | 1.439452741 | 0.905 | 0.907 | 1 | 2.64395E-05 | 0.994266526 | 0.978 | 0.976 | 0.642267425 | 0.002028142 | 5.28782E-05 |
| Rps27rt | 2.65824E-05 | -3.146407195 | 0.048 | 0.545 | 0.645739492 | 0.001521142 | -1.071814916 | 0.156 | 0.422 | 1 | 0.001521142 | 5.31641E-05 |
| Rbmxl1 | 2.68677E-05 | -2.860206804 | 0.048 | 0.571 | 0.652669065 | 0.000181772 | -1.060319277 | 0.311 | 0.63 | 1 | 0.000181772 | 5.37346E-05 |
| Snapc1 | 0.116836948 | -0.138481954 | 0.333 | 0.567 | 1 | 2.70544E-05 | -1.249813218 | 0.222 | 0.592 | 0.657206421 | 0.116836948 | 5.41081E-05 |
| Hs3st3a1 | 2.70822E-05 | -2.582268392 | 0.143 | 0.653 | 0.65788001 | 0.001038165 | -1.247664751 | 0.178 | 0.474 | 1 | 0.001038165 | 5.41636E-05 |
| Ftsj3 | 0.897666762 | 0.295260144 | 0.333 | 0.407 | 1 | 2.71137E-05 | -1.883742568 | 0.133 | 0.484 | 0.658646065 | 0.897666762 | 5.42267E-05 |
| Sdc2 | 2.74521E-05 | -1.79778733 | 0.476 | 0.881 | 0.666866038 | 6.89082E-05 | -0.255226714 | 0.489 | 0.869 | 1 | 6.89082E-05 | 5.49034E-05 |
| Abcg2 | 0.005096435 | -1.690800665 | 0.143 | 0.463 | 1 | 2.75162E-05 | -1.913862556 | 0.089 | 0.433 | 0.668424387 | 0.005096435 | 5.50317E-05 |
| Hadh | 0.000658884 | -1.470886475 | 0.143 | 0.582 | 1 | 2.7595E-05 | -1.568845787 | 0.4 | 0.63 | 0.67033797 | 0.000658884 | 5.51893E-05 |
| Arl6ip6 | 0.152256436 | -0.496153234 | 0.238 | 0.444 | 1 | 2.76553E-05 | -1.632282855 | 0.111 | 0.457 | 0.671803703 | 0.152256436 | 5.53099E-05 |
| Pa2g4 | 0.00519746 | -0.679865187 | 0.476 | 0.825 | 1 | 2.76626E-05 | -0.833028056 | 0.8 | 0.865 | 0.671980593 | 0.00519746 | 5.53245E-05 |
| Shisa2 | 0.009109721 | -3.424260315 | 0 | 0.254 | 1 | 2.79158E-05 | -1.718375848 | 0.133 | 0.495 | 0.678129776 | 0.009109721 | 5.58308E-05 |
| Smpd1 | 0.0006859 | -3.370622352 | 0 | 0.373 | 1 | 2.79547E-05 | -2.008089318 | 0.089 | 0.422 | 0.679075502 | 0.0006859 | 5.59086E-05 |
| Stc2 | 2.79725E-05 | -3.212940946 | 0.048 | 0.578 | 0.679508689 | 0.010178009 | -1.970369652 | 0.067 | 0.235 | 1 | 0.010178009 | 5.59443E-05 |
| Wdr83os | 2.81034E-05 | -1.814514714 | 0.238 | 0.746 | 0.682687753 | 0.033083546 | -0.477392086 | 0.511 | 0.612 | 1 | 0.033083546 | 5.6206E-05 |
| Tmf1 | 0.028590933 | -0.249060372 | 0.667 | 0.918 | 1 | 2.81296E-05 | -0.712987238 | 0.867 | 0.927 | 0.683323766 | 0.028590933 | 5.62584E-05 |
| Rgl1 | 0.51808165 | -0.197571249 | 0.571 | 0.743 | 1 | 2.81638E-05 | -0.901889915 | 0.444 | 0.813 | 0.684155257 | 0.51808165 | 5.63268E-05 |
| Ogfod3 | 0.002581377 | -1.921174822 | 0.048 | 0.399 | 1 | 2.82163E-05 | -2.516599759 | 0.067 | 0.384 | 0.685429537 | 0.002581377 | 5.64317E-05 |
| Phospho2 | 0.120800365 | -1.116305163 | 0.095 | 0.239 | 1 | 2.82389E-05 | -2.516599759 | 0.044 | 0.367 | 0.685978488 | 0.120800365 | 5.64769E-05 |
| Mfap1a | 0.074740313 | -0.849790188 | 0.143 | 0.354 | 1 | 2.84953E-05 | -2.476798751 | 0.044 | 0.367 | 0.692207852 | 0.074740313 | 5.69898E-05 |
| Rab34 | 0.005413534 | -0.998653574 | 0.238 | 0.593 | 1 | 2.86282E-05 | -0.96900411 | 0.222 | 0.619 | 0.695435576 | 0.005413534 | 5.72555E-05 |
| Vit | 0.784840569 | 0.338381413 | 0.333 | 0.328 | 1 | 2.86569E-05 | -1.041577686 | 0.178 | 0.567 | 0.696133149 | 0.784840569 | 5.7313E-05 |
| Med29 | 0.000427836 | -2.648156327 | 0.048 | 0.448 | 1 | 2.86742E-05 | -2.456478766 | 0.067 | 0.384 | 0.696552992 | 0.000427836 | 5.73475E-05 |
| Plpp5 | 8.02551E-05 | -1.675939484 | 0.19 | 0.709 | 1 | 2.86907E-05 | -0.776359032 | 0.244 | 0.668 | 0.696954483 | 8.02551E-05 | 5.73806E-05 |
| Pld3 | 2.87811E-05 | -3.90365706 | 0 | 0.493 | 0.699149919 | 0.001699511 | -0.197939378 | 0.2 | 0.509 | 1 | 0.001699511 | 5.75613E-05 |
| Urm1 | 0.228533577 | -0.233118828 | 0.19 | 0.34 | 1 | 2.8919E-05 | -1.597035333 | 0.044 | 0.381 | 0.702501005 | 0.228533577 | 5.78372E-05 |
| Celf1 | 0.03470084 | 0.723602924 | 0.857 | 0.78 | 1 | 2.89861E-05 | 0.916169708 | 1 | 0.952 | 0.7041295 | 0.03470084 | 5.79713E-05 |
| 5031425E22Rik | 2.89889E-05 | 1.584542 | 0.952 | 0.821 | 0.704199228 | 0.026364558 | 0.873026068 | 0.889 | 0.834 | 1 | 0.026364558 | 5.7977E-05 |
| Sult1c1 | 0.54776222 | 2.088809267 | 0.048 | 0.026 | 1 | 2.9057E-05 | 2.197645759 | 0.244 | 0.059 | 0.705853615 | 0.54776222 | 5.81132E-05 |
| Ckb | 2.90881E-05 | -2.945261409 | 0.333 | 0.746 | 0.706607485 | 0.000110677 | -2.599941064 | 0.311 | 0.581 | 1 | 0.000110677 | 5.81753E-05 |
| Myo9b | 2.91301E-05 | 1.434433454 | 0.905 | 0.802 | 0.707628373 | 0.013199091 | 0.744264754 | 0.911 | 0.917 | 1 | 0.013199091 | 5.82594E-05 |
| Cks1b | 0.01111008 | -1.826074119 | 0.095 | 0.381 | 1 | 2.9221E-05 | -2.526380779 | 0.067 | 0.391 | 0.7098375 | 0.01111008 | 5.84412E-05 |
| Rapgef6 | 3.00027E-05 | 1.041503552 | 1 | 0.888 | 0.728826785 | 0.022614613 | 0.577554395 | 0.978 | 0.945 | 1 | 0.022614613 | 6.00046E-05 |
| Il31ra | 3.00658E-05 | 3.368917186 | 0.286 | 0.052 | 0.730358432 | 0.725193026 | 0.946106992 | 0.067 | 0.055 | 1 | 0.725193026 | 6.01307E-05 |
| Akap13 | 0.062516753 | 1.109536433 | 0.952 | 0.832 | 1 | 3.00727E-05 | 0.928535891 | 0.956 | 0.869 | 0.730526367 | 0.062516753 | 6.01445E-05 |
| Tmem230 | 0.02507401 | -0.496153234 | 0.238 | 0.549 | 1 | 3.0124E-05 | -1.594912161 | 0.089 | 0.436 | 0.731771204 | 0.02507401 | 6.0247E-05 |
| Anp32b | 0.001365705 | -0.812626899 | 0.667 | 0.922 | 1 | 3.03497E-05 | -0.759256605 | 0.933 | 0.948 | 0.7372541 | 0.001365705 | 6.06984E-05 |
| Ndufs8 | 3.05266E-05 | -3.22709504 | 0.048 | 0.541 | 0.741552657 | 0.701481628 | -0.316927414 | 0.511 | 0.481 | 1 | 0.701481628 | 6.10523E-05 |
| Guk1 | 6.79197E-05 | -2.450349544 | 0.143 | 0.593 | 1 | 3.07039E-05 | -0.901889915 | 0.156 | 0.516 | 0.745859834 | 6.79197E-05 | 6.14069E-05 |
| Adh5 | 3.07446E-05 | -2.623908781 | 0.095 | 0.582 | 0.746847856 | 0.040297013 | -0.623314658 | 0.489 | 0.578 | 1 | 0.040297013 | 6.14883E-05 |
| Aktip | 0.068134298 | -0.849790188 | 0.143 | 0.358 | 1 | 3.10413E-05 | -0.889178863 | 0.111 | 0.478 | 0.754056321 | 0.068134298 | 6.20817E-05 |
| Gabpb1 | 0.236724415 | 1.051719948 | 0.476 | 0.466 | 1 | 3.10641E-05 | -1.112342723 | 0.244 | 0.633 | 0.754610117 | 0.236724415 | 6.21273E-05 |
| Bnip3l | 3.10985E-05 | -1.567019564 | 0.381 | 0.821 | 0.755445456 | 0.008060264 | -0.583146076 | 0.844 | 0.806 | 1 | 0.008060264 | 6.21961E-05 |
| Camsap2 | 3.13592E-05 | 1.599771186 | 0.857 | 0.869 | 0.761777692 | 0.412137502 | 0.476041431 | 0.956 | 0.941 | 1 | 0.412137502 | 6.27174E-05 |
| Ciao2b | 0.000570166 | -1.735619168 | 0.048 | 0.47 | 1 | 3.13702E-05 | -1.167783975 | 0.133 | 0.484 | 0.762044367 | 0.000570166 | 6.27394E-05 |
| Rab18 | 0.000154427 | -1.073973665 | 0.619 | 0.907 | 1 | 3.14129E-05 | -0.783469422 | 0.822 | 0.917 | 0.763082927 | 0.000154427 | 6.28249E-05 |

| | | | | | | | | | | | | |
|---|---|---|---|---|---|---|---|---|---|---|---|---|
| Gpn3 | 0.040756344 | -1.133583154 | 0.19 | 0.414 | 1 | 3.17262E-05 | -2.298780067 | 0.044 | 0.367 | 0.770693181 | 0.040756344 | 6.34514E-05 |
| BC031181 | 3.1834E-05 | -1.108967855 | 0.286 | 0.84 | 0.773312669 | 0.010718238 | -0.460380204 | 0.711 | 0.775 | 1 | 0.010718238 | 6.36671E-05 |
| Pcbp4 | 0.062529991 | -1.094412557 | 0.095 | 0.291 | 1 | 3.18668E-05 | -1.901889915 | 0.089 | 0.429 | 0.77410839 | 0.062529991 | 6.37326E-05 |
| Micos13 | 3.18858E-05 | -1.625788514 | 0.429 | 0.802 | 0.774570527 | 0.132930392 | -0.297613939 | 0.733 | 0.723 | 1 | 0.132930392 | 6.37706E-05 |
| Timm10b | 3.19867E-05 | -1.188724709 | 0.095 | 0.627 | 0.777021441 | 0.00353115 | -0.80878051 | 0.356 | 0.578 | 1 | 0.00353115 | 6.39724E-05 |
| Tmem254 | 0.000138615 | -1.827667377 | 0.333 | 0.72 | 1 | 3.21344E-05 | -1.298242851 | 0.644 | 0.737 | 0.780607805 | 0.000138615 | 6.42677E-05 |
| Prdx5 | 3.22486E-05 | -1.320581669 | 0.667 | 0.896 | 0.783383991 | 5.7831E-05 | -0.751749565 | 0.889 | 0.896 | 1 | 5.7831E-05 | 6.44962E-05 |
| Rimoc1 | 0.002118893 | -2.574155746 | 0.048 | 0.384 | 1 | 3.22831E-05 | -0.881712033 | 0.111 | 0.488 | 0.784222045 | 0.002118893 | 6.45652E-05 |
| Sertad4 | 0.205487585 | -0.179679569 | 0.19 | 0.343 | 1 | 3.25194E-05 | -1.286553765 | 0.133 | 0.502 | 0.78996018 | 0.205487585 | 6.50377E-05 |
| Utp14a | 3.25614E-05 | -2.921174822 | 0.048 | 0.563 | 0.790980722 | 0.498344175 | -0.124282336 | 0.489 | 0.519 | 1 | 0.498344175 | 6.51217E-05 |
| Mzt1 | 3.26106E-05 | -2.470886475 | 0.095 | 0.604 | 0.79217614 | 8.06182E-05 | -0.888950859 | 0.289 | 0.647 | 1 | 8.06182E-05 | 6.52201E-05 |
| Clu | 0.04879918 | -1.101521945 | 0.333 | 0.556 | 1 | 3.26352E-05 | -1.489641002 | 0.756 | 0.844 | 0.792773931 | 0.04879918 | 6.52693E-05 |
| Slc4a1ap | 0.171877304 | -0.155497931 | 0.238 | 0.437 | 1 | 3.26871E-05 | -1.662702251 | 0.111 | 0.464 | 0.794036024 | 0.171877304 | 6.53732E-05 |
| 2210016L21Rik | 0.003102119 | -1.76085646 | 0.095 | 0.444 | 1 | 3.2717E-05 | -2.040953952 | 0.089 | 0.426 | 0.794760768 | 0.003102119 | 6.54329E-05 |
| Arhgap18 | 0.952183966 | 0.866416846 | 0.048 | 0.052 | 1 | 3.27974E-05 | 1.812355603 | 0.378 | 0.128 | 0.796714273 | 0.952183966 | 6.55937E-05 |
| Slc25a46 | 0.170607425 | -0.171718283 | 0.143 | 0.306 | 1 | 3.30051E-05 | -1.450783161 | 0.044 | 0.381 | 0.801758791 | 0.170607425 | 6.6009E-05 |
| Ramac | 0.000547196 | -1.204790105 | 0.238 | 0.716 | 1 | 3.30823E-05 | -0.972279242 | 0.644 | 0.792 | 0.803634563 | 0.000547196 | 6.61635E-05 |
| Cdkn2d | 3.30931E-05 | -3.208871282 | 0.048 | 0.56 | 0.803896535 | 0.00159161 | -1.21739174 | 0.133 | 0.394 | 1 | 0.00159161 | 6.6185E-05 |
| Ghitm | 0.005685919 | -0.56830302 | 0.524 | 0.847 | 1 | 3.35828E-05 | -0.763400607 | 0.756 | 0.858 | 0.815793487 | 0.005685919 | 6.71645E-05 |
| Ei24 | 0.160375158 | 0.126283972 | 0.19 | 0.396 | 1 | 3.39327E-05 | -1.439324045 | 0.111 | 0.457 | 0.824292881 | 0.160375158 | 6.78642E-05 |
| Dnajc10 | 3.41758E-05 | -1.176727646 | 0.286 | 0.821 | 0.830197811 | 6.33196E-05 | -0.718289976 | 0.444 | 0.782 | 1 | 6.33196E-05 | 6.83504E-05 |
| Nabp1 | 0.324747609 | -1.063193826 | 0.048 | 0.119 | 1 | 3.45158E-05 | 1.425274829 | 0.378 | 0.128 | 0.838458752 | 0.324747609 | 6.90305E-05 |
| Ankrd17 | 0.004224488 | 0.763000349 | 1 | 0.951 | 1 | 3.46671E-05 | 0.872076968 | 1 | 0.993 | 0.842133343 | 0.004224488 | 6.9333E-05 |
| Map1lc3a | 0.000730783 | -1.151309323 | 0.619 | 0.858 | 1 | 3.48592E-05 | -0.760098052 | 0.867 | 0.886 | 0.846800124 | 0.000730783 | 6.97172E-05 |
| 2310022B05Rik | 0.204112491 | -0.292012517 | 0.19 | 0.347 | 1 | 3.49427E-05 | -1.502793959 | 0.089 | 0.433 | 0.848827262 | 0.204112491 | 6.98841E-05 |
| Pitpnc1 | 3.51585E-05 | 1.505772236 | 1 | 0.981 | 0.854070152 | 0.085847784 | 0.538861749 | 1 | 0.986 | 1 | 0.085847784 | 7.03158E-05 |
| Ubqln2 | 0.000240871 | -3.701267664 | 0 | 0.418 | 1 | 3.51731E-05 | -1.746239044 | 0.156 | 0.491 | 0.854425278 | 0.000240871 | 7.0345E-05 |
| Mtmr3 | 0.058593364 | 0.83955024 | 0.81 | 0.757 | 1 | 3.51741E-05 | 0.976579344 | 1 | 0.896 | 0.85444871 | 0.058593364 | 7.03469E-05 |
| Slc22a17 | 7.351E-05 | -3.955584852 | 0 | 0.463 | 1 | 3.52995E-05 | -1.056466952 | 0.089 | 0.45 | 0.857495421 | 7.351E-05 | 7.05978E-05 |
| Mrps23 | 0.007428147 | -2.054148687 | 0.048 | 0.328 | 1 | 3.53066E-05 | -2.754332726 | 0.022 | 0.332 | 0.857666749 | 0.007428147 | 7.06119E-05 |
| Iscu | 3.56194E-05 | -1.845864485 | 0.19 | 0.701 | 0.865266371 | 0.012786232 | -0.58163064 | 0.533 | 0.682 | 1 | 0.012786232 | 7.12375E-05 |
| Smdt1 | 0.005921313 | -0.889560717 | 0.762 | 0.847 | 1 | 3.64371E-05 | -0.988122312 | 0.756 | 0.81 | 0.885130862 | 0.005921313 | 7.28729E-05 |
| Tcaf1 | 3.66299E-05 | -1.622469684 | 0.333 | 0.817 | 0.889814121 | 0.00014299 | -0.805003819 | 0.4 | 0.827 | 1 | 0.00014299 | 7.32585E-05 |
| Runx1 | 0.136068923 | 0.708905305 | 0.952 | 0.978 | 1 | 3.66309E-05 | 1.177529522 | 1 | 0.986 | 0.889837035 | 0.136068923 | 7.32604E-05 |
| Isoc1 | 0.009297084 | -2.133583154 | 0.048 | 0.317 | 1 | 3.6914E-05 | -1.339295227 | 0.089 | 0.443 | 0.896715264 | 0.009297084 | 7.38267E-05 |
| Tmem160 | 8.38043E-05 | -2.420745831 | 0.048 | 0.534 | 1 | 3.70463E-05 | -2.053893008 | 0.089 | 0.415 | 0.899929489 | 8.38043E-05 | 7.40913E-05 |
| Large1 | 0.004729891 | 1.47959922 | 0.857 | 0.739 | 1 | 3.72913E-05 | 0.943001081 | 0.933 | 0.858 | 0.905880972 | 0.004729891 | 7.45813E-05 |
| Spats2 | 0.316110149 | 0.763039106 | 0.238 | 0.437 | 1 | 3.72918E-05 | -0.689879512 | 0.111 | 0.488 | 0.905891698 | 0.316110149 | 7.45822E-05 |
| Tenm2 | 0.063229431 | 2.194343681 | 0.429 | 0.284 | 1 | 3.7411E-05 | 1.849868581 | 0.689 | 0.343 | 0.908788363 | 0.063229431 | 7.48206E-05 |
| Rfx3 | 0.264655441 | 1.433859247 | 0.524 | 0.485 | 1 | 3.75214E-05 | 1.306026744 | 0.978 | 0.682 | 0.91147064 | 0.264655441 | 7.50415E-05 |
| Gm8797 | 0.004568994 | -2.221045996 | 0.048 | 0.351 | 1 | 3.76441E-05 | -2.476798751 | 0.044 | 0.36 | 0.914450682 | 0.004568994 | 7.52868E-05 |
| Olfml3 | 0.000193822 | -1.650928076 | 0.238 | 0.731 | 1 | 3.78412E-05 | -1.383016604 | 0.267 | 0.644 | 0.919238053 | 0.000193822 | 7.56809E-05 |
| Naxe | 3.78669E-05 | -1.911190733 | 0.19 | 0.69 | 0.91986212 | 0.004937762 | -0.638855509 | 0.356 | 0.571 | 1 | 0.004937762 | 7.57323E-05 |
| Htra1 | 0.001903327 | -1.686786575 | 0.381 | 0.687 | 1 | 3.78816E-05 | -0.733955632 | 0.644 | 0.948 | 0.920220152 | 0.001903327 | 7.57618E-05 |
| Cox20 | 3.80564E-05 | -2.785659851 | 0.048 | 0.56 | 0.924464993 | 0.000445703 | -1.061088509 | 0.156 | 0.464 | 1 | 0.000445703 | 7.61113E-05 |
| Brd4 | 3.81485E-05 | 1.098803236 | 1 | 0.899 | 0.926702707 | 0.000219055 | 0.748931526 | 1 | 0.997 | 1 | 0.000219055 | 7.62955E-05 |
| Dynlt3 | 0.001195187 | -1.015527393 | 0.333 | 0.728 | 1 | 3.82435E-05 | -1.091367713 | 0.756 | 0.796 | 0.929009985 | 0.001195187 | 7.64854E-05 |
| Cby1 | 0.022813103 | -1.865387043 | 0.048 | 0.272 | 1 | 3.83397E-05 | -3.960783604 | 0 | 0.287 | 0.931349129 | 0.022813103 | 7.6678E-05 |
| Wtip | 3.83738E-05 | -2.979670471 | 0.095 | 0.575 | 0.932176101 | 0.028077617 | -0.329571623 | 0.467 | 0.782 | 1 | 0.028077617 | 7.67461E-05 |
| Cmpk1 | 0.000733632 | -1.074129304 | 0.524 | 0.862 | 1 | 3.87011E-05 | -0.78479587 | 0.822 | 0.855 | 0.940127766 | 0.000733632 | 7.74008E-05 |
| Fam120aos | 0.001126026 | -3.196592952 | 0 | 0.351 | 1 | 3.89673E-05 | -1.738391182 | 0.067 | 0.391 | 0.946593289 | 0.001126026 | 7.79331E-05 |
| Dnajb1 | 3.89927E-05 | -2.51188616 | 0.524 | 0.791 | 0.947210009 | 0.013880276 | -0.821719566 | 0.822 | 0.789 | 1 | 0.013880276 | 7.79838E-05 |
| Msmo1 | 0.234778079 | -1.167006156 | 0.571 | 0.653 | 1 | 3.90106E-05 | -1.105307555 | 0.222 | 0.599 | 0.947644614 | 0.234778079 | 7.80196E-05 |
| Trio | 0.000194631 | 1.278470486 | 1 | 0.94 | 1 | 3.91815E-05 | 1.042923164 | 1 | 0.972 | 0.9517981 | 0.000194631 | 7.83616E-05 |
| Ctxnd1 | 0.532426384 | 0.503846766 | 0 | 0.019 | 1 | 3.92211E-05 | 5.301982419 | 0.133 | 0.017 | 0.952759555 | 0.532426384 | 7.84407E-05 |
| Med21 | 0.005779611 | -0.933558546 | 0.286 | 0.69 | 1 | 3.93591E-05 | -1.714519779 | 0.2 | 0.529 | 0.956110445 | 0.005779611 | 7.87166E-05 |
| BC004004 | 5.92406E-05 | -2.184209227 | 0.048 | 0.556 | 1 | 3.93791E-05 | -1.017367132 | 0.089 | 0.443 | 0.956595921 | 5.92406E-05 | 7.87566E-05 |
| Smn1 | 0.010535345 | -1.593014773 | 0.095 | 0.384 | 1 | 3.93913E-05 | -1.826482512 | 0.111 | 0.446 | 0.956893394 | 0.010535345 | 7.8781E-05 |
| Hat1 | 0.168014284 | -0.1627295 | 0.238 | 0.451 | 1 | 3.94611E-05 | -0.626255472 | 0.111 | 0.495 | 0.958589777 | 0.168014284 | 7.89207E-05 |
| Aph1a | 0.000556306 | -1.724259306 | 0.143 | 0.578 | 1 | 3.96307E-05 | -1.128755812 | 0.4 | 0.661 | 0.962709576 | 0.000556306 | 7.92599E-05 |
| Psmb5 | 3.96361E-05 | -1.552296312 | 0.476 | 0.817 | 0.962840755 | 4.01198E-05 | -0.702985846 | 0.733 | 0.827 | 0.974589028 | 4.01198E-05 | 7.92707E-05 |
| Scd2 | 3.98859E-05 | -1.813264312 | 0.333 | 0.813 | 0.968907189 | 0.00056188 | -1.532864813 | 0.578 | 0.682 | 1 | 0.00056188 | 7.97701E-05 |
| Msl3 | 0.008050474 | -1.558888989 | 0.143 | 0.444 | 1 | 4.00131E-05 | -1.34440815 | 0.111 | 0.46 | 0.971999062 | 0.008050474 | 8.00247E-05 |
| Rnf130 | 4.01324E-05 | -1.408145794 | 0.524 | 0.896 | 0.974895429 | 0.033493946 | -0.102188565 | 0.933 | 0.92 | 1 | 0.033493946 | 8.02631E-05 |
| Ubr5 | 4.01893E-05 | 1.129574301 | 1 | 0.989 | 0.976279546 | 0.349801545 | 0.267103895 | 1 | 1 | 1 | 0.349801545 | 8.03771E-05 |
| Sap30l | 0.002098696 | -2.648156327 | 0.048 | 0.384 | 1 | 4.03787E-05 | -0.969866708 | 0.133 | 0.491 | 0.98087929 | 0.002098696 | 8.07558E-05 |
| Crls1 | 0.015948754 | -1.486099569 | 0.095 | 0.366 | 1 | 4.04139E-05 | -1.504554417 | 0.067 | 0.412 | 0.981733658 | 0.015948754 | 8.08261E-05 |
| Cdc42ep4 | 0.079640101 | -0.569402216 | 0.333 | 0.593 | 1 | 4.05428E-05 | -0.985721505 | 0.289 | 0.72 | 0.984866844 | 0.079640101 | 8.10841E-05 |
| Gse1 | 0.010062339 | 1.196122291 | 0.667 | 0.563 | 1 | 4.06257E-05 | 1.193780532 | 0.911 | 0.578 | 0.986879641 | 0.010062339 | 8.12498E-05 |
| Rtn4 | 0.48264965 | 0.116054103 | 0.952 | 0.981 | 1 | 4.09961E-05 | 0.700022329 | 1 | 0.997 | 0.995877204 | 0.48264965 | 8.19905E-05 |
| Fam3c | 4.09964E-05 | -1.871578877 | 0.286 | 0.754 | 0.995883649 | 0.001235053 | -0.812093244 | 0.378 | 0.63 | 1 | 0.001235053 | 8.1991E-05 |
| Gadd45b | 0.010889612 | -0.854534339 | 0.762 | 0.858 | 1 | 4.1137E-05 | -0.950950733 | 0.867 | 0.934 | 0.999299129 | 0.010889612 | 8.22722E-05 |
| Spp1 | 0.106335315 | 1.844439029 | 0.143 | 0.317 | 1 | 4.11556E-05 | -2.630652 | 0.133 | 0.467 | 0.999750626 | 0.106335315 | 8.23094E-05 |
| Gm17477 | 4.12131E-05 | 4.351843673 | 0.143 | 0.011 | 1 | 0.974356719 | 1.576157382 | 0.067 | 0.069 | 1 | 0.974356719 | 8.24244E-05 |
| Nqo1 | 0.005347142 | -2.24509147 | 0.095 | 0.41 | 1 | 4.16947E-05 | -3.516599759 | 0.022 | 0.318 | 1 | 0.005347142 | 8.33877E-05 |
| Lzic | 0.945964828 | 0.642293536 | 0.286 | 0.347 | 1 | 4.16979E-05 | -3.425451871 | 0.022 | 0.318 | 1 | 0.945964828 | 8.3394E-05 |
| Zfp771 | 0.005713777 | -1.602352638 | 0.095 | 0.418 | 1 | 4.2159E-05 | -1.935837246 | 0.111 | 0.436 | 1 | 0.005713777 | 8.43163E-05 |
| Yeats4 | 0.004608188 | -1.392317423 | 0.143 | 0.496 | 1 | 4.21744E-05 | -1.513324627 | 0.133 | 0.471 | 1 | 0.004608188 | 8.4347E-05 |
| Cadm4 | 4.21942E-05 | 4.995699863 | 0.143 | 0.011 | 1 | 0.173348863 | 1.098110085 | 0.089 | 0.042 | 1 | 0.173348863 | 8.43867E-05 |
| Lamtor3 | 4.23312E-05 | -2.677903671 | 0.048 | 0.556 | 1 | 0.011521644 | -0.467170049 | 0.467 | 0.668 | 1 | 0.011521644 | 8.46605E-05 |
| Tcea1 | 0.542487341 | 0.137718867 | 0.905 | 0.937 | 1 | 4.30098E-05 | -0.597613838 | 0.867 | 0.941 | 1 | 0.542487341 | 8.60178E-05 |
| Ube2j2 | 0.08911422 | -0.52416761 | 0.429 | 0.694 | 1 | 4.3021E-05 | -0.86482782 | 0.689 | 0.817 | 1 | 0.08911422 | 8.60402E-05 |
| Bbln | 0.000103709 | -3.02666795 | 0.048 | 0.5 | 1 | 4.30311E-05 | -2.124282336 | 0.067 | 0.394 | 1 | 0.000103709 | 8.60604E-05 |
| Dyrk2 | 0.187410633 | -0.51887331 | 0.143 | 0.302 | 1 | 4.31304E-05 | -1.576489628 | 0.133 | 0.481 | 1 | 0.187410633 | 8.62589E-05 |
| Myh10 | 0.006688436 | -0.664097871 | 0.19 | 0.549 | 1 | 4.31583E-05 | -1.083640351 | 0.222 | 0.626 | 1 | 0.006688436 | 8.63147E-05 |
| Rab9 | 0.000689623 | -1.321210158 | 0.143 | 0.619 | 1 | 4.32321E-05 | -0.665449719 | 0.311 | 0.723 | 1 | 0.000689623 | 8.64624E-05 |
| Ndufb9 | 0.00012844 | -1.293294343 | 0.619 | 0.836 | 1 | 4.346E-05 | -0.707995209 | 0.756 | 0.837 | 1 | 0.00012844 | 8.69182E-05 |
| Sqle | 0.066409873 | -1.159885026 | 0.476 | 0.649 | 1 | 4.35021E-05 | -1.400774476 | 0.178 | 0.519 | 1 | 0.066409873 | 8.70024E-05 |
| Sgsm2 | 0.566689045 | 0.855610091 | 0.429 | 0.493 | 1 | 4.37707E-05 | 1.774911453 | 0.667 | 0.581 | 1 | 0.566689045 | 8.75395E-05 |
| Osgin2 | 0.430587942 | 1.793611538 | 0.286 | 0.463 | 1 | 4.37763E-05 | -0.410586521 | 0.111 | 0.484 | 1 | 0.430587942 | 8.75506E-05 |
| Irak1 | 0.103073303 | -0.527862093 | 0.238 | 0.463 | 1 | 4.37866E-05 | -1.206744496 | 0.178 | 0.543 | 1 | 0.103073303 | 8.75712E-05 |
| Mex3d | 0.028330767 | -2.184209227 | 0 | 0.19 | 1 | 4.41319E-05 | -2.008089318 | 0.044 | 0.36 | 1 | 0.028330767 | 8.82618E-05 |
| Bmyc | 0.030176208 | -1.586755783 | 0.095 | 0.317 | 1 | 4.41497E-05 | -3.960783604 | 0 | 0.284 | 1 | 0.030176208 | 8.82975E-05 |
| Dpm3 | 4.42895E-05 | -1.915478091 | 0.19 | 0.69 | 1 | 0.001113206 | -0.711206353 | 0.444 | 0.689 | 1 | 0.001113206 | 8.85771E-05 |
| Mmadhc | 0.039588533 | -0.601506234 | 0.238 | 0.526 | 1 | 4.43354E-05 | -0.805674599 | 0.267 | 0.644 | 1 | 0.039588533 | 8.86688E-05 |
| Col5a2 | 0.052202758 | -3.44776175 | 0.095 | 0.28 | 1 | 4.44125E-05 | -2.008805118 | 0.222 | 0.574 | 1 | 0.052202758 | 8.88231E-05 |
| Cdk4 | 9.30233E-05 | -2.16543202 | 0.19 | 0.668 | 1 | 4.44502E-05 | -0.947902312 | 0.533 | 0.727 | 1 | 9.30233E-05 | 8.88985E-05 |
| Hapln1 | 0.814540979 | 0.946035761 | 0.952 | 0.966 | 1 | 4.45769E-05 | -0.450917523 | 0.756 | 0.955 | 1 | 0.814540979 | 8.91519E-05 |

| | | | | | | | | | | | | |
|---|---|---|---|---|---|---|---|---|---|---|---|---|
| Gnpnat1 | 0.024368729 | -1.124970024 | 0.143 | 0.418 | 1 | 4.47047E-05 | -2.298780067 | 0.067 | 0.384 | 1 | 0.024368729 | 8.94074E-05 |
| Spg20 | 0.003187136 | -1.849790188 | 0.095 | 0.44 | 1 | 4.4782E-05 | -1.564854927 | 0.089 | 0.426 | 1 | 0.003187136 | 8.9562E-05 |
| Epyc | 0.000187337 | -2.033887457 | 0.333 | 0.743 | 1 | 4.48557E-05 | -1.696305781 | 0.289 | 0.619 | 1 | 0.000187337 | 8.97094E-05 |
| Ndufaf8 | 0.000204411 | -3.611630451 | 0 | 0.422 | 1 | 4.50703E-05 | -0.951133434 | 0.067 | 0.388 | 1 | 0.000204411 | 9.01386E-05 |
| Btg3 | 0.000426971 | -1.033587364 | 0.333 | 0.825 | 1 | 4.52388E-05 | -0.945744204 | 0.644 | 0.82 | 1 | 0.000426971 | 9.04755E-05 |
| Trappc2l | 4.53002E-05 | -2.292012517 | 0.095 | 0.597 | 1 | 0.000118178 | -1.510699157 | 0.133 | 0.45 | 1 | 0.000118178 | 9.05983E-05 |
| Lsm1 | 0.240894313 | 0.852272663 | 0.238 | 0.463 | 1 | 4.55304E-05 | -1.466674533 | 0.2 | 0.54 | 1 | 0.240894313 | 9.10587E-05 |
| Spag7 | 0.021134715 | -0.977279923 | 0.19 | 0.493 | 1 | 4.55981E-05 | -2.191396532 | 0.089 | 0.408 | 1 | 0.021134715 | 9.11941E-05 |
| Fbxo9 | 0.010877264 | -0.669636055 | 0.19 | 0.526 | 1 | 4.57075E-05 | -1.223818009 | 0.156 | 0.54 | 1 | 0.010877264 | 9.1413E-05 |
| Srxn1 | 0.028498479 | -0.596757175 | 0.143 | 0.448 | 1 | 4.59977E-05 | -2.062066939 | 0.089 | 0.426 | 1 | 0.028498479 | 9.19933E-05 |
| Snrpd3 | 4.65888E-05 | -1.606336152 | 0.381 | 0.806 | 1 | 0.000710186 | -0.649502753 | 0.822 | 0.837 | 1 | 0.000710186 | 9.31754E-05 |
| Abhd12 | 4.70675E-05 | -3.184209227 | 0.048 | 0.526 | 1 | 0.000145458 | -1.142353909 | 0.2 | 0.522 | 1 | 0.000145458 | 9.41328E-05 |
| Rpl7l1 | 0.0088663 | -0.715555612 | 0.381 | 0.784 | 1 | 4.72389E-05 | -0.868247071 | 0.511 | 0.789 | 1 | 0.0088663 | 9.44756E-05 |
| Pik3r1 | 0.006163241 | -1.0757625 | 0.81 | 0.903 | 1 | 4.73072E-05 | -0.56772825 | 0.889 | 0.972 | 1 | 0.006163241 | 9.46123E-05 |
| Poglut3 | 0.006165974 | -1.655351829 | 0.19 | 0.481 | 1 | 4.73294E-05 | -1.022184148 | 0.2 | 0.557 | 1 | 0.006165974 | 9.46566E-05 |
| Fhl1 | 0.090738163 | -1.164171474 | 0.095 | 0.276 | 1 | 4.74491E-05 | -1.971237961 | 0.089 | 0.419 | 1 | 0.090738163 | 9.48959E-05 |
| Tipin | 0.018588563 | -1.527862093 | 0.143 | 0.407 | 1 | 4.7713E-05 | -2.132844349 | 0.067 | 0.384 | 1 | 0.018588563 | 9.54238E-05 |
| Bloc1s4 | 0.002313572 | -2.326228232 | 0.048 | 0.381 | 1 | 4.79371E-05 | -2.115669206 | 0.111 | 0.422 | 1 | 0.002313572 | 9.58719E-05 |
| Myo1d | 0.000232192 | 1.341534573 | 1 | 0.978 | 1 | 4.82052E-05 | 0.782930326 | 1 | 0.986 | 1 | 0.000232192 | 9.6408E-05 |
| Noct | 0.099679221 | 1.183668453 | 0.524 | 0.489 | 1 | 4.82204E-05 | 0.967375609 | 0.844 | 0.564 | 1 | 0.099679221 | 9.64385E-05 |
| B3galnt2 | 0.270913622 | -0.381510668 | 0.19 | 0.317 | 1 | 4.84108E-05 | -1.938979233 | 0.067 | 0.401 | 1 | 0.270913622 | 9.68192E-05 |
| Mdh2 | 4.86017E-05 | -1.171172738 | 0.667 | 0.892 | 1 | 0.021081742 | -0.3672708 | 0.889 | 0.855 | 1 | 0.021081742 | 9.7201E-05 |
| Cdc34 | 0.045409494 | -0.785659851 | 0.095 | 0.317 | 1 | 4.87057E-05 | -2.071814916 | 0.133 | 0.443 | 1 | 0.045409494 | 9.7409E-05 |
| Eif3e | 0.001652775 | -0.831020384 | 0.762 | 0.925 | 1 | 4.8752E-05 | -0.516296193 | 0.889 | 0.934 | 1 | 0.001652775 | 9.75015E-05 |
| Socs6 | 0.020362963 | -0.849790188 | 0.143 | 0.44 | 1 | 4.90081E-05 | -0.709244837 | 0.133 | 0.509 | 1 | 0.020362963 | 9.80138E-05 |
| Trmt112 | 4.90737E-05 | -1.590335673 | 0.429 | 0.799 | 1 | 9.67408E-05 | -0.730003397 | 0.644 | 0.782 | 1 | 9.67408E-05 | 9.81449E-05 |
| Uxs1 | 4.91066E-05 | -1.370622352 | 0.19 | 0.743 | 1 | 0.005715989 | -0.131757456 | 0.289 | 0.619 | 1 | 0.005715989 | 9.82108E-05 |
| H2az1 | 0.002973624 | -1.334790246 | 0.81 | 0.866 | 1 | 4.95497E-05 | -0.946037018 | 0.933 | 0.931 | 1 | 0.002973624 | 9.9097E-05 |
| Tmem39a | 0.635529684 | 0.335902129 | 0.619 | 0.698 | 1 | 4.96903E-05 | -0.666336245 | 0.333 | 0.754 | 1 | 0.635529684 | 9.93781E-05 |
| Pcna | 5.01165E-05 | -2.512755201 | 0.19 | 0.649 | 1 | 0.01496332 | -0.404390255 | 0.711 | 0.761 | 1 | 0.01496332 | 0.00010023 |
| Fgfr2 | 5.02283E-05 | 0.900528366 | 0.952 | 0.959 | 1 | 0.000589682 | 0.762507054 | 1 | 0.997 | 1 | 0.000589682 | 0.000100454 |
| Akt1 | 5.02494E-05 | -1.949325862 | 0.19 | 0.672 | 1 | 0.031955248 | -0.319657207 | 0.711 | 0.827 | 1 | 0.031955248 | 0.000100496 |
| Prpf19 | 0.001192074 | -1.940938076 | 0.095 | 0.481 | 1 | 5.03119E-05 | -1.589945908 | 0.289 | 0.574 | 1 | 0.001192074 | 0.000100621 |
| Fars2 | 0.538719067 | 0.289107917 | 0.333 | 0.459 | 1 | 5.03605E-05 | 1.381105233 | 0.778 | 0.471 | 1 | 0.538719067 | 0.000100718 |
| Polr2j | 0.000497354 | -3.574155746 | 0 | 0.388 | 1 | 5.04999E-05 | -2.89183625 | 0.044 | 0.343 | 1 | 0.000497354 | 0.000100997 |
| Thoc2 | 5.06487E-05 | 0.882358389 | 1 | 0.925 | 1 | 0.020573582 | 0.621267302 | 0.978 | 0.938 | 1 | 0.020573582 | 0.000101295 |
| Gcsh | 0.000940123 | -2.602352638 | 0.048 | 0.418 | 1 | 5.08114E-05 | -2.911874003 | 0.044 | 0.343 | 1 | 0.000940123 | 0.00010162 |
| Ascc3 | 0.226830285 | 0.566243303 | 0.905 | 0.828 | 1 | 5.08595E-05 | 1.135762627 | 0.978 | 0.924 | 1 | 0.226830285 | 0.000101716 |
| Tmem14c | 5.10126E-05 | -1.548620654 | 0.381 | 0.787 | 1 | 0.014729997 | -0.669716472 | 0.622 | 0.692 | 1 | 0.014729997 | 0.000102023 |
| Rbm8a | 0.000720173 | -1.263904953 | 0.429 | 0.784 | 1 | 5.11954E-05 | -0.825941061 | 0.778 | 0.862 | 1 | 0.000720173 | 0.000102388 |
| Pla2g12a | 0.001785904 | -3.25696557 | 0 | 0.332 | 1 | 5.13341E-05 | -1.738391182 | 0.067 | 0.384 | 1 | 0.001785904 | 0.000102665 |
| Ddt | 0.000292561 | -1.946814643 | 0.095 | 0.56 | 1 | 5.16966E-05 | -1.768138526 | 0.133 | 0.457 | 1 | 0.000292561 | 0.00010339 |
| Nol7 | 5.23752E-05 | -1.952667369 | 0.381 | 0.743 | 1 | 0.008784273 | -0.745531042 | 0.756 | 0.74 | 1 | 0.008784273 | 0.000104748 |
| Pum1 | 0.002158521 | 0.719900938 | 0.952 | 0.955 | 1 | 5.24827E-05 | 0.580618521 | 1 | 0.986 | 1 | 0.002158521 | 0.000104963 |
| Csde1 | 0.001303036 | -0.637677809 | 0.905 | 0.981 | 1 | 5.29737E-05 | -0.419025601 | 1 | 0.997 | 1 | 0.001303036 | 0.000105945 |
| Polr3k | 0.042493745 | -1.280424543 | 0.048 | 0.254 | 1 | 5.30262E-05 | -4.085111739 | 0 | 0.28 | 1 | 0.042493745 | 0.00010605 |
| Tmx4 | 0.003809164 | -1.203427143 | 0.238 | 0.664 | 1 | 5.30599E-05 | -1.501802757 | 0.422 | 0.682 | 1 | 0.003809164 | 0.000106117 |
| Ahcyl2 | 0.00555097 | 1.15610238 | 0.857 | 0.631 | 1 | 5.34387E-05 | 1.1586397 | 0.956 | 0.851 | 1 | 0.00555097 | 0.000106875 |
| Necap2 | 5.38479E-05 | -2.184209227 | 0.048 | 0.56 | 1 | 0.067690812 | -0.448633093 | 0.467 | 0.616 | 1 | 0.067690812 | 0.000107693 |
| Btbd9 | 0.005827717 | 1.616701582 | 0.905 | 0.888 | 1 | 5.40183E-05 | 1.12679284 | 0.978 | 0.952 | 1 | 0.005827717 | 0.000108034 |
| Pam | 5.40266E-05 | -0.942028137 | 1 | 0.996 | 1 | 0.037153481 | -0.314237132 | 1 | 1 | 1 | 0.037153481 | 0.00010805 |
| Fbxo2 | 0.001542512 | -4.016226204 | 0.048 | 0.396 | 1 | 5.42855E-05 | -4.404390255 | 0 | 0.28 | 1 | 0.001542512 | 0.000108568 |
| Tcf12 | 0.034626782 | 0.592608915 | 0.952 | 0.959 | 1 | 5.46096E-05 | 0.671379772 | 1 | 0.997 | 1 | 0.034626782 | 0.000109216 |
| Alg2 | 0.003856445 | -2.337455488 | 0.048 | 0.358 | 1 | 5.4611E-05 | -1.564854927 | 0.111 | 0.443 | 1 | 0.003856445 | 0.000109219 |
| Aprt | 5.48784E-05 | -3.403043829 | 0.048 | 0.522 | 1 | 0.041120195 | -0.486852415 | 0.444 | 0.578 | 1 | 0.041120195 | 0.000109754 |
| Pccb | 0.095749049 | -0.473069621 | 0.143 | 0.366 | 1 | 5.5097E-05 | -2.071814916 | 0.111 | 0.426 | 1 | 0.095749049 | 0.000110191 |
| Rnf5 | 0.010124084 | -2.081115734 | 0.048 | 0.313 | 1 | 5.5157E-05 | -3.223818009 | 0.022 | 0.311 | 1 | 0.010124084 | 0.000110311 |
| Adipor1 | 0.00014543 | -0.914050387 | 0.762 | 0.959 | 1 | 5.52186E-05 | -0.541050723 | 0.978 | 0.962 | 1 | 0.00014543 | 0.000110434 |
| Pcdh17 | 0.725529959 | 0.392815454 | 0.286 | 0.272 | 1 | 5.52953E-05 | -0.161276543 | 0.533 | 0.201 | 1 | 0.725529959 | 0.000110588 |
| Mogs | 0.051184345 | -1.04047375 | 0.095 | 0.306 | 1 | 5.53565E-05 | -3.211745177 | 0.022 | 0.311 | 1 | 0.051184345 | 0.00011071 |
| Twist1 | 0.081277522 | -1.793833782 | 0.048 | 0.209 | 1 | 5.55688E-05 | -3.149817428 | 0.044 | 0.353 | 1 | 0.081277522 | 0.000111134 |
| Hgsnat | 0.009515438 | -1.272647192 | 0.19 | 0.493 | 1 | 5.58129E-05 | -1.638855509 | 0.156 | 0.491 | 1 | 0.009515438 | 0.000111623 |
| Pop5 | 0.007484696 | -1.818081329 | 0.048 | 0.347 | 1 | 5.5865E-05 | -2.115669206 | 0.089 | 0.405 | 1 | 0.007484696 | 0.000111727 |
| Acss1 | 0.532436372 | 1.088809267 | 0 | 0.019 | 1 | 5.58723E-05 | 1.91753784 | 0.2 | 0.042 | 1 | 0.532436372 | 0.000111741 |
| Ppp1r12b | 0.000137291 | 1.628507034 | 0.857 | 0.769 | 1 | 5.58965E-05 | 1.274033828 | 0.956 | 0.799 | 1 | 0.000137291 | 0.00011179 |
| Slc38a10 | 5.65885E-05 | -1.547331958 | 0.143 | 0.694 | 1 | 0.005631486 | -0.362442073 | 0.311 | 0.619 | 1 | 0.005631486 | 0.000113174 |
| Cr1l | 0.134673131 | -0.266849685 | 0.524 | 0.787 | 1 | 5.66805E-05 | -0.883627708 | 0.667 | 0.813 | 1 | 0.134673131 | 0.000113358 |
| Septin9 | 0.046730371 | 1.499453086 | 0.857 | 0.851 | 1 | 5.6809E-05 | 1.201945896 | 0.978 | 0.844 | 1 | 0.046730371 | 0.000113615 |
| Arsi | 0.024830378 | -2.054148687 | 0.095 | 0.332 | 1 | 5.7116E-05 | -1.448955808 | 0.222 | 0.599 | 1 | 0.024830378 | 0.000114229 |
| Sri | 5.71728E-05 | -1.525900577 | 0.333 | 0.772 | 1 | 0.028989136 | -0.338622485 | 0.8 | 0.789 | 1 | 0.028989136 | 0.000114342 |
| Coa3 | 5.73158E-05 | -3.90365706 | 0 | 0.47 | 1 | 0.000210134 | -1.4734319 | 0.244 | 0.516 | 1 | 0.000210134 | 0.000114628 |
| Casp8ap2 | 0.514030655 | 0.162809848 | 0.238 | 0.343 | 1 | 5.74918E-05 | -1.907344345 | 0.133 | 0.46 | 1 | 0.514030655 | 0.00011498 |
| Hint2 | 5.75291E-05 | -2.948280052 | 0.048 | 0.519 | 1 | 0.001957116 | -1.542995493 | 0.133 | 0.36 | 1 | 0.001957116 | 0.000115055 |
| Plod1 | 5.84977E-05 | -1.595688907 | 0.095 | 0.604 | 1 | 0.000462083 | -0.81335324 | 0.267 | 0.609 | 1 | 0.000462083 | 0.000116992 |
| Cdr2 | 0.086694145 | -1.133583154 | 0.19 | 0.369 | 1 | 5.85741E-05 | -1.048731303 | 0.2 | 0.599 | 1 | 0.086694145 | 0.000117145 |
| Zfp407 | 5.89758E-05 | 1.659124992 | 0.857 | 0.604 | 1 | 0.014481321 | 1.020107573 | 0.911 | 0.734 | 1 | 0.014481321 | 0.000117948 |
| Ttc4 | 0.035520827 | -0.926141075 | 0.048 | 0.269 | 1 | 5.91194E-05 | -2.435868487 | 0.044 | 0.349 | 1 | 0.035520827 | 0.000118235 |
| Slc26a2 | 5.91228E-05 | -1.756815672 | 0.524 | 0.843 | 1 | 0.000167284 | -0.950487788 | 0.622 | 0.827 | 1 | 0.000167284 | 0.000118242 |
| Cdc37 | 5.91643E-05 | -1.385843089 | 0.476 | 0.854 | 1 | 0.002416945 | -0.433572333 | 0.844 | 0.9 | 1 | 0.002416945 | 0.000118325 |
| Zmat2 | 5.97555E-05 | -2.031284579 | 0.143 | 0.649 | 1 | 0.005454674 | -0.973274548 | 0.444 | 0.595 | 1 | 0.005454674 | 0.000119507 |
| Mrps31 | 0.162057754 | -0.574155746 | 0.143 | 0.302 | 1 | 5.98341E-05 | -2.829996996 | 0.044 | 0.339 | 1 | 0.162057754 | 0.000119665 |
| Mt1 | 0.182972607 | -2.277211412 | 0.714 | 0.638 | 1 | 5.99283E-05 | -1.447539491 | 0.844 | 0.391 | 1 | 0.182972607 | 0.000119853 |
| B4galt7 | 0.001070309 | -1.841928071 | 0.095 | 0.489 | 1 | 6.03964E-05 | -2.316927414 | 0.044 | 0.349 | 1 | 0.001070309 | 0.000120789 |
| Aspn | 0.057467243 | -3.347586815 | 0.143 | 0.332 | 1 | 6.04177E-05 | -2.02527233 | 0.289 | 0.63 | 1 | 0.057467243 | 0.000120832 |
| Cebpb | 0.00058687 | -1.991381911 | 0.857 | 0.851 | 1 | 6.06116E-05 | -1.251578703 | 0.933 | 0.917 | 1 | 0.00058687 | 0.00012122 |
| Slc35g1 | 6.08006E-05 | -3.593014773 | 0.048 | 0.519 | 1 | 0.001844946 | -3.162417465 | 0.022 | 0.218 | 1 | 0.001844946 | 0.000121598 |
| Irx3 | 0.002855382 | -2.370622352 | 0.143 | 0.485 | 1 | 6.08783E-05 | -1.346674757 | 0.133 | 0.478 | 1 | 0.002855382 | 0.000121753 |
| Mapkapk3 | 0.818482987 | 0.425844254 | 0.048 | 0.06 | 1 | 6.10318E-05 | 1.55062229 | 0.422 | 0.159 | 1 | 0.818482987 | 0.00012206 |
| Card19 | 6.11194E-05 | -2.252227651 | 0.095 | 0.608 | 1 | 0.000192217 | -0.904749569 | 0.422 | 0.702 | 1 | 0.000192217 | 0.000122235 |
| Kctd10 | 0.109115985 | -0.718545655 | 0.143 | 0.328 | 1 | 6.11249E-05 | -1.098287127 | 0.178 | 0.557 | 1 | 0.109115985 | 0.000122246 |
| Lemd2 | 0.006538162 | -1.602352638 | 0.095 | 0.41 | 1 | 6.12188E-05 | -1.034784185 | 0.178 | 0.54 | 1 | 0.006538162 | 0.000122434 |
| Lrp1 | 0.007420781 | -0.408316082 | 0.857 | 0.996 | 1 | 6.12502E-05 | -0.514752331 | 0.911 | 1 | 1 | 0.007420781 | 0.000122497 |
| C1s1 | 0.08049829 | -1.249060372 | 0.048 | 0.213 | 1 | 6.12553E-05 | -1.130331381 | 0.111 | 0.46 | 1 | 0.08049829 | 0.000122507 |
| Rap2a | 0.101830605 | -0.65714511 | 0.143 | 0.351 | 1 | 6.15726E-05 | -2.421264074 | 0.044 | 0.356 | 1 | 0.101830605 | 0.000123141 |
| Rbmx | 0.783842348 | 0.866416846 | 0.429 | 0.481 | 1 | 6.2134E-05 | -1.172537504 | 0.178 | 0.55 | 1 | 0.783842348 | 0.000124264 |
| Col6a3 | 0.001692512 | 0.9250228 | 0.952 | 0.951 | 1 | 6.29606E-05 | 1.003322727 | 0.911 | 0.886 | 1 | 0.001692512 | 0.000125917 |
| Nrbp1 | 6.30311E-05 | -1.965956536 | 0.095 | 0.623 | 1 | 0.018529045 | -0.447051407 | 0.622 | 0.754 | 1 | 0.018529045 | 0.000126058 |
| Trib2 | 0.803188988 | 0.326606381 | 0.286 | 0.351 | 1 | 6.32481E-05 | -1.992492463 | 0.067 | 0.388 | 1 | 0.803188988 | 0.000126492 |

| | | | | | | | | | | | | |
|---|---|---|---|---|---|---|---|---|---|---|---|---|
| Tacc2 | 6.35611E-05 | 1.605059018 | 0.952 | 0.914 | 1 | 0.000593392 | 0.745247732 | 1 | 0.948 | 1 | 0.000593392 | 0.000127118 |
| Fgfr3 | 6.36716E-05 | -1.574155746 | 0.381 | 0.813 | 1 | 0.879897712 | 0.261829937 | 0.822 | 0.796 | 1 | 0.879897712 | 0.000127339 |
| Cct5 | 6.36746E-05 | -1.634217136 | 0.381 | 0.769 | 1 | 0.000381586 | -0.557718746 | 0.8 | 0.841 | 1 | 0.000381586 | 0.000127345 |
| Trappc3 | 6.37243E-05 | -1.163330498 | 0.143 | 0.683 | 1 | 6.77123E-05 | -0.652530446 | 0.378 | 0.72 | 1 | 6.77123E-05 | 0.000127444 |
| Aftph | 6.3801E-05 | 1.42865927 | 0.905 | 0.638 | 1 | 0.173352068 | 0.533798094 | 0.889 | 0.848 | 1 | 0.173352068 | 0.000127598 |
| Dguok | 0.258396011 | -0.548620654 | 0.19 | 0.31 | 1 | 6.38639E-05 | -1.674479418 | 0.089 | 0.405 | 1 | 0.258396011 | 0.000127724 |
| Fndc4 | 0.238160714 | -0.484657595 | 0.19 | 0.328 | 1 | 6.38694E-05 | -3.305612101 | 0.022 | 0.308 | 1 | 0.238160714 | 0.000127735 |
| Golga7 | 0.03301695 | -0.718545655 | 0.333 | 0.672 | 1 | 6.43014E-05 | -1.055136175 | 0.356 | 0.675 | 1 | 0.03301695 | 0.000128599 |
| Jup | 0.465428117 | 0.1445187 | 0.381 | 0.284 | 1 | 6.43841E-05 | 0.683072586 | 0.711 | 0.322 | 1 | 0.465428117 | 0.000128764 |
| Dusp10 | 0.507751878 | 0.804355877 | 0.143 | 0.22 | 1 | 6.46126E-05 | 1.098110085 | 0.4 | 0.142 | 1 | 0.507751878 | 0.000129221 |
| Hook1 | 0.632076134 | 1.673771768 | 0 | 0.011 | 1 | 6.50957E-05 | 1.750186782 | 0.222 | 0.052 | 1 | 0.632076134 | 0.000130187 |
| Clip3 | 0.063795847 | -1.233118828 | 0.048 | 0.228 | 1 | 6.51997E-05 | -1.412084647 | 0.067 | 0.391 | 1 | 0.063795847 | 0.000130395 |
| Pigs | 0.000106298 | -3.555046923 | 0 | 0.444 | 1 | 6.52597E-05 | -2.174908409 | 0.044 | 0.349 | 1 | 0.000106298 | 0.000130515 |
| Chmp5 | 6.54438E-05 | -1.611630451 | 0.286 | 0.72 | 1 | 0.05593335 | -0.50103135 | 0.711 | 0.744 | 1 | 0.05593335 | 0.000130883 |
| Keap1 | 0.065242745 | -0.979670471 | 0.143 | 0.358 | 1 | 6.55058E-05 | -1.65082815 | 0.156 | 0.484 | 1 | 0.065242745 | 0.000131007 |
| Tomm22 | 6.55862E-05 | -1.590931459 | 0.238 | 0.724 | 1 | 0.079493435 | -0.391695182 | 0.689 | 0.709 | 1 | 0.079493435 | 0.000131168 |
| Erg28 | 0.077468268 | -0.501866885 | 0.19 | 0.451 | 1 | 6.58624E-05 | -1.783627033 | 0.067 | 0.398 | 1 | 0.077468268 | 0.00013172 |
| Syvn1 | 0.330774752 | -0.171718283 | 0.19 | 0.31 | 1 | 6.59092E-05 | -1.849422495 | 0.089 | 0.412 | 1 | 0.330774752 | 0.000131814 |
| Mrtfa | 0.188266038 | 1.159198595 | 0.571 | 0.541 | 1 | 6.59762E-05 | 1.334177444 | 0.889 | 0.664 | 1 | 0.188266038 | 0.000131948 |
| Pink1 | 0.002045432 | -1.586755783 | 0.143 | 0.526 | 1 | 6.62614E-05 | -1.124282336 | 0.178 | 0.557 | 1 | 0.002045432 | 0.000132518 |
| Snapc5 | 0.020406007 | -0.839297815 | 0.095 | 0.384 | 1 | 6.65636E-05 | -2.223818009 | 0.089 | 0.391 | 1 | 0.020406007 | 0.000133123 |
| Rab14 | 0.046850916 | -0.415128737 | 0.857 | 0.966 | 1 | 6.70149E-05 | -0.55291278 | 0.956 | 0.976 | 1 | 0.046850916 | 0.000134025 |
| Zmat4 | 6.75064E-05 | 5.48112669 | 0.095 | 0.004 | 1 | 0.701092235 | 0.361144491 | 0.067 | 0.052 | 1 | 0.701092235 | 0.000135008 |
| Papss1 | 0.000153598 | -2.111330348 | 0.19 | 0.642 | 1 | 6.7745E-05 | -1.135687099 | 0.756 | 0.792 | 1 | 0.000153598 | 0.000135485 |
| Pdha1 | 0.187688391 | 0.113056813 | 0.381 | 0.631 | 1 | 6.77749E-05 | -0.826338882 | 0.733 | 0.796 | 1 | 0.187688391 | 0.000135545 |
| Prorsd1 | 0.002894957 | -1.962852853 | 0.095 | 0.444 | 1 | 6.78688E-05 | -2.271123724 | 0.067 | 0.374 | 1 | 0.002894957 | 0.000135733 |
| Dlg2 | 6.78835E-05 | 1.692450321 | 0.905 | 0.761 | 1 | 0.429717155 | 0.457607331 | 0.378 | 0.509 | 1 | 0.429717155 | 0.000135762 |
| Taf13 | 0.007617344 | -0.724507213 | 0.333 | 0.731 | 1 | 6.79758E-05 | -1.114275163 | 0.533 | 0.713 | 1 | 0.007617344 | 0.000135947 |
| Dock9 | 0.375113159 | -0.379339569 | 0.19 | 0.295 | 1 | 6.80273E-05 | 1.331998146 | 0.689 | 0.363 | 1 | 0.375113159 | 0.00013605 |
| Coq7 | 0.074492341 | -0.56326743 | 0.143 | 0.362 | 1 | 6.81164E-05 | -2.819427754 | 0.044 | 0.336 | 1 | 0.074492341 | 0.000136228 |
| Rwdd4a | 0.003013243 | -1.76085646 | 0.095 | 0.444 | 1 | 6.81239E-05 | -1.238539515 | 0.111 | 0.46 | 1 | 0.003013243 | 0.000136243 |
| Cln5 | 0.161060613 | -0.583616075 | 0.238 | 0.407 | 1 | 6.8278E-05 | -1.617051138 | 0.111 | 0.446 | 1 | 0.161060613 | 0.000136551 |
| Tmem175 | 0.529581574 | 1.049280903 | 0.238 | 0.343 | 1 | 6.84137E-05 | -1.346674757 | 0.044 | 0.349 | 1 | 0.529581574 | 0.000136823 |
| Rnf111 | 0.41345426 | 0.404311093 | 0.619 | 0.679 | 1 | 6.88381E-05 | 0.831687081 | 0.978 | 0.82 | 1 | 0.41345426 | 0.000137671 |
| Wdr74 | 0.024818629 | -1.359651234 | 0.095 | 0.343 | 1 | 6.90249E-05 | -1.867124496 | 0.111 | 0.426 | 1 | 0.024818629 | 0.000138045 |
| Enpp5 | 0.000938646 | -3.516052791 | 0 | 0.362 | 1 | 6.93482E-05 | -2.798054104 | 0.044 | 0.336 | 1 | 0.000938646 | 0.000138692 |
| Gm94 | 6.97312E-05 | 4.673771768 | 0.095 | 0.004 | 1 | 0.430487676 | 0.361144491 | 0 | 0.014 | 1 | 0.430487676 | 0.000139458 |
| Prelid3b | 0.004180713 | -1.735619168 | 0.095 | 0.429 | 1 | 7.02933E-05 | -0.549588171 | 0.111 | 0.471 | 1 | 0.004180713 | 0.000140582 |
| Nkd1 | 0.01324741 | -1.424260315 | 0.286 | 0.578 | 1 | 7.04669E-05 | -1.124282336 | 0.244 | 0.595 | 1 | 0.01324741 | 0.000140929 |
| Pnn | 0.287869385 | 0.409068541 | 0.905 | 0.896 | 1 | 7.05057E-05 | -0.738021213 | 0.978 | 0.934 | 1 | 0.287869385 | 0.000141006 |
| Mast2 | 0.000195115 | 1.694445336 | 0.857 | 0.78 | 1 | 7.05164E-05 | 1.068156863 | 0.911 | 0.83 | 1 | 0.000195115 | 0.000141028 |
| Rabggtb | 7.09046E-05 | -1.839297815 | 0.286 | 0.72 | 1 | 0.000136458 | -0.942444013 | 0.489 | 0.737 | 1 | 0.000136458 | 0.000141804 |
| 1110059G10Rik | 0.048738758 | -0.376854305 | 0.095 | 0.313 | 1 | 7.11139E-05 | -2.731964913 | 0.022 | 0.315 | 1 | 0.048738758 | 0.000142223 |
| Myd88 | 0.000688566 | -2.475975352 | 0.048 | 0.451 | 1 | 7.1188E-05 | -0.982263331 | 0.111 | 0.457 | 1 | 0.000688566 | 0.000142371 |
| Camta1 | 0.491520217 | 1.722212123 | 0.619 | 0.687 | 1 | 7.15048E-05 | 1.111166238 | 0.911 | 0.623 | 1 | 0.491520217 | 0.000143005 |
| Anapc5 | 7.60835E-05 | -1.174610423 | 0.476 | 0.899 | 1 | 7.17258E-05 | -0.688402304 | 0.756 | 0.913 | 1 | 7.60835E-05 | 0.000143446 |
| Pdk3 | 0.089847089 | -0.674956387 | 0.19 | 0.414 | 1 | 7.2758E-05 | -1.21001221 | 0.111 | 0.45 | 1 | 0.089847089 | 0.000145511 |
| Kpna4 | 0.43538729 | 0.340541024 | 0.857 | 0.884 | 1 | 7.29086E-05 | 0.662010971 | 0.978 | 0.958 | 1 | 0.43538729 | 0.000145812 |
| Gm57402 | 0.044716029 | -2.054148687 | 0 | 0.164 | 1 | 7.33572E-05 | -3.149817428 | 0.022 | 0.304 | 1 | 0.044716029 | 0.000146709 |
| Cln8 | 0.076496208 | -1.475975352 | 0.048 | 0.205 | 1 | 7.34128E-05 | -3.361321533 | 0.022 | 0.304 | 1 | 0.076496208 | 0.00014682 |
| Ecd | 0.609026583 | 0.157196242 | 0.19 | 0.254 | 1 | 7.347E-05 | -3.361321533 | 0.022 | 0.304 | 1 | 0.609026583 | 0.000146935 |
| Rest | 0.001195201 | -1.258783711 | 0.19 | 0.608 | 1 | 7.38595E-05 | -1.104706798 | 0.689 | 0.789 | 1 | 0.001195201 | 0.000147713 |
| Ddrgk1 | 7.46047E-05 | -2.163330498 | 0.238 | 0.653 | 1 | 0.062276851 | -0.42982147 | 0.622 | 0.692 | 1 | 0.062276851 | 0.000149204 |
| Gnptg | 0.000521889 | -1.793833782 | 0.048 | 0.474 | 1 | 7.49165E-05 | -1.776359032 | 0.089 | 0.401 | 1 | 0.000521889 | 0.000149827 |
| Lsm2 | 0.074190253 | -0.981580061 | 0.143 | 0.336 | 1 | 7.50648E-05 | -1.462604869 | 0.089 | 0.422 | 1 | 0.074190253 | 0.000150124 |
| Alkbh5 | 0.001288411 | -1.379796914 | 0.429 | 0.761 | 1 | 7.51159E-05 | -0.688418609 | 0.889 | 0.927 | 1 | 0.001288411 | 0.000150226 |
| Fam50a | 7.58037E-05 | -2.34303652 | 0.048 | 0.549 | 1 | 0.022608694 | -0.207302923 | 0.4 | 0.595 | 1 | 0.022608694 | 0.000151602 |
| Polr2c | 7.60669E-05 | -2.174225139 | 0.143 | 0.616 | 1 | 0.006568408 | -0.755048526 | 0.467 | 0.64 | 1 | 0.006568408 | 0.000152128 |
| Timm44 | 0.018721748 | -0.813494059 | 0.095 | 0.392 | 1 | 7.62139E-05 | -0.57996182 | 0.089 | 0.443 | 1 | 0.018721748 | 0.000152422 |
| Glrx5 | 7.62807E-05 | -2.169207064 | 0.143 | 0.616 | 1 | 0.065132469 | -0.529921137 | 0.556 | 0.63 | 1 | 0.065132469 | 0.000152556 |
| Hacd1 | 0.217109349 | 0.3012176 | 0.19 | 0.34 | 1 | 7.63798E-05 | -1.543996322 | 0.111 | 0.443 | 1 | 0.217109349 | 0.000152754 |
| Naga | 7.64555E-05 | -2.599246727 | 0.048 | 0.537 | 1 | 7.63882E-05 | -1.267539387 | 0.133 | 0.474 | 1 | 7.64555E-05 | 0.00015277 |
| Csgalnact2 | 0.030573372 | -0.72935241 | 0.381 | 0.709 | 1 | 7.68188E-05 | -0.792012297 | 0.178 | 0.567 | 1 | 0.030573372 | 0.000153632 |
| Pgm3 | 0.034084125 | -0.284297552 | 0.333 | 0.683 | 1 | 7.68468E-05 | -1.010824286 | 0.267 | 0.654 | 1 | 0.034084125 | 0.000153688 |
| Hjurp | 0.119865918 | 1.179007076 | 0.429 | 0.325 | 1 | 7.69212E-05 | -1.829996996 | 0.044 | 0.346 | 1 | 0.119865918 | 0.000153837 |
| Tmem159 | 0.200897264 | -0.243766072 | 0.19 | 0.366 | 1 | 7.71285E-05 | -1.504554417 | 0.067 | 0.388 | 1 | 0.200897264 | 0.000154251 |
| Rnps1 | 0.008491328 | -0.853475235 | 0.381 | 0.75 | 1 | 7.75877E-05 | -0.850905986 | 0.689 | 0.813 | 1 | 0.008491328 | 0.000155169 |
| Arsb | 0.270514552 | 0.981894063 | 0.143 | 0.261 | 1 | 7.76157E-05 | -3.223818009 | 0.067 | 0.381 | 1 | 0.270514552 | 0.000155225 |
| Nedd4 | 0.00010844 | -0.733575341 | 1 | 1 | 1 | 7.86514E-05 | -0.45413964 | 1 | 1 | 1 | 0.00010844 | 0.000157297 |
| Snap47 | 7.88772E-05 | -2.834022873 | 0.048 | 0.534 | 1 | 0.003636863 | -0.332524269 | 0.178 | 0.481 | 1 | 0.003636863 | 0.000157748 |
| Trim2 | 0.533335118 | 1.014515485 | 0.429 | 0.414 | 1 | 7.88815E-05 | 1.01701166 | 0.822 | 0.502 | 1 | 0.533335118 | 0.000157757 |
| Atn1 | 0.792003467 | 0.480000024 | 0.333 | 0.459 | 1 | 7.88853E-05 | -0.95892086 | 0.378 | 0.696 | 1 | 0.792003467 | 0.000157764 |
| Ldb1 | 0.873176776 | 0.122871103 | 0.429 | 0.526 | 1 | 7.88942E-05 | -0.790244559 | 0.533 | 0.772 | 1 | 0.873176776 | 0.000157782 |
| Nktr | 7.93218E-05 | 0.645613498 | 1 | 0.996 | 1 | 0.001271325 | 0.51682757 | 1 | 0.997 | 1 | 0.001271325 | 0.000158637 |
| Sf3b6 | 0.000271985 | -1.116305163 | 0.571 | 0.873 | 1 | 7.93916E-05 | -0.543435944 | 0.867 | 0.889 | 1 | 0.000271985 | 0.000158777 |
| Poglut2 | 0.001485572 | -1.548620654 | 0.143 | 0.545 | 1 | 7.94342E-05 | -0.949195629 | 0.156 | 0.536 | 1 | 0.001485572 | 0.000158862 |
| Tfb2m | 0.002316667 | -2.403043829 | 0.048 | 0.381 | 1 | 7.96174E-05 | -1.909384451 | 0.089 | 0.412 | 1 | 0.002316667 | 0.000159228 |
| Rab6a | 7.97242E-05 | -0.953889359 | 0.714 | 0.951 | 1 | 0.038999684 | -0.21578069 | 0.956 | 0.983 | 1 | 0.038999684 | 0.000159442 |
| Il16 | 7.99043E-05 | 2.055284554 | 0.81 | 0.799 | 1 | 0.000429527 | 0.964901365 | 0.822 | 0.862 | 1 | 0.000429527 | 0.000159802 |
| Exoc4 | 0.022038754 | 0.505034658 | 1 | 0.881 | 1 | 7.99866E-05 | 0.975599765 | 0.978 | 0.927 | 1 | 0.022038754 | 0.000159967 |
| Ebf1 | 0.000199145 | 1.096143777 | 0.952 | 0.97 | 1 | 8.00259E-05 | 0.887824267 | 0.978 | 0.965 | 1 | 0.000199145 | 0.000160045 |
| Cnot8 | 0.254899611 | -0.317183093 | 0.286 | 0.422 | 1 | 8.00345E-05 | -1.117827314 | 0.133 | 0.488 | 1 | 0.254899611 | 0.000160063 |
| Dusp3 | 0.047145713 | -1.120644099 | 0.048 | 0.25 | 1 | 8.01187E-05 | -1.299920989 | 0.111 | 0.446 | 1 | 0.047145713 | 0.000160231 |
| Arih1 | 0.000251934 | 1.383450036 | 1 | 0.989 | 1 | 8.07614E-05 | 0.630918572 | 1 | 0.997 | 1 | 0.000251934 | 0.000161516 |
| Nob1 | 0.063553605 | -0.255145134 | 0.19 | 0.474 | 1 | 8.11929E-05 | -0.870181055 | 0.178 | 0.561 | 1 | 0.063553605 | 0.000162379 |
| Fjx1 | 0.292663657 | -1.76085646 | 0.143 | 0.228 | 1 | 8.13621E-05 | -4.350350415 | 0 | 0.27 | 1 | 0.292663657 | 0.000162718 |
| Pphln1 | 0.12575814 | -0.489974659 | 0.333 | 0.549 | 1 | 8.15939E-05 | -0.764739949 | 0.222 | 0.647 | 1 | 0.12575814 | 0.000163181 |
| Rnf138 | 0.010127328 | -0.602352638 | 0.095 | 0.422 | 1 | 8.15985E-05 | -1.050281754 | 0.156 | 0.529 | 1 | 0.010127328 | 0.00016319 |
| Eea1 | 8.27148E-05 | 1.15327286 | 1 | 0.881 | 1 | 0.21268842 | 0.484853838 | 0.978 | 0.927 | 1 | 0.21268842 | 0.000165423 |
| Idh3g | 8.30395E-05 | -2.177977274 | 0.095 | 0.578 | 1 | 0.280452719 | -0.44937771 | 0.467 | 0.505 | 1 | 0.280452719 | 0.000166072 |
| Gprasp2 | 0.0393764 | -2.392317423 | 0 | 0.172 | 1 | 8.32333E-05 | -2.16492432 | 0.022 | 0.315 | 1 | 0.0393764 | 0.00016646 |
| Igfbp5 | 0.671486267 | -1.916191414 | 0.429 | 0.313 | 1 | 8.34151E-05 | -0.567022764 | 0.667 | 0.28 | 1 | 0.671486267 | 0.000166823 |
| Zfp560 | 0.058116787 | -0.637814383 | 0.143 | 0.396 | 1 | 8.39069E-05 | -1.008805118 | 0.067 | 0.401 | 1 | 0.058116787 | 0.000167807 |
| Tbc1d9 | 0.251580301 | -0.496153234 | 0 | 0.06 | 1 | 8.39278E-05 | 1.808603468 | 0.356 | 0.125 | 1 | 0.251580301 | 0.000167849 |
| Tsen34 | 8.39346E-05 | -1.197414766 | 0.19 | 0.743 | 1 | 0.000178982 | -0.765828365 | 0.556 | 0.782 | 1 | 0.000178982 | 0.000167862 |
| Arl6ip4 | 0.012075377 | -0.911190733 | 0.095 | 0.41 | 1 | 8.43185E-05 | -1.916840256 | 0.133 | 0.439 | 1 | 0.012075377 | 0.00016863 |
| Surf1 | 0.033172621 | -0.950719097 | 0.143 | 0.403 | 1 | 8.43659E-05 | -1.476798751 | 0.067 | 0.394 | 1 | 0.033172621 | 0.000168725 |
| Borcs8 | 0.016852948 | -1.611630451 | 0.095 | 0.347 | 1 | 8.43822E-05 | -3.11134328 | 0.022 | 0.301 | 1 | 0.016852948 | 0.000168757 |

| | | | | | | | | | | | | |
|---|---|---|---|---|---|---|---|---|---|---|---|---|
| Sbds | 8.44433E-05 | -1.607926482 | 0.571 | 0.84 | 1 | 0.000325717 | -0.899741448 | 0.822 | 0.83 | 1 | 0.000325717 | 0.000168879 |
| Mzt2 | 0.011110711 | -1.970084422 | 0.048 | 0.325 | 1 | 8.45575E-05 | -3.223818009 | 0.022 | 0.301 | 1 | 0.011110711 | 0.000169108 |
| Tpp1 | 0.000412287 | -1.611630451 | 0.143 | 0.601 | 1 | 8.53791E-05 | -1.030623229 | 0.244 | 0.602 | 1 | 0.000412287 | 0.000170751 |
| Atp5mpl | 8.54943E-05 | -1.581728965 | 0.333 | 0.75 | 1 | 0.777004375 | -0.182742066 | 0.711 | 0.571 | 1 | 0.777004375 | 0.000170981 |
| Ikzf5 | 0.408042108 | 0.514573173 | 0.19 | 0.328 | 1 | 8.63019E-05 | -1.04275245 | 0.089 | 0.398 | 1 | 0.408042108 | 0.000172596 |
| Rce1 | 0.012288683 | -1.337455488 | 0.095 | 0.384 | 1 | 8.6876E-05 | -1.375821103 | 0.089 | 0.412 | 1 | 0.012288683 | 0.000173744 |
| Tmem276 | 0.005284202 | -2.880817084 | 0 | 0.28 | 1 | 8.72116E-05 | -4.031172931 | 0 | 0.266 | 1 | 0.005284202 | 0.000174416 |
| Uap1l1 | 0.014285107 | -1.729950418 | 0.048 | 0.313 | 1 | 8.7433E-05 | -1.856086225 | 0.089 | 0.405 | 1 | 0.014285107 | 0.000174858 |
| Sh3pxd2a | 0.000699887 | 0.909516952 | 0.952 | 0.903 | 1 | 8.76005E-05 | 1.389306099 | 0.956 | 0.879 | 1 | 0.000699887 | 0.000175193 |
| Naa60 | 0.582564376 | 0.580662363 | 0.238 | 0.354 | 1 | 8.79762E-05 | -2.166593141 | 0.089 | 0.391 | 1 | 0.582564376 | 0.000175945 |
| Ciao2a | 0.000306811 | -3.516052791 | 0 | 0.407 | 1 | 8.83073E-05 | -1.776359032 | 0.156 | 0.464 | 1 | 0.000306811 | 0.000176607 |
| Tpbg | 0.001470365 | -4.033587364 | 0 | 0.343 | 1 | 8.87739E-05 | -2.617051138 | 0.067 | 0.363 | 1 | 0.001470365 | 0.00017754 |
| Abr | 0.957086794 | 0.61076197 | 0.333 | 0.407 | 1 | 8.87818E-05 | 1.144858345 | 0.8 | 0.533 | 1 | 0.957086794 | 0.000177556 |
| Abhd16a | 0.000129456 | -3.683780237 | 0 | 0.44 | 1 | 8.88119E-05 | -1.27534831 | 0.089 | 0.422 | 1 | 0.000129456 | 0.000177616 |
| Psmg2 | 0.001437372 | -2.208871282 | 0.095 | 0.451 | 1 | 8.90267E-05 | -2.425451871 | 0.044 | 0.339 | 1 | 0.001437372 | 0.000178046 |
| Tmcc1 | 0.022295802 | 0.80310136 | 0.905 | 0.769 | 1 | 8.90529E-05 | 0.950715812 | 1 | 0.927 | 1 | 0.022295802 | 0.000178098 |
| Mak16 | 0.019011112 | -1.060937853 | 0.19 | 0.478 | 1 | 8.92715E-05 | -1.258824459 | 0.222 | 0.557 | 1 | 0.019011112 | 0.000178535 |
| Cmbl | 0.003362101 | -3.146407195 | 0 | 0.302 | 1 | 8.94899E-05 | -1.938979233 | 0.067 | 0.384 | 1 | 0.003362101 | 0.000178972 |
| Arid1b | 0.000235501 | 1.524106916 | 0.905 | 0.918 | 1 | 8.95803E-05 | 1.018434707 | 1 | 0.969 | 1 | 0.000235501 | 0.000179152 |
| Pdlim5 | 0.032848635 | 0.840881754 | 1 | 0.888 | 1 | 9.01387E-05 | 1.110765386 | 0.956 | 0.865 | 1 | 0.032848635 | 0.000180269 |
| Slc7a6 | 0.184415198 | 0.728913322 | 0.429 | 0.34 | 1 | 9.07386E-05 | -2.901889915 | 0.022 | 0.308 | 1 | 0.184415198 | 0.000181469 |
| Leprotl1 | 9.08545E-05 | -1.975843691 | 0.095 | 0.586 | 1 | 0.016652173 | -0.62887142 | 0.422 | 0.599 | 1 | 0.016652173 | 0.000181701 |
| Rnf126 | 0.060914059 | -1.02666795 | 0.095 | 0.295 | 1 | 9.09224E-05 | -2.207698344 | 0.111 | 0.405 | 1 | 0.060914059 | 0.000181837 |
| Tmem97 | 0.000500613 | -2.365147222 | 0.095 | 0.507 | 1 | 9.11925E-05 | -2.80878051 | 0.022 | 0.308 | 1 | 0.000500613 | 0.000182377 |
| Adcy3 | 0.125503045 | -0.303508156 | 0.048 | 0.19 | 1 | 9.13885E-05 | -2.462604869 | 0.022 | 0.311 | 1 | 0.125503045 | 0.000182769 |
| Atic | 9.16407E-05 | -3.948280052 | 0 | 0.455 | 1 | 0.000973427 | -0.64570918 | 0.156 | 0.453 | 1 | 0.000973427 | 0.000183273 |
| Pbxip1 | 0.006770107 | -0.648156327 | 0.429 | 0.776 | 1 | 9.16637E-05 | -0.83350294 | 0.578 | 0.779 | 1 | 0.006770107 | 0.000183319 |
| Ppm1m | 0.013588332 | -1.20069735 | 0.143 | 0.448 | 1 | 9.18036E-05 | -1.526380779 | 0.067 | 0.381 | 1 | 0.013588332 | 0.000183599 |
| Thap11 | 0.052095308 | -1.555046923 | 0.048 | 0.228 | 1 | 9.21154E-05 | -2.526380779 | 0.067 | 0.356 | 1 | 0.052095308 | 0.000184222 |
| Tmem234 | 9.25613E-05 | -0.980352758 | 0.714 | 0.903 | 1 | 0.000104615 | -0.833273302 | 0.867 | 0.855 | 1 | 0.000104615 | 0.000185114 |
| Arpc3 | 0.000117387 | -0.933993683 | 0.762 | 0.922 | 1 | 9.324E-05 | -0.740735123 | 0.933 | 0.913 | 1 | 0.000117387 | 0.000186471 |
| Tiam2 | 9.37283E-05 | 1.267757684 | 0.952 | 0.966 | 1 | 0.000409443 | 0.90867171 | 1 | 0.938 | 1 | 0.000409443 | 0.000187448 |
| Nr2f2 | 9.39075E-05 | -3.326228232 | 0.048 | 0.526 | 1 | 0.090116455 | -0.870728983 | 0.556 | 0.561 | 1 | 0.090116455 | 0.000187806 |
| Cdk5rap3 | 0.014278018 | -1.424260315 | 0.095 | 0.373 | 1 | 9.44173E-05 | -2.134550671 | 0.067 | 0.367 | 1 | 0.014278018 | 0.000188826 |
| Rapgef2 | 9.44582E-05 | 1.262691295 | 1 | 0.866 | 1 | 0.003643528 | 0.653262198 | 1 | 0.979 | 1 | 0.003643528 | 0.000188907 |
| Mrps12 | 0.000382494 | -2.333722769 | 0.048 | 0.478 | 1 | 9.5375E-05 | -1.598213524 | 0.089 | 0.415 | 1 | 0.000382494 | 0.000190741 |
| Loxl3 | 0.00063057 | -1.290962903 | 0.333 | 0.757 | 1 | 9.56181E-05 | -1.152433376 | 0.244 | 0.633 | 1 | 0.00063057 | 0.000191227 |
| Eloa | 0.000406664 | -1.383678505 | 0.19 | 0.649 | 1 | 9.58638E-05 | -1.086079882 | 0.711 | 0.779 | 1 | 0.000406664 | 0.000191718 |
| Foxa3 | 0.031285171 | -0.107587946 | 0.143 | 0.392 | 1 | 9.60201E-05 | -2.938979233 | 0.044 | 0.332 | 1 | 0.031285171 | 0.000192031 |
| Mrps33 | 9.6622E-05 | -1.840801405 | 0.143 | 0.646 | 1 | 0.019102368 | -0.887950724 | 0.489 | 0.585 | 1 | 0.019102368 | 0.000193235 |
| Psmd2 | 0.001776656 | -0.945579459 | 0.429 | 0.754 | 1 | 9.68979E-05 | -0.680422001 | 0.756 | 0.862 | 1 | 0.001776656 | 0.000193786 |
| mt-Nd3 | 0.291034199 | 0.638776346 | 0.857 | 0.896 | 1 | 9.71859E-05 | 1.048261536 | 0.956 | 0.844 | 1 | 0.291034199 | 0.000194362 |
| Junos | 9.73988E-05 | 2.638147858 | 0.524 | 0.209 | 1 | 0.00746455 | 1.161119883 | 0.289 | 0.135 | 1 | 0.00746455 | 0.000194788 |
| Zbtb1 | 0.09949387 | -1.075166468 | 0.238 | 0.429 | 1 | 9.74332E-05 | -0.904245336 | 0.4 | 0.734 | 1 | 0.09949387 | 0.000194857 |
| Relch | 0.021438004 | 0.995699863 | 0.667 | 0.556 | 1 | 9.78171E-05 | 1.060142235 | 0.911 | 0.64 | 1 | 0.021438004 | 0.000195625 |
| Osbpl9 | 0.002101488 | 1.077850055 | 0.952 | 0.903 | 1 | 9.83042E-05 | 0.788183088 | 0.978 | 0.945 | 1 | 0.002101488 | 0.000196599 |
| Sccpdh | 0.030891537 | -0.984439715 | 0.048 | 0.276 | 1 | 9.8401E-05 | -1.112786697 | 0.044 | 0.353 | 1 | 0.030891537 | 0.000196792 |
| Psmc3 | 9.8418E-05 | -1.517572217 | 0.381 | 0.784 | 1 | 0.000449128 | -0.706873932 | 0.733 | 0.837 | 1 | 0.000449128 | 0.000196826 |
| Lman2 | 9.86077E-05 | -1.440443801 | 0.238 | 0.746 | 1 | 0.046170844 | -0.415991541 | 0.667 | 0.744 | 1 | 0.046170844 | 0.000197206 |
| Zfp91 | 0.002974477 | -1.006552589 | 0.667 | 0.806 | 1 | 9.89537E-05 | -0.648919192 | 0.911 | 0.913 | 1 | 0.002974477 | 0.000197898 |
| Sh3d19 | 0.013624642 | 0.842413803 | 0.952 | 0.97 | 1 | 9.89765E-05 | 1.088902587 | 0.978 | 0.962 | 1 | 0.013624642 | 0.000197943 |
| Nifk | 0.229223015 | -0.107587946 | 0.286 | 0.496 | 1 | 9.90774E-05 | -1.653210802 | 0.111 | 0.436 | 1 | 0.229223015 | 0.000198145 |
| Cd276 | 0.351308089 | -0.376854305 | 0.143 | 0.243 | 1 | 9.95202E-05 | -1.513324627 | 0.089 | 0.415 | 1 | 0.351308089 | 0.000199031 |
| Polr1g | 0.113002474 | -0.772484462 | 0.095 | 0.269 | 1 | 9.99112E-05 | -3.115669206 | 0.044 | 0.325 | 1 | 0.113002474 | 0.000199812 |
| Mrpl36 | 0.002837934 | -1.807354922 | 0.048 | 0.396 | 1 | 0.000100545 | -1.366170933 | 0.044 | 0.353 | 1 | 0.002837934 | 0.000201079 |
| Bub3 | 0.123754263 | -0.501486678 | 0.286 | 0.522 | 1 | 0.000100712 | -0.787557239 | 0.333 | 0.702 | 1 | 0.123754263 | 0.000201413 |
| Gpr155 | 0.647471572 | 0.55829455 | 0.048 | 0.078 | 1 | 0.000100835 | 1.471568481 | 0.333 | 0.111 | 1 | 0.647471572 | 0.000201661 |
| Ctdnep1 | 0.002169245 | -1.222392421 | 0.19 | 0.604 | 1 | 0.000101499 | -1.147869156 | 0.467 | 0.696 | 1 | 0.002169245 | 0.000202987 |
| Hsdl2 | 0.006696988 | -2.984439715 | 0 | 0.269 | 1 | 0.000101658 | -2.871516266 | 0.044 | 0.325 | 1 | 0.006696988 | 0.000203305 |
| Fdx1 | 0.000101785 | -2.611630451 | 0.095 | 0.545 | 1 | 0.408060054 | -0.312522235 | 0.644 | 0.609 | 1 | 0.408060054 | 0.00020356 |
| Polr2i | 0.007951128 | -2.146407195 | 0.048 | 0.325 | 1 | 0.000102796 | -1.147002412 | 0.044 | 0.339 | 1 | 0.007951128 | 0.000205582 |
| Rbm7 | 0.018467086 | -0.52393639 | 0.333 | 0.728 | 1 | 0.000103251 | -0.94858315 | 0.422 | 0.74 | 1 | 0.018467086 | 0.00020649 |
| Snhg4 | 0.298597761 | 0.43169698 | 0.286 | 0.493 | 1 | 0.000103523 | -1.656777417 | 0.133 | 0.46 | 1 | 0.298597761 | 0.000207035 |
| Myo10 | 0.016897895 | 1.202554964 | 0.905 | 0.851 | 1 | 0.000103799 | 0.920322966 | 0.978 | 0.972 | 1 | 0.016897895 | 0.000207588 |
| Tmem107 | 0.009842655 | -2.314912919 | 0.048 | 0.313 | 1 | 0.000103872 | -4.071814916 | 0 | 0.263 | 1 | 0.009842655 | 0.000207734 |
| Ndufs3 | 0.000104266 | -1.378549267 | 0.238 | 0.739 | 1 | 0.076467735 | -0.398721505 | 0.622 | 0.685 | 1 | 0.076467735 | 0.000208521 |
| Hnrnpf | 0.006176249 | -0.627094712 | 0.905 | 0.959 | 1 | 0.000104327 | -0.484144399 | 0.978 | 0.99 | 1 | 0.006176249 | 0.000208644 |
| Ppp3ca | 0.006846709 | 1.578852275 | 1 | 1 | 1 | 0.000104828 | 1.098320468 | 0.978 | 0.997 | 1 | 0.006846709 | 0.000209644 |
| Dbndd2 | 0.228798032 | -0.289702356 | 0.095 | 0.213 | 1 | 0.000104859 | -2.743192169 | 0.022 | 0.304 | 1 | 0.228798032 | 0.000209707 |
| Selenon | 0.257110556 | -0.430564892 | 0.143 | 0.261 | 1 | 0.000105089 | -1.719025857 | 0.067 | 0.367 | 1 | 0.257110556 | 0.000210166 |
| Mthfd2 | 0.314214272 | -0.464642261 | 0.333 | 0.474 | 1 | 0.000105128 | -1.690807588 | 0.133 | 0.46 | 1 | 0.314214272 | 0.000210245 |
| Fkrp | 0.014570072 | -2.683780237 | 0 | 0.228 | 1 | 0.000105165 | -1.638855509 | 0.067 | 0.367 | 1 | 0.014570072 | 0.00021032 |
| Fat4 | 0.001927532 | -2.519999976 | 0.19 | 0.545 | 1 | 0.000105459 | -1.141009422 | 0.267 | 0.64 | 1 | 0.001927532 | 0.000210908 |
| Rnft1 | 0.001637196 | -1.063193826 | 0.19 | 0.642 | 1 | 0.000105866 | -0.88939897 | 0.244 | 0.633 | 1 | 0.001637196 | 0.000211721 |
| Sin3b | 0.000106818 | -1.294895026 | 0.429 | 0.825 | 1 | 0.026596495 | -0.41070956 | 0.844 | 0.837 | 1 | 0.026596495 | 0.000213625 |
| Efcab14 | 0.00010793 | -1.174851173 | 0.286 | 0.776 | 1 | 0.045107187 | -0.401735802 | 0.8 | 0.789 | 1 | 0.045107187 | 0.000215849 |
| Samd4b | 0.148351688 | -0.131239214 | 0.619 | 0.825 | 1 | 0.000107957 | -0.687614821 | 0.644 | 0.824 | 1 | 0.148351688 | 0.000215903 |
| Yae1d1 | 0.008397301 | -1.38800443 | 0.143 | 0.451 | 1 | 0.000108139 | -1.163382156 | 0.111 | 0.443 | 1 | 0.008397301 | 0.000216266 |
| Cd320 | 0.020443846 | -1.196592952 | 0.095 | 0.358 | 1 | 0.000108149 | -2.187292133 | 0.044 | 0.336 | 1 | 0.020443846 | 0.000216287 |
| Trim27 | 0.002299904 | -2.564632972 | 0.048 | 0.381 | 1 | 0.000108359 | -1.059891749 | 0.178 | 0.522 | 1 | 0.002299904 | 0.000216706 |
| Phax | 0.00010906 | -1.840801405 | 0.143 | 0.642 | 1 | 0.002571882 | -0.588732029 | 0.422 | 0.702 | 1 | 0.002571882 | 0.000218108 |
| Sgms2 | 0.294277655 | 0.179383786 | 0.81 | 0.899 | 1 | 0.000109168 | 0.928282771 | 1 | 0.955 | 1 | 0.294277655 | 0.000218324 |
| Agr2 | 0.12097145 | -4.25696557 | 0 | 0.104 | 1 | 0.000109503 | 0.892995656 | 0.356 | 0.121 | 1 | 0.12097145 | 0.000218994 |
| Cav2 | 0.000109937 | -4.274595464 | 0 | 0.451 | 1 | 0.019598983 | -1.323353683 | 0.178 | 0.353 | 1 | 0.019598983 | 0.000219862 |
| Ddr2 | 0.031227898 | 0.876573461 | 0.857 | 0.91 | 1 | 0.000110388 | 0.73317855 | 0.933 | 0.896 | 1 | 0.031227898 | 0.000220764 |
| Mrpl45 | 0.244068229 | -0.264827688 | 0.143 | 0.28 | 1 | 0.000110958 | -2.194671664 | 0.044 | 0.336 | 1 | 0.244068229 | 0.000221904 |
| St3gal2 | 0.722808846 | -0.14385149 | 0.286 | 0.34 | 1 | 0.000111128 | -0.78103552 | 0.178 | 0.554 | 1 | 0.722808846 | 0.000222243 |
| Anp32e | 0.072168794 | -0.589262638 | 0.429 | 0.642 | 1 | 0.000111523 | -0.982601714 | 0.711 | 0.775 | 1 | 0.072168794 | 0.000223034 |
| Hibadh | 0.000111571 | -1.305588466 | 0.19 | 0.713 | 1 | 0.037564146 | -0.350558192 | 0.622 | 0.709 | 1 | 0.037564146 | 0.000223129 |
| Gtf3c4 | 0.094569663 | -1.20069735 | 0.095 | 0.254 | 1 | 0.000112928 | -1.662702251 | 0.111 | 0.433 | 1 | 0.094569663 | 0.000225844 |
| Stoml2 | 0.010876392 | -1.317183093 | 0.095 | 0.407 | 1 | 0.000113161 | -1.502793959 | 0.111 | 0.426 | 1 | 0.010876392 | 0.00022631 |
| Mphosph6 | 0.020668997 | -0.400575574 | 0.19 | 0.478 | 1 | 0.000114017 | -1.894356242 | 0.111 | 0.419 | 1 | 0.020668997 | 0.00022802 |
| Crlf2 | 0.096971422 | -0.555046923 | 0.19 | 0.396 | 1 | 0.000114067 | -1.983684006 | 0.044 | 0.336 | 1 | 0.096971422 | 0.00022812 |
| Matn1 | 0.007962839 | -3.332022881 | 0.19 | 0.485 | 1 | 0.000114087 | -3.605921619 | 0.133 | 0.439 | 1 | 0.007962839 | 0.000228161 |
| Srgap2 | 0.051814239 | 0.723647674 | 0.857 | 0.787 | 1 | 0.000114515 | 0.726141308 | 0.978 | 0.917 | 1 | 0.051814239 | 0.000229017 |
| Setd7 | 0.000114721 | -1.486099569 | 0.238 | 0.776 | 1 | 0.132715159 | -0.10158975 | 0.867 | 0.882 | 1 | 0.132715159 | 0.000229429 |
| Lmo4 | 0.001245233 | -1.106878831 | 0.857 | 0.933 | 1 | 0.000114836 | -0.593172693 | 0.933 | 0.927 | 1 | 0.001245233 | 0.00022966 |
| Jtb | 0.38231779 | 0.209103501 | 0.429 | 0.593 | 1 | 0.000114972 | -1.219804947 | 0.4 | 0.633 | 1 | 0.38231779 | 0.000229932 |

| | | | | | | | | | | | | |
|---|---|---|---|---|---|---|---|---|---|---|---|---|
| Prpf38a | 0.000115773 | -2.99156415 | 0.048 | 0.496 | 1 | 0.000170881 | -0.760534065 | 0.2 | 0.567 | 1 | 0.000170881 | 0.000231532 |
| Dr1 | 0.184515218 | -0.637814383 | 0.19 | 0.34 | 1 | 0.000115892 | -0.886783022 | 0.111 | 0.453 | 1 | 0.184515218 | 0.00023177 |
| Tmem276-zftraf1 | 0.018520509 | -1.801961663 | 0.048 | 0.284 | 1 | 0.000117707 | -1.674479418 | 0.089 | 0.408 | 1 | 0.018520509 | 0.0002354 |
| Dlg3 | 0.142172688 | -1.25696557 | 0.048 | 0.168 | 1 | 0.000117805 | -2.006226574 | 0.022 | 0.304 | 1 | 0.142172688 | 0.000235595 |
| Tor1aip1 | 0.096560172 | -0.304297592 | 0.762 | 0.944 | 1 | 0.000117841 | -0.484965306 | 0.956 | 0.979 | 1 | 0.096560172 | 0.000235667 |
| Gpr180 | 0.000118012 | -2.01272876 | 0.143 | 0.612 | 1 | 0.000346846 | -0.816498423 | 0.222 | 0.564 | 1 | 0.000346846 | 0.00023601 |
| Ndufb5 | 0.000118108 | -1.176332802 | 0.524 | 0.862 | 1 | 0.006699308 | -0.37679887 | 0.778 | 0.796 | 1 | 0.006699308 | 0.000236201 |
| Tbpl1 | 0.006999853 | -1.828728573 | 0.048 | 0.351 | 1 | 0.000118752 | -0.960783604 | 0.111 | 0.439 | 1 | 0.006999853 | 0.00023749 |
| Apmap | 0.01811051 | -0.606336152 | 0.095 | 0.392 | 1 | 0.000120279 | -1.462604869 | 0.111 | 0.436 | 1 | 0.01811051 | 0.000240544 |
| Ier5 | 0.000250574 | -1.754089772 | 0.714 | 0.881 | 1 | 0.000120321 | -1.149168296 | 0.8 | 0.799 | 1 | 0.000250574 | 0.000240627 |
| Traf7 | 0.051159065 | -0.811655059 | 0.095 | 0.325 | 1 | 0.000120591 | -1.674479418 | 0.133 | 0.439 | 1 | 0.051159065 | 0.000241166 |
| A2ml1 | 0.000120762 | 3.811275291 | 0.19 | 0.026 | 1 | 0.49622258 | -1.017367132 | 0 | 0.01 | 1 | 0.49622258 | 0.000241509 |
| Pdap1 | 0.000120949 | -1.366013098 | 0.476 | 0.825 | 1 | 0.007331698 | -0.564854927 | 0.778 | 0.81 | 1 | 0.007331698 | 0.000241884 |
| Pxdn | 0.038838853 | -2.44170545 | 0.095 | 0.295 | 1 | 0.000122382 | -0.168926251 | 0.133 | 0.498 | 1 | 0.038838853 | 0.000244749 |
| Entpd5 | 0.050629593 | -1.911190733 | 0 | 0.157 | 1 | 0.000122493 | 1.3379371 | 0.533 | 0.249 | 1 | 0.050629593 | 0.000244971 |
| Ppt2 | 0.08055997 | -0.8244791 | 0.095 | 0.28 | 1 | 0.000122903 | -1.970369652 | 0.089 | 0.388 | 1 | 0.08055997 | 0.000245792 |
| Banf1 | 0.00012297 | -1.603386457 | 0.19 | 0.694 | 1 | 0.006807595 | -0.827889333 | 0.533 | 0.64 | 1 | 0.006807595 | 0.000245925 |
| 2300009A05Rik | 0.008652685 | -2.107587946 | 0.048 | 0.321 | 1 | 0.000123111 | -2.731964913 | 0.044 | 0.322 | 1 | 0.008652685 | 0.000246207 |
| Rab3gap2 | 0.096131797 | 1.069142234 | 0.667 | 0.642 | 1 | 0.000123549 | 1.29179121 | 0.867 | 0.702 | 1 | 0.096131797 | 0.000247082 |
| Faf1 | 0.000193223 | 1.282041642 | 0.905 | 0.784 | 1 | 0.000123847 | 1.248088699 | 0.978 | 0.91 | 1 | 0.000193223 | 0.000247679 |
| Taf5l | 0.250119315 | -0.434752689 | 0.095 | 0.205 | 1 | 0.000124074 | -2.754332726 | 0.044 | 0.322 | 1 | 0.250119315 | 0.000248134 |
| Abhd17a | 0.000495256 | -1.818081329 | 0.143 | 0.56 | 1 | 0.000124305 | -1.336120979 | 0.222 | 0.543 | 1 | 0.000495256 | 0.000248595 |
| Gm44700 | 0.000124715 | 3.673771768 | 0.19 | 0.026 | 1 | 0.688289371 | 1.91753784 | 0.067 | 0.055 | 1 | 0.688289371 | 0.000249415 |
| Dnajc9 | 0.000874906 | -2.901137068 | 0.048 | 0.44 | 1 | 0.000124997 | -0.242926832 | 0.067 | 0.391 | 1 | 0.000874906 | 0.000249978 |
| Ywhab | 0.006982073 | -0.64543169 | 0.905 | 0.955 | 1 | 0.000125022 | -0.544362465 | 0.978 | 0.976 | 1 | 0.006982073 | 0.000250029 |
| Cuta | 0.000125162 | -1.330320987 | 0.19 | 0.705 | 1 | 0.001845051 | -1.194671664 | 0.422 | 0.581 | 1 | 0.001845051 | 0.000250309 |
| Smarca5 | 0.008058679 | -0.633283051 | 0.952 | 0.974 | 1 | 0.00012536 | -0.409684555 | 0.978 | 0.979 | 1 | 0.008058679 | 0.000250705 |
| Thap12 | 0.178082036 | -0.287057635 | 0.286 | 0.515 | 1 | 0.000126559 | -0.826985106 | 0.289 | 0.619 | 1 | 0.178082036 | 0.000253102 |
| Gpcpd1 | 0.000126824 | 1.530813814 | 0.857 | 0.56 | 1 | 0.082586196 | 0.183970822 | 0.822 | 0.55 | 1 | 0.082586196 | 0.000253632 |
| Cpeb1 | 0.651849243 | 0.129451251 | 0.143 | 0.19 | 1 | 0.000127665 | -2.72631835 | 0.044 | 0.329 | 1 | 0.651849243 | 0.000255313 |
| Mrpl51 | 0.000128728 | -2.15277672 | 0.048 | 0.53 | 1 | 0.035297109 | -0.762486858 | 0.311 | 0.467 | 1 | 0.035297109 | 0.00025744 |
| Slc25a11 | 0.000128799 | -3.005708332 | 0.048 | 0.493 | 1 | 0.107102586 | -0.264459994 | 0.267 | 0.398 | 1 | 0.107102586 | 0.000257581 |
| Nedd4l | 0.027614055 | 0.683430552 | 0.952 | 0.877 | 1 | 0.000129127 | 1.357931571 | 0.978 | 0.913 | 1 | 0.027614055 | 0.000258237 |
| Trnau1ap | 0.043299266 | -0.254335314 | 0.143 | 0.399 | 1 | 0.000129165 | -1.678383873 | 0.089 | 0.405 | 1 | 0.043299266 | 0.000258314 |
| Arl6 | 0.099705054 | -1.233118828 | 0.048 | 0.19 | 1 | 0.000129448 | -3.743192169 | 0 | 0.256 | 1 | 0.099705054 | 0.000258879 |
| Caml | 0.013131725 | -0.96430207 | 0.238 | 0.56 | 1 | 0.000130743 | -1.08775646 | 0.178 | 0.543 | 1 | 0.013131725 | 0.000261469 |
| Cgrrf1 | 0.003203982 | -1.348596045 | 0.19 | 0.556 | 1 | 0.000131122 | -0.698798049 | 0.222 | 0.616 | 1 | 0.003203982 | 0.000262227 |
| Plcb4 | 0.056411335 | 0.719575457 | 0.429 | 0.235 | 1 | 0.000131687 | 0.869841098 | 0.756 | 0.384 | 1 | 0.056411335 | 0.000263357 |
| Imp3 | 0.000131889 | -2.25696557 | 0.143 | 0.612 | 1 | 0.013878877 | -0.583207479 | 0.444 | 0.623 | 1 | 0.013878877 | 0.00026376 |
| Maob | 0.023614394 | -2.081115734 | 0.095 | 0.336 | 1 | 0.00013214 | -1.982263331 | 0.067 | 0.37 | 1 | 0.023614394 | 0.000264262 |
| Col8a1 | 0.956488923 | 0.551924759 | 0.762 | 0.772 | 1 | 0.000132814 | -1.15548088 | 0.422 | 0.796 | 1 | 0.956488923 | 0.00026561 |
| Tceal1 | 0.054035013 | -0.798716004 | 0.095 | 0.306 | 1 | 0.000133074 | -1.84048937 | 0.044 | 0.332 | 1 | 0.054035013 | 0.000266131 |
| Ano6 | 0.002046123 | 1.04174723 | 1 | 0.922 | 1 | 0.000133135 | 1.025902663 | 1 | 0.955 | 1 | 0.002046123 | 0.000266252 |
| Igdcc4 | 0.157499759 | -0.250500906 | 0.19 | 0.377 | 1 | 0.000133474 | -1.017367132 | 0.089 | 0.426 | 1 | 0.157499759 | 0.00026693 |
| Myh14 | 0.71454173 | 0.918884266 | 0.095 | 0.075 | 1 | 0.000133586 | 0.86810448 | 0.578 | 0.263 | 1 | 0.71454173 | 0.000267154 |
| Magi3 | 0.274671541 | 0.735172312 | 0.524 | 0.362 | 1 | 0.000134179 | 0.941265729 | 0.844 | 0.488 | 1 | 0.274671541 | 0.000268341 |
| Psmd13 | 0.123494449 | -0.63701577 | 0.333 | 0.541 | 1 | 0.000134761 | -1.204452685 | 0.511 | 0.685 | 1 | 0.123494449 | 0.000269504 |
| Memo1 | 0.203389456 | 0.825774861 | 0.619 | 0.59 | 1 | 0.000134929 | 1.116725764 | 0.911 | 0.696 | 1 | 0.203389456 | 0.000269841 |
| Spr | 0.000544056 | -3.535681598 | 0 | 0.384 | 1 | 0.000135089 | -2.184823878 | 0.067 | 0.356 | 1 | 0.000544056 | 0.00027016 |
| Timm22 | 0.001300888 | -1.989193245 | 0.048 | 0.429 | 1 | 0.000135184 | -1.197345798 | 0.089 | 0.401 | 1 | 0.001300888 | 0.000270349 |
| Dcaf13 | 0.578895468 | 0.281454345 | 0.286 | 0.396 | 1 | 0.000135322 | -0.979892427 | 0.111 | 0.453 | 1 | 0.578895468 | 0.000270626 |
| Rab12 | 0.000135801 | -1.323004331 | 0.333 | 0.787 | 1 | 0.007674653 | -0.509254962 | 0.622 | 0.834 | 1 | 0.007674653 | 0.000271583 |
| Lrba | 0.00046408 | 1.537950687 | 0.905 | 0.799 | 1 | 0.000136023 | 1.039593621 | 0.956 | 0.799 | 1 | 0.00046408 | 0.000272027 |
| Slc38a3 | 0.00947519 | -1.93109029 | 0.048 | 0.332 | 1 | 0.000136316 | -1.782901878 | 0.133 | 0.446 | 1 | 0.00947519 | 0.000272613 |
| Dact1 | 0.137293019 | -0.727676667 | 0.238 | 0.451 | 1 | 0.000136453 | -1.099829292 | 0.178 | 0.54 | 1 | 0.137293019 | 0.000272887 |
| Arl15 | 0.009550134 | 1.379710466 | 0.762 | 0.593 | 1 | 0.000136643 | 1.35226634 | 0.911 | 0.647 | 1 | 0.009550134 | 0.000273268 |
| Ppp3r1 | 0.000137423 | -1.45092398 | 0.143 | 0.672 | 1 | 0.000782703 | -0.690902256 | 0.644 | 0.782 | 1 | 0.000782703 | 0.000274827 |
| Tmem238 | 0.05529491 | -1.116305163 | 0.048 | 0.239 | 1 | 0.000138072 | -1.053893008 | 0.022 | 0.301 | 1 | 0.05529491 | 0.000276124 |
| Timm17b | 0.014516236 | -1.116305163 | 0.143 | 0.448 | 1 | 0.000138205 | -2.662702251 | 0.022 | 0.298 | 1 | 0.014516236 | 0.000276391 |
| Psme3 | 0.190811226 | -0.355975576 | 0.143 | 0.299 | 1 | 0.000138209 | -1.901889915 | 0.089 | 0.391 | 1 | 0.190811226 | 0.000276399 |
| Nr2f6 | 0.045514776 | -1.666078235 | 0.048 | 0.235 | 1 | 0.00013853 | -1.638855509 | 0.067 | 0.37 | 1 | 0.045514776 | 0.000277041 |
| Ccdc86 | 0.16946783 | -0.159118246 | 0.238 | 0.403 | 1 | 0.000138841 | -2.486852415 | 0.067 | 0.346 | 1 | 0.16946783 | 0.000277663 |
| Por | 0.130812746 | 0.698862749 | 0.667 | 0.53 | 1 | 0.00013993 | 1.10205727 | 0.844 | 0.606 | 1 | 0.130812746 | 0.000279841 |
| Ash2l | 0.170418885 | -0.280424543 | 0.143 | 0.321 | 1 | 0.000140713 | -1.589945908 | 0.089 | 0.405 | 1 | 0.170418885 | 0.000281406 |
| Ski | 0.000141301 | -1.181517631 | 0.524 | 0.91 | 1 | 0.005730884 | -0.324507869 | 0.933 | 0.986 | 1 | 0.005730884 | 0.000282581 |
| Serbp1 | 0.015007311 | -0.472584763 | 0.952 | 0.978 | 1 | 0.000141546 | -0.409796989 | 0.978 | 0.979 | 1 | 0.015007311 | 0.000283072 |
| Tmem115 | 0.007357894 | -2.184209227 | 0.048 | 0.328 | 1 | 0.000141564 | -0.716858021 | 0.133 | 0.474 | 1 | 0.007357894 | 0.000283109 |
| Fkbp3 | 0.000141731 | -1.469186186 | 0.333 | 0.784 | 1 | 0.000158824 | -1.084093246 | 0.556 | 0.734 | 1 | 0.000158824 | 0.000283442 |
| Mrpl24 | 0.000142593 | -1.540007523 | 0.286 | 0.72 | 1 | 0.335588155 | -0.168064028 | 0.689 | 0.696 | 1 | 0.335588155 | 0.000285167 |
| Imp4 | 0.803170433 | 0.60338244 | 0.238 | 0.235 | 1 | 0.000142596 | -2.674479418 | 0.044 | 0.318 | 1 | 0.803170433 | 0.000285172 |
| Trp53 | 0.175356192 | -0.62860353 | 0.476 | 0.612 | 1 | 0.000143718 | -0.921789472 | 0.733 | 0.81 | 1 | 0.175356192 | 0.000287416 |
| Cdc5l | 0.146529491 | -0.478875242 | 0.524 | 0.728 | 1 | 0.000145298 | -0.520109231 | 0.822 | 0.855 | 1 | 0.146529491 | 0.000290576 |
| Hsd11b1 | 0.000145414 | -4.46236492 | 0.048 | 0.493 | 1 | 0.16535115 | -1.776359032 | 0 | 0.042 | 1 | 0.16535115 | 0.000290807 |
| Mrpl50 | 0.010975782 | -1.295854583 | 0.19 | 0.481 | 1 | 0.000146124 | -1.322926154 | 0.156 | 0.495 | 1 | 0.010975782 | 0.000292227 |
| Cenpx | 0.0001463 | -3.841928071 | 0 | 0.437 | 1 | 0.000696368 | -0.265528261 | 0.156 | 0.464 | 1 | 0.000696368 | 0.000292578 |
| Ppp1r14b | 0.039726805 | -0.861802706 | 0.524 | 0.72 | 1 | 0.000146536 | -0.739239577 | 0.778 | 0.813 | 1 | 0.039726805 | 0.00029305 |
| Cycs | 0.000146731 | -1.411992786 | 0.619 | 0.881 | 1 | 0.000189111 | -0.835338722 | 0.778 | 0.817 | 1 | 0.000189111 | 0.000293441 |
| Hagh | 0.001045564 | -1.984439715 | 0.048 | 0.444 | 1 | 0.000147792 | -1.832627252 | 0.111 | 0.408 | 1 | 0.001045564 | 0.000295562 |
| Sfswap | 0.061273263 | 0.483526782 | 0.81 | 0.743 | 1 | 0.000148071 | 0.947693243 | 0.978 | 0.803 | 1 | 0.061273263 | 0.000296121 |
| Eaf1 | 0.072983565 | -0.940938076 | 0.143 | 0.351 | 1 | 0.00014876 | -1.530706705 | 0.089 | 0.384 | 1 | 0.072983565 | 0.000297498 |
| Mrpl12 | 0.000348457 | -2.420745831 | 0.095 | 0.504 | 1 | 0.000148909 | -1.264459994 | 0.2 | 0.509 | 1 | 0.000348457 | 0.000297796 |
| Nck2 | 0.146158537 | 0.851558887 | 0.619 | 0.664 | 1 | 0.000149009 | 1.167673679 | 0.889 | 0.779 | 1 | 0.146158537 | 0.000297996 |
| Fam104a | 0.014388684 | -0.673578924 | 0.714 | 0.922 | 1 | 0.000149259 | -0.608597688 | 0.867 | 0.941 | 1 | 0.014388684 | 0.000298495 |
| Gm36411 | 0.000149635 | 3.088809267 | 0.19 | 0.026 | 1 | 0.430487676 | 0.361144491 | 0 | 0.014 | 1 | 0.430487676 | 0.000299247 |
| Lrch3 | 0.011684284 | 1.04774661 | 0.905 | 0.881 | 1 | 0.000150244 | 0.973476011 | 0.978 | 0.945 | 1 | 0.011684284 | 0.000300464 |
| Lamtor1 | 0.00015068 | -1.461387816 | 0.381 | 0.81 | 1 | 0.000226913 | -0.814017159 | 0.644 | 0.803 | 1 | 0.000226913 | 0.000301338 |
| Slc48a1 | 0.021456075 | -1.242704877 | 0.143 | 0.403 | 1 | 0.000150837 | -1.124282336 | 0.133 | 0.467 | 1 | 0.021456075 | 0.000301651 |
| Tsnax | 0.038887062 | -0.873122692 | 0.238 | 0.489 | 1 | 0.000151763 | -1.167783975 | 0.133 | 0.481 | 1 | 0.038887062 | 0.000303504 |
| Sdhaf4 | 0.0012916 | -1.785659851 | 0.095 | 0.481 | 1 | 0.000151954 | -1.683249628 | 0.089 | 0.384 | 1 | 0.0012916 | 0.000303886 |
| Eif3d | 0.002537777 | -1.513226747 | 0.19 | 0.552 | 1 | 0.000152287 | -1.000191988 | 0.422 | 0.689 | 1 | 0.002537777 | 0.000304551 |
| Slc19a1 | 0.008113167 | -2.683780237 | 0 | 0.257 | 1 | 0.000152343 | -3.856086225 | 0 | 0.253 | 1 | 0.008113167 | 0.000304663 |
| B3glct | 0.00855621 | -0.675939484 | 0.238 | 0.623 | 1 | 0.000152453 | -1.299920989 | 0.156 | 0.474 | 1 | 0.00855621 | 0.000304883 |
| Pcyox1 | 0.080132521 | -0.752492987 | 0.143 | 0.351 | 1 | 0.000152617 | -1.486852415 | 0.067 | 0.37 | 1 | 0.080132521 | 0.000305212 |
| Gpr153 | 0.152165771 | -0.676725479 | 0.143 | 0.295 | 1 | 0.00015336 | -1.0583944 | 0.178 | 0.516 | 1 | 0.152165771 | 0.000306696 |
| Tmem176a | 0.000153849 | -1.778073356 | 0.571 | 0.772 | 1 | 0.172833477 | -0.510122195 | 0.778 | 0.685 | 1 | 0.172833477 | 0.000307674 |
| Mtss2 | 0.756282539 | 0.66124467 | 0.524 | 0.623 | 1 | 0.000153923 | -0.988627237 | 0.289 | 0.651 | 1 | 0.756282539 | 0.000307823 |
| Banp | 0.045157788 | 1.19333985 | 0.524 | 0.392 | 1 | 0.00015449 | 1.001248546 | 0.8 | 0.561 | 1 | 0.045157788 | 0.000308956 |
| Fxyd5 | 0.001391209 | -1.263401395 | 0.333 | 0.761 | 1 | 0.000154866 | -1.04620826 | 0.644 | 0.81 | 1 | 0.001391209 | 0.000309707 |

| | | | | | | | | | | | | |
|---|---|---|---|---|---|---|---|---|---|---|---|---|
| Anapc4 | 0.044925666 | -0.911190733 | 0.286 | 0.511 | 1 | 0.000155748 | -0.642857439 | 0.267 | 0.654 | 1 | 0.044925666 | 0.000311473 |
| Lypla2 | 0.008183775 | -1.424260315 | 0.095 | 0.403 | 1 | 0.000155757 | -1.199570463 | 0.111 | 0.446 | 1 | 0.008183775 | 0.00031149 |
| Stk19 | 0.836172951 | 0.332303486 | 0.381 | 0.53 | 1 | 0.000156212 | -0.983684006 | 0.178 | 0.526 | 1 | 0.836172951 | 0.0003124 |
| Ptprj | 0.082830418 | 1.641742913 | 0.81 | 0.716 | 1 | 0.000156568 | 0.963524045 | 0.889 | 0.547 | 1 | 0.082830418 | 0.000313112 |
| Derl2 | 0.004314905 | -1.496153234 | 0.095 | 0.433 | 1 | 0.000156738 | -1.785696897 | 0.089 | 0.391 | 1 | 0.004314905 | 0.000313452 |
| Nbn | 0.035745273 | -0.940938076 | 0.048 | 0.269 | 1 | 0.000156844 | -1.734779929 | 0.067 | 0.367 | 1 | 0.035745273 | 0.000313664 |
| Hycc1 | 0.263212987 | 0.939945881 | 0.381 | 0.31 | 1 | 0.000157552 | 0.963624753 | 0.689 | 0.349 | 1 | 0.263212987 | 0.00031508 |
| Hcfc1r1 | 0.000158129 | -1.619535649 | 0.429 | 0.784 | 1 | 0.00499521 | -0.786676323 | 0.6 | 0.689 | 1 | 0.00499521 | 0.000316232 |
| Sumo3 | 0.000282154 | -1.652657719 | 0.19 | 0.66 | 1 | 0.000158254 | -0.946284034 | 0.444 | 0.709 | 1 | 0.000282154 | 0.000316482 |
| Nudt9 | 0.000178837 | -1.958496448 | 0.143 | 0.593 | 1 | 0.000158388 | -0.9159651 | 0.133 | 0.481 | 1 | 0.000178837 | 0.00031675 |
| Zc3h15 | 0.224738862 | -0.100346825 | 0.857 | 0.925 | 1 | 0.000158402 | -0.488124073 | 0.867 | 0.945 | 1 | 0.224738862 | 0.000316778 |
| Cct7 | 0.036544588 | -0.553297141 | 0.476 | 0.761 | 1 | 0.000158561 | -0.754785621 | 0.756 | 0.82 | 1 | 0.036544588 | 0.000317096 |
| Mepce | 0.044663392 | -1.769171728 | 0.048 | 0.235 | 1 | 0.000158734 | -2.583713955 | 0.067 | 0.343 | 1 | 0.044663392 | 0.000317444 |
| Shisa4 | 0.000159317 | -1.482732718 | 0.095 | 0.59 | 1 | 0.004032998 | -1.316927414 | 0.133 | 0.356 | 1 | 0.004032998 | 0.000318609 |
| Sgk3 | 0.000159866 | 1.926752509 | 0.857 | 0.761 | 1 | 0.031699899 | 0.917848732 | 0.867 | 0.83 | 1 | 0.031699899 | 0.000319707 |
| Psma1 | 0.050392541 | -0.361223654 | 0.524 | 0.817 | 1 | 0.000160519 | -0.735596164 | 0.889 | 0.872 | 1 | 0.050392541 | 0.000321012 |
| Egln2 | 0.072447974 | -0.496153234 | 0.095 | 0.306 | 1 | 0.000160825 | -0.595786787 | 0.089 | 0.408 | 1 | 0.072447974 | 0.000321624 |
| Babam1 | 0.060241067 | -0.39661756 | 0.19 | 0.459 | 1 | 0.000160932 | -0.746915255 | 0.111 | 0.436 | 1 | 0.060241067 | 0.000321837 |
| Itgb1 | 0.00028221 | -0.745767124 | 0.952 | 0.996 | 1 | 0.00016125 | -0.533511616 | 1 | 0.99 | 1 | 0.00028221 | 0.000322474 |
| Klf13 | 0.054912118 | -0.243283807 | 0.667 | 0.825 | 1 | 0.000162747 | -0.683741609 | 0.956 | 0.965 | 1 | 0.054912118 | 0.000325467 |
| Fkbp2 | 0.00016283 | -1.781555453 | 0.19 | 0.638 | 1 | 0.000632299 | -1.16492432 | 0.511 | 0.654 | 1 | 0.000632299 | 0.000325634 |
| Smim3 | 0.000163686 | -1.843503926 | 0.286 | 0.728 | 1 | 0.001230384 | -0.624355939 | 0.311 | 0.675 | 1 | 0.001230384 | 0.000327346 |
| Sfxn3 | 0.431006367 | -0.252227651 | 0.143 | 0.22 | 1 | 0.000163725 | -2.456478766 | 0.067 | 0.343 | 1 | 0.431006367 | 0.000327423 |
| Upf3a | 0.000728231 | -1.478875242 | 0.095 | 0.537 | 1 | 0.000163905 | -1.11443455 | 0.244 | 0.578 | 1 | 0.000728231 | 0.000327784 |
| Setbp1 | 0.007560077 | 0.673771768 | 1 | 0.948 | 1 | 0.000164026 | 0.754997513 | 1 | 0.986 | 1 | 0.007560077 | 0.000328026 |
| Jarid2 | 0.02807258 | -0.516498503 | 1 | 0.978 | 1 | 0.000164189 | -0.528004522 | 0.978 | 0.986 | 1 | 0.02807258 | 0.000328351 |
| Bfar | 0.028443351 | -0.601022352 | 0.286 | 0.593 | 1 | 0.000164571 | -0.827889333 | 0.222 | 0.616 | 1 | 0.028443351 | 0.000329114 |
| Mapk3 | 0.00016489 | -1.84455354 | 0.143 | 0.604 | 1 | 0.292605829 | -0.1638107 | 0.489 | 0.554 | 1 | 0.292605829 | 0.000329752 |
| Cyc1 | 0.000166524 | -1.590931459 | 0.333 | 0.713 | 1 | 0.534773671 | -0.159687672 | 0.711 | 0.654 | 1 | 0.534773671 | 0.000333021 |
| Plet1 | 0.764289001 | 0.825774861 | 0.048 | 0.067 | 1 | 0.000166702 | 0.905465008 | 0.244 | 0.066 | 1 | 0.764289001 | 0.000333377 |
| Cfap68 | 0.002168336 | -1.828728573 | 0.048 | 0.407 | 1 | 0.000167341 | -3.11134328 | 0.022 | 0.284 | 1 | 0.002168336 | 0.000334655 |
| Mob3b | 0.530208549 | -1.300233024 | 0.143 | 0.194 | 1 | 0.000167453 | 1.121569058 | 0.489 | 0.215 | 1 | 0.530208549 | 0.000334879 |
| Mpnd | 0.020397608 | -2.475975352 | 0 | 0.209 | 1 | 0.000167523 | -2.946284034 | 0.022 | 0.284 | 1 | 0.020397608 | 0.000335018 |
| Macroh2a2 | 0.003790989 | -3.496153234 | 0 | 0.299 | 1 | 0.000167861 | -2.476798751 | 0.044 | 0.322 | 1 | 0.003790989 | 0.000335695 |
| Pknox2 | 0.000168002 | 2.255780951 | 0.571 | 0.265 | 1 | 0.656851839 | 0.909076261 | 0.289 | 0.394 | 1 | 0.656851839 | 0.000335976 |
| Lap3 | 0.005084204 | -1.970084422 | 0.143 | 0.444 | 1 | 0.000170241 | -1.198282917 | 0.178 | 0.516 | 1 | 0.005084204 | 0.000340454 |
| Chd4 | 0.092244392 | -0.379721537 | 0.905 | 0.922 | 1 | 0.000170244 | -0.711661817 | 1 | 0.99 | 1 | 0.092244392 | 0.000340459 |
| Tuft1 | 0.189198508 | -1.496153234 | 0.048 | 0.149 | 1 | 0.0001708 | 1.268035087 | 0.378 | 0.142 | 1 | 0.189198508 | 0.000341571 |
| Lpar4 | 0.024624938 | -1.818081329 | 0.095 | 0.336 | 1 | 0.000171029 | -2.361321533 | 0.067 | 0.349 | 1 | 0.024624938 | 0.000342028 |
| Cops6 | 0.000171217 | -1.834022873 | 0.143 | 0.619 | 1 | 0.005242617 | -0.758154437 | 0.489 | 0.637 | 1 | 0.005242617 | 0.000342404 |
| H1f4 | 0.014305754 | -2.788389644 | 0.19 | 0.44 | 1 | 0.000171584 | -2.500149238 | 0.111 | 0.408 | 1 | 0.014305754 | 0.000343139 |
| Yipf2 | 0.011826092 | -2.752492987 | 0 | 0.239 | 1 | 0.000171914 | -3.80878051 | 0 | 0.249 | 1 | 0.011826092 | 0.000343799 |
| Eif2b2 | 0.049194704 | -1.555046923 | 0.048 | 0.231 | 1 | 0.000172256 | -1.937513824 | 0.044 | 0.325 | 1 | 0.049194704 | 0.000344481 |
| Deptor | 0.745470241 | 0.434700605 | 0.333 | 0.451 | 1 | 0.000172611 | -1.223818009 | 0.089 | 0.415 | 1 | 0.745470241 | 0.000345193 |
| Eif2b5 | 0.010835272 | -1.774688733 | 0.095 | 0.366 | 1 | 0.000173799 | -1.472205639 | 0.156 | 0.46 | 1 | 0.010835272 | 0.000347568 |
| Pex3 | 0.197276543 | -0.420745831 | 0.286 | 0.466 | 1 | 0.000174113 | -1.63081819 | 0.111 | 0.405 | 1 | 0.197276543 | 0.000348196 |
| Mast4 | 0.000174698 | 1.278167086 | 1 | 0.929 | 1 | 0.021739027 | 1.032542739 | 0.978 | 0.886 | 1 | 0.021739027 | 0.000349366 |
| Prps1 | 0.010384587 | -1.574155746 | 0.048 | 0.332 | 1 | 0.000175619 | -1.747977231 | 0.089 | 0.381 | 1 | 0.010384587 | 0.000351207 |
| Sdf2l1 | 0.011110149 | -1.403043829 | 0.048 | 0.332 | 1 | 0.000175867 | -2.361321533 | 0.044 | 0.322 | 1 | 0.011110149 | 0.000351703 |
| Abi2 | 0.000176032 | -2.484080401 | 0.095 | 0.563 | 1 | 0.366788003 | 0.14918427 | 0.578 | 0.744 | 1 | 0.366788003 | 0.000352032 |
| Obsl1 | 0.912166461 | 0.503846766 | 0.286 | 0.31 | 1 | 0.000177146 | -1.206223949 | 0.089 | 0.405 | 1 | 0.912166461 | 0.00035426 |
| Hnrnpul2 | 0.335675825 | -0.157785503 | 0.571 | 0.78 | 1 | 0.000177168 | -0.667424661 | 0.867 | 0.913 | 1 | 0.335675825 | 0.000354306 |
| Zbtb7c | 0.000178119 | 2.094074583 | 0.81 | 0.56 | 1 | 0.132351037 | 1.39517612 | 0.422 | 0.388 | 1 | 0.132351037 | 0.000356207 |
| Nrf1 | 0.241679662 | 0.885717402 | 0.524 | 0.526 | 1 | 0.000178166 | 0.963701052 | 0.911 | 0.74 | 1 | 0.241679662 | 0.0003563 |
| Timm8a1 | 0.041529727 | -1.926141075 | 0.048 | 0.239 | 1 | 0.000178197 | -1.925736657 | 0.022 | 0.294 | 1 | 0.041529727 | 0.000356362 |
| Ltv1 | 0.990334012 | 0.206645757 | 0.381 | 0.451 | 1 | 0.000179395 | -0.727211383 | 0.111 | 0.45 | 1 | 0.990334012 | 0.000358757 |
| Zswim6 | 0.000180946 | 1.271545067 | 1 | 0.884 | 1 | 0.01359685 | 0.907451735 | 0.933 | 0.9 | 1 | 0.01359685 | 0.000361859 |
| Clcc1 | 0.000781262 | -1.940938076 | 0.095 | 0.5 | 1 | 0.000181226 | -0.782050145 | 0.156 | 0.512 | 1 | 0.000781262 | 0.000362419 |
| Fzd1 | 0.00148764 | -3.535681598 | 0.048 | 0.396 | 1 | 0.000181471 | -1.589945908 | 0.089 | 0.398 | 1 | 0.00148764 | 0.000362909 |
| Tada2b | 0.013379844 | -2.455511249 | 0 | 0.231 | 1 | 0.000181584 | -2.16492432 | 0.067 | 0.349 | 1 | 0.013379844 | 0.000363135 |
| Hectd2 | 0.000181833 | 1.3917364 | 0.905 | 0.735 | 1 | 0.230931893 | 0.625521564 | 0.6 | 0.616 | 1 | 0.230931893 | 0.000363633 |
| Map3k1 | 0.221512369 | 0.602110783 | 0.524 | 0.451 | 1 | 0.000183023 | 0.805063111 | 0.844 | 0.481 | 1 | 0.221512369 | 0.000366012 |
| Cux1 | 0.000183326 | 1.473957194 | 1 | 0.899 | 1 | 0.317339127 | 0.236455895 | 0.978 | 0.948 | 1 | 0.317339127 | 0.000366619 |
| Mrpl53 | 0.002121499 | -2.381510668 | 0.048 | 0.384 | 1 | 0.000183659 | -2.154870656 | 0.067 | 0.349 | 1 | 0.002121499 | 0.000367285 |
| Stub1 | 0.000183684 | -1.458678528 | 0.286 | 0.75 | 1 | 0.018098755 | -0.494465599 | 0.533 | 0.689 | 1 | 0.018098755 | 0.000367334 |
| Ccng1 | 0.050257602 | -0.772484462 | 0.19 | 0.448 | 1 | 0.000183735 | -1.103523776 | 0.111 | 0.439 | 1 | 0.050257602 | 0.000367437 |
| Pdzrn3 | 0.123900183 | 1.662042475 | 1 | 0.981 | 1 | 0.000184376 | 1.001388661 | 0.889 | 0.983 | 1 | 0.123900183 | 0.000368718 |
| Prmt1 | 0.000782755 | -1.56376414 | 0.286 | 0.687 | 1 | 0.000185475 | -1.204452685 | 0.644 | 0.74 | 1 | 0.000782755 | 0.000370915 |
| Pam16 | 0.000229782 | -2.060937853 | 0.095 | 0.545 | 1 | 0.000186242 | -1.026585662 | 0.089 | 0.405 | 1 | 0.000229782 | 0.00037245 |
| Pnkd | 0.000187334 | -1.365756597 | 0.095 | 0.593 | 1 | 0.015245593 | -0.589945908 | 0.244 | 0.471 | 1 | 0.015245593 | 0.000374632 |
| Orai1 | 0.065281104 | -0.715680321 | 0.381 | 0.612 | 1 | 0.000187376 | -0.634606036 | 0.2 | 0.536 | 1 | 0.065281104 | 0.000374716 |
| Acp2 | 0.020352259 | -1.359651234 | 0.095 | 0.354 | 1 | 0.000187927 | -1.421264074 | 0.089 | 0.398 | 1 | 0.020352259 | 0.000375819 |
| Copz1 | 0.000188732 | -1.708697869 | 0.19 | 0.649 | 1 | 0.002792058 | -0.659319611 | 0.378 | 0.657 | 1 | 0.002792058 | 0.000377429 |
| Gm56743 | 0.229423321 | -0.434752689 | 0.095 | 0.213 | 1 | 0.000189565 | -1.475356776 | 0.067 | 0.363 | 1 | 0.229423321 | 0.000379094 |
| Wdr5 | 0.5679245 | 0.157196242 | 0.19 | 0.272 | 1 | 0.000189933 | -1.199570463 | 0.133 | 0.443 | 1 | 0.5679245 | 0.000379831 |
| Tmem60 | 0.000941826 | -2.639111188 | 0.048 | 0.418 | 1 | 0.000190583 | -1.804193241 | 0.067 | 0.36 | 1 | 0.000941826 | 0.00038113 |
| Pip4k2c | 0.007846202 | -1.088728919 | 0.143 | 0.504 | 1 | 0.000191484 | -1.095369644 | 0.2 | 0.526 | 1 | 0.007846202 | 0.000382932 |
| Smim26 | 0.002796276 | -3.081115734 | 0 | 0.31 | 1 | 0.00019211 | -2.960783604 | 0.022 | 0.28 | 1 | 0.002796276 | 0.000384183 |
| Sdhd | 0.000192561 | -1.723444455 | 0.19 | 0.649 | 1 | 0.044689313 | -0.600151979 | 0.556 | 0.64 | 1 | 0.044689313 | 0.000385084 |
| Zfp469 | 0.004419667 | -1.948280052 | 0.143 | 0.448 | 1 | 0.000193436 | -1.042947501 | 0.333 | 0.713 | 1 | 0.004419667 | 0.000386835 |
| Rhbdf1 | 0.046483189 | -1.116305163 | 0.048 | 0.25 | 1 | 0.000193842 | -1.504554417 | 0.089 | 0.388 | 1 | 0.046483189 | 0.000387647 |
| Tubb2b | 0.000323219 | -2.679375058 | 0.095 | 0.545 | 1 | 0.000194103 | -1.26724029 | 0.133 | 0.446 | 1 | 0.000323219 | 0.000388168 |
| Rbm6 | 0.014268694 | 1.172829413 | 0.857 | 0.813 | 1 | 0.000194281 | 0.867912262 | 0.978 | 0.917 | 1 | 0.014268694 | 0.000388525 |
| Fyco1 | 0.049797571 | -1.314912919 | 0.095 | 0.306 | 1 | 0.000194953 | -1.184823878 | 0.044 | 0.332 | 1 | 0.049797571 | 0.000389869 |
| Ccn4 | 0.333445312 | 0.258734268 | 0.238 | 0.41 | 1 | 0.000195152 | -2.776359032 | 0.044 | 0.315 | 1 | 0.333445312 | 0.000390265 |
| Plrg1 | 0.782946477 | 0.316219763 | 0.19 | 0.243 | 1 | 0.000195168 | -2.328154669 | 0.067 | 0.339 | 1 | 0.782946477 | 0.000390299 |
| Eya3 | 0.000195168 | 1.687192283 | 0.714 | 0.448 | 1 | 0.887084816 | 0.431940269 | 0.578 | 0.564 | 1 | 0.887084816 | 0.000390299 |
| Scn1b | 0.045203568 | -1.120644099 | 0.19 | 0.425 | 1 | 0.000195586 | -1.199570463 | 0.222 | 0.554 | 1 | 0.045203568 | 0.000391134 |
| Cebpzos | 0.002753699 | -2.506137322 | 0.048 | 0.373 | 1 | 0.000195608 | -2.071814916 | 0.022 | 0.291 | 1 | 0.002753699 | 0.000391178 |
| Polr2f | 0.002210384 | -0.906172658 | 0.238 | 0.66 | 1 | 0.000195748 | -0.946977804 | 0.289 | 0.581 | 1 | 0.002210384 | 0.000391458 |
| Get3 | 0.00355719 | -2.25696557 | 0.048 | 0.362 | 1 | 0.000195789 | -1.495730567 | 0.089 | 0.398 | 1 | 0.00355719 | 0.00039154 |
| Rnf34 | 0.032389319 | -0.184209227 | 0.286 | 0.638 | 1 | 0.00019645 | -0.716858021 | 0.222 | 0.561 | 1 | 0.032389319 | 0.000392861 |
| Glrx2 | 0.000876597 | -1.655351829 | 0.095 | 0.504 | 1 | 0.000197429 | -1.08246216 | 0.156 | 0.478 | 1 | 0.000876597 | 0.00039482 |
| Il27 | 0.000197971 | 4.351843673 | 0.143 | 0.015 | 1 | 0.132760897 | -1.223818009 | 0 | 0.048 | 1 | 0.132760897 | 0.000395902 |
| Gxylt2 | 0.000199941 | -1.801007815 | 0.238 | 0.716 | 1 | 0.000956696 | -0.671124312 | 0.4 | 0.74 | 1 | 0.000956696 | 0.000399842 |
| Gmppb | 0.095827066 | -0.998653574 | 0.095 | 0.265 | 1 | 0.000200576 | -3.931637258 | 0 | 0.246 | 1 | 0.095827066 | 0.000401112 |
| Ints13 | 0.064274902 | -0.950719097 | 0.143 | 0.362 | 1 | 0.00020119 | -1.124282336 | 0.156 | 0.481 | 1 | 0.064274902 | 0.00040234 |
| Pon2 | 0.00020153 | -2.239117569 | 0.143 | 0.563 | 1 | 0.543233185 | 0.212930487 | 0.667 | 0.581 | 1 | 0.543233185 | 0.00040302 |

| | | | | | | | | | | | | |
|---|---|---|---|---|---|---|---|---|---|---|---|---|
| Dchs1 | 0.057733126 | -2.107587946 | 0 | 0.149 | 1 | 0.000202725 | -2.154870656 | 0.067 | 0.346 | 1 | 0.057733126 | 0.000405409 |
| Cpeb2 | 0.250811771 | -0.434752689 | 0.143 | 0.269 | 1 | 0.000203138 | 0.659740082 | 0.867 | 0.488 | 1 | 0.250811771 | 0.000406235 |
| Wdr77 | 0.256455863 | -0.586755783 | 0.19 | 0.31 | 1 | 0.000203908 | -1.375821103 | 0.067 | 0.36 | 1 | 0.256455863 | 0.000407774 |
| Josd2 | 0.085463422 | -0.363317551 | 0.238 | 0.485 | 1 | 0.000205056 | -1.951133434 | 0.089 | 0.374 | 1 | 0.085463422 | 0.00041007 |
| Trappc6a | 0.001361509 | -3.208871282 | 0 | 0.343 | 1 | 0.000205148 | -1.328154669 | 0.044 | 0.332 | 1 | 0.001361509 | 0.000410253 |
| 1700123O20Rik | 0.00245627 | -3.171718283 | 0 | 0.317 | 1 | 0.000205486 | -1.669443829 | 0.089 | 0.377 | 1 | 0.00245627 | 0.000410929 |
| Hspb11 | 0.012768893 | -1.574155746 | 0.095 | 0.377 | 1 | 0.000205563 | -2.061088509 | 0.022 | 0.291 | 1 | 0.012768893 | 0.000411083 |
| Hdac3 | 0.121297638 | -0.516052791 | 0.19 | 0.381 | 1 | 0.000205784 | -1.57996182 | 0.133 | 0.439 | 1 | 0.121297638 | 0.000411525 |
| Gsk3a | 0.127619176 | 0.547662999 | 0.238 | 0.504 | 1 | 0.000206778 | -1.07773975 | 0.378 | 0.668 | 1 | 0.127619176 | 0.000413514 |
| Mrpl30 | 0.00020679 | -1.718545655 | 0.19 | 0.649 | 1 | 0.159664162 | -0.450783161 | 0.467 | 0.533 | 1 | 0.159664162 | 0.000413537 |
| Pdgfa | 0.004219103 | -2.1627295 | 0.143 | 0.47 | 1 | 0.000206875 | -0.822110841 | 0.756 | 0.81 | 1 | 0.004219103 | 0.000413708 |
| Yaf2 | 0.000207123 | -0.453717967 | 0.143 | 0.705 | 1 | 0.040120246 | -0.232766966 | 0.644 | 0.785 | 1 | 0.040120246 | 0.000414203 |
| Mrpl23 | 0.000208487 | -1.619827604 | 0.19 | 0.672 | 1 | 0.118020469 | -0.671770131 | 0.556 | 0.571 | 1 | 0.118020469 | 0.000416931 |
| Commd8 | 0.00205901 | -3.159118246 | 0 | 0.325 | 1 | 0.000208555 | -2.124282336 | 0.067 | 0.346 | 1 | 0.00205901 | 0.000417066 |
| Tspan14 | 0.885280512 | 0.75177428 | 0.238 | 0.306 | 1 | 0.000209023 | 0.508574855 | 0.689 | 0.349 | 1 | 0.885280512 | 0.000418002 |
| Psmd8 | 0.001680717 | -1.083925268 | 0.476 | 0.799 | 1 | 0.000210189 | -0.781208432 | 0.844 | 0.834 | 1 | 0.001680717 | 0.000420334 |
| Gm21859 | 0.000212108 | 2.885275873 | 0.238 | 0.045 | 1 | 0.500277618 | 2.055041364 | 0.022 | 0.045 | 1 | 0.500277618 | 0.00042417 |
| Klhl28 | 0.180053708 | -0.56326743 | 0.19 | 0.351 | 1 | 0.000212404 | -1.798054104 | 0.111 | 0.408 | 1 | 0.180053708 | 0.000424764 |
| Hdhd2 | 0.00943834 | -1.159118246 | 0.143 | 0.451 | 1 | 0.000213936 | -1.58163064 | 0.111 | 0.408 | 1 | 0.00943834 | 0.000427826 |
| Timm13 | 0.000214233 | -1.363880774 | 0.381 | 0.757 | 1 | 0.156583711 | -0.172192321 | 0.644 | 0.671 | 1 | 0.156583711 | 0.000428421 |
| Ndufa11 | 0.000214461 | -1.25358293 | 0.429 | 0.802 | 1 | 0.486334409 | -0.197345798 | 0.711 | 0.654 | 1 | 0.486334409 | 0.000428875 |
| Yipf1 | 0.000215003 | -3.516052791 | 0 | 0.418 | 1 | 0.024184649 | -0.555332153 | 0.178 | 0.377 | 1 | 0.024184649 | 0.00042996 |
| Phf1 | 0.048768209 | -0.639111188 | 0.143 | 0.407 | 1 | 0.000215351 | -1.201450196 | 0.156 | 0.471 | 1 | 0.048768209 | 0.000430656 |
| Lxn | 0.000216944 | -3.785659851 | 0 | 0.422 | 1 | 0.000313753 | -1.776359032 | 0.044 | 0.318 | 1 | 0.000313753 | 0.000433841 |
| Ola1 | 0.000217049 | -1.137870513 | 0.19 | 0.675 | 1 | 0.021834761 | -0.835102823 | 0.6 | 0.651 | 1 | 0.021834761 | 0.000434051 |
| Sdad1 | 0.831484214 | 0.845252236 | 0.381 | 0.459 | 1 | 0.000217143 | -0.80878051 | 0.2 | 0.581 | 1 | 0.831484214 | 0.00043424 |
| Aldh9a1 | 0.271495399 | -0.295854583 | 0.143 | 0.269 | 1 | 0.000217294 | -2.626782676 | 0.044 | 0.308 | 1 | 0.271495399 | 0.000434541 |
| Cdc42ep5 | 0.290600796 | -0.880817084 | 0.095 | 0.19 | 1 | 0.000217356 | -3.271123724 | 0.022 | 0.277 | 1 | 0.290600796 | 0.000434666 |
| Isca1 | 0.156121199 | -0.280424543 | 0.238 | 0.481 | 1 | 0.000217745 | -1.124282336 | 0.178 | 0.505 | 1 | 0.156121199 | 0.000435442 |
| 2210016F16Rik | 0.129710701 | -0.507557997 | 0.143 | 0.336 | 1 | 0.000218253 | -1.656777417 | 0.089 | 0.384 | 1 | 0.129710701 | 0.000436458 |
| Sun2 | 0.445383355 | 0.569435108 | 0.476 | 0.496 | 1 | 0.000218321 | -1.322221713 | 0.2 | 0.509 | 1 | 0.445383355 | 0.000436594 |
| Efcab1 | 0.012758841 | -1.752492987 | 0.095 | 0.377 | 1 | 0.000218574 | -2.290932205 | 0.089 | 0.374 | 1 | 0.012758841 | 0.0004371 |
| Larp6 | 0.000218957 | -3.202745179 | 0.048 | 0.474 | 1 | 0.001283191 | -0.828573237 | 0.311 | 0.626 | 1 | 0.001283191 | 0.000437867 |
| Cachd1 | 0.241301848 | 1.614878079 | 0.238 | 0.149 | 1 | 0.00021946 | 1.278230854 | 0.6 | 0.308 | 1 | 0.241301848 | 0.000438872 |
| Hspbp1 | 0.125466136 | -0.970084422 | 0.095 | 0.239 | 1 | 0.000219953 | -2.975138897 | 0.022 | 0.277 | 1 | 0.125466136 | 0.000439857 |
| Phlda3 | 0.004113398 | -3.911190733 | 0 | 0.295 | 1 | 0.00022091 | -2.519867472 | 0.022 | 0.287 | 1 | 0.004113398 | 0.000441771 |
| Nfkbiz | 0.261486654 | 0.150209812 | 0.667 | 0.534 | 1 | 0.000221872 | 0.310263322 | 0.733 | 0.332 | 1 | 0.261486654 | 0.000443695 |
| Dynll2 | 0.000222143 | -1.870548749 | 0.143 | 0.593 | 1 | 0.084630472 | -0.313085348 | 0.689 | 0.671 | 1 | 0.084630472 | 0.000444236 |
| Sigmar1 | 0.022384137 | -1.599246727 | 0.095 | 0.332 | 1 | 0.000222414 | -3.72631835 | 0 | 0.242 | 1 | 0.022384137 | 0.000444779 |
| Ppp4r1 | 0.728655241 | 0.382217322 | 0.429 | 0.466 | 1 | 0.000222555 | 1.103433689 | 0.889 | 0.747 | 1 | 0.728655241 | 0.00044506 |
| Crip1 | 0.000222669 | -2.421486434 | 0.286 | 0.683 | 1 | 0.011342119 | -1.340117041 | 0.4 | 0.522 | 1 | 0.011342119 | 0.000445288 |
| Atp10a | 0.00022326 | 1.942452737 | 0.905 | 0.817 | 1 | 0.009089125 | 1.260352746 | 0.711 | 0.758 | 1 | 0.009089125 | 0.00044647 |
| Tbl1xr1 | 0.140167448 | -0.34415014 | 0.238 | 0.451 | 1 | 0.000223401 | 1.042367534 | 0.889 | 0.619 | 1 | 0.140167448 | 0.000446752 |
| Hic2 | 0.000223737 | 2.825774861 | 0.238 | 0.045 | 1 | 0.733884261 | 1.612683258 | 0.133 | 0.128 | 1 | 0.733884261 | 0.000447423 |
| Nsmce1 | 0.02885835 | -1.648156327 | 0.095 | 0.317 | 1 | 0.000223996 | -1.509572492 | 0.044 | 0.329 | 1 | 0.02885835 | 0.000447942 |
| Dab2 | 0.822527553 | -0.489974659 | 0.619 | 0.683 | 1 | 0.000224207 | -0.675381385 | 0.489 | 0.844 | 1 | 0.822527553 | 0.000448364 |
| 2310033P09Rik | 0.163315548 | -0.445169305 | 0.19 | 0.358 | 1 | 0.000225044 | -1.960783604 | 0.067 | 0.346 | 1 | 0.163315548 | 0.000450038 |
| Rbl2 | 0.00696769 | -1.811655059 | 0.143 | 0.448 | 1 | 0.000225057 | -0.569908155 | 0.2 | 0.581 | 1 | 0.00696769 | 0.000450064 |
| Uhmk1 | 0.943898524 | 0.586308926 | 0.286 | 0.31 | 1 | 0.000226106 | -0.78507625 | 0.111 | 0.429 | 1 | 0.943898524 | 0.000452161 |
| Tada3 | 0.877876783 | 0.271673324 | 0.19 | 0.216 | 1 | 0.000226895 | -3.824722054 | 0 | 0.242 | 1 | 0.877876783 | 0.000453738 |
| Npas3 | 0.018310556 | 1.254071732 | 0.905 | 0.821 | 1 | 0.00022727 | 1.014502255 | 0.867 | 0.875 | 1 | 0.018310556 | 0.000454488 |
| Psmc5 | 0.000227361 | -1.475975352 | 0.19 | 0.69 | 1 | 0.002408794 | -0.688324185 | 0.6 | 0.72 | 1 | 0.002408794 | 0.000454669 |
| Prkab1 | 0.135221474 | -1.184209227 | 0.048 | 0.172 | 1 | 0.000228028 | -1.901889915 | 0.044 | 0.318 | 1 | 0.135221474 | 0.000456005 |
| Acat1 | 0.000744717 | -1.010726407 | 0.333 | 0.78 | 1 | 0.000228505 | -0.528815708 | 0.644 | 0.81 | 1 | 0.000744717 | 0.000456958 |
| S100a4 | 0.006090556 | -3.06576777 | 0.238 | 0.549 | 1 | 0.000229182 | -0.908944672 | 0.133 | 0.443 | 1 | 0.006090556 | 0.000458312 |
| Terf2 | 0.115191387 | -0.522625445 | 0.19 | 0.399 | 1 | 0.000229187 | -1.113940392 | 0.2 | 0.54 | 1 | 0.115191387 | 0.000458322 |
| Mrps26 | 0.071053516 | -0.552296312 | 0.143 | 0.366 | 1 | 0.000229744 | -1.432404631 | 0.089 | 0.384 | 1 | 0.071053516 | 0.000459436 |
| Ufm1 | 0.000230787 | -1.6987824 | 0.19 | 0.646 | 1 | 0.001316708 | -0.848308874 | 0.489 | 0.668 | 1 | 0.001316708 | 0.00046152 |
| Mrps15 | 0.000231162 | -2.44170545 | 0.048 | 0.496 | 1 | 0.000436499 | -0.957385027 | 0.178 | 0.502 | 1 | 0.000436499 | 0.00046227 |
| Usp5 | 0.706900474 | 0.24081236 | 0.238 | 0.306 | 1 | 0.000231358 | -1.556393349 | 0.111 | 0.408 | 1 | 0.706900474 | 0.000462662 |
| Lmo7 | 0.970519166 | 0.240070424 | 0.81 | 0.772 | 1 | 0.000232789 | 0.671359166 | 0.911 | 0.619 | 1 | 0.970519166 | 0.000465523 |
| Pxdc1 | 0.008152293 | -1.706126396 | 0.476 | 0.701 | 1 | 0.000234225 | -0.985812398 | 0.222 | 0.578 | 1 | 0.008152293 | 0.000468394 |
| Fasn | 0.032815041 | -0.89470261 | 0.238 | 0.515 | 1 | 0.00023477 | -0.976436868 | 0.178 | 0.516 | 1 | 0.032815041 | 0.000469485 |
| Gsn | 0.000234992 | -1.143272211 | 0.952 | 0.985 | 1 | 0.001773641 | -0.442008283 | 1 | 0.997 | 1 | 0.001773641 | 0.000469929 |
| Spg21 | 0.000235285 | -1.617782678 | 0.095 | 0.556 | 1 | 0.003335603 | -0.751330238 | 0.444 | 0.664 | 1 | 0.003335603 | 0.000470514 |
| Sh3bgrl | 0.049098579 | -0.185422086 | 0.476 | 0.769 | 1 | 0.000235317 | -0.504171891 | 0.844 | 0.875 | 1 | 0.049098579 | 0.000470578 |
| C1qtnf4 | 0.010366173 | -2.865387043 | 0 | 0.246 | 1 | 0.000235854 | -2.526380779 | 0.022 | 0.284 | 1 | 0.010366173 | 0.000471653 |
| Mrpl2 | 0.000237579 | -3.709932525 | 0 | 0.418 | 1 | 0.000265469 | -1.486852415 | 0.067 | 0.353 | 1 | 0.000265469 | 0.000475101 |
| Ccdc115 | 0.077272063 | -0.707050016 | 0.143 | 0.34 | 1 | 0.000239001 | -2.787247349 | 0.044 | 0.304 | 1 | 0.077272063 | 0.000477945 |
| Timm21 | 0.0011579 | -3.337455488 | 0 | 0.351 | 1 | 0.000239498 | -2.174908409 | 0.044 | 0.315 | 1 | 0.0011579 | 0.000478939 |
| Spsb2 | 0.00207325 | -1.645379243 | 0.19 | 0.597 | 1 | 0.000240015 | -1.047661054 | 0.2 | 0.526 | 1 | 0.00207325 | 0.000479973 |
| Fam204a | 0.000241276 | -1.535681598 | 0.238 | 0.698 | 1 | 0.284976212 | -0.138266565 | 0.533 | 0.595 | 1 | 0.284976212 | 0.000482495 |
| Oaf | 0.000241588 | -1.36531344 | 0.429 | 0.858 | 1 | 0.174668282 | -0.344219105 | 0.622 | 0.813 | 1 | 0.174668282 | 0.000483118 |
| Ywhaz | 0.000241667 | -0.733920881 | 1 | 1 | 1 | 0.000758348 | -0.360538645 | 1 | 1 | 1 | 0.000758348 | 0.000483277 |
| Mpc2 | 0.000317883 | -0.975658267 | 0.762 | 0.922 | 1 | 0.000242253 | -0.564854927 | 0.933 | 0.886 | 1 | 0.000317883 | 0.000484448 |
| Pithd1 | 0.009143061 | -1.496153234 | 0.048 | 0.34 | 1 | 0.000242506 | -1.709244837 | 0.067 | 0.346 | 1 | 0.009143061 | 0.000484953 |
| Slc36a4 | 0.042719626 | -0.880817084 | 0.143 | 0.373 | 1 | 0.000242948 | -0.871516266 | 0.178 | 0.491 | 1 | 0.042719626 | 0.000485836 |
| Prrc2a | 0.01894321 | -0.532132531 | 0.476 | 0.761 | 1 | 0.000243098 | -0.623472476 | 0.867 | 0.875 | 1 | 0.01894321 | 0.000486138 |
| Impa2 | 0.000243139 | -3.962852853 | 0 | 0.418 | 1 | 0.046882142 | -0.571168701 | 0.244 | 0.439 | 1 | 0.046882142 | 0.000486218 |
| Apoe | 0.876372272 | -0.538278709 | 0.286 | 0.302 | 1 | 0.000245095 | -1.378038928 | 0.867 | 0.789 | 1 | 0.876372272 | 0.000490131 |
| Klhl21 | 0.087858582 | -0.651947906 | 0.381 | 0.623 | 1 | 0.000247954 | -0.86124793 | 0.333 | 0.716 | 1 | 0.087858582 | 0.000495847 |
| Apex1 | 0.002890864 | -1.899117901 | 0.143 | 0.485 | 1 | 0.000247989 | -2.456478766 | 0.067 | 0.332 | 1 | 0.002890864 | 0.000495916 |
| Tro | 0.708164804 | 0.634243403 | 0.143 | 0.187 | 1 | 0.000248474 | -3.282711698 | 0.022 | 0.273 | 1 | 0.708164804 | 0.000496886 |
| Crip2 | 0.000248929 | -1.672912426 | 0.429 | 0.799 | 1 | 0.002486007 | -1.034559332 | 0.6 | 0.768 | 1 | 0.002486007 | 0.000497795 |
| Trmt1 | 0.04962697 | -0.935037475 | 0.143 | 0.366 | 1 | 0.00024928 | -1.174908409 | 0.089 | 0.398 | 1 | 0.04962697 | 0.000498497 |
| Crispld2 | 0.000250154 | -2.597332068 | 0.286 | 0.664 | 1 | 0.029525454 | -1.167783975 | 0.111 | 0.273 | 1 | 0.029525454 | 0.000500246 |
| Fnip2 | 0.988897513 | 0.563347778 | 0.476 | 0.683 | 1 | 0.000250353 | 0.778768239 | 0.956 | 0.803 | 1 | 0.988897513 | 0.000500642 |
| Pigbos1 | 0.006831751 | -2.133583154 | 0.048 | 0.332 | 1 | 0.000251241 | -2.856086225 | 0.022 | 0.273 | 1 | 0.006831751 | 0.00050242 |
| Zfp36l2 | 0.000251251 | -3.254993328 | 0.238 | 0.604 | 1 | 0.941836302 | -0.633186759 | 0.867 | 0.675 | 1 | 0.941836302 | 0.000502438 |
| Urod | 0.009471552 | -2.666078235 | 0 | 0.25 | 1 | 0.000251678 | -2.513324627 | 0.044 | 0.304 | 1 | 0.009471552 | 0.000503293 |
| Dnajb14 | 0.000252697 | 0.543240205 | 0.095 | 0.556 | 1 | 0.591622194 | -0.269098889 | 0.622 | 0.574 | 1 | 0.591622194 | 0.000505331 |
| Lyar | 0.001054799 | -2.251040736 | 0.095 | 0.481 | 1 | 0.000253107 | -1.036819495 | 0.133 | 0.46 | 1 | 0.001054799 | 0.00050615 |
| Haus3 | 0.187146861 | -0.660128969 | 0.143 | 0.287 | 1 | 0.000254093 | -2.425451871 | 0.022 | 0.284 | 1 | 0.187146861 | 0.000508121 |
| Adprm | 0.026770933 | -2.455511249 | 0 | 0.194 | 1 | 0.000255027 | -3.792660845 | 0 | 0.239 | 1 | 0.026770933 | 0.000509989 |
| Prkra | 0.099316083 | -0.574155746 | 0.095 | 0.284 | 1 | 0.000255096 | -1.776359032 | 0.022 | 0.284 | 1 | 0.099316083 | 0.000510128 |
| Paxx | 0.060458871 | -1.081115734 | 0.095 | 0.295 | 1 | 0.000255905 | -3.792660845 | 0 | 0.239 | 1 | 0.060458871 | 0.000511745 |
| Ifnar2 | 0.000256013 | -1.885195525 | 0.19 | 0.631 | 1 | 0.000756639 | -0.451447079 | 0.667 | 0.768 | 1 | 0.000756639 | 0.000511961 |
| Pin4 | 0.029329569 | -0.561444694 | 0.095 | 0.343 | 1 | 0.000257654 | -2.390176396 | 0.044 | 0.304 | 1 | 0.029329569 | 0.000515242 |

| | | | | | | | | | | | | |
|---|---|---|---|---|---|---|---|---|---|---|---|---|
| Gfod1 | 0.000257931 | 1.572488432 | 0.81 | 0.522 | 1 | 0.405955298 | 0.281974278 | 0.311 | 0.246 | 1 | 0.405955298 | 0.000515796 |
| Chmp2a | 0.000260925 | -1.135923293 | 0.476 | 0.847 | 1 | 0.017926362 | -0.459466527 | 0.8 | 0.827 | 1 | 0.017926362 | 0.000521783 |
| Ppp1r8 | 0.263716335 | 0.983456267 | 0.286 | 0.205 | 1 | 0.000261669 | -1.372209849 | 0.067 | 0.356 | 1 | 0.263716335 | 0.00052327 |
| Rnf11 | 0.001303008 | -0.721529514 | 0.667 | 0.937 | 1 | 0.000261754 | -0.514608082 | 0.889 | 0.972 | 1 | 0.001303008 | 0.00052344 |
| Ehd3 | 0.000262524 | -1.440244493 | 0.19 | 0.668 | 1 | 0.000478164 | -0.976436868 | 0.156 | 0.467 | 1 | 0.000478164 | 0.000524979 |
| Eci1 | 0.06513306 | -0.728326676 | 0.143 | 0.381 | 1 | 0.000263095 | -1.414959497 | 0.067 | 0.356 | 1 | 0.06513306 | 0.000526121 |
| Klhl11 | 0.023860296 | -1.998653574 | 0.048 | 0.269 | 1 | 0.000263178 | -1.071814916 | 0.133 | 0.467 | 1 | 0.023860296 | 0.000526287 |
| Pnrc2 | 0.000534048 | -3.555046923 | 0 | 0.384 | 1 | 0.000264957 | -1.96900411 | 0.133 | 0.405 | 1 | 0.000534048 | 0.000529843 |
| Angptl1 | 0.023605153 | -2.984439715 | 0 | 0.201 | 1 | 0.000265519 | -4.361321533 | 0 | 0.239 | 1 | 0.023605153 | 0.000530967 |
| Flot1 | 0.000414731 | -2.427766259 | 0.048 | 0.474 | 1 | 0.000265658 | -1.427351404 | 0.111 | 0.415 | 1 | 0.000414731 | 0.000531245 |
| Malsu1 | 0.000266284 | -2.037723139 | 0.143 | 0.578 | 1 | 0.0096376 | -0.323353683 | 0.222 | 0.495 | 1 | 0.0096376 | 0.000532497 |
| Arfip2 | 0.00209136 | -3.159118246 | 0 | 0.325 | 1 | 0.000267476 | -0.806312255 | 0.089 | 0.384 | 1 | 0.00209136 | 0.000534881 |
| Rufy1 | 0.199482345 | -0.769171728 | 0.095 | 0.216 | 1 | 0.000268204 | -1.901889915 | 0.044 | 0.322 | 1 | 0.199482345 | 0.000536336 |
| Mrto4 | 0.011462662 | -2.292012517 | 0.048 | 0.306 | 1 | 0.000269576 | -2.124282336 | 0.067 | 0.339 | 1 | 0.011462662 | 0.00053908 |
| Insr | 0.367306495 | 0.709270449 | 0.524 | 0.522 | 1 | 0.000269784 | 1.123495404 | 0.956 | 0.657 | 1 | 0.367306495 | 0.000539495 |
| Ndufb6 | 0.000270363 | -1.363317551 | 0.381 | 0.765 | 1 | 0.001755002 | -0.8545842 | 0.6 | 0.699 | 1 | 0.001755002 | 0.000540652 |
| Cep63 | 0.75459511 | 0.812478039 | 0.286 | 0.287 | 1 | 0.000270374 | -1.72631835 | 0.067 | 0.349 | 1 | 0.75459511 | 0.000540675 |
| Mospd3 | 0.006654993 | -1.695462042 | 0.048 | 0.354 | 1 | 0.000271804 | -2.316927414 | 0.022 | 0.28 | 1 | 0.006654993 | 0.000543534 |
| Atg16l1 | 0.106819528 | -0.342530045 | 0.238 | 0.437 | 1 | 0.000272451 | -0.938415791 | 0.2 | 0.547 | 1 | 0.106819528 | 0.000544828 |
| Sfxn2 | 0.131323303 | -0.589262638 | 0.095 | 0.261 | 1 | 0.000274556 | -2.731964913 | 0.044 | 0.301 | 1 | 0.131323303 | 0.000549037 |
| Zfp275 | 0.01483334 | -1.522625445 | 0.048 | 0.313 | 1 | 0.000274972 | -1.116333583 | 0.089 | 0.405 | 1 | 0.01483334 | 0.000549868 |
| Cnpy3 | 0.079096477 | -0.505031385 | 0.238 | 0.463 | 1 | 0.000275196 | -1.326911502 | 0.089 | 0.381 | 1 | 0.079096477 | 0.000550316 |
| Cd14 | 0.043170033 | -2.222776884 | 0.143 | 0.358 | 1 | 0.000275643 | 1.262229465 | 0.467 | 0.208 | 1 | 0.043170033 | 0.00055121 |
| Mbd5 | 0.000277721 | 1.472137907 | 0.952 | 0.799 | 1 | 0.30761498 | 0.427815531 | 0.956 | 0.896 | 1 | 0.30761498 | 0.000555365 |
| Cox5a | 0.000278228 | -1.194124696 | 0.667 | 0.858 | 1 | 0.187364604 | -0.15082949 | 0.911 | 0.851 | 1 | 0.187364604 | 0.000556378 |
| Txndc9 | 0.000279641 | -1.648156327 | 0.238 | 0.66 | 1 | 0.001303907 | -0.671770131 | 0.489 | 0.72 | 1 | 0.001303907 | 0.000559203 |
| Nars | 0.914712652 | 0.173201454 | 0.762 | 0.836 | 1 | 0.000279931 | -0.702856509 | 0.822 | 0.903 | 1 | 0.914712652 | 0.000559784 |
| Pear1 | 0.179325941 | -0.834022873 | 0.095 | 0.224 | 1 | 0.000280292 | -1.003427941 | 0.089 | 0.401 | 1 | 0.179325941 | 0.000560506 |
| Spcs3 | 0.028393259 | -1.292012517 | 0.095 | 0.336 | 1 | 0.000281468 | -1.502793959 | 0.111 | 0.412 | 1 | 0.028393259 | 0.000562856 |
| Igfbp4 | 0.004829379 | -3.400667066 | 0.048 | 0.358 | 1 | 0.000281792 | -1.906890596 | 0.156 | 0.46 | 1 | 0.004829379 | 0.000563505 |
| Psmg4 | 0.037271558 | -0.891012851 | 0.143 | 0.399 | 1 | 0.000282385 | -1.307882274 | 0.111 | 0.405 | 1 | 0.037271558 | 0.000564691 |
| Klhdc3 | 0.7142076 | 0.179007076 | 0.286 | 0.354 | 1 | 0.000283528 | -1.199570463 | 0.089 | 0.401 | 1 | 0.7142076 | 0.000566975 |
| Anp32a | 0.036778903 | -0.744080747 | 0.714 | 0.843 | 1 | 0.000284129 | -0.816160041 | 0.8 | 0.81 | 1 | 0.036778903 | 0.000568178 |
| Klhl5 | 0.053606149 | -0.858723313 | 0.143 | 0.377 | 1 | 0.000284823 | -1.251039478 | 0.133 | 0.443 | 1 | 0.053606149 | 0.000569564 |
| Ube2q1 | 0.00028508 | -2.070389328 | 0.143 | 0.571 | 1 | 0.007865949 | -0.563166577 | 0.622 | 0.74 | 1 | 0.007865949 | 0.000570078 |
| Mtln | 0.000633756 | -3.465779585 | 0 | 0.377 | 1 | 0.000286197 | -3.124282336 | 0.022 | 0.27 | 1 | 0.000633756 | 0.000572312 |
| Slc27a3 | 0.021857904 | -2.535681598 | 0 | 0.205 | 1 | 0.000286875 | -2.989352756 | 0.022 | 0.27 | 1 | 0.021857904 | 0.000573669 |
| Shmt2 | 0.045800208 | -1.184209227 | 0.048 | 0.25 | 1 | 0.000286988 | -3.674479418 | 0 | 0.235 | 1 | 0.045800208 | 0.000573893 |
| Micu1 | 0.000287495 | 1.473548166 | 0.905 | 0.929 | 1 | 0.000783579 | 0.857928351 | 0.933 | 0.917 | 1 | 0.000783579 | 0.000574906 |
| Gcc1 | 0.055959236 | -1.516052791 | 0.048 | 0.224 | 1 | 0.00028803 | -1.638855509 | 0.044 | 0.322 | 1 | 0.055959236 | 0.000575976 |
| Txn2 | 0.000349389 | -1.244174016 | 0.238 | 0.746 | 1 | 0.00028803 | -0.954357334 | 0.6 | 0.737 | 1 | 0.000349389 | 0.000575978 |
| Exph5 | 0.004606304 | 3.088809267 | 0.095 | 0.011 | 1 | 0.000288532 | 1.387616703 | 0.222 | 0.059 | 1 | 0.004606304 | 0.000576981 |
| Mkrn1 | 0.000288655 | -2.593014773 | 0.095 | 0.507 | 1 | 0.350709302 | -0.192507559 | 0.689 | 0.706 | 1 | 0.350709302 | 0.000577227 |
| H2aj | 0.000289001 | -1.577316087 | 0.381 | 0.739 | 1 | 0.386057417 | -0.43081838 | 0.733 | 0.623 | 1 | 0.386057417 | 0.000577918 |
| Xrra1 | 0.57816106 | 1.351843673 | 0 | 0.015 | 1 | 0.000289308 | 4.683072586 | 0.067 | 0.003 | 1 | 0.57816106 | 0.000578531 |
| Rpp25l | 0.0216698 | -2.303508156 | 0 | 0.205 | 1 | 0.000289331 | -2.513324627 | 0.044 | 0.301 | 1 | 0.0216698 | 0.000578578 |
| Sprtn | 0.136617265 | -1.054148687 | 0.048 | 0.172 | 1 | 0.000290087 | -3.84048937 | 0 | 0.235 | 1 | 0.136617265 | 0.00058009 |
| Selenot | 0.00160488 | -0.98800633 | 0.381 | 0.743 | 1 | 0.000290325 | -0.54825296 | 0.822 | 0.882 | 1 | 0.00160488 | 0.000580567 |
| Arpc1a | 0.000290328 | -1.15201386 | 0.667 | 0.862 | 1 | 0.058106789 | -0.278959564 | 0.756 | 0.827 | 1 | 0.058106789 | 0.000580572 |
| Ap1s2 | 0.15417801 | 0.544488751 | 0.19 | 0.414 | 1 | 0.000290619 | -3.035745661 | 0.044 | 0.298 | 1 | 0.15417801 | 0.000581154 |
| Taf10 | 0.000290863 | -1.489458581 | 0.333 | 0.713 | 1 | 0.144224222 | -0.275864757 | 0.622 | 0.692 | 1 | 0.144224222 | 0.000581641 |
| Scara3 | 0.450365921 | -0.298762444 | 0.714 | 0.832 | 1 | 0.000291289 | -0.639349498 | 0.778 | 0.976 | 1 | 0.450365921 | 0.000582492 |
| Zfp267 | 0.409609815 | -0.348596045 | 0.095 | 0.168 | 1 | 0.000292692 | -3.824722054 | 0 | 0.235 | 1 | 0.409609815 | 0.000585299 |
| Sar1b | 0.000292813 | -1.213179981 | 0.524 | 0.813 | 1 | 0.000668191 | -0.628695642 | 0.711 | 0.813 | 1 | 0.000668191 | 0.000585541 |
| Srsf2 | 0.005104276 | -0.640819686 | 0.857 | 0.963 | 1 | 0.000293158 | -0.491178341 | 0.956 | 0.99 | 1 | 0.005104276 | 0.00058623 |
| Dnttip2 | 0.005922113 | -1.077969179 | 0.333 | 0.709 | 1 | 0.000294601 | -0.738391182 | 0.556 | 0.754 | 1 | 0.005922113 | 0.000589116 |
| Nae1 | 0.037650371 | -0.606336152 | 0.095 | 0.347 | 1 | 0.000295233 | -0.616487696 | 0.178 | 0.536 | 1 | 0.037650371 | 0.000590378 |
| Vamp8 | 0.000296107 | -1.287754085 | 0.571 | 0.821 | 1 | 0.001062583 | -0.776359032 | 0.778 | 0.779 | 1 | 0.001062583 | 0.000592126 |
| Dixdc1 | 0.258962312 | 0.570805189 | 0.667 | 0.698 | 1 | 0.000296839 | 0.911218535 | 0.822 | 0.709 | 1 | 0.258962312 | 0.00059359 |
| Phkg2 | 0.751249359 | 0.3917364 | 0.286 | 0.28 | 1 | 0.000297127 | -1.628513564 | 0.067 | 0.349 | 1 | 0.751249359 | 0.000594166 |
| Ccnd3 | 0.000299165 | -2.184209227 | 0.143 | 0.59 | 1 | 0.302786127 | 0.685614781 | 0.778 | 0.723 | 1 | 0.302786127 | 0.000598241 |
| Zw10 | 0.957253145 | 0.600522786 | 0.238 | 0.28 | 1 | 0.000299261 | -0.9740397 | 0.111 | 0.419 | 1 | 0.957253145 | 0.000598432 |
| Yrdc | 0.000873764 | -1.491693586 | 0.238 | 0.646 | 1 | 0.000299362 | -0.898127996 | 0.4 | 0.671 | 1 | 0.000873764 | 0.000598634 |
| H1f0 | 0.000300581 | -1.840254826 | 0.333 | 0.724 | 1 | 0.035490927 | -0.795273004 | 0.6 | 0.668 | 1 | 0.035490927 | 0.000601072 |
| Adgra2 | 0.049877467 | -2.054148687 | 0.048 | 0.228 | 1 | 0.000301042 | -0.894356242 | 0.111 | 0.415 | 1 | 0.049877467 | 0.000601993 |
| Taldo1 | 0.000301663 | -1.255838904 | 0.667 | 0.84 | 1 | 0.751906456 | -0.226487196 | 0.911 | 0.765 | 1 | 0.751906456 | 0.000603235 |
| Pbx1 | 0.000302627 | 1.208760777 | 1 | 0.944 | 1 | 0.005039515 | 0.855665505 | 0.978 | 0.958 | 1 | 0.005039515 | 0.000605163 |
| Pdgfra | 0.284021502 | -0.124184456 | 0.238 | 0.388 | 1 | 0.000303038 | -2.223818009 | 0.133 | 0.415 | 1 | 0.284021502 | 0.000605984 |
| Hpf1 | 0.000304165 | -1.314912919 | 0.095 | 0.575 | 1 | 0.004866402 | -0.697749198 | 0.267 | 0.502 | 1 | 0.004866402 | 0.000608237 |
| Rac1 | 0.000304596 | -0.731431371 | 0.952 | 0.981 | 1 | 0.001835779 | -0.37432142 | 1 | 0.993 | 1 | 0.001835779 | 0.0006091 |
| Chmp4b | 0.00030532 | -0.731953698 | 0.905 | 0.951 | 1 | 0.945413235 | 0.13345965 | 1 | 0.976 | 1 | 0.945413235 | 0.000610547 |
| Ppp1cc | 0.022249742 | -0.715555612 | 0.571 | 0.806 | 1 | 0.000305491 | -0.768844979 | 0.822 | 0.855 | 1 | 0.022249742 | 0.000610889 |
| Slc35f5 | 0.001297393 | -1.30974011 | 0.238 | 0.675 | 1 | 0.000305668 | -0.587630185 | 0.4 | 0.74 | 1 | 0.001297393 | 0.000611242 |
| Selenoh | 0.003843065 | -2.593014773 | 0.048 | 0.358 | 1 | 0.000305727 | -2.486852415 | 0.022 | 0.277 | 1 | 0.003843065 | 0.00061136 |
| Inpp5a | 0.125943995 | 1.364174475 | 0.667 | 0.69 | 1 | 0.000305744 | 0.878009476 | 0.867 | 0.754 | 1 | 0.125943995 | 0.000611395 |
| Myo1c | 0.484020604 | 0.609118809 | 0.333 | 0.552 | 1 | 0.000306485 | -0.804193241 | 0.267 | 0.592 | 1 | 0.484020604 | 0.000612876 |
| Rap2c | 0.175746578 | -0.438928365 | 0.238 | 0.399 | 1 | 0.000306609 | -0.916840256 | 0.111 | 0.419 | 1 | 0.175746578 | 0.000613124 |
| Eny2 | 0.000421251 | -0.929726684 | 0.524 | 0.873 | 1 | 0.000307687 | -0.598430455 | 0.844 | 0.893 | 1 | 0.000421251 | 0.00061528 |
| Plscr3 | 0.064201978 | -0.811655059 | 0.095 | 0.295 | 1 | 0.000307959 | -2.282711698 | 0.067 | 0.329 | 1 | 0.064201978 | 0.000615823 |
| 2810001A02Rik | 0.124564382 | -1.04047375 | 0.143 | 0.291 | 1 | 0.000308171 | -2.331877755 | 0.022 | 0.277 | 1 | 0.124564382 | 0.000616247 |
| Fam114a2 | 0.005485835 | -0.870548749 | 0.19 | 0.604 | 1 | 0.000308978 | -1.057894939 | 0.444 | 0.661 | 1 | 0.005485835 | 0.000617861 |
| Chpt1 | 0.000394973 | -1.161063891 | 0.238 | 0.694 | 1 | 0.000309051 | -0.951133434 | 0.622 | 0.779 | 1 | 0.000394973 | 0.000618007 |
| Meg3 | 0.000309431 | 1.133534926 | 0.905 | 0.761 | 1 | 0.277226956 | -0.108140125 | 0.711 | 0.879 | 1 | 0.277226956 | 0.000618766 |
| Stard3nl | 0.134551485 | 0.249745485 | 0.19 | 0.429 | 1 | 0.000309788 | -0.81719275 | 0.133 | 0.481 | 1 | 0.134551485 | 0.00061948 |
| Eif1ad | 0.051698611 | -1.02666795 | 0.095 | 0.306 | 1 | 0.000310056 | -1.404390255 | 0.067 | 0.349 | 1 | 0.051698611 | 0.000620016 |
| Cpe | 0.000311017 | -1.026981409 | 0.667 | 0.959 | 1 | 0.000613608 | -0.536855795 | 0.822 | 0.976 | 1 | 0.000613608 | 0.000621937 |
| Psmc2 | 0.000311821 | -1.370622352 | 0.286 | 0.746 | 1 | 0.028100305 | -0.563166577 | 0.711 | 0.737 | 1 | 0.028100305 | 0.000623545 |
| Cstb | 0.000312081 | 0.703665877 | 0.762 | 0.892 | 1 | 0.003388557 | -0.750748389 | 0.889 | 0.893 | 1 | 0.003388557 | 0.000624065 |
| Ring1 | 0.023033074 | -1.221045996 | 0.095 | 0.351 | 1 | 0.000312629 | -1.232535227 | 0.133 | 0.439 | 1 | 0.023033074 | 0.000625161 |
| Tspyl2 | 0.668956197 | 0.489347197 | 0.19 | 0.272 | 1 | 0.000313336 | -1.586388089 | 0.067 | 0.339 | 1 | 0.668956197 | 0.000626573 |
| Proser1 | 0.890575709 | 0.468657338 | 0.19 | 0.239 | 1 | 0.000313995 | -2.08246216 | 0.067 | 0.336 | 1 | 0.890575709 | 0.000627891 |
| Znrf3 | 0.009795419 | 1.170307381 | 0.714 | 0.586 | 1 | 0.000314412 | 0.871940973 | 0.889 | 0.675 | 1 | 0.009795419 | 0.000628726 |
| Dst | 0.000314928 | 1.150934311 | 1 | 0.959 | 1 | 0.00136204 | 0.848673437 | 0.978 | 0.972 | 1 | 0.00136204 | 0.000629757 |
| Tmub2 | 0.00244532 | -3.107587946 | 0 | 0.317 | 1 | 0.000315619 | -2.174908409 | 0.067 | 0.329 | 1 | 0.00244532 | 0.000631139 |
| Pigx | 0.00039754 | -2.818081329 | 0.048 | 0.451 | 1 | 0.000316299 | -1.223818009 | 0.178 | 0.471 | 1 | 0.00039754 | 0.000632498 |
| Ndufb3 | 0.000316802 | -1.117219708 | 0.476 | 0.847 | 1 | 0.000374822 | -0.933186045 | 0.667 | 0.761 | 1 | 0.000374822 | 0.000633504 |
| Asxl1 | 0.003149313 | 1.41597401 | 0.905 | 0.713 | 1 | 0.000316849 | 0.712157079 | 0.978 | 0.851 | 1 | 0.003149313 | 0.000633598 |
| Hcfc1 | 0.874042009 | 0.135806746 | 0.333 | 0.388 | 1 | 0.000317075 | -1.281428733 | 0.267 | 0.533 | 1 | 0.874042009 | 0.00063405 |

| | | | | | | | | | | | | |
|---|---|---|---|---|---|---|---|---|---|---|---|---|
| Vps13d | 0.034125494 | 1.049990018 | 0.667 | 0.586 | 1 | 0.000317513 | 1.141857981 | 0.956 | 0.83 | 1 | 0.034125494 | 0.000634926 |
| Mgat3 | 0.189306366 | 1.825774861 | 0.19 | 0.108 | 1 | 0.000317787 | 0.819134136 | 0.489 | 0.215 | 1 | 0.189306366 | 0.000635473 |
| Slc2a10 | 0.778421844 | 0.157196242 | 0.238 | 0.287 | 1 | 0.000318401 | -2.053893008 | 0.089 | 0.36 | 1 | 0.778421844 | 0.0006367 |
| Crim1 | 0.784024629 | -0.226988387 | 1 | 0.974 | 1 | 0.000318705 | -0.17398617 | 1 | 0.993 | 1 | 0.784024629 | 0.000637309 |
| H2-T22 | 0.001624991 | -3.564632972 | 0 | 0.336 | 1 | 0.000319214 | -1.034340211 | 0.111 | 0.401 | 1 | 0.001624991 | 0.000638326 |
| Pla2g7 | 0.002447418 | -2.499488946 | 0.143 | 0.481 | 1 | 0.000319823 | -2.723891613 | 0.067 | 0.329 | 1 | 0.002447418 | 0.000639545 |
| Gmppa | 0.00543958 | -1.709932525 | 0.095 | 0.418 | 1 | 0.000322399 | -1.206223949 | 0.111 | 0.415 | 1 | 0.00543958 | 0.000644695 |
| Ids | 0.059546355 | -1.475975352 | 0.048 | 0.22 | 1 | 0.000323083 | -1.147869156 | 0.133 | 0.45 | 1 | 0.059546355 | 0.000646061 |
| Tbc1d10b | 0.893175343 | 1.001346426 | 0.143 | 0.142 | 1 | 0.000324423 | -1.731964913 | 0.067 | 0.346 | 1 | 0.893175343 | 0.000648742 |
| 2310010J17Rik | 0.005107719 | -1.911190733 | 0.048 | 0.366 | 1 | 0.000326691 | -3.003427941 | 0.022 | 0.266 | 1 | 0.005107719 | 0.000653275 |
| Gga2 | 0.670151991 | 0.38254047 | 0.286 | 0.392 | 1 | 0.0003267 | -1.023519359 | 0.156 | 0.471 | 1 | 0.670151991 | 0.000653294 |
| Dnajc4 | 0.010006181 | -2.120644099 | 0.048 | 0.313 | 1 | 0.00032752 | -2.871516266 | 0.022 | 0.266 | 1 | 0.010006181 | 0.000654933 |
| Ilvbl | 0.011635706 | -2.555046923 | 0 | 0.239 | 1 | 0.000327567 | -2.989352756 | 0.022 | 0.266 | 1 | 0.011635706 | 0.000655027 |
| Cmc4 | 0.342790718 | 0.316219763 | 0.143 | 0.257 | 1 | 0.000327781 | -2.84048937 | 0.022 | 0.266 | 1 | 0.342790718 | 0.000655454 |
| Nfyb | 0.00073956 | -1.070389328 | 0.143 | 0.556 | 1 | 0.000327847 | -0.915695714 | 0.222 | 0.543 | 1 | 0.00073956 | 0.000655586 |
| Cmtm7 | 0.000327854 | -2.948280052 | 0.048 | 0.459 | 1 | 0.000791878 | -1.098287127 | 0.222 | 0.498 | 1 | 0.000791878 | 0.000655601 |
| Dyrk1a | 0.000329991 | 0.986892729 | 0.952 | 0.881 | 1 | 0.004765363 | 0.716060998 | 1 | 0.955 | 1 | 0.004765363 | 0.000659872 |
| Polr3d | 0.000331258 | -1.687684691 | 0.19 | 0.649 | 1 | 0.000715217 | -0.96829114 | 0.222 | 0.533 | 1 | 0.000715217 | 0.000662406 |
| Arpc5l | 0.001355523 | -1.167530486 | 0.143 | 0.593 | 1 | 0.000331827 | -1.189970239 | 0.444 | 0.644 | 1 | 0.001355523 | 0.000663544 |
| H2-DMb1 | 0.016627456 | -1.114130792 | 0.143 | 0.44 | 1 | 0.000333019 | -2.339295227 | 0.067 | 0.325 | 1 | 0.016627456 | 0.000665926 |
| Arhgef25 | 0.095316854 | 0.184836155 | 0.238 | 0.563 | 1 | 0.00033326 | -0.880828299 | 0.222 | 0.564 | 1 | 0.095316854 | 0.000666409 |
| Ppie | 0.109999725 | -0.461387816 | 0.143 | 0.336 | 1 | 0.000334583 | -1.697749198 | 0.067 | 0.346 | 1 | 0.109999725 | 0.000669054 |
| Osbpl5 | 0.524014737 | 0.487905222 | 0.19 | 0.272 | 1 | 0.000335859 | -1.395878755 | 0.089 | 0.391 | 1 | 0.524014737 | 0.000671605 |
| Txnip | 0.000336493 | -1.363138922 | 0.571 | 0.765 | 1 | 0.002703639 | -1.153173315 | 0.889 | 0.834 | 1 | 0.002703639 | 0.000672873 |
| Dgcr6 | 0.164386761 | -0.381510668 | 0.19 | 0.381 | 1 | 0.000337823 | -1.937513824 | 0.067 | 0.336 | 1 | 0.164386761 | 0.000675531 |
| Rbsn | 0.116708812 | -0.548620654 | 0.143 | 0.332 | 1 | 0.000337975 | -1.147869156 | 0.111 | 0.422 | 1 | 0.116708812 | 0.000675837 |
| Cdk17 | 0.000338039 | 2.170236866 | 0.762 | 0.694 | 1 | 0.200966674 | 0.568560705 | 0.978 | 0.824 | 1 | 0.200966674 | 0.000675965 |
| Ifngr2 | 0.085343643 | -0.556525852 | 0.19 | 0.44 | 1 | 0.000338444 | -1.181382743 | 0.4 | 0.619 | 1 | 0.085343643 | 0.000676773 |
| Gm11659 | 0.261628269 | 3.673771768 | 0.048 | 0.015 | 1 | 0.000339564 | 5.005000681 | 0.044 | 0 | 1 | 0.261628269 | 0.000679012 |
| Nmur1 | 0.492590603 | 0.673771768 | 0 | 0.022 | 1 | 0.000339564 | 5.490427508 | 0.044 | 0 | 1 | 0.492590603 | 0.000679012 |
| Gm35549 | 0.578156445 | 1.088809267 | 0 | 0.015 | 1 | 0.000339564 | 5.490427508 | 0.044 | 0 | 1 | 0.578156445 | 0.000679012 |
| E2f2 | 0.578159907 | 0.503846766 | 0 | 0.015 | 1 | 0.000339564 | 5.852997588 | 0.044 | 0 | 1 | 0.578159907 | 0.000679012 |
| Tmem179b | 0.000341613 | -1.672003069 | 0.143 | 0.582 | 1 | 0.001640419 | -1.174908409 | 0.267 | 0.505 | 1 | 0.001640419 | 0.00068311 |
| Ubxn2a | 0.000341775 | -2.02666795 | 0.143 | 0.567 | 1 | 0.627258915 | -0.221579537 | 0.622 | 0.595 | 1 | 0.627258915 | 0.000683433 |
| Gdf10 | 0.000355343 | -4.317750078 | 0.048 | 0.451 | 1 | 0.000343037 | -2.02198376 | 0.089 | 0.374 | 1 | 0.000355343 | 0.000685956 |
| Eif3l | 0.000344757 | -1.616247079 | 0.19 | 0.657 | 1 | 0.00807139 | -0.638855509 | 0.6 | 0.727 | 1 | 0.00807139 | 0.000689396 |
| Miga1 | 0.08445356 | -0.308080886 | 0.048 | 0.216 | 1 | 0.000344796 | -1.301820521 | 0.089 | 0.384 | 1 | 0.08445356 | 0.000689472 |
| Zeb1 | 0.002272297 | 1.373579602 | 0.857 | 0.802 | 1 | 0.000347227 | 1.221079328 | 0.867 | 0.896 | 1 | 0.002272297 | 0.000694334 |
| Magi2 | 0.649043475 | 0.899736919 | 0.571 | 0.75 | 1 | 0.000347518 | 1.083654102 | 0.778 | 0.699 | 1 | 0.649043475 | 0.000694915 |
| Gosr2 | 0.007929041 | -1.535681598 | 0.095 | 0.403 | 1 | 0.000348346 | -1.116333583 | 0.111 | 0.422 | 1 | 0.007929041 | 0.00069657 |
| Erlin2 | 0.040525162 | -0.379339569 | 0.048 | 0.261 | 1 | 0.000350755 | -2.174908409 | 0.044 | 0.304 | 1 | 0.040525162 | 0.000701387 |
| Tmem100 | 0.00041753 | -1.63000898 | 0.19 | 0.664 | 1 | 0.000351356 | -1.262550404 | 0.244 | 0.574 | 1 | 0.00041753 | 0.000702588 |
| Gtf2f1 | 0.000351415 | -1.744080747 | 0.238 | 0.649 | 1 | 0.301187828 | -0.252387162 | 0.6 | 0.633 | 1 | 0.301187828 | 0.000702707 |
| Golph3 | 0.059357551 | -0.430826986 | 0.81 | 0.966 | 1 | 0.000351659 | -0.353561143 | 0.889 | 0.986 | 1 | 0.059357551 | 0.000703195 |
| Pex13 | 0.14622558 | -0.337455488 | 0.333 | 0.56 | 1 | 0.000351896 | -1.017367132 | 0.311 | 0.606 | 1 | 0.14622558 | 0.000703669 |
| Slc27a1 | 0.070473418 | -1.081115734 | 0.048 | 0.224 | 1 | 0.000352447 | -1.659319611 | 0.067 | 0.336 | 1 | 0.070473418 | 0.00070477 |
| Tbcb | 0.000352567 | -1.616905393 | 0.19 | 0.634 | 1 | 0.051123669 | -0.465319254 | 0.467 | 0.574 | 1 | 0.051123669 | 0.00070501 |
| Ndufa12 | 0.000352586 | -1.540007523 | 0.333 | 0.687 | 1 | 0.101692114 | -0.326145944 | 0.622 | 0.637 | 1 | 0.101692114 | 0.000705048 |
| Sf3b2 | 0.004093424 | -0.73675025 | 0.857 | 0.922 | 1 | 0.000353749 | -0.486852415 | 0.933 | 0.955 | 1 | 0.004093424 | 0.000707374 |
| Dpm2 | 0.08558765 | -0.471905688 | 0.238 | 0.478 | 1 | 0.000354342 | -0.80878051 | 0.111 | 0.439 | 1 | 0.08558765 | 0.000708558 |
| Ppm1l | 0.492698493 | -0.326228232 | 0.095 | 0.153 | 1 | 0.000356749 | 0.669133395 | 0.778 | 0.401 | 1 | 0.492698493 | 0.00071337 |
| Ppip5k2 | 0.692478724 | -0.174225139 | 0.238 | 0.295 | 1 | 0.000356868 | -1.189052591 | 0.089 | 0.377 | 1 | 0.692478724 | 0.000713608 |
| Lratd2 | 0.022077175 | -2.769171728 | 0 | 0.205 | 1 | 0.000358046 | -2.247664751 | 0.089 | 0.353 | 1 | 0.022077175 | 0.000715963 |
| Bet1 | 0.021846861 | -0.905166945 | 0.286 | 0.575 | 1 | 0.000359582 | -0.876354822 | 0.178 | 0.509 | 1 | 0.021846861 | 0.000719034 |
| Vdac3 | 0.000360131 | -1.215840836 | 0.238 | 0.746 | 1 | 0.015067553 | -0.444788356 | 0.733 | 0.817 | 1 | 0.015067553 | 0.000720132 |
| Ppp1r12a | 0.000392146 | 0.974431246 | 1 | 0.925 | 1 | 0.000360484 | 0.63943387 | 0.978 | 0.993 | 1 | 0.000392146 | 0.000720837 |
| Ttc9c | 0.025295655 | -1.289702356 | 0.143 | 0.392 | 1 | 0.000360673 | -0.680332145 | 0.178 | 0.547 | 1 | 0.025295655 | 0.000721216 |
| Fry | 0.000361835 | 1.261896499 | 0.952 | 0.896 | 1 | 0.648486839 | 0.151307651 | 0.867 | 0.837 | 1 | 0.648486839 | 0.000723538 |
| Parp8 | 0.024418769 | 1.523455051 | 0.714 | 0.489 | 1 | 0.000362888 | 1.180118497 | 0.867 | 0.561 | 1 | 0.024418769 | 0.000725645 |
| Tbcel | 0.010314132 | -2.008052272 | 0.095 | 0.366 | 1 | 0.000363376 | -0.460763186 | 0.2 | 0.557 | 1 | 0.010314132 | 0.000726619 |
| Dctn3 | 0.000363719 | -2.280424543 | 0.143 | 0.59 | 1 | 0.109841734 | -0.821399997 | 0.578 | 0.619 | 1 | 0.109841734 | 0.000727307 |
| Epdr1 | 0.36299944 | -0.655351829 | 0.286 | 0.373 | 1 | 0.000363781 | -3.665655568 | 0.022 | 0.263 | 1 | 0.36299944 | 0.000727429 |
| Tex264 | 0.046929112 | -0.697787095 | 0.238 | 0.504 | 1 | 0.000365493 | -0.91748677 | 0.244 | 0.585 | 1 | 0.046929112 | 0.000730852 |
| Zfp330 | 0.000543576 | -1.584962501 | 0.143 | 0.59 | 1 | 0.000366624 | -0.589945908 | 0.2 | 0.561 | 1 | 0.000543576 | 0.000733113 |
| Daam2 | 0.52473612 | 0.490549944 | 0.333 | 0.485 | 1 | 0.000367552 | -0.825268633 | 0.133 | 0.446 | 1 | 0.52473612 | 0.00073497 |
| Rpf1 | 0.950782951 | 0.75490853 | 0.286 | 0.343 | 1 | 0.000367723 | -1.064161343 | 0.133 | 0.446 | 1 | 0.950782951 | 0.00073531 |
| Ube2e2 | 0.005130128 | 1.105203705 | 0.952 | 0.817 | 1 | 0.000369176 | 1.386500066 | 0.956 | 0.837 | 1 | 0.005130128 | 0.000738216 |
| Tmem203 | 0.091148719 | -1.081115734 | 0.095 | 0.257 | 1 | 0.000370611 | -3.620708162 | 0 | 0.228 | 1 | 0.091148719 | 0.000741086 |
| Dnal4 | 0.0419264 | -1.735619168 | 0.048 | 0.239 | 1 | 0.000371906 | -3.674479418 | 0 | 0.228 | 1 | 0.0419264 | 0.000743673 |
| Creld2 | 0.000372022 | -1.998653574 | 0.095 | 0.549 | 1 | 0.001898934 | -0.680332145 | 0.2 | 0.505 | 1 | 0.001898934 | 0.000743906 |
| Icmt | 0.137644435 | -0.599246727 | 0.048 | 0.183 | 1 | 0.000372145 | -3.0583944 | 0.022 | 0.263 | 1 | 0.137644435 | 0.000744152 |
| Gm26805 | 0.000372197 | 5.258734268 | 0.048 | 0 | 1 | 0.011555302 | 4.268035087 | 0.022 | 0 | 1 | 0.011555302 | 0.000744256 |
| Ptprn | 0.000372197 | 5.258734268 | 0.048 | 0 | 1 | 0.309824707 | 3.098110085 | 0.022 | 0.007 | 1 | 0.309824707 | 0.000744256 |
| Klhl30 | 0.000372197 | 4.673771768 | 0.048 | 0 | 1 | 0.496223879 | 0.683072586 | 0 | 0.01 | 1 | 0.496223879 | 0.000744256 |
| Mir205hg | 0.000372197 | 4.673771768 | 0.048 | 0 | 1 | 0.496223879 | 0.683072586 | 0 | 0.01 | 1 | 0.496223879 | 0.000744256 |
| Tbr1 | 0.000372197 | 5.258734268 | 0.048 | 0 | 1 | 0.29365335 | -0.776359032 | 0 | 0.024 | 1 | 0.29365335 | 0.000744256 |
| Rhov | 0.000372197 | 4.673771768 | 0.048 | 0 | 1 | 0.293637566 | -0.638855509 | 0 | 0.024 | 1 | 0.293637566 | 0.000744256 |
| Slco4a1 | 0.000372197 | 5.258734268 | 0.048 | 0 | 1 | 0.496223879 | 0.683072586 | 0 | 0.01 | 1 | 0.496223879 | 0.000744256 |
| Grhl3 | 0.000372197 | 5.258734268 | 0.048 | 0 | 1 | 0.260676661 | -1.124282336 | 0 | 0.028 | 1 | 0.260676661 | 0.000744256 |
| Gm43031 | 0.000372197 | 4.673771768 | 0.048 | 0 | 1 | 0.430487676 | 0.361144491 | 0 | 0.014 | 1 | 0.430487676 | 0.000744256 |
| Otop1 | 0.000372197 | 5.258734268 | 0.048 | 0 | 1 | 0.314689364 | 2.098110085 | 0.022 | 0.007 | 1 | 0.314689364 | 0.000744256 |
| 1700016F12Rik | 0.000372197 | 5.258734268 | 0.048 | 0 | 1 | 0.011555302 | 4.268035087 | 0.022 | 0 | 1 | 0.011555302 | 0.000744256 |
| Kpna7 | 0.000372197 | 5.258734268 | 0.048 | 0 | 1 | 0.496223879 | 0.683072586 | 0 | 0.01 | 1 | 0.496223879 | 0.000744256 |
| Zar1l | 0.000372197 | 5.995699863 | 0.048 | 0 | 1 | 0.030581005 | 3.683072586 | 0.044 | 0.007 | 1 | 0.030581005 | 0.000744256 |
| Tmem178b | 0.000372197 | 5.258734268 | 0.048 | 0 | 1 | 0.49622258 | -0.316927414 | 0 | 0.01 | 1 | 0.49622258 | 0.000744256 |
| 2310001H17Rik | 0.000372197 | 5.258734268 | 0.048 | 0 | 1 | 0.293647037 | -0.638855509 | 0 | 0.024 | 1 | 0.293647037 | 0.000744256 |
| Phldb3 | 0.000372197 | 4.673771768 | 0.048 | 0 | 1 | 0.430487676 | 0.361144491 | 0 | 0.014 | 1 | 0.430487676 | 0.000744256 |
| Cyp2b10 | 0.000372197 | 5.258734268 | 0.048 | 0 | 1 | 0.49622258 | 0.361144491 | 0 | 0.01 | 1 | 0.49622258 | 0.000744256 |
| Ntf5 | 0.000372197 | 4.673771768 | 0.048 | 0 | 1 | 0.260665169 | -0.776359032 | 0 | 0.028 | 1 | 0.260665169 | 0.000744256 |
| Klkb1 | 0.000372197 | 5.258734268 | 0.048 | 0 | 1 | 0.020284559 | 3.361144491 | 0.067 | 0.014 | 1 | 0.020284559 | 0.000744256 |
| Kcnj5 | 0.000372197 | 5.995699863 | 0.048 | 0 | 1 | 0.011555302 | 4.683072586 | 0.022 | 0 | 1 | 0.011555302 | 0.000744256 |
| Gm27188 | 0.000372197 | 4.673771768 | 0.048 | 0 | 1 | 0.675805699 | 0.683072586 | 0.022 | 0.014 | 1 | 0.675805699 | 0.000744256 |
| 4930429F24Rik | 0.000372197 | 4.673771768 | 0.048 | 0 | 1 | 0.376842245 | -0.316927414 | 0 | 0.017 | 1 | 0.376842245 | 0.000744256 |
| Mst1 | 0.000372197 | 5.258734268 | 0.048 | 0 | 1 | 0.430487676 | 0.361144491 | 0 | 0.014 | 1 | 0.430487676 | 0.000744256 |
| Gm57394 | 0.000372197 | 5.258734268 | 0.048 | 0 | 1 | 0.496223879 | 0.683072586 | 0 | 0.01 | 1 | 0.496223879 | 0.000744256 |
| Tns4 | 0.000372197 | 4.673771768 | 0.048 | 0 | 1 | 0.260653677 | -0.638855509 | 0 | 0.028 | 1 | 0.260653677 | 0.000744256 |
| Gm11940 | 0.000372197 | 5.258734268 | 0.048 | 0 | 1 | 0.129721052 | 3.268035087 | 0.022 | 0.003 | 1 | 0.129721052 | 0.000744256 |
| Cdhr18 | 0.000372197 | 7.48112669 | 0.048 | 0 | 1 | 0.129721052 | 4.005000681 | 0.022 | 0.003 | 1 | 0.129721052 | 0.000744256 |

| | | | | | | | | | | | | |
|---|---|---|---|---|---|---|---|---|---|---|---|---|
| Cdhr1 | 0.000372197 | 5.258734268 | 0.048 | 0 | 1 | 0.8202424 | 1.098110085 | 0.022 | 0.017 | 1 | 0.8202424 | 0.000744256 |
| 3110083C13Rik | 0.000372197 | 4.673771768 | 0.048 | 0 | 1 | 0.49622258 | 0.361144491 | 0 | 0.01 | 1 | 0.49622258 | 0.000744256 |
| Fgf9 | 0.000372197 | 4.673771768 | 0.048 | 0 | 1 | 0.030581005 | 4.098110085 | 0.044 | 0.007 | 1 | 0.030581005 | 0.000744256 |
| Pigz | 0.000372197 | 5.258734268 | 0.048 | 0 | 1 | 0.309824707 | 2.683072586 | 0.022 | 0.007 | 1 | 0.309824707 | 0.000744256 |
| Ovol1 | 0.000372197 | 4.673771768 | 0.048 | 0 | 1 | 0.496223879 | 0.683072586 | 0 | 0.01 | 1 | 0.496223879 | 0.000744256 |
| Gm36913 | 0.000372197 | 5.258734268 | 0.048 | 0 | 1 | 0.011555302 | 3.683072586 | 0.022 | 0 | 1 | 0.011555302 | 0.000744256 |
| Xlr5a | 0.000372197 | 5.258734268 | 0.048 | 0 | 1 | 0.011555302 | 4.683072586 | 0.022 | 0 | 1 | 0.011555302 | 0.000744256 |
| Tmem35a | 0.000372197 | 5.258734268 | 0.048 | 0 | 1 | 0.331919067 | -0.638855509 | 0 | 0.021 | 1 | 0.331919067 | 0.000744256 |
| Egfl6 | 0.000372197 | 5.673771768 | 0.048 | 0 | 1 | 0.184859981 | -2.486852415 | 0 | 0.038 | 1 | 0.184859981 | 0.000744256 |
| Acot1 | 0.015788644 | -2.818081329 | 0 | 0.224 | 1 | 0.000374149 | -3.691966845 | 0 | 0.228 | 1 | 0.015788644 | 0.000748158 |
| Etfrf1 | 0.049057183 | -1.555046923 | 0.048 | 0.231 | 1 | 0.000374205 | -3.709244837 | 0 | 0.228 | 1 | 0.049057183 | 0.00074827 |
| Colec12 | 0.000374812 | -1.571780939 | 0.571 | 0.929 | 1 | 0.005826418 | -0.18549316 | 0.756 | 0.976 | 1 | 0.005826418 | 0.000749483 |
| Thbs3 | 0.224094719 | -0.120644099 | 0.19 | 0.34 | 1 | 0.000375232 | -1.602329633 | 0.089 | 0.377 | 1 | 0.224094719 | 0.000750323 |
| Nelfe | 0.040227464 | -0.718545655 | 0.095 | 0.343 | 1 | 0.000377366 | -1.937513824 | 0.044 | 0.304 | 1 | 0.040227464 | 0.000754589 |
| Zfp146 | 0.791970393 | 0.247507013 | 0.286 | 0.358 | 1 | 0.000377898 | -0.795899219 | 0.178 | 0.512 | 1 | 0.791970393 | 0.000755654 |
| Tet1 | 0.014595565 | -1.901137068 | 0.048 | 0.31 | 1 | 0.000379224 | -2.674479418 | 0.022 | 0.27 | 1 | 0.014595565 | 0.000758304 |
| Akap12 | 0.027119671 | -3.574155746 | 0 | 0.194 | 1 | 0.000380775 | -2.597035333 | 0.044 | 0.308 | 1 | 0.027119671 | 0.000761405 |
| Pelo | 0.321801054 | -0.496153234 | 0.19 | 0.291 | 1 | 0.000382059 | -1.925736657 | 0.044 | 0.311 | 1 | 0.321801054 | 0.000763972 |
| Mrpl21 | 0.000382347 | -2.047327421 | 0.095 | 0.526 | 1 | 0.001840705 | -1.017367132 | 0.156 | 0.422 | 1 | 0.001840705 | 0.000764548 |
| Mboat7 | 0.115318144 | -0.558888989 | 0.095 | 0.272 | 1 | 0.0003828 | -1.113940392 | 0.067 | 0.346 | 1 | 0.115318144 | 0.000765454 |
| Pef1 | 0.002952165 | -1.002689088 | 0.095 | 0.481 | 1 | 0.00038282 | -0.508547486 | 0.133 | 0.467 | 1 | 0.002952165 | 0.000765493 |
| Ppp2r5b | 0.000385102 | -3.648156327 | 0 | 0.399 | 1 | 0.000575452 | -0.852980314 | 0.133 | 0.426 | 1 | 0.000575452 | 0.000770055 |
| Gm14164 | 0.000385514 | -3.392317423 | 0 | 0.396 | 1 | 0.001132837 | -1.233404058 | 0.133 | 0.394 | 1 | 0.001132837 | 0.00077088 |
| Manbal | 0.000385833 | -2.280424543 | 0.048 | 0.478 | 1 | 0.039226483 | -0.316927414 | 0.267 | 0.491 | 1 | 0.039226483 | 0.000771516 |
| Rtl3 | 0.037454481 | -0.98647886 | 0.048 | 0.265 | 1 | 0.000386012 | -2.050281754 | 0.022 | 0.273 | 1 | 0.037454481 | 0.000771875 |
| Cryab | 0.000386538 | -4.58046952 | 0.048 | 0.466 | 1 | 0.035702352 | 1.412983423 | 0.089 | 0.249 | 1 | 0.035702352 | 0.000772927 |
| Bola2 | 0.002312016 | -2.381510668 | 0.048 | 0.381 | 1 | 0.000386672 | -1.827889333 | 0.044 | 0.304 | 1 | 0.002312016 | 0.000773194 |
| Gab2 | 0.785290148 | 1.169427727 | 0.238 | 0.284 | 1 | 0.000388582 | 1.575431863 | 0.822 | 0.495 | 1 | 0.785290148 | 0.000777013 |
| Cd24a | 0.017009718 | -2.144277755 | 0.571 | 0.66 | 1 | 0.000390199 | -0.242022605 | 0.756 | 0.356 | 1 | 0.017009718 | 0.000780247 |
| Cfdp1 | 0.004627195 | -0.732569533 | 0.714 | 0.918 | 1 | 0.000392409 | -0.555714273 | 0.844 | 0.945 | 1 | 0.004627195 | 0.000784663 |
| Clns1a | 0.087986697 | -0.989193245 | 0.19 | 0.373 | 1 | 0.000395123 | -0.371375198 | 0.156 | 0.484 | 1 | 0.087986697 | 0.000790089 |
| Limk2 | 0.195391766 | 0.936806174 | 0.333 | 0.231 | 1 | 0.000395513 | 0.864519458 | 0.689 | 0.37 | 1 | 0.195391766 | 0.000790869 |
| Siva1 | 0.000395819 | -3.826074119 | 0 | 0.399 | 1 | 0.004933864 | -1.124282336 | 0.289 | 0.488 | 1 | 0.004933864 | 0.000791482 |
| Rfk | 0.000397657 | -2.399477214 | 0.095 | 0.5 | 1 | 0.317779903 | -0.307146393 | 0.578 | 0.574 | 1 | 0.317779903 | 0.000795155 |
| Emc8 | 0.015487259 | -0.90365706 | 0.143 | 0.455 | 1 | 0.000398852 | -0.527824196 | 0.244 | 0.578 | 1 | 0.015487259 | 0.000797544 |
| Spns1 | 0.006677075 | -1.599246727 | 0.048 | 0.354 | 1 | 0.00040134 | -1.509572492 | 0.067 | 0.343 | 1 | 0.006677075 | 0.000802519 |
| Wdtc1 | 0.486469059 | 0.728913322 | 0.143 | 0.243 | 1 | 0.00040223 | -2.017367132 | 0.089 | 0.353 | 1 | 0.486469059 | 0.000804299 |
| Mylk | 0.02357681 | -2.896083841 | 0.048 | 0.272 | 1 | 0.000402982 | -1.55589406 | 0.111 | 0.398 | 1 | 0.02357681 | 0.000805801 |
| Rilpl2 | 0.0006385 | -1.133583154 | 0.095 | 0.545 | 1 | 0.000403001 | -0.973755901 | 0.311 | 0.619 | 1 | 0.0006385 | 0.00080584 |
| Plekhj1 | 0.001814227 | -0.996079631 | 0.19 | 0.642 | 1 | 0.0004031 | -0.417387596 | 0.2 | 0.543 | 1 | 0.001814227 | 0.000806037 |
| Dhrs1 | 0.006290197 | -0.728326676 | 0.095 | 0.422 | 1 | 0.000404222 | -2.124282336 | 0.089 | 0.346 | 1 | 0.006290197 | 0.000808281 |
| Adam15 | 0.003423934 | -1.692550447 | 0.095 | 0.44 | 1 | 0.000404231 | -2.137106376 | 0.044 | 0.301 | 1 | 0.003423934 | 0.000808298 |
| E2f6 | 0.30441032 | -0.213753503 | 0.095 | 0.201 | 1 | 0.000405312 | -1.2318108 | 0.133 | 0.429 | 1 | 0.30441032 | 0.000810461 |
| Nus1 | 0.000405805 | -1.763633545 | 0.095 | 0.556 | 1 | 0.035891844 | -0.423549833 | 0.533 | 0.696 | 1 | 0.035891844 | 0.000811445 |
| Slc35e1 | 0.565946206 | 0.234660133 | 0.286 | 0.377 | 1 | 0.000406163 | -0.789415185 | 0.178 | 0.509 | 1 | 0.565946206 | 0.000812161 |
| Esyt1 | 0.904422864 | 0.237672653 | 0.333 | 0.41 | 1 | 0.000406272 | -1.486852415 | 0.089 | 0.377 | 1 | 0.904422864 | 0.000812379 |
| Mrpl27 | 0.000406503 | -2.377890352 | 0.048 | 0.474 | 1 | 0.000498649 | -1.216931624 | 0.178 | 0.478 | 1 | 0.000498649 | 0.00081284 |
| Eapp | 0.000407998 | -1.772484462 | 0.095 | 0.534 | 1 | 0.019356341 | -0.116333583 | 0.2 | 0.446 | 1 | 0.019356341 | 0.000815829 |
| Pcmtd2 | 1 | 0.561562264 | 0.286 | 0.358 | 1 | 0.000408166 | -1.249240606 | 0.156 | 0.471 | 1 | 1 | 0.000816166 |
| Slc25a5 | 0.028900809 | -0.675476933 | 0.762 | 0.806 | 1 | 0.000408182 | -0.916656625 | 0.889 | 0.903 | 1 | 0.028900809 | 0.000816197 |
| Phb2 | 0.073590849 | -0.626782676 | 0.476 | 0.668 | 1 | 0.000410897 | -0.785923897 | 0.711 | 0.775 | 1 | 0.073590849 | 0.000821624 |
| Cox17 | 0.000411681 | -1.747692001 | 0.19 | 0.616 | 1 | 0.320524756 | -0.316927414 | 0.511 | 0.516 | 1 | 0.320524756 | 0.000823193 |
| Kifbp | 0.054041079 | -0.530100566 | 0.238 | 0.552 | 1 | 0.000413691 | -0.527380222 | 0.222 | 0.612 | 1 | 0.054041079 | 0.000827211 |
| Fbh1 | 0.234779063 | -0.870548749 | 0.095 | 0.198 | 1 | 0.000413904 | -1.564854927 | 0.089 | 0.36 | 1 | 0.234779063 | 0.000827636 |
| Ggh | 0.000414834 | -1.460083979 | 0.19 | 0.66 | 1 | 0.007155686 | -0.683249628 | 0.489 | 0.637 | 1 | 0.007155686 | 0.000829496 |
| Wnk3 | 0.244291856 | -1.02666795 | 0.048 | 0.134 | 1 | 0.000415021 | -3.339295227 | 0.022 | 0.26 | 1 | 0.244291856 | 0.00082987 |
| Schip1 | 0.00041618 | 1.627297338 | 0.857 | 0.634 | 1 | 0.014280139 | 1.123598507 | 0.911 | 0.841 | 1 | 0.014280139 | 0.000832186 |
| Ccnk | 0.705352014 | 0.417432014 | 0.429 | 0.463 | 1 | 0.00041846 | -0.819427754 | 0.156 | 0.481 | 1 | 0.705352014 | 0.000836744 |
| Glt8d2 | 0.006715969 | -2.392317423 | 0.048 | 0.332 | 1 | 0.000419299 | -3.583713955 | 0 | 0.225 | 1 | 0.006715969 | 0.000838422 |
| Sec11a | 0.035329346 | -0.438437736 | 0.524 | 0.851 | 1 | 0.000420621 | -0.502793959 | 0.822 | 0.893 | 1 | 0.035329346 | 0.000841064 |
| Tshz2 | 0.029441431 | 0.934799843 | 1 | 0.966 | 1 | 0.000421107 | 0.984728286 | 0.933 | 0.855 | 1 | 0.029441431 | 0.000842036 |
| Hmgb3 | 0.028682332 | -2.593014773 | 0 | 0.19 | 1 | 0.000421437 | -3.271123724 | 0.022 | 0.26 | 1 | 0.028682332 | 0.000842695 |
| Gstm1 | 0.088323828 | 0.43169698 | 0.524 | 0.31 | 1 | 0.000421638 | 0.7219616 | 0.733 | 0.405 | 1 | 0.088323828 | 0.000843099 |
| Nr1d1 | 0.001736998 | -1.56326743 | 0.286 | 0.653 | 1 | 0.000421909 | -0.947770276 | 0.578 | 0.747 | 1 | 0.001736998 | 0.00084364 |
| Prcc | 0.066005699 | -0.870548749 | 0.143 | 0.358 | 1 | 0.000422081 | -0.80878051 | 0.222 | 0.557 | 1 | 0.066005699 | 0.000843983 |
| Thyn1 | 0.028283139 | -1.184209227 | 0.095 | 0.34 | 1 | 0.000422175 | -1.989352756 | 0.044 | 0.301 | 1 | 0.028283139 | 0.000844171 |
| U2af1l4 | 0.002587615 | -2.984439715 | 0 | 0.313 | 1 | 0.000426199 | -2.824722054 | 0.022 | 0.26 | 1 | 0.002587615 | 0.000852217 |
| Cdk5 | 0.034179007 | -0.672003069 | 0.095 | 0.354 | 1 | 0.000426483 | -1.552143876 | 0.067 | 0.332 | 1 | 0.034179007 | 0.000852784 |
| Exosc8 | 0.317003449 | -0.434752689 | 0.095 | 0.187 | 1 | 0.000426836 | -3.72631835 | 0 | 0.225 | 1 | 0.317003449 | 0.00085349 |
| Fmn1 | 0.000892763 | 2.321627282 | 0.857 | 0.877 | 1 | 0.000427517 | 1.382347163 | 0.933 | 0.934 | 1 | 0.000892763 | 0.000854851 |
| Trip6 | 0.02727441 | -1.403043829 | 0.095 | 0.336 | 1 | 0.000428305 | -0.991527127 | 0.111 | 0.408 | 1 | 0.02727441 | 0.000856426 |
| Zfp422 | 0.030684595 | -2.574155746 | 0 | 0.187 | 1 | 0.000430675 | -2.564854927 | 0.044 | 0.291 | 1 | 0.030684595 | 0.000861164 |
| Serp1 | 0.000430776 | -1.359651234 | 0.619 | 0.836 | 1 | 0.00061757 | -0.671358148 | 0.822 | 0.855 | 1 | 0.00061757 | 0.000861366 |
| Emc4 | 0.000614874 | -1.56872186 | 0.19 | 0.612 | 1 | 0.000430848 | -1.202284848 | 0.311 | 0.564 | 1 | 0.000614874 | 0.00086151 |
| Rmdn2 | 0.092922639 | -2.789752606 | 0.143 | 0.302 | 1 | 0.00043212 | -2.32442195 | 0.067 | 0.325 | 1 | 0.092922639 | 0.000864054 |
| Mospd1 | 0.115853992 | -0.496153234 | 0.143 | 0.325 | 1 | 0.000432513 | -1.223818009 | 0.111 | 0.401 | 1 | 0.115853992 | 0.00086484 |
| Lama4 | 0.510791773 | 0.311412624 | 0.476 | 0.638 | 1 | 0.000433965 | -1.089516918 | 0.244 | 0.585 | 1 | 0.510791773 | 0.000867742 |
| H2ax | 0.018919684 | -1.718545655 | 0.095 | 0.362 | 1 | 0.000436025 | -1.802354241 | 0.022 | 0.27 | 1 | 0.018919684 | 0.000871859 |
| Prss23 | 0.01705616 | -3.81004401 | 0 | 0.22 | 1 | 0.000436262 | -2.806106376 | 0.089 | 0.353 | 1 | 0.01705616 | 0.000872333 |
| Lrp5 | 0.167940394 | -0.648156327 | 0.143 | 0.31 | 1 | 0.000437101 | -0.953552034 | 0.111 | 0.419 | 1 | 0.167940394 | 0.00087401 |
| Plbd2 | 0.000437603 | -2.31869456 | 0.095 | 0.496 | 1 | 0.000479137 | -0.638855509 | 0.178 | 0.519 | 1 | 0.000479137 | 0.000875015 |
| Ccdc117 | 0.125364852 | -0.738041831 | 0.143 | 0.332 | 1 | 0.000438085 | -0.990699182 | 0.2 | 0.529 | 1 | 0.125364852 | 0.000875979 |
| Aip | 0.009169807 | -1.208871282 | 0.048 | 0.343 | 1 | 0.000438893 | -1.496836504 | 0.089 | 0.367 | 1 | 0.009169807 | 0.000877593 |
| 2310039H08Rik | 0.001910905 | -2.142145168 | 0.095 | 0.44 | 1 | 0.000439378 | -1.394929926 | 0.089 | 0.367 | 1 | 0.001910905 | 0.000878562 |
| Ensa | 0.019988583 | -0.776889641 | 0.333 | 0.668 | 1 | 0.000440501 | -0.75987091 | 0.622 | 0.803 | 1 | 0.019988583 | 0.000880808 |
| Adam9 | 0.020512982 | 1.448212067 | 0.619 | 0.563 | 1 | 0.00044098 | 0.98346752 | 0.8 | 0.554 | 1 | 0.020512982 | 0.000881765 |
| Ryk | 0.000441571 | -1.085220133 | 0.381 | 0.866 | 1 | 0.00234339 | -0.284471202 | 0.822 | 0.927 | 1 | 0.00234339 | 0.000882946 |
| Mfap4 | 0.125771626 | -2.087779465 | 0.048 | 0.179 | 1 | 0.000442959 | -5.276929346 | 0 | 0.225 | 1 | 0.125771626 | 0.000885722 |
| Gatad1 | 0.000443968 | -0.998653574 | 0.238 | 0.698 | 1 | 0.206502358 | -0.134382926 | 0.622 | 0.706 | 1 | 0.206502358 | 0.00088774 |
| Cbx1 | 0.002129263 | -1.018719197 | 0.571 | 0.854 | 1 | 0.000444926 | -0.428088412 | 0.844 | 0.903 | 1 | 0.002129263 | 0.000889653 |
| Phldb2 | 0.030188727 | -1.616905393 | 0.095 | 0.34 | 1 | 0.000446223 | -1.140049652 | 0.244 | 0.564 | 1 | 0.030188727 | 0.000892248 |
| Nt5c | 0.048008362 | -1.196592952 | 0.095 | 0.306 | 1 | 0.000447387 | -2.08246216 | 0.067 | 0.325 | 1 | 0.048008362 | 0.000894573 |
| Abcc1 | 0.031472373 | 1.374211486 | 0.333 | 0.168 | 1 | 0.000447614 | 0.736653626 | 0.733 | 0.388 | 1 | 0.031472373 | 0.000895028 |
| Vps28 | 0.000448221 | -1.142145168 | 0.238 | 0.713 | 1 | 0.09728995 | -0.679497493 | 0.578 | 0.585 | 1 | 0.09728995 | 0.00089624 |
| H2-T23 | 0.053997255 | -2.120644099 | 0.048 | 0.224 | 1 | 0.000448277 | 1.15951063 | 0.289 | 0.097 | 1 | 0.053997255 | 0.000896353 |
| Fcho2 | 0.00044921 | 0.945630024 | 0.952 | 0.903 | 1 | 0.640631348 | 0.566195713 | 1 | 0.969 | 1 | 0.640631348 | 0.000898217 |
| Churc1 | 0.003235601 | -1.467240542 | 0.19 | 0.541 | 1 | 0.000450337 | -1.142353909 | 0.289 | 0.554 | 1 | 0.003235601 | 0.000900471 |
| Slit2 | 0.038233289 | -0.354425124 | 0.381 | 0.172 | 1 | 0.000450876 | 0.516233565 | 0.667 | 0.308 | 1 | 0.038233289 | 0.000901548 |

| | | | | | | | | | | | | |
|---|---|---|---|---|---|---|---|---|---|---|---|---|
| Tafazzin | 0.005025634 | -1.155497931 | 0.143 | 0.504 | 1 | 0.000455312 | -0.909882705 | 0.133 | 0.443 | 1 | 0.005025634 | 0.000910416 |
| Ippk | 0.47288061 | 0.421853394 | 0.333 | 0.313 | 1 | 0.000455431 | -1.203059449 | 0.111 | 0.408 | 1 | 0.47288061 | 0.000910654 |
| Plpp1 | 0.8323746 | 0.745325029 | 0.429 | 0.526 | 1 | 0.000455902 | -0.518561275 | 0.222 | 0.592 | 1 | 0.8323746 | 0.000911597 |
| Gtf2i | 0.001720798 | -1.473433157 | 0.381 | 0.709 | 1 | 0.000456494 | -0.697749198 | 0.844 | 0.879 | 1 | 0.001720798 | 0.00091278 |
| Pin1 | 0.000715246 | -1.655351829 | 0.095 | 0.511 | 1 | 0.000458284 | -0.722919774 | 0.2 | 0.529 | 1 | 0.000715246 | 0.000916358 |
| Dnajc1 | 0.000459992 | 0.810921733 | 1 | 0.981 | 1 | 0.100921469 | 0.490427508 | 0.978 | 0.952 | 1 | 0.100921469 | 0.000919772 |
| Rpia | 0.551821935 | 0.728913322 | 0.143 | 0.213 | 1 | 0.000460456 | -2.098287127 | 0.044 | 0.298 | 1 | 0.551821935 | 0.000920701 |
| Sqstm1 | 0.002389478 | -0.568252462 | 0.857 | 0.97 | 1 | 0.000460813 | -0.571648889 | 0.956 | 0.986 | 1 | 0.002389478 | 0.000921414 |
| Ccn6 | 0.00112989 | -2.865387043 | 0.048 | 0.41 | 1 | 0.00046139 | -1.174908409 | 0.089 | 0.377 | 1 | 0.00112989 | 0.000922567 |
| Hspa13 | 0.010837481 | -0.861150051 | 0.19 | 0.541 | 1 | 0.000462646 | -0.987863138 | 0.2 | 0.509 | 1 | 0.010837481 | 0.000925079 |
| Nr1h2 | 0.029366326 | -0.93109029 | 0.143 | 0.414 | 1 | 0.000469018 | -0.987465646 | 0.133 | 0.439 | 1 | 0.029366326 | 0.000937817 |
| Utp3 | 0.000470055 | -2.022036502 | 0.19 | 0.582 | 1 | 0.022230922 | -0.571168701 | 0.6 | 0.689 | 1 | 0.022230922 | 0.000939888 |
| Zfp318 | 0.119916041 | -0.505031385 | 0.19 | 0.396 | 1 | 0.000470756 | -1.237492946 | 0.133 | 0.426 | 1 | 0.119916041 | 0.000941291 |
| Gpx3 | 0.000473777 | -2.772277639 | 0.238 | 0.646 | 1 | 0.010431913 | -1.727449432 | 0.067 | 0.246 | 1 | 0.010431913 | 0.000947329 |
| Cfap298 | 0.382432781 | 0.264380832 | 0.143 | 0.231 | 1 | 0.000476909 | -3.124282336 | 0.022 | 0.256 | 1 | 0.382432781 | 0.00095359 |
| C1ra | 0.059617807 | -2.054148687 | 0.048 | 0.216 | 1 | 0.000477868 | -0.645114499 | 0.111 | 0.405 | 1 | 0.059617807 | 0.000955508 |
| Atrnl1 | 0.542009927 | 0.96048517 | 0.476 | 0.448 | 1 | 0.000478162 | 0.837488214 | 0.911 | 0.505 | 1 | 0.542009927 | 0.000956095 |
| Irak2 | 0.94400642 | 0.822915207 | 0.19 | 0.231 | 1 | 0.000478694 | 1.484091551 | 0.533 | 0.277 | 1 | 0.94400642 | 0.000957159 |
| Asah1 | 0.008124781 | -1.086449179 | 0.238 | 0.601 | 1 | 0.000480996 | -0.86124793 | 0.556 | 0.716 | 1 | 0.008124781 | 0.00096176 |
| Ndufa3 | 0.000481621 | -1.724259306 | 0.19 | 0.597 | 1 | 0.157686132 | -0.541893779 | 0.378 | 0.457 | 1 | 0.157686132 | 0.000963011 |
| Mios | 0.306907771 | -0.295854583 | 0.143 | 0.246 | 1 | 0.000481847 | -3.11134328 | 0.022 | 0.256 | 1 | 0.306907771 | 0.000963463 |
| Srebf1 | 0.294172432 | -0.488115915 | 0.238 | 0.384 | 1 | 0.000482296 | -2.033134448 | 0.111 | 0.374 | 1 | 0.294172432 | 0.000964359 |
| Gna14 | 0.043489312 | 1.134406133 | 0.714 | 0.526 | 1 | 0.000482462 | 1.173865923 | 0.667 | 0.388 | 1 | 0.043489312 | 0.000964692 |
| Ambra1 | 0.003418211 | 0.636389546 | 0.857 | 0.862 | 1 | 0.000484007 | 0.852017829 | 0.956 | 0.882 | 1 | 0.003418211 | 0.00096778 |
| Orai3 | 0.018983831 | -2.392317423 | 0 | 0.213 | 1 | 0.000485311 | -2.824722054 | 0.022 | 0.256 | 1 | 0.018983831 | 0.000970386 |
| Cstf2t | 0.011775054 | -2.666078235 | 0 | 0.239 | 1 | 0.00048638 | -1.513324627 | 0.044 | 0.308 | 1 | 0.011775054 | 0.000972523 |
| Mrpl52 | 0.000486388 | -1.226232442 | 0.381 | 0.765 | 1 | 0.008033609 | -0.664850717 | 0.511 | 0.661 | 1 | 0.008033609 | 0.00097254 |
| Spsb3 | 0.273485928 | -0.133583154 | 0.095 | 0.213 | 1 | 0.000488023 | -1.329751454 | 0.067 | 0.329 | 1 | 0.273485928 | 0.000975807 |
| Nrp1 | 0.786955376 | -0.216999163 | 0.19 | 0.216 | 1 | 0.000489367 | -1.423842618 | 0.067 | 0.343 | 1 | 0.786955376 | 0.000978494 |
| Chrac1 | 0.000489406 | -3.555046923 | 0 | 0.388 | 1 | 0.005810367 | -0.912778231 | 0.089 | 0.294 | 1 | 0.005810367 | 0.000978572 |
| Tanc2 | 0.054704388 | 0.844049033 | 0.762 | 0.597 | 1 | 0.000489502 | 1.001148356 | 0.956 | 0.678 | 1 | 0.054704388 | 0.000978763 |
| Dip2c | 0.01759706 | 0.61090371 | 0.952 | 0.743 | 1 | 0.000490748 | 1.001096227 | 0.978 | 0.882 | 1 | 0.01759706 | 0.000981255 |
| Trim8 | 0.000492966 | -2.469186186 | 0.048 | 0.466 | 1 | 0.213003966 | -0.535634788 | 0.6 | 0.623 | 1 | 0.213003966 | 0.000985688 |
| Dhx58os | 0.000621222 | -1.71385396 | 0.381 | 0.761 | 1 | 0.000493994 | -0.552941606 | 0.289 | 0.651 | 1 | 0.000621222 | 0.000987744 |
| Meaf6 | 0.011750618 | -1.555046923 | 0.095 | 0.381 | 1 | 0.000496099 | -1.017367132 | 0.111 | 0.405 | 1 | 0.011750618 | 0.000991952 |
| Cd47 | 0.00049652 | -1.335217016 | 0.714 | 0.836 | 1 | 0.002768911 | -0.277633113 | 0.911 | 0.879 | 1 | 0.002768911 | 0.000992794 |
| Fbxl17 | 0.165612964 | 1.091264919 | 0.857 | 0.899 | 1 | 0.000496643 | 1.006085589 | 1 | 0.952 | 1 | 0.165612964 | 0.000993039 |
| Kdm6b | 0.43044207 | 0.562263452 | 0.714 | 0.843 | 1 | 0.000497484 | -0.340259918 | 0.911 | 0.91 | 1 | 0.43044207 | 0.00099472 |
| H13 | 0.000499828 | -1.082957081 | 0.619 | 0.914 | 1 | 0.034900143 | -0.289093206 | 0.889 | 0.948 | 1 | 0.034900143 | 0.000999406 |
| Snw1 | 0.000771963 | -1.251700254 | 0.429 | 0.795 | 1 | 0.000500169 | -0.621201644 | 0.8 | 0.844 | 1 | 0.000771963 | 0.001000087 |
| Trappc4 | 0.000501512 | -1.39661756 | 0.19 | 0.649 | 1 | 0.029099663 | -0.678114939 | 0.489 | 0.599 | 1 | 0.029099663 | 0.001002773 |
| Brd3 | 0.020405901 | -0.727107669 | 0.19 | 0.545 | 1 | 0.000501771 | -0.531608994 | 0.711 | 0.81 | 1 | 0.020405901 | 0.00100329 |
| Phf5a | 0.000501997 | -1.516052791 | 0.143 | 0.601 | 1 | 0.044510397 | -0.321928095 | 0.444 | 0.616 | 1 | 0.044510397 | 0.001003741 |
| Ier2 | 0.093353576 | -0.63448139 | 0.667 | 0.754 | 1 | 0.000502009 | -1.14761484 | 0.889 | 0.841 | 1 | 0.093353576 | 0.001003765 |
| Adsl | 0.447497511 | -0.289702356 | 0.143 | 0.216 | 1 | 0.000502337 | -1.428435729 | 0.044 | 0.298 | 1 | 0.447497511 | 0.001004422 |
| Tmem42 | 0.002310147 | -2.486099569 | 0.048 | 0.381 | 1 | 0.000503045 | -2.107004344 | 0.089 | 0.346 | 1 | 0.002310147 | 0.001005837 |
| Mrpl35 | 0.021345083 | -1.818081329 | 0.048 | 0.276 | 1 | 0.000504218 | -1.526380779 | 0.044 | 0.298 | 1 | 0.021345083 | 0.001008182 |
| Erc1 | 0.000505209 | 1.251433903 | 1 | 0.951 | 1 | 0.025394302 | 0.856655598 | 0.956 | 0.986 | 1 | 0.025394302 | 0.001010164 |
| Ndufa9 | 0.000505816 | -2.116305163 | 0.048 | 0.466 | 1 | 0.009843093 | -0.885770249 | 0.244 | 0.446 | 1 | 0.009843093 | 0.001011376 |
| Htra2 | 0.005840109 | -2.094412557 | 0.048 | 0.34 | 1 | 0.000506439 | -1.500149238 | 0.044 | 0.298 | 1 | 0.005840109 | 0.001012621 |
| Lias | 0.118176009 | -0.707050016 | 0.143 | 0.325 | 1 | 0.000506814 | -1.757500005 | 0.067 | 0.325 | 1 | 0.118176009 | 0.001013371 |
| Col25a1 | 0.521957617 | -0.299461323 | 0.286 | 0.216 | 1 | 0.000507124 | 1.256321666 | 0.622 | 0.325 | 1 | 0.521957617 | 0.001013991 |
| Gpatch8 | 0.000507739 | 0.91372256 | 0.905 | 0.884 | 1 | 0.365953294 | 0.450411829 | 1 | 0.958 | 1 | 0.365953294 | 0.001015221 |
| Mrtfb | 0.042707436 | 1.538612184 | 0.524 | 0.463 | 1 | 0.000508414 | 1.103712792 | 0.844 | 0.671 | 1 | 0.042707436 | 0.00101657 |
| Plekhb1 | 0.000513398 | -3.403043829 | 0.048 | 0.44 | 1 | 0.007076526 | -1.856086225 | 0.044 | 0.218 | 1 | 0.007076526 | 0.001026532 |
| Triqk | 0.029537486 | -1.984439715 | 0.048 | 0.257 | 1 | 0.000514093 | -2.174908409 | 0.044 | 0.294 | 1 | 0.029537486 | 0.001027921 |
| Usp24 | 0.000674019 | 1.073858925 | 0.952 | 0.869 | 1 | 0.000514334 | 0.794820944 | 1 | 0.931 | 1 | 0.000674019 | 0.001028403 |
| Eml3 | 0.32513182 | 0.65584986 | 0.143 | 0.246 | 1 | 0.000514605 | -0.80878051 | 0.067 | 0.339 | 1 | 0.32513182 | 0.001028945 |
| Psme4 | 0.00051604 | 0.86899539 | 1 | 0.892 | 1 | 0.095254489 | 0.452194373 | 1 | 0.986 | 1 | 0.095254489 | 0.001031814 |
| Lsm6 | 0.000516484 | -1.574155746 | 0.286 | 0.672 | 1 | 0.002620577 | -0.801363039 | 0.4 | 0.609 | 1 | 0.002620577 | 0.001032702 |
| Pxylp1 | 0.039901023 | -0.767663422 | 0.143 | 0.418 | 1 | 0.000517507 | -1.83350294 | 0.089 | 0.356 | 1 | 0.039901023 | 0.001034746 |
| Fbxo11 | 0.068373143 | 0.569435108 | 1 | 0.802 | 1 | 0.000521766 | 0.604121245 | 0.978 | 0.955 | 1 | 0.068373143 | 0.001043259 |
| Fryl | 0.010543872 | 1.137718867 | 0.857 | 0.687 | 1 | 0.00052225 | 0.850988862 | 0.933 | 0.716 | 1 | 0.010543872 | 0.001044226 |
| Spock2 | 0.004337528 | -0.460352512 | 0.238 | 0.597 | 1 | 0.000522643 | -1.124282336 | 0.111 | 0.419 | 1 | 0.004337528 | 0.001045012 |
| Uqcr11 | 0.000523163 | -1.0047901 | 0.619 | 0.869 | 1 | 0.003237507 | -0.730939908 | 0.733 | 0.768 | 1 | 0.003237507 | 0.001046052 |
| Aamdc | 0.000643874 | -2.674956387 | 0.048 | 0.433 | 1 | 0.000524242 | -0.330733213 | 0.111 | 0.429 | 1 | 0.000643874 | 0.001048209 |
| Mrpl17 | 0.000526449 | -1.461387816 | 0.238 | 0.701 | 1 | 0.003336298 | -0.893574646 | 0.4 | 0.606 | 1 | 0.003336298 | 0.001052621 |
| Wwox | 0.000527853 | 1.590113839 | 0.857 | 0.817 | 1 | 0.09271314 | 0.801056659 | 0.867 | 0.768 | 1 | 0.09271314 | 0.001055427 |
| Nol12 | 0.005987823 | -1.611630451 | 0.095 | 0.414 | 1 | 0.000528123 | -1.056466952 | 0.156 | 0.45 | 1 | 0.005987823 | 0.001055967 |
| Man2b2 | 0.022512472 | -0.865387043 | 0.048 | 0.295 | 1 | 0.000532259 | -1.628513564 | 0.111 | 0.377 | 1 | 0.022512472 | 0.001064236 |
| Timp1 | 0.000534028 | -0.421035016 | 0.429 | 0.828 | 1 | 0.001031093 | -2.072904977 | 0.311 | 0.616 | 1 | 0.001031093 | 0.00106777 |
| Fgf2 | 0.507097909 | -0.116543251 | 0.619 | 0.701 | 1 | 0.000537307 | -0.611548163 | 0.289 | 0.657 | 1 | 0.507097909 | 0.001074326 |
| Zfp53 | 0.832694375 | 0.503846766 | 0.19 | 0.183 | 1 | 0.000537417 | -3.299920989 | 0.044 | 0.287 | 1 | 0.832694375 | 0.001074546 |
| Rnf168 | 0.031040396 | -0.970084422 | 0.19 | 0.466 | 1 | 0.000537832 | -0.569314575 | 0.222 | 0.561 | 1 | 0.031040396 | 0.001075375 |
| Atp13a1 | 0.942866366 | 0.311201688 | 0.238 | 0.235 | 1 | 0.000538274 | -1.379937211 | 0.044 | 0.304 | 1 | 0.942866366 | 0.001076258 |
| Tmtc3 | 0.28266222 | -0.174225139 | 0.095 | 0.198 | 1 | 0.000541008 | -1.064161343 | 0.089 | 0.384 | 1 | 0.28266222 | 0.001081723 |
| Ndufaf4 | 0.028689627 | -0.98647886 | 0.143 | 0.41 | 1 | 0.000541942 | -1.105981519 | 0.178 | 0.484 | 1 | 0.028689627 | 0.001083591 |
| Ttc33 | 0.257399833 | -0.31112134 | 0.143 | 0.272 | 1 | 0.000543268 | -2.124282336 | 0.067 | 0.315 | 1 | 0.257399833 | 0.001086241 |
| Camk1d | 0.000543702 | 2.027547878 | 1 | 1 | 1 | 0.322481835 | 0.48072044 | 1 | 1 | 1 | 0.322481835 | 0.001087107 |
| Arhgap21 | 0.029674799 | 0.953091257 | 0.857 | 0.869 | 1 | 0.00054486 | 0.718975133 | 1 | 0.945 | 1 | 0.029674799 | 0.001089424 |
| Trip10 | 0.108133801 | -0.785659851 | 0.143 | 0.328 | 1 | 0.000545955 | -1.087445568 | 0.133 | 0.446 | 1 | 0.108133801 | 0.001091611 |
| Polr3gl | 0.004201882 | -2.24509147 | 0.048 | 0.354 | 1 | 0.000546622 | -1.316927414 | 0.111 | 0.412 | 1 | 0.004201882 | 0.001092945 |
| Tsen15 | 0.331491216 | -0.366642501 | 0.333 | 0.459 | 1 | 0.000549287 | -1.098287127 | 0.111 | 0.412 | 1 | 0.331491216 | 0.001098273 |
| Gm41307 | 0.000551057 | 4.121230745 | 0.19 | 0.034 | 1 | 0.548948369 | 0.180572246 | 0.022 | 0.042 | 1 | 0.548948369 | 0.00110181 |
| Ccnd1 | 0.000552556 | -3.754588405 | 0.095 | 0.485 | 1 | 0.000910492 | -0.332868958 | 0.689 | 0.322 | 1 | 0.000910492 | 0.001104807 |
| Nasp | 0.377042011 | 0.199285803 | 0.667 | 0.683 | 1 | 0.000557162 | -0.548125733 | 0.867 | 0.862 | 1 | 0.377042011 | 0.001114013 |
| Prpf38b | 0.134810375 | -0.248339565 | 0.714 | 0.847 | 1 | 0.000558823 | -0.600063026 | 0.889 | 0.913 | 1 | 0.134810375 | 0.001117334 |
| Dele1 | 0.000559335 | -3.348596045 | 0 | 0.381 | 1 | 0.017712402 | -0.497499659 | 0.133 | 0.339 | 1 | 0.017712402 | 0.001118357 |
| Gal3st4 | 0.041381787 | -1.891012851 | 0.048 | 0.25 | 1 | 0.000564031 | -2.683249628 | 0.044 | 0.287 | 1 | 0.041381787 | 0.001127743 |
| Ahsa1 | 0.000564771 | -1.805200037 | 0.095 | 0.537 | 1 | 0.184635746 | -0.300926935 | 0.467 | 0.564 | 1 | 0.184635746 | 0.001129223 |
| Glis2 | 0.775913997 | 1.733478014 | 0.286 | 0.317 | 1 | 0.000565078 | -0.313867614 | 0.244 | 0.609 | 1 | 0.775913997 | 0.001129837 |
| Grpel1 | 0.028695769 | -0.389922908 | 0.286 | 0.649 | 1 | 0.000565846 | -0.86582066 | 0.422 | 0.657 | 1 | 0.028695769 | 0.001131372 |
| Lyst | 0.127937981 | -1.326228232 | 0.095 | 0.243 | 1 | 0.000566669 | 0.769487338 | 0.778 | 0.446 | 1 | 0.127937981 | 0.001133018 |
| Orc6 | 0.084874294 | -0.81004401 | 0.19 | 0.396 | 1 | 0.000567291 | -0.964174193 | 0.133 | 0.429 | 1 | 0.084874294 | 0.00113426 |
| Bhlhb9 | 0.032394194 | -1.752492987 | 0.048 | 0.254 | 1 | 0.000567307 | -1.689481582 | 0.089 | 0.356 | 1 | 0.032394194 | 0.001134292 |
| Samd9l | 0.322949537 | -1.326228232 | 0.048 | 0.116 | 1 | 0.000568147 | -1.435868487 | 0.044 | 0.294 | 1 | 0.322949537 | 0.001135972 |
| Klc1 | 0.081612042 | 1.101387333 | 0.667 | 0.806 | 1 | 0.000569249 | 0.810006868 | 0.956 | 0.913 | 1 | 0.081612042 | 0.001138175 |
| Hmgcr | 0.444413408 | -0.641583673 | 0.619 | 0.731 | 1 | 0.00057094 | -0.979279947 | 0.467 | 0.696 | 1 | 0.444413408 | 0.001141554 |

| | | | | | | | | | | | | |
|---|---|---|---|---|---|---|---|---|---|---|---|---|
| Plin2 | 0.000977521 | -4.320581669 | 0 | 0.362 | 1 | 0.000574324 | -1.960783604 | 0.067 | 0.325 | 1 | 0.000977521 | 0.001148319 |
| Mrpl15 | 0.057494981 | -0.867601464 | 0.19 | 0.422 | 1 | 0.000574455 | -0.979892427 | 0.111 | 0.401 | 1 | 0.057494981 | 0.00114858 |
| Thap2 | 0.040535093 | -0.228301812 | 0.238 | 0.56 | 1 | 0.000577367 | -0.952013394 | 0.244 | 0.595 | 1 | 0.040535093 | 0.001154401 |
| Sh3glb1 | 0.007665405 | -0.451681648 | 0.952 | 0.985 | 1 | 0.000579637 | -0.479646131 | 0.978 | 0.993 | 1 | 0.007665405 | 0.001158937 |
| Gde1 | 0.000580998 | -2.648156327 | 0.048 | 0.437 | 1 | 0.494750292 | -0.389683756 | 0.556 | 0.543 | 1 | 0.494750292 | 0.001161658 |
| Wnt11 | 0.203516922 | -0.885195525 | 0.19 | 0.321 | 1 | 0.000582076 | -1.834203107 | 0.089 | 0.353 | 1 | 0.203516922 | 0.001163813 |
| Rhobtb3 | 0.057136283 | -0.556525852 | 0.19 | 0.463 | 1 | 0.000582513 | -1.112786697 | 0.2 | 0.512 | 1 | 0.057136283 | 0.001164686 |
| Bmi1 | 0.003232703 | -2.486099569 | 0.048 | 0.366 | 1 | 0.000582737 | -1.53415813 | 0.133 | 0.405 | 1 | 0.003232703 | 0.001165134 |
| Adprh | 0.00058293 | -1.955584852 | 0.095 | 0.511 | 1 | 0.000902752 | -0.713817566 | 0.111 | 0.401 | 1 | 0.000902752 | 0.001165519 |
| Ctnna1 | 0.00183951 | 0.890913928 | 0.952 | 0.989 | 1 | 0.000583106 | 0.629757967 | 1 | 0.986 | 1 | 0.00183951 | 0.001165872 |
| Mmgt2 | 0.064281908 | -0.899117901 | 0.095 | 0.306 | 1 | 0.000583169 | -1.180425414 | 0.067 | 0.343 | 1 | 0.064281908 | 0.001165997 |
| Tomm5 | 0.00058482 | -1.839297815 | 0.143 | 0.571 | 1 | 0.000651075 | -1.071814916 | 0.289 | 0.55 | 1 | 0.000651075 | 0.001169298 |
| Trappc1 | 0.073587872 | -0.818081329 | 0.095 | 0.287 | 1 | 0.000584841 | -2.255526869 | 0.022 | 0.26 | 1 | 0.073587872 | 0.00116934 |
| Pip4p1 | 0.000585132 | -1.830848625 | 0.143 | 0.556 | 1 | 0.070878407 | 0.328539826 | 0.2 | 0.415 | 1 | 0.070878407 | 0.001169922 |
| Rbck1 | 0.010250991 | -1.648156327 | 0.048 | 0.332 | 1 | 0.000586398 | -1.286553765 | 0.089 | 0.36 | 1 | 0.010250991 | 0.001172452 |
| Ino80e | 0.804044629 | 0.323274521 | 0.238 | 0.291 | 1 | 0.000586583 | -1.498257179 | 0.089 | 0.36 | 1 | 0.804044629 | 0.001172821 |
| Mfsd5 | 0.117675055 | -0.683780237 | 0.095 | 0.257 | 1 | 0.000588299 | -1.787247349 | 0.067 | 0.322 | 1 | 0.117675055 | 0.001176252 |
| Eng | 0.123890196 | -0.524497859 | 0.238 | 0.429 | 1 | 0.000590381 | -0.894112564 | 0.2 | 0.557 | 1 | 0.123890196 | 0.001180413 |
| Ube2n | 0.043212764 | -0.358238627 | 0.619 | 0.91 | 1 | 0.000591315 | -0.4832858 | 0.889 | 0.907 | 1 | 0.043212764 | 0.00118228 |
| Sumf1 | 0.075000581 | -0.619009982 | 0.095 | 0.287 | 1 | 0.000594794 | -2.162417465 | 0.067 | 0.311 | 1 | 0.075000581 | 0.001189235 |
| Baz1a | 0.011212699 | 1.744161096 | 0.667 | 0.433 | 1 | 0.00059778 | 0.390960146 | 0.889 | 0.533 | 1 | 0.011212699 | 0.001195202 |
| Usf1 | 0.120459619 | -1.133583154 | 0.048 | 0.179 | 1 | 0.000599249 | -2.137106376 | 0.067 | 0.311 | 1 | 0.120459619 | 0.001198139 |
| Faah | 0.000599559 | 3.673771768 | 0.19 | 0.034 | 1 | 0.331907298 | -0.316927414 | 0 | 0.021 | 1 | 0.331907298 | 0.001198759 |
| Stxbp5 | 0.000599618 | 1.921293174 | 0.762 | 0.616 | 1 | 0.011295083 | 0.698179479 | 0.889 | 0.716 | 1 | 0.011295083 | 0.001198876 |
| Cct4 | 0.276699766 | -0.296683957 | 0.667 | 0.813 | 1 | 0.000602561 | -0.657286214 | 0.889 | 0.851 | 1 | 0.276699766 | 0.001204759 |
| Aifm1 | 0.001213962 | -1.683780237 | 0.095 | 0.507 | 1 | 0.000603237 | -1.137106376 | 0.178 | 0.481 | 1 | 0.001213962 | 0.001206109 |
| Pole4 | 0.019194121 | -0.946814643 | 0.238 | 0.545 | 1 | 0.000605265 | -0.776359032 | 0.2 | 0.495 | 1 | 0.019194121 | 0.001210163 |
| Ets1 | 0.000606453 | 0.78322291 | 0.952 | 0.937 | 1 | 0.584906696 | 0.131601151 | 1 | 0.965 | 1 | 0.584906696 | 0.001212539 |
| Tmem250-ps | 0.675098175 | -0.184209227 | 0.143 | 0.175 | 1 | 0.000606978 | -3.506751973 | 0 | 0.215 | 1 | 0.675098175 | 0.001213587 |
| Cdkn2c | 0.244489115 | -1.184209227 | 0.095 | 0.201 | 1 | 0.000607633 | -3.085111739 | 0.022 | 0.249 | 1 | 0.244489115 | 0.001214897 |
| Mtif2 | 0.02244583 | -1.103835811 | 0.143 | 0.44 | 1 | 0.000608861 | -0.969866708 | 0.156 | 0.453 | 1 | 0.02244583 | 0.001217351 |
| Grip1 | 0.468123264 | 0.892412054 | 0.238 | 0.179 | 1 | 0.000610399 | 0.502500341 | 0.667 | 0.291 | 1 | 0.468123264 | 0.001220425 |
| Tle3 | 0.116858773 | -0.950719097 | 0.048 | 0.19 | 1 | 0.000612153 | 1.164343784 | 0.622 | 0.329 | 1 | 0.116858773 | 0.001223931 |
| Uqcc3 | 0.001778484 | -2.506137322 | 0.048 | 0.392 | 1 | 0.000612195 | -2.552143876 | 0.044 | 0.28 | 1 | 0.001778484 | 0.001224014 |
| Mrpl11 | 0.004699945 | -1.970084422 | 0.095 | 0.403 | 1 | 0.000612346 | -1.316927414 | 0.067 | 0.322 | 1 | 0.004699945 | 0.001224317 |
| Pde8a | 0.609286083 | 1.147598598 | 0.571 | 0.634 | 1 | 0.00061238 | 1.426610834 | 0.8 | 0.671 | 1 | 0.609286083 | 0.001224386 |
| Parp1 | 0.038668614 | -1.280424543 | 0.095 | 0.321 | 1 | 0.000612548 | -2.035745661 | 0.067 | 0.318 | 1 | 0.038668614 | 0.00122472 |
| Grhpr | 0.039452564 | -2.184209227 | 0 | 0.172 | 1 | 0.000614219 | -3.638855509 | 0 | 0.215 | 1 | 0.039452564 | 0.001228061 |
| Mia3 | 0.116467375 | -0.283392034 | 0.81 | 0.888 | 1 | 0.000615137 | -0.478315733 | 0.956 | 0.924 | 1 | 0.116467375 | 0.001229895 |
| Jpt2 | 0.056457379 | -0.862281133 | 0.095 | 0.302 | 1 | 0.000615184 | -1.638855509 | 0.067 | 0.329 | 1 | 0.056457379 | 0.00122999 |
| Atg101 | 0.000618554 | -3.434752689 | 0 | 0.377 | 1 | 0.002917974 | -1.357954682 | 0.267 | 0.467 | 1 | 0.002917974 | 0.001236726 |
| Cenpp | 0.184459849 | 1.223110359 | 0.095 | 0.239 | 1 | 0.000618809 | -2.124282336 | 0.022 | 0.26 | 1 | 0.184459849 | 0.001237236 |
| Etfdh | 0.128626901 | -0.926141075 | 0.095 | 0.246 | 1 | 0.000618968 | -1.188412673 | 0.111 | 0.394 | 1 | 0.128626901 | 0.001237553 |
| Phb | 0.125397849 | -0.648156327 | 0.19 | 0.377 | 1 | 0.000620831 | -1.095900535 | 0.089 | 0.37 | 1 | 0.125397849 | 0.001241276 |
| Gpalpp1 | 0.103507846 | -0.862281133 | 0.143 | 0.317 | 1 | 0.000621804 | -1.432404631 | 0.089 | 0.356 | 1 | 0.103507846 | 0.001243221 |
| Dhx33 | 0.857311318 | 0.406985227 | 0.19 | 0.231 | 1 | 0.000623095 | -3.003427941 | 0.022 | 0.249 | 1 | 0.857311318 | 0.001245801 |
| Cetn3 | 0.003284343 | -0.985391675 | 0.333 | 0.724 | 1 | 0.000625135 | -0.992492463 | 0.578 | 0.713 | 1 | 0.003284343 | 0.00124988 |
| Mtfr1 | 0.002080563 | -0.837190152 | 0.143 | 0.575 | 1 | 0.000625157 | -0.643656495 | 0.244 | 0.602 | 1 | 0.002080563 | 0.001249923 |
| Mxi1 | 0.672370695 | 0.444953077 | 0.286 | 0.261 | 1 | 0.000626066 | 0.106723216 | 0.822 | 0.394 | 1 | 0.672370695 | 0.001251741 |
| Scd1 | 0.000626143 | -3.03588648 | 0.286 | 0.638 | 1 | 0.001339502 | -1.472845557 | 0.2 | 0.495 | 1 | 0.001339502 | 0.001251895 |
| Grik3 | 0.229775799 | -0.326228232 | 0.048 | 0.149 | 1 | 0.000628646 | -2.674479418 | 0.022 | 0.249 | 1 | 0.229775799 | 0.001256897 |
| Flrt1 | 0.041859842 | -1.252227651 | 0.095 | 0.325 | 1 | 0.0006288 | -1.396654606 | 0.067 | 0.339 | 1 | 0.041859842 | 0.001257205 |
| Slc35b3 | 0.437474026 | 0.3012176 | 0.238 | 0.407 | 1 | 0.000629351 | -1.674479418 | 0.133 | 0.394 | 1 | 0.437474026 | 0.001258306 |
| Mrpl58 | 0.002244644 | -3.133583154 | 0 | 0.321 | 1 | 0.000630059 | -1.624355939 | 0.022 | 0.26 | 1 | 0.002244644 | 0.00125972 |
| Tmtc4 | 0.023226432 | -2.303508156 | 0 | 0.201 | 1 | 0.000631395 | -3.960783604 | 0 | 0.215 | 1 | 0.023226432 | 0.001262392 |
| 2310009B15Rik | 0.077005963 | -0.923163375 | 0.143 | 0.336 | 1 | 0.00063231 | -2.552143876 | 0.044 | 0.28 | 1 | 0.077005963 | 0.00126422 |
| Gm15867 | 0.002749132 | -3.555046923 | 0 | 0.313 | 1 | 0.000633434 | -1.765387915 | 0.044 | 0.291 | 1 | 0.002749132 | 0.001266467 |
| Aldh18a1 | 0.566183391 | 0.510025341 | 0.381 | 0.537 | 1 | 0.00063373 | -0.66055058 | 0.267 | 0.612 | 1 | 0.566183391 | 0.001267058 |
| Use1 | 0.000637093 | -1.098817736 | 0.524 | 0.832 | 1 | 0.037795474 | -0.431223546 | 0.711 | 0.699 | 1 | 0.037795474 | 0.001273779 |
| Commd4 | 0.004965842 | -2.120644099 | 0.048 | 0.347 | 1 | 0.000638586 | -1.276285429 | 0.089 | 0.36 | 1 | 0.004965842 | 0.001276763 |
| Pcolce2 | 0.033177601 | 0.978385277 | 0.095 | 0.336 | 1 | 0.000645578 | -2.373195633 | 0.067 | 0.318 | 1 | 0.033177601 | 0.00129074 |
| Cobl | 0.292445477 | 0.775217031 | 0.571 | 0.578 | 1 | 0.000647139 | 1.340558719 | 0.889 | 0.692 | 1 | 0.292445477 | 0.001293859 |
| Bckdha | 0.199650699 | -0.611630451 | 0.095 | 0.22 | 1 | 0.000652004 | -2.331877755 | 0.022 | 0.256 | 1 | 0.199650699 | 0.001303582 |
| Cdk6 | 0.049295567 | 1.760308518 | 0.714 | 0.683 | 1 | 0.000652247 | 0.669937184 | 0.778 | 0.412 | 1 | 0.049295567 | 0.001304069 |
| Wiz | 0.347107565 | -0.174225139 | 0.143 | 0.243 | 1 | 0.000652453 | -1.564854927 | 0.111 | 0.381 | 1 | 0.347107565 | 0.001304479 |
| Cyp26a1 | 0.268077408 | -2.940938076 | 0 | 0.056 | 1 | 0.000653174 | -4.461585767 | 0.022 | 0.246 | 1 | 0.268077408 | 0.001305921 |
| Abcf2 | 0.807582514 | 0.189572244 | 0.476 | 0.522 | 1 | 0.000654112 | -0.704944699 | 0.356 | 0.668 | 1 | 0.807582514 | 0.001307795 |
| Sipa1l1 | 0.058859232 | 0.781527948 | 0.81 | 0.761 | 1 | 0.000658227 | 0.79687582 | 1 | 0.827 | 1 | 0.058859232 | 0.00131602 |
| Rap1gds1 | 0.01459713 | 1.313369525 | 0.81 | 0.698 | 1 | 0.000658541 | 1.128085145 | 0.956 | 0.799 | 1 | 0.01459713 | 0.001316648 |
| Zfp511 | 0.077166283 | -0.828728573 | 0.048 | 0.22 | 1 | 0.000659431 | -2.255526869 | 0.022 | 0.256 | 1 | 0.077166283 | 0.001318428 |
| Pdzd2 | 0.360194516 | 0.323274521 | 0.524 | 0.489 | 1 | 0.000661709 | 1.239601075 | 0.733 | 0.512 | 1 | 0.360194516 | 0.00132298 |
| Mdfi | 0.128287227 | -0.368145621 | 0.571 | 0.836 | 1 | 0.000665423 | -0.676008507 | 0.4 | 0.799 | 1 | 0.128287227 | 0.001330404 |
| Lepr | 0.07934153 | -0.636083495 | 0.143 | 0.343 | 1 | 0.000665482 | -0.894112564 | 0.178 | 0.505 | 1 | 0.07934153 | 0.00133052 |
| Chmp6 | 0.028415202 | -2.280424543 | 0 | 0.19 | 1 | 0.000666982 | -2.124282336 | 0.022 | 0.256 | 1 | 0.028415202 | 0.001333519 |
| Farsa | 0.139520384 | -0.385121921 | 0.19 | 0.369 | 1 | 0.000667987 | -1.00994966 | 0.156 | 0.453 | 1 | 0.139520384 | 0.001335528 |
| Tpm1 | 0.019994377 | -0.958246219 | 1 | 0.97 | 1 | 0.000671267 | -0.979195641 | 0.933 | 0.952 | 1 | 0.019994377 | 0.001342083 |
| Zbtb38 | 0.00067504 | 1.212809979 | 0.81 | 0.884 | 1 | 0.423286052 | 0.748167614 | 0.844 | 0.837 | 1 | 0.423286052 | 0.001349625 |
| Far1 | 0.000677369 | 1.021285273 | 1 | 0.784 | 1 | 0.216639313 | 0.506118606 | 0.889 | 0.893 | 1 | 0.216639313 | 0.00135428 |
| 9130401M01Rik | 0.015916524 | -1.926141075 | 0.048 | 0.291 | 1 | 0.000678248 | -2.044847868 | 0.044 | 0.287 | 1 | 0.015916524 | 0.001356035 |
| Golim4 | 0.001391272 | -0.206025167 | 0.667 | 0.955 | 1 | 0.000678866 | -0.49397685 | 0.889 | 0.955 | 1 | 0.001391272 | 0.001357271 |
| Map2k3 | 0.051471294 | -0.29627501 | 0.81 | 0.918 | 1 | 0.000679344 | -0.562971103 | 0.844 | 0.9 | 1 | 0.051471294 | 0.001358227 |
| Get4 | 0.009952347 | -1.192476843 | 0.143 | 0.448 | 1 | 0.000679562 | -0.508547486 | 0.178 | 0.516 | 1 | 0.009952347 | 0.001358662 |
| Cdk8 | 0.000679565 | 0.760787414 | 0.952 | 0.948 | 1 | 0.265731238 | 0.345129211 | 0.978 | 0.945 | 1 | 0.265731238 | 0.001358669 |
| Esyt2 | 0.006700105 | 1.001825965 | 0.905 | 0.892 | 1 | 0.000682044 | 1.013112432 | 1 | 0.917 | 1 | 0.006700105 | 0.001363623 |
| Med10 | 0.000685568 | -1.295854583 | 0.048 | 0.466 | 1 | 0.004270835 | -0.559783938 | 0.133 | 0.381 | 1 | 0.004270835 | 0.001370666 |
| Clasp2 | 0.000686889 | 0.802681893 | 0.952 | 0.854 | 1 | 0.059443807 | 0.402964667 | 0.956 | 0.862 | 1 | 0.059443807 | 0.001373306 |
| Borcs7 | 0.015191262 | -0.998653574 | 0.095 | 0.377 | 1 | 0.000687963 | -2.154870656 | 0.089 | 0.332 | 1 | 0.015191262 | 0.001375453 |
| Acp1 | 0.000820814 | -1.617782678 | 0.095 | 0.53 | 1 | 0.000689772 | -1.076664316 | 0.356 | 0.606 | 1 | 0.000820814 | 0.001379068 |
| Septin5 | 0.094262028 | -1.233118828 | 0.048 | 0.194 | 1 | 0.000693077 | -3.545746104 | 0 | 0.211 | 1 | 0.094262028 | 0.001385674 |
| Nynrin | 0.086240508 | -1.326228232 | 0.048 | 0.209 | 1 | 0.000694095 | -2.071814916 | 0.089 | 0.339 | 1 | 0.086240508 | 0.001387707 |
| Arfgef1 | 0.122468872 | 0.66367589 | 0.81 | 0.851 | 1 | 0.000695241 | 0.712128619 | 1 | 0.941 | 1 | 0.122468872 | 0.00139 |
| Ccdc9 | 0.969961467 | 0.129451251 | 0.286 | 0.287 | 1 | 0.000696975 | -0.721317669 | 0.111 | 0.408 | 1 | 0.969961467 | 0.001393464 |
| Ak6 | 0.021338706 | -1.213753503 | 0.095 | 0.369 | 1 | 0.000698038 | -0.757500005 | 0.178 | 0.505 | 1 | 0.021338706 | 0.001395589 |
| Eif4e | 0.000700363 | 0.716207034 | 0.952 | 0.918 | 1 | 0.601033479 | 0.149690638 | 0.978 | 0.955 | 1 | 0.601033479 | 0.001400235 |
| Lonp1 | 0.59093481 | 0.150209812 | 0.19 | 0.257 | 1 | 0.000700744 | -1.271123724 | 0.067 | 0.318 | 1 | 0.59093481 | 0.001400998 |
| D030056L22Rik | 0.257582898 | -0.154047257 | 0.095 | 0.205 | 1 | 0.000701825 | -3.84048937 | 0 | 0.211 | 1 | 0.257582898 | 0.001403158 |
| Ephb3 | 0.322831128 | -0.496153234 | 0.143 | 0.239 | 1 | 0.000702517 | -1.434924623 | 0.156 | 0.422 | 1 | 0.322831128 | 0.00140454 |
| Fuz | 0.181779327 | -0.496153234 | 0.095 | 0.239 | 1 | 0.000704455 | -3.75987091 | 0 | 0.211 | 1 | 0.181779327 | 0.001408414 |

| | | | | | | | | | | | | |
|---|---|---|---|---|---|---|---|---|---|---|---|---|
| Hacd4 | 0.422167493 | -0.233118828 | 0.143 | 0.231 | 1 | 0.000705417 | -2.638855509 | 0.022 | 0.253 | 1 | 0.422167493 | 0.001410335 |
| Cep250 | 0.179565772 | -1.280424543 | 0.048 | 0.153 | 1 | 0.000706032 | -3.84048937 | 0 | 0.211 | 1 | 0.179565772 | 0.001411566 |
| 4931406C07Rik | 0.003400665 | -2.063193826 | 0.048 | 0.384 | 1 | 0.00070646 | -1.356455778 | 0.089 | 0.363 | 1 | 0.003400665 | 0.001412422 |
| Lsm4 | 0.000707093 | -1.455511249 | 0.19 | 0.619 | 1 | 0.293444828 | -0.223818009 | 0.489 | 0.505 | 1 | 0.293444828 | 0.001413685 |
| Gng12 | 0.049852873 | 0.732893173 | 0.857 | 0.903 | 1 | 0.000708274 | 0.926469836 | 1 | 0.952 | 1 | 0.049852873 | 0.001416046 |
| Slc35b1 | 0.000708358 | -1.192476843 | 0.238 | 0.69 | 1 | 0.090522232 | -0.562552141 | 0.511 | 0.595 | 1 | 0.090522232 | 0.001416214 |
| Ppp1r7 | 0.022896993 | -1.213753503 | 0.095 | 0.366 | 1 | 0.000709179 | -1.300439291 | 0.089 | 0.363 | 1 | 0.022896993 | 0.001417855 |
| Kcnq1ot1 | 0.00070925 | 1.287745176 | 0.952 | 0.929 | 1 | 0.007770889 | 0.656224685 | 1 | 0.99 | 1 | 0.007770889 | 0.001417998 |
| Gltp | 0.000709425 | -2.970084422 | 0.048 | 0.429 | 1 | 0.371644706 | 0.207987703 | 0.444 | 0.343 | 1 | 0.371644706 | 0.001418347 |
| Elapor1 | 0.186946218 | 1.866416846 | 0.143 | 0.067 | 1 | 0.000709749 | 1.894576691 | 0.178 | 0.045 | 1 | 0.186946218 | 0.001418994 |
| Rhoa | 0.017026482 | -0.553519453 | 0.905 | 0.989 | 1 | 0.000710727 | -0.405780805 | 0.978 | 0.997 | 1 | 0.017026482 | 0.001420948 |
| Gne | 0.210387943 | -0.735619168 | 0.095 | 0.213 | 1 | 0.000711798 | -0.731964913 | 0.022 | 0.256 | 1 | 0.210387943 | 0.00142309 |
| Galt | 0.016390081 | -2.370622352 | 0 | 0.22 | 1 | 0.000712428 | -2.776359032 | 0.022 | 0.246 | 1 | 0.016390081 | 0.001424348 |
| Tsr3 | 0.042577034 | -1.701267664 | 0.048 | 0.239 | 1 | 0.000712916 | -2.72631835 | 0.022 | 0.246 | 1 | 0.042577034 | 0.001425324 |
| Naglu | 0.004079775 | -2.818081329 | 0 | 0.291 | 1 | 0.000713099 | -2.792660845 | 0.022 | 0.246 | 1 | 0.004079775 | 0.001425689 |
| Cdc42se1 | 0.000718122 | -1.071186617 | 0.476 | 0.843 | 1 | 0.001355784 | -0.613818958 | 0.867 | 0.855 | 1 | 0.001355784 | 0.001435728 |
| Stk39 | 0.000719718 | 1.740678426 | 0.857 | 0.884 | 1 | 0.001283919 | 1.088088166 | 0.956 | 0.893 | 1 | 0.001283919 | 0.001438918 |
| Pycr1 | 0.077015258 | -0.718545655 | 0.095 | 0.299 | 1 | 0.000720467 | -1.815178281 | 0.044 | 0.287 | 1 | 0.077015258 | 0.001440415 |
| Rora | 0.003954924 | 0.950956041 | 1 | 0.996 | 1 | 0.000722024 | 0.952285567 | 1 | 0.986 | 1 | 0.003954924 | 0.001443527 |
| Lgals8 | 0.321300473 | 0.238385623 | 0.19 | 0.336 | 1 | 0.000722639 | -0.629810369 | 0.089 | 0.367 | 1 | 0.321300473 | 0.001444757 |
| Mrpl57 | 0.001718806 | -2.233118828 | 0.095 | 0.444 | 1 | 0.000726744 | -1.751555641 | 0.133 | 0.391 | 1 | 0.001718806 | 0.001452959 |
| Nomo1 | 0.061519992 | -0.729950418 | 0.143 | 0.373 | 1 | 0.000728508 | -1.292124023 | 0.133 | 0.412 | 1 | 0.061519992 | 0.001456486 |
| Snhg8 | 0.000730001 | -1.03170354 | 0.476 | 0.843 | 1 | 0.009223554 | -0.582517525 | 0.6 | 0.706 | 1 | 0.009223554 | 0.001459468 |
| Ddx39a | 0.991364558 | 0.518493542 | 0.238 | 0.246 | 1 | 0.000730446 | -1.553966611 | 0.089 | 0.353 | 1 | 0.991364558 | 0.001460358 |
| ENSMUSG00000121322 | 0.001488599 | -2.159118246 | 0.048 | 0.422 | 1 | 0.000731014 | -1.509572492 | 0.044 | 0.298 | 1 | 0.001488599 | 0.001461493 |
| Dcaf11 | 0.001139833 | -3.303508156 | 0 | 0.351 | 1 | 0.000732195 | -0.941418279 | 0.111 | 0.384 | 1 | 0.001139833 | 0.001463854 |
| Ankrd46 | 0.112659057 | 0.438555306 | 0.143 | 0.34 | 1 | 0.000734106 | -1.375821103 | 0.111 | 0.388 | 1 | 0.112659057 | 0.001467672 |
| Ust | 0.000734758 | 1.819212636 | 0.952 | 0.754 | 1 | 0.006521133 | 0.659368326 | 0.978 | 0.945 | 1 | 0.006521133 | 0.001468975 |
| Mob1a | 0.015281413 | -1.032820177 | 0.19 | 0.507 | 1 | 0.000735471 | -0.786954148 | 0.578 | 0.727 | 1 | 0.015281413 | 0.001470402 |
| A2m | 0.000737154 | -5.304938691 | 0.048 | 0.422 | 1 | 0.469733857 | -0.75987091 | 0.044 | 0.076 | 1 | 0.469733857 | 0.001473765 |
| Slc9a6 | 0.853925902 | 1.154397609 | 0.238 | 0.261 | 1 | 0.000737951 | -1.336826971 | 0.089 | 0.353 | 1 | 0.853925902 | 0.001475357 |
| Pfkl | 0.000740951 | -1.76585137 | 0.095 | 0.526 | 1 | 0.004048445 | -0.57996182 | 0.111 | 0.346 | 1 | 0.004048445 | 0.001481353 |
| Impa1 | 0.001155566 | -0.962852853 | 0.095 | 0.496 | 1 | 0.000741256 | -1.103523776 | 0.356 | 0.592 | 1 | 0.001155566 | 0.001481962 |
| Snrnp200 | 0.022579584 | -1.142145168 | 0.143 | 0.422 | 1 | 0.000746036 | -0.880828299 | 0.311 | 0.595 | 1 | 0.022579584 | 0.001491516 |
| Itpr1 | 0.862552407 | 0.19190276 | 0.952 | 0.854 | 1 | 0.000746714 | 0.889470819 | 1 | 0.972 | 1 | 0.862552407 | 0.001492871 |
| Dazap2 | 0.000747195 | -1.509450056 | 0.286 | 0.657 | 1 | 0.881653462 | -0.111066716 | 0.844 | 0.664 | 1 | 0.881653462 | 0.001493831 |
| Dpf2 | 0.051698657 | -1.054148687 | 0.095 | 0.306 | 1 | 0.000751322 | -0.526380779 | 0.2 | 0.512 | 1 | 0.051698657 | 0.00150208 |
| Gspt1 | 0.000755761 | -0.728529744 | 0.714 | 0.937 | 1 | 0.056292442 | -0.295865798 | 0.978 | 0.976 | 1 | 0.056292442 | 0.00151095 |
| Mrpl3 | 0.230648066 | -0.10233222 | 0.238 | 0.418 | 1 | 0.000757556 | -1.116333583 | 0.178 | 0.464 | 1 | 0.230648066 | 0.001514537 |
| Pth1r | 0.025482423 | -1.836285924 | 0.238 | 0.53 | 1 | 0.000764358 | -1.146197112 | 0.2 | 0.516 | 1 | 0.025482423 | 0.001528132 |
| Prpf8 | 0.00076488 | -1.213753503 | 0.19 | 0.657 | 1 | 0.04528759 | -0.509572492 | 0.622 | 0.73 | 1 | 0.04528759 | 0.001529175 |
| Arpp19 | 0.061851159 | -0.599246727 | 0.619 | 0.772 | 1 | 0.000766379 | -0.673902456 | 0.756 | 0.841 | 1 | 0.061851159 | 0.00153217 |
| Gm17024 | 0.000766448 | 2.936806174 | 0.19 | 0.034 | 1 | 0.661337779 | 0.290755163 | 0.044 | 0.062 | 1 | 0.661337779 | 0.001532309 |
| Rnf41 | 0.010976264 | -1.191298652 | 0.143 | 0.478 | 1 | 0.000766513 | -0.723552673 | 0.356 | 0.702 | 1 | 0.010976264 | 0.001532439 |
| Psma5 | 0.000767438 | -1.031100197 | 0.619 | 0.862 | 1 | 0.002462641 | -0.356663166 | 0.867 | 0.889 | 1 | 0.002462641 | 0.001534287 |
| Setd2 | 0.000768741 | 1.263683484 | 1 | 0.866 | 1 | 0.938824211 | 0.126737885 | 1 | 0.972 | 1 | 0.938824211 | 0.001536891 |
| Tcp11l2 | 0.000768798 | -1.676725479 | 0.238 | 0.642 | 1 | 0.158026901 | -0.455341443 | 0.822 | 0.744 | 1 | 0.158026901 | 0.001537004 |
| Camk2d | 0.000771001 | 1.650151537 | 1 | 0.974 | 1 | 0.001400695 | 1.042107513 | 1 | 0.986 | 1 | 0.001400695 | 0.001541409 |
| Dnmt1 | 0.730271622 | 0.223738847 | 0.286 | 0.384 | 1 | 0.000771993 | -0.960783604 | 0.178 | 0.491 | 1 | 0.730271622 | 0.00154339 |
| Ano1 | 0.547777007 | 2.295260144 | 0.048 | 0.026 | 1 | 0.000772766 | 2.188307894 | 0.2 | 0.055 | 1 | 0.547777007 | 0.001544934 |
| Atp6v0a2 | 0.003540317 | -1.834022873 | 0.143 | 0.463 | 1 | 0.000774396 | -0.805674599 | 0.289 | 0.644 | 1 | 0.003540317 | 0.001548191 |
| Sema5a | 0.066063972 | -1.811655059 | 0.143 | 0.343 | 1 | 0.000774718 | -0.870181055 | 0.156 | 0.478 | 1 | 0.066063972 | 0.001548836 |
| Fxr2 | 0.000777164 | -1.81004401 | 0.048 | 0.459 | 1 | 0.099372527 | -0.105845088 | 0.467 | 0.623 | 1 | 0.099372527 | 0.001553725 |
| Ndufv1 | 0.00078196 | -2.727107669 | 0.048 | 0.425 | 1 | 0.985519683 | -0.170086025 | 0.556 | 0.509 | 1 | 0.985519683 | 0.001563309 |
| Gpc4 | 0.029310988 | -2.150656668 | 0.095 | 0.313 | 1 | 0.000783363 | -0.150633663 | 0.156 | 0.453 | 1 | 0.029310988 | 0.001566112 |
| Zcchc10 | 0.05082931 | -1.998653574 | 0 | 0.157 | 1 | 0.000783978 | -3.620708162 | 0 | 0.208 | 1 | 0.05082931 | 0.001567342 |
| Zbtb41 | 0.021859878 | -1.348596045 | 0.095 | 0.351 | 1 | 0.000784276 | -0.4734319 | 0.156 | 0.495 | 1 | 0.021859878 | 0.001567936 |
| Dtd1 | 0.008458775 | -3.196592952 | 0 | 0.257 | 1 | 0.000784735 | -0.994999319 | 0.111 | 0.388 | 1 | 0.008458775 | 0.001568854 |
| Cdc73 | 0.000785358 | 1.055642403 | 0.905 | 0.802 | 1 | 0.566815078 | 0.311816779 | 0.956 | 0.872 | 1 | 0.566815078 | 0.001570099 |
| Htt | 0.00078582 | 1.569030935 | 0.762 | 0.649 | 1 | 0.178440746 | 0.656862707 | 0.822 | 0.782 | 1 | 0.178440746 | 0.001571023 |
| Slc3a2 | 0.430185287 | -0.167006156 | 0.857 | 0.858 | 1 | 0.000785893 | -0.449132749 | 0.978 | 0.934 | 1 | 0.430185287 | 0.001571169 |
| Nherf2 | 0.023305571 | -2.434752689 | 0 | 0.201 | 1 | 0.000785905 | -3.564854927 | 0 | 0.208 | 1 | 0.023305571 | 0.001571192 |
| Pck2 | 0.448159288 | 0.788414334 | 0.19 | 0.295 | 1 | 0.000785955 | -0.849422495 | 0.089 | 0.353 | 1 | 0.448159288 | 0.001571293 |
| Lgals3bp | 0.002095192 | -3.303508156 | 0 | 0.325 | 1 | 0.000785956 | -0.868723051 | 0.067 | 0.315 | 1 | 0.002095192 | 0.001571294 |
| Smim15 | 0.000787157 | -1.91868527 | 0.048 | 0.459 | 1 | 0.004735237 | -0.745164411 | 0.178 | 0.412 | 1 | 0.004735237 | 0.001573694 |
| Ang | 0.00474286 | -2.849790188 | 0.048 | 0.347 | 1 | 0.000787781 | -1.017367132 | 0.133 | 0.436 | 1 | 0.00474286 | 0.001574941 |
| Flywch2 | 0.599749006 | 0.866416846 | 0.095 | 0.142 | 1 | 0.000788063 | -3.526380779 | 0 | 0.208 | 1 | 0.599749006 | 0.001575504 |
| Uba3 | 0.828208793 | 0.296251347 | 0.524 | 0.642 | 1 | 0.000788177 | -0.416974591 | 0.311 | 0.696 | 1 | 0.828208793 | 0.001575732 |
| Ubxn6 | 0.000789052 | -1.602352638 | 0.19 | 0.619 | 1 | 0.007293251 | -0.761712257 | 0.467 | 0.626 | 1 | 0.007293251 | 0.001577481 |
| Zfp30 | 0.136444758 | -1.785659851 | 0 | 0.097 | 1 | 0.000789552 | -3.674479418 | 0 | 0.208 | 1 | 0.136444758 | 0.001578481 |
| Wfdc1 | 0.16360158 | -1.326228232 | 0 | 0.086 | 1 | 0.000791609 | -3.792660845 | 0 | 0.208 | 1 | 0.16360158 | 0.001582592 |
| Dusp23 | 0.126525826 | -0.807354922 | 0.048 | 0.187 | 1 | 0.000793268 | -3.80878051 | 0 | 0.208 | 1 | 0.126525826 | 0.001585907 |
| Tsr1 | 0.031173783 | -1.227765593 | 0.286 | 0.504 | 1 | 0.000797271 | -0.912778231 | 0.311 | 0.626 | 1 | 0.031173783 | 0.001593906 |
| 1110038F14Rik | 0.06631946 | -1.413691074 | 0.095 | 0.272 | 1 | 0.000797432 | -3.085111739 | 0.022 | 0.242 | 1 | 0.06631946 | 0.001594227 |
| Ubac1 | 0.08695652 | -1.067695219 | 0.095 | 0.272 | 1 | 0.000801819 | -2.071814916 | 0.067 | 0.311 | 1 | 0.08695652 | 0.001602995 |
| Mrpl34 | 0.000802177 | -3.268742738 | 0 | 0.366 | 1 | 0.018169618 | -0.571741313 | 0.089 | 0.26 | 1 | 0.018169618 | 0.001603711 |
| Ufl1 | 0.000803268 | -1.555046923 | 0.095 | 0.53 | 1 | 0.053768838 | -0.243717567 | 0.444 | 0.592 | 1 | 0.053768838 | 0.001605891 |
| Dennd4c | 0.56182332 | 0.55829455 | 0.381 | 0.373 | 1 | 0.000804201 | 0.611932636 | 0.956 | 0.727 | 1 | 0.56182332 | 0.001607755 |
| Rragc | 0.15875564 | 0.199840579 | 0.333 | 0.616 | 1 | 0.000804737 | -0.811692106 | 0.733 | 0.803 | 1 | 0.15875564 | 0.001608826 |
| Ppp4r3a | 0.95978459 | 0.425844254 | 0.571 | 0.701 | 1 | 0.000805728 | 0.763515579 | 0.956 | 0.865 | 1 | 0.95978459 | 0.001610806 |
| Pcbd2 | 0.086493397 | -0.361852142 | 0.19 | 0.433 | 1 | 0.000806292 | -1.076664316 | 0.089 | 0.363 | 1 | 0.086493397 | 0.001611934 |
| Srbd1 | 0.001072055 | -1.609316585 | 0.381 | 0.735 | 1 | 0.000807073 | -0.371693217 | 0.333 | 0.723 | 1 | 0.001072055 | 0.001613494 |
| Bbip1 | 0.027260235 | -0.98647886 | 0.19 | 0.463 | 1 | 0.000808308 | -0.711990213 | 0.222 | 0.533 | 1 | 0.027260235 | 0.001615963 |
| Bms1 | 0.097218442 | -0.807354922 | 0.286 | 0.466 | 1 | 0.000809154 | -0.761712257 | 0.244 | 0.564 | 1 | 0.097218442 | 0.001617654 |
| Ift57 | 0.185929015 | -0.564632972 | 0.238 | 0.396 | 1 | 0.000810205 | -1.387316742 | 0.111 | 0.384 | 1 | 0.185929015 | 0.001619754 |
| Mettl21a | 0.010915347 | -2.648156327 | 0 | 0.243 | 1 | 0.000810625 | -2.691966845 | 0.022 | 0.242 | 1 | 0.010915347 | 0.001620592 |
| Ing4 | 0.104968116 | -0.38923803 | 0.238 | 0.496 | 1 | 0.000810747 | -0.658395695 | 0.2 | 0.509 | 1 | 0.104968116 | 0.001620836 |
| Otub1 | 0.000811628 | -1.818081329 | 0.095 | 0.5 | 1 | 0.003996778 | -0.667424661 | 0.244 | 0.498 | 1 | 0.003996778 | 0.001622598 |
| Rasl11b | 0.34061799 | -1.067695219 | 0.095 | 0.175 | 1 | 0.000812231 | -4.545746104 | 0 | 0.208 | 1 | 0.34061799 | 0.001623802 |
| Rrp8 | 0.128796241 | -0.837190152 | 0.143 | 0.302 | 1 | 0.000812644 | -0.989352756 | 0.089 | 0.363 | 1 | 0.128796241 | 0.001624627 |
| Srsf1 | 0.12124674 | -0.448624864 | 0.476 | 0.724 | 1 | 0.000812842 | -0.703950537 | 0.8 | 0.813 | 1 | 0.12124674 | 0.001625023 |
| Map3k10 | 0.598005961 | 0.569435108 | 0.19 | 0.261 | 1 | 0.000814078 | -1.255526869 | 0.133 | 0.408 | 1 | 0.598005961 | 0.001627493 |
| Cnep1r1 | 0.000815222 | -1.191842552 | 0.238 | 0.731 | 1 | 0.002806329 | -0.671770131 | 0.467 | 0.779 | 1 | 0.002806329 | 0.001629778 |
| Etv5 | 0.012821255 | -3.146407195 | 0 | 0.235 | 1 | 0.000816334 | -0.871516266 | 0.133 | 0.419 | 1 | 0.012821255 | 0.001632002 |
| Hps5 | 0.162670585 | -0.40151636 | 0.143 | 0.321 | 1 | 0.000818553 | -0.997648897 | 0.156 | 0.433 | 1 | 0.162670585 | 0.001636437 |
| Gadd45a | 0.005373954 | -2.239117569 | 0.143 | 0.485 | 1 | 0.000819611 | -0.960281709 | 0.489 | 0.699 | 1 | 0.005373954 | 0.00163855 |
| Ctnnbl1 | 0.031341019 | -1.120644099 | 0.19 | 0.466 | 1 | 0.000822141 | -1.189466505 | 0.178 | 0.481 | 1 | 0.031341019 | 0.001643607 |
| Oip5os1 | 0.002431301 | -1.196592952 | 0.429 | 0.746 | 1 | 0.000823152 | -0.573938032 | 0.867 | 0.872 | 1 | 0.002431301 | 0.001645626 |

| | | | | | | | | | | | | |
|---|---|---|---|---|---|---|---|---|---|---|---|---|
| Sbno2 | 0.015794526 | 0.874958582 | 0.952 | 0.757 | 1 | 0.00082575 | 0.783925188 | 0.867 | 0.623 | 1 | 0.015794526 | 0.001650818 |
| Ccdc97 | 0.118622936 | -0.348596045 | 0.048 | 0.194 | 1 | 0.000828925 | -2.375821103 | 0.044 | 0.273 | 1 | 0.118622936 | 0.001657163 |
| Fam107a | 1 | 0.503846766 | 0.048 | 0.049 | 1 | 0.000832956 | -2.331877755 | 0.044 | 0.284 | 1 | 1 | 0.001665218 |
| Nabp2 | 0.018312179 | -1.252227651 | 0.143 | 0.414 | 1 | 0.000834219 | -0.589945908 | 0.133 | 0.443 | 1 | 0.018312179 | 0.001667741 |
| Decr1 | 0.009496355 | -1.763633545 | 0.095 | 0.373 | 1 | 0.000839762 | -1.294207337 | 0.067 | 0.325 | 1 | 0.009496355 | 0.001678819 |
| BC029722 | 0.00175687 | -3.159118246 | 0 | 0.332 | 1 | 0.000839973 | -2.331877755 | 0.044 | 0.273 | 1 | 0.00175687 | 0.00167924 |
| Pex14 | 0.646154374 | 0.607682577 | 0.429 | 0.459 | 1 | 0.000844486 | 1.1444488 | 0.778 | 0.588 | 1 | 0.646154374 | 0.001688258 |
| Gabarapl1 | 0.000844546 | -1.525900577 | 0.286 | 0.683 | 1 | 0.03030646 | -0.602329633 | 0.711 | 0.744 | 1 | 0.03030646 | 0.00168838 |
| Parl | 0.000846512 | -1.288160191 | 0.095 | 0.507 | 1 | 0.094344307 | 0.217409014 | 0.289 | 0.502 | 1 | 0.094344307 | 0.001692308 |
| Gosr1 | 0.775102936 | 0.731257262 | 0.333 | 0.343 | 1 | 0.000846678 | -1.316927414 | 0.133 | 0.408 | 1 | 0.775102936 | 0.00169264 |
| Chmp2b | 0.000847729 | -1.063193826 | 0.238 | 0.668 | 1 | 0.005795527 | -0.539319835 | 0.644 | 0.737 | 1 | 0.005795527 | 0.00169474 |
| Ypel3 | 0.00084841 | -1.093394064 | 0.524 | 0.847 | 1 | 0.000924517 | -0.705313785 | 0.778 | 0.848 | 1 | 0.000924517 | 0.001696101 |
| Ccdc59 | 0.031736878 | -0.744482444 | 0.429 | 0.713 | 1 | 0.000849618 | -0.663730177 | 0.578 | 0.734 | 1 | 0.031736878 | 0.001698515 |
| Mfsd10 | 0.009268036 | -0.616905393 | 0.095 | 0.403 | 1 | 0.000849983 | -1.276285429 | 0.089 | 0.349 | 1 | 0.009268036 | 0.001699244 |
| Tfpt | 0.070628805 | -1.008052272 | 0.048 | 0.224 | 1 | 0.000850172 | -2.331877755 | 0.044 | 0.273 | 1 | 0.070628805 | 0.001699621 |
| Ogfr | 0.02827575 | -0.77835505 | 0.19 | 0.489 | 1 | 0.000853651 | -1.112107622 | 0.222 | 0.512 | 1 | 0.02827575 | 0.001706574 |
| Arhgap17 | 0.000854873 | 1.236907906 | 0.905 | 0.784 | 1 | 0.002655171 | 1.085358178 | 0.933 | 0.834 | 1 | 0.002655171 | 0.001709015 |
| Phactr2 | 0.000859397 | -1.475975352 | 0.19 | 0.601 | 1 | 0.229864388 | -0.143182541 | 0.8 | 0.699 | 1 | 0.229864388 | 0.001718056 |
| Atp5md | 0.000860892 | -0.918386234 | 0.524 | 0.847 | 1 | 0.012901096 | -0.39313798 | 0.689 | 0.751 | 1 | 0.012901096 | 0.001721043 |
| Chsy1 | 0.032062764 | -0.669217943 | 0.333 | 0.601 | 1 | 0.000864546 | -0.614263521 | 0.4 | 0.775 | 1 | 0.032062764 | 0.001728344 |
| Heatr1 | 0.79707217 | 0.380990018 | 0.286 | 0.28 | 1 | 0.000865907 | -2.02527233 | 0.089 | 0.336 | 1 | 0.79707217 | 0.001731064 |
| Alx1 | 0.019588052 | -0.624627792 | 0.524 | 0.836 | 1 | 0.000866331 | -0.696775485 | 0.556 | 0.882 | 1 | 0.019588052 | 0.001731911 |
| Gna13 | 0.000866884 | -0.588213787 | 0.524 | 0.854 | 1 | 0.199014929 | -0.154496949 | 0.911 | 0.83 | 1 | 0.199014929 | 0.001733017 |
| Tex30 | 0.039623293 | -2.455511249 | 0 | 0.172 | 1 | 0.000867336 | -2.709244837 | 0.044 | 0.27 | 1 | 0.039623293 | 0.001733921 |
| Sntg1 | 0.000869248 | 4.410737362 | 0.095 | 0.007 | 1 | 0.309824707 | 2.683072586 | 0.022 | 0.007 | 1 | 0.309824707 | 0.00173774 |
| Gcnt7 | 0.000869248 | 4.410737362 | 0.095 | 0.007 | 1 | 0.430487676 | 0.361144491 | 0 | 0.014 | 1 | 0.430487676 | 0.00173774 |
| Slc5a5 | 0.000869248 | 4.410737362 | 0.095 | 0.007 | 1 | 0.424938789 | 1.683072586 | 0.067 | 0.042 | 1 | 0.424938789 | 0.00173774 |
| Gm36279 | 0.000869248 | 4.410737362 | 0.095 | 0.007 | 1 | 0.23197796 | -0.776359032 | 0 | 0.031 | 1 | 0.23197796 | 0.00173774 |
| Kctd16 | 0.000869279 | 5.761234609 | 0.095 | 0.007 | 1 | 0.232011406 | -1.709244837 | 0 | 0.031 | 1 | 0.232011406 | 0.001737803 |
| Zdbf2 | 0.018457153 | -1.666078235 | 0.095 | 0.366 | 1 | 0.000870183 | -1.115669206 | 0.156 | 0.446 | 1 | 0.018457153 | 0.001739608 |
| Fus | 0.000870258 | 0.83578865 | 1 | 0.974 | 1 | 0.071950566 | 0.338609925 | 1 | 1 | 1 | 0.071950566 | 0.001739759 |
| Pomgnt1 | 0.072535417 | -0.984439715 | 0.095 | 0.284 | 1 | 0.000871974 | -1.674479418 | 0.067 | 0.311 | 1 | 0.072535417 | 0.001743188 |
| Met | 0.163566442 | 1.562488434 | 0.524 | 0.425 | 1 | 0.000874656 | 1.284852705 | 0.933 | 0.689 | 1 | 0.163566442 | 0.001748547 |
| Paip2b | 0.023166591 | -1.133583154 | 0.095 | 0.369 | 1 | 0.000878364 | -1.89183625 | 0.089 | 0.336 | 1 | 0.023166591 | 0.001755957 |
| U2surp | 0.869708744 | 0.226312791 | 0.667 | 0.825 | 1 | 0.000880592 | -0.273061528 | 0.867 | 0.917 | 1 | 0.869708744 | 0.001760409 |
| Tbc1d10a | 0.313304807 | -0.167530486 | 0.524 | 0.701 | 1 | 0.000885811 | -0.350203569 | 0.4 | 0.792 | 1 | 0.313304807 | 0.001770837 |
| Mrps34 | 0.068570649 | -0.785659851 | 0.143 | 0.362 | 1 | 0.000886232 | -1.213091603 | 0.089 | 0.349 | 1 | 0.068570649 | 0.001771679 |
| Hadha | 0.167042655 | -0.2884262 | 0.571 | 0.802 | 1 | 0.000887942 | -0.450783161 | 0.889 | 0.824 | 1 | 0.167042655 | 0.001775096 |
| Agfg1 | 0.000893039 | 0.85029718 | 0.952 | 0.866 | 1 | 0.754773589 | 0.272788617 | 0.956 | 0.92 | 1 | 0.754773589 | 0.001785281 |
| Tspyl4 | 0.108421384 | -0.399477214 | 0.095 | 0.28 | 1 | 0.000893612 | -3.656777417 | 0 | 0.204 | 1 | 0.108421384 | 0.001786425 |
| Dnajc18 | 0.387500107 | -0.376854305 | 0.19 | 0.287 | 1 | 0.000894783 | -0.716858021 | 0.133 | 0.436 | 1 | 0.387500107 | 0.001788765 |
| Zkscan17 | 0.355495395 | -0.326228232 | 0.095 | 0.179 | 1 | 0.000895436 | -1.913862556 | 0.089 | 0.329 | 1 | 0.355495395 | 0.001790071 |
| Dgka | 0.768741946 | 0.926537838 | 0.238 | 0.328 | 1 | 0.000898644 | 0.962454986 | 0.711 | 0.436 | 1 | 0.768741946 | 0.00179648 |
| Irf9 | 0.014542766 | -1.20069735 | 0.095 | 0.396 | 1 | 0.000899436 | -1.178221143 | 0.111 | 0.388 | 1 | 0.014542766 | 0.001798063 |
| Smyd3 | 0.105453719 | 1.313875823 | 0.619 | 0.597 | 1 | 0.000900203 | 1.176112597 | 0.844 | 0.664 | 1 | 0.105453719 | 0.001799597 |
| Psmc6 | 0.00090186 | -0.768065791 | 0.762 | 0.881 | 1 | 0.008482972 | -0.265237937 | 0.889 | 0.913 | 1 | 0.008482972 | 0.001802907 |
| Tsfm | 0.010292545 | -2.752492987 | 0 | 0.246 | 1 | 0.000905143 | -1.552143876 | 0.067 | 0.311 | 1 | 0.010292545 | 0.001809467 |
| Ywhae | 0.000906839 | -0.602673557 | 1 | 0.993 | 1 | 0.013159554 | -0.271299331 | 1 | 0.997 | 1 | 0.013159554 | 0.001812855 |
| Spata5 | 0.835126793 | 0.417658728 | 0.619 | 0.687 | 1 | 0.000907891 | 1.287934644 | 0.8 | 0.647 | 1 | 0.835126793 | 0.001814958 |
| Nbea | 0.297173412 | 0.400753273 | 0.667 | 0.541 | 1 | 0.000910563 | 0.826920231 | 0.956 | 0.782 | 1 | 0.297173412 | 0.001820297 |
| Mcm4 | 0.148658493 | -0.616905393 | 0.143 | 0.321 | 1 | 0.00091181 | -2.500149238 | 0.022 | 0.246 | 1 | 0.148658493 | 0.001822789 |
| Slc39a13 | 0.706008501 | 0.23170131 | 0.619 | 0.679 | 1 | 0.000911881 | -0.467030974 | 0.333 | 0.682 | 1 | 0.706008501 | 0.001822931 |
| Mreg | 0.000911923 | 4.088809267 | 0.095 | 0.007 | 1 | 0.148044209 | -1.486852415 | 0 | 0.045 | 1 | 0.148044209 | 0.001823014 |
| Large2 | 0.000911923 | 4.088809267 | 0.095 | 0.007 | 1 | 0.952530085 | 1.223640968 | 0.022 | 0.024 | 1 | 0.952530085 | 0.001823014 |
| Fam110c | 0.000911923 | 4.088809267 | 0.095 | 0.007 | 1 | 0.18483667 | -1.223818009 | 0 | 0.038 | 1 | 0.18483667 | 0.001823014 |
| Cfap20 | 0.00091411 | -1.034047481 | 0.19 | 0.664 | 1 | 0.299810634 | -0.155176344 | 0.711 | 0.734 | 1 | 0.299810634 | 0.001827384 |
| Jmjd8 | 0.076939067 | -0.849790188 | 0.048 | 0.22 | 1 | 0.000916145 | -2.709244837 | 0.022 | 0.239 | 1 | 0.076939067 | 0.00183145 |
| Mrpl16 | 0.981958179 | 0.634243403 | 0.19 | 0.22 | 1 | 0.00091956 | -2.656777417 | 0.022 | 0.239 | 1 | 0.981958179 | 0.001838275 |
| Pld2 | 0.043862929 | 2.062337056 | 0.286 | 0.142 | 1 | 0.000923579 | -0.86124793 | 0.022 | 0.249 | 1 | 0.043862929 | 0.001846304 |
| Tor1a | 0.001517361 | -2.03588648 | 0.048 | 0.422 | 1 | 0.00092497 | -0.73945945 | 0.133 | 0.439 | 1 | 0.001517361 | 0.001849084 |
| Tmem259 | 0.156400985 | -0.103835811 | 0.286 | 0.493 | 1 | 0.00092644 | -0.673071224 | 0.222 | 0.557 | 1 | 0.156400985 | 0.001852021 |
| Cblb | 0.020217612 | 0.530514717 | 0.81 | 0.619 | 1 | 0.000927788 | 1.08393115 | 0.933 | 0.727 | 1 | 0.020217612 | 0.001854716 |
| Zwint | 0.000929725 | 0.879355901 | 0.81 | 0.746 | 1 | 0.768411403 | 0.205804282 | 0.8 | 0.765 | 1 | 0.768411403 | 0.001858585 |
| St6galnac6 | 0.480457209 | 0.418271035 | 0.286 | 0.399 | 1 | 0.000930368 | -0.960783604 | 0.111 | 0.401 | 1 | 0.480457209 | 0.00185987 |
| Ttc5 | 0.000932012 | -3.486099569 | 0 | 0.362 | 1 | 0.004197421 | -0.800743191 | 0.156 | 0.401 | 1 | 0.004197421 | 0.001863154 |
| Fance | 0.843785958 | 0.33137957 | 0.19 | 0.224 | 1 | 0.000933022 | -2.41846544 | 0.044 | 0.27 | 1 | 0.843785958 | 0.001865174 |
| Ahsa2 | 0.15409289 | -0.688798312 | 0.19 | 0.354 | 1 | 0.000935314 | -0.657551602 | 0.178 | 0.488 | 1 | 0.15409289 | 0.001869754 |
| Srgap3 | 0.043822351 | 1.351843673 | 0.333 | 0.172 | 1 | 0.000939212 | 2.098110085 | 0.711 | 0.453 | 1 | 0.043822351 | 0.001877541 |
| Vti1b | 0.003441605 | -0.625195179 | 0.286 | 0.75 | 1 | 0.000939495 | -0.787885286 | 0.511 | 0.699 | 1 | 0.003441605 | 0.001878107 |
| Chsy3 | 0.000939505 | 1.544720528 | 0.905 | 0.903 | 1 | 0.794394132 | 0.120899181 | 0.644 | 0.692 | 1 | 0.794394132 | 0.001878127 |
| Mrpl41 | 0.000940343 | -2.583616075 | 0.048 | 0.418 | 1 | 0.001607658 | -1.375821103 | 0.111 | 0.363 | 1 | 0.001607658 | 0.001879802 |
| Med30 | 0.000940371 | -2.593014773 | 0.048 | 0.418 | 1 | 0.001433646 | -0.7206496 | 0.089 | 0.339 | 1 | 0.001433646 | 0.001879857 |
| Ctps | 0.339938702 | -0.427766259 | 0.143 | 0.246 | 1 | 0.000941341 | -0.765387915 | 0.044 | 0.28 | 1 | 0.339938702 | 0.001881795 |
| Magohb | 0.048942616 | -1.01272876 | 0.095 | 0.31 | 1 | 0.000941516 | -1.865364039 | 0.089 | 0.329 | 1 | 0.048942616 | 0.001882145 |
| Lgmn | 0.000945983 | -1.965956536 | 0.19 | 0.571 | 1 | 0.423813559 | -0.391274755 | 0.444 | 0.46 | 1 | 0.423813559 | 0.001891072 |
| Pts | 0.001813732 | -3.268742738 | 0 | 0.332 | 1 | 0.000946468 | -2.41846544 | 0.044 | 0.27 | 1 | 0.001813732 | 0.00189204 |
| Plin3 | 0.000947185 | -3.611630451 | 0 | 0.362 | 1 | 0.102314444 | -0.686161223 | 0.378 | 0.453 | 1 | 0.102314444 | 0.001893473 |
| Med24 | 0.020204082 | -2.326228232 | 0 | 0.209 | 1 | 0.000948585 | -2.239759553 | 0.022 | 0.246 | 1 | 0.020204082 | 0.00189627 |
| Phykpl | 0.397495042 | -0.252227651 | 0.143 | 0.228 | 1 | 0.000950041 | -1.181997834 | 0.044 | 0.28 | 1 | 0.397495042 | 0.001899179 |
| Runx3 | 0.22929103 | -1.326228232 | 0.095 | 0.201 | 1 | 0.000955385 | -1.296749532 | 0.111 | 0.388 | 1 | 0.22929103 | 0.001909858 |
| Kif13b | 0.00507702 | 0.976375711 | 0.952 | 0.981 | 1 | 0.000956413 | 0.990788912 | 1 | 0.965 | 1 | 0.00507702 | 0.00191191 |
| Mical3 | 0.027778343 | 1.098363574 | 0.762 | 0.765 | 1 | 0.00095689 | 1.094074572 | 0.778 | 0.509 | 1 | 0.027778343 | 0.001912864 |
| Vdac1 | 0.000957847 | -1.197962383 | 0.571 | 0.787 | 1 | 0.02269993 | -0.328296228 | 0.844 | 0.827 | 1 | 0.02269993 | 0.001914777 |
| Usp37 | 0.000960248 | 1.455866737 | 0.762 | 0.567 | 1 | 0.102279185 | -0.239759553 | 0.778 | 0.827 | 1 | 0.102279185 | 0.001919573 |
| Tbc1d20 | 0.000962754 | -1.431762646 | 0.095 | 0.537 | 1 | 0.001508164 | -0.648133322 | 0.311 | 0.623 | 1 | 0.001508164 | 0.00192458 |
| Arsk | 0.105454464 | -0.690800665 | 0.095 | 0.276 | 1 | 0.000963137 | -0.394929926 | 0.089 | 0.363 | 1 | 0.105454464 | 0.001925346 |
| Tmem63b | 0.000966084 | -1.482732718 | 0.095 | 0.526 | 1 | 0.113612433 | 0.431847483 | 0.6 | 0.46 | 1 | 0.113612433 | 0.001931235 |
| Sbf2 | 0.107852587 | 0.715498893 | 0.667 | 0.612 | 1 | 0.000966333 | 0.78125298 | 0.978 | 0.824 | 1 | 0.107852587 | 0.001931732 |
| Commd6 | 0.004892753 | -1.379339569 | 0.143 | 0.478 | 1 | 0.000968167 | -0.796396786 | 0.111 | 0.388 | 1 | 0.004892753 | 0.001935396 |
| Crebl2 | 0.029943537 | -1.469186186 | 0.048 | 0.272 | 1 | 0.000969403 | -0.465319254 | 0.067 | 0.308 | 1 | 0.029943537 | 0.001937867 |
| Gas2l1 | 0.027334517 | -1.326228232 | 0.095 | 0.34 | 1 | 0.000971513 | -1.292124023 | 0.111 | 0.381 | 1 | 0.027334517 | 0.001942083 |
| Itpkc | 0.511779145 | -0.303508156 | 0.19 | 0.265 | 1 | 0.000973174 | -1.994999319 | 0.067 | 0.304 | 1 | 0.511779145 | 0.001945402 |
| Bpnt1 | 0.268222036 | -0.326228232 | 0.095 | 0.201 | 1 | 0.000981482 | -2.017367132 | 0.044 | 0.277 | 1 | 0.268222036 | 0.001962 |
| Ythdf1 | 0.016829214 | -0.838340168 | 0.19 | 0.522 | 1 | 0.000982561 | -0.700842396 | 0.511 | 0.716 | 1 | 0.016829214 | 0.001964156 |
| Cisd2 | 0.000986611 | -0.989193245 | 0.476 | 0.866 | 1 | 0.023177082 | -0.132127353 | 0.822 | 0.907 | 1 | 0.023177082 | 0.001972248 |
| Ssrp1 | 0.000986864 | -0.747692001 | 0.19 | 0.631 | 1 | 0.003440665 | -0.664520392 | 0.578 | 0.699 | 1 | 0.003440665 | 0.001972754 |
| Actr1b | 0.07979729 | -0.413691074 | 0.238 | 0.511 | 1 | 0.000987671 | -0.74848 | 0.178 | 0.481 | 1 | 0.07979729 | 0.001974366 |
| Wdr11 | 0.237856817 | 0.744854866 | 0.333 | 0.235 | 1 | 0.000988081 | -1.586388089 | 0.089 | 0.343 | 1 | 0.237856817 | 0.001975185 |

| | | | | | | | | | | | | |
|---|---|---|---|---|---|---|---|---|---|---|---|---|
| Tmem120a | 0.002991012 | -2.496153234 | 0.048 | 0.369 | 1 | 0.000992265 | -1.329751454 | 0.044 | 0.287 | 1 | 0.002991012 | 0.001983546 |
| Cryzl2 | 0.017612918 | -1.392317423 | 0.095 | 0.362 | 1 | 0.000993553 | -1.513324627 | 0.044 | 0.287 | 1 | 0.017612918 | 0.001986119 |
| Pitrm1 | 0.458830656 | 1.754390228 | 0.238 | 0.366 | 1 | 0.000993582 | -0.736466305 | 0.156 | 0.433 | 1 | 0.458830656 | 0.001986178 |
| Pycr2 | 0.011637698 | -1.455511249 | 0.095 | 0.384 | 1 | 0.000993792 | -1.105423309 | 0.089 | 0.356 | 1 | 0.011637698 | 0.001986596 |
| Alg3 | 0.077738549 | -1.303508156 | 0.048 | 0.205 | 1 | 0.000995672 | -3.404390255 | 0 | 0.201 | 1 | 0.077738549 | 0.001990352 |
| Fam149b | 0.39422188 | 0.552756367 | 0.143 | 0.243 | 1 | 0.000998027 | -1.071814916 | 0.067 | 0.329 | 1 | 0.39422188 | 0.001995058 |
| Ppa1 | 0.001000213 | -2.155497931 | 0.524 | 0.832 | 1 | 0.011479634 | -1.509048161 | 0.689 | 0.727 | 1 | 0.011479634 | 0.001999426 |
| Polr2b | 0.021827095 | -0.66121248 | 0.143 | 0.455 | 1 | 0.001000995 | -0.824722054 | 0.311 | 0.595 | 1 | 0.021827095 | 0.002000989 |
| Rchy1 | 0.090616466 | -0.822654058 | 0.19 | 0.392 | 1 | 0.001001938 | -1.116333583 | 0.156 | 0.429 | 1 | 0.090616466 | 0.002002872 |
| Rnaseh2c | 0.001002906 | -3.392317423 | 0 | 0.358 | 1 | 0.052201937 | -0.847442131 | 0.156 | 0.294 | 1 | 0.052201937 | 0.002004806 |
| Abhd14b | 0.081293766 | -0.849790188 | 0.048 | 0.216 | 1 | 0.001003258 | -1.47073275 | 0.022 | 0.246 | 1 | 0.081293766 | 0.002005509 |
| Tesk1 | 0.377872533 | 0.425844254 | 0.095 | 0.19 | 1 | 0.001006335 | -1.486852415 | 0.022 | 0.246 | 1 | 0.377872533 | 0.002011657 |
| Zkscan1 | 0.346868781 | 0.406985227 | 0.238 | 0.399 | 1 | 0.001006504 | -0.662702251 | 0.2 | 0.516 | 1 | 0.346868781 | 0.002011994 |
| Psma3 | 0.515155544 | 0.237600055 | 0.952 | 0.978 | 1 | 0.001006913 | -0.41846544 | 0.933 | 0.972 | 1 | 0.515155544 | 0.002012813 |
| Pwwp3b | 0.641574476 | -0.116305163 | 0.143 | 0.19 | 1 | 0.00100761 | -2.393743011 | 0.044 | 0.273 | 1 | 0.641574476 | 0.002014205 |
| Abt1 | 0.023323839 | -1.648156327 | 0.095 | 0.328 | 1 | 0.001009253 | -1.937513824 | 0.089 | 0.325 | 1 | 0.023323839 | 0.002017488 |
| Gnptab | 0.575646669 | 0.958928984 | 0.333 | 0.34 | 1 | 0.001010845 | -1.738391182 | 0.156 | 0.426 | 1 | 0.575646669 | 0.002020668 |
| Ogt | 0.001013604 | 0.936806174 | 1 | 0.903 | 1 | 0.2803173 | 0.303981063 | 1 | 0.965 | 1 | 0.2803173 | 0.00202618 |
| Aimp2 | 0.48229743 | 0.150209812 | 0.143 | 0.228 | 1 | 0.001017094 | -2.137106376 | 0.067 | 0.298 | 1 | 0.48229743 | 0.002033154 |
| Exoc5 | 0.179398261 | 0.722212123 | 0.667 | 0.683 | 1 | 0.00101734 | 0.798028693 | 0.933 | 0.817 | 1 | 0.179398261 | 0.002033645 |
| Zc3h7b | 0.014256319 | -1.07218261 | 0.095 | 0.399 | 1 | 0.001018322 | -0.526380779 | 0.2 | 0.54 | 1 | 0.014256319 | 0.002035606 |
| Zfp84 | 0.101674698 | -0.828728573 | 0.048 | 0.201 | 1 | 0.001018572 | -1.787247349 | 0.089 | 0.332 | 1 | 0.101674698 | 0.002036107 |
| Usp45 | 0.385564418 | 0.397647362 | 0.238 | 0.381 | 1 | 0.001018653 | -0.58541625 | 0.111 | 0.415 | 1 | 0.385564418 | 0.002036269 |
| Mrpl22 | 0.01255657 | -2.63000898 | 0 | 0.235 | 1 | 0.001020936 | -1.653210802 | 0.022 | 0.246 | 1 | 0.01255657 | 0.002040831 |
| Ufsp2 | 0.004333838 | -1.676725479 | 0.143 | 0.474 | 1 | 0.001021405 | -1.08246216 | 0.133 | 0.415 | 1 | 0.004333838 | 0.002041767 |
| Sirpa | 0.030264709 | -2.184209227 | 0 | 0.187 | 1 | 0.00102538 | -3.098287127 | 0.022 | 0.235 | 1 | 0.030264709 | 0.002049708 |
| Ddx47 | 0.001289556 | -2.098817736 | 0.048 | 0.429 | 1 | 0.001025584 | -0.537478488 | 0.222 | 0.547 | 1 | 0.001289556 | 0.002050116 |
| Slc35b4 | 0.01674886 | -2.574155746 | 0 | 0.22 | 1 | 0.001026112 | -0.901889915 | 0.067 | 0.308 | 1 | 0.01674886 | 0.002051172 |
| 2410002F23Rik | 0.171334504 | -0.540007523 | 0.286 | 0.448 | 1 | 0.001028713 | -1.094534992 | 0.156 | 0.439 | 1 | 0.171334504 | 0.002056367 |
| Lrrc8b | 0.818077498 | 1.029915578 | 0.238 | 0.246 | 1 | 0.001030007 | 1.908953994 | 0.533 | 0.332 | 1 | 0.818077498 | 0.002058954 |
| Abraxas2 | 0.094017548 | -0.660128969 | 0.143 | 0.328 | 1 | 0.001031386 | -0.901889915 | 0.178 | 0.464 | 1 | 0.094017548 | 0.002061708 |
| Cyp4a12b | 0.632076134 | -0.133583154 | 0 | 0.011 | 1 | 0.00103273 | 2.320502507 | 0.133 | 0.028 | 1 | 0.632076134 | 0.002064394 |
| Ado | 0.145682588 | -0.998653574 | 0.048 | 0.168 | 1 | 0.001032987 | -2.871516266 | 0.022 | 0.235 | 1 | 0.145682588 | 0.002064908 |
| Rptor | 0.001036243 | 1.20215074 | 0.81 | 0.713 | 1 | 0.002194853 | 1.157003775 | 0.889 | 0.806 | 1 | 0.002194853 | 0.002071411 |
| Slc25a33 | 0.061588228 | -1.184209227 | 0.048 | 0.231 | 1 | 0.001036267 | -2.328154669 | 0.044 | 0.273 | 1 | 0.061588228 | 0.00207146 |
| Rgs10 | 0.004822571 | -1.752492987 | 0.095 | 0.425 | 1 | 0.001037419 | -2.638855509 | 0.067 | 0.294 | 1 | 0.004822571 | 0.002073763 |
| Dlgap4 | 0.004622881 | 0.899775443 | 0.905 | 0.854 | 1 | 0.001040851 | 1.029652111 | 1 | 0.931 | 1 | 0.004622881 | 0.002080618 |
| Trim32 | 0.196379028 | -0.911190733 | 0 | 0.075 | 1 | 0.001043339 | -2.620708162 | 0.022 | 0.235 | 1 | 0.196379028 | 0.00208559 |
| Mxd4 | 0.018243464 | -1.164731853 | 0.476 | 0.698 | 1 | 0.001045747 | -0.538121086 | 0.778 | 0.865 | 1 | 0.018243464 | 0.002090401 |
| Tcf7l2 | 0.807169842 | 0.297743463 | 0.762 | 0.925 | 1 | 0.001046456 | 0.60596366 | 1 | 0.979 | 1 | 0.807169842 | 0.002091817 |
| Tenm3 | 0.001046819 | 1.167672599 | 1 | 0.996 | 1 | 0.012219822 | 0.505847364 | 1 | 1 | 1 | 0.012219822 | 0.002092542 |
| Akr7a5 | 0.009839825 | -1.509450056 | 0.048 | 0.336 | 1 | 0.001052179 | -1.264459994 | 0.067 | 0.308 | 1 | 0.009839825 | 0.002103251 |
| Coro1a | 0.001052327 | 1.536268244 | 0.095 | 0.007 | 1 | 0.532888979 | 0.595609745 | 0.067 | 0.045 | 1 | 0.532888979 | 0.002103546 |
| Triap1 | 0.042768624 | -1.233118828 | 0.048 | 0.254 | 1 | 0.001054442 | -1.7206496 | 0.089 | 0.332 | 1 | 0.042768624 | 0.002107771 |
| Mettl26 | 0.001055837 | -3.403043829 | 0 | 0.354 | 1 | 0.001117656 | -1.071814916 | 0.089 | 0.343 | 1 | 0.001117656 | 0.00211056 |
| Smpd2 | 0.048483206 | -1.818081329 | 0.048 | 0.231 | 1 | 0.001056336 | -2.286553765 | 0.022 | 0.242 | 1 | 0.048483206 | 0.002111556 |
| Fam168a | 0.001057773 | -0.932252349 | 0.476 | 0.862 | 1 | 0.321620362 | 0.207231808 | 0.844 | 0.858 | 1 | 0.321620362 | 0.002114427 |
| Wdr6 | 0.082836106 | -1.348596045 | 0.048 | 0.201 | 1 | 0.001058991 | -1.709244837 | 0.089 | 0.332 | 1 | 0.082836106 | 0.00211686 |
| Pfdn6 | 0.00106128 | -2.142145168 | 0.048 | 0.437 | 1 | 0.004024197 | -0.541893779 | 0.178 | 0.429 | 1 | 0.004024197 | 0.002121434 |
| Insig2 | 0.001065225 | -1.242704877 | 0.143 | 0.575 | 1 | 0.206418554 | -0.127449615 | 0.6 | 0.671 | 1 | 0.206418554 | 0.002129316 |
| Atxn2 | 0.001380339 | 1.110681447 | 0.952 | 0.884 | 1 | 0.001067004 | 0.669178145 | 1 | 0.979 | 1 | 0.001380339 | 0.00213287 |
| Wdr13 | 0.19196452 | 1.143257051 | 0.238 | 0.138 | 1 | 0.001067094 | -2.08246216 | 0.067 | 0.301 | 1 | 0.19196452 | 0.00213305 |
| Dysf | 0.001068106 | 1.713146006 | 0.619 | 0.388 | 1 | 0.302034008 | 0.8118059 | 0.311 | 0.266 | 1 | 0.302034008 | 0.002135071 |
| Ccndbp1 | 0.022089737 | -0.935037475 | 0.095 | 0.377 | 1 | 0.001070201 | -0.911874003 | 0.111 | 0.374 | 1 | 0.022089737 | 0.002139258 |
| Dexi | 0.052429369 | -1.385121921 | 0.095 | 0.287 | 1 | 0.0010705 | -1.201450196 | 0.156 | 0.426 | 1 | 0.052429369 | 0.002139853 |
| Fiz1 | 0.595578623 | 0.912959432 | 0.19 | 0.16 | 1 | 0.001071643 | -2.207698344 | 0.022 | 0.242 | 1 | 0.595578623 | 0.002142137 |
| Clp1 | 0.105849294 | -0.159118246 | 0.095 | 0.265 | 1 | 0.001071774 | -0.690385809 | 0.044 | 0.277 | 1 | 0.105849294 | 0.0021424 |
| Aff1 | 0.001072359 | 0.811514168 | 0.952 | 0.929 | 1 | 0.322919481 | 0.484491126 | 1 | 0.907 | 1 | 0.322919481 | 0.002143569 |
| Gtf3c6 | 0.00414957 | -2.955584852 | 0 | 0.291 | 1 | 0.001073229 | -1.271123724 | 0.089 | 0.336 | 1 | 0.00414957 | 0.002145305 |
| Sec16b | 0.256161184 | 1.05282997 | 0.476 | 0.433 | 1 | 0.001073682 | -0.977178042 | 0.089 | 0.353 | 1 | 0.256161184 | 0.002146211 |
| Copg1 | 0.005508564 | -0.732220592 | 0.333 | 0.776 | 1 | 0.001073733 | -0.644092157 | 0.467 | 0.799 | 1 | 0.005508564 | 0.002146313 |
| Zfp362 | 0.074747778 | -0.496153234 | 0.048 | 0.224 | 1 | 0.001077912 | -1.174908409 | 0.111 | 0.37 | 1 | 0.074747778 | 0.002154662 |
| Abca5 | 0.03895459 | -0.545396753 | 0.095 | 0.343 | 1 | 0.001078599 | -1.495730567 | 0.089 | 0.339 | 1 | 0.03895459 | 0.002156034 |
| Napa | 0.001083709 | -0.614709844 | 0.429 | 0.828 | 1 | 0.124105149 | 0.520257974 | 0.822 | 0.789 | 1 | 0.124105149 | 0.002166243 |
| Mettl16 | 0.160107196 | -0.332226973 | 0.286 | 0.541 | 1 | 0.001085082 | -0.80981433 | 0.356 | 0.616 | 1 | 0.160107196 | 0.002168987 |
| Pgk1 | 0.001087023 | -1.117095032 | 0.476 | 0.836 | 1 | 0.086322452 | -0.394658717 | 0.867 | 0.779 | 1 | 0.086322452 | 0.002172864 |
| Sptssa | 0.001203736 | -1.133583154 | 0.333 | 0.765 | 1 | 0.001087066 | -0.727860515 | 0.622 | 0.772 | 1 | 0.001203736 | 0.002172951 |
| Ccn1 | 0.001087801 | -2.539511764 | 0.81 | 0.84 | 1 | 0.273684983 | -1.735092369 | 0.756 | 0.616 | 1 | 0.273684983 | 0.002174418 |
| Slc25a24 | 0.005432709 | -1.745767124 | 0.143 | 0.459 | 1 | 0.001093571 | -0.603808562 | 0.178 | 0.488 | 1 | 0.005432709 | 0.002185947 |
| Pak1ip1 | 0.001100505 | -1.297249165 | 0.238 | 0.653 | 1 | 0.001480713 | -0.719025857 | 0.622 | 0.758 | 1 | 0.001480713 | 0.002199799 |
| Armcx1 | 0.008357938 | -3.081115734 | 0 | 0.257 | 1 | 0.001102911 | -1.911874003 | 0.089 | 0.329 | 1 | 0.008357938 | 0.002204605 |
| Zfp410 | 0.745526611 | 0.490549944 | 0.238 | 0.302 | 1 | 0.001104059 | -0.782590986 | 0.178 | 0.488 | 1 | 0.745526611 | 0.002206899 |
| Ankrd28 | 0.001104895 | 0.780995273 | 1 | 0.978 | 1 | 0.351722522 | 0.359625776 | 1 | 0.997 | 1 | 0.351722522 | 0.002208569 |
| Mrps18b | 0.022717056 | -1.015527393 | 0.095 | 0.354 | 1 | 0.001107274 | -1.392215541 | 0.067 | 0.315 | 1 | 0.022717056 | 0.002213323 |
| Pomp | 0.001107586 | -0.707050016 | 0.667 | 0.869 | 1 | 0.017418895 | -0.466987109 | 0.822 | 0.817 | 1 | 0.017418895 | 0.002213946 |
| Psmb3 | 0.00448151 | -0.749242452 | 0.667 | 0.866 | 1 | 0.001107745 | -0.619633529 | 0.867 | 0.858 | 1 | 0.00448151 | 0.002214264 |
| Ptgis | 0.001921806 | -1.246645819 | 0.619 | 0.862 | 1 | 0.001108348 | -0.520062909 | 0.533 | 0.882 | 1 | 0.001921806 | 0.002215468 |
| 1810009A15Rik | 0.001109976 | -2.184209227 | 0.095 | 0.463 | 1 | 0.008851414 | -0.827889333 | 0.111 | 0.311 | 1 | 0.008851414 | 0.00221872 |
| Steap3 | 0.041161288 | -1.060615282 | 0.238 | 0.5 | 1 | 0.001114796 | -0.868986172 | 0.378 | 0.689 | 1 | 0.041161288 | 0.00222835 |
| Rnf167 | 0.001116212 | -1.056621173 | 0.19 | 0.646 | 1 | 0.17281574 | -0.355401562 | 0.511 | 0.612 | 1 | 0.17281574 | 0.002231179 |
| Majin | 0.001292032 | -3.274595464 | 0.143 | 0.496 | 1 | 0.001117425 | -1.554597022 | 0.311 | 0.574 | 1 | 0.001292032 | 0.002233601 |
| Herpud2 | 0.878957081 | 0.485642171 | 0.476 | 0.604 | 1 | 0.001117749 | -0.636449011 | 0.6 | 0.806 | 1 | 0.878957081 | 0.002234248 |
| Immp2l | 0.249699166 | 1.029915578 | 0.476 | 0.455 | 1 | 0.001118691 | 1.053284886 | 0.667 | 0.398 | 1 | 0.249699166 | 0.002236131 |
| Kif1c | 0.196196906 | -0.530100566 | 0.19 | 0.34 | 1 | 0.001123163 | -0.543435944 | 0.178 | 0.488 | 1 | 0.196196906 | 0.002245064 |
| Ap4s1 | 0.881793412 | 0.127283415 | 0.429 | 0.459 | 1 | 0.00112396 | -0.198282917 | 0.2 | 0.55 | 1 | 0.881793412 | 0.002246656 |
| Rassf2 | 0.009001534 | -2.984439715 | 0 | 0.254 | 1 | 0.001126681 | -1.850905986 | 0.044 | 0.28 | 1 | 0.009001534 | 0.002252092 |
| Suclg2 | 0.001128498 | -2.683780237 | 0.048 | 0.41 | 1 | 0.932611101 | -0.173191936 | 0.511 | 0.457 | 1 | 0.932611101 | 0.002255722 |
| Lrrfip1 | 0.263111578 | 0.633986902 | 0.762 | 0.668 | 1 | 0.001128872 | 1.252980741 | 0.956 | 0.834 | 1 | 0.263111578 | 0.002256469 |
| Sipa1l2 | 0.001135715 | 2.825774861 | 0.476 | 0.228 | 1 | 0.185787605 | 1.193267318 | 0.333 | 0.242 | 1 | 0.185787605 | 0.00227014 |
| Srsf6 | 0.022451121 | -0.660980368 | 0.571 | 0.799 | 1 | 0.001136684 | -0.536422274 | 0.8 | 0.869 | 1 | 0.022451121 | 0.002272076 |
| Nol11 | 0.050378163 | -0.140009424 | 0.19 | 0.489 | 1 | 0.001141575 | -0.670564368 | 0.2 | 0.529 | 1 | 0.050378163 | 0.002281846 |
| Btd | 0.005053366 | -1.31869456 | 0.143 | 0.496 | 1 | 0.001142884 | -0.888469399 | 0.2 | 0.495 | 1 | 0.005053366 | 0.002284463 |
| Gtpbp4 | 0.274700693 | -0.273057573 | 0.714 | 0.802 | 1 | 0.001144956 | -0.560509457 | 0.778 | 0.824 | 1 | 0.274700693 | 0.0022886 |
| Phf23 | 0.00114656 | -1.208871282 | 0.048 | 0.448 | 1 | 0.197168673 | -0.50103135 | 0.6 | 0.609 | 1 | 0.197168673 | 0.002291805 |
| Ap1s1 | 0.001151966 | -3.292012517 | 0 | 0.351 | 1 | 0.002213773 | -0.544733331 | 0.156 | 0.429 | 1 | 0.002213773 | 0.002302606 |
| Braf | 0.095122473 | 0.715591943 | 0.905 | 0.869 | 1 | 0.001153931 | 0.70744149 | 1 | 0.965 | 1 | 0.095122473 | 0.002306531 |
| Sh3kbp1 | 0.001169183 | 1.592133299 | 0.81 | 0.619 | 1 | 0.065824097 | 1.745703574 | 0.356 | 0.26 | 1 | 0.065824097 | 0.002337 |
| Pou3f3 | 0.238329964 | -0.379339569 | 0.095 | 0.22 | 1 | 0.001171914 | -1.178221143 | 0.111 | 0.384 | 1 | 0.238329964 | 0.002342454 |

| | | | | | | | | | | | | |
|---|---|---|---|---|---|---|---|---|---|---|---|---|
| Rgs2 | 0.026995041 | -2.391816574 | 0.381 | 0.604 | 1 | 0.001174059 | -1.062881791 | 0.489 | 0.215 | 1 | 0.026995041 | 0.002346739 |
| Vps51 | 0.627693033 | 0.252307999 | 0.143 | 0.209 | 1 | 0.001174856 | -2.75987091 | 0.022 | 0.232 | 1 | 0.627693033 | 0.002348331 |
| Cntln | 0.00117519 | -2.320581669 | 0.048 | 0.437 | 1 | 0.014356807 | -0.513324627 | 0.267 | 0.481 | 1 | 0.014356807 | 0.002348999 |
| Bmp1 | 0.080923028 | -0.3935835 | 0.286 | 0.515 | 1 | 0.001175379 | -0.673071224 | 0.178 | 0.512 | 1 | 0.080923028 | 0.002349376 |
| Tmem185b | 0.005631533 | -2.769171728 | 0 | 0.276 | 1 | 0.001177877 | -2.674479418 | 0.022 | 0.232 | 1 | 0.005631533 | 0.002354366 |
| Dpp7 | 0.059794423 | -0.989193245 | 0.048 | 0.235 | 1 | 0.001179482 | -2.656777417 | 0.022 | 0.232 | 1 | 0.059794423 | 0.002357573 |
| Tlcd4 | 0.046658028 | -0.393972839 | 0.143 | 0.418 | 1 | 0.001180197 | -0.807497544 | 0.178 | 0.491 | 1 | 0.046658028 | 0.002359001 |
| Skic2 | 0.707202519 | 0.735172312 | 0.238 | 0.216 | 1 | 0.001183806 | -0.915186737 | 0.067 | 0.304 | 1 | 0.707202519 | 0.00236621 |
| Slc4a2 | 0.947988115 | 0.699766976 | 0.286 | 0.31 | 1 | 0.001184755 | -0.746239044 | 0.156 | 0.453 | 1 | 0.947988115 | 0.002368106 |
| Vps13a | 0.196916349 | -0.291124431 | 0.381 | 0.582 | 1 | 0.001190363 | 1.067607997 | 0.911 | 0.616 | 1 | 0.196916349 | 0.00237931 |
| Ncln | 0.216431438 | -0.666078235 | 0.143 | 0.261 | 1 | 0.001198908 | -1.989352756 | 0.067 | 0.294 | 1 | 0.216431438 | 0.002396378 |
| Kat5 | 0.418179524 | 0.673771768 | 0.286 | 0.224 | 1 | 0.001201506 | -1.463768802 | 0.089 | 0.339 | 1 | 0.418179524 | 0.002401569 |
| Acsf2 | 0.004581815 | -0.583616075 | 0.048 | 0.381 | 1 | 0.001206098 | -1.697749198 | 0.067 | 0.308 | 1 | 0.004581815 | 0.002410742 |
| Psmd12 | 0.007269981 | -0.877128897 | 0.571 | 0.832 | 1 | 0.001217074 | -0.563917199 | 0.911 | 0.9 | 1 | 0.007269981 | 0.002432666 |
| Rida | 0.024349289 | -1.834022873 | 0.048 | 0.269 | 1 | 0.001218892 | -2.404390255 | 0.044 | 0.263 | 1 | 0.024349289 | 0.002436298 |
| Glo1 | 0.146892583 | -0.363880774 | 0.524 | 0.757 | 1 | 0.001220217 | -0.681201799 | 0.511 | 0.758 | 1 | 0.146892583 | 0.002438945 |
| Nt5c3b | 0.001220323 | -1.745767124 | 0.095 | 0.504 | 1 | 0.064916943 | -0.152433376 | 0.244 | 0.446 | 1 | 0.064916943 | 0.002439158 |
| F2rl1 | 0.004520526 | 3.351843673 | 0.095 | 0.011 | 1 | 0.00122584 | 2.045642666 | 0.156 | 0.038 | 1 | 0.004520526 | 0.002450177 |
| Cacnb2 | 0.168837069 | 0.974925894 | 0.571 | 0.5 | 1 | 0.001226853 | 1.425850181 | 0.778 | 0.744 | 1 | 0.168837069 | 0.002452201 |
| Camk2n1 | 0.013637447 | -1.579569242 | 0.143 | 0.433 | 1 | 0.001227026 | -0.884611923 | 0.578 | 0.713 | 1 | 0.013637447 | 0.002452546 |
| Cog5 | 0.843222828 | 0.96048517 | 0.429 | 0.504 | 1 | 0.001227526 | 1.066099691 | 0.778 | 0.526 | 1 | 0.843222828 | 0.002453546 |
| Piezo1 | 0.00122809 | 1.207176615 | 0.952 | 0.937 | 1 | 0.494627385 | 0.635235928 | 0.867 | 0.844 | 1 | 0.494627385 | 0.002454671 |
| Ndufb8 | 0.019246952 | -0.708697869 | 0.476 | 0.78 | 1 | 0.001229343 | -0.757036935 | 0.778 | 0.789 | 1 | 0.019246952 | 0.002457174 |
| Tead1 | 0.001230811 | 1.002236264 | 0.952 | 0.966 | 1 | 0.024048385 | 0.49832887 | 1 | 0.969 | 1 | 0.024048385 | 0.002460107 |
| Usf2 | 0.050264129 | -0.601506234 | 0.238 | 0.526 | 1 | 0.001231663 | -0.941418279 | 0.378 | 0.606 | 1 | 0.050264129 | 0.002461809 |
| Tmem161a | 0.001237796 | -2.475975352 | 0.048 | 0.407 | 1 | 0.050006501 | -0.390927995 | 0.222 | 0.391 | 1 | 0.050006501 | 0.002474059 |
| Mrpl32 | 0.001238698 | -1.725399326 | 0.095 | 0.504 | 1 | 0.002354654 | -1.147002412 | 0.333 | 0.543 | 1 | 0.002354654 | 0.002475861 |
| Zhx2 | 0.001471133 | 1.372417213 | 0.857 | 0.728 | 1 | 0.001239135 | 0.792792953 | 0.956 | 0.806 | 1 | 0.001471133 | 0.002476734 |
| Srgn | 0.01752681 | -0.982273831 | 0.381 | 0.668 | 1 | 0.001249468 | -0.944756564 | 0.733 | 0.792 | 1 | 0.01752681 | 0.002497375 |
| Tent5a | 0.003650304 | -0.603528995 | 0.238 | 0.657 | 1 | 0.001252748 | -1.452086997 | 0.178 | 0.446 | 1 | 0.003650304 | 0.002503926 |
| Naa12 | 0.001254106 | -1.006713344 | 0.238 | 0.713 | 1 | 0.044746252 | -0.170537927 | 0.622 | 0.758 | 1 | 0.044746252 | 0.00250664 |
| Ramp2 | 0.001256535 | -3.801961663 | 0 | 0.351 | 1 | 0.001570516 | -2.071814916 | 0.044 | 0.263 | 1 | 0.001570516 | 0.002511491 |
| Mipol1 | 0.00125684 | -3.348596045 | 0 | 0.347 | 1 | 0.378907881 | -0.280401538 | 0.267 | 0.343 | 1 | 0.378907881 | 0.0025121 |
| Nudt16l1 | 0.013477211 | -2.516052791 | 0 | 0.231 | 1 | 0.001267443 | -1.390176396 | 0.044 | 0.28 | 1 | 0.013477211 | 0.002533279 |
| Gstt3 | 0.01007529 | -2.683780237 | 0 | 0.246 | 1 | 0.001267699 | -3.425451871 | 0 | 0.194 | 1 | 0.01007529 | 0.002533791 |
| Rnf25 | 0.118434416 | -0.61573485 | 0.238 | 0.44 | 1 | 0.001268092 | -0.556393349 | 0.133 | 0.419 | 1 | 0.118434416 | 0.002534575 |
| Ppp1r35 | 0.077166283 | -0.828728573 | 0.048 | 0.22 | 1 | 0.001273311 | -3.446210431 | 0 | 0.194 | 1 | 0.077166283 | 0.002545001 |
| 1300002E11Rik | 0.100897424 | 0.236366455 | 0.048 | 0.205 | 1 | 0.00127364 | -3.404390255 | 0 | 0.194 | 1 | 0.100897424 | 0.002545658 |
| Tmem80 | 0.553742143 | -0.133583154 | 0.143 | 0.201 | 1 | 0.001275307 | -1.960783604 | 0.044 | 0.27 | 1 | 0.553742143 | 0.002548987 |
| Pop4 | 0.006233679 | -1.636083495 | 0.048 | 0.358 | 1 | 0.001276421 | -1.975138897 | 0.044 | 0.27 | 1 | 0.006233679 | 0.002551212 |
| Atm | 0.601301886 | 0.468657338 | 0.143 | 0.205 | 1 | 0.001278377 | -1.937513824 | 0.044 | 0.27 | 1 | 0.601301886 | 0.002555121 |
| Lrmda | 0.27696789 | 0.863671469 | 0.524 | 0.496 | 1 | 0.001279836 | 1.118574188 | 0.844 | 0.505 | 1 | 0.27696789 | 0.002558034 |
| Poglut1 | 0.105661348 | -0.233118828 | 0.19 | 0.418 | 1 | 0.001281929 | -0.695439037 | 0.111 | 0.394 | 1 | 0.105661348 | 0.002562216 |
| Rbm4b | 0.145801245 | -0.530100566 | 0.143 | 0.325 | 1 | 0.001282649 | -0.71328035 | 0.133 | 0.419 | 1 | 0.145801245 | 0.002563653 |
| Irgm1 | 0.064662174 | -2.138726458 | 0.143 | 0.325 | 1 | 0.001282803 | -1.513324627 | 0.044 | 0.277 | 1 | 0.064662174 | 0.002563961 |
| Rnmt | 0.420644283 | 0.425844254 | 0.524 | 0.541 | 1 | 0.001282838 | -0.618677627 | 0.244 | 0.578 | 1 | 0.420644283 | 0.00256403 |
| Qars | 0.018318371 | -1.353028292 | 0.143 | 0.41 | 1 | 0.001285668 | -1.193916524 | 0.178 | 0.433 | 1 | 0.018318371 | 0.002569683 |
| Klf16 | 0.028075043 | -1.865387043 | 0.048 | 0.261 | 1 | 0.001286203 | -1.463768802 | 0.067 | 0.301 | 1 | 0.028075043 | 0.002570751 |
| Gm57315 | 0.066548289 | -1.107587946 | 0.095 | 0.287 | 1 | 0.001287027 | -1.931637258 | 0.044 | 0.27 | 1 | 0.066548289 | 0.002572397 |
| Sod1 | 0.001288261 | -1.077969179 | 0.619 | 0.795 | 1 | 0.116200525 | -0.282313659 | 0.778 | 0.74 | 1 | 0.116200525 | 0.002574863 |
| Pcgf2 | 0.136163569 | -1.063193826 | 0.095 | 0.231 | 1 | 0.001289279 | -1.2150478 | 0.111 | 0.37 | 1 | 0.136163569 | 0.002576896 |
| Bicd2 | 0.008290865 | -2.911190733 | 0 | 0.257 | 1 | 0.001290151 | -0.688486276 | 0.156 | 0.45 | 1 | 0.008290865 | 0.002578637 |
| Ppp6r1 | 0.014564388 | -2.593014773 | 0 | 0.228 | 1 | 0.001290441 | -1.372209849 | 0.089 | 0.329 | 1 | 0.014564388 | 0.002579217 |
| Fabp5 | 0.001291731 | -3.861503609 | 0.048 | 0.403 | 1 | 0.003585976 | -1.770385131 | 0.156 | 0.363 | 1 | 0.003585976 | 0.002581793 |
| Atp5k | 0.001292443 | -1.76585137 | 0.143 | 0.522 | 1 | 0.154578166 | -0.184823878 | 0.422 | 0.516 | 1 | 0.154578166 | 0.002583215 |
| Taf12 | 0.010875492 | -1.707050016 | 0.048 | 0.328 | 1 | 0.00129402 | -0.536095934 | 0.111 | 0.394 | 1 | 0.010875492 | 0.002586365 |
| Sde2 | 0.001294021 | -1.104670462 | 0.286 | 0.743 | 1 | 0.003272625 | -0.725639275 | 0.644 | 0.747 | 1 | 0.003272625 | 0.002586368 |
| Mrpl33 | 0.001295161 | -1.109130111 | 0.381 | 0.776 | 1 | 0.33774836 | 0.397101369 | 0.711 | 0.644 | 1 | 0.33774836 | 0.002588645 |
| Pir | 0.282159097 | -0.326228232 | 0.381 | 0.481 | 1 | 0.001300684 | 0.348377195 | 0.778 | 0.398 | 1 | 0.282159097 | 0.002599677 |
| Rreb1 | 0.675858266 | 1.057782371 | 0.429 | 0.459 | 1 | 0.001300779 | 1.146527215 | 0.733 | 0.46 | 1 | 0.675858266 | 0.002599867 |
| Antkmt | 0.029094106 | -0.709932525 | 0.143 | 0.422 | 1 | 0.001301962 | -1.539319835 | 0.178 | 0.415 | 1 | 0.029094106 | 0.00260223 |
| Cttnbp2 | 0.434190171 | -0.849790188 | 0.048 | 0.101 | 1 | 0.001305919 | 1.304560963 | 0.356 | 0.152 | 1 | 0.434190171 | 0.002610133 |
| Fez2 | 0.501519131 | -0.20069735 | 0.238 | 0.328 | 1 | 0.001308899 | -0.86124793 | 0.089 | 0.353 | 1 | 0.501519131 | 0.002616084 |
| Iqsec1 | 0.002744341 | 1.459894311 | 0.857 | 0.896 | 1 | 0.001309251 | 0.933152782 | 0.956 | 0.952 | 1 | 0.002744341 | 0.002616787 |
| Rbm33 | 0.005221085 | 1.286041614 | 0.762 | 0.776 | 1 | 0.001311684 | 0.954775858 | 0.956 | 0.848 | 1 | 0.005221085 | 0.002621648 |
| Hyi | 0.033856957 | -1.02666795 | 0.095 | 0.347 | 1 | 0.001316871 | -1.113940392 | 0.089 | 0.356 | 1 | 0.033856957 | 0.002632009 |
| Syne2 | 0.111165293 | 1.548240886 | 0.476 | 0.302 | 1 | 0.001319508 | 0.653050103 | 0.867 | 0.547 | 1 | 0.111165293 | 0.002637274 |
| Smim8 | 0.42926114 | 0.1445187 | 0.238 | 0.358 | 1 | 0.001322449 | -2.856086225 | 0.022 | 0.228 | 1 | 0.42926114 | 0.002643149 |
| Gpaa1 | 0.089894363 | -0.798716004 | 0.143 | 0.332 | 1 | 0.001322537 | -2.11134328 | 0.067 | 0.291 | 1 | 0.089894363 | 0.002643325 |
| Nicn1 | 0.213098298 | -0.516052791 | 0.095 | 0.216 | 1 | 0.00133136 | -2.792660845 | 0.022 | 0.228 | 1 | 0.213098298 | 0.002660947 |
| Rps6ka1 | 0.16331317 | -1.133583154 | 0.048 | 0.16 | 1 | 0.001332989 | 1.098110085 | 0.356 | 0.152 | 1 | 0.16331317 | 0.002664201 |
| Tm2d1 | 0.025878359 | -0.542753647 | 0.476 | 0.761 | 1 | 0.001336188 | -0.422579201 | 0.422 | 0.747 | 1 | 0.025878359 | 0.002670591 |
| Tbl2 | 0.052845391 | -1.475975352 | 0.048 | 0.228 | 1 | 0.001336531 | -2.620708162 | 0.022 | 0.228 | 1 | 0.052845391 | 0.002671275 |
| Tnfsf12 | 0.037640447 | -1.326228232 | 0.048 | 0.261 | 1 | 0.00133801 | -0.889817082 | 0.067 | 0.322 | 1 | 0.037640447 | 0.00267423 |
| H1f2 | 0.001338948 | -2.600489894 | 0.238 | 0.623 | 1 | 0.058633072 | -1.446210431 | 0.511 | 0.543 | 1 | 0.058633072 | 0.002676103 |
| Tmem242 | 0.004888766 | -1.574155746 | 0.095 | 0.425 | 1 | 0.001340436 | -0.96521585 | 0.133 | 0.401 | 1 | 0.004888766 | 0.002679075 |
| Cnn2 | 0.001344615 | -3.705606599 | 0.048 | 0.399 | 1 | 0.004593788 | -1.084332956 | 0.422 | 0.595 | 1 | 0.004593788 | 0.002687421 |
| Dek | 0.008853363 | -1.039665291 | 0.714 | 0.866 | 1 | 0.001344661 | -0.29224315 | 0.889 | 0.913 | 1 | 0.008853363 | 0.002687514 |
| AW112010 | 0.001344886 | -3.905813556 | 0.095 | 0.459 | 1 | 0.604926603 | -0.316927414 | 0.022 | 0.038 | 1 | 0.604926603 | 0.002687963 |
| Ephx2 | 0.00134751 | -3.977279923 | 0 | 0.347 | 1 | 0.182440693 | -1.316927414 | 0.022 | 0.076 | 1 | 0.182440693 | 0.002693205 |
| Fap | 0.190521225 | -0.920068884 | 0.667 | 0.694 | 1 | 0.001351289 | -0.503672882 | 0.711 | 0.938 | 1 | 0.190521225 | 0.002700751 |
| Ccdc25 | 0.012352797 | -0.93109029 | 0.238 | 0.556 | 1 | 0.001351795 | -0.511041881 | 0.311 | 0.682 | 1 | 0.012352797 | 0.002701763 |
| Denr | 0.001355935 | -1.042435266 | 0.286 | 0.694 | 1 | 0.01699586 | -0.772809965 | 0.6 | 0.661 | 1 | 0.01699586 | 0.002710032 |
| Ror1 | 0.09170692 | 0.908609282 | 0.619 | 0.53 | 1 | 0.00136184 | 1.653499471 | 0.667 | 0.554 | 1 | 0.09170692 | 0.002721825 |
| Snx19 | 0.003360575 | -3.120644099 | 0 | 0.302 | 1 | 0.001365822 | -0.796396786 | 0.111 | 0.377 | 1 | 0.003360575 | 0.002729778 |
| Kcnq5 | 0.001368482 | 1.55860328 | 0.952 | 0.772 | 1 | 0.553841113 | 0.361944729 | 0.889 | 0.907 | 1 | 0.553841113 | 0.002735091 |
| Cpq | 0.006084209 | -0.879987234 | 0.476 | 0.769 | 1 | 0.001369549 | -0.219292411 | 0.578 | 0.837 | 1 | 0.006084209 | 0.002737223 |
| Galnt7 | 0.103936974 | 1.849055685 | 0.476 | 0.425 | 1 | 0.001371369 | 1.45254082 | 0.711 | 0.405 | 1 | 0.103936974 | 0.002740858 |
| Tmx2 | 0.008883472 | -1.623908781 | 0.048 | 0.34 | 1 | 0.001373934 | -2.301820521 | 0.044 | 0.26 | 1 | 0.008883472 | 0.00274598 |
| Hspa14 | 0.164697462 | 0.586308926 | 0.238 | 0.474 | 1 | 0.001374419 | -1.155646507 | 0.156 | 0.419 | 1 | 0.164697462 | 0.002746948 |
| Tor2a | 0.003268661 | -2.292012517 | 0.048 | 0.366 | 1 | 0.001378568 | -1.162417465 | 0.089 | 0.329 | 1 | 0.003268661 | 0.002755236 |
| Zfp874b | 0.429159048 | 0.451379346 | 0.095 | 0.179 | 1 | 0.001379482 | -1.697749198 | 0.067 | 0.304 | 1 | 0.429159048 | 0.002757062 |
| Purg | 0.482625079 | -0.287057635 | 0.19 | 0.269 | 1 | 0.001385091 | -0.688183221 | 0.156 | 0.439 | 1 | 0.482625079 | 0.002768264 |
| Trerf1 | 0.471192559 | 1.16475812 | 0.19 | 0.149 | 1 | 0.001395904 | 1.254229287 | 0.444 | 0.215 | 1 | 0.471192559 | 0.002789859 |
| Tor1b | 0.116915704 | -0.392317423 | 0.238 | 0.474 | 1 | 0.001399988 | -1.02527233 | 0.156 | 0.429 | 1 | 0.116915704 | 0.002798017 |
| Calcoco1 | 0.145654346 | -0.589262638 | 0.095 | 0.254 | 1 | 0.001400642 | -1.530706705 | 0.111 | 0.356 | 1 | 0.145654346 | 0.002799323 |
| Slc31a1 | 0.158547004 | -0.26603345 | 0.429 | 0.728 | 1 | 0.001403373 | -0.633186759 | 0.622 | 0.754 | 1 | 0.158547004 | 0.002804777 |
| mt-Atp8 | 0.302604388 | 0.824263394 | 0.667 | 0.694 | 1 | 0.001407233 | 0.789536736 | 0.911 | 0.689 | 1 | 0.302604388 | 0.002812485 |

| | | | | | | | | | | | | |
|---|---|---|---|---|---|---|---|---|---|---|---|---|
| Gzf1 | 0.137933299 | -1.02666795 | 0.095 | 0.231 | 1 | 0.001409981 | -2.017367132 | 0.044 | 0.266 | 1 | 0.137933299 | 0.002817975 |
| Foxc2 | 0.018553934 | -1.399477214 | 0.048 | 0.302 | 1 | 0.001410923 | -0.814427073 | 0.156 | 0.426 | 1 | 0.018553934 | 0.002819855 |
| Tfap2b | 0.223580833 | -1.648156327 | 0.048 | 0.142 | 1 | 0.001411377 | -1.369015925 | 0.067 | 0.304 | 1 | 0.223580833 | 0.002820762 |
| Retreg2 | 0.016367921 | -1.03588648 | 0.143 | 0.444 | 1 | 0.001415261 | -0.791706997 | 0.178 | 0.471 | 1 | 0.016367921 | 0.00282852 |
| Rsbn1l | 0.936479769 | -0.253226401 | 0.524 | 0.478 | 1 | 0.001416499 | 0.779502349 | 0.889 | 0.675 | 1 | 0.936479769 | 0.002830992 |
| Baiap2l1 | 0.00363512 | 1.676313962 | 0.905 | 0.769 | 1 | 0.001417428 | 1.085366955 | 0.956 | 0.837 | 1 | 0.00363512 | 0.002832847 |
| Noc4l | 0.604075606 | 0.281454345 | 0.095 | 0.142 | 1 | 0.001418855 | -3.294207337 | 0 | 0.19 | 1 | 0.604075606 | 0.002835697 |
| Emd | 0.001500212 | -1.35864971 | 0.333 | 0.675 | 1 | 0.001420369 | -0.938165301 | 0.644 | 0.73 | 1 | 0.001500212 | 0.002838721 |
| 2900089D17Rik | 0.001442651 | 2.029915578 | 0.381 | 0.149 | 1 | 0.97821218 | 0.841770332 | 0.2 | 0.228 | 1 | 0.97821218 | 0.00288322 |
| Vangl1 | 0.104658972 | -1.348596045 | 0.048 | 0.187 | 1 | 0.001445538 | -1.776359032 | 0.044 | 0.273 | 1 | 0.104658972 | 0.002888987 |
| Pdcd4 | 0.00144561 | -0.780249255 | 0.857 | 0.985 | 1 | 0.052153483 | -0.137770818 | 0.978 | 0.976 | 1 | 0.052153483 | 0.00288913 |
| Ptgds | 0.005519941 | -3.555046923 | 0 | 0.28 | 1 | 0.001448438 | -3.743192169 | 0 | 0.19 | 1 | 0.005519941 | 0.002894778 |
| Cdca4 | 0.253600474 | -0.268742738 | 0.19 | 0.343 | 1 | 0.001453196 | -0.458701375 | 0.156 | 0.464 | 1 | 0.253600474 | 0.00290428 |
| Bcr | 0.001457002 | 1.955684184 | 0.762 | 0.623 | 1 | 0.004790288 | 0.969595345 | 0.8 | 0.685 | 1 | 0.004790288 | 0.00291188 |
| Iah1 | 0.045126939 | -1.264827688 | 0.048 | 0.25 | 1 | 0.001458172 | -1.983684006 | 0.067 | 0.294 | 1 | 0.045126939 | 0.002914218 |
| Ech1 | 0.001460188 | -1.321712751 | 0.238 | 0.642 | 1 | 0.22907901 | -0.34440815 | 0.578 | 0.609 | 1 | 0.22907901 | 0.002918244 |
| Ptpn12 | 0.978827942 | 0.265924647 | 0.619 | 0.679 | 1 | 0.001460332 | 1.126558515 | 0.889 | 0.772 | 1 | 0.978827942 | 0.002918532 |
| Adgrl2 | 0.495207759 | 1.390837804 | 0.095 | 0.164 | 1 | 0.001462316 | 1.771299769 | 0.444 | 0.218 | 1 | 0.495207759 | 0.002922494 |
| Fzd8 | 0.042350946 | -2.574155746 | 0 | 0.168 | 1 | 0.001463282 | -3.223818009 | 0.022 | 0.225 | 1 | 0.042350946 | 0.002924422 |
| Rab11b | 0.001466078 | -0.916191414 | 0.619 | 0.862 | 1 | 0.00732758 | -0.258455011 | 0.933 | 0.927 | 1 | 0.00732758 | 0.002930006 |
| Arl14ep | 0.008400259 | -1.960434252 | 0.048 | 0.34 | 1 | 0.001469204 | -0.834775719 | 0.178 | 0.478 | 1 | 0.008400259 | 0.002936249 |
| Ccdc47 | 0.009336851 | -0.718545655 | 0.524 | 0.825 | 1 | 0.001482617 | -0.486852415 | 0.933 | 0.91 | 1 | 0.009336851 | 0.002963035 |
| Zmiz1 | 0.001486377 | 1.389722056 | 0.905 | 0.948 | 1 | 0.001690164 | 0.805765981 | 1 | 0.976 | 1 | 0.001690164 | 0.002970544 |
| Rrp15 | 0.177460229 | -0.355975576 | 0.19 | 0.369 | 1 | 0.001486865 | -0.854892435 | 0.111 | 0.374 | 1 | 0.177460229 | 0.002971519 |
| Btbd7 | 0.059916806 | 0.88827628 | 0.762 | 0.799 | 1 | 0.001492454 | 0.589808792 | 1 | 0.948 | 1 | 0.059916806 | 0.00298268 |
| Utp20 | 0.223243548 | -0.24509147 | 0.143 | 0.299 | 1 | 0.001495838 | -0.920268444 | 0.111 | 0.36 | 1 | 0.223243548 | 0.002989438 |
| Slc25a32 | 0.196870769 | 0.129451251 | 0.095 | 0.235 | 1 | 0.001497039 | -0.476126009 | 0.067 | 0.322 | 1 | 0.196870769 | 0.002991836 |
| Tprkb | 0.145391081 | -0.260340297 | 0.143 | 0.321 | 1 | 0.001503443 | -1.184823878 | 0.089 | 0.336 | 1 | 0.145391081 | 0.003004626 |
| Pmf1 | 0.034549004 | -2.159118246 | 0 | 0.179 | 1 | 0.001505084 | -2.691966845 | 0.022 | 0.225 | 1 | 0.034549004 | 0.003007903 |
| Sec22a | 0.183048507 | 0.334969854 | 0.19 | 0.41 | 1 | 0.001508669 | -1.038026603 | 0.156 | 0.422 | 1 | 0.183048507 | 0.003015063 |
| Mrpl55 | 0.004967197 | -2.146407195 | 0.048 | 0.347 | 1 | 0.00151113 | -1.975138897 | 0.067 | 0.287 | 1 | 0.004967197 | 0.003019977 |
| Pcdhgc3 | 0.161456115 | -0.599246727 | 0.048 | 0.172 | 1 | 0.001513463 | -2.564854927 | 0.022 | 0.225 | 1 | 0.161456115 | 0.003024635 |
| Psph | 0.014823123 | -1.807354922 | 0.095 | 0.351 | 1 | 0.00151405 | -2.41846544 | 0.044 | 0.256 | 1 | 0.014823123 | 0.003025808 |
| Ube2v1 | 0.001514429 | -1.745767124 | 0.143 | 0.519 | 1 | 0.005163571 | -0.615396536 | 0.444 | 0.626 | 1 | 0.005163571 | 0.003026564 |
| Scaf8 | 0.003520469 | 0.782873205 | 1 | 0.922 | 1 | 0.001515207 | 0.626489058 | 1 | 0.983 | 1 | 0.003520469 | 0.003028119 |
| Hint3 | 0.064795146 | -0.233118828 | 0.143 | 0.351 | 1 | 0.001515305 | -2.41846544 | 0.044 | 0.256 | 1 | 0.064795146 | 0.003028315 |
| Mrps16 | 0.002164053 | -1.522625445 | 0.238 | 0.616 | 1 | 0.001516384 | -1.051993475 | 0.333 | 0.561 | 1 | 0.002164053 | 0.003030468 |
| Zfp408 | 0.613021963 | 0.323274521 | 0.095 | 0.146 | 1 | 0.001517627 | -2.255526869 | 0.022 | 0.232 | 1 | 0.613021963 | 0.00303295 |
| Mynn | 0.040092277 | -1.081115734 | 0.095 | 0.321 | 1 | 0.001518856 | -0.682576886 | 0.089 | 0.346 | 1 | 0.040092277 | 0.003035406 |
| Smpdl3a | 0.001522135 | -1.694298109 | 0.333 | 0.646 | 1 | 0.002544124 | -0.725392259 | 0.644 | 0.768 | 1 | 0.002544124 | 0.003041954 |
| Ppp5c | 0.119289909 | -0.303508156 | 0.143 | 0.351 | 1 | 0.00152641 | -0.486852415 | 0.111 | 0.388 | 1 | 0.119289909 | 0.003050491 |
| Zc3h11a | 0.691803706 | 0.110835573 | 0.286 | 0.373 | 1 | 0.001528387 | -0.802354241 | 0.156 | 0.436 | 1 | 0.691803706 | 0.003054439 |
| Mcm6 | 0.02893928 | -0.453084512 | 0.238 | 0.53 | 1 | 0.001534137 | -1.003988102 | 0.178 | 0.46 | 1 | 0.02893928 | 0.003065921 |
| Rab1b | 0.001539796 | -1.383059455 | 0.143 | 0.552 | 1 | 0.05719136 | -0.722919774 | 0.4 | 0.512 | 1 | 0.05719136 | 0.003077221 |
| Hsd17b12 | 0.001543089 | 0.95952625 | 0.905 | 0.825 | 1 | 0.051975825 | 1.046955997 | 0.867 | 0.723 | 1 | 0.051975825 | 0.003083797 |
| Ttc3 | 0.001543696 | -0.75296546 | 0.762 | 0.944 | 1 | 0.034836437 | -0.143252528 | 1 | 0.979 | 1 | 0.034836437 | 0.00308501 |
| Epb41l1 | 0.001543937 | 1.43600412 | 0.762 | 0.716 | 1 | 0.390936949 | 0.473208839 | 0.756 | 0.758 | 1 | 0.390936949 | 0.003085491 |
| Prkd1 | 0.001548162 | 1.579336966 | 0.714 | 0.582 | 1 | 0.001952409 | 1.180572246 | 0.778 | 0.633 | 1 | 0.001952409 | 0.003093927 |
| Efs | 0.014320977 | -2.413691074 | 0 | 0.228 | 1 | 0.001551454 | -2.11134328 | 0.044 | 0.263 | 1 | 0.014320977 | 0.003100501 |
| Trim24 | 0.194709664 | 1.010806755 | 0.476 | 0.418 | 1 | 0.001552944 | 0.843229811 | 0.733 | 0.439 | 1 | 0.194709664 | 0.003103477 |
| Rai1 | 0.290525879 | 0.550389352 | 0.429 | 0.373 | 1 | 0.001555976 | 1.441463656 | 0.867 | 0.799 | 1 | 0.290525879 | 0.00310953 |
| Med4 | 0.035776435 | -1.636083495 | 0.095 | 0.306 | 1 | 0.001559829 | -0.564854927 | 0.089 | 0.356 | 1 | 0.035776435 | 0.003117225 |
| Vps26b | 0.299864508 | -0.574155746 | 0.048 | 0.123 | 1 | 0.001563954 | -2.316927414 | 0.044 | 0.256 | 1 | 0.299864508 | 0.003125462 |
| Lrp10 | 0.003825595 | -0.978304929 | 0.095 | 0.444 | 1 | 0.001568507 | -1.055695251 | 0.356 | 0.578 | 1 | 0.003825595 | 0.003134554 |
| Id4 | 0.082856205 | -0.807354922 | 0.19 | 0.429 | 1 | 0.001571406 | -1.230215781 | 0.133 | 0.377 | 1 | 0.082856205 | 0.003140343 |
| 1110059E24Rik | 0.311641416 | 1.710297644 | 0.333 | 0.287 | 1 | 0.001571979 | -1.356455778 | 0.067 | 0.304 | 1 | 0.311641416 | 0.003141487 |
| Lsm8 | 0.001575268 | -1.91868527 | 0.095 | 0.47 | 1 | 0.129204703 | -0.500149238 | 0.422 | 0.502 | 1 | 0.129204703 | 0.003148055 |
| Pdcd7 | 0.172665297 | -0.40151636 | 0.143 | 0.302 | 1 | 0.00157683 | -1.462604869 | 0.133 | 0.384 | 1 | 0.172665297 | 0.003151173 |
| Nebl | 0.019806932 | 0.690942468 | 0.952 | 0.802 | 1 | 0.001576833 | 1.167143671 | 0.778 | 0.737 | 1 | 0.019806932 | 0.003151181 |
| Ube2e1 | 0.00964038 | -0.951937075 | 0.381 | 0.728 | 1 | 0.001578347 | -0.692661952 | 0.711 | 0.789 | 1 | 0.00964038 | 0.003154203 |
| R3hdm4 | 0.001590563 | -1.66251162 | 0.143 | 0.519 | 1 | 0.236259293 | -0.442458296 | 0.622 | 0.578 | 1 | 0.236259293 | 0.003178597 |
| Fhit | 0.95435574 | 1.147403921 | 0.333 | 0.392 | 1 | 0.001593652 | 1.116031993 | 0.844 | 0.471 | 1 | 0.95435574 | 0.003184764 |
| Rgs19 | 0.041846518 | -2.02666795 | 0 | 0.168 | 1 | 0.001595042 | -3.223818009 | 0 | 0.187 | 1 | 0.041846518 | 0.003187539 |
| Commd9 | 0.093353244 | -1.326228232 | 0.048 | 0.194 | 1 | 0.001600993 | -2.207698344 | 0.044 | 0.256 | 1 | 0.093353244 | 0.003199422 |
| Supt7l | 0.33598189 | -0.496153234 | 0.048 | 0.116 | 1 | 0.001602884 | -3.383016604 | 0 | 0.187 | 1 | 0.33598189 | 0.003203199 |
| Nfat5 | 0.00161059 | 0.725658509 | 1 | 0.993 | 1 | 0.001772021 | 0.633752829 | 1 | 1 | 1 | 0.001772021 | 0.003218586 |
| Elp2 | 0.375595553 | -0.233118828 | 0.19 | 0.295 | 1 | 0.001612971 | -1.286553765 | 0.089 | 0.339 | 1 | 0.375595553 | 0.00322334 |
| Hras | 0.001625102 | -2.692550447 | 0.048 | 0.396 | 1 | 0.209706994 | -0.605172382 | 0.511 | 0.516 | 1 | 0.209706994 | 0.003247563 |
| Nudc | 0.002406213 | -0.75918764 | 0.286 | 0.728 | 1 | 0.001627039 | -0.79913034 | 0.6 | 0.709 | 1 | 0.002406213 | 0.003251431 |
| Pde12 | 0.935815763 | 0.38426515 | 0.238 | 0.254 | 1 | 0.001629146 | -1.577454964 | 0.067 | 0.294 | 1 | 0.935815763 | 0.003255639 |
| Spry2 | 0.78565303 | 0.178605937 | 0.381 | 0.478 | 1 | 0.001629858 | -1.359238219 | 0.222 | 0.478 | 1 | 0.78565303 | 0.003257059 |
| Unkl | 0.084701309 | -0.774688733 | 0.143 | 0.347 | 1 | 0.001634256 | -0.81719275 | 0.178 | 0.474 | 1 | 0.084701309 | 0.003265842 |
| Snx29 | 0.004329485 | 1.716476219 | 0.619 | 0.437 | 1 | 0.001634411 | 1.196023842 | 0.733 | 0.564 | 1 | 0.004329485 | 0.00326615 |
| Hlf | 0.105145717 | -1.989193245 | 0.048 | 0.19 | 1 | 0.001634788 | 0.522607914 | 0.578 | 0.28 | 1 | 0.105145717 | 0.003266904 |
| Ergic2 | 0.07511309 | -0.416718766 | 0.476 | 0.799 | 1 | 0.001636177 | -0.512183705 | 0.711 | 0.851 | 1 | 0.07511309 | 0.003269677 |
| Ubqln4 | 0.993701179 | 0.281454345 | 0.19 | 0.194 | 1 | 0.001642488 | -0.616487696 | 0.067 | 0.315 | 1 | 0.993701179 | 0.003282278 |
| Arfgap1 | 0.145094071 | 0.516230491 | 0.762 | 0.735 | 1 | 0.001643621 | -0.459885368 | 0.378 | 0.751 | 1 | 0.145094071 | 0.00328454 |
| Dynlt2b | 0.005382112 | -2.233118828 | 0.048 | 0.343 | 1 | 0.001645214 | -1.983684006 | 0.089 | 0.311 | 1 | 0.005382112 | 0.00328772 |
| Tmem256 | 0.001649604 | -1.346887703 | 0.333 | 0.698 | 1 | 0.035719206 | -0.643220701 | 0.533 | 0.606 | 1 | 0.035719206 | 0.003296486 |
| Wdr33 | 0.001652943 | 0.8372705 | 1 | 0.903 | 1 | 0.015594617 | 0.520907008 | 1 | 0.958 | 1 | 0.015594617 | 0.003303154 |
| Rrp9 | 0.128338451 | -0.522625445 | 0.143 | 0.325 | 1 | 0.001662084 | -1.856086225 | 0.044 | 0.263 | 1 | 0.128338451 | 0.003321405 |
| Specc1 | 0.068611739 | -1.350074974 | 0.19 | 0.418 | 1 | 0.00166298 | -1.051830542 | 0.667 | 0.785 | 1 | 0.068611739 | 0.003323195 |
| Ing5 | 0.634373099 | 0.226312791 | 0.238 | 0.336 | 1 | 0.001666607 | -1.234465254 | 0.089 | 0.332 | 1 | 0.634373099 | 0.003330436 |
| Rrp36 | 0.989093147 | 0.753206235 | 0.238 | 0.291 | 1 | 0.001672422 | -1.101198723 | 0.067 | 0.304 | 1 | 0.989093147 | 0.003342048 |
| Skic8 | 0.001674126 | -0.882621581 | 0.286 | 0.746 | 1 | 0.200676419 | -0.241639287 | 0.711 | 0.747 | 1 | 0.200676419 | 0.003345449 |
| Wdfy2 | 0.053134341 | -0.741265732 | 0.095 | 0.306 | 1 | 0.001674319 | 0.919001762 | 0.644 | 0.346 | 1 | 0.053134341 | 0.003345835 |
| Ankrd44 | 0.049346327 | -1.360498134 | 0.19 | 0.455 | 1 | 0.001685584 | 1.165224281 | 0.667 | 0.374 | 1 | 0.049346327 | 0.003368326 |
| Polb | 0.001690156 | -1.542165632 | 0.095 | 0.5 | 1 | 0.931131899 | 0.157349289 | 0.6 | 0.595 | 1 | 0.931131899 | 0.003377455 |
| Ttf1 | 0.608777973 | 0.825774861 | 0.19 | 0.272 | 1 | 0.001691122 | -0.703950537 | 0.067 | 0.304 | 1 | 0.608777973 | 0.003379383 |
| Ncbp1 | 0.001695653 | -0.771013075 | 0.048 | 0.429 | 1 | 0.217409256 | -0.119120631 | 0.489 | 0.592 | 1 | 0.217409256 | 0.003388431 |
| Vta1 | 0.001698748 | -1.484657595 | 0.143 | 0.56 | 1 | 0.429880073 | -0.100116025 | 0.556 | 0.599 | 1 | 0.429880073 | 0.00339461 |
| Tuba4a | 0.251069117 | 1.425844254 | 0.095 | 0.041 | 1 | 0.001699806 | 1.254229287 | 0.4 | 0.19 | 1 | 0.251069117 | 0.003396723 |
| Dock2 | 0.632076134 | -1.880817084 | 0 | 0.011 | 1 | 0.001700597 | 1.808603468 | 0.111 | 0.021 | 1 | 0.632076134 | 0.003398302 |
| Crebzf | 0.652763051 | 0.222075798 | 0.333 | 0.433 | 1 | 0.001701407 | -0.686161223 | 0.222 | 0.505 | 1 | 0.652763051 | 0.00339992 |
| Micu3 | 0.001707144 | 1.691049759 | 0.667 | 0.481 | 1 | 0.006732813 | 0.930840235 | 0.733 | 0.678 | 1 | 0.006732813 | 0.003411374 |
| Slc39a14 | 0.073936375 | 1.280787511 | 0.857 | 0.884 | 1 | 0.001707702 | -0.195621118 | 0.311 | 0.678 | 1 | 0.073936375 | 0.003412487 |
| Itpa | 0.076963426 | -0.465779585 | 0.143 | 0.381 | 1 | 0.001712357 | -0.890918797 | 0.111 | 0.36 | 1 | 0.076963426 | 0.003421781 |
| Chpf2 | 0.040660962 | -0.672003069 | 0.095 | 0.343 | 1 | 0.001712643 | -0.947902312 | 0.178 | 0.457 | 1 | 0.040660962 | 0.003422352 |

| | | | | | | | | | | | | |
|---|---|---|---|---|---|---|---|---|---|---|---|---|
| Rock2 | 0.001713522 | 1.19592746 | 1 | 0.993 | 1 | 0.006275585 | 0.454069706 | 1 | 1 | 1 | 0.006275585 | 0.003424108 |
| 1600012H06Rik | 0.001715241 | -3.094412557 | 0 | 0.332 | 1 | 0.002144097 | -0.877642368 | 0.089 | 0.336 | 1 | 0.002144097 | 0.00342754 |
| Socs2 | 0.001721827 | -1.319996279 | 0.476 | 0.772 | 1 | 0.982297426 | -0.380724064 | 0.6 | 0.519 | 1 | 0.982297426 | 0.00344069 |
| Dnpep | 0.029988365 | -1.03588648 | 0.143 | 0.407 | 1 | 0.001727738 | -0.966020252 | 0.111 | 0.37 | 1 | 0.029988365 | 0.00345249 |
| Sppl2a | 0.00173064 | -0.589780755 | 0.762 | 0.914 | 1 | 0.012987755 | -0.301296762 | 0.889 | 0.862 | 1 | 0.012987755 | 0.003458286 |
| Zfp942 | 0.374834008 | -0.289702356 | 0.143 | 0.231 | 1 | 0.001732885 | -1.686161223 | 0.067 | 0.298 | 1 | 0.374834008 | 0.003462766 |
| Washc5 | 0.425022225 | -0.20069735 | 0.19 | 0.306 | 1 | 0.001746599 | -0.815733271 | 0.2 | 0.484 | 1 | 0.425022225 | 0.003490148 |
| Crtc2 | 0.425403209 | 0.521768674 | 0.143 | 0.254 | 1 | 0.00174792 | -1.259441919 | 0.089 | 0.322 | 1 | 0.425403209 | 0.003492784 |
| Ptprs | 0.002537173 | -1.34303652 | 0.095 | 0.481 | 1 | 0.001750168 | -1.02198376 | 0.378 | 0.585 | 1 | 0.002537173 | 0.003497272 |
| Coq2 | 0.092372791 | -0.804275529 | 0.238 | 0.422 | 1 | 0.001751429 | -0.84048937 | 0.111 | 0.37 | 1 | 0.092372791 | 0.00349979 |
| Irf2bp2 | 0.001757983 | -1.015527393 | 0.19 | 0.571 | 1 | 0.023826534 | -0.228143935 | 0.822 | 0.806 | 1 | 0.023826534 | 0.003512875 |
| Oaz2 | 0.096800019 | -0.352507557 | 0.286 | 0.586 | 1 | 0.00176342 | -0.650505868 | 0.467 | 0.702 | 1 | 0.096800019 | 0.00352373 |
| Terf2ip | 0.026741526 | -0.75918764 | 0.048 | 0.287 | 1 | 0.001770339 | -0.415417277 | 0.156 | 0.419 | 1 | 0.026741526 | 0.003537544 |
| Zfyve16 | 0.031613067 | -0.842803758 | 0.19 | 0.455 | 1 | 0.00177069 | -0.815178281 | 0.156 | 0.439 | 1 | 0.031613067 | 0.003538244 |
| Agpat2 | 0.001774225 | -3.303508156 | 0 | 0.332 | 1 | 0.006755613 | -1.21001221 | 0.067 | 0.253 | 1 | 0.006755613 | 0.003545301 |
| Samm50 | 0.001776898 | -2.403043829 | 0.048 | 0.392 | 1 | 0.038644485 | -0.573267167 | 0.222 | 0.398 | 1 | 0.038644485 | 0.003550639 |
| Btbd3 | 0.053193085 | -0.90449608 | 0.429 | 0.694 | 1 | 0.001785159 | -0.691210914 | 0.244 | 0.567 | 1 | 0.053193085 | 0.003567132 |
| Gdf5 | 0.3201691 | -0.763633545 | 0.048 | 0.123 | 1 | 0.001785199 | -3.938979233 | 0.022 | 0.218 | 1 | 0.3201691 | 0.003567211 |
| Smco4 | 0.006695819 | -3.02666795 | 0 | 0.269 | 1 | 0.00178765 | -0.989352756 | 0.111 | 0.367 | 1 | 0.006695819 | 0.003572105 |
| Pex11b | 0.044091414 | -1.063193826 | 0.048 | 0.254 | 1 | 0.001788207 | -1.824722054 | 0.022 | 0.228 | 1 | 0.044091414 | 0.003573216 |
| Cited2 | 0.042183532 | -0.862935564 | 0.143 | 0.373 | 1 | 0.001790049 | -0.181113558 | 0.467 | 0.211 | 1 | 0.042183532 | 0.003576895 |
| Stard4 | 0.095667856 | -0.280424543 | 0.286 | 0.541 | 1 | 0.001791741 | -0.938415791 | 0.2 | 0.46 | 1 | 0.095667856 | 0.003580272 |
| Podxl2 | 0.317881698 | -0.741265732 | 0.048 | 0.123 | 1 | 0.001796543 | -3.223818009 | 0 | 0.183 | 1 | 0.317881698 | 0.003589859 |
| Eif2b1 | 0.04420644 | -1.326228232 | 0.095 | 0.299 | 1 | 0.001804237 | -2.255526869 | 0.044 | 0.253 | 1 | 0.04420644 | 0.003605219 |
| Gramd1a | 0.968540663 | 0.271673324 | 0.333 | 0.377 | 1 | 0.001806094 | -0.593767619 | 0.222 | 0.543 | 1 | 0.968540663 | 0.003608927 |
| Gpsm1 | 0.729945572 | -0.262413135 | 0.333 | 0.403 | 1 | 0.00180924 | -0.644393692 | 0.222 | 0.529 | 1 | 0.729945572 | 0.003615207 |
| Wbp2 | 0.001815735 | -1.018105937 | 0.238 | 0.672 | 1 | 0.815106741 | 0.141699354 | 0.689 | 0.637 | 1 | 0.815106741 | 0.003628172 |
| Hebp2 | 0.005387847 | -3.120644099 | 0 | 0.28 | 1 | 0.001816116 | -3.404390255 | 0 | 0.183 | 1 | 0.005387847 | 0.003628935 |
| Acad9 | 0.196857648 | -0.911190733 | 0.048 | 0.149 | 1 | 0.00181644 | -3.316927414 | 0 | 0.183 | 1 | 0.196857648 | 0.00362958 |
| Ddb1 | 0.0254275 | -0.378316743 | 0.571 | 0.843 | 1 | 0.001816865 | -0.481377285 | 0.911 | 0.92 | 1 | 0.0254275 | 0.003630429 |
| Septin10 | 0.256160366 | -0.133583154 | 0.19 | 0.366 | 1 | 0.001820145 | -2.372209849 | 0.067 | 0.277 | 1 | 0.256160366 | 0.003636977 |
| Dtd2 | 0.04904551 | -1.341178574 | 0.095 | 0.291 | 1 | 0.00182941 | -3.404390255 | 0 | 0.183 | 1 | 0.04904551 | 0.003655474 |
| Maff | 0.001833081 | -2.380304905 | 0.238 | 0.586 | 1 | 0.44210698 | -0.716858021 | 0.6 | 0.557 | 1 | 0.44210698 | 0.003662802 |
| Cemip | 0.719970987 | -0.637814383 | 0.762 | 0.772 | 1 | 0.001838445 | -0.646298492 | 0.467 | 0.834 | 1 | 0.719970987 | 0.003673511 |
| Mpv17 | 0.034387784 | -0.337455488 | 0.095 | 0.358 | 1 | 0.001845944 | -1.135687099 | 0.067 | 0.301 | 1 | 0.034387784 | 0.003688481 |
| Ctdsp2 | 0.016069108 | -0.707422277 | 0.524 | 0.836 | 1 | 0.001848038 | -0.474978315 | 0.867 | 0.931 | 1 | 0.016069108 | 0.00369266 |
| Pprc1 | 0.250161671 | 0.918884266 | 0.476 | 0.437 | 1 | 0.001852782 | -0.638855509 | 0.267 | 0.571 | 1 | 0.250161671 | 0.003702131 |
| Sh3bp2 | 0.001852897 | 2.673771768 | 0.238 | 0.06 | 1 | 0.075960033 | -0.86124793 | 0.067 | 0.176 | 1 | 0.075960033 | 0.003702361 |
| Sec16a | 0.001854717 | 1.028999872 | 0.857 | 0.716 | 1 | 0.618449218 | 0.431326252 | 0.778 | 0.834 | 1 | 0.618449218 | 0.003705993 |
| Cdc123 | 0.00185753 | -0.951744831 | 0.333 | 0.817 | 1 | 0.026000182 | -0.483987082 | 0.733 | 0.754 | 1 | 0.026000182 | 0.003711611 |
| Hpse2 | 0.718638814 | 0.673771768 | 0.143 | 0.194 | 1 | 0.001858783 | -3.263346374 | 0.044 | 0.253 | 1 | 0.718638814 | 0.003714111 |
| Borcs6 | 0.002246946 | -3.292012517 | 0 | 0.321 | 1 | 0.001867919 | -0.657964332 | 0.111 | 0.37 | 1 | 0.002246946 | 0.003732348 |
| Ptbp2 | 0.348558371 | 0.536780656 | 0.81 | 0.784 | 1 | 0.001868562 | 0.714217689 | 0.956 | 0.886 | 1 | 0.348558371 | 0.003733632 |
| Atf1 | 0.020049997 | -0.319275472 | 0.143 | 0.481 | 1 | 0.001874884 | 0.873957206 | 0.844 | 0.661 | 1 | 0.020049997 | 0.003746253 |
| Elp3 | 0.125018826 | -0.133583154 | 0.19 | 0.403 | 1 | 0.001878274 | -0.71780685 | 0.133 | 0.401 | 1 | 0.125018826 | 0.003753021 |
| Strn4 | 0.581621203 | 0.944419358 | 0.333 | 0.302 | 1 | 0.001878404 | -0.486852415 | 0.156 | 0.471 | 1 | 0.581621203 | 0.003753279 |
| Psmc4 | 0.049578829 | -0.810663857 | 0.381 | 0.601 | 1 | 0.001884034 | -0.682576886 | 0.622 | 0.713 | 1 | 0.049578829 | 0.003764518 |
| S1pr2 | 0.197858677 | -0.611630451 | 0.095 | 0.22 | 1 | 0.001884927 | -1.116333583 | 0.111 | 0.36 | 1 | 0.197858677 | 0.003766301 |
| Mex3a | 0.689355774 | -0.811655059 | 0.143 | 0.168 | 1 | 0.001893261 | -1.486852415 | 0.067 | 0.294 | 1 | 0.689355774 | 0.003782938 |
| Etf1 | 0.059313206 | 0.422396953 | 1 | 0.94 | 1 | 0.001894601 | -0.401383073 | 1 | 0.986 | 1 | 0.059313206 | 0.003785613 |
| Efemp1 | 0.052912619 | 0.766881172 | 0.095 | 0.336 | 1 | 0.00189717 | -2.4734319 | 0.178 | 0.384 | 1 | 0.052912619 | 0.00379074 |
| Mmp16 | 0.013240113 | -1.444225442 | 0.143 | 0.455 | 1 | 0.001902893 | -0.107289265 | 0.244 | 0.512 | 1 | 0.013240113 | 0.003802165 |
| Pepd | 0.039803332 | 1.134612957 | 0.524 | 0.399 | 1 | 0.001905004 | -0.754332726 | 0.267 | 0.55 | 1 | 0.039803332 | 0.003806379 |
| Angptl2 | 0.043578732 | -1.159118246 | 0.143 | 0.377 | 1 | 0.001910345 | -1.154277076 | 0.156 | 0.419 | 1 | 0.043578732 | 0.00381704 |
| Fzd7 | 0.088461825 | -0.958496448 | 0.095 | 0.284 | 1 | 0.001912309 | -1.180425414 | 0.044 | 0.27 | 1 | 0.088461825 | 0.003820962 |
| Bst2 | 0.009211045 | -4.24509147 | 0.048 | 0.313 | 1 | 0.001916839 | 0.51561584 | 0.089 | 0.325 | 1 | 0.009211045 | 0.003830004 |
| Serpinb1a | 0.457281049 | 0.503846766 | 0 | 0.026 | 1 | 0.00192535 | 1.280974143 | 0.2 | 0.062 | 1 | 0.457281049 | 0.003846993 |
| Agpat3 | 0.00193164 | -1.386924164 | 0.143 | 0.56 | 1 | 0.016475323 | -0.573938032 | 0.578 | 0.713 | 1 | 0.016475323 | 0.003859548 |
| Lipa | 0.006759955 | -3.067695219 | 0 | 0.269 | 1 | 0.001937293 | -1.375821103 | 0.089 | 0.325 | 1 | 0.006759955 | 0.003870832 |
| Nup35 | 0.001943292 | -2.516052791 | 0.048 | 0.388 | 1 | 0.002452021 | -0.602329633 | 0.111 | 0.367 | 1 | 0.002452021 | 0.003882808 |
| Ammecr1 | 0.898251041 | 0.60338244 | 0.286 | 0.299 | 1 | 0.001949975 | -1.109485334 | 0.111 | 0.349 | 1 | 0.898251041 | 0.003896148 |
| Gtf3c1 | 0.569497066 | 0.1445187 | 0.524 | 0.485 | 1 | 0.001952253 | 0.67013353 | 0.933 | 0.713 | 1 | 0.569497066 | 0.003900694 |
| Mrpl40 | 0.007848952 | -2.25696557 | 0.048 | 0.325 | 1 | 0.001954742 | -2.071814916 | 0.022 | 0.225 | 1 | 0.007848952 | 0.003905664 |
| Bcl9l | 0.008036377 | -1.20069735 | 0.143 | 0.496 | 1 | 0.001956652 | -0.699397051 | 0.578 | 0.768 | 1 | 0.008036377 | 0.003909476 |
| Zbed3 | 0.013606377 | -2.63000898 | 0 | 0.231 | 1 | 0.001957559 | -0.989352756 | 0.044 | 0.27 | 1 | 0.013606377 | 0.003911286 |
| Ndufs5 | 0.013271598 | -0.686291456 | 0.571 | 0.873 | 1 | 0.001962636 | -0.477574602 | 0.667 | 0.751 | 1 | 0.013271598 | 0.003921419 |
| Gm53028 | 0.010102936 | -2.593014773 | 0 | 0.246 | 1 | 0.00196371 | -1.286553765 | 0.044 | 0.26 | 1 | 0.010102936 | 0.003923563 |
| Ing1 | 0.001965389 | -0.24822572 | 0.143 | 0.541 | 1 | 0.00202889 | -0.865364039 | 0.422 | 0.581 | 1 | 0.00202889 | 0.003926915 |
| Gns | 0.001971968 | -1.239972499 | 0.095 | 0.5 | 1 | 0.311153721 | -0.242196539 | 0.6 | 0.644 | 1 | 0.311153721 | 0.003940048 |
| Dis3 | 0.565099447 | 0.430915244 | 0.381 | 0.354 | 1 | 0.001981265 | -0.993388269 | 0.133 | 0.388 | 1 | 0.565099447 | 0.003958604 |
| Ptpn4 | 0.953854528 | 0.852741909 | 0.286 | 0.347 | 1 | 0.001983757 | 1.295620919 | 0.689 | 0.391 | 1 | 0.953854528 | 0.003963579 |
| Fmnl3 | 0.001984444 | 1.596695545 | 0.619 | 0.354 | 1 | 0.670146804 | 0.320974053 | 0.422 | 0.602 | 1 | 0.670146804 | 0.00396495 |
| Zmpste24 | 0.027095123 | -0.505551932 | 0.286 | 0.638 | 1 | 0.001986886 | -0.756550551 | 0.6 | 0.723 | 1 | 0.027095123 | 0.003969823 |
| Pcnx3 | 0.001987517 | -2.998653574 | 0 | 0.325 | 1 | 0.015667284 | 0.306696707 | 0.156 | 0.37 | 1 | 0.015667284 | 0.003971084 |
| Coq10a | 0.100670296 | -1.159118246 | 0.048 | 0.19 | 1 | 0.00199024 | -1.989352756 | 0.044 | 0.256 | 1 | 0.100670296 | 0.003976519 |
| Acot7 | 0.010814031 | -1.043219127 | 0.19 | 0.534 | 1 | 0.001991123 | -0.271123724 | 0.178 | 0.495 | 1 | 0.010814031 | 0.003978282 |
| Zxdc | 0.556322503 | 0.632744499 | 0.19 | 0.265 | 1 | 0.001991848 | -0.434622456 | 0.2 | 0.502 | 1 | 0.556322503 | 0.003979728 |
| Gm43331 | 0.285499161 | 1.866416846 | 0.095 | 0.045 | 1 | 0.00199284 | 2.254229287 | 0.244 | 0.1 | 1 | 0.285499161 | 0.003981709 |
| Apoh | 0.001995154 | 3.258734268 | 0.143 | 0.022 | 1 | 0.950034855 | 1.098110085 | 0.022 | 0.024 | 1 | 0.950034855 | 0.003986327 |
| Tmem30c | 0.001995154 | 3.258734268 | 0.143 | 0.022 | 1 | 0.430487676 | 0.361144491 | 0 | 0.014 | 1 | 0.430487676 | 0.003986327 |
| Ltbp3 | 0.060527491 | -0.567537867 | 0.476 | 0.754 | 1 | 0.001995273 | -0.463084249 | 0.444 | 0.789 | 1 | 0.060527491 | 0.003986565 |
| Irf1 | 0.459368605 | -0.633656758 | 0.238 | 0.321 | 1 | 0.001998871 | 0.741624769 | 0.511 | 0.253 | 1 | 0.459368605 | 0.003993746 |
| Msl1 | 0.147855054 | -0.545396753 | 0.286 | 0.511 | 1 | 0.00200097 | -0.582132919 | 0.8 | 0.858 | 1 | 0.147855054 | 0.003997936 |
| Zfp639 | 0.450323965 | -0.14822993 | 0.19 | 0.284 | 1 | 0.002004288 | -0.765387915 | 0.089 | 0.325 | 1 | 0.450323965 | 0.004004559 |
| Ebf3 | 0.00400868 | -3.303508156 | 0 | 0.295 | 1 | 0.002005177 | -0.91748677 | 0.133 | 0.412 | 1 | 0.00400868 | 0.004006334 |
| Mrpl37 | 0.133245388 | -0.970084422 | 0.095 | 0.235 | 1 | 0.002018965 | -2.017367132 | 0.044 | 0.256 | 1 | 0.133245388 | 0.004033854 |
| Lats2 | 0.002019132 | -1.718545655 | 0.095 | 0.485 | 1 | 0.017202376 | -0.195835597 | 0.444 | 0.696 | 1 | 0.017202376 | 0.004034188 |
| Hibch | 0.080658357 | -1.555046923 | 0.048 | 0.201 | 1 | 0.002023216 | -1.223818009 | 0.044 | 0.266 | 1 | 0.080658357 | 0.004042339 |
| Mrpl44 | 0.120857291 | -1.133583154 | 0.048 | 0.179 | 1 | 0.002026875 | -3.271123724 | 0 | 0.18 | 1 | 0.120857291 | 0.004049641 |
| Lcn4 | 0.472858494 | 1.351843673 | 0.048 | 0.022 | 1 | 0.00202803 | 3.683072586 | 0.089 | 0.014 | 1 | 0.472858494 | 0.004051948 |
| Tpd52l1 | 0.003676542 | -2.107587946 | 0.048 | 0.381 | 1 | 0.002032688 | -0.34440815 | 0.044 | 0.266 | 1 | 0.003676542 | 0.004061243 |
| Dgkz | 0.384078559 | 1.44596585 | 0.286 | 0.243 | 1 | 0.002033248 | -1.37003875 | 0.089 | 0.332 | 1 | 0.384078559 | 0.004062361 |
| Gm48024 | 0.268033526 | -1.081115734 | 0 | 0.056 | 1 | 0.002034522 | 4.268035087 | 0.067 | 0.007 | 1 | 0.268033526 | 0.004064906 |
| Aasdhppt | 0.04093555 | -0.950719097 | 0.143 | 0.388 | 1 | 0.0020383 | -0.998751454 | 0.111 | 0.37 | 1 | 0.04093555 | 0.004072446 |
| Rom1 | 0.119236694 | -1.370622352 | 0.048 | 0.179 | 1 | 0.002043312 | -3.425451871 | 0 | 0.18 | 1 | 0.119236694 | 0.004082449 |
| Gpatch2l | 0.186151071 | 1.258734268 | 0.333 | 0.235 | 1 | 0.002044686 | -1.614607962 | 0.111 | 0.346 | 1 | 0.186151071 | 0.004085192 |
| Cyb5r1 | 0.044745806 | -0.992984824 | 0.143 | 0.369 | 1 | 0.002045279 | -1.502793959 | 0.022 | 0.225 | 1 | 0.044745806 | 0.004086375 |
| L3hypdh | 0.085070393 | -0.911190733 | 0.048 | 0.213 | 1 | 0.002046745 | -3.486852415 | 0 | 0.18 | 1 | 0.085070393 | 0.004089301 |

| | | | | | | | | | | | | |
|---|---|---|---|---|---|---|---|---|---|---|---|---|
| Fem1b | 0.68487668 | 0.620660431 | 0.381 | 0.41 | 1 | 0.002052233 | -0.498257179 | 0.2 | 0.516 | 1 | 0.68487668 | 0.004100254 |
| Fam168b | 0.229698371 | -0.194124696 | 0.381 | 0.616 | 1 | 0.00205629 | -0.624058507 | 0.6 | 0.775 | 1 | 0.229698371 | 0.004108351 |
| Agbl3 | 0.002058157 | -4.530799377 | 0 | 0.328 | 1 | 0.675148376 | 0.45463112 | 0.2 | 0.242 | 1 | 0.675148376 | 0.004112078 |
| Mdm4 | 0.021492525 | 0.842830025 | 0.714 | 0.687 | 1 | 0.002060249 | 0.751552324 | 0.933 | 0.799 | 1 | 0.021492525 | 0.004116253 |
| Dcun1d2 | 0.59041679 | 0.425844254 | 0.095 | 0.149 | 1 | 0.002070419 | -1.459885368 | 0.022 | 0.225 | 1 | 0.59041679 | 0.004136551 |
| Dusp5 | 0.502357829 | -0.488115915 | 0.381 | 0.261 | 1 | 0.002072821 | 0.423205459 | 0.533 | 0.26 | 1 | 0.502357829 | 0.004141346 |
| Hsd17b4 | 0.002077029 | -3.268742738 | 0 | 0.325 | 1 | 0.167785788 | -0.657551602 | 0.378 | 0.467 | 1 | 0.167785788 | 0.004149744 |
| Slc13a5 | 0.111509261 | 0.42946457 | 0.095 | 0.284 | 1 | 0.002080276 | -4.751555641 | 0 | 0.18 | 1 | 0.111509261 | 0.004156224 |
| E130304I02Rik | 0.002086451 | 3.188344941 | 0.143 | 0.022 | 1 | 0.314689364 | 2.098110085 | 0.022 | 0.007 | 1 | 0.314689364 | 0.004168548 |
| Zfp651 | 0.092572112 | -1.098817736 | 0.048 | 0.205 | 1 | 0.002086461 | -0.718177961 | 0.178 | 0.481 | 1 | 0.092572112 | 0.004168568 |
| Max | 0.01671442 | -0.751410289 | 0.571 | 0.877 | 1 | 0.002090645 | -0.398422532 | 0.844 | 0.924 | 1 | 0.01671442 | 0.004176918 |
| Rbms1 | 0.002093902 | 1.165178956 | 1 | 0.974 | 1 | 0.030407917 | 0.73164909 | 1 | 0.986 | 1 | 0.030407917 | 0.004183419 |
| Cep350 | 0.355089039 | 0.614878079 | 0.762 | 0.75 | 1 | 0.002098364 | 0.850971002 | 0.978 | 0.855 | 1 | 0.355089039 | 0.004192324 |
| Foxn3 | 0.017609537 | 1.006733555 | 1 | 0.963 | 1 | 0.002103512 | 0.414018726 | 1 | 0.99 | 1 | 0.017609537 | 0.004202599 |
| Lcmt1 | 0.161329201 | -0.103835811 | 0.143 | 0.299 | 1 | 0.00210917 | -0.669443829 | 0.089 | 0.336 | 1 | 0.161329201 | 0.004213892 |
| Gtf2ird1 | 0.923924668 | -0.194124696 | 0.381 | 0.377 | 1 | 0.002109245 | 0.619619305 | 0.933 | 0.644 | 1 | 0.923924668 | 0.004214042 |
| Aak1 | 0.002109259 | 1.213054897 | 0.905 | 0.933 | 1 | 0.055967286 | 0.646864767 | 0.978 | 0.979 | 1 | 0.055967286 | 0.004214068 |
| Uhrf2 | 0.002112713 | 0.926921585 | 0.857 | 0.728 | 1 | 0.029363838 | 0.61050396 | 0.956 | 0.81 | 1 | 0.029363838 | 0.004220962 |
| Cntnap2 | 0.026542213 | -2.286230164 | 0.095 | 0.321 | 1 | 0.002113076 | -1.508547486 | 0.089 | 0.318 | 1 | 0.026542213 | 0.004221687 |
| Ercc1 | 0.02139252 | -1.818081329 | 0.048 | 0.276 | 1 | 0.002115264 | -1.361321533 | 0.111 | 0.343 | 1 | 0.02139252 | 0.004226054 |
| Nek9 | 0.127317407 | -0.239117569 | 0.333 | 0.56 | 1 | 0.002129187 | -0.191396532 | 0.333 | 0.671 | 1 | 0.127317407 | 0.00425384 |
| Wls | 0.002132714 | -0.671488831 | 0.714 | 0.854 | 1 | 0.078037471 | -0.134191613 | 0.933 | 0.958 | 1 | 0.078037471 | 0.004260879 |
| Tspan1 | 0.36506982 | 2.258734268 | 0.048 | 0.019 | 1 | 0.00213383 | 2.168499413 | 0.133 | 0.031 | 1 | 0.36506982 | 0.004263108 |
| Gfm1 | 0.121111064 | -0.752492987 | 0.095 | 0.254 | 1 | 0.002137198 | -0.871516266 | 0.111 | 0.363 | 1 | 0.121111064 | 0.004269828 |
| Habp4 | 0.089786496 | -0.865387043 | 0.095 | 0.272 | 1 | 0.00214051 | -1.174908409 | 0.111 | 0.349 | 1 | 0.089786496 | 0.004276438 |
| Cdk2ap2 | 0.002141868 | -3.359651234 | 0 | 0.325 | 1 | 0.060685091 | 0.28452321 | 0.578 | 0.367 | 1 | 0.060685091 | 0.004279148 |
| Emc1 | 0.029767864 | -1.233118828 | 0.048 | 0.276 | 1 | 0.002146852 | -1.57996182 | 0.067 | 0.287 | 1 | 0.029767864 | 0.004289095 |
| Snai1 | 0.043510308 | -1.732220592 | 0.095 | 0.313 | 1 | 0.002157883 | -1.191396532 | 0.133 | 0.377 | 1 | 0.043510308 | 0.00431111 |
| Brwd1 | 0.154215272 | 0.643609797 | 0.762 | 0.761 | 1 | 0.002158552 | 0.564461629 | 0.978 | 0.896 | 1 | 0.154215272 | 0.004312444 |
| Stx17 | 0.005746147 | -1.752492987 | 0.095 | 0.414 | 1 | 0.00216507 | -0.678695773 | 0.178 | 0.467 | 1 | 0.005746147 | 0.004325453 |
| Nudt3 | 0.002169632 | -1.297771786 | 0.143 | 0.56 | 1 | 0.024365012 | -0.531246535 | 0.467 | 0.623 | 1 | 0.024365012 | 0.004334557 |
| Slc35a1 | 0.003528395 | -1.901137068 | 0.048 | 0.384 | 1 | 0.002170176 | -1.330733213 | 0.222 | 0.45 | 1 | 0.003528395 | 0.004335643 |
| Chmp1a | 0.008777635 | -1.870548749 | 0.095 | 0.377 | 1 | 0.002171925 | -1.350350415 | 0.111 | 0.349 | 1 | 0.008777635 | 0.004339132 |
| Lmf1 | 0.03054433 | -1.718545655 | 0.048 | 0.257 | 1 | 0.002174458 | -2.526380779 | 0.022 | 0.215 | 1 | 0.03054433 | 0.004344188 |
| Gm56517 | 0.12938293 | -0.718545655 | 0.095 | 0.25 | 1 | 0.002180253 | -1.169370225 | 0.089 | 0.332 | 1 | 0.12938293 | 0.004355752 |
| Gm15581 | 0.00218039 | 3.199840579 | 0.238 | 0.063 | 1 | 0.054009537 | 2.576157382 | 0.089 | 0.031 | 1 | 0.054009537 | 0.004356025 |
| Cnot6l | 0.002187857 | 1.42675938 | 0.762 | 0.604 | 1 | 0.396297403 | -0.162012725 | 0.933 | 0.834 | 1 | 0.396297403 | 0.004370927 |
| Lcor | 0.009096945 | 1.137436235 | 1 | 0.854 | 1 | 0.002191441 | 0.689875472 | 1 | 0.958 | 1 | 0.009096945 | 0.004378079 |
| Plgrkt | 0.007096752 | -1.729950418 | 0.048 | 0.351 | 1 | 0.002191815 | -1.498257179 | 0.111 | 0.339 | 1 | 0.007096752 | 0.004378827 |
| Arl4a | 0.042381206 | -0.264827688 | 0.524 | 0.743 | 1 | 0.002192224 | -0.627267534 | 0.844 | 0.872 | 1 | 0.042381206 | 0.004379641 |
| Ifnar1 | 0.008716364 | -1.620848981 | 0.095 | 0.396 | 1 | 0.002195422 | -0.49218586 | 0.244 | 0.557 | 1 | 0.008716364 | 0.004386025 |
| Aurkaip1 | 0.002196184 | -1.159118246 | 0.286 | 0.698 | 1 | 0.003330988 | -0.850107372 | 0.467 | 0.637 | 1 | 0.003330988 | 0.004387544 |
| Dnajb4 | 0.035031928 | -1.085384066 | 0.476 | 0.72 | 1 | 0.002199465 | -0.557792495 | 0.8 | 0.841 | 1 | 0.035031928 | 0.004394093 |
| Zfp955b | 0.600822396 | 0.304537958 | 0.19 | 0.272 | 1 | 0.00219985 | -2.235790651 | 0.067 | 0.273 | 1 | 0.600822396 | 0.004394861 |
| Prkn | 0.346538371 | 0.737497365 | 0.571 | 0.597 | 1 | 0.002210785 | 1.380325904 | 0.8 | 0.554 | 1 | 0.346538371 | 0.004416683 |
| Brwd3 | 0.151904264 | -0.34415014 | 0.143 | 0.291 | 1 | 0.00221728 | 0.305401939 | 0.8 | 0.46 | 1 | 0.151904264 | 0.004429644 |
| Mul1 | 0.045793844 | -0.593014773 | 0.048 | 0.254 | 1 | 0.002218408 | -1.486852415 | 0.044 | 0.263 | 1 | 0.045793844 | 0.004431895 |
| Mpp1 | 0.015111558 | -1.506137322 | 0.095 | 0.369 | 1 | 0.002224596 | -0.889178863 | 0.178 | 0.453 | 1 | 0.015111558 | 0.004444244 |
| Rrp1 | 0.002226767 | -1.205373837 | 0.429 | 0.72 | 1 | 0.012303628 | -0.516236222 | 0.733 | 0.747 | 1 | 0.012303628 | 0.004448575 |
| Prxl2c | 0.089029688 | -0.672003069 | 0.238 | 0.489 | 1 | 0.002230835 | -0.718498846 | 0.533 | 0.696 | 1 | 0.089029688 | 0.004456693 |
| Efhd2 | 0.002242568 | -1.38923803 | 0.286 | 0.664 | 1 | 0.581676333 | -0.175608916 | 0.733 | 0.657 | 1 | 0.581676333 | 0.004480108 |
| Efnb2 | 0.632043426 | 0.579254169 | 0.333 | 0.302 | 1 | 0.002247698 | 0.829362028 | 0.667 | 0.356 | 1 | 0.632043426 | 0.004490344 |
| Fads1 | 0.027123924 | -1.280424543 | 0.143 | 0.392 | 1 | 0.002252427 | -0.686161223 | 0.111 | 0.374 | 1 | 0.027123924 | 0.004499781 |
| Smarcb1 | 0.002253496 | -1.059582573 | 0.143 | 0.56 | 1 | 0.006837442 | -0.942917183 | 0.444 | 0.588 | 1 | 0.006837442 | 0.004501914 |
| Slc30a1 | 0.135509259 | -0.718545655 | 0.095 | 0.246 | 1 | 0.002260904 | -2.174908409 | 0.067 | 0.28 | 1 | 0.135509259 | 0.004516695 |
| Lsm7 | 0.133750909 | -0.238765391 | 0.286 | 0.541 | 1 | 0.002261184 | -1.012533978 | 0.356 | 0.574 | 1 | 0.133750909 | 0.004517255 |
| Golga2 | 0.463985227 | -0.280424543 | 0.524 | 0.649 | 1 | 0.002264688 | -0.71285609 | 0.444 | 0.692 | 1 | 0.463985227 | 0.004524247 |
| Nkiras1 | 0.216529325 | -0.434752689 | 0.095 | 0.216 | 1 | 0.002272388 | -2.223818009 | 0.044 | 0.246 | 1 | 0.216529325 | 0.004539612 |
| Scx | 0.15405006 | -1.880817084 | 0 | 0.09 | 1 | 0.002277497 | -1.787247349 | 0.044 | 0.26 | 1 | 0.15405006 | 0.004549807 |
| Pfn2 | 0.012255312 | -1.298920886 | 0.095 | 0.399 | 1 | 0.002280957 | -1.223818009 | 0.111 | 0.353 | 1 | 0.012255312 | 0.00455671 |
| Pag1 | 0.260978576 | 2.243137413 | 0.19 | 0.123 | 1 | 0.002282849 | 0.7219616 | 0.533 | 0.273 | 1 | 0.260978576 | 0.004560486 |
| Vmac | 0.097257334 | -0.763633545 | 0.048 | 0.205 | 1 | 0.002286155 | -1.602329633 | 0.022 | 0.221 | 1 | 0.097257334 | 0.004567083 |
| Rnf215 | 0.002286688 | -2.116305163 | 0.095 | 0.433 | 1 | 0.003067524 | -0.721317669 | 0.156 | 0.422 | 1 | 0.003067524 | 0.004568148 |
| Lysmd2 | 0.420670965 | 0.214340149 | 0.048 | 0.108 | 1 | 0.002288903 | -3.247664751 | 0 | 0.176 | 1 | 0.420670965 | 0.004572568 |
| Pycrl | 0.137945102 | -1.098817736 | 0.095 | 0.231 | 1 | 0.002289172 | -3.223818009 | 0 | 0.176 | 1 | 0.137945102 | 0.004573104 |
| Cnot3 | 0.002292006 | -2.03588648 | 0.048 | 0.403 | 1 | 0.004525086 | -0.611110518 | 0.178 | 0.436 | 1 | 0.004525086 | 0.004578759 |
| Blvra | 0.002293616 | -2.545396753 | 0.048 | 0.381 | 1 | 0.004568628 | -1.394929926 | 0.111 | 0.322 | 1 | 0.004568628 | 0.004581971 |
| Mppe1 | 0.088710221 | -0.735619168 | 0.095 | 0.276 | 1 | 0.002295514 | -2.239759553 | 0.044 | 0.246 | 1 | 0.088710221 | 0.004585758 |
| Exosc9 | 0.860349166 | 0.812478039 | 0.286 | 0.332 | 1 | 0.002300419 | -1.0794281 | 0.044 | 0.263 | 1 | 0.860349166 | 0.004595547 |
| Hdac6 | 0.407863846 | 0.400753273 | 0.095 | 0.183 | 1 | 0.002301435 | -3.506751973 | 0 | 0.176 | 1 | 0.407863846 | 0.004597573 |
| Frzb | 0.00230417 | -2.735619168 | 0.048 | 0.403 | 1 | 0.008995251 | -3.564854927 | 0 | 0.135 | 1 | 0.008995251 | 0.00460303 |
| Kbtbd2 | 0.060660443 | -0.811655059 | 0.095 | 0.313 | 1 | 0.002306117 | -0.486852415 | 0.222 | 0.554 | 1 | 0.060660443 | 0.004606916 |
| Cinp | 0.0722456 | -1.413691074 | 0.048 | 0.209 | 1 | 0.002315627 | -1.334849322 | 0.022 | 0.221 | 1 | 0.0722456 | 0.004625893 |
| Kiz | 0.494677123 | 0.479393723 | 0.19 | 0.291 | 1 | 0.002317909 | -1.245034495 | 0.089 | 0.322 | 1 | 0.494677123 | 0.004630446 |
| Immt | 0.002319185 | -1.588323078 | 0.286 | 0.604 | 1 | 0.014939208 | -0.696029217 | 0.689 | 0.713 | 1 | 0.014939208 | 0.004632991 |
| Tead4 | 0.002319612 | 2.171271427 | 0.524 | 0.299 | 1 | 0.936095404 | 1.274267041 | 0.244 | 0.28 | 1 | 0.936095404 | 0.004633843 |
| Cmtm5 | 0.005772764 | -2.20069735 | 0.048 | 0.354 | 1 | 0.002320064 | -3.564854927 | 0 | 0.176 | 1 | 0.005772764 | 0.004634745 |
| Ptpn21 | 0.547290691 | 0.230828272 | 0.143 | 0.213 | 1 | 0.002320624 | -0.407529963 | 0.133 | 0.426 | 1 | 0.547290691 | 0.004635862 |
| Xpo5 | 0.287306007 | -0.221045996 | 0.143 | 0.276 | 1 | 0.002328075 | -0.799320181 | 0.156 | 0.419 | 1 | 0.287306007 | 0.00465073 |
| Scyl3 | 0.061982825 | -1.116305163 | 0.048 | 0.231 | 1 | 0.002331635 | -1.404390255 | 0.067 | 0.284 | 1 | 0.061982825 | 0.004657834 |
| Gnl3 | 0.498805947 | -0.249894541 | 0.762 | 0.866 | 1 | 0.002343625 | -0.652061048 | 0.778 | 0.834 | 1 | 0.498805947 | 0.004681757 |
| Srrt | 0.142964202 | -0.303508156 | 0.429 | 0.675 | 1 | 0.002351338 | -0.553225437 | 0.778 | 0.803 | 1 | 0.142964202 | 0.004697147 |
| Foxj2 | 0.027991533 | -1.834022873 | 0.048 | 0.261 | 1 | 0.002359602 | -0.81719275 | 0.178 | 0.457 | 1 | 0.027991533 | 0.004713636 |
| Fchsd2 | 0.003884287 | 1.025355986 | 0.952 | 0.925 | 1 | 0.00237001 | 1.476952099 | 0.978 | 0.938 | 1 | 0.003884287 | 0.004734403 |
| Plxnb2 | 0.002373229 | -2.288160191 | 0.095 | 0.429 | 1 | 0.467572819 | 0.109887253 | 0.689 | 0.557 | 1 | 0.467572819 | 0.004740825 |
| 4930453N24Rik | 0.005050331 | -1.377890352 | 0.143 | 0.493 | 1 | 0.002376206 | -0.443036183 | 0.2 | 0.495 | 1 | 0.005050331 | 0.004746765 |
| Atp9b | 0.004093195 | 1.19333985 | 0.905 | 0.75 | 1 | 0.002379399 | 0.695636048 | 0.978 | 0.813 | 1 | 0.004093195 | 0.004753137 |
| Heatr5a | 0.783892664 | 0.166601428 | 0.762 | 0.825 | 1 | 0.002381115 | 1.063248654 | 0.911 | 0.865 | 1 | 0.783892664 | 0.004756561 |
| Airn | 0.002387941 | 0.986229792 | 1 | 0.981 | 1 | 0.024265356 | 0.629158864 | 1 | 0.99 | 1 | 0.024265356 | 0.00477018 |
| Spon1 | 0.002391643 | -2.405713016 | 0.19 | 0.541 | 1 | 0.020995861 | -0.220986419 | 0.2 | 0.419 | 1 | 0.020995861 | 0.004777565 |
| Eif2d | 0.002395605 | -3.081115734 | 0 | 0.317 | 1 | 0.233363225 | 0.586857271 | 0.2 | 0.343 | 1 | 0.233363225 | 0.00478547 |
| Reps1 | 0.873446961 | 0.762308442 | 0.286 | 0.336 | 1 | 0.002411019 | 0.626929508 | 0.733 | 0.443 | 1 | 0.873446961 | 0.004816226 |
| Plxna1 | 0.017857793 | -2.535681598 | 0 | 0.216 | 1 | 0.002414189 | -1.416463087 | 0.089 | 0.311 | 1 | 0.017857793 | 0.00482255 |
| 3830406C13Rik | 0.010918105 | 0.379151019 | 0.095 | 0.425 | 1 | 0.002420819 | -0.394929926 | 0.111 | 0.384 | 1 | 0.010918105 | 0.004835777 |
| Mta1 | 0.002420945 | -0.779087197 | 0.143 | 0.549 | 1 | 0.082436696 | -0.217665782 | 0.378 | 0.633 | 1 | 0.082436696 | 0.004836029 |
| R3hdm1 | 0.172675432 | 0.623923218 | 0.714 | 0.743 | 1 | 0.002422254 | 0.887892378 | 1 | 0.83 | 1 | 0.172675432 | 0.00483864 |
| Atp6v1d | 0.002424065 | -0.880817084 | 0.381 | 0.728 | 1 | 0.097393065 | -0.51151447 | 0.8 | 0.709 | 1 | 0.097393065 | 0.004842253 |
| Cep20 | 0.00242824 | -3.146407195 | 0 | 0.317 | 1 | 0.021422466 | -0.452086997 | 0.133 | 0.308 | 1 | 0.021422466 | 0.004850583 |

| | | | | | | | | | | | | |
|---|---|---|---|---|---|---|---|---|---|---|---|---|
| Rcbtb2 | 0.063408329 | -1.370622352 | 0.095 | 0.276 | 1 | 0.002434077 | 0.177544553 | 0.111 | 0.377 | 1 | 0.063408329 | 0.00486223 |
| Nop14 | 0.022065315 | -0.617782678 | 0.381 | 0.728 | 1 | 0.002436784 | -0.531052219 | 0.489 | 0.723 | 1 | 0.022065315 | 0.00486763 |
| Hif1a | 0.771502729 | 0.520241204 | 0.952 | 0.981 | 1 | 0.002441071 | -0.410164893 | 1 | 0.983 | 1 | 0.771502729 | 0.004876184 |
| Gtf2b | 0.002441575 | -1.112824594 | 0.476 | 0.776 | 1 | 0.037155655 | -0.268974191 | 0.622 | 0.74 | 1 | 0.037155655 | 0.004877188 |
| Siae | 0.047630891 | -0.926141075 | 0.095 | 0.313 | 1 | 0.002441759 | -1.404390255 | 0.089 | 0.311 | 1 | 0.047630891 | 0.004877555 |
| Eef1e1 | 0.240176548 | -0.392317423 | 0.19 | 0.328 | 1 | 0.002442779 | -2.003427941 | 0.067 | 0.273 | 1 | 0.240176548 | 0.004879592 |
| Qsox2 | 0.093350721 | -1.303508156 | 0.048 | 0.194 | 1 | 0.00244717 | -2.620708162 | 0.022 | 0.211 | 1 | 0.093350721 | 0.004888352 |
| Orc5 | 0.449894359 | 0.999172054 | 0.143 | 0.25 | 1 | 0.002451609 | -2.564854927 | 0.022 | 0.211 | 1 | 0.449894359 | 0.004897208 |
| Mrps22 | 0.338898675 | -0.133583154 | 0.143 | 0.243 | 1 | 0.002452031 | -2.331877755 | 0.044 | 0.242 | 1 | 0.338898675 | 0.00489805 |
| Sorbs1 | 0.002873359 | 0.95580035 | 0.905 | 0.884 | 1 | 0.002460569 | 0.948704103 | 0.978 | 0.889 | 1 | 0.002873359 | 0.004915083 |
| Pkn2 | 0.817415021 | 0.715033733 | 0.905 | 0.91 | 1 | 0.002467603 | 0.523583037 | 1 | 0.983 | 1 | 0.817415021 | 0.004929118 |
| Ndufv2 | 0.083974301 | -0.316534836 | 0.524 | 0.776 | 1 | 0.002471692 | -0.69851843 | 0.756 | 0.772 | 1 | 0.083974301 | 0.004937275 |
| Prkcd | 0.286673723 | 1.300313372 | 0.19 | 0.123 | 1 | 0.00247321 | 1.00269052 | 0.556 | 0.298 | 1 | 0.286673723 | 0.004940304 |
| Rcbtb1 | 0.04300298 | -0.811655059 | 0.238 | 0.504 | 1 | 0.002473586 | -0.660071995 | 0.222 | 0.536 | 1 | 0.04300298 | 0.004941053 |
| Hmgn2 | 0.002473911 | -0.847060396 | 0.333 | 0.731 | 1 | 0.074197501 | -0.709244837 | 0.578 | 0.592 | 1 | 0.074197501 | 0.004941702 |
| Kif1b | 0.254036017 | 0.391432301 | 0.952 | 0.91 | 1 | 0.002482573 | 0.646802637 | 1 | 0.986 | 1 | 0.254036017 | 0.004958983 |
| Gja1 | 0.021398992 | -1.010132583 | 0.238 | 0.571 | 1 | 0.00249619 | -0.9912374 | 0.244 | 0.519 | 1 | 0.021398992 | 0.004986149 |
| Stk24 | 0.976575294 | 0.539550828 | 0.571 | 0.724 | 1 | 0.00249889 | 0.885757336 | 0.956 | 0.837 | 1 | 0.976575294 | 0.004991536 |
| Abtb2 | 0.002503167 | 1.372443442 | 0.952 | 0.892 | 1 | 0.847719347 | 0.794221674 | 0.867 | 0.907 | 1 | 0.847719347 | 0.005000068 |
| Bap1 | 0.072478029 | -1.413691074 | 0.048 | 0.209 | 1 | 0.002505878 | -0.82328008 | 0.111 | 0.36 | 1 | 0.072478029 | 0.005005477 |
| Cfap97 | 0.006226171 | -2.424260315 | 0.048 | 0.336 | 1 | 0.002513245 | -1.989352756 | 0.044 | 0.249 | 1 | 0.006226171 | 0.005020174 |
| Ror2 | 0.002515943 | 1.132277296 | 0.905 | 0.743 | 1 | 0.578163925 | 0.721014158 | 0.556 | 0.678 | 1 | 0.578163925 | 0.005025557 |
| Zcchc14 | 0.668391667 | 0.179007076 | 0.714 | 0.78 | 1 | 0.002523458 | -0.532095508 | 0.844 | 0.952 | 1 | 0.668391667 | 0.005040548 |
| Slc10a7 | 0.929515357 | 0.69025989 | 0.381 | 0.414 | 1 | 0.002525368 | 1.034147027 | 0.822 | 0.55 | 1 | 0.929515357 | 0.005044359 |
| Adgrl1 | 0.027185819 | -1.018719197 | 0.19 | 0.466 | 1 | 0.002526988 | -0.495491334 | 0.267 | 0.599 | 1 | 0.027185819 | 0.00504759 |
| Ptpn11 | 0.069835326 | -0.496153234 | 0.476 | 0.784 | 1 | 0.002527493 | -0.410036818 | 0.844 | 0.865 | 1 | 0.069835326 | 0.005048597 |
| Zbtb2 | 0.459199685 | -0.335217016 | 0.19 | 0.265 | 1 | 0.002531157 | -1.257765341 | 0.133 | 0.384 | 1 | 0.459199685 | 0.005055907 |
| Emg1 | 0.002531881 | -1.139296806 | 0.19 | 0.601 | 1 | 0.044248175 | -0.512478223 | 0.489 | 0.592 | 1 | 0.044248175 | 0.005057351 |
| Ier3 | 0.002538453 | -1.755174577 | 0.667 | 0.802 | 1 | 0.578298896 | 0.217098122 | 0.867 | 0.651 | 1 | 0.578298896 | 0.005070462 |
| Ralgapa1 | 0.00254332 | 0.820876062 | 0.905 | 0.84 | 1 | 0.004867551 | 0.598292166 | 1 | 0.927 | 1 | 0.004867551 | 0.005080172 |
| Rint1 | 0.995180058 | 0.285074661 | 0.381 | 0.444 | 1 | 0.002545467 | -0.574943745 | 0.244 | 0.547 | 1 | 0.995180058 | 0.005084454 |
| Smu1 | 0.008110577 | -0.862281133 | 0.19 | 0.526 | 1 | 0.002545695 | -0.80878051 | 0.444 | 0.647 | 1 | 0.008110577 | 0.005084909 |
| Map4k5 | 0.002550454 | 1.041250647 | 0.905 | 0.918 | 1 | 0.008071985 | 0.981492591 | 1 | 0.969 | 1 | 0.008071985 | 0.005094403 |
| Fam53a | 0.654589834 | 0.192645078 | 0.143 | 0.187 | 1 | 0.002551212 | -0.486852415 | 0.044 | 0.26 | 1 | 0.654589834 | 0.005095915 |
| Slc31a2 | 0.078592283 | -1.718545655 | 0 | 0.131 | 1 | 0.002552795 | -2.331877755 | 0.044 | 0.242 | 1 | 0.078592283 | 0.005099074 |
| Uqcc5 | 0.057966789 | -0.375471751 | 0.143 | 0.377 | 1 | 0.002556902 | -0.394929926 | 0.111 | 0.36 | 1 | 0.057966789 | 0.005107266 |
| Tmem101 | 0.016430416 | -2.370622352 | 0 | 0.22 | 1 | 0.002557524 | -3.174908409 | 0 | 0.173 | 1 | 0.016430416 | 0.005108508 |
| Angel2 | 0.201087844 | 0.275740694 | 0.333 | 0.597 | 1 | 0.00256028 | -0.496836504 | 0.267 | 0.578 | 1 | 0.201087844 | 0.005114004 |
| Hddc2 | 0.014596582 | -2.701267664 | 0 | 0.228 | 1 | 0.002572468 | -3.223818009 | 0 | 0.173 | 1 | 0.014596582 | 0.005138319 |
| Noc2l | 0.010153262 | -2.067695219 | 0.048 | 0.313 | 1 | 0.00257571 | -0.938415791 | 0.111 | 0.356 | 1 | 0.010153262 | 0.005144786 |
| Slc25a1 | 0.016930182 | -2.094412557 | 0.048 | 0.287 | 1 | 0.002577096 | -3.271123724 | 0 | 0.173 | 1 | 0.016930182 | 0.00514755 |
| Shfl | 0.218613795 | -0.940938076 | 0.048 | 0.142 | 1 | 0.002581599 | -3.361321533 | 0 | 0.173 | 1 | 0.218613795 | 0.005156534 |
| Gm42984 | 0.854728443 | 0.74303443 | 0.143 | 0.172 | 1 | 0.00258601 | -3.271123724 | 0 | 0.173 | 1 | 0.854728443 | 0.005165333 |
| Gatd3a | 0.012394426 | -2.146407195 | 0.048 | 0.302 | 1 | 0.002587836 | -0.664850717 | 0.089 | 0.332 | 1 | 0.012394426 | 0.005168974 |
| Tmem128 | 0.002590624 | -1.326228232 | 0.238 | 0.601 | 1 | 0.003200996 | -0.617322347 | 0.333 | 0.592 | 1 | 0.003200996 | 0.005174536 |
| Ube2z | 0.011259101 | -0.473069621 | 0.19 | 0.549 | 1 | 0.002595501 | -0.68721853 | 0.511 | 0.713 | 1 | 0.011259101 | 0.005184266 |
| Psma6 | 0.002598662 | -1.065313878 | 0.429 | 0.78 | 1 | 0.009232611 | -0.38012124 | 0.8 | 0.824 | 1 | 0.009232611 | 0.005190571 |
| Paqr6 | 0.113650949 | -1.683780237 | 0 | 0.108 | 1 | 0.002599888 | -3.506751973 | 0 | 0.173 | 1 | 0.113650949 | 0.005193016 |
| Evc2 | 0.178664642 | -0.633656758 | 0.143 | 0.284 | 1 | 0.002601066 | -0.979892427 | 0.133 | 0.391 | 1 | 0.178664642 | 0.005195366 |
| Tmem185a | 0.003132677 | -1.392317423 | 0.048 | 0.396 | 1 | 0.00260521 | -0.713817566 | 0.156 | 0.422 | 1 | 0.003132677 | 0.005203632 |
| Ppfibp1 | 0.121599361 | 0.49799404 | 0.857 | 0.955 | 1 | 0.002613854 | 0.901407185 | 0.978 | 0.924 | 1 | 0.121599361 | 0.005220876 |
| Dars | 0.355325125 | -0.228301812 | 0.476 | 0.631 | 1 | 0.002615096 | -1.056466952 | 0.489 | 0.637 | 1 | 0.355325125 | 0.005223354 |
| Sfxn1 | 0.002615526 | -2.167530486 | 0.048 | 0.396 | 1 | 0.433991408 | 0.439147003 | 0.267 | 0.391 | 1 | 0.433991408 | 0.005224211 |
| Srp72 | 0.002618644 | -0.672678646 | 0.571 | 0.896 | 1 | 0.003765545 | -0.556393349 | 0.867 | 0.855 | 1 | 0.003765545 | 0.005230431 |
| Ccdc107 | 0.018388879 | -1.427766259 | 0.048 | 0.302 | 1 | 0.002624628 | -0.816160041 | 0.133 | 0.384 | 1 | 0.018388879 | 0.005242367 |
| Tcf3 | 0.066549843 | -1.133583154 | 0.095 | 0.287 | 1 | 0.002626843 | -0.69105172 | 0.2 | 0.467 | 1 | 0.066549843 | 0.005246786 |
| Rab11a | 0.002630189 | -0.995507019 | 0.476 | 0.776 | 1 | 0.007971287 | -0.344599832 | 0.844 | 0.834 | 1 | 0.007971287 | 0.005253461 |
| Rad21 | 0.002637506 | -0.926141075 | 0.238 | 0.608 | 1 | 0.079117057 | -0.391889472 | 0.822 | 0.779 | 1 | 0.079117057 | 0.005268056 |
| Hilpda | 0.009104592 | -1.469186186 | 0.095 | 0.414 | 1 | 0.002642904 | -1.80878051 | 0.044 | 0.249 | 1 | 0.009104592 | 0.005278824 |
| Hsph1 | 0.002648305 | -1.513226747 | 0.476 | 0.757 | 1 | 0.035349389 | -0.738223904 | 0.933 | 0.799 | 1 | 0.035349389 | 0.005289596 |
| Clptm1 | 0.139018363 | -0.151654727 | 0.333 | 0.59 | 1 | 0.002650897 | -0.85858613 | 0.556 | 0.699 | 1 | 0.139018363 | 0.005294766 |
| Creb3l1 | 0.851638372 | 0.251707002 | 0.429 | 0.429 | 1 | 0.002666938 | -0.779270628 | 0.244 | 0.567 | 1 | 0.851638372 | 0.005326763 |
| Rai14 | 0.514474579 | 0.477374555 | 0.429 | 0.399 | 1 | 0.002671358 | 0.521242889 | 0.844 | 0.626 | 1 | 0.514474579 | 0.00533558 |
| Fzr1 | 0.032273335 | -2.107587946 | 0 | 0.183 | 1 | 0.002677114 | -2.017367132 | 0.044 | 0.242 | 1 | 0.032273335 | 0.00534706 |
| Pgls | 0.002688281 | -1.206646617 | 0.333 | 0.679 | 1 | 0.325622068 | -0.248340329 | 0.578 | 0.581 | 1 | 0.325622068 | 0.005369335 |
| Ddit3 | 0.127985133 | -0.905166945 | 0.238 | 0.422 | 1 | 0.002691951 | -0.557241743 | 0.133 | 0.408 | 1 | 0.127985133 | 0.005376656 |
| BC005624 | 0.00447293 | -0.681927064 | 0.286 | 0.728 | 1 | 0.002692805 | -0.683499424 | 0.489 | 0.671 | 1 | 0.00447293 | 0.00537836 |
| Emc2 | 0.035985405 | -0.266671388 | 0.238 | 0.575 | 1 | 0.002694571 | -0.91748677 | 0.333 | 0.554 | 1 | 0.035985405 | 0.005381882 |
| Gle1 | 0.570173819 | 0.225311267 | 0.238 | 0.34 | 1 | 0.002696156 | -0.829377415 | 0.133 | 0.398 | 1 | 0.570173819 | 0.005385044 |
| Usp1 | 0.002697243 | -1.940938076 | 0.143 | 0.504 | 1 | 0.012615536 | -0.862719317 | 0.4 | 0.557 | 1 | 0.012615536 | 0.005387212 |
| Ttc7 | 0.002709341 | 1.27436492 | 0.762 | 0.716 | 1 | 0.425107478 | 0.568586571 | 0.733 | 0.834 | 1 | 0.425107478 | 0.005411341 |
| Glb1 | 0.148473204 | -0.512094778 | 0.095 | 0.254 | 1 | 0.002711973 | -1.751555641 | 0.067 | 0.287 | 1 | 0.148473204 | 0.005416591 |
| Mpzl1 | 0.016102884 | -0.413691074 | 0.238 | 0.649 | 1 | 0.002713865 | -0.454430938 | 0.489 | 0.799 | 1 | 0.016102884 | 0.005420365 |
| Adcy6 | 0.809702103 | 0.793353383 | 0.238 | 0.239 | 1 | 0.002715773 | -0.448171947 | 0.156 | 0.429 | 1 | 0.809702103 | 0.005424171 |
| Uba5 | 0.152668631 | -0.541825092 | 0.429 | 0.642 | 1 | 0.002715788 | -0.696029217 | 0.556 | 0.72 | 1 | 0.152668631 | 0.005424201 |
| Clba1 | 0.162176307 | -1.392317423 | 0.048 | 0.16 | 1 | 0.002717265 | -2.486852415 | 0.044 | 0.239 | 1 | 0.162176307 | 0.005427146 |
| Nek7 | 0.07469828 | 0.608676739 | 0.905 | 0.769 | 1 | 0.002718734 | 0.638678467 | 1 | 0.834 | 1 | 0.07469828 | 0.005430077 |
| Casp3 | 0.002720699 | -1.733732059 | 0.143 | 0.522 | 1 | 0.012469152 | -0.867674199 | 0.4 | 0.554 | 1 | 0.012469152 | 0.005433996 |
| Ncoa2 | 0.825452481 | 0.408427201 | 0.571 | 0.735 | 1 | 0.002721324 | 1.021083885 | 1 | 0.851 | 1 | 0.825452481 | 0.005435243 |
| Zmat5 | 0.0071002 | -2.818081329 | 0 | 0.265 | 1 | 0.002724262 | -1.41846544 | 0.067 | 0.28 | 1 | 0.0071002 | 0.005441102 |
| Gigyf1 | 0.25546865 | -0.593014773 | 0.095 | 0.201 | 1 | 0.002725328 | -0.927425007 | 0.133 | 0.388 | 1 | 0.25546865 | 0.005443228 |
| Tmem222 | 0.002727767 | -2.107587946 | 0.095 | 0.425 | 1 | 0.045191525 | -0.124282336 | 0.156 | 0.336 | 1 | 0.045191525 | 0.005448094 |
| Aarsd1 | 0.004249336 | -3.094412557 | 0 | 0.291 | 1 | 0.002730028 | -1.500149238 | 0.044 | 0.249 | 1 | 0.004249336 | 0.005452604 |
| Srfbp1 | 0.002732726 | -2.564632972 | 0.048 | 0.373 | 1 | 0.059620982 | -0.316927414 | 0.2 | 0.391 | 1 | 0.059620982 | 0.005457985 |
| Gm26626 | 0.011747022 | 3.673771768 | 0.095 | 0.015 | 1 | 0.00273636 | 3.383512304 | 0.111 | 0.024 | 1 | 0.011747022 | 0.005465232 |
| Fn1 | 0.550811757 | 0.386920838 | 1 | 1 | 1 | 0.002738206 | -0.309227889 | 1 | 1 | 1 | 0.550811757 | 0.005468915 |
| Nosip | 0.107143129 | -1.133583154 | 0.048 | 0.187 | 1 | 0.002738406 | -2.124282336 | 0.022 | 0.215 | 1 | 0.107143129 | 0.005469314 |
| Zfp143 | 0.434917661 | 0.892412054 | 0.19 | 0.138 | 1 | 0.002740499 | -2.085111739 | 0.044 | 0.246 | 1 | 0.434917661 | 0.005473488 |
| Acbd3 | 0.402958529 | -0.184209227 | 0.81 | 0.925 | 1 | 0.00275259 | -0.39743004 | 0.933 | 0.972 | 1 | 0.402958529 | 0.005497604 |
| Syt17 | 0.239516787 | 1.847801167 | 0.143 | 0.075 | 1 | 0.00275305 | 2.875717664 | 0.178 | 0.059 | 1 | 0.239516787 | 0.005498521 |
| Tlcd1 | 0.002756513 | -2.496153234 | 0.048 | 0.373 | 1 | 0.003626705 | -1.722919774 | 0.067 | 0.27 | 1 | 0.003626705 | 0.005505429 |
| Ap3b1 | 0.002761433 | 0.801827051 | 0.952 | 0.78 | 1 | 0.126500725 | 0.482706471 | 0.933 | 0.896 | 1 | 0.126500725 | 0.005515241 |
| Rbm42 | 0.00276715 | -0.800714197 | 0.19 | 0.578 | 1 | 0.016971686 | -0.761712257 | 0.6 | 0.664 | 1 | 0.016971686 | 0.005526643 |
| Med17 | 0.094856312 | -1.067695219 | 0.095 | 0.265 | 1 | 0.002771735 | -0.960783604 | 0.111 | 0.353 | 1 | 0.094856312 | 0.005535788 |
| Ube2l3 | 0.103742876 | -0.300961474 | 0.619 | 0.877 | 1 | 0.002771739 | -0.430348045 | 0.889 | 0.903 | 1 | 0.103742876 | 0.005535796 |
| Tmem132a | 0.285499161 | -0.718545655 | 0 | 0.052 | 1 | 0.002772276 | -1.466674533 | 0.067 | 0.284 | 1 | 0.285499161 | 0.005536866 |
| Mall | 0.054533125 | -1.834022873 | 0.143 | 0.328 | 1 | 0.002773532 | -1.310903626 | 0.133 | 0.367 | 1 | 0.054533125 | 0.005539371 |
| Dhx30 | 0.096478026 | -0.709932525 | 0.19 | 0.403 | 1 | 0.002773732 | -0.815178281 | 0.2 | 0.481 | 1 | 0.096478026 | 0.00553977 |

| | | | | | | | | | | | | |
|---|---|---|---|---|---|---|---|---|---|---|---|---|
| Trappc5 | 0.055947476 | -0.619009982 | 0.095 | 0.306 | 1 | 0.002777024 | -1.589945908 | 0.111 | 0.325 | 1 | 0.055947476 | 0.005546336 |
| Atpif1 | 0.00277718 | -1.021108425 | 0.667 | 0.802 | 1 | 0.408492854 | -0.365964907 | 0.844 | 0.689 | 1 | 0.408492854 | 0.005546647 |
| Ccnq | 0.013586015 | -2.648156327 | 0 | 0.231 | 1 | 0.002777458 | -2.486852415 | 0.022 | 0.208 | 1 | 0.013586015 | 0.005547202 |
| Nit2 | 0.014422622 | -2.535681598 | 0 | 0.228 | 1 | 0.002777458 | -2.446210431 | 0.022 | 0.208 | 1 | 0.014422622 | 0.005547202 |
| Nudt2 | 0.002783209 | -3.02666795 | 0 | 0.31 | 1 | 0.033152647 | -1.271123724 | 0.067 | 0.197 | 1 | 0.033152647 | 0.005558672 |
| Rnh1 | 0.473100124 | -0.184209227 | 0.762 | 0.761 | 1 | 0.002790602 | -0.550799332 | 0.844 | 0.817 | 1 | 0.473100124 | 0.005573416 |
| Mrpl14 | 0.002794404 | -1.76085646 | 0.095 | 0.448 | 1 | 0.235626417 | -0.647317087 | 0.356 | 0.408 | 1 | 0.235626417 | 0.005581 |
| Utp14b | 0.949811658 | 0.79426706 | 0.381 | 0.511 | 1 | 0.002806379 | 1.197645759 | 0.711 | 0.467 | 1 | 0.949811658 | 0.005604882 |
| Cc2d1b | 0.637987403 | 0.569435108 | 0.095 | 0.146 | 1 | 0.002822743 | -1.361321533 | 0.044 | 0.256 | 1 | 0.637987403 | 0.005637518 |
| Csk | 0.106755015 | -0.392317423 | 0.048 | 0.201 | 1 | 0.00282299 | -0.811692106 | 0.089 | 0.318 | 1 | 0.106755015 | 0.005638011 |
| Hdac8 | 0.695751896 | 0.846517288 | 0.571 | 0.597 | 1 | 0.002824696 | 1.061134842 | 0.844 | 0.72 | 1 | 0.695751896 | 0.005641413 |
| Ctf1 | 0.020395986 | -2.455511249 | 0 | 0.209 | 1 | 0.002824963 | -1.736466305 | 0.067 | 0.277 | 1 | 0.020395986 | 0.005641947 |
| Cmas | 0.002826342 | -1.502816964 | 0.143 | 0.5 | 1 | 0.726126882 | 0.103846455 | 0.556 | 0.495 | 1 | 0.726126882 | 0.005644697 |
| Crnkl1 | 0.0185062 | -1.648156327 | 0.048 | 0.299 | 1 | 0.00283221 | -0.69105172 | 0.156 | 0.415 | 1 | 0.0185062 | 0.005656398 |
| Tmem237 | 0.602034298 | 0.459992477 | 0.286 | 0.437 | 1 | 0.002837507 | -0.387316742 | 0.133 | 0.408 | 1 | 0.602034298 | 0.005666962 |
| Ubxn1 | 0.002855547 | -1.280424543 | 0.333 | 0.657 | 1 | 0.228146445 | -0.345622847 | 0.8 | 0.706 | 1 | 0.228146445 | 0.005702941 |
| Suclg1 | 0.00285635 | -1.455511249 | 0.19 | 0.56 | 1 | 0.177401015 | -0.3076496 | 0.578 | 0.633 | 1 | 0.177401015 | 0.005704542 |
| Ccdc12 | 0.00285639 | -1.432427636 | 0.238 | 0.601 | 1 | 0.149691601 | 0.331850184 | 0.622 | 0.45 | 1 | 0.149691601 | 0.005704621 |
| Patj | 0.036968063 | 1.054861935 | 0.524 | 0.354 | 1 | 0.002864404 | 0.768093171 | 0.756 | 0.446 | 1 | 0.036968063 | 0.005720603 |
| Trappc2 | 0.026782779 | -1.593014773 | 0.048 | 0.265 | 1 | 0.002871371 | -1.931637258 | 0.067 | 0.27 | 1 | 0.026782779 | 0.005734497 |
| Tubgcp2 | 0.927917262 | 0.744854866 | 0.238 | 0.276 | 1 | 0.002871892 | -0.316927414 | 0.111 | 0.377 | 1 | 0.927917262 | 0.005735536 |
| Efnb1 | 0.002942844 | -2.849790188 | 0.048 | 0.369 | 1 | 0.002875366 | -0.977657422 | 0.156 | 0.405 | 1 | 0.002942844 | 0.005742464 |
| Ppp2r3d | 0.217985023 | -0.351763324 | 0.19 | 0.343 | 1 | 0.002877378 | -1.237492946 | 0.067 | 0.287 | 1 | 0.217985023 | 0.005746477 |
| Bcl2l1 | 0.345425833 | -0.450349544 | 0.286 | 0.425 | 1 | 0.002880569 | 1.274109601 | 0.644 | 0.478 | 1 | 0.345425833 | 0.00575284 |
| Csad | 0.002888889 | -1.473715845 | 0.19 | 0.552 | 1 | 0.563848756 | 0.498040692 | 0.622 | 0.675 | 1 | 0.563848756 | 0.005769432 |
| Ptgr2 | 0.582780861 | 0.316219763 | 0.095 | 0.146 | 1 | 0.002889612 | -1.80878051 | 0.022 | 0.215 | 1 | 0.582780861 | 0.005770873 |
| Pgap4 | 0.061235497 | -1.911190733 | 0 | 0.146 | 1 | 0.002889784 | -3.174908409 | 0 | 0.17 | 1 | 0.061235497 | 0.005771217 |
| Wdr89 | 0.002896316 | -0.79964408 | 0.238 | 0.675 | 1 | 0.014414441 | -0.669443829 | 0.556 | 0.654 | 1 | 0.014414441 | 0.005784242 |
| Kctd11 | 0.316572795 | -0.167530486 | 0.048 | 0.127 | 1 | 0.002898677 | -3.271123724 | 0 | 0.17 | 1 | 0.316572795 | 0.005788952 |
| Tusc1 | 0.094601638 | -1.752492987 | 0 | 0.119 | 1 | 0.002898894 | -3.271123724 | 0 | 0.17 | 1 | 0.094601638 | 0.005789384 |
| Klc4 | 0.259108505 | -0.911190733 | 0.048 | 0.131 | 1 | 0.002900955 | -1.638855509 | 0.067 | 0.277 | 1 | 0.259108505 | 0.005793495 |
| Ccdc167 | 0.654067372 | 1.786246497 | 0.143 | 0.119 | 1 | 0.002901274 | -1.280401538 | 0.022 | 0.215 | 1 | 0.654067372 | 0.005794131 |
| Gm2885 | 0.982550637 | 0.673771768 | 0.048 | 0.049 | 1 | 0.002901981 | 3.268035087 | 0.111 | 0.024 | 1 | 0.982550637 | 0.005795541 |
| Sike1 | 0.00290356 | -3.146407195 | 0 | 0.31 | 1 | 0.002991997 | -0.901889915 | 0.156 | 0.412 | 1 | 0.002991997 | 0.005798689 |
| Daam1 | 0.002907114 | 1.307154855 | 0.905 | 0.851 | 1 | 0.023103 | 0.984105457 | 0.956 | 0.869 | 1 | 0.023103 | 0.005805777 |
| Nt5dc2 | 0.012070448 | -1.891012851 | 0.048 | 0.321 | 1 | 0.002907276 | -1.638855509 | 0.022 | 0.215 | 1 | 0.012070448 | 0.005806099 |
| Ttc32 | 0.457599708 | -0.133583154 | 0.095 | 0.16 | 1 | 0.002907364 | -3.361321533 | 0 | 0.17 | 1 | 0.457599708 | 0.005806275 |
| Eda | 0.691216649 | 0.351843673 | 0.143 | 0.175 | 1 | 0.002910192 | 0.324702495 | 0.622 | 0.308 | 1 | 0.691216649 | 0.005811916 |
| Fpgs | 0.785751906 | 0.388369549 | 0.095 | 0.119 | 1 | 0.002910782 | -3.339295227 | 0 | 0.17 | 1 | 0.785751906 | 0.005813091 |
| Iars2 | 0.224166176 | -0.300233024 | 0.143 | 0.284 | 1 | 0.002912664 | -0.851516392 | 0.2 | 0.46 | 1 | 0.224166176 | 0.005816844 |
| Rufy3 | 0.601097673 | 0.491568436 | 0.429 | 0.425 | 1 | 0.002913096 | 0.704446237 | 0.8 | 0.526 | 1 | 0.601097673 | 0.005817705 |
| Exosc3 | 0.00946381 | -1.124970024 | 0.143 | 0.451 | 1 | 0.002914349 | -0.387316742 | 0.156 | 0.419 | 1 | 0.00946381 | 0.005820204 |
| Fbxo8 | 0.002915206 | -3.25696557 | 0 | 0.31 | 1 | 0.497977408 | 0.697572156 | 0.178 | 0.242 | 1 | 0.497977408 | 0.005821913 |
| Erbb2 | 0.559359973 | 0.951305743 | 0.143 | 0.104 | 1 | 0.002917916 | -1.204452685 | 0.022 | 0.215 | 1 | 0.559359973 | 0.005827318 |
| Rap1gap | 0.500771035 | -0.535681598 | 0.048 | 0.09 | 1 | 0.002928039 | -3.72631835 | 0 | 0.17 | 1 | 0.500771035 | 0.005847505 |
| Fbxw4 | 0.432065858 | 0.696491844 | 0.238 | 0.175 | 1 | 0.002928364 | -0.835656216 | 0.067 | 0.298 | 1 | 0.432065858 | 0.005848152 |
| Dctn5 | 0.013560474 | -2.63000898 | 0 | 0.231 | 1 | 0.002941892 | -1.749886821 | 0.089 | 0.298 | 1 | 0.013560474 | 0.005875129 |
| Maip1 | 0.044220452 | -1.120644099 | 0.095 | 0.313 | 1 | 0.00295075 | -0.789415185 | 0.089 | 0.318 | 1 | 0.044220452 | 0.005892793 |
| B4galt4 | 0.008032274 | -2.611630451 | 0 | 0.257 | 1 | 0.00295679 | -1.0583944 | 0.089 | 0.308 | 1 | 0.008032274 | 0.005904838 |
| Nup62 | 0.086873391 | -0.834022873 | 0.095 | 0.276 | 1 | 0.002957381 | -1.199570463 | 0.089 | 0.315 | 1 | 0.086873391 | 0.005906015 |
| Inpp4a | 0.061817979 | -0.295854583 | 0.143 | 0.354 | 1 | 0.002970298 | 1.103346734 | 0.667 | 0.505 | 1 | 0.061817979 | 0.005931774 |
| Pcgf5 | 0.574752671 | 0.745110549 | 0.429 | 0.608 | 1 | 0.002974295 | 0.992400644 | 0.822 | 0.751 | 1 | 0.574752671 | 0.005939743 |
| Map2k4 | 0.002976305 | 0.935624121 | 0.857 | 0.694 | 1 | 0.047826201 | 0.749841598 | 0.889 | 0.837 | 1 | 0.047826201 | 0.005943752 |
| Ptpn3 | 0.007391291 | 1.227329131 | 0.762 | 0.526 | 1 | 0.002979724 | 0.853424989 | 0.889 | 0.696 | 1 | 0.007391291 | 0.00595057 |
| Lmf2 | 0.149251912 | -0.555046923 | 0.095 | 0.243 | 1 | 0.002981905 | -2.071814916 | 0.044 | 0.239 | 1 | 0.149251912 | 0.005954918 |
| Pigc | 0.182857706 | -1.081115734 | 0.048 | 0.153 | 1 | 0.002981905 | -2.141355849 | 0.044 | 0.239 | 1 | 0.182857706 | 0.005954918 |
| Dnajb2 | 0.002988818 | -1.16543202 | 0.143 | 0.534 | 1 | 0.141086449 | -0.206223949 | 0.222 | 0.367 | 1 | 0.141086449 | 0.005968703 |
| Mars1 | 0.060046788 | -0.970084422 | 0.048 | 0.235 | 1 | 0.002989196 | -1.500149238 | 0.067 | 0.277 | 1 | 0.060046788 | 0.005969457 |
| Ccn5 | 0.00300488 | -4.050742085 | 0.048 | 0.366 | 1 | 0.08760086 | 0.161303912 | 0.089 | 0.211 | 1 | 0.08760086 | 0.006000731 |
| Crlf3 | 0.196304924 | -0.410950911 | 0.476 | 0.694 | 1 | 0.003006677 | 0.748660928 | 0.711 | 0.488 | 1 | 0.196304924 | 0.006004314 |
| Nutf2-ps1 | 0.046752116 | -0.984439715 | 0.095 | 0.313 | 1 | 0.003009218 | -2.017367132 | 0.044 | 0.239 | 1 | 0.046752116 | 0.00600938 |
| Klhl29 | 0.560907514 | 0.180034027 | 0.286 | 0.418 | 1 | 0.003011428 | -0.242360069 | 0.222 | 0.495 | 1 | 0.560907514 | 0.006013787 |
| Slc20a1 | 0.464824553 | 0.923000374 | 0.714 | 0.69 | 1 | 0.003015496 | 0.78228219 | 0.8 | 0.491 | 1 | 0.464824553 | 0.006021899 |
| Aldoc | 0.003024396 | -2.960434252 | 0.048 | 0.381 | 1 | 0.062440474 | -0.266886731 | 0.067 | 0.183 | 1 | 0.062440474 | 0.006039644 |
| Pias4 | 0.122175603 | -0.752492987 | 0.095 | 0.254 | 1 | 0.003036532 | -1.113940392 | 0.111 | 0.356 | 1 | 0.122175603 | 0.006063843 |
| Khsrp | 0.003036654 | -1.683780237 | 0.095 | 0.463 | 1 | 0.154113636 | -0.272073118 | 0.622 | 0.689 | 1 | 0.154113636 | 0.006064087 |
| Mgst2 | 0.003044221 | -3.880817084 | 0 | 0.31 | 1 | 0.052788904 | -0.638855509 | 0.044 | 0.159 | 1 | 0.052788904 | 0.006079174 |
| Ppp1r21 | 0.749989526 | 0.65387221 | 0.143 | 0.183 | 1 | 0.003048891 | -1.539319835 | 0.044 | 0.246 | 1 | 0.749989526 | 0.006088486 |
| Agpat1 | 0.204034537 | -0.818081329 | 0.143 | 0.261 | 1 | 0.003055055 | -1.526380779 | 0.089 | 0.304 | 1 | 0.204034537 | 0.006100776 |
| Hbegf | 0.395892036 | -0.471905688 | 0.619 | 0.437 | 1 | 0.003060729 | -0.245844316 | 0.667 | 0.325 | 1 | 0.395892036 | 0.00611209 |
| Stxbp4 | 0.27063573 | 1.114344359 | 0.286 | 0.213 | 1 | 0.003061333 | -0.880516264 | 0.111 | 0.356 | 1 | 0.27063573 | 0.006113294 |
| Klc2 | 0.028531207 | -2.413691074 | 0 | 0.19 | 1 | 0.003064339 | -1.518561275 | 0.044 | 0.253 | 1 | 0.028531207 | 0.006119287 |
| Ccdc112 | 0.009654158 | -2.926141075 | 0 | 0.25 | 1 | 0.003075942 | -1.356455778 | 0.067 | 0.287 | 1 | 0.009654158 | 0.006142424 |
| Prmt3 | 0.461235479 | 0.3917364 | 0.19 | 0.302 | 1 | 0.003087646 | -0.702985846 | 0.067 | 0.284 | 1 | 0.461235479 | 0.006165758 |
| Socs1 | 0.615120996 | -0.574155746 | 0.19 | 0.239 | 1 | 0.003089608 | -2.259441919 | 0.022 | 0.211 | 1 | 0.615120996 | 0.00616967 |
| Gpn1 | 0.055621886 | -1.535681598 | 0.048 | 0.224 | 1 | 0.003091558 | -2.620708162 | 0.022 | 0.204 | 1 | 0.055621886 | 0.006173558 |
| Ankmy2 | 0.075988584 | -1.666078235 | 0.048 | 0.205 | 1 | 0.003092288 | -2.638855509 | 0.022 | 0.204 | 1 | 0.075988584 | 0.006175013 |
| Znrd2 | 0.020458584 | -2.455511249 | 0 | 0.209 | 1 | 0.003096702 | -1.998751454 | 0.022 | 0.211 | 1 | 0.020458584 | 0.006183814 |
| Morn2 | 0.163452338 | -0.611630451 | 0.095 | 0.235 | 1 | 0.003098238 | -2.564854927 | 0.022 | 0.204 | 1 | 0.163452338 | 0.006186877 |
| Mpv17l2 | 0.108655481 | -0.496153234 | 0.143 | 0.336 | 1 | 0.00310468 | -2.506751973 | 0.022 | 0.204 | 1 | 0.108655481 | 0.006199721 |
| Trp53rka | 0.593100443 | -0.233118828 | 0.095 | 0.134 | 1 | 0.003105219 | -2.583713955 | 0.022 | 0.204 | 1 | 0.593100443 | 0.006200795 |
| Pum2 | 0.003107216 | 0.966553517 | 0.952 | 0.918 | 1 | 0.092628133 | 0.373918853 | 0.978 | 0.965 | 1 | 0.092628133 | 0.006204777 |
| Plscr1 | 0.04405835 | -1.683780237 | 0.238 | 0.459 | 1 | 0.003109006 | 0.104804738 | 0.644 | 0.343 | 1 | 0.04405835 | 0.006208346 |
| Med9 | 0.113012005 | -0.496153234 | 0.095 | 0.261 | 1 | 0.003112843 | -1.931637258 | 0.067 | 0.266 | 1 | 0.113012005 | 0.006215996 |
| St3gal4 | 0.003113502 | -0.32873074 | 0.333 | 0.728 | 1 | 0.933141557 | 0.199038994 | 0.578 | 0.63 | 1 | 0.933141557 | 0.00621731 |
| Tmem219 | 0.042453888 | -1.666078235 | 0.048 | 0.239 | 1 | 0.003124577 | -0.764386391 | 0.067 | 0.284 | 1 | 0.042453888 | 0.006239391 |
| Peak1 | 0.159840541 | 0.559815578 | 0.667 | 0.601 | 1 | 0.003131632 | 0.808603468 | 0.778 | 0.512 | 1 | 0.159840541 | 0.006253457 |
| Nxt1 | 0.23196064 | -1.054148687 | 0.048 | 0.138 | 1 | 0.003131825 | -1.47073275 | 0.044 | 0.246 | 1 | 0.23196064 | 0.006253841 |
| Wdr18 | 0.234178056 | -0.818081329 | 0.143 | 0.25 | 1 | 0.003136739 | -1.290932205 | 0.067 | 0.284 | 1 | 0.234178056 | 0.006263638 |
| Rad23a | 0.003137277 | -1.303508156 | 0.19 | 0.578 | 1 | 0.057950625 | -0.572688585 | 0.467 | 0.564 | 1 | 0.057950625 | 0.006264711 |
| Mrpl10 | 0.005375103 | -2.184209227 | 0.048 | 0.343 | 1 | 0.003140209 | -0.511305459 | 0.089 | 0.325 | 1 | 0.005375103 | 0.006270557 |
| Elovl1 | 0.003736972 | -1.59769126 | 0.19 | 0.56 | 1 | 0.003141551 | -0.594912161 | 0.156 | 0.426 | 1 | 0.003736972 | 0.006273234 |
| Nid1 | 0.720698583 | 0.986711079 | 0.19 | 0.254 | 1 | 0.003143185 | -2.298780067 | 0.044 | 0.242 | 1 | 0.720698583 | 0.006276491 |
| Slc35a4 | 0.059553486 | -0.683780237 | 0.143 | 0.377 | 1 | 0.003148399 | -1.199570463 | 0.089 | 0.304 | 1 | 0.059553486 | 0.006286886 |
| Trim37 | 0.668813089 | 0.360888812 | 0.286 | 0.369 | 1 | 0.003150948 | -0.936823705 | 0.156 | 0.422 | 1 | 0.668813089 | 0.006291967 |
| Fuom | 0.28406094 | 0.103916159 | 0.143 | 0.254 | 1 | 0.003155465 | -1.158229668 | 0.044 | 0.246 | 1 | 0.28406094 | 0.006300973 |
| Zim1 | 0.139378732 | -1.050254771 | 0.143 | 0.299 | 1 | 0.003167664 | -0.762486858 | 0.111 | 0.37 | 1 | 0.139378732 | 0.006325295 |
| Diablo | 0.00317154 | -1.901137068 | 0.095 | 0.422 | 1 | 0.034565533 | -0.11601472 | 0.244 | 0.46 | 1 | 0.034565533 | 0.006333021 |

| | | | | | | | | | | | | |
|---|---|---|---|---|---|---|---|---|---|---|---|---|
| Ube2b | 0.006031023 | -0.542229684 | 1 | 0.985 | 1 | 0.003179144 | -0.244239513 | 0.978 | 1 | 1 | 0.006031023 | 0.00634818 |
| Ndufv3 | 0.003182403 | -1.081115734 | 0.333 | 0.687 | 1 | 0.045247017 | -0.728741012 | 0.533 | 0.592 | 1 | 0.045247017 | 0.006354679 |
| Sec23ip | 0.057475601 | -0.724259306 | 0.286 | 0.541 | 1 | 0.003184136 | -0.508068901 | 0.333 | 0.671 | 1 | 0.057475601 | 0.006358133 |
| Ubfd1 | 0.039576611 | -0.807354922 | 0.238 | 0.493 | 1 | 0.003184415 | -0.213091603 | 0.244 | 0.557 | 1 | 0.039576611 | 0.00635869 |
| Phka2 | 0.272621701 | 1.345445902 | 0.476 | 0.481 | 1 | 0.003188126 | -2.154870656 | 0.067 | 0.27 | 1 | 0.272621701 | 0.006366088 |
| Zfp622 | 0.036547253 | -1.233118828 | 0.19 | 0.418 | 1 | 0.003193885 | -1.096905897 | 0.333 | 0.54 | 1 | 0.036547253 | 0.006377569 |
| Ndufs2 | 0.003200208 | -1.15277672 | 0.19 | 0.567 | 1 | 0.010635462 | -0.86124793 | 0.556 | 0.64 | 1 | 0.010635462 | 0.006390175 |
| Becn1 | 0.003202758 | -1.469186186 | 0.429 | 0.795 | 1 | 0.475046767 | 0.187970698 | 0.733 | 0.654 | 1 | 0.475046767 | 0.006395257 |
| Nphp1 | 0.07095166 | -0.60421298 | 0.095 | 0.291 | 1 | 0.003205018 | -1.564854927 | 0.067 | 0.28 | 1 | 0.07095166 | 0.006399764 |
| Ndst2 | 0.920524815 | 0.735172312 | 0.143 | 0.146 | 1 | 0.003213146 | -1.667424661 | 0.067 | 0.273 | 1 | 0.920524815 | 0.006415967 |
| Gm45159 | 0.003214203 | 2.429846185 | 0.333 | 0.127 | 1 | 0.132686876 | -1.404390255 | 0 | 0.048 | 1 | 0.132686876 | 0.006418075 |
| Smarcd1 | 0.855037849 | 0.503846766 | 0.286 | 0.287 | 1 | 0.003214593 | -0.893276784 | 0.133 | 0.374 | 1 | 0.855037849 | 0.006418852 |
| Fosl1 | 0.083150172 | -1.081115734 | 0.095 | 0.284 | 1 | 0.003224483 | -1.441623161 | 0.067 | 0.28 | 1 | 0.083150172 | 0.006438569 |
| Tmcc3 | 0.34394286 | 1.064962525 | 0.286 | 0.224 | 1 | 0.003228862 | 1.715133795 | 0.733 | 0.478 | 1 | 0.34394286 | 0.006447298 |
| Top3b | 0.371660661 | 1.075512444 | 0.333 | 0.284 | 1 | 0.003230274 | -0.560852997 | 0.156 | 0.419 | 1 | 0.371660661 | 0.006450113 |
| Psmb7 | 0.003232497 | -0.49061505 | 0.476 | 0.825 | 1 | 0.951797559 | 0.114788827 | 0.844 | 0.775 | 1 | 0.951797559 | 0.006454545 |
| Igf2r | 0.003234089 | -0.708331842 | 0.81 | 0.963 | 1 | 0.003334374 | -0.336502952 | 0.956 | 0.969 | 1 | 0.003334374 | 0.006457718 |
| Cops4 | 0.003244354 | -0.669217943 | 0.286 | 0.672 | 1 | 0.011587509 | -0.417905062 | 0.578 | 0.737 | 1 | 0.011587509 | 0.006478183 |
| Phf2 | 0.740043235 | 0.201903046 | 0.381 | 0.515 | 1 | 0.003246507 | -0.610234831 | 0.644 | 0.806 | 1 | 0.740043235 | 0.006482473 |
| Nanp | 0.039457724 | -2.25696557 | 0 | 0.172 | 1 | 0.003253621 | -3.247664751 | 0 | 0.166 | 1 | 0.039457724 | 0.006496656 |
| Rab31 | 0.003257943 | -0.942217629 | 0.286 | 0.739 | 1 | 0.100588866 | 0.532358189 | 0.822 | 0.83 | 1 | 0.100588866 | 0.006505272 |
| Zfp358 | 0.089324258 | -1.413691074 | 0 | 0.123 | 1 | 0.003266775 | -1.204452685 | 0.022 | 0.211 | 1 | 0.089324258 | 0.006522879 |
| Fstl4 | 0.719087874 | 1.55829455 | 0.048 | 0.034 | 1 | 0.003270594 | 1.992757085 | 0.178 | 0.059 | 1 | 0.719087874 | 0.006530491 |
| Map4k3 | 0.099319968 | 0.503846766 | 0.714 | 0.519 | 1 | 0.003290905 | 0.544068079 | 0.978 | 0.723 | 1 | 0.099319968 | 0.00657098 |
| Homer3 | 0.381840507 | -0.264827688 | 0.143 | 0.239 | 1 | 0.003295729 | -0.336826971 | 0.156 | 0.429 | 1 | 0.381840507 | 0.006580597 |
| Pde7a | 0.411781099 | 1.351843673 | 0.286 | 0.22 | 1 | 0.003299828 | 0.734787623 | 0.8 | 0.467 | 1 | 0.411781099 | 0.006588768 |
| Fam13b | 0.003309697 | 1.347727564 | 0.905 | 0.769 | 1 | 0.169970127 | 0.524883693 | 0.933 | 0.83 | 1 | 0.169970127 | 0.006608441 |
| Mtmr6 | 0.435959719 | 0.634243403 | 0.476 | 0.537 | 1 | 0.003312343 | -0.596476906 | 0.356 | 0.661 | 1 | 0.435959719 | 0.006613714 |
| Marchf7 | 0.003314765 | 1.059887658 | 0.905 | 0.847 | 1 | 0.070736055 | 0.65142696 | 0.978 | 0.92 | 1 | 0.070736055 | 0.006618543 |
| Fndc1 | 0.003322644 | -2.437363903 | 0.143 | 0.47 | 1 | 0.028728888 | -0.395584358 | 0.156 | 0.367 | 1 | 0.028728888 | 0.006634247 |
| Exosc4 | 0.149610248 | -0.555046923 | 0.095 | 0.243 | 1 | 0.003330749 | -2.141355849 | 0.044 | 0.235 | 1 | 0.149610248 | 0.006650405 |
| Gm49797 | 0.8002109 | 0.936806174 | 0.429 | 0.534 | 1 | 0.00333108 | 0.722094475 | 0.844 | 0.761 | 1 | 0.8002109 | 0.006651064 |
| Jade1 | 0.003333939 | -1.857609693 | 0.048 | 0.392 | 1 | 0.129460909 | -0.453466247 | 0.444 | 0.54 | 1 | 0.129460909 | 0.006656762 |
| Trp53inp2 | 0.007133828 | -1.471905688 | 0.095 | 0.425 | 1 | 0.003338699 | -0.16492432 | 0.178 | 0.464 | 1 | 0.007133828 | 0.00666625 |
| Nsa2 | 0.008098466 | -0.775422742 | 0.81 | 0.896 | 1 | 0.003343296 | -0.466201906 | 0.889 | 0.862 | 1 | 0.008098466 | 0.006675415 |
| Gtf2h3 | 0.032394038 | -1.769171728 | 0.048 | 0.254 | 1 | 0.003347304 | -2.035745661 | 0.044 | 0.235 | 1 | 0.032394038 | 0.006683404 |
| Mpdu1 | 0.006235995 | -1.870548749 | 0.095 | 0.392 | 1 | 0.003358974 | -1.02842232 | 0.111 | 0.339 | 1 | 0.006235995 | 0.006706665 |
| Pkd1 | 0.309750267 | 0.507494543 | 0.429 | 0.735 | 1 | 0.003361537 | -0.335786441 | 0.4 | 0.782 | 1 | 0.309750267 | 0.006711775 |
| Ssbp4 | 0.248151331 | -0.455511249 | 0.238 | 0.403 | 1 | 0.003377211 | -0.379937211 | 0.222 | 0.512 | 1 | 0.248151331 | 0.006743016 |
| Cyba | 0.003379566 | -2.99156415 | 0.048 | 0.362 | 1 | 0.606611455 | -0.266886731 | 0.533 | 0.488 | 1 | 0.606611455 | 0.006747711 |
| Spg7 | 0.004644276 | -1.38800443 | 0.095 | 0.433 | 1 | 0.003385379 | -1.255526869 | 0.156 | 0.394 | 1 | 0.004644276 | 0.006759297 |
| Vegfb | 0.154070699 | -0.574155746 | 0.095 | 0.25 | 1 | 0.003391686 | -1.667424661 | 0.044 | 0.242 | 1 | 0.154070699 | 0.006771868 |
| Bin1 | 0.704035501 | 0.810273768 | 0.238 | 0.351 | 1 | 0.0034028 | -1.008089318 | 0.089 | 0.325 | 1 | 0.704035501 | 0.00679402 |
| Cdh11 | 0.048075875 | -3.348596045 | 0.143 | 0.343 | 1 | 0.00340428 | -1.886783022 | 0.133 | 0.381 | 1 | 0.048075875 | 0.006796971 |
| Tpgs2 | 0.064635954 | -1.385121921 | 0.095 | 0.276 | 1 | 0.003409232 | -1.691966845 | 0.044 | 0.242 | 1 | 0.064635954 | 0.006806841 |
| Man2a2 | 0.144958013 | -0.586755783 | 0.143 | 0.299 | 1 | 0.003431619 | -1.184823878 | 0.111 | 0.343 | 1 | 0.144958013 | 0.006851462 |
| Ift52 | 0.088964407 | -0.729950418 | 0.143 | 0.347 | 1 | 0.003434283 | -1.02842232 | 0.111 | 0.329 | 1 | 0.088964407 | 0.006856772 |
| Ints14 | 0.673522725 | 0.410737362 | 0.19 | 0.261 | 1 | 0.003438931 | -1.017367132 | 0.089 | 0.318 | 1 | 0.673522725 | 0.006866035 |
| Tut7 | 0.045210178 | 1.007314149 | 0.905 | 0.918 | 1 | 0.003444796 | -0.337456227 | 0.933 | 0.972 | 1 | 0.045210178 | 0.006877725 |
| Fbxl12 | 0.830929531 | 0.614878079 | 0.286 | 0.291 | 1 | 0.003461173 | -1.174908409 | 0.156 | 0.381 | 1 | 0.830929531 | 0.006910366 |
| Loxl2 | 0.037531955 | -0.995672798 | 0.333 | 0.608 | 1 | 0.003470941 | -2.496836504 | 0.022 | 0.208 | 1 | 0.037531955 | 0.006929834 |
| Idh3a | 0.003478437 | -1.870548749 | 0.048 | 0.384 | 1 | 0.043009197 | -0.956337699 | 0.244 | 0.388 | 1 | 0.043009197 | 0.006944774 |
| Lyrm2 | 0.046081992 | -1.150656668 | 0.048 | 0.25 | 1 | 0.003486349 | -1.960783604 | 0.022 | 0.208 | 1 | 0.046081992 | 0.006960543 |
| Pop7 | 0.103838915 | -1.116305163 | 0.095 | 0.25 | 1 | 0.003509112 | -2.486852415 | 0.022 | 0.201 | 1 | 0.103838915 | 0.007005911 |
| 2510039O18Rik | 0.199480036 | -0.427766259 | 0.143 | 0.291 | 1 | 0.003513793 | -0.511943396 | 0.178 | 0.446 | 1 | 0.199480036 | 0.007015239 |
| Pik3cb | 0.084557399 | 1.636934999 | 0.476 | 0.388 | 1 | 0.003526772 | 0.935838657 | 0.733 | 0.516 | 1 | 0.084557399 | 0.007041107 |
| Nhp2 | 0.003535202 | -1.249060372 | 0.095 | 0.47 | 1 | 0.040334482 | 0.111530601 | 0.133 | 0.298 | 1 | 0.040334482 | 0.007057906 |
| Zfp322a | 0.639839548 | -0.434752689 | 0.143 | 0.175 | 1 | 0.003540347 | -1.765387915 | 0.067 | 0.273 | 1 | 0.639839548 | 0.00706816 |
| Galk1 | 0.003543807 | -2.280424543 | 0.048 | 0.362 | 1 | 0.159278889 | 0.490427508 | 0.2 | 0.346 | 1 | 0.159278889 | 0.007075055 |
| Abca9 | 0.037341667 | -2.303508156 | 0.048 | 0.243 | 1 | 0.003546552 | -0.776359032 | 0.067 | 0.287 | 1 | 0.037341667 | 0.007080525 |
| Edil3 | 0.786908047 | 0.356068259 | 0.619 | 0.735 | 1 | 0.003554537 | -0.307882274 | 0.533 | 0.869 | 1 | 0.786908047 | 0.007096439 |
| Mfsd1 | 0.00355826 | -2.303508156 | 0.048 | 0.362 | 1 | 0.385750212 | -0.343727473 | 0.356 | 0.391 | 1 | 0.385750212 | 0.00710386 |
| Tmem216 | 0.010253234 | -2.752492987 | 0 | 0.246 | 1 | 0.003559498 | -1.085111739 | 0.089 | 0.301 | 1 | 0.010253234 | 0.007106326 |
| Hipk2 | 0.015524942 | 0.970723197 | 0.952 | 0.802 | 1 | 0.003562722 | 0.680107128 | 0.956 | 0.747 | 1 | 0.015524942 | 0.007112752 |
| Plxnd1 | 0.402092763 | -0.385121921 | 0.048 | 0.108 | 1 | 0.003574703 | -0.647317087 | 0.089 | 0.315 | 1 | 0.402092763 | 0.007136627 |
| Slpi | 0.060258713 | 1.046035243 | 0.095 | 0.325 | 1 | 0.003588817 | -0.316927414 | 0.178 | 0.439 | 1 | 0.060258713 | 0.007164754 |
| Rnf38 | 0.305197419 | -0.199056992 | 0.333 | 0.504 | 1 | 0.003595584 | 0.993240968 | 0.867 | 0.678 | 1 | 0.305197419 | 0.00717824 |
| Asph | 0.003619251 | -1.30677587 | 0.333 | 0.679 | 1 | 0.505354346 | 0.309192412 | 0.822 | 0.799 | 1 | 0.505354346 | 0.007225403 |
| Rundc1 | 0.07854135 | -1.752492987 | 0 | 0.131 | 1 | 0.003619556 | -1.620708162 | 0.022 | 0.208 | 1 | 0.07854135 | 0.007226011 |
| Cryz | 0.138420052 | -0.940938076 | 0.048 | 0.172 | 1 | 0.003621116 | -3.044847868 | 0 | 0.163 | 1 | 0.138420052 | 0.00722912 |
| Scube2 | 0.070480969 | -0.535681598 | 0.048 | 0.228 | 1 | 0.003625022 | -2.683249628 | 0.022 | 0.204 | 1 | 0.070480969 | 0.007236904 |
| Endog | 0.030310792 | -2.184209227 | 0 | 0.187 | 1 | 0.003627472 | -3.124282336 | 0 | 0.163 | 1 | 0.030310792 | 0.007241785 |
| Mgrn1 | 0.00362765 | -1.709932525 | 0.095 | 0.437 | 1 | 0.475072117 | -0.194936889 | 0.533 | 0.561 | 1 | 0.475072117 | 0.007242141 |
| Slc27a4 | 0.07382541 | -1.752492987 | 0 | 0.134 | 1 | 0.003627683 | -0.952515988 | 0.044 | 0.249 | 1 | 0.07382541 | 0.007242206 |
| Acadsb | 0.003636743 | -1.169925001 | 0.333 | 0.705 | 1 | 0.045400873 | 0.319667855 | 0.467 | 0.754 | 1 | 0.045400873 | 0.007260259 |
| Unc45a | 0.003649365 | -3.133583154 | 0 | 0.299 | 1 | 0.939053682 | 0.15414412 | 0.444 | 0.467 | 1 | 0.939053682 | 0.007285413 |
| Ccdc8 | 0.065188054 | -1.880817084 | 0 | 0.142 | 1 | 0.003649913 | -3.247664751 | 0 | 0.163 | 1 | 0.065188054 | 0.007286504 |
| Asb4 | 0.069782925 | -3.107587946 | 0 | 0.138 | 1 | 0.003654548 | -1.693896872 | 0.044 | 0.239 | 1 | 0.069782925 | 0.00729574 |
| Gtf3a | 0.172830402 | -1.054148687 | 0.048 | 0.157 | 1 | 0.003655801 | -1.486852415 | 0.022 | 0.208 | 1 | 0.172830402 | 0.007298236 |
| Cactin | 0.498054776 | 0.451379346 | 0.048 | 0.097 | 1 | 0.003659159 | -3.294207337 | 0 | 0.163 | 1 | 0.498054776 | 0.007304928 |
| Chid1 | 0.033828741 | -1.599246727 | 0.095 | 0.31 | 1 | 0.003664669 | -1.029645462 | 0.111 | 0.329 | 1 | 0.033828741 | 0.007315908 |
| Ddx52 | 0.007105425 | -0.707050016 | 0.048 | 0.358 | 1 | 0.003671703 | -0.931637258 | 0.111 | 0.346 | 1 | 0.007105425 | 0.007329924 |
| Sec24b | 0.003672795 | 1.242219182 | 0.762 | 0.664 | 1 | 0.004012632 | 0.855253562 | 0.911 | 0.844 | 1 | 0.004012632 | 0.007332101 |
| Mycbp | 0.016085383 | -1.054148687 | 0.095 | 0.392 | 1 | 0.003674316 | -1.393743011 | 0.089 | 0.298 | 1 | 0.016085383 | 0.007335131 |
| D130009I18Rik | 0.035008289 | -0.701962771 | 0.476 | 0.754 | 1 | 0.003692076 | -0.997814335 | 0.756 | 0.72 | 1 | 0.035008289 | 0.007370521 |
| Twf1 | 0.003694762 | -0.964828697 | 0.19 | 0.593 | 1 | 0.054165383 | -0.585794265 | 0.6 | 0.647 | 1 | 0.054165383 | 0.007375873 |
| Rexo4 | 0.106090666 | -0.541241123 | 0.143 | 0.336 | 1 | 0.003699338 | -0.647844292 | 0.133 | 0.391 | 1 | 0.106090666 | 0.007384991 |
| Ipo7 | 0.003699992 | 1.178076605 | 0.905 | 0.858 | 1 | 0.068776638 | -0.159571086 | 0.911 | 0.917 | 1 | 0.068776638 | 0.007386294 |
| Cog2 | 0.778640404 | -0.233118828 | 0.286 | 0.302 | 1 | 0.003706804 | -0.584699739 | 0.156 | 0.422 | 1 | 0.778640404 | 0.007399867 |
| Nsmce2 | 0.29269571 | 0.424883418 | 0.857 | 0.78 | 1 | 0.003720067 | 0.48586672 | 0.978 | 0.82 | 1 | 0.29269571 | 0.007426295 |
| Metrn | 0.003721636 | -3.370622352 | 0 | 0.299 | 1 | 0.085201566 | -0.486852415 | 0.067 | 0.173 | 1 | 0.085201566 | 0.007429421 |
| Tmprss2 | 0.019692488 | 4.258734268 | 0.048 | 0.004 | 1 | 0.003726601 | 2.393565969 | 0.133 | 0.035 | 1 | 0.019692488 | 0.007439315 |
| Ift172 | 0.574078568 | 1.010806755 | 0.238 | 0.205 | 1 | 0.003728905 | -0.975138897 | 0.044 | 0.249 | 1 | 0.574078568 | 0.007443904 |
| Otud3 | 0.647051988 | 0.181918671 | 0.095 | 0.134 | 1 | 0.003729038 | -1.181997834 | 0.044 | 0.249 | 1 | 0.647051988 | 0.007444169 |
| Arfgef2 | 0.009269646 | 1.225311267 | 0.762 | 0.754 | 1 | 0.003748766 | 0.804799034 | 0.933 | 0.813 | 1 | 0.009269646 | 0.007483478 |
| Slc16a1 | 0.060244574 | -1.124970024 | 0.143 | 0.354 | 1 | 0.003760741 | -0.64605101 | 0.133 | 0.394 | 1 | 0.060244574 | 0.007507339 |
| Mrpl49 | 0.207125797 | -0.434752689 | 0.095 | 0.22 | 1 | 0.003770514 | -1.743192169 | 0.044 | 0.239 | 1 | 0.207125797 | 0.007526811 |
| Alkbh6 | 0.003785966 | -2.849790188 | 0 | 0.295 | 1 | 0.011271142 | -1.421264074 | 0.044 | 0.211 | 1 | 0.011271142 | 0.007557592 |

| | | | | | | | | | | | | |
|---|---|---|---|---|---|---|---|---|---|---|---|---|
| Dnm2 | 0.362776157 | 0.673771768 | 0.762 | 0.683 | 1 | 0.003791404 | 0.668044377 | 0.978 | 0.837 | 1 | 0.362776157 | 0.007568433 |
| Myo1b | 0.069906055 | -3.392317423 | 0 | 0.138 | 1 | 0.003797001 | -0.477919291 | 0.356 | 0.149 | 1 | 0.069906055 | 0.007579584 |
| Nhsl1 | 0.144964687 | -1.683780237 | 0 | 0.093 | 1 | 0.003798804 | 1.055041364 | 0.222 | 0.08 | 1 | 0.144964687 | 0.007583177 |
| Scaf1 | 0.151251988 | 0.753206235 | 0.476 | 0.418 | 1 | 0.003802548 | -0.492566066 | 0.222 | 0.516 | 1 | 0.151251988 | 0.007590636 |
| Cog6 | 0.357389714 | -0.496153234 | 0.19 | 0.28 | 1 | 0.003807698 | -0.735428768 | 0.111 | 0.346 | 1 | 0.357389714 | 0.007600898 |
| Arhgef7 | 0.210419927 | -0.34303652 | 0.238 | 0.437 | 1 | 0.00382028 | -1.053893008 | 0.133 | 0.367 | 1 | 0.210419927 | 0.007625965 |
| Sdhaf2 | 0.003823879 | -1.648156327 | 0.095 | 0.437 | 1 | 0.008445037 | -0.884611923 | 0.156 | 0.374 | 1 | 0.008445037 | 0.007633137 |
| Ebna1bp2 | 0.050995363 | -0.660128969 | 0.381 | 0.694 | 1 | 0.003837249 | -0.609709163 | 0.4 | 0.64 | 1 | 0.050995363 | 0.007659774 |
| Zfp703 | 0.053194958 | -0.544935607 | 0.286 | 0.601 | 1 | 0.003840075 | -0.525094764 | 0.644 | 0.785 | 1 | 0.053194958 | 0.007665403 |
| Smim12 | 0.021710607 | -2.326228232 | 0 | 0.205 | 1 | 0.003841494 | -1.691966845 | 0.044 | 0.239 | 1 | 0.021710607 | 0.007668231 |
| Zfp68 | 0.70918494 | 0.162809848 | 0.238 | 0.306 | 1 | 0.003849239 | -0.734335818 | 0.178 | 0.45 | 1 | 0.70918494 | 0.007683662 |
| Car6 | 0.003862306 | -4.564632972 | 0 | 0.299 | 1 | 0.376837034 | -0.124282336 | 0 | 0.017 | 1 | 0.376837034 | 0.007709694 |
| Slc1a4 | 0.003870214 | 1.908237021 | 0.619 | 0.459 | 1 | 0.512288033 | 0.262964097 | 0.4 | 0.595 | 1 | 0.512288033 | 0.007725449 |
| Grik5 | 0.312419766 | -0.752492987 | 0.048 | 0.119 | 1 | 0.00387556 | -0.879863608 | 0.067 | 0.284 | 1 | 0.312419766 | 0.007736101 |
| Soat1 | 0.003877431 | -1.254813899 | 0.19 | 0.59 | 1 | 0.409983852 | 0.133338574 | 0.511 | 0.612 | 1 | 0.409983852 | 0.007739828 |
| Gas8 | 0.267258192 | -0.555046923 | 0.095 | 0.198 | 1 | 0.003880838 | -1.47073275 | 0.044 | 0.239 | 1 | 0.267258192 | 0.007746616 |
| Psmd4 | 0.003883545 | -0.801313115 | 0.667 | 0.843 | 1 | 0.004421595 | -0.608049462 | 0.8 | 0.844 | 1 | 0.004421595 | 0.007752008 |
| Vps53 | 0.003886151 | -2.063193826 | 0.048 | 0.377 | 1 | 0.034270804 | -0.608158711 | 0.244 | 0.422 | 1 | 0.034270804 | 0.0077572 |
| Ubap2l | 0.003901926 | 0.625766395 | 1 | 0.944 | 1 | 0.931603753 | 0.327749371 | 0.978 | 0.976 | 1 | 0.931603753 | 0.007788626 |
| Lrrc47 | 0.488012175 | 0.192645078 | 0.143 | 0.228 | 1 | 0.003904146 | -1.255526869 | 0.111 | 0.339 | 1 | 0.488012175 | 0.007793049 |
| Cpne2 | 0.717475871 | 1.036341847 | 0.19 | 0.265 | 1 | 0.003918116 | -0.810916255 | 0.133 | 0.37 | 1 | 0.717475871 | 0.00782088 |
| Timm50 | 0.010137153 | -2.01272876 | 0.048 | 0.313 | 1 | 0.003921249 | -1.946284034 | 0.067 | 0.26 | 1 | 0.010137153 | 0.007827121 |
| Mbip | 0.364029746 | 0.926537838 | 0.333 | 0.276 | 1 | 0.003923389 | -0.204452685 | 0.089 | 0.329 | 1 | 0.364029746 | 0.007831385 |
| Mfsd14b | 0.003924283 | -1.314912919 | 0.143 | 0.507 | 1 | 0.188955174 | -0.250500052 | 0.6 | 0.685 | 1 | 0.188955174 | 0.007833166 |
| Ddx3y | 0.116498135 | -0.41151342 | 0.429 | 0.563 | 1 | 0.003934738 | -0.407216299 | 0.933 | 0.948 | 1 | 0.116498135 | 0.007853994 |
| 0610010K14Rik | 0.003937311 | -1.65714511 | 0.048 | 0.384 | 1 | 0.005902802 | -0.361321533 | 0.111 | 0.336 | 1 | 0.005902802 | 0.00785912 |
| As3mt | 0.519943428 | 0.119182916 | 0.095 | 0.153 | 1 | 0.003943079 | -1.003427941 | 0.044 | 0.246 | 1 | 0.519943428 | 0.00787061 |
| Alg1 | 0.008689912 | -2.067695219 | 0.048 | 0.321 | 1 | 0.00394457 | -0.702985846 | 0.089 | 0.318 | 1 | 0.008689912 | 0.00787358 |
| Apoo | 0.257347072 | -0.167530486 | 0.048 | 0.142 | 1 | 0.003947475 | -2.404390255 | 0.022 | 0.197 | 1 | 0.257347072 | 0.007879368 |
| Kin | 0.037519277 | -0.667696514 | 0.19 | 0.459 | 1 | 0.003950036 | -0.812622576 | 0.222 | 0.495 | 1 | 0.037519277 | 0.007884469 |
| Flrt2 | 0.931501385 | 0.480102412 | 0.762 | 0.776 | 1 | 0.003958873 | -0.339626462 | 0.733 | 0.941 | 1 | 0.931501385 | 0.007902074 |
| Drg2 | 0.062456469 | -1.20069735 | 0.095 | 0.28 | 1 | 0.003985346 | -0.627714951 | 0.111 | 0.346 | 1 | 0.062456469 | 0.00795481 |
| Abhd13 | 0.84124238 | 0.483947209 | 0.19 | 0.239 | 1 | 0.003988482 | -0.202284848 | 0.111 | 0.36 | 1 | 0.84124238 | 0.007961055 |
| Strn3 | 0.003989364 | 0.588238954 | 1 | 0.966 | 1 | 0.150057388 | 0.418060531 | 1 | 0.99 | 1 | 0.150057388 | 0.007962813 |
| Ppil1 | 0.112111576 | -1.326228232 | 0.048 | 0.183 | 1 | 0.003990051 | -2.271123724 | 0.022 | 0.197 | 1 | 0.112111576 | 0.007964181 |
| Ndufab1 | 0.003996435 | -0.816553859 | 0.476 | 0.784 | 1 | 0.009329975 | -0.504234493 | 0.756 | 0.765 | 1 | 0.009329975 | 0.007976898 |
| Thoc2l | 0.004003203 | 1.007125469 | 0.857 | 0.81 | 1 | 0.972535388 | 0.40799499 | 0.889 | 0.824 | 1 | 0.972535388 | 0.007990381 |
| Mrc2 | 0.180890245 | 0.175744904 | 0.476 | 0.769 | 1 | 0.004007052 | -0.122907686 | 0.467 | 0.851 | 1 | 0.180890245 | 0.007998048 |
| Gm56906 | 0.060055851 | -2.299686445 | 0.19 | 0.366 | 1 | 0.004009191 | -1.19663318 | 0.244 | 0.478 | 1 | 0.060055851 | 0.008002308 |
| Tecpr1 | 0.058525788 | -0.801961663 | 0.048 | 0.239 | 1 | 0.004011051 | -1.440309829 | 0.067 | 0.273 | 1 | 0.058525788 | 0.008006013 |
| Cnot9 | 0.024416769 | -1.849790188 | 0.048 | 0.269 | 1 | 0.004033332 | -1.089516918 | 0.089 | 0.304 | 1 | 0.024416769 | 0.008050396 |
| Slk | 0.004038389 | 0.894073189 | 1 | 0.929 | 1 | 0.044117127 | 0.427277596 | 1 | 0.955 | 1 | 0.044117127 | 0.00806047 |
| A430018G15Rik | 0.241188472 | -0.295854583 | 0.048 | 0.146 | 1 | 0.00403894 | -2.931637258 | 0 | 0.159 | 1 | 0.241188472 | 0.008061567 |
| H4c4 | 0.036745215 | -2.081115734 | 0.095 | 0.302 | 1 | 0.004042995 | -1.509572492 | 0.089 | 0.301 | 1 | 0.036745215 | 0.008069645 |
| Pacsin3 | 0.210526979 | -0.555046923 | 0.095 | 0.216 | 1 | 0.00404912 | -0.559783938 | 0.111 | 0.356 | 1 | 0.210526979 | 0.008081844 |
| Atp6v1c1 | 0.469541572 | 0.205002284 | 0.238 | 0.381 | 1 | 0.004051607 | -1.355846403 | 0.244 | 0.464 | 1 | 0.469541572 | 0.008086799 |
| Tmem165 | 0.026073433 | -0.53338614 | 0.524 | 0.81 | 1 | 0.004053071 | -0.475116307 | 0.733 | 0.858 | 1 | 0.026073433 | 0.008089714 |
| Dtx3 | 0.81573243 | 0.418514712 | 0.524 | 0.649 | 1 | 0.004053844 | -0.432404631 | 0.311 | 0.664 | 1 | 0.81573243 | 0.008091254 |
| L3mbtl3 | 0.004056228 | -3.535681598 | 0 | 0.295 | 1 | 0.01074608 | -0.854361544 | 0.156 | 0.367 | 1 | 0.01074608 | 0.008096004 |
| Ppp6r3 | 0.042968732 | 1.101192991 | 0.81 | 0.731 | 1 | 0.004056939 | 0.56933042 | 0.978 | 0.879 | 1 | 0.042968732 | 0.008097418 |
| Gnl3l | 0.015525405 | -0.546276713 | 0.19 | 0.526 | 1 | 0.004058156 | -0.251474352 | 0.2 | 0.522 | 1 | 0.015525405 | 0.008099842 |
| Mrpl48 | 0.293901547 | 0.866416846 | 0.714 | 0.802 | 1 | 0.004066664 | -0.602329633 | 0.644 | 0.768 | 1 | 0.293901547 | 0.008116791 |
| Mlx | 0.005797874 | -2.25696557 | 0.048 | 0.34 | 1 | 0.004067452 | -1.135687099 | 0.111 | 0.332 | 1 | 0.005797874 | 0.008118361 |
| Fam199x | 0.731164999 | 0.696491844 | 0.19 | 0.175 | 1 | 0.004068446 | -1.017367132 | 0.111 | 0.339 | 1 | 0.731164999 | 0.00812034 |
| Slc9a9 | 0.918194228 | 1.088809267 | 0.19 | 0.224 | 1 | 0.004074732 | 0.868287459 | 0.644 | 0.367 | 1 | 0.918194228 | 0.008132861 |
| Gm57390 | 0.069325635 | -1.785659851 | 0 | 0.138 | 1 | 0.00408072 | -3.149817428 | 0 | 0.159 | 1 | 0.069325635 | 0.008144787 |
| Slc22a15 | 0.412041545 | 0.795450324 | 0.19 | 0.336 | 1 | 0.00408097 | -0.846180482 | 0.067 | 0.277 | 1 | 0.412041545 | 0.008145285 |
| Sat2 | 0.028415202 | -2.25696557 | 0 | 0.19 | 1 | 0.004086687 | -3.316927414 | 0 | 0.159 | 1 | 0.028415202 | 0.008156673 |
| Dhx8 | 0.261366885 | -0.197713492 | 0.238 | 0.388 | 1 | 0.004088263 | -0.510407143 | 0.2 | 0.491 | 1 | 0.261366885 | 0.008159813 |
| Amotl1 | 0.262262696 | 0.48165102 | 0.81 | 0.806 | 1 | 0.00409028 | -0.50259822 | 0.778 | 0.931 | 1 | 0.262262696 | 0.00816383 |
| Spata21 | 0.004112952 | 3.995699863 | 0.095 | 0.011 | 1 | 0.833619716 | 0.361144491 | 0.022 | 0.028 | 1 | 0.833619716 | 0.008208987 |
| Hapln3 | 0.072482187 | -1.370622352 | 0.048 | 0.209 | 1 | 0.004120699 | -1.602329633 | 0.022 | 0.204 | 1 | 0.072482187 | 0.008224417 |
| Cbr3 | 0.125413951 | -1.326228232 | 0.048 | 0.175 | 1 | 0.004121638 | -3.564854927 | 0 | 0.159 | 1 | 0.125413951 | 0.008226289 |
| Abcf3 | 0.248293824 | -0.38923803 | 0.19 | 0.328 | 1 | 0.004123279 | -0.297818591 | 0.133 | 0.398 | 1 | 0.248293824 | 0.008229556 |
| Adat1 | 0.111943936 | 1.611487489 | 0.333 | 0.213 | 1 | 0.0041298 | -2.346674757 | 0.022 | 0.201 | 1 | 0.111943936 | 0.008242545 |
| Thoc6 | 0.987072735 | 0.631951592 | 0.286 | 0.343 | 1 | 0.004139719 | -0.280401538 | 0.156 | 0.419 | 1 | 0.987072735 | 0.008262301 |
| Tpst2 | 0.065271629 | -0.517848305 | 0.238 | 0.515 | 1 | 0.004141402 | -0.6640928 | 0.156 | 0.412 | 1 | 0.065271629 | 0.008265654 |
| Ppp1r2 | 0.004143842 | -0.65559931 | 0.667 | 0.851 | 1 | 0.157746263 | -0.304987801 | 0.911 | 0.865 | 1 | 0.157746263 | 0.008270512 |
| Rusc1 | 0.202127697 | -0.39661756 | 0.095 | 0.231 | 1 | 0.004149562 | -1.446210431 | 0.067 | 0.273 | 1 | 0.202127697 | 0.008281904 |
| Mcrs1 | 0.041190951 | -1.01272876 | 0.095 | 0.321 | 1 | 0.004153163 | -0.916840256 | 0.067 | 0.277 | 1 | 0.041190951 | 0.008289077 |
| Wdfy1 | 0.018273495 | 1.088809267 | 0.571 | 0.437 | 1 | 0.004161687 | 0.937336782 | 0.689 | 0.519 | 1 | 0.018273495 | 0.008306054 |
| Zdhhc4 | 0.004163836 | -2.896083841 | 0 | 0.291 | 1 | 0.004880612 | -1.346674757 | 0.067 | 0.27 | 1 | 0.004880612 | 0.008310335 |
| Ybx3 | 0.004179176 | -0.622161119 | 0.905 | 0.955 | 1 | 0.03545078 | -0.317721631 | 1 | 0.969 | 1 | 0.03545078 | 0.008340886 |
| B3galt6 | 0.00418011 | -2.940938076 | 0 | 0.291 | 1 | 0.004482625 | -1.089516918 | 0.044 | 0.235 | 1 | 0.004482625 | 0.008342746 |
| Rictor | 0.004187371 | 0.883225133 | 0.81 | 0.739 | 1 | 0.025388837 | 0.719122247 | 0.956 | 0.824 | 1 | 0.025388837 | 0.008357207 |
| Acvr1c | 0.004191879 | 4.159198595 | 0.095 | 0.011 | 1 | 0.945046771 | 0.595609745 | 0.022 | 0.024 | 1 | 0.945046771 | 0.008366187 |
| Kxd1 | 0.00419533 | -2.984439715 | 0 | 0.291 | 1 | 0.097247647 | 0.242499995 | 0.156 | 0.301 | 1 | 0.097247647 | 0.008373058 |
| Frg1 | 0.004199214 | -1.300961474 | 0.238 | 0.578 | 1 | 0.492069458 | 0.319774057 | 0.6 | 0.529 | 1 | 0.492069458 | 0.008380794 |
| Ube2i | 0.086928687 | -0.347289848 | 0.81 | 0.925 | 1 | 0.00420457 | -0.352292567 | 0.889 | 0.958 | 1 | 0.086928687 | 0.008391461 |
| Dolk | 0.063806775 | -0.93109029 | 0.048 | 0.231 | 1 | 0.004206804 | -1.72631835 | 0.044 | 0.235 | 1 | 0.063806775 | 0.008395911 |
| Herc4 | 0.004223324 | 0.574236094 | 0.905 | 0.69 | 1 | 0.018294958 | 0.89907668 | 0.911 | 0.723 | 1 | 0.018294958 | 0.008428811 |
| Gm20544 | 0.004230311 | 3.503846766 | 0.143 | 0.026 | 1 | 0.260665169 | -0.901889915 | 0 | 0.028 | 1 | 0.260665169 | 0.008442727 |
| Wsb2 | 0.004243067 | -0.948945655 | 0.19 | 0.616 | 1 | 0.008112432 | -0.293790915 | 0.422 | 0.699 | 1 | 0.008112432 | 0.008468131 |
| Mtus1 | 0.005478607 | 1.717463849 | 0.952 | 0.899 | 1 | 0.00424324 | 0.815452768 | 0.978 | 0.955 | 1 | 0.005478607 | 0.008468476 |
| Tgfb1i1 | 0.684829325 | 1.126877308 | 0.286 | 0.425 | 1 | 0.004254507 | -0.49236942 | 0.244 | 0.522 | 1 | 0.684829325 | 0.008490913 |
| Tgif1 | 0.004265688 | -1.235521318 | 0.476 | 0.72 | 1 | 0.600779431 | 0.223640968 | 0.8 | 0.72 | 1 | 0.600779431 | 0.00851318 |
| Dpp9 | 0.335335793 | -0.348596045 | 0.095 | 0.183 | 1 | 0.004269022 | -0.511305459 | 0.111 | 0.336 | 1 | 0.335335793 | 0.008519819 |
| Eif4a3 | 0.075910098 | -0.359071489 | 0.333 | 0.612 | 1 | 0.004277416 | -0.779634164 | 0.533 | 0.685 | 1 | 0.075910098 | 0.008536536 |
| Gga1 | 0.134668566 | -0.413691074 | 0.095 | 0.25 | 1 | 0.004277598 | -1.11134328 | 0.089 | 0.304 | 1 | 0.134668566 | 0.008536898 |
| Gart | 0.52853668 | 0.199840579 | 0.19 | 0.284 | 1 | 0.004279024 | -0.850905986 | 0.089 | 0.315 | 1 | 0.52853668 | 0.008539738 |
| Sgcb | 0.057837265 | -1.216999163 | 0.048 | 0.235 | 1 | 0.004288672 | -1.789415185 | 0.067 | 0.263 | 1 | 0.057837265 | 0.008558951 |
| Slc30a5 | 0.031197021 | -0.377508737 | 0.286 | 0.578 | 1 | 0.004295015 | -0.511099962 | 0.511 | 0.768 | 1 | 0.031197021 | 0.008571584 |
| Bpifb5 | 0.069730706 | -2.63000898 | 0 | 0.138 | 1 | 0.004301159 | -3.199570463 | 0.022 | 0.194 | 1 | 0.069730706 | 0.008583817 |
| Zmynd19 | 0.221686161 | -0.233118828 | 0.095 | 0.228 | 1 | 0.00431184 | -1.174908409 | 0.067 | 0.273 | 1 | 0.221686161 | 0.008605088 |
| Polr1a | 0.815897383 | 0.866416846 | 0.238 | 0.31 | 1 | 0.004319452 | -1.047320354 | 0.111 | 0.329 | 1 | 0.815897383 | 0.008620246 |
| Pom121 | 0.084475168 | -0.268742738 | 0.286 | 0.53 | 1 | 0.004326104 | -0.169925001 | 0.289 | 0.609 | 1 | 0.084475168 | 0.008633493 |
| Carhsp1 | 0.005764354 | -2.486099569 | 0.048 | 0.34 | 1 | 0.004333421 | -0.802354241 | 0.133 | 0.374 | 1 | 0.005764354 | 0.008648063 |
| Phlda1 | 0.044663418 | -0.61934263 | 0.81 | 0.869 | 1 | 0.004336006 | -1.191396532 | 0.756 | 0.744 | 1 | 0.044663418 | 0.008653211 |

| | | | | | | | | | | | | |
|---|---|---|---|---|---|---|---|---|---|---|---|---|
| Gm37612 | 0.004352726 | 3.673771768 | 0.095 | 0.011 | 1 | 0.492040936 | 2.268035087 | 0.022 | 0.01 | 1 | 0.492040936 | 0.008686505 |
| Blvrb | 0.083669264 | -1.067695219 | 0.095 | 0.272 | 1 | 0.00435419 | 0.84285303 | 0.533 | 0.301 | 1 | 0.083669264 | 0.00868942 |
| Acat2 | 0.295648816 | 0.503846766 | 0.19 | 0.336 | 1 | 0.00435718 | -1.137106376 | 0.044 | 0.235 | 1 | 0.295648816 | 0.008695376 |
| Epn1 | 0.004357988 | -1.648156327 | 0.095 | 0.429 | 1 | 0.01047364 | -0.716098508 | 0.267 | 0.495 | 1 | 0.01047364 | 0.008696985 |
| Lztr1 | 0.184718156 | -0.243766072 | 0.19 | 0.396 | 1 | 0.004360994 | -0.399389574 | 0.133 | 0.384 | 1 | 0.184718156 | 0.008702969 |
| Zfc3h1 | 0.004372959 | 0.79632495 | 0.952 | 0.881 | 1 | 0.92349937 | 0.327293712 | 1 | 0.958 | 1 | 0.92349937 | 0.008726795 |
| Uevld | 0.083682568 | -1.752492987 | 0 | 0.127 | 1 | 0.00438493 | -2.602329633 | 0.022 | 0.194 | 1 | 0.083682568 | 0.008750632 |
| Zfp24 | 0.13403744 | -0.358077098 | 0.286 | 0.504 | 1 | 0.004385989 | -0.372780649 | 0.289 | 0.626 | 1 | 0.13403744 | 0.008752742 |
| Apba3 | 0.268929139 | -0.326228232 | 0.095 | 0.201 | 1 | 0.00438742 | -1.138929112 | 0.067 | 0.266 | 1 | 0.268929139 | 0.008755591 |
| Eef2kmt | 0.246677786 | 0.264380832 | 0.095 | 0.22 | 1 | 0.004395416 | -2.545746104 | 0.022 | 0.194 | 1 | 0.246677786 | 0.008771512 |
| Frs2 | 0.67642026 | -0.146020258 | 0.571 | 0.69 | 1 | 0.004398932 | 0.717926418 | 0.911 | 0.865 | 1 | 0.67642026 | 0.008778513 |
| Ercc2 | 0.078540653 | -1.718545655 | 0 | 0.131 | 1 | 0.004400354 | -1.881712033 | 0.022 | 0.201 | 1 | 0.078540653 | 0.008781344 |
| Smim27 | 0.083456568 | -0.769171728 | 0.095 | 0.28 | 1 | 0.004409755 | -1.881712033 | 0.022 | 0.201 | 1 | 0.083456568 | 0.008800064 |
| Cdkal1 | 0.004411359 | 1.292522582 | 0.762 | 0.634 | 1 | 0.02572074 | 0.911060045 | 0.911 | 0.765 | 1 | 0.02572074 | 0.008803258 |
| Cables2 | 0.740122589 | 0.866416846 | 0.143 | 0.123 | 1 | 0.004427123 | -2.466674533 | 0.022 | 0.194 | 1 | 0.740122589 | 0.008834646 |
| Sppl2b | 0.011760587 | -2.04047375 | 0.048 | 0.306 | 1 | 0.004433415 | -1.37003875 | 0.044 | 0.235 | 1 | 0.011760587 | 0.008847175 |
| Gm29677 | 0.00443639 | 3.410737362 | 0.095 | 0.011 | 1 | 0.158797077 | 3.135584791 | 0.067 | 0.028 | 1 | 0.158797077 | 0.008853099 |
| Dkc1 | 0.004436783 | -2.506137322 | 0.048 | 0.351 | 1 | 0.006128872 | -1.144746438 | 0.089 | 0.291 | 1 | 0.006128872 | 0.00885388 |
| Sulf2 | 0.040420618 | -1.556794585 | 0.19 | 0.422 | 1 | 0.004445611 | -1.312173884 | 0.178 | 0.398 | 1 | 0.040420618 | 0.008871458 |
| Prps1l3 | 0.147764816 | -0.469186186 | 0.048 | 0.179 | 1 | 0.004455132 | -2.361321533 | 0.022 | 0.194 | 1 | 0.147764816 | 0.008890415 |
| Npepps | 0.007048171 | 1.055334562 | 0.905 | 0.847 | 1 | 0.004462377 | 0.609555563 | 1 | 0.938 | 1 | 0.007048171 | 0.008904841 |
| Jmjd1c | 0.019438803 | 0.573745465 | 1 | 0.989 | 1 | 0.004462499 | 0.681653309 | 1 | 0.99 | 1 | 0.019438803 | 0.008905084 |
| Unc119 | 0.004485334 | -2.896083841 | 0 | 0.287 | 1 | 0.020522123 | -1.242926832 | 0.067 | 0.218 | 1 | 0.020522123 | 0.00895055 |
| Lonrf1 | 0.059584363 | 1.218869808 | 0.476 | 0.362 | 1 | 0.004490026 | 0.979057414 | 0.622 | 0.398 | 1 | 0.059584363 | 0.008959892 |
| Pdcd6 | 0.004501575 | -0.841928071 | 0.571 | 0.832 | 1 | 0.061542615 | -0.149254864 | 0.822 | 0.837 | 1 | 0.061542615 | 0.008982885 |
| Gas6 | 0.587212015 | -0.5577923 | 0.19 | 0.265 | 1 | 0.004502273 | -0.100303473 | 0.533 | 0.256 | 1 | 0.587212015 | 0.008984275 |
| Fuca2 | 0.054139934 | -2.054148687 | 0 | 0.153 | 1 | 0.004504664 | -1.167783975 | 0.067 | 0.273 | 1 | 0.054139934 | 0.008989036 |
| Snap29 | 0.085025238 | -0.849790188 | 0.143 | 0.332 | 1 | 0.004511755 | -0.404390255 | 0.089 | 0.318 | 1 | 0.085025238 | 0.009003153 |
| Zfp280d | 0.440600215 | 0.803539643 | 0.667 | 0.795 | 1 | 0.004512384 | 0.637594956 | 0.978 | 0.913 | 1 | 0.440600215 | 0.009004406 |
| Avpi1 | 0.004534363 | -2.242704877 | 0.143 | 0.463 | 1 | 0.011455219 | 0.83979809 | 0.489 | 0.291 | 1 | 0.011455219 | 0.009048166 |
| Pex6 | 0.370280371 | 0.137718867 | 0.095 | 0.19 | 1 | 0.00453512 | -1.303988358 | 0.089 | 0.301 | 1 | 0.370280371 | 0.009049673 |
| Hpgd | 0.004536486 | -2.780606623 | 0.286 | 0.604 | 1 | 0.007246269 | -1.339864225 | 0.311 | 0.474 | 1 | 0.007246269 | 0.009052393 |
| Tmem205 | 0.004541631 | -2.337455488 | 0.048 | 0.351 | 1 | 0.11606786 | 0.385392038 | 0.444 | 0.294 | 1 | 0.11606786 | 0.009062636 |
| Macroh2a1 | 0.832137559 | 0.409068541 | 0.762 | 0.81 | 1 | 0.004542718 | -0.437312954 | 0.889 | 0.882 | 1 | 0.832137559 | 0.0090648 |
| Xdh | 0.715873069 | 1.995699863 | 0.048 | 0.034 | 1 | 0.004549878 | 1.402964667 | 0.133 | 0.035 | 1 | 0.715873069 | 0.009079055 |
| Hacd3 | 0.049931245 | -1.10233222 | 0.143 | 0.362 | 1 | 0.004549948 | -0.591934461 | 0.222 | 0.498 | 1 | 0.049931245 | 0.009079193 |
| Fam53b | 0.37898587 | -0.413691074 | 0.048 | 0.108 | 1 | 0.004552855 | -3.071814916 | 0 | 0.156 | 1 | 0.37898587 | 0.009084981 |
| Vamp2 | 0.072412491 | -0.445967477 | 0.286 | 0.515 | 1 | 0.004558337 | -0.508068901 | 0.244 | 0.554 | 1 | 0.072412491 | 0.009095896 |
| Snrnp25 | 0.019055493 | -2.475975352 | 0 | 0.213 | 1 | 0.004567042 | -3.044847868 | 0 | 0.156 | 1 | 0.019055493 | 0.009113225 |
| Nubp2 | 0.057558982 | -1.911190733 | 0 | 0.149 | 1 | 0.004567042 | -3.044847868 | 0 | 0.156 | 1 | 0.057558982 | 0.009113225 |
| Chmp7 | 0.509428832 | -0.133583154 | 0.048 | 0.09 | 1 | 0.004568468 | -0.975138897 | 0.067 | 0.273 | 1 | 0.509428832 | 0.009116066 |
| Polr2l | 0.004574999 | -1.413691074 | 0.143 | 0.496 | 1 | 0.006950188 | -1.174908409 | 0.089 | 0.287 | 1 | 0.006950188 | 0.009129067 |
| Rsph9 | 0.057606209 | -1.940938076 | 0 | 0.149 | 1 | 0.004579015 | -3.149817428 | 0 | 0.156 | 1 | 0.057606209 | 0.009137063 |
| Dhcr7 | 0.281111676 | -0.752492987 | 0.095 | 0.194 | 1 | 0.004579463 | -3.199570463 | 0 | 0.156 | 1 | 0.281111676 | 0.009137954 |
| Kras | 0.004587353 | 1.077326699 | 0.905 | 0.858 | 1 | 0.039162394 | 0.405199671 | 0.956 | 0.92 | 1 | 0.039162394 | 0.009153663 |
| Syt8 | 0.004587661 | -3.297771786 | 0.048 | 0.347 | 1 | 0.0464093 | -0.147002412 | 0.133 | 0.308 | 1 | 0.0464093 | 0.009154275 |
| Cspg4b | 0.681919718 | 0.55829455 | 0.048 | 0.075 | 1 | 0.00459011 | -3.223818009 | 0 | 0.156 | 1 | 0.681919718 | 0.009159151 |
| Ecm1 | 0.412425944 | 0.951305743 | 0.143 | 0.09 | 1 | 0.004590951 | -3.339295227 | 0 | 0.156 | 1 | 0.412425944 | 0.009160824 |
| Tcaim | 0.544173118 | 0.295260144 | 0.095 | 0.157 | 1 | 0.00459493 | -3.247664751 | 0 | 0.156 | 1 | 0.544173118 | 0.009168746 |
| Zbtb33 | 0.073336562 | -1.107587946 | 0.095 | 0.284 | 1 | 0.004601875 | -1.194671664 | 0.111 | 0.332 | 1 | 0.073336562 | 0.009182573 |
| Sult2b1 | 0.004606304 | 3.188344941 | 0.095 | 0.011 | 1 | 0.331922009 | -0.124282336 | 0 | 0.021 | 1 | 0.331922009 | 0.009191391 |
| Gm57013 | 0.004606702 | 3.258734268 | 0.095 | 0.011 | 1 | 0.293659664 | -0.316927414 | 0 | 0.024 | 1 | 0.293659664 | 0.009192183 |
| Tspo | 0.004607616 | -1.785659851 | 0.238 | 0.526 | 1 | 0.314107678 | -0.498257179 | 0.778 | 0.651 | 1 | 0.314107678 | 0.009194001 |
| Cyp7b1 | 0.442273654 | -0.940938076 | 0.048 | 0.097 | 1 | 0.004612739 | -3.446210431 | 0 | 0.156 | 1 | 0.442273654 | 0.009204201 |
| Itgb8 | 0.624447247 | -0.167530486 | 0.048 | 0.078 | 1 | 0.004612975 | -1.844665046 | 0.133 | 0.346 | 1 | 0.624447247 | 0.009204671 |
| Hdac1 | 0.004616447 | -0.980232377 | 0.476 | 0.765 | 1 | 0.021486755 | -0.177524357 | 0.756 | 0.813 | 1 | 0.021486755 | 0.009211582 |
| Bhlhe41 | 0.073253524 | -1.434752689 | 0.095 | 0.276 | 1 | 0.004620824 | -3.564854927 | 0 | 0.156 | 1 | 0.073253524 | 0.009220296 |
| Tbc1d5 | 0.121593528 | 0.554264673 | 0.714 | 0.612 | 1 | 0.00462696 | 0.815710355 | 0.889 | 0.64 | 1 | 0.121593528 | 0.00923251 |
| Tma16 | 0.025658857 | -2.04047375 | 0.048 | 0.265 | 1 | 0.00462825 | -1.526380779 | 0.089 | 0.291 | 1 | 0.025658857 | 0.00923508 |
| Igf1 | 0.04165848 | -1.863169218 | 0.048 | 0.254 | 1 | 0.004629605 | -1.704944699 | 0.089 | 0.308 | 1 | 0.04165848 | 0.009237777 |
| Fbln5 | 0.059646262 | -2.413691074 | 0.048 | 0.216 | 1 | 0.00462964 | -4.174908409 | 0 | 0.156 | 1 | 0.059646262 | 0.009237846 |
| Itprid2 | 0.118260942 | 1.339352729 | 0.429 | 0.272 | 1 | 0.004630148 | 0.422836814 | 0.711 | 0.415 | 1 | 0.118260942 | 0.009238858 |
| Pelp1 | 0.063540767 | -1.434752689 | 0.048 | 0.216 | 1 | 0.004634265 | -1.094534992 | 0.067 | 0.273 | 1 | 0.063540767 | 0.009247054 |
| Oard1 | 0.00463909 | -3.171718283 | 0 | 0.287 | 1 | 0.015110975 | -1.86124793 | 0.044 | 0.19 | 1 | 0.015110975 | 0.009256658 |
| Frmd4b | 0.732723397 | -0.13549022 | 0.571 | 0.668 | 1 | 0.004640554 | 0.912429442 | 0.867 | 0.657 | 1 | 0.732723397 | 0.009259574 |
| Mrpl43 | 0.004645493 | -1.558888989 | 0.095 | 0.444 | 1 | 0.058343804 | -0.509572492 | 0.311 | 0.446 | 1 | 0.058343804 | 0.009269406 |
| Chchd6 | 0.004908938 | -2.348596045 | 0.048 | 0.347 | 1 | 0.004649776 | -0.941418279 | 0.067 | 0.273 | 1 | 0.004908938 | 0.009277931 |
| Selenop | 0.039011413 | -1.175893959 | 0.143 | 0.381 | 1 | 0.004657559 | -1.091714473 | 0.178 | 0.408 | 1 | 0.039011413 | 0.009293424 |
| Fhdc1 | 0.080428034 | 1.33137957 | 0.333 | 0.201 | 1 | 0.004673617 | -2.255526869 | 0.022 | 0.197 | 1 | 0.080428034 | 0.009325392 |
| Sesn3 | 0.00467443 | 0.10210609 | 0.524 | 0.929 | 1 | 0.007401074 | -0.547475284 | 0.867 | 0.9 | 1 | 0.007401074 | 0.00932701 |
| Smc6 | 0.011649101 | -0.51736972 | 0.381 | 0.72 | 1 | 0.004677882 | -0.365094696 | 0.689 | 0.785 | 1 | 0.011649101 | 0.009333882 |
| Acvr1 | 0.00469144 | 1.694782261 | 0.952 | 0.884 | 1 | 0.270199084 | 0.416913047 | 0.933 | 0.882 | 1 | 0.270199084 | 0.009360871 |
| Utp6 | 0.425745991 | 1.026881518 | 0.333 | 0.299 | 1 | 0.004699279 | -0.568845787 | 0.156 | 0.412 | 1 | 0.425745991 | 0.009376475 |
| Rrp1b | 0.464800069 | 0.388369549 | 0.143 | 0.216 | 1 | 0.004712398 | -1.459885368 | 0.067 | 0.263 | 1 | 0.464800069 | 0.009402589 |
| Mrpl13 | 0.030951591 | -0.826074119 | 0.19 | 0.481 | 1 | 0.004727228 | -0.63081819 | 0.178 | 0.426 | 1 | 0.030951591 | 0.00943211 |
| Ccdc28b | 0.524228088 | 0.137718867 | 0.238 | 0.34 | 1 | 0.004728597 | -1.06586565 | 0.089 | 0.308 | 1 | 0.524228088 | 0.009434835 |
| Phf10 | 0.079623695 | -0.280424543 | 0.238 | 0.552 | 1 | 0.004738616 | -0.561549783 | 0.467 | 0.699 | 1 | 0.079623695 | 0.009454778 |
| Megf9 | 0.004745538 | -2.834022873 | 0.048 | 0.347 | 1 | 0.015182332 | 0.831595111 | 0.556 | 0.336 | 1 | 0.015182332 | 0.009468557 |
| Ahnak | 0.022293052 | -0.529199229 | 1 | 0.974 | 1 | 0.004776812 | -0.494508377 | 1 | 1 | 1 | 0.022293052 | 0.009530805 |
| Kpna6 | 0.99613683 | 0.995699863 | 0.286 | 0.354 | 1 | 0.004788958 | -0.638855509 | 0.2 | 0.491 | 1 | 0.99613683 | 0.009554981 |
| Htatip2 | 0.004791595 | -2.818081329 | 0 | 0.284 | 1 | 0.024256375 | -1.901889915 | 0.022 | 0.142 | 1 | 0.024256375 | 0.00956023 |
| Ro60 | 0.030776891 | -1.249060372 | 0.143 | 0.396 | 1 | 0.00480292 | -0.644854249 | 0.222 | 0.484 | 1 | 0.030776891 | 0.009582772 |
| Irak1bp1 | 0.016746661 | -2.159118246 | 0.048 | 0.287 | 1 | 0.004805787 | -1.184823878 | 0.111 | 0.322 | 1 | 0.016746661 | 0.009588479 |
| Idi1 | 0.08128935 | -1.617537092 | 0.524 | 0.623 | 1 | 0.004809782 | -1.188706441 | 0.289 | 0.481 | 1 | 0.08128935 | 0.009596431 |
| Coa8 | 0.004817041 | -0.90365706 | 0.095 | 0.466 | 1 | 0.135610641 | 0.151058965 | 0.289 | 0.481 | 1 | 0.135610641 | 0.009610879 |
| Glul | 0.004817137 | -1.757002419 | 0.81 | 0.821 | 1 | 0.015227415 | -0.837184225 | 0.889 | 0.799 | 1 | 0.015227415 | 0.009611069 |
| Sbspon | 0.004819004 | -4.376076782 | 0 | 0.287 | 1 | 0.086057273 | -0.255526869 | 0.067 | 0.176 | 1 | 0.086057273 | 0.009614785 |
| Neu1 | 0.008113325 | -2.718545655 | 0 | 0.257 | 1 | 0.004820444 | -1.290932205 | 0.067 | 0.27 | 1 | 0.008113325 | 0.009617651 |
| Zfp160 | 0.753626757 | 1.161321767 | 0.286 | 0.28 | 1 | 0.004824567 | -0.731964913 | 0.156 | 0.394 | 1 | 0.753626757 | 0.009625858 |
| Ubr7 | 0.235058138 | -0.337455488 | 0.19 | 0.332 | 1 | 0.004825164 | -0.252387162 | 0.133 | 0.377 | 1 | 0.235058138 | 0.009627045 |
| Zfp189 | 0.299864508 | -0.574155746 | 0.048 | 0.123 | 1 | 0.004826918 | -1.979892427 | 0.022 | 0.197 | 1 | 0.299864508 | 0.009630536 |
| Asrgl1 | 0.020829393 | -1.289702356 | 0.095 | 0.369 | 1 | 0.004835966 | -1.271123724 | 0.089 | 0.298 | 1 | 0.020829393 | 0.009648545 |
| Zfp667 | 0.43957004 | 0.129451251 | 0.048 | 0.104 | 1 | 0.004840719 | -2.743192169 | 0.022 | 0.19 | 1 | 0.43957004 | 0.009658005 |
| B9d2 | 0.050735123 | -1.940938076 | 0 | 0.157 | 1 | 0.004844805 | -1.979892427 | 0.022 | 0.197 | 1 | 0.050735123 | 0.009666138 |
| Rtf1 | 0.111089691 | -0.520985087 | 0.619 | 0.75 | 1 | 0.004850091 | -0.565653983 | 0.933 | 0.886 | 1 | 0.111089691 | 0.009676659 |
| Rrp7a | 0.118409332 | -0.704739856 | 0.143 | 0.313 | 1 | 0.004854045 | -0.628513564 | 0.111 | 0.349 | 1 | 0.118409332 | 0.009684529 |
| Smyd2 | 0.019597061 | -1.303508156 | 0.095 | 0.358 | 1 | 0.004870872 | -1.346674757 | 0.067 | 0.27 | 1 | 0.019597061 | 0.009718019 |
| Fam8a1 | 0.036500496 | -1.213753503 | 0.143 | 0.373 | 1 | 0.00488214 | -0.337989029 | 0.267 | 0.554 | 1 | 0.036500496 | 0.009740445 |

| | | | | | | | | | | | | |
|---|---|---|---|---|---|---|---|---|---|---|---|---|
| Ltbp2 | 0.994485153 | 1.367963338 | 0.238 | 0.276 | 1 | 0.004882481 | -1.427958726 | 0.156 | 0.384 | 1 | 0.994485153 | 0.009741124 |
| Bag4 | 0.857464343 | 0.236366455 | 0.143 | 0.164 | 1 | 0.004882755 | -1.960783604 | 0.022 | 0.197 | 1 | 0.857464343 | 0.009741669 |
| Erh | 0.076431165 | -0.526655322 | 0.476 | 0.765 | 1 | 0.004883853 | -0.510847896 | 0.644 | 0.779 | 1 | 0.076431165 | 0.009743854 |
| Ankra2 | 0.218482512 | 1.02467893 | 0.286 | 0.194 | 1 | 0.004890465 | -1.271123724 | 0.044 | 0.239 | 1 | 0.218482512 | 0.009757014 |
| Rprd1a | 0.004892857 | 1.742158743 | 0.619 | 0.478 | 1 | 0.852461143 | 0.492969703 | 0.422 | 0.581 | 1 | 0.852461143 | 0.009761775 |
| Dcps | 0.007100858 | -2.834022873 | 0 | 0.265 | 1 | 0.004905984 | -1.979892427 | 0.022 | 0.197 | 1 | 0.007100858 | 0.0097879 |
| Hyal2 | 0.128939306 | -1.107587946 | 0.048 | 0.175 | 1 | 0.004918227 | -1.47073275 | 0.044 | 0.232 | 1 | 0.128939306 | 0.009812265 |
| Smarcd3 | 0.193424663 | -0.548620654 | 0.048 | 0.16 | 1 | 0.004923288 | -2.564854927 | 0.022 | 0.19 | 1 | 0.193424663 | 0.009822337 |
| Mypop | 0.271422861 | 1.451379346 | 0.143 | 0.078 | 1 | 0.004927529 | -1.84048937 | 0.022 | 0.197 | 1 | 0.271422861 | 0.009830777 |
| Oxa1l | 0.199086232 | -0.574155746 | 0.095 | 0.22 | 1 | 0.00493429 | -0.450783161 | 0.133 | 0.367 | 1 | 0.199086232 | 0.009844232 |
| Obp2b | 0.661796616 | 0.803407048 | 0.238 | 0.231 | 1 | 0.004946502 | 5.905465008 | 0.089 | 0.017 | 1 | 0.661796616 | 0.009868536 |
| Ifrd2 | 0.022814738 | -1.849790188 | 0.048 | 0.272 | 1 | 0.004947559 | -1.681499846 | 0.067 | 0.26 | 1 | 0.022814738 | 0.00987064 |
| Fzd5 | 0.222553843 | -0.522625445 | 0.048 | 0.149 | 1 | 0.004948974 | -2.486852415 | 0.022 | 0.19 | 1 | 0.222553843 | 0.009873455 |
| Tmem138 | 0.021751126 | -2.455511249 | 0 | 0.205 | 1 | 0.004949337 | -2.564854927 | 0.022 | 0.19 | 1 | 0.021751126 | 0.009874178 |
| Dvl3 | 0.234560995 | -0.171718283 | 0.143 | 0.295 | 1 | 0.004954913 | -0.510699157 | 0.156 | 0.412 | 1 | 0.234560995 | 0.009885274 |
| Gstt1 | 0.083718251 | -1.880817084 | 0 | 0.127 | 1 | 0.00496038 | -2.425451871 | 0.022 | 0.19 | 1 | 0.083718251 | 0.009896155 |
| Ube2o | 0.355198765 | 0.733150315 | 0.476 | 0.44 | 1 | 0.004970522 | -0.321214772 | 0.267 | 0.606 | 1 | 0.355198765 | 0.009916339 |
| Msrb1 | 0.004971399 | -1.979670471 | 0.048 | 0.366 | 1 | 0.860410519 | 0.127256431 | 0.356 | 0.363 | 1 | 0.860410519 | 0.009918084 |
| Naaladl2 | 0.235653138 | 0.577163444 | 0.667 | 0.668 | 1 | 0.004971873 | 0.828758136 | 0.911 | 0.806 | 1 | 0.235653138 | 0.009919026 |
| Mrps36 | 0.025920019 | -0.785659851 | 0.095 | 0.369 | 1 | 0.004971903 | -1.286553765 | 0.067 | 0.263 | 1 | 0.025920019 | 0.009919087 |
| Zfp60 | 0.477138921 | 0.780686972 | 0.095 | 0.16 | 1 | 0.004973024 | -2.466674533 | 0.022 | 0.19 | 1 | 0.477138921 | 0.009921318 |
| Brpf1 | 0.236376695 | -0.31112134 | 0.143 | 0.269 | 1 | 0.00497659 | -0.960783604 | 0.222 | 0.439 | 1 | 0.236376695 | 0.009928414 |
| Mpc1 | 0.004984866 | -0.790509251 | 0.619 | 0.862 | 1 | 0.016501381 | -0.562552161 | 0.889 | 0.82 | 1 | 0.016501381 | 0.009944884 |
| Nlk | 0.077795337 | 1.508864841 | 0.571 | 0.493 | 1 | 0.00499676 | 0.663781248 | 0.867 | 0.64 | 1 | 0.077795337 | 0.009968551 |
| S100a16 | 0.005006843 | -1.216999163 | 0.381 | 0.672 | 1 | 0.012789552 | -0.889311848 | 0.667 | 0.709 | 1 | 0.012789552 | 0.009988617 |
| Ccz1 | 0.005008331 | -1.127128132 | 0.143 | 0.526 | 1 | 0.631405326 | 0.361144491 | 0.556 | 0.533 | 1 | 0.631405326 | 0.009991578 |
| Pnpla2 | 0.295175661 | -0.326228232 | 0.095 | 0.194 | 1 | 0.005009033 | -1.989352756 | 0.067 | 0.253 | 1 | 0.295175661 | 0.009992976 |
| Gm13832 | 0.005010028 | 3.493199522 | 0.19 | 0.049 | 1 | 0.617558746 | 1.435145073 | 0.022 | 0.038 | 1 | 0.617558746 | 0.009994955 |
| Slc35e3 | 0.065485969 | -0.6987824 | 0.286 | 0.552 | 1 | 0.00502461 | -0.598962782 | 0.178 | 0.446 | 1 | 0.065485969 | 0.010023973 |
| Has2 | 0.833798483 | -0.634276738 | 0.619 | 0.563 | 1 | 0.005036214 | -0.81719275 | 0.178 | 0.429 | 1 | 0.833798483 | 0.010047065 |
| Wdr20 | 0.827914277 | 0.866416846 | 0.333 | 0.388 | 1 | 0.005038206 | 1.03537433 | 0.733 | 0.55 | 1 | 0.827914277 | 0.010051029 |
| Ano10 | 0.34408071 | -0.314912919 | 0.19 | 0.31 | 1 | 0.005047721 | -0.409373663 | 0.133 | 0.377 | 1 | 0.34408071 | 0.010069962 |
| Rab5c | 0.005054054 | -1.25696557 | 0.238 | 0.575 | 1 | 0.480279536 | -0.137106376 | 0.689 | 0.682 | 1 | 0.480279536 | 0.010082564 |
| Selenoo | 0.942133083 | 0.55829455 | 0.143 | 0.16 | 1 | 0.005054775 | -1.404390255 | 0.022 | 0.197 | 1 | 0.942133083 | 0.010084 |
| Abl2 | 0.387662056 | 0.522528187 | 0.762 | 0.84 | 1 | 0.005055017 | 0.819973438 | 0.956 | 0.91 | 1 | 0.387662056 | 0.01008448 |
| Cfap410 | 0.086236464 | -0.828728573 | 0.048 | 0.213 | 1 | 0.005075194 | -1.425451871 | 0.022 | 0.197 | 1 | 0.086236464 | 0.010124629 |
| Bmp7 | 0.389619252 | -0.911190733 | 0.143 | 0.216 | 1 | 0.005083037 | -2.459885368 | 0.044 | 0.221 | 1 | 0.389619252 | 0.010140236 |
| Uchl5 | 0.500228768 | 0.451379346 | 0.19 | 0.291 | 1 | 0.005083579 | -0.258375231 | 0.133 | 0.384 | 1 | 0.500228768 | 0.010141314 |
| Gm56582 | 0.094305534 | -1.455511249 | 0 | 0.119 | 1 | 0.005084451 | -2.901889915 | 0 | 0.152 | 1 | 0.094305534 | 0.01014305 |
| Rab27a | 0.096909174 | -0.243766072 | 0.143 | 0.343 | 1 | 0.0050986 | -2.124282336 | 0.067 | 0.249 | 1 | 0.096909174 | 0.010171205 |
| Uros | 0.525462593 | 0.181918671 | 0.095 | 0.153 | 1 | 0.005100916 | -2.989352756 | 0 | 0.152 | 1 | 0.525462593 | 0.010175812 |
| Thnsl2 | 0.034503481 | -2.133583154 | 0 | 0.179 | 1 | 0.005100978 | -3.044847868 | 0 | 0.152 | 1 | 0.034503481 | 0.010175936 |
| Casp8 | 0.192182868 | -0.718545655 | 0.095 | 0.22 | 1 | 0.005102006 | -0.375821103 | 0.067 | 0.27 | 1 | 0.192182868 | 0.010177982 |
| Tmsb15b2 | 0.137073547 | -1.02666795 | 0.048 | 0.172 | 1 | 0.005115791 | -3.124282336 | 0 | 0.152 | 1 | 0.137073547 | 0.010205411 |
| Bola1 | 0.159590475 | -0.496153234 | 0.095 | 0.239 | 1 | 0.005115999 | -3.017367132 | 0 | 0.152 | 1 | 0.159590475 | 0.010205824 |
| Rmi1 | 0.120586929 | -1.370622352 | 0 | 0.104 | 1 | 0.005122708 | -3.124282336 | 0 | 0.152 | 1 | 0.120586929 | 0.010219174 |
| Slc25a14 | 0.33598189 | -0.496153234 | 0.048 | 0.116 | 1 | 0.005122812 | -3.149817428 | 0 | 0.152 | 1 | 0.33598189 | 0.010219381 |
| Cars | 0.005128194 | 0.97086801 | 0.714 | 0.504 | 1 | 0.435626243 | 0.261608818 | 0.467 | 0.692 | 1 | 0.435626243 | 0.010230089 |
| Gm14305 | 0.026765847 | -2.392317423 | 0 | 0.194 | 1 | 0.00512859 | -3.098287127 | 0 | 0.152 | 1 | 0.026765847 | 0.010230878 |
| Aig1 | 0.011619762 | 1.838786855 | 0.905 | 0.843 | 1 | 0.005130314 | 1.157956281 | 0.933 | 0.72 | 1 | 0.011619762 | 0.010234309 |
| Tmem131l | 0.228702366 | 0.775148788 | 0.429 | 0.351 | 1 | 0.005135132 | 1.192191329 | 0.867 | 0.64 | 1 | 0.228702366 | 0.010243894 |
| Cmtm3 | 0.376735683 | -0.218245551 | 0.286 | 0.388 | 1 | 0.005138926 | -0.514866791 | 0.111 | 0.349 | 1 | 0.376735683 | 0.010251443 |
| Cetn4 | 0.016889212 | -2.094412557 | 0.048 | 0.287 | 1 | 0.005139773 | -3.149817428 | 0 | 0.152 | 1 | 0.016889212 | 0.010253129 |
| Pkp4 | 0.014608806 | 0.764996528 | 0.857 | 0.489 | 1 | 0.005139835 | 1.013067962 | 0.889 | 0.72 | 1 | 0.014608806 | 0.010253253 |
| Dclre1a | 0.845912949 | 0.281454345 | 0.095 | 0.112 | 1 | 0.005145271 | -3.223818009 | 0 | 0.152 | 1 | 0.845912949 | 0.010264068 |
| Ar | 0.545975733 | -1.413691074 | 0.048 | 0.082 | 1 | 0.005155841 | 0.13763845 | 0.067 | 0.27 | 1 | 0.545975733 | 0.010285099 |
| Kank2 | 0.512804897 | 0.247507013 | 0.286 | 0.403 | 1 | 0.005157427 | -0.518561275 | 0.311 | 0.612 | 1 | 0.512804897 | 0.010288255 |
| Vps9d1 | 0.428753191 | -0.25696557 | 0.095 | 0.164 | 1 | 0.005157605 | -1.486852415 | 0.022 | 0.197 | 1 | 0.428753191 | 0.010288609 |
| Asb8 | 0.087091561 | -1.413691074 | 0.048 | 0.198 | 1 | 0.005158379 | -1.84048937 | 0.044 | 0.228 | 1 | 0.087091561 | 0.010290149 |
| Timm9 | 0.436869089 | 0.658174913 | 0.238 | 0.407 | 1 | 0.005177064 | -0.858821193 | 0.156 | 0.391 | 1 | 0.436869089 | 0.010327325 |
| Slc25a13 | 0.732234171 | 0.129451251 | 0.19 | 0.231 | 1 | 0.005181119 | -0.984352075 | 0.133 | 0.363 | 1 | 0.732234171 | 0.010335393 |
| Thbs2 | 0.173748441 | -2.502816964 | 0.048 | 0.157 | 1 | 0.005189369 | -4.953552034 | 0 | 0.152 | 1 | 0.173748441 | 0.010351808 |
| Ciao1 | 0.075565239 | -0.926141075 | 0.095 | 0.284 | 1 | 0.005191434 | -1.039393438 | 0.111 | 0.332 | 1 | 0.075565239 | 0.010355917 |
| Pim3 | 0.005248992 | -2.22214771 | 0.381 | 0.604 | 1 | 0.01023999 | 0.265987258 | 0.711 | 0.419 | 1 | 0.01023999 | 0.010470433 |
| Csf1 | 0.286296077 | -0.676725479 | 0.095 | 0.198 | 1 | 0.005250051 | -1.84048937 | 0.067 | 0.26 | 1 | 0.286296077 | 0.010472539 |
| Calml3 | 0.057003081 | -2.196592952 | 0.048 | 0.22 | 1 | 0.005250546 | -1.249240606 | 0.111 | 0.318 | 1 | 0.057003081 | 0.010473523 |
| Dnase1l1 | 0.02491957 | -2.434752689 | 0 | 0.198 | 1 | 0.005252044 | -1.75987091 | 0.044 | 0.228 | 1 | 0.02491957 | 0.010476505 |
| Bphl | 0.015573212 | -1.561444694 | 0.048 | 0.31 | 1 | 0.005267924 | -0.142093612 | 0.133 | 0.363 | 1 | 0.015573212 | 0.010508096 |
| Tcn2 | 0.005268734 | -2.911190733 | 0 | 0.28 | 1 | 0.059462945 | -0.745770713 | 0.2 | 0.336 | 1 | 0.059462945 | 0.010509708 |
| Vamp5 | 0.5470954 | 0.60829793 | 0.333 | 0.485 | 1 | 0.005269571 | -0.643821762 | 0.156 | 0.408 | 1 | 0.5470954 | 0.010511373 |
| Trim28 | 0.008409352 | -1.535681598 | 0.095 | 0.414 | 1 | 0.005277647 | -0.716334261 | 0.444 | 0.616 | 1 | 0.008409352 | 0.010527441 |
| Pdlim2 | 0.197875583 | -0.741265732 | 0.048 | 0.157 | 1 | 0.005278171 | -1.277756817 | 0.044 | 0.235 | 1 | 0.197875583 | 0.010528484 |
| Usp11 | 0.810823066 | 0.201283996 | 0.238 | 0.299 | 1 | 0.005281467 | -0.970369652 | 0.133 | 0.349 | 1 | 0.810823066 | 0.010535041 |
| Gphn | 0.005283232 | 2.730779973 | 1 | 1 | 1 | 0.038403254 | 0.878430348 | 1 | 1 | 1 | 0.038403254 | 0.010538552 |
| Fcf1 | 0.916616986 | 0.597420882 | 0.381 | 0.489 | 1 | 0.005283831 | -0.659319611 | 0.222 | 0.481 | 1 | 0.916616986 | 0.010539743 |
| Spg11 | 0.408194338 | -0.522625445 | 0.095 | 0.16 | 1 | 0.005284773 | -0.745770713 | 0.089 | 0.308 | 1 | 0.408194338 | 0.010541618 |
| Arhgef28 | 0.446232496 | 1.410737362 | 0.619 | 0.537 | 1 | 0.005298759 | 0.760414675 | 0.667 | 0.405 | 1 | 0.446232496 | 0.010569441 |
| Slc35f6 | 0.00530075 | -2.955584852 | 0 | 0.28 | 1 | 0.074402041 | -0.504554417 | 0.089 | 0.208 | 1 | 0.074402041 | 0.010573402 |
| Cyp51 | 0.108941298 | -0.609797992 | 0.619 | 0.754 | 1 | 0.005301954 | -0.896786637 | 0.533 | 0.657 | 1 | 0.108941298 | 0.010575797 |
| Taf15 | 0.005305325 | 1.102884452 | 0.81 | 0.784 | 1 | 0.329106116 | 0.394827618 | 0.911 | 0.792 | 1 | 0.329106116 | 0.010582504 |
| Ndrg3 | 0.005305811 | -1.20069735 | 0.143 | 0.522 | 1 | 0.071558632 | -0.509572492 | 0.489 | 0.619 | 1 | 0.071558632 | 0.010583471 |
| Nxn | 0.542399158 | 0.127097752 | 0.476 | 0.56 | 1 | 0.005308322 | 1.405087839 | 0.844 | 0.567 | 1 | 0.542399158 | 0.010588466 |
| Ccdc88a | 0.15492465 | -0.153727227 | 0.81 | 0.974 | 1 | 0.005309229 | -0.323459487 | 0.933 | 0.979 | 1 | 0.15492465 | 0.010590269 |
| Tnfaip1 | 0.573244547 | 0.281454345 | 0.286 | 0.407 | 1 | 0.0053196 | -0.762237924 | 0.333 | 0.567 | 1 | 0.573244547 | 0.010610902 |
| Dop1b | 0.629886513 | -0.478231326 | 0.19 | 0.243 | 1 | 0.005320767 | -0.986778812 | 0.089 | 0.298 | 1 | 0.629886513 | 0.010613223 |
| Snx16 | 0.415395258 | -0.785659851 | 0.095 | 0.157 | 1 | 0.005330829 | -1.609709163 | 0.044 | 0.228 | 1 | 0.415395258 | 0.010633239 |
| Stag1 | 0.005337054 | 0.796494939 | 1 | 0.922 | 1 | 0.01385009 | 0.505367528 | 1 | 0.993 | 1 | 0.01385009 | 0.010645624 |
| Mfsd11 | 0.051505928 | -0.658424663 | 0.095 | 0.325 | 1 | 0.00533855 | -0.830551133 | 0.178 | 0.439 | 1 | 0.051505928 | 0.010648599 |
| Lpcat1 | 0.932038121 | 0.503846766 | 0.286 | 0.358 | 1 | 0.005339681 | -0.918963428 | 0.133 | 0.363 | 1 | 0.932038121 | 0.010650851 |
| Msh3 | 0.065511861 | -2.370622352 | 0 | 0.142 | 1 | 0.005355908 | -2.071814916 | 0.022 | 0.194 | 1 | 0.065511861 | 0.01068313 |
| Ppp2r1a | 0.006303477 | -0.862281133 | 0.381 | 0.731 | 1 | 0.00536425 | -0.692794316 | 0.667 | 0.734 | 1 | 0.006303477 | 0.010699725 |
| Pgap2 | 0.005367035 | -1.81004401 | 0.095 | 0.414 | 1 | 0.226232547 | 0.53722172 | 0.578 | 0.426 | 1 | 0.226232547 | 0.010705264 |
| Xrcc5 | 0.7962719 | 0.444953077 | 0.238 | 0.306 | 1 | 0.005378863 | -0.316927414 | 0.156 | 0.388 | 1 | 0.7962719 | 0.010728793 |
| Prdm16os | 0.174092302 | -0.44170545 | 0.048 | 0.168 | 1 | 0.005380611 | -1.656777417 | 0.044 | 0.228 | 1 | 0.174092302 | 0.010732272 |
| Mrpl39 | 0.007603507 | -2.785659851 | 0 | 0.261 | 1 | 0.005380634 | -1.674479418 | 0.044 | 0.228 | 1 | 0.007603507 | 0.010732317 |
| Nol10 | 0.494678386 | 0.945851313 | 0.381 | 0.399 | 1 | 0.005383968 | -0.479198843 | 0.156 | 0.419 | 1 | 0.494678386 | 0.010738949 |
| Dok1 | 0.008201188 | -1.769171728 | 0.095 | 0.41 | 1 | 0.005386968 | -1.80878051 | 0.067 | 0.26 | 1 | 0.008201188 | 0.010744916 |
| Ttc8 | 0.042747421 | -1.31112134 | 0.048 | 0.254 | 1 | 0.005389317 | -1.53415813 | 0.044 | 0.228 | 1 | 0.042747421 | 0.010749589 |

| | | | | | | | | | | | | |
|---|---|---|---|---|---|---|---|---|---|---|---|---|
| Ethe1 | 0.100392236 | -1.718545655 | 0 | 0.116 | 1 | 0.005399932 | -2.017367132 | 0.044 | 0.221 | 1 | 0.100392236 | 0.010770705 |
| St13 | 0.0054231 | -0.726899442 | 0.81 | 0.94 | 1 | 0.012132648 | -0.35365052 | 0.956 | 0.962 | 1 | 0.012132648 | 0.01081679 |
| Kctd2 | 0.0670566 | -0.589262638 | 0.095 | 0.295 | 1 | 0.005437665 | -0.731964913 | 0.067 | 0.277 | 1 | 0.0670566 | 0.010845762 |
| Nfatc4 | 0.678872005 | 0.614878079 | 0.19 | 0.172 | 1 | 0.005438423 | -1.124282336 | 0.089 | 0.294 | 1 | 0.678872005 | 0.01084727 |
| Prxl2a | 0.005442219 | -3.403043829 | 0 | 0.28 | 1 | 0.098796921 | -0.792660845 | 0.111 | 0.218 | 1 | 0.098796921 | 0.01085482 |
| Ngrn | 0.094263842 | -1.208871282 | 0.048 | 0.194 | 1 | 0.005480579 | -1.881712033 | 0.044 | 0.221 | 1 | 0.094263842 | 0.010931122 |
| Map2k2 | 0.125256002 | -0.355025497 | 0.619 | 0.813 | 1 | 0.005482002 | -0.542995493 | 0.8 | 0.855 | 1 | 0.125256002 | 0.010933951 |
| Tmem184c | 0.061072516 | -1.20069735 | 0.048 | 0.231 | 1 | 0.005488388 | -0.348405645 | 0.089 | 0.318 | 1 | 0.061072516 | 0.010946654 |
| Lancl1 | 0.029196208 | -1.314912919 | 0.143 | 0.373 | 1 | 0.005506235 | -1.169370225 | 0.111 | 0.322 | 1 | 0.029196208 | 0.010982152 |
| Cyb561d2 | 0.030163102 | -1.849790188 | 0.048 | 0.257 | 1 | 0.005510747 | -2.583713955 | 0.022 | 0.187 | 1 | 0.030163102 | 0.010991125 |
| Prdm4 | 0.077682261 | 1.614878079 | 0.333 | 0.213 | 1 | 0.005524564 | -0.486852415 | 0.111 | 0.336 | 1 | 0.077682261 | 0.011018607 |
| Nid2 | 0.010231513 | -0.711421221 | 0.381 | 0.705 | 1 | 0.005538139 | -0.452336534 | 0.267 | 0.554 | 1 | 0.010231513 | 0.011045607 |
| Sdhb | 0.005538482 | -1.076249979 | 0.381 | 0.66 | 1 | 0.077647682 | -0.478608421 | 0.689 | 0.689 | 1 | 0.077647682 | 0.011046289 |
| Ppig | 0.747827127 | 0.241150585 | 0.905 | 0.862 | 1 | 0.0055416 | -0.382022442 | 0.889 | 0.934 | 1 | 0.747827127 | 0.011052491 |
| Ppan | 0.789513049 | 0.61199557 | 0.333 | 0.399 | 1 | 0.005547861 | -0.776359032 | 0.2 | 0.446 | 1 | 0.789513049 | 0.011064942 |
| Trmt10a | 0.363869798 | 1.276881615 | 0.286 | 0.246 | 1 | 0.005568765 | -2.425451871 | 0.022 | 0.187 | 1 | 0.363869798 | 0.011106519 |
| Cenpb | 0.081245149 | -0.426688414 | 0.238 | 0.481 | 1 | 0.005583095 | -0.727559734 | 0.667 | 0.713 | 1 | 0.081245149 | 0.011135019 |
| Med8 | 0.032111135 | -0.574155746 | 0.143 | 0.414 | 1 | 0.005583835 | -0.649123844 | 0.133 | 0.36 | 1 | 0.032111135 | 0.011136491 |
| Hace1 | 0.798078907 | 0.85064953 | 0.238 | 0.246 | 1 | 0.005590382 | -0.695439037 | 0.222 | 0.481 | 1 | 0.798078907 | 0.011149511 |
| Znhit3 | 0.007123207 | -2.769171728 | 0 | 0.265 | 1 | 0.005595838 | -2.383016604 | 0.022 | 0.187 | 1 | 0.007123207 | 0.011160362 |
| Dynlt1f | 0.032123562 | -1.849790188 | 0.048 | 0.254 | 1 | 0.005608872 | -2.404390255 | 0.022 | 0.187 | 1 | 0.032123562 | 0.011186285 |
| Epb41l4aos | 0.511896195 | -0.252227651 | 0.19 | 0.25 | 1 | 0.005608872 | -2.404390255 | 0.022 | 0.187 | 1 | 0.511896195 | 0.011186285 |
| Uvrag | 0.220382502 | 0.340541024 | 0.81 | 0.817 | 1 | 0.005629149 | 0.756409198 | 0.978 | 0.869 | 1 | 0.220382502 | 0.011226611 |
| Sypl | 0.022265395 | -0.563858093 | 0.81 | 0.903 | 1 | 0.005635996 | -0.406233955 | 0.889 | 0.9 | 1 | 0.022265395 | 0.011240227 |
| 2210408F21Rik | 0.005643475 | 2.025699223 | 0.619 | 0.437 | 1 | 0.08277806 | 1.037503321 | 0.489 | 0.415 | 1 | 0.08277806 | 0.011255101 |
| Nudt21 | 0.005650008 | -0.887622262 | 0.381 | 0.757 | 1 | 0.015735824 | -0.460331539 | 0.733 | 0.81 | 1 | 0.015735824 | 0.011268094 |
| Brd9 | 0.465734964 | 0.150209812 | 0.238 | 0.347 | 1 | 0.005659999 | -0.785696897 | 0.133 | 0.367 | 1 | 0.465734964 | 0.011287963 |
| Pet100 | 0.005667581 | -2.880817084 | 0 | 0.276 | 1 | 0.037433472 | -1.223818009 | 0.044 | 0.166 | 1 | 0.037433472 | 0.011303041 |
| Zbtb6 | 0.265663903 | -0.413691074 | 0.095 | 0.201 | 1 | 0.005697366 | -0.543996322 | 0.067 | 0.277 | 1 | 0.265663903 | 0.011362273 |
| Gatd1 | 0.071785987 | -1.555046923 | 0.048 | 0.209 | 1 | 0.005698523 | -0.925736657 | 0.022 | 0.194 | 1 | 0.071785987 | 0.011364574 |
| Sumf2 | 0.310867629 | -0.280424543 | 0.19 | 0.321 | 1 | 0.005700004 | -0.719025857 | 0.133 | 0.349 | 1 | 0.310867629 | 0.011367517 |
| Saal1 | 0.260503572 | 0.119182916 | 0.048 | 0.142 | 1 | 0.005703591 | -2.901889915 | 0 | 0.149 | 1 | 0.260503572 | 0.011374652 |
| Gm56860 | 0.113499821 | -1.455511249 | 0 | 0.108 | 1 | 0.005712034 | -2.960783604 | 0 | 0.149 | 1 | 0.113499821 | 0.01139144 |
| Vegfa | 0.809206077 | -0.149112853 | 0.429 | 0.526 | 1 | 0.005717863 | 0.485133209 | 0.667 | 0.377 | 1 | 0.809206077 | 0.011403033 |
| Kdm5a | 0.005725499 | 0.744426321 | 0.952 | 0.896 | 1 | 0.674811486 | 0.282110272 | 0.978 | 0.945 | 1 | 0.674811486 | 0.011418217 |
| Gpn2 | 0.05394162 | -1.849790188 | 0 | 0.153 | 1 | 0.005727978 | -2.989352756 | 0 | 0.149 | 1 | 0.05394162 | 0.011423147 |
| Rnaseh1 | 0.403622827 | -0.107587946 | 0.095 | 0.172 | 1 | 0.005741189 | -3.071814916 | 0 | 0.149 | 1 | 0.403622827 | 0.011449417 |
| Mrpl18 | 0.010224976 | -1.047327421 | 0.143 | 0.474 | 1 | 0.005752728 | -0.577454964 | 0.222 | 0.464 | 1 | 0.010224976 | 0.011472361 |
| Eln | 0.455000042 | -0.766800824 | 0.19 | 0.272 | 1 | 0.005754364 | 1.070888617 | 0.067 | 0.27 | 1 | 0.455000042 | 0.011475616 |
| Triobp | 0.023786607 | -1.413691074 | 0.048 | 0.287 | 1 | 0.005758716 | -0.648441558 | 0.156 | 0.384 | 1 | 0.023786607 | 0.011484269 |
| Snrk | 0.005765083 | -2.998653574 | 0 | 0.276 | 1 | 0.106146496 | -0.416463087 | 0.244 | 0.394 | 1 | 0.106146496 | 0.011496929 |
| Asf1a | 0.005765193 | -2.516052791 | 0.048 | 0.34 | 1 | 0.014276071 | -0.495264655 | 0.156 | 0.37 | 1 | 0.014276071 | 0.011497148 |
| Trim39 | 0.073183197 | 1.351843673 | 0.476 | 0.328 | 1 | 0.005772462 | -0.560381451 | 0.156 | 0.408 | 1 | 0.073183197 | 0.011511603 |
| Midn | 0.126633035 | -0.733403614 | 0.571 | 0.716 | 1 | 0.005775414 | -0.711851332 | 0.822 | 0.799 | 1 | 0.126633035 | 0.011517473 |
| Cdk7 | 0.753267329 | 0.427897913 | 0.429 | 0.616 | 1 | 0.005785193 | -0.461977747 | 0.422 | 0.727 | 1 | 0.753267329 | 0.011536917 |
| Vps4a | 0.005809517 | -1.355975576 | 0.143 | 0.489 | 1 | 0.016400165 | -0.296749532 | 0.2 | 0.457 | 1 | 0.016400165 | 0.011585283 |
| Cbx6 | 0.015248817 | -1.709932525 | 0.095 | 0.362 | 1 | 0.005810968 | -1.00994966 | 0.6 | 0.664 | 1 | 0.015248817 | 0.01158817 |
| Fubp3 | 0.163137349 | -0.595688907 | 0.19 | 0.354 | 1 | 0.00581222 | -0.268017813 | 0.2 | 0.484 | 1 | 0.163137349 | 0.011590659 |
| Gprasp1 | 0.24356639 | -0.380304905 | 0.381 | 0.56 | 1 | 0.005816731 | -0.678565219 | 0.356 | 0.647 | 1 | 0.24356639 | 0.011599627 |
| Tceal5 | 0.030469578 | -2.434752689 | 0 | 0.187 | 1 | 0.005824683 | -1.017367132 | 0.022 | 0.194 | 1 | 0.030469578 | 0.01161544 |
| Cyth2 | 0.005835247 | -1.032820177 | 0.143 | 0.526 | 1 | 0.019920466 | -0.709244837 | 0.4 | 0.55 | 1 | 0.019920466 | 0.011636444 |
| Dynlt2a2 | 0.005838434 | 2.673771768 | 0.19 | 0.049 | 1 | 0.392807003 | 0.567595369 | 0.044 | 0.083 | 1 | 0.392807003 | 0.011642781 |
| Parp9 | 0.066388661 | -2.268742738 | 0.048 | 0.209 | 1 | 0.005863725 | -0.667424661 | 0.044 | 0.235 | 1 | 0.066388661 | 0.011693066 |
| Surf2 | 0.009542349 | -2.801961663 | 0 | 0.25 | 1 | 0.00587338 | -2.089516918 | 0.044 | 0.218 | 1 | 0.009542349 | 0.011712263 |
| Acaa1a | 0.027787811 | -0.496153234 | 0.238 | 0.604 | 1 | 0.005897263 | -0.855347329 | 0.311 | 0.519 | 1 | 0.027787811 | 0.011759749 |
| Eps8l1 | 0.005900377 | 2.825774861 | 0.19 | 0.049 | 1 | 0.077394404 | -1.84048937 | 0 | 0.066 | 1 | 0.077394404 | 0.011765939 |
| Nfx1 | 0.005916513 | -1.133583154 | 0.238 | 0.597 | 1 | 0.046140887 | 0.603638119 | 0.822 | 0.74 | 1 | 0.046140887 | 0.011798021 |
| Tjap1 | 0.63276276 | 0.464318402 | 0.238 | 0.325 | 1 | 0.005927055 | -0.609709163 | 0.178 | 0.439 | 1 | 0.63276276 | 0.01181898 |
| Ngdn | 0.013255779 | -1.159118246 | 0.19 | 0.478 | 1 | 0.005936887 | -0.404390255 | 0.244 | 0.478 | 1 | 0.013255779 | 0.011838527 |
| Tmem150a | 0.021145929 | -1.926141075 | 0.048 | 0.276 | 1 | 0.005938636 | -1.404390255 | 0.089 | 0.291 | 1 | 0.021145929 | 0.011842005 |
| Ruvbl1 | 0.017336413 | -1.34415014 | 0.143 | 0.41 | 1 | 0.005940236 | -0.602329633 | 0.178 | 0.436 | 1 | 0.017336413 | 0.011845186 |
| Eno1b | 0.005940935 | -2.666078235 | 0 | 0.272 | 1 | 0.016967574 | -1.709244837 | 0.044 | 0.187 | 1 | 0.016967574 | 0.011846575 |
| Cisd3 | 0.005941152 | -2.648156327 | 0 | 0.272 | 1 | 0.013543127 | -1.103523776 | 0.044 | 0.201 | 1 | 0.013543127 | 0.011847007 |
| Ltn1 | 0.054534669 | -0.333147646 | 0.19 | 0.455 | 1 | 0.005954657 | -0.253875243 | 0.289 | 0.585 | 1 | 0.054534669 | 0.011873857 |
| Stip1 | 0.005969638 | -1.385121921 | 0.238 | 0.552 | 1 | 0.017395728 | -0.503340538 | 0.756 | 0.751 | 1 | 0.017395728 | 0.011903639 |
| N6amt1 | 0.005990756 | -1.901137068 | 0.048 | 0.358 | 1 | 0.008781639 | -1.638855509 | 0.067 | 0.242 | 1 | 0.008781639 | 0.011945623 |
| Zfp1 | 0.078771035 | -0.531342662 | 0.19 | 0.429 | 1 | 0.006000801 | -0.784053424 | 0.156 | 0.388 | 1 | 0.078771035 | 0.011965592 |
| Uba1 | 0.027098391 | -0.718545655 | 0.286 | 0.616 | 1 | 0.006003371 | -0.621080807 | 0.711 | 0.768 | 1 | 0.027098391 | 0.011970702 |
| Gm29418 | 0.015637018 | -2.611630451 | 0 | 0.224 | 1 | 0.006004095 | -0.709244837 | 0.111 | 0.332 | 1 | 0.015637018 | 0.011972142 |
| Tpgs1 | 0.021347323 | -1.834022873 | 0.048 | 0.276 | 1 | 0.006006431 | -1.743192169 | 0.067 | 0.249 | 1 | 0.021347323 | 0.011976784 |
| Sidt2 | 0.963361579 | 0.156496074 | 0.667 | 0.739 | 1 | 0.006026814 | -0.628513564 | 0.644 | 0.779 | 1 | 0.963361579 | 0.012017306 |
| Kars | 0.00603217 | -2.769171728 | 0 | 0.272 | 1 | 0.513987461 | 0.361144491 | 0.244 | 0.325 | 1 | 0.513987461 | 0.012027953 |
| Prrx1 | 0.006051805 | -0.951937075 | 0.333 | 0.683 | 1 | 0.218486919 | 0.836675563 | 0.778 | 0.858 | 1 | 0.218486919 | 0.012066986 |
| Ptrhd1 | 0.006054316 | -2.801961663 | 0 | 0.272 | 1 | 0.008897736 | -1.339295227 | 0.022 | 0.18 | 1 | 0.008897736 | 0.012071978 |
| Gm16133 | 0.006065574 | 1.685087081 | 0.476 | 0.265 | 1 | 0.014855396 | 1.301878542 | 0.556 | 0.519 | 1 | 0.014855396 | 0.012094356 |
| Cul3 | 0.006071824 | 0.625766395 | 0.952 | 0.929 | 1 | 0.656407007 | 0.103549825 | 1 | 0.979 | 1 | 0.656407007 | 0.012106782 |
| Nedd9 | 0.006075007 | 1.272922711 | 0.905 | 0.881 | 1 | 0.699113988 | 0.207473129 | 0.867 | 0.865 | 1 | 0.699113988 | 0.012113108 |
| Hsd17b11 | 0.406030059 | 0.586308926 | 0.286 | 0.463 | 1 | 0.006088334 | -0.597883728 | 0.222 | 0.481 | 1 | 0.406030059 | 0.012139599 |
| Ube2d-ps | 0.10446714 | -0.574155746 | 0.095 | 0.28 | 1 | 0.006101856 | -1.342462506 | 0.089 | 0.284 | 1 | 0.10446714 | 0.012166479 |
| Tm4sf1 | 0.00610378 | -4.215932317 | 0.238 | 0.507 | 1 | 0.038635234 | -1.764106772 | 0.2 | 0.381 | 1 | 0.038635234 | 0.012170305 |
| Sik2 | 0.187472779 | 0.929610672 | 0.476 | 0.422 | 1 | 0.006111671 | 0.929133913 | 0.889 | 0.671 | 1 | 0.187472779 | 0.01218599 |
| Larp4b | 0.00614147 | 0.870544731 | 0.905 | 0.847 | 1 | 0.031995721 | 0.662934941 | 0.978 | 0.903 | 1 | 0.031995721 | 0.012245223 |
| P4ha2 | 0.05639217 | -0.711295916 | 0.476 | 0.709 | 1 | 0.006160263 | -0.433610394 | 0.289 | 0.616 | 1 | 0.05639217 | 0.012282578 |
| Sp1 | 0.006165851 | -1.142921019 | 0.238 | 0.616 | 1 | 0.783100784 | 0.380888427 | 0.756 | 0.785 | 1 | 0.783100784 | 0.012293684 |
| Prr12 | 0.755837686 | 0.926537838 | 0.143 | 0.127 | 1 | 0.006173082 | -1.37003875 | 0.044 | 0.225 | 1 | 0.755837686 | 0.012308057 |
| Psen1 | 0.006196273 | 1.396237792 | 0.667 | 0.59 | 1 | 0.310750768 | 0.405433767 | 0.778 | 0.678 | 1 | 0.310750768 | 0.012354152 |
| Nsmce4a | 0.027678762 | -1.009924687 | 0.238 | 0.515 | 1 | 0.006200393 | -0.756796766 | 0.4 | 0.609 | 1 | 0.027678762 | 0.012362341 |
| 2310014F06Rik | 0.801462099 | 1.548240886 | 0.143 | 0.134 | 1 | 0.006216397 | -2.583713955 | 0.022 | 0.183 | 1 | 0.801462099 | 0.012394151 |
| Zfp532 | 0.006230354 | 1.65491274 | 0.571 | 0.347 | 1 | 0.538061935 | 0.283465127 | 0.356 | 0.471 | 1 | 0.538061935 | 0.01242189 |
| Men1 | 0.938674785 | 0.629377648 | 0.095 | 0.104 | 1 | 0.006245644 | -1.239759553 | 0.067 | 0.256 | 1 | 0.938674785 | 0.01245228 |
| Dnaaf2 | 0.524859929 | -0.25696557 | 0.095 | 0.149 | 1 | 0.006247487 | -2.404390255 | 0.022 | 0.183 | 1 | 0.524859929 | 0.012455943 |
| Akip1 | 0.065238833 | -1.880817084 | 0 | 0.142 | 1 | 0.006248274 | -2.466674533 | 0.022 | 0.183 | 1 | 0.065238833 | 0.012457507 |
| Acsl1 | 0.67144157 | -0.24438994 | 0.429 | 0.481 | 1 | 0.006260547 | 0.300602949 | 0.711 | 0.415 | 1 | 0.67144157 | 0.0124819 |
| Atf6 | 0.006270063 | 0.896164189 | 0.905 | 0.802 | 1 | 0.623202488 | 0.259264877 | 0.933 | 0.858 | 1 | 0.623202488 | 0.012500812 |
| Gm19951 | 0.006273428 | 1.809235185 | 0.952 | 0.97 | 1 | 0.785951235 | 0.933317931 | 0.978 | 0.99 | 1 | 0.785951235 | 0.0125075 |
| Ttc13 | 0.143205173 | -1.133583154 | 0.048 | 0.168 | 1 | 0.006278753 | -2.361321533 | 0.022 | 0.183 | 1 | 0.143205173 | 0.012518083 |
| Tacc1 | 0.380654641 | 0.167263143 | 0.762 | 0.884 | 1 | 0.006287515 | 0.483135016 | 0.978 | 0.872 | 1 | 0.380654641 | 0.012535498 |
| Nek8 | 0.280604124 | -0.695462042 | 0.095 | 0.187 | 1 | 0.006293858 | -2.425451871 | 0.022 | 0.183 | 1 | 0.280604124 | 0.012548103 |

| | | | | | | | | | | | | |
|---|---|---|---|---|---|---|---|---|---|---|---|---|
| Gm17586 | 0.501934276 | -0.233118828 | 0.095 | 0.153 | 1 | 0.006299484 | -1.294207337 | 0.022 | 0.19 | 1 | 0.501934276 | 0.012559285 |
| Snapc2 | 0.050450061 | -1.818081329 | 0 | 0.157 | 1 | 0.00630803 | -2.316927414 | 0.022 | 0.183 | 1 | 0.050450061 | 0.012576269 |
| Washc1 | 0.113628978 | -1.133583154 | 0.048 | 0.183 | 1 | 0.006308174 | -2.383016604 | 0.022 | 0.183 | 1 | 0.113628978 | 0.012576556 |
| Pus1 | 0.281404792 | -0.326228232 | 0.095 | 0.198 | 1 | 0.006322017 | -2.425451871 | 0.022 | 0.183 | 1 | 0.281404792 | 0.012604067 |
| Fhip2a | 0.012057812 | -1.413691074 | 0.143 | 0.429 | 1 | 0.006340028 | -0.61571901 | 0.378 | 0.696 | 1 | 0.012057812 | 0.01263986 |
| Pxmp4 | 0.018060732 | -1.623908781 | 0.095 | 0.343 | 1 | 0.006348094 | -2.223818009 | 0.022 | 0.183 | 1 | 0.018060732 | 0.01265589 |
| Lars | 0.006350559 | 1.221259563 | 0.762 | 0.646 | 1 | 0.053528623 | -0.129490627 | 0.622 | 0.747 | 1 | 0.053528623 | 0.012660789 |
| Pcif1 | 0.020037564 | -1.213753503 | 0.095 | 0.373 | 1 | 0.006353661 | -0.529921137 | 0.2 | 0.453 | 1 | 0.020037564 | 0.012666953 |
| Ak1 | 0.006357599 | -3.370622352 | 0 | 0.272 | 1 | 0.024137857 | -0.425451871 | 0.156 | 0.346 | 1 | 0.024137857 | 0.012674779 |
| Serhl | 0.010103427 | -2.63000898 | 0 | 0.246 | 1 | 0.00635825 | -1.404390255 | 0.022 | 0.19 | 1 | 0.010103427 | 0.012676072 |
| Hmbs | 0.050683621 | -1.970084422 | 0 | 0.157 | 1 | 0.006374668 | -2.84048937 | 0 | 0.145 | 1 | 0.050683621 | 0.0127087 |
| Tmem17 | 0.153900443 | -1.455511249 | 0 | 0.09 | 1 | 0.006374668 | -2.84048937 | 0 | 0.145 | 1 | 0.153900443 | 0.0127087 |
| Smagp | 0.006379534 | -3.648156327 | 0 | 0.272 | 1 | 0.687216983 | 1.567595369 | 0.022 | 0.035 | 1 | 0.687216983 | 0.01271837 |
| Hmgn3 | 0.072325752 | -0.67052414 | 0.429 | 0.623 | 1 | 0.006380385 | -0.89183625 | 0.644 | 0.685 | 1 | 0.072325752 | 0.01272006 |
| Tmsb15b1 | 0.036867867 | -2.159118246 | 0 | 0.175 | 1 | 0.006384197 | -2.871516266 | 0 | 0.145 | 1 | 0.036867867 | 0.012727635 |
| Ppp1r13l | 0.507699835 | -0.184209227 | 0.048 | 0.09 | 1 | 0.006386875 | -1.486852415 | 0.022 | 0.19 | 1 | 0.507699835 | 0.012732957 |
| Lef1 | 0.022912634 | -2.873122692 | 0.048 | 0.269 | 1 | 0.006392078 | -0.417531355 | 0.2 | 0.46 | 1 | 0.022912634 | 0.012743297 |
| Azin2 | 0.333592519 | 0.69394965 | 0.095 | 0.198 | 1 | 0.006393192 | -2.989352756 | 0 | 0.145 | 1 | 0.333592519 | 0.012745511 |
| Atp11a | 0.126602139 | 1.257125016 | 0.619 | 0.582 | 1 | 0.006396331 | 0.74386427 | 0.889 | 0.657 | 1 | 0.126602139 | 0.012751749 |
| Paf1 | 0.653879832 | 0.448058988 | 0.476 | 0.575 | 1 | 0.006396591 | -0.649635347 | 0.489 | 0.675 | 1 | 0.653879832 | 0.012752265 |
| Elac2 | 0.094474288 | -1.574155746 | 0 | 0.119 | 1 | 0.00640182 | -2.931637258 | 0 | 0.145 | 1 | 0.094474288 | 0.012762657 |
| Zfp513 | 0.285927079 | -0.133583154 | 0.048 | 0.134 | 1 | 0.006409681 | -3.071814916 | 0 | 0.145 | 1 | 0.285927079 | 0.012778279 |
| Alg6 | 0.072926247 | -1.348596045 | 0.048 | 0.209 | 1 | 0.006409939 | -2.960783604 | 0 | 0.145 | 1 | 0.072926247 | 0.012778791 |
| C1qtnf12 | 0.094472904 | -1.611630451 | 0 | 0.119 | 1 | 0.006409939 | -2.960783604 | 0 | 0.145 | 1 | 0.094472904 | 0.012778791 |
| Lin37 | 0.219747789 | -0.20069735 | 0.048 | 0.153 | 1 | 0.006409939 | -2.960783604 | 0 | 0.145 | 1 | 0.219747789 | 0.012778791 |
| Gdap2 | 0.232423369 | 1.451379346 | 0.333 | 0.287 | 1 | 0.00644724 | -0.486852415 | 0.067 | 0.273 | 1 | 0.232423369 | 0.012852914 |
| Ddah2 | 0.08143857 | -1.586755783 | 0.095 | 0.257 | 1 | 0.006460249 | -1.574315257 | 0.089 | 0.284 | 1 | 0.08143857 | 0.012878764 |
| Hgs | 0.337738228 | -0.361852142 | 0.143 | 0.239 | 1 | 0.006466086 | -1.02842232 | 0.111 | 0.318 | 1 | 0.337738228 | 0.012890362 |
| Trim12a | 0.303265167 | -0.548620654 | 0.048 | 0.127 | 1 | 0.006470632 | -3.545746104 | 0 | 0.145 | 1 | 0.303265167 | 0.012899394 |
| Pcdh18 | 0.145024626 | -1.940938076 | 0 | 0.093 | 1 | 0.006481287 | -3.886783022 | 0 | 0.145 | 1 | 0.145024626 | 0.012920567 |
| Znrf2 | 0.884996342 | 1.360888812 | 0.286 | 0.384 | 1 | 0.00649642 | 0.540114632 | 0.8 | 0.481 | 1 | 0.884996342 | 0.012950637 |
| Ercc3 | 0.902980621 | 1.380990018 | 0.143 | 0.146 | 1 | 0.00651993 | -0.743192169 | 0.044 | 0.232 | 1 | 0.902980621 | 0.012997351 |
| Boc | 0.641297041 | 0.84737109 | 0.429 | 0.444 | 1 | 0.006534686 | -0.349815708 | 0.378 | 0.702 | 1 | 0.641297041 | 0.01302667 |
| Zfp740 | 0.237619662 | -0.351763324 | 0.143 | 0.291 | 1 | 0.006555992 | -0.374122815 | 0.289 | 0.561 | 1 | 0.237619662 | 0.013069003 |
| Tfdp2 | 0.006558869 | -1.654155068 | 0.143 | 0.466 | 1 | 0.518805027 | 0.361144491 | 0.689 | 0.606 | 1 | 0.518805027 | 0.01307472 |
| Slc35d1 | 0.006565999 | -0.94420579 | 0.476 | 0.802 | 1 | 0.32640142 | -0.280750478 | 0.8 | 0.824 | 1 | 0.32640142 | 0.013088885 |
| Vps39 | 0.278000672 | -0.413691074 | 0.095 | 0.198 | 1 | 0.006585206 | -1.502793959 | 0.067 | 0.253 | 1 | 0.278000672 | 0.013127047 |
| Kirrel | 0.353723315 | 0.174539141 | 0.429 | 0.668 | 1 | 0.006594551 | -0.250500052 | 0.422 | 0.799 | 1 | 0.353723315 | 0.013145614 |
| Lynx1 | 0.093353244 | -1.348596045 | 0.048 | 0.194 | 1 | 0.00659895 | -1.674479418 | 0.044 | 0.221 | 1 | 0.093353244 | 0.013154355 |
| Zfyve1 | 0.634942321 | 1.443474148 | 0.381 | 0.399 | 1 | 0.006610292 | -0.407650584 | 0.356 | 0.671 | 1 | 0.634942321 | 0.013176887 |
| Atl3 | 0.397123362 | 0.500288947 | 0.571 | 0.701 | 1 | 0.006614923 | -0.525198727 | 0.733 | 0.834 | 1 | 0.397123362 | 0.01318609 |
| Arhgef12 | 0.006622034 | 0.966611134 | 0.952 | 0.985 | 1 | 0.014195087 | 0.432977236 | 1 | 0.993 | 1 | 0.014195087 | 0.013200217 |
| Aar2 | 0.208490502 | 0.185485286 | 0.095 | 0.235 | 1 | 0.006623982 | -1.331877755 | 0.067 | 0.26 | 1 | 0.208490502 | 0.013204087 |
| Clk3 | 0.383650769 | -0.361852142 | 0.143 | 0.228 | 1 | 0.006627236 | -1.041820176 | 0.089 | 0.298 | 1 | 0.383650769 | 0.013210552 |
| Zfp397 | 0.246360417 | -0.471905688 | 0.286 | 0.433 | 1 | 0.006632148 | -0.664850717 | 0.2 | 0.453 | 1 | 0.246360417 | 0.01322031 |
| Ccdc71 | 0.977857927 | 1.001346426 | 0.143 | 0.16 | 1 | 0.006634662 | -1.674479418 | 0.044 | 0.221 | 1 | 0.977857927 | 0.013225305 |
| Homer2 | 0.184899106 | -1.574155746 | 0 | 0.078 | 1 | 0.006641419 | -2.901889915 | 0.022 | 0.18 | 1 | 0.184899106 | 0.01323873 |
| Mtss1 | 0.577135075 | 0.339871031 | 0.095 | 0.149 | 1 | 0.006645358 | 0.124105294 | 0.356 | 0.163 | 1 | 0.577135075 | 0.013246555 |
| Rit1 | 0.116444095 | -0.813494059 | 0.19 | 0.373 | 1 | 0.006654727 | -0.146393037 | 0.222 | 0.498 | 1 | 0.116444095 | 0.013265169 |
| Nup54 | 0.114003081 | 0.140339568 | 0.238 | 0.493 | 1 | 0.006656232 | -0.257221168 | 0.311 | 0.585 | 1 | 0.114003081 | 0.013268158 |
| Nt5dc3 | 0.074908499 | 0.9861942 | 0.619 | 0.612 | 1 | 0.006661718 | 0.838830256 | 0.822 | 0.706 | 1 | 0.074908499 | 0.013279058 |
| Wrn | 0.122282838 | 1.648236676 | 0.381 | 0.299 | 1 | 0.00666297 | -0.371375198 | 0.2 | 0.464 | 1 | 0.122282838 | 0.013281545 |
| Ica1 | 0.006681544 | 1.213042178 | 0.714 | 0.653 | 1 | 0.161007705 | 0.576971162 | 0.289 | 0.478 | 1 | 0.161007705 | 0.013318444 |
| Prph | 0.367982232 | 2.088809267 | 0.048 | 0.019 | 1 | 0.006695593 | 4.268035087 | 0.044 | 0.003 | 1 | 0.367982232 | 0.013346354 |
| Gm2895 | 0.457292877 | 0.351843673 | 0 | 0.026 | 1 | 0.006695593 | 4.268035087 | 0.044 | 0.003 | 1 | 0.457292877 | 0.013346354 |
| Ccdc170 | 0.57816106 | 1.351843673 | 0 | 0.015 | 1 | 0.006695593 | 5.005000681 | 0.044 | 0.003 | 1 | 0.57816106 | 0.013346354 |
| Gm42912 | 0.632076134 | 1.088809267 | 0 | 0.011 | 1 | 0.006695593 | 4.490427508 | 0.044 | 0.003 | 1 | 0.632076134 | 0.013346354 |
| Acta2 | 0.006701618 | -3.58046952 | 0.048 | 0.34 | 1 | 0.155398027 | -2.863821874 | 0.022 | 0.083 | 1 | 0.155398027 | 0.013358324 |
| Nagk | 0.066309899 | -0.8244791 | 0.095 | 0.306 | 1 | 0.006704602 | -0.616487696 | 0.111 | 0.329 | 1 | 0.066309899 | 0.013364252 |
| Pea15a | 0.02073974 | -0.253018385 | 0.286 | 0.649 | 1 | 0.00670893 | -0.162893785 | 0.378 | 0.73 | 1 | 0.02073974 | 0.013372851 |
| Prg4 | 0.452309728 | 0.164526832 | 0.381 | 0.332 | 1 | 0.006721916 | -1.284717531 | 0.133 | 0.343 | 1 | 0.452309728 | 0.013398648 |
| Fli1 | 0.033222184 | 1.275492787 | 0.667 | 0.642 | 1 | 0.006730056 | -0.413788953 | 0.267 | 0.571 | 1 | 0.033222184 | 0.013414817 |
| Mkln1 | 0.006737868 | 0.879538973 | 0.857 | 0.933 | 1 | 0.030842965 | 0.565138149 | 1 | 0.962 | 1 | 0.030842965 | 0.013430336 |
| Cd2bp2 | 0.067763815 | -1.455511249 | 0.048 | 0.213 | 1 | 0.006773033 | -0.23119754 | 0.067 | 0.273 | 1 | 0.067763815 | 0.013500192 |
| Mkrn2 | 0.12053871 | -0.385121921 | 0.095 | 0.272 | 1 | 0.006788267 | -1.459885368 | 0.067 | 0.256 | 1 | 0.12053871 | 0.013530453 |
| Zdhhc3 | 0.00679476 | -0.950719097 | 0.19 | 0.563 | 1 | 0.062153935 | -0.435868487 | 0.444 | 0.581 | 1 | 0.062153935 | 0.013543352 |
| Sema4f | 0.396916958 | 0.214340149 | 0 | 0.034 | 1 | 0.006798693 | 3.683072586 | 0.044 | 0.003 | 1 | 0.396916958 | 0.013551164 |
| Gm44175 | 0.57816106 | 1.351843673 | 0 | 0.015 | 1 | 0.006798693 | 4.490427508 | 0.044 | 0.003 | 1 | 0.57816106 | 0.013551164 |
| Gm49164 | 0.63207476 | 1.351843673 | 0 | 0.011 | 1 | 0.006798693 | 4.005000681 | 0.044 | 0.003 | 1 | 0.63207476 | 0.013551164 |
| Dmwd | 0.138998215 | -0.63000898 | 0.095 | 0.246 | 1 | 0.006822737 | -0.865364039 | 0.089 | 0.291 | 1 | 0.138998215 | 0.013598925 |
| Ube2d1 | 0.006848405 | -1.101015292 | 0.143 | 0.507 | 1 | 0.012418191 | -0.195621118 | 0.2 | 0.446 | 1 | 0.012418191 | 0.01364991 |
| Ubxn8 | 0.011418507 | -1.672003069 | 0.095 | 0.366 | 1 | 0.006860271 | -0.404390255 | 0.133 | 0.377 | 1 | 0.011418507 | 0.01367348 |
| Rbm14 | 0.439446409 | -0.496153234 | 0.19 | 0.257 | 1 | 0.006909532 | -0.674479418 | 0.156 | 0.408 | 1 | 0.439446409 | 0.013771323 |
| Cenpt | 0.731194421 | 0.503846766 | 0.095 | 0.127 | 1 | 0.006929008 | -1.731964913 | 0.022 | 0.187 | 1 | 0.731194421 | 0.013810006 |
| Trip12 | 0.00693061 | 0.961855663 | 1 | 0.955 | 1 | 0.187752621 | 0.438379816 | 1 | 0.986 | 1 | 0.187752621 | 0.013813186 |
| Uqcrc2 | 0.006950089 | -0.553055625 | 0.571 | 0.817 | 1 | 0.014908192 | -0.266886731 | 0.867 | 0.889 | 1 | 0.014908192 | 0.013851874 |
| Carm1 | 0.942196206 | 0.569435108 | 0.333 | 0.407 | 1 | 0.006958041 | -0.386412514 | 0.311 | 0.612 | 1 | 0.942196206 | 0.013867667 |
| Ccdc66 | 0.569447138 | 0.295260144 | 0.19 | 0.257 | 1 | 0.006961306 | -0.845306386 | 0.111 | 0.325 | 1 | 0.569447138 | 0.013874151 |
| Mocs2 | 0.006969413 | -1.1627295 | 0.095 | 0.414 | 1 | 0.318075852 | -0.10791889 | 0.489 | 0.529 | 1 | 0.318075852 | 0.013890254 |
| Wbp4 | 0.006972282 | -1.117391121 | 0.381 | 0.694 | 1 | 0.034242047 | -0.420291252 | 0.689 | 0.799 | 1 | 0.034242047 | 0.013895952 |
| Uck1 | 0.217681657 | -0.289702356 | 0.095 | 0.228 | 1 | 0.006975792 | -1.57996182 | 0.089 | 0.273 | 1 | 0.217681657 | 0.013902923 |
| Dync2li1 | 0.117675055 | -0.683780237 | 0.095 | 0.257 | 1 | 0.007013341 | -0.679497493 | 0.111 | 0.325 | 1 | 0.117675055 | 0.013977495 |
| Dimt1 | 0.042707419 | -1.184209227 | 0.095 | 0.313 | 1 | 0.00703315 | -0.618096949 | 0.111 | 0.325 | 1 | 0.042707419 | 0.014016836 |
| Ecpas | 0.077662086 | 0.719575457 | 0.905 | 0.907 | 1 | 0.007038697 | 0.707547087 | 1 | 0.965 | 1 | 0.077662086 | 0.01402785 |
| D5Ertd579e | 0.882594411 | 0.33554186 | 0.619 | 0.709 | 1 | 0.007059503 | 0.443482767 | 0.889 | 0.754 | 1 | 0.882594411 | 0.014069168 |
| Ubxn7 | 0.00707192 | 1.453894944 | 0.762 | 0.634 | 1 | 0.012177601 | 0.755508422 | 0.933 | 0.913 | 1 | 0.012177601 | 0.014093827 |
| Ddx49 | 0.010608252 | -1.683780237 | 0.095 | 0.369 | 1 | 0.007077676 | -0.725012152 | 0.133 | 0.353 | 1 | 0.010608252 | 0.014105258 |
| Oxsr1 | 0.412013737 | 0.123574685 | 0.143 | 0.239 | 1 | 0.007079176 | -1.026585662 | 0.133 | 0.339 | 1 | 0.412013737 | 0.014108236 |
| Dus3l | 0.020286774 | -2.326228232 | 0 | 0.209 | 1 | 0.007090856 | -2.294207337 | 0.022 | 0.18 | 1 | 0.020286774 | 0.014131432 |
| Med22 | 0.168762571 | -0.870548749 | 0.095 | 0.22 | 1 | 0.00709307 | -0.754332726 | 0.133 | 0.353 | 1 | 0.168762571 | 0.014135827 |
| Csrnp1 | 0.007109174 | -1.713630165 | 0.19 | 0.496 | 1 | 0.145339478 | -0.136595597 | 0.422 | 0.512 | 1 | 0.145339478 | 0.014167807 |
| Traf3ip1 | 0.091443332 | -1.516052791 | 0.048 | 0.194 | 1 | 0.007112871 | -1.656777417 | 0.022 | 0.187 | 1 | 0.091443332 | 0.01417515 |
| Gm6980 | 0.578156445 | 1.088809267 | 0 | 0.015 | 1 | 0.007117306 | 2.490427508 | 0.044 | 0.003 | 1 | 0.578156445 | 0.014183955 |
| Nat9 | 0.11399785 | -1.107587946 | 0.048 | 0.183 | 1 | 0.007132386 | -2.871516266 | 0 | 0.142 | 1 | 0.11399785 | 0.014213901 |
| Ddx56 | 0.144276864 | -0.769171728 | 0.143 | 0.287 | 1 | 0.007133922 | -1.609709163 | 0.067 | 0.249 | 1 | 0.144276864 | 0.014216952 |
| Sdhaf3 | 0.100532627 | -1.718545655 | 0 | 0.116 | 1 | 0.007142412 | -2.871516266 | 0 | 0.142 | 1 | 0.100532627 | 0.014233811 |
| Zfp319 | 0.208844381 | -0.849790188 | 0 | 0.071 | 1 | 0.007151873 | -2.901889915 | 0 | 0.142 | 1 | 0.208844381 | 0.014252596 |
| Ggps1 | 0.007152266 | -0.962852853 | 0.333 | 0.69 | 1 | 0.168275969 | -0.131023914 | 0.6 | 0.692 | 1 | 0.168275969 | 0.014253378 |

| | | | | | | | | | | | | |
|---|---|---|---|---|---|---|---|---|---|---|---|---|
| Sephs1 | 0.186689851 | -0.361852142 | 0.095 | 0.228 | 1 | 0.007160431 | -0.57996182 | 0.111 | 0.343 | 1 | 0.186689851 | 0.01426959 |
| Hexdc | 0.222110654 | -0.648156327 | 0 | 0.067 | 1 | 0.007169305 | -3.098287127 | 0 | 0.142 | 1 | 0.222110654 | 0.014287211 |
| Zmat3 | 0.160581777 | 0.106087258 | 0.095 | 0.239 | 1 | 0.007169985 | -0.915186737 | 0.067 | 0.26 | 1 | 0.160581777 | 0.014288561 |
| Yes1 | 0.569208805 | 0.421632003 | 0.381 | 0.545 | 1 | 0.007181424 | 0.899883975 | 0.8 | 0.647 | 1 | 0.569208805 | 0.014311276 |
| H2-Q4 | 0.982439935 | 0.165430549 | 0.286 | 0.291 | 1 | 0.007189635 | -0.935837246 | 0.022 | 0.187 | 1 | 0.982439935 | 0.014327579 |
| Abhd8 | 0.007193028 | -1.623908781 | 0.048 | 0.351 | 1 | 0.009375055 | -0.819427754 | 0.089 | 0.28 | 1 | 0.009375055 | 0.014334316 |
| Vrk3 | 0.007199525 | -1.660128969 | 0.048 | 0.351 | 1 | 0.146465237 | -0.242926832 | 0.178 | 0.291 | 1 | 0.146465237 | 0.014347217 |
| Fas | 0.00847972 | -1.856049179 | 0.143 | 0.437 | 1 | 0.007200993 | -1.255526869 | 0.089 | 0.291 | 1 | 0.00847972 | 0.014350131 |
| Plekhh3 | 0.515369244 | 0.214340149 | 0.048 | 0.093 | 1 | 0.007204511 | -3.199570463 | 0 | 0.142 | 1 | 0.515369244 | 0.014357117 |
| Gm48302 | 0.007206392 | 2.773307441 | 0.238 | 0.075 | 1 | 0.315791203 | 2.135584791 | 0.089 | 0.055 | 1 | 0.315791203 | 0.014360852 |
| Gm29724 | 0.007207085 | 3.351843673 | 0.143 | 0.03 | 1 | 0.376837034 | -0.124282336 | 0 | 0.017 | 1 | 0.376837034 | 0.014362228 |
| Cnr1 | 0.007210102 | 3.580662363 | 0.143 | 0.03 | 1 | 0.293652719 | -2.709244837 | 0 | 0.024 | 1 | 0.293652719 | 0.014368219 |
| Hlcs | 0.831513269 | 0.441111011 | 0.143 | 0.127 | 1 | 0.007210298 | -3.361321533 | 0 | 0.142 | 1 | 0.831513269 | 0.014368608 |
| Mtmr12 | 0.007213148 | -0.187132623 | 0.143 | 0.541 | 1 | 0.025387279 | -0.443875343 | 0.511 | 0.678 | 1 | 0.025387279 | 0.014374267 |
| Acsl6 | 0.492590603 | 0.673771768 | 0 | 0.022 | 1 | 0.007218437 | -3.294207337 | 0 | 0.142 | 1 | 0.492590603 | 0.014384768 |
| Sart3 | 0.871234239 | 0.267779408 | 0.238 | 0.272 | 1 | 0.007229747 | -0.468930507 | 0.178 | 0.405 | 1 | 0.871234239 | 0.014407226 |
| Rigi | 0.885481183 | 1.227515538 | 0.19 | 0.224 | 1 | 0.007231435 | 0.298408736 | 0.089 | 0.311 | 1 | 0.885481183 | 0.014410576 |
| 2810013P06Rik | 0.195601318 | 1.421384606 | 0.333 | 0.239 | 1 | 0.007242683 | -1.743192169 | 0.044 | 0.218 | 1 | 0.195601318 | 0.01443291 |
| Gask1b | 0.4759797 | 1.351843673 | 0.048 | 0.022 | 1 | 0.007246389 | -4.339295227 | 0 | 0.142 | 1 | 0.4759797 | 0.014440268 |
| Saysd1 | 0.007250822 | -1.175893959 | 0.095 | 0.433 | 1 | 0.024848518 | 0.446033389 | 0.156 | 0.367 | 1 | 0.024848518 | 0.01444907 |
| Efna5 | 0.007254898 | 1.485207624 | 0.905 | 0.966 | 1 | 0.9377726 | 0.294147927 | 0.978 | 0.976 | 1 | 0.9377726 | 0.014457162 |
| Camkk2 | 0.048992807 | 0.989273593 | 0.571 | 0.444 | 1 | 0.007272029 | -0.588390442 | 0.289 | 0.571 | 1 | 0.048992807 | 0.014491176 |
| Sf3a2 | 0.37774394 | -0.14822993 | 0.19 | 0.299 | 1 | 0.007299551 | -0.80878051 | 0.133 | 0.343 | 1 | 0.37774394 | 0.014545818 |
| Mettl23 | 0.157526038 | -0.484657595 | 0.19 | 0.366 | 1 | 0.007302622 | -0.444039332 | 0.178 | 0.422 | 1 | 0.157526038 | 0.014551916 |
| Atf5 | 0.018984788 | -1.047327421 | 0.095 | 0.384 | 1 | 0.00732987 | -1.091367713 | 0.222 | 0.419 | 1 | 0.018984788 | 0.014606013 |
| Ass1 | 0.007335226 | -3.25696557 | 0 | 0.265 | 1 | 0.124358137 | -0.722919774 | 0.044 | 0.125 | 1 | 0.124358137 | 0.014616646 |
| Tkt | 0.007338624 | -0.880817084 | 0.476 | 0.754 | 1 | 0.673031164 | -0.251601166 | 0.8 | 0.651 | 1 | 0.673031164 | 0.014623394 |
| Csrp1 | 0.700852327 | -1.458942155 | 0.619 | 0.507 | 1 | 0.007354846 | -0.147646329 | 0.689 | 0.381 | 1 | 0.700852327 | 0.014655599 |
| Rab8b | 0.992860195 | 0.810273768 | 0.429 | 0.459 | 1 | 0.00736537 | 0.968133298 | 0.778 | 0.536 | 1 | 0.992860195 | 0.014676491 |
| Thrb | 0.007366651 | 1.450595912 | 0.857 | 0.81 | 1 | 0.026080232 | 0.835270231 | 0.911 | 0.82 | 1 | 0.026080232 | 0.014679035 |
| Ago2 | 0.113579341 | 0.836890893 | 0.762 | 0.754 | 1 | 0.007366713 | 0.704446237 | 1 | 0.934 | 1 | 0.113579341 | 0.014679158 |
| Socs4 | 0.23113785 | -0.496153234 | 0.333 | 0.481 | 1 | 0.007367588 | -0.41846544 | 0.311 | 0.612 | 1 | 0.23113785 | 0.014680895 |
| Itgb1bp1 | 0.090112208 | 0.59848364 | 0.143 | 0.354 | 1 | 0.007372935 | -0.397847409 | 0.111 | 0.339 | 1 | 0.090112208 | 0.01469151 |
| 1600020E01Rik | 0.926103036 | 0.214340149 | 0.524 | 0.631 | 1 | 0.007384431 | -0.195248857 | 0.4 | 0.678 | 1 | 0.926103036 | 0.014714332 |
| Gstz1 | 0.007385893 | -1.005708332 | 0.143 | 0.485 | 1 | 0.039003599 | -0.505588978 | 0.2 | 0.384 | 1 | 0.039003599 | 0.014717235 |
| Ap1m1 | 0.007406922 | -2.171718283 | 0.048 | 0.328 | 1 | 0.038323155 | -0.375821103 | 0.178 | 0.363 | 1 | 0.038323155 | 0.014758981 |
| Npm3 | 0.007421728 | -1.683780237 | 0.095 | 0.403 | 1 | 0.081743914 | -0.108934372 | 0.244 | 0.412 | 1 | 0.081743914 | 0.014788374 |
| Pantr1 | 0.0074409 | -2.593014773 | 0.143 | 0.444 | 1 | 0.00756797 | -0.785696897 | 0.111 | 0.318 | 1 | 0.00756797 | 0.014826432 |
| Rnf146 | 0.107774002 | -0.449610648 | 0.286 | 0.545 | 1 | 0.007463909 | -0.499452452 | 0.4 | 0.657 | 1 | 0.107774002 | 0.014872108 |
| Unc80 | 0.007478609 | 3.295260144 | 0.143 | 0.03 | 1 | 0.496223879 | 0.683072586 | 0 | 0.01 | 1 | 0.496223879 | 0.014901287 |
| Sphk1 | 0.278949404 | -0.51887331 | 0.143 | 0.265 | 1 | 0.007479571 | -2.174908409 | 0.044 | 0.211 | 1 | 0.278949404 | 0.014903198 |
| Pak1 | 0.052710366 | -0.580217499 | 0.429 | 0.668 | 1 | 0.007481434 | 0.647448676 | 0.889 | 0.73 | 1 | 0.052710366 | 0.014906896 |
| Plekhg5 | 0.247241099 | -1.008052272 | 0.048 | 0.138 | 1 | 0.007497745 | -1.066949161 | 0.089 | 0.291 | 1 | 0.247241099 | 0.014939274 |
| Gpr89 | 0.315920122 | 0.460778044 | 0.19 | 0.302 | 1 | 0.007513295 | -0.557935513 | 0.067 | 0.26 | 1 | 0.315920122 | 0.014970141 |
| Ly96 | 0.071304469 | -1.574155746 | 0.048 | 0.209 | 1 | 0.007519757 | -0.954357334 | 0.089 | 0.294 | 1 | 0.071304469 | 0.014982966 |
| Mcoln1 | 0.007532909 | -2.701267664 | 0 | 0.261 | 1 | 0.030722009 | -0.300439291 | 0.089 | 0.242 | 1 | 0.030722009 | 0.015009072 |
| Pkp1 | 0.173814074 | 2.351843673 | 0.048 | 0.011 | 1 | 0.007539359 | 1.583536913 | 0.133 | 0.038 | 1 | 0.173814074 | 0.015021877 |
| Epc2 | 0.025005486 | 0.98840737 | 0.762 | 0.735 | 1 | 0.007545268 | 0.568345405 | 1 | 0.862 | 1 | 0.025005486 | 0.015033606 |
| Ccdc32 | 0.059174048 | -1.516052791 | 0.048 | 0.22 | 1 | 0.007547159 | -1.47073275 | 0.044 | 0.218 | 1 | 0.059174048 | 0.015037359 |
| Utp23 | 0.131346189 | -1.413691074 | 0.048 | 0.172 | 1 | 0.007561603 | -1.681499846 | 0.067 | 0.246 | 1 | 0.131346189 | 0.015066029 |
| AU041133 | 0.007566992 | 2.756233928 | 0.19 | 0.052 | 1 | 0.132686876 | -1.486852415 | 0 | 0.048 | 1 | 0.132686876 | 0.015076725 |
| Smarcd2 | 0.093047555 | -1.434752689 | 0.048 | 0.194 | 1 | 0.007600449 | -1.367553487 | 0.133 | 0.322 | 1 | 0.093047555 | 0.015143131 |
| Rabgef1 | 0.087400101 | 0.611675647 | 0.762 | 0.705 | 1 | 0.007603386 | 0.798953298 | 0.822 | 0.73 | 1 | 0.087400101 | 0.01514896 |
| Gm49003 | 0.578159907 | 0.214340149 | 0 | 0.015 | 1 | 0.007605371 | 4.142504205 | 0.067 | 0.01 | 1 | 0.578159907 | 0.0151529 |
| Flywch1 | 0.012462861 | -2.496153234 | 0 | 0.235 | 1 | 0.007605602 | -0.954357334 | 0.089 | 0.287 | 1 | 0.012462861 | 0.015153359 |
| Pde4a | 0.570522686 | 0.405282932 | 0.19 | 0.276 | 1 | 0.007608205 | -0.432404631 | 0.111 | 0.322 | 1 | 0.570522686 | 0.015158524 |
| Zcchc9 | 0.07706048 | 0.100882099 | 0.095 | 0.302 | 1 | 0.007609306 | -0.98280991 | 0.156 | 0.367 | 1 | 0.07706048 | 0.015160711 |
| Ppa2 | 0.227701286 | 0.159198595 | 0.095 | 0.228 | 1 | 0.007609418 | -0.886783022 | 0.067 | 0.249 | 1 | 0.227701286 | 0.015160933 |
| Plekhm2 | 0.007611842 | 0.877600584 | 0.857 | 0.75 | 1 | 0.051766909 | 0.586048132 | 0.911 | 0.865 | 1 | 0.051766909 | 0.015165743 |
| Ccdc137 | 0.126330458 | -0.801961663 | 0.095 | 0.25 | 1 | 0.007611994 | -1.109485334 | 0.067 | 0.249 | 1 | 0.126330458 | 0.015166045 |
| Tead3 | 0.265380777 | -0.213753503 | 0.095 | 0.213 | 1 | 0.00761498 | -1.158229668 | 0.044 | 0.225 | 1 | 0.265380777 | 0.015171972 |
| Dlst | 0.023222006 | 0.810508104 | 0.762 | 0.649 | 1 | 0.007628538 | 0.849216848 | 0.889 | 0.74 | 1 | 0.023222006 | 0.015198882 |
| Cnot4 | 0.007635296 | 0.718271228 | 1 | 0.881 | 1 | 0.008819712 | 0.762585956 | 1 | 0.962 | 1 | 0.008819712 | 0.015212294 |
| Yars2 | 0.175232787 | -0.880817084 | 0.048 | 0.157 | 1 | 0.007643421 | -1.881712033 | 0.044 | 0.211 | 1 | 0.175232787 | 0.01522842 |
| Fh1 | 0.018062833 | -1.623908781 | 0.095 | 0.343 | 1 | 0.007644897 | -0.452086997 | 0.111 | 0.325 | 1 | 0.018062833 | 0.015231349 |
| Csnk1g2 | 0.059064445 | -0.129095763 | 0.381 | 0.679 | 1 | 0.007655278 | -0.493380458 | 0.644 | 0.737 | 1 | 0.059064445 | 0.015251952 |
| Akr1e1 | 0.007666019 | -2.801961663 | 0 | 0.261 | 1 | 0.010009975 | -1.486852415 | 0.067 | 0.239 | 1 | 0.010009975 | 0.015273269 |
| Zbed5 | 0.211504219 | -0.599246727 | 0.048 | 0.153 | 1 | 0.00766896 | -2.620708162 | 0.022 | 0.176 | 1 | 0.211504219 | 0.015279107 |
| Cacfd1 | 0.043360909 | -0.633656758 | 0.095 | 0.321 | 1 | 0.007669321 | -0.372209849 | 0.111 | 0.336 | 1 | 0.043360909 | 0.015279824 |
| Ccdc136 | 0.89891872 | 0.673771768 | 0.095 | 0.112 | 1 | 0.007671404 | -1.167783975 | 0.067 | 0.256 | 1 | 0.89891872 | 0.015283957 |
| Mis12 | 0.067010425 | 0.740885964 | 0.048 | 0.231 | 1 | 0.007676825 | -1.109485334 | 0.067 | 0.249 | 1 | 0.067010425 | 0.015294717 |
| Cdc40 | 0.007679906 | -0.896083841 | 0.19 | 0.541 | 1 | 0.15404026 | 0.749102457 | 0.689 | 0.609 | 1 | 0.15404026 | 0.015300831 |
| Dock10 | 0.113798245 | -2.326228232 | 0 | 0.108 | 1 | 0.007682547 | -2.446210431 | 0.044 | 0.215 | 1 | 0.113798245 | 0.015306072 |
| Fsip1 | 0.63207476 | 1.351843673 | 0 | 0.011 | 1 | 0.007687626 | 4.448607333 | 0.067 | 0.01 | 1 | 0.63207476 | 0.015316152 |
| Utp11 | 0.007718822 | -0.742874984 | 0.238 | 0.627 | 1 | 0.009932488 | -0.465604682 | 0.4 | 0.612 | 1 | 0.009932488 | 0.015378064 |
| Ptges | 0.544491988 | 0.503846766 | 0.095 | 0.157 | 1 | 0.007731082 | 0.982632868 | 0.244 | 0.1 | 1 | 0.544491988 | 0.015402394 |
| Cript | 0.007738164 | -0.206646617 | 0.095 | 0.437 | 1 | 0.274270323 | -0.207139268 | 0.378 | 0.453 | 1 | 0.274270323 | 0.015416449 |
| Ifi35 | 0.480454647 | 0.376091219 | 0.238 | 0.373 | 1 | 0.007745541 | 0.110182918 | 0.2 | 0.433 | 1 | 0.480454647 | 0.015431089 |
| Atp6ap2 | 0.542215849 | -0.148849911 | 0.81 | 0.843 | 1 | 0.007751956 | -0.529067363 | 0.978 | 0.924 | 1 | 0.542215849 | 0.01544382 |
| Rhog | 0.100033164 | -0.558888989 | 0.095 | 0.269 | 1 | 0.007759621 | -0.805213895 | 0.089 | 0.287 | 1 | 0.100033164 | 0.015459031 |
| Nup155 | 0.339578816 | 0.533290544 | 0.143 | 0.257 | 1 | 0.007765525 | -1.007242915 | 0.111 | 0.325 | 1 | 0.339578816 | 0.015470747 |
| Aagab | 0.140447233 | 1.304537958 | 0.286 | 0.179 | 1 | 0.007766317 | -1.624355939 | 0.067 | 0.246 | 1 | 0.140447233 | 0.015472319 |
| Lipo3 | 0.160753189 | -0.695462042 | 0.048 | 0.172 | 1 | 0.007769981 | -1.141355849 | 0.044 | 0.218 | 1 | 0.160753189 | 0.01547959 |
| Gm54771 | 0.457308648 | 0.673771768 | 0 | 0.026 | 1 | 0.007770303 | 3.268035087 | 0.067 | 0.01 | 1 | 0.457308648 | 0.015480228 |
| Adprs | 0.01561396 | -2.611630451 | 0 | 0.224 | 1 | 0.00778497 | -2.506751973 | 0.022 | 0.176 | 1 | 0.01561396 | 0.015509333 |
| Fam149a | 0.132757274 | -0.280424543 | 0.048 | 0.187 | 1 | 0.007807197 | -1.053893008 | 0.044 | 0.218 | 1 | 0.132757274 | 0.015553442 |
| D3Ertd751e | 0.481875957 | -0.496153234 | 0.143 | 0.209 | 1 | 0.007811227 | -1.674479418 | 0.022 | 0.183 | 1 | 0.481875957 | 0.01556144 |
| Mbtps1 | 0.007817294 | -1.049604762 | 0.286 | 0.642 | 1 | 0.012880454 | -0.459885368 | 0.511 | 0.73 | 1 | 0.012880454 | 0.015573479 |
| Psmc1 | 0.043707467 | -0.44309199 | 0.333 | 0.69 | 1 | 0.007818266 | -0.515072289 | 0.556 | 0.689 | 1 | 0.043707467 | 0.015575406 |
| Otud1 | 0.120787029 | -1.785659851 | 0 | 0.104 | 1 | 0.007828706 | -2.425451871 | 0.022 | 0.176 | 1 | 0.120787029 | 0.015596124 |
| Uckl1 | 0.030310792 | -2.184209227 | 0 | 0.187 | 1 | 0.007836087 | -1.589945908 | 0.022 | 0.183 | 1 | 0.030310792 | 0.01561077 |
| Prmt5 | 0.022510804 | -1.926141075 | 0.048 | 0.272 | 1 | 0.007845276 | -0.763183644 | 0.089 | 0.287 | 1 | 0.022510804 | 0.015629004 |
| Gm42679 | 0.439146102 | 0.569435108 | 0.143 | 0.09 | 1 | 0.007853399 | 3.683072586 | 0.067 | 0.01 | 1 | 0.439146102 | 0.015645123 |
| 4930505N22Rik | 0.924795272 | 1.351843673 | 0.048 | 0.045 | 1 | 0.007853399 | 3.490427508 | 0.067 | 0.01 | 1 | 0.924795272 | 0.015645123 |
| Sipa1l3 | 0.52060399 | 0.8372705 | 0.238 | 0.198 | 1 | 0.007877436 | 0.520801157 | 0.8 | 0.512 | 1 | 0.52060399 | 0.015692818 |
| Tox | 0.007883215 | -1.899117901 | 0.143 | 0.444 | 1 | 0.080735586 | -0.316927414 | 0.378 | 0.578 | 1 | 0.080735586 | 0.015704286 |
| Coprs | 0.016542951 | -2.455511249 | 0 | 0.22 | 1 | 0.007891323 | -2.404390255 | 0.022 | 0.176 | 1 | 0.016542951 | 0.015720374 |
| Zfp955a | 0.192427974 | 0.74303443 | 0.286 | 0.168 | 1 | 0.00791125 | -2.339295227 | 0.022 | 0.176 | 1 | 0.192427974 | 0.015759912 |

| | | | | | | | | | | | | |
|---|---|---|---|---|---|---|---|---|---|---|---|---|
| Epc1 | 0.007927794 | 1.107570195 | 0.905 | 0.847 | 1 | 0.20597485 | 0.594172082 | 0.933 | 0.945 | 1 | 0.20597485 | 0.015792738 |
| Eif4e3 | 0.876838649 | 0.271673324 | 0.143 | 0.168 | 1 | 0.007929704 | -1.098287127 | 0.044 | 0.225 | 1 | 0.876838649 | 0.015796528 |
| Ints12 | 0.031925424 | -1.834022873 | 0.048 | 0.254 | 1 | 0.00793383 | -1.223818009 | 0.089 | 0.284 | 1 | 0.031925424 | 0.015804715 |
| Fam219b | 0.122422819 | -0.998653574 | 0.048 | 0.179 | 1 | 0.007950468 | -1.017367132 | 0.022 | 0.183 | 1 | 0.122422819 | 0.015837725 |
| Samd12 | 0.949676521 | 1.639824436 | 0.095 | 0.108 | 1 | 0.007957162 | 2.312234891 | 0.2 | 0.08 | 1 | 0.949676521 | 0.015851007 |
| Pde3b | 0.007973434 | -1.774688733 | 0.095 | 0.418 | 1 | 0.53267647 | 1.158231078 | 0.422 | 0.505 | 1 | 0.53267647 | 0.015883291 |
| Nek4 | 0.720076804 | 1.188344941 | 0.095 | 0.078 | 1 | 0.007976321 | -2.871516266 | 0 | 0.138 | 1 | 0.720076804 | 0.015889021 |
| Trappc14 | 0.42890475 | -0.233118828 | 0.048 | 0.101 | 1 | 0.007986896 | -2.84048937 | 0 | 0.138 | 1 | 0.42890475 | 0.015910001 |
| Thop1 | 0.338320997 | -0.413691074 | 0.048 | 0.116 | 1 | 0.007986944 | -2.871516266 | 0 | 0.138 | 1 | 0.338320997 | 0.015910096 |
| Xab2 | 0.338320997 | -0.455511249 | 0.048 | 0.116 | 1 | 0.00799956 | -0.994999319 | 0.022 | 0.183 | 1 | 0.338320997 | 0.015935128 |
| Tpi1 | 0.008001076 | -0.929569262 | 0.619 | 0.776 | 1 | 0.046411225 | -0.348689976 | 0.844 | 0.803 | 1 | 0.046411225 | 0.015938136 |
| Timm29 | 0.896486021 | 0.780686972 | 0.143 | 0.168 | 1 | 0.008024972 | -0.681499846 | 0.089 | 0.287 | 1 | 0.896486021 | 0.015985543 |
| Fkbp1b | 0.030310792 | -2.184209227 | 0 | 0.187 | 1 | 0.008031857 | -2.960783604 | 0 | 0.138 | 1 | 0.030310792 | 0.015999203 |
| Nptxr | 0.163434053 | -1.081115734 | 0 | 0.086 | 1 | 0.008038979 | -3.124282336 | 0 | 0.138 | 1 | 0.163434053 | 0.016013333 |
| Rexo1 | 0.062814362 | -0.593014773 | 0.143 | 0.388 | 1 | 0.008050591 | -0.645114499 | 0.289 | 0.512 | 1 | 0.062814362 | 0.016036371 |
| Nr2f1 | 0.048074173 | -2.683780237 | 0 | 0.16 | 1 | 0.008057848 | -3.316927414 | 0 | 0.138 | 1 | 0.048074173 | 0.016050766 |
| Rxrb | 0.0177942 | -2.516052791 | 0 | 0.216 | 1 | 0.008057927 | -1.47073275 | 0.044 | 0.221 | 1 | 0.0177942 | 0.016050923 |
| Vps29 | 0.008059224 | -1.036721615 | 0.286 | 0.616 | 1 | 0.162836669 | -0.282980082 | 0.6 | 0.671 | 1 | 0.162836669 | 0.016053497 |
| Trnp1 | 0.323632587 | -0.496153234 | 0.048 | 0.123 | 1 | 0.008062945 | -3.294207337 | 0 | 0.138 | 1 | 0.323632587 | 0.016060879 |
| Fam25c | 0.907378967 | 0.936806174 | 0.048 | 0.056 | 1 | 0.008062945 | -3.361321533 | 0 | 0.138 | 1 | 0.907378967 | 0.016060879 |
| Gm26887 | 0.578234907 | 0.137718867 | 0.095 | 0.149 | 1 | 0.008063041 | -3.316927414 | 0 | 0.138 | 1 | 0.578234907 | 0.01606107 |
| Pgam5 | 0.056026058 | -0.984439715 | 0.095 | 0.302 | 1 | 0.00808112 | -0.979892427 | 0.089 | 0.291 | 1 | 0.056026058 | 0.016096936 |
| Erg | 0.016835294 | 1.628277614 | 0.952 | 0.854 | 1 | 0.008094533 | 1.476435806 | 0.867 | 0.92 | 1 | 0.016835294 | 0.016123544 |
| Kti12 | 0.045101473 | -1.718545655 | 0.048 | 0.235 | 1 | 0.008103209 | -0.557935513 | 0.089 | 0.287 | 1 | 0.045101473 | 0.016140755 |
| Cwc27 | 0.951176865 | 0.334634383 | 0.381 | 0.455 | 1 | 0.008108438 | 1.253821228 | 0.644 | 0.491 | 1 | 0.951176865 | 0.01615113 |
| Eif3j1 | 0.282975222 | -0.186050574 | 0.857 | 0.918 | 1 | 0.008126008 | -0.342206822 | 0.867 | 0.907 | 1 | 0.282975222 | 0.016185985 |
| Mtmr9 | 0.98109446 | 0.644024424 | 0.143 | 0.157 | 1 | 0.008139913 | -1.346674757 | 0.089 | 0.277 | 1 | 0.98109446 | 0.016213569 |
| Wdr48 | 0.00814287 | -1.150656668 | 0.095 | 0.429 | 1 | 0.948139255 | 0.462625809 | 0.422 | 0.502 | 1 | 0.948139255 | 0.016219435 |
| Syde1 | 0.451240353 | 0.726239188 | 0.095 | 0.175 | 1 | 0.008148985 | -1.421264074 | 0.044 | 0.221 | 1 | 0.451240353 | 0.016231563 |
| Trim23 | 0.852658563 | 0.106087258 | 0.238 | 0.22 | 1 | 0.008160098 | -1.372209849 | 0.089 | 0.28 | 1 | 0.852658563 | 0.016253609 |
| Zfp110 | 0.717732014 | 0.397647362 | 0.381 | 0.384 | 1 | 0.00816573 | -0.280401538 | 0.222 | 0.467 | 1 | 0.717732014 | 0.016264781 |
| Hspa1a | 0.688063512 | -2.61489078 | 0.619 | 0.496 | 1 | 0.008166448 | -1.457339135 | 0.844 | 0.567 | 1 | 0.688063512 | 0.016266206 |
| Mthfsd | 0.04179441 | -1.970084422 | 0 | 0.168 | 1 | 0.008166986 | -0.442458296 | 0.067 | 0.266 | 1 | 0.04179441 | 0.016267272 |
| Acd | 0.017305996 | -1.769171728 | 0.048 | 0.287 | 1 | 0.008183937 | -1.153428682 | 0.089 | 0.284 | 1 | 0.017305996 | 0.016300897 |
| F13a1 | 0.720806056 | 0.430863105 | 0.714 | 0.776 | 1 | 0.00819183 | -0.100334432 | 0.378 | 0.696 | 1 | 0.720806056 | 0.016316554 |
| Musk | 0.632076134 | 1.673771768 | 0 | 0.011 | 1 | 0.008196642 | 2.905465008 | 0.067 | 0.01 | 1 | 0.632076134 | 0.016326098 |
| Loxl4 | 0.387064705 | -0.545572788 | 0.667 | 0.735 | 1 | 0.008198995 | -0.656150761 | 0.489 | 0.779 | 1 | 0.387064705 | 0.016330767 |
| Ubac2 | 0.008223854 | -1.364363361 | 0.143 | 0.474 | 1 | 0.86449065 | 0.419053069 | 0.6 | 0.595 | 1 | 0.86449065 | 0.016380077 |
| Idh3b | 0.008235 | -0.89470261 | 0.238 | 0.601 | 1 | 0.022675697 | -0.700977221 | 0.444 | 0.574 | 1 | 0.022675697 | 0.016402185 |
| Styx | 0.473121451 | 0.901182264 | 0.333 | 0.31 | 1 | 0.008241748 | -0.322381844 | 0.178 | 0.436 | 1 | 0.473121451 | 0.01641557 |
| Ppp2r3a | 0.090065726 | 0.893552976 | 0.714 | 0.634 | 1 | 0.008251962 | 0.863246352 | 0.822 | 0.734 | 1 | 0.090065726 | 0.01643583 |
| Hdgfl2 | 0.21110769 | -0.275244304 | 0.19 | 0.358 | 1 | 0.008278025 | -0.334849322 | 0.133 | 0.381 | 1 | 0.21110769 | 0.016487524 |
| Utp18 | 0.0082781 | -0.917189474 | 0.143 | 0.507 | 1 | 0.313262839 | -0.305118112 | 0.556 | 0.602 | 1 | 0.313262839 | 0.016487673 |
| Rin1 | 0.070083823 | 3.258734268 | 0.095 | 0.026 | 1 | 0.008283909 | 3.168499413 | 0.067 | 0.01 | 1 | 0.070083823 | 0.016499194 |
| Kmt5a | 0.008312359 | -1.038181938 | 0.571 | 0.765 | 1 | 0.178355282 | -0.207814958 | 0.933 | 0.875 | 1 | 0.178355282 | 0.016555622 |
| Znhit6 | 0.959329931 | 0.433457438 | 0.333 | 0.407 | 1 | 0.008327733 | -0.464826109 | 0.156 | 0.377 | 1 | 0.959329931 | 0.016586116 |
| Leng1 | 0.013636181 | -2.735619168 | 0 | 0.231 | 1 | 0.008353711 | -1.513324627 | 0.067 | 0.249 | 1 | 0.013636181 | 0.016637637 |
| Faap20 | 0.018183749 | -1.469186186 | 0.048 | 0.302 | 1 | 0.008359078 | -1.691966845 | 0.044 | 0.215 | 1 | 0.018183749 | 0.016648282 |
| Xrcc6 | 0.364711061 | -0.133583154 | 0.143 | 0.246 | 1 | 0.008382224 | -1.998751454 | 0.044 | 0.208 | 1 | 0.364711061 | 0.016694186 |
| Ptbp3 | 0.469628467 | 0.516625371 | 0.905 | 0.925 | 1 | 0.008388628 | 0.764072308 | 1 | 0.945 | 1 | 0.469628467 | 0.016706886 |
| Ipo4 | 0.633432651 | 0.530813814 | 0.238 | 0.325 | 1 | 0.008389342 | -0.664850717 | 0.111 | 0.322 | 1 | 0.633432651 | 0.016708303 |
| Nkapd1 | 0.015237288 | -1.648156327 | 0.143 | 0.41 | 1 | 0.008422135 | -0.236585306 | 0.2 | 0.453 | 1 | 0.015237288 | 0.016773338 |
| Arsa | 0.010964251 | -2.648156327 | 0 | 0.243 | 1 | 0.008429587 | -1.960783604 | 0.044 | 0.208 | 1 | 0.010964251 | 0.016788116 |
| Pik3ip1 | 0.140839527 | -0.940938076 | 0.143 | 0.299 | 1 | 0.008443483 | -0.432404631 | 0.156 | 0.415 | 1 | 0.140839527 | 0.016815674 |
| 2610507I01Rik | 0.347895436 | -0.233118828 | 0.048 | 0.119 | 1 | 0.008443892 | -1.921789472 | 0.022 | 0.18 | 1 | 0.347895436 | 0.016816485 |
| Fgd4 | 0.054399745 | -2.303508156 | 0 | 0.153 | 1 | 0.008452681 | 0.377727375 | 0.511 | 0.27 | 1 | 0.054399745 | 0.016833915 |
| Pacs2 | 0.166735602 | -1.249060372 | 0.095 | 0.216 | 1 | 0.008453065 | -1.199570463 | 0.089 | 0.287 | 1 | 0.166735602 | 0.016834675 |
| Col3a1 | 0.062075194 | -5.346374805 | 0.095 | 0.261 | 1 | 0.008455489 | -1.901889915 | 0.178 | 0.381 | 1 | 0.062075194 | 0.016839484 |
| Prune1 | 0.739133098 | 0.258734268 | 0.143 | 0.183 | 1 | 0.008458995 | -0.847442131 | 0.067 | 0.246 | 1 | 0.739133098 | 0.016846435 |
| Taf2 | 0.730409433 | 0.910472026 | 0.333 | 0.34 | 1 | 0.008459983 | -0.602329633 | 0.178 | 0.415 | 1 | 0.730409433 | 0.016848395 |
| Capzb | 0.026545161 | -0.322358686 | 0.905 | 0.951 | 1 | 0.008468291 | -0.305211111 | 0.978 | 0.962 | 1 | 0.026545161 | 0.01686487 |
| Gm10762 | 0.194019947 | -0.249060372 | 0.095 | 0.224 | 1 | 0.008477933 | -1.502793959 | 0.044 | 0.215 | 1 | 0.194019947 | 0.01688399 |
| Abhd10 | 0.021969398 | -2.496153234 | 0 | 0.205 | 1 | 0.008491861 | -1.674479418 | 0.067 | 0.239 | 1 | 0.021969398 | 0.01691161 |
| Fgd1 | 0.05683149 | -0.896083841 | 0.048 | 0.239 | 1 | 0.008520286 | -1.827889333 | 0.089 | 0.263 | 1 | 0.05683149 | 0.016967977 |
| Aamp | 0.617325732 | 0.106087258 | 0.667 | 0.84 | 1 | 0.00856029 | -0.281895398 | 0.711 | 0.875 | 1 | 0.617325732 | 0.017047302 |
| Mdp1 | 0.09366015 | -1.303508156 | 0.048 | 0.194 | 1 | 0.008568679 | -1.881712033 | 0.044 | 0.208 | 1 | 0.09366015 | 0.017063937 |
| Gsk3b | 0.067915275 | 1.286077936 | 1 | 0.94 | 1 | 0.00857412 | 0.564994853 | 1 | 0.979 | 1 | 0.067915275 | 0.017074725 |
| Bmper | 0.084089072 | -0.26168798 | 0.143 | 0.362 | 1 | 0.00857886 | -2.012735683 | 0.089 | 0.284 | 1 | 0.084089072 | 0.017084123 |
| Picalm | 0.008582109 | 0.939480308 | 0.905 | 0.922 | 1 | 0.12278513 | 0.668089557 | 1 | 0.99 | 1 | 0.12278513 | 0.017090565 |
| Iqgap1 | 0.00858985 | 0.634312053 | 1 | 0.981 | 1 | 0.085795099 | 0.542728179 | 1 | 0.997 | 1 | 0.085795099 | 0.017105914 |
| Cgn | 0.173814074 | 1.866416846 | 0.048 | 0.011 | 1 | 0.008595281 | 1.250113179 | 0.2 | 0.076 | 1 | 0.173814074 | 0.017116684 |
| Ppp4c | 0.008618061 | -0.579208973 | 0.238 | 0.601 | 1 | 0.109842197 | -0.320202546 | 0.689 | 0.751 | 1 | 0.109842197 | 0.017161852 |
| Dbr1 | 0.059180385 | -1.555046923 | 0.048 | 0.22 | 1 | 0.008623791 | -1.237492946 | 0.089 | 0.273 | 1 | 0.059180385 | 0.017173213 |
| Slc29a3 | 0.008626254 | -2.146407195 | 0.048 | 0.321 | 1 | 0.13526975 | 0.320502507 | 0.156 | 0.291 | 1 | 0.13526975 | 0.017178096 |
| Mir100hg | 0.123204714 | 1.007141871 | 0.81 | 0.776 | 1 | 0.008630996 | 0.826513666 | 0.733 | 0.792 | 1 | 0.123204714 | 0.017187497 |
| Gm26911 | 0.417335663 | 1.171271427 | 0.095 | 0.052 | 1 | 0.008631943 | 2.82057611 | 0.156 | 0.055 | 1 | 0.417335663 | 0.017189377 |
| Ranbp3 | 0.411897018 | 0.121976131 | 0.238 | 0.366 | 1 | 0.008633387 | -0.63081819 | 0.2 | 0.436 | 1 | 0.411897018 | 0.017192239 |
| Zbtb21 | 0.059837426 | -0.960434252 | 0.19 | 0.399 | 1 | 0.008640014 | -0.163382156 | 0.178 | 0.439 | 1 | 0.059837426 | 0.017205379 |
| Wdr83 | 0.234341503 | 0.119182916 | 0.048 | 0.149 | 1 | 0.008644825 | -1.662702251 | 0.022 | 0.18 | 1 | 0.234341503 | 0.017214917 |
| Atxn3 | 0.911364858 | 0.434305833 | 0.238 | 0.239 | 1 | 0.008648798 | -1.202284848 | 0.089 | 0.284 | 1 | 0.911364858 | 0.017222794 |
| Gabpa | 0.40618058 | 0.794626163 | 0.333 | 0.276 | 1 | 0.008654227 | -0.147869156 | 0.178 | 0.422 | 1 | 0.40618058 | 0.017233559 |
| Pcsk7 | 0.917975739 | 0.72381245 | 0.19 | 0.224 | 1 | 0.008669357 | -0.219630212 | 0.156 | 0.388 | 1 | 0.917975739 | 0.017263557 |
| Eif2ak1 | 0.374391264 | -0.242704877 | 0.286 | 0.392 | 1 | 0.008733055 | -0.486852415 | 0.222 | 0.453 | 1 | 0.374391264 | 0.017389843 |
| Znrf1 | 0.180022275 | 0.635390676 | 0.667 | 0.649 | 1 | 0.008735788 | 0.530473461 | 0.956 | 0.668 | 1 | 0.180022275 | 0.017395261 |
| Ap2s1 | 0.008743739 | -0.864193254 | 0.286 | 0.642 | 1 | 0.020275324 | -0.623173886 | 0.578 | 0.664 | 1 | 0.020275324 | 0.017411026 |
| Mettl4 | 0.307714542 | 0.79930265 | 0.048 | 0.131 | 1 | 0.008744023 | -2.564854927 | 0.022 | 0.173 | 1 | 0.307714542 | 0.017411588 |
| Dqx1 | 0.008749952 | 0.735172312 | 0.143 | 0.03 | 1 | 0.293652719 | -0.901889915 | 0 | 0.024 | 1 | 0.293652719 | 0.017423343 |
| Bcl9 | 0.095320252 | -0.911190733 | 0.048 | 0.205 | 1 | 0.008763196 | -0.86124793 | 0.133 | 0.336 | 1 | 0.095320252 | 0.017449597 |
| Dhrs4 | 0.029767917 | -1.20069735 | 0.048 | 0.276 | 1 | 0.008764695 | -0.329751454 | 0.089 | 0.284 | 1 | 0.029767917 | 0.01745257 |
| Asnsd1 | 0.008783639 | -2.683780237 | 0 | 0.254 | 1 | 0.018620079 | -1.819427754 | 0.044 | 0.183 | 1 | 0.018620079 | 0.017490126 |
| Grsf1 | 0.011373559 | -1.059582573 | 0.238 | 0.575 | 1 | 0.008785547 | -0.98280991 | 0.489 | 0.612 | 1 | 0.011373559 | 0.017493908 |
| Zfp521 | 0.008790801 | 1.552101935 | 0.857 | 0.832 | 1 | 0.115960324 | 0.535404083 | 0.733 | 0.879 | 1 | 0.115960324 | 0.017504324 |
| Pigb | 0.036990147 | -2.208871282 | 0 | 0.175 | 1 | 0.008791893 | -2.404390255 | 0.022 | 0.173 | 1 | 0.036990147 | 0.017506488 |
| Cyp2j6 | 0.024220272 | -1.048694257 | 0.048 | 0.291 | 1 | 0.008792353 | -2.446210431 | 0.022 | 0.173 | 1 | 0.024220272 | 0.0175074 |
| Bcl11b | 0.492603827 | 0.214340149 | 0 | 0.022 | 1 | 0.008812112 | 0.78998779 | 0.133 | 0.038 | 1 | 0.492603827 | 0.01754657 |
| Pdhx | 0.794088438 | 0.842371372 | 0.286 | 0.347 | 1 | 0.008813117 | -0.107004344 | 0.133 | 0.377 | 1 | 0.794088438 | 0.017548562 |
| Coil | 0.64784005 | -0.326228232 | 0.095 | 0.127 | 1 | 0.008816211 | -2.361321533 | 0.022 | 0.173 | 1 | 0.64784005 | 0.017554696 |
| Izumo4 | 0.008830426 | -2.735619168 | 0 | 0.254 | 1 | 0.036119062 | -1.564854927 | 0.044 | 0.163 | 1 | 0.036119062 | 0.017582876 |

| | | | | | | | | | | | | |
|---|---|---|---|---|---|---|---|---|---|---|---|---|
| Usp15 | 0.108964571 | 1.033526831 | 0.667 | 0.623 | 1 | 0.008839176 | 0.632148651 | 0.978 | 0.803 | 1 | 0.108964571 | 0.017600221 |
| Pex2 | 0.283360511 | 0.304537958 | 0.19 | 0.347 | 1 | 0.008858804 | -0.141355849 | 0.156 | 0.412 | 1 | 0.283360511 | 0.01763913 |
| Kif21a | 0.008874641 | 0.747772349 | 0.905 | 0.873 | 1 | 0.077762663 | 0.548521915 | 1 | 0.92 | 1 | 0.077762663 | 0.017670523 |
| Nsun4 | 0.008885436 | 1.847801167 | 0.429 | 0.224 | 1 | 0.26909938 | 0.498648015 | 0.178 | 0.284 | 1 | 0.26909938 | 0.01769192 |
| Ube2k | 0.008914574 | 0.709071904 | 1 | 0.97 | 1 | 0.022422823 | 0.488621656 | 1 | 0.99 | 1 | 0.022422823 | 0.017749677 |
| Cic | 0.470240538 | 0.551781243 | 0.381 | 0.381 | 1 | 0.008920882 | -0.159187951 | 0.244 | 0.543 | 1 | 0.470240538 | 0.017762181 |
| Prpf3 | 0.636563469 | -0.206646617 | 0.333 | 0.388 | 1 | 0.008933468 | -0.242926832 | 0.222 | 0.536 | 1 | 0.636563469 | 0.01778713 |
| Unc5c | 0.008944377 | 1.160527103 | 0.952 | 0.922 | 1 | 0.138756252 | 0.865841545 | 0.8 | 0.896 | 1 | 0.138756252 | 0.017808752 |
| Anapc15 | 0.032365233 | -2.184209227 | 0 | 0.183 | 1 | 0.008947325 | -2.871516266 | 0 | 0.135 | 1 | 0.032365233 | 0.017814596 |
| Rnf141 | 0.423027883 | -0.413691074 | 0.048 | 0.101 | 1 | 0.008956448 | -2.901889915 | 0 | 0.135 | 1 | 0.423027883 | 0.017832679 |
| Etaa1 | 0.06537222 | -2.054148687 | 0 | 0.142 | 1 | 0.008972679 | -2.960783604 | 0 | 0.135 | 1 | 0.06537222 | 0.017864848 |
| Etfbkmt | 0.128164113 | -1.413691074 | 0 | 0.101 | 1 | 0.008973533 | -3.044847868 | 0 | 0.135 | 1 | 0.128164113 | 0.017866542 |
| Fam216a | 0.014603429 | -2.146407195 | 0.048 | 0.295 | 1 | 0.008977298 | -0.960783604 | 0.133 | 0.336 | 1 | 0.014603429 | 0.017874004 |
| Slc7a2 | 0.009002048 | -1.004300137 | 0.095 | 0.418 | 1 | 0.010820375 | 0.282695394 | 0.2 | 0.471 | 1 | 0.010820375 | 0.017923059 |
| Myl9 | 0.179963627 | -1.24509147 | 0.048 | 0.16 | 1 | 0.009005643 | -2.280401538 | 0.067 | 0.239 | 1 | 0.179963627 | 0.017930185 |
| Cebpa | 0.54961553 | 0.400753273 | 0.048 | 0.09 | 1 | 0.009007835 | -2.141355849 | 0.044 | 0.204 | 1 | 0.54961553 | 0.01793453 |
| Krt10 | 0.072926247 | -1.348596045 | 0.048 | 0.209 | 1 | 0.009008722 | -0.526380779 | 0.022 | 0.18 | 1 | 0.072926247 | 0.017936288 |
| Rassf1 | 0.026859992 | -0.422596627 | 0.286 | 0.582 | 1 | 0.0090098 | -0.346674757 | 0.6 | 0.73 | 1 | 0.026859992 | 0.017938423 |
| Rasd1 | 0.243674937 | -0.125634401 | 0.048 | 0.146 | 1 | 0.009012498 | -3.84048937 | 0 | 0.135 | 1 | 0.243674937 | 0.017943771 |
| Mme | 0.094875406 | -3.545396753 | 0 | 0.119 | 1 | 0.009026439 | -4.124282336 | 0 | 0.135 | 1 | 0.094875406 | 0.017971402 |
| Pcid2 | 0.05578928 | -1.496153234 | 0.048 | 0.224 | 1 | 0.009027424 | -0.731964913 | 0.067 | 0.26 | 1 | 0.05578928 | 0.017973353 |
| Sf3a3 | 0.009081392 | -0.626351957 | 0.143 | 0.478 | 1 | 0.086072085 | -0.241739917 | 0.356 | 0.533 | 1 | 0.086072085 | 0.018080312 |
| Alg9 | 0.180211683 | 1.179007076 | 0.286 | 0.19 | 1 | 0.009087178 | -0.776359032 | 0.067 | 0.253 | 1 | 0.180211683 | 0.01809178 |
| Rap1b | 0.044909924 | -0.450349544 | 0.857 | 0.993 | 1 | 0.009115377 | -0.406884111 | 0.956 | 0.986 | 1 | 0.044909924 | 0.018147664 |
| Echs1 | 0.009129618 | -1.486099569 | 0.095 | 0.396 | 1 | 0.986479897 | -0.211217973 | 0.467 | 0.422 | 1 | 0.986479897 | 0.018175885 |
| Slbp | 0.009133832 | -0.828728573 | 0.381 | 0.675 | 1 | 0.161810496 | -0.490054847 | 0.667 | 0.671 | 1 | 0.161810496 | 0.018184236 |
| Tcof1 | 0.061415149 | -1.184209227 | 0.095 | 0.291 | 1 | 0.009161271 | -0.986778812 | 0.133 | 0.339 | 1 | 0.061415149 | 0.018238613 |
| Twsg1 | 0.015684836 | -0.668359898 | 0.714 | 0.884 | 1 | 0.009164301 | -0.411796847 | 0.867 | 0.972 | 1 | 0.015684836 | 0.018244618 |
| Gon7 | 0.091703525 | -1.081115734 | 0.095 | 0.257 | 1 | 0.009179231 | -0.941418279 | 0.089 | 0.287 | 1 | 0.091703525 | 0.018274203 |
| 2700054A10Rik | 0.009180395 | 2.451379346 | 0.143 | 0.03 | 1 | 0.376843981 | -0.638855509 | 0 | 0.017 | 1 | 0.376843981 | 0.01827651 |
| Zfp385b | 0.020504221 | 1.383659205 | 0.905 | 0.914 | 1 | 0.00918679 | 1.286976651 | 0.844 | 0.872 | 1 | 0.020504221 | 0.018289184 |
| Gm56686 | 0.03937464 | -2.159118246 | 0 | 0.172 | 1 | 0.009190824 | -1.316927414 | 0.044 | 0.218 | 1 | 0.03937464 | 0.018297176 |
| Ghr | 0.049120072 | 1.245534771 | 0.714 | 0.78 | 1 | 0.009198586 | 1.163414124 | 0.756 | 0.723 | 1 | 0.049120072 | 0.018312558 |
| Gpm6b | 0.074408545 | -0.905166945 | 0.19 | 0.407 | 1 | 0.00920913 | -0.877642368 | 0.178 | 0.415 | 1 | 0.074408545 | 0.018333451 |
| Stac2 | 0.128253211 | -1.496153234 | 0 | 0.101 | 1 | 0.009215632 | -0.404390255 | 0.089 | 0.291 | 1 | 0.128253211 | 0.018346337 |
| Srpk2 | 0.574389288 | 0.164142592 | 0.81 | 0.892 | 1 | 0.009218301 | 0.701119042 | 1 | 0.913 | 1 | 0.574389288 | 0.018351626 |
| Pigm | 0.924411838 | 0.472137907 | 0.143 | 0.157 | 1 | 0.009220198 | -0.316927414 | 0.044 | 0.221 | 1 | 0.924411838 | 0.018355383 |
| Nckap1 | 0.082130095 | 0.92343895 | 0.905 | 0.97 | 1 | 0.009226252 | 0.477997932 | 1 | 0.976 | 1 | 0.082130095 | 0.01836738 |
| Erap1 | 0.247410437 | -0.527862093 | 0.143 | 0.261 | 1 | 0.009227172 | -0.316927414 | 0.178 | 0.401 | 1 | 0.247410437 | 0.018369203 |
| Sfrp2 | 0.009234153 | -6.926141075 | 0 | 0.254 | 1 | 0.128911026 | -3.931637258 | 0.022 | 0.087 | 1 | 0.128911026 | 0.018383036 |
| Elavl1 | 0.232108119 | -0.300233024 | 0.714 | 0.881 | 1 | 0.009269893 | -0.440851348 | 0.933 | 0.958 | 1 | 0.232108119 | 0.018453855 |
| Idh1 | 0.00928043 | -1.050254771 | 0.095 | 0.422 | 1 | 0.595860814 | -0.394929926 | 0.511 | 0.474 | 1 | 0.595860814 | 0.018474734 |
| Zcrb1 | 0.02897863 | -0.502467008 | 0.857 | 0.877 | 1 | 0.009281255 | -0.496362635 | 0.822 | 0.879 | 1 | 0.02897863 | 0.018476369 |
| Ccp110 | 0.338157101 | -0.93109029 | 0.095 | 0.172 | 1 | 0.009285459 | -1.094534992 | 0.044 | 0.218 | 1 | 0.338157101 | 0.018484697 |
| Rsrp1 | 0.199285942 | -0.29731554 | 0.952 | 0.959 | 1 | 0.00928584 | -0.350847688 | 0.978 | 0.993 | 1 | 0.199285942 | 0.018485453 |
| Clspn | 0.009293284 | 1.786246497 | 0.143 | 0.03 | 1 | 0.532520164 | -0.404390255 | 0.022 | 0.042 | 1 | 0.532520164 | 0.018500204 |
| Fmnl2 | 0.051612064 | 2.036686536 | 0.81 | 0.787 | 1 | 0.009293767 | 1.660524335 | 0.956 | 0.81 | 1 | 0.051612064 | 0.018501159 |
| Aida | 0.555261604 | 0.746622333 | 0.476 | 0.541 | 1 | 0.009297253 | -0.294207337 | 0.333 | 0.64 | 1 | 0.555261604 | 0.018508067 |
| Pfkm | 0.014422622 | -2.496153234 | 0 | 0.228 | 1 | 0.009341045 | -0.989352756 | 0.089 | 0.28 | 1 | 0.014422622 | 0.018594835 |
| Cnot10 | 0.195916451 | -0.30677587 | 0.286 | 0.5 | 1 | 0.009354185 | -0.223818009 | 0.333 | 0.606 | 1 | 0.195916451 | 0.018620869 |
| Car5b | 0.009383475 | -1.796548167 | 0.048 | 0.336 | 1 | 0.545035716 | 1.126679238 | 0.156 | 0.128 | 1 | 0.545035716 | 0.0186789 |
| Alas1 | 0.034223999 | 1.214340149 | 0.476 | 0.265 | 1 | 0.009385746 | 0.670016433 | 0.711 | 0.436 | 1 | 0.034223999 | 0.0186834 |
| Mpp7 | 0.202156435 | 0.444953077 | 0.524 | 0.351 | 1 | 0.009398782 | 0.708517603 | 0.8 | 0.536 | 1 | 0.202156435 | 0.018709226 |
| Kpnb1 | 0.146063231 | -0.469186186 | 0.381 | 0.593 | 1 | 0.009401533 | -0.522041844 | 0.667 | 0.765 | 1 | 0.146063231 | 0.018714676 |
| Pigyl | 0.009423284 | -2.701267664 | 0 | 0.25 | 1 | 0.020441408 | -0.884611923 | 0.067 | 0.218 | 1 | 0.020441408 | 0.01875777 |
| Upf1 | 0.534499982 | -0.126203624 | 0.333 | 0.451 | 1 | 0.009470631 | -0.516784921 | 0.578 | 0.74 | 1 | 0.534499982 | 0.01885157 |
| Chkb | 0.315806265 | 0.491568436 | 0.19 | 0.332 | 1 | 0.0094735 | -0.7206496 | 0.133 | 0.336 | 1 | 0.315806265 | 0.018857252 |
| Mecp2 | 0.785603596 | 0.829833078 | 0.619 | 0.735 | 1 | 0.009474156 | 0.991621651 | 0.956 | 0.824 | 1 | 0.785603596 | 0.018858552 |
| Pbrm1 | 0.343854161 | -0.254335314 | 0.952 | 0.903 | 1 | 0.009475487 | -0.397097763 | 0.956 | 0.948 | 1 | 0.343854161 | 0.018861188 |
| Pigf | 0.153954576 | -0.558888989 | 0.143 | 0.299 | 1 | 0.009483573 | -0.58227198 | 0.133 | 0.346 | 1 | 0.153954576 | 0.018877209 |
| Rcc1l | 0.049055654 | -1.516052791 | 0.048 | 0.231 | 1 | 0.009497577 | -0.764386391 | 0.089 | 0.287 | 1 | 0.049055654 | 0.018904949 |
| Vipas39 | 0.009499639 | -2.666078235 | 0 | 0.25 | 1 | 0.117587021 | 0.199989699 | 0.156 | 0.294 | 1 | 0.117587021 | 0.018909034 |
| Tns1 | 0.009504008 | 1.606612285 | 0.857 | 0.854 | 1 | 0.576815322 | 0.244813861 | 0.778 | 0.872 | 1 | 0.576815322 | 0.01891769 |
| Sap18b | 0.040249667 | -1.574155746 | 0.048 | 0.243 | 1 | 0.009549121 | -1.466674533 | 0.044 | 0.211 | 1 | 0.040249667 | 0.019007056 |
| Ppp1r15b | 0.009553472 | -0.73675025 | 0.238 | 0.631 | 1 | 0.009616442 | -0.505319164 | 0.689 | 0.806 | 1 | 0.009616442 | 0.019015675 |
| Tgif2 | 0.577246663 | 0.552756367 | 0.143 | 0.205 | 1 | 0.009568349 | -0.979892427 | 0.067 | 0.249 | 1 | 0.577246663 | 0.019045145 |
| Bmpr2 | 0.009588008 | 1.112921723 | 0.905 | 0.877 | 1 | 0.975392051 | 0.112015687 | 1 | 0.986 | 1 | 0.975392051 | 0.019084086 |
| 4930500G05Rik | 0.148844489 | -1.516052791 | 0.048 | 0.164 | 1 | 0.009602318 | -1.334849322 | 0.044 | 0.211 | 1 | 0.148844489 | 0.019112431 |
| Akt2 | 0.009616355 | -0.72091267 | 0.429 | 0.806 | 1 | 0.062074363 | 0.153001844 | 0.711 | 0.862 | 1 | 0.062074363 | 0.019140237 |
| B4galt1 | 0.009627437 | 1.173022271 | 0.905 | 0.72 | 1 | 0.047225301 | 0.482450788 | 0.911 | 0.685 | 1 | 0.047225301 | 0.019162186 |
| Fbxo33 | 0.009670292 | -1.888470657 | 0.095 | 0.388 | 1 | 0.296906536 | -0.255526869 | 0.4 | 0.439 | 1 | 0.296906536 | 0.01924707 |
| Ilf2 | 0.041506825 | -0.51736972 | 0.524 | 0.802 | 1 | 0.009682416 | -0.641384335 | 0.711 | 0.772 | 1 | 0.041506825 | 0.019271083 |
| Zmiz2 | 0.293573952 | -0.940938076 | 0.048 | 0.123 | 1 | 0.009693885 | -1.264459994 | 0.089 | 0.27 | 1 | 0.293573952 | 0.019293798 |
| Ppat | 0.014938737 | -1.241111619 | 0.095 | 0.392 | 1 | 0.009698916 | -0.648441558 | 0.111 | 0.315 | 1 | 0.014938737 | 0.019303762 |
| Pygo1 | 0.070820953 | -1.735619168 | 0.048 | 0.209 | 1 | 0.009704196 | -1.316927414 | 0.133 | 0.318 | 1 | 0.070820953 | 0.01931422 |
| Rab11fip2 | 0.038264793 | -1.19835341 | 0.19 | 0.429 | 1 | 0.009717529 | -0.30658547 | 0.2 | 0.464 | 1 | 0.038264793 | 0.019340627 |
| Klhl15 | 0.360444991 | 0.214340149 | 0.19 | 0.317 | 1 | 0.009719967 | -0.138929112 | 0.178 | 0.436 | 1 | 0.360444991 | 0.019345456 |
| Rbm45 | 0.468322253 | -0.264827688 | 0.095 | 0.153 | 1 | 0.009728213 | -2.526380779 | 0.022 | 0.17 | 1 | 0.468322253 | 0.019361787 |
| Tctn2 | 0.131079735 | -0.574155746 | 0.048 | 0.187 | 1 | 0.009741913 | -1.564854927 | 0.022 | 0.176 | 1 | 0.131079735 | 0.019388922 |
| Ccdc93 | 0.989684908 | 1.001346426 | 0.143 | 0.157 | 1 | 0.009747125 | -0.776359032 | 0.089 | 0.287 | 1 | 0.989684908 | 0.019399245 |
| Rpain | 0.043598735 | -1.116305163 | 0.048 | 0.254 | 1 | 0.009748969 | -0.387316742 | 0.111 | 0.318 | 1 | 0.043598735 | 0.019402895 |
| S1pr1 | 0.009763672 | -3.076656086 | 0.048 | 0.321 | 1 | 0.931973413 | 0.447856124 | 0.178 | 0.197 | 1 | 0.931973413 | 0.019432015 |
| Jam2 | 0.064249537 | -0.208391742 | 0.381 | 0.701 | 1 | 0.009787361 | 0.268035087 | 0.156 | 0.419 | 1 | 0.064249537 | 0.01947893 |
| Dctn1 | 0.013910567 | -0.535681598 | 0.143 | 0.493 | 1 | 0.009807689 | -0.745770713 | 0.422 | 0.578 | 1 | 0.013910567 | 0.019519187 |
| Dusp16 | 0.0282936 | 1.643009514 | 0.571 | 0.463 | 1 | 0.009815902 | 0.793956692 | 0.844 | 0.571 | 1 | 0.0282936 | 0.019535453 |
| Mtif3 | 0.050865994 | -2.054148687 | 0 | 0.157 | 1 | 0.009840598 | -2.361321533 | 0.022 | 0.17 | 1 | 0.050865994 | 0.019584359 |
| Ascc2 | 0.009858372 | -1.10233222 | 0.095 | 0.399 | 1 | 0.04871746 | 0.104804738 | 0.267 | 0.478 | 1 | 0.04871746 | 0.019619556 |
| Bloc1s6 | 0.076520618 | -0.870548749 | 0.048 | 0.22 | 1 | 0.009867033 | -1.141355849 | 0.067 | 0.242 | 1 | 0.076520618 | 0.019636707 |
| Zfp451 | 0.009869091 | 1.123024982 | 0.714 | 0.578 | 1 | 0.989926997 | 0.476319289 | 0.756 | 0.837 | 1 | 0.989926997 | 0.019640784 |
| Lppos | 0.604563988 | -0.154047257 | 0.143 | 0.19 | 1 | 0.009921349 | -2.339295227 | 0.022 | 0.17 | 1 | 0.604563988 | 0.019744265 |
| Ift22 | 0.099703161 | -1.303508156 | 0.048 | 0.19 | 1 | 0.009961586 | -1.294207337 | 0.022 | 0.176 | 1 | 0.099703161 | 0.019823938 |
| Erf | 0.00996628 | -1.12140844 | 0.333 | 0.668 | 1 | 0.013191229 | -0.448839089 | 0.689 | 0.837 | 1 | 0.013191229 | 0.019833233 |
| Usp16 | 0.017556317 | -0.683780237 | 0.619 | 0.869 | 1 | 0.009977896 | -0.388448711 | 0.889 | 0.865 | 1 | 0.017556317 | 0.019856233 |
| Gpc3 | 0.024939405 | -2.907428814 | 0.143 | 0.358 | 1 | 0.009983575 | -1.671770131 | 0.444 | 0.529 | 1 | 0.024939405 | 0.019867478 |
| Msl3l2 | 0.251958717 | -0.385121921 | 0.048 | 0.142 | 1 | 0.00999504 | -2.80878051 | 0 | 0.131 | 1 | 0.251958717 | 0.019890178 |
| Samd14 | 0.173820841 | -1.081115734 | 0 | 0.082 | 1 | 0.00999517 | -2.174908409 | 0.022 | 0.17 | 1 | 0.173820841 | 0.019890437 |
| Adam22 | 0.196462148 | -1.133583154 | 0 | 0.075 | 1 | 0.010004458 | -2.931637258 | 0 | 0.131 | 1 | 0.196462148 | 0.019908827 |
| Fam229b | 0.268852861 | -0.233118828 | 0.048 | 0.138 | 1 | 0.010004458 | -2.901889915 | 0 | 0.131 | 1 | 0.268852861 | 0.019908827 |

| | | | | | | | | | | | | |
|---|---|---|---|---|---|---|---|---|---|---|---|---|
| Zfp609 | 0.010020168 | 1.098724736 | 0.81 | 0.623 | 1 | 0.189571934 | 0.425813919 | 0.889 | 0.744 | 1 | 0.189571934 | 0.019939931 |
| Hspa2 | 0.136437434 | -0.732220592 | 0.095 | 0.257 | 1 | 0.010020785 | -0.616487696 | 0.089 | 0.287 | 1 | 0.136437434 | 0.019941155 |
| Zfp395 | 0.096796242 | -0.658424663 | 0.095 | 0.28 | 1 | 0.010035591 | -0.8698998 | 0.156 | 0.374 | 1 | 0.096796242 | 0.019970468 |
| Bex2 | 0.222258487 | -1.233118828 | 0 | 0.067 | 1 | 0.010050108 | -3.199570463 | 0 | 0.131 | 1 | 0.222258487 | 0.019999211 |
| Gucy1a2 | 0.878412101 | 0.735172312 | 0.048 | 0.041 | 1 | 0.010050108 | -3.223818009 | 0 | 0.131 | 1 | 0.878412101 | 0.019999211 |
| Rab4a | 0.034503481 | -2.159118246 | 0 | 0.179 | 1 | 0.010074842 | -0.432404631 | 0.044 | 0.218 | 1 | 0.034503481 | 0.020048182 |
| Cxcl14 | 0.113808989 | -3.146407195 | 0 | 0.108 | 1 | 0.010086676 | -3.824722054 | 0 | 0.131 | 1 | 0.113808989 | 0.020071612 |
| Srp54c | 0.191103305 | 1.106087258 | 0.333 | 0.261 | 1 | 0.010095739 | -0.736466305 | 0.089 | 0.287 | 1 | 0.191103305 | 0.020089553 |
| Gmpr2 | 0.010103777 | -2.648156327 | 0 | 0.246 | 1 | 0.020633909 | -0.794974711 | 0.067 | 0.218 | 1 | 0.020633909 | 0.020105467 |
| Stimate | 0.226552499 | -0.287057635 | 0.143 | 0.269 | 1 | 0.010163548 | -1.316927414 | 0.111 | 0.298 | 1 | 0.226552499 | 0.020223799 |
| Pmm2 | 0.648881892 | 0.123574685 | 0.19 | 0.25 | 1 | 0.010194602 | -0.852980314 | 0.111 | 0.304 | 1 | 0.648881892 | 0.020285274 |
| Cmc1 | 0.010205619 | -2.666078235 | 0 | 0.246 | 1 | 0.013753319 | -1.071814916 | 0.089 | 0.266 | 1 | 0.013753319 | 0.020307084 |
| Rrnad1 | 0.010207731 | -2.718545655 | 0 | 0.246 | 1 | 0.010807229 | -0.952515988 | 0.067 | 0.239 | 1 | 0.010807229 | 0.020311264 |
| Trim62 | 0.106901486 | -1.683780237 | 0 | 0.112 | 1 | 0.01020938 | -0.824722054 | 0.067 | 0.249 | 1 | 0.106901486 | 0.020314529 |
| Pltp | 0.115169446 | -1.950719097 | 0.095 | 0.235 | 1 | 0.010256593 | -1.486852415 | 0.089 | 0.27 | 1 | 0.115169446 | 0.020407989 |
| Zfp821 | 0.281924945 | 0.214340149 | 0.095 | 0.209 | 1 | 0.010262055 | -0.745770713 | 0.111 | 0.311 | 1 | 0.281924945 | 0.0204188 |
| Sirt1 | 0.010289102 | -1.542753647 | 0.333 | 0.59 | 1 | 0.151194226 | -0.330733213 | 0.778 | 0.768 | 1 | 0.151194226 | 0.020472338 |
| Cep170b | 0.508144266 | 0.673771768 | 0.143 | 0.235 | 1 | 0.010299487 | -0.583713955 | 0.133 | 0.349 | 1 | 0.508144266 | 0.020492894 |
| Prr13 | 0.010318681 | -1.525900577 | 0.238 | 0.519 | 1 | 0.843095362 | -0.401963518 | 0.667 | 0.543 | 1 | 0.843095362 | 0.020530887 |
| Atosa | 0.835585597 | 0.548240886 | 0.381 | 0.455 | 1 | 0.010319569 | 0.901252756 | 0.689 | 0.491 | 1 | 0.835585597 | 0.020532645 |
| Arl6ip5 | 0.010325765 | -0.611630451 | 0.952 | 0.996 | 1 | 0.038406655 | -0.14558453 | 1 | 1 | 1 | 0.038406655 | 0.020544908 |
| Rbm15b | 0.067357223 | -0.970084422 | 0.048 | 0.228 | 1 | 0.010327405 | -0.738391182 | 0.133 | 0.339 | 1 | 0.067357223 | 0.020548155 |
| Dock8 | 0.011463281 | 1.201818229 | 0.81 | 0.75 | 1 | 0.010344116 | 0.858106648 | 0.844 | 0.723 | 1 | 0.011463281 | 0.020581231 |
| Cep78 | 0.010347666 | 2.521768674 | 0.19 | 0.052 | 1 | 0.474803405 | -1.316927414 | 0.022 | 0.045 | 1 | 0.474803405 | 0.020588257 |
| Ppp1r10 | 0.010378407 | 0.949121496 | 0.905 | 0.694 | 1 | 0.581960705 | 0.11843007 | 0.933 | 0.806 | 1 | 0.581960705 | 0.020649102 |
| Wdr45 | 0.019087159 | -2.475975352 | 0 | 0.213 | 1 | 0.010382142 | -1.454430938 | 0.067 | 0.239 | 1 | 0.019087159 | 0.020656496 |
| Msantd3 | 0.041896711 | -2.107587946 | 0 | 0.168 | 1 | 0.010414172 | -0.502793959 | 0.067 | 0.249 | 1 | 0.041896711 | 0.02071989 |
| Itgbl1 | 0.063335503 | -4.0004205 | 0.048 | 0.213 | 1 | 0.010435682 | -0.715235274 | 0.178 | 0.384 | 1 | 0.063335503 | 0.02076246 |
| Rmc1 | 0.454693459 | 0.214340149 | 0.143 | 0.231 | 1 | 0.010482223 | -0.587016577 | 0.156 | 0.374 | 1 | 0.454693459 | 0.020854568 |
| Dvl2 | 0.0954683 | -0.249060372 | 0.095 | 0.291 | 1 | 0.010507375 | -0.89183625 | 0.111 | 0.308 | 1 | 0.0954683 | 0.020904346 |
| Tifa | 0.953237886 | 0.3917364 | 0.19 | 0.198 | 1 | 0.010510135 | -1.421264074 | 0.044 | 0.208 | 1 | 0.953237886 | 0.020909808 |
| Lrfn4 | 0.120477198 | -1.326228232 | 0 | 0.104 | 1 | 0.010511713 | -1.960783604 | 0.044 | 0.201 | 1 | 0.120477198 | 0.02091293 |
| Map2k1 | 0.365385659 | 0.30180299 | 0.238 | 0.41 | 1 | 0.010516973 | 0.951666517 | 0.667 | 0.488 | 1 | 0.365385659 | 0.02092334 |
| Cyhr1 | 0.373844248 | -0.522625445 | 0.095 | 0.168 | 1 | 0.01052567 | -1.174908409 | 0.089 | 0.27 | 1 | 0.373844248 | 0.020940551 |
| Usp47 | 0.013283035 | 0.728633703 | 1 | 0.899 | 1 | 0.010533863 | 0.460680165 | 1 | 0.986 | 1 | 0.013283035 | 0.020956765 |
| Map1a | 0.010544653 | -1.860206804 | 0.048 | 0.328 | 1 | 0.115593261 | 0.205663227 | 0.289 | 0.474 | 1 | 0.115593261 | 0.020978117 |
| Cldn10 | 0.010563638 | -4.074421082 | 0 | 0.246 | 1 | 0.106690443 | -1.602329633 | 0.022 | 0.093 | 1 | 0.106690443 | 0.021015685 |
| Tlr2 | 0.082047691 | -1.535681598 | 0.048 | 0.201 | 1 | 0.010582391 | -1.383016604 | 0.111 | 0.298 | 1 | 0.082047691 | 0.021052796 |
| Tsg101 | 0.010594532 | -0.87740242 | 0.476 | 0.739 | 1 | 0.185183255 | -0.168828775 | 0.778 | 0.837 | 1 | 0.185183255 | 0.02107682 |
| Dhx29 | 0.147661465 | -0.683780237 | 0.143 | 0.306 | 1 | 0.010612458 | -1.008805118 | 0.133 | 0.339 | 1 | 0.147661465 | 0.021112291 |
| Spaca6 | 0.010614898 | 1.43673257 | 0.524 | 0.354 | 1 | 0.613073079 | 0.618942249 | 0.422 | 0.602 | 1 | 0.613073079 | 0.021117119 |
| Cltb | 0.021307426 | -0.735619168 | 0.571 | 0.813 | 1 | 0.010621494 | -0.552698865 | 0.867 | 0.81 | 1 | 0.021307426 | 0.021130171 |
| Srp68 | 0.728104427 | 0.186989661 | 0.429 | 0.53 | 1 | 0.010621686 | -0.334000927 | 0.311 | 0.647 | 1 | 0.728104427 | 0.021130552 |
| Cers4 | 0.010632156 | 2.586308926 | 0.238 | 0.082 | 1 | 0.021019217 | -0.877642368 | 0.022 | 0.152 | 1 | 0.021019217 | 0.021151268 |
| Sap18 | 0.013844796 | -0.712809286 | 0.571 | 0.791 | 1 | 0.010639716 | -0.302806306 | 0.778 | 0.834 | 1 | 0.013844796 | 0.021166229 |
| Ivd | 0.010653749 | -1.796548167 | 0.048 | 0.328 | 1 | 0.064440921 | -0.327496656 | 0.2 | 0.349 | 1 | 0.064440921 | 0.021193996 |
| Rbm22 | 0.010660789 | -1.004300137 | 0.19 | 0.549 | 1 | 0.143380669 | -0.128398444 | 0.511 | 0.661 | 1 | 0.143380669 | 0.021207925 |
| Aars | 0.320510836 | 0.258734268 | 0.286 | 0.489 | 1 | 0.010665284 | -0.309233022 | 0.311 | 0.602 | 1 | 0.320510836 | 0.021216821 |
| Vars | 0.14729881 | -1.04504648 | 0.095 | 0.228 | 1 | 0.010689126 | -1.223818009 | 0.089 | 0.273 | 1 | 0.14729881 | 0.021263995 |
| Msh2 | 0.034838143 | -2.434752689 | 0 | 0.179 | 1 | 0.010690526 | -1.901889915 | 0.044 | 0.201 | 1 | 0.034838143 | 0.021266765 |
| Ssh3 | 0.268433862 | -0.535681598 | 0.048 | 0.131 | 1 | 0.010691165 | -1.960783604 | 0.044 | 0.201 | 1 | 0.268433862 | 0.021268028 |
| Chac2 | 0.194112985 | -1.107587946 | 0.048 | 0.149 | 1 | 0.010699808 | -0.691966845 | 0.044 | 0.215 | 1 | 0.194112985 | 0.021285129 |
| Stambp | 0.069842265 | -1.094412557 | 0.095 | 0.284 | 1 | 0.010705058 | -1.098287127 | 0.089 | 0.28 | 1 | 0.069842265 | 0.021295519 |
| Rpp30 | 0.020285617 | -2.348596045 | 0 | 0.209 | 1 | 0.010719391 | -1.298780067 | 0.044 | 0.208 | 1 | 0.020285617 | 0.021323877 |
| Polk | 0.283586062 | -0.709932525 | 0.19 | 0.302 | 1 | 0.010728379 | -0.929370651 | 0.111 | 0.308 | 1 | 0.283586062 | 0.021341659 |
| Stk3 | 0.633372828 | 0.149069792 | 0.571 | 0.776 | 1 | 0.010729838 | 0.913242358 | 0.844 | 0.782 | 1 | 0.633372828 | 0.021344547 |
| Tcf7 | 0.243396361 | -1.081115734 | 0.048 | 0.134 | 1 | 0.010729887 | -1.2318108 | 0.111 | 0.301 | 1 | 0.243396361 | 0.021344643 |
| Rdh14 | 0.020286113 | -2.370622352 | 0 | 0.209 | 1 | 0.010737695 | -1.316927414 | 0.089 | 0.266 | 1 | 0.020286113 | 0.021360091 |
| Nop2 | 0.377297767 | 0.96482055 | 0.286 | 0.239 | 1 | 0.010740015 | -0.834775719 | 0.111 | 0.311 | 1 | 0.377297767 | 0.021364683 |
| Bmp2k | 0.013059131 | 1.231934209 | 0.524 | 0.343 | 1 | 0.010788225 | 0.858015662 | 0.667 | 0.433 | 1 | 0.013059131 | 0.021460065 |
| Gnb5 | 0.16574497 | -0.828728573 | 0.048 | 0.168 | 1 | 0.010805432 | -1.57996182 | 0.067 | 0.235 | 1 | 0.16574497 | 0.021494107 |
| Zfp945 | 0.010826449 | -2.25696557 | 0.048 | 0.31 | 1 | 0.019533385 | -1.02842232 | 0.133 | 0.311 | 1 | 0.019533385 | 0.021535687 |
| Actr3 | 0.616666372 | 0.201740112 | 0.952 | 0.966 | 1 | 0.010829716 | -0.451064086 | 0.933 | 0.952 | 1 | 0.616666372 | 0.021542149 |
| Chd7 | 0.023005814 | 1.369618375 | 0.81 | 0.53 | 1 | 0.01083885 | 0.828503026 | 0.889 | 0.751 | 1 | 0.023005814 | 0.021560219 |
| Mtmr11 | 0.512686918 | 0.749499094 | 0.381 | 0.381 | 1 | 0.010863245 | -0.333606155 | 0.289 | 0.574 | 1 | 0.512686918 | 0.02160848 |
| Ttll4 | 0.709996255 | 1.206645757 | 0.238 | 0.224 | 1 | 0.010876766 | -0.129995987 | 0.2 | 0.471 | 1 | 0.709996255 | 0.021635228 |
| Esco1 | 0.572411473 | 0.46132215 | 0.381 | 0.496 | 1 | 0.01088887 | 0.529089129 | 0.867 | 0.675 | 1 | 0.572411473 | 0.021659173 |
| Specc1l | 0.211602025 | 0.893383893 | 0.571 | 0.634 | 1 | 0.010889958 | 0.992255453 | 0.822 | 0.716 | 1 | 0.211602025 | 0.021661325 |
| Synj1 | 0.835653478 | 0.73232395 | 0.333 | 0.358 | 1 | 0.010896119 | 0.695858741 | 0.711 | 0.498 | 1 | 0.835653478 | 0.021673513 |
| Appl2 | 0.923649013 | 0.16475812 | 0.476 | 0.582 | 1 | 0.010902415 | -0.260832852 | 0.378 | 0.699 | 1 | 0.923649013 | 0.021685968 |
| Scfd2 | 0.327177817 | 1.019767623 | 0.429 | 0.381 | 1 | 0.010934537 | 0.565075377 | 0.111 | 0.332 | 1 | 0.327177817 | 0.02174951 |
| Rps6ka3 | 0.22891889 | 0.720470707 | 0.905 | 0.873 | 1 | 0.010971114 | 0.78201676 | 0.978 | 0.9 | 1 | 0.22891889 | 0.021821863 |
| Dnlz | 0.021771112 | -1.196592952 | 0.095 | 0.354 | 1 | 0.0109722 | -1.089516918 | 0.067 | 0.246 | 1 | 0.021771112 | 0.021824011 |
| Ttll1 | 0.153212165 | -1.184209227 | 0.048 | 0.164 | 1 | 0.010979704 | -2.446210431 | 0.022 | 0.166 | 1 | 0.153212165 | 0.021838855 |
| C1qtnf2 | 0.196565371 | -1.718545655 | 0 | 0.075 | 1 | 0.010980028 | -2.564854927 | 0.022 | 0.166 | 1 | 0.196565371 | 0.021839496 |
| Abca1 | 0.530503007 | -0.326228232 | 0.286 | 0.216 | 1 | 0.010983864 | 0.211203864 | 0.556 | 0.298 | 1 | 0.530503007 | 0.021847082 |
| Mtmr2 | 0.010988493 | 1.330363945 | 0.619 | 0.504 | 1 | 0.528051509 | 0.741624769 | 0.422 | 0.612 | 1 | 0.528051509 | 0.021856239 |
| Gdpd5 | 0.021517614 | 2.493199522 | 0.238 | 0.093 | 1 | 0.011016562 | -0.432404631 | 0.111 | 0.322 | 1 | 0.021517614 | 0.021911759 |
| Neurl4 | 0.073771805 | -1.718545655 | 0 | 0.134 | 1 | 0.011020839 | -1.223818009 | 0.022 | 0.173 | 1 | 0.073771805 | 0.021920218 |
| Nde1 | 0.011024091 | -1.213753503 | 0.095 | 0.392 | 1 | 0.019440487 | -0.611548163 | 0.178 | 0.394 | 1 | 0.019440487 | 0.021926652 |
| Aspscr1 | 0.011052428 | -1.839297815 | 0.095 | 0.366 | 1 | 0.564445438 | 0.206258889 | 0.422 | 0.557 | 1 | 0.564445438 | 0.021982699 |
| Vps52 | 0.243545833 | -0.20069735 | 0.048 | 0.146 | 1 | 0.011055783 | -0.802354241 | 0.022 | 0.173 | 1 | 0.243545833 | 0.021989336 |
| Chd9 | 0.042110046 | 0.91692731 | 0.905 | 0.955 | 1 | 0.011091738 | 0.689053563 | 0.956 | 0.965 | 1 | 0.042110046 | 0.022060449 |
| Ogfrl1 | 0.274910725 | -0.695462042 | 0.048 | 0.134 | 1 | 0.011105635 | -1.061088509 | 0.022 | 0.173 | 1 | 0.274910725 | 0.022087935 |
| Exosc10 | 0.011107597 | -0.911190733 | 0.095 | 0.414 | 1 | 0.045455476 | 0.212442761 | 0.244 | 0.457 | 1 | 0.045455476 | 0.022091816 |
| Cbll1 | 0.01111562 | -1.063193826 | 0.095 | 0.414 | 1 | 0.040240404 | -0.504554417 | 0.311 | 0.488 | 1 | 0.040240404 | 0.022107682 |
| Smim31 | 0.011139193 | -3.133583154 | 0 | 0.243 | 1 | 0.077279075 | -0.749886821 | 0.044 | 0.142 | 1 | 0.077279075 | 0.022154304 |
| Hscb | 0.183588786 | -0.413691074 | 0.048 | 0.164 | 1 | 0.011149094 | -2.80878051 | 0 | 0.128 | 1 | 0.183588786 | 0.022173886 |
| Kif7 | 0.463956758 | 0.171271427 | 0.048 | 0.101 | 1 | 0.011149226 | -2.80878051 | 0 | 0.128 | 1 | 0.463956758 | 0.022174147 |
| Fastkd2 | 0.662029574 | 0.951305743 | 0.143 | 0.116 | 1 | 0.011159174 | -2.124282336 | 0.022 | 0.166 | 1 | 0.662029574 | 0.022193821 |
| Kctd15 | 0.93066253 | 0.651403955 | 0.19 | 0.205 | 1 | 0.011163724 | -0.790858602 | 0.067 | 0.246 | 1 | 0.93066253 | 0.02220282 |
| Zfp618 | 0.221237427 | 1.478755785 | 0.286 | 0.19 | 1 | 0.011163781 | -0.166102512 | 0.156 | 0.384 | 1 | 0.221237427 | 0.022202931 |
| Kansl1 | 0.014515667 | 1.197261706 | 1 | 0.966 | 1 | 0.011166675 | 0.561954826 | 1 | 0.993 | 1 | 0.014515667 | 0.022208655 |
| Traf4 | 0.011179244 | -2.955584852 | 0 | 0.243 | 1 | 0.011168954 | -1.095900535 | 0.133 | 0.329 | 1 | 0.011179244 | 0.022213162 |
| Ago1 | 0.17161385 | -0.60421298 | 0.095 | 0.239 | 1 | 0.011170246 | -1.017367132 | 0.089 | 0.277 | 1 | 0.17161385 | 0.022215717 |
| Sephs2 | 0.394078062 | -0.413691074 | 0.095 | 0.164 | 1 | 0.011179597 | -2.931637258 | 0 | 0.128 | 1 | 0.394078062 | 0.02223421 |
| Isoc2b | 0.536626714 | -0.355975576 | 0.095 | 0.142 | 1 | 0.011180037 | -2.84048937 | 0 | 0.128 | 1 | 0.536626714 | 0.022235081 |
| Rhbdd3 | 0.780296598 | 0.214340149 | 0.143 | 0.172 | 1 | 0.011186525 | -2.223818009 | 0.022 | 0.166 | 1 | 0.780296598 | 0.022247911 |

| | | | | | | | | | | | | |
|---|---|---|---|---|---|---|---|---|---|---|---|---|
| Zfp383 | 0.642127938 | 0.119182916 | 0.095 | 0.134 | 1 | 0.011186698 | -2.098287127 | 0.022 | 0.166 | 1 | 0.642127938 | 0.022248254 |
| Hspa4 | 0.547458483 | 0.106087258 | 0.857 | 0.937 | 1 | 0.011202289 | -0.301245792 | 0.956 | 0.958 | 1 | 0.547458483 | 0.022279086 |
| Atg13 | 0.333785787 | -0.12324121 | 0.19 | 0.302 | 1 | 0.011209531 | -1.174908409 | 0.156 | 0.346 | 1 | 0.333785787 | 0.022293408 |
| 2810004N23Rik | 0.172469175 | -0.392317423 | 0.238 | 0.399 | 1 | 0.011222828 | -0.522041844 | 0.156 | 0.388 | 1 | 0.172469175 | 0.022319705 |
| Tead2 | 0.576690908 | 0.247507013 | 0.048 | 0.086 | 1 | 0.011227852 | -3.199570463 | 0 | 0.128 | 1 | 0.576690908 | 0.022329639 |
| Zfp329 | 0.955699686 | 0.141276687 | 0.19 | 0.19 | 1 | 0.011230491 | -0.994999319 | 0.089 | 0.284 | 1 | 0.955699686 | 0.022334858 |
| Parn | 0.187157037 | -0.24509147 | 0.143 | 0.317 | 1 | 0.011248434 | -0.103523776 | 0.133 | 0.339 | 1 | 0.187157037 | 0.022370341 |
| Zfp367 | 0.21063528 | -0.970084422 | 0.095 | 0.224 | 1 | 0.011294516 | 0.259861155 | 0.067 | 0.246 | 1 | 0.21063528 | 0.022461466 |
| Psmd3 | 0.01130384 | -0.922079049 | 0.238 | 0.601 | 1 | 0.076414761 | -0.308314284 | 0.6 | 0.692 | 1 | 0.076414761 | 0.022479903 |
| Rpp21 | 0.366168683 | 0.106087258 | 0.143 | 0.25 | 1 | 0.011314127 | -0.746915255 | 0.067 | 0.246 | 1 | 0.366168683 | 0.022500245 |
| Lcn3 | 0.447399626 | -0.370622352 | 0.095 | 0.16 | 1 | 0.011321846 | 5.213587303 | 0.089 | 0.021 | 1 | 0.447399626 | 0.022515507 |
| Prickle2 | 0.784974239 | 1.376435661 | 0.286 | 0.313 | 1 | 0.01132863 | -0.890662659 | 0.2 | 0.433 | 1 | 0.784974239 | 0.022528922 |
| Btbd2 | 0.938469681 | 1.521768674 | 0.19 | 0.205 | 1 | 0.011334046 | 0.16649706 | 0.067 | 0.246 | 1 | 0.938469681 | 0.02253963 |
| Chst5 | 0.011338589 | -2.54926457 | 0.286 | 0.5 | 1 | 0.075087368 | -1.035745661 | 0.044 | 0.142 | 1 | 0.075087368 | 0.022548614 |
| Gna12 | 0.094939658 | 0.181918671 | 0.333 | 0.701 | 1 | 0.011349124 | -0.32765382 | 0.378 | 0.73 | 1 | 0.094939658 | 0.022569445 |
| Arl8a | 0.014655837 | -0.275763165 | 0.238 | 0.623 | 1 | 0.011374432 | -0.577454964 | 0.711 | 0.803 | 1 | 0.014655837 | 0.022619486 |
| Castor1 | 0.184469797 | -0.348596045 | 0.19 | 0.373 | 1 | 0.011394866 | -0.970369652 | 0.156 | 0.363 | 1 | 0.184469797 | 0.02265989 |
| Septin1 | 0.684716853 | 0.214340149 | 0.143 | 0.19 | 1 | 0.011411581 | -1.638855509 | 0.044 | 0.204 | 1 | 0.684716853 | 0.022692939 |
| Pofut1 | 0.859248283 | 0.199840579 | 0.238 | 0.276 | 1 | 0.011423365 | -0.200734396 | 0.156 | 0.401 | 1 | 0.859248283 | 0.022716236 |
| Mapk14 | 0.626424517 | 0.698575159 | 0.333 | 0.354 | 1 | 0.011431982 | 0.949466004 | 0.711 | 0.529 | 1 | 0.626424517 | 0.022733274 |
| Ash1l | 0.043312461 | 0.580454617 | 0.952 | 0.978 | 1 | 0.01143602 | 0.295780077 | 1 | 0.997 | 1 | 0.043312461 | 0.022741258 |
| 3010001F23Rik | 0.011437001 | 2.673771768 | 0.19 | 0.056 | 1 | 0.280275514 | 0.361144491 | 0.067 | 0.128 | 1 | 0.280275514 | 0.022743196 |
| Pgd | 0.030242343 | -0.785659851 | 0.286 | 0.541 | 1 | 0.011442631 | 0.460680165 | 0.822 | 0.55 | 1 | 0.030242343 | 0.022754327 |
| Zfp36 | 0.048176176 | -0.487142858 | 0.762 | 0.787 | 1 | 0.011446273 | -0.747561768 | 0.867 | 0.827 | 1 | 0.048176176 | 0.022761529 |
| Wdr35 | 0.011473472 | -0.17922582 | 0.143 | 0.496 | 1 | 0.299207689 | 0.161412667 | 0.378 | 0.616 | 1 | 0.299207689 | 0.022815303 |
| Letm2 | 0.711134535 | 1.334969854 | 0.333 | 0.373 | 1 | 0.011474894 | -0.301497373 | 0.289 | 0.571 | 1 | 0.711134535 | 0.022818115 |
| Cdc42bpa | 0.011497449 | 0.577387231 | 1 | 0.985 | 1 | 0.238504584 | -0.18891152 | 1 | 0.993 | 1 | 0.238504584 | 0.022862706 |
| Man1b1 | 0.05400374 | -1.067695219 | 0.095 | 0.302 | 1 | 0.011516777 | -0.124282336 | 0.222 | 0.467 | 1 | 0.05400374 | 0.022900918 |
| Mrgprf | 0.264234575 | -0.385121921 | 0.048 | 0.138 | 1 | 0.01155402 | -2.107004344 | 0.044 | 0.197 | 1 | 0.264234575 | 0.022974545 |
| Vrk1 | 0.065412796 | -2.159118246 | 0 | 0.142 | 1 | 0.011554887 | -2.071814916 | 0.044 | 0.197 | 1 | 0.065412796 | 0.022976259 |
| 4930586N03Rik | 0.082280431 | 3.088809267 | 0.048 | 0.007 | 1 | 0.011555302 | 4.268035087 | 0.022 | 0 | 1 | 0.082280431 | 0.02297708 |
| Zfp804b | 0.136517983 | -3.434752689 | 0 | 0.097 | 1 | 0.011555302 | 4.268035087 | 0.022 | 0 | 1 | 0.136517983 | 0.02297708 |
| Gm19557 | 0.171708932 | 2.673771768 | 0.048 | 0.011 | 1 | 0.011555302 | 4.268035087 | 0.022 | 0 | 1 | 0.171708932 | 0.02297708 |
| Mog | 0.208886118 | -0.849790188 | 0 | 0.071 | 1 | 0.011555302 | 4.683072586 | 0.022 | 0 | 1 | 0.208886118 | 0.02297708 |
| 4930425P05Rik | 0.261628269 | 2.936806174 | 0.048 | 0.015 | 1 | 0.011555302 | 3.683072586 | 0.022 | 0 | 1 | 0.261628269 | 0.02297708 |
| Trappc3l | 0.264194629 | 2.673771768 | 0.048 | 0.015 | 1 | 0.011555302 | 4.683072586 | 0.022 | 0 | 1 | 0.264194629 | 0.02297708 |
| Gm15551 | 0.425625368 | -0.326228232 | 0 | 0.03 | 1 | 0.011555302 | 4.268035087 | 0.022 | 0 | 1 | 0.425625368 | 0.02297708 |
| Gm29282 | 0.457292877 | 0.351843673 | 0 | 0.026 | 1 | 0.011555302 | 4.683072586 | 0.022 | 0 | 1 | 0.457292877 | 0.02297708 |
| 4933401D09Rik | 0.472861597 | 1.866416846 | 0.048 | 0.022 | 1 | 0.011555302 | 3.683072586 | 0.022 | 0 | 1 | 0.472861597 | 0.02297708 |
| Gm13710 | 0.479160274 | 0.735172312 | 0.048 | 0.022 | 1 | 0.011555302 | 4.683072586 | 0.022 | 0 | 1 | 0.479160274 | 0.02297708 |
| Ccnb1 | 0.492599419 | -0.718545655 | 0 | 0.022 | 1 | 0.011555302 | 3.683072586 | 0.022 | 0 | 1 | 0.492599419 | 0.02297708 |
| Gm42456 | 0.492608234 | 0.866416846 | 0 | 0.022 | 1 | 0.011555302 | 4.683072586 | 0.022 | 0 | 1 | 0.492608234 | 0.02297708 |
| Plaat1 | 0.492608234 | 0.866416846 | 0 | 0.022 | 1 | 0.011555302 | 3.683072586 | 0.022 | 0 | 1 | 0.492608234 | 0.02297708 |
| Tex56 | 0.532426384 | 0.866416846 | 0 | 0.019 | 1 | 0.011555302 | 4.268035087 | 0.022 | 0 | 1 | 0.532426384 | 0.02297708 |
| Gm30439 | 0.532426384 | 0.866416846 | 0 | 0.019 | 1 | 0.011555302 | 4.683072586 | 0.022 | 0 | 1 | 0.532426384 | 0.02297708 |
| Cxcl10 | 0.532435374 | -3.326228232 | 0 | 0.019 | 1 | 0.011555302 | 9.705440399 | 0.022 | 0 | 1 | 0.532435374 | 0.02297708 |
| Gm10827 | 0.578156445 | 1.088809267 | 0 | 0.015 | 1 | 0.011555302 | 4.268035087 | 0.022 | 0 | 1 | 0.578156445 | 0.02297708 |
| Mx1 | 0.578159907 | -0.233118828 | 0 | 0.015 | 1 | 0.011555302 | 7.727466706 | 0.022 | 0 | 1 | 0.578159907 | 0.02297708 |
| Gm39323 | 0.57816106 | 1.351843673 | 0 | 0.015 | 1 | 0.011555302 | 4.268035087 | 0.022 | 0 | 1 | 0.57816106 | 0.02297708 |
| Gm36278 | 0.63207476 | 1.351843673 | 0 | 0.011 | 1 | 0.011555302 | 4.268035087 | 0.022 | 0 | 1 | 0.63207476 | 0.02297708 |
| Gm14936 | 0.63207476 | 1.351843673 | 0 | 0.011 | 1 | 0.011555302 | 4.683072586 | 0.022 | 0 | 1 | 0.63207476 | 0.02297708 |
| Sp110 | 0.632076134 | 1.673771768 | 0 | 0.011 | 1 | 0.011555302 | 4.683072586 | 0.022 | 0 | 1 | 0.632076134 | 0.02297708 |
| Fam187b | 0.632076134 | 1.673771768 | 0 | 0.011 | 1 | 0.011555302 | 4.268035087 | 0.022 | 0 | 1 | 0.632076134 | 0.02297708 |
| Capn9 | 0.632076134 | 1.673771768 | 0 | 0.011 | 1 | 0.011555302 | 4.268035087 | 0.022 | 0 | 1 | 0.632076134 | 0.02297708 |
| Or8b54 | 0.632076134 | 1.673771768 | 0 | 0.011 | 1 | 0.011555302 | 4.683072586 | 0.022 | 0 | 1 | 0.632076134 | 0.02297708 |
| Gldc | 0.632076134 | 1.673771768 | 0 | 0.011 | 1 | 0.011555302 | 4.683072586 | 0.022 | 0 | 1 | 0.632076134 | 0.02297708 |
| Cxcr3 | 0.632076134 | 1.673771768 | 0 | 0.011 | 1 | 0.011555302 | 3.683072586 | 0.022 | 0 | 1 | 0.632076134 | 0.02297708 |
| Sord | 0.429992071 | -0.25696557 | 0.095 | 0.164 | 1 | 0.011567957 | -1.033134448 | 0.044 | 0.211 | 1 | 0.429992071 | 0.023002096 |
| Clip2 | 0.36228827 | 1.010806755 | 0.19 | 0.134 | 1 | 0.011586428 | -0.223818009 | 0.067 | 0.246 | 1 | 0.36228827 | 0.023038612 |
| Ric8a | 0.011635706 | -2.555046923 | 0 | 0.239 | 1 | 0.021429983 | -0.960783604 | 0.089 | 0.256 | 1 | 0.021429983 | 0.023136023 |
| Idh2 | 0.011636762 | -0.820545369 | 0.19 | 0.534 | 1 | 0.059558685 | -0.646428728 | 0.578 | 0.616 | 1 | 0.059558685 | 0.02313811 |
| Inafm1 | 0.011641124 | -2.171718283 | 0.048 | 0.306 | 1 | 0.012626552 | -0.931637258 | 0.089 | 0.263 | 1 | 0.012626552 | 0.023146732 |
| 2310011J03Rik | 0.011642165 | -2.171718283 | 0.048 | 0.306 | 1 | 0.015175452 | -1.564854927 | 0.089 | 0.249 | 1 | 0.015175452 | 0.02314879 |
| Bace1 | 0.028155053 | -1.203972482 | 0.095 | 0.34 | 1 | 0.011654049 | -0.430489973 | 0.222 | 0.495 | 1 | 0.028155053 | 0.023172282 |
| Zdhhc9 | 0.29825485 | -0.246793765 | 0.143 | 0.261 | 1 | 0.011668412 | -0.671770131 | 0.111 | 0.318 | 1 | 0.29825485 | 0.023200672 |
| B4galnt3 | 0.997578287 | 1.729114802 | 0.333 | 0.429 | 1 | 0.01166852 | -1.5585134 | 0.111 | 0.304 | 1 | 0.997578287 | 0.023200886 |
| Ift43 | 0.011682466 | -0.617782678 | 0.19 | 0.545 | 1 | 0.233539811 | 0.280974143 | 0.378 | 0.574 | 1 | 0.233539811 | 0.023228452 |
| Tra2a | 0.011687261 | 0.658529825 | 0.952 | 0.963 | 1 | 0.157687396 | 0.306352018 | 1 | 0.952 | 1 | 0.157687396 | 0.023237931 |
| Cpne8 | 0.011721157 | 1.092651333 | 1 | 0.985 | 1 | 0.019266189 | 0.489109858 | 1 | 0.983 | 1 | 0.019266189 | 0.023304929 |
| Trmt2a | 0.540775733 | 0.150209812 | 0.143 | 0.216 | 1 | 0.011727105 | -1.691966845 | 0.067 | 0.228 | 1 | 0.540775733 | 0.023316684 |
| Zfp954 | 0.380042247 | -0.208871282 | 0.095 | 0.175 | 1 | 0.011731733 | -1.466674533 | 0.044 | 0.204 | 1 | 0.380042247 | 0.023325833 |
| Arl2bp | 0.01706796 | -0.799845757 | 0.571 | 0.784 | 1 | 0.011774846 | -0.483880849 | 0.6 | 0.761 | 1 | 0.01706796 | 0.023411045 |
| Cc2d2a | 0.079893994 | -0.862281133 | 0.095 | 0.28 | 1 | 0.011783954 | -0.202756394 | 0.133 | 0.343 | 1 | 0.079893994 | 0.023429046 |
| Otx1 | 0.578156445 | 1.088809267 | 0 | 0.015 | 1 | 0.011793172 | 1.683072586 | 0.133 | 0.042 | 1 | 0.578156445 | 0.023447264 |
| Sqor | 0.011801714 | -2.701267664 | 0 | 0.239 | 1 | 0.013986686 | -0.714519779 | 0.133 | 0.329 | 1 | 0.013986686 | 0.023464147 |
| E2f5 | 0.043339008 | -0.970084422 | 0.095 | 0.332 | 1 | 0.011806298 | -0.544733331 | 0.156 | 0.388 | 1 | 0.043339008 | 0.023473208 |
| Ctso | 0.061181292 | -1.785659851 | 0 | 0.146 | 1 | 0.011825594 | -1.881712033 | 0.044 | 0.197 | 1 | 0.061181292 | 0.023511342 |
| Prr5l | 0.566849325 | -0.455511249 | 0.143 | 0.198 | 1 | 0.011837032 | 1.188307894 | 0.467 | 0.315 | 1 | 0.566849325 | 0.023533949 |
| Atp11c | 0.011850243 | 1.248886482 | 0.762 | 0.567 | 1 | 0.092301831 | 0.619346989 | 0.756 | 0.682 | 1 | 0.092301831 | 0.023560058 |
| Atf7 | 0.011852566 | 0.77802173 | 0.952 | 0.825 | 1 | 0.776677796 | 0.21350552 | 0.978 | 0.945 | 1 | 0.776677796 | 0.023564649 |
| Epm2aip1 | 0.011856831 | -1.535681598 | 0.095 | 0.381 | 1 | 0.04514837 | -0.365290435 | 0.222 | 0.412 | 1 | 0.04514837 | 0.023573078 |
| Tcf20 | 0.086688476 | 1.207686663 | 0.667 | 0.541 | 1 | 0.011880281 | 0.572962829 | 0.956 | 0.702 | 1 | 0.086688476 | 0.02361942 |
| Mapre3 | 0.011885366 | -0.888470657 | 0.238 | 0.601 | 1 | 0.986948518 | 0.931860742 | 0.533 | 0.651 | 1 | 0.986948518 | 0.023629469 |
| Cables1 | 0.21882248 | 1.991947728 | 0.238 | 0.164 | 1 | 0.01190759 | -1.84048937 | 0.022 | 0.17 | 1 | 0.21882248 | 0.023673388 |
| Gm16192 | 0.011916119 | 3.673771768 | 0.143 | 0.034 | 1 | 0.112236835 | 2.268035087 | 0.067 | 0.024 | 1 | 0.112236835 | 0.023690243 |
| AU022252 | 0.370324723 | -0.303508156 | 0.095 | 0.175 | 1 | 0.011926058 | -1.239759553 | 0.067 | 0.235 | 1 | 0.370324723 | 0.023709886 |
| Ntmt1 | 0.036264275 | -0.164171474 | 0.143 | 0.384 | 1 | 0.011926297 | -0.683249628 | 0.178 | 0.394 | 1 | 0.036264275 | 0.023710358 |
| Fanca | 0.011932339 | 3.673771768 | 0.095 | 0.015 | 1 | 0.492040936 | 2.683072586 | 0.022 | 0.01 | 1 | 0.492040936 | 0.023722298 |
| Foxj3 | 0.011942462 | 1.440152091 | 0.619 | 0.552 | 1 | 0.649562445 | 0.377887755 | 0.711 | 0.747 | 1 | 0.649562445 | 0.023742301 |
| Ccar2 | 0.584221981 | 0.55829455 | 0.048 | 0.086 | 1 | 0.011960318 | -1.881712033 | 0.044 | 0.197 | 1 | 0.584221981 | 0.023777586 |
| 1810030O07Rik | 0.208497431 | -0.672003069 | 0.19 | 0.325 | 1 | 0.011963605 | -0.412084647 | 0.2 | 0.405 | 1 | 0.208497431 | 0.023784082 |
| Atp2b1 | 0.018436079 | 0.576844718 | 1 | 0.985 | 1 | 0.011989945 | 0.439147003 | 0.978 | 0.99 | 1 | 0.018436079 | 0.023836131 |
| Eola1 | 0.077285621 | -0.807354922 | 0.048 | 0.22 | 1 | 0.012014995 | -1.638855509 | 0.067 | 0.228 | 1 | 0.077285621 | 0.023885631 |
| Zfp938 | 0.304044954 | -0.455511249 | 0.095 | 0.19 | 1 | 0.012020917 | -1.589945908 | 0.022 | 0.17 | 1 | 0.304044954 | 0.023897331 |
| Rnf14 | 0.867378184 | 0.177345942 | 0.429 | 0.556 | 1 | 0.012027267 | -0.267714439 | 0.333 | 0.64 | 1 | 0.867378184 | 0.02390988 |
| Mrps30 | 0.013853081 | -1.801961663 | 0.048 | 0.299 | 1 | 0.012038616 | -0.749886821 | 0.111 | 0.301 | 1 | 0.013853081 | 0.023932303 |
| Tmem50b | 0.012039205 | -1.548620654 | 0.048 | 0.325 | 1 | 0.039414903 | -0.665655568 | 0.244 | 0.408 | 1 | 0.039414903 | 0.023933468 |
| Tbca | 0.012049365 | -0.648156327 | 0.81 | 0.91 | 1 | 0.083225115 | -0.149817428 | 0.867 | 0.907 | 1 | 0.083225115 | 0.023953544 |

| | | | | | | | | | | | | |
|---|---|---|---|---|---|---|---|---|---|---|---|---|
| Naalad2 | 0.070631609 | -1.004300137 | 0.095 | 0.295 | 1 | 0.012050404 | -2.674479418 | 0.022 | 0.163 | 1 | 0.070631609 | 0.023955596 |
| Dusp22 | 0.039087936 | -1.280424543 | 0.095 | 0.317 | 1 | 0.012060297 | -0.352551324 | 0.044 | 0.204 | 1 | 0.039087936 | 0.023975142 |
| Tcf4 | 0.012073449 | 0.857584841 | 1 | 0.985 | 1 | 0.248985649 | 0.3760452 | 1 | 1 | 1 | 0.248985649 | 0.024001129 |
| 0610040F04Rik | 0.01208222 | 4.425844254 | 0.19 | 0.056 | 1 | 0.281437689 | 0.816339117 | 0.089 | 0.052 | 1 | 0.281437689 | 0.024018459 |
| Atp6v0b | 0.022448795 | -0.528574711 | 0.762 | 0.914 | 1 | 0.012085143 | -0.331765923 | 0.956 | 0.938 | 1 | 0.022448795 | 0.024024234 |
| Trappc6b | 0.135513205 | 0.129451251 | 0.333 | 0.653 | 1 | 0.012103183 | -0.323241188 | 0.6 | 0.772 | 1 | 0.135513205 | 0.024059878 |
| Cds2 | 0.060388551 | -1.094412557 | 0.095 | 0.295 | 1 | 0.012112586 | -0.2150478 | 0.156 | 0.384 | 1 | 0.060388551 | 0.024078458 |
| Spata1 | 0.012116642 | 1.56206138 | 0.619 | 0.41 | 1 | 0.346551626 | 0.605713703 | 0.244 | 0.374 | 1 | 0.346551626 | 0.024086471 |
| Ier5l | 0.27030822 | -0.512094778 | 0.095 | 0.198 | 1 | 0.012144952 | -1.754332726 | 0.044 | 0.201 | 1 | 0.27030822 | 0.024142405 |
| Phyhd1 | 0.686617231 | -0.348596045 | 0.143 | 0.172 | 1 | 0.012177943 | -0.598962782 | 0.067 | 0.253 | 1 | 0.686617231 | 0.024207584 |
| Gmfb | 0.022851864 | -0.804275529 | 0.286 | 0.627 | 1 | 0.012180952 | -0.477004629 | 0.556 | 0.74 | 1 | 0.022851864 | 0.024213528 |
| Klf9 | 0.012185798 | -0.967863182 | 0.905 | 0.937 | 1 | 0.123309384 | -0.178623107 | 0.978 | 0.965 | 1 | 0.123309384 | 0.024223103 |
| Pgrmc1 | 0.101407571 | -0.478513075 | 0.714 | 0.78 | 1 | 0.012198232 | -0.449682623 | 0.933 | 0.879 | 1 | 0.101407571 | 0.024247667 |
| Zfp512b | 0.994796597 | 0.644024424 | 0.143 | 0.153 | 1 | 0.012203286 | -0.928362126 | 0.089 | 0.27 | 1 | 0.994796597 | 0.024257652 |
| Kctd9 | 0.209426492 | 0.97333205 | 0.476 | 0.455 | 1 | 0.012226736 | 1.23169324 | 0.533 | 0.45 | 1 | 0.209426492 | 0.024303979 |
| Dzip1l | 0.012243485 | -1.807354922 | 0.048 | 0.321 | 1 | 0.016546803 | -1.152116544 | 0.156 | 0.336 | 1 | 0.016546803 | 0.024337067 |
| B230354K17Rik | 0.90887831 | 0.425844254 | 0.19 | 0.213 | 1 | 0.01224592 | -1.549588171 | 0.067 | 0.232 | 1 | 0.90887831 | 0.024341877 |
| Nfkb1 | 0.012269394 | 0.826806568 | 1 | 0.937 | 1 | 0.088969204 | 0.715546193 | 1 | 0.979 | 1 | 0.088969204 | 0.024388249 |
| Hexim1 | 0.28161781 | -0.312289041 | 0.429 | 0.575 | 1 | 0.012285349 | -0.62152563 | 0.822 | 0.789 | 1 | 0.28161781 | 0.024419769 |
| Rft1 | 0.892860555 | 0.55829455 | 0.19 | 0.216 | 1 | 0.012310467 | -2.271123724 | 0.022 | 0.163 | 1 | 0.892860555 | 0.024469387 |
| Nelfa | 0.078302441 | -0.120644099 | 0.095 | 0.302 | 1 | 0.012350888 | -0.378038928 | 0.244 | 0.516 | 1 | 0.078302441 | 0.024549232 |
| Mrpl46 | 0.155829047 | -0.940938076 | 0.048 | 0.164 | 1 | 0.012350971 | -1.686161223 | 0.044 | 0.197 | 1 | 0.155829047 | 0.024549396 |
| Mid1 | 0.489277083 | 0.510273035 | 0.667 | 0.608 | 1 | 0.012368768 | 0.28671965 | 0.867 | 0.536 | 1 | 0.489277083 | 0.024584549 |
| Tmem129 | 0.21528561 | -0.385121921 | 0.048 | 0.153 | 1 | 0.012381642 | -2.339295227 | 0.022 | 0.163 | 1 | 0.21528561 | 0.02460998 |
| Rnf216 | 0.01238418 | 1.051719948 | 0.714 | 0.806 | 1 | 0.712499973 | 0.1767844 | 0.978 | 0.907 | 1 | 0.712499973 | 0.024614993 |
| Zbtb14 | 0.876110828 | 1.451379346 | 0.095 | 0.093 | 1 | 0.012404556 | -0.901889915 | 0.022 | 0.17 | 1 | 0.876110828 | 0.02465524 |
| Uqcc4 | 0.133823936 | -0.593014773 | 0.095 | 0.25 | 1 | 0.012404781 | -1.094534992 | 0.089 | 0.27 | 1 | 0.133823936 | 0.024655684 |
| Tpmt | 0.315761485 | -0.611630451 | 0.048 | 0.119 | 1 | 0.012406023 | -2.602329633 | 0 | 0.125 | 1 | 0.315761485 | 0.024658136 |
| Caskin2 | 0.153858837 | -1.184209227 | 0 | 0.09 | 1 | 0.012419703 | -2.674479418 | 0 | 0.125 | 1 | 0.153858837 | 0.024685157 |
| Tbkbp1 | 0.966230998 | 1.258734268 | 0.048 | 0.052 | 1 | 0.012419728 | -2.709244837 | 0 | 0.125 | 1 | 0.966230998 | 0.024685206 |
| Tpm3-rs7 | 0.153703558 | -1.02666795 | 0 | 0.09 | 1 | 0.012432633 | -2.674479418 | 0 | 0.125 | 1 | 0.153703558 | 0.024710695 |
| Uqcrfs1 | 0.012433567 | -0.943612211 | 0.333 | 0.623 | 1 | 0.385397979 | -0.329454512 | 0.756 | 0.682 | 1 | 0.385397979 | 0.024712541 |
| Wrap73 | 0.618972991 | 0.673771768 | 0.048 | 0.082 | 1 | 0.012444517 | -2.743192169 | 0 | 0.125 | 1 | 0.618972991 | 0.024734167 |
| Dbt | 0.811968561 | 0.957564734 | 0.143 | 0.134 | 1 | 0.012444761 | -2.709244837 | 0 | 0.125 | 1 | 0.811968561 | 0.024734651 |
| Fbxo17 | 0.176107596 | -0.527862093 | 0.095 | 0.239 | 1 | 0.012447582 | -1.446210431 | 0.067 | 0.232 | 1 | 0.176107596 | 0.024740221 |
| Smndc1 | 0.033092022 | -0.538464039 | 0.429 | 0.765 | 1 | 0.012451078 | -0.559608758 | 0.756 | 0.785 | 1 | 0.033092022 | 0.024747127 |
| Zfp414 | 0.173743052 | -0.44170545 | 0.048 | 0.168 | 1 | 0.012455647 | -2.80878051 | 0 | 0.125 | 1 | 0.173743052 | 0.024756152 |
| Dgat2 | 0.544789083 | -0.516052791 | 0.095 | 0.142 | 1 | 0.012466513 | -2.989352756 | 0 | 0.125 | 1 | 0.544789083 | 0.024777613 |
| Igfals | 0.77384646 | 1.673771768 | 0.048 | 0.067 | 1 | 0.012466587 | -2.931637258 | 0 | 0.125 | 1 | 0.77384646 | 0.024777758 |
| Cep89 | 0.128254702 | -1.718545655 | 0 | 0.101 | 1 | 0.012475351 | -2.84048937 | 0 | 0.125 | 1 | 0.128254702 | 0.024795068 |
| C1rl | 0.19646186 | -1.081115734 | 0 | 0.075 | 1 | 0.012476723 | -2.80878051 | 0 | 0.125 | 1 | 0.19646186 | 0.024797777 |
| Kansl2 | 0.012483528 | -0.727107669 | 0.095 | 0.418 | 1 | 0.02564673 | -0.422461828 | 0.289 | 0.519 | 1 | 0.02564673 | 0.024811217 |
| Gm57068 | 0.012498529 | 3.351843673 | 0.095 | 0.015 | 1 | 0.492040936 | 2.683072586 | 0.022 | 0.01 | 1 | 0.492040936 | 0.024840846 |
| Gm57275 | 0.012498529 | 3.351843673 | 0.095 | 0.015 | 1 | 0.8202424 | 1.098110085 | 0.022 | 0.017 | 1 | 0.8202424 | 0.024840846 |
| Gm26583 | 0.012498529 | 3.351843673 | 0.095 | 0.015 | 1 | 0.030581005 | 3.683072586 | 0.044 | 0.007 | 1 | 0.030581005 | 0.024840846 |
| Rgs6 | 0.0125014 | 3.258734268 | 0.095 | 0.015 | 1 | 0.088895206 | -0.722919774 | 0.022 | 0.104 | 1 | 0.088895206 | 0.024846516 |
| Gtpbp10 | 0.755319062 | 0.425844254 | 0.095 | 0.123 | 1 | 0.012513182 | -2.071814916 | 0.022 | 0.163 | 1 | 0.755319062 | 0.024869784 |
| Satb1 | 0.251653515 | -0.718545655 | 0 | 0.06 | 1 | 0.012515906 | -3.425451871 | 0 | 0.125 | 1 | 0.251653515 | 0.024875164 |
| Furin | 0.012520619 | -0.836523934 | 0.429 | 0.757 | 1 | 0.046292711 | -0.162893785 | 0.778 | 0.855 | 1 | 0.046292711 | 0.024884473 |
| Cxcl12 | 0.039803207 | -3.648156327 | 0 | 0.172 | 1 | 0.012531825 | -3.691966845 | 0 | 0.125 | 1 | 0.039803207 | 0.024906604 |
| Pold2 | 0.100350957 | -1.25696557 | 0.048 | 0.19 | 1 | 0.012543851 | -2.071814916 | 0.022 | 0.163 | 1 | 0.100350957 | 0.024930354 |
| Rhno1 | 0.810571543 | 1.054861935 | 0.143 | 0.142 | 1 | 0.012543851 | -2.071814916 | 0.022 | 0.163 | 1 | 0.810571543 | 0.024930354 |
| Tdg | 0.022100514 | -0.557182667 | 0.143 | 0.433 | 1 | 0.012548355 | -0.299920989 | 0.244 | 0.547 | 1 | 0.022100514 | 0.024939249 |
| Pde5a | 0.945235824 | -0.611630451 | 0.095 | 0.097 | 1 | 0.012548638 | -4.638855509 | 0 | 0.125 | 1 | 0.945235824 | 0.024939808 |
| Golga3 | 0.776036396 | 0.210597747 | 0.524 | 0.679 | 1 | 0.01257659 | -0.226415121 | 0.467 | 0.775 | 1 | 0.776036396 | 0.02499501 |
| Mirg | 0.012580599 | 1.694530328 | 0.571 | 0.347 | 1 | 0.133313922 | -0.477663244 | 0.267 | 0.405 | 1 | 0.133313922 | 0.025002926 |
| Vkorc1l1 | 0.012583164 | 1.112656009 | 0.714 | 0.642 | 1 | 0.211240849 | 0.518925306 | 0.8 | 0.768 | 1 | 0.211240849 | 0.025007992 |
| Ubiad1 | 0.722967451 | 0.673771768 | 0.048 | 0.071 | 1 | 0.012603466 | -2.044847868 | 0.022 | 0.163 | 1 | 0.722967451 | 0.025048084 |
| Cdk16 | 0.012617377 | -2.01272876 | 0.048 | 0.302 | 1 | 0.923183923 | 0.892995656 | 0.333 | 0.398 | 1 | 0.923183923 | 0.025075556 |
| Gstk1 | 0.18217443 | -0.469186186 | 0.048 | 0.164 | 1 | 0.012632016 | -1.960783604 | 0.022 | 0.163 | 1 | 0.18217443 | 0.025104465 |
| Gps2 | 0.012648794 | -1.326228232 | 0.143 | 0.448 | 1 | 0.075958184 | -0.437942815 | 0.444 | 0.557 | 1 | 0.075958184 | 0.025137595 |
| Naa10 | 0.012653363 | -2.094412557 | 0.048 | 0.302 | 1 | 0.016242992 | -0.552143876 | 0.111 | 0.301 | 1 | 0.016242992 | 0.025146619 |
| Tirap | 0.096766013 | -0.807354922 | 0.048 | 0.205 | 1 | 0.012655409 | -1.280401538 | 0.044 | 0.208 | 1 | 0.096766013 | 0.025150658 |
| Pan2 | 0.094148668 | -0.926141075 | 0.095 | 0.269 | 1 | 0.01266425 | -1.153428682 | 0.067 | 0.239 | 1 | 0.094148668 | 0.025168116 |
| Prrx2 | 0.012674028 | -1.804275529 | 0.095 | 0.373 | 1 | 0.120701741 | -0.452538051 | 0.311 | 0.491 | 1 | 0.120701741 | 0.025187425 |
| Ube3a | 0.01277451 | 0.730095374 | 0.952 | 0.914 | 1 | 0.286262547 | 0.363221806 | 1 | 0.962 | 1 | 0.286262547 | 0.025385832 |
| Txndc16 | 0.012797825 | -2.984439715 | 0 | 0.235 | 1 | 0.074738723 | 0.203079645 | 0.111 | 0.253 | 1 | 0.074738723 | 0.025431866 |
| Stk11 | 0.014862895 | -1.063193826 | 0.238 | 0.545 | 1 | 0.012824269 | -0.534712054 | 0.711 | 0.779 | 1 | 0.014862895 | 0.025484076 |
| A630023A22Rik | 0.012829897 | 2.351843673 | 0.238 | 0.082 | 1 | 0.687197015 | 1.545569062 | 0.022 | 0.035 | 1 | 0.687197015 | 0.025495187 |
| Rnf114 | 0.03620418 | -1.089993886 | 0.143 | 0.392 | 1 | 0.012841858 | 0.125077133 | 0.222 | 0.491 | 1 | 0.03620418 | 0.025518803 |
| Chst1 | 0.340906217 | 1.747772349 | 0.095 | 0.049 | 1 | 0.012861189 | -1.597035333 | 0.044 | 0.204 | 1 | 0.340906217 | 0.025556968 |
| Kat2b | 0.550393288 | 1.430915244 | 0.19 | 0.302 | 1 | 0.012865969 | 0.571014079 | 0.711 | 0.408 | 1 | 0.550393288 | 0.025566405 |
| F3 | 0.012891532 | -2.820217073 | 0.571 | 0.642 | 1 | 0.558983944 | -0.656777417 | 0.844 | 0.668 | 1 | 0.558983944 | 0.025616872 |
| Notch2 | 0.01290427 | 0.95207293 | 0.905 | 0.91 | 1 | 0.036704261 | 0.423662054 | 0.978 | 0.969 | 1 | 0.036704261 | 0.025642019 |
| Tcfl5 | 0.11772097 | -0.981580061 | 0.143 | 0.302 | 1 | 0.012918347 | -0.454430938 | 0.133 | 0.336 | 1 | 0.11772097 | 0.02566981 |
| Dnajc19 | 0.073481172 | -0.556525852 | 0.286 | 0.541 | 1 | 0.012919077 | -0.369015925 | 0.356 | 0.571 | 1 | 0.073481172 | 0.025671252 |
| Taf4b | 0.769848962 | 0.15682769 | 0.524 | 0.563 | 1 | 0.012967619 | 1.540892575 | 0.644 | 0.491 | 1 | 0.769848962 | 0.025767078 |
| Plagl2 | 0.399527985 | 1.064962525 | 0.238 | 0.183 | 1 | 0.012969309 | -0.740138845 | 0.111 | 0.298 | 1 | 0.399527985 | 0.025770414 |
| Cbfa2t3 | 0.083931487 | -2.574155746 | 0 | 0.127 | 1 | 0.013008078 | -2.098287127 | 0.022 | 0.166 | 1 | 0.083931487 | 0.025846946 |
| Afap1l2 | 0.314986473 | 0.199840579 | 0.095 | 0.198 | 1 | 0.013025419 | -2.204452685 | 0.044 | 0.197 | 1 | 0.314986473 | 0.025881176 |
| Cdk12 | 0.013027602 | 1.173421495 | 0.952 | 0.881 | 1 | 0.438236726 | 0.447368909 | 0.978 | 0.955 | 1 | 0.438236726 | 0.025885486 |
| Sh3bgrl3 | 0.013038753 | -0.857150893 | 0.476 | 0.743 | 1 | 0.017771937 | -0.945516847 | 0.756 | 0.716 | 1 | 0.017771937 | 0.025907497 |
| Sec24d | 0.013048182 | 0.719930757 | 0.905 | 0.907 | 1 | 0.964122874 | 0.155675373 | 0.889 | 0.958 | 1 | 0.964122874 | 0.025926109 |
| Myrfl | 0.013094732 | 2.574236094 | 0.095 | 0.015 | 1 | 0.293647037 | -1.017367132 | 0 | 0.024 | 1 | 0.293647037 | 0.026017993 |
| Dhx15 | 0.971094084 | 0.178195585 | 0.714 | 0.769 | 1 | 0.013099932 | -0.324500634 | 0.911 | 0.875 | 1 | 0.971094084 | 0.026028255 |
| Ddx23 | 0.16820708 | -0.348596045 | 0.286 | 0.474 | 1 | 0.013101738 | -0.458283263 | 0.333 | 0.567 | 1 | 0.16820708 | 0.02603182 |
| Hccs | 0.164078102 | -0.512094778 | 0.095 | 0.235 | 1 | 0.013134358 | -0.991527127 | 0.133 | 0.332 | 1 | 0.164078102 | 0.026096205 |
| Hsd17b8 | 0.168577956 | -0.39661756 | 0.095 | 0.246 | 1 | 0.013166248 | -1.425451871 | 0.044 | 0.201 | 1 | 0.168577956 | 0.026159146 |
| Ctdspl | 0.544394241 | 0.101520319 | 0.857 | 0.97 | 1 | 0.013169768 | 0.575257435 | 0.978 | 1 | 1 | 0.544394241 | 0.026166093 |
| Setdb1 | 0.013205 | 1.620133804 | 0.524 | 0.354 | 1 | 0.219716423 | 0.127022777 | 0.311 | 0.488 | 1 | 0.219716423 | 0.026235627 |
| Atp6v0a1 | 0.524415732 | 1.143257051 | 0.333 | 0.493 | 1 | 0.013222095 | 1.045642666 | 0.667 | 0.495 | 1 | 0.524415732 | 0.026269365 |
| Nfatc1 | 0.39238923 | 0.675300856 | 0.857 | 0.944 | 1 | 0.013229688 | 0.728724692 | 1 | 0.979 | 1 | 0.39238923 | 0.026284352 |
| Ints3 | 0.100105518 | -0.496153234 | 0.048 | 0.205 | 1 | 0.01324448 | -0.802354241 | 0.089 | 0.277 | 1 | 0.100105518 | 0.026313543 |
| Trf | 0.048883421 | -3.645147577 | 0.048 | 0.231 | 1 | 0.013259012 | -1.977178042 | 0.044 | 0.201 | 1 | 0.048883421 | 0.026342222 |
| Tprg | 0.01329797 | 2.936806174 | 0.095 | 0.015 | 1 | 0.497611189 | 1.875717664 | 0.022 | 0.01 | 1 | 0.497611189 | 0.026419104 |
| Cant1 | 0.013308937 | -2.413691074 | 0 | 0.231 | 1 | 0.018514843 | -1.881712033 | 0.044 | 0.183 | 1 | 0.018514843 | 0.026440746 |
| Pan3 | 0.013366233 | 0.892889057 | 0.905 | 0.869 | 1 | 0.017439895 | 0.952282455 | 1 | 0.931 | 1 | 0.017439895 | 0.02655381 |
| Vegfd | 0.358377174 | 1.491568436 | 0.19 | 0.131 | 1 | 0.013369047 | 1.605070074 | 0.111 | 0.031 | 1 | 0.358377174 | 0.026559363 |

| | | | | | | | | | | | | |
|---|---|---|---|---|---|---|---|---|---|---|---|---|
| Slc35a5 | 0.017946357 | -2.818081329 | 0 | 0.216 | 1 | 0.013394457 | -0.161276543 | 0.044 | 0.201 | 1 | 0.017946357 | 0.026609503 |
| Kbtbd4 | 0.11339646 | -1.370622352 | 0 | 0.108 | 1 | 0.013457495 | -1.513324627 | 0.022 | 0.166 | 1 | 0.11339646 | 0.026733885 |
| Rapgef1 | 0.013458838 | 0.635218527 | 0.81 | 0.646 | 1 | 0.308780486 | 0.526796646 | 0.756 | 0.716 | 1 | 0.308780486 | 0.026736535 |
| Vdr | 0.092134859 | -0.381510668 | 0.095 | 0.287 | 1 | 0.013461684 | -0.504554417 | 0.044 | 0.201 | 1 | 0.092134859 | 0.026742152 |
| Slc2a13 | 0.013469154 | -1.619009982 | 0.048 | 0.321 | 1 | 0.440358689 | 1.045642666 | 0.311 | 0.304 | 1 | 0.440358689 | 0.026756891 |
| Atxn7l3 | 0.014865555 | -1.25696557 | 0.048 | 0.317 | 1 | 0.01347957 | -0.625049709 | 0.244 | 0.464 | 1 | 0.014865555 | 0.02677744 |
| Elp1 | 0.039650576 | 1.451379346 | 0.524 | 0.433 | 1 | 0.013494128 | -0.390176396 | 0.289 | 0.567 | 1 | 0.039650576 | 0.026806164 |
| Gm56635 | 0.01350285 | 2.673771768 | 0.095 | 0.015 | 1 | 0.206882538 | -0.901889915 | 0 | 0.035 | 1 | 0.206882538 | 0.026823374 |
| C630043F03Rik | 0.403134049 | 0.866416846 | 0.048 | 0.112 | 1 | 0.0135032 | -0.941418279 | 0.044 | 0.201 | 1 | 0.403134049 | 0.026824064 |
| Xpa | 0.211079621 | -0.801961663 | 0.095 | 0.213 | 1 | 0.013505601 | -0.614607962 | 0.133 | 0.346 | 1 | 0.211079621 | 0.026828801 |
| Thoc5 | 0.113254574 | -1.208871282 | 0.048 | 0.183 | 1 | 0.013522715 | -1.819427754 | 0.044 | 0.194 | 1 | 0.113254574 | 0.026862566 |
| St7 | 0.811303007 | 0.121546053 | 0.429 | 0.552 | 1 | 0.01352927 | 0.960950839 | 0.733 | 0.543 | 1 | 0.811303007 | 0.0268755 |
| Carmil1 | 0.013545682 | 1.843556763 | 0.81 | 0.806 | 1 | 0.220447731 | 0.606865932 | 0.956 | 0.931 | 1 | 0.220447731 | 0.026907878 |
| Fam43a | 0.188964423 | 0.881213848 | 0.048 | 0.164 | 1 | 0.013553654 | -2.901889915 | 0.022 | 0.159 | 1 | 0.188964423 | 0.026923607 |
| Lrrc1 | 0.675096949 | 0.137064436 | 0.476 | 0.631 | 1 | 0.013558131 | 1.081401627 | 0.867 | 0.817 | 1 | 0.675096949 | 0.026932439 |
| Gm26549 | 0.013561408 | -2.683780237 | 0 | 0.231 | 1 | 0.290850806 | -0.316927414 | 0.133 | 0.204 | 1 | 0.290850806 | 0.026938904 |
| Xpo6 | 0.745392981 | 1.020637764 | 0.381 | 0.466 | 1 | 0.01356734 | 0.83979809 | 0.756 | 0.699 | 1 | 0.745392981 | 0.026950607 |
| Dipk1a | 0.032775607 | -2.785659851 | 0 | 0.183 | 1 | 0.01356873 | -1.440309829 | 0.089 | 0.26 | 1 | 0.032775607 | 0.02695335 |
| Ndel1 | 0.078899902 | 0.811718811 | 0.857 | 0.914 | 1 | 0.013600855 | -0.166982774 | 1 | 0.958 | 1 | 0.078899902 | 0.027016727 |
| Tax1bp3 | 0.013630475 | -1.392317423 | 0.143 | 0.444 | 1 | 0.191409623 | -0.466674533 | 0.422 | 0.467 | 1 | 0.191409623 | 0.02707516 |
| Ctbs | 0.080647357 | -0.558888989 | 0.095 | 0.284 | 1 | 0.013641235 | -0.72631835 | 0.067 | 0.239 | 1 | 0.080647357 | 0.027096386 |
| Fra10ac1 | 0.013662106 | -2.849790188 | 0 | 0.231 | 1 | 0.035665468 | -0.743192169 | 0.067 | 0.204 | 1 | 0.035665468 | 0.027137559 |
| Mtfmt | 0.246951561 | -0.849790188 | 0.048 | 0.134 | 1 | 0.013679008 | -1.199570463 | 0.022 | 0.166 | 1 | 0.246951561 | 0.027170902 |
| Gm56955 | 0.013699654 | 2.957564734 | 0.286 | 0.119 | 1 | 0.247775233 | 0.905465008 | 0.044 | 0.104 | 1 | 0.247775233 | 0.027211628 |
| Rhob | 0.345117918 | -0.911190733 | 0.619 | 0.687 | 1 | 0.013700444 | -1.103259085 | 0.778 | 0.737 | 1 | 0.345117918 | 0.027213185 |
| Nom1 | 0.818328852 | 0.936806174 | 0.333 | 0.377 | 1 | 0.013708635 | -0.271123724 | 0.2 | 0.46 | 1 | 0.818328852 | 0.027229342 |
| Rnf6 | 0.172287801 | -0.169502729 | 0.333 | 0.563 | 1 | 0.013711306 | -0.284505936 | 0.444 | 0.661 | 1 | 0.172287801 | 0.027234611 |
| 5730455P16Rik | 0.024586599 | -1.548620654 | 0.048 | 0.284 | 1 | 0.01371191 | -0.686161223 | 0.111 | 0.311 | 1 | 0.024586599 | 0.027235803 |
| Tlk2 | 0.013713927 | 0.757207472 | 0.905 | 0.884 | 1 | 0.03108696 | 0.50309392 | 1 | 0.969 | 1 | 0.03108696 | 0.027239782 |
| Zbtb8os | 0.018017568 | -1.392317423 | 0.095 | 0.362 | 1 | 0.013719562 | -0.831500587 | 0.067 | 0.232 | 1 | 0.018017568 | 0.027250897 |
| Camk4 | 0.013753932 | -1.611630451 | 0.143 | 0.433 | 1 | 0.027938048 | 0.395724723 | 0.356 | 0.657 | 1 | 0.027938048 | 0.027318693 |
| Gm36220 | 0.092426668 | -0.785659851 | 0.143 | 0.328 | 1 | 0.01375505 | -0.754332726 | 0.044 | 0.201 | 1 | 0.092426668 | 0.027320898 |
| Rcan3 | 0.100336721 | -1.455511249 | 0 | 0.116 | 1 | 0.013758455 | -2.247664751 | 0.022 | 0.159 | 1 | 0.100336721 | 0.027327614 |
| Tug1 | 0.013764993 | -0.672003069 | 0.238 | 0.612 | 1 | 0.032289094 | -0.561234612 | 0.556 | 0.692 | 1 | 0.032289094 | 0.027340511 |
| Klhl26 | 0.199754554 | 1.308122295 | 0.286 | 0.205 | 1 | 0.013784515 | -0.446210431 | 0.089 | 0.287 | 1 | 0.199754554 | 0.027379017 |
| Gm19325 | 0.837550449 | 1.815790773 | 0.095 | 0.09 | 1 | 0.013799805 | -2.294207337 | 0.022 | 0.159 | 1 | 0.837550449 | 0.027409175 |
| Mttp | 0.275383697 | 1.281454345 | 0.19 | 0.119 | 1 | 0.01380081 | -2.404390255 | 0.022 | 0.159 | 1 | 0.275383697 | 0.027411158 |
| Lcmt2 | 0.173676192 | -0.911190733 | 0 | 0.082 | 1 | 0.013829556 | -2.526380779 | 0 | 0.121 | 1 | 0.173676192 | 0.027467855 |
| Pop1 | 0.078369064 | -1.718545655 | 0 | 0.131 | 1 | 0.01385664 | -0.776359032 | 0.022 | 0.166 | 1 | 0.078369064 | 0.027521274 |
| 9930104L06Rik | 0.538371401 | -0.133583154 | 0.048 | 0.086 | 1 | 0.0138725 | -2.674479418 | 0 | 0.121 | 1 | 0.538371401 | 0.027552553 |
| Hacl1 | 0.287079534 | -0.849790188 | 0.095 | 0.19 | 1 | 0.013877157 | -2.223818009 | 0.022 | 0.159 | 1 | 0.287079534 | 0.027561738 |
| C2cd2l | 0.128113406 | -1.280424543 | 0 | 0.101 | 1 | 0.01388525 | -2.674479418 | 0 | 0.121 | 1 | 0.128113406 | 0.027577701 |
| Cad | 0.153811633 | -1.184209227 | 0 | 0.09 | 1 | 0.013896809 | -2.80878051 | 0 | 0.121 | 1 | 0.153811633 | 0.027600498 |
| Ypel1 | 0.987342144 | 0.673771768 | 0.095 | 0.097 | 1 | 0.013907856 | -2.84048937 | 0 | 0.121 | 1 | 0.987342144 | 0.027622283 |
| Hsdl1 | 0.564286249 | 1.536268244 | 0.095 | 0.067 | 1 | 0.013908128 | -2.743192169 | 0 | 0.121 | 1 | 0.564286249 | 0.02762282 |
| Ccdc134 | 0.652151687 | 1.541321472 | 0.19 | 0.164 | 1 | 0.013916571 | -2.098287127 | 0.022 | 0.159 | 1 | 0.652151687 | 0.027639471 |
| Ntpcr | 0.081776521 | -1.516052791 | 0.048 | 0.201 | 1 | 0.013916891 | -2.80878051 | 0 | 0.121 | 1 | 0.081776521 | 0.027640103 |
| Acp5 | 0.236452955 | -0.849790188 | 0 | 0.063 | 1 | 0.013927863 | -8.388389776 | 0 | 0.121 | 1 | 0.236452955 | 0.027661741 |
| Pole2 | 0.749416568 | 0.410737362 | 0.095 | 0.127 | 1 | 0.013936251 | -3.124282336 | 0 | 0.121 | 1 | 0.749416568 | 0.027678283 |
| Osgep | 0.022902591 | -0.748760268 | 0.19 | 0.504 | 1 | 0.013949991 | -0.21001221 | 0.222 | 0.446 | 1 | 0.022902591 | 0.027705379 |
| Dut | 0.017840515 | -0.852923078 | 0.238 | 0.612 | 1 | 0.013952866 | -1.184823878 | 0.578 | 0.657 | 1 | 0.017840515 | 0.027711049 |
| Slc25a44 | 0.928600764 | 0.866416846 | 0.19 | 0.228 | 1 | 0.013975862 | -0.671770131 | 0.133 | 0.339 | 1 | 0.928600764 | 0.027756399 |
| Pxn | 0.241652655 | 0.969510339 | 0.476 | 0.425 | 1 | 0.013992642 | 1.015054346 | 0.711 | 0.547 | 1 | 0.241652655 | 0.027789489 |
| Crtc3 | 0.124486521 | 0.939583258 | 0.667 | 0.675 | 1 | 0.014008132 | 0.871585049 | 0.911 | 0.81 | 1 | 0.124486521 | 0.027820036 |
| Fgd6 | 0.014011861 | -1.298920886 | 0.143 | 0.444 | 1 | 0.102252841 | 0.242868641 | 0.778 | 0.599 | 1 | 0.102252841 | 0.027827389 |
| Cyrib | 0.619887831 | 1.054861935 | 0.381 | 0.377 | 1 | 0.014013677 | 0.656177359 | 0.8 | 0.529 | 1 | 0.619887831 | 0.02783097 |
| Thada | 0.160054718 | 0.830497271 | 0.524 | 0.44 | 1 | 0.014018906 | 0.843537258 | 0.622 | 0.512 | 1 | 0.160054718 | 0.027841283 |
| B930036N10Rik | 0.276981321 | 1.231253532 | 0.333 | 0.25 | 1 | 0.014041171 | -2.124282336 | 0.022 | 0.163 | 1 | 0.276981321 | 0.027885188 |
| Cep164 | 0.406296795 | -0.355975576 | 0.143 | 0.224 | 1 | 0.014074857 | -0.247664751 | 0.089 | 0.284 | 1 | 0.406296795 | 0.027951612 |
| Sec61a1 | 0.014093895 | -0.8244791 | 0.476 | 0.791 | 1 | 0.019606825 | -0.20320773 | 0.8 | 0.841 | 1 | 0.019606825 | 0.027989152 |
| B230219D22Rik | 0.051123318 | -0.483957937 | 0.619 | 0.832 | 1 | 0.014112048 | -0.376491968 | 0.8 | 0.875 | 1 | 0.051123318 | 0.028024945 |
| Crkl | 0.773996678 | 0.665597836 | 0.286 | 0.41 | 1 | 0.014116636 | -0.649635347 | 0.467 | 0.647 | 1 | 0.773996678 | 0.028033993 |
| Bop1 | 0.096743437 | -0.807354922 | 0.048 | 0.205 | 1 | 0.014135964 | -0.743192169 | 0.067 | 0.239 | 1 | 0.096743437 | 0.028072103 |
| Whamm | 0.847094159 | 0.281454345 | 0.143 | 0.168 | 1 | 0.014149369 | -1.181997834 | 0.067 | 0.235 | 1 | 0.847094159 | 0.028098534 |
| Lin54 | 0.790414444 | 1.276881615 | 0.19 | 0.194 | 1 | 0.014172853 | -0.994999319 | 0.133 | 0.322 | 1 | 0.790414444 | 0.028144835 |
| Nopchap1 | 0.058182105 | -1.26168798 | 0.143 | 0.343 | 1 | 0.014207737 | -0.499130745 | 0.133 | 0.346 | 1 | 0.058182105 | 0.028213614 |
| Cyp20a1 | 0.014212882 | -0.413691074 | 0.286 | 0.634 | 1 | 0.161882437 | 0.610172039 | 0.378 | 0.592 | 1 | 0.161882437 | 0.028223758 |
| Mrps5 | 0.014243816 | -1.683780237 | 0.048 | 0.313 | 1 | 0.046062909 | -0.638855509 | 0.156 | 0.315 | 1 | 0.046062909 | 0.028284745 |
| Tmub1 | 0.095793106 | -1.167530486 | 0.095 | 0.254 | 1 | 0.014246619 | -1.064161343 | 0.089 | 0.26 | 1 | 0.095793106 | 0.028290272 |
| Ism1 | 0.728384055 | 2.782674497 | 0.333 | 0.381 | 1 | 0.014281554 | -1.478815096 | 0.178 | 0.363 | 1 | 0.728384055 | 0.028359144 |
| Zdhhc5 | 0.014282778 | -0.911190733 | 0.286 | 0.619 | 1 | 0.081515288 | -0.196073019 | 0.644 | 0.751 | 1 | 0.081515288 | 0.028361558 |
| E130317F20Rik | 0.014288422 | 2.233199176 | 0.19 | 0.056 | 1 | 0.165324946 | -1.404390255 | 0 | 0.042 | 1 | 0.165324946 | 0.028372684 |
| Lars2 | 0.029373098 | 3.536628273 | 1 | 1 | 1 | 0.014324612 | 1.412916739 | 1 | 1 | 1 | 0.029373098 | 0.028444029 |
| Zfp617 | 0.014356312 | -2.475975352 | 0 | 0.228 | 1 | 0.097082401 | -0.204452685 | 0.089 | 0.204 | 1 | 0.097082401 | 0.028506519 |
| Ldah | 0.01438639 | -1.370622352 | 0.143 | 0.437 | 1 | 0.045264993 | -0.189466505 | 0.267 | 0.481 | 1 | 0.045264993 | 0.028565812 |
| Fnbp1 | 0.208639476 | 0.143257051 | 0.524 | 0.34 | 1 | 0.014422521 | 0.513147585 | 0.8 | 0.529 | 1 | 0.208639476 | 0.028637034 |
| Amz2 | 0.308082734 | -0.146407195 | 0.19 | 0.336 | 1 | 0.014441685 | -0.186343304 | 0.089 | 0.277 | 1 | 0.308082734 | 0.028674808 |
| Sdr39u1 | 0.014456736 | -2.496153234 | 0 | 0.228 | 1 | 0.034418418 | 0.42003818 | 0.067 | 0.208 | 1 | 0.034418418 | 0.028704475 |
| Polr3h | 0.05526447 | -0.911190733 | 0.143 | 0.358 | 1 | 0.014479086 | -0.686161223 | 0.111 | 0.301 | 1 | 0.05526447 | 0.028748527 |
| Alg10b | 0.099217749 | -0.555046923 | 0.048 | 0.205 | 1 | 0.014479628 | -1.033134448 | 0.067 | 0.235 | 1 | 0.099217749 | 0.028749596 |
| C1galt1 | 0.018378765 | 1.642293536 | 0.476 | 0.325 | 1 | 0.014505413 | 0.619569644 | 0.644 | 0.391 | 1 | 0.018378765 | 0.028800419 |
| Rnf113a2 | 0.339580841 | -0.326228232 | 0.143 | 0.239 | 1 | 0.014516795 | -1.352551324 | 0.067 | 0.228 | 1 | 0.339580841 | 0.028822852 |
| Cnst | 0.620967589 | -0.107587946 | 0.095 | 0.134 | 1 | 0.014518979 | -0.979892427 | 0.089 | 0.263 | 1 | 0.620967589 | 0.028827156 |
| Bpifa1 | 0.384108638 | 0.524024648 | 0.429 | 0.377 | 1 | 0.014529169 | 0.732204294 | 0.844 | 0.706 | 1 | 0.384108638 | 0.028847242 |
| Dnajc14 | 0.215161765 | -0.63000898 | 0.095 | 0.213 | 1 | 0.014530676 | -0.520211012 | 0.178 | 0.394 | 1 | 0.215161765 | 0.028850211 |
| Gm12592 | 0.014531867 | 2.55829455 | 0.143 | 0.034 | 1 | 0.032976507 | -2.404390255 | 0 | 0.093 | 1 | 0.032976507 | 0.028852558 |
| Ppp1r12c | 0.024141928 | -0.620848981 | 0.143 | 0.437 | 1 | 0.014532552 | -0.166102512 | 0.267 | 0.55 | 1 | 0.024141928 | 0.028853908 |
| Immp1l | 0.600152326 | 0.337949231 | 0.476 | 0.664 | 1 | 0.014537542 | -0.182626322 | 0.4 | 0.644 | 1 | 0.600152326 | 0.028863743 |
| Gale | 0.127589769 | -0.992984824 | 0.143 | 0.299 | 1 | 0.014538286 | -1.387316742 | 0.044 | 0.197 | 1 | 0.127589769 | 0.02886521 |
| Gm46430 | 0.014548163 | 2.233199176 | 0.19 | 0.056 | 1 | 0.070982579 | -1.602329633 | 0.022 | 0.107 | 1 | 0.070982579 | 0.028884676 |
| Nsd3 | 0.014552109 | 1.13423199 | 1 | 0.94 | 1 | 0.041695318 | 0.5232476 | 1 | 0.99 | 1 | 0.041695318 | 0.028892455 |
| Stard7 | 0.16223792 | 0.133203386 | 0.143 | 0.306 | 1 | 0.014557474 | -0.702358451 | 0.2 | 0.422 | 1 | 0.16223792 | 0.028903028 |
| Mvd | 0.014571092 | -2.735619168 | 0 | 0.228 | 1 | 0.087346514 | -1.526380779 | 0.022 | 0.1 | 1 | 0.087346514 | 0.028929867 |
| Iqce | 0.061286955 | -2.107587946 | 0 | 0.146 | 1 | 0.014572661 | -1.819427754 | 0.022 | 0.163 | 1 | 0.061286955 | 0.02893296 |
| Plpp3 | 0.085407584 | -0.387395929 | 0.952 | 0.929 | 1 | 0.014577296 | -0.312798501 | 0.956 | 0.972 | 1 | 0.085407584 | 0.028942095 |
| Fzd6 | 0.233121077 | -0.527862093 | 0.095 | 0.209 | 1 | 0.014597752 | -0.340774156 | 0.089 | 0.273 | 1 | 0.233121077 | 0.028982409 |
| Cldn12 | 0.380318428 | -0.44170545 | 0.095 | 0.168 | 1 | 0.014598401 | -1.124282336 | 0.067 | 0.235 | 1 | 0.380318428 | 0.028983688 |
| Prkcz | 0.018532244 | -0.840801405 | 0.095 | 0.366 | 1 | 0.014626133 | -0.513324627 | 0.133 | 0.332 | 1 | 0.018532244 | 0.029038342 |

| | | | | | | | | | | | | |
|---|---|---|---|---|---|---|---|---|---|---|---|---|
| Prkaa1 | 0.671681888 | 0.710297644 | 0.333 | 0.321 | 1 | 0.014634369 | -0.169370225 | 0.2 | 0.439 | 1 | 0.671681888 | 0.029054573 |
| Esd | 0.014634612 | -0.844076537 | 0.667 | 0.791 | 1 | 0.034882063 | -0.470173674 | 0.756 | 0.761 | 1 | 0.034882063 | 0.029055052 |
| Adgre5 | 0.213718691 | -0.849790188 | 0.048 | 0.149 | 1 | 0.014640576 | -1.476798751 | 0.067 | 0.225 | 1 | 0.213718691 | 0.029066806 |
| Mbd1 | 0.582209017 | 0.281454345 | 0.19 | 0.265 | 1 | 0.014694964 | -0.392215541 | 0.111 | 0.311 | 1 | 0.582209017 | 0.029173986 |
| Eya1 | 0.029002883 | 1.147304916 | 0.905 | 0.892 | 1 | 0.014713254 | 0.577560262 | 0.978 | 0.962 | 1 | 0.029002883 | 0.029210028 |
| Wdr82 | 0.277714266 | 0.708875569 | 0.476 | 0.451 | 1 | 0.014714748 | -0.602329633 | 0.356 | 0.561 | 1 | 0.277714266 | 0.029212973 |
| Spacdr | 0.014746127 | -0.785659851 | 0.286 | 0.642 | 1 | 0.561939338 | 0.151601492 | 0.778 | 0.709 | 1 | 0.561939338 | 0.029274806 |
| Prmt9 | 0.817049062 | 0.651403955 | 0.19 | 0.179 | 1 | 0.014749552 | 0.263533695 | 0.089 | 0.266 | 1 | 0.817049062 | 0.029281554 |
| Gpr27 | 0.248716669 | -0.818081329 | 0.048 | 0.134 | 1 | 0.014755868 | -1.141355849 | 0.044 | 0.197 | 1 | 0.248716669 | 0.029294001 |
| Actl6a | 0.014758844 | -0.596757175 | 0.143 | 0.451 | 1 | 0.021330969 | -0.491701815 | 0.422 | 0.578 | 1 | 0.021330969 | 0.029299864 |
| Acbd6 | 0.014770592 | -0.587895802 | 0.333 | 0.709 | 1 | 0.31346279 | 0.204609707 | 0.622 | 0.661 | 1 | 0.31346279 | 0.029323013 |
| Rev3l | 0.014775507 | 0.683687237 | 0.81 | 0.672 | 1 | 0.231619749 | 0.380145453 | 0.978 | 0.841 | 1 | 0.231619749 | 0.029332698 |
| Gfus | 0.059553997 | -1.455511249 | 0.048 | 0.22 | 1 | 0.014783806 | -0.738391182 | 0.044 | 0.204 | 1 | 0.059553997 | 0.029349051 |
| Ccng2 | 0.15985017 | -1.25696557 | 0.048 | 0.16 | 1 | 0.014850261 | -1.564854927 | 0.022 | 0.163 | 1 | 0.15985017 | 0.029479992 |
| Azin1 | 0.014851391 | -0.621029149 | 0.762 | 0.91 | 1 | 0.026883792 | -0.240691828 | 0.911 | 0.924 | 1 | 0.026883792 | 0.029482218 |
| Ly6c1 | 0.014908942 | -3.818081329 | 0 | 0.228 | 1 | 0.095598201 | -0.782590986 | 0.111 | 0.225 | 1 | 0.095598201 | 0.029595607 |
| Zc3h3 | 0.014912441 | 1.901182264 | 0.429 | 0.25 | 1 | 0.107665417 | -0.154870656 | 0.178 | 0.325 | 1 | 0.107665417 | 0.029602501 |
| Cdk2 | 0.104247861 | -1.04504648 | 0.048 | 0.198 | 1 | 0.014913097 | -1.137106376 | 0.111 | 0.291 | 1 | 0.104247861 | 0.029603794 |
| Zfp37 | 0.668203648 | 0.943378827 | 0.143 | 0.194 | 1 | 0.014916646 | -0.638855509 | 0.111 | 0.301 | 1 | 0.668203648 | 0.029610785 |
| Toe1 | 0.075827327 | -0.93109029 | 0.048 | 0.22 | 1 | 0.014930768 | -1.271123724 | 0.089 | 0.256 | 1 | 0.075827327 | 0.029638608 |
| Zfp212 | 0.527672711 | 0.983456267 | 0.238 | 0.209 | 1 | 0.015011242 | -0.798054104 | 0.111 | 0.304 | 1 | 0.527672711 | 0.029797147 |
| Mex3c | 0.015021312 | -0.790695441 | 0.238 | 0.56 | 1 | 0.21018128 | -0.372371741 | 0.778 | 0.765 | 1 | 0.21018128 | 0.029816985 |
| Cdkn2aip | 0.144654019 | -0.648156327 | 0.095 | 0.254 | 1 | 0.015042984 | -0.909882705 | 0.2 | 0.388 | 1 | 0.144654019 | 0.029859676 |
| Arid2 | 0.202708342 | 0.714413752 | 0.667 | 0.563 | 1 | 0.015045637 | 0.849310062 | 0.933 | 0.844 | 1 | 0.202708342 | 0.029864902 |
| Cep44 | 0.954974288 | 0.304537958 | 0.143 | 0.153 | 1 | 0.01505055 | -0.545746104 | 0.044 | 0.197 | 1 | 0.954974288 | 0.02987458 |
| Lrp12 | 0.075088522 | -0.648156327 | 0.143 | 0.362 | 1 | 0.015059974 | -0.651911662 | 0.244 | 0.457 | 1 | 0.075088522 | 0.029893145 |
| Zdhhc1 | 0.648731817 | 0.267779408 | 0.286 | 0.403 | 1 | 0.015069593 | -0.662702251 | 0.178 | 0.391 | 1 | 0.648731817 | 0.029912094 |
| Itfg1 | 0.192326244 | 0.410737362 | 0.381 | 0.679 | 1 | 0.015081639 | -0.425938036 | 0.489 | 0.626 | 1 | 0.192326244 | 0.029935822 |
| Ormdl2 | 0.144223614 | -0.672003069 | 0.048 | 0.179 | 1 | 0.015105847 | -0.72631835 | 0.067 | 0.235 | 1 | 0.144223614 | 0.029983508 |
| Slc35c2 | 0.039127832 | -2.054148687 | 0 | 0.172 | 1 | 0.015110688 | -1.798054104 | 0.044 | 0.19 | 1 | 0.039127832 | 0.029993044 |
| Ubtd2 | 0.948067976 | 0.754740609 | 0.571 | 0.754 | 1 | 0.015114804 | 1.063758581 | 0.756 | 0.834 | 1 | 0.948067976 | 0.030001151 |
| Cacna2d3 | 0.015142984 | 2.673771768 | 0.238 | 0.09 | 1 | 0.100403378 | 2.872897145 | 0.156 | 0.09 | 1 | 0.100403378 | 0.030056658 |
| Cfap418 | 0.137281565 | -0.619009982 | 0.095 | 0.246 | 1 | 0.015159301 | -0.547225033 | 0.111 | 0.298 | 1 | 0.137281565 | 0.030088798 |
| Morc4 | 0.209136364 | -0.636083495 | 0.095 | 0.224 | 1 | 0.015187353 | -0.702985846 | 0.067 | 0.242 | 1 | 0.209136364 | 0.03014405 |
| Agl | 0.767517499 | 0.808701348 | 0.095 | 0.127 | 1 | 0.015189872 | -1.339295227 | 0.022 | 0.163 | 1 | 0.767517499 | 0.030149012 |
| Slc27a6 | 0.015216136 | -1.729950418 | 0.048 | 0.31 | 1 | 0.054298824 | -0.308706907 | 0.156 | 0.315 | 1 | 0.054298824 | 0.030200742 |
| Porcn | 0.456301976 | -0.184209227 | 0.095 | 0.16 | 1 | 0.015217212 | -1.334849322 | 0.044 | 0.201 | 1 | 0.456301976 | 0.03020286 |
| R3hcc1 | 0.11964398 | -1.233118828 | 0.048 | 0.179 | 1 | 0.01522594 | -0.719025857 | 0.044 | 0.204 | 1 | 0.11964398 | 0.030220051 |
| Bahcc1 | 0.23330414 | 2.404311093 | 0.381 | 0.31 | 1 | 0.015248578 | -0.596151058 | 0.178 | 0.384 | 1 | 0.23330414 | 0.030264637 |
| Klhl2 | 0.047026809 | 1.046989091 | 0.381 | 0.209 | 1 | 0.015273565 | 0.963180505 | 0.556 | 0.363 | 1 | 0.047026809 | 0.030313848 |
| Six4 | 0.343098908 | 0.97333205 | 0.048 | 0.123 | 1 | 0.015273956 | -1.695439037 | 0.067 | 0.221 | 1 | 0.343098908 | 0.030314618 |
| Mboat2 | 0.081831262 | -0.926141075 | 0.048 | 0.216 | 1 | 0.015279043 | -0.252797076 | 0.222 | 0.478 | 1 | 0.081831262 | 0.030324637 |
| Pafah1b3 | 0.29690025 | -0.348596045 | 0.095 | 0.194 | 1 | 0.015324899 | -1.071814916 | 0.022 | 0.163 | 1 | 0.29690025 | 0.030414946 |
| Msx1 | 0.346221524 | -1.370622352 | 0.048 | 0.112 | 1 | 0.015329308 | -2.86124793 | 0.044 | 0.187 | 1 | 0.346221524 | 0.030423629 |
| Tjp2 | 0.015335649 | 1.330884054 | 0.524 | 0.31 | 1 | 0.015333574 | 0.460680165 | 0.8 | 0.509 | 1 | 0.015335649 | 0.03043203 |
| Sntg2 | 0.015341137 | 2.351843673 | 0.143 | 0.034 | 1 | 0.320927426 | -0.819427754 | 0.022 | 0.059 | 1 | 0.320927426 | 0.030446923 |
| Ascc1 | 0.108740261 | -0.849790188 | 0.095 | 0.261 | 1 | 0.015357464 | -1.153428682 | 0.089 | 0.256 | 1 | 0.108740261 | 0.030479076 |
| Fam171b | 0.865416309 | 0.866416846 | 0.143 | 0.138 | 1 | 0.015362958 | -1.667424661 | 0.044 | 0.194 | 1 | 0.865416309 | 0.030489896 |
| Casp6 | 0.40040211 | -0.413691074 | 0.048 | 0.104 | 1 | 0.015369215 | -2.223818009 | 0.022 | 0.156 | 1 | 0.40040211 | 0.030502217 |
| Gadd45g | 0.015389293 | -1.417726587 | 0.238 | 0.526 | 1 | 0.055490307 | -0.581034854 | 0.178 | 0.322 | 1 | 0.055490307 | 0.030541756 |
| Fem1a | 0.732130428 | 0.936806174 | 0.048 | 0.071 | 1 | 0.015433877 | -0.679497493 | 0.044 | 0.204 | 1 | 0.732130428 | 0.030629549 |
| Ero1a | 0.015448335 | -0.814514714 | 0.143 | 0.448 | 1 | 0.342591008 | 0.421580345 | 0.511 | 0.408 | 1 | 0.342591008 | 0.030658019 |
| Rdm1 | 0.156515116 | 0.619904896 | 0.19 | 0.351 | 1 | 0.015454733 | -0.674479418 | 0.156 | 0.367 | 1 | 0.156515116 | 0.030670618 |
| Ramp1 | 0.128477385 | -2.413691074 | 0 | 0.101 | 1 | 0.015460891 | -1.827889333 | 0.067 | 0.221 | 1 | 0.128477385 | 0.030682742 |
| Dcp2 | 0.086191719 | -0.39470797 | 0.143 | 0.354 | 1 | 0.015465941 | -0.1336202 | 0.156 | 0.367 | 1 | 0.086191719 | 0.030692687 |
| Cxcl16 | 0.425971597 | -0.280424543 | 0.048 | 0.101 | 1 | 0.015473396 | -2.638855509 | 0 | 0.118 | 1 | 0.425971597 | 0.030707365 |
| Ctla4 | 0.851548758 | 0.304537958 | 0.095 | 0.108 | 1 | 0.015473396 | -2.674479418 | 0 | 0.118 | 1 | 0.851548758 | 0.030707365 |
| Gm15706 | 0.323619271 | -0.370622352 | 0.095 | 0.187 | 1 | 0.015473516 | -2.602329633 | 0 | 0.118 | 1 | 0.323619271 | 0.030707603 |
| Ptgs2os | 0.01548532 | 1.747772349 | 0.19 | 0.056 | 1 | 0.184854061 | -2.071814916 | 0 | 0.038 | 1 | 0.184854061 | 0.030730845 |
| Adhfe1 | 0.086841095 | -1.133583154 | 0.048 | 0.209 | 1 | 0.015486542 | -2.709244837 | 0 | 0.118 | 1 | 0.086841095 | 0.03073325 |
| Mybl1 | 0.054407092 | -2.735619168 | 0 | 0.153 | 1 | 0.015498906 | -3.199570463 | 0 | 0.118 | 1 | 0.054407092 | 0.030757596 |
| Ndufa10 | 0.015508156 | -0.443336535 | 0.286 | 0.619 | 1 | 0.133944148 | -0.657964332 | 0.6 | 0.592 | 1 | 0.133944148 | 0.03077581 |
| Phgdh | 0.33127194 | -0.940938076 | 0.048 | 0.116 | 1 | 0.015509309 | -2.776359032 | 0 | 0.118 | 1 | 0.33127194 | 0.030778079 |
| Zfp948 | 0.852353818 | 0.995699863 | 0.333 | 0.478 | 1 | 0.015519041 | -0.489931808 | 0.333 | 0.592 | 1 | 0.852353818 | 0.030797242 |
| Cys1 | 0.222255538 | -0.970084422 | 0 | 0.067 | 1 | 0.01552035 | -2.871516266 | 0 | 0.118 | 1 | 0.222255538 | 0.030799819 |
| Snupn | 0.251331225 | -0.648156327 | 0.048 | 0.134 | 1 | 0.01552038 | -2.84048937 | 0 | 0.118 | 1 | 0.251331225 | 0.030799878 |
| Trabd2b | 0.040292016 | 1.09183696 | 0.762 | 0.765 | 1 | 0.015524271 | 0.921501971 | 0.822 | 0.907 | 1 | 0.040292016 | 0.030807539 |
| Lrp4 | 0.144967842 | -2.574155746 | 0 | 0.093 | 1 | 0.015530093 | -2.989352756 | 0 | 0.118 | 1 | 0.144967842 | 0.030819003 |
| Dhfr | 0.356892611 | -0.522625445 | 0.048 | 0.116 | 1 | 0.015530214 | -2.960783604 | 0 | 0.118 | 1 | 0.356892611 | 0.030819241 |
| Klhl12 | 0.640904793 | 0.410737362 | 0.238 | 0.325 | 1 | 0.015538259 | -0.494465599 | 0.156 | 0.377 | 1 | 0.640904793 | 0.030835081 |
| Klhl23 | 0.331269099 | -0.752492987 | 0.048 | 0.116 | 1 | 0.01554653 | -3.124282336 | 0 | 0.118 | 1 | 0.331269099 | 0.030851365 |
| Tmem38a | 0.400305588 | 0.673771768 | 0.048 | 0.112 | 1 | 0.015547665 | -2.098287127 | 0.022 | 0.156 | 1 | 0.400305588 | 0.030853601 |
| Map3k4 | 0.486348102 | -0.198767323 | 0.286 | 0.433 | 1 | 0.015554815 | -0.333229226 | 0.156 | 0.388 | 1 | 0.486348102 | 0.030867678 |
| Tesk2 | 0.226081172 | -2.233118828 | 0.048 | 0.138 | 1 | 0.015561731 | -3.199570463 | 0 | 0.118 | 1 | 0.226081172 | 0.030881295 |
| Stard10 | 0.074598193 | -0.798716004 | 0.095 | 0.299 | 1 | 0.015570586 | -0.551392667 | 0.2 | 0.412 | 1 | 0.074598193 | 0.030898729 |
| Usp14 | 0.07386359 | -0.577766999 | 0.429 | 0.69 | 1 | 0.015585604 | -0.338529668 | 0.6 | 0.73 | 1 | 0.07386359 | 0.030928297 |
| Alad | 0.015586004 | -2.593014773 | 0 | 0.224 | 1 | 0.060292083 | -0.316927414 | 0.044 | 0.156 | 1 | 0.060292083 | 0.030929085 |
| Ppih | 0.094520107 | -1.683780237 | 0 | 0.119 | 1 | 0.015590632 | -2.124282336 | 0.022 | 0.156 | 1 | 0.094520107 | 0.030938197 |
| Tmem67 | 0.03205914 | -2.02666795 | 0 | 0.183 | 1 | 0.015590866 | -2.124282336 | 0.022 | 0.156 | 1 | 0.03205914 | 0.030938656 |
| Ttc23l | 0.222366285 | -2.516052791 | 0 | 0.067 | 1 | 0.015596975 | -3.946284034 | 0 | 0.118 | 1 | 0.222366285 | 0.030950683 |
| Vhl | 0.987804091 | 0.544488751 | 0.19 | 0.213 | 1 | 0.015611129 | -0.811692106 | 0.133 | 0.318 | 1 | 0.987804091 | 0.030978551 |
| Crbn | 0.058462467 | -1.133583154 | 0.048 | 0.235 | 1 | 0.015619433 | -1.421264074 | 0.067 | 0.225 | 1 | 0.058462467 | 0.0309949 |
| Scgb1c1 | 0.202379273 | 1.79930265 | 0.095 | 0.037 | 1 | 0.015619871 | 0.663173029 | 0.333 | 0.173 | 1 | 0.202379273 | 0.030995762 |
| Phc1 | 0.407284178 | -0.434752689 | 0.095 | 0.168 | 1 | 0.015648782 | -0.736466305 | 0.111 | 0.284 | 1 | 0.407284178 | 0.031052679 |
| Fyn | 0.060573891 | 1.48112669 | 0.619 | 0.552 | 1 | 0.015652983 | 0.993615918 | 0.644 | 0.491 | 1 | 0.060573891 | 0.031060949 |
| Atxn7l1 | 0.170138821 | 0.808701348 | 0.524 | 0.425 | 1 | 0.015657036 | 1.028401114 | 0.667 | 0.474 | 1 | 0.170138821 | 0.031068928 |
| Chst2 | 0.015657281 | -2.133583154 | 0.048 | 0.306 | 1 | 0.023595722 | -2.339295227 | 0.022 | 0.142 | 1 | 0.023595722 | 0.031069411 |
| Nampt | 0.346916538 | 0.127578837 | 0.286 | 0.433 | 1 | 0.015665428 | 0.546533753 | 0.822 | 0.557 | 1 | 0.346916538 | 0.03108545 |
| Erbin | 0.015692412 | 0.935304148 | 0.952 | 0.888 | 1 | 0.028279603 | 0.47015992 | 0.978 | 0.965 | 1 | 0.028279603 | 0.031138572 |
| Pip5k1c | 0.755153506 | 0.281454345 | 0.19 | 0.235 | 1 | 0.015700547 | -0.638855509 | 0.089 | 0.27 | 1 | 0.755153506 | 0.031154586 |
| Dtx2 | 0.198208877 | -0.785659851 | 0.048 | 0.149 | 1 | 0.015702036 | -1.421264074 | 0.067 | 0.225 | 1 | 0.198208877 | 0.031157517 |
| 1110002L01Rik | 0.069909802 | -0.555046923 | 0.048 | 0.228 | 1 | 0.015713363 | -1.064161343 | 0.067 | 0.232 | 1 | 0.069909802 | 0.031179816 |
| Alg13 | 0.785635025 | 1.108987149 | 0.19 | 0.175 | 1 | 0.015743013 | -0.62568012 | 0.067 | 0.242 | 1 | 0.785635025 | 0.031238183 |
| Zfp593 | 0.015774614 | -1.984439715 | 0.048 | 0.291 | 1 | 0.019436553 | -2.044847868 | 0.022 | 0.149 | 1 | 0.019436553 | 0.031300389 |
| Dusp12 | 0.015779431 | -2.02666795 | 0.048 | 0.291 | 1 | 0.01802296 | 0.152557869 | 0.089 | 0.27 | 1 | 0.01802296 | 0.031309872 |
| Ormdl1 | 0.253055697 | -0.574155746 | 0.048 | 0.134 | 1 | 0.015789136 | -1.960783604 | 0.022 | 0.156 | 1 | 0.253055697 | 0.031328974 |
| Hdac9 | 0.48332231 | -0.648156327 | 0.048 | 0.097 | 1 | 0.015795265 | -3.519867472 | 0.044 | 0.183 | 1 | 0.48332231 | 0.03134104 |
| Galk2 | 0.668589314 | 0.351843673 | 0.19 | 0.261 | 1 | 0.015802167 | -0.705198004 | 0.133 | 0.339 | 1 | 0.668589314 | 0.031354626 |

| | | | | | | | | | | | | |
|---|---|---|---|---|---|---|---|---|---|---|---|---|
| Zfp644 | 0.015805537 | 0.643009514 | 0.81 | 0.72 | 1 | 0.097118613 | 0.779958681 | 0.844 | 0.862 | 1 | 0.097118613 | 0.031361259 |
| Cdkl5 | 0.086690371 | 1.516902919 | 0.429 | 0.328 | 1 | 0.015828949 | -0.49236942 | 0.222 | 0.429 | 1 | 0.086690371 | 0.031407341 |
| Arfrp1 | 0.028233392 | -1.184209227 | 0.095 | 0.34 | 1 | 0.015831678 | -0.16020191 | 0.133 | 0.343 | 1 | 0.028233392 | 0.031412713 |
| Tmem127 | 0.098739401 | -0.611630451 | 0.286 | 0.485 | 1 | 0.015837952 | -0.416463087 | 0.356 | 0.609 | 1 | 0.098739401 | 0.031425064 |
| Rgcc | 0.015838511 | -1.2619361 | 0.429 | 0.724 | 1 | 0.065268057 | -0.892768525 | 0.689 | 0.713 | 1 | 0.065268057 | 0.031426163 |
| Atpsckmt | 0.072702841 | -1.370622352 | 0.048 | 0.209 | 1 | 0.015842192 | -0.486852415 | 0.067 | 0.235 | 1 | 0.072702841 | 0.031433409 |
| Cbr1 | 0.01587764 | -1.486099569 | 0.095 | 0.366 | 1 | 0.163266177 | -0.524522833 | 0.356 | 0.439 | 1 | 0.163266177 | 0.031503181 |
| Wdr45b | 0.015884667 | -0.559847909 | 0.238 | 0.604 | 1 | 0.907597772 | 0.19513454 | 0.778 | 0.768 | 1 | 0.907597772 | 0.031517012 |
| Fgfr1 | 0.322263686 | -0.283923199 | 1 | 0.985 | 1 | 0.015886105 | -0.33199867 | 1 | 0.979 | 1 | 0.322263686 | 0.031519842 |
| Ddx54 | 0.548418071 | 0.113056813 | 0.238 | 0.336 | 1 | 0.015899799 | -0.776359032 | 0.178 | 0.37 | 1 | 0.548418071 | 0.031546794 |
| Anks1b | 0.015928177 | 3.826928556 | 0.381 | 0.213 | 1 | 0.612158998 | 0.767464773 | 0.156 | 0.208 | 1 | 0.612158998 | 0.031602647 |
| Htra3 | 0.090210896 | -0.255838904 | 0.286 | 0.534 | 1 | 0.015931209 | -0.204452685 | 0.222 | 0.453 | 1 | 0.090210896 | 0.031608615 |
| Bltp1 | 0.015936082 | 1.136983861 | 0.905 | 0.922 | 1 | 0.02121099 | 0.526347082 | 1 | 0.955 | 1 | 0.02121099 | 0.031618205 |
| Pdk1 | 0.062111824 | -1.120644099 | 0.095 | 0.291 | 1 | 0.01594202 | -1.085111739 | 0.089 | 0.26 | 1 | 0.062111824 | 0.031629893 |
| Zzz3 | 0.762265948 | 0.196238343 | 0.619 | 0.631 | 1 | 0.015959765 | 0.643749456 | 0.933 | 0.869 | 1 | 0.762265948 | 0.031664816 |
| Atg9a | 0.785234425 | 0.918884266 | 0.143 | 0.183 | 1 | 0.015987283 | -0.38467202 | 0.111 | 0.301 | 1 | 0.785234425 | 0.031718973 |
| Exosc2 | 0.206751149 | -0.940938076 | 0.048 | 0.146 | 1 | 0.015987304 | -1.506751973 | 0.044 | 0.194 | 1 | 0.206751149 | 0.031719014 |
| Srcap | 0.016010771 | 0.775268958 | 0.762 | 0.619 | 1 | 0.110840063 | -0.2831391 | 0.822 | 0.837 | 1 | 0.110840063 | 0.031765196 |
| Cep135 | 0.875583017 | 0.267779408 | 0.095 | 0.108 | 1 | 0.016016926 | -1.819427754 | 0.022 | 0.159 | 1 | 0.875583017 | 0.031777311 |
| Cep290 | 0.114548814 | -0.345421798 | 0.143 | 0.347 | 1 | 0.016027556 | -1.689481582 | 0.111 | 0.27 | 1 | 0.114548814 | 0.03179823 |
| Cfl2 | 0.016053732 | -0.541667893 | 0.476 | 0.806 | 1 | 0.233400661 | -0.132727404 | 0.778 | 0.789 | 1 | 0.233400661 | 0.031849742 |
| Vwa5a | 0.586738849 | 0.549650456 | 0.476 | 0.511 | 1 | 0.01607851 | -0.460763186 | 0.378 | 0.647 | 1 | 0.586738849 | 0.031898501 |
| Rnd2 | 0.155312849 | -0.970084422 | 0.048 | 0.164 | 1 | 0.016078785 | -1.446210431 | 0.044 | 0.194 | 1 | 0.155312849 | 0.031899042 |
| Smad3 | 0.428127199 | 0.644916905 | 0.714 | 0.761 | 1 | 0.016088084 | 0.774588732 | 0.889 | 0.83 | 1 | 0.428127199 | 0.031917342 |
| Camk2g | 0.465829155 | 0.420015175 | 0.286 | 0.466 | 1 | 0.01611284 | -0.108934372 | 0.333 | 0.606 | 1 | 0.465829155 | 0.031966056 |
| Sdhc | 0.016113819 | -0.752492987 | 0.286 | 0.631 | 1 | 0.32982679 | -0.23159536 | 0.578 | 0.585 | 1 | 0.32982679 | 0.031967982 |
| B4galt6 | 0.154264321 | -1.063193826 | 0.095 | 0.224 | 1 | 0.01613677 | -0.577454964 | 0.111 | 0.301 | 1 | 0.154264321 | 0.032013144 |
| Rmdn3 | 0.516049957 | 0.914779867 | 0.286 | 0.272 | 1 | 0.016150754 | -0.992492463 | 0.111 | 0.28 | 1 | 0.516049957 | 0.032040661 |
| Mdfic | 0.016174631 | 2.233199176 | 0.476 | 0.34 | 1 | 0.20476797 | 0.117320689 | 0.356 | 0.522 | 1 | 0.20476797 | 0.032087644 |
| Dnajc2 | 0.016182418 | -0.669636055 | 0.143 | 0.444 | 1 | 0.248907847 | -0.557935513 | 0.4 | 0.46 | 1 | 0.248907847 | 0.032102966 |
| Pvt1 | 0.040651046 | 0.651403955 | 0.571 | 0.388 | 1 | 0.016211159 | 1.483027596 | 0.511 | 0.356 | 1 | 0.040651046 | 0.032159516 |
| Zfp248 | 0.945384708 | 0.783000837 | 0.19 | 0.231 | 1 | 0.01621842 | -0.806312255 | 0.089 | 0.273 | 1 | 0.945384708 | 0.032173803 |
| Dhx16 | 0.709680063 | 0.230828272 | 0.19 | 0.243 | 1 | 0.016243839 | -0.432404631 | 0.133 | 0.325 | 1 | 0.709680063 | 0.032223816 |
| Chmp1b2 | 0.135228095 | -1.133583154 | 0.048 | 0.172 | 1 | 0.016247869 | -1.731964913 | 0.022 | 0.159 | 1 | 0.135228095 | 0.032231744 |
| Fdx2 | 0.046025315 | -1.01272876 | 0.095 | 0.313 | 1 | 0.016295871 | -0.542995493 | 0.156 | 0.353 | 1 | 0.046025315 | 0.032326187 |
| Smg8 | 0.435865045 | 0.808701348 | 0.095 | 0.168 | 1 | 0.016299307 | -1.881712033 | 0.044 | 0.187 | 1 | 0.435865045 | 0.032332947 |
| Caprin1 | 0.016343374 | 0.594653051 | 1 | 0.869 | 1 | 0.685700478 | 0.197235261 | 0.978 | 0.955 | 1 | 0.685700478 | 0.032419642 |
| Creb1 | 0.20858263 | 0.667773027 | 0.476 | 0.425 | 1 | 0.016394982 | 0.834449646 | 0.778 | 0.578 | 1 | 0.20858263 | 0.032521168 |
| Slc25a16 | 0.052439766 | -1.427766259 | 0.095 | 0.287 | 1 | 0.016418709 | -0.745770713 | 0.133 | 0.315 | 1 | 0.052439766 | 0.032567843 |
| Gstcd | 0.706028584 | 0.819622634 | 0.19 | 0.257 | 1 | 0.01642637 | -1.001425588 | 0.067 | 0.225 | 1 | 0.706028584 | 0.032582914 |
| Zfp574 | 0.032408879 | -2.233118828 | 0 | 0.183 | 1 | 0.016426742 | -0.624355939 | 0.089 | 0.27 | 1 | 0.032408879 | 0.032583645 |
| Trappc12 | 0.697099069 | 0.951305743 | 0.143 | 0.194 | 1 | 0.016438354 | -0.935837246 | 0.067 | 0.232 | 1 | 0.697099069 | 0.032606489 |
| Ndufaf3 | 0.016542951 | -2.496153234 | 0 | 0.22 | 1 | 0.024391361 | -1.84048937 | 0.022 | 0.142 | 1 | 0.024391361 | 0.032812233 |
| Ezh1 | 0.978832835 | 0.226312791 | 0.286 | 0.317 | 1 | 0.016557465 | -0.751555641 | 0.178 | 0.384 | 1 | 0.978832835 | 0.032840781 |
| Traf2 | 0.604250818 | 0.866416846 | 0.143 | 0.108 | 1 | 0.016567954 | -1.204452685 | 0.044 | 0.194 | 1 | 0.604250818 | 0.03286141 |
| Smap2 | 0.057994289 | -0.614094491 | 0.333 | 0.612 | 1 | 0.016576241 | 0.689767239 | 0.867 | 0.754 | 1 | 0.057994289 | 0.03287771 |
| Dgat1 | 0.69831848 | 0.198038337 | 0.19 | 0.25 | 1 | 0.016585446 | -1.286553765 | 0.067 | 0.228 | 1 | 0.69831848 | 0.032895814 |
| Coq10b | 0.144703461 | -0.367829137 | 0.81 | 0.862 | 1 | 0.016597876 | -0.292419712 | 0.889 | 0.903 | 1 | 0.144703461 | 0.032920263 |
| Ehd1 | 0.01660558 | -0.853087778 | 0.333 | 0.687 | 1 | 0.0524247 | -0.358528318 | 0.489 | 0.657 | 1 | 0.0524247 | 0.032935415 |
| Desi2 | 0.056836765 | 0.550389352 | 0.857 | 0.896 | 1 | 0.016631806 | 0.474685576 | 1 | 0.965 | 1 | 0.056836765 | 0.032986995 |
| Oser1 | 0.15534499 | -0.487691655 | 0.429 | 0.66 | 1 | 0.016638457 | -0.563238462 | 0.578 | 0.709 | 1 | 0.15534499 | 0.033000076 |
| Rae1 | 0.075948804 | -0.298920886 | 0.19 | 0.414 | 1 | 0.016638986 | -0.587456356 | 0.2 | 0.429 | 1 | 0.075948804 | 0.033001116 |
| Cyth1 | 0.016650407 | 1.257280671 | 0.667 | 0.504 | 1 | 0.072442594 | 1.556808814 | 0.867 | 0.713 | 1 | 0.072442594 | 0.033023578 |
| Uhrf1bp1 | 0.87885171 | 0.786246497 | 0.095 | 0.116 | 1 | 0.016691439 | -1.16492432 | 0.044 | 0.194 | 1 | 0.87885171 | 0.033104274 |
| Gramd2 | 0.852353775 | 0.311579803 | 0.524 | 0.646 | 1 | 0.016699367 | -0.866391233 | 0.156 | 0.353 | 1 | 0.852353775 | 0.033119865 |
| Afap1 | 0.666071703 | 0.249745485 | 0.238 | 0.321 | 1 | 0.01670214 | 0.574890934 | 0.711 | 0.474 | 1 | 0.666071703 | 0.033125318 |
| Zfp729a | 0.986121976 | 0.8372705 | 0.143 | 0.153 | 1 | 0.016737499 | -0.901889915 | 0.044 | 0.194 | 1 | 0.986121976 | 0.033194854 |
| Luzp1 | 0.016779196 | 0.876171413 | 0.952 | 0.959 | 1 | 0.243845279 | 0.511140217 | 1 | 0.983 | 1 | 0.243845279 | 0.033276851 |
| B3gnt2 | 0.016803157 | -1.22709504 | 0.476 | 0.664 | 1 | 0.28828561 | 0.352682913 | 0.867 | 0.692 | 1 | 0.28828561 | 0.033323968 |
| Huwe1 | 0.016808472 | 0.573252774 | 1 | 0.989 | 1 | 0.021125211 | 0.413045593 | 1 | 0.997 | 1 | 0.021125211 | 0.033334419 |
| Zdhhc20 | 0.016831936 | 0.839030958 | 0.905 | 0.776 | 1 | 0.667831028 | 0.242019497 | 0.956 | 0.896 | 1 | 0.667831028 | 0.033380558 |
| Klhl22 | 0.136436573 | -0.633656758 | 0.095 | 0.257 | 1 | 0.016873603 | -0.414959497 | 0.133 | 0.325 | 1 | 0.136436573 | 0.033462487 |
| Senp6 | 0.068438441 | 0.36444371 | 1 | 0.985 | 1 | 0.016929534 | -0.120589035 | 1 | 0.997 | 1 | 0.068438441 | 0.033572458 |
| Gm43305 | 0.016974715 | -2.326228232 | 0.048 | 0.287 | 1 | 0.107068573 | -0.575661682 | 0.178 | 0.308 | 1 | 0.107068573 | 0.033661288 |
| Tbc1d9b | 0.266658603 | 0.321255353 | 0.19 | 0.369 | 1 | 0.016983326 | -0.822455447 | 0.178 | 0.377 | 1 | 0.266658603 | 0.033678218 |
| Ift74 | 0.016996513 | -0.911190733 | 0.238 | 0.545 | 1 | 0.094787265 | 0.27368165 | 0.356 | 0.616 | 1 | 0.094787265 | 0.033704144 |
| Rab11fip3 | 0.017022732 | 1.744161096 | 0.619 | 0.552 | 1 | 0.173771977 | 0.316110738 | 0.4 | 0.616 | 1 | 0.173771977 | 0.033755691 |
| Srgap1 | 0.017030132 | 1.067202811 | 0.905 | 0.713 | 1 | 0.443608484 | 0.281541407 | 0.889 | 0.941 | 1 | 0.443608484 | 0.033770239 |
| Smim1 | 0.06518747 | -1.880817084 | 0 | 0.142 | 1 | 0.017033331 | -0.432404631 | 0.022 | 0.159 | 1 | 0.06518747 | 0.033776527 |
| Glrx3 | 0.017053234 | -0.550601018 | 0.429 | 0.769 | 1 | 0.062325489 | -0.245455659 | 0.822 | 0.803 | 1 | 0.062325489 | 0.033815656 |
| Mtch2 | 0.017059454 | -0.715106571 | 0.476 | 0.769 | 1 | 0.608855598 | 0.123645178 | 0.733 | 0.761 | 1 | 0.608855598 | 0.033827884 |
| Ten1 | 0.125188376 | -0.370622352 | 0.048 | 0.19 | 1 | 0.017074757 | -0.731964913 | 0.022 | 0.159 | 1 | 0.125188376 | 0.033857967 |
| Coa6 | 0.019271088 | -2.648156327 | 0 | 0.213 | 1 | 0.017106965 | -0.852980314 | 0.022 | 0.159 | 1 | 0.019271088 | 0.033921282 |
| Zeb2 | 0.299893699 | -0.238765391 | 0.095 | 0.201 | 1 | 0.017121969 | 0.207987703 | 0.289 | 0.135 | 1 | 0.299893699 | 0.033950777 |
| Arfgap2 | 0.179034815 | 0.877305162 | 0.429 | 0.362 | 1 | 0.017124569 | -0.564854927 | 0.133 | 0.349 | 1 | 0.179034815 | 0.033955887 |
| Msrb2 | 0.209667009 | -0.818081329 | 0.048 | 0.146 | 1 | 0.01714677 | -1.223818009 | 0.067 | 0.228 | 1 | 0.209667009 | 0.033999529 |
| 4930590J08Rik | 0.420968517 | 1.673771768 | 0.095 | 0.056 | 1 | 0.017215648 | -2.124282336 | 0.022 | 0.152 | 1 | 0.420968517 | 0.034134917 |
| Nol6 | 0.747755005 | 0.673771768 | 0.095 | 0.075 | 1 | 0.017222781 | -2.486852415 | 0 | 0.114 | 1 | 0.747755005 | 0.034148938 |
| Gm48678 | 0.313004724 | 2.088809267 | 0.095 | 0.049 | 1 | 0.017238354 | -2.638855509 | 0 | 0.114 | 1 | 0.313004724 | 0.034179547 |
| Zbtb49 | 0.587082947 | 0.55829455 | 0.095 | 0.063 | 1 | 0.017238354 | -2.564854927 | 0 | 0.114 | 1 | 0.587082947 | 0.034179547 |
| Sirt4 | 0.268100354 | -0.326228232 | 0 | 0.056 | 1 | 0.017238388 | -2.526380779 | 0 | 0.114 | 1 | 0.268100354 | 0.034179613 |
| Decr2 | 0.183905319 | -0.516052791 | 0.095 | 0.228 | 1 | 0.017244104 | -1.638855509 | 0.044 | 0.187 | 1 | 0.183905319 | 0.034190848 |
| Pex12 | 0.128113406 | -1.280424543 | 0 | 0.101 | 1 | 0.017252859 | -2.602329633 | 0 | 0.114 | 1 | 0.128113406 | 0.034208056 |
| Gm56815 | 0.285465396 | -0.326228232 | 0 | 0.052 | 1 | 0.017252859 | -2.638855509 | 0 | 0.114 | 1 | 0.285465396 | 0.034208056 |
| Dhrs13 | 0.291331722 | 0.832469514 | 0.048 | 0.134 | 1 | 0.017252859 | -2.602329633 | 0 | 0.114 | 1 | 0.291331722 | 0.034208056 |
| Zfp36l1 | 0.017257832 | -0.511308425 | 0.762 | 0.869 | 1 | 0.438161542 | -0.63979202 | 0.911 | 0.827 | 1 | 0.438161542 | 0.034217831 |
| Cmc2 | 0.350203145 | -0.159118246 | 0.095 | 0.183 | 1 | 0.017266293 | -2.638855509 | 0 | 0.114 | 1 | 0.350203145 | 0.034234462 |
| Alx3 | 0.172931902 | -0.984439715 | 0.095 | 0.224 | 1 | 0.017284684 | -1.624355939 | 0.044 | 0.19 | 1 | 0.172931902 | 0.034270607 |
| Elf1 | 0.017286509 | 0.7118049 | 0.952 | 0.922 | 1 | 0.903444779 | 0.29657249 | 1 | 0.965 | 1 | 0.903444779 | 0.034274196 |
| Tanc1 | 0.017298335 | 0.922290394 | 1 | 0.944 | 1 | 0.026374564 | 0.529826326 | 0.911 | 0.83 | 1 | 0.026374564 | 0.034297438 |
| Mast3 | 0.106613525 | -1.370622352 | 0 | 0.112 | 1 | 0.017300873 | -2.776359032 | 0 | 0.114 | 1 | 0.106613525 | 0.034302427 |
| C130050O18Rik | 0.872207635 | 0.586308926 | 0.048 | 0.056 | 1 | 0.017301714 | -2.84048937 | 0 | 0.114 | 1 | 0.872207635 | 0.034304079 |
| Foxd1 | 0.277513527 | -1.067695219 | 0.095 | 0.187 | 1 | 0.017303343 | -0.901889915 | 0.089 | 0.263 | 1 | 0.277513527 | 0.034307281 |
| Zfp825 | 0.299864508 | -0.574155746 | 0.048 | 0.123 | 1 | 0.017310254 | -2.80878051 | 0 | 0.114 | 1 | 0.299864508 | 0.034320864 |
| Il11 | 0.710182109 | 0.896164189 | 0.095 | 0.075 | 1 | 0.017320108 | -3.124282336 | 0 | 0.114 | 1 | 0.710182109 | 0.03434023 |
| Gcc2 | 0.017322568 | 0.434805122 | 0.238 | 0.537 | 1 | 0.04147245 | 0.670248546 | 0.778 | 0.599 | 1 | 0.04147245 | 0.034345066 |
| Nudt22 | 0.32017459 | -0.413691074 | 0.048 | 0.119 | 1 | 0.017329191 | -3.223818009 | 0 | 0.114 | 1 | 0.32017459 | 0.034358081 |
| Tfap2a | 0.938801699 | 0.793353383 | 0.19 | 0.209 | 1 | 0.017353368 | -0.71547679 | 0.089 | 0.26 | 1 | 0.938801699 | 0.034405597 |

| | | | | | | | | | | | | |
|---|---|---|---|---|---|---|---|---|---|---|---|---|
| Mkx | 0.637754157 | 2.881213848 | 0.19 | 0.153 | 1 | 0.017359682 | -3.506751973 | 0 | 0.114 | 1 | 0.637754157 | 0.034418005 |
| Zbed6 | 0.670026774 | 0.471333953 | 0.524 | 0.664 | 1 | 0.017369324 | 0.706447095 | 0.822 | 0.806 | 1 | 0.670026774 | 0.034436955 |
| Srebf2 | 0.348814191 | 0.756489566 | 0.667 | 0.754 | 1 | 0.017385165 | 0.69371983 | 0.867 | 0.737 | 1 | 0.348814191 | 0.034468087 |
| Ino80dos | 0.950410129 | 0.586308926 | 0.19 | 0.201 | 1 | 0.017390809 | -0.17873012 | 0.133 | 0.353 | 1 | 0.950410129 | 0.034479177 |
| B4galt2 | 0.597928198 | 0.214340149 | 0.095 | 0.142 | 1 | 0.017459228 | -2.044847868 | 0.022 | 0.152 | 1 | 0.597928198 | 0.03461363 |
| Pank4 | 0.720639083 | 0.351843673 | 0.095 | 0.127 | 1 | 0.017468188 | -1.506751973 | 0.044 | 0.19 | 1 | 0.720639083 | 0.034631239 |
| Plcl2 | 0.048072939 | -3.094412557 | 0 | 0.16 | 1 | 0.017514317 | -1.137106376 | 0.044 | 0.197 | 1 | 0.048072939 | 0.034721883 |
| Myg1 | 0.087955272 | -1.303508156 | 0.048 | 0.198 | 1 | 0.017524082 | -0.602329633 | 0.067 | 0.232 | 1 | 0.087955272 | 0.034741072 |
| Srf | 0.090234907 | -0.785659851 | 0.095 | 0.272 | 1 | 0.017525836 | -0.316927414 | 0.111 | 0.298 | 1 | 0.090234907 | 0.034744517 |
| Dennd4a | 0.412535161 | 3.09377552 | 0.429 | 0.44 | 1 | 0.017541595 | 0.354124064 | 0.822 | 0.533 | 1 | 0.412535161 | 0.034775482 |
| Ttyh3 | 0.267584172 | 1.226312791 | 0.381 | 0.306 | 1 | 0.017544613 | -0.168404889 | 0.133 | 0.339 | 1 | 0.267584172 | 0.034781413 |
| Usp6nl | 0.017544866 | 0.933203937 | 0.762 | 0.657 | 1 | 0.055470807 | 0.529650216 | 0.933 | 0.716 | 1 | 0.055470807 | 0.034781909 |
| Map7d3 | 0.412174704 | 1.060240115 | 0.333 | 0.306 | 1 | 0.01754819 | -1.328154669 | 0.067 | 0.225 | 1 | 0.412174704 | 0.034788441 |
| Nrd1 | 0.548620232 | 0.360749226 | 0.762 | 0.851 | 1 | 0.017552941 | -0.180637186 | 0.867 | 0.889 | 1 | 0.548620232 | 0.034797777 |
| Mtx2 | 0.017594529 | -1.660128969 | 0.048 | 0.302 | 1 | 0.907194399 | 0.75542343 | 0.333 | 0.398 | 1 | 0.907194399 | 0.03487949 |
| Pgghg | 0.061234946 | -1.849790188 | 0 | 0.146 | 1 | 0.017595421 | -1.901889915 | 0.022 | 0.152 | 1 | 0.061234946 | 0.034881243 |
| Hps1 | 0.081776276 | -1.475975352 | 0.048 | 0.201 | 1 | 0.017595421 | -1.901889915 | 0.022 | 0.152 | 1 | 0.081776276 | 0.034881243 |
| 1700061N14Rik | 0.86904731 | 0.489347197 | 0.19 | 0.183 | 1 | 0.017601807 | -1.728741012 | 0.089 | 0.242 | 1 | 0.86904731 | 0.03489379 |
| Mxd1 | 0.606031648 | 1.274600674 | 0.238 | 0.332 | 1 | 0.017611444 | 0.731616438 | 0.644 | 0.422 | 1 | 0.606031648 | 0.034912725 |
| Naa30 | 0.474872845 | -0.118786152 | 0.19 | 0.276 | 1 | 0.017619728 | -0.3932783 | 0.2 | 0.436 | 1 | 0.474872845 | 0.034929001 |
| Iqcn | 0.017622072 | 2.126283972 | 0.238 | 0.086 | 1 | 0.241450513 | -0.564854927 | 0.111 | 0.187 | 1 | 0.241450513 | 0.034933607 |
| Atxn2l | 0.333085379 | 0.482345697 | 0.571 | 0.649 | 1 | 0.017661607 | -0.418013539 | 0.644 | 0.81 | 1 | 0.333085379 | 0.035011283 |
| Orc2 | 0.081674388 | -0.179679569 | 0.143 | 0.362 | 1 | 0.01768873 | 0.407794585 | 0.2 | 0.429 | 1 | 0.081674388 | 0.035064568 |
| Zfp523 | 0.017688991 | -2.434752689 | 0 | 0.216 | 1 | 0.177060957 | -0.638855509 | 0.111 | 0.201 | 1 | 0.177060957 | 0.035065082 |
| Mybbp1a | 0.449631743 | 0.263752343 | 0.381 | 0.563 | 1 | 0.017706325 | -0.400140782 | 0.489 | 0.647 | 1 | 0.449631743 | 0.035099137 |
| Zbtb34 | 0.893446897 | 0.503846766 | 0.095 | 0.108 | 1 | 0.017736021 | -1.053893008 | 0.044 | 0.197 | 1 | 0.893446897 | 0.035157476 |
| Slc17a5 | 0.932740462 | 1.198686885 | 0.286 | 0.34 | 1 | 0.017816996 | -0.273338145 | 0.267 | 0.516 | 1 | 0.932740462 | 0.035316547 |
| Obp1b | 0.677100402 | 0.8372705 | 0.19 | 0.168 | 1 | 0.017828127 | -1.498257179 | 0.067 | 0.225 | 1 | 0.677100402 | 0.035338411 |
| Nsfl1c | 0.017830554 | -0.739856161 | 0.381 | 0.701 | 1 | 0.060300635 | -0.349348892 | 0.667 | 0.775 | 1 | 0.060300635 | 0.03534318 |
| Slc44a2 | 0.033803485 | -0.672540486 | 0.571 | 0.869 | 1 | 0.017889778 | -0.464309847 | 0.511 | 0.803 | 1 | 0.033803485 | 0.035459511 |
| Pias3 | 0.567930682 | 0.150209812 | 0.095 | 0.146 | 1 | 0.017896282 | -0.674479418 | 0.111 | 0.301 | 1 | 0.567930682 | 0.035472286 |
| Pcnp | 0.017926431 | -0.512831975 | 0.714 | 0.933 | 1 | 0.144829253 | -0.116333583 | 1 | 0.948 | 1 | 0.144829253 | 0.035531506 |
| Ets2 | 0.818983317 | -0.161734195 | 0.714 | 0.66 | 1 | 0.018028941 | -0.508745021 | 1 | 0.872 | 1 | 0.818983317 | 0.03573284 |
| Tnrc6c | 0.156362717 | -0.549316767 | 0.667 | 0.731 | 1 | 0.018033425 | 0.527565252 | 0.978 | 0.917 | 1 | 0.156362717 | 0.035741646 |
| Pank3 | 0.018057001 | -0.569084756 | 0.238 | 0.597 | 1 | 0.023791727 | -0.406121143 | 0.511 | 0.689 | 1 | 0.023791727 | 0.035787946 |
| St3gal6 | 0.018072797 | -0.496153234 | 0.381 | 0.743 | 1 | 0.08502914 | -0.2139391 | 0.756 | 0.834 | 1 | 0.08502914 | 0.035818968 |
| Gsr | 0.018078128 | -2.126203624 | 0.19 | 0.466 | 1 | 0.020385566 | 0.698502627 | 0.889 | 0.599 | 1 | 0.020385566 | 0.035829437 |
| Parp2 | 0.018092308 | -1.359651234 | 0.095 | 0.362 | 1 | 0.034725293 | -0.288913038 | 0.111 | 0.273 | 1 | 0.034725293 | 0.035857284 |
| Crcp | 0.055451982 | -1.574155746 | 0.048 | 0.224 | 1 | 0.018094956 | -1.334849322 | 0.067 | 0.221 | 1 | 0.055451982 | 0.035862485 |
| Mindy3 | 0.695929058 | 1.111529343 | 0.333 | 0.388 | 1 | 0.018228764 | 1.192232263 | 0.6 | 0.45 | 1 | 0.695929058 | 0.03612524 |
| Bach2 | 0.192548412 | 0.95415974 | 0.619 | 0.541 | 1 | 0.018271531 | 0.593574435 | 0.978 | 0.689 | 1 | 0.192548412 | 0.036209214 |
| Phldb1 | 0.018307781 | 0.696491844 | 0.857 | 0.862 | 1 | 0.339847115 | 0.333724023 | 0.822 | 0.91 | 1 | 0.339847115 | 0.036280388 |
| Mthfsl | 0.022970386 | -1.769171728 | 0.048 | 0.272 | 1 | 0.018370163 | -0.62568012 | 0.111 | 0.287 | 1 | 0.022970386 | 0.036402863 |
| Tgfb2 | 0.018370671 | -1.491801209 | 0.619 | 0.828 | 1 | 0.487744512 | 0.774401984 | 0.733 | 0.924 | 1 | 0.487744512 | 0.036403861 |
| Taf8 | 0.661102457 | 0.327996931 | 0.143 | 0.198 | 1 | 0.018374733 | -1.583713955 | 0.067 | 0.215 | 1 | 0.661102457 | 0.036411835 |
| Polr2h | 0.048737707 | -1.295854583 | 0.19 | 0.399 | 1 | 0.018384755 | -0.58163064 | 0.222 | 0.415 | 1 | 0.048737707 | 0.03643151 |
| Hmgcl | 0.2215174 | -0.437736548 | 0.19 | 0.336 | 1 | 0.018419952 | -0.37003875 | 0.2 | 0.426 | 1 | 0.2215174 | 0.036500609 |
| Eya4 | 0.08294284 | 0.923000374 | 0.714 | 0.705 | 1 | 0.018423844 | 0.800098241 | 0.867 | 0.761 | 1 | 0.08294284 | 0.036508249 |
| Eif2ak2 | 0.032453557 | -1.101015292 | 0.095 | 0.343 | 1 | 0.018446402 | -0.124282336 | 0.2 | 0.429 | 1 | 0.032453557 | 0.036552534 |
| Dtnbp1 | 0.118402663 | -0.455511249 | 0.286 | 0.507 | 1 | 0.018449916 | 1.000262762 | 0.733 | 0.623 | 1 | 0.118402663 | 0.036559432 |
| Fhod1 | 0.018490857 | 2.179007076 | 0.238 | 0.093 | 1 | 0.366628438 | -0.539319835 | 0.044 | 0.083 | 1 | 0.366628438 | 0.036639803 |
| Palm | 0.047943526 | -2.326228232 | 0 | 0.16 | 1 | 0.018527333 | -0.719025857 | 0.044 | 0.197 | 1 | 0.047943526 | 0.036711404 |
| Stk32b | 0.018537117 | 1.523409033 | 0.714 | 0.657 | 1 | 0.241418175 | 0.875717664 | 0.556 | 0.571 | 1 | 0.241418175 | 0.036730609 |
| Spire1 | 0.600775464 | 0.366424656 | 0.857 | 0.933 | 1 | 0.018552556 | 0.722703395 | 0.867 | 0.903 | 1 | 0.600775464 | 0.036760914 |
| 2810432F15Rik | 0.136109171 | -1.184209227 | 0 | 0.097 | 1 | 0.01856941 | 2.098110085 | 0.156 | 0.062 | 1 | 0.136109171 | 0.036793998 |
| Pdia5 | 0.067866888 | -0.626351957 | 0.19 | 0.407 | 1 | 0.018613314 | -0.129557278 | 0.244 | 0.491 | 1 | 0.067866888 | 0.036880173 |
| Yy1 | 0.092423501 | -0.504146025 | 0.714 | 0.776 | 1 | 0.018637624 | -0.377759558 | 0.956 | 0.92 | 1 | 0.092423501 | 0.036927888 |
| Hivep1 | 0.914096891 | 0.46732089 | 0.381 | 0.459 | 1 | 0.018643423 | 0.911583117 | 0.844 | 0.599 | 1 | 0.914096891 | 0.036939268 |
| Sesn2 | 0.035982939 | 1.327996931 | 0.476 | 0.302 | 1 | 0.018700887 | -0.786412697 | 0.111 | 0.291 | 1 | 0.035982939 | 0.037052051 |
| Rad17 | 0.284446461 | -0.488115915 | 0.286 | 0.418 | 1 | 0.018707128 | -0.174908409 | 0.222 | 0.464 | 1 | 0.284446461 | 0.037064299 |
| Amer1 | 0.840120592 | 0.351843673 | 0.095 | 0.112 | 1 | 0.018717232 | -2.425451871 | 0.022 | 0.149 | 1 | 0.840120592 | 0.037084128 |
| Vangl2 | 0.083678262 | -1.818081329 | 0 | 0.127 | 1 | 0.018746187 | -1.017367132 | 0.067 | 0.221 | 1 | 0.083678262 | 0.037140955 |
| Dhrs7b | 0.168025702 | -0.911190733 | 0.143 | 0.272 | 1 | 0.018752496 | -1.017367132 | 0.044 | 0.19 | 1 | 0.168025702 | 0.037153336 |
| Mrps6 | 0.018767141 | -0.677302673 | 0.524 | 0.836 | 1 | 0.414579282 | 0.104396348 | 0.8 | 0.799 | 1 | 0.414579282 | 0.037182077 |
| Wapl | 0.018779332 | 0.743504389 | 0.905 | 0.963 | 1 | 0.3668357 | 0.289345593 | 1 | 0.983 | 1 | 0.3668357 | 0.037206001 |
| Tmem164 | 0.061125086 | -0.639111188 | 0.238 | 0.515 | 1 | 0.01878379 | 0.665150678 | 0.889 | 0.789 | 1 | 0.061125086 | 0.037214749 |
| Xrn2 | 0.025935706 | 1.125444269 | 0.857 | 0.832 | 1 | 0.0187998 | 0.557114429 | 0.956 | 0.889 | 1 | 0.025935706 | 0.037246167 |
| Rilpl1 | 0.018806597 | -0.423089772 | 0.095 | 0.366 | 1 | 0.031063552 | -0.316927414 | 0.2 | 0.394 | 1 | 0.031063552 | 0.037259505 |
| Pfdn2 | 0.051436235 | -0.452337001 | 0.524 | 0.81 | 1 | 0.018812442 | -0.138757658 | 0.733 | 0.865 | 1 | 0.051436235 | 0.037270975 |
| Anks1 | 0.443385731 | 0.613075836 | 0.476 | 0.522 | 1 | 0.018864835 | 0.743193579 | 0.822 | 0.616 | 1 | 0.443385731 | 0.037373788 |
| Lrrc57 | 0.032646202 | -1.666078235 | 0.048 | 0.254 | 1 | 0.018891876 | -0.361321533 | 0.089 | 0.249 | 1 | 0.032646202 | 0.037426848 |
| Wdfy3 | 0.260016564 | -0.152198833 | 0.524 | 0.743 | 1 | 0.018898077 | 0.545569062 | 0.956 | 0.896 | 1 | 0.260016564 | 0.037439017 |
| Snhg5 | 0.129901144 | 0.125335143 | 0.19 | 0.418 | 1 | 0.018919479 | -0.21874702 | 0.089 | 0.273 | 1 | 0.129901144 | 0.03748101 |
| Steap1 | 0.01893698 | -1.636083495 | 0.19 | 0.459 | 1 | 0.266441689 | -0.442458296 | 0.111 | 0.183 | 1 | 0.266441689 | 0.037515352 |
| Tmem181a | 0.931057318 | 1.019767623 | 0.238 | 0.299 | 1 | 0.018950506 | 0.151352107 | 0.111 | 0.301 | 1 | 0.931057318 | 0.037541891 |
| Hdac7 | 0.13495007 | 2.295260144 | 0.143 | 0.063 | 1 | 0.019037456 | -0.618096949 | 0.022 | 0.156 | 1 | 0.13495007 | 0.037712487 |
| Prcp | 0.100534095 | -1.818081329 | 0 | 0.116 | 1 | 0.019071841 | -0.597035333 | 0.022 | 0.156 | 1 | 0.100534095 | 0.037779947 |
| Pmpca | 0.043769728 | -0.413691074 | 0.095 | 0.321 | 1 | 0.019073588 | -0.271123724 | 0.133 | 0.332 | 1 | 0.043769728 | 0.037783374 |
| A430005L14Rik | 0.025829079 | -1.381510668 | 0.095 | 0.34 | 1 | 0.019143551 | -0.746915255 | 0.089 | 0.256 | 1 | 0.025829079 | 0.037920626 |
| Larp4 | 0.458091824 | 0.311201688 | 0.905 | 0.899 | 1 | 0.019145128 | -0.327294129 | 1 | 0.952 | 1 | 0.458091824 | 0.037923721 |
| Trappc13 | 0.019162761 | -1.337455488 | 0.095 | 0.358 | 1 | 0.019516598 | -0.373758636 | 0.222 | 0.474 | 1 | 0.019516598 | 0.037958311 |
| Retsat | 0.163379068 | -1.02666795 | 0 | 0.086 | 1 | 0.019181919 | -2.404390255 | 0 | 0.111 | 1 | 0.163379068 | 0.037995892 |
| Cdpf1 | 0.522701208 | 0.150209812 | 0.095 | 0.153 | 1 | 0.019199257 | -2.446210431 | 0 | 0.111 | 1 | 0.522701208 | 0.038029903 |
| Pqlc3 | 0.019202759 | -2.611630451 | 0 | 0.213 | 1 | 0.029738827 | -0.80878051 | 0.067 | 0.204 | 1 | 0.029738827 | 0.038036772 |
| Hps3 | 0.396220964 | -0.496153234 | 0.048 | 0.104 | 1 | 0.019212182 | -2.223818009 | 0.022 | 0.149 | 1 | 0.396220964 | 0.038055256 |
| Zfp580 | 0.144697129 | -1.107587946 | 0.048 | 0.168 | 1 | 0.019212886 | -2.223818009 | 0.022 | 0.149 | 1 | 0.144697129 | 0.038056638 |
| Nrtn | 0.163569936 | -1.184209227 | 0 | 0.086 | 1 | 0.019215445 | -2.564854927 | 0 | 0.111 | 1 | 0.163569936 | 0.038061656 |
| Prr5 | 0.208952089 | -1.081115734 | 0 | 0.071 | 1 | 0.019215482 | -2.486852415 | 0 | 0.111 | 1 | 0.208952089 | 0.038061729 |
| Nagpa | 0.088612257 | -1.455511249 | 0 | 0.123 | 1 | 0.019230481 | -2.638855509 | 0 | 0.111 | 1 | 0.088612257 | 0.03809115 |
| Brf2 | 0.499824268 | 0.815790773 | 0.048 | 0.097 | 1 | 0.019230481 | -2.564854927 | 0 | 0.111 | 1 | 0.499824268 | 0.03809115 |
| Tapbpl | 0.044803144 | -2.133583154 | 0 | 0.164 | 1 | 0.019230593 | -2.638855509 | 0 | 0.111 | 1 | 0.044803144 | 0.03809137 |
| Gimap6 | 0.50008556 | 0.544488751 | 0.048 | 0.097 | 1 | 0.019230593 | -2.602329633 | 0 | 0.111 | 1 | 0.50008556 | 0.03809137 |
| Sfrp1 | 0.146611503 | -1.63000898 | 0.048 | 0.164 | 1 | 0.019241991 | 0.361144491 | 0.422 | 0.228 | 1 | 0.146611503 | 0.038113727 |
| Alg11 | 0.144697129 | -1.081115734 | 0.048 | 0.168 | 1 | 0.019246321 | -1.16492432 | 0.044 | 0.194 | 1 | 0.144697129 | 0.03812222 |
| Mbtd1 | 0.019251294 | 0.84737109 | 0.81 | 0.623 | 1 | 0.05636583 | 0.624668029 | 0.978 | 0.844 | 1 | 0.05636583 | 0.038131977 |
| Tmem204 | 0.032808168 | -2.801961663 | 0 | 0.183 | 1 | 0.019257468 | -2.743192169 | 0 | 0.111 | 1 | 0.032808168 | 0.038144086 |
| Zscan12 | 0.893461944 | 0.544488751 | 0.095 | 0.108 | 1 | 0.019257841 | -2.602329633 | 0 | 0.111 | 1 | 0.893461944 | 0.038144818 |
| Snx22 | 0.173729397 | -0.970084422 | 0 | 0.082 | 1 | 0.01926867 | -2.674479418 | 0 | 0.111 | 1 | 0.173729397 | 0.038166059 |

| | | | | | | | | | | | | |
|---|---|---|---|---|---|---|---|---|---|---|---|---|
| Ggct | 0.043222517 | -1.509450056 | 0.048 | 0.25 | 1 | 0.019269231 | -2.709244837 | 0 | 0.111 | 1 | 0.043222517 | 0.038167158 |
| Tyms | 0.828454643 | -0.133583154 | 0.095 | 0.112 | 1 | 0.019279091 | -2.776359032 | 0 | 0.111 | 1 | 0.828454643 | 0.038186499 |
| Nsg1 | 0.062579179 | -1.374591254 | 0.143 | 0.362 | 1 | 0.019291059 | -1.071814916 | 0.044 | 0.194 | 1 | 0.062579179 | 0.038209973 |
| Rab15 | 0.545989347 | 0.304537958 | 0.048 | 0.09 | 1 | 0.019297809 | -2.80878051 | 0 | 0.111 | 1 | 0.545989347 | 0.038223212 |
| Tmem62 | 0.729887791 | 1.088809267 | 0.048 | 0.071 | 1 | 0.019305096 | -2.84048937 | 0 | 0.111 | 1 | 0.729887791 | 0.038237505 |
| Car12 | 0.692474166 | 1.537994989 | 0.238 | 0.209 | 1 | 0.019328647 | -3.526380779 | 0 | 0.111 | 1 | 0.692474166 | 0.038283697 |
| Flrt3 | 0.341827059 | 0.351843673 | 0.048 | 0.123 | 1 | 0.019341736 | -3.339295227 | 0 | 0.111 | 1 | 0.341827059 | 0.038309369 |
| Arl5b | 0.019353133 | -0.729076323 | 0.19 | 0.507 | 1 | 0.181830134 | -0.451092652 | 0.667 | 0.657 | 1 | 0.181830134 | 0.038331721 |
| Tex14 | 0.019354666 | 1.406345593 | 0.857 | 0.672 | 1 | 0.677373591 | 0.444378353 | 0.667 | 0.675 | 1 | 0.677373591 | 0.03833473 |
| Arhgef18 | 0.595614272 | 0.291302131 | 0.143 | 0.216 | 1 | 0.019379007 | 0.770055888 | 0.533 | 0.349 | 1 | 0.595614272 | 0.038382469 |
| Bace2 | 0.044378066 | -2.04047375 | 0.048 | 0.235 | 1 | 0.019381588 | -2.098287127 | 0.022 | 0.149 | 1 | 0.044378066 | 0.03838753 |
| Coq5 | 0.264834949 | 0.614878079 | 0.095 | 0.216 | 1 | 0.01942859 | -1.167783975 | 0.067 | 0.218 | 1 | 0.264834949 | 0.038479709 |
| Gm27017 | 0.019435864 | 3.143257051 | 0.143 | 0.037 | 1 | 0.492711267 | -0.124282336 | 0.022 | 0.045 | 1 | 0.492711267 | 0.038493976 |
| Mad2l1bp | 0.18415515 | -0.940938076 | 0.048 | 0.153 | 1 | 0.019436809 | -2.149817428 | 0.022 | 0.149 | 1 | 0.18415515 | 0.038495828 |
| Gm45051 | 0.065014733 | -1.718545655 | 0 | 0.142 | 1 | 0.019437121 | -1.989352756 | 0.022 | 0.149 | 1 | 0.065014733 | 0.038496441 |
| Terf1 | 0.034078204 | -1.486099569 | 0.095 | 0.321 | 1 | 0.019449393 | -0.776359032 | 0.2 | 0.405 | 1 | 0.034078204 | 0.038520506 |
| Psmg1 | 0.019478453 | -0.926141075 | 0.048 | 0.302 | 1 | 0.269184567 | 0.361144491 | 0.2 | 0.322 | 1 | 0.269184567 | 0.038577496 |
| Gm5617 | 0.01947863 | -1.940938076 | 0.048 | 0.28 | 1 | 0.084091314 | -0.404390255 | 0.067 | 0.18 | 1 | 0.084091314 | 0.038577842 |
| Stx7 | 0.019493408 | -0.892575055 | 0.381 | 0.646 | 1 | 0.163003843 | -0.344112243 | 0.667 | 0.682 | 1 | 0.163003843 | 0.038606822 |
| Vill | 0.098214867 | -0.174225139 | 0.286 | 0.549 | 1 | 0.019536368 | -0.397492895 | 0.311 | 0.581 | 1 | 0.098214867 | 0.038691067 |
| Ptcd1 | 0.457614701 | 0.908237021 | 0.143 | 0.093 | 1 | 0.019543334 | -1.989352756 | 0.022 | 0.149 | 1 | 0.457614701 | 0.038704726 |
| Rab2b | 0.879748048 | 0.552756367 | 0.381 | 0.425 | 1 | 0.019571928 | -0.654797053 | 0.2 | 0.405 | 1 | 0.879748048 | 0.038760796 |
| Il17ra | 0.182233046 | -0.34303652 | 0.286 | 0.459 | 1 | 0.019573149 | -0.396654606 | 0.067 | 0.235 | 1 | 0.182233046 | 0.038763189 |
| Serpind1 | 0.425639612 | 0.503846766 | 0 | 0.03 | 1 | 0.019575332 | 3.683072586 | 0.067 | 0.014 | 1 | 0.425639612 | 0.03876747 |
| Colgalt2 | 0.019639414 | -1.403471231 | 0.381 | 0.634 | 1 | 0.335387668 | -0.238539515 | 0.2 | 0.298 | 1 | 0.335387668 | 0.038893121 |
| Tigd2 | 0.019643313 | -1.940938076 | 0.048 | 0.28 | 1 | 0.198916721 | -0.375821103 | 0.156 | 0.246 | 1 | 0.198916721 | 0.038900765 |
| Olfm1 | 0.22922422 | 0.137718867 | 0.143 | 0.306 | 1 | 0.019676468 | -0.316927414 | 0.067 | 0.218 | 1 | 0.22922422 | 0.038965772 |
| Vsig8 | 0.019692488 | 4.258734268 | 0.048 | 0.004 | 1 | 0.293637566 | -0.776359032 | 0 | 0.024 | 1 | 0.293637566 | 0.038997183 |
| Fermt1 | 0.019692488 | 4.258734268 | 0.048 | 0.004 | 1 | 0.331907298 | -0.316927414 | 0 | 0.021 | 1 | 0.331907298 | 0.038997183 |
| 9530097N15Rik | 0.019692488 | 4.258734268 | 0.048 | 0.004 | 1 | 0.496223879 | 0.683072586 | 0 | 0.01 | 1 | 0.496223879 | 0.038997183 |
| Gm43569 | 0.019692488 | 4.258734268 | 0.048 | 0.004 | 1 | 0.293637566 | -0.638855509 | 0 | 0.024 | 1 | 0.293637566 | 0.038997183 |
| Trp73 | 0.019692488 | 4.995699863 | 0.048 | 0.004 | 1 | 0.496223879 | 0.683072586 | 0 | 0.01 | 1 | 0.496223879 | 0.038997183 |
| Gm35113 | 0.019692488 | 4.258734268 | 0.048 | 0.004 | 1 | 0.496223879 | 0.683072586 | 0 | 0.01 | 1 | 0.496223879 | 0.038997183 |
| Gm15756 | 0.019692488 | 5.258734268 | 0.048 | 0.004 | 1 | 0.312251526 | 2.361144491 | 0.022 | 0.007 | 1 | 0.312251526 | 0.038997183 |
| Kcnj8 | 0.019692488 | 4.258734268 | 0.048 | 0.004 | 1 | 0.260665169 | -0.776359032 | 0 | 0.028 | 1 | 0.260665169 | 0.038997183 |
| Ptprh | 0.019692488 | 4.258734268 | 0.048 | 0.004 | 1 | 0.260653677 | -0.776359032 | 0 | 0.028 | 1 | 0.260653677 | 0.038997183 |
| Rsph6a | 0.019692488 | 4.258734268 | 0.048 | 0.004 | 1 | 0.309824707 | 2.683072586 | 0.022 | 0.007 | 1 | 0.309824707 | 0.038997183 |
| Hipk4 | 0.019692488 | 4.258734268 | 0.048 | 0.004 | 1 | 0.955025007 | 1.683072586 | 0.022 | 0.024 | 1 | 0.955025007 | 0.038997183 |
| Selenov | 0.019692488 | 5.258734268 | 0.048 | 0.004 | 1 | 0.314689364 | 2.098110085 | 0.022 | 0.007 | 1 | 0.314689364 | 0.038997183 |
| Gm44997 | 0.019692488 | 4.673771768 | 0.048 | 0.004 | 1 | 0.232482267 | 2.098110085 | 0.044 | 0.017 | 1 | 0.232482267 | 0.038997183 |
| Gm6410 | 0.019692488 | 4.258734268 | 0.048 | 0.004 | 1 | 0.312251526 | 2.361144491 | 0.022 | 0.007 | 1 | 0.312251526 | 0.038997183 |
| Best2 | 0.019692488 | 4.258734268 | 0.048 | 0.004 | 1 | 0.113529367 | 2.361144491 | 0.067 | 0.024 | 1 | 0.113529367 | 0.038997183 |
| Calb2 | 0.019692488 | 4.258734268 | 0.048 | 0.004 | 1 | 0.430487676 | 0.361144491 | 0 | 0.014 | 1 | 0.430487676 | 0.038997183 |
| Clec18a | 0.019692488 | 4.258734268 | 0.048 | 0.004 | 1 | 0.132894982 | 2.098110085 | 0.022 | 0.003 | 1 | 0.132894982 | 0.038997183 |
| 5730403I07Rik | 0.019692488 | 4.258734268 | 0.048 | 0.004 | 1 | 0.430483494 | -0.124282336 | 0 | 0.014 | 1 | 0.430483494 | 0.038997183 |
| Ngp | 0.019692488 | 4.258734268 | 0.048 | 0.004 | 1 | 0.94343746 | 0.852997588 | 0.089 | 0.093 | 1 | 0.94343746 | 0.038997183 |
| Stox1 | 0.019692488 | 4.258734268 | 0.048 | 0.004 | 1 | 0.492043882 | 2.361144491 | 0.022 | 0.01 | 1 | 0.492043882 | 0.038997183 |
| 1110002J07Rik | 0.019692488 | 4.258734268 | 0.048 | 0.004 | 1 | 0.675805699 | 0.683072586 | 0.022 | 0.014 | 1 | 0.675805699 | 0.038997183 |
| Havcr2 | 0.019692488 | 4.410737362 | 0.048 | 0.004 | 1 | 0.314689364 | 2.098110085 | 0.022 | 0.007 | 1 | 0.314689364 | 0.038997183 |
| Lrrc46 | 0.019692488 | 4.258734268 | 0.048 | 0.004 | 1 | 0.376837034 | -0.124282336 | 0 | 0.017 | 1 | 0.376837034 | 0.038997183 |
| Akap5 | 0.019692488 | 4.258734268 | 0.048 | 0.004 | 1 | 0.496223879 | 0.683072586 | 0 | 0.01 | 1 | 0.496223879 | 0.038997183 |
| Ttc9 | 0.019692488 | 5.258734268 | 0.048 | 0.004 | 1 | 0.293659664 | -0.316927414 | 0 | 0.024 | 1 | 0.293659664 | 0.038997183 |
| Ofcc1 | 0.019692488 | 4.258734268 | 0.048 | 0.004 | 1 | 0.496223879 | 0.683072586 | 0 | 0.01 | 1 | 0.496223879 | 0.038997183 |
| Fbxo43 | 0.019692488 | 4.258734268 | 0.048 | 0.004 | 1 | 0.031613022 | 3.490427508 | 0.044 | 0.007 | 1 | 0.031613022 | 0.038997183 |
| Parp10 | 0.019692488 | 4.258734268 | 0.048 | 0.004 | 1 | 0.18484555 | -1.404390255 | 0 | 0.038 | 1 | 0.18484555 | 0.038997183 |
| Kcnj4 | 0.019692488 | 4.258734268 | 0.048 | 0.004 | 1 | 0.030581005 | 3.42003818 | 0.044 | 0.007 | 1 | 0.030581005 | 0.038997183 |
| Wfikkn1 | 0.019692488 | 4.258734268 | 0.048 | 0.004 | 1 | 0.331907298 | -0.316927414 | 0 | 0.021 | 1 | 0.331907298 | 0.038997183 |
| Dgkk | 0.019692488 | 4.673771768 | 0.048 | 0.004 | 1 | 0.496223879 | 0.683072586 | 0 | 0.01 | 1 | 0.496223879 | 0.038997183 |
| Coasy | 0.184150046 | -0.970084422 | 0.048 | 0.153 | 1 | 0.019695517 | -1.84048937 | 0.022 | 0.149 | 1 | 0.184150046 | 0.039003121 |
| Lin7c | 0.128772643 | -0.383059455 | 0.286 | 0.537 | 1 | 0.019736346 | -0.536804237 | 0.622 | 0.696 | 1 | 0.128772643 | 0.03908317 |
| Smad9 | 0.236576657 | 1.181918671 | 0.19 | 0.112 | 1 | 0.019760537 | -1.881712033 | 0.022 | 0.152 | 1 | 0.236576657 | 0.039130596 |
| Cpsf2 | 0.019817507 | -1.465779585 | 0.048 | 0.299 | 1 | 0.288316272 | 0.46339455 | 0.467 | 0.388 | 1 | 0.288316272 | 0.03924228 |
| Zfp52 | 0.082739156 | 1.188344941 | 0.381 | 0.254 | 1 | 0.019856679 | -0.22981675 | 0.244 | 0.498 | 1 | 0.082739156 | 0.03931907 |
| Araf | 0.16255158 | -0.182365527 | 0.429 | 0.72 | 1 | 0.019858958 | -0.293790915 | 0.511 | 0.734 | 1 | 0.16255158 | 0.039323537 |
| Kremen1 | 0.019866266 | -0.723820597 | 0.19 | 0.507 | 1 | 0.426974579 | 0.263942332 | 0.6 | 0.547 | 1 | 0.426974579 | 0.039337864 |
| Fkbp4 | 0.01993794 | -1.034573149 | 0.143 | 0.433 | 1 | 0.129887902 | -0.545746104 | 0.578 | 0.585 | 1 | 0.129887902 | 0.039478359 |
| Lacc1 | 0.021858891 | -2.555046923 | 0 | 0.205 | 1 | 0.01996861 | -1.662702251 | 0.022 | 0.152 | 1 | 0.021858891 | 0.039538474 |
| Scyl1 | 0.238274412 | -0.259114036 | 0.286 | 0.451 | 1 | 0.019970752 | -0.178936411 | 0.311 | 0.574 | 1 | 0.238274412 | 0.039542674 |
| Ganab | 0.224286936 | -0.103350953 | 0.476 | 0.743 | 1 | 0.019979851 | -0.217977131 | 0.578 | 0.768 | 1 | 0.224286936 | 0.039560507 |
| Spred1 | 0.019983049 | 1.078060026 | 0.857 | 0.881 | 1 | 0.468101171 | 0.174241155 | 0.844 | 0.91 | 1 | 0.468101171 | 0.039566775 |
| Ccdc84 | 0.519943428 | 0.119182916 | 0.095 | 0.153 | 1 | 0.020003368 | -0.622735843 | 0.067 | 0.232 | 1 | 0.519943428 | 0.039606601 |
| Cyp4f17 | 0.612893855 | 0.811275291 | 0.238 | 0.22 | 1 | 0.020035347 | -1.298780067 | 0.067 | 0.218 | 1 | 0.612893855 | 0.03966928 |
| Renbp | 0.897672747 | 0.847801167 | 0.19 | 0.228 | 1 | 0.020051593 | -1.264459994 | 0.089 | 0.249 | 1 | 0.897672747 | 0.039701119 |
| Trim35 | 0.200947121 | 1.037147147 | 0.429 | 0.358 | 1 | 0.020061561 | -0.316927414 | 0.222 | 0.46 | 1 | 0.200947121 | 0.039720656 |
| Ipo9 | 0.020113057 | -0.695462042 | 0.048 | 0.302 | 1 | 0.490244751 | 0.28452321 | 0.356 | 0.474 | 1 | 0.490244751 | 0.039821579 |
| Hmgcll1 | 0.041723846 | 2.589707503 | 0.238 | 0.108 | 1 | 0.020142241 | -0.531940305 | 0.067 | 0.232 | 1 | 0.041723846 | 0.039878772 |
| Abcb10 | 0.065367452 | -1.940938076 | 0 | 0.142 | 1 | 0.02014326 | -0.743192169 | 0.067 | 0.225 | 1 | 0.065367452 | 0.039880769 |
| Gm17634 | 0.02019452 | 3.673771768 | 0.048 | 0.004 | 1 | 0.376837034 | -0.124282336 | 0 | 0.017 | 1 | 0.376837034 | 0.039981222 |
| Kif21b | 0.02019452 | 3.673771768 | 0.048 | 0.004 | 1 | 0.152163868 | 1.946106992 | 0.044 | 0.014 | 1 | 0.152163868 | 0.039981222 |
| Ccl21a | 0.02019452 | 3.673771768 | 0.048 | 0.004 | 1 | 0.496223879 | 0.683072586 | 0 | 0.01 | 1 | 0.496223879 | 0.039981222 |
| Gm57320 | 0.02019452 | 3.673771768 | 0.048 | 0.004 | 1 | 0.129721052 | 4.005000681 | 0.022 | 0.003 | 1 | 0.129721052 | 0.039981222 |
| Atp1a3 | 0.02019452 | 3.673771768 | 0.048 | 0.004 | 1 | 0.232008061 | -1.486852415 | 0 | 0.031 | 1 | 0.232008061 | 0.039981222 |
| Dmkn | 0.02019452 | 3.673771768 | 0.048 | 0.004 | 1 | 0.932582169 | 0.361144491 | 0.022 | 0.024 | 1 | 0.932582169 | 0.039981222 |
| Rab38 | 0.02019452 | 3.673771768 | 0.048 | 0.004 | 1 | 0.293656507 | -1.901889915 | 0 | 0.024 | 1 | 0.293656507 | 0.039981222 |
| 1700041G16Rik | 0.02019452 | 3.673771768 | 0.048 | 0.004 | 1 | 0.376837034 | -0.124282336 | 0 | 0.017 | 1 | 0.376837034 | 0.039981222 |
| Gm45370 | 0.02019452 | 3.673771768 | 0.048 | 0.004 | 1 | 0.541900769 | 1.098110085 | 0.044 | 0.028 | 1 | 0.541900769 | 0.039981222 |
| Card14 | 0.02019452 | 3.673771768 | 0.048 | 0.004 | 1 | 0.809042496 | 1.361144491 | 0.022 | 0.017 | 1 | 0.809042496 | 0.039981222 |
| Ttc6 | 0.02019452 | 3.673771768 | 0.048 | 0.004 | 1 | 0.426109575 | 2.268035087 | 0.044 | 0.024 | 1 | 0.426109575 | 0.039981222 |
| Gm35161 | 0.02019452 | 3.673771768 | 0.048 | 0.004 | 1 | 0.331914654 | -0.486852415 | 0 | 0.021 | 1 | 0.331914654 | 0.039981222 |
| Gm16549 | 0.02019452 | 3.673771768 | 0.048 | 0.004 | 1 | 0.331907298 | -0.486852415 | 0 | 0.021 | 1 | 0.331907298 | 0.039981222 |
| Gm16136 | 0.02019452 | 3.673771768 | 0.048 | 0.004 | 1 | 0.496223879 | 0.683072586 | 0 | 0.01 | 1 | 0.496223879 | 0.039981222 |
| Nol4 | 0.02019452 | 3.673771768 | 0.048 | 0.004 | 1 | 0.331919067 | -1.638855509 | 0 | 0.021 | 1 | 0.331919067 | 0.039981222 |
| Sh3rf2 | 0.02019452 | 3.673771768 | 0.048 | 0.004 | 1 | 0.293652719 | -1.486852415 | 0 | 0.024 | 1 | 0.293652719 | 0.039981222 |
| 2010001A14Rik | 0.020245643 | -2.326228232 | 0 | 0.209 | 1 | 0.077996475 | -0.667424661 | 0.044 | 0.142 | 1 | 0.077996475 | 0.0400814 |
| Grina | 0.020259686 | -1.274595464 | 0.143 | 0.433 | 1 | 0.774046547 | 0.148898195 | 0.533 | 0.522 | 1 | 0.774046547 | 0.040108917 |
| Cib2 | 0.061286863 | -1.970084422 | 0 | 0.146 | 1 | 0.020280182 | -1.589945908 | 0.022 | 0.152 | 1 | 0.061286863 | 0.040149078 |
| B9d1 | 0.020323803 | -2.370622352 | 0 | 0.209 | 1 | 0.035443514 | -0.749886821 | 0.044 | 0.17 | 1 | 0.035443514 | 0.040234549 |
| Pwwp2b | 0.163434053 | -1.02666795 | 0 | 0.086 | 1 | 0.020365481 | -1.184823878 | 0.044 | 0.187 | 1 | 0.163434053 | 0.040316209 |

| | | | | | | | | | | | | |
|---|---|---|---|---|---|---|---|---|---|---|---|---|
| Ube2j1 | 0.076182493 | -0.515111391 | 0.476 | 0.709 | 1 | 0.020405028 | -0.24890899 | 0.644 | 0.813 | 1 | 0.076182493 | 0.04039369 |
| Utp25 | 0.112336711 | 1.754390228 | 0.333 | 0.228 | 1 | 0.02041133 | -0.802354241 | 0.111 | 0.28 | 1 | 0.112336711 | 0.040406038 |
| 0610030E20Rik | 0.399361293 | -0.120644099 | 0.19 | 0.291 | 1 | 0.02041423 | -0.414959497 | 0.133 | 0.325 | 1 | 0.399361293 | 0.04041172 |
| Nup85 | 0.020431202 | -2.496153234 | 0 | 0.209 | 1 | 0.037271482 | -0.124282336 | 0.111 | 0.27 | 1 | 0.037271482 | 0.040444971 |
| Cast | 0.028178259 | 0.774700677 | 0.952 | 0.951 | 1 | 0.020434156 | 0.5315537 | 0.978 | 0.941 | 1 | 0.028178259 | 0.040450756 |
| Fbxo38 | 0.489263959 | 0.630703046 | 0.524 | 0.59 | 1 | 0.020468458 | -0.473046616 | 0.422 | 0.592 | 1 | 0.489263959 | 0.040517959 |
| Dnajc12 | 0.276089074 | 0.957564734 | 0.048 | 0.138 | 1 | 0.020488167 | -1.486852415 | 0.022 | 0.152 | 1 | 0.276089074 | 0.040556569 |
| Lrrc14 | 0.036911244 | -2.159118246 | 0 | 0.175 | 1 | 0.020509075 | -1.459885368 | 0.022 | 0.152 | 1 | 0.036911244 | 0.040597528 |
| Eif2a | 0.089444484 | -0.403672318 | 0.333 | 0.601 | 1 | 0.020553416 | -0.564854927 | 0.6 | 0.682 | 1 | 0.089444484 | 0.04068439 |
| Dkk3 | 0.020583781 | -2.063193826 | 0.095 | 0.347 | 1 | 0.525617855 | 0.377086035 | 0.6 | 0.606 | 1 | 0.525617855 | 0.04074387 |
| Ccni | 0.020606625 | -0.508246306 | 0.571 | 0.884 | 1 | 0.087668801 | -0.140357099 | 0.911 | 0.886 | 1 | 0.087668801 | 0.040788616 |
| Dedd2 | 0.740261153 | 1.578614535 | 0.143 | 0.134 | 1 | 0.020631763 | -0.83350294 | 0.089 | 0.253 | 1 | 0.740261153 | 0.040837857 |
| Rabep1 | 0.16694029 | 0.628257108 | 0.857 | 0.802 | 1 | 0.020643157 | 0.66848505 | 0.911 | 0.865 | 1 | 0.16694029 | 0.040860175 |
| Tfdp1 | 0.725218985 | 1.081754509 | 0.429 | 0.489 | 1 | 0.020701646 | -0.706873932 | 0.4 | 0.574 | 1 | 0.725218985 | 0.040974735 |
| P2ry12 | 0.020707684 | 2.673771768 | 0.048 | 0.004 | 1 | 0.376837034 | -0.124282336 | 0 | 0.017 | 1 | 0.376837034 | 0.040986559 |
| Slco1a5 | 0.020707684 | 2.936806174 | 0.048 | 0.004 | 1 | 0.129721052 | 4.005000681 | 0.022 | 0.003 | 1 | 0.129721052 | 0.040986559 |
| Gldn | 0.020707684 | 2.258734268 | 0.048 | 0.004 | 1 | 0.49622258 | -0.316927414 | 0 | 0.01 | 1 | 0.49622258 | 0.040986559 |
| Dnah8 | 0.020707684 | 3.258734268 | 0.048 | 0.004 | 1 | 0.765739892 | 0.776181991 | 0.022 | 0.031 | 1 | 0.765739892 | 0.040986559 |
| Asb11 | 0.020707684 | 3.088809267 | 0.048 | 0.004 | 1 | 0.430487676 | 0.361144491 | 0 | 0.014 | 1 | 0.430487676 | 0.040986559 |
| Cndp2 | 0.828357052 | 0.503846766 | 0.143 | 0.175 | 1 | 0.020796461 | -0.84048937 | 0.044 | 0.187 | 1 | 0.828357052 | 0.041160429 |
| Mindy1 | 0.073567843 | -0.240498358 | 0.19 | 0.463 | 1 | 0.0208429 | -0.737589462 | 0.333 | 0.509 | 1 | 0.073567843 | 0.041251373 |
| Gm17435 | 0.047705493 | 1.472137907 | 0.286 | 0.138 | 1 | 0.020853317 | -1.098287127 | 0.022 | 0.152 | 1 | 0.047705493 | 0.041271773 |
| Jak1 | 0.020856413 | -0.256455032 | 0.952 | 0.981 | 1 | 0.055237431 | -0.199982062 | 1 | 0.979 | 1 | 0.055237431 | 0.041277836 |
| Fbxo21 | 0.061571582 | -2.184209227 | 0 | 0.146 | 1 | 0.020866337 | -0.383016604 | 0.044 | 0.187 | 1 | 0.061571582 | 0.041297269 |
| Rbfa | 0.030430156 | -2.303508156 | 0 | 0.187 | 1 | 0.020882597 | -0.794974711 | 0.067 | 0.218 | 1 | 0.030430156 | 0.041329112 |
| Herc2 | 0.020913332 | 1.100732434 | 0.81 | 0.724 | 1 | 0.859912199 | 0.128659149 | 0.933 | 0.893 | 1 | 0.859912199 | 0.041389296 |
| Ppil3 | 0.30186248 | -0.361852142 | 0.143 | 0.25 | 1 | 0.02093487 | -0.329751454 | 0.111 | 0.291 | 1 | 0.30186248 | 0.041431472 |
| 2810002D19Rik | 0.08354242 | -1.718545655 | 0 | 0.127 | 1 | 0.020943776 | -1.199570463 | 0.022 | 0.152 | 1 | 0.08354242 | 0.04144891 |
| Ern1 | 0.790231977 | 0.718812732 | 0.571 | 0.571 | 1 | 0.020983151 | 0.606620526 | 0.867 | 0.678 | 1 | 0.790231977 | 0.041526009 |
| Lss | 0.654484788 | 0.74303443 | 0.19 | 0.272 | 1 | 0.020983505 | -1.039393438 | 0.022 | 0.152 | 1 | 0.654484788 | 0.041526702 |
| Fntb | 0.10354506 | -1.427766259 | 0.095 | 0.246 | 1 | 0.021009178 | -0.890918797 | 0.111 | 0.284 | 1 | 0.10354506 | 0.041576971 |
| Rap2b | 0.28963731 | -0.732220592 | 0.143 | 0.243 | 1 | 0.021028175 | -0.3932783 | 0.156 | 0.346 | 1 | 0.28963731 | 0.041614165 |
| Gm42726 | 0.021064888 | 2.48112669 | 0.143 | 0.037 | 1 | 0.485511539 | 1.683072586 | 0.067 | 0.045 | 1 | 0.485511539 | 0.041686046 |
| Psmg3 | 0.062015865 | -0.496153234 | 0.048 | 0.235 | 1 | 0.021065557 | -1.098287127 | 0.022 | 0.152 | 1 | 0.062015865 | 0.041687356 |
| Ing2 | 0.268628532 | 0.188344941 | 0.238 | 0.433 | 1 | 0.021080007 | -0.628129102 | 0.267 | 0.45 | 1 | 0.268628532 | 0.041715647 |
| C2cd2 | 0.639630098 | 1.291302131 | 0.19 | 0.175 | 1 | 0.021106544 | -0.612383297 | 0.067 | 0.232 | 1 | 0.639630098 | 0.041767602 |
| Id2 | 0.05478061 | -1.63391696 | 0.619 | 0.653 | 1 | 0.021139864 | -1.696699583 | 0.756 | 0.675 | 1 | 0.05478061 | 0.041832835 |
| Gpatch11 | 0.021152065 | -1.970084422 | 0.048 | 0.276 | 1 | 0.036448638 | -0.790858602 | 0.089 | 0.232 | 1 | 0.036448638 | 0.04185672 |
| Rp9 | 0.02118481 | -0.160218281 | 0.381 | 0.638 | 1 | 0.523774062 | -0.274699178 | 0.778 | 0.668 | 1 | 0.523774062 | 0.041920823 |
| Tmem236 | 0.853725152 | 2.393663848 | 0.048 | 0.041 | 1 | 0.021207722 | 2.020107573 | 0.067 | 0.014 | 1 | 0.853725152 | 0.041965677 |
| Dapp1 | 0.659316927 | 1.036341847 | 0.238 | 0.231 | 1 | 0.021213552 | 1.170003004 | 0.444 | 0.27 | 1 | 0.659316927 | 0.04197709 |
| Ilrun | 0.173677086 | 0.899507632 | 0.667 | 0.761 | 1 | 0.02121425 | 0.535978293 | 0.978 | 0.879 | 1 | 0.173677086 | 0.041978456 |
| Foxd2 | 0.088977675 | -1.752492987 | 0 | 0.123 | 1 | 0.021235214 | -0.486852415 | 0.022 | 0.152 | 1 | 0.088977675 | 0.042019494 |
| Ipmk | 0.347203033 | -0.152198833 | 0.143 | 0.261 | 1 | 0.02125556 | 0.863644832 | 0.444 | 0.27 | 1 | 0.347203033 | 0.042059321 |
| Ephb4 | 0.537002949 | 0.65584986 | 0.143 | 0.213 | 1 | 0.021257438 | -0.220497651 | 0.222 | 0.46 | 1 | 0.537002949 | 0.042062997 |
| 4930533B01Rik | 0.021265256 | 2.673771768 | 0.143 | 0.037 | 1 | 0.952529269 | 1.098110085 | 0.022 | 0.024 | 1 | 0.952529269 | 0.042078301 |
| Dcakd | 0.742440087 | 0.113056813 | 0.238 | 0.295 | 1 | 0.021302714 | -0.539319835 | 0.156 | 0.349 | 1 | 0.742440087 | 0.042151623 |
| Mrpl38 | 0.042811675 | -0.958496448 | 0.095 | 0.332 | 1 | 0.021342501 | -0.394929926 | 0.156 | 0.349 | 1 | 0.042811675 | 0.042229499 |
| Mbd6 | 0.552580113 | 0.108987149 | 0.143 | 0.213 | 1 | 0.021344335 | -0.151503805 | 0.111 | 0.311 | 1 | 0.552580113 | 0.042233089 |
| Lgi2 | 0.22219759 | -1.899117901 | 0.095 | 0.201 | 1 | 0.021353882 | -1.724774535 | 0.2 | 0.353 | 1 | 0.22219759 | 0.042251776 |
| Paox | 0.163567557 | -1.280424543 | 0 | 0.086 | 1 | 0.021377687 | -2.361321533 | 0 | 0.107 | 1 | 0.163567557 | 0.042298369 |
| Gm56524 | 0.464386198 | 2.214340149 | 0.095 | 0.06 | 1 | 0.021377687 | -2.361321533 | 0 | 0.107 | 1 | 0.464386198 | 0.042298369 |
| Ercc6l2 | 0.181483458 | -0.276840205 | 0.238 | 0.418 | 1 | 0.021379075 | -0.381057751 | 0.133 | 0.339 | 1 | 0.181483458 | 0.042301084 |
| Lrch4 | 0.145090655 | -1.02666795 | 0 | 0.093 | 1 | 0.02139571 | -2.404390255 | 0 | 0.107 | 1 | 0.145090655 | 0.042333644 |
| Zscan25 | 0.538021513 | 1.503846766 | 0.095 | 0.063 | 1 | 0.02139571 | -2.404390255 | 0 | 0.107 | 1 | 0.538021513 | 0.042333644 |
| Gm57463 | 0.106613525 | -1.496153234 | 0 | 0.112 | 1 | 0.021412495 | -2.564854927 | 0 | 0.107 | 1 | 0.106613525 | 0.042366496 |
| Mtg1 | 0.236422459 | -0.718545655 | 0 | 0.063 | 1 | 0.021412495 | -2.486852415 | 0 | 0.107 | 1 | 0.236422459 | 0.042366496 |
| Ncapd2 | 0.415139724 | -0.107587946 | 0.048 | 0.108 | 1 | 0.021412495 | -2.564854927 | 0 | 0.107 | 1 | 0.415139724 | 0.042366496 |
| Alkbh4 | 0.222378078 | -0.574155746 | 0 | 0.067 | 1 | 0.021412537 | -2.446210431 | 0 | 0.107 | 1 | 0.222378078 | 0.042366577 |
| Agpat5 | 0.021422745 | -0.994606741 | 0.143 | 0.429 | 1 | 0.300089112 | 0.343222583 | 0.267 | 0.363 | 1 | 0.300089112 | 0.042386556 |
| Gm14296 | 0.547845204 | 0.351843673 | 0.048 | 0.09 | 1 | 0.021428042 | -2.564854927 | 0 | 0.107 | 1 | 0.547845204 | 0.042396922 |
| Eno3 | 0.04802372 | -2.648156327 | 0 | 0.16 | 1 | 0.021428166 | -2.564854927 | 0 | 0.107 | 1 | 0.04802372 | 0.042397166 |
| Rnaseh2a | 0.709755848 | 0.780686972 | 0.095 | 0.134 | 1 | 0.021428166 | -2.674479418 | 0 | 0.107 | 1 | 0.709755848 | 0.042397166 |
| Sco1 | 0.127996876 | -1.233118828 | 0 | 0.101 | 1 | 0.021428208 | -2.486852415 | 0 | 0.107 | 1 | 0.127996876 | 0.042397247 |
| Snph | 0.805198946 | 0.918884266 | 0.095 | 0.082 | 1 | 0.021428208 | -2.486852415 | 0 | 0.107 | 1 | 0.805198946 | 0.042397247 |
| Mob3c | 0.156939835 | -0.735619168 | 0.095 | 0.235 | 1 | 0.021436686 | -1.064161343 | 0.067 | 0.221 | 1 | 0.156939835 | 0.042413841 |
| Foxk2 | 0.036675364 | 1.726239188 | 0.476 | 0.369 | 1 | 0.021438075 | -0.185921402 | 0.267 | 0.498 | 1 | 0.036675364 | 0.042416559 |
| Clstn3 | 0.396971401 | 0.351843673 | 0 | 0.034 | 1 | 0.021442348 | -2.564854927 | 0 | 0.107 | 1 | 0.396971401 | 0.042424922 |
| Zfp420 | 0.77384646 | 1.351843673 | 0.048 | 0.067 | 1 | 0.021442348 | -2.564854927 | 0 | 0.107 | 1 | 0.77384646 | 0.042424922 |
| Nckipsd | 0.163567557 | -1.233118828 | 0 | 0.086 | 1 | 0.02144268 | -2.674479418 | 0 | 0.107 | 1 | 0.163567557 | 0.042425571 |
| Gatad2b | 0.162926138 | 0.918884266 | 0.81 | 0.854 | 1 | 0.021449342 | 0.640168085 | 0.956 | 0.965 | 1 | 0.162926138 | 0.042438609 |
| Gtpbp6 | 0.697705209 | 0.188344941 | 0.143 | 0.183 | 1 | 0.021455828 | -2.638855509 | 0 | 0.107 | 1 | 0.697705209 | 0.042451304 |
| Osbpl7 | 0.912814742 | 1.472137907 | 0.048 | 0.056 | 1 | 0.021455828 | -2.638855509 | 0 | 0.107 | 1 | 0.912814742 | 0.042451304 |
| H2bc6 | 0.048015879 | -1.65714511 | 0.048 | 0.246 | 1 | 0.021479686 | -4.085111739 | 0 | 0.107 | 1 | 0.048015879 | 0.042497995 |
| Clcn3 | 0.040108146 | 1.232336578 | 0.619 | 0.481 | 1 | 0.021505472 | 0.469293295 | 0.822 | 0.578 | 1 | 0.040108146 | 0.042548459 |
| Gm56812 | 0.208885811 | -0.911190733 | 0 | 0.071 | 1 | 0.021507705 | -2.80878051 | 0 | 0.107 | 1 | 0.208885811 | 0.042552829 |
| Zfp646 | 0.929340808 | 0.214340149 | 0.19 | 0.183 | 1 | 0.02151967 | -0.583713955 | 0.156 | 0.346 | 1 | 0.929340808 | 0.042576244 |
| 2900060B14Rik | 0.816239924 | -0.574155746 | 0.143 | 0.16 | 1 | 0.021573917 | 2.03099589 | 0.2 | 0.097 | 1 | 0.816239924 | 0.0426824 |
| Aebp2 | 0.103100388 | 0.989799261 | 0.714 | 0.664 | 1 | 0.021576643 | 0.530635011 | 0.956 | 0.844 | 1 | 0.103100388 | 0.042687734 |
| Ticam1 | 0.196657642 | -0.718545655 | 0 | 0.075 | 1 | 0.02158521 | -1.539319835 | 0.044 | 0.18 | 1 | 0.196657642 | 0.042704499 |
| Dtnb | 0.822918396 | 0.739431194 | 0.524 | 0.679 | 1 | 0.021609374 | 0.649209134 | 0.889 | 0.758 | 1 | 0.822918396 | 0.042751783 |
| Ska2 | 0.245189675 | -1.208871282 | 0.048 | 0.134 | 1 | 0.021624345 | -2.044847868 | 0.022 | 0.145 | 1 | 0.245189675 | 0.042781078 |
| Med11 | 0.047392534 | -1.849790188 | 0 | 0.16 | 1 | 0.021624783 | -2.098287127 | 0.022 | 0.145 | 1 | 0.047392534 | 0.042781935 |
| Prpf39 | 0.021641893 | 0.64075671 | 0.857 | 0.754 | 1 | 0.03177504 | 0.637400728 | 0.933 | 0.875 | 1 | 0.03177504 | 0.042815415 |
| Mtbp | 0.021684451 | 2.55829455 | 0.238 | 0.093 | 1 | 0.118475508 | -1.674479418 | 0.022 | 0.09 | 1 | 0.118475508 | 0.042898687 |
| Zmym3 | 0.184642276 | -0.849790188 | 0 | 0.078 | 1 | 0.021686236 | -1.989352756 | 0.022 | 0.145 | 1 | 0.184642276 | 0.042902178 |
| Adamts10 | 0.036713981 | -0.481506458 | 0.143 | 0.422 | 1 | 0.021717849 | -0.375821103 | 0.311 | 0.567 | 1 | 0.036713981 | 0.042964033 |
| Sepsecs | 0.559357592 | 0.673771768 | 0.238 | 0.201 | 1 | 0.021719769 | -0.716858021 | 0.089 | 0.26 | 1 | 0.559357592 | 0.04296779 |
| Cept1 | 0.089581458 | -0.280424543 | 0.238 | 0.474 | 1 | 0.021721752 | 0.737998797 | 0.844 | 0.626 | 1 | 0.089581458 | 0.042971669 |
| Sdhaf1 | 0.106613525 | -1.455511249 | 0 | 0.112 | 1 | 0.02174795 | -2.044847868 | 0.022 | 0.145 | 1 | 0.106613525 | 0.043022927 |
| Pdik1l | 0.083814226 | -1.998653574 | 0 | 0.127 | 1 | 0.021748264 | -1.931637258 | 0.022 | 0.145 | 1 | 0.083814226 | 0.043023541 |
| Nr4a3 | 0.307599552 | -0.798716004 | 0.19 | 0.299 | 1 | 0.021773562 | -0.987687787 | 0.289 | 0.498 | 1 | 0.307599552 | 0.043073036 |
| Atxn7 | 0.736210669 | -0.180169948 | 0.714 | 0.72 | 1 | 0.021781481 | 0.837915631 | 0.933 | 0.848 | 1 | 0.736210669 | 0.043088529 |
| Tysnd1 | 0.057454773 | -1.818081329 | 0 | 0.149 | 1 | 0.021807881 | -1.989352756 | 0.022 | 0.145 | 1 | 0.057454773 | 0.043140178 |
| Adi1 | 0.088938003 | -1.718545655 | 0 | 0.123 | 1 | 0.021808196 | -1.901889915 | 0.022 | 0.145 | 1 | 0.088938003 | 0.043140794 |
| Gm44699 | 0.021811144 | 1.903253614 | 0.333 | 0.172 | 1 | 0.110297354 | 0.203079645 | 0.156 | 0.298 | 1 | 0.110297354 | 0.043146562 |
| Bbs7 | 0.021824955 | -2.496153234 | 0 | 0.205 | 1 | 0.071927864 | -0.140049652 | 0.089 | 0.208 | 1 | 0.071927864 | 0.043173582 |
| 1600014C10Rik | 0.10049151 | -1.574155746 | 0 | 0.116 | 1 | 0.021866962 | -1.901889915 | 0.022 | 0.145 | 1 | 0.10049151 | 0.043255761 |

| | | | | | | | | | | | | |
|---|---|---|---|---|---|---|---|---|---|---|---|---|
| Lmnb2 | 0.144846679 | -1.370622352 | 0 | 0.093 | 1 | 0.021867057 | -1.931637258 | 0.022 | 0.145 | 1 | 0.144846679 | 0.043255946 |
| Tmem38b | 0.021888438 | -2.455511249 | 0 | 0.205 | 1 | 0.610994898 | 0.828923453 | 0.289 | 0.26 | 1 | 0.610994898 | 0.043297772 |
| Rasa2 | 0.502743392 | 0.235478916 | 0.524 | 0.694 | 1 | 0.021906165 | 0.928448326 | 0.911 | 0.782 | 1 | 0.502743392 | 0.043332451 |
| Abracl | 0.055295009 | -1.169925001 | 0.238 | 0.437 | 1 | 0.02191914 | 0.390290837 | 0.667 | 0.405 | 1 | 0.055295009 | 0.043357832 |
| Dnaja4 | 0.843651338 | 0.335902129 | 0.19 | 0.228 | 1 | 0.021930801 | -1.211745177 | 0.133 | 0.294 | 1 | 0.843651338 | 0.043380641 |
| AW554918 | 0.054586443 | 1.359663177 | 0.476 | 0.358 | 1 | 0.02193581 | 1.074024467 | 0.578 | 0.439 | 1 | 0.054586443 | 0.04339044 |
| Fktn | 0.021948741 | -2.516052791 | 0 | 0.205 | 1 | 0.50150712 | 0.520801157 | 0.156 | 0.218 | 1 | 0.50150712 | 0.043415735 |
| Pim1 | 0.021965081 | 1.223110359 | 0.095 | 0.354 | 1 | 0.059751811 | 0.217600365 | 0.689 | 0.443 | 1 | 0.059751811 | 0.043447697 |
| Fetub | 0.196573996 | -1.849790188 | 0 | 0.075 | 1 | 0.021973347 | 0.875717664 | 0.267 | 0.131 | 1 | 0.196573996 | 0.043463867 |
| Cstf1 | 0.04179441 | -1.970084422 | 0 | 0.168 | 1 | 0.0219816 | -1.80878051 | 0.022 | 0.145 | 1 | 0.04179441 | 0.043480009 |
| Dnmt3a | 0.02205643 | 0.936806174 | 0.714 | 0.642 | 1 | 0.051267884 | 0.604271018 | 1 | 0.938 | 1 | 0.051267884 | 0.043626373 |
| Stxbp6 | 0.480727724 | 0.217324007 | 0.333 | 0.429 | 1 | 0.022093545 | 1.000812884 | 0.533 | 0.377 | 1 | 0.480727724 | 0.043698965 |
| Zfhx4 | 0.022095972 | -1.993501087 | 0.19 | 0.459 | 1 | 0.48622941 | -0.753860408 | 0.644 | 0.626 | 1 | 0.48622941 | 0.043703712 |
| Gorab | 0.02210403 | -0.978304929 | 0.143 | 0.407 | 1 | 0.252339449 | 0.215946576 | 0.267 | 0.422 | 1 | 0.252339449 | 0.043719471 |
| Dpysl2 | 0.124067974 | -0.659211516 | 0.286 | 0.47 | 1 | 0.022107693 | -0.213091603 | 0.2 | 0.405 | 1 | 0.124067974 | 0.043726636 |
| Ncoa7 | 0.290413704 | -0.886943187 | 0.143 | 0.243 | 1 | 0.022134768 | -0.586388089 | 0.089 | 0.249 | 1 | 0.290413704 | 0.043779589 |
| Hdac2 | 0.022145594 | -0.769171728 | 0.19 | 0.534 | 1 | 0.870836221 | 0.180572246 | 0.711 | 0.678 | 1 | 0.870836221 | 0.043800761 |
| Strip1 | 0.061588478 | -1.133583154 | 0.048 | 0.231 | 1 | 0.022152976 | -0.307882274 | 0.156 | 0.356 | 1 | 0.061588478 | 0.043815198 |
| Cd46 | 0.873114862 | 1.460778044 | 0.095 | 0.116 | 1 | 0.022161247 | -0.57996182 | 0.044 | 0.183 | 1 | 0.873114862 | 0.043831373 |
| Spata2 | 0.386523167 | -0.107587946 | 0.095 | 0.175 | 1 | 0.022190683 | -0.622735843 | 0.089 | 0.253 | 1 | 0.386523167 | 0.04388894 |
| Sertad3 | 0.022198797 | -3.01272876 | 0 | 0.205 | 1 | 0.030598456 | -0.404390255 | 0.111 | 0.277 | 1 | 0.030598456 | 0.043904807 |
| Yju2 | 0.437340914 | 0.912959432 | 0.238 | 0.187 | 1 | 0.022239434 | -1.638855509 | 0.067 | 0.208 | 1 | 0.437340914 | 0.043984276 |
| Zdhhc6 | 0.733765691 | 0.490549944 | 0.238 | 0.302 | 1 | 0.022250598 | -0.885770249 | 0.156 | 0.336 | 1 | 0.733765691 | 0.044006106 |
| Mllt6 | 0.022313033 | -1.303508156 | 0.095 | 0.351 | 1 | 0.04554822 | -0.299369042 | 0.267 | 0.481 | 1 | 0.04554822 | 0.044128195 |
| Cttnbp2nl | 0.309871788 | -0.186576242 | 0.381 | 0.582 | 1 | 0.022325123 | -0.169156793 | 0.511 | 0.83 | 1 | 0.309871788 | 0.044151835 |
| Rabep2 | 0.877036579 | 0.503846766 | 0.19 | 0.22 | 1 | 0.022329034 | -1.191396532 | 0.067 | 0.215 | 1 | 0.877036579 | 0.044159482 |
| Raver1 | 0.993227302 | 0.214340149 | 0.286 | 0.295 | 1 | 0.022342209 | -0.423126818 | 0.156 | 0.377 | 1 | 0.993227302 | 0.044185245 |
| Tlcd2 | 0.205052405 | -0.461387816 | 0.143 | 0.295 | 1 | 0.02235791 | -0.58227198 | 0.111 | 0.294 | 1 | 0.205052405 | 0.044215944 |
| Lamp2 | 0.02236385 | -0.423277122 | 0.905 | 0.996 | 1 | 0.038191177 | -0.208086092 | 1 | 0.997 | 1 | 0.038191177 | 0.044227557 |
| Nfs1 | 0.146169183 | -0.574155746 | 0.143 | 0.328 | 1 | 0.022418731 | -0.207698344 | 0.178 | 0.394 | 1 | 0.146169183 | 0.044334862 |
| Pde4dip | 0.022444011 | 0.589520431 | 0.857 | 0.951 | 1 | 0.067360938 | 0.631874078 | 0.978 | 0.969 | 1 | 0.067360938 | 0.044384288 |
| Tnpo3 | 0.139728467 | 0.655393238 | 0.619 | 0.601 | 1 | 0.022449539 | 0.592272489 | 0.889 | 0.74 | 1 | 0.139728467 | 0.044395097 |
| Tmem39b | 0.958984219 | 0.69025989 | 0.19 | 0.213 | 1 | 0.022457219 | -0.106248413 | 0.133 | 0.332 | 1 | 0.958984219 | 0.044410111 |
| Slc30a7 | 0.256565981 | -0.25696557 | 0.238 | 0.422 | 1 | 0.022459455 | -0.187940973 | 0.311 | 0.536 | 1 | 0.256565981 | 0.044414484 |
| Btg2 | 0.090904446 | -1.715795044 | 0.81 | 0.795 | 1 | 0.022483565 | -0.678178639 | 0.844 | 0.567 | 1 | 0.090904446 | 0.044461619 |
| Cdc23 | 0.300341544 | -0.184209227 | 0.19 | 0.332 | 1 | 0.022493245 | -0.282161996 | 0.178 | 0.384 | 1 | 0.300341544 | 0.044480544 |
| Foxd2os | 0.927974261 | 0.207072149 | 0.333 | 0.34 | 1 | 0.022518976 | -0.776359032 | 0.067 | 0.221 | 1 | 0.927974261 | 0.044530849 |
| Rbm12b2 | 0.022553018 | 2.133203386 | 0.238 | 0.093 | 1 | 0.138200208 | -0.57996182 | 0.067 | 0.152 | 1 | 0.138200208 | 0.044597397 |
| Trip4 | 1 | 0.403682604 | 0.333 | 0.403 | 1 | 0.022566009 | -0.40731365 | 0.244 | 0.474 | 1 | 1 | 0.044622793 |
| Atrn | 0.022584401 | -0.398856032 | 0.286 | 0.642 | 1 | 0.596918233 | 0.155622544 | 0.867 | 0.851 | 1 | 0.596918233 | 0.044658747 |
| Adgrg6 | 0.632120027 | 0.887299131 | 0.714 | 0.616 | 1 | 0.022643582 | -0.162935767 | 0.533 | 0.789 | 1 | 0.632120027 | 0.044774433 |
| Prkar2b | 0.041943764 | -2.107587946 | 0 | 0.168 | 1 | 0.022665786 | -0.75987091 | 0.067 | 0.215 | 1 | 0.041943764 | 0.044817833 |
| Acyp1 | 0.055855649 | -0.326228232 | 0.048 | 0.243 | 1 | 0.022670019 | -1.204452685 | 0.067 | 0.215 | 1 | 0.055855649 | 0.044826108 |
| Apcdd1 | 0.054384154 | -2.574155746 | 0 | 0.153 | 1 | 0.02271201 | -0.776359032 | 0.044 | 0.19 | 1 | 0.054384154 | 0.044908184 |
| Exosc5 | 0.10089491 | -0.326228232 | 0.048 | 0.205 | 1 | 0.022742425 | -1.84048937 | 0.044 | 0.176 | 1 | 0.10089491 | 0.044967633 |
| Gm20257 | 0.044669036 | -2.02666795 | 0 | 0.164 | 1 | 0.022768415 | -1.459885368 | 0.022 | 0.149 | 1 | 0.044669036 | 0.045018429 |
| Armt1 | 0.083636919 | -1.785659851 | 0 | 0.127 | 1 | 0.022768415 | -1.459885368 | 0.022 | 0.149 | 1 | 0.083636919 | 0.045018429 |
| Alpk1 | 0.022834997 | 0.766881172 | 0.667 | 0.493 | 1 | 0.88446455 | 0.903981516 | 0.289 | 0.381 | 1 | 0.88446455 | 0.045148557 |
| Apbb1 | 0.902093916 | 0.719575457 | 0.095 | 0.09 | 1 | 0.022837241 | -0.921789472 | 0.044 | 0.183 | 1 | 0.902093916 | 0.045152942 |
| Dctpp1 | 0.211877951 | -0.926141075 | 0.143 | 0.257 | 1 | 0.022876402 | -1.819427754 | 0.044 | 0.176 | 1 | 0.211877951 | 0.045229474 |
| Golgb1 | 0.539935841 | 0.112543916 | 0.905 | 0.989 | 1 | 0.022883113 | -0.197423915 | 0.911 | 0.958 | 1 | 0.539935841 | 0.045242588 |
| Dop1a | 0.02288879 | 0.944668291 | 0.714 | 0.556 | 1 | 0.149013402 | -0.213970223 | 0.4 | 0.574 | 1 | 0.149013402 | 0.045253683 |
| Ostf1 | 0.022928371 | -0.671056729 | 0.571 | 0.791 | 1 | 0.032052015 | -0.437221648 | 0.822 | 0.81 | 1 | 0.032052015 | 0.045331032 |
| Piezo2 | 0.547820704 | 0.524258226 | 0.667 | 0.705 | 1 | 0.022939892 | 0.638254082 | 0.933 | 0.889 | 1 | 0.547820704 | 0.045353545 |
| Nup98 | 0.125591683 | 1.156898257 | 0.857 | 0.873 | 1 | 0.0229736 | 0.582069267 | 1 | 0.952 | 1 | 0.125591683 | 0.045419414 |
| Blcap | 0.625323968 | 0.335902129 | 0.19 | 0.269 | 1 | 0.022996253 | -0.339295227 | 0.156 | 0.343 | 1 | 0.625323968 | 0.045463677 |
| Tmem248 | 0.130785852 | -0.455511249 | 0.524 | 0.743 | 1 | 0.022997688 | -0.230360855 | 0.667 | 0.844 | 1 | 0.130785852 | 0.045466483 |
| Fam207a | 0.036781406 | -1.785659851 | 0.048 | 0.246 | 1 | 0.023075988 | -0.463768802 | 0.156 | 0.353 | 1 | 0.036781406 | 0.045619474 |
| Mef2b | 0.242025191 | -0.672003069 | 0.095 | 0.198 | 1 | 0.023098295 | -0.979892427 | 0.067 | 0.215 | 1 | 0.242025191 | 0.045663058 |
| Adck2 | 0.153857942 | -1.233118828 | 0 | 0.09 | 1 | 0.023102631 | -0.960783604 | 0.022 | 0.149 | 1 | 0.153857942 | 0.045671531 |
| Tatdn3 | 0.880743585 | 2.946790262 | 0.143 | 0.142 | 1 | 0.023119982 | -0.754332726 | 0.044 | 0.183 | 1 | 0.880743585 | 0.04570543 |
| 2310022A10Rik | 0.100391649 | -1.535681598 | 0 | 0.116 | 1 | 0.023144197 | -1.754332726 | 0.044 | 0.176 | 1 | 0.100391649 | 0.045752741 |
| Uhrf1bp1l | 0.023237804 | 0.95841263 | 0.81 | 0.743 | 1 | 0.074318325 | 0.532969026 | 0.956 | 0.837 | 1 | 0.074318325 | 0.045935613 |
| Ep400 | 0.023272818 | 0.608348693 | 0.905 | 0.754 | 1 | 0.291463663 | 0.716987995 | 0.933 | 0.886 | 1 | 0.291463663 | 0.046004012 |
| Cep57 | 0.95046125 | 0.551375136 | 0.381 | 0.433 | 1 | 0.02328731 | -0.368589534 | 0.222 | 0.439 | 1 | 0.95046125 | 0.046032321 |
| Arhgap23 | 0.700815369 | 0.673771768 | 0.429 | 0.597 | 1 | 0.023297212 | 0.747202924 | 0.844 | 0.848 | 1 | 0.700815369 | 0.046051664 |
| Mrrf | 0.023301921 | -1.535681598 | 0.095 | 0.332 | 1 | 0.097398596 | -0.26178586 | 0.089 | 0.204 | 1 | 0.097398596 | 0.046060863 |
| Vps18 | 0.023306919 | -2.326228232 | 0 | 0.201 | 1 | 0.041146611 | -0.686161223 | 0.089 | 0.235 | 1 | 0.041146611 | 0.046070625 |
| Isg20 | 0.506172426 | -0.249060372 | 0.333 | 0.444 | 1 | 0.023312691 | -1.459885368 | 0.067 | 0.215 | 1 | 0.506172426 | 0.0460819 |
| Ankmy1 | 0.050952627 | -2.02666795 | 0.048 | 0.228 | 1 | 0.023321775 | -0.486852415 | 0.067 | 0.221 | 1 | 0.050952627 | 0.046099644 |
| Ptgfrn | 0.147622734 | -0.31112134 | 0.095 | 0.243 | 1 | 0.023334491 | -1.124282336 | 0.044 | 0.187 | 1 | 0.147622734 | 0.046124483 |
| Cebpz | 0.023348071 | -0.834022873 | 0.238 | 0.53 | 1 | 0.826547311 | -0.136355168 | 0.644 | 0.592 | 1 | 0.826547311 | 0.04615101 |
| Sertad1 | 0.02338067 | -1.399477214 | 0.286 | 0.549 | 1 | 0.024235467 | -0.44551162 | 0.556 | 0.699 | 1 | 0.024235467 | 0.046214684 |
| Pgm1 | 0.072702841 | -1.370622352 | 0.048 | 0.209 | 1 | 0.02341083 | -1.754332726 | 0.044 | 0.176 | 1 | 0.072702841 | 0.046273594 |
| Topors | 0.041891439 | -0.381912365 | 0.286 | 0.549 | 1 | 0.023439668 | -0.337853253 | 0.756 | 0.747 | 1 | 0.041891439 | 0.046329918 |
| Zer1 | 0.477945183 | 0.252307999 | 0.143 | 0.228 | 1 | 0.023472357 | -0.349348892 | 0.089 | 0.26 | 1 | 0.477945183 | 0.046393763 |
| Hmg20b | 0.023479163 | -0.548620654 | 0.333 | 0.679 | 1 | 0.299720805 | -0.457017186 | 0.578 | 0.567 | 1 | 0.299720805 | 0.046407055 |
| Daxx | 0.083469737 | -1.295854583 | 0.095 | 0.261 | 1 | 0.023530729 | -0.626782676 | 0.133 | 0.304 | 1 | 0.083469737 | 0.046507764 |
| Fam172a | 0.02353464 | 1.158660376 | 0.81 | 0.817 | 1 | 0.037907533 | 0.756782272 | 0.933 | 0.834 | 1 | 0.037907533 | 0.046515401 |
| Zfp503 | 0.023586239 | -2.107587946 | 0.048 | 0.28 | 1 | 0.141528692 | 0.256807831 | 0.489 | 0.343 | 1 | 0.141528692 | 0.046616168 |
| Ssx2ip | 0.152433181 | -0.574155746 | 0.095 | 0.25 | 1 | 0.023598416 | -1.421264074 | 0.2 | 0.346 | 1 | 0.152433181 | 0.046639947 |
| Msn | 0.219321694 | 0.581529628 | 0.952 | 0.944 | 1 | 0.023744224 | 0.403465429 | 1 | 0.945 | 1 | 0.219321694 | 0.04692466 |
| C430049B03Rik | 0.023748317 | -3.63000898 | 0 | 0.201 | 1 | 0.361627311 | 0.69556353 | 0.111 | 0.17 | 1 | 0.361627311 | 0.046932651 |
| Slc45a4 | 0.023790458 | -2.159118246 | 0.048 | 0.269 | 1 | 0.164707413 | 0.45174704 | 0.489 | 0.401 | 1 | 0.164707413 | 0.04701493 |
| Chd3 | 0.144578746 | -0.62268564 | 0.524 | 0.623 | 1 | 0.023808015 | -0.504554417 | 0.844 | 0.858 | 1 | 0.144578746 | 0.047049209 |
| Gcat | 0.354981605 | -0.496153234 | 0.048 | 0.112 | 1 | 0.023817692 | -2.316927414 | 0 | 0.104 | 1 | 0.354981605 | 0.047068102 |
| Arhgap29 | 0.145976991 | -0.786653101 | 0.524 | 0.739 | 1 | 0.023835219 | -0.479043492 | 0.689 | 0.91 | 1 | 0.145976991 | 0.047102321 |
| Gm1673 | 0.057694796 | -2.107587946 | 0 | 0.149 | 1 | 0.023853789 | -2.404390255 | 0 | 0.104 | 1 | 0.057694796 | 0.047138575 |
| Gm35162 | 0.065284654 | -1.880817084 | 0 | 0.142 | 1 | 0.023853789 | -2.404390255 | 0 | 0.104 | 1 | 0.065284654 | 0.047138575 |
| Coro1b | 0.499693471 | 0.503846766 | 0.048 | 0.097 | 1 | 0.023853789 | -2.404390255 | 0 | 0.104 | 1 | 0.499693471 | 0.047138575 |
| Zfp623 | 0.95783752 | 0.866416846 | 0.048 | 0.045 | 1 | 0.023853789 | -2.404390255 | 0 | 0.104 | 1 | 0.95783752 | 0.047138575 |
| Septin4 | 0.649593995 | 0.614878079 | 0.048 | 0.078 | 1 | 0.023869956 | -2.446210431 | 0 | 0.104 | 1 | 0.649593995 | 0.047170138 |
| Gins2 | 0.758302222 | 0.901182264 | 0.143 | 0.123 | 1 | 0.023884745 | -2.564854927 | 0 | 0.104 | 1 | 0.758302222 | 0.047199009 |
| Mat2b | 0.023906099 | -1.214971481 | 0.238 | 0.493 | 1 | 0.34680313 | 0.162021849 | 0.667 | 0.516 | 1 | 0.34680313 | 0.047240696 |
| Cyp26b1 | 0.023906493 | -1.196592952 | 0.238 | 0.519 | 1 | 0.027590925 | -0.736466305 | 0.244 | 0.474 | 1 | 0.027590925 | 0.047241465 |
| Gm11613 | 0.136525282 | 3.521768674 | 0.095 | 0.034 | 1 | 0.023909861 | -2.602329633 | 0 | 0.104 | 1 | 0.136525282 | 0.047248041 |
| Rwdd3 | 0.06544374 | -2.054148687 | 0 | 0.142 | 1 | 0.023910553 | -2.743192169 | 0 | 0.104 | 1 | 0.06544374 | 0.047249391 |
| Rnasel | 0.32890191 | -0.940938076 | 0.048 | 0.116 | 1 | 0.023922169 | -2.743192169 | 0 | 0.104 | 1 | 0.32890191 | 0.047272068 |

| | | | | | | | | | | | | |
|---|---|---|---|---|---|---|---|---|---|---|---|---|
| Gm43713 | 0.196423035 | -0.911190733 | 0 | 0.075 | 1 | 0.023932174 | -2.84048937 | 0 | 0.104 | 1 | 0.196423035 | 0.047291599 |
| Tmx3 | 0.023933904 | -0.763633545 | 0.286 | 0.604 | 1 | 0.646683885 | -0.174092636 | 0.778 | 0.685 | 1 | 0.646683885 | 0.047294977 |
| Man2c1 | 0.966410903 | 0.550389352 | 0.19 | 0.213 | 1 | 0.023934222 | -1.539319835 | 0.044 | 0.176 | 1 | 0.966410903 | 0.047295598 |
| Kctd17 | 0.523753665 | 0.638147858 | 0.286 | 0.243 | 1 | 0.023935518 | -0.792660845 | 0.089 | 0.242 | 1 | 0.523753665 | 0.047298127 |
| Vamp4 | 0.613543705 | 0.205002284 | 0.429 | 0.631 | 1 | 0.023935732 | -0.109122529 | 0.311 | 0.571 | 1 | 0.613543705 | 0.047298545 |
| Mrgbp | 0.338118358 | -0.695462042 | 0.095 | 0.172 | 1 | 0.023937317 | -1.298780067 | 0.067 | 0.211 | 1 | 0.338118358 | 0.04730164 |
| Atpaf1 | 0.813013432 | 1.586308926 | 0.19 | 0.194 | 1 | 0.023940303 | -0.746915255 | 0.067 | 0.218 | 1 | 0.813013432 | 0.047307467 |
| G0s2 | 0.245721732 | -2.780184721 | 0.048 | 0.134 | 1 | 0.023959107 | -3.383016604 | 0 | 0.104 | 1 | 0.245721732 | 0.047344175 |
| Nop53 | 0.023966791 | -0.770146852 | 0.429 | 0.724 | 1 | 0.142087403 | -0.568466181 | 0.778 | 0.696 | 1 | 0.142087403 | 0.047359174 |
| Tmem8b | 0.113338406 | -1.326228232 | 0 | 0.108 | 1 | 0.023972108 | -2.174908409 | 0.022 | 0.142 | 1 | 0.113338406 | 0.047369555 |
| Ptgfr | 0.08923057 | -3.303508156 | 0 | 0.123 | 1 | 0.023983143 | -4.137106376 | 0 | 0.104 | 1 | 0.08923057 | 0.047391095 |
| Mnt | 0.02398706 | -1.370622352 | 0.048 | 0.287 | 1 | 0.881835061 | 1.230560381 | 0.222 | 0.256 | 1 | 0.881835061 | 0.04739874 |
| Dusp14 | 0.556128413 | -0.718545655 | 0.333 | 0.396 | 1 | 0.023989557 | -0.638855509 | 0.111 | 0.28 | 1 | 0.556128413 | 0.047403615 |
| Xrcc1 | 0.337524906 | 0.458758877 | 0.095 | 0.187 | 1 | 0.024001787 | -0.557935513 | 0.089 | 0.256 | 1 | 0.337524906 | 0.047427488 |
| Pkig | 0.024003232 | -0.957665163 | 0.238 | 0.575 | 1 | 0.282267169 | 0.280526031 | 0.578 | 0.685 | 1 | 0.282267169 | 0.047430309 |
| Pias2 | 0.090500838 | 0.870953626 | 0.857 | 0.84 | 1 | 0.024008988 | 0.878596338 | 0.889 | 0.824 | 1 | 0.090500838 | 0.047441544 |
| Enpp1 | 0.054507248 | 0.44690587 | 0.667 | 0.896 | 1 | 0.024044075 | -0.289620068 | 0.511 | 0.82 | 1 | 0.054507248 | 0.047510032 |
| Pygo2 | 0.60438943 | 1.171271427 | 0.286 | 0.284 | 1 | 0.024049351 | -0.292124023 | 0.2 | 0.422 | 1 | 0.60438943 | 0.047520331 |
| 2410006H16Rik | 0.024054097 | -0.628118574 | 0.571 | 0.817 | 1 | 0.887257388 | -0.217977131 | 0.8 | 0.692 | 1 | 0.887257388 | 0.047529595 |
| Mecr | 0.387866204 | 0.963278385 | 0.286 | 0.213 | 1 | 0.024086115 | -0.731964913 | 0.089 | 0.249 | 1 | 0.387866204 | 0.04759209 |
| Bin3 | 0.024100765 | -0.880817084 | 0.095 | 0.377 | 1 | 0.044647073 | -0.609709163 | 0.156 | 0.318 | 1 | 0.044647073 | 0.047620683 |
| Lrif1 | 0.114115236 | -0.970084422 | 0.095 | 0.254 | 1 | 0.024110909 | -0.30740464 | 0.133 | 0.322 | 1 | 0.114115236 | 0.047640481 |
| Tbcc | 0.076499693 | -1.535681598 | 0.048 | 0.205 | 1 | 0.024117151 | -1.960783604 | 0.022 | 0.142 | 1 | 0.076499693 | 0.047652665 |
| Pdcd2 | 0.246961351 | -0.818081329 | 0.048 | 0.134 | 1 | 0.024186063 | -1.960783604 | 0.022 | 0.142 | 1 | 0.246961351 | 0.047787161 |
| Gcdh | 0.158916342 | 0.267779408 | 0.048 | 0.175 | 1 | 0.024186581 | -1.989352756 | 0.022 | 0.142 | 1 | 0.158916342 | 0.047788171 |
| 2310057M21Rik | 0.024219448 | -1.326228232 | 0.048 | 0.287 | 1 | 0.189674901 | 0.175277946 | 0.133 | 0.232 | 1 | 0.189674901 | 0.047852313 |
| Stk35 | 0.227005641 | 1.576910228 | 0.286 | 0.213 | 1 | 0.024236547 | -0.124282336 | 0.133 | 0.318 | 1 | 0.227005641 | 0.047885685 |
| Lag3 | 0.38847587 | 1.214340149 | 0.095 | 0.052 | 1 | 0.024349682 | 2.219125486 | 0.2 | 0.1 | 1 | 0.38847587 | 0.048106458 |
| Gm17484 | 0.806831419 | 2.405282932 | 0.143 | 0.134 | 1 | 0.024351579 | -2.174908409 | 0.067 | 0.201 | 1 | 0.806831419 | 0.048110159 |
| Gskip | 0.035110145 | -1.326228232 | 0.048 | 0.265 | 1 | 0.024406495 | -0.16020191 | 0.111 | 0.298 | 1 | 0.035110145 | 0.048217312 |
| Hsf2 | 0.365068931 | -0.558888989 | 0.143 | 0.228 | 1 | 0.024435198 | -0.437221648 | 0.178 | 0.394 | 1 | 0.365068931 | 0.048273317 |
| Gmeb2 | 0.024445343 | 1.513307095 | 0.524 | 0.396 | 1 | 0.171617169 | 0.270474132 | 0.244 | 0.412 | 1 | 0.171617169 | 0.048293111 |
| U2af2 | 0.615523258 | 0.255270413 | 0.524 | 0.612 | 1 | 0.024498029 | -0.542102276 | 0.6 | 0.682 | 1 | 0.615523258 | 0.048395905 |
| Sh3gl1 | 0.024511202 | -0.329444945 | 0.381 | 0.772 | 1 | 0.159772068 | -0.162058987 | 0.578 | 0.751 | 1 | 0.159772068 | 0.048421606 |
| Usp31 | 0.024528204 | -1.648156327 | 0.048 | 0.284 | 1 | 0.342608917 | 0.904428961 | 0.489 | 0.46 | 1 | 0.342608917 | 0.048454776 |
| Pfkfb3 | 0.024531619 | -1.970084422 | 0.095 | 0.332 | 1 | 0.043536157 | 0.203904749 | 0.222 | 0.446 | 1 | 0.043536157 | 0.048461438 |
| Ranbp9 | 0.049996158 | 0.822529202 | 0.81 | 0.701 | 1 | 0.024570341 | 0.588866416 | 1 | 0.869 | 1 | 0.049996158 | 0.048536979 |
| Adpgk | 0.027490417 | -1.25696557 | 0.095 | 0.34 | 1 | 0.024580332 | -0.761712257 | 0.111 | 0.277 | 1 | 0.027490417 | 0.048556472 |
| Tex2 | 0.927232572 | 0.866416846 | 0.238 | 0.265 | 1 | 0.024592876 | 1.128806883 | 0.667 | 0.502 | 1 | 0.927232572 | 0.048580943 |
| Aste1 | 0.156212318 | -0.593014773 | 0.095 | 0.239 | 1 | 0.024595813 | -0.847442131 | 0.089 | 0.246 | 1 | 0.156212318 | 0.048586672 |
| Fam234a | 0.024612582 | -0.953501538 | 0.143 | 0.422 | 1 | 0.127059478 | 0.215467036 | 0.222 | 0.377 | 1 | 0.127059478 | 0.048619384 |
| Cops5 | 0.049795191 | -0.440244493 | 0.333 | 0.657 | 1 | 0.0246154 | -0.423316758 | 0.6 | 0.723 | 1 | 0.049795191 | 0.048624881 |
| Ppp1r3c | 0.024628935 | -1.301424841 | 0.19 | 0.463 | 1 | 0.849179039 | 0.290755163 | 0.222 | 0.225 | 1 | 0.849179039 | 0.048651286 |
| Tmem212 | 0.024629562 | 2.417432014 | 0.238 | 0.097 | 1 | 0.568680838 | 1.576157382 | 0.111 | 0.152 | 1 | 0.568680838 | 0.048652508 |
| Stxbp1 | 0.16553817 | 0.144992102 | 0.286 | 0.556 | 1 | 0.024654284 | 0.86810448 | 0.622 | 0.626 | 1 | 0.16553817 | 0.048700733 |
| Grk6 | 0.897845826 | 0.365019062 | 0.286 | 0.332 | 1 | 0.024655264 | -0.931637258 | 0.089 | 0.253 | 1 | 0.897845826 | 0.048702646 |
| Fhip1b | 0.118162883 | 1.700738815 | 0.381 | 0.299 | 1 | 0.024706338 | -0.180865864 | 0.178 | 0.405 | 1 | 0.118162883 | 0.048802273 |
| Rlf | 0.685877911 | 0.299891593 | 0.667 | 0.862 | 1 | 0.024726576 | 0.558336358 | 1 | 0.91 | 1 | 0.685877911 | 0.048841749 |
| Dpysl3 | 0.663212331 | -1.233118828 | 0.095 | 0.123 | 1 | 0.024765802 | -0.141355849 | 0.244 | 0.111 | 1 | 0.663212331 | 0.04891826 |
| Rspry1 | 0.024800983 | 1.049280903 | 0.714 | 0.616 | 1 | 0.118739576 | -0.271798038 | 0.733 | 0.737 | 1 | 0.118739576 | 0.048986878 |
| Zswim7 | 0.275353193 | -0.213753503 | 0.095 | 0.209 | 1 | 0.024824607 | -1.223818009 | 0.044 | 0.18 | 1 | 0.275353193 | 0.049032953 |
| Smpd4 | 0.024839805 | -2.25696557 | 0 | 0.198 | 1 | 0.043120854 | -0.316927414 | 0.111 | 0.266 | 1 | 0.043120854 | 0.049062593 |
| Gm16599 | 0.26960409 | 2.195724471 | 0.19 | 0.123 | 1 | 0.024843579 | -1.486852415 | 0.044 | 0.18 | 1 | 0.26960409 | 0.049069955 |
| Col4a5 | 0.024847128 | -2.896083841 | 0.095 | 0.317 | 1 | 0.035960864 | -0.330733213 | 0.067 | 0.211 | 1 | 0.035960864 | 0.049076875 |
| Pold3 | 0.242258378 | -0.233118828 | 0.095 | 0.209 | 1 | 0.024853443 | -0.11339402 | 0.111 | 0.301 | 1 | 0.242258378 | 0.049089193 |
| Ddhd1 | 0.635832264 | 0.896164189 | 0.333 | 0.313 | 1 | 0.024862585 | 0.902271083 | 0.711 | 0.509 | 1 | 0.635832264 | 0.049107022 |
| Nexn | 0.47037125 | 1.8372705 | 0.095 | 0.06 | 1 | 0.024868935 | -1.614607962 | 0.022 | 0.145 | 1 | 0.47037125 | 0.049119406 |
| Ppm1h | 0.02731008 | 1.38426515 | 0.333 | 0.16 | 1 | 0.024892488 | 0.513147585 | 0.422 | 0.239 | 1 | 0.02731008 | 0.04916534 |
| Farsb | 0.024911899 | 1.451379346 | 0.524 | 0.399 | 1 | 0.039276521 | -0.468236737 | 0.267 | 0.488 | 1 | 0.039276521 | 0.049203195 |
| Cnnm3 | 0.024917022 | -2.370622352 | 0 | 0.198 | 1 | 0.035105732 | -0.638855509 | 0.089 | 0.235 | 1 | 0.035105732 | 0.049213187 |
| Ankrd39 | 0.024919411 | -2.413691074 | 0 | 0.198 | 1 | 0.042371877 | -0.754332726 | 0.044 | 0.163 | 1 | 0.042371877 | 0.049217844 |
| Dio2 | 0.216403129 | -2.752492987 | 0.048 | 0.149 | 1 | 0.024937004 | -3.060319277 | 0.067 | 0.204 | 1 | 0.216403129 | 0.049252153 |
| Srd5a3 | 0.128514269 | -1.081115734 | 0.048 | 0.175 | 1 | 0.024947133 | -0.656777417 | 0.067 | 0.218 | 1 | 0.128514269 | 0.049271906 |
| Cxxc5 | 0.082882348 | -0.713630165 | 0.286 | 0.515 | 1 | 0.024967502 | -0.453532342 | 0.644 | 0.785 | 1 | 0.082882348 | 0.049311627 |
| Cdc26 | 0.02501738 | -0.626351957 | 0.333 | 0.731 | 1 | 0.032201218 | -0.383016604 | 0.578 | 0.74 | 1 | 0.032201218 | 0.049408891 |
| Xpc | 0.025024813 | -2.475975352 | 0 | 0.198 | 1 | 0.249289475 | 0.256807831 | 0.133 | 0.225 | 1 | 0.249289475 | 0.049423385 |
| Tnrc18 | 0.02502965 | 0.615367211 | 0.952 | 0.903 | 1 | 0.984034875 | 0.212585779 | 0.956 | 0.965 | 1 | 0.984034875 | 0.049432816 |
| Nudcd1 | 0.025046352 | -1.648156327 | 0.095 | 0.325 | 1 | 0.480284311 | 0.268035087 | 0.178 | 0.242 | 1 | 0.480284311 | 0.049465384 |
| Txlna | 0.967338454 | 0.451379346 | 0.524 | 0.69 | 1 | 0.025047215 | -0.542347528 | 0.556 | 0.734 | 1 | 0.967338454 | 0.049467068 |
| Uggt2 | 0.497486969 | 0.77865834 | 0.524 | 0.604 | 1 | 0.025068209 | -0.181200765 | 0.356 | 0.702 | 1 | 0.497486969 | 0.049508002 |
| Syt14 | 0.025083263 | 4.258734268 | 0.095 | 0.019 | 1 | 0.49622258 | 0.361144491 | 0 | 0.01 | 1 | 0.49622258 | 0.049537355 |
| Gm31224 | 0.025083263 | 3.410737362 | 0.095 | 0.019 | 1 | 0.430487676 | 0.361144491 | 0 | 0.014 | 1 | 0.430487676 | 0.049537355 |
| Btrc | 0.620947322 | 0.97333205 | 0.429 | 0.496 | 1 | 0.025125877 | 0.833362265 | 0.844 | 0.747 | 1 | 0.620947322 | 0.049620445 |
| Zbtb39 | 0.083590408 | -1.718545655 | 0 | 0.127 | 1 | 0.0251404 | -0.622735843 | 0.089 | 0.249 | 1 | 0.083590408 | 0.04964876 |
| Niban2 | 0.50224774 | 0.993050284 | 0.714 | 0.888 | 1 | 0.025194445 | 0.552058681 | 1 | 0.927 | 1 | 0.50224774 | 0.049754131 |
| Nmd3 | 0.143443266 | -0.159118246 | 0.286 | 0.515 | 1 | 0.025198572 | -0.124282336 | 0.422 | 0.612 | 1 | 0.143443266 | 0.049762176 |
| Gls | 0.025211938 | 0.559247453 | 0.952 | 0.981 | 1 | 0.12284165 | 0.358475305 | 1 | 0.993 | 1 | 0.12284165 | 0.049788234 |
| Mcts1 | 0.672238034 | 0.267779408 | 0.381 | 0.522 | 1 | 0.025223975 | -0.336120979 | 0.289 | 0.526 | 1 | 0.672238034 | 0.049811701 |
| Sart1 | 0.224021379 | -0.203972482 | 0.19 | 0.373 | 1 | 0.025229862 | -0.394929926 | 0.311 | 0.522 | 1 | 0.224021379 | 0.049823179 |
| Tmed5 | 0.025263291 | -0.392317423 | 0.476 | 0.724 | 1 | 0.302452431 | 0.302648842 | 0.867 | 0.661 | 1 | 0.302452431 | 0.049888347 |
| Lrrc40 | 0.766830409 | 0.574236094 | 0.143 | 0.183 | 1 | 0.025338142 | -0.466674533 | 0.044 | 0.187 | 1 | 0.766830409 | 0.050034263 |
| Col4a4 | 0.025357125 | -3.25696557 | 0 | 0.198 | 1 | 0.901539577 | 1.098110085 | 0.067 | 0.076 | 1 | 0.901539577 | 0.050071266 |
| Ehd4 | 0.183425273 | -0.5423887 | 0.476 | 0.672 | 1 | 0.025389638 | 1.347023383 | 0.667 | 0.453 | 1 | 0.183425273 | 0.050134643 |
| Taf6 | 0.087673949 | -1.370622352 | 0.048 | 0.198 | 1 | 0.025486435 | -0.539319835 | 0.067 | 0.211 | 1 | 0.087673949 | 0.050323311 |
| Ncstn | 0.025487163 | -0.280424543 | 0.095 | 0.351 | 1 | 0.342385317 | -0.195621118 | 0.444 | 0.495 | 1 | 0.342385317 | 0.05032473 |
| Gm17178 | 0.93080287 | 1.295260144 | 0.048 | 0.045 | 1 | 0.025490999 | 1.78998779 | 0.178 | 0.08 | 1 | 0.93080287 | 0.050332207 |
| Tor3a | 0.025496096 | -1.10233222 | 0.095 | 0.347 | 1 | 0.202796456 | 0.164343784 | 0.222 | 0.339 | 1 | 0.202796456 | 0.05034214 |
| Nacc2 | 0.025526675 | -1.242704877 | 0.143 | 0.396 | 1 | 0.032386367 | -0.426980959 | 0.311 | 0.592 | 1 | 0.032386367 | 0.050401738 |
| Dcxr | 0.100573008 | -1.880817084 | 0 | 0.116 | 1 | 0.025552707 | -1.071814916 | 0.022 | 0.145 | 1 | 0.100573008 | 0.050452472 |
| Vapb | 0.025566076 | -0.462005011 | 0.333 | 0.687 | 1 | 0.066972425 | -0.45770607 | 0.622 | 0.72 | 1 | 0.066972425 | 0.050478528 |
| Tmem183a | 0.025585064 | -0.522625445 | 0.143 | 0.425 | 1 | 0.865003921 | 0.435145073 | 0.511 | 0.522 | 1 | 0.865003921 | 0.050515532 |
| Mettl2 | 0.920878041 | 0.351843673 | 0.19 | 0.213 | 1 | 0.025608264 | -0.573267167 | 0.067 | 0.218 | 1 | 0.920878041 | 0.050560745 |
| Smad5 | 0.025688803 | 1.03785808 | 0.667 | 0.597 | 1 | 0.711689176 | 0.400756347 | 0.533 | 0.709 | 1 | 0.711689176 | 0.050717692 |
| Acvr2a | 0.394753328 | 0.214340149 | 0.19 | 0.325 | 1 | 0.025694021 | 0.76922923 | 0.689 | 0.488 | 1 | 0.394753328 | 0.050727859 |
| Tnfrsf11b | 0.063110082 | -0.675939484 | 0.524 | 0.776 | 1 | 0.025728788 | -0.858300646 | 0.267 | 0.516 | 1 | 0.063110082 | 0.050795606 |
| Wnt5a | 0.356896239 | -0.623908781 | 0.048 | 0.116 | 1 | 0.025730591 | -1.620708162 | 0.044 | 0.176 | 1 | 0.356896239 | 0.050799119 |
| Smim20 | 0.300496132 | -0.233118828 | 0.095 | 0.194 | 1 | 0.025734627 | -1.223818009 | 0.022 | 0.145 | 1 | 0.300496132 | 0.050806983 |
| Babam2 | 0.351020148 | 1.236366455 | 0.619 | 0.668 | 1 | 0.025735024 | 0.947900274 | 0.844 | 0.64 | 1 | 0.351020148 | 0.050807757 |

| | | | | | | | | | | | | |
|---|---|---|---|---|---|---|---|---|---|---|---|---|
| Lrrc69 | 0.02575431 | 3.188344941 | 0.095 | 0.019 | 1 | 0.852144264 | 1.098110085 | 0.022 | 0.028 | 1 | 0.852144264 | 0.050845335 |
| Prex1 | 0.305211072 | -0.178671044 | 0.143 | 0.261 | 1 | 0.025768764 | 0.218404319 | 0.222 | 0.453 | 1 | 0.305211072 | 0.050873498 |
| Eif2b4 | 0.402650679 | 1.289107917 | 0.143 | 0.261 | 1 | 0.025774313 | -0.857495795 | 0.133 | 0.304 | 1 | 0.402650679 | 0.05088431 |
| Pik3r2 | 0.073826504 | -1.718545655 | 0 | 0.134 | 1 | 0.025779444 | -1.709244837 | 0.044 | 0.173 | 1 | 0.073826504 | 0.050894309 |
| Eid2 | 0.219216171 | -0.593014773 | 0.095 | 0.213 | 1 | 0.025832585 | -0.436666658 | 0.133 | 0.311 | 1 | 0.219216171 | 0.050997848 |
| Igf2 | 0.054476077 | -2.718545655 | 0 | 0.153 | 1 | 0.025856082 | -1.017367132 | 0.022 | 0.145 | 1 | 0.054476077 | 0.051043628 |
| Rbbp5 | 0.601731815 | 0.252307999 | 0.143 | 0.213 | 1 | 0.025864127 | 0.431533819 | 0.111 | 0.277 | 1 | 0.601731815 | 0.051059301 |
| Cep112 | 0.025881396 | 1.777960618 | 0.619 | 0.53 | 1 | 0.16084251 | 1.307563451 | 0.467 | 0.495 | 1 | 0.16084251 | 0.051092946 |
| Mitf | 0.025910882 | -1.724259306 | 0.095 | 0.351 | 1 | 0.921318668 | 0.262740787 | 0.2 | 0.221 | 1 | 0.921318668 | 0.051150391 |
| Coq9 | 0.097341608 | -1.280424543 | 0.095 | 0.25 | 1 | 0.025928736 | -1.686161223 | 0.044 | 0.173 | 1 | 0.097341608 | 0.051185172 |
| Zfand2b | 0.151384905 | -0.516052791 | 0.095 | 0.243 | 1 | 0.02593043 | -1.638855509 | 0.044 | 0.173 | 1 | 0.151384905 | 0.051188472 |
| Sim2 | 0.744804734 | 0.30251596 | 0.381 | 0.496 | 1 | 0.02597258 | -0.526380779 | 0.289 | 0.498 | 1 | 0.744804734 | 0.051270584 |
| Ddx6 | 0.203561492 | -0.149258223 | 0.952 | 0.978 | 1 | 0.025990811 | -0.302130412 | 1 | 0.993 | 1 | 0.203561492 | 0.051306099 |
| Extl1 | 0.185412637 | -0.633656758 | 0.143 | 0.284 | 1 | 0.026017862 | -0.731964913 | 0.089 | 0.253 | 1 | 0.185412637 | 0.051358795 |
| Glrx | 0.030537325 | -2.516052791 | 0 | 0.187 | 1 | 0.026091802 | -0.709244837 | 0.089 | 0.246 | 1 | 0.030537325 | 0.051502822 |
| Septin6 | 0.196947072 | -1.133583154 | 0.048 | 0.153 | 1 | 0.026151939 | -1.264459994 | 0.067 | 0.208 | 1 | 0.196947072 | 0.051619953 |
| Dock1 | 0.026176103 | 1.206514585 | 0.81 | 0.81 | 1 | 0.071290071 | 0.816988207 | 0.956 | 0.92 | 1 | 0.071290071 | 0.051667017 |
| Nufip1 | 0.307905317 | -0.355975576 | 0.238 | 0.373 | 1 | 0.026184525 | -0.172917111 | 0.178 | 0.384 | 1 | 0.307905317 | 0.051683421 |
| Mcmbp | 0.02622045 | -0.800714197 | 0.238 | 0.534 | 1 | 0.351433104 | 0.294619872 | 0.844 | 0.723 | 1 | 0.351433104 | 0.051753387 |
| Mcur1 | 0.055081449 | -0.923163375 | 0.095 | 0.317 | 1 | 0.026221082 | -0.508068901 | 0.156 | 0.332 | 1 | 0.055081449 | 0.051754619 |
| Dock6 | 0.321159089 | -0.480033568 | 0.143 | 0.243 | 1 | 0.026252216 | -0.472845557 | 0.2 | 0.412 | 1 | 0.321159089 | 0.051815252 |
| Crebbp | 0.026260255 | 0.489252906 | 1 | 0.974 | 1 | 0.168395164 | 0.28020376 | 1 | 1 | 1 | 0.168395164 | 0.051830909 |
| Epha7 | 0.02628748 | 0.934686122 | 0.619 | 0.463 | 1 | 0.416696147 | 1.428075499 | 0.333 | 0.339 | 1 | 0.416696147 | 0.051883929 |
| Usp50 | 0.026337238 | 1.243137413 | 0.048 | 0.287 | 1 | 0.032273352 | 0.302250802 | 0.111 | 0.273 | 1 | 0.032273352 | 0.051980827 |
| Ppp2r5d | 0.366279188 | 0.815790773 | 0.095 | 0.194 | 1 | 0.026337493 | -0.223818009 | 0.067 | 0.225 | 1 | 0.366279188 | 0.051981323 |
| Ctnnd1 | 0.026338243 | 0.712765899 | 0.857 | 0.81 | 1 | 0.55885995 | 0.166932118 | 0.956 | 0.903 | 1 | 0.55885995 | 0.051982783 |
| Csgalnact1 | 0.026373769 | 1.859338243 | 0.952 | 0.806 | 1 | 0.218293123 | 0.649597882 | 0.578 | 0.609 | 1 | 0.218293123 | 0.052051963 |
| Sulf1 | 0.026378395 | 2.226312791 | 0.381 | 0.213 | 1 | 0.768643062 | -1.484741669 | 0.222 | 0.239 | 1 | 0.768643062 | 0.05206097 |
| Bmpr1b | 0.026528643 | 0.797577969 | 0.857 | 0.716 | 1 | 0.440491705 | 0.72765668 | 0.889 | 0.792 | 1 | 0.440491705 | 0.052353518 |
| Cox5b-ps | 0.230647742 | -0.233118828 | 0.048 | 0.149 | 1 | 0.026528936 | -2.271123724 | 0 | 0.1 | 1 | 0.230647742 | 0.052354088 |
| Ndc1 | 0.278817629 | -0.849790188 | 0.048 | 0.127 | 1 | 0.02655525 | -2.361321533 | 0.022 | 0.138 | 1 | 0.278817629 | 0.052405319 |
| Chtop | 0.026558993 | -0.672003069 | 0.238 | 0.567 | 1 | 0.040367974 | -0.389827961 | 0.556 | 0.685 | 1 | 0.040367974 | 0.052412605 |
| Gstm4 | 0.496351051 | 0.400753273 | 0.048 | 0.097 | 1 | 0.026582654 | -2.526380779 | 0 | 0.1 | 1 | 0.496351051 | 0.052458671 |
| Ripk3 | 0.164991574 | -0.911190733 | 0.048 | 0.16 | 1 | 0.026582859 | -2.404390255 | 0 | 0.1 | 1 | 0.164991574 | 0.05245907 |
| Zfp579 | 0.844871807 | 0.503846766 | 0.095 | 0.082 | 1 | 0.026597702 | -2.486852415 | 0 | 0.1 | 1 | 0.844871807 | 0.052487966 |
| Lrwd1 | 0.974684278 | 1.719575457 | 0.095 | 0.097 | 1 | 0.026597702 | -2.486852415 | 0 | 0.1 | 1 | 0.974684278 | 0.052487966 |
| Naa80 | 0.520551663 | 0.304537958 | 0.048 | 0.093 | 1 | 0.026598163 | -2.564854927 | 0 | 0.1 | 1 | 0.520551663 | 0.052488863 |
| Adgrg1 | 0.396916958 | -0.233118828 | 0 | 0.034 | 1 | 0.026598214 | -2.446210431 | 0 | 0.1 | 1 | 0.396916958 | 0.052488963 |
| Rfwd3 | 0.913838422 | 1.102229783 | 0.238 | 0.257 | 1 | 0.026608039 | 0.352923984 | 0.178 | 0.408 | 1 | 0.913838422 | 0.05250809 |
| Gzmm | 0.815673353 | 1.214340149 | 0.048 | 0.037 | 1 | 0.026611319 | -2.526380779 | 0 | 0.1 | 1 | 0.815673353 | 0.052514476 |
| Hectd3 | 0.995710676 | 0.351843673 | 0.095 | 0.093 | 1 | 0.026612343 | -2.486852415 | 0 | 0.1 | 1 | 0.995710676 | 0.052516469 |
| Gpsm3 | 0.121648189 | -1.107587946 | 0.048 | 0.179 | 1 | 0.026624273 | -2.674479418 | 0 | 0.1 | 1 | 0.121648189 | 0.052539694 |
| Gm16536 | 0.196423035 | -0.911190733 | 0 | 0.075 | 1 | 0.026624273 | -2.674479418 | 0 | 0.1 | 1 | 0.196423035 | 0.052539694 |
| Cfap300 | 0.267936726 | -0.264827688 | 0.048 | 0.138 | 1 | 0.026624273 | -2.602329633 | 0 | 0.1 | 1 | 0.267936726 | 0.052539694 |
| Vnn1 | 0.352487813 | -1.054148687 | 0.048 | 0.112 | 1 | 0.026636051 | -2.871516266 | 0 | 0.1 | 1 | 0.352487813 | 0.052562622 |
| Cdk5r1 | 0.910775761 | 0.629377648 | 0.095 | 0.09 | 1 | 0.026638399 | -2.223818009 | 0.022 | 0.138 | 1 | 0.910775761 | 0.052567193 |
| Fez1 | 0.285527294 | -0.574155746 | 0 | 0.052 | 1 | 0.026645013 | -2.674479418 | 0 | 0.1 | 1 | 0.285527294 | 0.05258007 |
| Ppil4 | 0.042324094 | -0.613576095 | 0.381 | 0.679 | 1 | 0.026664715 | -0.458283263 | 0.733 | 0.806 | 1 | 0.042324094 | 0.052618424 |
| Fam184a | 0.486488953 | 0.698019314 | 0.286 | 0.243 | 1 | 0.026698063 | -0.388244818 | 0.222 | 0.446 | 1 | 0.486488953 | 0.05268334 |
| Rpa3 | 0.142258124 | -0.849790188 | 0.143 | 0.299 | 1 | 0.026707998 | -0.776359032 | 0.089 | 0.246 | 1 | 0.142258124 | 0.052702679 |
| Lbh | 0.026728188 | -1.280424543 | 0.143 | 0.422 | 1 | 0.085021175 | -0.583713955 | 0.044 | 0.142 | 1 | 0.085021175 | 0.05274198 |
| E130307A14Rik | 0.227803695 | -0.619009982 | 0.095 | 0.209 | 1 | 0.026772992 | -0.58541625 | 0.067 | 0.218 | 1 | 0.227803695 | 0.052829191 |
| Gm10778 | 0.026779265 | 3.088809267 | 0.095 | 0.019 | 1 | 0.496223879 | 0.683072586 | 0 | 0.01 | 1 | 0.496223879 | 0.052841401 |
| Gm57330 | 0.026779265 | 3.088809267 | 0.095 | 0.019 | 1 | 0.430487676 | 0.361144491 | 0 | 0.014 | 1 | 0.430487676 | 0.052841401 |
| Dnaaf9 | 0.663399533 | 0.253620137 | 0.429 | 0.646 | 1 | 0.026789567 | 0.597765274 | 0.956 | 0.813 | 1 | 0.663399533 | 0.052861454 |
| Med6 | 0.242368756 | -0.264827688 | 0.238 | 0.407 | 1 | 0.026794217 | -0.17942389 | 0.156 | 0.356 | 1 | 0.242368756 | 0.052870505 |
| Smad6 | 0.246946755 | 0.270049582 | 0.143 | 0.28 | 1 | 0.026816241 | -0.420215222 | 0.2 | 0.415 | 1 | 0.246946755 | 0.052913372 |
| Itpr2 | 0.026827513 | 0.481557495 | 0.905 | 0.918 | 1 | 0.08164264 | 0.504350636 | 0.911 | 0.841 | 1 | 0.08164264 | 0.052935311 |
| Dalrd3 | 0.115693904 | -0.695462042 | 0.048 | 0.194 | 1 | 0.026834876 | -1.071814916 | 0.067 | 0.215 | 1 | 0.115693904 | 0.052949641 |
| Cdkl3 | 0.689836949 | 0.378315884 | 0.238 | 0.213 | 1 | 0.026870868 | -1.328154669 | 0.089 | 0.239 | 1 | 0.689836949 | 0.053019692 |
| Fzd10 | 0.324806011 | -0.648156327 | 0 | 0.045 | 1 | 0.026872167 | -0.946284034 | 0.111 | 0.277 | 1 | 0.324806011 | 0.05302222 |
| Bag2 | 0.817534431 | 0.281454345 | 0.143 | 0.172 | 1 | 0.026878959 | 0.298408736 | 0.089 | 0.256 | 1 | 0.817534431 | 0.053035439 |
| Celsr2 | 0.822006883 | 0.97333205 | 0.048 | 0.037 | 1 | 0.026882356 | -1.960783604 | 0.022 | 0.138 | 1 | 0.822006883 | 0.053042051 |
| Art3 | 0.026962253 | -3.094412557 | 0 | 0.194 | 1 | 0.854457549 | 1.83507568 | 0.022 | 0.028 | 1 | 0.854457549 | 0.053197542 |
| Rab23 | 0.95814876 | 0.123574685 | 0.19 | 0.201 | 1 | 0.026966157 | -0.884611923 | 0.067 | 0.215 | 1 | 0.95814876 | 0.053205141 |
| Tom1l2 | 0.026983523 | 1.359390482 | 0.762 | 0.772 | 1 | 0.956622604 | 0.227661486 | 0.933 | 0.886 | 1 | 0.956622604 | 0.053238935 |
| Ankrd27 | 0.09695165 | -0.785659851 | 0.048 | 0.205 | 1 | 0.026986973 | -0.594912161 | 0.089 | 0.246 | 1 | 0.09695165 | 0.053245648 |
| Sco2 | 0.196858256 | -0.849790188 | 0.048 | 0.149 | 1 | 0.02704204 | -1.871516266 | 0.022 | 0.138 | 1 | 0.196858256 | 0.053352809 |
| Ankrd49 | 0.033569888 | 2.204286484 | 0.333 | 0.183 | 1 | 0.027084232 | -1.901889915 | 0.022 | 0.142 | 1 | 0.033569888 | 0.053434907 |
| Map3k5 | 0.027127729 | 0.636872185 | 0.952 | 0.896 | 1 | 0.377000933 | 0.207684091 | 0.933 | 0.896 | 1 | 0.377000933 | 0.053519545 |
| Kcnk3 | 0.027134954 | 3.503846766 | 0.095 | 0.019 | 1 | 0.545623751 | 0.197645759 | 0.044 | 0.028 | 1 | 0.545623751 | 0.053533601 |
| Kri1 | 0.163326056 | -0.413691074 | 0.143 | 0.31 | 1 | 0.027155631 | -1.0583944 | 0.089 | 0.242 | 1 | 0.163326056 | 0.053573834 |
| Rnpep | 0.027263158 | -0.849790188 | 0.095 | 0.362 | 1 | 0.063092541 | 0.487724988 | 0.622 | 0.439 | 1 | 0.063092541 | 0.053783037 |
| Mpg | 0.028288837 | -1.718545655 | 0.048 | 0.261 | 1 | 0.027270392 | -0.72631835 | 0.067 | 0.215 | 1 | 0.028288837 | 0.05379711 |
| Rnf40 | 0.274272408 | -0.184209227 | 0.143 | 0.272 | 1 | 0.027278375 | -0.438917938 | 0.156 | 0.36 | 1 | 0.274272408 | 0.053812639 |
| Spred2 | 0.027347411 | 0.371881426 | 0.19 | 0.507 | 1 | 0.572942561 | 0.251121704 | 0.867 | 0.751 | 1 | 0.572942561 | 0.053946942 |
| Sh2b1 | 0.063133213 | -1.516052791 | 0.048 | 0.216 | 1 | 0.027353511 | -0.979892427 | 0.133 | 0.294 | 1 | 0.063133213 | 0.053958808 |
| Tubgcp3 | 0.909659354 | 0.808701348 | 0.238 | 0.246 | 1 | 0.02736715 | -0.369015925 | 0.133 | 0.325 | 1 | 0.909659354 | 0.053985339 |
| Gm16153 | 0.196422748 | -0.970084422 | 0 | 0.075 | 1 | 0.027369445 | -1.686161223 | 0.022 | 0.142 | 1 | 0.196422748 | 0.053989803 |
| Rnf166 | 0.330210225 | 0.114804475 | 0.286 | 0.429 | 1 | 0.02741484 | -0.271123724 | 0.244 | 0.478 | 1 | 0.330210225 | 0.054078106 |
| Usp12 | 0.037419974 | -0.399477214 | 0.143 | 0.429 | 1 | 0.027449764 | -0.396991803 | 0.511 | 0.678 | 1 | 0.037419974 | 0.054146039 |
| Gss | 0.591472138 | 0.866416846 | 0.19 | 0.272 | 1 | 0.027462315 | -0.133215461 | 0.156 | 0.36 | 1 | 0.591472138 | 0.054170451 |
| Agrn | 0.769834545 | 0.115776315 | 0.238 | 0.291 | 1 | 0.027468274 | -0.58227198 | 0.133 | 0.322 | 1 | 0.769834545 | 0.054182042 |
| Ndufaf7 | 0.058804268 | -1.666078235 | 0.048 | 0.22 | 1 | 0.027474747 | -1.017367132 | 0.067 | 0.215 | 1 | 0.058804268 | 0.054194632 |
| Espnl | 0.027490621 | 2.866416846 | 0.095 | 0.019 | 1 | 0.293659664 | -0.316927414 | 0 | 0.024 | 1 | 0.293659664 | 0.054225508 |
| Dlx6 | 0.027490621 | 2.866416846 | 0.095 | 0.019 | 1 | 0.508845649 | 0.513147585 | 0.022 | 0.01 | 1 | 0.508845649 | 0.054225508 |
| Syt9 | 0.027490621 | 2.866416846 | 0.095 | 0.019 | 1 | 0.376843981 | -0.638855509 | 0 | 0.017 | 1 | 0.376843981 | 0.054225508 |
| Ppp1r1c | 0.027493336 | 4.737902105 | 0.095 | 0.019 | 1 | 0.077153538 | 3.335149283 | 0.044 | 0.01 | 1 | 0.077153538 | 0.054230788 |
| Nrcam | 0.027493336 | 3.574236094 | 0.095 | 0.019 | 1 | 0.353709246 | 3.142504205 | 0.067 | 0.038 | 1 | 0.353709246 | 0.054230788 |
| Slc1a1 | 0.027495436 | 1.699976838 | 0.619 | 0.481 | 1 | 0.18987006 | -0.101621738 | 0.2 | 0.311 | 1 | 0.18987006 | 0.054234872 |
| Gypc | 0.030620434 | -2.593014773 | 0 | 0.187 | 1 | 0.027524697 | -2.053893008 | 0.044 | 0.17 | 1 | 0.030620434 | 0.054291786 |
| Eepd1 | 0.365263682 | 2.016659481 | 0.143 | 0.09 | 1 | 0.027606745 | -1.877642368 | 0.044 | 0.173 | 1 | 0.365263682 | 0.054451357 |
| Fbxw8 | 0.027623224 | 0.750007354 | 0.762 | 0.709 | 1 | 0.673445856 | 0.321782059 | 0.533 | 0.727 | 1 | 0.673445856 | 0.054483406 |
| Marcks | 0.318708237 | -1.097057278 | 0.429 | 0.463 | 1 | 0.027699696 | -0.988376973 | 0.711 | 0.657 | 1 | 0.318708237 | 0.054632118 |
| Vsig10 | 0.517193716 | 1.661799126 | 0.286 | 0.269 | 1 | 0.027716194 | -0.295967755 | 0.178 | 0.391 | 1 | 0.517193716 | 0.054664201 |
| Coro1c | 0.798622024 | 1.351843673 | 0.333 | 0.392 | 1 | 0.027764606 | 0.632446513 | 0.622 | 0.436 | 1 | 0.798622024 | 0.054758339 |
| Nup93 | 0.29922756 | 1.441111011 | 0.429 | 0.414 | 1 | 0.027782324 | -1.223818009 | 0.133 | 0.308 | 1 | 0.29922756 | 0.054792791 |
| Zfp458 | 0.027811872 | 1.957564734 | 0.19 | 0.063 | 1 | 0.132416275 | -1.404390255 | 0.022 | 0.087 | 1 | 0.132416275 | 0.054850244 |

| | | | | | | | | | | | | |
|---|---|---|---|---|---|---|---|---|---|---|---|---|
| Map7d2 | 0.027842915 | -2.400904919 | 0.095 | 0.332 | 1 | 0.255031161 | -1.053893008 | 0.022 | 0.066 | 1 | 0.255031161 | 0.054910602 |
| Pex5 | 0.766041391 | 0.873080576 | 0.381 | 0.504 | 1 | 0.027844665 | -0.107359033 | 0.311 | 0.595 | 1 | 0.766041391 | 0.054914004 |
| Uchl1 | 0.027850303 | 2.351843673 | 0.095 | 0.019 | 1 | 0.119078256 | -2.743192169 | 0 | 0.052 | 1 | 0.119078256 | 0.054924967 |
| Jag1 | 0.074529607 | -1.399477214 | 0.048 | 0.216 | 1 | 0.027855461 | 1.038167545 | 0.467 | 0.294 | 1 | 0.074529607 | 0.054934996 |
| Klhdc1 | 0.250075901 | -0.44170545 | 0.048 | 0.142 | 1 | 0.027865338 | -0.620708162 | 0.044 | 0.183 | 1 | 0.250075901 | 0.054954198 |
| Spop | 0.19499082 | 0.314690674 | 0.905 | 0.925 | 1 | 0.027905546 | 0.823594933 | 0.978 | 0.917 | 1 | 0.19499082 | 0.055032372 |
| Ccm2 | 0.524544917 | 0.121230745 | 0.286 | 0.418 | 1 | 0.027917071 | -0.26178586 | 0.289 | 0.522 | 1 | 0.524544917 | 0.055054779 |
| Ints10 | 0.054092955 | -1.970084422 | 0 | 0.153 | 1 | 0.027934857 | -1.432404631 | 0.022 | 0.142 | 1 | 0.054092955 | 0.055089358 |
| Gm37422 | 0.027967396 | 3.325848464 | 0.143 | 0.041 | 1 | 0.366948101 | 0.361144491 | 0.022 | 0.055 | 1 | 0.366948101 | 0.055152616 |
| Helb | 0.047789162 | -2.303508156 | 0 | 0.16 | 1 | 0.027989676 | -0.381057751 | 0.067 | 0.215 | 1 | 0.047789162 | 0.05519593 |
| Hus1 | 0.966786142 | 0.544488751 | 0.095 | 0.101 | 1 | 0.027999988 | -1.404390255 | 0.022 | 0.142 | 1 | 0.966786142 | 0.055215977 |
| Exoc7 | 0.288343609 | -0.361852142 | 0.143 | 0.254 | 1 | 0.028005539 | -0.476126009 | 0.133 | 0.311 | 1 | 0.288343609 | 0.055226768 |
| Flnb | 0.028063885 | 1.520068385 | 0.857 | 0.869 | 1 | 0.916855952 | 0.594478663 | 0.956 | 0.907 | 1 | 0.916855952 | 0.055340189 |
| Ube3b | 0.02806728 | -1.818081329 | 0.048 | 0.261 | 1 | 0.203106971 | 0.264571232 | 0.222 | 0.353 | 1 | 0.203106971 | 0.055346788 |
| Sacm1l | 0.627886105 | 0.279693887 | 0.524 | 0.679 | 1 | 0.028072011 | 0.877627254 | 0.889 | 0.803 | 1 | 0.627886105 | 0.055355984 |
| Rab22a | 0.028075972 | -0.849790188 | 0.333 | 0.649 | 1 | 0.63199028 | 0.313050203 | 0.778 | 0.789 | 1 | 0.63199028 | 0.055363684 |
| Zfp932 | 0.899931007 | 0.441111011 | 0.143 | 0.134 | 1 | 0.028082732 | -1.375821103 | 0.022 | 0.142 | 1 | 0.899931007 | 0.055376823 |
| Lrrc42 | 0.253110612 | -0.392317423 | 0.095 | 0.205 | 1 | 0.028089137 | -0.630344009 | 0.178 | 0.36 | 1 | 0.253110612 | 0.055389275 |
| Wsb1 | 0.52905622 | -0.154047257 | 0.905 | 0.948 | 1 | 0.028110994 | -0.208346601 | 0.978 | 0.997 | 1 | 0.52905622 | 0.05543176 |
| Zfp949 | 0.965640838 | 1.223110359 | 0.19 | 0.224 | 1 | 0.028113178 | -0.108340792 | 0.178 | 0.388 | 1 | 0.965640838 | 0.055436005 |
| Pgs1 | 0.972913247 | 0.495434526 | 0.333 | 0.41 | 1 | 0.028153242 | -0.556393349 | 0.2 | 0.412 | 1 | 0.972913247 | 0.055513879 |
| Myo1g | 0.028216798 | 2.673771768 | 0.095 | 0.019 | 1 | 0.376837034 | -0.124282336 | 0 | 0.017 | 1 | 0.376837034 | 0.055637409 |
| Gm13375 | 0.11606501 | -0.672003069 | 0.048 | 0.194 | 1 | 0.028244181 | -1.316927414 | 0.022 | 0.142 | 1 | 0.11606501 | 0.055690628 |
| Ubald1 | 0.050968334 | -0.840801405 | 0.143 | 0.377 | 1 | 0.02825235 | -0.602329633 | 0.133 | 0.304 | 1 | 0.050968334 | 0.055706504 |
| Gpr146 | 0.028294424 | -1.818081329 | 0.048 | 0.261 | 1 | 0.308500297 | -0.564854927 | 0.089 | 0.145 | 1 | 0.308500297 | 0.055788273 |
| Brix1 | 0.058748875 | -0.619827604 | 0.429 | 0.675 | 1 | 0.028323905 | -0.349942471 | 0.533 | 0.709 | 1 | 0.058748875 | 0.055845566 |
| Naaa | 0.028330318 | -2.25696557 | 0 | 0.19 | 1 | 0.063600961 | -1.223818009 | 0.022 | 0.114 | 1 | 0.063600961 | 0.05585803 |
| Rab26os | 0.028374322 | -2.25696557 | 0 | 0.19 | 1 | 0.036608996 | -2.124282336 | 0 | 0.09 | 1 | 0.036608996 | 0.055943542 |
| Fbxo31 | 0.028374726 | -2.303508156 | 0 | 0.19 | 1 | 0.069127381 | -1.098287127 | 0.044 | 0.145 | 1 | 0.069127381 | 0.055944327 |
| Rmnd5a | 0.028376638 | 0.744325048 | 0.762 | 0.638 | 1 | 0.064616685 | 0.586319353 | 0.933 | 0.785 | 1 | 0.064616685 | 0.055948043 |
| Tgfa | 0.028984291 | -2.249060372 | 0.238 | 0.478 | 1 | 0.028378679 | -0.55491669 | 0.2 | 0.401 | 1 | 0.028984291 | 0.055952009 |
| Tesc | 0.028381469 | -2.791114281 | 0.048 | 0.261 | 1 | 0.735983802 | 1.163698427 | 0.089 | 0.08 | 1 | 0.735983802 | 0.05595743 |
| Slc25a53 | 0.252341893 | 1.569435108 | 0.19 | 0.112 | 1 | 0.028406264 | -0.597035333 | 0.044 | 0.176 | 1 | 0.252341893 | 0.056005613 |
| Fbxo6 | 0.028415202 | -2.25696557 | 0 | 0.19 | 1 | 0.083192695 | -1.039393438 | 0.089 | 0.194 | 1 | 0.083192695 | 0.05602298 |
| Edem2 | 0.028428653 | -1.752492987 | 0.048 | 0.261 | 1 | 0.06531284 | -0.124282336 | 0.089 | 0.221 | 1 | 0.06531284 | 0.056049119 |
| A330076H08Rik | 0.028431162 | 1.512308345 | 0.381 | 0.22 | 1 | 0.452226106 | 1.479980721 | 0.267 | 0.266 | 1 | 0.452226106 | 0.056053994 |
| Idnk | 0.067765718 | -1.516052791 | 0.048 | 0.213 | 1 | 0.028447581 | -0.84048937 | 0.089 | 0.235 | 1 | 0.067765718 | 0.056085898 |
| Prkaca | 0.086863709 | -0.455511249 | 0.095 | 0.295 | 1 | 0.028490004 | -0.282711698 | 0.111 | 0.28 | 1 | 0.086863709 | 0.056168327 |
| Adnp2 | 0.360014733 | 1.123574685 | 0.286 | 0.231 | 1 | 0.028520319 | -0.731964913 | 0.089 | 0.242 | 1 | 0.360014733 | 0.056227229 |
| Zfp142 | 0.060515205 | -0.413691074 | 0.143 | 0.373 | 1 | 0.028566577 | -0.486852415 | 0.267 | 0.481 | 1 | 0.060515205 | 0.056317105 |
| Cnot2 | 0.028581703 | 0.564650045 | 0.857 | 0.851 | 1 | 0.477946818 | 0.3368678 | 0.956 | 0.872 | 1 | 0.477946818 | 0.056346492 |
| 4933423P22Rik | 0.028587429 | 2.673771768 | 0.095 | 0.019 | 1 | 0.317138221 | 1.683072586 | 0.022 | 0.007 | 1 | 0.317138221 | 0.056357616 |
| Wnk2 | 0.428062885 | 1.129451251 | 0.19 | 0.142 | 1 | 0.028640437 | 0.728876276 | 0.178 | 0.381 | 1 | 0.428062885 | 0.0564606 |
| Ulk4 | 0.028679479 | 1.722026936 | 0.429 | 0.269 | 1 | 0.512670151 | 1.145107565 | 0.222 | 0.211 | 1 | 0.512670151 | 0.056536445 |
| Aff3 | 0.172043869 | 0.948770764 | 0.81 | 0.832 | 1 | 0.028693358 | 0.710716746 | 0.911 | 0.82 | 1 | 0.172043869 | 0.056563407 |
| Zc3h10 | 0.078679184 | -1.880817084 | 0 | 0.131 | 1 | 0.028697496 | -1.686161223 | 0.044 | 0.17 | 1 | 0.078679184 | 0.056571446 |
| Birc2 | 0.028727921 | -0.326228232 | 0.143 | 0.425 | 1 | 0.035639002 | -0.204452685 | 0.156 | 0.343 | 1 | 0.035639002 | 0.056630549 |
| Zfp770 | 0.590227247 | 0.119182916 | 0.095 | 0.142 | 1 | 0.02873075 | -1.017367132 | 0.022 | 0.142 | 1 | 0.590227247 | 0.056636044 |
| Apof | 0.028781675 | -2.785659851 | 0 | 0.19 | 1 | 0.49622258 | 0.361144491 | 0 | 0.01 | 1 | 0.49622258 | 0.056734964 |
| Psmb10 | 0.02879097 | -2.801961663 | 0 | 0.19 | 1 | 0.155745541 | 0.78260826 | 0.067 | 0.152 | 1 | 0.155745541 | 0.056753019 |
| Nfkbib | 0.128885929 | -0.509450056 | 0.143 | 0.325 | 1 | 0.028795299 | -0.392215541 | 0.133 | 0.315 | 1 | 0.128885929 | 0.056761428 |
| Stat1 | 0.028815614 | -1.777439344 | 0.048 | 0.272 | 1 | 0.677295203 | 0.867155651 | 0.222 | 0.287 | 1 | 0.677295203 | 0.056800888 |
| Ahi1 | 0.050931113 | -0.222392421 | 0.19 | 0.504 | 1 | 0.028815656 | -0.290932205 | 0.2 | 0.412 | 1 | 0.050931113 | 0.05680097 |
| Plpp2 | 0.380297304 | -0.326228232 | 0.048 | 0.108 | 1 | 0.028847516 | -0.549588171 | 0.022 | 0.142 | 1 | 0.380297304 | 0.056862854 |
| Prkag2 | 0.046529307 | 1.627078532 | 0.571 | 0.463 | 1 | 0.028851842 | 1.248025126 | 0.689 | 0.519 | 1 | 0.046529307 | 0.056871255 |
| Trim56 | 0.449110995 | 0.425844254 | 0.143 | 0.246 | 1 | 0.028895866 | -0.680675684 | 0.178 | 0.363 | 1 | 0.449110995 | 0.056956762 |
| Nle1 | 0.028980676 | 2.281454345 | 0.286 | 0.138 | 1 | 0.080069567 | -0.564854927 | 0.022 | 0.107 | 1 | 0.080069567 | 0.057121473 |
| Commd1 | 0.029012628 | -0.525383945 | 0.381 | 0.728 | 1 | 0.074181587 | -0.248540439 | 0.622 | 0.685 | 1 | 0.074181587 | 0.057183524 |
| Gm11837 | 0.236481694 | -0.849790188 | 0 | 0.063 | 1 | 0.029015055 | -0.247664751 | 0.022 | 0.142 | 1 | 0.236481694 | 0.057188237 |
| Gm56764 | 0.029024262 | 2.673771768 | 0.143 | 0.041 | 1 | 0.344256744 | -0.84048937 | 0.022 | 0.055 | 1 | 0.344256744 | 0.057206115 |
| Stambpl1 | 0.295441689 | -0.314912919 | 0.143 | 0.261 | 1 | 0.029040835 | -1.84048937 | 0.067 | 0.204 | 1 | 0.295441689 | 0.057238301 |
| Puf60 | 0.029073644 | -0.413691074 | 0.381 | 0.724 | 1 | 0.055324029 | -0.516599759 | 0.689 | 0.72 | 1 | 0.055324029 | 0.057302012 |
| Klc3 | 0.722967451 | 0.673771768 | 0.048 | 0.071 | 1 | 0.029195117 | -1.614607962 | 0.044 | 0.17 | 1 | 0.722967451 | 0.057537878 |
| Caap1 | 0.732637342 | 0.190688881 | 0.238 | 0.295 | 1 | 0.029246735 | -0.776359032 | 0.133 | 0.301 | 1 | 0.732637342 | 0.057638098 |
| Ak2 | 0.029251734 | -0.839297815 | 0.238 | 0.549 | 1 | 0.342810064 | -0.23159536 | 0.533 | 0.557 | 1 | 0.342810064 | 0.057647803 |
| Cstf3 | 0.98995193 | 0.712245915 | 0.571 | 0.687 | 1 | 0.029303886 | 0.753461914 | 0.911 | 0.813 | 1 | 0.98995193 | 0.057749054 |
| Angpt1 | 0.376595309 | -2.04047375 | 0.048 | 0.104 | 1 | 0.029340016 | -0.941418279 | 0.044 | 0.18 | 1 | 0.376595309 | 0.057819195 |
| Rfng | 0.611826876 | 0.673771768 | 0.095 | 0.142 | 1 | 0.029360131 | -1.589945908 | 0.044 | 0.17 | 1 | 0.611826876 | 0.057858244 |
| Pid1 | 0.258988 | 0.434746754 | 0.286 | 0.507 | 1 | 0.029381588 | 0.320502507 | 0.156 | 0.367 | 1 | 0.258988 | 0.057899898 |
| Mtrf1l | 0.075196364 | -1.683780237 | 0.048 | 0.205 | 1 | 0.029459912 | -0.918963428 | 0.067 | 0.211 | 1 | 0.075196364 | 0.058051938 |
| Fbxo7 | 0.636873983 | 0.242980197 | 0.429 | 0.586 | 1 | 0.029473118 | -0.54109546 | 0.244 | 0.464 | 1 | 0.636873983 | 0.058077571 |
| 9330020H09Rik | 0.163379068 | -0.970084422 | 0 | 0.086 | 1 | 0.029541516 | -2.223818009 | 0 | 0.097 | 1 | 0.163379068 | 0.058210332 |
| 1700037H04Rik | 0.113581507 | -1.574155746 | 0 | 0.108 | 1 | 0.029561464 | -2.271123724 | 0 | 0.097 | 1 | 0.113581507 | 0.058249048 |
| Zdhhc24 | 0.128212019 | -1.370622352 | 0 | 0.101 | 1 | 0.029561464 | -2.271123724 | 0 | 0.097 | 1 | 0.128212019 | 0.058249048 |
| Peg13 | 0.173820841 | -1.081115734 | 0 | 0.082 | 1 | 0.029579881 | -2.446210431 | 0 | 0.097 | 1 | 0.173820841 | 0.058284792 |
| Fbxo45 | 0.036793905 | -1.911190733 | 0.048 | 0.246 | 1 | 0.029594856 | -0.223818009 | 0.2 | 0.405 | 1 | 0.036793905 | 0.058313856 |
| 4933407K13Rik | 0.159808134 | 1.432763668 | 0.238 | 0.142 | 1 | 0.029598037 | -0.847442131 | 0.089 | 0.239 | 1 | 0.159808134 | 0.05832003 |
| Mlf1 | 0.029605995 | -2.522625445 | 0.048 | 0.265 | 1 | 0.073401694 | -0.528977891 | 0.089 | 0.215 | 1 | 0.073401694 | 0.058335475 |
| Wfs1 | 0.127996876 | -1.233118828 | 0 | 0.101 | 1 | 0.02961246 | -2.486852415 | 0 | 0.097 | 1 | 0.127996876 | 0.058348022 |
| Nat14 | 0.285465396 | -0.326228232 | 0 | 0.052 | 1 | 0.02961246 | -2.564854927 | 0 | 0.097 | 1 | 0.285465396 | 0.058348022 |
| Syce2 | 0.153935113 | -1.370622352 | 0 | 0.09 | 1 | 0.029612687 | -2.404390255 | 0 | 0.097 | 1 | 0.153935113 | 0.058348464 |
| Gm20033 | 0.88676233 | 1.629377648 | 0.095 | 0.093 | 1 | 0.029612687 | -2.404390255 | 0 | 0.097 | 1 | 0.88676233 | 0.058348464 |
| Maf | 0.029618354 | -1.191471131 | 0.381 | 0.619 | 1 | 0.088524884 | -0.282566818 | 0.644 | 0.886 | 1 | 0.088524884 | 0.058359461 |
| Qrsl1 | 0.196462148 | -1.233118828 | 0 | 0.075 | 1 | 0.029626507 | -2.638855509 | 0 | 0.097 | 1 | 0.196462148 | 0.058375285 |
| Slc39a6 | 0.097317141 | 0.160148049 | 0.238 | 0.552 | 1 | 0.029632716 | -0.355846403 | 0.378 | 0.55 | 1 | 0.097317141 | 0.058387334 |
| Srek1ip1 | 0.849026053 | 0.333921765 | 0.19 | 0.22 | 1 | 0.029634502 | -1.048731303 | 0.067 | 0.204 | 1 | 0.849026053 | 0.0583908 |
| Kif5a | 0.904268355 | 0.719575457 | 0.095 | 0.09 | 1 | 0.02963959 | -2.638855509 | 0 | 0.097 | 1 | 0.904268355 | 0.058400674 |
| Gm49708 | 0.200677602 | 1.351843673 | 0.19 | 0.104 | 1 | 0.029639988 | -2.638855509 | 0 | 0.097 | 1 | 0.200677602 | 0.058401447 |
| Akap17b | 0.324789396 | -0.413691074 | 0 | 0.045 | 1 | 0.029660923 | -2.709244837 | 0 | 0.097 | 1 | 0.324789396 | 0.058442076 |
| Anapc10 | 0.78923796 | 0.496233582 | 0.381 | 0.466 | 1 | 0.029678205 | 0.133733995 | 0.244 | 0.474 | 1 | 0.78923796 | 0.058475614 |
| Ndufb1 | 0.029684678 | -0.56376414 | 0.524 | 0.754 | 1 | 0.595842428 | -0.232535227 | 0.689 | 0.633 | 1 | 0.595842428 | 0.058488175 |
| Pcdhb22 | 0.236454006 | -1.081115734 | 0 | 0.063 | 1 | 0.029696604 | -3.044847868 | 0 | 0.097 | 1 | 0.236454006 | 0.05851132 |
| Moxd1 | 0.587441545 | 0.596327682 | 0.381 | 0.354 | 1 | 0.029719335 | -1.167783975 | 0.111 | 0.27 | 1 | 0.587441545 | 0.058555431 |
| Tmem231 | 0.291149844 | 0.673771768 | 0.048 | 0.134 | 1 | 0.029727857 | -2.174908409 | 0 | 0.097 | 1 | 0.291149844 | 0.058571969 |
| Rbm39 | 0.094408456 | 0.309230671 | 1 | 1 | 1 | 0.029741839 | 0.243208066 | 1 | 1 | 1 | 0.094408456 | 0.058599101 |
| Ube2u | 0.425631778 | -0.326228232 | 0 | 0.03 | 1 | 0.029782503 | 2.123645178 | 0.111 | 0.038 | 1 | 0.425631778 | 0.058678008 |
| Ddx20 | 0.039330597 | -2.159118246 | 0 | 0.172 | 1 | 0.02978709 | -0.606434031 | 0.089 | 0.249 | 1 | 0.039330597 | 0.05868691 |
| Egln1 | 0.029808906 | -0.832580898 | 0.286 | 0.563 | 1 | 0.037176694 | -0.370442923 | 0.556 | 0.664 | 1 | 0.037176694 | 0.058729241 |

| | | | | | | | | | | | | |
|---|---|---|---|---|---|---|---|---|---|---|---|---|
| Rpgrip1l | 0.029821048 | -1.984439715 | 0.048 | 0.257 | 1 | 0.169862618 | 0.605070074 | 0.133 | 0.249 | 1 | 0.169862618 | 0.058752802 |
| Ldlrap1 | 0.129288743 | -0.284649129 | 0.238 | 0.455 | 1 | 0.029847315 | 0.174523201 | 0.267 | 0.491 | 1 | 0.129288743 | 0.058803768 |
| Rpp38 | 0.085578898 | -1.496153234 | 0.048 | 0.198 | 1 | 0.02986554 | -2.044847868 | 0.022 | 0.135 | 1 | 0.085578898 | 0.05883913 |
| Setd5 | 0.029865711 | 0.620404838 | 1 | 0.933 | 1 | 0.175312694 | 0.439147003 | 0.978 | 0.993 | 1 | 0.175312694 | 0.058839461 |
| Helz | 0.562905634 | 0.448705212 | 0.952 | 0.899 | 1 | 0.029870678 | -0.198943341 | 0.911 | 0.948 | 1 | 0.562905634 | 0.0588491 |
| Prpf4b | 0.029920869 | 0.457357748 | 0.952 | 0.929 | 1 | 0.977940493 | 0.128836601 | 0.978 | 0.934 | 1 | 0.977940493 | 0.058946479 |
| Msh6 | 0.930536335 | 0.503846766 | 0.095 | 0.104 | 1 | 0.029956206 | -2.044847868 | 0.022 | 0.135 | 1 | 0.930536335 | 0.059015038 |
| Enpp4 | 0.356705238 | 0.162809848 | 0.095 | 0.183 | 1 | 0.030046209 | -2.017367132 | 0.022 | 0.135 | 1 | 0.356705238 | 0.059189644 |
| Nog | 0.030104684 | -3.445169305 | 0.048 | 0.254 | 1 | 0.086210295 | -2.071814916 | 0.022 | 0.1 | 1 | 0.086210295 | 0.059303076 |
| Gatc | 0.053823925 | -1.785659851 | 0 | 0.153 | 1 | 0.030135083 | -1.84048937 | 0.022 | 0.135 | 1 | 0.053823925 | 0.059362043 |
| LTO1 | 0.030218362 | -2.233118828 | 0 | 0.187 | 1 | 0.280515421 | 0.132875504 | 0.067 | 0.125 | 1 | 0.280515421 | 0.059523574 |
| Pih1d1 | 0.055451921 | -1.611630451 | 0.048 | 0.224 | 1 | 0.030219536 | -0.468930507 | 0.067 | 0.218 | 1 | 0.055451921 | 0.059525851 |
| Prim1 | 0.267226869 | -0.184209227 | 0.048 | 0.138 | 1 | 0.030272852 | -1.638855509 | 0.022 | 0.138 | 1 | 0.267226869 | 0.059629258 |
| Ppp2r2a | 0.228209652 | 0.401454121 | 0.952 | 0.866 | 1 | 0.030283698 | 0.494769055 | 0.978 | 0.924 | 1 | 0.228209652 | 0.059650294 |
| Mta2 | 0.031319965 | -1.242704877 | 0.095 | 0.343 | 1 | 0.030307246 | -0.571741313 | 0.444 | 0.571 | 1 | 0.031319965 | 0.059695962 |
| Ppp1r9b | 0.030431494 | -2.326228232 | 0 | 0.187 | 1 | 0.20193512 | 0.126124462 | 0.222 | 0.339 | 1 | 0.20193512 | 0.059936912 |
| Meltf | 0.465201684 | -0.543766101 | 0.81 | 0.724 | 1 | 0.030446487 | -0.639798139 | 0.489 | 0.706 | 1 | 0.465201684 | 0.059965986 |
| Ankhd1 | 0.052355458 | 0.630997744 | 0.952 | 0.828 | 1 | 0.030473313 | 0.439336346 | 0.956 | 0.893 | 1 | 0.052355458 | 0.060018004 |
| Vasp | 0.224607142 | -0.160055366 | 0.429 | 0.664 | 1 | 0.030480086 | -0.27904753 | 0.822 | 0.806 | 1 | 0.224607142 | 0.060031135 |
| Phf11d | 0.492599419 | -0.133583154 | 0 | 0.022 | 1 | 0.030581005 | 4.798549804 | 0.044 | 0.007 | 1 | 0.492599419 | 0.060226813 |
| Zbtb32 | 0.532436372 | 1.088809267 | 0 | 0.019 | 1 | 0.030581005 | 3.905465008 | 0.044 | 0.007 | 1 | 0.532436372 | 0.060226813 |
| Tmem275 | 0.57816106 | 0.351843673 | 0 | 0.015 | 1 | 0.030581005 | 3.42003818 | 0.044 | 0.007 | 1 | 0.57816106 | 0.060226813 |
| Ggt7 | 0.629723166 | 2.896164189 | 0.048 | 0.03 | 1 | 0.030581005 | 4.42003818 | 0.044 | 0.007 | 1 | 0.629723166 | 0.060226813 |
| Gm57096 | 0.632076134 | 1.673771768 | 0 | 0.011 | 1 | 0.030581005 | 3.683072586 | 0.044 | 0.007 | 1 | 0.632076134 | 0.060226813 |
| Syngr2 | 0.26382005 | -0.785659851 | 0.048 | 0.131 | 1 | 0.030601929 | 0.533325467 | 0.333 | 0.18 | 1 | 0.26382005 | 0.06026738 |
| Vps26a | 0.40584463 | 0.366343242 | 0.333 | 0.578 | 1 | 0.030630148 | -0.376469359 | 0.822 | 0.865 | 1 | 0.40584463 | 0.06032209 |
| Nr2c2ap | 0.127649013 | -1.184209227 | 0.048 | 0.175 | 1 | 0.030689566 | -1.017367132 | 0.067 | 0.204 | 1 | 0.127649013 | 0.060437283 |
| Src | 0.241080731 | -0.392317423 | 0.095 | 0.209 | 1 | 0.030720443 | -0.886783022 | 0.089 | 0.239 | 1 | 0.241080731 | 0.06049714 |
| Scand1 | 0.030731884 | -0.656617906 | 0.381 | 0.634 | 1 | 0.060753023 | -0.579149263 | 0.6 | 0.644 | 1 | 0.060753023 | 0.06051932 |
| Osgepl1 | 0.195766569 | -0.718545655 | 0.048 | 0.157 | 1 | 0.03078895 | -1.459885368 | 0.022 | 0.138 | 1 | 0.195766569 | 0.060629941 |
| Cbx4 | 0.092930612 | -0.429039038 | 0.143 | 0.347 | 1 | 0.030798314 | 0.206634542 | 0.644 | 0.398 | 1 | 0.092930612 | 0.060648092 |
| Txnrd3 | 0.036913874 | -2.303508156 | 0 | 0.175 | 1 | 0.030808467 | -0.979892427 | 0.044 | 0.173 | 1 | 0.036913874 | 0.060667772 |
| Pmm1 | 0.030816504 | -1.004300137 | 0.095 | 0.354 | 1 | 0.080320113 | -0.255526869 | 0.156 | 0.294 | 1 | 0.080320113 | 0.060683351 |
| Zbtb7a | 0.52646539 | 0.328943271 | 0.714 | 0.679 | 1 | 0.030839348 | -0.476975736 | 0.956 | 0.913 | 1 | 0.52646539 | 0.06072763 |
| Gins4 | 0.05399559 | -1.940938076 | 0 | 0.153 | 1 | 0.030841525 | -1.223818009 | 0.044 | 0.173 | 1 | 0.05399559 | 0.06073185 |
| Ddhd2 | 0.03087313 | -0.154047257 | 0.238 | 0.563 | 1 | 0.764411677 | 0.484292722 | 0.489 | 0.682 | 1 | 0.764411677 | 0.06079311 |
| Stk38l | 0.030900126 | 1.310190052 | 0.81 | 0.746 | 1 | 0.62243774 | 0.844863875 | 0.6 | 0.779 | 1 | 0.62243774 | 0.060845434 |
| Slc4a3 | 0.485133023 | 0.696491844 | 0.238 | 0.183 | 1 | 0.030924047 | -0.564854927 | 0.067 | 0.211 | 1 | 0.485133023 | 0.060891797 |
| Tmsb10 | 0.032569675 | -1.487951421 | 0.714 | 0.713 | 1 | 0.030962873 | -0.837635373 | 0.889 | 0.779 | 1 | 0.032569675 | 0.060967047 |
| Cog1 | 0.036821875 | -2.107587946 | 0 | 0.175 | 1 | 0.031096131 | -0.255526869 | 0.089 | 0.249 | 1 | 0.036821875 | 0.061225292 |
| Ddx24 | 0.031104424 | -0.488678113 | 0.667 | 0.873 | 1 | 0.424174781 | 0.493377203 | 0.889 | 0.848 | 1 | 0.424174781 | 0.061241364 |
| Acot8 | 0.036868724 | -2.184209227 | 0 | 0.175 | 1 | 0.031138623 | -1.039393438 | 0.044 | 0.173 | 1 | 0.036868724 | 0.061307632 |
| Nol9 | 0.191454721 | 1.503846766 | 0.333 | 0.261 | 1 | 0.031163855 | -0.477919291 | 0.156 | 0.346 | 1 | 0.191454721 | 0.061356525 |
| Matr3 | 0.076109231 | -0.356693513 | 0.857 | 0.974 | 1 | 0.031167635 | -0.13424521 | 1 | 0.986 | 1 | 0.076109231 | 0.061363849 |
| Dedd | 0.031186988 | -1.120644099 | 0.095 | 0.336 | 1 | 0.346975994 | 0.392395425 | 0.222 | 0.332 | 1 | 0.346975994 | 0.061401347 |
| Mllt11 | 0.154582393 | -0.732220592 | 0.143 | 0.291 | 1 | 0.031188707 | -0.326911502 | 0.156 | 0.339 | 1 | 0.154582393 | 0.061404678 |
| Npr2 | 0.08175365 | -0.484657595 | 0.143 | 0.358 | 1 | 0.031188916 | -0.182910066 | 0.244 | 0.505 | 1 | 0.08175365 | 0.061405083 |
| Zrsr2 | 0.675780542 | 0.195180571 | 0.476 | 0.683 | 1 | 0.031190349 | -0.435484024 | 0.578 | 0.716 | 1 | 0.675780542 | 0.061407861 |
| Hspb6 | 0.031228057 | -2.348596045 | 0.048 | 0.254 | 1 | 0.183590103 | -0.602329633 | 0.044 | 0.111 | 1 | 0.183590103 | 0.061480923 |
| Gm28153 | 0.091782708 | -1.611630451 | 0.048 | 0.194 | 1 | 0.031228227 | -0.486852415 | 0.067 | 0.215 | 1 | 0.091782708 | 0.061481253 |
| Prnd | 0.07959214 | 4.410737362 | 0.048 | 0.007 | 1 | 0.031264596 | 3.42003818 | 0.044 | 0.007 | 1 | 0.07959214 | 0.061551717 |
| Gm49083 | 0.080927838 | 3.258734268 | 0.048 | 0.007 | 1 | 0.031264596 | 3.683072586 | 0.044 | 0.007 | 1 | 0.080927838 | 0.061551717 |
| Tnfsf10 | 0.269367392 | 1.79930265 | 0.048 | 0.015 | 1 | 0.031264596 | 3.42003818 | 0.044 | 0.007 | 1 | 0.269367392 | 0.061551717 |
| Cd4 | 0.425629641 | -0.849790188 | 0 | 0.03 | 1 | 0.031264596 | 3.42003818 | 0.044 | 0.007 | 1 | 0.425629641 | 0.061551717 |
| Gm57367 | 0.532436372 | 1.088809267 | 0 | 0.019 | 1 | 0.031264596 | 3.098110085 | 0.044 | 0.007 | 1 | 0.532436372 | 0.061551717 |
| Nrxn3 | 0.492608234 | 0.866416846 | 0 | 0.022 | 1 | 0.031266129 | 3.168499413 | 0.044 | 0.007 | 1 | 0.492608234 | 0.061554688 |
| Usp10 | 0.031267688 | -1.564632972 | 0.143 | 0.362 | 1 | 0.714127346 | 0.19121949 | 0.711 | 0.595 | 1 | 0.714127346 | 0.061557707 |
| Rp2 | 0.117977257 | -1.370622352 | 0.048 | 0.179 | 1 | 0.031308244 | -0.709244837 | 0.089 | 0.239 | 1 | 0.117977257 | 0.061636281 |
| Runx1t1 | 0.475179585 | -0.637814383 | 0.095 | 0.153 | 1 | 0.031326108 | -2.65082815 | 0.044 | 0.163 | 1 | 0.475179585 | 0.061670892 |
| Cln3 | 0.920524815 | 0.735172312 | 0.143 | 0.146 | 1 | 0.031343985 | -1.486852415 | 0.067 | 0.197 | 1 | 0.920524815 | 0.061705524 |
| Has2os | 0.060441274 | 0.681633885 | 0.619 | 0.489 | 1 | 0.031351276 | 0.113540126 | 0.156 | 0.349 | 1 | 0.060441274 | 0.061719649 |
| Scrn3 | 0.171319218 | -0.548620654 | 0.048 | 0.168 | 1 | 0.031458785 | 0.635766871 | 0.044 | 0.18 | 1 | 0.171319218 | 0.061927914 |
| Uso1 | 0.41486694 | 0.155923463 | 0.762 | 0.862 | 1 | 0.031483189 | -0.281676567 | 0.889 | 0.934 | 1 | 0.41486694 | 0.061975186 |
| B4galt3 | 0.316053046 | -0.208871282 | 0.095 | 0.19 | 1 | 0.031493069 | -0.776359032 | 0.089 | 0.246 | 1 | 0.316053046 | 0.061994325 |
| Ppcdc | 0.850439119 | 1.923750021 | 0.095 | 0.09 | 1 | 0.031495787 | -1.199570463 | 0.022 | 0.138 | 1 | 0.850439119 | 0.06199959 |
| Hps4 | 0.129816799 | -0.379339569 | 0.095 | 0.269 | 1 | 0.031509084 | -0.316927414 | 0.067 | 0.211 | 1 | 0.129816799 | 0.062025345 |
| Nlgn1 | 0.246676543 | 1.649925026 | 0.143 | 0.075 | 1 | 0.031517666 | -4.350350415 | 0.022 | 0.131 | 1 | 0.246676543 | 0.062041969 |
| Gm10575 | 0.031519679 | 2.673771768 | 0.143 | 0.041 | 1 | 0.086186001 | -2.071814916 | 0 | 0.062 | 1 | 0.086186001 | 0.062045867 |
| Nrip1 | 0.51008503 | 0.773883252 | 0.857 | 0.854 | 1 | 0.031552088 | 0.479490015 | 1 | 0.824 | 1 | 0.51008503 | 0.062108642 |
| Fam136a | 0.229133793 | -0.326228232 | 0.143 | 0.28 | 1 | 0.031565633 | -0.709244837 | 0.044 | 0.173 | 1 | 0.229133793 | 0.062134878 |
| B3gnt5 | 0.63207476 | 1.351843673 | 0 | 0.011 | 1 | 0.031613022 | 3.683072586 | 0.044 | 0.007 | 1 | 0.63207476 | 0.06222666 |
| Arap1 | 0.913916989 | 1.107424945 | 0.19 | 0.216 | 1 | 0.031729166 | -0.339295227 | 0.111 | 0.284 | 1 | 0.913916989 | 0.062451592 |
| Zfp276 | 0.884607017 | 1.304537958 | 0.095 | 0.093 | 1 | 0.031741366 | -1.754332726 | 0.044 | 0.166 | 1 | 0.884607017 | 0.062475218 |
| Dnaja3 | 0.335898589 | 0.165977127 | 0.238 | 0.366 | 1 | 0.031757753 | 0.289294206 | 0.222 | 0.488 | 1 | 0.335898589 | 0.062506952 |
| Rxra | 0.031820358 | -1.849790188 | 0.048 | 0.254 | 1 | 0.372672364 | 0.42003818 | 0.222 | 0.343 | 1 | 0.372672364 | 0.06262818 |
| Dnmbp | 0.031821603 | 1.232336578 | 0.571 | 0.41 | 1 | 0.558515808 | 0.617919907 | 0.6 | 0.564 | 1 | 0.558515808 | 0.062630592 |
| Apc | 0.475815516 | 0.318467536 | 0.714 | 0.683 | 1 | 0.031833386 | 0.619932997 | 0.933 | 0.834 | 1 | 0.475815516 | 0.062653407 |
| Mvp | 0.206480346 | -0.413691074 | 0.143 | 0.299 | 1 | 0.031864354 | -0.737259213 | 0.178 | 0.353 | 1 | 0.206480346 | 0.062713372 |
| Ccnl2 | 0.435646905 | 0.531083351 | 0.81 | 0.91 | 1 | 0.031873771 | 0.377058981 | 1 | 0.955 | 1 | 0.435646905 | 0.062731604 |
| Enoph1 | 0.036911244 | -2.159118246 | 0 | 0.175 | 1 | 0.031890164 | -0.699397051 | 0.067 | 0.204 | 1 | 0.036911244 | 0.062763346 |
| Ogfod1 | 0.093693368 | -0.93109029 | 0.143 | 0.34 | 1 | 0.031922095 | -0.408425768 | 0.2 | 0.401 | 1 | 0.093693368 | 0.062825169 |
| Frmpd3 | 0.492590603 | 0.673771768 | 0 | 0.022 | 1 | 0.0319636 | 2.905465008 | 0.044 | 0.007 | 1 | 0.492590603 | 0.062905529 |
| Klhl41 | 0.63207476 | 1.351843673 | 0 | 0.011 | 1 | 0.031963989 | 2.683072586 | 0.044 | 0.007 | 1 | 0.63207476 | 0.062906282 |
| Gm56786 | 0.766624392 | 0.735172312 | 0.048 | 0.067 | 1 | 0.031963989 | 2.683072586 | 0.044 | 0.007 | 1 | 0.766624392 | 0.062906282 |
| Cbfa2t2 | 0.343745856 | 0.486144765 | 0.524 | 0.451 | 1 | 0.031995309 | 1.156443396 | 0.711 | 0.644 | 1 | 0.343745856 | 0.062966919 |
| Wdr53 | 0.122728228 | -0.701267664 | 0.095 | 0.254 | 1 | 0.032004354 | -0.515707278 | 0.089 | 0.246 | 1 | 0.122728228 | 0.062984429 |
| Slc15a4 | 0.224622546 | -0.496153234 | 0.095 | 0.213 | 1 | 0.032015897 | -0.330733213 | 0.111 | 0.284 | 1 | 0.224622546 | 0.063006777 |
| Mettl13 | 0.299350787 | 1.623145695 | 0.143 | 0.082 | 1 | 0.032019607 | -0.989352756 | 0.022 | 0.138 | 1 | 0.299350787 | 0.063013959 |
| Mgll | 0.032089326 | 1.610661724 | 0.667 | 0.597 | 1 | 0.182085274 | 0.323176641 | 0.533 | 0.654 | 1 | 0.182085274 | 0.063148928 |
| Spin4 | 0.100277091 | -1.455511249 | 0 | 0.116 | 1 | 0.032113045 | -1.662702251 | 0.044 | 0.166 | 1 | 0.100277091 | 0.063194843 |
| Sh3glb2 | 0.032223011 | -2.133583154 | 0 | 0.183 | 1 | 0.035207637 | -0.589945908 | 0.044 | 0.17 | 1 | 0.035207637 | 0.0634077 |
| Copa | 0.160511544 | 0.201376178 | 1 | 0.981 | 1 | 0.032246019 | -0.118023345 | 0.978 | 0.993 | 1 | 0.160511544 | 0.063452233 |
| Cpsf3 | 0.510931959 | 0.673771768 | 0.19 | 0.291 | 1 | 0.03228386 | -0.499130745 | 0.133 | 0.301 | 1 | 0.510931959 | 0.063525472 |
| Pip4k2b | 0.176592071 | -0.695462042 | 0.048 | 0.164 | 1 | 0.032295988 | -1.003427941 | 0.111 | 0.263 | 1 | 0.176592071 | 0.063548946 |
| 6820431F20Rik | 0.032305984 | 0.35834231 | 0.905 | 0.892 | 1 | 0.136669285 | 0.372921659 | 0.978 | 0.958 | 1 | 0.136669285 | 0.06356829 |
| Scin | 0.109401103 | -0.412574004 | 0.524 | 0.728 | 1 | 0.032306699 | 0.235329082 | 0.378 | 0.689 | 1 | 0.109401103 | 0.063569675 |
| Gm39556 | 0.271980818 | 2.351843673 | 0.048 | 0.015 | 1 | 0.032317506 | 2.197645759 | 0.044 | 0.007 | 1 | 0.271980818 | 0.063590592 |
| Snhg16 | 0.691807751 | 0.410737362 | 0.19 | 0.254 | 1 | 0.032341538 | -0.609709163 | 0.089 | 0.239 | 1 | 0.691807751 | 0.063637102 |

| | | | | | | | | | | | | |
|---|---|---|---|---|---|---|---|---|---|---|---|---|
| Cry1 | 0.075598877 | -1.073462162 | 0.19 | 0.388 | 1 | 0.032353447 | -0.316927414 | 0.378 | 0.585 | 1 | 0.075598877 | 0.063660148 |
| Cpsf4 | 0.032378014 | 1.791128718 | 0.333 | 0.175 | 1 | 0.153199211 | -0.142898014 | 0.111 | 0.215 | 1 | 0.153199211 | 0.063707692 |
| Emsy | 0.032415993 | -0.63701577 | 0.238 | 0.493 | 1 | 0.306967718 | 0.350364653 | 0.733 | 0.63 | 1 | 0.306967718 | 0.06378119 |
| Dnaaf10 | 0.042108512 | -2.392317423 | 0 | 0.168 | 1 | 0.032443158 | -1.255526869 | 0.067 | 0.201 | 1 | 0.042108512 | 0.063833757 |
| Trmt2b | 0.032449398 | -2.303508156 | 0 | 0.183 | 1 | 0.063051399 | -1.80878051 | 0.022 | 0.111 | 1 | 0.063051399 | 0.063845832 |
| Smarca2 | 0.05907312 | -0.197904135 | 0.619 | 0.802 | 1 | 0.032473631 | -0.316927414 | 0.889 | 0.844 | 1 | 0.05907312 | 0.063892726 |
| Edc3 | 0.197663857 | -0.648156327 | 0.095 | 0.22 | 1 | 0.032479441 | -0.86124793 | 0.111 | 0.27 | 1 | 0.197663857 | 0.063903967 |
| Chac1 | 0.746187038 | -0.438928365 | 0.19 | 0.157 | 1 | 0.032519598 | -2.583713955 | 0.022 | 0.131 | 1 | 0.746187038 | 0.063981673 |
| Maml3 | 0.048356584 | 1.038183194 | 0.667 | 0.537 | 1 | 0.032522128 | 1.013594266 | 0.844 | 0.595 | 1 | 0.048356584 | 0.063986568 |
| Cdc42bpb | 0.959853053 | 0.332303486 | 0.571 | 0.735 | 1 | 0.032565375 | 0.52614943 | 0.956 | 0.841 | 1 | 0.959853053 | 0.064070246 |
| Sanbr | 0.053733896 | 0.150209812 | 0.143 | 0.381 | 1 | 0.032587048 | -0.257370569 | 0.2 | 0.426 | 1 | 0.053733896 | 0.06411218 |
| Abhd4 | 0.104302643 | -1.434752689 | 0.048 | 0.187 | 1 | 0.032624391 | -0.901889915 | 0.089 | 0.232 | 1 | 0.104302643 | 0.064184431 |
| Rc3h1 | 0.175528944 | 0.451379346 | 0.762 | 0.873 | 1 | 0.03264383 | 0.55984082 | 1 | 0.955 | 1 | 0.175528944 | 0.06422204 |
| Gpr137 | 0.205567657 | -0.695462042 | 0.095 | 0.209 | 1 | 0.032666424 | -1.564854927 | 0.044 | 0.166 | 1 | 0.205567657 | 0.064265753 |
| Prim2 | 0.032679982 | 1.735172312 | 0.381 | 0.216 | 1 | 0.705155681 | 0.766904177 | 0.244 | 0.242 | 1 | 0.705155681 | 0.064291983 |
| Wdr3 | 0.188138062 | -0.264827688 | 0.095 | 0.239 | 1 | 0.032719338 | -0.674479418 | 0.133 | 0.304 | 1 | 0.188138062 | 0.064368121 |
| Lrrn4cl | 0.032791835 | -2.769171728 | 0 | 0.183 | 1 | 0.163242267 | -1.526380779 | 0.022 | 0.08 | 1 | 0.163242267 | 0.064508366 |
| Gm13483 | 0.032854578 | 1.762308442 | 0.333 | 0.172 | 1 | 0.528226585 | 1.0840349 | 0.222 | 0.201 | 1 | 0.528226585 | 0.064629733 |
| Zmat1 | 0.446361877 | -0.39661756 | 0.143 | 0.216 | 1 | 0.03288147 | -1.901889915 | 0.022 | 0.135 | 1 | 0.446361877 | 0.06468175 |
| Sgms1os1 | 0.144664875 | -1.133583154 | 0 | 0.093 | 1 | 0.032889024 | -2.174908409 | 0 | 0.093 | 1 | 0.144664875 | 0.06469636 |
| Efna4 | 0.544173668 | 0.214340149 | 0.048 | 0.09 | 1 | 0.032909546 | -2.223818009 | 0 | 0.093 | 1 | 0.544173668 | 0.064736054 |
| Pfdn1 | 0.032917625 | -0.870548749 | 0.286 | 0.567 | 1 | 0.577941425 | 0.205025289 | 0.578 | 0.606 | 1 | 0.577941425 | 0.064751679 |
| Zfyve19 | 0.196329837 | -0.849790188 | 0 | 0.075 | 1 | 0.03292843 | -2.316927414 | 0 | 0.093 | 1 | 0.196329837 | 0.064772579 |
| Ccdc174 | 0.423962399 | 0.292342661 | 0.333 | 0.507 | 1 | 0.032929841 | -0.149817428 | 0.289 | 0.578 | 1 | 0.423962399 | 0.064775307 |
| Rabgap1 | 0.455464798 | 0.150209812 | 0.714 | 0.675 | 1 | 0.032930177 | 0.850529332 | 0.911 | 0.827 | 1 | 0.455464798 | 0.064775957 |
| Hmga1 | 0.448506256 | -0.326228232 | 0.048 | 0.097 | 1 | 0.032945927 | -2.316927414 | 0 | 0.093 | 1 | 0.448506256 | 0.064806421 |
| Rccd1 | 0.650632629 | 0.673771768 | 0.095 | 0.134 | 1 | 0.032950615 | -2.361321533 | 0.022 | 0.131 | 1 | 0.650632629 | 0.064815487 |
| Ppp1r36 | 0.120716826 | -1.683780237 | 0 | 0.104 | 1 | 0.032988513 | -2.526380779 | 0 | 0.093 | 1 | 0.120716826 | 0.064888785 |
| Slc12a7 | 0.770713471 | 0.351843673 | 0.048 | 0.063 | 1 | 0.032989145 | -2.564854927 | 0 | 0.093 | 1 | 0.770713471 | 0.064890007 |
| Lcorl | 0.235360254 | 0.295905501 | 0.524 | 0.396 | 1 | 0.033009673 | 1.00816796 | 0.756 | 0.581 | 1 | 0.235360254 | 0.064929708 |
| Hyls1 | 0.65600418 | 2.325848464 | 0.048 | 0.078 | 1 | 0.033020748 | -2.871516266 | 0 | 0.093 | 1 | 0.65600418 | 0.064951126 |
| Ctdspl2 | 0.14461475 | -0.144987918 | 0.381 | 0.627 | 1 | 0.033024375 | 0.746637715 | 0.911 | 0.785 | 1 | 0.14461475 | 0.064958141 |
| Serping1 | 0.22231747 | -1.574155746 | 0 | 0.067 | 1 | 0.03302865 | -4.316927414 | 0 | 0.093 | 1 | 0.22231747 | 0.064966408 |
| Polr3c | 0.623717808 | 1.252307999 | 0.238 | 0.216 | 1 | 0.033030524 | -0.316927414 | 0.089 | 0.253 | 1 | 0.623717808 | 0.064970033 |
| Kptn | 0.033035853 | 2.069700444 | 0.381 | 0.209 | 1 | 0.198373399 | 1.150198597 | 0.178 | 0.308 | 1 | 0.198373399 | 0.064980338 |
| Rasl11a | 0.140497022 | -2.341178574 | 0.048 | 0.175 | 1 | 0.033037564 | -3.545746104 | 0 | 0.093 | 1 | 0.140497022 | 0.064983647 |
| Scap | 0.19637264 | -0.399477214 | 0.143 | 0.295 | 1 | 0.033086234 | -0.180865864 | 0.2 | 0.412 | 1 | 0.19637264 | 0.065077769 |
| Ttpal | 0.216750002 | -0.326228232 | 0.048 | 0.153 | 1 | 0.033235657 | -0.260343885 | 0.111 | 0.284 | 1 | 0.216750002 | 0.065366706 |
| Arnt | 0.033258534 | 1.206990693 | 0.714 | 0.75 | 1 | 0.067541294 | 0.659921622 | 0.844 | 0.813 | 1 | 0.067541294 | 0.065410938 |
| Gspt2 | 0.136163632 | -1.280424543 | 0 | 0.097 | 1 | 0.033263794 | -2.017367132 | 0.022 | 0.131 | 1 | 0.136163632 | 0.065421108 |
| Rasal2 | 0.586941755 | 0.632170863 | 0.476 | 0.481 | 1 | 0.033275196 | 0.735540006 | 0.733 | 0.54 | 1 | 0.586941755 | 0.065443154 |
| Lrp3 | 0.088896765 | -1.648156327 | 0 | 0.123 | 1 | 0.033278141 | -0.539319835 | 0.089 | 0.239 | 1 | 0.088896765 | 0.065448847 |
| Lrrk2 | 0.163271828 | -1.04504648 | 0.048 | 0.168 | 1 | 0.033304388 | -0.695439037 | 0.044 | 0.176 | 1 | 0.163271828 | 0.065499594 |
| Usp25 | 0.033818845 | 0.816849631 | 0.857 | 0.683 | 1 | 0.03331678 | 0.606256989 | 0.933 | 0.661 | 1 | 0.033818845 | 0.065523552 |
| Adamts9 | 0.105993173 | -1.993501087 | 0.19 | 0.351 | 1 | 0.033383723 | -0.375821103 | 0.156 | 0.339 | 1 | 0.105993173 | 0.065652974 |
| Zc3hav1l | 0.895445406 | 0.544488751 | 0.095 | 0.108 | 1 | 0.033451128 | -1.614607962 | 0.022 | 0.135 | 1 | 0.895445406 | 0.065783278 |
| Agtrap | 0.309921258 | -0.326228232 | 0.048 | 0.127 | 1 | 0.033468658 | -1.84048937 | 0.022 | 0.131 | 1 | 0.309921258 | 0.065817165 |
| E4f1 | 0.936855995 | 0.714413752 | 0.19 | 0.205 | 1 | 0.033494821 | -0.155646507 | 0.089 | 0.239 | 1 | 0.936855995 | 0.065867738 |
| Ophn1 | 0.036366993 | -1.014728337 | 0.238 | 0.53 | 1 | 0.033496163 | 0.63286114 | 0.667 | 0.498 | 1 | 0.036366993 | 0.065870332 |
| Fbxl14 | 0.244463041 | -0.167530486 | 0.048 | 0.146 | 1 | 0.033500284 | -0.794974711 | 0.089 | 0.239 | 1 | 0.244463041 | 0.065878298 |
| Cdc42ep3 | 0.03353984 | -0.937440632 | 0.667 | 0.806 | 1 | 0.049134688 | -0.459107425 | 0.911 | 0.9 | 1 | 0.049134688 | 0.065954759 |
| Nradd | 0.153811633 | -1.184209227 | 0 | 0.09 | 1 | 0.033567591 | -2.044847868 | 0.022 | 0.131 | 1 | 0.153811633 | 0.0660084 |
| Serpina3n | 0.033606613 | -2.347742127 | 0.286 | 0.478 | 1 | 0.230428389 | -2.316927414 | 0.022 | 0.069 | 1 | 0.230428389 | 0.066083822 |
| Mcm7 | 0.593914059 | -0.379339569 | 0.143 | 0.187 | 1 | 0.033657238 | -1.526380779 | 0.067 | 0.194 | 1 | 0.593914059 | 0.066181667 |
| Prr14 | 0.931200338 | 0.434305833 | 0.333 | 0.41 | 1 | 0.03365735 | -0.16492432 | 0.244 | 0.481 | 1 | 0.931200338 | 0.066181882 |
| Gm14966 | 0.943607145 | 0.786246497 | 0.095 | 0.108 | 1 | 0.033666661 | -1.901889915 | 0.022 | 0.131 | 1 | 0.943607145 | 0.066199878 |
| Sorbs2 | 0.555869016 | 0.173690479 | 0.905 | 0.914 | 1 | 0.033669312 | -0.446728361 | 0.933 | 0.958 | 1 | 0.555869016 | 0.066205001 |
| Med13 | 0.226251247 | 0.379013874 | 0.952 | 0.933 | 1 | 0.033678597 | 0.356153695 | 0.978 | 0.986 | 1 | 0.226251247 | 0.066222945 |
| Mtmr7 | 0.651717725 | 1.957564734 | 0.095 | 0.075 | 1 | 0.033725363 | 2.924080686 | 0.089 | 0.028 | 1 | 0.651717725 | 0.066313325 |
| Cutc | 0.79693741 | 1.088809267 | 0.095 | 0.119 | 1 | 0.033764248 | -1.776359032 | 0.022 | 0.131 | 1 | 0.79693741 | 0.066388472 |
| Cmklr1 | 0.130491872 | -0.116305163 | 0.143 | 0.34 | 1 | 0.033808302 | 0.20086966 | 0.356 | 0.637 | 1 | 0.130491872 | 0.066473602 |
| Col27a1 | 0.033817194 | 0.531990132 | 0.952 | 0.955 | 1 | 0.872739512 | 0.318183027 | 0.911 | 0.976 | 1 | 0.872739512 | 0.066490785 |
| Dlx5 | 0.794994777 | -0.413691074 | 0.143 | 0.16 | 1 | 0.033854988 | -0.397847409 | 0.044 | 0.176 | 1 | 0.794994777 | 0.066563816 |
| Usp22 | 0.834042029 | 1.185485286 | 0.381 | 0.481 | 1 | 0.033926746 | -0.624355939 | 0.4 | 0.554 | 1 | 0.834042029 | 0.066702467 |
| Fgf18 | 0.140363982 | 2.057513136 | 0.095 | 0.269 | 1 | 0.033953843 | -0.449641336 | 0.222 | 0.436 | 1 | 0.140363982 | 0.066754822 |
| 2510009E07Rik | 0.572453128 | 0.111529343 | 0.286 | 0.41 | 1 | 0.033961155 | 0.131793212 | 0.356 | 0.644 | 1 | 0.572453128 | 0.06676895 |
| Secisbp2l | 0.034019897 | -0.540671901 | 0.619 | 0.873 | 1 | 0.241403039 | -0.247430567 | 0.889 | 0.882 | 1 | 0.241403039 | 0.06688244 |
| Ice2 | 0.548335312 | -0.20069735 | 0.143 | 0.205 | 1 | 0.034053614 | -0.736466305 | 0.089 | 0.242 | 1 | 0.548335312 | 0.06694758 |
| Agps | 0.034105125 | 1.346305489 | 0.667 | 0.489 | 1 | 0.03533028 | 0.622772076 | 0.867 | 0.678 | 1 | 0.03533028 | 0.067047091 |
| Klhl42 | 0.050513042 | -1.818081329 | 0 | 0.157 | 1 | 0.034135643 | -0.979892427 | 0.067 | 0.201 | 1 | 0.050513042 | 0.067106043 |
| Sucla2 | 0.034149093 | -0.697787095 | 0.19 | 0.5 | 1 | 0.676308157 | 0.205663227 | 0.556 | 0.616 | 1 | 0.676308157 | 0.067132026 |
| Itch | 0.034167066 | 0.802458842 | 0.857 | 0.866 | 1 | 0.631379109 | 0.199229301 | 0.956 | 0.955 | 1 | 0.631379109 | 0.067166743 |
| Slc22a4 | 0.034338016 | 1.118710753 | 0.667 | 0.578 | 1 | 0.116532049 | 1.149892628 | 0.378 | 0.298 | 1 | 0.116532049 | 0.067496932 |
| Itih2 | 0.061234946 | -1.849790188 | 0 | 0.146 | 1 | 0.034372375 | -1.223818009 | 0.044 | 0.17 | 1 | 0.061234946 | 0.067563289 |
| R3hcc1l | 0.761742762 | 0.526930379 | 0.333 | 0.425 | 1 | 0.034379614 | 0.903462655 | 0.644 | 0.522 | 1 | 0.761742762 | 0.067577269 |
| Clec3a | 0.684849658 | 0.307319058 | 0.571 | 0.627 | 1 | 0.034389677 | -1.040953952 | 0.133 | 0.294 | 1 | 0.684849658 | 0.067596703 |
| Fig4 | 0.318899491 | 0.150209812 | 0.048 | 0.127 | 1 | 0.034422203 | -0.881712033 | 0.044 | 0.17 | 1 | 0.318899491 | 0.067659519 |
| Snrpn | 0.041988961 | -2.208871282 | 0 | 0.168 | 1 | 0.034434257 | -1.375821103 | 0.022 | 0.135 | 1 | 0.041988961 | 0.067682796 |
| Rab11fip5 | 0.770692315 | 0.841894526 | 0.333 | 0.388 | 1 | 0.034436291 | 0.753461914 | 0.644 | 0.682 | 1 | 0.770692315 | 0.067686724 |
| Ssh2 | 0.15597244 | 0.170155222 | 0.619 | 0.437 | 1 | 0.034450668 | 0.359757951 | 0.956 | 0.723 | 1 | 0.15597244 | 0.067714488 |
| Ap4b1 | 0.034455446 | -2.107587946 | 0 | 0.179 | 1 | 0.052084274 | -1.223818009 | 0.022 | 0.121 | 1 | 0.052084274 | 0.067723714 |
| Twf2 | 0.034504019 | -2.159118246 | 0 | 0.179 | 1 | 0.118185185 | -0.255526869 | 0.089 | 0.194 | 1 | 0.118185185 | 0.06781751 |
| Lefty1 | 0.034504557 | -2.107587946 | 0 | 0.179 | 1 | 0.034685193 | -1.316927414 | 0.022 | 0.135 | 1 | 0.034685193 | 0.067818549 |
| Plpbp | 0.034542739 | 0.578614535 | 0.19 | 0.507 | 1 | 0.2166981 | 0.162500673 | 0.333 | 0.533 | 1 | 0.2166981 | 0.067892278 |
| Acp6 | 0.034592015 | -2.184209227 | 0 | 0.179 | 1 | 0.492179896 | 0.704446237 | 0.133 | 0.197 | 1 | 0.492179896 | 0.067987422 |
| Mark3 | 0.604324356 | 0.255270413 | 0.81 | 0.914 | 1 | 0.034592695 | 0.50203044 | 0.978 | 0.976 | 1 | 0.604324356 | 0.067988735 |
| Eral1 | 0.753842929 | 0.710297644 | 0.095 | 0.131 | 1 | 0.03460011 | -1.124282336 | 0.044 | 0.17 | 1 | 0.753842929 | 0.068003052 |
| Sh2d4a | 0.228243709 | 0.719575457 | 0.19 | 0.104 | 1 | 0.034662678 | -1.404390255 | 0.022 | 0.135 | 1 | 0.228243709 | 0.068123854 |
| Mon1a | 0.034670484 | -2.233118828 | 0 | 0.179 | 1 | 0.035615764 | -0.792660845 | 0.089 | 0.228 | 1 | 0.035615764 | 0.068138925 |
| Angptl4 | 0.177015705 | -1.14385149 | 0.048 | 0.164 | 1 | 0.034682978 | -1.396654606 | 0.089 | 0.232 | 1 | 0.177015705 | 0.068163046 |
| Snx7 | 0.663063171 | 0.612889526 | 0.429 | 0.433 | 1 | 0.034687656 | 0.187812888 | 0.222 | 0.46 | 1 | 0.663063171 | 0.068172078 |
| Gm16066 | 0.213733922 | -0.44170545 | 0.048 | 0.153 | 1 | 0.034710227 | -0.466674533 | 0.067 | 0.208 | 1 | 0.213733922 | 0.068215654 |
| Mideas | 0.034720916 | -0.905166945 | 0.19 | 0.463 | 1 | 0.750253905 | 0.457897724 | 0.6 | 0.557 | 1 | 0.750253905 | 0.06823629 |
| Itgb3bp | 0.090206304 | -0.93109029 | 0.048 | 0.209 | 1 | 0.0347522 | -1.124282336 | 0.044 | 0.17 | 1 | 0.090206304 | 0.068296685 |
| Mpped2 | 0.125991195 | -0.9277927 | 0.286 | 0.5 | 1 | 0.034762914 | -0.496251113 | 0.422 | 0.651 | 1 | 0.125991195 | 0.068317367 |
| Galnt4 | 0.354981605 | -0.496153234 | 0.048 | 0.112 | 1 | 0.034802562 | -1.526380779 | 0.067 | 0.194 | 1 | 0.354981605 | 0.068393906 |
| Qpct | 0.357983845 | 0.247507013 | 0.048 | 0.119 | 1 | 0.034872286 | -1.709244837 | 0.044 | 0.163 | 1 | 0.357983845 | 0.068528497 |
| 4632427E13Rik | 0.034882131 | 1.838001163 | 0.571 | 0.515 | 1 | 0.108879705 | 0.747923731 | 0.644 | 0.561 | 1 | 0.108879705 | 0.068547499 |

| | | | | | | | | | | | | |
|---|---|---|---|---|---|---|---|---|---|---|---|---|
| Pdcd2l | 0.034892317 | -2.574155746 | 0 | 0.179 | 1 | 0.037926955 | -0.286553765 | 0.089 | 0.242 | 1 | 0.037926955 | 0.06856716 |
| 4930550C14Rik | 0.375155151 | 0.214340149 | 0.048 | 0.116 | 1 | 0.034896526 | -0.901889915 | 0.044 | 0.17 | 1 | 0.375155151 | 0.068575285 |
| Rnf128 | 0.034943329 | -2.880817084 | 0 | 0.179 | 1 | 0.240404756 | -0.432404631 | 0.022 | 0.069 | 1 | 0.240404756 | 0.068665621 |
| Tnks | 0.034952586 | 0.95495352 | 0.857 | 0.787 | 1 | 0.093424042 | 0.460146328 | 0.978 | 0.913 | 1 | 0.093424042 | 0.068683489 |
| Zfp386 | 0.034961317 | -1.666078235 | 0.048 | 0.25 | 1 | 0.03578668 | -1.003427941 | 0.111 | 0.26 | 1 | 0.03578668 | 0.068700341 |
| Rin2 | 0.035062416 | 2.139573715 | 0.952 | 0.918 | 1 | 0.500933197 | 0.534762925 | 0.911 | 0.948 | 1 | 0.500933197 | 0.068895459 |
| Corin | 0.035085677 | -3.413691074 | 0 | 0.179 | 1 | 0.306434953 | 0.5159626 | 0.289 | 0.232 | 1 | 0.306434953 | 0.068940349 |
| Gm34455 | 0.447243623 | 3.329817366 | 0.19 | 0.138 | 1 | 0.035139658 | -0.227660076 | 0.111 | 0.277 | 1 | 0.447243623 | 0.069044521 |
| Cd2ap | 0.035152376 | -0.131864637 | 0.762 | 0.918 | 1 | 0.081777499 | -0.152116544 | 0.911 | 0.917 | 1 | 0.081777499 | 0.069069063 |
| Rcc1 | 0.86259164 | 0.586308926 | 0.143 | 0.175 | 1 | 0.035156483 | -0.375821103 | 0.067 | 0.208 | 1 | 0.86259164 | 0.069076988 |
| Cdk9 | 0.035187261 | -0.83578333 | 0.19 | 0.47 | 1 | 0.395053234 | -0.192995086 | 0.622 | 0.616 | 1 | 0.395053234 | 0.069136379 |
| Klf2 | 0.200758813 | -1.373501983 | 0.571 | 0.698 | 1 | 0.035222569 | -1.277911612 | 0.711 | 0.72 | 1 | 0.200758813 | 0.069204509 |
| D130043K22Rik | 0.796672617 | 1.351843673 | 0.048 | 0.037 | 1 | 0.035266245 | 2.9310001 | 0.089 | 0.028 | 1 | 0.796672617 | 0.069288783 |
| Rngtt | 0.758606019 | 0.253439969 | 0.429 | 0.597 | 1 | 0.035279772 | 0.889710936 | 0.822 | 0.64 | 1 | 0.758606019 | 0.069314881 |
| Skap2 | 0.235693057 | 0.43127814 | 0.19 | 0.388 | 1 | 0.03529008 | 0.519096851 | 0.356 | 0.208 | 1 | 0.235693057 | 0.06933477 |
| Arel1 | 0.038938002 | 1.805875304 | 0.333 | 0.19 | 1 | 0.035347991 | -0.108340792 | 0.089 | 0.246 | 1 | 0.038938002 | 0.069446501 |
| Pik3r4 | 0.333559237 | 1.882358389 | 0.19 | 0.131 | 1 | 0.03537406 | -1.506751973 | 0.067 | 0.194 | 1 | 0.333559237 | 0.069496796 |
| Heca | 0.431689393 | 0.269017242 | 0.286 | 0.451 | 1 | 0.035402433 | 0.411267971 | 0.8 | 0.588 | 1 | 0.431689393 | 0.069551534 |
| Ubl7 | 0.065743333 | -0.984439715 | 0.095 | 0.291 | 1 | 0.035449354 | -0.776359032 | 0.089 | 0.228 | 1 | 0.065743333 | 0.06964205 |
| Rcan1 | 0.035451678 | -1.329327027 | 0.857 | 0.84 | 1 | 0.037192201 | -0.793261092 | 0.911 | 0.851 | 1 | 0.037192201 | 0.069646535 |
| Ints11 | 0.078524763 | -0.874664857 | 0.143 | 0.336 | 1 | 0.035452656 | 0.140545352 | 0.133 | 0.318 | 1 | 0.078524763 | 0.06964842 |
| Jmjd6 | 0.109856554 | -0.625788514 | 0.238 | 0.466 | 1 | 0.035458539 | -0.52728728 | 0.489 | 0.664 | 1 | 0.109856554 | 0.069659769 |
| Tspan2 | 0.48737261 | 0.451379346 | 0.19 | 0.284 | 1 | 0.035465264 | -0.638855509 | 0.067 | 0.211 | 1 | 0.48737261 | 0.069672743 |
| Tmpo | 0.035468211 | -1.150656668 | 0.19 | 0.459 | 1 | 0.198315986 | -0.385969058 | 0.667 | 0.654 | 1 | 0.198315986 | 0.069678428 |
| Pard6g | 0.216264544 | -1.392317423 | 0.048 | 0.142 | 1 | 0.035529697 | -1.342462506 | 0.089 | 0.225 | 1 | 0.216264544 | 0.069797034 |
| Mre11a | 0.471910916 | 0.376091219 | 0.095 | 0.16 | 1 | 0.035531361 | -0.518561275 | 0.067 | 0.211 | 1 | 0.471910916 | 0.069800244 |
| 2610021A01Rik | 0.196495505 | -1.233118828 | 0 | 0.075 | 1 | 0.035536643 | -0.695439037 | 0.022 | 0.135 | 1 | 0.196495505 | 0.069810433 |
| Wwtr1 | 0.035569732 | 0.601882069 | 1 | 0.996 | 1 | 0.28475393 | 0.268952346 | 1 | 1 | 1 | 0.28475393 | 0.069874258 |
| Uba2 | 0.244078827 | -0.205373837 | 0.286 | 0.489 | 1 | 0.035577388 | -0.525429447 | 0.578 | 0.664 | 1 | 0.244078827 | 0.069889026 |
| Hsd3b7 | 0.218149267 | -0.473069621 | 0.19 | 0.336 | 1 | 0.035626701 | -0.494120416 | 0.2 | 0.405 | 1 | 0.218149267 | 0.069984141 |
| Rgl2 | 0.957125073 | 0.351843673 | 0.143 | 0.142 | 1 | 0.035634777 | -0.867124496 | 0.067 | 0.204 | 1 | 0.957125073 | 0.069999717 |
| Sel1l | 0.308233048 | -0.180377366 | 0.571 | 0.705 | 1 | 0.035658831 | -0.405827918 | 0.756 | 0.779 | 1 | 0.308233048 | 0.07004611 |
| Ubxn4 | 0.413932696 | -0.155233318 | 1 | 0.993 | 1 | 0.035727961 | -0.15989159 | 1 | 0.993 | 1 | 0.413932696 | 0.070179434 |
| Gars | 0.035786397 | 1.39417517 | 0.667 | 0.59 | 1 | 0.206515326 | 0.188808396 | 0.467 | 0.685 | 1 | 0.206515326 | 0.070292128 |
| Ap3d1 | 0.790139557 | 0.285276056 | 0.619 | 0.769 | 1 | 0.035829318 | -0.124282336 | 0.689 | 0.837 | 1 | 0.790139557 | 0.070374897 |
| Glce | 0.035890153 | -0.989193245 | 0.19 | 0.448 | 1 | 0.044298148 | -0.277756817 | 0.244 | 0.46 | 1 | 0.044298148 | 0.070492202 |
| Atp6v1e1 | 0.03590304 | -0.254674971 | 0.619 | 0.78 | 1 | 0.448933881 | -0.359660399 | 0.8 | 0.768 | 1 | 0.448933881 | 0.070517051 |
| Rsph3b | 0.675684865 | 0.901182264 | 0.238 | 0.228 | 1 | 0.035909504 | -0.504554417 | 0.067 | 0.204 | 1 | 0.675684865 | 0.070529516 |
| Evc | 0.36148503 | 1.311201688 | 0.048 | 0.119 | 1 | 0.035912334 | -1.564854927 | 0.044 | 0.163 | 1 | 0.36148503 | 0.070534972 |
| G930009F23Rik | 0.036095473 | 2.258734268 | 0.238 | 0.097 | 1 | 0.205914278 | 1.91753784 | 0.111 | 0.062 | 1 | 0.205914278 | 0.070888062 |
| Agap1 | 0.036097056 | 1.195118169 | 0.714 | 0.552 | 1 | 0.141182002 | 0.555317039 | 0.889 | 0.751 | 1 | 0.141182002 | 0.070891114 |
| Vps26c | 0.835694907 | 0.24081236 | 0.143 | 0.164 | 1 | 0.036177944 | 0.388889482 | 0.111 | 0.284 | 1 | 0.835694907 | 0.071047044 |
| Arhgap28 | 0.34071055 | -0.660128969 | 0.048 | 0.119 | 1 | 0.036199478 | -2.404390255 | 0.022 | 0.128 | 1 | 0.34071055 | 0.071088555 |
| Kmt2e | 0.837549296 | 0.101841831 | 1 | 1 | 1 | 0.036269564 | 0.503441767 | 1 | 0.997 | 1 | 0.837549296 | 0.071223646 |
| Bzw2 | 0.523495215 | 1.010806755 | 0.619 | 0.761 | 1 | 0.036280146 | -0.593051819 | 0.689 | 0.671 | 1 | 0.523495215 | 0.071244043 |
| Cenpj | 0.295681943 | -0.849790188 | 0.048 | 0.123 | 1 | 0.036314875 | -2.174908409 | 0.022 | 0.128 | 1 | 0.295681943 | 0.07131098 |
| Znhit1 | 0.036318646 | -0.785659851 | 0.143 | 0.403 | 1 | 0.057951973 | -0.465319254 | 0.2 | 0.346 | 1 | 0.057951973 | 0.071318249 |
| Flii | 0.111848063 | -0.285586248 | 0.333 | 0.634 | 1 | 0.036326889 | -0.453193104 | 0.689 | 0.758 | 1 | 0.111848063 | 0.071334135 |
| Smarcal1 | 1 | 0.304537958 | 0.095 | 0.093 | 1 | 0.036436553 | -0.754332726 | 0.044 | 0.173 | 1 | 1 | 0.071545483 |
| Plcg1 | 0.970120955 | 0.894161836 | 0.238 | 0.254 | 1 | 0.036523025 | -0.141355849 | 0.178 | 0.367 | 1 | 0.970120955 | 0.071712119 |
| Abca2 | 0.376355449 | -0.611630451 | 0.048 | 0.108 | 1 | 0.036528188 | -1.486852415 | 0.044 | 0.163 | 1 | 0.376355449 | 0.071722068 |
| Drg1 | 0.036592882 | -0.251323423 | 0.286 | 0.657 | 1 | 0.891009444 | 0.168950326 | 0.667 | 0.74 | 1 | 0.891009444 | 0.071846726 |
| Otud5 | 0.348456176 | 0.36444371 | 0.286 | 0.541 | 1 | 0.036593289 | -0.347392695 | 0.467 | 0.657 | 1 | 0.348456176 | 0.071847509 |
| Hmgn2-ps | 0.128113406 | -1.280424543 | 0 | 0.101 | 1 | 0.036608996 | -2.124282336 | 0 | 0.09 | 1 | 0.128113406 | 0.071877774 |
| Hebp1 | 0.168377627 | 0.574236094 | 0.048 | 0.172 | 1 | 0.036608996 | -2.486852415 | 0 | 0.09 | 1 | 0.168377627 | 0.071877774 |
| Tmem14a | 0.100873252 | -1.326228232 | 0 | 0.116 | 1 | 0.036630047 | -2.174908409 | 0 | 0.09 | 1 | 0.100873252 | 0.071918334 |
| Tmem198b | 0.844871807 | 0.614878079 | 0.095 | 0.082 | 1 | 0.036630047 | -2.174908409 | 0 | 0.09 | 1 | 0.844871807 | 0.071918334 |
| Dgke | 1 | 0.464318402 | 0.095 | 0.097 | 1 | 0.036630047 | -2.223818009 | 0 | 0.09 | 1 | 1 | 0.071918334 |
| Kctd13 | 0.948863791 | 0.766881172 | 0.095 | 0.104 | 1 | 0.036682689 | -2.361321533 | 0 | 0.09 | 1 | 0.948863791 | 0.072019759 |
| Zfp994 | 0.078782564 | -1.970084422 | 0 | 0.131 | 1 | 0.03668311 | -2.404390255 | 0 | 0.09 | 1 | 0.078782564 | 0.07202057 |
| Catsper2 | 0.655943564 | 1.8372705 | 0.048 | 0.078 | 1 | 0.03668311 | -2.446210431 | 0 | 0.09 | 1 | 0.655943564 | 0.07202057 |
| Harbi1 | 0.944290482 | 0.719575457 | 0.095 | 0.093 | 1 | 0.036683321 | -2.564854927 | 0 | 0.09 | 1 | 0.944290482 | 0.072020976 |
| 5430402E10Rik | 0.570211706 | 1.351843673 | 0.048 | 0.026 | 1 | 0.03669673 | -2.526380779 | 0 | 0.09 | 1 | 0.570211706 | 0.072046811 |
| Lyrm1 | 0.390282351 | 0.889500459 | 0.095 | 0.187 | 1 | 0.036697432 | -2.526380779 | 0 | 0.09 | 1 | 0.390282351 | 0.072048163 |
| Drp2 | 0.415237051 | 0.192645078 | 0.048 | 0.108 | 1 | 0.036697432 | -2.486852415 | 0 | 0.09 | 1 | 0.415237051 | 0.072048163 |
| Fstl3 | 0.924795272 | 1.351843673 | 0.048 | 0.045 | 1 | 0.036697432 | -2.446210431 | 0 | 0.09 | 1 | 0.924795272 | 0.072048163 |
| Fam174c | 0.036707819 | -1.098817736 | 0.048 | 0.265 | 1 | 0.41758888 | 0.481438725 | 0.089 | 0.138 | 1 | 0.41758888 | 0.072068174 |
| Get1 | 0.173729397 | -0.970084422 | 0 | 0.082 | 1 | 0.036710773 | -2.526380779 | 0 | 0.09 | 1 | 0.173729397 | 0.072073865 |
| Gprc5b | 0.512935963 | -0.159118246 | 0.095 | 0.153 | 1 | 0.036711124 | -2.674479418 | 0 | 0.09 | 1 | 0.512935963 | 0.072074542 |
| Slc22a1 | 0.555949871 | 1.673771768 | 0.095 | 0.067 | 1 | 0.036722008 | -2.638855509 | 0 | 0.09 | 1 | 0.555949871 | 0.072095511 |
| Mpst | 0.083489388 | -1.611630451 | 0 | 0.127 | 1 | 0.0367317 | -0.531940305 | 0.044 | 0.173 | 1 | 0.083489388 | 0.072114183 |
| Ero1b | 0.451322061 | -0.295854583 | 0.048 | 0.101 | 1 | 0.036742725 | -3.071814916 | 0 | 0.09 | 1 | 0.451322061 | 0.072135423 |
| Adcy10 | 0.036761502 | 1.936806174 | 0.143 | 0.041 | 1 | 0.474343734 | 2.180572246 | 0.067 | 0.045 | 1 | 0.474343734 | 0.072171595 |
| Ints5 | 0.036773272 | -2.081115734 | 0 | 0.175 | 1 | 0.142702943 | -0.223818009 | 0.111 | 0.211 | 1 | 0.142702943 | 0.072194271 |
| Pstk | 0.036773843 | -2.054148687 | 0 | 0.175 | 1 | 0.054551831 | -0.539319835 | 0.044 | 0.159 | 1 | 0.054551831 | 0.072195371 |
| Cfh | 0.196587507 | -1.648156327 | 0 | 0.075 | 1 | 0.036782553 | -4.339295227 | 0 | 0.09 | 1 | 0.196587507 | 0.072212151 |
| Commd5 | 0.088894155 | -1.683780237 | 0 | 0.123 | 1 | 0.036791546 | -2.071814916 | 0 | 0.09 | 1 | 0.088894155 | 0.072229475 |
| Atad2b | 0.036818047 | 0.842015502 | 0.762 | 0.634 | 1 | 0.138860493 | 0.531272133 | 0.911 | 0.796 | 1 | 0.138860493 | 0.072280526 |
| Gm57383 | 0.036821875 | -2.107587946 | 0 | 0.175 | 1 | 0.038216693 | -1.404390255 | 0.022 | 0.131 | 1 | 0.038216693 | 0.0722879 |
| Slain2 | 0.036847737 | -0.896976874 | 0.429 | 0.664 | 1 | 0.632593908 | -0.134976937 | 0.911 | 0.817 | 1 | 0.632593908 | 0.072337719 |
| Pigu | 0.553138491 | 0.228986925 | 0.19 | 0.28 | 1 | 0.036849742 | -0.316927414 | 0.133 | 0.311 | 1 | 0.553138491 | 0.072341581 |
| Tcirg1 | 0.036868724 | -2.184209227 | 0 | 0.175 | 1 | 0.099012576 | -1.153428682 | 0.089 | 0.19 | 1 | 0.099012576 | 0.072378145 |
| G3bp1 | 0.036876688 | -0.155116316 | 0.81 | 0.918 | 1 | 0.10200521 | -0.211148817 | 0.978 | 0.903 | 1 | 0.10200521 | 0.072393486 |
| Mphosph8 | 0.036907878 | -0.685631033 | 0.333 | 0.627 | 1 | 0.366422844 | -0.131023914 | 0.6 | 0.612 | 1 | 0.366422844 | 0.072453564 |
| Llgl1 | 0.918439472 | 0.351843673 | 0.143 | 0.16 | 1 | 0.036935235 | -0.423842618 | 0.133 | 0.304 | 1 | 0.918439472 | 0.072506258 |
| Tm7sf2 | 0.03699375 | -2.280424543 | 0 | 0.175 | 1 | 0.051656525 | -1.674479418 | 0.022 | 0.118 | 1 | 0.051656525 | 0.072618962 |
| Atp8a2 | 0.037032819 | -2.413691074 | 0 | 0.175 | 1 | 0.063443099 | -0.805213895 | 0.089 | 0.211 | 1 | 0.063443099 | 0.072694207 |
| Exd2 | 0.472155519 | 2.736965594 | 0.095 | 0.16 | 1 | 0.037035753 | -1.960783604 | 0.022 | 0.128 | 1 | 0.472155519 | 0.072699859 |
| Zfp280c | 0.658631508 | 0.503846766 | 0.286 | 0.403 | 1 | 0.037088393 | -0.447324051 | 0.289 | 0.516 | 1 | 0.658631508 | 0.072801236 |
| Kif3b | 0.037106115 | -1.25696557 | 0.095 | 0.321 | 1 | 0.191156672 | 0.120136392 | 0.244 | 0.419 | 1 | 0.191156672 | 0.072835367 |
| Pafah1b2 | 0.037357125 | -0.673185122 | 0.381 | 0.701 | 1 | 0.037117279 | -0.214274237 | 0.689 | 0.855 | 1 | 0.037357125 | 0.072856866 |
| Bdh1 | 0.350395032 | -0.348596045 | 0.095 | 0.179 | 1 | 0.037151446 | -1.960783604 | 0.022 | 0.128 | 1 | 0.350395032 | 0.072922662 |
| Gm49980 | 0.153938468 | -1.455511249 | 0 | 0.09 | 1 | 0.037152503 | -1.84048937 | 0.022 | 0.128 | 1 | 0.153938468 | 0.072924697 |
| Zfp518a | 0.037163889 | -2.348596045 | 0 | 0.175 | 1 | 0.166571302 | -0.446210431 | 0.067 | 0.149 | 1 | 0.166571302 | 0.072946624 |
| Gm57146 | 0.037169368 | 1.442817333 | 0.476 | 0.366 | 1 | 0.63877295 | 0.505154794 | 0.244 | 0.332 | 1 | 0.63877295 | 0.072957174 |
| Slc10a6 | 0.037236918 | -2.648156327 | 0 | 0.175 | 1 | 0.854461673 | 1.545569062 | 0.022 | 0.028 | 1 | 0.854461673 | 0.073087248 |
| Gm35546 | 0.037241501 | -3.01272876 | 0 | 0.175 | 1 | 0.295787052 | -0.112786697 | 0.133 | 0.208 | 1 | 0.295787052 | 0.073096073 |
| Lrrc28 | 0.037263389 | -2.849790188 | 0 | 0.175 | 1 | 0.512029278 | 0.495445583 | 0.111 | 0.159 | 1 | 0.512029278 | 0.073138219 |

| | | | | | | | | | | | | |
|---|---|---|---|---|---|---|---|---|---|---|---|---|
| Crat | 0.08354242 | -1.718545655 | 0 | 0.127 | 1 | 0.037264545 | -1.84048937 | 0.022 | 0.128 | 1 | 0.08354242 | 0.073140443 |
| Pced1a | 0.722928975 | 0.464318402 | 0.143 | 0.116 | 1 | 0.037264545 | -1.84048937 | 0.022 | 0.128 | 1 | 0.722928975 | 0.073140443 |
| H1f10 | 0.037299724 | -3.403043829 | 0 | 0.175 | 1 | 0.041083518 | -0.664850717 | 0.089 | 0.235 | 1 | 0.041083518 | 0.073208179 |
| Prpf31 | 0.070448083 | -0.496153234 | 0.143 | 0.347 | 1 | 0.03730205 | -0.240691828 | 0.156 | 0.332 | 1 | 0.070448083 | 0.073212658 |
| Zfp317 | 0.744306335 | 0.351843673 | 0.286 | 0.272 | 1 | 0.03730325 | -0.178221143 | 0.133 | 0.298 | 1 | 0.744306335 | 0.073214968 |
| Ccser2 | 0.03731793 | -0.174851173 | 0.333 | 0.746 | 1 | 0.038345884 | -0.213304783 | 0.822 | 0.862 | 1 | 0.038345884 | 0.073243232 |
| Col4a3 | 0.037341744 | -3.171718283 | 0 | 0.175 | 1 | 0.638540729 | 0.290755163 | 0.044 | 0.031 | 1 | 0.638540729 | 0.073289083 |
| Jazf1 | 0.227505582 | 2.181173285 | 0.286 | 0.22 | 1 | 0.037357542 | 0.926838658 | 0.2 | 0.415 | 1 | 0.227505582 | 0.073319499 |
| Sgms1 | 0.840748659 | 0.737994211 | 0.714 | 0.806 | 1 | 0.037368819 | 0.590814078 | 0.978 | 0.862 | 1 | 0.840748659 | 0.073341209 |
| Lsmem1 | 0.513255622 | 1.625837291 | 0.19 | 0.157 | 1 | 0.037375393 | 1.361144491 | 0.378 | 0.232 | 1 | 0.513255622 | 0.073353865 |
| Spata24 | 0.041944343 | -2.159118246 | 0 | 0.168 | 1 | 0.037377861 | -1.80878051 | 0.022 | 0.128 | 1 | 0.041944343 | 0.073358617 |
| Fbxl15 | 0.073771805 | -1.718545655 | 0 | 0.134 | 1 | 0.037377861 | -1.80878051 | 0.022 | 0.128 | 1 | 0.073771805 | 0.073358617 |
| Pex1 | 1 | 0.866416846 | 0.143 | 0.153 | 1 | 0.037388133 | -0.901889915 | 0.044 | 0.166 | 1 | 1 | 0.073378394 |
| Acly | 0.037399647 | 0.555180595 | 0.905 | 0.877 | 1 | 0.05231344 | 0.799265604 | 0.822 | 0.785 | 1 | 0.05231344 | 0.07340056 |
| Tnpo2 | 0.92803331 | 0.258734268 | 0.286 | 0.328 | 1 | 0.037409404 | -0.103523776 | 0.244 | 0.471 | 1 | 0.92803331 | 0.073419345 |
| Igfbp2 | 0.037448082 | -4.550229907 | 0 | 0.175 | 1 | 0.064683465 | -0.435484024 | 0.111 | 0.253 | 1 | 0.064683465 | 0.073493805 |
| Tmem18 | 0.191380448 | -0.496153234 | 0.048 | 0.16 | 1 | 0.037456818 | -0.57996182 | 0.089 | 0.239 | 1 | 0.191380448 | 0.073510622 |
| Cipc | 0.03747872 | -1.455511249 | 0.048 | 0.261 | 1 | 0.074891727 | 0.180572246 | 0.067 | 0.183 | 1 | 0.074891727 | 0.073552785 |
| Tpra1 | 0.048172641 | -1.054148687 | 0.095 | 0.31 | 1 | 0.037547634 | -0.282711698 | 0.133 | 0.311 | 1 | 0.048172641 | 0.073685444 |
| Zfp423 | 0.756571613 | 0.214340149 | 0.095 | 0.127 | 1 | 0.037548908 | 0.199989699 | 0.089 | 0.249 | 1 | 0.756571613 | 0.073687896 |
| Tbc1d1 | 0.037559111 | 0.786820019 | 0.762 | 0.728 | 1 | 0.172474626 | 0.666264298 | 0.867 | 0.879 | 1 | 0.172474626 | 0.073707535 |
| Anxa2 | 0.437812808 | 0.117345178 | 1 | 0.996 | 1 | 0.03758235 | -0.214183184 | 0.978 | 1 | 1 | 0.437812808 | 0.073752267 |
| Zfp777 | 0.835914424 | 0.866416846 | 0.19 | 0.194 | 1 | 0.037584814 | -0.543996322 | 0.089 | 0.239 | 1 | 0.835914424 | 0.07375701 |
| Gpank1 | 0.601575449 | -0.184209227 | 0.048 | 0.078 | 1 | 0.037599815 | -1.80878051 | 0.022 | 0.128 | 1 | 0.601575449 | 0.073785883 |
| Ermp1 | 0.100530424 | -1.648156327 | 0 | 0.116 | 1 | 0.037646174 | -1.08246216 | 0.044 | 0.166 | 1 | 0.100530424 | 0.073875113 |
| Tnfrsf10b | 0.037660891 | -1.140009424 | 0.19 | 0.451 | 1 | 0.153478941 | -0.191396532 | 0.2 | 0.329 | 1 | 0.153478941 | 0.07390344 |
| Fhip2b | 0.805198946 | 0.918884266 | 0.095 | 0.082 | 1 | 0.037708429 | -1.709244837 | 0.022 | 0.128 | 1 | 0.805198946 | 0.073994933 |
| Grk5 | 0.037725737 | 1.881533049 | 0.667 | 0.72 | 1 | 0.204795008 | 0.781506545 | 0.8 | 0.723 | 1 | 0.204795008 | 0.074028242 |
| Smim14 | 0.141558113 | -0.277857143 | 0.952 | 0.981 | 1 | 0.037768269 | -0.332406146 | 1 | 0.976 | 1 | 0.141558113 | 0.074110097 |
| Cand2 | 0.13621393 | -1.280424543 | 0 | 0.097 | 1 | 0.037815453 | -1.638855509 | 0.022 | 0.128 | 1 | 0.13621393 | 0.074200897 |
| B4gat1 | 0.166002797 | -0.535681598 | 0.095 | 0.235 | 1 | 0.03783096 | -1.141355849 | 0.089 | 0.225 | 1 | 0.166002797 | 0.074230738 |
| Glb1l | 0.98939197 | 0.188344941 | 0.143 | 0.146 | 1 | 0.037835072 | -0.142898014 | 0.067 | 0.211 | 1 | 0.98939197 | 0.074238652 |
| Per1 | 0.037845154 | -0.403043829 | 0.095 | 0.347 | 1 | 0.889671303 | 0.21492375 | 0.378 | 0.37 | 1 | 0.889671303 | 0.074258053 |
| Jade2 | 0.932820644 | 0.410737362 | 0.238 | 0.287 | 1 | 0.037983816 | -0.35874759 | 0.156 | 0.329 | 1 | 0.932820644 | 0.074524862 |
| Armc10 | 0.262888295 | -0.911190733 | 0.048 | 0.131 | 1 | 0.037984213 | -1.149817428 | 0.044 | 0.166 | 1 | 0.262888295 | 0.074525625 |
| Gm12811 | 0.715863632 | 1.766881172 | 0.048 | 0.034 | 1 | 0.038035881 | 1.83507568 | 0.089 | 0.028 | 1 | 0.715863632 | 0.074625034 |
| Baiap2 | 0.038049235 | 1.251814854 | 0.762 | 0.653 | 1 | 0.061509246 | 0.815276445 | 0.867 | 0.74 | 1 | 0.061509246 | 0.074650726 |
| Tbl3 | 0.349149624 | -0.184209227 | 0.095 | 0.183 | 1 | 0.0381513 | -1.071814916 | 0.089 | 0.225 | 1 | 0.349149624 | 0.074847078 |
| Gripap1 | 0.038156087 | -0.426688414 | 0.238 | 0.549 | 1 | 0.556203909 | 0.280313417 | 0.578 | 0.612 | 1 | 0.556203909 | 0.074856287 |
| Il34 | 0.840474136 | 1.224464366 | 0.143 | 0.142 | 1 | 0.038192633 | -0.96900411 | 0.044 | 0.17 | 1 | 0.840474136 | 0.074926589 |
| Tgs1 | 0.10842816 | 1.115184682 | 0.571 | 0.545 | 1 | 0.038203842 | -0.326145944 | 0.378 | 0.564 | 1 | 0.10842816 | 0.07494815 |
| Dennd5b | 0.450593454 | -0.564632972 | 0.286 | 0.403 | 1 | 0.038214864 | -0.110778072 | 0.222 | 0.422 | 1 | 0.450593454 | 0.074969353 |
| Pcm1 | 0.038232105 | 0.603100911 | 0.952 | 0.896 | 1 | 0.608915497 | 0.250720502 | 0.956 | 0.958 | 1 | 0.608915497 | 0.075002515 |
| Trappc11 | 0.351296393 | -0.379339569 | 0.143 | 0.235 | 1 | 0.038241513 | -0.598962782 | 0.111 | 0.26 | 1 | 0.351296393 | 0.075020612 |
| Trmt13 | 0.469488176 | -0.337455488 | 0.19 | 0.28 | 1 | 0.038268835 | -0.191396532 | 0.067 | 0.208 | 1 | 0.469488176 | 0.075073167 |
| Stard3 | 0.235356878 | -0.174225139 | 0.095 | 0.224 | 1 | 0.038295037 | -1.486852415 | 0.067 | 0.19 | 1 | 0.235356878 | 0.075123565 |
| Sncaip | 0.467444599 | 1.548759188 | 0.238 | 0.187 | 1 | 0.038354169 | -1.806312255 | 0.089 | 0.225 | 1 | 0.467444599 | 0.075237296 |
| Slc25a15 | 0.128057475 | -1.370622352 | 0 | 0.101 | 1 | 0.038360281 | -0.972279242 | 0.044 | 0.166 | 1 | 0.128057475 | 0.075249051 |
| Gm14168 | 0.038404744 | 3.580662363 | 0.143 | 0.045 | 1 | 0.045429229 | -2.316927414 | 0 | 0.083 | 1 | 0.045429229 | 0.075334564 |
| Idua | 0.100277091 | -1.413691074 | 0 | 0.116 | 1 | 0.038410033 | -1.124282336 | 0.044 | 0.166 | 1 | 0.100277091 | 0.075344735 |
| Tnrc6a | 0.038415662 | 0.654662945 | 0.905 | 0.694 | 1 | 0.051595503 | 0.543735905 | 1 | 0.958 | 1 | 0.051595503 | 0.075355562 |
| Sh3bp1 | 0.593775495 | 2.60338244 | 0.095 | 0.067 | 1 | 0.038584106 | -1.223818009 | 0.022 | 0.131 | 1 | 0.593775495 | 0.07567948 |
| Btbd1 | 0.038594447 | -0.549975933 | 0.333 | 0.631 | 1 | 0.603564445 | 0.298660361 | 0.844 | 0.824 | 1 | 0.603564445 | 0.075699362 |
| Mn1 | 0.284208732 | -0.222392421 | 0.19 | 0.332 | 1 | 0.038620609 | -0.162654147 | 0.311 | 0.578 | 1 | 0.284208732 | 0.075749667 |
| Insig1 | 0.170026954 | -0.52393639 | 0.619 | 0.75 | 1 | 0.038629264 | -0.690385809 | 0.622 | 0.685 | 1 | 0.170026954 | 0.075766308 |
| Eif4h | 0.038891838 | -0.292856774 | 0.952 | 0.892 | 1 | 0.041204856 | -0.211745177 | 0.978 | 0.955 | 1 | 0.041204856 | 0.076271101 |
| Snhg17 | 0.646064891 | 0.410737362 | 0.095 | 0.142 | 1 | 0.038916199 | -1.191396532 | 0.022 | 0.131 | 1 | 0.646064891 | 0.076317928 |
| Slc18b1 | 0.430049488 | -0.44170545 | 0.095 | 0.157 | 1 | 0.038932539 | -0.141355849 | 0.067 | 0.211 | 1 | 0.430049488 | 0.076349336 |
| Ptk2 | 0.038937649 | 1.470916491 | 0.762 | 0.81 | 1 | 0.351612173 | 0.440769622 | 1 | 0.938 | 1 | 0.351612173 | 0.076359157 |
| Dpagt1 | 0.203121171 | -0.233118828 | 0.095 | 0.235 | 1 | 0.038945925 | -0.659319611 | 0.067 | 0.197 | 1 | 0.203121171 | 0.076375065 |
| Tbc1d15 | 0.088424713 | -0.326228232 | 0.381 | 0.694 | 1 | 0.03906841 | -0.479865985 | 0.778 | 0.799 | 1 | 0.088424713 | 0.07661048 |
| Gstp1 | 0.094421407 | -1.574155746 | 0 | 0.119 | 1 | 0.039080636 | -0.539319835 | 0.044 | 0.166 | 1 | 0.094421407 | 0.076633976 |
| Mad2l2 | 0.039182604 | -1.998653574 | 0 | 0.172 | 1 | 0.072816536 | -0.486852415 | 0.022 | 0.111 | 1 | 0.072816536 | 0.076829931 |
| Ints9 | 0.332481687 | 1.813174824 | 0.238 | 0.183 | 1 | 0.039193302 | -0.854892435 | 0.133 | 0.291 | 1 | 0.332481687 | 0.076850489 |
| Cptp | 0.039234728 | -2.159118246 | 0 | 0.172 | 1 | 0.388907488 | -0.124282336 | 0.044 | 0.083 | 1 | 0.388907488 | 0.076930093 |
| Tada2a | 0.51214224 | 0.710297644 | 0.238 | 0.328 | 1 | 0.039235224 | -0.194671664 | 0.133 | 0.322 | 1 | 0.51214224 | 0.076931045 |
| Fbl | 0.168424424 | -0.207583736 | 0.333 | 0.563 | 1 | 0.039333824 | -0.475881297 | 0.644 | 0.696 | 1 | 0.168424424 | 0.077120498 |
| Prkrip1 | 0.351731202 | -0.287057635 | 0.19 | 0.299 | 1 | 0.039368779 | -0.626782676 | 0.133 | 0.298 | 1 | 0.351731202 | 0.077187656 |
| Slc35g2 | 0.039376703 | -2.25696557 | 0 | 0.172 | 1 | 0.849488332 | 1.0259603 | 0.111 | 0.107 | 1 | 0.849488332 | 0.077202881 |
| Manea | 0.039376764 | -2.107587946 | 0 | 0.172 | 1 | 0.051704238 | -0.84048937 | 0.067 | 0.187 | 1 | 0.051704238 | 0.077202998 |
| Fam131a | 0.039417998 | -1.341178574 | 0.048 | 0.257 | 1 | 0.046589904 | -0.653210802 | 0.111 | 0.253 | 1 | 0.046589904 | 0.077282218 |
| Tmem9b | 0.039509596 | -0.567814219 | 0.238 | 0.522 | 1 | 1 | 0.553012046 | 0.511 | 0.536 | 1 | 1 | 0.077458184 |
| Cdt1 | 0.047893971 | -2.370622352 | 0 | 0.16 | 1 | 0.039549986 | -0.749886821 | 0.022 | 0.131 | 1 | 0.047893971 | 0.07753577 |
| Epn2 | 0.367411172 | 1.273488035 | 0.429 | 0.433 | 1 | 0.039581855 | 1.071637874 | 0.667 | 0.588 | 1 | 0.367411172 | 0.077596988 |
| Adamts5 | 0.495562783 | -0.683780237 | 0.048 | 0.09 | 1 | 0.039588124 | -2.72631835 | 0.022 | 0.125 | 1 | 0.495562783 | 0.077609029 |
| Pinx1 | 0.039613084 | -2.370622352 | 0 | 0.172 | 1 | 0.268861816 | 0.490427508 | 0.133 | 0.228 | 1 | 0.268861816 | 0.077656971 |
| Jcad | 0.061982825 | -1.116305163 | 0.048 | 0.231 | 1 | 0.039614209 | -0.587016577 | 0.133 | 0.298 | 1 | 0.061982825 | 0.077659132 |
| Nos1ap | 0.379563874 | 2.545583499 | 0.571 | 0.582 | 1 | 0.039642486 | 1.27650118 | 0.844 | 0.737 | 1 | 0.379563874 | 0.077713446 |
| Eeig1 | 0.086850167 | 0.944973494 | 0.619 | 0.53 | 1 | 0.039643228 | 0.356343505 | 0.644 | 0.426 | 1 | 0.086850167 | 0.077714871 |
| Gucd1 | 0.088737678 | -1.535681598 | 0 | 0.123 | 1 | 0.03970436 | -1.506751973 | 0.044 | 0.163 | 1 | 0.088737678 | 0.077832284 |
| Lima1 | 0.781730518 | -0.255145134 | 0.714 | 0.724 | 1 | 0.039749221 | -0.608390228 | 0.867 | 0.848 | 1 | 0.781730518 | 0.077918442 |
| Dnajb6 | 0.466221629 | -0.284440797 | 0.905 | 0.959 | 1 | 0.039823146 | -0.194502443 | 1 | 0.983 | 1 | 0.466221629 | 0.078060409 |
| Pbdc1 | 0.039837998 | -0.525383945 | 0.524 | 0.832 | 1 | 0.082238046 | 0.188307894 | 0.644 | 0.834 | 1 | 0.082238046 | 0.07808893 |
| Tmem232 | 0.80298679 | 1.171271427 | 0.048 | 0.037 | 1 | 0.039845535 | 4.490427508 | 0.067 | 0.017 | 1 | 0.80298679 | 0.078103404 |
| Edar | 0.57816106 | 1.351843673 | 0 | 0.015 | 1 | 0.039897053 | 2.268035087 | 0.111 | 0.042 | 1 | 0.57816106 | 0.078202331 |
| Cybc1 | 0.146344343 | -0.548620654 | 0.048 | 0.179 | 1 | 0.039909227 | -1.614607962 | 0.044 | 0.159 | 1 | 0.146344343 | 0.078225708 |
| Hmbox1 | 0.616397174 | 0.818720104 | 0.429 | 0.489 | 1 | 0.039915745 | 0.793255504 | 0.756 | 0.682 | 1 | 0.616397174 | 0.078238224 |
| B430010I23Rik | 0.039918039 | 2.130328536 | 0.429 | 0.287 | 1 | 0.73278161 | 0.223640968 | 0.289 | 0.339 | 1 | 0.73278161 | 0.078242628 |
| Pds5a | 0.826103338 | 0.35437694 | 0.571 | 0.709 | 1 | 0.039923898 | 0.452042608 | 0.956 | 0.903 | 1 | 0.826103338 | 0.078253878 |
| Ppp2r2d | 0.157557235 | -0.280424543 | 0.476 | 0.728 | 1 | 0.039950674 | -0.239759553 | 0.844 | 0.872 | 1 | 0.157557235 | 0.078305291 |
| Gtf3c3 | 0.240669743 | -0.40151636 | 0.19 | 0.332 | 1 | 0.039951431 | -0.286553765 | 0.089 | 0.239 | 1 | 0.240669743 | 0.078306746 |
| Ofd1 | 0.702003389 | 0.264380832 | 0.19 | 0.254 | 1 | 0.040016138 | -0.564854927 | 0.089 | 0.235 | 1 | 0.702003389 | 0.078430984 |
| Fibp | 0.040044879 | -0.531342662 | 0.19 | 0.481 | 1 | 0.183358447 | 0.350876156 | 0.222 | 0.37 | 1 | 0.183358447 | 0.078486167 |
| Irf3 | 0.26005996 | -0.512094778 | 0.143 | 0.257 | 1 | 0.040139822 | -0.187292133 | 0.133 | 0.304 | 1 | 0.26005996 | 0.078668439 |
| AA986860 | 0.040197747 | 2.179007076 | 0.19 | 0.071 | 1 | 0.81718262 | 1.304560963 | 0.044 | 0.055 | 1 | 0.81718262 | 0.078779636 |
| Ehbp1 | 0.041779399 | 0.502105432 | 0.857 | 0.701 | 1 | 0.040261324 | 0.794296785 | 0.911 | 0.865 | 1 | 0.041779399 | 0.078901674 |
| Zfp354c | 0.443509833 | -0.509450056 | 0.286 | 0.388 | 1 | 0.040293731 | -0.386600942 | 0.333 | 0.588 | 1 | 0.443509833 | 0.078963878 |
| Mrps35 | 0.09396339 | -1.303508156 | 0.048 | 0.194 | 1 | 0.040346154 | -0.931637258 | 0.089 | 0.221 | 1 | 0.09396339 | 0.079064497 |

| | | | | | | | | | | | | |
|---|---|---|---|---|---|---|---|---|---|---|---|---|
| Dse | 0.491568171 | 0.662466307 | 0.714 | 0.847 | 1 | 0.040430191 | 0.818215946 | 0.778 | 0.855 | 1 | 0.491568171 | 0.079225782 |
| Gnpda1 | 0.162745226 | -1.233118828 | 0.048 | 0.16 | 1 | 0.040544648 | -0.609709163 | 0.089 | 0.235 | 1 | 0.162745226 | 0.079445428 |
| Pdss2 | 0.040604692 | 1.565712022 | 0.524 | 0.325 | 1 | 0.430182332 | -0.438390926 | 0.356 | 0.412 | 1 | 0.430182332 | 0.079560642 |
| Nxph4 | 0.560128116 | 1.79930265 | 0.095 | 0.067 | 1 | 0.040685505 | -2.071814916 | 0.022 | 0.125 | 1 | 0.560128116 | 0.079715699 |
| 2810403D21Rik | 0.555258105 | 1.099780385 | 0.333 | 0.302 | 1 | 0.040695057 | 0.30164348 | 0.089 | 0.235 | 1 | 0.555258105 | 0.079734025 |
| Ston1 | 0.782842217 | 0.667345499 | 0.333 | 0.466 | 1 | 0.040698162 | -0.331475722 | 0.289 | 0.509 | 1 | 0.782842217 | 0.079739984 |
| Ptpdc1 | 0.222553843 | -0.522625445 | 0.048 | 0.149 | 1 | 0.040717886 | -0.463768802 | 0.044 | 0.17 | 1 | 0.222553843 | 0.079777826 |
| Lyplal1 | 0.065370176 | -2.02666795 | 0 | 0.142 | 1 | 0.040743447 | -2.124282336 | 0 | 0.087 | 1 | 0.065370176 | 0.079826865 |
| Gm5577 | 0.364537878 | 2.088809267 | 0.095 | 0.052 | 1 | 0.040748494 | 2.320502507 | 0.111 | 0.042 | 1 | 0.364537878 | 0.079836547 |
| Fpgt | 0.113338406 | -1.326228232 | 0 | 0.108 | 1 | 0.040764972 | -2.124282336 | 0 | 0.087 | 1 | 0.113338406 | 0.079868161 |
| Rogdi | 0.427439322 | -0.233118828 | 0.048 | 0.101 | 1 | 0.040764972 | -2.124282336 | 0 | 0.087 | 1 | 0.427439322 | 0.079868161 |
| Hyal1 | 0.466416324 | 1.351843673 | 0.143 | 0.101 | 1 | 0.040764972 | -2.124282336 | 0 | 0.087 | 1 | 0.466416324 | 0.079868161 |
| Cpt2 | 0.684003327 | 0.214340149 | 0.048 | 0.071 | 1 | 0.040784627 | -2.223818009 | 0 | 0.087 | 1 | 0.684003327 | 0.079905869 |
| Trp53cor1 | 0.163662481 | -1.818081329 | 0 | 0.086 | 1 | 0.040784705 | -2.174908409 | 0 | 0.087 | 1 | 0.163662481 | 0.079906019 |
| Atp23 | 0.304355016 | -0.233118828 | 0 | 0.049 | 1 | 0.040802413 | -2.271123724 | 0 | 0.087 | 1 | 0.304355016 | 0.079939989 |
| Nup37 | 0.926265024 | 1.088809267 | 0.095 | 0.097 | 1 | 0.040802647 | -2.316927414 | 0 | 0.087 | 1 | 0.926265024 | 0.079940438 |
| Mfsd8 | 0.13621393 | -1.280424543 | 0 | 0.097 | 1 | 0.040802725 | -2.223818009 | 0 | 0.087 | 1 | 0.13621393 | 0.079940588 |
| Lsm11 | 0.370790696 | -0.133583154 | 0 | 0.037 | 1 | 0.040818328 | -2.316927414 | 0 | 0.087 | 1 | 0.370790696 | 0.07997052 |
| Mfsd12 | 0.457281049 | 0.503846766 | 0 | 0.026 | 1 | 0.040819108 | -2.446210431 | 0 | 0.087 | 1 | 0.457281049 | 0.079972017 |
| Lipe | 0.568914119 | 0.129451251 | 0.048 | 0.086 | 1 | 0.040819108 | -2.271123724 | 0 | 0.087 | 1 | 0.568914119 | 0.079972017 |
| Tmem30b | 0.492608234 | 0.866416846 | 0 | 0.022 | 1 | 0.040832372 | -2.361321533 | 0 | 0.087 | 1 | 0.492608234 | 0.079997462 |
| Fgfbp3 | 0.106727131 | -1.455511249 | 0 | 0.112 | 1 | 0.040833152 | -2.486852415 | 0 | 0.087 | 1 | 0.106727131 | 0.079998958 |
| Gm973 | 0.34590997 | -0.320204445 | 0.19 | 0.31 | 1 | 0.040842447 | -0.986778812 | 0.089 | 0.232 | 1 | 0.34590997 | 0.080016789 |
| Gdf11 | 0.773866481 | 1.281454345 | 0.048 | 0.067 | 1 | 0.040845715 | -2.446210431 | 0 | 0.087 | 1 | 0.773866481 | 0.080023057 |
| Zfp280b | 0.346827513 | -0.233118828 | 0 | 0.041 | 1 | 0.040847276 | -2.361321533 | 0 | 0.087 | 1 | 0.346827513 | 0.080026051 |
| Nipsnap2 | 0.040868556 | -0.93109029 | 0.143 | 0.392 | 1 | 0.752849118 | 0.477168288 | 0.444 | 0.446 | 1 | 0.752849118 | 0.080066874 |
| Akna | 0.20895393 | -1.233118828 | 0 | 0.071 | 1 | 0.040874745 | -2.674479418 | 0 | 0.087 | 1 | 0.20895393 | 0.080078745 |
| Rnf152 | 0.3436979 | -1.107587946 | 0.048 | 0.112 | 1 | 0.040888559 | -2.743192169 | 0 | 0.087 | 1 | 0.3436979 | 0.080105245 |
| Smoc1 | 0.533956407 | 0.61076197 | 0.19 | 0.157 | 1 | 0.04091182 | -3.871516266 | 0 | 0.087 | 1 | 0.533956407 | 0.080149863 |
| Tmem135 | 0.040919453 | -1.073462162 | 0.143 | 0.384 | 1 | 0.336331821 | 0.330005979 | 0.556 | 0.488 | 1 | 0.336331821 | 0.080164505 |
| Col1a1 | 0.069993805 | -4.014787106 | 0.095 | 0.265 | 1 | 0.040947599 | -0.758362603 | 0.267 | 0.443 | 1 | 0.069993805 | 0.080218493 |
| Nxpe4 | 0.040971152 | -0.376854305 | 0.048 | 0.261 | 1 | 0.73420837 | 0.933050839 | 0.222 | 0.211 | 1 | 0.73420837 | 0.080263668 |
| Ddx19a | 0.056011253 | -1.370622352 | 0.095 | 0.284 | 1 | 0.040992694 | -0.278453266 | 0.2 | 0.388 | 1 | 0.056011253 | 0.080304987 |
| B930095G15Rik | 0.046733576 | 1.033314154 | 0.571 | 0.485 | 1 | 0.041065718 | 1.324217922 | 0.511 | 0.547 | 1 | 0.046733576 | 0.080445042 |
| Syt12 | 0.716588651 | -0.233118828 | 0.048 | 0.067 | 1 | 0.041090305 | -2.149817428 | 0.022 | 0.125 | 1 | 0.716588651 | 0.080492196 |
| Anxa8 | 0.995585553 | -0.649443873 | 0.81 | 0.653 | 1 | 0.041117099 | -0.75041058 | 0.644 | 0.353 | 1 | 0.995585553 | 0.080543581 |
| Fxyd4 | 0.396955847 | -0.233118828 | 0 | 0.034 | 1 | 0.041157171 | 2.335149283 | 0.111 | 0.042 | 1 | 0.396955847 | 0.08062043 |
| Bcor | 0.04116119 | -1.184209227 | 0.143 | 0.381 | 1 | 0.493539389 | 0.538151591 | 0.622 | 0.578 | 1 | 0.493539389 | 0.080628136 |
| Nostrin | 0.208981542 | -1.370622352 | 0 | 0.071 | 1 | 0.041162095 | -0.776359032 | 0.044 | 0.163 | 1 | 0.208981542 | 0.080629872 |
| Zfp35 | 0.073714373 | -1.683780237 | 0 | 0.134 | 1 | 0.041219568 | -1.901889915 | 0.022 | 0.125 | 1 | 0.073714373 | 0.080740083 |
| Slc11a2 | 0.941892706 | 0.277843091 | 0.333 | 0.396 | 1 | 0.041219593 | -0.373510942 | 0.222 | 0.429 | 1 | 0.941892706 | 0.080740131 |
| Sema4b | 0.313464368 | -0.303508156 | 0.048 | 0.127 | 1 | 0.04127811 | 1.045642666 | 0.156 | 0.066 | 1 | 0.313464368 | 0.080852338 |
| Gm43466 | 0.966871605 | 0.503846766 | 0.048 | 0.045 | 1 | 0.041378619 | 3.683072586 | 0.067 | 0.017 | 1 | 0.966871605 | 0.081045049 |
| Pogk | 0.137190686 | -0.735619168 | 0.095 | 0.246 | 1 | 0.041399976 | 0.299368294 | 0.156 | 0.343 | 1 | 0.137190686 | 0.081085994 |
| 1700025G04Rik | 0.041421205 | 0.733376128 | 0.952 | 0.966 | 1 | 0.603585588 | 0.640866026 | 0.867 | 0.958 | 1 | 0.603585588 | 0.081126693 |
| Ythdf3 | 0.268520972 | 0.479851285 | 0.762 | 0.787 | 1 | 0.041436343 | 0.650357011 | 0.978 | 0.931 | 1 | 0.268520972 | 0.081155715 |
| Ccdc92 | 0.041474359 | 2.48112669 | 0.143 | 0.045 | 1 | 0.476265473 | -0.486852415 | 0.022 | 0.045 | 1 | 0.476265473 | 0.081228596 |
| Rad18 | 0.768357285 | 0.195724471 | 0.143 | 0.175 | 1 | 0.041485231 | -1.564854927 | 0.022 | 0.128 | 1 | 0.768357285 | 0.081249437 |
| Rdh10 | 0.146193691 | -1.174225139 | 0.095 | 0.239 | 1 | 0.041495488 | -0.743192169 | 0.333 | 0.491 | 1 | 0.146193691 | 0.0812691 |
| Chd2 | 0.225934689 | 0.375218181 | 0.857 | 0.854 | 1 | 0.041500926 | 0.780428962 | 1 | 0.972 | 1 | 0.225934689 | 0.081279525 |
| Uba7 | 0.534722409 | -0.326228232 | 0.048 | 0.086 | 1 | 0.041542697 | -1.459885368 | 0.022 | 0.128 | 1 | 0.534722409 | 0.081359598 |
| Fubp1 | 0.625316366 | 0.269212786 | 0.905 | 0.951 | 1 | 0.041551084 | -0.185121781 | 0.978 | 0.962 | 1 | 0.625316366 | 0.081375675 |
| Tmcc2 | 0.768257326 | 0.150209812 | 0.048 | 0.063 | 1 | 0.041606363 | -1.931637258 | 0.022 | 0.125 | 1 | 0.768257326 | 0.081481637 |
| Tmem35b | 0.065075906 | -1.752492987 | 0 | 0.142 | 1 | 0.04160664 | -1.743192169 | 0.022 | 0.125 | 1 | 0.065075906 | 0.081482168 |
| Atxn1l | 0.584325334 | 0.825774861 | 0.143 | 0.209 | 1 | 0.041629513 | -0.237492946 | 0.178 | 0.37 | 1 | 0.584325334 | 0.08152601 |
| Eif4g3 | 0.041632539 | 0.423545138 | 0.952 | 0.918 | 1 | 0.045383339 | 0.45679673 | 0.978 | 0.948 | 1 | 0.045383339 | 0.081531811 |
| Mfn2 | 0.268791682 | -0.430564892 | 0.143 | 0.257 | 1 | 0.041657208 | 0.337297749 | 0.133 | 0.315 | 1 | 0.268791682 | 0.081579092 |
| Pou2f1 | 0.041691223 | 1.125871361 | 0.571 | 0.403 | 1 | 0.411760781 | 0.157981541 | 0.733 | 0.599 | 1 | 0.411760781 | 0.081644287 |
| Capza2 | 0.349946288 | -0.102133516 | 0.81 | 0.896 | 1 | 0.041726356 | -0.186735154 | 0.956 | 0.948 | 1 | 0.349946288 | 0.081711622 |
| Mafg | 0.169485039 | 0.746331335 | 0.714 | 0.806 | 1 | 0.041791396 | 0.378290475 | 0.956 | 0.927 | 1 | 0.169485039 | 0.081836271 |
| Cop1 | 0.097768496 | 0.866416846 | 0.762 | 0.858 | 1 | 0.041827589 | 0.673751958 | 0.978 | 0.934 | 1 | 0.097768496 | 0.081905632 |
| Zbtb11os1 | 0.113338406 | -1.370622352 | 0 | 0.108 | 1 | 0.041855683 | -1.709244837 | 0.022 | 0.125 | 1 | 0.113338406 | 0.081959467 |
| Eif5a2 | 0.727548938 | 0.79930265 | 0.048 | 0.071 | 1 | 0.041855906 | -1.674479418 | 0.022 | 0.125 | 1 | 0.727548938 | 0.081959894 |
| Gm15834 | 0.041877977 | 2.719575457 | 0.19 | 0.075 | 1 | 0.676121819 | 0.410054092 | 0.044 | 0.062 | 1 | 0.676121819 | 0.082002188 |
| Gorasp1 | 0.041896068 | -2.054148687 | 0 | 0.168 | 1 | 0.139606406 | -0.191396532 | 0.044 | 0.121 | 1 | 0.139606406 | 0.082036855 |
| Pigv | 0.041897355 | -2.02666795 | 0 | 0.168 | 1 | 0.6747639 | 1.042614973 | 0.156 | 0.142 | 1 | 0.6747639 | 0.082039321 |
| Cercam | 0.134046506 | -0.233118828 | 0.048 | 0.187 | 1 | 0.04192859 | -1.103523776 | 0.044 | 0.166 | 1 | 0.134046506 | 0.082099174 |
| Grpel2 | 0.065286989 | -1.998653574 | 0 | 0.142 | 1 | 0.041951832 | -0.664850717 | 0.133 | 0.284 | 1 | 0.065286989 | 0.082143707 |
| Ccl27a | 0.651055703 | 0.747772349 | 0.143 | 0.112 | 1 | 0.041978266 | -1.709244837 | 0.022 | 0.125 | 1 | 0.651055703 | 0.082194357 |
| Snhg15 | 0.041990313 | -2.081115734 | 0 | 0.168 | 1 | 0.096712156 | -0.901889915 | 0.022 | 0.1 | 1 | 0.096712156 | 0.08221744 |
| Ywhag | 0.041999898 | 0.826888481 | 0.762 | 0.78 | 1 | 0.084475893 | 0.648699542 | 0.911 | 0.9 | 1 | 0.084475893 | 0.082235805 |
| Cspp1 | 0.106211686 | 0.766881172 | 0.619 | 0.489 | 1 | 0.042057543 | 0.437093345 | 0.889 | 0.637 | 1 | 0.106211686 | 0.082346249 |
| Echdc2 | 0.042070782 | -1.801961663 | 0.048 | 0.239 | 1 | 0.217227874 | 0.207339155 | 0.133 | 0.232 | 1 | 0.217227874 | 0.082371613 |
| Cpt1a | 0.042106708 | -2.233118828 | 0 | 0.168 | 1 | 0.40271894 | -0.597035333 | 0.4 | 0.415 | 1 | 0.40271894 | 0.08244044 |
| Slc38a7 | 0.291544465 | 1.614878079 | 0.143 | 0.082 | 1 | 0.042110468 | -1.271123724 | 0.044 | 0.163 | 1 | 0.291544465 | 0.082447645 |
| Pcbp3 | 0.250894175 | 0.143257051 | 0.095 | 0.216 | 1 | 0.042196793 | -0.273858692 | 0.067 | 0.204 | 1 | 0.250894175 | 0.082613016 |
| Man1a | 0.55568231 | 0.211883103 | 0.762 | 0.869 | 1 | 0.04220057 | 0.590079383 | 0.978 | 0.924 | 1 | 0.55568231 | 0.082620251 |
| Med23 | 0.792862837 | 0.316219763 | 0.19 | 0.228 | 1 | 0.042224586 | -0.547225033 | 0.156 | 0.325 | 1 | 0.792862837 | 0.082666256 |
| Cracd | 0.669248332 | 0.85177007 | 0.143 | 0.205 | 1 | 0.042259576 | 0.193106599 | 0.111 | 0.28 | 1 | 0.669248332 | 0.08273328 |
| Anxa4 | 0.625062448 | 0.896599851 | 0.429 | 0.713 | 1 | 0.042357821 | 0.744862785 | 0.733 | 0.696 | 1 | 0.625062448 | 0.082921458 |
| Antxr2 | 0.042398312 | -1.111330348 | 0.095 | 0.336 | 1 | 0.587418147 | 0.742865671 | 0.422 | 0.519 | 1 | 0.587418147 | 0.082999008 |
| Upf2 | 0.042436695 | 0.712660781 | 0.714 | 0.675 | 1 | 0.287394619 | 0.324618615 | 0.956 | 0.779 | 1 | 0.287394619 | 0.083072518 |
| Lnpep | 0.691164852 | 0.484251286 | 0.857 | 0.951 | 1 | 0.042449781 | 0.427298865 | 0.978 | 0.983 | 1 | 0.691164852 | 0.083097578 |
| Vps50 | 0.042453888 | -1.718545655 | 0.048 | 0.239 | 1 | 0.894760074 | 0.501742821 | 0.222 | 0.256 | 1 | 0.894760074 | 0.083105444 |
| Ahnak2 | 0.042467705 | 0.918884266 | 0.286 | 0.127 | 1 | 0.313116139 | -0.674479418 | 0.044 | 0.09 | 1 | 0.313116139 | 0.083131903 |
| Zfp334 | 0.17385959 | -1.133583154 | 0 | 0.082 | 1 | 0.042516905 | -0.699397051 | 0.067 | 0.194 | 1 | 0.17385959 | 0.083226124 |
| Rnf185 | 0.042569563 | -1.017390137 | 0.143 | 0.369 | 1 | 0.918740181 | 0.656105539 | 0.333 | 0.426 | 1 | 0.918740181 | 0.083326959 |
| Afg3l2 | 0.158729562 | -0.283999997 | 0.238 | 0.429 | 1 | 0.042583348 | -0.213658143 | 0.311 | 0.557 | 1 | 0.158729562 | 0.083353355 |
| Fer | 0.959751266 | 0.368586937 | 0.381 | 0.478 | 1 | 0.04260335 | 1.169368769 | 0.578 | 0.474 | 1 | 0.959751266 | 0.083391654 |
| Tango2 | 0.655832576 | 0.351843673 | 0.19 | 0.153 | 1 | 0.042629457 | -1.674479418 | 0.044 | 0.159 | 1 | 0.655832576 | 0.083441644 |
| Ftsj1 | 0.048544603 | -0.463731756 | 0.048 | 0.25 | 1 | 0.042743854 | -0.819427754 | 0.111 | 0.249 | 1 | 0.048544603 | 0.083660671 |
| Mnat1 | 0.195840734 | 0.955914996 | 0.571 | 0.649 | 1 | 0.04281621 | 0.570304656 | 0.822 | 0.692 | 1 | 0.195840734 | 0.083799191 |
| Ncbp3 | 0.766299975 | 0.88827628 | 0.286 | 0.313 | 1 | 0.042880448 | -0.2150478 | 0.311 | 0.474 | 1 | 0.766299975 | 0.083922164 |
| Serpinb9 | 0.312039698 | -0.264827688 | 0.048 | 0.127 | 1 | 0.042912535 | -0.614607962 | 0.044 | 0.163 | 1 | 0.312039698 | 0.083983584 |
| Gba | 0.042918627 | -1.067695219 | 0.095 | 0.317 | 1 | 0.046890789 | -0.339295227 | 0.067 | 0.197 | 1 | 0.046890789 | 0.083995245 |
| 2500004C02Rik | 0.136258876 | -1.413691074 | 0 | 0.097 | 1 | 0.042943423 | -0.564854927 | 0.044 | 0.163 | 1 | 0.136258876 | 0.084042709 |
| Abcb8 | 0.050572964 | -1.849790188 | 0 | 0.157 | 1 | 0.042977895 | -0.802354241 | 0.044 | 0.163 | 1 | 0.050572964 | 0.084108691 |
| Irgq | 0.419591705 | 1.267779408 | 0.19 | 0.146 | 1 | 0.042985193 | -0.819427754 | 0.067 | 0.194 | 1 | 0.419591705 | 0.08412266 |

| | | | | | | | | | | | | |
|---|---|---|---|---|---|---|---|---|---|---|---|---|
| Rangrf | 0.280604124 | -0.695462042 | 0.095 | 0.187 | 1 | 0.043049444 | -0.802354241 | 0.044 | 0.163 | 1 | 0.280604124 | 0.084245634 |
| Mdga1 | 0.253731554 | 0.782147929 | 0.381 | 0.284 | 1 | 0.043147699 | -0.242926832 | 0.111 | 0.277 | 1 | 0.253731554 | 0.084433675 |
| Sgk1 | 0.043150155 | -0.820290663 | 0.714 | 0.724 | 1 | 0.494500589 | -0.570558897 | 0.911 | 0.682 | 1 | 0.494500589 | 0.084438374 |
| Cbx7 | 0.094635965 | -1.752492987 | 0 | 0.119 | 1 | 0.043159446 | -0.575661682 | 0.067 | 0.194 | 1 | 0.094635965 | 0.084456154 |
| Umps | 0.329441909 | -0.233118828 | 0.048 | 0.123 | 1 | 0.043172706 | -0.620708162 | 0.067 | 0.204 | 1 | 0.329441909 | 0.08448153 |
| Foxo3 | 0.586149925 | 0.262983741 | 0.714 | 0.821 | 1 | 0.043205847 | -0.101454406 | 0.911 | 0.913 | 1 | 0.586149925 | 0.084544949 |
| 1700017B05Rik | 0.418598017 | -0.611630451 | 0.048 | 0.101 | 1 | 0.04331848 | -0.662702251 | 0.111 | 0.263 | 1 | 0.418598017 | 0.084760469 |
| Susd6 | 0.043463661 | 0.936013702 | 0.81 | 0.795 | 1 | 0.192351507 | 0.670749308 | 0.956 | 0.844 | 1 | 0.192351507 | 0.085038231 |
| Dhx37 | 0.094179603 | 2.088809267 | 0.19 | 0.09 | 1 | 0.043502314 | -0.695439037 | 0.022 | 0.128 | 1 | 0.094179603 | 0.085112176 |
| Eps15l1 | 0.043640872 | 1.02857416 | 0.619 | 0.519 | 1 | 0.727877157 | 0.44987341 | 0.778 | 0.744 | 1 | 0.727877157 | 0.085377219 |
| Ndufaf5 | 0.231949841 | -0.911190733 | 0.048 | 0.138 | 1 | 0.043658074 | -0.638855509 | 0.022 | 0.128 | 1 | 0.231949841 | 0.08541012 |
| Zfp869 | 0.978338935 | 0.306989437 | 0.381 | 0.448 | 1 | 0.043697506 | -0.285494906 | 0.267 | 0.491 | 1 | 0.978338935 | 0.08548554 |
| Tle2 | 0.882731937 | 0.825774861 | 0.095 | 0.116 | 1 | 0.043717176 | -0.454430938 | 0.022 | 0.128 | 1 | 0.882731937 | 0.085523161 |
| Sacs | 0.050871609 | -2.348596045 | 0 | 0.157 | 1 | 0.043744363 | -0.223818009 | 0.022 | 0.128 | 1 | 0.050871609 | 0.085575157 |
| Sft2d3 | 0.065188443 | -1.785659851 | 0 | 0.142 | 1 | 0.04381154 | -1.589945908 | 0.044 | 0.156 | 1 | 0.065188443 | 0.085703629 |
| Edc4 | 0.69387462 | 0.38426515 | 0.19 | 0.272 | 1 | 0.043897512 | -0.695439037 | 0.111 | 0.26 | 1 | 0.69387462 | 0.085868032 |
| Rac3 | 0.173676192 | -0.911190733 | 0 | 0.082 | 1 | 0.043904203 | -0.618096949 | 0.044 | 0.166 | 1 | 0.173676192 | 0.085880826 |
| Ttc39b | 0.053603703 | 1.070430738 | 0.714 | 0.784 | 1 | 0.043983908 | 0.619275936 | 0.822 | 0.817 | 1 | 0.053603703 | 0.086033232 |
| Fam89a | 0.043984284 | -1.032206134 | 0.095 | 0.321 | 1 | 0.095952004 | -2.486852415 | 0 | 0.059 | 1 | 0.095952004 | 0.086033951 |
| Ninl | 0.885488583 | 0.388369549 | 0.095 | 0.108 | 1 | 0.044062648 | -0.776359032 | 0.044 | 0.166 | 1 | 0.885488583 | 0.086183778 |
| Blm | 0.044065338 | -1.785659851 | 0.143 | 0.354 | 1 | 0.29546512 | -0.124282336 | 0.022 | 0.062 | 1 | 0.29546512 | 0.086188922 |
| Kdm6a | 0.044070424 | 0.692183805 | 0.81 | 0.705 | 1 | 0.682100799 | 0.256372169 | 0.867 | 0.824 | 1 | 0.682100799 | 0.086198646 |
| Timm10 | 0.460886212 | -0.370622352 | 0.19 | 0.261 | 1 | 0.044081876 | -0.686161223 | 0.067 | 0.194 | 1 | 0.460886212 | 0.086220541 |
| Miga2 | 0.144717908 | -1.233118828 | 0 | 0.093 | 1 | 0.044152038 | -0.709244837 | 0.044 | 0.166 | 1 | 0.144717908 | 0.086354674 |
| Pdcd11 | 0.881125138 | 0.454603247 | 0.333 | 0.403 | 1 | 0.044253056 | -0.331877755 | 0.222 | 0.412 | 1 | 0.881125138 | 0.086547778 |
| A330023F24Rik | 0.553820882 | 0.59848364 | 0.19 | 0.146 | 1 | 0.044273886 | 1.622190344 | 0.333 | 0.235 | 1 | 0.553820882 | 0.086587595 |
| Cyb561 | 0.044275379 | 1.503846766 | 0.19 | 0.067 | 1 | 0.906750741 | 1.567595369 | 0.044 | 0.042 | 1 | 0.906750741 | 0.086590448 |
| Ifitm10 | 0.274576293 | 1.005393259 | 0.619 | 0.66 | 1 | 0.044310431 | 0.937047653 | 0.711 | 0.768 | 1 | 0.274576293 | 0.086657447 |
| Trit1 | 0.773607181 | 1.316219763 | 0.238 | 0.235 | 1 | 0.044455466 | -0.372780649 | 0.178 | 0.363 | 1 | 0.773607181 | 0.086934643 |
| Inpp4b | 0.111450466 | 1.476470327 | 0.286 | 0.172 | 1 | 0.044489161 | -1.334849322 | 0.044 | 0.159 | 1 | 0.111450466 | 0.086999037 |
| Zfp934 | 0.044545789 | 2.174965911 | 0.286 | 0.153 | 1 | 0.244038584 | -0.361321533 | 0.089 | 0.159 | 1 | 0.244038584 | 0.08710725 |
| Psmd10 | 0.11204789 | -0.25696557 | 0.048 | 0.198 | 1 | 0.044579041 | -1.486852415 | 0.044 | 0.156 | 1 | 0.11204789 | 0.087170791 |
| Tmem82 | 0.044590897 | 3.188344941 | 0.095 | 0.022 | 1 | 0.962383208 | 1.013221188 | 0.089 | 0.097 | 1 | 0.962383208 | 0.087193445 |
| Gm15991 | 0.044590897 | 3.188344941 | 0.095 | 0.022 | 1 | 0.787608894 | 0.513147585 | 0.044 | 0.055 | 1 | 0.787608894 | 0.087193445 |
| St6galnac4 | 0.886422617 | 0.162809848 | 0.286 | 0.28 | 1 | 0.044603125 | -0.831500587 | 0.089 | 0.221 | 1 | 0.886422617 | 0.087216811 |
| Gfer | 0.0446198 | -1.970084422 | 0 | 0.164 | 1 | 0.559546785 | 0.486675373 | 0.111 | 0.152 | 1 | 0.559546785 | 0.087248674 |
| Aptx | 0.762071023 | 0.376091219 | 0.143 | 0.183 | 1 | 0.044671617 | -0.158229668 | 0.111 | 0.266 | 1 | 0.762071023 | 0.087347681 |
| Anks3 | 0.87274741 | 0.659965968 | 0.286 | 0.291 | 1 | 0.044714166 | 0.170622585 | 0.2 | 0.433 | 1 | 0.87274741 | 0.087428975 |
| Gtf2h4 | 0.044760094 | -2.159118246 | 0 | 0.164 | 1 | 0.126035431 | -0.709244837 | 0.044 | 0.125 | 1 | 0.126035431 | 0.087516722 |
| Herc1 | 0.152281745 | 0.204585582 | 0.952 | 0.821 | 1 | 0.044763152 | 0.357965581 | 0.978 | 0.917 | 1 | 0.152281745 | 0.087522565 |
| 1810062G17Rik | 0.04476392 | -2.054148687 | 0 | 0.164 | 1 | 0.120422285 | -1.404390255 | 0.022 | 0.09 | 1 | 0.120422285 | 0.087524032 |
| Nectin2 | 0.044805331 | -2.208871282 | 0 | 0.164 | 1 | 0.11404908 | -0.952515988 | 0.133 | 0.232 | 1 | 0.11404908 | 0.087603143 |
| Suox | 0.190431735 | -0.574155746 | 0.095 | 0.224 | 1 | 0.044833748 | -1.459885368 | 0.044 | 0.156 | 1 | 0.190431735 | 0.087657431 |
| Abhd11 | 0.236219797 | -0.763633545 | 0.095 | 0.198 | 1 | 0.044834222 | -1.486852415 | 0.044 | 0.156 | 1 | 0.236219797 | 0.087658336 |
| Trappc9 | 0.775648636 | 0.735172312 | 0.429 | 0.526 | 1 | 0.044862281 | 1.131662645 | 0.711 | 0.578 | 1 | 0.775648636 | 0.087711939 |
| Bche | 0.044882091 | -2.280424543 | 0 | 0.164 | 1 | 0.050682869 | -2.044847868 | 0.022 | 0.118 | 1 | 0.050682869 | 0.08774978 |
| Plaat3 | 0.620410565 | 0.275222391 | 0.19 | 0.261 | 1 | 0.044897421 | 0.11747541 | 0.267 | 0.512 | 1 | 0.620410565 | 0.087779064 |
| Ppp2r1b | 0.948622969 | 1.107424945 | 0.19 | 0.216 | 1 | 0.044949829 | -0.414959497 | 0.156 | 0.318 | 1 | 0.948622969 | 0.087879171 |
| Otud7b | 0.096316801 | 0.528094312 | 0.857 | 0.735 | 1 | 0.044954347 | 0.534156621 | 0.956 | 0.896 | 1 | 0.096316801 | 0.087887801 |
| Slitrk6 | 0.044981929 | -2.752492987 | 0 | 0.164 | 1 | 0.119040864 | -2.361321533 | 0 | 0.052 | 1 | 0.119040864 | 0.087940484 |
| Bcl10 | 0.045014732 | -0.679375058 | 0.333 | 0.649 | 1 | 0.382703682 | -0.282980082 | 0.622 | 0.606 | 1 | 0.382703682 | 0.088003138 |
| Rbbp6 | 0.201401087 | -0.459569466 | 0.905 | 0.903 | 1 | 0.045071054 | -0.303717557 | 1 | 0.969 | 1 | 0.201401087 | 0.088110708 |
| Mettl5 | 0.210652454 | 0.252307999 | 0.095 | 0.235 | 1 | 0.045077936 | -0.486852415 | 0.111 | 0.256 | 1 | 0.210652454 | 0.088123852 |
| BC034090 | 0.991562202 | 0.629377648 | 0.095 | 0.097 | 1 | 0.045088054 | -1.404390255 | 0.044 | 0.156 | 1 | 0.991562202 | 0.088143175 |
| D16Ertd472e | 0.197011335 | -0.807354922 | 0.143 | 0.272 | 1 | 0.045095584 | -0.901889915 | 0.2 | 0.349 | 1 | 0.197011335 | 0.088157556 |
| 4930500A05Rik | 0.045104418 | 3.673771768 | 0.095 | 0.022 | 1 | 0.293659664 | -0.316927414 | 0 | 0.024 | 1 | 0.293659664 | 0.088174427 |
| Pcdhb14 | 0.373715976 | -0.648156327 | 0.048 | 0.108 | 1 | 0.045109775 | -2.044847868 | 0.022 | 0.121 | 1 | 0.373715976 | 0.088184659 |
| Lsamp | 0.457305494 | -0.849790188 | 0 | 0.026 | 1 | 0.045134375 | -0.337073987 | 0.2 | 0.388 | 1 | 0.457305494 | 0.088231638 |
| Knop1 | 0.045182596 | -1.116305163 | 0.143 | 0.381 | 1 | 0.346493909 | -0.513324627 | 0.4 | 0.429 | 1 | 0.346493909 | 0.088323725 |
| Fyttd1 | 0.640825757 | 0.155923463 | 0.857 | 0.877 | 1 | 0.04518365 | -0.250838223 | 0.933 | 0.924 | 1 | 0.640825757 | 0.088325738 |
| Bcl7a | 0.330083496 | -0.818081329 | 0.048 | 0.116 | 1 | 0.045246458 | 1.066401226 | 0.222 | 0.114 | 1 | 0.330083496 | 0.088445674 |
| Gm12743 | 0.938884995 | 0.908237021 | 0.095 | 0.097 | 1 | 0.045260229 | -2.017367132 | 0.022 | 0.121 | 1 | 0.938884995 | 0.08847197 |
| Gm11508 | 0.363561289 | 0.233199176 | 0.095 | 0.179 | 1 | 0.045274217 | 0.323991493 | 0.111 | 0.256 | 1 | 0.363561289 | 0.088498679 |
| Ccdc186 | 0.692635894 | 0.459850862 | 0.667 | 0.765 | 1 | 0.045325781 | -0.187644397 | 0.867 | 0.83 | 1 | 0.692635894 | 0.088597136 |
| Birc6 | 0.045334669 | 0.43406164 | 1 | 0.951 | 1 | 0.047016761 | 0.422308354 | 1 | 0.993 | 1 | 0.047016761 | 0.088614106 |
| Thtpa | 0.285527727 | -0.496153234 | 0 | 0.052 | 1 | 0.045339481 | -2.071814916 | 0 | 0.083 | 1 | 0.285527727 | 0.088623294 |
| Nudt14 | 0.313244997 | -0.233118828 | 0.048 | 0.127 | 1 | 0.045339481 | -2.017367132 | 0 | 0.083 | 1 | 0.313244997 | 0.088623294 |
| Fastkd5 | 0.218331105 | 2.410737362 | 0.095 | 0.041 | 1 | 0.045361416 | -2.124282336 | 0 | 0.083 | 1 | 0.218331105 | 0.088665175 |
| Acy1 | 0.208797419 | -0.718545655 | 0 | 0.071 | 1 | 0.045381446 | -2.124282336 | 0 | 0.083 | 1 | 0.208797419 | 0.088703416 |
| Chrna1os | 0.208921714 | -0.970084422 | 0 | 0.071 | 1 | 0.045381446 | -2.124282336 | 0 | 0.083 | 1 | 0.208921714 | 0.088703416 |
| Madd | 0.285465396 | -0.326228232 | 0 | 0.052 | 1 | 0.045399569 | -2.271123724 | 0 | 0.083 | 1 | 0.285465396 | 0.088738018 |
| Slc46a1 | 0.163379068 | -0.970084422 | 0 | 0.086 | 1 | 0.045415266 | -2.271123724 | 0 | 0.083 | 1 | 0.163379068 | 0.088767986 |
| Reno1 | 0.173676192 | -0.911190733 | 0 | 0.082 | 1 | 0.045415266 | -2.271123724 | 0 | 0.083 | 1 | 0.173676192 | 0.088767986 |
| Gm15411 | 0.342664981 | 1.586308926 | 0.095 | 0.049 | 1 | 0.045415266 | -2.361321533 | 0 | 0.083 | 1 | 0.342664981 | 0.088767986 |
| Pla2g6 | 0.799182193 | 0.614878079 | 0.095 | 0.078 | 1 | 0.045415266 | -2.361321533 | 0 | 0.083 | 1 | 0.799182193 | 0.088767986 |
| Zc3h8 | 0.236452955 | -0.849790188 | 0 | 0.063 | 1 | 0.045415786 | -2.361321533 | 0 | 0.083 | 1 | 0.236452955 | 0.088768979 |
| Vgll3 | 0.632076134 | 1.088809267 | 0 | 0.011 | 1 | 0.045415786 | -2.989352756 | 0 | 0.083 | 1 | 0.632076134 | 0.088768979 |
| Nap1l3 | 0.725258133 | 0.735172312 | 0.048 | 0.071 | 1 | 0.045416133 | -2.223818009 | 0 | 0.083 | 1 | 0.725258133 | 0.088769642 |
| Nudt6 | 0.13616383 | -1.326228232 | 0 | 0.097 | 1 | 0.045429229 | -2.316927414 | 0 | 0.083 | 1 | 0.13616383 | 0.088794644 |
| Rbm38 | 0.765801178 | 0.150209812 | 0.048 | 0.063 | 1 | 0.04543053 | -2.446210431 | 0 | 0.083 | 1 | 0.765801178 | 0.088797128 |
| Fstl5 | 0.54776222 | 2.088809267 | 0.048 | 0.026 | 1 | 0.045430791 | -3.124282336 | 0 | 0.083 | 1 | 0.54776222 | 0.088797624 |
| Hes6 | 0.236454357 | -0.785659851 | 0 | 0.063 | 1 | 0.045430964 | -2.271123724 | 0 | 0.083 | 1 | 0.236454357 | 0.088797955 |
| Galnt5 | 0.222228992 | -1.02666795 | 0 | 0.067 | 1 | 0.045441199 | -2.404390255 | 0 | 0.083 | 1 | 0.222228992 | 0.088817495 |
| Dffb | 0.701303834 | 1.088809267 | 0.095 | 0.075 | 1 | 0.0454425 | -2.446210431 | 0 | 0.083 | 1 | 0.701303834 | 0.088819978 |
| Map2k6 | 0.236480292 | -1.133583154 | 0 | 0.063 | 1 | 0.045452995 | -2.638855509 | 0 | 0.083 | 1 | 0.236480292 | 0.088840015 |
| Tmem144 | 0.492603827 | 0.214340149 | 0 | 0.022 | 1 | 0.045452995 | -2.446210431 | 0 | 0.083 | 1 | 0.492603827 | 0.088840015 |
| Kctd12b | 0.346873877 | -0.413691074 | 0 | 0.041 | 1 | 0.045455164 | -2.526380779 | 0 | 0.083 | 1 | 0.346873877 | 0.088844155 |
| Gm13889 | 0.128414409 | -4.120644099 | 0 | 0.101 | 1 | 0.045472426 | -3.446210431 | 0 | 0.083 | 1 | 0.128414409 | 0.08887711 |
| Bpifb1 | 0.396935107 | -0.233118828 | 0 | 0.034 | 1 | 0.045474594 | -3.223818009 | 0 | 0.083 | 1 | 0.396935107 | 0.08888125 |
| Slc16a7 | 0.045504608 | 2.740885964 | 0.143 | 0.045 | 1 | 0.197919408 | -0.191396532 | 0.022 | 0.076 | 1 | 0.197919408 | 0.088938547 |
| Zkscan6 | 0.427439322 | -0.233118828 | 0.048 | 0.101 | 1 | 0.045515023 | -1.960783604 | 0 | 0.083 | 1 | 0.427439322 | 0.08895843 |
| Gm20274 | 0.649186945 | 0.281454345 | 0.048 | 0.075 | 1 | 0.045515023 | -1.960783604 | 0 | 0.083 | 1 | 0.649186945 | 0.08895843 |
| Nlrc5 | 0.045591 | 1.440281637 | 0.476 | 0.313 | 1 | 0.575408978 | 1.057237804 | 0.244 | 0.232 | 1 | 0.575408978 | 0.089103461 |
| Atg3 | 0.093977292 | -0.623908781 | 0.524 | 0.746 | 1 | 0.045637456 | -0.399667845 | 0.756 | 0.782 | 1 | 0.093977292 | 0.089192136 |
| Zdhhc16 | 0.045642796 | -1.574155746 | 0.048 | 0.235 | 1 | 0.188020139 | -0.2150478 | 0.133 | 0.232 | 1 | 0.188020139 | 0.089202326 |
| Tmod2 | 0.173921884 | -1.370622352 | 0 | 0.082 | 1 | 0.045719696 | -1.103523776 | 0.044 | 0.159 | 1 | 0.173921884 | 0.089349102 |
| Steap2 | 0.555425832 | 0.495434526 | 0.524 | 0.537 | 1 | 0.045741012 | 0.468059695 | 0.178 | 0.381 | 1 | 0.555425832 | 0.089389784 |
| Inpp5b | 0.570969636 | 0.323274521 | 0.095 | 0.146 | 1 | 0.045758308 | -1.033134448 | 0.089 | 0.221 | 1 | 0.570969636 | 0.089422794 |

| | | | | | | | | | | | | |
|---|---|---|---|---|---|---|---|---|---|---|---|---|
| Itprip | 0.501787581 | -0.260340297 | 0.19 | 0.265 | 1 | 0.045804895 | -0.941418279 | 0.067 | 0.19 | 1 | 0.501787581 | 0.089511701 |
| Slc25a30 | 0.134101515 | 0.323274521 | 0.143 | 0.317 | 1 | 0.045873855 | -0.960783604 | 0.067 | 0.19 | 1 | 0.134101515 | 0.0896433 |
| Papolg | 0.255531932 | -0.516052791 | 0.095 | 0.201 | 1 | 0.045885951 | 0.374950291 | 0.089 | 0.242 | 1 | 0.255531932 | 0.089666381 |
| Cdk14 | 0.053244829 | 0.888669652 | 0.667 | 0.522 | 1 | 0.04588912 | 0.521609163 | 0.911 | 0.734 | 1 | 0.053244829 | 0.089672429 |
| Itga6 | 0.141487357 | 0.95883248 | 0.81 | 0.69 | 1 | 0.045929631 | 0.580261781 | 0.778 | 0.557 | 1 | 0.141487357 | 0.08974973 |
| Ccdc71l | 0.106727131 | -1.455511249 | 0 | 0.112 | 1 | 0.045995237 | -1.174908409 | 0.044 | 0.159 | 1 | 0.106727131 | 0.089874913 |
| Nmrk1 | 0.785773366 | 0.33137957 | 0.143 | 0.183 | 1 | 0.046017358 | -0.416463087 | 0.044 | 0.166 | 1 | 0.785773366 | 0.089917119 |
| Zfas1 | 0.04605442 | -0.574155746 | 0.429 | 0.735 | 1 | 0.071044996 | -0.564854927 | 0.556 | 0.623 | 1 | 0.071044996 | 0.08998783 |
| Kcna6 | 0.250359446 | 0.152721031 | 0.333 | 0.563 | 1 | 0.04606483 | 0.572790531 | 0.378 | 0.664 | 1 | 0.250359446 | 0.090007692 |
| Zfp219 | 0.832501289 | 0.866416846 | 0.095 | 0.116 | 1 | 0.046099885 | -0.916840256 | 0.111 | 0.253 | 1 | 0.832501289 | 0.090074571 |
| Scarb2 | 0.046104365 | -0.374591254 | 0.333 | 0.664 | 1 | 0.839407577 | 0.156618491 | 0.8 | 0.824 | 1 | 0.839407577 | 0.090083117 |
| Pate2 | 0.046139318 | 3.351843673 | 0.095 | 0.022 | 1 | 0.825853389 | 0.223640968 | 0.022 | 0.017 | 1 | 0.825853389 | 0.0901498 |
| Slc4a5 | 0.046140324 | 3.159198595 | 0.095 | 0.022 | 1 | 0.740122935 | -0.124282336 | 0.022 | 0.031 | 1 | 0.740122935 | 0.090151719 |
| Gm56862 | 0.222255538 | -1.02666795 | 0 | 0.067 | 1 | 0.046152082 | -1.776359032 | 0.022 | 0.121 | 1 | 0.222255538 | 0.090174149 |
| Tmem192 | 0.378843125 | 0.171271427 | 0.048 | 0.116 | 1 | 0.046154206 | -1.743192169 | 0.022 | 0.121 | 1 | 0.378843125 | 0.090178202 |
| Phf12 | 0.046215276 | -0.702604111 | 0.238 | 0.53 | 1 | 0.981013376 | 0.243251986 | 0.8 | 0.792 | 1 | 0.981013376 | 0.090294701 |
| Pald1 | 0.796434059 | 2.24081236 | 0.095 | 0.082 | 1 | 0.046246526 | -1.035745661 | 0.067 | 0.19 | 1 | 0.796434059 | 0.090354311 |
| Haus1 | 0.280793265 | -0.718545655 | 0.048 | 0.127 | 1 | 0.046295456 | -1.776359032 | 0.022 | 0.121 | 1 | 0.280793265 | 0.090447643 |
| Agk | 0.628118047 | 0.928344595 | 0.286 | 0.269 | 1 | 0.046313934 | -0.65082815 | 0.111 | 0.263 | 1 | 0.628118047 | 0.090482888 |
| Lmbrd2 | 0.438758354 | 0.540746349 | 0.286 | 0.444 | 1 | 0.04635881 | -0.500149238 | 0.267 | 0.429 | 1 | 0.438758354 | 0.090568481 |
| Hpn | 0.927794995 | 1.258734268 | 0.048 | 0.045 | 1 | 0.046419246 | -0.921789472 | 0.044 | 0.159 | 1 | 0.927794995 | 0.090683745 |
| Gpam | 0.591953669 | 0.524024648 | 0.095 | 0.149 | 1 | 0.046437712 | -1.674479418 | 0.022 | 0.121 | 1 | 0.591953669 | 0.090718963 |
| Mical1 | 0.813849549 | 1.223110359 | 0.238 | 0.228 | 1 | 0.046457591 | -0.440309829 | 0.133 | 0.301 | 1 | 0.813849549 | 0.090756875 |
| Atg4c | 0.716526111 | 0.60338244 | 0.095 | 0.134 | 1 | 0.046484602 | -1.039393438 | 0.067 | 0.19 | 1 | 0.716526111 | 0.090808386 |
| Gmip | 0.887106287 | 0.569435108 | 0.095 | 0.112 | 1 | 0.046539373 | -0.972279242 | 0.044 | 0.159 | 1 | 0.887106287 | 0.090912832 |
| Dusp19 | 0.053995916 | -1.880817084 | 0 | 0.153 | 1 | 0.046577008 | -1.674479418 | 0.022 | 0.121 | 1 | 0.053995916 | 0.090984599 |
| Nol3 | 0.153760172 | -1.081115734 | 0 | 0.09 | 1 | 0.046577008 | -1.674479418 | 0.022 | 0.121 | 1 | 0.153760172 | 0.090984599 |
| Tmem41a | 0.862687636 | 1.088809267 | 0.048 | 0.06 | 1 | 0.046577252 | -1.638855509 | 0.022 | 0.121 | 1 | 0.862687636 | 0.090985064 |
| Rhot2 | 0.354979527 | -0.535681598 | 0.048 | 0.112 | 1 | 0.04671516 | -1.638855509 | 0.022 | 0.121 | 1 | 0.354979527 | 0.091248013 |
| Pdxdc1 | 0.046775672 | 0.614156911 | 0.952 | 0.828 | 1 | 0.947321134 | 0.273764447 | 0.933 | 0.92 | 1 | 0.947321134 | 0.091363381 |
| Pten | 0.143222296 | -0.386210337 | 0.762 | 0.918 | 1 | 0.046783952 | -0.167480025 | 0.978 | 0.965 | 1 | 0.143222296 | 0.091379166 |
| Ell | 0.046788023 | 1.168777572 | 0.667 | 0.709 | 1 | 0.051407033 | 0.624178897 | 0.778 | 0.734 | 1 | 0.051407033 | 0.091386927 |
| Ell2 | 0.192707647 | 1.154886299 | 0.857 | 0.888 | 1 | 0.046809227 | 0.571214776 | 1 | 0.855 | 1 | 0.192707647 | 0.09142735 |
| 8430429K09Rik | 0.570707746 | 0.699766976 | 0.19 | 0.149 | 1 | 0.046835226 | -1.286553765 | 0.022 | 0.125 | 1 | 0.570707746 | 0.091476913 |
| Cgnl1 | 0.046837992 | -1.239972499 | 0.143 | 0.384 | 1 | 0.515138605 | 0.991538421 | 0.356 | 0.377 | 1 | 0.515138605 | 0.091482186 |
| Mir155hg | 0.046919172 | 2.38426515 | 0.19 | 0.078 | 1 | 0.165324946 | -1.404390255 | 0 | 0.042 | 1 | 0.165324946 | 0.091636936 |
| Wars1 | 0.797818007 | 1.323274521 | 0.143 | 0.172 | 1 | 0.046995568 | -0.84048937 | 0.067 | 0.19 | 1 | 0.797818007 | 0.091782553 |
| Pi4kb | 0.944240481 | 0.550389352 | 0.286 | 0.313 | 1 | 0.047016847 | -0.217794222 | 0.244 | 0.46 | 1 | 0.944240481 | 0.091823109 |
| Gm35608 | 0.047256943 | 2.566856564 | 0.19 | 0.075 | 1 | 0.244379289 | 1.498648015 | 0.022 | 0.069 | 1 | 0.244379289 | 0.092280667 |
| Cdh23 | 0.047344511 | 2.33273485 | 0.238 | 0.104 | 1 | 0.192017122 | 2.176112597 | 0.111 | 0.062 | 1 | 0.192017122 | 0.092447519 |
| Net1 | 0.944670948 | 0.103023126 | 0.333 | 0.34 | 1 | 0.047411521 | 0.185294656 | 0.711 | 0.457 | 1 | 0.944670948 | 0.09257519 |
| Ifrd1 | 0.448773793 | 0.491915098 | 1 | 0.974 | 1 | 0.047436893 | -0.110174938 | 1 | 0.993 | 1 | 0.448773793 | 0.092623527 |
| Map3k6 | 0.798263109 | 0.524024648 | 0.143 | 0.175 | 1 | 0.047528532 | -1.124282336 | 0.022 | 0.125 | 1 | 0.798263109 | 0.092798103 |
| Usf3 | 0.047561986 | 0.137718867 | 0.095 | 0.328 | 1 | 0.054165886 | 0.786166079 | 0.6 | 0.436 | 1 | 0.054165886 | 0.09286183 |
| Kif3a | 0.084384721 | -0.496153234 | 0.048 | 0.216 | 1 | 0.047597535 | -0.423842618 | 0.089 | 0.235 | 1 | 0.084384721 | 0.092929545 |
| Sirt3 | 0.073876282 | -1.849790188 | 0 | 0.134 | 1 | 0.047621933 | -0.564854927 | 0.044 | 0.159 | 1 | 0.073876282 | 0.092976017 |
| Tns3 | 0.231769028 | 1.181335882 | 0.667 | 0.728 | 1 | 0.047644785 | -0.405000661 | 0.8 | 0.896 | 1 | 0.231769028 | 0.093019545 |
| Pradc1 | 0.047658741 | -2.02666795 | 0 | 0.16 | 1 | 0.730163466 | 0.735540006 | 0.089 | 0.076 | 1 | 0.730163466 | 0.093046126 |
| Atp2a2 | 0.755262917 | 0.243699423 | 1 | 0.985 | 1 | 0.047690467 | -0.193940968 | 1 | 0.997 | 1 | 0.755262917 | 0.093106553 |
| Lyrm7 | 0.047711093 | 2.866416846 | 0.095 | 0.022 | 1 | 0.148059913 | -1.486852415 | 0 | 0.045 | 1 | 0.148059913 | 0.093145837 |
| Hhat | 0.146504399 | 1.328943271 | 0.381 | 0.302 | 1 | 0.047728642 | 0.559243999 | 0.2 | 0.412 | 1 | 0.146504399 | 0.09317926 |
| Mccc1 | 0.047748557 | -2.208871282 | 0 | 0.16 | 1 | 0.065676536 | 0.454253896 | 0.067 | 0.19 | 1 | 0.065676536 | 0.093217188 |
| Tmco4 | 0.04776409 | 1.192119984 | 0.524 | 0.418 | 1 | 0.88335538 | 0.508298185 | 0.4 | 0.498 | 1 | 0.88335538 | 0.093246772 |
| Nfkbil1 | 0.047786697 | -2.159118246 | 0 | 0.16 | 1 | 0.080883392 | -0.463768802 | 0.067 | 0.176 | 1 | 0.080883392 | 0.093289825 |
| Txnrd2 | 0.083489388 | -1.611630451 | 0 | 0.127 | 1 | 0.047810639 | -0.722919774 | 0.044 | 0.159 | 1 | 0.083489388 | 0.093335421 |
| Pfas | 0.255520517 | -0.233118828 | 0.048 | 0.142 | 1 | 0.047837294 | -1.053893008 | 0.022 | 0.125 | 1 | 0.255520517 | 0.093386181 |
| Tmie | 0.047846329 | -1.008052272 | 0.095 | 0.31 | 1 | 0.237333231 | -0.776359032 | 0.022 | 0.069 | 1 | 0.237333231 | 0.093403386 |
| Stx2 | 0.047858933 | -2.233118828 | 0 | 0.16 | 1 | 0.532593269 | 0.760240447 | 0.156 | 0.228 | 1 | 0.532593269 | 0.093427389 |
| Slc16a2 | 0.785901472 | 0.635351549 | 0.619 | 0.683 | 1 | 0.047907521 | 0.40239522 | 0.733 | 0.896 | 1 | 0.785901472 | 0.093519912 |
| Mb21d2 | 0.212636095 | 1.538612184 | 0.381 | 0.325 | 1 | 0.047917596 | 0.120830162 | 0.133 | 0.311 | 1 | 0.212636095 | 0.093539096 |
| Eefsec | 0.242577743 | -0.574155746 | 0.095 | 0.205 | 1 | 0.047957859 | -1.11134328 | 0.111 | 0.246 | 1 | 0.242577743 | 0.093615761 |
| Tgds | 0.063731289 | -1.496153234 | 0.048 | 0.216 | 1 | 0.047995037 | -0.602329633 | 0.089 | 0.232 | 1 | 0.063731289 | 0.093686551 |
| Rpa1 | 0.048032915 | -1.865387043 | 0.048 | 0.231 | 1 | 0.15737708 | -0.394929926 | 0.111 | 0.211 | 1 | 0.15737708 | 0.093758668 |
| Asah2 | 0.050950146 | -2.683780237 | 0 | 0.157 | 1 | 0.048104627 | -0.81335324 | 0.044 | 0.163 | 1 | 0.050950146 | 0.093895199 |
| Sec63 | 0.048128858 | -0.633049435 | 0.81 | 0.94 | 1 | 0.824837799 | 0.181925301 | 0.911 | 0.969 | 1 | 0.824837799 | 0.093941329 |
| Cks2 | 0.048129791 | -3.146407195 | 0 | 0.16 | 1 | 0.070167187 | 0.412983423 | 0.289 | 0.166 | 1 | 0.070167187 | 0.093943106 |
| Lasp1 | 0.048195845 | -0.350612391 | 0.143 | 0.414 | 1 | 0.920777952 | -0.284190627 | 0.622 | 0.533 | 1 | 0.920777952 | 0.09406885 |
| Gm16740 | 0.818080815 | 1.316219763 | 0.095 | 0.123 | 1 | 0.048204874 | -0.901889915 | 0.022 | 0.125 | 1 | 0.818080815 | 0.094086038 |
| Cep120 | 0.950782188 | 0.371208998 | 0.286 | 0.34 | 1 | 0.048232393 | -0.142898014 | 0.333 | 0.54 | 1 | 0.950782188 | 0.094138423 |
| Lrrd1 | 0.048257036 | 3.673771768 | 0.095 | 0.022 | 1 | 0.132717935 | -1.776359032 | 0 | 0.048 | 1 | 0.132717935 | 0.09418533 |
| Txlng | 0.882776042 | 0.119182916 | 0.381 | 0.433 | 1 | 0.048295168 | -0.316927414 | 0.289 | 0.45 | 1 | 0.882776042 | 0.094257913 |
| Uspl1 | 0.716554435 | 0.563347778 | 0.333 | 0.448 | 1 | 0.048360237 | -0.241739917 | 0.267 | 0.467 | 1 | 0.716554435 | 0.094381762 |
| Npas2 | 0.421799182 | 1.34285489 | 0.381 | 0.366 | 1 | 0.048436534 | 0.675079795 | 0.489 | 0.308 | 1 | 0.421799182 | 0.09452697 |
| Zfhx3 | 0.048521344 | 0.775752499 | 1 | 0.903 | 1 | 0.433921507 | 0.410831615 | 0.978 | 0.945 | 1 | 0.433921507 | 0.094688368 |
| Zbtb40 | 0.451614009 | -0.233118828 | 0.048 | 0.097 | 1 | 0.048526697 | -0.486852415 | 0.022 | 0.125 | 1 | 0.451614009 | 0.094698554 |
| Abcg1 | 0.324808024 | -0.413691074 | 0 | 0.045 | 1 | 0.048529631 | -0.286553765 | 0.022 | 0.125 | 1 | 0.324808024 | 0.094704137 |
| Med14 | 0.048697379 | 1.42946457 | 0.524 | 0.403 | 1 | 0.690780174 | 0.13875207 | 0.644 | 0.574 | 1 | 0.690780174 | 0.095023324 |
| Scrib | 0.822651628 | 1.521768674 | 0.143 | 0.146 | 1 | 0.048705326 | -1.204452685 | 0.067 | 0.187 | 1 | 0.822651628 | 0.095038443 |
| B430319H21Rik | 0.048800887 | 2.673771768 | 0.095 | 0.022 | 1 | 0.708987514 | 1.42003818 | 0.044 | 0.035 | 1 | 0.708987514 | 0.095220248 |
| Enox1 | 0.184495547 | -1.174225139 | 0.19 | 0.332 | 1 | 0.048865751 | -0.100937247 | 0.244 | 0.439 | 1 | 0.184495547 | 0.09534364 |
| Exosc7 | 0.048964627 | -0.752492987 | 0.048 | 0.25 | 1 | 0.081696177 | -0.881712033 | 0.067 | 0.173 | 1 | 0.081696177 | 0.095531719 |
| Nfil3 | 0.0490558 | -1.785659851 | 0.095 | 0.295 | 1 | 0.074275017 | 0.376826114 | 0.511 | 0.343 | 1 | 0.074275017 | 0.095705129 |
| Zfp655 | 0.049068154 | -0.321210158 | 0.286 | 0.616 | 1 | 0.713046676 | 0.368561963 | 0.511 | 0.664 | 1 | 0.713046676 | 0.095728624 |
| Scgb2b27 | 0.049100545 | 1.483088206 | 0.429 | 0.299 | 1 | 0.088879547 | -0.16492432 | 0.022 | 0.104 | 1 | 0.088879547 | 0.095790226 |
| Zfp512 | 0.049111295 | 1.514573173 | 0.333 | 0.187 | 1 | 0.492958541 | 0.551828053 | 0.178 | 0.253 | 1 | 0.492958541 | 0.095810671 |
| Erbb3 | 0.251681669 | -0.785659851 | 0 | 0.06 | 1 | 0.049115979 | 1.757073168 | 0.133 | 0.055 | 1 | 0.251681669 | 0.095819578 |
| Gtf2e2 | 0.049203283 | -0.31112134 | 0.19 | 0.47 | 1 | 0.079656254 | -0.375821103 | 0.378 | 0.536 | 1 | 0.079656254 | 0.095985602 |
| Tusc2 | 0.08614687 | -0.828728573 | 0.048 | 0.213 | 1 | 0.049365562 | -1.223818009 | 0.067 | 0.183 | 1 | 0.08614687 | 0.096294165 |
| Pon3 | 0.04938075 | -1.326228232 | 0.095 | 0.302 | 1 | 0.703792355 | 0.425914747 | 0.489 | 0.429 | 1 | 0.703792355 | 0.096323041 |
| Lrrc8d | 0.418727983 | 1.184625054 | 0.619 | 0.675 | 1 | 0.04939075 | 0.731394069 | 0.956 | 0.837 | 1 | 0.418727983 | 0.096342054 |
| Tmtc2 | 0.577710208 | -0.180377366 | 0.714 | 0.687 | 1 | 0.049422081 | 0.255534303 | 0.889 | 0.657 | 1 | 0.577710208 | 0.096401621 |
| Otor | 0.049422793 | -4.667667632 | 0.286 | 0.429 | 1 | 0.156858666 | -0.983965626 | 0.2 | 0.318 | 1 | 0.156858666 | 0.096402974 |
| Tcp11l1 | 0.199086232 | -0.593014773 | 0.095 | 0.22 | 1 | 0.049461404 | -0.690385809 | 0.133 | 0.273 | 1 | 0.199086232 | 0.096476378 |
| Cox19 | 0.049464202 | -0.421385465 | 0.19 | 0.466 | 1 | 0.29582872 | -0.140049652 | 0.356 | 0.443 | 1 | 0.29582872 | 0.096481697 |
| Macf1 | 0.179207761 | 0.651014616 | 1 | 0.978 | 1 | 0.049544998 | 0.473407708 | 1 | 0.986 | 1 | 0.179207761 | 0.09663529 |
| Tubgcp5 | 0.504101666 | 0.548240886 | 0.286 | 0.239 | 1 | 0.049579467 | -0.200734396 | 0.156 | 0.336 | 1 | 0.504101666 | 0.096700811 |
| Slc41a2 | 0.18358839 | -0.186197781 | 0.333 | 0.56 | 1 | 0.049669755 | -0.124282336 | 0.444 | 0.72 | 1 | 0.18358839 | 0.096872425 |
| Ssh1 | 0.049713459 | -0.760214834 | 0.143 | 0.392 | 1 | 0.910334898 | 0.387616703 | 0.422 | 0.585 | 1 | 0.910334898 | 0.096955491 |

| | | | | | | | | | | | | |
|---|---|---|---|---|---|---|---|---|---|---|---|---|
| Zfp101 | 0.049738042 | 1.420015175 | 0.333 | 0.187 | 1 | 0.900757632 | 0.694387899 | 0.244 | 0.291 | 1 | 0.900757632 | 0.097002211 |
| Maoa | 0.049773525 | -1.839297815 | 0.048 | 0.239 | 1 | 0.203709604 | -0.586388089 | 0.133 | 0.228 | 1 | 0.203709604 | 0.097069647 |
| Patz1 | 0.368180647 | -0.348596045 | 0.095 | 0.175 | 1 | 0.049775657 | -0.286553765 | 0.111 | 0.253 | 1 | 0.368180647 | 0.097073698 |
| Synj2 | 0.049858348 | -1.998653574 | 0.048 | 0.228 | 1 | 0.60114346 | 0.147019686 | 0.244 | 0.308 | 1 | 0.60114346 | 0.097230841 |
| Myof | 0.965364586 | -0.106838299 | 0.667 | 0.653 | 1 | 0.049875815 | 0.51932616 | 0.889 | 0.73 | 1 | 0.965364586 | 0.097264032 |
| Robo2 | 0.049881522 | -0.370622352 | 0.048 | 0.25 | 1 | 0.11259271 | -0.339607803 | 0.2 | 0.349 | 1 | 0.11259271 | 0.097274877 |
| Tafa4 | 0.049909318 | 2.503846766 | 0.095 | 0.022 | 1 | 0.079149465 | 3.005000681 | 0.044 | 0.01 | 1 | 0.079149465 | 0.097327696 |
| Myadml2os | 0.049909318 | 2.503846766 | 0.095 | 0.022 | 1 | 0.953500769 | 1.098110085 | 0.044 | 0.048 | 1 | 0.953500769 | 0.097327696 |
| Cpn1 | 0.049909318 | 2.503846766 | 0.095 | 0.022 | 1 | 0.496223879 | 0.683072586 | 0 | 0.01 | 1 | 0.496223879 | 0.097327696 |
| L1cam | 0.049909318 | 2.503846766 | 0.095 | 0.022 | 1 | 0.293637566 | -0.486852415 | 0 | 0.024 | 1 | 0.293637566 | 0.097327696 |
| Sms | 0.087591855 | 0.444953077 | 0.143 | 0.377 | 1 | 0.049922277 | 0.572041274 | 0.556 | 0.36 | 1 | 0.087591855 | 0.097352321 |
| Arpin | 0.18539133 | -0.326228232 | 0.048 | 0.164 | 1 | 0.049941202 | -1.103523776 | 0.067 | 0.187 | 1 | 0.18539133 | 0.097388281 |
| Tpk1 | 0.749491377 | 0.194302396 | 0.238 | 0.299 | 1 | 0.050053721 | -0.289960366 | 0.111 | 0.266 | 1 | 0.749491377 | 0.097602068 |
| Apobec3 | 0.164261037 | -0.118786152 | 0.095 | 0.25 | 1 | 0.050062193 | 0.900170708 | 0.422 | 0.266 | 1 | 0.164261037 | 0.097618162 |
| Umad1 | 0.374164369 | 0.311201688 | 0.143 | 0.261 | 1 | 0.050068776 | 1.162240423 | 0.511 | 0.398 | 1 | 0.374164369 | 0.097630669 |
| Fbxl6 | 0.050080534 | -1.008052272 | 0.048 | 0.246 | 1 | 0.112772416 | -0.17942389 | 0.111 | 0.221 | 1 | 0.112772416 | 0.097653008 |
| Pi4ka | 0.259297658 | 0.943378827 | 0.524 | 0.552 | 1 | 0.050123996 | 0.586368503 | 0.756 | 0.682 | 1 | 0.259297658 | 0.097735576 |
| Wdr7 | 0.050137433 | 1.861631783 | 0.429 | 0.313 | 1 | 0.552469966 | 0.71623945 | 0.422 | 0.478 | 1 | 0.552469966 | 0.097761104 |
| Safb | 0.128709611 | 0.385010537 | 0.81 | 0.713 | 1 | 0.050141543 | -0.41175096 | 0.889 | 0.848 | 1 | 0.128709611 | 0.097768912 |
| Urb2 | 0.928510588 | 1.171271427 | 0.095 | 0.112 | 1 | 0.050226271 | -0.425451871 | 0.067 | 0.194 | 1 | 0.928510588 | 0.097929863 |
| Zyx | 0.050247516 | -0.900219615 | 0.238 | 0.493 | 1 | 0.512024978 | -0.253362285 | 0.422 | 0.436 | 1 | 0.512024978 | 0.097970219 |
| Sfmbt1 | 0.05033024 | 0.818322557 | 0.238 | 0.608 | 1 | 0.236172781 | 0.337173929 | 0.867 | 0.768 | 1 | 0.236172781 | 0.098127347 |
| Slc44a1 | 0.050394064 | 0.964418405 | 0.857 | 0.884 | 1 | 0.898151063 | 0.551132672 | 0.956 | 0.903 | 1 | 0.898151063 | 0.098248566 |
| Plxna3 | 0.285622926 | -0.233118828 | 0 | 0.052 | 1 | 0.05045002 | -1.960783604 | 0 | 0.08 | 1 | 0.285622926 | 0.098354835 |
| Gm57095 | 0.050468368 | 2.351843673 | 0.095 | 0.022 | 1 | 0.232482267 | 2.098110085 | 0.044 | 0.017 | 1 | 0.232482267 | 0.09838968 |
| Gm16552 | 0.050468368 | 2.214340149 | 0.095 | 0.022 | 1 | 0.659638259 | 0.490427508 | 0.067 | 0.09 | 1 | 0.659638259 | 0.09838968 |
| Trpt1 | 0.106777151 | -1.611630451 | 0 | 0.112 | 1 | 0.050472291 | -2.017367132 | 0 | 0.08 | 1 | 0.106777151 | 0.098397129 |
| Gm32031 | 0.120415565 | -1.233118828 | 0 | 0.104 | 1 | 0.050472291 | -2.017367132 | 0 | 0.08 | 1 | 0.120415565 | 0.098397129 |
| Cfap46 | 0.532436372 | 1.088809267 | 0 | 0.019 | 1 | 0.050472291 | -2.017367132 | 0 | 0.08 | 1 | 0.532436372 | 0.098397129 |
| Gm15708 | 0.918802135 | 1.673771768 | 0.048 | 0.045 | 1 | 0.050472291 | -2.017367132 | 0 | 0.08 | 1 | 0.918802135 | 0.098397129 |
| Rfesd | 0.106726508 | -1.496153234 | 0 | 0.112 | 1 | 0.050492442 | -2.124282336 | 0 | 0.08 | 1 | 0.106726508 | 0.098435397 |
| Rbks | 0.173676192 | -0.911190733 | 0 | 0.082 | 1 | 0.050492442 | -2.174908409 | 0 | 0.08 | 1 | 0.173676192 | 0.098435397 |
| C230013L11Rik | 0.966230998 | 1.258734268 | 0.048 | 0.052 | 1 | 0.050492442 | -2.124282336 | 0 | 0.08 | 1 | 0.966230998 | 0.098435397 |
| Mier2 | 0.128057475 | -1.233118828 | 0 | 0.101 | 1 | 0.050492538 | -2.071814916 | 0 | 0.08 | 1 | 0.128057475 | 0.09843558 |
| Ralgapb | 0.749364832 | 0.515769934 | 0.333 | 0.489 | 1 | 0.050494278 | 0.814317119 | 0.689 | 0.571 | 1 | 0.749364832 | 0.098438884 |
| Prxl2b | 0.220191446 | -0.695462042 | 0.048 | 0.149 | 1 | 0.050510473 | -2.174908409 | 0 | 0.08 | 1 | 0.220191446 | 0.098469639 |
| Uqcc6 | 0.294625684 | -0.785659851 | 0.048 | 0.123 | 1 | 0.050510473 | -2.174908409 | 0 | 0.08 | 1 | 0.294625684 | 0.098469639 |
| Armc7 | 0.837532072 | 1.24081236 | 0.095 | 0.09 | 1 | 0.050510473 | -2.223818009 | 0 | 0.08 | 1 | 0.837532072 | 0.098469639 |
| Acot2 | 0.1363349 | -1.611630451 | 0 | 0.097 | 1 | 0.050526385 | -2.223818009 | 0 | 0.08 | 1 | 0.1363349 | 0.098499854 |
| Zkscan14 | 0.196422748 | -0.970084422 | 0 | 0.075 | 1 | 0.050526385 | -2.223818009 | 0 | 0.08 | 1 | 0.196422748 | 0.098499854 |
| Bbs1 | 0.324739547 | -0.133583154 | 0 | 0.045 | 1 | 0.050526963 | -2.271123724 | 0 | 0.08 | 1 | 0.324739547 | 0.098500952 |
| Gm45767 | 0.761856935 | 0.673771768 | 0.048 | 0.067 | 1 | 0.050526963 | -2.316927414 | 0 | 0.08 | 1 | 0.761856935 | 0.098500952 |
| Fam210b | 0.078592283 | -1.718545655 | 0 | 0.131 | 1 | 0.050527252 | -2.564854927 | 0 | 0.08 | 1 | 0.078592283 | 0.098501502 |
| Abcb7 | 0.95509539 | 0.441111011 | 0.19 | 0.201 | 1 | 0.050535525 | -0.124282336 | 0.133 | 0.287 | 1 | 0.95509539 | 0.098517211 |
| C030006K11Rik | 0.153760172 | -1.081115734 | 0 | 0.09 | 1 | 0.050541139 | -2.316927414 | 0 | 0.08 | 1 | 0.153760172 | 0.098527872 |
| 9230114K14Rik | 0.717182628 | 0.945851313 | 0.143 | 0.127 | 1 | 0.050561392 | -2.564854927 | 0 | 0.08 | 1 | 0.717182628 | 0.09856633 |
| Mtus2 | 0.760890285 | -0.326228232 | 0.048 | 0.063 | 1 | 0.050561392 | -2.404390255 | 0 | 0.08 | 1 | 0.760890285 | 0.09856633 |
| Sult5a1 | 0.71557581 | 0.620660431 | 0.19 | 0.172 | 1 | 0.050563418 | -2.446210431 | 0 | 0.08 | 1 | 0.71557581 | 0.098570176 |
| Il17rd | 0.222299448 | -1.326228232 | 0 | 0.067 | 1 | 0.050583672 | -2.776359032 | 0 | 0.08 | 1 | 0.222299448 | 0.098608636 |
| Reps2 | 0.236559138 | -1.785659851 | 0 | 0.063 | 1 | 0.0505938 | -2.743192169 | 0 | 0.08 | 1 | 0.236559138 | 0.098627867 |
| Rab39 | 0.113338406 | -1.370622352 | 0 | 0.108 | 1 | 0.050620809 | -1.901889915 | 0 | 0.08 | 1 | 0.113338406 | 0.098679152 |
| Thap1 | 0.050629901 | -1.880817084 | 0 | 0.157 | 1 | 0.257352214 | -0.237492946 | 0.089 | 0.159 | 1 | 0.257352214 | 0.098696415 |
| Smarca4 | 0.388339509 | 0.60338244 | 0.667 | 0.69 | 1 | 0.050655763 | 0.591814841 | 0.889 | 0.789 | 1 | 0.388339509 | 0.098745519 |
| Cnrip1 | 0.113789319 | -2.054148687 | 0 | 0.108 | 1 | 0.050677596 | -1.931637258 | 0.022 | 0.118 | 1 | 0.113789319 | 0.098786973 |
| Pygb | 0.09530482 | -0.911190733 | 0.048 | 0.205 | 1 | 0.050719813 | 1.084813263 | 0.444 | 0.287 | 1 | 0.09530482 | 0.098867127 |
| Gm14636 | 0.532435374 | 0.503846766 | 0 | 0.019 | 1 | 0.050760967 | 1.442064487 | 0.133 | 0.055 | 1 | 0.532435374 | 0.098945257 |
| Etv1 | 0.521979329 | -0.208871282 | 0.095 | 0.149 | 1 | 0.050864137 | -0.765387915 | 0.111 | 0.256 | 1 | 0.521979329 | 0.099141114 |
| Tlr3 | 0.05097769 | -2.280424543 | 0 | 0.157 | 1 | 0.187692533 | 1.83507568 | 0.044 | 0.111 | 1 | 0.187692533 | 0.099356655 |
| Dhrs11 | 0.05101093 | -2.280424543 | 0 | 0.157 | 1 | 0.904823699 | 1.03537433 | 0.111 | 0.125 | 1 | 0.904823699 | 0.099419745 |
| Fdps | 0.081328525 | -1.25696557 | 0.381 | 0.571 | 1 | 0.051049432 | -1.249813218 | 0.489 | 0.547 | 1 | 0.081328525 | 0.099492819 |
| Pdzd8 | 0.051052616 | -0.516052791 | 0.19 | 0.463 | 1 | 0.132296476 | 0.474303449 | 0.889 | 0.779 | 1 | 0.132296476 | 0.099498861 |
| Hr | 0.981869361 | 0.339871031 | 0.238 | 0.269 | 1 | 0.05106875 | -0.544733331 | 0.178 | 0.339 | 1 | 0.981869361 | 0.099529483 |
| Gm15270 | 0.339126856 | 0.98727124 | 0.333 | 0.302 | 1 | 0.051094617 | -1.017367132 | 0.044 | 0.156 | 1 | 0.339126856 | 0.099578574 |
| Atp6v1c2 | 0.051148394 | -2.496153234 | 0 | 0.157 | 1 | 0.685250124 | 0.776181991 | 0.022 | 0.035 | 1 | 0.685250124 | 0.09968063 |
| Zdhhc23 | 0.120716826 | -1.611630451 | 0 | 0.104 | 1 | 0.051175016 | -1.871516266 | 0.022 | 0.118 | 1 | 0.120716826 | 0.09973115 |
| Dipk1b | 0.518183263 | 0.24081236 | 0.095 | 0.16 | 1 | 0.051176011 | -2.071814916 | 0.022 | 0.118 | 1 | 0.518183263 | 0.099733039 |
| Arhgap6 | 0.05117674 | -2.727107669 | 0.048 | 0.224 | 1 | 0.50217242 | 0.740153681 | 0.311 | 0.439 | 1 | 0.50217242 | 0.09973442 |
| Rnd1 | 0.051185504 | -2.718545655 | 0 | 0.157 | 1 | 0.165339686 | -1.486852415 | 0 | 0.042 | 1 | 0.165339686 | 0.099751052 |
| Etnk2 | 0.051185812 | -2.752492987 | 0 | 0.157 | 1 | 0.132686876 | -1.486852415 | 0 | 0.048 | 1 | 0.132686876 | 0.099751636 |
| Setd1b | 0.051293271 | -0.582567986 | 0.143 | 0.407 | 1 | 0.138124946 | -0.442771347 | 0.6 | 0.654 | 1 | 0.138124946 | 0.099955542 |
| Bglap | 0.051323991 | -4.63092179 | 0.095 | 0.287 | 1 | 0.106886235 | -5.22981675 | 0 | 0.055 | 1 | 0.106886235 | 0.10001383 |
| Rrm1 | 0.117990253 | -1.911190733 | 0.048 | 0.179 | 1 | 0.051349236 | 0.154693614 | 0.067 | 0.194 | 1 | 0.117990253 | 0.100061728 |
| Rbm25 | 0.093704431 | 0.474402988 | 1 | 0.996 | 1 | 0.051379407 | 0.387561618 | 1 | 0.997 | 1 | 0.093704431 | 0.10011897 |
| Kctd6 | 0.314390552 | -0.20069735 | 0.048 | 0.127 | 1 | 0.051431852 | -0.381057751 | 0.089 | 0.228 | 1 | 0.314390552 | 0.100218469 |
| Litaf | 0.526191031 | 0.206063046 | 0.81 | 0.836 | 1 | 0.0514765 | 0.52277017 | 0.911 | 0.689 | 1 | 0.526191031 | 0.100303169 |
| Plekho2 | 0.44933556 | 0.712942365 | 0.333 | 0.295 | 1 | 0.051493191 | -0.340774156 | 0.156 | 0.318 | 1 | 0.44933556 | 0.100334833 |
| Stom | 0.158472799 | -0.763633545 | 0.048 | 0.172 | 1 | 0.051498367 | -1.871516266 | 0.022 | 0.118 | 1 | 0.158472799 | 0.100344653 |
| Arl13b | 0.17279563 | -0.553055625 | 0.476 | 0.716 | 1 | 0.051505571 | -0.283374854 | 0.711 | 0.789 | 1 | 0.17279563 | 0.100358319 |
| R74862 | 0.078679184 | -1.911190733 | 0 | 0.131 | 1 | 0.051523295 | -1.316927414 | 0.022 | 0.121 | 1 | 0.078679184 | 0.100391941 |
| Plxnb1 | 0.110928025 | -1.326228232 | 0.048 | 0.183 | 1 | 0.051616162 | -0.543996322 | 0.133 | 0.284 | 1 | 0.110928025 | 0.100568096 |
| Zfp105 | 0.208844381 | -0.785659851 | 0 | 0.071 | 1 | 0.051657192 | -1.638855509 | 0.022 | 0.118 | 1 | 0.208844381 | 0.100645919 |
| Gm35769 | 0.480458678 | -0.184209227 | 0.048 | 0.093 | 1 | 0.051657192 | -1.638855509 | 0.022 | 0.118 | 1 | 0.480458678 | 0.100645919 |
| Zfp868 | 0.745405831 | 0.59848364 | 0.143 | 0.187 | 1 | 0.051680532 | -0.296749532 | 0.067 | 0.194 | 1 | 0.745405831 | 0.100690187 |
| Zfp672 | 0.793208359 | 1.088809267 | 0.238 | 0.235 | 1 | 0.05169486 | -0.348405645 | 0.156 | 0.304 | 1 | 0.793208359 | 0.100717361 |
| Fam114a1 | 0.556133287 | 0.198238475 | 0.762 | 0.903 | 1 | 0.051788026 | -0.257292714 | 0.6 | 0.889 | 1 | 0.556133287 | 0.100894053 |
| 4833418N02Rik | 0.051891387 | 1.71684049 | 0.333 | 0.179 | 1 | 0.103160702 | -0.526380779 | 0.089 | 0.201 | 1 | 0.103160702 | 0.101090058 |
| Mbnl1 | 0.051912354 | 2.06896673 | 1 | 1 | 1 | 0.397624588 | 0.850119392 | 1 | 0.997 | 1 | 0.397624588 | 0.101129816 |
| Hs6st3 | 0.051984955 | -1.669636055 | 0.095 | 0.31 | 1 | 0.184864051 | -4.75987091 | 0 | 0.038 | 1 | 0.184864051 | 0.101267475 |
| P2rx4 | 0.229530297 | -0.701267664 | 0.238 | 0.369 | 1 | 0.052036435 | 0.463904066 | 0.178 | 0.356 | 1 | 0.229530297 | 0.101365079 |
| Plp2 | 0.054324436 | -0.772484462 | 0.286 | 0.541 | 1 | 0.052162398 | -0.845071783 | 0.689 | 0.647 | 1 | 0.054324436 | 0.10160388 |
| Bmp8a | 0.645782775 | -0.365756597 | 0.143 | 0.194 | 1 | 0.052391195 | -1.207698344 | 0.022 | 0.121 | 1 | 0.645782775 | 0.102037552 |
| Rnf139 | 0.079489653 | 0.281454345 | 0.095 | 0.284 | 1 | 0.052416595 | -0.178221143 | 0.267 | 0.474 | 1 | 0.079489653 | 0.102085691 |
| Osbpl11 | 0.170848294 | -0.608782089 | 0.286 | 0.478 | 1 | 0.052568143 | 0.86507042 | 0.778 | 0.668 | 1 | 0.170848294 | 0.102372877 |
| Fitm2 | 0.25648148 | -0.20069735 | 0.048 | 0.142 | 1 | 0.052571943 | -1.124282336 | 0.022 | 0.121 | 1 | 0.25648148 | 0.102380077 |
| Kansl1l | 0.18497571 | 1.371959034 | 0.619 | 0.582 | 1 | 0.052603831 | 0.845522467 | 0.889 | 0.862 | 1 | 0.18497571 | 0.102440498 |
| Gin1 | 0.793480127 | 0.451379346 | 0.143 | 0.127 | 1 | 0.052610186 | -0.749886821 | 0.044 | 0.156 | 1 | 0.793480127 | 0.10245254 |
| Thap3 | 0.218822624 | -0.543458949 | 0.095 | 0.224 | 1 | 0.052689037 | -0.776359032 | 0.089 | 0.218 | 1 | 0.218822624 | 0.102601939 |
| Fbxl9 | 0.930536335 | 0.544488751 | 0.095 | 0.104 | 1 | 0.052731672 | -1.089516918 | 0.022 | 0.121 | 1 | 0.930536335 | 0.102682714 |

| | | | | | | | | | | | | |
|---|---|---|---|---|---|---|---|---|---|---|---|---|
| Stard13 | 0.052909025 | 0.661307802 | 0.857 | 0.907 | 1 | 0.81856375 | 0.320318969 | 0.822 | 0.91 | 1 | 0.81856375 | 0.103018686 |
| Bmp5 | 0.844205895 | 1.346587947 | 0.381 | 0.466 | 1 | 0.052911553 | -0.234465254 | 0.556 | 0.727 | 1 | 0.844205895 | 0.103023473 |
| Mapkapk2 | 0.985604029 | 0.255270413 | 0.81 | 0.847 | 1 | 0.0529506 | 0.863514865 | 0.911 | 0.879 | 1 | 0.985604029 | 0.103097435 |
| Plekhb2 | 0.608959364 | 0.400753273 | 0.048 | 0.082 | 1 | 0.053041897 | -1.017367132 | 0.022 | 0.121 | 1 | 0.608959364 | 0.103270352 |
| Akap10 | 0.345846587 | 0.221259563 | 0.238 | 0.384 | 1 | 0.053050416 | 0.645812968 | 0.667 | 0.512 | 1 | 0.345846587 | 0.103286486 |
| Bltp2 | 0.053057282 | -0.858723313 | 0.429 | 0.619 | 1 | 0.400017226 | -0.250376221 | 0.8 | 0.72 | 1 | 0.400017226 | 0.103299488 |
| Tti1 | 0.293567806 | -0.818081329 | 0.048 | 0.123 | 1 | 0.053165059 | -0.695439037 | 0.022 | 0.121 | 1 | 0.293567806 | 0.103503594 |
| Pibf1 | 0.157404979 | 1.223738847 | 0.571 | 0.537 | 1 | 0.05325135 | 0.780369788 | 0.689 | 0.529 | 1 | 0.157404979 | 0.103666993 |
| Nudt18 | 0.139956182 | -0.970084422 | 0.048 | 0.179 | 1 | 0.053269193 | -0.709244837 | 0.022 | 0.121 | 1 | 0.139956182 | 0.10370078 |
| Prex2 | 0.053313983 | -1.839297815 | 0.048 | 0.235 | 1 | 0.278756995 | 0.545569062 | 0.2 | 0.325 | 1 | 0.278756995 | 0.103785584 |
| Taf5 | 0.186946898 | -0.683780237 | 0.095 | 0.224 | 1 | 0.053321291 | -0.486852415 | 0.089 | 0.225 | 1 | 0.186946898 | 0.103799422 |
| Srr | 0.05336799 | -0.584962501 | 0.095 | 0.306 | 1 | 0.778453909 | 0.603345394 | 0.289 | 0.304 | 1 | 0.778453909 | 0.103887837 |
| Lrrfip2 | 0.053454037 | 0.902317797 | 0.857 | 0.832 | 1 | 0.406236582 | 0.412351987 | 0.933 | 0.851 | 1 | 0.406236582 | 0.10405074 |
| Gm28875 | 0.208886118 | -0.849790188 | 0 | 0.071 | 1 | 0.053490613 | -0.486852415 | 0.022 | 0.121 | 1 | 0.208886118 | 0.104119979 |
| Sod2 | 0.066566237 | -0.516690165 | 0.667 | 0.817 | 1 | 0.053617961 | -0.426226056 | 0.733 | 0.72 | 1 | 0.066566237 | 0.104361037 |
| Eif2s3y | 0.563526322 | 1.201283996 | 0.381 | 0.369 | 1 | 0.053656642 | 0.605435175 | 0.8 | 0.657 | 1 | 0.563526322 | 0.104434249 |
| Pla2g15 | 0.446945662 | -0.413691074 | 0.048 | 0.097 | 1 | 0.0536806 | -0.674479418 | 0.022 | 0.121 | 1 | 0.446945662 | 0.104479593 |
| Krba1 | 0.43369756 | 1.137718867 | 0.143 | 0.093 | 1 | 0.053735677 | -0.454430938 | 0.022 | 0.121 | 1 | 0.43369756 | 0.104583832 |
| Sarnp | 0.053757324 | -0.294238118 | 0.952 | 0.922 | 1 | 0.074957449 | -0.151249383 | 0.911 | 0.948 | 1 | 0.074957449 | 0.104624799 |
| Gli3 | 0.053777116 | 1.275968806 | 0.667 | 0.668 | 1 | 0.162043113 | 1.081954317 | 0.578 | 0.595 | 1 | 0.162043113 | 0.104662254 |
| Id1 | 0.276065734 | -1.219698312 | 0.476 | 0.522 | 1 | 0.053835315 | 0.421265484 | 0.778 | 0.519 | 1 | 0.276065734 | 0.104772388 |
| Zcchc3 | 0.989591319 | 0.673771768 | 0.143 | 0.153 | 1 | 0.053910177 | -0.679497493 | 0.067 | 0.19 | 1 | 0.989591319 | 0.104914047 |
| Tent5c | 0.331261794 | -0.752492987 | 0.048 | 0.116 | 1 | 0.053911105 | -0.84048937 | 0.022 | 0.121 | 1 | 0.331261794 | 0.104915803 |
| 4930581F22Rik | 0.053922421 | -1.280424543 | 0.048 | 0.239 | 1 | 0.082699094 | -1.316927414 | 0.044 | 0.135 | 1 | 0.082699094 | 0.104937214 |
| Cul9 | 0.790350282 | 0.586308926 | 0.143 | 0.187 | 1 | 0.053932973 | -0.149817428 | 0.178 | 0.346 | 1 | 0.790350282 | 0.104957181 |
| Ubxn2b | 0.05399559 | -1.940938076 | 0 | 0.153 | 1 | 0.173777479 | 0.735540006 | 0.067 | 0.152 | 1 | 0.173777479 | 0.105075656 |
| St6galnac2 | 0.054064625 | 2.48112669 | 0.143 | 0.049 | 1 | 0.11902586 | -1.564854927 | 0 | 0.052 | 1 | 0.11902586 | 0.105206267 |
| Ncdn | 0.153703558 | -1.081115734 | 0 | 0.09 | 1 | 0.054080187 | -0.614607962 | 0.044 | 0.159 | 1 | 0.153703558 | 0.105235708 |
| Stradb | 0.972567341 | 0.866416846 | 0.095 | 0.097 | 1 | 0.054080187 | -0.589945908 | 0.044 | 0.159 | 1 | 0.972567341 | 0.105235708 |
| Wdr46 | 0.054113643 | 1.503846766 | 0.238 | 0.101 | 1 | 0.274649408 | -0.454430938 | 0.067 | 0.125 | 1 | 0.274649408 | 0.105298999 |
| Slc43a1 | 0.238508315 | -0.911190733 | 0.048 | 0.142 | 1 | 0.054171092 | -0.277756817 | 0.089 | 0.218 | 1 | 0.238508315 | 0.105407676 |
| Tbcd | 0.054184347 | -1.911190733 | 0.048 | 0.224 | 1 | 0.734135021 | 0.975253338 | 0.244 | 0.232 | 1 | 0.734135021 | 0.105432751 |
| 9430038I01Rik | 0.054222165 | -2.184209227 | 0 | 0.153 | 1 | 0.125098331 | -0.486852415 | 0.044 | 0.125 | 1 | 0.125098331 | 0.105504286 |
| Rab4b | 0.054225549 | -0.605451876 | 0.286 | 0.571 | 1 | 0.064633742 | -0.253733587 | 0.356 | 0.554 | 1 | 0.064633742 | 0.105510689 |
| 1110019D14Rik | 0.054280586 | -2.107587946 | 0 | 0.153 | 1 | 0.208976612 | -0.191396532 | 0.067 | 0.142 | 1 | 0.208976612 | 0.105614791 |
| Abcb1b | 0.054321798 | -2.326228232 | 0 | 0.153 | 1 | 0.146980783 | -1.446210431 | 0.044 | 0.118 | 1 | 0.146980783 | 0.105692737 |
| Hcfc2 | 0.783794318 | 0.233199176 | 0.381 | 0.381 | 1 | 0.054338424 | -0.110476536 | 0.311 | 0.554 | 1 | 0.783794318 | 0.105724184 |
| Kctd20 | 0.528930618 | -0.161417363 | 0.286 | 0.388 | 1 | 0.05438094 | -0.369568894 | 0.378 | 0.585 | 1 | 0.528930618 | 0.105804593 |
| Potegl | 0.862735204 | 1.214340149 | 0.048 | 0.06 | 1 | 0.054394997 | 2.426732833 | 0.156 | 0.073 | 1 | 0.862735204 | 0.105831179 |
| Itfg2 | 0.069521699 | -0.516052791 | 0.048 | 0.228 | 1 | 0.054404025 | 0.808603468 | 0.111 | 0.273 | 1 | 0.069521699 | 0.105848252 |
| Hsd17b7 | 0.054494309 | -1.120224772 | 0.429 | 0.664 | 1 | 0.07564427 | -0.442163598 | 0.333 | 0.509 | 1 | 0.07564427 | 0.106018987 |
| Cgref1 | 0.108738264 | -0.874664857 | 0.19 | 0.369 | 1 | 0.054529616 | -0.472845557 | 0.156 | 0.311 | 1 | 0.108738264 | 0.106085752 |
| Zfp148 | 0.519374017 | 0.173925881 | 0.857 | 0.866 | 1 | 0.054580171 | 0.43042448 | 0.956 | 0.955 | 1 | 0.519374017 | 0.106181347 |
| Itih5 | 0.054592363 | -4.486099569 | 0 | 0.153 | 1 | 0.184854431 | -2.743192169 | 0 | 0.038 | 1 | 0.184854431 | 0.106204401 |
| Klhdc8b | 0.164435561 | -1.02666795 | 0.048 | 0.16 | 1 | 0.054594818 | -0.155646507 | 0.089 | 0.228 | 1 | 0.164435561 | 0.106209043 |
| Ppm1g | 0.05463981 | -0.53267911 | 0.238 | 0.526 | 1 | 0.250085702 | -0.151503805 | 0.689 | 0.699 | 1 | 0.250085702 | 0.10629411 |
| Itgb7 | 0.054642531 | 1.541321472 | 0.286 | 0.142 | 1 | 0.385421976 | 0.496194451 | 0.244 | 0.356 | 1 | 0.385421976 | 0.106299255 |
| Ttc7b | 0.054753325 | 1.179007076 | 0.524 | 0.41 | 1 | 0.262271473 | 1.10471282 | 0.378 | 0.384 | 1 | 0.262271473 | 0.106508724 |
| D630003M21Rik | 0.513345575 | -0.194124696 | 0.095 | 0.157 | 1 | 0.054842133 | -1.033134448 | 0.089 | 0.215 | 1 | 0.513345575 | 0.106676607 |
| Ccdc91 | 0.756109111 | 0.572233741 | 0.19 | 0.231 | 1 | 0.054849978 | -0.459885368 | 0.089 | 0.218 | 1 | 0.756109111 | 0.106691435 |
| Fhl2 | 0.70391225 | 1.277112798 | 0.381 | 0.444 | 1 | 0.054881685 | 1.817522786 | 0.378 | 0.298 | 1 | 0.70391225 | 0.106751371 |
| Pacs1 | 0.054894961 | -0.771013075 | 0.095 | 0.313 | 1 | 0.326906128 | 0.935459748 | 0.556 | 0.478 | 1 | 0.326906128 | 0.106776465 |
| Wdr73 | 0.341753532 | -0.326228232 | 0.048 | 0.116 | 1 | 0.054923681 | -1.124282336 | 0.067 | 0.183 | 1 | 0.341753532 | 0.106830752 |
| Fbxo36 | 0.054927714 | -1.752492987 | 0.048 | 0.224 | 1 | 0.228137989 | -0.316927414 | 0.089 | 0.166 | 1 | 0.228137989 | 0.106838374 |
| Ncaph2 | 0.454435882 | 1.003920369 | 0.381 | 0.369 | 1 | 0.05494923 | -0.232535227 | 0.267 | 0.436 | 1 | 0.454435882 | 0.106879042 |
| Niban1 | 0.151937986 | -0.822654058 | 0.143 | 0.313 | 1 | 0.054961325 | 0.64125241 | 0.689 | 0.471 | 1 | 0.151937986 | 0.106901902 |
| Wbp1 | 0.055060506 | -1.133583154 | 0.048 | 0.239 | 1 | 0.183568948 | -0.234465254 | 0.089 | 0.176 | 1 | 0.183568948 | 0.107089353 |
| Mthfr | 0.108136371 | -0.704739856 | 0.095 | 0.261 | 1 | 0.055176445 | -0.350350415 | 0.156 | 0.315 | 1 | 0.108136371 | 0.107308451 |
| Gm44686 | 0.396971401 | 0.351843673 | 0 | 0.034 | 1 | 0.055190624 | 1.731982187 | 0.156 | 0.076 | 1 | 0.396971401 | 0.107335242 |
| Kdm5b | 0.082954366 | 0.425172452 | 0.905 | 0.877 | 1 | 0.0552678 | -0.337586884 | 1 | 0.993 | 1 | 0.082954366 | 0.10748107 |
| Ddx17 | 0.080998324 | -0.359651234 | 0.857 | 0.97 | 1 | 0.055288022 | -0.249813218 | 1 | 0.986 | 1 | 0.080998324 | 0.107519278 |
| Usp4 | 0.055458466 | 1.788414334 | 0.571 | 0.571 | 1 | 0.07706929 | 0.466939155 | 0.8 | 0.727 | 1 | 0.07706929 | 0.107841291 |
| Prkar2a | 0.055515754 | 0.12259758 | 0.19 | 0.463 | 1 | 0.323506627 | 0.330690695 | 0.578 | 0.654 | 1 | 0.323506627 | 0.107949508 |
| Stk40 | 0.055623406 | 0.739914124 | 0.714 | 0.687 | 1 | 0.839438611 | 0.245863706 | 0.867 | 0.796 | 1 | 0.839438611 | 0.108152848 |
| Sag | 0.237321829 | 1.119182916 | 0.19 | 0.108 | 1 | 0.055699751 | -1.08246216 | 0.044 | 0.152 | 1 | 0.237321829 | 0.108297039 |
| Nav1 | 0.468641747 | 0.163156608 | 0.619 | 0.664 | 1 | 0.05583185 | 1.244145605 | 0.578 | 0.588 | 1 | 0.468641747 | 0.108546505 |
| Efr3a | 0.794369353 | 0.395385099 | 0.619 | 0.649 | 1 | 0.055862648 | -0.295967795 | 0.756 | 0.789 | 1 | 0.794369353 | 0.10860466 |
| Il20rb | 0.117121182 | -1.496153234 | 0.048 | 0.179 | 1 | 0.056004844 | -0.515707278 | 0.111 | 0.253 | 1 | 0.117121182 | 0.108873145 |
| Abhd6 | 0.311307311 | -0.940938076 | 0.048 | 0.119 | 1 | 0.056113256 | 0.602152591 | 0.044 | 0.159 | 1 | 0.311307311 | 0.109077815 |
| Gm38910 | 0.136163632 | -1.280424543 | 0 | 0.097 | 1 | 0.056134646 | -1.901889915 | 0 | 0.076 | 1 | 0.136163632 | 0.109118193 |
| Lysmd1 | 0.268100354 | -0.326228232 | 0 | 0.056 | 1 | 0.056157169 | -1.960783604 | 0 | 0.076 | 1 | 0.268100354 | 0.10916071 |
| Cyp2d22 | 0.267917141 | -0.496153234 | 0 | 0.056 | 1 | 0.056177441 | -2.124282336 | 0 | 0.076 | 1 | 0.267917141 | 0.109198976 |
| Anks6 | 0.918519233 | 0.425844254 | 0.048 | 0.052 | 1 | 0.056177441 | -2.071814916 | 0 | 0.076 | 1 | 0.918519233 | 0.109198976 |
| Mcm5 | 0.732242151 | 1.780686972 | 0.048 | 0.071 | 1 | 0.056177548 | -2.017367132 | 0 | 0.076 | 1 | 0.732242151 | 0.109199179 |
| Gm56496 | 0.872207635 | 0.586308926 | 0.048 | 0.056 | 1 | 0.056177548 | -2.017367132 | 0 | 0.076 | 1 | 0.872207635 | 0.109199179 |
| Hip1r | 0.0561815 | 0.921911958 | 0.714 | 0.619 | 1 | 0.367961297 | 0.653546634 | 0.733 | 0.699 | 1 | 0.367961297 | 0.109206639 |
| D130020L05Rik | 0.163379068 | -0.970084422 | 0 | 0.086 | 1 | 0.056195461 | -2.124282336 | 0 | 0.076 | 1 | 0.163379068 | 0.109232992 |
| Sh2d4b | 0.531977913 | 1.188344941 | 0.095 | 0.063 | 1 | 0.056195461 | -2.174908409 | 0 | 0.076 | 1 | 0.531977913 | 0.109232992 |
| Creld1 | 0.113449548 | -1.496153234 | 0 | 0.108 | 1 | 0.05619589 | -2.071814916 | 0 | 0.076 | 1 | 0.113449548 | 0.109233802 |
| Eppk1 | 0.490506432 | 1.041503552 | 0.095 | 0.06 | 1 | 0.05619589 | 0.683072586 | 0.133 | 0.055 | 1 | 0.490506432 | 0.109233802 |
| Zfp930 | 0.765801178 | 0.150209812 | 0.048 | 0.063 | 1 | 0.05619589 | -2.071814916 | 0 | 0.076 | 1 | 0.765801178 | 0.109233802 |
| Slc6a9 | 0.773899393 | 1.614878079 | 0.048 | 0.067 | 1 | 0.05619589 | -2.071814916 | 0 | 0.076 | 1 | 0.773899393 | 0.109233802 |
| Dus4l | 0.163307215 | 1.55829455 | 0.19 | 0.101 | 1 | 0.05623601 | -2.316927414 | 0 | 0.076 | 1 | 0.163307215 | 0.109309531 |
| Gprc5a | 0.586595039 | -0.593014773 | 0.095 | 0.138 | 1 | 0.056250385 | -2.638855509 | 0 | 0.076 | 1 | 0.586595039 | 0.109336665 |
| Otoa | 0.213882822 | -1.555046923 | 0.048 | 0.142 | 1 | 0.056266585 | -2.743192169 | 0 | 0.076 | 1 | 0.213882822 | 0.109367242 |
| Vps33a | 0.638175594 | 1.150209812 | 0.143 | 0.19 | 1 | 0.056270214 | -0.11443455 | 0.156 | 0.336 | 1 | 0.638175594 | 0.109374091 |
| Speg | 0.500661815 | 1.281454345 | 0.095 | 0.06 | 1 | 0.056272949 | -0.941418279 | 0.067 | 0.183 | 1 | 0.500661815 | 0.109379254 |
| Sntb1 | 0.184940904 | -2.970084422 | 0 | 0.078 | 1 | 0.056278172 | -3.199570463 | 0 | 0.076 | 1 | 0.184940904 | 0.109389112 |
| Lpl | 0.136535001 | -2.648156327 | 0 | 0.097 | 1 | 0.05628461 | -4.044847868 | 0 | 0.076 | 1 | 0.136535001 | 0.109401262 |
| Dnajb5 | 0.268100354 | -0.326228232 | 0 | 0.056 | 1 | 0.056299845 | -1.84048937 | 0 | 0.076 | 1 | 0.268100354 | 0.109430018 |
| Insyn1 | 0.370840523 | 0.214340149 | 0 | 0.037 | 1 | 0.056299845 | -1.84048937 | 0 | 0.076 | 1 | 0.370840523 | 0.109430018 |
| Primpol | 0.376354176 | -0.496153234 | 0.048 | 0.108 | 1 | 0.056299845 | -1.84048937 | 0 | 0.076 | 1 | 0.376354176 | 0.109430018 |
| Rsl1 | 0.79666227 | 1.351843673 | 0.048 | 0.037 | 1 | 0.056299845 | -1.84048937 | 0 | 0.076 | 1 | 0.79666227 | 0.109430018 |
| Slc2a1 | 0.056306963 | -1.776261153 | 0.619 | 0.724 | 1 | 0.404282425 | -0.433536601 | 0.733 | 0.526 | 1 | 0.404282425 | 0.109443452 |
| Eri3 | 0.056414681 | 0.648632205 | 0.762 | 0.836 | 1 | 0.133687318 | 0.363206958 | 0.8 | 0.813 | 1 | 0.133687318 | 0.109646746 |
| Lamc1 | 0.056456343 | -0.484657595 | 0.19 | 0.414 | 1 | 0.310824805 | 0.139576703 | 0.778 | 0.633 | 1 | 0.310824805 | 0.109725368 |
| Stx12 | 0.204857791 | -0.19276109 | 0.476 | 0.765 | 1 | 0.056538786 | -0.227993685 | 0.667 | 0.761 | 1 | 0.204857791 | 0.109880937 |
| Dcaf6 | 0.094701202 | 1.068050707 | 0.81 | 0.694 | 1 | 0.056554343 | 0.844690768 | 0.911 | 0.827 | 1 | 0.094701202 | 0.109910293 |

| | | | | | | | | | | | | |
|---|---|---|---|---|---|---|---|---|---|---|---|---|
| Capn15 | 0.976187194 | 0.260990242 | 0.381 | 0.466 | 1 | 0.05656452 | -0.254271806 | 0.422 | 0.647 | 1 | 0.976187194 | 0.109929495 |
| Ralgds | 0.124817923 | -1.194124696 | 0.095 | 0.254 | 1 | 0.056571592 | -0.547881849 | 0.156 | 0.301 | 1 | 0.124817923 | 0.109942838 |
| Sec14l2 | 0.094305534 | -1.455511249 | 0 | 0.119 | 1 | 0.056611094 | -0.701591264 | 0.111 | 0.246 | 1 | 0.094305534 | 0.110017372 |
| Tomm70a | 0.056617769 | -0.34612779 | 0.381 | 0.716 | 1 | 0.85233539 | 0.144067376 | 0.778 | 0.754 | 1 | 0.85233539 | 0.110029967 |
| Phlpp1 | 0.249114489 | 0.763323353 | 0.524 | 0.448 | 1 | 0.056692068 | 0.64421665 | 0.844 | 0.664 | 1 | 0.249114489 | 0.110170144 |
| Zfr2 | 0.056694977 | 2.673771768 | 0.143 | 0.049 | 1 | 0.716733944 | 0.825091591 | 0.067 | 0.087 | 1 | 0.716733944 | 0.110175634 |
| Slmap | 0.076045021 | 0.435367028 | 0.857 | 0.769 | 1 | 0.056736316 | 0.700774588 | 0.956 | 0.886 | 1 | 0.076045021 | 0.110253623 |
| Zfp608 | 0.056761712 | 0.783618949 | 0.905 | 0.914 | 1 | 0.890696481 | 0.949262547 | 0.933 | 0.879 | 1 | 0.890696481 | 0.110301531 |
| Kcns1 | 0.057768344 | -2.133583154 | 0 | 0.149 | 1 | 0.056807713 | -0.276285429 | 0.067 | 0.183 | 1 | 0.057768344 | 0.11038831 |
| Gm49539 | 0.324789396 | -0.574155746 | 0 | 0.045 | 1 | 0.056837578 | 2.135584791 | 0.111 | 0.045 | 1 | 0.324789396 | 0.110444647 |
| Micu2 | 0.431235755 | 0.325371461 | 0.381 | 0.575 | 1 | 0.056888445 | 0.918404063 | 0.711 | 0.567 | 1 | 0.431235755 | 0.110540595 |
| Lig1 | 0.189199765 | -1.249060372 | 0.048 | 0.157 | 1 | 0.056906786 | -2.017367132 | 0.022 | 0.114 | 1 | 0.189199765 | 0.110575189 |
| Zfp652 | 0.056968335 | -0.569402216 | 0.286 | 0.563 | 1 | 0.913603109 | 0.138351933 | 0.8 | 0.737 | 1 | 0.913603109 | 0.110691279 |
| Smg9 | 0.656884481 | 0.281454345 | 0.095 | 0.134 | 1 | 0.057013337 | -0.526380779 | 0.067 | 0.194 | 1 | 0.656884481 | 0.110776153 |
| Dhps | 0.506377895 | 0.3012176 | 0.238 | 0.354 | 1 | 0.057079805 | 0.522080709 | 0.2 | 0.405 | 1 | 0.506377895 | 0.110901507 |
| Zfp111 | 0.618972991 | 0.673771768 | 0.048 | 0.082 | 1 | 0.057085922 | -1.709244837 | 0.022 | 0.114 | 1 | 0.618972991 | 0.110913041 |
| Narf | 0.622041944 | 0.38426515 | 0.143 | 0.201 | 1 | 0.057129602 | -0.84048937 | 0.044 | 0.156 | 1 | 0.622041944 | 0.110995413 |
| Zfp975 | 0.606026235 | 0.805875304 | 0.238 | 0.22 | 1 | 0.057201493 | -0.731964913 | 0.089 | 0.221 | 1 | 0.606026235 | 0.111130976 |
| Ovgp1 | 0.447355723 | 0.673771768 | 0.048 | 0.104 | 1 | 0.057271729 | -1.255526869 | 0.022 | 0.118 | 1 | 0.447355723 | 0.111263408 |
| Ercc6 | 0.334065944 | 0.849343332 | 0.286 | 0.205 | 1 | 0.057337909 | -0.101198723 | 0.133 | 0.28 | 1 | 0.334065944 | 0.111388183 |
| Arhgef17 | 0.141247073 | -0.936725825 | 0.19 | 0.354 | 1 | 0.057342247 | 0.34358612 | 0.222 | 0.446 | 1 | 0.141247073 | 0.11139636 |
| Cttn | 0.439643482 | -0.109970597 | 0.476 | 0.653 | 1 | 0.057357989 | -0.141355849 | 0.689 | 0.841 | 1 | 0.439643482 | 0.111426039 |
| Nin | 0.357252346 | 0.555590341 | 0.286 | 0.448 | 1 | 0.057365694 | 0.625877185 | 0.689 | 0.491 | 1 | 0.357252346 | 0.111440565 |
| Gm32282 | 0.057397102 | -1.818081329 | 0 | 0.149 | 1 | 0.132686876 | -1.404390255 | 0 | 0.048 | 1 | 0.132686876 | 0.111499777 |
| Vps33b | 0.057397189 | -1.785659851 | 0 | 0.149 | 1 | 0.972971072 | 1.163698427 | 0.111 | 0.121 | 1 | 0.972971072 | 0.11149994 |
| Zfp81 | 0.419192086 | 0.129451251 | 0.048 | 0.108 | 1 | 0.057433004 | -1.223818009 | 0.022 | 0.118 | 1 | 0.419192086 | 0.111567458 |
| Dcaf4 | 0.348717586 | -0.752492987 | 0.048 | 0.112 | 1 | 0.057445637 | -1.776359032 | 0.022 | 0.114 | 1 | 0.348717586 | 0.111591273 |
| Dnajc22 | 0.100442315 | -1.574155746 | 0 | 0.116 | 1 | 0.057445783 | -1.602329633 | 0.022 | 0.114 | 1 | 0.100442315 | 0.111591549 |
| Tmem126b | 0.057454773 | -1.818081329 | 0 | 0.149 | 1 | 0.559652674 | 0.41458375 | 0.111 | 0.149 | 1 | 0.559652674 | 0.111608495 |
| Pigl | 0.113583987 | -1.611630451 | 0 | 0.108 | 1 | 0.057455426 | -1.223818009 | 0.022 | 0.118 | 1 | 0.113583987 | 0.111609726 |
| Tep1 | 0.987543946 | 0.825774861 | 0.095 | 0.101 | 1 | 0.057516017 | -0.686161223 | 0.044 | 0.152 | 1 | 0.987543946 | 0.111723941 |
| Amn1 | 0.886080489 | 0.93423223 | 0.333 | 0.41 | 1 | 0.057540005 | -0.448633093 | 0.156 | 0.311 | 1 | 0.886080489 | 0.111769159 |
| Atoh8 | 0.057558982 | -1.911190733 | 0 | 0.149 | 1 | 0.090454814 | -1.404390255 | 0.044 | 0.131 | 1 | 0.090454814 | 0.111804927 |
| Gbe1 | 0.26504087 | 1.104406122 | 0.429 | 0.321 | 1 | 0.057579228 | 0.679025753 | 0.644 | 0.495 | 1 | 0.26504087 | 0.111843089 |
| Atpaf2 | 0.057607506 | -1.970084422 | 0 | 0.149 | 1 | 0.085715416 | -0.446210431 | 0.089 | 0.204 | 1 | 0.085715416 | 0.111896388 |
| Gm527 | 0.294238901 | 1.24081236 | 0.143 | 0.078 | 1 | 0.057621971 | -1.674479418 | 0.022 | 0.114 | 1 | 0.294238901 | 0.11192365 |
| Tarbp1 | 0.574604187 | 0.129451251 | 0.048 | 0.086 | 1 | 0.057621971 | -1.776359032 | 0.022 | 0.114 | 1 | 0.574604187 | 0.11192365 |
| Rbm43 | 0.093355136 | -1.348596045 | 0.048 | 0.194 | 1 | 0.057681483 | -0.531940305 | 0.067 | 0.183 | 1 | 0.093355136 | 0.112035812 |
| Luc7l3 | 0.132747151 | 0.661280823 | 1 | 0.884 | 1 | 0.057731492 | -0.216179633 | 0.978 | 0.972 | 1 | 0.132747151 | 0.112130059 |
| Ppm1f | 0.723619404 | 0.214340149 | 0.048 | 0.067 | 1 | 0.057808212 | -0.776359032 | 0.044 | 0.152 | 1 | 0.723619404 | 0.112274635 |
| Calhm2 | 0.057912882 | -2.370622352 | 0 | 0.149 | 1 | 0.12372564 | -0.161276543 | 0.089 | 0.194 | 1 | 0.12372564 | 0.112471863 |
| Slc25a48 | 0.05795305 | -2.455511249 | 0 | 0.149 | 1 | 0.165351478 | -1.404390255 | 0 | 0.042 | 1 | 0.165351478 | 0.112547545 |
| Fam13c | 0.057968374 | -2.555046923 | 0 | 0.149 | 1 | 0.753978972 | 0.311103809 | 0.156 | 0.176 | 1 | 0.753978972 | 0.112576416 |
| Tmem69 | 0.880422239 | 1.029915578 | 0.095 | 0.09 | 1 | 0.057969345 | -1.486852415 | 0.022 | 0.114 | 1 | 0.880422239 | 0.112578245 |
| Itga1 | 0.18392838 | 0.125335143 | 0.095 | 0.239 | 1 | 0.057990162 | -0.167031983 | 0.111 | 0.26 | 1 | 0.18392838 | 0.112617466 |
| Ugcg | 0.224865589 | -0.249060372 | 0.238 | 0.418 | 1 | 0.057990217 | -0.332752381 | 0.356 | 0.533 | 1 | 0.224865589 | 0.112617569 |
| Mafb | 0.058029508 | -1.907177671 | 0.143 | 0.347 | 1 | 0.763123741 | -0.781120407 | 0.356 | 0.398 | 1 | 0.763123741 | 0.112691592 |
| Zfp180 | 0.078679184 | -1.940938076 | 0 | 0.131 | 1 | 0.058073858 | -0.761712257 | 0.089 | 0.215 | 1 | 0.078679184 | 0.112775143 |
| Slurp1 | 0.058098432 | -5.162278587 | 0 | 0.149 | 1 | 0.148091611 | -2.526380779 | 0 | 0.045 | 1 | 0.148091611 | 0.112821436 |
| Fads3 | 0.211611036 | -0.455511249 | 0.333 | 0.53 | 1 | 0.058103856 | -0.241200088 | 0.289 | 0.519 | 1 | 0.211611036 | 0.112831654 |
| Dnajc24 | 0.518532604 | 0.367963338 | 0.143 | 0.228 | 1 | 0.058154415 | -0.83350294 | 0.089 | 0.218 | 1 | 0.518532604 | 0.112926895 |
| Cradd | 0.185079207 | -0.434752689 | 0.095 | 0.235 | 1 | 0.058167549 | 0.868375689 | 0.422 | 0.277 | 1 | 0.185079207 | 0.112951634 |
| Zfp783 | 0.798038625 | 0.214340149 | 0.143 | 0.168 | 1 | 0.058189052 | -0.695439037 | 0.044 | 0.152 | 1 | 0.798038625 | 0.112992138 |
| Tll1 | 0.058241206 | -0.718545655 | 0.048 | 0.239 | 1 | 0.160806945 | -0.314385219 | 0.378 | 0.55 | 1 | 0.160806945 | 0.113090374 |
| Rab39b | 0.286684167 | -0.361852142 | 0.095 | 0.201 | 1 | 0.058252039 | -0.303988358 | 0.111 | 0.253 | 1 | 0.286684167 | 0.113110778 |
| Cdr2l | 0.058392441 | -1.24509147 | 0.095 | 0.295 | 1 | 0.807412653 | 0.397670367 | 0.089 | 0.104 | 1 | 0.807412653 | 0.113375205 |
| Prep | 0.865777612 | 0.272892331 | 0.333 | 0.403 | 1 | 0.058396874 | -0.404390255 | 0.289 | 0.464 | 1 | 0.865777612 | 0.113383553 |
| 6330403L08Rik | 0.208980008 | -1.133583154 | 0 | 0.071 | 1 | 0.058410925 | -1.053893008 | 0.022 | 0.118 | 1 | 0.208980008 | 0.113410013 |
| Smox | 0.45309658 | 0.731874722 | 0.333 | 0.317 | 1 | 0.058414948 | -0.716858021 | 0.111 | 0.239 | 1 | 0.45309658 | 0.11341759 |
| Dcaf5 | 0.078032115 | -1.216999163 | 0.143 | 0.328 | 1 | 0.058506033 | 0.416997726 | 0.8 | 0.612 | 1 | 0.078032115 | 0.11358911 |
| Sars | 0.464413781 | 1.181290183 | 0.476 | 0.563 | 1 | 0.058528204 | -0.184091731 | 0.556 | 0.657 | 1 | 0.464413781 | 0.113630858 |
| Rab3il1 | 0.09813432 | -1.955584852 | 0.048 | 0.187 | 1 | 0.058616686 | -0.722919774 | 0.089 | 0.215 | 1 | 0.09813432 | 0.113797456 |
| Ddb2 | 0.910124359 | 0.458758877 | 0.19 | 0.183 | 1 | 0.058739028 | -0.124282336 | 0.067 | 0.187 | 1 | 0.910124359 | 0.114027783 |
| Adnp | 0.518561206 | 0.173201454 | 0.714 | 0.776 | 1 | 0.05874518 | 0.448797173 | 0.956 | 0.924 | 1 | 0.518561206 | 0.114039365 |
| Ap1g1 | 0.058835455 | 0.788273466 | 0.81 | 0.862 | 1 | 0.319928848 | 0.332842233 | 0.978 | 0.927 | 1 | 0.319928848 | 0.114209299 |
| Ehmt2 | 0.713807346 | 0.59848364 | 0.286 | 0.287 | 1 | 0.058860542 | -0.132502843 | 0.222 | 0.394 | 1 | 0.713807346 | 0.114256521 |
| Ccdc15 | 0.113568819 | 1.491568436 | 0.238 | 0.123 | 1 | 0.059101142 | -0.361321533 | 0.044 | 0.156 | 1 | 0.113568819 | 0.114709338 |
| 8030456M14Rik | 0.362165014 | 2.451379346 | 0.048 | 0.019 | 1 | 0.059122854 | 1.905465008 | 0.089 | 0.031 | 1 | 0.362165014 | 0.114750197 |
| Eif3j2 | 0.764355315 | 0.544488751 | 0.095 | 0.123 | 1 | 0.059146561 | -0.564854927 | 0.022 | 0.118 | 1 | 0.764355315 | 0.114794806 |
| Lpcat4 | 0.517211624 | 1.271673324 | 0.143 | 0.108 | 1 | 0.059180958 | -1.16492432 | 0.089 | 0.204 | 1 | 0.517211624 | 0.114859531 |
| Pcdh19 | 0.346886421 | -1.455511249 | 0 | 0.041 | 1 | 0.059310045 | -0.954357334 | 0.089 | 0.208 | 1 | 0.346886421 | 0.115102409 |
| Gm16279 | 0.059381233 | 2.673771768 | 0.143 | 0.049 | 1 | 0.086124205 | -1.709244837 | 0 | 0.062 | 1 | 0.086124205 | 0.115236334 |
| Zfp518b | 0.196422748 | -0.970084422 | 0 | 0.075 | 1 | 0.059392883 | -0.242926832 | 0.022 | 0.118 | 1 | 0.196422748 | 0.115258252 |
| Cox15 | 0.91729082 | 0.586308926 | 0.095 | 0.09 | 1 | 0.059426251 | -0.602329633 | 0.022 | 0.118 | 1 | 0.91729082 | 0.115321022 |
| Bbs5 | 0.639307266 | 1.388369549 | 0.143 | 0.123 | 1 | 0.059463517 | -1.539319835 | 0.044 | 0.145 | 1 | 0.639307266 | 0.115391125 |
| Kifc3 | 0.981247489 | 0.457453861 | 0.333 | 0.384 | 1 | 0.059489614 | -0.282711698 | 0.133 | 0.301 | 1 | 0.981247489 | 0.115440213 |
| Clcn6 | 0.93150888 | 0.474991904 | 0.238 | 0.287 | 1 | 0.059568624 | -0.145815497 | 0.178 | 0.349 | 1 | 0.93150888 | 0.115588826 |
| Ap1b1 | 0.10076252 | -0.834022873 | 0.095 | 0.265 | 1 | 0.059589376 | -0.423842618 | 0.133 | 0.284 | 1 | 0.10076252 | 0.115627858 |
| Srek1 | 0.059634462 | 0.656698254 | 0.952 | 0.869 | 1 | 0.811953872 | 0.321806343 | 0.978 | 0.924 | 1 | 0.811953872 | 0.115712654 |
| Tmem209 | 0.171239821 | -0.548620654 | 0.048 | 0.168 | 1 | 0.059785673 | -0.72631835 | 0.067 | 0.19 | 1 | 0.171239821 | 0.115997019 |
| Gm27003 | 0.059817347 | 1.521768674 | 0.238 | 0.108 | 1 | 0.127607098 | -0.124282336 | 0.044 | 0.128 | 1 | 0.127607098 | 0.116056578 |
| Chm | 0.8159712 | 0.806295853 | 0.429 | 0.672 | 1 | 0.059869309 | 0.712543705 | 0.778 | 0.754 | 1 | 0.8159712 | 0.116154285 |
| Ackr3 | 0.059895785 | 1.06829225 | 0.667 | 0.537 | 1 | 0.140133311 | -0.356663166 | 0.378 | 0.526 | 1 | 0.140133311 | 0.116204065 |
| Tmem140 | 0.102813066 | -0.632474705 | 0.333 | 0.549 | 1 | 0.06000763 | -0.36350578 | 0.311 | 0.536 | 1 | 0.102813066 | 0.116414345 |
| Tbc1d2b | 0.303919612 | -0.31112134 | 0.143 | 0.257 | 1 | 0.060096114 | -0.151503805 | 0.111 | 0.266 | 1 | 0.303919612 | 0.116580686 |
| Igf2bp3 | 0.794513304 | 1.525379928 | 0.238 | 0.228 | 1 | 0.060096767 | -1.235790651 | 0.067 | 0.183 | 1 | 0.794513304 | 0.116581912 |
| Gm16062 | 0.500020275 | 1.214340149 | 0.048 | 0.097 | 1 | 0.060174702 | -0.204452685 | 0.044 | 0.156 | 1 | 0.500020275 | 0.116728409 |
| Grb14 | 0.060337589 | -2.496153234 | 0.048 | 0.216 | 1 | 0.132745369 | -2.174908409 | 0 | 0.048 | 1 | 0.132745369 | 0.117034553 |
| Phkb | 0.567883803 | 0.787942788 | 0.333 | 0.347 | 1 | 0.060384219 | 0.715738775 | 0.667 | 0.484 | 1 | 0.567883803 | 0.117122183 |
| Capn1 | 0.060406278 | 2.036341847 | 0.19 | 0.078 | 1 | 0.490299592 | 0.361144491 | 0.067 | 0.104 | 1 | 0.490299592 | 0.117163638 |
| Zfp87 | 0.220184345 | -0.911190733 | 0.048 | 0.142 | 1 | 0.060443696 | -1.199570463 | 0.067 | 0.176 | 1 | 0.220184345 | 0.117233952 |
| Ankfy1 | 0.172611793 | 0.6858446 | 0.571 | 0.522 | 1 | 0.060466137 | 0.982632868 | 0.756 | 0.664 | 1 | 0.172611793 | 0.117276121 |
| Fggy | 0.616785892 | 1.503846766 | 0.095 | 0.149 | 1 | 0.060497621 | -0.946284034 | 0.067 | 0.183 | 1 | 0.616785892 | 0.117335281 |
| Dnhd1 | 0.160116854 | 1.586308926 | 0.238 | 0.138 | 1 | 0.060518939 | -1.539319835 | 0.044 | 0.145 | 1 | 0.160116854 | 0.117375336 |
| Tasp1 | 0.060554236 | 1.520738546 | 0.619 | 0.59 | 1 | 0.362774798 | 0.457774797 | 0.422 | 0.657 | 1 | 0.362774798 | 0.117441656 |
| Cbl | 0.749294612 | 0.346305489 | 0.429 | 0.466 | 1 | 0.060555705 | 0.650939965 | 0.889 | 0.678 | 1 | 0.749294612 | 0.117444416 |
| Pcbp2 | 0.171098027 | -0.27941248 | 0.952 | 0.978 | 1 | 0.060561427 | -0.19264746 | 0.978 | 0.997 | 1 | 0.171098027 | 0.117455167 |
| Nup50 | 0.065354722 | -0.741265732 | 0.333 | 0.575 | 1 | 0.060579429 | -0.257765341 | 0.578 | 0.72 | 1 | 0.065354722 | 0.117488991 |

| | | | | | | | | | | | | |
|---|---|---|---|---|---|---|---|---|---|---|---|---|
| Pspc1 | 0.060631998 | 0.785417124 | 0.667 | 0.578 | 1 | 0.14440575 | 0.401301618 | 0.844 | 0.668 | 1 | 0.14440575 | 0.117587756 |
| Zfp746 | 0.937280155 | 0.434305833 | 0.238 | 0.254 | 1 | 0.060642894 | -0.306873749 | 0.156 | 0.318 | 1 | 0.937280155 | 0.117608228 |
| Gch1 | 0.605903228 | -0.308080886 | 0.143 | 0.19 | 1 | 0.060660519 | -0.731964913 | 0.089 | 0.221 | 1 | 0.605903228 | 0.117641339 |
| Fbrs | 0.156891775 | 0.313024424 | 0.143 | 0.34 | 1 | 0.060729847 | 0.169448867 | 0.267 | 0.516 | 1 | 0.156891775 | 0.11777158 |
| B4galt5 | 0.060745374 | 1.577847348 | 0.476 | 0.384 | 1 | 0.715936668 | 0.548646266 | 0.333 | 0.436 | 1 | 0.715936668 | 0.117800747 |
| Aspg | 0.060779041 | 2.010806755 | 0.143 | 0.049 | 1 | 0.206882538 | -0.901889915 | 0 | 0.035 | 1 | 0.206882538 | 0.11786399 |
| Zcchc2 | 0.139513966 | -0.865387043 | 0.095 | 0.243 | 1 | 0.060837618 | -1.276285429 | 0.156 | 0.277 | 1 | 0.139513966 | 0.11797402 |
| Hprt | 0.060951225 | -0.512094778 | 0.429 | 0.687 | 1 | 0.273046339 | -0.237996945 | 0.711 | 0.706 | 1 | 0.273046339 | 0.118187397 |
| Acads | 0.083371142 | -1.535681598 | 0 | 0.127 | 1 | 0.061004042 | -0.21001221 | 0.044 | 0.156 | 1 | 0.083371142 | 0.118286591 |
| Wdyhv1 | 0.061062447 | -1.718545655 | 0 | 0.146 | 1 | 0.079806607 | -0.290932205 | 0.044 | 0.145 | 1 | 0.079806607 | 0.118396272 |
| Khnyn | 0.094099745 | 1.36444371 | 0.429 | 0.31 | 1 | 0.061095033 | -0.117827314 | 0.267 | 0.457 | 1 | 0.094099745 | 0.118457463 |
| Scaf11 | 0.061152472 | -0.459943496 | 0.905 | 0.951 | 1 | 0.077963602 | -0.176512295 | 1 | 0.979 | 1 | 0.077963602 | 0.11856532 |
| Gm46563 | 0.0611812 | -1.849790188 | 0 | 0.146 | 1 | 0.443620131 | -0.316927414 | 0.044 | 0.076 | 1 | 0.443620131 | 0.118619261 |
| Tradd | 0.061181292 | -1.785659851 | 0 | 0.146 | 1 | 0.07912331 | -1.486852415 | 0.022 | 0.104 | 1 | 0.07912331 | 0.118619433 |
| Arf3 | 0.939003306 | 0.625408746 | 0.333 | 0.474 | 1 | 0.061238163 | 0.670545488 | 0.756 | 0.626 | 1 | 0.939003306 | 0.118726214 |
| Zbtb24 | 0.577272477 | 0.129451251 | 0.143 | 0.209 | 1 | 0.061255531 | 0.174058939 | 0.089 | 0.211 | 1 | 0.577272477 | 0.118758821 |
| Ift122 | 0.061285487 | -1.940938076 | 0 | 0.146 | 1 | 0.066184324 | -1.039393438 | 0.067 | 0.176 | 1 | 0.066184324 | 0.118815064 |
| 2310061I04Rik | 0.360100913 | 0.171271427 | 0.048 | 0.119 | 1 | 0.061375641 | -1.124282336 | 0.044 | 0.149 | 1 | 0.360100913 | 0.118984313 |
| Zfp790 | 0.640268317 | 1.351843673 | 0.095 | 0.071 | 1 | 0.061379144 | -1.08246216 | 0.067 | 0.18 | 1 | 0.640268317 | 0.118990888 |
| Bcl2l11 | 0.190046903 | -0.726158839 | 0.095 | 0.235 | 1 | 0.061390274 | -0.391474367 | 0.422 | 0.253 | 1 | 0.190046903 | 0.119011783 |
| Otud4 | 0.061408106 | 0.786246497 | 0.619 | 0.53 | 1 | 0.628668601 | 0.197645759 | 0.778 | 0.744 | 1 | 0.628668601 | 0.119045256 |
| Pdpr | 0.758246592 | 0.503846766 | 0.143 | 0.19 | 1 | 0.061410309 | -0.251039478 | 0.111 | 0.263 | 1 | 0.758246592 | 0.119049392 |
| Ube2l6 | 0.061462558 | -2.303508156 | 0 | 0.146 | 1 | 0.120810184 | 0.723714571 | 0.022 | 0.093 | 1 | 0.120810184 | 0.119147469 |
| Sap30 | 0.061562404 | -2.326228232 | 0 | 0.146 | 1 | 0.083606745 | 0.503163496 | 0.489 | 0.329 | 1 | 0.083606745 | 0.119334878 |
| N4bp2l1 | 0.061571582 | -2.133583154 | 0 | 0.146 | 1 | 0.110110372 | 0.62996125 | 0.089 | 0.201 | 1 | 0.110110372 | 0.119352104 |
| Prr14l | 0.316323805 | 0.614878079 | 0.571 | 0.619 | 1 | 0.061581468 | 0.674027447 | 0.889 | 0.785 | 1 | 0.316323805 | 0.11937066 |
| Igtp | 0.061685586 | -2.648156327 | 0 | 0.146 | 1 | 0.754605206 | 1.825091591 | 0.067 | 0.083 | 1 | 0.754605206 | 0.119566061 |
| Slc6a6 | 0.061731518 | -0.515401118 | 0.857 | 0.948 | 1 | 0.228346508 | -0.111046509 | 0.978 | 0.913 | 1 | 0.228346508 | 0.119652256 |
| Rab30 | 0.061740218 | -1.683780237 | 0.048 | 0.228 | 1 | 0.704916698 | 1.238287743 | 0.178 | 0.235 | 1 | 0.704916698 | 0.119668582 |
| Mfsd2a | 0.061745507 | 2.029915578 | 0.19 | 0.075 | 1 | 0.164841739 | -1.404390255 | 0.022 | 0.08 | 1 | 0.164841739 | 0.119678506 |
| Olfm2 | 0.790349864 | 1.908237021 | 0.048 | 0.037 | 1 | 0.061823023 | 1.972579203 | 0.178 | 0.093 | 1 | 0.790349864 | 0.11982396 |
| Hspb1 | 0.061956442 | -2.366870217 | 0.762 | 0.72 | 1 | 0.349067235 | -1.122390277 | 0.956 | 0.737 | 1 | 0.349067235 | 0.120074284 |
| Mgst1 | 0.388649993 | -1.064996069 | 0.333 | 0.384 | 1 | 0.062070165 | 0.477603468 | 0.711 | 0.457 | 1 | 0.388649993 | 0.120287625 |
| Mmaa | 0.818724272 | 0.503846766 | 0.143 | 0.127 | 1 | 0.062135378 | 0.286182434 | 0.067 | 0.194 | 1 | 0.818724272 | 0.120409951 |
| Sox8 | 0.216716958 | -0.862281133 | 0.095 | 0.216 | 1 | 0.062271237 | -0.941418279 | 0.044 | 0.152 | 1 | 0.216716958 | 0.120664767 |
| Vps11 | 0.243478337 | -0.623908781 | 0.095 | 0.198 | 1 | 0.062287834 | -0.230215781 | 0.133 | 0.294 | 1 | 0.243478337 | 0.120695894 |
| Rhpn2 | 0.370806713 | -0.326228232 | 0 | 0.037 | 1 | 0.0624018 | 1.83507568 | 0.133 | 0.059 | 1 | 0.370806713 | 0.120909615 |
| Fam122a | 0.115715483 | -0.785659851 | 0.095 | 0.257 | 1 | 0.062421756 | -0.144746438 | 0.067 | 0.187 | 1 | 0.115715483 | 0.120947037 |
| Slc25a35 | 0.251790506 | -0.413691074 | 0 | 0.06 | 1 | 0.062460623 | -1.84048937 | 0 | 0.073 | 1 | 0.251790506 | 0.121019916 |
| Gm56976 | 0.97964157 | 2.188344941 | 0.048 | 0.049 | 1 | 0.062460623 | -1.84048937 | 0 | 0.073 | 1 | 0.97964157 | 0.121019916 |
| Dars2 | 0.222255538 | -1.081115734 | 0 | 0.067 | 1 | 0.062483303 | -1.960783604 | 0 | 0.073 | 1 | 0.222255538 | 0.121062443 |
| Ociad2 | 0.236481344 | -1.02666795 | 0 | 0.063 | 1 | 0.062483303 | -1.901889915 | 0 | 0.073 | 1 | 0.236481344 | 0.121062443 |
| D930015M05Rik | 0.275816289 | 1.908237021 | 0.095 | 0.045 | 1 | 0.062483303 | -1.901889915 | 0 | 0.073 | 1 | 0.275816289 | 0.121062443 |
| Vipr2 | 0.425629641 | -0.133583154 | 0 | 0.03 | 1 | 0.062483303 | -2.776359032 | 0 | 0.073 | 1 | 0.425629641 | 0.121062443 |
| Sema3b | 0.544576436 | 2.258734268 | 0.048 | 0.026 | 1 | 0.062483303 | -1.960783604 | 0 | 0.073 | 1 | 0.544576436 | 0.121062443 |
| Siah1b | 0.869523519 | 0.503846766 | 0.048 | 0.056 | 1 | 0.062483303 | -1.901889915 | 0 | 0.073 | 1 | 0.869523519 | 0.121062443 |
| A130010J15Rik | 0.985457491 | 0.766881172 | 0.048 | 0.049 | 1 | 0.062483303 | -1.901889915 | 0 | 0.073 | 1 | 0.985457491 | 0.121062443 |
| Tyrobp | 0.63207476 | 0.214340149 | 0 | 0.011 | 1 | 0.062496307 | -1.428435729 | 0.133 | 0.055 | 1 | 0.63207476 | 0.121086826 |
| Gm35612 | 0.425639612 | 0.503846766 | 0 | 0.03 | 1 | 0.062503597 | -2.017367132 | 0 | 0.073 | 1 | 0.425639612 | 0.121100494 |
| 4930558N01Rik | 0.818172757 | 1.936806174 | 0.048 | 0.063 | 1 | 0.062503597 | -2.017367132 | 0 | 0.073 | 1 | 0.818172757 | 0.121100494 |
| Noxred1 | 0.859875139 | 1.451379346 | 0.048 | 0.041 | 1 | 0.062503597 | -2.071814916 | 0 | 0.073 | 1 | 0.859875139 | 0.121100494 |
| Snhg9 | 0.144766727 | -1.233118828 | 0 | 0.093 | 1 | 0.062503716 | -1.960783604 | 0 | 0.073 | 1 | 0.144766727 | 0.121100717 |
| Hdac11 | 0.425614684 | 0.214340149 | 0 | 0.03 | 1 | 0.062503716 | -1.960783604 | 0 | 0.073 | 1 | 0.425614684 | 0.121100717 |
| Rlbp1 | 0.865447636 | 1.233199176 | 0.095 | 0.112 | 1 | 0.062503716 | -1.960783604 | 0 | 0.073 | 1 | 0.865447636 | 0.121100717 |
| Rps6ka6 | 0.457292877 | 0.351843673 | 0 | 0.026 | 1 | 0.062521503 | -2.071814916 | 0 | 0.073 | 1 | 0.457292877 | 0.121134068 |
| D030028A08Rik | 0.464386198 | 2.536268244 | 0.095 | 0.06 | 1 | 0.062521503 | -2.071814916 | 0 | 0.073 | 1 | 0.464386198 | 0.121134068 |
| 1700029H14Rik | 0.621091768 | 1.503846766 | 0.048 | 0.082 | 1 | 0.062521861 | -2.174908409 | 0 | 0.073 | 1 | 0.621091768 | 0.12113474 |
| Dhx34 | 0.699118062 | 1.150209812 | 0.095 | 0.075 | 1 | 0.062521861 | -2.271123724 | 0 | 0.073 | 1 | 0.699118062 | 0.12113474 |
| Dctd | 0.078487401 | -1.683780237 | 0 | 0.131 | 1 | 0.062521981 | -2.017367132 | 0 | 0.073 | 1 | 0.078487401 | 0.121134964 |
| Fam171a2 | 0.324767242 | -0.233118828 | 0 | 0.045 | 1 | 0.062521981 | -2.017367132 | 0 | 0.073 | 1 | 0.324767242 | 0.121134964 |
| Gm41409 | 0.251653515 | -0.718545655 | 0 | 0.06 | 1 | 0.062537023 | -2.124282336 | 0 | 0.073 | 1 | 0.251653515 | 0.121163167 |
| Twnk | 0.317974915 | -0.611630451 | 0.048 | 0.119 | 1 | 0.062537023 | -2.124282336 | 0 | 0.073 | 1 | 0.317974915 | 0.121163167 |
| Ptges3l | 0.173819828 | -1.133583154 | 0 | 0.082 | 1 | 0.062537739 | -2.174908409 | 0 | 0.073 | 1 | 0.173819828 | 0.12116451 |
| Gm14418 | 0.304385789 | -0.413691074 | 0 | 0.049 | 1 | 0.062537739 | -2.223818009 | 0 | 0.073 | 1 | 0.304385789 | 0.12116451 |
| Zfp229 | 0.075093029 | 2.673771768 | 0.095 | 0.026 | 1 | 0.062538217 | -2.071814916 | 0 | 0.073 | 1 | 0.075093029 | 0.121165405 |
| Ggnbp1 | 0.773169671 | 0.425844254 | 0.048 | 0.063 | 1 | 0.062538217 | -2.071814916 | 0 | 0.073 | 1 | 0.773169671 | 0.121165405 |
| Rab36 | 0.370840523 | 0.214340149 | 0 | 0.037 | 1 | 0.062550155 | -2.223818009 | 0 | 0.073 | 1 | 0.370840523 | 0.121187789 |
| Cip2a | 0.309681683 | 3.010806755 | 0.095 | 0.049 | 1 | 0.062551349 | -2.271123724 | 0 | 0.073 | 1 | 0.309681683 | 0.121190027 |
| Nmral1 | 0.196657642 | -0.718545655 | 0 | 0.075 | 1 | 0.062552066 | -2.271123724 | 0 | 0.073 | 1 | 0.196657642 | 0.12119137 |
| Epb41l3 | 0.409635495 | 0.673771768 | 0.095 | 0.052 | 1 | 0.062564482 | -2.486852415 | 0 | 0.073 | 1 | 0.409635495 | 0.121214649 |
| Exd1 | 0.651055703 | 0.710297644 | 0.143 | 0.112 | 1 | 0.062574272 | -2.486852415 | 0 | 0.073 | 1 | 0.651055703 | 0.121233004 |
| Hmga2 | 0.910111151 | 1.159198595 | 0.048 | 0.056 | 1 | 0.062575705 | -3.425451871 | 0 | 0.073 | 1 | 0.910111151 | 0.12123569 |
| Orc1 | 0.236454357 | -0.785659851 | 0 | 0.063 | 1 | 0.062580958 | -2.602329633 | 0 | 0.073 | 1 | 0.236454357 | 0.121245539 |
| Slc7a11 | 0.709200368 | 0.171271427 | 0.048 | 0.071 | 1 | 0.062591465 | -3.466674533 | 0 | 0.073 | 1 | 0.709200368 | 0.121265238 |
| Bdp1 | 0.677084559 | 0.228212324 | 0.952 | 0.884 | 1 | 0.062595907 | -0.227332494 | 0.911 | 0.931 | 1 | 0.677084559 | 0.121273567 |
| Fbln1 | 0.225213418 | -1.611630451 | 0.048 | 0.138 | 1 | 0.062600419 | -2.80878051 | 0 | 0.073 | 1 | 0.225213418 | 0.121282026 |
| Gm36640 | 0.396971401 | 0.351843673 | 0 | 0.034 | 1 | 0.062606389 | -3.199570463 | 0 | 0.073 | 1 | 0.396971401 | 0.121293218 |
| Oxnad1 | 0.393945924 | 0.444953077 | 0.095 | 0.183 | 1 | 0.062610759 | -1.174908409 | 0.044 | 0.149 | 1 | 0.393945924 | 0.121301411 |
| Gak | 0.112623782 | 1.109870882 | 0.524 | 0.56 | 1 | 0.062631103 | 0.482848984 | 0.844 | 0.702 | 1 | 0.112623782 | 0.121339552 |
| Echdc1 | 0.062682997 | -0.919752747 | 0.19 | 0.414 | 1 | 0.181990884 | 0.308677071 | 0.2 | 0.346 | 1 | 0.181990884 | 0.121436837 |
| Aox2 | 0.062711093 | 1.503846766 | 0.143 | 0.049 | 1 | 0.761677441 | -0.174908409 | 0.044 | 0.055 | 1 | 0.761677441 | 0.121489505 |
| Hdx | 0.062718995 | 1.936806174 | 0.143 | 0.049 | 1 | 0.815922253 | 0.946106992 | 0.044 | 0.038 | 1 | 0.815922253 | 0.121504319 |
| Mms22l | 0.062727106 | 2.325848464 | 0.143 | 0.049 | 1 | 0.679340743 | 0.361144491 | 0.022 | 0.035 | 1 | 0.679340743 | 0.121519522 |
| Zfp933 | 0.12811322 | -1.326228232 | 0 | 0.101 | 1 | 0.062738514 | -0.881712033 | 0.067 | 0.183 | 1 | 0.12811322 | 0.121540907 |
| Gem | 0.06274018 | -0.940938076 | 0.571 | 0.664 | 1 | 0.680061724 | 0.460680165 | 0.733 | 0.775 | 1 | 0.680061724 | 0.12154403 |
| Extl3 | 0.928110976 | 0.516230491 | 0.286 | 0.31 | 1 | 0.06275355 | -0.288168421 | 0.2 | 0.377 | 1 | 0.928110976 | 0.121569092 |
| Trp53inp1 | 0.062754562 | -1.175893959 | 0.19 | 0.396 | 1 | 0.505963197 | 0.419191777 | 0.533 | 0.443 | 1 | 0.505963197 | 0.121570989 |
| Cdk13 | 0.062829772 | 0.890395709 | 0.905 | 0.825 | 1 | 0.263607051 | 0.288941593 | 1 | 0.962 | 1 | 0.263607051 | 0.121711965 |
| Cramp1 | 0.062973232 | -0.504146025 | 0.143 | 0.392 | 1 | 0.283653692 | 0.476028129 | 0.667 | 0.626 | 1 | 0.283653692 | 0.121980836 |
| Vps45 | 0.658148175 | -0.154047257 | 0.143 | 0.183 | 1 | 0.06304563 | -0.412084647 | 0.089 | 0.225 | 1 | 0.658148175 | 0.122116509 |
| Slc25a26 | 0.173777275 | -1.233118828 | 0 | 0.082 | 1 | 0.063055205 | -1.84048937 | 0.022 | 0.111 | 1 | 0.173777275 | 0.12213445 |
| Ngef | 0.063088069 | 1.108331825 | 0.619 | 0.534 | 1 | 0.738530291 | 0.707555271 | 0.444 | 0.533 | 1 | 0.738530291 | 0.122196034 |
| Fxyd3 | 0.353378427 | -2.017163102 | 0.81 | 0.653 | 1 | 0.063117975 | -1.095459501 | 0.822 | 0.529 | 1 | 0.353378427 | 0.122252072 |
| Dnajc30 | 0.063129545 | -1.496153234 | 0.048 | 0.216 | 1 | 0.116618002 | -0.174908409 | 0.067 | 0.163 | 1 | 0.116618002 | 0.12227375 |
| Ahcyl1 | 0.300467436 | 0.343850882 | 0.81 | 0.825 | 1 | 0.063148674 | -0.283480931 | 0.933 | 0.903 | 1 | 0.300467436 | 0.122309592 |
| Ndufaf2 | 0.063155296 | -0.77835505 | 0.19 | 0.429 | 1 | 0.124577048 | 0.408710175 | 0.222 | 0.408 | 1 | 0.124577048 | 0.122322001 |
| Zfp120 | 0.332446545 | -0.648156327 | 0.048 | 0.116 | 1 | 0.063259204 | -1.743192169 | 0.022 | 0.111 | 1 | 0.332446545 | 0.122516682 |
| Ttc39c | 0.351231646 | -0.818081329 | 0.048 | 0.112 | 1 | 0.063260317 | -1.931637258 | 0.022 | 0.111 | 1 | 0.351231646 | 0.122518765 |

| | | | | | | | | | | | | |
|---|---|---|---|---|---|---|---|---|---|---|---|---|
| Gria3 | 0.672768253 | 0.285501178 | 0.333 | 0.437 | 1 | 0.063295989 | -0.192707477 | 0.244 | 0.443 | 1 | 0.672768253 | 0.122585595 |
| Pigo | 0.757115357 | 0.97333205 | 0.095 | 0.078 | 1 | 0.063462909 | -1.709244837 | 0.022 | 0.111 | 1 | 0.757115357 | 0.122898276 |
| Slc39a9 | 0.22542646 | 0.447703688 | 0.19 | 0.347 | 1 | 0.063484908 | 0.15951063 | 0.267 | 0.484 | 1 | 0.22542646 | 0.122939483 |
| Tmem161b | 0.877538247 | 1.146524765 | 0.238 | 0.28 | 1 | 0.063495409 | 0.460680165 | 0.156 | 0.325 | 1 | 0.877538247 | 0.122959151 |
| 8030462N17Rik | 0.063614424 | -0.870548749 | 0.238 | 0.47 | 1 | 0.076199807 | 0.869291395 | 0.778 | 0.661 | 1 | 0.076199807 | 0.123182052 |
| Cstf2 | 0.684716772 | 0.192645078 | 0.143 | 0.19 | 1 | 0.063628432 | -0.709244837 | 0.133 | 0.273 | 1 | 0.684716772 | 0.123208287 |
| Pak3 | 0.806431474 | 0.707560081 | 0.19 | 0.243 | 1 | 0.06363538 | -1.536095934 | 0.067 | 0.18 | 1 | 0.806431474 | 0.123221298 |
| Rassf3 | 0.065854506 | -0.901137068 | 0.095 | 0.291 | 1 | 0.063682346 | 0.41403944 | 0.689 | 0.467 | 1 | 0.065854506 | 0.123309251 |
| Srpx2 | 0.063727234 | -0.777439344 | 0.238 | 0.5 | 1 | 0.312483996 | 0.229899958 | 0.2 | 0.311 | 1 | 0.312483996 | 0.123393308 |
| Hmgxb3 | 0.372573399 | 1.334461595 | 0.381 | 0.332 | 1 | 0.06378015 | -0.26247963 | 0.2 | 0.394 | 1 | 0.372573399 | 0.123492392 |
| Smtn | 0.605951915 | 0.287190714 | 0.476 | 0.478 | 1 | 0.063854395 | -0.302028537 | 0.467 | 0.758 | 1 | 0.605951915 | 0.123631407 |
| Gemin6 | 0.222110654 | -0.648156327 | 0 | 0.067 | 1 | 0.063864138 | -1.602329633 | 0.022 | 0.111 | 1 | 0.222110654 | 0.123649647 |
| Wtap | 0.063898928 | -0.818081329 | 0.238 | 0.466 | 1 | 0.659585019 | 0.194325401 | 0.578 | 0.633 | 1 | 0.659585019 | 0.123714783 |
| Proser2 | 0.129582406 | 1.189386222 | 0.714 | 0.657 | 1 | 0.063902271 | 0.209141398 | 0.444 | 0.74 | 1 | 0.129582406 | 0.123721041 |
| Polg | 0.4671937 | 0.258734268 | 0.19 | 0.302 | 1 | 0.063931428 | 0.252022769 | 0.2 | 0.388 | 1 | 0.4671937 | 0.123775628 |
| Lpar1 | 0.080429539 | 0.955783848 | 0.429 | 0.881 | 1 | 0.063982541 | 0.742778832 | 0.8 | 0.882 | 1 | 0.080429539 | 0.123871317 |
| Nubpl | 0.752175119 | 1.018419939 | 0.143 | 0.198 | 1 | 0.064047435 | -0.342462506 | 0.044 | 0.149 | 1 | 0.752175119 | 0.123992796 |
| Kmt2b | 0.135077774 | -0.348596045 | 0.143 | 0.34 | 1 | 0.064118053 | 0.129271017 | 0.267 | 0.519 | 1 | 0.135077774 | 0.124124982 |
| Ripk1 | 0.064165887 | -0.332891963 | 0.429 | 0.716 | 1 | 0.185016168 | -0.198282917 | 0.733 | 0.761 | 1 | 0.185016168 | 0.124214513 |
| Nipsnap3b | 0.268929139 | -0.326228232 | 0.095 | 0.201 | 1 | 0.064244709 | -0.266886731 | 0.089 | 0.215 | 1 | 0.268929139 | 0.124362036 |
| 4930579G24Rik | 0.184816574 | -1.280424543 | 0 | 0.078 | 1 | 0.064258633 | -1.486852415 | 0.022 | 0.111 | 1 | 0.184816574 | 0.124388094 |
| Tbc1d7 | 0.307563163 | 0.901182264 | 0.048 | 0.131 | 1 | 0.064258633 | -1.486852415 | 0.022 | 0.111 | 1 | 0.307563163 | 0.124388094 |
| Fbxl2 | 0.064348412 | -0.830359658 | 0.333 | 0.59 | 1 | 0.440096402 | 0.658688428 | 0.444 | 0.505 | 1 | 0.440096402 | 0.124556105 |
| Pak4 | 0.186069285 | -0.849790188 | 0.048 | 0.153 | 1 | 0.064367102 | -0.667424661 | 0.044 | 0.149 | 1 | 0.186069285 | 0.124591081 |
| Lsg1 | 0.287860193 | 0.827068178 | 0.524 | 0.507 | 1 | 0.064371991 | -0.398721505 | 0.378 | 0.526 | 1 | 0.287860193 | 0.124600229 |
| Dennd6a | 0.064420356 | 1.48016777 | 0.524 | 0.474 | 1 | 0.334492379 | 0.813295164 | 0.556 | 0.578 | 1 | 0.334492379 | 0.124690729 |
| D330041H03Rik | 0.604519763 | 1.24081236 | 0.095 | 0.067 | 1 | 0.064422937 | -1.089516918 | 0.022 | 0.114 | 1 | 0.604519763 | 0.124695559 |
| Samd8 | 0.06446569 | -0.251040736 | 0.238 | 0.522 | 1 | 0.563441051 | -0.220251394 | 0.756 | 0.661 | 1 | 0.563441051 | 0.124775555 |
| Uimc1 | 0.06455047 | 0.993389702 | 0.667 | 0.746 | 1 | 0.743813312 | 0.480829864 | 0.689 | 0.709 | 1 | 0.743813312 | 0.124934177 |
| Unc93b1 | 0.064645231 | -1.20069735 | 0.048 | 0.228 | 1 | 0.311170295 | 0.849082538 | 0.267 | 0.208 | 1 | 0.311170295 | 0.125111457 |
| Lbr | 0.06464943 | -0.678744647 | 0.143 | 0.369 | 1 | 0.508282247 | 0.408897623 | 0.533 | 0.453 | 1 | 0.508282247 | 0.125119312 |
| Pdss1 | 0.064654618 | 1.388369549 | 0.333 | 0.187 | 1 | 0.532025183 | 0.361144491 | 0.111 | 0.156 | 1 | 0.532025183 | 0.125129016 |
| Hdac4 | 0.083734171 | 1.333696326 | 0.619 | 0.638 | 1 | 0.064734225 | 0.746266413 | 0.867 | 0.747 | 1 | 0.083734171 | 0.12527793 |
| 1700029J07Rik | 0.610916977 | 0.214340149 | 0.143 | 0.194 | 1 | 0.064847135 | -1.017367132 | 0.022 | 0.114 | 1 | 0.610916977 | 0.125489119 |
| Qtrt2 | 0.882628218 | 0.594044575 | 0.238 | 0.257 | 1 | 0.064947703 | -0.251832386 | 0.156 | 0.325 | 1 | 0.882628218 | 0.125677202 |
| Ppwd1 | 0.064994542 | -0.586755783 | 0.095 | 0.313 | 1 | 0.096723535 | 0.271258988 | 0.178 | 0.336 | 1 | 0.096723535 | 0.125764794 |
| Lsm14b | 0.065075906 | -1.785659851 | 0 | 0.142 | 1 | 0.298376327 | 0.122357632 | 0.089 | 0.156 | 1 | 0.298376327 | 0.125916938 |
| Arxes2 | 0.065076003 | -1.718545655 | 0 | 0.142 | 1 | 0.086715729 | -0.316927414 | 0.044 | 0.138 | 1 | 0.086715729 | 0.12591712 |
| Synpo2 | 0.268033526 | -1.081115734 | 0 | 0.056 | 1 | 0.065118506 | -0.901889915 | 0.022 | 0.114 | 1 | 0.268033526 | 0.125996592 |
| Rgp1 | 0.065133485 | -1.785659851 | 0 | 0.142 | 1 | 0.13901267 | -0.674479418 | 0.044 | 0.121 | 1 | 0.13901267 | 0.126024599 |
| Lsm10 | 0.065133874 | -1.752492987 | 0 | 0.142 | 1 | 0.125264548 | -0.486852415 | 0.067 | 0.156 | 1 | 0.125264548 | 0.126025327 |
| Srrd | 0.06518747 | -1.818081329 | 0 | 0.142 | 1 | 0.618980574 | 0.683072586 | 0.067 | 0.093 | 1 | 0.618980574 | 0.126125534 |
| Fbxo25 | 0.06518747 | -1.880817084 | 0 | 0.142 | 1 | 0.142171013 | 0.268035087 | 0.044 | 0.125 | 1 | 0.142171013 | 0.126125534 |
| Nipa1 | 0.065188054 | -1.880817084 | 0 | 0.142 | 1 | 0.086047953 | 0.567595369 | 0.044 | 0.138 | 1 | 0.086047953 | 0.126126625 |
| Dcun1d3 | 0.065213598 | 1.822294292 | 0.333 | 0.198 | 1 | 0.068852082 | 0.388451837 | 0.156 | 0.325 | 1 | 0.068852082 | 0.126174383 |
| Elmod2 | 0.06523786 | -1.880817084 | 0 | 0.142 | 1 | 0.177143538 | -0.294207337 | 0.089 | 0.17 | 1 | 0.177143538 | 0.126219742 |
| Cdkn2aipnl | 0.06523786 | -1.849790188 | 0 | 0.142 | 1 | 0.157449999 | 0.325520582 | 0.044 | 0.121 | 1 | 0.157449999 | 0.126219742 |
| C030037D09Rik | 0.065239417 | -1.911190733 | 0 | 0.142 | 1 | 0.55467343 | 0.304560963 | 0.044 | 0.069 | 1 | 0.55467343 | 0.126222652 |
| Ptrh2 | 0.124990956 | -0.648156327 | 0.095 | 0.254 | 1 | 0.065262214 | -0.564854927 | 0.089 | 0.208 | 1 | 0.124990956 | 0.126265271 |
| Gnpat | 0.542458836 | 1.747772349 | 0.19 | 0.168 | 1 | 0.065276451 | -0.849422495 | 0.089 | 0.211 | 1 | 0.542458836 | 0.126291886 |
| Prox1 | 0.163658913 | -1.496153234 | 0 | 0.086 | 1 | 0.065477102 | -0.80878051 | 0.022 | 0.114 | 1 | 0.163658913 | 0.126666952 |
| Trim26 | 0.065495829 | -1.01272876 | 0.095 | 0.291 | 1 | 0.212715947 | 0.311103809 | 0.244 | 0.391 | 1 | 0.212715947 | 0.126701954 |
| Numb | 0.294007391 | 0.605059018 | 0.667 | 0.563 | 1 | 0.065619772 | 0.369006609 | 0.956 | 0.758 | 1 | 0.294007391 | 0.126933589 |
| Gm15417 | 0.314941114 | -0.25696557 | 0.095 | 0.19 | 1 | 0.065625426 | -0.849422495 | 0.089 | 0.211 | 1 | 0.314941114 | 0.126944156 |
| Xaf1 | 0.065631966 | -2.785659851 | 0 | 0.142 | 1 | 0.731428882 | 0.223640968 | 0.089 | 0.104 | 1 | 0.731428882 | 0.126956377 |
| Dclk1 | 0.065640532 | -2.648156327 | 0 | 0.142 | 1 | 0.838319647 | 0.624178897 | 0.089 | 0.104 | 1 | 0.838319647 | 0.126972384 |
| Bglap2 | 0.06567392 | -4.133583154 | 0 | 0.142 | 1 | 0.106869533 | -3.339295227 | 0 | 0.055 | 1 | 0.106869533 | 0.127034776 |
| Pgpep1 | 0.233465583 | 0.55829455 | 0.048 | 0.149 | 1 | 0.065772882 | -0.367553487 | 0.044 | 0.152 | 1 | 0.233465583 | 0.127219692 |
| Arhgap5 | 0.309901723 | -0.151549701 | 0.714 | 0.869 | 1 | 0.065881738 | -0.194936889 | 1 | 0.938 | 1 | 0.309901723 | 0.127423073 |
| Mknk2 | 0.233550668 | -0.184209227 | 0.333 | 0.556 | 1 | 0.065945054 | -0.481408277 | 0.778 | 0.727 | 1 | 0.233550668 | 0.127541357 |
| Hmgb2 | 0.066032122 | -0.985391675 | 0.571 | 0.657 | 1 | 0.201324763 | -0.118116963 | 0.867 | 0.612 | 1 | 0.201324763 | 0.127704003 |
| Orc3 | 0.066060435 | 0.116823643 | 0.143 | 0.377 | 1 | 0.715814321 | 0.608304818 | 0.289 | 0.367 | 1 | 0.715814321 | 0.127756889 |
| Gpx2 | 0.457308648 | 0.673771768 | 0 | 0.026 | 1 | 0.066089674 | 1.304560963 | 0.089 | 0.031 | 1 | 0.457308648 | 0.127811503 |
| Tbc1d23 | 0.066094446 | 0.948631609 | 0.619 | 0.627 | 1 | 0.743356146 | 0.364250402 | 0.667 | 0.706 | 1 | 0.743356146 | 0.127820416 |
| Ap4m1 | 0.13475449 | -1.326228232 | 0.048 | 0.172 | 1 | 0.066104106 | -0.602329633 | 0.067 | 0.183 | 1 | 0.13475449 | 0.127838459 |
| Bmp3 | 0.285549799 | -0.911190733 | 0 | 0.052 | 1 | 0.06620858 | -0.831500587 | 0.067 | 0.183 | 1 | 0.285549799 | 0.128033584 |
| Cobll1 | 0.682720051 | 0.704007775 | 0.714 | 0.698 | 1 | 0.066219145 | 1.077351525 | 0.778 | 0.623 | 1 | 0.682720051 | 0.128053314 |
| Pals2 | 0.151256682 | 1.131877989 | 0.81 | 0.899 | 1 | 0.066386848 | 0.45745338 | 0.911 | 0.91 | 1 | 0.151256682 | 0.128366482 |
| Asap1 | 0.066437215 | -0.622771009 | 0.476 | 0.705 | 1 | 0.393657956 | 0.216819702 | 1 | 0.865 | 1 | 0.393657956 | 0.128460527 |
| Tmem184b | 0.066506504 | -0.628347846 | 0.19 | 0.451 | 1 | 0.95063124 | 0.47967288 | 0.422 | 0.554 | 1 | 0.95063124 | 0.128589894 |
| Tdp2 | 0.619749083 | 0.97333205 | 0.286 | 0.284 | 1 | 0.066623292 | -0.160624183 | 0.2 | 0.377 | 1 | 0.619749083 | 0.128807921 |
| Tle4 | 0.089074078 | 0.817212848 | 0.762 | 0.604 | 1 | 0.066624704 | 0.457652472 | 0.911 | 0.72 | 1 | 0.089074078 | 0.128810556 |
| Suz12 | 0.696583386 | 0.503846766 | 0.429 | 0.627 | 1 | 0.066630844 | 0.431390985 | 0.844 | 0.682 | 1 | 0.696583386 | 0.128822019 |
| Snapc4 | 0.466303684 | 0.295260144 | 0.095 | 0.172 | 1 | 0.066681016 | -1.404390255 | 0.044 | 0.142 | 1 | 0.466303684 | 0.128915674 |
| Stau1 | 0.066705156 | 0.443671251 | 0.905 | 0.78 | 1 | 0.930337717 | 0.228119073 | 0.978 | 0.958 | 1 | 0.930337717 | 0.128960735 |
| Zfp426 | 0.866981928 | 0.69394965 | 0.19 | 0.194 | 1 | 0.066972772 | -0.571741313 | 0.111 | 0.239 | 1 | 0.866981928 | 0.129460191 |
| Cemip2 | 0.067051354 | -0.718545655 | 0.429 | 0.724 | 1 | 0.87352908 | -0.205812221 | 0.733 | 0.609 | 1 | 0.87352908 | 0.129606824 |
| Rybp | 0.424305892 | -0.290729551 | 0.667 | 0.817 | 1 | 0.067056372 | 0.634520841 | 0.911 | 0.827 | 1 | 0.424305892 | 0.129616186 |
| Ino80b | 0.414281854 | -0.475975352 | 0.095 | 0.164 | 1 | 0.067066776 | -1.564854927 | 0.044 | 0.142 | 1 | 0.414281854 | 0.1296356 |
| Snx8 | 0.06715748 | 0.316219763 | 0.048 | 0.231 | 1 | 0.261571819 | 0.223640968 | 0.244 | 0.377 | 1 | 0.261571819 | 0.129804834 |
| Prx | 0.067172422 | 2.936806174 | 0.143 | 0.049 | 1 | 0.501086724 | 1.460680165 | 0.067 | 0.045 | 1 | 0.501086724 | 0.12983271 |
| Ubap1 | 0.067231905 | 0.788907478 | 0.286 | 0.593 | 1 | 0.521192144 | 0.263533695 | 0.711 | 0.664 | 1 | 0.521192144 | 0.129943681 |
| Zyg11b | 0.586908306 | 0.606940259 | 0.429 | 0.474 | 1 | 0.06729749 | 0.505676335 | 0.867 | 0.727 | 1 | 0.586908306 | 0.130066028 |
| Golga1 | 0.311278934 | 0.623145695 | 0.429 | 0.381 | 1 | 0.067332452 | 0.13875207 | 0.267 | 0.464 | 1 | 0.311278934 | 0.130131246 |
| B130055M24Rik | 0.067345096 | 1.866416846 | 0.286 | 0.16 | 1 | 0.095144309 | -0.191396532 | 0.089 | 0.204 | 1 | 0.095144309 | 0.13015483 |
| Dbn1 | 0.923277058 | 0.710297644 | 0.143 | 0.168 | 1 | 0.06735227 | -0.105423309 | 0.067 | 0.183 | 1 | 0.923277058 | 0.130168212 |
| Fzd3 | 0.067434517 | 0.335902129 | 0.048 | 0.231 | 1 | 0.11714813 | -0.265943485 | 0.156 | 0.287 | 1 | 0.11714813 | 0.13032162 |
| Clybl | 0.067597235 | 0.214340149 | 0.048 | 0.231 | 1 | 0.189942297 | 0.385392038 | 0.111 | 0.215 | 1 | 0.189942297 | 0.130625083 |
| Nhs | 0.584666608 | -0.27768438 | 0.238 | 0.284 | 1 | 0.067658214 | 0.180572246 | 0.756 | 0.471 | 1 | 0.584666608 | 0.130738793 |
| Ahctf1 | 0.85855359 | 0.1514905 | 0.619 | 0.757 | 1 | 0.067902237 | 0.274987848 | 0.956 | 0.879 | 1 | 0.85855359 | 0.13119376 |
| Dhx57 | 0.620616697 | 0.412506455 | 0.238 | 0.354 | 1 | 0.067918226 | -0.287383138 | 0.267 | 0.467 | 1 | 0.620616697 | 0.131223567 |
| Zscan21 | 0.746311264 | 0.316219763 | 0.095 | 0.123 | 1 | 0.067985166 | -0.776359032 | 0.067 | 0.176 | 1 | 0.746311264 | 0.13134835 |
| Slit3 | 0.480934137 | -2.399477214 | 0.048 | 0.093 | 1 | 0.068039932 | 1.809520921 | 0.267 | 0.18 | 1 | 0.480934137 | 0.131450432 |
| Cdc27 | 0.776432555 | 0.597668789 | 0.429 | 0.53 | 1 | 0.068198319 | 0.488958119 | 0.778 | 0.675 | 1 | 0.776432555 | 0.131745627 |
| Xpo7 | 0.687958858 | 0.405282932 | 0.476 | 0.653 | 1 | 0.068202367 | 0.470242387 | 0.978 | 0.779 | 1 | 0.687958858 | 0.131753171 |
| Prtg | 0.691128173 | -0.735619168 | 0.095 | 0.123 | 1 | 0.068209734 | -2.404390255 | 0.022 | 0.107 | 1 | 0.691128173 | 0.1317669 |
| Radil | 0.068218821 | 2.214340149 | 0.381 | 0.284 | 1 | 0.233372396 | 1.578735926 | 0.133 | 0.083 | 1 | 0.233372396 | 0.131783835 |
| Dhodh | 0.743880619 | 1.150209812 | 0.095 | 0.078 | 1 | 0.068224275 | -1.432404631 | 0.044 | 0.142 | 1 | 0.743880619 | 0.131793997 |

| | | | | | | | | | | | | |
|---|---|---|---|---|---|---|---|---|---|---|---|---|
| Api5 | 0.125389355 | -0.440244493 | 0.429 | 0.664 | 1 | 0.068224846 | -0.25301565 | 0.8 | 0.792 | 1 | 0.125389355 | 0.131795063 |
| St3gal1 | 0.068234959 | 1.668147218 | 0.476 | 0.354 | 1 | 0.723297449 | 0.585040503 | 0.2 | 0.256 | 1 | 0.723297449 | 0.131813909 |
| Med16 | 0.873345642 | 0.477374555 | 0.143 | 0.168 | 1 | 0.068325143 | -0.738391182 | 0.089 | 0.211 | 1 | 0.873345642 | 0.131981961 |
| Ppp2r3c | 0.209322295 | -0.555046923 | 0.095 | 0.216 | 1 | 0.068368492 | -0.638855509 | 0.067 | 0.176 | 1 | 0.209322295 | 0.132062734 |
| Tank | 0.068370817 | 1.819992509 | 0.571 | 0.504 | 1 | 0.400965089 | 0.165673369 | 0.578 | 0.495 | 1 | 0.400965089 | 0.132067065 |
| Nadk | 0.068419839 | -0.574155746 | 0.238 | 0.478 | 1 | 0.521167576 | 0.336269823 | 0.667 | 0.612 | 1 | 0.521167576 | 0.132158404 |
| Kcnk1 | 0.068504039 | -1.008052272 | 0.238 | 0.451 | 1 | 0.090473891 | -1.128240349 | 0.422 | 0.481 | 1 | 0.090473891 | 0.132315275 |
| Foxred1 | 0.068547514 | -0.849790188 | 0.048 | 0.228 | 1 | 0.123473775 | 0.147019686 | 0.067 | 0.166 | 1 | 0.123473775 | 0.132396267 |
| Itsn2 | 0.605554064 | 0.644559846 | 0.619 | 0.672 | 1 | 0.068558645 | 0.479141701 | 0.956 | 0.872 | 1 | 0.605554064 | 0.132417002 |
| Zfp748 | 0.184780696 | -1.133583154 | 0 | 0.078 | 1 | 0.068576925 | -0.416463087 | 0.067 | 0.183 | 1 | 0.184780696 | 0.132451055 |
| Gpd1l | 0.370826288 | -0.785659851 | 0 | 0.037 | 1 | 0.06860888 | -1.316927414 | 0.044 | 0.142 | 1 | 0.370826288 | 0.132510582 |
| Cs | 0.510787397 | -0.105201353 | 0.238 | 0.343 | 1 | 0.068616507 | 0.605070074 | 0.778 | 0.547 | 1 | 0.510787397 | 0.132524788 |
| Tdrd3 | 0.683096021 | 1.443474148 | 0.333 | 0.381 | 1 | 0.068640734 | 1.09378416 | 0.6 | 0.567 | 1 | 0.683096021 | 0.132569917 |
| Srsf10 | 0.1909605 | -0.192210884 | 0.81 | 0.918 | 1 | 0.068900616 | -0.210168948 | 0.911 | 0.976 | 1 | 0.1909605 | 0.133053937 |
| Clec16a | 0.195756901 | -0.602352638 | 0.19 | 0.351 | 1 | 0.068925435 | 1.190228287 | 0.6 | 0.505 | 1 | 0.195756901 | 0.133100154 |
| Pdhb | 0.069053756 | -0.56399588 | 0.333 | 0.634 | 1 | 0.111588016 | -0.351053198 | 0.689 | 0.685 | 1 | 0.111588016 | 0.13333909 |
| Slc2a8 | 0.069206809 | -1.648156327 | 0 | 0.138 | 1 | 0.078624934 | -1.602329633 | 0.022 | 0.104 | 1 | 0.078624934 | 0.133624036 |
| Gsdme | 0.069316732 | -0.911190733 | 0.143 | 0.358 | 1 | 0.295970857 | 0.879764497 | 0.244 | 0.388 | 1 | 0.295970857 | 0.133828655 |
| Ntrk2 | 0.372781629 | -1.828728573 | 0.048 | 0.112 | 1 | 0.069354564 | -1.109485334 | 0.067 | 0.173 | 1 | 0.372781629 | 0.133899073 |
| Zfp715 | 0.069379483 | -1.818081329 | 0 | 0.138 | 1 | 0.112460674 | -0.722919774 | 0.067 | 0.159 | 1 | 0.112460674 | 0.133945454 |
| Chst14 | 0.069380515 | -1.752492987 | 0 | 0.138 | 1 | 0.084587371 | -1.158229668 | 0.044 | 0.135 | 1 | 0.084587371 | 0.133947374 |
| Gm48226 | 0.069380515 | -1.752492987 | 0 | 0.138 | 1 | 0.080101789 | -0.602329633 | 0.022 | 0.107 | 1 | 0.080101789 | 0.133947374 |
| Arntl2 | 0.069405877 | 1.088809267 | 0.381 | 0.22 | 1 | 0.697276791 | 0.251226799 | 0.267 | 0.343 | 1 | 0.697276791 | 0.133994578 |
| D830050J10Rik | 0.069477594 | -1.880817084 | 0 | 0.138 | 1 | 0.830857135 | 0.180572246 | 0.044 | 0.052 | 1 | 0.830857135 | 0.134128051 |
| Gm13293 | 0.304385789 | -0.326228232 | 0 | 0.049 | 1 | 0.069504102 | -1.776359032 | 0 | 0.069 | 1 | 0.304385789 | 0.134177385 |
| Galnt3 | 0.16548778 | 5.258734268 | 0.048 | 0.011 | 1 | 0.069526834 | -2.017367132 | 0 | 0.069 | 1 | 0.16548778 | 0.134219688 |
| Vwa1 | 0.220184345 | -0.911190733 | 0.048 | 0.142 | 1 | 0.069526834 | -1.84048937 | 0 | 0.069 | 1 | 0.220184345 | 0.134219688 |
| 9930014A18Rik | 0.23647819 | -0.911190733 | 0 | 0.063 | 1 | 0.069526834 | -1.84048937 | 0 | 0.069 | 1 | 0.23647819 | 0.134219688 |
| Elp6 | 0.251683171 | -0.718545655 | 0 | 0.06 | 1 | 0.069526834 | -2.017367132 | 0 | 0.069 | 1 | 0.251683171 | 0.134219688 |
| Shld3 | 0.649186945 | 0.281454345 | 0.048 | 0.075 | 1 | 0.069526834 | -1.84048937 | 0 | 0.069 | 1 | 0.649186945 | 0.134219688 |
| Tlk1 | 0.069544368 | 0.72579398 | 0.762 | 0.657 | 1 | 0.925815 | 0.15951063 | 0.867 | 0.848 | 1 | 0.925815 | 0.134252317 |
| Thg1l | 0.184739689 | -1.081115734 | 0 | 0.078 | 1 | 0.06954704 | -2.017367132 | 0 | 0.069 | 1 | 0.184739689 | 0.13425729 |
| Zfp94 | 0.267953794 | -0.496153234 | 0 | 0.056 | 1 | 0.069547173 | -1.901889915 | 0 | 0.069 | 1 | 0.267953794 | 0.134257537 |
| Eid3 | 0.305657443 | 1.451379346 | 0.095 | 0.045 | 1 | 0.069547173 | -1.901889915 | 0 | 0.069 | 1 | 0.305657443 | 0.134257537 |
| Zfp709 | 0.25165389 | -0.648156327 | 0 | 0.06 | 1 | 0.069565253 | -1.960783604 | 0 | 0.069 | 1 | 0.25165389 | 0.134291182 |
| Rxfp3 | 0.63207476 | 1.351843673 | 0 | 0.011 | 1 | 0.069565253 | -1.960783604 | 0 | 0.069 | 1 | 0.63207476 | 0.134291182 |
| Usp35 | 0.864156366 | 0.214340149 | 0.048 | 0.056 | 1 | 0.069565253 | -1.960783604 | 0 | 0.069 | 1 | 0.864156366 | 0.134291182 |
| Osgin1 | 0.532436372 | -0.233118828 | 0 | 0.019 | 1 | 0.069579877 | -2.124282336 | 0 | 0.069 | 1 | 0.532436372 | 0.134318394 |
| 3222401L13Rik | 0.105292847 | 1.181918671 | 0.19 | 0.086 | 1 | 0.069580674 | -2.124282336 | 0 | 0.069 | 1 | 0.105292847 | 0.134319878 |
| Dnaaf4 | 0.267984803 | -0.849790188 | 0 | 0.056 | 1 | 0.069580674 | -2.124282336 | 0 | 0.069 | 1 | 0.267984803 | 0.134319878 |
| Rinl | 0.279052091 | 1.825774861 | 0.095 | 0.045 | 1 | 0.069581206 | -2.017367132 | 0 | 0.069 | 1 | 0.279052091 | 0.134320868 |
| Batf3 | 0.91015245 | 0.673771768 | 0.095 | 0.093 | 1 | 0.069592506 | -2.124282336 | 0 | 0.069 | 1 | 0.91015245 | 0.134341896 |
| Rab3a | 0.50481484 | 1.150209812 | 0.095 | 0.06 | 1 | 0.069605934 | -2.271123724 | 0 | 0.069 | 1 | 0.50481484 | 0.134366882 |
| D330023K18Rik | 0.094522876 | -1.611630451 | 0 | 0.119 | 1 | 0.069606186 | -1.84048937 | 0.022 | 0.107 | 1 | 0.094522876 | 0.134367352 |
| Klk8 | 0.144972469 | -2.611630451 | 0 | 0.093 | 1 | 0.069606731 | -2.638855509 | 0 | 0.069 | 1 | 0.144972469 | 0.134368366 |
| Gm17501 | 0.09733899 | -1.849790188 | 0.048 | 0.19 | 1 | 0.069606997 | -2.674479418 | 0 | 0.069 | 1 | 0.09733899 | 0.134368861 |
| Etv4 | 0.069629267 | -2.054148687 | 0 | 0.138 | 1 | 0.385735427 | -0.242926832 | 0.067 | 0.111 | 1 | 0.385735427 | 0.134410299 |
| Myo1f | 0.918798858 | 3.045740545 | 0.048 | 0.045 | 1 | 0.069631061 | -2.743192169 | 0 | 0.069 | 1 | 0.918798858 | 0.134413637 |
| Tspan11 | 0.466599481 | 1.673771768 | 0.048 | 0.022 | 1 | 0.06963957 | -2.931637258 | 0 | 0.069 | 1 | 0.466599481 | 0.13442947 |
| Alpl | 0.234246825 | -1.725399326 | 0.048 | 0.142 | 1 | 0.069647813 | -5.031172931 | 0 | 0.069 | 1 | 0.234246825 | 0.134444808 |
| Tctn3 | 0.23638565 | -0.648156327 | 0 | 0.063 | 1 | 0.069655657 | -1.709244837 | 0 | 0.069 | 1 | 0.23638565 | 0.134459403 |
| Tmem64 | 0.069658558 | -1.30677587 | 0.143 | 0.347 | 1 | 0.211765685 | -0.39313798 | 0.578 | 0.606 | 1 | 0.211765685 | 0.134464801 |
| Als2 | 0.70382437 | -0.341178574 | 0.19 | 0.22 | 1 | 0.069658767 | -0.539319835 | 0.133 | 0.263 | 1 | 0.70382437 | 0.134465191 |
| Scamp2 | 0.164480489 | -0.253018385 | 0.476 | 0.698 | 1 | 0.069690239 | -0.191396532 | 0.6 | 0.654 | 1 | 0.164480489 | 0.134523748 |
| Sass6 | 0.069728951 | -2.434752689 | 0 | 0.138 | 1 | 0.239535548 | -0.686161223 | 0.067 | 0.131 | 1 | 0.239535548 | 0.134595776 |
| Mx2 | 0.532426384 | 0.503846766 | 0 | 0.019 | 1 | 0.069757154 | 3.311103809 | 0.067 | 0.021 | 1 | 0.532426384 | 0.134648248 |
| Csnk2a1 | 0.328164865 | 0.500987112 | 0.714 | 0.81 | 1 | 0.069816033 | -0.260799401 | 0.889 | 0.882 | 1 | 0.328164865 | 0.134757787 |
| Hal | 0.58888737 | -2.101015292 | 0.095 | 0.134 | 1 | 0.069839119 | 1.683072586 | 0.133 | 0.062 | 1 | 0.58888737 | 0.134800735 |
| Wdr66 | 0.114495717 | 2.021695071 | 0.143 | 0.06 | 1 | 0.069840577 | -1.743192169 | 0.022 | 0.107 | 1 | 0.114495717 | 0.134803447 |
| Fat1 | 0.069872792 | 0.109482835 | 0.333 | 0.597 | 1 | 0.459268734 | -0.522041844 | 0.8 | 0.671 | 1 | 0.459268734 | 0.134863376 |
| Gm5165 | 0.973011511 | 0.747772349 | 0.095 | 0.101 | 1 | 0.069900638 | -0.731964913 | 0.044 | 0.149 | 1 | 0.973011511 | 0.134915177 |
| Trim68 | 0.337250672 | 1.503846766 | 0.095 | 0.049 | 1 | 0.069917707 | -1.286553765 | 0.022 | 0.111 | 1 | 0.337250672 | 0.134946928 |
| Fam83g | 0.957800805 | 1.010806755 | 0.048 | 0.052 | 1 | 0.069942118 | -1.255526869 | 0.022 | 0.111 | 1 | 0.957800805 | 0.134992336 |
| Eri2 | 0.069969604 | 2.726239188 | 0.19 | 0.082 | 1 | 0.490350464 | 0.875717664 | 0.044 | 0.076 | 1 | 0.490350464 | 0.135043463 |
| Slc37a3 | 0.354218996 | 0.276881615 | 0.238 | 0.403 | 1 | 0.070012262 | -0.16982312 | 0.444 | 0.595 | 1 | 0.354218996 | 0.135122808 |
| Arrdc3 | 0.593500837 | 0.276881615 | 0.238 | 0.336 | 1 | 0.07003875 | -0.140049652 | 0.6 | 0.398 | 1 | 0.593500837 | 0.135172074 |
| Tmeff1 | 0.915519387 | 1.866416846 | 0.048 | 0.056 | 1 | 0.0700772 | -1.776359032 | 0.022 | 0.107 | 1 | 0.915519387 | 0.135243586 |
| Slc7a14 | 0.070083823 | 3.843696769 | 0.095 | 0.026 | 1 | 0.809040217 | 1.460680165 | 0.022 | 0.017 | 1 | 0.809040217 | 0.135255903 |
| Papolb | 0.070083823 | 3.843696769 | 0.095 | 0.026 | 1 | 0.492040936 | 2.683072586 | 0.022 | 0.01 | 1 | 0.492040936 | 0.135255903 |
| Gm47167 | 0.070083823 | 2.995699863 | 0.095 | 0.026 | 1 | 0.184823348 | -1.223818009 | 0 | 0.038 | 1 | 0.184823348 | 0.135255903 |
| D130062J10Rik | 0.070083823 | 2.995699863 | 0.095 | 0.026 | 1 | 0.118986849 | -1.404390255 | 0 | 0.052 | 1 | 0.118986849 | 0.135255903 |
| Hsbp1l1 | 0.070083823 | 2.995699863 | 0.095 | 0.026 | 1 | 0.23197796 | -0.776359032 | 0 | 0.031 | 1 | 0.23197796 | 0.135255903 |
| Trpc1 | 0.070089724 | -1.399477214 | 0.048 | 0.22 | 1 | 0.18729549 | 0.396549827 | 0.178 | 0.298 | 1 | 0.18729549 | 0.135266879 |
| Khdrbs3 | 0.390719343 | -0.133583154 | 0.095 | 0.172 | 1 | 0.070114419 | 0.44987341 | 0.644 | 0.471 | 1 | 0.390719343 | 0.135312807 |
| Dpcd | 0.741073798 | 0.199840579 | 0.238 | 0.287 | 1 | 0.070171151 | 0.236816356 | 0.111 | 0.263 | 1 | 0.741073798 | 0.135418312 |
| Cdk5rap2 | 0.070211748 | -0.244614467 | 0.286 | 0.567 | 1 | 0.656114484 | 0.688021802 | 0.356 | 0.488 | 1 | 0.656114484 | 0.135493806 |
| Map3k20 | 0.070226908 | -0.252227651 | 0.143 | 0.388 | 1 | 0.243628427 | 0.29778243 | 0.622 | 0.46 | 1 | 0.243628427 | 0.135521998 |
| Slc25a12 | 0.070401997 | -0.211585666 | 0.095 | 0.306 | 1 | 0.685297566 | 0.375644061 | 0.289 | 0.37 | 1 | 0.685297566 | 0.135847552 |
| Alkbh1 | 0.522999689 | 0.389978802 | 0.619 | 0.713 | 1 | 0.070417775 | -0.114110531 | 0.4 | 0.64 | 1 | 0.522999689 | 0.135876887 |
| Pde7b | 0.079002895 | -2.341178574 | 0.048 | 0.209 | 1 | 0.070438562 | 0.771932518 | 0.356 | 0.242 | 1 | 0.079002895 | 0.135915533 |
| Acbd5 | 0.070466523 | 0.142849107 | 0.476 | 0.683 | 1 | 0.164803973 | -0.131624311 | 0.822 | 0.779 | 1 | 0.164803973 | 0.135967515 |
| Rab8a | 0.070611136 | -0.749439663 | 0.333 | 0.552 | 1 | 0.082163456 | -0.442864403 | 0.578 | 0.623 | 1 | 0.082163456 | 0.136236339 |
| Gm50013 | 0.359357379 | 1.503846766 | 0.143 | 0.09 | 1 | 0.070735938 | -0.513324627 | 0.044 | 0.145 | 1 | 0.359357379 | 0.136468303 |
| Kdm1b | 0.285622926 | -0.233118828 | 0 | 0.052 | 1 | 0.070759179 | -1.743192169 | 0.022 | 0.107 | 1 | 0.285622926 | 0.136511497 |
| Slc4a8 | 0.070791435 | 3.503846766 | 0.095 | 0.026 | 1 | 0.476265768 | -0.564854927 | 0.022 | 0.045 | 1 | 0.476265768 | 0.136571443 |
| Mtg2 | 0.630978014 | 0.503846766 | 0.095 | 0.138 | 1 | 0.070914407 | -0.404390255 | 0.044 | 0.145 | 1 | 0.630978014 | 0.136799962 |
| Usp48 | 0.163903459 | 1.072867723 | 0.524 | 0.511 | 1 | 0.070960027 | 0.384414271 | 0.8 | 0.654 | 1 | 0.163903459 | 0.136884728 |
| Gm57158 | 0.163568985 | -1.280424543 | 0 | 0.086 | 1 | 0.070982055 | -1.674479418 | 0.022 | 0.107 | 1 | 0.163568985 | 0.136925657 |
| Ptgr1 | 0.196379028 | -0.849790188 | 0 | 0.075 | 1 | 0.070982055 | -1.602329633 | 0.022 | 0.107 | 1 | 0.196379028 | 0.136925657 |
| Igbp1 | 0.070999409 | -0.501486678 | 0.333 | 0.604 | 1 | 0.268287846 | -0.179048139 | 0.6 | 0.671 | 1 | 0.268287846 | 0.136957902 |
| Fam171a1 | 0.203018746 | -0.970084422 | 0.048 | 0.153 | 1 | 0.071080217 | 0.142504205 | 0.111 | 0.249 | 1 | 0.203018746 | 0.137108036 |
| Fgl2 | 0.128470678 | -2.434752689 | 0 | 0.101 | 1 | 0.071113356 | -1.141355849 | 0.022 | 0.111 | 1 | 0.128470678 | 0.137169603 |
| BC051537 | 0.123201066 | 0.97333205 | 0.095 | 0.03 | 1 | 0.071134123 | 2.683072586 | 0.067 | 0.021 | 1 | 0.123201066 | 0.137208182 |
| Sphk2 | 0.23638565 | -0.648156327 | 0 | 0.063 | 1 | 0.071204501 | -1.602329633 | 0.022 | 0.107 | 1 | 0.23638565 | 0.137338922 |
| Wdr81 | 0.23638565 | -0.648156327 | 0 | 0.063 | 1 | 0.071204501 | -1.486852415 | 0.022 | 0.107 | 1 | 0.23638565 | 0.137338922 |
| Mfsd4a | 0.32202842 | 1.088809267 | 0.143 | 0.082 | 1 | 0.071275774 | -0.153428682 | 0.044 | 0.145 | 1 | 0.32202842 | 0.137471312 |
| Armc9 | 0.071307918 | -1.648156327 | 0.048 | 0.209 | 1 | 0.405535706 | 0.561082062 | 0.2 | 0.294 | 1 | 0.405535706 | 0.137531016 |
| Oca2 | 0.559404139 | -0.167530486 | 0.048 | 0.086 | 1 | 0.071371565 | 1.140545352 | 0.244 | 0.152 | 1 | 0.559404139 | 0.137649229 |

| | | | | | | | | | | | | |
|---|---|---|---|---|---|---|---|---|---|---|---|---|
| Ddx42 | 0.071371962 | 0.615780045 | 0.857 | 0.78 | 1 | 0.434862306 | 0.158074874 | 0.956 | 0.848 | 1 | 0.434862306 | 0.137649967 |
| Setd1a | 0.299849154 | 0.185485286 | 0.143 | 0.28 | 1 | 0.0713923 | -0.149817428 | 0.267 | 0.467 | 1 | 0.299849154 | 0.13768774 |
| Swap70 | 0.104726427 | -0.397690595 | 0.238 | 0.493 | 1 | 0.07139982 | 0.511067274 | 0.711 | 0.526 | 1 | 0.104726427 | 0.137701706 |
| Cap1 | 0.747222239 | 0.298346124 | 0.762 | 0.765 | 1 | 0.071456325 | 0.423921615 | 0.911 | 0.862 | 1 | 0.747222239 | 0.137806643 |
| Ankrd16 | 0.133573502 | 1.088809267 | 0.286 | 0.164 | 1 | 0.071496826 | -0.466674533 | 0.067 | 0.18 | 1 | 0.133573502 | 0.137881855 |
| F630206G17Rik | 0.071497464 | 2.825774861 | 0.095 | 0.026 | 1 | 0.293637566 | -0.638855509 | 0 | 0.024 | 1 | 0.293637566 | 0.137883041 |
| Ddr1 | 0.407881328 | 0.929610672 | 0.238 | 0.187 | 1 | 0.071499398 | 0.352155708 | 0.556 | 0.367 | 1 | 0.407881328 | 0.137886632 |
| Rnf103 | 0.071507871 | -0.593014773 | 0.333 | 0.604 | 1 | 0.103424503 | -0.26926098 | 0.489 | 0.633 | 1 | 0.103424503 | 0.137902366 |
| Abcc9 | 0.346862423 | -0.326228232 | 0 | 0.041 | 1 | 0.071553944 | -1.071814916 | 0.022 | 0.111 | 1 | 0.346862423 | 0.13798792 |
| Tet2 | 0.549240941 | 0.647299556 | 0.667 | 0.724 | 1 | 0.071558445 | 0.487647212 | 0.978 | 0.837 | 1 | 0.549240941 | 0.137996278 |
| Tefm | 0.663853825 | 1.171271427 | 0.143 | 0.116 | 1 | 0.071579224 | -0.979892427 | 0.022 | 0.111 | 1 | 0.663853825 | 0.138034864 |
| Iqcc | 0.752699207 | 1.029915578 | 0.095 | 0.078 | 1 | 0.071579224 | -0.979892427 | 0.022 | 0.111 | 1 | 0.752699207 | 0.138034864 |
| 2010004M13Rik | 0.156341059 | -0.880817084 | 0.048 | 0.164 | 1 | 0.071612345 | -0.979892427 | 0.022 | 0.111 | 1 | 0.156341059 | 0.138096362 |
| Fahd2a | 0.106672433 | -1.413691074 | 0 | 0.112 | 1 | 0.071642027 | -1.404390255 | 0.022 | 0.107 | 1 | 0.106672433 | 0.138151474 |
| Gm39090 | 0.363960255 | 0.825774861 | 0.238 | 0.172 | 1 | 0.071660272 | -0.466674533 | 0.067 | 0.18 | 1 | 0.363960255 | 0.138185349 |
| Dock11 | 0.93844607 | -0.259591039 | 0.286 | 0.321 | 1 | 0.07173121 | 0.259530755 | 0.622 | 0.433 | 1 | 0.93844607 | 0.138317054 |
| Agbl5 | 0.088793927 | -1.611630451 | 0 | 0.123 | 1 | 0.071797906 | -0.941418279 | 0.022 | 0.111 | 1 | 0.088793927 | 0.138440873 |
| Cenpa | 0.989456082 | -0.735619168 | 0.095 | 0.097 | 1 | 0.071832888 | -0.454430938 | 0.022 | 0.111 | 1 | 0.989456082 | 0.138505812 |
| Psme2 | 0.072005134 | -0.82721234 | 0.381 | 0.56 | 1 | 0.655957283 | -0.113674236 | 0.622 | 0.578 | 1 | 0.655957283 | 0.138825528 |
| Amd1 | 0.072077036 | -0.607926482 | 0.429 | 0.683 | 1 | 0.608963721 | -0.16492432 | 0.778 | 0.678 | 1 | 0.608963721 | 0.138958973 |
| Cdkn1b | 0.072182588 | -0.801961663 | 0.286 | 0.504 | 1 | 0.526197468 | -0.222600031 | 0.822 | 0.744 | 1 | 0.526197468 | 0.13915485 |
| Abca3 | 0.100392236 | -1.611630451 | 0 | 0.116 | 1 | 0.072240702 | -0.931637258 | 0.022 | 0.111 | 1 | 0.100392236 | 0.139262685 |
| Cabin1 | 0.072297205 | -0.436652222 | 0.19 | 0.463 | 1 | 0.234589563 | 0.182379002 | 0.4 | 0.522 | 1 | 0.234589563 | 0.139367525 |
| Slc10a3 | 0.237632205 | 0.150209812 | 0.095 | 0.213 | 1 | 0.072341217 | -0.564854927 | 0.089 | 0.204 | 1 | 0.237632205 | 0.139449183 |
| Nsun6 | 0.072366639 | 1.968515033 | 0.333 | 0.209 | 1 | 0.131966633 | -0.216400538 | 0.111 | 0.228 | 1 | 0.131966633 | 0.139496348 |
| Hfe | 0.42890475 | -0.184209227 | 0.048 | 0.101 | 1 | 0.072435306 | -0.709244837 | 0.067 | 0.18 | 1 | 0.42890475 | 0.139623739 |
| Tmem86a | 0.910091451 | 1.088809267 | 0.048 | 0.056 | 1 | 0.072452428 | -0.776359032 | 0.022 | 0.111 | 1 | 0.910091451 | 0.139655501 |
| AW011738 | 0.208922634 | -1.02666795 | 0 | 0.071 | 1 | 0.072454231 | -0.486852415 | 0.022 | 0.111 | 1 | 0.208922634 | 0.139658847 |
| Gm30025 | 0.144664875 | -1.133583154 | 0 | 0.093 | 1 | 0.072483686 | -0.819427754 | 0.022 | 0.111 | 1 | 0.144664875 | 0.139713487 |
| Wac | 0.950940804 | 0.105111079 | 0.905 | 0.955 | 1 | 0.072628797 | 0.292591043 | 1 | 0.983 | 1 | 0.950940804 | 0.139982652 |
| Tdrd7 | 0.072657859 | -0.701267664 | 0.048 | 0.224 | 1 | 0.216449787 | 0.264120037 | 0.178 | 0.287 | 1 | 0.216449787 | 0.140036553 |
| 1700037F03Rik | 0.072660337 | 2.773307441 | 0.19 | 0.086 | 1 | 0.119007856 | -1.638855509 | 0 | 0.052 | 1 | 0.119007856 | 0.14004115 |
| Grin2d | 0.106726508 | -1.648156327 | 0 | 0.112 | 1 | 0.072696534 | -0.361321533 | 0.022 | 0.111 | 1 | 0.106726508 | 0.140108282 |
| Nufip2 | 0.072698481 | 0.350357124 | 1 | 0.985 | 1 | 0.920585049 | 0.133851675 | 1 | 1 | 1 | 0.920585049 | 0.140111893 |
| Cul5 | 0.638243908 | 0.406549565 | 0.571 | 0.601 | 1 | 0.072743975 | -0.314922278 | 0.778 | 0.834 | 1 | 0.638243908 | 0.140196263 |
| Nav2 | 0.200944669 | 1.027170201 | 0.762 | 0.664 | 1 | 0.072816143 | 1.46584187 | 0.644 | 0.502 | 1 | 0.200944669 | 0.140330095 |
| Bbs2 | 0.420079637 | -0.574155746 | 0.048 | 0.101 | 1 | 0.072816536 | -0.486852415 | 0.022 | 0.111 | 1 | 0.420079637 | 0.140330825 |
| Rad9b | 0.441169573 | -0.413691074 | 0.095 | 0.16 | 1 | 0.072831188 | -0.37003875 | 0.089 | 0.215 | 1 | 0.441169573 | 0.140357995 |
| Nfya | 0.072841712 | 1.11189288 | 0.429 | 0.306 | 1 | 0.335813162 | 0.145107565 | 0.244 | 0.329 | 1 | 0.335813162 | 0.14037751 |
| Gm15879 | 0.681769711 | 0.150209812 | 0.048 | 0.071 | 1 | 0.072881195 | -0.513324627 | 0.044 | 0.149 | 1 | 0.681769711 | 0.140450721 |
| Abtb3 | 0.072904972 | -0.118786152 | 0.095 | 0.299 | 1 | 0.279607349 | 0.644937457 | 0.333 | 0.273 | 1 | 0.279607349 | 0.140494808 |
| Zfp971 | 0.584337797 | 0.719575457 | 0.048 | 0.086 | 1 | 0.072911718 | -0.280401538 | 0.022 | 0.111 | 1 | 0.584337797 | 0.140507318 |
| C78859 | 0.072925649 | 2.536268244 | 0.095 | 0.026 | 1 | 0.659788885 | -0.564854927 | 0.022 | 0.035 | 1 | 0.659788885 | 0.140533148 |
| Sdcbp2 | 0.072931551 | 2.79930265 | 0.095 | 0.026 | 1 | 0.450829449 | 1.005000681 | 0.022 | 0.048 | 1 | 0.450829449 | 0.14054409 |
| Cops3 | 0.074119539 | -0.569682269 | 0.333 | 0.597 | 1 | 0.072994055 | -0.148529882 | 0.578 | 0.685 | 1 | 0.074119539 | 0.140659978 |
| Gm17749 | 0.073100198 | 2.233199176 | 0.143 | 0.052 | 1 | 0.095952004 | -2.223818009 | 0 | 0.059 | 1 | 0.095952004 | 0.140856758 |
| Adam17 | 0.370061637 | 0.392815454 | 0.286 | 0.44 | 1 | 0.073130711 | 0.35394899 | 0.778 | 0.581 | 1 | 0.370061637 | 0.140913322 |
| Efna2 | 0.07318076 | 1.327996931 | 0.286 | 0.153 | 1 | 0.105720015 | -0.301820521 | 0.089 | 0.194 | 1 | 0.105720015 | 0.141006096 |
| Atrip | 0.07337342 | -0.75918764 | 0.095 | 0.302 | 1 | 0.105783245 | 0.216754582 | 0.133 | 0.273 | 1 | 0.105783245 | 0.141363181 |
| Naa15 | 0.073606351 | 0.773607035 | 0.857 | 0.858 | 1 | 0.528953797 | 0.167715553 | 0.933 | 0.931 | 1 | 0.528953797 | 0.141794807 |
| Smg5 | 0.192495069 | 1.462267662 | 0.333 | 0.261 | 1 | 0.073723945 | 0.219723065 | 0.222 | 0.426 | 1 | 0.192495069 | 0.142012669 |
| Ccdc68 | 0.555242546 | 0.766881172 | 0.048 | 0.09 | 1 | 0.073768342 | -0.240691828 | 0.111 | 0.253 | 1 | 0.555242546 | 0.142094917 |
| Zdhhc12 | 0.073772133 | -1.818081329 | 0 | 0.134 | 1 | 0.295418043 | -0.255526869 | 0.022 | 0.062 | 1 | 0.295418043 | 0.142101938 |
| Mettl14 | 0.073772242 | -1.683780237 | 0 | 0.134 | 1 | 0.202334862 | -0.749886821 | 0.067 | 0.135 | 1 | 0.202334862 | 0.142102141 |
| Grwd1 | 0.524313463 | 0.380990018 | 0.095 | 0.153 | 1 | 0.073793265 | -1.124282336 | 0.067 | 0.17 | 1 | 0.524313463 | 0.142141084 |
| Rfxank | 0.073826504 | -1.718545655 | 0 | 0.134 | 1 | 0.462387653 | 0.361144491 | 0.067 | 0.104 | 1 | 0.462387653 | 0.142202656 |
| Ankib1 | 0.073831689 | 1.140128044 | 0.667 | 0.646 | 1 | 0.673405033 | 0.194647431 | 0.778 | 0.779 | 1 | 0.673405033 | 0.142212259 |
| Add1 | 0.219192666 | -0.247474121 | 0.429 | 0.586 | 1 | 0.073872169 | 0.497465962 | 0.933 | 0.747 | 1 | 0.219192666 | 0.142287241 |
| Coa5 | 0.073875188 | -1.785659851 | 0 | 0.134 | 1 | 0.484255312 | 0.223640968 | 0.111 | 0.156 | 1 | 0.484255312 | 0.142292833 |
| Coq6 | 0.073875188 | -1.818081329 | 0 | 0.134 | 1 | 0.136819872 | -0.667424661 | 0.067 | 0.152 | 1 | 0.136819872 | 0.142292833 |
| Mapk7 | 0.073877376 | -1.752492987 | 0 | 0.134 | 1 | 0.896616346 | 0.989175714 | 0.133 | 0.152 | 1 | 0.896616346 | 0.142296886 |
| Pygl | 0.204867226 | -0.174225139 | 0.19 | 0.343 | 1 | 0.073907307 | 0.154693614 | 0.156 | 0.304 | 1 | 0.204867226 | 0.142352324 |
| Pisd | 0.377197164 | 0.150209812 | 0.095 | 0.187 | 1 | 0.073984618 | 0.348088338 | 0.133 | 0.301 | 1 | 0.377197164 | 0.142495512 |
| H4c9 | 0.07400812 | -2.02666795 | 0 | 0.134 | 1 | 0.331306855 | 0.290755163 | 0.022 | 0.059 | 1 | 0.331306855 | 0.142539039 |
| Adgrf5 | 0.074008449 | -2.107587946 | 0 | 0.134 | 1 | 0.086320512 | -1.199570463 | 0.022 | 0.104 | 1 | 0.086320512 | 0.142539647 |
| Ecsit | 0.25724614 | -0.167530486 | 0.048 | 0.142 | 1 | 0.074091038 | -0.463768802 | 0.067 | 0.18 | 1 | 0.25724614 | 0.142692594 |
| Cdyl2 | 0.074104408 | -2.02666795 | 0 | 0.134 | 1 | 0.539062701 | 0.640428249 | 0.133 | 0.187 | 1 | 0.539062701 | 0.142717352 |
| Lamb3 | 0.897785641 | 0.908237021 | 0.095 | 0.09 | 1 | 0.074196871 | 0.905465008 | 0.133 | 0.059 | 1 | 0.897785641 | 0.142888567 |
| Bckdhb | 0.683574248 | 0.586308926 | 0.333 | 0.354 | 1 | 0.074202416 | 0.401502229 | 0.178 | 0.339 | 1 | 0.683574248 | 0.142898834 |
| Mitd1 | 0.074248983 | -1.133583154 | 0.095 | 0.28 | 1 | 0.096889114 | -0.298780067 | 0.089 | 0.204 | 1 | 0.096889114 | 0.142985054 |
| Rbm48 | 0.089216195 | -0.865387043 | 0.095 | 0.272 | 1 | 0.07438909 | -0.242926832 | 0.067 | 0.18 | 1 | 0.089216195 | 0.143244444 |
| Nab1 | 0.074440602 | 0.744161096 | 0.667 | 0.593 | 1 | 0.17284946 | 0.361144491 | 0.911 | 0.785 | 1 | 0.17284946 | 0.1433398 |
| Glcci1 | 0.074460724 | 1.258734268 | 0.524 | 0.384 | 1 | 0.286782419 | 0.225184783 | 0.6 | 0.498 | 1 | 0.286782419 | 0.143377049 |
| Tatdn2 | 0.866710377 | 0.425844254 | 0.19 | 0.22 | 1 | 0.074479048 | -0.476126009 | 0.2 | 0.36 | 1 | 0.866710377 | 0.143410968 |
| Kif28 | 0.074648842 | 3.133203386 | 0.143 | 0.052 | 1 | 0.797698974 | 1.209141398 | 0.067 | 0.059 | 1 | 0.797698974 | 0.143725235 |
| Gm20712 | 0.251790506 | -0.413691074 | 0 | 0.06 | 1 | 0.074707021 | -1.346674757 | 0.044 | 0.138 | 1 | 0.251790506 | 0.143832903 |
| Crot | 0.627199284 | 0.586308926 | 0.095 | 0.146 | 1 | 0.074872767 | -0.437942815 | 0.089 | 0.208 | 1 | 0.627199284 | 0.144139603 |
| Zfand2a | 0.139522549 | -0.707050016 | 0.238 | 0.422 | 1 | 0.075019585 | -0.276929346 | 0.222 | 0.408 | 1 | 0.139522549 | 0.144411232 |
| Apobec1 | 0.075093029 | 3.48112669 | 0.095 | 0.026 | 1 | 0.846213537 | -0.174908409 | 0.044 | 0.052 | 1 | 0.846213537 | 0.144547095 |
| Gm56626 | 0.075093029 | 2.673771768 | 0.095 | 0.026 | 1 | 0.11902586 | -1.564854927 | 0 | 0.052 | 1 | 0.11902586 | 0.144547095 |
| Duoxa1 | 0.075118229 | 2.574236094 | 0.095 | 0.026 | 1 | 0.312251526 | 2.361144491 | 0.022 | 0.007 | 1 | 0.312251526 | 0.14459371 |
| Heatr5b | 0.8245231 | 0.586308926 | 0.19 | 0.187 | 1 | 0.075140368 | -0.520799747 | 0.156 | 0.298 | 1 | 0.8245231 | 0.144634661 |
| Supt20 | 0.163287325 | -0.326228232 | 0.333 | 0.552 | 1 | 0.075404693 | 0.480255703 | 0.689 | 0.498 | 1 | 0.163287325 | 0.145123518 |
| Cx3cl1 | 0.075455369 | -1.611630451 | 0.048 | 0.205 | 1 | 0.254184844 | -0.486852415 | 0.133 | 0.208 | 1 | 0.254184844 | 0.145217225 |
| Mrps27 | 0.346305727 | -0.599246727 | 0.095 | 0.172 | 1 | 0.075460992 | -0.169370225 | 0.067 | 0.18 | 1 | 0.346305727 | 0.145227623 |
| Nepro | 0.201498548 | 1.150209812 | 0.238 | 0.142 | 1 | 0.075476803 | -0.719025857 | 0.089 | 0.208 | 1 | 0.201498548 | 0.145256858 |
| Fancl | 0.969397411 | 1.177345942 | 0.19 | 0.209 | 1 | 0.075557807 | -0.124282336 | 0.089 | 0.215 | 1 | 0.969397411 | 0.145406632 |
| Dock4 | 0.917197562 | 1.044415148 | 0.286 | 0.332 | 1 | 0.075611684 | 0.792860732 | 0.622 | 0.54 | 1 | 0.917197562 | 0.145506242 |
| Arl4c | 0.907396046 | 0.815790773 | 0.048 | 0.056 | 1 | 0.075761031 | 0.553789569 | 0.156 | 0.073 | 1 | 0.907396046 | 0.145782329 |
| Gbf1 | 0.81873726 | 0.351843673 | 0.762 | 0.922 | 1 | 0.075822684 | 0.58182044 | 0.956 | 0.934 | 1 | 0.81873726 | 0.145896288 |
| Plekhg6 | 0.075844833 | 3.088809267 | 0.095 | 0.026 | 1 | 0.206882538 | -0.901889915 | 0 | 0.035 | 1 | 0.206882538 | 0.145937228 |
| Cadm2 | 0.075862932 | 2.673771768 | 0.095 | 0.026 | 1 | 0.116526752 | -1.361321533 | 0.022 | 0.093 | 1 | 0.116526752 | 0.14597068 |
| Cacng7 | 0.146121469 | -0.413691074 | 0.048 | 0.179 | 1 | 0.076010887 | 0.396191438 | 0.133 | 0.287 | 1 | 0.146121469 | 0.14624412 |
| Mcm3ap | 0.750408244 | 0.918884266 | 0.238 | 0.246 | 1 | 0.076036534 | 0.299743947 | 0.133 | 0.277 | 1 | 0.750408244 | 0.146291513 |
| Stim2 | 0.358483423 | 0.135525766 | 0.429 | 0.619 | 1 | 0.076103046 | 0.683072586 | 0.867 | 0.761 | 1 | 0.358483423 | 0.146414419 |
| Tbce | 0.458284749 | 0.766881172 | 0.476 | 0.556 | 1 | 0.076192244 | 0.223640968 | 0.267 | 0.484 | 1 | 0.458284749 | 0.146579229 |
| mt-Nd6 | 0.076245957 | 1.300962504 | 0.524 | 0.437 | 1 | 0.297267203 | 0.848750617 | 0.378 | 0.381 | 1 | 0.297267203 | 0.146678468 |
| Morc2a | 0.076272039 | 0.940710628 | 0.571 | 0.47 | 1 | 0.977940513 | 0.315701521 | 0.511 | 0.637 | 1 | 0.977940513 | 0.146726655 |

| | | | | | | | | | | | | |
|---|---|---|---|---|---|---|---|---|---|---|---|---|
| 1500004A13Rik | 0.394211774 | 1.503846766 | 0.095 | 0.052 | 1 | 0.076283162 | -0.989352756 | 0.044 | 0.142 | 1 | 0.394211774 | 0.146747203 |
| Fyb | 0.769092896 | 1.595769256 | 0.048 | 0.067 | 1 | 0.076302615 | -2.149817428 | 0.022 | 0.104 | 1 | 0.769092896 | 0.146783141 |
| Hdgfl3 | 0.076445781 | -0.94420579 | 0.238 | 0.437 | 1 | 0.162133871 | 0.590506467 | 0.4 | 0.571 | 1 | 0.162133871 | 0.147047605 |
| Ddx21 | 0.84584956 | 0.225891012 | 0.619 | 0.776 | 1 | 0.07654754 | -0.23468396 | 0.822 | 0.844 | 1 | 0.84584956 | 0.147235555 |
| 4930579K19Rik | 0.076591385 | 2.503846766 | 0.095 | 0.026 | 1 | 0.226015083 | 2.42003818 | 0.044 | 0.017 | 1 | 0.226015083 | 0.147316531 |
| Bvht | 0.076591385 | 2.503846766 | 0.095 | 0.026 | 1 | 0.148044209 | -1.486852415 | 0 | 0.045 | 1 | 0.148044209 | 0.147316531 |
| Pycard | 0.855723381 | 0.638147858 | 0.095 | 0.112 | 1 | 0.076595873 | 0.813656696 | 0.378 | 0.249 | 1 | 0.855723381 | 0.147324819 |
| Tmem117 | 0.076597033 | 0.768697025 | 0.619 | 0.478 | 1 | 0.152515372 | 1.344270673 | 0.311 | 0.256 | 1 | 0.152515372 | 0.14732696 |
| Lmx1a | 0.076611606 | 2.393663848 | 0.095 | 0.026 | 1 | 0.244524402 | 0.653325243 | 0.022 | 0.069 | 1 | 0.244524402 | 0.147353873 |
| Tmem241 | 0.762588632 | 0.775651382 | 0.19 | 0.179 | 1 | 0.076670033 | -0.255526869 | 0.089 | 0.218 | 1 | 0.762588632 | 0.147461772 |
| Klf11 | 0.117913785 | -0.849790188 | 0.095 | 0.254 | 1 | 0.076797766 | -0.16492432 | 0.156 | 0.315 | 1 | 0.117913785 | 0.147697635 |
| Gm12216 | 0.271131131 | 1.8372705 | 0.238 | 0.157 | 1 | 0.076798074 | 1.358290494 | 0.4 | 0.339 | 1 | 0.271131131 | 0.147698203 |
| Ube2q2 | 0.076882824 | 0.155413045 | 0.905 | 0.832 | 1 | 0.656538052 | 0.266765668 | 0.844 | 0.789 | 1 | 0.656538052 | 0.147854679 |
| Coq8b | 0.09325493 | 0.820320431 | 0.476 | 0.347 | 1 | 0.076923204 | -0.638855509 | 0.133 | 0.273 | 1 | 0.09325493 | 0.147929229 |
| Nop9 | 0.672400894 | 0.400753273 | 0.143 | 0.198 | 1 | 0.076967154 | -0.656777417 | 0.111 | 0.235 | 1 | 0.672400894 | 0.148010365 |
| Pnkp | 0.863468507 | 0.786246497 | 0.095 | 0.112 | 1 | 0.077095959 | -0.239759553 | 0.111 | 0.235 | 1 | 0.863468507 | 0.148248132 |
| Gm33906 | 0.457300763 | -0.718545655 | 0 | 0.026 | 1 | 0.077151385 | 3.268035087 | 0.044 | 0.01 | 1 | 0.457300763 | 0.148350434 |
| Gm15287 | 0.550953306 | 1.936806174 | 0.048 | 0.026 | 1 | 0.077151385 | 4.490427508 | 0.044 | 0.01 | 1 | 0.550953306 | 0.148350434 |
| Hap1 | 0.632076134 | 0.866416846 | 0 | 0.011 | 1 | 0.077151385 | 3.005000681 | 0.044 | 0.01 | 1 | 0.632076134 | 0.148350434 |
| Gm5444 | 0.492599419 | -0.233118828 | 0 | 0.022 | 1 | 0.077153 | 3.683072586 | 0.044 | 0.01 | 1 | 0.492599419 | 0.148353415 |
| Ruvbl2 | 0.110017456 | -0.672003069 | 0.048 | 0.198 | 1 | 0.077154459 | -0.506751973 | 0.089 | 0.208 | 1 | 0.110017456 | 0.148356107 |
| Seh1l | 0.1036112 | -0.337455488 | 0.286 | 0.552 | 1 | 0.077167667 | -0.141355849 | 0.289 | 0.522 | 1 | 0.1036112 | 0.148380484 |
| Zfp185 | 0.896682417 | 0.704145417 | 0.143 | 0.142 | 1 | 0.077304299 | -0.670564368 | 0.089 | 0.204 | 1 | 0.896682417 | 0.148632644 |
| Mutyh | 0.077343918 | 2.214340149 | 0.095 | 0.026 | 1 | 0.331922009 | -0.124282336 | 0 | 0.021 | 1 | 0.331922009 | 0.148705755 |
| Gm38882 | 0.094421823 | -1.611630451 | 0 | 0.119 | 1 | 0.077351591 | -1.709244837 | 0 | 0.066 | 1 | 0.094421823 | 0.148719913 |
| Gal3st1 | 0.229212234 | -0.741265732 | 0.048 | 0.146 | 1 | 0.077351591 | -1.709244837 | 0 | 0.066 | 1 | 0.229212234 | 0.148719913 |
| Pcyox1l | 0.285622926 | -0.233118828 | 0 | 0.052 | 1 | 0.077351591 | -1.709244837 | 0 | 0.066 | 1 | 0.285622926 | 0.148719913 |
| Tsku | 0.457281049 | 0.503846766 | 0 | 0.026 | 1 | 0.077351591 | -1.901889915 | 0 | 0.066 | 1 | 0.457281049 | 0.148719913 |
| Mon1b | 0.547761155 | 0.351843673 | 0.048 | 0.09 | 1 | 0.077351591 | -1.709244837 | 0 | 0.066 | 1 | 0.547761155 | 0.148719913 |
| Sun1 | 0.077370921 | -0.678744647 | 0.143 | 0.373 | 1 | 0.799595942 | 0.502500341 | 0.4 | 0.439 | 1 | 0.799595942 | 0.148755582 |
| Mfsd4b4 | 0.346802964 | -0.133583154 | 0 | 0.041 | 1 | 0.077374257 | -1.776359032 | 0 | 0.066 | 1 | 0.346802964 | 0.148761738 |
| Taf1c | 0.628535192 | 1.010806755 | 0.095 | 0.067 | 1 | 0.077374257 | -1.776359032 | 0 | 0.066 | 1 | 0.628535192 | 0.148761738 |
| C330024C12Rik | 0.741589569 | 1.214340149 | 0.048 | 0.034 | 1 | 0.077374257 | -1.776359032 | 0 | 0.066 | 1 | 0.741589569 | 0.148761738 |
| Ptpru | 0.817162882 | 0.304537958 | 0.095 | 0.078 | 1 | 0.077374257 | -1.901889915 | 0 | 0.066 | 1 | 0.817162882 | 0.148761738 |
| A130023E24Rik | 0.85370107 | 1.673771768 | 0.048 | 0.041 | 1 | 0.077374257 | -1.960783604 | 0 | 0.066 | 1 | 0.85370107 | 0.148761738 |
| Cdc7 | 0.985459372 | 1.258734268 | 0.048 | 0.049 | 1 | 0.077374257 | -1.84048937 | 0 | 0.066 | 1 | 0.985459372 | 0.148761738 |
| Lrrc27 | 0.186349485 | 1.918884266 | 0.143 | 0.071 | 1 | 0.077394256 | -1.960783604 | 0 | 0.066 | 1 | 0.186349485 | 0.148798642 |
| Nipal3 | 0.184642276 | -0.849790188 | 0 | 0.078 | 1 | 0.077394404 | -1.84048937 | 0 | 0.066 | 1 | 0.184642276 | 0.148798915 |
| Pafah2 | 0.267953794 | -0.496153234 | 0 | 0.056 | 1 | 0.077394404 | -1.84048937 | 0 | 0.066 | 1 | 0.267953794 | 0.148798915 |
| Lca5l | 0.509428832 | -0.184209227 | 0.048 | 0.09 | 1 | 0.077394404 | -1.84048937 | 0 | 0.066 | 1 | 0.509428832 | 0.148798915 |
| Intu | 0.769094408 | 0.918884266 | 0.048 | 0.067 | 1 | 0.077411589 | -2.071814916 | 0 | 0.066 | 1 | 0.769094408 | 0.148830624 |
| Rragb | 0.933800488 | 1.088809267 | 0.048 | 0.045 | 1 | 0.077411589 | -1.960783604 | 0 | 0.066 | 1 | 0.933800488 | 0.148830624 |
| 1700086O06Rik | 0.30672684 | 0.380990018 | 0.048 | 0.131 | 1 | 0.077412182 | -1.901889915 | 0 | 0.066 | 1 | 0.30672684 | 0.148831718 |
| C230035I16Rik | 0.425614684 | 0.214340149 | 0 | 0.03 | 1 | 0.077412182 | -1.901889915 | 0 | 0.066 | 1 | 0.425614684 | 0.148831718 |
| Prdm5 | 0.454194288 | 0.373966484 | 0.619 | 0.623 | 1 | 0.07741224 | 0.378388064 | 0.4 | 0.661 | 1 | 0.454194288 | 0.148831825 |
| Psmc3ip | 0.196495505 | -1.133583154 | 0 | 0.075 | 1 | 0.077427737 | -1.960783604 | 0 | 0.066 | 1 | 0.196495505 | 0.14886042 |
| Cenpo | 0.602420617 | 1.351843673 | 0.095 | 0.067 | 1 | 0.077438256 | -2.071814916 | 0 | 0.066 | 1 | 0.602420617 | 0.148879828 |
| Suv39h2 | 0.692869542 | 1.281454345 | 0.048 | 0.075 | 1 | 0.077439737 | -2.223818009 | 0 | 0.066 | 1 | 0.692869542 | 0.148882561 |
| Adra2a | 0.184817653 | -1.081115734 | 0 | 0.078 | 1 | 0.077440922 | -2.446210431 | 0 | 0.066 | 1 | 0.184817653 | 0.148884748 |
| Gm53610 | 0.479831566 | -0.20069735 | 0.048 | 0.097 | 1 | 0.077441219 | -2.017367132 | 0 | 0.066 | 1 | 0.479831566 | 0.148885295 |
| Itgb3 | 0.178637485 | 3.311201688 | 0.143 | 0.071 | 1 | 0.077452182 | -3.506751973 | 0 | 0.066 | 1 | 0.178637485 | 0.148905523 |
| Tnfrsf19 | 0.434472492 | -2.769171728 | 0.048 | 0.097 | 1 | 0.077452478 | -3.526380779 | 0 | 0.066 | 1 | 0.434472492 | 0.148906069 |
| Fabp7 | 0.106962556 | -1.970084422 | 0 | 0.112 | 1 | 0.077460478 | -2.404390255 | 0 | 0.066 | 1 | 0.106962556 | 0.14892083 |
| Bend7 | 0.39695844 | -0.413691074 | 0 | 0.034 | 1 | 0.077468626 | -2.526380779 | 0 | 0.066 | 1 | 0.39695844 | 0.148935864 |
| Mfap3l | 0.285582254 | -0.849790188 | 0 | 0.052 | 1 | 0.077470108 | -2.446210431 | 0 | 0.066 | 1 | 0.285582254 | 0.148938598 |
| Diaph3 | 0.442482843 | 2.524394144 | 0.095 | 0.056 | 1 | 0.077474404 | -2.871516266 | 0 | 0.066 | 1 | 0.442482843 | 0.148946525 |
| Rgs7bp | 0.915539771 | 2.700738815 | 0.048 | 0.056 | 1 | 0.077483885 | -3.098287167 | 0 | 0.066 | 1 | 0.915539771 | 0.148964018 |
| Recql | 0.237715905 | -0.818081329 | 0.048 | 0.138 | 1 | 0.077556531 | -0.539319835 | 0.044 | 0.142 | 1 | 0.237715905 | 0.149098046 |
| Lclat1 | 0.442108998 | 0.941967879 | 0.286 | 0.455 | 1 | 0.077567145 | -0.124282336 | 0.133 | 0.28 | 1 | 0.442108998 | 0.149117627 |
| Faxc | 0.215087908 | 2.179007076 | 0.143 | 0.075 | 1 | 0.077605573 | -1.80878051 | 0.022 | 0.104 | 1 | 0.215087908 | 0.149188521 |
| Usp20 | 0.97093274 | 0.586308926 | 0.095 | 0.101 | 1 | 0.077608121 | -1.901889915 | 0.022 | 0.104 | 1 | 0.97093274 | 0.149193222 |
| Bend3 | 0.911381619 | 0.304537958 | 0.095 | 0.086 | 1 | 0.077739682 | -0.776359032 | 0.044 | 0.145 | 1 | 0.911381619 | 0.149435906 |
| Oga | 0.07774968 | 0.465663572 | 0.857 | 0.787 | 1 | 0.913792413 | 0.11467393 | 0.933 | 0.886 | 1 | 0.913792413 | 0.149454348 |
| Blk | 0.578159907 | -0.326228232 | 0 | 0.015 | 1 | 0.077814953 | 4.448607333 | 0.044 | 0.01 | 1 | 0.578159907 | 0.149574739 |
| Casp12 | 0.077956711 | -1.413691074 | 0.048 | 0.213 | 1 | 0.087953887 | -0.300439291 | 0.156 | 0.301 | 1 | 0.087953887 | 0.149836174 |
| Col10a1 | 0.257936418 | -1.969283761 | 0.429 | 0.489 | 1 | 0.078028071 | -6.592586359 | 0.022 | 0.104 | 1 | 0.257936418 | 0.149967761 |
| Riiad1 | 0.096318818 | 2.304537958 | 0.19 | 0.09 | 1 | 0.078031747 | -0.181997834 | 0.044 | 0.142 | 1 | 0.096318818 | 0.149974541 |
| Vmp1 | 0.171260151 | -0.111128304 | 0.905 | 0.94 | 1 | 0.078040215 | 0.714642665 | 0.933 | 0.82 | 1 | 0.171260151 | 0.149990155 |
| Ipo8 | 0.339876698 | 0.61076197 | 0.19 | 0.325 | 1 | 0.078093904 | 0.124341627 | 0.289 | 0.46 | 1 | 0.339876698 | 0.15008915 |
| Cdc14b | 0.724599598 | 0.503846766 | 0.143 | 0.123 | 1 | 0.078115762 | 0.168499413 | 0.089 | 0.218 | 1 | 0.724599598 | 0.150129452 |
| Zfp946 | 0.173820587 | -1.233118828 | 0 | 0.082 | 1 | 0.078116048 | -1.638855509 | 0.022 | 0.104 | 1 | 0.173820587 | 0.15012998 |
| Elmod3 | 0.236753906 | -0.44170545 | 0.048 | 0.146 | 1 | 0.078119365 | -1.776359032 | 0.022 | 0.104 | 1 | 0.236753906 | 0.150136094 |
| Lpin2 | 0.309601234 | 1.002186912 | 0.667 | 0.75 | 1 | 0.078145258 | 0.858744586 | 0.956 | 0.855 | 1 | 0.309601234 | 0.150183834 |
| Spty2d1 | 0.349303562 | -0.226900305 | 0.619 | 0.694 | 1 | 0.078163816 | -0.464132339 | 0.778 | 0.754 | 1 | 0.349303562 | 0.15021805 |
| Cab39 | 0.59032252 | 0.347949757 | 0.857 | 0.847 | 1 | 0.078199183 | 0.557541704 | 1 | 0.948 | 1 | 0.59032252 | 0.150283254 |
| Zfp626 | 0.54961553 | 0.400753273 | 0.048 | 0.09 | 1 | 0.078239326 | -0.432404631 | 0.044 | 0.142 | 1 | 0.54961553 | 0.150357259 |
| Slc12a9 | 0.9034473 | 0.866416846 | 0.095 | 0.108 | 1 | 0.078348558 | -0.375821103 | 0.044 | 0.142 | 1 | 0.9034473 | 0.150558619 |
| Znhit2 | 0.078369064 | -1.611630451 | 0 | 0.131 | 1 | 0.244728837 | -0.404390255 | 0.044 | 0.1 | 1 | 0.244728837 | 0.150596419 |
| Tshz1 | 0.084623271 | -0.728326676 | 0.286 | 0.519 | 1 | 0.078406399 | 0.614468772 | 0.756 | 0.536 | 1 | 0.084623271 | 0.150665235 |
| Nup188 | 0.078419891 | 1.917042919 | 0.381 | 0.28 | 1 | 0.505233233 | 0.272788617 | 0.311 | 0.436 | 1 | 0.505233233 | 0.150690103 |
| Cdan1 | 0.451614009 | -0.233118828 | 0.048 | 0.097 | 1 | 0.07843757 | -0.346674757 | 0.044 | 0.142 | 1 | 0.451614009 | 0.150722687 |
| Rgma | 0.119860016 | -0.891012851 | 0.048 | 0.19 | 1 | 0.078444143 | -0.404390255 | 0.111 | 0.239 | 1 | 0.119860016 | 0.150734802 |
| Kmt5c | 0.078487401 | -1.683780237 | 0 | 0.131 | 1 | 0.880902002 | 0.975253338 | 0.133 | 0.135 | 1 | 0.880902002 | 0.150814529 |
| Aqp1 | 0.078540997 | -1.574155746 | 0.095 | 0.276 | 1 | 0.251649854 | -1.80878051 | 0.044 | 0.097 | 1 | 0.251649854 | 0.150913306 |
| Ret | 0.07854135 | -1.785659851 | 0 | 0.131 | 1 | 0.80624346 | 1.683072586 | 0.022 | 0.017 | 1 | 0.80624346 | 0.150913955 |
| Pla2g4a | 0.532423383 | 0.499139764 | 0.333 | 0.466 | 1 | 0.078573671 | 0.786577162 | 0.156 | 0.322 | 1 | 0.532423383 | 0.15097352 |
| Arl16 | 0.12676939 | -1.233118828 | 0.048 | 0.175 | 1 | 0.078636702 | -0.57996182 | 0.044 | 0.142 | 1 | 0.12676939 | 0.151089674 |
| Sh3bp5 | 0.078638808 | -2.133583154 | 0 | 0.131 | 1 | 0.362874876 | 0.859950348 | 0.067 | 0.121 | 1 | 0.362874876 | 0.151093554 |
| Gm13919 | 0.313004724 | 2.088809267 | 0.095 | 0.049 | 1 | 0.078708691 | -1.017367132 | 0.022 | 0.107 | 1 | 0.313004724 | 0.151222324 |
| Ndufaf1 | 0.684106853 | 1.247507013 | 0.143 | 0.127 | 1 | 0.078708691 | -1.017367132 | 0.022 | 0.107 | 1 | 0.684106853 | 0.151222324 |
| Acox3 | 0.078721534 | -2.326228232 | 0 | 0.131 | 1 | 0.256607397 | 0.193687745 | 0.089 | 0.163 | 1 | 0.256607397 | 0.151245988 |
| Ndp | 0.078793935 | -2.233118828 | 0 | 0.131 | 1 | 0.081280025 | -1.662702251 | 0.044 | 0.135 | 1 | 0.081280025 | 0.151379386 |
| Tyk2 | 0.153858613 | -1.413691074 | 0 | 0.09 | 1 | 0.078833297 | -0.286553765 | 0.044 | 0.142 | 1 | 0.153858613 | 0.151451905 |
| Rusf1 | 0.627437955 | 1.049280903 | 0.143 | 0.116 | 1 | 0.07887473 | -1.526380779 | 0.022 | 0.104 | 1 | 0.627437955 | 0.151528237 |
| Dock5 | 0.078920757 | -2.611630451 | 0 | 0.131 | 1 | 0.547615532 | 0.444667847 | 0.178 | 0.239 | 1 | 0.547615532 | 0.151613029 |
| Fnbp4 | 0.841076944 | 0.364334617 | 0.714 | 0.75 | 1 | 0.078977006 | 0.529182383 | 0.933 | 0.917 | 1 | 0.841076944 | 0.151716644 |
| Fabp3 | 0.079004883 | -3.81004401 | 0 | 0.131 | 1 | 0.30065951 | -0.776359032 | 0.044 | 0.09 | 1 | 0.30065951 | 0.151767994 |
| Cyp39a1 | 0.113688148 | -1.998653574 | 0 | 0.108 | 1 | 0.079023933 | -0.545746104 | 0.067 | 0.176 | 1 | 0.113688148 | 0.151803085 |

| | | | | | | | | | | | | |
|---|---|---|---|---|---|---|---|---|---|---|---|---|
| Wdr1 | 0.17281387 | -0.397545637 | 0.714 | 0.821 | 1 | 0.079148142 | -0.249302981 | 0.933 | 0.855 | 1 | 0.17281387 | 0.152031855 |
| Omp | 0.454217076 | 2.825774861 | 0.048 | 0.022 | 1 | 0.079149465 | 3.005000681 | 0.044 | 0.01 | 1 | 0.454217076 | 0.152034292 |
| Ccdc83 | 0.492607353 | 0.214340149 | 0 | 0.022 | 1 | 0.079149465 | 3.490427508 | 0.044 | 0.01 | 1 | 0.492607353 | 0.152034292 |
| Gm15893 | 0.532426384 | 0.866416846 | 0 | 0.019 | 1 | 0.079149465 | 2.683072586 | 0.044 | 0.01 | 1 | 0.532426384 | 0.152034292 |
| Gm11210 | 0.658161459 | 1.918884266 | 0.095 | 0.075 | 1 | 0.079227445 | -0.17942389 | 0.067 | 0.183 | 1 | 0.658161459 | 0.152177902 |
| Ndst1 | 0.079241873 | 1.12212934 | 0.619 | 0.653 | 1 | 0.080389833 | 0.904099465 | 0.778 | 0.779 | 1 | 0.080389833 | 0.152204471 |
| Tbck | 0.501669767 | 0.225311267 | 0.19 | 0.284 | 1 | 0.079297498 | 0.144652671 | 0.244 | 0.446 | 1 | 0.501669767 | 0.152306903 |
| Gm43113 | 0.532436372 | 1.088809267 | 0 | 0.019 | 1 | 0.079352256 | 1.683072586 | 0.067 | 0.021 | 1 | 0.532436372 | 0.152407731 |
| Smg7 | 0.079386692 | 0.866416846 | 0.619 | 0.593 | 1 | 0.353917594 | 0.308877182 | 0.911 | 0.737 | 1 | 0.353917594 | 0.152471136 |
| Gm16638 | 0.079430789 | 1.429846185 | 0.476 | 0.373 | 1 | 0.210043777 | -0.281428733 | 0.4 | 0.554 | 1 | 0.210043777 | 0.152552328 |
| A630072M18Rik | 0.228908726 | -0.413691074 | 0.048 | 0.149 | 1 | 0.079441656 | -0.72631835 | 0.089 | 0.204 | 1 | 0.228908726 | 0.152572335 |
| Gm26532 | 0.344407483 | -0.210308945 | 0.286 | 0.414 | 1 | 0.079531259 | -0.267714439 | 0.333 | 0.55 | 1 | 0.344407483 | 0.152737297 |
| Dclre1c | 0.079590133 | 2.460778044 | 0.238 | 0.119 | 1 | 0.120036976 | -1.446210431 | 0.022 | 0.09 | 1 | 0.120036976 | 0.152845677 |
| Kmo | 0.07959214 | 3.673771768 | 0.048 | 0.007 | 1 | 0.496223879 | 0.683072586 | 0 | 0.01 | 1 | 0.496223879 | 0.15284937 |
| Gpr158 | 0.07959214 | 3.673771768 | 0.048 | 0.007 | 1 | 0.08188835 | 2.098110085 | 0.044 | 0.01 | 1 | 0.08188835 | 0.15284937 |
| 1700064H15Rik | 0.07959214 | 3.673771768 | 0.048 | 0.007 | 1 | 0.496223879 | 0.683072586 | 0 | 0.01 | 1 | 0.496223879 | 0.15284937 |
| Coro2a | 0.07959214 | 3.936806174 | 0.048 | 0.007 | 1 | 0.49622258 | 0.361144491 | 0 | 0.01 | 1 | 0.49622258 | 0.15284937 |
| Lrp8os3 | 0.07959214 | 3.673771768 | 0.048 | 0.007 | 1 | 0.376842245 | -1.124282336 | 0 | 0.017 | 1 | 0.376842245 | 0.15284937 |
| Sema3a | 0.07959214 | 4.410737362 | 0.048 | 0.007 | 1 | 0.376842245 | -0.486852415 | 0 | 0.017 | 1 | 0.376842245 | 0.15284937 |
| Gm52829 | 0.07959214 | 4.410737362 | 0.048 | 0.007 | 1 | 0.658583583 | 1.946106992 | 0.022 | 0.014 | 1 | 0.658583583 | 0.15284937 |
| Gm43661 | 0.07959214 | 3.673771768 | 0.048 | 0.007 | 1 | 0.293647037 | -0.638855509 | 0 | 0.024 | 1 | 0.293647037 | 0.15284937 |
| Adap1 | 0.07959214 | 3.673771768 | 0.048 | 0.007 | 1 | 0.129721052 | 3.268035087 | 0.022 | 0.003 | 1 | 0.129721052 | 0.15284937 |
| Gm57292 | 0.07959214 | 4.673771768 | 0.048 | 0.007 | 1 | 0.112236835 | 2.490427508 | 0.067 | 0.024 | 1 | 0.112236835 | 0.15284937 |
| Tfpi2 | 0.07959214 | 3.673771768 | 0.048 | 0.007 | 1 | 0.496223879 | 0.683072586 | 0 | 0.01 | 1 | 0.496223879 | 0.15284937 |
| Il4i1 | 0.07959214 | 3.673771768 | 0.048 | 0.007 | 1 | 0.708994398 | 1.567595369 | 0.044 | 0.035 | 1 | 0.708994398 | 0.15284937 |
| Slc28a1 | 0.07959214 | 3.673771768 | 0.048 | 0.007 | 1 | 0.226024189 | 2.490427508 | 0.044 | 0.017 | 1 | 0.226024189 | 0.15284937 |
| Gm48796 | 0.07959214 | 3.673771768 | 0.048 | 0.007 | 1 | 0.496223879 | 0.683072586 | 0 | 0.01 | 1 | 0.496223879 | 0.15284937 |
| Mlip | 0.07959214 | 4.896164189 | 0.048 | 0.007 | 1 | 0.331907298 | -0.316927414 | 0 | 0.021 | 1 | 0.331907298 | 0.15284937 |
| Acpp | 0.07959214 | 3.673771768 | 0.048 | 0.007 | 1 | 0.17912725 | 0.545569062 | 0.089 | 0.042 | 1 | 0.17912725 | 0.15284937 |
| ENSMUSG00000121382 | 0.07959214 | 3.673771768 | 0.048 | 0.007 | 1 | 0.658587075 | 2.098110085 | 0.022 | 0.014 | 1 | 0.658587075 | 0.15284937 |
| Zmynd15 | 0.07959214 | 3.673771768 | 0.048 | 0.007 | 1 | 0.260684322 | -0.486852415 | 0 | 0.028 | 1 | 0.260684322 | 0.15284937 |
| Calm4 | 0.07959214 | 3.673771768 | 0.048 | 0.007 | 1 | 0.430486631 | -0.486852415 | 0 | 0.014 | 1 | 0.430486631 | 0.15284937 |
| Gm15327 | 0.07959214 | 3.673771768 | 0.048 | 0.007 | 1 | 0.496223879 | 0.683072586 | 0 | 0.01 | 1 | 0.496223879 | 0.15284937 |
| Gm45025 | 0.359208964 | 0.316219763 | 0.048 | 0.119 | 1 | 0.079592735 | -0.531940305 | 0.067 | 0.176 | 1 | 0.359208964 | 0.152850467 |
| Gm29707 | 0.079593353 | 4.258734268 | 0.048 | 0.007 | 1 | 0.129721052 | 3.268035087 | 0.022 | 0.003 | 1 | 0.129721052 | 0.152851604 |
| Rgs9 | 0.079593353 | 3.673771768 | 0.048 | 0.007 | 1 | 0.260653677 | -0.638855509 | 0 | 0.028 | 1 | 0.260653677 | 0.152851604 |
| Cfap47 | 0.079593353 | 3.673771768 | 0.048 | 0.007 | 1 | 0.506024628 | 1.098110085 | 0.022 | 0.01 | 1 | 0.506024628 | 0.152851604 |
| Ube2s | 0.07966715 | -1.017390137 | 0.095 | 0.291 | 1 | 0.653985599 | 0.683072586 | 0.378 | 0.405 | 1 | 0.653985599 | 0.152987445 |
| Pggt1b | 0.079703522 | -0.376854305 | 0.286 | 0.522 | 1 | 0.642361904 | 0.250899175 | 0.511 | 0.633 | 1 | 0.642361904 | 0.153054392 |
| Abce1 | 0.93354588 | 0.293999598 | 0.619 | 0.75 | 1 | 0.079805501 | -0.239172256 | 0.756 | 0.803 | 1 | 0.93354588 | 0.153242085 |
| Ephx4 | 0.457281049 | 0.503846766 | 0 | 0.026 | 1 | 0.079829167 | 2.683072586 | 0.044 | 0.01 | 1 | 0.457281049 | 0.153285639 |
| Cntfr | 0.079855508 | -1.091762979 | 0.095 | 0.284 | 1 | 0.197310678 | 0.318500154 | 0.133 | 0.228 | 1 | 0.197310678 | 0.153334114 |
| 2510002D24Rik | 0.114528621 | -1.107587946 | 0.095 | 0.254 | 1 | 0.079941627 | -0.651911662 | 0.133 | 0.263 | 1 | 0.114528621 | 0.153492591 |
| Abat | 0.307868314 | -0.20069735 | 0.143 | 0.261 | 1 | 0.080006669 | -0.268017813 | 0.133 | 0.273 | 1 | 0.307868314 | 0.153612271 |
| Pkn1 | 0.080093378 | -1.611630451 | 0.048 | 0.201 | 1 | 0.372940466 | 0.23638946 | 0.467 | 0.377 | 1 | 0.372940466 | 0.153771807 |
| Wrnip1 | 0.917868562 | 0.351843673 | 0.19 | 0.213 | 1 | 0.08016937 | -0.375821103 | 0.156 | 0.298 | 1 | 0.917868562 | 0.153911612 |
| Mrpl19 | 0.650633513 | -0.133583154 | 0.143 | 0.179 | 1 | 0.080216498 | -0.339295227 | 0.067 | 0.176 | 1 | 0.650633513 | 0.15399831 |
| Eno2 | 0.298673845 | -0.264827688 | 0.048 | 0.131 | 1 | 0.080235814 | -0.135089091 | 0.133 | 0.291 | 1 | 0.298673845 | 0.154033842 |
| Nmt1 | 0.136991606 | -0.160804623 | 0.286 | 0.515 | 1 | 0.080366688 | -0.279125381 | 0.333 | 0.484 | 1 | 0.136991606 | 0.154274571 |
| Zfp768 | 0.184739689 | -0.970084422 | 0 | 0.078 | 1 | 0.08041597 | -0.404390255 | 0.044 | 0.145 | 1 | 0.184739689 | 0.154365211 |
| Col7a1 | 0.3247899 | -0.574155746 | 0 | 0.045 | 1 | 0.080466083 | -0.454430938 | 0.022 | 0.107 | 1 | 0.3247899 | 0.154457375 |
| 4930447K03Rik | 0.359280535 | 3.451379346 | 0.048 | 0.019 | 1 | 0.080511939 | 1.460680165 | 0.044 | 0.01 | 1 | 0.359280535 | 0.154541705 |
| Sp2 | 0.333698685 | 1.165977127 | 0.333 | 0.291 | 1 | 0.080549753 | -0.208551253 | 0.178 | 0.332 | 1 | 0.333698685 | 0.154611243 |
| Spata5l1 | 0.304411431 | -0.413691074 | 0 | 0.049 | 1 | 0.080559135 | -0.124282336 | 0.022 | 0.107 | 1 | 0.304411431 | 0.154628496 |
| Zfp12 | 0.106776216 | -1.535681598 | 0 | 0.112 | 1 | 0.08060643 | -0.466674533 | 0.089 | 0.201 | 1 | 0.106776216 | 0.154715464 |
| Ccdc22 | 0.184693549 | -0.911190733 | 0 | 0.078 | 1 | 0.080614923 | -0.181997834 | 0.044 | 0.145 | 1 | 0.184693549 | 0.15473108 |
| Prps2 | 0.23638565 | -0.718545655 | 0 | 0.063 | 1 | 0.080664609 | -0.242926832 | 0.022 | 0.107 | 1 | 0.23638565 | 0.154822439 |
| Tagap1 | 0.963721266 | 1.171271427 | 0.095 | 0.108 | 1 | 0.080695267 | -0.352551324 | 0.022 | 0.107 | 1 | 0.963721266 | 0.154878808 |
| Ccdc102a | 0.120751051 | -1.683780237 | 0 | 0.104 | 1 | 0.080713009 | -0.23119754 | 0.111 | 0.239 | 1 | 0.120751051 | 0.154911429 |
| Dubr | 0.588263129 | 0.766881172 | 0.048 | 0.086 | 1 | 0.08078743 | -0.454430938 | 0.022 | 0.107 | 1 | 0.588263129 | 0.155048251 |
| Pkn3 | 0.080788462 | 2.773307441 | 0.286 | 0.179 | 1 | 0.711577542 | 0.776181991 | 0.067 | 0.087 | 1 | 0.711577542 | 0.155050148 |
| Kcmf1 | 0.08079614 | 0.480894316 | 0.905 | 0.944 | 1 | 0.704466426 | 0.243678721 | 1 | 0.972 | 1 | 0.704466426 | 0.155064264 |
| Zscan29 | 0.602342393 | 0.351843673 | 0.095 | 0.149 | 1 | 0.080802582 | -1.459885368 | 0.044 | 0.135 | 1 | 0.602342393 | 0.155076106 |
| Mlph | 0.440872629 | -0.666078235 | 0.143 | 0.209 | 1 | 0.080830559 | 1.635766871 | 0.022 | 0.107 | 1 | 0.440872629 | 0.155127538 |
| Washc4 | 0.767193655 | 0.32947586 | 0.476 | 0.575 | 1 | 0.080856309 | 0.534680746 | 0.756 | 0.716 | 1 | 0.767193655 | 0.155174876 |
| Pls3 | 0.86727629 | 0.416761149 | 0.571 | 0.593 | 1 | 0.080863037 | 0.542982814 | 0.8 | 0.664 | 1 | 0.86727629 | 0.155187244 |
| Spc24 | 0.080887687 | 1.719575457 | 0.143 | 0.052 | 1 | 0.936023974 | 1.875717664 | 0.022 | 0.021 | 1 | 0.936023974 | 0.155232555 |
| Sgk2 | 0.080927838 | 3.258734268 | 0.048 | 0.007 | 1 | 0.496223879 | 0.683072586 | 0 | 0.01 | 1 | 0.496223879 | 0.15530636 |
| Matk | 0.080927838 | 3.258734268 | 0.048 | 0.007 | 1 | 0.263501801 | -0.589945908 | 0.022 | 0.066 | 1 | 0.263501801 | 0.15530636 |
| Ccdc38 | 0.080927838 | 3.258734268 | 0.048 | 0.007 | 1 | 0.376837034 | -0.316927414 | 0 | 0.017 | 1 | 0.376837034 | 0.15530636 |
| Cdc20b | 0.080927838 | 3.258734268 | 0.048 | 0.007 | 1 | 0.232018094 | -0.638855509 | 0 | 0.031 | 1 | 0.232018094 | 0.15530636 |
| Isl1 | 0.080927838 | 3.258734268 | 0.048 | 0.007 | 1 | 0.238422127 | 1.223640968 | 0.067 | 0.031 | 1 | 0.238422127 | 0.15530636 |
| Cbs | 0.080927838 | 3.258734268 | 0.048 | 0.007 | 1 | 0.331907298 | -0.316927414 | 0 | 0.021 | 1 | 0.331907298 | 0.15530636 |
| Gm10138 | 0.08100846 | 1.756233928 | 0.286 | 0.157 | 1 | 0.820572505 | 1.765534746 | 0.133 | 0.135 | 1 | 0.820572505 | 0.155454549 |
| Chfr | 0.367813667 | 0.512308345 | 0.238 | 0.418 | 1 | 0.081055276 | 0.83237653 | 0.622 | 0.574 | 1 | 0.367813667 | 0.155540594 |
| Cenph | 0.285590043 | -1.02666795 | 0 | 0.052 | 1 | 0.081196116 | 2.197645759 | 0.044 | 0.01 | 1 | 0.285590043 | 0.155799423 |
| Snapc3 | 0.081208375 | 1.261958169 | 0.476 | 0.325 | 1 | 0.527503592 | 0.683072586 | 0.333 | 0.512 | 1 | 0.527503592 | 0.155821949 |
| Celf5 | 0.344880504 | 0.957564734 | 0.19 | 0.123 | 1 | 0.081281991 | -1.539319835 | 0.044 | 0.135 | 1 | 0.344880504 | 0.155957219 |
| Aen | 0.31289444 | 0.696491844 | 0.381 | 0.328 | 1 | 0.081394277 | -0.288168421 | 0.156 | 0.311 | 1 | 0.31289444 | 0.156163526 |
| Ccdc157 | 0.081401807 | -0.581728965 | 0.238 | 0.47 | 1 | 0.46466147 | 0.240850258 | 0.356 | 0.484 | 1 | 0.46466147 | 0.15617736 |
| Gm17276 | 0.532426384 | -0.133583154 | 0 | 0.019 | 1 | 0.08143648 | 2.576157382 | 0.089 | 0.035 | 1 | 0.532426384 | 0.156241059 |
| Slc26a11 | 0.081774238 | -1.455511249 | 0.048 | 0.201 | 1 | 0.796694917 | 0.720874619 | 0.222 | 0.287 | 1 | 0.796694917 | 0.156861451 |
| Nfkb2 | 0.475674363 | 0.619323984 | 0.333 | 0.291 | 1 | 0.081848849 | -0.17706975 | 0.2 | 0.363 | 1 | 0.475674363 | 0.156998463 |
| Rusc2 | 0.996398192 | 0.202980287 | 0.381 | 0.444 | 1 | 0.082004454 | -0.343727473 | 0.333 | 0.522 | 1 | 0.996398192 | 0.157284177 |
| Gm29128 | 0.082280431 | 3.088809267 | 0.048 | 0.007 | 1 | 0.496223879 | 0.683072586 | 0 | 0.01 | 1 | 0.496223879 | 0.157790792 |
| Gm19497 | 0.082280431 | 3.088809267 | 0.048 | 0.007 | 1 | 0.430487676 | 0.361144491 | 0 | 0.014 | 1 | 0.430487676 | 0.157790792 |
| Fam83c | 0.082280431 | 3.088809267 | 0.048 | 0.007 | 1 | 0.492040936 | 2.683072586 | 0.022 | 0.01 | 1 | 0.492040936 | 0.157790792 |
| Gm38505 | 0.082280431 | 3.088809267 | 0.048 | 0.007 | 1 | 0.227305413 | 2.460680165 | 0.044 | 0.017 | 1 | 0.227305413 | 0.157790792 |
| Fam184b | 0.082280431 | 2.936806174 | 0.048 | 0.007 | 1 | 0.496223879 | 0.683072586 | 0 | 0.01 | 1 | 0.496223879 | 0.157790792 |
| Epha1 | 0.082280431 | 3.088809267 | 0.048 | 0.007 | 1 | 0.687197015 | 1.545569062 | 0.022 | 0.035 | 1 | 0.687197015 | 0.157790792 |
| Camsap3 | 0.082280431 | 3.088809267 | 0.048 | 0.007 | 1 | 0.610038162 | 2.168499413 | 0.044 | 0.031 | 1 | 0.610038162 | 0.157790792 |
| Palm3 | 0.082280431 | 3.088809267 | 0.048 | 0.007 | 1 | 0.664308814 | 1.460680165 | 0.022 | 0.014 | 1 | 0.664308814 | 0.157790792 |
| A230103J11Rik | 0.082280431 | 3.088809267 | 0.048 | 0.007 | 1 | 0.430487676 | 0.361144491 | 0 | 0.014 | 1 | 0.430487676 | 0.157790792 |
| Reep6 | 0.082280431 | 3.088809267 | 0.048 | 0.007 | 1 | 0.496223879 | 0.683072586 | 0 | 0.01 | 1 | 0.496223879 | 0.157790792 |
| Mmp19 | 0.082280431 | 3.088809267 | 0.048 | 0.007 | 1 | 0.430487676 | 0.361144491 | 0 | 0.014 | 1 | 0.430487676 | 0.157790792 |
| Gm12059 | 0.082280431 | 2.936806174 | 0.048 | 0.007 | 1 | 0.430487676 | 0.361144491 | 0 | 0.014 | 1 | 0.430487676 | 0.157790792 |
| Gm12185 | 0.082280431 | 2.503846766 | 0.048 | 0.007 | 1 | 0.309826166 | 3.005000681 | 0.022 | 0.007 | 1 | 0.309826166 | 0.157790792 |
| Gm16045 | 0.082280431 | 3.088809267 | 0.048 | 0.007 | 1 | 0.936023974 | 1.460680165 | 0.022 | 0.021 | 1 | 0.936023974 | 0.157790792 |

| | | | | | | | | | | | | |
|---|---|---|---|---|---|---|---|---|---|---|---|---|
| Zfp750 | 0.082280431 | 3.088809267 | 0.048 | 0.007 | 1 | 0.430487676 | 0.361144491 | 0 | 0.014 | 1 | 0.430487676 | 0.157790792 |
| Scel | 0.082280431 | 2.936806174 | 0.048 | 0.007 | 1 | 0.496223879 | 0.683072586 | 0 | 0.01 | 1 | 0.496223879 | 0.157790792 |
| Amigo2 | 0.082280431 | 3.088809267 | 0.048 | 0.007 | 1 | 0.331907298 | -0.316927414 | 0 | 0.021 | 1 | 0.331907298 | 0.157790792 |
| Gm42477 | 0.082280431 | 3.088809267 | 0.048 | 0.007 | 1 | 0.331907298 | -0.776359032 | 0 | 0.021 | 1 | 0.331907298 | 0.157790792 |
| Muc13 | 0.082280431 | 3.088809267 | 0.048 | 0.007 | 1 | 0.496223879 | 0.683072586 | 0 | 0.01 | 1 | 0.496223879 | 0.157790792 |
| Egfl8 | 0.082280431 | 3.088809267 | 0.048 | 0.007 | 1 | 0.121443112 | 1.683072586 | 0.067 | 0.024 | 1 | 0.121443112 | 0.157790792 |
| Zfp64 | 0.220566813 | 0.805875304 | 0.476 | 0.41 | 1 | 0.082286949 | 0.201945896 | 0.267 | 0.453 | 1 | 0.220566813 | 0.157802757 |
| Mcoln2 | 0.496085409 | 1.109739925 | 0.381 | 0.586 | 1 | 0.082293945 | -0.138266565 | 0.244 | 0.443 | 1 | 0.496085409 | 0.157815597 |
| Tnfaip8 | 0.082328871 | -0.805481292 | 0.476 | 0.627 | 1 | 0.999330302 | 0.305139082 | 0.8 | 0.772 | 1 | 0.999330302 | 0.1578797 |
| Cep126 | 0.08233268 | -0.513855235 | 0.143 | 0.347 | 1 | 0.691964144 | 0.286182434 | 0.2 | 0.253 | 1 | 0.691964144 | 0.157886689 |
| Rab10os | 0.15520904 | -0.683780237 | 0.19 | 0.343 | 1 | 0.082339989 | -0.736466305 | 0.156 | 0.28 | 1 | 0.15520904 | 0.157900104 |
| Wdr19 | 0.748694478 | 1.351843673 | 0.095 | 0.082 | 1 | 0.082421841 | -1.098287127 | 0.044 | 0.138 | 1 | 0.748694478 | 0.158050322 |
| Pygm | 0.821731677 | 1.088809267 | 0.095 | 0.086 | 1 | 0.082443047 | -0.86124793 | 0.044 | 0.142 | 1 | 0.821731677 | 0.158089237 |
| Agfg2 | 0.082454157 | -1.196592952 | 0.095 | 0.272 | 1 | 0.088652272 | -0.515707278 | 0.089 | 0.204 | 1 | 0.088652272 | 0.158109626 |
| Myo9a | 0.08246371 | 1.038904361 | 0.81 | 0.843 | 1 | 0.315015376 | 0.368833757 | 0.978 | 0.927 | 1 | 0.315015376 | 0.158127157 |
| Cenpc1 | 0.082554319 | -0.866796614 | 0.19 | 0.422 | 1 | 0.244193757 | 0.361144491 | 0.667 | 0.543 | 1 | 0.244193757 | 0.158293422 |
| Mtx1 | 0.083573519 | -0.496153234 | 0.19 | 0.422 | 1 | 0.082631069 | 0.135009668 | 0.244 | 0.446 | 1 | 0.083573519 | 0.158434245 |
| Dgcr8 | 0.469944905 | 1.117378419 | 0.286 | 0.257 | 1 | 0.082636392 | -0.285218554 | 0.222 | 0.394 | 1 | 0.469944905 | 0.158444011 |
| Rnf170 | 0.187159565 | -0.66251162 | 0.143 | 0.28 | 1 | 0.082653467 | -0.589945908 | 0.133 | 0.26 | 1 | 0.187159565 | 0.158475338 |
| Adck1 | 0.588737763 | 0.119182916 | 0.095 | 0.142 | 1 | 0.082699094 | -1.346674757 | 0.044 | 0.135 | 1 | 0.588737763 | 0.158559048 |
| Map2k3os | 0.63712708 | 0.634243403 | 0.143 | 0.201 | 1 | 0.082731829 | 0.65559185 | 0.044 | 0.145 | 1 | 0.63712708 | 0.158619102 |
| Oasl2 | 0.082799564 | 1.586308926 | 0.095 | 0.026 | 1 | 0.854469184 | 1.928185084 | 0.022 | 0.028 | 1 | 0.854469184 | 0.15874336 |
| Zfp606 | 0.890671317 | 0.400753273 | 0.143 | 0.164 | 1 | 0.082839642 | -0.80878051 | 0.089 | 0.204 | 1 | 0.890671317 | 0.158816878 |
| Zgpat | 0.379945202 | -0.233118828 | 0.095 | 0.175 | 1 | 0.082861382 | 0.320502507 | 0.133 | 0.287 | 1 | 0.379945202 | 0.158856755 |
| Baz2a | 0.307548643 | 0.335902129 | 0.524 | 0.437 | 1 | 0.082916952 | 0.463258709 | 0.889 | 0.734 | 1 | 0.307548643 | 0.158958683 |
| Cep162 | 0.386857108 | -0.189726232 | 0.143 | 0.231 | 1 | 0.08293058 | -0.835656216 | 0.133 | 0.256 | 1 | 0.386857108 | 0.158983679 |
| Ptgs2 | 0.780604314 | 0.255919253 | 0.333 | 0.313 | 1 | 0.083016783 | -3.751555641 | 0.022 | 0.1 | 1 | 0.780604314 | 0.15914178 |
| Cage1 | 0.083119596 | 2.258734268 | 0.143 | 0.052 | 1 | 0.106819427 | -1.638855509 | 0 | 0.055 | 1 | 0.106819427 | 0.159330325 |
| Zfp444 | 0.083306174 | -1.496153234 | 0 | 0.127 | 1 | 0.164294518 | -0.124282336 | 0.133 | 0.239 | 1 | 0.164294518 | 0.159672429 |
| Pnpla6 | 0.181550846 | -1.159118246 | 0.048 | 0.153 | 1 | 0.083339691 | -0.142898014 | 0.089 | 0.211 | 1 | 0.181550846 | 0.159733878 |
| Rbbp8 | 0.083427521 | 1.979874895 | 0.571 | 0.534 | 1 | 0.097605781 | 0.258406386 | 0.822 | 0.564 | 1 | 0.097605781 | 0.159894891 |
| Nprl2 | 0.083432172 | -1.648156327 | 0 | 0.127 | 1 | 0.126330288 | -1.053893008 | 0.044 | 0.121 | 1 | 0.126330288 | 0.159903417 |
| Gm17251 | 0.083432172 | -1.574155746 | 0 | 0.127 | 1 | 0.107749217 | -1.602329633 | 0.022 | 0.093 | 1 | 0.107749217 | 0.159903417 |
| Kat6b | 0.194073586 | 0.841122825 | 0.667 | 0.743 | 1 | 0.083433852 | 0.65769374 | 0.978 | 0.896 | 1 | 0.194073586 | 0.159906497 |
| Ddit4 | 0.083488535 | -2.04504648 | 0.048 | 0.205 | 1 | 0.80456159 | -1.064161343 | 0.289 | 0.27 | 1 | 0.80456159 | 0.160006734 |
| ENSMUSG00000121487 | 0.083488896 | -1.648156327 | 0 | 0.127 | 1 | 0.331922009 | -0.124282336 | 0 | 0.021 | 1 | 0.331922009 | 0.160007396 |
| Cryl1 | 0.922237805 | 0.673771768 | 0.429 | 0.567 | 1 | 0.083538416 | 0.64654671 | 0.733 | 0.561 | 1 | 0.922237805 | 0.160098165 |
| Prelid3a | 0.08354242 | -1.818081329 | 0 | 0.127 | 1 | 0.643783629 | 0.390290837 | 0.089 | 0.118 | 1 | 0.643783629 | 0.160105505 |
| Prss36 | 0.083542913 | -1.648156327 | 0 | 0.127 | 1 | 0.327026632 | -0.819427754 | 0.044 | 0.087 | 1 | 0.327026632 | 0.160106407 |
| Cyp4f16 | 0.52641075 | 0.267779408 | 0.095 | 0.16 | 1 | 0.083571466 | -0.174908409 | 0.067 | 0.176 | 1 | 0.52641075 | 0.160158742 |
| Zcwpw2 | 0.083609233 | 1.171271427 | 0.19 | 0.078 | 1 | 0.56906854 | 0.531069493 | 0.044 | 0.069 | 1 | 0.56906854 | 0.160227962 |
| Atg16l2 | 0.91105258 | -0.112824594 | 0.429 | 0.437 | 1 | 0.083645509 | 0.105643758 | 0.378 | 0.609 | 1 | 0.91105258 | 0.160294447 |
| Eef1akmt4 | 0.468322398 | -0.233118828 | 0.095 | 0.153 | 1 | 0.083645515 | -1.223818009 | 0.044 | 0.135 | 1 | 0.468322398 | 0.160294458 |
| Rgsl1 | 0.08365005 | 2.351843673 | 0.048 | 0.007 | 1 | 0.496223879 | 0.683072586 | 0 | 0.01 | 1 | 0.496223879 | 0.16030277 |
| Nectin4 | 0.08365005 | 2.673771768 | 0.048 | 0.007 | 1 | 0.430486631 | -0.124282336 | 0 | 0.014 | 1 | 0.430486631 | 0.16030277 |
| Dennd2d | 0.08365005 | 2.673771768 | 0.048 | 0.007 | 1 | 0.309824707 | 2.683072586 | 0.022 | 0.007 | 1 | 0.309824707 | 0.16030277 |
| Prom1 | 0.08365005 | 2.351843673 | 0.048 | 0.007 | 1 | 0.096013784 | 0.683072586 | 0.111 | 0.048 | 1 | 0.096013784 | 0.16030277 |
| Gm13844 | 0.08365005 | 2.673771768 | 0.048 | 0.007 | 1 | 0.935071939 | 0.513147585 | 0.022 | 0.024 | 1 | 0.935071939 | 0.16030277 |
| Mypopos | 0.08365005 | 2.673771768 | 0.048 | 0.007 | 1 | 0.430487676 | 0.361144491 | 0 | 0.014 | 1 | 0.430487676 | 0.16030277 |
| 4930488L21Rik | 0.08365005 | 2.673771768 | 0.048 | 0.007 | 1 | 0.496223879 | -0.486852415 | 0 | 0.01 | 1 | 0.496223879 | 0.16030277 |
| Ly6c2 | 0.08365005 | 2.673771768 | 0.048 | 0.007 | 1 | 0.809040217 | 1.460680165 | 0.022 | 0.017 | 1 | 0.809040217 | 0.16030277 |
| Hgd | 0.08365005 | 2.673771768 | 0.048 | 0.007 | 1 | 0.376837034 | -0.124282336 | 0 | 0.017 | 1 | 0.376837034 | 0.16030277 |
| Ostm1 | 0.083680846 | -1.911190733 | 0 | 0.127 | 1 | 0.097206154 | -0.627714951 | 0.111 | 0.232 | 1 | 0.097206154 | 0.160359208 |
| Cacna1g | 0.350763915 | -0.599246727 | 0.095 | 0.183 | 1 | 0.083812774 | -0.594912161 | 0.067 | 0.173 | 1 | 0.350763915 | 0.160600966 |
| Tbx2 | 0.083832682 | -2.280424543 | 0 | 0.127 | 1 | 0.095978199 | -2.674479418 | 0 | 0.059 | 1 | 0.095978199 | 0.160637446 |
| 2700038G22Rik | 0.341550816 | 1.088809267 | 0.238 | 0.172 | 1 | 0.083863698 | 0.446033389 | 0.067 | 0.18 | 1 | 0.341550816 | 0.160694276 |
| A930007I19Rik | 0.083909404 | 2.761234609 | 0.19 | 0.09 | 1 | 0.181254321 | -1.446210431 | 0.022 | 0.076 | 1 | 0.181254321 | 0.160778019 |
| Dusp15 | 0.083945514 | -2.434752689 | 0 | 0.127 | 1 | 0.206909388 | -1.404390255 | 0 | 0.035 | 1 | 0.206909388 | 0.160844178 |
| H2-Aa | 0.083983042 | -4.067695219 | 0 | 0.127 | 1 | 0.184823348 | -1.316927414 | 0 | 0.038 | 1 | 0.184823348 | 0.160912932 |
| Smpd3 | 0.084034105 | -4.76085646 | 0 | 0.127 | 1 | 0.170704325 | -3.331877755 | 0.022 | 0.08 | 1 | 0.170704325 | 0.161006479 |
| Dbnl | 0.084041846 | -0.60421298 | 0.095 | 0.28 | 1 | 0.095426126 | -0.326387743 | 0.244 | 0.37 | 1 | 0.095426126 | 0.161020659 |
| Exoc3 | 0.084043 | -0.330143282 | 0.429 | 0.66 | 1 | 0.949725884 | 0.109469948 | 0.756 | 0.758 | 1 | 0.949725884 | 0.161022774 |
| Lcat | 0.400493027 | 1.526930379 | 0.143 | 0.093 | 1 | 0.084137149 | -1.017367132 | 0.044 | 0.138 | 1 | 0.400493027 | 0.161195238 |
| Ctns | 0.084166526 | -1.02666795 | 0.048 | 0.213 | 1 | 0.122447356 | -0.529921137 | 0.156 | 0.266 | 1 | 0.122447356 | 0.161249049 |
| Rerg | 0.994383901 | 0.710297644 | 0.238 | 0.261 | 1 | 0.084256661 | 1.161626536 | 0.489 | 0.436 | 1 | 0.994383901 | 0.161414138 |
| Gnpda2 | 0.150849549 | -0.361852142 | 0.095 | 0.243 | 1 | 0.084333105 | -0.333229226 | 0.111 | 0.239 | 1 | 0.150849549 | 0.161554137 |
| Mtfr2 | 0.437977203 | 1.908237021 | 0.143 | 0.097 | 1 | 0.084422507 | -0.709244837 | 0.044 | 0.138 | 1 | 0.437977203 | 0.161717853 |
| Brd7 | 0.270557124 | 0.700538677 | 0.571 | 0.563 | 1 | 0.084432102 | 0.255534303 | 0.867 | 0.657 | 1 | 0.270557124 | 0.161735424 |
| Cldn11 | 0.084468298 | -1.953964757 | 0.143 | 0.321 | 1 | 0.346649279 | -1.765387915 | 0.044 | 0.087 | 1 | 0.346649279 | 0.161801703 |
| Ranbp2 | 0.084527038 | 0.529302347 | 0.905 | 0.94 | 1 | 0.705479777 | 0.164953835 | 1 | 0.965 | 1 | 0.705479777 | 0.161909256 |
| Cmtm8 | 0.341696746 | -0.752492987 | 0.095 | 0.175 | 1 | 0.084544315 | -0.529921137 | 0.4 | 0.498 | 1 | 0.341696746 | 0.161940889 |
| Zxdb | 0.738393266 | 1.351843673 | 0.048 | 0.034 | 1 | 0.084587371 | -1.158229668 | 0.044 | 0.135 | 1 | 0.738393266 | 0.162019719 |
| Zscan26 | 0.091802933 | -0.66251162 | 0.238 | 0.455 | 1 | 0.08461693 | 0.132875504 | 0.289 | 0.533 | 1 | 0.091802933 | 0.162073835 |
| Flnc | 0.615132382 | 0.673771768 | 0.048 | 0.082 | 1 | 0.084777021 | -1.589945908 | 0.022 | 0.104 | 1 | 0.615132382 | 0.162366899 |
| Arhgap22 | 0.787207817 | 2.171271427 | 0.048 | 0.037 | 1 | 0.084892013 | -1.339295227 | 0.022 | 0.104 | 1 | 0.787207817 | 0.162577372 |
| Pde11a | 0.084962999 | 0.586308926 | 0.048 | 0.216 | 1 | 0.761401784 | 0.446033389 | 0.111 | 0.1 | 1 | 0.761401784 | 0.162707286 |
| Cert1 | 0.085183455 | 0.656698254 | 0.857 | 0.679 | 1 | 0.315872334 | 0.336171963 | 0.956 | 0.761 | 1 | 0.315872334 | 0.16311069 |
| Abca8b | 0.198157623 | -0.911190733 | 0.095 | 0.216 | 1 | 0.085195827 | -0.662702251 | 0.044 | 0.142 | 1 | 0.198157623 | 0.163133324 |
| Agt | 0.08530153 | -1.785659851 | 0.048 | 0.198 | 1 | 0.331922009 | -0.124282336 | 0 | 0.021 | 1 | 0.331922009 | 0.163326709 |
| Heyl | 0.943760954 | 0.451379346 | 0.048 | 0.052 | 1 | 0.08534846 | -2.339295227 | 0.022 | 0.1 | 1 | 0.943760954 | 0.16341256 |
| Fxr1 | 0.085415341 | 0.305085574 | 0.952 | 0.94 | 1 | 0.260987041 | 0.282627283 | 1 | 0.976 | 1 | 0.260987041 | 0.163534902 |
| Slc35e2 | 0.085484671 | 1.351843673 | 0.381 | 0.257 | 1 | 0.103764536 | 0.653325243 | 0.133 | 0.266 | 1 | 0.103764536 | 0.163661713 |
| Ncapg2 | 0.646768449 | 1.252307999 | 0.143 | 0.116 | 1 | 0.085497366 | 2.177837278 | 0.178 | 0.104 | 1 | 0.646768449 | 0.163684932 |
| Akr1b3 | 0.085595176 | -0.184209227 | 0.19 | 0.403 | 1 | 0.267004433 | 0.106425354 | 0.489 | 0.367 | 1 | 0.267004433 | 0.163863817 |
| Nt5m | 0.160361892 | -0.718545655 | 0.095 | 0.235 | 1 | 0.085688215 | -0.242926832 | 0.089 | 0.204 | 1 | 0.160361892 | 0.16403396 |
| Gm11973 | 0.400197582 | -0.159118246 | 0.095 | 0.172 | 1 | 0.085709513 | -0.305612101 | 0.133 | 0.273 | 1 | 0.400197582 | 0.164072905 |
| Zfp507 | 0.72984258 | 0.936806174 | 0.333 | 0.336 | 1 | 0.085712216 | -0.271123724 | 0.178 | 0.325 | 1 | 0.72984258 | 0.164077849 |
| Psme3ip1 | 0.085907838 | -0.75727805 | 0.19 | 0.381 | 1 | 0.186145832 | 0.123201066 | 0.289 | 0.443 | 1 | 0.186145832 | 0.164435519 |
| Fam222b | 0.086018711 | 1.056798872 | 0.571 | 0.593 | 1 | 0.157778966 | 0.852634051 | 0.733 | 0.647 | 1 | 0.157778966 | 0.164638203 |
| Irgm2 | 0.371977114 | -0.695462042 | 0.095 | 0.183 | 1 | 0.086033349 | -1.158229668 | 0.067 | 0.166 | 1 | 0.371977114 | 0.164664961 |
| Kbtbd7 | 0.834360754 | 0.629377648 | 0.095 | 0.116 | 1 | 0.086045891 | -0.404390255 | 0.044 | 0.138 | 1 | 0.834360754 | 0.164687887 |
| Gm29157 | 0.12053426 | -1.326228232 | 0 | 0.104 | 1 | 0.086101733 | -1.638855509 | 0 | 0.062 | 1 | 0.12053426 | 0.164789958 |
| Poli | 0.196521095 | -1.133583154 | 0 | 0.075 | 1 | 0.086101733 | -1.638855509 | 0 | 0.062 | 1 | 0.196521095 | 0.164789958 |
| 4933433G19Rik | 0.492608234 | 0.866416846 | 0 | 0.022 | 1 | 0.086101733 | -1.638855509 | 0 | 0.062 | 1 | 0.492608234 | 0.164789958 |
| Ddx28 | 0.644626546 | 0.150209812 | 0.048 | 0.075 | 1 | 0.086101733 | -1.638855509 | 0 | 0.062 | 1 | 0.644626546 | 0.164789958 |
| Zc2hc1c | 0.655561847 | 1.503846766 | 0.048 | 0.03 | 1 | 0.086101733 | -1.638855509 | 0 | 0.062 | 1 | 0.655561847 | 0.164789958 |
| Slc2a12 | 0.841369044 | 1.162809848 | 0.143 | 0.175 | 1 | 0.086101733 | -1.638855509 | 0 | 0.062 | 1 | 0.841369044 | 0.164789958 |
| Slx1b | 0.860101956 | 1.088809267 | 0.048 | 0.06 | 1 | 0.086101733 | -1.638855509 | 0 | 0.062 | 1 | 0.860101956 | 0.164789958 |

| | | | | | | | | | | | | |
|---|---|---|---|---|---|---|---|---|---|---|---|---|
| Mis18a | 0.100391649 | -1.535681598 | 0 | 0.116 | 1 | 0.086124205 | -1.709244837 | 0 | 0.062 | 1 | 0.100391649 | 0.164831031 |
| Gm56915 | 0.184642276 | -0.849790188 | 0 | 0.078 | 1 | 0.086124205 | -1.709244837 | 0 | 0.062 | 1 | 0.184642276 | 0.164831031 |
| Xrcc3 | 0.376794341 | 1.866416846 | 0.048 | 0.019 | 1 | 0.086124205 | -1.84048937 | 0 | 0.062 | 1 | 0.376794341 | 0.164831031 |
| Garin1a | 0.544919419 | 0.673771768 | 0.095 | 0.06 | 1 | 0.086124205 | -1.901889915 | 0 | 0.062 | 1 | 0.544919419 | 0.164831031 |
| Rtel1 | 0.557078677 | 1.150209812 | 0.095 | 0.063 | 1 | 0.086124205 | -1.776359032 | 0 | 0.062 | 1 | 0.557078677 | 0.164831031 |
| Ace | 0.573468511 | 1.214340149 | 0.048 | 0.026 | 1 | 0.086124205 | -1.709244837 | 0 | 0.062 | 1 | 0.573468511 | 0.164831031 |
| Acsf3 | 0.924795272 | 1.351843673 | 0.048 | 0.045 | 1 | 0.086124205 | -1.709244837 | 0 | 0.062 | 1 | 0.924795272 | 0.164831031 |
| A130051J06Rik | 0.153858613 | -1.326228232 | 0 | 0.09 | 1 | 0.086144033 | -1.776359032 | 0 | 0.062 | 1 | 0.153858613 | 0.164867271 |
| Morn4 | 0.196460997 | -1.02666795 | 0 | 0.075 | 1 | 0.086144033 | -1.776359032 | 0 | 0.062 | 1 | 0.196460997 | 0.164867271 |
| Rfx5 | 0.209094725 | -0.648156327 | 0 | 0.071 | 1 | 0.086144033 | -1.776359032 | 0 | 0.062 | 1 | 0.209094725 | 0.164867271 |
| Lzts3 | 0.324739547 | -0.326228232 | 0 | 0.045 | 1 | 0.086144033 | -1.776359032 | 0 | 0.062 | 1 | 0.324739547 | 0.164867271 |
| Arrdc2 | 0.332449589 | -0.785659851 | 0.048 | 0.116 | 1 | 0.086144033 | -2.174908409 | 0 | 0.062 | 1 | 0.332449589 | 0.164867271 |
| Tmem121 | 0.860084871 | 1.010806755 | 0.048 | 0.06 | 1 | 0.086144033 | -1.776359032 | 0 | 0.062 | 1 | 0.860084871 | 0.164867271 |
| Gba2 | 0.960849104 | 0.673771768 | 0.048 | 0.045 | 1 | 0.086144033 | -1.776359032 | 0 | 0.062 | 1 | 0.960849104 | 0.164867271 |
| Tmem220 | 0.982550637 | 0.673771768 | 0.048 | 0.049 | 1 | 0.086144033 | -1.776359032 | 0 | 0.062 | 1 | 0.982550637 | 0.164867271 |
| Bbof1 | 0.959951046 | 0.673771768 | 0.095 | 0.097 | 1 | 0.086160721 | -1.901889915 | 0 | 0.062 | 1 | 0.959951046 | 0.164897772 |
| Tnn | 0.109791647 | 1.010806755 | 0.143 | 0.056 | 1 | 0.086161217 | -2.017367132 | 0 | 0.062 | 1 | 0.109791647 | 0.164898678 |
| Dyrk1b | 0.285499161 | -0.648156327 | 0 | 0.052 | 1 | 0.086161382 | -1.84048937 | 0 | 0.062 | 1 | 0.285499161 | 0.16489898 |
| Gm56688 | 0.267985206 | -0.574155746 | 0 | 0.056 | 1 | 0.086174765 | -1.960783604 | 0 | 0.062 | 1 | 0.267985206 | 0.164923441 |
| Pik3c2g | 0.106939188 | -2.02666795 | 0 | 0.112 | 1 | 0.086196906 | -2.223818009 | 0 | 0.062 | 1 | 0.106939188 | 0.164963906 |
| Gm46440 | 0.65609845 | 2.451379346 | 0.048 | 0.078 | 1 | 0.086205994 | -2.316927414 | 0 | 0.062 | 1 | 0.65609845 | 0.164980514 |
| Pde1a | 0.216061841 | -0.273760812 | 0.048 | 0.153 | 1 | 0.086215908 | -3.316927414 | 0 | 0.062 | 1 | 0.216061841 | 0.164998632 |
| Cenpw | 0.153937349 | -1.535681598 | 0 | 0.09 | 1 | 0.086236561 | -1.564854927 | 0 | 0.062 | 1 | 0.153937349 | 0.165036378 |
| Hddc3 | 0.285622926 | -0.233118828 | 0 | 0.052 | 1 | 0.086236561 | -1.564854927 | 0 | 0.062 | 1 | 0.285622926 | 0.165036378 |
| Wdr5b | 0.3247899 | -0.326228232 | 0 | 0.045 | 1 | 0.086236561 | -1.564854927 | 0 | 0.062 | 1 | 0.3247899 | 0.165036378 |
| Kifc2 | 0.425639612 | 0.503846766 | 0 | 0.03 | 1 | 0.086236561 | -1.564854927 | 0 | 0.062 | 1 | 0.425639612 | 0.165036378 |
| Eps15 | 0.551808887 | 0.290443128 | 0.333 | 0.478 | 1 | 0.086243762 | 0.638678467 | 0.844 | 0.675 | 1 | 0.551808887 | 0.165049537 |
| Prpf4 | 0.194558353 | -0.44170545 | 0.143 | 0.28 | 1 | 0.086333881 | 0.313838777 | 0.156 | 0.315 | 1 | 0.194558353 | 0.165214224 |
| Rab5a | 0.216037967 | -0.253226401 | 0.81 | 0.933 | 1 | 0.086386844 | -0.222149189 | 0.933 | 0.976 | 1 | 0.216037967 | 0.165311002 |
| Tent2 | 0.933279376 | 0.619323984 | 0.476 | 0.668 | 1 | 0.086415729 | 0.63653 | 0.8 | 0.737 | 1 | 0.933279376 | 0.16536378 |
| Kcnk6 | 0.647471572 | 0.55829455 | 0.048 | 0.078 | 1 | 0.086449766 | -0.539319835 | 0.089 | 0.201 | 1 | 0.647471572 | 0.16542597 |
| Zfp788 | 0.52776362 | 1.304537958 | 0.048 | 0.093 | 1 | 0.086563852 | -0.776359032 | 0.044 | 0.138 | 1 | 0.52776362 | 0.165634403 |
| 9530077C05Rik | 0.319870046 | 1.908237021 | 0.095 | 0.049 | 1 | 0.086572146 | -1.053893008 | 0.022 | 0.104 | 1 | 0.319870046 | 0.165649555 |
| Mcc | 0.992320392 | 0.478972098 | 0.714 | 0.75 | 1 | 0.086577269 | 0.780652947 | 0.867 | 0.699 | 1 | 0.992320392 | 0.165658915 |
| Bcat2 | 0.478475937 | -0.219313028 | 0.19 | 0.269 | 1 | 0.086648546 | -0.526380779 | 0.089 | 0.208 | 1 | 0.478475937 | 0.165789122 |
| Gpat3 | 0.086665653 | 1.691919114 | 0.333 | 0.205 | 1 | 0.369772606 | 0.553789569 | 0.089 | 0.149 | 1 | 0.369772606 | 0.16582037 |
| Pcnx | 0.219022805 | 0.633184234 | 0.905 | 0.795 | 1 | 0.086666573 | 0.728504015 | 0.956 | 0.844 | 1 | 0.219022805 | 0.165822051 |
| Arid3a | 0.25165389 | -0.648156327 | 0 | 0.06 | 1 | 0.086757028 | -0.199570463 | 0.044 | 0.142 | 1 | 0.25165389 | 0.165987274 |
| Mtrfr | 0.086804928 | -1.611630451 | 0.048 | 0.198 | 1 | 0.215304569 | -0.149817428 | 0.089 | 0.163 | 1 | 0.215304569 | 0.16607476 |
| Pot1a | 0.487934712 | 0.866416846 | 0.19 | 0.142 | 1 | 0.086818473 | -0.459885368 | 0.044 | 0.142 | 1 | 0.487934712 | 0.166099499 |
| Parg | 0.086927059 | 1.162271429 | 0.619 | 0.53 | 1 | 0.12548777 | 0.620788308 | 0.8 | 0.737 | 1 | 0.12548777 | 0.166297805 |
| Ank | 0.125423145 | 1.126035852 | 0.952 | 0.97 | 1 | 0.086965117 | 0.727802709 | 0.978 | 0.972 | 1 | 0.125423145 | 0.166367302 |
| Itpr3 | 0.692144098 | -0.146962184 | 0.429 | 0.586 | 1 | 0.086984443 | 0.662543773 | 0.8 | 0.72 | 1 | 0.692144098 | 0.166402594 |
| Bak1 | 0.735703693 | 0.129451251 | 0.095 | 0.119 | 1 | 0.087033514 | -0.609709163 | 0.044 | 0.142 | 1 | 0.735703693 | 0.166492195 |
| Ppox | 0.232937926 | -0.133583154 | 0.048 | 0.149 | 1 | 0.087055775 | -1.174908409 | 0.067 | 0.163 | 1 | 0.232937926 | 0.166532841 |
| Zfp799 | 0.087121067 | 1.214340149 | 0.286 | 0.153 | 1 | 0.64644494 | 0.765534746 | 0.067 | 0.093 | 1 | 0.64644494 | 0.166652054 |
| Limch1 | 0.817690291 | 0.503846766 | 0.143 | 0.175 | 1 | 0.087216263 | 0.317891293 | 0.111 | 0.246 | 1 | 0.817690291 | 0.16682585 |
| Tbc1d16 | 0.844588485 | 1.146524765 | 0.19 | 0.201 | 1 | 0.087263688 | -0.220497651 | 0.2 | 0.353 | 1 | 0.844588485 | 0.166912425 |
| Cfap69 | 0.087274434 | 2.070775344 | 0.381 | 0.265 | 1 | 0.674387512 | 1.289158378 | 0.178 | 0.173 | 1 | 0.674387512 | 0.16693204 |
| Arxes1 | 0.153969332 | -1.496153234 | 0 | 0.09 | 1 | 0.087348588 | -1.486852415 | 0.022 | 0.1 | 1 | 0.153969332 | 0.167067401 |
| Trmt6 | 0.406172489 | -0.177084793 | 0.19 | 0.276 | 1 | 0.087368799 | -1.08246216 | 0.067 | 0.166 | 1 | 0.406172489 | 0.16710429 |
| Tspan5 | 0.087417086 | 1.142051288 | 0.714 | 0.634 | 1 | 0.23206216 | 0.222575068 | 0.8 | 0.619 | 1 | 0.23206216 | 0.167192425 |
| Vsir | 0.761718677 | 0.750939628 | 0.238 | 0.224 | 1 | 0.087511189 | -0.137957273 | 0.111 | 0.249 | 1 | 0.761718677 | 0.167364169 |
| Il17rc | 0.196462148 | -0.970084422 | 0 | 0.075 | 1 | 0.087626683 | -1.486852415 | 0.022 | 0.1 | 1 | 0.196462148 | 0.16757493 |
| Mlycd | 0.453165196 | -0.233118828 | 0.048 | 0.097 | 1 | 0.087626683 | -1.486852415 | 0.022 | 0.1 | 1 | 0.453165196 | 0.16757493 |
| Ogg1 | 0.094522876 | -1.611630451 | 0 | 0.119 | 1 | 0.087627306 | -1.564854927 | 0.022 | 0.1 | 1 | 0.094522876 | 0.167576068 |
| Ubr4 | 0.087813868 | 0.475715331 | 0.905 | 0.862 | 1 | 0.358903773 | 0.338918573 | 0.978 | 0.965 | 1 | 0.358903773 | 0.16791646 |
| F8 | 0.852959125 | 1.46732089 | 0.143 | 0.138 | 1 | 0.087819984 | -0.80878051 | 0.022 | 0.104 | 1 | 0.852959125 | 0.167927619 |
| Gm15728 | 0.859868645 | 1.451379346 | 0.048 | 0.041 | 1 | 0.087903446 | 1.846571318 | 0.133 | 0.066 | 1 | 0.859868645 | 0.168079876 |
| 4933431E20Rik | 0.63824517 | -0.133583154 | 0.048 | 0.075 | 1 | 0.087904608 | -1.446210431 | 0.022 | 0.1 | 1 | 0.63824517 | 0.168081996 |
| Pecr | 0.239965919 | 1.735172312 | 0.238 | 0.153 | 1 | 0.087954076 | -0.86124793 | 0.022 | 0.104 | 1 | 0.239965919 | 0.168172233 |
| Hdhd5 | 0.507332882 | 2.159198595 | 0.095 | 0.063 | 1 | 0.087954076 | -0.86124793 | 0.022 | 0.104 | 1 | 0.507332882 | 0.168172233 |
| Negr1 | 0.919565008 | 1.150209812 | 0.095 | 0.108 | 1 | 0.087982499 | -0.571741313 | 0.111 | 0.232 | 1 | 0.919565008 | 0.168224078 |
| Ubox5 | 0.087984283 | 1.843696769 | 0.19 | 0.086 | 1 | 0.731319906 | 1.320502507 | 0.156 | 0.142 | 1 | 0.731319906 | 0.168227332 |
| Virma | 0.140921302 | 0.628129102 | 0.619 | 0.593 | 1 | 0.087988555 | 0.114229751 | 0.6 | 0.72 | 1 | 0.140921302 | 0.168235123 |
| Brap | 0.907142017 | 0.698575159 | 0.333 | 0.396 | 1 | 0.088175663 | -0.105423309 | 0.4 | 0.644 | 1 | 0.907142017 | 0.168576379 |
| Plpp6 | 0.551474458 | 0.451379346 | 0.048 | 0.09 | 1 | 0.088180378 | -1.361321533 | 0.022 | 0.1 | 1 | 0.551474458 | 0.168584977 |
| Stxbp2 | 0.616977656 | 0.614878079 | 0.048 | 0.082 | 1 | 0.088223614 | -0.819427754 | 0.022 | 0.104 | 1 | 0.616977656 | 0.168663821 |
| B230217O12Rik | 0.770338347 | 1.041503552 | 0.095 | 0.082 | 1 | 0.088223614 | -0.819427754 | 0.022 | 0.104 | 1 | 0.770338347 | 0.168663821 |
| Rbl1 | 0.293648156 | 1.010806755 | 0.19 | 0.119 | 1 | 0.088262003 | -0.819427754 | 0.022 | 0.104 | 1 | 0.293648156 | 0.168733825 |
| Dph5 | 0.224378217 | -0.241836045 | 0.19 | 0.332 | 1 | 0.08828876 | -1.375821103 | 0.089 | 0.204 | 1 | 0.224378217 | 0.168782615 |
| Prmt6 | 0.153703558 | -1.02666795 | 0 | 0.09 | 1 | 0.088453665 | -1.316927414 | 0.022 | 0.1 | 1 | 0.153703558 | 0.169083278 |
| Ttl | 0.088484484 | -0.970084422 | 0.095 | 0.272 | 1 | 0.253769254 | 0.157003775 | 0.222 | 0.339 | 1 | 0.253769254 | 0.169139464 |
| Ak4 | 0.088565892 | -1.037723139 | 0.143 | 0.325 | 1 | 0.330045093 | 0.15951063 | 0.111 | 0.183 | 1 | 0.330045093 | 0.169287867 |
| Ccdc138 | 0.737733035 | 0.546915488 | 0.286 | 0.362 | 1 | 0.088614603 | -0.404390255 | 0.289 | 0.46 | 1 | 0.737733035 | 0.169376658 |
| Nsun3 | 0.088627461 | 1.673771768 | 0.143 | 0.052 | 1 | 0.866836726 | 1.152557869 | 0.067 | 0.062 | 1 | 0.866836726 | 0.169400096 |
| Nt5c3 | 0.248826707 | -0.337455488 | 0.333 | 0.493 | 1 | 0.088722471 | -0.170537927 | 0.489 | 0.654 | 1 | 0.248826707 | 0.169573266 |
| Mcts2 | 0.088737678 | -1.535681598 | 0 | 0.123 | 1 | 0.186096429 | -0.901889915 | 0.044 | 0.107 | 1 | 0.186096429 | 0.169600981 |
| Ints4 | 0.088740028 | -1.008052272 | 0.048 | 0.209 | 1 | 0.219632475 | 0.457004507 | 0.178 | 0.298 | 1 | 0.219632475 | 0.169605263 |
| Phf6 | 0.088894064 | 0.38254047 | 0.143 | 0.34 | 1 | 0.848261902 | 0.294507298 | 0.422 | 0.488 | 1 | 0.848261902 | 0.169885973 |
| Pdpn | 0.19848224 | -1.02666795 | 0.048 | 0.153 | 1 | 0.088903817 | 0.323176641 | 0.467 | 0.304 | 1 | 0.19848224 | 0.169903745 |
| Dcp1b | 0.558305772 | 1.569435108 | 0.143 | 0.108 | 1 | 0.088914926 | -0.486852415 | 0.022 | 0.104 | 1 | 0.558305772 | 0.169923988 |
| Paqr7 | 0.815913222 | 0.281454345 | 0.048 | 0.06 | 1 | 0.088976838 | -0.564854927 | 0.022 | 0.104 | 1 | 0.815913222 | 0.170036799 |
| Senp8 | 0.088983547 | -1.998653574 | 0 | 0.123 | 1 | 0.864452005 | 1.163698427 | 0.111 | 0.111 | 1 | 0.864452005 | 0.170049023 |
| Mlh1 | 0.513572074 | -0.133583154 | 0.048 | 0.093 | 1 | 0.089003372 | -0.602329633 | 0.022 | 0.104 | 1 | 0.513572074 | 0.170085143 |
| Zcchc4 | 0.089044489 | -1.818081329 | 0 | 0.123 | 1 | 0.902922038 | 1.075390009 | 0.156 | 0.163 | 1 | 0.902922038 | 0.170160057 |
| Lgr4 | 0.49276821 | 1.587184083 | 0.429 | 0.698 | 1 | 0.089044654 | 1.218566411 | 0.889 | 0.716 | 1 | 0.49276821 | 0.170160358 |
| Catspere2 | 0.089071632 | -1.849790188 | 0 | 0.123 | 1 | 0.856435785 | 0.830629775 | 0.133 | 0.131 | 1 | 0.856435785 | 0.170209509 |
| Gnao1 | 0.089118088 | -2.208871282 | 0 | 0.123 | 1 | 0.307651854 | -0.446210431 | 0.133 | 0.208 | 1 | 0.307651854 | 0.170294142 |
| Hmcn1 | 0.220154691 | -2.017390137 | 0.048 | 0.142 | 1 | 0.089119436 | -2.674479418 | 0.022 | 0.1 | 1 | 0.220154691 | 0.170296599 |
| Tacc3 | 0.089139619 | -2.326228232 | 0 | 0.123 | 1 | 0.379847138 | 0.402964667 | 0.044 | 0.087 | 1 | 0.379847138 | 0.170333366 |
| Mafk | 0.089141103 | 0.404311093 | 0.238 | 0.526 | 1 | 0.129437721 | -0.386771402 | 0.689 | 0.716 | 1 | 0.129437721 | 0.170336069 |
| Sharpin | 0.303726098 | 0.821328956 | 0.095 | 0.209 | 1 | 0.089146575 | -0.573267167 | 0.089 | 0.201 | 1 | 0.303726098 | 0.170346038 |
| Sgip1 | 0.089189858 | -2.849790188 | 0 | 0.123 | 1 | 0.648406793 | 0.91753784 | 0.067 | 0.052 | 1 | 0.648406793 | 0.170424885 |
| Plat | 0.089230178 | -2.818081329 | 0 | 0.123 | 1 | 0.553958026 | -0.526380779 | 0.089 | 0.118 | 1 | 0.553958026 | 0.170498332 |
| Kdm4c | 0.089263115 | 0.95495352 | 0.619 | 0.541 | 1 | 0.329465108 | 0.574193789 | 0.822 | 0.74 | 1 | 0.329465108 | 0.170558327 |
| Syne1 | 0.536957488 | 0.773105793 | 0.81 | 0.84 | 1 | 0.089292842 | 0.542727653 | 0.978 | 0.983 | 1 | 0.536957488 | 0.170612472 |
| Dnajc5 | 0.297641847 | 0.440520831 | 0.714 | 0.772 | 1 | 0.089668026 | -0.226462731 | 0.889 | 0.91 | 1 | 0.297641847 | 0.171295697 |

| | | | | | | | | | | | | |
|---|---|---|---|---|---|---|---|---|---|---|---|---|
| Prrc2c | 0.089698755 | -0.284440797 | 1 | 0.993 | 1 | 0.114572949 | -0.179728224 | 1 | 1 | 1 | 0.114572949 | 0.171351643 |
| Neu2 | 0.089751805 | 1.620660431 | 0.381 | 0.265 | 1 | 0.821620213 | 1.226214911 | 0.222 | 0.242 | 1 | 0.821620213 | 0.171448223 |
| Cyth3 | 0.933618113 | 0.551856811 | 0.714 | 0.903 | 1 | 0.089921894 | 0.488569562 | 0.933 | 0.855 | 1 | 0.933618113 | 0.171757841 |
| Hirip3 | 0.144874061 | -1.116305163 | 0.048 | 0.175 | 1 | 0.089929204 | -1.564854927 | 0.044 | 0.131 | 1 | 0.144874061 | 0.171771147 |
| Gtf2ird2 | 0.365969666 | -0.295854583 | 0.048 | 0.116 | 1 | 0.089931406 | -1.486852415 | 0.044 | 0.131 | 1 | 0.365969666 | 0.171775154 |
| Irs1 | 0.560492775 | -0.163712821 | 0.429 | 0.556 | 1 | 0.089940351 | 0.308593969 | 0.378 | 0.602 | 1 | 0.560492775 | 0.171791435 |
| Afg1l | 0.089973832 | 1.503846766 | 0.286 | 0.168 | 1 | 0.376426583 | 1.870158139 | 0.156 | 0.125 | 1 | 0.376426583 | 0.171852373 |
| Zbtb5 | 0.448810475 | 0.586308926 | 0.048 | 0.104 | 1 | 0.090165809 | -0.994999319 | 0.067 | 0.166 | 1 | 0.448810475 | 0.172201745 |
| Repin1 | 0.480458678 | -0.133583154 | 0.048 | 0.093 | 1 | 0.090310829 | -0.921789472 | 0.089 | 0.194 | 1 | 0.480458678 | 0.172465612 |
| Socs5 | 0.926689544 | 0.652076697 | 0.429 | 0.511 | 1 | 0.090321157 | 0.147019686 | 0.356 | 0.588 | 1 | 0.926689544 | 0.172484402 |
| Stn1 | 0.090405859 | -0.584962501 | 0.143 | 0.347 | 1 | 0.851544591 | 0.728876276 | 0.222 | 0.242 | 1 | 0.851544591 | 0.172638499 |
| Dhx36 | 0.090444999 | -0.24509147 | 0.571 | 0.78 | 1 | 0.198148392 | 0.328229869 | 0.911 | 0.744 | 1 | 0.198148392 | 0.1727097 |
| Khk | 0.33216244 | -0.648156327 | 0.095 | 0.175 | 1 | 0.090475577 | -0.877642368 | 0.067 | 0.166 | 1 | 0.33216244 | 0.172765324 |
| Pdp2 | 0.090500707 | -0.870548749 | 0.048 | 0.209 | 1 | 0.416727948 | 0.361144491 | 0.067 | 0.111 | 1 | 0.416727948 | 0.172811036 |
| Gm51425 | 0.090529511 | 1.60338244 | 0.143 | 0.052 | 1 | 0.908735329 | 1.683072586 | 0.067 | 0.076 | 1 | 0.908735329 | 0.17286343 |
| Gm10648 | 0.671803158 | 0.980749521 | 0.19 | 0.168 | 1 | 0.090648985 | 0.339928005 | 0.089 | 0.211 | 1 | 0.671803158 | 0.173080731 |
| Cav1 | 0.090714317 | 1.670794068 | 0.429 | 0.317 | 1 | 0.111674977 | 0.221684771 | 0.133 | 0.27 | 1 | 0.111674977 | 0.173199547 |
| Arhgap44 | 0.090763398 | 1.103916159 | 0.048 | 0.213 | 1 | 0.435037989 | 0.595609745 | 0.178 | 0.263 | 1 | 0.435037989 | 0.173288801 |
| Stk4 | 0.238433585 | -0.555046923 | 0.19 | 0.325 | 1 | 0.090781491 | 0.376017768 | 0.556 | 0.384 | 1 | 0.238433585 | 0.173321703 |
| Mtmr14 | 0.377672345 | -0.413691074 | 0.048 | 0.108 | 1 | 0.090790873 | 0.332575339 | 0.044 | 0.142 | 1 | 0.377672345 | 0.173338764 |
| Ric1 | 0.237097413 | 0.910006114 | 0.619 | 0.593 | 1 | 0.090912745 | 0.904540499 | 0.822 | 0.678 | 1 | 0.237097413 | 0.173560363 |
| Mark1 | 0.091035065 | -1.413691074 | 0.19 | 0.373 | 1 | 0.182969343 | 0.298408736 | 0.2 | 0.332 | 1 | 0.182969343 | 0.173782747 |
| Tufm | 0.091078341 | -0.828728573 | 0.048 | 0.209 | 1 | 0.113248595 | 0.381903051 | 0.089 | 0.201 | 1 | 0.113248595 | 0.173861418 |
| Bahd1 | 0.091099558 | 1.503846766 | 0.19 | 0.082 | 1 | 0.254231186 | -0.199570463 | 0.089 | 0.163 | 1 | 0.254231186 | 0.173899987 |
| Csf2ra | 0.157141827 | -0.60421298 | 0.143 | 0.295 | 1 | 0.091149169 | -1.124282336 | 0.044 | 0.135 | 1 | 0.157141827 | 0.173990167 |
| Slc36a1 | 0.798580785 | 0.97333205 | 0.095 | 0.082 | 1 | 0.091499198 | -1.316927414 | 0.044 | 0.131 | 1 | 0.798580785 | 0.174626293 |
| Trmt61b | 0.874914468 | 0.758163955 | 0.19 | 0.201 | 1 | 0.091530806 | -0.901889915 | 0.111 | 0.218 | 1 | 0.874914468 | 0.174683724 |
| Clk4 | 0.091565291 | 0.101192991 | 0.524 | 0.802 | 1 | 0.149186666 | 0.486465542 | 0.867 | 0.761 | 1 | 0.149186666 | 0.174746379 |
| Usp49 | 0.091682869 | 1.389978802 | 0.381 | 0.261 | 1 | 0.253846274 | 0.227190035 | 0.244 | 0.374 | 1 | 0.253846274 | 0.174959989 |
| Ccl25 | 0.091917615 | 1.484737943 | 0.381 | 0.261 | 1 | 0.547375821 | 0.471924922 | 0.289 | 0.405 | 1 | 0.547375821 | 0.175386382 |
| Nrip3 | 0.496264982 | -0.672003069 | 0.095 | 0.146 | 1 | 0.091949543 | -1.486852415 | 0.022 | 0.1 | 1 | 0.496264982 | 0.175444368 |
| Brf1 | 0.092253841 | -0.260340297 | 0.095 | 0.291 | 1 | 0.310531222 | 0.292431741 | 0.311 | 0.481 | 1 | 0.310531222 | 0.175996911 |
| Chchd4 | 0.107558988 | -0.807354922 | 0.048 | 0.198 | 1 | 0.092348899 | -0.557935513 | 0.133 | 0.26 | 1 | 0.107558988 | 0.176169479 |
| Ncor2 | 0.092352987 | 0.812241705 | 0.667 | 0.731 | 1 | 0.291353025 | 0.419428032 | 0.889 | 0.865 | 1 | 0.291353025 | 0.176176901 |
| Gprin3 | 0.497303221 | -1.280424543 | 0.048 | 0.09 | 1 | 0.092364369 | -1.255526869 | 0.111 | 0.218 | 1 | 0.497303221 | 0.176197562 |
| Phlpp2 | 0.340855263 | 0.781754449 | 0.429 | 0.384 | 1 | 0.09245323 | 0.133443159 | 0.378 | 0.602 | 1 | 0.340855263 | 0.176358861 |
| Mtm1 | 0.092541244 | -0.621684116 | 0.143 | 0.351 | 1 | 0.807694118 | 0.850182572 | 0.178 | 0.208 | 1 | 0.807694118 | 0.176518605 |
| Thap7 | 0.491006067 | 0.233199176 | 0.143 | 0.224 | 1 | 0.092633863 | -0.336826971 | 0.089 | 0.201 | 1 | 0.491006067 | 0.176686693 |
| Zc3h7a | 0.161509592 | 0.593661593 | 0.81 | 0.907 | 1 | 0.092647888 | 0.336079154 | 0.978 | 0.907 | 1 | 0.161509592 | 0.176712144 |
| Kif16b | 0.148198633 | 1.123574685 | 0.476 | 0.466 | 1 | 0.092665337 | 0.764209349 | 0.756 | 0.696 | 1 | 0.148198633 | 0.17674381 |
| Gm43329 | 0.248649751 | 1.472137907 | 0.143 | 0.075 | 1 | 0.092966415 | 1.875717664 | 0.2 | 0.128 | 1 | 0.248649751 | 0.177290076 |
| Zbtb43 | 0.259185888 | -0.44170545 | 0.143 | 0.265 | 1 | 0.093140125 | 0.207339155 | 0.222 | 0.391 | 1 | 0.259185888 | 0.177605167 |
| Epha3 | 0.52423707 | -1.054148687 | 0.048 | 0.09 | 1 | 0.093194521 | -3.263346374 | 0.022 | 0.097 | 1 | 0.52423707 | 0.177703824 |
| 4930523C07Rik | 0.790209539 | -0.11396621 | 0.952 | 0.896 | 1 | 0.093213269 | -0.24508048 | 0.956 | 0.934 | 1 | 0.790209539 | 0.177737825 |
| Akr1b10 | 0.093350721 | -1.303508156 | 0.048 | 0.194 | 1 | 0.167248448 | -0.181997834 | 0.067 | 0.149 | 1 | 0.167248448 | 0.177987085 |
| Haus8 | 0.093444408 | 0.803832308 | 0.524 | 0.437 | 1 | 0.481193084 | 0.494038762 | 0.267 | 0.391 | 1 | 0.481193084 | 0.178156959 |
| Marchf6 | 0.093460231 | 0.627870917 | 0.857 | 0.724 | 1 | 0.943800354 | 0.185572927 | 0.933 | 0.889 | 1 | 0.943800354 | 0.178185647 |
| Laptm4b | 0.104591675 | -0.818081329 | 0.143 | 0.343 | 1 | 0.093625662 | -0.134301092 | 0.267 | 0.45 | 1 | 0.104591675 | 0.178485559 |
| Phtf1 | 0.815220366 | 0.339871031 | 0.429 | 0.448 | 1 | 0.093764451 | 0.679470349 | 0.689 | 0.581 | 1 | 0.815220366 | 0.178737129 |
| Alox12 | 0.093820871 | -1.31441893 | 0.143 | 0.325 | 1 | 0.944742566 | -0.638855509 | 0.044 | 0.042 | 1 | 0.944742566 | 0.178839386 |
| Foxo4 | 0.113654256 | -1.752492987 | 0 | 0.108 | 1 | 0.093895549 | -0.589945908 | 0.067 | 0.166 | 1 | 0.113654256 | 0.178974723 |
| Rorc | 0.236343578 | -0.574155746 | 0 | 0.063 | 1 | 0.093973284 | -0.722919774 | 0.044 | 0.138 | 1 | 0.236343578 | 0.179115589 |
| Gm56830 | 0.886609917 | 0.889500459 | 0.238 | 0.254 | 1 | 0.094003214 | -0.390927995 | 0.156 | 0.298 | 1 | 0.886609917 | 0.179169823 |
| Ints1 | 0.922571476 | 0.908237021 | 0.095 | 0.112 | 1 | 0.094103651 | -1.124282336 | 0.044 | 0.131 | 1 | 0.922571476 | 0.179351805 |
| Uap1 | 0.492692713 | 0.405410137 | 0.952 | 0.914 | 1 | 0.094165166 | -0.288227102 | 0.822 | 0.913 | 1 | 0.492692713 | 0.179463253 |
| Kpna2 | 0.527721908 | 0.766881172 | 0.048 | 0.093 | 1 | 0.09425131 | -0.575661682 | 0.067 | 0.17 | 1 | 0.527721908 | 0.179619311 |
| Bicc1 | 0.602607843 | -0.267136792 | 0.667 | 0.769 | 1 | 0.094261927 | -0.96587437 | 0.6 | 0.855 | 1 | 0.602607843 | 0.179638542 |
| 9530026P05Rik | 0.094272887 | -0.598115645 | 0.19 | 0.403 | 1 | 0.18198815 | -0.316927414 | 0.311 | 0.446 | 1 | 0.18198815 | 0.179658396 |
| Klhl8 | 0.094305534 | -1.496153234 | 0 | 0.119 | 1 | 0.189303522 | 0.982632868 | 0.044 | 0.114 | 1 | 0.189303522 | 0.179717535 |
| Apeh | 0.094305534 | -1.455511249 | 0 | 0.119 | 1 | 0.15021953 | -0.57996182 | 0.067 | 0.149 | 1 | 0.15021953 | 0.179717535 |
| Mob2 | 0.937170368 | 0.544488751 | 0.333 | 0.407 | 1 | 0.094331799 | 0.582271945 | 0.756 | 0.592 | 1 | 0.937170368 | 0.179765109 |
| Rnf169 | 0.094351337 | -0.959553755 | 0.238 | 0.444 | 1 | 0.18920012 | 0.73692256 | 0.733 | 0.606 | 1 | 0.18920012 | 0.179800498 |
| Pdlim4 | 0.46806015 | 0.624528249 | 0.19 | 0.313 | 1 | 0.094355464 | -0.288913038 | 0.244 | 0.408 | 1 | 0.46806015 | 0.179807975 |
| Gpr137b | 0.561991567 | 2.233199176 | 0.143 | 0.116 | 1 | 0.094365047 | -0.286553765 | 0.067 | 0.17 | 1 | 0.561991567 | 0.179825331 |
| Oscp1 | 0.094365756 | -1.496153234 | 0 | 0.119 | 1 | 0.259955674 | 0.498648015 | 0.067 | 0.135 | 1 | 0.259955674 | 0.179826616 |
| Tmem170b | 0.73000611 | 0.889500459 | 0.19 | 0.179 | 1 | 0.0944046 | -0.21001221 | 0.111 | 0.239 | 1 | 0.73000611 | 0.179896972 |
| Jpt1 | 0.928087536 | 0.351843673 | 0.286 | 0.347 | 1 | 0.094407272 | 0.658126229 | 0.578 | 0.405 | 1 | 0.928087536 | 0.179901812 |
| 2310015A10Rik | 0.094472904 | -1.683780237 | 0 | 0.119 | 1 | 0.490252713 | 0.605070074 | 0.067 | 0.104 | 1 | 0.490252713 | 0.180020678 |
| Armcx6 | 0.094472904 | -1.611630451 | 0 | 0.119 | 1 | 0.306081442 | 0.223640968 | 0.067 | 0.125 | 1 | 0.306081442 | 0.180020678 |
| Snx33 | 0.094520107 | -1.752492987 | 0 | 0.119 | 1 | 0.676621251 | 1.098110085 | 0.2 | 0.201 | 1 | 0.676621251 | 0.180106164 |
| Zfp874a | 0.094520107 | -1.648156327 | 0 | 0.119 | 1 | 0.147388465 | 0.250113179 | 0.067 | 0.159 | 1 | 0.147388465 | 0.180106164 |
| Alkbh2 | 0.094521492 | -1.718545655 | 0 | 0.119 | 1 | 0.185254141 | -0.158229668 | 0.044 | 0.111 | 1 | 0.185254141 | 0.180108671 |
| Rexo5 | 0.094601638 | -1.785659851 | 0 | 0.119 | 1 | 0.622665392 | 0.859950348 | 0.089 | 0.125 | 1 | 0.622665392 | 0.180253806 |
| Nol4l | 0.094606483 | -1.880817084 | 0 | 0.119 | 1 | 0.687981382 | 1.152557869 | 0.2 | 0.194 | 1 | 0.687981382 | 0.180262579 |
| Dnai4 | 0.09464607 | -1.970084422 | 0 | 0.119 | 1 | 0.173002869 | 0.325520582 | 0.044 | 0.118 | 1 | 0.173002869 | 0.180334261 |
| Tmem119 | 0.094762057 | -2.348596045 | 0 | 0.119 | 1 | 0.11886609 | -1.564854927 | 0.022 | 0.09 | 1 | 0.11886609 | 0.180544266 |
| Tub | 0.094762799 | 2.586308926 | 0.143 | 0.056 | 1 | 0.162280518 | -0.316927414 | 0.022 | 0.083 | 1 | 0.162280518 | 0.18054561 |
| Lgals9 | 0.094763994 | -2.208871282 | 0 | 0.119 | 1 | 0.098299812 | -0.124282336 | 0.022 | 0.1 | 1 | 0.098299812 | 0.180547774 |
| A430033K04Rik | 0.722923749 | 0.673771768 | 0.048 | 0.071 | 1 | 0.094770207 | -1.124282336 | 0.022 | 0.1 | 1 | 0.722923749 | 0.180559022 |
| Zfp961 | 0.094787373 | 2.503846766 | 0.143 | 0.056 | 1 | 0.295520395 | -0.316927414 | 0.022 | 0.062 | 1 | 0.295520395 | 0.1805901 |
| Nlgn2 | 0.664219439 | 1.221259563 | 0.286 | 0.269 | 1 | 0.094817961 | 0.28452321 | 0.2 | 0.401 | 1 | 0.664219439 | 0.180645476 |
| Col24a1 | 0.094832088 | -3.81004401 | 0 | 0.119 | 1 | 0.625716651 | -0.237492946 | 0.067 | 0.09 | 1 | 0.625716651 | 0.18067105 |
| Dlc1 | 0.094912365 | 0.885797473 | 0.762 | 0.903 | 1 | 0.752328477 | 0.442064487 | 0.733 | 0.889 | 1 | 0.752328477 | 0.180816373 |
| Mcph1 | 0.094926044 | 1.69394965 | 0.286 | 0.16 | 1 | 0.172994493 | -0.298780067 | 0.089 | 0.173 | 1 | 0.172994493 | 0.180841134 |
| Abhd17b | 0.094931992 | -0.508431564 | 0.238 | 0.455 | 1 | 0.468938689 | 0.164497483 | 0.733 | 0.574 | 1 | 0.468938689 | 0.180851901 |
| Il1rap | 0.845989015 | 0.574236094 | 0.143 | 0.134 | 1 | 0.095156345 | -0.638855509 | 0.044 | 0.138 | 1 | 0.845989015 | 0.181257961 |
| Haus2 | 0.095188774 | -1.233118828 | 0.095 | 0.254 | 1 | 0.256291444 | 0.311103809 | 0.133 | 0.221 | 1 | 0.256291444 | 0.181316646 |
| Rita1 | 0.106726975 | -1.535681598 | 0 | 0.112 | 1 | 0.095193147 | -1.199570463 | 0.067 | 0.159 | 1 | 0.106726975 | 0.181324559 |
| Ube2r2 | 0.38352673 | 0.200742546 | 1 | 0.925 | 1 | 0.095284746 | -0.102587265 | 1 | 0.983 | 1 | 0.38352673 | 0.18149031 |
| Ppif | 0.095350587 | -1.752492987 | 0.048 | 0.201 | 1 | 0.098803546 | -0.242926832 | 0.022 | 0.1 | 1 | 0.098803546 | 0.181609439 |
| Ccsap | 0.517057607 | 0.129451251 | 0.048 | 0.093 | 1 | 0.095406092 | -1.255526869 | 0.022 | 0.1 | 1 | 0.517057607 | 0.181709861 |
| Tarbp2 | 0.268811897 | -0.718545655 | 0.095 | 0.19 | 1 | 0.095420856 | -0.743192169 | 0.044 | 0.135 | 1 | 0.268811897 | 0.181736572 |
| Tcaf2 | 0.509428832 | -0.233118828 | 0.048 | 0.09 | 1 | 0.095433623 | -1.871516266 | 0.022 | 0.097 | 1 | 0.509428832 | 0.181759669 |
| Cc2d1a | 0.209094725 | -0.648156327 | 0 | 0.071 | 1 | 0.095573869 | -0.255526869 | 0.044 | 0.135 | 1 | 0.209094725 | 0.182013373 |
| Smug1 | 0.58037501 | 0.451379346 | 0.048 | 0.086 | 1 | 0.095575105 | -0.286553765 | 0.044 | 0.135 | 1 | 0.58037501 | 0.18201561 |
| Gm28198 | 0.18415515 | -0.940938076 | 0.048 | 0.153 | 1 | 0.095601379 | 0.118287967 | 0.089 | 0.201 | 1 | 0.18415515 | 0.182063134 |
| Cd68 | 0.895445406 | 0.503846766 | 0.095 | 0.108 | 1 | 0.095757832 | -3.211745177 | 0.022 | 0.097 | 1 | 0.895445406 | 0.182346101 |
| Gm26944 | 0.095787749 | 2.036341847 | 0.19 | 0.09 | 1 | 0.559088601 | 1.612683258 | 0.089 | 0.069 | 1 | 0.559088601 | 0.182400205 |

| | | | | | | | | | | | | |
|---|---|---|---|---|---|---|---|---|---|---|---|---|
| Gm20300 | 0.208797419 | -0.718545655 | 0 | 0.071 | 1 | 0.095816427 | -0.223818009 | 0.044 | 0.135 | 1 | 0.208797419 | 0.182452066 |
| Fbxo44 | 0.208955157 | -0.970084422 | 0 | 0.071 | 1 | 0.095867515 | -1.564854927 | 0 | 0.059 | 1 | 0.208955157 | 0.18254445 |
| Comtd1 | 0.251580301 | -0.496153234 | 0 | 0.06 | 1 | 0.095867515 | -1.564854927 | 0 | 0.059 | 1 | 0.251580301 | 0.18254445 |
| B430105G09Rik | 0.267917141 | -0.413691074 | 0 | 0.056 | 1 | 0.095867515 | -1.564854927 | 0 | 0.059 | 1 | 0.267917141 | 0.18254445 |
| Gm57269 | 0.720638157 | 0.673771768 | 0.048 | 0.071 | 1 | 0.095867515 | -1.564854927 | 0 | 0.059 | 1 | 0.720638157 | 0.18254445 |
| Slc4a11 | 0.859875139 | 1.451379346 | 0.048 | 0.041 | 1 | 0.095867515 | -1.638855509 | 0 | 0.059 | 1 | 0.859875139 | 0.18254445 |
| Gm57084 | 0.860084871 | 1.150209812 | 0.048 | 0.06 | 1 | 0.095867515 | -1.564854927 | 0 | 0.059 | 1 | 0.860084871 | 0.18254445 |
| Kyat3 | 0.924111829 | 0.586308926 | 0.048 | 0.052 | 1 | 0.095867515 | -1.638855509 | 0 | 0.059 | 1 | 0.924111829 | 0.18254445 |
| Arntl | 0.29491387 | -0.557182667 | 0.19 | 0.302 | 1 | 0.095874742 | 1.195626591 | 0.622 | 0.398 | 1 | 0.29491387 | 0.182557518 |
| Fam221a | 0.106777151 | -1.574155746 | 0 | 0.112 | 1 | 0.095889652 | -1.638855509 | 0 | 0.059 | 1 | 0.106777151 | 0.18258448 |
| B630019K06Rik | 0.12067628 | -1.496153234 | 0 | 0.104 | 1 | 0.095889652 | -1.709244837 | 0 | 0.059 | 1 | 0.12067628 | 0.18258448 |
| Adrm1b | 0.209094725 | -0.648156327 | 0 | 0.071 | 1 | 0.095889652 | -1.638855509 | 0 | 0.059 | 1 | 0.209094725 | 0.18258448 |
| Carns1 | 0.359270744 | 2.673771768 | 0.048 | 0.019 | 1 | 0.095889652 | -1.709244837 | 0 | 0.059 | 1 | 0.359270744 | 0.18258448 |
| Dnah1 | 0.370840523 | 0.214340149 | 0 | 0.037 | 1 | 0.095889652 | -1.638855509 | 0 | 0.059 | 1 | 0.370840523 | 0.18258448 |
| Rab3d | 0.472861597 | 1.866416846 | 0.048 | 0.022 | 1 | 0.095889652 | -1.709244837 | 0 | 0.059 | 1 | 0.472861597 | 0.18258448 |
| 1700066M21Rik | 0.532436372 | 1.088809267 | 0 | 0.019 | 1 | 0.095889652 | -1.638855509 | 0 | 0.059 | 1 | 0.532436372 | 0.18258448 |
| Wdr76 | 0.585375147 | 1.188344941 | 0.095 | 0.067 | 1 | 0.095889652 | -1.709244837 | 0 | 0.059 | 1 | 0.585375147 | 0.18258448 |
| 0610009L18Rik | 0.58812323 | 0.97333205 | 0.048 | 0.086 | 1 | 0.095889652 | -1.709244837 | 0 | 0.059 | 1 | 0.58812323 | 0.18258448 |
| Osbp2 | 0.709494687 | 2.188344941 | 0.048 | 0.034 | 1 | 0.095889652 | -1.709244837 | 0 | 0.059 | 1 | 0.709494687 | 0.18258448 |
| Brd3os | 0.732158744 | 1.281454345 | 0.048 | 0.071 | 1 | 0.095889652 | -1.776359032 | 0 | 0.059 | 1 | 0.732158744 | 0.18258448 |
| Zfp354b | 0.578159907 | 0.866416846 | 0 | 0.015 | 1 | 0.095908838 | -1.84048937 | 0 | 0.059 | 1 | 0.578159907 | 0.182619171 |
| Zgrf1 | 0.604571771 | 1.586308926 | 0.095 | 0.067 | 1 | 0.095908838 | -1.776359032 | 0 | 0.059 | 1 | 0.604571771 | 0.182619171 |
| Gm10643 | 0.775775929 | 0.503846766 | 0.048 | 0.063 | 1 | 0.095908838 | -1.901889915 | 0 | 0.059 | 1 | 0.775775929 | 0.182619171 |
| Slc41a3 | 0.236581911 | -0.496153234 | 0 | 0.063 | 1 | 0.095909022 | -1.709244837 | 0 | 0.059 | 1 | 0.236581911 | 0.182619504 |
| Rasl10a | 0.267953794 | -0.496153234 | 0 | 0.056 | 1 | 0.095909022 | -1.709244837 | 0 | 0.059 | 1 | 0.267953794 | 0.182619504 |
| Zbtb8a | 0.396971401 | 0.351843673 | 0 | 0.034 | 1 | 0.095909022 | -1.709244837 | 0 | 0.059 | 1 | 0.396971401 | 0.182619504 |
| Dph1 | 0.425599726 | 0.351843673 | 0 | 0.03 | 1 | 0.095909022 | -1.709244837 | 0 | 0.059 | 1 | 0.425599726 | 0.182619504 |
| Olfml1 | 0.479147948 | 1.503846766 | 0.048 | 0.022 | 1 | 0.095909022 | -2.017367132 | 0 | 0.059 | 1 | 0.479147948 | 0.182619504 |
| 2310047K21Rik | 0.538371401 | -0.133583154 | 0.048 | 0.086 | 1 | 0.095909022 | -1.84048937 | 0 | 0.059 | 1 | 0.538371401 | 0.182619504 |
| 9330160F10Rik | 0.890779585 | 0.97333205 | 0.048 | 0.041 | 1 | 0.095909022 | -1.709244837 | 0 | 0.059 | 1 | 0.890779585 | 0.182619504 |
| Bik | 0.57816106 | 1.351843673 | 0 | 0.015 | 1 | 0.095925072 | -1.84048937 | 0 | 0.059 | 1 | 0.57816106 | 0.182648524 |
| Slc35f2 | 0.924795272 | 1.351843673 | 0.048 | 0.045 | 1 | 0.095925072 | -1.901889915 | 0 | 0.059 | 1 | 0.924795272 | 0.182648524 |
| Col19a1 | 0.492604708 | 0.351843673 | 0 | 0.022 | 1 | 0.095925625 | -2.071814916 | 0 | 0.059 | 1 | 0.492604708 | 0.182649524 |
| Pcx | 0.324927044 | 0.536268244 | 0.048 | 0.127 | 1 | 0.095925809 | -1.776359032 | 0 | 0.059 | 1 | 0.324927044 | 0.182649858 |
| Colq | 0.5651989 | -0.184209227 | 0.048 | 0.082 | 1 | 0.095925809 | -1.776359032 | 0 | 0.059 | 1 | 0.5651989 | 0.182649858 |
| Entpd3 | 0.271980818 | 2.351843673 | 0.048 | 0.015 | 1 | 0.095938354 | -1.901889915 | 0 | 0.059 | 1 | 0.271980818 | 0.182672539 |
| Snap23 | 0.589873606 | 0.477984703 | 0.524 | 0.694 | 1 | 0.095948412 | -0.158900533 | 0.733 | 0.834 | 1 | 0.589873606 | 0.182690726 |
| Ankrd37 | 0.267985206 | -0.574155746 | 0 | 0.056 | 1 | 0.095950529 | -2.017367132 | 0 | 0.059 | 1 | 0.267985206 | 0.182694553 |
| ENSMUSG00000121479 | 0.982551672 | 1.586308926 | 0.048 | 0.049 | 1 | 0.095950529 | -2.071814916 | 0 | 0.059 | 1 | 0.982551672 | 0.182694553 |
| Grin3a | 0.890779585 | 0.866416846 | 0.048 | 0.041 | 1 | 0.095951451 | -2.124282336 | 0 | 0.059 | 1 | 0.890779585 | 0.182696221 |
| Trib3 | 0.759553442 | 1.304537958 | 0.095 | 0.082 | 1 | 0.09595883 | -2.071814916 | 0 | 0.059 | 1 | 0.759553442 | 0.182709562 |
| Pms1 | 0.251707193 | -1.081115734 | 0 | 0.06 | 1 | 0.095961966 | -2.174908409 | 0 | 0.059 | 1 | 0.251707193 | 0.182715232 |
| Ankrd29 | 0.16360158 | -1.280424543 | 0 | 0.086 | 1 | 0.095964364 | -2.486852415 | 0 | 0.059 | 1 | 0.16360158 | 0.182719568 |
| Cdkl4 | 0.89926261 | 0.79930265 | 0.048 | 0.056 | 1 | 0.095975985 | -2.446210431 | 0 | 0.059 | 1 | 0.89926261 | 0.18274058 |
| Fgf13 | 0.396957792 | -0.718545655 | 0 | 0.034 | 1 | 0.09598115 | -3.098287127 | 0 | 0.059 | 1 | 0.396957792 | 0.182749919 |
| Itga2 | 0.37081917 | -0.413691074 | 0 | 0.037 | 1 | 0.095987606 | -3.149817428 | 0 | 0.059 | 1 | 0.37081917 | 0.182761592 |
| Gm57100 | 0.153703558 | -1.02666795 | 0 | 0.09 | 1 | 0.095992956 | -1.486852415 | 0 | 0.059 | 1 | 0.153703558 | 0.182771264 |
| 2010204K13Rik | 0.457281049 | 0.503846766 | 0 | 0.026 | 1 | 0.095992956 | -1.486852415 | 0 | 0.059 | 1 | 0.457281049 | 0.182771264 |
| Gtpbp3 | 0.775775929 | 0.503846766 | 0.048 | 0.063 | 1 | 0.095992956 | -1.486852415 | 0 | 0.059 | 1 | 0.775775929 | 0.182771264 |
| Ighmbp2 | 0.88770495 | 1.088809267 | 0.048 | 0.041 | 1 | 0.095992956 | -1.486852415 | 0 | 0.059 | 1 | 0.88770495 | 0.182771264 |
| Slc38a4 | 0.429313876 | -0.285586248 | 0.19 | 0.287 | 1 | 0.096075759 | -2.223818009 | 0.022 | 0.097 | 1 | 0.429313876 | 0.182920967 |
| Dnajc16 | 0.590245857 | 0.323274521 | 0.095 | 0.149 | 1 | 0.096127881 | -0.549588171 | 0.044 | 0.138 | 1 | 0.590245857 | 0.183015193 |
| Kdm4a | 0.096248962 | 1.518493542 | 0.381 | 0.261 | 1 | 0.74442996 | 0.91753784 | 0.289 | 0.367 | 1 | 0.74442996 | 0.18323406 |
| Eef1akmt3 | 0.106864405 | -1.648156327 | 0 | 0.112 | 1 | 0.096392095 | -1.743192169 | 0.022 | 0.097 | 1 | 0.106864405 | 0.183492753 |
| Cep192 | 0.341492332 | 0.430915244 | 0.19 | 0.287 | 1 | 0.096397917 | 0.247686442 | 0.133 | 0.277 | 1 | 0.341492332 | 0.183503275 |
| Slco2b1 | 0.161594235 | -1.133583154 | 0.048 | 0.16 | 1 | 0.096404834 | -0.979892427 | 0.022 | 0.1 | 1 | 0.161594235 | 0.183515776 |
| Mrps10 | 0.380985096 | 0.951305743 | 0.333 | 0.291 | 1 | 0.096559148 | 0.426732833 | 0.111 | 0.235 | 1 | 0.380985096 | 0.183794626 |
| Myc | 0.096676307 | -0.532132531 | 0.286 | 0.534 | 1 | 0.16961895 | -0.540415692 | 0.378 | 0.464 | 1 | 0.16961895 | 0.184006305 |
| Nup43 | 0.694721246 | 1.214340149 | 0.095 | 0.075 | 1 | 0.096704346 | -1.526380779 | 0.022 | 0.097 | 1 | 0.694721246 | 0.184056961 |
| Mocs3 | 0.136109171 | -1.233118828 | 0 | 0.097 | 1 | 0.096708292 | -1.486852415 | 0.022 | 0.097 | 1 | 0.136109171 | 0.18406409 |
| Gstt2 | 0.274721743 | -0.434752689 | 0.095 | 0.198 | 1 | 0.096745293 | -0.450783161 | 0.089 | 0.197 | 1 | 0.274721743 | 0.184130935 |
| Kdm2b | 0.09676128 | -0.811655059 | 0.095 | 0.28 | 1 | 0.58993808 | 0.410054092 | 0.244 | 0.322 | 1 | 0.58993808 | 0.184159814 |
| Prtn3 | 0.096777248 | 2.60964143 | 0.143 | 0.056 | 1 | 0.165804052 | 1.502500341 | 0.111 | 0.059 | 1 | 0.165804052 | 0.18418866 |
| Kat6a | 0.326902545 | -0.127846785 | 0.524 | 0.746 | 1 | 0.096868155 | 0.518143318 | 0.956 | 0.927 | 1 | 0.326902545 | 0.18435287 |
| Prkcb | 0.686435176 | -0.355975576 | 0.048 | 0.071 | 1 | 0.096920906 | -0.901889915 | 0.022 | 0.1 | 1 | 0.686435176 | 0.184448151 |
| Cep57l1 | 0.822906381 | 0.892412054 | 0.143 | 0.138 | 1 | 0.097019377 | -1.526380779 | 0.022 | 0.097 | 1 | 0.822906381 | 0.184625995 |
| Nsf | 0.763364819 | 1.037462387 | 0.429 | 0.556 | 1 | 0.097064067 | 0.797484376 | 0.778 | 0.692 | 1 | 0.763364819 | 0.184706701 |
| BC002059 | 0.396218801 | -0.785659851 | 0.048 | 0.104 | 1 | 0.097132163 | -0.743192169 | 0.022 | 0.1 | 1 | 0.396218801 | 0.184829669 |
| Snx1 | 0.097191416 | -0.226232442 | 0.429 | 0.743 | 1 | 0.957860362 | 0.170622585 | 0.667 | 0.675 | 1 | 0.957860362 | 0.18493666 |
| Slx4ip | 0.960960071 | 0.451379346 | 0.19 | 0.22 | 1 | 0.097255106 | 0.170622585 | 0.089 | 0.211 | 1 | 0.960960071 | 0.185051657 |
| Neto2 | 0.716179636 | -0.701267664 | 0.095 | 0.119 | 1 | 0.09726887 | -0.918963428 | 0.022 | 0.1 | 1 | 0.716179636 | 0.185076506 |
| Ap5s1 | 0.505969692 | -0.280424543 | 0.048 | 0.09 | 1 | 0.097332109 | -1.446210431 | 0.022 | 0.097 | 1 | 0.505969692 | 0.185190679 |
| Tnfrsf18 | 0.251619726 | -0.574155746 | 0 | 0.06 | 1 | 0.097332788 | -1.564854927 | 0.022 | 0.097 | 1 | 0.251619726 | 0.185191904 |
| Slc22a5 | 0.097396554 | 2.247507013 | 0.429 | 0.302 | 1 | 0.156950093 | 1.141363283 | 0.4 | 0.367 | 1 | 0.156950093 | 0.18530702 |
| Layn | 0.097468565 | -1.14385149 | 0.143 | 0.313 | 1 | 0.690995489 | 0.690452117 | 0.267 | 0.343 | 1 | 0.690995489 | 0.185437009 |
| Pld1 | 0.939610005 | 0.382217322 | 0.333 | 0.399 | 1 | 0.097495447 | 0.612683258 | 0.2 | 0.381 | 1 | 0.939610005 | 0.185485532 |
| Got2 | 0.097501717 | -0.542165632 | 0.286 | 0.534 | 1 | 0.370727959 | -0.396154105 | 0.578 | 0.54 | 1 | 0.370727959 | 0.185496849 |
| Gigyf2 | 0.269417402 | 0.611675647 | 0.762 | 0.743 | 1 | 0.097549182 | 0.656230974 | 0.933 | 0.837 | 1 | 0.269417402 | 0.185582522 |
| Wdr44 | 0.097619843 | 1.591309607 | 0.524 | 0.519 | 1 | 0.69677894 | 0.104034919 | 0.644 | 0.692 | 1 | 0.69677894 | 0.185710052 |
| Atp1b2 | 0.163483795 | -1.370622352 | 0 | 0.086 | 1 | 0.097642979 | -1.361321533 | 0.022 | 0.097 | 1 | 0.163483795 | 0.185751807 |
| Scmh1 | 0.98109162 | 0.303660139 | 0.571 | 0.66 | 1 | 0.09769086 | 0.581156049 | 0.889 | 0.844 | 1 | 0.98109162 | 0.185838216 |
| Smarce1 | 0.097704353 | -0.56399588 | 0.619 | 0.754 | 1 | 0.237862474 | -0.207839294 | 0.978 | 0.893 | 1 | 0.237862474 | 0.185862565 |
| Clock | 0.097743638 | -0.334162414 | 0.571 | 0.72 | 1 | 0.422218297 | 0.104899251 | 0.956 | 0.855 | 1 | 0.422218297 | 0.185933456 |
| Zranb3 | 0.520551663 | 0.351843673 | 0.048 | 0.093 | 1 | 0.097757209 | -0.387316742 | 0.022 | 0.1 | 1 | 0.520551663 | 0.185957946 |
| Gm56587 | 0.268011783 | -0.648156327 | 0 | 0.056 | 1 | 0.097783264 | -0.638855509 | 0.022 | 0.1 | 1 | 0.268011783 | 0.186004961 |
| Cchcr1 | 0.802845115 | 0.704145417 | 0.143 | 0.131 | 1 | 0.097950612 | -1.361321533 | 0.022 | 0.097 | 1 | 0.802845115 | 0.186306902 |
| Nsun5 | 0.507700057 | -0.133583154 | 0.048 | 0.09 | 1 | 0.097950726 | -1.316927414 | 0.022 | 0.097 | 1 | 0.507700057 | 0.186307107 |
| 2010320M18Rik | 0.766624392 | 0.673771768 | 0.048 | 0.067 | 1 | 0.097950726 | -1.316927414 | 0.022 | 0.097 | 1 | 0.766624392 | 0.186307107 |
| Stag2 | 0.442483591 | 0.46382492 | 0.81 | 0.858 | 1 | 0.098071646 | 0.421458855 | 1 | 0.965 | 1 | 0.442483591 | 0.186525245 |
| Spag1 | 0.098084745 | 1.432763668 | 0.238 | 0.123 | 1 | 0.350812009 | 0.294801996 | 0.111 | 0.183 | 1 | 0.350812009 | 0.186548873 |
| Map4k4 | 0.098088564 | 0.485838817 | 0.952 | 0.97 | 1 | 0.252368713 | 0.504793797 | 1 | 0.99 | 1 | 0.252368713 | 0.186555762 |
| Atad3a | 0.475181275 | 0.791128718 | 0.333 | 0.317 | 1 | 0.098123839 | 0.195806759 | 0.222 | 0.388 | 1 | 0.475181275 | 0.18661939 |
| Gm10371 | 0.996958033 | 0.638147858 | 0.19 | 0.213 | 1 | 0.098158676 | -0.549588171 | 0.111 | 0.228 | 1 | 0.996958033 | 0.186682226 |
| Cnot11 | 0.098206813 | -1.04504648 | 0.095 | 0.254 | 1 | 0.197080343 | 0.373217324 | 0.2 | 0.332 | 1 | 0.197080343 | 0.186769048 |
| Zfp809 | 0.869748023 | 0.97333205 | 0.143 | 0.146 | 1 | 0.098397154 | 0.316290255 | 0.133 | 0.27 | 1 | 0.869748023 | 0.187112308 |
| Bcl2 | 0.098501005 | 1.290914674 | 0.19 | 0.41 | 1 | 0.465763573 | -0.654797053 | 0.4 | 0.45 | 1 | 0.465763573 | 0.187299561 |
| Pcdhb17 | 0.324827658 | -0.718545655 | 0 | 0.045 | 1 | 0.098508331 | -0.387316742 | 0.022 | 0.1 | 1 | 0.324827658 | 0.18731277 |
| Eef1akmt1 | 0.144438142 | -0.335217016 | 0.19 | 0.407 | 1 | 0.098520479 | -0.142898014 | 0.2 | 0.356 | 1 | 0.144438142 | 0.187334673 |

| | | | | | | | | | | | | |
|---|---|---|---|---|---|---|---|---|---|---|---|---|
| Palmd | 0.262135677 | 1.277843091 | 0.048 | 0.142 | 1 | 0.098548181 | -0.725012152 | 0.089 | 0.197 | 1 | 0.262135677 | 0.187384618 |
| Ccdc149 | 0.208886118 | -0.849790188 | 0 | 0.071 | 1 | 0.098557863 | -0.518561275 | 0.022 | 0.1 | 1 | 0.208886118 | 0.187402073 |
| 1700008J07Rik | 0.222228992 | -0.849790188 | 0 | 0.067 | 1 | 0.098559731 | -1.223818009 | 0.022 | 0.097 | 1 | 0.222228992 | 0.187405442 |
| Reep4 | 0.768257326 | 0.214340149 | 0.048 | 0.063 | 1 | 0.098559731 | -1.223818009 | 0.022 | 0.097 | 1 | 0.768257326 | 0.187405442 |
| Ptp4a3 | 0.502505429 | -0.413691074 | 0.048 | 0.09 | 1 | 0.098640967 | -0.316927414 | 0.022 | 0.1 | 1 | 0.502505429 | 0.187551893 |
| Arhgap24 | 0.349927091 | -0.560283571 | 0.095 | 0.183 | 1 | 0.098683267 | -0.638855509 | 0.044 | 0.135 | 1 | 0.349927091 | 0.187628147 |
| Eeig2 | 0.098712373 | 0.380272081 | 0.81 | 0.754 | 1 | 0.780575704 | 0.116399005 | 0.578 | 0.785 | 1 | 0.780575704 | 0.187680614 |
| Gm12227 | 0.893497436 | 0.60338244 | 0.095 | 0.09 | 1 | 0.098778348 | 0.397670367 | 0.022 | 0.1 | 1 | 0.893497436 | 0.187799535 |
| Mosmo | 0.098822451 | -0.375859 | 0.476 | 0.739 | 1 | 0.186016477 | 0.113706941 | 0.533 | 0.813 | 1 | 0.186016477 | 0.187879025 |
| Smc4 | 0.098824556 | -0.614709844 | 0.333 | 0.593 | 1 | 0.439884008 | -0.569496309 | 0.667 | 0.585 | 1 | 0.439884008 | 0.187882819 |
| Tgm2 | 0.099102876 | 0.803692363 | 0.857 | 0.754 | 1 | 0.475269956 | -0.526929855 | 0.689 | 0.526 | 1 | 0.475269956 | 0.188384372 |
| Rsrc1 | 0.859047921 | 0.610529334 | 0.571 | 0.694 | 1 | 0.099124913 | 0.524056484 | 0.844 | 0.699 | 1 | 0.859047921 | 0.188424078 |
| Sh3rf1 | 0.200596758 | -0.505284246 | 0.476 | 0.634 | 1 | 0.099599189 | 0.486675373 | 0.911 | 0.73 | 1 | 0.200596758 | 0.18927838 |
| A930015D03Rik | 0.09969084 | -0.397690595 | 0.286 | 0.515 | 1 | 0.413407727 | 0.320502507 | 0.467 | 0.599 | 1 | 0.413407727 | 0.189443416 |
| Zfp758 | 0.099701742 | -1.303508156 | 0.048 | 0.19 | 1 | 0.114775816 | -0.564854927 | 0.089 | 0.19 | 1 | 0.114775816 | 0.189463046 |
| Traf6 | 0.858960965 | 0.625408746 | 0.19 | 0.231 | 1 | 0.099795351 | 0.147019686 | 0.2 | 0.343 | 1 | 0.858960965 | 0.189631589 |
| Zzef1 | 0.099810776 | 0.72926688 | 0.714 | 0.687 | 1 | 0.930631575 | 0.297917689 | 0.911 | 0.796 | 1 | 0.930631575 | 0.18965936 |
| Clk1 | 0.310805396 | 0.34619711 | 1 | 0.989 | 1 | 0.099858318 | 0.394184753 | 0.978 | 0.976 | 1 | 0.310805396 | 0.189744953 |
| Jade3 | 0.623343204 | 1.526930379 | 0.095 | 0.071 | 1 | 0.099903307 | -1.204452685 | 0.067 | 0.159 | 1 | 0.623343204 | 0.189825944 |
| ENSMUSG00000121485 | 0.71232227 | 0.918884266 | 0.095 | 0.075 | 1 | 0.099938874 | -1.695439037 | 0.044 | 0.131 | 1 | 0.71232227 | 0.18988997 |
| Kif19b | 0.592290112 | -0.154047257 | 0.095 | 0.142 | 1 | 0.100100596 | -0.44345982 | 0.111 | 0.232 | 1 | 0.592290112 | 0.190181063 |
| Cilk1 | 0.357693588 | 0.954323935 | 0.476 | 0.451 | 1 | 0.10013873 | 0.123645178 | 0.356 | 0.616 | 1 | 0.357693588 | 0.190249695 |
| Macir | 0.100184292 | -0.989193245 | 0.048 | 0.201 | 1 | 0.527484641 | 0.348918189 | 0.467 | 0.419 | 1 | 0.527484641 | 0.190331692 |
| Nif3l1 | 0.208945042 | -0.880817084 | 0.048 | 0.146 | 1 | 0.100232614 | 0.172110667 | 0.067 | 0.17 | 1 | 0.208945042 | 0.19041865 |
| Elk1 | 0.153741354 | -1.107587946 | 0.048 | 0.164 | 1 | 0.10028166 | -0.432404631 | 0.067 | 0.17 | 1 | 0.153741354 | 0.190506908 |
| Zfp654 | 0.165211692 | 0.629377648 | 0.571 | 0.485 | 1 | 0.100317195 | 1.057981973 | 0.778 | 0.612 | 1 | 0.165211692 | 0.190570851 |
| Nbas | 0.100335338 | 0.498329761 | 0.286 | 0.563 | 1 | 0.987418015 | 0.394396297 | 0.467 | 0.567 | 1 | 0.987418015 | 0.190603496 |
| Tet3 | 0.5646994 | 0.794937457 | 0.524 | 0.493 | 1 | 0.100404963 | 0.778831569 | 0.933 | 0.785 | 1 | 0.5646994 | 0.190728769 |
| Mtx3 | 0.100573155 | -1.752492987 | 0 | 0.116 | 1 | 0.127032989 | -0.454430938 | 0.044 | 0.128 | 1 | 0.127032989 | 0.191031351 |
| Nfe2l3 | 0.176025523 | 1.8372705 | 0.143 | 0.067 | 1 | 0.100624459 | 1.572041274 | 0.222 | 0.138 | 1 | 0.176025523 | 0.191123635 |
| A830082K12Rik | 0.10069752 | -2.02666795 | 0 | 0.116 | 1 | 0.1895925 | 1.045642666 | 0.044 | 0.114 | 1 | 0.1895925 | 0.191255049 |
| St8sia6 | 0.100700016 | -2.159118246 | 0 | 0.116 | 1 | 0.119061868 | -2.124282336 | 0 | 0.052 | 1 | 0.119061868 | 0.191259539 |
| Slu7 | 0.100786543 | -0.752492987 | 0.238 | 0.433 | 1 | 0.258456427 | -0.480157763 | 0.378 | 0.436 | 1 | 0.258456427 | 0.191415158 |
| Klk10 | 0.100817907 | -3.962852853 | 0 | 0.116 | 1 | 0.260665169 | -0.901889915 | 0 | 0.028 | 1 | 0.260665169 | 0.191471564 |
| Ide | 0.970866805 | 0.569435108 | 0.524 | 0.698 | 1 | 0.100827456 | -0.174092636 | 0.733 | 0.716 | 1 | 0.970866805 | 0.191488735 |
| Lifr | 0.224548622 | 1.131513718 | 0.238 | 0.451 | 1 | 0.100941628 | -0.649123844 | 0.133 | 0.253 | 1 | 0.224548622 | 0.191694044 |
| Gm43330 | 0.10098287 | 2.825774861 | 0.095 | 0.03 | 1 | 0.49148904 | 0.632446513 | 0.044 | 0.076 | 1 | 0.49148904 | 0.191768199 |
| 5031439G07Rik | 0.17048654 | 0.435782491 | 0.19 | 0.407 | 1 | 0.101302805 | 0.330556171 | 0.244 | 0.478 | 1 | 0.17048654 | 0.192343351 |
| Gm39469 | 0.101373035 | 1.195724471 | 0.238 | 0.116 | 1 | 0.533631826 | 0.277080226 | 0.089 | 0.128 | 1 | 0.533631826 | 0.192469577 |
| Zswim4 | 0.101417137 | 1.483947209 | 0.429 | 0.313 | 1 | 0.638432435 | 0.367870354 | 0.311 | 0.426 | 1 | 0.638432435 | 0.192548839 |
| Wdpcp | 0.804524518 | 1.811275291 | 0.238 | 0.243 | 1 | 0.101481366 | 1.592307122 | 0.333 | 0.239 | 1 | 0.804524518 | 0.192664264 |
| Creb5 | 0.382040565 | 1.359075242 | 0.286 | 0.231 | 1 | 0.101777908 | 1.195626591 | 0.289 | 0.211 | 1 | 0.382040565 | 0.193197073 |
| Klhl20 | 0.782048615 | 1.155923463 | 0.143 | 0.172 | 1 | 0.101940445 | -0.413788953 | 0.089 | 0.201 | 1 | 0.782048615 | 0.193489035 |
| Hira | 0.102000944 | -0.488115915 | 0.143 | 0.354 | 1 | 0.57768149 | 0.639351209 | 0.489 | 0.491 | 1 | 0.57768149 | 0.193597695 |
| Asap3 | 0.181361321 | 1.295260144 | 0.429 | 0.377 | 1 | 0.102029392 | 0.826436761 | 0.111 | 0.239 | 1 | 0.181361321 | 0.193648787 |
| Katnbl1 | 0.102156691 | 0.777607579 | 0.571 | 0.507 | 1 | 0.114722773 | 0.680094886 | 0.756 | 0.706 | 1 | 0.114722773 | 0.193877393 |
| Cyp2u1 | 0.928552379 | 0.214340149 | 0.095 | 0.104 | 1 | 0.102199601 | -0.144746438 | 0.044 | 0.135 | 1 | 0.928552379 | 0.193954444 |
| Pgap1 | 0.196462148 | -0.970084422 | 0 | 0.075 | 1 | 0.102509813 | -0.901889915 | 0.044 | 0.131 | 1 | 0.196462148 | 0.194511364 |
| Dhrs3 | 0.962201795 | 0.268630305 | 0.476 | 0.549 | 1 | 0.102547805 | -0.494465599 | 0.667 | 0.72 | 1 | 0.962201795 | 0.194579557 |
| Gm2415 | 0.173676192 | -0.911190733 | 0 | 0.082 | 1 | 0.10260097 | 0.259861155 | 0.067 | 0.173 | 1 | 0.173676192 | 0.19467498 |
| Ankdd1a | 0.102787795 | 2.673771768 | 0.095 | 0.03 | 1 | 0.331918331 | -0.776359032 | 0 | 0.021 | 1 | 0.331918331 | 0.19501026 |
| Gata3 | 0.102795046 | 3.214340149 | 0.095 | 0.03 | 1 | 0.232015706 | -2.709244837 | 0 | 0.031 | 1 | 0.232015706 | 0.19502327 |
| Mid2 | 0.157365934 | 0.877661603 | 0.619 | 0.653 | 1 | 0.102801428 | 0.846029563 | 0.711 | 0.675 | 1 | 0.157365934 | 0.195034722 |
| Aldh3a2 | 0.317975113 | -0.496153234 | 0.048 | 0.119 | 1 | 0.102907763 | -1.158229668 | 0.044 | 0.128 | 1 | 0.317975113 | 0.195225519 |
| Afg3l1 | 0.10293251 | -0.718545655 | 0.143 | 0.332 | 1 | 0.485719489 | 0.674510573 | 0.267 | 0.381 | 1 | 0.485719489 | 0.195269918 |
| Zfp74 | 0.264115824 | -0.413691074 | 0.048 | 0.138 | 1 | 0.103189889 | -0.249813218 | 0.089 | 0.201 | 1 | 0.264115824 | 0.195731624 |
| Fastkd1 | 0.789239895 | 1.518493542 | 0.095 | 0.127 | 1 | 0.103238009 | -0.361321533 | 0.067 | 0.166 | 1 | 0.789239895 | 0.195817932 |
| Tbc1d4 | 0.122365506 | 1.389563339 | 0.095 | 0.272 | 1 | 0.103280536 | 0.55654018 | 0.111 | 0.239 | 1 | 0.122365506 | 0.195894202 |
| Arhgef2 | 0.819123517 | 0.908237021 | 0.286 | 0.325 | 1 | 0.103280934 | -0.311280851 | 0.289 | 0.443 | 1 | 0.819123517 | 0.195894918 |
| Strbp | 0.528451931 | 0.340498699 | 0.476 | 0.646 | 1 | 0.103283587 | 0.44410594 | 0.933 | 0.83 | 1 | 0.528451931 | 0.195899675 |
| Sh3rf3 | 0.10332825 | -1.849790188 | 0.095 | 0.25 | 1 | 0.242963737 | 0.293505774 | 0.244 | 0.394 | 1 | 0.242963737 | 0.195979773 |
| Kank1 | 0.969555478 | -0.135797575 | 0.857 | 0.828 | 1 | 0.103432855 | 0.107674949 | 0.578 | 0.834 | 1 | 0.969555478 | 0.196167355 |
| Akap1 | 0.239532796 | -0.355975576 | 0.048 | 0.146 | 1 | 0.103462708 | -1.017367132 | 0.067 | 0.159 | 1 | 0.239532796 | 0.196220884 |
| Slc25a22 | 0.196857648 | -0.970084422 | 0.048 | 0.149 | 1 | 0.103529315 | -0.667424661 | 0.044 | 0.131 | 1 | 0.196857648 | 0.196340312 |
| Tmem116 | 0.512813368 | 1.214340149 | 0.143 | 0.104 | 1 | 0.103563134 | -0.667424661 | 0.044 | 0.131 | 1 | 0.512813368 | 0.196400945 |
| Supt3 | 0.103684946 | 1.291051989 | 0.524 | 0.485 | 1 | 0.174465663 | 1.080243605 | 0.533 | 0.467 | 1 | 0.174465663 | 0.196619324 |
| Mybl2 | 0.103693787 | 2.79930265 | 0.095 | 0.03 | 1 | 0.659789691 | -0.404390255 | 0.022 | 0.035 | 1 | 0.659789691 | 0.196635172 |
| Gm15787 | 0.764647844 | 1.832469514 | 0.095 | 0.131 | 1 | 0.103802609 | -0.638855509 | 0.044 | 0.131 | 1 | 0.764647844 | 0.196830237 |
| Cnot1 | 0.652310622 | 0.229321541 | 0.667 | 0.776 | 1 | 0.104019442 | 0.57096222 | 0.956 | 0.875 | 1 | 0.652310622 | 0.19721884 |
| Gm26688 | 0.104129536 | 1.79930265 | 0.143 | 0.056 | 1 | 0.418201984 | 1.567595369 | 0.067 | 0.042 | 1 | 0.418201984 | 0.197416111 |
| Plekhg3 | 0.162233321 | 1.589238257 | 0.333 | 0.243 | 1 | 0.104409385 | 0.375083682 | 0.711 | 0.581 | 1 | 0.162233321 | 0.197917451 |
| Bmp4 | 0.208982462 | -1.370622352 | 0 | 0.071 | 1 | 0.104545644 | -2.044847868 | 0.022 | 0.093 | 1 | 0.208982462 | 0.198161496 |
| Nek5 | 0.104615147 | 2.536268244 | 0.095 | 0.03 | 1 | 0.497611189 | 1.683072586 | 0.022 | 0.01 | 1 | 0.497611189 | 0.198285966 |
| Nsdhl | 0.71672318 | 0.166811779 | 0.286 | 0.366 | 1 | 0.104661008 | -0.110476536 | 0.111 | 0.228 | 1 | 0.71672318 | 0.19836809 |
| Ift88 | 0.104780383 | -0.180377366 | 0.19 | 0.422 | 1 | 0.5543186 | 0.274987848 | 0.267 | 0.388 | 1 | 0.5543186 | 0.198581837 |
| Psmd14 | 0.104894284 | 0.824809525 | 0.714 | 0.843 | 1 | 0.87227154 | 0.329165287 | 0.844 | 0.782 | 1 | 0.87227154 | 0.198785757 |
| Ankrd12 | 0.729817411 | 0.205374641 | 1 | 0.944 | 1 | 0.104903835 | 0.486380675 | 1 | 0.972 | 1 | 0.729817411 | 0.198802856 |
| Map2 | 0.104933057 | 1.701516758 | 0.524 | 0.53 | 1 | 0.260957314 | 0.487521777 | 0.244 | 0.388 | 1 | 0.260957314 | 0.198855168 |
| Cspg5 | 0.474473391 | 2.045740545 | 0.143 | 0.101 | 1 | 0.105015901 | -0.286553765 | 0.044 | 0.131 | 1 | 0.474473391 | 0.199003462 |
| Osbpl8 | 0.385526113 | 0.610609747 | 0.81 | 0.776 | 1 | 0.105018079 | 0.393238121 | 0.956 | 0.848 | 1 | 0.385526113 | 0.199007362 |
| Dnajc13 | 0.310748158 | 0.935419633 | 0.619 | 0.634 | 1 | 0.105084763 | 0.535867661 | 0.844 | 0.758 | 1 | 0.310748158 | 0.199126719 |
| Cox10 | 0.279950921 | -0.355975576 | 0.048 | 0.134 | 1 | 0.105224986 | -0.276285429 | 0.067 | 0.17 | 1 | 0.279950921 | 0.199377674 |
| Zadh2 | 0.803744045 | 0.214340149 | 0.19 | 0.216 | 1 | 0.105296556 | -0.147002412 | 0.156 | 0.291 | 1 | 0.803744045 | 0.199505748 |
| Rell1 | 0.105330084 | 0.123574685 | 0.238 | 0.459 | 1 | 0.272722208 | 0.219323672 | 0.844 | 0.678 | 1 | 0.272722208 | 0.199565741 |
| Fbxl18 | 0.374462733 | 2.088809267 | 0.143 | 0.093 | 1 | 0.105400846 | 0.435145073 | 0.044 | 0.135 | 1 | 0.374462733 | 0.199692354 |
| Zbtb48 | 0.105421242 | 1.908237021 | 0.143 | 0.056 | 1 | 0.140239497 | -0.941418279 | 0.044 | 0.118 | 1 | 0.140239497 | 0.199728845 |
| Ccdc141 | 0.627648035 | 1.79930265 | 0.095 | 0.071 | 1 | 0.105424234 | -0.792660845 | 0.067 | 0.163 | 1 | 0.627648035 | 0.1997342 |
| Gm15614 | 0.499272992 | -0.159118246 | 0.095 | 0.153 | 1 | 0.105451895 | 0.513147585 | 0.067 | 0.173 | 1 | 0.499272992 | 0.199783688 |
| Gclm | 0.105489721 | -0.270045397 | 0.333 | 0.556 | 1 | 0.579691057 | -0.193916524 | 0.778 | 0.585 | 1 | 0.579691057 | 0.19985136 |
| Ncoa6 | 0.541208157 | 1.077404504 | 0.429 | 0.504 | 1 | 0.105518229 | 0.867104534 | 0.889 | 0.789 | 1 | 0.541208157 | 0.199902361 |
| Ss18 | 0.159756904 | -0.378549267 | 0.381 | 0.679 | 1 | 0.105563695 | -0.218311533 | 0.711 | 0.772 | 1 | 0.159756904 | 0.199983696 |
| Pcgf3 | 0.147106889 | -0.519999976 | 0.238 | 0.429 | 1 | 0.105577499 | 0.730519749 | 0.6 | 0.619 | 1 | 0.147106889 | 0.20000839 |
| Gm10033 | 0.500606474 | -0.184209227 | 0.095 | 0.153 | 1 | 0.105801117 | -1.017367132 | 0.067 | 0.159 | 1 | 0.500606474 | 0.200408357 |
| Lsm14a | 0.105879746 | -0.611630451 | 0.429 | 0.646 | 1 | 0.325830792 | 0.176112597 | 0.889 | 0.896 | 1 | 0.325830792 | 0.200548972 |
| Nln | 0.696248478 | -0.355975576 | 0.095 | 0.119 | 1 | 0.10591359 | -1.11134328 | 0.089 | 0.187 | 1 | 0.696248478 | 0.200609492 |
| Tmc8 | 0.267917141 | -0.413691074 | 0 | 0.056 | 1 | 0.105978808 | 2.583536913 | 0.089 | 0.038 | 1 | 0.267917141 | 0.200726109 |
| Szt2 | 0.875010596 | 0.747772349 | 0.095 | 0.116 | 1 | 0.106083365 | -0.387316742 | 0.044 | 0.131 | 1 | 0.875010596 | 0.200913049 |
| Msto1 | 0.402633734 | 1.258734268 | 0.048 | 0.112 | 1 | 0.106083631 | -0.158229668 | 0.044 | 0.131 | 1 | 0.402633734 | 0.200913525 |

| | | | | | | | | | | | | |
|---|---|---|---|---|---|---|---|---|---|---|---|---|
| Snrnp48 | 0.106209453 | 0.684578523 | 0.762 | 0.75 | 1 | 0.884647003 | 0.209701777 | 0.778 | 0.782 | 1 | 0.884647003 | 0.201138458 |
| Appbp2 | 0.106266098 | 0.598705953 | 0.857 | 0.944 | 1 | 0.245013796 | -0.121403662 | 0.978 | 0.958 | 1 | 0.245013796 | 0.201239713 |
| Cuedc1 | 0.106288257 | -0.133583154 | 0.286 | 0.582 | 1 | 0.888219886 | 0.160119748 | 0.756 | 0.792 | 1 | 0.888219886 | 0.20127932 |
| Gata2 | 0.492606471 | -0.574155746 | 0 | 0.022 | 1 | 0.106356446 | -1.149817428 | 0.022 | 0.097 | 1 | 0.492606471 | 0.201401198 |
| Zdhhc2 | 0.106405553 | 2.001346426 | 0.19 | 0.09 | 1 | 0.897953988 | 1.757073168 | 0.089 | 0.1 | 1 | 0.897953988 | 0.201488964 |
| Mcm9 | 0.212861535 | 1.79930265 | 0.286 | 0.201 | 1 | 0.106489939 | 0.583536913 | 0.111 | 0.239 | 1 | 0.212861535 | 0.201639771 |
| Gm57071 | 0.106550248 | -1.326228232 | 0 | 0.112 | 1 | 0.933853613 | 1.098110085 | 0.044 | 0.048 | 1 | 0.933853613 | 0.20174754 |
| Zbtb22 | 0.106550248 | -1.326228232 | 0 | 0.112 | 1 | 0.486174415 | -0.153428682 | 0.111 | 0.152 | 1 | 0.486174415 | 0.20174754 |
| Armc5 | 0.106613525 | -1.370622352 | 0 | 0.112 | 1 | 0.138200208 | -0.57996182 | 0.067 | 0.152 | 1 | 0.138200208 | 0.201860606 |
| Trmt61a | 0.106672278 | -1.455511249 | 0 | 0.112 | 1 | 0.332124242 | -0.589945908 | 0.044 | 0.087 | 1 | 0.332124242 | 0.20196558 |
| Tk2 | 0.106672433 | -1.413691074 | 0 | 0.112 | 1 | 0.357544342 | -0.242926832 | 0.067 | 0.114 | 1 | 0.357544342 | 0.201965859 |
| Sh3bp5l | 0.106672433 | -1.413691074 | 0 | 0.112 | 1 | 0.722897503 | 0.486675373 | 0.133 | 0.163 | 1 | 0.722897503 | 0.201965859 |
| Mrm3 | 0.106672433 | -1.413691074 | 0 | 0.112 | 1 | 0.765203983 | 0.513147585 | 0.067 | 0.083 | 1 | 0.765203983 | 0.201965859 |
| Kcnip2 | 0.304430544 | -0.574155746 | 0 | 0.049 | 1 | 0.10669472 | -1.743192169 | 0.022 | 0.093 | 1 | 0.304430544 | 0.202005677 |
| Rbm19 | 0.106727131 | -1.455511249 | 0 | 0.112 | 1 | 0.748243998 | 0.390290837 | 0.111 | 0.135 | 1 | 0.748243998 | 0.202063581 |
| Tmem186 | 0.106727131 | -1.455511249 | 0 | 0.112 | 1 | 0.254588464 | -0.223818009 | 0.067 | 0.128 | 1 | 0.254588464 | 0.202063581 |
| H2-M3 | 0.106727131 | -1.455511249 | 0 | 0.112 | 1 | 0.12214448 | -1.404390255 | 0.044 | 0.121 | 1 | 0.12214448 | 0.202063581 |
| Mark2 | 0.278544685 | -0.111005306 | 0.429 | 0.638 | 1 | 0.106745188 | 0.61952677 | 0.978 | 0.83 | 1 | 0.278544685 | 0.202095841 |
| Stx11 | 0.106776216 | -1.574155746 | 0 | 0.112 | 1 | 0.268056706 | -0.776359032 | 0.067 | 0.125 | 1 | 0.268056706 | 0.202151271 |
| Yjefn3 | 0.106777774 | -1.496153234 | 0 | 0.112 | 1 | 0.381522305 | -0.387316742 | 0.067 | 0.111 | 1 | 0.381522305 | 0.202154055 |
| Dusp28 | 0.113542153 | -1.535681598 | 0 | 0.108 | 1 | 0.10677901 | -1.486852415 | 0 | 0.055 | 1 | 0.113542153 | 0.202156262 |
| Slc52a2 | 0.144606783 | -1.081115734 | 0 | 0.093 | 1 | 0.10677901 | -1.486852415 | 0 | 0.055 | 1 | 0.144606783 | 0.202156262 |
| Tkfc | 0.144664875 | -1.133583154 | 0 | 0.093 | 1 | 0.10677901 | -1.486852415 | 0 | 0.055 | 1 | 0.144664875 | 0.202156262 |
| Bora | 0.144907264 | -1.496153234 | 0 | 0.093 | 1 | 0.10677901 | -1.486852415 | 0 | 0.055 | 1 | 0.144907264 | 0.202156262 |
| 5730414N17Rik | 0.196329837 | -0.849790188 | 0 | 0.075 | 1 | 0.10677901 | -1.564854927 | 0 | 0.055 | 1 | 0.196329837 | 0.202156262 |
| Glra1 | 0.285527727 | -1.133583154 | 0 | 0.052 | 1 | 0.10677901 | -1.564854927 | 0 | 0.055 | 1 | 0.285527727 | 0.202156262 |
| Dnd1 | 0.359270744 | 2.673771768 | 0.048 | 0.019 | 1 | 0.10677901 | -1.486852415 | 0 | 0.055 | 1 | 0.359270744 | 0.202156262 |
| Asb1 | 0.555949871 | 1.673771768 | 0.095 | 0.067 | 1 | 0.10677901 | -1.486852415 | 0 | 0.055 | 1 | 0.555949871 | 0.202156262 |
| Gm50071 | 0.632076134 | 1.673771768 | 0 | 0.011 | 1 | 0.10677901 | -1.486852415 | 0 | 0.055 | 1 | 0.632076134 | 0.202156262 |
| Ciart | 0.837550449 | 1.815790773 | 0.095 | 0.09 | 1 | 0.10677901 | -1.486852415 | 0 | 0.055 | 1 | 0.837550449 | 0.202156262 |
| Ccne2 | 0.173893017 | -1.611630451 | 0 | 0.082 | 1 | 0.106800662 | -1.564854927 | 0 | 0.055 | 1 | 0.173893017 | 0.202194942 |
| Zfp213 | 0.184814956 | -1.133583154 | 0 | 0.078 | 1 | 0.106800662 | -1.638855509 | 0 | 0.055 | 1 | 0.184814956 | 0.202194942 |
| Fam217b | 0.251619726 | -0.574155746 | 0 | 0.06 | 1 | 0.106800662 | -1.564854927 | 0 | 0.055 | 1 | 0.251619726 | 0.202194942 |
| Neu3 | 0.532436372 | 1.088809267 | 0 | 0.019 | 1 | 0.106800662 | -1.564854927 | 0 | 0.055 | 1 | 0.532436372 | 0.202194942 |
| Gm9828 | 0.655561847 | 1.503846766 | 0.048 | 0.03 | 1 | 0.106800662 | -1.564854927 | 0 | 0.055 | 1 | 0.655561847 | 0.202194942 |
| Cep152 | 0.679768988 | 0.425844254 | 0.048 | 0.075 | 1 | 0.106800662 | -1.638855509 | 0 | 0.055 | 1 | 0.679768988 | 0.202194942 |
| Gm49041 | 0.725963137 | 0.281454345 | 0.048 | 0.067 | 1 | 0.106800662 | -1.638855509 | 0 | 0.055 | 1 | 0.725963137 | 0.202194942 |
| Gm17344 | 0.741589569 | 1.214340149 | 0.048 | 0.034 | 1 | 0.106800662 | -1.564854927 | 0 | 0.055 | 1 | 0.741589569 | 0.202194942 |
| Tincr | 0.815673353 | 1.214340149 | 0.048 | 0.037 | 1 | 0.106800662 | -1.564854927 | 0 | 0.055 | 1 | 0.815673353 | 0.202194942 |
| Rnf125 | 0.457281049 | 0.503846766 | 0 | 0.026 | 1 | 0.106819221 | -1.776359032 | 0 | 0.055 | 1 | 0.457281049 | 0.202228095 |
| 4930570G19Rik | 0.492608234 | 0.866416846 | 0 | 0.022 | 1 | 0.106819221 | -1.776359032 | 0 | 0.055 | 1 | 0.492608234 | 0.202228095 |
| Foxl1 | 0.492608234 | 0.866416846 | 0 | 0.022 | 1 | 0.106819221 | -1.901889915 | 0 | 0.055 | 1 | 0.492608234 | 0.202228095 |
| Ric3 | 0.929739496 | 0.766881172 | 0.048 | 0.052 | 1 | 0.106819221 | -1.709244837 | 0 | 0.055 | 1 | 0.929739496 | 0.202228095 |
| Gm48604 | 0.110202281 | 2.351843673 | 0.095 | 0.03 | 1 | 0.106819427 | -1.776359032 | 0 | 0.055 | 1 | 0.110202281 | 0.202228464 |
| Gm10941 | 0.30448834 | -0.133583154 | 0 | 0.049 | 1 | 0.106819427 | -1.638855509 | 0 | 0.055 | 1 | 0.30448834 | 0.202228464 |
| Adcy5 | 0.37081917 | -0.496153234 | 0 | 0.037 | 1 | 0.106819427 | -1.901889915 | 0 | 0.055 | 1 | 0.37081917 | 0.202228464 |
| Arl4d | 0.847205797 | 0.281454345 | 0.095 | 0.082 | 1 | 0.106819427 | -1.638855509 | 0 | 0.055 | 1 | 0.847205797 | 0.202228464 |
| Trpc4ap | 0.55160033 | 0.646971708 | 0.667 | 0.776 | 1 | 0.106826698 | 0.333623429 | 0.933 | 0.844 | 1 | 0.55160033 | 0.202241453 |
| Ccdc30 | 0.145389745 | 2.088809267 | 0.095 | 0.034 | 1 | 0.106835305 | -1.901889915 | 0 | 0.055 | 1 | 0.145389745 | 0.202256827 |
| Slc16a3 | 0.163434053 | -1.02666795 | 0 | 0.086 | 1 | 0.106835305 | -2.071814916 | 0 | 0.055 | 1 | 0.163434053 | 0.202256827 |
| Cbx8 | 0.292179707 | 1.673771768 | 0.095 | 0.045 | 1 | 0.106835511 | -1.709244837 | 0 | 0.055 | 1 | 0.292179707 | 0.202257195 |
| Spry1 | 0.517346031 | 0.866416846 | 0.095 | 0.06 | 1 | 0.106835511 | -2.271123724 | 0 | 0.055 | 1 | 0.517346031 | 0.202257195 |
| Lama3 | 0.173814074 | 1.214340149 | 0.048 | 0.011 | 1 | 0.106847058 | -1.84048937 | 0 | 0.055 | 1 | 0.173814074 | 0.202277822 |
| Ajuba | 0.304385789 | -0.326228232 | 0 | 0.049 | 1 | 0.106847058 | -1.84048937 | 0 | 0.055 | 1 | 0.304385789 | 0.202277822 |
| Kank4 | 0.304411431 | -0.574155746 | 0 | 0.049 | 1 | 0.106847058 | -2.017367132 | 0 | 0.055 | 1 | 0.304411431 | 0.202277822 |
| Gm48512 | 0.900104013 | 0.673771768 | 0.048 | 0.041 | 1 | 0.106848295 | -2.316927414 | 0 | 0.055 | 1 | 0.900104013 | 0.202280032 |
| Lbhd2 | 0.918806658 | 2.425844254 | 0.048 | 0.045 | 1 | 0.106848708 | -2.017367132 | 0 | 0.055 | 1 | 0.918806658 | 0.202280769 |
| Ccl19-ps2 | 0.251727085 | -1.02666795 | 0 | 0.06 | 1 | 0.106848914 | -1.960783604 | 0 | 0.055 | 1 | 0.251727085 | 0.202281137 |
| Scml2 | 0.304385789 | -0.574155746 | 0 | 0.049 | 1 | 0.10684912 | -1.776359032 | 0 | 0.055 | 1 | 0.304385789 | 0.202281506 |
| Slc44a3 | 0.106864405 | -1.648156327 | 0 | 0.112 | 1 | 0.179497566 | 2.435145073 | 0.089 | 0.045 | 1 | 0.179497566 | 0.202308809 |
| Il17rb | 0.106864405 | -1.683780237 | 0 | 0.112 | 1 | 0.941914357 | 1.155825583 | 0.111 | 0.125 | 1 | 0.941914357 | 0.202308809 |
| Rasgrp2 | 0.425614684 | 0.214340149 | 0 | 0.03 | 1 | 0.106865616 | -2.316927414 | 0 | 0.055 | 1 | 0.425614684 | 0.202310972 |
| Adora1 | 0.184945758 | -1.648156327 | 0 | 0.078 | 1 | 0.106867678 | -2.071814916 | 0 | 0.055 | 1 | 0.184945758 | 0.202314655 |
| Gm34728 | 0.57816106 | 1.351843673 | 0 | 0.015 | 1 | 0.106869946 | -2.071814916 | 0 | 0.055 | 1 | 0.57816106 | 0.202318706 |
| Gm19605 | 0.192751309 | 1.97333205 | 0.095 | 0.037 | 1 | 0.106894483 | -1.404390255 | 0 | 0.055 | 1 | 0.192751309 | 0.202362535 |
| Abca7 | 0.425599726 | 0.351843673 | 0 | 0.03 | 1 | 0.106894483 | -1.404390255 | 0 | 0.055 | 1 | 0.425599726 | 0.202362535 |
| Gm47675 | 0.551781566 | 1.747772349 | 0.095 | 0.067 | 1 | 0.106894483 | -1.404390255 | 0 | 0.055 | 1 | 0.551781566 | 0.202362535 |
| Gm8883 | 0.860094294 | 1.214340149 | 0.048 | 0.06 | 1 | 0.106894483 | -1.404390255 | 0 | 0.055 | 1 | 0.860094294 | 0.202362535 |
| Lca5 | 0.106929841 | -1.718545655 | 0 | 0.112 | 1 | 0.359996023 | -0.101198273 | 0.089 | 0.142 | 1 | 0.359996023 | 0.20242569 |
| Cenpv | 0.210056645 | 0.586308926 | 0.048 | 0.157 | 1 | 0.107007621 | -0.738391182 | 0.067 | 0.159 | 1 | 0.210056645 | 0.20256461 |
| Zfand1 | 0.603600115 | -0.280424543 | 0.048 | 0.078 | 1 | 0.107022462 | -0.86124793 | 0.022 | 0.097 | 1 | 0.603600115 | 0.202591116 |
| 4732440D04Rik | 0.107097349 | 0.995699863 | 0.381 | 0.25 | 1 | 0.138988599 | 0.158117395 | 0.222 | 0.377 | 1 | 0.138988599 | 0.202724856 |
| Tsc1 | 0.991440601 | 0.763039106 | 0.333 | 0.418 | 1 | 0.107162691 | 0.177373364 | 0.333 | 0.55 | 1 | 0.991440601 | 0.202841539 |
| Kctd18 | 0.10720225 | 1.936806174 | 0.238 | 0.127 | 1 | 0.198979532 | -0.294207337 | 0.089 | 0.173 | 1 | 0.198979532 | 0.202912177 |
| Dmac2 | 0.209667483 | -0.849790188 | 0.048 | 0.146 | 1 | 0.107294496 | -0.901889915 | 0.022 | 0.097 | 1 | 0.209667483 | 0.203076883 |
| Chml | 0.718367751 | 0.503846766 | 0.048 | 0.071 | 1 | 0.107399209 | -1.674479418 | 0.022 | 0.093 | 1 | 0.718367751 | 0.203263829 |
| Ccdc122 | 0.316728594 | 0.171271427 | 0.095 | 0.19 | 1 | 0.107519031 | -0.539319835 | 0.089 | 0.187 | 1 | 0.316728594 | 0.203477721 |
| Fam177a | 0.251790506 | -0.413691074 | 0 | 0.06 | 1 | 0.107665622 | -0.86124793 | 0.022 | 0.097 | 1 | 0.251790506 | 0.203739359 |
| D730003I15Rik | 0.324739547 | -0.233118828 | 0 | 0.045 | 1 | 0.107701798 | -0.776359032 | 0.022 | 0.097 | 1 | 0.324739547 | 0.203803919 |
| AW146154 | 0.304411431 | -0.413691074 | 0 | 0.049 | 1 | 0.107748603 | -1.486852415 | 0.022 | 0.093 | 1 | 0.304411431 | 0.203887444 |
| Mtdh | 0.935811478 | 0.191939258 | 1 | 0.974 | 1 | 0.107914935 | -0.124282336 | 0.978 | 0.965 | 1 | 0.935811478 | 0.204184237 |
| Homer1 | 0.848275034 | 0.376687744 | 0.476 | 0.496 | 1 | 0.107959225 | 0.453890999 | 0.756 | 0.616 | 1 | 0.848275034 | 0.204263255 |
| Msx2 | 0.597524069 | -0.280424543 | 0.048 | 0.078 | 1 | 0.107980563 | -0.856086225 | 0.089 | 0.19 | 1 | 0.597524069 | 0.204301324 |
| Eif4a2 | 0.108026469 | -0.172538762 | 0.857 | 0.978 | 1 | 0.168305084 | -0.154582376 | 0.978 | 0.952 | 1 | 0.168305084 | 0.20438322 |
| Exo5 | 0.163567557 | -1.233118828 | 0 | 0.086 | 1 | 0.108096119 | -1.404390255 | 0.022 | 0.093 | 1 | 0.163567557 | 0.204507468 |
| Phf7 | 0.451613788 | -0.326228232 | 0.048 | 0.097 | 1 | 0.108096858 | -1.526380779 | 0.022 | 0.093 | 1 | 0.451613788 | 0.204508785 |
| Zfp747 | 0.966230998 | 1.258734268 | 0.048 | 0.052 | 1 | 0.108097227 | -1.486852415 | 0.022 | 0.093 | 1 | 0.966230998 | 0.204509443 |
| Ogfod2 | 0.120586929 | -1.370622352 | 0 | 0.104 | 1 | 0.10809735 | -1.361321533 | 0.022 | 0.093 | 1 | 0.120586929 | 0.204509663 |
| Tdp1 | 0.896890958 | 0.316219763 | 0.143 | 0.131 | 1 | 0.10809735 | -1.361321533 | 0.022 | 0.093 | 1 | 0.896890958 | 0.204509663 |
| Faim | 0.928499002 | 0.503846766 | 0.095 | 0.104 | 1 | 0.10809735 | -1.361321533 | 0.022 | 0.093 | 1 | 0.928499002 | 0.204509663 |
| N4bp2l2 | 0.108233686 | 0.398681923 | 0.905 | 0.884 | 1 | 0.198552238 | 0.344656368 | 0.911 | 0.893 | 1 | 0.198552238 | 0.20475284 |
| Nadsyn1 | 0.108291308 | 2.503846766 | 0.095 | 0.03 | 1 | 0.555549391 | 0.91753784 | 0.022 | 0.042 | 1 | 0.555549391 | 0.204855609 |
| Sema3f | 0.108291308 | 2.503846766 | 0.095 | 0.03 | 1 | 0.23197796 | -1.017367132 | 0 | 0.031 | 1 | 0.23197796 | 0.204855609 |
| Gm12089 | 0.108291308 | 2.503846766 | 0.095 | 0.03 | 1 | 0.381591326 | 0.250113179 | 0.044 | 0.087 | 1 | 0.381591326 | 0.204855609 |
| Amhr2 | 0.108291308 | 2.503846766 | 0.095 | 0.03 | 1 | 0.184823348 | -1.223818009 | 0 | 0.038 | 1 | 0.184823348 | 0.204855609 |
| Ap2a2 | 0.659269495 | 0.372602233 | 0.381 | 0.496 | 1 | 0.108291787 | 0.844249384 | 0.822 | 0.74 | 1 | 0.659269495 | 0.204856462 |
| Atp6v1g2 | 0.267953794 | -0.496153234 | 0 | 0.056 | 1 | 0.108442485 | -1.404390255 | 0.022 | 0.093 | 1 | 0.267953794 | 0.205125198 |
| Trim46 | 0.196422748 | -0.970084422 | 0 | 0.075 | 1 | 0.108479699 | -0.526380779 | 0.022 | 0.097 | 1 | 0.196422748 | 0.205191553 |

| | | | | | | | | | | | | |
|---|---|---|---|---|---|---|---|---|---|---|---|---|
| Gm56980 | 0.57816106 | 1.351843673 | 0 | 0.015 | 1 | 0.108490762 | 2.83507568 | 0.067 | 0.024 | 1 | 0.57816106 | 0.205211279 |
| Gm14167 | 0.108582614 | -0.785659851 | 0.048 | 0.198 | 1 | 0.147363558 | 0.339118185 | 0.067 | 0.159 | 1 | 0.147363558 | 0.205375044 |
| Slco2a1 | 0.10871311 | 1.766881172 | 0.19 | 0.09 | 1 | 0.802729223 | 0.545569062 | 0.111 | 0.104 | 1 | 0.802729223 | 0.205607679 |
| Gm10353 | 0.268011783 | -0.648156327 | 0 | 0.056 | 1 | 0.10878658 | -1.526380779 | 0.022 | 0.093 | 1 | 0.268011783 | 0.205738641 |
| Kcng2 | 0.554963324 | 1.214340149 | 0.095 | 0.063 | 1 | 0.10878658 | -1.271123724 | 0.022 | 0.093 | 1 | 0.554963324 | 0.205738641 |
| Actn1 | 0.80218294 | 0.65058214 | 0.619 | 0.563 | 1 | 0.108791145 | -0.267051507 | 0.911 | 0.841 | 1 | 0.80218294 | 0.205746776 |
| Raph1 | 0.221326796 | 0.28243145 | 0.905 | 0.937 | 1 | 0.109015317 | 0.618360373 | 0.978 | 0.924 | 1 | 0.221326796 | 0.206146294 |
| Odf2 | 0.27164578 | 0.483088206 | 0.238 | 0.455 | 1 | 0.109138876 | 0.596915942 | 0.6 | 0.478 | 1 | 0.27164578 | 0.206366459 |
| Otulinl | 0.120634678 | -1.611630451 | 0 | 0.104 | 1 | 0.109216041 | -0.679497493 | 0.044 | 0.131 | 1 | 0.120634678 | 0.206503939 |
| Aph1c | 0.113543807 | -1.574155746 | 0 | 0.108 | 1 | 0.109228834 | -0.404390255 | 0.022 | 0.097 | 1 | 0.113543807 | 0.20652673 |
| Gm10635 | 0.163769036 | -2.984439715 | 0 | 0.086 | 1 | 0.109416163 | 0.599008321 | 0.022 | 0.097 | 1 | 0.163769036 | 0.20686043 |
| Mavs | 0.806554358 | 1.137718867 | 0.095 | 0.086 | 1 | 0.109467155 | -1.316927414 | 0.022 | 0.093 | 1 | 0.806554358 | 0.206951252 |
| Aph1b | 0.523771294 | 0.295260144 | 0.095 | 0.16 | 1 | 0.109514373 | -0.709244837 | 0.067 | 0.159 | 1 | 0.523771294 | 0.207035349 |
| Man2c1os | 0.10951716 | 2.516230491 | 0.143 | 0.06 | 1 | 0.112466333 | -1.513324627 | 0.044 | 0.125 | 1 | 0.112466333 | 0.207040312 |
| H6pd | 0.13013423 | -0.672003069 | 0.048 | 0.187 | 1 | 0.109587578 | 0.481438725 | 0.089 | 0.208 | 1 | 0.13013423 | 0.207165719 |
| Riox2 | 0.109662976 | 2.137718867 | 0.19 | 0.09 | 1 | 0.580578162 | -0.191396532 | 0.044 | 0.066 | 1 | 0.580578162 | 0.207299984 |
| Parp6 | 0.109807139 | 1.439306514 | 0.333 | 0.228 | 1 | 0.200849634 | 0.134635961 | 0.156 | 0.273 | 1 | 0.200849634 | 0.20755667 |
| Brcc3 | 0.139451892 | -0.75918764 | 0.143 | 0.299 | 1 | 0.109861254 | -0.331877755 | 0.244 | 0.405 | 1 | 0.139451892 | 0.207653012 |
| Cherp | 0.369145347 | 1.368426557 | 0.381 | 0.422 | 1 | 0.109892599 | -0.24890899 | 0.311 | 0.509 | 1 | 0.369145347 | 0.207708814 |
| Nfatc2ip | 0.549660257 | 0.400753273 | 0.048 | 0.09 | 1 | 0.110026549 | -0.989352756 | 0.044 | 0.128 | 1 | 0.549660257 | 0.207947257 |
| Fam161b | 0.110202281 | 2.351843673 | 0.095 | 0.03 | 1 | 0.936025227 | 1.683072586 | 0.022 | 0.021 | 1 | 0.936025227 | 0.208260019 |
| Pwwp2a | 0.733246311 | 0.582273413 | 0.286 | 0.396 | 1 | 0.11026512 | 0.469668948 | 0.667 | 0.533 | 1 | 0.733246311 | 0.208371844 |
| Cab39l | 0.324090984 | 0.252735528 | 0.381 | 0.675 | 1 | 0.110331386 | 1.011259672 | 0.822 | 0.706 | 1 | 0.324090984 | 0.208489756 |
| Ikbkb | 0.188128546 | -0.25696557 | 0.143 | 0.299 | 1 | 0.110364573 | 0.748660928 | 0.467 | 0.37 | 1 | 0.188128546 | 0.208548807 |
| Ccdc191 | 0.110547151 | -1.434752689 | 0.048 | 0.183 | 1 | 0.141165658 | -0.290932205 | 0.044 | 0.125 | 1 | 0.141165658 | 0.208873629 |
| Pcdh11x | 0.236553882 | -1.683780237 | 0 | 0.063 | 1 | 0.110569624 | -1.877642368 | 0.044 | 0.125 | 1 | 0.236553882 | 0.208913607 |
| Phc2 | 0.234678136 | -0.435526875 | 0.571 | 0.75 | 1 | 0.110628979 | -0.14614177 | 0.933 | 0.9 | 1 | 0.234678136 | 0.209019187 |
| Atp2b4 | 0.110655794 | 0.959041392 | 0.619 | 0.545 | 1 | 0.847860423 | 0.82057611 | 0.422 | 0.529 | 1 | 0.847860423 | 0.209066883 |
| Atf6b | 0.534563051 | 0.316219763 | 0.19 | 0.276 | 1 | 0.110881479 | 0.180572246 | 0.111 | 0.221 | 1 | 0.534563051 | 0.209468255 |
| Ankrd13b | 0.390395345 | 1.586308926 | 0.095 | 0.052 | 1 | 0.111197447 | -1.346674757 | 0.044 | 0.125 | 1 | 0.390395345 | 0.210030021 |
| Skp2 | 0.802298473 | 2.129451251 | 0.095 | 0.086 | 1 | 0.111293331 | -1.044847868 | 0.044 | 0.128 | 1 | 0.802298473 | 0.210200457 |
| Phtf2 | 0.964133605 | 0.516424832 | 0.476 | 0.623 | 1 | 0.111431042 | 0.727446324 | 0.933 | 0.848 | 1 | 0.964133605 | 0.210445207 |
| Tigit | 0.578159907 | 0.866416846 | 0 | 0.015 | 1 | 0.11161871 | 2.393565969 | 0.067 | 0.024 | 1 | 0.578159907 | 0.210778684 |
| L2hgdh | 0.882341453 | 0.351843673 | 0.143 | 0.164 | 1 | 0.111619486 | -0.597035333 | 0.067 | 0.163 | 1 | 0.882341453 | 0.210780062 |
| Taok2 | 0.561966789 | 0.420015175 | 0.238 | 0.332 | 1 | 0.111819747 | -0.101198723 | 0.289 | 0.502 | 1 | 0.561966789 | 0.211135838 |
| Slc35c1 | 0.153857942 | -1.233118828 | 0 | 0.09 | 1 | 0.111831202 | -1.316927414 | 0.044 | 0.125 | 1 | 0.153857942 | 0.211156187 |
| Snrnp70 | 0.111903842 | 0.360575965 | 1 | 0.974 | 1 | 0.294013664 | 0.156443396 | 0.978 | 0.983 | 1 | 0.294013664 | 0.211285214 |
| Cmtm4 | 0.80238217 | 0.673771768 | 0.381 | 0.489 | 1 | 0.11195594 | -0.450410419 | 0.444 | 0.581 | 1 | 0.80238217 | 0.211377747 |
| Ino80 | 0.112057236 | 0.89958371 | 0.857 | 0.828 | 1 | 0.246655617 | 0.40921115 | 0.978 | 0.882 | 1 | 0.246655617 | 0.211557648 |
| Zfp664 | 0.112069241 | -0.326228232 | 0.238 | 0.466 | 1 | 0.384905767 | -0.237492946 | 0.711 | 0.682 | 1 | 0.384905767 | 0.211578968 |
| Gm16156 | 0.112136063 | 2.214340149 | 0.095 | 0.03 | 1 | 0.950034855 | 0.946106992 | 0.022 | 0.024 | 1 | 0.950034855 | 0.211697629 |
| Gsap | 0.112152081 | 1.673771768 | 0.095 | 0.03 | 1 | 0.215278367 | -0.743192169 | 0.022 | 0.073 | 1 | 0.215278367 | 0.211726072 |
| Gm4631 | 0.690680012 | 0.79930265 | 0.048 | 0.075 | 1 | 0.112152225 | -0.749886821 | 0.067 | 0.159 | 1 | 0.690680012 | 0.211726328 |
| Eri1 | 0.804643245 | 0.455348249 | 0.286 | 0.295 | 1 | 0.112420377 | -0.124282336 | 0.133 | 0.253 | 1 | 0.804643245 | 0.212202412 |
| Adamts1 | 0.112719282 | -1.626962105 | 0.095 | 0.261 | 1 | 0.364386809 | -0.798054104 | 0.222 | 0.277 | 1 | 0.364386809 | 0.212732927 |
| Milr1 | 0.16548778 | 3.258734268 | 0.048 | 0.011 | 1 | 0.112882988 | 2.972579203 | 0.067 | 0.024 | 1 | 0.16548778 | 0.213023407 |
| Scml4 | 0.279442334 | -0.543458949 | 0.048 | 0.134 | 1 | 0.112950931 | -0.604504004 | 0.111 | 0.218 | 1 | 0.279442334 | 0.213143949 |
| Gm13986 | 0.939406941 | 1.376091219 | 0.143 | 0.153 | 1 | 0.112951944 | 0.15951063 | 0.089 | 0.201 | 1 | 0.939406941 | 0.213145747 |
| Gm13936 | 0.113106186 | 1.97333205 | 0.095 | 0.03 | 1 | 0.49622258 | -0.124282336 | 0 | 0.01 | 1 | 0.49622258 | 0.213419362 |
| Ralgps2 | 0.113125106 | -0.326228232 | 0.143 | 0.328 | 1 | 0.94151424 | 0.952679646 | 0.156 | 0.166 | 1 | 0.94151424 | 0.213452923 |
| Usp28 | 0.381146554 | -0.179679569 | 0.19 | 0.299 | 1 | 0.113153179 | 0.19121949 | 0.2 | 0.349 | 1 | 0.381146554 | 0.213502715 |
| Lrrc51 | 0.113396294 | -1.413691074 | 0 | 0.108 | 1 | 0.770549578 | 0.982632868 | 0.067 | 0.083 | 1 | 0.770549578 | 0.213933869 |
| 2510017J16Rik | 0.11339646 | -1.370622352 | 0 | 0.108 | 1 | 0.196831959 | 1.683072586 | 0.133 | 0.08 | 1 | 0.196831959 | 0.213934162 |
| Map3k11 | 0.11339646 | -1.370622352 | 0 | 0.108 | 1 | 0.168225263 | 0.313838777 | 0.089 | 0.183 | 1 | 0.168225263 | 0.213934162 |
| Tlcd5 | 0.113450209 | -1.413691074 | 0 | 0.108 | 1 | 0.427508837 | 0.683072586 | 0.044 | 0.08 | 1 | 0.427508837 | 0.214029469 |
| Amdhd2 | 0.113450209 | -1.413691074 | 0 | 0.108 | 1 | 0.121187145 | -1.174908409 | 0.022 | 0.09 | 1 | 0.121187145 | 0.214029469 |
| Pcdhga7 | 0.113542153 | -1.535681598 | 0 | 0.108 | 1 | 0.27524185 | -0.392215541 | 0.044 | 0.097 | 1 | 0.27524185 | 0.214192486 |
| Ago4 | 0.113543807 | -1.574155746 | 0 | 0.108 | 1 | 0.1762219 | -0.342462506 | 0.067 | 0.145 | 1 | 0.1762219 | 0.214195417 |
| Gm57286 | 0.113600123 | 1.223110359 | 0.476 | 0.392 | 1 | 0.516608167 | 0.645597881 | 0.244 | 0.339 | 1 | 0.516608167 | 0.214295259 |
| Gm49692 | 0.113623669 | -1.880817084 | 0 | 0.108 | 1 | 0.180634714 | -0.124282336 | 0.022 | 0.08 | 1 | 0.180634714 | 0.214337 |
| Grk2 | 0.113635912 | -0.463731756 | 0.095 | 0.276 | 1 | 0.913775294 | 0.352682913 | 0.378 | 0.405 | 1 | 0.913775294 | 0.214358704 |
| Fgfr4 | 0.11364632 | -1.648156327 | 0 | 0.108 | 1 | 0.406644621 | 0.361144491 | 0.022 | 0.052 | 1 | 0.406644621 | 0.214377154 |
| Sema4c | 0.113650949 | -1.683780237 | 0 | 0.108 | 1 | 0.279628641 | -0.960783604 | 0.089 | 0.145 | 1 | 0.279628641 | 0.214385361 |
| Gm57093 | 0.113682362 | -1.752492987 | 0 | 0.108 | 1 | 0.139882659 | -0.877642368 | 0.044 | 0.121 | 1 | 0.139882659 | 0.214441044 |
| Stard9 | 0.113685172 | -1.818081329 | 0 | 0.108 | 1 | 0.376157272 | 0.205025289 | 0.044 | 0.087 | 1 | 0.376157272 | 0.214446026 |
| Lin9 | 0.113685834 | -1.940938076 | 0 | 0.108 | 1 | 0.125749292 | -1.223818009 | 0.022 | 0.09 | 1 | 0.125749292 | 0.214447198 |
| Vcpkmt | 0.759464695 | 0.614878079 | 0.048 | 0.067 | 1 | 0.113733497 | -1.124282336 | 0.044 | 0.125 | 1 | 0.759464695 | 0.214531686 |
| Coch | 0.113743364 | -3.424260315 | 0 | 0.108 | 1 | 0.994463827 | 2.142504205 | 0.044 | 0.045 | 1 | 0.994463827 | 0.214549175 |
| Cdin1 | 0.163476132 | 1.311201688 | 0.238 | 0.134 | 1 | 0.113793219 | 0.339118185 | 0.044 | 0.128 | 1 | 0.163476132 | 0.214637541 |
| Afdn | 0.757944434 | 0.236366455 | 0.667 | 0.642 | 1 | 0.113806846 | 0.645902794 | 0.911 | 0.761 | 1 | 0.757944434 | 0.214661694 |
| Gm10131 | 0.113880063 | -1.233118828 | 0 | 0.108 | 1 | 0.357544208 | -0.191396532 | 0.067 | 0.114 | 1 | 0.357544208 | 0.214791457 |
| Zfp65 | 0.397614565 | -0.574155746 | 0.048 | 0.104 | 1 | 0.113959548 | -0.638855509 | 0.044 | 0.128 | 1 | 0.397614565 | 0.214932318 |
| Clpx | 0.113983295 | -0.517092571 | 0.381 | 0.623 | 1 | 0.378299723 | 0.232820394 | 0.8 | 0.73 | 1 | 0.378299723 | 0.214974398 |
| Atad2 | 0.400336094 | 1.235650655 | 0.286 | 0.228 | 1 | 0.114039185 | 0.667475731 | 0.089 | 0.208 | 1 | 0.400336094 | 0.215073435 |
| Zfp251 | 0.840503985 | 0.60338244 | 0.143 | 0.131 | 1 | 0.114073225 | 0.174058939 | 0.067 | 0.17 | 1 | 0.840503985 | 0.215133749 |
| Tbc1d13 | 0.114076195 | 2.121230745 | 0.238 | 0.134 | 1 | 0.215147725 | -0.213091603 | 0.089 | 0.17 | 1 | 0.215147725 | 0.215139012 |
| Gipc1 | 0.114715626 | -0.690800665 | 0.095 | 0.257 | 1 | 0.727183575 | -0.352551324 | 0.311 | 0.315 | 1 | 0.727183575 | 0.216271576 |
| Dcaf10 | 0.478895259 | 0.507859829 | 0.571 | 0.631 | 1 | 0.114784624 | 0.119696948 | 0.622 | 0.789 | 1 | 0.478895259 | 0.216393737 |
| Smchd1 | 0.114827696 | -0.339525055 | 0.762 | 0.91 | 1 | 0.457654001 | 0.43953239 | 0.933 | 0.91 | 1 | 0.457654001 | 0.216469992 |
| Zfp235 | 0.27086281 | -0.167530486 | 0.048 | 0.138 | 1 | 0.114858883 | -0.432404631 | 0.067 | 0.159 | 1 | 0.27086281 | 0.216525203 |
| Mga | 0.409385486 | 0.676683363 | 0.667 | 0.69 | 1 | 0.115021705 | -0.270732591 | 0.822 | 0.834 | 1 | 0.409385486 | 0.216813417 |
| 1500035N22Rik | 0.115075278 | 1.97333205 | 0.095 | 0.03 | 1 | 0.658583583 | 1.946106992 | 0.022 | 0.014 | 1 | 0.658583583 | 0.216908236 |
| Git1 | 0.642869784 | 0.351843673 | 0.095 | 0.142 | 1 | 0.115094395 | 0.327591932 | 0.089 | 0.204 | 1 | 0.642869784 | 0.216942071 |
| Zfp959 | 0.136109171 | -1.184209227 | 0 | 0.097 | 1 | 0.115138053 | -0.709244837 | 0.044 | 0.128 | 1 | 0.136109171 | 0.217019336 |
| Atf7ip | 0.775667083 | 0.279779713 | 0.571 | 0.642 | 1 | 0.115572442 | 0.499229083 | 0.911 | 0.803 | 1 | 0.775667083 | 0.217787894 |
| Tbc1d24 | 0.342363296 | 1.129451251 | 0.048 | 0.123 | 1 | 0.115616476 | -0.247664751 | 0.089 | 0.194 | 1 | 0.342363296 | 0.217865782 |
| Ahdc1 | 0.115647922 | 0.665778977 | 0.667 | 0.582 | 1 | 0.266495162 | 0.496100386 | 0.822 | 0.858 | 1 | 0.266495162 | 0.217921401 |
| Bicd1 | 0.473264418 | -0.522625445 | 0.048 | 0.097 | 1 | 0.115682969 | -2.271123724 | 0.022 | 0.09 | 1 | 0.473264418 | 0.217983389 |
| Pex16 | 0.671900318 | 0.673771768 | 0.095 | 0.142 | 1 | 0.115770894 | -0.674479418 | 0.044 | 0.128 | 1 | 0.671900318 | 0.218138889 |
| Dcaf1 | 0.375533927 | 0.991947728 | 0.429 | 0.418 | 1 | 0.115785935 | 0.67480497 | 0.667 | 0.557 | 1 | 0.375533927 | 0.218165488 |
| Plek | 0.116067137 | 1.503846766 | 0.095 | 0.03 | 1 | 0.18484555 | -1.564854927 | 0 | 0.038 | 1 | 0.18484555 | 0.218662694 |
| Gpr39 | 0.116074373 | 1.614878079 | 0.095 | 0.03 | 1 | 0.331919067 | -1.564854927 | 0 | 0.021 | 1 | 0.331919067 | 0.218675485 |
| Susd1 | 0.116074373 | 1.97333205 | 0.095 | 0.03 | 1 | 0.206897641 | -1.124282336 | 0 | 0.035 | 1 | 0.206897641 | 0.218675485 |
| E130308A19Rik | 0.821927541 | 1.36999102 | 0.19 | 0.179 | 1 | 0.116098114 | -0.343727473 | 0.156 | 0.273 | 1 | 0.821927541 | 0.218717456 |
| Arhgap42 | 0.296321252 | -0.575615225 | 0.524 | 0.701 | 1 | 0.116118167 | 0.563283665 | 0.711 | 0.699 | 1 | 0.296321252 | 0.218752905 |
| Ltbp4 | 0.9389077 | -0.413691074 | 0.095 | 0.097 | 1 | 0.11621431 | -0.238924902 | 0.4 | 0.246 | 1 | 0.9389077 | 0.218922854 |
| Shank1 | 0.827510585 | 0.137718867 | 0.143 | 0.164 | 1 | 0.116221404 | 0.211766867 | 0.067 | 0.17 | 1 | 0.827510585 | 0.218935393 |
| Plekha7 | 0.267810975 | 2.121230745 | 0.143 | 0.078 | 1 | 0.116234974 | 0.69556353 | 0.311 | 0.204 | 1 | 0.267810975 | 0.218959379 |

| | | | | | | | | | | | | |
|---|---|---|---|---|---|---|---|---|---|---|---|---|
| 2610037D02Rik | 1 | 0.442817333 | 0.286 | 0.336 | 1 | 0.116327752 | 1.068130928 | 0.644 | 0.536 | 1 | 1 | 0.219123357 |
| Gm9929 | 0.853709349 | 1.97333205 | 0.048 | 0.041 | 1 | 0.116401905 | -1.053893008 | 0.022 | 0.093 | 1 | 0.853709349 | 0.219254406 |
| Slc12a2 | 0.116438944 | -0.843707136 | 0.571 | 0.653 | 1 | 0.719800297 | -0.151148942 | 0.778 | 0.623 | 1 | 0.719800297 | 0.21931986 |
| Pdzrn4 | 0.648758467 | -2.712809286 | 0.095 | 0.131 | 1 | 0.116544995 | -4.635846759 | 0.022 | 0.09 | 1 | 0.648758467 | 0.219507254 |
| Dcbld2 | 0.116735256 | -0.309221807 | 0.286 | 0.519 | 1 | 0.503334605 | 0.33679463 | 0.333 | 0.446 | 1 | 0.503334605 | 0.219843393 |
| Scgn | 0.261628269 | 2.936806174 | 0.048 | 0.015 | 1 | 0.116778371 | 1.776181991 | 0.067 | 0.024 | 1 | 0.261628269 | 0.219919555 |
| Ttc14 | 0.56697974 | 0.272055647 | 0.714 | 0.784 | 1 | 0.116976702 | 0.506646019 | 0.867 | 0.754 | 1 | 0.56697974 | 0.220269855 |
| Gm12703 | 0.752561857 | -0.348596045 | 0.095 | 0.119 | 1 | 0.117004921 | -0.476126009 | 0.111 | 0.228 | 1 | 0.752561857 | 0.22031969 |
| Syn1 | 0.119520861 | 1.761234609 | 0.238 | 0.131 | 1 | 0.117091869 | -0.174908409 | 0.044 | 0.131 | 1 | 0.119520861 | 0.220473232 |
| Lrrc20 | 0.117273813 | -1.167530486 | 0.048 | 0.19 | 1 | 0.290720554 | 0.473619221 | 0.044 | 0.097 | 1 | 0.290720554 | 0.22079448 |
| Emb | 0.142110745 | -1.13764518 | 0.762 | 0.724 | 1 | 0.117363936 | 0.106425354 | 0.867 | 0.657 | 1 | 0.142110745 | 0.220953579 |
| Usp39 | 0.117364987 | -1.02666795 | 0.095 | 0.243 | 1 | 0.216581728 | 0.558744451 | 0.156 | 0.284 | 1 | 0.216581728 | 0.220955434 |
| Fam185a | 0.274517372 | 0.214340149 | 0.048 | 0.138 | 1 | 0.117442209 | -0.140049652 | 0.067 | 0.166 | 1 | 0.274517372 | 0.221091745 |
| Mss51 | 0.117652091 | 2.129451251 | 0.19 | 0.09 | 1 | 0.578603603 | 1.066401226 | 0.133 | 0.114 | 1 | 0.578603603 | 0.221462167 |
| Pakap.1 | 0.117739637 | -1.166951904 | 0.476 | 0.653 | 1 | 0.248542632 | 1.264571232 | 0.6 | 0.481 | 1 | 0.248542632 | 0.221616652 |
| Mbd4 | 0.47364177 | 1.258734268 | 0.048 | 0.101 | 1 | 0.117966663 | -0.86124793 | 0.022 | 0.093 | 1 | 0.47364177 | 0.222017193 |
| Stum | 0.118029954 | 1.936806174 | 0.095 | 0.03 | 1 | 0.496223879 | 0.683072586 | 0 | 0.01 | 1 | 0.496223879 | 0.222128837 |
| Ncam1 | 0.118113915 | -0.551941012 | 0.571 | 0.784 | 1 | 0.220079307 | -0.220251394 | 0.667 | 0.754 | 1 | 0.220079307 | 0.222276933 |
| Wipf2 | 0.229137587 | -0.159118246 | 0.143 | 0.299 | 1 | 0.118189673 | 0.620788308 | 0.533 | 0.415 | 1 | 0.229137587 | 0.222410547 |
| Eif2ak4 | 0.118239672 | 1.673771768 | 0.571 | 0.541 | 1 | 0.246034172 | 0.142779135 | 0.4 | 0.606 | 1 | 0.246034172 | 0.222498724 |
| Ldlrad2 | 0.595875107 | 1.281454345 | 0.095 | 0.067 | 1 | 0.118345556 | -0.819427754 | 0.022 | 0.093 | 1 | 0.595875107 | 0.222685441 |
| Platr25 | 0.810382062 | 0.951305743 | 0.143 | 0.187 | 1 | 0.118407569 | -0.352551324 | 0.044 | 0.131 | 1 | 0.810382062 | 0.222794785 |
| Plk4 | 0.865257151 | 0.747772349 | 0.095 | 0.086 | 1 | 0.118476039 | -1.776359032 | 0.022 | 0.09 | 1 | 0.865257151 | 0.222915507 |
| Pax3 | 0.346882058 | -1.611630451 | 0 | 0.041 | 1 | 0.118515941 | -1.158229668 | 0.044 | 0.128 | 1 | 0.346882058 | 0.222985854 |
| Garnl3 | 0.118535338 | -0.233118828 | 0.048 | 0.194 | 1 | 0.791825012 | 0.502500341 | 0.111 | 0.131 | 1 | 0.791825012 | 0.223020049 |
| Rttn | 0.11861554 | 2.511500339 | 0.19 | 0.093 | 1 | 0.910619191 | 0.205025289 | 0.089 | 0.083 | 1 | 0.910619191 | 0.223161433 |
| Haus7 | 0.982900121 | 0.393663848 | 0.19 | 0.201 | 1 | 0.118789529 | -0.276285429 | 0.089 | 0.19 | 1 | 0.982900121 | 0.223468106 |
| Mafa | 0.815913222 | 0.281454345 | 0.048 | 0.06 | 1 | 0.118863297 | -1.486852415 | 0.022 | 0.09 | 1 | 0.815913222 | 0.22359811 |
| Casp7 | 0.987346235 | 0.825774861 | 0.095 | 0.097 | 1 | 0.118868883 | -1.638855509 | 0.022 | 0.09 | 1 | 0.987346235 | 0.223607955 |
| Taok1 | 0.473224827 | 0.619181881 | 0.81 | 0.847 | 1 | 0.118911151 | -0.185844139 | 1 | 0.976 | 1 | 0.473224827 | 0.223682441 |
| B830012L14Rik | 0.191903244 | 1.48112669 | 0.429 | 0.336 | 1 | 0.118938852 | 0.408653417 | 0.289 | 0.491 | 1 | 0.191903244 | 0.223731254 |
| Polr1has | 0.136337276 | -1.611630451 | 0 | 0.097 | 1 | 0.118986849 | -1.404390255 | 0 | 0.052 | 1 | 0.136337276 | 0.223815828 |
| Gm19412 | 0.184739689 | -0.970084422 | 0 | 0.078 | 1 | 0.118986849 | -1.486852415 | 0 | 0.052 | 1 | 0.184739689 | 0.223815828 |
| Gm43138 | 0.267953794 | -0.496153234 | 0 | 0.056 | 1 | 0.118986849 | -1.404390255 | 0 | 0.052 | 1 | 0.267953794 | 0.223815828 |
| Tinagl1 | 0.310184716 | -0.998653574 | 0.048 | 0.119 | 1 | 0.118986849 | -1.404390255 | 0 | 0.052 | 1 | 0.310184716 | 0.223815828 |
| 6530402F18Rik | 0.492599419 | 0.503846766 | 0 | 0.022 | 1 | 0.118986849 | -1.404390255 | 0 | 0.052 | 1 | 0.492599419 | 0.223815828 |
| Pbx4 | 0.532436372 | 1.088809267 | 0 | 0.019 | 1 | 0.118986849 | -1.404390255 | 0 | 0.052 | 1 | 0.532436372 | 0.223815828 |
| Ccne1 | 0.723385652 | 0.629377648 | 0.095 | 0.075 | 1 | 0.118986849 | -1.404390255 | 0 | 0.052 | 1 | 0.723385652 | 0.223815828 |
| Fndc10 | 0.97964157 | 1.451379346 | 0.048 | 0.049 | 1 | 0.118986849 | -1.404390255 | 0 | 0.052 | 1 | 0.97964157 | 0.223815828 |
| Smpdl3b | 0.136297285 | -1.413691074 | 0 | 0.097 | 1 | 0.119007856 | -1.486852415 | 0 | 0.052 | 1 | 0.136297285 | 0.223852841 |
| Mdm1 | 0.144848782 | -1.413691074 | 0 | 0.093 | 1 | 0.119007856 | -1.709244837 | 0 | 0.052 | 1 | 0.144848782 | 0.223852841 |
| 5730409E04Rik | 0.304385789 | -0.326228232 | 0 | 0.049 | 1 | 0.119007856 | -1.564854927 | 0 | 0.052 | 1 | 0.304385789 | 0.223852841 |
| Zfp526 | 0.396916958 | 0.214340149 | 0 | 0.034 | 1 | 0.119007856 | -1.709244837 | 0 | 0.052 | 1 | 0.396916958 | 0.223852841 |
| Itgb1bp2 | 0.420968517 | 1.673771768 | 0.095 | 0.056 | 1 | 0.119007856 | -1.486852415 | 0 | 0.052 | 1 | 0.420968517 | 0.223852841 |
| Gm45435 | 0.492608234 | 0.866416846 | 0 | 0.022 | 1 | 0.119007856 | -1.486852415 | 0 | 0.052 | 1 | 0.492608234 | 0.223852841 |
| Limk1 | 0.532436372 | 1.088809267 | 0 | 0.019 | 1 | 0.119007856 | -1.486852415 | 0 | 0.052 | 1 | 0.532436372 | 0.223852841 |
| Hps6 | 0.632076134 | 1.673771768 | 0 | 0.011 | 1 | 0.119007856 | -1.486852415 | 0 | 0.052 | 1 | 0.632076134 | 0.223852841 |
| Gli1 | 0.695160218 | 0.735172312 | 0.095 | 0.071 | 1 | 0.119007856 | -1.486852415 | 0 | 0.052 | 1 | 0.695160218 | 0.223852841 |
| Fam83e | 0.872207635 | 0.586308926 | 0.048 | 0.056 | 1 | 0.119007856 | -1.638855509 | 0 | 0.052 | 1 | 0.872207635 | 0.223852841 |
| Il17re | 0.277250164 | 1.673771768 | 0.048 | 0.015 | 1 | 0.119025629 | -1.638855509 | 0 | 0.052 | 1 | 0.277250164 | 0.223884158 |
| Cracr2a | 0.425639612 | 0.503846766 | 0 | 0.03 | 1 | 0.119025629 | -1.709244837 | 0 | 0.052 | 1 | 0.425639612 | 0.223884158 |
| A430105J06Rik | 0.81881728 | 1.088809267 | 0.048 | 0.037 | 1 | 0.11902586 | -1.564854927 | 0 | 0.052 | 1 | 0.81881728 | 0.223884565 |
| B3galnt1 | 0.236343578 | -0.574155746 | 0 | 0.063 | 1 | 0.119040171 | -1.901889915 | 0 | 0.052 | 1 | 0.236343578 | 0.22390978 |
| Tprn | 0.324739547 | -0.133583154 | 0 | 0.045 | 1 | 0.119040171 | -1.709244837 | 0 | 0.052 | 1 | 0.324739547 | 0.22390978 |
| Gm19265 | 0.552820087 | 0.918884266 | 0.095 | 0.063 | 1 | 0.119040171 | -1.709244837 | 0 | 0.052 | 1 | 0.552820087 | 0.22390978 |
| 5430427G11Rik | 0.924791296 | 1.351843673 | 0.048 | 0.045 | 1 | 0.119040171 | -1.776359032 | 0 | 0.052 | 1 | 0.924791296 | 0.22390978 |
| Rrm2 | 0.28888789 | 0.425844254 | 0.095 | 0.045 | 1 | 0.119040864 | -1.960783604 | 0 | 0.052 | 1 | 0.28888789 | 0.223911 |
| Il10 | 0.409489432 | 1.908237021 | 0.095 | 0.056 | 1 | 0.119040864 | -2.124282336 | 0 | 0.052 | 1 | 0.409489432 | 0.223911 |
| Disc1 | 0.454224194 | 3.088809267 | 0.048 | 0.022 | 1 | 0.119040864 | -1.776359032 | 0 | 0.052 | 1 | 0.454224194 | 0.223911 |
| Cd248 | 0.578156445 | 1.088809267 | 0 | 0.015 | 1 | 0.119040864 | -1.776359032 | 0 | 0.052 | 1 | 0.578156445 | 0.223911 |
| Zfp239 | 0.304411431 | -0.413691074 | 0 | 0.049 | 1 | 0.119051482 | -1.776359032 | 0 | 0.052 | 1 | 0.304411431 | 0.223929708 |
| Adamts4 | 0.12844124 | -1.970084422 | 0 | 0.101 | 1 | 0.119052866 | -1.901889915 | 0 | 0.052 | 1 | 0.12844124 | 0.223932148 |
| Gm31045 | 0.425599726 | 0.351843673 | 0 | 0.03 | 1 | 0.11905379 | -1.709244837 | 0 | 0.052 | 1 | 0.425599726 | 0.223933775 |
| Foxj1 | 0.645818514 | 1.010806755 | 0.048 | 0.03 | 1 | 0.119063022 | -2.017367132 | 0 | 0.052 | 1 | 0.645818514 | 0.223950041 |
| Rtn4rl1 | 0.952180751 | 0.866416846 | 0.048 | 0.052 | 1 | 0.119063253 | -2.446210431 | 0 | 0.052 | 1 | 0.952180751 | 0.223950448 |
| Mcm2 | 0.251682795 | -0.849790188 | 0 | 0.06 | 1 | 0.119067869 | -2.071814916 | 0 | 0.052 | 1 | 0.251682795 | 0.223958581 |
| Gm17167 | 0.802987008 | 1.088809267 | 0.048 | 0.037 | 1 | 0.119071332 | -2.174908409 | 0 | 0.052 | 1 | 0.802987008 | 0.223964681 |
| Dlk1 | 0.457292877 | 0.351843673 | 0 | 0.026 | 1 | 0.119072255 | -3.223818009 | 0 | 0.052 | 1 | 0.457292877 | 0.223966308 |
| Rnf150 | 0.155994996 | 3.182785415 | 0.19 | 0.101 | 1 | 0.119077563 | -2.526380779 | 0 | 0.052 | 1 | 0.155994996 | 0.223975661 |
| Ccn3 | 0.37081917 | -0.648156327 | 0 | 0.037 | 1 | 0.119077794 | -2.80878051 | 0 | 0.052 | 1 | 0.37081917 | 0.223976067 |
| Angptl7 | 1 | -0.772484462 | 0.095 | 0.093 | 1 | 0.119084488 | -2.989352756 | 0 | 0.052 | 1 | 1 | 0.22398786 |
| Mdga2 | 0.370835778 | -2.475975352 | 0 | 0.037 | 1 | 0.119086565 | -3.404390255 | 0 | 0.052 | 1 | 0.370835778 | 0.22399152 |
| Tvp23a | 0.285465396 | -0.326228232 | 0 | 0.052 | 1 | 0.119091873 | -1.316927414 | 0 | 0.052 | 1 | 0.285465396 | 0.224000872 |
| Lrfn3 | 0.425639612 | 0.503846766 | 0 | 0.03 | 1 | 0.119091873 | -1.316927414 | 0 | 0.052 | 1 | 0.425639612 | 0.224000872 |
| Syne3 | 0.119122258 | 1.235198754 | 0.333 | 0.239 | 1 | 0.828763617 | 0.376826114 | 0.311 | 0.339 | 1 | 0.828763617 | 0.224054404 |
| Prpsap2 | 0.119202709 | -0.399477214 | 0.095 | 0.272 | 1 | 0.493180522 | 1.329435632 | 0.244 | 0.232 | 1 | 0.493180522 | 0.224196132 |
| Cdk5rap1 | 0.119240584 | -1.208871282 | 0.048 | 0.179 | 1 | 0.617897618 | 0.429315994 | 0.111 | 0.145 | 1 | 0.617897618 | 0.224262851 |
| Spry4 | 0.136258876 | -1.455511249 | 0 | 0.097 | 1 | 0.119256664 | -1.446210431 | 0.022 | 0.09 | 1 | 0.136258876 | 0.224291176 |
| Mtcl1 | 0.495558723 | -0.683780237 | 0.048 | 0.09 | 1 | 0.119278911 | -0.618096949 | 0.067 | 0.159 | 1 | 0.495558723 | 0.224330363 |
| Uck2 | 0.11946509 | -0.642373975 | 0.286 | 0.552 | 1 | 0.824761412 | 0.195957409 | 0.533 | 0.488 | 1 | 0.824761412 | 0.224658273 |
| Kdm2a | 0.119568198 | 0.387800551 | 0.857 | 0.772 | 1 | 0.945441471 | 0.187377424 | 0.978 | 0.962 | 1 | 0.945441471 | 0.224839842 |
| Sec22c | 0.196423035 | -0.911190733 | 0 | 0.075 | 1 | 0.119573867 | -0.486852415 | 0.022 | 0.093 | 1 | 0.196423035 | 0.224849824 |
| Zkscan5 | 0.176282202 | 1.775651382 | 0.19 | 0.104 | 1 | 0.119602904 | -0.316927414 | 0.022 | 0.093 | 1 | 0.176282202 | 0.224900954 |
| Zfp976 | 0.822949222 | 0.503846766 | 0.095 | 0.116 | 1 | 0.119826854 | -0.575661682 | 0.067 | 0.156 | 1 | 0.822949222 | 0.225295233 |
| Vopp1 | 0.119859707 | -0.849790188 | 0.048 | 0.19 | 1 | 0.283404896 | 0.205025289 | 0.178 | 0.284 | 1 | 0.283404896 | 0.225353064 |
| Gmpr | 0.954841465 | 0.97333205 | 0.048 | 0.045 | 1 | 0.11998422 | -0.446210431 | 0.022 | 0.093 | 1 | 0.954841465 | 0.225572227 |
| Poc1b | 0.779344758 | 0.233199176 | 0.143 | 0.175 | 1 | 0.119994939 | -0.223818009 | 0.156 | 0.294 | 1 | 0.779344758 | 0.225591093 |
| Fdxacb1 | 0.222110654 | -0.718545655 | 0 | 0.067 | 1 | 0.120036174 | -1.404390255 | 0.022 | 0.09 | 1 | 0.222110654 | 0.225663664 |
| Ptges2 | 0.153760172 | -1.081115734 | 0 | 0.09 | 1 | 0.120285556 | 0.361144491 | 0.067 | 0.163 | 1 | 0.153760172 | 0.226102498 |
| Inf2 | 0.120308841 | 0.829422638 | 0.429 | 0.347 | 1 | 0.54373537 | 0.808603468 | 0.378 | 0.401 | 1 | 0.54373537 | 0.226143464 |
| Mapk1ip1 | 0.120360233 | 1.578614535 | 0.238 | 0.123 | 1 | 0.847548993 | 0.875717664 | 0.133 | 0.131 | 1 | 0.847548993 | 0.22623388 |
| Hgh1 | 0.120415565 | -1.233118828 | 0 | 0.104 | 1 | 0.132686876 | -1.404390255 | 0 | 0.048 | 1 | 0.132686876 | 0.226331221 |
| Agrp | 0.608959364 | 0.400753273 | 0.048 | 0.082 | 1 | 0.120422285 | -1.316927414 | 0.022 | 0.09 | 1 | 0.608959364 | 0.226343044 |
| Atmin | 0.120477198 | -1.280424543 | 0 | 0.104 | 1 | 0.412123272 | 0.15951063 | 0.089 | 0.135 | 1 | 0.412123272 | 0.226439641 |
| Mks1 | 0.120477198 | -1.280424543 | 0 | 0.104 | 1 | 0.761447202 | 0.683072586 | 0.022 | 0.031 | 1 | 0.761447202 | 0.226439641 |
| Gm56692 | 0.120477198 | -1.370622352 | 0 | 0.104 | 1 | 0.148464731 | -1.223818009 | 0.022 | 0.083 | 1 | 0.148464731 | 0.226439641 |
| Nf2 | 0.277097423 | -0.271086678 | 0.19 | 0.328 | 1 | 0.120500203 | -0.238924902 | 0.422 | 0.571 | 1 | 0.277097423 | 0.226480107 |
| Frmd8 | 0.120528032 | -0.637814383 | 0.286 | 0.504 | 1 | 0.799012945 | 0.358570548 | 0.467 | 0.63 | 1 | 0.799012945 | 0.226529057 |

| | | | | | | | | | | | | |
|---|---|---|---|---|---|---|---|---|---|---|---|---|
| Ppp1r16a | 0.120534085 | -1.370622352 | 0 | 0.104 | 1 | 0.350979033 | -0.271123724 | 0.044 | 0.087 | 1 | 0.350979033 | 0.226539704 |
| Tubg1 | 0.12053426 | -1.326228232 | 0 | 0.104 | 1 | 0.162178563 | -0.486852415 | 0.022 | 0.083 | 1 | 0.162178563 | 0.226540013 |
| Rad1 | 0.120586227 | -1.413691074 | 0 | 0.104 | 1 | 0.201605048 | -0.638855509 | 0.044 | 0.107 | 1 | 0.201605048 | 0.226631415 |
| Gm6712 | 0.120586227 | -1.535681598 | 0 | 0.104 | 1 | 0.496037494 | -0.404390255 | 0.089 | 0.125 | 1 | 0.496037494 | 0.226631415 |
| Isoc2a | 0.120586753 | -1.455511249 | 0 | 0.104 | 1 | 0.529300652 | -0.486852415 | 0.022 | 0.042 | 1 | 0.529300652 | 0.226632342 |
| Ipp | 0.120586929 | -1.370622352 | 0 | 0.104 | 1 | 0.125633133 | -1.053893008 | 0.044 | 0.121 | 1 | 0.125633133 | 0.22663265 |
| Masp2 | 0.862687636 | 1.088809267 | 0.048 | 0.06 | 1 | 0.120661177 | -0.174908409 | 0.022 | 0.093 | 1 | 0.862687636 | 0.226763234 |
| Csdc2 | 0.12067628 | -1.496153234 | 0 | 0.104 | 1 | 0.939498715 | 0.320502507 | 0.089 | 0.093 | 1 | 0.939498715 | 0.226789795 |
| Gm4924 | 0.120678913 | -1.648156327 | 0 | 0.104 | 1 | 0.406646829 | 0.545569062 | 0.022 | 0.052 | 1 | 0.406646829 | 0.226794425 |
| Gm56952 | 0.236422109 | -0.785659851 | 0 | 0.063 | 1 | 0.120741715 | -0.271123724 | 0.022 | 0.093 | 1 | 0.236422109 | 0.226904869 |
| Zfp169 | 0.120751051 | -1.718545655 | 0 | 0.104 | 1 | 0.439561776 | 1.439147003 | 0.156 | 0.131 | 1 | 0.439561776 | 0.226921285 |
| Mamdc2 | 0.120756316 | -2.184209227 | 0 | 0.104 | 1 | 0.369103384 | -0.518561275 | 0.067 | 0.111 | 1 | 0.369103384 | 0.226930544 |
| Nup88 | 0.602048181 | 0.34285489 | 0.286 | 0.41 | 1 | 0.120769079 | 0.238287743 | 0.244 | 0.457 | 1 | 0.602048181 | 0.226952988 |
| Lonrf3 | 0.120784396 | -1.683780237 | 0 | 0.104 | 1 | 0.141615262 | -0.653210802 | 0.067 | 0.156 | 1 | 0.141615262 | 0.226979922 |
| Olfml2b | 0.120787029 | -1.911190733 | 0 | 0.104 | 1 | 0.631004768 | -0.776359032 | 0.044 | 0.062 | 1 | 0.631004768 | 0.226984551 |
| Lfng | 0.120788082 | -1.818081329 | 0 | 0.104 | 1 | 0.132718194 | -1.564854927 | 0 | 0.048 | 1 | 0.132718194 | 0.226986403 |
| Xndc1 | 0.923594926 | 0.634243403 | 0.143 | 0.16 | 1 | 0.120795873 | -0.352551324 | 0.089 | 0.197 | 1 | 0.923594926 | 0.227000103 |
| Sap25 | 0.709465802 | 1.936806174 | 0.048 | 0.034 | 1 | 0.120805947 | -1.271123724 | 0.022 | 0.09 | 1 | 0.709465802 | 0.227017818 |
| Tbc1d25 | 0.556910165 | 1.029915578 | 0.048 | 0.09 | 1 | 0.120810184 | 0.320502507 | 0.022 | 0.093 | 1 | 0.556910165 | 0.227025268 |
| Gm56730 | 0.590073855 | 0.847801167 | 0.19 | 0.287 | 1 | 0.120887091 | -0.743192169 | 0.089 | 0.187 | 1 | 0.590073855 | 0.227160494 |
| Lman1l | 0.30448834 | -0.133583154 | 0 | 0.049 | 1 | 0.120890055 | -0.316927414 | 0.022 | 0.093 | 1 | 0.30448834 | 0.227165705 |
| Atp8b1 | 0.120908462 | -2.081115734 | 0 | 0.104 | 1 | 0.762464433 | 0.361144491 | 0.089 | 0.104 | 1 | 0.762464433 | 0.227198068 |
| Unc79 | 0.905918164 | 1.351843673 | 0.095 | 0.093 | 1 | 0.120948267 | 0.42003818 | 0.022 | 0.093 | 1 | 0.905918164 | 0.22726805 |
| Car2 | 0.120957239 | -3.303508156 | 0 | 0.104 | 1 | 0.831324582 | -0.255526869 | 0.022 | 0.028 | 1 | 0.831324582 | 0.227283825 |
| Ncald | 0.723619404 | 0.214340149 | 0.048 | 0.067 | 1 | 0.120960712 | 0.587915353 | 0.022 | 0.093 | 1 | 0.723619404 | 0.22728993 |
| Nmnat3 | 0.121076692 | 2.901182264 | 0.19 | 0.093 | 1 | 0.211155048 | 0.115388077 | 0.067 | 0.142 | 1 | 0.211155048 | 0.227493818 |
| Thsd7a | 0.548705461 | 0.951305743 | 0.095 | 0.063 | 1 | 0.12109855 | -0.14832781 | 0.089 | 0.194 | 1 | 0.548705461 | 0.227532241 |
| Retreg3 | 0.121174619 | -0.589262638 | 0.095 | 0.269 | 1 | 0.699272729 | 0.105643758 | 0.4 | 0.429 | 1 | 0.699272729 | 0.227665949 |
| Gemin8 | 0.184739419 | -1.184209227 | 0 | 0.078 | 1 | 0.121187145 | -1.223818009 | 0.022 | 0.09 | 1 | 0.184739419 | 0.227687966 |
| Fbxw7 | 0.121236518 | 1.335603032 | 0.667 | 0.511 | 1 | 0.216385408 | 0.468808046 | 0.8 | 0.609 | 1 | 0.216385408 | 0.227774743 |
| Pcp4 | 0.121251915 | -2.434752689 | 0.048 | 0.183 | 1 | 0.496223879 | -0.124282336 | 0 | 0.01 | 1 | 0.496223879 | 0.227801803 |
| Macrod2 | 0.511567719 | 0.895205269 | 0.429 | 0.455 | 1 | 0.121384666 | 1.903381693 | 0.378 | 0.287 | 1 | 0.511567719 | 0.228035096 |
| Atxn7l2 | 0.324739547 | -0.133583154 | 0 | 0.045 | 1 | 0.121565863 | -1.124282336 | 0.022 | 0.09 | 1 | 0.324739547 | 0.228353467 |
| H2ac6 | 0.12176749 | -1.940938076 | 0.048 | 0.175 | 1 | 0.852797828 | 1.03537433 | 0.067 | 0.076 | 1 | 0.852797828 | 0.228707659 |
| Svep1 | 0.121804205 | -0.107587946 | 0.143 | 0.336 | 1 | 0.405456751 | 0.454937568 | 0.422 | 0.623 | 1 | 0.405456751 | 0.228772146 |
| Nup133 | 0.91673339 | 0.281454345 | 0.143 | 0.16 | 1 | 0.121867646 | -1.071814916 | 0.089 | 0.18 | 1 | 0.91673339 | 0.22888357 |
| Dram2 | 0.555581302 | 0.572233741 | 0.286 | 0.246 | 1 | 0.121910415 | 0.740297455 | 0.533 | 0.401 | 1 | 0.555581302 | 0.228958681 |
| Sec61a2 | 0.121978885 | 0.892412054 | 0.571 | 0.504 | 1 | 0.267675402 | 0.582722061 | 0.311 | 0.529 | 1 | 0.267675402 | 0.229078921 |
| Pitpnb | 0.867224661 | 0.265189291 | 0.619 | 0.776 | 1 | 0.122014386 | -0.109485334 | 0.822 | 0.824 | 1 | 0.867224661 | 0.229141261 |
| Pex7 | 0.271806594 | 0.365019062 | 0.19 | 0.336 | 1 | 0.122085746 | -0.110476536 | 0.156 | 0.277 | 1 | 0.271806594 | 0.229266562 |
| Myl3 | 0.122142305 | 1.673771768 | 0.095 | 0.03 | 1 | 0.661444614 | 1.683072586 | 0.022 | 0.014 | 1 | 0.661444614 | 0.229365867 |
| Gfra1 | 0.12214588 | 1.595114824 | 0.476 | 0.396 | 1 | 0.915749255 | 1.340811489 | 0.333 | 0.415 | 1 | 0.915749255 | 0.229372145 |
| Eef2k | 0.499040442 | -0.385121921 | 0.095 | 0.146 | 1 | 0.122213189 | 0.484292722 | 0.111 | 0.235 | 1 | 0.499040442 | 0.229490315 |
| Sema4d | 0.122435219 | 2.289107917 | 0.238 | 0.131 | 1 | 0.679860749 | -0.47073275 | 0.2 | 0.163 | 1 | 0.679860749 | 0.229880055 |
| Ifit3 | 0.162327706 | -2.979670471 | 0.048 | 0.164 | 1 | 0.122791556 | 2.633031904 | 0.044 | 0.125 | 1 | 0.162327706 | 0.230505345 |
| Txndc11 | 0.122920786 | -0.506137322 | 0.333 | 0.556 | 1 | 0.229156587 | 0.179410188 | 0.489 | 0.661 | 1 | 0.229156587 | 0.230732053 |
| Itga5 | 0.694477517 | 1.171271427 | 0.333 | 0.444 | 1 | 0.123233192 | 0.230105631 | 0.289 | 0.481 | 1 | 0.694477517 | 0.231279965 |
| Otulin | 0.996639607 | 1.646056389 | 0.476 | 0.649 | 1 | 0.123234642 | 0.850367332 | 0.689 | 0.664 | 1 | 0.996639607 | 0.231282507 |
| Rabepk | 0.804744138 | 0.625408746 | 0.238 | 0.235 | 1 | 0.12346034 | 0.37431988 | 0.156 | 0.277 | 1 | 0.804744138 | 0.231678224 |
| Nkiras2 | 0.788998424 | 0.786246497 | 0.095 | 0.127 | 1 | 0.123538707 | -1.223818009 | 0.044 | 0.121 | 1 | 0.788998424 | 0.231815603 |
| Ccnl1 | 0.123541375 | 0.391455529 | 1 | 0.955 | 1 | 0.948806561 | 0.228932031 | 1 | 0.969 | 1 | 0.948806561 | 0.231820279 |
| Metap1 | 0.517256898 | 0.254232876 | 0.19 | 0.287 | 1 | 0.123645096 | 0.390290837 | 0.289 | 0.509 | 1 | 0.517256898 | 0.232002083 |
| Csnk1d | 0.750918173 | 0.231934209 | 0.571 | 0.608 | 1 | 0.12378505 | 0.550414842 | 0.867 | 0.761 | 1 | 0.750918173 | 0.232247362 |
| Ppp2r5e | 0.12399887 | 0.511884085 | 0.714 | 0.616 | 1 | 0.740204117 | 0.543228924 | 0.822 | 0.806 | 1 | 0.740204117 | 0.232622021 |
| Cdc25a | 0.618392893 | 0.648236676 | 0.095 | 0.138 | 1 | 0.124280283 | -0.612383297 | 0.133 | 0.242 | 1 | 0.618392893 | 0.233114977 |
| Egr1 | 0.219243423 | -1.995321714 | 0.619 | 0.597 | 1 | 0.124291026 | -1.154784424 | 0.733 | 0.675 | 1 | 0.219243423 | 0.233133792 |
| Ptprt | 0.124398683 | -2.727107669 | 0.048 | 0.172 | 1 | 0.789464486 | 0.635766871 | 0.133 | 0.159 | 1 | 0.789464486 | 0.233322333 |
| Enpp2 | 0.340807445 | -0.564157341 | 0.857 | 0.907 | 1 | 0.124410897 | -0.940391815 | 0.556 | 0.668 | 1 | 0.340807445 | 0.233343722 |
| Mtf1 | 0.978174576 | 0.894161836 | 0.238 | 0.291 | 1 | 0.124421397 | 0.463627449 | 0.222 | 0.398 | 1 | 0.978174576 | 0.233362111 |
| Hey1 | 0.192656654 | 0.735172312 | 0.19 | 0.101 | 1 | 0.124484698 | -0.446210431 | 0.044 | 0.128 | 1 | 0.192656654 | 0.233472956 |
| Dpy19l3 | 0.744421685 | 0.281454345 | 0.143 | 0.175 | 1 | 0.124516426 | 0.513147585 | 0.111 | 0.239 | 1 | 0.744421685 | 0.233528511 |
| Mcam | 0.770356896 | 1.316219763 | 0.095 | 0.082 | 1 | 0.124751184 | -0.342462506 | 0.044 | 0.128 | 1 | 0.770356896 | 0.233939509 |
| Carnmt1 | 0.124883162 | -0.422596627 | 0.381 | 0.608 | 1 | 0.224769107 | -0.27272761 | 0.6 | 0.671 | 1 | 0.224769107 | 0.234170519 |
| Scly | 0.551267885 | 1.236366455 | 0.19 | 0.153 | 1 | 0.124923168 | -0.502793959 | 0.089 | 0.19 | 1 | 0.551267885 | 0.234240537 |
| Snx32 | 0.860084871 | 1.010806755 | 0.048 | 0.06 | 1 | 0.124934666 | -1.124282336 | 0.044 | 0.121 | 1 | 0.860084871 | 0.234260662 |
| Gm49774 | 0.124967068 | 1.79930265 | 0.143 | 0.06 | 1 | 0.765728546 | 1.42003818 | 0.022 | 0.031 | 1 | 0.765728546 | 0.234317368 |
| Sgsm3 | 0.145669871 | -0.623908781 | 0.143 | 0.299 | 1 | 0.125102281 | 0.891659208 | 0.378 | 0.298 | 1 | 0.145669871 | 0.234553981 |
| Zmym5 | 0.827107746 | 0.642623069 | 0.667 | 0.81 | 1 | 0.125177397 | -0.155124972 | 0.956 | 0.875 | 1 | 0.827107746 | 0.234685413 |
| Ube2g1 | 0.880810941 | 0.363916505 | 0.714 | 0.784 | 1 | 0.125331887 | 0.769487338 | 0.978 | 0.913 | 1 | 0.880810941 | 0.234955692 |
| Smurf1 | 0.125408772 | -0.611630451 | 0.143 | 0.332 | 1 | 0.901357016 | 0.854058783 | 0.4 | 0.484 | 1 | 0.901357016 | 0.235090183 |
| Ptprz1 | 0.57816106 | 1.351843673 | 0 | 0.015 | 1 | 0.125439407 | 1.098110085 | 0.111 | 0.052 | 1 | 0.57816106 | 0.235143769 |
| Hbp1 | 0.125484599 | -0.516784567 | 0.619 | 0.739 | 1 | 0.508062818 | -0.126341858 | 0.889 | 0.837 | 1 | 0.508062818 | 0.235222813 |
| Fndc3a | 0.125508666 | 0.257457143 | 1 | 0.985 | 1 | 0.17137505 | 0.703618291 | 0.978 | 0.931 | 1 | 0.17137505 | 0.235264906 |
| Wnk4 | 0.200374297 | -0.51887331 | 0.286 | 0.451 | 1 | 0.125614104 | -0.398721505 | 0.156 | 0.277 | 1 | 0.200374297 | 0.235449306 |
| Acer3 | 0.751972437 | 0.656535063 | 0.571 | 0.646 | 1 | 0.125859499 | 0.86188981 | 0.911 | 0.747 | 1 | 0.751972437 | 0.235878385 |
| Pcgf6 | 0.848865644 | 0.704145417 | 0.19 | 0.231 | 1 | 0.126020665 | -0.158229668 | 0.111 | 0.221 | 1 | 0.848865644 | 0.236160122 |
| Brd8 | 0.126222165 | 0.926958388 | 0.762 | 0.769 | 1 | 0.245268684 | 0.180572246 | 0.778 | 0.792 | 1 | 0.245268684 | 0.236512296 |
| Phf20l1 | 0.823267727 | 0.303660139 | 0.762 | 0.784 | 1 | 0.126267535 | 0.517211517 | 0.956 | 0.772 | 1 | 0.823267727 | 0.236591579 |
| Tmem201 | 0.897445293 | 0.586308926 | 0.095 | 0.108 | 1 | 0.126354798 | -0.709244837 | 0.044 | 0.125 | 1 | 0.897445293 | 0.236744062 |
| Tiparp | 0.647158803 | 0.424557856 | 0.857 | 0.821 | 1 | 0.12641102 | -0.536789986 | 0.978 | 0.882 | 1 | 0.647158803 | 0.236842293 |
| 2010007H06Rik | 0.126520325 | 1.738866796 | 0.286 | 0.183 | 1 | 0.952491713 | 1.163698427 | 0.111 | 0.118 | 1 | 0.952491713 | 0.237033258 |
| Pgm2 | 0.126599243 | -0.774688733 | 0.143 | 0.317 | 1 | 0.143512504 | 0.189533113 | 0.267 | 0.446 | 1 | 0.143512504 | 0.237171117 |
| Ncs1 | 0.126605675 | 1.621304348 | 0.333 | 0.243 | 1 | 0.283395008 | 0.866518727 | 0.422 | 0.415 | 1 | 0.283395008 | 0.237182353 |
| Celf4 | 0.697541628 | 1.478755785 | 0.238 | 0.231 | 1 | 0.12677798 | -1.017367132 | 0.067 | 0.152 | 1 | 0.697541628 | 0.237483304 |
| Cep68 | 0.570033342 | 0.96105372 | 0.19 | 0.164 | 1 | 0.126812362 | 1.683072586 | 0.244 | 0.183 | 1 | 0.570033342 | 0.237543349 |
| 2810402E24Rik | 0.127012032 | 0.327996931 | 0.048 | 0.19 | 1 | 0.168165958 | -0.153428682 | 0.044 | 0.114 | 1 | 0.168165958 | 0.237892007 |
| Tcerg1 | 0.127068137 | 0.665837586 | 0.81 | 0.802 | 1 | 0.827798473 | 0.219844399 | 0.933 | 0.875 | 1 | 0.827798473 | 0.237989962 |
| Tapt1 | 0.127152987 | -0.636083495 | 0.286 | 0.489 | 1 | 0.982872876 | 0.27368165 | 0.644 | 0.723 | 1 | 0.982872876 | 0.238138093 |
| Atp11b | 0.127252424 | 0.612781138 | 0.762 | 0.679 | 1 | 0.209162686 | 0.336964679 | 0.933 | 0.806 | 1 | 0.209162686 | 0.238311669 |
| Utp4 | 0.813718142 | 0.658174913 | 0.381 | 0.418 | 1 | 0.127466078 | 0.132057417 | 0.333 | 0.55 | 1 | 0.813718142 | 0.238684556 |
| Pign | 0.12748058 | 1.295260144 | 0.476 | 0.429 | 1 | 0.624733886 | 0.989499588 | 0.422 | 0.443 | 1 | 0.624733886 | 0.238709862 |
| Ercc5 | 0.645052131 | 1.017424634 | 0.333 | 0.336 | 1 | 0.127567523 | 0.175730983 | 0.222 | 0.401 | 1 | 0.645052131 | 0.238861573 |
| Gm33989 | 0.127649249 | -0.445169305 | 0.143 | 0.306 | 1 | 0.236584971 | 0.380509816 | 0.267 | 0.429 | 1 | 0.236584971 | 0.239004167 |
| Nedd1 | 0.64373078 | 0.766881172 | 0.143 | 0.213 | 1 | 0.127742776 | -0.404390255 | 0.133 | 0.249 | 1 | 0.64373078 | 0.239167335 |
| Rian | 0.12782344 | 1.36385369 | 0.571 | 0.541 | 1 | 0.364520474 | 0.482728189 | 0.533 | 0.685 | 1 | 0.364520474 | 0.239308048 |
| Hapstr1 | 0.905018798 | 0.451379346 | 0.619 | 0.806 | 1 | 0.127872236 | 0.438835776 | 0.933 | 0.92 | 1 | 0.905018798 | 0.239393163 |
| Lin52 | 0.127880857 | 0.113056813 | 0.19 | 0.433 | 1 | 0.622118426 | 0.750186782 | 0.4 | 0.405 | 1 | 0.622118426 | 0.2394082 |
| Cklf | 0.178192398 | 0.550389352 | 0.048 | 0.168 | 1 | 0.127910257 | -0.286553765 | 0.044 | 0.128 | 1 | 0.178192398 | 0.23945948 |

| | | | | | | | | | | | | |
|---|---|---|---|---|---|---|---|---|---|---|---|---|
| Nudt8 | 0.127996876 | -1.233118828 | 0 | 0.101 | 1 | 0.22632047 | 0.595609745 | 0.044 | 0.107 | 1 | 0.22632047 | 0.239610551 |
| Shroom4 | 0.128021412 | 1.790972979 | 0.524 | 0.478 | 1 | 0.966936945 | 0.701784846 | 0.444 | 0.529 | 1 | 0.966936945 | 0.239653342 |
| 4930430F08Rik | 0.128057475 | -1.280424543 | 0 | 0.101 | 1 | 0.164309241 | -1.223818009 | 0.022 | 0.08 | 1 | 0.164309241 | 0.239716233 |
| Faap100 | 0.128113406 | -1.280424543 | 0 | 0.101 | 1 | 0.419119453 | 0.188307894 | 0.111 | 0.166 | 1 | 0.419119453 | 0.239813768 |
| Zkscan8 | 0.128113406 | -1.280424543 | 0 | 0.101 | 1 | 0.388907488 | -0.124282336 | 0.044 | 0.083 | 1 | 0.388907488 | 0.239813768 |
| Med1 | 0.26741386 | 0.680724528 | 0.476 | 0.474 | 1 | 0.128142794 | -0.242926832 | 0.533 | 0.644 | 1 | 0.26741386 | 0.239865013 |
| Rfc4 | 0.128164113 | -1.370622352 | 0 | 0.101 | 1 | 0.346432578 | -0.564854927 | 0.022 | 0.055 | 1 | 0.346432578 | 0.239902186 |
| Efcab15 | 0.128211833 | -1.496153234 | 0 | 0.101 | 1 | 0.132703959 | -1.486852415 | 0 | 0.048 | 1 | 0.132703959 | 0.239985391 |
| Vps37d | 0.128212019 | -1.370622352 | 0 | 0.101 | 1 | 0.300659003 | -0.731964913 | 0.044 | 0.09 | 1 | 0.300659003 | 0.239985716 |
| Mkln1os | 0.128254143 | -1.683780237 | 0 | 0.101 | 1 | 0.238126817 | -0.589945908 | 0.022 | 0.069 | 1 | 0.238126817 | 0.240059161 |
| Tspan32 | 0.128254889 | -1.718545655 | 0 | 0.101 | 1 | 0.133714345 | -1.223818009 | 0.022 | 0.087 | 1 | 0.133714345 | 0.240060461 |
| Rfc5 | 0.128319187 | -1.535681598 | 0 | 0.101 | 1 | 0.430238982 | 0.223640968 | 0.067 | 0.107 | 1 | 0.430238982 | 0.24017256 |
| Ccno | 0.128327573 | -1.880817084 | 0 | 0.101 | 1 | 0.184806696 | -1.017367132 | 0 | 0.038 | 1 | 0.184806696 | 0.24018718 |
| Smim5 | 0.128357762 | -1.818081329 | 0 | 0.101 | 1 | 0.132717935 | -2.174908409 | 0 | 0.048 | 1 | 0.132717935 | 0.24023981 |
| Megf8 | 0.383720343 | -0.116305163 | 0.143 | 0.243 | 1 | 0.128370071 | -0.108340792 | 0.133 | 0.249 | 1 | 0.383720343 | 0.240261267 |
| Rffl | 0.128374347 | -1.648156327 | 0 | 0.101 | 1 | 0.644895223 | 0.589963182 | 0.156 | 0.128 | 1 | 0.644895223 | 0.240268721 |
| Haus4 | 0.128404161 | -1.818081329 | 0 | 0.101 | 1 | 0.418161012 | 0.875717664 | 0.044 | 0.083 | 1 | 0.418161012 | 0.240320694 |
| Chchd3 | 0.128444906 | -0.696686221 | 0.429 | 0.597 | 1 | 0.209692106 | 0.467723442 | 0.889 | 0.713 | 1 | 0.209692106 | 0.240391718 |
| Thbd | 0.128446643 | -2.208871282 | 0 | 0.101 | 1 | 0.206909388 | -1.486852415 | 0 | 0.035 | 1 | 0.206909388 | 0.240394746 |
| Tnc | 0.128537373 | -4.634567262 | 0 | 0.101 | 1 | 0.173809289 | -0.672408068 | 0.044 | 0.111 | 1 | 0.173809289 | 0.240552889 |
| Kif13a | 0.128542959 | 1.414499281 | 0.619 | 0.586 | 1 | 0.371639651 | 0.395158524 | 0.844 | 0.668 | 1 | 0.371639651 | 0.240562626 |
| Klf6 | 0.128543817 | -0.212551201 | 0.81 | 0.903 | 1 | 0.64597197 | -0.302382866 | 0.956 | 0.882 | 1 | 0.64597197 | 0.240564121 |
| Dhx40 | 0.12896498 | -0.755539863 | 0.381 | 0.619 | 1 | 0.532122096 | 0.29688995 | 0.778 | 0.671 | 1 | 0.532122096 | 0.241297994 |
| Smcr8 | 0.333182369 | -0.184209227 | 0.095 | 0.187 | 1 | 0.129296835 | -0.446210431 | 0.089 | 0.187 | 1 | 0.333182369 | 0.241875999 |
| Obi1 | 0.129361521 | -1.081115734 | 0.048 | 0.175 | 1 | 0.215745785 | -0.53415813 | 0.111 | 0.194 | 1 | 0.215745785 | 0.24198864 |
| Pcsk5 | 0.129375173 | -1.752492987 | 0.143 | 0.302 | 1 | 0.323812145 | 0.391438414 | 0.2 | 0.308 | 1 | 0.323812145 | 0.24201241 |
| Bsdc1 | 0.129557275 | -0.115282338 | 0.286 | 0.549 | 1 | 0.175003107 | -0.258824459 | 0.556 | 0.602 | 1 | 0.175003107 | 0.242329462 |
| Gm43445 | 0.129719906 | 1.766881172 | 0.19 | 0.097 | 1 | 0.907786668 | 1.152557869 | 0.067 | 0.066 | 1 | 0.907786668 | 0.242612559 |
| Cdh15 | 0.167544534 | 2.936806174 | 0.048 | 0.011 | 1 | 0.129721052 | 3.268035087 | 0.022 | 0.003 | 1 | 0.167544534 | 0.242614552 |
| Ctcflos | 0.17843975 | 2.410737362 | 0.095 | 0.037 | 1 | 0.129721052 | 3.683072586 | 0.022 | 0.003 | 1 | 0.17843975 | 0.242614552 |
| Gdnf | 0.324789396 | -1.280424543 | 0 | 0.045 | 1 | 0.129721052 | 3.098110085 | 0.022 | 0.003 | 1 | 0.324789396 | 0.242614552 |
| 4930565N06Rik | 0.367982232 | 2.088809267 | 0.048 | 0.019 | 1 | 0.129721052 | 3.268035087 | 0.022 | 0.003 | 1 | 0.367982232 | 0.242614552 |
| AA467197 | 0.39694807 | -0.413691074 | 0 | 0.034 | 1 | 0.129721052 | 3.683072586 | 0.022 | 0.003 | 1 | 0.39694807 | 0.242614552 |
| Ncapg | 0.39694807 | -0.413691074 | 0 | 0.034 | 1 | 0.129721052 | 3.268035087 | 0.022 | 0.003 | 1 | 0.39694807 | 0.242614552 |
| Phtf1os | 0.454205211 | 3.188344941 | 0.048 | 0.022 | 1 | 0.129721052 | 3.268035087 | 0.022 | 0.003 | 1 | 0.454205211 | 0.242614552 |
| Bsx | 0.457281049 | 0.503846766 | 0 | 0.026 | 1 | 0.129721052 | 3.268035087 | 0.022 | 0.003 | 1 | 0.457281049 | 0.242614552 |
| Aurkb | 0.492604708 | -0.233118828 | 0 | 0.022 | 1 | 0.129721052 | 4.005000681 | 0.022 | 0.003 | 1 | 0.492604708 | 0.242614552 |
| Gm14009 | 0.532432377 | 0.673771768 | 0 | 0.019 | 1 | 0.129721052 | 3.683072586 | 0.022 | 0.003 | 1 | 0.532432377 | 0.242614552 |
| Rsad2 | 0.532432377 | 0.673771768 | 0 | 0.019 | 1 | 0.129721052 | 7.075390009 | 0.022 | 0.003 | 1 | 0.532432377 | 0.242614552 |
| Mrc1 | 0.532436372 | 1.088809267 | 0 | 0.019 | 1 | 0.129721052 | 3.098110085 | 0.022 | 0.003 | 1 | 0.532436372 | 0.242614552 |
| Pax6os1 | 0.578156445 | 1.088809267 | 0 | 0.015 | 1 | 0.129721052 | 3.268035087 | 0.022 | 0.003 | 1 | 0.578156445 | 0.242614552 |
| Gm30003 | 0.57816106 | 1.351843673 | 0 | 0.015 | 1 | 0.129721052 | 3.268035087 | 0.022 | 0.003 | 1 | 0.57816106 | 0.242614552 |
| Prss32 | 0.57816106 | 1.351843673 | 0 | 0.015 | 1 | 0.129721052 | 3.683072586 | 0.022 | 0.003 | 1 | 0.57816106 | 0.242614552 |
| Gm57468 | 0.63207476 | 1.351843673 | 0 | 0.011 | 1 | 0.129721052 | 3.683072586 | 0.022 | 0.003 | 1 | 0.63207476 | 0.242614552 |
| Gfy | 0.63207476 | 1.351843673 | 0 | 0.011 | 1 | 0.129721052 | 3.268035087 | 0.022 | 0.003 | 1 | 0.63207476 | 0.242614552 |
| Atp2c2 | 0.63207476 | 0.503846766 | 0 | 0.011 | 1 | 0.129721052 | 3.683072586 | 0.022 | 0.003 | 1 | 0.63207476 | 0.242614552 |
| H3c11 | 0.63207476 | 1.088809267 | 0 | 0.011 | 1 | 0.129721052 | 4.005000681 | 0.022 | 0.003 | 1 | 0.63207476 | 0.242614552 |
| Gm49975 | 0.63207476 | 1.351843673 | 0 | 0.011 | 1 | 0.129721052 | 3.268035087 | 0.022 | 0.003 | 1 | 0.63207476 | 0.242614552 |
| Brinp2 | 0.632076134 | 1.673771768 | 0 | 0.011 | 1 | 0.129721052 | 3.683072586 | 0.022 | 0.003 | 1 | 0.632076134 | 0.242614552 |
| Dkk1 | 0.632076134 | 1.673771768 | 0 | 0.011 | 1 | 0.129721052 | 3.683072586 | 0.022 | 0.003 | 1 | 0.632076134 | 0.242614552 |
| 4930428N03Rik | 0.71266839 | 1.79930265 | 0.048 | 0.034 | 1 | 0.129721052 | 3.268035087 | 0.022 | 0.003 | 1 | 0.71266839 | 0.242614552 |
| E330011O21Rik | 0.91021959 | 0.181918671 | 0.095 | 0.104 | 1 | 0.129721052 | 4.268035087 | 0.022 | 0.003 | 1 | 0.91021959 | 0.242614552 |
| Slc9a1 | 0.372417428 | 1.332303486 | 0.429 | 0.485 | 1 | 0.12974603 | 0.831395333 | 0.667 | 0.585 | 1 | 0.372417428 | 0.242658028 |
| Spidr | 0.129858091 | 1.239368944 | 0.476 | 0.392 | 1 | 0.996897107 | 0.987610544 | 0.311 | 0.363 | 1 | 0.996897107 | 0.242853059 |
| Ddi2 | 0.129871821 | 0.908237021 | 0.714 | 0.761 | 1 | 0.282215109 | 0.188537167 | 0.933 | 0.938 | 1 | 0.282215109 | 0.242876952 |
| Mysm1 | 0.130037647 | 0.405282932 | 0.667 | 0.575 | 1 | 0.39335634 | 0.36462506 | 0.733 | 0.664 | 1 | 0.39335634 | 0.243165504 |
| Pola1 | 0.986736894 | 1.454937166 | 0.19 | 0.231 | 1 | 0.130147935 | 0.405087839 | 0.178 | 0.311 | 1 | 0.986736894 | 0.243357386 |
| Dbf4 | 0.347456154 | -0.880817084 | 0.048 | 0.112 | 1 | 0.13044828 | -0.819427754 | 0.022 | 0.09 | 1 | 0.347456154 | 0.243879807 |
| Prkab2 | 0.130510347 | -0.899117901 | 0.095 | 0.246 | 1 | 0.167628036 | -0.137106376 | 0.244 | 0.391 | 1 | 0.167628036 | 0.243987744 |
| Lactb | 0.130515912 | -0.837190152 | 0.095 | 0.246 | 1 | 0.198474323 | 0.199989699 | 0.156 | 0.266 | 1 | 0.198474323 | 0.24399742 |
| Fxn | 0.300961054 | -0.133583154 | 0.048 | 0.131 | 1 | 0.130565713 | 0.335149283 | 0.067 | 0.159 | 1 | 0.300961054 | 0.244084021 |
| Odad1 | 0.130620547 | 1.179007076 | 0.286 | 0.172 | 1 | 0.479905576 | 1.251733275 | 0.222 | 0.208 | 1 | 0.479905576 | 0.244179366 |
| Gm28791 | 0.251683171 | -0.718545655 | 0 | 0.06 | 1 | 0.130926054 | -0.776359032 | 0.022 | 0.09 | 1 | 0.251683171 | 0.244710476 |
| Tec | 0.604518835 | 0.97333205 | 0.095 | 0.067 | 1 | 0.131025139 | 0.320502507 | 0.133 | 0.066 | 1 | 0.604518835 | 0.244882691 |
| Hmcn2 | 0.875318873 | 0.936806174 | 0.048 | 0.041 | 1 | 0.131112429 | -1.84048937 | 0.022 | 0.087 | 1 | 0.875318873 | 0.245034389 |
| Zfp445 | 0.69797801 | 0.457834368 | 0.381 | 0.537 | 1 | 0.13123546 | -0.158943343 | 0.422 | 0.612 | 1 | 0.69797801 | 0.245248174 |
| 2810414N06Rik | 0.454205211 | 2.451379346 | 0.048 | 0.022 | 1 | 0.131300581 | 2.683072586 | 0.022 | 0.003 | 1 | 0.454205211 | 0.24536132 |
| Adarb2 | 0.457300763 | -0.849790188 | 0 | 0.026 | 1 | 0.131300581 | 2.683072586 | 0.022 | 0.003 | 1 | 0.457300763 | 0.24536132 |
| Dmbt1 | 0.460380341 | 1.171271427 | 0.048 | 0.022 | 1 | 0.131300581 | 2.683072586 | 0.022 | 0.003 | 1 | 0.460380341 | 0.24536132 |
| Gm43623 | 0.532426384 | 0.866416846 | 0 | 0.019 | 1 | 0.131300581 | 2.683072586 | 0.022 | 0.003 | 1 | 0.532426384 | 0.24536132 |
| Fndc7 | 0.578156445 | 0.503846766 | 0 | 0.015 | 1 | 0.131300581 | 2.683072586 | 0.022 | 0.003 | 1 | 0.578156445 | 0.24536132 |
| Foxs1 | 0.578159907 | 0.673771768 | 0 | 0.015 | 1 | 0.131300581 | 2.683072586 | 0.022 | 0.003 | 1 | 0.578159907 | 0.24536132 |
| AU023762 | 0.578159907 | 0.673771768 | 0 | 0.015 | 1 | 0.131300581 | 2.683072586 | 0.022 | 0.003 | 1 | 0.578159907 | 0.24536132 |
| Ren1 | 0.57816106 | 1.351843673 | 0 | 0.015 | 1 | 0.131300581 | 2.683072586 | 0.022 | 0.003 | 1 | 0.57816106 | 0.24536132 |
| Zfp981 | 0.63207476 | 1.088809267 | 0 | 0.011 | 1 | 0.131300581 | 2.683072586 | 0.022 | 0.003 | 1 | 0.63207476 | 0.24536132 |
| Tpte | 0.63207476 | 0.866416846 | 0 | 0.011 | 1 | 0.131300581 | 2.683072586 | 0.022 | 0.003 | 1 | 0.63207476 | 0.24536132 |
| Gm44036 | 0.632076134 | 1.673771768 | 0 | 0.011 | 1 | 0.131300581 | 2.683072586 | 0.022 | 0.003 | 1 | 0.632076134 | 0.24536132 |
| Gm5082 | 0.632076134 | 1.673771768 | 0 | 0.011 | 1 | 0.131300581 | 2.683072586 | 0.022 | 0.003 | 1 | 0.632076134 | 0.24536132 |
| Gm48860 | 0.632076134 | 1.673771768 | 0 | 0.011 | 1 | 0.131300581 | 2.683072586 | 0.022 | 0.003 | 1 | 0.632076134 | 0.24536132 |
| Nxt2 | 0.401793043 | -0.326228232 | 0.048 | 0.104 | 1 | 0.131345279 | -0.731964913 | 0.022 | 0.09 | 1 | 0.401793043 | 0.245438976 |
| Entrep1 | 0.758437281 | -0.455511249 | 0.048 | 0.063 | 1 | 0.131548236 | -1.602329633 | 0.022 | 0.087 | 1 | 0.758437281 | 0.245791534 |
| Ttyh2 | 0.68243969 | 1.171271427 | 0.095 | 0.138 | 1 | 0.131631657 | -0.889817082 | 0.089 | 0.183 | 1 | 0.68243969 | 0.24593642 |
| Tmem260 | 0.491020952 | 0.914779867 | 0.286 | 0.246 | 1 | 0.131716032 | 0.427815531 | 0.156 | 0.304 | 1 | 0.491020952 | 0.24608295 |
| Prpf40b | 0.131983727 | -0.413691074 | 0.048 | 0.187 | 1 | 0.468301844 | 0.223640968 | 0.178 | 0.249 | 1 | 0.468301844 | 0.24654775 |
| Serac1 | 0.292155863 | 1.586308926 | 0.095 | 0.045 | 1 | 0.131984923 | -1.709244837 | 0.022 | 0.087 | 1 | 0.292155863 | 0.246549826 |
| Zfp113 | 0.721275807 | 0.150209812 | 0.048 | 0.067 | 1 | 0.1319855 | -1.564854927 | 0.022 | 0.087 | 1 | 0.721275807 | 0.246550828 |
| Ercc8 | 0.759396864 | 0.896164189 | 0.143 | 0.127 | 1 | 0.132080803 | -0.802354241 | 0.067 | 0.152 | 1 | 0.759396864 | 0.246716267 |
| Patl1 | 0.132082422 | 1.171271427 | 0.429 | 0.336 | 1 | 0.632165501 | 0.496194451 | 0.333 | 0.46 | 1 | 0.632165501 | 0.246719078 |
| Exog | 0.267953794 | -0.496153234 | 0 | 0.056 | 1 | 0.132178104 | -0.638855509 | 0.022 | 0.09 | 1 | 0.267953794 | 0.246885157 |
| Wdr43 | 0.132220735 | 0.698862749 | 0.667 | 0.705 | 1 | 0.944803726 | 0.218046951 | 0.889 | 0.789 | 1 | 0.944803726 | 0.246959148 |
| Mgat5 | 0.805962826 | 0.673771768 | 0.476 | 0.549 | 1 | 0.132272242 | 0.611581544 | 0.822 | 0.765 | 1 | 0.805962826 | 0.247048538 |
| Eif3b | 0.132366924 | -0.538278709 | 0.333 | 0.552 | 1 | 0.678256719 | -0.109485334 | 0.733 | 0.668 | 1 | 0.678256719 | 0.247212845 |
| 1500015L24Rik | 0.347190422 | 1.351843673 | 0.143 | 0.086 | 1 | 0.13262998 | -0.316927414 | 0.022 | 0.09 | 1 | 0.347190422 | 0.247669248 |
| 2310034G01Rik | 0.268100354 | -0.326228232 | 0 | 0.056 | 1 | 0.132666686 | -1.316927414 | 0 | 0.048 | 1 | 0.268100354 | 0.247732923 |
| Gm4013 | 0.285622926 | -0.233118828 | 0 | 0.052 | 1 | 0.132666686 | -1.404390255 | 0 | 0.048 | 1 | 0.285622926 | 0.247732923 |
| Gpr45 | 0.324767242 | -0.233118828 | 0 | 0.045 | 1 | 0.132666686 | -1.404390255 | 0 | 0.048 | 1 | 0.324767242 | 0.247732923 |
| Lrrc75a | 0.457300763 | -0.133583154 | 0 | 0.026 | 1 | 0.132666686 | -1.316927414 | 0 | 0.048 | 1 | 0.457300763 | 0.247732923 |
| Lrrc8dos | 0.457308648 | 0.673771768 | 0 | 0.026 | 1 | 0.132666686 | -1.316927414 | 0 | 0.048 | 1 | 0.457308648 | 0.247732923 |

| | | | | | | | | | | | | |
|---|---|---|---|---|---|---|---|---|---|---|---|---|
| Ada | 0.492599419 | 0.503846766 | 0 | 0.022 | 1 | 0.132666686 | -1.316927414 | 0 | 0.048 | 1 | 0.492599419 | 0.247732923 |
| Sbsn | 0.570964864 | -0.184209227 | 0.048 | 0.082 | 1 | 0.132666686 | -1.316927414 | 0 | 0.048 | 1 | 0.570964864 | 0.247732923 |
| Gm56948 | 0.57816106 | 1.351843673 | 0 | 0.015 | 1 | 0.132666686 | -1.316927414 | 0 | 0.048 | 1 | 0.57816106 | 0.247732923 |
| Rpusd1 | 0.96060963 | 1.088809267 | 0.048 | 0.052 | 1 | 0.132666686 | -1.564854927 | 0 | 0.048 | 1 | 0.96060963 | 0.247732923 |
| Cdkn2b | 0.153811856 | -1.280424543 | 0 | 0.09 | 1 | 0.132686876 | -1.564854927 | 0 | 0.048 | 1 | 0.153811856 | 0.247767944 |
| Lrrc17 | 0.173988472 | -2.496153234 | 0 | 0.082 | 1 | 0.132686876 | -1.638855509 | 0 | 0.048 | 1 | 0.173988472 | 0.247767944 |
| Hrct1 | 0.222110654 | -0.648156327 | 0 | 0.067 | 1 | 0.132686876 | -1.564854927 | 0 | 0.048 | 1 | 0.222110654 | 0.247767944 |
| Mterf1a | 0.251619726 | -0.574155746 | 0 | 0.06 | 1 | 0.132686876 | -1.404390255 | 0 | 0.048 | 1 | 0.251619726 | 0.247767944 |
| Smim36 | 0.346827513 | -0.233118828 | 0 | 0.041 | 1 | 0.132686876 | -1.404390255 | 0 | 0.048 | 1 | 0.346827513 | 0.247767944 |
| Mbnl3 | 0.425301468 | 0.673771768 | 0.095 | 0.052 | 1 | 0.132686876 | -1.486852415 | 0 | 0.048 | 1 | 0.425301468 | 0.247767944 |
| Sirt5 | 0.4540883 | -0.633656758 | 0.143 | 0.209 | 1 | 0.132686876 | -1.638855509 | 0 | 0.048 | 1 | 0.4540883 | 0.247767944 |
| Car11 | 0.550953306 | 1.936806174 | 0.048 | 0.026 | 1 | 0.132686876 | -1.486852415 | 0 | 0.048 | 1 | 0.550953306 | 0.247767944 |
| Gm34934 | 0.715873069 | 1.673771768 | 0.048 | 0.034 | 1 | 0.132686876 | -1.404390255 | 0 | 0.048 | 1 | 0.715873069 | 0.247767944 |
| Rab17 | 0.960611849 | 1.088809267 | 0.048 | 0.052 | 1 | 0.132686876 | -1.404390255 | 0 | 0.048 | 1 | 0.960611849 | 0.247767944 |
| 2010310C07Rik | 0.251706443 | -0.911190733 | 0 | 0.06 | 1 | 0.132696228 | -0.242926832 | 0.022 | 0.09 | 1 | 0.251706443 | 0.247784167 |
| Npff | 0.251580301 | -0.496153234 | 0 | 0.06 | 1 | 0.1327037 | -1.564854927 | 0 | 0.048 | 1 | 0.251580301 | 0.247797128 |
| Sox13 | 0.57816106 | 1.351843673 | 0 | 0.015 | 1 | 0.1327037 | -1.564854927 | 0 | 0.048 | 1 | 0.57816106 | 0.247797128 |
| Shkbp1 | 0.629700127 | 2.088809267 | 0.048 | 0.03 | 1 | 0.1327037 | -1.564854927 | 0 | 0.048 | 1 | 0.629700127 | 0.247797128 |
| Fxyd7 | 0.960849104 | 0.673771768 | 0.048 | 0.045 | 1 | 0.1327037 | -1.564854927 | 0 | 0.048 | 1 | 0.960849104 | 0.247797128 |
| Gm14327 | 0.251790506 | -0.413691074 | 0 | 0.06 | 1 | 0.132703959 | -1.486852415 | 0 | 0.048 | 1 | 0.251790506 | 0.247797577 |
| Eme1 | 0.285499161 | -0.413691074 | 0 | 0.052 | 1 | 0.132703959 | -1.776359032 | 0 | 0.048 | 1 | 0.285499161 | 0.247797577 |
| Opn1mw | 0.576921108 | 1.351843673 | 0.095 | 0.067 | 1 | 0.132703959 | -1.776359032 | 0 | 0.048 | 1 | 0.576921108 | 0.247797577 |
| Ccdc18 | 0.912928517 | -0.535681598 | 0.048 | 0.052 | 1 | 0.132703959 | -2.124282336 | 0 | 0.048 | 1 | 0.912928517 | 0.247797577 |
| Coro2b | 0.434294571 | 1.48112669 | 0.143 | 0.101 | 1 | 0.132717159 | -1.638855509 | 0 | 0.048 | 1 | 0.434294571 | 0.247820473 |
| Pram1 | 0.438359353 | 1.586308926 | 0.095 | 0.056 | 1 | 0.132717159 | -1.638855509 | 0 | 0.048 | 1 | 0.438359353 | 0.247820473 |
| Gm31282 | 0.457308648 | 0.673771768 | 0 | 0.026 | 1 | 0.132717159 | -1.638855509 | 0 | 0.048 | 1 | 0.457308648 | 0.247820473 |
| Lama1 | 0.222155241 | -0.718545655 | 0 | 0.067 | 1 | 0.132717935 | -1.84048937 | 0 | 0.048 | 1 | 0.222155241 | 0.24782182 |
| Gpatch3 | 0.304355016 | -0.233118828 | 0 | 0.049 | 1 | 0.132717935 | -1.84048937 | 0 | 0.048 | 1 | 0.304355016 | 0.24782182 |
| Lrrn3 | 0.396948718 | -0.496153234 | 0 | 0.034 | 1 | 0.132717935 | -1.776359032 | 0 | 0.048 | 1 | 0.396948718 | 0.24782182 |
| Gm29292 | 0.924795272 | 1.351843673 | 0.048 | 0.045 | 1 | 0.132717935 | -1.709244837 | 0 | 0.048 | 1 | 0.924795272 | 0.24782182 |
| Gm34312 | 0.63207476 | 1.351843673 | 0 | 0.011 | 1 | 0.132718194 | -1.564854927 | 0 | 0.048 | 1 | 0.63207476 | 0.247822269 |
| Fgf1 | 0.905776468 | -0.413691074 | 0.143 | 0.149 | 1 | 0.132718194 | -1.776359032 | 0 | 0.048 | 1 | 0.905776468 | 0.247822269 |
| Ccnj | 0.969041235 | 1.351843673 | 0.048 | 0.052 | 1 | 0.132718194 | -1.564854927 | 0 | 0.048 | 1 | 0.969041235 | 0.247822269 |
| Col26a1 | 0.425624656 | -0.648156327 | 0 | 0.03 | 1 | 0.132727253 | -2.124282336 | 0 | 0.048 | 1 | 0.425624656 | 0.247837981 |
| Dtl | 0.630729388 | -0.167530486 | 0.048 | 0.078 | 1 | 0.132729323 | -1.901889915 | 0 | 0.048 | 1 | 0.630729388 | 0.247841573 |
| Gm44148 | 0.264691916 | 2.451379346 | 0.095 | 0.045 | 1 | 0.132738123 | -2.174908409 | 0 | 0.048 | 1 | 0.264691916 | 0.247856836 |
| Gm17382 | 0.921796548 | 1.908237021 | 0.048 | 0.045 | 1 | 0.132739158 | -1.709244837 | 0 | 0.048 | 1 | 0.921796548 | 0.247858632 |
| Ptk2b | 0.719072865 | 1.766881172 | 0.048 | 0.034 | 1 | 0.132743816 | -2.271123724 | 0 | 0.048 | 1 | 0.719072865 | 0.247866712 |
| Cdh20 | 0.623370066 | -1.849790188 | 0.048 | 0.075 | 1 | 0.132745369 | -3.294207337 | 0 | 0.048 | 1 | 0.623370066 | 0.247869406 |
| Mapk10 | 0.359280535 | 3.188344941 | 0.048 | 0.019 | 1 | 0.132746405 | -2.124282336 | 0 | 0.048 | 1 | 0.359280535 | 0.247871201 |
| Rnf180 | 0.85372741 | 2.281454345 | 0.048 | 0.041 | 1 | 0.132748993 | -3.044847868 | 0 | 0.048 | 1 | 0.85372741 | 0.24787569 |
| A230056P14Rik | 0.285499161 | -0.648156327 | 0 | 0.052 | 1 | 0.132760897 | -1.223818009 | 0 | 0.048 | 1 | 0.285499161 | 0.247896339 |
| Gm13199 | 0.359270744 | 2.673771768 | 0.048 | 0.019 | 1 | 0.132760897 | -1.223818009 | 0 | 0.048 | 1 | 0.359270744 | 0.247896339 |
| C330011M18Rik | 0.396971401 | 0.351843673 | 0 | 0.034 | 1 | 0.132760897 | -1.223818009 | 0 | 0.048 | 1 | 0.396971401 | 0.247896339 |
| Srsf12 | 0.425599726 | 0.351843673 | 0 | 0.03 | 1 | 0.132760897 | -1.223818009 | 0 | 0.048 | 1 | 0.425599726 | 0.247896339 |
| 1700030C12Rik | 0.425599726 | 0.351843673 | 0 | 0.03 | 1 | 0.132760897 | -1.223818009 | 0 | 0.048 | 1 | 0.425599726 | 0.247896339 |
| Neurl2 | 0.472861597 | 1.866416846 | 0.048 | 0.022 | 1 | 0.132760897 | -1.223818009 | 0 | 0.048 | 1 | 0.472861597 | 0.247896339 |
| Dmrta1 | 0.532436372 | 1.088809267 | 0 | 0.019 | 1 | 0.132760897 | -1.223818009 | 0 | 0.048 | 1 | 0.532436372 | 0.247896339 |
| Jrk | 0.532436372 | 1.088809267 | 0 | 0.019 | 1 | 0.132760897 | -1.223818009 | 0 | 0.048 | 1 | 0.532436372 | 0.247896339 |
| Gm43774 | 0.636138263 | 1.79930265 | 0.048 | 0.03 | 1 | 0.132760897 | -1.223818009 | 0 | 0.048 | 1 | 0.636138263 | 0.247896339 |
| 1700109K24Rik | 0.71266839 | 1.79930265 | 0.048 | 0.034 | 1 | 0.132760897 | -1.223818009 | 0 | 0.048 | 1 | 0.71266839 | 0.247896339 |
| Polm | 0.81881728 | 0.97333205 | 0.048 | 0.037 | 1 | 0.132760897 | -1.223818009 | 0 | 0.048 | 1 | 0.81881728 | 0.247896339 |
| Gm56912 | 0.856787907 | 1.55829455 | 0.048 | 0.041 | 1 | 0.132760897 | -1.223818009 | 0 | 0.048 | 1 | 0.856787907 | 0.247896339 |
| Gm32342 | 0.425614684 | 0.214340149 | 0 | 0.03 | 1 | 0.132852687 | -1.486852415 | 0.022 | 0.087 | 1 | 0.425614684 | 0.248055538 |
| Cd164l2 | 0.132862182 | 1.673771768 | 0.143 | 0.06 | 1 | 0.426859946 | 1.235613609 | 0.111 | 0.08 | 1 | 0.426859946 | 0.248072005 |
| Styk1 | 0.173814074 | 2.088809267 | 0.048 | 0.011 | 1 | 0.132894982 | 1.361144491 | 0.022 | 0.003 | 1 | 0.173814074 | 0.248128887 |
| Gm44767 | 0.578156445 | 0.866416846 | 0 | 0.015 | 1 | 0.132894982 | 0.460680165 | 0.022 | 0.003 | 1 | 0.578156445 | 0.248128887 |
| Pcsk2 | 0.632076134 | 0.866416846 | 0 | 0.011 | 1 | 0.132894982 | 0.875717664 | 0.022 | 0.003 | 1 | 0.632076134 | 0.248128887 |
| Npas1 | 0.632076134 | 1.673771768 | 0 | 0.011 | 1 | 0.132894982 | 2.098110085 | 0.022 | 0.003 | 1 | 0.632076134 | 0.248128887 |
| 1700029J03Rik | 0.864156366 | 0.281454345 | 0.048 | 0.056 | 1 | 0.132894982 | 2.098110085 | 0.022 | 0.003 | 1 | 0.864156366 | 0.248128887 |
| Rcor3 | 0.320139494 | -0.246793765 | 0.286 | 0.451 | 1 | 0.132938783 | 0.266507617 | 0.333 | 0.578 | 1 | 0.320139494 | 0.248204846 |
| Gm40438 | 0.132994635 | 1.464318402 | 0.143 | 0.06 | 1 | 0.65758319 | 1.460680165 | 0.067 | 0.052 | 1 | 0.65758319 | 0.248301698 |
| Spa17 | 0.204539388 | -1.107587946 | 0.048 | 0.146 | 1 | 0.13308291 | -0.271123724 | 0.022 | 0.09 | 1 | 0.204539388 | 0.24845476 |
| Myo18a | 0.942093974 | 0.492706209 | 0.476 | 0.578 | 1 | 0.133102482 | 0.721418898 | 0.911 | 0.879 | 1 | 0.942093974 | 0.248488693 |
| Rcor1 | 0.986527375 | -0.156947147 | 0.571 | 0.638 | 1 | 0.133175507 | 0.473619221 | 0.956 | 0.83 | 1 | 0.986527375 | 0.248615299 |
| Zfp628 | 0.304355016 | -0.496153234 | 0 | 0.049 | 1 | 0.133284611 | -1.361321533 | 0.022 | 0.087 | 1 | 0.304355016 | 0.248804434 |
| Tmem191 | 0.370840523 | 0.214340149 | 0 | 0.037 | 1 | 0.133410996 | -0.486852415 | 0.022 | 0.09 | 1 | 0.370840523 | 0.249023499 |
| Xylt2 | 0.818482987 | 0.281454345 | 0.048 | 0.06 | 1 | 0.133456565 | -0.124282336 | 0.022 | 0.09 | 1 | 0.818482987 | 0.249102476 |
| Pierce2 | 0.207486144 | -0.911190733 | 0.048 | 0.146 | 1 | 0.133498485 | -0.174908409 | 0.022 | 0.09 | 1 | 0.207486144 | 0.249175125 |
| Ankrd11 | 0.133621465 | 0.580144247 | 1 | 0.989 | 1 | 0.29505372 | 0.224253014 | 1 | 1 | 1 | 0.29505372 | 0.249388233 |
| Tbc1d19 | 0.133766864 | -0.984439715 | 0.095 | 0.243 | 1 | 0.214202744 | 0.598539076 | 0.178 | 0.291 | 1 | 0.214202744 | 0.249640155 |
| Napb | 0.133804359 | -1.434752689 | 0.048 | 0.172 | 1 | 0.165524045 | -0.346674757 | 0.044 | 0.118 | 1 | 0.165524045 | 0.249705112 |
| ENSMUSG00000121477 | 0.133804359 | -1.392317423 | 0.048 | 0.172 | 1 | 0.227469845 | 0.268035087 | 0.111 | 0.197 | 1 | 0.227469845 | 0.249705112 |
| Rsf1 | 0.397370533 | -0.247125686 | 0.762 | 0.694 | 1 | 0.133811893 | 0.271592906 | 0.978 | 0.882 | 1 | 0.397370533 | 0.249718164 |
| Paqr3 | 0.252195327 | -0.611630451 | 0.048 | 0.134 | 1 | 0.134141294 | -1.223818009 | 0.022 | 0.087 | 1 | 0.252195327 | 0.250288701 |
| Prkaa2 | 0.134191399 | 1.316219763 | 0.381 | 0.261 | 1 | 0.559598162 | 0.305767734 | 0.489 | 0.588 | 1 | 0.559598162 | 0.250375466 |
| Tfcp2 | 0.153362905 | -0.672003069 | 0.143 | 0.302 | 1 | 0.134348917 | 0.298408736 | 0.178 | 0.318 | 1 | 0.153362905 | 0.250648202 |
| Rragd | 0.3247899 | -0.496153234 | 0 | 0.045 | 1 | 0.134566316 | -1.174908409 | 0.022 | 0.087 | 1 | 0.3247899 | 0.251024538 |
| Donson | 0.54437166 | 1.60338244 | 0.095 | 0.063 | 1 | 0.134566316 | -1.174908409 | 0.022 | 0.087 | 1 | 0.54437166 | 0.251024538 |
| Acaca | 0.134629236 | 0.889875818 | 0.619 | 0.657 | 1 | 0.248051829 | 1.016947233 | 0.556 | 0.626 | 1 | 0.248051829 | 0.251133441 |
| Garem1 | 0.239006996 | 2.756233928 | 0.524 | 0.537 | 1 | 0.134948593 | 0.371713733 | 0.689 | 0.561 | 1 | 0.239006996 | 0.251686063 |
| Tle1 | 0.134986251 | 0.689136077 | 0.333 | 0.616 | 1 | 0.385025585 | 0.610827702 | 0.756 | 0.727 | 1 | 0.385025585 | 0.251751214 |
| Rtca | 0.135098124 | -0.555046923 | 0.095 | 0.25 | 1 | 0.381951797 | 0.172110667 | 0.2 | 0.291 | 1 | 0.381951797 | 0.251944745 |
| Zdhhc13 | 0.215927545 | -0.355975576 | 0.048 | 0.153 | 1 | 0.135204586 | -0.84048937 | 0.044 | 0.121 | 1 | 0.215927545 | 0.252128892 |
| Rmnd5b | 0.135228095 | -1.133583154 | 0.048 | 0.172 | 1 | 0.283949573 | 0.189533113 | 0.156 | 0.249 | 1 | 0.283949573 | 0.252169553 |
| Atr | 0.821721051 | 0.866416846 | 0.143 | 0.183 | 1 | 0.13526075 | -0.245034495 | 0.133 | 0.253 | 1 | 0.821721051 | 0.25222603 |
| Birc3 | 0.913126977 | 0.933014594 | 0.238 | 0.25 | 1 | 0.135552057 | 0.210584815 | 0.133 | 0.26 | 1 | 0.913126977 | 0.252729753 |
| Grk4 | 0.424147373 | -0.233118828 | 0.143 | 0.22 | 1 | 0.135601808 | 0.283665739 | 0.133 | 0.26 | 1 | 0.424147373 | 0.252815766 |
| Phf8 | 0.135607809 | 1.527693508 | 0.524 | 0.53 | 1 | 0.882533437 | 0.10428866 | 0.667 | 0.654 | 1 | 0.882533437 | 0.252826141 |
| Larp1 | 0.969077482 | 0.166948094 | 0.667 | 0.776 | 1 | 0.13567491 | 0.467294543 | 0.978 | 0.931 | 1 | 0.969077482 | 0.252942138 |
| Asns | 0.135770456 | -0.192476843 | 0.19 | 0.396 | 1 | 0.13984596 | 0.407438144 | 0.111 | 0.218 | 1 | 0.13984596 | 0.253107296 |
| Sh3d21 | 0.172526215 | -1.249060372 | 0.048 | 0.164 | 1 | 0.135830376 | -0.47073275 | 0.111 | 0.211 | 1 | 0.172526215 | 0.253210861 |
| Adgra3 | 0.135967242 | 1.143775726 | 0.762 | 0.746 | 1 | 0.595928559 | 0.305060944 | 0.622 | 0.875 | 1 | 0.595928559 | 0.253447393 |
| 4833411C07Rik | 0.13616383 | -1.370622352 | 0 | 0.097 | 1 | 0.165306929 | -1.124282336 | 0 | 0.042 | 1 | 0.165306929 | 0.253787072 |
| Efcab2 | 0.889048931 | 0.472137907 | 0.143 | 0.134 | 1 | 0.136172824 | -0.749886821 | 0.044 | 0.125 | 1 | 0.889048931 | 0.253802609 |
| Parp11 | 0.136213138 | -1.326228232 | 0 | 0.097 | 1 | 0.14169169 | 0.309614191 | 0.067 | 0.156 | 1 | 0.14169169 | 0.253872256 |
| Gm28154 | 0.136213732 | -1.455511249 | 0 | 0.097 | 1 | 0.734338054 | 0.393565969 | 0.089 | 0.111 | 1 | 0.734338054 | 0.253873283 |
| Gm4890 | 0.13621393 | -1.280424543 | 0 | 0.097 | 1 | 0.532520164 | -0.316927414 | 0.022 | 0.042 | 1 | 0.532520164 | 0.253873625 |

| | | | | | | | | | | | | |
|---|---|---|---|---|---|---|---|---|---|---|---|---|
| Pxmp2 | 0.136259668 | -1.326228232 | 0 | 0.097 | 1 | 0.260665169 | -1.017367132 | 0 | 0.028 | 1 | 0.260665169 | 0.253952639 |
| Ccdc39 | 0.136297285 | -1.413691074 | 0 | 0.097 | 1 | 0.987030734 | 0.846571318 | 0.089 | 0.097 | 1 | 0.987030734 | 0.254017621 |
| 2810029C07Rik | 0.136297285 | -1.413691074 | 0 | 0.097 | 1 | 0.765728546 | 1.223640968 | 0.022 | 0.031 | 1 | 0.765728546 | 0.254017621 |
| Zfp566 | 0.136299265 | -1.455511249 | 0 | 0.097 | 1 | 0.390770626 | -0.16492432 | 0.067 | 0.111 | 1 | 0.390770626 | 0.254021041 |
| Cited1 | 0.136300849 | -1.683780237 | 0 | 0.097 | 1 | 0.18483667 | -2.743192169 | 0 | 0.038 | 1 | 0.18483667 | 0.254023777 |
| Kif3c | 0.877539888 | 0.316219763 | 0.095 | 0.108 | 1 | 0.136333464 | 0.122357632 | 0.067 | 0.163 | 1 | 0.877539888 | 0.254080115 |
| Fut10 | 0.1363349 | -1.496153234 | 0 | 0.097 | 1 | 0.345836529 | 0.460680165 | 0.089 | 0.149 | 1 | 0.345836529 | 0.254082595 |
| Tsga10 | 0.136421007 | -1.648156327 | 0 | 0.097 | 1 | 0.876421435 | 0.460680165 | 0.089 | 0.1 | 1 | 0.876421435 | 0.254231322 |
| Nusap1 | 0.136427736 | -2.370622352 | 0 | 0.097 | 1 | 0.967465715 | 1.235613609 | 0.067 | 0.073 | 1 | 0.967465715 | 0.254242945 |
| Akap6 | 0.136445747 | -1.849790188 | 0 | 0.097 | 1 | 0.328390395 | -0.316927414 | 0.044 | 0.09 | 1 | 0.328390395 | 0.254274053 |
| Rsrc2 | 0.5535034 | 0.183734524 | 1 | 0.955 | 1 | 0.136482644 | -0.140925001 | 0.933 | 0.962 | 1 | 0.5535034 | 0.254337776 |
| Adamtsl1 | 0.136490079 | -3.76085646 | 0 | 0.097 | 1 | 0.572534372 | -1.425451871 | 0.067 | 0.09 | 1 | 0.572534372 | 0.254350617 |
| Camk2b | 0.136564485 | -1.081115734 | 0 | 0.097 | 1 | 0.61665314 | 0.765534746 | 0.067 | 0.093 | 1 | 0.61665314 | 0.254479112 |
| Pecam1 | 0.883248729 | 1.60338244 | 0.095 | 0.112 | 1 | 0.136706982 | 2.234482527 | 0.178 | 0.1 | 1 | 0.883248729 | 0.254725165 |
| Brms1 | 0.395367815 | 0.477374555 | 0.19 | 0.313 | 1 | 0.136997112 | 0.290755163 | 0.156 | 0.291 | 1 | 0.395367815 | 0.255226014 |
| Scgb1b27 | 0.137098296 | 1.574236094 | 0.286 | 0.19 | 1 | 0.661337779 | 0.481438725 | 0.044 | 0.062 | 1 | 0.661337779 | 0.25540065 |
| Fbf1 | 0.277533773 | 1.162809848 | 0.238 | 0.16 | 1 | 0.137210615 | 0.503748887 | 0.089 | 0.19 | 1 | 0.277533773 | 0.255594476 |
| Plekhf2 | 0.137495933 | -0.392317423 | 0.143 | 0.336 | 1 | 0.758770963 | -0.226563386 | 0.6 | 0.516 | 1 | 0.758770963 | 0.256086735 |
| Satb2 | 0.455311072 | -1.295854583 | 0.048 | 0.097 | 1 | 0.137671406 | -1.709244837 | 0.022 | 0.087 | 1 | 0.455311072 | 0.256389396 |
| Ampd3 | 0.307426853 | -1.124970024 | 0.143 | 0.235 | 1 | 0.137692858 | -0.486852415 | 0.044 | 0.121 | 1 | 0.307426853 | 0.256426393 |
| Ppp3cc | 0.137709892 | 1.33273485 | 0.476 | 0.437 | 1 | 0.965140572 | 1.704767657 | 0.222 | 0.256 | 1 | 0.965140572 | 0.256455769 |
| Gm10125 | 0.324140258 | -1.233118828 | 0.048 | 0.116 | 1 | 0.13771339 | 1.765534746 | 0.178 | 0.111 | 1 | 0.324140258 | 0.256461802 |
| Poln | 0.256356158 | -0.20069735 | 0.048 | 0.142 | 1 | 0.137767878 | 0.653325243 | 0.067 | 0.156 | 1 | 0.256356158 | 0.256555767 |
| Nacc1 | 0.137976795 | 1.281454345 | 0.381 | 0.295 | 1 | 0.774134163 | 0.436911999 | 0.333 | 0.453 | 1 | 0.774134163 | 0.256915994 |
| Them4 | 0.812721728 | 0.258734268 | 0.143 | 0.172 | 1 | 0.13798617 | -0.609709163 | 0.044 | 0.121 | 1 | 0.812721728 | 0.256932156 |
| Tcea3 | 0.550788759 | 0.791128718 | 0.19 | 0.287 | 1 | 0.138016955 | 0.272788617 | 0.133 | 0.26 | 1 | 0.550788759 | 0.25698523 |
| Slco3a1 | 0.761037572 | 1.400753273 | 0.333 | 0.362 | 1 | 0.138093772 | 0.643544222 | 0.289 | 0.197 | 1 | 0.761037572 | 0.257117654 |
| Gm47283 | 0.138102447 | 1.039012567 | 0.952 | 0.918 | 1 | 0.217756608 | 0.437960088 | 0.978 | 0.913 | 1 | 0.217756608 | 0.257132608 |
| Mir22hg | 0.229719858 | 0.951305743 | 0.524 | 0.504 | 1 | 0.138374491 | 0.658126229 | 0.667 | 0.602 | 1 | 0.229719858 | 0.257601483 |
| Mta3 | 0.138570703 | -0.496153234 | 0.143 | 0.317 | 1 | 0.162167745 | -0.328900055 | 0.378 | 0.491 | 1 | 0.162167745 | 0.257939566 |
| Garin4 | 0.138662267 | 2.143257051 | 0.19 | 0.097 | 1 | 0.90731367 | 1.921232323 | 0.067 | 0.076 | 1 | 0.90731367 | 0.258097311 |
| Gm56844 | 0.138709139 | 2.536268244 | 0.095 | 0.034 | 1 | 0.497228696 | 0.268035087 | 0.022 | 0.045 | 1 | 0.497228696 | 0.258178053 |
| Cep112it | 0.138709139 | 2.536268244 | 0.095 | 0.034 | 1 | 0.376842245 | -0.486852415 | 0 | 0.017 | 1 | 0.376842245 | 0.258178053 |
| Mroh5 | 0.138709139 | 2.896164189 | 0.095 | 0.034 | 1 | 0.765720982 | 0.946106992 | 0.022 | 0.031 | 1 | 0.765720982 | 0.258178053 |
| Fads2 | 0.191357273 | 0.539916021 | 0.19 | 0.399 | 1 | 0.13871837 | 0.259861155 | 0.133 | 0.27 | 1 | 0.191357273 | 0.258193954 |
| 4921504A21Rik | 0.138867088 | 2.088809267 | 0.143 | 0.063 | 1 | 0.804442995 | 1.64125241 | 0.067 | 0.083 | 1 | 0.804442995 | 0.258450108 |
| Fnip1 | 0.300491995 | 0.445224031 | 0.905 | 0.929 | 1 | 0.138903908 | 0.41651902 | 1 | 0.979 | 1 | 0.300491995 | 0.25851352 |
| Ank1 | 0.138908705 | 1.267779408 | 0.19 | 0.093 | 1 | 0.575602616 | 0.583536913 | 0.044 | 0.069 | 1 | 0.575602616 | 0.258521781 |
| Scamp1 | 0.138925828 | -0.6082636 | 0.238 | 0.455 | 1 | 0.67220577 | 0.149094014 | 0.511 | 0.536 | 1 | 0.67220577 | 0.258551271 |
| Krcc1 | 0.139083811 | -0.436652222 | 0.476 | 0.679 | 1 | 0.347823641 | -0.127957994 | 0.644 | 0.644 | 1 | 0.347823641 | 0.258823315 |
| C2cd3 | 0.139279067 | -0.75918764 | 0.095 | 0.243 | 1 | 0.565950649 | 0.404213213 | 0.311 | 0.28 | 1 | 0.565950649 | 0.259159475 |
| Plcb1 | 0.139350697 | 1.005466943 | 0.762 | 0.84 | 1 | 0.139611179 | 0.45864102 | 0.867 | 0.945 | 1 | 0.139611179 | 0.259282776 |
| Usp8 | 0.139413535 | -0.188724709 | 0.429 | 0.672 | 1 | 0.195634714 | -0.254514807 | 0.778 | 0.74 | 1 | 0.195634714 | 0.259390936 |
| Gtf2f2 | 0.139434563 | 0.121546053 | 0.524 | 0.813 | 1 | 0.209739274 | 0.392746489 | 0.844 | 0.709 | 1 | 0.209739274 | 0.259427128 |
| Pigq | 0.139509445 | -0.496153234 | 0.048 | 0.183 | 1 | 0.184281752 | 0.242499995 | 0.111 | 0.208 | 1 | 0.184281752 | 0.259556004 |
| Natd1 | 0.910742352 | 0.786246497 | 0.095 | 0.112 | 1 | 0.139556739 | -1.199570463 | 0.067 | 0.145 | 1 | 0.910742352 | 0.259637394 |
| Pip5k1b | 0.303013622 | 0.214340149 | 0.048 | 0.131 | 1 | 0.139625464 | -1.271123724 | 0.111 | 0.197 | 1 | 0.303013622 | 0.259755658 |
| Gm56829 | 0.52656069 | 0.908237021 | 0.143 | 0.101 | 1 | 0.139772973 | -0.638855509 | 0.044 | 0.121 | 1 | 0.52656069 | 0.260009463 |
| 5430405H02Rik | 0.803353239 | 2.029915578 | 0.095 | 0.123 | 1 | 0.139775463 | -0.659319611 | 0.067 | 0.152 | 1 | 0.803353239 | 0.260013746 |
| Pbx2 | 0.352380585 | -0.348596045 | 0.095 | 0.179 | 1 | 0.139815308 | -0.107004344 | 0.111 | 0.218 | 1 | 0.352380585 | 0.260082295 |
| Zfyve27 | 0.416069023 | -0.233118828 | 0.143 | 0.224 | 1 | 0.140284925 | -0.365290435 | 0.133 | 0.242 | 1 | 0.416069023 | 0.26088999 |
| Cyb5b | 0.140317778 | -0.138996651 | 0.286 | 0.571 | 1 | 0.518713686 | 0.310029176 | 0.622 | 0.571 | 1 | 0.518713686 | 0.260946478 |
| Mmp28 | 0.140434402 | 2.483440556 | 0.286 | 0.183 | 1 | 0.197393091 | 1.361144491 | 0.156 | 0.1 | 1 | 0.197393091 | 0.261146983 |
| Tyw5 | 0.190061613 | -0.574155746 | 0.048 | 0.16 | 1 | 0.140444905 | -0.446210431 | 0.089 | 0.187 | 1 | 0.190061613 | 0.261165038 |
| Vps41 | 0.815824284 | 0.473149969 | 0.476 | 0.578 | 1 | 0.140485174 | 0.678785228 | 0.622 | 0.602 | 1 | 0.815824284 | 0.261234263 |
| Sp7 | 0.140497338 | -0.574155746 | 0.048 | 0.183 | 1 | 0.146441616 | -0.638855509 | 0.044 | 0.121 | 1 | 0.146441616 | 0.261255175 |
| Krt6a | 0.544576436 | 2.258734268 | 0.048 | 0.026 | 1 | 0.140501527 | -1.776359032 | 0.022 | 0.087 | 1 | 0.544576436 | 0.261262375 |
| Macrod1 | 0.140641757 | -0.326228232 | 0.095 | 0.261 | 1 | 0.141727073 | 0.157003775 | 0.156 | 0.291 | 1 | 0.141727073 | 0.26150341 |
| Gm48321 | 0.629700127 | 2.088809267 | 0.048 | 0.03 | 1 | 0.140643088 | 2.142504205 | 0.089 | 0.042 | 1 | 0.629700127 | 0.261505698 |
| Tab1 | 0.208921714 | -0.970084422 | 0 | 0.071 | 1 | 0.1407561 | 2.435145073 | 0.111 | 0.059 | 1 | 0.208921714 | 0.26169992 |
| Vwa8 | 0.243892182 | -0.601506234 | 0.238 | 0.407 | 1 | 0.140858072 | 1.098110085 | 0.511 | 0.45 | 1 | 0.243892182 | 0.261875148 |
| H2-DMa | 0.308765546 | 0.503846766 | 0.095 | 0.205 | 1 | 0.14089662 | -0.191396532 | 0.044 | 0.125 | 1 | 0.308765546 | 0.261941383 |
| Itpk1 | 0.141011073 | -0.370622352 | 0.048 | 0.183 | 1 | 0.889691957 | 1.168499413 | 0.156 | 0.159 | 1 | 0.889691957 | 0.262138024 |
| Syn3 | 0.159207799 | -1.02812743 | 0.381 | 0.556 | 1 | 0.141011383 | 0.342648148 | 0.133 | 0.266 | 1 | 0.159207799 | 0.262138555 |
| Arid3b | 0.324822624 | -0.718545655 | 0 | 0.045 | 1 | 0.14112379 | 1.476621709 | 0.111 | 0.055 | 1 | 0.324822624 | 0.262331656 |
| Id3 | 0.141262391 | -0.398905737 | 0.571 | 0.604 | 1 | 0.829454518 | -0.745662235 | 0.556 | 0.453 | 1 | 0.829454518 | 0.262569718 |
| Nt5dc1 | 0.65694726 | -0.93177161 | 0.524 | 0.5 | 1 | 0.141278044 | -0.295123044 | 0.244 | 0.346 | 1 | 0.65694726 | 0.262596603 |
| Fhad1 | 0.141456117 | 4.281454345 | 0.143 | 0.063 | 1 | 0.231991339 | -1.017367132 | 0 | 0.031 | 1 | 0.231991339 | 0.2629024 |
| Arhgef11 | 0.310325254 | 0.614878079 | 0.524 | 0.496 | 1 | 0.141669231 | 0.467272842 | 0.844 | 0.727 | 1 | 0.310325254 | 0.26326829 |
| Abl1 | 0.141671771 | 1.44596585 | 0.429 | 0.366 | 1 | 0.720888719 | 0.102654011 | 0.6 | 0.54 | 1 | 0.720888719 | 0.263272651 |
| Nrbp2 | 0.965708259 | 0.376091219 | 0.429 | 0.496 | 1 | 0.14173718 | -0.153428682 | 0.2 | 0.346 | 1 | 0.965708259 | 0.263384932 |
| Tinf2 | 0.317975113 | -0.496153234 | 0.048 | 0.119 | 1 | 0.141769654 | -0.242926832 | 0.044 | 0.121 | 1 | 0.317975113 | 0.263440673 |
| Map3k2 | 0.38307373 | 0.189481965 | 0.81 | 0.91 | 1 | 0.141796723 | -0.106071357 | 0.933 | 0.938 | 1 | 0.38307373 | 0.263487135 |
| Pptc7 | 0.927487841 | 0.586308926 | 0.286 | 0.317 | 1 | 0.141810527 | 0.566463399 | 0.711 | 0.644 | 1 | 0.927487841 | 0.263510828 |
| Notum | 0.94876154 | -0.503766418 | 0.333 | 0.313 | 1 | 0.142018517 | -0.805401832 | 0.556 | 0.689 | 1 | 0.94876154 | 0.263867775 |
| Nphp4 | 0.142030847 | 2.451379346 | 0.095 | 0.034 | 1 | 0.293637566 | -0.486852415 | 0 | 0.024 | 1 | 0.293637566 | 0.263888932 |
| Arhgef9 | 0.676835518 | -0.355975576 | 0.095 | 0.123 | 1 | 0.142049687 | -0.124282336 | 0.067 | 0.159 | 1 | 0.676835518 | 0.26392126 |
| Acaa2 | 0.142119435 | -0.785659851 | 0.048 | 0.179 | 1 | 0.804194846 | -0.564854927 | 0.244 | 0.249 | 1 | 0.804194846 | 0.264040936 |
| Sumo1 | 0.142266367 | -0.200235466 | 0.857 | 0.944 | 1 | 0.183652836 | -0.149660212 | 0.978 | 0.948 | 1 | 0.183652836 | 0.264293014 |
| Plscr2 | 0.142341108 | 1.271673324 | 0.143 | 0.063 | 1 | 0.658583583 | 1.946106992 | 0.022 | 0.014 | 1 | 0.658583583 | 0.264421226 |
| Gm44649 | 0.142689378 | 1.775651382 | 0.238 | 0.138 | 1 | 0.506779734 | 1.122357632 | 0.2 | 0.18 | 1 | 0.506779734 | 0.265018497 |
| Metap1d | 0.557776852 | 0.582273413 | 0.619 | 0.683 | 1 | 0.142722573 | 0.237762076 | 0.356 | 0.592 | 1 | 0.557776852 | 0.265075414 |
| 0610040J01Rik | 0.519873629 | 0.85614851 | 0.143 | 0.228 | 1 | 0.14288053 | -0.281974815 | 0.244 | 0.405 | 1 | 0.519873629 | 0.265346215 |
| Gm42439 | 0.7536435 | 0.735172312 | 0.19 | 0.187 | 1 | 0.142949216 | 1.730378301 | 0.222 | 0.163 | 1 | 0.7536435 | 0.265463954 |
| Kifap3 | 0.142960555 | -0.445663463 | 0.333 | 0.59 | 1 | 0.291277278 | 0.3895729 | 0.444 | 0.633 | 1 | 0.291277278 | 0.265483391 |
| Zfp202 | 0.143137003 | 2.451379346 | 0.095 | 0.034 | 1 | 0.165324946 | -1.223818009 | 0 | 0.042 | 1 | 0.165324946 | 0.265785804 |
| Sppl3 | 0.143425517 | 0.376091219 | 0.857 | 0.772 | 1 | 0.375542263 | 0.371870898 | 0.978 | 0.92 | 1 | 0.375542263 | 0.266280155 |
| Agpat4 | 0.143639374 | 0.838265805 | 0.667 | 0.84 | 1 | 0.213293851 | 0.921091676 | 0.644 | 0.813 | 1 | 0.213293851 | 0.266646478 |
| Kdm4b | 0.420461201 | 0.79930265 | 0.238 | 0.179 | 1 | 0.143664346 | -0.251832386 | 0.156 | 0.27 | 1 | 0.420461201 | 0.266689248 |
| Slc44a4 | 0.532434375 | 0.214340149 | 0 | 0.019 | 1 | 0.144068362 | 2.946106992 | 0.044 | 0.014 | 1 | 0.532434375 | 0.267381032 |
| Wdr62 | 0.709465802 | 1.936806174 | 0.048 | 0.034 | 1 | 0.144068362 | 3.361144491 | 0.044 | 0.014 | 1 | 0.709465802 | 0.267381032 |
| Coq3 | 0.144201507 | -1.159118246 | 0.048 | 0.168 | 1 | 0.491958747 | 0.532130688 | 0.2 | 0.284 | 1 | 0.491958747 | 0.267608939 |
| Gm37768 | 0.144257061 | 2.258734268 | 0.095 | 0.034 | 1 | 0.700757366 | 0.995016592 | 0.067 | 0.09 | 1 | 0.700757366 | 0.267704022 |
| Heatr6 | 0.910652234 | 1.161321767 | 0.238 | 0.269 | 1 | 0.144628609 | -0.290932205 | 0.156 | 0.28 | 1 | 0.910652234 | 0.268339784 |
| D17H6S53E | 0.144664875 | -1.133583154 | 0 | 0.093 | 1 | 0.483989161 | -0.316927414 | 0.044 | 0.073 | 1 | 0.483989161 | 0.268401823 |
| Spring1 | 0.144707548 | -0.623908781 | 0.048 | 0.179 | 1 | 0.197971501 | -0.286553765 | 0.067 | 0.142 | 1 | 0.197971501 | 0.268474821 |
| Lrrc32 | 0.144717908 | -1.233118828 | 0 | 0.093 | 1 | 0.971332021 | 0.513147585 | 0.067 | 0.069 | 1 | 0.971332021 | 0.268492543 |
| 2210408I21Rik | 0.549556067 | 1.075693767 | 0.476 | 0.582 | 1 | 0.144748629 | 0.652748049 | 0.667 | 0.609 | 1 | 0.549556067 | 0.268545093 |

| | | | | | | | | | | | | |
|---|---|---|---|---|---|---|---|---|---|---|---|---|
| Calr3 | 0.578473027 | 0.400753273 | 0.048 | 0.086 | 1 | 0.144764714 | -0.731964913 | 0.022 | 0.087 | 1 | 0.578473027 | 0.268572606 |
| Foxred2 | 0.144765886 | -1.280424543 | 0 | 0.093 | 1 | 0.308655594 | 1.197645759 | 0.067 | 0.125 | 1 | 0.308655594 | 0.26857461 |
| Relt | 0.144810913 | -1.280424543 | 0 | 0.093 | 1 | 0.148025014 | -1.223818009 | 0 | 0.045 | 1 | 0.148025014 | 0.268651625 |
| Fdxr | 0.144849834 | -1.648156327 | 0 | 0.093 | 1 | 0.896390428 | 0.78998779 | 0.067 | 0.062 | 1 | 0.896390428 | 0.268718194 |
| Caprin2 | 0.144884545 | -1.648156327 | 0 | 0.093 | 1 | 0.916482138 | 0.966865552 | 0.111 | 0.125 | 1 | 0.916482138 | 0.268777559 |
| Zfp780b | 0.144886859 | -1.370622352 | 0 | 0.093 | 1 | 0.416233084 | 0.337297749 | 0.111 | 0.173 | 1 | 0.416233084 | 0.268781517 |
| Me3 | 0.144940078 | -1.683780237 | 0 | 0.093 | 1 | 0.596051504 | 0.721140627 | 0.178 | 0.228 | 1 | 0.596051504 | 0.26887253 |
| Usp18 | 0.144943443 | -1.818081329 | 0 | 0.093 | 1 | 0.692503623 | 0.250113179 | 0.067 | 0.087 | 1 | 0.692503623 | 0.268878285 |
| Gm26771 | 0.144947229 | -2.611630451 | 0 | 0.093 | 1 | 0.371676476 | 0.723714571 | 0.111 | 0.073 | 1 | 0.371676476 | 0.268884759 |
| Gjb6 | 0.145012008 | -1.911190733 | 0 | 0.093 | 1 | 0.496223879 | -0.486852415 | 0 | 0.01 | 1 | 0.496223879 | 0.268995533 |
| Plppr3 | 0.454205211 | 2.451379346 | 0.048 | 0.022 | 1 | 0.145064168 | 2.683072586 | 0.044 | 0.014 | 1 | 0.454205211 | 0.269084723 |
| Gm45518 | 0.492599419 | 0.351843673 | 0 | 0.022 | 1 | 0.145064168 | 3.098110085 | 0.044 | 0.014 | 1 | 0.492599419 | 0.269084723 |
| Bdh2 | 0.852271211 | 0.79930265 | 0.048 | 0.06 | 1 | 0.145079347 | -1.674479418 | 0.022 | 0.083 | 1 | 0.852271211 | 0.269110677 |
| Rras2 | 0.467411946 | 0.958152909 | 0.619 | 0.799 | 1 | 0.145221486 | 1.187916044 | 0.911 | 0.855 | 1 | 0.467411946 | 0.269353692 |
| Kirrel3 | 0.38220648 | -0.828728573 | 0.048 | 0.112 | 1 | 0.145252972 | -1.031172931 | 0.044 | 0.121 | 1 | 0.38220648 | 0.269407518 |
| Rnf144b | 0.257157307 | 1.536268244 | 0.238 | 0.168 | 1 | 0.145309586 | 0.327591932 | 0.111 | 0.221 | 1 | 0.257157307 | 0.269504296 |
| Gm15283 | 0.396416287 | 1.242219182 | 0.333 | 0.317 | 1 | 0.145371061 | -0.271123724 | 0.222 | 0.36 | 1 | 0.396416287 | 0.269609377 |
| Tgfb1 | 0.145471325 | 0.239875241 | 0.429 | 0.791 | 1 | 0.977263956 | 0.320955401 | 0.8 | 0.855 | 1 | 0.977263956 | 0.269780744 |
| Slc13a4 | 0.396968161 | -1.02666795 | 0 | 0.034 | 1 | 0.145545882 | 2.168499413 | 0.089 | 0.042 | 1 | 0.396968161 | 0.269908161 |
| Rab7 | 0.145569212 | -0.433756697 | 0.857 | 0.896 | 1 | 0.287798668 | 0.301718213 | 0.978 | 0.958 | 1 | 0.287798668 | 0.269948028 |
| Tpp2 | 0.14557702 | 0.52481654 | 0.905 | 0.836 | 1 | 0.394556402 | 0.251558041 | 0.978 | 0.893 | 1 | 0.394556402 | 0.269961372 |
| Dip2b | 0.752279229 | 0.289107917 | 0.524 | 0.504 | 1 | 0.14566332 | 0.450124983 | 0.844 | 0.671 | 1 | 0.752279229 | 0.270108838 |
| Ralb | 0.149098577 | -0.190414377 | 0.286 | 0.541 | 1 | 0.14566983 | -0.297818591 | 0.444 | 0.567 | 1 | 0.149098577 | 0.27011996 |
| Neb | 0.145675762 | -1.741265732 | 0.048 | 0.172 | 1 | 0.659788885 | -0.404390255 | 0.022 | 0.035 | 1 | 0.659788885 | 0.270130096 |
| Zfp106 | 0.206423898 | -0.328480684 | 0.81 | 0.884 | 1 | 0.145684094 | -0.157270748 | 0.911 | 0.844 | 1 | 0.206423898 | 0.270144333 |
| Ulk1 | 0.145724484 | -0.355975576 | 0.238 | 0.459 | 1 | 0.186677883 | -0.426551905 | 0.556 | 0.606 | 1 | 0.186677883 | 0.270213343 |
| Cpne1 | 0.145781287 | -0.377508737 | 0.381 | 0.623 | 1 | 0.780360901 | 0.260839585 | 0.844 | 0.841 | 1 | 0.780360901 | 0.27031039 |
| Zfp287 | 0.210775815 | 1.97333205 | 0.143 | 0.075 | 1 | 0.145807283 | -0.602329633 | 0.022 | 0.087 | 1 | 0.210775815 | 0.270354802 |
| A330040F15Rik | 0.285569273 | -1.413691074 | 0 | 0.052 | 1 | 0.146062155 | 2.513147585 | 0.044 | 0.014 | 1 | 0.285569273 | 0.270790158 |
| Irag1 | 0.373857158 | 2.088809267 | 0.048 | 0.019 | 1 | 0.146062155 | 2.460680165 | 0.044 | 0.014 | 1 | 0.373857158 | 0.270790158 |
| Ahrr | 0.632076134 | 1.088809267 | 0 | 0.011 | 1 | 0.146062155 | 3.197645759 | 0.044 | 0.014 | 1 | 0.632076134 | 0.270790158 |
| Mtrr | 0.163484033 | -1.081115734 | 0 | 0.086 | 1 | 0.146145964 | -0.589945908 | 0.022 | 0.087 | 1 | 0.163484033 | 0.270933286 |
| Zfp607a | 0.324767242 | -0.233118828 | 0 | 0.045 | 1 | 0.14618424 | -0.589945908 | 0.022 | 0.087 | 1 | 0.324767242 | 0.270998648 |
| Adck5 | 0.553287629 | 0.503846766 | 0.048 | 0.09 | 1 | 0.14618424 | -0.589945908 | 0.022 | 0.087 | 1 | 0.553287629 | 0.270998648 |
| Zfp335 | 0.857593936 | 0.503846766 | 0.095 | 0.112 | 1 | 0.146409092 | -0.394929926 | 0.111 | 0.215 | 1 | 0.857593936 | 0.271382562 |
| Dmtn | 0.146468912 | 2.351843673 | 0.095 | 0.034 | 1 | 0.833619716 | 0.223640968 | 0.022 | 0.028 | 1 | 0.833619716 | 0.271484682 |
| Arrdc4 | 0.146479897 | -0.690800665 | 0.095 | 0.25 | 1 | 0.884540137 | 0.424338318 | 0.444 | 0.426 | 1 | 0.884540137 | 0.271503434 |
| Pcdhb16 | 0.492599419 | 0.503846766 | 0 | 0.022 | 1 | 0.14653572 | -1.404390255 | 0.022 | 0.083 | 1 | 0.492599419 | 0.271598723 |
| Ighm | 0.57816106 | -2.184209227 | 0 | 0.015 | 1 | 0.146541346 | -1.564854927 | 0.022 | 0.083 | 1 | 0.57816106 | 0.271608326 |
| Cdc37l1 | 0.146545074 | -0.391519693 | 0.476 | 0.728 | 1 | 0.880627576 | 0.337650789 | 0.867 | 0.844 | 1 | 0.880627576 | 0.27161469 |
| Iws1 | 0.513962103 | 0.283418532 | 0.619 | 0.649 | 1 | 0.146744897 | -0.194515874 | 0.711 | 0.73 | 1 | 0.513962103 | 0.271955729 |
| Tcf7l1 | 0.249631455 | 1.232671972 | 0.524 | 0.53 | 1 | 0.146805122 | 0.802261941 | 0.8 | 0.789 | 1 | 0.249631455 | 0.2720585 |
| Clasrp | 0.52367678 | 0.947946731 | 0.286 | 0.257 | 1 | 0.147406012 | -0.124282336 | 0.133 | 0.253 | 1 | 0.52367678 | 0.273083492 |
| Csnk1g1 | 0.14751427 | 1.012706288 | 0.667 | 0.642 | 1 | 0.216811087 | 0.64054797 | 0.911 | 0.82 | 1 | 0.216811087 | 0.273268079 |
| Nsmaf | 0.147596663 | 1.61076197 | 0.381 | 0.313 | 1 | 0.649225805 | 0.789159845 | 0.311 | 0.433 | 1 | 0.649225805 | 0.273408552 |
| Dcdc2b | 0.147630084 | 2.673771768 | 0.095 | 0.034 | 1 | 0.96679366 | 1.852997588 | 0.044 | 0.045 | 1 | 0.96679366 | 0.273465526 |
| Gm36862 | 0.147674063 | 2.159198595 | 0.095 | 0.034 | 1 | 0.293647037 | -0.638855509 | 0 | 0.024 | 1 | 0.293647037 | 0.273540497 |
| Gm4285 | 0.293485583 | -0.44170545 | 0.048 | 0.131 | 1 | 0.147677932 | 0.147019686 | 0.022 | 0.087 | 1 | 0.293485583 | 0.273547092 |
| Dzip3 | 0.85699981 | 0.556076725 | 0.333 | 0.422 | 1 | 0.147718213 | 0.636205981 | 0.222 | 0.405 | 1 | 0.85699981 | 0.273615756 |
| Gm41609 | 0.14775242 | -1.849790188 | 0.048 | 0.164 | 1 | 0.802463629 | 0.320502507 | 0.067 | 0.08 | 1 | 0.802463629 | 0.273674063 |
| Zfp688 | 0.285499161 | -0.413691074 | 0 | 0.052 | 1 | 0.148025014 | -1.223818009 | 0 | 0.045 | 1 | 0.285499161 | 0.274138624 |
| H4c8 | 0.285566676 | -0.718545655 | 0 | 0.052 | 1 | 0.148025014 | -1.316927414 | 0 | 0.045 | 1 | 0.285566676 | 0.274138624 |
| Wasf1 | 0.3247899 | -0.574155746 | 0 | 0.045 | 1 | 0.148025014 | -1.404390255 | 0 | 0.045 | 1 | 0.3247899 | 0.274138624 |
| Inca1 | 0.339088057 | 1.825774861 | 0.095 | 0.049 | 1 | 0.148025014 | -1.404390255 | 0 | 0.045 | 1 | 0.339088057 | 0.274138624 |
| Mas1 | 0.39694807 | -0.133583154 | 0 | 0.034 | 1 | 0.148025014 | -1.223818009 | 0 | 0.045 | 1 | 0.39694807 | 0.274138624 |
| Phf11c | 0.425599726 | 0.351843673 | 0 | 0.03 | 1 | 0.148025014 | -1.223818009 | 0 | 0.045 | 1 | 0.425599726 | 0.274138624 |
| Gm14403 | 0.425639612 | 0.503846766 | 0 | 0.03 | 1 | 0.148025014 | -1.223818009 | 0 | 0.045 | 1 | 0.425639612 | 0.274138624 |
| Trim21 | 0.425639612 | 0.503846766 | 0 | 0.03 | 1 | 0.148025014 | -1.223818009 | 0 | 0.045 | 1 | 0.425639612 | 0.274138624 |
| Gm56656 | 0.457308648 | 0.673771768 | 0 | 0.026 | 1 | 0.148025014 | -1.223818009 | 0 | 0.045 | 1 | 0.457308648 | 0.274138624 |
| Pcdhb2 | 0.532426384 | 0.866416846 | 0 | 0.019 | 1 | 0.148025014 | -1.316927414 | 0 | 0.045 | 1 | 0.532426384 | 0.274138624 |
| Tnfrsf25 | 0.578156445 | 0.866416846 | 0 | 0.015 | 1 | 0.148025014 | -1.223818009 | 0 | 0.045 | 1 | 0.578156445 | 0.274138624 |
| Tnfrsf14 | 0.578156445 | 1.088809267 | 0 | 0.015 | 1 | 0.148025014 | -1.404390255 | 0 | 0.045 | 1 | 0.578156445 | 0.274138624 |
| Lor | 0.57816106 | 1.351843673 | 0 | 0.015 | 1 | 0.148025014 | -1.223818009 | 0 | 0.045 | 1 | 0.57816106 | 0.274138624 |
| 9330133O14Rik | 0.616977656 | 0.614878079 | 0.048 | 0.082 | 1 | 0.148025014 | -1.223818009 | 0 | 0.045 | 1 | 0.616977656 | 0.274138624 |
| Gm20635 | 0.646855374 | 1.472137907 | 0.095 | 0.071 | 1 | 0.148025014 | -1.223818009 | 0 | 0.045 | 1 | 0.646855374 | 0.274138624 |
| Gm56573 | 0.71266839 | 1.79930265 | 0.048 | 0.034 | 1 | 0.148025014 | -1.223818009 | 0 | 0.045 | 1 | 0.71266839 | 0.274138624 |
| D430020J02Rik | 0.718367751 | 0.503846766 | 0.048 | 0.071 | 1 | 0.148025014 | -1.223818009 | 0 | 0.045 | 1 | 0.718367751 | 0.274138624 |
| B230303O12Rik | 0.787193017 | 1.673771768 | 0.048 | 0.037 | 1 | 0.148025014 | -1.316927414 | 0 | 0.045 | 1 | 0.787193017 | 0.274138624 |
| Prox2 | 0.813144372 | 0.936806174 | 0.048 | 0.063 | 1 | 0.148025014 | -1.316927414 | 0 | 0.045 | 1 | 0.813144372 | 0.274138624 |
| Fam234b | 0.869523519 | 0.503846766 | 0.048 | 0.056 | 1 | 0.148025014 | -1.316927414 | 0 | 0.045 | 1 | 0.869523519 | 0.274138624 |
| Camkmt | 0.665649757 | 1.419350492 | 0.286 | 0.287 | 1 | 0.148044053 | 0.65505821 | 0.156 | 0.291 | 1 | 0.665649757 | 0.274171065 |
| Ube2t | 0.16352901 | -1.233118828 | 0 | 0.086 | 1 | 0.148044209 | -1.316927414 | 0 | 0.045 | 1 | 0.16352901 | 0.274171331 |
| Cenps | 0.208921714 | -0.970084422 | 0 | 0.071 | 1 | 0.148044209 | -1.316927414 | 0 | 0.045 | 1 | 0.208921714 | 0.274171331 |
| Gm49463 | 0.236343578 | -0.574155746 | 0 | 0.063 | 1 | 0.148044209 | -1.316927414 | 0 | 0.045 | 1 | 0.236343578 | 0.274171331 |
| Cox6b2 | 0.267985206 | -0.785659851 | 0 | 0.056 | 1 | 0.148044209 | -1.564854927 | 0 | 0.045 | 1 | 0.267985206 | 0.274171331 |
| Chek2 | 0.324739547 | -0.133583154 | 0 | 0.045 | 1 | 0.148044209 | -1.316927414 | 0 | 0.045 | 1 | 0.324739547 | 0.274171331 |
| Smim24 | 0.324767242 | -0.233118828 | 0 | 0.045 | 1 | 0.148044209 | -1.486852415 | 0 | 0.045 | 1 | 0.324767242 | 0.274171331 |
| Gm10432 | 0.346846605 | -0.413691074 | 0 | 0.041 | 1 | 0.148044209 | -1.316927414 | 0 | 0.045 | 1 | 0.346846605 | 0.274171331 |
| A630052C17Rik | 0.396971401 | 0.351843673 | 0 | 0.034 | 1 | 0.148044209 | -1.316927414 | 0 | 0.045 | 1 | 0.396971401 | 0.274171331 |
| Dhrs9 | 0.45730234 | -1.752492987 | 0 | 0.026 | 1 | 0.148044209 | -1.564854927 | 0 | 0.045 | 1 | 0.45730234 | 0.274171331 |
| Gm11457 | 0.532436372 | 1.088809267 | 0 | 0.019 | 1 | 0.148044209 | -1.316927414 | 0 | 0.045 | 1 | 0.532436372 | 0.274171331 |
| Aard | 0.532436372 | 1.088809267 | 0 | 0.019 | 1 | 0.148044209 | -1.316927414 | 0 | 0.045 | 1 | 0.532436372 | 0.274171331 |
| Adm | 0.790349864 | 1.55829455 | 0.048 | 0.037 | 1 | 0.148044209 | -1.316927414 | 0 | 0.045 | 1 | 0.790349864 | 0.274171331 |
| Egln3 | 0.861474289 | 0.281454345 | 0.048 | 0.056 | 1 | 0.148044209 | -1.404390255 | 0 | 0.045 | 1 | 0.861474289 | 0.274171331 |
| Cldn34c1 | 0.864156685 | 0.214340149 | 0.048 | 0.056 | 1 | 0.148044209 | -1.316927414 | 0 | 0.045 | 1 | 0.864156685 | 0.274171331 |
| Gm17690 | 0.173819828 | -1.133583154 | 0 | 0.082 | 1 | 0.148059913 | -1.486852415 | 0 | 0.045 | 1 | 0.173819828 | 0.274198089 |
| Fbln2 | 0.324767242 | -0.233118828 | 0 | 0.045 | 1 | 0.148059913 | -1.486852415 | 0 | 0.045 | 1 | 0.324767242 | 0.274198089 |
| Hcls1 | 0.492590603 | 0.673771768 | 0 | 0.022 | 1 | 0.148059913 | -1.564854927 | 0 | 0.045 | 1 | 0.492590603 | 0.274198089 |
| Gdf15 | 0.659453916 | -3.101015292 | 0.048 | 0.071 | 1 | 0.148059913 | -1.84048937 | 0 | 0.045 | 1 | 0.659453916 | 0.274198089 |
| Gm10382 | 0.904673267 | 0.936806174 | 0.048 | 0.056 | 1 | 0.148059913 | -1.84048937 | 0 | 0.045 | 1 | 0.904673267 | 0.274198089 |
| Tent5b | 0.169616506 | 2.258734268 | 0.048 | 0.011 | 1 | 0.148060204 | -1.404390255 | 0 | 0.045 | 1 | 0.169616506 | 0.274198585 |
| Mfsd3 | 0.196329837 | -0.785659851 | 0 | 0.075 | 1 | 0.148060204 | -1.404390255 | 0 | 0.045 | 1 | 0.196329837 | 0.274198585 |
| 4933427G23Rik | 0.370840523 | 0.214340149 | 0 | 0.037 | 1 | 0.148060204 | -1.404390255 | 0 | 0.045 | 1 | 0.370840523 | 0.274198585 |
| Zfp14 | 0.425614684 | 0.214340149 | 0 | 0.03 | 1 | 0.148060204 | -1.404390255 | 0 | 0.045 | 1 | 0.425614684 | 0.274198585 |
| Aoc2 | 0.457281049 | 0.503846766 | 0 | 0.026 | 1 | 0.148060204 | -1.638855509 | 0 | 0.045 | 1 | 0.457281049 | 0.274198585 |
| 4632404H12Rik | 0.479147948 | 1.351843673 | 0.048 | 0.022 | 1 | 0.148060204 | -1.564854927 | 0 | 0.045 | 1 | 0.479147948 | 0.274198585 |
| Ano7 | 0.54776222 | 2.088809267 | 0.048 | 0.026 | 1 | 0.148060204 | -1.564854927 | 0 | 0.045 | 1 | 0.54776222 | 0.274198585 |
| Gm15559 | 0.815691551 | 1.088809267 | 0.048 | 0.063 | 1 | 0.148060204 | -1.404390255 | 0 | 0.045 | 1 | 0.815691551 | 0.274198585 |

| | | | | | | | | | | | | |
|---|---|---|---|---|---|---|---|---|---|---|---|---|
| Gm49700 | 0.918794646 | 1.866416846 | 0.048 | 0.045 | 1 | 0.148060204 | -1.404390255 | 0 | 0.045 | 1 | 0.918794646 | 0.274198585 |
| 4930481B07Rik | 0.492590603 | 0.673771768 | 0 | 0.022 | 1 | 0.148064631 | 2.683072586 | 0.044 | 0.014 | 1 | 0.492590603 | 0.274206126 |
| Gm42744 | 0.57816106 | 1.351843673 | 0 | 0.015 | 1 | 0.148064631 | 2.683072586 | 0.044 | 0.014 | 1 | 0.57816106 | 0.274206126 |
| Enc1 | 0.826079082 | 0.936806174 | 0.095 | 0.086 | 1 | 0.148072128 | -1.709244837 | 0 | 0.045 | 1 | 0.826079082 | 0.2742189 |
| Cops7b | 0.436441527 | -0.133583154 | 0.048 | 0.104 | 1 | 0.148072696 | -0.440309829 | 0.067 | 0.152 | 1 | 0.436441527 | 0.274219869 |
| Gm17225 | 0.209040746 | -1.785659851 | 0 | 0.071 | 1 | 0.148073 | -1.709244837 | 0 | 0.045 | 1 | 0.209040746 | 0.274220387 |
| Shank3 | 0.268011783 | -0.648156327 | 0 | 0.056 | 1 | 0.148073 | -1.638855509 | 0 | 0.045 | 1 | 0.268011783 | 0.274220387 |
| Acad10 | 0.346265118 | 1.425844254 | 0.095 | 0.049 | 1 | 0.148073 | -1.84048937 | 0 | 0.045 | 1 | 0.346265118 | 0.274220387 |
| AU040972 | 0.532426384 | 0.866416846 | 0 | 0.019 | 1 | 0.148073 | -2.124282336 | 0 | 0.045 | 1 | 0.532426384 | 0.274220387 |
| Hlx | 0.209065702 | -1.880817084 | 0.048 | 0.142 | 1 | 0.148073291 | -2.316927414 | 0 | 0.045 | 1 | 0.209065702 | 0.274220882 |
| Arhgef37 | 0.346827513 | -0.413691074 | 0 | 0.041 | 1 | 0.148073291 | -1.84048937 | 0 | 0.045 | 1 | 0.346827513 | 0.274220882 |
| Gm34894 | 0.359270744 | 3.673771768 | 0.048 | 0.019 | 1 | 0.148073291 | -1.486852415 | 0 | 0.045 | 1 | 0.359270744 | 0.274220882 |
| Gdpd4 | 0.396963624 | -0.970084422 | 0 | 0.034 | 1 | 0.148073883 | 2.490427508 | 0.044 | 0.014 | 1 | 0.396963624 | 0.274221891 |
| Slc14a2 | 0.396964273 | -0.911190733 | 0 | 0.034 | 1 | 0.148075733 | 2.320502507 | 0.044 | 0.014 | 1 | 0.396964273 | 0.274225043 |
| Zfp612 | 0.285527294 | -0.574155746 | 0 | 0.052 | 1 | 0.148082596 | -1.709244837 | 0 | 0.045 | 1 | 0.285527294 | 0.274236738 |
| Trpv2 | 0.304471562 | -1.940938076 | 0 | 0.049 | 1 | 0.148082596 | -1.960783604 | 0 | 0.045 | 1 | 0.304471562 | 0.274236738 |
| Phex | 0.629723166 | 6.42865927 | 0.048 | 0.03 | 1 | 0.148083469 | -1.776359032 | 0 | 0.045 | 1 | 0.629723166 | 0.274238224 |
| Rarres2 | 0.324820107 | -0.648156327 | 0 | 0.045 | 1 | 0.14808376 | -1.564854927 | 0 | 0.045 | 1 | 0.324820107 | 0.274238719 |
| Slc1a3 | 0.396962328 | -1.574155746 | 0 | 0.034 | 1 | 0.14809481 | -2.071814916 | 0 | 0.045 | 1 | 0.396962328 | 0.274257547 |
| Ppcs | 0.208955157 | -0.970084422 | 0 | 0.071 | 1 | 0.148108186 | -1.124282336 | 0 | 0.045 | 1 | 0.208955157 | 0.274280337 |
| Zfp324 | 0.370840523 | 0.214340149 | 0 | 0.037 | 1 | 0.148108186 | -1.124282336 | 0 | 0.045 | 1 | 0.370840523 | 0.274280337 |
| Amigo1 | 0.396971401 | 0.351843673 | 0 | 0.034 | 1 | 0.148108186 | -1.124282336 | 0 | 0.045 | 1 | 0.396971401 | 0.274280337 |
| Panx2 | 0.57816106 | 1.351843673 | 0 | 0.015 | 1 | 0.148108186 | -1.124282336 | 0 | 0.045 | 1 | 0.57816106 | 0.274280337 |
| 4833413G10Rik | 0.63207476 | 1.351843673 | 0 | 0.011 | 1 | 0.148108186 | -1.124282336 | 0 | 0.045 | 1 | 0.63207476 | 0.274280337 |
| Nr2c2 | 0.336971239 | 0.219393377 | 0.333 | 0.5 | 1 | 0.148221386 | 0.374200644 | 0.867 | 0.72 | 1 | 0.336971239 | 0.274473192 |
| Actr5 | 0.473894763 | -0.496153234 | 0.048 | 0.093 | 1 | 0.148268205 | -0.84048937 | 0.044 | 0.118 | 1 | 0.473894763 | 0.27455295 |
| Lpin3 | 0.285527727 | -0.718545655 | 0 | 0.052 | 1 | 0.148464731 | -1.271123724 | 0.022 | 0.083 | 1 | 0.285527727 | 0.274887686 |
| Fendrr | 0.148481904 | 0.33748838 | 0.095 | 0.257 | 1 | 0.940134787 | 0.617781126 | 0.222 | 0.26 | 1 | 0.940134787 | 0.274916933 |
| Gm16268 | 0.773910509 | 2.789248985 | 0.048 | 0.067 | 1 | 0.148680117 | 1.841770332 | 0.156 | 0.093 | 1 | 0.773910509 | 0.275254456 |
| Dnah12 | 0.820819496 | -1.098817736 | 0.095 | 0.112 | 1 | 0.148701437 | 3.875717664 | 0.089 | 0.042 | 1 | 0.820819496 | 0.275290756 |
| Pomt2 | 0.306024127 | -0.348596045 | 0.095 | 0.19 | 1 | 0.148711211 | 0.495445583 | 0.156 | 0.304 | 1 | 0.306024127 | 0.275307398 |
| Tgfbi | 0.267806461 | -2.02666795 | 0.048 | 0.127 | 1 | 0.148713764 | -0.223818009 | 0.067 | 0.156 | 1 | 0.267806461 | 0.275311745 |
| Parva | 0.148717175 | 0.956897591 | 0.81 | 0.78 | 1 | 0.314747774 | 0.619949667 | 0.911 | 0.834 | 1 | 0.314747774 | 0.275317551 |
| Gm26827 | 0.148777478 | 2.214340149 | 0.095 | 0.034 | 1 | 0.293579277 | -0.271123724 | 0.022 | 0.062 | 1 | 0.293579277 | 0.275420218 |
| Gm6169 | 0.285622926 | -0.233118828 | 0 | 0.052 | 1 | 0.14894071 | -1.271123724 | 0.022 | 0.083 | 1 | 0.285622926 | 0.275698086 |
| Rad9a | 0.927798188 | 1.258734268 | 0.048 | 0.045 | 1 | 0.14894071 | -1.174908409 | 0.022 | 0.083 | 1 | 0.927798188 | 0.275698086 |
| N4bp2os | 0.222228992 | -0.849790188 | 0 | 0.067 | 1 | 0.148940868 | -1.124282336 | 0.022 | 0.083 | 1 | 0.222228992 | 0.275698354 |
| Mkks | 0.236581911 | -0.496153234 | 0 | 0.063 | 1 | 0.148940868 | -1.124282336 | 0.022 | 0.083 | 1 | 0.236581911 | 0.275698354 |
| Slc38a2 | 0.148957476 | -0.16192778 | 1 | 0.996 | 1 | 0.318314557 | 0.282205398 | 1 | 0.993 | 1 | 0.318314557 | 0.275726622 |
| Gm31763 | 0.47224907 | -0.455511249 | 0.048 | 0.093 | 1 | 0.149096311 | 0.154693614 | 0.089 | 0.187 | 1 | 0.47224907 | 0.275962912 |
| Ahcy | 0.5690434 | -0.184209227 | 0.048 | 0.082 | 1 | 0.149414151 | -1.071814916 | 0.022 | 0.083 | 1 | 0.5690434 | 0.276503714 |
| Mtres1 | 0.149415877 | -0.413691074 | 0.095 | 0.243 | 1 | 0.504473494 | 0.946106992 | 0.133 | 0.194 | 1 | 0.504473494 | 0.27650665 |
| Mapk13 | 0.184739689 | -0.970084422 | 0 | 0.078 | 1 | 0.14949605 | -0.80878051 | 0.044 | 0.118 | 1 | 0.184739689 | 0.276643032 |
| Slc66a2 | 0.149582561 | 0.336330673 | 0.333 | 0.604 | 1 | 0.695379096 | 0.207110863 | 0.578 | 0.578 | 1 | 0.695379096 | 0.27679018 |
| Ubr1 | 0.14964745 | 0.69025989 | 0.667 | 0.709 | 1 | 0.5936447 | 0.209656738 | 0.778 | 0.744 | 1 | 0.5936447 | 0.276900541 |
| Gpt2 | 0.424500903 | -0.413691074 | 0.048 | 0.101 | 1 | 0.149688721 | -1.191396532 | 0.044 | 0.114 | 1 | 0.424500903 | 0.276970729 |
| Dync1i2 | 0.149696623 | 0.271673324 | 0.524 | 0.817 | 1 | 0.589483784 | 0.354081099 | 0.889 | 0.872 | 1 | 0.589483784 | 0.276984167 |
| Topbp1 | 0.540857659 | 0.803407048 | 0.286 | 0.25 | 1 | 0.149809964 | -0.114883638 | 0.2 | 0.322 | 1 | 0.540857659 | 0.277176903 |
| 1810034E14Rik | 0.149931669 | 1.97333205 | 0.095 | 0.034 | 1 | 0.376842245 | -0.486852415 | 0 | 0.017 | 1 | 0.376842245 | 0.277383832 |
| Capn10 | 0.149944962 | -1.280424543 | 0.048 | 0.164 | 1 | 0.265809947 | 0.245667274 | 0.089 | 0.163 | 1 | 0.265809947 | 0.277406432 |
| 8430426J06Rik | 0.149960708 | 2.866416846 | 0.095 | 0.034 | 1 | 0.932074328 | 1.765534746 | 0.044 | 0.048 | 1 | 0.932074328 | 0.277433203 |
| A4galt | 0.149998109 | 0.964551164 | 0.667 | 0.601 | 1 | 0.21063849 | 0.408617001 | 0.644 | 0.716 | 1 | 0.21063849 | 0.277496785 |
| Gm17058 | 0.532436372 | 1.088809267 | 0 | 0.019 | 1 | 0.150103609 | 2.098110085 | 0.044 | 0.014 | 1 | 0.532436372 | 0.277676125 |
| Golph3l | 0.969967987 | 0.793353383 | 0.19 | 0.22 | 1 | 0.150259841 | -0.213091603 | 0.133 | 0.249 | 1 | 0.969967987 | 0.277941661 |
| Zscan22 | 0.239223183 | 1.351843673 | 0.19 | 0.112 | 1 | 0.150324538 | -0.776359032 | 0.044 | 0.118 | 1 | 0.239223183 | 0.278051609 |
| Wasl | 0.845997866 | 0.199169714 | 0.619 | 0.75 | 1 | 0.150436053 | -0.154870656 | 0.844 | 0.927 | 1 | 0.845997866 | 0.2782411 |
| Bbs9 | 0.150490067 | -0.297771786 | 0.238 | 0.444 | 1 | 0.756080536 | 0.983274494 | 0.578 | 0.647 | 1 | 0.756080536 | 0.278332873 |
| Gm16124 | 0.423798862 | -0.370622352 | 0.095 | 0.164 | 1 | 0.150508735 | 0.290755163 | 0.067 | 0.159 | 1 | 0.423798862 | 0.278364591 |
| 9530059O14Rik | 0.382757625 | 2.340348034 | 0.143 | 0.09 | 1 | 0.150736116 | -2.375821103 | 0.022 | 0.083 | 1 | 0.382757625 | 0.278750855 |
| Casp2 | 0.196462148 | -0.970084422 | 0 | 0.075 | 1 | 0.150761541 | -0.743192169 | 0.044 | 0.118 | 1 | 0.196462148 | 0.27879404 |
| Mvk | 0.333630039 | -0.648156327 | 0.048 | 0.116 | 1 | 0.150761541 | -0.776359032 | 0.044 | 0.118 | 1 | 0.333630039 | 0.27879404 |
| Zfp33b | 0.260424153 | 0.410737362 | 0.048 | 0.142 | 1 | 0.15078769 | 0.327591932 | 0.111 | 0.221 | 1 | 0.260424153 | 0.278838452 |
| Synpo | 0.206307347 | 0.2329026 | 0.762 | 0.776 | 1 | 0.150811781 | 0.126549271 | 0.489 | 0.754 | 1 | 0.206307347 | 0.278879368 |
| Alms1 | 0.151053117 | -1.434752689 | 0.048 | 0.164 | 1 | 0.504831466 | 0.643544222 | 0.178 | 0.142 | 1 | 0.504831466 | 0.27928919 |
| Cfap53 | 0.151109321 | 2.088809267 | 0.095 | 0.034 | 1 | 0.211019767 | 1.098110085 | 0.089 | 0.045 | 1 | 0.211019767 | 0.279384615 |
| Osmr | 0.409490936 | 0.861775463 | 0.667 | 0.772 | 1 | 0.151145749 | -0.256216 | 0.422 | 0.619 | 1 | 0.409490936 | 0.27944646 |
| 1700102P08Rik | 0.151305415 | 2.045740545 | 0.19 | 0.101 | 1 | 0.344811716 | 0.180572246 | 0.111 | 0.176 | 1 | 0.344811716 | 0.279717501 |
| Fhip1a | 0.344337239 | 1.516728058 | 0.429 | 0.437 | 1 | 0.151336121 | 0.573639345 | 0.711 | 0.571 | 1 | 0.344337239 | 0.279769621 |
| Mcm3 | 0.291375153 | -0.496153234 | 0.048 | 0.131 | 1 | 0.151374685 | -1.158229668 | 0.044 | 0.114 | 1 | 0.291375153 | 0.279835075 |
| 4930594M22Rik | 0.994183949 | 1.010806755 | 0.048 | 0.049 | 1 | 0.151455972 | 0.313838777 | 0.044 | 0.118 | 1 | 0.994183949 | 0.279973033 |
| Trmo | 0.932575038 | 0.586308926 | 0.095 | 0.104 | 1 | 0.15159229 | -0.709244837 | 0.044 | 0.118 | 1 | 0.932575038 | 0.280204358 |
| Flna | 0.642402589 | 0.287713335 | 0.524 | 0.571 | 1 | 0.151694158 | -0.282347182 | 0.844 | 0.827 | 1 | 0.642402589 | 0.280377198 |
| Pbx3 | 0.151783017 | 1.214340149 | 0.524 | 0.425 | 1 | 0.200916058 | 0.263533695 | 0.844 | 0.602 | 1 | 0.200916058 | 0.28052795 |
| Slc25a40 | 0.537725187 | 1.001346426 | 0.095 | 0.16 | 1 | 0.151900967 | -0.264459994 | 0.067 | 0.152 | 1 | 0.537725187 | 0.28072803 |
| Klhdc4 | 0.786941173 | 0.376091219 | 0.143 | 0.179 | 1 | 0.151921325 | 0.132875504 | 0.111 | 0.208 | 1 | 0.786941173 | 0.280762561 |
| Bcorl1 | 0.391936228 | -0.496153234 | 0.095 | 0.172 | 1 | 0.151937145 | -0.216400538 | 0.222 | 0.332 | 1 | 0.391936228 | 0.280789395 |
| Per2 | 0.629490495 | 0.183805781 | 0.286 | 0.418 | 1 | 0.152079567 | -0.424221427 | 0.356 | 0.478 | 1 | 0.629490495 | 0.28103094 |
| Oas1b | 0.167544534 | 2.936806174 | 0.048 | 0.011 | 1 | 0.152163868 | 1.946106992 | 0.044 | 0.014 | 1 | 0.167544534 | 0.281173894 |
| Iqcg | 0.492590603 | 0.673771768 | 0 | 0.022 | 1 | 0.152163868 | 1.946106992 | 0.044 | 0.014 | 1 | 0.492590603 | 0.281173894 |
| H3c7 | 0.492599419 | -0.133583154 | 0 | 0.022 | 1 | 0.152163868 | 1.946106992 | 0.044 | 0.014 | 1 | 0.492599419 | 0.281173894 |
| Gm30092 | 0.57816106 | 1.351843673 | 0 | 0.015 | 1 | 0.152163868 | 1.946106992 | 0.044 | 0.014 | 1 | 0.57816106 | 0.281173894 |
| Zfat | 0.64125812 | 1.281454345 | 0.143 | 0.201 | 1 | 0.152196752 | 0.576157382 | 0.133 | 0.263 | 1 | 0.64125812 | 0.281229653 |
| Snx31 | 0.15230504 | 2.536268244 | 0.095 | 0.034 | 1 | 0.331914654 | -0.638855509 | 0 | 0.021 | 1 | 0.331914654 | 0.281413255 |
| Zfp871 | 0.152422051 | 0.766881172 | 0.667 | 0.649 | 1 | 0.89216828 | 0.108378421 | 0.756 | 0.789 | 1 | 0.89216828 | 0.28161162 |
| Fam151b | 0.677501769 | 0.451379346 | 0.048 | 0.075 | 1 | 0.152611688 | -0.486852415 | 0.044 | 0.118 | 1 | 0.677501769 | 0.281933048 |
| Acox1 | 0.15283965 | 0.235962961 | 0.429 | 0.765 | 1 | 0.215850059 | 0.363869129 | 0.778 | 0.737 | 1 | 0.215850059 | 0.282319341 |
| Nubp1 | 0.582871457 | 0.766881172 | 0.286 | 0.265 | 1 | 0.152949065 | -0.427351404 | 0.156 | 0.26 | 1 | 0.582871457 | 0.282504714 |
| Il18 | 0.866918641 | 0.528094312 | 0.333 | 0.396 | 1 | 0.153051237 | 0.278088752 | 0.178 | 0.311 | 1 | 0.866918641 | 0.282677792 |
| Star | 0.865330016 | 2.199840579 | 0.048 | 0.06 | 1 | 0.153059952 | -1.017367132 | 0.044 | 0.114 | 1 | 0.865330016 | 0.282692555 |
| Kcnip3 | 0.532436372 | 1.088809267 | 0 | 0.019 | 1 | 0.153195762 | 1.513147585 | 0.044 | 0.014 | 1 | 0.532436372 | 0.282922583 |
| Acvr1b | 0.594387605 | 0.829422638 | 0.381 | 0.381 | 1 | 0.153351189 | 0.254229287 | 0.178 | 0.301 | 1 | 0.594387605 | 0.28318579 |
| Lipg | 0.153352184 | -2.880817084 | 0.048 | 0.16 | 1 | 0.823042771 | 0.875717664 | 0.022 | 0.017 | 1 | 0.823042771 | 0.283187476 |
| Zfp763 | 0.153464994 | 1.97333205 | 0.095 | 0.034 | 1 | 0.539321288 | -0.271123724 | 0.044 | 0.069 | 1 | 0.539321288 | 0.283378484 |
| Fbxl4 | 0.593100443 | -0.44170545 | 0.095 | 0.134 | 1 | 0.153509905 | -0.352551324 | 0.044 | 0.121 | 1 | 0.593100443 | 0.283454519 |
| Adam32 | 0.330910872 | 2.462267662 | 0.143 | 0.09 | 1 | 0.153609031 | 2.581192972 | 0.156 | 0.093 | 1 | 0.330910872 | 0.283622328 |
| Phf21a | 0.388188849 | 0.647682539 | 0.81 | 0.832 | 1 | 0.153671646 | 0.326521095 | 1 | 0.924 | 1 | 0.388188849 | 0.283728317 |
| Aldh4a1 | 0.153693413 | 2.405282932 | 0.238 | 0.149 | 1 | 0.899977549 | 1.203079645 | 0.133 | 0.149 | 1 | 0.899977549 | 0.283765161 |
| Dhx35 | 0.153760172 | -1.133583154 | 0 | 0.09 | 1 | 0.84251778 | 0.995016592 | 0.089 | 0.083 | 1 | 0.84251778 | 0.283878154 |

| | | | | | | | | | | | | |
|---|---|---|---|---|---|---|---|---|---|---|---|---|
| Tamm41 | 0.153760172 | -1.081115734 | 0 | 0.09 | 1 | 0.358519478 | 0.205025289 | 0.067 | 0.118 | 1 | 0.358519478 | 0.283878154 |
| Mapkap1 | 0.315626964 | -0.226900305 | 0.619 | 0.735 | 1 | 0.153784276 | 0.566802897 | 0.911 | 0.782 | 1 | 0.315626964 | 0.283918949 |
| Gyg | 0.153811633 | -1.184209227 | 0 | 0.09 | 1 | 0.904257497 | 0.933050839 | 0.089 | 0.1 | 1 | 0.904257497 | 0.283965247 |
| 2900005J15Rik | 0.153811856 | -1.133583154 | 0 | 0.09 | 1 | 0.821680938 | 1.361144491 | 0.044 | 0.038 | 1 | 0.821680938 | 0.283965625 |
| Abhd3 | 0.153811856 | -1.413691074 | 0 | 0.09 | 1 | 0.42546306 | 1.142504205 | 0.044 | 0.08 | 1 | 0.42546306 | 0.283965625 |
| Trub1 | 0.153811856 | -1.233118828 | 0 | 0.09 | 1 | 0.901730233 | 1.020107573 | 0.089 | 0.1 | 1 | 0.901730233 | 0.283965625 |
| Tbk1 | 0.707847934 | 0.939062147 | 0.381 | 0.433 | 1 | 0.153845869 | 0.522389737 | 0.711 | 0.619 | 1 | 0.707847934 | 0.284023187 |
| Tmem94 | 0.153858613 | -1.370622352 | 0 | 0.09 | 1 | 0.629422286 | 0.744473131 | 0.089 | 0.121 | 1 | 0.629422286 | 0.284044753 |
| Serp2 | 0.153899101 | -1.326228232 | 0 | 0.09 | 1 | 0.379809495 | 0.361144491 | 0.044 | 0.087 | 1 | 0.379809495 | 0.284113269 |
| Gng3 | 0.153899101 | -1.326228232 | 0 | 0.09 | 1 | 0.217102799 | 1.5342092 | 0.2 | 0.145 | 1 | 0.217102799 | 0.284113269 |
| Cln6 | 0.153901338 | -1.233118828 | 0 | 0.09 | 1 | 0.430238982 | 0.446033389 | 0.067 | 0.107 | 1 | 0.430238982 | 0.284117054 |
| 2810433D01Rik | 0.153901338 | -1.233118828 | 0 | 0.09 | 1 | 0.241195354 | -0.375821103 | 0.022 | 0.069 | 1 | 0.241195354 | 0.284117054 |
| Dgkq | 0.942764118 | 0.908237021 | 0.095 | 0.104 | 1 | 0.153902241 | -0.191396532 | 0.044 | 0.118 | 1 | 0.942764118 | 0.284118582 |
| Dcaf15 | 0.153935113 | -1.326228232 | 0 | 0.09 | 1 | 0.274873309 | -0.686161223 | 0.044 | 0.093 | 1 | 0.274873309 | 0.284174206 |
| Wdr27 | 0.153935113 | -1.370622352 | 0 | 0.09 | 1 | 0.298262476 | 0.223640968 | 0.022 | 0.062 | 1 | 0.298262476 | 0.284174206 |
| Dtwd1 | 0.153937349 | -1.370622352 | 0 | 0.09 | 1 | 0.716733944 | 0.995016592 | 0.067 | 0.087 | 1 | 0.716733944 | 0.284177991 |
| H2-K2 | 0.153938468 | -1.455511249 | 0 | 0.09 | 1 | 0.184806696 | -1.017367132 | 0 | 0.038 | 1 | 0.184806696 | 0.284179883 |
| Gm30489 | 0.153939139 | -1.535681598 | 0 | 0.09 | 1 | 0.512490525 | 2.098110085 | 0.044 | 0.028 | 1 | 0.512490525 | 0.284181019 |
| Cdc42ep1 | 0.280793265 | -0.683780237 | 0.048 | 0.127 | 1 | 0.153954499 | -0.264459994 | 0.067 | 0.156 | 1 | 0.280793265 | 0.284207011 |
| 1810024B03Rik | 0.153999523 | -1.535681598 | 0 | 0.09 | 1 | 0.345581787 | 0.91753784 | 0.044 | 0.09 | 1 | 0.345581787 | 0.284283192 |
| Basp1 | 0.154001088 | -1.752492987 | 0 | 0.09 | 1 | 0.602156559 | -0.677674758 | 0.133 | 0.156 | 1 | 0.602156559 | 0.284285841 |
| Slc24a3 | 0.154029935 | -2.208871282 | 0 | 0.09 | 1 | 0.926675192 | 1.318075769 | 0.089 | 0.09 | 1 | 0.926675192 | 0.28433465 |
| C1qtnf3 | 0.154062358 | -1.648156327 | 0 | 0.09 | 1 | 0.293637566 | -0.486852415 | 0 | 0.024 | 1 | 0.293637566 | 0.284389506 |
| H2-Eb1 | 0.15407242 | -3.24509147 | 0 | 0.09 | 1 | 0.312564649 | -0.564854927 | 0.022 | 0.059 | 1 | 0.312564649 | 0.284406529 |
| H1f5 | 0.15410238 | -4.159118246 | 0 | 0.09 | 1 | 0.944008652 | 0.567595369 | 0.022 | 0.021 | 1 | 0.944008652 | 0.284457216 |
| Dync2i2 | 0.632404643 | 0.316219763 | 0.095 | 0.138 | 1 | 0.154200303 | -0.518561275 | 0.044 | 0.118 | 1 | 0.632404643 | 0.284622873 |
| Klhdc10 | 0.154339482 | -0.239516599 | 0.238 | 0.463 | 1 | 0.591016403 | 0.282401376 | 0.711 | 0.644 | 1 | 0.591016403 | 0.284858289 |
| Muc5ac | 0.532432377 | -2.648156327 | 0 | 0.019 | 1 | 0.15484014 | 2.83507568 | 0.067 | 0.028 | 1 | 0.532432377 | 0.285704812 |
| Ptar1 | 0.155761428 | -0.633656758 | 0.286 | 0.455 | 1 | 0.377168011 | 0.297879598 | 0.356 | 0.547 | 1 | 0.377168011 | 0.287261234 |
| Lgals3 | 0.156384131 | -0.716790551 | 0.905 | 0.881 | 1 | 0.38381491 | -0.51597613 | 1 | 0.907 | 1 | 0.38381491 | 0.288312266 |
| Galc | 0.451614009 | -0.326228232 | 0.048 | 0.097 | 1 | 0.156413554 | -0.802354241 | 0.067 | 0.145 | 1 | 0.451614009 | 0.288361909 |
| Gm50012 | 0.905419792 | 0.673771768 | 0.095 | 0.108 | 1 | 0.156517176 | 1.635766871 | 0.244 | 0.183 | 1 | 0.905419792 | 0.288536725 |
| Cdh19 | 0.156585144 | -0.381510668 | 0.048 | 0.175 | 1 | 0.950035371 | 0.982632868 | 0.022 | 0.024 | 1 | 0.950035371 | 0.28865138 |
| Rbm41 | 0.691398392 | 0.882358389 | 0.238 | 0.22 | 1 | 0.156640507 | -0.309695845 | 0.222 | 0.356 | 1 | 0.691398392 | 0.288744766 |
| Ttc23 | 0.172426011 | -0.984439715 | 0.095 | 0.224 | 1 | 0.156775574 | -0.246538086 | 0.178 | 0.298 | 1 | 0.172426011 | 0.288972567 |
| Ehbp1l1 | 0.156794444 | 1.019546605 | 0.619 | 0.653 | 1 | 0.795672459 | 0.572199369 | 0.578 | 0.747 | 1 | 0.795672459 | 0.28900439 |
| Cidea | 0.17867807 | -0.741265732 | 0.095 | 0.235 | 1 | 0.156935141 | -0.468930507 | 0.067 | 0.149 | 1 | 0.17867807 | 0.289241644 |
| Gm47644 | 0.411436112 | 2.010806755 | 0.095 | 0.056 | 1 | 0.157005796 | 1.583536913 | 0.089 | 0.042 | 1 | 0.411436112 | 0.289360772 |
| Foxn2 | 0.226189217 | -0.65714511 | 0.19 | 0.336 | 1 | 0.157010506 | 0.743735368 | 0.622 | 0.536 | 1 | 0.226189217 | 0.289368713 |
| Prrg2 | 0.692363104 | 1.351843673 | 0.095 | 0.138 | 1 | 0.157017007 | -0.294207337 | 0.089 | 0.18 | 1 | 0.692363104 | 0.289379673 |
| Endod1 | 0.157191811 | -0.303508156 | 0.143 | 0.31 | 1 | 0.716977474 | -0.242926832 | 0.422 | 0.419 | 1 | 0.716977474 | 0.289674356 |
| Gm26632 | 0.264194629 | 2.673771768 | 0.048 | 0.015 | 1 | 0.157205906 | 2.557541704 | 0.067 | 0.028 | 1 | 0.264194629 | 0.289698115 |
| Asap2 | 0.157212521 | -0.653694511 | 0.524 | 0.739 | 1 | 0.982963986 | 0.147649546 | 0.444 | 0.512 | 1 | 0.982963986 | 0.289709266 |
| Il1r1 | 0.499340035 | 0.512308345 | 0.571 | 0.56 | 1 | 0.157272368 | -0.277756817 | 0.178 | 0.304 | 1 | 0.499340035 | 0.289810138 |
| Brca2 | 0.715153984 | 1.6858446 | 0.095 | 0.134 | 1 | 0.157294941 | -2.545746104 | 0.022 | 0.08 | 1 | 0.715153984 | 0.289848184 |
| Mtmr1 | 0.188669664 | -0.157169975 | 0.19 | 0.366 | 1 | 0.157560777 | 0.182379002 | 0.289 | 0.498 | 1 | 0.188669664 | 0.290296155 |
| Snx9 | 0.157597771 | -0.11172372 | 0.524 | 0.84 | 1 | 0.176739371 | 0.347847669 | 0.956 | 0.896 | 1 | 0.176739371 | 0.290358485 |
| Mllt1 | 0.318480978 | -0.522625445 | 0.095 | 0.179 | 1 | 0.157625604 | -0.425451871 | 0.111 | 0.208 | 1 | 0.318480978 | 0.290405377 |
| Yod1 | 0.194531079 | 0.233199176 | 0.143 | 0.299 | 1 | 0.157673371 | -0.593372739 | 0.222 | 0.353 | 1 | 0.194531079 | 0.29048585 |
| Washc2 | 0.157875829 | -0.38923803 | 0.286 | 0.5 | 1 | 0.185773496 | -0.547493767 | 0.667 | 0.647 | 1 | 0.185773496 | 0.290826881 |
| Rarres1 | 0.158179596 | -1.171718283 | 0.095 | 0.239 | 1 | 0.53953886 | 0.517263693 | 0.178 | 0.246 | 1 | 0.53953886 | 0.291338408 |
| Sdc1 | 0.158209031 | -1.133583154 | 0.143 | 0.28 | 1 | 0.462799559 | -0.448171947 | 0.644 | 0.578 | 1 | 0.462799559 | 0.291387964 |
| Ccny | 0.158219423 | 0.968408171 | 0.81 | 0.832 | 1 | 0.525690116 | 0.296374236 | 0.933 | 0.927 | 1 | 0.525690116 | 0.291405461 |
| Gm37245 | 0.158241154 | 1.586308926 | 0.095 | 0.034 | 1 | 0.309824707 | 3.098110085 | 0.022 | 0.007 | 1 | 0.309824707 | 0.291442044 |
| Taf1d | 0.494633623 | 0.180392817 | 0.81 | 0.832 | 1 | 0.158430242 | 0.273959921 | 0.933 | 0.772 | 1 | 0.494633623 | 0.291760342 |
| Zpbp | 0.686300024 | 0.673771768 | 0.048 | 0.075 | 1 | 0.158487111 | -1.017367132 | 0.022 | 0.083 | 1 | 0.686300024 | 0.291856058 |
| Cdon | 0.158564076 | 0.160148049 | 0.238 | 0.44 | 1 | 0.208462593 | -0.394371499 | 0.356 | 0.453 | 1 | 0.208462593 | 0.291985585 |
| Gm43062 | 0.370840523 | 0.214340149 | 0 | 0.037 | 1 | 0.158671292 | 1.83507568 | 0.111 | 0.059 | 1 | 0.370840523 | 0.292166005 |
| Zfp708 | 0.23638565 | -0.648156327 | 0 | 0.063 | 1 | 0.159024463 | -0.776359032 | 0.022 | 0.083 | 1 | 0.23638565 | 0.292760147 |
| Podnl1 | 0.387852734 | -0.159118246 | 0.048 | 0.112 | 1 | 0.159068352 | -0.776359032 | 0.022 | 0.083 | 1 | 0.387852734 | 0.292833964 |
| Mnd1 | 0.805641202 | 0.735172312 | 0.048 | 0.063 | 1 | 0.15909804 | -0.743192169 | 0.022 | 0.083 | 1 | 0.805641202 | 0.292883893 |
| Serpinc1 | 0.443831812 | -0.718545655 | 0.048 | 0.097 | 1 | 0.159289859 | -0.472205639 | 0.067 | 0.152 | 1 | 0.443831812 | 0.293206459 |
| Cyb5d2 | 0.159311467 | 0.574236094 | 0.048 | 0.175 | 1 | 0.476083899 | 0.42003818 | 0.089 | 0.131 | 1 | 0.476083899 | 0.293242791 |
| Mcm10 | 0.574700194 | 0.304537958 | 0.048 | 0.086 | 1 | 0.159333386 | -1.199570463 | 0.044 | 0.114 | 1 | 0.574700194 | 0.293279644 |
| Plau | 0.159451508 | 1.425844254 | 0.095 | 0.034 | 1 | 0.206897641 | -1.124282336 | 0 | 0.035 | 1 | 0.206897641 | 0.293478233 |
| Luzp2 | 0.159535499 | -2.140592213 | 0.238 | 0.369 | 1 | 0.622401213 | 0.705509975 | 0.244 | 0.239 | 1 | 0.622401213 | 0.293619422 |
| Snx11 | 0.159887486 | 1.451379346 | 0.238 | 0.142 | 1 | 0.182163919 | -0.513324627 | 0.089 | 0.17 | 1 | 0.182163919 | 0.294210964 |
| Spx | 0.222155241 | -0.718545655 | 0 | 0.067 | 1 | 0.160101688 | -0.686161223 | 0.022 | 0.083 | 1 | 0.222155241 | 0.294570825 |
| Shld1 | 0.251790506 | -0.413691074 | 0 | 0.06 | 1 | 0.160101688 | -0.686161223 | 0.022 | 0.083 | 1 | 0.251790506 | 0.294570825 |
| Gtf3c5 | 0.160125552 | -0.718545655 | 0.048 | 0.172 | 1 | 0.215558436 | 0.125077133 | 0.067 | 0.142 | 1 | 0.215558436 | 0.294610911 |
| Med15 | 0.16014543 | 0.761234609 | 0.571 | 0.578 | 1 | 0.510892073 | 0.395091823 | 0.689 | 0.664 | 1 | 0.510892073 | 0.294644302 |
| Zfyve26 | 0.944545221 | 0.69394965 | 0.19 | 0.205 | 1 | 0.16016 | -0.557935513 | 0.111 | 0.208 | 1 | 0.944545221 | 0.294668774 |
| Gnai1 | 0.669410013 | -0.695462042 | 0.238 | 0.276 | 1 | 0.160284854 | -0.142567689 | 0.422 | 0.554 | 1 | 0.669410013 | 0.294878474 |
| Tchh | 0.160447383 | -1.769171728 | 0.048 | 0.16 | 1 | 0.165324946 | -1.404390255 | 0 | 0.042 | 1 | 0.165324946 | 0.295151403 |
| Fut4 | 0.16044765 | -1.496153234 | 0.048 | 0.16 | 1 | 0.260684322 | -0.486852415 | 0 | 0.028 | 1 | 0.260684322 | 0.295151851 |
| Tceanc | 0.222277822 | -1.02666795 | 0 | 0.067 | 1 | 0.160614508 | -0.638855509 | 0.022 | 0.083 | 1 | 0.222277822 | 0.295431995 |
| P3h2 | 0.160768949 | 1.280899526 | 0.81 | 0.769 | 1 | 0.906219837 | 1.236678491 | 0.489 | 0.647 | 1 | 0.906219837 | 0.295691243 |
| Antxr1 | 0.384179717 | -0.24312022 | 0.571 | 0.709 | 1 | 0.16092322 | 0.27364651 | 0.8 | 0.92 | 1 | 0.384179717 | 0.295950157 |
| Eva1a | 0.41268394 | -0.970084422 | 0.048 | 0.101 | 1 | 0.160923746 | 0.299743947 | 0.111 | 0.221 | 1 | 0.41268394 | 0.29595104 |
| Pi4k2a | 0.498325238 | 0.382308954 | 0.762 | 0.791 | 1 | 0.160995143 | -0.124282336 | 0.667 | 0.727 | 1 | 0.498325238 | 0.29607085 |
| Uchl3 | 0.161050925 | 0.136337637 | 0.238 | 0.444 | 1 | 0.33620585 | 0.265837531 | 0.533 | 0.412 | 1 | 0.33620585 | 0.29616445 |
| Mbnl2 | 0.161091171 | 0.339576943 | 1 | 1 | 1 | 0.184870859 | 0.522470364 | 1 | 1 | 1 | 0.184870859 | 0.296231977 |
| Ythdf2 | 0.516025702 | 0.385526799 | 0.571 | 0.709 | 1 | 0.161178629 | -0.288306736 | 0.844 | 0.765 | 1 | 0.516025702 | 0.296378708 |
| Vps54 | 0.16132265 | 0.997661379 | 0.571 | 0.541 | 1 | 0.273729369 | 0.929428251 | 0.8 | 0.772 | 1 | 0.273729369 | 0.296620302 |
| Cp | 0.271433665 | 0.597133308 | 0.429 | 0.631 | 1 | 0.161440971 | -1.103523776 | 0.111 | 0.197 | 1 | 0.271433665 | 0.296818756 |
| Mex3b | 0.906165334 | 0.281454345 | 0.095 | 0.104 | 1 | 0.161622295 | -1.564854927 | 0.022 | 0.08 | 1 | 0.906165334 | 0.297122823 |
| 4930447C04Rik | 0.675315714 | 0.129451251 | 0.048 | 0.075 | 1 | 0.161627855 | -1.674479418 | 0.022 | 0.08 | 1 | 0.675315714 | 0.297132147 |
| Spaca1 | 0.618972991 | 0.673771768 | 0.048 | 0.082 | 1 | 0.161632386 | -0.539319835 | 0.022 | 0.083 | 1 | 0.618972991 | 0.297139743 |
| Kif26a | 0.492590603 | 0.673771768 | 0 | 0.022 | 1 | 0.161734214 | -0.361321533 | 0.022 | 0.083 | 1 | 0.492590603 | 0.297310471 |
| Tbc1d31 | 0.659969575 | 1.43673257 | 0.095 | 0.071 | 1 | 0.161734214 | -0.361321533 | 0.022 | 0.083 | 1 | 0.659969575 | 0.297310471 |
| Gm11906 | 0.209079391 | -2.184209227 | 0 | 0.071 | 1 | 0.161784225 | -0.446210431 | 0.022 | 0.083 | 1 | 0.209079391 | 0.297394314 |
| Mvb12b | 0.16178786 | -0.208871282 | 0.143 | 0.31 | 1 | 0.332579085 | 0.214303103 | 0.222 | 0.336 | 1 | 0.332579085 | 0.297400408 |
| Plcl1 | 0.161815006 | -1.114130792 | 0.095 | 0.235 | 1 | 0.204661483 | -0.486852415 | 0.133 | 0.225 | 1 | 0.204661483 | 0.297445916 |
| Strn | 0.161857278 | 1.120921452 | 0.524 | 0.549 | 1 | 0.86869617 | 0.191932108 | 0.756 | 0.685 | 1 | 0.86869617 | 0.297516777 |
| Gm42951 | 0.878411976 | 0.866416846 | 0.048 | 0.041 | 1 | 0.161993786 | 2.320502507 | 0.067 | 0.028 | 1 | 0.878411976 | 0.297745586 |
| Adal | 0.162011393 | -0.574155746 | 0.048 | 0.172 | 1 | 0.17298457 | -0.638855509 | 0.067 | 0.142 | 1 | 0.17298457 | 0.297775095 |
| 6430548M08Rik | 0.162042719 | 1.896164189 | 0.143 | 0.067 | 1 | 0.786204898 | 1.027720758 | 0.111 | 0.135 | 1 | 0.786204898 | 0.297827595 |
| Ece2 | 0.285527294 | -0.648156327 | 0 | 0.052 | 1 | 0.162137416 | -0.486852415 | 0.022 | 0.083 | 1 | 0.285527294 | 0.297986291 |

| | | | | | | | | | | | | |
|---|---|---|---|---|---|---|---|---|---|---|---|---|
| Oma1 | 0.162177112 | -1.303508156 | 0.048 | 0.16 | 1 | 0.82963452 | 0.816339117 | 0.133 | 0.156 | 1 | 0.82963452 | 0.298052808 |
| Endov | 0.349001577 | 1.281454345 | 0.19 | 0.127 | 1 | 0.162178563 | -0.486852415 | 0.022 | 0.083 | 1 | 0.349001577 | 0.29805524 |
| B3gntl1 | 0.465575833 | 0.214340149 | 0.048 | 0.101 | 1 | 0.162305718 | -0.361321533 | 0.022 | 0.083 | 1 | 0.465575833 | 0.29826829 |
| Apod | 0.530023253 | -0.935037475 | 0.095 | 0.06 | 1 | 0.16241918 | 0.435145073 | 0.022 | 0.083 | 1 | 0.530023253 | 0.29845837 |
| Sec23a | 0.39490154 | -0.15658498 | 0.524 | 0.795 | 1 | 0.162419727 | -0.277257268 | 0.689 | 0.848 | 1 | 0.39490154 | 0.298459287 |
| Arhgef10 | 0.786957402 | 0.658821426 | 0.19 | 0.235 | 1 | 0.162625007 | 0.13875207 | 0.2 | 0.356 | 1 | 0.786957402 | 0.29880312 |
| Gm17354 | 0.954841465 | 0.97333205 | 0.048 | 0.045 | 1 | 0.162684771 | -0.432404631 | 0.022 | 0.083 | 1 | 0.954841465 | 0.298903208 |
| Gm26728 | 0.793501451 | 1.351843673 | 0.048 | 0.037 | 1 | 0.162704087 | -1.526380779 | 0.022 | 0.08 | 1 | 0.793501451 | 0.298935555 |
| Brca1 | 0.808181144 | 1.054861935 | 0.048 | 0.063 | 1 | 0.162796963 | 2.005000681 | 0.067 | 0.028 | 1 | 0.808181144 | 0.299091076 |
| Herc3 | 0.802987008 | 1.171271427 | 0.048 | 0.037 | 1 | 0.162956957 | 0.42003818 | 0.022 | 0.083 | 1 | 0.802987008 | 0.299358945 |
| Sash1 | 0.163168669 | 1.305593858 | 0.905 | 0.948 | 1 | 0.221452229 | 0.449290232 | 1 | 0.986 | 1 | 0.221452229 | 0.299713323 |
| Axin1 | 0.163191278 | -0.255145134 | 0.19 | 0.388 | 1 | 0.206958456 | -0.12066202 | 0.578 | 0.647 | 1 | 0.206958456 | 0.299751163 |
| Sgtb | 0.600272659 | 1.410737362 | 0.095 | 0.067 | 1 | 0.163240237 | -1.361321533 | 0.022 | 0.08 | 1 | 0.600272659 | 0.2998331 |
| Ctc1 | 0.184739689 | -1.081115734 | 0 | 0.078 | 1 | 0.16324176 | -1.446210431 | 0.022 | 0.08 | 1 | 0.184739689 | 0.299835648 |
| Mmachc | 0.236343578 | -0.574155746 | 0 | 0.063 | 1 | 0.163274605 | 0.250113179 | 0.022 | 0.083 | 1 | 0.236343578 | 0.299890613 |
| C1qtnf9 | 0.862687636 | 1.088809267 | 0.048 | 0.06 | 1 | 0.163274768 | 0.197645759 | 0.022 | 0.083 | 1 | 0.862687636 | 0.299890886 |
| Dmd | 0.858791894 | -0.184209227 | 0.048 | 0.056 | 1 | 0.163296506 | 2.350876156 | 0.267 | 0.187 | 1 | 0.858791894 | 0.299927264 |
| Pex10 | 0.520499169 | 0.351843673 | 0.048 | 0.093 | 1 | 0.163311774 | -0.124282336 | 0.022 | 0.083 | 1 | 0.520499169 | 0.299952812 |
| Snx13 | 0.225091393 | 0.798132106 | 0.571 | 0.563 | 1 | 0.163314639 | 0.5265681 | 0.778 | 0.671 | 1 | 0.225091393 | 0.299957607 |
| Eml4 | 0.439608212 | -0.545989438 | 0.667 | 0.713 | 1 | 0.163315362 | 0.505534401 | 0.956 | 0.851 | 1 | 0.439608212 | 0.299958817 |
| Fech | 0.601162448 | 1.199840579 | 0.286 | 0.31 | 1 | 0.163365832 | 0.274267041 | 0.2 | 0.339 | 1 | 0.601162448 | 0.300043269 |
| Fhl3 | 0.813344052 | 0.150209812 | 0.048 | 0.06 | 1 | 0.163374367 | -0.223818009 | 0.022 | 0.083 | 1 | 0.813344052 | 0.30005755 |
| Il2 | 0.163379068 | -0.970084422 | 0 | 0.086 | 1 | 0.667598499 | 0.223640968 | 0.022 | 0.035 | 1 | 0.667598499 | 0.300065417 |
| Copg2 | 0.163388344 | -0.184209227 | 0.143 | 0.31 | 1 | 0.762190598 | -0.289620068 | 0.4 | 0.336 | 1 | 0.762190598 | 0.300080938 |
| Pnma8b | 0.800649369 | 0.503846766 | 0.048 | 0.063 | 1 | 0.163399792 | -0.271123724 | 0.022 | 0.083 | 1 | 0.800649369 | 0.300100093 |
| Zfp956 | 0.163434053 | -1.081115734 | 0 | 0.086 | 1 | 0.496490519 | -0.16492432 | 0.067 | 0.1 | 1 | 0.496490519 | 0.300157417 |
| Scamp5 | 0.163434053 | -1.02666795 | 0 | 0.086 | 1 | 0.183618787 | -1.017367132 | 0.022 | 0.076 | 1 | 0.183618787 | 0.300157417 |
| Rdh5 | 0.163434053 | -1.133583154 | 0 | 0.086 | 1 | 0.420164314 | -0.271123724 | 0.044 | 0.08 | 1 | 0.420164314 | 0.300157417 |
| Zbtb12 | 0.163434053 | -1.02666795 | 0 | 0.086 | 1 | 0.208370172 | -0.86124793 | 0.044 | 0.104 | 1 | 0.208370172 | 0.300157417 |
| Ap2a1 | 0.163444289 | -0.531342662 | 0.286 | 0.459 | 1 | 0.402932245 | 0.531660882 | 0.356 | 0.554 | 1 | 0.402932245 | 0.300174542 |
| Entpd6 | 0.163483795 | -1.133583154 | 0 | 0.086 | 1 | 0.199734354 | 0.304560963 | 0.022 | 0.076 | 1 | 0.199734354 | 0.300240639 |
| Ficd | 0.163484033 | -1.184209227 | 0 | 0.086 | 1 | 0.186612308 | 0.19121949 | 0.067 | 0.145 | 1 | 0.186612308 | 0.300241037 |
| Rhobtb2 | 0.163484033 | -1.280424543 | 0 | 0.086 | 1 | 0.439307027 | -0.486852415 | 0.044 | 0.076 | 1 | 0.439307027 | 0.300241037 |
| Mro | 0.163484033 | -1.081115734 | 0 | 0.086 | 1 | 0.331922009 | -0.124282336 | 0 | 0.021 | 1 | 0.331922009 | 0.300241037 |
| Usp33 | 0.163514466 | 0.679246898 | 0.571 | 0.53 | 1 | 0.4447314 | 0.597048842 | 0.733 | 0.734 | 1 | 0.4447314 | 0.300291952 |
| Mrnip | 0.16352901 | -1.280424543 | 0 | 0.086 | 1 | 0.803858895 | 1.583536913 | 0.089 | 0.111 | 1 | 0.803858895 | 0.300316282 |
| Catsperd | 0.163567557 | -1.233118828 | 0 | 0.086 | 1 | 0.69751685 | 1.567595369 | 0.067 | 0.055 | 1 | 0.69751685 | 0.300380768 |
| Abcd1 | 0.163568985 | -1.413691074 | 0 | 0.086 | 1 | 0.384912978 | 0.426732833 | 0.067 | 0.118 | 1 | 0.384912978 | 0.300383156 |
| BC049715 | 0.16356946 | -1.455511249 | 0 | 0.086 | 1 | 0.972325608 | 1.595609745 | 0.044 | 0.045 | 1 | 0.972325608 | 0.300383952 |
| Hook2 | 0.163606338 | -1.233118828 | 0 | 0.086 | 1 | 0.559680974 | 1.03099589 | 0.133 | 0.104 | 1 | 0.559680974 | 0.300445643 |
| Adra2c | 0.163630367 | -1.370622352 | 0 | 0.086 | 1 | 0.496223879 | -0.316927414 | 0 | 0.01 | 1 | 0.496223879 | 0.300485836 |
| Lypd1 | 0.163638217 | -1.752492987 | 0 | 0.086 | 1 | 0.552268171 | 0.42003818 | 0.022 | 0.042 | 1 | 0.552268171 | 0.300498968 |
| Sergef | 0.163662244 | -1.574155746 | 0 | 0.086 | 1 | 0.885240007 | 0.612683258 | 0.089 | 0.1 | 1 | 0.885240007 | 0.300539157 |
| Mmp23 | 0.163662481 | -1.455511249 | 0 | 0.086 | 1 | 0.744358033 | 0.223640968 | 0.022 | 0.031 | 1 | 0.744358033 | 0.300539555 |
| Ggta1 | 0.351234556 | -0.752492987 | 0.048 | 0.112 | 1 | 0.163691309 | -0.149817428 | 0.044 | 0.118 | 1 | 0.351234556 | 0.300587773 |
| Trpc3 | 0.16370054 | -1.683780237 | 0 | 0.086 | 1 | 0.672954772 | 0.825091591 | 0.089 | 0.073 | 1 | 0.672954772 | 0.300603213 |
| Ksr1 | 0.163713146 | -2.02666795 | 0 | 0.086 | 1 | 0.751936505 | 1.726141308 | 0.089 | 0.08 | 1 | 0.751936505 | 0.300624298 |
| Xlr3b | 0.163722898 | -1.718545655 | 0 | 0.086 | 1 | 0.446742942 | 0.15951063 | 0.022 | 0.048 | 1 | 0.446742942 | 0.300640608 |
| Trmt5 | 0.163800426 | -0.911190733 | 0 | 0.086 | 1 | 0.328123368 | -0.316927414 | 0.022 | 0.059 | 1 | 0.328123368 | 0.300770273 |
| Wdr41 | 0.613448612 | 0.673771768 | 0.19 | 0.291 | 1 | 0.163849922 | 0.449275402 | 0.156 | 0.294 | 1 | 0.613448612 | 0.300853048 |
| Dmac2l | 0.741239269 | 0.162809848 | 0.143 | 0.175 | 1 | 0.163866635 | -0.960783604 | 0.044 | 0.114 | 1 | 0.741239269 | 0.300880996 |
| Ganc | 0.174775826 | -0.25696557 | 0.048 | 0.168 | 1 | 0.163927821 | -0.413788953 | 0.111 | 0.204 | 1 | 0.174775826 | 0.300983311 |
| Gm12236 | 0.675067981 | -0.280424543 | 0.048 | 0.071 | 1 | 0.164309241 | -1.223818009 | 0.022 | 0.08 | 1 | 0.675067981 | 0.301620955 |
| Barx2 | 0.20983778 | -0.763633545 | 0.048 | 0.153 | 1 | 0.164310937 | -1.743192169 | 0.022 | 0.08 | 1 | 0.20983778 | 0.30162379 |
| Hipk1 | 0.307470846 | 0.413133202 | 0.905 | 0.948 | 1 | 0.164334698 | -0.124282336 | 0.956 | 0.952 | 1 | 0.307470846 | 0.301663503 |
| Sox12 | 0.268011783 | -0.648156327 | 0 | 0.056 | 1 | 0.16484123 | -1.223818009 | 0.022 | 0.08 | 1 | 0.268011783 | 0.302509828 |
| Tmem104 | 0.542240288 | 1.425844254 | 0.095 | 0.063 | 1 | 0.16484123 | -1.223818009 | 0.022 | 0.08 | 1 | 0.542240288 | 0.302509828 |
| Dnajc17 | 0.938674785 | 0.629377648 | 0.095 | 0.104 | 1 | 0.16484123 | -1.174908409 | 0.022 | 0.08 | 1 | 0.938674785 | 0.302509828 |
| Fen1 | 0.196495505 | -1.133583154 | 0 | 0.075 | 1 | 0.164841909 | -1.124282336 | 0.022 | 0.08 | 1 | 0.196495505 | 0.302510963 |
| Wdr70 | 0.164908296 | 0.590171566 | 0.667 | 0.683 | 1 | 0.913222766 | 0.374819428 | 0.733 | 0.734 | 1 | 0.913222766 | 0.302621846 |
| Nat8f1 | 0.173729397 | -1.02666795 | 0 | 0.082 | 1 | 0.165306929 | -1.124282336 | 0 | 0.042 | 1 | 0.173729397 | 0.303287477 |
| Pcdhgc5 | 0.222378078 | -0.574155746 | 0 | 0.067 | 1 | 0.165306929 | -1.124282336 | 0 | 0.042 | 1 | 0.222378078 | 0.303287477 |
| Rad54b | 0.268010172 | -0.718545655 | 0 | 0.056 | 1 | 0.165306929 | -1.316927414 | 0 | 0.042 | 1 | 0.268010172 | 0.303287477 |
| Gm9887 | 0.344460627 | 1.503846766 | 0.095 | 0.049 | 1 | 0.165306929 | -1.124282336 | 0 | 0.042 | 1 | 0.344460627 | 0.303287477 |
| Zfp28 | 0.425599726 | 0.351843673 | 0 | 0.03 | 1 | 0.165306929 | -1.124282336 | 0 | 0.042 | 1 | 0.425599726 | 0.303287477 |
| Asgr1 | 0.425599726 | 0.214340149 | 0 | 0.03 | 1 | 0.165306929 | -1.124282336 | 0 | 0.042 | 1 | 0.425599726 | 0.303287477 |
| Gm56698 | 0.466343532 | 1.673771768 | 0.095 | 0.06 | 1 | 0.165306929 | -1.223818009 | 0 | 0.042 | 1 | 0.466343532 | 0.303287477 |
| 1700037C18Rik | 0.532426384 | 0.866416846 | 0 | 0.019 | 1 | 0.165306929 | -1.223818009 | 0 | 0.042 | 1 | 0.532426384 | 0.303287477 |
| Ttpa | 0.57816106 | 1.351843673 | 0 | 0.015 | 1 | 0.165306929 | -1.223818009 | 0 | 0.042 | 1 | 0.57816106 | 0.303287477 |
| Fchsd1 | 0.57816106 | 1.351843673 | 0 | 0.015 | 1 | 0.165306929 | -1.124282336 | 0 | 0.042 | 1 | 0.57816106 | 0.303287477 |
| Faim2 | 0.632076134 | 0.866416846 | 0 | 0.011 | 1 | 0.165306929 | -1.316927414 | 0 | 0.042 | 1 | 0.632076134 | 0.303287477 |
| A830008E24Rik | 0.656051514 | 2.708537186 | 0.048 | 0.078 | 1 | 0.165306929 | -1.124282336 | 0 | 0.042 | 1 | 0.656051514 | 0.303287477 |
| Zfp870 | 0.71266839 | 1.79930265 | 0.048 | 0.034 | 1 | 0.165306929 | -1.124282336 | 0 | 0.042 | 1 | 0.71266839 | 0.303287477 |
| Inhca | 0.904673267 | 0.936806174 | 0.048 | 0.056 | 1 | 0.165306929 | -1.124282336 | 0 | 0.042 | 1 | 0.904673267 | 0.303287477 |
| Spns2 | 0.184926613 | -1.574155746 | 0 | 0.078 | 1 | 0.165324946 | -1.223818009 | 0 | 0.042 | 1 | 0.184926613 | 0.303317554 |
| Garin5a | 0.267953794 | -0.496153234 | 0 | 0.056 | 1 | 0.165324946 | -1.223818009 | 0 | 0.042 | 1 | 0.267953794 | 0.303317554 |
| Egr2 | 0.297035631 | 0.400753273 | 0.143 | 0.078 | 1 | 0.165324946 | -2.526380779 | 0 | 0.042 | 1 | 0.297035631 | 0.303317554 |
| Gpr19 | 0.324739547 | -0.133583154 | 0 | 0.045 | 1 | 0.165324946 | -1.223818009 | 0 | 0.042 | 1 | 0.324739547 | 0.303317554 |
| Gdpgp1 | 0.346827513 | -0.133583154 | 0 | 0.041 | 1 | 0.165324946 | -1.223818009 | 0 | 0.042 | 1 | 0.346827513 | 0.303317554 |
| C230034O21Rik | 0.376794341 | 1.866416846 | 0.048 | 0.019 | 1 | 0.165324946 | -1.223818009 | 0 | 0.042 | 1 | 0.376794341 | 0.303317554 |
| Tyw3 | 0.38281368 | 1.766881172 | 0.095 | 0.052 | 1 | 0.165324946 | -1.223818009 | 0 | 0.042 | 1 | 0.38281368 | 0.303317554 |
| Tulp2 | 0.39694807 | -0.133583154 | 0 | 0.034 | 1 | 0.165324946 | -1.223818009 | 0 | 0.042 | 1 | 0.39694807 | 0.303317554 |
| Sytl4 | 0.425599726 | 0.351843673 | 0 | 0.03 | 1 | 0.165324946 | -1.223818009 | 0 | 0.042 | 1 | 0.425599726 | 0.303317554 |
| Irf8 | 0.466502439 | 0.815790773 | 0.095 | 0.056 | 1 | 0.165324946 | -1.486852415 | 0 | 0.042 | 1 | 0.466502439 | 0.303317554 |
| Frat2 | 0.492590603 | 0.673771768 | 0 | 0.022 | 1 | 0.165324946 | -1.223818009 | 0 | 0.042 | 1 | 0.492590603 | 0.303317554 |
| Saxo2 | 0.492608234 | 0.866416846 | 0 | 0.022 | 1 | 0.165324946 | -1.316927414 | 0 | 0.042 | 1 | 0.492608234 | 0.303317554 |
| Prss53 | 0.536308039 | 0.866416846 | 0.095 | 0.06 | 1 | 0.165324946 | -1.316927414 | 0 | 0.042 | 1 | 0.536308039 | 0.303317554 |
| Gm43112 | 0.57816106 | 1.351843673 | 0 | 0.015 | 1 | 0.165324946 | -1.404390255 | 0 | 0.042 | 1 | 0.57816106 | 0.303317554 |
| Zfp54 | 0.57816106 | 1.351843673 | 0 | 0.015 | 1 | 0.165324946 | -1.223818009 | 0 | 0.042 | 1 | 0.57816106 | 0.303317554 |
| Psd | 0.629700127 | 2.088809267 | 0.048 | 0.03 | 1 | 0.165324946 | -1.316927414 | 0 | 0.042 | 1 | 0.629700127 | 0.303317554 |
| Zfp114 | 0.63207476 | 1.351843673 | 0 | 0.011 | 1 | 0.165324946 | -1.223818009 | 0 | 0.042 | 1 | 0.63207476 | 0.303317554 |
| Aldh3b1 | 0.632917461 | 1.936806174 | 0.048 | 0.03 | 1 | 0.165324946 | -1.223818009 | 0 | 0.042 | 1 | 0.632917461 | 0.303317554 |
| Gm56938 | 0.686271924 | 0.614878079 | 0.048 | 0.075 | 1 | 0.165324946 | -1.223818009 | 0 | 0.042 | 1 | 0.686271924 | 0.303317554 |
| Dennd11 | 0.89926261 | 0.79930265 | 0.048 | 0.056 | 1 | 0.165324946 | -1.638855509 | 0 | 0.042 | 1 | 0.89926261 | 0.303317554 |
| Gm15862 | 0.918789496 | 1.55829455 | 0.048 | 0.045 | 1 | 0.165324946 | -1.316927414 | 0 | 0.042 | 1 | 0.918789496 | 0.303317554 |
| Gm26881 | 0.918798858 | 1.766881172 | 0.048 | 0.045 | 1 | 0.165324946 | -1.223818009 | 0 | 0.042 | 1 | 0.918798858 | 0.303317554 |
| Elfn1 | 0.266771227 | 2.451379346 | 0.048 | 0.015 | 1 | 0.165339358 | -1.564854927 | 0 | 0.042 | 1 | 0.266771227 | 0.303341613 |
| Lpar6 | 0.41354048 | 1.088809267 | 0.095 | 0.052 | 1 | 0.165339358 | -1.404390255 | 0 | 0.042 | 1 | 0.41354048 | 0.303341613 |

| | | | | | | | | | | | | |
|---|---|---|---|---|---|---|---|---|---|---|---|---|
| Gm26555 | 0.479147948 | 1.503846766 | 0.048 | 0.022 | 1 | 0.165339358 | -1.404390255 | 0 | 0.042 | 1 | 0.479147948 | 0.303341613 |
| Arhgap4 | 0.492590603 | 0.503846766 | 0 | 0.022 | 1 | 0.165339358 | -1.404390255 | 0 | 0.042 | 1 | 0.492590603 | 0.303341613 |
| Abraxas1 | 0.185001031 | -0.785659851 | 0 | 0.078 | 1 | 0.165339686 | -1.316927414 | 0 | 0.042 | 1 | 0.185001031 | 0.30334216 |
| Rnft2 | 0.304355016 | -0.326228232 | 0 | 0.049 | 1 | 0.165339686 | -1.316927414 | 0 | 0.042 | 1 | 0.304355016 | 0.30334216 |
| Chaf1a | 0.304431942 | -0.648156327 | 0 | 0.049 | 1 | 0.165339686 | -1.316927414 | 0 | 0.042 | 1 | 0.304431942 | 0.30334216 |
| Gm47469 | 0.346802964 | -0.133583154 | 0 | 0.041 | 1 | 0.165339686 | -1.316927414 | 0 | 0.042 | 1 | 0.346802964 | 0.30334216 |
| Nudt17 | 0.57672979 | 1.088809267 | 0.048 | 0.026 | 1 | 0.165339686 | -1.316927414 | 0 | 0.042 | 1 | 0.57672979 | 0.30334216 |
| Gpt | 0.57816106 | 1.351843673 | 0 | 0.015 | 1 | 0.165339686 | -1.486852415 | 0 | 0.042 | 1 | 0.57816106 | 0.30334216 |
| Olfml2a | 0.532435374 | 0.503846766 | 0 | 0.019 | 1 | 0.16535115 | -1.564854927 | 0 | 0.042 | 1 | 0.532435374 | 0.303361297 |
| Chaf1b | 0.904667709 | 0.866416846 | 0.048 | 0.056 | 1 | 0.16535115 | -1.638855509 | 0 | 0.042 | 1 | 0.904667709 | 0.303361297 |
| Lypd6 | 0.970926602 | 0.281454345 | 0.048 | 0.049 | 1 | 0.16535115 | -1.901889915 | 0 | 0.042 | 1 | 0.970926602 | 0.303361297 |
| Gm27253 | 0.186194818 | 2.088809267 | 0.095 | 0.037 | 1 | 0.165351478 | -1.404390255 | 0 | 0.042 | 1 | 0.186194818 | 0.303361844 |
| Klhl3 | 0.547777007 | 2.088809267 | 0.048 | 0.026 | 1 | 0.165351478 | -1.404390255 | 0 | 0.042 | 1 | 0.547777007 | 0.303361844 |
| Lingo1 | 0.638246407 | -0.184209227 | 0.048 | 0.075 | 1 | 0.165359339 | -1.709244837 | 0 | 0.042 | 1 | 0.638246407 | 0.303374966 |
| Rbpms2 | 0.793508072 | 1.451379346 | 0.048 | 0.037 | 1 | 0.165359994 | -1.84048937 | 0 | 0.042 | 1 | 0.793508072 | 0.30337606 |
| Dok5 | 0.623342453 | 0.521768674 | 0.143 | 0.112 | 1 | 0.165360321 | -2.071814916 | 0 | 0.042 | 1 | 0.623342453 | 0.303376607 |
| Krt16 | 0.952887277 | -2.796548167 | 0.095 | 0.093 | 1 | 0.165360321 | -5.065120263 | 0 | 0.042 | 1 | 0.952887277 | 0.303376607 |
| Pabpc4l | 0.23647819 | -0.911190733 | 0 | 0.063 | 1 | 0.165360649 | -1.486852415 | 0 | 0.042 | 1 | 0.23647819 | 0.303377153 |
| Axdnd1 | 0.346860242 | -0.574155746 | 0 | 0.041 | 1 | 0.165360649 | -1.960783604 | 0 | 0.042 | 1 | 0.346860242 | 0.303377153 |
| Dnaaf3 | 0.492599419 | 0.351843673 | 0 | 0.022 | 1 | 0.165360649 | -1.486852415 | 0 | 0.042 | 1 | 0.492599419 | 0.303377153 |
| Gm1604b | 0.457292877 | 0.351843673 | 0 | 0.026 | 1 | 0.165364252 | -1.709244837 | 0 | 0.042 | 1 | 0.457292877 | 0.303383168 |
| Gngt2 | 0.285551097 | -0.911190733 | 0 | 0.052 | 1 | 0.165368837 | -1.960783604 | 0 | 0.042 | 1 | 0.285551097 | 0.303390822 |
| Shisal1 | 0.304469698 | -1.184209227 | 0 | 0.049 | 1 | 0.165369165 | -2.271123724 | 0 | 0.042 | 1 | 0.304469698 | 0.303391369 |
| Slc25a23 | 0.492590603 | 0.673771768 | 0 | 0.022 | 1 | 0.165370849 | -1.071814916 | 0.022 | 0.08 | 1 | 0.492590603 | 0.303394181 |
| Panx3 | 0.221028022 | -3.067695219 | 0.048 | 0.142 | 1 | 0.165377353 | -4.931637258 | 0 | 0.042 | 1 | 0.221028022 | 0.303405037 |
| 9530077C14Rik | 0.236343578 | -0.574155746 | 0 | 0.063 | 1 | 0.16537899 | -1.017367132 | 0 | 0.042 | 1 | 0.236343578 | 0.30340777 |
| Tstd3 | 0.285622926 | -0.233118828 | 0 | 0.052 | 1 | 0.16537899 | -1.017367132 | 0 | 0.042 | 1 | 0.285622926 | 0.30340777 |
| Pcdhgb7 | 0.346827513 | -0.133583154 | 0 | 0.041 | 1 | 0.16537899 | -1.017367132 | 0 | 0.042 | 1 | 0.346827513 | 0.30340777 |
| Gm10734 | 0.346846605 | -0.326228232 | 0 | 0.041 | 1 | 0.16537899 | -1.017367132 | 0 | 0.042 | 1 | 0.346846605 | 0.30340777 |
| Clic6 | 0.359286409 | 3.021695071 | 0.048 | 0.019 | 1 | 0.16537899 | -1.017367132 | 0 | 0.042 | 1 | 0.359286409 | 0.30340777 |
| Tmem198 | 0.370819763 | -0.233118828 | 0 | 0.037 | 1 | 0.16537899 | -1.017367132 | 0 | 0.042 | 1 | 0.370819763 | 0.30340777 |
| Lrrc75b | 0.425639612 | 0.503846766 | 0 | 0.03 | 1 | 0.16537899 | -1.017367132 | 0 | 0.042 | 1 | 0.425639612 | 0.30340777 |
| Acvrl1 | 0.457281049 | 0.503846766 | 0 | 0.026 | 1 | 0.16537899 | -1.017367132 | 0 | 0.042 | 1 | 0.457281049 | 0.30340777 |
| Gm29491 | 0.492590603 | 0.673771768 | 0 | 0.022 | 1 | 0.16537899 | -1.017367132 | 0 | 0.042 | 1 | 0.492590603 | 0.30340777 |
| Gm10135 | 0.492608234 | 0.866416846 | 0 | 0.022 | 1 | 0.16537899 | -1.017367132 | 0 | 0.042 | 1 | 0.492608234 | 0.30340777 |
| Moap1 | 0.492608234 | 0.866416846 | 0 | 0.022 | 1 | 0.16537899 | -1.017367132 | 0 | 0.042 | 1 | 0.492608234 | 0.30340777 |
| Gm36266 | 0.54776222 | 2.088809267 | 0.048 | 0.026 | 1 | 0.16537899 | -1.017367132 | 0 | 0.042 | 1 | 0.54776222 | 0.30340777 |
| Arhgef33 | 0.63207476 | 1.351843673 | 0 | 0.011 | 1 | 0.16537899 | -1.017367132 | 0 | 0.042 | 1 | 0.63207476 | 0.30340777 |
| Cnksr1 | 0.784037261 | 1.79930265 | 0.048 | 0.037 | 1 | 0.16537899 | -1.017367132 | 0 | 0.042 | 1 | 0.784037261 | 0.30340777 |
| S100a7a | 0.95783752 | 0.866416846 | 0.048 | 0.045 | 1 | 0.16537899 | -1.017367132 | 0 | 0.042 | 1 | 0.95783752 | 0.30340777 |
| Msra | 0.970090906 | 1.188344941 | 0.238 | 0.291 | 1 | 0.165452931 | 0.571362199 | 0.511 | 0.384 | 1 | 0.970090906 | 0.30353119 |
| Bclaf3 | 0.777677906 | 1.235650655 | 0.333 | 0.373 | 1 | 0.165464546 | -0.429627547 | 0.378 | 0.509 | 1 | 0.777677906 | 0.303550576 |
| Gm16150 | 0.16548778 | 3.258734268 | 0.048 | 0.011 | 1 | 0.331922009 | -0.124282336 | 0 | 0.021 | 1 | 0.331922009 | 0.303589355 |
| Oip5 | 0.16548778 | 3.258734268 | 0.048 | 0.011 | 1 | 0.49622258 | 0.361144491 | 0 | 0.01 | 1 | 0.49622258 | 0.303589355 |
| Itpka | 0.16548778 | 3.258734268 | 0.048 | 0.011 | 1 | 0.331914654 | -0.638855509 | 0 | 0.021 | 1 | 0.331914654 | 0.303589355 |
| Miat | 0.16548778 | 3.673771768 | 0.048 | 0.011 | 1 | 0.955025729 | 1.361144491 | 0.022 | 0.024 | 1 | 0.955025729 | 0.303589355 |
| Tmem130 | 0.16548778 | 4.258734268 | 0.048 | 0.011 | 1 | 0.49622258 | 0.361144491 | 0 | 0.01 | 1 | 0.49622258 | 0.303589355 |
| Trim66 | 0.16548778 | 3.258734268 | 0.048 | 0.011 | 1 | 0.29365335 | -0.776359032 | 0 | 0.024 | 1 | 0.29365335 | 0.303589355 |
| Tgfbr3l | 0.16548778 | 3.673771768 | 0.048 | 0.011 | 1 | 0.430487676 | 0.361144491 | 0 | 0.014 | 1 | 0.430487676 | 0.303589355 |
| Gm45820 | 0.16548778 | 3.673771768 | 0.048 | 0.011 | 1 | 0.496223879 | 0.683072586 | 0 | 0.01 | 1 | 0.496223879 | 0.303589355 |
| Spata2l | 0.16548778 | 3.258734268 | 0.048 | 0.011 | 1 | 0.293659664 | -0.316927414 | 0 | 0.024 | 1 | 0.293659664 | 0.303589355 |
| Gm26631 | 0.16548778 | 3.258734268 | 0.048 | 0.011 | 1 | 0.941347173 | 1.098110085 | 0.022 | 0.021 | 1 | 0.941347173 | 0.303589355 |
| Gm35721 | 0.16548778 | 3.258734268 | 0.048 | 0.011 | 1 | 0.317138221 | 1.683072586 | 0.022 | 0.007 | 1 | 0.317138221 | 0.303589355 |
| Fhl4 | 0.16548778 | 3.258734268 | 0.048 | 0.011 | 1 | 0.941346407 | 0.946106992 | 0.022 | 0.021 | 1 | 0.941346407 | 0.303589355 |
| Rundc3a | 0.16548778 | 3.258734268 | 0.048 | 0.011 | 1 | 0.8202424 | 1.098110085 | 0.022 | 0.017 | 1 | 0.8202424 | 0.303589355 |
| 4930544I03Rik | 0.16548778 | 3.258734268 | 0.048 | 0.011 | 1 | 0.496223879 | -0.638855509 | 0 | 0.01 | 1 | 0.496223879 | 0.303589355 |
| Gm48250 | 0.16548778 | 3.258734268 | 0.048 | 0.011 | 1 | 0.312251526 | 2.361144491 | 0.022 | 0.007 | 1 | 0.312251526 | 0.303589355 |
| Marveld2 | 0.16548778 | 4.258734268 | 0.048 | 0.011 | 1 | 0.8202424 | 1.098110085 | 0.022 | 0.017 | 1 | 0.8202424 | 0.303589355 |
| Gm41335 | 0.16548778 | 3.258734268 | 0.048 | 0.011 | 1 | 0.496223879 | -1.316927414 | 0 | 0.01 | 1 | 0.496223879 | 0.303589355 |
| Gm41396 | 0.16548778 | 3.258734268 | 0.048 | 0.011 | 1 | 0.293637566 | -0.486852415 | 0 | 0.024 | 1 | 0.293637566 | 0.303589355 |
| Csnka2ip | 0.16548778 | 3.258734268 | 0.048 | 0.011 | 1 | 0.496223879 | -2.017367132 | 0 | 0.01 | 1 | 0.496223879 | 0.303589355 |
| Gng5c | 0.16548778 | 3.258734268 | 0.048 | 0.011 | 1 | 0.555515361 | 0.567595369 | 0.022 | 0.042 | 1 | 0.555515361 | 0.303589355 |
| Gm17315 | 0.16548778 | 3.258734268 | 0.048 | 0.011 | 1 | 0.293637566 | -0.486852415 | 0 | 0.024 | 1 | 0.293637566 | 0.303589355 |
| 1700061H18Rik | 0.16548778 | 4.258734268 | 0.048 | 0.011 | 1 | 0.314689364 | 1.460680165 | 0.022 | 0.007 | 1 | 0.314689364 | 0.303589355 |
| Rnf165 | 0.16548778 | 3.258734268 | 0.048 | 0.011 | 1 | 0.494823351 | 1.683072586 | 0.022 | 0.01 | 1 | 0.494823351 | 0.303589355 |
| Gm17227 | 0.16548778 | 3.258734268 | 0.048 | 0.011 | 1 | 0.309824707 | 2.683072586 | 0.022 | 0.007 | 1 | 0.309824707 | 0.303589355 |
| Gm9895 | 0.16548778 | 3.258734268 | 0.048 | 0.011 | 1 | 0.49622258 | 0.361144491 | 0 | 0.01 | 1 | 0.49622258 | 0.303589355 |
| Iqca | 0.165491632 | 3.088809267 | 0.048 | 0.011 | 1 | 0.496223879 | 0.683072586 | 0 | 0.01 | 1 | 0.496223879 | 0.303595783 |
| Plpp7 | 0.165491632 | 3.351843673 | 0.048 | 0.011 | 1 | 0.331216021 | 0.513147585 | 0.022 | 0.059 | 1 | 0.331216021 | 0.303595783 |
| Jph2 | 0.165491632 | 4.133203386 | 0.048 | 0.011 | 1 | 0.50040834 | 1.683072586 | 0.022 | 0.01 | 1 | 0.50040834 | 0.303595783 |
| Tnfaip8l1 | 0.165491632 | 4.159198595 | 0.048 | 0.011 | 1 | 0.492043882 | 2.683072586 | 0.022 | 0.01 | 1 | 0.492043882 | 0.303595783 |
| St8sia5 | 0.165491632 | 3.936806174 | 0.048 | 0.011 | 1 | 0.317138221 | 1.683072586 | 0.022 | 0.007 | 1 | 0.317138221 | 0.303595783 |
| Zfp957 | 0.165492915 | 3.258734268 | 0.048 | 0.011 | 1 | 0.675805699 | 0.875717664 | 0.022 | 0.014 | 1 | 0.675805699 | 0.303597926 |
| Ermard | 0.165536654 | 1.323274521 | 0.238 | 0.146 | 1 | 0.449203111 | 0.607784459 | 0.111 | 0.17 | 1 | 0.449203111 | 0.303670923 |
| Ninj2 | 0.544557277 | 0.794626163 | 0.571 | 0.608 | 1 | 0.165541586 | 0.458441693 | 0.333 | 0.543 | 1 | 0.544557277 | 0.303679155 |
| Atp1a2 | 0.196592969 | -2.611630451 | 0 | 0.075 | 1 | 0.165804868 | 0.223640968 | 0.089 | 0.18 | 1 | 0.196592969 | 0.304118483 |
| Gm6225 | 0.165848734 | 1.297904866 | 0.333 | 0.239 | 1 | 0.874668828 | 0.470078863 | 0.178 | 0.201 | 1 | 0.874668828 | 0.304191666 |
| Dolpp1 | 0.236343578 | -0.648156327 | 0 | 0.063 | 1 | 0.165897576 | -1.223818009 | 0.022 | 0.08 | 1 | 0.236343578 | 0.304273147 |
| Ttll12 | 0.810632857 | 0.735172312 | 0.048 | 0.063 | 1 | 0.165897576 | -1.017367132 | 0.022 | 0.08 | 1 | 0.810632857 | 0.304273147 |
| E230016M11Rik | 0.881937898 | 1.171271427 | 0.19 | 0.198 | 1 | 0.166001322 | 0.771609261 | 0.2 | 0.332 | 1 | 0.881937898 | 0.304446204 |
| Mterf4 | 0.242832356 | -0.233118828 | 0.048 | 0.146 | 1 | 0.166083211 | -0.286553765 | 0.067 | 0.145 | 1 | 0.242832356 | 0.304582788 |
| Axin2 | 0.458438075 | 0.150209812 | 0.19 | 0.299 | 1 | 0.166198243 | -0.153428682 | 0.2 | 0.343 | 1 | 0.458438075 | 0.304774631 |
| Cyb561a3 | 0.401793043 | -0.326228232 | 0.048 | 0.104 | 1 | 0.166421908 | -0.960783604 | 0.022 | 0.08 | 1 | 0.401793043 | 0.305147564 |
| Gm56555 | 0.825533132 | 0.123574685 | 0.143 | 0.175 | 1 | 0.166440008 | -0.286553765 | 0.044 | 0.118 | 1 | 0.825533132 | 0.30517774 |
| Sox4 | 0.166588696 | -0.978839425 | 0.381 | 0.5 | 1 | 0.731236156 | -0.845103888 | 0.756 | 0.592 | 1 | 0.731236156 | 0.305425599 |
| Plag1 | 0.4673821 | 0.281454345 | 0.048 | 0.101 | 1 | 0.166608388 | -0.383016604 | 0.044 | 0.118 | 1 | 0.4673821 | 0.305458421 |
| Tom1l1 | 0.20895393 | -1.133583154 | 0 | 0.071 | 1 | 0.166628448 | -0.709244837 | 0.044 | 0.114 | 1 | 0.20895393 | 0.305491857 |
| Cplx2 | 0.236481694 | -0.849790188 | 0 | 0.063 | 1 | 0.166723258 | -0.518561275 | 0.044 | 0.114 | 1 | 0.236481694 | 0.305649872 |
| Gm16351 | 0.869730687 | -1.034573149 | 0.143 | 0.16 | 1 | 0.166852708 | -0.647317087 | 0.178 | 0.294 | 1 | 0.869730687 | 0.305865591 |
| Ankzf1 | 0.556479065 | 0.766881172 | 0.095 | 0.149 | 1 | 0.1669645 | -1.053893008 | 0.044 | 0.111 | 1 | 0.556479065 | 0.306051856 |
| Tsc22d2 | 0.167073143 | 0.708919711 | 0.857 | 0.881 | 1 | 0.215336201 | 0.539093058 | 0.956 | 0.882 | 1 | 0.215336201 | 0.306232852 |
| Hectd2os | 0.854703644 | 2.362570079 | 0.19 | 0.205 | 1 | 0.167140595 | -0.609709163 | 0.044 | 0.118 | 1 | 0.854703644 | 0.306345211 |
| Pde4c | 0.726653001 | 2.109870882 | 0.095 | 0.078 | 1 | 0.167164655 | 1.098110085 | 0.089 | 0.042 | 1 | 0.726653001 | 0.306385288 |
| Fah | 0.95783752 | 0.866416846 | 0.048 | 0.045 | 1 | 0.167336193 | -0.901889915 | 0.022 | 0.08 | 1 | 0.95783752 | 0.306670984 |
| Nsmf | 0.960298344 | 0.405282932 | 0.286 | 0.321 | 1 | 0.167349198 | 0.464237984 | 0.2 | 0.353 | 1 | 0.960298344 | 0.306692642 |
| Ppp1r13b | 0.48516078 | 0.146524765 | 0.286 | 0.429 | 1 | 0.167380501 | 0.696011642 | 0.756 | 0.671 | 1 | 0.48516078 | 0.30674477 |
| Arih2 | 0.189356878 | 0.939062147 | 0.619 | 0.75 | 1 | 0.167477424 | 0.591150097 | 0.889 | 0.796 | 1 | 0.189356878 | 0.30690616 |
| Gm30455 | 0.167544534 | 2.936806174 | 0.048 | 0.011 | 1 | 0.293659664 | -0.316927414 | 0 | 0.024 | 1 | 0.293659664 | 0.307017897 |

| | | | | | | | | | | | | |
|---|---|---|---|---|---|---|---|---|---|---|---|---|
| Chrnb2 | 0.167544534 | 2.936806174 | 0.048 | 0.011 | 1 | 0.50040834 | 1.683072586 | 0.022 | 0.01 | 1 | 0.50040834 | 0.307017897 |
| Gm128 | 0.167544534 | 2.936806174 | 0.048 | 0.011 | 1 | 0.42222537 | 1.460680165 | 0.067 | 0.042 | 1 | 0.42222537 | 0.307017897 |
| 9130230L23Rik | 0.167544534 | 2.936806174 | 0.048 | 0.011 | 1 | 0.887849977 | 2.055041364 | 0.044 | 0.042 | 1 | 0.887849977 | 0.307017897 |
| Art4 | 0.167544534 | 2.936806174 | 0.048 | 0.011 | 1 | 0.496223879 | 0.683072586 | 0 | 0.01 | 1 | 0.496223879 | 0.307017897 |
| Cblc | 0.167544534 | 2.936806174 | 0.048 | 0.011 | 1 | 0.430483494 | -0.124282336 | 0 | 0.014 | 1 | 0.430483494 | 0.307017897 |
| Gm57038 | 0.167544534 | 2.936806174 | 0.048 | 0.011 | 1 | 0.765720982 | 0.946106992 | 0.022 | 0.031 | 1 | 0.765720982 | 0.307017897 |
| Jag2 | 0.167544534 | 2.936806174 | 0.048 | 0.011 | 1 | 0.293637566 | -0.638855509 | 0 | 0.024 | 1 | 0.293637566 | 0.307017897 |
| Amph | 0.167544534 | 2.936806174 | 0.048 | 0.011 | 1 | 0.314689364 | 1.513147585 | 0.022 | 0.007 | 1 | 0.314689364 | 0.307017897 |
| Gm27010 | 0.167544534 | 2.936806174 | 0.048 | 0.011 | 1 | 0.617514334 | 1.098110085 | 0.022 | 0.038 | 1 | 0.617514334 | 0.307017897 |
| 2600014E21Rik | 0.167601721 | 0.36999102 | 0.048 | 0.172 | 1 | 0.260665169 | -2.223818009 | 0 | 0.028 | 1 | 0.260665169 | 0.307113106 |
| Gm57325 | 0.268043994 | -0.849790188 | 0 | 0.056 | 1 | 0.167651965 | 1.83507568 | 0.067 | 0.028 | 1 | 0.268043994 | 0.307196748 |
| Gm6277 | 0.677300419 | -0.280424543 | 0.048 | 0.071 | 1 | 0.167651965 | 2.320502507 | 0.067 | 0.028 | 1 | 0.677300419 | 0.307196748 |
| Lrrc4 | 0.167667581 | 1.631951592 | 0.286 | 0.194 | 1 | 0.393093379 | 1.481438725 | 0.222 | 0.197 | 1 | 0.393093379 | 0.307222745 |
| Dpp8 | 0.167718862 | 0.181918671 | 0.333 | 0.578 | 1 | 0.409287903 | 0.122632405 | 0.711 | 0.775 | 1 | 0.409287903 | 0.307308108 |
| Prmt2 | 0.167724974 | -0.834022873 | 0.095 | 0.228 | 1 | 0.434265722 | 0.439147003 | 0.178 | 0.249 | 1 | 0.434265722 | 0.307318282 |
| Ercc4 | 0.312426684 | -0.683780237 | 0.048 | 0.119 | 1 | 0.168100468 | -0.316927414 | 0.089 | 0.176 | 1 | 0.312426684 | 0.307943168 |
| Gm43948 | 0.7027822 | 1.410737362 | 0.095 | 0.078 | 1 | 0.168100796 | 1.545569062 | 0.111 | 0.059 | 1 | 0.7027822 | 0.307943714 |
| Rrp12 | 0.720688542 | 0.815790773 | 0.19 | 0.179 | 1 | 0.168145256 | 0.290755163 | 0.089 | 0.183 | 1 | 0.720688542 | 0.308017685 |
| Shroom2 | 0.168187411 | -0.695462042 | 0.095 | 0.231 | 1 | 0.919793098 | -0.699397051 | 0.356 | 0.315 | 1 | 0.919793098 | 0.308087817 |
| 1700085D22Rik | 0.849946817 | 1.041503552 | 0.095 | 0.086 | 1 | 0.168202754 | -0.191396532 | 0.044 | 0.114 | 1 | 0.849946817 | 0.308113342 |
| Ndrg4 | 0.333055137 | -0.133583154 | 0.048 | 0.123 | 1 | 0.168207004 | -0.255526869 | 0.044 | 0.114 | 1 | 0.333055137 | 0.308120412 |
| Parp4 | 0.525686837 | -0.203972482 | 0.238 | 0.321 | 1 | 0.168352939 | 0.699313227 | 0.6 | 0.471 | 1 | 0.525686837 | 0.308363165 |
| Nbeal1 | 0.168666842 | 0.658396542 | 0.905 | 0.937 | 1 | 0.788015576 | 0.337685518 | 0.911 | 0.931 | 1 | 0.788015576 | 0.30888518 |
| Rc3h2 | 0.413827893 | 0.54955974 | 0.667 | 0.81 | 1 | 0.168785088 | 0.254835589 | 0.956 | 0.931 | 1 | 0.413827893 | 0.309081771 |
| Wdr47 | 0.168910077 | -0.880817084 | 0.143 | 0.306 | 1 | 0.361857031 | 0.971418081 | 0.378 | 0.381 | 1 | 0.361857031 | 0.30928954 |
| Tmem19 | 0.586671517 | 1.351843673 | 0.238 | 0.231 | 1 | 0.169042337 | -0.178221143 | 0.156 | 0.273 | 1 | 0.586671517 | 0.309509363 |
| Marchf8 | 0.276618401 | 0.487071053 | 0.238 | 0.437 | 1 | 0.16944799 | 0.205750809 | 0.356 | 0.543 | 1 | 0.276618401 | 0.310183359 |
| Capza1 | 0.169469854 | 0.167664688 | 0.619 | 0.84 | 1 | 0.67124251 | 0.216472282 | 0.889 | 0.896 | 1 | 0.67124251 | 0.310219676 |
| Runx2 | 0.594734867 | -0.553691643 | 0.762 | 0.799 | 1 | 0.169500451 | -0.475075247 | 0.889 | 0.844 | 1 | 0.594734867 | 0.310270499 |
| Wdhd1 | 0.983128998 | 1.247507013 | 0.095 | 0.097 | 1 | 0.169577374 | 0.268035087 | 0.044 | 0.118 | 1 | 0.983128998 | 0.310398262 |
| Gm16759 | 0.867636519 | 1.041503552 | 0.095 | 0.09 | 1 | 0.169583424 | -1.432404631 | 0.022 | 0.08 | 1 | 0.867636519 | 0.310408311 |
| Slc30a10 | 0.169616506 | 2.673771768 | 0.048 | 0.011 | 1 | 0.49622258 | -0.124282336 | 0 | 0.01 | 1 | 0.49622258 | 0.310463253 |
| Ptgir | 0.169616506 | 2.451379346 | 0.048 | 0.011 | 1 | 0.496223879 | 0.683072586 | 0 | 0.01 | 1 | 0.496223879 | 0.310463253 |
| Cfap161 | 0.169616506 | 2.673771768 | 0.048 | 0.011 | 1 | 0.661444614 | 1.875717664 | 0.022 | 0.014 | 1 | 0.661444614 | 0.310463253 |
| Hs3st5 | 0.169616506 | 2.673771768 | 0.048 | 0.011 | 1 | 0.539099877 | -0.695439037 | 0.022 | 0.042 | 1 | 0.539099877 | 0.310463253 |
| Gm47561 | 0.169616506 | 2.673771768 | 0.048 | 0.011 | 1 | 0.814636613 | 1.098110085 | 0.022 | 0.017 | 1 | 0.814636613 | 0.310463253 |
| Pde8b | 0.169616506 | 2.673771768 | 0.048 | 0.011 | 1 | 0.331907298 | -0.486852415 | 0 | 0.021 | 1 | 0.331907298 | 0.310463253 |
| Gm49267 | 0.169617801 | 2.673771768 | 0.048 | 0.011 | 1 | 0.326732519 | 2.197645759 | 0.044 | 0.021 | 1 | 0.326732519 | 0.310465404 |
| Cars2 | 0.33424206 | -0.535681598 | 0.095 | 0.183 | 1 | 0.169687974 | -0.271123724 | 0.089 | 0.176 | 1 | 0.33424206 | 0.310581939 |
| Man1c1 | 0.815160268 | 0.771952162 | 0.238 | 0.276 | 1 | 0.169697651 | 0.540114632 | 0.244 | 0.412 | 1 | 0.815160268 | 0.31059801 |
| Gm42047 | 0.773914775 | -0.470886475 | 0.19 | 0.22 | 1 | 0.169822957 | -1.085111739 | 0.067 | 0.142 | 1 | 0.773914775 | 0.310806078 |
| Wwp2 | 0.169978601 | 0.388063861 | 0.905 | 0.933 | 1 | 0.287830112 | 0.225070369 | 0.978 | 0.965 | 1 | 0.287830112 | 0.311064478 |
| Zfp692 | 0.458898129 | -0.107587946 | 0.095 | 0.16 | 1 | 0.170324615 | 0.277080226 | 0.067 | 0.149 | 1 | 0.458898129 | 0.311638755 |
| 5930430L01Rik | 0.170656898 | -0.574155746 | 0.048 | 0.168 | 1 | 0.241227006 | 1.383512304 | 0.2 | 0.152 | 1 | 0.241227006 | 0.312190019 |
| Nemf | 0.170693701 | 0.671160554 | 0.857 | 0.78 | 1 | 0.200040338 | 0.579979093 | 0.911 | 0.806 | 1 | 0.200040338 | 0.312251062 |
| Ago3 | 0.170791931 | 0.608480307 | 0.905 | 0.784 | 1 | 0.593648574 | 0.202385407 | 0.911 | 0.907 | 1 | 0.593648574 | 0.312413978 |
| Sav1 | 0.671022218 | 0.333572571 | 0.714 | 0.84 | 1 | 0.170912618 | 0.416506489 | 0.778 | 0.889 | 1 | 0.671022218 | 0.312614113 |
| Sirt2 | 0.920403228 | 0.237952706 | 0.524 | 0.608 | 1 | 0.170943563 | -0.110241397 | 0.533 | 0.678 | 1 | 0.920403228 | 0.312665424 |
| Gpi1 | 0.171035068 | -0.381510668 | 0.571 | 0.705 | 1 | 0.769022939 | -0.121248276 | 0.889 | 0.765 | 1 | 0.769022939 | 0.312817141 |
| Arhgap31 | 0.174868991 | 0.915918807 | 0.81 | 0.795 | 1 | 0.171043851 | 0.322897022 | 0.844 | 0.855 | 1 | 0.174868991 | 0.312831704 |
| Tmsb4x | 0.171132355 | -0.901767479 | 0.857 | 0.799 | 1 | 0.646870032 | -0.923997695 | 0.956 | 0.841 | 1 | 0.646870032 | 0.312978428 |
| Hpcal1 | 0.171353047 | -0.989193245 | 0.048 | 0.164 | 1 | 0.198970429 | 0.9310001 | 0.222 | 0.145 | 1 | 0.198970429 | 0.313344228 |
| Ptprf | 0.171582629 | -0.180888869 | 0.714 | 0.828 | 1 | 0.758097947 | 0.167372748 | 0.933 | 0.907 | 1 | 0.758097947 | 0.313724659 |
| Kctd8 | 0.171707632 | 2.258734268 | 0.048 | 0.011 | 1 | 0.671505935 | -1.044847868 | 0.022 | 0.035 | 1 | 0.671505935 | 0.313931752 |
| Gm14471 | 0.171708932 | 2.673771768 | 0.048 | 0.011 | 1 | 0.759306107 | 0.460680165 | 0.022 | 0.031 | 1 | 0.759306107 | 0.313933907 |
| Snap25 | 0.171708932 | 2.673771768 | 0.048 | 0.011 | 1 | 0.376842245 | -0.638855509 | 0 | 0.017 | 1 | 0.376842245 | 0.313933907 |
| Vtcn1 | 0.171708932 | 2.673771768 | 0.048 | 0.011 | 1 | 0.496223879 | 0.683072586 | 0 | 0.01 | 1 | 0.496223879 | 0.313933907 |
| C77080 | 0.171708932 | 2.673771768 | 0.048 | 0.011 | 1 | 0.978440959 | -0.564854927 | 0.067 | 0.066 | 1 | 0.978440959 | 0.313933907 |
| Gm44957 | 0.171708932 | 2.673771768 | 0.048 | 0.011 | 1 | 0.496223879 | 0.683072586 | 0 | 0.01 | 1 | 0.496223879 | 0.313933907 |
| Pirb | 0.171708932 | 2.673771768 | 0.048 | 0.011 | 1 | 0.935071939 | 0.361144491 | 0.022 | 0.024 | 1 | 0.935071939 | 0.313933907 |
| Gm3854 | 0.171708932 | 2.673771768 | 0.048 | 0.011 | 1 | 0.376837034 | -0.124282336 | 0 | 0.017 | 1 | 0.376837034 | 0.313933907 |
| Zfp108 | 0.171708932 | 2.673771768 | 0.048 | 0.011 | 1 | 0.331914654 | -0.638855509 | 0 | 0.021 | 1 | 0.331914654 | 0.313933907 |
| Sdr42e1 | 0.171708932 | 2.673771768 | 0.048 | 0.011 | 1 | 0.314689364 | 2.098110085 | 0.022 | 0.007 | 1 | 0.314689364 | 0.313933907 |
| Gm26537 | 0.171708932 | 2.673771768 | 0.048 | 0.011 | 1 | 0.496223879 | 0.683072586 | 0 | 0.01 | 1 | 0.496223879 | 0.313933907 |
| Gm32742 | 0.171708932 | 2.673771768 | 0.048 | 0.011 | 1 | 0.496223879 | 0.683072586 | 0 | 0.01 | 1 | 0.496223879 | 0.313933907 |
| Hey2 | 0.171708932 | 2.673771768 | 0.048 | 0.011 | 1 | 0.49622258 | -0.124282336 | 0 | 0.01 | 1 | 0.49622258 | 0.313933907 |
| Gm16364.1 | 0.171708932 | 2.673771768 | 0.048 | 0.011 | 1 | 0.910549015 | 1.765534746 | 0.044 | 0.042 | 1 | 0.910549015 | 0.313933907 |
| Hcn2 | 0.171708932 | 2.673771768 | 0.048 | 0.011 | 1 | 0.658583583 | 2.361144491 | 0.022 | 0.014 | 1 | 0.658583583 | 0.313933907 |
| Tjp3 | 0.171708932 | 2.673771768 | 0.048 | 0.011 | 1 | 0.936023974 | 2.197645759 | 0.022 | 0.021 | 1 | 0.936023974 | 0.313933907 |
| Lratd1 | 0.171708932 | 2.673771768 | 0.048 | 0.011 | 1 | 0.496223879 | 0.683072586 | 0 | 0.01 | 1 | 0.496223879 | 0.313933907 |
| 5730507C01Rik | 0.171708932 | 2.673771768 | 0.048 | 0.011 | 1 | 0.293637566 | -0.486852415 | 0 | 0.024 | 1 | 0.293637566 | 0.313933907 |
| Rec8 | 0.171708932 | 2.673771768 | 0.048 | 0.011 | 1 | 0.331907298 | -0.486852415 | 0 | 0.021 | 1 | 0.331907298 | 0.313933907 |
| Dsg3 | 0.171708932 | 2.673771768 | 0.048 | 0.011 | 1 | 0.496223879 | 0.683072586 | 0 | 0.01 | 1 | 0.496223879 | 0.313933907 |
| Psmf1 | 0.273135709 | -0.184209227 | 0.143 | 0.272 | 1 | 0.172280597 | 0.294801996 | 0.133 | 0.242 | 1 | 0.273135709 | 0.314880591 |
| Tox4 | 0.385623871 | 0.377530316 | 0.429 | 0.627 | 1 | 0.172358385 | -0.254514807 | 0.756 | 0.744 | 1 | 0.385623871 | 0.315009356 |
| Gm19409 | 0.915528848 | 1.673771768 | 0.048 | 0.056 | 1 | 0.172603358 | 0.486675373 | 0.067 | 0.152 | 1 | 0.915528848 | 0.315414796 |
| Rab3gap1 | 0.188997269 | 0.498329761 | 0.714 | 0.806 | 1 | 0.172729273 | 0.455609175 | 0.867 | 0.799 | 1 | 0.188997269 | 0.315623144 |
| Pacsin2 | 0.366053536 | 0.643788901 | 0.762 | 0.84 | 1 | 0.173000593 | 0.595609745 | 0.956 | 0.848 | 1 | 0.366053536 | 0.31607198 |
| Lmbrd1 | 0.173011943 | -0.27289873 | 0.571 | 0.772 | 1 | 0.754033384 | 0.124827424 | 0.889 | 0.81 | 1 | 0.754033384 | 0.316090753 |
| A430110L20Rik | 0.173124678 | 1.79930265 | 0.143 | 0.067 | 1 | 0.494929772 | 0.683072586 | 0.044 | 0.076 | 1 | 0.494929772 | 0.316277202 |
| Ints6 | 0.173154671 | 0.827002165 | 0.762 | 0.66 | 1 | 0.645470551 | 0.712922514 | 0.867 | 0.817 | 1 | 0.645470551 | 0.316326801 |
| Spryd4 | 0.173676192 | -0.911190733 | 0 | 0.082 | 1 | 0.227164429 | -0.84048937 | 0.022 | 0.069 | 1 | 0.227164429 | 0.317188965 |
| Gm16867 | 0.173676192 | -0.911190733 | 0 | 0.082 | 1 | 0.553874252 | 0.460680165 | 0.022 | 0.042 | 1 | 0.553874252 | 0.317188965 |
| Pcdhgb5 | 0.173676192 | -0.911190733 | 0 | 0.082 | 1 | 0.199734918 | 0.513147585 | 0.022 | 0.076 | 1 | 0.199734918 | 0.317188965 |
| Ss18l1 | 0.173729397 | -0.970084422 | 0 | 0.082 | 1 | 0.387530512 | 1.750186782 | 0.089 | 0.059 | 1 | 0.387530512 | 0.31727689 |
| Zfp428 | 0.173729397 | -1.081115734 | 0 | 0.082 | 1 | 0.250725289 | -0.901889915 | 0.022 | 0.066 | 1 | 0.250725289 | 0.31727689 |
| Katnb1 | 0.173729397 | -0.970084422 | 0 | 0.082 | 1 | 0.29730293 | -0.124282336 | 0.022 | 0.062 | 1 | 0.29730293 | 0.31727689 |
| Top1mt | 0.173729397 | -0.970084422 | 0 | 0.082 | 1 | 0.199155372 | -0.223818009 | 0.022 | 0.076 | 1 | 0.199155372 | 0.31727689 |
| Il13ra1 | 0.173752816 | 1.458758877 | 0.381 | 0.306 | 1 | 0.178723585 | 0.245667274 | 0.533 | 0.381 | 1 | 0.178723585 | 0.317315591 |
| E2f1 | 0.173777275 | -1.081115734 | 0 | 0.082 | 1 | 0.180634714 | -0.124282336 | 0.022 | 0.08 | 1 | 0.180634714 | 0.317356008 |
| Nlrc3 | 0.173777275 | -1.081115734 | 0 | 0.082 | 1 | 0.683686674 | 1.946106992 | 0.111 | 0.097 | 1 | 0.683686674 | 0.317356008 |
| Mrm1 | 0.173777528 | -1.02666795 | 0 | 0.082 | 1 | 0.35815185 | 0.197645759 | 0.044 | 0.087 | 1 | 0.35815185 | 0.317356427 |
| Gtf2h2 | 0.516074765 | 0.214340149 | 0.095 | 0.153 | 1 | 0.173797782 | 0.162815775 | 0.156 | 0.277 | 1 | 0.516074765 | 0.317389895 |
| Mir181a-1hg | 0.173814074 | 2.351843673 | 0.048 | 0.011 | 1 | 0.430486631 | -0.124282336 | 0 | 0.014 | 1 | 0.430486631 | 0.317416816 |
| Gm42705 | 0.173814074 | 2.351843673 | 0.048 | 0.011 | 1 | 0.317138221 | 1.683072586 | 0.022 | 0.007 | 1 | 0.317138221 | 0.317416816 |
| Slc23a4 | 0.173814074 | 2.351843673 | 0.048 | 0.011 | 1 | 0.823042771 | 0.875717664 | 0.022 | 0.017 | 1 | 0.823042771 | 0.317416816 |
| Dscaml1 | 0.173814074 | 0.425844254 | 0.048 | 0.011 | 1 | 0.331914654 | -1.223818009 | 0 | 0.021 | 1 | 0.331914654 | 0.317416816 |
| Trim16 | 0.173814074 | 2.088809267 | 0.048 | 0.011 | 1 | 0.80624346 | 2.42003818 | 0.022 | 0.017 | 1 | 0.80624346 | 0.317416816 |

| | | | | | | | | | | | | |
|---|---|---|---|---|---|---|---|---|---|---|---|---|
| B4galnt2 | 0.173814074 | 2.351843673 | 0.048 | 0.011 | 1 | 0.496223879 | 0.683072586 | 0 | 0.01 | 1 | 0.496223879 | 0.317416816 |
| Gm49313 | 0.173814074 | 2.351843673 | 0.048 | 0.011 | 1 | 0.496223879 | 0.683072586 | 0 | 0.01 | 1 | 0.496223879 | 0.317416816 |
| Tmem53 | 0.173819828 | -1.233118828 | 0 | 0.082 | 1 | 0.587491145 | 1.039216396 | 0.044 | 0.069 | 1 | 0.587491145 | 0.317426323 |
| Gm50240 | 0.173820587 | -1.233118828 | 0 | 0.082 | 1 | 0.183590103 | -0.674479418 | 0.044 | 0.111 | 1 | 0.183590103 | 0.317427578 |
| Lnx2 | 0.173857057 | -1.184209227 | 0 | 0.082 | 1 | 0.987769776 | 1.154693614 | 0.156 | 0.17 | 1 | 0.987769776 | 0.317487838 |
| Nudt7 | 0.173857057 | -1.184209227 | 0 | 0.082 | 1 | 0.47918822 | -0.223818009 | 0.022 | 0.045 | 1 | 0.47918822 | 0.317487838 |
| Vstm4 | 0.173858577 | -1.280424543 | 0 | 0.082 | 1 | 0.270417291 | -0.316927414 | 0.044 | 0.097 | 1 | 0.270417291 | 0.317490349 |
| D130040H23Rik | 0.173891498 | -1.326228232 | 0 | 0.082 | 1 | 0.66805079 | 1.410054092 | 0.089 | 0.076 | 1 | 0.66805079 | 0.317544743 |
| Twist2 | 0.173915554 | -1.280424543 | 0 | 0.082 | 1 | 0.269484004 | 0.15951063 | 0.022 | 0.066 | 1 | 0.269484004 | 0.317584488 |
| Rasa4 | 0.173919352 | -1.413691074 | 0 | 0.082 | 1 | 0.978881707 | 0.995016592 | 0.067 | 0.069 | 1 | 0.978881707 | 0.317590763 |
| Adcy7 | 0.173972016 | -2.208871282 | 0 | 0.082 | 1 | 0.812151577 | 1.320502507 | 0.067 | 0.08 | 1 | 0.812151577 | 0.317677769 |
| Psmb8 | 0.173984674 | -1.940938076 | 0 | 0.082 | 1 | 0.617529321 | 2.982632868 | 0.022 | 0.038 | 1 | 0.617529321 | 0.317698682 |
| Nptx1 | 0.1740229 | -2.701267664 | 0 | 0.082 | 1 | 0.184823348 | -1.486852415 | 0 | 0.038 | 1 | 0.184823348 | 0.31776183 |
| Slc4a7 | 0.174213377 | 0.969575029 | 0.762 | 0.795 | 1 | 0.21391805 | 0.365921632 | 0.956 | 0.824 | 1 | 0.21391805 | 0.318076453 |
| Rhof | 0.490548052 | 1.195724471 | 0.143 | 0.101 | 1 | 0.174495363 | -0.486852415 | 0.067 | 0.145 | 1 | 0.490548052 | 0.318542094 |
| Slc25a28 | 0.174735547 | -0.530100566 | 0.19 | 0.34 | 1 | 0.269451931 | 0.209141398 | 0.222 | 0.343 | 1 | 0.269451931 | 0.318938583 |
| Klhl18 | 0.689651289 | 0.750939628 | 0.238 | 0.228 | 1 | 0.175054949 | 0.387616703 | 0.156 | 0.287 | 1 | 0.689651289 | 0.319465663 |
| Cadm1 | 0.823191661 | -0.206733075 | 0.667 | 0.649 | 1 | 0.175164391 | 0.483184717 | 0.956 | 0.955 | 1 | 0.823191661 | 0.319646218 |
| Slf2 | 0.175169384 | -0.266671388 | 0.286 | 0.489 | 1 | 0.390880473 | 0.131138886 | 0.756 | 0.671 | 1 | 0.390880473 | 0.319654456 |
| Csrp2 | 0.175207359 | -1.297249165 | 0.238 | 0.388 | 1 | 0.861415846 | -0.33895372 | 0.711 | 0.623 | 1 | 0.861415846 | 0.319717099 |
| Frmd8os | 0.295039357 | 1.780686972 | 0.143 | 0.086 | 1 | 0.1753169 | -1.017367132 | 0.022 | 0.08 | 1 | 0.295039357 | 0.319897784 |
| L3mbtl2 | 0.815673353 | 1.214340149 | 0.048 | 0.037 | 1 | 0.175317073 | -0.776359032 | 0.022 | 0.08 | 1 | 0.815673353 | 0.31989807 |
| Trim44 | 0.175387774 | 1.122957489 | 0.857 | 0.832 | 1 | 0.906455814 | 0.193810502 | 0.956 | 0.882 | 1 | 0.906455814 | 0.320014676 |
| Zfp57 | 0.732774634 | 0.503846766 | 0.095 | 0.131 | 1 | 0.175574907 | -0.486852415 | 0.067 | 0.142 | 1 | 0.732774634 | 0.320323266 |
| Kif5c | 0.784047982 | 2.780686972 | 0.048 | 0.037 | 1 | 0.175923906 | -0.731964913 | 0.022 | 0.08 | 1 | 0.784047982 | 0.320898591 |
| Fmo1 | 0.175947671 | 2.088809267 | 0.048 | 0.011 | 1 | 0.206897641 | -1.316927414 | 0 | 0.035 | 1 | 0.206897641 | 0.32093776 |
| Ifi203 | 0.175947671 | 2.088809267 | 0.048 | 0.011 | 1 | 0.376842245 | -0.486852415 | 0 | 0.017 | 1 | 0.376842245 | 0.32093776 |
| Lypd6b | 0.175947671 | 1.351843673 | 0.048 | 0.011 | 1 | 0.49622258 | -0.124282336 | 0 | 0.01 | 1 | 0.49622258 | 0.32093776 |
| Gm57242 | 0.175947671 | 1.79930265 | 0.048 | 0.011 | 1 | 0.675805699 | 0.223640968 | 0.022 | 0.014 | 1 | 0.675805699 | 0.32093776 |
| Gm36198 | 0.3247899 | -0.326228232 | 0 | 0.045 | 1 | 0.176113295 | 1.098110085 | 0.067 | 0.028 | 1 | 0.3247899 | 0.321210698 |
| 1700036A12Rik | 0.231954566 | -0.970084422 | 0.048 | 0.138 | 1 | 0.176268393 | 0.152557869 | 0.133 | 0.239 | 1 | 0.231954566 | 0.32146624 |
| Gm57198 | 0.611709251 | 1.119182916 | 0.095 | 0.149 | 1 | 0.176471068 | 0.339118185 | 0.089 | 0.187 | 1 | 0.611709251 | 0.321800098 |
| Nadk2 | 0.197654463 | 0.607682577 | 0.048 | 0.16 | 1 | 0.17655367 | 0.402964667 | 0.133 | 0.249 | 1 | 0.197654463 | 0.321936141 |
| Stx1a | 0.474362221 | 1.258734268 | 0.143 | 0.101 | 1 | 0.176818963 | -0.459885368 | 0.067 | 0.142 | 1 | 0.474362221 | 0.32237298 |
| Heg1 | 0.177072761 | 0.621683257 | 0.095 | 0.25 | 1 | 0.196456287 | 0.711086962 | 0.067 | 0.145 | 1 | 0.196456287 | 0.322790759 |
| Efhb | 0.244244748 | 1.756233928 | 0.143 | 0.075 | 1 | 0.177096005 | -0.486852415 | 0.022 | 0.08 | 1 | 0.244244748 | 0.322829014 |
| Hspa12a | 0.1773735 | 1.811275291 | 0.19 | 0.101 | 1 | 0.505129828 | -0.174908409 | 0.067 | 0.1 | 1 | 0.505129828 | 0.323285641 |
| Togaram1 | 0.586935438 | 0.180034027 | 0.476 | 0.612 | 1 | 0.1776436 | -0.138266565 | 0.756 | 0.775 | 1 | 0.586935438 | 0.323729951 |
| Isy1 | 0.522994241 | 0.324362936 | 0.333 | 0.481 | 1 | 0.177822513 | -0.275507486 | 0.622 | 0.651 | 1 | 0.522994241 | 0.324024179 |
| B230307C23Rik | 0.177951106 | -0.648156327 | 0.048 | 0.164 | 1 | 0.525383761 | 0.705792663 | 0.111 | 0.159 | 1 | 0.525383761 | 0.324235616 |
| Plekha5 | 0.178010466 | 0.822182874 | 0.762 | 0.69 | 1 | 0.246313186 | 0.383283469 | 0.978 | 0.834 | 1 | 0.246313186 | 0.324333206 |
| Gm57184 | 0.562574042 | 1.044415148 | 0.143 | 0.22 | 1 | 0.178087334 | -1.044847868 | 0.044 | 0.111 | 1 | 0.562574042 | 0.324459569 |
| AI661453 | 0.178096747 | 1.503846766 | 0.048 | 0.011 | 1 | 0.658583583 | 2.946106992 | 0.022 | 0.014 | 1 | 0.658583583 | 0.324475043 |
| Smarcad1 | 0.178146107 | -0.287754085 | 0.333 | 0.556 | 1 | 0.502918462 | 0.511424964 | 0.622 | 0.692 | 1 | 0.502918462 | 0.324556178 |
| Gm49602 | 0.97964157 | 1.451379346 | 0.048 | 0.049 | 1 | 0.178199306 | -0.539319835 | 0.022 | 0.08 | 1 | 0.97964157 | 0.324643619 |
| Gm48693 | 0.178454209 | 2.55829455 | 0.095 | 0.037 | 1 | 0.590623425 | -0.564854927 | 0.022 | 0.038 | 1 | 0.590623425 | 0.325062513 |
| Gm46367 | 0.178454209 | 3.143257051 | 0.095 | 0.037 | 1 | 0.796264725 | 0.723714571 | 0.089 | 0.08 | 1 | 0.796264725 | 0.325062513 |
| Gm56796 | 0.178827814 | 2.493199522 | 0.19 | 0.112 | 1 | 0.919697584 | 0.795547315 | 0.089 | 0.097 | 1 | 0.919697584 | 0.325676241 |
| Gm56741 | 0.423027883 | -0.413691074 | 0.048 | 0.101 | 1 | 0.178831395 | -0.316927414 | 0.022 | 0.08 | 1 | 0.423027883 | 0.325682122 |
| Steap4 | 0.994855961 | 2.3693311 | 0.143 | 0.157 | 1 | 0.178868887 | -2.271123724 | 0.022 | 0.076 | 1 | 0.994855961 | 0.325743696 |
| Nsd2 | 0.790085489 | 0.441919017 | 0.429 | 0.515 | 1 | 0.178887439 | 0.626041641 | 0.711 | 0.633 | 1 | 0.790085489 | 0.325774163 |
| Kcnd2 | 0.179073029 | 3.341928204 | 0.143 | 0.067 | 1 | 0.52451989 | 0.516065068 | 0.178 | 0.235 | 1 | 0.52451989 | 0.326078908 |
| Ppil6 | 0.185434129 | -0.940938076 | 0.048 | 0.153 | 1 | 0.17927688 | -0.191396532 | 0.067 | 0.145 | 1 | 0.185434129 | 0.32641356 |
| St6gal1 | 0.368858354 | -0.308080886 | 0.048 | 0.116 | 1 | 0.179352638 | -0.486852415 | 0.044 | 0.114 | 1 | 0.368858354 | 0.326537908 |
| Rad51d | 0.655975202 | 1.029915578 | 0.048 | 0.078 | 1 | 0.179459978 | -0.271123724 | 0.022 | 0.08 | 1 | 0.655975202 | 0.326714073 |
| Rnf227 | 0.852115916 | 0.995699863 | 0.095 | 0.086 | 1 | 0.1794632 | -1.602329633 | 0.022 | 0.076 | 1 | 0.852115916 | 0.32671936 |
| 1810053B23Rik | 0.17977986 | -1.718545655 | 0.048 | 0.164 | 1 | 0.184854431 | -2.361321533 | 0 | 0.038 | 1 | 0.184854431 | 0.327238921 |
| 1700123M08Rik | 0.179905876 | 1.503846766 | 0.143 | 0.067 | 1 | 0.659788885 | -0.404390255 | 0.022 | 0.035 | 1 | 0.659788885 | 0.327445628 |
| Nek6 | 0.180067797 | 0.985715774 | 0.381 | 0.276 | 1 | 0.610482144 | 0.75994135 | 0.267 | 0.374 | 1 | 0.610482144 | 0.327711183 |
| Morrbid | 0.180550128 | 2.978626349 | 0.143 | 0.071 | 1 | 0.42228109 | -1.223818009 | 0.022 | 0.048 | 1 | 0.42228109 | 0.328501908 |
| Irf7 | 0.222302397 | -1.280424543 | 0 | 0.067 | 1 | 0.180602462 | 1.098110085 | 0.022 | 0.08 | 1 | 0.222302397 | 0.328587674 |
| Mcat | 0.823729147 | 0.586308926 | 0.048 | 0.06 | 1 | 0.180634714 | 0.197645759 | 0.022 | 0.08 | 1 | 0.823729147 | 0.328640528 |
| Myo5c | 0.370807306 | -0.574155746 | 0 | 0.037 | 1 | 0.180657647 | -1.602329633 | 0.022 | 0.076 | 1 | 0.370807306 | 0.328678109 |
| Gm14066 | 0.997092023 | 0.614878079 | 0.048 | 0.049 | 1 | 0.180665561 | 0.268035087 | 0.022 | 0.08 | 1 | 0.997092023 | 0.328691077 |
| Ccdc62 | 0.18072612 | -0.984439715 | 0.095 | 0.22 | 1 | 0.969311524 | 0.58153456 | 0.178 | 0.197 | 1 | 0.969311524 | 0.32879031 |
| Cers5 | 0.180769067 | -0.133583154 | 0.333 | 0.586 | 1 | 0.334880705 | 0.344104588 | 0.8 | 0.758 | 1 | 0.334880705 | 0.328860678 |
| C79798 | 0.181001241 | 2.295260144 | 0.095 | 0.037 | 1 | 0.690919606 | 1.410054092 | 0.044 | 0.062 | 1 | 0.690919606 | 0.329241034 |
| Crym | 0.681938179 | 0.673771768 | 0.048 | 0.075 | 1 | 0.18100761 | 1.683072586 | 0.133 | 0.08 | 1 | 0.681938179 | 0.329251465 |
| Tsc2 | 0.181058809 | -0.807354922 | 0.095 | 0.216 | 1 | 0.265426778 | 0.180572246 | 0.111 | 0.19 | 1 | 0.265426778 | 0.329335326 |
| Nhej1 | 0.622562199 | 0.936806174 | 0.19 | 0.16 | 1 | 0.181066897 | 1.435145073 | 0.089 | 0.176 | 1 | 0.622562199 | 0.329348572 |
| H2-K1 | 0.181452376 | -0.979670471 | 0.667 | 0.694 | 1 | 0.85431914 | -0.180564811 | 0.867 | 0.713 | 1 | 0.85431914 | 0.329979788 |
| Jmy | 0.243293275 | 0.374804821 | 0.381 | 0.575 | 1 | 0.181791753 | -0.287690193 | 0.8 | 0.796 | 1 | 0.243293275 | 0.330535264 |
| Gramd4 | 0.287503371 | -0.891012851 | 0.095 | 0.183 | 1 | 0.181823685 | 0.498648015 | 0.111 | 0.221 | 1 | 0.287503371 | 0.330587517 |
| Plb1 | 0.621155057 | 2.420015175 | 0.048 | 0.082 | 1 | 0.181847577 | -1.316927414 | 0.022 | 0.076 | 1 | 0.621155057 | 0.330626613 |
| Ntn4 | 0.49826337 | 0.490549944 | 0.19 | 0.142 | 1 | 0.181957195 | -0.638855509 | 0.067 | 0.142 | 1 | 0.49826337 | 0.330805969 |
| Slc8a2 | 0.208922634 | -1.081115734 | 0 | 0.071 | 1 | 0.182120253 | 1.958707029 | 0.133 | 0.08 | 1 | 0.208922634 | 0.33107272 |
| Ebf4 | 0.918805176 | 3.744161096 | 0.048 | 0.045 | 1 | 0.182125579 | -0.124282336 | 0.044 | 0.114 | 1 | 0.918805176 | 0.331081431 |
| Ankrd52 | 0.182284576 | -0.413691074 | 0.19 | 0.373 | 1 | 0.824656392 | 0.503748887 | 0.356 | 0.484 | 1 | 0.824656392 | 0.331341485 |
| Chn2 | 0.182305299 | 2.451379346 | 0.095 | 0.037 | 1 | 0.490132999 | -0.454430938 | 0.044 | 0.073 | 1 | 0.490132999 | 0.331375377 |
| Pitpnm2 | 0.182339744 | -0.648156327 | 0.143 | 0.291 | 1 | 0.780852046 | 0.850800032 | 0.244 | 0.315 | 1 | 0.780852046 | 0.331431706 |
| Thap4 | 0.182365513 | -0.469186186 | 0.048 | 0.164 | 1 | 0.609385284 | 0.841770332 | 0.178 | 0.239 | 1 | 0.609385284 | 0.331473846 |
| Fbxl5 | 0.531257053 | 0.564761146 | 0.476 | 0.698 | 1 | 0.182402809 | 0.270291061 | 0.844 | 0.699 | 1 | 0.531257053 | 0.331534833 |
| Cand1 | 0.357756551 | 0.130873327 | 0.381 | 0.649 | 1 | 0.182414803 | -0.208346601 | 0.889 | 0.851 | 1 | 0.357756551 | 0.331554445 |
| Gemin2 | 0.896684646 | 0.808701348 | 0.143 | 0.142 | 1 | 0.182611361 | -0.124282336 | 0.067 | 0.145 | 1 | 0.896684646 | 0.331875814 |
| Bud23 | 0.182681763 | -0.449610648 | 0.19 | 0.354 | 1 | 0.195486921 | 0.2793504 | 0.2 | 0.339 | 1 | 0.195486921 | 0.331990899 |
| Lrrcc1 | 0.18273483 | -0.14822993 | 0.095 | 0.228 | 1 | 0.453165067 | 0.390290837 | 0.133 | 0.201 | 1 | 0.453165067 | 0.332077642 |
| Alg12 | 0.867345526 | 0.629377648 | 0.095 | 0.112 | 1 | 0.183030918 | -1.071814916 | 0.022 | 0.076 | 1 | 0.867345526 | 0.332561518 |
| Bivm | 0.517057607 | 0.258734268 | 0.048 | 0.093 | 1 | 0.183070193 | -0.638855509 | 0.044 | 0.111 | 1 | 0.517057607 | 0.332625691 |
| Pamr1 | 0.183255484 | -0.849790188 | 0.095 | 0.228 | 1 | 0.985252187 | -0.365290435 | 0.178 | 0.183 | 1 | 0.985252187 | 0.332928396 |
| Tex10 | 0.384563185 | 0.728913322 | 0.381 | 0.351 | 1 | 0.18329271 | 0.581192972 | 0.489 | 0.394 | 1 | 0.384563185 | 0.332989202 |
| Usp21 | 0.539532325 | -0.264827688 | 0.095 | 0.142 | 1 | 0.1834317 | -0.271123724 | 0.089 | 0.173 | 1 | 0.539532325 | 0.333216211 |
| Wdcp | 0.701849009 | 0.673771768 | 0.19 | 0.168 | 1 | 0.183445087 | 0.323176641 | 0.111 | 0.204 | 1 | 0.701849009 | 0.333238073 |
| Sdha | 0.560340859 | 0.39267739 | 0.619 | 0.728 | 1 | 0.183504631 | -0.161022883 | 0.778 | 0.765 | 1 | 0.560340859 | 0.333335313 |
| Col5a3 | 0.183584264 | 1.735172312 | 0.095 | 0.037 | 1 | 0.878773794 | 1.250113179 | 0.044 | 0.052 | 1 | 0.878773794 | 0.333465345 |
| Tmem95 | 0.183586088 | 2.188344941 | 0.095 | 0.037 | 1 | 0.480781131 | -0.124282336 | 0.022 | 0.045 | 1 | 0.480781131 | 0.333468324 |
| Jmjd4 | 0.184739419 | -1.02666795 | 0 | 0.078 | 1 | 0.183618604 | -1.071814916 | 0.022 | 0.076 | 1 | 0.184739419 | 0.333521416 |
| Pdzk1ip1 | 0.196598718 | -1.535681598 | 0 | 0.075 | 1 | 0.183618604 | -1.124282336 | 0.022 | 0.076 | 1 | 0.196598718 | 0.333521416 |
| D2hgdh | 0.208886118 | -0.849790188 | 0 | 0.071 | 1 | 0.183618787 | -1.017367132 | 0.022 | 0.076 | 1 | 0.208886118 | 0.333521716 |

| | | | | | | | | | | | | |
|---|---|---|---|---|---|---|---|---|---|---|---|---|
| Mpzl3 | 0.675317969 | 0.719575457 | 0.095 | 0.071 | 1 | 0.183618787 | -1.017367132 | 0.022 | 0.076 | 1 | 0.675317969 | 0.333521716 |
| Wars2 | 0.951437208 | 0.989273593 | 0.286 | 0.325 | 1 | 0.183629705 | 0.388451837 | 0.178 | 0.301 | 1 | 0.951437208 | 0.333539541 |
| Als2cl | 0.694510638 | 0.96105372 | 0.19 | 0.175 | 1 | 0.184440561 | -0.662702251 | 0.067 | 0.142 | 1 | 0.694510638 | 0.334862801 |
| Gm38115 | 0.184642276 | -0.911190733 | 0 | 0.078 | 1 | 0.698591406 | 0.426732833 | 0.089 | 0.114 | 1 | 0.698591406 | 0.335191782 |
| Gm7694 | 0.184642276 | -0.849790188 | 0 | 0.078 | 1 | 0.441257002 | 0.429315994 | 0.111 | 0.173 | 1 | 0.441257002 | 0.335191782 |
| Six5 | 0.184642276 | -0.849790188 | 0 | 0.078 | 1 | 0.265549958 | -0.446210431 | 0.044 | 0.097 | 1 | 0.265549958 | 0.335191782 |
| Camk2n2 | 0.184642276 | -0.849790188 | 0 | 0.078 | 1 | 0.227164429 | -0.901889915 | 0.022 | 0.069 | 1 | 0.227164429 | 0.335191782 |
| Gm10130 | 0.184693549 | -0.911190733 | 0 | 0.078 | 1 | 0.201928208 | -1.223818009 | 0.022 | 0.073 | 1 | 0.201928208 | 0.335275391 |
| Gm57457 | 0.184693549 | -0.911190733 | 0 | 0.078 | 1 | 0.313790052 | -0.486852415 | 0.022 | 0.059 | 1 | 0.313790052 | 0.335275391 |
| Dclre1b | 0.184693549 | -0.911190733 | 0 | 0.078 | 1 | 0.203242543 | -1.271123724 | 0.022 | 0.073 | 1 | 0.203242543 | 0.335275391 |
| Nek3 | 0.184693549 | -1.02666795 | 0 | 0.078 | 1 | 0.845607744 | 0.632446513 | 0.067 | 0.076 | 1 | 0.845607744 | 0.335275391 |
| Ak5 | 0.20900547 | -1.413691074 | 0 | 0.071 | 1 | 0.184699526 | 0.71623945 | 0.067 | 0.149 | 1 | 0.20900547 | 0.335285138 |
| Itga3 | 0.184739419 | -1.133583154 | 0 | 0.078 | 1 | 0.474802224 | -0.564854927 | 0.022 | 0.045 | 1 | 0.474802224 | 0.335350185 |
| Ddx51 | 0.184739689 | -0.970084422 | 0 | 0.078 | 1 | 0.328724919 | -0.686161223 | 0.044 | 0.087 | 1 | 0.328724919 | 0.335350625 |
| 4732491K20Rik | 0.184739689 | -0.970084422 | 0 | 0.078 | 1 | 0.347517921 | -0.486852415 | 0.022 | 0.055 | 1 | 0.347517921 | 0.335350625 |
| Zfp995 | 0.184739689 | -0.970084422 | 0 | 0.078 | 1 | 0.354652542 | 0.223640968 | 0.044 | 0.087 | 1 | 0.354652542 | 0.335350625 |
| Parm1 | 0.451322275 | -0.522625445 | 0.048 | 0.101 | 1 | 0.184774853 | 0.42003818 | 0.111 | 0.059 | 1 | 0.451322275 | 0.335407959 |
| Accs | 0.184779887 | -1.081115734 | 0 | 0.078 | 1 | 0.477727467 | -0.316927414 | 0.022 | 0.045 | 1 | 0.477727467 | 0.335416167 |
| 9330179D12Rik | 0.184779887 | -1.081115734 | 0 | 0.078 | 1 | 0.640731026 | 2.402964667 | 0.067 | 0.052 | 1 | 0.640731026 | 0.335416167 |
| ENSMUSG00000121502 | 0.184779887 | -1.081115734 | 0 | 0.078 | 1 | 0.663415888 | 0.91753784 | 0.067 | 0.09 | 1 | 0.663415888 | 0.335416167 |
| Map3k13 | 0.184779887 | -1.133583154 | 0 | 0.078 | 1 | 0.664307093 | 0.808603468 | 0.022 | 0.014 | 1 | 0.664307093 | 0.335416167 |
| Fam72a | 0.184780966 | -1.02666795 | 0 | 0.078 | 1 | 0.34534552 | -0.638855509 | 0.022 | 0.055 | 1 | 0.34534552 | 0.335417927 |
| Usp27x | 0.525779634 | 0.503846766 | 0.048 | 0.093 | 1 | 0.184787784 | -0.901889915 | 0.022 | 0.076 | 1 | 0.525779634 | 0.335429043 |
| Bud13 | 0.184801014 | -0.970084422 | 0.095 | 0.213 | 1 | 0.213494124 | -0.147002412 | 0.089 | 0.166 | 1 | 0.213494124 | 0.335450614 |
| Dennd6b | 0.196329837 | -0.785659851 | 0 | 0.075 | 1 | 0.184806696 | -1.124282336 | 0 | 0.038 | 1 | 0.196329837 | 0.335459877 |
| Catip | 0.236343578 | -0.574155746 | 0 | 0.063 | 1 | 0.184806696 | -1.486852415 | 0 | 0.038 | 1 | 0.236343578 | 0.335459877 |
| Gm57182 | 0.251619726 | -0.574155746 | 0 | 0.06 | 1 | 0.184806696 | -1.124282336 | 0 | 0.038 | 1 | 0.251619726 | 0.335459877 |
| Hemk1 | 0.267917141 | -0.413691074 | 0 | 0.056 | 1 | 0.184806696 | -1.124282336 | 0 | 0.038 | 1 | 0.267917141 | 0.335459877 |
| H2ac18 | 0.268100354 | -0.326228232 | 0 | 0.056 | 1 | 0.184806696 | -1.017367132 | 0 | 0.038 | 1 | 0.268100354 | 0.335459877 |
| Gm50069 | 0.285622926 | -0.233118828 | 0 | 0.052 | 1 | 0.184806696 | -1.017367132 | 0 | 0.038 | 1 | 0.285622926 | 0.335459877 |
| Dhtkd1 | 0.30448834 | -0.133583154 | 0 | 0.049 | 1 | 0.184806696 | -1.124282336 | 0 | 0.038 | 1 | 0.30448834 | 0.335459877 |
| Gm47962 | 0.370769338 | -0.233118828 | 0 | 0.037 | 1 | 0.184806696 | -1.223818009 | 0 | 0.038 | 1 | 0.370769338 | 0.335459877 |
| 4930452G13Rik | 0.370840523 | 0.214340149 | 0 | 0.037 | 1 | 0.184806696 | -1.017367132 | 0 | 0.038 | 1 | 0.370840523 | 0.335459877 |
| Mzf1 | 0.492608234 | 0.866416846 | 0 | 0.022 | 1 | 0.184806696 | -1.124282336 | 0 | 0.038 | 1 | 0.492608234 | 0.335459877 |
| Gm38134 | 0.532435374 | 0.503846766 | 0 | 0.019 | 1 | 0.184806696 | -1.017367132 | 0 | 0.038 | 1 | 0.532435374 | 0.335459877 |
| Plxdc1 | 0.711492968 | 0.304537958 | 0.048 | 0.071 | 1 | 0.184806696 | -1.017367132 | 0 | 0.038 | 1 | 0.711492968 | 0.335459877 |
| Zfp119b | 0.773866629 | 1.536268244 | 0.048 | 0.067 | 1 | 0.184806696 | -1.017367132 | 0 | 0.038 | 1 | 0.773866629 | 0.335459877 |
| 9630013D21Rik | 0.799824033 | 1.258734268 | 0.048 | 0.037 | 1 | 0.184806696 | -1.017367132 | 0 | 0.038 | 1 | 0.799824033 | 0.335459877 |
| Urah | 0.86009419 | 1.088809267 | 0.048 | 0.06 | 1 | 0.184806696 | -1.017367132 | 0 | 0.038 | 1 | 0.86009419 | 0.335459877 |
| Gm36899 | 0.184814956 | -1.184209227 | 0 | 0.078 | 1 | 0.549343282 | 0.683072586 | 0.044 | 0.028 | 1 | 0.549343282 | 0.335473343 |
| Paqr8 | 0.184816574 | -1.184209227 | 0 | 0.078 | 1 | 0.687919128 | 0.905465008 | 0.044 | 0.062 | 1 | 0.687919128 | 0.335475982 |
| Gm15446 | 0.184816574 | -1.233118828 | 0 | 0.078 | 1 | 0.220154 | 0.304560963 | 0.022 | 0.073 | 1 | 0.220154 | 0.335475982 |
| Slc26a7 | 0.208922941 | -0.911190733 | 0 | 0.071 | 1 | 0.184823348 | -1.404390255 | 0 | 0.038 | 1 | 0.208922941 | 0.335487027 |
| Zfp607b | 0.23638565 | -0.648156327 | 0 | 0.063 | 1 | 0.184823348 | -1.124282336 | 0 | 0.038 | 1 | 0.23638565 | 0.335487027 |
| Nudt16 | 0.251619726 | -0.574155746 | 0 | 0.06 | 1 | 0.184823348 | -1.124282336 | 0 | 0.038 | 1 | 0.251619726 | 0.335487027 |
| Cd200r1 | 0.425614684 | -0.574155746 | 0 | 0.03 | 1 | 0.184823348 | -1.223818009 | 0 | 0.038 | 1 | 0.425614684 | 0.335487027 |
| Stk32c | 0.492608234 | 0.866416846 | 0 | 0.022 | 1 | 0.184823348 | -1.404390255 | 0 | 0.038 | 1 | 0.492608234 | 0.335487027 |
| Gm4681 | 0.532435374 | 0.503846766 | 0 | 0.019 | 1 | 0.184823348 | -1.124282336 | 0 | 0.038 | 1 | 0.532435374 | 0.335487027 |
| Gpr35 | 0.532436372 | 1.088809267 | 0 | 0.019 | 1 | 0.184823348 | -1.124282336 | 0 | 0.038 | 1 | 0.532436372 | 0.335487027 |
| Dph2 | 0.570211706 | 1.503846766 | 0.048 | 0.026 | 1 | 0.184823348 | -1.124282336 | 0 | 0.038 | 1 | 0.570211706 | 0.335487027 |
| Tedc1 | 0.573468511 | 1.351843673 | 0.048 | 0.026 | 1 | 0.184823348 | -1.124282336 | 0 | 0.038 | 1 | 0.573468511 | 0.335487027 |
| Gm47163 | 0.57816106 | 1.351843673 | 0 | 0.015 | 1 | 0.184823348 | -1.124282336 | 0 | 0.038 | 1 | 0.57816106 | 0.335487027 |
| Emc9 | 0.715863632 | 1.97333205 | 0.048 | 0.034 | 1 | 0.184823348 | -1.223818009 | 0 | 0.038 | 1 | 0.715863632 | 0.335487027 |
| Pmel | 0.982553176 | 1.425844254 | 0.048 | 0.049 | 1 | 0.184823348 | -1.124282336 | 0 | 0.038 | 1 | 0.982553176 | 0.335487027 |
| Stard6 | 0.346827513 | -0.233118828 | 0 | 0.041 | 1 | 0.1848363 | -1.316927414 | 0 | 0.038 | 1 | 0.346827513 | 0.335508142 |
| 1700028E10Rik | 0.492606471 | -0.326228232 | 0 | 0.022 | 1 | 0.1848363 | -1.404390255 | 0 | 0.038 | 1 | 0.492606471 | 0.335508142 |
| Ccdc189 | 0.267917141 | -0.413691074 | 0 | 0.056 | 1 | 0.18483667 | -1.223818009 | 0 | 0.038 | 1 | 0.267917141 | 0.335508745 |
| Carmil2 | 0.454205211 | 2.451379346 | 0.048 | 0.022 | 1 | 0.18483667 | -1.223818009 | 0 | 0.038 | 1 | 0.454205211 | 0.335508745 |
| Zfp677 | 0.457281049 | 0.503846766 | 0 | 0.026 | 1 | 0.18483667 | -1.486852415 | 0 | 0.038 | 1 | 0.457281049 | 0.335508745 |
| Zfp712 | 0.457308648 | 0.673771768 | 0 | 0.026 | 1 | 0.18483667 | -1.223818009 | 0 | 0.038 | 1 | 0.457308648 | 0.335508745 |
| Arg2 | 0.463491968 | 1.936806174 | 0.048 | 0.022 | 1 | 0.18483667 | -1.486852415 | 0 | 0.038 | 1 | 0.463491968 | 0.335508745 |
| Fgd5 | 0.492590603 | 0.351843673 | 0 | 0.022 | 1 | 0.18483667 | -1.404390255 | 0 | 0.038 | 1 | 0.492590603 | 0.335508745 |
| Gm34189 | 0.613101537 | 0.638147858 | 0.048 | 0.082 | 1 | 0.18483667 | -1.901889915 | 0 | 0.038 | 1 | 0.613101537 | 0.335508745 |
| Prrg4 | 0.709487466 | 2.088809267 | 0.048 | 0.034 | 1 | 0.18483667 | -1.404390255 | 0 | 0.038 | 1 | 0.709487466 | 0.335508745 |
| C030014I23Rik | 0.818140453 | 1.088809267 | 0.048 | 0.063 | 1 | 0.18483667 | -1.404390255 | 0 | 0.038 | 1 | 0.818140453 | 0.335508745 |
| 4933424G06Rik | 0.988367256 | 1.171271427 | 0.048 | 0.049 | 1 | 0.18483667 | -1.404390255 | 0 | 0.038 | 1 | 0.988367256 | 0.335508745 |
| Slc2a4 | 0.184844626 | -1.233118828 | 0 | 0.078 | 1 | 0.260653677 | -0.638855509 | 0 | 0.028 | 1 | 0.260653677 | 0.335521717 |
| Gm56606 | 0.251619726 | -0.574155746 | 0 | 0.06 | 1 | 0.18484555 | -1.404390255 | 0 | 0.038 | 1 | 0.251619726 | 0.335523223 |
| Pak6 | 0.373857158 | 2.088809267 | 0.048 | 0.019 | 1 | 0.18484555 | -1.564854927 | 0 | 0.038 | 1 | 0.373857158 | 0.335523223 |
| Sstr2 | 0.430431493 | -0.295854583 | 0.048 | 0.104 | 1 | 0.18484555 | -1.404390255 | 0 | 0.038 | 1 | 0.430431493 | 0.335523223 |
| Nrgn | 0.693076093 | 1.852741909 | 0.048 | 0.075 | 1 | 0.18484555 | -1.709244837 | 0 | 0.038 | 1 | 0.693076093 | 0.335523223 |
| Ccdc148 | 0.425637475 | -0.718545655 | 0 | 0.03 | 1 | 0.18484666 | -2.271123724 | 0 | 0.038 | 1 | 0.425637475 | 0.335525033 |
| Tbx3os1 | 0.600957465 | 0.171271427 | 0.048 | 0.082 | 1 | 0.18484666 | -1.638855509 | 0 | 0.038 | 1 | 0.600957465 | 0.335525033 |
| Trac | 0.744838936 | 0.866416846 | 0.048 | 0.034 | 1 | 0.18484666 | -1.486852415 | 0 | 0.038 | 1 | 0.744838936 | 0.335525033 |
| Pdlim3 | 0.878412101 | 0.400753273 | 0.048 | 0.041 | 1 | 0.18484666 | -1.960783604 | 0 | 0.038 | 1 | 0.878412101 | 0.335525033 |
| Emid1 | 0.184850021 | -1.133583154 | 0 | 0.078 | 1 | 0.955025729 | 1.361144491 | 0.022 | 0.024 | 1 | 0.955025729 | 0.335530511 |
| Tenm1 | 0.277250164 | 0.97333205 | 0.048 | 0.015 | 1 | 0.184853321 | -3.017367132 | 0 | 0.038 | 1 | 0.277250164 | 0.335535891 |
| 9630014M24Rik | 0.304430544 | -0.911190733 | 0 | 0.049 | 1 | 0.184853321 | -1.84048937 | 0 | 0.038 | 1 | 0.304430544 | 0.335535891 |
| Tmprss7 | 0.532426384 | 0.673771768 | 0 | 0.019 | 1 | 0.184853321 | -1.709244837 | 0 | 0.038 | 1 | 0.532426384 | 0.335535891 |
| Gab3 | 0.285570571 | -0.970084422 | 0 | 0.052 | 1 | 0.184854801 | -1.709244837 | 0 | 0.038 | 1 | 0.285570571 | 0.335538304 |
| Slc2a3 | 0.991275314 | 0.79930265 | 0.048 | 0.049 | 1 | 0.184858871 | -2.017367132 | 0 | 0.038 | 1 | 0.991275314 | 0.335544939 |
| Fabp4 | 0.285571003 | -1.535681598 | 0 | 0.052 | 1 | 0.184859611 | -2.271123724 | 0 | 0.038 | 1 | 0.285571003 | 0.335546146 |
| Mamstr | 0.304385789 | -0.326228232 | 0 | 0.049 | 1 | 0.184867751 | -0.901889915 | 0 | 0.038 | 1 | 0.304385789 | 0.335559416 |
| Il3ra | 0.304411431 | -0.413691074 | 0 | 0.049 | 1 | 0.184867751 | -0.901889915 | 0 | 0.038 | 1 | 0.304411431 | 0.335559416 |
| Fkbpl | 0.304430544 | -0.574155746 | 0 | 0.049 | 1 | 0.184867751 | -0.901889915 | 0 | 0.038 | 1 | 0.304430544 | 0.335559416 |
| Gm12840 | 0.396971401 | 0.351843673 | 0 | 0.034 | 1 | 0.184867751 | -0.901889915 | 0 | 0.038 | 1 | 0.396971401 | 0.335559416 |
| Gm57238 | 0.425599726 | 0.351843673 | 0 | 0.03 | 1 | 0.184867751 | -0.901889915 | 0 | 0.038 | 1 | 0.425599726 | 0.335559416 |
| 9330111N05Rik | 0.457281049 | 0.214340149 | 0 | 0.026 | 1 | 0.184867751 | -0.901889915 | 0 | 0.038 | 1 | 0.457281049 | 0.335559416 |
| Slc19a3 | 0.457292877 | 2.503846766 | 0.048 | 0.022 | 1 | 0.184867751 | -0.901889915 | 0 | 0.038 | 1 | 0.457292877 | 0.335559416 |
| Vsig10l | 0.63207476 | 1.088809267 | 0 | 0.011 | 1 | 0.184867751 | -0.901889915 | 0 | 0.038 | 1 | 0.63207476 | 0.335559416 |
| Gm32687 | 0.632917461 | 1.936806174 | 0.048 | 0.03 | 1 | 0.184867751 | -0.901889915 | 0 | 0.038 | 1 | 0.632917461 | 0.335559416 |
| Gm19426 | 0.784037261 | 1.79930265 | 0.048 | 0.037 | 1 | 0.184867751 | -0.901889915 | 0 | 0.038 | 1 | 0.784037261 | 0.335559416 |
| 2210406O10Rik | 0.813128768 | 0.79930265 | 0.048 | 0.063 | 1 | 0.184867751 | -0.901889915 | 0 | 0.038 | 1 | 0.813128768 | 0.335559416 |
| Enkd1 | 0.85370107 | 1.673771768 | 0.048 | 0.041 | 1 | 0.184867751 | -0.901889915 | 0 | 0.038 | 1 | 0.85370107 | 0.335559416 |
| Itga8 | 0.184876992 | -1.611630451 | 0 | 0.078 | 1 | 0.421413347 | -0.589945908 | 0.044 | 0.08 | 1 | 0.421413347 | 0.335574481 |
| A630001G21Rik | 0.184887779 | -1.280424543 | 0 | 0.078 | 1 | 0.295444118 | -0.124282336 | 0.022 | 0.062 | 1 | 0.295444118 | 0.335592068 |
| Vcan | 0.18491792 | -2.955584852 | 0.048 | 0.149 | 1 | 0.634491566 | 1.632446513 | 0.044 | 0.066 | 1 | 0.634491566 | 0.335641203 |
| Greb1l | 0.184932006 | -1.785659851 | 0 | 0.078 | 1 | 0.828866812 | 0.185572927 | 0.111 | 0.128 | 1 | 0.828866812 | 0.335664165 |

| | | | | | | | | | | | | |
|---|---|---|---|---|---|---|---|---|---|---|---|---|
| Slc12a8 | 0.184946028 | 1.054861935 | 0.143 | 0.067 | 1 | 0.615755764 | 0.683072586 | 0.022 | 0.038 | 1 | 0.615755764 | 0.335687022 |
| 1700024B18Rik | 0.184954926 | -2.785659851 | 0 | 0.078 | 1 | 0.542493488 | -0.686161223 | 0.067 | 0.093 | 1 | 0.542493488 | 0.335701527 |
| Cyfip1 | 0.18540297 | 1.130085543 | 0.667 | 0.653 | 1 | 0.304801162 | 0.40973726 | 0.889 | 0.83 | 1 | 0.304801162 | 0.336431678 |
| Kat2a | 0.185434714 | -0.880817084 | 0.048 | 0.153 | 1 | 0.382795245 | 0.172110667 | 0.111 | 0.173 | 1 | 0.382795245 | 0.336483395 |
| 6030443J06Rik | 0.961702232 | 1.316219763 | 0.095 | 0.108 | 1 | 0.185687841 | 0.495445583 | 0.044 | 0.114 | 1 | 0.961702232 | 0.336895707 |
| Taok3 | 0.729460358 | 0.911271081 | 0.381 | 0.526 | 1 | 0.18599685 | 0.430530554 | 0.822 | 0.675 | 1 | 0.729460358 | 0.337398871 |
| Casp9 | 0.764232894 | 0.735172312 | 0.048 | 0.067 | 1 | 0.186106628 | -0.486852415 | 0.044 | 0.111 | 1 | 0.764232894 | 0.337577578 |
| Tut1 | 0.678477115 | 0.565247311 | 0.143 | 0.205 | 1 | 0.186185209 | 0.143913775 | 0.133 | 0.246 | 1 | 0.678477115 | 0.337705486 |
| Zfp935 | 0.635543451 | 0.503846766 | 0.19 | 0.153 | 1 | 0.186254439 | 0.15951063 | 0.044 | 0.114 | 1 | 0.635543451 | 0.337818161 |
| 1190005I06Rik | 0.876931167 | 0.901182264 | 0.095 | 0.116 | 1 | 0.186259388 | -0.280401538 | 0.044 | 0.111 | 1 | 0.876931167 | 0.337826216 |
| Nudt13 | 0.315042054 | -0.427766259 | 0.143 | 0.25 | 1 | 0.18632404 | -0.531940305 | 0.156 | 0.253 | 1 | 0.315042054 | 0.337931432 |
| Dhcr24 | 0.300639533 | -0.587895802 | 0.238 | 0.369 | 1 | 0.186576192 | -0.468930507 | 0.067 | 0.142 | 1 | 0.300639533 | 0.338341708 |
| Tgfbr1 | 0.186582121 | 0.962276861 | 0.619 | 0.575 | 1 | 0.273791651 | 0.148218934 | 0.844 | 0.716 | 1 | 0.273791651 | 0.338351354 |
| Gm42659 | 0.540194551 | -0.133583154 | 0.048 | 0.086 | 1 | 0.186689996 | -0.242926832 | 0.044 | 0.111 | 1 | 0.540194551 | 0.338526837 |
| Gabpb2 | 0.854830352 | 0.763323353 | 0.333 | 0.429 | 1 | 0.186945828 | -0.468520592 | 0.556 | 0.588 | 1 | 0.854830352 | 0.338942914 |
| Orc4 | 0.187050717 | 1.031323772 | 0.524 | 0.53 | 1 | 0.246981517 | 0.607028395 | 0.711 | 0.678 | 1 | 0.246981517 | 0.339113464 |
| Zfp472 | 0.222227681 | -0.911190733 | 0 | 0.067 | 1 | 0.187103018 | -0.204452685 | 0.044 | 0.111 | 1 | 0.222227681 | 0.339198497 |
| Pcdhga9 | 0.370840523 | 0.214340149 | 0 | 0.037 | 1 | 0.187375724 | 2.005000681 | 0.089 | 0.045 | 1 | 0.370840523 | 0.339641787 |
| Ephb2 | 0.904863907 | 0.566477755 | 0.333 | 0.369 | 1 | 0.187398918 | 1.210319589 | 0.711 | 0.664 | 1 | 0.904863907 | 0.339679481 |
| Edem3 | 0.789981864 | 0.921911958 | 0.524 | 0.608 | 1 | 0.187494025 | -0.149817428 | 0.711 | 0.751 | 1 | 0.789981864 | 0.339834041 |
| Eaf2 | 0.82911125 | 0.576910228 | 0.19 | 0.187 | 1 | 0.187600735 | 0.563333342 | 0.089 | 0.183 | 1 | 0.82911125 | 0.340007434 |
| Tmc6 | 0.373773585 | 0.46732089 | 0.333 | 0.261 | 1 | 0.187713484 | 0.548771494 | 0.2 | 0.349 | 1 | 0.373773585 | 0.340190615 |
| Mlh3 | 0.523365791 | 0.129451251 | 0.095 | 0.157 | 1 | 0.187718726 | -1.280401538 | 0.067 | 0.135 | 1 | 0.523365791 | 0.340199132 |
| Zfp689 | 0.187751816 | 1.60338244 | 0.143 | 0.067 | 1 | 0.327170273 | 0.147019686 | 0.044 | 0.09 | 1 | 0.327170273 | 0.340252888 |
| Gm42939 | 0.709465802 | 1.936806174 | 0.048 | 0.034 | 1 | 0.187819301 | 1.42003818 | 0.156 | 0.097 | 1 | 0.709465802 | 0.340362512 |
| Evi5l | 0.837704711 | 0.250560337 | 0.286 | 0.287 | 1 | 0.187987696 | 0.258574758 | 0.111 | 0.215 | 1 | 0.837704711 | 0.340636018 |
| Pdk4 | 0.200906655 | -2.438928365 | 0.143 | 0.246 | 1 | 0.188144761 | -0.804767448 | 0.156 | 0.26 | 1 | 0.200906655 | 0.340891071 |
| Zc3h6 | 0.188327122 | -1.167530486 | 0.095 | 0.209 | 1 | 0.915554665 | 0.55062229 | 0.111 | 0.121 | 1 | 0.915554665 | 0.34118714 |
| Zfp93 | 0.188735947 | 2.214340149 | 0.095 | 0.037 | 1 | 0.671398648 | 1.020107573 | 0.067 | 0.052 | 1 | 0.671398648 | 0.341850637 |
| Actr8 | 0.188737801 | -0.718545655 | 0.095 | 0.224 | 1 | 0.206826099 | 0.197645759 | 0.2 | 0.336 | 1 | 0.206826099 | 0.341853645 |
| Tmem202 | 0.558208169 | 1.049280903 | 0.143 | 0.108 | 1 | 0.188775958 | 2.268035087 | 0.111 | 0.062 | 1 | 0.558208169 | 0.341915553 |
| Mindy4 | 0.188838462 | -0.586755783 | 0.143 | 0.295 | 1 | 0.455566462 | 1.064162754 | 0.222 | 0.304 | 1 | 0.455566462 | 0.342016958 |
| Gcg | 0.209005777 | -1.413691074 | 0 | 0.071 | 1 | 0.188874972 | 1.744473131 | 0.089 | 0.045 | 1 | 0.209005777 | 0.342076189 |
| Zfp711 | 0.96988282 | -0.233118828 | 0.048 | 0.045 | 1 | 0.189005656 | -0.474468691 | 0.067 | 0.142 | 1 | 0.96988282 | 0.342288175 |
| Alg8 | 0.189116536 | 1.79930265 | 0.19 | 0.104 | 1 | 0.973671689 | 1.525531309 | 0.089 | 0.093 | 1 | 0.973671689 | 0.342468008 |
| Cep83 | 0.189533279 | -0.60421298 | 0.381 | 0.556 | 1 | 0.692877814 | 0.152827255 | 0.711 | 0.699 | 1 | 0.692877814 | 0.343143695 |
| Dcun1d4 | 0.422232953 | 1.469413269 | 0.286 | 0.235 | 1 | 0.189937026 | 0.268035087 | 0.111 | 0.204 | 1 | 0.422232953 | 0.343797978 |
| Bri3 | 0.190399927 | 0.10560451 | 0.857 | 0.925 | 1 | 0.438735287 | 0.167469285 | 0.933 | 0.955 | 1 | 0.438735287 | 0.344547722 |
| Mbd2 | 0.190510362 | -0.183650152 | 0.905 | 0.974 | 1 | 0.644657912 | 0.320257421 | 1 | 0.976 | 1 | 0.644657912 | 0.344726526 |
| Pik3c3 | 0.538092925 | 0.74303443 | 0.238 | 0.194 | 1 | 0.190582394 | 0.256807831 | 0.111 | 0.208 | 1 | 0.538092925 | 0.344843139 |
| Rbm28 | 0.190644812 | -0.418674481 | 0.333 | 0.534 | 1 | 0.63742724 | 0.272984303 | 0.644 | 0.599 | 1 | 0.63742724 | 0.34494418 |
| Gm17106 | 0.190737407 | 1.393663848 | 0.19 | 0.104 | 1 | 0.384116335 | 0.946106992 | 0.2 | 0.163 | 1 | 0.384116335 | 0.345094056 |
| Parp12 | 0.393091661 | 0.825774861 | 0.238 | 0.172 | 1 | 0.191076795 | 0.229899958 | 0.133 | 0.232 | 1 | 0.393091661 | 0.345643248 |
| Gm20275 | 0.19125008 | 2.107424945 | 0.19 | 0.112 | 1 | 0.391795143 | 2.393565969 | 0.222 | 0.194 | 1 | 0.391795143 | 0.345923567 |
| Nemp2 | 0.191414462 | 2.088809267 | 0.095 | 0.037 | 1 | 0.406607361 | 0.683072586 | 0.022 | 0.052 | 1 | 0.406607361 | 0.346189428 |
| Ankrd2 | 0.191414462 | 2.088809267 | 0.095 | 0.037 | 1 | 0.272266345 | 1.946106992 | 0.089 | 0.052 | 1 | 0.272266345 | 0.346189428 |
| Gm56935 | 0.191434794 | 2.780686972 | 0.095 | 0.037 | 1 | 0.402446709 | 2.304560963 | 0.067 | 0.042 | 1 | 0.402446709 | 0.346222308 |
| Hnrnpl | 0.606110827 | -0.105201353 | 0.857 | 0.951 | 1 | 0.19144491 | -0.186182594 | 0.978 | 0.979 | 1 | 0.606110827 | 0.346238666 |
| Scaf4 | 0.224926763 | 0.668477468 | 0.524 | 0.496 | 1 | 0.191553839 | 0.330056813 | 0.889 | 0.751 | 1 | 0.224926763 | 0.346414805 |
| Chdh | 0.265217017 | -0.355975576 | 0.048 | 0.138 | 1 | 0.191896705 | 0.172110667 | 0.222 | 0.36 | 1 | 0.265217017 | 0.346969064 |
| Bysl | 0.192116301 | -0.496153234 | 0.048 | 0.16 | 1 | 0.425924593 | 0.245667274 | 0.133 | 0.194 | 1 | 0.425924593 | 0.347323929 |
| Prr16 | 0.492599419 | -1.233118828 | 0 | 0.022 | 1 | 0.192466129 | -0.346674757 | 0.044 | 0.111 | 1 | 0.492599419 | 0.347889047 |
| Trim11 | 0.34847756 | 1.298732336 | 0.286 | 0.239 | 1 | 0.192741256 | 0.152557869 | 0.156 | 0.263 | 1 | 0.34847756 | 0.348333321 |
| Rab20 | 0.193420756 | -1.303508156 | 0.048 | 0.149 | 1 | 0.947237337 | 0.852997588 | 0.156 | 0.163 | 1 | 0.947237337 | 0.349429924 |
| Agbl4 | 0.19345468 | 2.181918671 | 0.143 | 0.071 | 1 | 0.855991671 | 1.055041364 | 0.067 | 0.062 | 1 | 0.855991671 | 0.349484646 |
| Traf3 | 0.349848155 | -0.353028292 | 0.381 | 0.549 | 1 | 0.193481245 | 0.229706968 | 0.911 | 0.72 | 1 | 0.349848155 | 0.349527497 |
| Zfp866 | 0.193609092 | 1.171271427 | 0.238 | 0.146 | 1 | 0.245996883 | 0.549216839 | 0.133 | 0.221 | 1 | 0.245996883 | 0.349733703 |
| Sema6d | 0.871465156 | 0.710297644 | 0.095 | 0.112 | 1 | 0.193613906 | 0.454253896 | 0.067 | 0.142 | 1 | 0.871465156 | 0.349741467 |
| Cfap70 | 0.362152637 | 1.425844254 | 0.048 | 0.119 | 1 | 0.193986349 | -0.859185464 | 0.156 | 0.253 | 1 | 0.362152637 | 0.350341995 |
| Gm44710 | 0.194115887 | 1.97333205 | 0.095 | 0.037 | 1 | 0.295444359 | -0.255526869 | 0.022 | 0.062 | 1 | 0.295444359 | 0.350550796 |
| Ctnnal1 | 0.908176802 | 0.832469514 | 0.571 | 0.619 | 1 | 0.19433811 | -0.137603714 | 0.489 | 0.637 | 1 | 0.908176802 | 0.35090892 |
| Igf2bp1 | 0.936805841 | 1.010806755 | 0.048 | 0.045 | 1 | 0.194811426 | 1.744473131 | 0.089 | 0.045 | 1 | 0.936805841 | 0.351671361 |
| Gm15892 | 0.578510387 | 0.400753273 | 0.048 | 0.086 | 1 | 0.195259355 | -0.155646507 | 0.089 | 0.173 | 1 | 0.578510387 | 0.352392494 |
| Cebpg | 0.195400369 | -0.247297763 | 0.429 | 0.675 | 1 | 0.575643393 | 0.107858062 | 0.8 | 0.779 | 1 | 0.575643393 | 0.352619434 |
| Zfp867 | 0.195490682 | -0.970084422 | 0.048 | 0.149 | 1 | 0.750558379 | 0.313838777 | 0.067 | 0.083 | 1 | 0.750558379 | 0.352764757 |
| Capn11 | 0.324767242 | -0.233118828 | 0 | 0.045 | 1 | 0.1958708 | -0.589945908 | 0.022 | 0.076 | 1 | 0.324767242 | 0.35337623 |
| Mrs2 | 0.323619406 | -0.326228232 | 0.095 | 0.187 | 1 | 0.19591137 | 0.253084745 | 0.133 | 0.242 | 1 | 0.323619406 | 0.353441475 |
| Sh3pxd2b | 0.196031875 | -0.704739856 | 0.143 | 0.28 | 1 | 0.269656553 | 0.216930746 | 0.333 | 0.491 | 1 | 0.269656553 | 0.353635254 |
| Sh2b3 | 0.518406365 | 1.115776315 | 0.19 | 0.153 | 1 | 0.196136844 | 0.380509816 | 0.111 | 0.211 | 1 | 0.518406365 | 0.353804026 |
| Clcf1 | 0.891519311 | 1.289107917 | 0.095 | 0.108 | 1 | 0.196139846 | 0.410054092 | 0.111 | 0.218 | 1 | 0.891519311 | 0.353808852 |
| Trp53bp2 | 0.196209496 | 1.041503552 | 0.524 | 0.437 | 1 | 0.354543893 | 0.303900539 | 0.756 | 0.661 | 1 | 0.354543893 | 0.353920825 |
| Ipo13 | 0.196251598 | 2.021695071 | 0.143 | 0.071 | 1 | 0.337895195 | 0.470078863 | 0.089 | 0.152 | 1 | 0.337895195 | 0.353988506 |
| 9430015G10Rik | 0.196329837 | -0.785659851 | 0 | 0.075 | 1 | 0.53630117 | 0.846571318 | 0.044 | 0.073 | 1 | 0.53630117 | 0.354114269 |
| Gpbp1l1 | 0.995247125 | 0.400753273 | 0.429 | 0.47 | 1 | 0.196363169 | 0.364946061 | 0.844 | 0.647 | 1 | 0.995247125 | 0.354167843 |
| Sirt6 | 0.196379028 | -0.849790188 | 0 | 0.075 | 1 | 0.295444359 | -0.255526869 | 0.022 | 0.062 | 1 | 0.295444359 | 0.354193334 |
| A330048O09Rik | 0.196379028 | -0.849790188 | 0 | 0.075 | 1 | 0.765720982 | 1.361144491 | 0.022 | 0.031 | 1 | 0.765720982 | 0.354193334 |
| 2410022M11Rik | 0.196379028 | -0.849790188 | 0 | 0.075 | 1 | 0.835943453 | 0.513147585 | 0.022 | 0.028 | 1 | 0.835943453 | 0.354193334 |
| Pyroxd2 | 0.196379028 | -0.849790188 | 0 | 0.075 | 1 | 0.358152023 | 0.410054092 | 0.044 | 0.087 | 1 | 0.358152023 | 0.354193334 |
| Uprt | 0.196379028 | -0.849790188 | 0 | 0.075 | 1 | 0.787621874 | 1.020107573 | 0.044 | 0.055 | 1 | 0.787621874 | 0.354193334 |
| P2ry2 | 0.196422748 | -0.970084422 | 0 | 0.075 | 1 | 0.293659664 | -0.316927414 | 0 | 0.024 | 1 | 0.293659664 | 0.354263599 |
| 4930512B01Rik | 0.196423035 | -1.184209227 | 0 | 0.075 | 1 | 0.213147645 | 1.841770332 | 0.178 | 0.128 | 1 | 0.213147645 | 0.354264062 |
| Rdh12 | 0.196423035 | -0.911190733 | 0 | 0.075 | 1 | 0.268666992 | -0.124282336 | 0.022 | 0.066 | 1 | 0.268666992 | 0.354264062 |
| Fbxw17 | 0.196423035 | -0.911190733 | 0 | 0.075 | 1 | 0.356253 | 0.647448676 | 0.067 | 0.118 | 1 | 0.356253 | 0.354264062 |
| Bcdin3d | 0.251580301 | -0.496153234 | 0 | 0.06 | 1 | 0.19643899 | -0.539319835 | 0.022 | 0.076 | 1 | 0.251580301 | 0.354289704 |
| Arrb1 | 0.196460997 | -1.02666795 | 0 | 0.075 | 1 | 0.89527588 | 0.490427508 | 0.067 | 0.073 | 1 | 0.89527588 | 0.354325071 |
| Lrch2 | 0.196460997 | -1.02666795 | 0 | 0.075 | 1 | 0.628262013 | 0.776181991 | 0.089 | 0.121 | 1 | 0.628262013 | 0.354325071 |
| Gm14399 | 0.196462148 | -0.970084422 | 0 | 0.075 | 1 | 0.252512009 | -0.638855509 | 0.044 | 0.097 | 1 | 0.252512009 | 0.35432692 |
| Cdc6 | 0.196495505 | -1.184209227 | 0 | 0.075 | 1 | 0.521501458 | 1.152557869 | 0.089 | 0.066 | 1 | 0.521501458 | 0.354380526 |
| Etaa1os | 0.196496367 | -1.184209227 | 0 | 0.075 | 1 | 0.232018094 | -0.638855509 | 0 | 0.031 | 1 | 0.232018094 | 0.354381912 |
| 4932422M17Rik | 0.196523971 | -1.184209227 | 0 | 0.075 | 1 | 0.459383005 | -0.174908409 | 0.044 | 0.076 | 1 | 0.459383005 | 0.35442627 |
| Fam13a | 0.196523971 | -1.184209227 | 0 | 0.075 | 1 | 0.847509275 | 0.513147585 | 0.022 | 0.028 | 1 | 0.847509275 | 0.35442627 |
| Cenpk | 0.196547259 | -1.496153234 | 0 | 0.075 | 1 | 0.232000895 | -1.124282336 | 0 | 0.031 | 1 | 0.232000895 | 0.354463693 |
| Lamb1 | 0.196630625 | -2.303508156 | 0 | 0.075 | 1 | 0.33989087 | -1.271123724 | 0.022 | 0.055 | 1 | 0.33989087 | 0.354597647 |
| Ccdc73 | 0.196657642 | -0.718545655 | 0 | 0.075 | 1 | 0.617514334 | 1.098110085 | 0.022 | 0.038 | 1 | 0.617514334 | 0.354641056 |
| Gm14295 | 0.196657642 | -0.718545655 | 0 | 0.075 | 1 | 0.400430002 | -0.539319835 | 0.044 | 0.08 | 1 | 0.400430002 | 0.354641056 |
| Atp6v0e2 | 0.196657642 | -0.718545655 | 0 | 0.075 | 1 | 0.936023974 | 1.460680165 | 0.022 | 0.021 | 1 | 0.936023974 | 0.354641056 |
| Trim13 | 0.196657642 | -0.718545655 | 0 | 0.075 | 1 | 0.287259788 | -0.564854927 | 0.044 | 0.093 | 1 | 0.287259788 | 0.354641056 |
| Pde1b | 0.196657642 | -0.718545655 | 0 | 0.075 | 1 | 0.670048992 | 1.361144491 | 0.022 | 0.014 | 1 | 0.670048992 | 0.354641056 |

| | | | | | | | | | | | | |
|---|---|---|---|---|---|---|---|---|---|---|---|---|
| Gm56489 | 0.196657642 | -0.718545655 | 0 | 0.075 | 1 | 0.387952149 | -0.316927414 | 0.022 | 0.052 | 1 | 0.387952149 | 0.354641056 |
| Usb1 | 0.196857648 | -0.970084422 | 0.048 | 0.149 | 1 | 0.211288067 | -0.871516266 | 0.067 | 0.131 | 1 | 0.211288067 | 0.354962363 |
| Hspg2 | 0.196886779 | 0.399725786 | 0.905 | 0.944 | 1 | 0.699385671 | 0.245058553 | 0.711 | 0.934 | 1 | 0.699385671 | 0.355009155 |
| Dis3l2 | 0.196963847 | 1.183024609 | 0.524 | 0.526 | 1 | 0.205027761 | 1.00302754 | 0.622 | 0.578 | 1 | 0.205027761 | 0.355132937 |
| 1700030K09Rik | 0.787619621 | 0.464318402 | 0.095 | 0.119 | 1 | 0.197094406 | -0.223818009 | 0.022 | 0.076 | 1 | 0.787619621 | 0.355342608 |
| Acadl | 0.473284583 | -0.452760638 | 0.524 | 0.534 | 1 | 0.197188555 | -0.340386387 | 0.778 | 0.754 | 1 | 0.473284583 | 0.355493784 |
| Lekr1 | 0.339402421 | 0.74303443 | 0.048 | 0.123 | 1 | 0.197201178 | -0.486852415 | 0.022 | 0.076 | 1 | 0.339402421 | 0.355514051 |
| Gm56560 | 0.67730218 | -0.280424543 | 0.048 | 0.071 | 1 | 0.197716916 | -0.432404631 | 0.022 | 0.076 | 1 | 0.67730218 | 0.356341853 |
| 2900052L18Rik | 0.603600115 | -0.133583154 | 0.048 | 0.078 | 1 | 0.197784427 | -0.271123724 | 0.022 | 0.076 | 1 | 0.603600115 | 0.356450175 |
| D430041D05Rik | 0.727607937 | 2.644024424 | 0.048 | 0.071 | 1 | 0.197851739 | 2.899883975 | 0.089 | 0.045 | 1 | 0.727607937 | 0.356558167 |
| Yap1 | 0.248940353 | 0.351843673 | 0.667 | 0.739 | 1 | 0.198164142 | -0.131136007 | 0.867 | 0.841 | 1 | 0.248940353 | 0.357059257 |
| Gm11713 | 0.198347037 | 2.088809267 | 0.143 | 0.071 | 1 | 0.844407538 | 0.719598462 | 0.089 | 0.104 | 1 | 0.844407538 | 0.357352527 |
| Nlgn3 | 0.532426384 | 0.866416846 | 0 | 0.019 | 1 | 0.198471982 | -0.223818009 | 0.022 | 0.076 | 1 | 0.532426384 | 0.357552837 |
| Zfp808 | 0.378181658 | -0.469186186 | 0.048 | 0.112 | 1 | 0.198547044 | -0.404390255 | 0.022 | 0.076 | 1 | 0.378181658 | 0.35767316 |
| Kctd1 | 0.718452047 | 0.222560656 | 0.238 | 0.321 | 1 | 0.198614466 | 0.544590632 | 0.556 | 0.45 | 1 | 0.718452047 | 0.357781225 |
| Csnk1g3 | 0.537273001 | 0.435782491 | 0.571 | 0.649 | 1 | 0.198752792 | 0.343704043 | 0.978 | 0.844 | 1 | 0.537273001 | 0.358002912 |
| Naa25 | 0.198753416 | 1.010806755 | 0.429 | 0.418 | 1 | 0.736879012 | 0.282788854 | 0.444 | 0.478 | 1 | 0.736879012 | 0.358003911 |
| Ifih1 | 0.73978548 | -0.280424543 | 0.095 | 0.119 | 1 | 0.198771353 | 1.632446513 | 0.067 | 0.142 | 1 | 0.73978548 | 0.358032655 |
| Galnt18 | 0.198886707 | 0.971822954 | 0.762 | 0.757 | 1 | 0.200598792 | 0.656600375 | 0.867 | 0.747 | 1 | 0.200598792 | 0.358217491 |
| Fam76b | 0.199030737 | -0.233118828 | 0.238 | 0.425 | 1 | 0.712962167 | 0.505019704 | 0.489 | 0.512 | 1 | 0.712962167 | 0.35844824 |
| Zc3h18 | 0.543366974 | 0.562569205 | 0.429 | 0.638 | 1 | 0.199176385 | -0.142093612 | 0.756 | 0.751 | 1 | 0.543366974 | 0.358681537 |
| Gas2l3 | 0.236481344 | -1.326228232 | 0 | 0.063 | 1 | 0.199269727 | -1.404390255 | 0.022 | 0.073 | 1 | 0.236481344 | 0.35883103 |
| Dusp4 | 0.7319334 | 1.010806755 | 0.048 | 0.034 | 1 | 0.199292044 | -1.960783604 | 0.022 | 0.073 | 1 | 0.7319334 | 0.35886677 |
| Hormad2 | 0.199372751 | 0.811275291 | 0.048 | 0.16 | 1 | 0.90156084 | 1.776181991 | 0.067 | 0.076 | 1 | 0.90156084 | 0.358996009 |
| Snx30 | 0.199467167 | 0.483947209 | 0.048 | 0.16 | 1 | 0.426177664 | 1.301201951 | 0.2 | 0.176 | 1 | 0.426177664 | 0.359147184 |
| Safb2 | 0.19953475 | 0.594413976 | 0.619 | 0.638 | 1 | 0.906155711 | 0.17894116 | 0.889 | 0.81 | 1 | 0.906155711 | 0.359255383 |
| Rtkn | 0.199716858 | -0.427766259 | 0.143 | 0.291 | 1 | 0.312262586 | 0.231861474 | 0.267 | 0.408 | 1 | 0.312262586 | 0.359546893 |
| Ak8 | 0.446392013 | 2.295260144 | 0.095 | 0.056 | 1 | 0.199819727 | 0.268035087 | 0.022 | 0.076 | 1 | 0.446392013 | 0.359711531 |
| Plekhh2 | 0.199851113 | 1.07635978 | 0.762 | 0.731 | 1 | 0.244712527 | 1.098110085 | 0.6 | 0.699 | 1 | 0.244712527 | 0.359761758 |
| Gm9967 | 0.960849104 | 0.766881172 | 0.048 | 0.045 | 1 | 0.199857141 | 0.223640968 | 0.022 | 0.076 | 1 | 0.960849104 | 0.359771406 |
| Crppa | 0.910409289 | 0.731874722 | 0.238 | 0.276 | 1 | 0.199899347 | 1.474485964 | 0.267 | 0.215 | 1 | 0.910409289 | 0.359838946 |
| Trp53i11 | 0.42752413 | -0.167530486 | 0.048 | 0.104 | 1 | 0.199917672 | -0.316927414 | 0.022 | 0.076 | 1 | 0.42752413 | 0.359868268 |
| Arhgap35 | 0.507049597 | 0.313875823 | 0.333 | 0.522 | 1 | 0.200225806 | 0.572041274 | 0.889 | 0.754 | 1 | 0.507049597 | 0.360361239 |
| Tmem132e | 0.492599419 | 0.503846766 | 0 | 0.022 | 1 | 0.200603908 | -1.316927414 | 0.022 | 0.073 | 1 | 0.492599419 | 0.360965888 |
| Cyb5r4 | 0.784481066 | 0.417432014 | 0.19 | 0.243 | 1 | 0.200987626 | 0.481438725 | 0.444 | 0.308 | 1 | 0.784481066 | 0.361579227 |
| Osbpl2 | 0.201054475 | -1.04047375 | 0.143 | 0.257 | 1 | 0.273826642 | 0.149094014 | 0.2 | 0.308 | 1 | 0.273826642 | 0.361686048 |
| Vps37c | 0.991489423 | 0.623145695 | 0.143 | 0.16 | 1 | 0.201191591 | 0.246973471 | 0.133 | 0.235 | 1 | 0.991489423 | 0.361905126 |
| Nup160 | 0.536763072 | 0.171271427 | 0.238 | 0.328 | 1 | 0.201396667 | 0.16601415 | 0.222 | 0.36 | 1 | 0.536763072 | 0.362232716 |
| Dido1 | 0.201445154 | 0.72986633 | 0.667 | 0.724 | 1 | 0.370284156 | 0.309965767 | 0.778 | 0.872 | 1 | 0.370284156 | 0.362310158 |
| Arhgap32 | 0.676771446 | 0.146729243 | 0.619 | 0.597 | 1 | 0.201662852 | 0.436954095 | 0.911 | 0.817 | 1 | 0.676771446 | 0.362657798 |
| Mtmr4 | 0.236422459 | -0.718545655 | 0 | 0.063 | 1 | 0.201772833 | -0.255526869 | 0.044 | 0.107 | 1 | 0.236422459 | 0.362833389 |
| Fzd4 | 0.370840523 | 0.214340149 | 0 | 0.037 | 1 | 0.201928208 | -1.404390255 | 0.022 | 0.073 | 1 | 0.370840523 | 0.363081416 |
| Postn | 0.202065973 | -0.877681471 | 0.238 | 0.392 | 1 | 0.447416547 | -2.189052591 | 0.067 | 0.104 | 1 | 0.447416547 | 0.363301289 |
| Prkcg | 0.287079255 | 1.080681386 | 0.381 | 0.328 | 1 | 0.2021409 | -0.466674533 | 0.2 | 0.332 | 1 | 0.287079255 | 0.363420856 |
| St3gal5 | 0.20257742 | -0.923163375 | 0.095 | 0.216 | 1 | 0.287241349 | 0.165224281 | 0.222 | 0.149 | 1 | 0.287241349 | 0.364117228 |
| Zswim9 | 0.28385815 | 2.451379346 | 0.095 | 0.045 | 1 | 0.202627037 | 0.132875504 | 0.044 | 0.111 | 1 | 0.28385815 | 0.364196358 |
| Smad2 | 0.447522532 | 0.621130288 | 0.333 | 0.56 | 1 | 0.202642998 | 0.617919907 | 0.778 | 0.657 | 1 | 0.447522532 | 0.364221812 |
| Brms1l | 0.220598509 | -0.582567986 | 0.238 | 0.384 | 1 | 0.202644403 | 0.117218908 | 0.289 | 0.457 | 1 | 0.220598509 | 0.364224052 |
| Nanos1 | 0.267953794 | -0.496153234 | 0 | 0.056 | 1 | 0.202665386 | 0.132875504 | 0.044 | 0.111 | 1 | 0.267953794 | 0.364257513 |
| Nmnat1 | 0.843963442 | 1.48112669 | 0.095 | 0.09 | 1 | 0.202826426 | -0.979892427 | 0.044 | 0.104 | 1 | 0.843963442 | 0.364514293 |
| Rapgefl1 | 0.658782751 | 1.351843673 | 0.048 | 0.03 | 1 | 0.203094074 | 0.168499413 | 0.044 | 0.111 | 1 | 0.658782751 | 0.364940945 |
| Elac1 | 0.856482079 | 0.815790773 | 0.095 | 0.086 | 1 | 0.203242543 | -1.071814916 | 0.022 | 0.073 | 1 | 0.856482079 | 0.365177555 |
| Gm19705 | 0.971771371 | 2.43673257 | 0.095 | 0.108 | 1 | 0.20365668 | 0.7849522 | 0.044 | 0.111 | 1 | 0.971771371 | 0.365837316 |
| Camp | 0.492608234 | 0.866416846 | 0 | 0.022 | 1 | 0.203895946 | -1.124282336 | 0.022 | 0.073 | 1 | 0.492608234 | 0.366218335 |
| Psat1 | 0.229775799 | -0.264827688 | 0.048 | 0.149 | 1 | 0.203896143 | -1.124282336 | 0.022 | 0.073 | 1 | 0.229775799 | 0.366218649 |
| H2ac20 | 0.684003327 | -0.280424543 | 0.048 | 0.071 | 1 | 0.203896143 | -0.960783604 | 0.022 | 0.073 | 1 | 0.684003327 | 0.366218649 |
| Prickle1 | 0.203898361 | 0.641196164 | 0.667 | 0.649 | 1 | 0.931391168 | 0.451470688 | 0.467 | 0.54 | 1 | 0.931391168 | 0.366222181 |
| Chtf8 | 0.341130224 | 0.129451251 | 0.048 | 0.123 | 1 | 0.203936031 | -0.901889915 | 0.044 | 0.104 | 1 | 0.341130224 | 0.366282157 |
| A230057D06Rik | 0.297213793 | -0.891012851 | 0.19 | 0.306 | 1 | 0.204068046 | -0.1336202 | 0.133 | 0.225 | 1 | 0.297213793 | 0.366492325 |
| Nfatc2 | 0.719618489 | 1.438767402 | 0.571 | 0.608 | 1 | 0.204081468 | 0.822074894 | 0.444 | 0.651 | 1 | 0.719618489 | 0.366513691 |
| Stx3 | 0.204316679 | -0.24509147 | 0.143 | 0.306 | 1 | 0.734400902 | 0.243787541 | 0.533 | 0.554 | 1 | 0.734400902 | 0.366888052 |
| Cul2 | 0.28831684 | 1.252928647 | 0.429 | 0.384 | 1 | 0.204328528 | 0.198636961 | 0.356 | 0.536 | 1 | 0.28831684 | 0.366906909 |
| Dtna | 0.760891107 | -0.133583154 | 0.048 | 0.063 | 1 | 0.204528194 | 1.651363726 | 0.067 | 0.145 | 1 | 0.760891107 | 0.367224606 |
| Dynlt1c | 0.248716669 | -0.818081329 | 0.048 | 0.134 | 1 | 0.204546826 | -0.901889915 | 0.022 | 0.073 | 1 | 0.248716669 | 0.367254247 |
| Zfp629 | 0.204670238 | 1.673771768 | 0.143 | 0.071 | 1 | 0.392315274 | 0.147019686 | 0.111 | 0.166 | 1 | 0.392315274 | 0.36745057 |
| Arl10 | 0.807768462 | 0.400753273 | 0.143 | 0.168 | 1 | 0.20471994 | -0.531940305 | 0.111 | 0.19 | 1 | 0.807768462 | 0.367529626 |
| Appl1 | 0.895917778 | 0.38689062 | 0.667 | 0.75 | 1 | 0.205043136 | 0.591043448 | 0.889 | 0.841 | 1 | 0.895917778 | 0.368043584 |
| Gnb4 | 0.936695708 | 0.287123095 | 0.714 | 0.791 | 1 | 0.205125083 | -0.185682881 | 0.622 | 0.886 | 1 | 0.936695708 | 0.368173866 |
| Mus81 | 0.315761485 | -0.611630451 | 0.048 | 0.119 | 1 | 0.20519517 | -0.84048937 | 0.022 | 0.073 | 1 | 0.315761485 | 0.368285283 |
| Gm16159 | 0.425639612 | 0.503846766 | 0 | 0.03 | 1 | 0.205548014 | -0.16492432 | 0.044 | 0.107 | 1 | 0.425639612 | 0.368846042 |
| Iffo1 | 0.307653794 | -0.385121921 | 0.048 | 0.127 | 1 | 0.205551003 | -0.316927414 | 0.067 | 0.135 | 1 | 0.307653794 | 0.368850792 |
| Ppp2r5a | 0.230364717 | -0.235647654 | 0.619 | 0.769 | 1 | 0.205655556 | -0.112027184 | 0.933 | 0.924 | 1 | 0.230364717 | 0.369016905 |
| Tbc1d22b | 0.206114225 | 1.979874895 | 0.238 | 0.157 | 1 | 0.252192919 | 0.361144491 | 0.111 | 0.201 | 1 | 0.252192919 | 0.369745376 |
| Gm26542 | 0.706651951 | 0.936806174 | 0.143 | 0.123 | 1 | 0.206533821 | 1.180572246 | 0.222 | 0.166 | 1 | 0.706651951 | 0.370411423 |
| Gm37529 | 0.206554098 | 1.957564734 | 0.143 | 0.075 | 1 | 0.337816978 | 0.470078863 | 0.089 | 0.152 | 1 | 0.337816978 | 0.370443601 |
| Lmln | 0.880619091 | 0.410737362 | 0.095 | 0.086 | 1 | 0.206623939 | 0.95724755 | 0.044 | 0.111 | 1 | 0.880619091 | 0.370554426 |
| ENSMUSG00000121307 | 0.206654721 | 1.007015176 | 0.381 | 0.31 | 1 | 0.763136946 | 0.479539192 | 0.244 | 0.315 | 1 | 0.763136946 | 0.370603268 |
| Sec14l1 | 0.427236768 | 0.547099964 | 0.619 | 0.81 | 1 | 0.206779836 | 0.251406496 | 0.933 | 0.827 | 1 | 0.427236768 | 0.370801771 |
| Ccdc181 | 0.251790506 | -0.413691074 | 0 | 0.06 | 1 | 0.206882538 | -0.901889915 | 0 | 0.035 | 1 | 0.251790506 | 0.370964691 |
| Trmt12 | 0.268100354 | -0.326228232 | 0 | 0.056 | 1 | 0.206882538 | -0.901889915 | 0 | 0.035 | 1 | 0.268100354 | 0.370964691 |
| Sting1 | 0.285571003 | -0.648156327 | 0 | 0.052 | 1 | 0.206882538 | -0.901889915 | 0 | 0.035 | 1 | 0.285571003 | 0.370964691 |
| Nthl1 | 0.304355016 | -0.233118828 | 0 | 0.049 | 1 | 0.206882538 | -0.901889915 | 0 | 0.035 | 1 | 0.304355016 | 0.370964691 |
| Rnf113a1 | 0.359270744 | 2.673771768 | 0.048 | 0.019 | 1 | 0.206882538 | -0.901889915 | 0 | 0.035 | 1 | 0.359270744 | 0.370964691 |
| Tmem79 | 0.370769338 | -0.849790188 | 0 | 0.037 | 1 | 0.206882538 | -0.901889915 | 0 | 0.035 | 1 | 0.370769338 | 0.370964691 |
| Tmem169 | 0.373857158 | 2.088809267 | 0.048 | 0.019 | 1 | 0.206882538 | -0.901889915 | 0 | 0.035 | 1 | 0.373857158 | 0.370964691 |
| Gm43566 | 0.396971401 | 0.351843673 | 0 | 0.034 | 1 | 0.206882538 | -0.901889915 | 0 | 0.035 | 1 | 0.396971401 | 0.370964691 |
| Naif1 | 0.425639612 | 0.503846766 | 0 | 0.03 | 1 | 0.206882538 | -0.901889915 | 0 | 0.035 | 1 | 0.425639612 | 0.370964691 |
| 1810059H22Rik | 0.425639612 | 0.503846766 | 0 | 0.03 | 1 | 0.206882538 | -0.901889915 | 0 | 0.035 | 1 | 0.425639612 | 0.370964691 |
| Irx3os | 0.425639612 | 0.503846766 | 0 | 0.03 | 1 | 0.206882538 | -0.901889915 | 0 | 0.035 | 1 | 0.425639612 | 0.370964691 |
| Teddm2 | 0.429310057 | 0.614878079 | 0.095 | 0.052 | 1 | 0.206882538 | -1.017367132 | 0 | 0.035 | 1 | 0.429310057 | 0.370964691 |
| 1700022I11Rik | 0.457281049 | 0.503846766 | 0 | 0.026 | 1 | 0.206882538 | -0.901889915 | 0 | 0.035 | 1 | 0.457281049 | 0.370964691 |
| 4930522L14Rik | 0.457281049 | 0.351843673 | 0 | 0.026 | 1 | 0.206882538 | -0.901889915 | 0 | 0.035 | 1 | 0.457281049 | 0.370964691 |
| Snn | 0.457292877 | 0.351843673 | 0 | 0.026 | 1 | 0.206882538 | -0.901889915 | 0 | 0.035 | 1 | 0.457292877 | 0.370964691 |
| Fbxl16 | 0.457308648 | 0.673771768 | 0 | 0.026 | 1 | 0.206882538 | -0.901889915 | 0 | 0.035 | 1 | 0.457308648 | 0.370964691 |
| Zkscan7 | 0.511386191 | 1.866416846 | 0.095 | 0.063 | 1 | 0.206882538 | -0.901889915 | 0 | 0.035 | 1 | 0.511386191 | 0.370964691 |
| Cacna2d2 | 0.532426384 | 0.866416846 | 0 | 0.019 | 1 | 0.206882538 | -0.901889915 | 0 | 0.035 | 1 | 0.532426384 | 0.370964691 |
| Faap24 | 0.532436372 | 1.088809267 | 0 | 0.019 | 1 | 0.206882538 | -0.901889915 | 0 | 0.035 | 1 | 0.532436372 | 0.370964691 |
| Kcna1 | 0.578156445 | 1.088809267 | 0 | 0.015 | 1 | 0.206882538 | -0.901889915 | 0 | 0.035 | 1 | 0.578156445 | 0.370964691 |

| | | | | | | | | | | | | |
|---|---|---|---|---|---|---|---|---|---|---|---|---|
| Taf9 | 0.578156445 | 1.088809267 | 0 | 0.015 | 1 | 0.206882538 | -0.901889915 | 0 | 0.035 | 1 | 0.578156445 | 0.370964691 |
| Vash2 | 0.63207476 | 1.351843673 | 0 | 0.011 | 1 | 0.206882538 | -0.901889915 | 0 | 0.035 | 1 | 0.63207476 | 0.370964691 |
| Gm28379 | 0.63207476 | 1.351843673 | 0 | 0.011 | 1 | 0.206882538 | -0.901889915 | 0 | 0.035 | 1 | 0.63207476 | 0.370964691 |
| Upb1 | 0.632076134 | 1.673771768 | 0 | 0.011 | 1 | 0.206882538 | -0.901889915 | 0 | 0.035 | 1 | 0.632076134 | 0.370964691 |
| Phf2os1 | 0.632076134 | 1.673771768 | 0 | 0.011 | 1 | 0.206882538 | -0.901889915 | 0 | 0.035 | 1 | 0.632076134 | 0.370964691 |
| Gm10638 | 0.907384734 | 1.010806755 | 0.048 | 0.056 | 1 | 0.206882538 | -0.901889915 | 0 | 0.035 | 1 | 0.907384734 | 0.370964691 |
| Rad51c | 0.251619726 | -0.648156327 | 0 | 0.06 | 1 | 0.206897641 | -1.017367132 | 0 | 0.035 | 1 | 0.251619726 | 0.370988649 |
| Gm47093 | 0.261628269 | 2.936806174 | 0.048 | 0.015 | 1 | 0.206897641 | -1.223818009 | 0 | 0.035 | 1 | 0.261628269 | 0.370988649 |
| Kif18a | 0.304452918 | -0.718545655 | 0 | 0.049 | 1 | 0.206897641 | -1.017367132 | 0 | 0.035 | 1 | 0.304452918 | 0.370988649 |
| 4930434J08Rik | 0.30448834 | -0.133583154 | 0 | 0.049 | 1 | 0.206897641 | -1.017367132 | 0 | 0.035 | 1 | 0.30448834 | 0.370988649 |
| 1700007L15Rik | 0.324739547 | -0.133583154 | 0 | 0.045 | 1 | 0.206897641 | -1.124282336 | 0 | 0.035 | 1 | 0.324739547 | 0.370988649 |
| Cyria | 0.392357017 | 2.38426515 | 0.095 | 0.052 | 1 | 0.206897641 | -1.017367132 | 0 | 0.035 | 1 | 0.392357017 | 0.370988649 |
| Ccdc159 | 0.396971401 | 0.351843673 | 0 | 0.034 | 1 | 0.206897641 | -1.017367132 | 0 | 0.035 | 1 | 0.396971401 | 0.370988649 |
| A930033M14Rik | 0.396971401 | 0.351843673 | 0 | 0.034 | 1 | 0.206897641 | -1.124282336 | 0 | 0.035 | 1 | 0.396971401 | 0.370988649 |
| 1700080G18Rik | 0.457308648 | 0.673771768 | 0 | 0.026 | 1 | 0.206897641 | -1.124282336 | 0 | 0.035 | 1 | 0.457308648 | 0.370988649 |
| Txnl4b | 0.457308648 | 0.673771768 | 0 | 0.026 | 1 | 0.206897641 | -1.017367132 | 0 | 0.035 | 1 | 0.457308648 | 0.370988649 |
| Gm39529 | 0.57816106 | 0.673771768 | 0 | 0.015 | 1 | 0.206897641 | -1.124282336 | 0 | 0.035 | 1 | 0.57816106 | 0.370988649 |
| Bmp15 | 0.768257486 | 0.281454345 | 0.048 | 0.063 | 1 | 0.206897641 | -1.124282336 | 0 | 0.035 | 1 | 0.768257486 | 0.370988649 |
| 5031415H12Rik | 0.790349864 | 1.55829455 | 0.048 | 0.037 | 1 | 0.206897641 | -1.017367132 | 0 | 0.035 | 1 | 0.790349864 | 0.370988649 |
| Gm17455 | 0.79666227 | 1.258734268 | 0.048 | 0.037 | 1 | 0.206897641 | -1.017367132 | 0 | 0.035 | 1 | 0.79666227 | 0.370988649 |
| Spon2 | 0.891145213 | 0.400753273 | 0.048 | 0.056 | 1 | 0.206897641 | -1.017367132 | 0 | 0.035 | 1 | 0.891145213 | 0.370988649 |
| 9530056K15Rik | 0.918789496 | 1.55829455 | 0.048 | 0.045 | 1 | 0.206897641 | -1.017367132 | 0 | 0.035 | 1 | 0.918789496 | 0.370988649 |
| Gm46212 | 0.930795883 | 1.171271427 | 0.048 | 0.045 | 1 | 0.206897641 | -1.017367132 | 0 | 0.035 | 1 | 0.930795883 | 0.370988649 |
| Pip5kl1 | 0.930798611 | 1.171271427 | 0.048 | 0.045 | 1 | 0.206897641 | -1.017367132 | 0 | 0.035 | 1 | 0.930798611 | 0.370988649 |
| Gm57455 | 0.966230998 | 1.258734268 | 0.048 | 0.052 | 1 | 0.206897641 | -1.017367132 | 0 | 0.035 | 1 | 0.966230998 | 0.370988649 |
| Gm13274 | 0.457292877 | 0.351843673 | 0 | 0.026 | 1 | 0.206908969 | -1.316927414 | 0 | 0.035 | 1 | 0.457292877 | 0.371006616 |
| Gm48768 | 0.457292877 | 0.214340149 | 0 | 0.026 | 1 | 0.206908969 | -1.223818009 | 0 | 0.035 | 1 | 0.457292877 | 0.371006616 |
| Gm48163 | 0.457308648 | 0.673771768 | 0 | 0.026 | 1 | 0.206908969 | -1.316927414 | 0 | 0.035 | 1 | 0.457308648 | 0.371006616 |
| Lrrc4b | 0.57816106 | 1.351843673 | 0 | 0.015 | 1 | 0.206908969 | -1.223818009 | 0 | 0.035 | 1 | 0.57816106 | 0.371006616 |
| Adgrb2 | 0.359270744 | 2.673771768 | 0.048 | 0.019 | 1 | 0.206909388 | -1.404390255 | 0 | 0.035 | 1 | 0.359270744 | 0.371007282 |
| 4930403P22Rik | 0.373857158 | 2.088809267 | 0.048 | 0.019 | 1 | 0.206909388 | -1.124282336 | 0 | 0.035 | 1 | 0.373857158 | 0.371007282 |
| Kdm8 | 0.454205211 | 3.451379346 | 0.048 | 0.022 | 1 | 0.206909388 | -1.316927414 | 0 | 0.035 | 1 | 0.454205211 | 0.371007282 |
| Hopx | 0.573468511 | 1.351843673 | 0.048 | 0.026 | 1 | 0.206909388 | -1.404390255 | 0 | 0.035 | 1 | 0.573468511 | 0.371007282 |
| Efcab11 | 0.629713293 | 3.83423644 | 0.048 | 0.03 | 1 | 0.206909388 | -1.404390255 | 0 | 0.035 | 1 | 0.629713293 | 0.371007282 |
| Nlrp5-ps | 0.382765313 | 0.97333205 | 0.048 | 0.019 | 1 | 0.20691652 | -1.316927414 | 0 | 0.035 | 1 | 0.382765313 | 0.371018594 |
| 4933404O12Rik | 0.492590603 | 0.673771768 | 0 | 0.022 | 1 | 0.20691652 | -1.486852415 | 0 | 0.035 | 1 | 0.492590603 | 0.371018594 |
| She | 0.492608234 | 0.866416846 | 0 | 0.022 | 1 | 0.206917779 | -1.404390255 | 0 | 0.035 | 1 | 0.492608234 | 0.37102059 |
| Fam81a | 0.930798611 | 1.171271427 | 0.048 | 0.045 | 1 | 0.206917779 | -1.404390255 | 0 | 0.035 | 1 | 0.930798611 | 0.37102059 |
| Upk1b | 0.981928216 | -0.785659851 | 0.048 | 0.045 | 1 | 0.206917779 | -1.564854927 | 0 | 0.035 | 1 | 0.981928216 | 0.37102059 |
| 1700016P03Rik | 0.251724834 | -0.970084422 | 0 | 0.06 | 1 | 0.206918198 | -1.84048937 | 0 | 0.035 | 1 | 0.251724834 | 0.371021256 |
| Gm16287 | 0.963421211 | 1.171271427 | 0.048 | 0.052 | 1 | 0.206918198 | -1.223818009 | 0 | 0.035 | 1 | 0.963421211 | 0.371021256 |
| Gm49575 | 0.268029902 | -0.911190733 | 0 | 0.056 | 1 | 0.206920296 | -1.638855509 | 0 | 0.035 | 1 | 0.268029902 | 0.371024583 |
| 4930527F14Rik | 0.346827513 | -0.133583154 | 0 | 0.041 | 1 | 0.206923652 | -1.84048937 | 0 | 0.035 | 1 | 0.346827513 | 0.371029906 |
| Tnfsf11 | 0.773956881 | 1.699766976 | 0.048 | 0.067 | 1 | 0.206924071 | -2.446210431 | 0 | 0.035 | 1 | 0.773956881 | 0.371030572 |
| Cdk18 | 0.567009484 | 1.673771768 | 0.048 | 0.026 | 1 | 0.206924491 | -1.316927414 | 0 | 0.035 | 1 | 0.567009484 | 0.371031237 |
| H1f3 | 0.555576977 | -1.159118246 | 0.048 | 0.082 | 1 | 0.206926589 | -1.776359032 | 0 | 0.035 | 1 | 0.555576977 | 0.371034564 |
| Ndnf | 0.304470164 | -1.535681598 | 0 | 0.049 | 1 | 0.206928267 | -2.404390255 | 0 | 0.035 | 1 | 0.304470164 | 0.371037226 |
| Col14a1 | 0.533093785 | -1.233118828 | 0.048 | 0.086 | 1 | 0.206928267 | -2.486852415 | 0 | 0.035 | 1 | 0.533093785 | 0.371037226 |
| Ppp1r3f | 0.251580301 | -0.496153234 | 0 | 0.06 | 1 | 0.206932881 | -0.776359032 | 0 | 0.035 | 1 | 0.251580301 | 0.371044545 |
| Gm7628 | 0.267953794 | -0.496153234 | 0 | 0.056 | 1 | 0.206932881 | -0.776359032 | 0 | 0.035 | 1 | 0.267953794 | 0.371044545 |
| Dhh | 0.285622926 | -0.233118828 | 0 | 0.052 | 1 | 0.206932881 | -0.776359032 | 0 | 0.035 | 1 | 0.285622926 | 0.371044545 |
| Tmem44 | 0.346827513 | -0.133583154 | 0 | 0.041 | 1 | 0.206932881 | -0.776359032 | 0 | 0.035 | 1 | 0.346827513 | 0.371044545 |
| Gm57027 | 0.346860242 | -0.496153234 | 0 | 0.041 | 1 | 0.206932881 | -0.776359032 | 0 | 0.035 | 1 | 0.346860242 | 0.371044545 |
| Ptp4a1.1 | 0.370840523 | 0.214340149 | 0 | 0.037 | 1 | 0.206932881 | -0.776359032 | 0 | 0.035 | 1 | 0.370840523 | 0.371044545 |
| Gpr3 | 0.370840523 | 0.214340149 | 0 | 0.037 | 1 | 0.206932881 | -0.776359032 | 0 | 0.035 | 1 | 0.370840523 | 0.371044545 |
| Gm16091 | 0.39694807 | -0.574155746 | 0 | 0.034 | 1 | 0.206932881 | -0.776359032 | 0 | 0.035 | 1 | 0.39694807 | 0.371044545 |
| Aldh5a1 | 0.425614684 | 0.214340149 | 0 | 0.03 | 1 | 0.206932881 | -0.776359032 | 0 | 0.035 | 1 | 0.425614684 | 0.371044545 |
| Ccdc17 | 0.425639612 | 0.503846766 | 0 | 0.03 | 1 | 0.206932881 | -0.776359032 | 0 | 0.035 | 1 | 0.425639612 | 0.371044545 |
| Gm16152 | 0.457281049 | 0.503846766 | 0 | 0.026 | 1 | 0.206932881 | -0.776359032 | 0 | 0.035 | 1 | 0.457281049 | 0.371044545 |
| Gm16008 | 0.457281049 | 0.503846766 | 0 | 0.026 | 1 | 0.206932881 | -0.776359032 | 0 | 0.035 | 1 | 0.457281049 | 0.371044545 |
| Ovca2 | 0.492608234 | 0.866416846 | 0 | 0.022 | 1 | 0.206932881 | -0.776359032 | 0 | 0.035 | 1 | 0.492608234 | 0.371044545 |
| Gm48696 | 0.532436372 | 1.088809267 | 0 | 0.019 | 1 | 0.206932881 | -0.776359032 | 0 | 0.035 | 1 | 0.532436372 | 0.371044545 |
| 9330151L19Rik | 0.544576436 | 2.258734268 | 0.048 | 0.026 | 1 | 0.206932881 | -0.776359032 | 0 | 0.035 | 1 | 0.544576436 | 0.371044545 |
| Ghrl | 0.578156445 | 1.088809267 | 0 | 0.015 | 1 | 0.206932881 | -0.776359032 | 0 | 0.035 | 1 | 0.578156445 | 0.371044545 |
| Gm42067 | 0.609665403 | 0.150209812 | 0.048 | 0.078 | 1 | 0.206932881 | -0.776359032 | 0 | 0.035 | 1 | 0.609665403 | 0.371044545 |
| Hcar2 | 0.63207476 | 1.351843673 | 0 | 0.011 | 1 | 0.206932881 | -0.776359032 | 0 | 0.035 | 1 | 0.63207476 | 0.371044545 |
| 2310068J16Rik | 0.709465802 | 1.936806174 | 0.048 | 0.034 | 1 | 0.206932881 | -0.776359032 | 0 | 0.035 | 1 | 0.709465802 | 0.371044545 |
| Tnfrsf23 | 0.580185564 | 1.460778044 | 0.381 | 0.437 | 1 | 0.207095915 | 0.700350578 | 0.178 | 0.325 | 1 | 0.580185564 | 0.371303111 |
| Cog7 | 0.20713214 | -0.308080886 | 0.095 | 0.231 | 1 | 0.487686338 | 0.88398528 | 0.156 | 0.218 | 1 | 0.487686338 | 0.371360556 |
| Elmo2 | 0.207486144 | -0.911190733 | 0.048 | 0.146 | 1 | 0.573403356 | 0.250113179 | 0.156 | 0.208 | 1 | 0.573403356 | 0.371921788 |
| Suv39h1 | 0.234447954 | -0.785659851 | 0.048 | 0.138 | 1 | 0.207582905 | -0.638855509 | 0.044 | 0.107 | 1 | 0.234447954 | 0.372075147 |
| Kpna3 | 0.207727933 | 1.254819218 | 0.524 | 0.47 | 1 | 0.376436541 | 0.283141979 | 0.822 | 0.706 | 1 | 0.376436541 | 0.372304972 |
| Oxr1 | 0.20777768 | -0.450475659 | 0.619 | 0.728 | 1 | 0.415307649 | 0.137197442 | 0.867 | 0.716 | 1 | 0.415307649 | 0.372383796 |
| Maea | 0.317069191 | -0.194124696 | 0.333 | 0.511 | 1 | 0.207861077 | 0.413771422 | 0.711 | 0.644 | 1 | 0.317069191 | 0.372515927 |
| Zc3h14 | 0.20798728 | -0.500599139 | 0.429 | 0.634 | 1 | 0.23624862 | -0.223818009 | 0.844 | 0.82 | 1 | 0.23624862 | 0.372715852 |
| Zranb1 | 0.208083508 | 0.299293151 | 0.905 | 0.836 | 1 | 0.614679263 | 0.224390614 | 0.933 | 0.907 | 1 | 0.614679263 | 0.37286827 |
| Ino80d | 0.668987707 | 0.666277231 | 0.476 | 0.631 | 1 | 0.20835984 | 0.368963996 | 0.956 | 0.837 | 1 | 0.668987707 | 0.373305857 |
| Mamld1 | 0.401478924 | 1.125871361 | 0.381 | 0.336 | 1 | 0.208368522 | -0.691966845 | 0.089 | 0.163 | 1 | 0.401478924 | 0.373319603 |
| Rab5b | 0.20847458 | 0.705480627 | 0.476 | 0.455 | 1 | 0.42470146 | 0.400808709 | 0.6 | 0.571 | 1 | 0.42470146 | 0.373487509 |
| Gm10509 | 0.208797419 | -0.718545655 | 0 | 0.071 | 1 | 0.406553529 | 1.005000681 | 0.022 | 0.052 | 1 | 0.406553529 | 0.373998476 |
| Rab29 | 0.208844381 | -1.02666795 | 0 | 0.071 | 1 | 0.458286526 | -0.223818009 | 0.044 | 0.076 | 1 | 0.458286526 | 0.374072786 |
| Arrdc1 | 0.208844381 | -0.849790188 | 0 | 0.071 | 1 | 0.390923314 | 0.410054092 | 0.044 | 0.083 | 1 | 0.390923314 | 0.374072786 |
| Cfap141 | 0.208844381 | -0.849790188 | 0 | 0.071 | 1 | 0.606692213 | 0.728876276 | 0.067 | 0.097 | 1 | 0.606692213 | 0.374072786 |
| 1500002C15Rik | 0.208844381 | -0.785659851 | 0 | 0.071 | 1 | 0.386564371 | -0.404390255 | 0.022 | 0.052 | 1 | 0.386564371 | 0.374072786 |
| Gng7 | 0.208844381 | -0.849790188 | 0 | 0.071 | 1 | 0.50681717 | 0.197645759 | 0.044 | 0.073 | 1 | 0.50681717 | 0.374072786 |
| Rgs11 | 0.208844381 | -0.785659851 | 0 | 0.071 | 1 | 0.928508635 | 0.595609745 | 0.044 | 0.048 | 1 | 0.928508635 | 0.374072786 |
| G530011O06Rik | 0.208844381 | -0.785659851 | 0 | 0.071 | 1 | 0.376837034 | -0.316927414 | 0 | 0.017 | 1 | 0.376837034 | 0.374072786 |
| Slf1 | 0.528180846 | 0.487071053 | 0.238 | 0.34 | 1 | 0.208852227 | 0.648802684 | 0.222 | 0.36 | 1 | 0.528180846 | 0.374085202 |
| Bcap29 | 0.294367627 | 0.918884266 | 0.286 | 0.216 | 1 | 0.208856364 | 0.168499413 | 0.089 | 0.176 | 1 | 0.294367627 | 0.374091747 |
| Entpd2 | 0.208885811 | -0.911190733 | 0 | 0.071 | 1 | 0.299154247 | 0.757073168 | 0.022 | 0.062 | 1 | 0.299154247 | 0.37413834 |
| Ttf2 | 0.208885811 | -0.911190733 | 0 | 0.071 | 1 | 0.405392777 | 0.223640968 | 0.022 | 0.052 | 1 | 0.405392777 | 0.37413834 |
| Glp2r | 0.208885811 | -1.02666795 | 0 | 0.071 | 1 | 0.555549391 | 0.776181991 | 0.022 | 0.042 | 1 | 0.555549391 | 0.37413834 |
| Bok | 0.208886118 | -1.133583154 | 0 | 0.071 | 1 | 0.659789959 | -0.223818009 | 0.022 | 0.035 | 1 | 0.659789959 | 0.374138826 |
| Sh2d3c | 0.208886118 | -0.849790188 | 0 | 0.071 | 1 | 0.497256285 | 0.223640968 | 0.022 | 0.045 | 1 | 0.497256285 | 0.374138826 |
| Ccdc61 | 0.208886118 | -0.849790188 | 0 | 0.071 | 1 | 0.439307211 | -0.432404631 | 0.044 | 0.076 | 1 | 0.439307211 | 0.374138826 |
| Gm16973 | 0.208886118 | -0.970084422 | 0 | 0.071 | 1 | 0.745237189 | 1.460680165 | 0.089 | 0.08 | 1 | 0.745237189 | 0.374138826 |
| H2-D1 | 0.208903087 | -0.44170545 | 0.762 | 0.746 | 1 | 0.225129116 | -0.259323437 | 0.911 | 0.806 | 1 | 0.225129116 | 0.374165674 |
| 1110017D15Rik | 0.208921714 | -1.184209227 | 0 | 0.071 | 1 | 0.76524345 | 1.223640968 | 0.067 | 0.083 | 1 | 0.76524345 | 0.374195145 |

| | | | | | | | | | | | | |
|---|---|---|---|---|---|---|---|---|---|---|---|---|
| Ppp1r18os | 0.208921714 | -0.970084422 | 0 | 0.071 | 1 | 0.376837034 | -0.124282336 | 0 | 0.017 | 1 | 0.376837034 | 0.374195145 |
| Ptpn6 | 0.208922634 | -1.02666795 | 0 | 0.071 | 1 | 0.753274427 | 1.605070074 | 0.044 | 0.059 | 1 | 0.753274427 | 0.374196601 |
| Def8 | 0.208945673 | -0.818081329 | 0.048 | 0.146 | 1 | 0.467299899 | 0.383512304 | 0.111 | 0.166 | 1 | 0.467299899 | 0.374233051 |
| Eid2b | 0.20895393 | -1.184209227 | 0 | 0.071 | 1 | 0.390923314 | 0.410054092 | 0.044 | 0.083 | 1 | 0.390923314 | 0.374246116 |
| H4c16 | 0.208980008 | -1.370622352 | 0 | 0.071 | 1 | 0.940059902 | 0.460680165 | 0.022 | 0.024 | 1 | 0.940059902 | 0.374287372 |
| Vegfc | 0.208980008 | -1.184209227 | 0 | 0.071 | 1 | 0.238825064 | -1.404390255 | 0.067 | 0.125 | 1 | 0.238825064 | 0.374287372 |
| Pola2 | 0.208980008 | -1.133583154 | 0 | 0.071 | 1 | 0.530270552 | -0.375821103 | 0.044 | 0.069 | 1 | 0.530270552 | 0.374287372 |
| Lix1 | 0.208996267 | -1.280424543 | 0 | 0.071 | 1 | 0.87573411 | 1.522607914 | 0.089 | 0.087 | 1 | 0.87573411 | 0.374313095 |
| Amy1 | 0.209006524 | -0.818081329 | 0.095 | 0.213 | 1 | 0.664502484 | 1.084813263 | 0.178 | 0.232 | 1 | 0.664502484 | 0.37432932 |
| Kctd14 | 0.209010071 | -1.184209227 | 0 | 0.071 | 1 | 0.239643796 | -0.819427754 | 0.022 | 0.069 | 1 | 0.239643796 | 0.374334933 |
| 2810454H06Rik | 0.209021114 | -1.280424543 | 0 | 0.071 | 1 | 0.776241121 | 0.15951063 | 0.044 | 0.055 | 1 | 0.776241121 | 0.374352403 |
| Gria4 | 0.209066203 | -2.849790188 | 0 | 0.071 | 1 | 0.824696063 | 0.9310001 | 0.067 | 0.08 | 1 | 0.824696063 | 0.374423728 |
| Cpa6 | 0.209075097 | -2.370622352 | 0 | 0.071 | 1 | 0.669547031 | -1.124282336 | 0.022 | 0.035 | 1 | 0.669547031 | 0.374437798 |
| Itga11 | 0.209078923 | -0.67052414 | 0.19 | 0.347 | 1 | 0.322679469 | 0.264282952 | 0.4 | 0.554 | 1 | 0.322679469 | 0.37444385 |
| Iqank1 | 0.314726171 | 2.673771768 | 0.095 | 0.049 | 1 | 0.20909216 | 2.946106992 | 0.067 | 0.031 | 1 | 0.314726171 | 0.374464788 |
| A430090L17Rik | 0.632076134 | 1.673771768 | 0 | 0.011 | 1 | 0.20909216 | 2.531069493 | 0.067 | 0.031 | 1 | 0.632076134 | 0.374464788 |
| Ubqln1 | 0.514287034 | -0.243766072 | 0.476 | 0.578 | 1 | 0.20929019 | -0.203833635 | 0.778 | 0.768 | 1 | 0.514287034 | 0.374777996 |
| Ypel2 | 0.53371476 | 2.367440528 | 0.286 | 0.433 | 1 | 0.209453388 | 0.424669629 | 0.711 | 0.599 | 1 | 0.53371476 | 0.375036055 |
| Rara | 0.825695327 | 0.43673257 | 0.143 | 0.172 | 1 | 0.209477322 | -0.172537504 | 0.244 | 0.349 | 1 | 0.825695327 | 0.375073895 |
| Gm4221 | 0.799819908 | 0.576910228 | 0.143 | 0.175 | 1 | 0.209522356 | 0.361144491 | 0.111 | 0.204 | 1 | 0.799819908 | 0.375145095 |
| Ranbp6 | 0.331266055 | -0.718545655 | 0.048 | 0.116 | 1 | 0.209534344 | -0.564854927 | 0.089 | 0.163 | 1 | 0.331266055 | 0.375164046 |
| Ptn | 0.798782752 | -2.078144168 | 0.381 | 0.34 | 1 | 0.209599683 | -0.13575056 | 0.578 | 0.394 | 1 | 0.798782752 | 0.375267339 |
| Gtpbp8 | 0.209929406 | -0.555046923 | 0.095 | 0.216 | 1 | 0.918179297 | 0.636778934 | 0.2 | 0.235 | 1 | 0.918179297 | 0.375788457 |
| Sin3a | 0.210514277 | 1.137718867 | 0.381 | 0.302 | 1 | 0.382712503 | 0.426732833 | 0.533 | 0.467 | 1 | 0.382712503 | 0.376712293 |
| Cnbd2 | 0.234275769 | -0.548620654 | 0.048 | 0.146 | 1 | 0.210541817 | 0.395495996 | 0.111 | 0.204 | 1 | 0.234275769 | 0.376755778 |
| Ggnbp2 | 0.210766621 | 0.559259887 | 0.952 | 0.944 | 1 | 0.651246302 | 0.172976241 | 1 | 0.976 | 1 | 0.651246302 | 0.377110674 |
| Map3k7 | 0.764350751 | 0.325848464 | 0.381 | 0.519 | 1 | 0.211788088 | 0.659740082 | 0.689 | 0.661 | 1 | 0.764350751 | 0.378721981 |
| Gm19522 | 0.330083699 | -1.02666795 | 0.048 | 0.116 | 1 | 0.212459545 | -0.124282336 | 0.067 | 0.138 | 1 | 0.330083699 | 0.379780032 |
| Itpkb | 0.212790412 | 1.186505941 | 0.524 | 0.519 | 1 | 0.856468714 | 0.867912262 | 0.356 | 0.408 | 1 | 0.856468714 | 0.380301065 |
| Cpsf1 | 0.317975113 | -0.496153234 | 0.048 | 0.119 | 1 | 0.212837769 | -0.181997834 | 0.067 | 0.142 | 1 | 0.317975113 | 0.380375621 |
| Usp54 | 0.246115851 | 0.918884266 | 0.381 | 0.321 | 1 | 0.212900031 | 0.479383604 | 0.356 | 0.505 | 1 | 0.246115851 | 0.380473639 |
| Gm14326 | 0.727019032 | 0.918884266 | 0.19 | 0.183 | 1 | 0.213096434 | -0.124282336 | 0.111 | 0.197 | 1 | 0.727019032 | 0.380782778 |
| 5530601H04Rik | 0.213299386 | -0.30605035 | 0.19 | 0.358 | 1 | 0.273781019 | 0.424338318 | 0.244 | 0.408 | 1 | 0.273781019 | 0.381102144 |
| Nprl3 | 0.72081399 | 1.396931562 | 0.19 | 0.179 | 1 | 0.213626727 | 0.591150097 | 0.111 | 0.208 | 1 | 0.72081399 | 0.381617075 |
| Gm1976 | 0.91745856 | 0.843696769 | 0.143 | 0.164 | 1 | 0.213690415 | -0.16492432 | 0.089 | 0.166 | 1 | 0.91745856 | 0.381717236 |
| Chst15 | 0.869600423 | 0.214340149 | 0.095 | 0.108 | 1 | 0.213753953 | -0.416463087 | 0.089 | 0.166 | 1 | 0.869600423 | 0.381817153 |
| Podn | 0.324767242 | -0.326228232 | 0 | 0.045 | 1 | 0.213828533 | -1.053893008 | 0.022 | 0.073 | 1 | 0.324767242 | 0.381934424 |
| Icam1 | 0.213922328 | -0.445663463 | 0.238 | 0.392 | 1 | 0.954590084 | -0.313242367 | 0.578 | 0.457 | 1 | 0.954590084 | 0.382081894 |
| Zbtb9 | 0.935729463 | 0.995699863 | 0.095 | 0.093 | 1 | 0.214553048 | -0.346674757 | 0.044 | 0.107 | 1 | 0.935729463 | 0.383073086 |
| Ttll5 | 0.214598608 | 1.608676739 | 0.524 | 0.496 | 1 | 0.264206597 | 0.568705561 | 0.578 | 0.609 | 1 | 0.264206597 | 0.383144653 |
| Spindoc | 0.214690822 | -0.451759114 | 0.238 | 0.41 | 1 | 0.808213162 | 0.431289315 | 0.489 | 0.55 | 1 | 0.808213162 | 0.383289494 |
| Atl1 | 0.214763692 | 0.575739685 | 0.381 | 0.276 | 1 | 0.437435438 | -0.316927414 | 0.156 | 0.208 | 1 | 0.437435438 | 0.383403941 |
| Slc35a3 | 0.214772446 | -0.133583154 | 0.333 | 0.575 | 1 | 0.411165617 | 0.500869255 | 0.622 | 0.633 | 1 | 0.411165617 | 0.383417688 |
| 6030407O03Rik | 0.215017736 | -0.271086678 | 0.048 | 0.153 | 1 | 0.728501877 | 1.0259603 | 0.067 | 0.087 | 1 | 0.728501877 | 0.383802845 |
| Son | 0.215178239 | 0.226825688 | 1 | 1 | 1 | 0.44433048 | 0.181886026 | 1 | 1 | 1 | 0.44433048 | 0.384054803 |
| Ltbp1 | 0.329239744 | -0.117641611 | 0.095 | 0.194 | 1 | 0.215184699 | -1.081213614 | 0.156 | 0.232 | 1 | 0.329239744 | 0.384064943 |
| Gm14963 | 0.997890744 | 0.464318402 | 0.095 | 0.097 | 1 | 0.215222398 | -0.638855509 | 0.022 | 0.073 | 1 | 0.997890744 | 0.384124115 |
| Dgki | 0.215255621 | -1.741265732 | 0.048 | 0.149 | 1 | 0.802675537 | 0.821778857 | 0.111 | 0.1 | 1 | 0.802675537 | 0.38417626 |
| Vrk2 | 0.215330123 | -0.807354922 | 0.048 | 0.149 | 1 | 0.993240072 | 1.492741374 | 0.178 | 0.194 | 1 | 0.993240072 | 0.384293184 |
| Nceh1 | 0.215450091 | -1.159118246 | 0.048 | 0.142 | 1 | 0.246381442 | 0.193687745 | 0.067 | 0.135 | 1 | 0.246381442 | 0.38448144 |
| Ylpm1 | 0.499424609 | 0.355645243 | 0.429 | 0.649 | 1 | 0.215534179 | 0.304113546 | 0.933 | 0.761 | 1 | 0.499424609 | 0.384613376 |
| Map3k3 | 0.215875162 | 1.125567817 | 0.619 | 0.735 | 1 | 0.929134332 | 0.102515265 | 0.889 | 0.872 | 1 | 0.929134332 | 0.385148238 |
| Polr2a | 0.920922581 | 0.271353755 | 0.762 | 0.713 | 1 | 0.215884013 | 0.298929095 | 0.778 | 0.744 | 1 | 0.920922581 | 0.385162119 |
| Flvcr2 | 0.546675006 | 0.614878079 | 0.095 | 0.149 | 1 | 0.21593394 | -1.071814916 | 0.044 | 0.104 | 1 | 0.546675006 | 0.385240413 |
| Atp13a3 | 0.216125406 | 0.365782864 | 0.905 | 0.731 | 1 | 0.432085488 | 0.321398312 | 0.933 | 0.882 | 1 | 0.432085488 | 0.385540621 |
| Clk2 | 0.644080819 | 0.54665985 | 0.286 | 0.399 | 1 | 0.216216476 | 0.35429082 | 0.311 | 0.491 | 1 | 0.644080819 | 0.385683387 |
| Stat5b | 0.815875223 | 1.529630792 | 0.333 | 0.422 | 1 | 0.216363114 | 0.851110478 | 0.778 | 0.73 | 1 | 0.815875223 | 0.38591323 |
| Frrs1 | 0.285549799 | -0.648156327 | 0 | 0.052 | 1 | 0.216607348 | -1.975138897 | 0.067 | 0.128 | 1 | 0.285549799 | 0.386295952 |
| Gm41724 | 0.856526182 | 1.65584986 | 0.143 | 0.138 | 1 | 0.217353875 | -0.486852415 | 0.022 | 0.073 | 1 | 0.856526182 | 0.387465043 |
| Rb1 | 0.546498304 | 0.434984908 | 0.429 | 0.366 | 1 | 0.217518744 | 0.413506107 | 0.822 | 0.606 | 1 | 0.546498304 | 0.387723084 |
| Dapk2 | 0.217559055 | -2.506137322 | 0.048 | 0.138 | 1 | 0.659605288 | -0.800743191 | 0.267 | 0.215 | 1 | 0.659605288 | 0.387786167 |
| Tafa5 | 0.217587991 | 0.873190074 | 0.571 | 0.53 | 1 | 0.30035239 | 0.685287007 | 0.422 | 0.692 | 1 | 0.30035239 | 0.387831447 |
| Gm20705 | 0.264194629 | 2.673771768 | 0.048 | 0.015 | 1 | 0.217596841 | 1.946106992 | 0.067 | 0.031 | 1 | 0.264194629 | 0.387845296 |
| Gm52870 | 0.362165014 | 2.451379346 | 0.048 | 0.019 | 1 | 0.217596841 | 2.168499413 | 0.067 | 0.031 | 1 | 0.362165014 | 0.387845296 |
| A930005H10Rik | 0.281778562 | -0.611630451 | 0.048 | 0.127 | 1 | 0.217740381 | -0.674479418 | 0.044 | 0.104 | 1 | 0.281778562 | 0.388069889 |
| Prr7 | 0.217847785 | -1.02666795 | 0.095 | 0.201 | 1 | 0.269074952 | -0.342462506 | 0.089 | 0.156 | 1 | 0.269074952 | 0.388237912 |
| Gm16541 | 0.74032607 | -1.807354922 | 0.048 | 0.067 | 1 | 0.218015682 | -0.731964913 | 0.022 | 0.073 | 1 | 0.74032607 | 0.388500526 |
| Tmem86b | 0.969041235 | 1.351843673 | 0.048 | 0.052 | 1 | 0.218047177 | -0.432404631 | 0.022 | 0.073 | 1 | 0.969041235 | 0.388549783 |
| Endou | 0.333997812 | -0.34415014 | 0.095 | 0.19 | 1 | 0.218203592 | -0.746915255 | 0.089 | 0.163 | 1 | 0.333997812 | 0.388794377 |
| Zfp687 | 0.981011824 | 0.766881172 | 0.095 | 0.097 | 1 | 0.218236187 | 0.632446513 | 0.089 | 0.173 | 1 | 0.981011824 | 0.388845341 |
| Gm26621 | 0.218331105 | 2.410737362 | 0.095 | 0.041 | 1 | 0.344255941 | -0.776359032 | 0.022 | 0.055 | 1 | 0.344255941 | 0.388993738 |
| Thegl | 0.218389693 | 3.766881172 | 0.095 | 0.041 | 1 | 0.530765204 | 1.905465008 | 0.044 | 0.028 | 1 | 0.530765204 | 0.389085328 |
| Opa3 | 0.867957995 | 0.33878752 | 0.238 | 0.287 | 1 | 0.218611542 | 0.463904066 | 0.2 | 0.343 | 1 | 0.867957995 | 0.389432077 |
| Rhot1 | 0.218635359 | 0.570678275 | 0.524 | 0.481 | 1 | 0.96632251 | 0.405652752 | 0.578 | 0.682 | 1 | 0.96632251 | 0.389469298 |
| Ddx59 | 0.638247026 | -0.184209227 | 0.048 | 0.075 | 1 | 0.218818634 | -0.316927414 | 0.022 | 0.073 | 1 | 0.638247026 | 0.389755674 |
| Prorp | 0.218977193 | 1.65491274 | 0.238 | 0.16 | 1 | 0.945336997 | 0.525531309 | 0.111 | 0.111 | 1 | 0.945336997 | 0.390003375 |
| Pusl1 | 0.478820484 | -0.184209227 | 0.048 | 0.093 | 1 | 0.219428341 | -0.316927414 | 0.022 | 0.073 | 1 | 0.478820484 | 0.390707884 |
| Hao1 | 0.569042745 | -0.133583154 | 0.048 | 0.082 | 1 | 0.219539452 | 2.393565969 | 0.067 | 0.031 | 1 | 0.569042745 | 0.390881333 |
| Gmps | 0.360486758 | 0.393119948 | 0.381 | 0.657 | 1 | 0.219640027 | -0.132289914 | 0.733 | 0.754 | 1 | 0.360486758 | 0.391038312 |
| Mob1b | 0.222750729 | 0.979510624 | 0.524 | 0.463 | 1 | 0.219661731 | 0.555064974 | 0.867 | 0.706 | 1 | 0.222750729 | 0.391072186 |
| Adamts17 | 0.841315127 | 1.68887866 | 0.19 | 0.19 | 1 | 0.219662542 | -0.277181235 | 0.111 | 0.201 | 1 | 0.841315127 | 0.391073452 |
| Crlf1 | 0.544173668 | 0.258734268 | 0.048 | 0.09 | 1 | 0.220014852 | 1.071637874 | 0.2 | 0.135 | 1 | 0.544173668 | 0.391623169 |
| Golga4 | 0.220048718 | 0.360158941 | 0.952 | 0.877 | 1 | 0.601685993 | 0.183022337 | 0.978 | 0.948 | 1 | 0.601685993 | 0.391675998 |
| 6330418K02Rik | 0.236343578 | -0.648156327 | 0 | 0.063 | 1 | 0.220115784 | -0.255526869 | 0.022 | 0.073 | 1 | 0.236343578 | 0.39178061 |
| Tsen54 | 0.285622926 | -0.233118828 | 0 | 0.052 | 1 | 0.220115784 | -0.255526869 | 0.022 | 0.073 | 1 | 0.285622926 | 0.39178061 |
| Mpzl2 | 0.241410881 | -0.303508156 | 0.048 | 0.146 | 1 | 0.220123383 | 0.132875504 | 0.289 | 0.194 | 1 | 0.241410881 | 0.391792462 |
| Mettl15 | 0.550390599 | 0.999172054 | 0.143 | 0.224 | 1 | 0.220169204 | 0.261608818 | 0.089 | 0.176 | 1 | 0.550390599 | 0.39186393 |
| Resf1 | 0.967112564 | -0.22439541 | 0.81 | 0.716 | 1 | 0.220338486 | -0.790572892 | 0.778 | 0.651 | 1 | 0.967112564 | 0.392127924 |
| L3mbtl1 | 0.220502945 | 1.766881172 | 0.143 | 0.075 | 1 | 0.909554942 | 0.553789569 | 0.111 | 0.125 | 1 | 0.909554942 | 0.392384341 |
| 4732471J01Rik | 0.22080705 | 0.468657338 | 0.048 | 0.153 | 1 | 0.570542158 | 0.442064487 | 0.067 | 0.097 | 1 | 0.570542158 | 0.392858347 |
| CJ186046Rik | 0.713704917 | 0.673771768 | 0.143 | 0.123 | 1 | 0.220825069 | 0.481438725 | 0.067 | 0.138 | 1 | 0.713704917 | 0.392886427 |
| Gm50232 | 0.988367841 | 1.171271427 | 0.048 | 0.049 | 1 | 0.220915594 | 0.304560963 | 0.022 | 0.073 | 1 | 0.988367841 | 0.393027489 |
| Gm57271 | 0.39694807 | -0.233118828 | 0 | 0.034 | 1 | 0.220945781 | 0.513147585 | 0.022 | 0.073 | 1 | 0.39694807 | 0.393074524 |
| Fam220a.1 | 0.388537847 | 2.311201688 | 0.095 | 0.052 | 1 | 0.220971941 | 0.197645759 | 0.022 | 0.073 | 1 | 0.388537847 | 0.393115284 |
| Map3k12 | 0.220981978 | -0.623908781 | 0.048 | 0.149 | 1 | 0.615996506 | 0.683072586 | 0.178 | 0.239 | 1 | 0.615996506 | 0.393130921 |
| Uaca | 0.271340768 | 0.302323537 | 0.571 | 0.81 | 1 | 0.221045783 | -0.228269794 | 0.733 | 0.792 | 1 | 0.271340768 | 0.393230328 |
| Rab40c | 0.221057314 | -0.233118828 | 0.143 | 0.299 | 1 | 0.425539757 | 0.816823975 | 0.378 | 0.339 | 1 | 0.425539757 | 0.393248292 |
| Cxcl1 | 0.221095836 | 0.250560337 | 0.048 | 0.153 | 1 | 0.293647037 | -1.017367132 | 0 | 0.024 | 1 | 0.293647037 | 0.393308303 |

| | | | | | | | | | | | | |
|---|---|---|---|---|---|---|---|---|---|---|---|---|
| Pou5f1 | 0.221193344 | 2.295260144 | 0.095 | 0.041 | 1 | 0.244269117 | -0.124282336 | 0.022 | 0.069 | 1 | 0.244269117 | 0.393460193 |
| Gm56904 | 0.221193344 | 2.295260144 | 0.095 | 0.041 | 1 | 0.231991339 | -0.901889915 | 0 | 0.031 | 1 | 0.231991339 | 0.393460193 |
| Ppm1k | 0.670178094 | 1.195724471 | 0.143 | 0.127 | 1 | 0.221449577 | -0.316927414 | 0.044 | 0.104 | 1 | 0.670178094 | 0.393859239 |
| Plscr4 | 0.221574552 | -0.752492987 | 0.143 | 0.272 | 1 | 0.386660672 | 0.977815852 | 0.178 | 0.273 | 1 | 0.386660672 | 0.394053822 |
| 2310002F09Rik | 0.221641087 | 1.97333205 | 0.143 | 0.075 | 1 | 0.871906337 | 0.683072586 | 0.044 | 0.052 | 1 | 0.871906337 | 0.394157403 |
| Gm12940 | 0.221722614 | 0.60529203 | 0.143 | 0.302 | 1 | 0.837218829 | 0.528349991 | 0.356 | 0.363 | 1 | 0.837218829 | 0.394284311 |
| Tmco6 | 0.22173579 | -0.785659851 | 0.048 | 0.142 | 1 | 0.722093254 | 0.875717664 | 0.044 | 0.059 | 1 | 0.722093254 | 0.394304819 |
| Zfp385a | 0.356947467 | 0.289107917 | 0.143 | 0.25 | 1 | 0.221848276 | -0.223818009 | 0.067 | 0.138 | 1 | 0.356947467 | 0.394479894 |
| Mthfd1l | 0.221855397 | 1.09434745 | 0.524 | 0.563 | 1 | 0.334008048 | 0.60576686 | 0.378 | 0.547 | 1 | 0.334008048 | 0.394490977 |
| Pdlim7 | 0.221906514 | 0.780686972 | 0.524 | 0.478 | 1 | 0.422812852 | 0.831595111 | 0.489 | 0.478 | 1 | 0.422812852 | 0.394570527 |
| Trim34a | 0.222110654 | -0.648156327 | 0 | 0.067 | 1 | 0.261911033 | -0.242926832 | 0.044 | 0.1 | 1 | 0.261911033 | 0.394888165 |
| Pigw | 0.222110654 | -0.648156327 | 0 | 0.067 | 1 | 0.534128516 | -0.124282336 | 0.022 | 0.042 | 1 | 0.534128516 | 0.394888165 |
| Ttc39a | 0.222155241 | -0.718545655 | 0 | 0.067 | 1 | 0.338332548 | 1.726141308 | 0.133 | 0.097 | 1 | 0.338332548 | 0.39495753 |
| Aldh16a1 | 0.222155241 | -0.718545655 | 0 | 0.067 | 1 | 0.943054607 | 0.998574412 | 0.111 | 0.125 | 1 | 0.943054607 | 0.39495753 |
| Parp16 | 0.222155241 | -0.718545655 | 0 | 0.067 | 1 | 0.441223708 | -0.191396532 | 0.022 | 0.048 | 1 | 0.441223708 | 0.39495753 |
| 4930556M19Rik | 0.222155241 | -0.785659851 | 0 | 0.067 | 1 | 0.346432578 | -0.564854927 | 0.022 | 0.055 | 1 | 0.346432578 | 0.39495753 |
| Rwdd2b | 0.222155241 | -0.718545655 | 0 | 0.067 | 1 | 0.347517921 | -0.486852415 | 0.022 | 0.055 | 1 | 0.347517921 | 0.39495753 |
| Ephx3 | 0.222155241 | -0.785659851 | 0 | 0.067 | 1 | 0.351068509 | -0.242926832 | 0.044 | 0.087 | 1 | 0.351068509 | 0.39495753 |
| Cd38 | 0.222194249 | -0.970084422 | 0 | 0.067 | 1 | 0.496223879 | 0.683072586 | 0 | 0.01 | 1 | 0.496223879 | 0.395018214 |
| Ung | 0.222194249 | -0.849790188 | 0 | 0.067 | 1 | 0.417050545 | 0.78998779 | 0.044 | 0.083 | 1 | 0.417050545 | 0.395018214 |
| Mfsd2b | 0.222194249 | -0.970084422 | 0 | 0.067 | 1 | 0.345543073 | 0.223640968 | 0.044 | 0.09 | 1 | 0.345543073 | 0.395018214 |
| Gm45509 | 0.222194577 | -0.785659851 | 0 | 0.067 | 1 | 0.35815185 | 0.460680165 | 0.044 | 0.087 | 1 | 0.35815185 | 0.395018724 |
| Zfp563 | 0.222194577 | -0.785659851 | 0 | 0.067 | 1 | 0.259198647 | -0.242926832 | 0.044 | 0.1 | 1 | 0.259198647 | 0.395018724 |
| Ttc30b | 0.222228664 | -0.970084422 | 0 | 0.067 | 1 | 0.575239232 | -0.375821103 | 0.044 | 0.066 | 1 | 0.575239232 | 0.395071749 |
| Mrm2 | 0.222228664 | -0.970084422 | 0 | 0.067 | 1 | 0.936452267 | 0.78998779 | 0.067 | 0.073 | 1 | 0.936452267 | 0.395071749 |
| Zfp846 | 0.222255538 | -0.970084422 | 0 | 0.067 | 1 | 0.621894124 | 0.410054092 | 0.044 | 0.066 | 1 | 0.621894124 | 0.395113551 |
| Zfp595 | 0.222257504 | -1.280424543 | 0 | 0.067 | 1 | 0.628865315 | 0.735540006 | 0.044 | 0.066 | 1 | 0.628865315 | 0.39511661 |
| Dtwd2 | 0.222257504 | -1.133583154 | 0 | 0.067 | 1 | 0.286196506 | -0.16492432 | 0.044 | 0.097 | 1 | 0.286196506 | 0.39511661 |
| Ncoa3 | 0.883683893 | 0.421308492 | 0.714 | 0.84 | 1 | 0.22227397 | 0.65173462 | 0.978 | 0.913 | 1 | 0.883683893 | 0.395142223 |
| Cdc25b | 0.222294533 | -1.081115734 | 0 | 0.067 | 1 | 0.295752136 | 0.13875207 | 0.044 | 0.093 | 1 | 0.295752136 | 0.395174207 |
| Klf8 | 0.222345974 | -1.818081329 | 0 | 0.067 | 1 | 0.833127239 | 0.7849522 | 0.089 | 0.083 | 1 | 0.833127239 | 0.395254215 |
| Nalcn | 0.222348922 | -2.233118828 | 0 | 0.067 | 1 | 0.581464179 | 1.135584791 | 0.067 | 0.048 | 1 | 0.581464179 | 0.395258801 |
| Mpi | 0.222378078 | -0.574155746 | 0 | 0.067 | 1 | 0.577907507 | -0.255526869 | 0.044 | 0.066 | 1 | 0.577907507 | 0.395304147 |
| Gm28501 | 0.457308648 | 0.673771768 | 0 | 0.026 | 1 | 0.222453122 | 1.683072586 | 0.067 | 0.031 | 1 | 0.457308648 | 0.395420852 |
| Hmgn5 | 0.222799809 | -0.509450056 | 0.238 | 0.399 | 1 | 0.50797962 | 0.20029048 | 0.489 | 0.384 | 1 | 0.50797962 | 0.395959864 |
| Gpr173 | 0.757613212 | 0.926537838 | 0.095 | 0.131 | 1 | 0.222937544 | 0.576157382 | 0.067 | 0.142 | 1 | 0.757613212 | 0.396173939 |
| Zfp112 | 0.223140192 | 1.97333205 | 0.143 | 0.071 | 1 | 0.339365053 | -0.264459994 | 0.067 | 0.118 | 1 | 0.339365053 | 0.396488838 |
| Unc13d | 0.716066226 | 0.451379346 | 0.048 | 0.071 | 1 | 0.223777564 | 1.561082062 | 0.156 | 0.104 | 1 | 0.716066226 | 0.39747873 |
| Gm13684 | 0.224057062 | 1.825774861 | 0.095 | 0.041 | 1 | 0.966814799 | 0.78998779 | 0.067 | 0.069 | 1 | 0.966814799 | 0.397912557 |
| Gm5820 | 0.224077604 | 2.188344941 | 0.095 | 0.041 | 1 | 0.293654613 | -1.124282336 | 0 | 0.024 | 1 | 0.293654613 | 0.397944436 |
| Panx1 | 0.719080203 | 1.55829455 | 0.048 | 0.034 | 1 | 0.224259682 | -1.124282336 | 0.022 | 0.069 | 1 | 0.719080203 | 0.39822696 |
| Shld2 | 0.935530611 | 1.072867723 | 0.19 | 0.22 | 1 | 0.224389985 | -0.213091603 | 0.156 | 0.253 | 1 | 0.935530611 | 0.398429105 |
| Tnfrsf21 | 0.277970221 | 1.076593317 | 0.571 | 0.541 | 1 | 0.224541123 | 0.313139118 | 0.844 | 0.765 | 1 | 0.277970221 | 0.39866353 |
| Tap1 | 0.97924348 | 1.010806755 | 0.095 | 0.101 | 1 | 0.224679417 | 0.361144491 | 0.044 | 0.107 | 1 | 0.97924348 | 0.398877993 |
| Shq1 | 1 | 0.258734268 | 0.095 | 0.093 | 1 | 0.22475833 | 0.256807831 | 0.044 | 0.107 | 1 | 1 | 0.399000354 |
| Nsrp1 | 0.225368651 | -0.418674481 | 0.333 | 0.507 | 1 | 0.557322855 | 0.356779299 | 0.467 | 0.581 | 1 | 0.557322855 | 0.399946274 |
| Dancr | 0.225384405 | 1.780686972 | 0.238 | 0.16 | 1 | 0.559546785 | 0.486675373 | 0.111 | 0.152 | 1 | 0.559546785 | 0.399970679 |
| Mycbp2 | 0.646941892 | 0.470488169 | 0.952 | 0.907 | 1 | 0.225398967 | 0.41677336 | 1 | 0.979 | 1 | 0.646941892 | 0.399993239 |
| Acvr2b | 0.544576436 | 2.258734268 | 0.048 | 0.026 | 1 | 0.225716988 | -1.017367132 | 0.022 | 0.069 | 1 | 0.544576436 | 0.400485817 |
| Cyb5rl | 0.856787907 | 1.55829455 | 0.048 | 0.041 | 1 | 0.225717833 | -1.124282336 | 0.022 | 0.069 | 1 | 0.856787907 | 0.400487125 |
| Lztfl1 | 0.225895413 | -0.184209227 | 0.524 | 0.754 | 1 | 0.965640225 | 0.227066442 | 0.756 | 0.772 | 1 | 0.965640225 | 0.400762088 |
| Ly6a | 0.225994772 | -2.525008096 | 0.429 | 0.451 | 1 | 0.99045146 | -0.164719842 | 0.844 | 0.612 | 1 | 0.99045146 | 0.400915907 |
| 4931403E22Rik | 0.457308648 | 0.673771768 | 0 | 0.026 | 1 | 0.226015083 | 2.42003818 | 0.044 | 0.017 | 1 | 0.457308648 | 0.400947348 |
| Gm57092 | 0.632076134 | 1.673771768 | 0 | 0.011 | 1 | 0.226015083 | 2.683072586 | 0.044 | 0.017 | 1 | 0.632076134 | 0.400947348 |
| Icam2 | 0.632076134 | 1.088809267 | 0 | 0.011 | 1 | 0.226015083 | 2.905465008 | 0.044 | 0.017 | 1 | 0.632076134 | 0.400947348 |
| Gm28874 | 0.632917461 | 1.936806174 | 0.048 | 0.03 | 1 | 0.226015083 | 2.683072586 | 0.044 | 0.017 | 1 | 0.632917461 | 0.400947348 |
| 4933406J09Rik | 0.532432377 | 0.673771768 | 0 | 0.019 | 1 | 0.226021153 | 3.78260826 | 0.044 | 0.017 | 1 | 0.532432377 | 0.400956745 |
| Doc2b | 0.532436372 | 1.088809267 | 0 | 0.019 | 1 | 0.226021153 | 2.875717664 | 0.044 | 0.017 | 1 | 0.532436372 | 0.400956745 |
| Rhebl1 | 0.226041553 | 2.536268244 | 0.143 | 0.075 | 1 | 0.89612222 | 1.567595369 | 0.089 | 0.09 | 1 | 0.89612222 | 0.400988322 |
| Gm12353 | 0.226058557 | -1.303508156 | 0.048 | 0.138 | 1 | 0.927779015 | 0.599656578 | 0.133 | 0.142 | 1 | 0.927779015 | 0.401014642 |
| Gm54798 | 0.304385789 | -0.326228232 | 0 | 0.049 | 1 | 0.226441946 | -0.960783604 | 0.022 | 0.069 | 1 | 0.304385789 | 0.401607937 |
| Gm37083 | 0.356859608 | 1.503846766 | 0.143 | 0.086 | 1 | 0.226441946 | -0.960783604 | 0.022 | 0.069 | 1 | 0.356859608 | 0.401607937 |
| Lmo1 | 0.532426384 | 0.866416846 | 0 | 0.019 | 1 | 0.226442157 | -0.901889915 | 0.022 | 0.069 | 1 | 0.532426384 | 0.401608264 |
| Clstn2 | 0.676753434 | 0.895465378 | 0.19 | 0.269 | 1 | 0.226598841 | 1.025046623 | 0.511 | 0.519 | 1 | 0.676753434 | 0.401850647 |
| Gopc | 0.226609809 | -0.648156327 | 0.19 | 0.325 | 1 | 0.617206378 | 0.790828767 | 0.333 | 0.488 | 1 | 0.617206378 | 0.401867613 |
| Ctps2 | 0.736697267 | 1.36298423 | 0.286 | 0.317 | 1 | 0.226775409 | 0.168499413 | 0.289 | 0.457 | 1 | 0.736697267 | 0.402123732 |
| Rars | 0.928132467 | 0.214340149 | 0.429 | 0.496 | 1 | 0.226853543 | -0.339295227 | 0.511 | 0.557 | 1 | 0.928132467 | 0.402244556 |
| Itgav | 0.226958332 | 0.39796114 | 0.905 | 0.854 | 1 | 0.642808571 | 0.169613034 | 0.911 | 0.844 | 1 | 0.642808571 | 0.40240658 |
| Paxip1 | 0.695257963 | 0.574236094 | 0.143 | 0.194 | 1 | 0.227052684 | 0.197645759 | 0.178 | 0.287 | 1 | 0.695257963 | 0.402552446 |
| Casp4 | 0.715685897 | 0.251707002 | 0.333 | 0.377 | 1 | 0.227119746 | -0.481116046 | 0.178 | 0.277 | 1 | 0.715685897 | 0.402656113 |
| Pard6a | 0.493128106 | 0.304537958 | 0.048 | 0.097 | 1 | 0.227164429 | -0.84048937 | 0.022 | 0.069 | 1 | 0.493128106 | 0.40272518 |
| Zfp704 | 0.22947706 | 0.251889887 | 0.381 | 0.619 | 1 | 0.22728303 | 0.306567958 | 0.756 | 0.775 | 1 | 0.22947706 | 0.402908484 |
| Il12rb1 | 0.709487466 | 2.866416846 | 0.048 | 0.034 | 1 | 0.227305413 | 2.875717664 | 0.044 | 0.017 | 1 | 0.709487466 | 0.402943075 |
| Fcsk | 0.57816106 | 1.351843673 | 0 | 0.015 | 1 | 0.227884427 | -0.776359032 | 0.022 | 0.069 | 1 | 0.57816106 | 0.403837543 |
| Pomt1 | 0.22823149 | -0.385121921 | 0.048 | 0.149 | 1 | 0.680289378 | 0.523873991 | 0.133 | 0.17 | 1 | 0.680289378 | 0.404373368 |
| Asb13 | 0.33871995 | 1.351843673 | 0.143 | 0.086 | 1 | 0.228250717 | 0.223640968 | 0.111 | 0.197 | 1 | 0.33871995 | 0.404403044 |
| Gm57385 | 0.396916958 | 0.214340149 | 0 | 0.034 | 1 | 0.228596086 | 2.197645759 | 0.044 | 0.017 | 1 | 0.396916958 | 0.404936001 |
| Gm41790 | 0.228996129 | 1.108987149 | 0.238 | 0.149 | 1 | 0.354619089 | 1.172110667 | 0.244 | 0.211 | 1 | 0.354619089 | 0.40555303 |
| Ppp6c | 0.229407389 | 0.840065519 | 0.524 | 0.534 | 1 | 0.808507967 | 0.500869255 | 0.733 | 0.716 | 1 | 0.808507967 | 0.406187027 |
| Prrg1 | 0.629863438 | 0.710297644 | 0.143 | 0.19 | 1 | 0.229416525 | -0.204452685 | 0.089 | 0.163 | 1 | 0.629863438 | 0.406201108 |
| 4933406C10Rik | 0.229517907 | 2.329817366 | 0.19 | 0.112 | 1 | 0.856198973 | 1.361144491 | 0.089 | 0.107 | 1 | 0.856198973 | 0.406357345 |
| Arid1a | 0.78729641 | 0.342664246 | 0.571 | 0.668 | 1 | 0.229528821 | 0.359515248 | 0.978 | 0.862 | 1 | 0.78729641 | 0.406374163 |
| Dym | 0.22973574 | -0.655351829 | 0.238 | 0.396 | 1 | 0.608043104 | 0.495987034 | 0.533 | 0.54 | 1 | 0.608043104 | 0.40669297 |
| Tnip2 | 0.229791063 | 0.712942365 | 0.524 | 0.481 | 1 | 0.828511221 | 0.603073264 | 0.356 | 0.422 | 1 | 0.828511221 | 0.406778193 |
| Kdm7a | 0.23025057 | 1.236654735 | 0.667 | 0.604 | 1 | 0.812117355 | -0.144746438 | 0.956 | 0.754 | 1 | 0.812117355 | 0.407485814 |
| Zranb2 | 0.734148292 | 0.33022986 | 0.714 | 0.806 | 1 | 0.230257872 | -0.237824867 | 0.8 | 0.772 | 1 | 0.734148292 | 0.407497056 |
| D11Wsu47e | 0.454714319 | -0.133583154 | 0.048 | 0.097 | 1 | 0.230258389 | -0.486852415 | 0.089 | 0.159 | 1 | 0.454714319 | 0.407497852 |
| Usp32 | 0.230471982 | 0.680113295 | 0.714 | 0.612 | 1 | 0.527636622 | 0.165079352 | 0.978 | 0.83 | 1 | 0.527636622 | 0.407826629 |
| Adra1b | 0.311354441 | 1.673771768 | 0.19 | 0.131 | 1 | 0.230596398 | -1.174908409 | 0.044 | 0.1 | 1 | 0.311354441 | 0.408018096 |
| Rpa2 | 0.230636651 | -0.880817084 | 0.095 | 0.205 | 1 | 0.314311382 | 0.513147585 | 0.111 | 0.183 | 1 | 0.314311382 | 0.408080037 |
| Nfia | 0.316358669 | 0.619149756 | 0.952 | 0.907 | 1 | 0.230694663 | 0.46046594 | 1 | 0.962 | 1 | 0.316358669 | 0.408169298 |
| Maco1 | 0.848720474 | 0.154806826 | 0.619 | 0.664 | 1 | 0.230699454 | 0.593752596 | 0.844 | 0.744 | 1 | 0.848720474 | 0.40817667 |
| Bcl2l2 | 0.230933361 | -0.370622352 | 0.095 | 0.213 | 1 | 0.405856968 | 0.347469554 | 0.2 | 0.301 | 1 | 0.405856968 | 0.408536505 |
| D030045P18Rik | 0.285527294 | -0.849790188 | 0 | 0.052 | 1 | 0.231081483 | 0.58546179 | 0.178 | 0.298 | 1 | 0.285527294 | 0.408764315 |
| Gk5 | 0.231268168 | 0.746622333 | 0.476 | 0.425 | 1 | 0.841713712 | 0.662854307 | 0.6 | 0.595 | 1 | 0.841713712 | 0.40905137 |
| Dck | 0.979245522 | 0.97333205 | 0.095 | 0.101 | 1 | 0.231797866 | -0.153428682 | 0.067 | 0.135 | 1 | 0.979245522 | 0.409865481 |
| Stat5a | 0.231956895 | 1.284729477 | 0.333 | 0.257 | 1 | 0.662878802 | 0.430306516 | 0.267 | 0.363 | 1 | 0.662878802 | 0.410109788 |
| Gm12204 | 0.261628269 | 2.936806174 | 0.048 | 0.015 | 1 | 0.23197796 | -0.776359032 | 0 | 0.031 | 1 | 0.261628269 | 0.410142146 |

| | | | | | | | | | | | | |
|---|---|---|---|---|---|---|---|---|---|---|---|---|
| Kcnk7 | 0.261628269 | 2.936806174 | 0.048 | 0.015 | 1 | 0.23197796 | -0.776359032 | 0 | 0.031 | 1 | 0.261628269 | 0.410142146 |
| Tubg2 | 0.285499161 | -0.413691074 | 0 | 0.052 | 1 | 0.23197796 | -0.776359032 | 0 | 0.031 | 1 | 0.285499161 | 0.410142146 |
| Slc5a6 | 0.285549799 | -0.785659851 | 0 | 0.052 | 1 | 0.23197796 | -0.776359032 | 0 | 0.031 | 1 | 0.285549799 | 0.410142146 |
| K230015D01Rik | 0.304355016 | -0.233118828 | 0 | 0.049 | 1 | 0.23197796 | -0.776359032 | 0 | 0.031 | 1 | 0.304355016 | 0.410142146 |
| Il4ra | 0.3247899 | -0.496153234 | 0 | 0.045 | 1 | 0.23197796 | -0.776359032 | 0 | 0.031 | 1 | 0.3247899 | 0.410142146 |
| Zfp772 | 0.346802964 | -0.133583154 | 0 | 0.041 | 1 | 0.23197796 | -0.776359032 | 0 | 0.031 | 1 | 0.346802964 | 0.410142146 |
| Fbxl12os | 0.379775085 | 1.351843673 | 0.048 | 0.019 | 1 | 0.23197796 | -0.776359032 | 0 | 0.031 | 1 | 0.379775085 | 0.410142146 |
| Gm15927 | 0.425614684 | 0.214340149 | 0 | 0.03 | 1 | 0.23197796 | -0.776359032 | 0 | 0.031 | 1 | 0.425614684 | 0.410142146 |
| Gm29170 | 0.457292877 | 2.258734268 | 0.048 | 0.022 | 1 | 0.23197796 | -0.776359032 | 0 | 0.031 | 1 | 0.457292877 | 0.410142146 |
| Gm56791 | 0.457308648 | 0.673771768 | 0 | 0.026 | 1 | 0.23197796 | -0.776359032 | 0 | 0.031 | 1 | 0.457308648 | 0.410142146 |
| Gm56589 | 0.457308648 | 0.673771768 | 0 | 0.026 | 1 | 0.23197796 | -0.776359032 | 0 | 0.031 | 1 | 0.457308648 | 0.410142146 |
| Gpr37l1 | 0.492599419 | 0.351843673 | 0 | 0.022 | 1 | 0.23197796 | -0.776359032 | 0 | 0.031 | 1 | 0.492599419 | 0.410142146 |
| Ccdc113 | 0.492599419 | 0.503846766 | 0 | 0.022 | 1 | 0.23197796 | -0.776359032 | 0 | 0.031 | 1 | 0.492599419 | 0.410142146 |
| Gm56795 | 0.492608234 | 0.866416846 | 0 | 0.022 | 1 | 0.23197796 | -1.017367132 | 0 | 0.031 | 1 | 0.492608234 | 0.410142146 |
| Zfp128 | 0.532426384 | 0.866416846 | 0 | 0.019 | 1 | 0.23197796 | -0.776359032 | 0 | 0.031 | 1 | 0.532426384 | 0.410142146 |
| Vwa3a | 0.532426384 | 0.673771768 | 0 | 0.019 | 1 | 0.23197796 | -0.776359032 | 0 | 0.031 | 1 | 0.532426384 | 0.410142146 |
| Arhgap45 | 0.532426384 | 0.866416846 | 0 | 0.019 | 1 | 0.23197796 | -0.776359032 | 0 | 0.031 | 1 | 0.532426384 | 0.410142146 |
| Gm14634 | 0.532426384 | 0.866416846 | 0 | 0.019 | 1 | 0.23197796 | -0.776359032 | 0 | 0.031 | 1 | 0.532426384 | 0.410142146 |
| Sp140 | 0.54776222 | 2.088809267 | 0.048 | 0.026 | 1 | 0.23197796 | -0.776359032 | 0 | 0.031 | 1 | 0.54776222 | 0.410142146 |
| Adam11 | 0.578156445 | 1.088809267 | 0 | 0.015 | 1 | 0.23197796 | -0.776359032 | 0 | 0.031 | 1 | 0.578156445 | 0.410142146 |
| Ldhd | 0.578159907 | 0.673771768 | 0 | 0.015 | 1 | 0.23197796 | -0.776359032 | 0 | 0.031 | 1 | 0.578159907 | 0.410142146 |
| Gm14455 | 0.57816106 | 1.351843673 | 0 | 0.015 | 1 | 0.23197796 | -0.776359032 | 0 | 0.031 | 1 | 0.57816106 | 0.410142146 |
| Mettl18 | 0.63207476 | 1.088809267 | 0 | 0.011 | 1 | 0.23197796 | -1.124282336 | 0 | 0.031 | 1 | 0.63207476 | 0.410142146 |
| Gm47204 | 0.632076134 | 1.673771768 | 0 | 0.011 | 1 | 0.23197796 | -0.776359032 | 0 | 0.031 | 1 | 0.632076134 | 0.410142146 |
| Zwilch | 0.645432064 | 0.544488751 | 0.048 | 0.078 | 1 | 0.23197796 | -0.901889915 | 0 | 0.031 | 1 | 0.645432064 | 0.410142146 |
| Serinc2 | 0.662059486 | 1.214340149 | 0.048 | 0.03 | 1 | 0.23197796 | -0.776359032 | 0 | 0.031 | 1 | 0.662059486 | 0.410142146 |
| Ssc4d | 0.686300024 | 0.673771768 | 0.048 | 0.075 | 1 | 0.23197796 | -0.776359032 | 0 | 0.031 | 1 | 0.686300024 | 0.410142146 |
| Btbd19 | 0.744838936 | 1.088809267 | 0.048 | 0.034 | 1 | 0.23197796 | -0.776359032 | 0 | 0.031 | 1 | 0.744838936 | 0.410142146 |
| Bco2 | 0.862957731 | 1.351843673 | 0.048 | 0.041 | 1 | 0.23197796 | -0.776359032 | 0 | 0.031 | 1 | 0.862957731 | 0.410142146 |
| Tcea2 | 0.969041235 | 1.351843673 | 0.048 | 0.052 | 1 | 0.23197796 | -0.776359032 | 0 | 0.031 | 1 | 0.969041235 | 0.410142146 |
| Gm5817 | 0.985459372 | 1.425844254 | 0.048 | 0.049 | 1 | 0.23197796 | -0.776359032 | 0 | 0.031 | 1 | 0.985459372 | 0.410142146 |
| B230208H11Rik | 0.268010172 | -0.718545655 | 0 | 0.056 | 1 | 0.231991339 | -0.901889915 | 0 | 0.031 | 1 | 0.268010172 | 0.410162696 |
| Gm48855 | 0.274597491 | 2.088809267 | 0.048 | 0.015 | 1 | 0.231991339 | -0.901889915 | 0 | 0.031 | 1 | 0.274597491 | 0.410162696 |
| Nme4 | 0.346827513 | -0.133583154 | 0 | 0.041 | 1 | 0.231991339 | -0.901889915 | 0 | 0.031 | 1 | 0.346827513 | 0.410162696 |
| Rwdd2a | 0.359270744 | 3.088809267 | 0.048 | 0.019 | 1 | 0.231991339 | -0.901889915 | 0 | 0.031 | 1 | 0.359270744 | 0.410162696 |
| Xylb | 0.359270744 | 2.673771768 | 0.048 | 0.019 | 1 | 0.231991339 | -1.124282336 | 0 | 0.031 | 1 | 0.359270744 | 0.410162696 |
| 2610029K11Rik | 0.396916958 | 0.214340149 | 0 | 0.034 | 1 | 0.231991339 | -1.316927414 | 0 | 0.031 | 1 | 0.396916958 | 0.410162696 |
| P2ry1 | 0.425614684 | 0.214340149 | 0 | 0.03 | 1 | 0.231991339 | -1.124282336 | 0 | 0.031 | 1 | 0.425614684 | 0.410162696 |
| Prss46 | 0.454205211 | 2.451379346 | 0.048 | 0.022 | 1 | 0.231991339 | -1.017367132 | 0 | 0.031 | 1 | 0.454205211 | 0.410162696 |
| Gm45250 | 0.457281049 | 0.503846766 | 0 | 0.026 | 1 | 0.231991339 | -0.901889915 | 0 | 0.031 | 1 | 0.457281049 | 0.410162696 |
| Gm715 | 0.457281049 | 0.351843673 | 0 | 0.026 | 1 | 0.231991339 | -0.901889915 | 0 | 0.031 | 1 | 0.457281049 | 0.410162696 |
| Fbxw10 | 0.4759797 | 1.673771768 | 0.048 | 0.022 | 1 | 0.231991339 | -1.017367132 | 0 | 0.031 | 1 | 0.4759797 | 0.410162696 |
| Tmem25 | 0.532436372 | 1.088809267 | 0 | 0.019 | 1 | 0.231991339 | -0.901889915 | 0 | 0.031 | 1 | 0.532436372 | 0.410162696 |
| Ly6g5b | 0.532436372 | 1.088809267 | 0 | 0.019 | 1 | 0.231991339 | -1.017367132 | 0 | 0.031 | 1 | 0.532436372 | 0.410162696 |
| Ppp1r14a | 0.547771462 | 2.214340149 | 0.048 | 0.026 | 1 | 0.231991339 | -0.901889915 | 0 | 0.031 | 1 | 0.547771462 | 0.410162696 |
| Slc25a18 | 0.578159907 | 0.866416846 | 0 | 0.015 | 1 | 0.231991339 | -0.901889915 | 0 | 0.031 | 1 | 0.578159907 | 0.410162696 |
| C2cd4b | 0.632076134 | 1.673771768 | 0 | 0.011 | 1 | 0.231991339 | -1.017367132 | 0 | 0.031 | 1 | 0.632076134 | 0.410162696 |
| Gm31834 | 0.658782751 | 1.351843673 | 0.048 | 0.03 | 1 | 0.231991339 | -1.017367132 | 0 | 0.031 | 1 | 0.658782751 | 0.410162696 |
| Samd13 | 0.741589569 | 1.088809267 | 0.048 | 0.034 | 1 | 0.231991339 | -1.017367132 | 0 | 0.031 | 1 | 0.741589569 | 0.410162696 |
| Gm29994 | 0.927798188 | 1.258734268 | 0.048 | 0.045 | 1 | 0.231991339 | -1.124282336 | 0 | 0.031 | 1 | 0.927798188 | 0.410162696 |
| Peg12 | 0.25165389 | -0.785659851 | 0 | 0.06 | 1 | 0.232000895 | -1.124282336 | 0 | 0.031 | 1 | 0.25165389 | 0.410177374 |
| Lhfpl2 | 0.304476689 | -1.611630451 | 0 | 0.049 | 1 | 0.232000895 | -1.486852415 | 0 | 0.031 | 1 | 0.304476689 | 0.410177374 |
| Gm49077 | 0.324767242 | -0.233118828 | 0 | 0.045 | 1 | 0.232000895 | -1.223818009 | 0 | 0.031 | 1 | 0.324767242 | 0.410177374 |
| Cd80 | 0.901972854 | 0.866416846 | 0.048 | 0.056 | 1 | 0.232000895 | -1.316927414 | 0 | 0.031 | 1 | 0.901972854 | 0.410177374 |
| Gpr135 | 0.304385789 | -0.326228232 | 0 | 0.049 | 1 | 0.232001372 | -1.017367132 | 0 | 0.031 | 1 | 0.304385789 | 0.410178108 |
| Insl6 | 0.457281049 | 0.503846766 | 0 | 0.026 | 1 | 0.232001372 | -1.017367132 | 0 | 0.031 | 1 | 0.457281049 | 0.410178108 |
| Gm46218 | 0.519007718 | 0.351843673 | 0.048 | 0.093 | 1 | 0.232001372 | -1.316927414 | 0 | 0.031 | 1 | 0.519007718 | 0.410178108 |
| Ano4 | 0.532432377 | 0.673771768 | 0 | 0.019 | 1 | 0.232001372 | -1.017367132 | 0 | 0.031 | 1 | 0.532432377 | 0.410178108 |
| Nrg4 | 0.57816106 | 1.351843673 | 0 | 0.015 | 1 | 0.232001372 | -1.486852415 | 0 | 0.031 | 1 | 0.57816106 | 0.410178108 |
| Efna3 | 0.85370107 | 1.673771768 | 0.048 | 0.041 | 1 | 0.232001372 | -1.017367132 | 0 | 0.031 | 1 | 0.85370107 | 0.410178108 |
| Gnmt | 0.921792415 | 1.451379346 | 0.048 | 0.045 | 1 | 0.232001372 | -1.017367132 | 0 | 0.031 | 1 | 0.921792415 | 0.410178108 |
| Acsm3 | 0.492590603 | 0.673771768 | 0 | 0.022 | 1 | 0.232006628 | -1.223818009 | 0 | 0.031 | 1 | 0.492590603 | 0.410186181 |
| Egflam | 0.673116577 | 0.214340149 | 0.048 | 0.075 | 1 | 0.232006628 | -1.223818009 | 0 | 0.031 | 1 | 0.673116577 | 0.410186181 |
| Kif15 | 0.790367465 | 1.60338244 | 0.048 | 0.037 | 1 | 0.232008061 | -1.486852415 | 0 | 0.031 | 1 | 0.790367465 | 0.410188382 |
| Shox2 | 0.915524318 | 1.425844254 | 0.048 | 0.056 | 1 | 0.232008061 | -1.486852415 | 0 | 0.031 | 1 | 0.915524318 | 0.410188382 |
| Tnfaip3 | 0.266774723 | 2.451379346 | 0.048 | 0.015 | 1 | 0.232008539 | -1.124282336 | 0 | 0.031 | 1 | 0.266774723 | 0.410189116 |
| Mns1 | 0.304431942 | -1.02666795 | 0 | 0.049 | 1 | 0.232008539 | -1.124282336 | 0 | 0.031 | 1 | 0.304431942 | 0.410189116 |
| Bend5 | 0.492608234 | 0.866416846 | 0 | 0.022 | 1 | 0.232008539 | -1.404390255 | 0 | 0.031 | 1 | 0.492608234 | 0.410189116 |
| Pcdhb12 | 0.578156445 | 1.088809267 | 0 | 0.015 | 1 | 0.232008539 | -1.404390255 | 0 | 0.031 | 1 | 0.578156445 | 0.410189116 |
| 1700034P13Rik | 0.918802447 | 3.513307095 | 0.048 | 0.045 | 1 | 0.232008539 | -1.564854927 | 0 | 0.031 | 1 | 0.918802447 | 0.410189116 |
| Pced1b | 0.457292877 | 2.258734268 | 0.048 | 0.022 | 1 | 0.232011406 | -1.404390255 | 0 | 0.031 | 1 | 0.457292877 | 0.410193519 |
| Ppargc1b | 0.457300763 | 0.214340149 | 0 | 0.026 | 1 | 0.232014272 | -1.709244837 | 0 | 0.031 | 1 | 0.457300763 | 0.410197922 |
| St6galnac3 | 0.945819522 | 0.214340149 | 0.048 | 0.045 | 1 | 0.232014272 | -2.271123724 | 0 | 0.031 | 1 | 0.945819522 | 0.410197922 |
| Mst1r | 0.632076134 | 1.673771768 | 0 | 0.011 | 1 | 0.232015228 | -2.271123724 | 0 | 0.031 | 1 | 0.632076134 | 0.41019939 |
| Pik3ap1 | 0.532432377 | 0.503846766 | 0 | 0.019 | 1 | 0.232015706 | -2.486852415 | 0 | 0.031 | 1 | 0.532432377 | 0.410200124 |
| Syk | 0.578159907 | -0.911190733 | 0 | 0.015 | 1 | 0.232015706 | -2.017367132 | 0 | 0.031 | 1 | 0.578159907 | 0.410200124 |
| Serpina1d | 0.252436477 | -2.492140171 | 0.048 | 0.134 | 1 | 0.232017139 | -4.137106376 | 0 | 0.031 | 1 | 0.252436477 | 0.410202325 |
| Thnsl1 | 0.267917141 | -0.413691074 | 0 | 0.056 | 1 | 0.232018094 | -0.638855509 | 0 | 0.031 | 1 | 0.267917141 | 0.410203793 |
| H3c15 | 0.267917141 | -0.648156327 | 0 | 0.056 | 1 | 0.232018094 | -0.638855509 | 0 | 0.031 | 1 | 0.267917141 | 0.410203793 |
| Gm56974 | 0.267985206 | -0.574155746 | 0 | 0.056 | 1 | 0.232018094 | -0.638855509 | 0 | 0.031 | 1 | 0.267985206 | 0.410203793 |
| Gm54310 | 0.28555153 | -0.574155746 | 0 | 0.052 | 1 | 0.232018094 | -0.638855509 | 0 | 0.031 | 1 | 0.28555153 | 0.410203793 |
| Gm57388 | 0.304355016 | -0.233118828 | 0 | 0.049 | 1 | 0.232018094 | -0.638855509 | 0 | 0.031 | 1 | 0.304355016 | 0.410203793 |
| 5033430I15Rik | 0.304355016 | -0.233118828 | 0 | 0.049 | 1 | 0.232018094 | -0.638855509 | 0 | 0.031 | 1 | 0.304355016 | 0.410203793 |
| H3c6 | 0.324767242 | -0.496153234 | 0 | 0.045 | 1 | 0.232018094 | -0.638855509 | 0 | 0.031 | 1 | 0.324767242 | 0.410203793 |
| Dnase1l2 | 0.373857158 | 2.088809267 | 0.048 | 0.019 | 1 | 0.232018094 | -0.638855509 | 0 | 0.031 | 1 | 0.373857158 | 0.410203793 |
| Pcdh1 | 0.396971401 | 0.351843673 | 0 | 0.034 | 1 | 0.232018094 | -0.638855509 | 0 | 0.031 | 1 | 0.396971401 | 0.410203793 |
| Fancf | 0.425599726 | 0.351843673 | 0 | 0.03 | 1 | 0.232018094 | -0.638855509 | 0 | 0.031 | 1 | 0.425599726 | 0.410203793 |
| Ccdc74a | 0.425639612 | 0.503846766 | 0 | 0.03 | 1 | 0.232018094 | -0.638855509 | 0 | 0.031 | 1 | 0.425639612 | 0.410203793 |
| Samd15 | 0.457281049 | 0.503846766 | 0 | 0.026 | 1 | 0.232018094 | -0.638855509 | 0 | 0.031 | 1 | 0.457281049 | 0.410203793 |
| 1700031C06Rik | 0.522328464 | 0.710297644 | 0.048 | 0.093 | 1 | 0.232018094 | -0.638855509 | 0 | 0.031 | 1 | 0.522328464 | 0.410203793 |
| St8sia2 | 0.532426384 | 0.503846766 | 0 | 0.019 | 1 | 0.232018094 | -0.638855509 | 0 | 0.031 | 1 | 0.532426384 | 0.410203793 |
| Epn3 | 0.532426384 | 0.866416846 | 0 | 0.019 | 1 | 0.232018094 | -0.638855509 | 0 | 0.031 | 1 | 0.532426384 | 0.410203793 |
| Gm50321 | 0.532432377 | 0.673771768 | 0 | 0.019 | 1 | 0.232018094 | -0.638855509 | 0 | 0.031 | 1 | 0.532432377 | 0.410203793 |
| Gm13166 | 0.550944111 | 1.936806174 | 0.048 | 0.026 | 1 | 0.232018094 | -0.638855509 | 0 | 0.031 | 1 | 0.550944111 | 0.410203793 |
| Gm15764 | 0.578156445 | 1.088809267 | 0 | 0.015 | 1 | 0.232018094 | -0.638855509 | 0 | 0.031 | 1 | 0.578156445 | 0.410203793 |
| Tlcd3b | 0.57816106 | 1.351843673 | 0 | 0.015 | 1 | 0.232018094 | -0.638855509 | 0 | 0.031 | 1 | 0.57816106 | 0.410203793 |
| Fam241b | 0.57816106 | 1.351843673 | 0 | 0.015 | 1 | 0.232018094 | -0.638855509 | 0 | 0.031 | 1 | 0.57816106 | 0.410203793 |
| B3galt4 | 0.57816106 | 1.351843673 | 0 | 0.015 | 1 | 0.232018094 | -0.638855509 | 0 | 0.031 | 1 | 0.57816106 | 0.410203793 |

| | | | | | | | | | | | | |
|---|---|---|---|---|---|---|---|---|---|---|---|---|
| Gm11175 | 0.629700127 | 2.088809267 | 0.048 | 0.03 | 1 | 0.232018094 | -0.638855509 | 0 | 0.031 | 1 | 0.629700127 | 0.410203793 |
| Gm42670 | 0.63207476 | 1.351843673 | 0 | 0.011 | 1 | 0.232018094 | -0.638855509 | 0 | 0.031 | 1 | 0.63207476 | 0.410203793 |
| Gm47814 | 0.632076134 | 1.673771768 | 0 | 0.011 | 1 | 0.232018094 | -0.638855509 | 0 | 0.031 | 1 | 0.632076134 | 0.410203793 |
| Selenok-ps1 | 0.636138263 | 1.79930265 | 0.048 | 0.03 | 1 | 0.232018094 | -0.638855509 | 0 | 0.031 | 1 | 0.636138263 | 0.410203793 |
| 2810032G03Rik | 0.773911991 | 1.351843673 | 0.048 | 0.067 | 1 | 0.232018094 | -0.638855509 | 0 | 0.031 | 1 | 0.773911991 | 0.410203793 |
| Pbld1 | 0.866047111 | 1.258734268 | 0.048 | 0.041 | 1 | 0.232018094 | -0.638855509 | 0 | 0.031 | 1 | 0.866047111 | 0.410203793 |
| 4930458D05Rik | 0.907384734 | 1.010806755 | 0.048 | 0.056 | 1 | 0.232018094 | -0.638855509 | 0 | 0.031 | 1 | 0.907384734 | 0.410203793 |
| A430046D13Rik | 0.921792415 | 1.451379346 | 0.048 | 0.045 | 1 | 0.232018094 | -0.638855509 | 0 | 0.031 | 1 | 0.921792415 | 0.410203793 |
| Gtpbp2 | 0.897440948 | 0.617918533 | 0.381 | 0.399 | 1 | 0.232456899 | 0.311816779 | 0.311 | 0.484 | 1 | 0.897440948 | 0.410877588 |
| Slc24a4 | 0.304385789 | -0.413691074 | 0 | 0.049 | 1 | 0.232482267 | 2.905465008 | 0.044 | 0.017 | 1 | 0.304385789 | 0.41091653 |
| ENSMUSG00000121456 | 0.709478182 | 2.214340149 | 0.048 | 0.034 | 1 | 0.232482267 | 2.42003818 | 0.044 | 0.017 | 1 | 0.709478182 | 0.41091653 |
| Pdgfc | 0.393852562 | 0.857078981 | 0.143 | 0.25 | 1 | 0.232755555 | 0.544658557 | 0.289 | 0.46 | 1 | 0.393852562 | 0.411335961 |
| Gm16093 | 0.233229031 | 1.491568436 | 0.238 | 0.146 | 1 | 0.249522611 | -0.264459994 | 0.067 | 0.131 | 1 | 0.249522611 | 0.412062282 |
| Rpusd4 | 0.23327272 | 0.644024424 | 0.048 | 0.149 | 1 | 0.441784911 | 0.15951063 | 0.089 | 0.131 | 1 | 0.441784911 | 0.412129278 |
| Nars2 | 0.975005461 | 0.464318402 | 0.19 | 0.198 | 1 | 0.233328778 | 0.741966275 | 0.111 | 0.201 | 1 | 0.975005461 | 0.412215237 |
| Rhbdf2 | 0.61241669 | 0.576910228 | 0.238 | 0.205 | 1 | 0.233338233 | -0.387316742 | 0.044 | 0.104 | 1 | 0.61241669 | 0.412229735 |
| Eogt | 0.307198578 | 0.673771768 | 0.143 | 0.246 | 1 | 0.233597566 | -0.116902806 | 0.222 | 0.343 | 1 | 0.307198578 | 0.412627309 |
| Gm32191 | 0.578156445 | 1.088809267 | 0 | 0.015 | 1 | 0.233802347 | 2.197645759 | 0.044 | 0.017 | 1 | 0.578156445 | 0.412941157 |
| Rnf32 | 0.915528315 | 1.351843673 | 0.048 | 0.056 | 1 | 0.233802347 | 2.197645759 | 0.044 | 0.017 | 1 | 0.915528315 | 0.412941157 |
| Bcl2l13 | 0.233886496 | -0.589262638 | 0.143 | 0.265 | 1 | 0.485361078 | -0.161952754 | 0.333 | 0.391 | 1 | 0.485361078 | 0.413070099 |
| Sugp1 | 0.619228268 | 0.866416846 | 0.238 | 0.216 | 1 | 0.234026335 | -0.16020191 | 0.178 | 0.273 | 1 | 0.619228268 | 0.413284345 |
| Etv3 | 0.234138885 | -0.413691074 | 0.143 | 0.28 | 1 | 0.916057846 | 0.186057785 | 0.356 | 0.374 | 1 | 0.916057846 | 0.413456753 |
| Dicer1 | 0.25933708 | 0.208885719 | 0.286 | 0.526 | 1 | 0.234467852 | 0.632024981 | 0.756 | 0.754 | 1 | 0.25933708 | 0.41396053 |
| Nipsnap1 | 0.882598156 | 1.295260144 | 0.095 | 0.09 | 1 | 0.234578975 | -0.674479418 | 0.067 | 0.128 | 1 | 0.882598156 | 0.414130655 |
| 4930549G23Rik | 0.679706589 | 0.503846766 | 0.048 | 0.075 | 1 | 0.234612023 | -0.387316742 | 0.044 | 0.104 | 1 | 0.679706589 | 0.414181245 |
| Mccc2 | 0.234782186 | -0.870548749 | 0.095 | 0.198 | 1 | 0.309572181 | 0.342035668 | 0.111 | 0.176 | 1 | 0.309572181 | 0.414441698 |
| Kdm3b | 0.235236173 | 1.094833055 | 0.476 | 0.466 | 1 | 0.410722381 | 0.266908421 | 0.756 | 0.633 | 1 | 0.410722381 | 0.415136289 |
| B230217C12Rik | 0.777549848 | 0.840881754 | 0.143 | 0.183 | 1 | 0.235512376 | 0.531069493 | 0.111 | 0.194 | 1 | 0.777549848 | 0.415558673 |
| Prkg2 | 0.285559366 | -0.987541619 | 0.095 | 0.205 | 1 | 0.235832354 | -2.620708162 | 0.022 | 0.069 | 1 | 0.285559366 | 0.416047809 |
| AU015336 | 0.23601705 | 1.351843673 | 0.286 | 0.201 | 1 | 0.949740962 | 0.638678467 | 0.067 | 0.073 | 1 | 0.949740962 | 0.416330051 |
| Runx2os1 | 0.350370971 | -0.133583154 | 0.048 | 0.119 | 1 | 0.236026534 | -0.191396532 | 0.067 | 0.135 | 1 | 0.350370971 | 0.416344543 |
| Clcn5 | 0.308254016 | 2.195231326 | 0.381 | 0.354 | 1 | 0.236189806 | -0.356943093 | 0.333 | 0.446 | 1 | 0.308254016 | 0.416593987 |
| Coq8a | 0.236343578 | -0.574155746 | 0 | 0.063 | 1 | 0.250724386 | -0.960783604 | 0.022 | 0.066 | 1 | 0.250724386 | 0.416828868 |
| Tbx6 | 0.236343578 | -0.718545655 | 0 | 0.063 | 1 | 0.513841138 | 0.320502507 | 0.067 | 0.104 | 1 | 0.513841138 | 0.416828868 |
| Pex11g | 0.236343578 | -0.574155746 | 0 | 0.063 | 1 | 0.32914342 | -0.191396532 | 0.022 | 0.059 | 1 | 0.32914342 | 0.416828868 |
| BC051226 | 0.236343578 | -0.574155746 | 0 | 0.063 | 1 | 0.849821852 | 1.098110085 | 0.022 | 0.028 | 1 | 0.849821852 | 0.416828868 |
| Aldh1l1 | 0.23638565 | -0.785659851 | 0 | 0.063 | 1 | 0.936025227 | 1.683072586 | 0.022 | 0.021 | 1 | 0.936025227 | 0.416893125 |
| Nrde2 | 0.23638565 | -0.648156327 | 0 | 0.063 | 1 | 0.670869263 | 0.320502507 | 0.067 | 0.087 | 1 | 0.670869263 | 0.416893125 |
| Pcdhgb4 | 0.23638565 | -0.648156327 | 0 | 0.063 | 1 | 0.366250765 | 2.197645759 | 0.089 | 0.059 | 1 | 0.366250765 | 0.416893125 |
| Iqck | 0.236422109 | -0.785659851 | 0 | 0.063 | 1 | 0.603818627 | 0.304560963 | 0.044 | 0.066 | 1 | 0.603818627 | 0.416948804 |
| Zfp963 | 0.236422109 | -0.849790188 | 0 | 0.063 | 1 | 0.616633976 | 0.290755163 | 0.089 | 0.121 | 1 | 0.616633976 | 0.416948804 |
| Inka1 | 0.236422109 | -0.911190733 | 0 | 0.063 | 1 | 0.273281063 | 0.361144491 | 0.044 | 0.097 | 1 | 0.273281063 | 0.416948804 |
| Ppp1r14c | 0.236422109 | -0.970084422 | 0 | 0.063 | 1 | 0.309616342 | -0.901889915 | 0.022 | 0.059 | 1 | 0.309616342 | 0.416948804 |
| 2610027K06Rik | 0.236422109 | -0.785659851 | 0 | 0.063 | 1 | 0.530911177 | -0.316927414 | 0.022 | 0.042 | 1 | 0.530911177 | 0.416948804 |
| Gm49735 | 0.236422109 | -0.785659851 | 0 | 0.063 | 1 | 0.427678952 | -0.387316742 | 0.044 | 0.08 | 1 | 0.427678952 | 0.416948804 |
| Tcte2 | 0.236422109 | -0.785659851 | 0 | 0.063 | 1 | 0.552232538 | 0.361144491 | 0.022 | 0.042 | 1 | 0.552232538 | 0.416948804 |
| Psmb9 | 0.236422109 | -0.911190733 | 0 | 0.063 | 1 | 0.936023974 | 1.460680165 | 0.022 | 0.021 | 1 | 0.936023974 | 0.416948804 |
| 9030612E09Rik | 0.236422459 | -0.718545655 | 0 | 0.063 | 1 | 0.326732519 | 1.875717664 | 0.044 | 0.021 | 1 | 0.326732519 | 0.416949339 |
| Tnfsf13os | 0.236422459 | -1.370622352 | 0 | 0.063 | 1 | 0.363486871 | 0.15951063 | 0.022 | 0.055 | 1 | 0.363486871 | 0.416949339 |
| Oxld1 | 0.236422459 | -0.718545655 | 0 | 0.063 | 1 | 0.500262731 | 0.683072586 | 0.022 | 0.045 | 1 | 0.500262731 | 0.416949339 |
| Tspan15 | 0.57816106 | 1.351843673 | 0 | 0.015 | 1 | 0.236447234 | 2.098110085 | 0.044 | 0.017 | 1 | 0.57816106 | 0.416987174 |
| Gm15788 | 0.236452955 | -1.02666795 | 0 | 0.063 | 1 | 0.885646453 | 1.098110085 | 0.044 | 0.052 | 1 | 0.885646453 | 0.416995909 |
| Gm16001 | 0.236452955 | -0.849790188 | 0 | 0.063 | 1 | 0.633916264 | 0.410054092 | 0.067 | 0.09 | 1 | 0.633916264 | 0.416995909 |
| Slc24a5 | 0.236454357 | -0.785659851 | 0 | 0.063 | 1 | 0.356316838 | -0.124282336 | 0.044 | 0.087 | 1 | 0.356316838 | 0.41699805 |
| Tmem59l | 0.236454357 | -0.785659851 | 0 | 0.063 | 1 | 0.260653677 | -0.638855509 | 0 | 0.028 | 1 | 0.260653677 | 0.41699805 |
| 6820445E23Rik | 0.236454357 | -0.785659851 | 0 | 0.063 | 1 | 0.667598499 | 0.223640968 | 0.022 | 0.035 | 1 | 0.667598499 | 0.41699805 |
| Mybph | 0.236480292 | -1.233118828 | 0 | 0.063 | 1 | 0.765741153 | 1.852997588 | 0.022 | 0.031 | 1 | 0.765741153 | 0.417037656 |
| Zbp1 | 0.23650132 | -1.081115734 | 0 | 0.063 | 1 | 0.769054632 | 0.905465008 | 0.089 | 0.107 | 1 | 0.769054632 | 0.417069766 |
| Dclk2 | 0.236520594 | -1.133583154 | 0 | 0.063 | 1 | 0.623740403 | -0.223818009 | 0.089 | 0.114 | 1 | 0.623740403 | 0.417099196 |
| E2f8 | 0.236522346 | -1.785659851 | 0 | 0.063 | 1 | 0.496223879 | 0.683072586 | 0 | 0.01 | 1 | 0.496223879 | 0.417101872 |
| Pou3f1 | 0.236537063 | -1.413691074 | 0 | 0.063 | 1 | 0.750173272 | 1.739656115 | 0.044 | 0.059 | 1 | 0.750173272 | 0.417124344 |
| Kcne4 | 0.236538465 | -1.718545655 | 0 | 0.063 | 1 | 0.376842245 | -0.776359032 | 0 | 0.017 | 1 | 0.376842245 | 0.417126484 |
| Lin7a | 0.23656124 | -1.611630451 | 0 | 0.063 | 1 | 0.421610828 | 0.203079645 | 0.156 | 0.114 | 1 | 0.421610828 | 0.417161259 |
| Mettl27 | 0.794695257 | 0.673771768 | 0.095 | 0.078 | 1 | 0.236568359 | -0.686161223 | 0.022 | 0.069 | 1 | 0.794695257 | 0.41717213 |
| Sfrp4 | 0.23657105 | -2.555046923 | 0 | 0.063 | 1 | 0.601352461 | -2.37003875 | 0.022 | 0.038 | 1 | 0.601352461 | 0.417176238 |
| Dzip1 | 0.500772156 | -0.785659851 | 0.048 | 0.09 | 1 | 0.236577744 | -0.686161223 | 0.022 | 0.069 | 1 | 0.500772156 | 0.417186459 |
| Nlrc4 | 0.236581911 | -0.496153234 | 0 | 0.063 | 1 | 0.753258193 | 1.320502507 | 0.044 | 0.059 | 1 | 0.753258193 | 0.417192821 |
| Rbm12b1 | 0.800793775 | 0.97333205 | 0.095 | 0.082 | 1 | 0.2366751 | -0.387316742 | 0.067 | 0.131 | 1 | 0.800793775 | 0.417335097 |
| Eml6 | 0.236883106 | 1.786246497 | 0.238 | 0.153 | 1 | 0.383038878 | 0.274987848 | 0.178 | 0.253 | 1 | 0.383038878 | 0.417652606 |
| Ppfibp2 | 0.725232502 | -0.648156327 | 0.095 | 0.119 | 1 | 0.237090392 | -1.709244837 | 0.067 | 0.125 | 1 | 0.725232502 | 0.417968929 |
| Pakap | 0.525089518 | -0.377890352 | 0.19 | 0.261 | 1 | 0.237103427 | -0.219965684 | 0.133 | 0.228 | 1 | 0.525089518 | 0.417988819 |
| Fam133b | 0.922821386 | 0.387671752 | 0.619 | 0.765 | 1 | 0.237206879 | -0.293468441 | 0.756 | 0.765 | 1 | 0.922821386 | 0.418146655 |
| Gm30054 | 0.237249394 | 1.97333205 | 0.095 | 0.041 | 1 | 0.708995259 | 1.683072586 | 0.044 | 0.035 | 1 | 0.708995259 | 0.418211514 |
| Cpsf6 | 0.237425502 | 0.886045652 | 0.667 | 0.754 | 1 | 0.411812068 | 0.383628505 | 0.933 | 0.931 | 1 | 0.411812068 | 0.418480136 |
| Ptch2 | 0.274597491 | 2.088809267 | 0.048 | 0.015 | 1 | 0.237772641 | 1.545569062 | 0.044 | 0.017 | 1 | 0.274597491 | 0.419009454 |
| Adssl1 | 0.808137022 | 0.55829455 | 0.048 | 0.063 | 1 | 0.238118274 | -0.84048937 | 0.022 | 0.069 | 1 | 0.808137022 | 0.419536235 |
| Gm43696 | 0.238361805 | 1.623145695 | 0.143 | 0.075 | 1 | 0.406873805 | 0.15951063 | 0.067 | 0.111 | 1 | 0.406873805 | 0.419907259 |
| Riok1 | 0.238427176 | 0.468096741 | 0.571 | 0.549 | 1 | 0.728396745 | 0.115388077 | 0.511 | 0.592 | 1 | 0.728396745 | 0.420006833 |
| Fbxl20 | 0.238458875 | 0.948879006 | 0.476 | 0.466 | 1 | 0.277843783 | 0.305705505 | 0.756 | 0.64 | 1 | 0.277843783 | 0.420055114 |
| Rpgrip1 | 0.409927861 | 0.620660431 | 0.095 | 0.179 | 1 | 0.239092287 | -0.371375198 | 0.111 | 0.19 | 1 | 0.409927861 | 0.421019452 |
| Gm11149 | 0.396916958 | 0.214340149 | 0 | 0.034 | 1 | 0.239110666 | 1.683072586 | 0.044 | 0.017 | 1 | 0.396916958 | 0.421047422 |
| Kdm3a | 0.24103613 | 0.59792801 | 0.429 | 0.608 | 1 | 0.239124368 | 0.567595369 | 0.867 | 0.713 | 1 | 0.24103613 | 0.421068272 |
| Slc38a9 | 0.239517437 | 1.205622932 | 0.286 | 0.22 | 1 | 0.824312855 | 0.502007027 | 0.267 | 0.318 | 1 | 0.824312855 | 0.421666271 |
| Tstd2 | 0.266315219 | -0.496153234 | 0.143 | 0.257 | 1 | 0.240251302 | 0.118287967 | 0.222 | 0.349 | 1 | 0.266315219 | 0.422781916 |
| Gm17057 | 0.240320909 | 2.351843673 | 0.095 | 0.041 | 1 | 0.613823544 | 1.78998779 | 0.044 | 0.031 | 1 | 0.613823544 | 0.422887678 |
| Slc2a9 | 0.323332113 | 3.316219763 | 0.095 | 0.049 | 1 | 0.240441658 | 0.776181991 | 0.044 | 0.017 | 1 | 0.323332113 | 0.423071126 |
| 1700112J16Rik | 0.598185743 | 1.088809267 | 0.095 | 0.067 | 1 | 0.240526101 | -0.446210431 | 0.022 | 0.069 | 1 | 0.598185743 | 0.423199396 |
| Nek11 | 0.446328101 | 1.425844254 | 0.095 | 0.056 | 1 | 0.240867571 | 1.808603468 | 0.089 | 0.048 | 1 | 0.446328101 | 0.423717955 |
| Chic2 | 0.69658508 | 0.327181619 | 0.667 | 0.817 | 1 | 0.241042294 | -0.186746423 | 0.956 | 0.865 | 1 | 0.69658508 | 0.4239832 |
| Cyp2ab1 | 0.532435374 | 0.503846766 | 0 | 0.019 | 1 | 0.241249968 | -0.223818009 | 0.022 | 0.069 | 1 | 0.532435374 | 0.424298389 |
| Hells | 0.241622411 | -1.303508156 | 0.048 | 0.134 | 1 | 0.5788741 | 1.162240423 | 0.089 | 0.128 | 1 | 0.5788741 | 0.424863433 |
| 4933417E11Rik | 0.241799439 | 1.586308926 | 0.095 | 0.041 | 1 | 0.496223879 | 0.683072586 | 0 | 0.01 | 1 | 0.496223879 | 0.425131909 |
| Myoz3 | 0.24181731 | 1.977777955 | 0.238 | 0.16 | 1 | 0.837955762 | 1.00269052 | 0.2 | 0.221 | 1 | 0.837955762 | 0.425159008 |
| Smurf2 | 0.24182975 | -0.408097057 | 0.857 | 0.918 | 1 | 0.543779513 | 0.237595139 | 1 | 0.979 | 1 | 0.543779513 | 0.425177872 |
| Sv2c | 0.241865236 | 1.489347197 | 0.095 | 0.041 | 1 | 0.430483494 | -0.901889915 | 0 | 0.014 | 1 | 0.430483494 | 0.425231679 |
| Gdpd1 | 0.75843612 | -0.184209227 | 0.048 | 0.063 | 1 | 0.241949037 | -1.053893008 | 0.044 | 0.097 | 1 | 0.75843612 | 0.425358738 |
| Ptprv | 0.877649952 | 0.258734268 | 0.048 | 0.056 | 1 | 0.242065788 | -0.223818009 | 0.022 | 0.069 | 1 | 0.877649952 | 0.42553573 |
| Dstyk | 0.242121359 | -0.197241792 | 0.333 | 0.545 | 1 | 0.352323256 | 0.582094938 | 0.622 | 0.654 | 1 | 0.352323256 | 0.425619966 |

| | | | | | | | | | | | | |
|---|---|---|---|---|---|---|---|---|---|---|---|---|
| Syf2 | 0.242420416 | -0.315580988 | 0.429 | 0.582 | 1 | 0.244953661 | -0.199570463 | 0.711 | 0.675 | 1 | 0.244953661 | 0.426073175 |
| Zfpm1 | 0.284778828 | -0.555046923 | 0.095 | 0.194 | 1 | 0.242464017 | 0.336269823 | 0.156 | 0.263 | 1 | 0.284778828 | 0.426139234 |
| Elk3 | 0.242518548 | 0.264380832 | 0.286 | 0.507 | 1 | 0.40416777 | 0.942388704 | 0.444 | 0.685 | 1 | 0.40416777 | 0.426221849 |
| Pank1 | 0.947285389 | 0.614878079 | 0.429 | 0.481 | 1 | 0.24263513 | 1.15951063 | 0.444 | 0.446 | 1 | 0.947285389 | 0.426398454 |
| Marchf9 | 0.304385789 | -0.496153234 | 0 | 0.049 | 1 | 0.242751375 | 0.304560963 | 0.022 | 0.069 | 1 | 0.304385789 | 0.426574521 |
| Pex26 | 0.492608234 | 0.866416846 | 0 | 0.022 | 1 | 0.242751375 | -0.255526869 | 0.022 | 0.069 | 1 | 0.492608234 | 0.426574521 |
| Gm12279 | 0.251619726 | -0.718545655 | 0 | 0.06 | 1 | 0.242859454 | 0.268035087 | 0.022 | 0.069 | 1 | 0.251619726 | 0.426738194 |
| 4930509H03Rik | 0.954841465 | 0.97333205 | 0.048 | 0.045 | 1 | 0.243511394 | -0.191396532 | 0.022 | 0.069 | 1 | 0.954841465 | 0.427724988 |
| Ift140 | 0.472245713 | -0.535681598 | 0.048 | 0.093 | 1 | 0.243668048 | 0.393565969 | 0.044 | 0.104 | 1 | 0.472245713 | 0.427961979 |
| Ranbp10 | 0.278890791 | 0.879355901 | 0.524 | 0.463 | 1 | 0.243759072 | 0.5448155 | 0.667 | 0.612 | 1 | 0.278890791 | 0.428099659 |
| Dhdds | 0.604168713 | 0.16475812 | 0.286 | 0.392 | 1 | 0.243791525 | 0.240850258 | 0.178 | 0.291 | 1 | 0.604168713 | 0.428148742 |
| AI597479 | 0.424500903 | -0.370622352 | 0.048 | 0.101 | 1 | 0.244058723 | -0.486852415 | 0.044 | 0.1 | 1 | 0.424500903 | 0.428552786 |
| Top3a | 0.296732346 | -0.818081329 | 0.048 | 0.123 | 1 | 0.244163475 | 0.435145073 | 0.067 | 0.138 | 1 | 0.296732346 | 0.428711147 |
| Glmn | 0.318784719 | -0.289702356 | 0.095 | 0.194 | 1 | 0.244178123 | 0.567595369 | 0.111 | 0.194 | 1 | 0.318784719 | 0.42873329 |
| 4930539J05Rik | 0.81881728 | 1.088809267 | 0.048 | 0.037 | 1 | 0.244269117 | -0.124282336 | 0.022 | 0.069 | 1 | 0.81881728 | 0.428870832 |
| Apex2 | 0.537193032 | 0.965426851 | 0.381 | 0.425 | 1 | 0.244327647 | 0.656272527 | 0.222 | 0.353 | 1 | 0.537193032 | 0.428959295 |
| Pla2g4b | 0.655561847 | 1.503846766 | 0.048 | 0.03 | 1 | 0.244346435 | 0.15951063 | 0.022 | 0.069 | 1 | 0.655561847 | 0.42898769 |
| Arl5c | 0.659925739 | 1.029915578 | 0.095 | 0.071 | 1 | 0.244346435 | 1.329435632 | 0.022 | 0.069 | 1 | 0.659925739 | 0.42898769 |
| 4930511M06Rik | 0.866047111 | 1.258734268 | 0.048 | 0.041 | 1 | 0.244346435 | 0.481438725 | 0.022 | 0.069 | 1 | 0.866047111 | 0.42898769 |
| Gm56494 | 0.790349864 | 1.766881172 | 0.048 | 0.037 | 1 | 0.244378645 | 0.42003818 | 0.022 | 0.069 | 1 | 0.790349864 | 0.429036368 |
| Sema7a | 0.444393494 | 2.489347197 | 0.095 | 0.056 | 1 | 0.244488775 | 1.435145073 | 0.022 | 0.069 | 1 | 0.444393494 | 0.429202788 |
| Fam20a | 0.319264877 | 3.113056813 | 0.19 | 0.127 | 1 | 0.244505087 | 1.098110085 | 0.022 | 0.069 | 1 | 0.319264877 | 0.429227436 |
| Sptbn1 | 0.480255991 | 0.543483738 | 0.952 | 0.981 | 1 | 0.2447286 | 0.330746538 | 0.978 | 1 | 1 | 0.480255991 | 0.429565113 |
| Mettl17 | 0.748287671 | 1.088809267 | 0.095 | 0.078 | 1 | 0.244728837 | -0.404390255 | 0.044 | 0.1 | 1 | 0.748287671 | 0.42956547 |
| Epha5 | 0.244907832 | 0.866416846 | 0.095 | 0.041 | 1 | 0.551245041 | 0.290755163 | 0.044 | 0.028 | 1 | 0.551245041 | 0.429835818 |
| Nup210l | 0.245029214 | 2.166811779 | 0.476 | 0.451 | 1 | 0.467670358 | 0.688719149 | 0.6 | 0.519 | 1 | 0.467670358 | 0.430019112 |
| Wdr91 | 0.398543057 | 0.171271427 | 0.048 | 0.112 | 1 | 0.24520415 | -0.468930507 | 0.067 | 0.135 | 1 | 0.398543057 | 0.430283225 |
| Wdr55 | 0.732130428 | 0.936806174 | 0.048 | 0.071 | 1 | 0.245345382 | -0.404390255 | 0.044 | 0.1 | 1 | 0.732130428 | 0.430496407 |
| Zfp182 | 0.568655103 | 0.698019314 | 0.19 | 0.299 | 1 | 0.245351723 | 0.232411177 | 0.178 | 0.304 | 1 | 0.568655103 | 0.430505978 |
| Osbpl10 | 0.245368361 | -0.233118828 | 0.048 | 0.146 | 1 | 0.643703883 | 0.832216025 | 0.267 | 0.346 | 1 | 0.643703883 | 0.43053109 |
| Tbc1d32 | 0.245442427 | 1.072867723 | 0.286 | 0.198 | 1 | 0.448218239 | 0.946106992 | 0.111 | 0.17 | 1 | 0.448218239 | 0.43064287 |
| Mertk | 0.245616179 | -0.413691074 | 0.333 | 0.537 | 1 | 0.338991723 | 0.330499174 | 0.489 | 0.696 | 1 | 0.338991723 | 0.43090505 |
| Katnip | 0.245620323 | 1.552756367 | 0.429 | 0.396 | 1 | 0.297622466 | -0.130158902 | 0.289 | 0.412 | 1 | 0.297622466 | 0.430911304 |
| Tbx15 | 0.425614684 | 0.214340149 | 0 | 0.03 | 1 | 0.245815295 | 1.536231198 | 0.111 | 0.066 | 1 | 0.425614684 | 0.431205431 |
| Dglucy | 0.246208913 | 0.989273593 | 0.19 | 0.108 | 1 | 0.294407898 | -0.941418279 | 0.044 | 0.09 | 1 | 0.294407898 | 0.431798998 |
| Cox18 | 0.522046706 | 0.789248985 | 0.286 | 0.265 | 1 | 0.246301504 | 0.285480221 | 0.2 | 0.322 | 1 | 0.522046706 | 0.431938577 |
| Hs6st1 | 0.246725824 | -0.911190733 | 0.095 | 0.194 | 1 | 0.562181374 | -0.135687099 | 0.267 | 0.311 | 1 | 0.562181374 | 0.432578016 |
| Zfp583 | 0.769036344 | 0.866416846 | 0.048 | 0.067 | 1 | 0.246788259 | -0.124282336 | 0.044 | 0.1 | 1 | 0.769036344 | 0.432672073 |
| Exoc2 | 0.418805339 | 0.940417427 | 0.333 | 0.295 | 1 | 0.246892937 | -0.151503805 | 0.267 | 0.381 | 1 | 0.418805339 | 0.432829751 |
| Scaper | 0.247023494 | 0.917358411 | 0.714 | 0.694 | 1 | 0.871580619 | 0.487724988 | 0.844 | 0.761 | 1 | 0.871580619 | 0.433026381 |
| Borcs5 | 0.607730376 | 0.638147858 | 0.143 | 0.108 | 1 | 0.247127792 | -0.346674757 | 0.067 | 0.131 | 1 | 0.607730376 | 0.433183439 |
| Pou6f1 | 0.338038591 | -0.611630451 | 0.095 | 0.179 | 1 | 0.247212515 | 0.361144491 | 0.111 | 0.197 | 1 | 0.338038591 | 0.433311003 |
| Mettl8 | 0.329147987 | 1.815790773 | 0.143 | 0.086 | 1 | 0.247414633 | 0.180572246 | 0.044 | 0.1 | 1 | 0.329147987 | 0.433615266 |
| Gm26879 | 0.409489432 | 1.908237021 | 0.095 | 0.056 | 1 | 0.247415536 | 0.13875207 | 0.044 | 0.1 | 1 | 0.409489432 | 0.433616625 |
| Rpap1 | 0.251619726 | -0.574155746 | 0 | 0.06 | 1 | 0.24745508 | 0.595609745 | 0.044 | 0.104 | 1 | 0.251619726 | 0.433676144 |
| Gm20045 | 0.247489943 | 0.97333205 | 0.143 | 0.071 | 1 | 0.990790529 | 0.71623945 | 0.089 | 0.093 | 1 | 0.990790529 | 0.433728614 |
| Tnfsf9 | 0.285551097 | -0.849790188 | 0 | 0.052 | 1 | 0.247497294 | -1.776359032 | 0.022 | 0.066 | 1 | 0.285551097 | 0.433739678 |
| Gys1 | 0.464091254 | 0.281454345 | 0.048 | 0.101 | 1 | 0.247758284 | 0.618942249 | 0.044 | 0.104 | 1 | 0.464091254 | 0.434132401 |
| Chst8 | 0.247948576 | 0.866416846 | 0.095 | 0.041 | 1 | 0.449526036 | 1.050804371 | 0.022 | 0.048 | 1 | 0.449526036 | 0.434418656 |
| Frk | 0.913296674 | 0.572233741 | 0.143 | 0.168 | 1 | 0.248155811 | 1.036182336 | 0.289 | 0.221 | 1 | 0.913296674 | 0.434730316 |
| Nmi | 0.543922905 | -0.252227651 | 0.143 | 0.198 | 1 | 0.248262158 | -0.454430938 | 0.133 | 0.204 | 1 | 0.543922905 | 0.434890217 |
| Mllt3 | 0.248293846 | 0.538901824 | 0.905 | 0.91 | 1 | 0.871943027 | 0.538834228 | 0.911 | 0.862 | 1 | 0.871943027 | 0.434937858 |
| AI467606 | 0.267985206 | -0.574155746 | 0 | 0.056 | 1 | 0.248302399 | -1.174908409 | 0.022 | 0.066 | 1 | 0.267985206 | 0.434950717 |
| Adcy9 | 0.436809255 | 0.771952162 | 0.143 | 0.25 | 1 | 0.24844641 | 0.219788642 | 0.311 | 0.481 | 1 | 0.436809255 | 0.435167201 |
| Myh9 | 0.845669646 | 0.940417427 | 0.762 | 0.772 | 1 | 0.248594065 | 0.234612085 | 0.978 | 0.9 | 1 | 0.845669646 | 0.435389122 |
| Ldlrad3 | 0.24924619 | 0.629377648 | 0.762 | 0.825 | 1 | 0.695918694 | 0.493172885 | 0.889 | 0.869 | 1 | 0.695918694 | 0.436368717 |
| Ppp3cb | 0.32488127 | 0.550389352 | 0.714 | 0.612 | 1 | 0.249357306 | 0.397364679 | 0.844 | 0.803 | 1 | 0.32488127 | 0.436535546 |
| Dock3 | 0.249512081 | 1.425844254 | 0.095 | 0.041 | 1 | 0.949212876 | 0.473619221 | 0.067 | 0.066 | 1 | 0.949212876 | 0.436767883 |
| Zfp335os | 0.619461129 | 0.719575457 | 0.19 | 0.16 | 1 | 0.249882304 | -0.237492946 | 0.067 | 0.128 | 1 | 0.619461129 | 0.437323442 |
| Swt1 | 0.631117102 | 0.773529548 | 0.524 | 0.642 | 1 | 0.250124656 | 0.480726107 | 0.889 | 0.865 | 1 | 0.631117102 | 0.437686968 |
| Brd1 | 0.250655807 | -0.22480356 | 0.905 | 0.866 | 1 | 0.5177425 | 0.185361279 | 0.978 | 0.934 | 1 | 0.5177425 | 0.438483281 |
| Qtrt1 | 0.301398764 | 1.103023126 | 0.333 | 0.269 | 1 | 0.250719017 | 0.1454158 | 0.244 | 0.384 | 1 | 0.301398764 | 0.438578009 |
| 1700048O20Rik | 0.800649369 | 0.614878079 | 0.048 | 0.063 | 1 | 0.250724386 | -0.960783604 | 0.022 | 0.066 | 1 | 0.800649369 | 0.438586055 |
| Dok4 | 0.710539439 | -0.295854583 | 0.095 | 0.119 | 1 | 0.250725289 | -0.901889915 | 0.022 | 0.066 | 1 | 0.710539439 | 0.438587408 |
| Gm13778 | 0.251107945 | 1.673771768 | 0.095 | 0.041 | 1 | 0.928515555 | 0.83507568 | 0.044 | 0.048 | 1 | 0.928515555 | 0.439160689 |
| Zfp398 | 0.831234516 | 0.534220415 | 0.238 | 0.299 | 1 | 0.251166148 | 0.458441693 | 0.244 | 0.408 | 1 | 0.831234516 | 0.439247862 |
| Spag4 | 0.251580301 | -0.496153234 | 0 | 0.06 | 1 | 0.299191224 | 0.361144491 | 0.022 | 0.062 | 1 | 0.299191224 | 0.439867954 |
| Hk1os | 0.251580301 | -0.496153234 | 0 | 0.06 | 1 | 0.331922009 | -0.124282336 | 0 | 0.021 | 1 | 0.331922009 | 0.439867954 |
| Pcdhga11 | 0.251580301 | -0.496153234 | 0 | 0.06 | 1 | 0.553874252 | 0.460680165 | 0.022 | 0.042 | 1 | 0.553874252 | 0.439867954 |
| Klhdc8a | 0.251619726 | -0.718545655 | 0 | 0.06 | 1 | 0.835943453 | 0.513147585 | 0.022 | 0.028 | 1 | 0.835943453 | 0.439926966 |
| Gm36975 | 0.251619726 | -0.718545655 | 0 | 0.06 | 1 | 0.737261441 | 1.280974143 | 0.111 | 0.1 | 1 | 0.737261441 | 0.439926966 |
| Gm56816 | 0.251619726 | -0.574155746 | 0 | 0.06 | 1 | 0.553402891 | 0.250113179 | 0.044 | 0.069 | 1 | 0.553402891 | 0.439926966 |
| 4833419F23Rik | 0.251619726 | -0.648156327 | 0 | 0.06 | 1 | 0.658583583 | 2.946106992 | 0.022 | 0.014 | 1 | 0.658583583 | 0.439926966 |
| Banf2 | 0.251653515 | -0.785659851 | 0 | 0.06 | 1 | 0.293659664 | -0.316927414 | 0 | 0.024 | 1 | 0.293659664 | 0.439977538 |
| Exoc3l2 | 0.251653515 | -0.970084422 | 0 | 0.06 | 1 | 0.494823351 | 1.946106992 | 0.022 | 0.01 | 1 | 0.494823351 | 0.439977538 |
| Fut2 | 0.251653515 | -0.718545655 | 0 | 0.06 | 1 | 0.430487676 | 0.361144491 | 0 | 0.014 | 1 | 0.430487676 | 0.439977538 |
| Gm12992 | 0.25165389 | -0.648156327 | 0 | 0.06 | 1 | 0.8040091 | 0.683072586 | 0.044 | 0.055 | 1 | 0.8040091 | 0.4399781 |
| D530017H19Rik | 0.25165389 | -0.648156327 | 0 | 0.06 | 1 | 0.994464236 | 0.223640968 | 0.044 | 0.045 | 1 | 0.994464236 | 0.4399781 |
| Tcap | 0.251682795 | -0.911190733 | 0 | 0.06 | 1 | 0.49622258 | 0.361144491 | 0 | 0.01 | 1 | 0.49622258 | 0.440021361 |
| Nyap1 | 0.251683171 | -0.718545655 | 0 | 0.06 | 1 | 0.698813251 | 0.473619221 | 0.067 | 0.087 | 1 | 0.698813251 | 0.440021923 |
| Gm5113 | 0.251683171 | -0.718545655 | 0 | 0.06 | 1 | 0.723627069 | 0.683072586 | 0.044 | 0.059 | 1 | 0.723627069 | 0.440021923 |
| Ubn2 | 0.251702354 | 0.773529548 | 1 | 0.94 | 1 | 0.476101445 | 0.140909798 | 0.978 | 0.948 | 1 | 0.476101445 | 0.440050633 |
| Rbpjl | 0.251704191 | -0.849790188 | 0 | 0.06 | 1 | 0.344700089 | 0.320502507 | 0.044 | 0.09 | 1 | 0.344700089 | 0.440053382 |
| Nr6a1os | 0.251706443 | -0.911190733 | 0 | 0.06 | 1 | 0.846219861 | 0.42003818 | 0.044 | 0.052 | 1 | 0.846219861 | 0.440056752 |
| Garin2 | 0.251707569 | -1.081115734 | 0 | 0.06 | 1 | 0.576705638 | 0.628624802 | 0.089 | 0.125 | 1 | 0.576705638 | 0.440058437 |
| Rhbdl2 | 0.25172671 | -1.081115734 | 0 | 0.06 | 1 | 0.529301833 | -0.404390255 | 0.022 | 0.042 | 1 | 0.529301833 | 0.440087084 |
| Gm47889 | 0.251727085 | -1.326228232 | 0 | 0.06 | 1 | 0.423610863 | -1.223818009 | 0.022 | 0.048 | 1 | 0.423610863 | 0.440087645 |
| Enkur | 0.251741347 | -1.184209227 | 0 | 0.06 | 1 | 0.329242657 | 0.250113179 | 0.022 | 0.059 | 1 | 0.329242657 | 0.440108988 |
| Azgp1 | 0.251741347 | -1.133583154 | 0 | 0.06 | 1 | 0.331922009 | -0.124282336 | 0 | 0.021 | 1 | 0.331922009 | 0.440108988 |
| Cgas | 0.251768367 | -1.683780237 | 0 | 0.06 | 1 | 0.260665169 | -0.901889915 | 0 | 0.028 | 1 | 0.260665169 | 0.440149423 |
| Gm33887 | 0.251790506 | -0.413691074 | 0 | 0.06 | 1 | 0.311583783 | -0.638855509 | 0.022 | 0.059 | 1 | 0.311583783 | 0.440182553 |
| Mansc1 | 0.251790506 | -0.413691074 | 0 | 0.06 | 1 | 0.509579374 | 1.683072586 | 0.067 | 0.045 | 1 | 0.509579374 | 0.440182553 |
| Mettl22 | 0.251790506 | -0.413691074 | 0 | 0.06 | 1 | 0.34534552 | -0.638855509 | 0.022 | 0.055 | 1 | 0.34534552 | 0.440182553 |
| Pcdhgb6 | 0.251790506 | -0.413691074 | 0 | 0.06 | 1 | 0.666728162 | 0.91753784 | 0.067 | 0.052 | 1 | 0.666728162 | 0.440182553 |
| Usp53 | 0.252179527 | -0.326228232 | 0.476 | 0.616 | 1 | 0.789410664 | -0.162976049 | 0.933 | 0.817 | 1 | 0.789410664 | 0.440764541 |
| Neil1 | 0.370840523 | 0.214340149 | 0 | 0.037 | 1 | 0.252329345 | -0.84048937 | 0.022 | 0.066 | 1 | 0.370840523 | 0.440988591 |
| Bid | 0.425599726 | 0.351843673 | 0 | 0.03 | 1 | 0.252329345 | -0.84048937 | 0.022 | 0.066 | 1 | 0.425599726 | 0.440988591 |
| 9030025P20Rik | 0.457308648 | 0.673771768 | 0 | 0.026 | 1 | 0.252329345 | -0.776359032 | 0.022 | 0.066 | 1 | 0.457308648 | 0.440988591 |

| | | | | | | | | | | | | |
|---|---|---|---|---|---|---|---|---|---|---|---|---|
| Gm12349 | 0.252420701 | -0.392317423 | 0.095 | 0.205 | 1 | 0.553874252 | 0.460680165 | 0.022 | 0.042 | 1 | 0.553874252 | 0.441125192 |
| Zfp653 | 0.252832023 | -0.326228232 | 0.048 | 0.142 | 1 | 0.344908855 | 0.231376616 | 0.156 | 0.239 | 1 | 0.344908855 | 0.441740015 |
| Wfdc18 | 0.253011496 | 0.97333205 | 0.381 | 0.306 | 1 | 0.522406469 | 1.081036572 | 0.2 | 0.183 | 1 | 0.522406469 | 0.442008175 |
| Ubap2 | 0.253061688 | 0.911931505 | 0.714 | 0.847 | 1 | 0.300530207 | 0.492531415 | 0.933 | 0.879 | 1 | 0.300530207 | 0.442083158 |
| Rabl6 | 0.571880257 | 0.463204782 | 0.381 | 0.642 | 1 | 0.253089363 | 0.303428993 | 0.8 | 0.734 | 1 | 0.571880257 | 0.442124501 |
| Fbp1 | 0.616977656 | 1.188344941 | 0.048 | 0.082 | 1 | 0.253128217 | -0.709244837 | 0.022 | 0.066 | 1 | 0.616977656 | 0.44218254 |
| Ccnb1ip1 | 0.646752221 | 0.214340149 | 0.048 | 0.075 | 1 | 0.253128217 | -0.901889915 | 0.022 | 0.066 | 1 | 0.646752221 | 0.44218254 |
| Cdh13 | 0.745868181 | -1.425763906 | 0.095 | 0.116 | 1 | 0.253380115 | -1.989352756 | 0.022 | 0.066 | 1 | 0.745868181 | 0.442558748 |
| Mib1 | 0.631327652 | -0.118786152 | 0.714 | 0.743 | 1 | 0.25353325 | 0.317423114 | 0.956 | 0.917 | 1 | 0.631327652 | 0.442787392 |
| Fam219a | 0.355214225 | 0.207797303 | 0.238 | 0.392 | 1 | 0.253804324 | 0.106015284 | 0.333 | 0.495 | 1 | 0.355214225 | 0.443192014 |
| Mknk1 | 0.253838848 | -0.648156327 | 0.095 | 0.201 | 1 | 0.400849931 | 0.765534746 | 0.178 | 0.273 | 1 | 0.400849931 | 0.443243535 |
| Il1rl2 | 0.254108703 | 1.55829455 | 0.095 | 0.041 | 1 | 0.376837034 | -0.124282336 | 0 | 0.017 | 1 | 0.376837034 | 0.443646173 |
| Lpin1 | 0.728490158 | 0.434305833 | 0.333 | 0.366 | 1 | 0.254197648 | 0.11747541 | 0.178 | 0.284 | 1 | 0.728490158 | 0.443778852 |
| Hexim2 | 0.304355016 | -0.326228232 | 0 | 0.049 | 1 | 0.254225598 | -0.638855509 | 0.022 | 0.066 | 1 | 0.304355016 | 0.443820541 |
| Fahd1 | 0.375036024 | -0.574155746 | 0.048 | 0.108 | 1 | 0.254225598 | -0.638855509 | 0.022 | 0.066 | 1 | 0.375036024 | 0.443820541 |
| Tyro3 | 0.632076134 | 1.673771768 | 0 | 0.011 | 1 | 0.254225598 | -0.638855509 | 0.022 | 0.066 | 1 | 0.632076134 | 0.443820541 |
| 2610306M01Rik | 0.926908894 | 0.673771768 | 0.048 | 0.052 | 1 | 0.254225598 | -0.638855509 | 0.022 | 0.066 | 1 | 0.926908894 | 0.443820541 |
| Il6st | 0.654454132 | 0.289464888 | 0.762 | 0.896 | 1 | 0.254256564 | -0.127893589 | 0.933 | 0.92 | 1 | 0.654454132 | 0.443866727 |
| Rap1a | 0.901090159 | 0.149004049 | 0.905 | 0.955 | 1 | 0.254410658 | 0.365185657 | 1 | 0.976 | 1 | 0.901090159 | 0.444096533 |
| Stk17b | 0.254679952 | -0.611630451 | 0.19 | 0.306 | 1 | 0.3517846 | -0.402415313 | 0.556 | 0.398 | 1 | 0.3517846 | 0.444498026 |
| Lemd3 | 0.25558775 | 0.143950821 | 0.286 | 0.519 | 1 | 0.28727708 | 0.552784557 | 0.733 | 0.651 | 1 | 0.28727708 | 0.445850402 |
| Zswim3 | 0.255760662 | 1.233199176 | 0.143 | 0.078 | 1 | 0.885635102 | 1.320502507 | 0.044 | 0.052 | 1 | 0.885635102 | 0.446107807 |
| Gm26533 | 0.255790428 | 1.150209812 | 0.095 | 0.041 | 1 | 0.420998748 | 1.683072586 | 0.044 | 0.024 | 1 | 0.420998748 | 0.446152113 |
| Lamc2 | 0.256170163 | 1.275222391 | 0.143 | 0.075 | 1 | 0.315817184 | -1.960783604 | 0.022 | 0.059 | 1 | 0.315817184 | 0.446717173 |
| Gm12762 | 0.257297491 | 1.451379346 | 0.095 | 0.041 | 1 | 0.734040487 | 0.875717664 | 0.044 | 0.035 | 1 | 0.734040487 | 0.448392983 |
| Msrb3 | 0.656421479 | 0.436331543 | 0.619 | 0.832 | 1 | 0.257524865 | 0.332201535 | 0.822 | 0.879 | 1 | 0.656421479 | 0.448730674 |
| Odf2l | 0.257536234 | -0.425763906 | 0.143 | 0.28 | 1 | 0.7529532 | 0.634709565 | 0.178 | 0.215 | 1 | 0.7529532 | 0.448747556 |
| Acad11 | 0.946436164 | 0.719575457 | 0.095 | 0.093 | 1 | 0.258060852 | 0.527794361 | 0.067 | 0.131 | 1 | 0.946436164 | 0.449526301 |
| Rgl3 | 0.675326052 | 0.351843673 | 0.048 | 0.075 | 1 | 0.25817855 | -0.181997834 | 0.089 | 0.156 | 1 | 0.675326052 | 0.449700936 |
| Tmem199 | 0.258209282 | -0.133583154 | 0.048 | 0.142 | 1 | 0.324978288 | -0.191396532 | 0.067 | 0.121 | 1 | 0.324978288 | 0.449746531 |
| Ripor1 | 0.258677028 | -0.512094778 | 0.143 | 0.257 | 1 | 0.375409987 | 0.111166238 | 0.356 | 0.512 | 1 | 0.375409987 | 0.450440252 |
| Sympk | 0.258677132 | -0.496153234 | 0.143 | 0.257 | 1 | 0.326815587 | 0.140545352 | 0.2 | 0.304 | 1 | 0.326815587 | 0.450440405 |
| Gapvd1 | 0.258720437 | 0.711386059 | 0.762 | 0.772 | 1 | 0.435264547 | 0.340038858 | 0.978 | 0.882 | 1 | 0.435264547 | 0.45050461 |
| Ric8b | 0.259030862 | -0.560283571 | 0.286 | 0.429 | 1 | 0.293789009 | 0.364946061 | 0.644 | 0.609 | 1 | 0.293789009 | 0.450964736 |
| Plxna4 | 0.259276152 | 0.125335143 | 0.048 | 0.142 | 1 | 0.922589644 | 1.324618615 | 0.333 | 0.436 | 1 | 0.922589644 | 0.45132818 |
| Ptx3 | 0.35697083 | -1.469186186 | 0.095 | 0.183 | 1 | 0.259373927 | -0.746915255 | 0.044 | 0.097 | 1 | 0.35697083 | 0.45147302 |
| Zc2hc1a | 0.259377905 | -0.107587946 | 0.143 | 0.276 | 1 | 0.34843912 | 0.289158378 | 0.289 | 0.388 | 1 | 0.34843912 | 0.451478913 |
| Chek1 | 0.497542969 | 0.8372705 | 0.19 | 0.138 | 1 | 0.259506075 | 0.15951063 | 0.133 | 0.221 | 1 | 0.497542969 | 0.451668746 |
| Gm38190 | 0.259700754 | 2.088809267 | 0.143 | 0.082 | 1 | 0.797823945 | 1.0259603 | 0.089 | 0.111 | 1 | 0.797823945 | 0.451957026 |
| Gnb1l | 0.296731385 | -0.752492987 | 0.048 | 0.123 | 1 | 0.259903385 | 0.76880246 | 0.067 | 0.135 | 1 | 0.296731385 | 0.452257001 |
| Eif2b3 | 0.740466229 | 0.100882099 | 0.381 | 0.474 | 1 | 0.26013393 | 0.238972621 | 0.289 | 0.439 | 1 | 0.740466229 | 0.452598198 |
| Ythdc2 | 0.260528278 | 0.807038299 | 0.381 | 0.306 | 1 | 0.531419861 | 0.534680746 | 0.2 | 0.291 | 1 | 0.531419861 | 0.453181572 |
| Apba2 | 0.266771227 | 2.79930265 | 0.048 | 0.015 | 1 | 0.260653677 | -0.901889915 | 0 | 0.028 | 1 | 0.266771227 | 0.453367015 |
| C130036L24Rik | 0.268100354 | -0.326228232 | 0 | 0.056 | 1 | 0.260653677 | -0.638855509 | 0 | 0.028 | 1 | 0.268100354 | 0.453367015 |
| Ccl19 | 0.3247899 | -0.326228232 | 0 | 0.045 | 1 | 0.260653677 | -0.638855509 | 0 | 0.028 | 1 | 0.3247899 | 0.453367015 |
| AV099323 | 0.346846605 | -0.326228232 | 0 | 0.041 | 1 | 0.260653677 | -0.638855509 | 0 | 0.028 | 1 | 0.346846605 | 0.453367015 |
| Vwa3b | 0.359270744 | 3.088809267 | 0.048 | 0.019 | 1 | 0.260653677 | -0.638855509 | 0 | 0.028 | 1 | 0.359270744 | 0.453367015 |
| Arhgap11a | 0.396948718 | -0.413691074 | 0 | 0.034 | 1 | 0.260653677 | -0.776359032 | 0 | 0.028 | 1 | 0.396948718 | 0.453367015 |
| Sfrp5 | 0.396957792 | -1.574155746 | 0 | 0.034 | 1 | 0.260653677 | -1.564854927 | 0 | 0.028 | 1 | 0.396957792 | 0.453367015 |
| Plac8 | 0.39695844 | -0.849790188 | 0 | 0.034 | 1 | 0.260653677 | -0.901889915 | 0 | 0.028 | 1 | 0.39695844 | 0.453367015 |
| Stk35os1 | 0.396971401 | 0.351843673 | 0 | 0.034 | 1 | 0.260653677 | -0.638855509 | 0 | 0.028 | 1 | 0.396971401 | 0.453367015 |
| Gm43625 | 0.396971401 | 0.351843673 | 0 | 0.034 | 1 | 0.260653677 | -0.776359032 | 0 | 0.028 | 1 | 0.396971401 | 0.453367015 |
| Tigd5 | 0.396971401 | 0.351843673 | 0 | 0.034 | 1 | 0.260653677 | -0.638855509 | 0 | 0.028 | 1 | 0.396971401 | 0.453367015 |
| Gm47123 | 0.425624656 | -0.133583154 | 0 | 0.03 | 1 | 0.260653677 | -0.638855509 | 0 | 0.028 | 1 | 0.425624656 | 0.453367015 |
| Sema6c | 0.425639612 | 0.503846766 | 0 | 0.03 | 1 | 0.260653677 | -0.638855509 | 0 | 0.028 | 1 | 0.425639612 | 0.453367015 |
| Gm56817 | 0.425639612 | 0.503846766 | 0 | 0.03 | 1 | 0.260653677 | -0.638855509 | 0 | 0.028 | 1 | 0.425639612 | 0.453367015 |
| Gm47664 | 0.442336143 | 1.503846766 | 0.095 | 0.056 | 1 | 0.260653677 | -0.638855509 | 0 | 0.028 | 1 | 0.442336143 | 0.453367015 |
| Pgbd1 | 0.457281049 | 0.503846766 | 0 | 0.026 | 1 | 0.260653677 | -0.638855509 | 0 | 0.028 | 1 | 0.457281049 | 0.453367015 |
| 1700102H20Rik | 0.457281049 | 0.503846766 | 0 | 0.026 | 1 | 0.260653677 | -0.638855509 | 0 | 0.028 | 1 | 0.457281049 | 0.453367015 |
| Irag2 | 0.457292877 | -0.233118828 | 0 | 0.026 | 1 | 0.260653677 | -0.638855509 | 0 | 0.028 | 1 | 0.457292877 | 0.453367015 |
| Cct6b | 0.457292877 | 0.214340149 | 0 | 0.026 | 1 | 0.260653677 | -0.776359032 | 0 | 0.028 | 1 | 0.457292877 | 0.453367015 |
| Serpinb5 | 0.492608234 | 0.866416846 | 0 | 0.022 | 1 | 0.260653677 | -0.638855509 | 0 | 0.028 | 1 | 0.492608234 | 0.453367015 |
| Coa4 | 0.492608234 | 0.866416846 | 0 | 0.022 | 1 | 0.260653677 | -0.638855509 | 0 | 0.028 | 1 | 0.492608234 | 0.453367015 |
| Llgl2 | 0.492608234 | 0.866416846 | 0 | 0.022 | 1 | 0.260653677 | -0.638855509 | 0 | 0.028 | 1 | 0.492608234 | 0.453367015 |
| Gm42788 | 0.50322484 | 1.825774861 | 0.095 | 0.063 | 1 | 0.260653677 | -0.638855509 | 0 | 0.028 | 1 | 0.50322484 | 0.453367015 |
| Rpusd3 | 0.532426384 | 0.866416846 | 0 | 0.019 | 1 | 0.260653677 | -0.638855509 | 0 | 0.028 | 1 | 0.532426384 | 0.453367015 |
| Cxcr4 | 0.532432377 | 0.351843673 | 0 | 0.019 | 1 | 0.260653677 | -0.638855509 | 0 | 0.028 | 1 | 0.532432377 | 0.453367015 |
| Gm12963 | 0.532436372 | 1.088809267 | 0 | 0.019 | 1 | 0.260653677 | -0.638855509 | 0 | 0.028 | 1 | 0.532436372 | 0.453367015 |
| Gm48529 | 0.532436372 | 1.088809267 | 0 | 0.019 | 1 | 0.260653677 | -0.638855509 | 0 | 0.028 | 1 | 0.532436372 | 0.453367015 |
| Gm45464 | 0.544576436 | 2.673771768 | 0.048 | 0.026 | 1 | 0.260653677 | -0.638855509 | 0 | 0.028 | 1 | 0.544576436 | 0.453367015 |
| Gm20548 | 0.550953306 | 1.936806174 | 0.048 | 0.026 | 1 | 0.260653677 | -0.638855509 | 0 | 0.028 | 1 | 0.550953306 | 0.453367015 |
| Gm42702 | 0.57816106 | 1.351843673 | 0 | 0.015 | 1 | 0.260653677 | -0.638855509 | 0 | 0.028 | 1 | 0.57816106 | 0.453367015 |
| Tigar | 0.63207476 | 1.088809267 | 0 | 0.011 | 1 | 0.260653677 | -0.776359032 | 0 | 0.028 | 1 | 0.63207476 | 0.453367015 |
| Syngr3 | 0.63207476 | 1.351843673 | 0 | 0.011 | 1 | 0.260653677 | -0.776359032 | 0 | 0.028 | 1 | 0.63207476 | 0.453367015 |
| ENSMUSG00000121439 | 0.632076134 | 1.673771768 | 0 | 0.011 | 1 | 0.260653677 | -0.776359032 | 0 | 0.028 | 1 | 0.632076134 | 0.453367015 |
| Zfp941 | 0.632076134 | 1.673771768 | 0 | 0.011 | 1 | 0.260653677 | -0.901889915 | 0 | 0.028 | 1 | 0.632076134 | 0.453367015 |
| Pla2g12b | 0.719072865 | 1.351843673 | 0.048 | 0.034 | 1 | 0.260653677 | -0.776359032 | 0 | 0.028 | 1 | 0.719072865 | 0.453367015 |
| Cpz | 0.865329917 | 2.150209812 | 0.048 | 0.06 | 1 | 0.260653677 | -0.776359032 | 0 | 0.028 | 1 | 0.865329917 | 0.453367015 |
| 9130004C02Rik | 0.95783752 | 0.766881172 | 0.048 | 0.045 | 1 | 0.260653677 | -0.776359032 | 0 | 0.028 | 1 | 0.95783752 | 0.453367015 |
| Fbp2 | 0.264194629 | 2.673771768 | 0.048 | 0.015 | 1 | 0.260665169 | -1.124282336 | 0 | 0.028 | 1 | 0.264194629 | 0.453384008 |
| Tbc1d2 | 0.362165014 | 2.451379346 | 0.048 | 0.019 | 1 | 0.260665169 | -0.901889915 | 0 | 0.028 | 1 | 0.362165014 | 0.453384008 |
| Arhgap40 | 0.365063958 | 2.258734268 | 0.048 | 0.019 | 1 | 0.260665169 | -0.776359032 | 0 | 0.028 | 1 | 0.365063958 | 0.453384008 |
| Gm2350 | 0.425614684 | -0.233118828 | 0 | 0.03 | 1 | 0.260665169 | -0.901889915 | 0 | 0.028 | 1 | 0.425614684 | 0.453384008 |
| Fam174b | 0.457292877 | 0.351843673 | 0 | 0.026 | 1 | 0.260665169 | -0.776359032 | 0 | 0.028 | 1 | 0.457292877 | 0.453384008 |
| Myoc | 0.457308648 | 0.673771768 | 0 | 0.026 | 1 | 0.260665169 | -0.776359032 | 0 | 0.028 | 1 | 0.457308648 | 0.453384008 |
| Gm27042 | 0.472861597 | 1.866416846 | 0.048 | 0.022 | 1 | 0.260665169 | -0.776359032 | 0 | 0.028 | 1 | 0.472861597 | 0.453384008 |
| Myo7a | 0.492608234 | 0.866416846 | 0 | 0.022 | 1 | 0.260665169 | -0.776359032 | 0 | 0.028 | 1 | 0.492608234 | 0.453384008 |
| Gm30978 | 0.492608234 | 0.866416846 | 0 | 0.022 | 1 | 0.260665169 | -0.776359032 | 0 | 0.028 | 1 | 0.492608234 | 0.453384008 |
| Reep2 | 0.492608234 | 0.866416846 | 0 | 0.022 | 1 | 0.260665169 | -0.776359032 | 0 | 0.028 | 1 | 0.492608234 | 0.453384008 |
| Krtcap3 | 0.532426384 | 0.866416846 | 0 | 0.019 | 1 | 0.260665169 | -0.776359032 | 0 | 0.028 | 1 | 0.532426384 | 0.453384008 |
| Itpripl1 | 0.532436372 | 1.088809267 | 0 | 0.019 | 1 | 0.260665169 | -0.901889915 | 0 | 0.028 | 1 | 0.532436372 | 0.453384008 |
| Gstm6 | 0.532436372 | 1.088809267 | 0 | 0.019 | 1 | 0.260665169 | -1.017367132 | 0 | 0.028 | 1 | 0.532436372 | 0.453384008 |
| Nppc | 0.578156445 | 0.866416846 | 0 | 0.015 | 1 | 0.260665169 | -1.017367132 | 0 | 0.028 | 1 | 0.578156445 | 0.453384008 |
| Kbtbd8 | 0.578156445 | 1.088809267 | 0 | 0.015 | 1 | 0.260665169 | -0.901889915 | 0 | 0.028 | 1 | 0.578156445 | 0.453384008 |
| C030005K06Rik | 0.578156445 | 1.088809267 | 0 | 0.015 | 1 | 0.260665169 | -0.776359032 | 0 | 0.028 | 1 | 0.578156445 | 0.453384008 |
| Ildr1 | 0.57816106 | 1.351843673 | 0 | 0.015 | 1 | 0.260665169 | -0.776359032 | 0 | 0.028 | 1 | 0.57816106 | 0.453384008 |
| Prrt1 | 0.629700127 | 2.088809267 | 0.048 | 0.03 | 1 | 0.260665169 | -0.901889915 | 0 | 0.028 | 1 | 0.629700127 | 0.453384008 |
| Gm48008 | 0.632076134 | 1.673771768 | 0 | 0.011 | 1 | 0.260665169 | -0.776359032 | 0 | 0.028 | 1 | 0.632076134 | 0.453384008 |

| | | | | | | | | | | | | |
|---|---|---|---|---|---|---|---|---|---|---|---|---|
| Cdc20 | 0.749869817 | -0.326228232 | 0.048 | 0.067 | 1 | 0.260665169 | -1.017367132 | 0 | 0.028 | 1 | 0.749869817 | 0.453384008 |
| Tmem40 | 0.985457491 | 0.766881172 | 0.048 | 0.049 | 1 | 0.260665169 | -0.901889915 | 0 | 0.028 | 1 | 0.985457491 | 0.453384008 |
| 6430550D23Rik | 0.550953306 | 1.936806174 | 0.048 | 0.026 | 1 | 0.260672831 | -1.017367132 | 0 | 0.028 | 1 | 0.550953306 | 0.453395337 |
| Rgs20 | 0.753530714 | -0.611630451 | 0.048 | 0.063 | 1 | 0.260672831 | -1.124282336 | 0 | 0.028 | 1 | 0.753530714 | 0.453395337 |
| Ablim3 | 0.274597491 | 2.088809267 | 0.048 | 0.015 | 1 | 0.260673378 | -0.901889915 | 0 | 0.028 | 1 | 0.274597491 | 0.453396146 |
| Rnf43 | 0.454224194 | 2.503846766 | 0.048 | 0.022 | 1 | 0.260673378 | -1.124282336 | 0 | 0.028 | 1 | 0.454224194 | 0.453396146 |
| Odf3l2 | 0.578156445 | 1.088809267 | 0 | 0.015 | 1 | 0.260673378 | -0.901889915 | 0 | 0.028 | 1 | 0.578156445 | 0.453396146 |
| Arhgef4 | 0.579997314 | 0.503846766 | 0.048 | 0.026 | 1 | 0.260673378 | -0.901889915 | 0 | 0.028 | 1 | 0.579997314 | 0.453396146 |
| Csmd3 | 0.63207476 | 1.088809267 | 0 | 0.011 | 1 | 0.260673378 | -1.223818009 | 0 | 0.028 | 1 | 0.63207476 | 0.453396146 |
| Trim69 | 0.64258579 | 1.55829455 | 0.048 | 0.03 | 1 | 0.260673378 | -1.124282336 | 0 | 0.028 | 1 | 0.64258579 | 0.453396146 |
| Kif20b | 0.7608916 | -0.535681598 | 0.048 | 0.063 | 1 | 0.260673378 | -1.223818009 | 0 | 0.028 | 1 | 0.7608916 | 0.453396146 |
| Gm16083 | 0.266771227 | 2.673771768 | 0.048 | 0.015 | 1 | 0.260676661 | -1.316927414 | 0 | 0.028 | 1 | 0.266771227 | 0.453401001 |
| 2010009K17Rik | 0.567008887 | 1.041503552 | 0.048 | 0.026 | 1 | 0.260676661 | -1.486852415 | 0 | 0.028 | 1 | 0.567008887 | 0.453401001 |
| Adamts18 | 0.761116934 | -0.455511249 | 0.048 | 0.034 | 1 | 0.260676661 | -1.124282336 | 0 | 0.028 | 1 | 0.761116934 | 0.453401001 |
| Smco2 | 0.261628269 | 2.936806174 | 0.048 | 0.015 | 1 | 0.260678303 | -1.223818009 | 0 | 0.028 | 1 | 0.261628269 | 0.453403428 |
| 9330154J02Rik | 0.396962328 | -0.496153234 | 0 | 0.034 | 1 | 0.260678303 | -1.316927414 | 0 | 0.028 | 1 | 0.396962328 | 0.453403428 |
| Gm42997 | 0.532435374 | 0.503846766 | 0 | 0.019 | 1 | 0.260678303 | -1.223818009 | 0 | 0.028 | 1 | 0.532435374 | 0.453403428 |
| Folr2 | 0.699759084 | 0.316219763 | 0.095 | 0.071 | 1 | 0.260678303 | -1.564854927 | 0 | 0.028 | 1 | 0.699759084 | 0.453403428 |
| 1700003E16Rik | 0.454217076 | 3.258734268 | 0.048 | 0.022 | 1 | 0.26067885 | -1.017367132 | 0 | 0.028 | 1 | 0.454217076 | 0.453404237 |
| Hoxb9 | 0.578156445 | 1.088809267 | 0 | 0.015 | 1 | 0.26067885 | -1.017367132 | 0 | 0.028 | 1 | 0.578156445 | 0.453404237 |
| 8030442B05Rik | 0.57816106 | 0.673771768 | 0 | 0.015 | 1 | 0.26067885 | -1.017367132 | 0 | 0.028 | 1 | 0.57816106 | 0.453404237 |
| Pnpla3 | 0.850006903 | 1.188344941 | 0.095 | 0.119 | 1 | 0.260679945 | -1.486852415 | 0 | 0.028 | 1 | 0.850006903 | 0.453405856 |
| Nipal1 | 0.50319368 | -0.133583154 | 0.048 | 0.093 | 1 | 0.260681586 | -1.709244837 | 0 | 0.028 | 1 | 0.50319368 | 0.453408283 |
| Gm15726 | 0.677501769 | -0.133583154 | 0.048 | 0.075 | 1 | 0.260681586 | -2.316927414 | 0 | 0.028 | 1 | 0.677501769 | 0.453408283 |
| Ibsp | 0.784076328 | 2.8832843 | 0.048 | 0.037 | 1 | 0.260682133 | -3.72631835 | 0 | 0.028 | 1 | 0.784076328 | 0.453409092 |
| Aloxe3 | 0.269367392 | 1.55829455 | 0.048 | 0.015 | 1 | 0.260684322 | -0.486852415 | 0 | 0.028 | 1 | 0.269367392 | 0.453412329 |
| Poll | 0.324739547 | -0.133583154 | 0 | 0.045 | 1 | 0.260684322 | -0.486852415 | 0 | 0.028 | 1 | 0.324739547 | 0.453412329 |
| Gins1 | 0.346879876 | -0.718545655 | 0 | 0.041 | 1 | 0.260684322 | -0.486852415 | 0 | 0.028 | 1 | 0.346879876 | 0.453412329 |
| Neil2 | 0.396916958 | 0.214340149 | 0 | 0.034 | 1 | 0.260684322 | -0.486852415 | 0 | 0.028 | 1 | 0.396916958 | 0.453412329 |
| Mrap | 0.39694807 | -0.133583154 | 0 | 0.034 | 1 | 0.260684322 | -0.486852415 | 0 | 0.028 | 1 | 0.39694807 | 0.453412329 |
| Mcrip2 | 0.425639612 | 0.503846766 | 0 | 0.03 | 1 | 0.260684322 | -0.486852415 | 0 | 0.028 | 1 | 0.425639612 | 0.453412329 |
| Gm11627 | 0.492590603 | 0.673771768 | 0 | 0.022 | 1 | 0.260684322 | -0.486852415 | 0 | 0.028 | 1 | 0.492590603 | 0.453412329 |
| Gm50146 | 0.492590603 | 0.673771768 | 0 | 0.022 | 1 | 0.260684322 | -0.486852415 | 0 | 0.028 | 1 | 0.492590603 | 0.453412329 |
| Gm16861 | 0.492599419 | 0.351843673 | 0 | 0.022 | 1 | 0.260684322 | -0.486852415 | 0 | 0.028 | 1 | 0.492599419 | 0.453412329 |
| Shd | 0.492608234 | 0.866416846 | 0 | 0.022 | 1 | 0.260684322 | -0.486852415 | 0 | 0.028 | 1 | 0.492608234 | 0.453412329 |
| D330050G23Rik | 0.498606083 | 1.351843673 | 0.095 | 0.06 | 1 | 0.260684322 | -0.486852415 | 0 | 0.028 | 1 | 0.498606083 | 0.453412329 |
| Gm42595 | 0.532436372 | 1.088809267 | 0 | 0.019 | 1 | 0.260684322 | -0.486852415 | 0 | 0.028 | 1 | 0.532436372 | 0.453412329 |
| Amt | 0.532436372 | 1.088809267 | 0 | 0.019 | 1 | 0.260684322 | -0.486852415 | 0 | 0.028 | 1 | 0.532436372 | 0.453412329 |
| 1700018L02Rik | 0.532436372 | 1.088809267 | 0 | 0.019 | 1 | 0.260684322 | -0.486852415 | 0 | 0.028 | 1 | 0.532436372 | 0.453412329 |
| Zfp977 | 0.578156445 | 0.866416846 | 0 | 0.015 | 1 | 0.260684322 | -0.486852415 | 0 | 0.028 | 1 | 0.578156445 | 0.453412329 |
| 4933427D14Rik | 0.578156445 | 1.088809267 | 0 | 0.015 | 1 | 0.260684322 | -0.486852415 | 0 | 0.028 | 1 | 0.578156445 | 0.453412329 |
| Gm20939 | 0.578156445 | 1.088809267 | 0 | 0.015 | 1 | 0.260684322 | -0.486852415 | 0 | 0.028 | 1 | 0.578156445 | 0.453412329 |
| 1700064N11Rik | 0.632917461 | 1.936806174 | 0.048 | 0.03 | 1 | 0.260684322 | -0.486852415 | 0 | 0.028 | 1 | 0.632917461 | 0.453412329 |
| Zfp953 | 0.636128533 | 1.673771768 | 0.048 | 0.03 | 1 | 0.260684322 | -0.486852415 | 0 | 0.028 | 1 | 0.636128533 | 0.453412329 |
| Cacnb3 | 0.828388925 | 0.766881172 | 0.048 | 0.037 | 1 | 0.260684322 | -0.486852415 | 0 | 0.028 | 1 | 0.828388925 | 0.453412329 |
| Ect2 | 0.831581629 | -0.413691074 | 0.048 | 0.037 | 1 | 0.260684322 | -0.486852415 | 0 | 0.028 | 1 | 0.831581629 | 0.453412329 |
| Dync2h1 | 0.497855674 | 0.863297504 | 0.571 | 0.627 | 1 | 0.261081297 | 0.193488134 | 0.6 | 0.713 | 1 | 0.497855674 | 0.45399915 |
| 2610307P16Rik | 0.426970251 | 0.438555306 | 0.19 | 0.127 | 1 | 0.261552077 | -1.105423309 | 0.044 | 0.097 | 1 | 0.426970251 | 0.454694664 |
| Gm28535 | 0.261628269 | 2.936806174 | 0.048 | 0.015 | 1 | 0.555574908 | 0.513147585 | 0.022 | 0.042 | 1 | 0.555574908 | 0.454807186 |
| Ppp1r16b | 0.261628269 | 2.936806174 | 0.048 | 0.015 | 1 | 0.936026228 | 1.683072586 | 0.022 | 0.021 | 1 | 0.936026228 | 0.454807186 |
| Gm43646 | 0.261628269 | 2.936806174 | 0.048 | 0.015 | 1 | 0.331922009 | -0.124282336 | 0 | 0.021 | 1 | 0.331922009 | 0.454807186 |
| Dmrta2 | 0.261628269 | 3.351843673 | 0.048 | 0.015 | 1 | 0.430487676 | 0.361144491 | 0 | 0.014 | 1 | 0.430487676 | 0.454807186 |
| Slc26a1 | 0.261628269 | 2.936806174 | 0.048 | 0.015 | 1 | 0.951997643 | 0.875717664 | 0.022 | 0.021 | 1 | 0.951997643 | 0.454807186 |
| Svip | 0.261628269 | 2.936806174 | 0.048 | 0.015 | 1 | 0.331919067 | -0.638855509 | 0 | 0.021 | 1 | 0.331919067 | 0.454807186 |
| Rsf1os1 | 0.261628269 | 2.936806174 | 0.048 | 0.015 | 1 | 0.299276035 | 0.15951063 | 0.022 | 0.062 | 1 | 0.299276035 | 0.454807186 |
| Tex12 | 0.261628269 | 2.936806174 | 0.048 | 0.015 | 1 | 0.376842245 | -0.316927414 | 0 | 0.017 | 1 | 0.376842245 | 0.454807186 |
| Gm47095 | 0.261628269 | 2.936806174 | 0.048 | 0.015 | 1 | 0.512490525 | 2.098110085 | 0.044 | 0.028 | 1 | 0.512490525 | 0.454807186 |
| Enpp3 | 0.261628269 | 2.936806174 | 0.048 | 0.015 | 1 | 0.49622258 | 0.361144491 | 0 | 0.01 | 1 | 0.49622258 | 0.454807186 |
| Gm15608 | 0.261628269 | 2.936806174 | 0.048 | 0.015 | 1 | 0.49622258 | 0.361144491 | 0 | 0.01 | 1 | 0.49622258 | 0.454807186 |
| Arhgap27os2 | 0.261628269 | 2.936806174 | 0.048 | 0.015 | 1 | 0.293659664 | -0.316927414 | 0 | 0.024 | 1 | 0.293659664 | 0.454807186 |
| Zfp85 | 0.261628269 | 2.936806174 | 0.048 | 0.015 | 1 | 0.935071939 | 0.513147585 | 0.022 | 0.024 | 1 | 0.935071939 | 0.454807186 |
| 9930004E17Rik | 0.261628269 | 2.936806174 | 0.048 | 0.015 | 1 | 0.331922009 | -0.124282336 | 0 | 0.021 | 1 | 0.331922009 | 0.454807186 |
| Espl1 | 0.261628269 | 3.936806174 | 0.048 | 0.015 | 1 | 0.496223879 | 0.683072586 | 0 | 0.01 | 1 | 0.496223879 | 0.454807186 |
| 2600002D14Rik | 0.261628269 | 2.936806174 | 0.048 | 0.015 | 1 | 0.661444614 | 1.683072586 | 0.022 | 0.014 | 1 | 0.661444614 | 0.454807186 |
| Prdm9 | 0.261628269 | 3.351843673 | 0.048 | 0.015 | 1 | 0.854461673 | 1.361144491 | 0.022 | 0.028 | 1 | 0.854461673 | 0.454807186 |
| C2 | 0.261628269 | 2.936806174 | 0.048 | 0.015 | 1 | 0.932582169 | 0.223640968 | 0.022 | 0.024 | 1 | 0.932582169 | 0.454807186 |
| Gm49893 | 0.261628269 | 3.351843673 | 0.048 | 0.015 | 1 | 0.8202424 | 1.098110085 | 0.022 | 0.017 | 1 | 0.8202424 | 0.454807186 |
| Gm50462 | 0.261628269 | 2.936806174 | 0.048 | 0.015 | 1 | 0.823042771 | 0.875717664 | 0.022 | 0.017 | 1 | 0.823042771 | 0.454807186 |
| Doc2g | 0.261628269 | 3.351843673 | 0.048 | 0.015 | 1 | 0.80624346 | 2.42003818 | 0.022 | 0.017 | 1 | 0.80624346 | 0.454807186 |
| Gm50186 | 0.261628269 | 2.936806174 | 0.048 | 0.015 | 1 | 0.497611189 | 1.683072586 | 0.022 | 0.01 | 1 | 0.497611189 | 0.454807186 |
| Rnf138rt1 | 0.261628269 | 2.936806174 | 0.048 | 0.015 | 1 | 0.376837034 | -0.316927414 | 0 | 0.017 | 1 | 0.376837034 | 0.454807186 |
| Ryr1 | 0.261635242 | 3.410737362 | 0.048 | 0.015 | 1 | 0.658587075 | 2.098110085 | 0.022 | 0.014 | 1 | 0.658587075 | 0.454817484 |
| Ankle1 | 0.261635242 | 3.088809267 | 0.048 | 0.015 | 1 | 0.309824707 | 3.098110085 | 0.022 | 0.007 | 1 | 0.309824707 | 0.454817484 |
| Dnah5 | 0.261635242 | 3.088809267 | 0.048 | 0.015 | 1 | 0.309824707 | 3.42003818 | 0.022 | 0.007 | 1 | 0.309824707 | 0.454817484 |
| A430093F15Rik | 0.261635242 | 2.673771768 | 0.048 | 0.015 | 1 | 0.954659411 | 0.683072586 | 0.022 | 0.021 | 1 | 0.954659411 | 0.454817484 |
| Ksr2 | 0.261637567 | 4.451379346 | 0.048 | 0.015 | 1 | 0.448135556 | 0.13875207 | 0.022 | 0.048 | 1 | 0.448135556 | 0.454820917 |
| Tsnaxip1 | 0.261637567 | 3.188344941 | 0.048 | 0.015 | 1 | 0.376837034 | -0.486852415 | 0 | 0.017 | 1 | 0.376837034 | 0.454820917 |
| Agtpbp1 | 0.860847157 | 0.345300827 | 0.333 | 0.444 | 1 | 0.26193218 | 0.849082538 | 0.711 | 0.64 | 1 | 0.860847157 | 0.455255894 |
| Dcbld1 | 0.649718423 | 0.97333205 | 0.048 | 0.078 | 1 | 0.261953688 | -1.158229668 | 0.044 | 0.093 | 1 | 0.649718423 | 0.455287642 |
| Zfp263 | 0.869880516 | 0.673771768 | 0.143 | 0.179 | 1 | 0.262313874 | 0.111530601 | 0.156 | 0.246 | 1 | 0.869880516 | 0.455819179 |
| Mocs1 | 0.262943924 | 1.889500459 | 0.143 | 0.078 | 1 | 0.331216021 | 0.513147585 | 0.022 | 0.059 | 1 | 0.331216021 | 0.456748341 |
| Slx4 | 0.307564996 | 0.60338244 | 0.048 | 0.131 | 1 | 0.263195653 | -0.124282336 | 0.111 | 0.19 | 1 | 0.307564996 | 0.457119354 |
| Usp3 | 0.416838424 | 0.70304956 | 0.619 | 0.619 | 1 | 0.263228415 | 0.674852079 | 0.867 | 0.81 | 1 | 0.416838424 | 0.457167631 |
| Mtpap | 0.499815295 | 0.146034136 | 0.286 | 0.425 | 1 | 0.263312796 | 0.309614191 | 0.267 | 0.419 | 1 | 0.499815295 | 0.457291964 |
| Zbtb26 | 0.966230998 | 1.258734268 | 0.048 | 0.052 | 1 | 0.263404078 | -1.017367132 | 0.044 | 0.093 | 1 | 0.966230998 | 0.457426447 |
| Jak2 | 0.382340071 | 0.534822416 | 0.714 | 0.735 | 1 | 0.263923372 | -0.435211432 | 0.867 | 0.737 | 1 | 0.382340071 | 0.458191198 |
| Brsk1 | 0.47389253 | -0.413691074 | 0.048 | 0.093 | 1 | 0.264021389 | 0.180572246 | 0.111 | 0.19 | 1 | 0.47389253 | 0.458335484 |
| Eva1c | 0.264180212 | -0.574155746 | 0.048 | 0.138 | 1 | 0.952284602 | 0.728876276 | 0.067 | 0.066 | 1 | 0.952284602 | 0.458569239 |
| Gm26930 | 0.264194629 | 2.673771768 | 0.048 | 0.015 | 1 | 0.506024628 | 0.513147585 | 0.022 | 0.01 | 1 | 0.506024628 | 0.458590456 |
| Kcnc4 | 0.264194629 | 2.673771768 | 0.048 | 0.015 | 1 | 0.936023974 | 1.460680165 | 0.022 | 0.021 | 1 | 0.936023974 | 0.458590456 |
| Mroh7 | 0.264194629 | 2.673771768 | 0.048 | 0.015 | 1 | 0.835943453 | 0.513147585 | 0.022 | 0.028 | 1 | 0.835943453 | 0.458590456 |
| Gm10399 | 0.264194629 | 2.673771768 | 0.048 | 0.015 | 1 | 0.293647037 | -0.901889915 | 0 | 0.024 | 1 | 0.293647037 | 0.458590456 |
| A930016O22Rik | 0.264194629 | 2.673771768 | 0.048 | 0.015 | 1 | 0.530754573 | 1.361144491 | 0.044 | 0.028 | 1 | 0.530754573 | 0.458590456 |
| Oacyl | 0.264194629 | 2.673771768 | 0.048 | 0.015 | 1 | 0.496223879 | 0.683072586 | 0 | 0.01 | 1 | 0.496223879 | 0.458590456 |
| Cpne5 | 0.26419812 | 2.673771768 | 0.048 | 0.015 | 1 | 0.293647037 | -0.776359032 | 0 | 0.024 | 1 | 0.293647037 | 0.458595594 |
| Kmt2c | 0.73251451 | 0.374211486 | 0.667 | 0.731 | 1 | 0.264668785 | 0.124985791 | 1 | 0.983 | 1 | 0.73251451 | 0.459288005 |
| Scube1 | 0.265032455 | 0.751113856 | 0.333 | 0.582 | 1 | 0.671911477 | 0.287587884 | 0.311 | 0.308 | 1 | 0.671911477 | 0.459822708 |
| Pitpnm1 | 0.681919718 | 0.55829455 | 0.048 | 0.075 | 1 | 0.26550515 | -0.486852415 | 0.044 | 0.097 | 1 | 0.681919718 | 0.460517315 |

| | | | | | | | | | | | | |
|---|---|---|---|---|---|---|---|---|---|---|---|---|
| Car13 | 0.655561847 | 1.503846766 | 0.048 | 0.03 | 1 | 0.265621111 | 0.223640968 | 0.044 | 0.1 | 1 | 0.655561847 | 0.460687647 |
| Rilp | 0.425599726 | 0.351843673 | 0 | 0.03 | 1 | 0.266028511 | -0.432404631 | 0.022 | 0.066 | 1 | 0.425599726 | 0.461285853 |
| Lmbr1l | 0.266071874 | 0.214340149 | 0.238 | 0.425 | 1 | 0.738114103 | 0.477168288 | 0.356 | 0.45 | 1 | 0.738114103 | 0.461349506 |
| Galnt15 | 0.274434227 | 0.786246497 | 0.048 | 0.138 | 1 | 0.266168508 | -0.802354241 | 0.089 | 0.152 | 1 | 0.274434227 | 0.461491342 |
| Gm26672 | 0.266256853 | 2.188344941 | 0.095 | 0.045 | 1 | 0.833619716 | 0.361144491 | 0.022 | 0.028 | 1 | 0.833619716 | 0.461620994 |
| Dock7 | 0.349035387 | 0.186505941 | 0.905 | 0.836 | 1 | 0.26627278 | 0.12759685 | 0.867 | 0.872 | 1 | 0.349035387 | 0.461644367 |
| Rbm20 | 1 | 0.388369549 | 0.095 | 0.097 | 1 | 0.266302981 | 0.256807831 | 0.044 | 0.1 | 1 | 1 | 0.461688684 |
| Cdh4 | 0.266771227 | 1.258734268 | 0.048 | 0.015 | 1 | 0.76573422 | 2.42003818 | 0.022 | 0.031 | 1 | 0.76573422 | 0.462375566 |
| Gm42531 | 0.266771227 | 2.088809267 | 0.048 | 0.015 | 1 | 0.331922009 | -0.124282336 | 0 | 0.021 | 1 | 0.331922009 | 0.462375566 |
| Gpr15lg | 0.266771227 | 2.258734268 | 0.048 | 0.015 | 1 | 0.376837034 | -0.316927414 | 0 | 0.017 | 1 | 0.376837034 | 0.462375566 |
| Ccdc116 | 0.266771227 | 2.451379346 | 0.048 | 0.015 | 1 | 0.744358033 | 0.223640968 | 0.022 | 0.031 | 1 | 0.744358033 | 0.462375566 |
| 1700105P06Rik | 0.266771227 | 2.451379346 | 0.048 | 0.015 | 1 | 0.496223879 | 0.683072586 | 0 | 0.01 | 1 | 0.496223879 | 0.462375566 |
| Slc29a4 | 0.266774723 | 2.451379346 | 0.048 | 0.015 | 1 | 0.430487676 | 0.361144491 | 0 | 0.014 | 1 | 0.430487676 | 0.462380693 |
| Gm39185 | 0.266774723 | 2.451379346 | 0.048 | 0.015 | 1 | 0.672920453 | 1.098110085 | 0.022 | 0.014 | 1 | 0.672920453 | 0.462380693 |
| Mid1ip1 | 0.267034891 | -1.20069735 | 0.048 | 0.134 | 1 | 0.654208266 | 0.318500154 | 0.2 | 0.228 | 1 | 0.654208266 | 0.462762148 |
| Zfp369 | 0.26726639 | 1.580662363 | 0.238 | 0.168 | 1 | 0.505119533 | 0.205025289 | 0.2 | 0.27 | 1 | 0.505119533 | 0.463101456 |
| Helz2 | 0.462928379 | 0.989273593 | 0.19 | 0.142 | 1 | 0.267321065 | 0.254229287 | 0.111 | 0.19 | 1 | 0.462928379 | 0.463181578 |
| Bnip1 | 0.26757288 | 0.33137957 | 0.095 | 0.216 | 1 | 0.404580895 | 0.250113179 | 0.133 | 0.197 | 1 | 0.404580895 | 0.463550513 |
| Rfc1 | 0.33466177 | -0.174967075 | 0.429 | 0.597 | 1 | 0.267593038 | -0.41954248 | 0.6 | 0.574 | 1 | 0.33466177 | 0.463580041 |
| Cep83os | 0.583579863 | 1.311201688 | 0.143 | 0.112 | 1 | 0.267668198 | -0.316927414 | 0.044 | 0.097 | 1 | 0.583579863 | 0.463690131 |
| Samd5 | 0.979005859 | 1.278470486 | 0.19 | 0.22 | 1 | 0.267860387 | 0.400672855 | 0.178 | 0.294 | 1 | 0.979005859 | 0.463971587 |
| 2700099C18Rik | 0.267881873 | 3.052283391 | 0.095 | 0.045 | 1 | 0.658587075 | 2.42003818 | 0.022 | 0.014 | 1 | 0.658587075 | 0.464003048 |
| Gm30835 | 0.267902502 | 5.133203386 | 0.095 | 0.045 | 1 | 0.430483494 | -0.124282336 | 0 | 0.014 | 1 | 0.430483494 | 0.464033254 |
| Haus5 | 0.267917141 | -0.413691074 | 0 | 0.056 | 1 | 0.687208248 | 1.098110085 | 0.022 | 0.035 | 1 | 0.687208248 | 0.464054688 |
| Gm12258 | 0.267917141 | -0.413691074 | 0 | 0.056 | 1 | 0.854461673 | 1.683072586 | 0.022 | 0.028 | 1 | 0.854461673 | 0.464054688 |
| Otub2 | 0.267917141 | -0.413691074 | 0 | 0.056 | 1 | 0.472922228 | 2.197645759 | 0.067 | 0.045 | 1 | 0.472922228 | 0.464054688 |
| Klf7 | 0.26793413 | 1.277191173 | 0.81 | 0.795 | 1 | 0.52829848 | 0.237187499 | 0.889 | 0.824 | 1 | 0.52829848 | 0.464079561 |
| 9530052E02Rik | 0.267936726 | -0.264827688 | 0.048 | 0.138 | 1 | 0.917004392 | 1.683072586 | 0.089 | 0.1 | 1 | 0.917004392 | 0.464083363 |
| Daglb | 0.489836719 | 0.214340149 | 0.048 | 0.097 | 1 | 0.267937827 | -0.346674757 | 0.067 | 0.128 | 1 | 0.489836719 | 0.464084976 |
| Gm14221 | 0.267953794 | -0.849790188 | 0 | 0.056 | 1 | 0.734553436 | 1.045642666 | 0.067 | 0.055 | 1 | 0.734553436 | 0.464108352 |
| Abtb1 | 0.267953794 | -0.496153234 | 0 | 0.056 | 1 | 0.655534067 | 0.15951063 | 0.044 | 0.062 | 1 | 0.655534067 | 0.464108352 |
| Slc22a18 | 0.267953794 | -0.496153234 | 0 | 0.056 | 1 | 0.908650103 | 1.875717664 | 0.044 | 0.042 | 1 | 0.908650103 | 0.464108352 |
| Miip | 0.267984803 | -0.648156327 | 0 | 0.056 | 1 | 0.430487676 | 0.361144491 | 0 | 0.014 | 1 | 0.430487676 | 0.464153752 |
| Vmn2r29 | 0.267985206 | -0.574155746 | 0 | 0.056 | 1 | 0.484094948 | 1.513147585 | 0.067 | 0.045 | 1 | 0.484094948 | 0.464154341 |
| Mboat1 | 0.267985206 | -0.718545655 | 0 | 0.056 | 1 | 0.62519394 | 1.320502507 | 0.044 | 0.031 | 1 | 0.62519394 | 0.464154341 |
| Tmem267 | 0.267985206 | -0.785659851 | 0 | 0.056 | 1 | 0.95885962 | 1.402964667 | 0.044 | 0.048 | 1 | 0.95885962 | 0.464154341 |
| Gm45871 | 0.267985206 | -0.574155746 | 0 | 0.056 | 1 | 0.299223847 | 0.612683258 | 0.022 | 0.062 | 1 | 0.299223847 | 0.464154341 |
| Bcan | 0.268010172 | -0.718545655 | 0 | 0.056 | 1 | 0.376842245 | -0.316927414 | 0 | 0.017 | 1 | 0.376842245 | 0.464190892 |
| Nsun7 | 0.26801138 | -0.970084422 | 0 | 0.056 | 1 | 0.976334347 | 1.576157382 | 0.067 | 0.073 | 1 | 0.976334347 | 0.46419266 |
| Gm35330 | 0.26801138 | -0.911190733 | 0 | 0.056 | 1 | 0.430486631 | -0.776359032 | 0 | 0.014 | 1 | 0.430486631 | 0.46419266 |
| Cmya5 | 0.268032318 | -0.970084422 | 0 | 0.056 | 1 | 0.627214989 | 1.78260826 | 0.067 | 0.052 | 1 | 0.627214989 | 0.464223312 |
| Foxa2 | 0.268033526 | -0.911190733 | 0 | 0.056 | 1 | 0.376837034 | -0.316927414 | 0 | 0.017 | 1 | 0.376837034 | 0.46422508 |
| Pthlh | 0.268033526 | -1.535681598 | 0 | 0.056 | 1 | 0.27499811 | -1.931637258 | 0.022 | 0.062 | 1 | 0.27499811 | 0.46422508 |
| Cd300a | 0.268033526 | -0.911190733 | 0 | 0.056 | 1 | 0.508055107 | 1.050804371 | 0.044 | 0.073 | 1 | 0.508055107 | 0.46422508 |
| Stpg2 | 0.26804802 | -1.081115734 | 0 | 0.056 | 1 | 0.331914654 | -0.638855509 | 0 | 0.021 | 1 | 0.331914654 | 0.464246299 |
| Rassf4 | 0.26804802 | -0.970084422 | 0 | 0.056 | 1 | 0.496223879 | 0.683072586 | 0 | 0.01 | 1 | 0.496223879 | 0.464246299 |
| Mapt | 0.268058487 | -0.970084422 | 0 | 0.056 | 1 | 0.522794177 | 0.473619221 | 0.044 | 0.073 | 1 | 0.522794177 | 0.464261622 |
| Fbn2 | 0.26806976 | -1.718545655 | 0 | 0.056 | 1 | 0.307711004 | -2.709244837 | 0.022 | 0.059 | 1 | 0.307711004 | 0.464278123 |
| Reg4 | 0.268086667 | -1.818081329 | 0 | 0.056 | 1 | 0.608532443 | -0.316927414 | 0.022 | 0.038 | 1 | 0.608532443 | 0.464302873 |
| A530072M11Rik | 0.268100354 | -0.326228232 | 0 | 0.056 | 1 | 0.313790052 | -0.486852415 | 0.022 | 0.059 | 1 | 0.313790052 | 0.464322907 |
| Noxo1 | 0.268100354 | -0.326228232 | 0 | 0.056 | 1 | 0.955025007 | 1.513147585 | 0.022 | 0.024 | 1 | 0.955025007 | 0.464322907 |
| Acrbp | 0.71232227 | 0.815790773 | 0.095 | 0.075 | 1 | 0.268336899 | -0.57996182 | 0.067 | 0.125 | 1 | 0.71232227 | 0.464669107 |
| Msantd2 | 0.268372904 | 0.572685643 | 0.524 | 0.507 | 1 | 0.89134798 | 0.256062445 | 0.644 | 0.765 | 1 | 0.89134798 | 0.464721793 |
| Sh2b2 | 0.65600418 | 1.8372705 | 0.095 | 0.075 | 1 | 0.268436206 | 0.495445583 | 0.044 | 0.1 | 1 | 0.65600418 | 0.464814415 |
| Brat1 | 0.357279452 | 2.088809267 | 0.095 | 0.052 | 1 | 0.268609517 | -0.191396532 | 0.022 | 0.066 | 1 | 0.357279452 | 0.465067961 |
| Tle5 | 0.351483369 | -0.444622933 | 0.524 | 0.53 | 1 | 0.268734455 | -0.556393349 | 0.756 | 0.633 | 1 | 0.351483369 | 0.465250702 |
| Zfp141 | 0.268957372 | 1.503846766 | 0.143 | 0.078 | 1 | 0.523788862 | 0.495445583 | 0.067 | 0.1 | 1 | 0.523788862 | 0.465576676 |
| Parp3 | 0.269100723 | 0.365019062 | 0.143 | 0.276 | 1 | 0.636578445 | 0.826436761 | 0.222 | 0.304 | 1 | 0.636578445 | 0.465786247 |
| Gm3235 | 0.269369726 | 2.258734268 | 0.048 | 0.015 | 1 | 0.945292241 | 0.966865552 | 0.067 | 0.073 | 1 | 0.945292241 | 0.466179402 |
| Sox2ot | 0.270191239 | 2.156713331 | 0.19 | 0.123 | 1 | 0.40365515 | 0.142504205 | 0.111 | 0.166 | 1 | 0.40365515 | 0.467379172 |
| Tmem91 | 0.396971401 | 0.351843673 | 0 | 0.034 | 1 | 0.270360214 | 0.223640968 | 0.022 | 0.066 | 1 | 0.396971401 | 0.467625782 |
| Tlcd3a | 0.346860242 | -0.496153234 | 0 | 0.041 | 1 | 0.270390574 | 0.15951063 | 0.022 | 0.066 | 1 | 0.346860242 | 0.467670086 |
| Pkd2l2 | 0.337250672 | 1.673771768 | 0.095 | 0.049 | 1 | 0.270391259 | 1.039216396 | 0.022 | 0.066 | 1 | 0.337250672 | 0.467671085 |
| Gm11476 | 0.800649369 | 0.614878079 | 0.048 | 0.063 | 1 | 0.270391259 | 0.42003818 | 0.022 | 0.066 | 1 | 0.800649369 | 0.467671085 |
| Megf6 | 0.849665028 | 0.735172312 | 0.048 | 0.06 | 1 | 0.270417964 | 0.846571318 | 0.022 | 0.066 | 1 | 0.849665028 | 0.467710053 |
| Btbd16 | 0.649670552 | 1.918884266 | 0.095 | 0.075 | 1 | 0.270478896 | 0.750186782 | 0.022 | 0.066 | 1 | 0.649670552 | 0.467798959 |
| Eya2 | 0.270488805 | -0.124970024 | 0.571 | 0.418 | 1 | 0.55481867 | 0.221713519 | 0.911 | 0.824 | 1 | 0.55481867 | 0.467813417 |
| Mpp2 | 0.576537487 | 0.304537958 | 0.048 | 0.086 | 1 | 0.270567643 | 0.180572246 | 0.044 | 0.097 | 1 | 0.576537487 | 0.467928437 |
| Agap3 | 0.615480733 | 0.129451251 | 0.19 | 0.265 | 1 | 0.270892568 | 1.018256778 | 0.511 | 0.529 | 1 | 0.615480733 | 0.468402353 |
| Lpgat1 | 0.270971826 | -0.413691074 | 0.476 | 0.593 | 1 | 0.298201583 | -0.128614757 | 0.911 | 0.765 | 1 | 0.298201583 | 0.468517921 |
| Ghdc | 0.545142445 | 0.530813814 | 0.095 | 0.149 | 1 | 0.271232002 | 0.235613609 | 0.089 | 0.166 | 1 | 0.545142445 | 0.468897205 |
| Oit3 | 0.866839888 | 0.425844254 | 0.048 | 0.056 | 1 | 0.271488546 | 2.557541704 | 0.067 | 0.035 | 1 | 0.866839888 | 0.469271061 |
| Gtf2h1 | 0.724847368 | 0.204286484 | 0.524 | 0.563 | 1 | 0.271722077 | 0.534078243 | 0.711 | 0.73 | 1 | 0.724847368 | 0.469611267 |
| Stard5 | 0.567293655 | 0.849343332 | 0.238 | 0.216 | 1 | 0.271820987 | 0.571179706 | 0.178 | 0.294 | 1 | 0.567293655 | 0.469755326 |
| Dtx3l | 0.271824334 | -1.392317423 | 0.048 | 0.127 | 1 | 0.277092276 | 0.78260826 | 0.067 | 0.128 | 1 | 0.277092276 | 0.4697602 |
| Stmn2 | 0.271976146 | 2.088809267 | 0.048 | 0.015 | 1 | 0.492040936 | 2.268035087 | 0.022 | 0.01 | 1 | 0.492040936 | 0.469981268 |
| 9930111J21Rik2 | 0.27197965 | 1.673771768 | 0.048 | 0.015 | 1 | 0.492040936 | 2.683072586 | 0.022 | 0.01 | 1 | 0.492040936 | 0.46998637 |
| Gm17359 | 0.271980818 | 2.351843673 | 0.048 | 0.015 | 1 | 0.852141123 | 0.946106992 | 0.022 | 0.028 | 1 | 0.852141123 | 0.469988071 |
| Cyb561d1 | 0.271980818 | 2.351843673 | 0.048 | 0.015 | 1 | 0.430487676 | 0.361144491 | 0 | 0.014 | 1 | 0.430487676 | 0.469988071 |
| Gm40323 | 0.271980818 | 2.351843673 | 0.048 | 0.015 | 1 | 0.496223879 | 0.683072586 | 0 | 0.01 | 1 | 0.496223879 | 0.469988071 |
| Mm2pr | 0.271980818 | 2.351843673 | 0.048 | 0.015 | 1 | 0.849821852 | 0.683072586 | 0.022 | 0.028 | 1 | 0.849821852 | 0.469988071 |
| Proser3 | 0.271980818 | 2.351843673 | 0.048 | 0.015 | 1 | 0.36692495 | 0.683072586 | 0.022 | 0.055 | 1 | 0.36692495 | 0.469988071 |
| Tmem147os | 0.271980818 | 2.351843673 | 0.048 | 0.015 | 1 | 0.936025227 | 1.683072586 | 0.022 | 0.021 | 1 | 0.936025227 | 0.469988071 |
| Gm56994 | 0.271980818 | 2.351843673 | 0.048 | 0.015 | 1 | 0.496223879 | 0.683072586 | 0 | 0.01 | 1 | 0.496223879 | 0.469988071 |
| Gm47071 | 0.271980818 | 2.351843673 | 0.048 | 0.015 | 1 | 0.41758974 | 1.946106992 | 0.044 | 0.024 | 1 | 0.41758974 | 0.469988071 |
| Gm41291 | 0.271980818 | 2.351843673 | 0.048 | 0.015 | 1 | 0.293659664 | -0.316927414 | 0 | 0.024 | 1 | 0.293659664 | 0.469988071 |
| Dgkg | 0.271980818 | 2.351843673 | 0.048 | 0.015 | 1 | 0.806246349 | 1.875717664 | 0.022 | 0.017 | 1 | 0.806246349 | 0.469988071 |
| Abcc10 | 0.271980818 | 2.351843673 | 0.048 | 0.015 | 1 | 0.430244927 | -0.316927414 | 0.022 | 0.048 | 1 | 0.430244927 | 0.469988071 |
| Gm30541 | 0.271980818 | 2.351843673 | 0.048 | 0.015 | 1 | 0.309824707 | 3.42003818 | 0.022 | 0.007 | 1 | 0.309824707 | 0.469988071 |
| 3110056K07Rik | 0.921052351 | 0.932304781 | 0.238 | 0.291 | 1 | 0.272037402 | 0.326587269 | 0.267 | 0.394 | 1 | 0.921052351 | 0.470070456 |
| Mrps11 | 0.272201045 | -0.763633545 | 0.095 | 0.187 | 1 | 0.433335622 | -0.255526869 | 0.089 | 0.131 | 1 | 0.433335622 | 0.470308681 |
| Csnk2a2 | 0.885576487 | 0.126020346 | 0.667 | 0.709 | 1 | 0.272607714 | 0.100469358 | 0.822 | 0.82 | 1 | 0.885576487 | 0.470900463 |
| Bmpr1a | 0.544900374 | 0.595114824 | 0.667 | 0.817 | 1 | 0.272621997 | 0.260973059 | 0.978 | 0.931 | 1 | 0.544900374 | 0.47092124 |
| Rubcn | 0.272840777 | -0.392317423 | 0.19 | 0.317 | 1 | 0.594307992 | 0.956091081 | 0.333 | 0.384 | 1 | 0.594307992 | 0.471239465 |
| Vps13c | 0.909117424 | 0.159198595 | 0.333 | 0.388 | 1 | 0.273631282 | 0.289158378 | 0.556 | 0.443 | 1 | 0.909117424 | 0.472388486 |
| Lsp1 | 0.530294352 | -0.314912919 | 0.286 | 0.366 | 1 | 0.273996239 | -0.242196539 | 0.378 | 0.543 | 1 | 0.530294352 | 0.472918539 |
| Synj2bp | 0.515930779 | 0.217622733 | 0.476 | 0.746 | 1 | 0.274447782 | 0.58153456 | 0.8 | 0.803 | 1 | 0.515930779 | 0.473573979 |
| Depdc7 | 0.274597491 | 1.673771768 | 0.048 | 0.015 | 1 | 0.88973484 | 1.683072586 | 0.044 | 0.042 | 1 | 0.88973484 | 0.4737912 |

| | | | | | | | | | | | | |
|---|---|---|---|---|---|---|---|---|---|---|---|---|
| Cd101 | 0.274597491 | 1.866416846 | 0.048 | 0.015 | 1 | 0.331907298 | -0.486852415 | 0 | 0.021 | 1 | 0.331907298 | 0.4737912 |
| Gm49890 | 0.274597491 | 2.088809267 | 0.048 | 0.015 | 1 | 0.45092657 | 0.339118185 | 0.067 | 0.042 | 1 | 0.45092657 | 0.4737912 |
| Gm48089 | 0.274597491 | 2.088809267 | 0.048 | 0.015 | 1 | 0.496223879 | -1.017367132 | 0 | 0.01 | 1 | 0.496223879 | 0.4737912 |
| Abhd15 | 0.274597491 | 1.936806174 | 0.048 | 0.015 | 1 | 0.54059033 | -0.316927414 | 0.044 | 0.069 | 1 | 0.54059033 | 0.4737912 |
| Macc1 | 0.274597491 | 1.351843673 | 0.048 | 0.015 | 1 | 0.293637566 | -0.486852415 | 0 | 0.024 | 1 | 0.293637566 | 0.4737912 |
| Sox7 | 0.274597491 | 2.088809267 | 0.048 | 0.015 | 1 | 0.293657138 | -1.564854927 | 0 | 0.024 | 1 | 0.293657138 | 0.4737912 |
| Zfp7 | 0.274597491 | 2.088809267 | 0.048 | 0.015 | 1 | 0.833619716 | 0.361144491 | 0.022 | 0.028 | 1 | 0.833619716 | 0.4737912 |
| Gm53058 | 0.274597491 | 1.673771768 | 0.048 | 0.015 | 1 | 0.779059159 | 1.250113179 | 0.067 | 0.059 | 1 | 0.779059159 | 0.4737912 |
| Tff2 | 0.274597491 | 1.866416846 | 0.048 | 0.015 | 1 | 0.552216857 | 0.361144491 | 0.022 | 0.042 | 1 | 0.552216857 | 0.4737912 |
| Pex11a | 0.274829216 | -1.107587946 | 0.048 | 0.127 | 1 | 0.926733543 | 0.757073168 | 0.044 | 0.048 | 1 | 0.926733543 | 0.474127334 |
| Cox11 | 0.635431581 | 0.292949984 | 0.238 | 0.328 | 1 | 0.275354555 | 0.264571232 | 0.222 | 0.346 | 1 | 0.635431581 | 0.474888979 |
| Zfp992 | 0.930336124 | 0.901182264 | 0.143 | 0.142 | 1 | 0.275361588 | -0.486852415 | 0.111 | 0.176 | 1 | 0.930336124 | 0.474899172 |
| Spast | 0.382762725 | 0.673771768 | 0.238 | 0.381 | 1 | 0.275760087 | 0.491646516 | 0.711 | 0.595 | 1 | 0.382762725 | 0.475476549 |
| Timmdc1 | 0.298556386 | 1.074734082 | 0.333 | 0.284 | 1 | 0.275811005 | 0.361144491 | 0.156 | 0.242 | 1 | 0.298556386 | 0.4755503 |
| Gm57301 | 0.275836128 | 1.908237021 | 0.095 | 0.045 | 1 | 0.955023924 | 1.268035087 | 0.022 | 0.024 | 1 | 0.955023924 | 0.475586687 |
| Atp6v1h | 0.512942236 | 0.325249463 | 0.571 | 0.757 | 1 | 0.275865469 | 0.239386772 | 0.889 | 0.796 | 1 | 0.512942236 | 0.475629181 |
| Gm31718 | 0.915524318 | 1.425844254 | 0.048 | 0.056 | 1 | 0.275911127 | 1.859950348 | 0.089 | 0.052 | 1 | 0.915524318 | 0.475695304 |
| Xkr6 | 0.276025748 | -0.522625445 | 0.048 | 0.134 | 1 | 0.622561365 | 1.573448095 | 0.111 | 0.097 | 1 | 0.622561365 | 0.475861282 |
| Megf11 | 0.324831685 | -1.413691074 | 0 | 0.045 | 1 | 0.276944981 | 2.180572246 | 0.067 | 0.035 | 1 | 0.324831685 | 0.47719144 |
| Uqcrc1 | 0.27702351 | -0.431762646 | 0.429 | 0.545 | 1 | 0.476965504 | -0.360254845 | 0.733 | 0.619 | 1 | 0.476965504 | 0.477304995 |
| Tnni1 | 0.277250164 | 1.503846766 | 0.048 | 0.015 | 1 | 0.503208953 | 0.361144491 | 0.022 | 0.01 | 1 | 0.503208953 | 0.477632674 |
| Lgals7 | 0.277250164 | 1.351843673 | 0.048 | 0.015 | 1 | 0.523426772 | 1.304560963 | 0.044 | 0.028 | 1 | 0.523426772 | 0.477632674 |
| Dpys | 0.277250164 | 1.866416846 | 0.048 | 0.015 | 1 | 0.376842245 | -0.316927414 | 0 | 0.017 | 1 | 0.376842245 | 0.477632674 |
| Zbtb7b | 0.802317603 | 0.638147858 | 0.143 | 0.127 | 1 | 0.277261639 | 0.405087839 | 0.156 | 0.263 | 1 | 0.802317603 | 0.477649261 |
| Sgcd | 0.277551876 | 1.290556552 | 0.619 | 0.78 | 1 | 0.345993148 | 0.945562681 | 0.778 | 0.931 | 1 | 0.345993148 | 0.478068709 |
| Leng9 | 0.729839742 | 0.866416846 | 0.048 | 0.071 | 1 | 0.27768208 | -1.017367132 | 0.022 | 0.062 | 1 | 0.729839742 | 0.478256822 |
| Atp2c1 | 0.927952809 | 0.489347197 | 0.333 | 0.414 | 1 | 0.277753692 | 0.632446513 | 0.711 | 0.588 | 1 | 0.927952809 | 0.47836027 |
| Tmprss5 | 0.324739547 | -0.133583154 | 0 | 0.045 | 1 | 0.278574089 | -0.84048937 | 0.022 | 0.062 | 1 | 0.324739547 | 0.479544654 |
| 4931414P19Rik | 0.853725905 | 2.171271427 | 0.048 | 0.041 | 1 | 0.278574089 | -0.84048937 | 0.022 | 0.062 | 1 | 0.853725905 | 0.479544654 |
| 9330188P03Rik | 0.922502659 | -0.666078235 | 0.238 | 0.231 | 1 | 0.278613263 | 0.394094913 | 0.178 | 0.284 | 1 | 0.922502659 | 0.479601176 |
| Ikbkg | 0.946237143 | 0.380990018 | 0.143 | 0.153 | 1 | 0.278628239 | 0.525531309 | 0.089 | 0.166 | 1 | 0.946237143 | 0.479622783 |
| Gm48882 | 0.278974785 | -0.413691074 | 0.048 | 0.134 | 1 | 0.941346407 | 1.098110085 | 0.022 | 0.021 | 1 | 0.941346407 | 0.480122639 |
| Gm49959 | 0.279805041 | -0.752492987 | 0.048 | 0.127 | 1 | 0.948654232 | 0.659225844 | 0.156 | 0.166 | 1 | 0.948654232 | 0.481319221 |
| Gm32618 | 0.279864362 | -1.774688733 | 0.095 | 0.179 | 1 | 0.677689825 | 0.812059027 | 0.156 | 0.208 | 1 | 0.677689825 | 0.481404663 |
| Slfn2 | 0.279915475 | 0.866416846 | 0.048 | 0.015 | 1 | 0.293652719 | -1.316927414 | 0 | 0.024 | 1 | 0.293652719 | 0.481478277 |
| 9930021J03Rik | 0.551215181 | 0.128122191 | 0.619 | 0.545 | 1 | 0.279999372 | 0.212651278 | 0.844 | 0.713 | 1 | 0.551215181 | 0.481599096 |
| Gm20470 | 0.550953306 | 1.936806174 | 0.048 | 0.026 | 1 | 0.280232364 | 1.776181991 | 0.067 | 0.035 | 1 | 0.550953306 | 0.481934551 |
| Zfp524 | 0.606700221 | 1.150209812 | 0.095 | 0.067 | 1 | 0.28035123 | -0.84048937 | 0.022 | 0.062 | 1 | 0.606700221 | 0.482105648 |
| Taco1 | 0.818482987 | 0.425844254 | 0.048 | 0.06 | 1 | 0.28035123 | -0.709244837 | 0.022 | 0.062 | 1 | 0.818482987 | 0.482105648 |
| Zfp940 | 0.96060963 | 1.088809267 | 0.048 | 0.052 | 1 | 0.28035123 | -0.84048937 | 0.022 | 0.062 | 1 | 0.96060963 | 0.482105648 |
| Jpx | 0.370269495 | 1.79930265 | 0.143 | 0.093 | 1 | 0.280771714 | 0.361144491 | 0.089 | 0.159 | 1 | 0.370269495 | 0.482710672 |
| Malt1 | 0.281045442 | -0.413691074 | 0.048 | 0.134 | 1 | 0.90573131 | 1.039216396 | 0.133 | 0.145 | 1 | 0.90573131 | 0.483104344 |
| Rprd2 | 0.974079907 | 0.459328099 | 0.524 | 0.653 | 1 | 0.281527463 | -0.168604667 | 0.778 | 0.806 | 1 | 0.974079907 | 0.483797213 |
| Ndfip2 | 0.38705199 | 0.512948973 | 0.81 | 0.825 | 1 | 0.281821755 | 0.467918331 | 0.889 | 0.872 | 1 | 0.38705199 | 0.484220008 |
| Ireb2 | 0.282313258 | 0.444953077 | 0.571 | 0.578 | 1 | 0.698246532 | 0.431236387 | 0.711 | 0.72 | 1 | 0.698246532 | 0.484925741 |
| Tomm40 | 0.282645085 | 0.462267662 | 0.238 | 0.448 | 1 | 0.791952465 | 0.228694195 | 0.533 | 0.581 | 1 | 0.791952465 | 0.485401926 |
| Erich1 | 0.282938574 | -0.295854583 | 0.048 | 0.134 | 1 | 0.403624826 | 0.863644832 | 0.089 | 0.138 | 1 | 0.403624826 | 0.485822911 |
| Ip6k1 | 0.283031318 | 0.800728943 | 0.571 | 0.627 | 1 | 0.915101211 | 0.310263322 | 0.822 | 0.806 | 1 | 0.915101211 | 0.485955908 |
| Mycbpap | 0.943759773 | 0.673771768 | 0.048 | 0.052 | 1 | 0.283542478 | 2.045642666 | 0.067 | 0.035 | 1 | 0.943759773 | 0.486688619 |
| Rtp4 | 0.283691798 | -1.586755783 | 0.048 | 0.127 | 1 | 0.627496876 | 0.600610426 | 0.067 | 0.093 | 1 | 0.627496876 | 0.486902559 |
| Tars2 | 0.728003448 | 0.103916159 | 0.19 | 0.243 | 1 | 0.284083343 | 0.564131513 | 0.2 | 0.315 | 1 | 0.728003448 | 0.48746334 |
| Morc3 | 0.28484499 | 0.962276861 | 0.381 | 0.332 | 1 | 0.506048984 | -0.32442195 | 0.356 | 0.381 | 1 | 0.506048984 | 0.488553312 |
| C430014B12Rik | 0.285465396 | -0.326228232 | 0 | 0.052 | 1 | 0.926730234 | 0.595609745 | 0.044 | 0.048 | 1 | 0.926730234 | 0.4894403 |
| Bicdl1 | 0.285465396 | -0.326228232 | 0 | 0.052 | 1 | 0.5539024 | 0.744473131 | 0.022 | 0.042 | 1 | 0.5539024 | 0.4894403 |
| D6Ertd527e | 0.285465396 | -0.326228232 | 0 | 0.052 | 1 | 0.612016574 | 0.361144491 | 0.044 | 0.066 | 1 | 0.612016574 | 0.4894403 |
| Zfp433 | 0.285465396 | -0.326228232 | 0 | 0.052 | 1 | 0.763583976 | 0.808603468 | 0.022 | 0.031 | 1 | 0.763583976 | 0.4894403 |
| Gm29684 | 0.285465396 | -0.326228232 | 0 | 0.052 | 1 | 0.825853389 | 0.683072586 | 0.022 | 0.017 | 1 | 0.825853389 | 0.4894403 |
| Srd5a1 | 0.285465396 | -0.326228232 | 0 | 0.052 | 1 | 0.555534078 | 1.197645759 | 0.022 | 0.042 | 1 | 0.555534078 | 0.4894403 |
| Pus7l | 0.285465396 | -0.326228232 | 0 | 0.052 | 1 | 0.293556836 | -0.375821103 | 0.022 | 0.062 | 1 | 0.293556836 | 0.4894403 |
| Pcdhga2 | 0.285465396 | -0.326228232 | 0 | 0.052 | 1 | 0.595927847 | -0.124282336 | 0.022 | 0.038 | 1 | 0.595927847 | 0.4894403 |
| Gnasas1 | 0.285499161 | -0.574155746 | 0 | 0.052 | 1 | 0.687918987 | 1.152557869 | 0.044 | 0.062 | 1 | 0.687918987 | 0.489488551 |
| Gm42869 | 0.285499161 | -0.413691074 | 0 | 0.052 | 1 | 0.541900769 | 1.098110085 | 0.044 | 0.028 | 1 | 0.541900769 | 0.489488551 |
| Sars2 | 0.285499161 | -0.413691074 | 0 | 0.052 | 1 | 0.552216857 | 0.361144491 | 0.022 | 0.042 | 1 | 0.552216857 | 0.489488551 |
| Exoc8 | 0.285499161 | -0.496153234 | 0 | 0.052 | 1 | 0.30653433 | 0.361144491 | 0.067 | 0.125 | 1 | 0.30653433 | 0.489488551 |
| Mmd | 0.285499161 | -0.413691074 | 0 | 0.052 | 1 | 0.555571506 | 1.361144491 | 0.022 | 0.042 | 1 | 0.555571506 | 0.489488551 |
| Ccdc120 | 0.285499161 | -0.413691074 | 0 | 0.052 | 1 | 0.75170438 | 1.020107573 | 0.044 | 0.059 | 1 | 0.75170438 | 0.489488551 |
| Zc4h2 | 0.285499161 | -0.496153234 | 0 | 0.052 | 1 | 0.29730293 | -0.124282336 | 0.022 | 0.062 | 1 | 0.29730293 | 0.489488551 |
| A330041J22Rik | 0.285499161 | 1.866416846 | 0.095 | 0.045 | 1 | 0.990772207 | 0.875717664 | 0.044 | 0.045 | 1 | 0.990772207 | 0.489488551 |
| Gm15337 | 0.285527294 | -0.574155746 | 0 | 0.052 | 1 | 0.56661406 | 1.28452321 | 0.067 | 0.048 | 1 | 0.56661406 | 0.489528753 |
| Chit1 | 0.285527727 | -0.911190733 | 0 | 0.052 | 1 | 0.49622258 | -1.223818009 | 0 | 0.01 | 1 | 0.49622258 | 0.489529371 |
| Lrp11 | 0.285527727 | -0.496153234 | 0 | 0.052 | 1 | 0.667598499 | 0.223640968 | 0.022 | 0.035 | 1 | 0.667598499 | 0.489529371 |
| BC024063 | 0.285527727 | -0.496153234 | 0 | 0.052 | 1 | 0.552232538 | 0.361144491 | 0.022 | 0.042 | 1 | 0.552232538 | 0.489529371 |
| 4930471E19Rik | 0.285527727 | -1.184209227 | 0 | 0.052 | 1 | 0.314689364 | 1.460680165 | 0.022 | 0.007 | 1 | 0.314689364 | 0.489529371 |
| 4933412O06Rik | 0.285527727 | -0.648156327 | 0 | 0.052 | 1 | 0.309824707 | 2.683072586 | 0.022 | 0.007 | 1 | 0.309824707 | 0.489529371 |
| Asxl3 | 0.285551097 | -1.02666795 | 0 | 0.052 | 1 | 0.293652719 | -1.017367132 | 0 | 0.024 | 1 | 0.293652719 | 0.489562765 |
| Nox1 | 0.285551097 | -0.970084422 | 0 | 0.052 | 1 | 0.854464766 | 2.78998779 | 0.022 | 0.028 | 1 | 0.854464766 | 0.489562765 |
| Erich6 | 0.28555153 | -0.574155746 | 0 | 0.052 | 1 | 0.579492135 | 0.42003818 | 0.044 | 0.069 | 1 | 0.579492135 | 0.489563384 |
| Cenpl | 0.285566676 | -0.718545655 | 0 | 0.052 | 1 | 0.29730293 | -0.124282336 | 0.022 | 0.062 | 1 | 0.29730293 | 0.489585026 |
| Mok | 0.285566676 | -0.718545655 | 0 | 0.052 | 1 | 0.697563751 | 0.361144491 | 0.067 | 0.087 | 1 | 0.697563751 | 0.489585026 |
| Gm9968 | 0.285566676 | -0.718545655 | 0 | 0.052 | 1 | 0.67587272 | 1.268035087 | 0.089 | 0.076 | 1 | 0.67587272 | 0.489585026 |
| Nfkbie | 0.285566676 | -0.718545655 | 0 | 0.052 | 1 | 0.450829449 | 0.683072586 | 0.022 | 0.048 | 1 | 0.450829449 | 0.489585026 |
| 4833422C13Rik | 0.285569273 | -0.849790188 | 0 | 0.052 | 1 | 0.8202424 | 1.098110085 | 0.022 | 0.017 | 1 | 0.8202424 | 0.489588736 |
| Rims1 | 0.285570571 | -1.574155746 | 0 | 0.052 | 1 | 0.726320704 | 1.039216396 | 0.044 | 0.035 | 1 | 0.726320704 | 0.489590591 |
| Ptpn22 | 0.285570571 | -1.326228232 | 0 | 0.052 | 1 | 0.552245367 | 0.513147585 | 0.022 | 0.042 | 1 | 0.552245367 | 0.489590591 |
| Dnai3 | 0.285571003 | -0.648156327 | 0 | 0.052 | 1 | 0.309824707 | 3.42003818 | 0.022 | 0.007 | 1 | 0.309824707 | 0.489591209 |
| Tspyl5 | 0.285571003 | -0.648156327 | 0 | 0.052 | 1 | 0.747015076 | 1.683072586 | 0.067 | 0.055 | 1 | 0.747015076 | 0.489591209 |
| Efcab7 | 0.285582254 | -1.02666795 | 0 | 0.052 | 1 | 0.683283723 | 0.595609745 | 0.022 | 0.035 | 1 | 0.683283723 | 0.489607284 |
| C730034F03Rik | 0.285582254 | -0.849790188 | 0 | 0.052 | 1 | 0.555549391 | 1.098110085 | 0.022 | 0.042 | 1 | 0.555549391 | 0.489607284 |
| Ccnf | 0.285582254 | -0.911190733 | 0 | 0.052 | 1 | 0.331914654 | -0.486852415 | 0 | 0.021 | 1 | 0.331914654 | 0.489607284 |
| Card6 | 0.285583552 | -0.849790188 | 0 | 0.052 | 1 | 0.935071939 | 0.513147585 | 0.022 | 0.024 | 1 | 0.935071939 | 0.489609139 |
| Ube2c | 0.285586581 | -3.094412557 | 0 | 0.052 | 1 | 0.326732519 | 2.460680165 | 0.044 | 0.021 | 1 | 0.326732519 | 0.489613467 |
| Bmf | 0.285592639 | -0.970084422 | 0 | 0.052 | 1 | 0.568055486 | -0.242926832 | 0.067 | 0.093 | 1 | 0.568055486 | 0.489622122 |
| Edn2 | 0.285598264 | -1.911190733 | 0 | 0.052 | 1 | 0.293657138 | -3.916840256 | 0 | 0.024 | 1 | 0.293657138 | 0.489630159 |
| Gjb2 | 0.285603456 | -2.496153234 | 0 | 0.052 | 1 | 0.740122935 | -0.901889915 | 0.022 | 0.031 | 1 | 0.740122935 | 0.489637578 |
| Gm6710 | 0.285622926 | -0.233118828 | 0 | 0.052 | 1 | 0.387952149 | -0.316927414 | 0.022 | 0.052 | 1 | 0.387952149 | 0.489665395 |
| Gm41349 | 0.285622926 | -0.233118828 | 0 | 0.052 | 1 | 0.49622258 | 0.361144491 | 0 | 0.01 | 1 | 0.49622258 | 0.489665395 |
| Apaf1 | 0.285712604 | 0.523212091 | 0.429 | 0.347 | 1 | 0.504114406 | 0.283375633 | 0.4 | 0.512 | 1 | 0.504114406 | 0.489793516 |
| Gm47920 | 0.396971401 | 0.351843673 | 0 | 0.034 | 1 | 0.286020697 | 1.605070074 | 0.089 | 0.052 | 1 | 0.396971401 | 0.490233554 |

| | | | | | | | | | | | | |
|---|---|---|---|---|---|---|---|---|---|---|---|---|
| S1pr3 | 0.286145715 | -0.785659851 | 0.048 | 0.131 | 1 | 0.826709378 | -0.316927414 | 0.022 | 0.028 | 1 | 0.826709378 | 0.490412059 |
| Mapk8ip3 | 0.286311646 | 0.652076697 | 0.571 | 0.563 | 1 | 0.440746935 | 0.277080226 | 0.533 | 0.661 | 1 | 0.440746935 | 0.490648933 |
| Zbtb16 | 0.286976389 | 0.473632153 | 0.381 | 0.328 | 1 | 0.915840324 | 0.901252756 | 0.289 | 0.349 | 1 | 0.915840324 | 0.49159733 |
| Stard8 | 0.287438383 | -0.413691074 | 0.19 | 0.325 | 1 | 0.289804321 | 1.715738775 | 0.311 | 0.301 | 1 | 0.289804321 | 0.492255942 |
| 3300002I08Rik | 0.518540524 | 0.351843673 | 0.095 | 0.16 | 1 | 0.287882736 | 0.559690171 | 0.089 | 0.159 | 1 | 0.518540524 | 0.492889002 |
| Zfr | 0.477586743 | 0.187307901 | 0.952 | 0.996 | 1 | 0.288392284 | 0.133893338 | 1 | 0.997 | 1 | 0.477586743 | 0.493614459 |
| Fzd10os | 0.288828982 | 1.766881172 | 0.095 | 0.045 | 1 | 0.416574156 | -0.300439291 | 0.133 | 0.187 | 1 | 0.416574156 | 0.494235784 |
| Pax9 | 0.288852374 | 2.233199176 | 0.095 | 0.045 | 1 | 0.657860717 | -1.316927414 | 0.022 | 0.035 | 1 | 0.657860717 | 0.494269054 |
| Cacna1e | 0.728465456 | -0.763633545 | 0.048 | 0.067 | 1 | 0.28905589 | -1.189052591 | 0.111 | 0.173 | 1 | 0.728465456 | 0.494558472 |
| Dcaf12 | 0.389924296 | 0.825774861 | 0.429 | 0.448 | 1 | 0.289378085 | -0.162125687 | 0.511 | 0.561 | 1 | 0.389924296 | 0.495016494 |
| Cacnb1 | 0.290025048 | 2.351843673 | 0.143 | 0.086 | 1 | 0.783371065 | 0.426732833 | 0.089 | 0.107 | 1 | 0.783371065 | 0.495935568 |
| Ehhadh | 0.290488785 | 1.673771768 | 0.095 | 0.045 | 1 | 0.936023974 | 1.875717664 | 0.022 | 0.021 | 1 | 0.936023974 | 0.496593836 |
| Bcar3 | 0.290516227 | 0.901182264 | 0.143 | 0.078 | 1 | 0.364036192 | 0.519573854 | 0.089 | 0.149 | 1 | 0.364036192 | 0.496632776 |
| Tcf24 | 0.425614684 | 0.214340149 | 0 | 0.03 | 1 | 0.290736539 | -0.539319835 | 0.022 | 0.062 | 1 | 0.425614684 | 0.496945343 |
| Gm57450 | 0.368034131 | 1.79930265 | 0.143 | 0.09 | 1 | 0.291438471 | 0.223640968 | 0.044 | 0.097 | 1 | 0.368034131 | 0.49794056 |
| 4933406I18Rik | 0.291910064 | 0.969510339 | 0.143 | 0.276 | 1 | 0.764744919 | 0.9205719 | 0.2 | 0.249 | 1 | 0.764744919 | 0.498608642 |
| Fbxo42 | 0.291937954 | 1.510833197 | 0.429 | 0.44 | 1 | 0.399853624 | 0.574138215 | 0.689 | 0.682 | 1 | 0.399853624 | 0.498648139 |
| Ldlr | 0.642785689 | -0.276929922 | 0.667 | 0.705 | 1 | 0.292193223 | -0.189669787 | 0.556 | 0.609 | 1 | 0.642785689 | 0.499009566 |
| Zfp39 | 0.729839742 | 0.866416846 | 0.048 | 0.071 | 1 | 0.292584143 | -0.361321533 | 0.044 | 0.093 | 1 | 0.729839742 | 0.499562805 |
| Lrp2bp | 0.370806713 | -0.233118828 | 0 | 0.037 | 1 | 0.292659668 | -0.316927414 | 0.044 | 0.093 | 1 | 0.370806713 | 0.499669655 |
| Arhgap10 | 0.797990587 | 0.752843339 | 0.524 | 0.653 | 1 | 0.293062407 | 0.245098276 | 0.711 | 0.654 | 1 | 0.797990587 | 0.500239239 |
| Sos2 | 0.293099109 | 0.991343458 | 0.571 | 0.59 | 1 | 0.492942299 | 0.268035087 | 0.889 | 0.792 | 1 | 0.492942299 | 0.50029113 |
| Stx5a | 0.293413595 | 0.479393723 | 0.667 | 0.728 | 1 | 0.574398077 | 0.131957856 | 0.8 | 0.841 | 1 | 0.574398077 | 0.500735652 |
| Efnb3 | 0.71266839 | 1.79930265 | 0.048 | 0.034 | 1 | 0.293485399 | 1.91753784 | 0.089 | 0.052 | 1 | 0.71266839 | 0.500837119 |
| Spc25 | 0.324843766 | -1.880817084 | 0 | 0.045 | 1 | 0.293637566 | -0.486852415 | 0 | 0.024 | 1 | 0.324843766 | 0.501052112 |
| Acox2 | 0.346846605 | -0.326228232 | 0 | 0.041 | 1 | 0.293637566 | -0.486852415 | 0 | 0.024 | 1 | 0.346846605 | 0.501052112 |
| Zfp184 | 0.359270744 | 2.673771768 | 0.048 | 0.019 | 1 | 0.293637566 | -0.486852415 | 0 | 0.024 | 1 | 0.359270744 | 0.501052112 |
| Cdc42ep2 | 0.359270744 | 2.673771768 | 0.048 | 0.019 | 1 | 0.293637566 | -0.486852415 | 0 | 0.024 | 1 | 0.359270744 | 0.501052112 |
| Tedc2 | 0.370840523 | 0.214340149 | 0 | 0.037 | 1 | 0.293637566 | -0.486852415 | 0 | 0.024 | 1 | 0.370840523 | 0.501052112 |
| Gm30934 | 0.376794341 | 1.866416846 | 0.048 | 0.019 | 1 | 0.293637566 | -0.638855509 | 0 | 0.024 | 1 | 0.376794341 | 0.501052112 |
| Gm57065 | 0.396955847 | -0.574155746 | 0 | 0.034 | 1 | 0.293637566 | -0.486852415 | 0 | 0.024 | 1 | 0.396955847 | 0.501052112 |
| Gm20492 | 0.457308648 | 0.673771768 | 0 | 0.026 | 1 | 0.293637566 | -0.486852415 | 0 | 0.024 | 1 | 0.457308648 | 0.501052112 |
| Tes3-ps | 0.492590603 | 0.673771768 | 0 | 0.022 | 1 | 0.293637566 | -0.486852415 | 0 | 0.024 | 1 | 0.492590603 | 0.501052112 |
| Gpr21 | 0.492599419 | 0.503846766 | 0 | 0.022 | 1 | 0.293637566 | -0.486852415 | 0 | 0.024 | 1 | 0.492599419 | 0.501052112 |
| Gm13134 | 0.492608234 | 0.866416846 | 0 | 0.022 | 1 | 0.293637566 | -0.486852415 | 0 | 0.024 | 1 | 0.492608234 | 0.501052112 |
| Gm16573 | 0.492608234 | 0.866416846 | 0 | 0.022 | 1 | 0.293637566 | -0.486852415 | 0 | 0.024 | 1 | 0.492608234 | 0.501052112 |
| 4921515G04Rik | 0.492608234 | 0.866416846 | 0 | 0.022 | 1 | 0.293637566 | -0.486852415 | 0 | 0.024 | 1 | 0.492608234 | 0.501052112 |
| 4930444P10Rik | 0.532426384 | 0.673771768 | 0 | 0.019 | 1 | 0.293637566 | -0.486852415 | 0 | 0.024 | 1 | 0.532426384 | 0.501052112 |
| Pnp2 | 0.532426384 | 0.866416846 | 0 | 0.019 | 1 | 0.293637566 | -0.486852415 | 0 | 0.024 | 1 | 0.532426384 | 0.501052112 |
| Gm9920 | 0.532426384 | 0.866416846 | 0 | 0.019 | 1 | 0.293637566 | -0.638855509 | 0 | 0.024 | 1 | 0.532426384 | 0.501052112 |
| Gm40528 | 0.532432377 | -1.133583154 | 0 | 0.019 | 1 | 0.293637566 | -0.776359032 | 0 | 0.024 | 1 | 0.532432377 | 0.501052112 |
| Gm15860 | 0.532436372 | 1.088809267 | 0 | 0.019 | 1 | 0.293637566 | -0.486852415 | 0 | 0.024 | 1 | 0.532436372 | 0.501052112 |
| Gm57243 | 0.532436372 | 1.088809267 | 0 | 0.019 | 1 | 0.293637566 | -0.776359032 | 0 | 0.024 | 1 | 0.532436372 | 0.501052112 |
| 6820408C15Rik | 0.550953306 | 1.936806174 | 0.048 | 0.026 | 1 | 0.293637566 | -0.486852415 | 0 | 0.024 | 1 | 0.550953306 | 0.501052112 |
| Pctp | 0.578156445 | 1.088809267 | 0 | 0.015 | 1 | 0.293637566 | -0.776359032 | 0 | 0.024 | 1 | 0.578156445 | 0.501052112 |
| A330102I10Rik | 0.578156445 | 1.088809267 | 0 | 0.015 | 1 | 0.293637566 | -0.486852415 | 0 | 0.024 | 1 | 0.578156445 | 0.501052112 |
| Gm49801 | 0.578156445 | 1.088809267 | 0 | 0.015 | 1 | 0.293637566 | -0.776359032 | 0 | 0.024 | 1 | 0.578156445 | 0.501052112 |
| I830077J02Rik | 0.632076134 | 1.673771768 | 0 | 0.011 | 1 | 0.293637566 | -0.486852415 | 0 | 0.024 | 1 | 0.632076134 | 0.501052112 |
| Gm17745 | 0.632076134 | 1.673771768 | 0 | 0.011 | 1 | 0.293637566 | -0.486852415 | 0 | 0.024 | 1 | 0.632076134 | 0.501052112 |
| Kcns3 | 0.632076134 | 1.673771768 | 0 | 0.011 | 1 | 0.293637566 | -0.486852415 | 0 | 0.024 | 1 | 0.632076134 | 0.501052112 |
| Zfp41 | 0.632076134 | 1.673771768 | 0 | 0.011 | 1 | 0.293637566 | -0.638855509 | 0 | 0.024 | 1 | 0.632076134 | 0.501052112 |
| P2rx6 | 0.632076134 | 1.673771768 | 0 | 0.011 | 1 | 0.293637566 | -0.486852415 | 0 | 0.024 | 1 | 0.632076134 | 0.501052112 |
| Aif1 | 0.632076134 | 1.673771768 | 0 | 0.011 | 1 | 0.293637566 | -0.486852415 | 0 | 0.024 | 1 | 0.632076134 | 0.501052112 |
| 9130008F23Rik | 0.632076134 | 1.673771768 | 0 | 0.011 | 1 | 0.293637566 | -0.638855509 | 0 | 0.024 | 1 | 0.632076134 | 0.501052112 |
| 3830408C21Rik | 0.854880747 | 0.866416846 | 0.048 | 0.06 | 1 | 0.293637566 | -0.486852415 | 0 | 0.024 | 1 | 0.854880747 | 0.501052112 |
| Stil | 0.858792447 | -0.574155746 | 0.048 | 0.056 | 1 | 0.293637566 | -0.776359032 | 0 | 0.024 | 1 | 0.858792447 | 0.501052112 |
| Gm57006 | 0.954841465 | 0.97333205 | 0.048 | 0.045 | 1 | 0.293637566 | -0.486852415 | 0 | 0.024 | 1 | 0.954841465 | 0.501052112 |
| Tex26 | 0.979644203 | 2.503846766 | 0.048 | 0.049 | 1 | 0.293637566 | -0.901889915 | 0 | 0.024 | 1 | 0.979644203 | 0.501052112 |
| Reep1 | 0.324739547 | -0.233118828 | 0 | 0.045 | 1 | 0.293647037 | -1.124282336 | 0 | 0.024 | 1 | 0.324739547 | 0.501065491 |
| Ankrd61 | 0.373857158 | 2.088809267 | 0.048 | 0.019 | 1 | 0.293647037 | -0.638855509 | 0 | 0.024 | 1 | 0.373857158 | 0.501065491 |
| Napepld | 0.396916958 | 0.214340149 | 0 | 0.034 | 1 | 0.293647037 | -0.638855509 | 0 | 0.024 | 1 | 0.396916958 | 0.501065491 |
| 1700022H01Rik | 0.396916958 | 0.214340149 | 0 | 0.034 | 1 | 0.293647037 | -0.638855509 | 0 | 0.024 | 1 | 0.396916958 | 0.501065491 |
| A230108P19Rik | 0.39695844 | -0.133583154 | 0 | 0.034 | 1 | 0.293647037 | -0.638855509 | 0 | 0.024 | 1 | 0.39695844 | 0.501065491 |
| Agtr1a | 0.39695844 | -0.133583154 | 0 | 0.034 | 1 | 0.293647037 | -0.776359032 | 0 | 0.024 | 1 | 0.39695844 | 0.501065491 |
| Ttbk1 | 0.425599726 | 0.214340149 | 0 | 0.03 | 1 | 0.293647037 | -1.124282336 | 0 | 0.024 | 1 | 0.425599726 | 0.501065491 |
| A630014C17Rik | 0.457292877 | 2.258734268 | 0.048 | 0.022 | 1 | 0.293647037 | -0.901889915 | 0 | 0.024 | 1 | 0.457292877 | 0.501065491 |
| 4732465J04Rik | 0.457292877 | 0.351843673 | 0 | 0.026 | 1 | 0.293647037 | -0.638855509 | 0 | 0.024 | 1 | 0.457292877 | 0.501065491 |
| Gm9970 | 0.457300763 | 0.214340149 | 0 | 0.026 | 1 | 0.293647037 | -0.638855509 | 0 | 0.024 | 1 | 0.457300763 | 0.501065491 |
| Zfp660 | 0.492590603 | 0.503846766 | 0 | 0.022 | 1 | 0.293647037 | -0.638855509 | 0 | 0.024 | 1 | 0.492590603 | 0.501065491 |
| Tbx4 | 0.492599419 | 0.351843673 | 0 | 0.022 | 1 | 0.293647037 | -0.638855509 | 0 | 0.024 | 1 | 0.492599419 | 0.501065491 |
| Gm56897 | 0.492599419 | 0.503846766 | 0 | 0.022 | 1 | 0.293647037 | -0.638855509 | 0 | 0.024 | 1 | 0.492599419 | 0.501065491 |
| Gm10190 | 0.492599419 | 0.503846766 | 0 | 0.022 | 1 | 0.293647037 | -0.638855509 | 0 | 0.024 | 1 | 0.492599419 | 0.501065491 |
| Gm56649 | 0.532426384 | 0.866416846 | 0 | 0.019 | 1 | 0.293647037 | -0.901889915 | 0 | 0.024 | 1 | 0.532426384 | 0.501065491 |
| Gja4 | 0.532432377 | 0.673771768 | 0 | 0.019 | 1 | 0.293647037 | -0.901889915 | 0 | 0.024 | 1 | 0.532432377 | 0.501065491 |
| Adamts7 | 0.578156445 | 1.088809267 | 0 | 0.015 | 1 | 0.293647037 | -0.638855509 | 0 | 0.024 | 1 | 0.578156445 | 0.501065491 |
| Tmem255a | 0.578156445 | 1.088809267 | 0 | 0.015 | 1 | 0.293647037 | -0.638855509 | 0 | 0.024 | 1 | 0.578156445 | 0.501065491 |
| Gask1a | 0.578159907 | 0.503846766 | 0 | 0.015 | 1 | 0.293647037 | -0.901889915 | 0 | 0.024 | 1 | 0.578159907 | 0.501065491 |
| Prodh | 0.57816106 | 1.351843673 | 0 | 0.015 | 1 | 0.293647037 | -0.638855509 | 0 | 0.024 | 1 | 0.57816106 | 0.501065491 |
| Mir133a-1hg | 0.57816106 | 1.351843673 | 0 | 0.015 | 1 | 0.293647037 | -0.638855509 | 0 | 0.024 | 1 | 0.57816106 | 0.501065491 |
| Nat2 | 0.629700127 | 2.088809267 | 0.048 | 0.03 | 1 | 0.293647037 | -0.638855509 | 0 | 0.024 | 1 | 0.629700127 | 0.501065491 |
| Ccdc187 | 0.632076134 | 1.673771768 | 0 | 0.011 | 1 | 0.293647037 | -0.901889915 | 0 | 0.024 | 1 | 0.632076134 | 0.501065491 |
| Stxbp5l | 0.632076134 | 1.088809267 | 0 | 0.011 | 1 | 0.293647037 | -0.776359032 | 0 | 0.024 | 1 | 0.632076134 | 0.501065491 |
| H4c14 | 0.653365386 | 1.150209812 | 0.095 | 0.071 | 1 | 0.293647037 | -0.901889915 | 0 | 0.024 | 1 | 0.653365386 | 0.501065491 |
| Gm56813 | 0.741589569 | 1.214340149 | 0.048 | 0.034 | 1 | 0.293647037 | -0.638855509 | 0 | 0.024 | 1 | 0.741589569 | 0.501065491 |
| Kif9 | 0.818482987 | 0.281454345 | 0.048 | 0.06 | 1 | 0.293647037 | -0.638855509 | 0 | 0.024 | 1 | 0.818482987 | 0.501065491 |
| Vil1 | 0.865290804 | 1.171271427 | 0.048 | 0.06 | 1 | 0.293647037 | -0.776359032 | 0 | 0.024 | 1 | 0.865290804 | 0.501065491 |
| Pole | 0.910096909 | 1.281454345 | 0.048 | 0.056 | 1 | 0.293647037 | -0.776359032 | 0 | 0.024 | 1 | 0.910096909 | 0.501065491 |
| Pkmyt1 | 0.933600076 | 1.304537958 | 0.095 | 0.093 | 1 | 0.293647037 | -0.776359032 | 0 | 0.024 | 1 | 0.933600076 | 0.501065491 |
| Gm2449 | 0.346868423 | -0.648156327 | 0 | 0.041 | 1 | 0.293652719 | -0.901889915 | 0 | 0.024 | 1 | 0.346868423 | 0.501073519 |
| Hsf5 | 0.425599726 | 0.351843673 | 0 | 0.03 | 1 | 0.293652719 | -1.017367132 | 0 | 0.024 | 1 | 0.425599726 | 0.501073519 |
| Btk | 0.457300763 | -0.326228232 | 0 | 0.026 | 1 | 0.293652719 | -1.486852415 | 0 | 0.024 | 1 | 0.457300763 | 0.501073519 |
| Pcdhb21 | 0.492599419 | 0.351843673 | 0 | 0.022 | 1 | 0.293652719 | -1.017367132 | 0 | 0.024 | 1 | 0.492599419 | 0.501073519 |
| Tube1 | 0.715883852 | 1.673771768 | 0.048 | 0.034 | 1 | 0.293652719 | -1.017367132 | 0 | 0.024 | 1 | 0.715883852 | 0.501073519 |
| Gm30085 | 0.954990531 | 0.866416846 | 0.048 | 0.052 | 1 | 0.293652719 | -0.901889915 | 0 | 0.024 | 1 | 0.954990531 | 0.501073519 |
| Tmem145 | 0.362165014 | 2.451379346 | 0.048 | 0.019 | 1 | 0.29365335 | -0.776359032 | 0 | 0.024 | 1 | 0.362165014 | 0.50107441 |
| Gm31728 | 0.370840523 | 0.214340149 | 0 | 0.037 | 1 | 0.29365335 | -1.223818009 | 0 | 0.024 | 1 | 0.370840523 | 0.50107441 |
| Cd1d1 | 0.396916958 | 0.214340149 | 0 | 0.034 | 1 | 0.29365335 | -1.017367132 | 0 | 0.024 | 1 | 0.396916958 | 0.50107441 |
| Gm45397 | 0.486438116 | 1.188344941 | 0.095 | 0.06 | 1 | 0.29365335 | -0.776359032 | 0 | 0.024 | 1 | 0.486438116 | 0.50107441 |

| | | | | | | | | | | | | |
|---|---|---|---|---|---|---|---|---|---|---|---|---|
| Gm57236 | 0.492590603 | 0.673771768 | 0 | 0.022 | 1 | 0.29365335 | -0.776359032 | 0 | 0.024 | 1 | 0.492590603 | 0.50107441 |
| Unc13a | 0.532436372 | 1.088809267 | 0 | 0.019 | 1 | 0.29365335 | -0.776359032 | 0 | 0.024 | 1 | 0.532436372 | 0.50107441 |
| Dlec1 | 0.578159907 | 0.866416846 | 0 | 0.015 | 1 | 0.29365335 | -0.776359032 | 0 | 0.024 | 1 | 0.578159907 | 0.50107441 |
| Sptssb | 0.57816106 | 1.351843673 | 0 | 0.015 | 1 | 0.29365335 | -1.124282336 | 0 | 0.024 | 1 | 0.57816106 | 0.50107441 |
| A730036I17Rik | 0.629733978 | 3.936806174 | 0.048 | 0.03 | 1 | 0.29365335 | -0.776359032 | 0 | 0.024 | 1 | 0.629733978 | 0.50107441 |
| Srl | 0.632076134 | 1.673771768 | 0 | 0.011 | 1 | 0.29365335 | -1.017367132 | 0 | 0.024 | 1 | 0.632076134 | 0.50107441 |
| Pcdhb10 | 0.632076134 | 1.673771768 | 0 | 0.011 | 1 | 0.29365335 | -1.223818009 | 0 | 0.024 | 1 | 0.632076134 | 0.50107441 |
| Lurap1 | 0.751343528 | 0.866416846 | 0.048 | 0.034 | 1 | 0.29365335 | -0.776359032 | 0 | 0.024 | 1 | 0.751343528 | 0.50107441 |
| Tox2 | 0.532436372 | 1.088809267 | 0 | 0.019 | 1 | 0.293654613 | -1.404390255 | 0 | 0.024 | 1 | 0.532436372 | 0.501076194 |
| Gm29865 | 0.457300763 | -0.413691074 | 0 | 0.026 | 1 | 0.293656507 | -1.316927414 | 0 | 0.024 | 1 | 0.457300763 | 0.50107887 |
| Slc26a8 | 0.492590603 | 0.673771768 | 0 | 0.022 | 1 | 0.293656507 | -1.223818009 | 0 | 0.024 | 1 | 0.492590603 | 0.50107887 |
| Adgrf2 | 0.492590603 | 0.673771768 | 0 | 0.022 | 1 | 0.293656507 | -1.223818009 | 0 | 0.024 | 1 | 0.492590603 | 0.50107887 |
| Gm43154 | 0.518280777 | -1.516052791 | 0.048 | 0.086 | 1 | 0.293656507 | -2.124282336 | 0 | 0.024 | 1 | 0.518280777 | 0.50107887 |
| Reln | 0.63207476 | 1.351843673 | 0 | 0.011 | 1 | 0.293656507 | -1.316927414 | 0 | 0.024 | 1 | 0.63207476 | 0.50107887 |
| St8sia4 | 0.532434375 | 0.214340149 | 0 | 0.019 | 1 | 0.293657138 | -1.223818009 | 0 | 0.024 | 1 | 0.532434375 | 0.501079762 |
| Notch4 | 0.324739547 | -0.133583154 | 0 | 0.045 | 1 | 0.293659664 | -0.316927414 | 0 | 0.024 | 1 | 0.324739547 | 0.501083329 |
| Tslp | 0.324767242 | -0.413691074 | 0 | 0.045 | 1 | 0.293659664 | -0.316927414 | 0 | 0.024 | 1 | 0.324767242 | 0.501083329 |
| Bfsp2 | 0.346827513 | -0.233118828 | 0 | 0.041 | 1 | 0.293659664 | -0.316927414 | 0 | 0.024 | 1 | 0.346827513 | 0.501083329 |
| Pllp | 0.346847151 | -0.233118828 | 0 | 0.041 | 1 | 0.293659664 | -0.316927414 | 0 | 0.024 | 1 | 0.346847151 | 0.501083329 |
| Gm43260 | 0.359270744 | 2.673771768 | 0.048 | 0.019 | 1 | 0.293659664 | -0.316927414 | 0 | 0.024 | 1 | 0.359270744 | 0.501083329 |
| C230038L03Rik | 0.359270744 | 2.673771768 | 0.048 | 0.019 | 1 | 0.293659664 | -0.316927414 | 0 | 0.024 | 1 | 0.359270744 | 0.501083329 |
| Pld4 | 0.359280535 | 3.088809267 | 0.048 | 0.019 | 1 | 0.293659664 | -0.316927414 | 0 | 0.024 | 1 | 0.359280535 | 0.501083329 |
| 9330136K24Rik | 0.370790696 | -0.326228232 | 0 | 0.037 | 1 | 0.293659664 | -0.316927414 | 0 | 0.024 | 1 | 0.370790696 | 0.501083329 |
| Fadd | 0.425599726 | 0.351843673 | 0 | 0.03 | 1 | 0.293659664 | -0.316927414 | 0 | 0.024 | 1 | 0.425599726 | 0.501083329 |
| Atp6v0d2 | 0.425624656 | -0.574155746 | 0 | 0.03 | 1 | 0.293659664 | -4.996407513 | 0 | 0.024 | 1 | 0.425624656 | 0.501083329 |
| Cdca8 | 0.425636051 | -2.348596045 | 0 | 0.03 | 1 | 0.293659664 | -0.316927414 | 0 | 0.024 | 1 | 0.425636051 | 0.501083329 |
| Lmtk3 | 0.425639612 | 0.503846766 | 0 | 0.03 | 1 | 0.293659664 | -0.316927414 | 0 | 0.024 | 1 | 0.425639612 | 0.501083329 |
| Gm56728 | 0.425639612 | 0.503846766 | 0 | 0.03 | 1 | 0.293659664 | -0.316927414 | 0 | 0.024 | 1 | 0.425639612 | 0.501083329 |
| Zfp973 | 0.454205211 | 2.451379346 | 0.048 | 0.022 | 1 | 0.293659664 | -0.316927414 | 0 | 0.024 | 1 | 0.454205211 | 0.501083329 |
| Gm12781 | 0.457281049 | 0.503846766 | 0 | 0.026 | 1 | 0.293659664 | -0.316927414 | 0 | 0.024 | 1 | 0.457281049 | 0.501083329 |
| Dlk2 | 0.457292877 | 0.351843673 | 0 | 0.026 | 1 | 0.293659664 | -0.316927414 | 0 | 0.024 | 1 | 0.457292877 | 0.501083329 |
| Cmtr2 | 0.457308648 | 0.673771768 | 0 | 0.026 | 1 | 0.293659664 | -0.316927414 | 0 | 0.024 | 1 | 0.457308648 | 0.501083329 |
| Gm56859 | 0.457308648 | 0.673771768 | 0 | 0.026 | 1 | 0.293659664 | -0.316927414 | 0 | 0.024 | 1 | 0.457308648 | 0.501083329 |
| Hhipl1 | 0.457308648 | 0.673771768 | 0 | 0.026 | 1 | 0.293659664 | -0.316927414 | 0 | 0.024 | 1 | 0.457308648 | 0.501083329 |
| Cdk20 | 0.457308648 | 0.673771768 | 0 | 0.026 | 1 | 0.293659664 | -0.316927414 | 0 | 0.024 | 1 | 0.457308648 | 0.501083329 |
| D630039A03Rik | 0.472861597 | 1.866416846 | 0.048 | 0.022 | 1 | 0.293659664 | -0.316927414 | 0 | 0.024 | 1 | 0.472861597 | 0.501083329 |
| Cerox1 | 0.4759797 | 1.503846766 | 0.048 | 0.022 | 1 | 0.293659664 | -0.316927414 | 0 | 0.024 | 1 | 0.4759797 | 0.501083329 |
| Prrg3 | 0.492590603 | 0.673771768 | 0 | 0.022 | 1 | 0.293659664 | -0.316927414 | 0 | 0.024 | 1 | 0.492590603 | 0.501083329 |
| Gm2814 | 0.492608234 | 0.866416846 | 0 | 0.022 | 1 | 0.293659664 | -0.316927414 | 0 | 0.024 | 1 | 0.492608234 | 0.501083329 |
| Zfp3 | 0.492608234 | 0.866416846 | 0 | 0.022 | 1 | 0.293659664 | -0.316927414 | 0 | 0.024 | 1 | 0.492608234 | 0.501083329 |
| 9530062K07Rik | 0.532432377 | 0.503846766 | 0 | 0.019 | 1 | 0.293659664 | -0.316927414 | 0 | 0.024 | 1 | 0.532432377 | 0.501083329 |
| Fgf20 | 0.532436372 | 1.088809267 | 0 | 0.019 | 1 | 0.293659664 | -0.316927414 | 0 | 0.024 | 1 | 0.532436372 | 0.501083329 |
| Cxcr5 | 0.532436372 | 1.088809267 | 0 | 0.019 | 1 | 0.293659664 | -0.316927414 | 0 | 0.024 | 1 | 0.532436372 | 0.501083329 |
| Gm12111 | 0.532436372 | 1.088809267 | 0 | 0.019 | 1 | 0.293659664 | -0.316927414 | 0 | 0.024 | 1 | 0.532436372 | 0.501083329 |
| Med9os | 0.532436372 | 1.088809267 | 0 | 0.019 | 1 | 0.293659664 | -0.316927414 | 0 | 0.024 | 1 | 0.532436372 | 0.501083329 |
| C030010C08Rik | 0.532436372 | 1.088809267 | 0 | 0.019 | 1 | 0.293659664 | -0.316927414 | 0 | 0.024 | 1 | 0.532436372 | 0.501083329 |
| A730090N16Rik | 0.550953306 | 1.936806174 | 0.048 | 0.026 | 1 | 0.293659664 | -0.316927414 | 0 | 0.024 | 1 | 0.550953306 | 0.501083329 |
| 4930563E22Rik | 0.567009484 | 1.673771768 | 0.048 | 0.026 | 1 | 0.293659664 | -0.316927414 | 0 | 0.024 | 1 | 0.567009484 | 0.501083329 |
| Gm37233 | 0.570211706 | 1.214340149 | 0.048 | 0.026 | 1 | 0.293659664 | -0.316927414 | 0 | 0.024 | 1 | 0.570211706 | 0.501083329 |
| Gm50250 | 0.578159907 | 0.866416846 | 0 | 0.015 | 1 | 0.293659664 | -0.316927414 | 0 | 0.024 | 1 | 0.578159907 | 0.501083329 |
| Gm56824 | 0.57816106 | 1.351843673 | 0 | 0.015 | 1 | 0.293659664 | -0.316927414 | 0 | 0.024 | 1 | 0.57816106 | 0.501083329 |
| 5033425G24Rik | 0.57816106 | 1.351843673 | 0 | 0.015 | 1 | 0.293659664 | -0.316927414 | 0 | 0.024 | 1 | 0.57816106 | 0.501083329 |
| Nnt | 0.57816106 | 1.351843673 | 0 | 0.015 | 1 | 0.293659664 | -0.316927414 | 0 | 0.024 | 1 | 0.57816106 | 0.501083329 |
| Gm5608 | 0.629713293 | 2.673771768 | 0.048 | 0.03 | 1 | 0.293659664 | -0.316927414 | 0 | 0.024 | 1 | 0.629713293 | 0.501083329 |
| 9130221H12Rik | 0.632076134 | 1.673771768 | 0 | 0.011 | 1 | 0.293659664 | -0.316927414 | 0 | 0.024 | 1 | 0.632076134 | 0.501083329 |
| 1500026H17Rik | 0.632076134 | 1.673771768 | 0 | 0.011 | 1 | 0.293659664 | -0.316927414 | 0 | 0.024 | 1 | 0.632076134 | 0.501083329 |
| Mapk8ip2 | 0.632076134 | 1.673771768 | 0 | 0.011 | 1 | 0.293659664 | -0.316927414 | 0 | 0.024 | 1 | 0.632076134 | 0.501083329 |
| Ctsw | 0.770713471 | 0.351843673 | 0.048 | 0.063 | 1 | 0.293659664 | -0.316927414 | 0 | 0.024 | 1 | 0.770713471 | 0.501083329 |
| Odad4 | 0.985459372 | 1.258734268 | 0.048 | 0.049 | 1 | 0.293659664 | -0.316927414 | 0 | 0.024 | 1 | 0.985459372 | 0.501083329 |
| Thsd1 | 0.293851038 | 1.586308926 | 0.095 | 0.045 | 1 | 0.3304229 | -0.686161223 | 0.044 | 0.087 | 1 | 0.3304229 | 0.501353643 |
| A830082N09Rik | 0.294097203 | 1.55829455 | 0.143 | 0.082 | 1 | 0.54460932 | 1.946106992 | 0.067 | 0.048 | 1 | 0.54460932 | 0.501701241 |
| 1700109H08Rik | 0.91718805 | 0.625408746 | 0.238 | 0.265 | 1 | 0.294190302 | 0.132875504 | 0.2 | 0.301 | 1 | 0.91718805 | 0.501832671 |
| Abhd2 | 0.294279944 | 0.693642971 | 0.857 | 0.944 | 1 | 0.923163619 | 0.313530607 | 0.978 | 0.962 | 1 | 0.923163619 | 0.501959203 |
| 2700097O09Rik | 0.416576078 | 0.866416846 | 0.286 | 0.243 | 1 | 0.294467977 | 0.42003818 | 0.178 | 0.287 | 1 | 0.416576078 | 0.502224564 |
| Coq4 | 0.710123498 | 0.97333205 | 0.095 | 0.075 | 1 | 0.29451315 | 0.638678467 | 0.044 | 0.097 | 1 | 0.710123498 | 0.502288305 |
| Zfp61 | 0.954992271 | 0.936806174 | 0.048 | 0.052 | 1 | 0.294560165 | 0.595609745 | 0.044 | 0.097 | 1 | 0.954992271 | 0.502354639 |
| Zfp952 | 0.294625684 | -0.818081329 | 0.048 | 0.123 | 1 | 0.498180161 | 0.500869255 | 0.111 | 0.163 | 1 | 0.498180161 | 0.502447075 |
| Usp2 | 0.294752202 | -0.61573485 | 0.19 | 0.325 | 1 | 0.780857624 | 0.548423059 | 0.178 | 0.215 | 1 | 0.780857624 | 0.502625544 |
| 4933439C10Rik | 0.993863205 | 0.951305743 | 0.095 | 0.104 | 1 | 0.294834829 | -0.316927414 | 0.044 | 0.093 | 1 | 0.993863205 | 0.502742082 |
| Zmym6 | 0.51320005 | 0.738866796 | 0.238 | 0.187 | 1 | 0.295054627 | -0.251039478 | 0.111 | 0.18 | 1 | 0.51320005 | 0.503052022 |
| Lat2 | 0.542240288 | 1.425844254 | 0.095 | 0.063 | 1 | 0.295149029 | -0.174908409 | 0.067 | 0.125 | 1 | 0.542240288 | 0.503185109 |
| Gm16853 | 0.457281049 | 0.503846766 | 0 | 0.026 | 1 | 0.295331356 | 0.808603468 | 0.044 | 0.097 | 1 | 0.457281049 | 0.503442102 |
| C920021L13Rik | 0.295522534 | 1.889500459 | 0.143 | 0.082 | 1 | 0.322427593 | 0.638678467 | 0.044 | 0.093 | 1 | 0.322427593 | 0.5037115 |
| Lrrk1 | 0.810179215 | 0.168916006 | 0.571 | 0.642 | 1 | 0.2958738 | -0.215329274 | 0.622 | 0.754 | 1 | 0.810179215 | 0.504206294 |
| Rabif | 0.883586874 | 0.487905222 | 0.238 | 0.254 | 1 | 0.295953427 | 0.634162986 | 0.178 | 0.284 | 1 | 0.883586874 | 0.504318423 |
| Zfp964 | 0.457281049 | 0.503846766 | 0 | 0.026 | 1 | 0.29596827 | 1.268035087 | 0.067 | 0.035 | 1 | 0.457281049 | 0.504339324 |
| Med27 | 0.661564184 | 0.655235816 | 0.333 | 0.489 | 1 | 0.296093494 | 0.261245921 | 0.333 | 0.526 | 1 | 0.661564184 | 0.504515631 |
| Wdr90 | 0.36450864 | 1.936806174 | 0.095 | 0.052 | 1 | 0.296352763 | -0.271123724 | 0.044 | 0.093 | 1 | 0.36450864 | 0.504880566 |
| Tmem87a | 0.547271157 | 0.773607035 | 0.524 | 0.582 | 1 | 0.296362522 | 0.511561334 | 0.867 | 0.73 | 1 | 0.547271157 | 0.5048943 |
| Papln | 0.709500188 | 1.995699863 | 0.048 | 0.034 | 1 | 0.29637443 | -0.191396532 | 0.022 | 0.062 | 1 | 0.709500188 | 0.504911057 |
| Pnpla7 | 0.296691399 | -0.295854583 | 0.048 | 0.131 | 1 | 0.304563689 | 0.424338318 | 0.089 | 0.159 | 1 | 0.304563689 | 0.505357011 |
| Atf2 | 0.296786837 | -0.174702142 | 0.619 | 0.754 | 1 | 0.693009816 | 0.160347218 | 0.933 | 0.858 | 1 | 0.693009816 | 0.505491247 |
| Smad4 | 0.29680603 | 0.380990018 | 0.905 | 0.892 | 1 | 0.607132141 | 0.274267041 | 1 | 0.969 | 1 | 0.607132141 | 0.505518241 |
| Ino80c | 0.852936434 | 0.48112669 | 0.19 | 0.183 | 1 | 0.297014369 | -0.237492946 | 0.089 | 0.152 | 1 | 0.852936434 | 0.505811203 |
| Il1rapl1 | 0.297216488 | -0.208871282 | 0.048 | 0.131 | 1 | 0.802406533 | 1.402964667 | 0.067 | 0.059 | 1 | 0.802406533 | 0.506095335 |
| Syt2 | 0.459388879 | -0.14822993 | 0.048 | 0.101 | 1 | 0.297521903 | 1.402964667 | 0.133 | 0.09 | 1 | 0.459388879 | 0.506524523 |
| Slc39a10 | 0.297640109 | -0.445967477 | 0.238 | 0.358 | 1 | 0.365449701 | 0.143913775 | 0.556 | 0.457 | 1 | 0.365449701 | 0.506690584 |
| Zfp951 | 0.297653035 | -0.233118828 | 0.048 | 0.131 | 1 | 0.603701633 | 1.313838777 | 0.089 | 0.069 | 1 | 0.603701633 | 0.506708741 |
| Lgals4 | 0.297782951 | -0.233118828 | 0.048 | 0.131 | 1 | 0.612663423 | 0.887892378 | 0.111 | 0.156 | 1 | 0.612663423 | 0.506891216 |
| Cacul1 | 0.297825941 | 1.075512444 | 0.429 | 0.444 | 1 | 0.42541309 | 0.231759701 | 0.8 | 0.668 | 1 | 0.42541309 | 0.50695159 |
| Nek10 | 0.411201324 | -0.970084422 | 0.048 | 0.101 | 1 | 0.298202733 | 0.417178526 | 0.067 | 0.121 | 1 | 0.411201324 | 0.507480597 |
| Gm43727 | 0.532436372 | 1.088809267 | 0 | 0.019 | 1 | 0.298232319 | 1.304560963 | 0.067 | 0.035 | 1 | 0.532436372 | 0.507522121 |
| 2900009J06Rik | 0.727548938 | 0.79930265 | 0.048 | 0.071 | 1 | 0.298356681 | 0.313838777 | 0.022 | 0.062 | 1 | 0.727548938 | 0.507696654 |
| Glis1 | 0.298439082 | -0.170577362 | 0.095 | 0.198 | 1 | 0.887908276 | 1.345024826 | 0.244 | 0.298 | 1 | 0.887908276 | 0.507812278 |
| Pp2d1 | 0.346827513 | -0.133583154 | 0 | 0.041 | 1 | 0.299191224 | 0.361144491 | 0.022 | 0.062 | 1 | 0.346827513 | 0.50886706 |
| Gm26724 | 0.544576436 | 2.258734268 | 0.048 | 0.026 | 1 | 0.299191224 | 0.361144491 | 0.022 | 0.062 | 1 | 0.544576436 | 0.50886706 |
| Gm16845 | 0.818185674 | 1.281454345 | 0.048 | 0.063 | 1 | 0.299191224 | 1.361144491 | 0.022 | 0.062 | 1 | 0.818185674 | 0.50886706 |
| 4930556J24Rik | 0.346860242 | -0.496153234 | 0 | 0.041 | 1 | 0.299223847 | 0.290755163 | 0.022 | 0.062 | 1 | 0.346860242 | 0.508912784 |

| | | | | | | | | | | | | |
|---|---|---|---|---|---|---|---|---|---|---|---|---|
| Gm15956 | 0.57816106 | 1.351843673 | 0 | 0.015 | 1 | 0.299223847 | 0.290755163 | 0.022 | 0.062 | 1 | 0.57816106 | 0.508912784 |
| Slc7a8 | 0.655561847 | 1.503846766 | 0.048 | 0.03 | 1 | 0.299252842 | 0.744473131 | 0.022 | 0.062 | 1 | 0.655561847 | 0.50895342 |
| Hes1 | 0.299364793 | -0.50364777 | 0.571 | 0.358 | 1 | 0.324261498 | -0.374194615 | 0.756 | 0.529 | 1 | 0.324261498 | 0.509110306 |
| Cry2 | 0.300032316 | 0.664929891 | 0.81 | 0.802 | 1 | 0.566553652 | 0.115581222 | 0.711 | 0.851 | 1 | 0.566553652 | 0.510045242 |
| Galnt11 | 0.300067867 | -0.277318632 | 0.381 | 0.556 | 1 | 0.94413617 | 0.783236748 | 0.444 | 0.588 | 1 | 0.94413617 | 0.510095009 |
| Npm2 | 0.324739547 | -0.133583154 | 0 | 0.045 | 1 | 0.300659678 | -0.686161223 | 0.044 | 0.09 | 1 | 0.324739547 | 0.510923114 |
| Zfp775 | 0.300722381 | 1.936806174 | 0.143 | 0.082 | 1 | 0.346923896 | 1.442064487 | 0.089 | 0.055 | 1 | 0.346923896 | 0.511010811 |
| Dnal1 | 0.301303058 | 0.673771768 | 0.095 | 0.194 | 1 | 0.334881245 | 0.346037599 | 0.156 | 0.235 | 1 | 0.334881245 | 0.511822583 |
| Mad1l1 | 0.717289702 | 0.791128718 | 0.19 | 0.265 | 1 | 0.301361415 | 1.189664023 | 0.444 | 0.491 | 1 | 0.717289702 | 0.511904128 |
| Ccdc171 | 0.301365897 | 1.115776315 | 0.19 | 0.123 | 1 | 0.632134322 | 0.618942249 | 0.067 | 0.093 | 1 | 0.632134322 | 0.51191039 |
| Nckap5 | 0.301848266 | 1.43372068 | 0.286 | 0.216 | 1 | 0.447772154 | 0.499850762 | 0.222 | 0.308 | 1 | 0.447772154 | 0.512584156 |
| Arv1 | 0.301945417 | 1.400753273 | 0.143 | 0.082 | 1 | 0.486312983 | -0.255526869 | 0.044 | 0.073 | 1 | 0.486312983 | 0.512719799 |
| Ap4e1 | 0.302225294 | 1.553477534 | 0.286 | 0.224 | 1 | 0.996818444 | 0.558376839 | 0.289 | 0.339 | 1 | 0.996818444 | 0.51311046 |
| Col4a6 | 0.302345175 | 1.150209812 | 0.095 | 0.045 | 1 | 0.430483494 | -0.638855509 | 0 | 0.014 | 1 | 0.430483494 | 0.513277745 |
| Gm57189 | 0.355410306 | 1.088809267 | 0.095 | 0.049 | 1 | 0.302547716 | 1.481438725 | 0.156 | 0.114 | 1 | 0.355410306 | 0.513560311 |
| Pcdhb7 | 0.304462706 | -0.849790188 | 0 | 0.049 | 1 | 0.302694877 | -1.743192169 | 0.022 | 0.059 | 1 | 0.304462706 | 0.513765565 |
| Rftn1 | 0.565993325 | 0.351843673 | 0.143 | 0.101 | 1 | 0.302851537 | -0.450783161 | 0.067 | 0.125 | 1 | 0.565993325 | 0.513984021 |
| Trpm3 | 0.303232183 | -1.341178574 | 0.048 | 0.127 | 1 | 0.32624829 | -0.486852415 | 0.178 | 0.249 | 1 | 0.32624829 | 0.514514609 |
| Ccdc57 | 0.303303393 | -0.522625445 | 0.048 | 0.127 | 1 | 0.830484947 | 0.15951063 | 0.111 | 0.125 | 1 | 0.830484947 | 0.514613838 |
| Anapc1 | 0.303392875 | -0.385121921 | 0.381 | 0.526 | 1 | 0.608303767 | 0.194786105 | 0.533 | 0.702 | 1 | 0.608303767 | 0.514738513 |
| Tef | 0.303511245 | -0.192961701 | 0.333 | 0.522 | 1 | 0.569980243 | 0.199140006 | 0.533 | 0.651 | 1 | 0.569980243 | 0.514903414 |
| Dlg5 | 0.303606304 | 0.890869889 | 0.333 | 0.265 | 1 | 0.460892209 | 0.447624012 | 0.489 | 0.474 | 1 | 0.460892209 | 0.51503582 |
| Atp6v0d1 | 0.304015422 | 0.135147982 | 0.429 | 0.653 | 1 | 0.457166852 | 0.269770138 | 0.8 | 0.803 | 1 | 0.457166852 | 0.515605467 |
| Creb3l2 | 0.539217129 | 0.897812042 | 0.952 | 0.993 | 1 | 0.304264492 | 0.685975394 | 0.978 | 0.99 | 1 | 0.539217129 | 0.515952103 |
| Gm43149 | 0.304355016 | -0.233118828 | 0 | 0.049 | 1 | 0.730158045 | 0.982632868 | 0.044 | 0.035 | 1 | 0.730158045 | 0.516078056 |
| Zik1 | 0.304355016 | -0.233118828 | 0 | 0.049 | 1 | 0.681309135 | 0.460680165 | 0.022 | 0.035 | 1 | 0.681309135 | 0.516078056 |
| E530011L22Rik | 0.304355016 | -0.233118828 | 0 | 0.049 | 1 | 0.331250628 | 0.757073168 | 0.022 | 0.059 | 1 | 0.331250628 | 0.516078056 |
| Gm57395 | 0.304355016 | -0.233118828 | 0 | 0.049 | 1 | 0.420889035 | 1.875717664 | 0.067 | 0.042 | 1 | 0.420889035 | 0.516078056 |
| Ranbp3l | 0.304355016 | -0.574155746 | 0 | 0.049 | 1 | 0.527689453 | -0.564854927 | 0.022 | 0.042 | 1 | 0.527689453 | 0.516078056 |
| Gm49085 | 0.304355016 | -0.233118828 | 0 | 0.049 | 1 | 0.685243348 | 0.875717664 | 0.022 | 0.035 | 1 | 0.685243348 | 0.516078056 |
| Ap2b1 | 0.810195259 | 0.437573547 | 0.619 | 0.795 | 1 | 0.304359064 | 0.118287967 | 0.844 | 0.834 | 1 | 0.810195259 | 0.516083688 |
| Asphd2 | 0.304385789 | -0.326228232 | 0 | 0.049 | 1 | 0.376837034 | -0.124282336 | 0 | 0.017 | 1 | 0.376837034 | 0.51612087 |
| Tmc4 | 0.304385789 | -0.326228232 | 0 | 0.049 | 1 | 0.613121994 | 0.513147585 | 0.067 | 0.093 | 1 | 0.613121994 | 0.51612087 |
| Tmem143 | 0.304385789 | -0.413691074 | 0 | 0.049 | 1 | 0.416068746 | 0.735540006 | 0.044 | 0.083 | 1 | 0.416068746 | 0.51612087 |
| Dna2 | 0.304385789 | -0.326228232 | 0 | 0.049 | 1 | 0.715891525 | 1.513147585 | 0.067 | 0.055 | 1 | 0.715891525 | 0.51612087 |
| Shroom1 | 0.304385789 | -0.413691074 | 0 | 0.049 | 1 | 0.387952149 | -0.316927414 | 0.022 | 0.052 | 1 | 0.387952149 | 0.51612087 |
| Gm12248 | 0.304385789 | -0.326228232 | 0 | 0.049 | 1 | 0.549167674 | 0.76880246 | 0.089 | 0.128 | 1 | 0.549167674 | 0.51612087 |
| Snhg10 | 0.304385789 | -0.326228232 | 0 | 0.049 | 1 | 0.687197015 | 1.223640968 | 0.022 | 0.035 | 1 | 0.687197015 | 0.51612087 |
| Cabyr | 0.304385789 | -0.413691074 | 0 | 0.049 | 1 | 0.712836275 | 0.197645759 | 0.044 | 0.059 | 1 | 0.712836275 | 0.51612087 |
| Med20 | 0.636651566 | 0.644024424 | 0.286 | 0.284 | 1 | 0.304399043 | 0.276225594 | 0.244 | 0.398 | 1 | 0.636651566 | 0.516139309 |
| Asf1b | 0.304410965 | -0.718545655 | 0 | 0.049 | 1 | 0.76573443 | 1.304560963 | 0.022 | 0.031 | 1 | 0.76573443 | 0.516155894 |
| Ybey | 0.304410965 | -0.496153234 | 0 | 0.049 | 1 | 0.313790052 | -0.486852415 | 0.022 | 0.059 | 1 | 0.313790052 | 0.516155894 |
| Cyp2j9 | 0.304411431 | -0.413691074 | 0 | 0.049 | 1 | 0.430487676 | 0.361144491 | 0 | 0.014 | 1 | 0.430487676 | 0.516156543 |
| C030034L19Rik | 0.304430544 | -0.574155746 | 0 | 0.049 | 1 | 0.764217967 | 0.709067795 | 0.133 | 0.125 | 1 | 0.764217967 | 0.516183132 |
| Map3k21 | 0.304430544 | -0.574155746 | 0 | 0.049 | 1 | 0.726758375 | 1.03099589 | 0.067 | 0.055 | 1 | 0.726758375 | 0.516183132 |
| Gm53 | 0.304430544 | -0.574155746 | 0 | 0.049 | 1 | 0.952529269 | 1.361144491 | 0.022 | 0.024 | 1 | 0.952529269 | 0.516183132 |
| Mastl | 0.304431942 | -0.911190733 | 0 | 0.049 | 1 | 0.714378739 | 0.223640968 | 0.044 | 0.059 | 1 | 0.714378739 | 0.516185077 |
| 2610206C17Rik | 0.304431942 | -0.785659851 | 0 | 0.049 | 1 | 0.75717489 | 0.435145073 | 0.022 | 0.031 | 1 | 0.75717489 | 0.516185077 |
| 1700030J22Rik | 0.304432409 | -0.496153234 | 0 | 0.049 | 1 | 0.331922009 | -0.124282336 | 0 | 0.021 | 1 | 0.331922009 | 0.516185726 |
| Gm36033 | 0.304432409 | -0.496153234 | 0 | 0.049 | 1 | 0.965278751 | 0.960606562 | 0.089 | 0.09 | 1 | 0.965278751 | 0.516185726 |
| Slc16a6 | 0.304432409 | -0.496153234 | 0 | 0.049 | 1 | 0.858194269 | -0.181997834 | 0.044 | 0.052 | 1 | 0.858194269 | 0.516185726 |
| Rcor2 | 0.304444528 | -0.648156327 | 0 | 0.049 | 1 | 0.617556294 | 1.005000681 | 0.022 | 0.038 | 1 | 0.617556294 | 0.516202586 |
| Trim17 | 0.304449189 | -0.574155746 | 0 | 0.049 | 1 | 0.406607361 | 0.513147585 | 0.022 | 0.052 | 1 | 0.406607361 | 0.51620907 |
| Ttll10 | 0.304452918 | -0.785659851 | 0 | 0.049 | 1 | 0.450515412 | 0.683072586 | 0.067 | 0.111 | 1 | 0.450515412 | 0.516214257 |
| Prss35 | 0.30445991 | -1.280424543 | 0 | 0.049 | 1 | 0.996309481 | 0.481438725 | 0.044 | 0.045 | 1 | 0.996309481 | 0.516223983 |
| Gpr137c | 0.304460376 | -1.370622352 | 0 | 0.049 | 1 | 0.930306355 | 0.320502507 | 0.044 | 0.048 | 1 | 0.930306355 | 0.516224631 |
| E130114P18Rik | 0.304466901 | -1.233118828 | 0 | 0.049 | 1 | 0.476265768 | -0.404390255 | 0.022 | 0.045 | 1 | 0.476265768 | 0.516233708 |
| Eda2r | 0.304475291 | -1.496153234 | 0 | 0.049 | 1 | 0.823047007 | 0.683072586 | 0.022 | 0.017 | 1 | 0.823047007 | 0.516245379 |
| Ifi44 | 0.304477155 | -1.998653574 | 0 | 0.049 | 1 | 0.80624346 | 5.005000681 | 0.022 | 0.017 | 1 | 0.80624346 | 0.516247972 |
| D130017N08Rik | 0.30448834 | -0.133583154 | 0 | 0.049 | 1 | 0.885635102 | 1.513147585 | 0.044 | 0.052 | 1 | 0.885635102 | 0.516263531 |
| Atad5 | 0.606804665 | 0.766881172 | 0.095 | 0.067 | 1 | 0.304676439 | -1.602329633 | 0.022 | 0.059 | 1 | 0.606804665 | 0.516525145 |
| Stx6 | 0.766995324 | 0.599224814 | 0.381 | 0.485 | 1 | 0.304722696 | 0.515127949 | 0.6 | 0.509 | 1 | 0.766995324 | 0.516589471 |
| 2900076A07Rik | 0.416600843 | -0.208871282 | 0.095 | 0.168 | 1 | 0.305013483 | 0.223640968 | 0.133 | 0.218 | 1 | 0.416600843 | 0.516993741 |
| Zfp710 | 0.579393515 | 0.843696769 | 0.19 | 0.157 | 1 | 0.305671831 | 0.632446513 | 0.4 | 0.336 | 1 | 0.579393515 | 0.517908394 |
| Ripk2 | 0.668728113 | 2.153514956 | 0.238 | 0.25 | 1 | 0.305751961 | 1.050804371 | 0.2 | 0.298 | 1 | 0.668728113 | 0.51801966 |
| Trp53rkb | 0.527502744 | 0.55829455 | 0.048 | 0.093 | 1 | 0.305961239 | -0.602329633 | 0.067 | 0.118 | 1 | 0.527502744 | 0.518310198 |
| Fip1l1 | 0.377567013 | 0.214340149 | 0.524 | 0.705 | 1 | 0.30615816 | 0.65806858 | 0.844 | 0.754 | 1 | 0.377567013 | 0.518583501 |
| Gm15494 | 0.365063958 | 2.258734268 | 0.048 | 0.019 | 1 | 0.306315797 | 1.098110085 | 0.067 | 0.035 | 1 | 0.365063958 | 0.518802226 |
| Zfp277 | 0.306649556 | 0.351843673 | 0.19 | 0.332 | 1 | 0.778943637 | 0.385392038 | 0.378 | 0.471 | 1 | 0.778943637 | 0.519265161 |
| Gm28809 | 0.765961976 | 0.866416846 | 0.095 | 0.078 | 1 | 0.306650846 | 1.325520582 | 0.156 | 0.114 | 1 | 0.765961976 | 0.519266951 |
| 2700069I18Rik | 0.425614684 | 0.214340149 | 0 | 0.03 | 1 | 0.306650922 | -1.124282336 | 0.022 | 0.059 | 1 | 0.425614684 | 0.519267056 |
| Zfp346 | 0.59712679 | 0.943378827 | 0.238 | 0.209 | 1 | 0.307203528 | 0.578735926 | 0.133 | 0.215 | 1 | 0.59712679 | 0.520033048 |
| Siah1a | 0.555633177 | 0.433457438 | 0.286 | 0.455 | 1 | 0.307325437 | 0.415300261 | 0.667 | 0.606 | 1 | 0.555633177 | 0.520201949 |
| Pou2f2 | 0.307536988 | 1.614878079 | 0.095 | 0.045 | 1 | 0.434761384 | 2.202446745 | 0.089 | 0.059 | 1 | 0.434761384 | 0.520494977 |
| Ppp4r1l-ps | 0.89512561 | 0.460778044 | 0.381 | 0.44 | 1 | 0.307861146 | 0.242499995 | 0.311 | 0.464 | 1 | 0.89512561 | 0.520943806 |
| Serpine1 | 0.499856364 | 0.183805781 | 0.429 | 0.351 | 1 | 0.307892823 | -1.11739595 | 0.156 | 0.228 | 1 | 0.499856364 | 0.520987656 |
| Supt5 | 0.805153552 | 0.742484518 | 0.571 | 0.608 | 1 | 0.308032521 | -0.238768351 | 0.778 | 0.727 | 1 | 0.805153552 | 0.521181008 |
| Irx5 | 0.308222433 | 1.952925839 | 0.048 | 0.131 | 1 | 0.877720746 | 0.450411829 | 0.067 | 0.062 | 1 | 0.877720746 | 0.521443798 |
| Pigg | 0.953154927 | 0.472137907 | 0.143 | 0.153 | 1 | 0.308500406 | -0.539319835 | 0.089 | 0.145 | 1 | 0.953154927 | 0.521828311 |
| St3gal3 | 0.446162164 | 1.2926816 | 0.286 | 0.422 | 1 | 0.308593636 | 0.205025289 | 0.289 | 0.415 | 1 | 0.446162164 | 0.52195724 |
| Nova1 | 0.50066303 | 1.258734268 | 0.095 | 0.06 | 1 | 0.308629768 | -1.017367132 | 0.022 | 0.059 | 1 | 0.50066303 | 0.522007203 |
| Ppme1 | 0.309002384 | 0.45953787 | 0.762 | 0.754 | 1 | 0.588780279 | 0.189083745 | 0.733 | 0.727 | 1 | 0.588780279 | 0.522522295 |
| Rnpc3 | 0.686455162 | 0.40230874 | 0.524 | 0.627 | 1 | 0.309005133 | 0.21896448 | 0.356 | 0.564 | 1 | 0.686455162 | 0.522526093 |
| Akap11 | 0.30951379 | -0.291333442 | 0.571 | 0.784 | 1 | 0.522050803 | 0.18646596 | 0.622 | 0.744 | 1 | 0.522050803 | 0.523228794 |
| Il15 | 0.57816106 | 1.351843673 | 0 | 0.015 | 1 | 0.309616342 | -0.960783604 | 0.022 | 0.059 | 1 | 0.57816106 | 0.523370405 |
| Cacng5 | 0.359280535 | 3.451379346 | 0.048 | 0.019 | 1 | 0.309824707 | 2.683072586 | 0.022 | 0.007 | 1 | 0.359280535 | 0.523658065 |
| Slc5a10 | 0.367982232 | 2.088809267 | 0.048 | 0.019 | 1 | 0.309824707 | 3.098110085 | 0.022 | 0.007 | 1 | 0.367982232 | 0.523658065 |
| Cd27 | 0.457281049 | 0.503846766 | 0 | 0.026 | 1 | 0.309824707 | 3.098110085 | 0.022 | 0.007 | 1 | 0.457281049 | 0.523658065 |
| Gm56633 | 0.492608234 | 0.866416846 | 0 | 0.022 | 1 | 0.309824707 | 2.683072586 | 0.022 | 0.007 | 1 | 0.492608234 | 0.523658065 |
| Wdr38 | 0.532436372 | 1.088809267 | 0 | 0.019 | 1 | 0.309824707 | 2.683072586 | 0.022 | 0.007 | 1 | 0.532436372 | 0.523658065 |
| Map3k15 | 0.550953306 | 1.936806174 | 0.048 | 0.026 | 1 | 0.309824707 | 3.098110085 | 0.022 | 0.007 | 1 | 0.550953306 | 0.523658065 |
| Bcl2l14 | 0.578159907 | 0.866416846 | 0 | 0.015 | 1 | 0.309824707 | 3.098110085 | 0.022 | 0.007 | 1 | 0.578159907 | 0.523658065 |
| Gm31816 | 0.57816106 | 1.351843673 | 0 | 0.015 | 1 | 0.309824707 | 3.098110085 | 0.022 | 0.007 | 1 | 0.57816106 | 0.523658065 |
| Gm28731 | 0.57816106 | 1.351843673 | 0 | 0.015 | 1 | 0.309824707 | 3.098110085 | 0.022 | 0.007 | 1 | 0.57816106 | 0.523658065 |
| Gm17163 | 0.57816106 | 1.351843673 | 0 | 0.015 | 1 | 0.309824707 | 2.683072586 | 0.022 | 0.007 | 1 | 0.57816106 | 0.523658065 |
| Gm30655 | 0.57816106 | 1.351843673 | 0 | 0.015 | 1 | 0.309824707 | 3.098110085 | 0.022 | 0.007 | 1 | 0.57816106 | 0.523658065 |
| Abcg5 | 0.57816106 | 1.351843673 | 0 | 0.015 | 1 | 0.309824707 | 2.683072586 | 0.022 | 0.007 | 1 | 0.57816106 | 0.523658065 |
| Pla2g4c | 0.63207476 | 1.351843673 | 0 | 0.011 | 1 | 0.309824707 | 2.683072586 | 0.022 | 0.007 | 1 | 0.63207476 | 0.523658065 |

| | | | | | | | | | | | | |
|---|---|---|---|---|---|---|---|---|---|---|---|---|
| Gm40630 | 0.632076134 | 0.673771768 | 0 | 0.011 | 1 | 0.309824707 | 4.798549804 | 0.022 | 0.007 | 1 | 0.632076134 | 0.523658065 |
| Gm50020 | 0.632076134 | 1.673771768 | 0 | 0.011 | 1 | 0.309824707 | 3.098110085 | 0.022 | 0.007 | 1 | 0.632076134 | 0.523658065 |
| Gm15675 | 0.668622608 | 0.766881172 | 0.048 | 0.03 | 1 | 0.309824707 | 3.905465008 | 0.022 | 0.007 | 1 | 0.668622608 | 0.523658065 |
| Soat2 | 0.859868645 | 1.451379346 | 0.048 | 0.041 | 1 | 0.309824707 | 3.098110085 | 0.022 | 0.007 | 1 | 0.859868645 | 0.523658065 |
| Igdcc3 | 0.457281049 | 0.503846766 | 0 | 0.026 | 1 | 0.309826166 | 2.946106992 | 0.022 | 0.007 | 1 | 0.457281049 | 0.523660079 |
| Cartpt | 0.532426384 | 0.866416846 | 0 | 0.019 | 1 | 0.309826166 | 3.005000681 | 0.022 | 0.007 | 1 | 0.532426384 | 0.523660079 |
| Gm9195 | 0.532426384 | 0.866416846 | 0 | 0.019 | 1 | 0.309826166 | 3.268035087 | 0.022 | 0.007 | 1 | 0.532426384 | 0.523660079 |
| Xpnpep1 | 0.309862832 | 0.230828272 | 0.238 | 0.422 | 1 | 0.354066704 | 1.005000681 | 0.222 | 0.356 | 1 | 0.354066704 | 0.52371069 |
| Gm13008 | 0.492608234 | 0.866416846 | 0 | 0.022 | 1 | 0.30986553 | 0.982632868 | 0.067 | 0.035 | 1 | 0.492608234 | 0.523714413 |
| Kat7 | 0.310005568 | 0.949745229 | 0.476 | 0.403 | 1 | 0.634230113 | -0.204452685 | 0.578 | 0.578 | 1 | 0.634230113 | 0.523907684 |
| F8a | 0.444313791 | 1.425844254 | 0.095 | 0.056 | 1 | 0.310601014 | -0.901889915 | 0.022 | 0.059 | 1 | 0.444313791 | 0.524729038 |
| Zfp382 | 0.380297304 | -0.413691074 | 0.048 | 0.108 | 1 | 0.310653768 | 0.274987848 | 0.111 | 0.187 | 1 | 0.380297304 | 0.524801772 |
| Map1s | 0.656324705 | 0.393663848 | 0.143 | 0.209 | 1 | 0.310838623 | 0.212752651 | 0.2 | 0.318 | 1 | 0.656324705 | 0.525056596 |
| Zdhhc18 | 0.311156236 | -0.326228232 | 0.286 | 0.433 | 1 | 0.822750897 | 0.500869255 | 0.333 | 0.457 | 1 | 0.822750897 | 0.525494269 |
| Gm26903 | 0.311315359 | 2.843696769 | 0.095 | 0.049 | 1 | 0.497228696 | 0.268035087 | 0.022 | 0.045 | 1 | 0.497228696 | 0.525713466 |
| Zfp944 | 0.311336192 | -0.348596045 | 0.095 | 0.19 | 1 | 0.760900808 | 0.739656115 | 0.222 | 0.221 | 1 | 0.760900808 | 0.525742159 |
| Rangap1 | 0.311491181 | 0.448805403 | 0.238 | 0.437 | 1 | 0.326494889 | 0.483566594 | 0.6 | 0.54 | 1 | 0.326494889 | 0.525955607 |
| Adat2 | 0.370840523 | 0.214340149 | 0 | 0.037 | 1 | 0.311583783 | -0.638855509 | 0.022 | 0.059 | 1 | 0.370840523 | 0.526083113 |
| Gm45570 | 0.532436372 | 1.088809267 | 0 | 0.019 | 1 | 0.311583783 | -0.638855509 | 0.022 | 0.059 | 1 | 0.532436372 | 0.526083113 |
| Phaf1 | 0.317745467 | -0.463731756 | 0.143 | 0.243 | 1 | 0.311896766 | 0.129474256 | 0.289 | 0.408 | 1 | 0.317745467 | 0.526513939 |
| Slc16a12 | 0.63207476 | 1.351843673 | 0 | 0.011 | 1 | 0.312251526 | 2.268035087 | 0.022 | 0.007 | 1 | 0.63207476 | 0.527002037 |
| Hnrnpm | 0.533456253 | -0.174548287 | 0.857 | 0.948 | 1 | 0.312298406 | 0.219125486 | 1 | 0.983 | 1 | 0.533456253 | 0.527066518 |
| Abhd18 | 0.31242451 | -0.880817084 | 0.048 | 0.119 | 1 | 0.985882424 | 0.741175541 | 0.133 | 0.145 | 1 | 0.985882424 | 0.527239945 |
| Gm10184 | 0.370840523 | 0.214340149 | 0 | 0.037 | 1 | 0.312564649 | -0.564854927 | 0.022 | 0.059 | 1 | 0.370840523 | 0.527432638 |
| Gm21781 | 0.312656267 | 1.258734268 | 0.095 | 0.045 | 1 | 0.693836065 | 1.531069493 | 0.044 | 0.062 | 1 | 0.693836065 | 0.527558593 |
| Gm41231 | 0.312656267 | 1.258734268 | 0.095 | 0.045 | 1 | 0.5539024 | 0.595609745 | 0.022 | 0.042 | 1 | 0.5539024 | 0.527558593 |
| Enah | 0.313535871 | -0.263212515 | 0.857 | 0.937 | 1 | 0.994010333 | -0.207036557 | 0.978 | 0.855 | 1 | 0.994010333 | 0.528767 |
| Pou2af3 | 0.372024717 | 0.995699863 | 0.143 | 0.261 | 1 | 0.313902101 | 0.154693614 | 0.111 | 0.18 | 1 | 0.372024717 | 0.529269673 |
| Gm31323 | 0.893852518 | 0.673771768 | 0.048 | 0.056 | 1 | 0.314019223 | -0.549588171 | 0.044 | 0.09 | 1 | 0.893852518 | 0.529430374 |
| Mief2 | 0.355458734 | 0.918884266 | 0.143 | 0.082 | 1 | 0.314209039 | -0.316927414 | 0.089 | 0.149 | 1 | 0.355458734 | 0.529690758 |
| Pfkfb4 | 0.44731972 | 0.892412054 | 0.286 | 0.246 | 1 | 0.314481055 | 0.195806759 | 0.178 | 0.263 | 1 | 0.44731972 | 0.530063776 |
| Mon2 | 0.314518526 | 0.21668409 | 0.857 | 0.787 | 1 | 0.840750818 | 0.250113179 | 0.911 | 0.917 | 1 | 0.840750818 | 0.530115149 |
| Mgmt | 0.314653379 | -0.970084422 | 0.048 | 0.119 | 1 | 0.584775482 | 1.658410532 | 0.089 | 0.128 | 1 | 0.584775482 | 0.53030001 |
| Ucp3 | 0.457292877 | 0.351843673 | 0 | 0.026 | 1 | 0.314689364 | 2.098110085 | 0.022 | 0.007 | 1 | 0.457292877 | 0.530349332 |
| Plekhg4 | 0.457292877 | 0.351843673 | 0 | 0.026 | 1 | 0.314689364 | 2.098110085 | 0.022 | 0.007 | 1 | 0.457292877 | 0.530349332 |
| Gm11827 | 0.492590603 | 0.673771768 | 0 | 0.022 | 1 | 0.314689364 | 2.098110085 | 0.022 | 0.007 | 1 | 0.492590603 | 0.530349332 |
| Popdc2 | 0.532426384 | 0.866416846 | 0 | 0.019 | 1 | 0.314689364 | 2.098110085 | 0.022 | 0.007 | 1 | 0.532426384 | 0.530349332 |
| Dscam | 0.532426384 | 0.866416846 | 0 | 0.019 | 1 | 0.314689364 | 1.946106992 | 0.022 | 0.007 | 1 | 0.532426384 | 0.530349332 |
| 4930483P17Rik | 0.57816106 | 1.351843673 | 0 | 0.015 | 1 | 0.314689364 | 2.098110085 | 0.022 | 0.007 | 1 | 0.57816106 | 0.530349332 |
| Duox1 | 0.629700127 | 2.088809267 | 0.048 | 0.03 | 1 | 0.314689364 | 1.946106992 | 0.022 | 0.007 | 1 | 0.629700127 | 0.530349332 |
| Txlnb | 0.63207476 | 1.351843673 | 0 | 0.011 | 1 | 0.314689364 | 1.683072586 | 0.022 | 0.007 | 1 | 0.63207476 | 0.530349332 |
| Gm57150 | 0.632076134 | 1.673771768 | 0 | 0.011 | 1 | 0.314689364 | 2.098110085 | 0.022 | 0.007 | 1 | 0.632076134 | 0.530349332 |
| Ddx4 | 0.632076134 | 1.673771768 | 0 | 0.011 | 1 | 0.314689364 | 1.946106992 | 0.022 | 0.007 | 1 | 0.632076134 | 0.530349332 |
| Nkrf | 0.357465589 | -0.413691074 | 0.048 | 0.112 | 1 | 0.314943745 | -0.223818009 | 0.089 | 0.149 | 1 | 0.357465589 | 0.530697928 |
| Spsb1 | 0.3151034 | -0.233118828 | 0.476 | 0.657 | 1 | 0.325000585 | 0.205515335 | 0.489 | 0.706 | 1 | 0.325000585 | 0.530916647 |
| Tmem29 | 0.330326942 | 1.24081236 | 0.381 | 0.354 | 1 | 0.315387032 | -0.330733213 | 0.178 | 0.253 | 1 | 0.330326942 | 0.531305084 |
| Ccdc162 | 0.316186439 | 0.866416846 | 0.095 | 0.045 | 1 | 0.739875423 | 0.757073168 | 0.044 | 0.035 | 1 | 0.739875423 | 0.532399013 |
| Bcl2l12 | 0.316572795 | -0.133583154 | 0.048 | 0.127 | 1 | 0.734538905 | 0.460680165 | 0.067 | 0.083 | 1 | 0.734538905 | 0.532927255 |
| Esr1 | 0.830750915 | 0.435367028 | 0.381 | 0.399 | 1 | 0.317128201 | 0.305371238 | 0.444 | 0.644 | 1 | 0.830750915 | 0.533686106 |
| Insc | 0.42563249 | -1.184209227 | 0 | 0.03 | 1 | 0.317138221 | 0.683072586 | 0.022 | 0.007 | 1 | 0.42563249 | 0.533699791 |
| Gm57409 | 0.532426384 | 0.866416846 | 0 | 0.019 | 1 | 0.317138221 | 1.683072586 | 0.022 | 0.007 | 1 | 0.532426384 | 0.533699791 |
| Gm48202 | 0.632076134 | 1.673771768 | 0 | 0.011 | 1 | 0.317138221 | 1.361144491 | 0.022 | 0.007 | 1 | 0.632076134 | 0.533699791 |
| Gm43150 | 0.324807521 | -0.648156327 | 0 | 0.045 | 1 | 0.317556327 | 2.683072586 | 0.044 | 0.021 | 1 | 0.324807521 | 0.534270633 |
| Rab7b | 0.574866695 | 0.60338244 | 0.143 | 0.104 | 1 | 0.317564659 | 2.490427508 | 0.044 | 0.021 | 1 | 0.574866695 | 0.534282006 |
| Rasgef1c | 0.853730569 | 2.304537958 | 0.048 | 0.041 | 1 | 0.317569659 | 2.442064487 | 0.044 | 0.021 | 1 | 0.853730569 | 0.534288829 |
| 9830144P21Rik | 0.815691551 | 1.088809267 | 0.048 | 0.063 | 1 | 0.317570214 | 2.531069493 | 0.044 | 0.021 | 1 | 0.815691551 | 0.534289587 |
| Sim1 | 0.469719345 | 1.673771768 | 0.048 | 0.022 | 1 | 0.317571325 | 2.531069493 | 0.044 | 0.021 | 1 | 0.469719345 | 0.534291104 |
| Spink5 | 0.532436372 | 1.088809267 | 0 | 0.019 | 1 | 0.317571325 | 2.361144491 | 0.044 | 0.021 | 1 | 0.532436372 | 0.534291104 |
| Zc3h12a | 0.317975113 | -0.496153234 | 0.048 | 0.119 | 1 | 0.384110637 | 0.495445583 | 0.067 | 0.114 | 1 | 0.384110637 | 0.534842054 |
| Gm35853 | 0.318156593 | 3.936806174 | 0.095 | 0.049 | 1 | 0.661444614 | 1.683072586 | 0.022 | 0.014 | 1 | 0.661444614 | 0.535089569 |
| Nr1d2 | 0.318259894 | -0.319996279 | 0.619 | 0.739 | 1 | 0.402793211 | 0.297641549 | 0.867 | 0.844 | 1 | 0.402793211 | 0.535230428 |
| Cyp4v3 | 0.318988624 | 1.460778044 | 0.19 | 0.131 | 1 | 0.368592259 | 0.235613609 | 0.089 | 0.138 | 1 | 0.368592259 | 0.536223506 |
| Garin5b | 0.7319334 | 1.088809267 | 0.048 | 0.034 | 1 | 0.319089091 | 2.531069493 | 0.044 | 0.021 | 1 | 0.7319334 | 0.536360335 |
| Ankrd35 | 0.885488583 | 0.425844254 | 0.095 | 0.108 | 1 | 0.319179369 | 0.244499572 | 0.156 | 0.235 | 1 | 0.885488583 | 0.536483268 |
| Hspa1b | 0.593751866 | -1.690574732 | 0.619 | 0.552 | 1 | 0.31972131 | -1.088698367 | 0.822 | 0.585 | 1 | 0.593751866 | 0.537220904 |
| Efl1 | 0.794085439 | -0.557182667 | 0.286 | 0.299 | 1 | 0.320030616 | 0.935838657 | 0.489 | 0.457 | 1 | 0.794085439 | 0.537641637 |
| Flt3l | 0.88770495 | 1.088809267 | 0.048 | 0.041 | 1 | 0.320769182 | 0.402964667 | 0.044 | 0.093 | 1 | 0.88770495 | 0.538645495 |
| Synrg | 0.320853982 | -0.364363361 | 0.19 | 0.302 | 1 | 0.342930951 | 0.342035668 | 0.267 | 0.349 | 1 | 0.342930951 | 0.538760686 |
| Armh3 | 0.853002536 | 0.586308926 | 0.286 | 0.317 | 1 | 0.321032896 | 1.137793787 | 0.444 | 0.457 | 1 | 0.853002536 | 0.539003672 |
| Coro7 | 0.393411718 | 0.214340149 | 0.095 | 0.175 | 1 | 0.321145449 | 0.439147003 | 0.133 | 0.218 | 1 | 0.393411718 | 0.539156499 |
| Fbxl8 | 0.608931651 | 0.304537958 | 0.048 | 0.082 | 1 | 0.321598342 | 0.595609745 | 0.044 | 0.093 | 1 | 0.608931651 | 0.539771191 |
| A230072E10Rik | 0.321607841 | 1.866416846 | 0.095 | 0.049 | 1 | 0.491587946 | 1.268035087 | 0.044 | 0.076 | 1 | 0.491587946 | 0.539784078 |
| Cfap45 | 0.321613947 | 2.088809267 | 0.095 | 0.049 | 1 | 0.895371025 | 1.098110085 | 0.133 | 0.156 | 1 | 0.895371025 | 0.539792363 |
| Slc16a10 | 0.617464636 | -0.20069735 | 0.19 | 0.25 | 1 | 0.322248874 | 0.496496344 | 0.156 | 0.249 | 1 | 0.617464636 | 0.540653411 |
| 1700113A16Rik | 0.560093825 | 1.60338244 | 0.095 | 0.067 | 1 | 0.322344171 | 0.852997588 | 0.044 | 0.093 | 1 | 0.560093825 | 0.540782578 |
| Ppm1d | 0.480165811 | 0.48263028 | 0.19 | 0.313 | 1 | 0.322474581 | -0.476126009 | 0.444 | 0.502 | 1 | 0.480165811 | 0.540959307 |
| Trim59 | 0.565819678 | -0.348596045 | 0.095 | 0.142 | 1 | 0.322502532 | 0.188307894 | 0.089 | 0.149 | 1 | 0.565819678 | 0.540997181 |
| C4b | 0.540615335 | -0.184209227 | 0.048 | 0.09 | 1 | 0.322772054 | -0.578192727 | 0.156 | 0.225 | 1 | 0.540615335 | 0.541362309 |
| Gm49454 | 0.784072994 | 3.79930265 | 0.048 | 0.037 | 1 | 0.323064501 | 0.857101986 | 0.089 | 0.052 | 1 | 0.784072994 | 0.541758331 |
| Atf3 | 0.809340047 | -0.17537266 | 0.857 | 0.825 | 1 | 0.323504563 | -0.214701423 | 0.933 | 0.841 | 1 | 0.809340047 | 0.542353923 |
| Rsbn1 | 0.323544083 | -0.31112134 | 0.476 | 0.631 | 1 | 0.970268143 | 0.186814311 | 0.867 | 0.785 | 1 | 0.970268143 | 0.542407392 |
| Gm20342 | 0.323694993 | 1.60338244 | 0.19 | 0.123 | 1 | 0.325460655 | 0.563333342 | 0.178 | 0.284 | 1 | 0.325460655 | 0.542611537 |
| Tmod1 | 0.373857158 | 2.088809267 | 0.048 | 0.019 | 1 | 0.324011243 | -0.686161223 | 0.022 | 0.059 | 1 | 0.373857158 | 0.5430392 |
| Phkg1 | 0.324822624 | -0.785659851 | 0 | 0.045 | 1 | 0.32402804 | -0.271123724 | 0.022 | 0.059 | 1 | 0.324822624 | 0.543061909 |
| Acad8 | 0.324611142 | -0.303508156 | 0.095 | 0.187 | 1 | 0.372958831 | 0.24012909 | 0.156 | 0.235 | 1 | 0.372958831 | 0.54384989 |
| Gm28375 | 0.324739547 | -0.133583154 | 0 | 0.045 | 1 | 0.44805663 | 0.513147585 | 0.022 | 0.048 | 1 | 0.44805663 | 0.54402332 |
| Wnt2b | 0.324739547 | -0.233118828 | 0 | 0.045 | 1 | 0.708987514 | 1.42003818 | 0.044 | 0.035 | 1 | 0.708987514 | 0.54402332 |
| Stbd1 | 0.324739547 | -0.133583154 | 0 | 0.045 | 1 | 0.328724221 | -0.776359032 | 0.044 | 0.087 | 1 | 0.328724221 | 0.54402332 |
| Gm57314 | 0.324739547 | -0.233118828 | 0 | 0.045 | 1 | 0.983977613 | 0.298408736 | 0.089 | 0.09 | 1 | 0.983977613 | 0.54402332 |
| Gm47863 | 0.324739547 | -0.133583154 | 0 | 0.045 | 1 | 0.555534078 | 0.875717664 | 0.022 | 0.042 | 1 | 0.555534078 | 0.54402332 |
| Cdc45 | 0.324739547 | -0.326228232 | 0 | 0.045 | 1 | 0.430487676 | 0.361144491 | 0 | 0.014 | 1 | 0.430487676 | 0.54402332 |
| Gm36457 | 0.324739547 | -0.133583154 | 0 | 0.045 | 1 | 0.496223879 | 0.683072586 | 0 | 0.01 | 1 | 0.496223879 | 0.54402332 |
| Lst1 | 0.324739547 | -0.133583154 | 0 | 0.045 | 1 | 0.932582169 | 0.361144491 | 0.022 | 0.024 | 1 | 0.932582169 | 0.54402332 |
| Ribc1 | 0.324739547 | -0.326228232 | 0 | 0.045 | 1 | 0.376837034 | -0.124282336 | 0 | 0.017 | 1 | 0.376837034 | 0.54402332 |
| Gdap1 | 0.324767242 | -0.413691074 | 0 | 0.045 | 1 | 0.430487676 | 0.361144491 | 0 | 0.014 | 1 | 0.430487676 | 0.544060723 |
| Inka2 | 0.324767242 | -0.233118828 | 0 | 0.045 | 1 | 0.968326985 | 1.268035087 | 0.067 | 0.069 | 1 | 0.968326985 | 0.544060723 |
| Spaar | 0.324767242 | -0.413691074 | 0 | 0.045 | 1 | 0.430487676 | 0.361144491 | 0 | 0.014 | 1 | 0.430487676 | 0.544060723 |
| Med18 | 0.324767242 | -0.233118828 | 0 | 0.045 | 1 | 0.430707223 | -0.686161223 | 0.044 | 0.076 | 1 | 0.430707223 | 0.544060723 |

| | | | | | | | | | | | | |
|---|---|---|---|---|---|---|---|---|---|---|---|---|
| Nupr1l | 0.324767242 | -0.233118828 | 0 | 0.045 | 1 | 0.555549391 | 0.776181991 | 0.022 | 0.042 | 1 | 0.555549391 | 0.544060723 |
| Gm14547 | 0.324767242 | -0.233118828 | 0 | 0.045 | 1 | 0.49622258 | 0.361144491 | 0 | 0.01 | 1 | 0.49622258 | 0.544060723 |
| Ankrd33b | 0.324767242 | -0.233118828 | 0 | 0.045 | 1 | 0.983394262 | 0.966865552 | 0.044 | 0.045 | 1 | 0.983394262 | 0.544060723 |
| Cpt1b | 0.324767242 | -0.233118828 | 0 | 0.045 | 1 | 0.782759021 | 0.744473131 | 0.044 | 0.055 | 1 | 0.782759021 | 0.544060723 |
| 9330159F19Rik | 0.324789396 | -0.413691074 | 0 | 0.045 | 1 | 0.376842245 | -0.776359032 | 0 | 0.017 | 1 | 0.376842245 | 0.544090641 |
| Adcy1 | 0.324789396 | -0.496153234 | 0 | 0.045 | 1 | 0.932582169 | 0.361144491 | 0.022 | 0.024 | 1 | 0.932582169 | 0.544090641 |
| 1700030N03Rik | 0.324789396 | -0.496153234 | 0 | 0.045 | 1 | 0.376837034 | -0.124282336 | 0 | 0.017 | 1 | 0.376837034 | 0.544090641 |
| Il18r1 | 0.3247899 | -0.718545655 | 0 | 0.045 | 1 | 0.376837034 | -0.316927414 | 0 | 0.017 | 1 | 0.376837034 | 0.544091321 |
| Gm20646 | 0.3247899 | -0.574155746 | 0 | 0.045 | 1 | 0.686456799 | 0.683072586 | 0.044 | 0.062 | 1 | 0.686456799 | 0.544091321 |
| Foxm1 | 0.3247899 | -0.496153234 | 0 | 0.045 | 1 | 0.806246349 | 2.197645759 | 0.022 | 0.017 | 1 | 0.806246349 | 0.544091321 |
| Rtn2 | 0.3247899 | -0.326228232 | 0 | 0.045 | 1 | 0.450803673 | 1.946106992 | 0.022 | 0.048 | 1 | 0.450803673 | 0.544091321 |
| ENSMUSG00000121374 | 0.3247899 | -0.326228232 | 0 | 0.045 | 1 | 0.687208248 | 1.098110085 | 0.022 | 0.035 | 1 | 0.687208248 | 0.544091321 |
| Gm15839 | 0.3247899 | -0.326228232 | 0 | 0.045 | 1 | 0.376842245 | -0.638855509 | 0 | 0.017 | 1 | 0.376842245 | 0.544091321 |
| Heph | 0.3247899 | -0.718545655 | 0 | 0.045 | 1 | 0.672920453 | 0.875717664 | 0.022 | 0.014 | 1 | 0.672920453 | 0.544091321 |
| Pcdhb20 | 0.324806011 | -0.496153234 | 0 | 0.045 | 1 | 0.530911177 | -0.316927414 | 0.022 | 0.042 | 1 | 0.530911177 | 0.544113077 |
| Slc45a3 | 0.324807521 | -0.849790188 | 0 | 0.045 | 1 | 0.385364644 | -0.486852415 | 0.022 | 0.052 | 1 | 0.385364644 | 0.544115116 |
| Gm27241 | 0.324808024 | -0.413691074 | 0 | 0.045 | 1 | 0.44949712 | 0.361144491 | 0.022 | 0.048 | 1 | 0.44949712 | 0.544115796 |
| Gm46515 | 0.324808024 | -1.184209227 | 0 | 0.045 | 1 | 0.683278676 | 0.683072586 | 0.022 | 0.035 | 1 | 0.683278676 | 0.544115796 |
| Slc45a1 | 0.324817086 | -0.574155746 | 0 | 0.045 | 1 | 0.495731057 | 0.180572246 | 0.022 | 0.045 | 1 | 0.495731057 | 0.544128033 |
| Drc3 | 0.324820107 | -0.648156327 | 0 | 0.045 | 1 | 0.365828548 | 0.15951063 | 0.022 | 0.055 | 1 | 0.365828548 | 0.544132112 |
| Nexmif | 0.324820107 | -1.081115734 | 0 | 0.045 | 1 | 0.761454895 | 0.757073168 | 0.022 | 0.031 | 1 | 0.761454895 | 0.544132112 |
| Kcnq1 | 0.324821114 | -0.970084422 | 0 | 0.045 | 1 | 0.687223223 | 1.460680165 | 0.022 | 0.035 | 1 | 0.687223223 | 0.544133471 |
| Kntc1 | 0.32482212 | -0.496153234 | 0 | 0.045 | 1 | 0.331907298 | -0.316927414 | 0 | 0.021 | 1 | 0.331907298 | 0.544134831 |
| E330013P04Rik | 0.324822624 | -0.718545655 | 0 | 0.045 | 1 | 0.646587902 | 0.460680165 | 0.133 | 0.114 | 1 | 0.646587902 | 0.544135511 |
| Adad1 | 0.324830678 | -1.326228232 | 0 | 0.045 | 1 | 0.37684485 | -1.404390255 | 0 | 0.017 | 1 | 0.37684485 | 0.544146387 |
| Nhsl2 | 0.324831685 | -1.818081329 | 0 | 0.045 | 1 | 0.611693765 | 1.078210528 | 0.089 | 0.121 | 1 | 0.611693765 | 0.544147746 |
| A430010J10Rik | 0.324837726 | -1.326228232 | 0 | 0.045 | 1 | 0.972285017 | 0.205025289 | 0.067 | 0.066 | 1 | 0.972285017 | 0.544155903 |
| Adam2 | 0.324838229 | -1.940938076 | 0 | 0.045 | 1 | 0.675807368 | 0.683072586 | 0.022 | 0.014 | 1 | 0.675807368 | 0.544156583 |
| Lnpk | 0.385617816 | 1.069700444 | 0.381 | 0.392 | 1 | 0.324869602 | 0.597681095 | 0.244 | 0.384 | 1 | 0.385617816 | 0.544198946 |
| Setd6 | 0.425971378 | -0.326228232 | 0.048 | 0.101 | 1 | 0.325328037 | -0.819427754 | 0.044 | 0.087 | 1 | 0.425971378 | 0.544817742 |
| Alkbh8 | 0.328143659 | 0.304537958 | 0.143 | 0.261 | 1 | 0.325754698 | 0.162240423 | 0.178 | 0.277 | 1 | 0.328143659 | 0.545393272 |
| Clec2d | 0.492590603 | 0.673771768 | 0 | 0.022 | 1 | 0.32605595 | -0.316927414 | 0.022 | 0.059 | 1 | 0.492590603 | 0.545799418 |
| Lig4 | 0.982550637 | 0.673771768 | 0.048 | 0.049 | 1 | 0.32607682 | -0.255526869 | 0.022 | 0.059 | 1 | 0.982550637 | 0.545827547 |
| Rbak | 0.326288826 | 1.029915578 | 0.19 | 0.123 | 1 | 0.510669598 | 0.280974143 | 0.156 | 0.208 | 1 | 0.510669598 | 0.546113254 |
| Epg5 | 0.326407924 | 1.418957869 | 0.333 | 0.291 | 1 | 0.865470984 | 1.149094014 | 0.267 | 0.329 | 1 | 0.865470984 | 0.546273715 |
| Gm12536 | 0.882081196 | 0.97333205 | 0.143 | 0.146 | 1 | 0.326440253 | 1.288793647 | 0.222 | 0.18 | 1 | 0.882081196 | 0.546317267 |
| Ap5z1 | 0.3264482 | 1.521768674 | 0.19 | 0.131 | 1 | 0.798775062 | 0.707320132 | 0.156 | 0.152 | 1 | 0.798775062 | 0.546327973 |
| Slc7a7 | 0.928363519 | 0.625408746 | 0.19 | 0.198 | 1 | 0.326451366 | 0.614592848 | 0.178 | 0.291 | 1 | 0.928363519 | 0.546332238 |
| Rhobtb1 | 0.326517313 | -1.054148687 | 0.048 | 0.116 | 1 | 0.605063558 | 0.587915353 | 0.067 | 0.093 | 1 | 0.605063558 | 0.54642107 |
| 0610040B10Rik | 0.368211282 | 1.936806174 | 0.095 | 0.052 | 1 | 0.326732519 | 1.875717664 | 0.044 | 0.021 | 1 | 0.368211282 | 0.5467109 |
| Gm43560 | 0.75704869 | 0.55829455 | 0.048 | 0.067 | 1 | 0.326732519 | 2.197645759 | 0.044 | 0.021 | 1 | 0.75704869 | 0.5467109 |
| Gm34471 | 0.346861878 | -0.648156327 | 0 | 0.041 | 1 | 0.326759174 | 1.683072586 | 0.044 | 0.021 | 1 | 0.346861878 | 0.54674679 |
| Wrap53 | 0.651192687 | 1.351843673 | 0.095 | 0.071 | 1 | 0.327019473 | -0.709244837 | 0.067 | 0.114 | 1 | 0.651192687 | 0.54709721 |
| Zmynd11 | 0.327126052 | 0.269921949 | 0.905 | 0.922 | 1 | 0.731087598 | 0.155492308 | 1 | 0.965 | 1 | 0.731087598 | 0.547240651 |
| Cwf19l1 | 0.358902491 | 0.129451251 | 0.048 | 0.119 | 1 | 0.327170273 | 0.410054092 | 0.044 | 0.09 | 1 | 0.358902491 | 0.547300159 |
| Aqr | 0.457884572 | -0.142145168 | 0.286 | 0.388 | 1 | 0.327407622 | 0.356779299 | 0.378 | 0.578 | 1 | 0.457884572 | 0.547619492 |
| Gfpt2 | 0.327514271 | 0.76624855 | 0.905 | 0.869 | 1 | 0.674497975 | 0.907531413 | 0.756 | 0.851 | 1 | 0.674497975 | 0.547762944 |
| Slc23a3 | 0.675315714 | 0.351843673 | 0.048 | 0.075 | 1 | 0.327663102 | 0.19121949 | 0.067 | 0.118 | 1 | 0.675315714 | 0.547963096 |
| Gm14286 | 0.425599726 | 0.214340149 | 0 | 0.03 | 1 | 0.328150343 | -0.124282336 | 0.022 | 0.059 | 1 | 0.425599726 | 0.548618039 |
| Ceacam1 | 0.328155668 | 0.519966432 | 0.238 | 0.157 | 1 | 0.642268334 | -0.719025857 | 0.133 | 0.152 | 1 | 0.642268334 | 0.548625194 |
| D17H6S56E-5 | 0.565195739 | -0.531342662 | 0.048 | 0.086 | 1 | 0.328222602 | -0.242926832 | 0.022 | 0.059 | 1 | 0.565195739 | 0.548715127 |
| Gm11696 | 0.910091451 | 1.088809267 | 0.048 | 0.056 | 1 | 0.328292114 | 2.42003818 | 0.044 | 0.021 | 1 | 0.910091451 | 0.548808516 |
| Arnt2 | 0.328421883 | -0.385121921 | 0.048 | 0.123 | 1 | 0.43497292 | 1.264759955 | 0.289 | 0.273 | 1 | 0.43497292 | 0.548982832 |
| Chordc1 | 0.328709816 | -0.103835811 | 0.381 | 0.56 | 1 | 0.381991812 | 0.265915035 | 0.733 | 0.692 | 1 | 0.381991812 | 0.549369488 |
| AW046200 | 0.328712778 | 1.304537958 | 0.143 | 0.082 | 1 | 0.992454287 | 1.618942249 | 0.067 | 0.069 | 1 | 0.992454287 | 0.549373466 |
| Letmd1 | 0.352467199 | 1.351843673 | 0.143 | 0.086 | 1 | 0.328991315 | -0.280401538 | 0.067 | 0.118 | 1 | 0.352467199 | 0.549747345 |
| Nudcd3 | 0.808560195 | 0.261388722 | 0.524 | 0.701 | 1 | 0.329763233 | 0.5427207 | 0.8 | 0.772 | 1 | 0.808560195 | 0.550782676 |
| Dusp29 | 0.457292877 | 0.351843673 | 0 | 0.026 | 1 | 0.329852568 | 2.320502507 | 0.044 | 0.021 | 1 | 0.457292877 | 0.550902419 |
| Iqch | 0.979646835 | 3.60338244 | 0.048 | 0.049 | 1 | 0.329858674 | 0.946106992 | 0.044 | 0.021 | 1 | 0.979646835 | 0.550910603 |
| Mgat4a | 0.638558711 | 0.102045094 | 0.619 | 0.821 | 1 | 0.330126767 | -0.16372756 | 0.822 | 0.775 | 1 | 0.638558711 | 0.551269851 |
| Farp2 | 0.507313755 | 0.843052853 | 0.286 | 0.459 | 1 | 0.330458367 | 0.770339422 | 0.511 | 0.64 | 1 | 0.507313755 | 0.551714002 |
| Smap1 | 0.3309695 | -0.262866171 | 0.619 | 0.784 | 1 | 0.872863672 | 0.187771011 | 0.867 | 0.841 | 1 | 0.872863672 | 0.552398191 |
| Abcb6 | 0.738393266 | 1.351843673 | 0.048 | 0.034 | 1 | 0.331306855 | 0.612683258 | 0.022 | 0.059 | 1 | 0.738393266 | 0.552849477 |
| Tarsl2 | 0.856803385 | 1.586308926 | 0.048 | 0.041 | 1 | 0.331307872 | 0.905465008 | 0.022 | 0.059 | 1 | 0.856803385 | 0.552850838 |
| Poc1a | 0.88770495 | 1.088809267 | 0.048 | 0.041 | 1 | 0.331328477 | 0.223640968 | 0.022 | 0.059 | 1 | 0.88770495 | 0.552878394 |
| Gm34086 | 0.81881728 | 1.088809267 | 0.048 | 0.037 | 1 | 0.331364848 | 0.304560963 | 0.022 | 0.059 | 1 | 0.81881728 | 0.552927034 |
| Tnik | 0.563271619 | -1.054148687 | 0.048 | 0.082 | 1 | 0.331380361 | 1.254229287 | 0.022 | 0.059 | 1 | 0.563271619 | 0.552947779 |
| Spata48 | 0.492599419 | 0.503846766 | 0 | 0.022 | 1 | 0.331402225 | 1.020107573 | 0.044 | 0.021 | 1 | 0.492599419 | 0.552977016 |
| Atp2a1 | 0.57816106 | 1.351843673 | 0 | 0.015 | 1 | 0.331402225 | 1.513147585 | 0.044 | 0.021 | 1 | 0.57816106 | 0.552977016 |
| Sema3e | 0.957372496 | 1.39417517 | 0.19 | 0.19 | 1 | 0.331720853 | 2.190032575 | 0.089 | 0.055 | 1 | 0.957372496 | 0.553402981 |
| Fam227a | 0.575302194 | 1.766881172 | 0.143 | 0.116 | 1 | 0.331792267 | 1.683072586 | 0.156 | 0.114 | 1 | 0.575302194 | 0.553498426 |
| A530046M15Rik | 0.331864774 | 1.526930379 | 0.143 | 0.086 | 1 | 0.376842245 | -0.316927414 | 0 | 0.017 | 1 | 0.376842245 | 0.55359532 |
| D430018E03Rik | 0.379775085 | 1.673771768 | 0.048 | 0.019 | 1 | 0.331907298 | -0.316927414 | 0 | 0.021 | 1 | 0.379775085 | 0.553652142 |
| Gpc2 | 0.38281368 | 1.766881172 | 0.095 | 0.052 | 1 | 0.331907298 | -0.316927414 | 0 | 0.021 | 1 | 0.38281368 | 0.553652142 |
| Fcrla | 0.396916958 | -0.496153234 | 0 | 0.034 | 1 | 0.331907298 | -0.316927414 | 0 | 0.021 | 1 | 0.396916958 | 0.553652142 |
| Zfp658 | 0.396916958 | 0.214340149 | 0 | 0.034 | 1 | 0.331907298 | -0.316927414 | 0 | 0.021 | 1 | 0.396916958 | 0.553652142 |
| Gm3830 | 0.396916958 | 0.214340149 | 0 | 0.034 | 1 | 0.331907298 | -0.486852415 | 0 | 0.021 | 1 | 0.396916958 | 0.553652142 |
| Gm57009 | 0.396916958 | 0.214340149 | 0 | 0.034 | 1 | 0.331907298 | -0.316927414 | 0 | 0.021 | 1 | 0.396916958 | 0.553652142 |
| Gm49179 | 0.396916958 | 0.214340149 | 0 | 0.034 | 1 | 0.331907298 | -0.316927414 | 0 | 0.021 | 1 | 0.396916958 | 0.553652142 |
| Gm57349 | 0.396935107 | -0.233118828 | 0 | 0.034 | 1 | 0.331907298 | -0.316927414 | 0 | 0.021 | 1 | 0.396935107 | 0.553652142 |
| Gm30786 | 0.425599726 | 0.214340149 | 0 | 0.03 | 1 | 0.331907298 | -0.316927414 | 0 | 0.021 | 1 | 0.425599726 | 0.553652142 |
| Katnal2 | 0.425599726 | 0.351843673 | 0 | 0.03 | 1 | 0.331907298 | -0.486852415 | 0 | 0.021 | 1 | 0.425599726 | 0.553652142 |
| D930032P07Rik | 0.425599726 | 0.351843673 | 0 | 0.03 | 1 | 0.331907298 | -0.316927414 | 0 | 0.021 | 1 | 0.425599726 | 0.553652142 |
| Styxl1 | 0.425614684 | 0.214340149 | 0 | 0.03 | 1 | 0.331907298 | -0.901889915 | 0 | 0.021 | 1 | 0.425614684 | 0.553652142 |
| B130024G19Rik | 0.425614684 | -0.133583154 | 0 | 0.03 | 1 | 0.331907298 | -0.316927414 | 0 | 0.021 | 1 | 0.425614684 | 0.553652142 |
| Capn3 | 0.425639612 | 0.503846766 | 0 | 0.03 | 1 | 0.331907298 | -0.316927414 | 0 | 0.021 | 1 | 0.425639612 | 0.553652142 |
| Mfsd7a | 0.457281049 | 0.503846766 | 0 | 0.026 | 1 | 0.331907298 | -0.316927414 | 0 | 0.021 | 1 | 0.457281049 | 0.553652142 |
| Zfp174 | 0.457281049 | 0.503846766 | 0 | 0.026 | 1 | 0.331907298 | -0.316927414 | 0 | 0.021 | 1 | 0.457281049 | 0.553652142 |
| Atf7ip2 | 0.457292877 | -0.133583154 | 0 | 0.026 | 1 | 0.331907298 | -0.316927414 | 0 | 0.021 | 1 | 0.457292877 | 0.553652142 |
| Gm41760 | 0.457292877 | 0.351843673 | 0 | 0.026 | 1 | 0.331907298 | -0.316927414 | 0 | 0.021 | 1 | 0.457292877 | 0.553652142 |
| Abca8a | 0.457304706 | -0.413691074 | 0 | 0.026 | 1 | 0.331907298 | -0.316927414 | 0 | 0.021 | 1 | 0.457304706 | 0.553652142 |
| Myef2l | 0.472861597 | 1.866416846 | 0.048 | 0.022 | 1 | 0.331907298 | -0.486852415 | 0 | 0.021 | 1 | 0.472861597 | 0.553652142 |
| Zc3h12d | 0.492590603 | 0.503846766 | 0 | 0.022 | 1 | 0.331907298 | -0.316927414 | 0 | 0.021 | 1 | 0.492590603 | 0.553652142 |
| Aplp1 | 0.492599419 | 0.351843673 | 0 | 0.022 | 1 | 0.331907298 | -0.316927414 | 0 | 0.021 | 1 | 0.492599419 | 0.553652142 |
| Slc35f1 | 0.492599419 | -0.496153234 | 0 | 0.022 | 1 | 0.331907298 | -0.316927414 | 0 | 0.021 | 1 | 0.492599419 | 0.553652142 |
| Pla2g10os | 0.492604708 | -0.133583154 | 0 | 0.022 | 1 | 0.331907298 | -0.316927414 | 0 | 0.021 | 1 | 0.492604708 | 0.553652142 |
| Nudt12 | 0.492608234 | 0.866416846 | 0 | 0.022 | 1 | 0.331907298 | -0.316927414 | 0 | 0.021 | 1 | 0.492608234 | 0.553652142 |
| D330050I16Rik | 0.492608234 | 0.866416846 | 0 | 0.022 | 1 | 0.331907298 | -0.316927414 | 0 | 0.021 | 1 | 0.492608234 | 0.553652142 |

| | | | | | | | | | | | | |
|---|---|---|---|---|---|---|---|---|---|---|---|---|
| Myorg | 0.532436372 | 1.088809267 | 0 | 0.019 | 1 | 0.331907298 | -0.316927414 | 0 | 0.021 | 1 | 0.532436372 | 0.553652142 |
| 4930417H01Rik | 0.544576436 | 2.258734268 | 0.048 | 0.026 | 1 | 0.331907298 | -0.486852415 | 0 | 0.021 | 1 | 0.544576436 | 0.553652142 |
| Gm26812 | 0.544576436 | 2.258734268 | 0.048 | 0.026 | 1 | 0.331907298 | -0.316927414 | 0 | 0.021 | 1 | 0.544576436 | 0.553652142 |
| Plppr2 | 0.544576436 | 2.673771768 | 0.048 | 0.026 | 1 | 0.331907298 | -0.316927414 | 0 | 0.021 | 1 | 0.544576436 | 0.553652142 |
| Gm26660 | 0.544576436 | 2.673771768 | 0.048 | 0.026 | 1 | 0.331907298 | -0.316927414 | 0 | 0.021 | 1 | 0.544576436 | 0.553652142 |
| Traf1 | 0.570211706 | 1.351843673 | 0.048 | 0.026 | 1 | 0.331907298 | -0.486852415 | 0 | 0.021 | 1 | 0.570211706 | 0.553652142 |
| Necab3 | 0.57672979 | 0.673771768 | 0.048 | 0.026 | 1 | 0.331907298 | -0.316927414 | 0 | 0.021 | 1 | 0.57672979 | 0.553652142 |
| Pgap6 | 0.578156445 | 1.088809267 | 0 | 0.015 | 1 | 0.331907298 | -0.316927414 | 0 | 0.021 | 1 | 0.578156445 | 0.553652142 |
| Gm16189 | 0.578156445 | 0.866416846 | 0 | 0.015 | 1 | 0.331907298 | -0.486852415 | 0 | 0.021 | 1 | 0.578156445 | 0.553652142 |
| Dpysl5 | 0.578159907 | 0.866416846 | 0 | 0.015 | 1 | 0.331907298 | -0.486852415 | 0 | 0.021 | 1 | 0.578159907 | 0.553652142 |
| Mmp11 | 0.578159907 | 0.866416846 | 0 | 0.015 | 1 | 0.331907298 | -0.316927414 | 0 | 0.021 | 1 | 0.578159907 | 0.553652142 |
| Gm43714 | 0.57816106 | 1.351843673 | 0 | 0.015 | 1 | 0.331907298 | -0.316927414 | 0 | 0.021 | 1 | 0.57816106 | 0.553652142 |
| Gm57291 | 0.57816106 | 1.351843673 | 0 | 0.015 | 1 | 0.331907298 | -0.486852415 | 0 | 0.021 | 1 | 0.57816106 | 0.553652142 |
| 1700028J19Rik | 0.57816106 | 1.351843673 | 0 | 0.015 | 1 | 0.331907298 | -0.486852415 | 0 | 0.021 | 1 | 0.57816106 | 0.553652142 |
| Pdf | 0.57816106 | 1.351843673 | 0 | 0.015 | 1 | 0.331907298 | -0.316927414 | 0 | 0.021 | 1 | 0.57816106 | 0.553652142 |
| Mettl24 | 0.57816106 | 1.351843673 | 0 | 0.015 | 1 | 0.331907298 | -0.316927414 | 0 | 0.021 | 1 | 0.57816106 | 0.553652142 |
| 2810021J22Rik | 0.57816106 | 1.351843673 | 0 | 0.015 | 1 | 0.331907298 | -0.316927414 | 0 | 0.021 | 1 | 0.57816106 | 0.553652142 |
| Fgf2os | 0.612659924 | 2.46732089 | 0.095 | 0.071 | 1 | 0.331907298 | -0.316927414 | 0 | 0.021 | 1 | 0.612659924 | 0.553652142 |
| Kcnn1 | 0.629700127 | 2.088809267 | 0.048 | 0.03 | 1 | 0.331907298 | -0.316927414 | 0 | 0.021 | 1 | 0.629700127 | 0.553652142 |
| Gm57069 | 0.63207476 | 1.351843673 | 0 | 0.011 | 1 | 0.331907298 | -0.486852415 | 0 | 0.021 | 1 | 0.63207476 | 0.553652142 |
| Mfap5 | 0.63207476 | 0.351843673 | 0 | 0.011 | 1 | 0.331907298 | -1.017367132 | 0 | 0.021 | 1 | 0.63207476 | 0.553652142 |
| Rem2 | 0.63207476 | 1.351843673 | 0 | 0.011 | 1 | 0.331907298 | -0.316927414 | 0 | 0.021 | 1 | 0.63207476 | 0.553652142 |
| Tbc1d10c | 0.63207476 | 1.351843673 | 0 | 0.011 | 1 | 0.331907298 | -0.316927414 | 0 | 0.021 | 1 | 0.63207476 | 0.553652142 |
| Dtx4 | 0.63207476 | 1.351843673 | 0 | 0.011 | 1 | 0.331907298 | -0.486852415 | 0 | 0.021 | 1 | 0.63207476 | 0.553652142 |
| Emx2os | 0.63207476 | 1.088809267 | 0 | 0.011 | 1 | 0.331907298 | -0.316927414 | 0 | 0.021 | 1 | 0.63207476 | 0.553652142 |
| Mccc1os | 0.632076134 | 1.673771768 | 0 | 0.011 | 1 | 0.331907298 | -0.486852415 | 0 | 0.021 | 1 | 0.632076134 | 0.553652142 |
| Mycn | 0.632076134 | 1.673771768 | 0 | 0.011 | 1 | 0.331907298 | -0.316927414 | 0 | 0.021 | 1 | 0.632076134 | 0.553652142 |
| 3300002A11Rik | 0.632076134 | 1.088809267 | 0 | 0.011 | 1 | 0.331907298 | -0.316927414 | 0 | 0.021 | 1 | 0.632076134 | 0.553652142 |
| Cd83 | 0.632076134 | 1.673771768 | 0 | 0.011 | 1 | 0.331907298 | -0.316927414 | 0 | 0.021 | 1 | 0.632076134 | 0.553652142 |
| Spef2 | 0.632076134 | 1.088809267 | 0 | 0.011 | 1 | 0.331907298 | -0.638855509 | 0 | 0.021 | 1 | 0.632076134 | 0.553652142 |
| Gm20468 | 0.632076134 | 1.673771768 | 0 | 0.011 | 1 | 0.331907298 | -1.124282336 | 0 | 0.021 | 1 | 0.632076134 | 0.553652142 |
| Crb3 | 0.632076134 | 1.673771768 | 0 | 0.011 | 1 | 0.331907298 | -0.316927414 | 0 | 0.021 | 1 | 0.632076134 | 0.553652142 |
| Sema4g | 0.632076134 | 1.673771768 | 0 | 0.011 | 1 | 0.331907298 | -0.316927414 | 0 | 0.021 | 1 | 0.632076134 | 0.553652142 |
| Kcnu1 | 0.632917461 | 1.936806174 | 0.048 | 0.03 | 1 | 0.331907298 | -0.316927414 | 0 | 0.021 | 1 | 0.632917461 | 0.553652142 |
| Adam1a | 0.658782751 | 1.351843673 | 0.048 | 0.03 | 1 | 0.331907298 | -0.316927414 | 0 | 0.021 | 1 | 0.658782751 | 0.553652142 |
| Tnnc1 | 0.896555897 | 0.673771768 | 0.048 | 0.056 | 1 | 0.331907298 | -0.316927414 | 0 | 0.021 | 1 | 0.896555897 | 0.553652142 |
| 1500009L16Rik | 0.954841465 | 0.97333205 | 0.048 | 0.045 | 1 | 0.331907298 | -0.316927414 | 0 | 0.021 | 1 | 0.954841465 | 0.553652142 |
| H2bc9 | 0.96386048 | 0.351843673 | 0.048 | 0.045 | 1 | 0.331907298 | -0.316927414 | 0 | 0.021 | 1 | 0.96386048 | 0.553652142 |
| Igf1os | 0.370807306 | -0.133583154 | 0 | 0.037 | 1 | 0.331914654 | -0.486852415 | 0 | 0.021 | 1 | 0.370807306 | 0.55366197 |
| A330015K06Rik | 0.370831626 | -1.081115734 | 0 | 0.037 | 1 | 0.331914654 | -0.776359032 | 0 | 0.021 | 1 | 0.370831626 | 0.55366197 |
| Rspo1 | 0.396957792 | -0.718545655 | 0 | 0.034 | 1 | 0.331914654 | -0.486852415 | 0 | 0.021 | 1 | 0.396957792 | 0.55366197 |
| Nkain1 | 0.396957792 | -0.785659851 | 0 | 0.034 | 1 | 0.331914654 | -1.017367132 | 0 | 0.021 | 1 | 0.396957792 | 0.55366197 |
| Hsd17b14 | 0.396971401 | 0.351843673 | 0 | 0.034 | 1 | 0.331914654 | -2.602329633 | 0 | 0.021 | 1 | 0.396971401 | 0.55366197 |
| Gm16833 | 0.396971401 | 0.351843673 | 0 | 0.034 | 1 | 0.331914654 | -0.486852415 | 0 | 0.021 | 1 | 0.396971401 | 0.55366197 |
| Them6 | 0.425599726 | 0.351843673 | 0 | 0.03 | 1 | 0.331914654 | -0.486852415 | 0 | 0.021 | 1 | 0.425599726 | 0.55366197 |
| Opn3 | 0.425614684 | 0.214340149 | 0 | 0.03 | 1 | 0.331914654 | -1.017367132 | 0 | 0.021 | 1 | 0.425614684 | 0.55366197 |
| Gm11767 | 0.457308648 | 0.673771768 | 0 | 0.026 | 1 | 0.331914654 | -0.486852415 | 0 | 0.021 | 1 | 0.457308648 | 0.55366197 |
| Scn8a | 0.457308648 | 0.673771768 | 0 | 0.026 | 1 | 0.331914654 | -0.638855509 | 0 | 0.021 | 1 | 0.457308648 | 0.55366197 |
| Nrg2 | 0.492608234 | 0.866416846 | 0 | 0.022 | 1 | 0.331914654 | -0.638855509 | 0 | 0.021 | 1 | 0.492608234 | 0.55366197 |
| Vcam1 | 0.532426384 | 0.866416846 | 0 | 0.019 | 1 | 0.331914654 | -1.223818009 | 0 | 0.021 | 1 | 0.532426384 | 0.55366197 |
| Fbxo24 | 0.532426384 | 0.673771768 | 0 | 0.019 | 1 | 0.331914654 | -0.486852415 | 0 | 0.021 | 1 | 0.532426384 | 0.55366197 |
| Gm44837 | 0.532432377 | 0.673771768 | 0 | 0.019 | 1 | 0.331914654 | -0.486852415 | 0 | 0.021 | 1 | 0.532432377 | 0.55366197 |
| 2900041M22Rik | 0.532436372 | 1.088809267 | 0 | 0.019 | 1 | 0.331914654 | -0.486852415 | 0 | 0.021 | 1 | 0.532436372 | 0.55366197 |
| Cenpe | 0.55415519 | 1.989273593 | 0.048 | 0.026 | 1 | 0.331914654 | -0.486852415 | 0 | 0.021 | 1 | 0.55415519 | 0.55366197 |
| Ppp1r42 | 0.578156445 | 0.866416846 | 0 | 0.015 | 1 | 0.331914654 | -0.901889915 | 0 | 0.021 | 1 | 0.578156445 | 0.55366197 |
| B230319C09Rik | 0.578156445 | 1.088809267 | 0 | 0.015 | 1 | 0.331914654 | -0.486852415 | 0 | 0.021 | 1 | 0.578156445 | 0.55366197 |
| Slc17a7 | 0.578156445 | 1.088809267 | 0 | 0.015 | 1 | 0.331914654 | -0.486852415 | 0 | 0.021 | 1 | 0.578156445 | 0.55366197 |
| Mtnr1a | 0.578156445 | 0.866416846 | 0 | 0.015 | 1 | 0.331914654 | -0.486852415 | 0 | 0.021 | 1 | 0.578156445 | 0.55366197 |
| Dnah7a | 0.578159907 | 0.214340149 | 0 | 0.015 | 1 | 0.331914654 | -0.486852415 | 0 | 0.021 | 1 | 0.578159907 | 0.55366197 |
| Pnpla1 | 0.57816106 | 1.351843673 | 0 | 0.015 | 1 | 0.331914654 | -0.486852415 | 0 | 0.021 | 1 | 0.57816106 | 0.55366197 |
| Frs3 | 0.57816106 | 1.351843673 | 0 | 0.015 | 1 | 0.331914654 | -0.486852415 | 0 | 0.021 | 1 | 0.57816106 | 0.55366197 |
| Frmpd4 | 0.579997897 | -0.648156327 | 0.048 | 0.026 | 1 | 0.331914654 | -1.84048937 | 0 | 0.021 | 1 | 0.579997897 | 0.55366197 |
| Frmd3 | 0.63207476 | 1.351843673 | 0 | 0.011 | 1 | 0.331914654 | -0.638855509 | 0 | 0.021 | 1 | 0.63207476 | 0.55366197 |
| Stmn3 | 0.632076134 | 1.673771768 | 0 | 0.011 | 1 | 0.331914654 | -0.638855509 | 0 | 0.021 | 1 | 0.632076134 | 0.55366197 |
| 1110032F04Rik | 0.632076134 | 1.673771768 | 0 | 0.011 | 1 | 0.331914654 | -0.486852415 | 0 | 0.021 | 1 | 0.632076134 | 0.55366197 |
| Ppm1j | 0.632076134 | 1.673771768 | 0 | 0.011 | 1 | 0.331914654 | -0.486852415 | 0 | 0.021 | 1 | 0.632076134 | 0.55366197 |
| Abcb4 | 0.632076134 | 1.673771768 | 0 | 0.011 | 1 | 0.331914654 | -0.638855509 | 0 | 0.021 | 1 | 0.632076134 | 0.55366197 |
| Gm26787 | 0.632076134 | 1.673771768 | 0 | 0.011 | 1 | 0.331914654 | -0.486852415 | 0 | 0.021 | 1 | 0.632076134 | 0.55366197 |
| Tecta | 0.632076134 | 0.351843673 | 0 | 0.011 | 1 | 0.331914654 | -0.486852415 | 0 | 0.021 | 1 | 0.632076134 | 0.55366197 |
| 4933428G20Rik | 0.632076134 | 1.673771768 | 0 | 0.011 | 1 | 0.331914654 | -0.486852415 | 0 | 0.021 | 1 | 0.632076134 | 0.55366197 |
| Bex1 | 0.632076134 | 1.673771768 | 0 | 0.011 | 1 | 0.331914654 | -0.486852415 | 0 | 0.021 | 1 | 0.632076134 | 0.55366197 |
| Zcchc18 | 0.632076134 | 1.673771768 | 0 | 0.011 | 1 | 0.331914654 | -0.486852415 | 0 | 0.021 | 1 | 0.632076134 | 0.55366197 |
| Kif6 | 0.658782751 | 1.214340149 | 0.048 | 0.03 | 1 | 0.331914654 | -0.486852415 | 0 | 0.021 | 1 | 0.658782751 | 0.55366197 |
| Gm3455 | 0.784067278 | 3.258734268 | 0.048 | 0.037 | 1 | 0.331914654 | -0.776359032 | 0 | 0.021 | 1 | 0.784067278 | 0.55366197 |
| 9430037G07Rik | 0.79666227 | 1.351843673 | 0.048 | 0.037 | 1 | 0.331914654 | -0.638855509 | 0 | 0.021 | 1 | 0.79666227 | 0.55366197 |
| Gulo | 0.904678224 | 1.137718867 | 0.048 | 0.056 | 1 | 0.331914654 | -0.901889915 | 0 | 0.021 | 1 | 0.904678224 | 0.55366197 |
| Il1r2 | 0.979645995 | 2.38426515 | 0.048 | 0.049 | 1 | 0.331914654 | -1.709244837 | 0 | 0.021 | 1 | 0.979645995 | 0.55366197 |
| Lipt2 | 0.985457491 | 0.766881172 | 0.048 | 0.049 | 1 | 0.331914654 | -0.638855509 | 0 | 0.021 | 1 | 0.985457491 | 0.55366197 |
| Anln | 0.985461775 | 1.819622634 | 0.048 | 0.049 | 1 | 0.331914654 | -0.638855509 | 0 | 0.021 | 1 | 0.985461775 | 0.55366197 |
| Knl1 | 0.370833405 | -1.535681598 | 0 | 0.037 | 1 | 0.331918331 | -0.776359032 | 0 | 0.021 | 1 | 0.370833405 | 0.553666884 |
| Ripor2 | 0.396957792 | -2.054148687 | 0 | 0.034 | 1 | 0.331918331 | -1.486852415 | 0 | 0.021 | 1 | 0.396957792 | 0.553666884 |
| 2810407A14Rik | 0.532432377 | 0.673771768 | 0 | 0.019 | 1 | 0.331918331 | -1.223818009 | 0 | 0.021 | 1 | 0.532432377 | 0.553666884 |
| Gtf2a1l | 0.532432377 | 0.503846766 | 0 | 0.019 | 1 | 0.331918331 | -1.017367132 | 0 | 0.021 | 1 | 0.532432377 | 0.553666884 |
| Gm48898 | 0.57816106 | 1.351843673 | 0 | 0.015 | 1 | 0.331918331 | -0.901889915 | 0 | 0.021 | 1 | 0.57816106 | 0.553666884 |
| Tpx2 | 0.613087907 | 0.866416846 | 0.048 | 0.082 | 1 | 0.331918331 | -0.901889915 | 0 | 0.021 | 1 | 0.613087907 | 0.553666884 |
| Ryr2 | 0.63207476 | 1.351843673 | 0 | 0.011 | 1 | 0.331918331 | -0.901889915 | 0 | 0.021 | 1 | 0.63207476 | 0.553666884 |
| Rgs16 | 0.841862066 | 0.503846766 | 0.048 | 0.06 | 1 | 0.331918331 | -0.776359032 | 0 | 0.021 | 1 | 0.841862066 | 0.553666884 |
| Gm44127 | 0.869134278 | 0.735172312 | 0.048 | 0.041 | 1 | 0.331918331 | -0.901889915 | 0 | 0.021 | 1 | 0.869134278 | 0.553666884 |
| Ckap2 | 0.933804498 | 0.281454345 | 0.048 | 0.045 | 1 | 0.331918331 | -0.776359032 | 0 | 0.021 | 1 | 0.933804498 | 0.553666884 |
| 4933412E12Rik | 0.425599726 | 0.351843673 | 0 | 0.03 | 1 | 0.331919067 | -0.638855509 | 0 | 0.021 | 1 | 0.425599726 | 0.553667867 |
| Kcnj2 | 0.457292877 | -1.940938076 | 0 | 0.026 | 1 | 0.331919067 | -1.124282336 | 0 | 0.021 | 1 | 0.457292877 | 0.553667867 |
| Tex15 | 0.463493535 | 1.747772349 | 0.048 | 0.022 | 1 | 0.331919067 | -1.017367132 | 0 | 0.021 | 1 | 0.463493535 | 0.553667867 |
| Gm36371 | 0.544576436 | 2.258734268 | 0.048 | 0.026 | 1 | 0.331919067 | -0.638855509 | 0 | 0.021 | 1 | 0.544576436 | 0.553667867 |
| Tfap4 | 0.544576436 | 2.258734268 | 0.048 | 0.026 | 1 | 0.331919067 | -0.638855509 | 0 | 0.021 | 1 | 0.544576436 | 0.553667867 |
| Gpr183 | 0.63207476 | 1.351843673 | 0 | 0.011 | 1 | 0.331919067 | -1.960783604 | 0 | 0.021 | 1 | 0.63207476 | 0.553667867 |
| 1700109G14Rik | 0.632076134 | 1.673771768 | 0 | 0.011 | 1 | 0.331919067 | -1.017367132 | 0 | 0.021 | 1 | 0.632076134 | 0.553667867 |
| Gm11290 | 0.862957731 | 1.503846766 | 0.048 | 0.041 | 1 | 0.331919067 | -1.638855509 | 0 | 0.021 | 1 | 0.862957731 | 0.553667867 |
| Agbl2 | 0.9001036 | 0.586308926 | 0.048 | 0.041 | 1 | 0.331920538 | -1.223818009 | 0 | 0.021 | 1 | 0.9001036 | 0.553669832 |

| | | | | | | | | | | | | |
|---|---|---|---|---|---|---|---|---|---|---|---|---|
| Txndc8 | 0.532432377 | 0.351843673 | 0 | 0.019 | 1 | 0.331921273 | -1.564854927 | 0 | 0.021 | 1 | 0.532432377 | 0.553670815 |
| Oscar | 0.632076134 | 1.673771768 | 0 | 0.011 | 1 | 0.331921273 | -2.486852415 | 0 | 0.021 | 1 | 0.632076134 | 0.553670815 |
| Fst | 0.958517695 | 0.843696769 | 0.095 | 0.101 | 1 | 0.331921273 | -3.691966845 | 0 | 0.021 | 1 | 0.958517695 | 0.553670815 |
| Phf19 | 0.359280535 | 2.866416846 | 0.048 | 0.019 | 1 | 0.331922009 | -0.124282336 | 0 | 0.021 | 1 | 0.359280535 | 0.553671798 |
| Gm13056 | 0.370840523 | 0.214340149 | 0 | 0.037 | 1 | 0.331922009 | -0.124282336 | 0 | 0.021 | 1 | 0.370840523 | 0.553671798 |
| Ffar4 | 0.376794341 | 1.673771768 | 0.048 | 0.019 | 1 | 0.331922009 | -0.124282336 | 0 | 0.021 | 1 | 0.376794341 | 0.553671798 |
| Gm57475 | 0.396971401 | 0.351843673 | 0 | 0.034 | 1 | 0.331922009 | -0.124282336 | 0 | 0.021 | 1 | 0.396971401 | 0.553671798 |
| BB218582 | 0.425614684 | 0.214340149 | 0 | 0.03 | 1 | 0.331922009 | -0.124282336 | 0 | 0.021 | 1 | 0.425614684 | 0.553671798 |
| Gm57411 | 0.425639612 | 0.503846766 | 0 | 0.03 | 1 | 0.331922009 | -0.124282336 | 0 | 0.021 | 1 | 0.425639612 | 0.553671798 |
| Hhla1 | 0.425639612 | 0.503846766 | 0 | 0.03 | 1 | 0.331922009 | -0.124282336 | 0 | 0.021 | 1 | 0.425639612 | 0.553671798 |
| Elovl2 | 0.457292877 | 0.214340149 | 0 | 0.026 | 1 | 0.331922009 | -0.124282336 | 0 | 0.021 | 1 | 0.457292877 | 0.553671798 |
| Cfap126 | 0.457308648 | 0.673771768 | 0 | 0.026 | 1 | 0.331922009 | -0.124282336 | 0 | 0.021 | 1 | 0.457308648 | 0.553671798 |
| Zfp691 | 0.457308648 | 0.673771768 | 0 | 0.026 | 1 | 0.331922009 | -0.124282336 | 0 | 0.021 | 1 | 0.457308648 | 0.553671798 |
| Opn1sw | 0.492590603 | 0.673771768 | 0 | 0.022 | 1 | 0.331922009 | -0.124282336 | 0 | 0.021 | 1 | 0.492590603 | 0.553671798 |
| Pianp | 0.492590603 | 0.673771768 | 0 | 0.022 | 1 | 0.331922009 | -0.124282336 | 0 | 0.021 | 1 | 0.492590603 | 0.553671798 |
| Gm43598 | 0.492599419 | 0.351843673 | 0 | 0.022 | 1 | 0.331922009 | -0.124282336 | 0 | 0.021 | 1 | 0.492599419 | 0.553671798 |
| 4932443L11Rik | 0.492608234 | 0.866416846 | 0 | 0.022 | 1 | 0.331922009 | -0.124282336 | 0 | 0.021 | 1 | 0.492608234 | 0.553671798 |
| Fgl1 | 0.492608234 | 0.866416846 | 0 | 0.022 | 1 | 0.331922009 | -0.124282336 | 0 | 0.021 | 1 | 0.492608234 | 0.553671798 |
| Gpr87 | 0.532426384 | 0.866416846 | 0 | 0.019 | 1 | 0.331922009 | -0.124282336 | 0 | 0.021 | 1 | 0.532426384 | 0.553671798 |
| C530020B09Rik | 0.532426384 | 0.866416846 | 0 | 0.019 | 1 | 0.331922009 | -0.124282336 | 0 | 0.021 | 1 | 0.532426384 | 0.553671798 |
| Muc4 | 0.532426384 | -0.413691074 | 0 | 0.019 | 1 | 0.331922009 | -0.124282336 | 0 | 0.021 | 1 | 0.532426384 | 0.553671798 |
| 1010001B22Rik | 0.532436372 | 1.088809267 | 0 | 0.019 | 1 | 0.331922009 | -0.124282336 | 0 | 0.021 | 1 | 0.532436372 | 0.553671798 |
| Gm57397 | 0.544576436 | 2.258734268 | 0.048 | 0.026 | 1 | 0.331922009 | -0.124282336 | 0 | 0.021 | 1 | 0.544576436 | 0.553671798 |
| Plp1 | 0.54776222 | 2.088809267 | 0.048 | 0.026 | 1 | 0.331922009 | -0.124282336 | 0 | 0.021 | 1 | 0.54776222 | 0.553671798 |
| Stag3 | 0.570211706 | 1.503846766 | 0.048 | 0.026 | 1 | 0.331922009 | -0.124282336 | 0 | 0.021 | 1 | 0.570211706 | 0.553671798 |
| ENSMUSG00000121346 | 0.578156445 | 1.088809267 | 0 | 0.015 | 1 | 0.331922009 | -0.124282336 | 0 | 0.021 | 1 | 0.578156445 | 0.553671798 |
| 5930403N24Rik | 0.578156445 | 1.088809267 | 0 | 0.015 | 1 | 0.331922009 | -0.124282336 | 0 | 0.021 | 1 | 0.578156445 | 0.553671798 |
| Gm6403 | 0.578159907 | 0.503846766 | 0 | 0.015 | 1 | 0.331922009 | -0.124282336 | 0 | 0.021 | 1 | 0.578159907 | 0.553671798 |
| 2410004P03Rik | 0.57816106 | 1.351843673 | 0 | 0.015 | 1 | 0.331922009 | -0.124282336 | 0 | 0.021 | 1 | 0.57816106 | 0.553671798 |
| Cideb | 0.57816106 | 1.351843673 | 0 | 0.015 | 1 | 0.331922009 | -0.124282336 | 0 | 0.021 | 1 | 0.57816106 | 0.553671798 |
| Lrrc24 | 0.57816106 | 1.351843673 | 0 | 0.015 | 1 | 0.331922009 | -0.124282336 | 0 | 0.021 | 1 | 0.57816106 | 0.553671798 |
| Gm49584 | 0.57816106 | 1.351843673 | 0 | 0.015 | 1 | 0.331922009 | -0.124282336 | 0 | 0.021 | 1 | 0.57816106 | 0.553671798 |
| Gm50340 | 0.57816106 | 1.351843673 | 0 | 0.015 | 1 | 0.331922009 | -0.124282336 | 0 | 0.021 | 1 | 0.57816106 | 0.553671798 |
| Gm3625 | 0.629700127 | 2.088809267 | 0.048 | 0.03 | 1 | 0.331922009 | -0.124282336 | 0 | 0.021 | 1 | 0.629700127 | 0.553671798 |
| Calr4 | 0.63207476 | 1.351843673 | 0 | 0.011 | 1 | 0.331922009 | -0.124282336 | 0 | 0.021 | 1 | 0.63207476 | 0.553671798 |
| Gm42982 | 0.63207476 | 1.351843673 | 0 | 0.011 | 1 | 0.331922009 | -0.124282336 | 0 | 0.021 | 1 | 0.63207476 | 0.553671798 |
| Gm26826 | 0.63207476 | 1.351843673 | 0 | 0.011 | 1 | 0.331922009 | -0.124282336 | 0 | 0.021 | 1 | 0.63207476 | 0.553671798 |
| Gm28151 | 0.632076134 | 1.673771768 | 0 | 0.011 | 1 | 0.331922009 | -0.124282336 | 0 | 0.021 | 1 | 0.632076134 | 0.553671798 |
| C8g | 0.632076134 | 1.673771768 | 0 | 0.011 | 1 | 0.331922009 | -0.124282336 | 0 | 0.021 | 1 | 0.632076134 | 0.553671798 |
| Gm15747 | 0.632076134 | 1.673771768 | 0 | 0.011 | 1 | 0.331922009 | -0.124282336 | 0 | 0.021 | 1 | 0.632076134 | 0.553671798 |
| Gm15402 | 0.632076134 | 1.088809267 | 0 | 0.011 | 1 | 0.331922009 | -0.124282336 | 0 | 0.021 | 1 | 0.632076134 | 0.553671798 |
| Ido2 | 0.632076134 | 1.673771768 | 0 | 0.011 | 1 | 0.331922009 | -0.124282336 | 0 | 0.021 | 1 | 0.632076134 | 0.553671798 |
| Gm3604 | 0.632076134 | 1.673771768 | 0 | 0.011 | 1 | 0.331922009 | -0.124282336 | 0 | 0.021 | 1 | 0.632076134 | 0.553671798 |
| Nudt15 | 0.632076134 | 1.673771768 | 0 | 0.011 | 1 | 0.331922009 | -0.124282336 | 0 | 0.021 | 1 | 0.632076134 | 0.553671798 |
| Gm10053 | 0.632076134 | 1.673771768 | 0 | 0.011 | 1 | 0.331922009 | -0.124282336 | 0 | 0.021 | 1 | 0.632076134 | 0.553671798 |
| Gm29650 | 0.632076134 | 1.673771768 | 0 | 0.011 | 1 | 0.331922009 | -0.124282336 | 0 | 0.021 | 1 | 0.632076134 | 0.553671798 |
| Prelid2 | 0.890779585 | 0.673771768 | 0.048 | 0.041 | 1 | 0.331922009 | -0.124282336 | 0 | 0.021 | 1 | 0.890779585 | 0.553671798 |
| Gm47512 | 0.985459372 | 1.258734268 | 0.048 | 0.049 | 1 | 0.331922009 | -0.124282336 | 0 | 0.021 | 1 | 0.985459372 | 0.553671798 |
| Gm16070 | 0.332083509 | 0.333147579 | 0.571 | 0.53 | 1 | 0.392096202 | 0.717838004 | 0.533 | 0.585 | 1 | 0.392096202 | 0.553887561 |
| Ccdc6 | 0.855100193 | 0.629377648 | 0.381 | 0.448 | 1 | 0.332211647 | 0.494942989 | 0.622 | 0.623 | 1 | 0.855100193 | 0.554058715 |
| Upp2 | 0.446096507 | 1.334356246 | 0.476 | 0.545 | 1 | 0.332252692 | 0.628995913 | 0.289 | 0.474 | 1 | 0.446096507 | 0.554113532 |
| Srrm1 | 0.332782185 | 0.267779408 | 0.952 | 0.985 | 1 | 0.754832199 | 0.166685083 | 1 | 0.983 | 1 | 0.754832199 | 0.554820387 |
| Cplane1 | 0.332929014 | 0.186106468 | 0.286 | 0.466 | 1 | 0.487143883 | 0.381182245 | 0.667 | 0.599 | 1 | 0.487143883 | 0.5550163 |
| Adamtsl3 | 0.333298594 | 2.361320951 | 0.714 | 0.698 | 1 | 0.441055901 | 0.338408139 | 0.711 | 0.789 | 1 | 0.441055901 | 0.555509235 |
| Myo15a | 0.333372685 | 1.766881172 | 0.143 | 0.09 | 1 | 0.648063697 | 0.470078863 | 0.089 | 0.118 | 1 | 0.648063697 | 0.555608023 |
| Aldh1l2 | 0.454581689 | 0.614878079 | 0.143 | 0.093 | 1 | 0.333548206 | -0.294207337 | 0.044 | 0.09 | 1 | 0.454581689 | 0.555842006 |
| Ssbp2 | 0.333637312 | 0.417873543 | 0.714 | 0.601 | 1 | 0.594718372 | 0.411476167 | 0.978 | 0.848 | 1 | 0.594718372 | 0.555960768 |
| Setd4 | 0.333710188 | 2.088809267 | 0.095 | 0.049 | 1 | 0.490615597 | 1.290755163 | 0.089 | 0.062 | 1 | 0.490615597 | 0.556057887 |
| Insyn2a | 0.333815942 | 1.351843673 | 0.095 | 0.049 | 1 | 0.410749809 | 1.013221188 | 0.089 | 0.059 | 1 | 0.410749809 | 0.556198801 |
| Lrrc73 | 0.813128768 | 0.936806174 | 0.048 | 0.063 | 1 | 0.333826679 | -0.539319835 | 0.044 | 0.087 | 1 | 0.813128768 | 0.556213106 |
| Zc3h4 | 0.903697012 | 0.203451833 | 0.333 | 0.328 | 1 | 0.334514255 | 0.506749814 | 0.311 | 0.474 | 1 | 0.903697012 | 0.557128723 |
| Gas7 | 0.826374436 | 2.190989662 | 0.19 | 0.201 | 1 | 0.33452477 | 1.161119883 | 0.244 | 0.204 | 1 | 0.826374436 | 0.557142718 |
| Nol8 | 0.334701814 | 0.146034136 | 0.238 | 0.369 | 1 | 0.391466806 | 0.606256989 | 0.2 | 0.291 | 1 | 0.391466806 | 0.557378324 |
| Trpm4 | 0.334719623 | 2.258734268 | 0.143 | 0.09 | 1 | 0.911378283 | 0.647448676 | 0.111 | 0.111 | 1 | 0.911378283 | 0.557402019 |
| Hmg20a | 0.768766556 | 0.866416846 | 0.476 | 0.601 | 1 | 0.334786097 | 0.727015934 | 0.6 | 0.633 | 1 | 0.768766556 | 0.557490464 |
| Snx14 | 0.884388328 | 0.736823938 | 0.429 | 0.496 | 1 | 0.33489041 | 0.297181433 | 0.533 | 0.602 | 1 | 0.884388328 | 0.557629233 |
| Nrxn2 | 0.608897862 | 0.400753273 | 0.048 | 0.082 | 1 | 0.334940602 | 0.177837278 | 0.178 | 0.27 | 1 | 0.608897862 | 0.557695997 |
| Psme1 | 0.511189854 | -0.24509147 | 0.429 | 0.507 | 1 | 0.335232715 | 0.196572724 | 0.711 | 0.543 | 1 | 0.511189854 | 0.558084457 |
| Trp53bp1 | 0.482715412 | 0.766881172 | 0.429 | 0.422 | 1 | 0.3360099 | 0.217949855 | 0.356 | 0.512 | 1 | 0.482715412 | 0.559117148 |
| Trim36 | 0.532426384 | 0.866416846 | 0 | 0.019 | 1 | 0.336119631 | 1.304560963 | 0.044 | 0.021 | 1 | 0.532426384 | 0.559262856 |
| Gm26779 | 0.532426384 | 0.866416846 | 0 | 0.019 | 1 | 0.336122404 | 1.460680165 | 0.044 | 0.021 | 1 | 0.532426384 | 0.559266538 |
| Gm49173 | 0.578156445 | 1.088809267 | 0 | 0.015 | 1 | 0.336122404 | 1.460680165 | 0.044 | 0.021 | 1 | 0.578156445 | 0.559266538 |
| B3gnt6 | 0.632076134 | 1.673771768 | 0 | 0.011 | 1 | 0.336122404 | 1.460680165 | 0.044 | 0.021 | 1 | 0.632076134 | 0.559266538 |
| C030013E06Rik | 0.863644336 | 0.119182916 | 0.095 | 0.108 | 1 | 0.336366361 | -0.425451871 | 0.067 | 0.118 | 1 | 0.863644336 | 0.559590393 |
| Prmt7 | 0.336954338 | 1.631951592 | 0.238 | 0.187 | 1 | 0.451827276 | 0.669266787 | 0.156 | 0.232 | 1 | 0.451827276 | 0.56037045 |
| Snx4 | 0.896313488 | 0.268515289 | 0.524 | 0.679 | 1 | 0.337064118 | 0.398558453 | 0.867 | 0.799 | 1 | 0.896313488 | 0.560516017 |
| Acbd4 | 0.337152681 | -0.574155746 | 0.048 | 0.116 | 1 | 0.506676772 | 0.361144491 | 0.067 | 0.1 | 1 | 0.506676772 | 0.560633432 |
| Peli2 | 0.969048738 | 0.97333205 | 0.19 | 0.201 | 1 | 0.337272339 | 0.492129803 | 0.222 | 0.329 | 1 | 0.969048738 | 0.560792048 |
| Vti1a | 0.957987795 | 0.150209812 | 0.524 | 0.586 | 1 | 0.337306035 | 0.789620365 | 0.822 | 0.675 | 1 | 0.957987795 | 0.560836709 |
| Gm15987 | 0.337419573 | -2.701267664 | 0.048 | 0.112 | 1 | 0.492040936 | 2.683072586 | 0.022 | 0.01 | 1 | 0.492040936 | 0.560987178 |
| Polh | 0.337932486 | 0.181918671 | 0.048 | 0.123 | 1 | 0.71338542 | 0.578735926 | 0.089 | 0.111 | 1 | 0.71338542 | 0.561666607 |
| Tcf25 | 0.419847053 | 0.285754482 | 0.905 | 0.966 | 1 | 0.3380751 | 0.120362892 | 0.956 | 0.983 | 1 | 0.419847053 | 0.561855426 |
| Iffo2 | 0.338820412 | 1.386195178 | 0.524 | 0.526 | 1 | 0.699238952 | -0.344177448 | 0.733 | 0.609 | 1 | 0.699238952 | 0.562841552 |
| Zbtb46 | 0.33986077 | 1.58450443 | 0.381 | 0.347 | 1 | 0.67431613 | 0.337780498 | 0.333 | 0.422 | 1 | 0.67431613 | 0.564216197 |
| Slc8b1 | 0.516390116 | 0.620660431 | 0.238 | 0.198 | 1 | 0.340108098 | 0.316750372 | 0.244 | 0.346 | 1 | 0.516390116 | 0.564542677 |
| Tnni2 | 0.340168473 | 0.351843673 | 0.048 | 0.123 | 1 | 0.381723291 | 0.238287743 | 0.067 | 0.114 | 1 | 0.381723291 | 0.564622356 |
| Gm48742 | 0.340317491 | 0.281454345 | 0.048 | 0.123 | 1 | 0.406553529 | 0.268035087 | 0.022 | 0.052 | 1 | 0.406553529 | 0.564818987 |
| Ptpn23 | 0.607996904 | 0.54907602 | 0.381 | 0.444 | 1 | 0.340446046 | 0.14309754 | 0.311 | 0.464 | 1 | 0.607996904 | 0.564988582 |
| Wdr4 | 0.665138226 | 0.388369549 | 0.095 | 0.134 | 1 | 0.340655515 | 0.65559185 | 0.089 | 0.152 | 1 | 0.665138226 | 0.565264851 |
| Cdkl1 | 0.34092612 | 1.410737362 | 0.095 | 0.049 | 1 | 0.993422097 | 0.624178897 | 0.089 | 0.093 | 1 | 0.993422097 | 0.565621621 |
| Gpr68 | 0.949860134 | 0.304537958 | 0.095 | 0.09 | 1 | 0.340980098 | -1.071814916 | 0.022 | 0.055 | 1 | 0.949860134 | 0.565692768 |
| Enpep | 0.346827513 | -0.133583154 | 0 | 0.041 | 1 | 0.340984118 | -1.316927414 | 0.022 | 0.055 | 1 | 0.346827513 | 0.565698068 |
| Rufy4 | 0.968254366 | 1.847801167 | 0.143 | 0.168 | 1 | 0.341296882 | 0.429315994 | 0.067 | 0.121 | 1 | 0.968254366 | 0.566110202 |
| Bbx | 0.379982021 | 0.227227833 | 0.905 | 0.888 | 1 | 0.341378301 | 0.47891352 | 0.978 | 0.917 | 1 | 0.379982021 | 0.566217458 |
| Mms19 | 0.673872656 | 0.714413752 | 0.143 | 0.194 | 1 | 0.342111434 | 0.58546179 | 0.156 | 0.246 | 1 | 0.673872656 | 0.567182634 |
| Fat3 | 0.57073199 | -1.150656668 | 0.095 | 0.142 | 1 | 0.342248819 | 0.908632287 | 0.2 | 0.159 | 1 | 0.57073199 | 0.567363383 |
| Zfp943 | 0.796905186 | 0.657469955 | 0.238 | 0.243 | 1 | 0.342275571 | 0.605070074 | 0.156 | 0.246 | 1 | 0.796905186 | 0.567398576 |
| Gm9975 | 0.342664981 | 1.586308926 | 0.095 | 0.049 | 1 | 0.585819504 | 1.098110085 | 0.067 | 0.048 | 1 | 0.585819504 | 0.567910673 |

| | | | | | | | | | | | | |
|---|---|---|---|---|---|---|---|---|---|---|---|---|
| C5ar2 | 0.532436372 | 1.088809267 | 0 | 0.019 | 1 | 0.343164642 | -0.84048937 | 0.022 | 0.055 | 1 | 0.532436372 | 0.568567313 |
| Gm16835 | 0.803145338 | 0.673771768 | 0.048 | 0.063 | 1 | 0.34316625 | -1.174908409 | 0.022 | 0.055 | 1 | 0.803145338 | 0.568569425 |
| Tnk2 | 0.63171544 | -0.175893959 | 0.286 | 0.358 | 1 | 0.34363901 | 0.145107565 | 0.222 | 0.325 | 1 | 0.63171544 | 0.569190251 |
| Ttll3 | 0.343776369 | -1.337455488 | 0.143 | 0.22 | 1 | 0.546112432 | 0.464237984 | 0.222 | 0.318 | 1 | 0.546112432 | 0.569370546 |
| Cnksr2 | 0.370832219 | -0.911190733 | 0 | 0.037 | 1 | 0.344000254 | 0.435145073 | 0.044 | 0.09 | 1 | 0.370832219 | 0.569664333 |
| Zdhhc17 | 0.67012315 | 0.757295123 | 0.333 | 0.354 | 1 | 0.344178086 | 0.41458375 | 0.289 | 0.471 | 1 | 0.67012315 | 0.569897617 |
| Bend4 | 0.57816106 | 1.351843673 | 0 | 0.015 | 1 | 0.344255941 | -0.84048937 | 0.022 | 0.055 | 1 | 0.57816106 | 0.569999729 |
| Gm50431 | 0.640268317 | 1.351843673 | 0.095 | 0.071 | 1 | 0.344257012 | -0.709244837 | 0.022 | 0.055 | 1 | 0.640268317 | 0.570001134 |
| Kcnrg | 0.532426384 | 0.866416846 | 0 | 0.019 | 1 | 0.344430448 | 1.361144491 | 0.111 | 0.073 | 1 | 0.532426384 | 0.570228562 |
| Hykk | 0.344460627 | 1.503846766 | 0.095 | 0.049 | 1 | 0.587470466 | 1.361144491 | 0.044 | 0.069 | 1 | 0.587470466 | 0.57026813 |
| Gm30963 | 0.344460627 | 1.503846766 | 0.095 | 0.049 | 1 | 0.615717475 | 0.567595369 | 0.022 | 0.038 | 1 | 0.615717475 | 0.57026813 |
| Gm15983 | 0.344460627 | 1.088809267 | 0.095 | 0.049 | 1 | 0.84962781 | 0.435145073 | 0.044 | 0.052 | 1 | 0.84962781 | 0.57026813 |
| Adarb1 | 0.344628195 | 0.516230491 | 0.19 | 0.299 | 1 | 0.600309267 | 0.511306238 | 0.289 | 0.384 | 1 | 0.600309267 | 0.570487797 |
| Ttc28 | 0.344778514 | 1.351843673 | 0.857 | 0.903 | 1 | 0.805068027 | 0.346497715 | 0.933 | 0.917 | 1 | 0.805068027 | 0.570684804 |
| Hnmt | 0.425599726 | 0.351843673 | 0 | 0.03 | 1 | 0.345345252 | -0.709244837 | 0.022 | 0.055 | 1 | 0.425599726 | 0.571427161 |
| Ubxn11 | 0.770713471 | 0.150209812 | 0.048 | 0.063 | 1 | 0.34534552 | -0.638855509 | 0.022 | 0.055 | 1 | 0.770713471 | 0.571427511 |
| Ythdc1 | 0.345569482 | 0.245195001 | 1 | 0.978 | 1 | 0.946061093 | 0.219816724 | 0.956 | 0.986 | 1 | 0.946061093 | 0.571720697 |
| Atg7 | 0.346401071 | 1.417432014 | 0.286 | 0.235 | 1 | 0.73812522 | 1.057468101 | 0.2 | 0.263 | 1 | 0.73812522 | 0.57280844 |
| Gm17036 | 0.79666227 | 1.351843673 | 0.048 | 0.037 | 1 | 0.346427229 | 0.446033389 | 0.044 | 0.09 | 1 | 0.79666227 | 0.572842633 |
| Gm13427 | 0.346802964 | -0.133583154 | 0 | 0.041 | 1 | 0.592392472 | -0.223818009 | 0.022 | 0.038 | 1 | 0.592392472 | 0.573333632 |
| Ttc38 | 0.346802964 | -0.133583154 | 0 | 0.041 | 1 | 0.821674021 | 0.982632868 | 0.044 | 0.038 | 1 | 0.821674021 | 0.573333632 |
| 1700056N10Rik | 0.346802964 | -0.326228232 | 0 | 0.041 | 1 | 0.366897007 | 0.513147585 | 0.022 | 0.055 | 1 | 0.366897007 | 0.573333632 |
| Slc1a2 | 0.346827513 | -0.233118828 | 0 | 0.041 | 1 | 0.473939177 | 0.960606562 | 0.089 | 0.062 | 1 | 0.473939177 | 0.573365703 |
| A530013C23Rik | 0.346827513 | -0.233118828 | 0 | 0.041 | 1 | 0.595909655 | 1.567595369 | 0.089 | 0.069 | 1 | 0.595909655 | 0.573365703 |
| Sv2a | 0.346827513 | -0.133583154 | 0 | 0.041 | 1 | 0.630232557 | 0.513147585 | 0.044 | 0.066 | 1 | 0.630232557 | 0.573365703 |
| Arsj | 0.346827513 | -0.233118828 | 0 | 0.041 | 1 | 0.974172726 | -0.404390255 | 0.044 | 0.045 | 1 | 0.974172726 | 0.573365703 |
| D830036C21Rik | 0.346827513 | -0.133583154 | 0 | 0.041 | 1 | 0.855379145 | 1.188307894 | 0.067 | 0.08 | 1 | 0.855379145 | 0.573365703 |
| Upk1a | 0.346827513 | -0.233118828 | 0 | 0.041 | 1 | 0.911606948 | 1.039216396 | 0.067 | 0.076 | 1 | 0.911606948 | 0.573365703 |
| Gas2 | 0.346827513 | -0.133583154 | 0 | 0.041 | 1 | 0.426266324 | -0.776359032 | 0.022 | 0.048 | 1 | 0.426266324 | 0.573365703 |
| Gm45494 | 0.346827513 | -0.133583154 | 0 | 0.041 | 1 | 0.936025227 | 1.683072586 | 0.022 | 0.021 | 1 | 0.936025227 | 0.573365703 |
| Gm39302 | 0.346827513 | -0.133583154 | 0 | 0.041 | 1 | 0.496223879 | 0.683072586 | 0 | 0.01 | 1 | 0.496223879 | 0.573365703 |
| 4833445I07Rik | 0.346827513 | -0.233118828 | 0 | 0.041 | 1 | 0.49622258 | 0.361144491 | 0 | 0.01 | 1 | 0.49622258 | 0.573365703 |
| Stac3 | 0.346827513 | -0.133583154 | 0 | 0.041 | 1 | 0.81549849 | 1.098110085 | 0.044 | 0.055 | 1 | 0.81549849 | 0.573365703 |
| Hs1bp3 | 0.346827513 | -0.326228232 | 0 | 0.041 | 1 | 0.423581872 | 1.776181991 | 0.067 | 0.042 | 1 | 0.423581872 | 0.573365703 |
| Rslcan18 | 0.346827513 | -0.133583154 | 0 | 0.041 | 1 | 0.977857988 | 0.180572246 | 0.044 | 0.045 | 1 | 0.977857988 | 0.573365703 |
| Sall2 | 0.346827513 | -0.133583154 | 0 | 0.041 | 1 | 0.61754131 | 1.197645759 | 0.022 | 0.038 | 1 | 0.61754131 | 0.573365703 |
| Gm56696 | 0.346827513 | -0.133583154 | 0 | 0.041 | 1 | 0.500221785 | 0.875717664 | 0.022 | 0.045 | 1 | 0.500221785 | 0.573365703 |
| Gm49741 | 0.346827513 | -0.133583154 | 0 | 0.041 | 1 | 0.936025978 | 1.513147585 | 0.022 | 0.021 | 1 | 0.936025978 | 0.573365703 |
| Zfp300 | 0.346827513 | -0.233118828 | 0 | 0.041 | 1 | 0.937567971 | 0.683072586 | 0.022 | 0.024 | 1 | 0.937567971 | 0.573365703 |
| Gm15246 | 0.346827513 | -0.133583154 | 0 | 0.041 | 1 | 0.951997392 | 0.513147585 | 0.022 | 0.021 | 1 | 0.951997392 | 0.573365703 |
| Gm16751 | 0.346846605 | -0.326228232 | 0 | 0.041 | 1 | 0.617557928 | 1.180572246 | 0.022 | 0.038 | 1 | 0.617557928 | 0.573390643 |
| Muc1 | 0.346846605 | -0.326228232 | 0 | 0.041 | 1 | 0.53630117 | 1.039216396 | 0.044 | 0.073 | 1 | 0.53630117 | 0.573390643 |
| Actr3b | 0.346846605 | -0.496153234 | 0 | 0.041 | 1 | 0.833619716 | 0.361144491 | 0.022 | 0.028 | 1 | 0.833619716 | 0.573390643 |
| Ano8 | 0.346846605 | -0.496153234 | 0 | 0.041 | 1 | 0.635186993 | -0.124282336 | 0.044 | 0.062 | 1 | 0.635186993 | 0.573390643 |
| Scn2b | 0.346846605 | -0.326228232 | 0 | 0.041 | 1 | 0.386564371 | -0.404390255 | 0.022 | 0.052 | 1 | 0.386564371 | 0.573390643 |
| Ptger4 | 0.346846605 | -0.326228232 | 0 | 0.041 | 1 | 0.437141879 | -0.539319835 | 0.022 | 0.048 | 1 | 0.437141879 | 0.573390643 |
| Nrm | 0.346846605 | -0.413691074 | 0 | 0.041 | 1 | 0.347517921 | -0.486852415 | 0.022 | 0.055 | 1 | 0.347517921 | 0.573390643 |
| Zbtb45 | 0.346847151 | -0.233118828 | 0 | 0.041 | 1 | 0.449441826 | 0.268035087 | 0.022 | 0.048 | 1 | 0.449441826 | 0.573391356 |
| Zfp873 | 0.346847151 | -0.233118828 | 0 | 0.041 | 1 | 0.936023974 | 2.197645759 | 0.022 | 0.021 | 1 | 0.936023974 | 0.573391356 |
| Amdhd1 | 0.346847151 | -0.496153234 | 0 | 0.041 | 1 | 0.744358033 | 0.223640968 | 0.022 | 0.031 | 1 | 0.744358033 | 0.573391356 |
| Arrb2 | 0.346847151 | -0.233118828 | 0 | 0.041 | 1 | 0.366924684 | 0.435145073 | 0.022 | 0.055 | 1 | 0.366924684 | 0.573391356 |
| Efcab5 | 0.346847151 | -0.413691074 | 0 | 0.041 | 1 | 0.839072605 | 0.513147585 | 0.044 | 0.038 | 1 | 0.839072605 | 0.573391356 |
| Gm26590 | 0.346847151 | -0.233118828 | 0 | 0.041 | 1 | 0.658583583 | 1.946106992 | 0.022 | 0.014 | 1 | 0.658583583 | 0.573391356 |
| Ptpn18 | 0.346860242 | -0.413691074 | 0 | 0.041 | 1 | 0.49874463 | 0.595609745 | 0.022 | 0.045 | 1 | 0.49874463 | 0.573408456 |
| Efhd1 | 0.346860242 | -0.574155746 | 0 | 0.041 | 1 | 0.687217233 | 1.82057611 | 0.022 | 0.035 | 1 | 0.687217233 | 0.573408456 |
| Gper1 | 0.346862423 | -0.326228232 | 0 | 0.041 | 1 | 0.376842245 | -0.776359032 | 0 | 0.017 | 1 | 0.376842245 | 0.573411306 |
| Cenpn | 0.346862423 | -0.326228232 | 0 | 0.041 | 1 | 0.823042771 | 0.683072586 | 0.022 | 0.017 | 1 | 0.823042771 | 0.573411306 |
| Tfpi | 0.346871695 | -0.648156327 | 0 | 0.041 | 1 | 0.49622258 | 0.361144491 | 0 | 0.01 | 1 | 0.49622258 | 0.573423418 |
| Fbxo5 | 0.346871695 | -1.081115734 | 0 | 0.041 | 1 | 0.852141123 | 0.946106992 | 0.022 | 0.028 | 1 | 0.852141123 | 0.573423418 |
| Aipl1 | 0.346873877 | -1.413691074 | 0 | 0.041 | 1 | 0.496223879 | 0.683072586 | 0 | 0.01 | 1 | 0.496223879 | 0.573426267 |
| Rims2 | 0.346879331 | -0.849790188 | 0 | 0.041 | 1 | 0.987932665 | 1.772339924 | 0.067 | 0.069 | 1 | 0.987932665 | 0.573433392 |
| Gm33302 | 0.346889693 | -2.348596045 | 0 | 0.041 | 1 | 0.451804285 | -1.960783604 | 0.044 | 0.073 | 1 | 0.451804285 | 0.573446927 |
| Baz1b | 0.347258927 | -0.358238627 | 0.667 | 0.787 | 1 | 0.61659301 | -0.100361456 | 0.911 | 0.879 | 1 | 0.61659301 | 0.573929092 |
| Gabrr2 | 0.915506323 | 1.995699863 | 0.048 | 0.056 | 1 | 0.347459145 | -0.316927414 | 0.044 | 0.087 | 1 | 0.915506323 | 0.574190432 |
| Gm43773 | 0.57816106 | 1.351843673 | 0 | 0.015 | 1 | 0.347502232 | 1.905465008 | 0.067 | 0.038 | 1 | 0.57816106 | 0.574246663 |
| Tmem151a | 0.472861597 | 1.866416846 | 0.048 | 0.022 | 1 | 0.347517921 | -0.486852415 | 0.022 | 0.055 | 1 | 0.472861597 | 0.574267136 |
| Gm11423 | 0.729839742 | 0.866416846 | 0.048 | 0.071 | 1 | 0.347517921 | -0.564854927 | 0.022 | 0.055 | 1 | 0.729839742 | 0.574267136 |
| Arhgef40 | 0.768103519 | 0.207453763 | 0.429 | 0.433 | 1 | 0.347733663 | 0.160347218 | 0.378 | 0.616 | 1 | 0.768103519 | 0.574548625 |
| Kcnk5 | 0.506942016 | 0.144718008 | 0.524 | 0.679 | 1 | 0.348162299 | 0.188505105 | 0.444 | 0.664 | 1 | 0.506942016 | 0.575107611 |
| Rbm4 | 0.417788375 | 0.697487837 | 0.429 | 0.433 | 1 | 0.348283506 | 0.129136981 | 0.4 | 0.571 | 1 | 0.417788375 | 0.575265611 |
| Mospd2 | 0.34867308 | 0.228461257 | 0.333 | 0.556 | 1 | 0.817165084 | 0.599008321 | 0.489 | 0.522 | 1 | 0.817165084 | 0.575773243 |
| Fblim1 | 0.72996115 | 1.214340149 | 0.048 | 0.071 | 1 | 0.348814051 | 0.73821414 | 0.067 | 0.118 | 1 | 0.72996115 | 0.575956869 |
| Abcc5 | 0.569983718 | 0.334969854 | 0.524 | 0.552 | 1 | 0.349072406 | -0.154496949 | 0.578 | 0.754 | 1 | 0.569983718 | 0.576293267 |
| Orai2 | 0.493771861 | 0.840881754 | 0.095 | 0.157 | 1 | 0.349161817 | 0.972579203 | 0.044 | 0.09 | 1 | 0.493771861 | 0.576409659 |
| Tamalin | 0.585768777 | -0.434752689 | 0.19 | 0.25 | 1 | 0.34931331 | 0.645597881 | 0.356 | 0.284 | 1 | 0.585768777 | 0.576606831 |
| Strada | 0.349498509 | -0.268742738 | 0.143 | 0.243 | 1 | 0.9927721 | 1.007507537 | 0.2 | 0.232 | 1 | 0.9927721 | 0.57684781 |
| Galnt2 | 0.734535691 | 0.291901132 | 0.476 | 0.649 | 1 | 0.349595036 | 0.513147585 | 0.844 | 0.765 | 1 | 0.734535691 | 0.576973383 |
| Mark4 | 0.758587877 | 0.201283996 | 0.238 | 0.291 | 1 | 0.349805006 | 0.811396683 | 0.267 | 0.422 | 1 | 0.758587877 | 0.57724647 |
| Ryr3 | 0.709465802 | 1.936806174 | 0.048 | 0.034 | 1 | 0.349937208 | 0.536231198 | 0.044 | 0.09 | 1 | 0.709465802 | 0.577418366 |
| Dnah17 | 0.349944798 | 1.464318402 | 0.095 | 0.049 | 1 | 0.377151335 | 1.168499413 | 0.111 | 0.076 | 1 | 0.377151335 | 0.577428234 |
| Cir1 | 0.604020961 | 0.194799963 | 0.333 | 0.463 | 1 | 0.350224789 | -0.280401538 | 0.467 | 0.502 | 1 | 0.604020961 | 0.577792176 |
| Rhbdd2 | 0.739610706 | 0.252307999 | 0.19 | 0.239 | 1 | 0.350367213 | 0.188307894 | 0.111 | 0.176 | 1 | 0.739610706 | 0.577977242 |
| Glyr1 | 0.351477746 | 0.44169029 | 0.81 | 0.765 | 1 | 0.745587881 | 0.358909487 | 0.911 | 0.841 | 1 | 0.745587881 | 0.579418886 |
| Gab1 | 0.351698568 | 0.461468164 | 0.762 | 0.735 | 1 | 0.50833252 | 0.189798815 | 0.889 | 0.803 | 1 | 0.50833252 | 0.579705253 |
| 2010109A12Rik | 0.991820392 | 1.247507013 | 0.095 | 0.104 | 1 | 0.351785887 | 0.41458375 | 0.067 | 0.118 | 1 | 0.991820392 | 0.579818463 |
| Rassf8 | 0.858663532 | 0.815790773 | 0.238 | 0.246 | 1 | 0.352068373 | 0.202640669 | 0.156 | 0.235 | 1 | 0.858663532 | 0.580184607 |
| Cyp3a13 | 0.449497156 | 0.406985227 | 0.095 | 0.172 | 1 | 0.352167208 | -0.901889915 | 0.022 | 0.055 | 1 | 0.449497156 | 0.580312674 |
| Hivep3 | 0.352418212 | 0.780686972 | 0.524 | 0.534 | 1 | 0.961436763 | -0.30705717 | 0.578 | 0.484 | 1 | 0.961436763 | 0.580637828 |
| Ttc17 | 0.608522948 | 0.147225953 | 0.333 | 0.485 | 1 | 0.352490154 | 0.199681069 | 0.689 | 0.609 | 1 | 0.608522948 | 0.580731 |
| Fbxo22 | 0.353189447 | -0.30677587 | 0.286 | 0.422 | 1 | 0.593335526 | 0.742778832 | 0.267 | 0.356 | 1 | 0.593335526 | 0.581636109 |
| Col6a2 | 0.353429668 | 0.184471146 | 0.952 | 0.966 | 1 | 0.370082805 | 0.377420031 | 0.8 | 0.958 | 1 | 0.370082805 | 0.581946806 |
| Prune2 | 0.52600739 | 0.532759458 | 0.238 | 0.332 | 1 | 0.353699949 | 0.843868798 | 0.222 | 0.187 | 1 | 0.52600739 | 0.582296245 |
| Serpini1 | 0.549843626 | 0.281454345 | 0.048 | 0.09 | 1 | 0.354323056 | 0.42003818 | 0.067 | 0.121 | 1 | 0.549843626 | 0.583101284 |
| Mettl3 | 0.758739157 | 0.699766976 | 0.19 | 0.175 | 1 | 0.355434351 | 0.502500341 | 0.111 | 0.176 | 1 | 0.758739157 | 0.584535125 |
| Gm19744 | 0.355460892 | 1.451379346 | 0.095 | 0.049 | 1 | 0.616659915 | 2.055041364 | 0.067 | 0.052 | 1 | 0.616659915 | 0.584569339 |
| Xrn1 | 0.401547489 | 0.853592805 | 0.667 | 0.761 | 1 | 0.355831252 | 0.298574668 | 0.933 | 0.865 | 1 | 0.401547489 | 0.585046624 |
| Cul4a | 0.980242969 | 0.550389352 | 0.381 | 0.53 | 1 | 0.35583474 | -0.139812034 | 0.644 | 0.709 | 1 | 0.980242969 | 0.585051118 |
| Cpsf4l | 0.356106009 | 2.325848464 | 0.143 | 0.093 | 1 | 1 | 1.304560963 | 0.089 | 0.097 | 1 | 1 | 0.585400528 |

| | | | | | | | | | | | | |
|---|---|---|---|---|---|---|---|---|---|---|---|---|
| Lmbr1 | 0.742570094 | -0.154047257 | 0.143 | 0.172 | 1 | 0.356106212 | 1.815176122 | 0.111 | 0.176 | 1 | 0.742570094 | 0.58540079 |
| Ptcd3 | 0.356192838 | 1.24875018 | 0.381 | 0.362 | 1 | 0.410065739 | 0.361144491 | 0.289 | 0.436 | 1 | 0.410065739 | 0.585512338 |
| Ephb1 | 0.532432377 | -0.496153234 | 0 | 0.019 | 1 | 0.356245003 | 1.587915353 | 0.067 | 0.038 | 1 | 0.532432377 | 0.585579504 |
| Adipor2 | 0.833819105 | 0.508569176 | 0.381 | 0.601 | 1 | 0.357241519 | -0.161838341 | 0.778 | 0.706 | 1 | 0.833819105 | 0.586861534 |
| Fam120a | 0.357457255 | 0.46661386 | 0.81 | 0.858 | 1 | 0.753234698 | 0.139722275 | 0.933 | 0.924 | 1 | 0.753234698 | 0.58713882 |
| Tpcn2 | 0.357540693 | 1.704145417 | 0.19 | 0.127 | 1 | 0.642402593 | 1.669775764 | 0.178 | 0.242 | 1 | 0.642402593 | 0.587246039 |
| Thsd4 | 0.909362077 | 0.435782491 | 0.476 | 0.608 | 1 | 0.35776774 | -0.453064102 | 0.778 | 0.699 | 1 | 0.909362077 | 0.587537725 |
| Cx3cr1 | 0.357995917 | 1.811275291 | 0.19 | 0.127 | 1 | 0.643499894 | 1.141502681 | 0.178 | 0.163 | 1 | 0.643499894 | 0.587830758 |
| Mfhas1 | 0.371017874 | 1.285501178 | 0.238 | 0.172 | 1 | 0.358250191 | 1.347888395 | 0.356 | 0.294 | 1 | 0.371017874 | 0.588157183 |
| Lgi4 | 0.358961275 | 1.351843673 | 0.095 | 0.049 | 1 | 0.540019204 | 1.42003818 | 0.044 | 0.073 | 1 | 0.540019204 | 0.589069353 |
| Sytl2 | 0.726643819 | 1.79930265 | 0.095 | 0.078 | 1 | 0.358965173 | -0.271123724 | 0.022 | 0.055 | 1 | 0.726643819 | 0.589074351 |
| Kdm5c | 0.433385782 | 0.43623586 | 0.667 | 0.701 | 1 | 0.359020829 | 0.245863706 | 0.889 | 0.806 | 1 | 0.433385782 | 0.589145702 |
| Lrrc71 | 0.359106848 | 1.029915578 | 0.095 | 0.049 | 1 | 0.362913332 | 0.966865552 | 0.067 | 0.121 | 1 | 0.362913332 | 0.589255968 |
| Cdc42se2 | 0.652471641 | 0.31892905 | 0.333 | 0.5 | 1 | 0.35918727 | 0.774046246 | 0.711 | 0.682 | 1 | 0.652471641 | 0.589359045 |
| Zfp983 | 0.991688689 | 1.623145695 | 0.095 | 0.101 | 1 | 0.35925801 | -0.86124793 | 0.044 | 0.083 | 1 | 0.991688689 | 0.589449702 |
| Platr1 | 0.359270744 | 2.673771768 | 0.048 | 0.019 | 1 | 0.376837034 | -0.124282336 | 0 | 0.017 | 1 | 0.376837034 | 0.589466021 |
| 4930532G15Rik | 0.359270744 | 2.673771768 | 0.048 | 0.019 | 1 | 0.450869096 | 0.83507568 | 0.022 | 0.048 | 1 | 0.450869096 | 0.589466021 |
| Gm56773 | 0.359270744 | 2.673771768 | 0.048 | 0.019 | 1 | 0.430487676 | 0.361144491 | 0 | 0.014 | 1 | 0.430487676 | 0.589466021 |
| Baat | 0.359270744 | 2.673771768 | 0.048 | 0.019 | 1 | 0.430487676 | 0.361144491 | 0 | 0.014 | 1 | 0.430487676 | 0.589466021 |
| Gm15672 | 0.359270744 | 2.673771768 | 0.048 | 0.019 | 1 | 0.492040936 | 2.683072586 | 0.022 | 0.01 | 1 | 0.492040936 | 0.589466021 |
| Aass | 0.359270744 | 2.673771768 | 0.048 | 0.019 | 1 | 0.61754131 | 0.875717664 | 0.022 | 0.038 | 1 | 0.61754131 | 0.589466021 |
| Gm44386 | 0.359270744 | 2.673771768 | 0.048 | 0.019 | 1 | 0.937567971 | 0.683072586 | 0.022 | 0.024 | 1 | 0.937567971 | 0.589466021 |
| 1700126G02Rik | 0.359270744 | 2.673771768 | 0.048 | 0.019 | 1 | 0.936023974 | 1.460680165 | 0.022 | 0.021 | 1 | 0.936023974 | 0.589466021 |
| Alpk3 | 0.359270744 | 4.673771768 | 0.048 | 0.019 | 1 | 0.496774618 | 2.005000681 | 0.067 | 0.045 | 1 | 0.496774618 | 0.589466021 |
| Adora2a | 0.359270744 | 2.673771768 | 0.048 | 0.019 | 1 | 0.854461673 | 1.361144491 | 0.022 | 0.028 | 1 | 0.854461673 | 0.589466021 |
| A930001M01Rik | 0.359270744 | 2.673771768 | 0.048 | 0.019 | 1 | 0.500262731 | 0.683072586 | 0.022 | 0.045 | 1 | 0.500262731 | 0.589466021 |
| Gm49719 | 0.359270744 | 2.673771768 | 0.048 | 0.019 | 1 | 0.496223879 | 0.683072586 | 0 | 0.01 | 1 | 0.496223879 | 0.589466021 |
| Cacna1h | 0.359270744 | 2.673771768 | 0.048 | 0.019 | 1 | 0.496223879 | 0.683072586 | 0 | 0.01 | 1 | 0.496223879 | 0.589466021 |
| Rasgef1a | 0.359280535 | 3.673771768 | 0.048 | 0.019 | 1 | 0.496223879 | -0.124282336 | 0 | 0.01 | 1 | 0.496223879 | 0.589478567 |
| Fer1l5 | 0.35928543 | 4.673771768 | 0.048 | 0.019 | 1 | 0.664307093 | 1.460680165 | 0.022 | 0.014 | 1 | 0.664307093 | 0.58948484 |
| Zdhhc19 | 0.359286409 | 2.825774861 | 0.048 | 0.019 | 1 | 0.854466386 | 1.683072586 | 0.022 | 0.028 | 1 | 0.854466386 | 0.589486095 |
| Fem1c | 0.359434759 | 0.104928932 | 0.238 | 0.422 | 1 | 0.50920136 | 0.281710024 | 0.622 | 0.547 | 1 | 0.50920136 | 0.589676172 |
| Spice1 | 0.359688565 | 1.400753273 | 0.238 | 0.187 | 1 | 0.51763746 | 1.020107573 | 0.111 | 0.152 | 1 | 0.51763746 | 0.590001266 |
| Rnd3 | 0.578658502 | -0.155379078 | 0.571 | 0.604 | 1 | 0.359735103 | -0.396654606 | 0.756 | 0.63 | 1 | 0.578658502 | 0.590060862 |
| 1700001L05Rik | 0.360257067 | 1.815790773 | 0.143 | 0.093 | 1 | 0.815597676 | 1.506194824 | 0.111 | 0.135 | 1 | 0.815597676 | 0.59072898 |
| Gm10516 | 0.360277575 | 0.464318402 | 0.048 | 0.119 | 1 | 0.635710125 | 1.180572246 | 0.133 | 0.18 | 1 | 0.635710125 | 0.590755219 |
| Slc9a5 | 0.360900122 | 2.586308926 | 0.095 | 0.052 | 1 | 0.512490525 | 1.83507568 | 0.044 | 0.028 | 1 | 0.512490525 | 0.591551346 |
| Nat10 | 0.552590029 | -0.260340297 | 0.19 | 0.25 | 1 | 0.361539319 | 0.337297749 | 0.244 | 0.37 | 1 | 0.552590029 | 0.592367959 |
| Etnk1 | 0.361754793 | 0.541814616 | 0.81 | 0.851 | 1 | 0.926278091 | 0.303466215 | 0.933 | 0.858 | 1 | 0.926278091 | 0.592643055 |
| Proca1 | 0.361850071 | 0.638147858 | 0.048 | 0.119 | 1 | 0.958146231 | 1.344270673 | 0.111 | 0.125 | 1 | 0.958146231 | 0.592764668 |
| B3galt5 | 0.361958682 | 0.79930265 | 0.048 | 0.119 | 1 | 0.430486631 | -0.776359032 | 0 | 0.014 | 1 | 0.430486631 | 0.592903276 |
| Cda | 0.361962798 | 1.119182916 | 0.048 | 0.119 | 1 | 0.555534078 | 1.683072586 | 0.022 | 0.042 | 1 | 0.555534078 | 0.592908528 |
| Il33 | 0.362014731 | 1.918884266 | 0.048 | 0.119 | 1 | 0.84982823 | 0.595609745 | 0.022 | 0.028 | 1 | 0.84982823 | 0.592974796 |
| Cfap157 | 0.362165014 | 2.451379346 | 0.048 | 0.019 | 1 | 0.793019324 | 1.329435632 | 0.067 | 0.059 | 1 | 0.793019324 | 0.593166531 |
| Chn1os3 | 0.362165014 | 2.451379346 | 0.048 | 0.019 | 1 | 0.492043882 | 2.361144491 | 0.022 | 0.01 | 1 | 0.492043882 | 0.593166531 |
| Gpr157 | 0.362165014 | 2.451379346 | 0.048 | 0.019 | 1 | 0.448078778 | 0.180572246 | 0.022 | 0.048 | 1 | 0.448078778 | 0.593166531 |
| Ckm | 0.362165014 | 2.451379346 | 0.048 | 0.019 | 1 | 0.49622258 | 0.361144491 | 0 | 0.01 | 1 | 0.49622258 | 0.593166531 |
| Brme1 | 0.362165014 | 2.451379346 | 0.048 | 0.019 | 1 | 0.37684485 | -0.776359032 | 0 | 0.017 | 1 | 0.37684485 | 0.593166531 |
| Gm56868 | 0.362165014 | 2.451379346 | 0.048 | 0.019 | 1 | 0.496223879 | 0.683072586 | 0 | 0.01 | 1 | 0.496223879 | 0.593166531 |
| Gm57047 | 0.362165014 | 2.451379346 | 0.048 | 0.019 | 1 | 0.661444614 | 1.683072586 | 0.022 | 0.014 | 1 | 0.661444614 | 0.593166531 |
| Gm20324 | 0.362165014 | 2.451379346 | 0.048 | 0.019 | 1 | 0.49622258 | 0.361144491 | 0 | 0.01 | 1 | 0.49622258 | 0.593166531 |
| Cdhr4 | 0.570813494 | -0.233118828 | 0.143 | 0.198 | 1 | 0.362358473 | -0.124282336 | 0.022 | 0.055 | 1 | 0.570813494 | 0.593413284 |
| Dync2i1 | 0.390404322 | 0.18995599 | 0.238 | 0.369 | 1 | 0.362445687 | 0.562442881 | 0.244 | 0.367 | 1 | 0.390404322 | 0.593524497 |
| Impact | 0.362714454 | 0.460424486 | 0.571 | 0.638 | 1 | 0.466889118 | 0.305914971 | 0.644 | 0.678 | 1 | 0.466889118 | 0.593867132 |
| Cep70 | 0.362873392 | 1.204286484 | 0.238 | 0.179 | 1 | 0.997623797 | 0.624178897 | 0.111 | 0.118 | 1 | 0.997623797 | 0.594069686 |
| Pomk | 0.725234921 | 0.735172312 | 0.048 | 0.071 | 1 | 0.362938926 | -0.638855509 | 0.044 | 0.083 | 1 | 0.725234921 | 0.594153188 |
| Clic5 | 0.363036767 | -0.940938076 | 0.048 | 0.108 | 1 | 0.833269 | 0.683072586 | 0.044 | 0.038 | 1 | 0.833269 | 0.59427784 |
| Lrp6 | 0.623708364 | 0.606657572 | 0.905 | 0.955 | 1 | 0.363130136 | 0.156974819 | 0.978 | 0.986 | 1 | 0.623708364 | 0.594396776 |
| Lrriq1 | 0.555466509 | -0.129455269 | 0.333 | 0.325 | 1 | 0.363836594 | -0.714958488 | 0.178 | 0.239 | 1 | 0.555466509 | 0.595296121 |
| Sirt7 | 0.835162534 | 0.372602233 | 0.143 | 0.175 | 1 | 0.364582776 | 0.921232323 | 0.222 | 0.183 | 1 | 0.835162534 | 0.596244951 |
| Nnat | 0.365063958 | 2.088809267 | 0.048 | 0.019 | 1 | 0.955025007 | 1.83507568 | 0.022 | 0.024 | 1 | 0.955025007 | 0.596856223 |
| Chrna7 | 0.365063958 | 2.088809267 | 0.048 | 0.019 | 1 | 0.687231957 | 0.757073168 | 0.022 | 0.035 | 1 | 0.687231957 | 0.596856223 |
| Spata32 | 0.365063958 | 2.258734268 | 0.048 | 0.019 | 1 | 0.705167793 | 2.304560963 | 0.044 | 0.035 | 1 | 0.705167793 | 0.596856223 |
| Gm46404 | 0.36506982 | 2.258734268 | 0.048 | 0.019 | 1 | 0.376837034 | -0.124282336 | 0 | 0.017 | 1 | 0.376837034 | 0.596863667 |
| T2 | 0.36506982 | 2.258734268 | 0.048 | 0.019 | 1 | 0.938685475 | 1.513147585 | 0.022 | 0.021 | 1 | 0.938685475 | 0.596863667 |
| Agmo | 0.365168617 | -0.718545655 | 0.095 | 0.179 | 1 | 0.759310848 | 0.567595369 | 0.022 | 0.031 | 1 | 0.759310848 | 0.596989115 |
| Wasf3 | 0.384797718 | 0.103916159 | 0.143 | 0.257 | 1 | 0.365574713 | -0.330733213 | 0.178 | 0.239 | 1 | 0.384797718 | 0.597504556 |
| Rsph10b | 0.73311071 | 0.951305743 | 0.095 | 0.078 | 1 | 0.365768653 | 0.361144491 | 0.022 | 0.055 | 1 | 0.73311071 | 0.597750599 |
| Tor4a | 0.396916958 | 0.214340149 | 0 | 0.034 | 1 | 0.365792613 | 0.290755163 | 0.022 | 0.055 | 1 | 0.396916958 | 0.59778099 |
| Ralbp1 | 0.366016112 | 0.590782403 | 0.81 | 0.817 | 1 | 0.443024989 | -0.151148942 | 0.956 | 0.913 | 1 | 0.443024989 | 0.59806443 |
| Tnip1 | 0.366030945 | 0.79930265 | 0.238 | 0.172 | 1 | 0.602529638 | 0.562778352 | 0.178 | 0.232 | 1 | 0.602529638 | 0.598083238 |
| Scd4 | 0.629700127 | 2.088809267 | 0.048 | 0.03 | 1 | 0.366897007 | 1.098110085 | 0.022 | 0.055 | 1 | 0.629700127 | 0.5991806 |
| A530041M06Rik | 0.396971401 | 0.351843673 | 0 | 0.034 | 1 | 0.366924684 | 0.757073168 | 0.022 | 0.055 | 1 | 0.396971401 | 0.599215645 |
| Gbp7 | 0.473994575 | 1.503846766 | 0.048 | 0.101 | 1 | 0.36692495 | 2.82057611 | 0.022 | 0.055 | 1 | 0.473994575 | 0.599215982 |
| Gm10848 | 0.457281049 | 0.503846766 | 0 | 0.026 | 1 | 0.366948101 | 0.361144491 | 0.022 | 0.055 | 1 | 0.457281049 | 0.599245293 |
| Pot1b | 0.8548874 | 0.866416846 | 0.048 | 0.06 | 1 | 0.366948899 | 1.612683258 | 0.022 | 0.055 | 1 | 0.8548874 | 0.599246304 |
| 4930599N23Rik | 0.984248518 | 0.451379346 | 0.143 | 0.149 | 1 | 0.36706181 | 0.426732833 | 0.111 | 0.17 | 1 | 0.984248518 | 0.599389247 |
| Gm19967 | 0.655561847 | 1.503846766 | 0.048 | 0.03 | 1 | 0.367249804 | 2.135584791 | 0.089 | 0.059 | 1 | 0.655561847 | 0.59962719 |
| Usp40 | 0.966006241 | 0.84692067 | 0.286 | 0.328 | 1 | 0.367611066 | 0.423205459 | 0.244 | 0.377 | 1 | 0.966006241 | 0.600084236 |
| Ngb | 0.367982232 | 2.088809267 | 0.048 | 0.019 | 1 | 0.430487676 | 0.361144491 | 0 | 0.014 | 1 | 0.430487676 | 0.60055354 |
| Ccdc173 | 0.368189581 | 1.088809267 | 0.143 | 0.09 | 1 | 0.398090882 | -0.158229668 | 0.044 | 0.083 | 1 | 0.398090882 | 0.600815594 |
| Epb41l2 | 0.567413628 | 0.227515538 | 0.81 | 0.787 | 1 | 0.368402877 | 0.375777761 | 0.889 | 0.886 | 1 | 0.567413628 | 0.601085074 |
| Dnah7b | 0.368415209 | -1.02666795 | 0.048 | 0.108 | 1 | 0.580549202 | 1.068362742 | 0.133 | 0.114 | 1 | 0.580549202 | 0.601100651 |
| Dmap1 | 0.852115916 | 0.995699863 | 0.095 | 0.086 | 1 | 0.3685076 | -0.404390255 | 0.067 | 0.111 | 1 | 0.852115916 | 0.601217349 |
| Pou2af1 | 0.368806266 | 1.532415918 | 0.333 | 0.295 | 1 | 0.855212557 | 0.481438725 | 0.156 | 0.176 | 1 | 0.855212557 | 0.60159447 |
| Rtn3 | 0.368824752 | 0.526389335 | 0.952 | 0.978 | 1 | 0.552760744 | 0.380362407 | 1 | 0.983 | 1 | 0.552760744 | 0.601617807 |
| Rasgef1b | 0.369836493 | 1.019767623 | 0.238 | 0.179 | 1 | 0.687918564 | 1.15951063 | 0.044 | 0.062 | 1 | 0.687918564 | 0.602893954 |
| Ttc19 | 0.729011441 | 0.177916741 | 0.619 | 0.649 | 1 | 0.370640917 | 0.226071794 | 0.667 | 0.758 | 1 | 0.729011441 | 0.603907144 |
| Ikbke | 0.370709917 | 1.155923463 | 0.238 | 0.194 | 1 | 0.512071901 | 0.710039634 | 0.267 | 0.249 | 1 | 0.512071901 | 0.603993992 |
| Ccdc169 | 0.370790696 | -0.413691074 | 0 | 0.037 | 1 | 0.936023974 | 1.460680165 | 0.022 | 0.021 | 1 | 0.936023974 | 0.604095652 |
| Btbd8 | 0.370790696 | -0.233118828 | 0 | 0.037 | 1 | 0.448112847 | 0.290755163 | 0.022 | 0.048 | 1 | 0.448112847 | 0.604095652 |
| Cntn4 | 0.370790696 | -1.535681598 | 0 | 0.037 | 1 | 0.802396304 | 0.335149283 | 0.067 | 0.059 | 1 | 0.802396304 | 0.604095652 |
| Trip13 | 0.370790696 | -0.496153234 | 0 | 0.037 | 1 | 0.49622258 | 0.361144491 | 0 | 0.01 | 1 | 0.49622258 | 0.604095652 |
| Gm35867 | 0.370790696 | -0.133583154 | 0 | 0.037 | 1 | 0.555561299 | 0.683072586 | 0.022 | 0.042 | 1 | 0.555561299 | 0.604095652 |
| Spata17 | 0.370806713 | -0.326228232 | 0 | 0.037 | 1 | 0.935071939 | 0.513147585 | 0.022 | 0.024 | 1 | 0.935071939 | 0.604115808 |
| 4930405A21Rik | 0.370806713 | -0.326228232 | 0 | 0.037 | 1 | 0.951997643 | 0.875717664 | 0.022 | 0.021 | 1 | 0.951997643 | 0.604115808 |
| Gm11464 | 0.370806713 | -0.233118828 | 0 | 0.037 | 1 | 0.406607361 | 0.83507568 | 0.022 | 0.052 | 1 | 0.406607361 | 0.604115808 |
| Gm33508 | 0.370806713 | -0.233118828 | 0 | 0.037 | 1 | 0.430486631 | -0.124282336 | 0 | 0.014 | 1 | 0.430486631 | 0.604115808 |

| | | | | | | | | | | | | |
|---|---|---|---|---|---|---|---|---|---|---|---|---|
| Gm29966 | 0.370806713 | -0.326228232 | 0 | 0.037 | 1 | 0.835209112 | 0.683072586 | 0.044 | 0.038 | 1 | 0.835209112 | 0.604115808 |
| Gm53065 | 0.370807306 | -0.133583154 | 0 | 0.037 | 1 | 0.921934145 | 0.612683258 | 0.044 | 0.042 | 1 | 0.921934145 | 0.604116554 |
| S100a3 | 0.370807306 | -0.133583154 | 0 | 0.037 | 1 | 0.376837034 | -0.124282336 | 0 | 0.017 | 1 | 0.376837034 | 0.604116554 |
| Fyb2 | 0.370807306 | -0.133583154 | 0 | 0.037 | 1 | 0.389604052 | 0.612683258 | 0.089 | 0.149 | 1 | 0.389604052 | 0.604116554 |
| Tmprss9 | 0.370807306 | -0.133583154 | 0 | 0.037 | 1 | 0.651436295 | 1.531069493 | 0.067 | 0.052 | 1 | 0.651436295 | 0.604116554 |
| Sgo1 | 0.370807306 | -0.133583154 | 0 | 0.037 | 1 | 0.847509429 | 0.683072586 | 0.022 | 0.028 | 1 | 0.847509429 | 0.604116554 |
| Abcc3 | 0.37081739 | -0.326228232 | 0 | 0.037 | 1 | 0.811836522 | 1.361144491 | 0.022 | 0.017 | 1 | 0.811836522 | 0.604129244 |
| Sh3bgr | 0.37081739 | -0.326228232 | 0 | 0.037 | 1 | 0.430487676 | 0.361144491 | 0 | 0.014 | 1 | 0.430487676 | 0.604129244 |
| Lpxn | 0.37081739 | -0.326228232 | 0 | 0.037 | 1 | 0.952530085 | 1.223640968 | 0.022 | 0.024 | 1 | 0.952530085 | 0.604129244 |
| Ctss | 0.37081917 | -1.326228232 | 0 | 0.037 | 1 | 0.907791348 | 1.442064487 | 0.067 | 0.066 | 1 | 0.907791348 | 0.604131483 |
| Kdm4d | 0.370819763 | -0.233118828 | 0 | 0.037 | 1 | 0.959990133 | 0.361144491 | 0.022 | 0.021 | 1 | 0.959990133 | 0.604132229 |
| Grin1 | 0.370822729 | -0.413691074 | 0 | 0.037 | 1 | 0.430487676 | 0.361144491 | 0 | 0.014 | 1 | 0.430487676 | 0.604135961 |
| 4930417O13Rik | 0.370826288 | -0.785659851 | 0 | 0.037 | 1 | 0.492040936 | 3.268035087 | 0.022 | 0.01 | 1 | 0.492040936 | 0.60414044 |
| Dnah2 | 0.370826288 | -1.326228232 | 0 | 0.037 | 1 | 0.854466828 | 2.045642666 | 0.022 | 0.028 | 1 | 0.854466828 | 0.60414044 |
| Ppfia2 | 0.370828067 | -1.683780237 | 0 | 0.037 | 1 | 0.489743413 | -0.57996182 | 0.022 | 0.045 | 1 | 0.489743413 | 0.604142679 |
| Rarb | 0.37082866 | -0.849790188 | 0 | 0.037 | 1 | 0.498258387 | 0.998574412 | 0.067 | 0.045 | 1 | 0.498258387 | 0.604143425 |
| 4930435F18Rik | 0.370831626 | -0.970084422 | 0 | 0.037 | 1 | 0.636636737 | 1.098110085 | 0.044 | 0.031 | 1 | 0.636636737 | 0.604147157 |
| Cenpm | 0.370833405 | -1.02666795 | 0 | 0.037 | 1 | 0.658583583 | 2.361144491 | 0.022 | 0.014 | 1 | 0.658583583 | 0.604149396 |
| Ccna2 | 0.370833998 | -1.535681598 | 0 | 0.037 | 1 | 0.430487676 | 0.361144491 | 0 | 0.014 | 1 | 0.430487676 | 0.604150143 |
| Ctnna3 | 0.370834592 | -1.574155746 | 0 | 0.037 | 1 | 0.430483494 | -0.124282336 | 0 | 0.014 | 1 | 0.430483494 | 0.604150889 |
| Pclaf | 0.370836964 | -2.701267664 | 0 | 0.037 | 1 | 0.430486631 | -0.901889915 | 0 | 0.014 | 1 | 0.430486631 | 0.604153874 |
| Anxa9 | 0.370840523 | 0.214340149 | 0 | 0.037 | 1 | 0.426109575 | 2.005000681 | 0.044 | 0.024 | 1 | 0.426109575 | 0.604158352 |
| Ccdc60 | 0.370840523 | 0.214340149 | 0 | 0.037 | 1 | 0.877093314 | 0.825091591 | 0.067 | 0.076 | 1 | 0.877093314 | 0.604158352 |
| 9530018F02Rik | 0.370840523 | 0.214340149 | 0 | 0.037 | 1 | 0.661727509 | -0.124282336 | 0.022 | 0.035 | 1 | 0.661727509 | 0.604158352 |
| Gm20036 | 0.370840523 | 0.214340149 | 0 | 0.037 | 1 | 0.430487676 | 0.361144491 | 0 | 0.014 | 1 | 0.430487676 | 0.604158352 |
| Gm34961 | 0.370840523 | 0.214340149 | 0 | 0.037 | 1 | 0.796722331 | 1.42003818 | 0.044 | 0.038 | 1 | 0.796722331 | 0.604158352 |
| Gm17259 | 0.370840523 | 0.214340149 | 0 | 0.037 | 1 | 0.809040217 | 1.460680165 | 0.022 | 0.017 | 1 | 0.809040217 | 0.604158352 |
| Gm31814 | 0.370840523 | 0.214340149 | 0 | 0.037 | 1 | 0.49622258 | 0.361144491 | 0 | 0.01 | 1 | 0.49622258 | 0.604158352 |
| 4931408D14Rik | 0.370840523 | 0.214340149 | 0 | 0.037 | 1 | 0.552232538 | 0.361144491 | 0.022 | 0.042 | 1 | 0.552232538 | 0.604158352 |
| Gm26737 | 0.370910053 | 0.351843673 | 0.048 | 0.019 | 1 | 0.936025227 | 1.683072586 | 0.022 | 0.021 | 1 | 0.936025227 | 0.604245838 |
| Simc1 | 0.600219513 | 0.124307948 | 0.286 | 0.373 | 1 | 0.371187502 | 0.19182452 | 0.356 | 0.467 | 1 | 0.600219513 | 0.604594843 |
| Plch2 | 0.502808926 | 2.24081236 | 0.095 | 0.06 | 1 | 0.371940032 | 0.361144491 | 0.067 | 0.118 | 1 | 0.502808926 | 0.605540677 |
| Pstpip1 | 0.371965888 | -0.308080886 | 0.143 | 0.231 | 1 | 0.952238683 | 0.557541704 | 0.222 | 0.26 | 1 | 0.952238683 | 0.605573154 |
| Kdm5d | 0.488678646 | -0.133583154 | 0.143 | 0.224 | 1 | 0.372338173 | 0.392853351 | 0.311 | 0.439 | 1 | 0.488678646 | 0.606040631 |
| Zfp27 | 0.959285803 | 0.503846766 | 0.095 | 0.093 | 1 | 0.372409158 | 0.125077133 | 0.089 | 0.142 | 1 | 0.959285803 | 0.606129735 |
| Gm14744 | 0.372444438 | 0.162809848 | 0.095 | 0.19 | 1 | 0.631986825 | 0.635319455 | 0.333 | 0.346 | 1 | 0.631986825 | 0.606174017 |
| Erp27 | 0.480452211 | 0.351843673 | 0.095 | 0.168 | 1 | 0.372861415 | 1.545569062 | 0.178 | 0.142 | 1 | 0.480452211 | 0.606697195 |
| Kdm1a | 0.560010467 | 0.252070538 | 0.952 | 0.907 | 1 | 0.37369177 | 0.147241793 | 0.978 | 0.958 | 1 | 0.560010467 | 0.607738001 |
| Gm14440 | 0.373857158 | 2.088809267 | 0.048 | 0.019 | 1 | 0.662778712 | 0.290755163 | 0.044 | 0.062 | 1 | 0.662778712 | 0.607945142 |
| Lrriq3 | 0.373857158 | 2.088809267 | 0.048 | 0.019 | 1 | 1 | 0.875717664 | 0.044 | 0.045 | 1 | 1 | 0.607945142 |
| Gm43597 | 0.373857158 | 2.088809267 | 0.048 | 0.019 | 1 | 0.617529321 | 0.982632868 | 0.022 | 0.038 | 1 | 0.617529321 | 0.607945142 |
| ENSMUSG00000121482 | 0.373857158 | 2.088809267 | 0.048 | 0.019 | 1 | 0.49622258 | 0.361144491 | 0 | 0.01 | 1 | 0.49622258 | 0.607945142 |
| Gm15635 | 0.373857158 | 2.088809267 | 0.048 | 0.019 | 1 | 0.496223879 | 0.683072586 | 0 | 0.01 | 1 | 0.496223879 | 0.607945142 |
| Gm12714 | 0.373857158 | 2.088809267 | 0.048 | 0.019 | 1 | 0.376837034 | -0.316927414 | 0 | 0.017 | 1 | 0.376837034 | 0.607945142 |
| Gm56546 | 0.373857158 | 2.088809267 | 0.048 | 0.019 | 1 | 0.687197015 | 0.808603468 | 0.022 | 0.035 | 1 | 0.687197015 | 0.607945142 |
| Slc25a47 | 0.951653578 | 1.947946731 | 0.095 | 0.108 | 1 | 0.374303971 | 0.538682677 | 0.111 | 0.18 | 1 | 0.951653578 | 0.60850448 |
| Tmem41b | 0.569654191 | 0.132752615 | 0.381 | 0.597 | 1 | 0.37437677 | 0.427931582 | 0.556 | 0.689 | 1 | 0.569654191 | 0.608595574 |
| Rpgr | 0.603113968 | 1.595769256 | 0.143 | 0.119 | 1 | 0.375154988 | 0.513147585 | 0.044 | 0.087 | 1 | 0.603113968 | 0.609568712 |
| Ppp2r2b | 0.881505681 | -0.264827688 | 0.048 | 0.041 | 1 | 0.375325346 | 1.209141398 | 0.133 | 0.093 | 1 | 0.881505681 | 0.609781577 |
| Arfgef3 | 0.9521828 | 0.866416846 | 0.048 | 0.052 | 1 | 0.37653208 | -0.375821103 | 0.044 | 0.083 | 1 | 0.9521828 | 0.611287753 |
| Whrn | 0.376574297 | 0.351843673 | 0.19 | 0.336 | 1 | 0.900832882 | 0.366306196 | 0.4 | 0.453 | 1 | 0.900832882 | 0.611340393 |
| Pla2r1 | 0.376794341 | 0.673771768 | 0.048 | 0.019 | 1 | 0.430483494 | -0.124282336 | 0 | 0.014 | 1 | 0.430483494 | 0.611614707 |
| Mettl5os | 0.376794341 | 1.866416846 | 0.048 | 0.019 | 1 | 0.811838207 | 1.268035087 | 0.022 | 0.017 | 1 | 0.811838207 | 0.611614707 |
| Fabp12 | 0.376794341 | 1.866416846 | 0.048 | 0.019 | 1 | 0.430486631 | -0.486852415 | 0 | 0.014 | 1 | 0.430486631 | 0.611614707 |
| Gm21411 | 0.376794341 | 1.866416846 | 0.048 | 0.019 | 1 | 0.935071939 | 0.513147585 | 0.022 | 0.024 | 1 | 0.935071939 | 0.611614707 |
| Epo | 0.376794341 | 1.866416846 | 0.048 | 0.019 | 1 | 0.788320053 | 1.242499995 | 0.067 | 0.059 | 1 | 0.788320053 | 0.611614707 |
| Trhde | 0.376794341 | 1.214340149 | 0.048 | 0.019 | 1 | 0.728253477 | 0.928185084 | 0.044 | 0.035 | 1 | 0.728253477 | 0.611614707 |
| Itgae | 0.376794341 | 1.088809267 | 0.048 | 0.019 | 1 | 0.430483494 | -0.124282336 | 0 | 0.014 | 1 | 0.430483494 | 0.611614707 |
| Engase | 0.376794341 | 1.866416846 | 0.048 | 0.019 | 1 | 0.753258193 | 1.513147585 | 0.044 | 0.059 | 1 | 0.753258193 | 0.611614707 |
| Adcy4 | 0.376794341 | 1.866416846 | 0.048 | 0.019 | 1 | 0.933318841 | 0.460680165 | 0.044 | 0.042 | 1 | 0.933318841 | 0.611614707 |
| Tspyl3 | 0.396916958 | 0.214340149 | 0 | 0.034 | 1 | 0.376837034 | -0.124282336 | 0 | 0.017 | 1 | 0.396916958 | 0.611667918 |
| Gm34552 | 0.396971401 | 0.351843673 | 0 | 0.034 | 1 | 0.376837034 | -0.124282336 | 0 | 0.017 | 1 | 0.396971401 | 0.611667918 |
| Gm57028 | 0.425599726 | 0.351843673 | 0 | 0.03 | 1 | 0.376837034 | -0.124282336 | 0 | 0.017 | 1 | 0.425599726 | 0.611667918 |
| Cfap91 | 0.425599726 | 0.214340149 | 0 | 0.03 | 1 | 0.376837034 | -0.124282336 | 0 | 0.017 | 1 | 0.425599726 | 0.611667918 |
| 5830428M24Rik | 0.454205211 | 2.451379346 | 0.048 | 0.022 | 1 | 0.376837034 | -0.124282336 | 0 | 0.017 | 1 | 0.454205211 | 0.611667918 |
| Gm57273 | 0.457281049 | 0.503846766 | 0 | 0.026 | 1 | 0.376837034 | -0.124282336 | 0 | 0.017 | 1 | 0.457281049 | 0.611667918 |
| Gm960 | 0.457281049 | 0.351843673 | 0 | 0.026 | 1 | 0.376837034 | -0.124282336 | 0 | 0.017 | 1 | 0.457281049 | 0.611667918 |
| Gm16174 | 0.457292877 | 2.258734268 | 0.048 | 0.022 | 1 | 0.376837034 | -0.316927414 | 0 | 0.017 | 1 | 0.457292877 | 0.611667918 |
| Gm15853 | 0.457292877 | 0.351843673 | 0 | 0.026 | 1 | 0.376837034 | -0.124282336 | 0 | 0.017 | 1 | 0.457292877 | 0.611667918 |
| Rsf1os2 | 0.457292877 | 0.351843673 | 0 | 0.026 | 1 | 0.376837034 | -0.124282336 | 0 | 0.017 | 1 | 0.457292877 | 0.611667918 |
| Kiss1r | 0.457292877 | 0.351843673 | 0 | 0.026 | 1 | 0.376837034 | -0.316927414 | 0 | 0.017 | 1 | 0.457292877 | 0.611667918 |
| A930001C03Rik | 0.457292877 | 0.214340149 | 0 | 0.026 | 1 | 0.376837034 | -0.486852415 | 0 | 0.017 | 1 | 0.457292877 | 0.611667918 |
| Foxq1 | 0.469724015 | 0.866416846 | 0.048 | 0.022 | 1 | 0.376837034 | -0.316927414 | 0 | 0.017 | 1 | 0.469724015 | 0.611667918 |
| Lrrc56 | 0.492590603 | 0.673771768 | 0 | 0.022 | 1 | 0.376837034 | -0.124282336 | 0 | 0.017 | 1 | 0.492590603 | 0.611667918 |
| Ap1g2 | 0.492590603 | 0.673771768 | 0 | 0.022 | 1 | 0.376837034 | -0.124282336 | 0 | 0.017 | 1 | 0.492590603 | 0.611667918 |
| Cdkn3 | 0.492604708 | -0.233118828 | 0 | 0.022 | 1 | 0.376837034 | -1.223818009 | 0 | 0.017 | 1 | 0.492604708 | 0.611667918 |
| Gm56689 | 0.492608234 | 0.866416846 | 0 | 0.022 | 1 | 0.376837034 | -0.124282336 | 0 | 0.017 | 1 | 0.492608234 | 0.611667918 |
| Gm37634 | 0.532426384 | 0.866416846 | 0 | 0.019 | 1 | 0.376837034 | -0.124282336 | 0 | 0.017 | 1 | 0.532426384 | 0.611667918 |
| Gm19391 | 0.532426384 | 0.866416846 | 0 | 0.019 | 1 | 0.376837034 | -0.124282336 | 0 | 0.017 | 1 | 0.532426384 | 0.611667918 |
| Gm11542 | 0.532426384 | 0.503846766 | 0 | 0.019 | 1 | 0.376837034 | -0.124282336 | 0 | 0.017 | 1 | 0.532426384 | 0.611667918 |
| Cdca7l | 0.532426384 | 0.866416846 | 0 | 0.019 | 1 | 0.376837034 | -0.776359032 | 0 | 0.017 | 1 | 0.532426384 | 0.611667918 |
| Rem1 | 0.532432377 | 0.673771768 | 0 | 0.019 | 1 | 0.376837034 | -0.124282336 | 0 | 0.017 | 1 | 0.532432377 | 0.611667918 |
| Gabrp | 0.532432377 | 0.214340149 | 0 | 0.019 | 1 | 0.376837034 | -0.124282336 | 0 | 0.017 | 1 | 0.532432377 | 0.611667918 |
| Gm56516 | 0.532436372 | 1.088809267 | 0 | 0.019 | 1 | 0.376837034 | -0.124282336 | 0 | 0.017 | 1 | 0.532436372 | 0.611667918 |
| Cfap20dc | 0.570211706 | 1.503846766 | 0.048 | 0.026 | 1 | 0.376837034 | -0.124282336 | 0 | 0.017 | 1 | 0.570211706 | 0.611667918 |
| Vstm2b | 0.578156445 | 1.088809267 | 0 | 0.015 | 1 | 0.376837034 | -0.124282336 | 0 | 0.017 | 1 | 0.578156445 | 0.611667918 |
| 6030468B19Rik | 0.578156445 | 1.088809267 | 0 | 0.015 | 1 | 0.376837034 | -0.124282336 | 0 | 0.017 | 1 | 0.578156445 | 0.611667918 |
| Gm10457 | 0.578156445 | 1.088809267 | 0 | 0.015 | 1 | 0.376837034 | -0.901889915 | 0 | 0.017 | 1 | 0.578156445 | 0.611667918 |
| Cbln2 | 0.578156445 | 1.088809267 | 0 | 0.015 | 1 | 0.376837034 | -0.124282336 | 0 | 0.017 | 1 | 0.578156445 | 0.611667918 |
| Wdr95 | 0.578159907 | 0.503846766 | 0 | 0.015 | 1 | 0.376837034 | -0.124282336 | 0 | 0.017 | 1 | 0.578159907 | 0.611667918 |
| Aldh1a2 | 0.578159907 | 0.503846766 | 0 | 0.015 | 1 | 0.376837034 | -0.316927414 | 0 | 0.017 | 1 | 0.578159907 | 0.611667918 |
| B3gat2 | 0.57816106 | 1.351843673 | 0 | 0.015 | 1 | 0.376837034 | -0.486852415 | 0 | 0.017 | 1 | 0.57816106 | 0.611667918 |
| Fcgr3 | 0.57816106 | 1.351843673 | 0 | 0.015 | 1 | 0.376837034 | -0.776359032 | 0 | 0.017 | 1 | 0.57816106 | 0.611667918 |
| Gm57465 | 0.57816106 | 1.351843673 | 0 | 0.015 | 1 | 0.376837034 | -0.124282336 | 0 | 0.017 | 1 | 0.57816106 | 0.611667918 |
| Ptk6 | 0.57816106 | 1.351843673 | 0 | 0.015 | 1 | 0.376837034 | -0.124282336 | 0 | 0.017 | 1 | 0.57816106 | 0.611667918 |
| Gm12503 | 0.57816106 | 1.351843673 | 0 | 0.015 | 1 | 0.376837034 | -0.124282336 | 0 | 0.017 | 1 | 0.57816106 | 0.611667918 |
| Gm31597 | 0.57816106 | 1.351843673 | 0 | 0.015 | 1 | 0.376837034 | -0.124282336 | 0 | 0.017 | 1 | 0.57816106 | 0.611667918 |
| 2810408A11Rik | 0.57816106 | 1.351843673 | 0 | 0.015 | 1 | 0.376837034 | -0.316927414 | 0 | 0.017 | 1 | 0.57816106 | 0.611667918 |

| | | | | | | | | | | | | |
|---|---|---|---|---|---|---|---|---|---|---|---|---|
| Gm6034 | 0.57816106 | 0.673771768 | 0 | 0.015 | 1 | 0.376837034 | -0.316927414 | 0 | 0.017 | 1 | 0.57816106 | 0.611667918 |
| Gm11110 | 0.57816106 | 1.351843673 | 0 | 0.015 | 1 | 0.376837034 | -0.316927414 | 0 | 0.017 | 1 | 0.57816106 | 0.611667918 |
| 4930444A19Rik | 0.63207476 | 1.351843673 | 0 | 0.011 | 1 | 0.376837034 | -0.124282336 | 0 | 0.017 | 1 | 0.63207476 | 0.611667918 |
| 4930505A04Rik | 0.63207476 | 1.351843673 | 0 | 0.011 | 1 | 0.376837034 | -0.316927414 | 0 | 0.017 | 1 | 0.63207476 | 0.611667918 |
| Klhl10 | 0.63207476 | 1.351843673 | 0 | 0.011 | 1 | 0.376837034 | -0.124282336 | 0 | 0.017 | 1 | 0.63207476 | 0.611667918 |
| B930094E09Rik | 0.63207476 | 1.351843673 | 0 | 0.011 | 1 | 0.376837034 | -0.124282336 | 0 | 0.017 | 1 | 0.63207476 | 0.611667918 |
| Fam110a | 0.632076134 | 1.673771768 | 0 | 0.011 | 1 | 0.376837034 | -0.124282336 | 0 | 0.017 | 1 | 0.632076134 | 0.611667918 |
| Cavin4 | 0.632076134 | 1.673771768 | 0 | 0.011 | 1 | 0.376837034 | -0.124282336 | 0 | 0.017 | 1 | 0.632076134 | 0.611667918 |
| Zfp786 | 0.632076134 | 0.866416846 | 0 | 0.011 | 1 | 0.376837034 | -0.124282336 | 0 | 0.017 | 1 | 0.632076134 | 0.611667918 |
| Gm45442 | 0.632076134 | 1.673771768 | 0 | 0.011 | 1 | 0.376837034 | -0.124282336 | 0 | 0.017 | 1 | 0.632076134 | 0.611667918 |
| Rassf7 | 0.632076134 | 1.673771768 | 0 | 0.011 | 1 | 0.376837034 | -0.124282336 | 0 | 0.017 | 1 | 0.632076134 | 0.611667918 |
| Nbeal2 | 0.632076134 | 1.673771768 | 0 | 0.011 | 1 | 0.376837034 | -0.124282336 | 0 | 0.017 | 1 | 0.632076134 | 0.611667918 |
| Raet1e | 0.632076134 | 1.673771768 | 0 | 0.011 | 1 | 0.376837034 | -0.124282336 | 0 | 0.017 | 1 | 0.632076134 | 0.611667918 |
| Gm47819 | 0.632076134 | 1.673771768 | 0 | 0.011 | 1 | 0.376837034 | -0.124282336 | 0 | 0.017 | 1 | 0.632076134 | 0.611667918 |
| Serpinb6c | 0.632076134 | 1.673771768 | 0 | 0.011 | 1 | 0.376837034 | -0.316927414 | 0 | 0.017 | 1 | 0.632076134 | 0.611667918 |
| Ltb4r1 | 0.632076134 | 1.673771768 | 0 | 0.011 | 1 | 0.376837034 | -0.316927414 | 0 | 0.017 | 1 | 0.632076134 | 0.611667918 |
| Gm2629 | 0.632076134 | 1.673771768 | 0 | 0.011 | 1 | 0.376837034 | -0.316927414 | 0 | 0.017 | 1 | 0.632076134 | 0.611667918 |
| Mki67 | 0.642596287 | 0.844883684 | 0.048 | 0.03 | 1 | 0.376837034 | -1.017367132 | 0 | 0.017 | 1 | 0.642596287 | 0.611667918 |
| Gm33037 | 0.709478182 | 2.410737362 | 0.048 | 0.034 | 1 | 0.376837034 | -0.124282336 | 0 | 0.017 | 1 | 0.709478182 | 0.611667918 |
| Gm12829 | 0.784037261 | 1.79930265 | 0.048 | 0.037 | 1 | 0.376837034 | -0.124282336 | 0 | 0.017 | 1 | 0.784037261 | 0.611667918 |
| Tafa3 | 0.784065849 | 2.088809267 | 0.048 | 0.037 | 1 | 0.376837034 | -0.124282336 | 0 | 0.017 | 1 | 0.784065849 | 0.611667918 |
| Cish | 0.982550637 | 1.351843673 | 0.048 | 0.049 | 1 | 0.376837034 | -0.124282336 | 0 | 0.017 | 1 | 0.982550637 | 0.611667918 |
| Hpse | 0.425599726 | 0.214340149 | 0 | 0.03 | 1 | 0.376842245 | -0.316927414 | 0 | 0.017 | 1 | 0.425599726 | 0.611674412 |
| Gm11963 | 0.457281049 | 0.503846766 | 0 | 0.026 | 1 | 0.376842245 | -0.316927414 | 0 | 0.017 | 1 | 0.457281049 | 0.611674412 |
| 9130023H24Rik | 0.457308648 | 0.673771768 | 0 | 0.026 | 1 | 0.376842245 | -0.486852415 | 0 | 0.017 | 1 | 0.457308648 | 0.611674412 |
| Aspa | 0.492604708 | 0.351843673 | 0 | 0.022 | 1 | 0.376842245 | -0.486852415 | 0 | 0.017 | 1 | 0.492604708 | 0.611674412 |
| Lrrn1 | 0.492606471 | -0.648156327 | 0 | 0.022 | 1 | 0.376842245 | -0.316927414 | 0 | 0.017 | 1 | 0.492606471 | 0.611674412 |
| Meig1 | 0.532426384 | 0.866416846 | 0 | 0.019 | 1 | 0.376842245 | -0.901889915 | 0 | 0.017 | 1 | 0.532426384 | 0.611674412 |
| Gm34917 | 0.532432377 | 0.673771768 | 0 | 0.019 | 1 | 0.376842245 | -0.486852415 | 0 | 0.017 | 1 | 0.532432377 | 0.611674412 |
| Emx2 | 0.532432377 | 0.214340149 | 0 | 0.019 | 1 | 0.376842245 | -0.316927414 | 0 | 0.017 | 1 | 0.532432377 | 0.611674412 |
| Igfbp6 | 0.532434375 | 0.351843673 | 0 | 0.019 | 1 | 0.376842245 | -2.776359032 | 0 | 0.017 | 1 | 0.532434375 | 0.611674412 |
| Gm2237 | 0.532436372 | 1.088809267 | 0 | 0.019 | 1 | 0.376842245 | -0.316927414 | 0 | 0.017 | 1 | 0.532436372 | 0.611674412 |
| Elapor2 | 0.578156445 | 0.866416846 | 0 | 0.015 | 1 | 0.376842245 | -0.486852415 | 0 | 0.017 | 1 | 0.578156445 | 0.611674412 |
| Syt3 | 0.578156445 | 1.088809267 | 0 | 0.015 | 1 | 0.376842245 | -0.316927414 | 0 | 0.017 | 1 | 0.578156445 | 0.611674412 |
| Gm31356 | 0.578156445 | 1.088809267 | 0 | 0.015 | 1 | 0.376842245 | -1.017367132 | 0 | 0.017 | 1 | 0.578156445 | 0.611674412 |
| Scn9a | 0.578159907 | 0.673771768 | 0 | 0.015 | 1 | 0.376842245 | -0.776359032 | 0 | 0.017 | 1 | 0.578159907 | 0.611674412 |
| Cxxc4 | 0.578159907 | 0.866416846 | 0 | 0.015 | 1 | 0.376842245 | -0.486852415 | 0 | 0.017 | 1 | 0.578159907 | 0.611674412 |
| Gm26608 | 0.57816106 | 1.351843673 | 0 | 0.015 | 1 | 0.376842245 | -0.316927414 | 0 | 0.017 | 1 | 0.57816106 | 0.611674412 |
| BC051019 | 0.57816106 | 1.351843673 | 0 | 0.015 | 1 | 0.376842245 | -0.776359032 | 0 | 0.017 | 1 | 0.57816106 | 0.611674412 |
| 5033403F01Rik | 0.57816106 | 1.351843673 | 0 | 0.015 | 1 | 0.376842245 | -0.638855509 | 0 | 0.017 | 1 | 0.57816106 | 0.611674412 |
| Thbs4 | 0.57816106 | 1.351843673 | 0 | 0.015 | 1 | 0.376842245 | -0.901889915 | 0 | 0.017 | 1 | 0.57816106 | 0.611674412 |
| Cacna1b | 0.63207476 | 0.673771768 | 0 | 0.011 | 1 | 0.376842245 | -0.486852415 | 0 | 0.017 | 1 | 0.63207476 | 0.611674412 |
| Gm12472 | 0.63207476 | 1.351843673 | 0 | 0.011 | 1 | 0.376842245 | -0.638855509 | 0 | 0.017 | 1 | 0.63207476 | 0.611674412 |
| Tm6sf1 | 0.63207476 | 1.351843673 | 0 | 0.011 | 1 | 0.376842245 | -0.486852415 | 0 | 0.017 | 1 | 0.63207476 | 0.611674412 |
| D6Ertd474e | 0.632076134 | 1.673771768 | 0 | 0.011 | 1 | 0.376842245 | -0.776359032 | 0 | 0.017 | 1 | 0.632076134 | 0.611674412 |
| Apobr | 0.632076134 | 1.673771768 | 0 | 0.011 | 1 | 0.376842245 | -0.316927414 | 0 | 0.017 | 1 | 0.632076134 | 0.611674412 |
| Syt7 | 0.632076134 | 1.673771768 | 0 | 0.011 | 1 | 0.376842245 | -0.486852415 | 0 | 0.017 | 1 | 0.632076134 | 0.611674412 |
| Mis18bp1 | 0.728718166 | 0.719575457 | 0.048 | 0.034 | 1 | 0.376842245 | -0.776359032 | 0 | 0.017 | 1 | 0.728718166 | 0.611674412 |
| Lypd3 | 0.75134323 | 0.425844254 | 0.048 | 0.034 | 1 | 0.376842245 | -0.316927414 | 0 | 0.017 | 1 | 0.75134323 | 0.611674412 |
| Kif11 | 0.847081853 | 0.648236676 | 0.048 | 0.06 | 1 | 0.376842245 | -0.901889915 | 0 | 0.017 | 1 | 0.847081853 | 0.611674412 |
| 4930481A15Rik | 0.859875139 | 1.451379346 | 0.048 | 0.041 | 1 | 0.376842245 | -0.486852415 | 0 | 0.017 | 1 | 0.859875139 | 0.611674412 |
| Gm4107 | 0.866047663 | 1.304537958 | 0.048 | 0.041 | 1 | 0.376842245 | -0.901889915 | 0 | 0.017 | 1 | 0.866047663 | 0.611674412 |
| Crybb3 | 0.425639612 | 0.503846766 | 0 | 0.03 | 1 | 0.376843981 | -0.776359032 | 0 | 0.017 | 1 | 0.425639612 | 0.611676577 |
| Nckap1l | 0.492608234 | 0.866416846 | 0 | 0.022 | 1 | 0.376843981 | -0.638855509 | 0 | 0.017 | 1 | 0.492608234 | 0.611676577 |
| Gm5627 | 0.532436372 | 1.088809267 | 0 | 0.019 | 1 | 0.376843981 | -0.638855509 | 0 | 0.017 | 1 | 0.532436372 | 0.611676577 |
| Sult6b1 | 0.57816106 | 1.351843673 | 0 | 0.015 | 1 | 0.376843981 | -0.901889915 | 0 | 0.017 | 1 | 0.57816106 | 0.611676577 |
| Cdk15 | 0.853427036 | -0.184209227 | 0.048 | 0.056 | 1 | 0.376843981 | -0.638855509 | 0 | 0.017 | 1 | 0.853427036 | 0.611676577 |
| Gm48395 | 0.479147948 | 1.503846766 | 0.048 | 0.022 | 1 | 0.37684485 | -0.486852415 | 0 | 0.017 | 1 | 0.479147948 | 0.611677659 |
| Ildr2 | 0.578159907 | 0.214340149 | 0 | 0.015 | 1 | 0.37684485 | -2.361321533 | 0 | 0.017 | 1 | 0.578159907 | 0.611677659 |
| Mab21l2 | 0.632076134 | 0.503846766 | 0 | 0.011 | 1 | 0.37684485 | -1.124282336 | 0 | 0.017 | 1 | 0.632076134 | 0.611677659 |
| Rorb | 0.632076134 | 1.673771768 | 0 | 0.011 | 1 | 0.37684485 | -0.776359032 | 0 | 0.017 | 1 | 0.632076134 | 0.611677659 |
| Gm42209 | 0.636152163 | 2.595769256 | 0.048 | 0.03 | 1 | 0.37684485 | -1.486852415 | 0 | 0.017 | 1 | 0.636152163 | 0.611677659 |
| 1600002D24Rik | 0.818213398 | 1.351843673 | 0.048 | 0.063 | 1 | 0.37684485 | -0.776359032 | 0 | 0.017 | 1 | 0.818213398 | 0.611677659 |
| Gabbr2 | 0.96386048 | 0.425844254 | 0.048 | 0.045 | 1 | 0.37684485 | -1.564854927 | 0 | 0.017 | 1 | 0.96386048 | 0.611677659 |
| St7l | 0.464433632 | 0.11189288 | 0.333 | 0.493 | 1 | 0.376857795 | 0.188622378 | 0.444 | 0.654 | 1 | 0.464433632 | 0.611693792 |
| Gm4258 | 0.553626241 | 1.474462959 | 0.048 | 0.09 | 1 | 0.376954513 | -2.480157763 | 0.022 | 0.052 | 1 | 0.553626241 | 0.611814321 |
| Ddit4l | 0.377037742 | -0.145246072 | 0.238 | 0.358 | 1 | 0.561438612 | -0.838464534 | 0.178 | 0.232 | 1 | 0.561438612 | 0.611918025 |
| Arfip1 | 0.73115786 | 0.434632272 | 0.524 | 0.769 | 1 | 0.377896961 | 0.419127675 | 0.844 | 0.824 | 1 | 0.73115786 | 0.612987809 |
| Avl9 | 0.738258597 | 1.543711227 | 0.429 | 0.481 | 1 | 0.378102995 | 0.631542286 | 0.822 | 0.699 | 1 | 0.738258597 | 0.613244115 |
| Mb | 0.9521828 | 0.866416846 | 0.048 | 0.052 | 1 | 0.378141783 | -1.271123724 | 0.022 | 0.052 | 1 | 0.9521828 | 0.613292357 |
| Fbxw5 | 0.378157686 | -0.413691074 | 0.095 | 0.168 | 1 | 0.877084894 | 0.605070074 | 0.111 | 0.125 | 1 | 0.877084894 | 0.613312137 |
| Pard3b | 0.378297303 | 0.725766405 | 0.714 | 0.694 | 1 | 0.42363316 | 0.217949855 | 0.933 | 0.841 | 1 | 0.42363316 | 0.613485757 |
| Tnfrsf22 | 0.379111609 | 1.380990018 | 0.19 | 0.134 | 1 | 0.574287311 | 0.304560963 | 0.044 | 0.069 | 1 | 0.574287311 | 0.614497605 |
| Gm28050 | 0.379238344 | 1.60338244 | 0.095 | 0.052 | 1 | 0.541961436 | 2.03099589 | 0.089 | 0.069 | 1 | 0.541961436 | 0.614654966 |
| Bbox1 | 0.379775085 | 1.673771768 | 0.048 | 0.019 | 1 | 0.49622258 | 0.361144491 | 0 | 0.01 | 1 | 0.49622258 | 0.615321055 |
| Zfp850 | 0.379775085 | 1.673771768 | 0.048 | 0.019 | 1 | 0.823042771 | 0.875717664 | 0.022 | 0.017 | 1 | 0.823042771 | 0.615321055 |
| Rad51ap2 | 0.379775085 | 1.673771768 | 0.048 | 0.019 | 1 | 0.49622258 | 0.361144491 | 0 | 0.01 | 1 | 0.49622258 | 0.615321055 |
| Gm41541 | 0.379775085 | 1.503846766 | 0.048 | 0.019 | 1 | 0.496223879 | 0.683072586 | 0 | 0.01 | 1 | 0.496223879 | 0.615321055 |
| Six3os1 | 0.379775085 | 1.503846766 | 0.048 | 0.019 | 1 | 0.658587075 | 2.42003818 | 0.022 | 0.014 | 1 | 0.658587075 | 0.615321055 |
| Rnf157 | 0.8052569 | 1.544488751 | 0.095 | 0.082 | 1 | 0.379809495 | 0.583536913 | 0.044 | 0.087 | 1 | 0.8052569 | 0.615363738 |
| Sh3bgrl2 | 0.945819313 | 0.451379346 | 0.048 | 0.045 | 1 | 0.379933229 | 0.857101986 | 0.044 | 0.087 | 1 | 0.945819313 | 0.615517199 |
| Ppp4r3b | 0.915669704 | 0.398637884 | 0.667 | 0.728 | 1 | 0.380013268 | 0.173211541 | 0.889 | 0.862 | 1 | 0.915669704 | 0.615616453 |
| Ipo11 | 0.899629996 | 0.789248985 | 0.19 | 0.22 | 1 | 0.380222166 | 0.672503344 | 0.2 | 0.315 | 1 | 0.899629996 | 0.615875436 |
| Nck1 | 0.380557835 | -0.254813899 | 0.429 | 0.619 | 1 | 0.637931148 | 0.127472145 | 0.756 | 0.723 | 1 | 0.637931148 | 0.616291404 |
| Gm16193 | 0.529861656 | 1.193778827 | 0.143 | 0.216 | 1 | 0.380775965 | 0.586857271 | 0.178 | 0.277 | 1 | 0.529861656 | 0.616561595 |
| Ap3m1 | 0.381629619 | 0.228212324 | 0.333 | 0.519 | 1 | 0.490189558 | 0.391841289 | 0.289 | 0.405 | 1 | 0.490189558 | 0.617618072 |
| Pgap3 | 0.614943447 | 0.503846766 | 0.048 | 0.082 | 1 | 0.382317713 | 0.325520582 | 0.067 | 0.118 | 1 | 0.614943447 | 0.618468593 |
| Maml1 | 0.634413366 | 0.866416846 | 0.286 | 0.418 | 1 | 0.382322782 | 0.195232552 | 0.467 | 0.637 | 1 | 0.634413366 | 0.618474854 |
| Man2a1 | 0.382391642 | 1.041033542 | 0.476 | 0.638 | 1 | 0.909288098 | 0.815176122 | 0.533 | 0.512 | 1 | 0.909288098 | 0.618559916 |
| Nme5 | 0.58246925 | 0.544488751 | 0.048 | 0.086 | 1 | 0.382514898 | 1.600610426 | 0.133 | 0.1 | 1 | 0.58246925 | 0.618712148 |
| Elf2 | 0.967115477 | 0.116960307 | 0.857 | 0.825 | 1 | 0.382814923 | 0.196290479 | 0.978 | 0.945 | 1 | 0.967115477 | 0.619082581 |
| Vwce | 0.382971897 | 1.472137907 | 0.095 | 0.052 | 1 | 0.652620086 | -0.271123724 | 0.044 | 0.062 | 1 | 0.652620086 | 0.61927632 |
| Sfxn5 | 0.383198025 | 0.95952625 | 0.286 | 0.231 | 1 | 0.989367415 | 0.316110738 | 0.444 | 0.516 | 1 | 0.989367415 | 0.619555323 |
| Kcnc3 | 0.383313506 | 0.866416846 | 0.143 | 0.086 | 1 | 0.792971235 | 0.683072586 | 0.089 | 0.107 | 1 | 0.792971235 | 0.619697768 |
| Epha4 | 0.738451119 | 1.157933628 | 0.333 | 0.354 | 1 | 0.383664775 | -1.514866791 | 0.356 | 0.353 | 1 | 0.738451119 | 0.62013089 |
| Ctcf | 0.420396107 | -0.133583154 | 0.524 | 0.705 | 1 | 0.383690455 | -0.154076938 | 0.911 | 0.813 | 1 | 0.420396107 | 0.620162545 |
| Slc25a51 | 0.383695238 | 0.29423295 | 0.429 | 0.619 | 1 | 0.72833866 | 0.238287743 | 0.733 | 0.678 | 1 | 0.72833866 | 0.62016844 |
| Snx27 | 0.977141342 | 0.224088126 | 0.476 | 0.578 | 1 | 0.383727565 | 0.520519573 | 0.733 | 0.709 | 1 | 0.977141342 | 0.620208286 |

| | | | | | | | | | | | | |
|---|---|---|---|---|---|---|---|---|---|---|---|---|
| Trmt11 | 0.383916613 | 0.943378827 | 0.238 | 0.187 | 1 | 0.468021799 | 0.513147585 | 0.2 | 0.287 | 1 | 0.468021799 | 0.62044126 |
| A730017L22Rik | 0.396916958 | 0.214340149 | 0 | 0.034 | 1 | 0.384163146 | -0.638855509 | 0.022 | 0.052 | 1 | 0.396916958 | 0.620744969 |
| 1700120C14Rik | 0.532892507 | -0.326228232 | 0.048 | 0.086 | 1 | 0.384163146 | -0.709244837 | 0.022 | 0.052 | 1 | 0.532892507 | 0.620744969 |
| Ehd2 | 0.819397536 | 0.37881072 | 0.429 | 0.556 | 1 | 0.385099488 | 0.161535466 | 0.489 | 0.737 | 1 | 0.819397536 | 0.62189736 |
| Inpp1 | 0.541380192 | 0.783954685 | 0.238 | 0.336 | 1 | 0.385237246 | 0.4725056 | 0.244 | 0.381 | 1 | 0.541380192 | 0.622066756 |
| Gm44777 | 0.492590603 | 0.673771768 | 0 | 0.022 | 1 | 0.385364644 | -0.486852415 | 0.022 | 0.052 | 1 | 0.492590603 | 0.622223379 |
| Slc6a15 | 0.532426384 | 0.866416846 | 0 | 0.019 | 1 | 0.385364644 | -0.638855509 | 0.022 | 0.052 | 1 | 0.532426384 | 0.622223379 |
| A330069E16Rik | 0.642465438 | 2.866416846 | 0.095 | 0.071 | 1 | 0.385364644 | -0.84048937 | 0.022 | 0.052 | 1 | 0.642465438 | 0.622223379 |
| Rnf122 | 0.679706589 | 0.351843673 | 0.048 | 0.075 | 1 | 0.385364644 | -0.486852415 | 0.022 | 0.052 | 1 | 0.679706589 | 0.622223379 |
| 1500015A07Rik | 0.866839783 | 0.351843673 | 0.048 | 0.056 | 1 | 0.385364644 | -0.564854927 | 0.022 | 0.052 | 1 | 0.866839783 | 0.622223379 |
| Coa7 | 0.982550637 | 1.351843673 | 0.048 | 0.049 | 1 | 0.385364644 | -0.638855509 | 0.022 | 0.052 | 1 | 0.982550637 | 0.622223379 |
| Chd6 | 0.385664091 | 0.291302131 | 0.762 | 0.761 | 1 | 0.55782043 | 0.309883476 | 0.956 | 0.865 | 1 | 0.55782043 | 0.622591391 |
| Gm11714 | 0.738393266 | 1.351843673 | 0.048 | 0.034 | 1 | 0.386007187 | -0.223818009 | 0.044 | 0.083 | 1 | 0.738393266 | 0.623012826 |
| 1700093J21Rik | 0.904673267 | 0.936806174 | 0.048 | 0.056 | 1 | 0.386007187 | -0.223818009 | 0.044 | 0.083 | 1 | 0.904673267 | 0.623012826 |
| Sertad2 | 0.386465613 | 0.67989787 | 0.714 | 0.929 | 1 | 0.804552552 | 0.257740106 | 0.889 | 0.927 | 1 | 0.804552552 | 0.623575556 |
| Gm19531 | 0.4759797 | 1.673771768 | 0.048 | 0.022 | 1 | 0.386564371 | -0.404390255 | 0.022 | 0.052 | 1 | 0.4759797 | 0.62369673 |
| Bloc1s3 | 0.862687636 | 1.088809267 | 0.048 | 0.06 | 1 | 0.386564371 | -0.404390255 | 0.022 | 0.052 | 1 | 0.862687636 | 0.62369673 |
| Fign | 0.997404848 | 0.599224814 | 0.286 | 0.332 | 1 | 0.386620448 | 0.603552247 | 0.244 | 0.374 | 1 | 0.997404848 | 0.623765526 |
| Adam12 | 0.604413131 | 0.195039593 | 0.333 | 0.306 | 1 | 0.386637563 | 1.026176191 | 0.378 | 0.377 | 1 | 0.604413131 | 0.623786521 |
| Stk36 | 0.784047982 | 2.088809267 | 0.048 | 0.037 | 1 | 0.386986849 | -0.174908409 | 0.044 | 0.083 | 1 | 0.784047982 | 0.624214876 |
| Pip5k1a | 0.387446998 | 0.516902919 | 0.571 | 0.575 | 1 | 0.47864745 | 0.165224281 | 0.867 | 0.855 | 1 | 0.47864745 | 0.624778819 |
| Gm56783 | 0.396916958 | 0.214340149 | 0 | 0.034 | 1 | 0.387952149 | -0.316927414 | 0.022 | 0.052 | 1 | 0.396916958 | 0.625397429 |
| Rps6ka2 | 0.388023589 | 0.146524765 | 0.19 | 0.321 | 1 | 0.483462442 | 1.207734577 | 0.133 | 0.194 | 1 | 0.483462442 | 0.625484872 |
| Ess2 | 0.659925739 | 1.029915578 | 0.095 | 0.071 | 1 | 0.388171626 | 0.242499995 | 0.067 | 0.114 | 1 | 0.659925739 | 0.625666041 |
| Lax1 | 0.388538974 | 2.010806755 | 0.095 | 0.052 | 1 | 0.687208248 | 1.683072586 | 0.022 | 0.035 | 1 | 0.687208248 | 0.626115413 |
| Inpp5e | 0.585271001 | 2.16475812 | 0.143 | 0.112 | 1 | 0.388777658 | 0.498648015 | 0.089 | 0.145 | 1 | 0.585271001 | 0.626407249 |
| Gm19710 | 0.388882903 | 1.351843673 | 0.143 | 0.093 | 1 | 0.554757415 | 0.750186782 | 0.089 | 0.128 | 1 | 0.554757415 | 0.626535894 |
| Chrnb1 | 0.457300763 | 0.214340149 | 0 | 0.026 | 1 | 0.388907488 | -0.223818009 | 0.044 | 0.083 | 1 | 0.457300763 | 0.626565942 |
| Sugp2 | 0.42612811 | 1.002836044 | 0.381 | 0.328 | 1 | 0.389536404 | 0.982632868 | 0.333 | 0.346 | 1 | 0.42612811 | 0.627334198 |
| Abitram | 0.389576681 | -0.469186186 | 0.095 | 0.164 | 1 | 0.573422546 | 0.390290837 | 0.089 | 0.121 | 1 | 0.573422546 | 0.627383371 |
| Rsph3a | 0.390077384 | -0.370622352 | 0.095 | 0.172 | 1 | 0.447540323 | 0.189533113 | 0.156 | 0.225 | 1 | 0.447540323 | 0.627994402 |
| Cdca7 | 0.390395345 | 1.586308926 | 0.095 | 0.052 | 1 | 0.8202424 | 1.098110085 | 0.022 | 0.017 | 1 | 0.8202424 | 0.628382165 |
| Gm47271 | 0.390395345 | 1.586308926 | 0.095 | 0.052 | 1 | 0.85645734 | 0.946106992 | 0.044 | 0.052 | 1 | 0.85645734 | 0.628382165 |
| Zfp760 | 0.390395345 | 1.586308926 | 0.095 | 0.052 | 1 | 0.497965827 | 0.808603468 | 0.089 | 0.062 | 1 | 0.497965827 | 0.628382165 |
| Ntng1 | 0.532432377 | 0.351843673 | 0 | 0.019 | 1 | 0.390418979 | -0.918963428 | 0.022 | 0.052 | 1 | 0.532432377 | 0.628410979 |
| Tctn1 | 0.679687267 | 0.503846766 | 0.048 | 0.075 | 1 | 0.390658019 | 0.473619221 | 0.067 | 0.114 | 1 | 0.679687267 | 0.62870235 |
| Apbb3 | 0.934604576 | 0.586308926 | 0.095 | 0.104 | 1 | 0.391277064 | 0.513147585 | 0.067 | 0.118 | 1 | 0.934604576 | 0.629456387 |
| Atg12 | 0.391641861 | 0.474991904 | 0.286 | 0.474 | 1 | 0.543752271 | 0.255466413 | 0.556 | 0.484 | 1 | 0.543752271 | 0.629900374 |
| Fras1 | 0.39202236 | -0.970084422 | 0.048 | 0.104 | 1 | 0.449883068 | 0.972579203 | 0.089 | 0.142 | 1 | 0.449883068 | 0.63036319 |
| Deaf1 | 0.392462685 | 0.544488751 | 0.238 | 0.388 | 1 | 0.528303236 | 0.369412107 | 0.356 | 0.529 | 1 | 0.528303236 | 0.630898411 |
| Bend6 | 0.39261705 | 1.298732336 | 0.238 | 0.187 | 1 | 0.40541487 | 0.250113179 | 0.022 | 0.052 | 1 | 0.40541487 | 0.631085952 |
| Gm3734 | 0.784047982 | 2.896164189 | 0.048 | 0.037 | 1 | 0.392807003 | 0.304560963 | 0.044 | 0.083 | 1 | 0.784047982 | 0.631316665 |
| Nfib | 0.393244042 | 0.418161569 | 0.857 | 0.724 | 1 | 0.88685218 | 0.166706132 | 0.978 | 0.858 | 1 | 0.88685218 | 0.631847208 |
| Piga | 0.546029484 | 0.304537958 | 0.048 | 0.09 | 1 | 0.393364398 | 0.320502507 | 0.067 | 0.111 | 1 | 0.546029484 | 0.631993247 |
| Snrnp35 | 0.393413069 | -0.648156327 | 0.048 | 0.104 | 1 | 0.936873444 | 0.223640968 | 0.089 | 0.093 | 1 | 0.936873444 | 0.632052295 |
| Surf6 | 0.802661763 | 0.233199176 | 0.19 | 0.228 | 1 | 0.393438732 | 0.470078863 | 0.178 | 0.277 | 1 | 0.802661763 | 0.632083428 |
| Elp4 | 0.393962676 | 0.214340149 | 0.286 | 0.466 | 1 | 0.645519831 | 0.627790151 | 0.4 | 0.401 | 1 | 0.645519831 | 0.632718762 |
| Atp6v0a4 | 0.395842768 | 1.454093731 | 0.333 | 0.269 | 1 | 0.631001441 | 0.119134868 | 0.778 | 0.827 | 1 | 0.631001441 | 0.634994039 |
| F420014N23Rik | 0.396088017 | 1.914779867 | 0.19 | 0.134 | 1 | 0.770533339 | 0.683072586 | 0.133 | 0.163 | 1 | 0.770533339 | 0.635290316 |
| Gmcl1 | 0.396145718 | -0.133583154 | 0.143 | 0.239 | 1 | 0.667698683 | 0.664213559 | 0.289 | 0.381 | 1 | 0.667698683 | 0.635360006 |
| Csf1r | 0.557344141 | 1.55829455 | 0.048 | 0.026 | 1 | 0.396178213 | -0.877642368 | 0.067 | 0.038 | 1 | 0.557344141 | 0.63539925 |
| Mcmdc2 | 0.396333048 | 2.021695071 | 0.143 | 0.093 | 1 | 0.735368111 | 0.558744451 | 0.156 | 0.19 | 1 | 0.735368111 | 0.63558621 |
| Skil | 0.539287477 | 0.14517171 | 0.905 | 0.851 | 1 | 0.396402898 | -0.231989504 | 0.978 | 0.903 | 1 | 0.539287477 | 0.635670539 |
| Bcat1 | 0.396712358 | 0.430453507 | 0.524 | 0.504 | 1 | 0.680145449 | 0.886059608 | 0.267 | 0.26 | 1 | 0.680145449 | 0.636044021 |
| Gm14023 | 0.396916958 | 0.214340149 | 0 | 0.034 | 1 | 0.496223879 | 0.683072586 | 0 | 0.01 | 1 | 0.496223879 | 0.636290845 |
| Gm20457 | 0.396916958 | 0.214340149 | 0 | 0.034 | 1 | 0.852141123 | 0.946106992 | 0.022 | 0.028 | 1 | 0.852141123 | 0.636290845 |
| Zfp365 | 0.396916958 | 0.214340149 | 0 | 0.034 | 1 | 0.40531543 | 0.180572246 | 0.022 | 0.052 | 1 | 0.40531543 | 0.636290845 |
| Gm12002 | 0.396916958 | 0.214340149 | 0 | 0.034 | 1 | 0.480781131 | -0.124282336 | 0.022 | 0.045 | 1 | 0.480781131 | 0.636290845 |
| Shpk | 0.396916958 | 0.214340149 | 0 | 0.034 | 1 | 0.429570554 | 1.513147585 | 0.044 | 0.024 | 1 | 0.429570554 | 0.636290845 |
| Gm5141 | 0.396916958 | 0.214340149 | 0 | 0.034 | 1 | 0.951997643 | 0.875717664 | 0.022 | 0.021 | 1 | 0.951997643 | 0.636290845 |
| Gm20508 | 0.396916958 | 0.214340149 | 0 | 0.034 | 1 | 0.759306107 | 0.567595369 | 0.022 | 0.031 | 1 | 0.759306107 | 0.636290845 |
| Ypel4 | 0.396935107 | -0.233118828 | 0 | 0.034 | 1 | 0.476265768 | -0.564854927 | 0.022 | 0.045 | 1 | 0.476265768 | 0.636312735 |
| Bub1b | 0.396935107 | -0.233118828 | 0 | 0.034 | 1 | 0.954659411 | 0.683072586 | 0.022 | 0.021 | 1 | 0.954659411 | 0.636312735 |
| Amz1 | 0.396935107 | -0.133583154 | 0 | 0.034 | 1 | 0.765728546 | 1.545569062 | 0.022 | 0.031 | 1 | 0.765728546 | 0.636312735 |
| Gm15545 | 0.39694807 | -0.133583154 | 0 | 0.034 | 1 | 0.658583583 | 2.361144491 | 0.022 | 0.014 | 1 | 0.658583583 | 0.63632837 |
| Lif | 0.39694807 | -0.413691074 | 0 | 0.034 | 1 | 0.677376317 | -0.941418279 | 0.022 | 0.035 | 1 | 0.677376317 | 0.63632837 |
| A330033J07Rik | 0.39694807 | -0.233118828 | 0 | 0.034 | 1 | 0.811836522 | 1.808603468 | 0.022 | 0.017 | 1 | 0.811836522 | 0.63632837 |
| Hs6st2 | 0.39694807 | -0.133583154 | 0 | 0.034 | 1 | 0.404151886 | 0.268035087 | 0.022 | 0.052 | 1 | 0.404151886 | 0.63632837 |
| Mansc4 | 0.396948718 | -0.326228232 | 0 | 0.034 | 1 | 0.811836522 | 1.098110085 | 0.022 | 0.017 | 1 | 0.811836522 | 0.636329152 |
| Gm56874 | 0.396948718 | -0.326228232 | 0 | 0.034 | 1 | 0.80624346 | 1.683072586 | 0.022 | 0.017 | 1 | 0.80624346 | 0.636329152 |
| Polq | 0.396948718 | -1.326228232 | 0 | 0.034 | 1 | 0.496223879 | 0.683072586 | 0 | 0.01 | 1 | 0.496223879 | 0.636329152 |
| Kif20a | 0.396948718 | -0.574155746 | 0 | 0.034 | 1 | 0.658583583 | 2.683072586 | 0.022 | 0.014 | 1 | 0.658583583 | 0.636329152 |
| Gm53018 | 0.396955847 | -0.233118828 | 0 | 0.034 | 1 | 0.415886571 | 3.061584209 | 0.044 | 0.024 | 1 | 0.415886571 | 0.63633775 |
| Aurka | 0.39695844 | -0.133583154 | 0 | 0.034 | 1 | 0.936023974 | 1.460680165 | 0.022 | 0.021 | 1 | 0.936023974 | 0.636340877 |
| Ablim2 | 0.39695844 | -0.574155746 | 0 | 0.034 | 1 | 0.496223879 | 0.683072586 | 0 | 0.01 | 1 | 0.496223879 | 0.636340877 |
| Plcd4 | 0.396962328 | -0.574155746 | 0 | 0.034 | 1 | 0.720559127 | 1.098110085 | 0.044 | 0.059 | 1 | 0.720559127 | 0.636345567 |
| Gm28729 | 0.396963624 | -0.574155746 | 0 | 0.034 | 1 | 0.955025729 | 1.361144491 | 0.022 | 0.024 | 1 | 0.955025729 | 0.63634713 |
| Fgd3 | 0.396964273 | -0.970084422 | 0 | 0.034 | 1 | 0.826709901 | -0.564854927 | 0.022 | 0.028 | 1 | 0.826709901 | 0.636347911 |
| A430035B10Rik | 0.396964921 | -0.233118828 | 0 | 0.034 | 1 | 0.846807438 | 0.361144491 | 0.044 | 0.038 | 1 | 0.846807438 | 0.636348693 |
| Pcdhb9 | 0.396966217 | -1.233118828 | 0 | 0.034 | 1 | 0.545663876 | -0.124282336 | 0.022 | 0.042 | 1 | 0.545663876 | 0.636350256 |
| Iigp1 | 0.396970105 | -1.818081329 | 0 | 0.034 | 1 | 0.823048066 | -0.722919774 | 0.022 | 0.017 | 1 | 0.823048066 | 0.636354946 |
| Serinc4 | 0.396971401 | 0.351843673 | 0 | 0.034 | 1 | 0.952529269 | 1.098110085 | 0.022 | 0.024 | 1 | 0.952529269 | 0.636356509 |
| Gm14443 | 0.396971401 | 0.351843673 | 0 | 0.034 | 1 | 0.661727509 | -0.404390255 | 0.022 | 0.035 | 1 | 0.661727509 | 0.636356509 |
| 4921511C10Rik | 0.396971401 | 0.351843673 | 0 | 0.034 | 1 | 0.858160219 | 0.91753784 | 0.044 | 0.052 | 1 | 0.858160219 | 0.636356509 |
| Gm57313 | 0.396971401 | 0.351843673 | 0 | 0.034 | 1 | 0.496223879 | 0.683072586 | 0 | 0.01 | 1 | 0.496223879 | 0.636356509 |
| Plxna4os1 | 0.396971401 | 0.351843673 | 0 | 0.034 | 1 | 0.49622258 | 0.361144491 | 0 | 0.01 | 1 | 0.49622258 | 0.636356509 |
| Gm39167 | 0.396971401 | 0.351843673 | 0 | 0.034 | 1 | 0.806246349 | 1.875717664 | 0.022 | 0.017 | 1 | 0.806246349 | 0.636356509 |
| Mc1r | 0.396971401 | 0.351843673 | 0 | 0.034 | 1 | 0.658583583 | 2.361144491 | 0.022 | 0.014 | 1 | 0.658583583 | 0.636356509 |
| Swsap1 | 0.396971401 | 0.351843673 | 0 | 0.034 | 1 | 0.430487676 | 0.361144491 | 0 | 0.014 | 1 | 0.430487676 | 0.636356509 |
| Dppa5a | 0.396971401 | 0.351843673 | 0 | 0.034 | 1 | 0.724349113 | 1.223640968 | 0.044 | 0.035 | 1 | 0.724349113 | 0.636356509 |
| Pcdhga1 | 0.396971401 | 0.351843673 | 0 | 0.034 | 1 | 0.91244769 | 1.180572246 | 0.044 | 0.042 | 1 | 0.91244769 | 0.636356509 |
| Itga9 | 0.397631735 | 0.39876472 | 0.238 | 0.403 | 1 | 0.396996466 | 0.666198768 | 0.444 | 0.502 | 1 | 0.397631735 | 0.636386738 |
| Snhg14 | 0.397131576 | 0.129451251 | 0.048 | 0.112 | 1 | 0.569189849 | -0.280401538 | 0.067 | 0.093 | 1 | 0.569189849 | 0.636549663 |
| Phf3 | 0.464909838 | 0.353821322 | 0.905 | 0.862 | 1 | 0.397371395 | -0.115802105 | 1 | 0.941 | 1 | 0.464909838 | 0.636838764 |
| Cacna1d | 0.562827214 | 0.298732336 | 0.19 | 0.146 | 1 | 0.3979307 | 0.898240681 | 0.356 | 0.36 | 1 | 0.562827214 | 0.637512558 |
| Wnt5b | 0.667616617 | -0.858723313 | 0.143 | 0.183 | 1 | 0.398096164 | -0.21001221 | 0.044 | 0.083 | 1 | 0.667616617 | 0.637711772 |
| 2700049A03Rik | 0.926048451 | 0.400753273 | 0.19 | 0.216 | 1 | 0.398116446 | 0.275897205 | 0.156 | 0.225 | 1 | 0.926048451 | 0.637736187 |
| Slc7a6os | 0.987384934 | 0.503846766 | 0.333 | 0.369 | 1 | 0.398585925 | 0.477958156 | 0.244 | 0.367 | 1 | 0.987384934 | 0.638301111 |

| | | | | | | | | | | | | |
|---|---|---|---|---|---|---|---|---|---|---|---|---|
| Gm8098 | 0.860084871 | 1.010806755 | 0.048 | 0.06 | 1 | 0.399079543 | -0.375821103 | 0.022 | 0.052 | 1 | 0.860084871 | 0.638894605 |
| Prc1 | 0.703633643 | -0.413691074 | 0.095 | 0.075 | 1 | 0.399094795 | -0.174908409 | 0.022 | 0.052 | 1 | 0.703633643 | 0.638912934 |
| Dusp1 | 0.399724395 | -0.604485856 | 0.905 | 0.787 | 1 | 0.491425101 | -0.495328197 | 0.978 | 0.848 | 1 | 0.491425101 | 0.639669198 |
| Dus1l | 0.399797063 | 0.214340149 | 0.048 | 0.112 | 1 | 0.412058378 | 0.361144491 | 0.044 | 0.083 | 1 | 0.412058378 | 0.639756434 |
| Ddx60 | 0.40001236 | 1.889500459 | 0.095 | 0.052 | 1 | 0.809042781 | 1.595609745 | 0.022 | 0.017 | 1 | 0.809042781 | 0.640014831 |
| Gm43111 | 0.547771462 | 2.214340149 | 0.048 | 0.026 | 1 | 0.400333085 | -0.124282336 | 0.022 | 0.052 | 1 | 0.547771462 | 0.640399591 |
| Dab1 | 0.472860821 | 1.214340149 | 0.048 | 0.022 | 1 | 0.400764863 | 1.140545352 | 0.089 | 0.059 | 1 | 0.472860821 | 0.64091725 |
| Wdr36 | 0.832782273 | 0.780686972 | 0.286 | 0.407 | 1 | 0.401549428 | 0.307563451 | 0.267 | 0.391 | 1 | 0.832782273 | 0.641856913 |
| Dmxl2 | 0.433661669 | -0.25696557 | 0.095 | 0.164 | 1 | 0.401596096 | 0.925642889 | 0.2 | 0.159 | 1 | 0.433661669 | 0.641912768 |
| Dnah10 | 0.899259517 | 0.735172312 | 0.048 | 0.056 | 1 | 0.401607549 | -0.223818009 | 0.022 | 0.052 | 1 | 0.899259517 | 0.641926475 |
| Vac14 | 0.401792827 | -0.370622352 | 0.048 | 0.104 | 1 | 0.989950919 | 1.076415014 | 0.156 | 0.176 | 1 | 0.989950919 | 0.642148179 |
| Gm16794 | 0.425599726 | 0.351843673 | 0 | 0.03 | 1 | 0.402813251 | 0.290755163 | 0.022 | 0.052 | 1 | 0.425599726 | 0.643367987 |
| Nup205 | 0.678846206 | 0.819622634 | 0.238 | 0.228 | 1 | 0.402814232 | 0.804378882 | 0.178 | 0.273 | 1 | 0.678846206 | 0.643369159 |
| Pemt | 0.732158744 | 1.281454345 | 0.048 | 0.071 | 1 | 0.402834833 | 0.223640968 | 0.022 | 0.052 | 1 | 0.732158744 | 0.643393763 |
| Zfp791 | 0.492599419 | 0.351843673 | 0 | 0.022 | 1 | 0.402852816 | 0.15951063 | 0.022 | 0.052 | 1 | 0.492599419 | 0.643415241 |
| A3galt2 | 0.403319253 | 2.536268244 | 0.19 | 0.142 | 1 | 0.658155837 | 0.880753254 | 0.289 | 0.311 | 1 | 0.658155837 | 0.643972086 |
| Dpf3 | 0.403534817 | 0.936806174 | 0.286 | 0.228 | 1 | 0.430486631 | -0.124282336 | 0 | 0.014 | 1 | 0.430486631 | 0.644229286 |
| Septin2 | 0.403595971 | 1.088809267 | 0.19 | 0.134 | 1 | 0.809088424 | 1.229354619 | 0.178 | 0.19 | 1 | 0.809088424 | 0.644302235 |
| Gm56599 | 0.403849766 | 1.258734268 | 0.095 | 0.052 | 1 | 0.658587075 | 3.268035087 | 0.022 | 0.014 | 1 | 0.658587075 | 0.644604899 |
| Slc6a17 | 0.762799336 | 0.600522786 | 0.238 | 0.235 | 1 | 0.404597642 | 0.744473131 | 0.378 | 0.374 | 1 | 0.762799336 | 0.645496033 |
| Tom1 | 0.405079815 | 0.791128718 | 0.429 | 0.418 | 1 | 0.54418616 | 0.243650928 | 0.644 | 0.574 | 1 | 0.54418616 | 0.646069974 |
| 2500002B13Rik | 0.744838936 | 1.088809267 | 0.048 | 0.034 | 1 | 0.40531543 | 0.180572246 | 0.022 | 0.052 | 1 | 0.744838936 | 0.646350262 |
| Duxbl1 | 0.610955243 | 0.451379346 | 0.048 | 0.082 | 1 | 0.405340571 | 0.435145073 | 0.022 | 0.052 | 1 | 0.610955243 | 0.646380163 |
| Vamp1 | 0.71454173 | 0.918884266 | 0.095 | 0.075 | 1 | 0.405405205 | 0.361144491 | 0.022 | 0.052 | 1 | 0.71454173 | 0.646457029 |
| Kctd21 | 0.454205211 | 2.451379346 | 0.048 | 0.022 | 1 | 0.406607361 | 0.513147585 | 0.022 | 0.052 | 1 | 0.454205211 | 0.647885176 |
| Gpr161 | 0.492599419 | 0.503846766 | 0 | 0.022 | 1 | 0.406607361 | 0.513147585 | 0.022 | 0.052 | 1 | 0.492599419 | 0.647885176 |
| Cd59a | 0.632076134 | 1.673771768 | 0 | 0.011 | 1 | 0.406607361 | 0.83507568 | 0.022 | 0.052 | 1 | 0.632076134 | 0.647885176 |
| Chmp4c | 0.527414465 | -0.683780237 | 0.048 | 0.086 | 1 | 0.406628062 | 0.435145073 | 0.022 | 0.052 | 1 | 0.527414465 | 0.647909743 |
| Sinhcaf | 0.454439722 | 1.908237021 | 0.095 | 0.06 | 1 | 0.406786438 | 0.147019686 | 0.089 | 0.055 | 1 | 0.454439722 | 0.64809767 |
| Spata6 | 0.474783852 | 1.077404504 | 0.429 | 0.504 | 1 | 0.406790348 | 0.454253896 | 0.6 | 0.637 | 1 | 0.474783852 | 0.648102309 |
| Sorcs2 | 0.571891384 | 1.36999102 | 0.143 | 0.108 | 1 | 0.407610769 | 2.425274829 | 0.133 | 0.104 | 1 | 0.571891384 | 0.649074999 |
| Junb | 0.435877318 | -1.084148519 | 0.857 | 0.791 | 1 | 0.407821949 | -0.368653689 | 0.933 | 0.775 | 1 | 0.435877318 | 0.649325156 |
| 6430590A07Rik | 0.408078912 | 1.388369549 | 0.143 | 0.093 | 1 | 0.753432733 | 1.098110085 | 0.089 | 0.114 | 1 | 0.753432733 | 0.649629426 |
| Abcd2 | 0.98545851 | 1.503846766 | 0.048 | 0.049 | 1 | 0.408084204 | 0.972579203 | 0.044 | 0.083 | 1 | 0.98545851 | 0.64963569 |
| Vav2 | 0.538356911 | -0.348596045 | 0.095 | 0.146 | 1 | 0.408453705 | 1.477958156 | 0.222 | 0.201 | 1 | 0.538356911 | 0.650072981 |
| Zfx | 0.718531869 | 0.242980197 | 0.571 | 0.597 | 1 | 0.408611868 | 0.28671965 | 0.889 | 0.83 | 1 | 0.718531869 | 0.650260078 |
| Medag | 0.408672939 | -0.194497534 | 0.333 | 0.429 | 1 | 0.428457216 | 0.221071608 | 0.156 | 0.225 | 1 | 0.428457216 | 0.650332306 |
| Wipi2 | 0.896747598 | 0.195039593 | 0.476 | 0.552 | 1 | 0.408883314 | -0.200413405 | 0.711 | 0.689 | 1 | 0.896747598 | 0.650581063 |
| Mef2d | 0.957219257 | 0.358517679 | 0.619 | 0.776 | 1 | 0.40901013 | 0.345129211 | 0.978 | 0.931 | 1 | 0.957219257 | 0.650730974 |
| Ccdc85a | 0.425631778 | -0.326228232 | 0 | 0.03 | 1 | 0.409200081 | 0.889523464 | 0.044 | 0.083 | 1 | 0.425631778 | 0.650955455 |
| Shc2 | 0.409489432 | 1.908237021 | 0.095 | 0.056 | 1 | 0.864970277 | 0.402964667 | 0.089 | 0.1 | 1 | 0.864970277 | 0.651297269 |
| Guf1 | 0.655344583 | 0.565247311 | 0.19 | 0.164 | 1 | 0.40961351 | 0.513147585 | 0.089 | 0.142 | 1 | 0.655344583 | 0.651443792 |
| Cacng4 | 0.409639871 | 1.063718286 | 0.286 | 0.22 | 1 | 0.44785419 | 0.170622585 | 0.089 | 0.135 | 1 | 0.44785419 | 0.651474918 |
| Ihh | 0.40967441 | -0.370622352 | 0.095 | 0.052 | 1 | 0.430486631 | -2.223818009 | 0 | 0.014 | 1 | 0.430486631 | 0.651515698 |
| Stam2 | 0.615678746 | 0.727352808 | 0.381 | 0.407 | 1 | 0.409819121 | 0.356712245 | 0.644 | 0.564 | 1 | 0.615678746 | 0.65168653 |
| Zfp592 | 0.885752009 | 0.359075242 | 0.381 | 0.444 | 1 | 0.409864126 | 0.472413794 | 0.578 | 0.692 | 1 | 0.885752009 | 0.65173965 |
| Tmem132b | 0.979647018 | 2.957564734 | 0.048 | 0.049 | 1 | 0.410365863 | 3.027026987 | 0.067 | 0.042 | 1 | 0.979647018 | 0.652331585 |
| Gmeb1 | 0.410983009 | 0.762705496 | 0.381 | 0.354 | 1 | 0.865442935 | 1.163698427 | 0.289 | 0.36 | 1 | 0.865442935 | 0.653058984 |
| Rnf31 | 0.778157927 | 0.673771768 | 0.095 | 0.127 | 1 | 0.411903963 | 0.304560963 | 0.089 | 0.142 | 1 | 0.778157927 | 0.654143052 |
| Brpf3 | 0.411962358 | -0.463731756 | 0.143 | 0.22 | 1 | 0.524166859 | 0.264929176 | 0.244 | 0.318 | 1 | 0.524166859 | 0.654211731 |
| Adcy2 | 0.765213207 | 0.286748645 | 0.429 | 0.392 | 1 | 0.412221018 | 0.603050503 | 0.778 | 0.775 | 1 | 0.765213207 | 0.654515868 |
| Notch3 | 0.898198211 | 0.247507013 | 0.095 | 0.086 | 1 | 0.412234484 | 0.486675373 | 0.111 | 0.076 | 1 | 0.898198211 | 0.654531698 |
| Akirin2 | 0.900585185 | -0.186050574 | 0.714 | 0.664 | 1 | 0.412619102 | -0.151806765 | 0.844 | 0.799 | 1 | 0.900585185 | 0.654983681 |
| Impg2 | 0.492604708 | -0.133583154 | 0 | 0.022 | 1 | 0.412959518 | 1.595609745 | 0.067 | 0.042 | 1 | 0.492604708 | 0.655383472 |
| Gm9456 | 0.413308526 | 1.825774861 | 0.095 | 0.056 | 1 | 0.705150377 | 2.03099589 | 0.044 | 0.035 | 1 | 0.705150377 | 0.655793114 |
| Xrcc4 | 0.521769357 | 0.439306514 | 0.238 | 0.373 | 1 | 0.413621072 | 0.503748887 | 0.222 | 0.318 | 1 | 0.521769357 | 0.656159753 |
| Tnpo1 | 0.707372854 | 0.286369835 | 0.905 | 0.918 | 1 | 0.413631539 | 0.119393644 | 0.978 | 0.958 | 1 | 0.707372854 | 0.656172028 |
| Gpx6 | 0.532436372 | 1.088809267 | 0 | 0.019 | 1 | 0.414183647 | 2.005000681 | 0.044 | 0.024 | 1 | 0.532436372 | 0.6568192 |
| Urgcp | 0.807618172 | 0.754390228 | 0.333 | 0.369 | 1 | 0.414278596 | 0.516969296 | 0.378 | 0.592 | 1 | 0.807618172 | 0.656930437 |
| Cbfb | 0.904570377 | 0.399223374 | 0.667 | 0.832 | 1 | 0.414462271 | 0.176361531 | 0.933 | 0.858 | 1 | 0.904570377 | 0.657145568 |
| Larp1b | 0.414881128 | -0.202008295 | 0.429 | 0.53 | 1 | 0.734559138 | 0.29626164 | 0.8 | 0.675 | 1 | 0.734559138 | 0.657635905 |
| Rhbdd1 | 0.48335528 | 0.741790191 | 0.238 | 0.347 | 1 | 0.415305401 | 0.174731367 | 0.222 | 0.311 | 1 | 0.48335528 | 0.658132226 |
| Ist1 | 0.415451154 | 0.497183036 | 0.714 | 0.78 | 1 | 0.601439952 | 0.283665739 | 0.889 | 0.907 | 1 | 0.601439952 | 0.658302646 |
| Helq | 0.9019727 | 0.918884266 | 0.048 | 0.056 | 1 | 0.415655652 | 0.393565969 | 0.067 | 0.111 | 1 | 0.9019727 | 0.658541683 |
| Garre1 | 0.415987261 | 0.19151222 | 0.667 | 0.843 | 1 | 0.488893216 | 0.105763658 | 0.911 | 0.834 | 1 | 0.488893216 | 0.658929121 |
| AU040320 | 0.868329134 | 0.503846766 | 0.19 | 0.228 | 1 | 0.416570851 | 0.5265681 | 0.178 | 0.263 | 1 | 0.868329134 | 0.659610428 |
| Mfn1 | 0.504651622 | 0.614878079 | 0.381 | 0.399 | 1 | 0.416774687 | 0.635766871 | 0.489 | 0.471 | 1 | 0.504651622 | 0.659848235 |
| 9330159M07Rik | 0.417110049 | -0.611630451 | 0.048 | 0.101 | 1 | 0.722093786 | 0.905465008 | 0.044 | 0.059 | 1 | 0.722093786 | 0.660239305 |
| Zfyve28 | 0.57816106 | 0.214340149 | 0 | 0.015 | 1 | 0.417584497 | 1.83507568 | 0.044 | 0.024 | 1 | 0.57816106 | 0.660792182 |
| Pfkfb1 | 0.89388747 | 0.766881172 | 0.048 | 0.041 | 1 | 0.417584497 | 1.83507568 | 0.044 | 0.024 | 1 | 0.89388747 | 0.660792182 |
| Sh3yl1 | 0.640373682 | -0.133583154 | 0.048 | 0.075 | 1 | 0.418161533 | 1.410054092 | 0.044 | 0.083 | 1 | 0.640373682 | 0.661463999 |
| Gm38973 | 0.57816106 | 1.351843673 | 0 | 0.015 | 1 | 0.418201984 | 1.982632868 | 0.067 | 0.042 | 1 | 0.57816106 | 0.661511069 |
| Ccdc85c | 0.960849104 | 0.766881172 | 0.048 | 0.045 | 1 | 0.418227757 | 0.583536913 | 0.044 | 0.083 | 1 | 0.960849104 | 0.661541058 |
| Pde6h | 0.429160329 | -0.544935607 | 0.333 | 0.257 | 1 | 0.418255225 | -0.776359032 | 0.044 | 0.08 | 1 | 0.429160329 | 0.661573017 |
| 4930517O19Rik | 0.41901514 | 1.726239188 | 0.19 | 0.149 | 1 | 0.532745561 | 0.649125254 | 0.156 | 0.221 | 1 | 0.532745561 | 0.662456593 |
| Slc25a21 | 0.419133801 | 1.866416846 | 0.095 | 0.056 | 1 | 0.55196236 | 1.998574412 | 0.067 | 0.048 | 1 | 0.55196236 | 0.662594458 |
| Zfp286 | 0.926908894 | 0.673771768 | 0.048 | 0.052 | 1 | 0.419507562 | 0.329435632 | 0.067 | 0.107 | 1 | 0.926908894 | 0.663028529 |
| Brd8dc | 0.419746114 | 0.586308926 | 0.095 | 0.168 | 1 | 0.506901159 | 0.449275402 | 0.156 | 0.218 | 1 | 0.506901159 | 0.663305428 |
| H3c14 | 0.420080955 | -0.611630451 | 0.048 | 0.101 | 1 | 0.534128516 | -0.124282336 | 0.022 | 0.042 | 1 | 0.534128516 | 0.663693901 |
| Anxa6 | 0.439272444 | 0.970972777 | 0.476 | 0.593 | 1 | 0.420111968 | 0.243058421 | 0.4 | 0.599 | 1 | 0.439272444 | 0.66372987 |
| Micall1 | 0.597787524 | 0.575037814 | 0.476 | 0.496 | 1 | 0.420128822 | 1.007507537 | 0.378 | 0.401 | 1 | 0.597787524 | 0.663749416 |
| Tex9 | 0.451614009 | -0.233118828 | 0.048 | 0.097 | 1 | 0.42018605 | 0.578735926 | 0.067 | 0.111 | 1 | 0.451614009 | 0.663815784 |
| Map3k8 | 0.496677265 | 0.199285803 | 0.476 | 0.451 | 1 | 0.420393595 | 0.201203578 | 0.222 | 0.315 | 1 | 0.496677265 | 0.664056415 |
| Neurl1a | 0.862687636 | 1.088809267 | 0.048 | 0.06 | 1 | 0.420889035 | 1.946106992 | 0.067 | 0.042 | 1 | 0.862687636 | 0.66463049 |
| Zfp119a | 0.621091768 | 1.503846766 | 0.048 | 0.082 | 1 | 0.420951019 | 0.397670367 | 0.067 | 0.111 | 1 | 0.621091768 | 0.664702278 |
| Slc16a9 | 0.966230998 | 1.258734268 | 0.048 | 0.052 | 1 | 0.421213137 | -0.174908409 | 0.044 | 0.08 | 1 | 0.966230998 | 0.665005767 |
| Elf4 | 0.48984713 | 0.944843493 | 0.524 | 0.575 | 1 | 0.421711499 | 0.788254823 | 0.533 | 0.647 | 1 | 0.48984713 | 0.665582409 |
| Rars2 | 0.604188833 | 0.786246497 | 0.286 | 0.243 | 1 | 0.422047585 | 1.412424996 | 0.289 | 0.298 | 1 | 0.604188833 | 0.665971006 |
| Timp3 | 0.734562444 | -0.283377964 | 1 | 0.948 | 1 | 0.422106142 | 0.119264351 | 0.978 | 0.924 | 1 | 0.734562444 | 0.666038689 |
| Gm20682 | 0.991275714 | 1.088809267 | 0.048 | 0.049 | 1 | 0.42222537 | 1.776181991 | 0.067 | 0.042 | 1 | 0.991275714 | 0.666176477 |
| Zfp827 | 0.422399734 | 1.414579428 | 0.429 | 0.392 | 1 | 0.824313454 | 1.041120107 | 0.4 | 0.46 | 1 | 0.824313454 | 0.666377932 |
| Wnt9a | 0.422597127 | 0.840881754 | 0.048 | 0.108 | 1 | 0.900112624 | 0.776181991 | 0.067 | 0.076 | 1 | 0.900112624 | 0.666605923 |
| Srsf11 | 0.586170324 | 0.387509522 | 1 | 1 | 1 | 0.423314948 | 0.14141387 | 1 | 0.997 | 1 | 0.586170324 | 0.66743435 |
| Spats2l | 0.42504554 | 0.167003997 | 0.714 | 0.91 | 1 | 0.423418179 | 0.208237824 | 0.911 | 0.927 | 1 | 0.42504554 | 0.667553404 |
| Optn | 0.423487187 | 0.652710152 | 0.524 | 0.571 | 1 | 0.954459411 | 0.522895461 | 0.422 | 0.519 | 1 | 0.954459411 | 0.667632976 |
| A230087F16Rik | 0.425599726 | 0.214340149 | 0 | 0.03 | 1 | 0.423581872 | 1.583536913 | 0.067 | 0.042 | 1 | 0.425599726 | 0.667742142 |
| Rnf10 | 0.423588518 | -0.195805669 | 0.81 | 0.91 | 1 | 0.792371161 | 0.139186028 | 1 | 0.965 | 1 | 0.792371161 | 0.667749804 |
| Pwp2 | 0.424077859 | 1.673771768 | 0.143 | 0.097 | 1 | 0.549586684 | 0.460680165 | 0.089 | 0.125 | 1 | 0.549586684 | 0.668313688 |

| | | | | | | | | | | | | |
|---|---|---|---|---|---|---|---|---|---|---|---|---|
| Vps37a | 0.856821949 | 0.16421667 | 0.524 | 0.698 | 1 | 0.424256335 | 0.49570245 | 0.844 | 0.82 | 1 | 0.856821949 | 0.668519232 |
| 0610009E02Rik | 0.424500903 | -0.370622352 | 0.048 | 0.101 | 1 | 0.473334511 | 0.361144491 | 0.089 | 0.135 | 1 | 0.473334511 | 0.668800789 |
| Gng2 | 0.42554795 | 1.267779408 | 0.048 | 0.108 | 1 | 0.48614806 | 0.946106992 | 0.089 | 0.128 | 1 | 0.48614806 | 0.670004842 |
| Ifi204 | 0.425599726 | 0.351843673 | 0 | 0.03 | 1 | 0.932582169 | 0.223640968 | 0.022 | 0.024 | 1 | 0.932582169 | 0.670064325 |
| Cutal | 0.425599726 | 0.351843673 | 0 | 0.03 | 1 | 0.496223879 | 0.683072586 | 0 | 0.01 | 1 | 0.496223879 | 0.670064325 |
| BB557941 | 0.425599726 | 0.351843673 | 0 | 0.03 | 1 | 0.49622258 | 0.361144491 | 0 | 0.01 | 1 | 0.49622258 | 0.670064325 |
| 6430573P05Rik | 0.425599726 | 0.351843673 | 0 | 0.03 | 1 | 0.80624346 | 2.098110085 | 0.022 | 0.017 | 1 | 0.80624346 | 0.670064325 |
| Rrh | 0.425599726 | 0.351843673 | 0 | 0.03 | 1 | 0.825853389 | 0.683072586 | 0.022 | 0.017 | 1 | 0.825853389 | 0.670064325 |
| Gm12724 | 0.425599726 | 0.351843673 | 0 | 0.03 | 1 | 0.430487676 | 0.361144491 | 0 | 0.014 | 1 | 0.430487676 | 0.670064325 |
| Rab3b | 0.425599726 | 0.214340149 | 0 | 0.03 | 1 | 0.496223879 | 0.683072586 | 0 | 0.01 | 1 | 0.496223879 | 0.670064325 |
| Arhgef16 | 0.425599726 | 0.351843673 | 0 | 0.03 | 1 | 0.679341763 | 0.435145073 | 0.022 | 0.035 | 1 | 0.679341763 | 0.670064325 |
| Gm26810 | 0.425599726 | 0.351843673 | 0 | 0.03 | 1 | 0.627053233 | 1.361144491 | 0.044 | 0.031 | 1 | 0.627053233 | 0.670064325 |
| 6530437J22Rik | 0.425599726 | 0.351843673 | 0 | 0.03 | 1 | 0.957324717 | 0.513147585 | 0.022 | 0.021 | 1 | 0.957324717 | 0.670064325 |
| Mmp15 | 0.425599726 | 0.351843673 | 0 | 0.03 | 1 | 0.543165705 | -0.124282336 | 0.044 | 0.069 | 1 | 0.543165705 | 0.670064325 |
| Gm56848 | 0.425599726 | 0.351843673 | 0 | 0.03 | 1 | 0.952529269 | 1.098110085 | 0.022 | 0.024 | 1 | 0.952529269 | 0.670064325 |
| Nsg2 | 0.425599726 | 0.214340149 | 0 | 0.03 | 1 | 0.492040936 | 2.683072586 | 0.022 | 0.01 | 1 | 0.492040936 | 0.670064325 |
| AA465934 | 0.425599726 | 0.351843673 | 0 | 0.03 | 1 | 0.49622258 | 0.361144491 | 0 | 0.01 | 1 | 0.49622258 | 0.670064325 |
| Krt222 | 0.425599726 | 0.351843673 | 0 | 0.03 | 1 | 0.496223879 | 0.683072586 | 0 | 0.01 | 1 | 0.496223879 | 0.670064325 |
| Ism2 | 0.425599726 | 0.351843673 | 0 | 0.03 | 1 | 0.916239979 | 0.683072586 | 0.044 | 0.042 | 1 | 0.916239979 | 0.670064325 |
| Rtl6 | 0.425599726 | 0.351843673 | 0 | 0.03 | 1 | 0.548936906 | 0.180572246 | 0.022 | 0.042 | 1 | 0.548936906 | 0.670064325 |
| Hunk | 0.425599726 | -0.133583154 | 0 | 0.03 | 1 | 0.450885236 | 1.460680165 | 0.022 | 0.048 | 1 | 0.450885236 | 0.670064325 |
| Gm49958 | 0.425599726 | 0.351843673 | 0 | 0.03 | 1 | 0.819747895 | 1.304560963 | 0.044 | 0.038 | 1 | 0.819747895 | 0.670064325 |
| Gm50295 | 0.425599726 | 0.351843673 | 0 | 0.03 | 1 | 0.761441218 | 0.567595369 | 0.022 | 0.031 | 1 | 0.761441218 | 0.670064325 |
| Pcdhb5 | 0.425599726 | 0.351843673 | 0 | 0.03 | 1 | 0.811836522 | 1.098110085 | 0.022 | 0.017 | 1 | 0.811836522 | 0.670064325 |
| Gm33732 | 0.425599726 | 0.351843673 | 0 | 0.03 | 1 | 0.430486631 | -0.486852415 | 0 | 0.014 | 1 | 0.430486631 | 0.670064325 |
| Batf2 | 0.425599726 | 0.214340149 | 0 | 0.03 | 1 | 0.765728546 | 1.223640968 | 0.022 | 0.031 | 1 | 0.765728546 | 0.670064325 |
| Sec31b | 0.425599726 | 0.351843673 | 0 | 0.03 | 1 | 0.835943453 | 0.513147585 | 0.022 | 0.028 | 1 | 0.835943453 | 0.670064325 |
| Kcnd1 | 0.425599726 | 0.351843673 | 0 | 0.03 | 1 | 0.535080001 | 0.982632868 | 0.044 | 0.073 | 1 | 0.535080001 | 0.670064325 |
| Fancb | 0.425599726 | 0.351843673 | 0 | 0.03 | 1 | 0.49622258 | 0.361144491 | 0 | 0.01 | 1 | 0.49622258 | 0.670064325 |
| Gm12655 | 0.425614684 | 0.214340149 | 0 | 0.03 | 1 | 0.430487676 | -0.901889915 | 0 | 0.014 | 1 | 0.430487676 | 0.670081509 |
| Kctd7 | 0.425614684 | 0.214340149 | 0 | 0.03 | 1 | 0.910545146 | 0.875717664 | 0.044 | 0.042 | 1 | 0.910545146 | 0.670081509 |
| A730049H05Rik | 0.425614684 | 0.214340149 | 0 | 0.03 | 1 | 0.930092617 | -0.124282336 | 0.022 | 0.024 | 1 | 0.930092617 | 0.670081509 |
| Slc5a11 | 0.425614684 | 0.214340149 | 0 | 0.03 | 1 | 0.430483494 | -0.124282336 | 0 | 0.014 | 1 | 0.430483494 | 0.670081509 |
| Gm15910 | 0.425614684 | 0.214340149 | 0 | 0.03 | 1 | 0.835943453 | 0.513147585 | 0.022 | 0.028 | 1 | 0.835943453 | 0.670081509 |
| C130046K22Rik | 0.425614684 | 0.214340149 | 0 | 0.03 | 1 | 0.500244121 | 1.098110085 | 0.022 | 0.045 | 1 | 0.500244121 | 0.670081509 |
| Krt76 | 0.425614684 | 0.214340149 | 0 | 0.03 | 1 | 0.430483494 | -0.124282336 | 0 | 0.014 | 1 | 0.430483494 | 0.670081509 |
| Arvcf | 0.425614684 | -0.133583154 | 0 | 0.03 | 1 | 0.962655732 | 0.223640968 | 0.022 | 0.021 | 1 | 0.962655732 | 0.670081509 |
| Gpr156 | 0.425614684 | 0.214340149 | 0 | 0.03 | 1 | 0.95502578 | 2.235613609 | 0.022 | 0.024 | 1 | 0.95502578 | 0.670081509 |
| Pmaip1 | 0.425614684 | 0.214340149 | 0 | 0.03 | 1 | 0.516127799 | 2.03099589 | 0.044 | 0.028 | 1 | 0.516127799 | 0.670081509 |
| Gdpd2 | 0.425614684 | 0.214340149 | 0 | 0.03 | 1 | 0.496223879 | 0.683072586 | 0 | 0.01 | 1 | 0.496223879 | 0.670081509 |
| Gm32816 | 0.425624656 | -0.911190733 | 0 | 0.03 | 1 | 0.430483494 | -0.901889915 | 0 | 0.014 | 1 | 0.430483494 | 0.670092964 |
| Odad2 | 0.425624656 | -0.133583154 | 0 | 0.03 | 1 | 0.952529269 | 1.098110085 | 0.022 | 0.024 | 1 | 0.952529269 | 0.670092964 |
| Gm12511 | 0.425625368 | -0.413691074 | 0 | 0.03 | 1 | 0.496223879 | -0.124282336 | 0 | 0.01 | 1 | 0.496223879 | 0.670093782 |
| Mlxipl | 0.425625368 | -0.133583154 | 0 | 0.03 | 1 | 0.726291198 | 1.683072586 | 0.044 | 0.035 | 1 | 0.726291198 | 0.670093782 |
| Ddc | 0.425625368 | -1.133583154 | 0 | 0.03 | 1 | 0.430487676 | 0.361144491 | 0 | 0.014 | 1 | 0.430487676 | 0.670093782 |
| Kif14 | 0.425629641 | -0.233118828 | 0 | 0.03 | 1 | 0.430487676 | 0.361144491 | 0 | 0.014 | 1 | 0.430487676 | 0.670098691 |
| Nudt10 | 0.425629641 | -0.133583154 | 0 | 0.03 | 1 | 0.492040936 | 2.268035087 | 0.022 | 0.01 | 1 | 0.492040936 | 0.670098691 |
| Gm32585 | 0.42563249 | -0.648156327 | 0 | 0.03 | 1 | 0.496223879 | 0.683072586 | 0 | 0.01 | 1 | 0.496223879 | 0.670101963 |
| Gm31812 | 0.425633914 | -0.413691074 | 0 | 0.03 | 1 | 0.658587075 | 2.513147585 | 0.022 | 0.014 | 1 | 0.658587075 | 0.6701036 |
| Pappa | 0.425635339 | -0.718545655 | 0 | 0.03 | 1 | 0.98452369 | 0.571564271 | 0.133 | 0.138 | 1 | 0.98452369 | 0.670105236 |
| Mmp9 | 0.425636051 | -0.970084422 | 0 | 0.03 | 1 | 0.735977919 | -4.312411933 | 0.044 | 0.055 | 1 | 0.735977919 | 0.670106054 |
| Tll2 | 0.425636051 | -0.648156327 | 0 | 0.03 | 1 | 0.555577743 | 1.605070074 | 0.022 | 0.042 | 1 | 0.555577743 | 0.670106054 |
| Esco2 | 0.425636763 | -1.184209227 | 0 | 0.03 | 1 | 0.496223879 | 0.683072586 | 0 | 0.01 | 1 | 0.496223879 | 0.670106872 |
| Gm47200 | 0.425638188 | -1.648156327 | 0 | 0.03 | 1 | 0.496223879 | 0.683072586 | 0 | 0.01 | 1 | 0.496223879 | 0.670108508 |
| H2bc21 | 0.425639612 | 0.503846766 | 0 | 0.03 | 1 | 0.932582169 | 0.361144491 | 0.022 | 0.024 | 1 | 0.932582169 | 0.670110145 |
| Dok7 | 0.425639612 | 0.503846766 | 0 | 0.03 | 1 | 0.685254641 | 0.683072586 | 0.022 | 0.035 | 1 | 0.685254641 | 0.670110145 |
| Vamp9 | 0.425639612 | 0.503846766 | 0 | 0.03 | 1 | 0.494823351 | 1.946106992 | 0.022 | 0.01 | 1 | 0.494823351 | 0.670110145 |
| Cldn15 | 0.425639612 | 0.503846766 | 0 | 0.03 | 1 | 0.87875596 | 1.020107573 | 0.044 | 0.052 | 1 | 0.87875596 | 0.670110145 |
| Tril | 0.425639612 | 0.503846766 | 0 | 0.03 | 1 | 0.430487676 | 0.361144491 | 0 | 0.014 | 1 | 0.430487676 | 0.670110145 |
| A430078I02Rik | 0.425639612 | 0.503846766 | 0 | 0.03 | 1 | 0.430487676 | 0.361144491 | 0 | 0.014 | 1 | 0.430487676 | 0.670110145 |
| Gm45238 | 0.425639612 | 0.503846766 | 0 | 0.03 | 1 | 0.47918822 | -0.223818009 | 0.022 | 0.045 | 1 | 0.47918822 | 0.670110145 |
| Hyal3 | 0.425639612 | 0.503846766 | 0 | 0.03 | 1 | 0.477727467 | -0.316927414 | 0.022 | 0.045 | 1 | 0.477727467 | 0.670110145 |
| 4930533K18Rik | 0.425639612 | 0.503846766 | 0 | 0.03 | 1 | 0.716679936 | 1.268035087 | 0.044 | 0.035 | 1 | 0.716679936 | 0.670110145 |
| Zfp647 | 0.425639612 | 0.503846766 | 0 | 0.03 | 1 | 0.744358033 | 0.223640968 | 0.022 | 0.031 | 1 | 0.744358033 | 0.670110145 |
| Gm56727 | 0.425639612 | 0.503846766 | 0 | 0.03 | 1 | 0.496223879 | 0.683072586 | 0 | 0.01 | 1 | 0.496223879 | 0.670110145 |
| Eme2 | 0.425639612 | 0.503846766 | 0 | 0.03 | 1 | 0.692345886 | 0.946106992 | 0.044 | 0.062 | 1 | 0.692345886 | 0.670110145 |
| Gm4566 | 0.425639612 | 0.503846766 | 0 | 0.03 | 1 | 0.806246349 | 2.460680165 | 0.022 | 0.017 | 1 | 0.806246349 | 0.670110145 |
| Gm26511 | 0.492590603 | 0.673771768 | 0 | 0.022 | 1 | 0.426109575 | 2.005000681 | 0.044 | 0.024 | 1 | 0.492590603 | 0.67064978 |
| A430102K17Rik | 0.492608234 | 0.866416846 | 0 | 0.022 | 1 | 0.426109575 | 2.005000681 | 0.044 | 0.024 | 1 | 0.492608234 | 0.67064978 |
| Gm15852 | 0.57816106 | 1.351843673 | 0 | 0.015 | 1 | 0.426109575 | 1.683072586 | 0.044 | 0.024 | 1 | 0.57816106 | 0.67064978 |
| Mmp13 | 0.426621893 | -3.792668909 | 0.095 | 0.153 | 1 | 0.923182359 | -0.864736916 | 0.044 | 0.048 | 1 | 0.923182359 | 0.671237547 |
| ENSMUSG00000121500 | 0.7957781 | 1.451379346 | 0.095 | 0.086 | 1 | 0.4275083 | 0.250113179 | 0.044 | 0.08 | 1 | 0.7957781 | 0.672253253 |
| Kcnab1 | 0.813343641 | 0.214340149 | 0.048 | 0.06 | 1 | 0.427594344 | -0.776359032 | 0.022 | 0.048 | 1 | 0.813343641 | 0.672351765 |
| Gm49066 | 0.770713471 | 0.351843673 | 0.048 | 0.063 | 1 | 0.427629025 | 1.683072586 | 0.067 | 0.042 | 1 | 0.770713471 | 0.672391467 |
| Zfp473 | 0.532432377 | 0.214340149 | 0 | 0.019 | 1 | 0.427844671 | 1.83507568 | 0.044 | 0.024 | 1 | 0.532432377 | 0.67263828 |
| Snhg11 | 0.632076134 | 1.673771768 | 0 | 0.011 | 1 | 0.427865475 | 1.361144491 | 0.044 | 0.024 | 1 | 0.632076134 | 0.672662085 |
| Gdpd3 | 0.532436372 | 1.088809267 | 0 | 0.019 | 1 | 0.4285282 | 0.304560963 | 0.044 | 0.08 | 1 | 0.532436372 | 0.673419982 |
| Senp5 | 0.511710222 | 0.663787679 | 0.571 | 0.698 | 1 | 0.428905036 | 0.443402173 | 0.822 | 0.869 | 1 | 0.511710222 | 0.673850542 |
| Rph3al | 0.492608234 | 0.866416846 | 0 | 0.022 | 1 | 0.428920253 | -0.486852415 | 0.022 | 0.048 | 1 | 0.492608234 | 0.673867922 |
| Shmt1 | 0.532436372 | 1.088809267 | 0 | 0.019 | 1 | 0.428920253 | -0.564854927 | 0.022 | 0.048 | 1 | 0.532436372 | 0.673867922 |
| Gm26982 | 0.629700127 | 2.088809267 | 0.048 | 0.03 | 1 | 0.428920253 | -0.404390255 | 0.022 | 0.048 | 1 | 0.629700127 | 0.673867922 |
| Tnfaip2 | 0.761826274 | 0.614878079 | 0.048 | 0.067 | 1 | 0.428920253 | -0.486852415 | 0.022 | 0.048 | 1 | 0.761826274 | 0.673867922 |
| Gp1bb | 0.918789496 | 1.55829455 | 0.048 | 0.045 | 1 | 0.428920253 | -0.404390255 | 0.022 | 0.048 | 1 | 0.918789496 | 0.673867922 |
| Tia1 | 0.429093565 | -0.380676016 | 0.286 | 0.388 | 1 | 0.705029547 | -0.178936411 | 0.489 | 0.488 | 1 | 0.705029547 | 0.674065843 |
| Ptger3 | 0.42931619 | -0.8244791 | 0.095 | 0.164 | 1 | 0.528899417 | 0.536231198 | 0.044 | 0.073 | 1 | 0.528899417 | 0.674319989 |
| Dmtf1 | 0.429471246 | -0.220237537 | 0.333 | 0.466 | 1 | 0.459239509 | 0.62187243 | 0.578 | 0.592 | 1 | 0.459239509 | 0.674496941 |
| Gemin5 | 0.861762365 | 0.866416846 | 0.238 | 0.265 | 1 | 0.429856858 | -0.16301473 | 0.2 | 0.28 | 1 | 0.861762365 | 0.674936797 |
| Grb2 | 0.429981511 | 0.181918671 | 0.619 | 0.795 | 1 | 0.630123029 | 0.202698987 | 0.956 | 0.886 | 1 | 0.630123029 | 0.675078922 |
| Dzank1 | 0.94656814 | 0.400753273 | 0.048 | 0.052 | 1 | 0.430327413 | 1.320502507 | 0.067 | 0.042 | 1 | 0.94656814 | 0.675473144 |
| Gm26839 | 0.532432377 | 0.673771768 | 0 | 0.019 | 1 | 0.430483494 | -0.316927414 | 0 | 0.014 | 1 | 0.532432377 | 0.675650949 |
| Pebp4 | 0.532436372 | 1.088809267 | 0 | 0.019 | 1 | 0.430483494 | -0.124282336 | 0 | 0.014 | 1 | 0.532436372 | 0.675650949 |
| 4833408A19Rik | 0.532436372 | 1.088809267 | 0 | 0.019 | 1 | 0.430483494 | -0.124282336 | 0 | 0.014 | 1 | 0.532436372 | 0.675650949 |
| Pi15 | 0.578159907 | 0.214340149 | 0 | 0.015 | 1 | 0.430483494 | -0.316927414 | 0 | 0.014 | 1 | 0.578159907 | 0.675650949 |
| Tmeff2 | 0.578159907 | 0.503846766 | 0 | 0.015 | 1 | 0.430483494 | -1.404390255 | 0 | 0.014 | 1 | 0.578159907 | 0.675650949 |
| Caln1 | 0.578159907 | 0.866416846 | 0 | 0.015 | 1 | 0.430483494 | -1.638855509 | 0 | 0.014 | 1 | 0.578159907 | 0.675650949 |
| Flacc1 | 0.632076134 | 0.866416846 | 0 | 0.011 | 1 | 0.430483494 | -0.638855509 | 0 | 0.014 | 1 | 0.632076134 | 0.675650949 |
| D430040L24Rik | 0.632076134 | 1.673771768 | 0 | 0.011 | 1 | 0.430483494 | -0.124282336 | 0 | 0.014 | 1 | 0.632076134 | 0.675650949 |
| E2f7 | 0.632076134 | -0.574155746 | 0 | 0.011 | 1 | 0.430483494 | -0.486852415 | 0 | 0.014 | 1 | 0.632076134 | 0.675650949 |

| | | | | | | | | | | | |
|---|---|---|---|---|---|---|---|---|---|---|---|
| Cntnap1 | 0.632076134 | 1.673771768 | 0 | 0.011 | 1 | 0.430483494 | -0.124282336 | 0 | 0.014 | 1 | 0.632076134 | 0.675650949 |
| Gm16675 | 0.71266839 | 1.79930265 | 0.048 | 0.034 | 1 | 0.430483494 | -0.316927414 | 0 | 0.014 | 1 | 0.71266839 | 0.675650949 |
| Mtarc1 | 0.784065849 | 2.351843673 | 0.048 | 0.037 | 1 | 0.430483494 | -0.124282336 | 0 | 0.014 | 1 | 0.784065849 | 0.675650949 |
| Gm26671 | 0.875317586 | 0.936806174 | 0.048 | 0.041 | 1 | 0.430483494 | -0.316927414 | 0 | 0.014 | 1 | 0.875317586 | 0.675650949 |
| Clca2 | 0.90740681 | 0.673771768 | 0.095 | 0.108 | 1 | 0.430483494 | -0.316927414 | 0 | 0.014 | 1 | 0.90740681 | 0.675650949 |
| Ggt5 | 0.457281049 | 0.503846766 | 0 | 0.026 | 1 | 0.430486631 | -0.124282336 | 0 | 0.014 | 1 | 0.457281049 | 0.675654522 |
| 9430065F17Rik | 0.457281049 | 0.503846766 | 0 | 0.026 | 1 | 0.430486631 | -0.124282336 | 0 | 0.014 | 1 | 0.457281049 | 0.675654522 |
| Gm50373 | 0.463486483 | 1.351843673 | 0.048 | 0.022 | 1 | 0.430486631 | -0.486852415 | 0 | 0.014 | 1 | 0.463486483 | 0.675654522 |
| Gm57036 | 0.492608234 | 0.866416846 | 0 | 0.022 | 1 | 0.430486631 | -0.124282336 | 0 | 0.014 | 1 | 0.492608234 | 0.675654522 |
| Adgrv1 | 0.527414914 | -0.880817084 | 0.048 | 0.086 | 1 | 0.430486631 | -1.017367132 | 0 | 0.014 | 1 | 0.527414914 | 0.675654522 |
| Sp100 | 0.532434375 | -0.574155746 | 0 | 0.019 | 1 | 0.430486631 | -1.223818009 | 0 | 0.014 | 1 | 0.532434375 | 0.675654522 |
| Ankrd45 | 0.578156445 | 1.088809267 | 0 | 0.015 | 1 | 0.430486631 | -0.486852415 | 0 | 0.014 | 1 | 0.578156445 | 0.675654522 |
| Gm12979 | 0.578156445 | 0.866416846 | 0 | 0.015 | 1 | 0.430486631 | -0.316927414 | 0 | 0.014 | 1 | 0.578156445 | 0.675654522 |
| Tent5d | 0.578156445 | 1.088809267 | 0 | 0.015 | 1 | 0.430486631 | -0.124282336 | 0 | 0.014 | 1 | 0.578156445 | 0.675654522 |
| Gm16897 | 0.578159907 | 0.866416846 | 0 | 0.015 | 1 | 0.430486631 | -0.124282336 | 0 | 0.014 | 1 | 0.578159907 | 0.675654522 |
| Lingo2 | 0.578159907 | 0.673771768 | 0 | 0.015 | 1 | 0.430486631 | -0.486852415 | 0 | 0.014 | 1 | 0.578159907 | 0.675654522 |
| Dpp4 | 0.57816106 | 1.351843673 | 0 | 0.015 | 1 | 0.430486631 | -1.124282336 | 0 | 0.014 | 1 | 0.57816106 | 0.675654522 |
| Guca1a | 0.57816106 | 1.351843673 | 0 | 0.015 | 1 | 0.430486631 | -0.124282336 | 0 | 0.014 | 1 | 0.57816106 | 0.675654522 |
| C130060C02Rik | 0.57816106 | 1.351843673 | 0 | 0.015 | 1 | 0.430486631 | -0.124282336 | 0 | 0.014 | 1 | 0.57816106 | 0.675654522 |
| Gm56646 | 0.57816106 | 1.351843673 | 0 | 0.015 | 1 | 0.430486631 | -0.776359032 | 0 | 0.014 | 1 | 0.57816106 | 0.675654522 |
| Rasgrp1 | 0.63207476 | 0.503846766 | 0 | 0.011 | 1 | 0.430486631 | -0.316927414 | 0 | 0.014 | 1 | 0.63207476 | 0.675654522 |
| Usp51 | 0.63207476 | 1.351843673 | 0 | 0.011 | 1 | 0.430486631 | -0.638855509 | 0 | 0.014 | 1 | 0.63207476 | 0.675654522 |
| Gm20597 | 0.632076134 | 1.673771768 | 0 | 0.011 | 1 | 0.430486631 | -0.486852415 | 0 | 0.014 | 1 | 0.632076134 | 0.675654522 |
| Gm56752 | 0.632076134 | 1.673771768 | 0 | 0.011 | 1 | 0.430486631 | -0.316927414 | 0 | 0.014 | 1 | 0.632076134 | 0.675654522 |
| Ptx4 | 0.632076134 | 1.673771768 | 0 | 0.011 | 1 | 0.430486631 | -0.124282336 | 0 | 0.014 | 1 | 0.632076134 | 0.675654522 |
| Btla | 0.787219796 | 1.756233928 | 0.048 | 0.037 | 1 | 0.430486631 | -0.124282336 | 0 | 0.014 | 1 | 0.787219796 | 0.675654522 |
| H2-Q10 | 0.936805112 | 0.735172312 | 0.048 | 0.045 | 1 | 0.430486631 | -0.316927414 | 0 | 0.014 | 1 | 0.936805112 | 0.675654522 |
| Cdk1 | 0.972894124 | -1.280424543 | 0.048 | 0.045 | 1 | 0.430486631 | -0.124282336 | 0 | 0.014 | 1 | 0.972894124 | 0.675654522 |
| Ttc39aos1 | 0.457281049 | 0.503846766 | 0 | 0.026 | 1 | 0.430487676 | 0.361144491 | 0 | 0.014 | 1 | 0.457281049 | 0.675655713 |
| Zfp853 | 0.457281049 | 0.351843673 | 0 | 0.026 | 1 | 0.430487676 | 0.361144491 | 0 | 0.014 | 1 | 0.457281049 | 0.675655713 |
| Gm56588 | 0.457281049 | 0.503846766 | 0 | 0.026 | 1 | 0.430487676 | 0.361144491 | 0 | 0.014 | 1 | 0.457281049 | 0.675655713 |
| Gm49492 | 0.457281049 | 0.503846766 | 0 | 0.026 | 1 | 0.430487676 | 0.361144491 | 0 | 0.014 | 1 | 0.457281049 | 0.675655713 |
| Shcbp1 | 0.457304706 | -0.326228232 | 0 | 0.026 | 1 | 0.430487676 | 0.361144491 | 0 | 0.014 | 1 | 0.457304706 | 0.675655713 |
| C230012O17Rik | 0.457308648 | 0.673771768 | 0 | 0.026 | 1 | 0.430487676 | 0.361144491 | 0 | 0.014 | 1 | 0.457308648 | 0.675655713 |
| 4933408J17Rik | 0.457308648 | 0.673771768 | 0 | 0.026 | 1 | 0.430487676 | 0.361144491 | 0 | 0.014 | 1 | 0.457308648 | 0.675655713 |
| Adap2 | 0.457308648 | 0.673771768 | 0 | 0.026 | 1 | 0.430487676 | 0.361144491 | 0 | 0.014 | 1 | 0.457308648 | 0.675655713 |
| Tomm20l | 0.457308648 | 0.673771768 | 0 | 0.026 | 1 | 0.430487676 | -0.316927414 | 0 | 0.014 | 1 | 0.457308648 | 0.675655713 |
| 5730424H11Rik | 0.457308648 | 0.673771768 | 0 | 0.026 | 1 | 0.430487676 | 0.361144491 | 0 | 0.014 | 1 | 0.457308648 | 0.675655713 |
| 5033406O09Rik | 0.4759797 | 1.673771768 | 0.048 | 0.022 | 1 | 0.430487676 | 0.361144491 | 0 | 0.014 | 1 | 0.4759797 | 0.675655713 |
| Gm28411 | 0.492590603 | 0.673771768 | 0 | 0.022 | 1 | 0.430487676 | 0.361144491 | 0 | 0.014 | 1 | 0.492590603 | 0.675655713 |
| 4933431K23Rik | 0.492590603 | 0.214340149 | 0 | 0.022 | 1 | 0.430487676 | -0.776359032 | 0 | 0.014 | 1 | 0.492590603 | 0.675655713 |
| Gm15908 | 0.492599419 | -0.133583154 | 0 | 0.022 | 1 | 0.430487676 | 0.361144491 | 0 | 0.014 | 1 | 0.492599419 | 0.675655713 |
| Bub1 | 0.492606471 | -0.970084422 | 0 | 0.022 | 1 | 0.430487676 | 0.361144491 | 0 | 0.014 | 1 | 0.492606471 | 0.675655713 |
| Jsrp1 | 0.492608234 | 0.866416846 | 0 | 0.022 | 1 | 0.430487676 | 0.361144491 | 0 | 0.014 | 1 | 0.492608234 | 0.675655713 |
| Gm48420 | 0.492608234 | 0.866416846 | 0 | 0.022 | 1 | 0.430487676 | 0.361144491 | 0 | 0.014 | 1 | 0.492608234 | 0.675655713 |
| Psma8 | 0.492608234 | 0.866416846 | 0 | 0.022 | 1 | 0.430487676 | 0.361144491 | 0 | 0.014 | 1 | 0.492608234 | 0.675655713 |
| Fam178b | 0.532426384 | -0.133583154 | 0 | 0.019 | 1 | 0.430487676 | 0.361144491 | 0 | 0.014 | 1 | 0.532426384 | 0.675655713 |
| G730003C15Rik | 0.532426384 | 0.866416846 | 0 | 0.019 | 1 | 0.430487676 | 0.361144491 | 0 | 0.014 | 1 | 0.532426384 | 0.675655713 |
| Armh1 | 0.532426384 | 0.866416846 | 0 | 0.019 | 1 | 0.430487676 | 0.361144491 | 0 | 0.014 | 1 | 0.532426384 | 0.675655713 |
| Gm6598 | 0.532426384 | 0.503846766 | 0 | 0.019 | 1 | 0.430487676 | 0.361144491 | 0 | 0.014 | 1 | 0.532426384 | 0.675655713 |
| 1110035H17Rik | 0.532426384 | 0.866416846 | 0 | 0.019 | 1 | 0.430487676 | 0.361144491 | 0 | 0.014 | 1 | 0.532426384 | 0.675655713 |
| BC050972 | 0.532426384 | 0.866416846 | 0 | 0.019 | 1 | 0.430487676 | 0.361144491 | 0 | 0.014 | 1 | 0.532426384 | 0.675655713 |
| Gm57374 | 0.532426384 | 0.503846766 | 0 | 0.019 | 1 | 0.430487676 | 0.361144491 | 0 | 0.014 | 1 | 0.532426384 | 0.675655713 |
| Sox30 | 0.532426384 | 0.866416846 | 0 | 0.019 | 1 | 0.430487676 | 0.361144491 | 0 | 0.014 | 1 | 0.532426384 | 0.675655713 |
| Tbx21 | 0.532426384 | 0.866416846 | 0 | 0.019 | 1 | 0.430487676 | 0.361144491 | 0 | 0.014 | 1 | 0.532426384 | 0.675655713 |
| Gm47484 | 0.532426384 | 0.673771768 | 0 | 0.019 | 1 | 0.430487676 | 0.361144491 | 0 | 0.014 | 1 | 0.532426384 | 0.675655713 |
| Gm56767 | 0.532426384 | 0.866416846 | 0 | 0.019 | 1 | 0.430487676 | 0.361144491 | 0 | 0.014 | 1 | 0.532426384 | 0.675655713 |
| A930007A09Rik | 0.532426384 | 0.866416846 | 0 | 0.019 | 1 | 0.430487676 | 0.361144491 | 0 | 0.014 | 1 | 0.532426384 | 0.675655713 |
| Gm32261 | 0.532426384 | 0.866416846 | 0 | 0.019 | 1 | 0.430487676 | 0.361144491 | 0 | 0.014 | 1 | 0.532426384 | 0.675655713 |
| Gm29481 | 0.532432377 | 0.673771768 | 0 | 0.019 | 1 | 0.430487676 | 0.361144491 | 0 | 0.014 | 1 | 0.532432377 | 0.675655713 |
| Cfap206 | 0.532432377 | 0.351843673 | 0 | 0.019 | 1 | 0.430487676 | 0.361144491 | 0 | 0.014 | 1 | 0.532432377 | 0.675655713 |
| Wnt10b | 0.532432377 | 0.503846766 | 0 | 0.019 | 1 | 0.430487676 | 0.361144491 | 0 | 0.014 | 1 | 0.532432377 | 0.675655713 |
| 2810459M11Rik | 0.532436372 | 1.088809267 | 0 | 0.019 | 1 | 0.430487676 | 0.361144491 | 0 | 0.014 | 1 | 0.532436372 | 0.675655713 |
| Nhlrc1 | 0.532436372 | 1.088809267 | 0 | 0.019 | 1 | 0.430487676 | 0.361144491 | 0 | 0.014 | 1 | 0.532436372 | 0.675655713 |
| Gm36445 | 0.532436372 | 1.088809267 | 0 | 0.019 | 1 | 0.430487676 | 0.361144491 | 0 | 0.014 | 1 | 0.532436372 | 0.675655713 |
| Pdxp | 0.532436372 | 1.088809267 | 0 | 0.019 | 1 | 0.430487676 | 0.361144491 | 0 | 0.014 | 1 | 0.532436372 | 0.675655713 |
| Gp5 | 0.532436372 | 1.088809267 | 0 | 0.019 | 1 | 0.430487676 | 0.361144491 | 0 | 0.014 | 1 | 0.532436372 | 0.675655713 |
| Gm26691 | 0.54776222 | 2.088809267 | 0.048 | 0.026 | 1 | 0.430487676 | 0.361144491 | 0 | 0.014 | 1 | 0.54776222 | 0.675655713 |
| Gm9908 | 0.567009484 | 1.673771768 | 0.048 | 0.026 | 1 | 0.430487676 | 0.361144491 | 0 | 0.014 | 1 | 0.567009484 | 0.675655713 |
| Gm38250 | 0.567009484 | 1.673771768 | 0.048 | 0.026 | 1 | 0.430487676 | 0.361144491 | 0 | 0.014 | 1 | 0.567009484 | 0.675655713 |
| B930059L03Rik | 0.578156445 | 1.088809267 | 0 | 0.015 | 1 | 0.430487676 | 0.361144491 | 0 | 0.014 | 1 | 0.578156445 | 0.675655713 |
| Gm30881 | 0.578156445 | 1.088809267 | 0 | 0.015 | 1 | 0.430487676 | 0.361144491 | 0 | 0.014 | 1 | 0.578156445 | 0.675655713 |
| Dlgap5 | 0.578159907 | 0.214340149 | 0 | 0.015 | 1 | 0.430487676 | 0.361144491 | 0 | 0.014 | 1 | 0.578159907 | 0.675655713 |
| Ltb | 0.578159907 | -0.233118828 | 0 | 0.015 | 1 | 0.430487676 | 0.361144491 | 0 | 0.014 | 1 | 0.578159907 | 0.675655713 |
| P2rx3 | 0.57816106 | 1.351843673 | 0 | 0.015 | 1 | 0.430487676 | 0.361144491 | 0 | 0.014 | 1 | 0.57816106 | 0.675655713 |
| 4631405J19Rik | 0.57816106 | -0.326228232 | 0 | 0.015 | 1 | 0.430487676 | 0.361144491 | 0 | 0.014 | 1 | 0.57816106 | 0.675655713 |
| Frmd5 | 0.57816106 | 1.351843673 | 0 | 0.015 | 1 | 0.430487676 | -1.709244837 | 0 | 0.014 | 1 | 0.57816106 | 0.675655713 |
| Rab42 | 0.57816106 | 1.351843673 | 0 | 0.015 | 1 | 0.430487676 | 0.361144491 | 0 | 0.014 | 1 | 0.57816106 | 0.675655713 |
| Parvaos | 0.57816106 | 1.351843673 | 0 | 0.015 | 1 | 0.430487676 | 0.361144491 | 0 | 0.014 | 1 | 0.57816106 | 0.675655713 |
| 5830454E08Rik | 0.57816106 | 1.351843673 | 0 | 0.015 | 1 | 0.430487676 | 0.361144491 | 0 | 0.014 | 1 | 0.57816106 | 0.675655713 |
| Coro6 | 0.57816106 | 1.351843673 | 0 | 0.015 | 1 | 0.430487676 | 0.361144491 | 0 | 0.014 | 1 | 0.57816106 | 0.675655713 |
| Pgf | 0.57816106 | 1.351843673 | 0 | 0.015 | 1 | 0.430487676 | 0.361144491 | 0 | 0.014 | 1 | 0.57816106 | 0.675655713 |
| BC037039 | 0.57816106 | 1.351843673 | 0 | 0.015 | 1 | 0.430487676 | 0.361144491 | 0 | 0.014 | 1 | 0.57816106 | 0.675655713 |
| Cyp2c55 | 0.57816106 | 1.351843673 | 0 | 0.015 | 1 | 0.430487676 | 0.361144491 | 0 | 0.014 | 1 | 0.57816106 | 0.675655713 |
| Abcc2 | 0.57816106 | 1.351843673 | 0 | 0.015 | 1 | 0.430487676 | 0.361144491 | 0 | 0.014 | 1 | 0.57816106 | 0.675655713 |
| Gm21060 | 0.57816106 | 1.351843673 | 0 | 0.015 | 1 | 0.430487676 | 0.361144491 | 0 | 0.014 | 1 | 0.57816106 | 0.675655713 |
| Gm26594 | 0.63207476 | 1.351843673 | 0 | 0.011 | 1 | 0.430487676 | 0.361144491 | 0 | 0.014 | 1 | 0.63207476 | 0.675655713 |
| Pcdh15 | 0.63207476 | 1.351843673 | 0 | 0.011 | 1 | 0.430487676 | 0.361144491 | 0 | 0.014 | 1 | 0.63207476 | 0.675655713 |
| Gm49291 | 0.63207476 | 1.351843673 | 0 | 0.011 | 1 | 0.430487676 | 0.361144491 | 0 | 0.014 | 1 | 0.63207476 | 0.675655713 |
| Ogdhl | 0.63207476 | 1.351843673 | 0 | 0.011 | 1 | 0.430487676 | 0.361144491 | 0 | 0.014 | 1 | 0.63207476 | 0.675655713 |
| Dynlt2a3 | 0.63207476 | 1.351843673 | 0 | 0.011 | 1 | 0.430487676 | 0.361144491 | 0 | 0.014 | 1 | 0.63207476 | 0.675655713 |
| Gm21860 | 0.63207476 | 1.351843673 | 0 | 0.011 | 1 | 0.430487676 | 0.361144491 | 0 | 0.014 | 1 | 0.63207476 | 0.675655713 |
| Gm14302 | 0.632076134 | 1.673771768 | 0 | 0.011 | 1 | 0.430487676 | 0.361144491 | 0 | 0.014 | 1 | 0.632076134 | 0.675655713 |
| Otud7a | 0.632076134 | 1.673771768 | 0 | 0.011 | 1 | 0.430487676 | 0.361144491 | 0 | 0.014 | 1 | 0.632076134 | 0.675655713 |
| Gm10629 | 0.632076134 | 1.673771768 | 0 | 0.011 | 1 | 0.430487676 | 0.361144491 | 0 | 0.014 | 1 | 0.632076134 | 0.675655713 |
| Gm57003 | 0.632076134 | 1.673771768 | 0 | 0.011 | 1 | 0.430487676 | 0.361144491 | 0 | 0.014 | 1 | 0.632076134 | 0.675655713 |
| Nacad | 0.632076134 | 1.673771768 | 0 | 0.011 | 1 | 0.430487676 | 0.361144491 | 0 | 0.014 | 1 | 0.632076134 | 0.675655713 |
| Ccdc69 | 0.632076134 | 0.673771768 | 0 | 0.011 | 1 | 0.430487676 | 0.361144491 | 0 | 0.014 | 1 | 0.632076134 | 0.675655713 |

| | | | | | | | | | | | | |
|---|---|---|---|---|---|---|---|---|---|---|---|---|
| Hic1 | 0.632076134 | 1.673771768 | 0 | 0.011 | 1 | 0.430487676 | 0.361144491 | 0 | 0.014 | 1 | 0.632076134 | 0.675655713 |
| 1700003D09Rik | 0.632076134 | 1.673771768 | 0 | 0.011 | 1 | 0.430487676 | 0.361144491 | 0 | 0.014 | 1 | 0.632076134 | 0.675655713 |
| Gm26531 | 0.632076134 | 1.673771768 | 0 | 0.011 | 1 | 0.430487676 | 0.361144491 | 0 | 0.014 | 1 | 0.632076134 | 0.675655713 |
| Gm47882 | 0.632076134 | 1.673771768 | 0 | 0.011 | 1 | 0.430487676 | 0.361144491 | 0 | 0.014 | 1 | 0.632076134 | 0.675655713 |
| Gm5083 | 0.632076134 | 1.673771768 | 0 | 0.011 | 1 | 0.430487676 | 0.361144491 | 0 | 0.014 | 1 | 0.632076134 | 0.675655713 |
| Gm48342 | 0.632076134 | 1.673771768 | 0 | 0.011 | 1 | 0.430487676 | 0.361144491 | 0 | 0.014 | 1 | 0.632076134 | 0.675655713 |
| Sult4a1 | 0.632076134 | 1.673771768 | 0 | 0.011 | 1 | 0.430487676 | 0.361144491 | 0 | 0.014 | 1 | 0.632076134 | 0.675655713 |
| Plin4 | 0.632076134 | 1.673771768 | 0 | 0.011 | 1 | 0.430487676 | 0.361144491 | 0 | 0.014 | 1 | 0.632076134 | 0.675655713 |
| Gm17949 | 0.632076134 | 1.673771768 | 0 | 0.011 | 1 | 0.430487676 | 0.361144491 | 0 | 0.014 | 1 | 0.632076134 | 0.675655713 |
| Prob1 | 0.632076134 | 1.673771768 | 0 | 0.011 | 1 | 0.430487676 | 0.361144491 | 0 | 0.014 | 1 | 0.632076134 | 0.675655713 |
| Gm46210 | 0.655561847 | 1.503846766 | 0.048 | 0.03 | 1 | 0.430487676 | 0.361144491 | 0 | 0.014 | 1 | 0.655561847 | 0.675655713 |
| Lrrc15 | 0.657226866 | -1.752492987 | 0.048 | 0.071 | 1 | 0.430487676 | 0.361144491 | 0 | 0.014 | 1 | 0.657226866 | 0.675655713 |
| Gm15832 | 0.709465802 | 2.351843673 | 0.048 | 0.034 | 1 | 0.430487676 | 0.361144491 | 0 | 0.014 | 1 | 0.709465802 | 0.675655713 |
| AV039307 | 0.709465802 | 1.936806174 | 0.048 | 0.034 | 1 | 0.430487676 | 0.361144491 | 0 | 0.014 | 1 | 0.709465802 | 0.675655713 |
| Ica1l | 0.709493655 | 2.295260144 | 0.048 | 0.034 | 1 | 0.430487676 | 0.361144491 | 0 | 0.014 | 1 | 0.709493655 | 0.675655713 |
| Eldr | 0.71268473 | 1.866416846 | 0.048 | 0.034 | 1 | 0.430487676 | 0.361144491 | 0 | 0.014 | 1 | 0.71268473 | 0.675655713 |
| Iqgap3 | 0.790364455 | 1.780686972 | 0.048 | 0.037 | 1 | 0.430487676 | 0.361144491 | 0 | 0.014 | 1 | 0.790364455 | 0.675655713 |
| 1110006O24Rik | 0.823729147 | 0.586308926 | 0.048 | 0.06 | 1 | 0.430487676 | 0.361144491 | 0 | 0.014 | 1 | 0.823729147 | 0.675655713 |
| H2ac8 | 0.973832371 | -1.392317423 | 0.048 | 0.049 | 1 | 0.430487676 | 0.361144491 | 0 | 0.014 | 1 | 0.973832371 | 0.675655713 |
| Gm13594 | 0.97964157 | 1.451379346 | 0.048 | 0.049 | 1 | 0.430487676 | 0.361144491 | 0 | 0.014 | 1 | 0.97964157 | 0.675655713 |
| Dcp1a | 0.431055463 | 0.823667198 | 0.381 | 0.362 | 1 | 0.482960489 | 0.368030877 | 0.289 | 0.419 | 1 | 0.482960489 | 0.676302114 |
| Tmtc1 | 0.54673676 | -0.140436825 | 0.19 | 0.265 | 1 | 0.431151287 | 0.271258988 | 0.133 | 0.194 | 1 | 0.54673676 | 0.676411142 |
| Gm15890 | 0.460380341 | 2.088809267 | 0.048 | 0.022 | 1 | 0.431300474 | 1.361144491 | 0.044 | 0.024 | 1 | 0.460380341 | 0.676580849 |
| Wdr37 | 0.573998405 | 1.053619838 | 0.286 | 0.44 | 1 | 0.432041016 | 0.622531044 | 0.489 | 0.547 | 1 | 0.573998405 | 0.677422592 |
| Tbc1d12 | 0.432198624 | -0.105838164 | 0.238 | 0.362 | 1 | 0.696058556 | 0.389081368 | 0.4 | 0.522 | 1 | 0.696058556 | 0.677601598 |
| Rnf24 | 0.746747611 | 0.429846185 | 0.286 | 0.384 | 1 | 0.432614356 | 0.693268353 | 0.556 | 0.54 | 1 | 0.746747611 | 0.678073531 |
| Ppip5k1 | 0.615156256 | 1.300313372 | 0.095 | 0.149 | 1 | 0.432826676 | 0.683072586 | 0.111 | 0.176 | 1 | 0.615156256 | 0.67831442 |
| Gm42829 | 0.492608234 | 0.866416846 | 0 | 0.022 | 1 | 0.433034806 | 1.180572246 | 0.067 | 0.042 | 1 | 0.492608234 | 0.678550469 |
| Ppm1e | 0.560558468 | 1.010806755 | 0.048 | 0.026 | 1 | 0.433055612 | 1.612683258 | 0.044 | 0.024 | 1 | 0.560558468 | 0.678574061 |
| Mettl25 | 0.433158539 | -0.512094778 | 0.143 | 0.216 | 1 | 0.552315691 | 1.603638119 | 0.222 | 0.218 | 1 | 0.552315691 | 0.678690758 |
| Sned1 | 0.434245694 | 0.55829455 | 0.524 | 0.59 | 1 | 0.900707644 | 0.518591723 | 0.578 | 0.737 | 1 | 0.900707644 | 0.679922065 |
| Eml5 | 0.434446494 | -1.233118828 | 0.048 | 0.097 | 1 | 0.675789584 | 0.891659208 | 0.133 | 0.176 | 1 | 0.675789584 | 0.680149231 |
| Hkdc1 | 0.63207476 | 1.351843673 | 0 | 0.011 | 1 | 0.434781808 | 1.304560963 | 0.044 | 0.024 | 1 | 0.63207476 | 0.680528396 |
| Me2 | 0.43485406 | -0.295854583 | 0.143 | 0.224 | 1 | 0.59336337 | 0.600610426 | 0.133 | 0.173 | 1 | 0.59336337 | 0.680610066 |
| Sestd1 | 0.507610573 | 0.683294542 | 0.619 | 0.698 | 1 | 0.434992184 | 0.44378164 | 0.667 | 0.63 | 1 | 0.507610573 | 0.680766168 |
| Pja2 | 0.434995809 | 0.764537232 | 0.714 | 0.828 | 1 | 0.635332341 | 0.252722031 | 0.844 | 0.875 | 1 | 0.635332341 | 0.680770265 |
| Cracdl | 0.790558847 | 1.106087258 | 0.143 | 0.183 | 1 | 0.435083952 | 1.093403083 | 0.378 | 0.398 | 1 | 0.790558847 | 0.680869858 |
| Evl | 0.476217045 | 0.561878888 | 0.381 | 0.354 | 1 | 0.435871309 | -0.101704488 | 0.422 | 0.478 | 1 | 0.476217045 | 0.68175882 |
| Ptp4a1 | 0.436396472 | 2.171271427 | 0.095 | 0.056 | 1 | 0.908650103 | 1.197645759 | 0.044 | 0.042 | 1 | 0.908650103 | 0.682351063 |
| A730081D07Rik | 0.808167817 | 1.041503552 | 0.048 | 0.063 | 1 | 0.43647399 | 0.402964667 | 0.067 | 0.107 | 1 | 0.808167817 | 0.682438436 |
| D630045J12Rik | 0.437365373 | 1.178076605 | 0.429 | 0.53 | 1 | 0.906026784 | 0.292757516 | 0.533 | 0.619 | 1 | 0.906026784 | 0.683442276 |
| Fbxo30 | 0.45029692 | 0.252814297 | 0.476 | 0.66 | 1 | 0.438157328 | 0.204418988 | 0.8 | 0.81 | 1 | 0.45029692 | 0.684332812 |
| 4930537H20Rik | 0.492599419 | 0.503846766 | 0 | 0.022 | 1 | 0.438286667 | 1.268035087 | 0.044 | 0.024 | 1 | 0.492599419 | 0.684478132 |
| AB124611 | 0.499982599 | 0.908237021 | 0.048 | 0.097 | 1 | 0.438979434 | 0.953720176 | 0.156 | 0.121 | 1 | 0.499982599 | 0.685255924 |
| Pip4k2a | 0.884179452 | 0.43673257 | 0.19 | 0.224 | 1 | 0.439097146 | 0.342035668 | 0.222 | 0.287 | 1 | 0.884179452 | 0.685387988 |
| Acacb | 0.439142082 | -0.940938076 | 0.048 | 0.097 | 1 | 0.545676244 | -0.174908409 | 0.044 | 0.069 | 1 | 0.545676244 | 0.685438396 |
| Mpv17l | 0.849221195 | 1.896164189 | 0.19 | 0.246 | 1 | 0.439655097 | 0.865936643 | 0.133 | 0.197 | 1 | 0.849221195 | 0.68601359 |
| Rnls | 0.439729607 | 0.150209812 | 0.048 | 0.104 | 1 | 0.781449948 | 0.223640968 | 0.067 | 0.083 | 1 | 0.781449948 | 0.686097086 |
| Aff2 | 0.439820429 | 0.181918671 | 0.048 | 0.104 | 1 | 0.778334035 | 1.021352886 | 0.156 | 0.19 | 1 | 0.778334035 | 0.686198848 |
| Phyhipl | 0.440125527 | 0.586308926 | 0.19 | 0.131 | 1 | 0.658587075 | 2.098110085 | 0.022 | 0.014 | 1 | 0.658587075 | 0.686540575 |
| Mapk9 | 0.440269822 | 1.632744499 | 0.381 | 0.343 | 1 | 0.980738015 | 1.376411248 | 0.289 | 0.325 | 1 | 0.980738015 | 0.686702128 |
| Cog3 | 0.606078456 | 0.470488169 | 0.333 | 0.448 | 1 | 0.440775926 | 0.123201066 | 0.444 | 0.592 | 1 | 0.606078456 | 0.687268435 |
| Ndor1 | 0.440999049 | 0.171271427 | 0.048 | 0.104 | 1 | 0.552334987 | 0.277080226 | 0.111 | 0.149 | 1 | 0.552334987 | 0.687517937 |
| Riok3 | 0.441159547 | -0.238452272 | 0.857 | 0.866 | 1 | 0.572274777 | 0.206923573 | 1 | 0.955 | 1 | 0.572274777 | 0.687697348 |
| Gm28403 | 0.58558921 | 1.928344595 | 0.143 | 0.108 | 1 | 0.442004264 | 0.476621709 | 0.067 | 0.107 | 1 | 0.58558921 | 0.688640758 |
| Drosha | 1 | 0.59848364 | 0.286 | 0.31 | 1 | 0.442151149 | 0.297418894 | 0.244 | 0.356 | 1 | 1 | 0.688804659 |
| Gm26633 | 0.442599701 | 1.258734268 | 0.143 | 0.097 | 1 | 0.558010091 | 1.383512304 | 0.089 | 0.069 | 1 | 0.558010091 | 0.689304907 |
| Rapgef4 | 0.4427004 | 1.705022702 | 0.238 | 0.194 | 1 | 0.478965896 | 1.129474256 | 0.178 | 0.152 | 1 | 0.478965896 | 0.689417156 |
| Dync1li2 | 0.442919299 | 0.195724471 | 0.333 | 0.522 | 1 | 0.707299423 | 0.581986461 | 0.667 | 0.664 | 1 | 0.707299423 | 0.689661093 |
| Xpnpep3 | 0.442989081 | 0.374211486 | 0.238 | 0.332 | 1 | 0.704359987 | 1.036709541 | 0.333 | 0.381 | 1 | 0.704359987 | 0.689738836 |
| Tshz3 | 0.444079808 | -0.103835811 | 0.095 | 0.16 | 1 | 0.720735837 | 0.458578642 | 0.267 | 0.322 | 1 | 0.720735837 | 0.69095274 |
| Cbwd1 | 0.444686076 | 0.859990576 | 0.429 | 0.466 | 1 | 0.595542154 | 0.531069493 | 0.2 | 0.277 | 1 | 0.595542154 | 0.691626445 |
| Tox3 | 0.445084207 | -0.623908781 | 0.048 | 0.101 | 1 | 0.587315134 | 1.084435149 | 0.133 | 0.17 | 1 | 0.587315134 | 0.692068463 |
| Zc3hav1 | 0.48697118 | 0.348641241 | 0.524 | 0.687 | 1 | 0.445373334 | 0.468808046 | 0.756 | 0.623 | 1 | 0.48697118 | 0.692389261 |
| Gm34343 | 0.445384922 | -0.455511249 | 0.048 | 0.097 | 1 | 0.601043236 | 0.933050839 | 0.067 | 0.097 | 1 | 0.601043236 | 0.692402115 |
| Tubb3 | 0.445392698 | -0.413691074 | 0.048 | 0.097 | 1 | 0.723609176 | 0.435145073 | 0.044 | 0.059 | 1 | 0.723609176 | 0.692410741 |
| Ctu1 | 0.87495898 | 0.673771768 | 0.048 | 0.056 | 1 | 0.446200608 | -0.255526869 | 0.044 | 0.076 | 1 | 0.87495898 | 0.693306233 |
| Hook3 | 0.696153516 | 0.660376174 | 0.857 | 0.933 | 1 | 0.446296834 | 0.338362016 | 0.933 | 0.969 | 1 | 0.696153516 | 0.693412804 |
| Atp7a | 0.446378653 | 0.759501642 | 0.476 | 0.511 | 1 | 0.601140158 | 0.13763845 | 0.578 | 0.671 | 1 | 0.601140158 | 0.693503404 |
| Cnnm4 | 0.558636825 | 0.762080186 | 0.524 | 0.582 | 1 | 0.446493615 | 0.230325281 | 0.378 | 0.509 | 1 | 0.558636825 | 0.693630682 |
| Mras | 0.446708495 | -0.150441805 | 0.762 | 0.858 | 1 | 0.532024518 | 0.151109276 | 0.756 | 0.817 | 1 | 0.532024518 | 0.693868511 |
| Cep295 | 0.446765009 | -0.955584852 | 0.143 | 0.201 | 1 | 0.849259297 | 0.280974143 | 0.2 | 0.225 | 1 | 0.849259297 | 0.693931044 |
| Fbxo46 | 0.446996615 | 0.351843673 | 0.048 | 0.104 | 1 | 0.451173727 | 0.435145073 | 0.067 | 0.107 | 1 | 0.451173727 | 0.694187256 |
| Naa20 | 0.447131091 | -0.20069735 | 0.476 | 0.623 | 1 | 0.555693326 | 0.21492375 | 0.622 | 0.716 | 1 | 0.555693326 | 0.694335969 |
| Enox2 | 0.490961154 | 0.351843673 | 0.143 | 0.209 | 1 | 0.447159019 | 1.310518418 | 0.267 | 0.235 | 1 | 0.490961154 | 0.694366849 |
| Traf3ip2 | 0.447980827 | 0.995699863 | 0.143 | 0.093 | 1 | 0.489253147 | 1.28714391 | 0.156 | 0.131 | 1 | 0.489253147 | 0.695274833 |
| Dhdh | 0.787201474 | 1.866416846 | 0.048 | 0.037 | 1 | 0.448078778 | 0.361144491 | 0.022 | 0.048 | 1 | 0.787201474 | 0.695382964 |
| Espn | 0.492608234 | 0.866416846 | 0 | 0.022 | 1 | 0.448123066 | 0.15951063 | 0.022 | 0.048 | 1 | 0.492608234 | 0.69543185 |
| Zfp931 | 0.932575038 | 0.586308926 | 0.095 | 0.104 | 1 | 0.448278105 | 0.132875504 | 0.067 | 0.107 | 1 | 0.932575038 | 0.695602951 |
| Zfp467 | 0.44850537 | -0.413691074 | 0.048 | 0.097 | 1 | 0.567721894 | -0.191396532 | 0.111 | 0.142 | 1 | 0.567721894 | 0.695853673 |
| Rnf182 | 0.448912265 | 0.710297644 | 0.048 | 0.104 | 1 | 0.869939979 | 0.605070074 | 0.067 | 0.076 | 1 | 0.869939979 | 0.696302308 |
| Gm52988 | 0.448986599 | 1.775651382 | 0.048 | 0.104 | 1 | 0.567731669 | 1.541053581 | 0.156 | 0.138 | 1 | 0.567731669 | 0.696384232 |
| Manba | 0.877837412 | 0.503846766 | 0.143 | 0.172 | 1 | 0.44937362 | 0.979790118 | 0.156 | 0.228 | 1 | 0.877837412 | 0.69681059 |
| 2310058D17Rik | 0.991275314 | 1.010806755 | 0.048 | 0.049 | 1 | 0.449441826 | 0.268035087 | 0.022 | 0.048 | 1 | 0.991275314 | 0.696885697 |
| Cbarp | 0.532426384 | 0.866416846 | 0 | 0.019 | 1 | 0.449463946 | 0.513147585 | 0.022 | 0.048 | 1 | 0.532426384 | 0.696910053 |
| Npr3 | 0.850661302 | 1.719575457 | 0.238 | 0.228 | 1 | 0.449482377 | 0.435145073 | 0.022 | 0.048 | 1 | 0.850661302 | 0.696930346 |
| Gm33046 | 0.71268473 | 1.866416846 | 0.048 | 0.034 | 1 | 0.449510445 | 0.42003818 | 0.022 | 0.048 | 1 | 0.71268473 | 0.69696125 |
| Disp1 | 0.44958778 | 0.474991904 | 0.238 | 0.396 | 1 | 0.800839807 | 1.009300818 | 0.2 | 0.253 | 1 | 0.800839807 | 0.697046388 |
| Ldb2 | 0.449988126 | -0.676725479 | 0.19 | 0.265 | 1 | 0.534322725 | -1.094534992 | 0.133 | 0.173 | 1 | 0.534322725 | 0.697486939 |
| Gm7072 | 0.450482861 | 0.102229783 | 0.19 | 0.299 | 1 | 0.834871817 | 0.953720176 | 0.133 | 0.156 | 1 | 0.834871817 | 0.698030914 |
| Npc1 | 0.472796914 | 0.676210813 | 0.524 | 0.724 | 1 | 0.450492701 | 0.475392663 | 0.756 | 0.792 | 1 | 0.472796914 | 0.698041728 |
| Plcb3 | 0.662224133 | 0.569435108 | 0.095 | 0.142 | 1 | 0.450514927 | 0.361144491 | 0.133 | 0.194 | 1 | 0.662224133 | 0.698066155 |
| Gm57387 | 0.748090146 | 0.97333205 | 0.048 | 0.034 | 1 | 0.450829449 | 0.683072586 | 0.022 | 0.048 | 1 | 0.748090146 | 0.698411706 |
| Crocc | 0.853709349 | 2.55829455 | 0.048 | 0.041 | 1 | 0.450829449 | 0.683072586 | 0.022 | 0.048 | 1 | 0.853709349 | 0.698411706 |
| 2810405F17Rik | 0.87495898 | 0.673771768 | 0.048 | 0.056 | 1 | 0.450829449 | 0.683072586 | 0.022 | 0.048 | 1 | 0.87495898 | 0.698411706 |
| Gm46447 | 0.632076134 | 1.673771768 | 0 | 0.011 | 1 | 0.450869096 | 0.513147585 | 0.022 | 0.048 | 1 | 0.632076134 | 0.69845525 |
| Clhc1 | 0.719087874 | 1.536268244 | 0.048 | 0.034 | 1 | 0.450885519 | 1.15951063 | 0.022 | 0.048 | 1 | 0.719087874 | 0.698473287 |
| Rsad1 | 0.710218793 | 1.021695071 | 0.143 | 0.123 | 1 | 0.450898049 | -0.204452685 | 0.067 | 0.104 | 1 | 0.710218793 | 0.698487047 |

| | | | | | | | | | | | | |
|---|---|---|---|---|---|---|---|---|---|---|---|---|
| Rab11fip4 | 0.450947838 | 0.606264948 | 0.857 | 0.873 | 1 | 0.726352955 | 0.18821074 | 0.822 | 0.917 | 1 | 0.726352955 | 0.698541723 |
| Fam193b | 0.451025174 | 0.250560337 | 0.19 | 0.284 | 1 | 0.78997051 | 0.322541652 | 0.422 | 0.54 | 1 | 0.78997051 | 0.69862664 |
| Mov10 | 0.545952774 | 0.258734268 | 0.048 | 0.09 | 1 | 0.451411102 | 1.268035087 | 0.178 | 0.152 | 1 | 0.545952774 | 0.69905022 |
| Iqsec2 | 0.451542738 | -0.40151636 | 0.143 | 0.22 | 1 | 0.88069205 | 1.411770564 | 0.289 | 0.322 | 1 | 0.88069205 | 0.699194632 |
| Dmpk | 0.451614009 | -0.370622352 | 0.048 | 0.097 | 1 | 0.524823439 | 0.776181991 | 0.133 | 0.19 | 1 | 0.524823439 | 0.699272805 |
| Lrrc49 | 0.452621023 | -0.122087516 | 0.381 | 0.571 | 1 | 0.679909683 | 0.622951594 | 0.422 | 0.574 | 1 | 0.679909683 | 0.700376256 |
| Rbfox1 | 0.452648798 | -2.107587946 | 0.095 | 0.146 | 1 | 0.866938871 | 1.368199249 | 0.111 | 0.111 | 1 | 0.866938871 | 0.700406662 |
| Ube4b | 0.589367063 | 0.415037499 | 0.667 | 0.806 | 1 | 0.452737456 | 0.404771424 | 0.933 | 0.855 | 1 | 0.589367063 | 0.700503708 |
| Gm48027 | 0.736180094 | 1.088809267 | 0.143 | 0.127 | 1 | 0.452808049 | 0.966865552 | 0.2 | 0.173 | 1 | 0.736180094 | 0.700580969 |
| Itga10 | 0.826356658 | 0.127028589 | 0.667 | 0.791 | 1 | 0.453356016 | 0.337541592 | 0.689 | 0.872 | 1 | 0.826356658 | 0.701180355 |
| Ttc12 | 0.790341526 | 1.55829455 | 0.048 | 0.037 | 1 | 0.453384396 | 1.045642666 | 0.044 | 0.08 | 1 | 0.790341526 | 0.701211382 |
| Cd74 | 0.45417235 | -3.033587364 | 0.095 | 0.157 | 1 | 0.627509477 | 0.446033389 | 0.044 | 0.066 | 1 | 0.627509477 | 0.702072177 |
| 2610203C22Rik | 0.454205211 | 2.866416846 | 0.048 | 0.022 | 1 | 0.49622258 | 0.361144491 | 0 | 0.01 | 1 | 0.49622258 | 0.702108048 |
| Platr22 | 0.454205211 | 3.188344941 | 0.048 | 0.022 | 1 | 0.852141123 | 0.946106992 | 0.022 | 0.028 | 1 | 0.852141123 | 0.702108048 |
| A230083N12Rik | 0.454205211 | 2.451379346 | 0.048 | 0.022 | 1 | 0.496223879 | 0.683072586 | 0 | 0.01 | 1 | 0.496223879 | 0.702108048 |
| Abcc6 | 0.454205211 | 2.451379346 | 0.048 | 0.022 | 1 | 0.658583583 | 2.683072586 | 0.022 | 0.014 | 1 | 0.658583583 | 0.702108048 |
| Hspb2 | 0.454205211 | 2.451379346 | 0.048 | 0.022 | 1 | 0.930092697 | 0.223640968 | 0.022 | 0.024 | 1 | 0.930092697 | 0.702108048 |
| Pam16l | 0.454205211 | 2.451379346 | 0.048 | 0.022 | 1 | 0.765720982 | 0.946106992 | 0.022 | 0.031 | 1 | 0.765720982 | 0.702108048 |
| Camk2a | 0.454205211 | 2.451379346 | 0.048 | 0.022 | 1 | 0.944572107 | 0.875717664 | 0.044 | 0.048 | 1 | 0.944572107 | 0.702108048 |
| Cabp1 | 0.454217076 | 3.536268244 | 0.048 | 0.022 | 1 | 0.661444614 | 1.683072586 | 0.022 | 0.014 | 1 | 0.661444614 | 0.702121 |
| Gm17231 | 0.454217076 | 2.995699863 | 0.048 | 0.022 | 1 | 0.516141367 | 2.460680165 | 0.044 | 0.028 | 1 | 0.516141367 | 0.702121 |
| Itgam | 0.454231312 | 3.673771768 | 0.048 | 0.022 | 1 | 0.500308819 | 2.282534657 | 0.022 | 0.045 | 1 | 0.500308819 | 0.70213654 |
| Tmem240 | 0.775775929 | 0.503846766 | 0.048 | 0.063 | 1 | 0.454440842 | 0.846571318 | 0.044 | 0.08 | 1 | 0.775775929 | 0.702365205 |
| Usp46 | 0.753618857 | 0.866416846 | 0.095 | 0.123 | 1 | 0.45514157 | -0.852980314 | 0.089 | 0.125 | 1 | 0.753618857 | 0.703129291 |
| Llph | 0.455456487 | -0.108836492 | 0.476 | 0.619 | 1 | 0.934555145 | -0.207698344 | 0.622 | 0.522 | 1 | 0.934555145 | 0.703472363 |
| D6Wsu163e | 0.790211603 | 0.735172312 | 0.095 | 0.078 | 1 | 0.45562865 | -0.16492432 | 0.067 | 0.104 | 1 | 0.790211603 | 0.703659833 |
| Hif1an | 0.45610684 | 0.780686972 | 0.095 | 0.164 | 1 | 0.937205207 | 0.744473131 | 0.178 | 0.194 | 1 | 0.937205207 | 0.704180231 |
| Erlin1 | 0.456263184 | 1.150209812 | 0.238 | 0.198 | 1 | 0.660169028 | 0.485133209 | 0.244 | 0.322 | 1 | 0.660169028 | 0.704350275 |
| Tbccd1 | 0.945499728 | 0.451379346 | 0.095 | 0.09 | 1 | 0.456314463 | 0.64125241 | 0.111 | 0.173 | 1 | 0.945499728 | 0.704406037 |
| Btbd10 | 0.942382672 | 0.54665985 | 0.286 | 0.351 | 1 | 0.456410649 | 0.152003093 | 0.622 | 0.543 | 1 | 0.942382672 | 0.704510617 |
| Cnnm2 | 0.754971741 | 0.241925981 | 0.333 | 0.433 | 1 | 0.456872201 | 1.033979748 | 0.244 | 0.332 | 1 | 0.754971741 | 0.705012194 |
| Cfap43 | 0.803443033 | 0.79930265 | 0.238 | 0.22 | 1 | 0.456937414 | 0.970354538 | 0.156 | 0.121 | 1 | 0.803443033 | 0.705083027 |
| 4930556G22Rik | 0.457281049 | 0.503846766 | 0 | 0.026 | 1 | 0.492040936 | 2.268035087 | 0.022 | 0.01 | 1 | 0.492040936 | 0.70545614 |
| BC055324 | 0.457281049 | 0.503846766 | 0 | 0.026 | 1 | 0.492040936 | 2.268035087 | 0.022 | 0.01 | 1 | 0.492040936 | 0.70545614 |
| Gm13270 | 0.457281049 | 0.351843673 | 0 | 0.026 | 1 | 0.496223879 | 0.683072586 | 0 | 0.01 | 1 | 0.496223879 | 0.70545614 |
| Hspa12b | 0.457281049 | 0.503846766 | 0 | 0.026 | 1 | 0.610057126 | 1.905465008 | 0.044 | 0.031 | 1 | 0.610057126 | 0.70545614 |
| Bcl2l15 | 0.457281049 | 0.351843673 | 0 | 0.026 | 1 | 0.955023924 | 1.683072586 | 0.022 | 0.024 | 1 | 0.955023924 | 0.70545614 |
| Htr6 | 0.457281049 | 0.214340149 | 0 | 0.026 | 1 | 0.936023974 | 2.197645759 | 0.022 | 0.021 | 1 | 0.936023974 | 0.70545614 |
| Xrcc2 | 0.457281049 | 0.503846766 | 0 | 0.026 | 1 | 0.952529269 | 1.098110085 | 0.022 | 0.024 | 1 | 0.952529269 | 0.70545614 |
| Iqcd | 0.457281049 | 0.503846766 | 0 | 0.026 | 1 | 0.762072975 | 2.066401226 | 0.067 | 0.059 | 1 | 0.762072975 | 0.70545614 |
| Gm15929 | 0.457281049 | 0.503846766 | 0 | 0.026 | 1 | 0.661444614 | 1.683072586 | 0.022 | 0.014 | 1 | 0.661444614 | 0.70545614 |
| A030001D20Rik | 0.457281049 | 0.503846766 | 0 | 0.026 | 1 | 0.8202424 | 1.098110085 | 0.022 | 0.017 | 1 | 0.8202424 | 0.70545614 |
| Gm32389 | 0.457281049 | 0.214340149 | 0 | 0.026 | 1 | 0.814636613 | 1.098110085 | 0.022 | 0.017 | 1 | 0.814636613 | 0.70545614 |
| Gm57050 | 0.457281049 | 0.503846766 | 0 | 0.026 | 1 | 0.744358033 | 0.223640968 | 0.022 | 0.031 | 1 | 0.744358033 | 0.70545614 |
| Prima1 | 0.457281049 | 0.503846766 | 0 | 0.026 | 1 | 0.492043882 | 2.683072586 | 0.022 | 0.01 | 1 | 0.492043882 | 0.70545614 |
| Begain | 0.457281049 | 0.503846766 | 0 | 0.026 | 1 | 0.80624346 | 1.683072586 | 0.022 | 0.017 | 1 | 0.80624346 | 0.70545614 |
| Gm47460 | 0.457281049 | 0.503846766 | 0 | 0.026 | 1 | 0.705167793 | 1.78998779 | 0.044 | 0.035 | 1 | 0.705167793 | 0.70545614 |
| Hk3 | 0.457281049 | 0.503846766 | 0 | 0.026 | 1 | 0.80624346 | 2.098110085 | 0.022 | 0.017 | 1 | 0.80624346 | 0.70545614 |
| BC037032 | 0.457281049 | 0.503846766 | 0 | 0.026 | 1 | 0.496223879 | 0.683072586 | 0 | 0.01 | 1 | 0.496223879 | 0.70545614 |
| Nme3 | 0.457281049 | 0.503846766 | 0 | 0.026 | 1 | 0.835943453 | 0.513147585 | 0.022 | 0.028 | 1 | 0.835943453 | 0.70545614 |
| Tesmin | 0.457281049 | 0.503846766 | 0 | 0.026 | 1 | 0.476265768 | -0.404390255 | 0.022 | 0.045 | 1 | 0.476265768 | 0.70545614 |
| Gm14964 | 0.457281049 | 0.503846766 | 0 | 0.026 | 1 | 0.509367939 | -0.459885368 | 0.044 | 0.073 | 1 | 0.509367939 | 0.70545614 |
| Gm14798 | 0.457281049 | 0.351843673 | 0 | 0.026 | 1 | 0.57694994 | 1.242499995 | 0.067 | 0.048 | 1 | 0.57694994 | 0.70545614 |
| Tsx | 0.457281049 | 0.503846766 | 0 | 0.026 | 1 | 0.672920453 | 1.098110085 | 0.022 | 0.014 | 1 | 0.672920453 | 0.70545614 |
| Syt6 | 0.457292877 | 2.258734268 | 0.048 | 0.022 | 1 | 0.496223879 | -0.776359032 | 0 | 0.01 | 1 | 0.496223879 | 0.705468979 |
| 2200002D01Rik | 0.457292877 | 2.258734268 | 0.048 | 0.022 | 1 | 0.872783394 | 0.683072586 | 0.067 | 0.076 | 1 | 0.872783394 | 0.705468979 |
| Gm39363 | 0.457292877 | 2.258734268 | 0.048 | 0.022 | 1 | 0.808235119 | 1.180572246 | 0.044 | 0.038 | 1 | 0.808235119 | 0.705468979 |
| Gng13 | 0.457292877 | 2.258734268 | 0.048 | 0.022 | 1 | 0.932582169 | 0.361144491 | 0.022 | 0.024 | 1 | 0.932582169 | 0.705468979 |
| Gm26620 | 0.457292877 | 0.351843673 | 0 | 0.026 | 1 | 0.496223879 | 0.683072586 | 0 | 0.01 | 1 | 0.496223879 | 0.705468979 |
| Gm4673 | 0.457292877 | 0.214340149 | 0 | 0.026 | 1 | 0.796730175 | 1.78998779 | 0.044 | 0.038 | 1 | 0.796730175 | 0.705468979 |
| Grtp1 | 0.457292877 | 0.351843673 | 0 | 0.026 | 1 | 0.496223879 | 0.683072586 | 0 | 0.01 | 1 | 0.496223879 | 0.705468979 |
| Gm45894 | 0.457292877 | 0.351843673 | 0 | 0.026 | 1 | 0.966790692 | 1.460680165 | 0.044 | 0.045 | 1 | 0.966790692 | 0.705468979 |
| Slc26a6 | 0.457292877 | 0.351843673 | 0 | 0.026 | 1 | 0.552232538 | 0.361144491 | 0.022 | 0.042 | 1 | 0.552232538 | 0.705468979 |
| Klhl40 | 0.457292877 | -0.233118828 | 0 | 0.026 | 1 | 0.500288783 | 0.513147585 | 0.022 | 0.045 | 1 | 0.500288783 | 0.705468979 |
| Stab2 | 0.457292877 | 0.351843673 | 0 | 0.026 | 1 | 0.955023924 | 1.683072586 | 0.022 | 0.024 | 1 | 0.955023924 | 0.705468979 |
| Gm11630 | 0.457292877 | 0.351843673 | 0 | 0.026 | 1 | 0.664308814 | 1.460680165 | 0.022 | 0.014 | 1 | 0.664308814 | 0.705468979 |
| Tepsin | 0.457292877 | 0.351843673 | 0 | 0.026 | 1 | 0.497209727 | 0.268035087 | 0.022 | 0.045 | 1 | 0.497209727 | 0.705468979 |
| Gm38604 | 0.457292877 | 0.214340149 | 0 | 0.026 | 1 | 0.828665129 | 0.513147585 | 0.022 | 0.017 | 1 | 0.828665129 | 0.705468979 |
| Sema3g | 0.457292877 | 0.214340149 | 0 | 0.026 | 1 | 0.49622258 | 0.361144491 | 0 | 0.01 | 1 | 0.49622258 | 0.705468979 |
| Wnt7b | 0.457292877 | 0.351843673 | 0 | 0.026 | 1 | 0.49622258 | 0.361144491 | 0 | 0.01 | 1 | 0.49622258 | 0.705468979 |
| Gm33696 | 0.457292877 | 0.214340149 | 0 | 0.026 | 1 | 0.76573422 | 1.098110085 | 0.022 | 0.031 | 1 | 0.76573422 | 0.705468979 |
| Ccdc14 | 0.457292877 | 0.214340149 | 0 | 0.026 | 1 | 0.817437668 | 0.946106992 | 0.022 | 0.017 | 1 | 0.817437668 | 0.705468979 |
| Dnajc28 | 0.457292877 | 0.351843673 | 0 | 0.026 | 1 | 0.796736449 | 2.045642666 | 0.044 | 0.038 | 1 | 0.796736449 | 0.705468979 |
| Cldn14 | 0.457292877 | -0.648156327 | 0 | 0.026 | 1 | 0.664308814 | 1.460680165 | 0.022 | 0.014 | 1 | 0.664308814 | 0.705468979 |
| Gm50334 | 0.457292877 | 0.214340149 | 0 | 0.026 | 1 | 0.637304676 | 0.928185084 | 0.044 | 0.066 | 1 | 0.637304676 | 0.705468979 |
| Tk1 | 0.457299974 | -0.496153234 | 0 | 0.026 | 1 | 0.955023924 | 1.268035087 | 0.022 | 0.024 | 1 | 0.955023924 | 0.705476682 |
| Col13a1 | 0.457300763 | -1.611630451 | 0 | 0.026 | 1 | 0.49622258 | -0.638855509 | 0 | 0.01 | 1 | 0.49622258 | 0.705477538 |
| Slfn8 | 0.457300763 | -0.413691074 | 0 | 0.026 | 1 | 0.494823351 | 1.946106992 | 0.022 | 0.01 | 1 | 0.494823351 | 0.705477538 |
| Gm44577 | 0.457300763 | -0.785659851 | 0 | 0.026 | 1 | 0.914346156 | 1.460680165 | 0.044 | 0.042 | 1 | 0.914346156 | 0.705477538 |
| Trim30a | 0.45730234 | -0.133583154 | 0 | 0.026 | 1 | 0.936026061 | 1.683072586 | 0.022 | 0.021 | 1 | 0.936026061 | 0.70547925 |
| Igsf11 | 0.457304706 | -0.326228232 | 0 | 0.026 | 1 | 0.661444614 | 1.683072586 | 0.022 | 0.014 | 1 | 0.661444614 | 0.705481817 |
| Cdr1os | 0.457304706 | -1.326228232 | 0 | 0.026 | 1 | 0.496223879 | 0.683072586 | 0 | 0.01 | 1 | 0.496223879 | 0.705481817 |
| Ifit1 | 0.457305494 | -3.01272876 | 0 | 0.026 | 1 | 0.854469332 | 1.772339924 | 0.022 | 0.028 | 1 | 0.854469332 | 0.705482673 |
| Knstrn | 0.457306283 | -1.081115734 | 0 | 0.026 | 1 | 0.496223879 | 0.683072586 | 0 | 0.01 | 1 | 0.496223879 | 0.705483529 |
| Gpnmb | 0.457306283 | -0.496153234 | 0 | 0.026 | 1 | 0.944008652 | 0.946106992 | 0.022 | 0.021 | 1 | 0.944008652 | 0.705483529 |
| Cenpf | 0.45730786 | -2.718545655 | 0 | 0.026 | 1 | 0.496223879 | 0.683072586 | 0 | 0.01 | 1 | 0.496223879 | 0.705485241 |
| Bnipl | 0.457308648 | 0.673771768 | 0 | 0.026 | 1 | 0.814636613 | 1.098110085 | 0.022 | 0.017 | 1 | 0.814636613 | 0.705486096 |
| Mup5 | 0.457308648 | 0.673771768 | 0 | 0.026 | 1 | 0.796722331 | 1.683072586 | 0.044 | 0.038 | 1 | 0.796722331 | 0.705486096 |
| Plekhn1 | 0.457308648 | 0.673771768 | 0 | 0.026 | 1 | 0.61382235 | 1.683072586 | 0.044 | 0.031 | 1 | 0.61382235 | 0.705486096 |
| Gm42918 | 0.457308648 | 0.673771768 | 0 | 0.026 | 1 | 0.496223879 | 0.683072586 | 0 | 0.01 | 1 | 0.496223879 | 0.705486096 |
| Irf5 | 0.457308648 | 0.673771768 | 0 | 0.026 | 1 | 0.496223879 | 0.683072586 | 0 | 0.01 | 1 | 0.496223879 | 0.705486096 |
| Gm32856 | 0.457308648 | 0.673771768 | 0 | 0.026 | 1 | 0.496223879 | 0.683072586 | 0 | 0.01 | 1 | 0.496223879 | 0.705486096 |
| Ngfr | 0.457308648 | 0.673771768 | 0 | 0.026 | 1 | 0.994464399 | 0.42003818 | 0.044 | 0.045 | 1 | 0.994464399 | 0.705486096 |
| Anxa11os | 0.457308648 | 0.673771768 | 0 | 0.026 | 1 | 0.659895574 | 0.42003818 | 0.044 | 0.062 | 1 | 0.659895574 | 0.705486096 |
| Gm49189 | 0.457308648 | 0.673771768 | 0 | 0.026 | 1 | 0.856449138 | 0.595609745 | 0.044 | 0.052 | 1 | 0.856449138 | 0.705486096 |
| Gm56765 | 0.457308648 | 0.673771768 | 0 | 0.026 | 1 | 0.477727467 | -0.404390255 | 0.022 | 0.045 | 1 | 0.477727467 | 0.705486096 |
| Fam83h | 0.457308648 | 0.673771768 | 0 | 0.026 | 1 | 0.854457549 | 1.098110085 | 0.022 | 0.028 | 1 | 0.854457549 | 0.705486096 |
| Gm56721 | 0.457308648 | 0.673771768 | 0 | 0.026 | 1 | 0.800333063 | 1.28452321 | 0.067 | 0.083 | 1 | 0.800333063 | 0.705486096 |

| | | | | | | | | | | | | |
|---|---|---|---|---|---|---|---|---|---|---|---|---|
| Gm4544 | 0.457308648 | 0.673771768 | 0 | 0.026 | 1 | 0.496223879 | 0.683072586 | 0 | 0.01 | 1 | 0.496223879 | 0.705486096 |
| Kcnj6 | 0.457308648 | 0.673771768 | 0 | 0.026 | 1 | 0.936025227 | 2.683072586 | 0.022 | 0.021 | 1 | 0.936025227 | 0.705486096 |
| H2-Q7 | 0.457308648 | 0.673771768 | 0 | 0.026 | 1 | 0.833619716 | -0.124282336 | 0.022 | 0.028 | 1 | 0.833619716 | 0.705486096 |
| Gm50214 | 0.457308648 | 0.673771768 | 0 | 0.026 | 1 | 0.849825041 | 0.808603468 | 0.022 | 0.028 | 1 | 0.849825041 | 0.705486096 |
| Gm36608 | 0.457308648 | 0.673771768 | 0 | 0.026 | 1 | 0.496223879 | -0.124282336 | 0 | 0.01 | 1 | 0.496223879 | 0.705486096 |
| Ercc6l | 0.457308648 | 0.673771768 | 0 | 0.026 | 1 | 0.496223879 | 0.683072586 | 0 | 0.01 | 1 | 0.496223879 | 0.705486096 |
| Vav3 | 0.69380915 | 0.730698836 | 0.714 | 0.735 | 1 | 0.457636078 | 0.298180534 | 0.889 | 0.74 | 1 | 0.69380915 | 0.705841376 |
| Uvssa | 0.481977374 | 1.140339568 | 0.333 | 0.284 | 1 | 0.45792458 | 0.583536913 | 0.2 | 0.284 | 1 | 0.481977374 | 0.706154239 |
| Glipr2 | 0.458425959 | 2.010806755 | 0.095 | 0.06 | 1 | 0.889417434 | 0.361144491 | 0.067 | 0.073 | 1 | 0.889417434 | 0.706697559 |
| Zfp62 | 0.459873703 | 0.129451251 | 0.19 | 0.284 | 1 | 0.526986319 | 0.390290837 | 0.311 | 0.439 | 1 | 0.526986319 | 0.708263584 |
| 4933400C23Rik | 0.460380341 | 1.936806174 | 0.048 | 0.022 | 1 | 0.492043882 | 2.460680165 | 0.022 | 0.01 | 1 | 0.492043882 | 0.708810624 |
| Pcdhgb1 | 0.460380341 | 1.673771768 | 0.048 | 0.022 | 1 | 0.527061766 | 1.513147585 | 0.044 | 0.028 | 1 | 0.527061766 | 0.708810624 |
| Cplane2 | 0.460388202 | 2.088809267 | 0.048 | 0.022 | 1 | 0.990772207 | 1.197645759 | 0.044 | 0.045 | 1 | 0.990772207 | 0.708819108 |
| Nlrp12 | 0.460388202 | 2.214340149 | 0.048 | 0.022 | 1 | 0.658583583 | 2.361144491 | 0.022 | 0.014 | 1 | 0.658583583 | 0.708819108 |
| Akap8 | 0.460951874 | 0.365649472 | 0.286 | 0.433 | 1 | 0.812379855 | 0.154018827 | 0.6 | 0.602 | 1 | 0.812379855 | 0.709427118 |
| Rpap2 | 0.802845045 | 0.673771768 | 0.143 | 0.131 | 1 | 0.462336667 | 0.91753784 | 0.156 | 0.125 | 1 | 0.802845045 | 0.710918141 |
| Taf3 | 0.927921485 | 0.882358389 | 0.381 | 0.47 | 1 | 0.463238226 | 0.257255248 | 0.489 | 0.581 | 1 | 0.927921485 | 0.711886797 |
| Bcl6 | 0.753498986 | 0.678123792 | 0.667 | 0.716 | 1 | 0.463730406 | 0.581124379 | 0.933 | 0.851 | 1 | 0.753498986 | 0.712414922 |
| Nap1l4 | 0.464017091 | -0.294519373 | 0.571 | 0.668 | 1 | 0.788124073 | 0.198177041 | 0.844 | 0.782 | 1 | 0.788124073 | 0.712722321 |
| Tmem65 | 0.732821657 | 0.205002284 | 0.667 | 0.869 | 1 | 0.4644182 | 0.422308354 | 0.956 | 0.945 | 1 | 0.732821657 | 0.713152136 |
| Ip6k2 | 0.464923546 | -0.252227651 | 0.238 | 0.332 | 1 | 0.85600592 | 0.180572246 | 0.422 | 0.412 | 1 | 0.85600592 | 0.713693188 |
| Tent4a | 0.573946129 | 0.55829455 | 0.143 | 0.209 | 1 | 0.465488414 | 0.172110667 | 0.267 | 0.37 | 1 | 0.573946129 | 0.714297364 |
| 6330562C20Rik | 0.465520368 | 0.214340149 | 0.048 | 0.101 | 1 | 0.68327161 | 0.567595369 | 0.022 | 0.035 | 1 | 0.68327161 | 0.714331523 |
| Spata6l | 0.466284412 | 0.79930265 | 0.143 | 0.097 | 1 | 0.977371521 | 1.098110085 | 0.067 | 0.069 | 1 | 0.977371521 | 0.715147671 |
| Acer2 | 0.466597919 | 1.673771768 | 0.048 | 0.022 | 1 | 0.6175503 | 1.098110085 | 0.022 | 0.038 | 1 | 0.6175503 | 0.71548222 |
| Gm48877 | 0.466597919 | 1.97333205 | 0.048 | 0.022 | 1 | 0.49622258 | 0.361144491 | 0 | 0.01 | 1 | 0.49622258 | 0.71548222 |
| Gm15336 | 0.466597919 | 1.55829455 | 0.048 | 0.022 | 1 | 0.763594148 | 1.098110085 | 0.022 | 0.031 | 1 | 0.763594148 | 0.71548222 |
| Smc2 | 0.704703697 | -0.295854583 | 0.143 | 0.175 | 1 | 0.467869724 | -0.84048937 | 0.089 | 0.125 | 1 | 0.704703697 | 0.71683737 |
| Gm11655 | 0.468377087 | 1.43673257 | 0.143 | 0.104 | 1 | 0.788320053 | 1.242499995 | 0.067 | 0.059 | 1 | 0.788320053 | 0.717377078 |
| Tmem88 | 0.468600625 | 2.281454345 | 0.143 | 0.104 | 1 | 0.848389101 | 0.83507568 | 0.067 | 0.08 | 1 | 0.848389101 | 0.717614705 |
| Skic3 | 0.46877781 | 0.136337637 | 0.429 | 0.623 | 1 | 0.654919156 | 0.614263428 | 0.467 | 0.585 | 1 | 0.654919156 | 0.717802985 |
| Slc38a1 | 0.574073587 | 0.791454788 | 0.571 | 0.694 | 1 | 0.469014595 | 0.531069493 | 0.156 | 0.218 | 1 | 0.574073587 | 0.7180545 |
| Eml1 | 0.469056166 | 0.75177428 | 0.143 | 0.235 | 1 | 0.604631538 | 1.518685768 | 0.356 | 0.443 | 1 | 0.604631538 | 0.718098645 |
| Arf4os | 0.647471572 | 0.55829455 | 0.048 | 0.078 | 1 | 0.470232105 | 1.393565969 | 0.089 | 0.062 | 1 | 0.647471572 | 0.719345978 |
| Stat2 | 0.565856089 | 0.284001027 | 0.333 | 0.455 | 1 | 0.470711296 | 0.556963817 | 0.222 | 0.311 | 1 | 0.565856089 | 0.719853467 |
| Tnnt3 | 0.509612404 | -0.510652803 | 0.095 | 0.149 | 1 | 0.470812361 | -0.16492432 | 0.044 | 0.076 | 1 | 0.509612404 | 0.719960443 |
| Fam110b | 0.707509367 | 1.771619091 | 0.429 | 0.5 | 1 | 0.470830883 | 1.20029048 | 0.378 | 0.419 | 1 | 0.707509367 | 0.719980045 |
| Msi1 | 0.868416371 | 0.451379346 | 0.143 | 0.168 | 1 | 0.471251375 | 0.213587303 | 0.111 | 0.163 | 1 | 0.868416371 | 0.720424892 |
| Zcchc24 | 0.740592845 | 0.885869208 | 0.429 | 0.451 | 1 | 0.471836886 | 0.285856364 | 0.356 | 0.509 | 1 | 0.740592845 | 0.721043725 |
| Sprr1a | 0.47224907 | -0.496153234 | 0.048 | 0.093 | 1 | 0.626642759 | -1.819427754 | 0.044 | 0.062 | 1 | 0.626642759 | 0.721478955 |
| Dazap1 | 0.551791752 | 0.697230741 | 0.476 | 0.56 | 1 | 0.4723483 | 0.271507279 | 0.8 | 0.727 | 1 | 0.551791752 | 0.721583684 |
| Gm56549 | 0.472861597 | 1.866416846 | 0.048 | 0.022 | 1 | 0.494823351 | 1.946106992 | 0.022 | 0.01 | 1 | 0.494823351 | 0.722125104 |
| Zfp13 | 0.860084871 | 1.010806755 | 0.048 | 0.06 | 1 | 0.473337706 | -0.776359032 | 0.022 | 0.045 | 1 | 0.860084871 | 0.722626828 |
| Nfyc | 0.979152117 | 0.498133115 | 0.429 | 0.537 | 1 | 0.473552105 | 0.671359166 | 0.578 | 0.609 | 1 | 0.979152117 | 0.722852614 |
| Rabggta | 0.588025216 | 0.673771768 | 0.048 | 0.086 | 1 | 0.473829746 | 1.268035087 | 0.089 | 0.062 | 1 | 0.588025216 | 0.723144864 |
| Gm17092 | 0.474136073 | -0.44170545 | 0.095 | 0.149 | 1 | 0.563709837 | 0.320502507 | 0.067 | 0.097 | 1 | 0.563709837 | 0.723467131 |
| Mgme1 | 0.474362221 | 1.258734268 | 0.143 | 0.101 | 1 | 0.968776602 | 0.91753784 | 0.133 | 0.149 | 1 | 0.968776602 | 0.723704925 |
| Gca | 0.4759797 | 1.673771768 | 0.048 | 0.022 | 1 | 0.496223879 | 0.683072586 | 0 | 0.01 | 1 | 0.496223879 | 0.725402725 |
| Tex45 | 0.4759797 | 1.673771768 | 0.048 | 0.022 | 1 | 0.955025729 | 1.361144491 | 0.022 | 0.024 | 1 | 0.955025729 | 0.725402725 |
| Gm3055 | 0.4759797 | 1.673771768 | 0.048 | 0.022 | 1 | 0.80624346 | 1.683072586 | 0.022 | 0.017 | 1 | 0.80624346 | 0.725402725 |
| Gm28529 | 0.4759797 | 1.673771768 | 0.048 | 0.022 | 1 | 0.687197015 | 1.223640968 | 0.022 | 0.035 | 1 | 0.687197015 | 0.725402725 |
| Cep55 | 0.476005214 | 1.171271427 | 0.048 | 0.022 | 1 | 0.496223879 | 0.683072586 | 0 | 0.01 | 1 | 0.496223879 | 0.725429464 |
| 9630028H03Rik | 0.557715318 | 1.271673324 | 0.143 | 0.104 | 1 | 0.476020317 | 0.274987848 | 0.111 | 0.163 | 1 | 0.557715318 | 0.725445292 |
| Galm | 0.476101154 | 1.586308926 | 0.143 | 0.101 | 1 | 0.550164758 | -0.421264074 | 0.067 | 0.093 | 1 | 0.550164758 | 0.725529999 |
| Exoc6 | 0.513486849 | 1.735172312 | 0.095 | 0.063 | 1 | 0.476265768 | -0.404390255 | 0.022 | 0.045 | 1 | 0.513486849 | 0.725702454 |
| Vcpip1 | 0.85524413 | 0.147485174 | 0.429 | 0.567 | 1 | 0.476665687 | 0.218908229 | 0.822 | 0.72 | 1 | 0.85524413 | 0.726121197 |
| Gcnt1 | 0.477049761 | 0.457834368 | 0.476 | 0.437 | 1 | 0.823339156 | 0.59721185 | 0.267 | 0.322 | 1 | 0.823339156 | 0.726523048 |
| Xpo1 | 0.711894092 | 0.162809848 | 0.381 | 0.53 | 1 | 0.477316251 | 0.237155774 | 0.733 | 0.571 | 1 | 0.711894092 | 0.726801699 |
| Zfp384 | 0.477412554 | 1.368917186 | 0.286 | 0.254 | 1 | 0.949562338 | 0.51748652 | 0.356 | 0.415 | 1 | 0.949562338 | 0.726902361 |
| Best1 | 0.532436372 | 1.088809267 | 0 | 0.019 | 1 | 0.477727467 | -0.486852415 | 0.022 | 0.045 | 1 | 0.532436372 | 0.727231401 |
| Tspan32os | 0.71266839 | 1.79930265 | 0.048 | 0.034 | 1 | 0.477727467 | -0.316927414 | 0.022 | 0.045 | 1 | 0.71266839 | 0.727231401 |
| Stoml3 | 0.815913357 | 0.214340149 | 0.048 | 0.06 | 1 | 0.477727467 | -0.404390255 | 0.022 | 0.045 | 1 | 0.815913357 | 0.727231401 |
| Fbxo10 | 0.478602046 | 2.029915578 | 0.19 | 0.149 | 1 | 0.749001424 | 0.91753784 | 0.156 | 0.145 | 1 | 0.749001424 | 0.728144174 |
| Mapk8 | 0.904799482 | 1.025244138 | 0.333 | 0.466 | 1 | 0.479017621 | 0.496273766 | 0.711 | 0.64 | 1 | 0.904799482 | 0.728577361 |
| Prdm14 | 0.479147948 | 1.351843673 | 0.048 | 0.022 | 1 | 0.936023974 | 1.875717664 | 0.022 | 0.021 | 1 | 0.936023974 | 0.72871314 |
| Nr1h3 | 0.532426384 | 0.866416846 | 0 | 0.019 | 1 | 0.47918822 | -0.316927414 | 0.022 | 0.045 | 1 | 0.532426384 | 0.728755089 |
| Gm6994 | 0.578159907 | 0.866416846 | 0 | 0.015 | 1 | 0.47918822 | -0.223818009 | 0.022 | 0.045 | 1 | 0.578159907 | 0.728755089 |
| Gm19500 | 0.933802207 | 1.088809267 | 0.048 | 0.045 | 1 | 0.47918822 | -0.223818009 | 0.022 | 0.045 | 1 | 0.933802207 | 0.728755089 |
| Gm56901 | 0.985457491 | 0.766881172 | 0.048 | 0.049 | 1 | 0.47918822 | -0.223818009 | 0.022 | 0.045 | 1 | 0.985457491 | 0.728755089 |
| Ednra | 0.653499629 | -0.718545655 | 0.048 | 0.075 | 1 | 0.479291871 | -2.638855509 | 0.022 | 0.045 | 1 | 0.653499629 | 0.728863044 |
| Gm26749 | 0.907384734 | 1.010806755 | 0.048 | 0.056 | 1 | 0.480205094 | 0.531069493 | 0.067 | 0.107 | 1 | 0.907384734 | 0.729813256 |
| Nqo2 | 0.516070086 | 1.129451251 | 0.143 | 0.104 | 1 | 0.480316473 | 0.426732833 | 0.089 | 0.131 | 1 | 0.516070086 | 0.729929032 |
| Fes | 0.976796229 | 1.041503552 | 0.095 | 0.097 | 1 | 0.48072475 | 0.862042727 | 0.089 | 0.135 | 1 | 0.976796229 | 0.730353214 |
| Dip2a | 0.57472459 | 1.47583239 | 0.19 | 0.164 | 1 | 0.480769489 | 0.107760255 | 0.222 | 0.301 | 1 | 0.57472459 | 0.730399676 |
| Ssc5d | 0.492608234 | 0.866416846 | 0 | 0.022 | 1 | 0.480781131 | -0.124282336 | 0.022 | 0.045 | 1 | 0.492608234 | 0.730411766 |
| N4bp3 | 0.532436372 | 1.088809267 | 0 | 0.019 | 1 | 0.480781131 | -0.124282336 | 0.022 | 0.045 | 1 | 0.532436372 | 0.730411766 |
| Fam118b | 0.540137564 | 0.97333205 | 0.19 | 0.287 | 1 | 0.480841718 | 0.616983396 | 0.222 | 0.318 | 1 | 0.540137564 | 0.730474678 |
| Kcnb1 | 0.761157642 | 2.65387221 | 0.19 | 0.183 | 1 | 0.480993544 | 0.375288674 | 0.2 | 0.284 | 1 | 0.761157642 | 0.730632299 |
| Samhd1 | 0.481155159 | 0.290814239 | 0.667 | 0.627 | 1 | 0.952124337 | 0.148840888 | 0.578 | 0.54 | 1 | 0.952124337 | 0.730800031 |
| Abcd3 | 0.561837473 | 1.366343242 | 0.476 | 0.593 | 1 | 0.481264583 | 0.346037599 | 0.778 | 0.682 | 1 | 0.561837473 | 0.730913567 |
| Soga1 | 0.674589658 | 0.761234609 | 0.381 | 0.414 | 1 | 0.481521003 | 0.549806055 | 0.511 | 0.561 | 1 | 0.674589658 | 0.73117953 |
| Rgs12 | 0.680302384 | -0.326228232 | 0.095 | 0.123 | 1 | 0.481851616 | 0.583536913 | 0.2 | 0.159 | 1 | 0.680302384 | 0.731522253 |
| Gm16310 | 0.800830966 | 1.137718867 | 0.095 | 0.082 | 1 | 0.483387894 | 0.397670367 | 0.067 | 0.104 | 1 | 0.800830966 | 0.733111932 |
| Arc | 0.924132133 | -0.295854583 | 0.048 | 0.052 | 1 | 0.483744351 | -0.638855509 | 0.022 | 0.045 | 1 | 0.924132133 | 0.733480105 |
| Slc6a12 | 0.483789678 | -0.469186186 | 0.238 | 0.321 | 1 | 0.754642706 | 1.868628239 | 0.089 | 0.08 | 1 | 0.754642706 | 0.733526903 |
| Fut8 | 0.936791145 | 0.746122612 | 0.571 | 0.698 | 1 | 0.484069546 | 0.216930746 | 0.956 | 0.9 | 1 | 0.936791145 | 0.733815767 |
| Ssu72 | 0.484487967 | -0.145312447 | 0.667 | 0.802 | 1 | 0.985084213 | 0.16020009 | 0.844 | 0.744 | 1 | 0.985084213 | 0.734247344 |
| Gm26910 | 0.910091451 | 1.088809267 | 0.048 | 0.056 | 1 | 0.484564742 | 0.268035087 | 0.044 | 0.076 | 1 | 0.910091451 | 0.734326495 |
| Lins1 | 0.485471145 | 0.995699863 | 0.143 | 0.097 | 1 | 0.945357883 | 1.397670367 | 0.111 | 0.118 | 1 | 0.945357883 | 0.735260058 |
| Osbp | 0.485651666 | 0.110835573 | 0.381 | 0.59 | 1 | 0.920803215 | 0.148150768 | 0.844 | 0.796 | 1 | 0.920803215 | 0.735445791 |
| Setx | 0.676353606 | 0.351843673 | 0.286 | 0.41 | 1 | 0.485918228 | 0.269323783 | 0.622 | 0.581 | 1 | 0.676353606 | 0.735719932 |
| Lhfp | 0.957590314 | 0.438258425 | 0.476 | 0.627 | 1 | 0.486888665 | 0.605847074 | 0.533 | 0.827 | 1 | 0.957590314 | 0.736716758 |
| Grem1 | 0.969882907 | -0.184209227 | 0.048 | 0.045 | 1 | 0.488240656 | -0.602329633 | 0.022 | 0.045 | 1 | 0.969882907 | 0.738102374 |
| Unk | 0.488330584 | 1.205622932 | 0.286 | 0.272 | 1 | 0.740931612 | 0.812355603 | 0.222 | 0.298 | 1 | 0.740931612 | 0.738194409 |
| Gm43328 | 0.834600729 | 1.088809267 | 0.095 | 0.086 | 1 | 0.48855557 | 0.286182434 | 0.156 | 0.215 | 1 | 0.834600729 | 0.738424595 |
| Sgf29 | 0.88361085 | 0.536268244 | 0.143 | 0.134 | 1 | 0.488577384 | 0.118287967 | 0.156 | 0.211 | 1 | 0.88361085 | 0.738446908 |
| Sema4a | 0.524619333 | 0.912959432 | 0.143 | 0.104 | 1 | 0.488695534 | -1.187292133 | 0.067 | 0.1 | 1 | 0.524619333 | 0.738567743 |
| Wwc2 | 0.489582017 | 0.276777858 | 0.714 | 0.903 | 1 | 0.772585937 | 0.103076338 | 0.867 | 0.907 | 1 | 0.772585937 | 0.739473483 |

| | | | | | | | | | | | | |
|---|---|---|---|---|---|---|---|---|---|---|---|---|
| Dchs2 | 0.74272003 | -0.522625445 | 0.048 | 0.067 | 1 | 0.48974149 | 2.235613609 | 0.067 | 0.045 | 1 | 0.74272003 | 0.739636253 |
| Gm39929 | 0.490354231 | 1.503846766 | 0.095 | 0.06 | 1 | 0.687216983 | 0.982632868 | 0.022 | 0.035 | 1 | 0.687216983 | 0.74026119 |
| 9130410C08Rik | 0.515250719 | 0.710297644 | 0.095 | 0.06 | 1 | 0.490400683 | 0.638678467 | 0.044 | 0.076 | 1 | 0.515250719 | 0.740308536 |
| Slc22a21 | 0.86181432 | 0.728913322 | 0.286 | 0.325 | 1 | 0.490617541 | 0.577277922 | 0.156 | 0.221 | 1 | 0.86181432 | 0.740529511 |
| Fam117a | 0.490914698 | 1.396931562 | 0.19 | 0.142 | 1 | 0.784300845 | -0.213091603 | 0.133 | 0.145 | 1 | 0.784300845 | 0.740832155 |
| Gm42941 | 0.659977312 | 0.569435108 | 0.143 | 0.112 | 1 | 0.490947215 | 1.339118185 | 0.067 | 0.107 | 1 | 0.659977312 | 0.740865262 |
| Gm36447 | 0.921792415 | 1.451379346 | 0.048 | 0.045 | 1 | 0.491296045 | -0.446210431 | 0.044 | 0.073 | 1 | 0.921792415 | 0.741220286 |
| Gm47438 | 0.492599419 | 0.351843673 | 0 | 0.022 | 1 | 0.492040936 | 3.005000681 | 0.022 | 0.01 | 1 | 0.492599419 | 0.741977589 |
| Slc47a1 | 0.492599419 | 0.351843673 | 0 | 0.022 | 1 | 0.492040936 | 3.005000681 | 0.022 | 0.01 | 1 | 0.492599419 | 0.741977589 |
| Gm47270 | 0.492608234 | 0.866416846 | 0 | 0.022 | 1 | 0.492040936 | 3.005000681 | 0.022 | 0.01 | 1 | 0.492608234 | 0.741977589 |
| Ager | 0.492608234 | 0.866416846 | 0 | 0.022 | 1 | 0.492040936 | 2.683072586 | 0.022 | 0.01 | 1 | 0.492608234 | 0.741977589 |
| Gm44448 | 0.532426384 | 0.866416846 | 0 | 0.019 | 1 | 0.492040936 | 2.268035087 | 0.022 | 0.01 | 1 | 0.532426384 | 0.741977589 |
| Gm34121 | 0.532436372 | 1.088809267 | 0 | 0.019 | 1 | 0.492040936 | 2.683072586 | 0.022 | 0.01 | 1 | 0.532436372 | 0.741977589 |
| Fam217a | 0.578156445 | 1.088809267 | 0 | 0.015 | 1 | 0.492040936 | 2.268035087 | 0.022 | 0.01 | 1 | 0.578156445 | 0.741977589 |
| Gm49864 | 0.578156445 | 1.088809267 | 0 | 0.015 | 1 | 0.492040936 | 2.683072586 | 0.022 | 0.01 | 1 | 0.578156445 | 0.741977589 |
| Gm16183 | 0.578159907 | 0.503846766 | 0 | 0.015 | 1 | 0.492040936 | 2.268035087 | 0.022 | 0.01 | 1 | 0.578159907 | 0.741977589 |
| Icam5 | 0.57816106 | 1.351843673 | 0 | 0.015 | 1 | 0.492040936 | 2.683072586 | 0.022 | 0.01 | 1 | 0.57816106 | 0.741977589 |
| Gm16068 | 0.57816106 | 1.351843673 | 0 | 0.015 | 1 | 0.492040936 | 2.268035087 | 0.022 | 0.01 | 1 | 0.57816106 | 0.741977589 |
| Gm6345 | 0.63207476 | 1.351843673 | 0 | 0.011 | 1 | 0.492040936 | 2.268035087 | 0.022 | 0.01 | 1 | 0.63207476 | 0.741977589 |
| Aldh3b3 | 0.63207476 | 1.351843673 | 0 | 0.011 | 1 | 0.492040936 | 2.683072586 | 0.022 | 0.01 | 1 | 0.63207476 | 0.741977589 |
| Klf1 | 0.632076134 | 1.673771768 | 0 | 0.011 | 1 | 0.492040936 | 2.683072586 | 0.022 | 0.01 | 1 | 0.632076134 | 0.741977589 |
| Odad3 | 0.632076134 | 1.673771768 | 0 | 0.011 | 1 | 0.492040936 | 2.683072586 | 0.022 | 0.01 | 1 | 0.632076134 | 0.741977589 |
| Acot4 | 0.632076134 | 1.673771768 | 0 | 0.011 | 1 | 0.492040936 | 2.268035087 | 0.022 | 0.01 | 1 | 0.632076134 | 0.741977589 |
| Gm6270 | 0.632076134 | 1.673771768 | 0 | 0.011 | 1 | 0.492040936 | 2.268035087 | 0.022 | 0.01 | 1 | 0.632076134 | 0.741977589 |
| Gm50209 | 0.632076134 | 1.673771768 | 0 | 0.011 | 1 | 0.492040936 | 3.005000681 | 0.022 | 0.01 | 1 | 0.632076134 | 0.741977589 |
| Gm48677 | 0.815691551 | 1.088809267 | 0.048 | 0.063 | 1 | 0.492040936 | 2.683072586 | 0.022 | 0.01 | 1 | 0.815691551 | 0.741977589 |
| Fam124a | 0.492608234 | 0.866416846 | 0 | 0.022 | 1 | 0.492043882 | 2.361144491 | 0.022 | 0.01 | 1 | 0.492608234 | 0.741980582 |
| Cfap276 | 0.57816106 | 1.351843673 | 0 | 0.015 | 1 | 0.492043882 | 2.683072586 | 0.022 | 0.01 | 1 | 0.57816106 | 0.741980582 |
| Slc6a4 | 0.57816106 | 1.351843673 | 0 | 0.015 | 1 | 0.492043882 | 2.098110085 | 0.022 | 0.01 | 1 | 0.57816106 | 0.741980582 |
| Sidt1 | 0.632076134 | 1.673771768 | 0 | 0.011 | 1 | 0.492043882 | 2.268035087 | 0.022 | 0.01 | 1 | 0.632076134 | 0.741980582 |
| Gm47507 | 0.658816973 | 0.258734268 | 0.048 | 0.03 | 1 | 0.492043882 | 2.683072586 | 0.022 | 0.01 | 1 | 0.658816973 | 0.741980582 |
| Agap2 | 0.71266839 | 1.79930265 | 0.048 | 0.034 | 1 | 0.492043882 | 2.098110085 | 0.022 | 0.01 | 1 | 0.71266839 | 0.741980582 |
| Tsix | 0.780815746 | 1.503846766 | 0.143 | 0.138 | 1 | 0.492043882 | 2.946106992 | 0.022 | 0.01 | 1 | 0.780815746 | 0.741980582 |
| Gm29266 | 0.815671306 | -0.213753503 | 0.048 | 0.037 | 1 | 0.492043882 | 3.946106992 | 0.022 | 0.01 | 1 | 0.815671306 | 0.741980582 |
| Gm33424 | 0.81881728 | 1.088809267 | 0.048 | 0.037 | 1 | 0.492043882 | 2.361144491 | 0.022 | 0.01 | 1 | 0.81881728 | 0.741980582 |
| Abcb1a | 0.972893889 | 0.281454345 | 0.048 | 0.045 | 1 | 0.492043882 | 2.946106992 | 0.022 | 0.01 | 1 | 0.972893889 | 0.741980582 |
| Rtl4 | 0.979647676 | 3.560561157 | 0.048 | 0.049 | 1 | 0.492044864 | 3.168499413 | 0.022 | 0.01 | 1 | 0.979647676 | 0.74198158 |
| Gm11638 | 0.492461073 | 1.936806174 | 0.095 | 0.06 | 1 | 0.607952753 | 1.536231198 | 0.089 | 0.073 | 1 | 0.607952753 | 0.742404238 |
| Sap30bp | 0.54534445 | 0.651403955 | 0.333 | 0.493 | 1 | 0.492494923 | 0.513147585 | 0.489 | 0.557 | 1 | 0.54534445 | 0.742438596 |
| Hhipl2 | 0.492590603 | 0.673771768 | 0 | 0.022 | 1 | 0.545586636 | 0.946106992 | 0.044 | 0.028 | 1 | 0.545586636 | 0.742535704 |
| Gm13524 | 0.492590603 | 0.673771768 | 0 | 0.022 | 1 | 0.765720982 | 1.683072586 | 0.022 | 0.031 | 1 | 0.765720982 | 0.742535704 |
| Kcnn3 | 0.492590603 | 0.673771768 | 0 | 0.022 | 1 | 0.812078309 | 1.098110085 | 0.044 | 0.038 | 1 | 0.812078309 | 0.742535704 |
| Gm6260 | 0.492590603 | 0.673771768 | 0 | 0.022 | 1 | 0.49622258 | -1.017367132 | 0 | 0.01 | 1 | 0.49622258 | 0.742535704 |
| C530005A16Rik | 0.492590603 | 0.673771768 | 0 | 0.022 | 1 | 0.496223879 | 0.683072586 | 0 | 0.01 | 1 | 0.496223879 | 0.742535704 |
| Wdr86 | 0.492590603 | 0.673771768 | 0 | 0.022 | 1 | 0.670048992 | 1.361144491 | 0.022 | 0.014 | 1 | 0.670048992 | 0.742535704 |
| 2310040G24Rik | 0.492590603 | 0.673771768 | 0 | 0.022 | 1 | 0.935071939 | 0.223640968 | 0.022 | 0.024 | 1 | 0.935071939 | 0.742535704 |
| Phox2a | 0.492590603 | 0.351843673 | 0 | 0.022 | 1 | 0.811836522 | 1.361144491 | 0.022 | 0.017 | 1 | 0.811836522 | 0.742535704 |
| Gm56847 | 0.492590603 | 0.673771768 | 0 | 0.022 | 1 | 0.496223879 | 0.683072586 | 0 | 0.01 | 1 | 0.496223879 | 0.742535704 |
| Nrn1l | 0.492590603 | 0.673771768 | 0 | 0.022 | 1 | 0.496223879 | 0.683072586 | 0 | 0.01 | 1 | 0.496223879 | 0.742535704 |
| A330074K22Rik | 0.492590603 | 0.673771768 | 0 | 0.022 | 1 | 0.496223879 | 0.683072586 | 0 | 0.01 | 1 | 0.496223879 | 0.742535704 |
| Thy1 | 0.492590603 | 0.503846766 | 0 | 0.022 | 1 | 0.49622258 | -2.271123724 | 0 | 0.01 | 1 | 0.49622258 | 0.742535704 |
| Mtfp1 | 0.492590603 | 0.673771768 | 0 | 0.022 | 1 | 0.8202424 | 1.098110085 | 0.022 | 0.017 | 1 | 0.8202424 | 0.742535704 |
| Dnah9 | 0.492590603 | 0.673771768 | 0 | 0.022 | 1 | 0.80624346 | 2.098110085 | 0.022 | 0.017 | 1 | 0.80624346 | 0.742535704 |
| Taco1os | 0.492590603 | 0.673771768 | 0 | 0.022 | 1 | 0.672920453 | 0.875717664 | 0.022 | 0.014 | 1 | 0.672920453 | 0.742535704 |
| Ccdc40 | 0.492590603 | 0.673771768 | 0 | 0.022 | 1 | 0.687216983 | 2.442064487 | 0.022 | 0.035 | 1 | 0.687216983 | 0.742535704 |
| Plek2 | 0.492590603 | 0.503846766 | 0 | 0.022 | 1 | 0.835943453 | 0.513147585 | 0.022 | 0.028 | 1 | 0.835943453 | 0.742535704 |
| Zbtb42 | 0.492590603 | 0.673771768 | 0 | 0.022 | 1 | 0.937567971 | 0.683072586 | 0.022 | 0.024 | 1 | 0.937567971 | 0.742535704 |
| Shc3 | 0.492590603 | 0.673771768 | 0 | 0.022 | 1 | 0.496223879 | -0.316927414 | 0 | 0.01 | 1 | 0.496223879 | 0.742535704 |
| Etohd2 | 0.492590603 | 0.673771768 | 0 | 0.022 | 1 | 0.80624346 | 1.683072586 | 0.022 | 0.017 | 1 | 0.80624346 | 0.742535704 |
| Gm5089 | 0.492590603 | 0.673771768 | 0 | 0.022 | 1 | 0.49622258 | 0.361144491 | 0 | 0.01 | 1 | 0.49622258 | 0.742535704 |
| Rmi2 | 0.492590603 | 0.673771768 | 0 | 0.022 | 1 | 0.936025978 | 1.513147585 | 0.022 | 0.021 | 1 | 0.936025978 | 0.742535704 |
| Nrros | 0.492590603 | -0.133583154 | 0 | 0.022 | 1 | 0.672920453 | 0.875717664 | 0.022 | 0.014 | 1 | 0.672920453 | 0.742535704 |
| Gm20522 | 0.492590603 | 0.673771768 | 0 | 0.022 | 1 | 0.496223879 | 0.683072586 | 0 | 0.01 | 1 | 0.496223879 | 0.742535704 |
| Fsd1 | 0.492590603 | 0.503846766 | 0 | 0.022 | 1 | 0.849825041 | 0.808603468 | 0.022 | 0.028 | 1 | 0.849825041 | 0.742535704 |
| Dner | 0.492599419 | 0.351843673 | 0 | 0.022 | 1 | 0.49622258 | 0.361144491 | 0 | 0.01 | 1 | 0.49622258 | 0.74254465 |
| Spi1 | 0.492599419 | -0.133583154 | 0 | 0.022 | 1 | 0.925115145 | -1.545746104 | 0.022 | 0.024 | 1 | 0.925115145 | 0.74254465 |
| Mir670hg | 0.492599419 | 0.351843673 | 0 | 0.022 | 1 | 0.761447202 | 0.683072586 | 0.022 | 0.031 | 1 | 0.761447202 | 0.74254465 |
| Eqtn | 0.492599419 | 0.503846766 | 0 | 0.022 | 1 | 0.617551118 | 1.005000681 | 0.022 | 0.038 | 1 | 0.617551118 | 0.74254465 |
| Cxcl2 | 0.492599419 | -0.413691074 | 0 | 0.022 | 1 | 0.496223879 | 0.683072586 | 0 | 0.01 | 1 | 0.496223879 | 0.74254465 |
| Zscan2 | 0.492599419 | 0.351843673 | 0 | 0.022 | 1 | 0.951997643 | 0.875717664 | 0.022 | 0.021 | 1 | 0.951997643 | 0.74254465 |
| Gm56858 | 0.492599419 | 0.503846766 | 0 | 0.022 | 1 | 0.936023974 | 1.460680165 | 0.022 | 0.021 | 1 | 0.936023974 | 0.74254465 |
| Calml4 | 0.492599419 | 0.503846766 | 0 | 0.022 | 1 | 0.835943453 | 0.513147585 | 0.022 | 0.028 | 1 | 0.835943453 | 0.74254465 |
| Klhl31 | 0.492599419 | 0.503846766 | 0 | 0.022 | 1 | 0.944009091 | 0.776181991 | 0.022 | 0.021 | 1 | 0.944009091 | 0.74254465 |
| Ddo | 0.492599419 | 0.351843673 | 0 | 0.022 | 1 | 0.954659411 | 0.361144491 | 0.022 | 0.021 | 1 | 0.954659411 | 0.74254465 |
| BC025920 | 0.492599419 | 0.503846766 | 0 | 0.022 | 1 | 0.835943453 | 0.513147585 | 0.022 | 0.028 | 1 | 0.835943453 | 0.74254465 |
| Kcnh4 | 0.492599419 | 0.503846766 | 0 | 0.022 | 1 | 0.494823351 | 2.098110085 | 0.022 | 0.01 | 1 | 0.494823351 | 0.74254465 |
| 4930595D18Rik | 0.492599419 | 0.503846766 | 0 | 0.022 | 1 | 0.80624346 | 1.683072586 | 0.022 | 0.017 | 1 | 0.80624346 | 0.74254465 |
| Cep72 | 0.492599419 | 0.503846766 | 0 | 0.022 | 1 | 0.496223879 | 0.683072586 | 0 | 0.01 | 1 | 0.496223879 | 0.74254465 |
| Gm43434 | 0.492599419 | -0.233118828 | 0 | 0.022 | 1 | 0.496223879 | -0.124282336 | 0 | 0.01 | 1 | 0.496223879 | 0.74254465 |
| AW822252 | 0.492599419 | 0.503846766 | 0 | 0.022 | 1 | 0.675805699 | 0.223640968 | 0.022 | 0.014 | 1 | 0.675805699 | 0.74254465 |
| Gm15261 | 0.492599419 | 0.503846766 | 0 | 0.022 | 1 | 0.817164199 | 1.531069493 | 0.044 | 0.055 | 1 | 0.817164199 | 0.74254465 |
| Gm15492 | 0.492603827 | 0.214340149 | 0 | 0.022 | 1 | 0.92760325 | -0.124282336 | 0.022 | 0.024 | 1 | 0.92760325 | 0.742549123 |
| 9330104G04Rik | 0.492604708 | -0.133583154 | 0 | 0.022 | 1 | 0.955025007 | 2.098110085 | 0.022 | 0.024 | 1 | 0.955025007 | 0.742550018 |
| Trim71 | 0.492604708 | 0.351843673 | 0 | 0.022 | 1 | 0.496223879 | -0.124282336 | 0 | 0.01 | 1 | 0.496223879 | 0.742550018 |
| Vwc2 | 0.492604708 | -0.233118828 | 0 | 0.022 | 1 | 0.496223879 | 0.683072586 | 0 | 0.01 | 1 | 0.496223879 | 0.742550018 |
| Kcnb2 | 0.492606471 | -1.370622352 | 0 | 0.022 | 1 | 0.496223879 | -0.316927414 | 0 | 0.01 | 1 | 0.496223879 | 0.742551807 |
| H2ac24 | 0.492606471 | -0.413691074 | 0 | 0.022 | 1 | 0.496223879 | 0.683072586 | 0 | 0.01 | 1 | 0.496223879 | 0.742551807 |
| Inha | 0.492608234 | 0.866416846 | 0 | 0.022 | 1 | 0.765728546 | 1.223640968 | 0.022 | 0.031 | 1 | 0.765728546 | 0.742553596 |
| Rgs5 | 0.492608234 | 0.866416846 | 0 | 0.022 | 1 | 0.635830063 | 1.03099589 | 0.044 | 0.066 | 1 | 0.635830063 | 0.742553596 |
| Gm45902 | 0.492608234 | 0.866416846 | 0 | 0.022 | 1 | 0.63804111 | -0.124282336 | 0.044 | 0.062 | 1 | 0.63804111 | 0.742553596 |
| Gm13703 | 0.492608234 | 0.866416846 | 0 | 0.022 | 1 | 0.528913413 | 1.683072586 | 0.044 | 0.028 | 1 | 0.528913413 | 0.742553596 |
| Gm57446 | 0.492608234 | 0.866416846 | 0 | 0.022 | 1 | 0.821674021 | 0.982632868 | 0.044 | 0.038 | 1 | 0.821674021 | 0.742553596 |
| Fgf7 | 0.492608234 | 0.866416846 | 0 | 0.022 | 1 | 0.937567614 | -0.316927414 | 0.022 | 0.024 | 1 | 0.937567614 | 0.742553596 |
| Gm16098 | 0.492608234 | 0.866416846 | 0 | 0.022 | 1 | 0.658583583 | 2.361144491 | 0.022 | 0.014 | 1 | 0.658583583 | 0.742553596 |
| Gm15441 | 0.492608234 | 0.866416846 | 0 | 0.022 | 1 | 0.746512944 | 0.361144491 | 0.022 | 0.031 | 1 | 0.746512944 | 0.742553596 |
| Gm43063 | 0.492608234 | 0.866416846 | 0 | 0.022 | 1 | 0.496223879 | 0.683072586 | 0 | 0.01 | 1 | 0.496223879 | 0.742553596 |
| Gm43728 | 0.492608234 | 0.866416846 | 0 | 0.022 | 1 | 0.833619716 | 0.361144491 | 0.022 | 0.028 | 1 | 0.833619716 | 0.742553596 |
| 4930557J02Rik | 0.492608234 | 0.866416846 | 0 | 0.022 | 1 | 0.936023974 | 2.460680165 | 0.022 | 0.021 | 1 | 0.936023974 | 0.742553596 |

| | | | | | | | | | | | | |
|---|---|---|---|---|---|---|---|---|---|---|---|---|
| Wdr54 | 0.492608234 | 0.866416846 | 0 | 0.022 | 1 | 0.744358033 | 0.223640968 | 0.022 | 0.031 | 1 | 0.744358033 | 0.742553596 |
| Ano2 | 0.492608234 | 0.866416846 | 0 | 0.022 | 1 | 0.664308814 | 1.460680165 | 0.022 | 0.014 | 1 | 0.664308814 | 0.742553596 |
| Gm57192 | 0.492608234 | 0.866416846 | 0 | 0.022 | 1 | 0.9741699 | 1.180572246 | 0.044 | 0.045 | 1 | 0.9741699 | 0.742553596 |
| Vstm5 | 0.492608234 | 0.866416846 | 0 | 0.022 | 1 | 0.496223879 | 0.683072586 | 0 | 0.01 | 1 | 0.496223879 | 0.742553596 |
| D830035M03Rik | 0.492608234 | 0.866416846 | 0 | 0.022 | 1 | 0.930092697 | 0.223640968 | 0.022 | 0.024 | 1 | 0.930092697 | 0.742553596 |
| Tgm4 | 0.492608234 | 0.866416846 | 0 | 0.022 | 1 | 0.823042771 | 0.875717664 | 0.022 | 0.017 | 1 | 0.823042771 | 0.742553596 |
| Pfn4 | 0.492608234 | 0.866416846 | 0 | 0.022 | 1 | 0.496223879 | -0.124282336 | 0 | 0.01 | 1 | 0.496223879 | 0.742553596 |
| D630036H23Rik | 0.492608234 | 0.866416846 | 0 | 0.022 | 1 | 0.503208953 | 1.361144491 | 0.022 | 0.01 | 1 | 0.503208953 | 0.742553596 |
| Gm56760 | 0.492608234 | 0.866416846 | 0 | 0.022 | 1 | 0.833619716 | 0.361144491 | 0.022 | 0.028 | 1 | 0.833619716 | 0.742553596 |
| Gm41293 | 0.492608234 | 0.866416846 | 0 | 0.022 | 1 | 0.765720982 | 1.361144491 | 0.022 | 0.031 | 1 | 0.765720982 | 0.742553596 |
| Zfp474 | 0.492608234 | 0.866416846 | 0 | 0.022 | 1 | 0.496223879 | 0.683072586 | 0 | 0.01 | 1 | 0.496223879 | 0.742553596 |
| Taf6l | 0.492608234 | 0.866416846 | 0 | 0.022 | 1 | 0.532520164 | -0.223818009 | 0.022 | 0.042 | 1 | 0.532520164 | 0.742553596 |
| Fbxl19 | 0.59310178 | -0.167530486 | 0.095 | 0.134 | 1 | 0.492653492 | 0.454253896 | 0.133 | 0.194 | 1 | 0.59310178 | 0.742599521 |
| Snap91 | 0.63207476 | 1.088809267 | 0 | 0.011 | 1 | 0.492711267 | 0.15951063 | 0.022 | 0.045 | 1 | 0.63207476 | 0.742658141 |
| Wdr26 | 0.492718633 | 0.420705531 | 1 | 0.933 | 1 | 0.699844836 | 0.207768056 | 1 | 0.99 | 1 | 0.699844836 | 0.742665615 |
| Npnt | 0.492818314 | -0.133583154 | 0.095 | 0.153 | 1 | 0.974515471 | -1.191396532 | 0.378 | 0.318 | 1 | 0.974515471 | 0.742766738 |
| Eed | 0.493065689 | 0.207072149 | 0.476 | 0.403 | 1 | 0.501698098 | 0.170622585 | 0.644 | 0.54 | 1 | 0.501698098 | 0.743017604 |
| Hectd4 | 0.493536134 | 0.108987149 | 0.238 | 0.354 | 1 | 0.997897662 | -0.124282336 | 0.6 | 0.571 | 1 | 0.997897662 | 0.743494352 |
| Grik4 | 0.720240271 | 1.714413752 | 0.19 | 0.172 | 1 | 0.493928983 | 1.474964391 | 0.133 | 0.107 | 1 | 0.720240271 | 0.743892126 |
| Klhl33 | 0.956430899 | 2.123574685 | 0.095 | 0.101 | 1 | 0.494241994 | 0.15951063 | 0.022 | 0.045 | 1 | 0.956430899 | 0.744208839 |
| Ntm | 0.592981051 | -0.413691074 | 0.048 | 0.082 | 1 | 0.494248627 | -0.316927414 | 0.022 | 0.045 | 1 | 0.592981051 | 0.744215549 |
| Esrra | 0.629700127 | 2.088809267 | 0.048 | 0.03 | 1 | 0.494966204 | 0.982632868 | 0.044 | 0.076 | 1 | 0.629700127 | 0.744940865 |
| A330035P11Rik | 0.54776222 | 2.088809267 | 0.048 | 0.026 | 1 | 0.495714932 | 0.180572246 | 0.022 | 0.045 | 1 | 0.54776222 | 0.745696571 |
| Sgsm1 | 0.532426384 | 0.866416846 | 0 | 0.019 | 1 | 0.49622258 | 0.361144491 | 0 | 0.01 | 1 | 0.532426384 | 0.746208311 |
| Sox5os4 | 0.532426384 | 0.866416846 | 0 | 0.019 | 1 | 0.49622258 | 0.361144491 | 0 | 0.01 | 1 | 0.532426384 | 0.746208311 |
| Cdkn2a | 0.532432377 | 0.673771768 | 0 | 0.019 | 1 | 0.49622258 | 0.361144491 | 0 | 0.01 | 1 | 0.532432377 | 0.746208311 |
| Ankfn1 | 0.532432377 | 0.673771768 | 0 | 0.019 | 1 | 0.49622258 | 0.361144491 | 0 | 0.01 | 1 | 0.532432377 | 0.746208311 |
| Slc39a2 | 0.532432377 | 0.214340149 | 0 | 0.019 | 1 | 0.49622258 | 0.361144491 | 0 | 0.01 | 1 | 0.532432377 | 0.746208311 |
| Iqub | 0.532434375 | 0.351843673 | 0 | 0.019 | 1 | 0.49622258 | 0.361144491 | 0 | 0.01 | 1 | 0.532434375 | 0.746208311 |
| Fam186a | 0.532436372 | 1.088809267 | 0 | 0.019 | 1 | 0.49622258 | 0.361144491 | 0 | 0.01 | 1 | 0.532436372 | 0.746208311 |
| Serpinf2 | 0.544589441 | 2.503846766 | 0.048 | 0.026 | 1 | 0.49622258 | 0.361144491 | 0 | 0.01 | 1 | 0.544589441 | 0.746208311 |
| Gm17655 | 0.550953306 | 1.936806174 | 0.048 | 0.026 | 1 | 0.49622258 | 0.361144491 | 0 | 0.01 | 1 | 0.550953306 | 0.746208311 |
| Srrm3 | 0.578156445 | 1.088809267 | 0 | 0.015 | 1 | 0.49622258 | 0.361144491 | 0 | 0.01 | 1 | 0.578156445 | 0.746208311 |
| Gm29571 | 0.578156445 | 1.088809267 | 0 | 0.015 | 1 | 0.49622258 | 0.361144491 | 0 | 0.01 | 1 | 0.578156445 | 0.746208311 |
| Mfsd9 | 0.578159907 | 0.866416846 | 0 | 0.015 | 1 | 0.49622258 | 0.361144491 | 0 | 0.01 | 1 | 0.578159907 | 0.746208311 |
| Ptchd4 | 0.578159907 | 0.503846766 | 0 | 0.015 | 1 | 0.49622258 | -1.017367132 | 0 | 0.01 | 1 | 0.578159907 | 0.746208311 |
| Gm36201 | 0.578159907 | 0.673771768 | 0 | 0.015 | 1 | 0.49622258 | 0.361144491 | 0 | 0.01 | 1 | 0.578159907 | 0.746208311 |
| 1700016C15Rik | 0.57816106 | 1.351843673 | 0 | 0.015 | 1 | 0.49622258 | 0.361144491 | 0 | 0.01 | 1 | 0.57816106 | 0.746208311 |
| Nsl1 | 0.57816106 | 1.351843673 | 0 | 0.015 | 1 | 0.49622258 | 0.361144491 | 0 | 0.01 | 1 | 0.57816106 | 0.746208311 |
| Gm57214 | 0.57816106 | 1.351843673 | 0 | 0.015 | 1 | 0.49622258 | 0.361144491 | 0 | 0.01 | 1 | 0.57816106 | 0.746208311 |
| Gm36246 | 0.57816106 | 1.351843673 | 0 | 0.015 | 1 | 0.49622258 | 0.361144491 | 0 | 0.01 | 1 | 0.57816106 | 0.746208311 |
| Lpar3 | 0.631905465 | 2.351843673 | 0.095 | 0.071 | 1 | 0.49622258 | 0.361144491 | 0 | 0.01 | 1 | 0.631905465 | 0.746208311 |
| 5730419F03Rik | 0.63207476 | 1.351843673 | 0 | 0.011 | 1 | 0.49622258 | -0.124282336 | 0 | 0.01 | 1 | 0.63207476 | 0.746208311 |
| Slc39a12 | 0.63207476 | 1.088809267 | 0 | 0.011 | 1 | 0.49622258 | -0.316927414 | 0 | 0.01 | 1 | 0.63207476 | 0.746208311 |
| Plppr1 | 0.63207476 | 1.088809267 | 0 | 0.011 | 1 | 0.49622258 | 0.361144491 | 0 | 0.01 | 1 | 0.63207476 | 0.746208311 |
| Syne4 | 0.63207476 | 1.351843673 | 0 | 0.011 | 1 | 0.49622258 | 0.361144491 | 0 | 0.01 | 1 | 0.63207476 | 0.746208311 |
| Entrep2 | 0.63207476 | 1.088809267 | 0 | 0.011 | 1 | 0.49622258 | 0.361144491 | 0 | 0.01 | 1 | 0.63207476 | 0.746208311 |
| Sez6l2 | 0.63207476 | 1.351843673 | 0 | 0.011 | 1 | 0.49622258 | 0.361144491 | 0 | 0.01 | 1 | 0.63207476 | 0.746208311 |
| Gm16104 | 0.63207476 | 1.351843673 | 0 | 0.011 | 1 | 0.49622258 | 0.361144491 | 0 | 0.01 | 1 | 0.63207476 | 0.746208311 |
| 6430562O15Rik | 0.63207476 | 1.088809267 | 0 | 0.011 | 1 | 0.49622258 | 0.361144491 | 0 | 0.01 | 1 | 0.63207476 | 0.746208311 |
| D430040D24Rik | 0.632076134 | 1.673771768 | 0 | 0.011 | 1 | 0.49622258 | 0.361144491 | 0 | 0.01 | 1 | 0.632076134 | 0.746208311 |
| Gm37748 | 0.632076134 | 1.673771768 | 0 | 0.011 | 1 | 0.49622258 | 0.361144491 | 0 | 0.01 | 1 | 0.632076134 | 0.746208311 |
| Snx20 | 0.632076134 | 1.673771768 | 0 | 0.011 | 1 | 0.49622258 | 0.361144491 | 0 | 0.01 | 1 | 0.632076134 | 0.746208311 |
| Dusp13 | 0.632076134 | 1.673771768 | 0 | 0.011 | 1 | 0.49622258 | 0.361144491 | 0 | 0.01 | 1 | 0.632076134 | 0.746208311 |
| Gm10101 | 0.632076134 | 1.673771768 | 0 | 0.011 | 1 | 0.49622258 | 0.361144491 | 0 | 0.01 | 1 | 0.632076134 | 0.746208311 |
| Gm36329 | 0.632076134 | 1.673771768 | 0 | 0.011 | 1 | 0.49622258 | 0.361144491 | 0 | 0.01 | 1 | 0.632076134 | 0.746208311 |
| Gm19696 | 0.632076134 | 1.673771768 | 0 | 0.011 | 1 | 0.49622258 | 0.361144491 | 0 | 0.01 | 1 | 0.632076134 | 0.746208311 |
| Cybb | 0.632076134 | -1.133583154 | 0 | 0.011 | 1 | 0.49622258 | -0.124282336 | 0 | 0.01 | 1 | 0.632076134 | 0.746208311 |
| Necab1 | 0.636128533 | 1.451379346 | 0.048 | 0.03 | 1 | 0.49622258 | -1.124282336 | 0 | 0.01 | 1 | 0.636128533 | 0.746208311 |
| Dmp1 | 0.70948781 | 6.48112669 | 0.048 | 0.034 | 1 | 0.49622258 | -0.316927414 | 0 | 0.01 | 1 | 0.70948781 | 0.746208311 |
| Trim38 | 0.70948781 | 2.55829455 | 0.048 | 0.034 | 1 | 0.49622258 | 0.361144491 | 0 | 0.01 | 1 | 0.70948781 | 0.746208311 |
| Dnajc6 | 0.761580634 | 1.088809267 | 0.095 | 0.078 | 1 | 0.49622258 | 0.361144491 | 0 | 0.01 | 1 | 0.761580634 | 0.746208311 |
| Ch25h | 0.812495066 | 0.673771768 | 0.048 | 0.037 | 1 | 0.49622258 | -0.901889915 | 0 | 0.01 | 1 | 0.812495066 | 0.746208311 |
| Igsf6 | 0.532426384 | 0.866416846 | 0 | 0.019 | 1 | 0.496223879 | 0.683072586 | 0 | 0.01 | 1 | 0.532426384 | 0.74620962 |
| Gm56533 | 0.532426384 | 0.866416846 | 0 | 0.019 | 1 | 0.496223879 | 0.683072586 | 0 | 0.01 | 1 | 0.532426384 | 0.74620962 |
| Gm10037 | 0.532426384 | 0.866416846 | 0 | 0.019 | 1 | 0.496223879 | 0.683072586 | 0 | 0.01 | 1 | 0.532426384 | 0.74620962 |
| Rgs22 | 0.532426384 | 0.673771768 | 0 | 0.019 | 1 | 0.496223879 | -0.124282336 | 0 | 0.01 | 1 | 0.532426384 | 0.74620962 |
| Awat2 | 0.532426384 | 0.214340149 | 0 | 0.019 | 1 | 0.496223879 | 0.683072586 | 0 | 0.01 | 1 | 0.532426384 | 0.74620962 |
| Spag6 | 0.532432377 | 0.673771768 | 0 | 0.019 | 1 | 0.496223879 | 0.683072586 | 0 | 0.01 | 1 | 0.532432377 | 0.74620962 |
| Itgal | 0.532432377 | 0.351843673 | 0 | 0.019 | 1 | 0.496223879 | 0.683072586 | 0 | 0.01 | 1 | 0.532432377 | 0.74620962 |
| Pcyt1b | 0.532432377 | 0.673771768 | 0 | 0.019 | 1 | 0.496223879 | 0.683072586 | 0 | 0.01 | 1 | 0.532432377 | 0.74620962 |
| Gm2447 | 0.532435374 | -0.648156327 | 0 | 0.019 | 1 | 0.496223879 | 0.683072586 | 0 | 0.01 | 1 | 0.532435374 | 0.74620962 |
| Gm57180 | 0.532436372 | 1.088809267 | 0 | 0.019 | 1 | 0.496223879 | -0.124282336 | 0 | 0.01 | 1 | 0.532436372 | 0.74620962 |
| Gm9821 | 0.532436372 | 1.088809267 | 0 | 0.019 | 1 | 0.496223879 | 0.683072586 | 0 | 0.01 | 1 | 0.532436372 | 0.74620962 |
| Ubxn10 | 0.532436372 | 1.088809267 | 0 | 0.019 | 1 | 0.496223879 | 0.683072586 | 0 | 0.01 | 1 | 0.532436372 | 0.74620962 |
| Gm10244 | 0.532436372 | 1.088809267 | 0 | 0.019 | 1 | 0.496223879 | 0.683072586 | 0 | 0.01 | 1 | 0.532436372 | 0.74620962 |
| Mlkl | 0.532436372 | 1.088809267 | 0 | 0.019 | 1 | 0.496223879 | 0.683072586 | 0 | 0.01 | 1 | 0.532436372 | 0.74620962 |
| Abcg4 | 0.532436372 | 1.088809267 | 0 | 0.019 | 1 | 0.496223879 | 0.683072586 | 0 | 0.01 | 1 | 0.532436372 | 0.74620962 |
| Gm15915 | 0.532436372 | 1.088809267 | 0 | 0.019 | 1 | 0.496223879 | 0.683072586 | 0 | 0.01 | 1 | 0.532436372 | 0.74620962 |
| 1700093K21Rik | 0.532436372 | 1.088809267 | 0 | 0.019 | 1 | 0.496223879 | 0.683072586 | 0 | 0.01 | 1 | 0.532436372 | 0.74620962 |
| Gm51903 | 0.532436372 | 1.088809267 | 0 | 0.019 | 1 | 0.496223879 | 0.683072586 | 0 | 0.01 | 1 | 0.532436372 | 0.74620962 |
| Gm32934 | 0.532436372 | 1.088809267 | 0 | 0.019 | 1 | 0.496223879 | -0.316927414 | 0 | 0.01 | 1 | 0.532436372 | 0.74620962 |
| Gm48952 | 0.532436372 | 1.088809267 | 0 | 0.019 | 1 | 0.496223879 | 0.683072586 | 0 | 0.01 | 1 | 0.532436372 | 0.74620962 |
| Gm16618 | 0.532436372 | 1.088809267 | 0 | 0.019 | 1 | 0.496223879 | 0.683072586 | 0 | 0.01 | 1 | 0.532436372 | 0.74620962 |
| 1700047M11Rik | 0.536546698 | -0.184209227 | 0.048 | 0.086 | 1 | 0.496223879 | 0.683072586 | 0 | 0.01 | 1 | 0.536546698 | 0.74620962 |
| F730043M19Rik | 0.544576436 | 2.258734268 | 0.048 | 0.026 | 1 | 0.496223879 | 0.683072586 | 0 | 0.01 | 1 | 0.544576436 | 0.74620962 |
| Gm32036 | 0.544576436 | 2.258734268 | 0.048 | 0.026 | 1 | 0.496223879 | 0.683072586 | 0 | 0.01 | 1 | 0.544576436 | 0.74620962 |
| Ddx43 | 0.550953306 | 1.936806174 | 0.048 | 0.026 | 1 | 0.496223879 | 0.683072586 | 0 | 0.01 | 1 | 0.550953306 | 0.74620962 |
| Zfp947 | 0.554143605 | 1.79930265 | 0.048 | 0.026 | 1 | 0.496223879 | -0.124282336 | 0 | 0.01 | 1 | 0.554143605 | 0.74620962 |
| Creg2 | 0.578156445 | 0.866416846 | 0 | 0.015 | 1 | 0.496223879 | 0.683072586 | 0 | 0.01 | 1 | 0.578156445 | 0.74620962 |
| Acbd7 | 0.578156445 | 1.088809267 | 0 | 0.015 | 1 | 0.496223879 | 0.683072586 | 0 | 0.01 | 1 | 0.578156445 | 0.74620962 |
| Pm20d2 | 0.578156445 | 1.088809267 | 0 | 0.015 | 1 | 0.496223879 | 0.683072586 | 0 | 0.01 | 1 | 0.578156445 | 0.74620962 |
| Gm2164 | 0.578156445 | 1.088809267 | 0 | 0.015 | 1 | 0.496223879 | 0.683072586 | 0 | 0.01 | 1 | 0.578156445 | 0.74620962 |
| Ptpro | 0.578156445 | 1.088809267 | 0 | 0.015 | 1 | 0.496223879 | 0.683072586 | 0 | 0.01 | 1 | 0.578156445 | 0.74620962 |
| Gm11739 | 0.578156445 | 1.088809267 | 0 | 0.015 | 1 | 0.496223879 | 0.683072586 | 0 | 0.01 | 1 | 0.578156445 | 0.74620962 |
| H3c1 | 0.578156445 | 1.088809267 | 0 | 0.015 | 1 | 0.496223879 | 0.683072586 | 0 | 0.01 | 1 | 0.578156445 | 0.74620962 |
| Depdc1b | 0.578156445 | 0.503846766 | 0 | 0.015 | 1 | 0.496223879 | 0.683072586 | 0 | 0.01 | 1 | 0.578156445 | 0.74620962 |
| Acy3 | 0.578156445 | 1.088809267 | 0 | 0.015 | 1 | 0.496223879 | 0.683072586 | 0 | 0.01 | 1 | 0.578156445 | 0.74620962 |

| | | | | | | | | | | | | |
|---|---|---|---|---|---|---|---|---|---|---|---|---|
| Was | 0.578156445 | 1.088809267 | 0 | 0.015 | 1 | 0.496223879 | 0.683072586 | 0 | 0.01 | 1 | 0.578156445 | 0.74620962 |
| Ppef1 | 0.578156445 | 1.088809267 | 0 | 0.015 | 1 | 0.496223879 | -0.486852415 | 0 | 0.01 | 1 | 0.578156445 | 0.74620962 |
| Sphkap | 0.578159907 | 0.866416846 | 0 | 0.015 | 1 | 0.496223879 | -0.486852415 | 0 | 0.01 | 1 | 0.578159907 | 0.74620962 |
| Hmgcs2 | 0.578159907 | 0.351843673 | 0 | 0.015 | 1 | 0.496223879 | -0.638855509 | 0 | 0.01 | 1 | 0.578159907 | 0.74620962 |
| Lpar5 | 0.578159907 | 0.673771768 | 0 | 0.015 | 1 | 0.496223879 | 0.683072586 | 0 | 0.01 | 1 | 0.578159907 | 0.74620962 |
| Gm57103 | 0.57816106 | 1.351843673 | 0 | 0.015 | 1 | 0.496223879 | 0.683072586 | 0 | 0.01 | 1 | 0.57816106 | 0.74620962 |
| Atad3aos | 0.57816106 | 1.351843673 | 0 | 0.015 | 1 | 0.496223879 | 0.683072586 | 0 | 0.01 | 1 | 0.57816106 | 0.74620962 |
| Spaca7b | 0.57816106 | 1.351843673 | 0 | 0.015 | 1 | 0.496223879 | 0.683072586 | 0 | 0.01 | 1 | 0.57816106 | 0.74620962 |
| Acaa1b | 0.57816106 | 1.351843673 | 0 | 0.015 | 1 | 0.496223879 | 0.683072586 | 0 | 0.01 | 1 | 0.57816106 | 0.74620962 |
| Gm48765 | 0.57816106 | 1.351843673 | 0 | 0.015 | 1 | 0.496223879 | 0.683072586 | 0 | 0.01 | 1 | 0.57816106 | 0.74620962 |
| Gm36738 | 0.57816106 | 1.351843673 | 0 | 0.015 | 1 | 0.496223879 | 0.683072586 | 0 | 0.01 | 1 | 0.57816106 | 0.74620962 |
| Ciita | 0.57816106 | 1.351843673 | 0 | 0.015 | 1 | 0.496223879 | 0.683072586 | 0 | 0.01 | 1 | 0.57816106 | 0.74620962 |
| Gm36419 | 0.57816106 | 1.351843673 | 0 | 0.015 | 1 | 0.496223879 | 0.683072586 | 0 | 0.01 | 1 | 0.57816106 | 0.74620962 |
| A730056A06Rik | 0.629723166 | 3.258734268 | 0.048 | 0.03 | 1 | 0.496223879 | 0.683072586 | 0 | 0.01 | 1 | 0.629723166 | 0.74620962 |
| Gm13381 | 0.63207476 | 1.351843673 | 0 | 0.011 | 1 | 0.496223879 | 0.683072586 | 0 | 0.01 | 1 | 0.63207476 | 0.74620962 |
| 8030451A03Rik | 0.63207476 | 1.351843673 | 0 | 0.011 | 1 | 0.496223879 | 0.683072586 | 0 | 0.01 | 1 | 0.63207476 | 0.74620962 |
| Spink2 | 0.63207476 | 1.351843673 | 0 | 0.011 | 1 | 0.496223879 | 0.683072586 | 0 | 0.01 | 1 | 0.63207476 | 0.74620962 |
| Lmo3 | 0.63207476 | 1.351843673 | 0 | 0.011 | 1 | 0.496223879 | 0.683072586 | 0 | 0.01 | 1 | 0.63207476 | 0.74620962 |
| Zfp575 | 0.63207476 | 1.351843673 | 0 | 0.011 | 1 | 0.496223879 | 0.683072586 | 0 | 0.01 | 1 | 0.63207476 | 0.74620962 |
| B4galnt4 | 0.63207476 | 1.351843673 | 0 | 0.011 | 1 | 0.496223879 | 0.683072586 | 0 | 0.01 | 1 | 0.63207476 | 0.74620962 |
| Gsta2 | 0.63207476 | 1.088809267 | 0 | 0.011 | 1 | 0.496223879 | 0.683072586 | 0 | 0.01 | 1 | 0.63207476 | 0.74620962 |
| Gm57061 | 0.63207476 | 0.673771768 | 0 | 0.011 | 1 | 0.496223879 | 0.683072586 | 0 | 0.01 | 1 | 0.63207476 | 0.74620962 |
| Abi3 | 0.63207476 | 1.351843673 | 0 | 0.011 | 1 | 0.496223879 | 0.683072586 | 0 | 0.01 | 1 | 0.63207476 | 0.74620962 |
| Dio3os | 0.63207476 | 1.351843673 | 0 | 0.011 | 1 | 0.496223879 | 0.683072586 | 0 | 0.01 | 1 | 0.63207476 | 0.74620962 |
| Zfp85os | 0.63207476 | 1.351843673 | 0 | 0.011 | 1 | 0.496223879 | 0.683072586 | 0 | 0.01 | 1 | 0.63207476 | 0.74620962 |
| Gm47467 | 0.63207476 | 1.351843673 | 0 | 0.011 | 1 | 0.496223879 | 0.683072586 | 0 | 0.01 | 1 | 0.63207476 | 0.74620962 |
| Gm56602 | 0.63207476 | 1.088809267 | 0 | 0.011 | 1 | 0.496223879 | 0.683072586 | 0 | 0.01 | 1 | 0.63207476 | 0.74620962 |
| Trem3 | 0.63207476 | 0.503846766 | 0 | 0.011 | 1 | 0.496223879 | 0.683072586 | 0 | 0.01 | 1 | 0.63207476 | 0.74620962 |
| Gm30984 | 0.63207476 | 1.351843673 | 0 | 0.011 | 1 | 0.496223879 | 0.683072586 | 0 | 0.01 | 1 | 0.63207476 | 0.74620962 |
| Skida1 | 0.632076134 | 1.673771768 | 0 | 0.011 | 1 | 0.496223879 | 0.683072586 | 0 | 0.01 | 1 | 0.632076134 | 0.74620962 |
| Ttc30a1 | 0.632076134 | 1.673771768 | 0 | 0.011 | 1 | 0.496223879 | 0.683072586 | 0 | 0.01 | 1 | 0.632076134 | 0.74620962 |
| Zfp661 | 0.632076134 | 1.673771768 | 0 | 0.011 | 1 | 0.496223879 | 0.683072586 | 0 | 0.01 | 1 | 0.632076134 | 0.74620962 |
| Myh7b | 0.632076134 | 1.673771768 | 0 | 0.011 | 1 | 0.496223879 | 0.683072586 | 0 | 0.01 | 1 | 0.632076134 | 0.74620962 |
| 4933429H19Rik | 0.632076134 | 1.673771768 | 0 | 0.011 | 1 | 0.496223879 | 0.683072586 | 0 | 0.01 | 1 | 0.632076134 | 0.74620962 |
| 4930589L23Rik | 0.632076134 | 1.673771768 | 0 | 0.011 | 1 | 0.496223879 | 0.683072586 | 0 | 0.01 | 1 | 0.632076134 | 0.74620962 |
| Gm42701 | 0.632076134 | 1.673771768 | 0 | 0.011 | 1 | 0.496223879 | 0.683072586 | 0 | 0.01 | 1 | 0.632076134 | 0.74620962 |
| Gm43017 | 0.632076134 | 1.673771768 | 0 | 0.011 | 1 | 0.496223879 | 0.683072586 | 0 | 0.01 | 1 | 0.632076134 | 0.74620962 |
| Tmbim7 | 0.632076134 | 1.673771768 | 0 | 0.011 | 1 | 0.496223879 | -0.776359032 | 0 | 0.01 | 1 | 0.632076134 | 0.74620962 |
| Gm15997 | 0.632076134 | 1.088809267 | 0 | 0.011 | 1 | 0.496223879 | 0.683072586 | 0 | 0.01 | 1 | 0.632076134 | 0.74620962 |
| Gm33318 | 0.632076134 | 1.673771768 | 0 | 0.011 | 1 | 0.496223879 | 0.683072586 | 0 | 0.01 | 1 | 0.632076134 | 0.74620962 |
| Ccdc142os | 0.632076134 | 1.673771768 | 0 | 0.011 | 1 | 0.496223879 | 0.683072586 | 0 | 0.01 | 1 | 0.632076134 | 0.74620962 |
| Nfkbid | 0.632076134 | 1.673771768 | 0 | 0.011 | 1 | 0.496223879 | 0.683072586 | 0 | 0.01 | 1 | 0.632076134 | 0.74620962 |
| Klk14 | 0.632076134 | 1.673771768 | 0 | 0.011 | 1 | 0.496223879 | 0.683072586 | 0 | 0.01 | 1 | 0.632076134 | 0.74620962 |
| 2310016G11Rik | 0.632076134 | 1.673771768 | 0 | 0.011 | 1 | 0.496223879 | 0.683072586 | 0 | 0.01 | 1 | 0.632076134 | 0.74620962 |
| Sec1 | 0.632076134 | 1.673771768 | 0 | 0.011 | 1 | 0.496223879 | 0.683072586 | 0 | 0.01 | 1 | 0.632076134 | 0.74620962 |
| Tpbgl | 0.632076134 | 1.673771768 | 0 | 0.011 | 1 | 0.496223879 | 0.683072586 | 0 | 0.01 | 1 | 0.632076134 | 0.74620962 |
| Gm45502 | 0.632076134 | 1.673771768 | 0 | 0.011 | 1 | 0.496223879 | 0.683072586 | 0 | 0.01 | 1 | 0.632076134 | 0.74620962 |
| Proscos | 0.632076134 | 1.673771768 | 0 | 0.011 | 1 | 0.496223879 | 0.683072586 | 0 | 0.01 | 1 | 0.632076134 | 0.74620962 |
| Angptl6 | 0.632076134 | 1.673771768 | 0 | 0.011 | 1 | 0.496223879 | 0.683072586 | 0 | 0.01 | 1 | 0.632076134 | 0.74620962 |
| Stra6 | 0.632076134 | 1.673771768 | 0 | 0.011 | 1 | 0.496223879 | 0.683072586 | 0 | 0.01 | 1 | 0.632076134 | 0.74620962 |
| Fmnl1 | 0.632076134 | 1.673771768 | 0 | 0.011 | 1 | 0.496223879 | 0.683072586 | 0 | 0.01 | 1 | 0.632076134 | 0.74620962 |
| 2210039B01Rik | 0.632076134 | 1.673771768 | 0 | 0.011 | 1 | 0.496223879 | 0.683072586 | 0 | 0.01 | 1 | 0.632076134 | 0.74620962 |
| Tmem63c | 0.632076134 | 1.673771768 | 0 | 0.011 | 1 | 0.496223879 | 0.683072586 | 0 | 0.01 | 1 | 0.632076134 | 0.74620962 |
| Gsc | 0.632076134 | 1.673771768 | 0 | 0.011 | 1 | 0.496223879 | 0.683072586 | 0 | 0.01 | 1 | 0.632076134 | 0.74620962 |
| D030051J21Rik | 0.632076134 | 1.673771768 | 0 | 0.011 | 1 | 0.496223879 | 0.683072586 | 0 | 0.01 | 1 | 0.632076134 | 0.74620962 |
| Gm50019 | 0.632076134 | 1.673771768 | 0 | 0.011 | 1 | 0.496223879 | 0.683072586 | 0 | 0.01 | 1 | 0.632076134 | 0.74620962 |
| Hspe1-rs1 | 0.632076134 | 1.673771768 | 0 | 0.011 | 1 | 0.496223879 | 0.683072586 | 0 | 0.01 | 1 | 0.632076134 | 0.74620962 |
| Gm50203 | 0.632076134 | 1.673771768 | 0 | 0.011 | 1 | 0.496223879 | 0.683072586 | 0 | 0.01 | 1 | 0.632076134 | 0.74620962 |
| Gm1141 | 0.632076134 | 1.673771768 | 0 | 0.011 | 1 | 0.496223879 | 0.683072586 | 0 | 0.01 | 1 | 0.632076134 | 0.74620962 |
| Atp2a3 | 0.715863632 | 1.55829455 | 0.048 | 0.034 | 1 | 0.496223879 | 0.683072586 | 0 | 0.01 | 1 | 0.715863632 | 0.74620962 |
| Xk | 0.738393266 | 1.351843673 | 0.048 | 0.034 | 1 | 0.496223879 | 0.683072586 | 0 | 0.01 | 1 | 0.738393266 | 0.74620962 |
| Cenpu | 0.748090146 | 0.586308926 | 0.048 | 0.034 | 1 | 0.496223879 | 0.683072586 | 0 | 0.01 | 1 | 0.748090146 | 0.74620962 |
| Apobec2 | 0.773953771 | 2.870808615 | 0.048 | 0.067 | 1 | 0.496223879 | 0.683072586 | 0 | 0.01 | 1 | 0.773953771 | 0.74620962 |
| Tmem108 | 0.787193017 | 1.97333205 | 0.048 | 0.037 | 1 | 0.496223879 | 0.683072586 | 0 | 0.01 | 1 | 0.787193017 | 0.74620962 |
| Skap1 | 0.79666227 | 1.351843673 | 0.048 | 0.037 | 1 | 0.496223879 | 0.683072586 | 0 | 0.01 | 1 | 0.79666227 | 0.74620962 |
| B530045E10Rik | 0.825197354 | 0.866416846 | 0.048 | 0.037 | 1 | 0.496223879 | -0.124282336 | 0 | 0.01 | 1 | 0.825197354 | 0.74620962 |
| Mkrn2os | 0.875318873 | 0.79930265 | 0.048 | 0.041 | 1 | 0.496223879 | -0.316927414 | 0 | 0.01 | 1 | 0.875318873 | 0.74620962 |
| Amd2 | 0.496414193 | 0.400753273 | 0.048 | 0.097 | 1 | 0.502638178 | 0.268035087 | 0.067 | 0.1 | 1 | 0.502638178 | 0.746401335 |
| Adamtsl4 | 0.49643143 | 0.808701348 | 0.19 | 0.142 | 1 | 0.74409818 | 0.410054092 | 0.156 | 0.194 | 1 | 0.74409818 | 0.746418696 |
| Sorbs2os | 0.496498663 | -0.25696557 | 0.095 | 0.153 | 1 | 0.71357271 | 0.296049463 | 0.156 | 0.19 | 1 | 0.71357271 | 0.746486404 |
| 4921536K21Rik | 0.496521286 | 1.281454345 | 0.095 | 0.06 | 1 | 0.986421662 | 0.513147585 | 0.067 | 0.069 | 1 | 0.986421662 | 0.746509185 |
| Mylip | 0.496608955 | -0.88788336 | 0.333 | 0.366 | 1 | 0.71312409 | -0.23952291 | 0.756 | 0.557 | 1 | 0.71312409 | 0.746597456 |
| Gm9725 | 0.773910509 | 1.410737362 | 0.048 | 0.067 | 1 | 0.497228696 | 0.268035087 | 0.022 | 0.045 | 1 | 0.773910509 | 0.747221016 |
| S100a5 | 0.957847914 | 0.451379346 | 0.048 | 0.045 | 1 | 0.497228696 | 0.683072586 | 0.022 | 0.045 | 1 | 0.957847914 | 0.747221016 |
| Gm56492 | 0.822006883 | 0.866416846 | 0.048 | 0.037 | 1 | 0.497244215 | 0.435145073 | 0.022 | 0.045 | 1 | 0.822006883 | 0.747236621 |
| Bpifc | 0.682763067 | 1.762308442 | 0.19 | 0.175 | 1 | 0.497256285 | 0.290755163 | 0.022 | 0.045 | 1 | 0.682763067 | 0.747248757 |
| Ptpre | 0.497284318 | 0.473344678 | 0.19 | 0.299 | 1 | 0.954799583 | 0.598539076 | 0.2 | 0.204 | 1 | 0.954799583 | 0.747276943 |
| Rasa3 | 0.601458541 | -0.254335314 | 0.476 | 0.56 | 1 | 0.497478668 | 0.244066439 | 0.444 | 0.581 | 1 | 0.601458541 | 0.747472311 |
| Zfp457 | 0.532426384 | 0.866416846 | 0 | 0.019 | 1 | 0.497611189 | 1.683072586 | 0.022 | 0.01 | 1 | 0.532426384 | 0.747605483 |
| 4930524B15Rik | 0.63207476 | 1.351843673 | 0 | 0.011 | 1 | 0.497611189 | 1.683072586 | 0.022 | 0.01 | 1 | 0.63207476 | 0.747605483 |
| Myrip | 0.616977656 | 0.614878079 | 0.048 | 0.082 | 1 | 0.497612165 | 0.776181991 | 0.022 | 0.01 | 1 | 0.616977656 | 0.747606463 |
| Phf14 | 0.498734991 | 0.100882099 | 0.429 | 0.575 | 1 | 0.739568151 | 0.134868635 | 0.756 | 0.74 | 1 | 0.739568151 | 0.748733391 |
| Slc17a9 | 0.655435954 | 0.780686972 | 0.095 | 0.134 | 1 | 0.499001871 | 0.125077133 | 0.089 | 0.128 | 1 | 0.655435954 | 0.749000875 |
| Galns | 0.499761607 | 1.451379346 | 0.048 | 0.097 | 1 | 0.500262731 | 1.005000681 | 0.022 | 0.045 | 1 | 0.500262731 | 0.74976155 |
| Tnfrsf1b | 0.499881461 | 1.088809267 | 0.048 | 0.097 | 1 | 0.861113159 | 1.256807831 | 0.111 | 0.125 | 1 | 0.861113159 | 0.749881447 |
| Map2k5 | 0.797835865 | 0.320816777 | 0.381 | 0.429 | 1 | 0.500108293 | 0.295002134 | 0.689 | 0.661 | 1 | 0.797835865 | 0.750108281 |
| Tmem37 | 0.603600115 | -0.280424543 | 0.048 | 0.078 | 1 | 0.500262731 | 0.683072586 | 0.022 | 0.045 | 1 | 0.603600115 | 0.750262662 |
| Pbld2 | 0.655561847 | 1.503846766 | 0.048 | 0.03 | 1 | 0.500277618 | 0.595609745 | 0.022 | 0.045 | 1 | 0.655561847 | 0.750277541 |
| Foxo6 | 0.578156445 | 1.088809267 | 0 | 0.015 | 1 | 0.500299088 | 0.946106992 | 0.022 | 0.045 | 1 | 0.578156445 | 0.750298998 |
| Gbp6 | 0.754700198 | 0.351843673 | 0.048 | 0.067 | 1 | 0.500316833 | 0.972579203 | 0.022 | 0.045 | 1 | 0.754700198 | 0.750316733 |
| Col18a1 | 0.532426384 | 0.866416846 | 0 | 0.019 | 1 | 0.50040834 | 1.683072586 | 0.022 | 0.01 | 1 | 0.532426384 | 0.750408173 |
| 6530401F13Rik | 0.632076134 | 1.088809267 | 0 | 0.011 | 1 | 0.50040834 | 1.683072586 | 0.022 | 0.01 | 1 | 0.632076134 | 0.750408173 |
| Myo18b | 0.68198598 | 0.108987149 | 0.048 | 0.075 | 1 | 0.50040834 | 1.683072586 | 0.022 | 0.01 | 1 | 0.68198598 | 0.750408173 |
| Plekhg1 | 0.950973449 | 0.576910228 | 0.095 | 0.104 | 1 | 0.50099047 | -0.619490184 | 0.2 | 0.239 | 1 | 0.950973449 | 0.750989489 |
| Kansl3 | 0.502332242 | 0.412104198 | 0.333 | 0.526 | 1 | 0.913260084 | 0.313183453 | 0.578 | 0.702 | 1 | 0.913260084 | 0.752326803 |
| Prkx | 0.695162779 | 0.699766976 | 0.19 | 0.269 | 1 | 0.502860632 | 0.131898399 | 0.533 | 0.439 | 1 | 0.695162779 | 0.752852449 |
| Actr2 | 0.684677749 | 0.446433524 | 0.857 | 0.832 | 1 | 0.502966214 | 0.257812704 | 0.933 | 0.927 | 1 | 0.684677749 | 0.752957415 |
| 9530034E10Rik | 0.63207476 | 1.351843673 | 0 | 0.011 | 1 | 0.503208953 | 1.361144491 | 0.022 | 0.01 | 1 | 0.63207476 | 0.753198655 |

| | | | | | | | | | | | | |
|---|---|---|---|---|---|---|---|---|---|---|---|---|
| Gm47772 | 0.63207476 | 1.088809267 | 0 | 0.011 | 1 | 0.503208953 | 1.098110085 | 0.022 | 0.01 | 1 | 0.63207476 | 0.753198655 |
| Gm52054 | 0.63207476 | 1.351843673 | 0 | 0.011 | 1 | 0.503208953 | 1.361144491 | 0.022 | 0.01 | 1 | 0.63207476 | 0.753198655 |
| Susd2 | 0.632076134 | 1.673771768 | 0 | 0.011 | 1 | 0.503208953 | 1.361144491 | 0.022 | 0.01 | 1 | 0.632076134 | 0.753198655 |
| Plcxd2 | 0.632076134 | 1.673771768 | 0 | 0.011 | 1 | 0.503208953 | 1.361144491 | 0.022 | 0.01 | 1 | 0.632076134 | 0.753198655 |
| Tonsl | 0.918802135 | 1.673771768 | 0.048 | 0.045 | 1 | 0.503208953 | 1.361144491 | 0.022 | 0.01 | 1 | 0.918802135 | 0.753198655 |
| Polr1b | 0.5276228 | 1.188344941 | 0.048 | 0.093 | 1 | 0.503239693 | 1.361144491 | 0.156 | 0.131 | 1 | 0.5276228 | 0.753229197 |
| Cask | 0.944069583 | 0.227515538 | 0.714 | 0.769 | 1 | 0.503504925 | 0.213045852 | 0.911 | 0.858 | 1 | 0.944069583 | 0.75349264 |
| Mrpl1 | 0.504089804 | 1.243783927 | 0.286 | 0.269 | 1 | 0.698177679 | 0.671484612 | 0.222 | 0.291 | 1 | 0.698177679 | 0.754073078 |
| Ttll7 | 0.504503973 | 0.760483401 | 0.286 | 0.239 | 1 | 0.948533234 | 0.995016592 | 0.356 | 0.478 | 1 | 0.948533234 | 0.754483687 |
| Wdr59 | 0.676660768 | 0.761827944 | 0.333 | 0.336 | 1 | 0.504793593 | 0.485771146 | 0.2 | 0.273 | 1 | 0.676660768 | 0.754770615 |
| Tshr | 0.57816106 | 1.351843673 | 0 | 0.015 | 1 | 0.5054846 | -1.404390255 | 0.044 | 0.069 | 1 | 0.57816106 | 0.75545452 |
| D430001F17Rik | 0.532426384 | 0.503846766 | 0 | 0.019 | 1 | 0.506024628 | 1.098110085 | 0.022 | 0.01 | 1 | 0.532426384 | 0.755988332 |
| Carf | 0.506607967 | 0.323274521 | 0.095 | 0.164 | 1 | 0.624823637 | 1.152557869 | 0.089 | 0.125 | 1 | 0.624823637 | 0.756564302 |
| Eps8 | 0.507273564 | 0.283825249 | 0.762 | 0.784 | 1 | 0.534463741 | 0.402820836 | 0.867 | 0.896 | 1 | 0.534463741 | 0.75722066 |
| Tnfsf13 | 0.507699835 | -0.184209227 | 0.048 | 0.09 | 1 | 0.721650485 | 0.205025289 | 0.089 | 0.107 | 1 | 0.721650485 | 0.757640547 |
| Dpm1 | 0.507750205 | 0.290643516 | 0.905 | 0.881 | 1 | 0.8255651 | 0.15325164 | 0.911 | 0.858 | 1 | 0.8255651 | 0.757690139 |
| Tfip11 | 0.628554427 | 0.472137907 | 0.095 | 0.146 | 1 | 0.509836081 | 0.180572246 | 0.089 | 0.128 | 1 | 0.628554427 | 0.759739332 |
| Fam167a | 0.526374442 | 0.510833197 | 0.333 | 0.287 | 1 | 0.510376539 | 1.27368165 | 0.178 | 0.152 | 1 | 0.526374442 | 0.760268866 |
| Anxa11 | 0.511061273 | 0.745267359 | 0.714 | 0.795 | 1 | 0.819143374 | 0.328843999 | 0.911 | 0.848 | 1 | 0.819143374 | 0.760938921 |
| Prkg1 | 0.51123443 | -0.473400283 | 0.667 | 0.743 | 1 | 0.595297374 | 0.201083622 | 0.933 | 0.945 | 1 | 0.595297374 | 0.761108217 |
| Cep43 | 0.5114152 | 0.683825432 | 0.238 | 0.358 | 1 | 0.986680784 | 0.4513423 | 0.333 | 0.353 | 1 | 0.986680784 | 0.761284893 |
| Btaf1 | 0.512251819 | 0.487636539 | 0.857 | 0.929 | 1 | 0.969368132 | 0.193073029 | 1 | 0.931 | 1 | 0.969368132 | 0.762101712 |
| Sptb | 0.632076134 | 1.673771768 | 0 | 0.011 | 1 | 0.512490525 | 2.320502507 | 0.044 | 0.028 | 1 | 0.632076134 | 0.762334512 |
| Best3 | 0.570211706 | 1.503846766 | 0.048 | 0.026 | 1 | 0.512513901 | 1.905465008 | 0.044 | 0.028 | 1 | 0.570211706 | 0.762357303 |
| Tnfsf13b | 0.896995455 | 0.673771768 | 0.048 | 0.041 | 1 | 0.512513901 | 2.098110085 | 0.044 | 0.028 | 1 | 0.896995455 | 0.762357303 |
| Pabpc1l | 0.636128533 | 1.79930265 | 0.048 | 0.03 | 1 | 0.512915392 | 1.442064487 | 0.089 | 0.066 | 1 | 0.636128533 | 0.762748585 |
| Trmt9b | 0.513158301 | 1.029915578 | 0.095 | 0.06 | 1 | 0.52275247 | 0.205025289 | 0.044 | 0.073 | 1 | 0.52275247 | 0.76298516 |
| Nup107 | 0.611828793 | 0.649925026 | 0.19 | 0.261 | 1 | 0.513870746 | 0.361144491 | 0.222 | 0.308 | 1 | 0.611828793 | 0.763678349 |
| Igsf9b | 0.787201474 | 1.866416846 | 0.048 | 0.037 | 1 | 0.514004217 | 0.946106992 | 0.067 | 0.045 | 1 | 0.787201474 | 0.763808098 |
| Gm57200 | 0.514089901 | 1.64135029 | 0.143 | 0.108 | 1 | 0.723482561 | 0.699560709 | 0.178 | 0.17 | 1 | 0.723482561 | 0.763891376 |
| Cstpp1 | 0.79271747 | 0.645574876 | 0.286 | 0.31 | 1 | 0.514167028 | 0.172699402 | 0.289 | 0.394 | 1 | 0.79271747 | 0.763966323 |
| Stpg4 | 0.638246407 | -0.413691074 | 0.048 | 0.075 | 1 | 0.514179096 | 1.683072586 | 0.089 | 0.066 | 1 | 0.638246407 | 0.763978049 |
| Gm4876 | 0.514392257 | 0.79930265 | 0.095 | 0.164 | 1 | 0.946634112 | 0.435145073 | 0.089 | 0.09 | 1 | 0.946634112 | 0.76418512 |
| Mcub | 0.578396207 | 1.951305743 | 0.19 | 0.16 | 1 | 0.514532769 | 1.15951063 | 0.178 | 0.156 | 1 | 0.578396207 | 0.764321567 |
| 5330438D12Rik | 0.789446691 | 0.243783927 | 0.238 | 0.28 | 1 | 0.514794581 | 0.577719586 | 0.244 | 0.349 | 1 | 0.789446691 | 0.764575701 |
| Gli2 | 0.515623145 | -0.560283571 | 0.238 | 0.325 | 1 | 0.903712879 | 0.83507568 | 0.378 | 0.505 | 1 | 0.903712879 | 0.765379062 |
| Klrg2 | 0.660317509 | 1.866416846 | 0.095 | 0.075 | 1 | 0.516774949 | 1.258574758 | 0.089 | 0.135 | 1 | 0.660317509 | 0.76649355 |
| Arfgap3 | 0.516804198 | -0.108559131 | 0.667 | 0.862 | 1 | 0.66654908 | 0.101833171 | 0.889 | 0.907 | 1 | 0.66654908 | 0.766521817 |
| Supv3l1 | 0.516893784 | 0.264380832 | 0.143 | 0.216 | 1 | 0.57378629 | 0.614901084 | 0.222 | 0.294 | 1 | 0.57378629 | 0.766608384 |
| Crtc1 | 0.583228011 | 1.001346426 | 0.095 | 0.153 | 1 | 0.516930435 | 0.99005034 | 0.267 | 0.246 | 1 | 0.583228011 | 0.766643795 |
| Cd59b | 0.517057607 | 0.214340149 | 0.048 | 0.093 | 1 | 0.733079378 | 1.329435632 | 0.156 | 0.152 | 1 | 0.733079378 | 0.766766645 |
| Lrpprc | 0.7394969 | 0.664782984 | 0.333 | 0.354 | 1 | 0.517784353 | 0.391841289 | 0.578 | 0.512 | 1 | 0.7394969 | 0.76746807 |
| Adgrg3 | 0.632076134 | 1.673771768 | 0 | 0.011 | 1 | 0.517959493 | 2.242499995 | 0.044 | 0.028 | 1 | 0.632076134 | 0.76763695 |
| Dnm3os | 0.770233275 | 0.311201688 | 0.143 | 0.127 | 1 | 0.518144996 | -0.638855509 | 0.089 | 0.121 | 1 | 0.770233275 | 0.767815755 |
| Gm46412 | 0.518826147 | 0.304537958 | 0.048 | 0.093 | 1 | 0.804016776 | 0.744473131 | 0.044 | 0.055 | 1 | 0.804016776 | 0.768471723 |
| Chaserr | 0.666567231 | 0.214340149 | 0.81 | 0.825 | 1 | 0.518847898 | 0.266886901 | 0.933 | 0.893 | 1 | 0.666567231 | 0.768492655 |
| Stim1 | 0.519099697 | 0.414209554 | 0.333 | 0.507 | 1 | 0.576531786 | -0.439324045 | 0.756 | 0.633 | 1 | 0.576531786 | 0.768734898 |
| Flt1 | 0.725502576 | 1.150209812 | 0.048 | 0.034 | 1 | 0.519781311 | 2.055041364 | 0.044 | 0.028 | 1 | 0.725502576 | 0.769390011 |
| Evi5 | 0.545719712 | 0.257280671 | 0.429 | 0.552 | 1 | 0.519784081 | 0.37112858 | 0.778 | 0.723 | 1 | 0.545719712 | 0.769392672 |
| Acyp2 | 0.90639346 | 1.151545022 | 0.429 | 0.496 | 1 | 0.52010642 | 0.600610426 | 0.289 | 0.381 | 1 | 0.90639346 | 0.769702152 |
| Man2b1 | 0.520292302 | 0.472137907 | 0.238 | 0.396 | 1 | 0.759766328 | -0.191396532 | 0.511 | 0.502 | 1 | 0.759766328 | 0.769880524 |
| Ifit3b | 0.520549194 | -1.574155746 | 0.048 | 0.09 | 1 | 0.636641849 | 3.174925683 | 0.044 | 0.031 | 1 | 0.636641849 | 0.770126924 |
| Trmt1l | 0.671491058 | 0.481904213 | 0.476 | 0.522 | 1 | 0.520628754 | 0.34917185 | 0.556 | 0.574 | 1 | 0.671491058 | 0.770203208 |
| Dot1l | 0.867980108 | 0.178716239 | 0.571 | 0.698 | 1 | 0.520982832 | 0.430152258 | 0.844 | 0.824 | 1 | 0.867980108 | 0.770542553 |
| 0610043K17Rik | 0.649078652 | 0.694731386 | 0.381 | 0.396 | 1 | 0.521227393 | 0.902385615 | 0.289 | 0.284 | 1 | 0.649078652 | 0.77077679 |
| Slfn9 | 0.526139068 | 0.926537838 | 0.048 | 0.093 | 1 | 0.52124944 | 1.605070074 | 0.067 | 0.045 | 1 | 0.526139068 | 0.770797901 |
| Pcdhga12 | 0.768522056 | 1.316219763 | 0.143 | 0.138 | 1 | 0.521332548 | 1.683072586 | 0.111 | 0.087 | 1 | 0.768522056 | 0.770877471 |
| Rin3 | 0.521436269 | 0.866416846 | 0.095 | 0.16 | 1 | 0.547342988 | 0.456003678 | 0.156 | 0.211 | 1 | 0.547342988 | 0.770976755 |
| Zfp839 | 0.750334736 | 0.673771768 | 0.095 | 0.131 | 1 | 0.521740419 | 0.435145073 | 0.067 | 0.1 | 1 | 0.750334736 | 0.771267773 |
| Spopl | 0.74941989 | 0.52282987 | 0.286 | 0.381 | 1 | 0.521798073 | 0.464649067 | 0.333 | 0.491 | 1 | 0.74941989 | 0.771322917 |
| Svopl | 0.818172884 | 2.614878079 | 0.048 | 0.063 | 1 | 0.522026725 | 0.423510372 | 0.244 | 0.325 | 1 | 0.818172884 | 0.771541548 |
| Gm38551 | 0.896555897 | 0.673771768 | 0.048 | 0.056 | 1 | 0.522791571 | 0.435145073 | 0.044 | 0.073 | 1 | 0.896555897 | 0.772272115 |
| Mto1 | 0.653872794 | 0.59848364 | 0.19 | 0.16 | 1 | 0.522852487 | 0.683072586 | 0.133 | 0.187 | 1 | 0.653872794 | 0.772330251 |
| Ptcra | 0.709478182 | 2.536268244 | 0.048 | 0.034 | 1 | 0.523426772 | 1.42003818 | 0.044 | 0.028 | 1 | 0.709478182 | 0.772877958 |
| Dgkd | 0.976514995 | 0.85064953 | 0.333 | 0.399 | 1 | 0.523459943 | 0.110182918 | 0.533 | 0.446 | 1 | 0.976514995 | 0.772909574 |
| Armc8 | 0.523594766 | 0.123574685 | 0.286 | 0.384 | 1 | 0.808869315 | 0.531526114 | 0.467 | 0.602 | 1 | 0.808869315 | 0.773038053 |
| Ntn1 | 0.523765324 | 2.155923463 | 0.095 | 0.063 | 1 | 0.55778165 | 0.723714571 | 0.089 | 0.066 | 1 | 0.55778165 | 0.773200533 |
| Cyb5d1 | 0.524054884 | 0.451379346 | 0.048 | 0.093 | 1 | 0.61127372 | 1.235613609 | 0.067 | 0.097 | 1 | 0.61127372 | 0.773476246 |
| Elovl6 | 0.757061473 | 0.64357153 | 0.619 | 0.772 | 1 | 0.524334867 | 0.136584233 | 0.533 | 0.564 | 1 | 0.757061473 | 0.773742682 |
| Kazn | 0.524805208 | -0.484657595 | 0.143 | 0.213 | 1 | 0.757837792 | 0.563773658 | 0.178 | 0.208 | 1 | 0.757837792 | 0.774189909 |
| Zfp697 | 0.524989558 | -0.194124696 | 0.095 | 0.153 | 1 | 0.670511082 | 1.390290837 | 0.133 | 0.125 | 1 | 0.670511082 | 0.77436508 |
| Hdac5 | 0.955028872 | 0.185485286 | 0.238 | 0.265 | 1 | 0.525992256 | 0.325520582 | 0.289 | 0.388 | 1 | 0.955028872 | 0.775316659 |
| Prss12 | 0.532435374 | -0.849790188 | 0 | 0.019 | 1 | 0.526080233 | -0.901889915 | 0.022 | 0.042 | 1 | 0.532435374 | 0.775400054 |
| Slc4a4 | 0.526195972 | 0.492626228 | 0.524 | 0.582 | 1 | 0.886037946 | -0.144634171 | 0.689 | 0.664 | 1 | 0.886037946 | 0.775509743 |
| Ube2cbp | 0.707125428 | 1.815790773 | 0.095 | 0.078 | 1 | 0.526466235 | 0.683072586 | 0.044 | 0.073 | 1 | 0.707125428 | 0.775765773 |
| Rab43 | 0.526540059 | 0.357403198 | 0.286 | 0.44 | 1 | 0.689690849 | 0.423205459 | 0.667 | 0.654 | 1 | 0.689690849 | 0.775835684 |
| Vstm2l | 0.57816106 | 1.351843673 | 0 | 0.015 | 1 | 0.527061766 | 1.513147585 | 0.044 | 0.028 | 1 | 0.57816106 | 0.776329427 |
| Tcp11 | 0.614796124 | 1.55829455 | 0.095 | 0.071 | 1 | 0.527061766 | 1.513147585 | 0.044 | 0.028 | 1 | 0.614796124 | 0.776329427 |
| B230216N24Rik | 0.873611511 | 0.214340149 | 0.143 | 0.127 | 1 | 0.527645442 | 0.99005034 | 0.244 | 0.242 | 1 | 0.873611511 | 0.776881171 |
| 1600010M07Rik | 0.527893858 | 0.446702859 | 0.19 | 0.272 | 1 | 0.957477628 | 0.941990884 | 0.244 | 0.294 | 1 | 0.957477628 | 0.777115791 |
| Nr4a1 | 0.527962349 | 0.32771201 | 0.714 | 0.634 | 1 | 0.901983057 | -0.382515755 | 0.867 | 0.723 | 1 | 0.901983057 | 0.777180456 |
| Slc20a2 | 0.528440943 | 0.720565979 | 0.524 | 0.537 | 1 | 0.887361266 | 0.251356346 | 0.289 | 0.343 | 1 | 0.887361266 | 0.777632056 |
| Gm43768 | 0.532436372 | 1.088809267 | 0 | 0.019 | 1 | 0.528913413 | 1.683072586 | 0.044 | 0.028 | 1 | 0.532436372 | 0.778077428 |
| Mterf1b | 0.532436372 | 1.088809267 | 0 | 0.019 | 1 | 0.528913413 | 2.168499413 | 0.044 | 0.028 | 1 | 0.532436372 | 0.778077428 |
| Evi2b | 0.573468511 | 1.351843673 | 0.048 | 0.026 | 1 | 0.528913413 | 1.905465008 | 0.044 | 0.028 | 1 | 0.573468511 | 0.778077428 |
| Gm10785 | 0.529242006 | -0.849790188 | 0.048 | 0.086 | 1 | 0.917404477 | 1.410054092 | 0.067 | 0.076 | 1 | 0.917404477 | 0.778386911 |
| Pacrg | 0.824249521 | 1.2970512 | 0.238 | 0.243 | 1 | 0.529409773 | 0.788941221 | 0.156 | 0.218 | 1 | 0.824249521 | 0.778544838 |
| Uqcc1 | 0.529586278 | 0.719575457 | 0.524 | 0.493 | 1 | 0.683317612 | 0.483027596 | 0.422 | 0.567 | 1 | 0.683317612 | 0.77871093 |
| Mtf2 | 0.97877029 | 0.357403198 | 0.381 | 0.481 | 1 | 0.530048276 | 0.166281588 | 0.711 | 0.633 | 1 | 0.97877029 | 0.779145377 |
| Ptpra | 0.530350549 | 0.112494797 | 0.905 | 0.94 | 1 | 0.602122331 | 0.312234891 | 0.956 | 0.958 | 1 | 0.602122331 | 0.779429393 |
| Zfp78 | 0.578156445 | 1.088809267 | 0 | 0.015 | 1 | 0.530911177 | -0.316927414 | 0.022 | 0.042 | 1 | 0.578156445 | 0.779955677 |
| Gm26759 | 0.985459372 | 1.258734268 | 0.048 | 0.049 | 1 | 0.532293594 | 0.875717664 | 0.067 | 0.1 | 1 | 0.985459372 | 0.781250717 |
| A830018L16Rik | 0.532426384 | 0.866416846 | 0 | 0.019 | 1 | 0.947540957 | 0.91753784 | 0.022 | 0.024 | 1 | 0.947540957 | 0.781374914 |
| B3galt2 | 0.532426384 | 0.673771768 | 0 | 0.019 | 1 | 0.6175503 | 0.776181991 | 0.022 | 0.038 | 1 | 0.6175503 | 0.781374914 |
| Fam78b | 0.532426384 | 0.866416846 | 0 | 0.019 | 1 | 0.849825041 | 1.098110085 | 0.022 | 0.028 | 1 | 0.849825041 | 0.781374914 |
| Mal | 0.532426384 | 0.866416846 | 0 | 0.019 | 1 | 0.842877623 | 0.361144491 | 0.022 | 0.028 | 1 | 0.842877623 | 0.781374914 |
| 5430402O13Rik | 0.532426384 | 0.866416846 | 0 | 0.019 | 1 | 0.936023974 | 1.460680165 | 0.022 | 0.021 | 1 | 0.936023974 | 0.781374914 |
| Gm29443 | 0.532426384 | 0.503846766 | 0 | 0.019 | 1 | 0.811836522 | 0.946106992 | 0.022 | 0.017 | 1 | 0.811836522 | 0.781374914 |

| | | | | | | | | | | | | |
|---|---|---|---|---|---|---|---|---|---|---|---|---|
| Cd79a | 0.532426384 | -1.496153234 | 0 | 0.019 | 1 | 0.874225478 | 0.776181991 | 0.067 | 0.076 | 1 | 0.874225478 | 0.781374914 |
| Pgbd5 | 0.532426384 | 0.673771768 | 0 | 0.019 | 1 | 0.823042771 | 0.875717664 | 0.022 | 0.017 | 1 | 0.823042771 | 0.781374914 |
| Slc6a20a | 0.532426384 | 0.673771768 | 0 | 0.019 | 1 | 0.806247793 | 1.513147585 | 0.022 | 0.017 | 1 | 0.806247793 | 0.781374914 |
| Cfap54 | 0.532426384 | 0.866416846 | 0 | 0.019 | 1 | 0.854466828 | 1.304560963 | 0.022 | 0.028 | 1 | 0.854466828 | 0.781374914 |
| 9130230N09Rik | 0.532426384 | 0.866416846 | 0 | 0.019 | 1 | 0.835943453 | 0.513147585 | 0.022 | 0.028 | 1 | 0.835943453 | 0.781374914 |
| D630024D03Rik | 0.532426384 | 0.866416846 | 0 | 0.019 | 1 | 0.854457549 | 1.513147585 | 0.022 | 0.028 | 1 | 0.854457549 | 0.781374914 |
| Marchf10 | 0.532426384 | 0.866416846 | 0 | 0.019 | 1 | 0.658583583 | 2.946106992 | 0.022 | 0.014 | 1 | 0.658583583 | 0.781374914 |
| Gm6566 | 0.532426384 | 0.866416846 | 0 | 0.019 | 1 | 0.80624346 | 2.098110085 | 0.022 | 0.017 | 1 | 0.80624346 | 0.781374914 |
| 1700023H06Rik | 0.532426384 | 0.866416846 | 0 | 0.019 | 1 | 0.80624346 | 1.683072586 | 0.022 | 0.017 | 1 | 0.80624346 | 0.781374914 |
| Gpr18 | 0.532426384 | 0.503846766 | 0 | 0.019 | 1 | 0.672920453 | 0.875717664 | 0.022 | 0.014 | 1 | 0.672920453 | 0.781374914 |
| Gm15564 | 0.532426384 | 0.866416846 | 0 | 0.019 | 1 | 0.611930508 | 1.808603468 | 0.044 | 0.031 | 1 | 0.611930508 | 0.781374914 |
| Rsph1 | 0.532426384 | 0.503846766 | 0 | 0.019 | 1 | 0.972326404 | 1.402964667 | 0.044 | 0.045 | 1 | 0.972326404 | 0.781374914 |
| Gm56488 | 0.532426384 | 0.866416846 | 0 | 0.019 | 1 | 0.959990133 | -0.124282336 | 0.022 | 0.021 | 1 | 0.959990133 | 0.781374914 |
| Gm39526 | 0.532426384 | 0.866416846 | 0 | 0.019 | 1 | 0.80624346 | 1.683072586 | 0.022 | 0.017 | 1 | 0.80624346 | 0.781374914 |
| Gm9530 | 0.532432377 | 0.673771768 | 0 | 0.019 | 1 | 0.936023974 | 2.197645759 | 0.022 | 0.021 | 1 | 0.936023974 | 0.781380518 |
| C1ql3 | 0.532432377 | 0.673771768 | 0 | 0.019 | 1 | 0.849821852 | 0.808603468 | 0.022 | 0.028 | 1 | 0.849821852 | 0.781380518 |
| Kcnd3 | 0.532432377 | 0.503846766 | 0 | 0.019 | 1 | 0.658583583 | 1.946106992 | 0.022 | 0.014 | 1 | 0.658583583 | 0.781380518 |
| Rhd | 0.532432377 | 0.673771768 | 0 | 0.019 | 1 | 0.80624346 | 2.683072586 | 0.022 | 0.017 | 1 | 0.80624346 | 0.781380518 |
| Tnfrsf9 | 0.532432377 | 0.351843673 | 0 | 0.019 | 1 | 0.664307093 | 1.460680165 | 0.022 | 0.014 | 1 | 0.664307093 | 0.781380518 |
| Chd5 | 0.532432377 | 0.673771768 | 0 | 0.019 | 1 | 0.799108542 | 0.567595369 | 0.044 | 0.055 | 1 | 0.799108542 | 0.781380518 |
| Gm20559 | 0.532432377 | 0.214340149 | 0 | 0.019 | 1 | 0.732110016 | 1.098110085 | 0.044 | 0.035 | 1 | 0.732110016 | 0.781380518 |
| Atp8b3 | 0.532432377 | 0.503846766 | 0 | 0.019 | 1 | 0.972326404 | 1.765534746 | 0.044 | 0.045 | 1 | 0.972326404 | 0.781380518 |
| Aatk | 0.532432377 | 0.673771768 | 0 | 0.019 | 1 | 0.955025007 | 1.83507568 | 0.022 | 0.024 | 1 | 0.955025007 | 0.781380518 |
| Serpinb6b | 0.532432377 | 0.503846766 | 0 | 0.019 | 1 | 0.936023974 | 2.197645759 | 0.022 | 0.021 | 1 | 0.936023974 | 0.781380518 |
| Rpl3l | 0.532432377 | 0.673771768 | 0 | 0.019 | 1 | 0.672920453 | 0.875717664 | 0.022 | 0.014 | 1 | 0.672920453 | 0.781380518 |
| Hs3st6 | 0.532432377 | 0.503846766 | 0 | 0.019 | 1 | 0.759313649 | 0.290755163 | 0.022 | 0.031 | 1 | 0.759313649 | 0.781380518 |
| Gm38825 | 0.532434375 | 0.351843673 | 0 | 0.019 | 1 | 0.951997643 | 0.875717664 | 0.022 | 0.021 | 1 | 0.951997643 | 0.781382386 |
| Gm28055 | 0.532435374 | 0.503846766 | 0 | 0.019 | 1 | 0.734040487 | 0.875717664 | 0.044 | 0.035 | 1 | 0.734040487 | 0.78138332 |
| Cfap299 | 0.532435374 | -0.849790188 | 0 | 0.019 | 1 | 0.95502578 | 2.42003818 | 0.022 | 0.024 | 1 | 0.95502578 | 0.78138332 |
| H2ac4 | 0.532435374 | -1.455511249 | 0 | 0.019 | 1 | 0.854457549 | 1.098110085 | 0.022 | 0.028 | 1 | 0.854457549 | 0.78138332 |
| Gm42658 | 0.532436372 | 1.088809267 | 0 | 0.019 | 1 | 0.854738731 | 0.513147585 | 0.044 | 0.052 | 1 | 0.854738731 | 0.781384254 |
| Morn3 | 0.532436372 | 1.088809267 | 0 | 0.019 | 1 | 0.854457549 | 1.83507568 | 0.022 | 0.028 | 1 | 0.854457549 | 0.781384254 |
| Gm17135 | 0.532436372 | 1.088809267 | 0 | 0.019 | 1 | 0.936023974 | 1.875717664 | 0.022 | 0.021 | 1 | 0.936023974 | 0.781384254 |
| Gm43915 | 0.532436372 | 1.088809267 | 0 | 0.019 | 1 | 0.8202424 | 1.098110085 | 0.022 | 0.017 | 1 | 0.8202424 | 0.781384254 |
| 4930557K07Rik | 0.532436372 | 1.088809267 | 0 | 0.019 | 1 | 0.937567971 | 0.683072586 | 0.022 | 0.024 | 1 | 0.937567971 | 0.781384254 |
| Gm12758 | 0.532436372 | 1.088809267 | 0 | 0.019 | 1 | 0.80624346 | 2.098110085 | 0.022 | 0.017 | 1 | 0.80624346 | 0.781384254 |
| Aqp11 | 0.532436372 | 1.088809267 | 0 | 0.019 | 1 | 0.687216983 | 0.982632868 | 0.022 | 0.035 | 1 | 0.687216983 | 0.781384254 |
| Gm44759 | 0.532436372 | 1.088809267 | 0 | 0.019 | 1 | 0.854457549 | 1.098110085 | 0.022 | 0.028 | 1 | 0.854457549 | 0.781384254 |
| Gm57017 | 0.532436372 | 1.088809267 | 0 | 0.019 | 1 | 0.765720982 | 1.361144491 | 0.022 | 0.031 | 1 | 0.765720982 | 0.781384254 |
| Iho1 | 0.532436372 | 1.088809267 | 0 | 0.019 | 1 | 0.941347173 | 1.223640968 | 0.022 | 0.021 | 1 | 0.941347173 | 0.781384254 |
| Lgr5 | 0.532436372 | 1.088809267 | 0 | 0.019 | 1 | 0.669552268 | -0.805213895 | 0.022 | 0.035 | 1 | 0.669552268 | 0.781384254 |
| Efcab9 | 0.532436372 | 1.088809267 | 0 | 0.019 | 1 | 0.955023924 | 1.683072586 | 0.022 | 0.024 | 1 | 0.955023924 | 0.781384254 |
| Gm56531 | 0.532436372 | 1.088809267 | 0 | 0.019 | 1 | 0.536315951 | 1.197645759 | 0.044 | 0.028 | 1 | 0.536315951 | 0.781384254 |
| Gm54621 | 0.532436372 | 1.088809267 | 0 | 0.019 | 1 | 0.595927847 | -0.124282336 | 0.022 | 0.038 | 1 | 0.595927847 | 0.781384254 |
| Mapk12 | 0.532436372 | 1.088809267 | 0 | 0.019 | 1 | 0.765728546 | 1.223640968 | 0.022 | 0.031 | 1 | 0.765728546 | 0.781384254 |
| A230028O05Rik | 0.532436372 | 1.088809267 | 0 | 0.019 | 1 | 0.746512944 | 0.361144491 | 0.022 | 0.031 | 1 | 0.746512944 | 0.781384254 |
| Gm16278 | 0.532436372 | 1.088809267 | 0 | 0.019 | 1 | 0.947540055 | 0.361144491 | 0.022 | 0.024 | 1 | 0.947540055 | 0.781384254 |
| Rps6ka4 | 0.532436372 | 1.088809267 | 0 | 0.019 | 1 | 0.667598499 | 0.223640968 | 0.022 | 0.035 | 1 | 0.667598499 | 0.781384254 |
| Arhgap27 | 0.57816106 | 0.673771768 | 0 | 0.015 | 1 | 0.532520164 | -0.223818009 | 0.022 | 0.042 | 1 | 0.57816106 | 0.781462603 |
| Gm3331 | 0.918519165 | 0.214340149 | 0.048 | 0.052 | 1 | 0.532520164 | -0.316927414 | 0.022 | 0.042 | 1 | 0.918519165 | 0.781462603 |
| Mtor | 0.53261515 | 1.123574685 | 0.333 | 0.325 | 1 | 0.949417597 | 0.517062635 | 0.311 | 0.384 | 1 | 0.949417597 | 0.781551402 |
| Bloc1s2 | 0.967831287 | 0.673771768 | 0.19 | 0.209 | 1 | 0.533031162 | 0.470078863 | 0.111 | 0.152 | 1 | 0.967831287 | 0.781940104 |
| Kcnj15 | 0.533258563 | -0.326228232 | 0.048 | 0.09 | 1 | 0.848250833 | -0.738754079 | 0.422 | 0.36 | 1 | 0.848250833 | 0.782152431 |
| F5 | 0.533306163 | -0.540758536 | 0.381 | 0.444 | 1 | 0.981311799 | 0.25186615 | 0.089 | 0.09 | 1 | 0.981311799 | 0.782196862 |
| Cacna1c | 0.533630613 | 0.348412767 | 0.381 | 0.34 | 1 | 0.599516535 | 0.161374979 | 0.378 | 0.453 | 1 | 0.599516535 | 0.782499595 |
| Gdf9 | 0.784047982 | 2.410737362 | 0.048 | 0.037 | 1 | 0.534128516 | -0.124282336 | 0.022 | 0.042 | 1 | 0.784047982 | 0.78296376 |
| Pcdhgc4 | 0.813144372 | 0.936806174 | 0.048 | 0.063 | 1 | 0.534128516 | -0.316927414 | 0.022 | 0.042 | 1 | 0.813144372 | 0.78296376 |
| Pde1c | 0.534162725 | 1.107071473 | 0.333 | 0.474 | 1 | 0.861275722 | -0.819427754 | 0.311 | 0.256 | 1 | 0.861275722 | 0.782995633 |
| 4930593A02Rik | 0.800634017 | 0.503846766 | 0.048 | 0.063 | 1 | 0.53446475 | 1.304560963 | 0.044 | 0.028 | 1 | 0.800634017 | 0.783276931 |
| Myt1l | 0.632076134 | 1.673771768 | 0 | 0.011 | 1 | 0.534474944 | 0.982632868 | 0.044 | 0.028 | 1 | 0.632076134 | 0.783286422 |
| 3110009E18Rik | 0.534722409 | -0.455511249 | 0.048 | 0.086 | 1 | 0.827714954 | 0.852997588 | 0.089 | 0.083 | 1 | 0.827714954 | 0.783516763 |
| Prag1 | 0.535271925 | 0.304537958 | 0.095 | 0.149 | 1 | 0.592392185 | -0.316927414 | 0.022 | 0.038 | 1 | 0.592392185 | 0.784027816 |
| Ptcd2 | 0.727523208 | 0.79930265 | 0.048 | 0.071 | 1 | 0.535371089 | 0.960606562 | 0.067 | 0.1 | 1 | 0.727523208 | 0.784119975 |
| Krit1 | 0.645440866 | 1.027254596 | 0.524 | 0.701 | 1 | 0.536228488 | 0.311255615 | 0.756 | 0.734 | 1 | 0.645440866 | 0.784915985 |
| Sfxn4 | 0.68623671 | 0.281454345 | 0.048 | 0.071 | 1 | 0.53630117 | 1.039216396 | 0.044 | 0.073 | 1 | 0.68623671 | 0.784983395 |
| Tmem106a | 0.860084871 | 1.010806755 | 0.048 | 0.06 | 1 | 0.536918291 | 1.808603468 | 0.089 | 0.069 | 1 | 0.860084871 | 0.785555331 |
| Lpcat3 | 0.537514909 | 0.319422195 | 0.333 | 0.481 | 1 | 0.833478114 | 0.526796646 | 0.467 | 0.467 | 1 | 0.833478114 | 0.786107541 |
| Wbp1l | 0.537639977 | 0.60964143 | 0.381 | 0.34 | 1 | 0.689257478 | 0.185294656 | 0.711 | 0.675 | 1 | 0.689257478 | 0.786223209 |
| Sptlc2 | 0.537928102 | 0.407531048 | 0.762 | 0.877 | 1 | 0.76274891 | 0.439830907 | 0.867 | 0.82 | 1 | 0.76274891 | 0.786489561 |
| Gm49417 | 0.960020107 | 0.133203386 | 0.19 | 0.205 | 1 | 0.538154599 | 0.509740983 | 0.289 | 0.273 | 1 | 0.960020107 | 0.786698826 |
| Foxp2 | 0.538821657 | 0.23435011 | 0.238 | 0.351 | 1 | 0.943125744 | 0.235861516 | 0.4 | 0.422 | 1 | 0.943125744 | 0.787314536 |
| Nectin3 | 0.657128966 | 1.017148282 | 0.381 | 0.418 | 1 | 0.538892447 | 0.201945896 | 0.4 | 0.564 | 1 | 0.657128966 | 0.787379825 |
| Thumpd3 | 0.539438567 | 0.157196242 | 0.19 | 0.265 | 1 | 0.583560939 | 0.444285727 | 0.2 | 0.27 | 1 | 0.583560939 | 0.787883166 |
| Vldlr | 0.888915917 | 1.548240886 | 0.095 | 0.093 | 1 | 0.542905243 | 0.808603468 | 0.067 | 0.097 | 1 | 0.888915917 | 0.791064383 |
| Aaas | 0.543133336 | 1.320134813 | 0.095 | 0.16 | 1 | 0.804360874 | 1.442064487 | 0.067 | 0.083 | 1 | 0.804360874 | 0.791272852 |
| Nmb | 0.57816106 | 1.351843673 | 0 | 0.015 | 1 | 0.543134781 | -0.280401538 | 0.044 | 0.069 | 1 | 0.57816106 | 0.791274171 |
| Mapre2 | 0.543372759 | 1.015560285 | 0.619 | 0.765 | 1 | 0.818126772 | 0.428755397 | 0.8 | 0.772 | 1 | 0.818126772 | 0.791491563 |
| Lrrc75aos2 | 0.544334756 | 1.214340149 | 0.095 | 0.063 | 1 | 0.707742902 | 0.536231198 | 0.067 | 0.087 | 1 | 0.707742902 | 0.792369185 |
| 4930534D22Rik | 0.544576436 | 2.258734268 | 0.048 | 0.026 | 1 | 0.852141123 | 0.946106992 | 0.022 | 0.028 | 1 | 0.852141123 | 0.792589378 |
| 1700001P01Rik | 0.544576436 | 2.258734268 | 0.048 | 0.026 | 1 | 0.705150377 | 2.03099589 | 0.044 | 0.035 | 1 | 0.705150377 | 0.792589378 |
| Zfp811 | 0.544576436 | 2.258734268 | 0.048 | 0.026 | 1 | 0.685254641 | 0.683072586 | 0.022 | 0.035 | 1 | 0.685254641 | 0.792589378 |
| Pask | 0.544589441 | 2.503846766 | 0.048 | 0.026 | 1 | 0.940060108 | 0.15951063 | 0.022 | 0.024 | 1 | 0.940060108 | 0.792601223 |
| Gm30173 | 0.544589441 | 3.503846766 | 0.048 | 0.026 | 1 | 0.806246349 | 1.875717664 | 0.022 | 0.017 | 1 | 0.806246349 | 0.792601223 |
| Appbp2os | 0.544589441 | 2.503846766 | 0.048 | 0.026 | 1 | 0.765720982 | 1.361144491 | 0.022 | 0.031 | 1 | 0.765720982 | 0.792601223 |
| Tph2 | 0.544606159 | 3.936806174 | 0.048 | 0.026 | 1 | 0.968638537 | 1.513147585 | 0.044 | 0.045 | 1 | 0.968638537 | 0.79261645 |
| Zfp385c | 0.545728451 | 1.243783927 | 0.143 | 0.228 | 1 | 0.903835924 | 0.701219933 | 0.267 | 0.291 | 1 | 0.903835924 | 0.79363736 |
| Dnaaf5 | 0.802985311 | 0.918884266 | 0.095 | 0.082 | 1 | 0.545989229 | 1.429315994 | 0.111 | 0.09 | 1 | 0.802985311 | 0.79387422 |
| B230206L02Rik | 0.547187991 | 1.811275291 | 0.143 | 0.112 | 1 | 0.689902534 | 0.448607333 | 0.2 | 0.187 | 1 | 0.689902534 | 0.794961284 |
| Morn1 | 0.823255524 | 0.458758877 | 0.143 | 0.183 | 1 | 0.547263716 | 1.082168542 | 0.222 | 0.204 | 1 | 0.823255524 | 0.795029857 |
| Csmd1 | 0.57816106 | 1.351843673 | 0 | 0.015 | 1 | 0.547298125 | -1.486852415 | 0.022 | 0.042 | 1 | 0.57816106 | 0.795061012 |
| Pdzk1 | 0.54776222 | 2.088809267 | 0.048 | 0.026 | 1 | 0.950033996 | 0.946106992 | 0.022 | 0.024 | 1 | 0.950033996 | 0.79548099 |
| 2410080I02Rik | 0.54776222 | 2.088809267 | 0.048 | 0.026 | 1 | 0.707090231 | 1.683072586 | 0.044 | 0.035 | 1 | 0.707090231 | 0.79548099 |
| Rec114 | 0.54780329 | 0.304537958 | 0.048 | 0.09 | 1 | 0.925916521 | 1.098110085 | 0.133 | 0.138 | 1 | 0.925916521 | 0.795518136 |
| Arhgap39 | 0.710830798 | 0.171271427 | 0.143 | 0.187 | 1 | 0.548589176 | 0.907778873 | 0.156 | 0.215 | 1 | 0.710830798 | 0.796228268 |
| Kif1a | 0.787193017 | 1.673771768 | 0.048 | 0.037 | 1 | 0.549596879 | 0.723714571 | 0.044 | 0.069 | 1 | 0.787193017 | 0.797137029 |
| Fam161a | 0.54961553 | 0.400753273 | 0.048 | 0.09 | 1 | 0.649175351 | 0.470078863 | 0.089 | 0.118 | 1 | 0.649175351 | 0.797153829 |
| Trim12c | 0.549880372 | 0.323274521 | 0.095 | 0.157 | 1 | 0.578286168 | 0.298408736 | 0.089 | 0.118 | 1 | 0.578286168 | 0.79739232 |
| Gm16576 | 0.837997597 | 0.747772349 | 0.19 | 0.19 | 1 | 0.550326131 | 1.361144491 | 0.133 | 0.114 | 1 | 0.837997597 | 0.797793412 |

| | | | | | | | | | | | | |
|---|---|---|---|---|---|---|---|---|---|---|---|---|
| A130014A01Rik | 0.994183949 | 1.010806755 | 0.048 | 0.049 | 1 | 0.5504649 | 1.894576691 | 0.067 | 0.048 | 1 | 0.994183949 | 0.797918194 |
| Stk11ip | 0.550710951 | 1.281454345 | 0.095 | 0.063 | 1 | 0.96831838 | 0.42003818 | 0.067 | 0.069 | 1 | 0.96831838 | 0.798139351 |
| BC065397 | 0.717949775 | 1.304537958 | 0.095 | 0.078 | 1 | 0.550755195 | 0.587915353 | 0.067 | 0.1 | 1 | 0.717949775 | 0.798179105 |
| Cse1l | 0.550842388 | 0.60338244 | 0.381 | 0.396 | 1 | 0.878608963 | 0.446874365 | 0.467 | 0.519 | 1 | 0.878608963 | 0.79825744 |
| Alox5ap | 0.550944111 | 1.55829455 | 0.048 | 0.026 | 1 | 0.941347173 | 0.757073168 | 0.022 | 0.021 | 1 | 0.941347173 | 0.798348809 |
| Gng4 | 0.550953306 | 1.936806174 | 0.048 | 0.026 | 1 | 0.617514334 | 1.098110085 | 0.022 | 0.038 | 1 | 0.617514334 | 0.798357066 |
| Bc1 | 0.692878573 | -1.107587946 | 0.095 | 0.123 | 1 | 0.551292198 | -0.597035333 | 0.067 | 0.093 | 1 | 0.692878573 | 0.798661308 |
| Zfp326 | 0.663259527 | 0.222560656 | 0.952 | 0.922 | 1 | 0.551716726 | 0.39151814 | 0.956 | 0.917 | 1 | 0.663259527 | 0.799042106 |
| Odf3l1 | 0.985460078 | 1.351843673 | 0.048 | 0.049 | 1 | 0.551724542 | 0.857101986 | 0.067 | 0.1 | 1 | 0.985460078 | 0.799049114 |
| Myo3b | 0.942814629 | 0.673771768 | 0.048 | 0.045 | 1 | 0.552269311 | 0.490427508 | 0.022 | 0.042 | 1 | 0.942814629 | 0.79953723 |
| Ears2 | 0.553143166 | 1.129451251 | 0.143 | 0.108 | 1 | 0.846409385 | 0.739656115 | 0.133 | 0.131 | 1 | 0.846409385 | 0.80031897 |
| Brip1os | 0.597424133 | 0.796454287 | 0.238 | 0.343 | 1 | 0.553367057 | 0.551828053 | 0.267 | 0.388 | 1 | 0.597424133 | 0.800519014 |
| Syt11 | 0.553534138 | 0.673771768 | 0.048 | 0.09 | 1 | 0.742935934 | 0.361144491 | 0.089 | 0.107 | 1 | 0.742935934 | 0.800668234 |
| Catsperg1 | 0.632076134 | 1.673771768 | 0 | 0.011 | 1 | 0.553874252 | 0.460680165 | 0.022 | 0.042 | 1 | 0.632076134 | 0.800971817 |
| Cd274 | 0.790349864 | 1.673771768 | 0.048 | 0.037 | 1 | 0.553912635 | 0.683072586 | 0.022 | 0.042 | 1 | 0.790349864 | 0.801006062 |
| Man1a2 | 0.553962797 | 0.417731608 | 0.81 | 0.802 | 1 | 0.632052658 | 0.114598208 | 0.956 | 0.817 | 1 | 0.632052658 | 0.801050813 |
| Adgb | 0.554143605 | 1.79930265 | 0.048 | 0.026 | 1 | 0.716677627 | 1.197645759 | 0.044 | 0.035 | 1 | 0.716677627 | 0.801212075 |
| Vmo1 | 0.632076134 | 1.673771768 | 0 | 0.011 | 1 | 0.555515361 | 1.304560963 | 0.022 | 0.042 | 1 | 0.632076134 | 0.802433406 |
| Gm41505 | 0.875318873 | 0.866416846 | 0.048 | 0.041 | 1 | 0.555547514 | 1.719598462 | 0.089 | 0.069 | 1 | 0.875318873 | 0.802461988 |
| Tektip1 | 0.632076134 | 1.673771768 | 0 | 0.011 | 1 | 0.555549391 | 0.776181991 | 0.022 | 0.042 | 1 | 0.632076134 | 0.802463656 |
| Gm11747 | 0.632076134 | 1.673771768 | 0 | 0.011 | 1 | 0.555549391 | 0.776181991 | 0.022 | 0.042 | 1 | 0.632076134 | 0.802463656 |
| Gm41664 | 0.930801273 | 1.171271427 | 0.048 | 0.045 | 1 | 0.555576609 | 0.757073168 | 0.022 | 0.042 | 1 | 0.930801273 | 0.80248785 |
| Gm48521 | 0.712049 | 0.786246497 | 0.095 | 0.131 | 1 | 0.555579161 | 1.098110085 | 0.022 | 0.042 | 1 | 0.712049 | 0.802490117 |
| Slc23a2 | 0.672449074 | 0.315529973 | 0.714 | 0.705 | 1 | 0.555708286 | 0.246139592 | 0.844 | 0.737 | 1 | 0.672449074 | 0.802604873 |
| Tspan18 | 0.556386688 | 0.357196907 | 0.667 | 0.728 | 1 | 0.875337366 | 0.752075374 | 0.467 | 0.612 | 1 | 0.875337366 | 0.80320723 |
| Stx18 | 0.913852471 | 0.631951592 | 0.429 | 0.567 | 1 | 0.556607101 | 0.639129239 | 0.422 | 0.651 | 1 | 0.913852471 | 0.803402737 |
| Gtdc1 | 0.556759819 | 1.111351836 | 0.381 | 0.388 | 1 | 0.830609504 | 0.194572925 | 0.467 | 0.439 | 1 | 0.830609504 | 0.803538142 |
| Gm42937 | 0.557019275 | 1.503846766 | 0.048 | 0.09 | 1 | 0.875329055 | 0.875717664 | 0.044 | 0.052 | 1 | 0.875329055 | 0.803768077 |
| Eipr1 | 0.622582013 | -0.361852142 | 0.19 | 0.228 | 1 | 0.557419955 | 0.361144491 | 0.156 | 0.211 | 1 | 0.622582013 | 0.804122904 |
| Ntng2 | 0.775497172 | 0.673771768 | 0.143 | 0.127 | 1 | 0.558399175 | 0.478714088 | 0.111 | 0.149 | 1 | 0.775497172 | 0.804988711 |
| Plxna2 | 0.667036695 | 0.171271427 | 0.524 | 0.575 | 1 | 0.559845102 | 0.140701962 | 0.689 | 0.792 | 1 | 0.667036695 | 0.806263666 |
| Aatf | 0.659000822 | 0.555774558 | 0.286 | 0.377 | 1 | 0.559989533 | 0.875717664 | 0.378 | 0.408 | 1 | 0.659000822 | 0.806390789 |
| Dph7 | 0.560004491 | 1.029915578 | 0.095 | 0.157 | 1 | 0.938080073 | 0.963180505 | 0.133 | 0.152 | 1 | 0.938080073 | 0.806403952 |
| Katnal1 | 0.560042188 | 0.777607579 | 0.143 | 0.216 | 1 | 0.633697079 | 0.490427508 | 0.111 | 0.149 | 1 | 0.633697079 | 0.806437124 |
| Cfp | 0.560128368 | 1.60338244 | 0.095 | 0.067 | 1 | 0.980254379 | 0.683072586 | 0.089 | 0.093 | 1 | 0.980254379 | 0.806512947 |
| Nipbl | 0.661078338 | 0.341130542 | 1 | 0.978 | 1 | 0.560255526 | 0.196748002 | 1 | 0.997 | 1 | 0.661078338 | 0.806624798 |
| B230369F24Rik | 0.671166976 | 1.383094607 | 0.19 | 0.175 | 1 | 0.560763548 | 0.223640968 | 0.311 | 0.284 | 1 | 0.671166976 | 0.807071339 |
| Gpr107 | 0.895380531 | 0.902396143 | 0.381 | 0.515 | 1 | 0.562130464 | 0.272984303 | 0.533 | 0.685 | 1 | 0.895380531 | 0.808270269 |
| Gm15738 | 0.915506323 | 1.258734268 | 0.048 | 0.056 | 1 | 0.562656292 | 0.460680165 | 0.067 | 0.097 | 1 | 0.915506323 | 0.808730481 |
| Cep85 | 0.85610838 | 0.492442003 | 0.429 | 0.489 | 1 | 0.562672733 | 0.300602949 | 0.311 | 0.394 | 1 | 0.85610838 | 0.808744862 |
| Zfp950 | 0.562686477 | 0.730852863 | 0.429 | 0.466 | 1 | 0.580700207 | 0.513147585 | 0.356 | 0.484 | 1 | 0.580700207 | 0.808756882 |
| Lama2 | 0.562747943 | 0.284280389 | 0.905 | 0.892 | 1 | 0.852581896 | 0.292113505 | 0.6 | 0.737 | 1 | 0.852581896 | 0.808810638 |
| Cdo1 | 0.710262591 | 0.274821253 | 0.619 | 0.552 | 1 | 0.563256888 | -0.124282336 | 0.778 | 0.654 | 1 | 0.710262591 | 0.809255454 |
| Optc | 0.563778058 | 1.258734268 | 0.048 | 0.026 | 1 | 0.847506963 | 0.567595369 | 0.022 | 0.028 | 1 | 0.847506963 | 0.809710417 |
| Gm56962 | 0.563778058 | 1.258734268 | 0.048 | 0.026 | 1 | 0.61382235 | 1.683072586 | 0.044 | 0.031 | 1 | 0.61382235 | 0.809710417 |
| Phrf1 | 0.797916753 | 0.59848364 | 0.476 | 0.545 | 1 | 0.563887806 | 0.277728483 | 0.711 | 0.754 | 1 | 0.797916753 | 0.809806154 |
| Igf2bp2 | 0.564160192 | 0.384675065 | 0.333 | 0.414 | 1 | 0.610536615 | 0.576157382 | 0.822 | 0.744 | 1 | 0.610536615 | 0.810043662 |
| Ttll11 | 0.564286249 | 1.536268244 | 0.095 | 0.067 | 1 | 0.791458206 | 0.205025289 | 0.067 | 0.08 | 1 | 0.791458206 | 0.810153527 |
| Tmed8 | 0.994896639 | 1.064962525 | 0.143 | 0.16 | 1 | 0.564829091 | 0.744473131 | 0.133 | 0.18 | 1 | 0.994896639 | 0.81062628 |
| Oxsm | 0.723619226 | 0.150209812 | 0.048 | 0.067 | 1 | 0.565034102 | 2.235613609 | 0.067 | 0.048 | 1 | 0.723619226 | 0.810804668 |
| Il15ra | 0.97968872 | 1.902040755 | 0.238 | 0.295 | 1 | 0.565108654 | 0.757073168 | 0.2 | 0.28 | 1 | 0.97968872 | 0.810869518 |
| Gm47990 | 0.565783556 | -0.593014773 | 0.095 | 0.142 | 1 | 0.847506963 | 0.683072586 | 0.022 | 0.028 | 1 | 0.847506963 | 0.811456079 |
| 4933405D12Rik | 0.566587257 | 1.107424945 | 0.143 | 0.104 | 1 | 0.964958869 | 1.250113179 | 0.178 | 0.204 | 1 | 0.964958869 | 0.812153395 |
| Gm42917 | 0.915524318 | 1.425844254 | 0.048 | 0.056 | 1 | 0.566926975 | 1.098110085 | 0.067 | 0.1 | 1 | 0.915524318 | 0.812447755 |
| Gm26788 | 0.567009484 | 1.673771768 | 0.048 | 0.026 | 1 | 0.687197015 | 0.808603468 | 0.022 | 0.035 | 1 | 0.687197015 | 0.812519213 |
| Gm29340 | 0.567009484 | 1.673771768 | 0.048 | 0.026 | 1 | 0.811838207 | 1.268035087 | 0.022 | 0.017 | 1 | 0.811838207 | 0.812519213 |
| Tmem131 | 0.568083894 | 0.378588528 | 0.762 | 0.739 | 1 | 0.929599712 | 0.228299335 | 0.911 | 0.869 | 1 | 0.929599712 | 0.813448477 |
| Or2k2 | 0.924111829 | 0.586308926 | 0.048 | 0.052 | 1 | 0.56825396 | 0.857101986 | 0.067 | 0.097 | 1 | 0.924111829 | 0.813595357 |
| Rad51b | 0.57110659 | -0.532132531 | 0.143 | 0.198 | 1 | 0.569305016 | 1.610316244 | 0.089 | 0.069 | 1 | 0.57110659 | 0.81450183 |
| Zfp974 | 0.773169671 | 0.150209812 | 0.048 | 0.063 | 1 | 0.570110509 | -0.181997834 | 0.089 | 0.118 | 1 | 0.773169671 | 0.815195025 |
| Gm28229 | 0.570211706 | 1.503846766 | 0.048 | 0.026 | 1 | 0.81549849 | 1.098110085 | 0.044 | 0.055 | 1 | 0.81549849 | 0.815282023 |
| Zfp438 | 0.614950399 | 1.351843673 | 0.19 | 0.164 | 1 | 0.570483023 | -0.346674757 | 0.067 | 0.093 | 1 | 0.614950399 | 0.815515166 |
| Sdccag8 | 0.813754627 | 1.3917364 | 0.381 | 0.44 | 1 | 0.573148033 | 0.733537653 | 0.444 | 0.509 | 1 | 0.813754627 | 0.817797398 |
| Fank1 | 0.573468511 | 1.351843673 | 0.048 | 0.026 | 1 | 0.854464619 | 1.223640968 | 0.022 | 0.028 | 1 | 0.854464619 | 0.818070889 |
| AI480526 | 0.573631039 | 0.569435108 | 0.238 | 0.209 | 1 | 0.598359769 | 0.506749814 | 0.2 | 0.266 | 1 | 0.598359769 | 0.818209509 |
| Trim25 | 0.582031041 | 0.335902129 | 0.143 | 0.216 | 1 | 0.574119461 | 0.463627449 | 0.378 | 0.318 | 1 | 0.582031041 | 0.818625766 |
| Kcnj13 | 0.632076134 | 1.673771768 | 0 | 0.011 | 1 | 0.57431066 | 0.361144491 | 0.044 | 0.069 | 1 | 0.632076134 | 0.818788586 |
| Tmem265 | 0.720689518 | 0.388369549 | 0.095 | 0.127 | 1 | 0.574485043 | 0.520134015 | 0.2 | 0.266 | 1 | 0.720689518 | 0.818937021 |
| Alkbh3 | 0.575369104 | 0.569435108 | 0.19 | 0.265 | 1 | 0.906355952 | 0.446033389 | 0.222 | 0.253 | 1 | 0.906355952 | 0.819688602 |
| Gm20632 | 0.578156445 | 1.088809267 | 0 | 0.015 | 1 | 0.835943453 | 0.513147585 | 0.022 | 0.028 | 1 | 0.835943453 | 0.822048015 |
| Slc9b1 | 0.578156445 | 1.088809267 | 0 | 0.015 | 1 | 0.677376317 | 0.180572246 | 0.022 | 0.035 | 1 | 0.677376317 | 0.822048015 |
| Gpr63 | 0.578156445 | 1.088809267 | 0 | 0.015 | 1 | 0.809040217 | 1.460680165 | 0.022 | 0.017 | 1 | 0.809040217 | 0.822048015 |
| Gm52698 | 0.578156445 | 1.088809267 | 0 | 0.015 | 1 | 0.809040217 | 1.460680165 | 0.022 | 0.017 | 1 | 0.809040217 | 0.822048015 |
| Lrrc19 | 0.578156445 | 0.673771768 | 0 | 0.015 | 1 | 0.687197015 | 0.808603468 | 0.022 | 0.035 | 1 | 0.687197015 | 0.822048015 |
| Zmynd12 | 0.578156445 | 1.088809267 | 0 | 0.015 | 1 | 0.828665129 | 0.223640968 | 0.022 | 0.017 | 1 | 0.828665129 | 0.822048015 |
| Dpf1 | 0.578156445 | 0.866416846 | 0 | 0.015 | 1 | 0.954659411 | 0.683072586 | 0.022 | 0.021 | 1 | 0.954659411 | 0.822048015 |
| Bean1 | 0.578156445 | 1.088809267 | 0 | 0.015 | 1 | 0.954659411 | 0.683072586 | 0.022 | 0.021 | 1 | 0.954659411 | 0.822048015 |
| Apc2 | 0.578156445 | 1.088809267 | 0 | 0.015 | 1 | 0.670048992 | 1.361144491 | 0.022 | 0.014 | 1 | 0.670048992 | 0.822048015 |
| Plxnc1 | 0.578156445 | 0.866416846 | 0 | 0.015 | 1 | 0.735986925 | 1.020107573 | 0.044 | 0.035 | 1 | 0.735986925 | 0.822048015 |
| Evi2a | 0.578156445 | 1.088809267 | 0 | 0.015 | 1 | 0.955025007 | 1.83507568 | 0.022 | 0.024 | 1 | 0.955025007 | 0.822048015 |
| Tcam1 | 0.578156445 | 0.866416846 | 0 | 0.015 | 1 | 0.672920453 | 1.098110085 | 0.022 | 0.014 | 1 | 0.672920453 | 0.822048015 |
| Angel1 | 0.578156445 | 0.351843673 | 0 | 0.015 | 1 | 0.938685475 | 1.683072586 | 0.022 | 0.021 | 1 | 0.938685475 | 0.822048015 |
| 9430024F10Rik | 0.578156445 | 1.088809267 | 0 | 0.015 | 1 | 0.661444614 | 1.875717664 | 0.022 | 0.014 | 1 | 0.661444614 | 0.822048015 |
| Gm47802 | 0.578156445 | 1.088809267 | 0 | 0.015 | 1 | 0.807293078 | 0.683072586 | 0.044 | 0.055 | 1 | 0.807293078 | 0.822048015 |
| Tex43 | 0.578156445 | 1.088809267 | 0 | 0.015 | 1 | 0.809040217 | 1.460680165 | 0.022 | 0.017 | 1 | 0.809040217 | 0.822048015 |
| Ncf2 | 0.578159907 | 0.866416846 | 0 | 0.015 | 1 | 0.735980034 | 0.42003818 | 0.044 | 0.035 | 1 | 0.735980034 | 0.822050936 |
| Gm42567 | 0.578159907 | 0.503846766 | 0 | 0.015 | 1 | 0.837136013 | 0.435145073 | 0.044 | 0.038 | 1 | 0.837136013 | 0.822050936 |
| Cpne9 | 0.578159907 | 0.214340149 | 0 | 0.015 | 1 | 0.617606099 | 1.583536913 | 0.044 | 0.031 | 1 | 0.617606099 | 0.822050936 |
| Niban3 | 0.578159907 | 0.214340149 | 0 | 0.015 | 1 | 0.955023924 | 2.005000681 | 0.022 | 0.024 | 1 | 0.955023924 | 0.822050936 |
| Asic2 | 0.578159907 | 0.866416846 | 0 | 0.015 | 1 | 0.675805699 | -1.361321533 | 0.022 | 0.014 | 1 | 0.675805699 | 0.822050936 |
| Gm56614 | 0.578159907 | 0.351843673 | 0 | 0.015 | 1 | 0.722460992 | 1.545569062 | 0.044 | 0.035 | 1 | 0.722460992 | 0.822050936 |
| Gm29570 | 0.57816106 | 1.351843673 | 0 | 0.015 | 1 | 0.664307093 | 1.268035087 | 0.022 | 0.014 | 1 | 0.664307093 | 0.822051909 |
| 9430060I03Rik | 0.57816106 | 1.351843673 | 0 | 0.015 | 1 | 0.937567971 | 0.683072586 | 0.022 | 0.024 | 1 | 0.937567971 | 0.822051909 |
| Gm37362 | 0.57816106 | 1.351843673 | 0 | 0.015 | 1 | 0.951997643 | 0.875717664 | 0.022 | 0.021 | 1 | 0.951997643 | 0.822051909 |
| Myocos | 0.57816106 | 1.351843673 | 0 | 0.015 | 1 | 0.708987514 | 1.42003818 | 0.044 | 0.035 | 1 | 0.708987514 | 0.822051909 |
| Gm13421 | 0.57816106 | 1.351843673 | 0 | 0.015 | 1 | 0.823042771 | 0.361144491 | 0.022 | 0.017 | 1 | 0.823042771 | 0.822051909 |
| Fer1l4 | 0.57816106 | 1.351843673 | 0 | 0.015 | 1 | 0.664308814 | 1.460680165 | 0.022 | 0.014 | 1 | 0.664308814 | 0.822051909 |
| Bglap3 | 0.57816106 | 1.351843673 | 0 | 0.015 | 1 | 0.806246349 | 2.460680165 | 0.022 | 0.017 | 1 | 0.806246349 | 0.822051909 |

| | | | | | | | | | | | | |
|---|---|---|---|---|---|---|---|---|---|---|---|---|
| Gm42938 | 0.57816106 | 1.351843673 | 0 | 0.015 | 1 | 0.787608894 | 0.513147585 | 0.044 | 0.055 | 1 | 0.787608894 | 0.822051909 |
| Gm12473 | 0.57816106 | 1.351843673 | 0 | 0.015 | 1 | 0.817155885 | 1.605070074 | 0.044 | 0.055 | 1 | 0.817155885 | 0.822051909 |
| A230083G16Rik | 0.57816106 | 1.351843673 | 0 | 0.015 | 1 | 0.952529269 | 1.098110085 | 0.022 | 0.024 | 1 | 0.952529269 | 0.822051909 |
| Ttll13 | 0.57816106 | 1.351843673 | 0 | 0.015 | 1 | 0.615717475 | 0.567595369 | 0.022 | 0.038 | 1 | 0.615717475 | 0.822051909 |
| Gm57240 | 0.57816106 | 1.351843673 | 0 | 0.015 | 1 | 0.658583583 | 2.361144491 | 0.022 | 0.014 | 1 | 0.658583583 | 0.822051909 |
| Ttc41 | 0.57816106 | 1.351843673 | 0 | 0.015 | 1 | 0.68327161 | 0.567595369 | 0.022 | 0.035 | 1 | 0.68327161 | 0.822051909 |
| Adora2b | 0.57816106 | 1.351843673 | 0 | 0.015 | 1 | 0.645291449 | 1.320502507 | 0.067 | 0.052 | 1 | 0.645291449 | 0.822051909 |
| Slc10a1 | 0.57816106 | 1.351843673 | 0 | 0.015 | 1 | 0.950033996 | 0.946106992 | 0.022 | 0.024 | 1 | 0.950033996 | 0.822051909 |
| 1700087M22Rik | 0.57816106 | 1.351843673 | 0 | 0.015 | 1 | 0.988929908 | 0.460680165 | 0.044 | 0.045 | 1 | 0.988929908 | 0.822051909 |
| Gm49067 | 0.57816106 | 1.351843673 | 0 | 0.015 | 1 | 0.6175503 | 1.098110085 | 0.022 | 0.038 | 1 | 0.6175503 | 0.822051909 |
| 1700085D07Rik | 0.57816106 | 1.351843673 | 0 | 0.015 | 1 | 0.914340513 | 0.776181991 | 0.044 | 0.042 | 1 | 0.914340513 | 0.822051909 |
| Prr18 | 0.57816106 | 1.351843673 | 0 | 0.015 | 1 | 0.687208248 | 1.098110085 | 0.022 | 0.035 | 1 | 0.687208248 | 0.822051909 |
| Gm41604 | 0.57816106 | 1.351843673 | 0 | 0.015 | 1 | 0.880479454 | 1.020107573 | 0.044 | 0.052 | 1 | 0.880479454 | 0.822051909 |
| C3 | 0.57816106 | 1.351843673 | 0 | 0.015 | 1 | 0.935071939 | 0.361144491 | 0.022 | 0.024 | 1 | 0.935071939 | 0.822051909 |
| B3galt1 | 0.578673017 | 1.379858049 | 0.429 | 0.466 | 1 | 0.81501834 | 0.69361608 | 0.333 | 0.353 | 1 | 0.81501834 | 0.822483573 |
| Nutf2 | 0.579405938 | 1.059061924 | 0.19 | 0.164 | 1 | 0.989218448 | 0.875717664 | 0.133 | 0.149 | 1 | 0.989218448 | 0.823100635 |
| Trmt10b | 0.893706501 | 0.351843673 | 0.095 | 0.086 | 1 | 0.579518083 | 0.42003818 | 0.044 | 0.069 | 1 | 0.893706501 | 0.823194957 |
| Ankrd9 | 0.616977656 | 0.614878079 | 0.048 | 0.082 | 1 | 0.580578162 | -0.255526869 | 0.044 | 0.066 | 1 | 0.616977656 | 0.824085321 |
| Gm13830 | 0.96060963 | 1.088809267 | 0.048 | 0.052 | 1 | 0.580578162 | -0.255526869 | 0.044 | 0.066 | 1 | 0.96060963 | 0.824085321 |
| Qser1 | 0.580621563 | 0.583573959 | 0.476 | 0.496 | 1 | 0.738356165 | 0.328229869 | 0.822 | 0.74 | 1 | 0.738356165 | 0.824121727 |
| Zbtb37 | 0.601061141 | 0.726239188 | 0.095 | 0.149 | 1 | 0.581048748 | 0.416286045 | 0.156 | 0.204 | 1 | 0.601061141 | 0.824479849 |
| Zbtb25 | 0.937887387 | 1.843696769 | 0.095 | 0.093 | 1 | 0.581819108 | 0.64125241 | 0.133 | 0.18 | 1 | 0.937887387 | 0.825124742 |
| Ankrd42 | 0.665273929 | 0.896164189 | 0.143 | 0.205 | 1 | 0.582720775 | 0.702181409 | 0.178 | 0.235 | 1 | 0.665273929 | 0.825878049 |
| Ptprn2 | 0.867618878 | 3.157933628 | 0.143 | 0.134 | 1 | 0.58302378 | 0.769784219 | 0.067 | 0.097 | 1 | 0.867618878 | 0.826130832 |
| Bnc2 | 0.585363919 | 0.587357016 | 0.476 | 0.496 | 1 | 0.798026494 | 0.235736293 | 0.578 | 0.706 | 1 | 0.798026494 | 0.82807692 |
| Trim3 | 0.845618438 | 1.487905222 | 0.238 | 0.261 | 1 | 0.585589935 | 0.750186782 | 0.311 | 0.332 | 1 | 0.845618438 | 0.828264298 |
| Sfmbt2 | 0.771497544 | 1.673771768 | 0.048 | 0.067 | 1 | 0.585984541 | 0.383512304 | 0.111 | 0.149 | 1 | 0.771497544 | 0.8285912 |
| Gm9856 | 0.95431196 | 1.866416846 | 0.095 | 0.101 | 1 | 0.586163086 | 1.250113179 | 0.044 | 0.069 | 1 | 0.95431196 | 0.828739009 |
| Zdhhc14 | 0.846148946 | 0.400753273 | 0.095 | 0.116 | 1 | 0.586844458 | 0.45359074 | 0.244 | 0.208 | 1 | 0.846148946 | 0.829302498 |
| Rcan2 | 0.607047505 | 0.380990018 | 0.048 | 0.082 | 1 | 0.587088564 | -0.84048937 | 0.022 | 0.038 | 1 | 0.607047505 | 0.829504146 |
| Rtl5 | 0.862687636 | 1.088809267 | 0.048 | 0.06 | 1 | 0.58735262 | 0.583536913 | 0.067 | 0.093 | 1 | 0.862687636 | 0.829722139 |
| Cyp4f13 | 0.858282446 | 0.936806174 | 0.143 | 0.134 | 1 | 0.587741594 | 0.905465008 | 0.089 | 0.125 | 1 | 0.858282446 | 0.830043006 |
| Polr3b | 0.588799393 | 1.39876472 | 0.286 | 0.287 | 1 | 0.813858908 | 1.065977465 | 0.333 | 0.377 | 1 | 0.813858908 | 0.830914061 |
| Gm20735 | 0.632076134 | 1.673771768 | 0 | 0.011 | 1 | 0.590623425 | -0.404390255 | 0.022 | 0.038 | 1 | 0.632076134 | 0.83241082 |
| Slc9b2 | 0.632076134 | 1.673771768 | 0 | 0.011 | 1 | 0.590647301 | -2.686161223 | 0.022 | 0.038 | 1 | 0.632076134 | 0.832430368 |
| Prkar1b | 0.938731921 | 0.866416846 | 0.19 | 0.194 | 1 | 0.591260519 | 0.494411022 | 0.244 | 0.325 | 1 | 0.938731921 | 0.832932037 |
| Gm16023 | 0.965217224 | 0.65584986 | 0.19 | 0.216 | 1 | 0.591471371 | 1.071637874 | 0.178 | 0.256 | 1 | 0.965217224 | 0.833104359 |
| Mbtps2 | 0.839640595 | 0.525379928 | 0.333 | 0.354 | 1 | 0.593159371 | 0.984899475 | 0.289 | 0.308 | 1 | 0.839640595 | 0.834480703 |
| Rad23b | 0.718544958 | 0.157933628 | 0.81 | 0.858 | 1 | 0.593535022 | 0.208710807 | 0.933 | 0.938 | 1 | 0.718544958 | 0.834786222 |
| Cd200r3 | 0.709465802 | 1.936806174 | 0.048 | 0.034 | 1 | 0.594160317 | -0.124282336 | 0.022 | 0.038 | 1 | 0.709465802 | 0.835294152 |
| Kank3 | 0.957802897 | 1.010806755 | 0.048 | 0.052 | 1 | 0.594160317 | -0.223818009 | 0.022 | 0.038 | 1 | 0.957802897 | 0.835294152 |
| Med12l | 0.595488463 | -0.118786152 | 0.095 | 0.142 | 1 | 0.613866297 | 0.591150097 | 0.156 | 0.201 | 1 | 0.613866297 | 0.836370417 |
| Dnajc27 | 0.595747023 | 0.843696769 | 0.19 | 0.16 | 1 | 0.996985601 | 0.91753784 | 0.156 | 0.176 | 1 | 0.996985601 | 0.836579531 |
| Lcn11 | 0.595875107 | 1.281454345 | 0.095 | 0.067 | 1 | 0.969029253 | 1.076415014 | 0.133 | 0.152 | 1 | 0.969029253 | 0.836683071 |
| Trim65 | 0.924795272 | 1.351843673 | 0.048 | 0.045 | 1 | 0.595927847 | -0.124282336 | 0.022 | 0.038 | 1 | 0.924795272 | 0.836725695 |
| Gm26885 | 0.629700127 | 2.088809267 | 0.048 | 0.03 | 1 | 0.596402572 | 0.612683258 | 0.067 | 0.048 | 1 | 0.629700127 | 0.837109116 |
| Slc12a1 | 0.597569344 | 0.405999443 | 0.238 | 0.201 | 1 | 0.66366529 | -1.602329633 | 0.022 | 0.035 | 1 | 0.66366529 | 0.838049567 |
| Eif5b | 0.59830676 | 0.230537377 | 1 | 0.955 | 1 | 0.916376747 | 0.122445278 | 1 | 0.99 | 1 | 0.916376747 | 0.838642541 |
| Vps4b | 0.789730937 | 0.262496187 | 0.667 | 0.698 | 1 | 0.598747718 | 0.259861155 | 0.778 | 0.754 | 1 | 0.789730937 | 0.838996606 |
| Rassf5 | 0.598748902 | 1.819622634 | 0.143 | 0.116 | 1 | 0.904884521 | 1.268035087 | 0.133 | 0.156 | 1 | 0.904884521 | 0.838997556 |
| Avil | 0.600178437 | 0.812155252 | 0.381 | 0.377 | 1 | 0.692214578 | 0.543142325 | 0.489 | 0.609 | 1 | 0.692214578 | 0.840142717 |
| Rfx1 | 0.668937304 | 0.805875304 | 0.238 | 0.22 | 1 | 0.600336529 | 0.390290837 | 0.267 | 0.349 | 1 | 0.668937304 | 0.84026911 |
| Zfp282 | 0.600421099 | 2.221259563 | 0.19 | 0.272 | 1 | 0.65706463 | 0.896476224 | 0.289 | 0.322 | 1 | 0.65706463 | 0.840336702 |
| Kcnq3 | 0.746672225 | -0.275244304 | 0.095 | 0.123 | 1 | 0.601916784 | 1.657537494 | 0.111 | 0.093 | 1 | 0.746672225 | 0.841529753 |
| Focad | 0.602135826 | 0.586308926 | 0.333 | 0.317 | 1 | 0.779214645 | 0.831171225 | 0.244 | 0.318 | 1 | 0.779214645 | 0.841704099 |
| Ppfia4 | 0.602370482 | 1.214340149 | 0.095 | 0.067 | 1 | 0.691369024 | 1.605070074 | 0.067 | 0.055 | 1 | 0.691369024 | 0.841890766 |
| Gm30906 | 0.71266839 | 1.79930265 | 0.048 | 0.034 | 1 | 0.603746724 | 0.180572246 | 0.067 | 0.093 | 1 | 0.71266839 | 0.842983341 |
| Fam228b | 0.813144372 | 0.936806174 | 0.048 | 0.063 | 1 | 0.603859309 | 0.304560963 | 0.044 | 0.066 | 1 | 0.813144372 | 0.843072553 |
| Eps8l2 | 0.612961901 | 0.151393804 | 0.81 | 0.769 | 1 | 0.604038378 | -0.10018666 | 0.911 | 0.875 | 1 | 0.612961901 | 0.843214394 |
| Nsd1 | 0.766533535 | 0.278186631 | 0.762 | 0.776 | 1 | 0.604398353 | 0.462213316 | 0.978 | 0.948 | 1 | 0.766533535 | 0.843499337 |
| Ralgps1 | 0.604598232 | 1.295260144 | 0.095 | 0.067 | 1 | 0.664789858 | 1.28452321 | 0.089 | 0.121 | 1 | 0.664789858 | 0.843657442 |
| Cacna2d1 | 0.685078016 | 0.638147858 | 0.524 | 0.634 | 1 | 0.604613586 | 0.371593441 | 0.533 | 0.754 | 1 | 0.685078016 | 0.843669583 |
| Usp13 | 0.761665256 | 1.188344941 | 0.095 | 0.082 | 1 | 0.6063133 | 0.612683258 | 0.111 | 0.156 | 1 | 0.761665256 | 0.845010782 |
| Evpl | 0.818482987 | 0.281454345 | 0.048 | 0.06 | 1 | 0.606594188 | 0.42003818 | 0.044 | 0.066 | 1 | 0.818482987 | 0.845231867 |
| Irak3 | 0.607178966 | 0.541321472 | 0.286 | 0.414 | 1 | 0.798558074 | 1.404356558 | 0.244 | 0.256 | 1 | 0.798558074 | 0.845691635 |
| Rfx2 | 0.795284687 | 0.417432014 | 0.143 | 0.183 | 1 | 0.60776227 | 0.794863412 | 0.244 | 0.339 | 1 | 0.795284687 | 0.846149563 |
| Bfsp1 | 0.871904734 | 1.766881172 | 0.095 | 0.093 | 1 | 0.607882956 | 1.770535427 | 0.067 | 0.097 | 1 | 0.871904734 | 0.846244224 |
| Ifi27l2a | 0.700071195 | -0.413691074 | 0.048 | 0.071 | 1 | 0.609308987 | 1.209141398 | 0.044 | 0.066 | 1 | 0.700071195 | 0.847360532 |
| Clip1 | 0.892158331 | 0.399149388 | 0.762 | 0.907 | 1 | 0.609465299 | 0.328186653 | 0.956 | 0.941 | 1 | 0.892158331 | 0.847482647 |
| Fastkd3 | 0.609665403 | 0.150209812 | 0.048 | 0.078 | 1 | 0.695723289 | 0.946106992 | 0.067 | 0.09 | 1 | 0.695723289 | 0.847638902 |
| 4933424G05Rik | 0.61051998 | 1.97333205 | 0.095 | 0.071 | 1 | 0.867052953 | 0.460680165 | 0.067 | 0.076 | 1 | 0.867052953 | 0.848305314 |
| Gen1 | 0.853720939 | 2.574236094 | 0.048 | 0.041 | 1 | 0.612112936 | 0.361144491 | 0.022 | 0.038 | 1 | 0.853720939 | 0.849543626 |
| 1110046J04Rik | 0.612256266 | 0.989273593 | 0.143 | 0.116 | 1 | 0.994516495 | 0.709067795 | 0.133 | 0.142 | 1 | 0.994516495 | 0.849654797 |
| Gm17056 | 0.612985035 | 0.503846766 | 0.048 | 0.082 | 1 | 0.821108387 | 0.995016592 | 0.133 | 0.131 | 1 | 0.821108387 | 0.850219417 |
| Obp1a | 0.927006322 | 0.524024648 | 0.19 | 0.213 | 1 | 0.613330903 | 0.193687745 | 0.178 | 0.228 | 1 | 0.927006322 | 0.85048701 |
| Dmc1 | 0.802267666 | 1.188344941 | 0.095 | 0.086 | 1 | 0.613489384 | 1.045642666 | 0.089 | 0.125 | 1 | 0.802267666 | 0.850609544 |
| Gm43387 | 0.963421211 | 1.171271427 | 0.048 | 0.052 | 1 | 0.61382235 | 1.683072586 | 0.044 | 0.031 | 1 | 0.963421211 | 0.850866822 |
| H2bc18 | 0.963860445 | 0.425844254 | 0.048 | 0.045 | 1 | 0.613907968 | 0.545569062 | 0.022 | 0.038 | 1 | 0.963860445 | 0.850932943 |
| Ctnnd2 | 0.614087257 | 1.43600412 | 0.286 | 0.291 | 1 | 0.683160613 | 0.118287967 | 0.111 | 0.131 | 1 | 0.683160613 | 0.851071355 |
| Zmym4 | 0.614631578 | 0.402785238 | 0.667 | 0.642 | 1 | 0.778313044 | 0.24060296 | 0.867 | 0.824 | 1 | 0.778313044 | 0.851491179 |
| Gm35019 | 0.615153735 | 0.503846766 | 0.048 | 0.082 | 1 | 0.924650924 | 1.280974143 | 0.067 | 0.066 | 1 | 0.924650924 | 0.851893352 |
| 2610005L07Rik | 0.615392131 | 0.538612184 | 0.429 | 0.578 | 1 | 0.64897774 | 0.698393385 | 0.467 | 0.581 | 1 | 0.64897774 | 0.852076787 |
| 9530046B11Rik | 0.71266839 | 1.79930265 | 0.048 | 0.034 | 1 | 0.615717475 | 0.567595369 | 0.022 | 0.038 | 1 | 0.71266839 | 0.852326941 |
| Marcksl1 | 0.616335275 | -0.729076323 | 0.333 | 0.347 | 1 | 0.863733543 | -0.370794286 | 0.733 | 0.543 | 1 | 0.863733543 | 0.852801378 |
| Gm13708 | 0.859875139 | 1.451379346 | 0.048 | 0.041 | 1 | 0.617514334 | 1.098110085 | 0.022 | 0.038 | 1 | 0.859875139 | 0.853704715 |
| Pcgf1 | 0.774811077 | 0.629377648 | 0.095 | 0.078 | 1 | 0.61754131 | 1.197645759 | 0.022 | 0.038 | 1 | 0.774811077 | 0.85372535 |
| Gmnn | 0.915723736 | -0.326228232 | 0.048 | 0.052 | 1 | 0.6175503 | 0.776181991 | 0.022 | 0.038 | 1 | 0.915723736 | 0.853732227 |
| Galnt17 | 0.853725604 | 4.126283972 | 0.048 | 0.041 | 1 | 0.617567735 | 2.586857271 | 0.022 | 0.038 | 1 | 0.853725604 | 0.853745563 |
| Gm49915 | 0.787193017 | 1.673771768 | 0.048 | 0.037 | 1 | 0.617599877 | 1.42003818 | 0.044 | 0.031 | 1 | 0.787193017 | 0.853770146 |
| Mcf2l | 0.954990531 | 0.936806174 | 0.048 | 0.052 | 1 | 0.617612024 | 1.683072586 | 0.044 | 0.031 | 1 | 0.954990531 | 0.853779436 |
| Firre | 0.617735028 | 0.134612957 | 0.19 | 0.265 | 1 | 0.80860755 | 0.804751143 | 0.267 | 0.294 | 1 | 0.80860755 | 0.853873491 |
| Nphp3 | 0.61780811 | 0.36999102 | 0.143 | 0.213 | 1 | 0.809448152 | 0.697572156 | 0.178 | 0.208 | 1 | 0.809448152 | 0.853929359 |
| 5430416N02Rik | 0.6196134 | 0.472137907 | 0.286 | 0.25 | 1 | 0.865934136 | 1.168499413 | 0.178 | 0.218 | 1 | 0.865934136 | 0.855306034 |
| Hecw2 | 0.619875963 | 0.807038299 | 0.19 | 0.164 | 1 | 0.945413423 | 1.011953442 | 0.2 | 0.215 | 1 | 0.945413423 | 0.855505716 |
| Slc12a6 | 0.94508272 | 0.456541051 | 0.381 | 0.451 | 1 | 0.620915023 | -0.32320456 | 0.844 | 0.682 | 1 | 0.94508272 | 0.85629458 |
| C030013C21Rik | 0.620967462 | 0.735172312 | 0.048 | 0.082 | 1 | 0.87267348 | 0.857101986 | 0.111 | 0.128 | 1 | 0.87267348 | 0.856334335 |
| Scrn2 | 0.620967462 | 0.735172312 | 0.048 | 0.082 | 1 | 0.790860727 | 1.180572246 | 0.044 | 0.055 | 1 | 0.790860727 | 0.856334335 |
| Chn1 | 0.621146607 | 1.258734268 | 0.095 | 0.071 | 1 | 0.779106606 | -0.174908409 | 0.067 | 0.08 | 1 | 0.779106606 | 0.856470107 |

| | | | | | | | | | | | | |
|---|---|---|---|---|---|---|---|---|---|---|---|---|
| Rap1gap2 | 0.621256531 | 2.126283972 | 0.048 | 0.082 | 1 | 0.698578207 | 0.647448676 | 0.111 | 0.145 | 1 | 0.698578207 | 0.856553384 |
| Gm36037 | 0.945820148 | 0.351843673 | 0.048 | 0.045 | 1 | 0.621393986 | 1.683072586 | 0.044 | 0.031 | 1 | 0.945820148 | 0.856657486 |
| Zfp638 | 0.661299552 | 0.356093146 | 0.762 | 0.869 | 1 | 0.621418948 | 0.153996106 | 0.933 | 0.913 | 1 | 0.661299552 | 0.856676387 |
| Sap130 | 0.73471339 | 0.510833197 | 0.333 | 0.433 | 1 | 0.621793012 | 0.421057393 | 0.511 | 0.464 | 1 | 0.73471339 | 0.856959474 |
| Herc6 | 0.801495832 | 1.595769256 | 0.143 | 0.179 | 1 | 0.621941274 | 1.750186782 | 0.111 | 0.145 | 1 | 0.801495832 | 0.8570716 |
| Ubr2 | 0.789369151 | 0.52377381 | 0.619 | 0.802 | 1 | 0.622042773 | 0.23182203 | 0.956 | 0.903 | 1 | 0.789369151 | 0.857148334 |
| Cep295nl | 0.851769955 | 0.464318402 | 0.095 | 0.112 | 1 | 0.622573937 | 0.235613609 | 0.111 | 0.145 | 1 | 0.851769955 | 0.857549567 |
| Gm15458 | 0.921792415 | 1.451379346 | 0.048 | 0.045 | 1 | 0.622660035 | 1.513147585 | 0.067 | 0.052 | 1 | 0.921792415 | 0.85761455 |
| Rab28 | 0.623476614 | 0.158597597 | 0.476 | 0.683 | 1 | 0.874565685 | 0.481438725 | 0.622 | 0.619 | 1 | 0.874565685 | 0.85823014 |
| Ccdc152 | 0.79666227 | 0.866416846 | 0.048 | 0.037 | 1 | 0.624204867 | 1.683072586 | 0.067 | 0.052 | 1 | 0.79666227 | 0.858778018 |
| Zmiz1os1 | 0.834791327 | 1.088809267 | 0.095 | 0.119 | 1 | 0.624807729 | 1.397670367 | 0.111 | 0.097 | 1 | 0.834791327 | 0.859230759 |
| Polr3e | 0.700714988 | 1.495434526 | 0.238 | 0.254 | 1 | 0.625032773 | 0.385392038 | 0.311 | 0.412 | 1 | 0.700714988 | 0.859399579 |
| Kmt2d | 0.911122233 | 0.912959432 | 0.333 | 0.451 | 1 | 0.625342628 | 0.308789086 | 0.733 | 0.74 | 1 | 0.911122233 | 0.859631854 |
| Ube3c | 0.626431838 | 0.805502466 | 0.476 | 0.627 | 1 | 0.668556468 | 0.330338445 | 0.8 | 0.74 | 1 | 0.668556468 | 0.860446829 |
| Tert | 0.787193017 | 1.673771768 | 0.048 | 0.037 | 1 | 0.627053233 | 1.683072586 | 0.044 | 0.031 | 1 | 0.787193017 | 0.860910709 |
| 6330411D24Rik | 0.969049905 | 2.258734268 | 0.048 | 0.052 | 1 | 0.627093866 | 1.15951063 | 0.044 | 0.031 | 1 | 0.969049905 | 0.860941016 |
| Numbl | 0.628441415 | 0.905587443 | 0.286 | 0.276 | 1 | 0.894233186 | 0.952993378 | 0.356 | 0.457 | 1 | 0.894233186 | 0.861944218 |
| Ncaph | 0.629700127 | 2.088809267 | 0.048 | 0.03 | 1 | 0.814637721 | 1.098110085 | 0.022 | 0.017 | 1 | 0.814637721 | 0.862878004 |
| Gm43378 | 0.629700127 | 2.088809267 | 0.048 | 0.03 | 1 | 0.849821852 | 0.808603468 | 0.022 | 0.028 | 1 | 0.849821852 | 0.862878004 |
| Dpysl4 | 0.629700127 | 2.503846766 | 0.048 | 0.03 | 1 | 0.847019251 | 0.361144491 | 0.067 | 0.076 | 1 | 0.847019251 | 0.862878004 |
| Hsh2d | 0.629700127 | 3.088809267 | 0.048 | 0.03 | 1 | 0.687208248 | 1.42003818 | 0.022 | 0.035 | 1 | 0.687208248 | 0.862878004 |
| 9530082P21Rik | 0.629700127 | 2.088809267 | 0.048 | 0.03 | 1 | 0.823042771 | 0.875717664 | 0.022 | 0.017 | 1 | 0.823042771 | 0.862878004 |
| Spdef | 0.629700127 | 2.825774861 | 0.048 | 0.03 | 1 | 0.885639832 | 1.242499995 | 0.044 | 0.052 | 1 | 0.885639832 | 0.862878004 |
| Gm56999 | 0.629713293 | 2.673771768 | 0.048 | 0.03 | 1 | 0.685250124 | 0.776181991 | 0.022 | 0.035 | 1 | 0.685250124 | 0.862887754 |
| Hdc | 0.629722696 | 3.673771768 | 0.048 | 0.03 | 1 | 0.817437668 | 0.180572246 | 0.022 | 0.017 | 1 | 0.817437668 | 0.862894718 |
| Syt16 | 0.629722696 | 2.088809267 | 0.048 | 0.03 | 1 | 0.941347785 | 0.982632868 | 0.022 | 0.021 | 1 | 0.941347785 | 0.862894718 |
| Gm31305 | 0.629728337 | 2.088809267 | 0.048 | 0.03 | 1 | 0.998154737 | 0.513147585 | 0.044 | 0.045 | 1 | 0.998154737 | 0.862898896 |
| Galnt16 | 0.629729747 | 3.503846766 | 0.048 | 0.03 | 1 | 0.927603499 | -0.124282336 | 0.022 | 0.024 | 1 | 0.927603499 | 0.86289994 |
| Ppara | 0.629729747 | 2.866416846 | 0.048 | 0.03 | 1 | 0.726319687 | 1.03099589 | 0.044 | 0.035 | 1 | 0.726319687 | 0.86289994 |
| Gpr176 | 0.629733038 | 2.866416846 | 0.048 | 0.03 | 1 | 0.653980676 | -1.564854927 | 0.022 | 0.035 | 1 | 0.653980676 | 0.862902377 |
| Gm30371 | 0.629735859 | 3.503846766 | 0.048 | 0.03 | 1 | 0.664309387 | 1.460680165 | 0.022 | 0.014 | 1 | 0.664309387 | 0.862904466 |
| Gbp3 | 0.629741405 | -0.911190733 | 0.048 | 0.075 | 1 | 0.821680938 | 3.206634542 | 0.044 | 0.038 | 1 | 0.821680938 | 0.862908573 |
| Sobp | 0.71188729 | 0.379677881 | 0.81 | 0.903 | 1 | 0.63032464 | 0.384255162 | 0.911 | 0.958 | 1 | 0.71188729 | 0.863340128 |
| Ranbp17 | 0.631124973 | -0.828728573 | 0.095 | 0.127 | 1 | 0.875239178 | 1.158231078 | 0.178 | 0.208 | 1 | 0.875239178 | 0.863931215 |
| Gm26797 | 0.631863623 | -0.370622352 | 0.048 | 0.075 | 1 | 0.946780555 | 0.460680165 | 0.133 | 0.145 | 1 | 0.946780555 | 0.864475608 |
| Mettl21c | 0.63207476 | 1.351843673 | 0 | 0.011 | 1 | 0.670048992 | 1.361144491 | 0.022 | 0.014 | 1 | 0.670048992 | 0.864631018 |
| Inava | 0.63207476 | 1.088809267 | 0 | 0.011 | 1 | 0.80624346 | 2.098110085 | 0.022 | 0.017 | 1 | 0.80624346 | 0.864631018 |
| Gm14320 | 0.63207476 | 1.351843673 | 0 | 0.011 | 1 | 0.806246349 | 1.875717664 | 0.022 | 0.017 | 1 | 0.806246349 | 0.864631018 |
| Gm11266 | 0.63207476 | 1.351843673 | 0 | 0.011 | 1 | 0.658583583 | 2.683072586 | 0.022 | 0.014 | 1 | 0.658583583 | 0.864631018 |
| Drc1 | 0.63207476 | 1.351843673 | 0 | 0.011 | 1 | 0.941346407 | 1.361144491 | 0.022 | 0.021 | 1 | 0.941346407 | 0.864631018 |
| Muc5b | 0.63207476 | 1.351843673 | 0 | 0.011 | 1 | 0.730173512 | 1.197645759 | 0.044 | 0.035 | 1 | 0.730173512 | 0.864631018 |
| 4921516A02Rik | 0.63207476 | 1.351843673 | 0 | 0.011 | 1 | 0.761447202 | 0.683072586 | 0.022 | 0.031 | 1 | 0.761447202 | 0.864631018 |
| Slc26a10 | 0.63207476 | 1.351843673 | 0 | 0.011 | 1 | 0.957324717 | 0.513147585 | 0.022 | 0.021 | 1 | 0.957324717 | 0.864631018 |
| Gm40437 | 0.63207476 | 0.673771768 | 0 | 0.011 | 1 | 0.854464766 | 1.42003818 | 0.022 | 0.028 | 1 | 0.854464766 | 0.864631018 |
| Tex49 | 0.63207476 | -0.911190733 | 0 | 0.011 | 1 | 0.937567971 | 0.683072586 | 0.022 | 0.024 | 1 | 0.937567971 | 0.864631018 |
| Myh11 | 0.63207476 | 1.351843673 | 0 | 0.011 | 1 | 0.809040217 | 1.460680165 | 0.022 | 0.017 | 1 | 0.809040217 | 0.864631018 |
| Liph | 0.63207476 | 1.088809267 | 0 | 0.011 | 1 | 0.658587075 | 2.098110085 | 0.022 | 0.014 | 1 | 0.658587075 | 0.864631018 |
| Pcdhb19 | 0.63207476 | 1.351843673 | 0 | 0.011 | 1 | 0.765738001 | 1.567595369 | 0.022 | 0.031 | 1 | 0.765738001 | 0.864631018 |
| A030005L19Rik | 0.632076134 | 1.673771768 | 0 | 0.011 | 1 | 0.661444614 | 1.875717664 | 0.022 | 0.014 | 1 | 0.661444614 | 0.864632029 |
| Dnajb3 | 0.632076134 | 1.673771768 | 0 | 0.011 | 1 | 0.658588239 | 1.875717664 | 0.022 | 0.014 | 1 | 0.658588239 | 0.864632029 |
| Gm17090 | 0.632076134 | 1.673771768 | 0 | 0.011 | 1 | 0.658583583 | 2.361144491 | 0.022 | 0.014 | 1 | 0.658583583 | 0.864632029 |
| Fam209 | 0.632076134 | 1.673771768 | 0 | 0.011 | 1 | 0.951997643 | 0.875717664 | 0.022 | 0.021 | 1 | 0.951997643 | 0.864632029 |
| A730011C13Rik | 0.632076134 | 1.673771768 | 0 | 0.011 | 1 | 0.951997643 | 0.875717664 | 0.022 | 0.021 | 1 | 0.951997643 | 0.864632029 |
| Ermap | 0.632076134 | 1.673771768 | 0 | 0.011 | 1 | 0.658583583 | 1.946106992 | 0.022 | 0.014 | 1 | 0.658583583 | 0.864632029 |
| Oas1a | 0.632076134 | 0.351843673 | 0 | 0.011 | 1 | 0.658583583 | 2.361144491 | 0.022 | 0.014 | 1 | 0.658583583 | 0.864632029 |
| Zfp11 | 0.632076134 | 1.673771768 | 0 | 0.011 | 1 | 0.687197015 | 1.223640968 | 0.022 | 0.035 | 1 | 0.687197015 | 0.864632029 |
| Gm57270 | 0.632076134 | 1.673771768 | 0 | 0.011 | 1 | 0.80624346 | 2.098110085 | 0.022 | 0.017 | 1 | 0.80624346 | 0.864632029 |
| Gnb3 | 0.632076134 | 1.673771768 | 0 | 0.011 | 1 | 0.806246349 | 2.460680165 | 0.022 | 0.017 | 1 | 0.806246349 | 0.864632029 |
| Zfp109 | 0.632076134 | 1.673771768 | 0 | 0.011 | 1 | 0.661444614 | 1.683072586 | 0.022 | 0.014 | 1 | 0.661444614 | 0.864632029 |
| Gm10974 | 0.632076134 | 1.673771768 | 0 | 0.011 | 1 | 0.944008652 | 0.946106992 | 0.022 | 0.021 | 1 | 0.944008652 | 0.864632029 |
| Gm15489 | 0.632076134 | 1.673771768 | 0 | 0.011 | 1 | 0.957324717 | 0.513147585 | 0.022 | 0.021 | 1 | 0.957324717 | 0.864632029 |
| 1500002F19Rik | 0.632076134 | 1.673771768 | 0 | 0.011 | 1 | 0.670048992 | 1.361144491 | 0.022 | 0.014 | 1 | 0.670048992 | 0.864632029 |
| Pkd1l3 | 0.632076134 | 1.673771768 | 0 | 0.011 | 1 | 0.687208248 | 1.098110085 | 0.022 | 0.035 | 1 | 0.687208248 | 0.864632029 |
| Filip1 | 0.632076134 | -0.648156327 | 0 | 0.011 | 1 | 0.930092617 | -0.124282336 | 0.022 | 0.024 | 1 | 0.930092617 | 0.864632029 |
| Sycp3 | 0.632076134 | 1.673771768 | 0 | 0.011 | 1 | 0.952529269 | 1.098110085 | 0.022 | 0.024 | 1 | 0.952529269 | 0.864632029 |
| Syt1 | 0.632076134 | 1.673771768 | 0 | 0.011 | 1 | 0.636631625 | 0.91753784 | 0.044 | 0.031 | 1 | 0.636631625 | 0.864632029 |
| Myl6b | 0.632076134 | 1.673771768 | 0 | 0.011 | 1 | 0.670048992 | 1.361144491 | 0.022 | 0.014 | 1 | 0.670048992 | 0.864632029 |
| Dtnbos | 0.632076134 | 1.673771768 | 0 | 0.011 | 1 | 0.852141123 | 0.946106992 | 0.022 | 0.028 | 1 | 0.852141123 | 0.864632029 |
| Stab1 | 0.632076134 | 1.673771768 | 0 | 0.011 | 1 | 0.954659411 | 0.683072586 | 0.022 | 0.021 | 1 | 0.954659411 | 0.864632029 |
| Gm49521 | 0.632076134 | 1.088809267 | 0 | 0.011 | 1 | 0.852141123 | 0.946106992 | 0.022 | 0.028 | 1 | 0.852141123 | 0.864632029 |
| Car15 | 0.632076134 | 1.673771768 | 0 | 0.011 | 1 | 0.80624346 | 2.098110085 | 0.022 | 0.017 | 1 | 0.80624346 | 0.864632029 |
| Fgf12 | 0.632076134 | 0.866416846 | 0 | 0.011 | 1 | 0.806246349 | 2.005000681 | 0.022 | 0.017 | 1 | 0.806246349 | 0.864632029 |
| Dynlt1b | 0.632076134 | 1.673771768 | 0 | 0.011 | 1 | 0.937567971 | 0.683072586 | 0.022 | 0.024 | 1 | 0.937567971 | 0.864632029 |
| Gm3435 | 0.632076134 | 1.673771768 | 0 | 0.011 | 1 | 0.854457549 | 1.098110085 | 0.022 | 0.028 | 1 | 0.854457549 | 0.864632029 |
| Gm35550 | 0.632076134 | 1.673771768 | 0 | 0.011 | 1 | 0.957324717 | 0.223640968 | 0.022 | 0.021 | 1 | 0.957324717 | 0.864632029 |
| Gm50163 | 0.632076134 | 1.673771768 | 0 | 0.011 | 1 | 0.648138259 | 0.808603468 | 0.044 | 0.031 | 1 | 0.648138259 | 0.864632029 |
| Ska3 | 0.901963435 | 0.866416846 | 0.048 | 0.056 | 1 | 0.632805446 | 1.683072586 | 0.044 | 0.031 | 1 | 0.901963435 | 0.865168159 |
| Gm7160 | 0.632917461 | 1.936806174 | 0.048 | 0.03 | 1 | 0.966792311 | 1.776181991 | 0.044 | 0.045 | 1 | 0.966792311 | 0.86525041 |
| Gm57272 | 0.632917461 | 1.936806174 | 0.048 | 0.03 | 1 | 0.8202424 | 1.098110085 | 0.022 | 0.017 | 1 | 0.8202424 | 0.86525041 |
| Paqr4 | 0.632917461 | 1.936806174 | 0.048 | 0.03 | 1 | 0.738006063 | -0.316927414 | 0.022 | 0.031 | 1 | 0.738006063 | 0.86525041 |
| Zbtb44 | 0.893340273 | 0.44596585 | 0.333 | 0.407 | 1 | 0.633331305 | 0.237008191 | 0.667 | 0.612 | 1 | 0.893340273 | 0.865554068 |
| Sbk1 | 0.944804785 | 0.951305743 | 0.095 | 0.104 | 1 | 0.635186993 | -0.124282336 | 0.044 | 0.062 | 1 | 0.944804785 | 0.86691147 |
| Gm41496 | 0.635907859 | 0.759501642 | 0.095 | 0.134 | 1 | 0.673304101 | 0.326928776 | 0.178 | 0.163 | 1 | 0.673304101 | 0.867436913 |
| Gm15345 | 0.636138263 | 1.79930265 | 0.048 | 0.03 | 1 | 0.955025007 | 1.683072586 | 0.022 | 0.024 | 1 | 0.955025007 | 0.867604637 |
| Ccdc82 | 0.637477459 | 0.531997807 | 0.333 | 0.444 | 1 | 0.882718204 | 0.944369851 | 0.4 | 0.481 | 1 | 0.882718204 | 0.868577408 |
| Cpped1 | 0.637675903 | -0.233118828 | 0.143 | 0.179 | 1 | 0.856275989 | 0.683072586 | 0.178 | 0.211 | 1 | 0.856275989 | 0.868721248 |
| Phactr4 | 0.958118453 | 0.614878079 | 0.429 | 0.597 | 1 | 0.637812914 | 0.182998983 | 0.711 | 0.609 | 1 | 0.958118453 | 0.868820514 |
| Zcwpw1 | 0.638246407 | -0.133583154 | 0.048 | 0.075 | 1 | 0.745472714 | 0.966865552 | 0.044 | 0.059 | 1 | 0.745472714 | 0.869134338 |
| Lct | 0.639363014 | 1.673771768 | 0.048 | 0.03 | 1 | 0.854464619 | 1.808603468 | 0.022 | 0.028 | 1 | 0.854464619 | 0.869940964 |
| Zfp26 | 0.641751941 | 0.323274521 | 0.19 | 0.265 | 1 | 0.802443855 | 0.928185084 | 0.244 | 0.336 | 1 | 0.802443855 | 0.871658328 |
| Popdc3 | 0.64258579 | 1.451379346 | 0.048 | 0.03 | 1 | 0.806248082 | 2.567595369 | 0.022 | 0.017 | 1 | 0.806248082 | 0.872255082 |
| Map6 | 0.989455055 | 0.766881172 | 0.095 | 0.097 | 1 | 0.643731325 | 0.583536913 | 0.133 | 0.176 | 1 | 0.989455055 | 0.873072631 |
| Slc39a11 | 0.64423977 | 2.088809267 | 0.381 | 0.466 | 1 | 0.994238426 | 1.802654202 | 0.244 | 0.304 | 1 | 0.994238426 | 0.873434659 |
| Fxyd2 | 0.716022952 | -2.496153234 | 0.286 | 0.295 | 1 | 0.646343508 | -0.81233833 | 0.067 | 0.09 | 1 | 0.716022952 | 0.874927086 |
| Gm47761 | 0.647471572 | 0.55829455 | 0.048 | 0.078 | 1 | 0.806246349 | 2.197645759 | 0.022 | 0.017 | 1 | 0.806246349 | 0.875723707 |
| Taf1b | 0.648558969 | 0.773307441 | 0.286 | 0.28 | 1 | 0.940347997 | 0.875717664 | 0.267 | 0.294 | 1 | 0.940347997 | 0.876489201 |
| Igsf3 | 0.696051287 | 0.180392817 | 0.476 | 0.653 | 1 | 0.650511893 | 0.2633947 | 0.8 | 0.893 | 1 | 0.696051287 | 0.877858063 |
| Rhbdl3 | 0.651794844 | 1.171271427 | 0.048 | 0.078 | 1 | 0.670164513 | 1.383512304 | 0.133 | 0.17 | 1 | 0.670164513 | 0.878753169 |

| | | | | | | | | | | | | |
|---|---|---|---|---|---|---|---|---|---|---|---|---|
| Pla1a | 0.732240497 | 1.029915578 | 0.048 | 0.071 | 1 | 0.652015384 | 0.268035087 | 0.044 | 0.031 | 1 | 0.732240497 | 0.878906707 |
| Tmem120b | 0.810311249 | 1.460778044 | 0.143 | 0.134 | 1 | 0.653055117 | 1.296049463 | 0.178 | 0.173 | 1 | 0.810311249 | 0.879629248 |
| Sae1 | 0.993161481 | 0.35682708 | 0.429 | 0.601 | 1 | 0.653312065 | 0.389765169 | 0.733 | 0.751 | 1 | 0.993161481 | 0.879807476 |
| Ccdc125 | 0.653956972 | 1.129451251 | 0.048 | 0.078 | 1 | 0.996109072 | 1.76880246 | 0.089 | 0.097 | 1 | 0.996109072 | 0.880254223 |
| a | 0.732873402 | 2.604350367 | 0.143 | 0.19 | 1 | 0.654813855 | 1.308677071 | 0.222 | 0.218 | 1 | 0.732873402 | 0.880846526 |
| Capn5 | 0.654912325 | 0.55829455 | 0.095 | 0.067 | 1 | 0.956899512 | 1.460680165 | 0.067 | 0.066 | 1 | 0.956899512 | 0.880914497 |
| Opa1 | 0.894986195 | 0.614878079 | 0.429 | 0.549 | 1 | 0.655150992 | 0.247253525 | 0.422 | 0.509 | 1 | 0.894986195 | 0.881079162 |
| 4930473A02Rik | 0.655561847 | 1.503846766 | 0.048 | 0.03 | 1 | 0.933853613 | 1.361144491 | 0.044 | 0.048 | 1 | 0.933853613 | 0.881362359 |
| Ttc27 | 0.656430882 | 0.273841161 | 0.238 | 0.205 | 1 | 0.815967896 | 1.013221188 | 0.178 | 0.187 | 1 | 0.815967896 | 0.881960261 |
| Camsap1 | 0.657198843 | 0.570678275 | 0.333 | 0.433 | 1 | 0.981572599 | 0.308437441 | 0.711 | 0.696 | 1 | 0.981572599 | 0.882487367 |
| Yme1l1 | 0.817905233 | 0.271386648 | 0.81 | 0.918 | 1 | 0.65803506 | 0.26217804 | 0.956 | 0.91 | 1 | 0.817905233 | 0.88305998 |
| Cap2 | 0.8753172 | 0.936806174 | 0.048 | 0.041 | 1 | 0.658583583 | 2.361144491 | 0.022 | 0.014 | 1 | 0.8753172 | 0.883434831 |
| Gm56530 | 0.79666227 | 1.258734268 | 0.048 | 0.037 | 1 | 0.658587075 | 2.098110085 | 0.022 | 0.014 | 1 | 0.79666227 | 0.883437215 |
| Ubash3b | 0.732460135 | 1.984111888 | 0.143 | 0.187 | 1 | 0.661264412 | -0.169370225 | 0.089 | 0.111 | 1 | 0.732460135 | 0.885258202 |
| Abhd1 | 0.985457491 | 0.766881172 | 0.048 | 0.049 | 1 | 0.661727242 | -0.223818009 | 0.022 | 0.035 | 1 | 0.985457491 | 0.885571541 |
| Nkx6-1 | 0.822006883 | 0.97333205 | 0.048 | 0.037 | 1 | 0.661727509 | -0.316927414 | 0.022 | 0.035 | 1 | 0.822006883 | 0.885571722 |
| Pdp1 | 0.864168997 | 0.410737362 | 0.143 | 0.131 | 1 | 0.662479745 | 1.313838777 | 0.178 | 0.17 | 1 | 0.864168997 | 0.886080077 |
| Socs3 | 0.663193941 | -0.553763957 | 0.524 | 0.526 | 1 | 0.681753966 | -0.996072354 | 0.6 | 0.509 | 1 | 0.681753966 | 0.886561679 |
| Elovl7 | 0.775444022 | 0.631460963 | 0.238 | 0.243 | 1 | 0.663807556 | 2.091878132 | 0.156 | 0.149 | 1 | 0.775444022 | 0.88697464 |
| Gm5535 | 0.665338999 | 0.97333205 | 0.048 | 0.03 | 1 | 0.664307093 | 1.268035087 | 0.022 | 0.014 | 1 | 0.665338999 | 0.887310272 |
| Sgo2a | 0.872226982 | 0.735172312 | 0.048 | 0.041 | 1 | 0.664308814 | 1.223640968 | 0.022 | 0.014 | 1 | 0.872226982 | 0.887311428 |
| Fam117b | 0.665050379 | -0.144987918 | 0.476 | 0.537 | 1 | 0.74572548 | 0.204238436 | 0.778 | 0.72 | 1 | 0.74572548 | 0.887808751 |
| Abca4 | 0.689265129 | 0.719575457 | 0.19 | 0.164 | 1 | 0.665158488 | 1.921232323 | 0.111 | 0.097 | 1 | 0.689265129 | 0.887881162 |
| Prpf6 | 0.814377619 | 0.381856242 | 0.571 | 0.679 | 1 | 0.66594445 | 0.408973016 | 0.756 | 0.706 | 1 | 0.814377619 | 0.88840689 |
| Mroh1 | 0.688016328 | 1.062337056 | 0.19 | 0.168 | 1 | 0.666891388 | 0.583536913 | 0.156 | 0.208 | 1 | 0.688016328 | 0.889038653 |
| Cpsf7 | 0.949935408 | 0.275740694 | 0.476 | 0.578 | 1 | 0.666932624 | 0.180572246 | 0.667 | 0.751 | 1 | 0.949935408 | 0.889066123 |
| Clmn | 0.732838253 | 0.425844254 | 0.238 | 0.313 | 1 | 0.667416532 | 0.160096972 | 0.733 | 0.675 | 1 | 0.732838253 | 0.889388237 |
| Kics2 | 0.784037261 | 1.79930265 | 0.048 | 0.037 | 1 | 0.667598499 | 0.223640968 | 0.022 | 0.035 | 1 | 0.784037261 | 0.889509242 |
| Tekt2 | 0.790349864 | 1.55829455 | 0.048 | 0.037 | 1 | 0.667598499 | 0.223640968 | 0.022 | 0.035 | 1 | 0.790349864 | 0.889509242 |
| Ldhc | 0.668622608 | 0.425844254 | 0.048 | 0.03 | 1 | 0.728223757 | 1.304560963 | 0.044 | 0.035 | 1 | 0.728223757 | 0.890189024 |
| Cdh24 | 0.668740738 | 0.908237021 | 0.095 | 0.071 | 1 | 0.982370621 | 1.066401226 | 0.111 | 0.121 | 1 | 0.982370621 | 0.890267301 |
| P4ha3 | 0.669061053 | 0.742158743 | 0.238 | 0.34 | 1 | 0.797057047 | 0.695145418 | 0.222 | 0.225 | 1 | 0.797057047 | 0.890479414 |
| Ngf | 0.67003977 | 0.697487837 | 0.286 | 0.254 | 1 | 0.799050423 | 2.914398132 | 0.067 | 0.083 | 1 | 0.799050423 | 0.891126247 |
| Psrc1 | 0.712689836 | 1.908237021 | 0.048 | 0.034 | 1 | 0.670048992 | 1.361144491 | 0.022 | 0.014 | 1 | 0.712689836 | 0.891132332 |
| Gm17529 | 0.670916455 | 0.766881172 | 0.095 | 0.071 | 1 | 0.862871659 | 0.65559185 | 0.111 | 0.131 | 1 | 0.862871659 | 0.89170402 |
| Rhoq | 0.890125122 | 0.192645078 | 0.762 | 0.888 | 1 | 0.67172611 | 0.394599788 | 0.911 | 0.945 | 1 | 0.890125122 | 0.892236253 |
| Dach1 | 0.671910312 | 0.150209812 | 0.048 | 0.03 | 1 | 0.812275607 | -0.972279242 | 0.044 | 0.052 | 1 | 0.812275607 | 0.892357157 |
| Gsg1l | 0.960849104 | 0.766881172 | 0.048 | 0.045 | 1 | 0.671941816 | 1.761075098 | 0.089 | 0.076 | 1 | 0.960849104 | 0.892377828 |
| Trak2 | 0.672114731 | 0.690778193 | 0.429 | 0.526 | 1 | 0.713214018 | 0.134451933 | 0.644 | 0.692 | 1 | 0.713214018 | 0.89249125 |
| Top1 | 0.786046489 | 0.27985756 | 0.952 | 0.996 | 1 | 0.672349738 | 0.247686442 | 1 | 0.997 | 1 | 0.786046489 | 0.892645306 |
| Ptk7 | 0.673290211 | 1.054861935 | 0.333 | 0.354 | 1 | 0.8202387 | 0.380773298 | 0.578 | 0.581 | 1 | 0.8202387 | 0.893260714 |
| Pou2f3 | 0.673613205 | 2.008190807 | 0.143 | 0.119 | 1 | 0.945614069 | 1.03537433 | 0.133 | 0.152 | 1 | 0.945614069 | 0.89347166 |
| Cep170 | 0.673728014 | 0.448351654 | 0.381 | 0.537 | 1 | 0.875286109 | 0.35842499 | 0.578 | 0.678 | 1 | 0.875286109 | 0.893546591 |
| Ifit2 | 0.965750989 | 1.316219763 | 0.095 | 0.108 | 1 | 0.674692979 | -0.674479418 | 0.067 | 0.083 | 1 | 0.965750989 | 0.894175342 |
| Nkd2 | 0.675065029 | -0.133583154 | 0.048 | 0.071 | 1 | 0.916247337 | 0.410054092 | 0.044 | 0.042 | 1 | 0.916247337 | 0.894417265 |
| Gatad2a | 0.763301198 | 0.157196242 | 0.571 | 0.709 | 1 | 0.676848279 | 0.44683824 | 0.956 | 0.872 | 1 | 0.763301198 | 0.895572965 |
| Obp2a | 0.961651492 | 0.518493542 | 0.286 | 0.354 | 1 | 0.677410737 | 2.721546734 | 0.133 | 0.176 | 1 | 0.961651492 | 0.895936167 |
| Foxk1 | 0.677920941 | 0.883694837 | 0.381 | 0.414 | 1 | 0.889761104 | 0.44663239 | 0.489 | 0.606 | 1 | 0.889761104 | 0.89626508 |
| Slc25a34 | 0.679687267 | 0.503846766 | 0.048 | 0.075 | 1 | 0.92662154 | 1.137104217 | 0.178 | 0.194 | 1 | 0.92662154 | 0.897399753 |
| Apoa1 | 0.715863632 | 1.673771768 | 0.048 | 0.034 | 1 | 0.680719295 | 0.683072586 | 0.067 | 0.052 | 1 | 0.715863632 | 0.898059831 |
| Fan1 | 0.681679263 | 0.710297644 | 0.143 | 0.116 | 1 | 0.859171847 | 0.875717664 | 0.111 | 0.107 | 1 | 0.859171847 | 0.898671908 |
| Raver2 | 0.997092023 | 0.936806174 | 0.048 | 0.049 | 1 | 0.682012502 | 0.402964667 | 0.044 | 0.062 | 1 | 0.997092023 | 0.898883951 |
| Cubn | 0.996598342 | 1.876264632 | 0.143 | 0.16 | 1 | 0.684123528 | 0.309614191 | 0.089 | 0.114 | 1 | 0.996598342 | 0.900222055 |
| Pitpna | 0.684309493 | 0.284782363 | 1 | 0.963 | 1 | 0.921741486 | 0.293444251 | 0.978 | 0.972 | 1 | 0.921741486 | 0.900339504 |
| 6720427I07Rik | 0.801266525 | 0.586308926 | 0.286 | 0.284 | 1 | 0.685132987 | 0.88398528 | 0.244 | 0.349 | 1 | 0.801266525 | 0.900858764 |
| Rbm44 | 0.748090146 | 0.673771768 | 0.048 | 0.034 | 1 | 0.685255394 | 0.83507568 | 0.022 | 0.035 | 1 | 0.748090146 | 0.900935833 |
| Wdr25 | 0.68623671 | 0.281454345 | 0.048 | 0.071 | 1 | 0.689388459 | 1.618942249 | 0.044 | 0.062 | 1 | 0.689388459 | 0.901552598 |
| Kantr | 0.686349108 | 1.513307095 | 0.143 | 0.209 | 1 | 0.790143706 | 0.298408736 | 0.178 | 0.208 | 1 | 0.790143706 | 0.901623118 |
| Racgap1 | 0.841862533 | 0.316219763 | 0.048 | 0.06 | 1 | 0.687208248 | 1.098110085 | 0.022 | 0.035 | 1 | 0.841862533 | 0.90216132 |
| Ccnb2 | 0.790737419 | -0.303508156 | 0.095 | 0.116 | 1 | 0.689069581 | 1.04835705 | 0.178 | 0.17 | 1 | 0.790737419 | 0.903322275 |
| Tulp3 | 0.690154719 | 1.303822158 | 0.524 | 0.623 | 1 | 0.986546726 | 0.631120507 | 0.4 | 0.547 | 1 | 0.986546726 | 0.903995902 |
| Adamts6 | 0.690589038 | -0.167530486 | 0.143 | 0.179 | 1 | 0.994688728 | 0.268035087 | 0.222 | 0.242 | 1 | 0.994688728 | 0.904264856 |
| Ccdc77 | 0.691033822 | 0.214340149 | 0.19 | 0.243 | 1 | 0.753221188 | 0.478714088 | 0.178 | 0.215 | 1 | 0.753221188 | 0.904539901 |
| Tasor2 | 0.691176346 | 0.586308926 | 0.429 | 0.586 | 1 | 0.969432821 | 0.239237823 | 0.733 | 0.723 | 1 | 0.969432821 | 0.90462795 |
| Prickle3 | 0.845661131 | 0.832469514 | 0.143 | 0.138 | 1 | 0.691994674 | 0.713446235 | 0.111 | 0.145 | 1 | 0.845661131 | 0.905132719 |
| Scn2a | 0.693038715 | 1.351843673 | 0.048 | 0.075 | 1 | 0.807293078 | 1.098110085 | 0.044 | 0.055 | 1 | 0.807293078 | 0.90577477 |
| Gm56807 | 0.693099156 | 1.054861935 | 0.048 | 0.075 | 1 | 0.859106422 | 1.223640968 | 0.067 | 0.062 | 1 | 0.859106422 | 0.905811872 |
| Ube2ql1 | 0.982550637 | 0.673771768 | 0.048 | 0.049 | 1 | 0.693884671 | 1.329435632 | 0.044 | 0.062 | 1 | 0.982550637 | 0.906293405 |
| Gon4l | 0.695956883 | 0.535007698 | 0.429 | 0.5 | 1 | 0.888169824 | 0.370050045 | 0.467 | 0.55 | 1 | 0.888169824 | 0.907557783 |
| Prdm10 | 0.716603467 | 0.195724471 | 0.143 | 0.183 | 1 | 0.696852845 | 0.765534746 | 0.178 | 0.225 | 1 | 0.716603467 | 0.908101802 |
| Ap5m1 | 0.841097142 | 0.503846766 | 0.143 | 0.179 | 1 | 0.697114399 | 0.493248027 | 0.156 | 0.197 | 1 | 0.841097142 | 0.908260313 |
| Apba1 | 0.69773418 | 0.385526799 | 0.143 | 0.194 | 1 | 0.69830137 | 0.19121949 | 0.244 | 0.308 | 1 | 0.69830137 | 0.908635374 |
| Aven | 0.958697163 | 0.474991904 | 0.381 | 0.425 | 1 | 0.698231735 | 0.981555828 | 0.444 | 0.564 | 1 | 0.958697163 | 0.908935914 |
| Zfand6 | 0.942999385 | 0.254031649 | 0.905 | 0.877 | 1 | 0.698563628 | 0.318631064 | 0.933 | 0.934 | 1 | 0.942999385 | 0.909136113 |
| Gm3336 | 0.766007512 | 1.088809267 | 0.095 | 0.078 | 1 | 0.699054884 | -0.549588171 | 0.044 | 0.059 | 1 | 0.766007512 | 0.909432037 |
| Mpp4 | 0.706916727 | 0.171271427 | 0.048 | 0.071 | 1 | 0.699621145 | 0.83507568 | 0.067 | 0.09 | 1 | 0.706916727 | 0.909772544 |
| Gm13391 | 0.784047982 | 2.410737362 | 0.048 | 0.037 | 1 | 0.700106927 | 0.402964667 | 0.067 | 0.087 | 1 | 0.784047982 | 0.910064145 |
| Lrrc23 | 0.701523688 | 0.536268244 | 0.143 | 0.116 | 1 | 0.891166471 | 0.78260826 | 0.133 | 0.156 | 1 | 0.891166471 | 0.910911891 |
| Cwf19l2 | 0.848637347 | 0.719575457 | 0.429 | 0.578 | 1 | 0.703439099 | 0.482375236 | 0.511 | 0.54 | 1 | 0.848637347 | 0.912051632 |
| Plekha4 | 0.729258811 | 0.410737362 | 0.095 | 0.131 | 1 | 0.705132895 | 0.946106992 | 0.111 | 0.097 | 1 | 0.729258811 | 0.91305339 |
| 5330439K02Rik | 0.969051492 | 1.766881172 | 0.048 | 0.052 | 1 | 0.705167793 | 1.982632868 | 0.044 | 0.035 | 1 | 0.969051492 | 0.91307397 |
| Ccnt2 | 0.705351167 | 0.7077191 | 0.524 | 0.653 | 1 | 0.954719218 | 0.152557869 | 0.733 | 0.706 | 1 | 0.954719218 | 0.913182065 |
| Fam111a | 0.706917983 | 0.258734268 | 0.048 | 0.071 | 1 | 0.937567971 | 0.683072586 | 0.022 | 0.024 | 1 | 0.937567971 | 0.914102931 |
| B930046K06Rik | 0.856787907 | 1.55829455 | 0.048 | 0.041 | 1 | 0.707072702 | 1.683072586 | 0.044 | 0.035 | 1 | 0.856787907 | 0.914193598 |
| Bbs10 | 0.97964157 | 1.451379346 | 0.048 | 0.049 | 1 | 0.707072702 | 1.683072586 | 0.044 | 0.035 | 1 | 0.97964157 | 0.914193598 |
| 1700084C06Rik | 0.707904491 | 0.918884266 | 0.095 | 0.075 | 1 | 0.843197492 | 0.935838657 | 0.089 | 0.104 | 1 | 0.843197492 | 0.914680214 |
| Gm42797 | 0.709465802 | 1.936806174 | 0.048 | 0.034 | 1 | 0.914274488 | 0.290755163 | 0.044 | 0.048 | 1 | 0.914274488 | 0.91558988 |
| Qrich2 | 0.709465802 | 1.936806174 | 0.048 | 0.034 | 1 | 0.951997643 | 0.875717664 | 0.022 | 0.021 | 1 | 0.951997643 | 0.91558988 |
| Rskr | 0.709487466 | 2.410737362 | 0.048 | 0.034 | 1 | 0.945046897 | 0.683072586 | 0.022 | 0.024 | 1 | 0.945046897 | 0.915602468 |
| Phka1 | 0.710147179 | 0.929028823 | 0.381 | 0.41 | 1 | 0.991678408 | 0.481438725 | 0.444 | 0.588 | 1 | 0.991678408 | 0.915985342 |
| Grb10 | 0.710830594 | 0.439744646 | 0.81 | 0.91 | 1 | 0.996684797 | 0.139155452 | 0.933 | 0.962 | 1 | 0.996684797 | 0.916381054 |
| Aspm | 0.849993358 | 0.614878079 | 0.095 | 0.086 | 1 | 0.711324756 | 0.197645759 | 0.044 | 0.059 | 1 | 0.849993358 | 0.916666604 |
| Zfp937 | 0.71266839 | 1.79930265 | 0.048 | 0.034 | 1 | 0.809040217 | 1.460680165 | 0.022 | 0.017 | 1 | 0.809040217 | 0.917440546 |
| Gm17494 | 0.71266839 | 1.79930265 | 0.048 | 0.034 | 1 | 0.955023924 | 1.683072586 | 0.022 | 0.024 | 1 | 0.955023924 | 0.917440546 |
| Arl9 | 0.71266839 | 1.79930265 | 0.048 | 0.034 | 1 | 0.933837904 | 0.966865552 | 0.067 | 0.066 | 1 | 0.933837904 | 0.917440546 |
| Rnf225 | 0.71266839 | 1.79930265 | 0.048 | 0.034 | 1 | 0.938685475 | 1.268035087 | 0.022 | 0.021 | 1 | 0.938685475 | 0.917440546 |
| Gm13523 | 0.918789496 | 1.55829455 | 0.048 | 0.045 | 1 | 0.712786874 | 1.180572246 | 0.067 | 0.055 | 1 | 0.918789496 | 0.91750862 |
| Gm41556 | 1 | 0.425844254 | 0.095 | 0.097 | 1 | 0.713019976 | 0.498648015 | 0.067 | 0.087 | 1 | 1 | 0.917642466 |

| | | | | | | | | | | | | |
|---|---|---|---|---|---|---|---|---|---|---|---|---|
| Brip1 | 0.754274416 | 2.27643627 | 0.143 | 0.127 | 1 | 0.713026434 | 0.265220071 | 0.111 | 0.138 | 1 | 0.754274416 | 0.917646172 |
| Zc3h12b | 0.806484677 | 0.230828272 | 0.238 | 0.299 | 1 | 0.715295056 | 0.607784459 | 0.311 | 0.412 | 1 | 0.806484677 | 0.918943095 |
| Gm15265 | 0.715863632 | 1.673771768 | 0.048 | 0.034 | 1 | 0.765728546 | 1.545569062 | 0.022 | 0.031 | 1 | 0.765728546 | 0.919266524 |
| 2810036E18Rik | 0.715873069 | 1.673771768 | 0.048 | 0.034 | 1 | 0.840564713 | 0.361144491 | 0.022 | 0.028 | 1 | 0.840564713 | 0.919271887 |
| Gm34045 | 0.715873069 | 1.673771768 | 0.048 | 0.034 | 1 | 0.916236913 | 0.683072586 | 0.044 | 0.042 | 1 | 0.916236913 | 0.919271887 |
| Spink10 | 0.722278262 | 1.258734268 | 0.048 | 0.034 | 1 | 0.846824149 | 0.15951063 | 0.044 | 0.038 | 1 | 0.846824149 | 0.922870636 |
| Cntn3 | 0.722283873 | 1.351843673 | 0.048 | 0.034 | 1 | 0.809040217 | 1.460680165 | 0.022 | 0.017 | 1 | 0.809040217 | 0.922873753 |
| Aars2 | 0.722290144 | 1.451379346 | 0.048 | 0.034 | 1 | 0.85645734 | 1.098110085 | 0.044 | 0.052 | 1 | 0.85645734 | 0.922877236 |
| Mlxip | 0.99881878 | 0.271673324 | 0.381 | 0.474 | 1 | 0.724040936 | 0.319525402 | 0.711 | 0.651 | 1 | 0.99881878 | 0.923846595 |
| Mau2 | 0.724498068 | 0.456541051 | 0.333 | 0.433 | 1 | 0.972248218 | 0.101931796 | 0.711 | 0.702 | 1 | 0.972248218 | 0.924098685 |
| Pde10a | 0.962661243 | 2.161872729 | 0.095 | 0.101 | 1 | 0.725792323 | 0.221071608 | 0.289 | 0.336 | 1 | 0.962661243 | 0.92481015 |
| Napg | 0.728065791 | 0.255919253 | 0.333 | 0.429 | 1 | 0.918135808 | 0.207339155 | 0.467 | 0.498 | 1 | 0.918135808 | 0.926051786 |
| Parp14 | 0.728808364 | 1.159198595 | 0.095 | 0.078 | 1 | 0.939091268 | 0.818002166 | 0.089 | 0.097 | 1 | 0.939091268 | 0.926455096 |
| Dnm3 | 0.729963459 | 0.684419012 | 0.238 | 0.235 | 1 | 0.733241449 | 0.244951474 | 0.178 | 0.215 | 1 | 0.733241449 | 0.927080266 |
| Aacs | 0.974349698 | 0.873796376 | 0.381 | 0.418 | 1 | 0.730634282 | 0.352923984 | 0.289 | 0.377 | 1 | 0.974349698 | 0.92744211 |
| Pcsk6 | 0.989839407 | 0.598174149 | 0.524 | 0.578 | 1 | 0.730880564 | 0.872733806 | 0.178 | 0.221 | 1 | 0.989839407 | 0.927574729 |
| Gm29107 | 0.784037261 | 1.79930265 | 0.048 | 0.037 | 1 | 0.732103032 | 0.982632868 | 0.044 | 0.035 | 1 | 0.784037261 | 0.928231214 |
| Fanci | 0.732298208 | 1.936806174 | 0.048 | 0.071 | 1 | 0.983407978 | 1.332575339 | 0.067 | 0.069 | 1 | 0.983407978 | 0.92833575 |
| 1700027I24Rik | 0.869136842 | 1.171271427 | 0.048 | 0.041 | 1 | 0.733004449 | 0.313838777 | 0.044 | 0.059 | 1 | 0.869136842 | 0.928713376 |
| Lrrc45 | 0.8806889 | 0.376091219 | 0.143 | 0.164 | 1 | 0.73370418 | 0.697572156 | 0.2 | 0.249 | 1 | 0.8806889 | 0.929086537 |
| Slc25a37 | 0.766935858 | 0.503846766 | 0.238 | 0.295 | 1 | 0.735150911 | 0.798549804 | 0.2 | 0.249 | 1 | 0.766935858 | 0.92985496 |
| Usp36 | 0.737297331 | 0.181918671 | 0.524 | 0.582 | 1 | 0.996602644 | 0.376826114 | 0.667 | 0.779 | 1 | 0.996602644 | 0.930987307 |
| Atg2b | 0.737596622 | 0.544488751 | 0.333 | 0.347 | 1 | 0.796028903 | 0.801717083 | 0.422 | 0.478 | 1 | 0.796028903 | 0.931144467 |
| Gm9801 | 0.793383319 | 0.766881172 | 0.143 | 0.179 | 1 | 0.737913988 | 0.180572246 | 0.111 | 0.138 | 1 | 0.793383319 | 0.931310923 |
| Klhl36 | 0.738393266 | 1.351843673 | 0.048 | 0.034 | 1 | 0.765720982 | 1.361144491 | 0.022 | 0.031 | 1 | 0.765720982 | 0.931561917 |
| Zfp72 | 0.738393266 | 1.351843673 | 0.048 | 0.034 | 1 | 0.810571259 | 0.946106992 | 0.044 | 0.055 | 1 | 0.810571259 | 0.931561917 |
| Bicral | 0.738558108 | 0.188344941 | 0.476 | 0.537 | 1 | 0.822109029 | 0.134635961 | 0.511 | 0.592 | 1 | 0.822109029 | 0.931648137 |
| Gm50399 | 0.799824033 | 1.258734268 | 0.048 | 0.037 | 1 | 0.740122935 | -0.124282336 | 0.022 | 0.031 | 1 | 0.799824033 | 0.932463911 |
| Cfap61 | 0.982554806 | -0.469186186 | 0.048 | 0.049 | 1 | 0.740823546 | 1.28452321 | 0.067 | 0.055 | 1 | 0.982554806 | 0.932827566 |
| Tbc1d22a | 0.741311421 | 0.985715774 | 0.286 | 0.291 | 1 | 0.779645545 | 0.657537494 | 0.356 | 0.363 | 1 | 0.779645545 | 0.933080219 |
| Gm14410 | 0.741559901 | 1.181918671 | 0.143 | 0.127 | 1 | 0.979599398 | 1.142504205 | 0.156 | 0.17 | 1 | 0.979599398 | 0.933208715 |
| Dus2 | 0.741595385 | 0.614878079 | 0.095 | 0.131 | 1 | 0.959069489 | 1.460680165 | 0.089 | 0.1 | 1 | 0.959069489 | 0.933227055 |
| Zfp442 | 0.771430007 | 0.936806174 | 0.048 | 0.067 | 1 | 0.742354298 | 0.846571318 | 0.044 | 0.059 | 1 | 0.771430007 | 0.933618692 |
| Trrap | 0.75355063 | 0.284893429 | 0.571 | 0.716 | 1 | 0.742904181 | 0.218404319 | 0.844 | 0.855 | 1 | 0.75355063 | 0.93390174 |
| Vars2 | 0.890779585 | 0.866416846 | 0.048 | 0.041 | 1 | 0.744358033 | 0.223640968 | 0.022 | 0.031 | 1 | 0.890779585 | 0.934647185 |
| Clec2l | 0.921792415 | 1.451379346 | 0.048 | 0.045 | 1 | 0.744358033 | 0.223640968 | 0.022 | 0.031 | 1 | 0.921792415 | 0.934647185 |
| Fam120c | 0.860911702 | 0.116823643 | 0.286 | 0.328 | 1 | 0.745172329 | 1.068362742 | 0.333 | 0.37 | 1 | 0.860911702 | 0.935062858 |
| Plcg2 | 0.746550832 | 0.683825432 | 0.143 | 0.19 | 1 | 0.899083515 | 0.367949669 | 0.467 | 0.522 | 1 | 0.899083515 | 0.935763519 |
| Gm16272 | 0.979644203 | 2.393663848 | 0.048 | 0.049 | 1 | 0.747037062 | 0.808603468 | 0.044 | 0.059 | 1 | 0.979644203 | 0.936009752 |
| Zfp990 | 0.747755005 | 0.55829455 | 0.095 | 0.075 | 1 | 0.962754392 | 0.95724755 | 0.111 | 0.114 | 1 | 0.962754392 | 0.936372462 |
| Mmp25 | 0.748089844 | 0.586308926 | 0.048 | 0.034 | 1 | 0.81552994 | 1.15951063 | 0.044 | 0.055 | 1 | 0.81552994 | 0.936541273 |
| Dhx58 | 0.748287671 | 1.088809267 | 0.095 | 0.078 | 1 | 0.884070782 | 2.335149283 | 0.044 | 0.042 | 1 | 0.884070782 | 0.936640904 |
| 9430091E24Rik | 0.815642277 | 1.010806755 | 0.048 | 0.063 | 1 | 0.749978658 | 0.975253338 | 0.133 | 0.125 | 1 | 0.815642277 | 0.937489328 |
| Gm15271 | 0.750471577 | 0.327996931 | 0.19 | 0.228 | 1 | 0.80624346 | 2.098110085 | 0.022 | 0.017 | 1 | 0.80624346 | 0.937735566 |
| Cntd1 | 0.751342336 | 0.766881172 | 0.048 | 0.034 | 1 | 0.825853389 | 0.223640968 | 0.022 | 0.017 | 1 | 0.825853389 | 0.938169366 |
| Mtmr10 | 0.751868899 | 1.150209812 | 0.333 | 0.377 | 1 | 0.805767792 | 1.254229287 | 0.311 | 0.374 | 1 | 0.805767792 | 0.938430957 |
| Smad1 | 0.752130152 | 0.10560451 | 0.571 | 0.668 | 1 | 0.900388006 | 0.160304847 | 0.756 | 0.702 | 1 | 0.900388006 | 0.938560538 |
| Tjp1 | 0.953485612 | 0.569976355 | 0.619 | 0.701 | 1 | 0.752783911 | 0.156355226 | 0.911 | 0.81 | 1 | 0.953485612 | 0.938884205 |
| Tpcn1 | 0.752956978 | 0.403682604 | 0.286 | 0.381 | 1 | 0.76579707 | 0.403436213 | 0.511 | 0.606 | 1 | 0.76579707 | 0.938969745 |
| Taf1a | 0.995747381 | 0.366343242 | 0.238 | 0.257 | 1 | 0.753267283 | 0.970354538 | 0.222 | 0.298 | 1 | 0.995747381 | 0.939122966 |
| Gm49519 | 0.754598222 | 0.586308926 | 0.048 | 0.034 | 1 | 0.941347173 | 1.098110085 | 0.022 | 0.021 | 1 | 0.941347173 | 0.939777967 |
| Ankrd13c | 0.754673499 | 0.981576044 | 0.524 | 0.646 | 1 | 0.867410558 | 0.154230172 | 0.778 | 0.744 | 1 | 0.867410558 | 0.939814908 |
| Arhgef5 | 0.983096692 | 0.547239362 | 0.286 | 0.336 | 1 | 0.755021402 | 0.446033389 | 0.311 | 0.367 | 1 | 0.983096692 | 0.939985487 |
| Cstdc2 | 0.936805841 | 0.55829455 | 0.048 | 0.045 | 1 | 0.755040184 | 0.361144491 | 0.022 | 0.031 | 1 | 0.936805841 | 0.939994688 |
| Itga2b | 0.755319062 | 0.425844254 | 0.095 | 0.123 | 1 | 0.994517834 | 1.024109504 | 0.133 | 0.142 | 1 | 0.994517834 | 0.940131239 |
| Acot9 | 0.987320853 | 0.589049089 | 0.429 | 0.563 | 1 | 0.755339202 | 0.158545939 | 0.644 | 0.633 | 1 | 0.987320853 | 0.940141094 |
| Ccnyl1 | 0.757145666 | 0.673771768 | 0.238 | 0.243 | 1 | 0.850023027 | 0.56307281 | 0.333 | 0.377 | 1 | 0.850023027 | 0.941021772 |
| Pcdh9 | 0.758640579 | 0.264380832 | 0.524 | 0.705 | 1 | 0.81187006 | 0.935838657 | 0.711 | 0.685 | 1 | 0.81187006 | 0.94174563 |
| Gm29825 | 0.853720939 | 2.088809267 | 0.048 | 0.041 | 1 | 0.759306107 | 0.460680165 | 0.022 | 0.031 | 1 | 0.853720939 | 0.94206645 |
| Pml | 0.760890285 | -0.233118828 | 0.048 | 0.063 | 1 | 0.917074352 | 1.15951063 | 0.133 | 0.138 | 1 | 0.917074352 | 0.942826544 |
| Pikfyve | 0.761507699 | 0.613650775 | 0.476 | 0.634 | 1 | 0.892480488 | 0.573773944 | 0.6 | 0.671 | 1 | 0.892480488 | 0.943121422 |
| Gm41764 | 0.810898642 | 1.041503552 | 0.095 | 0.086 | 1 | 0.762368136 | 1.567595369 | 0.111 | 0.107 | 1 | 0.810898642 | 0.943531097 |
| D030016M11Rik | 0.921792415 | 1.451379346 | 0.048 | 0.045 | 1 | 0.76358991 | 0.982632868 | 0.022 | 0.031 | 1 | 0.921792415 | 0.944110269 |
| 2410018L13Rik | 0.908000182 | 0.524024648 | 0.19 | 0.198 | 1 | 0.7640668 | 0.503748887 | 0.133 | 0.159 | 1 | 0.908000182 | 0.944335525 |
| Top2a | 0.874955349 | -1.977279923 | 0.048 | 0.056 | 1 | 0.765739892 | 1.098110085 | 0.022 | 0.031 | 1 | 0.874955349 | 0.945122202 |
| R3hdml | 0.778245281 | -1.660128969 | 0.048 | 0.063 | 1 | 0.765741783 | 2.972579203 | 0.022 | 0.031 | 1 | 0.778245281 | 0.945123088 |
| Gm14858 | 0.765801178 | 0.150209812 | 0.048 | 0.063 | 1 | 0.861032947 | 1.232411177 | 0.089 | 0.087 | 1 | 0.861032947 | 0.945150912 |
| Qrich1 | 0.896418 | 0.390317821 | 0.524 | 0.619 | 1 | 0.766298245 | 0.290196737 | 0.889 | 0.817 | 1 | 0.896418 | 0.94538349 |
| Plekhm3 | 1 | 0.673771768 | 0.19 | 0.198 | 1 | 0.766715882 | 0.468059695 | 0.111 | 0.135 | 1 | 1 | 0.94557852 |
| Bicra | 0.766904494 | 0.595769256 | 0.143 | 0.179 | 1 | 0.941360245 | 0.556808814 | 0.311 | 0.353 | 1 | 0.941360245 | 0.945666485 |
| Ndufs1 | 0.766919899 | 0.606657572 | 0.333 | 0.455 | 1 | 0.919945991 | 0.141699354 | 0.556 | 0.602 | 1 | 0.919945991 | 0.945673667 |
| Mc5r | 0.985459372 | 1.258734268 | 0.048 | 0.049 | 1 | 0.767925091 | 2.073019104 | 0.067 | 0.083 | 1 | 0.985459372 | 0.946141237 |
| Zglp1 | 0.768257486 | 0.281454345 | 0.048 | 0.063 | 1 | 0.873031295 | 0.624178897 | 0.067 | 0.062 | 1 | 0.873031295 | 0.946295407 |
| Naa16 | 0.786868859 | 0.586308926 | 0.286 | 0.377 | 1 | 0.768374773 | 0.48487986 | 0.378 | 0.509 | 1 | 0.786868859 | 0.946349754 |
| Lockd | 0.76912025 | 1.524024648 | 0.048 | 0.067 | 1 | 0.804392553 | 1.361144491 | 0.044 | 0.038 | 1 | 0.804392553 | 0.946694541 |
| Khdc4 | 0.932028545 | 0.390077891 | 0.762 | 0.825 | 1 | 0.769846157 | 0.386615179 | 0.933 | 0.875 | 1 | 0.932028545 | 0.947029209 |
| Crnde | 0.991562202 | 0.544488751 | 0.095 | 0.097 | 1 | 0.770638477 | 0.776181991 | 0.089 | 0.08 | 1 | 0.991562202 | 0.947393292 |
| Ncoa4 | 0.770795398 | 0.438555306 | 0.238 | 0.243 | 1 | 0.779595701 | 0.117218908 | 0.489 | 0.498 | 1 | 0.779595701 | 0.94746525 |
| Gm20732 | 0.839460965 | 1.079223218 | 0.286 | 0.321 | 1 | 0.772218582 | 1.013221188 | 0.333 | 0.391 | 1 | 0.839460965 | 0.948115626 |
| Ank2 | 0.772329995 | 0.766881172 | 0.238 | 0.295 | 1 | 0.811306841 | 0.429731577 | 0.378 | 0.394 | 1 | 0.811306841 | 0.948166369 |
| Ptpn14 | 0.77418288 | 0.14902679 | 0.619 | 0.746 | 1 | 0.97928517 | 0.248025126 | 0.867 | 0.886 | 1 | 0.97928517 | 0.949006628 |
| Kif17 | 0.94813989 | 0.214340149 | 0.095 | 0.101 | 1 | 0.776494187 | 0.460680165 | 0.111 | 0.135 | 1 | 0.94813989 | 0.950045152 |
| Slc39a8 | 0.909152259 | -0.437074001 | 0.429 | 0.493 | 1 | 0.777078636 | 0.51748652 | 0.2 | 0.246 | 1 | 0.909152259 | 0.950306066 |
| E130102H24Rik | 0.938403825 | 0.521768674 | 0.19 | 0.22 | 1 | 0.77883916 | 0.838723457 | 0.2 | 0.201 | 1 | 0.938403825 | 0.951087883 |
| Grhl1 | 0.780325865 | 0.503846766 | 0.095 | 0.119 | 1 | 0.91862292 | 0.210584815 | 0.222 | 0.235 | 1 | 0.91862292 | 0.951743275 |
| Cep128 | 0.814564233 | 2.032225739 | 0.095 | 0.123 | 1 | 0.783450032 | 1.223640968 | 0.111 | 0.104 | 1 | 0.814564233 | 0.953106111 |
| Gpd2 | 0.872983749 | 1.04884286 | 0.524 | 0.683 | 1 | 0.783810924 | 0.429931715 | 0.844 | 0.806 | 1 | 0.872983749 | 0.953262284 |
| Dok3 | 0.78405632 | 1.97333205 | 0.048 | 0.037 | 1 | 0.806586479 | 0.473619221 | 0.067 | 0.08 | 1 | 0.806586479 | 0.953368327 |
| Mast1 | 0.784071327 | 3.281454345 | 0.048 | 0.037 | 1 | 0.941347632 | 1.098110085 | 0.022 | 0.021 | 1 | 0.941347632 | 0.953374808 |
| A530017D24Rik | 0.985459372 | 1.258734268 | 0.048 | 0.049 | 1 | 0.787241092 | 0.460680165 | 0.067 | 0.08 | 1 | 0.985459372 | 0.954733647 |
| Gm56507 | 0.966230998 | 1.258734268 | 0.048 | 0.052 | 1 | 0.787621983 | 1.393565969 | 0.044 | 0.055 | 1 | 0.966230998 | 0.954895578 |
| Kif23 | 0.788193244 | -0.233118828 | 0.048 | 0.063 | 1 | 0.854464766 | 1.567595369 | 0.022 | 0.028 | 1 | 0.854464766 | 0.955137898 |
| Armc6 | 0.89388747 | 0.866416846 | 0.048 | 0.041 | 1 | 0.788320053 | 1.242499995 | 0.067 | 0.059 | 1 | 0.89388747 | 0.9551916 |
| Ube2h | 0.789597298 | 0.229606906 | 1 | 0.896 | 1 | 0.892619246 | 0.361144491 | 0.978 | 0.976 | 1 | 0.892619246 | 0.955730703 |
| Gnal | 0.930798611 | 1.088809267 | 0.048 | 0.045 | 1 | 0.790082458 | 0.223640968 | 0.067 | 0.08 | 1 | 0.930798611 | 0.955934626 |
| Gm56933 | 0.790349864 | 1.55829455 | 0.048 | 0.037 | 1 | 0.80624346 | 2.098110085 | 0.022 | 0.017 | 1 | 0.80624346 | 0.956046821 |
| B430212C06Rik | 0.790349864 | 1.55829455 | 0.048 | 0.037 | 1 | 0.936023974 | 1.460680165 | 0.022 | 0.021 | 1 | 0.936023974 | 0.956046821 |
| Gm30551 | 0.793501451 | 1.088809267 | 0.048 | 0.037 | 1 | 0.936025978 | 2.098110085 | 0.022 | 0.021 | 1 | 0.936025978 | 0.957358349 |

| | | | | | | | | | | | | |
|---|---|---|---|---|---|---|---|---|---|---|---|---|
| Ube4bos1 | 0.793508072 | 1.451379346 | 0.048 | 0.037 | 1 | 0.95528377 | 1.005000681 | 0.044 | 0.048 | 1 | 0.95528377 | 0.957361084 |
| Isg15 | 0.795673155 | -0.216999163 | 0.048 | 0.063 | 1 | 0.945322224 | 2.139930261 | 0.067 | 0.073 | 1 | 0.945322224 | 0.95825054 |
| Ccrl2 | 0.79666227 | 1.171271427 | 0.048 | 0.037 | 1 | 0.954659411 | 0.683072586 | 0.022 | 0.021 | 1 | 0.954659411 | 0.958653768 |
| Hk1 | 0.799093237 | 0.193084589 | 0.524 | 0.799 | 1 | 0.851388495 | 0.41152668 | 0.8 | 0.806 | 1 | 0.851388495 | 0.959636473 |
| Plcd1 | 0.800098621 | 0.586308926 | 0.476 | 0.59 | 1 | 0.916921192 | 0.537874671 | 0.489 | 0.616 | 1 | 0.916921192 | 0.960039439 |
| Phc3 | 0.800338373 | 0.254819218 | 0.429 | 0.507 | 1 | 0.833998194 | 0.243540525 | 0.644 | 0.647 | 1 | 0.833998194 | 0.960135235 |
| Stx1b | 0.856787907 | 1.55829455 | 0.048 | 0.041 | 1 | 0.801305703 | 0.242499995 | 0.089 | 0.104 | 1 | 0.856787907 | 0.960520576 |
| Rps6kb1 | 0.843163818 | -0.121262883 | 0.667 | 0.761 | 1 | 0.801332748 | 0.135321165 | 0.933 | 0.903 | 1 | 0.843163818 | 0.960531323 |
| Wasf2 | 0.934523682 | 0.250981993 | 0.81 | 0.929 | 1 | 0.802332215 | 0.147236943 | 0.978 | 0.945 | 1 | 0.934523682 | 0.960927447 |
| Dagla | 0.915506323 | 1.673771768 | 0.048 | 0.056 | 1 | 0.802369204 | 0.905465008 | 0.067 | 0.059 | 1 | 0.915506323 | 0.960942069 |
| Itgb6 | 0.969051544 | 2.088809267 | 0.048 | 0.052 | 1 | 0.802385429 | 0.587915353 | 0.067 | 0.059 | 1 | 0.969051544 | 0.960948481 |
| Slco1b2 | 0.802987008 | 1.088809267 | 0.048 | 0.037 | 1 | 0.883933301 | 1.632446513 | 0.044 | 0.052 | 1 | 0.883933301 | 0.961185881 |
| Gm37982 | 0.803145338 | 0.258734268 | 0.048 | 0.063 | 1 | 0.854457549 | 1.098110085 | 0.022 | 0.028 | 1 | 0.854457549 | 0.961248242 |
| Arid4a | 0.804014191 | 0.331051871 | 0.667 | 0.616 | 1 | 0.936519685 | -0.110383432 | 0.911 | 0.796 | 1 | 0.936519685 | 0.961589563 |
| Pafah1b1 | 0.971760551 | 0.232984697 | 1 | 0.993 | 1 | 0.804241132 | 0.192521401 | 1 | 1 | 1 | 0.971760551 | 0.961678466 |
| Gm43707 | 0.933802207 | 1.088809267 | 0.048 | 0.045 | 1 | 0.805650661 | 0.808603468 | 0.044 | 0.055 | 1 | 0.933802207 | 0.962228335 |
| Bsn | 0.957800805 | 0.79930265 | 0.048 | 0.052 | 1 | 0.80618568 | 0.982632868 | 0.111 | 0.131 | 1 | 0.957800805 | 0.962436009 |
| Gm34655 | 0.815673353 | 1.214340149 | 0.048 | 0.037 | 1 | 0.80624346 | 2.42003818 | 0.022 | 0.017 | 1 | 0.815673353 | 0.962458403 |
| Clvs1 | 0.921315205 | 0.214340149 | 0.048 | 0.052 | 1 | 0.80624346 | 2.42003818 | 0.022 | 0.017 | 1 | 0.921315205 | 0.962458403 |
| Gm11520 | 0.853728162 | 2.010806755 | 0.048 | 0.041 | 1 | 0.806246349 | 2.268035087 | 0.022 | 0.017 | 1 | 0.853728162 | 0.962459523 |
| Srcin1 | 0.901975479 | 0.815790773 | 0.048 | 0.056 | 1 | 0.807507114 | 0.595609745 | 0.111 | 0.1 | 1 | 0.901975479 | 0.962946489 |
| Rev1 | 0.821684539 | 0.434700605 | 0.571 | 0.616 | 1 | 0.808041649 | 0.203842025 | 0.8 | 0.82 | 1 | 0.821684539 | 0.963151992 |
| Gm32673 | 0.809630777 | 0.760483401 | 0.19 | 0.172 | 1 | 0.808066156 | 0.572041274 | 0.178 | 0.218 | 1 | 0.809630777 | 0.963161399 |
| Nxpe3 | 0.808137022 | 0.673771768 | 0.048 | 0.063 | 1 | 0.863715585 | 1.15951063 | 0.067 | 0.062 | 1 | 0.863715585 | 0.963188598 |
| Slc25a42 | 0.8081653 | 0.351843673 | 0.095 | 0.078 | 1 | 0.97601511 | 1.242499995 | 0.044 | 0.045 | 1 | 0.97601511 | 0.963199448 |
| Pcyt1a | 0.931051727 | 0.327996931 | 0.476 | 0.716 | 1 | 0.808308897 | 0.359113956 | 0.822 | 0.799 | 1 | 0.931051727 | 0.963254521 |
| Tnrc6b | 0.82382811 | 0.136830782 | 0.952 | 0.974 | 1 | 0.809079636 | 0.193992761 | 0.978 | 0.979 | 1 | 0.82382811 | 0.963549415 |
| Bmx | 0.915506323 | 2.48112669 | 0.048 | 0.056 | 1 | 0.814636613 | 1.098110085 | 0.022 | 0.017 | 1 | 0.915506323 | 0.965640415 |
| Sytl5 | 0.815692705 | 1.400753273 | 0.048 | 0.063 | 1 | 0.949334008 | 0.180572246 | 0.022 | 0.021 | 1 | 0.949334008 | 0.966030821 |
| Nfkbia | 0.936444915 | -0.502305461 | 0.762 | 0.608 | 1 | 0.818191742 | -0.170162697 | 0.911 | 0.727 | 1 | 0.936444915 | 0.966945757 |
| Klhl7 | 0.907098174 | 0.702530761 | 0.429 | 0.567 | 1 | 0.818374731 | 0.382677653 | 0.6 | 0.571 | 1 | 0.907098174 | 0.967012262 |
| Msh4 | 0.81881728 | 1.088809267 | 0.048 | 0.037 | 1 | 0.953500769 | 1.320502507 | 0.044 | 0.048 | 1 | 0.953500769 | 0.967172822 |
| Gm29681 | 0.81881728 | 0.866416846 | 0.048 | 0.037 | 1 | 0.875329398 | 0.966865552 | 0.044 | 0.052 | 1 | 0.875329398 | 0.967172822 |
| 4933433G15Rik | 0.81881728 | 1.088809267 | 0.048 | 0.037 | 1 | 0.944573962 | 0.757073168 | 0.044 | 0.048 | 1 | 0.944573962 | 0.967172822 |
| Tsbp1 | 0.81881728 | 1.088809267 | 0.048 | 0.037 | 1 | 0.891628083 | 1.098110085 | 0.044 | 0.042 | 1 | 0.891628083 | 0.967172822 |
| Relb | 0.819419798 | 0.503846766 | 0.286 | 0.366 | 1 | 0.888869039 | 0.652826299 | 0.356 | 0.443 | 1 | 0.888869039 | 0.967390791 |
| Sync | 0.85370107 | 1.673771768 | 0.048 | 0.041 | 1 | 0.819747895 | 1.304560963 | 0.044 | 0.038 | 1 | 0.85370107 | 0.967509179 |
| Prdm1 | 0.930795883 | 1.088809267 | 0.048 | 0.045 | 1 | 0.819806573 | -2.098287127 | 0.022 | 0.028 | 1 | 0.930795883 | 0.967530329 |
| Prdm15 | 0.822713745 | 0.866416846 | 0.143 | 0.134 | 1 | 0.96115863 | 0.614901084 | 0.222 | 0.263 | 1 | 0.96115863 | 0.968569584 |
| Megf10 | 0.869134278 | 0.451379346 | 0.048 | 0.041 | 1 | 0.823042771 | 0.875717664 | 0.022 | 0.017 | 1 | 0.869134278 | 0.968686139 |
| 1700010K24Rik | 0.918519165 | 0.150209812 | 0.048 | 0.052 | 1 | 0.824407303 | -0.316927414 | 0.022 | 0.028 | 1 | 0.918519165 | 0.969167205 |
| Gm35188 | 0.836649017 | 0.673771768 | 0.095 | 0.119 | 1 | 0.826502332 | 1.075390009 | 0.133 | 0.166 | 1 | 0.836649017 | 0.969898559 |
| Casc1 | 0.82847504 | 0.181918671 | 0.095 | 0.112 | 1 | 0.9091447 | 1.683072586 | 0.156 | 0.173 | 1 | 0.9091447 | 0.970579188 |
| A930037H05Rik | 0.921315205 | 0.351843673 | 0.048 | 0.052 | 1 | 0.829012818 | -0.124282336 | 0.022 | 0.028 | 1 | 0.921315205 | 0.970763383 |
| Dand5 | 0.933336741 | 0.726239188 | 0.143 | 0.138 | 1 | 0.834675064 | 1.259861155 | 0.111 | 0.111 | 1 | 0.933336741 | 0.972667665 |
| Inhba | 0.905585182 | 1.170493065 | 0.19 | 0.209 | 1 | 0.838252226 | -0.375821103 | 0.022 | 0.028 | 1 | 0.905585182 | 0.973837658 |
| Gm15952 | 0.995710668 | 0.214340149 | 0.095 | 0.093 | 1 | 0.838524879 | 0.683072586 | 0.067 | 0.076 | 1 | 0.995710668 | 0.973925785 |
| H4c3 | 0.960849104 | 0.766881172 | 0.048 | 0.045 | 1 | 0.839081256 | 0.683072586 | 0.044 | 0.038 | 1 | 0.960849104 | 0.974105158 |
| Fsd1l | 0.99278889 | -0.252227651 | 0.143 | 0.138 | 1 | 0.839806956 | 0.653325243 | 0.178 | 0.211 | 1 | 0.99278889 | 0.974338189 |
| Zfp516 | 0.841418873 | 0.969637566 | 0.476 | 0.638 | 1 | 0.931033739 | 0.124679806 | 0.889 | 0.827 | 1 | 0.931033739 | 0.974852026 |
| Stk10 | 0.969626561 | 0.95986141 | 0.333 | 0.422 | 1 | 0.841650599 | 0.561936149 | 0.511 | 0.592 | 1 | 0.969626561 | 0.974925467 |
| Atcayos | 0.960849104 | 0.766881172 | 0.048 | 0.045 | 1 | 0.842081174 | 1.197645759 | 0.067 | 0.062 | 1 | 0.960849104 | 0.975061644 |
| Slc8a3 | 0.842281576 | 0.777373546 | 0.429 | 0.474 | 1 | 0.966350178 | 0.247443992 | 0.267 | 0.301 | 1 | 0.966350178 | 0.975124899 |
| Itsn1 | 0.945401399 | 0.255628358 | 0.381 | 0.463 | 1 | 0.842862706 | 0.356643079 | 0.689 | 0.723 | 1 | 0.945401399 | 0.975307871 |
| Gdi2 | 0.873912259 | 0.237586494 | 0.857 | 0.94 | 1 | 0.843975653 | 0.144432886 | 0.978 | 0.972 | 1 | 0.873912259 | 0.975656403 |
| E2f3 | 0.993887193 | 0.644024424 | 0.19 | 0.209 | 1 | 0.845280172 | 0.206258889 | 0.311 | 0.349 | 1 | 0.993887193 | 0.976061775 |
| Mdn1 | 0.845336906 | 0.463061151 | 0.381 | 0.459 | 1 | 0.938220936 | -0.124282336 | 0.756 | 0.633 | 1 | 0.938220936 | 0.976079327 |
| Lrp8 | 0.898588431 | 0.227515538 | 0.238 | 0.272 | 1 | 0.848378723 | 0.376017768 | 0.289 | 0.349 | 1 | 0.898588431 | 0.977010988 |
| Polg2 | 0.902692883 | -0.184209227 | 0.19 | 0.201 | 1 | 0.848536221 | 0.83698527 | 0.244 | 0.287 | 1 | 0.902692883 | 0.977058724 |
| Gm48956 | 0.849665028 | 0.400753273 | 0.048 | 0.06 | 1 | 0.944008652 | 0.683072586 | 0.022 | 0.021 | 1 | 0.944008652 | 0.977399396 |
| Pcdhga5 | 0.852101758 | 0.866416846 | 0.095 | 0.086 | 1 | 0.978944224 | 1.223640968 | 0.089 | 0.093 | 1 | 0.978944224 | 0.97812611 |
| Gm38537 | 0.853716123 | 3.503846766 | 0.048 | 0.041 | 1 | 0.852141123 | 0.946106992 | 0.022 | 0.028 | 1 | 0.853716123 | 0.978137752 |
| C330018D20Rik | 0.89388747 | 0.866416846 | 0.048 | 0.041 | 1 | 0.853050792 | 0.875717664 | 0.044 | 0.052 | 1 | 0.89388747 | 0.97840593 |
| Dcaf8 | 0.85367296 | 0.315401677 | 0.667 | 0.769 | 1 | 0.861011546 | 0.252022769 | 0.867 | 0.848 | 1 | 0.861011546 | 0.978588397 |
| Abca6 | 0.85370107 | 2.673771768 | 0.048 | 0.041 | 1 | 0.955025007 | 1.513147585 | 0.022 | 0.024 | 1 | 0.955025007 | 0.978596623 |
| Zfp341 | 0.853715972 | 2.673771768 | 0.048 | 0.041 | 1 | 0.994464236 | 0.361144491 | 0.044 | 0.045 | 1 | 0.994464236 | 0.978600983 |
| Gm42743 | 0.976738154 | -0.413691074 | 0.048 | 0.049 | 1 | 0.854466975 | 0.982632868 | 0.022 | 0.028 | 1 | 0.976738154 | 0.978820139 |
| Il18bp | 0.85825884 | 0.400753273 | 0.095 | 0.082 | 1 | 0.97274657 | 1.476621709 | 0.089 | 0.097 | 1 | 0.97274657 | 0.979909443 |
| Crebrf | 0.858854892 | 0.936806174 | 0.571 | 0.787 | 1 | 0.887970433 | 0.451835508 | 0.844 | 0.785 | 1 | 0.887970433 | 0.980078059 |
| Mocos | 0.896995562 | 0.766881172 | 0.048 | 0.041 | 1 | 0.859055851 | 1.750186782 | 0.067 | 0.062 | 1 | 0.896995562 | 0.980134747 |
| 4930578M01Rik | 0.899259517 | 0.735172312 | 0.048 | 0.056 | 1 | 0.859862723 | 0.683072586 | 0.044 | 0.052 | 1 | 0.899259517 | 0.980361544 |
| Pcdhga8 | 0.859868645 | 1.351843673 | 0.048 | 0.041 | 1 | 0.908650103 | 1.197645759 | 0.044 | 0.042 | 1 | 0.908650103 | 0.980363203 |
| 4833438C02Rik | 0.862687636 | 1.088809267 | 0.048 | 0.06 | 1 | 0.935071939 | 0.513147585 | 0.022 | 0.024 | 1 | 0.935071939 | 0.981145315 |
| Cpt1c | 0.863529756 | 0.719575457 | 0.143 | 0.179 | 1 | 0.935868697 | 1.435145073 | 0.156 | 0.163 | 1 | 0.935868697 | 0.981375872 |
| Msantd1 | 0.865290804 | 1.171271427 | 0.048 | 0.06 | 1 | 0.985236045 | 0.361144491 | 0.044 | 0.045 | 1 | 0.985236045 | 0.981853432 |
| Rps6kc1 | 0.92224816 | 0.830497271 | 0.238 | 0.243 | 1 | 0.86664451 | 0.920111783 | 0.333 | 0.433 | 1 | 0.92224816 | 0.982216313 |
| Gm41077 | 0.931971818 | 0.451379346 | 0.143 | 0.16 | 1 | 0.871315737 | 1.201945896 | 0.133 | 0.163 | 1 | 0.931971818 | 0.98344036 |
| Pcdhga6 | 0.872207635 | 0.586308926 | 0.048 | 0.056 | 1 | 0.882429463 | 0.91753784 | 0.089 | 0.087 | 1 | 0.882429463 | 0.983669112 |
| Epm2a | 0.873124156 | 1.199840579 | 0.095 | 0.116 | 1 | 0.910467444 | 1.062920657 | 0.133 | 0.152 | 1 | 0.910467444 | 0.98390252 |
| Gm44899 | 0.873657428 | 1.214340149 | 0.143 | 0.142 | 1 | 0.983690836 | 0.739656115 | 0.133 | 0.145 | 1 | 0.983690836 | 0.984037555 |
| Gria1 | 0.891009781 | 1.486144765 | 0.095 | 0.093 | 1 | 0.877055665 | 1.78260826 | 0.044 | 0.052 | 1 | 0.891009781 | 0.984884691 |
| Ddx11 | 0.923542405 | 0.536268244 | 0.095 | 0.108 | 1 | 0.885126399 | 0.498648015 | 0.089 | 0.087 | 1 | 0.923542405 | 0.986804056 |
| Dnmt3b | 0.921315271 | 0.351843673 | 0.048 | 0.052 | 1 | 0.887846181 | 1.197645759 | 0.044 | 0.042 | 1 | 0.921315271 | 0.987421521 |
| Gm5544 | 0.889737763 | 0.629377648 | 0.143 | 0.142 | 1 | 0.997962872 | 0.907778873 | 0.156 | 0.17 | 1 | 0.997962872 | 0.987842239 |
| Aox1 | 0.951535679 | 0.648236676 | 0.095 | 0.097 | 1 | 0.890469138 | 0.329435632 | 0.089 | 0.087 | 1 | 0.951535679 | 0.98800299 |
| Mir17hg | 0.890779585 | 0.673771768 | 0.048 | 0.041 | 1 | 1 | 0.757073168 | 0.044 | 0.045 | 1 | 1 | 0.988070901 |
| Flot2 | 0.893226358 | 0.339871031 | 0.238 | 0.284 | 1 | 0.913174352 | 0.262229465 | 0.356 | 0.336 | 1 | 0.913174352 | 0.988599389 |
| Fam222a | 0.988369564 | 0.866416846 | 0.048 | 0.049 | 1 | 0.898606929 | 1.545569062 | 0.067 | 0.066 | 1 | 0.988369564 | 0.989719445 |
| Mtcp1 | 0.987342144 | 0.673771768 | 0.095 | 0.097 | 1 | 0.901661335 | 0.955152132 | 0.133 | 0.156 | 1 | 0.987342144 | 0.990329507 |
| 2310008N11Rik | 0.936799984 | 1.195724471 | 0.095 | 0.097 | 1 | 0.905396226 | 0.595609745 | 0.111 | 0.111 | 1 | 0.936799984 | 0.991050126 |
| Gm35417 | 0.912810205 | 1.60338244 | 0.048 | 0.056 | 1 | 0.909349255 | 0.557541704 | 0.089 | 0.1 | 1 | 0.912810205 | 0.991782442 |
| E230029C05Rik | 0.940954861 | 0.503846766 | 0.048 | 0.052 | 1 | 0.916236913 | 0.83507568 | 0.044 | 0.042 | 1 | 0.940954861 | 0.992983745 |
| Pde4b | 0.969370808 | 0.645625462 | 0.667 | 0.746 | 1 | 0.921076162 | -0.220284156 | 0.867 | 0.765 | 1 | 0.969370808 | 0.993771028 |
| Igflr1 | 0.921792415 | 1.451379346 | 0.048 | 0.045 | 1 | 0.992618127 | 1.361144491 | 0.044 | 0.045 | 1 | 0.992618127 | 0.993883574 |
| Col15a1 | 0.979645135 | 2.171271427 | 0.048 | 0.049 | 1 | 0.922702524 | 1.041526557 | 0.133 | 0.138 | 1 | 0.979645135 | 0.9940251 |
| Ptgs1 | 0.925354349 | 0.271673324 | 0.333 | 0.358 | 1 | 0.966337792 | 1.301201951 | 0.156 | 0.183 | 1 | 0.966337792 | 0.994428027 |
| Gng8 | 0.927794995 | 1.088809267 | 0.048 | 0.045 | 1 | 0.954659411 | -0.223818009 | 0.022 | 0.021 | 1 | 0.954659411 | 0.994786437 |
| Rpp40 | 0.927798188 | 1.258734268 | 0.048 | 0.045 | 1 | 0.933318841 | 0.460680165 | 0.044 | 0.042 | 1 | 0.933318841 | 0.994786898 |

| | | | | | | | | | | | | |
|---|---|---|---|---|---|---|---|---|---|---|---|---|
| Rab27b | 0.928774292 | 0.425844254 | 0.381 | 0.399 | 1 | 0.96012074 | 0.156324699 | 0.578 | 0.498 | 1 | 0.96012074 | 0.994926898 |
| Gm43848 | 0.987128902 | 0.400753273 | 0.095 | 0.093 | 1 | 0.929312066 | 0.683072586 | 0.089 | 0.09 | 1 | 0.987128902 | 0.995003216 |
| Gm34102 | 0.930798611 | 1.171271427 | 0.048 | 0.045 | 1 | 0.951712893 | 0.91753784 | 0.044 | 0.048 | 1 | 0.951712893 | 0.995211168 |
| Gm57404 | 0.98545851 | 1.171271427 | 0.048 | 0.049 | 1 | 0.933318841 | 0.460680165 | 0.044 | 0.042 | 1 | 0.98545851 | 0.995553623 |
| Gm17103 | 0.933804626 | 1.088809267 | 0.048 | 0.045 | 1 | 0.979705049 | 0.946106992 | 0.044 | 0.045 | 1 | 0.979705049 | 0.995618172 |
| Gm1123 | 0.966235059 | 1.425844254 | 0.048 | 0.052 | 1 | 0.933812514 | 1.335149283 | 0.111 | 0.125 | 1 | 0.966235059 | 0.995619217 |
| Bdkrb2 | 0.939808708 | 0.936806174 | 0.048 | 0.045 | 1 | 0.936025978 | 2.513147585 | 0.022 | 0.021 | 1 | 0.939808708 | 0.995907325 |
| Rarg | 0.973730687 | 0.159463821 | 0.762 | 0.836 | 1 | 0.984700272 | -0.100002142 | 0.978 | 0.931 | 1 | 0.984700272 | 0.999309923 |
| Usp38 | 0.977822034 | 0.645574876 | 0.286 | 0.328 | 1 | 0.977268151 | 0.504735345 | 0.511 | 0.55 | 1 | 0.977822034 | 0.999483263 |
