## Supplementary File 2 for "Loss of PRDM16 Drives Nasal Septal Deviation through Dysregulated TGFβ2 Signaling"

**Cell cluster composition: Spatial seq 4-wk_WT & cKO cart_14clusters**

| Group | Cluster | Cell_Count | Percentage |
|---|---|---|---|
| CLW1430_cKO1 | 0 | 1388 | 16.51791 |
| CLW1432_17_cKO2 | 0 | 1695 | 19.808344 |
| CLW1434_WT2 | 0 | 1623 | 17.42351 |
| CLW1435_WT1 | 0 | 2189 | 22.189559 |
| CLW1430_cKO1 | 1 | 1621 | 19.29073 |
| CLW1432_17_cKO2 | 1 | 1593 | 18.616338 |
| CLW1434_WT2 | 1 | 1864 | 20.010735 |
| CLW1435_WT1 | 1 | 1684 | 17.070451 |
| CLW1430_cKO1 | 2 | 2062 | 24.538855 |
| CLW1432_17_cKO2 | 2 | 1789 | 20.90686 |
| CLW1434_WT2 | 2 | 1091 | 11.712292 |
| CLW1435_WT1 | 2 | 1663 | 16.857577 |
| CLW1430_cKO1 | 3 | 986 | 11.733905 |
| CLW1432_17_cKO2 | 3 | 1053 | 12.305715 |
| CLW1434_WT2 | 3 | 1993 | 21.395598 |
| CLW1435_WT1 | 3 | 980 | 9.9341105 |
| CLW1430_cKO1 | 4 | 753 | 8.9610853 |
| CLW1432_17_cKO2 | 4 | 691 | 8.07526 |
| CLW1434_WT2 | 4 | 853 | 9.1572732 |
| CLW1435_WT1 | 4 | 1321 | 13.390775 |
| CLW1430_cKO1 | 5 | 385 | 4.581697 |
| CLW1432_17_cKO2 | 5 | 414 | 4.8381442 |
| CLW1434_WT2 | 5 | 481 | 5.1637144 |
| CLW1435_WT1 | 5 | 367 | 3.720223 |
| CLW1430_cKO1 | 6 | 376 | 4.4745924 |
| CLW1432_17_cKO2 | 6 | 419 | 4.8965759 |
| CLW1434_WT2 | 6 | 412 | 4.4229737 |
| CLW1435_WT1 | 6 | 576 | 5.8388241 |
| CLW1430_cKO1 | 7 | 208 | 2.4753064 |
| CLW1432_17_cKO2 | 7 | 202 | 2.3606404 |
| CLW1434_WT2 | 7 | 186 | 1.9967794 |
| CLW1435_WT1 | 7 | 370 | 3.7506336 |
| CLW1430_cKO1 | 8 | 233 | 2.7728192 |
| CLW1432_17_cKO2 | 8 | 177 | 2.0684819 |
| CLW1434_WT2 | 8 | 207 | 2.2222222 |
| CLW1435_WT1 | 8 | 240 | 2.4328434 |
| CLW1430_cKO1 | 9 | 173 | 2.0587885 |
| CLW1432_17_cKO2 | 9 | 196 | 2.2905224 |
| CLW1434_WT2 | 9 | 204 | 2.1900161 |
| CLW1435_WT1 | 9 | 191 | 1.9361379 |
| CLW1430_cKO1 | 10 | 95 | 1.1305486 |
| CLW1432_17_cKO2 | 10 | 176 | 2.0567956 |
| CLW1434_WT2 | 10 | 204 | 2.1900161 |

| | | | |
|---|---|---|---|
| CLW1435_WT1 | 10 | 210 | 2.128738 |
| CLW1430_cKO1 | 11 | 44 | 0.5236225 |
| CLW1432_17_cKO2 | 11 | 78 | 0.9115344 |
| CLW1434_WT2 | 11 | 64 | 0.6870639 |
| CLW1435_WT1 | 11 | 29 | 0.2939686 |
| CLW1430_cKO1 | 12 | 79 | 0.9401404 |
| CLW1432_17_cKO2 | 12 | 74 | 0.8647891 |
| CLW1434_WT2 | 12 | 13 | 0.1395598 |
| CLW1435_WT1 | 12 | 44 | 0.4460213 |
| CLW1430_cKO1 | 13 | 0 | 0 |
| CLW1432_17_cKO2 | 13 | 0 | 0 |
| CLW1434_WT2 | 13 | 120 | 1.2882448 |
| CLW1435_WT1 | 13 | 1 | 0.0101368 |

**Spatial seq: cluster 0**

| | cKO_p_val | cKO_avg_log2FC | cKO_pct.1 | cKO_pct.2 | cKO_p_val_adj | WT_p_val | WT_avg_log2FC | WT_pct.1 | WT_pct.2 | WT_p_val_adj | max_pval | minimump_p_val |
|---|---|---|---|---|---|---|---|---|---|---|---|---|
| Hbb-bs | 0 | 4.238093823 | 0.651 | 0.051 | 0 | 0 | 4.111206606 | 0.536 | 0.047 | 0 | 0 | 0 |
| Hba-a2 | 0 | 4.014988234 | 0.555 | 0.043 | 0 | 0 | 3.717578197 | 0.672 | 0.072 | 0 | 0 | 0 |
| S100a9 | 0 | 4.991388932 | 0.331 | 0.013 | 0 | 0 | 4.325292389 | 0.423 | 0.027 | 0 | 0 | 0 |
| S100a8 | 0 | 4.751836754 | 0.242 | 0.01 | 0 | 0 | 4.127223467 | 0.261 | 0.016 | 0 | 0 | 0 |
| Hbb-bt | 0 | 4.733021471 | 0.202 | 0.008 | 0 | 0 | 4.619146367 | 0.22 | 0.009 | 0 | 0 | 0 |
| Ngp | 1.9285E-270 | 4.661677303 | 0.113 | 0.005 | 2.8879E-266 | 0 | 4.79219211 | 0.241 | 0.01 | 0 | 1.9285E-270 | 0 |
| Camp | 5.6922E-185 | 4.424861745 | 0.08 | 0.004 | 8.524E-181 | 0 | 4.714213158 | 0.143 | 0.006 | 0 | 5.6922E-185 | 0 |
| Car2 | 2.3158E-260 | 4.356989632 | 0.111 | 0.005 | 3.468E-256 | 9.7935E-305 | 4.527324547 | 0.112 | 0.005 | 1.4666E-300 | 2.3158E-260 | 1.9587E-304 |
| Spp1 | 2.3766E-243 | 2.69407408 | 0.205 | 0.038 | 3.559E-239 | 1.28857E-96 | 1.97818733 | 0.137 | 0.046 | 1.92963E-92 | 1.28857E-96 | 4.7533E-243 |
| Ppbp | 5.6654E-179 | 6.639058401 | 0.064 | 0.001 | 8.4839E-175 | 8.36608E-98 | 6.051723668 | 0.032 | 0.001 | 1.25282E-93 | 8.36608E-98 | 1.1331E-178 |
| Obp2a | 4.8023E-131 | -4.561338678 | 0.014 | 0.19 | 7.1915E-127 | 7.8934E-173 | -5.677057486 | 0.007 | 0.189 | 1.182E-168 | 4.8023E-131 | 1.5787E-172 |
| Acp5 | 2.1397E-160 | 3.246554812 | 0.099 | 0.012 | 3.2042E-156 | 5.50379E-65 | 2.556878462 | 0.056 | 0.012 | 8.24192E-61 | 5.50379E-65 | 4.2794E-160 |
| Slc4a1 | 8.5006E-124 | 4.745973605 | 0.05 | 0.002 | 1.273E-119 | 2.212E-143 | 4.346912432 | 0.055 | 0.003 | 3.3125E-139 | 8.5006E-124 | 4.4241E-143 |
| Col1a1 | 2.54768E-50 | -0.694950376 | 0.467 | 0.585 | 3.81515E-46 | 3.1786E-140 | -0.889898253 | 0.437 | 0.665 | 4.76E-136 | 2.54768E-50 | 6.3572E-140 |
| Lyz2 | 6.15495E-54 | 3.263398322 | 0.03 | 0.003 | 9.21704E-50 | 3.8659E-124 | 3.651767013 | 0.055 | 0.004 | 5.7892E-120 | 6.15495E-54 | 7.7318E-124 |
| Mpo | 2.0001E-123 | 5.124485228 | 0.048 | 0.001 | 2.9951E-119 | 4.1695E-117 | 4.308989948 | 0.045 | 0.002 | 6.2438E-113 | 4.1695E-117 | 4.0002E-123 |
| Ltf | 8.95439E-71 | 3.768548241 | 0.033 | 0.002 | 1.34092E-66 | 2.1762E-122 | 3.374714131 | 0.058 | 0.005 | 3.2589E-118 | 8.95439E-71 | 4.3524E-122 |
| Ctsk | 2.4568E-116 | 3.202710396 | 0.075 | 0.009 | 3.6791E-112 | 2.90521E-30 | 2.021293488 | 0.033 | 0.009 | 4.35056E-26 | 2.90521E-30 | 4.9136E-116 |
| Obp2b | 2.751E-106 | -6.304633782 | 0.003 | 0.144 | 4.1196E-102 | 6.6293E-108 | -6.961055033 | 0.002 | 0.119 | 9.9274E-104 | 2.751E-106 | 1.3259E-107 |
| Sparc | 2.7562E-106 | -1.370603515 | 0.175 | 0.372 | 4.1273E-102 | 1.20511E-69 | -1.206088112 | 0.148 | 0.281 | 1.80465E-65 | 1.20511E-69 | 5.5123E-106 |
| Col1a2 | 7.6512E-31 | -0.645570605 | 0.361 | 0.447 | 1.14577E-26 | 2.2654E-105 | -0.913885902 | 0.335 | 0.524 | 3.3924E-101 | 7.6512E-31 | 4.5308E-105 |
| Elane | 2.0283E-104 | 5.158973605 | 0.04 | 0.001 | 3.0374E-100 | 1.78559E-72 | 3.861733044 | 0.03 | 0.002 | 2.67393E-68 | 1.78559E-72 | 4.0567E-104 |
| Tmsb4x | 1.1052E-103 | 2.101164557 | 0.102 | 0.022 | 1.6551E-99 | 5.261E-103 | 1.86187175 | 0.101 | 0.025 | 7.8783E-99 | 5.261E-103 | 2.2105E-103 |
| Pf4 | 1.5451E-100 | 5.236378108 | 0.039 | 0.001 | 2.31381E-96 | 1.54888E-48 | 4.440152699 | 0.018 | 0.001 | 2.31945E-44 | 1.54888E-48 | 3.0902E-100 |
| Hist1h2ap | 4.97876E-96 | 4.107095091 | 0.043 | 0.002 | 7.45569E-92 | 4.7796E-100 | 3.957538143 | 0.041 | 0.002 | 7.15747E-96 | 4.97876E-96 | 9.5592E-100 |
| Hist1h1e | 1.1216E-99 | 3.338130605 | 0.056 | 0.005 | 1.67962E-95 | 8.4568E-100 | 3.018098565 | 0.054 | 0.006 | 1.26641E-95 | 1.1216E-99 | 1.6914E-99 |
| Ighm | 1.09418E-93 | 3.466421012 | 0.054 | 0.005 | 1.63853E-89 | 8.11948E-58 | 2.76949148 | 0.036 | 0.005 | 1.21589E-53 | 8.11948E-58 | 2.18835E-93 |
| Cd24a | 7.00867E-92 | 2.846749773 | 0.057 | 0.006 | 1.04955E-87 | 1.5578E-91 | 3.120085746 | 0.045 | 0.004 | 2.3328E-87 | 1.5578E-91 | 1.40173E-91 |
| Mgp | 5.21628E-90 | -5.454811065 | 0.004 | 0.126 | 7.81138E-86 | 3.18062E-34 | -5.13843772 | 0.001 | 0.04 | 4.76298E-30 | 3.18062E-34 | 1.04326E-89 |
| Alas2 | 2.62106E-73 | 4.049434523 | 0.033 | 0.002 | 3.92504E-69 | 1.86614E-80 | 3.660402238 | 0.036 | 0.003 | 2.79455E-76 | 2.62106E-73 | 3.73229E-80 |
| Rrm2 | 1.21018E-71 | 3.930338125 | 0.033 | 0.002 | 1.81225E-67 | 3.11624E-75 | 3.391913402 | 0.035 | 0.003 | 4.66656E-71 | 1.21018E-71 | 6.23247E-75 |
| Tmcc2 | 1.14577E-18 | 2.920310665 | 0.012 | 0.002 | 1.7158E-14 | 1.12004E-74 | 4.5962719 | 0.027 | 0.001 | 1.67725E-70 | 1.14577E-18 | 2.24007E-74 |
| Mmp9 | 1.26356E-74 | 2.986981705 | 0.048 | 0.006 | 1.89219E-70 | 3.45159E-28 | 2.431885083 | 0.021 | 0.004 | 5.16876E-24 | 3.45159E-28 | 2.52713E-74 |
| Lcn2 | 1.31037E-40 | 3.562606341 | 0.019 | 0.001 | 1.96228E-36 | 7.14531E-67 | 3.727516434 | 0.029 | 0.002 | 1.07001E-62 | 1.31037E-40 | 1.42906E-66 |
| Scgb1c1 | 3.8035E-45 | -4.909729111 | 0.003 | 0.067 | 5.69574E-41 | 1.08033E-65 | -4.646902083 | 0.005 | 0.082 | 1.6178E-61 | 3.8035E-45 | 2.16067E-65 |
| Col3a1 | 1.35823E-42 | -1.80043329 | 0.043 | 0.129 | 2.03395E-38 | 4.79735E-63 | -1.819632343 | 0.052 | 0.154 | 7.18404E-59 | 1.35823E-42 | 9.59471E-63 |
| Mmp13 | 4.15872E-61 | 2.359766717 | 0.059 | 0.012 | 6.22769E-57 | 2.57656E-17 | 1.415699655 | 0.032 | 0.013 | 3.8584E-13 | 2.57656E-17 | 8.31745E-61 |
| Fn1 | 4.60724E-60 | -5.481787779 | 0.002 | 0.085 | 6.89934E-56 | 2.76362E-16 | -2.916045298 | 0.003 | 0.024 | 4.13852E-12 | 2.76362E-16 | 9.21448E-60 |
| Obp1a | 5.76835E-38 | -6.769290296 | 0 | 0.052 | 8.6381E-34 | 8.71378E-60 | -6.410755366 | 0.001 | 0.068 | 1.30489E-55 | 5.76835E-38 | 1.74276E-59 |
| Hist1h1a | 1.94353E-56 | 3.617747895 | 0.029 | 0.002 | 2.91044E-52 | 1.26365E-52 | 3.645181501 | 0.023 | 0.002 | 1.89232E-48 | 1.26365E-52 | 3.88707E-56 |
| Bpgm | 3.23541E-29 | 2.392681339 | 0.024 | 0.004 | 4.84503E-25 | 2.27049E-56 | 2.796804888 | 0.034 | 0.005 | 3.40006E-52 | 3.23541E-29 | 4.54098E-56 |
| Gpx1 | 9.59524E-55 | 2.961193319 | 0.034 | 0.004 | 1.43689E-50 | 4.14889E-51 | 2.824540888 | 0.031 | 0.004 | 6.21297E-47 | 4.14889E-51 | 1.91905E-54 |
| Col2a1 | 4.55009E-51 | -7.98832077 | 0 | 0.069 | 6.81376E-47 | 8.22507E-15 | -6.260153628 | 0 | 0.016 | 1.2317E-10 | 8.22507E-15 | 9.10018E-51 |
| Ccn2 | 2.32186E-49 | -6.349347335 | 0.001 | 0.068 | 3.47698E-45 | 2.43028E-24 | -4.117973617 | 0.002 | 0.031 | 3.63934E-20 | 2.43028E-24 | 4.64371E-49 |
| Lcn3 | 1.27348E-45 | -5.865334992 | 0.001 | 0.065 | 1.90703E-41 | 9.94518E-49 | -7.441961234 | 0 | 0.055 | 1.48929E-44 | 1.27348E-45 | 1.98904E-48 |
| Col11a2 | 2.9757E-46 | -5.08844535 | 0.002 | 0.067 | 4.45611E-42 | 2.01481E-28 | -4.881081626 | 0.001 | 0.034 | 3.01717E-24 | 2.01481E-28 | 5.9514E-46 |
| Tubb1 | 4.33325E-46 | 4.848360823 | 0.018 | 0.001 | 6.48905E-42 | 2.56168E-29 | 4.663386096 | 0.01 | 0 | 3.83612E-25 | 2.56168E-29 | 8.66651E-46 |
| Lcn11 | 2.79258E-23 | -4.820665942 | 0.001 | 0.034 | 4.18189E-19 | 2.06667E-45 | -5.567198079 | 0.002 | 0.054 | 3.09483E-41 | 2.79258E-23 | 4.13333E-45 |
| Prtn3 | 5.63913E-30 | 3.896113955 | 0.014 | 0.001 | 8.44459E-26 | 4.08429E-45 | 4.448714712 | 0.017 | 0.001 | 6.11623E-41 | 5.63913E-30 | 8.16858E-45 |
| Bpifb6 | 1.97633E-22 | -3.416675936 | 0.005 | 0.039 | 2.95955E-18 | 2.73259E-42 | -3.876472373 | 0.005 | 0.056 | 4.09206E-38 | 1.97633E-22 | 5.46519E-42 |
| Bpifb4 | 2.48002E-15 | -2.657530107 | 0.006 | 0.032 | 3.71383E-11 | 8.51611E-41 | -4.108928478 | 0.004 | 0.053 | 1.27529E-36 | 2.48002E-15 | 1.70322E-40 |
| Hist1h1b | 3.1079E-33 | 3.277204122 | 0.018 | 0.002 | 4.65408E-29 | 7.69588E-40 | 2.686214026 | 0.024 | 0.003 | 1.15246E-35 | 3.1079E-33 | 1.53918E-39 |
| Igkc | 4.39853E-39 | 4.246639804 | 0.019 | 0.002 | 6.58679E-35 | 2.50681E-22 | 3.818664322 | 0.012 | 0.001 | 3.75394E-18 | 2.50681E-22 | 8.79705E-39 |
| Col27a1 | 1.80578E-36 | -6.024467936 | 0.001 | 0.051 | 2.70416E-32 | 1.49474E-09 | -4.76347766 | 0 | 0.01 | 2.23838E-05 | 1.49474E-09 | 3.61156E-36 |
| Col9a3 | 6.18309E-35 | -7.384299934 | 0 | 0.047 | 9.25918E-31 | 7.7454E-15 | -4.444017821 | 0.001 | 0.017 | 1.15987E-10 | 7.7454E-15 | 1.23662E-34 |
| Ctsg | 1.17027E-32 | 4.274625578 | 0.014 | 0.001 | 1.75247E-28 | 4.0316E-34 | 4.224303123 | 0.013 | 0.001 | 6.03732E-30 | 1.17027E-32 | 8.0632E-34 |
| Coro1a | 9.64294E-14 | 2.209817282 | 0.012 | 0.002 | 1.44403E-09 | 5.01818E-34 | 3.288149605 | 0.016 | 0.002 | 7.51473E-30 | 9.64294E-14 | 1.00364E-33 |
| Slc25a37 | 6.40188E-19 | 2.894654475 | 0.012 | 0.002 | 9.58682E-15 | 1.11203E-32 | 2.855190198 | 0.018 | 0.002 | 1.66527E-28 | 6.40188E-19 | 2.22407E-32 |
| Car1 | 3.08404E-27 | 3.791777295 | 0.013 | 0.001 | 4.61836E-23 | 1.32114E-31 | 3.804858522 | 0.013 | 0.001 | 1.97841E-27 | 3.08404E-27 | 2.64229E-31 |
| Vmo1 | 6.99102E-16 | -3.228882176 | 0.003 | 0.027 | 1.04691E-11 | 1.94291E-31 | -3.39468296 | 0.004 | 0.044 | 2.90951E-27 | 6.99102E-16 | 3.88583E-31 |
| Col9a2 | 3.31911E-31 | -5.295277487 | 0.001 | 0.044 | 4.97036E-27 | 1.17295E-12 | -4.512252556 | 0.001 | 0.014 | 1.7565E-08 | 1.17295E-12 | 6.63822E-31 |
| App | 7.6741E-25 | -3.008048323 | 0.006 | 0.045 | 1.1492E-20 | 5.3139E-31 | -3.166228786 | 0.005 | 0.045 | 7.95757E-27 | 7.6741E-25 | 1.06278E-30 |
| Tln1 | 7.36056E-31 | 2.089368923 | 0.033 | 0.008 | 1.10224E-26 | 1.80247E-07 | 1.120015659 | 0.016 | 0.007 | 0.002699197 | 1.80247E-07 | 1.47211E-30 |
| Epb41 | 2.37295E-28 | 3.542257695 | 0.014 | 0.001 | 3.55349E-24 | 8.71152E-31 | 3.110845073 | 0.015 | 0.001 | 1.30455E-26 | 2.37295E-28 | 1.7423E-30 |
| Prdx2 | 1.7244E-14 | 2.106158581 | 0.014 | 0.003 | 2.58229E-10 | 3.30549E-30 | 2.646545107 | 0.019 | 0.003 | 4.94998E-26 | 1.7244E-14 | 6.61099E-30 |
| Itm2b | 4.49128E-29 | -2.7071579 | 0.01 | 0.059 | 6.72569E-25 | 6.40369E-27 | -2.196798796 | 0.015 | 0.056 | 9.58953E-23 | 6.40369E-27 | 8.98256E-29 |
| Igfbp5 | 1.08698E-23 | -3.641353363 | 0.007 | 0.046 | 1.62775E-19 | 3.67572E-28 | -3.285307607 | 0.006 | 0.042 | 5.50439E-24 | 1.08698E-23 | 7.35144E-28 |
| Alad | 3.79499E-24 | 3.137122054 | 0.014 | 0.001 | 5.683E-20 | 3.9344E-28 | 2.921111591 | 0.015 | 0.002 | 5.89177E-24 | 3.79499E-24 | 7.8688E-28 |
| Obp1b | 2.01979E-17 | -2.697136923 | 0.006 | 0.034 | 3.02463E-13 | 1.25147E-27 | -4.274092819 | 0.002 | 0.034 | 1.87408E-23 | 2.01979E-17 | 2.50295E-27 |
| Ucp2 | 1.13102E-25 | 2.040521043 | 0.027 | 0.006 | 1.6937E-21 | 1.33663E-22 | 1.636460991 | 0.023 | 0.006 | 2.0016E-18 | 1.33663E-22 | 2.26204E-25 |
| Gpx3 | 8.03715E-11 | -1.157476117 | 0.023 | 0.05 | 1.20356E-06 | 4.89214E-25 | -1.802184659 | 0.017 | 0.057 | 7.32598E-21 | 8.03715E-11 | 9.78428E-25 |
| Mki67 | 6.72499E-23 | 3.244289499 | 0.013 | 0.001 | 1.00707E-18 | 7.91159E-22 | 3.181234401 | 0.011 | 0.001 | 1.18476E-17 | 7.91159E-22 | 1.345E-22 |
| Blvrb | 3.67964E-10 | 1.727770682 | 0.012 | 0.003 | 5.51027E-06 | 2.38006E-22 | 1.977362068 | 0.021 | 0.005 | 3.56414E-18 | 3.67964E-10 | 4.76012E-22 |
| Bpifb3 | 3.5013E-22 | -7.176224815 | 0 | 0.03 | 5.2432E-18 | 3.59844E-17 | -6.821580614 | 0 | 0.018 | 5.38866E-13 | 3.59844E-17 | 7.00261E-22 |
| Gpx6 | 3.87572E-07 | -1.603556742 | 0.008 | 0.023 | 0.005803889 | 3.57712E-22 | -3.052385276 | 0.004 | 0.033 | 5.35674E-18 | 3.87572E-07 | 7.15424E-22 |
| Bgn | 1.44862E-19 | -1.79133441 | 0.016 | 0.054 | 2.16931E-15 | 4.51778E-22 | -1.71578595 | 0.016 | 0.053 | 6.76538E-18 | 1.44862E-19 | 9.03556E-22 |
| Arf5 | 0.30577761 | 0.179334058 | 0.013 | 0.011 | 1 | 3.44451E-21 | 1.470741018 | 0.029 | 0.009 | 5.15816E-17 | 0.30577761 | 6.88903E-21 |
| Bpifa1 | 1.58019E-20 | -3.611070796 | 0.003 | 0.033 | 2.36633E-16 | 9.5853E-15 | -4.128241953 | 0.001 | 0.018 | 1.4354E-10 | 9.5853E-15 | 3.16037E-20 |
| Stmn1 | 6.77116E-17 | 2.318387557 | 0.013 | 0.002 | 1.01398E-12 | 3.19023E-19 | 2.0113094 | 0.017 | 0.004 | 4.77736E-15 | 6.77116E-17 | 6.38045E-19 |
| Ccdc80 | 3.53542E-19 | -3.649890044 | 0.002 | 0.03 | 5.29429E-15 | 3.44612E-07 | -2.318611486 | 0.002 | 0.011 | 0.005160561 | 3.44612E-07 | 7.07084E-19 |
| Arhgdib | 9.72223E-12 | 2.022190279 | 0.012 | 0.003 | 1.4559E-07 | 1.04679E-18 | 2.0113094 | 0.015 | 0.003 | 1.56757E-14 | 9.72223E-12 | 2.09358E-18 |
| Gsn | 1.95955E-18 | -3.548529329 | 0.003 | 0.03 | 2.93443E-14 | 1.03241E-13 | -3.076153441 | 0.003 | 0.02 | 1.54604E-09 | 1.03241E-13 | 3.9191E-18 |
| Hist1h2ae | 1.20057E-16 | 3.000363917 | 0.01 | 0.001 | 1.79786E-12 | 2.29351E-18 | 2.682686652 | 0.011 | 0.002 | 3.43453E-14 | 1.20057E-16 | 4.58702E-18 |
| Wfdc18 | 8.79283E-13 | -2.822177409 | 0.003 | 0.022 | 1.31673E-08 | 1.4191E-17 | -3.24921815 | 0.003 | 0.026 | 2.1251E-13 | 8.79283E-13 | 2.83819E-17 |
| Lcn4 | 1.55078E-17 | -4.952107713 | 0.001 | 0.025 | 2.32229E-13 | 7.36574E-17 | -3.76347766 | 0.001 | 0.02 | 1.10302E-12 | 7.36574E-17 | 3.10156E-17 |
| Cxcl12 | 1.59341E-16 | 1.765898663 | 0.022 | 0.006 | 2.38614E-12 | 6.3724E-09 | 1.203954478 | 0.013 | 0.005 | 9.54267E-05 | 6.3724E-09 | 3.18683E-16 |
| Mup5 | 1.1746E-11 | -2.977609777 | 0.003 | 0.021 | 1.75897E-07 | 2.04827E-16 | -3.985466699 | 0.001 | 0.02 | 3.06728E-12 | 1.1746E-11 | 4.09653E-16 |
| Dcn | 4.58112E-16 | -1.772784014 | 0.013 | 0.044 | 6.86023E-12 | 1.85495E-11 | -1.785345634 | 0.007 | 0.025 | 2.77778E-07 | 1.85495E-11 | 9.16224E-16 |
| Mup4 | 7.5696E-08 | -2.473567272 | 0.003 | 0.015 | 0.001133548 | 3.02497E-15 | -4.151920949 | 0.001 | 0.019 | 4.52989E-11 | 7.5696E-08 | 6.04994E-15 |
| mt-Atp8 | 0.001280771 | -0.854095241 | 0.013 | 0.022 | 1 | 4.52564E-15 | -2.506266261 | 0.004 | 0.025 | 6.77714E-11 | 0.001280771 | 9.05128E-15 |
| Tubb5 | 3.62367E-06 | 1.250459267 | 0.012 | 0.005 | 0.054264406 | 5.03078E-15 | 1.717578197 | 0.016 | 0.004 | 7.53359E-11 | 3.62367E-06 | 1.00616E-14 |
| Igfbp7 | 1.92337E-08 | -1.994282849 | 0.005 | 0.019 | 0.000288025 | 2.2734E-14 | -2.088226274 | 0.007 | 0.028 | 3.40441E-10 | 1.92337E-08 | 4.5468E-14 |
| Serinc3 | 9.05112E-12 | 1.294024286 | 0.023 | 0.009 | 1.35541E-07 | 2.73683E-14 | 1.346493591 | 0.021 | 0.007 | 4.09841E-10 | 9.05112E-12 | 5.47366E-14 |
| Tnc | 5.89915E-14 | 1.69635773 | 0.022 | 0.007 | 8.83398E-10 | 0.205551637 | 0.4778772 | 0.013 | 0.01 | 1 | 0.205551637 | 1.17983E-13 |
| Timp2 | 1.47399E-09 | -1.560104022 | 0.01 | 0.03 | 2.2073E-05 | 7.37734E-14 | -2.148561937 | 0.006 | 0.026 | 1.10476E-09 | 1.47399E-09 | 1.47547E-13 |
| Ptms | 9.40179E-09 | -1.570278913 | 0.009 | 0.027 | 0.000140792 | 7.63593E-14 | -1.932507765 | 0.009 | 0.031 | 1.14348E-09 | 9.40179E-09 | 1.52719E-13 |

| | | | | | | | | | | | | |
|---|---|---|---|---|---|---|---|---|---|---|---|---|
| Serpinh1 | 4.17159E-11 | -1.640007279 | 0.011 | 0.033 | 6.24696E-07 | 3.11116E-13 | -1.666250094 | 0.011 | 0.033 | 4.65896E-09 | 4.17159E-11 | 6.22231E-13 |
| Dmbt1 | 6.65116E-13 | -3.645628018 | 0.001 | 0.019 | 9.96011E-09 | 3.11491E-11 | -3.732851696 | 0.001 | 0.013 | 4.66458E-07 | 3.11491E-11 | 1.33023E-12 |
| Smoc2 | 6.91335E-13 | -5.748574319 | 0 | 0.017 | 1.03527E-08 | 4.6412E-09 | -3.63254679 | 0.001 | 0.011 | 6.95019E-05 | 4.6412E-09 | 1.38267E-12 |
| Apoe | 1.31177E-05 | -1.03591147 | 0.015 | 0.029 | 0.196438217 | 9.81755E-13 | -1.484843834 | 0.017 | 0.041 | 1.47018E-08 | 1.31177E-05 | 1.96351E-12 |
| mt-Nd3 | 3.98703E-12 | -3.401734527 | 0.002 | 0.019 | 5.97058E-08 | 5.88191E-11 | -3.246078443 | 0.002 | 0.014 | 8.80816E-07 | 5.88191E-11 | 7.97407E-12 |
| Postn | 5.42211E-12 | -2.731784661 | 0.003 | 0.021 | 8.11961E-08 | 7.98073E-09 | -1.805826543 | 0.005 | 0.018 | 0.000119511 | 7.98073E-09 | 1.08442E-11 |
| Maf | 2.38583E-07 | -2.811563735 | 0.001 | 0.011 | 0.003572774 | 2.72255E-11 | -3.846671595 | 0.001 | 0.013 | 4.07702E-07 | 2.38583E-07 | 5.4451E-11 |
| Sost | 3.43225E-07 | -1.939628774 | 0.005 | 0.017 | 0.0051398 | 4.43287E-11 | -2.369512384 | 0.004 | 0.02 | 6.63822E-07 | 3.43225E-07 | 8.86574E-11 |
| Ahnak | 1.25326E-10 | -1.876004734 | 0.007 | 0.026 | 1.87675E-06 | 2.35335E-05 | -1.063367287 | 0.01 | 0.02 | 0.352414806 | 2.35335E-05 | 2.50651E-10 |
| H3f3a | 1.20841E-07 | 1.141719766 | 0.016 | 0.006 | 0.001809589 | 1.43022E-10 | 1.1478114 | 0.018 | 0.007 | 2.14175E-06 | 1.20841E-07 | 2.86043E-10 |
| Col11a1 | 3.30168E-10 | -3.163611819 | 0.002 | 0.016 | 4.94427E-06 | 7.51613E-08 | -2.91152274 | 0.001 | 0.01 | 0.001125541 | 7.51613E-08 | 6.60336E-10 |
| mt-Nd1 | 0.000472317 | -1.324475774 | 0.007 | 0.015 | 1 | 3.79454E-10 | -2.675191127 | 0.002 | 0.015 | 5.68232E-06 | 0.000472317 | 7.58907E-10 |
| Vim | 9.11088E-05 | -0.787675862 | 0.023 | 0.038 | 1 | 6.65826E-10 | -1.308040151 | 0.014 | 0.033 | 9.97074E-06 | 9.11088E-05 | 1.33165E-09 |
| Atpif1 | 9.59076E-10 | 1.619273749 | 0.014 | 0.004 | 1.43622E-05 | 0.000188047 | 0.904394196 | 0.01 | 0.005 | 1 | 0.000188047 | 1.91815E-09 |
| Sfrp2 | 2.1246E-07 | -2.865334992 | 0.001 | 0.011 | 0.003181588 | 1.88814E-09 | -3.097215057 | 0.001 | 0.012 | 2.82749E-05 | 2.1246E-07 | 3.77628E-09 |
| mt-Cytb | 0.008273914 | -0.637066004 | 0.014 | 0.021 | 1 | 2.71402E-09 | -1.91152274 | 0.006 | 0.019 | 4.06424E-05 | 0.008273914 | 5.42804E-09 |
| Col12a1 | 1.80105E-06 | -1.49956248 | 0.007 | 0.019 | 0.02697072 | 4.1247E-09 | -3.205921316 | 0.001 | 0.011 | 6.17674E-05 | 1.80105E-06 | 8.2494E-09 |
| Mt1 | 4.5636E-09 | 1.131294786 | 0.023 | 0.01 | 6.83399E-05 | 0.04139059 | 0.263288617 | 0.019 | 0.015 | 1 | 0.04139059 | 9.12719E-09 |
| Igf2 | 7.30831E-09 | -2.938235539 | 0.002 | 0.014 | 0.000109442 | 3.55541E-08 | -2.558546209 | 0.002 | 0.012 | 0.000532422 | 3.55541E-08 | 1.46166E-08 |
| Dbp | 1.13931E-08 | -2.129271364 | 0.004 | 0.017 | 0.000170611 | 1.29459E-07 | -2.39468296 | 0.002 | 0.012 | 0.001938647 | 1.29459E-07 | 2.27861E-08 |
| mt-Nd4l | 7.16141E-06 | -1.724528845 | 0.005 | 0.015 | 0.107242175 | 1.14242E-08 | -2.755403538 | 0.002 | 0.012 | 0.000171077 | 7.16141E-06 | 2.28484E-08 |
| Dmp1 | 0.001465004 | -0.819657417 | 0.01 | 0.019 | 1 | 1.29135E-08 | -1.604490505 | 0.007 | 0.02 | 0.00019338 | 0.001465004 | 2.5827E-08 |
| Tpm1 | 1.0063E-05 | -1.128678028 | 0.012 | 0.025 | 0.150692741 | 6.35898E-08 | -1.292471348 | 0.012 | 0.027 | 0.000952258 | 1.0063E-05 | 1.2718E-07 |
| mt-Nd6 | 2.07102E-07 | -2.656259569 | 0.002 | 0.013 | 0.003101352 | 6.91295E-08 | -2.860815778 | 0.001 | 0.01 | 0.001035214 | 2.07102E-07 | 1.38259E-07 |
| Sdc4 | 1.06544E-07 | -2.975388537 | 0.001 | 0.012 | 0.001595492 | 1.60416E-05 | -2.052820938 | 0.003 | 0.01 | 0.240223094 | 1.60416E-05 | 2.13087E-07 |
| Mxra8 | 1.45602E-07 | -2.151639177 | 0.003 | 0.015 | 0.002180391 | 1.23067E-06 | -1.902960726 | 0.003 | 0.012 | 0.018429256 | 1.23067E-06 | 2.91204E-07 |
| Serpinf1 | 3.34078E-05 | -1.057517 | 0.012 | 0.024 | 0.500281593 | 1.70972E-07 | -1.199999522 | 0.01 | 0.024 | 0.002560303 | 3.34078E-05 | 3.41944E-07 |
| Mmp2 | 4.5618E-07 | -1.5301508 | 0.006 | 0.02 | 0.00683129 | 2.70972E-07 | -1.619946753 | 0.006 | 0.017 | 0.004057805 | 4.5618E-07 | 5.41944E-07 |
| Ibsp | 4.80412E-07 | 0.760644677 | 0.04 | 0.024 | 0.007194164 | 0.030276078 | 0.571829759 | 0.029 | 0.023 | 1 | 0.030276078 | 9.60823E-07 |
| Nisch | 0.000309044 | -1.158182023 | 0.007 | 0.015 | 1 | 7.19332E-07 | -1.76713283 | 0.004 | 0.014 | 0.010772002 | 0.000309044 | 1.43866E-06 |
| Arglu1 | 7.93764E-06 | -1.723720386 | 0.005 | 0.014 | 0.118866184 | 7.77107E-07 | -1.749502936 | 0.004 | 0.014 | 0.011637172 | 7.93764E-06 | 1.55421E-06 |
| Maged1 | 1.24596E-06 | -2.20668054 | 0.003 | 0.013 | 0.018658247 | 7.79656E-05 | -1.396194427 | 0.004 | 0.011 | 1 | 7.79656E-05 | 2.49192E-06 |
| Lrp1 | 2.62314E-05 | -1.219277894 | 0.008 | 0.018 | 0.392815173 | 2.18411E-06 | -1.442148318 | 0.005 | 0.015 | 0.032706984 | 2.62314E-05 | 4.36821E-06 |
| mt-Nd5 | 0.022265497 | -0.850370553 | 0.008 | 0.013 | 1 | 2.75596E-06 | -1.822912128 | 0.004 | 0.014 | 0.041270451 | 0.022265497 | 5.51191E-06 |
| mt-Co3 | 0.334577843 | -0.359659481 | 0.015 | 0.017 | 1 | 2.75732E-06 | -1.414955355 | 0.008 | 0.018 | 0.041290825 | 0.334577843 | 5.51463E-06 |
| mt-Atp6 | 0.000114348 | -1.334055123 | 0.007 | 0.016 | 1 | 3.29606E-06 | -1.521870559 | 0.006 | 0.016 | 0.04935846 | 0.000114348 | 6.5921E-06 |
| Tpm2 | 0.067549273 | -0.774569528 | 0.006 | 0.01 | 1 | 3.6526E-06 | -1.709168072 | 0.005 | 0.015 | 0.054697727 | 0.067549273 | 7.30519E-06 |
| Col6a3 | 0.017209854 | -0.752543222 | 0.007 | 0.013 | 1 | 5.37746E-06 | -1.743578102 | 0.003 | 0.012 | 0.080527427 | 0.017209854 | 1.07549E-05 |
| Selenom | 0.001016918 | -1.244748581 | 0.004 | 0.011 | 1 | 6.38263E-06 | -1.581147638 | 0.004 | 0.012 | 0.095579921 | 0.001016918 | 1.27652E-05 |
| Tsc22d1 | 0.000765504 | -1.100564992 | 0.007 | 0.015 | 1 | 7.26495E-06 | -2.03308472 | 0.002 | 0.01 | 0.10879265 | 0.000765504 | 1.45299E-05 |
| Kctd12 | 0.00174629 | -0.864199458 | 0.01 | 0.018 | 1 | 1.00452E-05 | -1.111450494 | 0.009 | 0.02 | 0.150426579 | 0.00174629 | 2.00903E-05 |
| Cald1 | 0.001565468 | -1.102729576 | 0.006 | 0.012 | 1 | 1.21904E-05 | -1.778767531 | 0.003 | 0.01 | 0.182551499 | 0.001565468 | 2.43807E-05 |
| Jund | 1.47188E-05 | -1.708856687 | 0.004 | 0.014 | 0.220414091 | 6.7613E-05 | -1.343040173 | 0.005 | 0.013 | 1 | 6.7613E-05 | 2.94374E-05 |
| Fth1 | 1.56793E-05 | -1.194707899 | 0.01 | 0.022 | 0.234797107 | 0.004819904 | -0.785157206 | 0.015 | 0.022 | 1 | 0.004819904 | 3.13583E-05 |
| Htra1 | 1.58791E-05 | -1.806991006 | 0.004 | 0.012 | 0.237790097 | 1.69978E-05 | -1.704897634 | 0.003 | 0.011 | 0.254541636 | 1.69978E-05 | 3.1758E-05 |
| Col16a1 | 2.03479E-05 | -2.019535641 | 0.002 | 0.01 | 0.304710352 | 9.62059E-05 | -1.448122219 | 0.004 | 0.011 | 1 | 9.62059E-05 | 4.06955E-05 |
| Nucb1 | 6.65859E-05 | -1.401962531 | 0.006 | 0.015 | 0.997124261 | 0.000980133 | -0.955523736 | 0.004 | 0.01 | 1 | 0.000980133 | 0.000133167 |
| Akr1a1 | 8.79602E-05 | -1.189967296 | 0.008 | 0.018 | 1 | 0.002579158 | -0.83570974 | 0.01 | 0.017 | 1 | 0.002579158 | 0.000175913 |
| mt-Co2 | 0.057226083 | -0.470589572 | 0.03 | 0.037 | 1 | 0.000143474 | -0.908414011 | 0.025 | 0.038 | 1 | 0.057226083 | 0.000286928 |
| Id3 | 0.029327655 | -0.629686474 | 0.009 | 0.014 | 1 | 0.000303976 | -1.14573431 | 0.006 | 0.013 | 1 | 0.029327655 | 0.000607859 |
| Igfbp4 | 0.073977867 | -0.561514971 | 0.009 | 0.013 | 1 | 0.001027979 | -0.786856351 | 0.011 | 0.019 | 1 | 0.073977867 | 0.002054901 |
| Ptprs | 0.310059353 | -0.322751093 | 0.008 | 0.01 | 1 | 0.001071988 | -1.167493753 | 0.004 | 0.01 | 1 | 0.310059353 | 0.002142826 |
| Rsrp1 | 0.001082401 | -0.785630014 | 0.016 | 0.026 | 1 | 0.141461969 | -0.540675779 | 0.015 | 0.018 | 1 | 0.141461969 | 0.002163631 |
| Pcolce | 0.006433687 | -0.821848964 | 0.007 | 0.013 | 1 | 0.003127983 | -0.738039919 | 0.006 | 0.011 | 1 | 0.006433687 | 0.006246183 |
| Sdc3 | 0.026184817 | -0.791236934 | 0.006 | 0.01 | 1 | 0.006420913 | -0.9886906 | 0.005 | 0.01 | 1 | 0.026184817 | 0.012800599 |
| Cst3 | 0.009645206 | -0.684513166 | 0.015 | 0.022 | 1 | 0.092959313 | -0.611923118 | 0.008 | 0.011 | 1 | 0.092959313 | 0.019197383 |
| Ddx17 | 0.011216318 | -0.801696706 | 0.008 | 0.014 | 1 | 0.045902063 | -0.660067853 | 0.007 | 0.01 | 1 | 0.045902063 | 0.022306831 |
| Srsf6 | 0.019893018 | -0.806991006 | 0.007 | 0.012 | 1 | 0.085582158 | -0.73592453 | 0.007 | 0.01 | 1 | 0.085582158 | 0.039390304 |
| mt-Co1 | 0.593099868 | -0.211140189 | 0.017 | 0.019 | 1 | 0.03367347 | -0.714515637 | 0.013 | 0.017 | 1 | 0.593099868 | 0.066213038 |
| Oaz1 | 0.07648577 | -0.689533424 | 0.008 | 0.012 | 1 | 0.621005853 | -0.474117427 | 0.012 | 0.012 | 1 | 0.621005853 | 0.147121467 |

Spatial seq: cluster 1

| | cKO_p_val | cKO_avg_log2FC | cKO_pct.1 | cKO_pct.2 | cKO_p_val_adj | WT_p_val | WT_avg_log2FC | WT_pct.1 | WT_pct.2 | WT_p_val_adj | max_pval | minimump_p_val |
|---|---|---|---|---|---|---|---|---|---|---|---|---|
| Col1a2 | 0 | 1.833365465 | 0.833 | 0.338 | 0 | 0 | 1.647182652 | 0.817 | 0.412 | 0 | 0 | 0 |
| Sparc | 0 | 1.471377685 | 0.692 | 0.253 | 0 | 0 | 2.018878796 | 0.632 | 0.169 | 0 | 0 | 0 |
| Col1a1 | 0 | 1.797477428 | 0.895 | 0.486 | 0 | 0 | 1.689195407 | 0.894 | 0.558 | 0 | 0 | 0 |
| Col3a1 | 0 | 2.944374719 | 0.361 | 0.056 | 0 | 0 | 2.748357892 | 0.402 | 0.073 | 0 | 0 | 0 |
| Gpx3 | 1.8246E-136 | 2.248573124 | 0.128 | 0.026 | 2.7323E-132 | 7.9272E-214 | 2.568651146 | 0.151 | 0.026 | 1.1871E-209 | 1.8246E-136 | 1.5854E-213 |
| Col11a2 | 5.73827E-57 | 1.635406138 | 0.112 | 0.042 | 8.59306E-53 | 7.1761E-194 | 3.369854198 | 0.1 | 0.01 | 1.0746E-189 | 5.73827E-57 | 1.4352E-193 |
| App | 2.54462E-72 | 2.010840156 | 0.092 | 0.025 | 3.81057E-68 | 2.2003E-183 | 2.836135316 | 0.119 | 0.018 | 3.295E-179 | 2.54462E-72 | 4.4007E-183 |
| Obp2a | 1.0232E-112 | -3.25286289 | 0.028 | 0.188 | 1.5323E-108 | 7.1035E-131 | -3.669111743 | 0.021 | 0.183 | 1.0637E-126 | 1.0232E-112 | 1.4207E-130 |
| Obp2b | 2.36376E-85 | -3.387039072 | 0.018 | 0.142 | 3.53973E-81 | 1.34943E-88 | -4.37959097 | 0.007 | 0.116 | 2.02077E-84 | 2.36376E-85 | 2.69886E-88 |
| Mgp | 9.2115E-77 | -3.569074848 | 0.014 | 0.124 | 1.37942E-72 | 1.49231E-10 | -1.445538043 | 0.015 | 0.036 | 2.23473E-06 | 1.49231E-10 | 1.8423E-76 |
| Bgn | 6.86719E-12 | 0.706955128 | 0.07 | 0.041 | 1.02836E-07 | 2.87784E-72 | 1.541660675 | 0.102 | 0.032 | 4.30957E-68 | 6.86719E-12 | 5.75569E-72 |
| Hbb-bs | 3.51254E-62 | -2.023040506 | 0.066 | 0.182 | 5.26003E-58 | 5.49595E-50 | -1.879369057 | 0.068 | 0.162 | 8.23019E-46 | 5.49595E-50 | 7.02508E-62 |
| Serpinf1 | 5.93197E-26 | 1.487154486 | 0.046 | 0.016 | 8.88312E-22 | 1.54776E-57 | 1.971870361 | 0.057 | 0.013 | 2.31778E-53 | 5.93197E-26 | 3.09553E-57 |
| Lcn3 | 3.21245E-50 | -8.265373743 | 0 | 0.065 | 4.81065E-46 | 5.07619E-42 | -5.501336926 | 0.002 | 0.053 | 7.60159E-38 | 5.07619E-42 | 6.42491E-50 |
| Col2a1 | 1.07533E-48 | -4.7287068 | 0.003 | 0.069 | 1.6103E-44 | 4.30734E-12 | -4.117963385 | 0.001 | 0.015 | 6.45025E-08 | 4.30734E-12 | 2.15065E-48 |
| Hba-a2 | 2.3452E-45 | -1.646111662 | 0.06 | 0.154 | 3.51193E-41 | 2.48021E-34 | -1.16025343 | 0.123 | 0.207 | 3.71411E-30 | 2.48021E-34 | 4.69039E-45 |
| S100a9 | 2.41614E-40 | -2.602912676 | 0.017 | 0.083 | 3.61817E-36 | 3.26822E-45 | -1.92710411 | 0.041 | 0.12 | 4.89416E-41 | 2.41614E-40 | 6.53644E-45 |
| Obp1a | 1.04239E-28 | -3.225358065 | 0.007 | 0.051 | 1.56098E-24 | 7.02266E-43 | -3.492501522 | 0.008 | 0.066 | 1.05164E-38 | 1.04239E-28 | 1.40453E-42 |
| Ccn2 | 2.58888E-42 | -3.523483135 | 0.006 | 0.068 | 3.87685E-38 | 5.40844E-17 | -2.812443047 | 0.005 | 0.03 | 8.09914E-13 | 5.40844E-17 | 5.17776E-42 |
| Dcn | 8.65439E-09 | 0.664185288 | 0.056 | 0.034 | 0.0001296 | 3.64487E-42 | 1.695122364 | 0.051 | 0.014 | 5.45819E-38 | 8.65439E-09 | 7.28974E-42 |
| Fn1 | 1.12543E-41 | -2.674259016 | 0.015 | 0.083 | 1.68533E-37 | 2.27465E-08 | -1.633165046 | 0.008 | 0.023 | 0.000340629 | 2.27465E-08 | 2.25086E-41 |
| Lcn11 | 1.95541E-21 | -3.615383676 | 0.003 | 0.034 | 2.92822E-17 | 2.32084E-39 | -4.539514191 | 0.003 | 0.052 | 3.47546E-35 | 1.95541E-21 | 4.64169E-39 |
| Postn | 2.94281E-37 | 1.937666435 | 0.044 | 0.011 | 4.40687E-33 | 2.32085E-32 | 1.893068793 | 0.037 | 0.01 | 3.47547E-28 | 2.32085E-32 | 5.88563E-37 |
| Hbb-bt | 6.60772E-27 | -2.597912232 | 0.008 | 0.051 | 9.89506E-23 | 2.11549E-35 | -2.698518784 | 0.01 | 0.06 | 3.16794E-31 | 6.60772E-27 | 4.23098E-35 |
| Vim | 1.39322E-17 | 0.91941346 | 0.06 | 0.029 | 2.08635E-13 | 3.14785E-35 | 1.325614513 | 0.061 | 0.022 | 4.71391E-31 | 1.39322E-17 | 6.2957E-35 |
| Ngp | 3.57311E-17 | -2.997406118 | 0.004 | 0.029 | 5.35073E-13 | 4.79332E-35 | -2.596005538 | 0.013 | 0.066 | 7.17799E-31 | 3.57311E-17 | 9.58663E-35 |
| Serpinh1 | 2.25765E-10 | 0.784227984 | 0.046 | 0.025 | 3.38083E-06 | 9.6352E-35 | 1.355361857 | 0.06 | 0.022 | 1.44287E-30 | 2.25765E-10 | 1.92704E-34 |
| S100a8 | 4.42887E-34 | -2.895896297 | 0.009 | 0.062 | 6.63224E-30 | 4.97226E-33 | -1.99493153 | 0.02 | 0.075 | 7.44595E-29 | 4.97226E-33 | 8.85775E-34 |
| Scgb1c1 | 2.13685E-33 | -2.951099221 | 0.012 | 0.066 | 3.19993E-29 | 3.04538E-28 | -2.216274374 | 0.026 | 0.076 | 4.56045E-24 | 3.04538E-28 | 4.2737E-33 |
| Col6a3 | 1.1079E-06 | 0.979386491 | 0.02 | 0.01 | 0.01659085 | 8.13399E-31 | 2.039888784 | 0.027 | 0.006 | 1.21807E-26 | 1.1079E-06 | 1.6268E-30 |
| Sgms2 | 3.46053E-29 | 2.152604065 | 0.031 | 0.007 | 5.18214E-25 | 1.79986E-20 | 2.035588647 | 0.019 | 0.004 | 2.69529E-16 | 1.79986E-20 | 6.92106E-29 |
| Tnn | 3.76018E-27 | 2.540176682 | 0.021 | 0.003 | 5.63086E-23 | 1.45759E-21 | 2.20759596 | 0.018 | 0.004 | 2.18275E-17 | 1.45759E-21 | 7.52035E-27 |
| Aspn | 4.75775E-26 | 2.325388721 | 0.022 | 0.004 | 7.12472E-22 | 3.69005E-20 | 2.164515439 | 0.016 | 0.003 | 5.52586E-16 | 3.69005E-20 | 9.51549E-26 |
| Alpl | 2.31412E-11 | 1.56051713 | 0.018 | 0.006 | 3.4654E-07 | 2.3409E-23 | 2.139424458 | 0.019 | 0.004 | 3.5055E-19 | 2.31412E-11 | 4.6818E-23 |
| Camp | 6.6296E-12 | -2.506771 | 0.003 | 0.021 | 9.92783E-08 | 3.15909E-22 | -2.563233086 | 0.007 | 0.039 | 4.73074E-18 | 6.6296E-12 | 6.31818E-22 |
| Thbs2 | 3.18087E-06 | 1.273447792 | 0.012 | 0.005 | 0.047633558 | 4.96677E-22 | 1.998068608 | 0.019 | 0.004 | 7.43774E-18 | 3.18087E-06 | 9.93354E-22 |
| Sfrp2 | 7.72303E-13 | 1.490233879 | 0.021 | 0.007 | 1.15652E-08 | 1.48198E-20 | 1.688763049 | 0.024 | 0.007 | 2.21926E-16 | 7.72303E-13 | 2.96396E-20 |
| Kctd12 | 9.87227E-05 | 0.713850251 | 0.024 | 0.014 | 1 | 2.29914E-19 | 1.243761117 | 0.036 | 0.014 | 3.44296E-15 | 9.87227E-05 | 4.59828E-19 |
| Ibsp | 7.02531E-19 | 1.184032872 | 0.05 | 0.022 | 1.05204E-14 | 5.16932E-09 | 0.714118623 | 0.038 | 0.021 | 7.74105E-05 | 5.16932E-09 | 1.40506E-18 |
| Col9a3 | 1.5308E-18 | -1.812365491 | 0.012 | 0.045 | 2.29237E-14 | 1.63955E-08 | -2.173992137 | 0.004 | 0.016 | 0.000245523 | 1.63955E-08 | 3.0616E-18 |
| Car2 | 6.79142E-15 | -2.391716451 | 0.005 | 0.029 | 1.01702E-10 | 4.40849E-18 | -2.534488966 | 0.005 | 0.031 | 6.60172E-14 | 6.79142E-15 | 8.81698E-18 |
| Mmp2 | 1.68948E-12 | 1.124584407 | 0.032 | 0.014 | 2.52999E-08 | 1.0138E-17 | 1.352273106 | 0.03 | 0.011 | 1.51817E-13 | 1.68948E-12 | 2.0276E-17 |
| Vmo1 | 2.18189E-15 | -2.80343418 | 0.004 | 0.027 | 3.26738E-11 | 1.14178E-17 | -2.053471066 | 0.012 | 0.041 | 1.70982E-13 | 2.18189E-15 | 2.28356E-17 |
| Bpifb3 | 1.38251E-17 | -3.311688936 | 0.003 | 0.029 | 2.07031E-13 | 5.1669E-10 | -2.844383509 | 0.003 | 0.017 | 7.73743E-06 | 5.1669E-10 | 2.76502E-17 |
| Bpifb6 | 3.80081E-17 | -2.460063747 | 0.009 | 0.038 | 5.69172E-13 | 8.16174E-11 | -1.325518897 | 0.026 | 0.051 | 1.22222E-06 | 8.16174E-11 | 7.60163E-17 |
| Col9a2 | 5.28286E-17 | -1.860869507 | 0.012 | 0.042 | 7.91109E-13 | 6.62628E-05 | -1.406768473 | 0.005 | 0.013 | 0.992285206 | 6.62628E-05 | 1.05657E-16 |
| Ptms | 5.31105E-05 | 0.722689856 | 0.033 | 0.021 | 0.795329909 | 6.16022E-16 | 1.240962484 | 0.046 | 0.022 | 9.22492E-12 | 5.31105E-05 | 1.23204E-15 |
| Ptn | 7.78898E-16 | 2.125139183 | 0.015 | 0.003 | 1.1664E-11 | 1.13666E-11 | 1.642998632 | 0.015 | 0.005 | 1.70215E-07 | 1.13666E-11 | 1.5578E-15 |
| Obp1b | 1.40266E-13 | -2.090257031 | 0.009 | 0.033 | 2.10049E-09 | 2.82007E-15 | -2.335355125 | 0.008 | 0.032 | 4.22305E-11 | 1.40266E-13 | 5.64014E-15 |
| Col4a1 | 4.21183E-15 | 1.719715725 | 0.02 | 0.006 | 6.30722E-11 | 7.68931E-11 | 1.163946777 | 0.023 | 0.01 | 1.15147E-06 | 7.68931E-11 | 8.42367E-15 |
| Lcn4 | 6.33174E-14 | -2.764960032 | 0.003 | 0.024 | 9.48177E-10 | 1.87744E-13 | -3.166015514 | 0.002 | 0.02 | 2.81147E-09 | 1.87744E-13 | 1.26635E-13 |
| Itm2b | 7.30045E-14 | -1.355033598 | 0.024 | 0.056 | 1.09324E-09 | 0.000145651 | -0.706247396 | 0.036 | 0.051 | 1 | 0.000145651 | 1.46009E-13 |
| Acp5 | 4.46921E-13 | -2.053877597 | 0.009 | 0.032 | 6.69264E-09 | 0.00127227 | -0.956334525 | 0.014 | 0.023 | 1 | 0.00127227 | 8.93841E-13 |
| Apoe | 0.308139162 | 0.235775849 | 0.029 | 0.026 | 1 | 6.08734E-13 | 1.040990499 | 0.057 | 0.032 | 9.11579E-09 | 0.308139162 | 1.21747E-12 |
| Lrp1 | 3.35537E-08 | 0.966371307 | 0.028 | 0.014 | 0.000502466 | 2.5379E-12 | 1.196649326 | 0.025 | 0.01 | 3.8005E-08 | 3.35537E-08 | 5.0758E-12 |
| Fmod | 0.109962885 | 0.264124127 | 0.017 | 0.014 | 1 | 8.35513E-12 | 1.537973834 | 0.016 | 0.005 | 1.25118E-07 | 0.109962885 | 1.67103E-11 |
| Bpifa1 | 1.79554E-11 | -1.790455259 | 0.01 | 0.032 | 2.68882E-07 | 4.67773E-07 | -2.002349503 | 0.005 | 0.017 | 0.007004904 | 4.67773E-07 | 3.59108E-11 |
| Pcolce | 5.14083E-11 | 1.261629277 | 0.023 | 0.009 | 7.69839E-07 | 1.09727E-08 | 1.001920934 | 0.019 | 0.008 | 0.000164317 | 1.09727E-08 | 1.02817E-10 |
| Tmsb4x | 5.97154E-11 | -1.298114407 | 0.017 | 0.041 | 8.94239E-07 | 8.03003E-10 | -1.091316545 | 0.022 | 0.044 | 1.2025E-05 | 8.03003E-10 | 1.19431E-10 |
| Dmbt1 | 2.84746E-10 | -2.462397262 | 0.003 | 0.019 | 4.26407E-06 | 2.2431E-08 | -2.577566437 | 0.002 | 0.013 | 0.000335904 | 2.2431E-08 | 5.69492E-10 |
| Col16a1 | 1.02661E-05 | 1.039986502 | 0.015 | 0.007 | 0.153734261 | 2.96669E-10 | 1.239729363 | 0.019 | 0.007 | 4.44262E-06 | 1.02661E-05 | 5.93338E-10 |
| Bpifb4 | 3.13254E-10 | -1.842029425 | 0.011 | 0.031 | 4.69098E-06 | 3.26879E-09 | -1.37237163 | 0.025 | 0.048 | 4.89502E-05 | 3.26879E-09 | 6.26508E-10 |
| Tnfrsf19 | 0.000312258 | 1.074874959 | 0.01 | 0.005 | 1 | 3.38697E-10 | 1.611045485 | 0.014 | 0.005 | 5.07199E-06 | 0.000312258 | 6.77394E-10 |
| Col5a1 | 5.56509E-10 | 1.31716917 | 0.021 | 0.008 | 8.33372E-06 | 4.58455E-10 | 1.461352553 | 0.015 | 0.005 | 6.86536E-06 | 5.56509E-10 | 9.1691E-10 |
| Timp2 | 0.001305582 | 0.432933216 | 0.034 | 0.024 | 1 | 9.6171E-10 | 0.79512855 | 0.036 | 0.019 | 1.44016E-05 | 0.001305582 | 1.92342E-09 |
| Slc4a1 | 2.45117E-07 | -2.338058197 | 0.002 | 0.013 | 0.003670623 | 1.13978E-09 | -2.469384785 | 0.003 | 0.015 | 1.70682E-05 | 2.45117E-07 | 2.27956E-09 |
| Col5a2 | 1.59384E-05 | 1.197498939 | 0.011 | 0.005 | 0.23867761 | 2.75165E-09 | 1.432206207 | 0.014 | 0.005 | 4.1206E-05 | 1.59384E-05 | 5.5033E-09 |
| Mepe | 2.85393E-09 | -2.260981974 | 0.004 | 0.019 | 4.27376E-05 | 0.000547779 | -1.326893547 | 0.005 | 0.012 | 1 | 0.000547779 | 5.70787E-09 |
| Hist1h1e | 5.31336E-09 | -2.213285232 | 0.003 | 0.017 | 7.95675E-05 | 8.56913E-08 | -1.546466952 | 0.006 | 0.018 | 0.001283228 | 8.56913E-08 | 1.06267E-08 |
| Ctsk | 5.6628E-09 | -1.699289253 | 0.008 | 0.024 | 8.48004E-05 | 0.017136377 | -0.767466138 | 0.009 | 0.014 | 1 | 0.017136377 | 1.13256E-08 |
| Lum | 7.01674E-09 | 1.343662892 | 0.016 | 0.006 | 0.000105076 | 4.58671E-07 | 1.307151903 | 0.011 | 0.004 | 0.006868602 | 4.58671E-07 | 1.40335E-08 |
| Mup5 | 2.46281E-05 | -1.382597806 | 0.009 | 0.02 | 0.368805505 | 8.99406E-09 | -1.901602811 | 0.005 | 0.018 | 0.000134686 | 2.46281E-05 | 1.79881E-08 |
| Col12a1 | 1.1421E-06 | 0.791320677 | 0.027 | 0.014 | 0.017102978 | 9.82266E-09 | 1.116704381 | 0.017 | 0.007 | 0.000147094 | 1.1421E-06 | 1.96453E-08 |
| Cd24a | 1.06644E-08 | -1.913999211 | 0.004 | 0.018 | 0.000159699 | 2.56165E-08 | -2.241397326 | 0.003 | 0.015 | 0.000383606 | 2.56165E-08 | 2.13288E-08 |
| Gsn | 1.76209E-08 | -1.494876504 | 0.011 | 0.028 | 0.000263872 | 1.13605E-05 | -1.320007161 | 0.008 | 0.019 | 0.170123597 | 1.13605E-05 | 3.52417E-08 |
| Dmp1 | 5.37168E-08 | -1.819906614 | 0.006 | 0.02 | 0.000804409 | 9.07465E-05 | -1.06668288 | 0.01 | 0.019 | 1 | 9.07465E-05 | 1.07434E-07 |
| Spp1 | 0.06518869 | -0.426682898 | 0.062 | 0.07 | 1 | 7.71008E-08 | 0.291676485 | 0.084 | 0.059 | 0.001154584 | 0.06518869 | 1.54202E-07 |
| Ighm | 7.90529E-08 | -2.112883335 | 0.004 | 0.016 | 0.001183817 | 5.76271E-05 | -1.334506731 | 0.005 | 0.013 | 0.862965583 | 5.76271E-05 | 1.58106E-07 |
| Sost | 1.00843E-07 | -1.914657225 | 0.004 | 0.017 | 0.001510131 | 1.1104E-05 | -1.457510685 | 0.008 | 0.019 | 0.166282328 | 1.1104E-05 | 2.01687E-07 |
| Wfdc18 | 1.74082E-07 | -1.611774886 | 0.007 | 0.021 | 0.002606871 | 1.62197E-05 | -1.171777231 | 0.012 | 0.023 | 0.242889491 | 1.62197E-05 | 3.48163E-07 |
| Ccdc80 | 1.86219E-07 | -1.256684399 | 0.012 | 0.028 | 0.002788636 | 0.060470797 | 0.366016868 | 0.012 | 0.009 | 1 | 0.060470797 | 3.72439E-07 |
| Gpx6 | 2.17072E-05 | -1.328735804 | 0.011 | 0.022 | 0.325065023 | 2.88744E-07 | -1.309882944 | 0.014 | 0.03 | 0.004323939 | 2.17072E-05 | 5.77488E-07 |
| Ahnak | 0.001113794 | 0.516376774 | 0.03 | 0.021 | 1 | 3.47128E-07 | 0.785195871 | 0.028 | 0.016 | 0.005198244 | 0.001113794 | 6.94256E-07 |
| Mpo | 4.04288E-07 | -2.48839247 | 0.002 | 0.012 | 0.006054207 | 5.89856E-06 | -1.689845241 | 0.004 | 0.012 | 0.088330948 | 5.89856E-06 | 8.08575E-07 |
| Hist1h2ap | 4.05167E-07 | -2.43592505 | 0.002 | 0.012 | 0.006067383 | 1.86993E-06 | -1.955732775 | 0.003 | 0.012 | 0.028002247 | 1.86993E-06 | 8.10335E-07 |
| Mup4 | 5.72839E-06 | -1.799594159 | 0.005 | 0.015 | 0.085782667 | 8.14185E-07 | -1.725645962 | 0.006 | 0.017 | 0.012192422 | 5.72839E-06 | 1.62837E-06 |
| mt-Nd1 | 8.24981E-07 | -1.892114656 | 0.004 | 0.015 | 0.012354088 | 1.02822E-06 | -2.113240975 | 0.004 | 0.014 | 0.015397641 | 1.02822E-06 | 1.64996E-06 |
| Tnc | 0.019323706 | 0.556001649 | 0.013 | 0.009 | 1 | 1.24383E-06 | 0.929389243 | 0.019 | 0.009 | 0.018626323 | 0.019323706 | 2.48765E-06 |
| mt-Nd3 | 2.5288E-05 | -1.326219745 | 0.008 | 0.018 | 0.3786881 | 1.42624E-06 | -2.079744063 | 0.004 | 0.014 | 0.02135796 | 2.5288E-05 | 2.85248E-06 |
| Thbs1 | 2.55158E-06 | -1.067293218 | 0.016 | 0.032 | 0.03820994 | 0.695578981 | -0.278186985 | 0.01 | 0.01 | 1 | 0.695578981 | 5.10316E-06 |
| Ptprs | 0.002236103 | 0.729198965 | 0.015 | 0.009 | 1 | 3.83822E-06 | 1.018118161 | 0.016 | 0.007 | 0.057477297 | 0.002236103 | 7.67642E-06 |
| Mxra8 | 0.099891557 | 0.348757054 | 0.016 | 0.012 | 1 | 4.91132E-06 | 0.893948423 | 0.017 | 0.009 | 0.073546981 | 0.099891557 | 9.82261E-06 |
| Rsrp1 | 1.16186E-05 | -1.083550738 | 0.014 | 0.027 | 0.173988877 | 0.034776518 | -0.592086706 | 0.014 | 0.019 | 1 | 0.034776518 | 2.32371E-05 |
| Col11a1 | 0.171081611 | -0.369747974 | 0.011 | 0.014 | 1 | 1.99657E-05 | 0.828222769 | 0.014 | 0.007 | 0.298985748 | 0.171081611 | 3.99309E-05 |
| mt-Co1 | 2.33515E-05 | -1.185465337 | 0.009 | 0.02 | 0.349689265 | 0.000583589 | -1.054953587 | 0.01 | 0.018 | 1 | 0.000583589 | 4.67025E-05 |
| Oaz1 | 2.44298E-05 | -1.596916927 | 0.004 | 0.013 | 0.365837 | 0.001628033 | -0.961661667 | 0.007 | 0.013 | 1 | 0.001628033 | 4.88591E-05 |

| | | | | | | | | | | | | |
|---|---|---|---|---|---|---|---|---|---|---|---|---|
| Gpx1 | 2.72222E-05 | -1.774915229 | 0.003 | 0.011 | 0.407652236 | 0.000254184 | -1.328181092 | 0.004 | 0.011 | 1 | 0.000254184 | 5.44436E-05 |
| mt-Nd5 | 3.18507E-05 | -1.597326842 | 0.005 | 0.013 | 0.476964321 | 0.239524389 | -0.343658429 | 0.01 | 0.012 | 1 | 0.239524389 | 6.37004E-05 |
| Ucp2 | 3.34708E-05 | -1.6935069 | 0.003 | 0.011 | 0.501225405 | 0.000145805 | -1.45303258 | 0.004 | 0.011 | 1 | 0.000145805 | 6.69405E-05 |
| Selenop | 0.039805145 | 0.526254306 | 0.01 | 0.007 | 1 | 3.63675E-05 | 1.044267225 | 0.012 | 0.005 | 0.544602764 | 0.039805145 | 7.27336E-05 |
| Cald1 | 0.555433375 | 0.124584407 | 0.012 | 0.011 | 1 | 3.68873E-05 | 0.978269666 | 0.015 | 0.007 | 0.552387269 | 0.555433375 | 7.37732E-05 |
| Tpm1 | 0.293888993 | -0.216970279 | 0.02 | 0.023 | 1 | 3.75007E-05 | 0.456996148 | 0.034 | 0.022 | 0.561572906 | 0.293888993 | 7.5E-05 |
| Col6a2 | 0.002814376 | 0.609124036 | 0.016 | 0.01 | 1 | 7.35296E-05 | 0.998561922 | 0.011 | 0.005 | 1 | 0.002814376 | 0.000147054 |
| mt-Atp8 | 7.78644E-05 | -1.002965643 | 0.012 | 0.023 | 1 | 0.000584968 | -1.005133824 | 0.013 | 0.022 | 1 | 0.000584968 | 0.000155723 |
| Serpine2 | 0.00011623 | -0.96951916 | 0.009 | 0.019 | 1 | 0.288931517 | 0.232533862 | 0.01 | 0.008 | 1 | 0.288931517 | 0.000232447 |
| Mmp13 | 0.706931269 | -0.114681833 | 0.02 | 0.021 | 1 | 0.000172732 | 0.60508803 | 0.024 | 0.015 | 1 | 0.706931269 | 0.000345435 |
| Ppic | 0.007758701 | 0.748646727 | 0.01 | 0.006 | 1 | 0.000310697 | 0.977961035 | 0.01 | 0.005 | 1 | 0.007758701 | 0.000621297 |
| Igfbp4 | 0.339612412 | -0.25796273 | 0.01 | 0.012 | 1 | 0.000443847 | 0.543695526 | 0.025 | 0.016 | 1 | 0.339612412 | 0.000887496 |
| Col6a1 | 0.091606898 | 0.454122035 | 0.012 | 0.009 | 1 | 0.000539517 | 0.778677114 | 0.012 | 0.006 | 1 | 0.091606898 | 0.001078742 |
| mt-Co3 | 0.00064917 | -1.035135649 | 0.01 | 0.019 | 1 | 0.610366107 | -0.286271523 | 0.015 | 0.016 | 1 | 0.610366107 | 0.001297919 |
| mt-Nd4l | 0.000949488 | -0.91625401 | 0.007 | 0.014 | 1 | 0.027052765 | -0.707279132 | 0.007 | 0.011 | 1 | 0.027052765 | 0.001898073 |
| Tpm2 | 0.001006108 | -1.418003142 | 0.004 | 0.01 | 1 | 0.220970678 | -0.545073717 | 0.011 | 0.013 | 1 | 0.220970678 | 0.002011205 |
| Mt1 | 0.001150171 | -0.955092089 | 0.007 | 0.014 | 1 | 0.001580596 | -0.938578054 | 0.01 | 0.017 | 1 | 0.001580596 | 0.002299018 |
| Akr1a1 | 0.013127726 | -0.693976604 | 0.011 | 0.017 | 1 | 0.001333947 | -0.789492444 | 0.01 | 0.017 | 1 | 0.013127726 | 0.002666115 |
| Arf5 | 0.181843037 | -0.454839911 | 0.009 | 0.011 | 1 | 0.001583786 | -0.972083858 | 0.008 | 0.014 | 1 | 0.181843037 | 0.003165064 |
| Arglu1 | 0.001884329 | -0.930910706 | 0.007 | 0.014 | 1 | 0.10197639 | -0.510829503 | 0.009 | 0.013 | 1 | 0.10197639 | 0.003765107 |
| Igfbp5 | 0.002043386 | -1.541617409 | 0.03 | 0.041 | 1 | 0.634933884 | -0.413116565 | 0.037 | 0.035 | 1 | 0.634933884 | 0.004082596 |
| Selenom | 0.002078655 | -1.216312925 | 0.005 | 0.01 | 1 | 0.411071196 | -0.275613042 | 0.009 | 0.011 | 1 | 0.411071196 | 0.00415299 |
| Gja1 | 0.004547903 | 0.677344734 | 0.013 | 0.008 | 1 | 0.002088972 | 0.675143439 | 0.015 | 0.009 | 1 | 0.004547903 | 0.00417358 |
| Cst3 | 0.002221156 | -0.793495943 | 0.014 | 0.023 | 1 | 0.014202043 | 0.43047324 | 0.014 | 0.009 | 1 | 0.014202043 | 0.004437379 |
| mt-Cytb | 0.002929303 | -0.890793144 | 0.013 | 0.021 | 1 | 0.066297286 | -0.617598789 | 0.013 | 0.017 | 1 | 0.066297286 | 0.005850026 |
| Maf | 0.003700344 | -1.005791687 | 0.005 | 0.01 | 1 | 0.454902673 | -0.334506731 | 0.01 | 0.011 | 1 | 0.454902673 | 0.007386996 |
| Timp3 | 0.090385593 | 0.201100614 | 0.027 | 0.022 | 1 | 0.004830314 | 0.564954331 | 0.013 | 0.008 | 1 | 0.090385593 | 0.009637296 |
| Nucb1 | 0.045682945 | -0.63813959 | 0.01 | 0.014 | 1 | 0.00526481 | 0.5345624 | 0.013 | 0.008 | 1 | 0.045682945 | 0.010501902 |
| Maged1 | 0.394010384 | 0.133095906 | 0.012 | 0.01 | 1 | 0.014406436 | 0.432206207 | 0.014 | 0.009 | 1 | 0.394010384 | 0.028605326 |
| Serinc3 | 0.017201602 | -0.76736842 | 0.007 | 0.012 | 1 | 0.566466091 | -0.248846132 | 0.009 | 0.01 | 1 | 0.566466091 | 0.034107308 |
| Dbp | 0.017880096 | -0.692484075 | 0.01 | 0.016 | 1 | 0.412694494 | -0.430431151 | 0.009 | 0.01 | 1 | 0.412694494 | 0.035440495 |
| Jund | 0.354736743 | -0.305528413 | 0.01 | 0.012 | 1 | 0.019259234 | -0.715236179 | 0.008 | 0.012 | 1 | 0.354736743 | 0.03814755 |
| Sdc3 | 0.31344583 | -0.277825484 | 0.008 | 0.01 | 1 | 0.044894894 | 0.461352553 | 0.012 | 0.008 | 1 | 0.31344583 | 0.087774236 |
| mt-Atp6 | 0.179875317 | -0.429975784 | 0.012 | 0.015 | 1 | 0.045099025 | -0.745098325 | 0.011 | 0.015 | 1 | 0.179875317 | 0.088164129 |
| mt-Co2 | 0.086043666 | -0.394678036 | 0.03 | 0.037 | 1 | 0.955156133 | -0.252376333 | 0.036 | 0.035 | 1 | 0.955156133 | 0.164683819 |
| Fth1 | 0.15588414 | -0.315407669 | 0.017 | 0.021 | 1 | 0.932272469 | -0.193999276 | 0.02 | 0.02 | 1 | 0.932272469 | 0.287468415 |
| Cnn2 | 0.383435993 | 0.260068763 | 0.01 | 0.009 | 1 | 0.167123113 | 0.286528752 | 0.013 | 0.011 | 1 | 0.383435993 | 0.306316091 |
| Srsf6 | 0.201798267 | -0.299782906 | 0.009 | 0.012 | 1 | 0.366358679 | -0.492843758 | 0.008 | 0.01 | 1 | 0.366358679 | 0.362873994 |
| Htra1 | 0.26492878 | -0.414391889 | 0.009 | 0.011 | 1 | 0.210063425 | 0.332069536 | 0.011 | 0.009 | 1 | 0.26492878 | 0.376000208 |
| Mbd2 | 0.249346974 | -0.341551082 | 0.009 | 0.012 | 1 | 0.344966568 | 0.174189876 | 0.011 | 0.01 | 1 | 0.344966568 | 0.436520034 |
| Ddx17 | 0.297890945 | -0.33341781 | 0.011 | 0.013 | 1 | 0.365574542 | 0.148469597 | 0.011 | 0.009 | 1 | 0.365574542 | 0.507042875 |

**Spatial seq: cluster 2**

| | cKO_p_val | cKO_avg_log2FC | cKO_pct.1 | cKO_pct.2 | cKO_p_val_adj | WT_p_val | WT_avg_log2FC | WT_pct.1 | WT_pct.2 | WT_p_val_adj | max_pval | minimump_p_val |
|---|---|---|---|---|---|---|---|---|---|---|---|---|
| Mgp | 0 | 4.242137329 | 0.373 | 0.024 | 0 | 0 | 3.950983373 | 0.157 | 0.011 | 0 | 0 | 0 |
| Obp2b | 0 | 2.931195854 | 0.343 | 0.052 | 0 | 0 | 3.378590064 | 0.399 | 0.045 | 0 | 0 | 0 |
| Fn1 | 0 | 4.920264226 | 0.272 | 0.01 | 0 | 4.453E-298 | 4.74630574 | 0.111 | 0.005 | 6.6683E-294 | 4.453E-298 | 0 |
| Col2a1 | 0 | 5.500606948 | 0.23 | 0.006 | 0 | 1.2385E-234 | 5.55038553 | 0.077 | 0.002 | 1.8547E-230 | 1.2385E-234 | 0 |
| Lcn3 | 0 | 6.260250246 | 0.22 | 0.004 | 0 | 0 | 7.492857383 | 0.289 | 0.003 | 0 | 0 | 0 |
| Ccn2 | 0 | 5.143187734 | 0.223 | 0.007 | 0 | 0 | 5.401883007 | 0.15 | 0.004 | 0 | 0 | 0 |
| Obp2a | 2.416E-257 | 1.701219445 | 0.338 | 0.105 | 3.618E-253 | 0 | 1.960220474 | 0.413 | 0.11 | 0 | 2.416E-257 | 0 |
| Col1a1 | 2.4242E-268 | -1.584074878 | 0.351 | 0.626 | 3.6302E-264 | 4.03541E-41 | -0.655065609 | 0.528 | 0.636 | 6.04302E-37 | 4.03541E-41 | 4.8483E-268 |
| Col1a2 | 1.3553E-191 | -1.511510414 | 0.24 | 0.488 | 2.0296E-187 | 4.76754E-56 | -0.763047771 | 0.357 | 0.509 | 7.1394E-52 | 4.76754E-56 | 2.7106E-191 |
| Hbb-bs | 5.2702E-152 | -3.54400916 | 0.024 | 0.2 | 7.8921E-148 | 3.99939E-63 | -2.385702486 | 0.041 | 0.162 | 5.98909E-59 | 3.99939E-63 | 1.054E-151 |
| Col9a2 | 4.4058E-147 | 2.544401241 | 0.105 | 0.016 | 6.5976E-143 | 2.62798E-37 | 2.000031368 | 0.036 | 0.007 | 3.9354E-33 | 2.62798E-37 | 8.8115E-147 |
| Col9a3 | 8.5782E-147 | 2.45030792 | 0.11 | 0.018 | 1.2846E-142 | 8.22746E-28 | 1.440059267 | 0.036 | 0.01 | 1.23206E-23 | 8.22746E-28 | 1.7156E-146 |
| Lcn11 | 1.91343E-21 | 1.017067725 | 0.05 | 0.021 | 2.86536E-17 | 7.6672E-142 | 2.106753353 | 0.135 | 0.028 | 1.1482E-137 | 1.91343E-21 | 1.5334E-141 |
| Hba-a2 | 1.0224E-131 | -3.58677633 | 0.018 | 0.171 | 1.5311E-127 | 4.30563E-89 | -2.331944661 | 0.053 | 0.214 | 6.44769E-85 | 4.30563E-89 | 2.0449E-131 |
| Obp1a | 4.47312E-32 | 1.042784721 | 0.077 | 0.033 | 6.6985E-28 | 1.5169E-129 | 1.853712642 | 0.153 | 0.038 | 2.2716E-125 | 4.47312E-32 | 3.0339E-129 |
| Col11a2 | 1.5911E-119 | 1.834117913 | 0.13 | 0.033 | 2.3827E-115 | 3.64932E-10 | 0.836282255 | 0.045 | 0.024 | 5.46485E-06 | 3.64932E-10 | 3.1823E-119 |
| Lcn4 | 3.14507E-86 | 2.473230509 | 0.059 | 0.009 | 4.70975E-82 | 1.33025E-94 | 2.48577819 | 0.063 | 0.009 | 1.99205E-90 | 3.14507E-86 | 2.66051E-94 |
| Ccdc80 | 6.77217E-79 | 2.300126597 | 0.067 | 0.013 | 1.01413E-74 | 6.09563E-18 | 1.766041958 | 0.024 | 0.007 | 9.1282E-14 | 6.09563E-18 | 1.35443E-78 |
| Col27a1 | 1.16315E-72 | 1.550934701 | 0.093 | 0.027 | 1.74181E-68 | 7.99991E-07 | 0.87594102 | 0.016 | 0.007 | 0.011979859 | 7.99991E-07 | 2.32629E-72 |
| S100a9 | 3.38958E-70 | -4.064699678 | 0.006 | 0.09 | 5.0759E-66 | 1.76701E-58 | -2.857823396 | 0.018 | 0.12 | 2.6461E-54 | 1.76701E-58 | 6.77917E-70 |
| Thbs1 | 6.22694E-69 | 1.942339314 | 0.07 | 0.016 | 9.32484E-65 | 9.48312E-05 | 1.008696229 | 0.017 | 0.009 | 1 | 9.48312E-05 | 1.24539E-68 |
| Cnmd | 1.86056E-68 | 2.775334954 | 0.044 | 0.006 | 2.78618E-64 | 3.16656E-13 | 2.218828734 | 0.011 | 0.002 | 4.74192E-09 | 3.16656E-13 | 3.72111E-68 |
| Dmbt1 | 2.04635E-50 | 1.985751113 | 0.043 | 0.008 | 3.0644E-46 | 1.805E-55 | 2.268713337 | 0.039 | 0.006 | 2.70299E-51 | 2.04635E-50 | 3.61E-55 |
| Itm2b | 8.92335E-52 | 1.405778385 | 0.097 | 0.036 | 1.33627E-47 | 2.99975E-30 | 1.211276503 | 0.091 | 0.041 | 4.49213E-26 | 2.99975E-30 | 1.78467E-51 |
| Smoc2 | 2.34622E-50 | 2.507385481 | 0.038 | 0.006 | 3.51346E-46 | 4.68939E-12 | 1.313346333 | 0.02 | 0.007 | 7.02236E-08 | 4.68939E-12 | 4.69244E-50 |
| Acan | 4.92836E-50 | 1.51467256 | 0.063 | 0.018 | 7.38022E-46 | 0.000613367 | 0.537906591 | 0.011 | 0.006 | 1 | 0.000613367 | 9.85672E-50 |
| S100a8 | 2.32476E-49 | -3.61907852 | 0.005 | 0.065 | 3.48133E-45 | 1.30097E-31 | -2.354906511 | 0.014 | 0.073 | 1.94821E-27 | 1.30097E-31 | 4.64952E-49 |
| Cytl1 | 4.62362E-49 | 2.022249694 | 0.044 | 0.009 | 6.92386E-45 | 4.29028E-17 | 2.179045241 | 0.013 | 0.002 | 6.42469E-13 | 4.29028E-17 | 9.24723E-49 |
| Obp1b | 2.60883E-29 | 1.032713751 | 0.055 | 0.021 | 3.90672E-25 | 1.34559E-48 | 1.606409534 | 0.07 | 0.021 | 2.01503E-44 | 2.60883E-29 | 2.69119E-48 |
| A2ml1 | 3.55835E-09 | 1.159570031 | 0.015 | 0.005 | 5.32862E-05 | 4.43117E-44 | 2.543590803 | 0.028 | 0.004 | 6.63567E-40 | 3.55835E-09 | 8.86233E-44 |
| Hbb-bt | 4.37775E-41 | -3.584422831 | 0.004 | 0.054 | 6.55569E-37 | 5.38053E-29 | -2.95002958 | 0.008 | 0.058 | 8.05734E-25 | 5.38053E-29 | 8.75551E-41 |
| Spp1 | 2.3926E-40 | -2.132912125 | 0.021 | 0.082 | 3.58292E-36 | 7.0531E-19 | -1.65218145 | 0.026 | 0.07 | 1.0562E-14 | 7.0531E-19 | 4.7852E-40 |
| Ngp | 2.06808E-24 | -3.574215674 | 0.002 | 0.031 | 3.09695E-20 | 1.02255E-32 | -3.154938292 | 0.008 | 0.064 | 1.53127E-28 | 2.06808E-24 | 2.04511E-32 |
| Maged2 | 1.56758E-32 | 2.112081105 | 0.03 | 0.007 | 2.34745E-28 | 0.004714181 | 0.787884844 | 0.011 | 0.006 | 1 | 0.004714181 | 3.13516E-32 |
| Mup5 | 2.72077E-13 | 0.852770679 | 0.032 | 0.014 | 4.07435E-09 | 8.2269E-28 | 1.470181335 | 0.04 | 0.012 | 1.23198E-23 | 2.72077E-13 | 1.64538E-27 |
| Wfdc18 | 1.01165E-27 | 1.339391068 | 0.04 | 0.013 | 1.51494E-23 | 9.67708E-09 | 0.591615252 | 0.036 | 0.019 | 0.000144914 | 9.67708E-09 | 2.0233E-27 |
| Bpifa1 | 1.75651E-27 | 1.089941818 | 0.053 | 0.02 | 2.63037E-23 | 7.3589E-05 | 0.383735661 | 0.023 | 0.013 | 1 | 7.3589E-05 | 3.51301E-27 |
| Six2 | 9.6885E-27 | 2.030287014 | 0.025 | 0.006 | 1.45085E-22 | 9.90258E-05 | 1.049834924 | 0.013 | 0.007 | 1 | 9.90258E-05 | 1.9377E-26 |
| Col3a1 | 1.04686E-26 | -1.137888206 | 0.066 | 0.127 | 1.56767E-22 | 0.023587735 | -0.301826144 | 0.121 | 0.136 | 1 | 0.023587735 | 2.09372E-26 |
| Maf | 4.86852E-26 | 2.177128402 | 0.024 | 0.005 | 7.2906E-22 | 1.54183E-12 | 1.388753735 | 0.024 | 0.009 | 2.30889E-08 | 1.54183E-12 | 9.73703E-26 |
| Auts2 | 1.46531E-25 | 1.966032472 | 0.025 | 0.006 | 2.1943E-21 | 0.000405244 | 0.816331531 | 0.011 | 0.005 | 1 | 0.000405244 | 2.93062E-25 |
| Car2 | 1.71666E-25 | -3.92623435 | 0.002 | 0.031 | 2.5707E-21 | 6.89681E-17 | -3.204978975 | 0.003 | 0.03 | 1.0328E-12 | 6.89681E-17 | 3.43332E-25 |
| Cdkn1c | 1.76292E-24 | 2.19936362 | 0.021 | 0.004 | 2.63998E-20 | 1.65601E-11 | 1.471411179 | 0.017 | 0.005 | 2.47987E-07 | 1.65601E-11 | 3.52585E-24 |
| Dbp | 6.4943E-23 | 1.563111891 | 0.032 | 0.01 | 9.72521E-19 | 0.000596628 | 0.793972174 | 0.016 | 0.009 | 1 | 0.000596628 | 1.29886E-22 |
| Foxo3 | 5.0771E-22 | 2.160916456 | 0.017 | 0.003 | 7.60296E-18 | 3.72942E-07 | 1.197869115 | 0.012 | 0.005 | 0.005584811 | 3.72942E-07 | 1.01542E-21 |
| Wwp2 | 5.85155E-22 | 1.699508001 | 0.026 | 0.007 | 8.76269E-18 | 1.00677E-05 | 1.207146929 | 0.012 | 0.005 | 0.150764194 | 1.00677E-05 | 1.17031E-21 |
| Fmod | 6.18611E-22 | 1.589465489 | 0.031 | 0.01 | 9.2637E-18 | 0.002204713 | 0.911247889 | 0.012 | 0.006 | 1 | 0.002204713 | 1.23722E-21 |
| Gsn | 9.04737E-22 | 1.142351362 | 0.046 | 0.019 | 1.35484E-17 | 1.06336E-09 | 0.898308833 | 0.031 | 0.014 | 1.59238E-05 | 1.06336E-09 | 1.80947E-21 |
| Igfbp7 | 7.26119E-21 | 1.52283388 | 0.033 | 0.011 | 1.08736E-16 | 2.1772E-10 | 0.867572157 | 0.041 | 0.021 | 3.26035E-06 | 2.1772E-10 | 1.45224E-20 |
| Scgb1c1 | 0.001063434 | -0.755688166 | 0.045 | 0.059 | 1 | 2.32824E-19 | -1.89492498 | 0.028 | 0.074 | 3.48654E-15 | 0.001063434 | 4.65648E-19 |
| Camp | 3.95606E-17 | -3.384750486 | 0.002 | 0.023 | 5.92419E-13 | 2.70091E-19 | -2.986951746 | 0.005 | 0.038 | 4.04461E-15 | 3.95606E-17 | 5.40182E-19 |
| Htra1 | 3.10134E-18 | 1.724183886 | 0.024 | 0.007 | 4.64425E-14 | 2.44568E-09 | 1.374746877 | 0.02 | 0.008 | 3.66241E-05 | 2.44568E-09 | 6.20268E-18 |
| Dcn | 9.72872E-18 | 0.903574079 | 0.062 | 0.032 | 1.45688E-13 | 0.317654421 | 0.181079457 | 0.024 | 0.021 | 1 | 0.317654421 | 1.94574E-17 |
| Zfp36l2 | 9.83503E-18 | 1.784954609 | 0.021 | 0.006 | 1.4728E-13 | 5.90059E-06 | 1.063311156 | 0.016 | 0.007 | 0.088361363 | 5.90059E-06 | 1.96701E-17 |
| Bpifb6 | 2.1899E-17 | -2.106518713 | 0.011 | 0.039 | 3.27937E-13 | 1.40834E-13 | -1.703727181 | 0.019 | 0.051 | 2.10899E-09 | 1.40834E-13 | 4.37979E-17 |
| mt-Nd4l | 2.29407E-17 | 1.42486041 | 0.026 | 0.009 | 3.43536E-13 | 0.000936443 | 0.618608974 | 0.016 | 0.009 | 1 | 0.000936443 | 4.58813E-17 |
| Sdc4 | 6.28441E-17 | 1.608554862 | 0.022 | 0.006 | 9.41091E-13 | 3.82145E-05 | 0.880774174 | 0.015 | 0.007 | 0.572261532 | 3.82145E-05 | 1.25688E-16 |
| Col11a1 | 6.5207E-17 | 1.392857093 | 0.027 | 0.009 | 9.76475E-13 | 0.130002731 | 0.397057039 | 0.011 | 0.008 | 1 | 0.130002731 | 1.30414E-16 |
| Acp5 | 2.90275E-16 | -2.139637988 | 0.009 | 0.033 | 4.34687E-12 | 0.009579194 | -0.755350531 | 0.015 | 0.022 | 1 | 0.009579194 | 5.80551E-16 |
| Mup4 | 1.78649E-09 | 0.871949986 | 0.022 | 0.01 | 2.67526E-05 | 6.19354E-16 | 1.043885658 | 0.033 | 0.012 | 9.27483E-12 | 1.78649E-09 | 1.23871E-15 |
| Gpx3 | 3.04417E-15 | -1.211022621 | 0.022 | 0.052 | 4.55864E-11 | 0.000701056 | -0.529552706 | 0.036 | 0.051 | 1 | 0.000701056 | 6.08833E-15 |
| Timp2 | 8.63017E-15 | 0.943023867 | 0.044 | 0.021 | 1.29237E-10 | 0.001422377 | 0.564408097 | 0.03 | 0.021 | 1 | 0.001422377 | 1.72603E-14 |
| Bpifb3 | 1.12748E-14 | 0.453060162 | 0.041 | 0.019 | 1.68841E-10 | 5.61369E-05 | -1.799486163 | 0.006 | 0.016 | 0.840650434 | 5.61369E-05 | 2.25497E-14 |
| Ptms | 2.60448E-14 | 1.035640248 | 0.04 | 0.019 | 3.90021E-10 | 1.94118E-07 | 0.695679667 | 0.041 | 0.024 | 0.002906918 | 1.94118E-07 | 5.20897E-14 |
| Dmp1 | 7.7478E-14 | -2.692179011 | 0.003 | 0.021 | 1.16023E-09 | 9.88393E-05 | -1.098218975 | 0.008 | 0.019 | 1 | 9.88393E-05 | 1.54956E-13 |
| Sparc | 1.59602E-13 | -0.375314031 | 0.293 | 0.349 | 2.39004E-09 | 1.90433E-11 | -0.45018988 | 0.206 | 0.262 | 2.85174E-07 | 1.90433E-11 | 3.19205E-13 |
| App | 2.58891E-13 | 0.751920954 | 0.058 | 0.032 | 3.87689E-09 | 0.007716998 | 0.317805981 | 0.046 | 0.035 | 1 | 0.007716998 | 5.17782E-13 |
| Mepe | 6.83379E-13 | -2.702232676 | 0.003 | 0.02 | 1.02336E-08 | 2.88251E-05 | -2.035053973 | 0.003 | 0.012 | 0.431656589 | 2.88251E-05 | 1.36676E-12 |
| Mpo | 2.26001E-12 | -5.675690888 | 0 | 0.013 | 3.38436E-08 | 9.10567E-07 | -2.532143718 | 0.002 | 0.012 | 0.013635742 | 9.10567E-07 | 4.52001E-12 |
| mt-Nd3 | 2.29832E-12 | 0.948215881 | 0.029 | 0.013 | 3.44174E-08 | 5.26808E-07 | 0.95442403 | 0.021 | 0.01 | 0.007888949 | 5.26808E-07 | 4.59664E-12 |
| Bpifb4 | 4.37785E-12 | -1.757643324 | 0.011 | 0.032 | 6.55583E-08 | 3.45353E-12 | -1.6836987 | 0.019 | 0.048 | 5.17166E-08 | 4.37785E-12 | 6.90706E-12 |
| Ibsp | 6.33405E-12 | -1.402672394 | 0.011 | 0.032 | 9.48525E-08 | 0.138318497 | -0.34961868 | 0.02 | 0.025 | 1 | 0.138318497 | 1.26681E-11 |
| Gpx6 | 9.57391E-12 | -2.062822391 | 0.006 | 0.024 | 1.43369E-07 | 0.000119641 | -1.064964232 | 0.016 | 0.029 | 1 | 0.000119641 | 1.91478E-11 |
| Ctsk | 2.8202E-11 | -1.865015608 | 0.008 | 0.025 | 4.22325E-07 | 8.11028E-05 | -1.335843301 | 0.005 | 0.015 | 1 | 8.11028E-05 | 5.6404E-11 |
| Arglu1 | 4.92823E-11 | 1.151916458 | 0.023 | 0.01 | 7.38003E-07 | 3.38805E-05 | 0.761936392 | 0.02 | 0.011 | 0.507360897 | 3.38805E-05 | 9.85647E-11 |
| Slc4a1 | 4.95421E-11 | -3.370250916 | 0.001 | 0.013 | 7.41893E-07 | 1.69215E-07 | -2.366133767 | 0.003 | 0.015 | 0.002533991 | 1.69215E-07 | 9.90841E-11 |
| Nucb1 | 6.02508E-11 | 1.108289526 | 0.024 | 0.01 | 9.02256E-07 | 0.170096337 | 0.28985798 | 0.011 | 0.009 | 1 | 0.170096337 | 1.20502E-10 |
| Bgn | 1.87144E-10 | 0.66953967 | 0.066 | 0.041 | 2.80248E-06 | 0.031126561 | 0.247073113 | 0.053 | 0.044 | 1 | 0.031126561 | 3.74287E-10 |
| Vmo1 | 0.001854274 | -0.678410856 | 0.016 | 0.024 | 1 | 2.59569E-10 | -1.616514785 | 0.015 | 0.039 | 3.88705E-06 | 0.001854274 | 5.19138E-10 |
| Ighm | 1.00884E-09 | -2.238371941 | 0.004 | 0.017 | 1.51074E-05 | 0.000967688 | -1.182064583 | 0.005 | 0.012 | 1 | 0.000967688 | 2.01768E-09 |
| Mmp13 | 1.58882E-09 | -1.520323982 | 0.009 | 0.024 | 2.37926E-05 | 0.001041361 | -0.954133978 | 0.009 | 0.018 | 1 | 0.001041361 | 3.17764E-09 |
| Rarg | 2.53144E-09 | 1.526244508 | 0.011 | 0.003 | 3.79083E-05 | 2.30131E-09 | 1.698371263 | 0.012 | 0.003 | 3.44621E-05 | 2.53144E-09 | 4.60261E-09 |
| Col12a1 | 3.8847E-09 | 1.025189146 | 0.028 | 0.014 | 5.81734E-05 | 0.422034319 | 0.208648954 | 0.011 | 0.009 | 1 | 0.422034319 | 7.7694E-09 |
| Jund | 6.10203E-09 | 1.099827947 | 0.021 | 0.009 | 9.13779E-05 | 4.28819E-06 | 0.936376874 | 0.02 | 0.01 | 0.064215608 | 4.28819E-06 | 1.22041E-08 |
| Rsrp1 | 6.7161E-09 | 0.730494058 | 0.037 | 0.021 | 0.000100574 | 0.001645739 | 0.598542389 | 0.025 | 0.016 | 1 | 0.001645739 | 1.34322E-08 |
| Sost | 7.31078E-09 | -1.979173316 | 0.005 | 0.017 | 0.000109479 | 1.44842E-06 | -1.804441045 | 0.006 | 0.019 | 0.021690103 | 1.44842E-06 | 1.46216E-08 |
| mt-Nd2 | 7.66169E-09 | 1.079842838 | 0.018 | 0.008 | 0.000114734 | 5.38317E-07 | 1.084527642 | 0.015 | 0.006 | 0.008061302 | 5.38317E-07 | 1.53234E-08 |
| Nisch | 7.74038E-09 | 1.027119734 | 0.023 | 0.011 | 0.000115912 | 2.21416E-05 | 0.865241132 | 0.021 | 0.011 | 0.331569984 | 2.21416E-05 | 1.54808E-08 |
| mt-Co1 | 1.82325E-08 | 0.748874078 | 0.029 | 0.015 | 0.000273032 | 0.007527744 | 0.454917226 | 0.023 | 0.015 | 1 | 0.007527744 | 3.64651E-08 |
| Oaz1 | 1.99867E-08 | 0.957355491 | 0.02 | 0.009 | 0.000299301 | 8.67785E-05 | 0.564206025 | 0.02 | 0.011 | 1 | 8.67785E-05 | 3.99735E-08 |
| Col16a1 | 2.23521E-08 | 1.236737891 | 0.016 | 0.007 | 0.000334723 | 0.000562504 | 0.86856149 | 0.015 | 0.008 | 1 | 0.000562504 | 4.47042E-08 |
| Ahnak | 2.52582E-08 | 0.825928515 | 0.035 | 0.019 | 0.000378241 | 0.005526151 | 0.468740015 | 0.025 | 0.017 | 1 | 0.005526151 | 5.05163E-08 |
| Hist1h2ap | 3.23691E-08 | -2.384750486 | 0.002 | 0.012 | 0.000484727 | 8.14005E-06 | -2.20122684 | 0.002 | 0.011 | 0.121897219 | 8.14005E-06 | 6.47382E-08 |
| Srsf6 | 5.20291E-08 | 0.97583923 | 0.02 | 0.009 | 0.000779135 | 0.094485645 | 0.482721631 | 0.012 | 0.009 | 1 | 0.094485645 | 1.04058E-07 |
| mt-Atp8 | 1.45093E-07 | 0.70215758 | 0.031 | 0.018 | 0.002172769 | 8.14195E-08 | 0.674944573 | 0.034 | 0.018 | 0.001219257 | 1.45093E-07 | 1.62839E-07 |
| Cd24a | 9.34583E-08 | -1.353762793 | 0.006 | 0.018 | 0.001399538 | 0.000167018 | -1.020554404 | 0.005 | 0.014 | 1 | 0.000167018 | 1.86917E-07 |

Spatial seq: cluster 3

| | cKO_p_val | cKO_avg_log2FC | cKO_pct.1 | cKO_pct.2 | cKO_p_val_adj | WT_p_val | WT_avg_log2FC | WT_pct.1 | WT_pct.2 | WT_p_val_adj | max_pval | minimump_p_val |
|---|---|---|---|---|---|---|---|---|---|---|---|---|
| Hba-a2 | 1.73339E-57 | -2.811584036 | 0.022 | 0.152 | 2.59575E-53 | 2.5203E-142 | -3.407360538 | 0.022 | 0.222 | 3.7742E-138 | 1.73339E-57 | 5.0407E-142 |
| Obp2a | 3.01251E-95 | -8.332549156 | 0 | 0.179 | 4.51123E-91 | 2.7986E-139 | -10.15537717 | 0 | 0.181 | 4.1908E-135 | 3.01251E-95 | 5.5971E-139 |
| Sparc | 3.44263E-42 | -0.509461199 | 0.179 | 0.357 | 5.15534E-38 | 2.8469E-121 | -1.374849578 | 0.074 | 0.287 | 4.2632E-117 | 3.44263E-42 | 5.6938E-121 |
| Col3a1 | 3.14831E-64 | -5.992775597 | 0.002 | 0.129 | 4.7146E-60 | 3.986E-114 | -5.806039736 | 0.003 | 0.158 | 5.969E-110 | 3.14831E-64 | 7.972E-114 |
| Hbb-bs | 1.1738E-65 | -2.778915138 | 0.03 | 0.178 | 1.75776E-61 | 2.5322E-112 | -4.074863634 | 0.01 | 0.169 | 3.7919E-108 | 1.1738E-65 | 5.0643E-112 |
| S100a9 | 1.04769E-38 | -5.736844275 | 0.001 | 0.08 | 1.56892E-34 | 1.24339E-88 | -6.091978977 | 0.002 | 0.124 | 1.86198E-84 | 1.04769E-38 | 2.48678E-88 |
| Obp2b | 1.10706E-68 | -7.92868044 | 0 | 0.134 | 1.65783E-64 | 8.58189E-83 | -9.335553497 | 0 | 0.113 | 1.28514E-78 | 1.10706E-68 | 1.71638E-82 |
| Mgp | 6.54813E-60 | -8.345335311 | 0 | 0.117 | 9.80583E-56 | 2.21962E-27 | -7.033154567 | 0 | 0.038 | 3.32388E-23 | 2.21962E-27 | 1.30963E-59 |
| Scgb1c1 | 1.93756E-31 | -7.540100441 | 0 | 0.063 | 2.9015E-27 | 8.79277E-57 | -8.547020363 | 0 | 0.079 | 1.31672E-52 | 1.93756E-31 | 1.75855E-56 |
| Spp1 | 2.07308E-38 | -6.546441968 | 0 | 0.078 | 3.10444E-34 | 1.45175E-53 | -7.08378064 | 0 | 0.076 | 2.174E-49 | 2.07308E-38 | 2.90351E-53 |
| S100a8 | 2.32359E-25 | -3.889401789 | 0.004 | 0.058 | 3.47958E-21 | 5.81858E-52 | -5.366086435 | 0.002 | 0.076 | 8.71333E-48 | 2.32359E-25 | 1.16372E-51 |
| Ngp | 2.88656E-13 | -4.370175439 | 0.001 | 0.028 | 4.32262E-09 | 1.31641E-46 | -6.894337067 | 0 | 0.066 | 1.97133E-42 | 2.88656E-13 | 2.63282E-46 |
| Obp1a | 2.47773E-24 | -7.069637059 | 0 | 0.049 | 3.7104E-20 | 2.53673E-46 | -8.300728133 | 0 | 0.065 | 3.79876E-42 | 2.47773E-24 | 5.07347E-46 |
| Col1a2 | 7.34111E-45 | -0.728904216 | 0.282 | 0.452 | 1.09933E-40 | 1.35723E-06 | -0.380585268 | 0.499 | 0.484 | 0.020324491 | 1.35723E-06 | 1.46822E-44 |
| Itm2b | 7.28718E-28 | -6.095815712 | 0 | 0.057 | 1.09125E-23 | 1.32447E-40 | -7.694204074 | 0 | 0.057 | 1.9834E-36 | 7.28718E-28 | 2.64894E-40 |
| Fn1 | 3.66929E-39 | -6.593135203 | 0 | 0.079 | 5.49476E-35 | 2.29534E-17 | -6.331451433 | 0 | 0.024 | 3.43727E-13 | 2.29534E-17 | 7.33857E-39 |
| Bpifb6 | 7.84969E-19 | -6.650189892 | 0 | 0.037 | 1.17549E-14 | 6.92155E-39 | -7.711983991 | 0 | 0.055 | 1.0365E-34 | 7.84969E-19 | 1.38431E-38 |
| Hbb-bt | 5.3987E-20 | -3.343879758 | 0.004 | 0.048 | 8.08456E-16 | 3.60802E-38 | -3.968411802 | 0.003 | 0.06 | 5.40301E-34 | 5.3987E-20 | 7.21604E-38 |
| Lcn3 | 4.56836E-30 | -7.490533226 | 0 | 0.06 | 6.84113E-26 | 6.6715E-37 | -8.007673597 | 0 | 0.052 | 9.99058E-33 | 4.56836E-30 | 1.3343E-36 |
| Bpifb4 | 9.85767E-16 | -6.325806146 | 0 | 0.031 | 1.47619E-11 | 1.32689E-36 | -7.692925656 | 0 | 0.051 | 1.98702E-32 | 9.85767E-16 | 2.65378E-36 |
| Lcn11 | 3.05872E-16 | -6.450517548 | 0 | 0.032 | 4.58044E-12 | 1.8255E-36 | -7.945691726 | 0 | 0.051 | 2.73368E-32 | 3.05872E-16 | 3.65099E-36 |
| Gpx3 | 5.4545E-18 | -2.055683619 | 0.008 | 0.05 | 8.16812E-14 | 4.05381E-36 | -3.420107772 | 0.003 | 0.058 | 6.07057E-32 | 5.4545E-18 | 8.10761E-36 |
| Col2a1 | 4.44444E-32 | -7.287199141 | 0 | 0.064 | 6.65554E-28 | 2.44243E-11 | -5.824837331 | 0 | 0.015 | 3.65754E-07 | 2.44243E-11 | 8.88887E-32 |
| Ccn2 | 1.03362E-31 | -7.235806528 | 0 | 0.064 | 1.54785E-27 | 1.3893E-21 | -6.870786917 | 0 | 0.03 | 2.08047E-17 | 1.3893E-21 | 2.06724E-31 |
| Col11a2 | 3.82562E-31 | -7.202730915 | 0 | 0.062 | 5.72886E-27 | 4.53633E-23 | -6.777375977 | 0 | 0.032 | 6.79315E-19 | 4.53633E-23 | 7.65124E-31 |
| App | 1.42326E-21 | -6.592113826 | 0 | 0.043 | 2.13133E-17 | 3.97162E-31 | -7.160704617 | 0 | 0.044 | 5.94749E-27 | 1.42326E-21 | 7.94323E-31 |
| Vmo1 | 3.38796E-13 | -5.880133512 | 0 | 0.025 | 5.07347E-09 | 2.39526E-30 | -7.313262486 | 0 | 0.043 | 3.5869E-26 | 3.38796E-13 | 4.79052E-30 |
| Apoe | 5.31021E-15 | -5.062280108 | 0 | 0.03 | 7.95204E-11 | 6.80839E-30 | -5.647891902 | 0.001 | 0.043 | 1.01956E-25 | 5.31021E-15 | 1.36168E-29 |
| Igfbp5 | 9.69527E-22 | -6.548549631 | 0 | 0.044 | 1.45187E-17 | 1.17401E-29 | -7.408242686 | 0 | 0.042 | 1.75808E-25 | 9.69527E-22 | 2.34802E-29 |
| Camp | 2.7063E-10 | -4.481736278 | 0 | 0.02 | 4.05269E-06 | 7.63494E-26 | -4.621615164 | 0.001 | 0.039 | 1.14333E-21 | 2.7063E-10 | 1.52699E-25 |
| Col27a1 | 9.54322E-24 | -6.911587662 | 0 | 0.047 | 1.4291E-19 | 7.82419E-08 | -5.334734017 | 0 | 0.01 | 0.001171673 | 7.82419E-08 | 1.90864E-23 |
| Ptms | 2.36434E-13 | -4.87342329 | 0 | 0.027 | 3.5406E-09 | 1.09409E-22 | -6.755498127 | 0 | 0.031 | 1.63839E-18 | 2.36434E-13 | 2.18817E-22 |
| Col9a3 | 1.36812E-21 | -5.094373737 | 0 | 0.044 | 2.04877E-17 | 3.57453E-12 | -6.020979853 | 0 | 0.016 | 5.35286E-08 | 3.57453E-12 | 2.73625E-21 |
| Ibsp | 1.13968E-15 | -6.062280108 | 0 | 0.031 | 1.70667E-11 | 2.4482E-20 | -5.558998852 | 0 | 0.029 | 3.66618E-16 | 1.13968E-15 | 4.8964E-20 |
| Car2 | 2.34068E-11 | -2.82207641 | 0.003 | 0.027 | 3.50517E-07 | 1.38163E-19 | -3.703121423 | 0.002 | 0.03 | 2.06899E-15 | 2.34068E-11 | 2.76326E-19 |
| Col9a2 | 2.59182E-19 | -4.594155858 | 0.001 | 0.041 | 3.88126E-15 | 1.62978E-09 | -4.076936259 | 0.001 | 0.014 | 2.44059E-05 | 1.62978E-09 | 5.18365E-19 |
| Col1a1 | 8.49338E-19 | -0.679900382 | 0.536 | 0.567 | 1.27188E-14 | 1.36343E-15 | -0.648678481 | 0.679 | 0.609 | 2.04173E-11 | 1.36343E-15 | 1.69868E-18 |
| Acp5 | 5.90204E-16 | -6.213397841 | 0 | 0.031 | 8.83831E-12 | 3.30909E-18 | -6.478186807 | 0 | 0.025 | 4.95537E-14 | 5.90204E-16 | 6.61819E-18 |
| Bgn | 4.95207E-11 | -1.15280569 | 0.018 | 0.051 | 7.41572E-07 | 5.19759E-18 | -1.123437927 | 0.015 | 0.051 | 7.78339E-14 | 4.95207E-11 | 1.03952E-17 |
| Gpx6 | 6.2132E-11 | -4.216052294 | 0.001 | 0.023 | 9.30426E-07 | 5.94775E-18 | -3.344241102 | 0.003 | 0.031 | 8.90675E-14 | 6.2132E-11 | 1.18955E-17 |
| Tmsb4x | 3.29337E-07 | -0.72529079 | 0.016 | 0.039 | 0.004931829 | 1.69023E-16 | -1.16105688 | 0.012 | 0.045 | 2.53112E-12 | 3.29337E-07 | 3.38046E-16 |
| Dmp1 | 2.16113E-10 | -5.45501894 | 0 | 0.019 | 3.2363E-06 | 3.48709E-15 | -6.033154567 | 0 | 0.02 | 5.22192E-11 | 2.16113E-10 | 6.97418E-15 |
| Sost | 5.60246E-09 | -5.221346594 | 0 | 0.016 | 8.38968E-05 | 1.10977E-14 | -6.168084147 | 0 | 0.02 | 1.66189E-10 | 5.60246E-09 | 2.21955E-14 |
| Gsn | 5.4957E-14 | -5.038482537 | 0 | 0.028 | 8.22981E-10 | 1.11033E-14 | -6.264180737 | 0 | 0.02 | 1.66272E-10 | 5.4957E-14 | 2.22067E-14 |
| Mmp13 | 2.43302E-11 | -4.04745269 | 0.001 | 0.023 | 3.64345E-07 | 1.79579E-14 | -5.954253739 | 0 | 0.019 | 2.68919E-10 | 2.43302E-11 | 3.59158E-14 |
| Bpifb3 | 3.06537E-14 | -6.475103186 | 0 | 0.028 | 4.59039E-10 | 4.31113E-13 | -6.386264317 | 0 | 0.017 | 6.45592E-09 | 4.31113E-13 | 6.13073E-14 |
| Vim | 1.18605E-06 | -0.423822681 | 0.016 | 0.038 | 0.017761059 | 1.73873E-13 | -1.216339316 | 0.008 | 0.033 | 2.60375E-09 | 1.18605E-06 | 3.47747E-13 |
| Serpinf1 | 1.9762E-06 | -1.012210269 | 0.007 | 0.024 | 0.029593643 | 1.03209E-12 | -1.991947317 | 0.004 | 0.025 | 1.54556E-08 | 1.9762E-06 | 2.06418E-12 |
| Serpinh1 | 0.001294869 | -0.226812989 | 0.018 | 0.031 | 1 | 1.07418E-12 | -1.161669308 | 0.009 | 0.033 | 1.60858E-08 | 0.001294869 | 2.14836E-12 |
| Ctsk | 1.67759E-12 | -5.776047879 | 0 | 0.024 | 2.51219E-08 | 4.32954E-12 | -5.723299305 | 0 | 0.016 | 6.48348E-08 | 4.32954E-12 | 3.35519E-12 |
| Postn | 9.74015E-11 | -5.395375993 | 0 | 0.02 | 1.45859E-06 | 1.83002E-12 | -3.84339315 | 0.001 | 0.018 | 2.74046E-08 | 9.74015E-11 | 3.66005E-12 |
| Dcn | 1.33954E-07 | -0.898181148 | 0.017 | 0.041 | 0.002005959 | 2.45181E-12 | -1.640837144 | 0.004 | 0.024 | 3.67159E-08 | 1.33954E-07 | 4.90362E-12 |
| Mepe | 1.06186E-09 | -5.357408143 | 0 | 0.018 | 1.59014E-05 | 1.25327E-09 | -5.386264317 | 0 | 0.012 | 1.87678E-05 | 1.25327E-09 | 2.12373E-09 |
| Slc4a1 | 6.48939E-06 | -3.019360383 | 0.001 | 0.012 | 0.097178604 | 1.09188E-09 | -2.922608113 | 0.001 | 0.015 | 1.63509E-05 | 6.48939E-06 | 2.18375E-09 |
| Mpo | 1.70027E-06 | -4.571532949 | 0 | 0.011 | 0.025461525 | 2.13967E-09 | -4.274473492 | 0 | 0.013 | 3.20416E-05 | 1.70027E-06 | 4.27935E-09 |
| Hist1h2ap | 5.45888E-06 | -3.512293745 | 0 | 0.011 | 0.081746726 | 2.45492E-09 | -5.182730923 | 0 | 0.012 | 3.67625E-05 | 5.45888E-06 | 4.90985E-09 |
| Tnc | 5.86813E-06 | -3.537980389 | 0 | 0.011 | 0.087875304 | 4.70215E-09 | -3.743198862 | 0.001 | 0.013 | 7.04147E-05 | 5.86813E-06 | 9.4043E-09 |
| Acta1 | 0.008926395 | 1.275307489 | 0.013 | 0.007 | 1 | 5.87695E-09 | 1.537874759 | 0.027 | 0.013 | 8.80074E-05 | 0.008926395 | 1.17539E-08 |
| Bpifa1 | 0.001445504 | 1.132414418 | 0.038 | 0.026 | 1 | 7.39351E-09 | 1.740071416 | 0.026 | 0.012 | 0.000110718 | 0.001445504 | 1.4787E-08 |
| Hist1h1e | 4.47296E-07 | -2.47436429 | 0.002 | 0.016 | 0.006698262 | 1.00229E-08 | -1.796548287 | 0.004 | 0.018 | 0.000150092 | 4.47296E-07 | 2.00457E-08 |
| Timp2 | 0.000232758 | -0.336346528 | 0.014 | 0.028 | 1 | 1.13758E-07 | -1.002414316 | 0.009 | 0.024 | 0.001703526 | 0.000232758 | 2.27516E-07 |
| Cd24a | 0.015266177 | -0.249604854 | 0.009 | 0.016 | 1 | 1.80382E-07 | -1.812936517 | 0.003 | 0.014 | 0.002701224 | 0.015266177 | 3.60764E-07 |
| Serpine2 | 3.06489E-06 | -1.844083517 | 0.004 | 0.019 | 0.045896696 | 5.16418E-07 | -3.360729225 | 0.001 | 0.01 | 0.007733353 | 3.06489E-06 | 1.03283E-06 |
| Igfbp4 | 0.014184905 | -0.757946073 | 0.006 | 0.013 | 1 | 5.23413E-07 | -1.169174208 | 0.006 | 0.02 | 0.007838106 | 0.014184905 | 1.04683E-06 |
| Thbs1 | 6.35975E-07 | -1.082785763 | 0.011 | 0.031 | 0.009523726 | 0.000135655 | -1.055874644 | 0.004 | 0.011 | 1 | 0.000135655 | 1.27195E-06 |
| Col6a3 | 0.005341729 | -0.698445061 | 0.005 | 0.012 | 1 | 7.87753E-07 | -1.678905185 | 0.002 | 0.011 | 0.011796608 | 0.005341729 | 1.57551E-06 |
| Mmp2 | 0.000584658 | -0.829029171 | 0.008 | 0.018 | 1 | 8.13209E-07 | -1.325963807 | 0.005 | 0.017 | 0.012177804 | 0.000584658 | 1.62642E-06 |
| Ahnak | 7.16998E-06 | -1.242805022 | 0.009 | 0.025 | 0.107370455 | 9.56753E-07 | -0.797077231 | 0.007 | 0.02 | 0.014327375 | 7.16998E-06 | 1.9135E-06 |
| Kctd12 | 0.000196496 | -1.019360383 | 0.006 | 0.017 | 1 | 1.31111E-06 | -0.949979084 | 0.007 | 0.02 | 0.019633933 | 0.000196496 | 2.62223E-06 |
| Gpx1 | 0.020557576 | -0.413991672 | 0.005 | 0.01 | 1 | 1.35664E-06 | -2.285178192 | 0.002 | 0.011 | 0.020315717 | 0.020557576 | 2.71328E-06 |
| Serinc3 | 0.062731968 | -0.128589453 | 0.007 | 0.012 | 1 | 1.51858E-06 | -1.612267992 | 0.002 | 0.012 | 0.022740663 | 0.062731968 | 3.03715E-06 |
| Gja1 | 0.001020378 | -1.55963927 | 0.002 | 0.01 | 1 | 5.21476E-06 | -1.422790193 | 0.002 | 0.011 | 0.078091019 | 0.001020378 | 1.04295E-05 |
| Tsc22d1 | 2.9864E-05 | -1.28613073 | 0.003 | 0.015 | 0.447212967 | 6.24487E-06 | -1.758488733 | 0.001 | 0.01 | 0.093516978 | 2.9864E-05 | 1.24897E-05 |
| Col12a1 | 8.63137E-06 | -1.386605784 | 0.005 | 0.018 | 0.129254815 | 0.010461657 | -0.494822121 | 0.005 | 0.01 | 1 | 0.010461657 | 1.72627E-05 |
| Ighm | 1.36823E-05 | -1.845580347 | 0.003 | 0.016 | 0.20489293 | 9.84872E-06 | -1.787429642 | 0.003 | 0.013 | 0.147484564 | 1.36823E-05 | 1.96973E-05 |
| Lrp1 | 1.17688E-05 | -1.64691476 | 0.005 | 0.018 | 0.176238373 | 1.43062E-05 | -1.006832021 | 0.005 | 0.015 | 0.214234637 | 1.43062E-05 | 2.35375E-05 |
| Ptprs | 0.000584777 | -1.546441968 | 0.003 | 0.011 | 1 | 1.37959E-05 | -1.649298723 | 0.002 | 0.01 | 0.206593301 | 0.000584777 | 2.75916E-05 |
| Arf5 | 0.020753244 | -0.290477135 | 0.006 | 0.012 | 1 | 1.3987E-05 | -1.048138995 | 0.005 | 0.015 | 0.209455284 | 0.020753244 | 2.79738E-05 |
| Igfbp7 | 4.69016E-05 | -1.194678643 | 0.005 | 0.017 | 0.702351941 | 1.56037E-05 | -0.481302889 | 0.012 | 0.026 | 0.233665341 | 4.69016E-05 | 3.12071E-05 |
| Id3 | 0.00163479 | -1.142395252 | 0.005 | 0.014 | 1 | 1.58387E-05 | -0.710418013 | 0.004 | 0.013 | 0.237184018 | 0.00163479 | 3.16771E-05 |
| Tln1 | 4.24991E-05 | -2.014721488 | 0.003 | 0.014 | 0.636424012 | 1.98022E-05 | -1.615150501 | 0.002 | 0.01 | 0.296538463 | 4.24991E-05 | 3.96041E-05 |
| Wfdc18 | 0.699982337 | 0.660843561 | 0.02 | 0.019 | 1 | 2.07022E-05 | 1.338884128 | 0.031 | 0.019 | 0.310014731 | 0.699982337 | 4.14039E-05 |
| Tpm1 | 0.001377932 | -0.557727149 | 0.013 | 0.024 | 1 | 3.28849E-05 | -0.244270249 | 0.013 | 0.026 | 0.4924516 | 0.001377932 | 6.57687E-05 |
| Col16a1 | 0.002834126 | -0.829029171 | 0.003 | 0.01 | 1 | 3.94032E-05 | -1.336750362 | 0.003 | 0.011 | 0.590062581 | 0.002834126 | 7.88048E-05 |
| Pcolce | 0.000260139 | -1.276488148 | 0.003 | 0.013 | 1 | 4.28411E-05 | -0.752154167 | 0.003 | 0.012 | 0.641544915 | 0.000260139 | 8.56803E-05 |
| mt-Co3 | 0.615502044 | 0.661296456 | 0.018 | 0.017 | 1 | 4.31921E-05 | -0.623763631 | 0.007 | 0.018 | 0.646801239 | 0.615502044 | 8.63823E-05 |
| Selenom | 0.083291724 | -0.166064158 | 0.006 | 0.01 | 1 | 4.98449E-05 | -1.14583134 | 0.003 | 0.012 | 0.746426757 | 0.083291724 | 9.96872E-05 |
| Mxra8 | 0.12813108 | -0.17123379 | 0.009 | 0.013 | 1 | 5.43402E-05 | -1.24639655 | 0.003 | 0.011 | 0.813745064 | 0.12813108 | 0.000108678 |
| Sfrp2 | 0.000186293 | -1.892355106 | 0.002 | 0.011 | 1 | 8.44547E-05 | -1.175426123 | 0.003 | 0.011 | 1 | 0.000186293 | 0.000168902 |
| Sdc3 | 8.44893E-05 | -2.477317607 | 0.001 | 0.01 | 1 | 0.003930098 | -0.714837726 | 0.004 | 0.01 | 1 | 0.003930098 | 0.000168972 |
| Mt1 | 0.005784117 | -0.638144551 | 0.006 | 0.013 | 1 | 0.00011366 | -0.501591761 | 0.007 | 0.017 | 1 | 0.005784117 | 0.000227307 |
| mt-Co2 | 0.086612008 | 0.142307071 | 0.028 | 0.036 | 1 | 0.000167561 | -0.296338638 | 0.023 | 0.037 | 1 | 0.086612008 | 0.000335095 |
| Mbd2 | 0.120224396 | -0.290477135 | 0.008 | 0.012 | 1 | 0.000230323 | -0.85234125 | 0.004 | 0.011 | 1 | 0.120224396 | 0.000460593 |
| Htra1 | 0.000242134 | -1.543626952 | 0.003 | 0.012 | 1 | 0.000445626 | -0.965757349 | 0.004 | 0.01 | 1 | 0.000445626 | 0.00048421 |
| Obp1b | 0.000268203 | 1.14330493 | 0.041 | 0.027 | 1 | 0.811423698 | 0.317623114 | 0.027 | 0.028 | 1 | 0.811423698 | 0.000536334 |
| Maged1 | 0.012947369 | -0.620442549 | 0.005 | 0.011 | 1 | 0.000495105 | -0.715645732 | 0.004 | 0.011 | 1 | 0.012947369 | 0.000989964 |

| | | | | | | | | | | | | |
|---|---|---|---|---|---|---|---|---|---|---|---|---|
| Maf | 0.002803692 | -1.404713858 | 0.003 | 0.01 | 1 | 0.000536898 | -0.835409671 | 0.005 | 0.012 | 1 | 0.002803692 | 0.001073507 |
| Cald1 | 0.021293291 | -0.689304407 | 0.006 | 0.012 | 1 | 0.000555018 | -1.103571386 | 0.003 | 0.01 | 1 | 0.021293291 | 0.001109728 |
| Fth1 | 0.000705037 | -0.643763623 | 0.01 | 0.022 | 1 | 0.0041337 | 0.158823386 | 0.013 | 0.022 | 1 | 0.0041337 | 0.001409578 |
| Ccdc80 | 0.00149419 | -0.686584906 | 0.015 | 0.026 | 1 | 0.00074166 | -0.809714056 | 0.004 | 0.01 | 1 | 0.00149419 | 0.00148277 |
| Cst3 | 0.010503618 | -0.28613073 | 0.013 | 0.022 | 1 | 0.002015255 | -0.666436968 | 0.005 | 0.011 | 1 | 0.010503618 | 0.004026449 |
| Arglu1 | 0.099995682 | -0.219442883 | 0.009 | 0.013 | 1 | 0.002422006 | -0.398864354 | 0.006 | 0.013 | 1 | 0.099995682 | 0.004838146 |
| Igf2 | 0.01596638 | -0.655836456 | 0.006 | 0.013 | 1 | 0.002777936 | -0.789262567 | 0.005 | 0.011 | 1 | 0.01596638 | 0.005548155 |
| Mup4 | 0.002845339 | 1.190689064 | 0.02 | 0.012 | 1 | 0.28436372 | 0.772025984 | 0.017 | 0.015 | 1 | 0.28436372 | 0.005682581 |
| Ddx17 | 0.012212031 | -0.443631162 | 0.007 | 0.014 | 1 | 0.002848759 | -0.489243966 | 0.005 | 0.011 | 1 | 0.012212031 | 0.005689402 |
| mt-Nd3 | 0.126326942 | 0.817772584 | 0.02 | 0.016 | 1 | 0.003134303 | 1.108577937 | 0.017 | 0.011 | 1 | 0.126326942 | 0.006258781 |
| Timp3 | 0.016917806 | -0.154651749 | 0.016 | 0.024 | 1 | 0.0057877 | -0.580174362 | 0.004 | 0.01 | 1 | 0.016917806 | 0.011541903 |
| Jund | 0.180699854 | -0.227079316 | 0.009 | 0.012 | 1 | 0.009894134 | -0.354065495 | 0.007 | 0.012 | 1 | 0.180699854 | 0.019690375 |
| Dbp | 0.010496021 | -0.415470601 | 0.008 | 0.016 | 1 | 0.269239367 | 0.304331814 | 0.008 | 0.01 | 1 | 0.269239367 | 0.020881876 |
| mt-Co1 | 0.459241372 | 0.267339224 | 0.016 | 0.019 | 1 | 0.03086182 | -0.125625752 | 0.012 | 0.017 | 1 | 0.459241372 | 0.060771189 |
| Oaz1 | 0.036794716 | -0.553087281 | 0.007 | 0.012 | 1 | 0.424639497 | 0.434450983 | 0.011 | 0.013 | 1 | 0.424639497 | 0.072235581 |
| mt-Nd1 | 0.040204661 | 0.904577411 | 0.018 | 0.013 | 1 | 0.091061548 | 0.950199871 | 0.015 | 0.012 | 1 | 0.091061548 | 0.078792907 |
| mt-Cytb | 0.770491204 | 0.290826487 | 0.019 | 0.02 | 1 | 0.047104549 | 0.106775694 | 0.012 | 0.017 | 1 | 0.770491204 | 0.091990259 |
| Mup5 | 0.052928813 | 0.866909155 | 0.023 | 0.017 | 1 | 0.285000322 | 0.713787988 | 0.018 | 0.016 | 1 | 0.285000322 | 0.103056167 |
| Srsf6 | 0.062498817 | -0.399118395 | 0.007 | 0.012 | 1 | 0.094570324 | 0.172263285 | 0.007 | 0.01 | 1 | 0.094570324 | 0.121091532 |
| mt-Atp6 | 0.463249404 | 0.42395157 | 0.013 | 0.015 | 1 | 0.164830474 | 0.171831578 | 0.011 | 0.015 | 1 | 0.463249404 | 0.302491863 |
| mt-Nd4l | 0.85699774 | 0.408619917 | 0.012 | 0.013 | 1 | 0.544914549 | 0.59136079 | 0.011 | 0.01 | 1 | 0.85699774 | 0.792897232 |
| Dmbt1 | 0.791596155 | 0.446388959 | 0.015 | 0.016 | 1 | 0.754022062 | 0.524628209 | 0.01 | 0.011 | 1 | 0.791596155 | 0.939494854 |
| mt-Nd5 | 0.841332952 | 0.527456146 | 0.011 | 0.012 | 1 | 0.868777971 | 0.383615899 | 0.011 | 0.012 | 1 | 0.868777971 | 0.974824768 |

Spatial seq: cluster 4

| | cKO_p_val | cKO_avg_log2FC | cKO_pct.1 | cKO_pct.2 | cKO_p_val_adj | WT_p_val | WT_avg_log2FC | WT_pct.1 | WT_pct.2 | WT_p_val_adj | max_pval | minimump_p_val |
|---|---|---|---|---|---|---|---|---|---|---|---|---|
| Scgb1c1 | 0 | 5.48998285 | 0.49 | 0.015 | 0 | 0 | 5.98681356 | 0.495 | 0.012 | 0 | 0 | 0 |
| Bpifb6 | 0 | 5.7711934 | 0.305 | 0.007 | 0 | 0 | 5.22395975 | 0.32 | 0.011 | 0 | 0 | 0 |
| Bpifb4 | 0 | 5.38908816 | 0.239 | 0.007 | 0 | 0 | 5.08887115 | 0.29 | 0.012 | 0 | 0 | 0 |
| Vmo1 | 0 | 4.78752157 | 0.185 | 0.007 | 0 | 0 | 5.20350826 | 0.253 | 0.008 | 0 | 0 | 0 |
| Gpx6 | 0 | 4.91390051 | 0.163 | 0.007 | 0 | 0 | 4.4549422 | 0.164 | 0.009 | 0 | 0 | 0 |
| Bpifb3 | 2.15E-159 | 3.88982437 | 0.128 | 0.015 | 3.22E-155 | 5.05E-252 | 4.9755252 | 0.097 | 0.004 | 7.55E-248 | 2.15E-159 | 1.01E-251 |
| Apoe | 2.69E-28 | 1.71651523 | 0.071 | 0.022 | 4.03E-24 | 1.05E-75 | 1.8529967 | 0.107 | 0.028 | 1.57E-71 | 2.69E-28 | 2.09E-75 |
| Inmt | 5.56E-10 | 2.47141772 | 0.01 | 0.002 | 8.33E-06 | 3.75E-69 | 3.53396682 | 0.034 | 0.003 | 5.62E-65 | 5.56E-10 | 7.50E-69 |
| Selenom | 2.54E-25 | 2.14384307 | 0.035 | 0.007 | 3.80E-21 | 7.73E-69 | 2.87094398 | 0.046 | 0.006 | 1.16E-64 | 2.54E-25 | 1.55E-68 |
| Ptgds | 2.88E-66 | 4.52514971 | 0.03 | 0.001 | 4.32E-62 | 1.44E-56 | 3.81089449 | 0.026 | 0.002 | 2.16E-52 | 1.44E-56 | 5.77E-66 |
| Ugt2a1 | 2.76E-08 | 1.75935143 | 0.015 | 0.004 | 0.00041311 | 1.77E-59 | 2.93326849 | 0.036 | 0.004 | 2.65E-55 | 2.76E-08 | 3.53E-59 |
| Ces1d | 1.01E-45 | 3.49776382 | 0.028 | 0.002 | 1.51E-41 | 4.38E-53 | 3.15106614 | 0.029 | 0.003 | 6.56E-49 | 1.01E-45 | 8.76E-53 |
| Akr1a1 | 2.20E-11 | 1.40707808 | 0.037 | 0.014 | 3.30E-07 | 6.69E-53 | 2.19389585 | 0.055 | 0.011 | 1.00E-48 | 2.20E-11 | 1.34E-52 |
| Acsm4 | 4.28E-20 | 3.13015815 | 0.015 | 0.002 | 6.41E-16 | 2.65E-51 | 3.22593199 | 0.027 | 0.002 | 3.98E-47 | 4.28E-20 | 5.31E-51 |
| Apod | 3.01E-23 | 2.44376138 | 0.027 | 0.005 | 4.51E-19 | 1.90E-43 | 2.80203393 | 0.032 | 0.004 | 2.85E-39 | 3.01E-23 | 3.80E-43 |
| Chil6 | 2.76E-35 | 3.28811051 | 0.025 | 0.002 | 4.14E-31 | 6.86E-43 | 2.99146674 | 0.027 | 0.003 | 1.03E-38 | 2.76E-35 | 1.37E-42 |
| Cyp2f2 | 6.73E-11 | 1.9197231 | 0.017 | 0.004 | 1.01E-06 | 1.75E-42 | 2.40103603 | 0.034 | 0.005 | 2.62E-38 | 6.73E-11 | 3.50E-42 |
| Muc5b | 9.30E-15 | 1.93084934 | 0.022 | 0.005 | 1.39E-10 | 5.03E-42 | 2.89099872 | 0.024 | 0.002 | 7.53E-38 | 9.30E-15 | 1.01E-41 |
| Obp2b | 1.85E-19 | -1.9662895 | 0.046 | 0.125 | 2.78E-15 | 5.42E-38 | -2.9585459 | 0.02 | 0.105 | 8.12E-34 | 1.85E-19 | 1.08E-37 |
| Obp2a | 1.11E-18 | -1.5235423 | 0.08 | 0.165 | 1.67E-14 | 2.25E-34 | -2.0436671 | 0.069 | 0.164 | 3.37E-30 | 1.11E-18 | 4.50E-34 |
| Slc22a20 | 2.57E-13 | 2.87307301 | 0.01 | 0.001 | 3.85E-09 | 4.47E-33 | 3.19001242 | 0.018 | 0.002 | 6.69E-29 | 2.57E-13 | 8.94E-33 |
| Mt1 | 2.11E-10 | 1.4417337 | 0.03 | 0.011 | 3.16E-06 | 1.11E-28 | 1.6634666 | 0.043 | 0.012 | 1.66E-24 | 2.11E-10 | 2.22E-28 |
| Hba-a2 | 3.97E-12 | -1.1828368 | 0.078 | 0.142 | 5.95E-08 | 4.53E-27 | -1.3956433 | 0.111 | 0.201 | 6.79E-23 | 3.97E-12 | 9.07E-27 |
| Cyb5a | 2.25E-05 | 1.49070906 | 0.011 | 0.004 | 0.33733405 | 3.58E-25 | 2.62658308 | 0.02 | 0.003 | 5.37E-21 | 2.25E-05 | 7.17E-25 |
| Col1a1 | 2.29E-23 | 0.26480016 | 0.703 | 0.551 | 3.42E-19 | 0.73151662 | -0.1331381 | 0.629 | 0.619 | 1 | 0.73151662 | 4.57E-23 |
| Hbb-bs | 3.14E-22 | -1.7835144 | 0.075 | 0.168 | 4.70E-18 | 2.10E-20 | -1.4364092 | 0.082 | 0.152 | 3.15E-16 | 2.10E-20 | 6.27E-22 |
| Lcn3 | 7.90E-11 | -2.2082195 | 0.017 | 0.056 | 1.18E-06 | 7.29E-22 | -4.0169279 | 0.004 | 0.049 | 1.09E-17 | 7.90E-11 | 1.46E-21 |
| Spp1 | 0.00022372 | 0.41253636 | 0.092 | 0.066 | 1 | 3.52E-21 | 0.70223017 | 0.111 | 0.058 | 5.27E-17 | 0.00022372 | 7.04E-21 |
| S100a9 | 6.60E-15 | -2.2032724 | 0.021 | 0.075 | 9.88E-11 | 5.11E-21 | -1.5115828 | 0.048 | 0.113 | 7.65E-17 | 6.60E-15 | 1.02E-20 |
| Furin | 2.65E-10 | 2.04228539 | 0.015 | 0.004 | 3.96E-06 | 8.32E-21 | 2.16026508 | 0.02 | 0.004 | 1.25E-16 | 2.65E-10 | 1.66E-20 |
| Aldh2 | 0.0008587 | 0.87420409 | 0.019 | 0.01 | 1 | 4.98E-20 | 1.92107741 | 0.026 | 0.007 | 7.46E-16 | 0.0008587 | 9.96E-20 |
| Igfbp7 | 9.51E-05 | 0.89906822 | 0.028 | 0.015 | 1 | 1.08E-19 | 1.27459775 | 0.052 | 0.02 | 1.62E-15 | 9.51E-05 | 2.17E-19 |
| Ugdh | 0.00230394 | 1.13610742 | 0.012 | 0.005 | 1 | 1.63E-19 | 2.29443031 | 0.017 | 0.003 | 2.45E-15 | 0.00230394 | 3.27E-19 |
| Col1a2 | 4.28E-18 | 0.23348371 | 0.575 | 0.418 | 6.41E-14 | 0.06180123 | -0.2123844 | 0.482 | 0.487 | 1 | 0.06180123 | 8.56E-18 |
| Igfbp4 | 5.11E-07 | 1.30459863 | 0.026 | 0.011 | 0.00764546 | 4.83E-18 | 1.41471871 | 0.04 | 0.015 | 7.23E-14 | 5.11E-07 | 9.65E-18 |
| Fn1 | 8.74E-17 | -2.3713989 | 0.017 | 0.075 | 1.31E-12 | 0.00145059 | -1.0854913 | 0.011 | 0.021 | 1 | 0.00145059 | 1.75E-16 |
| Tagln | 1.08E-06 | 1.6648017 | 0.012 | 0.003 | 0.01611919 | 1.43E-16 | 2.12808467 | 0.017 | 0.003 | 2.14E-12 | 1.08E-06 | 2.86E-16 |
| Map1b | 1.09E-05 | 1.61825911 | 0.012 | 0.004 | 0.16250843 | 2.24E-16 | 1.89723067 | 0.018 | 0.004 | 3.35E-12 | 1.09E-05 | 4.48E-16 |
| Cldn11 | 2.15E-05 | 1.48371699 | 0.012 | 0.004 | 0.32172442 | 2.73E-16 | 2.3155433 | 0.015 | 0.003 | 4.09E-12 | 2.15E-05 | 5.46E-16 |
| Dmp1 | 0.07907631 | 0.4356677 | 0.023 | 0.017 | 1 | 2.51E-15 | 1.30642191 | 0.038 | 0.015 | 3.76E-11 | 0.07907631 | 5.03E-15 |
| Fth1 | 0.13406408 | 0.32731196 | 0.026 | 0.02 | 1 | 6.62E-15 | 1.09179941 | 0.043 | 0.018 | 9.91E-11 | 0.13406408 | 1.32E-14 |
| Myl9 | 0.00060059 | 1.23798703 | 0.012 | 0.005 | 1 | 1.39E-14 | 1.93367267 | 0.017 | 0.004 | 2.08E-10 | 0.00060059 | 2.77E-14 |
| Igf2 | 2.23E-14 | 1.72074207 | 0.033 | 0.01 | 3.34E-10 | 1.35E-11 | 1.51563136 | 0.023 | 0.008 | 2.02E-07 | 1.35E-11 | 4.46E-14 |
| Ngp | 7.47E-05 | -1.5581939 | 0.009 | 0.026 | 1 | 2.95E-14 | -1.6750816 | 0.021 | 0.06 | 4.42E-10 | 7.47E-05 | 5.90E-14 |
| Hbb-bt | 4.06E-10 | -2.0104851 | 0.011 | 0.046 | 6.08E-06 | 4.50E-14 | -1.7084275 | 0.017 | 0.055 | 6.74E-10 | 4.06E-10 | 9.00E-14 |
| Id3 | 0.37670299 | 0.3053198 | 0.015 | 0.013 | 1 | 5.82E-14 | 1.44405804 | 0.029 | 0.01 | 8.72E-10 | 0.37670299 | 1.16E-13 |
| Wls | 9.44E-05 | 1.60458417 | 0.01 | 0.003 | 1 | 1.68E-13 | 2.50818838 | 0.011 | 0.002 | 2.52E-09 | 9.44E-05 | 3.37E-13 |
| Acta2 | 3.15E-13 | 1.98267054 | 0.021 | 0.005 | 4.72E-09 | 6.38E-12 | 1.54060986 | 0.02 | 0.006 | 9.56E-08 | 6.38E-12 | 6.30E-13 |
| Gja1 | 0.01570042 | 0.79758281 | 0.015 | 0.008 | 1 | 9.17E-13 | 1.50333898 | 0.024 | 0.008 | 1.37E-08 | 0.01570042 | 1.83E-12 |
| Flna | 0.06481513 | 0.65117373 | 0.012 | 0.008 | 1 | 3.91E-12 | 1.65241777 | 0.019 | 0.006 | 5.86E-08 | 0.06481513 | 7.82E-12 |
| Obp1a | 0.00064067 | -1.0616518 | 0.026 | 0.044 | 1 | 1.77E-11 | -1.6797294 | 0.024 | 0.059 | 2.65E-07 | 0.00064067 | 3.54E-11 |
| S100a8 | 1.19E-08 | -1.5180351 | 0.02 | 0.055 | 0.00017829 | 6.14E-11 | -1.1877605 | 0.032 | 0.069 | 9.19E-07 | 1.19E-08 | 1.23E-10 |
| Slc6a6 | 0.00858327 | 0.8280515 | 0.012 | 0.006 | 1 | 9.45E-11 | 1.62975036 | 0.016 | 0.005 | 1.42E-06 | 0.00858327 | 1.89E-10 |
| Camp | 0.01411587 | -0.9621161 | 0.01 | 0.019 | 1 | 9.73E-11 | -1.8597572 | 0.01 | 0.036 | 1.46E-06 | 0.01411587 | 1.95E-10 |
| Cd81 | 0.01268895 | 0.86761858 | 0.012 | 0.006 | 1 | 1.18E-10 | 1.62881808 | 0.015 | 0.004 | 1.77E-06 | 0.01268895 | 2.37E-10 |
| mt-Atp8 | 0.1242157 | 0.28731101 | 0.026 | 0.02 | 1 | 1.42E-10 | 0.90575474 | 0.039 | 0.018 | 2.13E-06 | 0.1242157 | 2.85E-10 |
| Marcks | 7.65E-06 | 1.4401136 | 0.016 | 0.006 | 0.11457749 | 1.70E-10 | 1.32921824 | 0.023 | 0.009 | 2.55E-06 | 7.65E-06 | 3.40E-10 |
| Ctsk | 0.10186801 | 0.28731101 | 0.027 | 0.02 | 1 | 3.64E-10 | 1.15383881 | 0.028 | 0.012 | 5.45E-06 | 0.10186801 | 7.28E-10 |
| Col2a1 | 6.16E-10 | -1.693327 | 0.021 | 0.06 | 9.22E-06 | 8.14E-05 | -2.0960747 | 0.004 | 0.014 | 1 | 8.14E-05 | 1.23E-09 |
| mt-Co2 | 0.05991768 | 0.2511163 | 0.044 | 0.035 | 1 | 6.35E-10 | 0.65193284 | 0.058 | 0.032 | 9.51E-06 | 0.05991768 | 1.27E-09 |
| Col11a2 | 1.04E-09 | -1.7022943 | 0.02 | 0.058 | 1.55E-05 | 4.13E-09 | -2.0448494 | 0.008 | 0.03 | 6.19E-05 | 4.13E-09 | 2.07E-09 |
| mt-Cytb | 0.00020835 | 0.75101432 | 0.033 | 0.018 | 1 | 1.25E-09 | 0.83376425 | 0.032 | 0.014 | 1.87E-05 | 0.00020835 | 2.50E-09 |
| Cd164 | 0.00011148 | 1.51307687 | 0.011 | 0.004 | 1 | 2.24E-09 | 1.70013969 | 0.012 | 0.003 | 3.36E-05 | 0.00011148 | 4.49E-09 |
| Clu | 9.13E-06 | 1.35786943 | 0.017 | 0.007 | 0.13676283 | 2.26E-09 | 1.5525825 | 0.013 | 0.004 | 3.39E-05 | 9.13E-06 | 4.52E-09 |
| Maged1 | 0.00010625 | 1.17322687 | 0.021 | 0.01 | 1 | 3.69E-09 | 1.37343897 | 0.022 | 0.008 | 5.52E-05 | 0.00010625 | 7.37E-09 |
| Selenow | 0.00075505 | 1.11527391 | 0.016 | 0.008 | 1 | 3.18E-08 | 1.34418335 | 0.017 | 0.006 | 0.00047657 | 0.00075505 | 6.36E-08 |
| mt-Atp6 | 0.12203817 | 0.34381994 | 0.019 | 0.014 | 1 | 3.20E-08 | 0.9005058 | 0.028 | 0.013 | 0.00047974 | 0.12203817 | 6.41E-08 |
| Lcn11 | 0.0005901 | -1.4581929 | 0.014 | 0.029 | 1 | 4.61E-08 | -1.6684468 | 0.021 | 0.046 | 0.0006905 | 0.0005901 | 9.22E-08 |
| Ctnnb1 | 0.2377375 | 0.36672034 | 0.012 | 0.009 | 1 | 6.71E-08 | 1.43710528 | 0.016 | 0.006 | 0.00100444 | 0.2377375 | 1.34E-07 |
| Tnc | 0.9904531 | 0.14021181 | 0.01 | 0.01 | 1 | 1.21E-07 | 1.14770161 | 0.022 | 0.01 | 0.00180462 | 0.9904531 | 2.41E-07 |
| Lcn4 | 1.46E-05 | -2.2643839 | 0.005 | 0.022 | 0.21845908 | 2.68E-07 | -2.33782 | 0.003 | 0.018 | 0.00401574 | 1.46E-05 | 5.36E-07 |
| Fau | 0.1482357 | 0.67072653 | 0.01 | 0.006 | 1 | 3.10E-07 | 1.34418335 | 0.016 | 0.006 | 0.004638 | 0.1482357 | 6.19E-07 |
| Ptprs | 3.82E-07 | 1.38497205 | 0.023 | 0.009 | 0.00571559 | 0.00097254 | 0.90707846 | 0.015 | 0.008 | 1 | 0.00097254 | 7.63E-07 |
| Cald1 | 0.00106401 | 0.93486687 | 0.019 | 0.01 | 1 | 3.84E-07 | 1.22713992 | 0.018 | 0.008 | 0.00575639 | 0.00106401 | 7.69E-07 |
| Mepe | 0.01395725 | 0.54569049 | 0.024 | 0.015 | 1 | 4.72E-07 | 1.05273661 | 0.021 | 0.009 | 0.0070753 | 0.01395725 | 9.45E-07 |
| Col4a1 | 0.10391927 | 0.59672595 | 0.012 | 0.008 | 1 | 4.76E-07 | 1.00826198 | 0.023 | 0.011 | 0.00712712 | 0.10391927 | 9.52E-07 |
| Sdc4 | 0.18412513 | 0.40769213 | 0.013 | 0.01 | 1 | 5.60E-07 | 1.01342369 | 0.017 | 0.007 | 0.00838873 | 0.18412513 | 1.12E-06 |
| Cnn2 | 0.03954499 | 0.69899038 | 0.014 | 0.009 | 1 | 5.65E-07 | 1.15204457 | 0.022 | 0.01 | 0.00845574 | 0.03954499 | 1.13E-06 |
| Tpm1 | 0.25461637 | 0.20838332 | 0.027 | 0.022 | 1 | 6.25E-07 | 0.66550038 | 0.04 | 0.022 | 0.00935995 | 0.25461637 | 1.25E-06 |
| Tgfbr2 | 0.00630619 | 1.08457712 | 0.01 | 0.004 | 1 | 1.23E-06 | 1.48649331 | 0.011 | 0.003 | 0.01843848 | 0.00630619 | 2.46E-06 |
| Aqp1 | 0.02938552 | 0.92311369 | 0.01 | 0.005 | 1 | 1.30E-06 | 1.36722746 | 0.014 | 0.005 | 0.01942552 | 0.02938552 | 2.59E-06 |
| Ptms | 0.47907181 | 0.13651733 | 0.026 | 0.023 | 1 | 1.47E-06 | 0.6573974 | 0.042 | 0.025 | 0.02201837 | 0.47907181 | 2.94E-06 |
| Arglu1 | 0.01737699 | 0.63206491 | 0.019 | 0.012 | 1 | 1.69E-06 | 0.9538142 | 0.023 | 0.011 | 0.02528358 | 0.01737699 | 3.38E-06 |
| Rsrp1 | 0.05785226 | 0.2792575 | 0.032 | 0.024 | 1 | 2.31E-06 | 0.78240513 | 0.03 | 0.016 | 0.03451981 | 0.05785226 | 4.61E-06 |
| Nisch | 0.00342923 | 0.80571774 | 0.022 | 0.013 | 1 | 2.31E-06 | 0.89857835 | 0.023 | 0.011 | 0.0345341 | 0.00342923 | 4.61E-06 |
| Mgp | 2.39E-06 | -0.9559488 | 0.069 | 0.107 | 0.03580727 | 0.33661881 | -0.3061408 | 0.029 | 0.033 | 1 | 0.33661881 | 4.78E-06 |
| mt-Nd5 | 0.00014192 | 0.89668557 | 0.022 | 0.011 | 1 | 2.71E-06 | 0.66944051 | 0.022 | 0.01 | 0.04058245 | 0.00014192 | 5.42E-06 |
| Tpm2 | 0.00332223 | 0.98082919 | 0.016 | 0.008 | 1 | 2.80E-06 | 0.67873216 | 0.023 | 0.011 | 0.04195992 | 0.00332223 | 5.60E-06 |
| Fgfr1 | 0.00131773 | 1.07511679 | 0.013 | 0.006 | 1 | 2.84E-06 | 1.33535178 | 0.013 | 0.005 | 0.04245845 | 0.00131773 | 5.67E-06 |
| Car2 | 0.00062239 | -1.1487012 | 0.011 | 0.026 | 1 | 3.36E-06 | -1.2924057 | 0.011 | 0.028 | 0.05025179 | 0.00062239 | 6.71E-06 |
| Cst3 | 0.02450748 | 0.42561403 | 0.029 | 0.02 | 1 | 3.72E-06 | 1.03175033 | 0.02 | 0.009 | 0.05576258 | 0.02450748 | 7.45E-06 |
| Sfrp2 | 3.85E-06 | 1.27128589 | 0.021 | 0.008 | 0.05761033 | 7.14E-05 | 0.85491986 | 0.018 | 0.009 | 1 | 7.14E-05 | 7.69E-06 |
| Tmem59 | 0.12525979 | 0.56763304 | 0.01 | 0.007 | 1 | 4.21E-06 | 1.31780635 | 0.012 | 0.005 | 0.0630462 | 0.12525979 | 8.42E-06 |
| Ptov1 | 0.0232479 | 0.71373942 | 0.019 | 0.012 | 1 | 5.12E-06 | 1.2127325 | 0.014 | 0.006 | 0.07663871 | 0.0232479 | 1.02E-05 |
| Id2 | 0.0062842 | 1.03734344 | 0.011 | 0.005 | 1 | 5.19E-06 | 1.50818838 | 0.011 | 0.004 | 0.07770322 | 0.0062842 | 1.04E-05 |
| Surf4 | 9.98E-05 | 1.25568903 | 0.015 | 0.006 | 1 | 8.58E-06 | 1.30465498 | 0.011 | 0.004 | 0.128529 | 9.98E-05 | 1.72E-05 |

| | | | | | | | | | | | | |
|---|---|---|---|---|---|---|---|---|---|---|---|---|
| Tle5 | 0.00193059 | 1.31013682 | 0.01 | 0.004 | 1 | 9.25E-06 | 1.48649331 | 0.01 | 0.004 | 0.13854327 | 0.00193059 | 1.85E-05 |
| Actn1 | 0.00540354 | 1.01057653 | 0.012 | 0.006 | 1 | 1.06E-05 | 1.2306544 | 0.012 | 0.005 | 0.15945476 | 0.00540354 | 2.13E-05 |
| Jund | 0.6349553 | 0.10368594 | 0.013 | 0.012 | 1 | 1.36E-05 | 0.91595788 | 0.021 | 0.01 | 0.20307347 | 0.6349553 | 2.71E-05 |
| Mmp2 | 0.00969779 | 0.63955389 | 0.026 | 0.016 | 1 | 1.51E-05 | 0.89026111 | 0.025 | 0.013 | 0.22574471 | 0.00969779 | 3.01E-05 |
| Dcn | 0.0002561 | 0.61825911 | 0.056 | 0.037 | 1 | 1.68E-05 | 0.71665842 | 0.034 | 0.019 | 0.25156735 | 0.0002561 | 3.36E-05 |
| Gpx3 | 1.82E-05 | 0.61623994 | 0.068 | 0.043 | 0.27246188 | 2.66E-05 | 0.48354085 | 0.068 | 0.047 | 0.39798783 | 2.66E-05 | 3.64E-05 |
| Ndfip1 | 0.09427921 | 0.67072653 | 0.01 | 0.007 | 1 | 2.02E-05 | 1.28579596 | 0.011 | 0.004 | 0.30296528 | 0.09427921 | 4.05E-05 |
| Lamp2 | 0.04465511 | 0.75004898 | 0.012 | 0.007 | 1 | 2.03E-05 | 1.22511622 | 0.011 | 0.004 | 0.304715 | 0.04465511 | 4.07E-05 |
| Ank | 0.40381665 | 0.42561403 | 0.01 | 0.008 | 1 | 2.25E-05 | 1.16791865 | 0.012 | 0.005 | 0.33744677 | 0.40381665 | 4.51E-05 |
| Ccn2 | 2.72E-05 | -1.0027034 | 0.032 | 0.058 | 0.4072999 | 0.00028882 | -1.1415017 | 0.014 | 0.027 | 1 | 0.00028882 | 5.44E-05 |
| Cd63 | 8.48E-05 | 1.26910955 | 0.016 | 0.007 | 1 | 2.83E-05 | 1.14045659 | 0.014 | 0.006 | 0.42417172 | 8.48E-05 | 5.67E-05 |
| Tln1 | 0.09009637 | -0.6476349 | 0.008 | 0.013 | 1 | 2.86E-05 | 1.09660644 | 0.017 | 0.008 | 0.4289444 | 0.09009637 | 5.73E-05 |
| Ahnak | 0.56670523 | 0.11936756 | 0.025 | 0.023 | 1 | 2.87E-05 | 0.75758478 | 0.029 | 0.017 | 0.4304343 | 0.56670523 | 5.75E-05 |
| Cpxm1 | 0.1424133 | 0.60543507 | 0.01 | 0.007 | 1 | 3.54E-05 | 1.1416494 | 0.015 | 0.007 | 0.53074399 | 0.1424133 | 7.09E-05 |
| Acp5 | 0.0037482 | 0.49300087 | 0.039 | 0.026 | 1 | 3.69E-05 | 0.48707539 | 0.033 | 0.02 | 0.55273692 | 0.0037482 | 7.38E-05 |
| Maf | 0.45531926 | -0.2278282 | 0.008 | 0.01 | 1 | 3.82E-05 | 0.95996642 | 0.019 | 0.01 | 0.57273375 | 0.45531926 | 7.65E-05 |
| Aebp1 | 0.01272665 | 1.06571809 | 0.01 | 0.005 | 1 | 5.08E-05 | 1.36189894 | 0.01 | 0.004 | 0.76139616 | 0.01272665 | 0.00010169 |
| Irf2bp2 | 5.33E-05 | 1.2497642 | 0.018 | 0.008 | 0.79831291 | 0.00454998 | 0.66276542 | 0.015 | 0.009 | 1 | 0.00454998 | 0.00010662 |
| Col5a1 | 0.03400835 | 0.77484947 | 0.016 | 0.01 | 1 | 6.27E-05 | 1.11446139 | 0.014 | 0.006 | 0.93965083 | 0.03400835 | 0.00012549 |
| Vim | 0.19726778 | -0.3209376 | 0.029 | 0.036 | 1 | 6.63E-05 | 0.58574936 | 0.043 | 0.028 | 0.99223376 | 0.19726778 | 0.00013251 |
| S100a4 | 0.00357587 | 0.97310183 | 0.012 | 0.006 | 1 | 6.75E-05 | 1.09820411 | 0.011 | 0.005 | 1 | 0.00357587 | 0.00013498 |
| Tnfrsf19 | 7.20E-05 | 1.42561403 | 0.013 | 0.005 | 1 | 0.00023473 | 1.1456183 | 0.012 | 0.006 | 1 | 0.00023473 | 0.00014408 |
| Pcolce | 0.00154641 | 0.87307301 | 0.02 | 0.011 | 1 | 8.93E-05 | 0.89403887 | 0.018 | 0.009 | 1 | 0.00154641 | 0.00017861 |
| Arf6 | 0.01643299 | 1.02872388 | 0.01 | 0.005 | 1 | 0.00012527 | 1.36275794 | 0.01 | 0.004 | 1 | 0.01643299 | 0.00025052 |
| Eid1 | 0.006343 | 1.06806203 | 0.01 | 0.005 | 1 | 0.00014803 | 1.05688734 | 0.011 | 0.004 | 1 | 0.006343 | 0.00029605 |
| Col6a2 | 0.22045571 | 0.38330323 | 0.014 | 0.01 | 1 | 0.00016134 | 1.10779766 | 0.012 | 0.005 | 1 | 0.22045571 | 0.00032266 |
| Ddx17 | 0.00201247 | 0.79625741 | 0.021 | 0.012 | 1 | 0.00018919 | 0.93109412 | 0.017 | 0.009 | 1 | 0.00201247 | 0.00037834 |
| Atox1 | 0.26918314 | -0.4692037 | 0.009 | 0.012 | 1 | 0.00019945 | 0.96762 | 0.012 | 0.005 | 1 | 0.26918314 | 0.00039886 |
| Slc4a1 | 0.00098303 | -2.0419915 | 0.002 | 0.011 | 1 | 0.00024541 | -1.4977609 | 0.005 | 0.014 | 1 | 0.00098303 | 0.00049077 |
| Serinc3 | 0.53746415 | 0.20064767 | 0.013 | 0.011 | 1 | 0.00031068 | 0.86328334 | 0.017 | 0.009 | 1 | 0.53746415 | 0.00062125 |
| Col12a1 | 0.21895865 | 0.39854579 | 0.021 | 0.016 | 1 | 0.00032313 | 1.0050947 | 0.016 | 0.008 | 1 | 0.21895865 | 0.00064616 |
| Mrc2 | 0.25316354 | 0.49961462 | 0.01 | 0.007 | 1 | 0.00044628 | 0.99186754 | 0.013 | 0.007 | 1 | 0.25316354 | 0.00089236 |
| Serpinf1 | 0.22930215 | 0.27993658 | 0.026 | 0.021 | 1 | 0.00047417 | 0.60104799 | 0.032 | 0.02 | 1 | 0.22930215 | 0.00094812 |
| Col9a2 | 0.00048443 | -1.0790064 | 0.02 | 0.038 | 1 | 0.13169915 | -0.5785729 | 0.008 | 0.012 | 1 | 0.13169915 | 0.00096863 |
| Col6a1 | 0.10624942 | 0.52837361 | 0.013 | 0.009 | 1 | 0.00052894 | 0.88582591 | 0.013 | 0.007 | 1 | 0.10624942 | 0.00105759 |
| Mmp13 | 0.05335837 | 0.43887013 | 0.028 | 0.02 | 1 | 0.00054863 | 0.72098003 | 0.025 | 0.015 | 1 | 0.05335837 | 0.00109695 |
| Lmna | 0.00268128 | 0.89163546 | 0.016 | 0.008 | 1 | 0.00056941 | 0.96762 | 0.012 | 0.006 | 1 | 0.00268128 | 0.00113851 |
| Mpo | 0.00186013 | -1.9922385 | 0.002 | 0.011 | 1 | 0.00069357 | -1.5348803 | 0.004 | 0.012 | 1 | 0.00186013 | 0.00138666 |
| Dmbt1 | 0.00199062 | -1.4943327 | 0.006 | 0.017 | 1 | 0.00073611 | -1.6063714 | 0.004 | 0.012 | 1 | 0.00199062 | 0.00147168 |
| Alpl | 0.00099163 | 1.06806203 | 0.016 | 0.008 | 1 | 0.01016619 | 0.79129722 | 0.011 | 0.006 | 1 | 0.01016619 | 0.00198227 |
| Kctd12 | 0.00569332 | 0.72517432 | 0.025 | 0.015 | 1 | 0.00111445 | 0.65972146 | 0.027 | 0.017 | 1 | 0.00569332 | 0.00222766 |
| Eef1a1 | 0.04363741 | 0.58315531 | 0.018 | 0.012 | 1 | 0.00113015 | 0.83210503 | 0.014 | 0.008 | 1 | 0.04363741 | 0.00225903 |
| Ibsp | 0.00119853 | 0.63834336 | 0.04 | 0.026 | 1 | 0.00309569 | 0.47823516 | 0.034 | 0.023 | 1 | 0.00309569 | 0.00239563 |
| Obp1b | 0.0013889 | -1.2420889 | 0.015 | 0.03 | 1 | 0.10114444 | -0.7446613 | 0.023 | 0.028 | 1 | 0.10114444 | 0.00277587 |
| Hist1h2ap | 0.0230216 | -1.106881 | 0.004 | 0.01 | 1 | 0.00163692 | -1.4323106 | 0.004 | 0.011 | 1 | 0.0230216 | 0.00327115 |
| Mbd2 | 0.2173241 | 0.4174401 | 0.015 | 0.011 | 1 | 0.00174104 | 0.86931792 | 0.016 | 0.009 | 1 | 0.2173241 | 0.00347904 |
| Cd24a | 0.00263066 | -1.2442374 | 0.006 | 0.016 | 1 | 0.00378641 | -1.1792214 | 0.006 | 0.013 | 1 | 0.00378641 | 0.0052544 |
| Pabpn1 | 0.00313486 | 1.00338103 | 0.015 | 0.008 | 1 | 0.0041044 | 0.96762 | 0.011 | 0.006 | 1 | 0.0041044 | 0.00625989 |
| Col6a3 | 0.00397935 | 0.81598948 | 0.019 | 0.011 | 1 | 0.09230979 | 0.43988237 | 0.013 | 0.01 | 1 | 0.09230979 | 0.00794287 |
| Mup5 | 0.0040385 | -1.3632208 | 0.008 | 0.019 | 1 | 0.07299524 | -0.8026975 | 0.011 | 0.017 | 1 | 0.07299524 | 0.00806069 |
| Col9a3 | 0.01432508 | -0.6925674 | 0.027 | 0.04 | 1 | 0.00424636 | -1.2590598 | 0.007 | 0.014 | 1 | 0.01432508 | 0.0084747 |
| Itm2b | 0.32334908 | -0.4018971 | 0.045 | 0.051 | 1 | 0.00499428 | 0.15734464 | 0.061 | 0.047 | 1 | 0.32334908 | 0.00996361 |
| Tagln2 | 0.01001168 | 1.01057653 | 0.012 | 0.006 | 1 | 0.00507046 | 0.92497566 | 0.011 | 0.006 | 1 | 0.01001168 | 0.0101152 |
| Sost | 0.15899682 | 0.29045445 | 0.019 | 0.014 | 1 | 0.00545417 | 0.34810027 | 0.024 | 0.016 | 1 | 0.15899682 | 0.01087859 |
| Maged2 | 0.3627441 | 0.20556555 | 0.015 | 0.012 | 1 | 0.00549498 | 0.68073885 | 0.011 | 0.006 | 1 | 0.3627441 | 0.01095976 |
| Thbs1 | 0.00697522 | -0.8454898 | 0.017 | 0.03 | 1 | 0.04398388 | 0.44977169 | 0.014 | 0.01 | 1 | 0.04398388 | 0.01390179 |
| Nucb1 | 0.1011641 | 0.41231721 | 0.018 | 0.013 | 1 | 0.0113158 | 0.62776999 | 0.014 | 0.008 | 1 | 0.1011641 | 0.02250356 |
| Selenop | 0.03151432 | 0.77536007 | 0.012 | 0.007 | 1 | 0.01241665 | 0.86070479 | 0.011 | 0.006 | 1 | 0.03151432 | 0.02467912 |
| mt-Nd4l | 0.7286713 | -0.2029754 | 0.012 | 0.013 | 1 | 0.0124798 | 0.54808111 | 0.015 | 0.009 | 1 | 0.7286713 | 0.02480385 |
| Timp3 | 0.33342762 | 0.14654142 | 0.027 | 0.023 | 1 | 0.01363885 | 0.67388879 | 0.013 | 0.008 | 1 | 0.33342762 | 0.02709168 |
| Dbp | 0.02844324 | 0.5873651 | 0.021 | 0.014 | 1 | 0.01795469 | 0.50049399 | 0.015 | 0.009 | 1 | 0.02844324 | 0.035587 |
| Itgb5 | 0.10395721 | 0.79625741 | 0.01 | 0.006 | 1 | 0.01882036 | 0.80468143 | 0.01 | 0.006 | 1 | 0.10395721 | 0.03728652 |
| Tmsb4x | 0.51602877 | -0.2334717 | 0.033 | 0.036 | 1 | 0.01889028 | 0.3146807 | 0.049 | 0.039 | 1 | 0.51602877 | 0.03742373 |
| Mup4 | 0.11444851 | -0.9667034 | 0.008 | 0.013 | 1 | 0.02032151 | -1.0715184 | 0.01 | 0.016 | 1 | 0.11444851 | 0.04023005 |
| Oaz1 | 0.25857422 | 0.3053198 | 0.015 | 0.011 | 1 | 0.02084019 | 0.40520007 | 0.017 | 0.012 | 1 | 0.25857422 | 0.04124607 |
| Mxra8 | 0.73885937 | 0.24873627 | 0.014 | 0.013 | 1 | 0.04711699 | 0.50001445 | 0.014 | 0.01 | 1 | 0.73885937 | 0.09201398 |
| Nupr1 | 0.04924807 | 0.72517432 | 0.01 | 0.006 | 1 | 0.06296355 | 0.54135524 | 0.01 | 0.007 | 1 | 0.06296355 | 0.09607077 |
| Sgms2 | 0.06752604 | 0.56962434 | 0.017 | 0.011 | 1 | 0.05751645 | 0.63320096 | 0.01 | 0.007 | 1 | 0.06752604 | 0.11172475 |
| Wfdc18 | 0.06663076 | -0.7251004 | 0.012 | 0.019 | 1 | 0.24743525 | -0.5295665 | 0.018 | 0.022 | 1 | 0.24743525 | 0.12882187 |
| mt-Nd3 | 0.45201628 | -0.359163 | 0.014 | 0.016 | 1 | 0.08564217 | 0.10702305 | 0.016 | 0.011 | 1 | 0.45201628 | 0.16394975 |
| Rcn3 | 0.10121328 | 0.64011855 | 0.012 | 0.008 | 1 | 0.08715313 | 0.49118195 | 0.01 | 0.007 | 1 | 0.10121328 | 0.16671059 |
| Serpinh1 | 0.10259111 | 0.29785849 | 0.036 | 0.028 | 1 | 0.2842589 | 0.11908164 | 0.033 | 0.029 | 1 | 0.2842589 | 0.19465729 |
| Tpt1 | 0.39155555 | 0.4181195 | 0.014 | 0.011 | 1 | 0.12528099 | 0.5134441 | 0.01 | 0.007 | 1 | 0.39155555 | 0.23486666 |
| Ighm | 0.5811585 | -0.3151431 | 0.012 | 0.014 | 1 | 0.16814243 | -0.4210771 | 0.008 | 0.012 | 1 | 0.5811585 | 0.30801299 |
| Hist1h1e | 0.17119255 | -0.4447507 | 0.01 | 0.015 | 1 | 0.18475031 | -0.4099004 | 0.012 | 0.016 | 1 | 0.18475031 | 0.31307822 |
| Arf5 | 0.26202236 | 0.34219803 | 0.014 | 0.011 | 1 | 0.18268871 | 0.26718028 | 0.016 | 0.013 | 1 | 0.26202236 | 0.33200226 |
| Tsc22d1 | 0.19507373 | -0.5129854 | 0.01 | 0.014 | 1 | 0.43831533 | 0.16966377 | 0.01 | 0.008 | 1 | 0.43831533 | 0.3520937 |
| Htra1 | 0.72397756 | 0.16574691 | 0.012 | 0.011 | 1 | 0.19794369 | 0.30465498 | 0.012 | 0.009 | 1 | 0.72397756 | 0.35670567 |
| Ucp2 | 0.2022904 | 0.44376138 | 0.013 | 0.01 | 1 | 0.21842497 | 0.3337479 | 0.012 | 0.009 | 1 | 0.21842497 | 0.36365939 |
| Igfbp5 | 0.32640201 | -1.2116944 | 0.035 | 0.039 | 1 | 0.21085034 | -0.2854754 | 0.04 | 0.034 | 1 | 0.32640201 | 0.37724281 |
| Postn | 0.33614955 | -0.1442416 | 0.015 | 0.018 | 1 | 0.27307368 | 0.25664961 | 0.018 | 0.015 | 1 | 0.33614955 | 0.47157813 |
| Sdc3 | 0.46680617 | 0.28425818 | 0.011 | 0.009 | 1 | 0.27754445 | 0.31933156 | 0.011 | 0.009 | 1 | 0.46680617 | 0.47805798 |
| Bpifa1 | 0.38059963 | -0.3842521 | 0.024 | 0.028 | 1 | 0.62666575 | -0.570626 | 0.013 | 0.015 | 1 | 0.62666575 | 0.61634318 |
| Col3a1 | 0.46410612 | -0.1131255 | 0.121 | 0.113 | 1 | 0.4541536 | -0.2388674 | 0.143 | 0.133 | 1 | 0.46410612 | 0.70205171 |

Spatial seq: cluster 5

| | cKO_p_val | cKO_avg_log2FC | cKO_pct.1 | cKO_pct.2 | cKO_p_val_adj | WT_p_val | WT_avg_log2FC | WT_pct.1 | WT_pct.2 | WT_p_val_adj | max_pval | minimump_p_val |
|---|---|---|---|---|---|---|---|---|---|---|---|---|
| Dmp1 | 0 | 5.30470245 | 0.235 | 0.006 | 0 | 0 | 4.38579309 | 0.184 | 0.01 | 0 | 0 | 0 |
| Sost | 0 | 5.22894809 | 0.191 | 0.006 | 0 | 0 | 5.46732298 | 0.23 | 0.007 | 0 | 0 | 0 |
| Itm2b | 2.37E-262 | 3.24436348 | 0.309 | 0.037 | 3.55E-258 | 0 | 3.51010497 | 0.338 | 0.035 | 0 | 2.37E-262 | 0 |
| Mepe | 8.31E-282 | 4.48208657 | 0.17 | 0.008 | 1.24E-277 | 4.31E-210 | 4.47195815 | 0.116 | 0.006 | 6.46E-206 | 4.31E-210 | 1.66E-281 |
| Gsn | 1.20E-46 | 2.5076769 | 0.101 | 0.021 | 1.80E-42 | 5.85E-92 | 3.5107774 | 0.104 | 0.013 | 8.76E-88 | 1.20E-46 | 1.17E-91 |
| Col1a1 | 3.43E-57 | 0.72914298 | 0.792 | 0.552 | 5.14E-53 | 1.38E-52 | 0.71282512 | 0.814 | 0.611 | 2.07E-48 | 1.38E-52 | 6.86E-57 |
| Col1a2 | 1.35E-48 | 0.72727787 | 0.687 | 0.419 | 2.02E-44 | 1.56E-42 | 0.65664757 | 0.697 | 0.477 | 2.33E-38 | 1.56E-42 | 2.70E-48 |
| Dkk1 | 1.10E-38 | 3.83567682 | 0.029 | 0.002 | 1.64E-34 | 2.18E-33 | 3.56540065 | 0.026 | 0.002 | 3.27E-29 | 2.18E-33 | 2.20E-38 |
| Ccn4 | 5.51E-09 | 2.53082224 | 0.016 | 0.003 | 8.25E-05 | 9.56E-33 | 3.17542185 | 0.032 | 0.003 | 1.43E-28 | 5.51E-09 | 1.91E-32 |
| Obp2a | 4.60E-30 | -4.1619753 | 0.015 | 0.165 | 6.89E-26 | 6.11E-31 | -4.4646197 | 0.014 | 0.16 | 9.16E-27 | 4.60E-30 | 1.22E-30 |
| Ibsp | 1.60E-07 | 1.10160312 | 0.056 | 0.025 | 0.00239975 | 4.99E-27 | 1.76253923 | 0.08 | 0.022 | 7.48E-23 | 1.60E-07 | 9.99E-27 |
| Obp2b | 3.58E-25 | -5.8765467 | 0.003 | 0.124 | 5.36E-21 | 3.25E-21 | -5.7622411 | 0.002 | 0.1 | 4.87E-17 | 3.25E-21 | 7.16E-25 |
| Sparc | 3.11E-12 | 0.45139144 | 0.452 | 0.33 | 4.66E-08 | 2.42E-24 | 0.66755864 | 0.408 | 0.247 | 3.63E-20 | 3.11E-12 | 4.85E-24 |
| Bambi | 2.56E-22 | 2.89071818 | 0.026 | 0.003 | 3.84E-18 | 1.22E-10 | 2.7337164 | 0.013 | 0.002 | 1.82E-06 | 1.22E-10 | 5.13E-22 |
| Bmp3 | 8.87E-21 | 3.14553208 | 0.024 | 0.003 | 1.33E-16 | 1.77E-11 | 2.58615921 | 0.017 | 0.003 | 2.66E-07 | 1.77E-11 | 1.77E-20 |
| Ackr3 | 2.60E-20 | 3.62197012 | 0.016 | 0.001 | 3.89E-16 | 1.77E-17 | 3.38353004 | 0.015 | 0.002 | 2.65E-13 | 1.77E-17 | 5.20E-20 |
| Mgp | 1.64E-18 | -3.5511747 | 0.011 | 0.108 | 2.46E-14 | 4.91E-05 | -2.0314103 | 0.008 | 0.033 | 0.73577192 | 4.91E-05 | 3.28E-18 |
| Ndrg1 | 5.20E-14 | 2.59401606 | 0.023 | 0.004 | 7.79E-10 | 1.80E-15 | 2.91059416 | 0.018 | 0.002 | 2.70E-11 | 5.20E-14 | 3.61E-15 |
| Obp1a | 7.99E-09 | -4.0149697 | 0.003 | 0.045 | 0.00011959 | 2.13E-12 | -5.3123582 | 0.001 | 0.057 | 3.19E-08 | 7.99E-09 | 4.26E-12 |
| Fn1 | 3.74E-12 | -3.1202295 | 0.009 | 0.073 | 5.60E-08 | 0.0004535 | -2.3340282 | 0.004 | 0.021 | 1 | 0.0004535 | 7.48E-12 |
| Lcn3 | 7.26E-12 | -6.0237666 | 0 | 0.056 | 1.09E-07 | 1.86E-10 | -6.0201432 | 0 | 0.046 | 2.78E-06 | 1.86E-10 | 1.45E-11 |
| Ccn2 | 1.64E-11 | -4.1814591 | 0.003 | 0.059 | 2.45E-07 | 1.38E-05 | -3.2937643 | 0.002 | 0.026 | 0.20646316 | 1.38E-05 | 3.27E-11 |
| Id3 | 3.44E-05 | 1.28651504 | 0.029 | 0.012 | 0.51461455 | 1.88E-11 | 1.67926861 | 0.037 | 0.011 | 2.81E-07 | 3.44E-05 | 3.76E-11 |
| Scgb1c1 | 5.87E-11 | -4.0701526 | 0.004 | 0.058 | 8.79E-07 | 4.17E-11 | -3.0929654 | 0.012 | 0.069 | 6.24E-07 | 5.87E-11 | 8.33E-11 |
| Wif1 | 5.96E-11 | 2.35728598 | 0.021 | 0.004 | 8.92E-07 | 4.98E-11 | 2.98669714 | 0.012 | 0.002 | 7.46E-07 | 5.96E-11 | 9.96E-11 |
| Pcsk6 | 1.27E-08 | 2.4968749 | 0.014 | 0.002 | 0.00019086 | 1.03E-10 | 2.78207942 | 0.014 | 0.002 | 1.54E-06 | 1.27E-08 | 2.06E-10 |
| Satb2 | 8.97E-06 | 2.20067363 | 0.011 | 0.003 | 0.13429857 | 1.84E-10 | 2.34669327 | 0.018 | 0.003 | 2.76E-06 | 8.97E-06 | 3.68E-10 |
| Ptgis | 0.00054978 | 1.27589288 | 0.02 | 0.008 | 1 | 2.45E-10 | 2.12672759 | 0.022 | 0.005 | 3.66E-06 | 0.00054978 | 4.89E-10 |
| Ank | 1.04E-05 | 1.62032039 | 0.021 | 0.007 | 0.1550027 | 3.95E-10 | 2.09787273 | 0.021 | 0.005 | 5.91E-06 | 1.04E-05 | 7.89E-10 |
| Lcn11 | 8.89E-07 | -4.9837509 | 0 | 0.029 | 0.01331053 | 2.03E-09 | -4.3710503 | 0.002 | 0.045 | 3.05E-05 | 8.89E-07 | 4.07E-09 |
| Fat1 | 7.45E-05 | 1.75321466 | 0.015 | 0.005 | 1 | 2.26E-09 | 2.03422551 | 0.02 | 0.005 | 3.38E-05 | 7.45E-05 | 4.51E-09 |
| Col2a1 | 5.49E-09 | -2.3483234 | 0.01 | 0.059 | 8.22E-05 | 0.00646871 | -2.2429762 | 0.002 | 0.013 | 1 | 0.00646871 | 1.10E-08 |
| Sdc3 | 3.91E-08 | 1.83228623 | 0.028 | 0.008 | 0.0005859 | 1.01E-08 | 1.6527964 | 0.027 | 0.008 | 0.00015095 | 3.91E-08 | 2.02E-08 |
| Col11a2 | 1.07E-08 | -2.5552929 | 0.01 | 0.057 | 0.00016077 | 0.00496268 | -1.3060846 | 0.012 | 0.028 | 1 | 0.00496268 | 2.15E-08 |
| Gja1 | 0.00054903 | 1.28608865 | 0.02 | 0.008 | 1 | 4.42E-08 | 1.66699028 | 0.028 | 0.009 | 0.00066181 | 0.00054903 | 8.84E-08 |
| mt-Atp6 | 2.00E-07 | 1.41102772 | 0.036 | 0.014 | 0.00298955 | 0.25355711 | 0.33261809 | 0.019 | 0.014 | 1 | 0.25355711 | 3.99E-07 |
| Tpt1 | 0.20296778 | 0.48019616 | 0.016 | 0.011 | 1 | 2.33E-07 | 1.60633709 | 0.022 | 0.007 | 0.00349239 | 0.20296778 | 4.66E-07 |
| Kazald1 | 3.22E-07 | 2.44005677 | 0.013 | 0.002 | 0.0048293 | 2.77E-06 | 2.13870023 | 0.012 | 0.003 | 0.04154645 | 2.77E-06 | 6.45E-07 |
| Cadm1 | 0.00516345 | 1.48019616 | 0.013 | 0.005 | 1 | 4.62E-07 | 2.1862286 | 0.013 | 0.003 | 0.00691215 | 0.00516345 | 9.23E-07 |
| App | 4.18E-06 | -2.305679 | 0.008 | 0.039 | 0.06260914 | 0.00015516 | -1.5677212 | 0.013 | 0.038 | 1 | 0.00015516 | 8.36E-06 |
| Hbb-bs | 0.92877445 | -0.3787809 | 0.166 | 0.16 | 1 | 5.82E-06 | -1.128236 | 0.094 | 0.147 | 0.08711121 | 0.92877445 | 1.16E-05 |
| Cdkn1a | 0.00286643 | 1.32025525 | 0.011 | 0.004 | 1 | 7.19E-06 | 1.95302943 | 0.013 | 0.003 | 0.10770376 | 0.00286643 | 1.44E-05 |
| Col9a2 | 7.86E-06 | -2.1192037 | 0.008 | 0.038 | 0.11764542 | 0.02567716 | -1.6691317 | 0.004 | 0.012 | 1 | 0.02567716 | 1.57E-05 |
| Cd81 | 0.10192323 | 0.82721524 | 0.011 | 0.006 | 1 | 1.03E-05 | 1.7337164 | 0.017 | 0.005 | 0.15382197 | 0.10192323 | 2.05E-05 |
| Ngp | 0.55744493 | 0.10018788 | 0.028 | 0.024 | 1 | 1.34E-05 | -1.4249551 | 0.022 | 0.057 | 0.20079978 | 0.55744493 | 2.68E-05 |
| Fkbp10 | 0.00055143 | 1.44509236 | 0.016 | 0.006 | 1 | 1.95E-05 | 1.95302943 | 0.013 | 0.004 | 0.29130225 | 0.00055143 | 3.89E-05 |
| Bpifb6 | 0.0002062 | -1.9977094 | 0.01 | 0.034 | 1 | 2.39E-05 | -1.7997831 | 0.017 | 0.048 | 0.35725666 | 0.0002062 | 4.77E-05 |
| Hba-a2 | 0.02652172 | -0.5843586 | 0.113 | 0.137 | 1 | 2.56E-05 | -0.8505345 | 0.142 | 0.193 | 0.38278875 | 0.02652172 | 5.11E-05 |
| Htra1 | 0.34961593 | -0.4543808 | 0.008 | 0.011 | 1 | 6.51E-05 | 1.35886799 | 0.022 | 0.009 | 0.97443066 | 0.34961593 | 0.00013014 |
| Ctnnb1 | 0.48122073 | 0.3563245 | 0.011 | 0.009 | 1 | 6.52E-05 | 1.47995981 | 0.018 | 0.006 | 0.97678899 | 0.48122073 | 0.00013045 |
| Ptprs | 0.03121012 | 0.88751575 | 0.018 | 0.01 | 1 | 9.70E-05 | 1.34223363 | 0.021 | 0.008 | 1 | 0.03121012 | 0.00019391 |
| Pmp22 | 0.00032802 | 1.75321466 | 0.011 | 0.003 | 1 | 0.00010925 | 1.58615921 | 0.011 | 0.003 | 1 | 0.00032802 | 0.0002185 |
| Cd74 | 0.83443849 | 0.22269994 | 0.01 | 0.009 | 1 | 0.00013399 | 1.7337164 | 0.014 | 0.005 | 1 | 0.83443849 | 0.00026796 |
| Obp1b | 0.81592247 | -0.152488 | 0.03 | 0.028 | 1 | 0.00016838 | -2.1768687 | 0.007 | 0.029 | 1 | 0.81592247 | 0.00033672 |
| Jund | 0.00138773 | 1.23968729 | 0.024 | 0.011 | 1 | 0.00017086 | 1.15036315 | 0.025 | 0.011 | 1 | 0.00138773 | 0.00034169 |
| Col9a3 | 0.00017739 | -1.3838985 | 0.014 | 0.04 | 1 | 0.17053391 | -0.8305471 | 0.008 | 0.014 | 1 | 0.17053391 | 0.00035474 |
| Bpifb4 | 0.00020182 | -2.2617357 | 0.006 | 0.028 | 1 | 0.16812762 | -0.6651917 | 0.034 | 0.044 | 1 | 0.16812762 | 0.0004036 |
| Hbb-bt | 0.01024842 | -0.7852053 | 0.025 | 0.044 | 1 | 0.00021045 | -1.0922542 | 0.024 | 0.052 | 1 | 0.01024842 | 0.00042085 |
| Vmo1 | 0.09254036 | -0.6721896 | 0.014 | 0.023 | 1 | 0.00022731 | -1.7223484 | 0.013 | 0.037 | 1 | 0.09254036 | 0.00045456 |
| mt-Co2 | 0.00026739 | 0.72402294 | 0.059 | 0.034 | 1 | 0.0135962 | 0.34116923 | 0.051 | 0.034 | 1 | 0.0135962 | 0.0005347 |
| Nisch | 0.50934718 | 0.30632829 | 0.016 | 0.013 | 1 | 0.00027148 | 1.17112171 | 0.026 | 0.012 | 1 | 0.50934718 | 0.00054289 |
| Ppic | 0.0903199 | 0.91863827 | 0.011 | 0.006 | 1 | 0.00027756 | 1.47332671 | 0.015 | 0.006 | 1 | 0.0903199 | 0.00055505 |
| Lcn4 | 0.00027771 | -2.3657264 | 0.003 | 0.021 | 1 | 0.00099971 | -2.0285506 | 0.002 | 0.017 | 1 | 0.00099971 | 0.00055534 |
| Mrc2 | 0.00045455 | 1.5179982 | 0.018 | 0.007 | 1 | 0.00027971 | 1.389762 | 0.018 | 0.007 | 1 | 0.00045455 | 0.00055934 |
| Ddx17 | 0.21312037 | 0.7801817 | 0.018 | 0.012 | 1 | 0.00042252 | 1.19023053 | 0.021 | 0.009 | 1 | 0.21312037 | 0.00084487 |
| S100a8 | 0.00045024 | -1.1137832 | 0.025 | 0.053 | 1 | 0.00068828 | -0.8457958 | 0.037 | 0.066 | 1 | 0.00068828 | 0.00090028 |
| Cfh | 0.00047884 | 1.42564 | 0.019 | 0.007 | 1 | 0.00381808 | 1.5081567 | 0.011 | 0.004 | 1 | 0.00381808 | 0.00095746 |
| Ctdsp2 | 0.06043938 | 0.91496573 | 0.013 | 0.007 | 1 | 0.00054307 | 1.34669327 | 0.015 | 0.006 | 1 | 0.06043938 | 0.00108584 |
| Serpinh1 | 0.55573157 | 0.13016865 | 0.033 | 0.029 | 1 | 0.00060115 | 0.75659662 | 0.048 | 0.028 | 1 | 0.55573157 | 0.00120193 |
| Bpifb3 | 0.00070083 | -1.9927397 | 0.006 | 0.025 | 1 | 0.0135062 | -1.5722702 | 0.005 | 0.015 | 1 | 0.0135062 | 0.00140117 |
| Mxra8 | 0.28635617 | -0.4627227 | 0.009 | 0.013 | 1 | 0.00111433 | 1.19384179 | 0.021 | 0.01 | 1 | 0.28635617 | 0.00222743 |
| Igfbp5 | 0.00116485 | -2.1611373 | 0.018 | 0.04 | 1 | 0.00130268 | -1.59296 | 0.015 | 0.036 | 1 | 0.00130268 | 0.00232834 |
| Thbs1 | 0.00122855 | -1.5114886 | 0.01 | 0.03 | 1 | 0.82525131 | -0.1306285 | 0.009 | 0.01 | 1 | 0.82525131 | 0.0024556 |
| Tubb5 | 0.04246781 | 1.06019241 | 0.011 | 0.006 | 1 | 0.00124634 | 1.28731473 | 0.015 | 0.006 | 1 | 0.04246781 | 0.00249112 |
| Lmna | 0.45537961 | 0.40271741 | 0.011 | 0.009 | 1 | 0.00171351 | 1.37526243 | 0.015 | 0.006 | 1 | 0.45537961 | 0.00342409 |
| Bgn | 0.00226587 | 0.63210557 | 0.069 | 0.046 | 1 | 0.01523078 | 0.42598218 | 0.062 | 0.045 | 1 | 0.01523078 | 0.00452661 |
| Ahnak | 0.66205963 | 0.15306575 | 0.025 | 0.023 | 1 | 0.00230543 | 0.79442781 | 0.032 | 0.018 | 1 | 0.66205963 | 0.00460554 |
| Adamts10 | 0.31087183 | 0.60121156 | 0.01 | 0.007 | 1 | 0.00260456 | 1.2157326 | 0.014 | 0.006 | 1 | 0.31087183 | 0.00520235 |
| Pabpn1 | 0.19687375 | 0.58815541 | 0.013 | 0.008 | 1 | 0.00388411 | 1.3226478 | 0.014 | 0.006 | 1 | 0.19687375 | 0.00775314 |
| Vim | 0.11470252 | -0.4947129 | 0.025 | 0.036 | 1 | 0.00390847 | 0.66408221 | 0.046 | 0.029 | 1 | 0.11470252 | 0.00780166 |
| Col3a1 | 0.44065446 | -0.2694748 | 0.106 | 0.114 | 1 | 0.00408572 | 0.15617137 | 0.17 | 0.132 | 1 | 0.44065446 | 0.00815474 |
| Serpinf1 | 0.00434604 | 0.78854773 | 0.036 | 0.021 | 1 | 0.16247075 | 0.41267639 | 0.028 | 0.021 | 1 | 0.16247075 | 0.00867319 |
| S100a9 | 0.04607569 | -0.7398254 | 0.054 | 0.072 | 1 | 0.00464336 | -0.7685001 | 0.078 | 0.107 | 1 | 0.04607569 | 0.00926516 |
| Lox | 0.00951538 | 1.23663913 | 0.014 | 0.006 | 1 | 0.00653493 | 1.11222802 | 0.013 | 0.006 | 1 | 0.00951538 | 0.01302716 |
| Ptov1 | 0.71796082 | 0.2296527 | 0.014 | 0.012 | 1 | 0.00686275 | 1.14591115 | 0.014 | 0.006 | 1 | 0.71796082 | 0.0136784 |
| Lamp2 | 0.3842703 | 0.55390585 | 0.01 | 0.007 | 1 | 0.00722728 | 1.36090713 | 0.012 | 0.005 | 1 | 0.3842703 | 0.01440232 |
| Akr1a1 | 0.23964648 | 0.48807259 | 0.021 | 0.016 | 1 | 0.00772293 | 0.9747245 | 0.027 | 0.015 | 1 | 0.23964648 | 0.01538622 |
| Sgms2 | 0.00967085 | 0.95351331 | 0.021 | 0.011 | 1 | 0.18694594 | 0.74485696 | 0.011 | 0.007 | 1 | 0.18694594 | 0.01924818 |
| Mup4 | 0.04543348 | -1.4743211 | 0.005 | 0.013 | 1 | 0.01000344 | -1.3073109 | 0.005 | 0.016 | 1 | 0.04543348 | 0.01990681 |
| Cnn2 | 0.13934829 | 0.83894453 | 0.014 | 0.009 | 1 | 0.0104977 | 0.9036414 | 0.02 | 0.011 | 1 | 0.13934829 | 0.02088521 |
| Fkbp9 | 0.27691877 | 0.60121156 | 0.01 | 0.007 | 1 | 0.01081471 | 1.20708721 | 0.012 | 0.005 | 1 | 0.27691877 | 0.02151246 |
| Tmem59 | 0.02264177 | 1.01624906 | 0.014 | 0.007 | 1 | 0.01191307 | 1.12540341 | 0.012 | 0.005 | 1 | 0.02264177 | 0.02368421 |
| Maf | 0.01424873 | -1.9927397 | 0.001 | 0.01 | 1 | 0.32531738 | 0.44709517 | 0.014 | 0.011 | 1 | 0.32531738 | 0.02829443 |
| Dmbt1 | 0.02531685 | -0.8799151 | 0.006 | 0.016 | 1 | 0.01573782 | -2.057697 | 0.002 | 0.011 | 1 | 0.02531685 | 0.03122796 |
| Wfdc18 | 0.0167359 | -1.2542799 | 0.008 | 0.019 | 1 | 0.8312103 | -0.3258113 | 0.022 | 0.021 | 1 | 0.8312103 | 0.03319171 |
| Tsc22d1 | 0.03380482 | -1.2467853 | 0.005 | 0.014 | 1 | 0.01945713 | 1.01911862 | 0.015 | 0.008 | 1 | 0.03380482 | 0.03853568 |

| | | | | | | | | | | | | |
|---|---|---|---|---|---|---|---|---|---|---|---|---|
| Eef1a1 | 0.17907203 | 0.5502746 | 0.018 | 0.012 | 1 | 0.02095494 | 0.88395903 | 0.015 | 0.008 | 1 | 0.17907203 | 0.04147078 |
| mt-Co3 | 0.07009843 | 0.52148437 | 0.025 | 0.017 | 1 | 0.02148659 | 0.62268509 | 0.026 | 0.016 | 1 | 0.07009843 | 0.0425115 |
| Apoe | 0.59993863 | -0.219478 | 0.024 | 0.027 | 1 | 0.0227417 | -0.893172 | 0.022 | 0.037 | 1 | 0.59993863 | 0.04496621 |
| Car2 | 0.19903021 | -0.4238172 | 0.018 | 0.025 | 1 | 0.02688563 | -0.8049834 | 0.014 | 0.027 | 1 | 0.19903021 | 0.05304842 |
| Maged2 | 0.02789219 | -1.4627227 | 0.004 | 0.012 | 1 | 0.05101119 | 0.94230302 | 0.012 | 0.006 | 1 | 0.05101119 | 0.0550064 |
| Acp5 | 0.37277422 | -0.4500689 | 0.023 | 0.028 | 1 | 0.02969386 | -1.1417612 | 0.011 | 0.022 | 1 | 0.37277422 | 0.05850599 |
| Pcolce | 0.67478511 | -0.1139497 | 0.01 | 0.012 | 1 | 0.03192469 | 0.7337164 | 0.018 | 0.01 | 1 | 0.67478511 | 0.06283019 |
| Timp3 | 0.38440064 | -0.2954592 | 0.019 | 0.024 | 1 | 0.03368485 | 1.05045182 | 0.015 | 0.008 | 1 | 0.38440064 | 0.06623503 |
| Gpx6 | 0.03849611 | -1.1429495 | 0.01 | 0.02 | 1 | 0.81193159 | -0.1387336 | 0.028 | 0.027 | 1 | 0.81193159 | 0.07551028 |
| Tln1 | 0.043328 | 0.80358818 | 0.02 | 0.012 | 1 | 0.0812141 | 0.77704383 | 0.014 | 0.008 | 1 | 0.0812141 | 0.08477869 |
| Col6a1 | 0.09605897 | -1.0281451 | 0.004 | 0.01 | 1 | 0.0503977 | 0.83920953 | 0.013 | 0.007 | 1 | 0.09605897 | 0.09825546 |
| Tnc | 0.23398802 | 0.60121156 | 0.014 | 0.01 | 1 | 0.05391689 | 0.74765559 | 0.018 | 0.011 | 1 | 0.23398802 | 0.10492676 |
| Arglu1 | 0.5476413 | 0.22483568 | 0.015 | 0.013 | 1 | 0.06095645 | 0.58498102 | 0.019 | 0.012 | 1 | 0.5476413 | 0.11819721 |
| Mup5 | 0.1507674 | -0.4506577 | 0.011 | 0.018 | 1 | 0.06525127 | -1.0383317 | 0.008 | 0.016 | 1 | 0.1507674 | 0.12624481 |
| Ighm | 0.69671315 | -0.2358142 | 0.013 | 0.014 | 1 | 0.06628314 | 0.87056727 | 0.018 | 0.011 | 1 | 0.69671315 | 0.12817283 |
| Cst3 | 0.66159414 | -0.1057415 | 0.019 | 0.021 | 1 | 0.0685825 | 0.74765559 | 0.017 | 0.01 | 1 | 0.66159414 | 0.13246143 |
| Sdc4 | 0.07388637 | -1.1375563 | 0.004 | 0.01 | 1 | 0.25257627 | 0.37002578 | 0.012 | 0.008 | 1 | 0.25257627 | 0.14231355 |
| Camp | 0.2407312 | -0.5152742 | 0.013 | 0.018 | 1 | 0.07602029 | -0.6014678 | 0.022 | 0.033 | 1 | 0.2407312 | 0.14626149 |
| Mpo | 0.15944442 | -0.7492857 | 0.005 | 0.01 | 1 | 0.08170736 | -0.9443555 | 0.005 | 0.011 | 1 | 0.15944442 | 0.15673862 |
| Mbd2 | 0.08609313 | 0.7215058 | 0.018 | 0.011 | 1 | 0.8208846 | 0.14137437 | 0.011 | 0.01 | 1 | 0.8208846 | 0.16477424 |
| Sfrp2 | 0.08654899 | -1.0281451 | 0.004 | 0.01 | 1 | 0.58447169 | 0.23537625 | 0.012 | 0.01 | 1 | 0.58447169 | 0.16560726 |
| mt-Nd3 | 0.08787701 | -0.706217 | 0.009 | 0.017 | 1 | 0.18711903 | -1.0205838 | 0.007 | 0.012 | 1 | 0.18711903 | 0.16803165 |
| Tagln2 | 0.21312529 | 0.79385664 | 0.01 | 0.006 | 1 | 0.08959424 | 0.87344116 | 0.011 | 0.006 | 1 | 0.21312529 | 0.17116135 |
| mt-Co1 | 0.24293918 | 0.35111621 | 0.024 | 0.018 | 1 | 0.09692397 | 0.54273693 | 0.024 | 0.016 | 1 | 0.24293918 | 0.18445369 |
| mt-Cytb | 0.09750701 | 0.49419331 | 0.028 | 0.019 | 1 | 0.77880867 | -0.3053834 | 0.015 | 0.017 | 1 | 0.77880867 | 0.1855064 |
| Maged1 | 0.3950579 | 0.3995777 | 0.014 | 0.011 | 1 | 0.09947117 | 0.63418072 | 0.015 | 0.01 | 1 | 0.3950579 | 0.18904782 |
| mt-Nd6 | 0.11220807 | -1.1853848 | 0.005 | 0.011 | 1 | 0.09962134 | 0.71899489 | 0.013 | 0.008 | 1 | 0.11220807 | 0.18931826 |
| Tmed9 | 0.44293248 | 0.43128656 | 0.011 | 0.009 | 1 | 0.10008795 | 0.80612489 | 0.011 | 0.006 | 1 | 0.44293248 | 0.1901583 |
| Col5a1 | 0.10093018 | 0.75321466 | 0.016 | 0.01 | 1 | 0.20316862 | 0.70080178 | 0.011 | 0.007 | 1 | 0.20316862 | 0.19167346 |
| mt-Nd5 | 0.84543032 | 0.23126195 | 0.013 | 0.012 | 1 | 0.10306929 | -1.1306285 | 0.006 | 0.012 | 1 | 0.84543032 | 0.1955153 |
| Cd24a | 0.10841528 | -0.8064811 | 0.009 | 0.016 | 1 | 0.14767913 | -0.7528424 | 0.007 | 0.013 | 1 | 0.14767913 | 0.2050767 |
| Slc4a1 | 0.22346492 | -0.5280715 | 0.006 | 0.011 | 1 | 0.12005992 | -0.7068562 | 0.007 | 0.013 | 1 | 0.22346492 | 0.22570546 |
| Col12a1 | 0.49525089 | -0.1363088 | 0.014 | 0.017 | 1 | 0.12091226 | 0.6588624 | 0.014 | 0.009 | 1 | 0.49525089 | 0.22720474 |
| Tpm2 | 0.1384418 | 0.9321848 | 0.014 | 0.009 | 1 | 0.3259358 | 0.22470275 | 0.017 | 0.013 | 1 | 0.3259358 | 0.25771746 |
| Ucp2 | 0.14112724 | 0.67229466 | 0.015 | 0.01 | 1 | 0.69181417 | -0.2168956 | 0.008 | 0.01 | 1 | 0.69181417 | 0.26233759 |
| Col11a1 | 0.6834334 | 0.16209993 | 0.015 | 0.013 | 1 | 0.14209729 | 0.559687 | 0.013 | 0.008 | 1 | 0.6834334 | 0.26400294 |
| Tmsb10 | 0.20378924 | 0.73991783 | 0.01 | 0.006 | 1 | 0.15300487 | 0.58980699 | 0.012 | 0.007 | 1 | 0.20378924 | 0.28259926 |
| Postn | 0.15484772 | -0.5638964 | 0.011 | 0.018 | 1 | 0.97908323 | 0.10264197 | 0.015 | 0.015 | 1 | 0.97908323 | 0.28571762 |
| Igfbp7 | 0.27059064 | -0.4379268 | 0.011 | 0.016 | 1 | 0.1684321 | 0.38825527 | 0.031 | 0.023 | 1 | 0.27059064 | 0.30849483 |
| Bpgm | 0.51809922 | 0.44201297 | 0.01 | 0.008 | 1 | 0.17992761 | -0.6319331 | 0.006 | 0.011 | 1 | 0.51809922 | 0.32748128 |
| Rsrp1 | 0.19479629 | -0.4978732 | 0.018 | 0.025 | 1 | 0.1913079 | 0.24151104 | 0.024 | 0.017 | 1 | 0.19479629 | 0.34601708 |
| Selenom | 0.85005582 | 0.13306273 | 0.01 | 0.009 | 1 | 0.19593378 | -0.6462173 | 0.006 | 0.01 | 1 | 0.85005582 | 0.35347752 |
| Tpm1 | 0.21375005 | -0.2911795 | 0.016 | 0.023 | 1 | 0.32758831 | 0.29314381 | 0.029 | 0.024 | 1 | 0.32758831 | 0.38181102 |
| Hist1h2ap | 0.50271046 | -0.1943179 | 0.008 | 0.01 | 1 | 0.21816704 | -0.5733384 | 0.006 | 0.01 | 1 | 0.50271046 | 0.38873722 |
| Ptms | 0.22180576 | 0.31669743 | 0.03 | 0.023 | 1 | 0.24223434 | 0.16028015 | 0.033 | 0.026 | 1 | 0.24223434 | 0.39441373 |
| Dcn | 0.48268376 | -0.2207901 | 0.034 | 0.039 | 1 | 0.23272283 | -0.4658481 | 0.015 | 0.021 | 1 | 0.48268376 | 0.41128574 |
| Igfbp4 | 0.23964946 | -0.5479549 | 0.008 | 0.012 | 1 | 0.7588027 | 0.19469018 | 0.019 | 0.017 | 1 | 0.7588027 | 0.42186706 |
| Kctd12 | 0.26072757 | -0.4431826 | 0.011 | 0.016 | 1 | 0.46059848 | 0.30900098 | 0.021 | 0.018 | 1 | 0.46059848 | 0.45347627 |
| Col6a3 | 0.26678971 | -0.4910925 | 0.008 | 0.012 | 1 | 0.60861969 | -0.1731742 | 0.008 | 0.01 | 1 | 0.60861969 | 0.46240267 |
| Igf2 | 0.41527148 | -0.3413029 | 0.009 | 0.012 | 1 | 0.31172931 | 0.47995981 | 0.013 | 0.009 | 1 | 0.41527148 | 0.52628345 |
| Hist1h1e | 0.62975335 | -0.103258 | 0.013 | 0.015 | 1 | 0.34327113 | -0.3731988 | 0.012 | 0.016 | 1 | 0.62975335 | 0.5687072 |
| Timp2 | 0.34598208 | 0.24095535 | 0.031 | 0.026 | 1 | 0.41502345 | 0.20952522 | 0.026 | 0.022 | 1 | 0.41502345 | 0.57226056 |
| Dbp | 0.39522823 | -0.356703 | 0.011 | 0.015 | 1 | 0.3688379 | 0.3048731 | 0.013 | 0.01 | 1 | 0.39522823 | 0.6016344 |
| Serinc3 | 0.53863861 | 0.31580934 | 0.014 | 0.011 | 1 | 0.40632762 | 0.37526243 | 0.013 | 0.01 | 1 | 0.53863861 | 0.6475531 |
| Arf5 | 0.42700842 | 0.3762452 | 0.014 | 0.011 | 1 | 0.77159635 | 0.12897128 | 0.014 | 0.013 | 1 | 0.77159635 | 0.67168065 |
| Oaz1 | 0.53612848 | 0.43817295 | 0.014 | 0.011 | 1 | 0.43654535 | -0.4088227 | 0.009 | 0.012 | 1 | 0.53612848 | 0.68251886 |
| Ctsk | 0.63992889 | -0.2542799 | 0.019 | 0.021 | 1 | 0.6651875 | -0.2253533 | 0.012 | 0.014 | 1 | 0.6651875 | 0.87034879 |
| mt-Nd4l | 0.70239364 | -0.1272037 | 0.011 | 0.013 | 1 | 0.8836825 | 0.10025538 | 0.011 | 0.01 | 1 | 0.8836825 | 0.91143045 |

Spatial seq: cluster 6

| | cKO_p_val | cKO_avg_log2FC | cKO_pct.1 | cKO_pct.2 | cKO_p_val_adj | WT_p_val | WT_avg_log2FC | WT_pct.1 | WT_pct.2 | WT_p_val_adj | max_pval | minimump_p_val |
|---|---|---|---|---|---|---|---|---|---|---|---|---|
| Obp2a | 0 | 3.33015812 | 1 | 0.117 | 0 | 0 | 3.50736091 | 1 | 0.107 | 0 | 0 | 0 |
| Hba-a2 | 3.73E-20 | -2.2206054 | 0.026 | 0.142 | 5.59E-16 | 4.44E-48 | -3.7543366 | 0.012 | 0.201 | 6.65E-44 | 3.73E-20 | 8.88E-48 |
| Col1a1 | 1.72E-20 | -0.8518848 | 0.447 | 0.569 | 2.58E-16 | 6.26E-41 | -1.0731368 | 0.465 | 0.629 | 9.38E-37 | 1.72E-20 | 1.25E-40 |
| Sparc | 2.46E-15 | -0.5507873 | 0.189 | 0.343 | 3.68E-11 | 4.30E-37 | -1.5039426 | 0.077 | 0.264 | 6.43E-33 | 2.46E-15 | 8.59E-37 |
| Col1a2 | 4.37E-33 | -1.1111908 | 0.223 | 0.442 | 6.54E-29 | 2.75E-36 | -1.0550079 | 0.309 | 0.496 | 4.11E-32 | 4.37E-33 | 5.50E-36 |
| Hbb-bs | 2.08E-24 | -2.4381417 | 0.03 | 0.167 | 3.11E-20 | 5.89E-34 | -3.4703653 | 0.011 | 0.152 | 8.82E-30 | 2.08E-24 | 1.18E-33 |
| S100a9 | 4.34E-14 | -4.26248 | 0.004 | 0.074 | 6.49E-10 | 7.00E-27 | -4.5589019 | 0.003 | 0.111 | 1.05E-22 | 4.34E-14 | 1.40E-26 |
| Obp2b | 5.16E-26 | -7.4547207 | 0 | 0.124 | 7.73E-22 | 9.78E-26 | -7.5795299 | 0 | 0.101 | 1.46E-21 | 9.78E-26 | 1.03E-25 |
| Mgp | 3.13E-20 | -4.2829745 | 0.006 | 0.108 | 4.68E-16 | 3.67E-09 | -5.277131 | 0 | 0.034 | 5.50E-05 | 3.67E-09 | 6.25E-20 |
| Scgb1c1 | 2.60E-12 | -6.0657362 | 0 | 0.058 | 3.90E-08 | 5.61E-18 | -6.7909968 | 0 | 0.071 | 8.40E-14 | 2.60E-12 | 1.12E-17 |
| Col3a1 | 1.87E-14 | -1.8699539 | 0.029 | 0.118 | 2.80E-10 | 1.05E-16 | -1.2927129 | 0.045 | 0.139 | 1.57E-12 | 1.87E-14 | 2.09E-16 |
| Spp1 | 1.43E-13 | -3.7469823 | 0.004 | 0.072 | 2.14E-09 | 3.50E-16 | -4.3258048 | 0.002 | 0.067 | 5.23E-12 | 1.43E-13 | 6.99E-16 |
| S100a8 | 8.17E-09 | -2.5864399 | 0.008 | 0.054 | 0.00012237 | 7.19E-16 | -3.8741663 | 0.003 | 0.068 | 1.08E-11 | 8.17E-09 | 1.44E-15 |
| Obp1a | 1.02E-09 | -5.5952728 | 0 | 0.045 | 1.53E-05 | 2.25E-14 | -5.543865 | 0.001 | 0.058 | 3.36E-10 | 1.02E-09 | 4.49E-14 |
| Ngp | 1.51E-05 | -3.4839479 | 0.001 | 0.025 | 0.22604293 | 4.40E-14 | -3.5488948 | 0.002 | 0.059 | 6.59E-10 | 1.51E-05 | 8.81E-14 |
| Hbb-bt | 2.46E-08 | -2.8791915 | 0.004 | 0.045 | 0.00036858 | 6.96E-13 | -4.2250127 | 0.002 | 0.054 | 1.04E-08 | 2.46E-08 | 1.39E-12 |
| Bpifb6 | 1.06E-07 | -5.1758256 | 0 | 0.034 | 0.0015843 | 3.85E-12 | -4.9546976 | 0.001 | 0.049 | 5.76E-08 | 1.06E-07 | 7.69E-12 |
| Lcn3 | 8.24E-12 | -6.0161689 | 0 | 0.056 | 1.23E-07 | 5.02E-12 | -6.25165 | 0 | 0.046 | 7.51E-08 | 8.24E-12 | 1.00E-11 |
| Fn1 | 5.06E-12 | -2.7886515 | 0.009 | 0.073 | 7.58E-08 | 1.19E-05 | -3.5721378 | 0.001 | 0.021 | 0.17750917 | 1.19E-05 | 1.01E-11 |
| Itm2b | 8.63E-10 | -3.618566 | 0.004 | 0.052 | 1.29E-05 | 1.05E-11 | -3.6111319 | 0.003 | 0.051 | 1.58E-07 | 8.63E-10 | 2.11E-11 |
| Bpifb4 | 4.36E-06 | -3.848982 | 0.001 | 0.028 | 0.0652951 | 1.86E-11 | -4.9356225 | 0.001 | 0.046 | 2.78E-07 | 4.36E-06 | 3.71E-11 |
| Lcn11 | 2.82E-06 | -3.9738973 | 0.001 | 0.029 | 0.04222676 | 2.01E-11 | -5.1885943 | 0.001 | 0.046 | 3.02E-07 | 2.82E-06 | 4.03E-11 |
| Obp1b | 0.06528507 | 0.703147 | 0.039 | 0.028 | 1 | 2.41E-10 | 1.44880989 | 0.06 | 0.026 | 3.61E-06 | 0.06528507 | 4.82E-10 |
| Gpx3 | 5.73E-06 | -1.6048371 | 0.013 | 0.047 | 0.08574003 | 3.30E-10 | -2.764577 | 0.007 | 0.051 | 4.94E-06 | 5.73E-06 | 6.60E-10 |
| Col2a1 | 8.32E-10 | -2.9978868 | 0.008 | 0.059 | 1.25E-05 | 0.00083756 | -3.0641372 | 0.001 | 0.013 | 1 | 0.00083756 | 1.66E-09 |
| App | 6.13E-05 | -1.637737 | 0.011 | 0.039 | 0.91799283 | 8.55E-10 | -3.8160145 | 0.001 | 0.039 | 1.28E-05 | 6.13E-05 | 1.71E-09 |
| Vmo1 | 1.26E-05 | -4.4057692 | 0 | 0.023 | 0.18937777 | 1.41E-09 | -4.5555739 | 0.001 | 0.038 | 2.10E-05 | 1.26E-05 | 2.81E-09 |
| Col11a2 | 5.28E-09 | -2.3943432 | 0.009 | 0.057 | 7.90E-05 | 2.27E-07 | -3.4315567 | 0.001 | 0.028 | 0.00339318 | 2.27E-07 | 1.06E-08 |
| Camp | 0.00029327 | -3.007372 | 0.001 | 0.019 | 1 | 7.76E-09 | -3.6068509 | 0.001 | 0.035 | 0.00011617 | 0.00029327 | 1.55E-08 |
| Ccn2 | 2.02E-08 | -2.2888712 | 0.011 | 0.058 | 0.00030284 | 1.27E-05 | -2.7837616 | 0.004 | 0.026 | 0.19039267 | 1.27E-05 | 4.04E-08 |
| Mup4 | 0.20610803 | 0.76081234 | 0.018 | 0.012 | 1 | 2.92E-08 | 1.52687164 | 0.036 | 0.014 | 0.0004372 | 0.20610803 | 5.84E-08 |
| Car2 | 0.00074307 | -1.3477121 | 0.006 | 0.025 | 1 | 1.21E-06 | -2.763135 | 0.002 | 0.027 | 0.01818975 | 0.00074307 | 2.43E-06 |
| Acan | 0.00053877 | 1.1303858 | 0.048 | 0.027 | 1 | 1.80E-06 | 1.67908735 | 0.018 | 0.006 | 0.02697567 | 0.00053877 | 3.60E-06 |
| Mup5 | 1.17E-05 | 1.59913133 | 0.038 | 0.017 | 0.17556991 | 1.91E-06 | 1.43127368 | 0.034 | 0.015 | 0.02857053 | 1.17E-05 | 3.82E-06 |
| Gpx6 | 0.00011085 | -3.3301822 | 0.001 | 0.021 | 1 | 1.96E-06 | -2.7398652 | 0.003 | 0.028 | 0.02936322 | 0.00011085 | 3.92E-06 |
| Tmsb4x | 0.00116076 | -0.9132761 | 0.015 | 0.037 | 1 | 2.35E-06 | -1.4680069 | 0.011 | 0.041 | 0.03521487 | 0.00116076 | 4.70E-06 |
| Apoe | 4.29E-05 | -1.9911036 | 0.004 | 0.028 | 0.64200353 | 2.36E-06 | -1.7586891 | 0.009 | 0.038 | 0.03530638 | 4.29E-05 | 4.72E-06 |
| Igfbp5 | 0.000692 | -1.8873122 | 0.016 | 0.04 | 1 | 1.25E-05 | -1.7233501 | 0.01 | 0.036 | 0.1874272 | 0.000692 | 2.50E-05 |
| Dmp1 | 0.00013949 | -3.9806547 | 0 | 0.018 | 1 | 1.84E-05 | -4.277131 | 0 | 0.018 | 0.2757045 | 0.00013949 | 3.68E-05 |
| Acp5 | 0.00103931 | -1.0063535 | 0.009 | 0.028 | 1 | 5.14E-05 | -2.3883119 | 0.003 | 0.022 | 0.77024594 | 0.00103931 | 0.00010287 |
| Bgn | 0.00195629 | -0.7852992 | 0.024 | 0.048 | 1 | 0.00011328 | -0.7706769 | 0.02 | 0.047 | 1 | 0.00195629 | 0.00022655 |
| Serpinh1 | 0.05272878 | -0.2420473 | 0.018 | 0.03 | 1 | 0.00013235 | -1.5570876 | 0.009 | 0.03 | 1 | 0.05272878 | 0.00026469 |
| Slc4a1 | 0.02310485 | -1.5449961 | 0.003 | 0.011 | 1 | 0.00021444 | -3.7803443 | 0 | 0.014 | 1 | 0.02310485 | 0.00042884 |
| Ibsp | 0.00023877 | -1.5671145 | 0.006 | 0.028 | 1 | 0.00068365 | -1.3180587 | 0.008 | 0.025 | 1 | 0.00068365 | 0.00047748 |
| Mmp13 | 0.01545428 | -1.1380409 | 0.009 | 0.021 | 1 | 0.00026409 | -1.5917666 | 0.002 | 0.017 | 1 | 0.01545428 | 0.00052811 |
| Serpinf1 | 0.00981231 | -1.2466822 | 0.009 | 0.023 | 1 | 0.00027609 | -1.193099 | 0.005 | 0.022 | 1 | 0.00981231 | 0.00055211 |
| Kctd12 | 0.00455745 | -1.7886515 | 0.004 | 0.017 | 1 | 0.00030023 | -1.7162139 | 0.003 | 0.019 | 1 | 0.00455745 | 0.00060037 |
| Ckm | 0.000532 | 1.87446912 | 0.01 | 0.003 | 1 | 0.00325365 | 1.85066398 | 0.01 | 0.004 | 1 | 0.00325365 | 0.00106371 |
| Sost | 0.00141144 | -2.741688 | 0.001 | 0.015 | 1 | 0.0005802 | -2.400977 | 0.003 | 0.017 | 1 | 0.00141144 | 0.00116006 |
| Ighm | 0.0114425 | -1.7256875 | 0.004 | 0.015 | 1 | 0.0005897 | -3.5589019 | 0 | 0.012 | 1 | 0.0114425 | 0.00117906 |
| Thbs1 | 0.00061186 | -1.5038909 | 0.009 | 0.03 | 1 | 0.10148758 | -0.3621353 | 0.005 | 0.01 | 1 | 0.10148758 | 0.00122335 |
| mt-Nd3 | 0.24231266 | 0.26727286 | 0.021 | 0.016 | 1 | 0.00061298 | 1.25363324 | 0.023 | 0.011 | 1 | 0.24231266 | 0.00122559 |
| Dcn | 0.00361215 | -0.7160014 | 0.019 | 0.039 | 1 | 0.00077551 | -1.3519395 | 0.006 | 0.022 | 1 | 0.00361215 | 0.00155041 |
| Dmbt1 | 0.20632455 | 0.79232331 | 0.021 | 0.016 | 1 | 0.00089209 | 1.23798238 | 0.021 | 0.01 | 1 | 0.20632455 | 0.00178339 |
| Mepe | 0.10447434 | -0.848982 | 0.009 | 0.016 | 1 | 0.00095126 | -3.6302407 | 0 | 0.011 | 1 | 0.10447434 | 0.00190161 |
| Vim | 0.00187413 | -0.8705431 | 0.015 | 0.036 | 1 | 0.0009678 | -0.7685917 | 0.012 | 0.031 | 1 | 0.00187413 | 0.00193467 |
| Hist1h1e | 0.00137502 | -2.6084215 | 0.001 | 0.015 | 1 | 0.00099752 | -1.7700433 | 0.003 | 0.016 | 1 | 0.00137502 | 0.00199405 |
| Ctsk | 0.01381112 | -1.1026857 | 0.009 | 0.022 | 1 | 0.00140033 | -1.6251697 | 0.002 | 0.014 | 1 | 0.01381112 | 0.0027987 |
| mt-Nd1 | 0.43940017 | 0.6221588 | 0.016 | 0.013 | 1 | 0.00142368 | 1.34850017 | 0.023 | 0.012 | 1 | 0.43940017 | 0.00284534 |
| Gpx1 | 0.03568927 | -1.4701421 | 0.003 | 0.01 | 1 | 0.00144735 | -3.3823132 | 0 | 0.01 | 1 | 0.03568927 | 0.00289261 |
| Ddx17 | 0.04758581 | -0.8766176 | 0.005 | 0.013 | 1 | 0.00144738 | -3.4267073 | 0 | 0.01 | 1 | 0.04758581 | 0.00289266 |
| Acta1 | 0.00628737 | 1.32691581 | 0.016 | 0.008 | 1 | 0.00154972 | 1.01102923 | 0.027 | 0.015 | 1 | 0.00628737 | 0.00309704 |
| Col27a1 | 0.00171611 | 0.91816866 | 0.063 | 0.041 | 1 | 0.00356528 | 1.69245457 | 0.016 | 0.008 | 1 | 0.00356528 | 0.00342928 |
| Mmp2 | 0.86280318 | 0.15595028 | 0.016 | 0.017 | 1 | 0.0017517 | -1.2236154 | 0.003 | 0.015 | 1 | 0.86280318 | 0.00350034 |
| Timp2 | 0.00178238 | -1.3360492 | 0.009 | 0.027 | 1 | 0.01789324 | -0.7680729 | 0.011 | 0.022 | 1 | 0.01789324 | 0.00356158 |
| Mpo | 0.0123358 | -2.0888534 | 0.001 | 0.01 | 1 | 0.00234554 | -1.9266337 | 0.001 | 0.011 | 1 | 0.0123358 | 0.00468559 |
| Mt1 | 0.02707865 | -0.8308139 | 0.004 | 0.013 | 1 | 0.00285562 | -1.3693741 | 0.004 | 0.016 | 1 | 0.02707865 | 0.00570309 |
| Id3 | 0.04718401 | -1.1004814 | 0.005 | 0.013 | 1 | 0.00311911 | -2.1490261 | 0.002 | 0.013 | 1 | 0.04718401 | 0.00622849 |
| Lcn4 | 0.00435554 | 0.9702657 | 0.034 | 0.019 | 1 | 0.04649697 | 0.55704546 | 0.024 | 0.016 | 1 | 0.04649697 | 0.0086921 |
| Igfbp4 | 0.06754053 | -0.7697024 | 0.005 | 0.012 | 1 | 0.00505132 | -0.7767109 | 0.006 | 0.018 | 1 | 0.06754053 | 0.01007713 |
| Postn | 0.00540666 | -1.5802246 | 0.005 | 0.018 | 1 | 0.01596459 | -1.0688137 | 0.006 | 0.016 | 1 | 0.01596459 | 0.01078409 |
| mt-Cytb | 0.69305581 | 0.15990829 | 0.018 | 0.02 | 1 | 0.00541473 | 1.12140946 | 0.027 | 0.016 | 1 | 0.69305581 | 0.01080014 |
| Cd24a | 0.19573027 | -0.4667234 | 0.01 | 0.016 | 1 | 0.00629787 | -0.9843492 | 0.003 | 0.013 | 1 | 0.19573027 | 0.01255608 |
| Bpifa1 | 0.00780314 | 0.81327976 | 0.043 | 0.027 | 1 | 0.00727615 | 1.03645702 | 0.024 | 0.014 | 1 | 0.00780314 | 0.01449937 |
| Ahnak | 0.01410774 | -0.8095034 | 0.01 | 0.023 | 1 | 0.00754717 | -1.1509876 | 0.007 | 0.019 | 1 | 0.01410774 | 0.01503738 |
| Mxra8 | 0.16954957 | -0.6542252 | 0.008 | 0.013 | 1 | 0.00862037 | -1.4768308 | 0.002 | 0.011 | 1 | 0.16954957 | 0.01716643 |
| Lrp1 | 0.04314386 | -0.6032412 | 0.008 | 0.017 | 1 | 0.01054185 | -1.4866499 | 0.004 | 0.014 | 1 | 0.04314386 | 0.02097258 |
| Akr1a1 | 0.01079226 | -1.5022221 | 0.005 | 0.017 | 1 | 0.01126037 | -0.682215 | 0.006 | 0.016 | 1 | 0.01126037 | 0.02146804 |
| Tnc | 0.01268488 | -2.0636161 | 0.001 | 0.01 | 1 | 0.01411828 | -1.565535 | 0.003 | 0.011 | 1 | 0.01411828 | 0.02520885 |
| mt-Atp8 | 0.52455187 | -0.155477 | 0.018 | 0.021 | 1 | 0.01283688 | 0.99399191 | 0.031 | 0.02 | 1 | 0.52455187 | 0.02550898 |
| Htra1 | 0.20447426 | -0.4467831 | 0.006 | 0.011 | 1 | 0.01332917 | -1.8197185 | 0.002 | 0.01 | 1 | 0.20447426 | 0.02648067 |
| Nisch | 0.37700788 | -0.1964832 | 0.01 | 0.014 | 1 | 0.01503984 | -1.001223 | 0.004 | 0.013 | 1 | 0.37700788 | 0.02985347 |
| Fth1 | 0.11730277 | -0.2311503 | 0.013 | 0.021 | 1 | 0.01903603 | -0.7680729 | 0.01 | 0.021 | 1 | 0.11730277 | 0.03770969 |
| Pcolce | 0.07776682 | -0.9833488 | 0.005 | 0.012 | 1 | 0.01937374 | -1.5787104 | 0.003 | 0.011 | 1 | 0.07776682 | 0.03837213 |
| mt-Nd2 | 0.69374343 | 0.23535085 | 0.009 | 0.01 | 1 | 0.01964927 | 1.00625209 | 0.013 | 0.007 | 1 | 0.69374343 | 0.03891245 |
| Serinc3 | 0.0370677 | -0.7203144 | 0.004 | 0.012 | 1 | 0.02213619 | -0.8988887 | 0.003 | 0.011 | 1 | 0.0370677 | 0.04378236 |
| Tpm2 | 0.02326985 | 1.13861693 | 0.016 | 0.009 | 1 | 0.48041788 | 0.61768681 | 0.015 | 0.013 | 1 | 0.48041788 | 0.04599822 |
| Ptms | 0.8696466 | 0.19691145 | 0.023 | 0.024 | 1 | 0.02346722 | -0.3570464 | 0.015 | 0.027 | 1 | 0.8696466 | 0.04638374 |
| Hist1h2ap | 0.03319835 | -1.0205474 | 0.003 | 0.01 | 1 | 0.02415738 | -0.8048452 | 0.003 | 0.01 | 1 | 0.03319835 | 0.04773118 |
| Col6a3 | 0.67478075 | -0.106352 | 0.01 | 0.012 | 1 | 0.02463985 | -1.1047792 | 0.003 | 0.01 | 1 | 0.67478075 | 0.04867258 |

| | | | | | | | | | | | | |
|---|---|---|---|---|---|---|---|---|---|---|---|---|
| Sfrp2 | 0.34693873 | 0.35481998 | 0.006 | 0.01 | 1 | 0.02502422 | -0.6764963 | 0.003 | 0.01 | 1 | 0.34693873 | 0.04942223 |
| Tpm1 | 0.02714757 | -0.9833488 | 0.011 | 0.023 | 1 | 0.0347781 | -0.3037034 | 0.014 | 0.025 | 1 | 0.0347781 | 0.05355815 |
| Wfdc18 | 0.0276561 | 0.89431393 | 0.029 | 0.018 | 1 | 0.17141095 | 0.42128959 | 0.027 | 0.021 | 1 | 0.17141095 | 0.05454733 |
| Gsn | 0.02785947 | 0.79604082 | 0.036 | 0.024 | 1 | 0.15878352 | 0.55321927 | 0.022 | 0.016 | 1 | 0.15878352 | 0.05494279 |
| Igf2 | 0.25719307 | -0.3337053 | 0.008 | 0.012 | 1 | 0.0290441 | -1.3972635 | 0.003 | 0.01 | 1 | 0.25719307 | 0.05724464 |
| mt-Nd6 | 0.04627038 | 1.33885542 | 0.018 | 0.01 | 1 | 0.13525905 | 0.85811978 | 0.012 | 0.008 | 1 | 0.13525905 | 0.09039981 |
| Selenom | 0.09443346 | -0.4701421 | 0.004 | 0.01 | 1 | 0.04668993 | -0.8777241 | 0.004 | 0.011 | 1 | 0.09443346 | 0.0911999 |
| A2ml1 | 0.2068636 | 0.85639 | 0.011 | 0.007 | 1 | 0.0475115 | 1.23917518 | 0.012 | 0.007 | 1 | 0.2068636 | 0.09276565 |
| Ucp2 | 0.28489903 | -0.4955274 | 0.006 | 0.01 | 1 | 0.06903845 | -0.4484024 | 0.004 | 0.01 | 1 | 0.28489903 | 0.13331059 |
| mt-Co1 | 0.89303395 | 0.43016702 | 0.019 | 0.018 | 1 | 0.07900317 | 0.92426264 | 0.023 | 0.016 | 1 | 0.89303395 | 0.15176484 |
| Dbp | 0.08416933 | -0.3491054 | 0.008 | 0.015 | 1 | 0.94842754 | 0.51614878 | 0.01 | 0.01 | 1 | 0.94842754 | 0.16125419 |
| mt-Atp6 | 0.08769044 | -0.587588 | 0.008 | 0.015 | 1 | 0.41376347 | 0.72871812 | 0.017 | 0.014 | 1 | 0.41376347 | 0.16769127 |
| Mbd2 | 0.08881536 | -0.6617197 | 0.005 | 0.012 | 1 | 0.21731459 | -0.4267073 | 0.006 | 0.01 | 1 | 0.21731459 | 0.16974255 |
| Arglu1 | 0.3202001 | -0.1651871 | 0.009 | 0.013 | 1 | 0.12046899 | 0.75519033 | 0.017 | 0.012 | 1 | 0.3202001 | 0.22642521 |
| Sdc4 | 0.66559855 | 0.38735394 | 0.011 | 0.01 | 1 | 0.17033148 | 0.61768681 | 0.012 | 0.008 | 1 | 0.66559855 | 0.31165015 |
| mt-Co3 | 0.21095168 | -0.3911908 | 0.011 | 0.017 | 1 | 0.18303673 | 0.82497866 | 0.021 | 0.016 | 1 | 0.21095168 | 0.33257101 |
| Bpifb3 | 0.21191719 | -0.5634039 | 0.018 | 0.025 | 1 | 0.21486459 | -1.0100497 | 0.01 | 0.015 | 1 | 0.21486459 | 0.37892549 |
| Srsf6 | 0.28725867 | -0.1567255 | 0.008 | 0.012 | 1 | 0.64933131 | 0.23775311 | 0.008 | 0.01 | 1 | 0.64933131 | 0.49199979 |
| Cst3 | 0.5159201 | 0.14806668 | 0.018 | 0.021 | 1 | 0.30311445 | -0.2033431 | 0.007 | 0.01 | 1 | 0.5159201 | 0.51435054 |
| Jund | 0.41716997 | -0.2053994 | 0.009 | 0.012 | 1 | 0.50513131 | 0.4446221 | 0.009 | 0.011 | 1 | 0.50513131 | 0.66030915 |
| mt-Co2 | 0.55760376 | 0.17801767 | 0.031 | 0.036 | 1 | 0.74237405 | 0.59496673 | 0.036 | 0.035 | 1 | 0.74237405 | 0.80428556 |
| mt-Nd5 | 0.65386763 | 0.35222986 | 0.01 | 0.012 | 1 | 0.66680411 | 0.66551533 | 0.013 | 0.012 | 1 | 0.66680411 | 0.88019239 |
| Oaz1 | 0.75566922 | 0.74673715 | 0.013 | 0.011 | 1 | 0.74471482 | 0.41083524 | 0.011 | 0.012 | 1 | 0.75566922 | 0.93482948 |

Spatial seq: cluster 7

| | cKO_p_val | cKO_avg_log2FC | cKO_pct.1 | cKO_pct.2 | cKO_p_val_adj | WT_p_val | WT_avg_log2FC | WT_pct.1 | WT_pct.2 | WT_p_val_adj | max_pval | minimump_p_val |
|---|---|---|---|---|---|---|---|---|---|---|---|---|
| Obp1a | 0 | 5.64116662 | 0.89 | 0.022 | 0 | 0 | 5.37918391 | 0.96 | 0.028 | 0 | 0 | 0 |
| Col1a1 | 0.57553603 | -0.2060753 | 0.59 | 0.563 | 1 | 1.68E-27 | -1.0770525 | 0.414 | 0.626 | 2.52E-23 | 0.57553603 | 3.36E-27 |
| Col1a2 | 0.00627256 | -0.2981329 | 0.373 | 0.433 | 1 | 3.93E-25 | -1.1201288 | 0.273 | 0.493 | 5.89E-21 | 0.00627256 | 7.86E-25 |
| Hba-a2 | 0.00011444 | -1.1902398 | 0.073 | 0.138 | 1 | 2.60E-21 | -2.5332812 | 0.034 | 0.196 | 3.90E-17 | 0.00011444 | 5.21E-21 |
| Obp2a | 0.03389951 | 0.38590717 | 0.193 | 0.157 | 1 | 1.70E-16 | 0.9676259 | 0.273 | 0.15 | 2.55E-12 | 0.03389951 | 3.40E-16 |
| Hbb-bs | 1.89E-06 | -1.4488531 | 0.078 | 0.162 | 0.02830964 | 7.17E-14 | -2.1896845 | 0.034 | 0.148 | 1.07E-09 | 1.89E-06 | 1.43E-13 |
| Obp2b | 6.24E-13 | -4.8796604 | 0.005 | 0.121 | 9.35E-09 | 1.81E-13 | -4.7150155 | 0.005 | 0.098 | 2.71E-09 | 6.24E-13 | 3.62E-13 |
| S100a9 | 0.00088194 | -1.3560984 | 0.029 | 0.072 | 1 | 8.80E-12 | -2.6922889 | 0.018 | 0.108 | 1.32E-07 | 0.00088194 | 1.76E-11 |
| Mgp | 5.91E-09 | -2.7068998 | 0.017 | 0.105 | 8.84E-05 | 1.22E-05 | -4.413846 | 0 | 0.033 | 0.18341849 | 1.22E-05 | 1.18E-08 |
| Obp1b | 0.10921355 | 0.66736057 | 0.041 | 0.028 | 1 | 1.35E-08 | 1.53740264 | 0.067 | 0.027 | 0.00020174 | 0.10921355 | 2.69E-08 |
| Scgb1c1 | 5.76E-06 | -3.4893649 | 0.005 | 0.057 | 0.08619064 | 1.83E-07 | -2.7521153 | 0.013 | 0.069 | 0.00274419 | 5.76E-06 | 3.67E-07 |
| Lcn11 | 0.00050326 | -3.9868646 | 0 | 0.029 | 1 | 3.62E-07 | -5.3263832 | 0 | 0.045 | 0.00542798 | 0.00050326 | 7.25E-07 |
| S100a8 | 0.00555632 | -1.2722668 | 0.022 | 0.053 | 1 | 4.14E-07 | -2.3295999 | 0.013 | 0.066 | 0.00619905 | 0.00555632 | 8.28E-07 |
| Mup4 | 0.72335063 | 0.54846714 | 0.015 | 0.013 | 1 | 4.14E-07 | 1.59200308 | 0.041 | 0.015 | 0.00620008 | 0.72335063 | 8.28E-07 |
| Lcn3 | 1.23E-06 | -5.0268803 | 0 | 0.054 | 0.01843478 | 8.93E-07 | -4.3873364 | 0.002 | 0.045 | 0.01337935 | 1.23E-06 | 1.79E-06 |
| Ngp | 0.10522913 | -0.8969576 | 0.012 | 0.025 | 1 | 1.01E-06 | -2.6856098 | 0.009 | 0.057 | 0.01513633 | 0.10522913 | 2.02E-06 |
| Hbb-bt | 0.00877836 | -1.2060331 | 0.017 | 0.044 | 1 | 1.29E-05 | -1.9396903 | 0.011 | 0.052 | 0.19281461 | 0.00877836 | 2.58E-05 |
| Ccn2 | 1.41E-05 | -2.7682243 | 0.007 | 0.057 | 0.21146138 | 0.00097385 | -2.2446785 | 0.004 | 0.026 | 1 | 0.00097385 | 2.82E-05 |
| Fn1 | 2.20E-05 | -1.9523924 | 0.017 | 0.071 | 0.3293913 | 0.00062605 | -3.7121429 | 0 | 0.021 | 1 | 0.00062605 | 4.40E-05 |
| Bpifb6 | 0.03590119 | -1.3673566 | 0.015 | 0.033 | 1 | 2.28E-05 | -2.2777273 | 0.009 | 0.047 | 0.3410552 | 0.03590119 | 4.55E-05 |
| Spp1 | 0.40907895 | -0.2477505 | 0.059 | 0.069 | 1 | 3.53E-05 | -1.5448445 | 0.022 | 0.065 | 0.52892846 | 0.40907895 | 7.06E-05 |
| mt-Nd6 | 0.42311244 | 0.50851501 | 0.015 | 0.011 | 1 | 3.70E-05 | 1.88082286 | 0.023 | 0.008 | 0.554591 | 0.42311244 | 7.41E-05 |
| Camp | 0.20880433 | -0.6828584 | 0.01 | 0.018 | 1 | 8.16E-05 | -2.7435659 | 0.004 | 0.034 | 1 | 0.20880433 | 0.00016324 |
| Bpifb4 | 0.02846971 | -1.5303605 | 0.01 | 0.027 | 1 | 0.00012199 | -2.2585678 | 0.011 | 0.044 | 1 | 0.02846971 | 0.00024396 |
| Col9a2 | 0.59914494 | -0.2009894 | 0.032 | 0.037 | 1 | 0.00050749 | 1.51974134 | 0.027 | 0.011 | 1 | 0.59914494 | 0.00101472 |
| Mepe | 0.00062892 | 1.36033026 | 0.037 | 0.015 | 1 | 0.11019054 | -0.7478469 | 0.004 | 0.011 | 1 | 0.11019054 | 0.00125744 |
| Col11a2 | 0.0006311 | -1.7296793 | 0.017 | 0.056 | 1 | 0.01621101 | -1.3361642 | 0.011 | 0.028 | 1 | 0.01621101 | 0.00126181 |
| Car2 | 0.0522119 | -1.046912 | 0.01 | 0.025 | 1 | 0.00076375 | -2.31777 | 0.004 | 0.027 | 1 | 0.0522119 | 0.00152691 |
| mt-Co1 | 0.84697227 | 0.50217348 | 0.02 | 0.018 | 1 | 0.00080452 | 1.40427901 | 0.034 | 0.016 | 1 | 0.84697227 | 0.0016084 |
| Gpx6 | 0.06406767 | -1.0083443 | 0.007 | 0.02 | 1 | 0.00145489 | -2.2008523 | 0.005 | 0.028 | 1 | 0.06406767 | 0.00290767 |
| Acta1 | 0.32495624 | 1.14743649 | 0.012 | 0.008 | 1 | 0.00283925 | 1.27238515 | 0.031 | 0.015 | 1 | 0.32495624 | 0.00567043 |
| Postn | 0.10532298 | -0.9176019 | 0.007 | 0.018 | 1 | 0.00292861 | -3.2378465 | 0 | 0.016 | 1 | 0.10532298 | 0.00584865 |
| Gpx3 | 0.38928577 | -0.2213299 | 0.037 | 0.046 | 1 | 0.00449856 | -0.8007992 | 0.023 | 0.05 | 1 | 0.38928577 | 0.00897689 |
| Sost | 0.01087549 | 1.2251458 | 0.029 | 0.014 | 1 | 0.01442446 | -1.9564335 | 0.004 | 0.017 | 1 | 0.01442446 | 0.0216327 |
| Col11a1 | 0.82578367 | 0.22952908 | 0.015 | 0.013 | 1 | 0.01226208 | 1.53819664 | 0.018 | 0.008 | 1 | 0.82578367 | 0.0243738 |
| Mup5 | 0.90710491 | 0.1871648 | 0.017 | 0.018 | 1 | 0.01441717 | 1.024907 | 0.029 | 0.016 | 1 | 0.90710491 | 0.02862649 |
| Igfbp7 | 0.15458303 | -0.7942195 | 0.007 | 0.016 | 1 | 0.02217038 | -0.5721396 | 0.009 | 0.024 | 1 | 0.15458303 | 0.04384924 |
| Col2a1 | 0.0253987 | -0.9996886 | 0.032 | 0.057 | 1 | 0.42477196 | 0.66383583 | 0.016 | 0.012 | 1 | 0.42477196 | 0.05015231 |
| Wfdc18 | 0.8246877 | 0.46185603 | 0.017 | 0.019 | 1 | 0.03051013 | 0.89342611 | 0.034 | 0.021 | 1 | 0.8246877 | 0.06008939 |
| Vim | 0.21199326 | 0.36007539 | 0.046 | 0.035 | 1 | 0.03175505 | -1.0073147 | 0.014 | 0.03 | 1 | 0.21199326 | 0.06250172 |
| Ighm | 0.23834001 | -0.7363989 | 0.007 | 0.014 | 1 | 0.03243839 | -1.6889532 | 0.002 | 0.011 | 1 | 0.23834001 | 0.06382452 |
| Apoe | 0.73836667 | 0.24457296 | 0.029 | 0.027 | 1 | 0.03270223 | -0.5904906 | 0.02 | 0.037 | 1 | 0.73836667 | 0.06433502 |
| Mpo | 0.30649668 | -0.5062388 | 0.005 | 0.01 | 1 | 0.03790972 | -1.6551649 | 0.002 | 0.011 | 1 | 0.30649668 | 0.07438229 |
| mt-Co3 | 0.67193128 | 0.5980979 | 0.02 | 0.017 | 1 | 0.03806934 | 1.12597185 | 0.027 | 0.016 | 1 | 0.67193128 | 0.07468941 |
| Hist1h2ap | 0.04156902 | -2.0572539 | 0 | 0.01 | 1 | 0.12313711 | -0.9638131 | 0.004 | 0.01 | 1 | 0.12313711 | 0.08141006 |
| Akr1a1 | 0.58747511 | 0.35524138 | 0.02 | 0.016 | 1 | 0.04361994 | -1.3091052 | 0.005 | 0.016 | 1 | 0.58747511 | 0.08533719 |
| Igfbp5 | 0.06640181 | -1.7545194 | 0.022 | 0.039 | 1 | 0.04464399 | -1.1867312 | 0.02 | 0.036 | 1 | 0.06640181 | 0.0872949 |
| Gja1 | 0.45807046 | 1.10624481 | 0.012 | 0.009 | 1 | 0.04838382 | -1.5851174 | 0.002 | 0.01 | 1 | 0.45807046 | 0.09442665 |
| Col6a3 | 0.91250683 | 0.27616981 | 0.012 | 0.012 | 1 | 0.04923534 | -1.5851174 | 0.002 | 0.01 | 1 | 0.91250683 | 0.09604657 |
| Ahnak | 0.11849952 | 0.64204125 | 0.034 | 0.022 | 1 | 0.04946852 | -1.1512964 | 0.007 | 0.018 | 1 | 0.11849952 | 0.09648991 |
| Acp5 | 0.05582594 | -0.7310257 | 0.012 | 0.028 | 1 | 0.085977 | -1.0336014 | 0.011 | 0.021 | 1 | 0.085977 | 0.10853535 |
| mt-Nd2 | 0.0562971 | 1.35778357 | 0.02 | 0.01 | 1 | 0.28792424 | 1.09910113 | 0.011 | 0.007 | 1 | 0.28792424 | 0.10942483 |
| Igfbp4 | 0.32756375 | 0.8255084 | 0.017 | 0.012 | 1 | 0.05953558 | -1.075662 | 0.007 | 0.018 | 1 | 0.32756375 | 0.11552668 |
| Dmp1 | 0.24732605 | -0.3837548 | 0.01 | 0.017 | 1 | 0.06327201 | -1.075662 | 0.007 | 0.018 | 1 | 0.24732605 | 0.12254067 |
| Ptms | 0.06367089 | -0.585502 | 0.01 | 0.024 | 1 | 0.54052 | 0.4422913 | 0.031 | 0.026 | 1 | 0.54052 | 0.1232878 |
| Vmo1 | 0.53554471 | 0.32469678 | 0.027 | 0.022 | 1 | 0.06489376 | -0.7635771 | 0.022 | 0.036 | 1 | 0.53554471 | 0.12557632 |
| Dmbt1 | 0.06914788 | -1.4936019 | 0.005 | 0.016 | 1 | 0.19682762 | 1.06030972 | 0.016 | 0.011 | 1 | 0.19682762 | 0.13351432 |
| Sdf4 | 0.07072082 | 1.38086719 | 0.015 | 0.007 | 1 | 0.07067811 | 1.63297486 | 0.011 | 0.005 | 1 | 0.07072082 | 0.13636082 |
| Ibsp | 0.76759769 | 0.24760066 | 0.029 | 0.027 | 1 | 0.07153966 | -0.2080406 | 0.013 | 0.025 | 1 | 0.76759769 | 0.13796139 |
| Bpifa1 | 0.62611117 | 0.19611415 | 0.032 | 0.028 | 1 | 0.07307326 | 0.78582635 | 0.023 | 0.014 | 1 | 0.62611117 | 0.14080682 |
| Serpinh1 | 0.07773492 | -0.8771723 | 0.015 | 0.029 | 1 | 0.29466054 | -0.2366285 | 0.022 | 0.029 | 1 | 0.29466054 | 0.14942711 |
| Mmp13 | 0.37833745 | -0.3454216 | 0.015 | 0.021 | 1 | 0.08203364 | -0.9958419 | 0.007 | 0.017 | 1 | 0.37833745 | 0.15733777 |
| Thbs1 | 0.08529335 | -0.8827211 | 0.015 | 0.029 | 1 | 0.78087224 | 0.12509634 | 0.009 | 0.01 | 1 | 0.78087224 | 0.16331174 |
| Id3 | 0.09231627 | 0.92228197 | 0.022 | 0.013 | 1 | 0.2911146 | -0.5369501 | 0.007 | 0.012 | 1 | 0.2911146 | 0.17611025 |
| Ctsk | 0.10615642 | -0.9760036 | 0.01 | 0.021 | 1 | 0.09399482 | -1.0888838 | 0.005 | 0.014 | 1 | 0.10615642 | 0.17915462 |
| Hist1h1e | 0.4154884 | -0.2796463 | 0.01 | 0.015 | 1 | 0.10118109 | -0.6391221 | 0.007 | 0.016 | 1 | 0.4154884 | 0.19212457 |
| Slc4a1 | 0.1035936 | -1.1487523 | 0.002 | 0.011 | 1 | 0.10874598 | -0.89985 | 0.005 | 0.013 | 1 | 0.10874598 | 0.19645557 |
| Rsrp1 | 0.10392911 | 0.75644252 | 0.037 | 0.024 | 1 | 0.95663352 | 0.26226439 | 0.018 | 0.018 | 1 | 0.95663352 | 0.19705695 |
| Wwp2 | 0.20762684 | -0.3722956 | 0.005 | 0.012 | 1 | 0.10906573 | 1.14707103 | 0.011 | 0.006 | 1 | 0.20762684 | 0.20623612 |
| Ddx17 | 0.7300363 | 0.34151852 | 0.015 | 0.013 | 1 | 0.10976543 | 0.82561994 | 0.016 | 0.009 | 1 | 0.7300363 | 0.2074824 |
| Col9a3 | 0.11026281 | 0.64058123 | 0.054 | 0.038 | 1 | 0.87323079 | -0.198769 | 0.014 | 0.014 | 1 | 0.87323079 | 0.20836773 |
| mt-Atp8 | 0.11458503 | -0.7093306 | 0.01 | 0.021 | 1 | 0.86368704 | 0.23155617 | 0.022 | 0.021 | 1 | 0.86368704 | 0.21604033 |
| Smoc2 | 0.12546123 | -0.9868646 | 0.005 | 0.014 | 1 | 0.13585687 | 1.30343358 | 0.014 | 0.008 | 1 | 0.13585687 | 0.23518193 |
| Mmp2 | 0.44721122 | 0.58017599 | 0.022 | 0.017 | 1 | 0.13582098 | -0.3603305 | 0.007 | 0.015 | 1 | 0.44721122 | 0.25319462 |
| mt-Co2 | 0.14217391 | 0.45135769 | 0.049 | 0.035 | 1 | 0.74372811 | 0.16977008 | 0.032 | 0.035 | 1 | 0.74372811 | 0.2641344 |
| Ucp2 | 0.9690354 | 0.49376124 | 0.01 | 0.01 | 1 | 0.14322953 | -1.0356038 | 0.004 | 0.01 | 1 | 0.9690354 | 0.26594436 |
| Eef1a1 | 0.18610193 | 0.8304431 | 0.02 | 0.012 | 1 | 0.50559067 | 0.83497984 | 0.011 | 0.008 | 1 | 0.50559067 | 0.33756993 |
| Fth1 | 0.64359193 | 0.15316585 | 0.017 | 0.02 | 1 | 0.18678299 | -0.4033005 | 0.013 | 0.021 | 1 | 0.64359193 | 0.3386781 |
| Itm2b | 0.19890071 | 0.50217348 | 0.063 | 0.05 | 1 | 0.49220143 | 0.37078885 | 0.054 | 0.048 | 1 | 0.49220143 | 0.35823992 |
| Tpm1 | 0.92065949 | 0.14703669 | 0.022 | 0.023 | 1 | 0.20793398 | -0.2251479 | 0.016 | 0.025 | 1 | 0.92065949 | 0.37263142 |
| Maged1 | 0.2079499 | 0.78047465 | 0.017 | 0.011 | 1 | 0.518442 | -0.284563 | 0.007 | 0.01 | 1 | 0.518442 | 0.37265664 |
| Rrbp1 | 0.93639441 | 0.28195216 | 0.01 | 0.009 | 1 | 0.21081209 | 0.96790219 | 0.011 | 0.006 | 1 | 0.93639441 | 0.37718244 |
| mt-Nd1 | 0.80656648 | 0.63462378 | 0.015 | 0.013 | 1 | 0.22866282 | 0.61119435 | 0.018 | 0.012 | 1 | 0.80656648 | 0.40503896 |
| Pcolce | 0.90343476 | 0.27616981 | 0.012 | 0.012 | 1 | 0.23921038 | -0.7154254 | 0.005 | 0.011 | 1 | 0.90343476 | 0.42119916 |
| Timp2 | 0.24965208 | -0.3467605 | 0.017 | 0.026 | 1 | 0.35164555 | -0.2625367 | 0.016 | 0.022 | 1 | 0.35164555 | 0.436978 |
| Serinc3 | 0.42120796 | -0.3302724 | 0.007 | 0.012 | 1 | 0.25477184 | -0.6345054 | 0.005 | 0.01 | 1 | 0.42120796 | 0.44463499 |
| Cd24a | 0.33411843 | -0.2347921 | 0.01 | 0.016 | 1 | 0.2573384 | -0.3491032 | 0.007 | 0.013 | 1 | 0.33411843 | 0.44845375 |
| Nisch | 0.85421715 | 0.45854655 | 0.015 | 0.014 | 1 | 0.2609428 | -0.3659115 | 0.007 | 0.013 | 1 | 0.85421715 | 0.45379445 |
| Col12a1 | 0.26328135 | -0.2894274 | 0.01 | 0.017 | 1 | 0.68661259 | 0.34972724 | 0.011 | 0.009 | 1 | 0.68661259 | 0.45724563 |
| Oaz1 | 0.28110884 | 0.92228197 | 0.017 | 0.011 | 1 | 0.47499733 | -0.1089914 | 0.009 | 0.012 | 1 | 0.47499733 | 0.4831955 |

| | | | | | | | | | | | | |
|---|---|---|---|---|---|---|---|---|---|---|---|---|
| mt-Nd4l | 0.7243542 | 0.56835056 | 0.015 | 0.013 | 1 | 0.31148251 | 0.29922133 | 0.014 | 0.01 | 1 | 0.7243542 | 0.52594367 |
| Htra1 | 0.48386508 | -0.2869883 | 0.007 | 0.011 | 1 | 0.31677231 | -0.5339786 | 0.005 | 0.01 | 1 | 0.48386508 | 0.53319992 |
| Gpx1 | 0.31993407 | -0.4808534 | 0.005 | 0.01 | 1 | 0.54813512 | -0.1667265 | 0.007 | 0.01 | 1 | 0.54813512 | 0.53751033 |
| Arglu1 | 0.32809852 | -0.2071946 | 0.007 | 0.013 | 1 | 0.89122462 | 0.41058244 | 0.013 | 0.012 | 1 | 0.89122462 | 0.54854841 |
| Maf | 0.33402616 | -0.4019021 | 0.005 | 0.01 | 1 | 0.68659936 | 0.23304426 | 0.009 | 0.011 | 1 | 0.68659936 | 0.55647884 |
| mt-Nd3 | 0.34530655 | 0.79369026 | 0.022 | 0.016 | 1 | 0.86937683 | 0.13993485 | 0.013 | 0.012 | 1 | 0.86937683 | 0.57137649 |
| Mt1 | 0.36479859 | 0.58687065 | 0.017 | 0.012 | 1 | 0.57219135 | -0.3091052 | 0.013 | 0.016 | 1 | 0.57219135 | 0.59651917 |
| Tnc | 0.9959505 | 0.2732873 | 0.01 | 0.01 | 1 | 0.38767009 | -0.3736889 | 0.007 | 0.011 | 1 | 0.9959505 | 0.62505208 |
| Dbp | 0.3911402 | -0.3874025 | 0.01 | 0.015 | 1 | 0.52588407 | 0.50114965 | 0.013 | 0.01 | 1 | 0.52588407 | 0.62928975 |
| mt-Cytb | 0.47399704 | 0.50022784 | 0.024 | 0.019 | 1 | 0.52820395 | 0.52677546 | 0.02 | 0.016 | 1 | 0.52820395 | 0.72332088 |
| Col27a1 | 0.47448763 | 0.23576829 | 0.049 | 0.042 | 1 | 0.47705016 | 0.66676318 | 0.011 | 0.008 | 1 | 0.47705016 | 0.72383674 |
| Mbd2 | 0.77357973 | 0.32756896 | 0.01 | 0.011 | 1 | 0.51850077 | -0.2631893 | 0.007 | 0.01 | 1 | 0.77357973 | 0.76815849 |
| Lcn4 | 0.94266081 | 0.47353343 | 0.02 | 0.02 | 1 | 0.5240515 | 0.40856327 | 0.02 | 0.016 | 1 | 0.94266081 | 0.77347303 |
| Srsf6 | 0.54459141 | 0.6555834 | 0.015 | 0.011 | 1 | 0.56938325 | -0.3332368 | 0.007 | 0.01 | 1 | 0.56938325 | 0.79260302 |
| mt-Nd5 | 0.59348102 | 0.29692837 | 0.015 | 0.012 | 1 | 0.82725142 | 0.10143295 | 0.011 | 0.012 | 1 | 0.82725142 | 0.83474232 |
| Lrp1 | 0.92840769 | 0.21094219 | 0.017 | 0.016 | 1 | 0.92802562 | 0.24717458 | 0.013 | 0.013 | 1 | 0.92840769 | 0.99481969 |

**Spatial seq: cluster 8**

| | cKO_p_val | cKO_avg_log2FC | cKO_pct.1 | cKO_pct.2 | cKO_p_val_adj | WT_p_val | WT_avg_log2FC | WT_pct.1 | WT_pct.2 | WT_p_val_adj | max_pval | minimump_p_val |
|---|---|---|---|---|---|---|---|---|---|---|---|---|
| Igfbp5 | 0 | 6.51563574 | 0.644 | 0.024 | 0 | 0 | 5.50439113 | 0.555 | 0.023 | 0 | 0 | 0 |
| Slpi | 0 | 7.27366295 | 0.195 | 0.001 | 0 | 1.32E-271 | 6.23162204 | 0.103 | 0.001 | 1.98E-267 | 1.32E-271 | 0 |
| Obp2b | 8.41E-85 | 1.73602581 | 0.429 | 0.11 | 1.26E-80 | 3.49E-266 | 2.42022952 | 0.586 | 0.084 | 5.23E-262 | 8.41E-85 | 6.98E-266 |
| Pthlh | 1.52E-263 | 6.920026 | 0.098 | 0.001 | 2.27E-259 | 9.86E-125 | 6.00214019 | 0.049 | 0.001 | 1.48E-120 | 9.86E-125 | 3.04E-263 |
| Parm1 | 9.51E-155 | 6.920026 | 0.056 | 0 | 1.42E-150 | 4.35E-43 | 5.04124001 | 0.022 | 0.001 | 6.51E-39 | 4.35E-43 | 1.90E-154 |
| Gdf6 | 2.42E-141 | 6.68870045 | 0.054 | 0 | 3.62E-137 | 1.04E-77 | 6.28224811 | 0.027 | 0 | 1.55E-73 | 1.04E-77 | 4.83E-141 |
| Obp2a | 3.69E-52 | 1.30358527 | 0.434 | 0.151 | 5.53E-48 | 1.47E-137 | 1.6167398 | 0.597 | 0.143 | 2.20E-133 | 3.69E-52 | 2.94E-137 |
| Postn | 1.38E-109 | 3.53669736 | 0.161 | 0.014 | 2.06E-105 | 1.24E-12 | 1.90472769 | 0.056 | 0.014 | 1.85E-08 | 1.24E-12 | 2.75E-109 |
| Lrrc17 | 4.43E-73 | 5.3350635 | 0.041 | 0.001 | 6.63E-69 | 6.75E-15 | 3.80420082 | 0.016 | 0.001 | 1.01E-10 | 6.75E-15 | 8.86E-73 |
| Sfrp1 | 2.92E-67 | 4.77318461 | 0.049 | 0.002 | 4.38E-63 | 2.94E-15 | 3.23388509 | 0.022 | 0.002 | 4.40E-11 | 2.94E-15 | 5.85E-67 |
| Mfap4 | 8.75E-64 | 4.88255129 | 0.044 | 0.002 | 1.31E-59 | 1.48E-07 | 2.74530713 | 0.016 | 0.002 | 0.00222128 | 1.48E-07 | 1.75E-63 |
| Tsc22d1 | 1.61E-59 | 3.22200083 | 0.105 | 0.011 | 2.42E-55 | 2.76E-54 | 3.38916332 | 0.074 | 0.007 | 4.14E-50 | 2.76E-54 | 3.23E-59 |
| Kctd12 | 6.69E-55 | 2.99483459 | 0.112 | 0.014 | 1.00E-50 | 9.55E-11 | 1.7261983 | 0.058 | 0.017 | 1.43E-06 | 9.55E-11 | 1.34E-54 |
| Mgp | 5.23E-47 | 1.61559318 | 0.32 | 0.098 | 7.84E-43 | 5.24E-32 | 2.1154025 | 0.13 | 0.03 | 7.85E-28 | 5.24E-32 | 1.05E-46 |
| Gpha2 | 7.32E-34 | 3.89896438 | 0.039 | 0.003 | 1.10E-29 | 3.38E-44 | 4.63017142 | 0.027 | 0.001 | 5.06E-40 | 7.32E-34 | 6.75E-44 |
| Ogn | 3.55E-42 | 3.41449796 | 0.066 | 0.006 | 5.31E-38 | 0.00072207 | 2.00669368 | 0.013 | 0.004 | 1 | 0.00072207 | 7.10E-42 |
| Tpm1 | 1.45E-36 | 2.41223136 | 0.115 | 0.02 | 2.17E-32 | 3.00E-06 | 1.24815101 | 0.058 | 0.024 | 0.04491413 | 3.00E-06 | 2.90E-36 |
| Eln | 5.84E-30 | 4.17752222 | 0.029 | 0.002 | 8.74E-26 | 6.88E-18 | 3.42233018 | 0.022 | 0.002 | 1.03E-13 | 6.88E-18 | 1.17E-29 |
| Col1a2 | 1.57E-28 | -1.9651723 | 0.185 | 0.438 | 2.35E-24 | 1.63E-16 | -0.9971998 | 0.3 | 0.491 | 2.44E-12 | 1.63E-16 | 3.13E-28 |
| Col1a1 | 5.11E-28 | -1.5007367 | 0.327 | 0.569 | 7.65E-24 | 6.01E-16 | -0.8629094 | 0.445 | 0.624 | 9.01E-12 | 6.01E-16 | 1.02E-27 |
| Igsf3 | 1.21E-27 | 3.9911091 | 0.029 | 0.002 | 1.82E-23 | 1.10E-08 | 3.38916332 | 0.011 | 0.001 | 0.00016453 | 1.10E-08 | 2.43E-27 |
| Serpine2 | 5.87E-24 | 2.3672439 | 0.08 | 0.015 | 8.79E-20 | 1.62E-17 | 2.55266205 | 0.045 | 0.008 | 2.42E-13 | 1.62E-17 | 1.17E-23 |
| Hbb-bs | 5.73E-17 | -3.8169044 | 0.01 | 0.164 | 8.58E-13 | 1.26E-10 | -2.4099711 | 0.04 | 0.147 | 1.89E-06 | 1.26E-10 | 1.15E-16 |
| Smoc2 | 0.00025024 | 1.40906408 | 0.034 | 0.013 | 1 | 8.68E-17 | 2.3408003 | 0.045 | 0.008 | 1.30E-12 | 0.00025024 | 1.74E-16 |
| Fndc1 | 4.12E-09 | 2.63462378 | 0.022 | 0.004 | 6.17E-05 | 2.69E-16 | 3.29400608 | 0.022 | 0.002 | 4.03E-12 | 4.12E-09 | 5.39E-16 |
| Sparc | 2.93E-15 | -1.296505 | 0.161 | 0.34 | 4.39E-11 | 3.44E-07 | -0.8435846 | 0.152 | 0.257 | 0.00514824 | 3.44E-07 | 5.86E-15 |
| Hba-a2 | 5.94E-14 | -3.3722956 | 0.01 | 0.139 | 8.90E-10 | 9.62E-14 | -2.1654255 | 0.056 | 0.194 | 1.44E-09 | 9.62E-14 | 1.19E-13 |
| Chad | 1.28E-12 | 1.74724134 | 0.088 | 0.028 | 1.92E-08 | 7.81E-12 | 2.80420082 | 0.022 | 0.003 | 1.17E-07 | 7.81E-12 | 2.56E-12 |
| S100a9 | 1.45E-07 | -3.2731913 | 0.005 | 0.072 | 0.00216878 | 5.37E-11 | -3.1494414 | 0.011 | 0.108 | 8.04E-07 | 1.45E-07 | 1.07E-10 |
| Dbp | 0.42544469 | 0.48262069 | 0.02 | 0.015 | 1 | 1.52E-09 | 1.97175491 | 0.038 | 0.009 | 2.28E-05 | 0.42544469 | 3.05E-09 |
| Col3a1 | 8.00E-09 | -2.2617474 | 0.024 | 0.116 | 0.00011978 | 4.29E-05 | -1.0633489 | 0.069 | 0.135 | 0.64315658 | 4.29E-05 | 1.60E-08 |
| Mxra8 | 0.00011747 | 1.48706659 | 0.034 | 0.012 | 1 | 5.31E-08 | 1.98165949 | 0.036 | 0.01 | 0.00079528 | 0.00011747 | 1.06E-07 |
| Maf | 0.00022601 | 1.68162126 | 0.027 | 0.009 | 1 | 2.05E-07 | 1.79090399 | 0.036 | 0.01 | 0.00307486 | 0.00022601 | 4.11E-07 |
| Bgn | 0.00238304 | 0.742833 | 0.078 | 0.046 | 1 | 6.32E-07 | 1.07527254 | 0.094 | 0.044 | 0.00945731 | 0.00238304 | 1.26E-06 |
| Scgb1c1 | 0.00010096 | -2.486179 | 0.012 | 0.057 | 1 | 6.45E-07 | -3.2797217 | 0.009 | 0.068 | 0.00966197 | 0.00010096 | 1.29E-06 |
| Thbs2 | 1.33E-06 | 2.3350635 | 0.024 | 0.006 | 0.01992037 | 0.02193175 | 1.31234772 | 0.016 | 0.007 | 1 | 0.02193175 | 2.66E-06 |
| Ngp | 0.00356245 | -2.4946592 | 0.002 | 0.025 | 1 | 4.61E-06 | -2.9495731 | 0.007 | 0.057 | 0.06900057 | 0.00356245 | 9.22E-06 |
| Spp1 | 4.71E-06 | -2.4936019 | 0.012 | 0.07 | 0.07051454 | 0.00110862 | -1.4300332 | 0.027 | 0.065 | 1 | 0.00110862 | 9.42E-06 |
| Hbb-bt | 4.37E-05 | -2.6307208 | 0.002 | 0.044 | 0.65423656 | 6.14E-06 | -3.0384987 | 0.004 | 0.052 | 0.09188529 | 4.37E-05 | 1.23E-05 |
| Col11a2 | 6.60E-06 | -3.1514364 | 0.005 | 0.056 | 0.09876824 | 0.00288726 | -2.2450427 | 0.004 | 0.028 | 1 | 0.00288726 | 1.32E-05 |
| Bpifb6 | 0.0004757 | -3.1845728 | 0.002 | 0.034 | 1 | 7.53E-06 | -3.7681836 | 0.002 | 0.047 | 0.11278873 | 0.0004757 | 1.51E-05 |
| Gas1 | 0.36224513 | 0.90210409 | 0.01 | 0.006 | 1 | 8.14E-06 | 2.32307413 | 0.016 | 0.003 | 0.12184039 | 0.36224513 | 1.63E-05 |
| Bicc1 | 0.00104362 | 2.27616981 | 0.012 | 0.003 | 1 | 1.05E-05 | 2.5240929 | 0.013 | 0.002 | 0.15790962 | 0.00104362 | 2.11E-05 |
| Col2a1 | 1.20E-05 | -2.8197546 | 0.007 | 0.058 | 0.18018474 | 0.04781672 | -1.8776232 | 0.002 | 0.013 | 1 | 0.04781672 | 2.41E-05 |
| S100a8 | 1.41E-05 | -3.0195532 | 0.005 | 0.053 | 0.21053976 | 2.16E-05 | -1.8353744 | 0.016 | 0.066 | 0.32352654 | 2.16E-05 | 2.81E-05 |
| Timp2 | 0.09673013 | 0.59597785 | 0.039 | 0.026 | 1 | 1.59E-05 | 1.29155252 | 0.051 | 0.021 | 0.23828809 | 0.09673013 | 3.18E-05 |
| Map2k3 | 0.02115538 | 1.7990106 | 0.01 | 0.003 | 1 | 3.87E-05 | 2.48227272 | 0.011 | 0.002 | 0.57887592 | 0.02115538 | 7.73E-05 |
| Mup4 | 0.09166685 | 0.94274607 | 0.022 | 0.013 | 1 | 8.33E-05 | 1.35002492 | 0.038 | 0.015 | 1 | 0.09166685 | 0.00016661 |
| Vmo1 | 0.00194307 | -3.4164806 | 0 | 0.023 | 1 | 0.00010726 | -3.3690599 | 0.002 | 0.037 | 1 | 0.00194307 | 0.0002145 |
| Selenow | 0.04592451 | 1.19551214 | 0.017 | 0.008 | 1 | 0.00010918 | 1.85518475 | 0.022 | 0.007 | 1 | 0.04592451 | 0.00021836 |
| Lcn11 | 0.00400077 | -2.3973866 | 0.005 | 0.029 | 1 | 0.00011444 | -2.9999302 | 0.007 | 0.044 | 1 | 0.00400077 | 0.00022886 |
| Obp1a | 0.00011998 | -3.0180833 | 0.005 | 0.044 | 1 | 0.00043376 | -2.1815357 | 0.018 | 0.056 | 1 | 0.00043376 | 0.00023995 |
| Camp | 0.01696125 | -2.0180833 | 0.002 | 0.018 | 1 | 0.00022906 | -3.0074415 | 0.002 | 0.034 | 1 | 0.01696125 | 0.00045806 |
| Cald1 | 0.0002701 | 1.55168744 | 0.029 | 0.01 | 1 | 0.58742733 | 0.54786106 | 0.011 | 0.009 | 1 | 0.58742733 | 0.00054012 |
| Bpifb4 | 0.00062992 | -3.8621532 | 0 | 0.028 | 1 | 0.00029789 | -2.1590164 | 0.009 | 0.044 | 1 | 0.00062992 | 0.00059569 |
| Amelx | 0.2270514 | 1.04966128 | 0.01 | 0.005 | 1 | 0.00037042 | 1.45370357 | 0.018 | 0.005 | 1 | 0.2270514 | 0.0007407 |
| Gpx3 | 0.00045132 | -2.0083443 | 0.01 | 0.046 | 1 | 0.00400165 | -1.2532495 | 0.02 | 0.05 | 1 | 0.00400165 | 0.00090243 |
| Ccn2 | 0.00052614 | -1.5917267 | 0.017 | 0.057 | 1 | 0.05405473 | -1.3319359 | 0.011 | 0.026 | 1 | 0.05405473 | 0.00105201 |
| Sfi1 | 0.00079664 | 1.90879874 | 0.017 | 0.005 | 1 | 0.00300018 | 2.03866607 | 0.013 | 0.004 | 1 | 0.00300018 | 0.00159265 |
| Itm2b | 0.00083587 | -1.8175835 | 0.015 | 0.051 | 1 | 0.0531749 | -0.9276031 | 0.029 | 0.049 | 1 | 0.0531749 | 0.00167103 |
| Thbs1 | 0.00129364 | -2.7038555 | 0.002 | 0.029 | 1 | 0.09073696 | -1.3921964 | 0.002 | 0.01 | 1 | 0.09073696 | 0.0025856 |
| Car2 | 0.00356835 | -2.379182 | 0.002 | 0.025 | 1 | 0.00141371 | -2.5823802 | 0.002 | 0.027 | 1 | 0.00356835 | 0.00282541 |
| Acp5 | 0.00168709 | -2.7470855 | 0.002 | 0.028 | 1 | 0.03281526 | -1.2017979 | 0.007 | 0.021 | 1 | 0.03281526 | 0.00337134 |
| Ibsp | 0.00191805 | -2.5956738 | 0.002 | 0.028 | 1 | 0.01429648 | -1.6108367 | 0.007 | 0.025 | 1 | 0.01429648 | 0.00383242 |
| Obp1b | 0.30833602 | 0.72862127 | 0.037 | 0.028 | 1 | 0.00212893 | 0.86940795 | 0.051 | 0.027 | 1 | 0.30833602 | 0.00425334 |
| Maged2 | 0.97715544 | 0.13212344 | 0.012 | 0.012 | 1 | 0.0024883 | 1.61305933 | 0.018 | 0.006 | 1 | 0.97715544 | 0.00497041 |
| Gpx6 | 0.00342042 | -3.3444166 | 0 | 0.02 | 1 | 0.03720302 | -1.287969 | 0.011 | 0.027 | 1 | 0.03720302 | 0.00682915 |
| Auts2 | 0.0040945 | 1.31876168 | 0.024 | 0.01 | 1 | 0.15712625 | 0.951758 | 0.011 | 0.006 | 1 | 0.15712625 | 0.00817224 |
| Nisch | 0.85880844 | 0.25977537 | 0.015 | 0.014 | 1 | 0.00537585 | 1.11232311 | 0.027 | 0.012 | 1 | 0.85880844 | 0.01072279 |
| Hist1h1e | 0.03858736 | -1.6191328 | 0.002 | 0.015 | 1 | 0.00681562 | -2.9237196 | 0 | 0.016 | 1 | 0.03858736 | 0.01358479 |
| Dmp1 | 0.00686279 | -2.991366 | 0 | 0.018 | 1 | 0.03519604 | -1.4975494 | 0.004 | 0.018 | 1 | 0.03519604 | 0.01367848 |
| Mlf2 | 0.16110067 | 1.2140481 | 0.01 | 0.005 | 1 | 0.00826713 | 1.73095183 | 0.013 | 0.005 | 1 | 0.16110067 | 0.01646592 |
| Cytl1 | 0.42117917 | 0.61168697 | 0.022 | 0.017 | 1 | 0.00848001 | 1.49839239 | 0.011 | 0.004 | 1 | 0.42117917 | 0.01688812 |
| Cd24a | 0.0098131 | -2.8448456 | 0 | 0.016 | 1 | 0.1233242 | -1.0426825 | 0.004 | 0.013 | 1 | 0.1233242 | 0.01952991 |
| Ahnak | 0.01029031 | 0.91627386 | 0.041 | 0.022 | 1 | 0.49669847 | 0.46237316 | 0.022 | 0.018 | 1 | 0.49669847 | 0.02047473 |
| Sost | 0.01308289 | -2.7576936 | 0 | 0.015 | 1 | 0.19506155 | -0.8888214 | 0.009 | 0.017 | 1 | 0.19506155 | 0.02599463 |
| Acan | 0.44048347 | -0.1788159 | 0.022 | 0.028 | 1 | 0.01338262 | 1.22727563 | 0.016 | 0.006 | 1 | 0.44048347 | 0.02658615 |
| Ski | 0.17035302 | 0.94274607 | 0.015 | 0.008 | 1 | 0.01809208 | 1.57040363 | 0.016 | 0.006 | 1 | 0.17035302 | 0.03585683 |
| mt-Co2 | 0.02112675 | -1.1037284 | 0.015 | 0.036 | 1 | 0.87302623 | 0.18859904 | 0.034 | 0.035 | 1 | 0.87302623 | 0.04180717 |
| Mmp13 | 0.0224306 | -1.5837997 | 0.005 | 0.021 | 1 | 0.20775715 | -0.6726129 | 0.009 | 0.017 | 1 | 0.20775715 | 0.04435808 |
| Ywhag | 0.13206053 | 1.2140481 | 0.01 | 0.005 | 1 | 0.02355071 | 1.53118232 | 0.011 | 0.004 | 1 | 0.13206053 | 0.04654679 |
| Mpo | 0.04156936 | -2.10788 | 0 | 0.01 | 1 | 0.02581327 | -2.3387571 | 0 | 0.011 | 1 | 0.04156936 | 0.05096022 |
| Jund | 0.69415968 | 0.17855901 | 0.01 | 0.012 | 1 | 0.02678823 | 0.96595189 | 0.022 | 0.011 | 1 | 0.69415968 | 0.05285886 |
| Mepe | 0.0280265 | -1.8889382 | 0.002 | 0.016 | 1 | 0.02849548 | -2.4437267 | 0 | 0.011 | 1 | 0.02849548 | 0.05526751 |
| mt-Nd2 | 0.27696942 | -0.7093306 | 0.005 | 0.01 | 1 | 0.02899198 | 1.27646318 | 0.016 | 0.007 | 1 | 0.27696942 | 0.05714342 |
| Hist1h2ap | 0.12601026 | -1.0486408 | 0.002 | 0.01 | 1 | 0.03146524 | -2.2401933 | 0 | 0.01 | 1 | 0.12601026 | 0.06194041 |
| Col9a2 | 0.03416704 | -0.9503387 | 0.017 | 0.037 | 1 | 0.70858705 | 0.1090554 | 0.013 | 0.011 | 1 | 0.70858705 | 0.0671667 |
| Slc4a1 | 0.03426989 | -2.1567896 | 0 | 0.011 | 1 | 0.04228525 | -1.5881166 | 0.002 | 0.013 | 1 | 0.04228525 | 0.06736536 |
| Tnc | 0.04219427 | -2.082789 | 0 | 0.01 | 1 | 0.18309752 | -0.8006612 | 0.004 | 0.011 | 1 | 0.18309752 | 0.08260818 |
| Ighm | 0.04329416 | -1.1567896 | 0.002 | 0.014 | 1 | 0.06827758 | -1.3657242 | 0.002 | 0.011 | 1 | 0.06827758 | 0.08471393 |
| Gpx1 | 0.0434739 | -2.082789 | 0 | 0.01 | 1 | 0.52050794 | -0.1730791 | 0.007 | 0.01 | 1 | 0.52050794 | 0.08505782 |
| Six2 | 0.36681371 | 0.65056532 | 0.015 | 0.01 | 1 | 0.04371469 | 1.11303891 | 0.016 | 0.007 | 1 | 0.36681371 | 0.0855184 |
| Mmp2 | 0.05251144 | -1.3326394 | 0.005 | 0.017 | 1 | 0.52521153 | -0.2642789 | 0.011 | 0.015 | 1 | 0.52521153 | 0.10226543 |
| Tmsb4x | 0.19928796 | -0.2107284 | 0.024 | 0.036 | 1 | 0.05586706 | -0.7035938 | 0.022 | 0.04 | 1 | 0.19928796 | 0.108613 |
| Acta1 | 0.32937788 | 0.76737899 | 0.012 | 0.008 | 1 | 0.05991637 | -0.8072339 | 0.004 | 0.016 | 1 | 0.32937788 | 0.11624278 |
| Htra1 | 0.45230716 | 0.74260646 | 0.015 | 0.011 | 1 | 0.06098008 | 1.15650256 | 0.018 | 0.009 | 1 | 0.45230716 | 0.11824159 |

| | | | | | | | | | | | | |
|---|---|---|---|---|---|---|---|---|---|---|---|---|
| Wfdc18 | 0.62687251 | 0.21329453 | 0.022 | 0.019 | 1 | 0.06906148 | 0.5562733 | 0.034 | 0.021 | 1 | 0.62687251 | 0.13335347 |
| Igf2 | 0.07569252 | -1.3860357 | 0.002 | 0.012 | 1 | 0.25748611 | -0.6332045 | 0.004 | 0.01 | 1 | 0.25748611 | 0.14565569 |
| Ptov1 | 0.07639983 | 0.94963246 | 0.022 | 0.012 | 1 | 0.07857161 | 1.52566532 | 0.013 | 0.007 | 1 | 0.07857161 | 0.14696273 |
| Pcolce | 0.08033378 | -1.3373618 | 0.002 | 0.012 | 1 | 0.08658041 | -0.8137767 | 0.002 | 0.011 | 1 | 0.08658041 | 0.15421405 |
| Maged1 | 0.43871893 | 0.58017599 | 0.015 | 0.011 | 1 | 0.08209489 | 0.91523213 | 0.018 | 0.01 | 1 | 0.43871893 | 0.15745021 |
| Dcn | 0.08224856 | -0.4808534 | 0.022 | 0.039 | 1 | 0.42311392 | -0.1654255 | 0.016 | 0.021 | 1 | 0.42311392 | 0.15773229 |
| Atp1b1 | 0.13931167 | 1.14919695 | 0.01 | 0.005 | 1 | 0.08897048 | 1.18529098 | 0.011 | 0.005 | 1 | 0.13931167 | 0.17002522 |
| Serinc3 | 0.20299275 | -0.7523993 | 0.005 | 0.012 | 1 | 0.09078372 | -1.3250822 | 0.002 | 0.01 | 1 | 0.20299275 | 0.17332576 |
| Ccdc80 | 0.09106975 | -0.9065225 | 0.012 | 0.025 | 1 | 0.64400957 | 0.45056386 | 0.011 | 0.009 | 1 | 0.64400957 | 0.17384581 |
| mt-Co1 | 0.09177715 | -1.1161476 | 0.007 | 0.019 | 1 | 0.10119295 | -0.8999334 | 0.007 | 0.017 | 1 | 0.10119295 | 0.17513126 |
| Ddx17 | 0.09263796 | 0.87563188 | 0.022 | 0.013 | 1 | 0.73710277 | 0.38166878 | 0.011 | 0.01 | 1 | 0.73710277 | 0.17669413 |
| Fth1 | 0.54434691 | 0.54365012 | 0.024 | 0.02 | 1 | 0.10228991 | 0.52126685 | 0.031 | 0.02 | 1 | 0.54434691 | 0.1941166 |
| Ucp2 | 0.1203755 | -1.1243681 | 0.002 | 0.01 | 1 | 0.10789137 | -1.3043236 | 0.002 | 0.01 | 1 | 0.1203755 | 0.20414219 |
| mt-Nd1 | 0.13049448 | -1.1406699 | 0.005 | 0.014 | 1 | 0.13687623 | 0.70266279 | 0.02 | 0.012 | 1 | 0.13687623 | 0.24396014 |
| Mup5 | 0.31908192 | 0.31639965 | 0.024 | 0.018 | 1 | 0.14825227 | 0.30496819 | 0.025 | 0.016 | 1 | 0.31908192 | 0.27452581 |
| mt-Nd4l | 0.15037443 | -1.0515176 | 0.005 | 0.013 | 1 | 0.79889691 | -0.2489105 | 0.009 | 0.01 | 1 | 0.79889691 | 0.2781364 |
| Igfbp7 | 0.15458303 | -0.7942195 | 0.007 | 0.016 | 1 | 0.28278389 | 0.35061008 | 0.031 | 0.023 | 1 | 0.28278389 | 0.28527015 |
| Bpifb3 | 0.32882481 | -0.9958534 | 0.017 | 0.024 | 1 | 0.15505249 | -1.1090876 | 0.007 | 0.015 | 1 | 0.32882481 | 0.28606371 |
| Lcn3 | 0.84029486 | -0.2987701 | 0.056 | 0.053 | 1 | 0.16952448 | -0.283262 | 0.058 | 0.043 | 1 | 0.84029486 | 0.31031042 |
| Ctdsp2 | 0.51278295 | 0.70279528 | 0.01 | 0.007 | 1 | 0.17408825 | 1.21923832 | 0.011 | 0.006 | 1 | 0.51278295 | 0.31786978 |
| Col9a3 | 0.44707313 | -0.2855229 | 0.032 | 0.039 | 1 | 0.22764089 | 0.42233018 | 0.02 | 0.013 | 1 | 0.44707313 | 0.4034614 |
| Id3 | 0.72920581 | 0.38753092 | 0.015 | 0.013 | 1 | 0.2446477 | 0.81627365 | 0.018 | 0.012 | 1 | 0.72920581 | 0.4294429 |
| App | 0.24492465 | 0.30531615 | 0.049 | 0.038 | 1 | 0.7026443 | 0.13933877 | 0.04 | 0.037 | 1 | 0.7026443 | 0.42986121 |
| Igfbp4 | 0.33028295 | 0.64856297 | 0.017 | 0.012 | 1 | 0.2491715 | 0.53949759 | 0.025 | 0.017 | 1 | 0.33028295 | 0.43625656 |
| mt-Cytb | 0.27567694 | -0.4160023 | 0.012 | 0.02 | 1 | 0.53854786 | 0.41953697 | 0.02 | 0.016 | 1 | 0.53854786 | 0.4753561 |
| Timp3 | 0.63652439 | 0.18193374 | 0.027 | 0.023 | 1 | 0.27839663 | 0.73708662 | 0.013 | 0.009 | 1 | 0.63652439 | 0.47928858 |
| Arglu1 | 0.71816517 | 0.48957345 | 0.015 | 0.013 | 1 | 0.30091415 | -0.2945331 | 0.007 | 0.012 | 1 | 0.71816517 | 0.51127898 |
| Dmbt1 | 0.53141114 | -0.4808534 | 0.012 | 0.016 | 1 | 0.30365706 | 0.5146942 | 0.016 | 0.011 | 1 | 0.53141114 | 0.51510651 |
| Fn1 | 0.33519839 | -0.4619495 | 0.059 | 0.07 | 1 | 0.31121497 | -0.561705 | 0.013 | 0.02 | 1 | 0.33519839 | 0.52557518 |
| Akr1a1 | 0.34791821 | 0.51233335 | 0.022 | 0.016 | 1 | 0.7432403 | 0.20594149 | 0.018 | 0.016 | 1 | 0.7432403 | 0.57478934 |
| Mt1 | 0.36097558 | -0.4397236 | 0.007 | 0.012 | 1 | 0.45846667 | 0.18840091 | 0.011 | 0.016 | 1 | 0.45846667 | 0.59164778 |
| Cst3 | 0.36534723 | -0.4274372 | 0.015 | 0.021 | 1 | 0.44370278 | -0.379021 | 0.007 | 0.01 | 1 | 0.44370278 | 0.59721586 |
| mt-Nd5 | 0.39692217 | -0.2009894 | 0.007 | 0.012 | 1 | 0.57382902 | -0.1756213 | 0.009 | 0.012 | 1 | 0.57382902 | 0.63629714 |
| mt-Atp6 | 0.4028752 | -0.1989151 | 0.01 | 0.015 | 1 | 0.51685968 | 0.40275238 | 0.018 | 0.014 | 1 | 0.51685968 | 0.64344197 |
| mt-Co3 | 0.98101969 | 0.16959814 | 0.017 | 0.017 | 1 | 0.50363574 | 0.33684228 | 0.02 | 0.016 | 1 | 0.98101969 | 0.75362252 |
| Selenom | 0.93678372 | 0.24760066 | 0.01 | 0.009 | 1 | 0.50454676 | 0.53830676 | 0.013 | 0.01 | 1 | 0.93678372 | 0.75452609 |

Spatial seq: cluster 9

| | cKO_p_val | cKO_avg_log2FC | cKO_pct.1 | cKO_pct.2 | cKO_p_val_adj | WT_p_val | WT_avg_log2FC | WT_pct.1 | WT_pct.2 | WT_p_val_adj | max_pval | minimump_p_val |
|---|---|---|---|---|---|---|---|---|---|---|---|---|
| Obp2b | 0 | 3.34039358 | 1 | 0.099 | 0 | 0 | 3.75737606 | 1 | 0.077 | 0 | 0 | 0 |
| Hba-a2 | 1.89E-09 | -2.2132674 | 0.03 | 0.139 | 2.83E-05 | 3.02E-17 | -3.0313461 | 0.025 | 0.195 | 4.52E-13 | 1.89E-09 | 6.04E-17 |
| Obp2a | 5.39E-17 | -6.7136292 | 0 | 0.161 | 8.08E-13 | 5.33E-17 | -6.0301861 | 0.003 | 0.156 | 7.98E-13 | 5.39E-17 | 1.07E-16 |
| Hbb-bs | 2.87E-10 | -1.9551745 | 0.041 | 0.163 | 4.30E-06 | 6.36E-14 | -3.1049602 | 0.013 | 0.147 | 9.52E-10 | 2.87E-10 | 1.27E-13 |
| Sparc | 1.15E-06 | -0.4471604 | 0.201 | 0.339 | 0.01717922 | 7.79E-14 | -1.294447 | 0.086 | 0.258 | 1.17E-09 | 1.15E-06 | 1.56E-13 |
| Col1a2 | 3.23E-13 | -0.9742498 | 0.247 | 0.436 | 4.83E-09 | 2.87E-09 | -0.7080208 | 0.354 | 0.49 | 4.30E-05 | 2.87E-09 | 6.45E-13 |
| Mgp | 4.52E-11 | -5.7261096 | 0 | 0.106 | 6.77E-07 | 0.00024242 | -3.9081956 | 0 | 0.033 | 1 | 0.00024242 | 9.04E-11 |
| S100a9 | 6.09E-06 | -2.3074884 | 0.011 | 0.072 | 0.09119715 | 1.71E-10 | -3.4535549 | 0.008 | 0.107 | 2.56E-06 | 6.09E-06 | 3.42E-10 |
| Col3a1 | 4.75E-06 | -1.2932195 | 0.038 | 0.115 | 0.07115347 | 1.11E-08 | -1.4906234 | 0.035 | 0.136 | 0.00016581 | 4.75E-06 | 2.21E-08 |
| Col1a1 | 1.69E-06 | -0.719344 | 0.499 | 0.565 | 0.02529437 | 1.17E-08 | -0.8347522 | 0.559 | 0.621 | 0.00017477 | 1.69E-06 | 2.33E-08 |
| Scgb1c1 | 2.53E-06 | -4.9208748 | 0 | 0.057 | 0.03787709 | 7.60E-08 | -5.4220614 | 0 | 0.068 | 0.00113871 | 2.53E-06 | 1.52E-07 |
| Fn1 | 3.14E-07 | -3.9739095 | 0.003 | 0.071 | 0.00470949 | 0.0041009 | -3.2064925 | 0 | 0.02 | 1 | 0.0041009 | 6.29E-07 |
| mt-Nd3 | 4.79E-07 | 1.95550487 | 0.049 | 0.015 | 0.00717584 | 0.00279094 | 1.61494102 | 0.028 | 0.011 | 1 | 0.00279094 | 9.58E-07 |
| S100a8 | 4.75E-05 | -2.8639805 | 0.005 | 0.053 | 0.71086999 | 1.17E-06 | -3.2443323 | 0.005 | 0.066 | 0.01754707 | 4.75E-05 | 2.34E-06 |
| Obp1a | 4.02E-05 | -4.4504114 | 0 | 0.044 | 0.6022667 | 1.31E-06 | -5.1757692 | 0 | 0.056 | 0.01963632 | 4.02E-05 | 2.62E-06 |
| Col2a1 | 1.97E-06 | -4.6679735 | 0 | 0.058 | 0.02956452 | 0.02378842 | -2.6998784 | 0 | 0.013 | 1 | 0.02378842 | 3.95E-06 |
| Ccn2 | 2.28E-06 | -4.6165809 | 0 | 0.057 | 0.03407573 | 0.001234 | -3.745828 | 0 | 0.026 | 1 | 0.001234 | 4.55E-06 |
| Lcn3 | 4.31E-06 | -4.8713076 | 0 | 0.054 | 0.06450363 | 1.73E-05 | -4.8827146 | 0 | 0.045 | 0.2597089 | 1.73E-05 | 8.61E-06 |
| Itm2b | 2.46E-05 | -3.47659 | 0.003 | 0.051 | 0.36881457 | 6.27E-06 | -4.5692451 | 0 | 0.049 | 0.09389796 | 2.46E-05 | 1.25E-05 |
| Col11a2 | 8.46E-06 | -3.5821663 | 0.003 | 0.056 | 0.12671848 | 0.00081132 | -3.652417 | 0 | 0.028 | 1 | 0.00081132 | 1.69E-05 |
| Lcn11 | 0.00097933 | -3.8312919 | 0 | 0.029 | 1 | 1.96E-05 | -4.8207328 | 0 | 0.044 | 0.29293261 | 0.00097933 | 3.91E-05 |
| Ngp | 0.01699808 | -1.7509498 | 0.005 | 0.025 | 1 | 2.39E-05 | -2.7671517 | 0.008 | 0.057 | 0.35838125 | 0.01699808 | 4.79E-05 |
| Spp1 | 0.00978817 | -1.1071503 | 0.035 | 0.069 | 1 | 2.64E-05 | -2.1465812 | 0.013 | 0.065 | 0.39597743 | 0.00978817 | 5.29E-05 |
| Hbb-bt | 0.00084297 | -2.0581857 | 0.008 | 0.044 | 1 | 2.78E-05 | -2.8560774 | 0.005 | 0.052 | 0.41592148 | 0.00084297 | 5.55E-05 |
| Gsn | 0.09622004 | 0.88058274 | 0.038 | 0.024 | 1 | 3.25E-05 | 1.64997255 | 0.043 | 0.016 | 0.48714609 | 0.09622004 | 6.51E-05 |
| Gpx3 | 0.0145566 | -0.8438605 | 0.019 | 0.046 | 1 | 4.18E-05 | -2.8178674 | 0.005 | 0.05 | 0.62592754 | 0.0145566 | 8.36E-05 |
| App | 0.00011867 | -3.9728882 | 0 | 0.039 | 1 | 8.55E-05 | -4.0357457 | 0 | 0.038 | 1 | 0.00011867 | 0.00017098 |
| Camp | 0.06788761 | -1.2731294 | 0.005 | 0.018 | 1 | 0.00020739 | -3.827159 | 0 | 0.034 | 1 | 0.06788761 | 0.00041473 |
| Bpifb6 | 0.13380349 | -0.6912614 | 0.019 | 0.033 | 1 | 0.00021981 | -2.5832335 | 0.008 | 0.047 | 1 | 0.13380349 | 0.00043957 |
| Bgn | 0.3011357 | -0.1764753 | 0.035 | 0.047 | 1 | 0.00028628 | -1.7107316 | 0.008 | 0.046 | 1 | 0.3011357 | 0.00057248 |
| Mup4 | 0.00055092 | 1.90567372 | 0.033 | 0.012 | 1 | 0.04031619 | 1.26693455 | 0.028 | 0.015 | 1 | 0.04031619 | 0.00110154 |
| Igfbp5 | 0.00193578 | -2.6042332 | 0.008 | 0.04 | 1 | 0.00103647 | -2.6952019 | 0.005 | 0.036 | 1 | 0.00193578 | 0.00207186 |
| Car2 | 0.0063931 | -2.2236093 | 0.003 | 0.025 | 1 | 0.00104276 | -3.4028299 | 0 | 0.027 | 1 | 0.0063931 | 0.00208443 |
| mt-Co3 | 0.00141345 | 1.60218287 | 0.038 | 0.016 | 1 | 0.50073496 | 0.87909369 | 0.02 | 0.016 | 1 | 0.50073496 | 0.0028249 |
| Apoe | 0.00388422 | -2.4430544 | 0.003 | 0.027 | 1 | 0.00193608 | -1.7824443 | 0.008 | 0.037 | 1 | 0.00388422 | 0.00386841 |
| Tmsb4x | 0.03960965 | -0.7035734 | 0.016 | 0.037 | 1 | 0.00239564 | -1.307561 | 0.01 | 0.041 | 1 | 0.03960965 | 0.00478554 |
| Bpifb4 | 0.10813129 | -0.6892729 | 0.014 | 0.027 | 1 | 0.00242884 | -1.7529174 | 0.013 | 0.044 | 1 | 0.10813129 | 0.00485178 |
| Gpx6 | 0.04301783 | -1.5968266 | 0.005 | 0.02 | 1 | 0.00247892 | -2.6998784 | 0.003 | 0.027 | 1 | 0.04301783 | 0.0049517 |
| mt-Nd1 | 0.33998631 | 0.63265522 | 0.019 | 0.013 | 1 | 0.00525019 | 1.11684473 | 0.028 | 0.012 | 1 | 0.33998631 | 0.01047282 |
| Wfdc18 | 0.10531424 | 1.10597237 | 0.03 | 0.018 | 1 | 0.00773289 | 0.73869464 | 0.041 | 0.021 | 1 | 0.10531424 | 0.01540598 |
| Serpinf1 | 0.26816932 | -0.5243141 | 0.014 | 0.022 | 1 | 0.0086658 | -2.1732492 | 0.003 | 0.022 | 1 | 0.26816932 | 0.0172565 |
| Lcn4 | 0.33104415 | 0.7321909 | 0.027 | 0.02 | 1 | 0.00867862 | 1.30309582 | 0.033 | 0.016 | 1 | 0.33104415 | 0.01728192 |
| Col9a3 | 0.01122692 | -1.2450407 | 0.014 | 0.039 | 1 | 0.01269642 | 1.42719474 | 0.028 | 0.013 | 1 | 0.01269642 | 0.0223278 |
| Vmo1 | 0.12965688 | -0.9255279 | 0.011 | 0.023 | 1 | 0.01176429 | -1.5949969 | 0.013 | 0.036 | 1 | 0.12965688 | 0.02339018 |
| Obp1b | 0.61593352 | 0.75916961 | 0.033 | 0.028 | 1 | 0.01207075 | 1.05182928 | 0.048 | 0.027 | 1 | 0.61593352 | 0.0239958 |
| Ibsp | 0.02446802 | -1.1092766 | 0.008 | 0.027 | 1 | 0.01220859 | -1.4284153 | 0.005 | 0.025 | 1 | 0.02446802 | 0.02426814 |
| Dcn | 0.01270134 | -0.9187547 | 0.014 | 0.039 | 1 | 0.02567586 | -1.1698823 | 0.005 | 0.021 | 1 | 0.02567586 | 0.02524136 |
| Ctsk | 0.01306569 | -2.15322 | 0.003 | 0.021 | 1 | 0.05689368 | -1.5933223 | 0.003 | 0.014 | 1 | 0.05689368 | 0.02596067 |
| Igfbp7 | 0.01315897 | -2.654022 | 0 | 0.016 | 1 | 0.26628469 | -0.3941996 | 0.015 | 0.024 | 1 | 0.26628469 | 0.02614479 |
| Hist1h1e | 0.01846646 | -2.4693657 | 0 | 0.015 | 1 | 0.08475692 | -1.1472336 | 0.005 | 0.016 | 1 | 0.08475692 | 0.03659191 |
| mt-Atp6 | 0.85675455 | 0.18424897 | 0.014 | 0.015 | 1 | 0.02071244 | 1.38292309 | 0.028 | 0.014 | 1 | 0.85675455 | 0.04099587 |
| Slc4a1 | 0.0448814 | -2.0012169 | 0 | 0.011 | 1 | 0.02127287 | -2.4114089 | 0 | 0.013 | 1 | 0.0448814 | 0.0420932 |
| Cd24a | 0.10977922 | -1.0943263 | 0.005 | 0.016 | 1 | 0.02408451 | -2.4563213 | 0 | 0.013 | 1 | 0.10977922 | 0.04758895 |
| Vim | 0.02434687 | -0.6613669 | 0.014 | 0.036 | 1 | 0.04568914 | -0.8689879 | 0.013 | 0.03 | 1 | 0.04568914 | 0.04810097 |
| Serpinh1 | 0.13788031 | -0.7215996 | 0.016 | 0.029 | 1 | 0.02494715 | -0.6548275 | 0.01 | 0.029 | 1 | 0.13788031 | 0.04927194 |
| Mxra8 | 0.02633266 | -2.3548538 | 0 | 0.013 | 1 | 0.9926188 | 0.72072809 | 0.01 | 0.01 | 1 | 0.9926188 | 0.05197191 |
| Acan | 0.03497468 | 0.94961707 | 0.046 | 0.028 | 1 | 0.02665597 | 1.6889416 | 0.015 | 0.006 | 1 | 0.03497468 | 0.05260141 |
| mt-Nd6 | 0.28126801 | 0.99878312 | 0.016 | 0.01 | 1 | 0.02836055 | 1.63539975 | 0.018 | 0.008 | 1 | 0.28126801 | 0.05591678 |
| Mt1 | 0.03086728 | -2.3037796 | 0 | 0.012 | 1 | 0.39241849 | 0.19654522 | 0.01 | 0.016 | 1 | 0.39241849 | 0.06078177 |
| Dmp1 | 0.03106865 | -1.8312919 | 0.003 | 0.017 | 1 | 0.0592319 | -1.3151281 | 0.005 | 0.018 | 1 | 0.0592319 | 0.06117204 |
| Ighm | 0.06069466 | -1.5915128 | 0.003 | 0.014 | 1 | 0.03250244 | -2.1899666 | 0 | 0.011 | 1 | 0.06069466 | 0.06394848 |
| Serinc3 | 0.03621326 | -2.1958643 | 0 | 0.012 | 1 | 0.61339208 | 0.89915931 | 0.008 | 0.01 | 1 | 0.61339208 | 0.07111511 |
| Mpo | 0.16258191 | -0.943992 | 0.003 | 0.01 | 1 | 0.03639868 | -2.1563358 | 0 | 0.011 | 1 | 0.16258191 | 0.0714725 |
| Mmp13 | 0.03667551 | -1.428227 | 0.005 | 0.021 | 1 | 0.16462996 | -0.4901915 | 0.008 | 0.017 | 1 | 0.16462996 | 0.07200593 |
| Thbs1 | 0.03914127 | -0.9523073 | 0.011 | 0.029 | 1 | 0.61255808 | 0.40165965 | 0.008 | 0.01 | 1 | 0.61255808 | 0.0767505 |
| Mepe | 0.04140403 | -1.7333655 | 0.003 | 0.016 | 1 | 0.03976342 | -2.2613054 | 0 | 0.011 | 1 | 0.04140403 | 0.0779457 |
| Gpx1 | 0.05572991 | -1.9272163 | 0 | 0.01 | 1 | 0.04749865 | -2.0133778 | 0 | 0.01 | 1 | 0.05572991 | 0.09274117 |
| Col9a1 | 0.21299241 | -0.6991883 | 0.005 | 0.013 | 1 | 0.04842079 | 1.88708648 | 0.01 | 0.004 | 1 | 0.21299241 | 0.094497 |
| Htra1 | 0.31097928 | -0.1314156 | 0.005 | 0.011 | 1 | 0.04998719 | -2.0504672 | 0 | 0.01 | 1 | 0.31097928 | 0.09747566 |
| Tpm1 | 0.05939356 | -0.5897372 | 0.008 | 0.023 | 1 | 0.12749378 | -0.629489 | 0.013 | 0.025 | 1 | 0.12749378 | 0.11525952 |
| Acp5 | 0.09682524 | -0.9958636 | 0.014 | 0.028 | 1 | 0.06114731 | -0.7533459 | 0.008 | 0.021 | 1 | 0.09682524 | 0.11855563 |
| Arf5 | 0.12523882 | -1.1314156 | 0.003 | 0.011 | 1 | 0.06290114 | -1.4945045 | 0.003 | 0.013 | 1 | 0.12523882 | 0.12184573 |
| Lrp1 | 0.0909523 | -1.0741484 | 0.005 | 0.017 | 1 | 0.06291405 | -1.4563213 | 0.003 | 0.013 | 1 | 0.0909523 | 0.12186992 |
| Timp2 | 0.06450185 | -0.7854882 | 0.011 | 0.026 | 1 | 0.56404652 | -0.2260769 | 0.018 | 0.022 | 1 | 0.56404652 | 0.12484321 |
| Nisch | 0.06750535 | -1.4222531 | 0.003 | 0.014 | 1 | 0.3878451 | 0.13973887 | 0.008 | 0.013 | 1 | 0.3878451 | 0.13045372 |
| Sost | 0.14153744 | -1.0065503 | 0.005 | 0.015 | 1 | 0.06815943 | -1.4507832 | 0.005 | 0.017 | 1 | 0.14153744 | 0.13167316 |
| Col11a1 | 0.07115457 | -1.4518783 | 0.003 | 0.014 | 1 | 0.35220906 | 1.07973156 | 0.013 | 0.008 | 1 | 0.35220906 | 0.13724616 |
| Cst3 | 0.08211229 | -1.0905644 | 0.008 | 0.021 | 1 | 0.29790095 | -0.1965997 | 0.005 | 0.01 | 1 | 0.29790095 | 0.15748216 |
| Ptms | 0.10412811 | -0.6557203 | 0.011 | 0.024 | 1 | 0.08400631 | -0.6132907 | 0.013 | 0.027 | 1 | 0.10412811 | 0.16095555 |
| Jund | 0.5106969 | 0.33413173 | 0.008 | 0.012 | 1 | 0.09407799 | -1.3650533 | 0.003 | 0.012 | 1 | 0.5106969 | 0.1793053 |
| Postn | 0.15706515 | -0.7620292 | 0.008 | 0.018 | 1 | 0.09670495 | -0.7184342 | 0.005 | 0.015 | 1 | 0.15706515 | 0.18405805 |
| Acta1 | 0.52926651 | 0.69189443 | 0.011 | 0.008 | 1 | 0.09726962 | 1.30309582 | 0.025 | 0.015 | 1 | 0.52926651 | 0.18507786 |
| Kctd12 | 0.09809822 | -1.0639526 | 0.005 | 0.016 | 1 | 0.11897551 | -0.6142819 | 0.008 | 0.018 | 1 | 0.11897551 | 0.18657319 |
| Mbd2 | 0.11547455 | -1.1314156 | 0.003 | 0.011 | 1 | 0.95307216 | 0.51269096 | 0.01 | 0.01 | 1 | 0.95307216 | 0.21761472 |
| Pcolce | 0.54058549 | 0.43174253 | 0.008 | 0.012 | 1 | 0.11997216 | -1.2228312 | 0.003 | 0.011 | 1 | 0.54058549 | 0.225551 |
| mt-Nd4l | 0.74871117 | 0.34962391 | 0.011 | 0.013 | 1 | 0.12562392 | 1.10620377 | 0.018 | 0.01 | 1 | 0.74871117 | 0.23546648 |
| Maged1 | 0.13046914 | -1.1092766 | 0.003 | 0.011 | 1 | 0.64875094 | 0.22108741 | 0.008 | 0.01 | 1 | 0.64875094 | 0.24391608 |
| Dbp | 0.53164916 | 0.26456814 | 0.011 | 0.015 | 1 | 0.13291865 | -1.2097751 | 0.003 | 0.01 | 1 | 0.53164916 | 0.24816994 |
| Dmbt1 | 0.6415836 | 0.42886002 | 0.019 | 0.016 | 1 | 0.16664963 | 1.12864116 | 0.018 | 0.011 | 1 | 0.6415836 | 0.30552716 |
| Mup5 | 0.16709999 | 1.07936064 | 0.027 | 0.018 | 1 | 0.34472521 | -0.6508078 | 0.01 | 0.016 | 1 | 0.34472521 | 0.30627757 |
| Cfl1 | 0.17303141 | 1.21265147 | 0.011 | 0.005 | 1 | 0.41087203 | 1.05851507 | 0.01 | 0.007 | 1 | 0.41087203 | 0.31612295 |
| Sfrp2 | 0.17425126 | -0.8930681 | 0.003 | 0.01 | 1 | 0.63884938 | 0.4630602 | 0.008 | 0.01 | 1 | 0.63884938 | 0.31813902 |
| mt-Nd5 | 0.75131331 | 0.45250109 | 0.014 | 0.012 | 1 | 0.21202384 | -0.5882867 | 0.005 | 0.012 | 1 | 0.75131331 | 0.37909357 |
| Srsf6 | 0.27276543 | 0.1121246 | 0.005 | 0.012 | 1 | 0.36105422 | -0.1563358 | 0.005 | 0.01 | 1 | 0.36105422 | 0.47112988 |
| Fth1 | 0.34275078 | 0.69922284 | 0.027 | 0.02 | 1 | 0.27390127 | -0.2260769 | 0.013 | 0.021 | 1 | 0.34275078 | 0.47278064 |
| mt-Co1 | 0.77047255 | 0.23010867 | 0.016 | 0.018 | 1 | 0.31374098 | 0.80151075 | 0.023 | 0.016 | 1 | 0.77047255 | 0.52904856 |

| | | | | | | | | | | | | |
|---|---|---|---|---|---|---|---|---|---|---|---|---|
| Col6a3 | 0.5308957 | 0.16151262 | 0.008 | 0.012 | 1 | 0.32197474 | -0.487309 | 0.005 | 0.01 | 1 | 0.5308957 | 0.54028175 |
| Tpm2 | 0.33850376 | 1.29423901 | 0.014 | 0.009 | 1 | 0.65517182 | 0.79739659 | 0.015 | 0.013 | 1 | 0.65517182 | 0.56242272 |
| Akr1a1 | 0.98379832 | 0.33581811 | 0.016 | 0.016 | 1 | 0.34918592 | -0.4771747 | 0.01 | 0.016 | 1 | 0.98379832 | 0.57644103 |
| Tnc | 0.39375255 | -0.3252807 | 0.005 | 0.01 | 1 | 0.52062587 | 0.13196151 | 0.008 | 0.011 | 1 | 0.52062587 | 0.63246403 |
| Cd63 | 0.44073956 | 0.83528439 | 0.011 | 0.007 | 1 | 0.41727438 | 1.31285038 | 0.01 | 0.007 | 1 | 0.44073956 | 0.66043085 |
| mt-Nd2 | 0.51300804 | 0.89817918 | 0.014 | 0.01 | 1 | 0.46048334 | 1.29856616 | 0.01 | 0.007 | 1 | 0.51300804 | 0.70892178 |
| Arglu1 | 0.53231187 | 0.64514617 | 0.016 | 0.013 | 1 | 0.55427458 | 0.58289997 | 0.015 | 0.012 | 1 | 0.55427458 | 0.78126781 |
| Oaz1 | 0.54562201 | 0.31404749 | 0.008 | 0.012 | 1 | 0.94016203 | 0.57158465 | 0.013 | 0.012 | 1 | 0.94016203 | 0.79354064 |
| mt-Co2 | 0.55267735 | 0.60693041 | 0.041 | 0.035 | 1 | 0.64671772 | 0.37102038 | 0.03 | 0.035 | 1 | 0.64671772 | 0.79990245 |
| Ucp2 | 0.72329523 | 0.37793609 | 0.008 | 0.01 | 1 | 0.68664158 | -0.1078954 | 0.008 | 0.01 | 1 | 0.72329523 | 0.9018065 |

Spatial seq: cluster 10

| | cKO_p_val | cKO_avg_log2FC | cKO_pct.1 | cKO_pct.2 | cKO_p_val_adj | WT_p_val | WT_avg_log2FC | WT_pct.1 | WT_pct.2 | WT_p_val_adj | max_pval | minimump_p_val |
|---|---|---|---|---|---|---|---|---|---|---|---|---|
| Lcn11 | 0 | 6.33851332 | 0.9 | 0.014 | 0 | 0 | 5.64619884 | 0.944 | 0.023 | 0 | 0 | 0 |
| Obp2a | 0.00021346 | 0.65905863 | 0.236 | 0.157 | 1 | 6.52E-21 | 1.14369919 | 0.314 | 0.15 | 9.76E-17 | 0.00021346 | 1.30E-20 |
| Hba-a2 | 7.73E-05 | -1.5102172 | 0.055 | 0.138 | 1 | 7.20E-14 | -2.2395853 | 0.048 | 0.194 | 1.08E-09 | 7.73E-05 | 1.44E-13 |
| Col1a1 | 0.11052049 | -0.365841 | 0.557 | 0.564 | 1 | 5.74E-12 | -0.8638895 | 0.505 | 0.623 | 8.60E-08 | 0.11052049 | 1.15E-11 |
| Col1a2 | 0.05141272 | -0.2953081 | 0.38 | 0.432 | 1 | 8.31E-11 | -0.7837102 | 0.341 | 0.49 | 1.24E-06 | 0.05141272 | 1.66E-10 |
| Obp2b | 4.04E-09 | -4.8556301 | 0.004 | 0.12 | 6.04E-05 | 1.94E-10 | -4.6940506 | 0.005 | 0.098 | 2.91E-06 | 4.04E-09 | 3.88E-10 |
| Hbb-bs | 0.00068882 | -1.2460011 | 0.089 | 0.161 | 1 | 1.08E-09 | -2.170668 | 0.041 | 0.147 | 1.62E-05 | 0.00068882 | 2.16E-09 |
| Mup5 | 0.1379772 | 1.25889441 | 0.03 | 0.018 | 1 | 1.15E-09 | 1.96491246 | 0.053 | 0.015 | 1.72E-05 | 0.1379772 | 2.30E-09 |
| Sparc | 0.00069054 | -0.3286192 | 0.229 | 0.338 | 1 | 8.45E-08 | -0.8335884 | 0.138 | 0.257 | 0.00126499 | 0.00069054 | 1.69E-07 |
| Scgb1c1 | 0.00047828 | -2.8799676 | 0.007 | 0.056 | 1 | 1.08E-07 | -4.4905917 | 0.002 | 0.068 | 0.00162198 | 0.00047828 | 2.17E-07 |
| S100a9 | 0.00070603 | -1.8536637 | 0.018 | 0.072 | 1 | 2.93E-07 | -1.9322816 | 0.029 | 0.107 | 0.00438068 | 0.00070603 | 5.85E-07 |
| Col3a1 | 0.01280127 | -0.8393949 | 0.066 | 0.114 | 1 | 2.44E-06 | -1.2469281 | 0.056 | 0.136 | 0.03656938 | 0.01280127 | 4.88E-06 |
| Lcn3 | 0.0002534 | -3.4163862 | 0.004 | 0.054 | 1 | 1.08E-05 | -4.9519527 | 0 | 0.045 | 0.16138894 | 0.0002534 | 2.16E-05 |
| Obp1a | 0.00045046 | -3.9965868 | 0 | 0.044 | 1 | 1.65E-05 | -2.9197182 | 0.007 | 0.056 | 0.24640182 | 0.00045046 | 3.29E-05 |
| Mgp | 2.03E-05 | -2.2680355 | 0.026 | 0.105 | 0.30471993 | 0.00050211 | -2.9754117 | 0.002 | 0.033 | 1 | 0.00050211 | 4.07E-05 |
| Bpifb4 | 0.04383026 | -1.6628695 | 0.007 | 0.027 | 1 | 3.46E-05 | -3.6359252 | 0.002 | 0.044 | 0.51866148 | 0.04383026 | 6.93E-05 |
| Spp1 | 0.01935201 | -1.143002 | 0.033 | 0.069 | 1 | 3.48E-05 | -1.8512904 | 0.014 | 0.065 | 0.52116652 | 0.01935201 | 6.96E-05 |
| Hbb-bt | 0.00130328 | -2.6082082 | 0.004 | 0.044 | 1 | 4.33E-05 | -2.5088807 | 0.007 | 0.052 | 0.64841851 | 0.00130328 | 8.66E-05 |
| Bpifb6 | 0.17865032 | -0.9823293 | 0.018 | 0.033 | 1 | 5.27E-05 | -2.6524715 | 0.005 | 0.047 | 0.78927071 | 0.17865032 | 0.00010541 |
| Ngp | 0.0087512 | -2.8884292 | 0 | 0.025 | 1 | 8.51E-05 | -2.2491975 | 0.012 | 0.057 | 1 | 0.0087512 | 0.00017024 |
| Tacc1 | 0.05973589 | 1.83593639 | 0.011 | 0.004 | 1 | 8.56E-05 | 2.24361232 | 0.014 | 0.003 | 1 | 0.05973589 | 0.00017117 |
| S100a8 | 0.01272038 | -1.1818749 | 0.018 | 0.052 | 1 | 0.00014988 | -1.7221911 | 0.019 | 0.065 | 1 | 0.01272038 | 0.00029974 |
| Obp1b | 0.61617059 | 0.67299782 | 0.033 | 0.028 | 1 | 0.0001543 | 1.1552939 | 0.058 | 0.027 | 1 | 0.61617059 | 0.00030857 |
| Col27a1 | 0.0853777 | 0.58690885 | 0.063 | 0.041 | 1 | 0.00025472 | 1.37994996 | 0.024 | 0.008 | 1 | 0.0853777 | 0.00050937 |
| Ccn2 | 0.00044677 | -2.5751754 | 0.007 | 0.057 | 1 | 0.00753214 | -1.8082662 | 0.005 | 0.026 | 1 | 0.00753214 | 0.00089335 |
| Gpx3 | 0.33655512 | -0.2507436 | 0.033 | 0.046 | 1 | 0.00044966 | -1.4648797 | 0.012 | 0.05 | 1 | 0.33655512 | 0.00089912 |
| Col11a2 | 0.06385534 | -0.7956572 | 0.03 | 0.055 | 1 | 0.00060305 | -3.7216551 | 0 | 0.028 | 1 | 0.06385534 | 0.00120574 |
| mt-Atp8 | 0.79764501 | 0.32240903 | 0.018 | 0.021 | 1 | 0.00076181 | 1.57695965 | 0.043 | 0.02 | 1 | 0.79764501 | 0.00152305 |
| Car2 | 0.02611038 | -1.7697847 | 0.004 | 0.025 | 1 | 0.00078407 | -3.472068 | 0 | 0.027 | 1 | 0.02611038 | 0.00156753 |
| Camp | 0.39356689 | -0.3998351 | 0.011 | 0.018 | 1 | 0.00117527 | -2.3071536 | 0.005 | 0.034 | 1 | 0.39356689 | 0.00234916 |
| Wfdc18 | 0.97552129 | 0.15562602 | 0.018 | 0.019 | 1 | 0.00413757 | 1.27995417 | 0.041 | 0.021 | 1 | 0.97552129 | 0.00825801 |
| Gpx6 | 0.28869652 | -0.7244241 | 0.011 | 0.02 | 1 | 0.00498201 | -2.1818176 | 0.005 | 0.027 | 1 | 0.28869652 | 0.0099392 |
| Sost | 0.96707758 | 0.19492658 | 0.015 | 0.014 | 1 | 0.00737779 | -3.1123633 | 0 | 0.017 | 1 | 0.96707758 | 0.01470115 |
| Rab34 | 0.02970226 | 2.0618178 | 0.011 | 0.003 | 1 | 0.00966898 | 1.71226966 | 0.012 | 0.004 | 1 | 0.02970226 | 0.01924447 |
| Bpifa1 | 0.1875539 | 0.89038418 | 0.041 | 0.028 | 1 | 0.01257247 | 0.90330891 | 0.029 | 0.014 | 1 | 0.1875539 | 0.02498687 |
| Slc4a1 | 0.26501139 | -0.5393549 | 0.004 | 0.011 | 1 | 0.01832222 | -2.480647 | 0 | 0.013 | 1 | 0.26501139 | 0.03630873 |
| Foxc1 | 0.27817979 | 1.23021533 | 0.011 | 0.006 | 1 | 0.01985669 | 1.84699476 | 0.01 | 0.003 | 1 | 0.27817979 | 0.03931909 |
| Igfbp5 | 0.03619184 | -2.1504086 | 0.015 | 0.039 | 1 | 0.0211765 | -1.3415745 | 0.014 | 0.036 | 1 | 0.03619184 | 0.04190455 |
| Ibsp | 0.21073668 | -0.655452 | 0.015 | 0.027 | 1 | 0.02235749 | -1.4976534 | 0.007 | 0.025 | 1 | 0.21073668 | 0.04421512 |
| Fn1 | 0.03382665 | -0.7004255 | 0.037 | 0.07 | 1 | 0.02605959 | -1.2658377 | 0.005 | 0.02 | 1 | 0.03382665 | 0.05144007 |
| Aldoa | 0.02697187 | 2.24402113 | 0.011 | 0.003 | 1 | 0.13920341 | 1.72098688 | 0.01 | 0.005 | 1 | 0.13920341 | 0.05321625 |
| Dmp1 | 0.02861734 | -2.3819686 | 0 | 0.017 | 1 | 0.0489654 | -1.3843661 | 0.005 | 0.018 | 1 | 0.0489654 | 0.05641572 |
| Vim | 0.03040658 | -1.3886945 | 0.011 | 0.035 | 1 | 0.06802511 | -0.7432061 | 0.014 | 0.03 | 1 | 0.06802511 | 0.05988861 |
| Apoe | 0.0469273 | -1.401314 | 0.007 | 0.027 | 1 | 0.0306304 | -0.9966134 | 0.017 | 0.037 | 1 | 0.0469273 | 0.06032257 |
| Serpinh1 | 0.75390744 | 0.10036538 | 0.026 | 0.029 | 1 | 0.03864716 | -0.9191172 | 0.012 | 0.029 | 1 | 0.75390744 | 0.07580071 |
| Ahnak | 0.11424644 | 0.91839855 | 0.037 | 0.023 | 1 | 0.04011625 | -1.4595854 | 0.005 | 0.018 | 1 | 0.11424644 | 0.07862318 |
| Col11a1 | 0.05326151 | -2.0039064 | 0 | 0.014 | 1 | 0.41105003 | 1.0104935 | 0.012 | 0.008 | 1 | 0.41105003 | 0.10368624 |
| Tmsb4x | 0.81135148 | 0.28954947 | 0.033 | 0.036 | 1 | 0.05967363 | -0.3644908 | 0.022 | 0.04 | 1 | 0.81135148 | 0.11578632 |
| Mmp13 | 0.25936319 | -0.555385 | 0.011 | 0.021 | 1 | 0.06052255 | -1.3050083 | 0.005 | 0.017 | 1 | 0.25936319 | 0.11738212 |
| mt-Nd3 | 0.20396826 | 1.17187135 | 0.026 | 0.016 | 1 | 0.0605258 | 0.84890435 | 0.022 | 0.012 | 1 | 0.20396826 | 0.11738824 |
| Bgn | 0.10067157 | -0.7367776 | 0.026 | 0.047 | 1 | 0.06470517 | -0.5451669 | 0.027 | 0.046 | 1 | 0.10067157 | 0.12522358 |
| Vmo1 | 0.09249864 | -1.2154105 | 0.007 | 0.023 | 1 | 0.06857874 | -0.6541579 | 0.019 | 0.036 | 1 | 0.09249864 | 0.13245443 |
| mt-Co1 | 0.06988611 | -1.5149708 | 0.004 | 0.019 | 1 | 0.63581706 | 0.62158573 | 0.019 | 0.016 | 1 | 0.63581706 | 0.13488814 |
| Id3 | 0.42783518 | -0.2519364 | 0.007 | 0.013 | 1 | 0.07030302 | -1.4401679 | 0.002 | 0.012 | 1 | 0.42783518 | 0.13566353 |
| Hist1h1e | 0.97193477 | 0.32975101 | 0.015 | 0.015 | 1 | 0.07123734 | -1.2164717 | 0.005 | 0.016 | 1 | 0.97193477 | 0.13739992 |
| Mt1 | 0.19798166 | -0.8434417 | 0.004 | 0.012 | 1 | 0.07571382 | -1.3092957 | 0.005 | 0.016 | 1 | 0.19798166 | 0.14569506 |
| Timp2 | 0.12040725 | -0.5977972 | 0.011 | 0.026 | 1 | 0.08644722 | -0.7168219 | 0.01 | 0.022 | 1 | 0.12040725 | 0.16542132 |
| Tnc | 0.1000125 | -1.4733917 | 0 | 0.01 | 1 | 0.09162332 | -1.2790131 | 0.002 | 0.011 | 1 | 0.1000125 | 0.1748518 |
| Mpo | 0.30391718 | -0.4901674 | 0.004 | 0.01 | 1 | 0.09663843 | -1.2187526 | 0.002 | 0.011 | 1 | 0.30391718 | 0.18393787 |
| Mepe | 0.1085397 | -0.6897452 | 0.004 | 0.016 | 1 | 0.10600862 | -1.3242019 | 0.002 | 0.011 | 1 | 0.1085397 | 0.20077941 |
| Serinc3 | 0.61225473 | 0.60817746 | 0.015 | 0.011 | 1 | 0.11190779 | -1.2118989 | 0.002 | 0.01 | 1 | 0.61225473 | 0.21129222 |
| Calm3 | 0.11427669 | 1.56363907 | 0.015 | 0.007 | 1 | 0.54942324 | 0.60510617 | 0.01 | 0.007 | 1 | 0.54942324 | 0.21549421 |
| Hist1h2ap | 0.68482986 | 0.15438392 | 0.007 | 0.01 | 1 | 0.1165495 | -1.1197052 | 0.002 | 0.01 | 1 | 0.68482986 | 0.21951522 |
| Col6a3 | 0.21956704 | -0.7279645 | 0.004 | 0.012 | 1 | 0.11927363 | -0.5565471 | 0.002 | 0.01 | 1 | 0.21956704 | 0.22432107 |
| Pura | 0.16666344 | 1.72723013 | 0.011 | 0.005 | 1 | 0.12052986 | 1.94775774 | 0.01 | 0.004 | 1 | 0.16666344 | 0.22653228 |
| Maged1 | 0.25661618 | -0.655452 | 0.004 | 0.011 | 1 | 0.12247048 | -0.5991914 | 0.002 | 0.01 | 1 | 0.25661618 | 0.22994194 |
| Gpx1 | 0.31382242 | -0.4649301 | 0.004 | 0.01 | 1 | 0.12807578 | -1.0750822 | 0.002 | 0.01 | 1 | 0.31382242 | 0.23974815 |
| Col9a2 | 0.53907184 | -0.1868817 | 0.03 | 0.037 | 1 | 0.13116011 | 1.02661316 | 0.019 | 0.011 | 1 | 0.53907184 | 0.24511725 |
| Lcn4 | 0.1324963 | -0.7594427 | 0.007 | 0.02 | 1 | 0.64339421 | 0.19690662 | 0.019 | 0.016 | 1 | 0.64339421 | 0.24743733 |
| Lrp1 | 0.81937319 | 0.12683759 | 0.015 | 0.017 | 1 | 0.13669228 | -0.9350587 | 0.005 | 0.013 | 1 | 0.81937319 | 0.25469978 |
| Mup4 | 0.80075326 | -0.1950905 | 0.011 | 0.013 | 1 | 0.13993831 | 0.879003 | 0.024 | 0.015 | 1 | 0.80075326 | 0.26029389 |
| Acan | 0.80371325 | -0.1096158 | 0.026 | 0.028 | 1 | 0.14165248 | 1.48604478 | 0.012 | 0.006 | 1 | 0.80371325 | 0.26323953 |
| mt-Co3 | 0.50141032 | 0.60461085 | 0.022 | 0.017 | 1 | 0.14653655 | -0.7110007 | 0.007 | 0.016 | 1 | 0.50141032 | 0.27160015 |
| mt-Nd2 | 0.17448699 | 0.92209304 | 0.018 | 0.01 | 1 | 0.52119406 | 1.2293281 | 0.01 | 0.007 | 1 | 0.52119406 | 0.31852827 |
| Tpm1 | 0.95497895 | 0.28624937 | 0.022 | 0.023 | 1 | 0.19293558 | -0.1757253 | 0.014 | 0.025 | 1 | 0.95497895 | 0.34864703 |
| mt-Nd1 | 0.19737277 | 1.24402113 | 0.022 | 0.013 | 1 | 0.20406989 | 0.38686938 | 0.019 | 0.012 | 1 | 0.20406989 | 0.35578954 |
| Kctd12 | 0.50282567 | -0.1899655 | 0.011 | 0.016 | 1 | 0.20149686 | -0.4165166 | 0.01 | 0.018 | 1 | 0.50282567 | 0.36239274 |
| Acta1 | 0.20231355 | 1.37677634 | 0.015 | 0.008 | 1 | 0.78364923 | 0.4445631 | 0.017 | 0.015 | 1 | 0.78364923 | 0.36369633 |
| Jund | 0.21055373 | -0.8237235 | 0.004 | 0.012 | 1 | 0.4229299 | -0.4224659 | 0.007 | 0.011 | 1 | 0.4229299 | 0.37677458 |
| Igf2 | 0.21281184 | -0.7766383 | 0.004 | 0.012 | 1 | 0.61650947 | 0.23181765 | 0.007 | 0.01 | 1 | 0.61650947 | 0.3803348 |
| Serpinf1 | 0.21976053 | -0.6628695 | 0.011 | 0.022 | 1 | 0.32428871 | -0.2221673 | 0.014 | 0.022 | 1 | 0.32428871 | 0.39122637 |
| mt-Atp6 | 0.31640212 | -0.1848222 | 0.007 | 0.015 | 1 | 0.22568834 | -0.2391204 | 0.007 | 0.014 | 1 | 0.31640212 | 0.40044145 |
| Nucb1 | 0.44530521 | 1.0557176 | 0.018 | 0.013 | 1 | 0.2274033 | 0.96318778 | 0.014 | 0.009 | 1 | 0.44530521 | 0.40309434 |
| mt-Co2 | 0.24218563 | -0.1682393 | 0.022 | 0.036 | 1 | 0.47609819 | 0.66044534 | 0.041 | 0.035 | 1 | 0.47609819 | 0.42571738 |
| Pcolce | 0.28211624 | 1.11519115 | 0.018 | 0.011 | 1 | 0.2600386 | -0.7005935 | 0.005 | 0.01 | 1 | 0.28211624 | 0.45245713 |
| Clta | 0.72054578 | 0.53506991 | 0.011 | 0.009 | 1 | 0.26621806 | 1.2735279 | 0.012 | 0.007 | 1 | 0.72054578 | 0.46156407 |
| Mxra8 | 0.78697895 | 0.44620998 | 0.015 | 0.013 | 1 | 0.27022258 | -0.5991914 | 0.005 | 0.01 | 1 | 0.78697895 | 0.46742492 |
| Oaz1 | 0.27523051 | 1.196268 | 0.018 | 0.011 | 1 | 0.97111466 | 0.50234659 | 0.012 | 0.012 | 1 | 0.97111466 | 0.47470919 |
| Fth1 | 0.27728863 | -0.5830162 | 0.011 | 0.02 | 1 | 0.85044466 | 0.17387565 | 0.022 | 0.02 | 1 | 0.85044466 | 0.47768828 |
| Ptgis | 0.29491151 | 0.9996024 | 0.015 | 0.009 | 1 | 0.32882214 | 0.99138467 | 0.01 | 0.006 | 1 | 0.32882214 | 0.50285023 |
| Sfrp2 | 0.32058024 | -0.4392434 | 0.004 | 0.01 | 1 | 0.57699276 | 0.39382214 | 0.007 | 0.01 | 1 | 0.57699276 | 0.53838879 |
| Postn | 0.40110219 | -0.3082046 | 0.011 | 0.018 | 1 | 0.34922075 | -0.4611275 | 0.01 | 0.015 | 1 | 0.40110219 | 0.57648636 |
| App | 0.3636978 | 0.68602568 | 0.048 | 0.038 | 1 | 0.55039705 | -0.1719634 | 0.031 | 0.037 | 1 | 0.55039705 | 0.59511951 |
| Igfbp7 | 0.51119574 | -0.1848222 | 0.011 | 0.016 | 1 | 0.36448828 | -0.2906244 | 0.017 | 0.024 | 1 | 0.51119574 | 0.59612485 |
| Cst3 | 0.56757256 | 0.55214343 | 0.026 | 0.021 | 1 | 0.38679472 | 1.24067902 | 0.014 | 0.01 | 1 | 0.56757256 | 0.62397928 |
| Ddx17 | 0.39883172 | 0.72206843 | 0.018 | 0.013 | 1 | 0.99431135 | 0.49485206 | 0.01 | 0.01 | 1 | 0.99431135 | 0.63859669 |

| | | | | | | | | | | | | |
|---|---|---|---|---|---|---|---|---|---|---|---|---|
| Maged2 | 0.48258239 | 0.14356095 | 0.007 | 0.012 | 1 | 0.42054396 | 0.94775774 | 0.01 | 0.006 | 1 | 0.48258239 | 0.66423069 |
| mt-Nd4l | 0.42667738 | -0.4421202 | 0.007 | 0.013 | 1 | 0.92918142 | 0.36133428 | 0.01 | 0.01 | 1 | 0.92918142 | 0.67130117 |
| Akr1a1 | 0.43258708 | 0.59194443 | 0.022 | 0.016 | 1 | 0.86861756 | 0.47554653 | 0.017 | 0.016 | 1 | 0.86861756 | 0.67804258 |
| Ighm | 0.67482985 | 0.20029975 | 0.011 | 0.014 | 1 | 0.43870465 | -0.2391204 | 0.007 | 0.011 | 1 | 0.67482985 | 0.68494753 |
| Gsn | 0.60565155 | 0.52124942 | 0.03 | 0.025 | 1 | 0.45833093 | -0.3802965 | 0.012 | 0.017 | 1 | 0.60565155 | 0.70659462 |
| Srsf6 | 0.52092595 | -0.1848222 | 0.007 | 0.012 | 1 | 0.57893194 | 0.80190687 | 0.012 | 0.009 | 1 | 0.57893194 | 0.77048806 |
| Dbp | 0.60735795 | -0.1053877 | 0.011 | 0.015 | 1 | 0.66315957 | 0.33242159 | 0.012 | 0.01 | 1 | 0.66315957 | 0.84583222 |
| Arglu1 | 0.75412369 | 0.67144236 | 0.015 | 0.013 | 1 | 0.63555041 | 0.51366191 | 0.014 | 0.012 | 1 | 0.75412369 | 0.8671765 |
| Mmp2 | 0.8641441 | 0.51819609 | 0.018 | 0.017 | 1 | 0.71113222 | 0.44826992 | 0.017 | 0.015 | 1 | 0.8641441 | 0.9165554 |
| Eef1a1 | 0.72346971 | 0.40840795 | 0.015 | 0.012 | 1 | 0.75351116 | 0.66104434 | 0.01 | 0.008 | 1 | 0.75351116 | 0.923531 |
| Ptms | 0.8788245 | 0.219947 | 0.022 | 0.024 | 1 | 0.75678742 | 0.24181904 | 0.029 | 0.027 | 1 | 0.8788245 | 0.94084764 |
| Ptprs | 0.87024038 | 0.49324974 | 0.011 | 0.01 | 1 | 0.86728191 | 0.63609798 | 0.01 | 0.009 | 1 | 0.87024038 | 0.98238591 |

Spatial seq: cluster 11

| | cKO_p_val | cKO_avg_log2FC | cKO_pct.1 | cKO_pct.2 | cKO_p_val_adj | WT_p_val | WT_avg_log2FC | WT_pct.1 | WT_pct.2 | WT_p_val_adj | max_pval | minimump_p_val |
|---|---|---|---|---|---|---|---|---|---|---|---|---|
| Col9a3 | 4.61E-157 | 4.09571797 | 0.5 | 0.035 | 6.91E-153 | 0 | 6.67504373 | 0.667 | 0.01 | 0 | 4.61E-157 | 0 |
| Col9a2 | 1.83E-181 | 4.35846219 | 0.516 | 0.033 | 2.74E-177 | 3.04E-176 | 5.62225024 | 0.323 | 0.01 | 4.56E-172 | 3.04E-176 | 3.66E-181 |
| Col1a1 | 8.34E-13 | -1.9795245 | 0.27 | 0.566 | 1.25E-08 | 0.03308073 | -0.8092162 | 0.645 | 0.62 | 1 | 0.03308073 | 1.67E-12 |
| Col1a2 | 1.46E-09 | -1.6828392 | 0.156 | 0.433 | 2.19E-05 | 0.02951454 | -0.7388162 | 0.441 | 0.487 | 1 | 0.02951454 | 2.92E-09 |
| Col9a1 | 1.26E-07 | 2.70270347 | 0.066 | 0.012 | 0.00188611 | 0.00687844 | 3.14198512 | 0.022 | 0.004 | 1 | 0.00687844 | 2.52E-07 |
| Tsc22d1 | 0.77748077 | 0.77479509 | 0.016 | 0.013 | 1 | 1.15E-06 | 2.865227 | 0.054 | 0.008 | 0.01716883 | 0.77748077 | 2.29E-06 |
| Hbb-bs | 1.41E-06 | -5.0454894 | 0 | 0.161 | 0.02108773 | 0.00023152 | -3.5832987 | 0.011 | 0.145 | 1 | 0.00023152 | 2.82E-06 |
| Obp2a | 5.31E-06 | -4.0952639 | 0.008 | 0.159 | 0.07954027 | 4.23E-05 | -4.9208589 | 0 | 0.154 | 0.63373872 | 4.23E-05 | 1.06E-05 |
| Wfdc18 | 6.57E-06 | 2.1162295 | 0.074 | 0.018 | 0.09831121 | 0.00025558 | 2.20915614 | 0.075 | 0.021 | 1 | 0.00025558 | 1.31E-05 |
| Hba-a2 | 4.03E-05 | -3.0158563 | 0.008 | 0.137 | 0.6036437 | 1.14E-05 | -3.339836 | 0.011 | 0.192 | 0.170473 | 4.03E-05 | 2.28E-05 |
| Acan | 3.58E-05 | 1.70416246 | 0.09 | 0.028 | 0.53559649 | 0.60184277 | 1.13424947 | 0.011 | 0.006 | 1 | 0.60184277 | 7.15E-05 |
| Obp2b | 5.15E-05 | -4.6917997 | 0 | 0.119 | 0.77082434 | 0.0016809 | -4.1010353 | 0 | 0.096 | 1 | 0.0016809 | 0.00010295 |
| Col3a1 | 7.39E-05 | -4.079893 | 0 | 0.114 | 1 | 0.00014906 | -4.0352469 | 0 | 0.134 | 1 | 0.00014906 | 0.00014778 |
| Kctd3 | 8.05E-05 | 4.30134091 | 0.016 | 0.002 | 1 | 0.00609716 | 3.98070422 | 0.011 | 0.001 | 1 | 0.00609716 | 0.00016096 |
| Mgp | 0.00017056 | -4.10805 | 0 | 0.104 | 1 | 0.07728252 | -1.7986363 | 0 | 0.032 | 1 | 0.07728252 | 0.00034109 |
| Sorbs1 | 0.00021106 | 3.73946202 | 0.016 | 0.002 | 1 | 0.00476444 | 4.15758198 | 0.011 | 0.001 | 1 | 0.00476444 | 0.00042207 |
| S100a9 | 0.00222237 | -3.502329 | 0 | 0.071 | 1 | 0.00091696 | -3.6681373 | 0 | 0.106 | 1 | 0.00222237 | 0.00183309 |
| Sparc | 0.00599039 | -0.5485997 | 0.205 | 0.337 | 1 | 0.00093423 | -0.9389681 | 0.097 | 0.255 | 1 | 0.00599039 | 0.00186758 |
| mt-Atp6 | 0.09637172 | 1.30650261 | 0.033 | 0.015 | 1 | 0.00135428 | 1.93967695 | 0.054 | 0.014 | 1 | 0.09637172 | 0.00270672 |
| Fn1 | 0.00240054 | -3.3568706 | 0 | 0.07 | 1 | 0.16708041 | -1.0969332 | 0 | 0.02 | 1 | 0.16708041 | 0.00479532 |
| Dnajc10 | 0.00307357 | 3.17009637 | 0.016 | 0.003 | 1 | 0.05140309 | 4.01817892 | 0.011 | 0.002 | 1 | 0.05140309 | 0.0061377 |
| Mb | 0.00409308 | 2.96573787 | 0.016 | 0.003 | 1 | 0.01484589 | 2.28882651 | 0.022 | 0.004 | 1 | 0.01484589 | 0.0081694 |
| Timp2 | 0.28738973 | 1.25382745 | 0.041 | 0.026 | 1 | 0.00430544 | 2.21190914 | 0.065 | 0.022 | 1 | 0.28738973 | 0.00859234 |
| Col2a1 | 0.0066362 | -3.0499139 | 0 | 0.057 | 1 | 0.27654416 | -0.5903191 | 0 | 0.013 | 1 | 0.27654416 | 0.01322837 |
| Ccn2 | 0.00697257 | -2.9985212 | 0 | 0.056 | 1 | 0.11988989 | -1.6362687 | 0 | 0.025 | 1 | 0.11988989 | 0.01389653 |
| Maf | 0.00722168 | 2.39903758 | 0.033 | 0.009 | 1 | 0.30891486 | 1.82316294 | 0.022 | 0.011 | 1 | 0.30891486 | 0.0143912 |
| Scgb1c1 | 0.00723426 | -3.3028152 | 0 | 0.056 | 1 | 0.00965441 | -3.3125021 | 0 | 0.067 | 1 | 0.00965441 | 0.01441618 |
| Col11a2 | 0.00752888 | -2.9654456 | 0 | 0.055 | 1 | 0.1069534 | -1.5428577 | 0 | 0.027 | 1 | 0.1069534 | 0.01500107 |
| Spp1 | 0.00806803 | -2.3091567 | 0.008 | 0.069 | 1 | 0.01159451 | -2.8502375 | 0 | 0.064 | 1 | 0.01159451 | 0.01607096 |
| Rexo2 | 0.00831574 | 2.26320578 | 0.025 | 0.006 | 1 | 0.26243201 | 2.17334929 | 0.011 | 0.004 | 1 | 0.26243201 | 0.01656234 |
| Lcn3 | 0.00871068 | -3.2532479 | 0 | 0.053 | 1 | 0.03861717 | -2.7731554 | 0 | 0.044 | 1 | 0.03861717 | 0.01734549 |
| S100a8 | 0.00956235 | -2.8338187 | 0 | 0.052 | 1 | 0.03425708 | -1.7208022 | 0.011 | 0.065 | 1 | 0.03425708 | 0.01903326 |
| Itm2b | 0.0109543 | -2.859971 | 0 | 0.05 | 1 | 0.0296779 | -2.4596858 | 0 | 0.048 | 1 | 0.0296779 | 0.02178861 |
| Lsm14b | 0.01144453 | 2.81101528 | 0.016 | 0.003 | 1 | 0.03478718 | 4.22171231 | 0.011 | 0.002 | 1 | 0.03478718 | 0.02275808 |
| Gnai2 | 0.01144453 | 2.81101528 | 0.016 | 0.003 | 1 | 0.31701865 | 2.23820044 | 0.011 | 0.004 | 1 | 0.31701865 | 0.02275808 |
| Nupr1 | 0.0115695 | 2.58513387 | 0.025 | 0.006 | 1 | 0.66283645 | 1.44273919 | 0.011 | 0.007 | 1 | 0.66283645 | 0.02300515 |
| Isyna1 | 0.01241776 | 3.15449952 | 0.016 | 0.003 | 1 | 0.08206257 | 3.09618143 | 0.011 | 0.002 | 1 | 0.08206257 | 0.02468132 |
| Irs1 | 0.02253045 | 2.69365833 | 0.016 | 0.004 | 1 | 0.01598441 | 3.92625643 | 0.011 | 0.001 | 1 | 0.02253045 | 0.03171331 |
| Ccdc80 | 0.58660513 | 0.60090119 | 0.033 | 0.025 | 1 | 0.01829186 | 2.47169057 | 0.032 | 0.009 | 1 | 0.58660513 | 0.03624913 |
| Ngp | 0.07991492 | -1.7241942 | 0 | 0.025 | 1 | 0.01875665 | -2.6609307 | 0 | 0.056 | 1 | 0.07991492 | 0.03716148 |
| Obp1a | 0.01909491 | -2.8323518 | 0 | 0.043 | 1 | 0.01986587 | -3.0662099 | 0 | 0.055 | 1 | 0.01986587 | 0.0378252 |
| Bpifa1 | 0.04518212 | 1.32733612 | 0.057 | 0.028 | 1 | 0.02001471 | 1.96346751 | 0.043 | 0.014 | 1 | 0.04518212 | 0.03962882 |
| Stk25 | 0.02258224 | 2.58513387 | 0.016 | 0.004 | 1 | 0.19387702 | 2.57261948 | 0.011 | 0.003 | 1 | 0.19387702 | 0.04465452 |
| Dmbt1 | 0.02476377 | 2.3186189 | 0.041 | 0.016 | 1 | 0.31503337 | -0.4063189 | 0 | 0.011 | 1 | 0.31503337 | 0.0489143 |
| Hbb-bt | 0.05720738 | -1.4439733 | 0.008 | 0.043 | 1 | 0.02513193 | -2.3342711 | 0 | 0.051 | 1 | 0.05720738 | 0.04963224 |
| Igfbp5 | 0.02550644 | -3.312317 | 0 | 0.039 | 1 | 0.06529922 | -2.1737244 | 0 | 0.035 | 1 | 0.06529922 | 0.05036231 |
| App | 0.02803675 | -2.3548285 | 0 | 0.038 | 1 | 0.05862163 | -1.9261864 | 0 | 0.037 | 1 | 0.05862163 | 0.05528745 |
| mt-Nd3 | 0.02895075 | 1.7336564 | 0.041 | 0.016 | 1 | 0.39187756 | 0.98533566 | 0.022 | 0.012 | 1 | 0.39187756 | 0.05706334 |
| Snrnp70 | 0.03190699 | 2.52373333 | 0.016 | 0.004 | 1 | 0.42064199 | 2.60789495 | 0.011 | 0.005 | 1 | 0.42064199 | 0.06279593 |
| Thbs1 | 0.05719243 | -1.9329633 | 0 | 0.029 | 1 | 0.03271817 | 2.2415208 | 0.032 | 0.01 | 1 | 0.05719243 | 0.06436587 |
| Bpifb6 | 0.04129086 | -2.4129046 | 0 | 0.033 | 1 | 0.03352287 | -2.4774658 | 0 | 0.046 | 1 | 0.04129086 | 0.06592195 |
| Bpifb4 | 0.06450601 | -2.0885209 | 0 | 0.027 | 1 | 0.03945132 | -2.4584074 | 0 | 0.044 | 1 | 0.06450601 | 0.07734624 |
| mt-Co3 | 0.03948261 | 1.34493018 | 0.041 | 0.017 | 1 | 0.6861363 | 0.72307461 | 0.011 | 0.016 | 1 | 0.6861363 | 0.07740635 |
| Lcn11 | 0.05991079 | -2.2132323 | 0 | 0.028 | 1 | 0.03984473 | -2.7111735 | 0 | 0.043 | 1 | 0.05991079 | 0.07810185 |
| Tcf4 | 0.04308647 | 2.46483964 | 0.016 | 0.004 | 1 | 0.07583607 | 3.83984168 | 0.011 | 0.002 | 1 | 0.07583607 | 0.0843165 |
| Bgn | 0.04321472 | -1.5830477 | 0.008 | 0.047 | 1 | 0.66944885 | 0.98843158 | 0.054 | 0.045 | 1 | 0.66944885 | 0.08456194 |
| Wwp2 | 0.60674091 | 0.97941281 | 0.016 | 0.011 | 1 | 0.04376671 | 2.28882651 | 0.022 | 0.006 | 1 | 0.60674091 | 0.0856179 |
| Tcap | 0.04510606 | 2.66575233 | 0.016 | 0.004 | 1 | 0.15990366 | 2.77425334 | 0.011 | 0.003 | 1 | 0.15990366 | 0.08817756 |
| Tmsb4x | 0.09627032 | -1.2706825 | 0.008 | 0.036 | 1 | 0.04882105 | -2.0158236 | 0 | 0.04 | 1 | 0.09627032 | 0.09525861 |
| Apoe | 0.06662965 | -1.8279421 | 0 | 0.027 | 1 | 0.05937671 | -2.0018507 | 0 | 0.037 | 1 | 0.06662965 | 0.11522782 |
| mt-Nd1 | 0.05956009 | 1.64926421 | 0.033 | 0.013 | 1 | 0.88200309 | 0.36373132 | 0.011 | 0.012 | 1 | 0.88200309 | 0.11557277 |
| Vmo1 | 0.09377263 | -1.6428482 | 0 | 0.023 | 1 | 0.06206957 | -2.0787442 | 0 | 0.036 | 1 | 0.09377263 | 0.12028651 |
| Acp5 | 0.06244749 | -1.9761126 | 0 | 0.028 | 1 | 0.15621461 | -1.2436686 | 0 | 0.021 | 1 | 0.15621461 | 0.12099529 |
| Ugt2a1 | 0.06500983 | 2.15449952 | 0.016 | 0.005 | 1 | 0.71711845 | 1.3061045 | 0.011 | 0.007 | 1 | 0.71711845 | 0.12579337 |
| Camp | 0.13491461 | -1.2488562 | 0 | 0.018 | 1 | 0.07413389 | -1.7175998 | 0 | 0.033 | 1 | 0.13491461 | 0.14277194 |
| Car2 | 0.07990896 | -1.6089806 | 0 | 0.025 | 1 | 0.11450633 | -1.2932707 | 0 | 0.026 | 1 | 0.11450633 | 0.15343248 |
| Gpx3 | 0.1273232 | -0.5584157 | 0.016 | 0.046 | 1 | 0.08566996 | -1.294704 | 0.011 | 0.049 | 1 | 0.1273232 | 0.16400058 |
| Ifitm10 | 0.11022055 | 1.6329624 | 0.025 | 0.01 | 1 | 0.09225745 | 3.53818598 | 0.011 | 0.002 | 1 | 0.11022055 | 0.17600346 |
| Serpinf1 | 0.09738502 | -1.5096897 | 0 | 0.022 | 1 | 0.99961227 | 0.52463945 | 0.022 | 0.021 | 1 | 0.99961227 | 0.1852862 |
| Ctsk | 0.1040807 | -1.5387626 | 0 | 0.021 | 1 | 0.25901129 | -0.4887811 | 0 | 0.014 | 1 | 0.25901129 | 0.19732861 |
| Vim | 0.10549221 | -1.2289261 | 0.008 | 0.035 | 1 | 0.65771088 | 0.7504066 | 0.022 | 0.03 | 1 | 0.65771088 | 0.19985582 |
| Mmp13 | 0.10658251 | -1.4030568 | 0 | 0.021 | 1 | 0.21163192 | -0.7197355 | 0 | 0.017 | 1 | 0.21163192 | 0.20180518 |
| Gpx6 | 0.11341812 | -1.5707843 | 0 | 0.02 | 1 | 0.10800267 | -1.5926517 | 0 | 0.027 | 1 | 0.11341812 | 0.20434076 |
| mt-Atp8 | 0.76961151 | 0.47207121 | 0.025 | 0.021 | 1 | 0.12358831 | 1.53721414 | 0.043 | 0.02 | 1 | 0.76961151 | 0.23190255 |
| Ibsp | 0.19897605 | -0.8220415 | 0.008 | 0.027 | 1 | 0.12674926 | -1.3272847 | 0 | 0.024 | 1 | 0.19897605 | 0.23743315 |
| Dmp1 | 0.1437404 | -1.2177337 | 0 | 0.017 | 1 | 0.19951549 | -0.7986363 | 0 | 0.017 | 1 | 0.19951549 | 0.2668195 |
| Col12a1 | 0.14729985 | -1.1249238 | 0 | 0.017 | 1 | 0.87364777 | 1.11890151 | 0.011 | 0.009 | 1 | 0.87364777 | 0.27290245 |
| Kctd12 | 0.15548303 | -1.0410513 | 0 | 0.016 | 1 | 0.1914406 | -0.842418 | 0 | 0.018 | 1 | 0.1914406 | 0.28679108 |
| Dcn | 0.20387477 | -0.7263018 | 0.016 | 0.039 | 1 | 0.15620863 | -1.0704001 | 0 | 0.021 | 1 | 0.20387477 | 0.28801613 |
| Mepe | 0.16016234 | -1.1201229 | 0 | 0.016 | 1 | 0.32229498 | -0.1517461 | 0 | 0.01 | 1 | 0.32229498 | 0.29467271 |
| Cd24a | 0.16255831 | -1.0712133 | 0 | 0.016 | 1 | 0.27755492 | -0.3467621 | 0 | 0.013 | 1 | 0.27755492 | 0.29869141 |
| Serf2 | 0.17183188 | 1.76292099 | 0.016 | 0.006 | 1 | 0.49003306 | 1.72694762 | 0.011 | 0.005 | 1 | 0.49003306 | 0.31413756 |
| Dbp | 0.17428537 | -0.9573934 | 0 | 0.015 | 1 | 0.33232202 | -0.1067586 | 0 | 0.01 | 1 | 0.33232202 | 0.31819535 |
| Col8a1 | 0.17604315 | 1.83567733 | 0.016 | 0.006 | 1 | 0.51582968 | 2.43321642 | 0.011 | 0.006 | 1 | 0.51582968 | 0.32109512 |
| Hist1h1e | 0.17870024 | -0.8513061 | 0 | 0.015 | 1 | 0.22147125 | -0.631739 | 0 | 0.016 | 1 | 0.22147125 | 0.32546671 |
| Sost | 0.17960187 | -0.9840613 | 0 | 0.015 | 1 | 0.20798696 | -0.9335659 | 0 | 0.017 | 1 | 0.20798696 | 0.32694691 |
| Smoc2 | 0.77399982 | 1.20782902 | 0.016 | 0.013 | 1 | 0.18112504 | 1.61505474 | 0.022 | 0.009 | 1 | 0.77399982 | 0.32944379 |
| Acta1 | 0.29481511 | 1.28426739 | 0.016 | 0.008 | 1 | 0.18443982 | 1.21843718 | 0.032 | 0.015 | 1 | 0.29481511 | 0.3348616 |
| mt-Nd5 | 0.18723957 | 1.57264293 | 0.025 | 0.012 | 1 | 0.92402009 | 0.51121893 | 0.011 | 0.012 | 1 | 0.92402009 | 0.33942048 |
| Nucb1 | 0.63020101 | 0.77479509 | 0.008 | 0.013 | 1 | 0.19593355 | 2.11128833 | 0.022 | 0.009 | 1 | 0.63020101 | 0.35347714 |
| Igfbp4 | 0.70401111 | 0.38077537 | 0.008 | 0.012 | 1 | 0.19678086 | -0.7986363 | 0 | 0.018 | 1 | 0.70401111 | 0.35483901 |
| Atox1 | 0.19683673 | 1.73946202 | 0.025 | 0.012 | 1 | 0.56474669 | 1.5212726 | 0.011 | 0.006 | 1 | 0.56474669 | 0.35492876 |
| Gsn | 0.23653248 | -0.8011973 | 0.008 | 0.025 | 1 | 0.20799074 | -1.0296625 | 0 | 0.017 | 1 | 0.23653248 | 0.37272134 |
| Mup5 | 0.21270497 | 1.06794949 | 0.033 | 0.018 | 1 | 0.68195728 | -0.139035 | 0.011 | 0.016 | 1 | 0.68195728 | 0.38016654 |
| Mt1 | 0.21790264 | -0.68572 | 0 | 0.012 | 1 | 0.22540001 | -0.7239975 | 0 | 0.016 | 1 | 0.22540001 | 0.38832372 |
| Htra1 | 0.55406764 | 1.06430171 | 0.016 | 0.011 | 1 | 0.22819513 | 1.65877612 | 0.022 | 0.009 | 1 | 0.55406764 | 0.40431725 |

| | | | | | | | | | | | | |
|---|---|---|---|---|---|---|---|---|---|---|---|---|
| Srsf6 | 0.61180721 | 0.97941281 | 0.016 | 0.011 | 1 | 0.22821232 | 1.95322348 | 0.022 | 0.009 | 1 | 0.61180721 | 0.40434377 |
| Tpt1 | 0.23070555 | -0.6461917 | 0 | 0.012 | 1 | 0.7117158 | 1.35921584 | 0.011 | 0.007 | 1 | 0.7117158 | 0.40818605 |
| Chad | 0.77210277 | 0.61417355 | 0.025 | 0.029 | 1 | 0.23456188 | 2.45232524 | 0.011 | 0.003 | 1 | 0.77210277 | 0.41410448 |
| Mmp2 | 0.44396372 | -0.148692 | 0.008 | 0.017 | 1 | 0.23683289 | -0.5809509 | 0 | 0.015 | 1 | 0.44396372 | 0.41757597 |
| Id3 | 0.23743339 | 1.6624396 | 0.025 | 0.013 | 1 | 0.92091671 | 1.3294685 | 0.011 | 0.012 | 1 | 0.92091671 | 0.41849217 |
| Bpifb3 | 0.25027271 | -0.6484163 | 0.008 | 0.024 | 1 | 0.23768493 | -1.1517461 | 0 | 0.015 | 1 | 0.25027271 | 0.41887574 |
| Mup4 | 0.24073829 | 1.2961976 | 0.025 | 0.013 | 1 | 0.62940813 | 0.92625643 | 0.022 | 0.015 | 1 | 0.62940813 | 0.42352165 |
| Tpm1 | 0.86841922 | 1.02832241 | 0.025 | 0.023 | 1 | 0.24608458 | 0.98905156 | 0.043 | 0.024 | 1 | 0.86841922 | 0.43161154 |
| mt-Nd6 | 0.25095926 | -0.7616689 | 0 | 0.011 | 1 | 0.77054606 | 0.96689842 | 0.011 | 0.008 | 1 | 0.77054606 | 0.43893798 |
| Slc4a1 | 0.25229778 | -0.3831573 | 0 | 0.011 | 1 | 0.26759788 | -0.3018496 | 0 | 0.013 | 1 | 0.26759788 | 0.44094139 |
| Mbd2 | 0.59022263 | 1.07894849 | 0.016 | 0.011 | 1 | 0.25488636 | 1.60077052 | 0.022 | 0.01 | 1 | 0.59022263 | 0.44480567 |
| Cytl1 | 0.51296603 | 0.76439992 | 0.025 | 0.017 | 1 | 0.25981908 | 2.74254448 | 0.011 | 0.004 | 1 | 0.51296603 | 0.4521322 |
| Ptprs | 0.2632844 | -0.3670376 | 0 | 0.01 | 1 | 0.85055448 | 1.19732816 | 0.011 | 0.009 | 1 | 0.85055448 | 0.45725013 |
| Lrp1 | 0.48249747 | 0.96403759 | 0.025 | 0.016 | 1 | 0.26662781 | -0.3522791 | 0 | 0.013 | 1 | 0.48249747 | 0.46216523 |
| Arf5 | 0.55914648 | 1.07894849 | 0.016 | 0.011 | 1 | 0.2666284 | -0.3903184 | 0 | 0.013 | 1 | 0.55914648 | 0.4621661 |
| Serinc3 | 0.73120439 | 0.42921573 | 0.008 | 0.012 | 1 | 0.2703376 | 2.30958507 | 0.022 | 0.01 | 1 | 0.73120439 | 0.46759278 |
| Tnc | 0.27191501 | -0.3091567 | 0 | 0.01 | 1 | 0.99389182 | 1.49131937 | 0.011 | 0.011 | 1 | 0.99389182 | 0.46989225 |
| Jund | 0.70754883 | 0.3405115 | 0.008 | 0.012 | 1 | 0.29999346 | -0.2613706 | 0 | 0.011 | 1 | 0.70754883 | 0.50999084 |
| Lcn4 | 0.3124971 | 1.22256379 | 0.033 | 0.02 | 1 | 0.67212551 | 0.47494355 | 0.011 | 0.016 | 1 | 0.67212551 | 0.52733976 |
| Cst3 | 0.73364284 | 0.52749525 | 0.016 | 0.021 | 1 | 0.32352647 | -0.1067586 | 0 | 0.01 | 1 | 0.73364284 | 0.54238356 |
| Per1 | 0.81077539 | 0.50878299 | 0.008 | 0.01 | 1 | 0.32399424 | 2.14198512 | 0.011 | 0.004 | 1 | 0.81077539 | 0.54301622 |
| Fth1 | 0.34144722 | -0.4265795 | 0.008 | 0.02 | 1 | 0.51648053 | 0.45232524 | 0.011 | 0.02 | 1 | 0.51648053 | 0.56630824 |
| Arglu1 | 0.65641917 | 0.22605278 | 0.008 | 0.013 | 1 | 0.38715973 | 1.99744748 | 0.022 | 0.012 | 1 | 0.65641917 | 0.6244268 |
| mt-Co1 | 0.4012923 | -0.3507358 | 0.008 | 0.018 | 1 | 0.70726341 | 0.64772093 | 0.022 | 0.016 | 1 | 0.70726341 | 0.6415491 |
| Oaz1 | 0.73052547 | 0.32079327 | 0.008 | 0.012 | 1 | 0.40973964 | 1.81324747 | 0.022 | 0.012 | 1 | 0.73052547 | 0.65159271 |
| Nisch | 0.60372337 | 0.19580649 | 0.008 | 0.014 | 1 | 0.42160052 | 1.65323794 | 0.022 | 0.012 | 1 | 0.60372337 | 0.66545404 |
| Postn | 0.42254171 | -0.153399 | 0.008 | 0.018 | 1 | 0.73308392 | 0.97148569 | 0.011 | 0.015 | 1 | 0.73308392 | 0.66654193 |
| Cald1 | 0.55344332 | 1.13141591 | 0.016 | 0.011 | 1 | 0.8412015 | 1.22171231 | 0.011 | 0.009 | 1 | 0.8412015 | 0.80058713 |
| Col6a3 | 0.62162808 | 1.02832241 | 0.016 | 0.012 | 1 | 0.93874388 | 1.03009224 | 0.011 | 0.01 | 1 | 0.93874388 | 0.85683469 |
| Mmp9 | 0.77869606 | 1.20782902 | 0.016 | 0.014 | 1 | 0.70068983 | 1.45232524 | 0.011 | 0.007 | 1 | 0.77869606 | 0.91041342 |
| mt-Nd4l | 0.72028034 | 0.72211478 | 0.016 | 0.013 | 1 | 0.94503263 | 1.29456288 | 0.011 | 0.01 | 1 | 0.94503263 | 0.92175691 |
| Pcolce | 0.727223 | 0.43627049 | 0.008 | 0.012 | 1 | 0.9637403 | 1.47820387 | 0.011 | 0.01 | 1 | 0.9637403 | 0.92559271 |
| Maged1 | 0.79195385 | 1.10120129 | 0.008 | 0.011 | 1 | 0.93065875 | 0.98765698 | 0.011 | 0.01 | 1 | 0.93065875 | 0.9567168 |
| Serpine2 | 0.96312725 | 0.47933921 | 0.016 | 0.017 | 1 | 0.80307752 | 1.2802645 | 0.011 | 0.008 | 1 | 0.96312725 | 0.96122154 |
| mt-Co2 | 0.89238763 | 0.47518158 | 0.033 | 0.035 | 1 | 0.90044382 | 0.57785612 | 0.032 | 0.035 | 1 | 0.90044382 | 0.98841958 |

**Spatial seq: cluster 12**

| | cKO_p_val | cKO_avg_log2FC | cKO_pct.1 | cKO_pct.2 | cKO_p_val_adj | WT_p_val | WT_avg_log2FC | WT_pct.1 | WT_pct.2 | WT_p_val_adj | max_pval | minimump_p_val |
|---|---|---|---|---|---|---|---|---|---|---|---|---|
| Col27a1 | 0 | 5.1548959 | 0.967 | 0.033 | 0 | 0 | 7.48699277 | 0.789 | 0.006 | 0 | 0 | 0 |
| Col10a1 | 6.20E-12 | 2.90790151 | 0.059 | 0.008 | 9.28E-08 | 0.00718628 | 3.68969152 | 0.018 | 0.002 | 1 | 0.00718628 | 1.24E-11 |
| Col1a2 | 1.16E-10 | -1.5254185 | 0.163 | 0.434 | 1.73E-06 | 0.18057147 | -0.4459191 | 0.404 | 0.487 | 1 | 0.18057147 | 2.31E-10 |
| Col1a1 | 1.60E-10 | -1.4415842 | 0.34 | 0.566 | 2.39E-06 | 0.00700677 | -0.7493775 | 0.421 | 0.621 | 1 | 0.00700677 | 3.20E-10 |
| Obp2a | 2.54E-07 | -4.4245729 | 0.007 | 0.159 | 0.00380656 | 0.00136273 | -4.2118716 | 0 | 0.154 | 1 | 0.00136273 | 5.08E-07 |
| Zdhhc3 | 3.91E-07 | 3.77938677 | 0.02 | 0.002 | 0.0058615 | 0.0001109 | 5.06820314 | 0.018 | 0.001 | 1 | 0.0001109 | 7.83E-07 |
| Hbb-bs | 4.48E-06 | -3.0516037 | 0.026 | 0.162 | 0.06706978 | 0.80578468 | -0.5500355 | 0.14 | 0.144 | 1 | 0.80578468 | 8.96E-06 |
| Obp2b | 5.67E-06 | -5.0211087 | 0 | 0.119 | 0.08491089 | 0.0140068 | -3.392048 | 0 | 0.096 | 1 | 0.0140068 | 1.13E-05 |
| F13a1 | 0.01549779 | 2.16467692 | 0.02 | 0.005 | 1 | 2.30E-05 | 5.33123755 | 0.018 | 0.001 | 0.3439149 | 0.01549779 | 4.59E-05 |
| Mapre2 | 0.00063675 | 3.27688643 | 0.013 | 0.002 | 1 | 0.00016975 | 4.93069962 | 0.018 | 0.001 | 1 | 0.00063675 | 0.00033948 |
| Mgp | 0.00021924 | -2.8511837 | 0.013 | 0.104 | 1 | 0.16703926 | -1.089649 | 0 | 0.032 | 1 | 0.16703926 | 0.00043844 |
| Col11a1 | 0.1686494 | 1.17650236 | 0.026 | 0.013 | 1 | 0.0002669 | 2.86656928 | 0.053 | 0.008 | 1 | 0.1686494 | 0.00053373 |
| Hba-a2 | 0.00040411 | -1.9279724 | 0.039 | 0.137 | 1 | 0.14909385 | -1.044497 | 0.123 | 0.191 | 1 | 0.14909385 | 0.00080805 |
| Fn1 | 0.00066746 | -3.6861796 | 0 | 0.07 | 1 | 0.27986298 | -0.3879459 | 0 | 0.02 | 1 | 0.27986298 | 0.00133448 |
| Atxn10 | 0.00100947 | 3.10131486 | 0.02 | 0.004 | 1 | 0.0430548 | 3.41285131 | 0.018 | 0.003 | 1 | 0.0430548 | 0.00201791 |
| Plec | 0.00510966 | 3.47560602 | 0.02 | 0.004 | 1 | 0.00108706 | 3.34573712 | 0.035 | 0.005 | 1 | 0.00510966 | 0.00217293 |
| Sdf4 | 0.38794941 | 1.35312201 | 0.013 | 0.007 | 1 | 0.00198807 | 3.12960369 | 0.035 | 0.005 | 1 | 0.38794941 | 0.00397218 |
| Spp1 | 0.0022854 | -2.6384657 | 0.007 | 0.069 | 1 | 0.15061487 | -1.1402751 | 0.018 | 0.064 | 1 | 0.15061487 | 0.00456558 |
| Col2a1 | 0.0023442 | -3.3792229 | 0 | 0.057 | 1 | 0.3947477 | 0.11866821 | 0 | 0.013 | 1 | 0.3947477 | 0.0046829 |
| Ccn2 | 0.00249162 | -3.3278303 | 0 | 0.056 | 1 | 0.223832 | -0.9272814 | 0 | 0.025 | 1 | 0.223832 | 0.00497704 |
| Lcn3 | 0.00327842 | -3.582557 | 0 | 0.054 | 1 | 0.10573782 | -2.0641681 | 0 | 0.044 | 1 | 0.10573782 | 0.00654609 |
| Prpf6 | 0.00520726 | 3.04242117 | 0.013 | 0.002 | 1 | 0.01025747 | 4.48324064 | 0.018 | 0.002 | 1 | 0.01025747 | 0.01038741 |
| Cytl1 | 0.00565238 | 1.45295728 | 0.046 | 0.017 | 1 | 0.08380098 | 3.45153178 | 0.018 | 0.004 | 1 | 0.08380098 | 0.01127281 |
| mt-Nd1 | 0.46662389 | 0.30365334 | 0.007 | 0.013 | 1 | 0.00574647 | 2.67588572 | 0.053 | 0.012 | 1 | 0.46662389 | 0.01145993 |
| Cd24a | 0.68436463 | 0.94144353 | 0.02 | 0.016 | 1 | 0.00624688 | 2.37890398 | 0.053 | 0.012 | 1 | 0.68436463 | 0.01245474 |
| Acan | 0.07068618 | 1.14760341 | 0.052 | 0.028 | 1 | 0.00680086 | 2.43593492 | 0.035 | 0.006 | 1 | 0.07068618 | 0.01355546 |
| Hnrnpul1 | 0.00693749 | 2.45745867 | 0.02 | 0.005 | 1 | 0.20890576 | 3.10472902 | 0.018 | 0.005 | 1 | 0.20890576 | 0.01382686 |
| Scgb1c1 | 0.00780698 | -2.6310646 | 0.007 | 0.056 | 1 | 0.04295444 | -2.6035148 | 0 | 0.067 | 1 | 0.04295444 | 0.01555302 |
| Obp1a | 0.00861354 | -3.1616608 | 0 | 0.043 | 1 | 0.22002432 | -0.7705806 | 0.018 | 0.055 | 1 | 0.22002432 | 0.01715289 |
| Fgfr3 | 0.01178032 | 2.36757317 | 0.026 | 0.008 | 1 | 0.20081524 | 3.12960369 | 0.018 | 0.005 | 1 | 0.20081524 | 0.02342187 |
| G0s2 | 0.01870159 | 2.05146631 | 0.02 | 0.005 | 1 | 0.01283102 | 4.36038389 | 0.018 | 0.002 | 1 | 0.01870159 | 0.0254974 |
| Clk1 | 0.07414132 | 2.55056808 | 0.013 | 0.004 | 1 | 0.01447129 | 3.80516873 | 0.018 | 0.002 | 1 | 0.07414132 | 0.02873316 |
| Rab11fip4 | 0.06717764 | 2.13553058 | 0.013 | 0.004 | 1 | 0.01448304 | 3.74627505 | 0.018 | 0.002 | 1 | 0.06717764 | 0.02875632 |
| Bpifa1 | 0.38393931 | 0.70375906 | 0.039 | 0.028 | 1 | 0.01585252 | 1.98413888 | 0.053 | 0.014 | 1 | 0.38393931 | 0.03145374 |
| Bgn | 0.01809325 | -1.9123568 | 0.007 | 0.047 | 1 | 0.31072154 | -0.4833129 | 0.018 | 0.045 | 1 | 0.31072154 | 0.03585913 |
| Aqp1 | 0.01974289 | 2.19442427 | 0.02 | 0.005 | 1 | 0.27091406 | 2.44761673 | 0.018 | 0.006 | 1 | 0.27091406 | 0.03909599 |
| Dmbt1 | 0.02098568 | 1.56129448 | 0.039 | 0.016 | 1 | 0.43198052 | 0.30266839 | 0 | 0.011 | 1 | 0.43198052 | 0.04153097 |
| Col11a2 | 0.02211687 | -1.7071131 | 0.013 | 0.055 | 1 | 0.20736993 | -0.8338704 | 0 | 0.027 | 1 | 0.20736993 | 0.04374458 |
| Hbb-bt | 0.02553206 | -1.7732823 | 0.007 | 0.043 | 1 | 0.93327498 | 0.70223071 | 0.053 | 0.051 | 1 | 0.93327498 | 0.05041223 |
| Camk1 | 0.12478199 | 2.04242117 | 0.013 | 0.005 | 1 | 0.02892742 | 3.55724122 | 0.018 | 0.003 | 1 | 0.12478199 | 0.05701804 |
| Mxra8 | 0.02906564 | 1.77938677 | 0.033 | 0.013 | 1 | 0.4423766 | 0.68969152 | 0 | 0.01 | 1 | 0.4423766 | 0.05728647 |
| mt-Co1 | 0.63956795 | 0.65423164 | 0.013 | 0.018 | 1 | 0.03005767 | 2.36776342 | 0.053 | 0.016 | 1 | 0.63956795 | 0.05921187 |
| Khdc4 | 0.08315532 | 2.1548959 | 0.013 | 0.004 | 1 | 0.03109592 | 3.53215024 | 0.018 | 0.003 | 1 | 0.08315532 | 0.06122489 |
| Crebzf | 0.03171626 | 2.60946177 | 0.013 | 0.003 | 1 | 0.16730868 | 2.71770589 | 0.018 | 0.005 | 1 | 0.16730868 | 0.06242659 |
| mt-Cytb | 0.24154746 | -0.7980421 | 0.007 | 0.02 | 1 | 0.03266373 | 1.80516873 | 0.053 | 0.016 | 1 | 0.24154746 | 0.06426054 |
| Lcn11 | 0.03495151 | -2.5425413 | 0 | 0.028 | 1 | 0.10792924 | -2.0021862 | 0 | 0.043 | 1 | 0.10792924 | 0.06868141 |
| Acp5 | 0.0367656 | -2.3054216 | 0 | 0.028 | 1 | 0.26743112 | -0.5346813 | 0 | 0.021 | 1 | 0.26743112 | 0.07217949 |
| Bpifb4 | 0.0382493 | -2.4178299 | 0 | 0.027 | 1 | 0.10722939 | -1.7494201 | 0 | 0.044 | 1 | 0.10722939 | 0.07503558 |
| Timp2 | 0.04210842 | -2.0971302 | 0 | 0.026 | 1 | 0.25795336 | -0.4300477 | 0 | 0.022 | 1 | 0.25795336 | 0.08244371 |
| Fkbp8 | 0.04298379 | 2.43361193 | 0.013 | 0.003 | 1 | 0.07507295 | 3.18067787 | 0.018 | 0.004 | 1 | 0.07507295 | 0.08411998 |
| Rsrp1 | 0.23195453 | 0.91829986 | 0.039 | 0.024 | 1 | 0.04458673 | 2.06820314 | 0.053 | 0.018 | 1 | 0.23195453 | 0.08718549 |
| S100a8 | 0.07300471 | -0.8353231 | 0.02 | 0.052 | 1 | 0.04716237 | -2.0128808 | 0 | 0.065 | 1 | 0.07300471 | 0.09210045 |
| Car2 | 0.04965313 | -1.9382897 | 0 | 0.025 | 1 | 0.21705074 | -0.5842834 | 0 | 0.026 | 1 | 0.21705074 | 0.09684083 |
| Bpifb3 | 0.04995014 | -2.567127 | 0 | 0.025 | 1 | 0.35573206 | -0.4427588 | 0 | 0.015 | 1 | 0.35573206 | 0.09740526 |
| Obp1b | 0.43081817 | 0.48936792 | 0.039 | 0.028 | 1 | 0.05060643 | 1.58461766 | 0.07 | 0.028 | 1 | 0.43081817 | 0.09865185 |
| Timp3 | 0.06371647 | 1.20111892 | 0.046 | 0.023 | 1 | 0.47262912 | 1.89028535 | 0.018 | 0.009 | 1 | 0.47262912 | 0.12337314 |
| Cpxm1 | 0.06945202 | 2.14711855 | 0.02 | 0.007 | 1 | 0.37954749 | 2.18067787 | 0.018 | 0.007 | 1 | 0.37954749 | 0.13408046 |
| Gpx6 | 0.075996 | -1.9000933 | 0 | 0.02 | 1 | 0.20872705 | -0.8836644 | 0 | 0.027 | 1 | 0.20872705 | 0.14621661 |
| mt-Atp8 | 0.50113217 | -0.2758957 | 0.013 | 0.021 | 1 | 0.08387431 | 2.02089743 | 0.053 | 0.02 | 1 | 0.50113217 | 0.16071372 |
| Col9a3 | 0.08560545 | 0.94844503 | 0.065 | 0.039 | 1 | 0.78649924 | 1.51566212 | 0.018 | 0.014 | 1 | 0.78649924 | 0.1638826 |
| Camp | 0.09378709 | -1.5781652 | 0 | 0.018 | 1 | 0.16249564 | -1.0086125 | 0 | 0.033 | 1 | 0.16249564 | 0.17877816 |
| mt-Nd5 | 0.37087322 | 0.91520062 | 0.02 | 0.012 | 1 | 0.10114159 | 2.2302599 | 0.035 | 0.012 | 1 | 0.37087322 | 0.19205355 |
| Dmp1 | 0.10126221 | -1.5470427 | 0 | 0.017 | 1 | 0.99009997 | 0.9143978 | 0.018 | 0.017 | 1 | 0.99009997 | 0.19227038 |
| App | 0.10660948 | -1.0950824 | 0.013 | 0.038 | 1 | 0.43699068 | -0.2153483 | 0.018 | 0.037 | 1 | 0.43699068 | 0.20185338 |
| mt-Nd3 | 0.10775775 | 0.98064498 | 0.033 | 0.016 | 1 | 0.68318046 | 1.69432297 | 0.018 | 0.012 | 1 | 0.68318046 | 0.20390378 |
| Kctd12 | 0.11134777 | -1.3703604 | 0 | 0.016 | 1 | 0.98133335 | 0.87049498 | 0.018 | 0.018 | 1 | 0.98133335 | 0.21029722 |
| Igfbp7 | 0.11268331 | -1.3652715 | 0 | 0.016 | 1 | 0.23964714 | -0.5957107 | 0 | 0.024 | 1 | 0.23964714 | 0.21266909 |
| Mepe | 0.11540901 | -1.4494319 | 0 | 0.016 | 1 | 0.43886996 | 0.55724122 | 0 | 0.01 | 1 | 0.43886996 | 0.21749879 |
| Per1 | 0.73616159 | 1.19442427 | 0.013 | 0.01 | 1 | 0.11695708 | 2.85097242 | 0.018 | 0.004 | 1 | 0.73616159 | 0.2202352 |
| Ibsp | 0.11712092 | -1.1513506 | 0.007 | 0.027 | 1 | 0.23233966 | -0.6182974 | 0 | 0.024 | 1 | 0.23233966 | 0.22052454 |
| Tmsb4x | 0.12333518 | -1.0128613 | 0.013 | 0.036 | 1 | 0.87089263 | 0.69838772 | 0.035 | 0.04 | 1 | 0.87089263 | 0.2314588 |
| Arglu1 | 0.13093779 | 1.73499265 | 0.026 | 0.013 | 1 | 0.70115453 | 1.36776342 | 0.018 | 0.012 | 1 | 0.70115453 | 0.24473088 |
| Sost | 0.13252099 | -1.3133704 | 0 | 0.015 | 1 | 0.32472603 | -0.2245786 | 0 | 0.017 | 1 | 0.32472603 | 0.24748017 |
| Col9a2 | 0.13476753 | 0.91520062 | 0.059 | 0.036 | 1 | 0.41435937 | 0.27119016 | 0 | 0.012 | 1 | 0.41435937 | 0.25137277 |
| Ighm | 0.13742807 | -1.3080761 | 0 | 0.014 | 1 | 0.42083706 | 0.62858 | 0 | 0.011 | 1 | 0.42083706 | 0.25596966 |
| Irf2bp2 | 0.13909244 | 1.59947768 | 0.02 | 0.009 | 1 | 0.51186938 | 1.74627505 | 0.018 | 0.009 | 1 | 0.51186938 | 0.25883817 |
| Apoe | 0.29396285 | -0.5663881 | 0.013 | 0.027 | 1 | 0.14031062 | -1.2928633 | 0 | 0.037 | 1 | 0.29396285 | 0.26093416 |
| Mup5 | 0.16632387 | 0.96468987 | 0.033 | 0.018 | 1 | 0.33492871 | -0.433236 | 0 | 0.016 | 1 | 0.33492871 | 0.30498411 |
| Bpifb6 | 0.16752394 | -1.1533201 | 0.013 | 0.033 | 1 | 0.68290074 | -0.1809893 | 0.035 | 0.046 | 1 | 0.68290074 | 0.3069836 |
| Itm2b | 0.16917818 | -0.8615811 | 0.026 | 0.05 | 1 | 0.27705076 | -0.7494201 | 0.018 | 0.048 | 1 | 0.27705076 | 0.3097351 |
| Jund | 0.17361266 | -0.9954003 | 0 | 0.012 | 1 | 0.66079594 | 1.4534933 | 0.018 | 0.011 | 1 | 0.66079594 | 0.31708397 |
| Pcolce | 0.1791235 | -0.9000933 | 0 | 0.012 | 1 | 0.43886925 | 0.58923134 | 0 | 0.01 | 1 | 0.43886925 | 0.32616177 |
| Sh3bgrl | 0.32133497 | 1.51885922 | 0.013 | 0.007 | 1 | 0.18347165 | 2.67588572 | 0.018 | 0.005 | 1 | 0.32133497 | 0.33328145 |
| Arf5 | 0.19196663 | -0.8499699 | 0 | 0.011 | 1 | 0.76445941 | 1.32404204 | 0.018 | 0.013 | 1 | 0.76445941 | 0.34708208 |
| Serpinf1 | 0.19239571 | -0.8353231 | 0.007 | 0.022 | 1 | 0.47886094 | 1.23362675 | 0.035 | 0.021 | 1 | 0.47886094 | 0.34777531 |
| Maged1 | 0.19565008 | -0.8279435 | 0 | 0.011 | 1 | 0.45072821 | 0.68969152 | 0 | 0.01 | 1 | 0.45072821 | 0.35302121 |
| Slc4a1 | 0.19942204 | -0.7124663 | 0 | 0.011 | 1 | 0.38590356 | 0.40713766 | 0 | 0.013 | 1 | 0.38590356 | 0.35907493 |
| Mmp2 | 0.30923038 | -0.4780011 | 0.007 | 0.017 | 1 | 0.2031278 | 1.7224283 | 0.035 | 0.015 | 1 | 0.30923038 | 0.36499469 |
| Ptms | 0.75226323 | 0.36660524 | 0.02 | 0.024 | 1 | 0.21146486 | -0.8119926 | 0 | 0.027 | 1 | 0.75226323 | 0.37821234 |
| Ucp2 | 0.21258525 | -0.6882188 | 0 | 0.01 | 1 | 0.53450536 | 1.69664428 | 0.018 | 0.01 | 1 | 0.53450536 | 0.37997801 |
| Hist1h2ap | 0.21673153 | -0.6129307 | 0 | 0.01 | 1 | 0.44711905 | 0.76077462 | 0 | 0.01 | 1 | 0.44711905 | 0.38649051 |
| Tpm1 | 0.76680269 | 0.69901335 | 0.026 | 0.023 | 1 | 0.23182909 | -0.6350083 | 0 | 0.024 | 1 | 0.76680269 | 0.40991345 |
| Gpx3 | 0.24997753 | -0.5640211 | 0.026 | 0.046 | 1 | 0.26808581 | -0.5857167 | 0.018 | 0.049 | 1 | 0.26808581 | 0.43746629 |
| Ccdc80 | 0.26287985 | 0.76337445 | 0.039 | 0.025 | 1 | 0.50380989 | 1.83554238 | 0.018 | 0.009 | 1 | 0.50380989 | 0.45665389 |
| Ahnak | 0.42625979 | 0.13914183 | 0.013 | 0.023 | 1 | 0.30406753 | -0.1644576 | 0 | 0.018 | 1 | 0.42625979 | 0.515678 |
| Fau | 0.33472651 | 1.44548603 | 0.013 | 0.007 | 1 | 0.36319178 | 2.17096271 | 0.018 | 0.007 | 1 | 0.36319178 | 0.55741118 |

| | | | | | | | | | | | | |
|---|---|---|---|---|---|---|---|---|---|---|---|---|
| Srsf6 | 0.33641962 | 1.90024116 | 0.02 | 0.011 | 1 | 0.45932276 | 0.64193839 | 0 | 0.01 | 1 | 0.45932276 | 0.55966108 |
| Acta1 | 0.47718251 | 0.95495833 | 0.013 | 0.008 | 1 | 0.34554187 | -0.4075303 | 0 | 0.015 | 1 | 0.47718251 | 0.57168456 |
| Tpt1 | 0.35034136 | 1.04467715 | 0.02 | 0.011 | 1 | 0.36982216 | 2.66221078 | 0.018 | 0.007 | 1 | 0.36982216 | 0.57794366 |
| mt-Co3 | 0.36636739 | 1.24244578 | 0.026 | 0.017 | 1 | 0.93674797 | 0.84323678 | 0.018 | 0.016 | 1 | 0.93674797 | 0.59850971 |
| Lrp1 | 0.74517329 | 0.63472852 | 0.013 | 0.017 | 1 | 0.38493987 | 0.35670823 | 0 | 0.013 | 1 | 0.74517329 | 0.62170103 |
| Nisch | 0.44740254 | -0.1335026 | 0.007 | 0.014 | 1 | 0.39674554 | 0.34573712 | 0 | 0.012 | 1 | 0.44740254 | 0.63608406 |
| Igfbp5 | 0.39693634 | -1.3154827 | 0.026 | 0.039 | 1 | 0.4679281 | -0.4631783 | 0.018 | 0.035 | 1 | 0.4679281 | 0.63631422 |
| Cnn2 | 0.58840287 | 1.06972852 | 0.013 | 0.009 | 1 | 0.42413517 | 0.57635004 | 0 | 0.011 | 1 | 0.58840287 | 0.6683797 |
| Tnc | 0.67254049 | 0.96346983 | 0.013 | 0.01 | 1 | 0.42635341 | 0.60222868 | 0 | 0.011 | 1 | 0.67254049 | 0.67092959 |
| Mpo | 0.68129193 | 0.34475854 | 0.007 | 0.01 | 1 | 0.43084236 | 0.66221078 | 0 | 0.011 | 1 | 0.68129193 | 0.67605958 |
| Vmo1 | 0.43398226 | -0.3804846 | 0.013 | 0.022 | 1 | 0.45250593 | -0.368092 | 0.018 | 0.036 | 1 | 0.45250593 | 0.67962392 |
| Sdc4 | 0.68247147 | 0.88053349 | 0.007 | 0.01 | 1 | 0.43472717 | 2.39013124 | 0.018 | 0.008 | 1 | 0.68247147 | 0.68046663 |
| Fth1 | 0.52395548 | -0.1670322 | 0.013 | 0.02 | 1 | 0.43641068 | 1.16131254 | 0.035 | 0.02 | 1 | 0.52395548 | 0.68236708 |
| Bpgm | 0.4772925 | 1.66044569 | 0.013 | 0.008 | 1 | 0.43655234 | 0.70363071 | 0 | 0.011 | 1 | 0.4772925 | 0.68252674 |
| Mup4 | 0.44033931 | 1.23506625 | 0.02 | 0.013 | 1 | 0.89982548 | 0.62858 | 0.018 | 0.015 | 1 | 0.89982548 | 0.68677991 |
| Selenom | 0.62902089 | 1.36153425 | 0.013 | 0.009 | 1 | 0.44120242 | 0.68969152 | 0 | 0.01 | 1 | 0.62902089 | 0.68774527 |
| Sfrp2 | 0.64698611 | 0.98930984 | 0.013 | 0.009 | 1 | 0.4483182 | 0.66221078 | 0 | 0.01 | 1 | 0.64698611 | 0.69564719 |
| Dbp | 0.62204976 | 0.72953822 | 0.02 | 0.015 | 1 | 0.44831894 | 0.60222868 | 0 | 0.01 | 1 | 0.62204976 | 0.695648 |
| Mbd2 | 0.83160943 | 0.74963942 | 0.013 | 0.011 | 1 | 0.45072812 | 0.71065114 | 0 | 0.01 | 1 | 0.83160943 | 0.6983004 |
| Gpx1 | 0.6952055 | 0.36999583 | 0.007 | 0.01 | 1 | 0.45560717 | 0.80516873 | 0 | 0.01 | 1 | 0.6952055 | 0.70363644 |
| Nucb1 | 0.98203662 | 0.44548603 | 0.013 | 0.013 | 1 | 0.48461001 | 2.39766491 | 0.018 | 0.009 | 1 | 0.98203662 | 0.73437315 |
| Cst3 | 0.50143399 | 0.19818619 | 0.013 | 0.021 | 1 | 0.59015392 | 1.60877152 | 0.018 | 0.01 | 1 | 0.59015392 | 0.75143193 |
| Gsn | 0.52437957 | 0.69192393 | 0.033 | 0.025 | 1 | 0.96300716 | 0.6827721 | 0.018 | 0.017 | 1 | 0.96300716 | 0.77378521 |
| mt-Co2 | 0.54623677 | 0.14587252 | 0.026 | 0.036 | 1 | 0.99065136 | 0.69838772 | 0.035 | 0.035 | 1 | 0.99065136 | 0.79409893 |
| Col6a3 | 0.86641896 | 0.69901335 | 0.013 | 0.012 | 1 | 0.56394555 | 1.73907954 | 0.018 | 0.01 | 1 | 0.86641896 | 0.80985651 |
| mt-Nd4l | 0.97278236 | 0.39280571 | 0.013 | 0.013 | 1 | 0.57719577 | 1.41285131 | 0.018 | 0.01 | 1 | 0.97278236 | 0.82123658 |
| Serpinh1 | 0.79249716 | 0.34198146 | 0.033 | 0.029 | 1 | 0.60490772 | 0.1493399 | 0.018 | 0.029 | 1 | 0.79249716 | 0.84390209 |
| Postn | 0.659689 | 0.10696143 | 0.013 | 0.018 | 1 | 0.8870388 | 1.09092322 | 0.018 | 0.015 | 1 | 0.8870388 | 0.88418843 |
| Oaz1 | 0.84938853 | 1.00459971 | 0.013 | 0.011 | 1 | 0.71100644 | 1.77542139 | 0.018 | 0.012 | 1 | 0.84938853 | 0.91648272 |
| Akr1a1 | 0.71954828 | 1.19944234 | 0.02 | 0.016 | 1 | 0.92476059 | 1.00642694 | 0.018 | 0.016 | 1 | 0.92476059 | 0.92134683 |
| Dcn | 0.72121284 | 0.1752307 | 0.033 | 0.038 | 1 | 0.85206745 | 0.64193839 | 0.018 | 0.021 | 1 | 0.85206745 | 0.92227772 |
| Ctsk | 0.89763174 | 0.14276215 | 0.02 | 0.021 | 1 | 0.78939722 | 1.22522431 | 0.018 | 0.013 | 1 | 0.89763174 | 0.95564647 |
| Wfdc18 | 0.93907111 | 0.17205645 | 0.02 | 0.019 | 1 | 0.85568886 | 0.77787704 | 0.018 | 0.021 | 1 | 0.93907111 | 0.97917429 |

**Spatial seq: cluster 13**

| | WT_p_val | WT_avg_log2FC | WT_pct.1 | WT_pct.2 | WT_p_val_adj |
|---|---|---|---|---|---|
| Sparc | 1.72E-10 | -5.4117 | 0 | 0.256 | 2.58E-06 |
| Col1a2 | 7.80E-09 | 0.10338509 | 0.992 | 0.484 | 0.00011673 |
| Hba-a2 | 9.64E-08 | -5.3070836 | 0 | 0.192 | 0.00144418 |
| Wdfy3 | 2.54E-07 | 6.29932157 | 0.017 | 0.001 | 0.00379637 |
| Obp2a | 2.96E-06 | -5.3026813 | 0 | 0.154 | 0.04429092 |
| Hbb-bs | 6.29E-06 | -4.9654142 | 0 | 0.145 | 0.09415766 |
| Col3a1 | 1.50E-05 | -4.4170692 | 0 | 0.135 | 0.22406276 |
| S100a9 | 0.00015443 | -4.0499597 | 0 | 0.106 | 1 |
| Obp2b | 0.00033583 | -4.4828576 | 0 | 0.096 | 1 |
| Ambn | 0.00099651 | 4.85186259 | 0.017 | 0.002 | 1 |
| Scgb1c1 | 0.00313552 | -3.6943245 | 0 | 0.067 | 1 |
| S100a8 | 0.00377044 | -3.1036905 | 0 | 0.065 | 1 |
| Spp1 | 0.00395914 | -3.2320599 | 0 | 0.064 | 1 |
| Ngp | 0.00729924 | -3.0427531 | 0 | 0.056 | 1 |
| Obp1a | 0.0078518 | -3.4480323 | 0 | 0.055 | 1 |
| Hbb-bt | 0.01058189 | -2.7160935 | 0 | 0.051 | 1 |
| Gpx3 | 0.01210746 | -2.6779584 | 0 | 0.049 | 1 |
| Itm2b | 0.01306444 | -2.8415082 | 0 | 0.048 | 1 |
| Bpifb6 | 0.01524382 | -2.8592881 | 0 | 0.046 | 1 |
| Bgn | 0.01610252 | -2.5756598 | 0 | 0.046 | 1 |
| Lcn3 | 0.01823191 | -3.1549777 | 0 | 0.044 | 1 |
| Bpifb4 | 0.01873133 | -2.8402298 | 0 | 0.044 | 1 |
| Lcn11 | 0.01896782 | -3.0929959 | 0 | 0.044 | 1 |
| Tmsb4x | 0.02451859 | -2.397646 | 0 | 0.04 | 1 |
| App | 0.03088293 | -2.3080087 | 0 | 0.037 | 1 |
| Apoe | 0.03138514 | -2.383673 | 0 | 0.037 | 1 |
| Vmo1 | 0.03318916 | -2.4605666 | 0 | 0.036 | 1 |
| mt-Co2 | 0.03523897 | -2.397646 | 0 | 0.035 | 1 |
| Igfbp5 | 0.03537875 | -2.5555468 | 0 | 0.035 | 1 |
| Camp | 0.04150444 | -2.0994221 | 0 | 0.033 | 1 |
| Mgp | 0.04373298 | -2.1804587 | 0 | 0.033 | 1 |
| Vim | 0.05412409 | -1.9627733 | 0 | 0.03 | 1 |
| Serpinh1 | 0.05658497 | -1.9438524 | 0 | 0.029 | 1 |
| Obp1b | 0.0621297 | -2.3245599 | 0 | 0.028 | 1 |
| Col11a2 | 0.06574967 | -1.9246801 | 0 | 0.027 | 1 |
| Gpx6 | 0.06655854 | -1.974474 | 0 | 0.027 | 1 |
| Ptms | 0.06820346 | -1.9028023 | 0 | 0.027 | 1 |
| Car2 | 0.0716137 | -1.675093 | 0 | 0.026 | 1 |
| Ccn2 | 0.07585056 | -2.018091 | 0 | 0.025 | 1 |
| Tpm1 | 0.08098189 | -1.725818 | 0 | 0.025 | 1 |
| Ibsp | 0.08131441 | -1.7091071 | 0 | 0.025 | 1 |
| Igfbp7 | 0.08613756 | -1.6865204 | 0 | 0.024 | 1 |
| Timp2 | 0.09873768 | -1.5208574 | 0 | 0.022 | 1 |
| Serpinf1 | 0.10207655 | -1.4488713 | 0 | 0.022 | 1 |
| Wfdc18 | 0.10424258 | -1.9028023 | 0 | 0.021 | 1 |
| Dcn | 0.10554088 | -1.4522225 | 0 | 0.021 | 1 |
| Acp5 | 0.10554592 | -1.6254909 | 0 | 0.021 | 1 |
| mt-Atp8 | 0.10959758 | -1.6693452 | 0 | 0.021 | 1 |
| Fth1 | 0.11097609 | -1.5208574 | 0 | 0.021 | 1 |
| Fn1 | 0.11476385 | -1.4787556 | 0 | 0.02 | 1 |
| Ahnak | 0.13362251 | -1.2552673 | 0 | 0.018 | 1 |
| Kctd12 | 0.13591809 | -1.2242404 | 0 | 0.018 | 1 |
| Rsrp1 | 0.13826197 | -1.3588899 | 0 | 0.018 | 1 |
| Igfbp4 | 0.14063963 | -1.1804587 | 0 | 0.018 | 1 |
| Dmp1 | 0.14306852 | -1.1804587 | 0 | 0.017 | 1 |
| Sost | 0.15063926 | -1.3153883 | 0 | 0.017 | 1 |
| Gsn | 0.15064266 | -1.4114849 | 0 | 0.017 | 1 |
| mt-Cytb | 0.15326114 | -1.2968682 | 0 | 0.017 | 1 |
| mt-Co1 | 0.15326174 | -1.3263873 | 0 | 0.017 | 1 |
| Mmp13 | 0.15391777 | -1.1015579 | 0 | 0.017 | 1 |
| Lcn4 | 0.15392759 | -1.49834 | 0 | 0.017 | 1 |
| mt-Co3 | 0.15728517 | -1.2514252 | 0 | 0.016 | 1 |
| Mup5 | 0.15934612 | -1.5240457 | 0 | 0.016 | 1 |
| Akr1a1 | 0.16002905 | -1.0886957 | 0 | 0.016 | 1 |
| Hist1h1e | 0.16282918 | -1.0135614 | 0 | 0.016 | 1 |
| Ltf | 0.16283175 | -1.1185309 | 0 | 0.016 | 1 |
| Mt1 | 0.16641178 | -1.1058199 | 0 | 0.016 | 1 |
| Mup4 | 0.16787653 | -1.46555 | 0 | 0.015 | 1 |
| Postn | 0.1700788 | -1.0044592 | 0 | 0.015 | 1 |
| Mmp2 | 0.1769135 | -0.9627733 | 0 | 0.015 | 1 |
| Bpifb3 | 0.17770058 | -1.5335684 | 0 | 0.015 | 1 |
| Bpifa1 | 0.18165119 | -1.4421454 | 0 | 0.015 | 1 |
| mt-Atp6 | 0.18406399 | -1.0494066 | 0 | 0.014 | 1 |
| Lyz2 | 0.18733933 | -0.8247997 | 0 | 0.014 | 1 |
| Col9a3 | 0.19584577 | -1.168284 | 0 | 0.014 | 1 |
| Ctsk | 0.19759236 | -0.8706034 | 0 | 0.014 | 1 |
| Lrp1 | 0.20478218 | -0.7341014 | 0 | 0.013 | 1 |
| Arf5 | 0.20478274 | -0.7721408 | 0 | 0.013 | 1 |
| Slc4a1 | 0.20570102 | -0.683672 | 0 | 0.013 | 1 |
| Tpm2 | 0.20943534 | -0.9814492 | 0 | 0.013 | 1 |
| Col2a1 | 0.21420716 | -0.9721415 | 0 | 0.013 | 1 |
| Cd24a | 0.21517182 | -0.7285844 | 0 | 0.013 | 1 |
| Nisch | 0.2161469 | -0.7450726 | 0 | 0.012 | 1 |
| mt-Nd1 | 0.21615173 | -1.0226065 | 0 | 0.012 | 1 |
| Oaz1 | 0.21811388 | -0.9101318 | 0 | 0.012 | 1 |
| Col4a1 | 0.21910098 | -0.7395974 | 0 | 0.012 | 1 |
| Id3 | 0.22412993 | -0.6490457 | 0 | 0.012 | 1 |
| Arglu1 | 0.22413185 | -0.7285844 | 0 | 0.012 | 1 |
| mt-Nd3 | 0.2272188 | -0.9906973 | 0 | 0.012 | 1 |
| mt-Nd5 | 0.22825667 | -0.8756041 | 0 | 0.012 | 1 |
| Col9a2 | 0.23353427 | -0.8196195 | 0 | 0.012 | 1 |
| Jund | 0.23676869 | -0.6431929 | 0 | 0.011 | 1 |
| Ighm | 0.24005563 | -0.4622297 | 0 | 0.011 | 1 |
| Cnn2 | 0.24340174 | -0.5144596 | 0 | 0.011 | 1 |
| Tnc | 0.24566196 | -0.488581 | 0 | 0.011 | 1 |
| Mpo | 0.25025958 | -0.4285989 | 0 | 0.011 | 1 |

| | | | | | |
|---|---|---|---|---|---|
| Maf | 0.25026198 | -0.5771954 | 0 | 0.011 | 1 |
| Dmbt1 | 0.25143032 | -0.7881413 | 0 | 0.011 | 1 |
| Bpgm | 0.25615328 | -0.387179 | 0 | 0.011 | 1 |
| Marcks | 0.25855914 | -0.4622297 | 0 | 0.01 | 1 |
| Pcolce | 0.25855914 | -0.5015783 | 0 | 0.01 | 1 |
| Mepe | 0.25855988 | -0.5335684 | 0 | 0.01 | 1 |
| Cst3 | 0.25977215 | -0.488581 | 0 | 0.01 | 1 |
| Selenom | 0.26099023 | -0.4011182 | 0 | 0.01 | 1 |
| Mxra8 | 0.26221684 | -0.4011182 | 0 | 0.01 | 1 |
| Serinc3 | 0.26345035 | -0.4217776 | 0 | 0.01 | 1 |
| Thbs1 | 0.26345141 | -0.488581 | 0 | 0.01 | 1 |
| Bpifb9a | 0.26403542 | 2.96433732 | 0.017 | 0.008 | 1 |
| mt-Nd4l | 0.26469444 | -0.683672 | 0 | 0.01 | 1 |
| Hist1h2ap | 0.26719246 | -0.3300351 | 0 | 0.01 | 1 |
| Gja1 | 0.26719269 | -0.3588899 | 0 | 0.01 | 1 |
| Col6a3 | 0.26845488 | -0.3588899 | 0 | 0.01 | 1 |
| Sfrp2 | 0.26845596 | -0.4285989 | 0 | 0.01 | 1 |
| Dbp | 0.26845674 | -0.488581 | 0 | 0.01 | 1 |
| Mbd2 | 0.27100178 | -0.3801585 | 0 | 0.01 | 1 |
| Maged1 | 0.27100188 | -0.4011182 | 0 | 0.01 | 1 |
| Gpx1 | 0.27618263 | -0.2856409 | 0 | 0.01 | 1 |
| Igf2 | 0.27618319 | -0.3227303 | 0 | 0.01 | 1 |
| Ddx17 | 0.2761833 | -0.3300351 | 0 | 0.01 | 1 |
| Rrm2 | 0.27881956 | -0.270534 | 0 | 0.01 | 1 |
| Ucp2 | 0.27882072 | -0.4011182 | 0 | 0.01 | 1 |
| Srsf6 | 0.2801518 | -0.4488713 | 0 | 0.01 | 1 |
| Acta1 | 0.53323632 | 1.10292435 | 0.008 | 0.015 | 1 |
| Col1a1 | 0.91351898 | -0.8136947 | 1 | 0.618 | 1 |
